# asteRIa enables robust interaction modeling between chromatin modifications and epigenetic readers

**DOI:** 10.1101/2024.03.15.585146

**Authors:** Mara Stadler, Saulius Lukauskas, Till Bartke, Christian L. Müller

## Abstract

Chromatin, the nucleoprotein complex consisting of DNA and histone proteins, plays a crucial role in regulating gene expression by controlling access to DNA. Chromatin modifications are key players in this regulation, as they help to orchestrate DNA transcription, replication, and repair. These modifications recruit epigenetic “reader” proteins, which mediate downstream events. Most modifications occur in distinctive combinations within a nucleosome, suggesting that epigenetic information can be encoded in combinatorial chromatin modifications. A detailed understanding of how multiple modifications cooperate in recruiting such proteins has, however, remained largely elusive. Here, we integrate nucleosome affinity purification data with high-throughput quantitative proteomics and hierarchical interaction modeling to estimate combinatorial effects of chromatin modifications on protein recruitment. This is facilitated by the computational workflow `asteRIa` which combines hierarchical interaction modeling, stability-based model selection, and replicate-consistency checks for **a st**able **e**stimation of **R**obust **I**nter**a**ctions among chromatin modifications. `asteRIa` identifies several epigenetic reader candidates responding to specific *interactions* between chromatin modifications. For the polycomb protein CBX8, we independently validate our results using genome-wide ChIP-Seq and bisulphite datasets. We provide the first quantitative framework for identifying cooperative effects of chromatin modifications on protein binding.

## Introduction

Eukaryotic cells store the genetic material in the nucleus where it is packaged into chromatin, a nucleo-protein complex made up primarily of DNA and histone proteins. Both DNA and histones carry chemical modifications that can either directly affect chromatin structure or recruit so-called epigenetic reader proteins that mediate downstream events. As these modifications are involved in the regulation of all DNA-templated processes, such as transcription, DNA replication, or DNA repair, they play central roles in controlling chromatin function [1]. The basic repeating unit of chromatin is the nucleosome, which coordinates 147 bp of DNA wrapped around an octamer consisting of two copies each of the core histones H2A, H2B, H3, and H4 [2]. Nucleosomes are folded into higher-order structures to form chromatin. Since DNA and histone modifications show extensive overlap in the genome [3] and decorate histones and nucleosomes in specific combinations [4, 5, 6, 7, 8, 9, 10], it is likely that these modifications act in a concerted manner. This is supported by the observation that most chromatin regulators contain multiple modification binding domains or are part of multi-subunit complexes harbouring multiple such domains, and are therefore likely to read out multiple chromatin modifications [11]. Indeed, the idea that combinations of histone modifications may form a ‘histone code’ that together with DNA modifications could store epigenetic information in the chromatin template, thereby expanding the genetic information encoded in the DNA sequence, has been around for over two decades [12, 13, 14].

To date, the functions and readers of a host of *individual* chromatin modifications have been described (see, e.g., [15, 16] and references therein for an overview). Moreover, epigenetic regulators that read the modification status of more than one epigenetic mark on histones or the DNA have been described using functional and structural studies [17, 18, 19, 20, 21, 22, 23]. Several DNA repair factors were also found to recognize dual histone modification signatures, ranging from individual interactions [24, 25, 26, 27, 28, 29] to combinatorial ones [30, 31]. One prime example is the ubiquitin ligase UHRF1, an essential player in DNA methylation maintenance, that recognizes a triple modification signature on histone H3 [32, 33, 34] and the DNA [35, 36, 37].

The gap in knowledge about the combinatorial nature of factors that read multiple DNA and histone modifications can be partially attributed to the fact that one of the most prevailing high-throughput technology to study histone modifications and their readers is chromatin immunoprecipitation followed by deep sequencing (ChIP-seq). Here, antibodies are used to detect the localization of specific modifications *or* chromatin-binding proteins at a genome-wide scale [38]. Despite its groundbreaking influence on our understanding of the histone code through community efforts such as the NIH Roadmap Epigenomics Mapping Consortium [39], ENCODE [40], and ChIP-Atlas [41], ChIP-seq alone can only probe a single modification or reader protein in each experiment, thus making it difficult to assess combinatorial synergies or antagonistic effects on epigenetic readers. However, careful integration of multiple genome-wide ChIP-seq experiments of individual modifications enabled the application of *multivariate* statistical analysis techniques to uncover chromatin states and interactions. For example, using hidden Markov modeling techniques, the `ChromHMM` method [42, 43] revealed cell-type specific discrete chromatin states that characterize the combinatorial presence or absence of modifications on the genome. Alternatively, sparse partial correlation estimation techniques were proposed to learn multivariate association networks between histone modifications [44]. The latter framework was extended in [45, 46] to include both histone modifications and a small set of chromatin modifiers. Using linear regression and sparse partial correlation estimation, the studies derived *de novo* high-confidence backbones of “chromatin signaling networks” from ChIP-Seq data. There, the inferred network edges are to be interpreted as additive (or main) effects between histone modifications on chromatin modifiers and vice versa. The analysis of the derived chromatin signaling networks revealed both histone-protein interactions known from literature and several novel hypothetical interactions. To show the power of the network approach, the authors were also able to experimentally verify the statistically hypothesized interactions between H4K20me1 and members of the polycomb repressive complexes 1 and 2 (PRC1 and PRC2, respectively) [46]. Nevertheless, none of these ChIP-Seq-based computational approaches allow the statistical estimation of how *multiple* histone modifications co-operate in recruiting epigenetic regulators.

In this contribution, we present a statistical interaction modeling approach, termed `asteRIa`, that tackles this challenge. Rather than considering genome-wide ChIP-Seq data, `asteRIa` uses novel nucleosome affinity purification data with high-throughput quantitative proteomics, as provided in the Modification Atlas of Regulation by Chromatin States (MARCS), to make robust and reproducible predictions of combinatorial effects of chromatin modifications on chromatin-interacting proteins. The MARCS data, available at https://marcs.helmholtz-muenchen.de comprises a collection of Stable Isotope Labeling with Amino acids in Cell culture (SILAC) nucleosome affinity purification (SNAP) experiments [47] that probe the binding of proteins from HeLa S3 nuclear extracts to a library of semi-synthetic di-nucleosomes (referred to as nucleosomes throughout the manuscript) incorporating biologically meaningful combinations of chromatin modifications representing promoter, enhancer and heterochromatin modification states. Each affinity purification measures the relative abundances of nuclear proteins on a modified nucleosome in relation to an unmodified control nucleosome using the SILAC labelling and quantitative proteomics as a read out. This allows the high-throughput identification of proteins that are either recruited or excluded by the modification(s), and also indicates the relative extent of the recruitment or exclusion. Collectively, the MARCS data set catalogs the binding responses of 1915 nuclear proteins to nucleosomes carrying 55 different modification signatures. The constructive nature of these data, paired with an appropriate statistical model, thus enables the direct analysis of combinatorial effects of different modification features on the nucleosome binding of the measured proteins. At its core, `asteRIa` uses a linear regression model with pairwise (or “two-way”) interactions among chromatin modifications to predict the binding affinities of each protein. Regression models with pairwise interactions have a long tradition in statistics and experimental design [48, 49, 50] but are notoriously difficult to estimate in the presence of noisy, scarce data and/or incomplete experimental designs, and are prone to misinterpretation [51, 52]. As we will show, the `asteRIa` framework incorporates several model and design principles that (i) guard against common pitfalls and (ii) take the properties of the MARCS data (and biological data in general) into account. Firstly, we posit that our framework should work in the underdetermined regime, i.e., the number of features *q* (here the chromatin modifications) and pairwise interactions exceeds the number of measurements *n*. We achieve this by including sparsity-inducing penalization of the model coefficients [53, 54, 55]. Secondly, we assume that the underlying interaction model obeys the so-called “strong hierarchy” principle [53, 50, 56], i.e., interactions among features are only included in the model if both features are present as main effects. Thirdly, we embrace the principle of statistical “stability” [57, 58, 59] for model selection, implying that interactions are only included when they are reproducibly identified across subsets of the data. To respect the ubiquitous measurement variability of biological systems, we also require replicate consistency [60] of our combinatorial models. This means that models with interactions need to be (at least partially) consistent across available technical or biological replicates, further ensuring the general robustness and validity of the resulting models. While these design principles and the underlying computational workflow, available at https://github.com/marastadler/asteRIa.git, are general, we illustrate the framework to detect novel combinatorial interactions between chromatin modifications on epigenetic reader recruitment.

On the MARCS data, we show that considering interaction effects between chromatin modifications can consistently improve the predictive performance of the binding profiles of a subset of proteins. `asteRIa` not only recovers known binding patterns, such as, e.g, the well-known H3K27me3-CBX8 pairing, but also identifies novel interaction effects between chromatin modifications on the binding behavior of proteins not yet implicated as epigenetic readers (e.g., ACTL8). Our analysis also allows to define and quantify the extent of distinct modes of apparent chromatin modification interactions, ranging from synergistic and antagonistic to competitive effects. Our post-hoc model analysis shows that proteins belonging to the same protein complexes do read combinatorial chromatin modification signatures in a similar fashion, thus allowing the delineation of a protein complex - chromatin modification interaction network.

Independent confirmation of the identified combinatorial interactions is challenging due to the uniqueness of the MARCS data and the accompanying statistical analysis. Nevertheless, we provide a validation workflow on ENCODE ChIP-Seq, ChIP-Atlas ChIP-Seq, and WGBS (Whole Genome Bisulfite Sequencing) data that demonstrates that our findings are not limited to a specific cell type or experimental setup. Specifically, we show that one of the found combinatorial interactions for CBX8 are consistent with these orthogonal datasets. The latter analysis also illustrates how to validate other interactions found in this study, thus inviting the generation of new ChIP-Seq data collections for previously understudied proteins.

## Materials and Methods

### The Modification Atlas of Regulation by Chromatin States dataset

The Modification Atlas of Regulation by Chromatin States (MARCS), as introduced in [61], builds on two experimental components: (i) a designed library of engineered di-nucleosomes (referred to as nucleosomes throughout the manuscript) comprising combinatorial chromatin modifications and (ii) nucleosome affinity purifications coupled to high-throughput quantitative proteomics measurements employing SILAC labeling (SNAP) [47]. The modified nucleosomes were assembled from a biotinylated DNA containing two 601 nucleosome positioning sequences [62] and histone octamers containing semi-synthetic site-specifically modified histones H3.1 and H4 prepared by native chemical ligation [63]. Some nucleosomes were also assembled using CpG-methylated DNA (5mC) or the histone variant H2A.Z. The complete library design matrix comprises *n*_total_ = 55 modified nucleosomes with *q* = 12 possible chromatin modifications (see left panel of Fig. 1 for a conceptual picture). The available modifications include six lysine residues on the tails of histone H3 (K4, K9, K14, K18, K23 and K27) and five on histone H4 (K5, K8, K12 and K20) as well as the variant histone H2A.Z and CpG methylated (5mC) DNA on both DNA strands (symmetric methylation), respectively. The lysines are modified with acetylation (ac) or mono-, di-, or tri-methylation (me1, me2, me3). H3-5ac denotes that multiple acetylations (K9, K14, K18, K23, K27) on the tails of histone H3 are present. H4-4ac denotes that multiple acetylations (K8, K5, K12, K16) on the tails of histone H4 are present. For our computational analysis, we do not consider engineered nucleosomes that include subsets of acetylations (namely, not all five acetylations on H3 or not all four acetylations on H4) since building their mathematical products would result in perfectly collinear (thus fully redundant, and therefore not distinguishable) pair-wise interaction features (see **Interaction modeling strategy** for further clarification). Our analysis thus excludes 22 nucleosomes from the initial nucleosome library and considers a subset of *n* = 33 nucleosomes, resulting in the design matrix *L* ∈ {0, 1 }^33×12^. The (transposed) design matrix *L* with the available combinatorial modifications is shown in the top panel (Step 1) of Fig. 2). Note that the design pattern in *L* does not follow any particular statistical experimental design guideline [50] but is driven by biological expertise about common modification co-occurrences.

**Fig 1.**
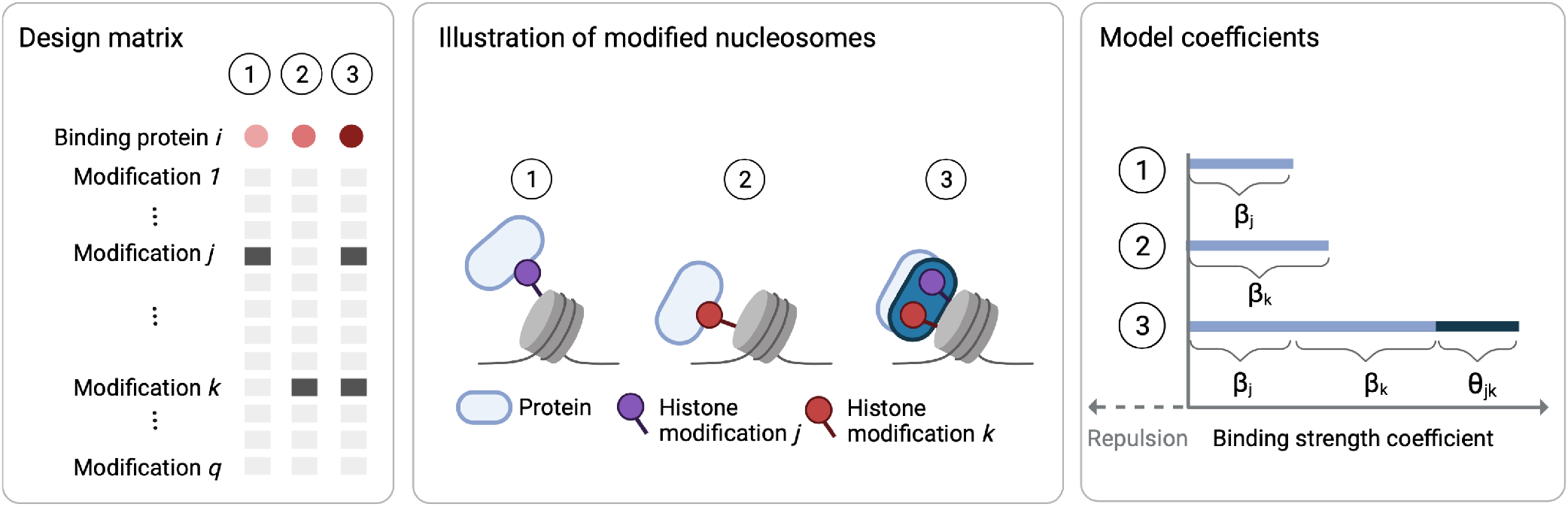
Left: Three exemplary columns of the design matrix *L*. Dark gray boxes indicate that a modification has been installed on the respective nucleosome. Above the design, the binding behavior of an exemplary protein to the modified nucleosomes is shown by color. The shade of red indicates the strength of the binding effect. Center: Illustration of two *individual* binding effects of chromatin modifications *j* and *k* on a protein *P*_*i*_ (1 and 2). Synergistic *combinatorial* effect of the modifications *j* and *k* on protein *P*_*i*_ (dark blue) compared to expected binding effect under independence of modification *j* and *k* (light blue) (3). Right: Model coefficients/estimated binding strength of protein *P*_*i*_ for the three scenarios. Light blue bar in scenario 3 shows the binding strength under independence of modification *j* and *k, β*_*j*_ + *β*_*k*_. Dark blue shows the additional combinatorial effect *θ*_*jk*_ that goes beyond additive combinatorial effects (created with BioRender.com).

**Fig 2.**
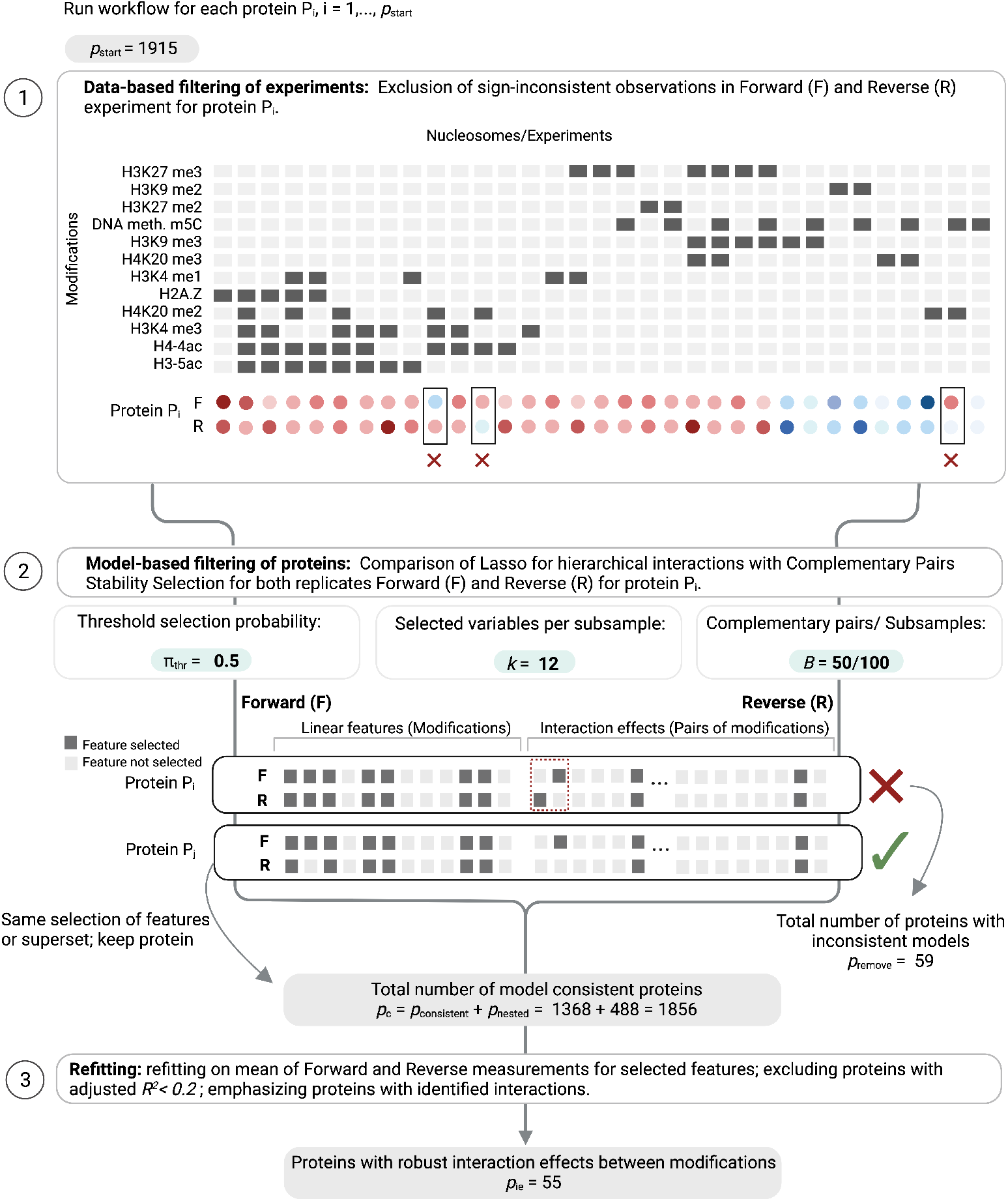
Graphical representation of the `asteRIa` workflow for the robust detection of hierarchical interactions (created with BioRender.com). Step 1: Design matrix and measured binding behavior for a protein (two replicates F and R). Removal of observations with different signs in the replicates (sign consistency). Step 2: Hierarchical interaction modeling with default complementary pairs stability selection (CPSS) parameters. Comparison of selected features for each replicate (nested model consistency): The first example shows a protein prediction model that gets filtered out since the selected features from forward and reverse replicate are neither identical nor nested. The second example shows a “consistent” protein model where the selected features learned from reverse replicate is a nested subset of the features learned from the forward replicate. Step 3: Least-squares refitting on averaged replicate data for final prediction model building. The intersection of two selected feature sets is used for refitting. Models with adjusted *R*^2^ *<* 0.2 are discarded.

For each modified nucleosome in MARCS, SNAP experiments are provided in two experimental ‘label-swap’ replicates of the nucleosome affinity purification process, a ‘forward’ (F) and ‘reverse’ (R) nucleosome pull-downs. Nucleosomes are immobilized on streptavidin beads and incubated with nuclear extracts from HeLa S3 cells cultured either in isotopically light or heavy-labelled SILAC media. In the ‘forward’ experiments the heavy extracts are incubated with the modified and the light extracts with the unmodified nucleosome, in the ‘reverse’ experiments the extracts are exchanged. Bound proteins are eluted from the beads and identified and quantified by mass spectrometry. For each SNAP experiment the relative abundance of a given protein on the modified nucleosome is determined in relation to the unmodified nucleosome by measuring the ratios between the heavy and the light peptides (H/L ratios) identified for that particular protein [47]. The H/L ratios indicate binding preferences to the modified or the unmodified nucleosomes and allow the unbiased identification of proteins that are either recruited or excluded by the modification(s) present on the modified nucleosomes. In addition, the SILAC enrichment ratios also indicate a relative “strength” of the recruitment or exclusion of a given protein by the modifications. In total, the MARCS dataset comprises the binding behavior of *p* = 1915 proteins in the forward (F) and reverse (R) experiment. For our analysis, we consider the protein measurement matrices *P* ^*F*^, *P* ^*R*^ ∈ ℝ^33×1915^ that correspond to the subset of *n* = 33 nucleosomes, described above.

### Interaction modeling strategy

We aim at predicting the binding profile of each protein captured in MARCS (*P*_*i*_)_1≤*i*≤1915_ (either from the forward or reverse experiment) from the combinations of nucleosome modifications (*L*_*j*_)_1≤*j*≤12_. Given the binary design matrix *L*, the baseline model of uncovering (joint) additive effects of the modifications on a binding profile *Y* = *P*_*i*_ ∈ ℝ^*n*^, *i* = 1, …, *p*, is the linear model

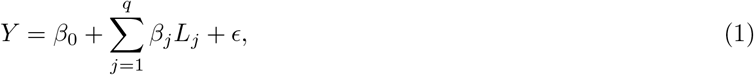

where *β*_0_ ∈ ℝ^*n*^ is a protein-specific (constant) intercept, *β*_*j*_ is the effect of modification *j* on the binding profile *Y* = *P*_*i*_ of protein *i*, and *ϵ* models the technical and biological noise component. In [61], a simplified version of this baseline model was investigated through “feature effect estimates” via pairwise comparisons of the enrichments of individual proteins on nucleosomes differing by a single modification feature. This, however, only allowed robust prediction of the effects of individual modifications or blocks of modifications and did not provide any information on combinatorial effects. Here, we extend the baseline model by including all pairwise interactions between modifications. For each protein binding profile *Y* = *P*_*i*_, *i* = 1, …, *p*, the core model in `asteRIa` thus reads

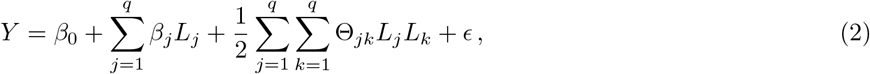

where Θ_*jk*_ models interaction effects between epigenetic readers that cannot be captured by linear additive effects. Robustly and reproducibly estimating non-zero entries in the interaction matrix Θ from replicated data is at the heart of the `asteRIa` workflow. The sign of the interaction coefficients also allows a characterization of epigenetic reader interplay. For example, when 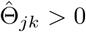 we interpret the two modifications *j* and *k* to have a synergistic binding effect if both *β*_*j*_ *>* 0 and *β*_*k*_ *>* 0 (see Fig. 1 for illustration).

To guarantee identifiability and interpretability of individual interaction models, we first need to ensure that the interaction design matrix *L*_*j*_*L*_*k*_ has no co-linear columns. In the concrete example of the MARCS data, we group modifications of the complete design matrix to a set of *n* = 33 non-redundant nucleosomes (see top panel (Step 1) of Fig. 2). Secondly, to enable estimation in the present underdetermined regime (*q*(*q* +1)*/*2 *> n*) with *q*(*q* +1)*/*2 = 78, we perform regularized maximum-likelihood estimation with *𝓁*_1_-norm (lasso) penalization [64] on the linear and interaction coefficients, respectively. Given the log-likelihood function of the model 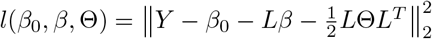, the (all-pairs) lasso problem reads

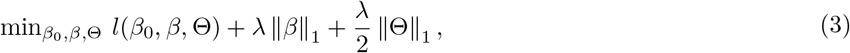

where *λ >* 0 is a tuning parameter and controls the sparsity levels of the coefficients *β* and Θ, respectively. To further ease model interpretability, we follow the statistical principle of hierarchy (also known as marginality or heredity) and allow the presence of an interaction in the model *only if* the associated linear (main) effects are in the model as well [see 53, and references therein]. In mathematical terms, this so-called strong hierarchy principle can be expressed as

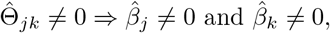

implying that interaction effects are only present if both linear effects enter the model. This hierarchy can be achieved by adding a constraint on the interaction effects Θ_*j*_ ∈ ℝ^*q*^ and a symmetry constraint on Θ. The corresponding optimization problem with hierarchical interactions thus reads

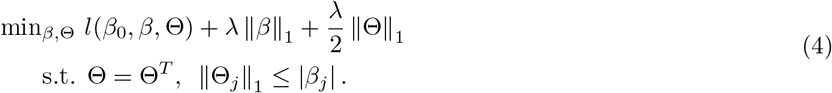

To solve the non-convex optimization problem in (4), we follow Bien, Taylor, and Tibshirani [53] who proposed a convex relaxation of the problem and provide an efficient implementation in the corresponding R package `hierNet` [65] (v1.9). In `asteRIa`, we use `hierNet` to model each protein binding profile *Y* = *P*_*i*_, *i* = 1, …, *p* with hierarchical interactions. Apart from reducing the number of spurious interaction effects, a major advantage of the strong hierarchy constraint is the so called “practical sparsity”. The strong hierarchy constraint favors models that “reuse” measured variables. In the context of the MARCS data, this becomes important when generating hypotheses for follow-up functional analysis (where experiments are complex and costly). Concretely, our models assumes that a protein or protein complex must have a domain capable of recognizing a particular chromatin modification. Thus, if there exists a response of a protein to an interaction effect between two modifications, a (possibly small) linear effect to both modifications is expected.

### Stability-based model selection for hierarchical interactions

One of the core challenges in high-dimensional penalized regression is determining a suitable regularization parameter *λ* that trades off sparsity (i.e., interpretability) of the model coefficients and out-of-sample predictive performance of the model [66, 67]. Standard procedures for (hierarchical) interaction models include cross-validation [53] and Information Criteria, including the Aikake (AIC) and the extended Bayesian Information Criterion (BIC) [55]. However, it has been observed that, both in simulation and practice, cross-validation and Information criteria tend to select more predictors (and interactions) than necessary [55].

To address this shortcoming, we follow the principle of stability [57] in `asteRIa` and introduce stability selection [58] for the identification of a reproducible set of predictive features *and* interactions. Stability selection has been proven useful across several scientific applications, ranging from network learning [68, 69] to data-driven partial differential equation identification [70, 71]. In the regression context, stability selection repeatedly learns sparse regression models from subsamples of the data of fixed size (e.g., *n*_*s*_ = ⌊ *n/*2 ⌋), records the frequency of all selected predictors across the models, and selects the most frequent predictors to fit the final regression model. Here, we use a variant of stability selection, the so-called complementary pairs stability selection (CPSS) [59] which draws *B* subsamples as complementary pairs { (*A*_2*h*−1_, *A*_2*h*_) : *h* = 1, …, *B* }, with *A*_2*h*−1_ ∩ *A*_2*h*_ = ∅ of samples { 1, …*n*} of size ⌊*n/*2⌋. Drawing complementary pairs is particularly beneficial when dealing with unbalanced experimental designs, as the resulting random splits ensure that individual subsamples are independent of each other. After applying a variable selection procedure *S* (e.g., using the first *k* predictors that enter the penalized model), each feature *j* in the model gets an individual estimated selection probability 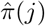, given by

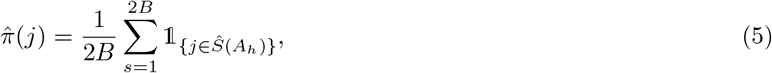

and the final selection set is given by 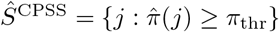, for a threshold *π*_thr_ defining the minimum selection frequency. In our workflow we use the corresponding R package `stabs` [72] (v0.6-4) that provides an efficient implementation of CPSS. The CPSS procedure includes the following hyperparameters: The set of regularization parameters Λ, a threshold *π*_thr_ ∈ [0, 1], the number of predictors *k* that first enter the sparse model, and the number of complementary splits *B*. In `asteRIa`, we set as default parameters Λ to be the internal *λ*-path in Bien and Tibshirani [65], *π*_thr_ = 0.5, *k* = 12, and *B* = 50, resulting in 100 subsamples. For the MARCS data, this means that chromatin modifications (as main or interaction effects) are part of the pairwise interaction model 2 for protein binding profile *i, Y* = *P*_*i*_, if it is among the *k* = 12 selected modifications in at least 50 subsamples. While these default values may need to be tuned in other scenarios, we verified in a realistic semi-synthetic simulation scenario (see Supplementary information and Fig. S1 and S2 for details) that hierarchical interaction modeling with stability selection greatly outperforms cross-validation, particularly in terms of false positive rate.

### Replicate consistency

Biological datasets typically include replicated measurements (replicates) to probe different sources of variability in the underlying experimental procedure or study object [73]. The MARCS dataset, for example, comprises two technical replicates of the SILAC-based protein binding affinities. Replicate consistency, i.e., assessing how consistent two or multiple replicated measurements are in terms of direction or size, is an important property to evaluate experimental protocols and downstream analysis quality (see, e.g., [74] for a discussion in the context of RNA sequencing data).

In `asteRIa`, we propose and include two replicate-consistency mechanisms: (i) data sign-consistency and (ii) nested model consistency. While there are alternative ways of performing filtering, data sign-consistency can be considered as a data filtering step that ensures that replicated measurements agree on the direction, i.e., the sign of the measured unit, and removes experiments where sign consistency does not hold. In MARCS, we perform data sign consistency for each protein *P*_*i*_ separately using the forward and reverse replicates (see Fig. 2, Step 1) and remove nucleosomes (experiments) where measured protein binding affinities disagree in sign. Although this reduction in sample size (for each protein *n*_*i*_ ≤ *n* samples are available) decreases the power for subsequent hierarchical interaction modeling, the filtering increases the chance of estimating pairs of consistent interaction models. In a second post-hoc step, nested model consistency further ensures that only pairs of consistent interaction models are considered for downstream analysis. Nested model consistency deems estimated interaction models valid only if they comprise the same set of features (main and interaction coefficients) across replicates *or* one model comprises a nested subset of main and interaction effects of the other model (see Fig. 2, Step 2, for illustration).

### The asteRIa workflow

The `asteRIa` workflow incorporates the described model and design principles as illustrated in Figure 2 on the MARCS data. `asteRIa` comprises three main steps: Step (1) uses sign consistency to filter pairs of forward and reverse experiments for each protein (*P*_*i*_)_1≤*i*≤*p*=1915_. Step (2) comprises model estimation using the hierarchical interaction model, CPSS-based model selection, and the post-hoc nested model consistency filter. Step (3) performs least-squares “refitting” to estimate main and interaction effect sizes on the selected model coefficients from averaged replicate data. The resulting signed model coefficients are then used for functional categorization and downstream analysis.

On the MARCS data, the experiment filtering step (1) removes on average 11 experiments across all proteins. In step (2), using the internal *λ*-path in Bien and Tibshirani [65], and CPSS parameters *π*_thr_ = 0.5, *k* = 12, and *B* = 50, `asteRIa` learns *p*_consistent_ = 1368 fully consistent regression models across forward and reverse replicates, as well as *p*_*nested*_ = 488 models that obey the nested model consistency criterion. Only *p*_remove_ = 59 models are inconsistent across replicates. Among all *p*_c_ = 1856 consistent models, `asteRIa` identifies 58 models that include robust interaction coefficients. The refitting estimation process in step (3) uses the averaged binding affinities as outcome and performs least-squares refitting on the *intersection* of the per-replicate selected features. The refit coefficients are the final effect sizes. For downstream analysis, `asteRIa` removes poorly-performing prediction models with adjusted *R*^2^ below 0.2 (three out of 58).

## Results

### Enhanced Predictive Performance of Protein Binding through Chromatin Modification interaction

We first quantify the overall predictive performance of `asteRIa` models for all proteins included in the MARCS dataset and then assess the degree to which hierarchical interaction modeling improves overall predictive performance of protein binding affinities. For a majority of the p=1915 protein binding profiles, `asteRIa` deems main effects models (i.e., the baseline linear model in 1) to be sufficient for robust prediction. For more than 200 proteins, main effects models achieve adjusted *R*^2^ *>* 0.8, and for more than 500 proteins, main effects models achieve adjusted *R*^2^ *>* 0.5 (see Figure S4 for a list of top protein binding models and associated coefficients). The top-six protein binding models achieve near-perfect predictive performance and include the protein ING5, a dimeric, bivalent reader of histone H3K4 me3 [75], with an *R*^2^ = 0.99, the methyl–lysine histone-binding protein L3MBTL3 (*R*^2^ = 0.99), SMARCC2 (*R*^2^ = 0.99) which is part of the chromatin remodeling complex SNF/SWI, the histone acetyltransferase KAT7 (*R*^2^ = 0.98), the YAF2 protein (*R*^2^ = 0.98), and the histone lysine demethylase KDM2B (*R*^2^ = 0.98).

However, `asteRIa` also identifies a set of *p*_ie_ = 55 models that comprise stable interaction effects among modifications with enhanced predictive performance. This provides statistical evidence that cooperative effects between chromatin modifications may play a crucial role in the binding of specific reader proteins and thus in controlling chromatin function. Figure 3a shows the modification design matrix (left panel) and binding profiles (both the ‘forward’ and the ‘reverse’ experiments) of the 55 proteins explained by interaction models. The proteins are sorted by data density (i.e., in terms of number of experiments removed due to sign consistency filtering step (1) in `asteRIa`, Fig. 3a, gray boxes). Figure 3c shows the corresponding predictive performance of the models in terms of adjusted *R*^2^ both for main effects (light blue) and interaction models (dark blue), respectively. While the light blue segment denotes the proportion of variance explained by all selected main effects combined, the dark blue portion represents the additional explained variance attributed solely to one interaction. We observe that the inclusion of robust interaction among modifications can boost the performance of up to 0.5 (e.g., for proteins CDKAL1 and PEX11B). For others, such as, e.g., RFC3, the binding behavior can only be sufficiently described by taking into account interaction effects. While the improvement is less dramatic for proteins with well-performing main effects models, `asteRIa` still provides evidence for stable interactions among modifications. Figure 3b illustrates the stabilities (inclusion probabilities) 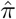 of all model coefficients for the protein CBX8. On both forward and reverse experimental data, `asteRIa` estimates a high selection probability (≈ 0.7) of an interaction effect between DNA methylation m5C and H3K9me3 while all other interaction effects emit a low inclusion probability. For detailed model inspection, we provide similar stability plots for all other proteins in the Supplementary Material. To illustrate the improvement in binding prediction, Figure 3c (right panel) shows predicted vs. observed binding profiles for the protein RNF2. Comparison of the fits of both the main effect (light gray) and interaction model (dark blue) visually and quantitatively (*R*^2^ = 0.76 vs. *R*^2^ = 0.9) confirm the enhanced predictive performance of the interaction model.

**Fig 3.**
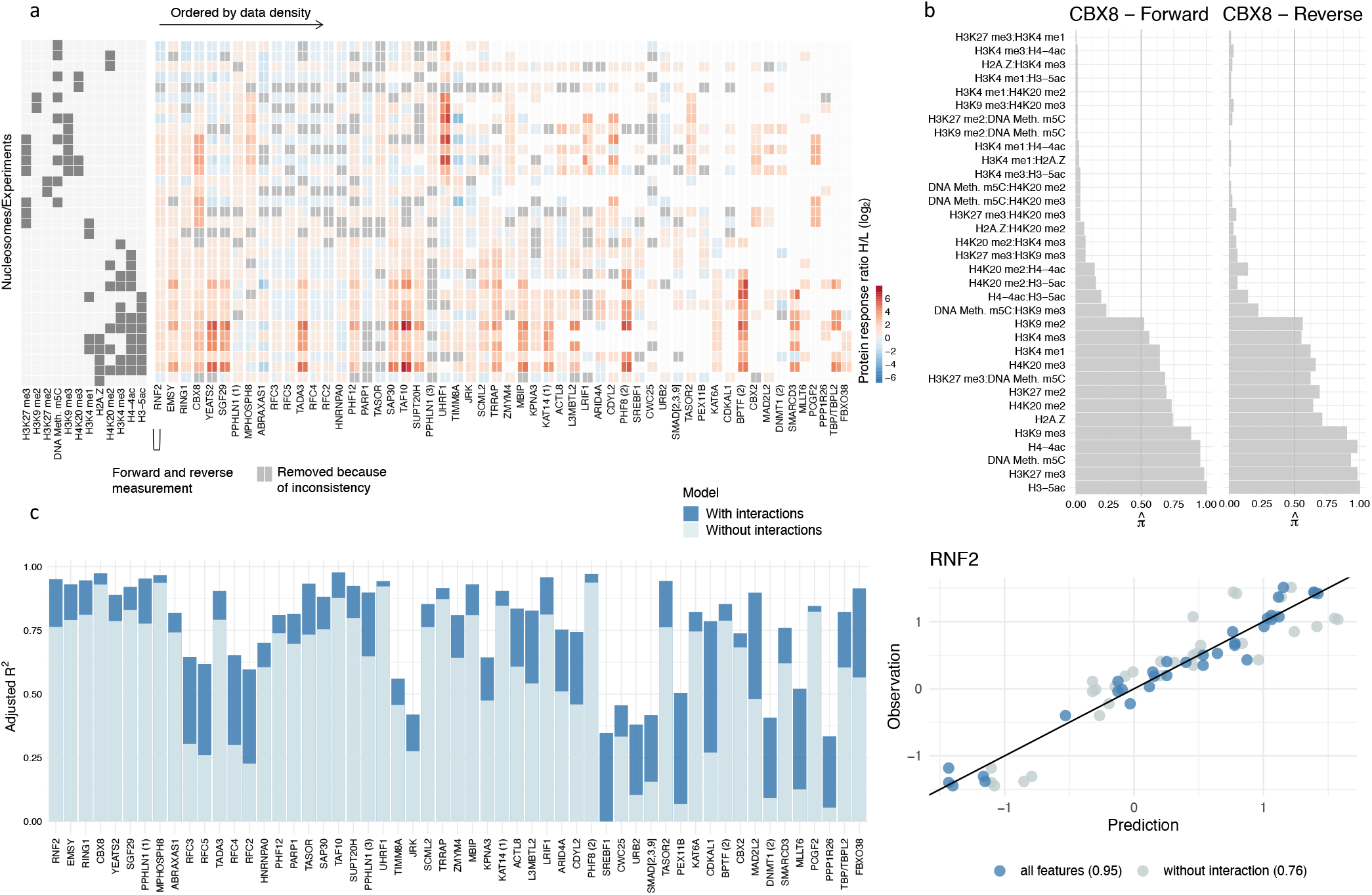
**a**, Observed protein binding profiles (forward and reverse experiment) for the *p*_ie_ = 55 proteins for which interactions between modifications have been detected. Proteins are arranged from left to right based on data density (number of non-zero measurements), with proteins with the highest data density being on the left. **b**, Stability plots for CBX8 of the hierarchical interaction model with Complementary pairs stability selection (CPSS). Vertical lines show the threshold for the selection probability threshold *π*_thr_ = 0.5. Stability plots for all proteins are provided in Extended Fig. 3b. **c**, Adjusted *R*^2^ for all *p*_*ie*_ = 55 proteins of the main effect (light blue) and interaction model (dark blue) (left panel). Scatter plot of observed vs. predicted values for the protein RNF2 (right panel). Scatter plots for all proteins are provided in Extended Fig. 3c.

### Modes of Chromatin Modification Interactions

To categorize the interaction effects uncovered in `asteRIa`, we establish potential modes of chromatin modification interactions. This is achieved by contrasting the effects of individual chromatin modifications (modification *j* and *k*) on the binding behavior of specific proteins, represented by the linear model coefficients 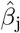 and 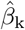 with the combinatorial effects identified during our analysis, represented by 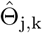 for the corresponding pair (see Fig. 4a and b). We define three major modes: synergistic combinatorial behavior, antagonistic combinatorial behavior, and conflicting combinatorial behavior. We further divide these into two sub-modes each of which describes the direction of the combinatorial effect, either towards binding (b, Θ_*j,k*_ *>* 0) or towards repulsion (r, Θ_*j,k*_ *<* 0). The direction and strength of the combinatorial effect is color-coded in Fig. 4b

**Fig 4.**
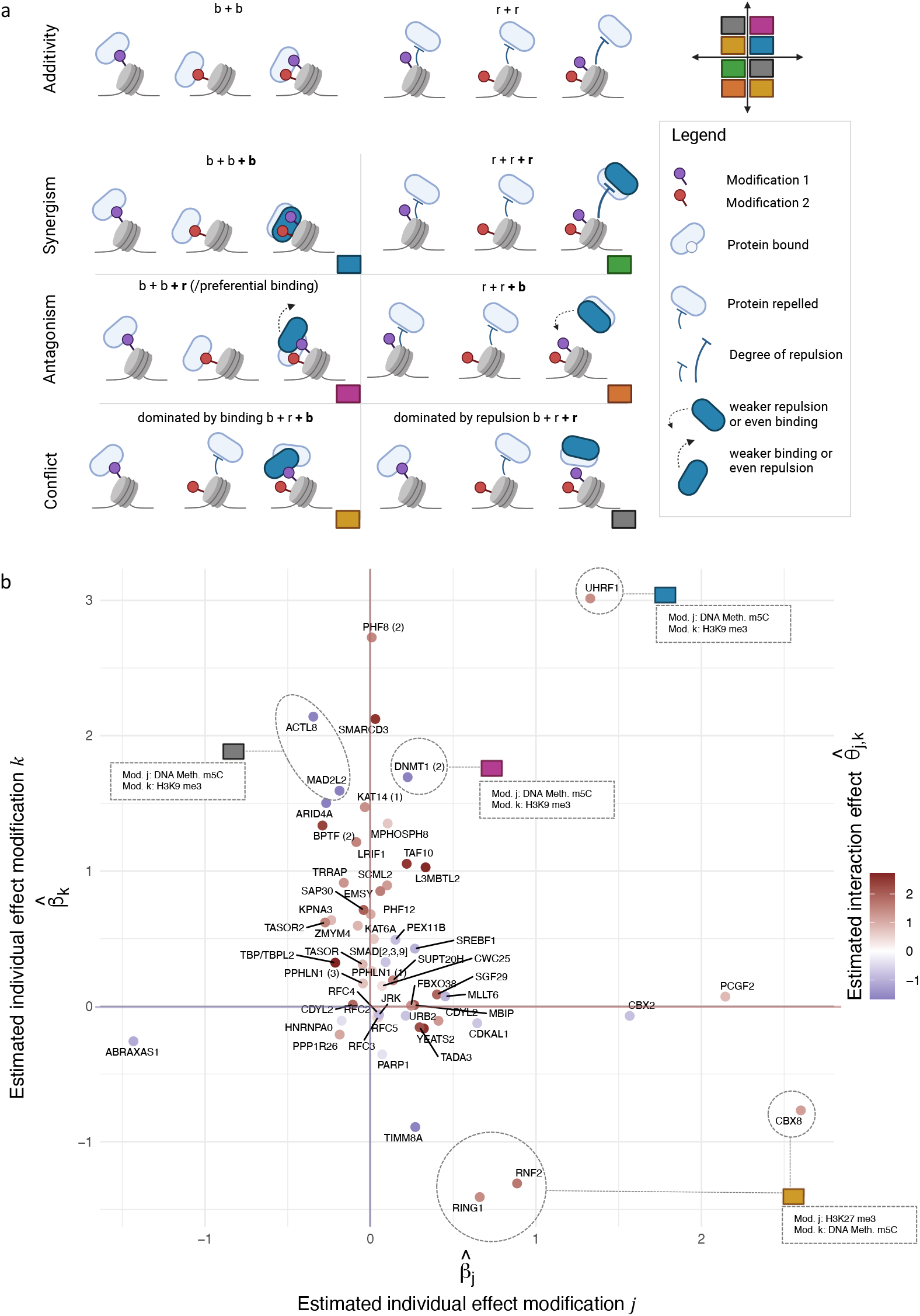
**a**, Six combinations of combinatorial interaction effects of two modifications on the binding behavior of chromatin-associated proteins (created with BioRender.com). The top row illustrates what would be expected under a purely additive dependence of the effects in a scenario where a protein shows *individual* binding effects to two distinct chromatin modifications (overall *additive* effect is the sum of both individual binding effects, b + b) (left) and where a protein is repelled by two distinct chromatin modifications (right). The rows below shows different modes of deviations due to (directional) interaction effects. **b**, Scatter plot of protein binding effects with unspecific linear effects *β*_*j*_ and *β*_*k*_ on the *x*- and *y*-axis and corresponding interaction effect Θ_*j,k*_ represented by color. For some example proteins detailed information is provided.

The ‘Synergy b+b+b’ category (shown in blue in Fig. 4) includes proteins that bind to two modifications individually and exhibit particularly strong binding, i.e., stronger than the sum of the two individual effects when both modifications are present. For example, we uncover that UHRF1 (Fig. 4b, 1st quadrant) responds in a synergistic way to an interaction effect between DNA methylation m5C and H3K9me3. UHRF1 is a RING-type E3 ubiquitin ligase that plays an essential role in DNA methylation by mediating the recruitment of the maintenance DNA methyltransferase DNMT1 [76]. UHRF1 is known to bind to H3K9me3 via a tandem tudor domain and to recognize hemi-methylated DNA via a SRA domain. Our analysis therefore validates previously known binding behaviors and, additionally, unveils that there is a true synergistic effect between H3K9me3 and DNA methylation in the recruitment of UHRF1.

For the maintenance DNA methyltransferase DNMT1 [77], we identify an individual binding effect to H3K9me3 and a modest individual binding to DNA methylation. Furthermore, we also identify an interaction effect between DNA methylation m5C and H3K9me3. In this case, however, the addition of DNA methylation m5C leads to a reduction in binding of DNMT1 to H3K9me3. We define this behavior as ‘Antagonism b+b+r’ or preferential binding (pink category in Fig. 4). UHRF1 and DNMT1 were found to interact with each other (see references in [76]), and binding of DNMT1 to H3K9me3 is likely mediated through UHRF1 (see above). Both UHRF1 and DNMT1 are flexible multi-domain proteins, that consist of several different domains and can change their shape or structure. They are involved in a complex network of interactions, both within themselves (intra-molecular) and with each other (inter-molecular). This network helps control their function through allosteric regulation events involving conformational rearrangements of autoinhibitory domains (changes in the structure of certain domains within the proteins) in both molecules [78, 76]. The antagonistic effect of DNA methylation on the recruitment of DNMT1 to H3K9me3 indicates that while symmetric DNA methylation stimulates binding of UHRF1 to the doubly modified nucleosomes (see above), it disrupts the interaction with DNMT1. This suggests a mechanism within DNMT1 that senses symmetrically methylated DNA (the end product of the DNA methylation reaction) and triggers the release from chromatin upon completion of its enzymatic reaction. Apart from the catalytic domain of DNMT1, which is responsible for the main activity of the protein, this observed behavior could involve a CXXC domain that has a special ability to bind to certain DNA sequences, specifically sequences that contain un-methylated CpG nucleotides, and could contribute to sensing the DNA methylation status.

Two proteins, MAD2L2 and ACTL8, exhibit a similar behavior with respect to DNA methylation m5C and H3K9me3. However, for these proteins, DNA methylation m5C exhibits a slight repulsive effect on its own. These proteins belong to the category ‘Conflict, dominated by repulsion b+r+r’ (grey category in Fig. 4).

Proteins in the ‘Conflict, dominated by binding b+r+b’ category (yellow category in Fig. 4) are repelled by one modification and bind to another modification if they are considered individually. In combination, these modifications show a stronger binding effect on the protein than expected under additivity. The chromodomain-containing protein CBX8, which is a component of the polycomb repressive complex 1 (PRC1) [79], also falls into this category. Our analysis reveals that DNA methylation m5C enhances the binding of CBX8 to H3K27me3, while DNA methylation m5C itself exhibits a slight repulsive effect on CBX8. The association of CBX8 with both DNA and H3K27me3 has been investigated in Connelly, Weaver, Alpsoy, Gu, Musselman, and Dykhuizen [80]. Here, the authors identified a dual interaction mechanism for the CBX8 chromodomain, where the engagement of both DNA and H3K27me3 mediates the association of CBX8 with chromatin. Similar binding behaviors are observed for the PRC1 subunits RNF2 and RING1. However, in contrast to CBX8, RNF2, and RING1 are shared among multiple complexes, including the canonical polycomb repressive complex 1 (PCR1) and various non-canonical versions of the complex (ncPRC) [79]. The nucleosome binding profiles of these shared subunits reflect a superposition of the binding profiles of all the complexes they are associated with. This introduces additional complexity to the interpretation of combinatorial effects.

### Chromatin modification interaction in the recruitment of proteins and complexes

Our analysis suggests that proteins within the same protein complex tend to exhibit similar binding patterns not only to individual chromatin modifications, but also with regards to interaction effects of modifications.

Our analysis reveals seven distinct combinations of chromatin modifications demonstrating a robust combinatorial effect on the shortlisted 55 proteins (see Fig. 5a). While six of the discovered interactions affect multiple proteins, H2A.Z incorporation appears to interact solely with H4K20 me2, influencing only the protein FBXO38. Given FBXO38’s notably low data density (see Fig. 3a, last column), we did not investigate this interaction further. Notably, TAF10 and TBP, which are both part of the Transcription Factor II D (TFIID) complex, respond similarly to the combination of H4K20me2 and H3K4me3. Similarly, members of the PRC1 complex, such as CBX8, RNF2, RING1, CBX2, and PCGF2, are found to respond to the combination of H3K27me3 and DNA methylation m5C (Fig. 5a). In addition to examining the effect sizes obtained from `asteRIa`, our approach allows for the interpretation of protein-specific selection probabilities for each individual chromatin modification and each interaction between chromatin modification combinations. The selection probability indicates how stable a feature is in predicting a proteins binding profile across subsamples.

**Fig 5.**
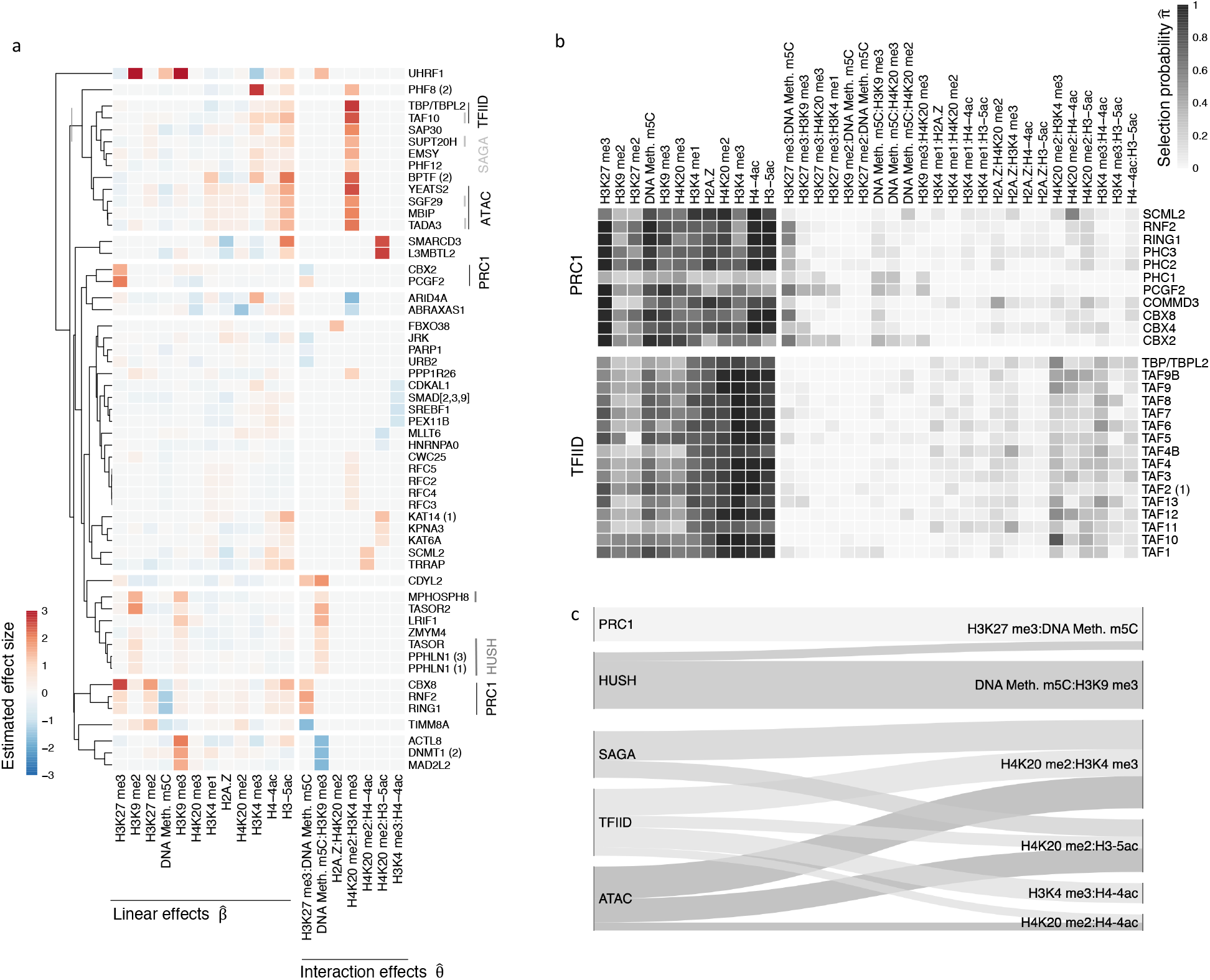
**a**, Clustered representation of robustly estimated linear and interaction coefficients for *p*_*I*_ = 55 proteins for which interaction coefficients have been identified. Proteins of the PRC1 and TFIID complex are highlighted. **b**, Selection probabilities 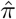 for all proteins in the TFIID and PRC1 complex. Selection probability plots for all protein complexes are provided in Extended Fig. 5b. **c**, Sankey diagram of mean selection probabilities (*>* .2) for interaction effects for proteins within a complex.

We observe that proteins belonging to the same complex show similar modification selection probability patterns (see Fig. 5b for an illustration using the TFIID and PRC1 complex, respectively). These similarities in selection probability patterns justify the exploration of mean selection probabilities over proteins within the same complexes, leading to a more general analysis of how entire complexes respond to interaction effects between chromatin modifications (see Fig. 5c).

One major discovery is that the Ada-Two-A-containing (ATAC), Spt-Ada-Gcn5 acetyltransferase (SAGA), and TFIID complexes exhibit multiple combinations of co-operative chromatin modifications that stimulate their binding (Fig. 5c). Notably, our analysis reveals several interactions where H4K20me2 is involved, particularly in conjunction with H3K4me3, H3-5ac, and H4-4ac.

SAGA is a highly conserved transcriptional co-activator with four distinct functional modules. Its enzymatic functions, including histone acetylation and deubiquitination modules, play crucial roles in chromatin structure and gene expression [81]. The ATAC complex, which shares subunits with SAGA, also exhibits histone acetyltransferase activity [81]. TFIID, another essential transcription factor, is also a histone acetyltransferase, but additionally recognizes core promoter sequences, recruits the transcription pre-initiation complex, and interacts with SAGA subunits. TFIID contributes to transcription initiation and gene expression by collaborating with cofactors, gene-specific regulators, and chromatin modifications associated with active genomic regions [82]. As such ATAC, SAGA, and TFIID are all protein complexes that possess activities that are intricately involved in the process of transcription initiation and that thereby contribute to the regulation of chromatin structure and gene expression.

H4K20me2 is a pervasive modification found on 80% of all histone H4 proteins, marking nearly every nucleosome throughout the genome. Since newly incorporated histone H4 is unmodified at K20 (H4K20me0), the H4K20me2 modification serves as a marker of not yet replicated ‘old’ chromatin, while H4K20me0 marks newly replicated chromatin during the cell cycle. This modification is used by the DNA repair machinery to determine between different DNA repair pathways in different cell cycle phases [83]. The synergistic effect between H4K20me2 and active modifications in recruiting protein complexes associated with transcriptional initiation is therefore surprising and hints to a so far unknown possible function of this modification in the context of promoter regulation.

In contrast, members of the repressive PRC1 and HUSH complexes show a response to an interaction effect between H3K27me3 and DNA methylation m5C and an interaction effect between DNA methylation m5C and H3K9me3, respectively.

The human silencing hub (HUSH) complex is well-established for its role in transcriptionally repressing long interspersed element-1 retrotransposons (L1s) and retroviruses through the modification of histone H3 lysine 9 trimethylation (H3K9me3) [84]. Our analysis not only confirms H3K9me3 to be an important binding determinant, in line with previous findings, but it also reveals the involvement of DNA methylation m5C in this regulatory process. Furthermore, our analysis uncovers a previously unreported synergistic interaction between these two modifications, indicating a more complex interplay between H3K9me3 and DNA methylation m5C than previously known.

As a last example, we find that for several members of the PRC1 complex, there is an increased likelihood of responding to an interaction between H3K27me3 and DNA methylation m5C, as previously discussed for CBX8 and the subunits RNF2 and RING1. The PRC1 complex is known to be capable of recognizing H3K27me3 and facilitating transcriptional repression [79], while there are no known associations between the PRC1 complex and methylated DNA. Our results suggest a distinct behavior of DNA methylation and H3K27me3 on regulating the recruitment of the PRC1 complex, with DNA methylation m5C having minimal or even a slightly repulsive effect and H3K27me3 having a binding effect on their own. However, in combination, our analysis reveals an interaction between these two modifications that enhances binding.

### Validation of the effects of H3K27me3 and DNA methylation on the binding of CBX8 with ChIP-seq and WGBS data

To validate and compare our findings with orthogonal data sources, we leverage publicly accessible ChIP-seq and WGBS (Whole Genome Bisulfite Sequencing) datasets from the ENCODE project (https://www.encodeproject.org)[40, 85, 86, 87] and ChIP-Atlas (https://chip-atlas.org) [88, 89, 90]. Specifically, we design a validation workflow that compares partial correlations from modification co-occurrence patterns with `asteRIa`’s linear and interaction coefficients.

Given the unique design of the MARCS data, our ability to independently validate our discoveries hinges on the availability of ChIP-seq/WGBS experiments that encompass chromatin modifications for which we have identified interaction effects *and* are available in the same cell type. After a comprehensive web search, we have identified only the trio of H3K27me3 (ChIP-seq), methylated DNA (WGBS), and the CBX8 protein (ChIP-seq) as the only adequate data set.

As previously described, `asteRIa` reveals a modest interaction effect between H3K27me3 and methylated DNA concerning the binding of CBX8 in the nucleosome binding data. This interaction effect is categorized as ‘conflict, dominated by binding b+r+b’ (see Fig. 4a). We detect a slight repulsive effect of methylated DNA on CBX8 and a recruitment to H3K27me3. Notably, we identify an additional positive interaction effect on CBX8 binding when methylated DNA and H3K27me3 co-occur. Consequently, our results indicate a subtle enhancing effect on CBX8 binding when methylated DNA co-occurs with H3K27me3 (see Fig. 4b, lower right corner), resulting in improved predictive accuracy (see Fig. 3c).

For this combination, we found matching ChIP-seq and WGBS experiments in A549 (human lung carcinoma epithelial cells), K562 (human myelogenous leukemia cells), and H1 cells (human embryonic stem cells) on ENCODE. Additionally, we use mES cell (mouse embryonic stem cells) data from ChIP-Atlas. For these four cell types, we perform the following analysis workflow: (1) We calculate averages of WGBS data and averages of fold-change values to a reference genome in the ChIP-seq data within consecutive genome bins of 1000 base pairs (bp) with no spacing between bins. We accomplish this by utilizing the ‘bins’ mode within deeptools on the Galaxy web platform [91], and we ensure the exclusion of blacklisted regions (hg38 for A549, K562, and H1 cells and mm9 for mES cells) during these calculations. (2) We then conduct a genome-wide analysis of the behavior of H3K27me3 and methylated DNA in CBX8 peak regions. We observe increased H3K27me3 fold-changes and simultaneously decreased DNA methylation values (decreased in K562, A549 and mES; unaffected in H1) in CBX8-bound regions across all cell types under investigation (see Fig. 6a and Fig. S3). This substantiates the (linear) dependencies identified in the `asteRIa` workflow. We compute Kendall’s partial correlations [92] (package version v1.1) of the genome-wide co-occurrence patterns between CBX8, methylated DNA, H3K27me3, and the “interaction” between methylated DNA and H3K27me3 (i.e., the product of WGBS and H3K23me3 ChIP-seq values, denoted by H3K27me3:WGBS). We use this rank-based correlation coefficient to account for the fact that WGBS and ChIP-seq data are measured and interpreted on different scales. The resulting partial correlations patterns are shown in Fig. 6b. The interpretation of the partial correlation coefficients aligns with the coefficients in `asteRIa`’s interaction model. Specifically, the first column of each partial correlation matrix (CBX8) can be understood as follows. The partial correlation between CBX8 and H3K27me3, as well as between CBX8 and the WGBS abundances, reflects the individual (linear) effects of these modifications on CBX8 binding (after conditioning on all other effects). We observe that they are (moderately) positive for CBX8 and H3K27me3 across all cell types, and negative for CBX8 and WGBS (b+r pattern). Furthermore, the partial correlation between CBX8 and the product of WGBS and H3K27me3 ChIP-seq values represents the additional combinatorial interaction effect, complementing the individual effects. This partial correlation is positive across all cell types, leading to the b+r+b pattern observed in `asteRIa`, Furthermore, it tends to be larger in magnitude than the negative partial correlation between the CBX8 and WGBS data, which also aligns with the `asteRIa` results on the CBX8 nucleosome binding data.

**Fig 6.**
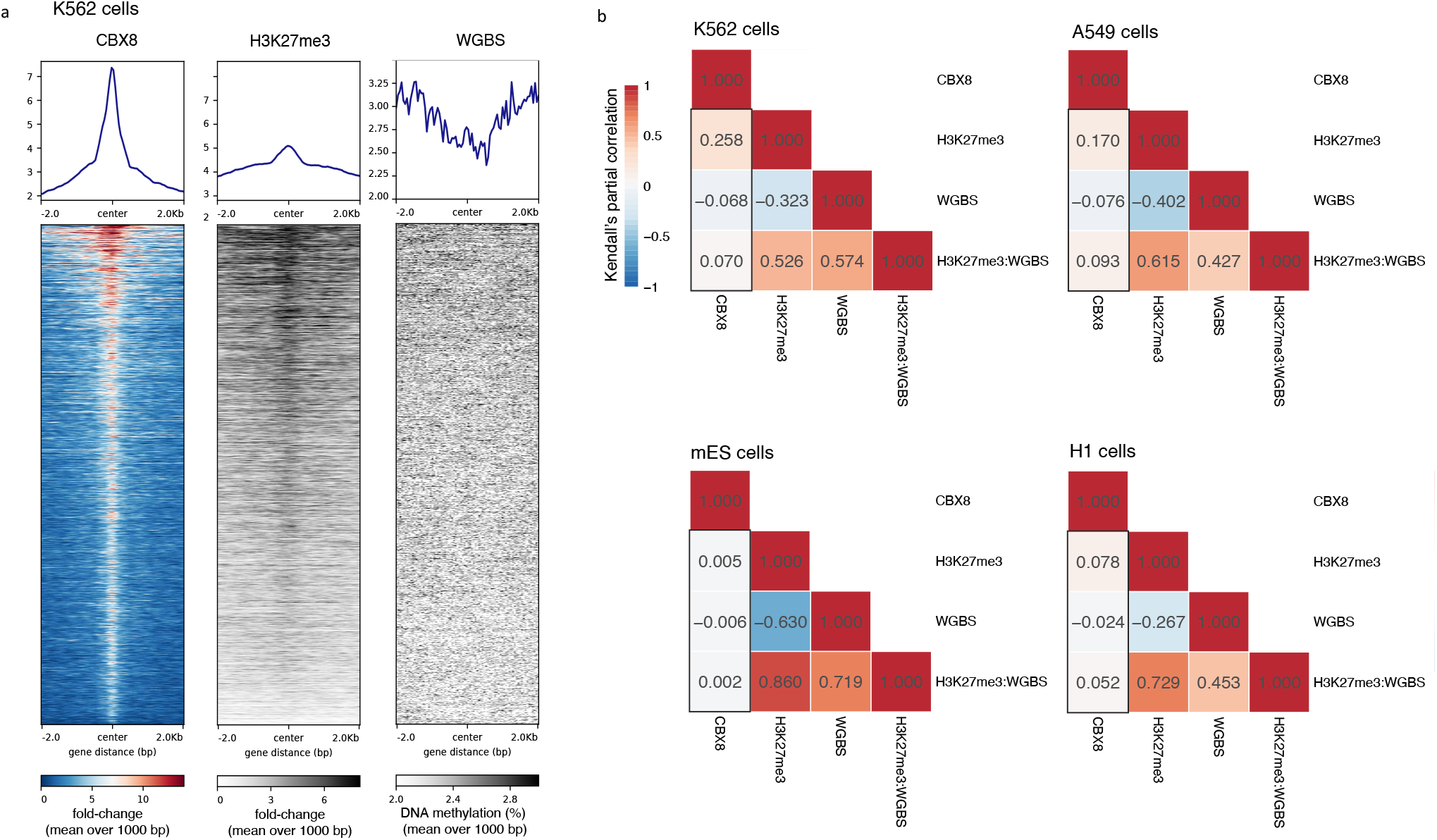
**a**, Heatmap for score distributions across CBX8 IDR thresholded peaks in K562 cells created with deeptools on the Galaxy web platform. **b**, Kendall’s partial correlation between CBX8, H3K27me3, WGBS data, and the product between H3K27me3 and WGBS for A549, K562, H1, and mES cells. The first column in each partial correlation plots recapitulates the main and interaction effects, derived by `asteRIa`. For ENCODE and ChIP-Atlas identifier see caption of Fig. S3.

In summary, this analysis provides evidence that `asteRIa`’s estimated main and interaction effects can be recapitulated using other high-throughput experimental data. Moreover, this type of analysis provides a recipe for further validation and invites to perform new ChIP-Seq experiments for other candidate proteins that show evidence of combinatorial interaction effects.

## Discussion

While many functions and readers of individual chromatin modifications have been described [15, 16], the understanding of how multiple modifications cooperate in recruiting epigenetic regulators has remained largely elusive. To gain insights into these cooperative effects, we have introduced `asteRIa`, a workflow for the robust statistical detection of interaction effects, and applied the workflow to the recently published MARCS nucleosome binding dataset. The MARCS data comprise a library of semi-synthetic nucleosomes followed by nucleosome affinity purification with high-throughput quantitative proteomics measurements. Despite MARCS’ novelty and uniqueness to probe the binding behavior of proteins to combinatorial chromatin modifications at a large scale, the imbalanced design matrix and the low sample size pose considerable challenges for consistent statistical interaction estimation. `asteRIa` presents a first step toward identifying robust combinatorial effects between chromatin modifications and is tailored specifically to address these challenges. At its core, `asteRIa` combines the lasso for hierarchical interactions [53] with the complementary pairs stability selection (CPSS) concept [59], and incorporates replicate consistency mechanisms to minimize the identification of spurious interaction effects. We also confirm in a realistic synthetic simulation scenario that combining the interaction model with CPSS reduces the number of spurious effects considerably and leads to more robust results compared to the standard cross-validation procedure (see Supplementary information and Fig. S1 and S2).

By employing `asteRIa` in conjunction with the MARCS dataset, our study provides the first quantitative framework for the identification of cooperative effects of chromatin modifications on protein binding. We identify a list of 55 epigenetic reader candidates that likely respond to combinatorial modification effects. For the set of 55 proteins we confirmed that interactions enhance predictive performance of protein binding.

To evaluate the validity of `asteRIa`’s data consistency checks, we performed a sensitivity analysis, comparing `asteRIa`’s sign-consistency checks to distance-based consistency filtering and no data filtering. Our analysis demonstrates that requiring data sign-consistency results in the largest number of replicate consistent models and gives the largest set of robustly identified proteins responding to chromatin modification interactions (see Suppl. Fig. S5).

For the 55 proteins identified, we observed consistent responses to these combinations across multiple proteins within the same protein complex, further substantiating the robustness of our findings. The derived candidate set also allowed for a quantitative categorization of different modes of potential chromatin modification interactions.

While our analysis is naturally limited to combinations of chromatin modifications that co-occur in at least one MARCS experiment, we were able to both recapitulate established effects of chromatin modifications on protein binding behavior and discover novel interaction effects between chromatin modifications, potentially promising candidates for future functional analyses. An intriguing finding of our analysis is the discovery of several combinations of cooperative chromatin modifications that elicit responses of the ATAC, SAGA, and TFIID complexes. In particular, we identified several interactions involving the H4K20me2 modification, especially in combination with H3K4me3, H3-5ac, and H4-4ac. Another intriguing finding from our analysis is the similar binding profile observed for the proteins DNMT1, MAD2L2, and ACTL8 - all exhibiting a repulsive combinatorial effect in response to DNA methylation m5c and H3K9me3. The function of ACTL8 has not been extensively studied. However, its analogous behavior to MAD2L2 and especially DNMT1 provides an initial hint to a potential function of ACTL8.

We demonstrated the generalizability of our findings beyond a specific cell type or experimental setup by comparing the interaction effect of H3K27me3 and methylated DNA on CBX8, as identified by `asteRIa`, using publicly available ChIP-seq and WGBS data from K562, A549, H1, and mES cells sourced from ENCODE and ChIP-Atlas. Our analysis revealed that, even with the modest improvement in predictive accuracy observed for CBX8 when considering the identified interaction effect between H3K27me3 and methylated DNA, similar patterns are consistently observed in ChIP-seq and WGBS experiments across these diverse cell types.

However, it is important to note that the majority of combinatorial chromatin modification interaction effects identified by `asteRIa`, particularly those characterized by strong interaction effect sizes, are not present in publicly available ChIP-seq datasets. Consequently, we posit that our study serves as a first unbiased attempt to identify chromatin regulators that respond to more than one modification and thereby act as a hypothesis generator, suggesting specific combinations of proteins and chromatin modifications worthy of further investigation in future biological experiments. In particular, we recommend focusing on proteins that exhibit relatively poor predictive accuracy when considering individual chromatin modification effects alone. For instance, proteins like RFC2, RFC3, RFC4, and RFC5 show a substantial enhancement in predictive accuracy when considering the identified interaction effect between H4K20me2 and H3K4me3.

Moreover, `asteRIa` functions as a versatile tool that can be readily updated whenever new nucleosome affinity purification experiments become available. As tools are developed to conduct a greater number of experiments with additional combinations of modifications, our workflow can be conveniently extended to explore more and higher-order interaction effects between chromatin modifications, allowing a more comprehensive understanding of the combinatorial complexity of chromatin modifications.

Even though our statistical workflow has been specifically designed and optimized for the MARCS dataset, its methodology and approach can be broadly applied in scenarios where robust assessment of hierarchical interactions is required, particularly in data-scarce regimes with high levels of noise.

In conclusion, our study provides compelling evidence that large-scale SILAC nucleosome affinity purification data, when combined with `asteRIa`, is a potent resource for generating hypotheses related to epigenetic reader candidates.

## Data availability

The `asteRIa` workflow, the processed data, and the code for reproducing all figures and results are available at https://figshare.com/articles/software/asteRIa/25003103 and (partly, without large files) at https://github.com/marastadler/asteRIa.git. The MARCS data is available at https://marcs.helmholtz-muenchen.de. Mass spectrometry data for MARCS was submitted to the PRIDE database (https://www.ebi.ac.uk/pride/) (accession number: PXD018966).

ENCODE K562 identifier: ENCFF405HIO, ENCFF687ZGN, ENCFF522HZT, ENCFF459XNY; ENCODE A549 identifier: ENCFF702IOJ, ENCFF081CPV, ENCFF723WVM, ENCFF552VXR; ENCODE H1 identifier: ENCFF345VHG, ENCFF284JDC, ENCFF975NYJ, ENCFF483UZG; ChIP-Atlas mES identifier: SRX426373, SRX006968, DRX001152, SRX5090173.05.

## Acknowledgments

We greatly acknowledge Roberto Olayo Alarcon for valuable discussion on the data validation. We acknowledge the ENCODE Consortium and the ENCODE production laboratorys generating the particular datasets used in this study.

## Author Contributions

M.S. developed asteRIa, conducted the analysis on the MARCS data, ChIP-Seq and bisulfite data and conducted the analysis on synthetic data. C.L.M. and T.B. supervised the work. M.S. and C.L.M. conceived the statistical workflow. S.L. and T.B. analyzed the results and provided feedback. M.S., C.L.M. and T.B. wrote the manuscript. All authors read and approved the final manuscript.

