## Supplementary material for "asteRIa enables robust interaction modeling between chromatin modifications and epigenetic readers"

### Supplementary information

#### Synthetic data generation and performance comparison

We construct a realistic synthetic data scenario to demonstrate that employing complementary pairs stability selection yields significantly more robust and accurate outcomes compared to the default implementation of cross-validation in the Lasso for hierarchical interactions (hiernet). To achieve this, we employ an asymmetric Laplace distribution to model the non-zero coefficient estimates derived from actual data across all proteins (see Fig. S1a and b).

In order to maintain consistent sparsity levels for both proteins and features (chromatin modifications or interactions between chromatin modifications), we opt for the simplest approach: retaining the same sparsity pattern as observed in the estimated coefficients from the real data. Additionally, we model the distribution of estimated intercepts for all proteins using a Laplace distribution. Introducing a normally distributed error term, akin to the noise inherent in the actual data, further enhances the fidelity of our synthetic data. Furthermore, we introduce variations in this noise level to illustrate how the quality of outcomes responds to differing degrees of noise. Using the simulated intercept, the product of the simulated coefficients, and the true experimental design matrix containing interaction terms, along with the simulated error term, we generate a synthetic dataset representing protein binding, $P_{syn} = I_{syn} + \Theta_{syn}(L, L_{int}) + E_{syn}$, with $P_{syn},\ I_{syn},\ E_{syn} \sim (p, n)$, $\Theta_{syn} \sim (p, q^2/2)$, $(L, L_{int}) \sim (q^2/2, n)$ (see Fig. S1c).

We compare the following two approaches using the `hierNet` model: one employing 5-fold cross-validation with the 1-standard error (1se) rule, and the other utilizing complementary pairs stability selection (CPSS). These experiments were conducted on a subset of 58 synthetic proteins known to exhibit interactions. Across both experiment sets, we conduct 20 replicates for each configuration to ensure robustness and reliability of our findings (see Fig. S2).

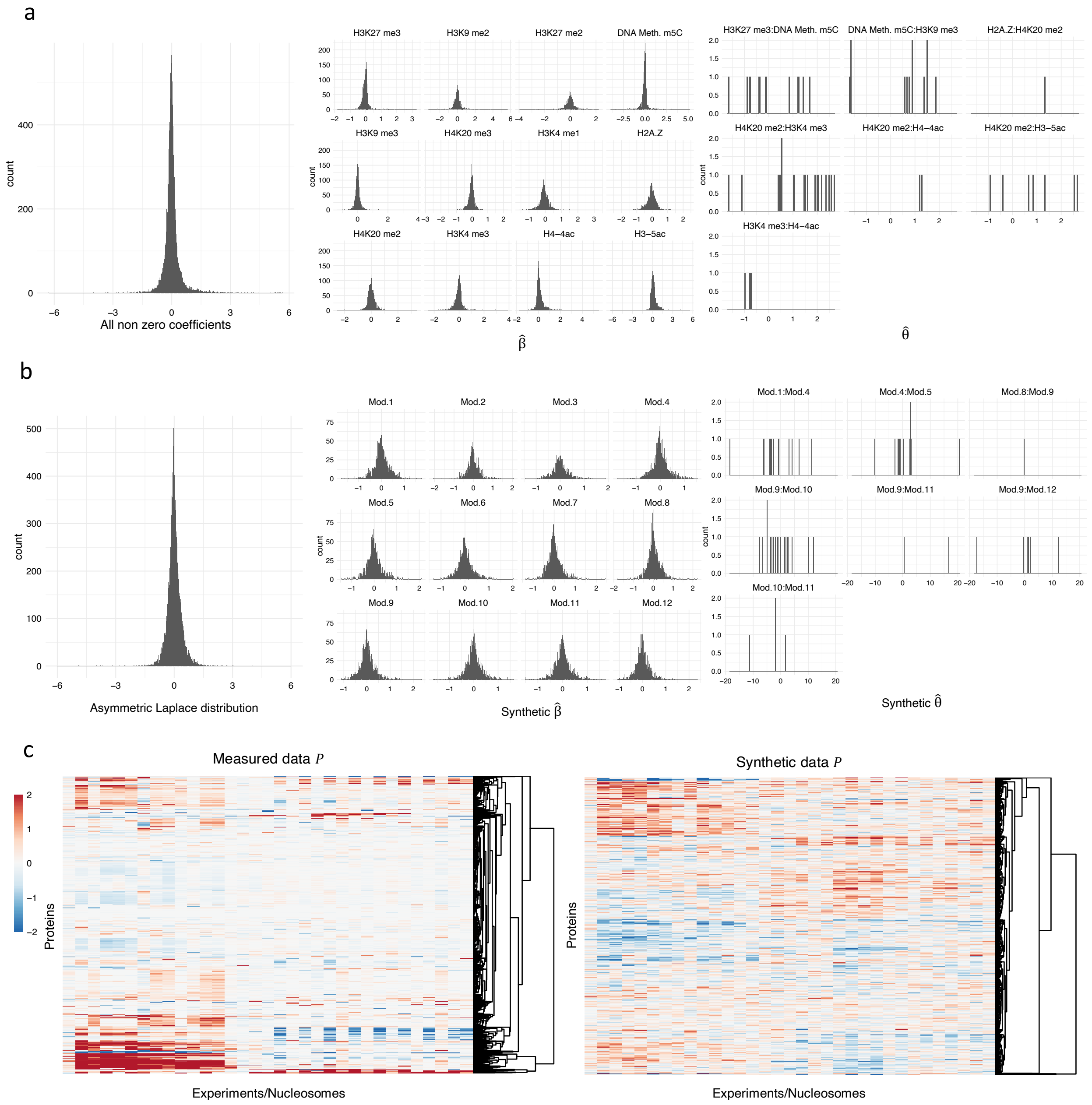


**Fig S1.** **a**, Joint and individual distributions of all non zero estimated coefficients $\hat{\beta}$ and $th\hat{e}ta$ in the statistical workflow. **b**, Asymmetric Laplace distribution fitted to the joint distribution estimated coefficients in **a**. **c**, Left: clustered heatmap of proteins binding measures $P$(mean of forward and reverse experiment); right: clustered heatmap of synthetic protein binding measures $P$.

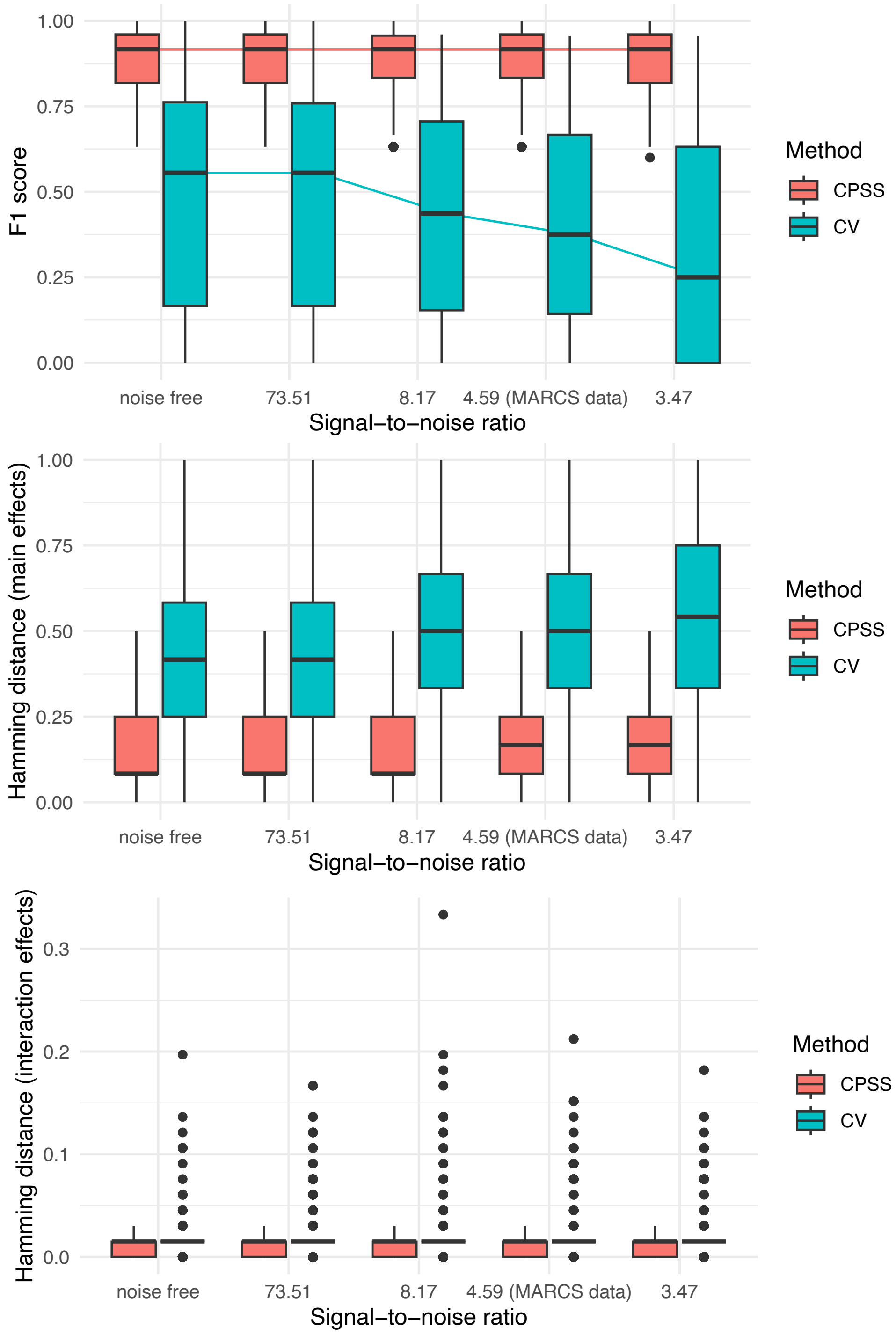


**Fig S2.** **a**, F1 score for five signal-to-noise ratios (SNR) for hiernet with CPSS and 5-fold cross-validation (1se rule). SNR of 4.59 corresponds to the SNR observed in the MARCS data. Noise free corresponds to 0% noise, SNR = 73.51 corresponds to 25% of the noise observed for the MARCS data; SNR = 8.17 corresponds to 75% of the noise observed for the MARCS data and SNR = 3.47 corresponds to 125% of the noise observed for the MARCS data. **b**, Hamming distance main effects for hiernet with CPSS and 5-fold cross-validation (1se rule) for five SNRs. **c**, Hamming distance interaction effects for hiernet with CPSS and 5-fold cross-validation (1se rule) for five SNRs.

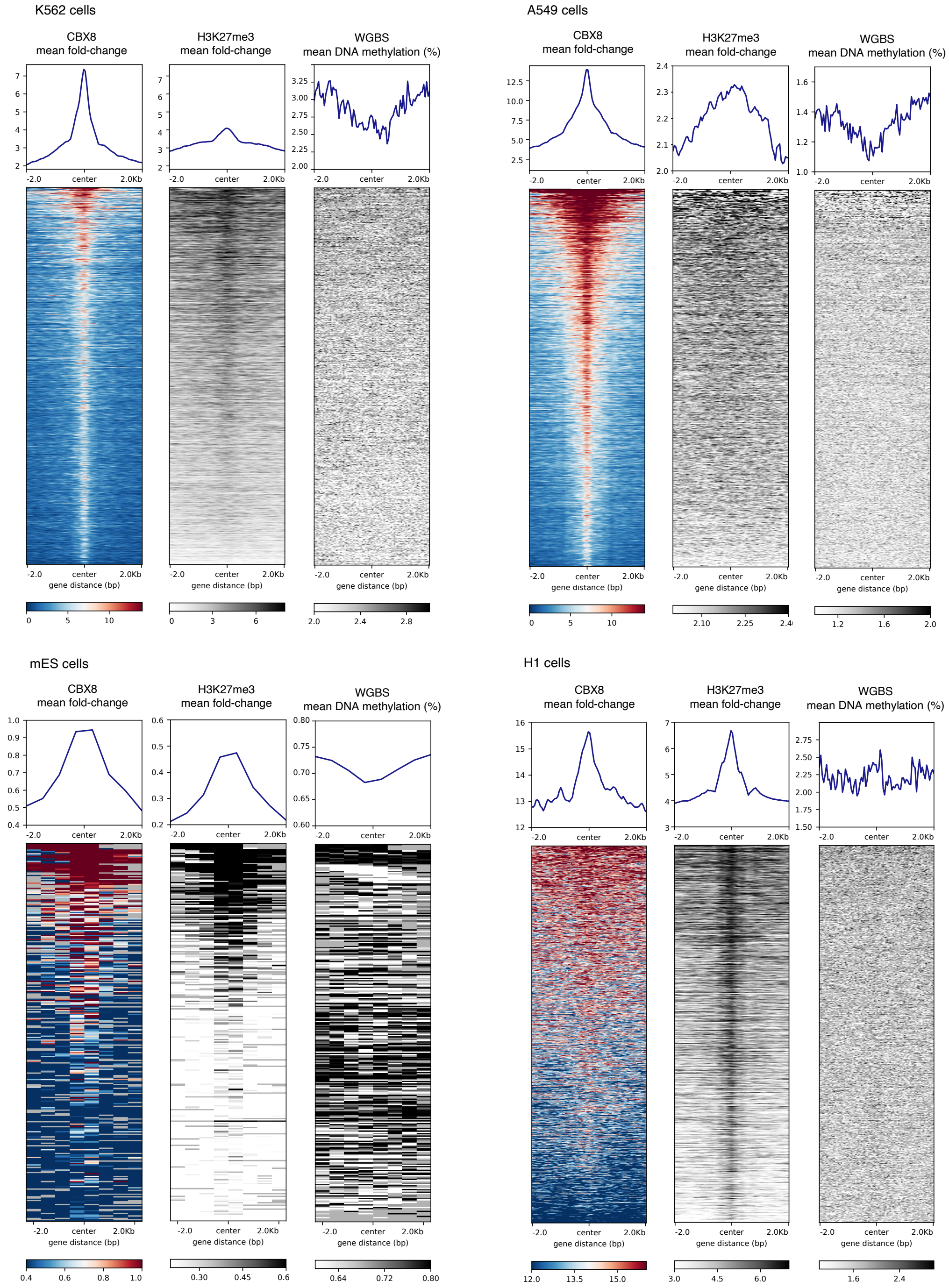


**Fig S3.** Heatmaps of score distributions across CBX8 IDR thresholded peaks in K562, A549, H1 and mES cells. mES heatmaps are based on 500 bp bins for visualization purposes because of missing values. Fold-changes in mES ChIP-Atlas experiments are scaled between 0 and 1 while mean fold-changes in ENCODE experiments represent raw values. ENCODE K562 identifier: ENCFF405HIO, ENCFF687ZGN, ENCFF522HZT, ENCFF459XNY; ENCODE A549 identifier: ENCFF702IOJ, ENCFF081CPV, ENCFF723WVM, ENCFF552VXR; ENCODE H1 identifier: ENCFF345VHG, ENCFF284JDC, ENCFF975NYJ, ENCFF483UZG; ChIP-Atlas mES identifier: SRX426373, SRX006968, DRX001152, SRX5090173.05.

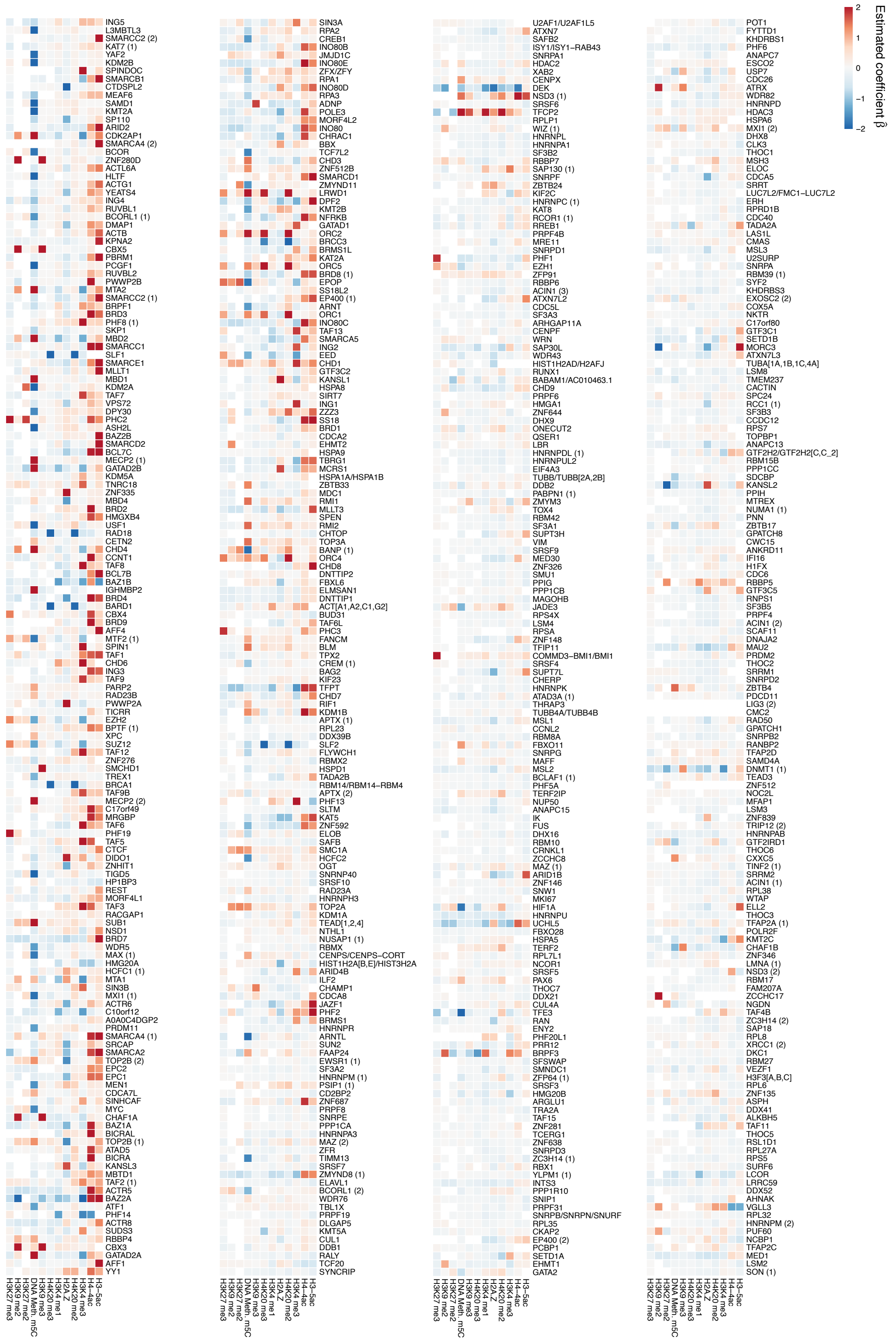


**Fig S4.** Heatmap of estimated main effects in the linear model. The proteins are ordered in descending order of predictive performance ($R^2$). A full list of all proteins is provided in Extended Fig. S4.

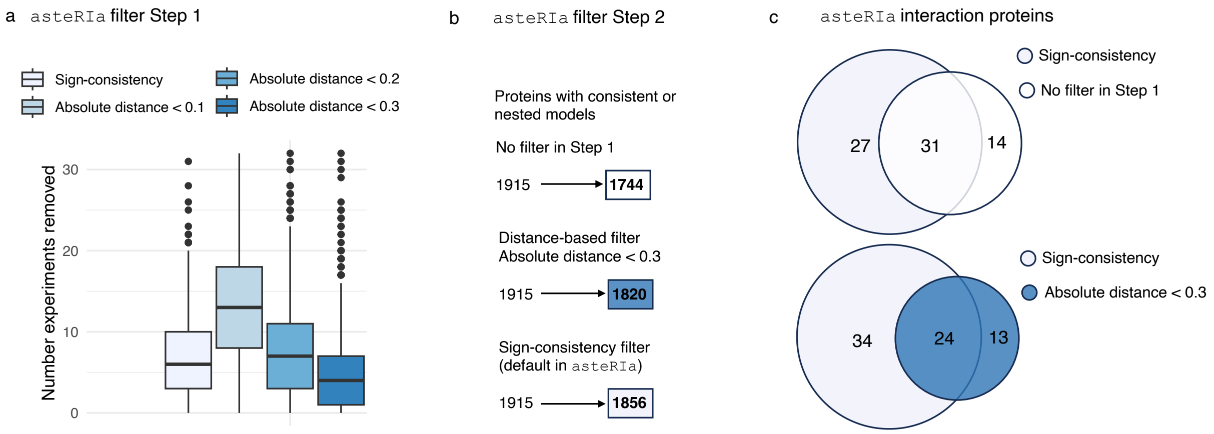


**Fig S5.** Sensitivity analysis of filter step 1 in asteRIa. **a**, Boxplots showing the number of experiments removed from the total of 1915 proteins for different filters: sign-consistency (default in asteRIa) and three distance-based filters. Retaining experiments where the absolute distance is $< 0.1$ or $< 0.2$ removes more experiments than requiring sign-consistency. **b**, Number of proteins filtered out in the model-based filtering in step 2 in asteRIa for the cases: no filter in step 1, distance-based filter ($< 0.3$) in step 1, and sign-consistency filter in step 1. **c**, Venn diagrams representing the number and overlap of proteins with identified interactions by asteRIa. The upper Venn diagram compares sign-consistency to no filter in step 1, the lower Venn diagram compares sign-consistency to the distance-based filter ($< 0.3$) in step 1.

### Overview Extended Figures

- **Extended Fig. 3b** Extension of Fig. 3b. Full list of stability plots for all 1915 proteins.
- **Extended Fig. 3c** Extended Fig. 3c. Extension of Fig. 3c. Full list of scatter plots for all 55 proteins with robust interaction effects.
- **Extended Fig. 5b** Extension of Fig. 5b. Selection probability heatmaps and model coefficients for all 1915 protein complexes.
- **Extended Fig. S4** Extension of Fig. S4. Heatmap of estimated main effects in the linear model for all 1915 proteins.
