## Extended Material for "asteRIa enables robust interaction modeling between chromatin modifications and epigenetic readers"

**Extended Fig. 3b.** Extension of Fig. 3b. Full list of stability plots for all 1915 proteins.

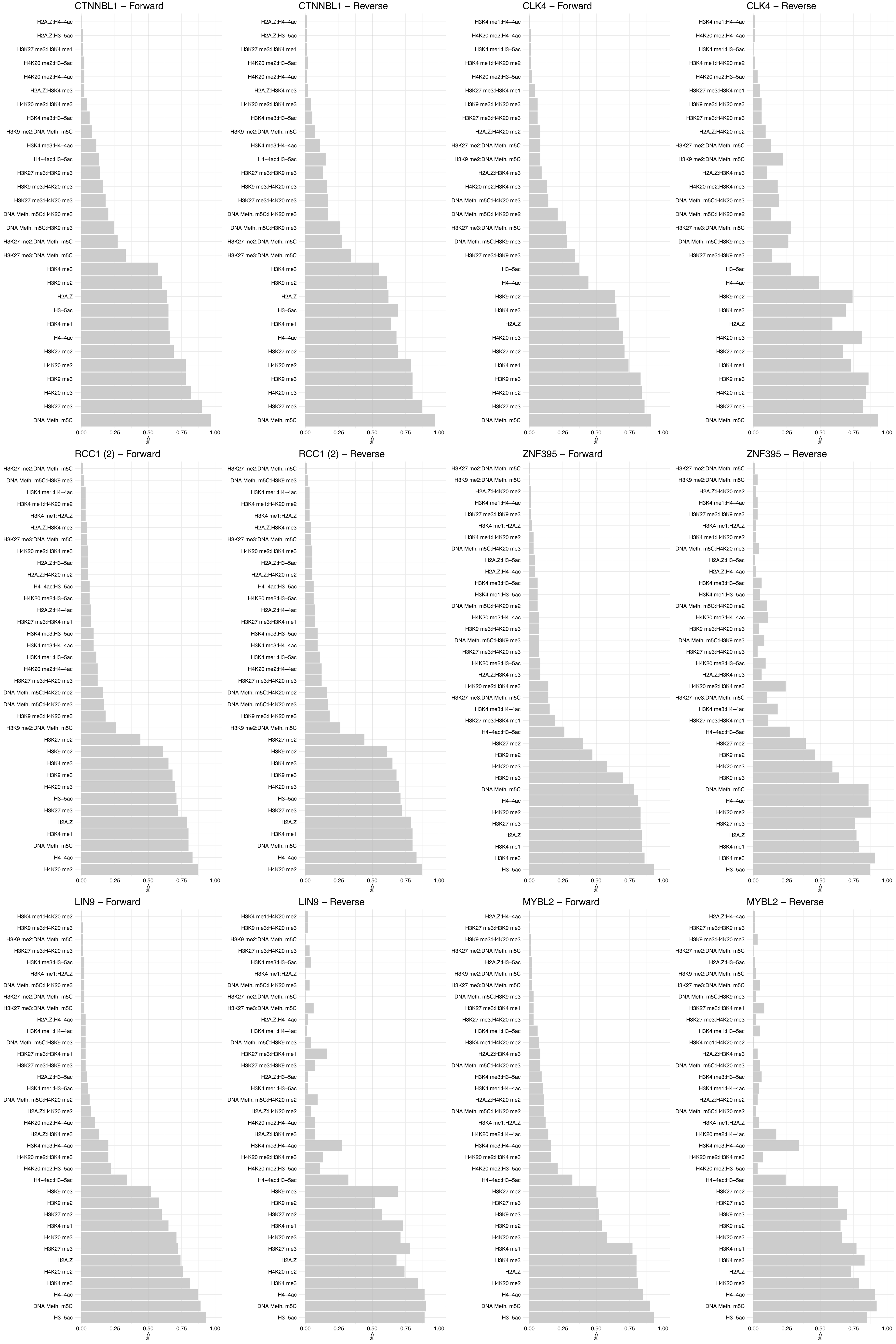
CTNNBL1 – Forward
H2A.Z:H4–4ac
H2A.Z:H3–5ac
H3K27 me3:H3K4 me1
H4K20 me2:H3–5ac
H4K20 me2:H4–4ac
H2A.Z:H3K4 me3
H4K20 me2:H3K4 me3
H3K4 me3:H3–5ac
H3K9 me2:DNA Meth. m5C
H3K4 me3:H4–4ac
H4–4ac:H3–5ac
H3K27 me3:H3K9 me3
H3K9 me3:H4K20 me3
H3K27 me3:H4K20 me3
DNA Meth. m5C:H4K20 me3
DNA Meth. m5C:H3K9 me3
H3K27 me2:DNA Meth. m5C
H3K27 me3:DNA Meth. m5C
H3K4 me3
H3K9 me2
H2A.Z
H3–5ac
H3K4 me1
H4–4ac
H3K27 me2
H4K20 me2
H3K9 me3
H4K20 me3
H3K27 me3
DNA Meth. m5C
0.00 0.25 0.50 0.75 1.00
π̂
CTNNBL1 – Reverse
H2A.Z:H4–4ac
H2A.Z:H3–5ac
H3K27 me3:H3K4 me1
H4K20 me2:H3–5ac
H4K20 me2:H4–4ac
H2A.Z:H3K4 me3
H4K20 me2:H3K4 me3
H3K4 me3:H3–5ac
H3K9 me2:DNA Meth. m5C
H3K4 me3:H4–4ac
H4–4ac:H3–5ac
H3K27 me3:H3K9 me3
H3K9 me3:H4K20 me3
H3K27 me3:H4K20 me3
DNA Meth. m5C:H4K20 me3
DNA Meth. m5C:H3K9 me3
H3K27 me2:DNA Meth. m5C
H3K27 me3:DNA Meth. m5C
H3K4 me3
H3K9 me2
H2A.Z
H3–5ac
H3K4 me1
H4–4ac
H3K27 me2
H4K20 me2
H3K9 me3
H4K20 me3
H3K27 me3
DNA Meth. m5C
0.00 0.25 0.50 0.75 1.00
π̂
CLK4 – Forward
H3K4 me1:H4–4ac
H4K20 me2:H4–4ac
H3K4 me1:H3–5ac
H3K4 me1:H4K20 me2
H4K20 me2:H3–5ac
H3K27 me3:H3K4 me1
H3K9 me3:H4K20 me3
H3K27 me3:H4K20 me3
H2A.Z:H4K20 me2
H3K27 me2:DNA Meth. m5C
H3K9 me2:DNA Meth. m5C
H2A.Z:H3K4 me3
H4K20 me2:H3K4 me3
DNA Meth. m5C:H4K20 me3
DNA Meth. m5C:H4K20 me2
H3K27 me3:DNA Meth. m5C
DNA Meth. m5C:H3K9 me3
H3K27 me3:H3K9 me3
H3–5ac
H4–4ac
H3K9 me2
H3K4 me3
H2A.Z
H4K20 me3
H3K27 me2
H3K4 me1
H3K9 me3
H4K20 me2
H3K27 me3
DNA Meth. m5C
0.00 0.25 0.50 0.75 1.00
π̂
CLK4 – Reverse
H3K4 me1:H4–4ac
H4K20 me2:H4–4ac
H3K4 me1:H3–5ac
H3K4 me1:H4K20 me2
H4K20 me2:H3–5ac
H3K27 me3:H3K4 me1
H3K9 me3:H4K20 me3
H3K27 me3:H4K20 me3
H2A.Z:H4K20 me2
H3K27 me2:DNA Meth. m5C
H3K9 me2:DNA Meth. m5C
H2A.Z:H3K4 me3
H4K20 me2:H3K4 me3
DNA Meth. m5C:H4K20 me3
DNA Meth. m5C:H4K20 me2
H3K27 me3:DNA Meth. m5C
DNA Meth. m5C:H3K9 me3
H3K27 me3:H3K9 me3
H3–5ac
H4–4ac
H3K9 me2
H3K4 me3
H2A.Z
H4K20 me3
H3K27 me2
H3K4 me1
H3K9 me3
H4K20 me2
H3K27 me3
DNA Meth. m5C
0.00 0.25 0.50 0.75 1.00
π̂
RCC1 (2) – Forward
H3K27 me2:DNA Meth. m5C
DNA Meth. m5C:H3K9 me3
H3K4 me1:H4–4ac
H3K4 me1:H4K20 me2
H3K4 me1:H2A.Z
H2A.Z:H3K4 me3
H3K27 me3:DNA Meth. m5C
H4K20 me2:H3K4 me3
H2A.Z:H3–5ac
H2A.Z:H4K20 me2
H4–4ac:H3–5ac
H4K20 me2:H3–5ac
H2A.Z:H4–4ac
H3K27 me3:H3K4 me1
H3K4 me3:H3–5ac
H3K4 me3:H4–4ac
H3K4 me1:H3–5ac
H4K20 me2:H4–4ac
H3K27 me3:H4K20 me3
DNA Meth. m5C:H4K20 me2
DNA Meth. m5C:H4K20 me3
H3K9 me3:H4K20 me3
H3K9 me2:DNA Meth. m5C
H3K27 me2
H3K9 me2
H3K4 me3
H3K9 me3
H4K20 me3
H3–5ac
H3K27 me3
H2A.Z
H3K4 me1
DNA Meth. m5C
H4–4ac
H4K20 me2
0.00 0.25 0.50 0.75 1.00
π̂
RCC1 (2) – Reverse
H3K27 me2:DNA Meth. m5C
DNA Meth. m5C:H3K9 me3
H3K4 me1:H4–4ac
H3K4 me1:H4K20 me2
H3K4 me1:H2A.Z
H2A.Z:H3K4 me3
H3K27 me3:DNA Meth. m5C
H4K20 me2:H3K4 me3
H2A.Z:H3–5ac
H2A.Z:H4K20 me2
H4–4ac:H3–5ac
H4K20 me2:H3–5ac
H2A.Z:H4–4ac
H3K27 me3:H3K4 me1
H3K4 me3:H3–5ac
H3K4 me3:H4–4ac
H3K4 me1:H3–5ac
H4K20 me2:H4–4ac
H3K27 me3:H4K20 me3
DNA Meth. m5C:H4K20 me2
DNA Meth. m5C:H4K20 me3
H3K9 me3:H4K20 me3
H3K9 me2:DNA Meth. m5C
H3K27 me2
H3K9 me2
H3K4 me3
H3K9 me3
H4K20 me3
H3–5ac
H3K27 me3
H2A.Z
H3K4 me1
DNA Meth. m5C
H4–4ac
H4K20 me2
0.00 0.25 0.50 0.75 1.00
π̂
ZNF395 – Forward
H3K27 me2:DNA Meth. m5C
H3K9 me2:DNA Meth. m5C
H2A.Z:H4K20 me2
H3K4 me1:H4–4ac
H3K27 me3:H3K9 me3
H3K4 me1:H2A.Z
H3K4 me1:H4K20 me2
DNA Meth. m5C:H4K20 me3
H2A.Z:H3–5ac
H2A.Z:H4–4ac
H3K4 me3:H3–5ac
H3K4 me1:H3–5ac
DNA Meth. m5C:H4K20 me2
H4K20 me2:H4–4ac
H3K9 me3:H4K20 me3
DNA Meth. m5C:H3K9 me3
H3K27 me3:H4K20 me3
H4K20 me2:H3–5ac
H2A.Z:H3K4 me3
H4K20 me2:H3K4 me3
H3K27 me3:DNA Meth. m5C
H3K4 me3:H4–4ac
H3K27 me3:H3K4 me1
H4–4ac:H3–5ac
H3K27 me2
H3K9 me2
H4K20 me3
H3K9 me3
DNA Meth. m5C
H4–4ac
H4K20 me2
H3K27 me3
H2A.Z
H3K4 me1
H3K4 me3
H3–5ac
0.00 0.25 0.50 0.75 1.00
π̂
ZNF395 – Reverse
H3K27 me2:DNA Meth. m5C
H3K9 me2:DNA Meth. m5C
H2A.Z:H4K20 me2
H3K4 me1:H4–4ac
H3K27 me3:H3K9 me3
H3K4 me1:H2A.Z
H3K4 me1:H4K20 me2
DNA Meth. m5C:H4K20 me3
H2A.Z:H3–5ac
H2A.Z:H4–4ac
H3K4 me3:H3–5ac
H3K4 me1:H3–5ac
DNA Meth. m5C:H4K20 me2
H4K20 me2:H4–4ac
H3K9 me3:H4K20 me3
DNA Meth. m5C:H3K9 me3
H3K27 me3:H4K20 me3
H4K20 me2:H3–5ac
H2A.Z:H3K4 me3
H4K20 me2:H3K4 me3
H3K27 me3:DNA Meth. m5C
H3K4 me3:H4–4ac
H3K27 me3:H3K4 me1
H4–4ac:H3–5ac
H3K27 me2
H3K9 me2
H4K20 me3
H3K9 me3
DNA Meth. m5C
H4–4ac
H4K20 me2
H3K27 me3
H2A.Z
H3K4 me1
H3K4 me3
H3–5ac
0.00 0.25 0.50 0.75 1.00
π̂
LIN9 – Forward
H3K4 me1:H4K20 me2
H3K9 me3:H4K20 me3
H3K9 me2:DNA Meth. m5C
H3K27 me3:H4K20 me3
H3K4 me3:H3–5ac
H3K4 me1:H2A.Z
DNA Meth. m5C:H4K20 me3
H3K27 me2:DNA Meth. m5C
H3K27 me3:DNA Meth. m5C
H2A.Z:H4–4ac
H3K4 me1:H4–4ac
DNA Meth. m5C:H3K9 me3
H3K27 me3:H3K4 me1
H3K27 me3:H3K9 me3
H2A.Z:H3–5ac
H3K4 me1:H3–5ac
DNA Meth. m5C:H4K20 me2
H2A.Z:H4K20 me2
H4K20 me2:H4–4ac
H2A.Z:H3K4 me3
H3K4 me3:H4–4ac
H4K20 me2:H3K4 me3
H4K20 me2:H3–5ac
H4–4ac:H3–5ac
H3K9 me3
H3K9 me2
H3K27 me2
H3K4 me1
H4K20 me3
H3K27 me3
H2A.Z
H4K20 me2
H3K4 me3
H4–4ac
DNA Meth. m5C
H3–5ac
0.00 0.25 0.50 0.75 1.00
π̂
LIN9 – Reverse
H3K4 me1:H4K20 me2
H3K9 me3:H4K20 me3
H3K9 me2:DNA Meth. m5C
H3K27 me3:H4K20 me3
H3K4 me3:H3–5ac
H3K4 me1:H2A.Z
DNA Meth. m5C:H4K20 me3
H3K27 me2:DNA Meth. m5C
H3K27 me3:DNA Meth. m5C
H2A.Z:H4–4ac
H3K4 me1:H4–4ac
DNA Meth. m5C:H3K9 me3
H3K27 me3:H3K4 me1
H3K27 me3:H3K9 me3
H2A.Z:H3–5ac
H3K4 me1:H3–5ac
DNA Meth. m5C:H4K20 me2
H2A.Z:H4K20 me2
H4K20 me2:H4–4ac
H2A.Z:H3K4 me3
H3K4 me3:H4–4ac
H4K20 me2:H3K4 me3
H4K20 me2:H3–5ac
H4–4ac:H3–5ac
H3K9 me3
H3K9 me2
H3K27 me2
H3K4 me1
H4K20 me3
H3K27 me3
H2A.Z
H4K20 me2
H3K4 me3
H4–4ac
DNA Meth. m5C
H3–5ac
0.00 0.25 0.50 0.75 1.00
π̂
MYBL2 – Forward
H2A.Z:H4–4ac
H3K27 me3:H3K9 me3
H3K9 me3:H4K20 me3
H3K27 me2:DNA Meth. m5C
H2A.Z:H3–5ac
H3K9 me2:DNA Meth. m5C
H3K27 me3:DNA Meth. m5C
DNA Meth. m5C:H3K9 me3
H3K27 me3:H3K4 me1
H3K27 me3:H4K20 me3
H3K4 me1:H3–5ac
H3K4 me1:H4K20 me2
H2A.Z:H3K4 me3
DNA Meth. m5C:H4K20 me3
H3K4 me3:H3–5ac
H3K4 me1:H4–4ac
H2A.Z:H4K20 me2
DNA Meth. m5C:H4K20 me2
H3K4 me1:H2A.Z
H4K20 me2:H4–4ac
H3K4 me3:H4–4ac
H4K20 me2:H3K4 me3
H4K20 me2:H3–5ac
H4–4ac:H3–5ac
H3K27 me2
H3K27 me3
H3K9 me3
H3K9 me2
H4K20 me3
H3K4 me1
H3K4 me3
H2A.Z
H4K20 me2
H4–4ac
DNA Meth. m5C
H3–5ac
0.00 0.25 0.50 0.75 1.00
π̂
MYBL2 – Reverse
H2A.Z:H4–4ac
H3K27 me3:H3K9 me3
H3K9 me3:H4K20 me3
H3K27 me2:DNA Meth. m5C
H2A.Z:H3–5ac
H3K9 me2:DNA Meth. m5C
H3K27 me3:DNA Meth. m5C
DNA Meth. m5C:H3K9 me3
H3K27 me3:H3K4 me1
H3K27 me3:H4K20 me3
H3K4 me1:H3–5ac
H3K4 me1:H4K20 me2
H2A.Z:H3K4 me3
DNA Meth. m5C:H4K20 me3
H3K4 me3:H3–5ac
H3K4 me1:H4–4ac
H2A.Z:H4K20 me2
DNA Meth. m5C:H4K20 me2
H3K4 me1:H2A.Z
H4K20 me2:H4–4ac
H3K4 me3:H4–4ac
H4K20 me2:H3K4 me3
H4K20 me2:H3–5ac
H4–4ac:H3–5ac
H3K27 me2
H3K27 me3
H3K9 me3
H3K9 me2
H4K20 me3
H3K4 me1
H3K4 me3
H2A.Z
H4K20 me2
H4–4ac
DNA Meth. m5C
H3–5ac
0.00 0.25 0.50 0.75 1.00
π̂

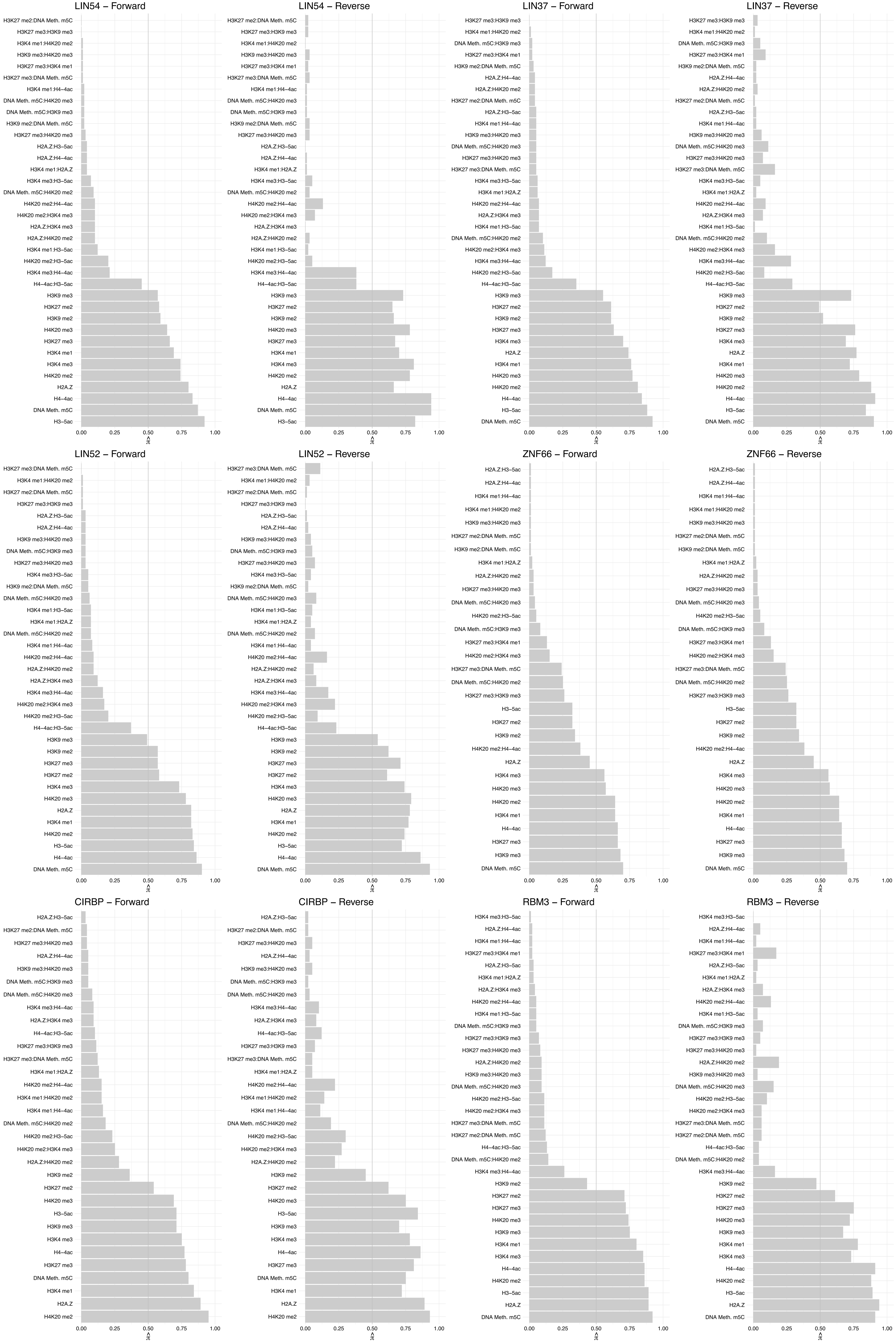
LIN54 – Forward
H3K27 me2:DNA Meth. m5C
H3K27 me3:H3K9 me3
H3K4 me1:H4K20 me2
H3K9 me3:H4K20 me3
H3K27 me3:H3K4 me1
H3K27 me3:DNA Meth. m5C
H3K4 me1:H4–4ac
DNA Meth. m5C:H4K20 me3
DNA Meth. m5C:H3K9 me3
H3K9 me2:DNA Meth. m5C
H3K27 me3:H4K20 me3
H2A.Z:H3–5ac
H2A.Z:H4–4ac
H3K4 me1:H2A.Z
H3K4 me3:H3–5ac
DNA Meth. m5C:H4K20 me2
H4K20 me2:H4–4ac
H4K20 me2:H3K4 me3
H2A.Z:H3K4 me3
H2A.Z:H4K20 me2
H3K4 me1:H3–5ac
H4K20 me2:H3–5ac
H3K4 me3:H4–4ac
H4–4ac:H3–5ac
H3K9 me3
H3K27 me2
H3K9 me2
H4K20 me3
H3K27 me3
H3K4 me1
H3K4 me3
H4K20 me2
H2A.Z
H4–4ac
DNA Meth. m5C
H3–5ac
0.00 0.25 0.50 0.75 1.00
$\hat{\pi}$
LIN54 – Reverse
H3K27 me2:DNA Meth. m5C
H3K27 me3:H3K9 me3
H3K4 me1:H4K20 me2
H3K9 me3:H4K20 me3
H3K27 me3:H3K4 me1
H3K27 me3:DNA Meth. m5C
H3K4 me1:H4–4ac
DNA Meth. m5C:H4K20 me3
DNA Meth. m5C:H3K9 me3
H3K9 me2:DNA Meth. m5C
H3K27 me3:H4K20 me3
H2A.Z:H3–5ac
H2A.Z:H4–4ac
H3K4 me1:H2A.Z
H3K4 me3:H3–5ac
DNA Meth. m5C:H4K20 me2
H4K20 me2:H4–4ac
H4K20 me2:H3K4 me3
H2A.Z:H3K4 me3
H2A.Z:H4K20 me2
H3K4 me1:H3–5ac
H4K20 me2:H3–5ac
H3K4 me3:H4–4ac
H4–4ac:H3–5ac
H3K9 me3
H3K27 me2
H3K9 me2
H4K20 me3
H3K27 me3
H3K4 me1
H3K4 me3
H4K20 me2
H2A.Z
H4–4ac
DNA Meth. m5C
H3–5ac
0.00 0.25 0.50 0.75 1.00
$\hat{\pi}$
LIN37 – Forward
H3K27 me3:H3K9 me3
H3K4 me1:H4K20 me2
DNA Meth. m5C:H3K9 me3
H3K27 me3:H3K4 me1
H3K9 me2:DNA Meth. m5C
H2A.Z:H4–4ac
H2A.Z:H4K20 me2
H3K27 me2:DNA Meth. m5C
H2A.Z:H3–5ac
H3K4 me1:H4–4ac
H3K9 me3:H4K20 me3
DNA Meth. m5C:H4K20 me3
H3K27 me3:H4K20 me3
H3K27 me3:DNA Meth. m5C
H3K4 me3:H3–5ac
H3K4 me1:H2A.Z
H4K20 me2:H4–4ac
H2A.Z:H3K4 me3
H3K4 me1:H3–5ac
DNA Meth. m5C:H4K20 me2
H4K20 me2:H3K4 me3
H3K4 me3:H4–4ac
H4K20 me2:H3–5ac
H4–4ac:H3–5ac
H3K9 me3
H3K27 me2
H3K9 me2
H3K27 me3
H3K4 me3
H2A.Z
H3K4 me1
H4K20 me3
H4K20 me2
H4–4ac
H3–5ac
DNA Meth. m5C
0.00 0.25 0.50 0.75 1.00
$\hat{\pi}$
LIN37 – Reverse
H3K27 me3:H3K9 me3
H3K4 me1:H4K20 me2
DNA Meth. m5C:H3K9 me3
H3K27 me3:H3K4 me1
H3K9 me2:DNA Meth. m5C
H2A.Z:H4–4ac
H2A.Z:H4K20 me2
H3K27 me2:DNA Meth. m5C
H2A.Z:H3–5ac
H3K4 me1:H4–4ac
H3K9 me3:H4K20 me3
DNA Meth. m5C:H4K20 me3
H3K27 me3:H4K20 me3
H3K27 me3:DNA Meth. m5C
H3K4 me3:H3–5ac
H3K4 me1:H2A.Z
H4K20 me2:H4–4ac
H2A.Z:H3K4 me3
H3K4 me1:H3–5ac
DNA Meth. m5C:H4K20 me2
H4K20 me2:H3K4 me3
H3K4 me3:H4–4ac
H4K20 me2:H3–5ac
H4–4ac:H3–5ac
H3K9 me3
H3K27 me2
H3K9 me2
H3K27 me3
H3K4 me3
H2A.Z
H3K4 me1
H4K20 me3
H4K20 me2
H4–4ac
H3–5ac
DNA Meth. m5C
0.00 0.25 0.50 0.75 1.00
$\hat{\pi}$
LIN52 – Forward
H3K27 me3:DNA Meth. m5C
H3K4 me1:H4K20 me2
H3K27 me2:DNA Meth. m5C
H3K27 me3:H3K9 me3
H2A.Z:H3–5ac
H2A.Z:H4–4ac
H3K9 me3:H4K20 me3
DNA Meth. m5C:H3K9 me3
H3K27 me3:H4K20 me3
H3K4 me3:H3–5ac
H3K9 me2:DNA Meth. m5C
DNA Meth. m5C:H4K20 me3
H3K4 me1:H3–5ac
H3K4 me1:H2A.Z
DNA Meth. m5C:H4K20 me2
H3K4 me1:H4–4ac
H4K20 me2:H4–4ac
H2A.Z:H4K20 me2
H2A.Z:H3K4 me3
H3K4 me3:H4–4ac
H4K20 me2:H3K4 me3
H4K20 me2:H3–5ac
H4–4ac:H3–5ac
H3K9 me3
H3K9 me2
H3K27 me3
H3K27 me2
H3K4 me3
H4K20 me3
H2A.Z
H3K4 me1
H4K20 me2
H3–5ac
H4–4ac
DNA Meth. m5C
0.00 0.25 0.50 0.75 1.00
$\hat{\pi}$
LIN52 – Reverse
H3K27 me3:DNA Meth. m5C
H3K4 me1:H4K20 me2
H3K27 me2:DNA Meth. m5C
H3K27 me3:H3K9 me3
H2A.Z:H3–5ac
H2A.Z:H4–4ac
H3K9 me3:H4K20 me3
DNA Meth. m5C:H3K9 me3
H3K27 me3:H4K20 me3
H3K4 me3:H3–5ac
H3K9 me2:DNA Meth. m5C
DNA Meth. m5C:H4K20 me3
H3K4 me1:H3–5ac
H3K4 me1:H2A.Z
DNA Meth. m5C:H4K20 me2
H3K4 me1:H4–4ac
H4K20 me2:H4–4ac
H2A.Z:H4K20 me2
H2A.Z:H3K4 me3
H3K4 me3:H4–4ac
H4K20 me2:H3K4 me3
H4K20 me2:H3–5ac
H4–4ac:H3–5ac
H3K9 me3
H3K9 me2
H3K27 me3
H3K27 me2
H3K4 me3
H4K20 me3
H2A.Z
H3K4 me1
H4K20 me2
H3–5ac
H4–4ac
DNA Meth. m5C
0.00 0.25 0.50 0.75 1.00
$\hat{\pi}$
ZNF66 – Forward
H2A.Z:H3–5ac
H2A.Z:H4–4ac
H3K4 me1:H4–4ac
H3K4 me1:H4K20 me2
H3K9 me3:H4K20 me3
H3K27 me2:DNA Meth. m5C
H3K9 me2:DNA Meth. m5C
H3K4 me1:H2A.Z
H2A.Z:H4K20 me2
H3K27 me3:H4K20 me3
DNA Meth. m5C:H4K20 me3
H4K20 me2:H3–5ac
DNA Meth. m5C:H3K9 me3
H3K27 me3:H3K4 me1
H4K20 me2:H3K4 me3
H3K27 me3:DNA Meth. m5C
DNA Meth. m5C:H4K20 me2
H3K27 me3:H3K9 me3
H3–5ac
H3K27 me2
H3K9 me2
H4K20 me2:H4–4ac
H2A.Z
H3K4 me3
H4K20 me3
H4K20 me2
H3K4 me1
H4–4ac
H3K27 me3
H3K9 me3
DNA Meth. m5C
0.00 0.25 0.50 0.75 1.00
$\hat{\pi}$
ZNF66 – Reverse
H2A.Z:H3–5ac
H2A.Z:H4–4ac
H3K4 me1:H4–4ac
H3K4 me1:H4K20 me2
H3K9 me3:H4K20 me3
H3K27 me2:DNA Meth. m5C
H3K9 me2:DNA Meth. m5C
H3K4 me1:H2A.Z
H2A.Z:H4K20 me2
H3K27 me3:H4K20 me3
DNA Meth. m5C:H4K20 me3
H4K20 me2:H3–5ac
DNA Meth. m5C:H3K9 me3
H3K27 me3:H3K4 me1
H4K20 me2:H3K4 me3
H3K27 me3:DNA Meth. m5C
DNA Meth. m5C:H4K20 me2
H3K27 me3:H3K9 me3
H3–5ac
H3K27 me2
H3K9 me2
H4K20 me2:H4–4ac
H2A.Z
H3K4 me3
H4K20 me3
H4K20 me2
H3K4 me1
H4–4ac
H3K27 me3
H3K9 me3
DNA Meth. m5C
0.00 0.25 0.50 0.75 1.00
$\hat{\pi}$
CIRBP – Forward
H2A.Z:H3–5ac
H3K27 me2:DNA Meth. m5C
H3K27 me3:H4K20 me3
H2A.Z:H4–4ac
H3K9 me3:H4K20 me3
DNA Meth. m5C:H3K9 me3
DNA Meth. m5C:H4K20 me3
H3K4 me3:H4–4ac
H2A.Z:H3K4 me3
H4–4ac:H3–5ac
H3K27 me3:H3K9 me3
H3K27 me3:DNA Meth. m5C
H3K4 me1:H2A.Z
H4K20 me2:H4–4ac
H3K4 me1:H4K20 me2
H3K4 me1:H4–4ac
DNA Meth. m5C:H4K20 me2
H4K20 me2:H3–5ac
H4K20 me2:H3K4 me3
H2A.Z:H4K20 me2
H3K9 me2
H3K27 me2
H4K20 me3
H3–5ac
H3K9 me3
H3K4 me3
H4–4ac
H3K27 me3
DNA Meth. m5C
H3K4 me1
H2A.Z
H4K20 me2
0.00 0.25 0.50 0.75 1.00
$\hat{\pi}$
CIRBP – Reverse
H2A.Z:H3–5ac
H3K27 me2:DNA Meth. m5C
H3K27 me3:H4K20 me3
H2A.Z:H4–4ac
H3K9 me3:H4K20 me3
DNA Meth. m5C:H3K9 me3
DNA Meth. m5C:H4K20 me3
H3K4 me3:H4–4ac
H2A.Z:H3K4 me3
H4–4ac:H3–5ac
H3K27 me3:H3K9 me3
H3K27 me3:DNA Meth. m5C
H3K4 me1:H2A.Z
H4K20 me2:H4–4ac
H3K4 me1:H4K20 me2
H3K4 me1:H4–4ac
DNA Meth. m5C:H4K20 me2
H4K20 me2:H3–5ac
H4K20 me2:H3K4 me3
H2A.Z:H4K20 me2
H3K9 me2
H3K27 me2
H4K20 me3
H3–5ac
H3K9 me3
H3K4 me3
H4–4ac
H3K27 me3
DNA Meth. m5C
H3K4 me1
H2A.Z
H4K20 me2
0.00 0.25 0.50 0.75 1.00
$\hat{\pi}$
RBM3 – Forward
H3K4 me3:H3–5ac
H2A.Z:H4–4ac
H3K4 me1:H4–4ac
H3K27 me3:H3K4 me1
H2A.Z:H3–5ac
H3K4 me1:H2A.Z
H2A.Z:H3K4 me3
H4K20 me2:H4–4ac
H3K4 me1:H3–5ac
DNA Meth. m5C:H3K9 me3
H3K27 me3:H3K9 me3
H3K27 me3:H4K20 me3
H2A.Z:H4K20 me2
H3K9 me3:H4K20 me3
DNA Meth. m5C:H4K20 me3
H4K20 me2:H3–5ac
H4K20 me2:H3K4 me3
H3K27 me3:DNA Meth. m5C
H3K27 me2:DNA Meth. m5C
H4–4ac:H3–5ac
DNA Meth. m5C:H4K20 me2
H3K4 me3:H4–4ac
H3K9 me2
H3K27 me2
H3K27 me3
H4K20 me3
H3K9 me3
H3K4 me1
H3K4 me3
H4–4ac
H4K20 me2
H3–5ac
H2A.Z
DNA Meth. m5C
0.00 0.25 0.50 0.75 1.00
$\hat{\pi}$
RBM3 – Reverse
H3K4 me3:H3–5ac
H2A.Z:H4–4ac
H3K4 me1:H4–4ac
H3K27 me3:H3K4 me1
H2A.Z:H3–5ac
H3K4 me1:H2A.Z
H2A.Z:H3K4 me3
H4K20 me2:H4–4ac
H3K4 me1:H3–5ac
DNA Meth. m5C:H3K9 me3
H3K27 me3:H3K9 me3
H3K27 me3:H4K20 me3
H2A.Z:H4K20 me2
H3K9 me3:H4K20 me3
DNA Meth. m5C:H4K20 me3
H4K20 me2:H3–5ac
H4K20 me2:H3K4 me3
H3K27 me3:DNA Meth. m5C
H3K27 me2:DNA Meth. m5C
H4–4ac:H3–5ac
DNA Meth. m5C:H4K20 me2
H3K4 me3:H4–4ac
H3K9 me2
H3K27 me2
H3K27 me3
H4K20 me3
H3K9 me3
H3K4 me1
H3K4 me3
H4–4ac
H4K20 me2
H3–5ac
H2A.Z
DNA Meth. m5C
0.00 0.25 0.50 0.75 1.00
$\hat{\pi}$

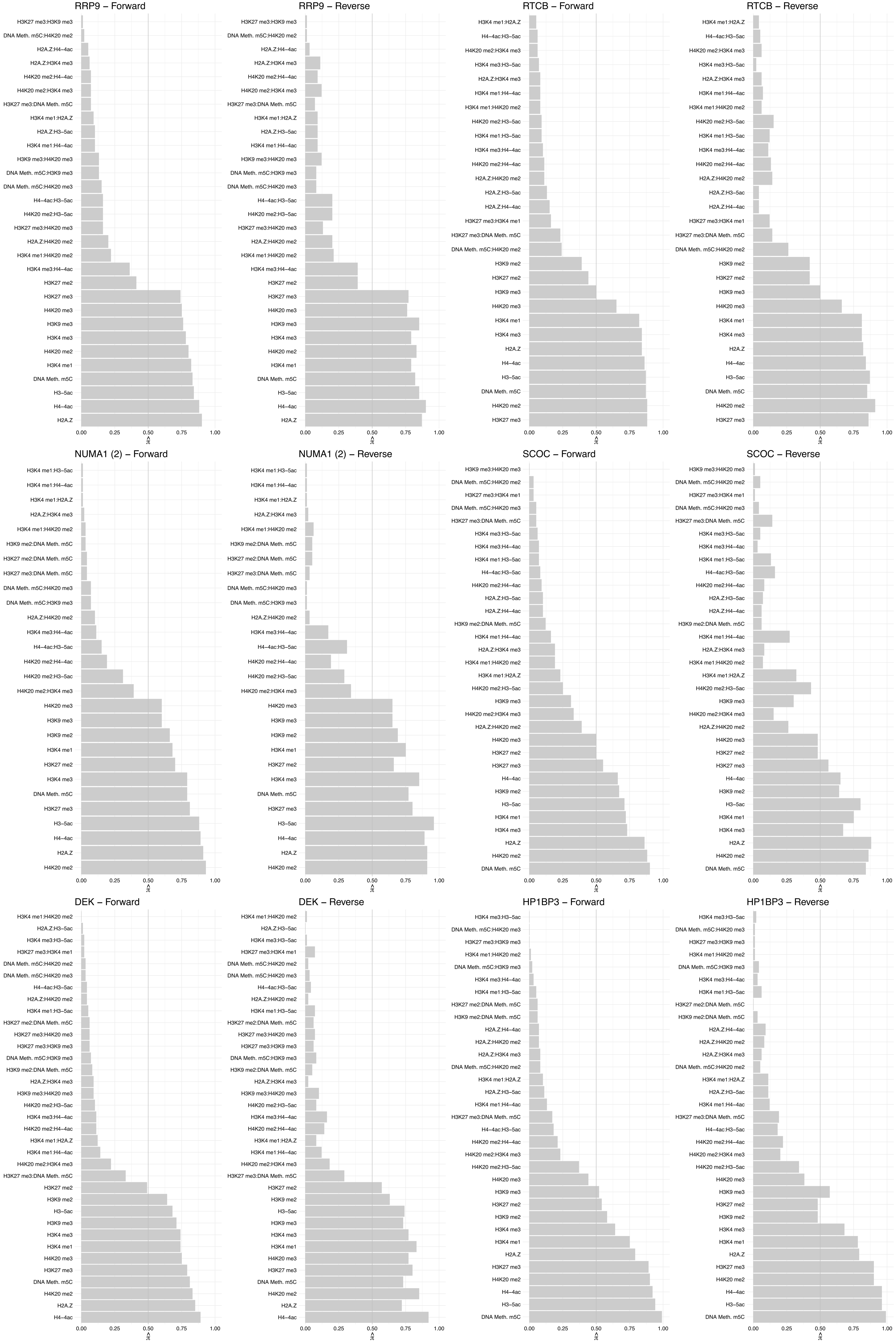
RRP9 – Forward
H3K27 me3:H3K9 me3
DNA Meth. m5C:H4K20 me2
H2A.Z:H4−4ac
H2A.Z:H3K4 me3
H4K20 me2:H4−4ac
H4K20 me2:H3K4 me3
H3K27 me3:DNA Meth. m5C
H3K4 me1:H2A.Z
H2A.Z:H3−5ac
H3K4 me1:H4−4ac
H3K9 me3:H4K20 me3
DNA Meth. m5C:H3K9 me3
DNA Meth. m5C:H4K20 me3
H4−4ac:H3−5ac
H4K20 me2:H3−5ac
H3K27 me3:H4K20 me3
H2A.Z:H4K20 me2
H3K4 me1:H4K20 me2
H3K4 me3:H4−4ac
H3K27 me2
H3K27 me3
H4K20 me3
H3K9 me3
H3K4 me3
H4K20 me2
H3K4 me1
DNA Meth. m5C
H3−5ac
H4−4ac
H2A.Z
0.00 0.25 0.50 0.75 1.00
π̂
RRP9 – Reverse
H3K27 me3:H3K9 me3
DNA Meth. m5C:H4K20 me2
H2A.Z:H4−4ac
H2A.Z:H3K4 me3
H4K20 me2:H4−4ac
H4K20 me2:H3K4 me3
H3K27 me3:DNA Meth. m5C
H3K4 me1:H2A.Z
H2A.Z:H3−5ac
H3K4 me1:H4−4ac
H3K9 me3:H4K20 me3
DNA Meth. m5C:H3K9 me3
DNA Meth. m5C:H4K20 me3
H4−4ac:H3−5ac
H4K20 me2:H3−5ac
H3K27 me3:H4K20 me3
H2A.Z:H4K20 me2
H3K4 me1:H4K20 me2
H3K4 me3:H4−4ac
H3K27 me2
H3K27 me3
H4K20 me3
H3K9 me3
H3K4 me3
H4K20 me2
H3K4 me1
DNA Meth. m5C
H3−5ac
H4−4ac
H2A.Z
0.00 0.25 0.50 0.75 1.00
π̂
RTCB – Forward
H3K4 me1:H2A.Z
H4−4ac:H3−5ac
H4K20 me2:H3K4 me3
H3K4 me3:H3−5ac
H2A.Z:H3K4 me3
H3K4 me1:H4−4ac
H3K4 me1:H4K20 me2
H4K20 me2:H3−5ac
H3K4 me1:H3−5ac
H3K4 me3:H4−4ac
H4K20 me2:H4−4ac
H2A.Z:H4K20 me2
H2A.Z:H3−5ac
H2A.Z:H4−4ac
H3K27 me3:H3K4 me1
H3K27 me3:DNA Meth. m5C
DNA Meth. m5C:H4K20 me2
H3K9 me2
H3K27 me2
H3K9 me3
H4K20 me3
H3K4 me1
H3K4 me3
H2A.Z
H4−4ac
H3−5ac
DNA Meth. m5C
H4K20 me2
H3K27 me3
0.00 0.25 0.50 0.75 1.00
π̂
RTCB – Reverse
H3K4 me1:H2A.Z
H4−4ac:H3−5ac
H4K20 me2:H3K4 me3
H3K4 me3:H3−5ac
H2A.Z:H3K4 me3
H3K4 me1:H4−4ac
H3K4 me1:H4K20 me2
H4K20 me2:H3−5ac
H3K4 me1:H3−5ac
H3K4 me3:H4−4ac
H4K20 me2:H4−4ac
H2A.Z:H4K20 me2
H2A.Z:H3−5ac
H2A.Z:H4−4ac
H3K27 me3:H3K4 me1
H3K27 me3:DNA Meth. m5C
DNA Meth. m5C:H4K20 me2
H3K9 me2
H3K27 me2
H3K9 me3
H4K20 me3
H3K4 me1
H3K4 me3
H2A.Z
H4−4ac
H3−5ac
DNA Meth. m5C
H4K20 me2
H3K27 me3
0.00 0.25 0.50 0.75 1.00
π̂
NUMA1 (2) – Forward
H3K4 me1:H3−5ac
H3K4 me1:H4−4ac
H3K4 me1:H2A.Z
H2A.Z:H3K4 me3
H3K4 me1:H4K20 me2
H3K9 me2:DNA Meth. m5C
H3K27 me2:DNA Meth. m5C
H3K27 me3:DNA Meth. m5C
DNA Meth. m5C:H4K20 me3
DNA Meth. m5C:H3K9 me3
H2A.Z:H4K20 me2
H3K4 me3:H4−4ac
H4−4ac:H3−5ac
H4K20 me2:H4−4ac
H4K20 me2:H3−5ac
H4K20 me2:H3K4 me3
H4K20 me3
H3K9 me3
H3K9 me2
H3K4 me1
H3K27 me2
H3K4 me3
DNA Meth. m5C
H3K27 me3
H3−5ac
H4−4ac
H2A.Z
H4K20 me2
0.00 0.25 0.50 0.75 1.00
π̂
NUMA1 (2) – Reverse
H3K4 me1:H3−5ac
H3K4 me1:H4−4ac
H3K4 me1:H2A.Z
H2A.Z:H3K4 me3
H3K4 me1:H4K20 me2
H3K9 me2:DNA Meth. m5C
H3K27 me2:DNA Meth. m5C
H3K27 me3:DNA Meth. m5C
DNA Meth. m5C:H4K20 me3
DNA Meth. m5C:H3K9 me3
H2A.Z:H4K20 me2
H3K4 me3:H4−4ac
H4−4ac:H3−5ac
H4K20 me2:H4−4ac
H4K20 me2:H3−5ac
H4K20 me2:H3K4 me3
H4K20 me3
H3K9 me3
H3K9 me2
H3K4 me1
H3K27 me2
H3K4 me3
DNA Meth. m5C
H3K27 me3
H3−5ac
H4−4ac
H2A.Z
H4K20 me2
0.00 0.25 0.50 0.75 1.00
π̂
SCOC – Forward
H3K9 me3:H4K20 me3
DNA Meth. m5C:H4K20 me2
H3K27 me3:H3K4 me1
DNA Meth. m5C:H4K20 me3
H3K27 me3:DNA Meth. m5C
H3K4 me3:H3−5ac
H3K4 me3:H4−4ac
H3K4 me1:H3−5ac
H4−4ac:H3−5ac
H4K20 me2:H4−4ac
H2A.Z:H3−5ac
H2A.Z:H4−4ac
H3K9 me2:DNA Meth. m5C
H3K4 me1:H4−4ac
H2A.Z:H3K4 me3
H3K4 me1:H4K20 me2
H3K4 me1:H2A.Z
H4K20 me2:H3−5ac
H3K9 me3
H4K20 me2:H3K4 me3
H2A.Z:H4K20 me2
H4K20 me3
H3K27 me2
H3K27 me3
H4−4ac
H3K9 me2
H3−5ac
H3K4 me1
H3K4 me3
H2A.Z
H4K20 me2
DNA Meth. m5C
0.00 0.25 0.50 0.75 1.00
π̂
SCOC – Reverse
H3K9 me3:H4K20 me3
DNA Meth. m5C:H4K20 me2
H3K27 me3:H3K4 me1
DNA Meth. m5C:H4K20 me3
H3K27 me3:DNA Meth. m5C
H3K4 me3:H3−5ac
H3K4 me3:H4−4ac
H3K4 me1:H3−5ac
H4−4ac:H3−5ac
H4K20 me2:H4−4ac
H2A.Z:H3−5ac
H2A.Z:H4−4ac
H3K9 me2:DNA Meth. m5C
H3K4 me1:H4−4ac
H2A.Z:H3K4 me3
H3K4 me1:H4K20 me2
H3K4 me1:H2A.Z
H4K20 me2:H3−5ac
H3K9 me3
H4K20 me2:H3K4 me3
H2A.Z:H4K20 me2
H4K20 me3
H3K27 me2
H3K27 me3
H4−4ac
H3K9 me2
H3−5ac
H3K4 me1
H3K4 me3
H2A.Z
H4K20 me2
DNA Meth. m5C
0.00 0.25 0.50 0.75 1.00
π̂
DEK – Forward
H3K4 me1:H4K20 me2
H2A.Z:H3−5ac
H3K4 me3:H3−5ac
H3K27 me3:H3K4 me1
DNA Meth. m5C:H4K20 me2
DNA Meth. m5C:H4K20 me3
H4−4ac:H3−5ac
H2A.Z:H4K20 me2
H3K4 me1:H3−5ac
H3K27 me2:DNA Meth. m5C
H3K27 me3:H4K20 me3
H3K27 me3:H3K9 me3
DNA Meth. m5C:H3K9 me3
H3K9 me2:DNA Meth. m5C
H2A.Z:H3K4 me3
H3K9 me3:H4K20 me3
H4K20 me2:H3−5ac
H3K4 me3:H4−4ac
H4K20 me2:H4−4ac
H3K4 me1:H2A.Z
H3K4 me1:H4−4ac
H4K20 me2:H3K4 me3
H3K27 me3:DNA Meth. m5C
H3K27 me2
H3K9 me2
H3−5ac
H3K9 me3
H3K4 me3
H3K4 me1
H4K20 me3
H3K27 me3
DNA Meth. m5C
H4K20 me2
H2A.Z
H4−4ac
0.00 0.25 0.50 0.75 1.00
π̂
DEK – Reverse
H3K4 me1:H4K20 me2
H2A.Z:H3−5ac
H3K4 me3:H3−5ac
H3K27 me3:H3K4 me1
DNA Meth. m5C:H4K20 me2
DNA Meth. m5C:H4K20 me3
H4−4ac:H3−5ac
H2A.Z:H4K20 me2
H3K4 me1:H3−5ac
H3K27 me2:DNA Meth. m5C
H3K27 me3:H4K20 me3
H3K27 me3:H3K9 me3
DNA Meth. m5C:H3K9 me3
H3K9 me2:DNA Meth. m5C
H2A.Z:H3K4 me3
H3K9 me3:H4K20 me3
H4K20 me2:H3−5ac
H3K4 me3:H4−4ac
H4K20 me2:H4−4ac
H3K4 me1:H2A.Z
H3K4 me1:H4−4ac
H4K20 me2:H3K4 me3
H3K27 me3:DNA Meth. m5C
H3K27 me2
H3K9 me2
H3−5ac
H3K9 me3
H3K4 me3
H3K4 me1
H4K20 me3
H3K27 me3
DNA Meth. m5C
H4K20 me2
H2A.Z
H4−4ac
0.00 0.25 0.50 0.75 1.00
π̂
HP1BP3 – Forward
H3K4 me3:H3−5ac
DNA Meth. m5C:H4K20 me3
H3K27 me3:H3K9 me3
H3K4 me1:H4K20 me2
DNA Meth. m5C:H3K9 me3
H3K4 me3:H4−4ac
H3K4 me1:H3−5ac
H3K27 me2:DNA Meth. m5C
H3K9 me2:DNA Meth. m5C
H2A.Z:H4−4ac
H2A.Z:H4K20 me2
H2A.Z:H3K4 me3
DNA Meth. m5C:H4K20 me2
H3K4 me1:H2A.Z
H2A.Z:H3−5ac
H3K4 me1:H4−4ac
H3K27 me3:DNA Meth. m5C
H4−4ac:H3−5ac
H4K20 me2:H4−4ac
H4K20 me2:H3K4 me3
H4K20 me2:H3−5ac
H4K20 me3
H3K9 me3
H3K27 me2
H3K9 me2
H3K4 me3
H3K4 me1
H2A.Z
H3K27 me3
H4K20 me2
H4−4ac
H3−5ac
DNA Meth. m5C
0.00 0.25 0.50 0.75 1.00
π̂
HP1BP3 – Reverse
H3K4 me3:H3−5ac
DNA Meth. m5C:H4K20 me3
H3K27 me3:H3K9 me3
H3K4 me1:H4K20 me2
DNA Meth. m5C:H3K9 me3
H3K4 me3:H4−4ac
H3K4 me1:H3−5ac
H3K27 me2:DNA Meth. m5C
H3K9 me2:DNA Meth. m5C
H2A.Z:H4−4ac
H2A.Z:H4K20 me2
H2A.Z:H3K4 me3
DNA Meth. m5C:H4K20 me2
H3K4 me1:H2A.Z
H2A.Z:H3−5ac
H3K4 me1:H4−4ac
H3K27 me3:DNA Meth. m5C
H4−4ac:H3−5ac
H4K20 me2:H4−4ac
H4K20 me2:H3K4 me3
H4K20 me2:H3−5ac
H4K20 me3
H3K9 me3
H3K27 me2
H3K9 me2
H3K4 me3
H3K4 me1
H2A.Z
H3K27 me3
H4K20 me2
H4−4ac
H3−5ac
DNA Meth. m5C
0.00 0.25 0.50 0.75 1.00
π̂

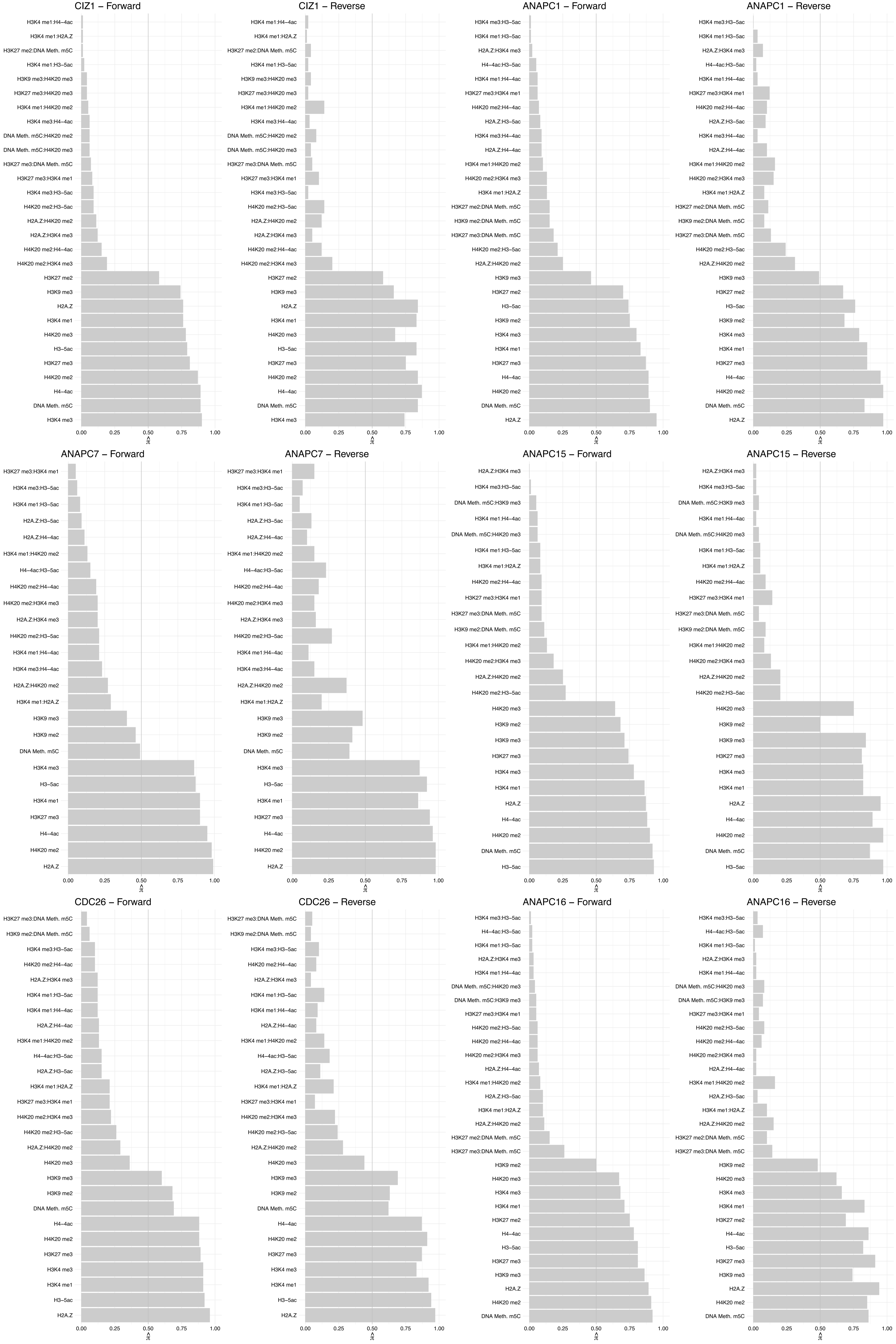
CIZ1 – Forward
H3K4 me1:H4-4ac
H3K4 me1:H2A.Z
H3K27 me2:DNA Meth. m5C
H3K4 me1:H3-5ac
H3K9 me3:H4K20 me3
H3K27 me3:H4K20 me3
H3K4 me1:H4K20 me2
H3K4 me3:H4-4ac
DNA Meth. m5C:H4K20 me2
DNA Meth. m5C:H4K20 me3
H3K27 me3:DNA Meth. m5C
H3K27 me3:H3K4 me1
H3K4 me3:H3-5ac
H4K20 me2:H3-5ac
H2A.Z:H4K20 me2
H2A.Z:H3K4 me3
H4K20 me2:H4-4ac
H4K20 me2:H3K4 me3
H3K27 me2
H3K9 me3
H2A.Z
H3K4 me1
H4K20 me3
H3-5ac
H3K27 me3
H4K20 me2
H4-4ac
DNA Meth. m5C
H3K4 me3
0.00 0.25 0.50 0.75 1.00
$\hat{\pi}$
CIZ1 – Reverse
H3K4 me1:H4-4ac
H3K4 me1:H2A.Z
H3K27 me2:DNA Meth. m5C
H3K4 me1:H3-5ac
H3K9 me3:H4K20 me3
H3K27 me3:H4K20 me3
H3K4 me1:H4K20 me2
H3K4 me3:H4-4ac
DNA Meth. m5C:H4K20 me2
DNA Meth. m5C:H4K20 me3
H3K27 me3:DNA Meth. m5C
H3K27 me3:H3K4 me1
H3K4 me3:H3-5ac
H4K20 me2:H3-5ac
H2A.Z:H4K20 me2
H2A.Z:H3K4 me3
H4K20 me2:H4-4ac
H4K20 me2:H3K4 me3
H3K27 me2
H3K9 me3
H2A.Z
H3K4 me1
H4K20 me3
H3-5ac
H3K27 me3
H4K20 me2
H4-4ac
DNA Meth. m5C
H3K4 me3
0.00 0.25 0.50 0.75 1.00
$\hat{\pi}$
ANAPC1 – Forward
H3K4 me3:H3-5ac
H3K4 me1:H3-5ac
H2A.Z:H3K4 me3
H4-4ac:H3-5ac
H3K4 me1:H4-4ac
H3K27 me3:H3K4 me1
H4K20 me2:H4-4ac
H2A.Z:H3-5ac
H3K4 me3:H4-4ac
H2A.Z:H4-4ac
H3K4 me1:H4K20 me2
H4K20 me2:H3K4 me3
H3K4 me1:H2A.Z
H3K27 me2:DNA Meth. m5C
H3K9 me2:DNA Meth. m5C
H3K27 me3:DNA Meth. m5C
H4K20 me2:H3-5ac
H2A.Z:H4K20 me2
H3K9 me3
H3K27 me2
H3-5ac
H3K9 me2
H3K4 me3
H3K4 me1
H3K27 me3
H4-4ac
H4K20 me2
DNA Meth. m5C
H2A.Z
0.00 0.25 0.50 0.75 1.00
$\hat{\pi}$
ANAPC1 – Reverse
H3K4 me3:H3-5ac
H3K4 me1:H3-5ac
H2A.Z:H3K4 me3
H4-4ac:H3-5ac
H3K4 me1:H4-4ac
H3K27 me3:H3K4 me1
H4K20 me2:H4-4ac
H2A.Z:H3-5ac
H3K4 me3:H4-4ac
H2A.Z:H4-4ac
H3K4 me1:H4K20 me2
H4K20 me2:H3K4 me3
H3K4 me1:H2A.Z
H3K27 me2:DNA Meth. m5C
H3K9 me2:DNA Meth. m5C
H3K27 me3:DNA Meth. m5C
H4K20 me2:H3-5ac
H2A.Z:H4K20 me2
H3K9 me3
H3K27 me2
H3-5ac
H3K9 me2
H3K4 me3
H3K4 me1
H3K27 me3
H4-4ac
H4K20 me2
DNA Meth. m5C
H2A.Z
0.00 0.25 0.50 0.75 1.00
$\hat{\pi}$
ANAPC7 – Forward
H3K27 me3:H3K4 me1
H3K4 me3:H3-5ac
H3K4 me1:H3-5ac
H2A.Z:H3-5ac
H2A.Z:H4-4ac
H3K4 me1:H4K20 me2
H4-4ac:H3-5ac
H4K20 me2:H4-4ac
H4K20 me2:H3K4 me3
H2A.Z:H3K4 me3
H4K20 me2:H3-5ac
H3K4 me1:H4-4ac
H3K4 me3:H4-4ac
H2A.Z:H4K20 me2
H3K4 me1:H2A.Z
H3K9 me3
H3K9 me2
DNA Meth. m5C
H3K4 me3
H3-5ac
H3K4 me1
H3K27 me3
H4-4ac
H4K20 me2
H2A.Z
0.00 0.25 0.50 0.75 1.00
$\hat{\pi}$
ANAPC7 – Reverse
H3K27 me3:H3K4 me1
H3K4 me3:H3-5ac
H3K4 me1:H3-5ac
H2A.Z:H3-5ac
H2A.Z:H4-4ac
H3K4 me1:H4K20 me2
H4-4ac:H3-5ac
H4K20 me2:H4-4ac
H4K20 me2:H3K4 me3
H2A.Z:H3K4 me3
H4K20 me2:H3-5ac
H3K4 me1:H4-4ac
H3K4 me3:H4-4ac
H2A.Z:H4K20 me2
H3K4 me1:H2A.Z
H3K9 me3
H3K9 me2
DNA Meth. m5C
H3K4 me3
H3-5ac
H3K4 me1
H3K27 me3
H4-4ac
H4K20 me2
H2A.Z
0.00 0.25 0.50 0.75 1.00
$\hat{\pi}$
ANAPC15 – Forward
H2A.Z:H3K4 me3
H3K4 me3:H3-5ac
DNA Meth. m5C:H3K9 me3
H3K4 me1:H4-4ac
DNA Meth. m5C:H4K20 me3
H3K4 me1:H3-5ac
H3K4 me1:H2A.Z
H4K20 me2:H4-4ac
H3K27 me3:H3K4 me1
H3K27 me3:DNA Meth. m5C
H3K9 me2:DNA Meth. m5C
H3K4 me1:H4K20 me2
H4K20 me2:H3K4 me3
H2A.Z:H4K20 me2
H4K20 me2:H3-5ac
H4K20 me3
H3K9 me2
H3K9 me3
H3K27 me3
H3K4 me3
H3K4 me1
H2A.Z
H4-4ac
H4K20 me2
DNA Meth. m5C
H3-5ac
0.00 0.25 0.50 0.75 1.00
$\hat{\pi}$
ANAPC15 – Reverse
H2A.Z:H3K4 me3
H3K4 me3:H3-5ac
DNA Meth. m5C:H3K9 me3
H3K4 me1:H4-4ac
DNA Meth. m5C:H4K20 me3
H3K4 me1:H3-5ac
H3K4 me1:H2A.Z
H4K20 me2:H4-4ac
H3K27 me3:H3K4 me1
H3K27 me3:DNA Meth. m5C
H3K9 me2:DNA Meth. m5C
H3K4 me1:H4K20 me2
H4K20 me2:H3K4 me3
H2A.Z:H4K20 me2
H4K20 me2:H3-5ac
H4K20 me3
H3K9 me2
H3K9 me3
H3K27 me3
H3K4 me3
H3K4 me1
H2A.Z
H4-4ac
H4K20 me2
DNA Meth. m5C
H3-5ac
0.00 0.25 0.50 0.75 1.00
$\hat{\pi}$
CDC26 – Forward
H3K27 me3:DNA Meth. m5C
H3K9 me2:DNA Meth. m5C
H3K4 me3:H3-5ac
H4K20 me2:H4-4ac
H2A.Z:H3K4 me3
H3K4 me1:H3-5ac
H3K4 me1:H4-4ac
H2A.Z:H4-4ac
H3K4 me1:H4K20 me2
H4-4ac:H3-5ac
H2A.Z:H3-5ac
H3K4 me1:H2A.Z
H3K27 me3:H3K4 me1
H4K20 me2:H3K4 me3
H4K20 me2:H3-5ac
H2A.Z:H4K20 me2
H4K20 me3
H3K9 me3
H3K9 me2
DNA Meth. m5C
H4-4ac
H4K20 me2
H3K27 me3
H3K4 me3
H3K4 me1
H3-5ac
H2A.Z
0.00 0.25 0.50 0.75 1.00
$\hat{\pi}$
CDC26 – Reverse
H3K27 me3:DNA Meth. m5C
H3K9 me2:DNA Meth. m5C
H3K4 me3:H3-5ac
H4K20 me2:H4-4ac
H2A.Z:H3K4 me3
H3K4 me1:H3-5ac
H3K4 me1:H4-4ac
H2A.Z:H4-4ac
H3K4 me1:H4K20 me2
H4-4ac:H3-5ac
H2A.Z:H3-5ac
H3K4 me1:H2A.Z
H3K27 me3:H3K4 me1
H4K20 me2:H3K4 me3
H4K20 me2:H3-5ac
H2A.Z:H4K20 me2
H4K20 me3
H3K9 me3
H3K9 me2
DNA Meth. m5C
H4-4ac
H4K20 me2
H3K27 me3
H3K4 me3
H3K4 me1
H3-5ac
H2A.Z
0.00 0.25 0.50 0.75 1.00
$\hat{\pi}$
ANAPC16 – Forward
H3K4 me3:H3-5ac
H4-4ac:H3-5ac
H3K4 me1:H3-5ac
H2A.Z:H3K4 me3
H3K4 me1:H4-4ac
DNA Meth. m5C:H4K20 me3
DNA Meth. m5C:H3K9 me3
H3K27 me3:H3K4 me1
H4K20 me2:H3-5ac
H4K20 me2:H4-4ac
H4K20 me2:H3K4 me3
H2A.Z:H4-4ac
H3K4 me1:H4K20 me2
H2A.Z:H3-5ac
H3K4 me1:H2A.Z
H2A.Z:H4K20 me2
H3K27 me2:DNA Meth. m5C
H3K27 me3:DNA Meth. m5C
H3K9 me2
H4K20 me3
H3K4 me3
H3K4 me1
H3K27 me2
H4-4ac
H3-5ac
H3K27 me3
H3K9 me3
H2A.Z
H4K20 me2
DNA Meth. m5C
0.00 0.25 0.50 0.75 1.00
$\hat{\pi}$
ANAPC16 – Reverse
H3K4 me3:H3-5ac
H4-4ac:H3-5ac
H3K4 me1:H3-5ac
H2A.Z:H3K4 me3
H3K4 me1:H4-4ac
DNA Meth. m5C:H4K20 me3
DNA Meth. m5C:H3K9 me3
H3K27 me3:H3K4 me1
H4K20 me2:H3-5ac
H4K20 me2:H4-4ac
H4K20 me2:H3K4 me3
H2A.Z:H4-4ac
H3K4 me1:H4K20 me2
H2A.Z:H3-5ac
H3K4 me1:H2A.Z
H2A.Z:H4K20 me2
H3K27 me2:DNA Meth. m5C
H3K27 me3:DNA Meth. m5C
H3K9 me2
H4K20 me3
H3K4 me3
H3K4 me1
H3K27 me2
H4-4ac
H3-5ac
H3K27 me3
H3K9 me3
H2A.Z
H4K20 me2
DNA Meth. m5C
0.00 0.25 0.50 0.75 1.00
$\hat{\pi}$

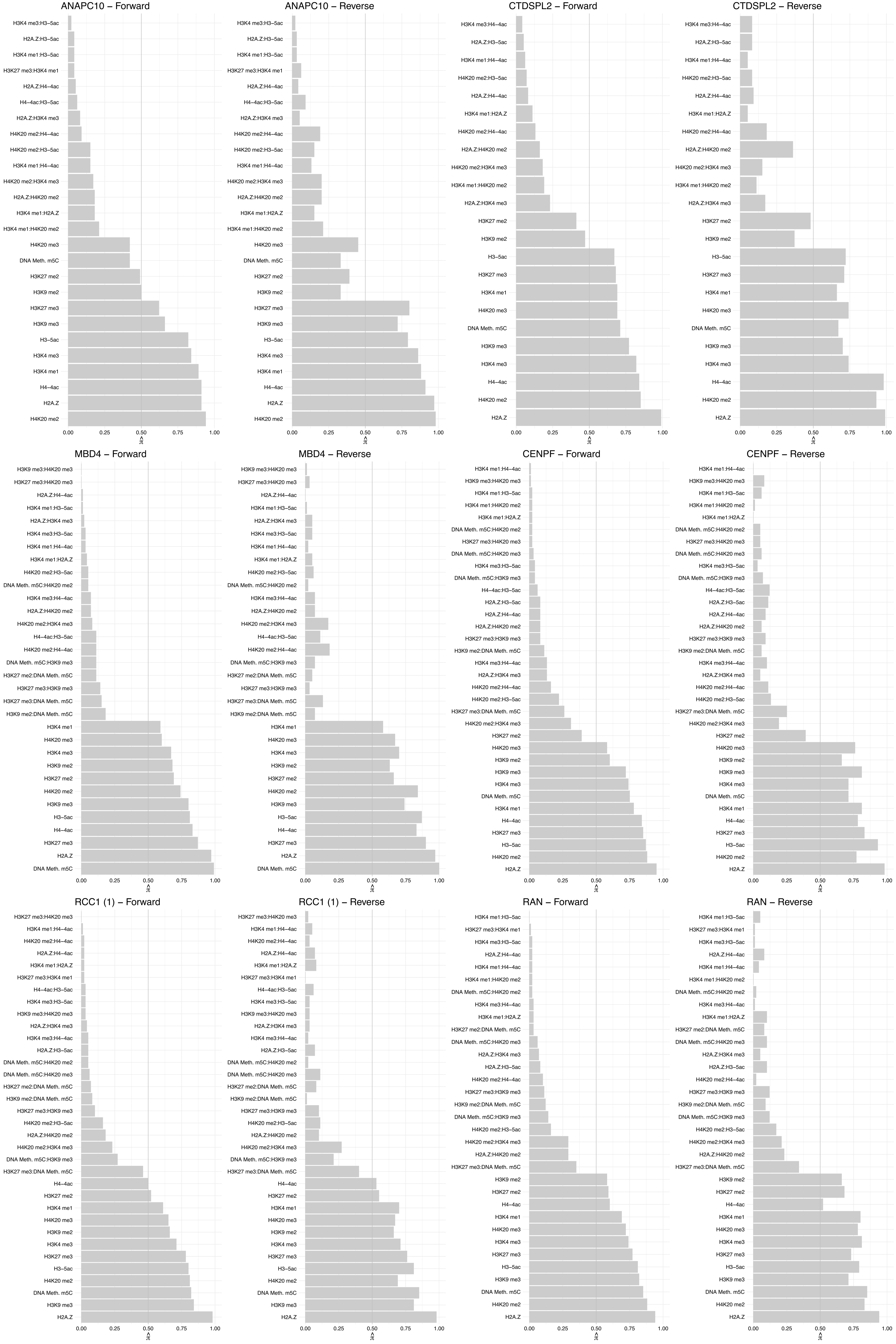

ANAPC10 – Forward
ANAPC10 – Reverse
H3K4 me3:H3–5ac
H2A.Z:H3–5ac
H3K4 me1:H3–5ac
H3K27 me3:H3K4 me1
H2A.Z:H4–4ac
H4–4ac:H3–5ac
H2A.Z:H3K4 me3
H4K20 me2:H4–4ac
H4K20 me2:H3–5ac
H3K4 me1:H4–4ac
H4K20 me2:H3K4 me3
H2A.Z:H4K20 me2
H3K4 me1:H2A.Z
H3K4 me1:H4K20 me2
H4K20 me3
DNA Meth. m5C
H3K27 me2
H3K9 me2
H3K27 me3
H3K9 me3
H3–5ac
H3K4 me3
H3K4 me1
H4–4ac
H2A.Z
H4K20 me2
CTDSPL2 – Forward
CTDSPL2 – Reverse
H3K4 me3:H4–4ac
H4K20 me2:H3–5ac
H3K4 me1:H2A.Z
H2A.Z:H4K20 me2
H4K20 me2:H3K4 me3
MBD4 – Forward
MBD4 – Reverse
H3K9 me3:H4K20 me3
H3K27 me3:H4K20 me3
DNA Meth. m5C:H4K20 me2
H3K4 me3:H4–4ac
DNA Meth. m5C:H3K9 me3
H3K27 me2:DNA Meth. m5C
H3K27 me3:H3K9 me3
H3K27 me3:DNA Meth. m5C
H3K9 me2:DNA Meth. m5C
CENPF – Forward
CENPF – Reverse
DNA Meth. m5C:H4K20 me3
H3K27 me3:H4K20 me3
RCC1 (1) – Forward
RCC1 (1) – Reverse
RAN – Forward
RAN – Reverse
0.00
0.25
0.50
0.75
1.00
$\hat{\pi}$

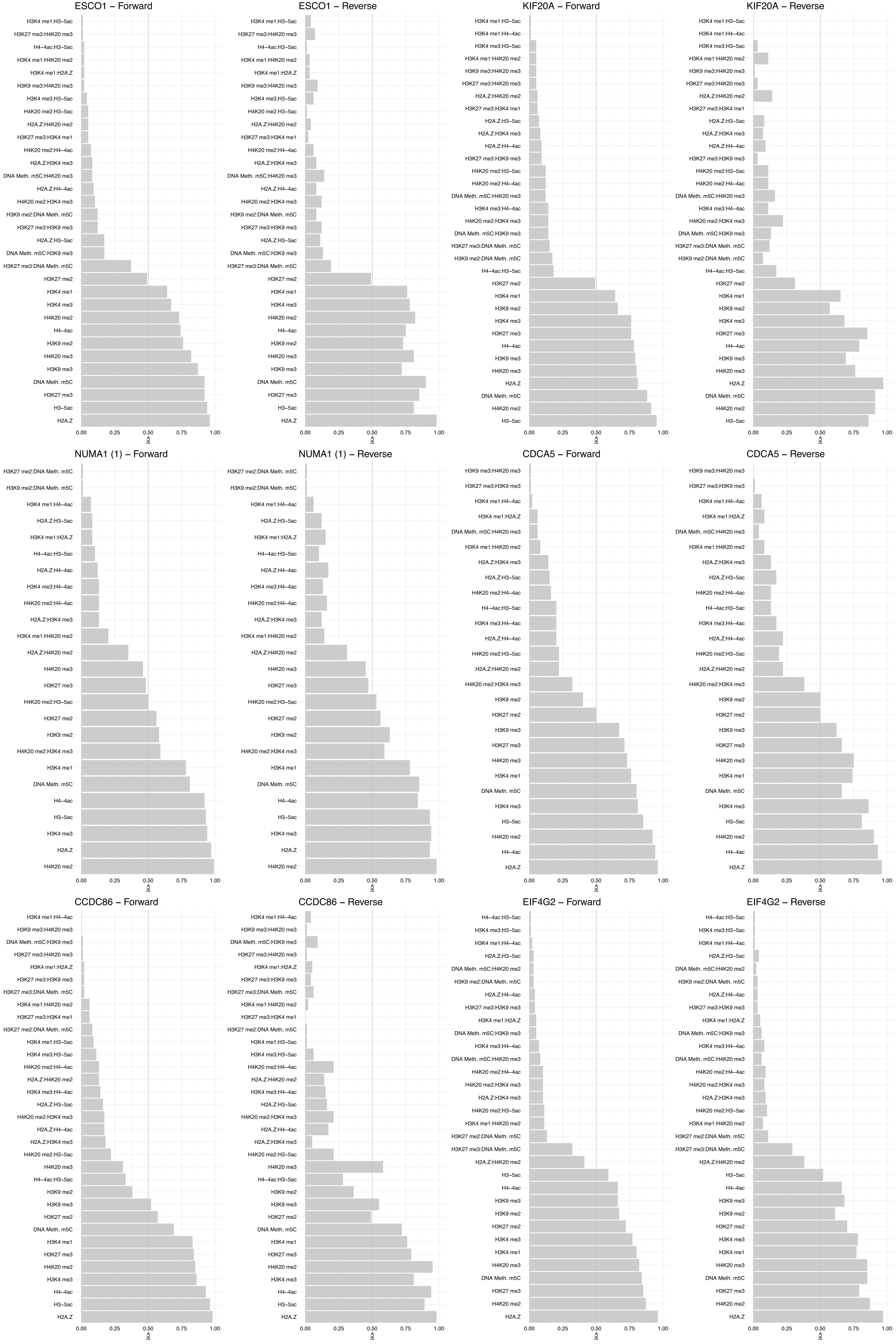

ESCO1 – Forward
ESCO1 – Reverse
H3K4 me1:H3−5ac
H3K27 me3:H4K20 me3
H4−4ac:H3−5ac
H3K4 me1:H4K20 me2
H3K4 me1:H2A.Z
H3K9 me3:H4K20 me3
H3K4 me3:H3−5ac
H4K20 me2:H3−5ac
H2A.Z:H4K20 me2
H3K27 me3:H3K4 me1
H4K20 me2:H4−4ac
H2A.Z:H3K4 me3
DNA Meth. m5C:H4K20 me3
H2A.Z:H4−4ac
H4K20 me2:H3K4 me3
H3K9 me2:DNA Meth. m5C
H3K27 me3:H3K9 me3
H2A.Z:H3−5ac
DNA Meth. m5C:H3K9 me3
H3K27 me3:DNA Meth. m5C
H3K27 me2
H3K4 me1
H3K4 me3
H4K20 me2
H4−4ac
H3K9 me2
H4K20 me3
H3K9 me3
DNA Meth. m5C
H3K27 me3
H3−5ac
H2A.Z
KIF20A – Forward
KIF20A – Reverse
H3K4 me1:H3−5ac
H3K4 me1:H4−4ac
H3K4 me3:H3−5ac
H3K4 me1:H4K20 me2
H3K9 me3:H4K20 me3
H3K27 me3:H4K20 me3
H2A.Z:H4K20 me2
H3K27 me3:H3K4 me1
H2A.Z:H3−5ac
H2A.Z:H3K4 me3
H2A.Z:H4−4ac
H3K27 me3:H3K9 me3
H4K20 me2:H3−5ac
H4K20 me2:H4−4ac
DNA Meth. m5C:H4K20 me3
H3K4 me3:H4−4ac
H4K20 me2:H3K4 me3
DNA Meth. m5C:H3K9 me3
H3K27 me3:DNA Meth. m5C
H3K9 me2:DNA Meth. m5C
H4−4ac:H3−5ac
H3K27 me2
H3K4 me1
H3K9 me2
H3K4 me3
H3K27 me3
H4−4ac
H3K9 me3
H4K20 me3
H2A.Z
DNA Meth. m5C
H4K20 me2
H3−5ac
NUMA1 (1) – Forward
NUMA1 (1) – Reverse
H3K27 me2:DNA Meth. m5C
H3K9 me2:DNA Meth. m5C
H3K4 me1:H4−4ac
H2A.Z:H3−5ac
H3K4 me1:H2A.Z
H4−4ac:H3−5ac
H2A.Z:H4−4ac
H3K4 me3:H4−4ac
H4K20 me2:H4−4ac
H2A.Z:H3K4 me3
H3K4 me1:H4K20 me2
H2A.Z:H4K20 me2
H4K20 me3
H3K27 me3
H4K20 me2:H3−5ac
H3K27 me2
H3K9 me2
H4K20 me2:H3K4 me3
H3K4 me1
DNA Meth. m5C
H4−4ac
H3−5ac
H3K4 me3
H2A.Z
H4K20 me2
CDCA5 – Forward
CDCA5 – Reverse
H3K9 me3:H4K20 me3
H3K27 me3:H3K9 me3
H3K4 me1:H4−4ac
H3K4 me1:H2A.Z
DNA Meth. m5C:H4K20 me3
H3K4 me1:H4K20 me2
H2A.Z:H3K4 me3
H2A.Z:H3−5ac
H4K20 me2:H4−4ac
H4−4ac:H3−5ac
H3K4 me3:H4−4ac
H2A.Z:H4−4ac
H4K20 me2:H3−5ac
H2A.Z:H4K20 me2
H4K20 me2:H3K4 me3
H3K9 me2
H3K27 me2
H3K9 me3
H3K27 me3
H4K20 me3
H3K4 me1
DNA Meth. m5C
H3K4 me3
H3−5ac
H4K20 me2
H4−4ac
H2A.Z
CCDC86 – Forward
CCDC86 – Reverse
H3K4 me1:H4−4ac
H3K9 me3:H4K20 me3
DNA Meth. m5C:H3K9 me3
H3K27 me3:H4K20 me3
H3K4 me1:H2A.Z
H3K27 me3:H3K9 me3
H3K27 me3:DNA Meth. m5C
H3K4 me1:H4K20 me2
H3K27 me3:H3K4 me1
H3K27 me2:DNA Meth. m5C
H3K4 me1:H3−5ac
H3K4 me3:H3−5ac
H4K20 me2:H4−4ac
H2A.Z:H4K20 me2
H3K4 me3:H4−4ac
H2A.Z:H3−5ac
H4K20 me2:H3K4 me3
H2A.Z:H4−4ac
H2A.Z:H3K4 me3
H4K20 me2:H3−5ac
H4K20 me3
H4−4ac:H3−5ac
H3K9 me2
H3K9 me3
H3K27 me2
DNA Meth. m5C
H3K4 me1
H3K27 me3
H4K20 me2
H3K4 me3
H4−4ac
H3−5ac
H2A.Z
EIF4G2 – Forward
EIF4G2 – Reverse
H4−4ac:H3−5ac
H3K4 me3:H3−5ac
H3K4 me1:H4−4ac
H2A.Z:H3−5ac
DNA Meth. m5C:H4K20 me2
H3K9 me2:DNA Meth. m5C
H2A.Z:H4−4ac
H3K27 me3:H3K9 me3
H3K4 me1:H2A.Z
DNA Meth. m5C:H3K9 me3
H3K4 me3:H4−4ac
DNA Meth. m5C:H4K20 me3
H4K20 me2:H4−4ac
H4K20 me2:H3K4 me3
H2A.Z:H3K4 me3
H4K20 me2:H3−5ac
H3K4 me1:H4K20 me2
H3K27 me2:DNA Meth. m5C
H3K27 me3:DNA Meth. m5C
H2A.Z:H4K20 me2
H3−5ac
H4−4ac
H3K9 me3
H3K9 me2
H3K27 me2
H3K4 me3
H3K4 me1
H4K20 me3
DNA Meth. m5C
H3K27 me3
H4K20 me2
H2A.Z
0.00
0.25
0.50
0.75
1.00
π̂

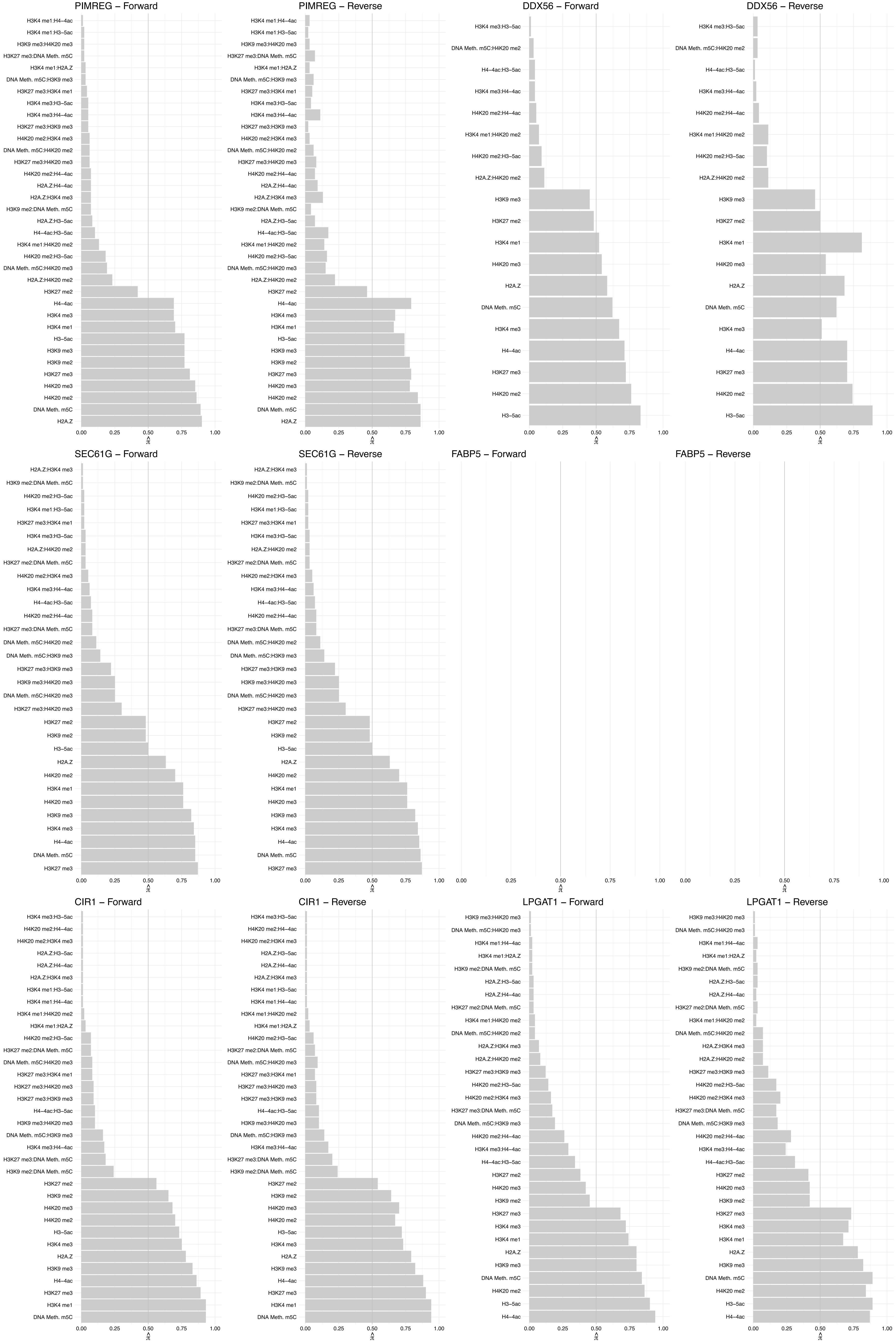

PIMREG – Forward
H3K4 me1:H4–4ac
H3K4 me1:H3–5ac
H3K9 me3:H4K20 me3
H3K27 me3:DNA Meth. m5C
H3K4 me1:H2A.Z
DNA Meth. m5C:H3K9 me3
H3K27 me3:H3K4 me1
H3K4 me3:H3–5ac
H3K4 me3:H4–4ac
H3K27 me3:H3K9 me3
H4K20 me2:H3K4 me3
DNA Meth. m5C:H4K20 me2
H3K27 me3:H4K20 me3
H4K20 me2:H4–4ac
H2A.Z:H4–4ac
H2A.Z:H3K4 me3
H3K9 me2:DNA Meth. m5C
H2A.Z:H3–5ac
H4–4ac:H3–5ac
H3K4 me1:H4K20 me2
H4K20 me2:H3–5ac
DNA Meth. m5C:H4K20 me3
H2A.Z:H4K20 me2
H3K27 me2
H4–4ac
H3K4 me3
H3K4 me1
H3–5ac
H3K9 me3
H3K9 me2
H3K27 me3
H4K20 me3
H4K20 me2
DNA Meth. m5C
H2A.Z
0.00 0.25 0.50 0.75 1.00
π̂
PIMREG – Reverse
DDX56 – Forward
H3K4 me3:H3–5ac
DNA Meth. m5C:H4K20 me2
H4–4ac:H3–5ac
H3K4 me3:H4–4ac
H4K20 me2:H4–4ac
H3K4 me1:H4K20 me2
H4K20 me2:H3–5ac
H2A.Z:H4K20 me2
H3K9 me3
H3K27 me2
H3K4 me1
H4K20 me3
H2A.Z
DNA Meth. m5C
H3K4 me3
H4–4ac
H3K27 me3
H4K20 me2
H3–5ac
DDX56 – Reverse
SEC61G – Forward
H2A.Z:H3K4 me3
H3K9 me2:DNA Meth. m5C
H4K20 me2:H3–5ac
H3K4 me1:H3–5ac
H3K27 me3:H3K4 me1
H3K4 me3:H3–5ac
H2A.Z:H4K20 me2
H3K27 me2:DNA Meth. m5C
H4K20 me2:H3K4 me3
H3K4 me3:H4–4ac
H4–4ac:H3–5ac
H4K20 me2:H4–4ac
H3K27 me3:DNA Meth. m5C
DNA Meth. m5C:H4K20 me2
DNA Meth. m5C:H3K9 me3
H3K27 me3:H3K9 me3
H3K9 me3:H4K20 me3
DNA Meth. m5C:H4K20 me3
H3K27 me3:H4K20 me3
H3K27 me2
H3K9 me2
H3–5ac
H2A.Z
H4K20 me2
H3K4 me1
H4K20 me3
H3K9 me3
H3K4 me3
H4–4ac
DNA Meth. m5C
H3K27 me3
SEC61G – Reverse
FABP5 – Forward
FABP5 – Reverse
CIR1 – Forward
H3K4 me3:H3–5ac
H4K20 me2:H4–4ac
H4K20 me2:H3K4 me3
H2A.Z:H3–5ac
H2A.Z:H4–4ac
H2A.Z:H3K4 me3
H3K4 me1:H3–5ac
H3K4 me1:H4–4ac
H3K4 me1:H4K20 me2
H3K4 me1:H2A.Z
H4K20 me2:H3–5ac
H3K27 me2:DNA Meth. m5C
DNA Meth. m5C:H4K20 me3
H3K27 me3:H3K4 me1
H3K27 me3:H4K20 me3
H3K27 me3:H3K9 me3
H4–4ac:H3–5ac
H3K9 me3:H4K20 me3
DNA Meth. m5C:H3K9 me3
H3K4 me3:H4–4ac
H3K27 me3:DNA Meth. m5C
H3K9 me2:DNA Meth. m5C
H3K27 me2
H3K9 me2
H4K20 me3
H4K20 me2
H3–5ac
H3K4 me3
H2A.Z
H3K9 me3
H4–4ac
H3K27 me3
H3K4 me1
DNA Meth. m5C
CIR1 – Reverse
LPGAT1 – Forward
H3K9 me3:H4K20 me3
DNA Meth. m5C:H4K20 me3
H3K4 me1:H4–4ac
H3K4 me1:H2A.Z
H3K9 me2:DNA Meth. m5C
H2A.Z:H3–5ac
H2A.Z:H4–4ac
H3K27 me2:DNA Meth. m5C
H3K4 me1:H4K20 me2
DNA Meth. m5C:H4K20 me2
H2A.Z:H3K4 me3
H2A.Z:H4K20 me2
H3K27 me3:H3K9 me3
H4K20 me2:H3–5ac
H4K20 me2:H3K4 me3
H3K27 me3:DNA Meth. m5C
DNA Meth. m5C:H3K9 me3
H4K20 me2:H4–4ac
H3K4 me3:H4–4ac
H4–4ac:H3–5ac
H3K27 me2
H4K20 me3
H3K9 me2
H3K27 me3
H3K4 me3
H3K4 me1
H2A.Z
H3K9 me3
DNA Meth. m5C
H4K20 me2
H3–5ac
H4–4ac
LPGAT1 – Reverse

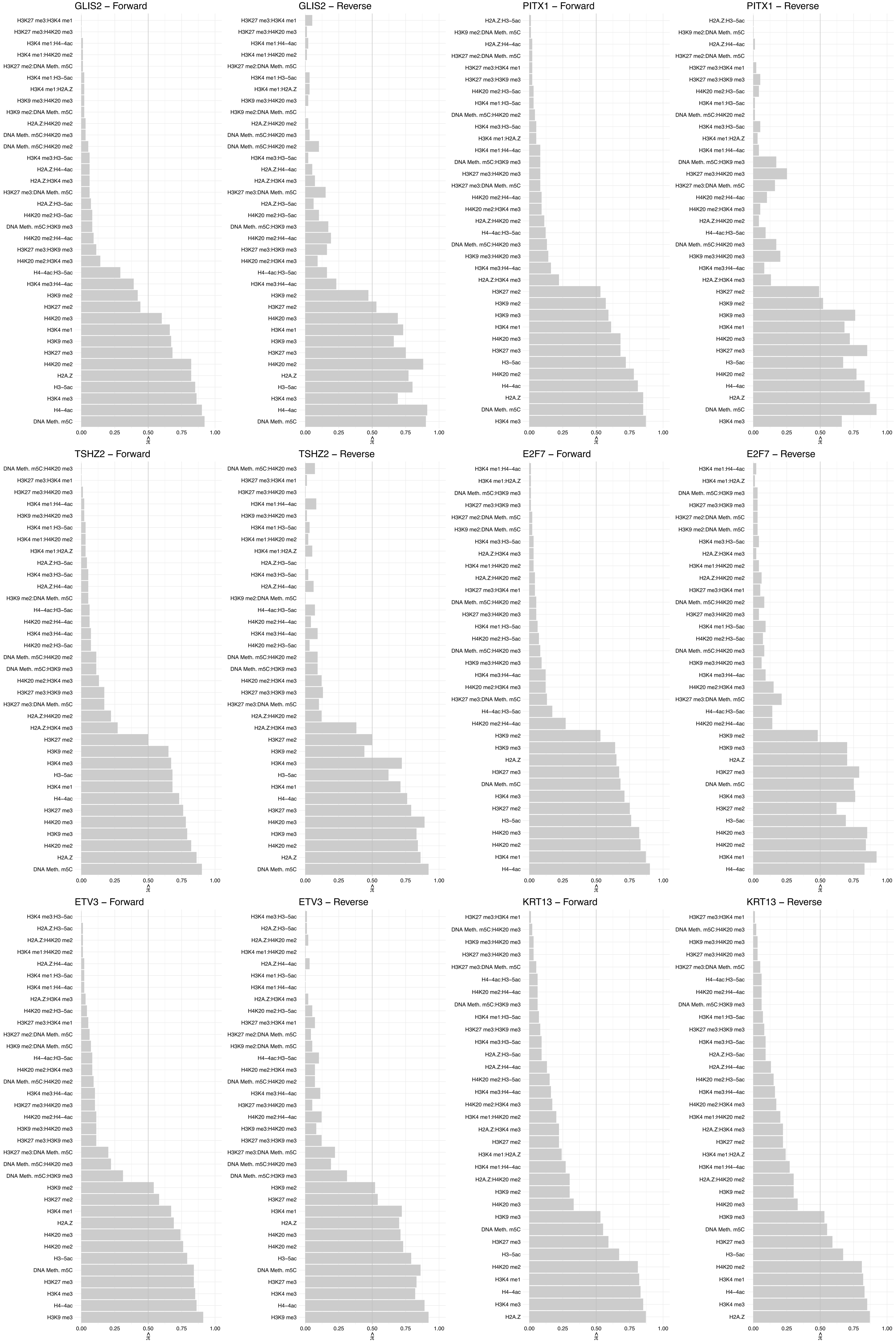
GLIS2 – Forward
H3K27 me3:H3K4 me1
H3K27 me3:H4K20 me3
H3K4 me1:H4–4ac
H3K4 me1:H4K20 me2
H3K27 me2:DNA Meth. m5C
H3K4 me1:H3–5ac
H3K4 me1:H2A.Z
H3K9 me3:H4K20 me3
H3K9 me2:DNA Meth. m5C
H2A.Z:H4K20 me2
DNA Meth. m5C:H4K20 me3
DNA Meth. m5C:H4K20 me2
H3K4 me3:H3–5ac
H2A.Z:H4–4ac
H2A.Z:H3K4 me3
H3K27 me3:DNA Meth. m5C
H2A.Z:H3–5ac
H4K20 me2:H3–5ac
DNA Meth. m5C:H3K9 me3
H4K20 me2:H4–4ac
H3K27 me3:H3K9 me3
H4K20 me2:H3K4 me3
H4–4ac:H3–5ac
H3K4 me3:H4–4ac
H3K9 me2
H3K27 me2
H4K20 me3
H3K4 me1
H3K9 me3
H3K27 me3
H4K20 me2
H2A.Z
H3–5ac
H3K4 me3
H4–4ac
DNA Meth. m5C
0.00 0.25 0.50 0.75 1.00
$\hat{\pi}$
GLIS2 – Reverse
H3K27 me3:H3K4 me1
H3K27 me3:H4K20 me3
H3K4 me1:H4–4ac
H3K4 me1:H4K20 me2
H3K27 me2:DNA Meth. m5C
H3K4 me1:H3–5ac
H3K4 me1:H2A.Z
H3K9 me3:H4K20 me3
H3K9 me2:DNA Meth. m5C
H2A.Z:H4K20 me2
DNA Meth. m5C:H4K20 me3
DNA Meth. m5C:H4K20 me2
H3K4 me3:H3–5ac
H2A.Z:H4–4ac
H2A.Z:H3K4 me3
H3K27 me3:DNA Meth. m5C
H2A.Z:H3–5ac
H4K20 me2:H3–5ac
DNA Meth. m5C:H3K9 me3
H4K20 me2:H4–4ac
H3K27 me3:H3K9 me3
H4K20 me2:H3K4 me3
H4–4ac:H3–5ac
H3K4 me3:H4–4ac
H3K9 me2
H3K27 me2
H4K20 me3
H3K4 me1
H3K9 me3
H3K27 me3
H4K20 me2
H2A.Z
H3–5ac
H3K4 me3
H4–4ac
DNA Meth. m5C
0.00 0.25 0.50 0.75 1.00
$\hat{\pi}$
PITX1 – Forward
H2A.Z:H3–5ac
H3K9 me2:DNA Meth. m5C
H2A.Z:H4–4ac
H3K27 me2:DNA Meth. m5C
H3K27 me3:H3K4 me1
H3K27 me3:H3K9 me3
H4K20 me2:H3–5ac
H3K4 me1:H3–5ac
DNA Meth. m5C:H4K20 me2
H3K4 me3:H3–5ac
H3K4 me1:H2A.Z
H3K4 me1:H4–4ac
DNA Meth. m5C:H3K9 me3
H3K27 me3:H4K20 me3
H3K27 me3:DNA Meth. m5C
H4K20 me2:H4–4ac
H4K20 me2:H3K4 me3
H2A.Z:H4K20 me2
H4–4ac:H3–5ac
DNA Meth. m5C:H4K20 me3
H3K9 me3:H4K20 me3
H3K4 me3:H4–4ac
H2A.Z:H3K4 me3
H3K27 me2
H3K9 me2
H3K9 me3
H3K4 me1
H4K20 me3
H3K27 me3
H3–5ac
H4K20 me2
H4–4ac
H2A.Z
DNA Meth. m5C
H3K4 me3
0.00 0.25 0.50 0.75 1.00
$\hat{\pi}$
PITX1 – Reverse
H2A.Z:H3–5ac
H3K9 me2:DNA Meth. m5C
H2A.Z:H4–4ac
H3K27 me2:DNA Meth. m5C
H3K27 me3:H3K4 me1
H3K27 me3:H3K9 me3
H4K20 me2:H3–5ac
H3K4 me1:H3–5ac
DNA Meth. m5C:H4K20 me2
H3K4 me3:H3–5ac
H3K4 me1:H2A.Z
H3K4 me1:H4–4ac
DNA Meth. m5C:H3K9 me3
H3K27 me3:H4K20 me3
H3K27 me3:DNA Meth. m5C
H4K20 me2:H4–4ac
H4K20 me2:H3K4 me3
H2A.Z:H4K20 me2
H4–4ac:H3–5ac
DNA Meth. m5C:H4K20 me3
H3K9 me3:H4K20 me3
H3K4 me3:H4–4ac
H2A.Z:H3K4 me3
H3K27 me2
H3K9 me2
H3K9 me3
H3K4 me1
H4K20 me3
H3K27 me3
H3–5ac
H4K20 me2
H4–4ac
H2A.Z
DNA Meth. m5C
H3K4 me3
0.00 0.25 0.50 0.75 1.00
$\hat{\pi}$
TSHZ2 – Forward
DNA Meth. m5C:H4K20 me3
H3K27 me3:H3K4 me1
H3K27 me3:H4K20 me3
H3K4 me1:H4–4ac
H3K9 me3:H4K20 me3
H3K4 me1:H3–5ac
H3K4 me1:H4K20 me2
H3K4 me1:H2A.Z
H2A.Z:H3–5ac
H3K4 me3:H3–5ac
H2A.Z:H4–4ac
H3K9 me2:DNA Meth. m5C
H4–4ac:H3–5ac
H4K20 me2:H4–4ac
H3K4 me3:H4–4ac
H4K20 me2:H3–5ac
DNA Meth. m5C:H4K20 me2
DNA Meth. m5C:H3K9 me3
H4K20 me2:H3K4 me3
H3K27 me3:H3K9 me3
H3K27 me3:DNA Meth. m5C
H2A.Z:H4K20 me2
H2A.Z:H3K4 me3
H3K27 me2
H3K9 me2
H3K4 me3
H3–5ac
H3K4 me1
H4–4ac
H3K27 me3
H4K20 me3
H3K9 me3
H4K20 me2
H2A.Z
DNA Meth. m5C
0.00 0.25 0.50 0.75 1.00
$\hat{\pi}$
TSHZ2 – Reverse
DNA Meth. m5C:H4K20 me3
H3K27 me3:H3K4 me1
H3K27 me3:H4K20 me3
H3K4 me1:H4–4ac
H3K9 me3:H4K20 me3
H3K4 me1:H3–5ac
H3K4 me1:H4K20 me2
H3K4 me1:H2A.Z
H2A.Z:H3–5ac
H3K4 me3:H3–5ac
H2A.Z:H4–4ac
H3K9 me2:DNA Meth. m5C
H4–4ac:H3–5ac
H4K20 me2:H4–4ac
H3K4 me3:H4–4ac
H4K20 me2:H3–5ac
DNA Meth. m5C:H4K20 me2
DNA Meth. m5C:H3K9 me3
H4K20 me2:H3K4 me3
H3K27 me3:H3K9 me3
H3K27 me3:DNA Meth. m5C
H2A.Z:H4K20 me2
H2A.Z:H3K4 me3
H3K27 me2
H3K9 me2
H3K4 me3
H3–5ac
H3K4 me1
H4–4ac
H3K27 me3
H4K20 me3
H3K9 me3
H4K20 me2
H2A.Z
DNA Meth. m5C
0.00 0.25 0.50 0.75 1.00
$\hat{\pi}$
E2F7 – Forward
H3K4 me1:H4–4ac
H3K4 me1:H2A.Z
DNA Meth. m5C:H3K9 me3
H3K27 me3:H3K9 me3
H3K27 me2:DNA Meth. m5C
H3K9 me2:DNA Meth. m5C
H3K4 me3:H3–5ac
H2A.Z:H3K4 me3
H3K4 me1:H4K20 me2
H2A.Z:H4K20 me2
H3K27 me3:H3K4 me1
DNA Meth. m5C:H4K20 me2
H3K27 me3:H4K20 me3
H3K4 me1:H3–5ac
H4K20 me2:H3–5ac
DNA Meth. m5C:H4K20 me3
H3K9 me3:H4K20 me3
H3K4 me3:H4–4ac
H4K20 me2:H3K4 me3
H3K27 me3:DNA Meth. m5C
H4–4ac:H3–5ac
H4K20 me2:H4–4ac
H3K9 me2
H3K9 me3
H2A.Z
H3K27 me3
DNA Meth. m5C
H3K4 me3
H3K27 me2
H3–5ac
H4K20 me3
H4K20 me2
H3K4 me1
H4–4ac
0.00 0.25 0.50 0.75 1.00
$\hat{\pi}$
E2F7 – Reverse
H3K4 me1:H4–4ac
H3K4 me1:H2A.Z
DNA Meth. m5C:H3K9 me3
H3K27 me3:H3K9 me3
H3K27 me2:DNA Meth. m5C
H3K9 me2:DNA Meth. m5C
H3K4 me3:H3–5ac
H2A.Z:H3K4 me3
H3K4 me1:H4K20 me2
H2A.Z:H4K20 me2
H3K27 me3:H3K4 me1
DNA Meth. m5C:H4K20 me2
H3K27 me3:H4K20 me3
H3K4 me1:H3–5ac
H4K20 me2:H3–5ac
DNA Meth. m5C:H4K20 me3
H3K9 me3:H4K20 me3
H3K4 me3:H4–4ac
H4K20 me2:H3K4 me3
H3K27 me3:DNA Meth. m5C
H4–4ac:H3–5ac
H4K20 me2:H4–4ac
H3K9 me2
H3K9 me3
H2A.Z
H3K27 me3
DNA Meth. m5C
H3K4 me3
H3K27 me2
H3–5ac
H4K20 me3
H4K20 me2
H3K4 me1
H4–4ac
0.00 0.25 0.50 0.75 1.00
$\hat{\pi}$
ETV3 – Forward
H3K4 me3:H3–5ac
H2A.Z:H3–5ac
H2A.Z:H4K20 me2
H3K4 me1:H4K20 me2
H2A.Z:H4–4ac
H3K4 me1:H3–5ac
H3K4 me1:H4–4ac
H2A.Z:H3K4 me3
H4K20 me2:H3–5ac
H3K27 me3:H3K4 me1
H3K27 me2:DNA Meth. m5C
H3K9 me2:DNA Meth. m5C
H4–4ac:H3–5ac
H4K20 me2:H3K4 me3
DNA Meth. m5C:H4K20 me2
H3K4 me3:H4–4ac
H3K27 me3:H4K20 me3
H4K20 me2:H4–4ac
H3K9 me3:H4K20 me3
H3K27 me3:H3K9 me3
H3K27 me3:DNA Meth. m5C
DNA Meth. m5C:H4K20 me3
DNA Meth. m5C:H3K9 me3
H3K9 me2
H3K27 me2
H3K4 me1
H2A.Z
H4K20 me3
H4K20 me2
H3–5ac
DNA Meth. m5C
H3K27 me3
H3K4 me3
H4–4ac
H3K9 me3
0.00 0.25 0.50 0.75 1.00
$\hat{\pi}$
ETV3 – Reverse
H3K4 me3:H3–5ac
H2A.Z:H3–5ac
H2A.Z:H4K20 me2
H3K4 me1:H4K20 me2
H2A.Z:H4–4ac
H3K4 me1:H3–5ac
H3K4 me1:H4–4ac
H2A.Z:H3K4 me3
H4K20 me2:H3–5ac
H3K27 me3:H3K4 me1
H3K27 me2:DNA Meth. m5C
H3K9 me2:DNA Meth. m5C
H4–4ac:H3–5ac
H4K20 me2:H3K4 me3
DNA Meth. m5C:H4K20 me2
H3K4 me3:H4–4ac
H3K27 me3:H4K20 me3
H4K20 me2:H4–4ac
H3K9 me3:H4K20 me3
H3K27 me3:H3K9 me3
H3K27 me3:DNA Meth. m5C
DNA Meth. m5C:H4K20 me3
DNA Meth. m5C:H3K9 me3
H3K9 me2
H3K27 me2
H3K4 me1
H2A.Z
H4K20 me3
H4K20 me2
H3–5ac
DNA Meth. m5C
H3K27 me3
H3K4 me3
H4–4ac
H3K9 me3
0.00 0.25 0.50 0.75 1.00
$\hat{\pi}$
KRT13 – Forward
H3K27 me3:H3K4 me1
DNA Meth. m5C:H4K20 me3
H3K9 me3:H4K20 me3
H3K27 me3:H4K20 me3
H3K27 me3:DNA Meth. m5C
H4–4ac:H3–5ac
H4K20 me2:H4–4ac
DNA Meth. m5C:H3K9 me3
H3K4 me1:H3–5ac
H3K27 me3:H3K9 me3
H3K4 me3:H3–5ac
H2A.Z:H3–5ac
H2A.Z:H4–4ac
H4K20 me2:H3–5ac
H3K4 me3:H4–4ac
H4K20 me2:H3K4 me3
H3K4 me1:H4K20 me2
H2A.Z:H3K4 me3
H3K27 me2
H3K4 me1:H2A.Z
H3K4 me1:H4–4ac
H2A.Z:H4K20 me2
H3K9 me2
H4K20 me3
H3K9 me3
DNA Meth. m5C
H3K27 me3
H3–5ac
H4K20 me2
H3K4 me1
H4–4ac
H3K4 me3
H2A.Z
0.00 0.25 0.50 0.75 1.00
$\hat{\pi}$
KRT13 – Reverse
H3K27 me3:H3K4 me1
DNA Meth. m5C:H4K20 me3
H3K9 me3:H4K20 me3
H3K27 me3:H4K20 me3
H3K27 me3:DNA Meth. m5C
H4–4ac:H3–5ac
H4K20 me2:H4–4ac
DNA Meth. m5C:H3K9 me3
H3K4 me1:H3–5ac
H3K27 me3:H3K9 me3
H3K4 me3:H3–5ac
H2A.Z:H3–5ac
H2A.Z:H4–4ac
H4K20 me2:H3–5ac
H3K4 me3:H4–4ac
H4K20 me2:H3K4 me3
H3K4 me1:H4K20 me2
H2A.Z:H3K4 me3
H3K27 me2
H3K4 me1:H2A.Z
H3K4 me1:H4–4ac
H2A.Z:H4K20 me2
H3K9 me2
H4K20 me3
H3K9 me3
DNA Meth. m5C
H3K27 me3
H3–5ac
H4K20 me2
H3K4 me1
H4–4ac
H3K4 me3
H2A.Z
0.00 0.25 0.50 0.75 1.00
$\hat{\pi}$

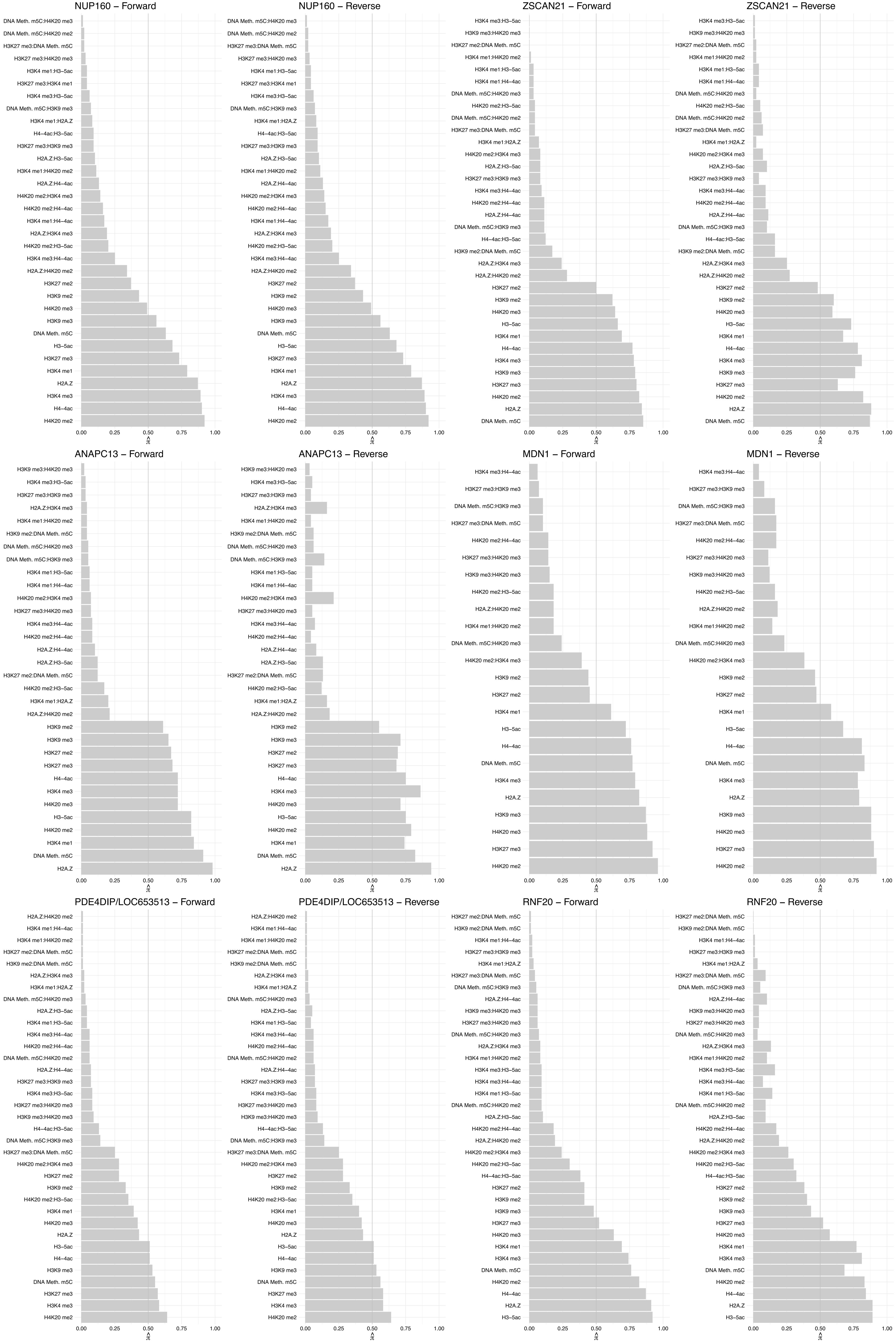
NUP160 − Forward
DNA Meth. m5C:H4K20 me3
DNA Meth. m5C:H4K20 me2
H3K27 me3:DNA Meth. m5C
H3K27 me3:H4K20 me3
H3K4 me1:H3−5ac
H3K27 me3:H3K4 me1
H3K4 me3:H3−5ac
DNA Meth. m5C:H3K9 me3
H3K4 me1:H2A.Z
H4−4ac:H3−5ac
H3K27 me3:H3K9 me3
H2A.Z:H3−5ac
H3K4 me1:H4K20 me2
H2A.Z:H4−4ac
H4K20 me2:H3K4 me3
H4K20 me2:H4−4ac
H3K4 me1:H4−4ac
H2A.Z:H3K4 me3
H4K20 me2:H3−5ac
H3K4 me3:H4−4ac
H2A.Z:H4K20 me2
H3K27 me2
H3K9 me2
H4K20 me3
H3K9 me3
DNA Meth. m5C
H3−5ac
H3K27 me3
H3K4 me1
H2A.Z
H3K4 me3
H4−4ac
H4K20 me2
0.00 0.25 0.50 0.75 1.00
π̂
NUP160 − Reverse
DNA Meth. m5C:H4K20 me3
DNA Meth. m5C:H4K20 me2
H3K27 me3:DNA Meth. m5C
H3K27 me3:H4K20 me3
H3K4 me1:H3−5ac
H3K27 me3:H3K4 me1
H3K4 me3:H3−5ac
DNA Meth. m5C:H3K9 me3
H3K4 me1:H2A.Z
H4−4ac:H3−5ac
H3K27 me3:H3K9 me3
H2A.Z:H3−5ac
H3K4 me1:H4K20 me2
H2A.Z:H4−4ac
H4K20 me2:H3K4 me3
H4K20 me2:H4−4ac
H3K4 me1:H4−4ac
H2A.Z:H3K4 me3
H4K20 me2:H3−5ac
H3K4 me3:H4−4ac
H2A.Z:H4K20 me2
H3K27 me2
H3K9 me2
H4K20 me3
H3K9 me3
DNA Meth. m5C
H3−5ac
H3K27 me3
H3K4 me1
H2A.Z
H3K4 me3
H4−4ac
H4K20 me2
0.00 0.25 0.50 0.75 1.00
π̂
ZSCAN21 − Forward
H3K4 me3:H3−5ac
H3K9 me3:H4K20 me3
H3K27 me2:DNA Meth. m5C
H3K4 me1:H4K20 me2
H3K4 me1:H3−5ac
H3K4 me1:H4−4ac
DNA Meth. m5C:H4K20 me3
H4K20 me2:H3−5ac
DNA Meth. m5C:H4K20 me2
H3K27 me3:DNA Meth. m5C
H3K4 me1:H2A.Z
H4K20 me2:H3K4 me3
H2A.Z:H3−5ac
H3K27 me3:H3K9 me3
H3K4 me3:H4−4ac
H4K20 me2:H4−4ac
H2A.Z:H4−4ac
DNA Meth. m5C:H3K9 me3
H4−4ac:H3−5ac
H3K9 me2:DNA Meth. m5C
H2A.Z:H3K4 me3
H2A.Z:H4K20 me2
H3K27 me2
H3K9 me2
H4K20 me3
H3−5ac
H3K4 me1
H4−4ac
H3K4 me3
H3K9 me3
H3K27 me3
H4K20 me2
H2A.Z
DNA Meth. m5C
0.00 0.25 0.50 0.75 1.00
π̂
ZSCAN21 − Reverse
H3K4 me3:H3−5ac
H3K9 me3:H4K20 me3
H3K27 me2:DNA Meth. m5C
H3K4 me1:H4K20 me2
H3K4 me1:H3−5ac
H3K4 me1:H4−4ac
DNA Meth. m5C:H4K20 me3
H4K20 me2:H3−5ac
DNA Meth. m5C:H4K20 me2
H3K27 me3:DNA Meth. m5C
H3K4 me1:H2A.Z
H4K20 me2:H3K4 me3
H2A.Z:H3−5ac
H3K27 me3:H3K9 me3
H3K4 me3:H4−4ac
H4K20 me2:H4−4ac
H2A.Z:H4−4ac
DNA Meth. m5C:H3K9 me3
H4−4ac:H3−5ac
H3K9 me2:DNA Meth. m5C
H2A.Z:H3K4 me3
H2A.Z:H4K20 me2
H3K27 me2
H3K9 me2
H4K20 me3
H3−5ac
H3K4 me1
H4−4ac
H3K4 me3
H3K9 me3
H3K27 me3
H4K20 me2
H2A.Z
DNA Meth. m5C
0.00 0.25 0.50 0.75 1.00
π̂
ANAPC13 − Forward
H3K9 me3:H4K20 me3
H3K4 me3:H3−5ac
H3K27 me3:H3K9 me3
H2A.Z:H3K4 me3
H3K4 me1:H4K20 me2
H3K9 me2:DNA Meth. m5C
DNA Meth. m5C:H4K20 me3
DNA Meth. m5C:H3K9 me3
H3K4 me1:H3−5ac
H3K4 me1:H4−4ac
H4K20 me2:H3K4 me3
H3K27 me3:H4K20 me3
H3K4 me3:H4−4ac
H4K20 me2:H4−4ac
H2A.Z:H4−4ac
H2A.Z:H3−5ac
H3K27 me2:DNA Meth. m5C
H4K20 me2:H3−5ac
H3K4 me1:H2A.Z
H2A.Z:H4K20 me2
H3K9 me2
H3K9 me3
H3K27 me2
H3K27 me3
H4−4ac
H3K4 me3
H4K20 me3
H3−5ac
H4K20 me2
H3K4 me1
DNA Meth. m5C
H2A.Z
0.00 0.25 0.50 0.75 1.00
π̂
ANAPC13 − Reverse
H3K9 me3:H4K20 me3
H3K4 me3:H3−5ac
H3K27 me3:H3K9 me3
H2A.Z:H3K4 me3
H3K4 me1:H4K20 me2
H3K9 me2:DNA Meth. m5C
DNA Meth. m5C:H4K20 me3
DNA Meth. m5C:H3K9 me3
H3K4 me1:H3−5ac
H3K4 me1:H4−4ac
H4K20 me2:H3K4 me3
H3K27 me3:H4K20 me3
H3K4 me3:H4−4ac
H4K20 me2:H4−4ac
H2A.Z:H4−4ac
H2A.Z:H3−5ac
H3K27 me2:DNA Meth. m5C
H4K20 me2:H3−5ac
H3K4 me1:H2A.Z
H2A.Z:H4K20 me2
H3K9 me2
H3K9 me3
H3K27 me2
H3K27 me3
H4−4ac
H3K4 me3
H4K20 me3
H3−5ac
H4K20 me2
H3K4 me1
DNA Meth. m5C
H2A.Z
0.00 0.25 0.50 0.75 1.00
π̂
MDN1 − Forward
H3K4 me3:H4−4ac
H3K27 me3:H3K9 me3
DNA Meth. m5C:H3K9 me3
H3K27 me3:DNA Meth. m5C
H4K20 me2:H4−4ac
H3K27 me3:H4K20 me3
H3K9 me3:H4K20 me3
H4K20 me2:H3−5ac
H2A.Z:H4K20 me2
H3K4 me1:H4K20 me2
DNA Meth. m5C:H4K20 me3
H4K20 me2:H3K4 me3
H3K9 me2
H3K27 me2
H3K4 me1
H3−5ac
H4−4ac
DNA Meth. m5C
H3K4 me3
H2A.Z
H3K9 me3
H4K20 me3
H3K27 me3
H4K20 me2
0.00 0.25 0.50 0.75 1.00
π̂
MDN1 − Reverse
H3K4 me3:H4−4ac
H3K27 me3:H3K9 me3
DNA Meth. m5C:H3K9 me3
H3K27 me3:DNA Meth. m5C
H4K20 me2:H4−4ac
H3K27 me3:H4K20 me3
H3K9 me3:H4K20 me3
H4K20 me2:H3−5ac
H2A.Z:H4K20 me2
H3K4 me1:H4K20 me2
DNA Meth. m5C:H4K20 me3
H4K20 me2:H3K4 me3
H3K9 me2
H3K27 me2
H3K4 me1
H3−5ac
H4−4ac
DNA Meth. m5C
H3K4 me3
H2A.Z
H3K9 me3
H4K20 me3
H3K27 me3
H4K20 me2
0.00 0.25 0.50 0.75 1.00
π̂
PDE4DIP/LOC653513 − Forward
H2A.Z:H4K20 me2
H3K4 me1:H4−4ac
H3K4 me1:H4K20 me2
H3K27 me2:DNA Meth. m5C
H3K9 me2:DNA Meth. m5C
H2A.Z:H3K4 me3
H3K4 me1:H2A.Z
DNA Meth. m5C:H4K20 me3
H2A.Z:H3−5ac
H3K4 me1:H3−5ac
H3K4 me3:H4−4ac
H4K20 me2:H4−4ac
DNA Meth. m5C:H4K20 me2
H2A.Z:H4−4ac
H3K27 me3:H3K9 me3
H3K4 me3:H3−5ac
H3K27 me3:H4K20 me3
H3K9 me3:H4K20 me3
H4−4ac:H3−5ac
DNA Meth. m5C:H3K9 me3
H3K27 me3:DNA Meth. m5C
H4K20 me2:H3K4 me3
H3K27 me2
H3K9 me2
H4K20 me2:H3−5ac
H3K4 me1
H4K20 me3
H2A.Z
H3−5ac
H4−4ac
H3K9 me3
DNA Meth. m5C
H3K27 me3
H3K4 me3
H4K20 me2
0.00 0.25 0.50 0.75 1.00
π̂
PDE4DIP/LOC653513 − Reverse
H2A.Z:H4K20 me2
H3K4 me1:H4−4ac
H3K4 me1:H4K20 me2
H3K27 me2:DNA Meth. m5C
H3K9 me2:DNA Meth. m5C
H2A.Z:H3K4 me3
H3K4 me1:H2A.Z
DNA Meth. m5C:H4K20 me3
H2A.Z:H3−5ac
H3K4 me1:H3−5ac
H3K4 me3:H4−4ac
H4K20 me2:H4−4ac
DNA Meth. m5C:H4K20 me2
H2A.Z:H4−4ac
H3K27 me3:H3K9 me3
H3K4 me3:H3−5ac
H3K27 me3:H4K20 me3
H3K9 me3:H4K20 me3
H4−4ac:H3−5ac
DNA Meth. m5C:H3K9 me3
H3K27 me3:DNA Meth. m5C
H4K20 me2:H3K4 me3
H3K27 me2
H3K9 me2
H4K20 me2:H3−5ac
H3K4 me1
H4K20 me3
H2A.Z
H3−5ac
H4−4ac
H3K9 me3
DNA Meth. m5C
H3K27 me3
H3K4 me3
H4K20 me2
0.00 0.25 0.50 0.75 1.00
π̂
RNF20 − Forward
H3K27 me2:DNA Meth. m5C
H3K9 me2:DNA Meth. m5C
H3K4 me1:H4−4ac
H3K27 me3:H3K9 me3
H3K4 me1:H2A.Z
H3K27 me3:DNA Meth. m5C
DNA Meth. m5C:H3K9 me3
H2A.Z:H4−4ac
H3K9 me3:H4K20 me3
H3K27 me3:H4K20 me3
DNA Meth. m5C:H4K20 me3
H2A.Z:H3K4 me3
H3K4 me1:H4K20 me2
H3K4 me3:H3−5ac
H3K4 me3:H4−4ac
H3K4 me1:H3−5ac
DNA Meth. m5C:H4K20 me2
H2A.Z:H3−5ac
H4K20 me2:H4−4ac
H2A.Z:H4K20 me2
H4K20 me2:H3K4 me3
H4K20 me2:H3−5ac
H4−4ac:H3−5ac
H3K27 me2
H3K9 me2
H3K9 me3
H3K27 me3
H4K20 me3
H3K4 me1
H3K4 me3
DNA Meth. m5C
H4K20 me2
H4−4ac
H2A.Z
H3−5ac
0.00 0.25 0.50 0.75 1.00
π̂
RNF20 − Reverse
H3K27 me2:DNA Meth. m5C
H3K9 me2:DNA Meth. m5C
H3K4 me1:H4−4ac
H3K27 me3:H3K9 me3
H3K4 me1:H2A.Z
H3K27 me3:DNA Meth. m5C
DNA Meth. m5C:H3K9 me3
H2A.Z:H4−4ac
H3K9 me3:H4K20 me3
H3K27 me3:H4K20 me3
DNA Meth. m5C:H4K20 me3
H2A.Z:H3K4 me3
H3K4 me1:H4K20 me2
H3K4 me3:H3−5ac
H3K4 me3:H4−4ac
H3K4 me1:H3−5ac
DNA Meth. m5C:H4K20 me2
H2A.Z:H3−5ac
H4K20 me2:H4−4ac
H2A.Z:H4K20 me2
H4K20 me2:H3K4 me3
H4K20 me2:H3−5ac
H4−4ac:H3−5ac
H3K27 me2
H3K9 me2
H3K9 me3
H3K27 me3
H4K20 me3
H3K4 me1
H3K4 me3
DNA Meth. m5C
H4K20 me2
H4−4ac
H2A.Z
H3−5ac
0.00 0.25 0.50 0.75 1.00
π̂

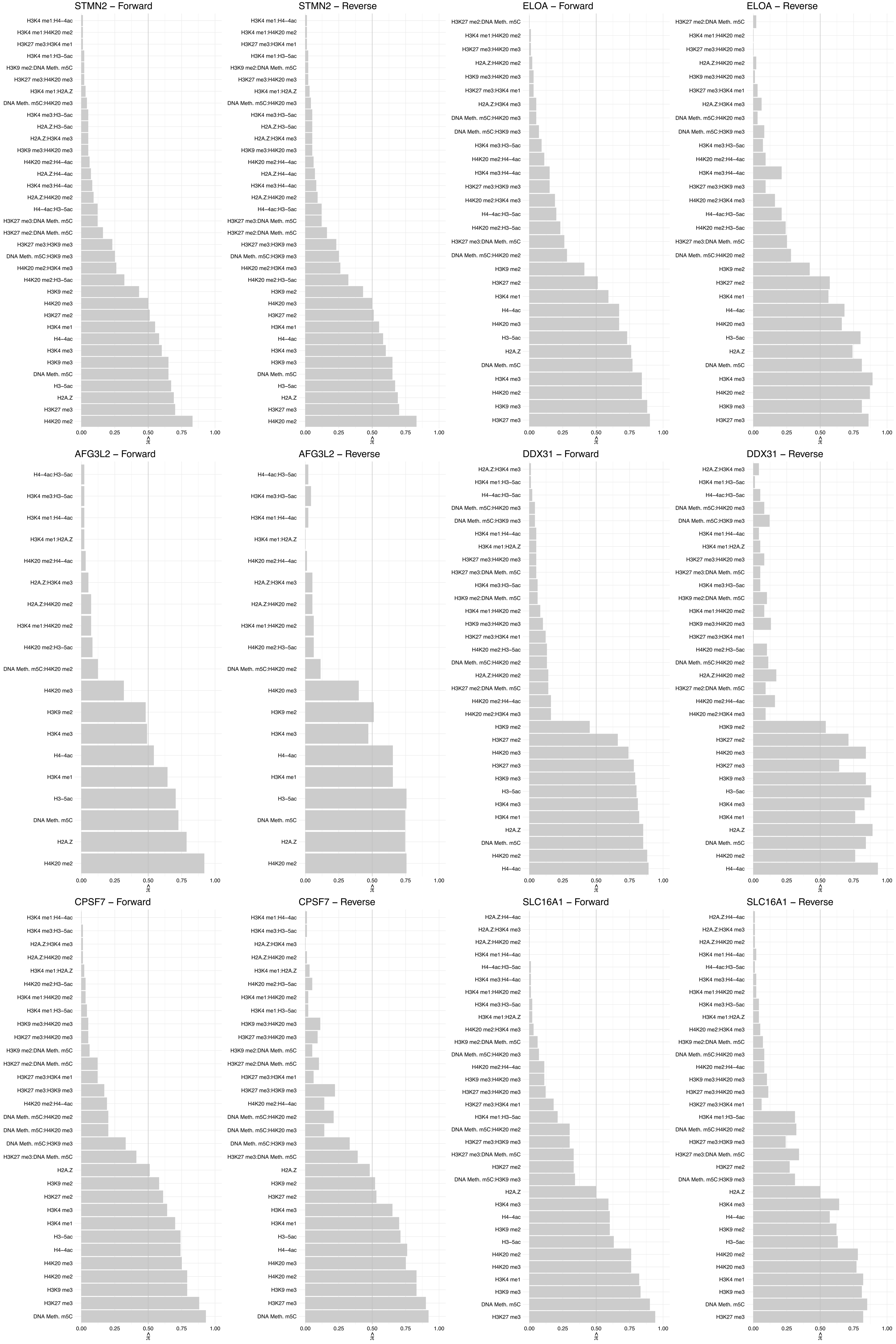
STMN2 – Forward
H3K4 me1:H4–4ac
H3K4 me1:H4K20 me2
H3K27 me3:H3K4 me1
H3K4 me1:H3–5ac
H3K9 me2:DNA Meth. m5C
H3K27 me3:H4K20 me3
H3K4 me1:H2A.Z
DNA Meth. m5C:H4K20 me3
H3K4 me3:H3–5ac
H2A.Z:H3–5ac
H2A.Z:H3K4 me3
H3K9 me3:H4K20 me3
H4K20 me2:H4–4ac
H2A.Z:H4–4ac
H3K4 me3:H4–4ac
H2A.Z:H4K20 me2
H4–4ac:H3–5ac
H3K27 me3:DNA Meth. m5C
H3K27 me2:DNA Meth. m5C
H3K27 me3:H3K9 me3
DNA Meth. m5C:H3K9 me3
H4K20 me2:H3K4 me3
H4K20 me2:H3–5ac
H3K9 me2
H4K20 me3
H3K27 me2
H3K4 me1
H4–4ac
H3K4 me3
H3K9 me3
DNA Meth. m5C
H3–5ac
H2A.Z
H3K27 me3
H4K20 me2
0.00 0.25 0.50 0.75 1.00
π̂
STMN2 – Reverse
H3K4 me1:H4–4ac
H3K4 me1:H4K20 me2
H3K27 me3:H3K4 me1
H3K4 me1:H3–5ac
H3K9 me2:DNA Meth. m5C
H3K27 me3:H4K20 me3
H3K4 me1:H2A.Z
DNA Meth. m5C:H4K20 me3
H3K4 me3:H3–5ac
H2A.Z:H3–5ac
H2A.Z:H3K4 me3
H3K9 me3:H4K20 me3
H4K20 me2:H4–4ac
H2A.Z:H4–4ac
H3K4 me3:H4–4ac
H2A.Z:H4K20 me2
H4–4ac:H3–5ac
H3K27 me3:DNA Meth. m5C
H3K27 me2:DNA Meth. m5C
H3K27 me3:H3K9 me3
DNA Meth. m5C:H3K9 me3
H4K20 me2:H3K4 me3
H4K20 me2:H3–5ac
H3K9 me2
H4K20 me3
H3K27 me2
H3K4 me1
H4–4ac
H3K4 me3
H3K9 me3
DNA Meth. m5C
H3–5ac
H2A.Z
H3K27 me3
H4K20 me2
0.00 0.25 0.50 0.75 1.00
π̂
ELOA – Forward
H3K27 me2:DNA Meth. m5C
H3K4 me1:H4K20 me2
H3K27 me3:H4K20 me3
H2A.Z:H4K20 me2
H3K9 me3:H4K20 me3
H3K27 me3:H3K4 me1
H2A.Z:H3K4 me3
DNA Meth. m5C:H4K20 me3
DNA Meth. m5C:H3K9 me3
H3K4 me3:H3–5ac
H4K20 me2:H4–4ac
H3K4 me3:H4–4ac
H3K27 me3:H3K9 me3
H4K20 me2:H3K4 me3
H4–4ac:H3–5ac
H4K20 me2:H3–5ac
H3K27 me3:DNA Meth. m5C
DNA Meth. m5C:H4K20 me2
H3K9 me2
H3K27 me2
H3K4 me1
H4–4ac
H4K20 me3
H3–5ac
H2A.Z
DNA Meth. m5C
H3K4 me3
H4K20 me2
H3K9 me3
H3K27 me3
0.00 0.25 0.50 0.75 1.00
π̂
ELOA – Reverse
H3K27 me2:DNA Meth. m5C
H3K4 me1:H4K20 me2
H3K27 me3:H4K20 me3
H2A.Z:H4K20 me2
H3K9 me3:H4K20 me3
H3K27 me3:H3K4 me1
H2A.Z:H3K4 me3
DNA Meth. m5C:H4K20 me3
DNA Meth. m5C:H3K9 me3
H3K4 me3:H3–5ac
H4K20 me2:H4–4ac
H3K4 me3:H4–4ac
H3K27 me3:H3K9 me3
H4K20 me2:H3K4 me3
H4–4ac:H3–5ac
H4K20 me2:H3–5ac
H3K27 me3:DNA Meth. m5C
DNA Meth. m5C:H4K20 me2
H3K9 me2
H3K27 me2
H3K4 me1
H4–4ac
H4K20 me3
H3–5ac
H2A.Z
DNA Meth. m5C
H3K4 me3
H4K20 me2
H3K9 me3
H3K27 me3
0.00 0.25 0.50 0.75 1.00
π̂
AFG3L2 – Forward
H4–4ac:H3–5ac
H3K4 me3:H3–5ac
H3K4 me1:H4–4ac
H3K4 me1:H2A.Z
H4K20 me2:H4–4ac
H2A.Z:H3K4 me3
H2A.Z:H4K20 me2
H3K4 me1:H4K20 me2
H4K20 me2:H3–5ac
DNA Meth. m5C:H4K20 me2
H4K20 me3
H3K9 me2
H3K4 me3
H4–4ac
H3K4 me1
H3–5ac
DNA Meth. m5C
H2A.Z
H4K20 me2
0.00 0.25 0.50 0.75 1.00
π̂
AFG3L2 – Reverse
H4–4ac:H3–5ac
H3K4 me3:H3–5ac
H3K4 me1:H4–4ac
H3K4 me1:H2A.Z
H4K20 me2:H4–4ac
H2A.Z:H3K4 me3
H2A.Z:H4K20 me2
H3K4 me1:H4K20 me2
H4K20 me2:H3–5ac
DNA Meth. m5C:H4K20 me2
H4K20 me3
H3K9 me2
H3K4 me3
H4–4ac
H3K4 me1
H3–5ac
DNA Meth. m5C
H2A.Z
H4K20 me2
0.00 0.25 0.50 0.75 1.00
π̂
DDX31 – Forward
H2A.Z:H3K4 me3
H3K4 me1:H3–5ac
H4–4ac:H3–5ac
DNA Meth. m5C:H4K20 me3
DNA Meth. m5C:H3K9 me3
H3K4 me1:H4–4ac
H3K4 me1:H2A.Z
H3K27 me3:H4K20 me3
H3K27 me3:DNA Meth. m5C
H3K4 me3:H3–5ac
H3K9 me2:DNA Meth. m5C
H3K4 me1:H4K20 me2
H3K9 me3:H4K20 me3
H3K27 me3:H3K4 me1
H4K20 me2:H3–5ac
DNA Meth. m5C:H4K20 me2
H2A.Z:H4K20 me2
H3K27 me2:DNA Meth. m5C
H4K20 me2:H4–4ac
H4K20 me2:H3K4 me3
H3K9 me2
H3K27 me2
H4K20 me3
H3K27 me3
H3K9 me3
H3–5ac
H3K4 me3
H3K4 me1
H2A.Z
DNA Meth. m5C
H4K20 me2
H4–4ac
0.00 0.25 0.50 0.75 1.00
π̂
DDX31 – Reverse
H2A.Z:H3K4 me3
H3K4 me1:H3–5ac
H4–4ac:H3–5ac
DNA Meth. m5C:H4K20 me3
DNA Meth. m5C:H3K9 me3
H3K4 me1:H4–4ac
H3K4 me1:H2A.Z
H3K27 me3:H4K20 me3
H3K27 me3:DNA Meth. m5C
H3K4 me3:H3–5ac
H3K9 me2:DNA Meth. m5C
H3K4 me1:H4K20 me2
H3K9 me3:H4K20 me3
H3K27 me3:H3K4 me1
H4K20 me2:H3–5ac
DNA Meth. m5C:H4K20 me2
H2A.Z:H4K20 me2
H3K27 me2:DNA Meth. m5C
H4K20 me2:H4–4ac
H4K20 me2:H3K4 me3
H3K9 me2
H3K27 me2
H4K20 me3
H3K27 me3
H3K9 me3
H3–5ac
H3K4 me3
H3K4 me1
H2A.Z
DNA Meth. m5C
H4K20 me2
H4–4ac
0.00 0.25 0.50 0.75 1.00
π̂
CPSF7 – Forward
H3K4 me1:H4–4ac
H3K4 me3:H3–5ac
H2A.Z:H3K4 me3
H2A.Z:H4K20 me2
H3K4 me1:H2A.Z
H4K20 me2:H3–5ac
H3K4 me1:H4K20 me2
H3K4 me1:H3–5ac
H3K9 me3:H4K20 me3
H3K27 me3:H4K20 me3
H3K9 me2:DNA Meth. m5C
H3K27 me2:DNA Meth. m5C
H3K27 me3:H3K4 me1
H3K27 me3:H3K9 me3
H4K20 me2:H4–4ac
DNA Meth. m5C:H4K20 me2
DNA Meth. m5C:H4K20 me3
DNA Meth. m5C:H3K9 me3
H3K27 me3:DNA Meth. m5C
H2A.Z
H3K9 me2
H3K27 me2
H3K4 me3
H3K4 me1
H3–5ac
H4–4ac
H4K20 me3
H4K20 me2
H3K9 me3
H3K27 me3
DNA Meth. m5C
0.00 0.25 0.50 0.75 1.00
π̂
CPSF7 – Reverse
H3K4 me1:H4–4ac
H3K4 me3:H3–5ac
H2A.Z:H3K4 me3
H2A.Z:H4K20 me2
H3K4 me1:H2A.Z
H4K20 me2:H3–5ac
H3K4 me1:H4K20 me2
H3K4 me1:H3–5ac
H3K9 me3:H4K20 me3
H3K27 me3:H4K20 me3
H3K9 me2:DNA Meth. m5C
H3K27 me2:DNA Meth. m5C
H3K27 me3:H3K4 me1
H3K27 me3:H3K9 me3
H4K20 me2:H4–4ac
DNA Meth. m5C:H4K20 me2
DNA Meth. m5C:H4K20 me3
DNA Meth. m5C:H3K9 me3
H3K27 me3:DNA Meth. m5C
H2A.Z
H3K9 me2
H3K27 me2
H3K4 me3
H3K4 me1
H3–5ac
H4–4ac
H4K20 me3
H4K20 me2
H3K9 me3
H3K27 me3
DNA Meth. m5C
0.00 0.25 0.50 0.75 1.00
π̂
SLC16A1 – Forward
H2A.Z:H4–4ac
H2A.Z:H3K4 me3
H2A.Z:H4K20 me2
H3K4 me1:H4–4ac
H4–4ac:H3–5ac
H3K4 me3:H4–4ac
H3K4 me1:H4K20 me2
H3K4 me3:H3–5ac
H3K4 me1:H2A.Z
H4K20 me2:H3K4 me3
H3K9 me2:DNA Meth. m5C
DNA Meth. m5C:H4K20 me3
H4K20 me2:H4–4ac
H3K9 me3:H4K20 me3
H3K27 me3:H4K20 me3
H3K27 me3:H3K4 me1
H3K4 me1:H3–5ac
DNA Meth. m5C:H4K20 me2
H3K27 me3:H3K9 me3
H3K27 me3:DNA Meth. m5C
H3K27 me2
DNA Meth. m5C:H3K9 me3
H2A.Z
H3K4 me3
H4–4ac
H3K9 me2
H3–5ac
H4K20 me2
H4K20 me3
H3K4 me1
H3K9 me3
DNA Meth. m5C
H3K27 me3
0.00 0.25 0.50 0.75 1.00
π̂
SLC16A1 – Reverse
H2A.Z:H4–4ac
H2A.Z:H3K4 me3
H2A.Z:H4K20 me2
H3K4 me1:H4–4ac
H4–4ac:H3–5ac
H3K4 me3:H4–4ac
H3K4 me1:H4K20 me2
H3K4 me3:H3–5ac
H3K4 me1:H2A.Z
H4K20 me2:H3K4 me3
H3K9 me2:DNA Meth. m5C
DNA Meth. m5C:H4K20 me3
H4K20 me2:H4–4ac
H3K9 me3:H4K20 me3
H3K27 me3:H4K20 me3
H3K27 me3:H3K4 me1
H3K4 me1:H3–5ac
DNA Meth. m5C:H4K20 me2
H3K27 me3:H3K9 me3
H3K27 me3:DNA Meth. m5C
H3K27 me2
DNA Meth. m5C:H3K9 me3
H2A.Z
H3K4 me3
H4–4ac
H3K9 me2
H3–5ac
H4K20 me2
H4K20 me3
H3K4 me1
H3K9 me3
DNA Meth. m5C
H3K27 me3
0.00 0.25 0.50 0.75 1.00
π̂

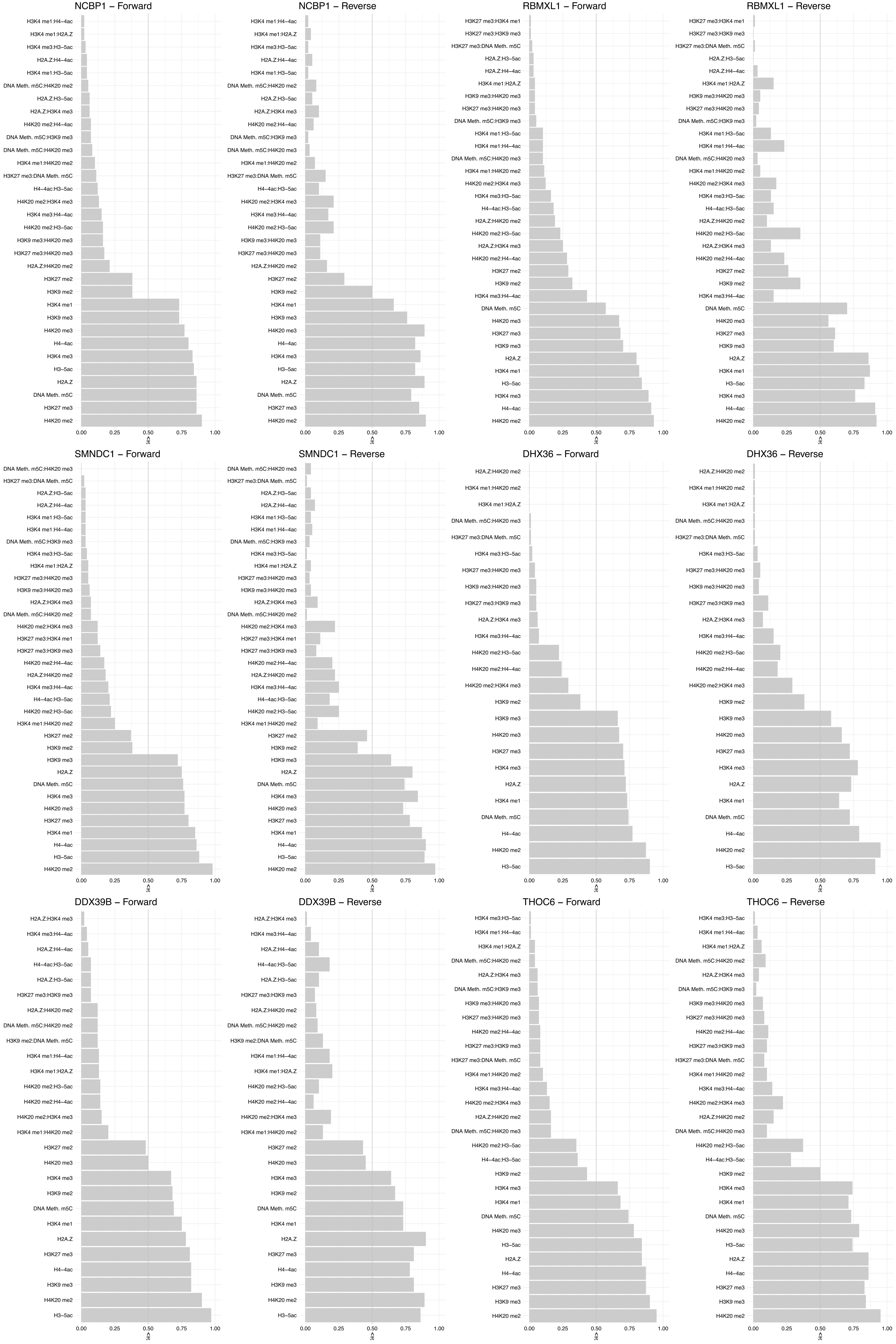

NCBP1 – Forward
NCBP1 – Reverse
RBMXL1 – Forward
RBMXL1 – Reverse
SMNDC1 – Forward
SMNDC1 – Reverse
DHX36 – Forward
DHX36 – Reverse
DDX39B – Forward
DDX39B – Reverse
THOC6 – Forward
THOC6 – Reverse

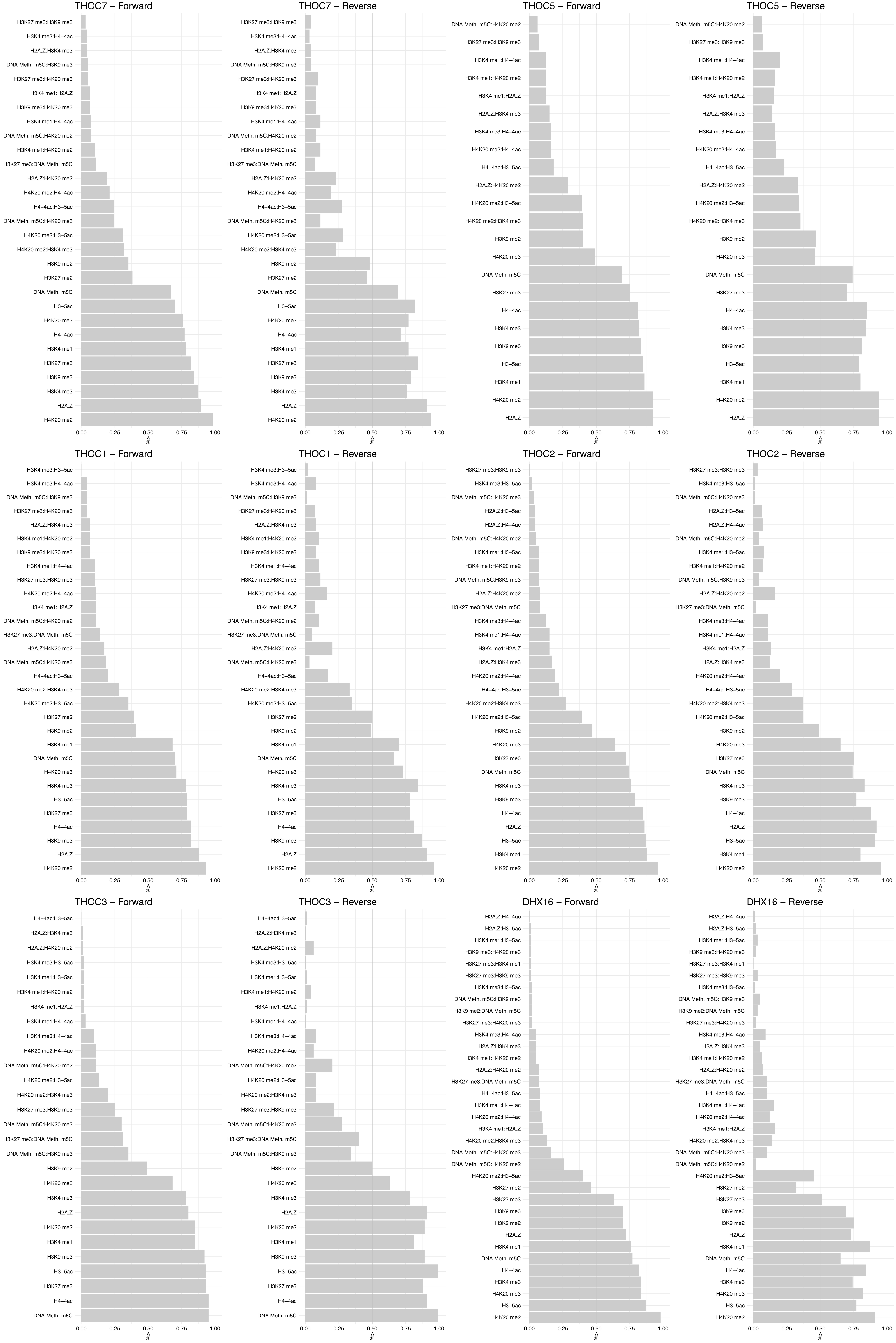

THOC7 – Forward
H3K27 me3:H3K9 me3
H3K4 me3:H4–4ac
H2A.Z:H3K4 me3
DNA Meth. m5C:H3K9 me3
H3K27 me3:H4K20 me3
H3K4 me1:H2A.Z
H3K9 me3:H4K20 me3
H3K4 me1:H4–4ac
DNA Meth. m5C:H4K20 me2
H3K4 me1:H4K20 me2
H3K27 me3:DNA Meth. m5C
H2A.Z:H4K20 me2
H4K20 me2:H4–4ac
H4–4ac:H3–5ac
DNA Meth. m5C:H4K20 me3
H4K20 me2:H3–5ac
H4K20 me2:H3K4 me3
H3K9 me2
H3K27 me2
DNA Meth. m5C
H3–5ac
H4K20 me3
H4–4ac
H3K4 me1
H3K27 me3
H3K9 me3
H3K4 me3
H2A.Z
H4K20 me2
0.00 0.25 0.50 0.75 1.00
π̂
THOC7 – Reverse
H3K27 me3:H3K9 me3
H3K4 me3:H4–4ac
H2A.Z:H3K4 me3
DNA Meth. m5C:H3K9 me3
H3K27 me3:H4K20 me3
H3K4 me1:H2A.Z
H3K9 me3:H4K20 me3
H3K4 me1:H4–4ac
DNA Meth. m5C:H4K20 me2
H3K4 me1:H4K20 me2
H3K27 me3:DNA Meth. m5C
H2A.Z:H4K20 me2
H4K20 me2:H4–4ac
H4–4ac:H3–5ac
DNA Meth. m5C:H4K20 me3
H4K20 me2:H3–5ac
H4K20 me2:H3K4 me3
H3K9 me2
H3K27 me2
DNA Meth. m5C
H3–5ac
H4K20 me3
H4–4ac
H3K4 me1
H3K27 me3
H3K9 me3
H3K4 me3
H2A.Z
H4K20 me2
0.00 0.25 0.50 0.75 1.00
π̂
THOC5 – Forward
DNA Meth. m5C:H4K20 me2
H3K27 me3:H3K9 me3
H3K4 me1:H4–4ac
H3K4 me1:H4K20 me2
H3K4 me1:H2A.Z
H2A.Z:H3K4 me3
H3K4 me3:H4–4ac
H4K20 me2:H4–4ac
H4–4ac:H3–5ac
H2A.Z:H4K20 me2
H4K20 me2:H3–5ac
H4K20 me2:H3K4 me3
H3K9 me2
H4K20 me3
DNA Meth. m5C
H3K27 me3
H4–4ac
H3K4 me3
H3K9 me3
H3–5ac
H3K4 me1
H4K20 me2
H2A.Z
0.00 0.25 0.50 0.75 1.00
π̂
THOC5 – Reverse
DNA Meth. m5C:H4K20 me2
H3K27 me3:H3K9 me3
H3K4 me1:H4–4ac
H3K4 me1:H4K20 me2
H3K4 me1:H2A.Z
H2A.Z:H3K4 me3
H3K4 me3:H4–4ac
H4K20 me2:H4–4ac
H4–4ac:H3–5ac
H2A.Z:H4K20 me2
H4K20 me2:H3–5ac
H4K20 me2:H3K4 me3
H3K9 me2
H4K20 me3
DNA Meth. m5C
H3K27 me3
H4–4ac
H3K4 me3
H3K9 me3
H3–5ac
H3K4 me1
H4K20 me2
H2A.Z
0.00 0.25 0.50 0.75 1.00
π̂
THOC1 – Forward
H3K4 me3:H3–5ac
H3K4 me3:H4–4ac
DNA Meth. m5C:H3K9 me3
H3K27 me3:H4K20 me3
H2A.Z:H3K4 me3
H3K4 me1:H4K20 me2
H3K9 me3:H4K20 me3
H3K4 me1:H4–4ac
H3K27 me3:H3K9 me3
H4K20 me2:H4–4ac
H3K4 me1:H2A.Z
DNA Meth. m5C:H4K20 me2
H3K27 me3:DNA Meth. m5C
H2A.Z:H4K20 me2
DNA Meth. m5C:H4K20 me3
H4–4ac:H3–5ac
H4K20 me2:H3K4 me3
H4K20 me2:H3–5ac
H3K27 me2
H3K9 me2
H3K4 me1
DNA Meth. m5C
H4K20 me3
H3K4 me3
H3–5ac
H3K27 me3
H4–4ac
H3K9 me3
H2A.Z
H4K20 me2
0.00 0.25 0.50 0.75 1.00
π̂
THOC1 – Reverse
H3K4 me3:H3–5ac
H3K4 me3:H4–4ac
DNA Meth. m5C:H3K9 me3
H3K27 me3:H4K20 me3
H2A.Z:H3K4 me3
H3K4 me1:H4K20 me2
H3K9 me3:H4K20 me3
H3K4 me1:H4–4ac
H3K27 me3:H3K9 me3
H4K20 me2:H4–4ac
H3K4 me1:H2A.Z
DNA Meth. m5C:H4K20 me2
H3K27 me3:DNA Meth. m5C
H2A.Z:H4K20 me2
DNA Meth. m5C:H4K20 me3
H4–4ac:H3–5ac
H4K20 me2:H3K4 me3
H4K20 me2:H3–5ac
H3K27 me2
H3K9 me2
H3K4 me1
DNA Meth. m5C
H4K20 me3
H3K4 me3
H3–5ac
H3K27 me3
H4–4ac
H3K9 me3
H2A.Z
H4K20 me2
0.00 0.25 0.50 0.75 1.00
π̂
THOC2 – Forward
H3K27 me3:H3K9 me3
H3K4 me3:H3–5ac
DNA Meth. m5C:H4K20 me3
H2A.Z:H3–5ac
H2A.Z:H4–4ac
DNA Meth. m5C:H4K20 me2
H3K4 me1:H3–5ac
H3K4 me1:H4K20 me2
DNA Meth. m5C:H3K9 me3
H2A.Z:H4K20 me2
H3K27 me3:DNA Meth. m5C
H3K4 me3:H4–4ac
H3K4 me1:H4–4ac
H3K4 me1:H2A.Z
H2A.Z:H3K4 me3
H4K20 me2:H4–4ac
H4–4ac:H3–5ac
H4K20 me2:H3K4 me3
H4K20 me2:H3–5ac
H3K9 me2
H4K20 me3
H3K27 me3
DNA Meth. m5C
H3K4 me3
H3K9 me3
H4–4ac
H2A.Z
H3–5ac
H3K4 me1
H4K20 me2
0.00 0.25 0.50 0.75 1.00
π̂
THOC2 – Reverse
H3K27 me3:H3K9 me3
H3K4 me3:H3–5ac
DNA Meth. m5C:H4K20 me3
H2A.Z:H3–5ac
H2A.Z:H4–4ac
DNA Meth. m5C:H4K20 me2
H3K4 me1:H3–5ac
H3K4 me1:H4K20 me2
DNA Meth. m5C:H3K9 me3
H2A.Z:H4K20 me2
H3K27 me3:DNA Meth. m5C
H3K4 me3:H4–4ac
H3K4 me1:H4–4ac
H3K4 me1:H2A.Z
H2A.Z:H3K4 me3
H4K20 me2:H4–4ac
H4–4ac:H3–5ac
H4K20 me2:H3K4 me3
H4K20 me2:H3–5ac
H3K9 me2
H4K20 me3
H3K27 me3
DNA Meth. m5C
H3K4 me3
H3K9 me3
H4–4ac
H2A.Z
H3–5ac
H3K4 me1
H4K20 me2
0.00 0.25 0.50 0.75 1.00
π̂
THOC3 – Forward
H4–4ac:H3–5ac
H2A.Z:H3K4 me3
H2A.Z:H4K20 me2
H3K4 me3:H3–5ac
H3K4 me1:H3–5ac
H3K4 me1:H4K20 me2
H3K4 me1:H2A.Z
H3K4 me1:H4–4ac
H3K4 me3:H4–4ac
H4K20 me2:H4–4ac
DNA Meth. m5C:H4K20 me2
H4K20 me2:H3–5ac
H4K20 me2:H3K4 me3
H3K27 me3:H3K9 me3
DNA Meth. m5C:H4K20 me3
H3K27 me3:DNA Meth. m5C
DNA Meth. m5C:H3K9 me3
H3K9 me2
H4K20 me3
H3K4 me3
H2A.Z
H4K20 me2
H3K4 me1
H3K9 me3
H3–5ac
H3K27 me3
H4–4ac
DNA Meth. m5C
0.00 0.25 0.50 0.75 1.00
π̂
THOC3 – Reverse
H4–4ac:H3–5ac
H2A.Z:H3K4 me3
H2A.Z:H4K20 me2
H3K4 me3:H3–5ac
H3K4 me1:H3–5ac
H3K4 me1:H4K20 me2
H3K4 me1:H2A.Z
H3K4 me1:H4–4ac
H3K4 me3:H4–4ac
H4K20 me2:H4–4ac
DNA Meth. m5C:H4K20 me2
H4K20 me2:H3–5ac
H4K20 me2:H3K4 me3
H3K27 me3:H3K9 me3
DNA Meth. m5C:H4K20 me3
H3K27 me3:DNA Meth. m5C
DNA Meth. m5C:H3K9 me3
H3K9 me2
H4K20 me3
H3K4 me3
H2A.Z
H4K20 me2
H3K4 me1
H3K9 me3
H3–5ac
H3K27 me3
H4–4ac
DNA Meth. m5C
0.00 0.25 0.50 0.75 1.00
π̂
DHX16 – Forward
H2A.Z:H4–4ac
H2A.Z:H3–5ac
H3K4 me1:H3–5ac
H3K9 me3:H4K20 me3
H3K27 me3:H3K4 me1
H3K27 me3:H3K9 me3
H3K4 me3:H3–5ac
DNA Meth. m5C:H3K9 me3
H3K9 me2:DNA Meth. m5C
H3K27 me3:H4K20 me3
H3K4 me3:H4–4ac
H2A.Z:H3K4 me3
H3K4 me1:H4K20 me2
H2A.Z:H4K20 me2
H3K27 me3:DNA Meth. m5C
H4–4ac:H3–5ac
H3K4 me1:H4–4ac
H4K20 me2:H4–4ac
H3K4 me1:H2A.Z
H4K20 me2:H3K4 me3
DNA Meth. m5C:H4K20 me3
DNA Meth. m5C:H4K20 me2
H4K20 me2:H3–5ac
H3K27 me2
H3K27 me3
H3K9 me3
H3K9 me2
H2A.Z
H3K4 me1
DNA Meth. m5C
H4–4ac
H3K4 me3
H4K20 me3
H3–5ac
H4K20 me2
0.00 0.25 0.50 0.75 1.00
π̂
DHX16 – Reverse
H2A.Z:H4–4ac
H2A.Z:H3–5ac
H3K4 me1:H3–5ac
H3K9 me3:H4K20 me3
H3K27 me3:H3K4 me1
H3K27 me3:H3K9 me3
H3K4 me3:H3–5ac
DNA Meth. m5C:H3K9 me3
H3K9 me2:DNA Meth. m5C
H3K27 me3:H4K20 me3
H3K4 me3:H4–4ac
H2A.Z:H3K4 me3
H3K4 me1:H4K20 me2
H2A.Z:H4K20 me2
H3K27 me3:DNA Meth. m5C
H4–4ac:H3–5ac
H3K4 me1:H4–4ac
H4K20 me2:H4–4ac
H3K4 me1:H2A.Z
H4K20 me2:H3K4 me3
DNA Meth. m5C:H4K20 me3
DNA Meth. m5C:H4K20 me2
H4K20 me2:H3–5ac
H3K27 me2
H3K27 me3
H3K9 me3
H3K9 me2
H2A.Z
H3K4 me1
DNA Meth. m5C
H4–4ac
H3K4 me3
H4K20 me3
H3–5ac
H4K20 me2
0.00 0.25 0.50 0.75 1.00
π̂

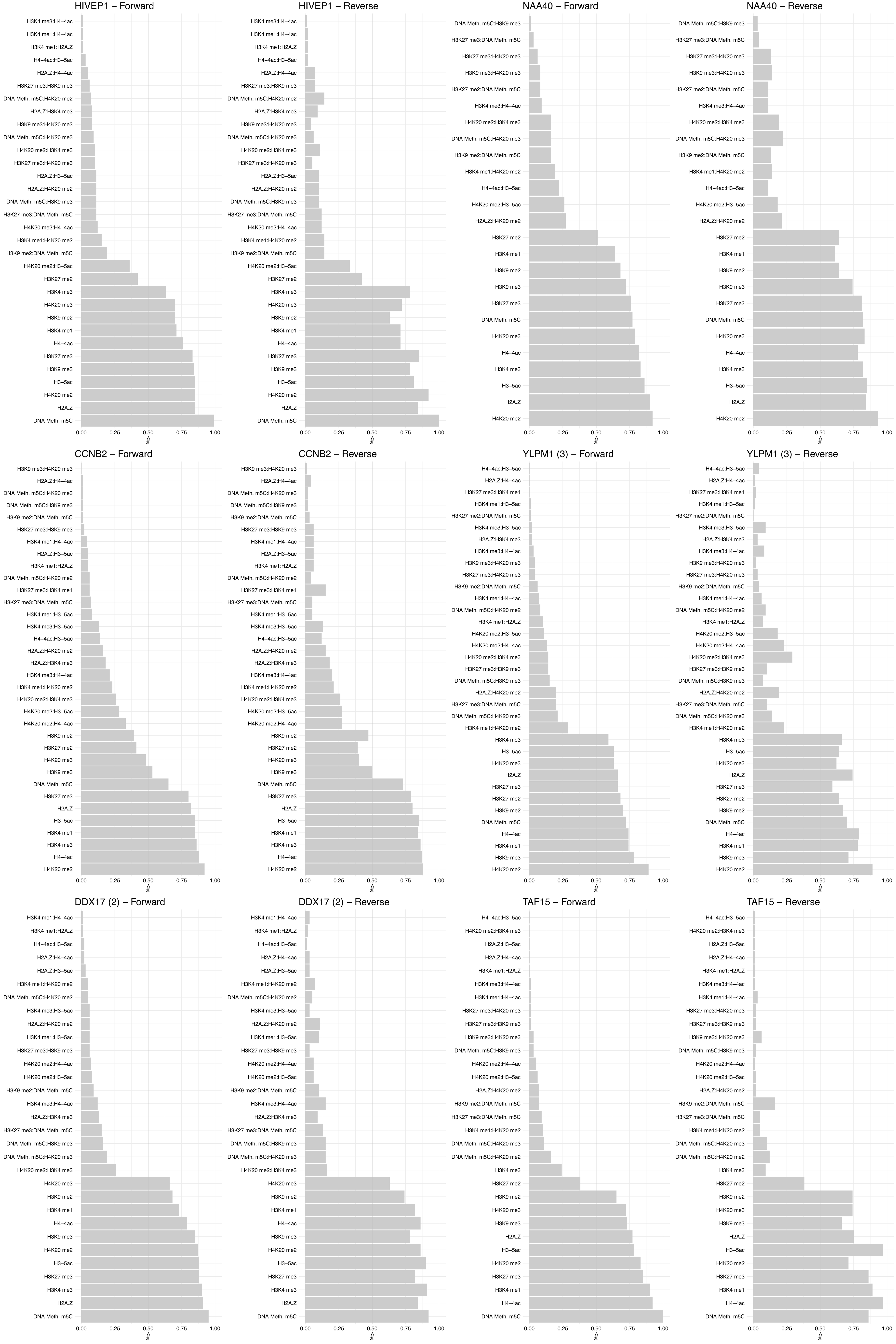
HIVEP1 − Forward
H3K4 me3:H4−4ac
H3K4 me1:H4−4ac
H3K4 me1:H2A.Z
H4−4ac:H3−5ac
H2A.Z:H4−4ac
H3K27 me3:H3K9 me3
DNA Meth. m5C:H4K20 me2
H2A.Z:H3K4 me3
H3K9 me3:H4K20 me3
DNA Meth. m5C:H4K20 me3
H4K20 me2:H3K4 me3
H3K27 me3:H4K20 me3
H2A.Z:H3−5ac
H2A.Z:H4K20 me2
DNA Meth. m5C:H3K9 me3
H3K27 me3:DNA Meth. m5C
H4K20 me2:H4−4ac
H3K4 me1:H4K20 me2
H3K9 me2:DNA Meth. m5C
H4K20 me2:H3−5ac
H3K27 me2
H3K4 me3
H4K20 me3
H3K9 me2
H3K4 me1
H4−4ac
H3K27 me3
H3K9 me3
H3−5ac
H4K20 me2
H2A.Z
DNA Meth. m5C
0.00 0.25 0.50 0.75 1.00
$\hat{\pi}$
HIVEP1 − Reverse
H3K4 me3:H4−4ac
H3K4 me1:H4−4ac
H3K4 me1:H2A.Z
H4−4ac:H3−5ac
H2A.Z:H4−4ac
H3K27 me3:H3K9 me3
DNA Meth. m5C:H4K20 me2
H2A.Z:H3K4 me3
H3K9 me3:H4K20 me3
DNA Meth. m5C:H4K20 me3
H4K20 me2:H3K4 me3
H3K27 me3:H4K20 me3
H2A.Z:H3−5ac
H2A.Z:H4K20 me2
DNA Meth. m5C:H3K9 me3
H3K27 me3:DNA Meth. m5C
H4K20 me2:H4−4ac
H3K4 me1:H4K20 me2
H3K9 me2:DNA Meth. m5C
H4K20 me2:H3−5ac
H3K27 me2
H3K4 me3
H4K20 me3
H3K9 me2
H3K4 me1
H4−4ac
H3K27 me3
H3K9 me3
H3−5ac
H4K20 me2
H2A.Z
DNA Meth. m5C
0.00 0.25 0.50 0.75 1.00
$\hat{\pi}$
NAA40 − Forward
DNA Meth. m5C:H3K9 me3
H3K27 me3:DNA Meth. m5C
H3K27 me3:H4K20 me3
H3K9 me3:H4K20 me3
H3K27 me2:DNA Meth. m5C
H3K4 me3:H4−4ac
H4K20 me2:H3K4 me3
DNA Meth. m5C:H4K20 me3
H3K9 me2:DNA Meth. m5C
H3K4 me1:H4K20 me2
H4−4ac:H3−5ac
H4K20 me2:H3−5ac
H2A.Z:H4K20 me2
H3K27 me2
H3K4 me1
H3K9 me2
H3K9 me3
H3K27 me3
DNA Meth. m5C
H4K20 me3
H4−4ac
H3K4 me3
H3−5ac
H2A.Z
H4K20 me2
0.00 0.25 0.50 0.75 1.00
$\hat{\pi}$
NAA40 − Reverse
DNA Meth. m5C:H3K9 me3
H3K27 me3:DNA Meth. m5C
H3K27 me3:H4K20 me3
H3K9 me3:H4K20 me3
H3K27 me2:DNA Meth. m5C
H3K4 me3:H4−4ac
H4K20 me2:H3K4 me3
DNA Meth. m5C:H4K20 me3
H3K9 me2:DNA Meth. m5C
H3K4 me1:H4K20 me2
H4−4ac:H3−5ac
H4K20 me2:H3−5ac
H2A.Z:H4K20 me2
H3K27 me2
H3K4 me1
H3K9 me2
H3K9 me3
H3K27 me3
DNA Meth. m5C
H4K20 me3
H4−4ac
H3K4 me3
H3−5ac
H2A.Z
H4K20 me2
0.00 0.25 0.50 0.75 1.00
$\hat{\pi}$
CCNB2 − Forward
H3K9 me3:H4K20 me3
H2A.Z:H4−4ac
DNA Meth. m5C:H4K20 me3
DNA Meth. m5C:H3K9 me3
H3K9 me2:DNA Meth. m5C
H3K27 me3:H3K9 me3
H3K4 me1:H4−4ac
H2A.Z:H3−5ac
H3K4 me1:H2A.Z
DNA Meth. m5C:H4K20 me2
H3K27 me3:H3K4 me1
H3K27 me3:DNA Meth. m5C
H3K4 me1:H3−5ac
H3K4 me3:H3−5ac
H4−4ac:H3−5ac
H2A.Z:H4K20 me2
H2A.Z:H3K4 me3
H3K4 me3:H4−4ac
H3K4 me1:H4K20 me2
H4K20 me2:H3K4 me3
H4K20 me2:H3−5ac
H4K20 me2:H4−4ac
H3K9 me2
H3K27 me2
H4K20 me3
H3K9 me3
DNA Meth. m5C
H3K27 me3
H2A.Z
H3−5ac
H3K4 me1
H3K4 me3
H4−4ac
H4K20 me2
0.00 0.25 0.50 0.75 1.00
$\hat{\pi}$
CCNB2 − Reverse
H3K9 me3:H4K20 me3
H2A.Z:H4−4ac
DNA Meth. m5C:H4K20 me3
DNA Meth. m5C:H3K9 me3
H3K9 me2:DNA Meth. m5C
H3K27 me3:H3K9 me3
H3K4 me1:H4−4ac
H2A.Z:H3−5ac
H3K4 me1:H2A.Z
DNA Meth. m5C:H4K20 me2
H3K27 me3:H3K4 me1
H3K27 me3:DNA Meth. m5C
H3K4 me1:H3−5ac
H3K4 me3:H3−5ac
H4−4ac:H3−5ac
H2A.Z:H4K20 me2
H2A.Z:H3K4 me3
H3K4 me3:H4−4ac
H3K4 me1:H4K20 me2
H4K20 me2:H3K4 me3
H4K20 me2:H3−5ac
H4K20 me2:H4−4ac
H3K9 me2
H3K27 me2
H4K20 me3
H3K9 me3
DNA Meth. m5C
H3K27 me3
H2A.Z
H3−5ac
H3K4 me1
H3K4 me3
H4−4ac
H4K20 me2
0.00 0.25 0.50 0.75 1.00
$\hat{\pi}$
YLPM1 (3) − Forward
H4−4ac:H3−5ac
H2A.Z:H4−4ac
H3K27 me3:H3K4 me1
H3K4 me1:H3−5ac
H3K27 me2:DNA Meth. m5C
H3K4 me3:H3−5ac
H2A.Z:H3K4 me3
H3K4 me3:H4−4ac
H3K9 me3:H4K20 me3
H3K27 me3:H4K20 me3
H3K9 me2:DNA Meth. m5C
H3K4 me1:H4−4ac
DNA Meth. m5C:H4K20 me2
H3K4 me1:H2A.Z
H4K20 me2:H3−5ac
H4K20 me2:H4−4ac
H4K20 me2:H3K4 me3
H3K27 me3:H3K9 me3
DNA Meth. m5C:H3K9 me3
H2A.Z:H4K20 me2
H3K27 me3:DNA Meth. m5C
DNA Meth. m5C:H4K20 me3
H3K4 me1:H4K20 me2
H3K4 me3
H3−5ac
H4K20 me3
H2A.Z
H3K27 me3
H3K27 me2
H3K9 me2
DNA Meth. m5C
H4−4ac
H3K4 me1
H3K9 me3
H4K20 me2
0.00 0.25 0.50 0.75 1.00
$\hat{\pi}$
YLPM1 (3) − Reverse
H4−4ac:H3−5ac
H2A.Z:H4−4ac
H3K27 me3:H3K4 me1
H3K4 me1:H3−5ac
H3K27 me2:DNA Meth. m5C
H3K4 me3:H3−5ac
H2A.Z:H3K4 me3
H3K4 me3:H4−4ac
H3K9 me3:H4K20 me3
H3K27 me3:H4K20 me3
H3K9 me2:DNA Meth. m5C
H3K4 me1:H4−4ac
DNA Meth. m5C:H4K20 me2
H3K4 me1:H2A.Z
H4K20 me2:H3−5ac
H4K20 me2:H4−4ac
H4K20 me2:H3K4 me3
H3K27 me3:H3K9 me3
DNA Meth. m5C:H3K9 me3
H2A.Z:H4K20 me2
H3K27 me3:DNA Meth. m5C
DNA Meth. m5C:H4K20 me3
H3K4 me1:H4K20 me2
H3K4 me3
H3−5ac
H4K20 me3
H2A.Z
H3K27 me3
H3K27 me2
H3K9 me2
DNA Meth. m5C
H4−4ac
H3K4 me1
H3K9 me3
H4K20 me2
0.00 0.25 0.50 0.75 1.00
$\hat{\pi}$
DDX17 (2) − Forward
H3K4 me1:H4−4ac
H3K4 me1:H2A.Z
H4−4ac:H3−5ac
H2A.Z:H4−4ac
H2A.Z:H3−5ac
H3K4 me1:H4K20 me2
DNA Meth. m5C:H4K20 me2
H3K4 me3:H3−5ac
H2A.Z:H4K20 me2
H3K4 me1:H3−5ac
H3K27 me3:H3K9 me3
H4K20 me2:H4−4ac
H4K20 me2:H3−5ac
H3K9 me2:DNA Meth. m5C
H3K4 me3:H4−4ac
H2A.Z:H3K4 me3
H3K27 me3:DNA Meth. m5C
DNA Meth. m5C:H3K9 me3
DNA Meth. m5C:H4K20 me3
H4K20 me2:H3K4 me3
H4K20 me3
H3K9 me2
H3K4 me1
H4−4ac
H3K9 me3
H4K20 me2
H3−5ac
H3K27 me3
H3K4 me3
H2A.Z
DNA Meth. m5C
0.00 0.25 0.50 0.75 1.00
$\hat{\pi}$
DDX17 (2) − Reverse
H3K4 me1:H4−4ac
H3K4 me1:H2A.Z
H4−4ac:H3−5ac
H2A.Z:H4−4ac
H2A.Z:H3−5ac
H3K4 me1:H4K20 me2
DNA Meth. m5C:H4K20 me2
H3K4 me3:H3−5ac
H2A.Z:H4K20 me2
H3K4 me1:H3−5ac
H3K27 me3:H3K9 me3
H4K20 me2:H4−4ac
H4K20 me2:H3−5ac
H3K9 me2:DNA Meth. m5C
H3K4 me3:H4−4ac
H2A.Z:H3K4 me3
H3K27 me3:DNA Meth. m5C
DNA Meth. m5C:H3K9 me3
DNA Meth. m5C:H4K20 me3
H4K20 me2:H3K4 me3
H4K20 me3
H3K9 me2
H3K4 me1
H4−4ac
H3K9 me3
H4K20 me2
H3−5ac
H3K27 me3
H3K4 me3
H2A.Z
DNA Meth. m5C
0.00 0.25 0.50 0.75 1.00
$\hat{\pi}$
TAF15 − Forward
H4−4ac:H3−5ac
H4K20 me2:H3K4 me3
H2A.Z:H3−5ac
H2A.Z:H4−4ac
H3K4 me1:H2A.Z
H3K4 me3:H4−4ac
H3K4 me1:H4−4ac
H3K27 me3:H4K20 me3
H3K27 me3:H3K9 me3
H3K9 me3:H4K20 me3
DNA Meth. m5C:H3K9 me3
H4K20 me2:H4−4ac
H4K20 me2:H3−5ac
H2A.Z:H4K20 me2
H3K9 me2:DNA Meth. m5C
H3K27 me3:DNA Meth. m5C
H3K4 me1:H4K20 me2
DNA Meth. m5C:H4K20 me3
DNA Meth. m5C:H4K20 me2
H3K4 me3
H3K27 me2
H3K9 me2
H4K20 me3
H3K9 me3
H2A.Z
H3−5ac
H4K20 me2
H3K27 me3
H3K4 me1
H4−4ac
DNA Meth. m5C
0.00 0.25 0.50 0.75 1.00
$\hat{\pi}$
TAF15 − Reverse
H4−4ac:H3−5ac
H4K20 me2:H3K4 me3
H2A.Z:H3−5ac
H2A.Z:H4−4ac
H3K4 me1:H2A.Z
H3K4 me3:H4−4ac
H3K4 me1:H4−4ac
H3K27 me3:H4K20 me3
H3K27 me3:H3K9 me3
H3K9 me3:H4K20 me3
DNA Meth. m5C:H3K9 me3
H4K20 me2:H4−4ac
H4K20 me2:H3−5ac
H2A.Z:H4K20 me2
H3K9 me2:DNA Meth. m5C
H3K27 me3:DNA Meth. m5C
H3K4 me1:H4K20 me2
DNA Meth. m5C:H4K20 me3
DNA Meth. m5C:H4K20 me2
H3K4 me3
H3K27 me2
H3K9 me2
H4K20 me3
H3K9 me3
H2A.Z
H3−5ac
H4K20 me2
H3K27 me3
H3K4 me1
H4−4ac
DNA Meth. m5C
0.00 0.25 0.50 0.75 1.00
$\hat{\pi}$

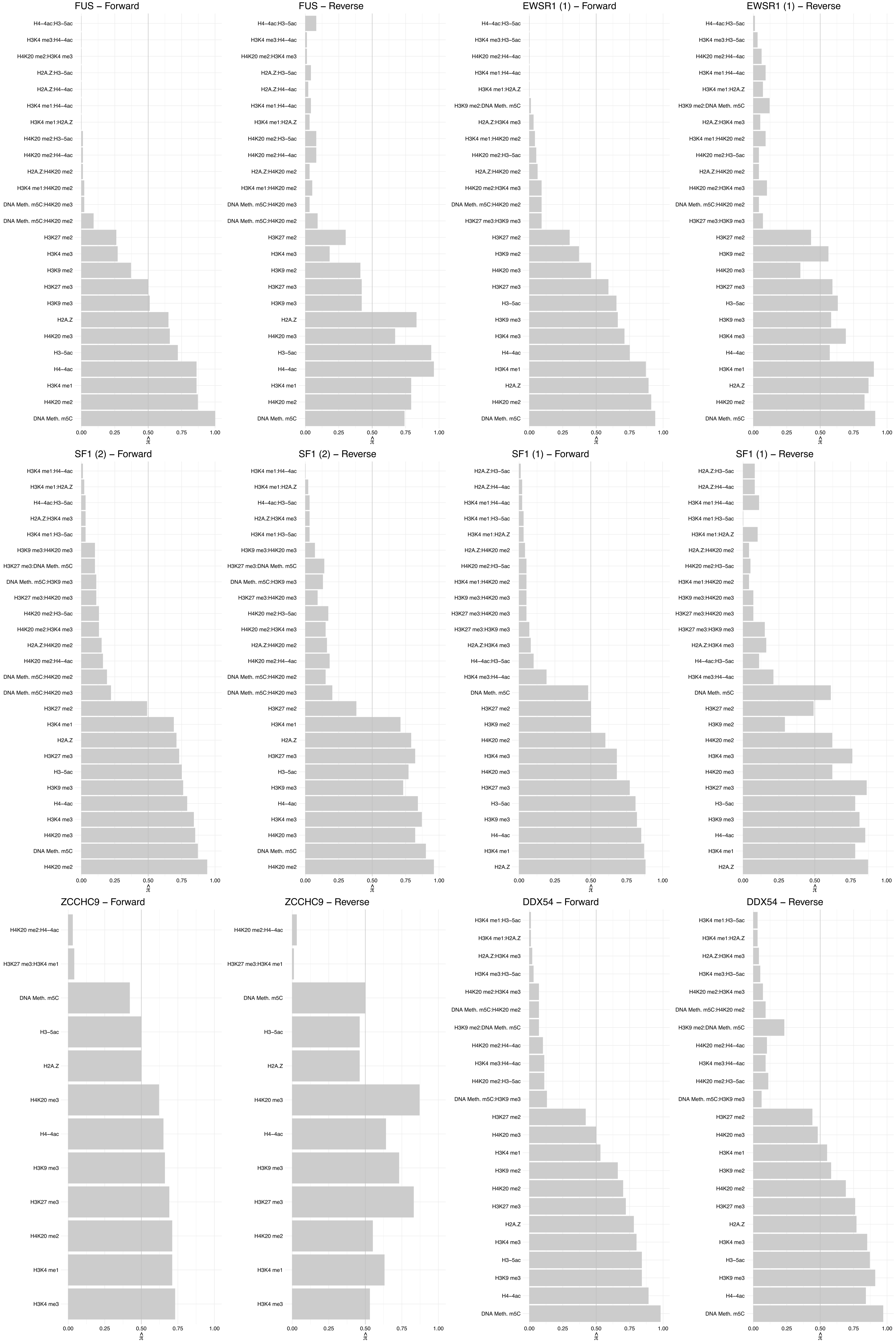
FUS – Forward
H4−4ac:H3−5ac
H3K4 me3:H4−4ac
H4K20 me2:H3K4 me3
H2A.Z:H3−5ac
H2A.Z:H4−4ac
H3K4 me1:H4−4ac
H3K4 me1:H2A.Z
H4K20 me2:H3−5ac
H4K20 me2:H4−4ac
H2A.Z:H4K20 me2
H3K4 me1:H4K20 me2
DNA Meth. m5C:H4K20 me3
DNA Meth. m5C:H4K20 me2
H3K27 me2
H3K4 me3
H3K9 me2
H3K27 me3
H3K9 me3
H2A.Z
H4K20 me3
H3−5ac
H4−4ac
H3K4 me1
H4K20 me2
DNA Meth. m5C
0.00 0.25 0.50 0.75 1.00
π̂
FUS – Reverse
H4−4ac:H3−5ac
H3K4 me3:H4−4ac
H4K20 me2:H3K4 me3
H2A.Z:H3−5ac
H2A.Z:H4−4ac
H3K4 me1:H4−4ac
H3K4 me1:H2A.Z
H4K20 me2:H3−5ac
H4K20 me2:H4−4ac
H2A.Z:H4K20 me2
H3K4 me1:H4K20 me2
DNA Meth. m5C:H4K20 me3
DNA Meth. m5C:H4K20 me2
H3K27 me2
H3K4 me3
H3K9 me2
H3K27 me3
H3K9 me3
H2A.Z
H4K20 me3
H3−5ac
H4−4ac
H3K4 me1
H4K20 me2
DNA Meth. m5C
0.00 0.25 0.50 0.75 1.00
π̂
EWSR1 (1) – Forward
H4−4ac:H3−5ac
H3K4 me3:H3−5ac
H4K20 me2:H4−4ac
H3K4 me1:H4−4ac
H3K4 me1:H2A.Z
H3K9 me2:DNA Meth. m5C
H2A.Z:H3K4 me3
H3K4 me1:H4K20 me2
H4K20 me2:H3−5ac
H2A.Z:H4K20 me2
H4K20 me2:H3K4 me3
DNA Meth. m5C:H4K20 me2
H3K27 me3:H3K9 me3
H3K27 me2
H3K9 me2
H4K20 me3
H3K27 me3
H3−5ac
H3K9 me3
H3K4 me3
H4−4ac
H3K4 me1
H2A.Z
H4K20 me2
DNA Meth. m5C
0.00 0.25 0.50 0.75 1.00
π̂
EWSR1 (1) – Reverse
H4−4ac:H3−5ac
H3K4 me3:H3−5ac
H4K20 me2:H4−4ac
H3K4 me1:H4−4ac
H3K4 me1:H2A.Z
H3K9 me2:DNA Meth. m5C
H2A.Z:H3K4 me3
H3K4 me1:H4K20 me2
H4K20 me2:H3−5ac
H2A.Z:H4K20 me2
H4K20 me2:H3K4 me3
DNA Meth. m5C:H4K20 me2
H3K27 me3:H3K9 me3
H3K27 me2
H3K9 me2
H4K20 me3
H3K27 me3
H3−5ac
H3K9 me3
H3K4 me3
H4−4ac
H3K4 me1
H2A.Z
H4K20 me2
DNA Meth. m5C
0.00 0.25 0.50 0.75 1.00
π̂
SF1 (2) – Forward
H3K4 me1:H4−4ac
H3K4 me1:H2A.Z
H4−4ac:H3−5ac
H2A.Z:H3K4 me3
H3K4 me1:H3−5ac
H3K9 me3:H4K20 me3
H3K27 me3:DNA Meth. m5C
DNA Meth. m5C:H3K9 me3
H3K27 me3:H4K20 me3
H4K20 me2:H3−5ac
H4K20 me2:H3K4 me3
H2A.Z:H4K20 me2
H4K20 me2:H4−4ac
DNA Meth. m5C:H4K20 me2
DNA Meth. m5C:H4K20 me3
H3K27 me2
H3K4 me1
H2A.Z
H3K27 me3
H3−5ac
H3K9 me3
H4−4ac
H3K4 me3
H4K20 me3
DNA Meth. m5C
H4K20 me2
0.00 0.25 0.50 0.75 1.00
π̂
SF1 (2) – Reverse
H3K4 me1:H4−4ac
H3K4 me1:H2A.Z
H4−4ac:H3−5ac
H2A.Z:H3K4 me3
H3K4 me1:H3−5ac
H3K9 me3:H4K20 me3
H3K27 me3:DNA Meth. m5C
DNA Meth. m5C:H3K9 me3
H3K27 me3:H4K20 me3
H4K20 me2:H3−5ac
H4K20 me2:H3K4 me3
H2A.Z:H4K20 me2
H4K20 me2:H4−4ac
DNA Meth. m5C:H4K20 me2
DNA Meth. m5C:H4K20 me3
H3K27 me2
H3K4 me1
H2A.Z
H3K27 me3
H3−5ac
H3K9 me3
H4−4ac
H3K4 me3
H4K20 me3
DNA Meth. m5C
H4K20 me2
0.00 0.25 0.50 0.75 1.00
π̂
SF1 (1) – Forward
H2A.Z:H3−5ac
H2A.Z:H4−4ac
H3K4 me1:H4−4ac
H3K4 me1:H3−5ac
H3K4 me1:H2A.Z
H2A.Z:H4K20 me2
H4K20 me2:H3−5ac
H3K4 me1:H4K20 me2
H3K9 me3:H4K20 me3
H3K27 me3:H4K20 me3
H3K27 me3:H3K9 me3
H2A.Z:H3K4 me3
H4−4ac:H3−5ac
H3K4 me3:H4−4ac
DNA Meth. m5C
H3K27 me2
H3K9 me2
H4K20 me2
H3K4 me3
H4K20 me3
H3K27 me3
H3−5ac
H3K9 me3
H4−4ac
H3K4 me1
H2A.Z
0.00 0.25 0.50 0.75 1.00
π̂
SF1 (1) – Reverse
H2A.Z:H3−5ac
H2A.Z:H4−4ac
H3K4 me1:H4−4ac
H3K4 me1:H3−5ac
H3K4 me1:H2A.Z
H2A.Z:H4K20 me2
H4K20 me2:H3−5ac
H3K4 me1:H4K20 me2
H3K9 me3:H4K20 me3
H3K27 me3:H4K20 me3
H3K27 me3:H3K9 me3
H2A.Z:H3K4 me3
H4−4ac:H3−5ac
H3K4 me3:H4−4ac
DNA Meth. m5C
H3K27 me2
H3K9 me2
H4K20 me2
H3K4 me3
H4K20 me3
H3K27 me3
H3−5ac
H3K9 me3
H4−4ac
H3K4 me1
H2A.Z
0.00 0.25 0.50 0.75 1.00
π̂
ZCCHC9 – Forward
H4K20 me2:H4−4ac
H3K27 me3:H3K4 me1
DNA Meth. m5C
H3−5ac
H2A.Z
H4K20 me3
H4−4ac
H3K9 me3
H3K27 me3
H4K20 me2
H3K4 me1
H3K4 me3
0.00 0.25 0.50 0.75 1.00
π̂
ZCCHC9 – Reverse
H4K20 me2:H4−4ac
H3K27 me3:H3K4 me1
DNA Meth. m5C
H3−5ac
H2A.Z
H4K20 me3
H4−4ac
H3K9 me3
H3K27 me3
H4K20 me2
H3K4 me1
H3K4 me3
0.00 0.25 0.50 0.75 1.00
π̂
DDX54 – Forward
H3K4 me1:H3−5ac
H3K4 me1:H2A.Z
H2A.Z:H3K4 me3
H3K4 me3:H3−5ac
H4K20 me2:H3K4 me3
DNA Meth. m5C:H4K20 me2
H3K9 me2:DNA Meth. m5C
H4K20 me2:H4−4ac
H3K4 me3:H4−4ac
H4K20 me2:H3−5ac
DNA Meth. m5C:H3K9 me3
H3K27 me2
H4K20 me3
H3K4 me1
H3K9 me2
H4K20 me2
H3K27 me3
H2A.Z
H3K4 me3
H3−5ac
H3K9 me3
H4−4ac
DNA Meth. m5C
0.00 0.25 0.50 0.75 1.00
π̂
DDX54 – Reverse
H3K4 me1:H3−5ac
H3K4 me1:H2A.Z
H2A.Z:H3K4 me3
H3K4 me3:H3−5ac
H4K20 me2:H3K4 me3
DNA Meth. m5C:H4K20 me2
H3K9 me2:DNA Meth. m5C
H4K20 me2:H4−4ac
H3K4 me3:H4−4ac
H4K20 me2:H3−5ac
DNA Meth. m5C:H3K9 me3
H3K27 me2
H4K20 me3
H3K4 me1
H3K9 me2
H4K20 me2
H3K27 me3
H2A.Z
H3K4 me3
H3−5ac
H3K9 me3
H4−4ac
DNA Meth. m5C
0.00 0.25 0.50 0.75 1.00
π̂

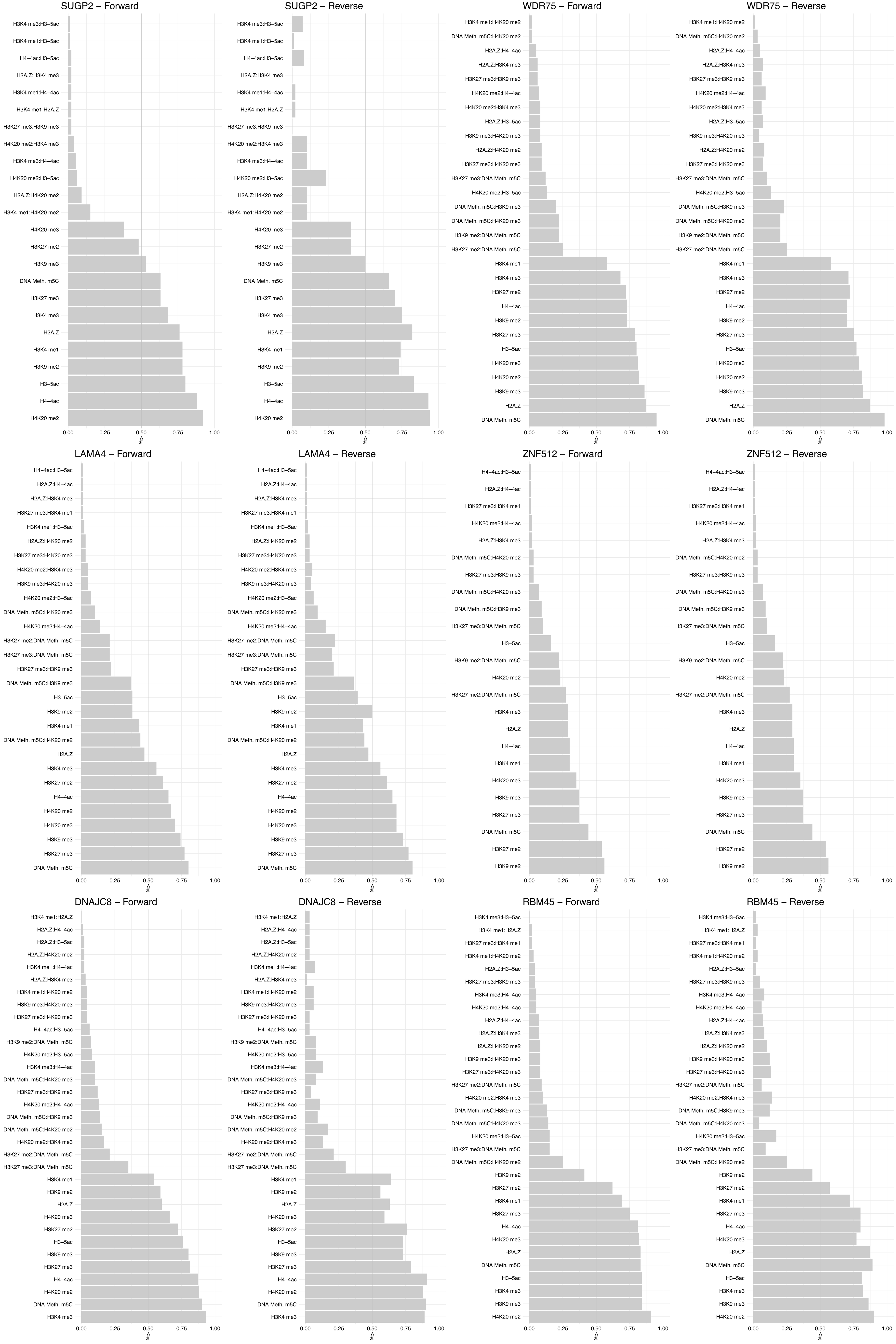
SUGP2 − Forward
H3K4 me3:H3−5ac
H3K4 me1:H3−5ac
H4−4ac:H3−5ac
H2A.Z:H3K4 me3
H3K4 me1:H4−4ac
H3K4 me1:H2A.Z
H3K27 me3:H3K9 me3
H4K20 me2:H3K4 me3
H3K4 me3:H4−4ac
H4K20 me2:H3−5ac
H2A.Z:H4K20 me2
H3K4 me1:H4K20 me2
H4K20 me3
H3K27 me2
H3K9 me3
DNA Meth. m5C
H3K27 me3
H3K4 me3
H2A.Z
H3K4 me1
H3K9 me2
H3−5ac
H4−4ac
H4K20 me2
0.00 0.25 0.50 0.75 1.00
$\hat{\pi}$
SUGP2 − Reverse
H3K4 me3:H3−5ac
H3K4 me1:H3−5ac
H4−4ac:H3−5ac
H2A.Z:H3K4 me3
H3K4 me1:H4−4ac
H3K4 me1:H2A.Z
H3K27 me3:H3K9 me3
H4K20 me2:H3K4 me3
H3K4 me3:H4−4ac
H4K20 me2:H3−5ac
H2A.Z:H4K20 me2
H3K4 me1:H4K20 me2
H4K20 me3
H3K27 me2
H3K9 me3
DNA Meth. m5C
H3K27 me3
H3K4 me3
H2A.Z
H3K4 me1
H3K9 me2
H3−5ac
H4−4ac
H4K20 me2
0.00 0.25 0.50 0.75 1.00
$\hat{\pi}$
WDR75 − Forward
H3K4 me1:H4K20 me2
DNA Meth. m5C:H4K20 me2
H2A.Z:H4−4ac
H2A.Z:H3K4 me3
H3K27 me3:H3K9 me3
H4K20 me2:H4−4ac
H4K20 me2:H3K4 me3
H2A.Z:H3−5ac
H3K9 me3:H4K20 me3
H2A.Z:H4K20 me2
H3K27 me3:H4K20 me3
H3K27 me3:DNA Meth. m5C
H4K20 me2:H3−5ac
DNA Meth. m5C:H3K9 me3
DNA Meth. m5C:H4K20 me3
H3K9 me2:DNA Meth. m5C
H3K27 me2:DNA Meth. m5C
H3K4 me1
H3K4 me3
H3K27 me2
H4−4ac
H3K9 me2
H3K27 me3
H3−5ac
H4K20 me3
H4K20 me2
H3K9 me3
H2A.Z
DNA Meth. m5C
0.00 0.25 0.50 0.75 1.00
$\hat{\pi}$
WDR75 − Reverse
H3K4 me1:H4K20 me2
DNA Meth. m5C:H4K20 me2
H2A.Z:H4−4ac
H2A.Z:H3K4 me3
H3K27 me3:H3K9 me3
H4K20 me2:H4−4ac
H4K20 me2:H3K4 me3
H2A.Z:H3−5ac
H3K9 me3:H4K20 me3
H2A.Z:H4K20 me2
H3K27 me3:H4K20 me3
H3K27 me3:DNA Meth. m5C
H4K20 me2:H3−5ac
DNA Meth. m5C:H3K9 me3
DNA Meth. m5C:H4K20 me3
H3K9 me2:DNA Meth. m5C
H3K27 me2:DNA Meth. m5C
H3K4 me1
H3K4 me3
H3K27 me2
H4−4ac
H3K9 me2
H3K27 me3
H3−5ac
H4K20 me3
H4K20 me2
H3K9 me3
H2A.Z
DNA Meth. m5C
0.00 0.25 0.50 0.75 1.00
$\hat{\pi}$
LAMA4 − Forward
H4−4ac:H3−5ac
H2A.Z:H4−4ac
H2A.Z:H3K4 me3
H3K27 me3:H3K4 me1
H3K4 me1:H3−5ac
H2A.Z:H4K20 me2
H3K27 me3:H4K20 me3
H4K20 me2:H3K4 me3
H3K9 me3:H4K20 me3
H4K20 me2:H3−5ac
DNA Meth. m5C:H4K20 me3
H4K20 me2:H4−4ac
H3K27 me2:DNA Meth. m5C
H3K27 me3:DNA Meth. m5C
H3K27 me3:H3K9 me3
DNA Meth. m5C:H3K9 me3
H3−5ac
H3K9 me2
H3K4 me1
DNA Meth. m5C:H4K20 me2
H2A.Z
H3K4 me3
H3K27 me2
H4−4ac
H4K20 me2
H4K20 me3
H3K9 me3
H3K27 me3
DNA Meth. m5C
0.00 0.25 0.50 0.75 1.00
$\hat{\pi}$
LAMA4 − Reverse
H4−4ac:H3−5ac
H2A.Z:H4−4ac
H2A.Z:H3K4 me3
H3K27 me3:H3K4 me1
H3K4 me1:H3−5ac
H2A.Z:H4K20 me2
H3K27 me3:H4K20 me3
H4K20 me2:H3K4 me3
H3K9 me3:H4K20 me3
H4K20 me2:H3−5ac
DNA Meth. m5C:H4K20 me3
H4K20 me2:H4−4ac
H3K27 me2:DNA Meth. m5C
H3K27 me3:DNA Meth. m5C
H3K27 me3:H3K9 me3
DNA Meth. m5C:H3K9 me3
H3−5ac
H3K9 me2
H3K4 me1
DNA Meth. m5C:H4K20 me2
H2A.Z
H3K4 me3
H3K27 me2
H4−4ac
H4K20 me2
H4K20 me3
H3K9 me3
H3K27 me3
DNA Meth. m5C
0.00 0.25 0.50 0.75 1.00
$\hat{\pi}$
ZNF512 − Forward
H4−4ac:H3−5ac
H2A.Z:H4−4ac
H3K27 me3:H3K4 me1
H4K20 me2:H4−4ac
H2A.Z:H3K4 me3
DNA Meth. m5C:H4K20 me2
H3K27 me3:H3K9 me3
DNA Meth. m5C:H4K20 me3
DNA Meth. m5C:H3K9 me3
H3K27 me3:DNA Meth. m5C
H3−5ac
H3K9 me2:DNA Meth. m5C
H4K20 me2
H3K27 me2:DNA Meth. m5C
H3K4 me3
H2A.Z
H4−4ac
H3K4 me1
H4K20 me3
H3K9 me3
H3K27 me3
DNA Meth. m5C
H3K27 me2
H3K9 me2
0.00 0.25 0.50 0.75 1.00
$\hat{\pi}$
ZNF512 − Reverse
H4−4ac:H3−5ac
H2A.Z:H4−4ac
H3K27 me3:H3K4 me1
H4K20 me2:H4−4ac
H2A.Z:H3K4 me3
DNA Meth. m5C:H4K20 me2
H3K27 me3:H3K9 me3
DNA Meth. m5C:H4K20 me3
DNA Meth. m5C:H3K9 me3
H3K27 me3:DNA Meth. m5C
H3−5ac
H3K9 me2:DNA Meth. m5C
H4K20 me2
H3K27 me2:DNA Meth. m5C
H3K4 me3
H2A.Z
H4−4ac
H3K4 me1
H4K20 me3
H3K9 me3
H3K27 me3
DNA Meth. m5C
H3K27 me2
H3K9 me2
0.00 0.25 0.50 0.75 1.00
$\hat{\pi}$
DNAJC8 − Forward
H3K4 me1:H2A.Z
H2A.Z:H4−4ac
H2A.Z:H3−5ac
H2A.Z:H4K20 me2
H3K4 me1:H4−4ac
H2A.Z:H3K4 me3
H3K4 me1:H4K20 me2
H3K9 me3:H4K20 me3
H3K27 me3:H4K20 me3
H4−4ac:H3−5ac
H3K9 me2:DNA Meth. m5C
H4K20 me2:H3−5ac
H3K4 me3:H4−4ac
DNA Meth. m5C:H4K20 me3
H3K27 me3:H3K9 me3
H4K20 me2:H4−4ac
DNA Meth. m5C:H3K9 me3
DNA Meth. m5C:H4K20 me2
H4K20 me2:H3K4 me3
H3K27 me2:DNA Meth. m5C
H3K27 me3:DNA Meth. m5C
H3K4 me1
H3K9 me2
H2A.Z
H4K20 me3
H3K27 me2
H3−5ac
H3K9 me3
H3K27 me3
H4−4ac
H4K20 me2
DNA Meth. m5C
H3K4 me3
0.00 0.25 0.50 0.75 1.00
$\hat{\pi}$
DNAJC8 − Reverse
H3K4 me1:H2A.Z
H2A.Z:H4−4ac
H2A.Z:H3−5ac
H2A.Z:H4K20 me2
H3K4 me1:H4−4ac
H2A.Z:H3K4 me3
H3K4 me1:H4K20 me2
H3K9 me3:H4K20 me3
H3K27 me3:H4K20 me3
H4−4ac:H3−5ac
H3K9 me2:DNA Meth. m5C
H4K20 me2:H3−5ac
H3K4 me3:H4−4ac
DNA Meth. m5C:H4K20 me3
H3K27 me3:H3K9 me3
H4K20 me2:H4−4ac
DNA Meth. m5C:H3K9 me3
DNA Meth. m5C:H4K20 me2
H4K20 me2:H3K4 me3
H3K27 me2:DNA Meth. m5C
H3K27 me3:DNA Meth. m5C
H3K4 me1
H3K9 me2
H2A.Z
H4K20 me3
H3K27 me2
H3−5ac
H3K9 me3
H3K27 me3
H4−4ac
H4K20 me2
DNA Meth. m5C
H3K4 me3
0.00 0.25 0.50 0.75 1.00
$\hat{\pi}$
RBM45 − Forward
H3K4 me3:H3−5ac
H3K4 me1:H2A.Z
H3K27 me3:H3K4 me1
H3K4 me1:H4K20 me2
H2A.Z:H3−5ac
H3K27 me3:H3K9 me3
H3K4 me3:H4−4ac
H4K20 me2:H4−4ac
H2A.Z:H4−4ac
H2A.Z:H3K4 me3
H2A.Z:H4K20 me2
H3K9 me3:H4K20 me3
H3K27 me3:H4K20 me3
H3K27 me2:DNA Meth. m5C
H4K20 me2:H3K4 me3
DNA Meth. m5C:H3K9 me3
DNA Meth. m5C:H4K20 me3
H4K20 me2:H3−5ac
H3K27 me3:DNA Meth. m5C
DNA Meth. m5C:H4K20 me2
H3K9 me2
H3K27 me2
H3K4 me1
H3K27 me3
H4−4ac
H4K20 me3
H2A.Z
DNA Meth. m5C
H3−5ac
H3K4 me3
H3K9 me3
H4K20 me2
0.00 0.25 0.50 0.75 1.00
$\hat{\pi}$
RBM45 − Reverse
H3K4 me3:H3−5ac
H3K4 me1:H2A.Z
H3K27 me3:H3K4 me1
H3K4 me1:H4K20 me2
H2A.Z:H3−5ac
H3K27 me3:H3K9 me3
H3K4 me3:H4−4ac
H4K20 me2:H4−4ac
H2A.Z:H4−4ac
H2A.Z:H3K4 me3
H2A.Z:H4K20 me2
H3K9 me3:H4K20 me3
H3K27 me3:H4K20 me3
H3K27 me2:DNA Meth. m5C
H4K20 me2:H3K4 me3
DNA Meth. m5C:H3K9 me3
DNA Meth. m5C:H4K20 me3
H4K20 me2:H3−5ac
H3K27 me3:DNA Meth. m5C
DNA Meth. m5C:H4K20 me2
H3K9 me2
H3K27 me2
H3K4 me1
H3K27 me3
H4−4ac
H4K20 me3
H2A.Z
DNA Meth. m5C
H3−5ac
H3K4 me3
H3K9 me3
H4K20 me2
0.00 0.25 0.50 0.75 1.00
$\hat{\pi}$

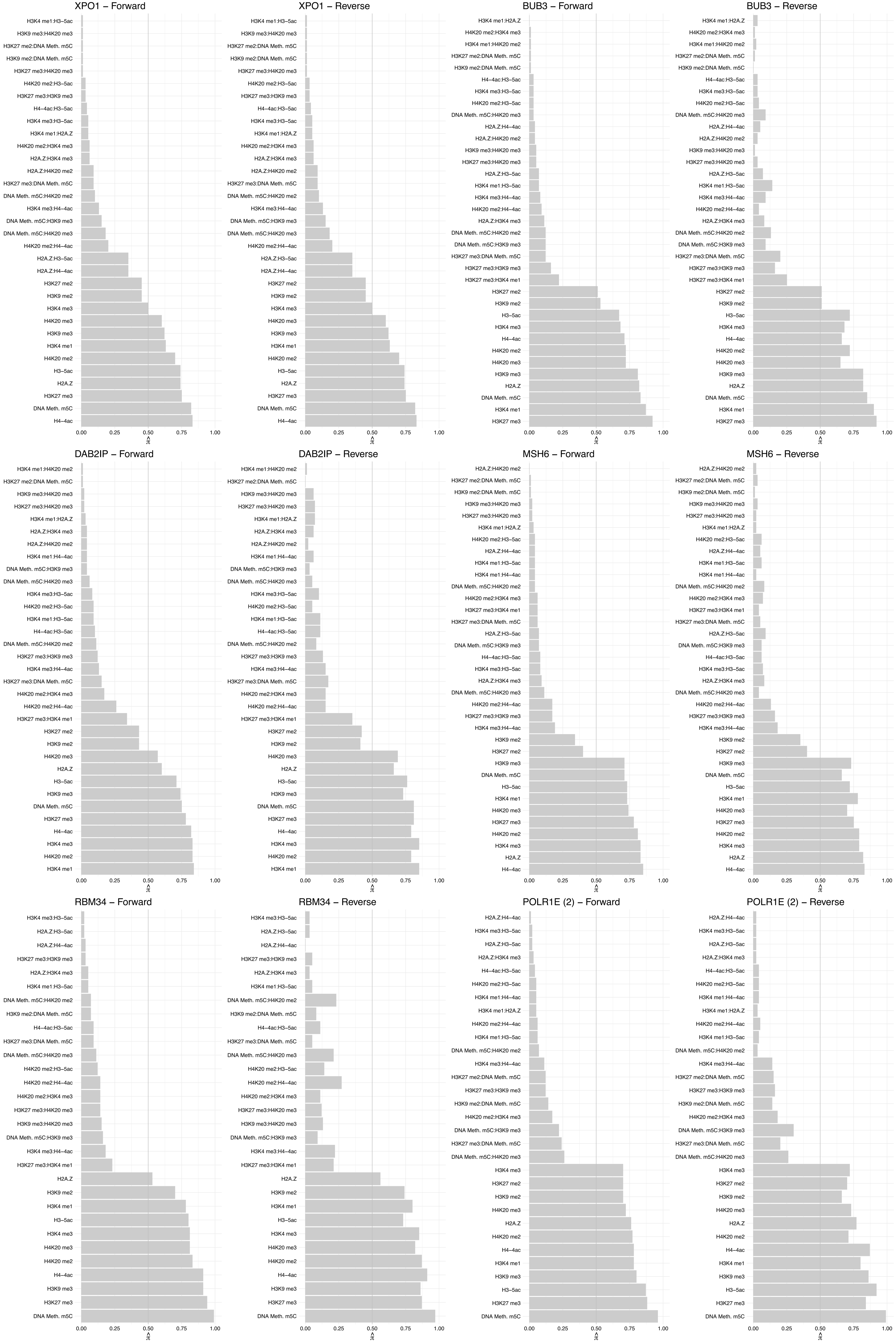
XPO1 – Forward
H3K4 me1:H3–5ac
H3K9 me3:H4K20 me3
H3K27 me2:DNA Meth. m5C
H3K9 me2:DNA Meth. m5C
H3K27 me3:H4K20 me3
H4K20 me2:H3–5ac
H3K27 me3:H3K9 me3
H4–4ac:H3–5ac
H3K4 me3:H3–5ac
H3K4 me1:H2A.Z
H4K20 me2:H3K4 me3
H2A.Z:H3K4 me3
H2A.Z:H4K20 me2
H3K27 me3:DNA Meth. m5C
DNA Meth. m5C:H4K20 me2
H3K4 me3:H4–4ac
DNA Meth. m5C:H3K9 me3
DNA Meth. m5C:H4K20 me3
H4K20 me2:H4–4ac
H2A.Z:H3–5ac
H2A.Z:H4–4ac
H3K27 me2
H3K9 me2
H3K4 me3
H4K20 me3
H3K9 me3
H3K4 me1
H4K20 me2
H3–5ac
H2A.Z
H3K27 me3
DNA Meth. m5C
H4–4ac
0.00 0.25 0.50 0.75 1.00
π̂
XPO1 – Reverse
H3K4 me1:H3–5ac
H3K9 me3:H4K20 me3
H3K27 me2:DNA Meth. m5C
H3K9 me2:DNA Meth. m5C
H3K27 me3:H4K20 me3
H4K20 me2:H3–5ac
H3K27 me3:H3K9 me3
H4–4ac:H3–5ac
H3K4 me3:H3–5ac
H3K4 me1:H2A.Z
H4K20 me2:H3K4 me3
H2A.Z:H3K4 me3
H2A.Z:H4K20 me2
H3K27 me3:DNA Meth. m5C
DNA Meth. m5C:H4K20 me2
H3K4 me3:H4–4ac
DNA Meth. m5C:H3K9 me3
DNA Meth. m5C:H4K20 me3
H4K20 me2:H4–4ac
H2A.Z:H3–5ac
H2A.Z:H4–4ac
H3K27 me2
H3K9 me2
H3K4 me3
H4K20 me3
H3K9 me3
H3K4 me1
H4K20 me2
H3–5ac
H2A.Z
H3K27 me3
DNA Meth. m5C
H4–4ac
0.00 0.25 0.50 0.75 1.00
π̂
BUB3 – Forward
H3K4 me1:H3–5ac
H4K20 me2:H3K4 me3
H3K4 me1:H4K20 me2
H3K27 me2:DNA Meth. m5C
H3K9 me2:DNA Meth. m5C
H4–4ac:H3–5ac
H3K4 me3:H3–5ac
H4K20 me2:H3–5ac
DNA Meth. m5C:H4K20 me3
H2A.Z:H4–4ac
H2A.Z:H4K20 me2
H3K9 me3:H4K20 me3
H3K27 me3:H4K20 me3
H2A.Z:H3–5ac
H3K4 me1:H3–5ac
H3K4 me3:H4–4ac
H4K20 me2:H4–4ac
H2A.Z:H3K4 me3
DNA Meth. m5C:H4K20 me2
DNA Meth. m5C:H3K9 me3
H3K27 me3:DNA Meth. m5C
H3K27 me3:H3K9 me3
H3K27 me3:H3K4 me1
H3K27 me2
H3K9 me2
H3–5ac
H3K4 me3
H4–4ac
H4K20 me2
H4K20 me3
H3K9 me3
H2A.Z
DNA Meth. m5C
H3K4 me1
H3K27 me3
0.00 0.25 0.50 0.75 1.00
π̂
BUB3 – Reverse
H3K4 me1:H3–5ac
H4K20 me2:H3K4 me3
H3K4 me1:H4K20 me2
H3K27 me2:DNA Meth. m5C
H3K9 me2:DNA Meth. m5C
H4–4ac:H3–5ac
H3K4 me3:H3–5ac
H4K20 me2:H3–5ac
DNA Meth. m5C:H4K20 me3
H2A.Z:H4–4ac
H2A.Z:H4K20 me2
H3K9 me3:H4K20 me3
H3K27 me3:H4K20 me3
H2A.Z:H3–5ac
H3K4 me1:H3–5ac
H3K4 me3:H4–4ac
H4K20 me2:H4–4ac
H2A.Z:H3K4 me3
DNA Meth. m5C:H4K20 me2
DNA Meth. m5C:H3K9 me3
H3K27 me3:DNA Meth. m5C
H3K27 me3:H3K9 me3
H3K27 me3:H3K4 me1
H3K27 me2
H3K9 me2
H3–5ac
H3K4 me3
H4–4ac
H4K20 me2
H4K20 me3
H3K9 me3
H2A.Z
DNA Meth. m5C
H3K4 me1
H3K27 me3
0.00 0.25 0.50 0.75 1.00
π̂
DAB2IP – Forward
H3K4 me1:H4K20 me2
H3K27 me2:DNA Meth. m5C
H3K9 me3:H4K20 me3
H3K27 me3:H4K20 me3
H3K4 me1:H2A.Z
H2A.Z:H3K4 me3
H2A.Z:H4K20 me2
H3K4 me1:H4–4ac
DNA Meth. m5C:H3K9 me3
DNA Meth. m5C:H4K20 me3
H3K4 me3:H3–5ac
H4K20 me2:H3–5ac
H3K4 me1:H3–5ac
H4–4ac:H3–5ac
DNA Meth. m5C:H4K20 me2
H3K27 me3:H3K9 me3
H3K4 me3:H4–4ac
H3K27 me3:DNA Meth. m5C
H4K20 me2:H3K4 me3
H4K20 me2:H4–4ac
H3K27 me3:H3K4 me1
H3K27 me2
H3K9 me2
H4K20 me3
H2A.Z
H3–5ac
H3K9 me3
DNA Meth. m5C
H3K27 me3
H4–4ac
H3K4 me3
H4K20 me2
H3K4 me1
0.00 0.25 0.50 0.75 1.00
π̂
DAB2IP – Reverse
H3K4 me1:H4K20 me2
H3K27 me2:DNA Meth. m5C
H3K9 me3:H4K20 me3
H3K27 me3:H4K20 me3
H3K4 me1:H2A.Z
H2A.Z:H3K4 me3
H2A.Z:H4K20 me2
H3K4 me1:H4–4ac
DNA Meth. m5C:H3K9 me3
DNA Meth. m5C:H4K20 me3
H3K4 me3:H3–5ac
H4K20 me2:H3–5ac
H3K4 me1:H3–5ac
H4–4ac:H3–5ac
DNA Meth. m5C:H4K20 me2
H3K27 me3:H3K9 me3
H3K4 me3:H4–4ac
H3K27 me3:DNA Meth. m5C
H4K20 me2:H3K4 me3
H4K20 me2:H4–4ac
H3K27 me3:H3K4 me1
H3K27 me2
H3K9 me2
H4K20 me3
H2A.Z
H3–5ac
H3K9 me3
DNA Meth. m5C
H3K27 me3
H4–4ac
H3K4 me3
H4K20 me2
H3K4 me1
0.00 0.25 0.50 0.75 1.00
π̂
MSH6 – Forward
H2A.Z:H4K20 me2
H3K27 me2:DNA Meth. m5C
H3K9 me2:DNA Meth. m5C
H3K9 me3:H4K20 me3
H3K27 me3:H4K20 me3
H3K4 me1:H2A.Z
H4K20 me2:H3–5ac
H2A.Z:H4–4ac
H3K4 me1:H3–5ac
H3K4 me1:H4–4ac
DNA Meth. m5C:H4K20 me2
H4K20 me2:H3K4 me3
H3K27 me3:H3K4 me1
H3K27 me3:DNA Meth. m5C
H2A.Z:H3–5ac
DNA Meth. m5C:H3K9 me3
H4–4ac:H3–5ac
H3K4 me3:H3–5ac
H2A.Z:H3K4 me3
DNA Meth. m5C:H4K20 me3
H4K20 me2:H4–4ac
H3K27 me3:H3K9 me3
H3K4 me3:H4–4ac
H3K9 me2
H3K27 me2
H3K9 me3
DNA Meth. m5C
H3–5ac
H3K4 me1
H4K20 me3
H3K27 me3
H4K20 me2
H3K4 me3
H2A.Z
H4–4ac
0.00 0.25 0.50 0.75 1.00
π̂
MSH6 – Reverse
H2A.Z:H4K20 me2
H3K27 me2:DNA Meth. m5C
H3K9 me2:DNA Meth. m5C
H3K9 me3:H4K20 me3
H3K27 me3:H4K20 me3
H3K4 me1:H2A.Z
H4K20 me2:H3–5ac
H2A.Z:H4–4ac
H3K4 me1:H3–5ac
H3K4 me1:H4–4ac
DNA Meth. m5C:H4K20 me2
H4K20 me2:H3K4 me3
H3K27 me3:H3K4 me1
H3K27 me3:DNA Meth. m5C
H2A.Z:H3–5ac
DNA Meth. m5C:H3K9 me3
H4–4ac:H3–5ac
H3K4 me3:H3–5ac
H2A.Z:H3K4 me3
DNA Meth. m5C:H4K20 me3
H4K20 me2:H4–4ac
H3K27 me3:H3K9 me3
H3K4 me3:H4–4ac
H3K9 me2
H3K27 me2
H3K9 me3
DNA Meth. m5C
H3–5ac
H3K4 me1
H4K20 me3
H3K27 me3
H4K20 me2
H3K4 me3
H2A.Z
H4–4ac
0.00 0.25 0.50 0.75 1.00
π̂
RBM34 – Forward
H3K4 me3:H3–5ac
H2A.Z:H3–5ac
H2A.Z:H4–4ac
H3K27 me3:H3K9 me3
H2A.Z:H3K4 me3
H3K4 me1:H3–5ac
DNA Meth. m5C:H4K20 me2
H3K9 me2:DNA Meth. m5C
H4–4ac:H3–5ac
H3K27 me3:DNA Meth. m5C
DNA Meth. m5C:H4K20 me3
H4K20 me2:H3–5ac
H4K20 me2:H4–4ac
H4K20 me2:H3K4 me3
H3K27 me3:H4K20 me3
H3K9 me3:H4K20 me3
DNA Meth. m5C:H3K9 me3
H3K4 me3:H4–4ac
H3K27 me3:H3K4 me1
H2A.Z
H3K9 me2
H3K4 me1
H3–5ac
H3K4 me3
H4K20 me3
H4K20 me2
H4–4ac
H3K9 me3
H3K27 me3
DNA Meth. m5C
0.00 0.25 0.50 0.75 1.00
π̂
RBM34 – Reverse
H3K4 me3:H3–5ac
H2A.Z:H3–5ac
H2A.Z:H4–4ac
H3K27 me3:H3K9 me3
H2A.Z:H3K4 me3
H3K4 me1:H3–5ac
DNA Meth. m5C:H4K20 me2
H3K9 me2:DNA Meth. m5C
H4–4ac:H3–5ac
H3K27 me3:DNA Meth. m5C
DNA Meth. m5C:H4K20 me3
H4K20 me2:H3–5ac
H4K20 me2:H4–4ac
H4K20 me2:H3K4 me3
H3K27 me3:H4K20 me3
H3K9 me3:H4K20 me3
DNA Meth. m5C:H3K9 me3
H3K4 me3:H4–4ac
H3K27 me3:H3K4 me1
H2A.Z
H3K9 me2
H3K4 me1
H3–5ac
H3K4 me3
H4K20 me3
H4K20 me2
H4–4ac
H3K9 me3
H3K27 me3
DNA Meth. m5C
0.00 0.25 0.50 0.75 1.00
π̂
POLR1E (2) – Forward
H2A.Z:H4–4ac
H3K4 me3:H3–5ac
H2A.Z:H3–5ac
H2A.Z:H3K4 me3
H4–4ac:H3–5ac
H4K20 me2:H3–5ac
H3K4 me1:H4–4ac
H3K4 me1:H2A.Z
H4K20 me2:H4–4ac
H3K4 me1:H3–5ac
DNA Meth. m5C:H4K20 me2
H3K4 me3:H4–4ac
H3K27 me2:DNA Meth. m5C
H3K27 me3:H3K9 me3
H3K9 me2:DNA Meth. m5C
H4K20 me2:H3K4 me3
DNA Meth. m5C:H3K9 me3
H3K27 me3:DNA Meth. m5C
DNA Meth. m5C:H4K20 me3
H3K4 me3
H3K27 me2
H3K9 me2
H4K20 me3
H2A.Z
H4K20 me2
H4–4ac
H3K4 me1
H3K9 me3
H3–5ac
H3K27 me3
DNA Meth. m5C
0.00 0.25 0.50 0.75 1.00
π̂
POLR1E (2) – Reverse
H2A.Z:H4–4ac
H3K4 me3:H3–5ac
H2A.Z:H3–5ac
H2A.Z:H3K4 me3
H4–4ac:H3–5ac
H4K20 me2:H3–5ac
H3K4 me1:H4–4ac
H3K4 me1:H2A.Z
H4K20 me2:H4–4ac
H3K4 me1:H3–5ac
DNA Meth. m5C:H4K20 me2
H3K4 me3:H4–4ac
H3K27 me2:DNA Meth. m5C
H3K27 me3:H3K9 me3
H3K9 me2:DNA Meth. m5C
H4K20 me2:H3K4 me3
DNA Meth. m5C:H3K9 me3
H3K27 me3:DNA Meth. m5C
DNA Meth. m5C:H4K20 me3
H3K4 me3
H3K27 me2
H3K9 me2
H4K20 me3
H2A.Z
H4K20 me2
H4–4ac
H3K4 me1
H3K9 me3
H3–5ac
H3K27 me3
DNA Meth. m5C
0.00 0.25 0.50 0.75 1.00
π̂

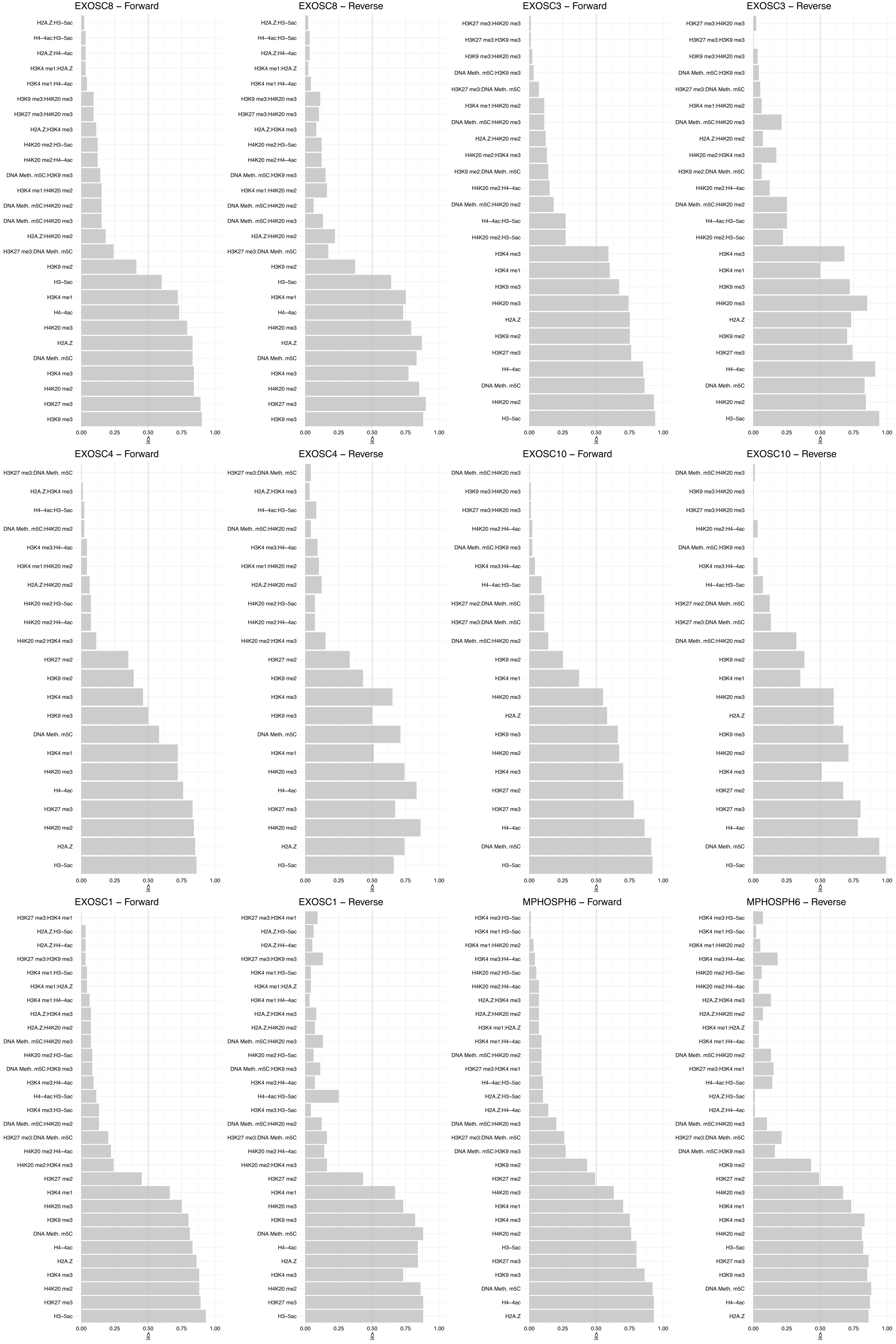
EXOSC8 – Forward
H2A.Z:H3−5ac
H4−4ac:H3−5ac
H2A.Z:H4−4ac
H3K4 me1:H2A.Z
H3K4 me1:H4−4ac
H3K9 me3:H4K20 me3
H3K27 me3:H4K20 me3
H2A.Z:H3K4 me3
H4K20 me2:H3−5ac
H4K20 me2:H4−4ac
DNA Meth. m5C:H3K9 me3
H3K4 me1:H4K20 me2
DNA Meth. m5C:H4K20 me2
DNA Meth. m5C:H4K20 me3
H2A.Z:H4K20 me2
H3K27 me3:DNA Meth. m5C
H3K9 me2
H3−5ac
H3K4 me1
H4−4ac
H4K20 me3
H2A.Z
DNA Meth. m5C
H3K4 me3
H4K20 me2
H3K27 me3
H3K9 me3
0.00
0.25
0.50
0.75
1.00
$\hat{\pi}$
EXOSC8 – Reverse
H2A.Z:H3−5ac
H4−4ac:H3−5ac
H2A.Z:H4−4ac
H3K4 me1:H2A.Z
H3K4 me1:H4−4ac
H3K9 me3:H4K20 me3
H3K27 me3:H4K20 me3
H2A.Z:H3K4 me3
H4K20 me2:H3−5ac
H4K20 me2:H4−4ac
DNA Meth. m5C:H3K9 me3
H3K4 me1:H4K20 me2
DNA Meth. m5C:H4K20 me2
DNA Meth. m5C:H4K20 me3
H2A.Z:H4K20 me2
H3K27 me3:DNA Meth. m5C
H3K9 me2
H3−5ac
H3K4 me1
H4−4ac
H4K20 me3
H2A.Z
DNA Meth. m5C
H3K4 me3
H4K20 me2
H3K27 me3
H3K9 me3
0.00
0.25
0.50
0.75
1.00
$\hat{\pi}$
EXOSC3 – Forward
H3K27 me3:H4K20 me3
H3K27 me3:H3K9 me3
H3K9 me3:H4K20 me3
DNA Meth. m5C:H3K9 me3
H3K27 me3:DNA Meth. m5C
H3K4 me1:H4K20 me2
DNA Meth. m5C:H4K20 me3
H2A.Z:H4K20 me2
H4K20 me2:H3K4 me3
H3K9 me2:DNA Meth. m5C
H4K20 me2:H4−4ac
DNA Meth. m5C:H4K20 me2
H4−4ac:H3−5ac
H4K20 me2:H3−5ac
H3K4 me3
H3K4 me1
H3K9 me3
H4K20 me3
H2A.Z
H3K9 me2
H3K27 me3
H4−4ac
DNA Meth. m5C
H4K20 me2
H3−5ac
0.00
0.25
0.50
0.75
1.00
$\hat{\pi}$
EXOSC3 – Reverse
H3K27 me3:H4K20 me3
H3K27 me3:H3K9 me3
H3K9 me3:H4K20 me3
DNA Meth. m5C:H3K9 me3
H3K27 me3:DNA Meth. m5C
H3K4 me1:H4K20 me2
DNA Meth. m5C:H4K20 me3
H2A.Z:H4K20 me2
H4K20 me2:H3K4 me3
H3K9 me2:DNA Meth. m5C
H4K20 me2:H4−4ac
DNA Meth. m5C:H4K20 me2
H4−4ac:H3−5ac
H4K20 me2:H3−5ac
H3K4 me3
H3K4 me1
H3K9 me3
H4K20 me3
H2A.Z
H3K9 me2
H3K27 me3
H4−4ac
DNA Meth. m5C
H4K20 me2
H3−5ac
0.00
0.25
0.50
0.75
1.00
$\hat{\pi}$
EXOSC4 – Forward
H3K27 me3:DNA Meth. m5C
H2A.Z:H3K4 me3
H4−4ac:H3−5ac
DNA Meth. m5C:H4K20 me2
H3K4 me3:H4−4ac
H3K4 me1:H4K20 me2
H2A.Z:H4K20 me2
H4K20 me2:H3−5ac
H4K20 me2:H4−4ac
H4K20 me2:H3K4 me3
H3K27 me2
H3K9 me2
H3K4 me3
H3K9 me3
DNA Meth. m5C
H3K4 me1
H4K20 me3
H4−4ac
H3K27 me3
H4K20 me2
H2A.Z
H3−5ac
0.00
0.25
0.50
0.75
1.00
$\hat{\pi}$
EXOSC4 – Reverse
H3K27 me3:DNA Meth. m5C
H2A.Z:H3K4 me3
H4−4ac:H3−5ac
DNA Meth. m5C:H4K20 me2
H3K4 me3:H4−4ac
H3K4 me1:H4K20 me2
H2A.Z:H4K20 me2
H4K20 me2:H3−5ac
H4K20 me2:H4−4ac
H4K20 me2:H3K4 me3
H3K27 me2
H3K9 me2
H3K4 me3
H3K9 me3
DNA Meth. m5C
H3K4 me1
H4K20 me3
H4−4ac
H3K27 me3
H4K20 me2
H2A.Z
H3−5ac
0.00
0.25
0.50
0.75
1.00
$\hat{\pi}$
EXOSC10 – Forward
DNA Meth. m5C:H4K20 me3
H3K9 me3:H4K20 me3
H3K27 me3:H4K20 me3
H4K20 me2:H4−4ac
DNA Meth. m5C:H3K9 me3
H3K4 me3:H4−4ac
H4−4ac:H3−5ac
H3K27 me2:DNA Meth. m5C
H3K27 me3:DNA Meth. m5C
DNA Meth. m5C:H4K20 me2
H3K9 me2
H3K4 me1
H4K20 me3
H2A.Z
H3K9 me3
H4K20 me2
H3K4 me3
H3K27 me2
H3K27 me3
H4−4ac
DNA Meth. m5C
H3−5ac
0.00
0.25
0.50
0.75
1.00
$\hat{\pi}$
EXOSC10 – Reverse
DNA Meth. m5C:H4K20 me3
H3K9 me3:H4K20 me3
H3K27 me3:H4K20 me3
H4K20 me2:H4−4ac
DNA Meth. m5C:H3K9 me3
H3K4 me3:H4−4ac
H4−4ac:H3−5ac
H3K27 me2:DNA Meth. m5C
H3K27 me3:DNA Meth. m5C
DNA Meth. m5C:H4K20 me2
H3K9 me2
H3K4 me1
H4K20 me3
H2A.Z
H3K9 me3
H4K20 me2
H3K4 me3
H3K27 me2
H3K27 me3
H4−4ac
DNA Meth. m5C
H3−5ac
0.00
0.25
0.50
0.75
1.00
$\hat{\pi}$
EXOSC1 – Forward
H3K27 me3:H3K4 me1
H2A.Z:H3−5ac
H2A.Z:H4−4ac
H3K27 me3:H3K9 me3
H3K4 me1:H3−5ac
H3K4 me1:H2A.Z
H3K4 me1:H4−4ac
H2A.Z:H3K4 me3
H2A.Z:H4K20 me2
DNA Meth. m5C:H4K20 me3
H4K20 me2:H3−5ac
DNA Meth. m5C:H3K9 me3
H3K4 me3:H4−4ac
H4−4ac:H3−5ac
H3K4 me3:H3−5ac
DNA Meth. m5C:H4K20 me2
H3K27 me3:DNA Meth. m5C
H4K20 me2:H4−4ac
H4K20 me2:H3K4 me3
H3K27 me2
H3K4 me1
H4K20 me3
H3K9 me3
DNA Meth. m5C
H4−4ac
H2A.Z
H3K4 me3
H4K20 me2
H3K27 me3
H3−5ac
0.00
0.25
0.50
0.75
1.00
$\hat{\pi}$
EXOSC1 – Reverse
H3K27 me3:H3K4 me1
H2A.Z:H3−5ac
H2A.Z:H4−4ac
H3K27 me3:H3K9 me3
H3K4 me1:H3−5ac
H3K4 me1:H2A.Z
H3K4 me1:H4−4ac
H2A.Z:H3K4 me3
H2A.Z:H4K20 me2
DNA Meth. m5C:H4K20 me3
H4K20 me2:H3−5ac
DNA Meth. m5C:H3K9 me3
H3K4 me3:H4−4ac
H4−4ac:H3−5ac
H3K4 me3:H3−5ac
DNA Meth. m5C:H4K20 me2
H3K27 me3:DNA Meth. m5C
H4K20 me2:H4−4ac
H4K20 me2:H3K4 me3
H3K27 me2
H3K4 me1
H4K20 me3
H3K9 me3
DNA Meth. m5C
H4−4ac
H2A.Z
H3K4 me3
H4K20 me2
H3K27 me3
H3−5ac
0.00
0.25
0.50
0.75
1.00
$\hat{\pi}$
MPHOSPH6 – Forward
H3K4 me3:H3−5ac
H3K4 me1:H3−5ac
H3K4 me1:H4K20 me2
H3K4 me3:H4−4ac
H4K20 me2:H3−5ac
H4K20 me2:H4−4ac
H2A.Z:H3K4 me3
H2A.Z:H4K20 me2
H3K4 me1:H2A.Z
H3K4 me1:H4−4ac
DNA Meth. m5C:H4K20 me2
H3K27 me3:H3K4 me1
H4−4ac:H3−5ac
H2A.Z:H3−5ac
H2A.Z:H4−4ac
DNA Meth. m5C:H4K20 me3
H3K27 me3:DNA Meth. m5C
DNA Meth. m5C:H3K9 me3
H3K9 me2
H3K27 me2
H4K20 me3
H3K4 me1
H3K4 me3
H4K20 me2
H3−5ac
H3K27 me3
H3K9 me3
DNA Meth. m5C
H4−4ac
H2A.Z
0.00
0.25
0.50
0.75
1.00
$\hat{\pi}$
MPHOSPH6 – Reverse
H3K4 me3:H3−5ac
H3K4 me1:H3−5ac
H3K4 me1:H4K20 me2
H3K4 me3:H4−4ac
H4K20 me2:H3−5ac
H4K20 me2:H4−4ac
H2A.Z:H3K4 me3
H2A.Z:H4K20 me2
H3K4 me1:H2A.Z
H3K4 me1:H4−4ac
DNA Meth. m5C:H4K20 me2
H3K27 me3:H3K4 me1
H4−4ac:H3−5ac
H2A.Z:H3−5ac
H2A.Z:H4−4ac
DNA Meth. m5C:H4K20 me3
H3K27 me3:DNA Meth. m5C
DNA Meth. m5C:H3K9 me3
H3K9 me2
H3K27 me2
H4K20 me3
H3K4 me1
H3K4 me3
H4K20 me2
H3−5ac
H3K27 me3
H3K9 me3
DNA Meth. m5C
H4−4ac
H2A.Z
0.00
0.25
0.50
0.75
1.00
$\hat{\pi}$

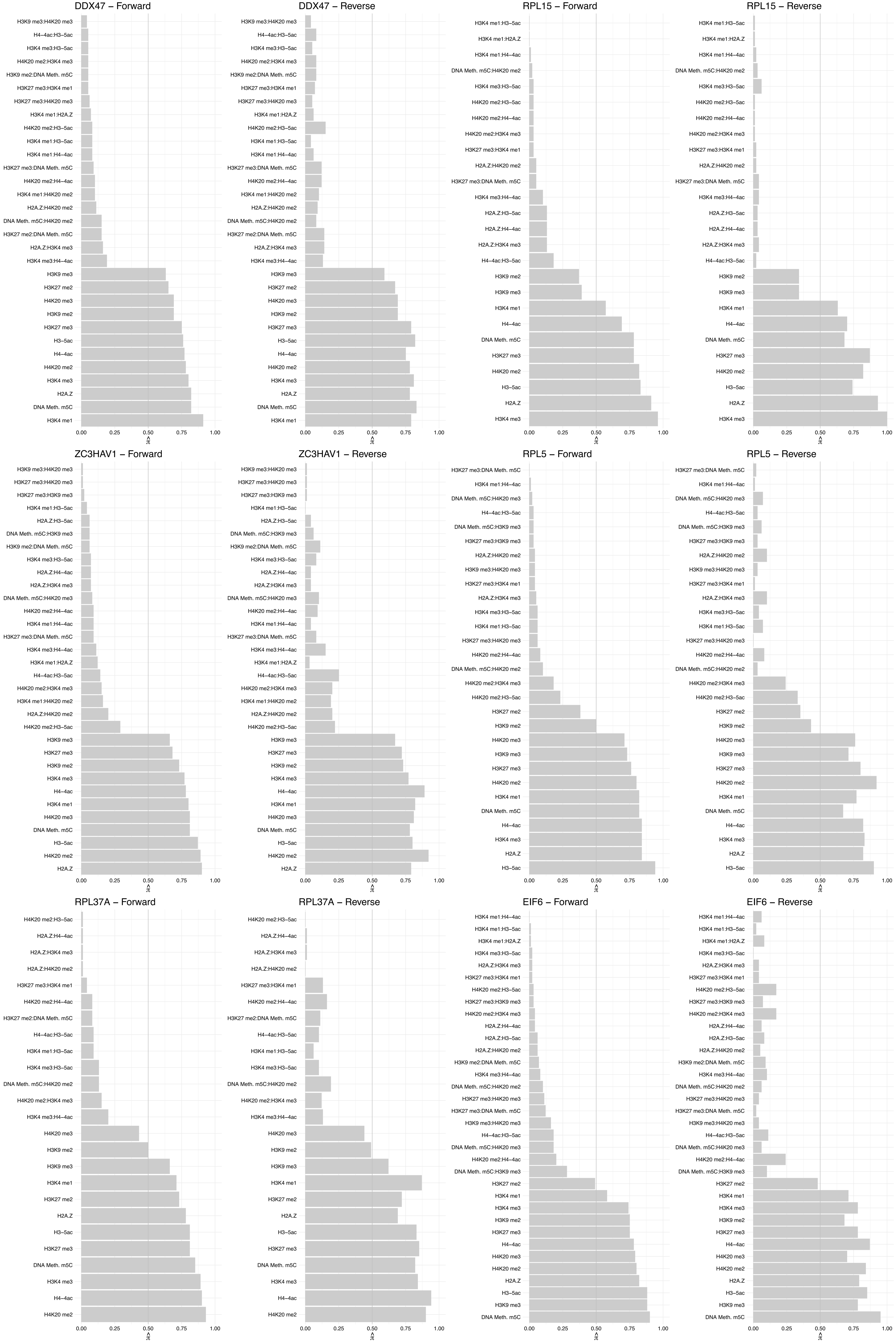
DDX47 – Forward
DDX47 – Reverse
RPL15 – Forward
RPL15 – Reverse
ZC3HAV1 – Forward
ZC3HAV1 – Reverse
RPL5 – Forward
RPL5 – Reverse
RPL37A – Forward
RPL37A – Reverse
EIF6 – Forward
EIF6 – Reverse
H3K9 me3:H4K20 me3
H4–4ac:H3–5ac
H3K4 me3:H3–5ac
H4K20 me2:H3K4 me3
H3K9 me2:DNA Meth. m5C
H3K27 me3:H3K4 me1
H3K27 me3:H4K20 me3
H3K4 me1:H2A.Z
H4K20 me2:H3–5ac
H3K4 me1:H3–5ac
H3K4 me1:H4–4ac
H3K27 me3:DNA Meth. m5C
H4K20 me2:H4–4ac
H3K4 me1:H4K20 me2
H2A.Z:H4K20 me2
DNA Meth. m5C:H4K20 me2
H3K27 me2:DNA Meth. m5C
H2A.Z:H3K4 me3
H3K4 me3:H4–4ac
H3K9 me3
H3K27 me2
H4K20 me3
H3K9 me2
H3K27 me3
H3–5ac
H4–4ac
H4K20 me2
H3K4 me3
H2A.Z
DNA Meth. m5C
H3K4 me1
0.00
0.25
0.50
0.75
1.00
$\hat{\pi}$

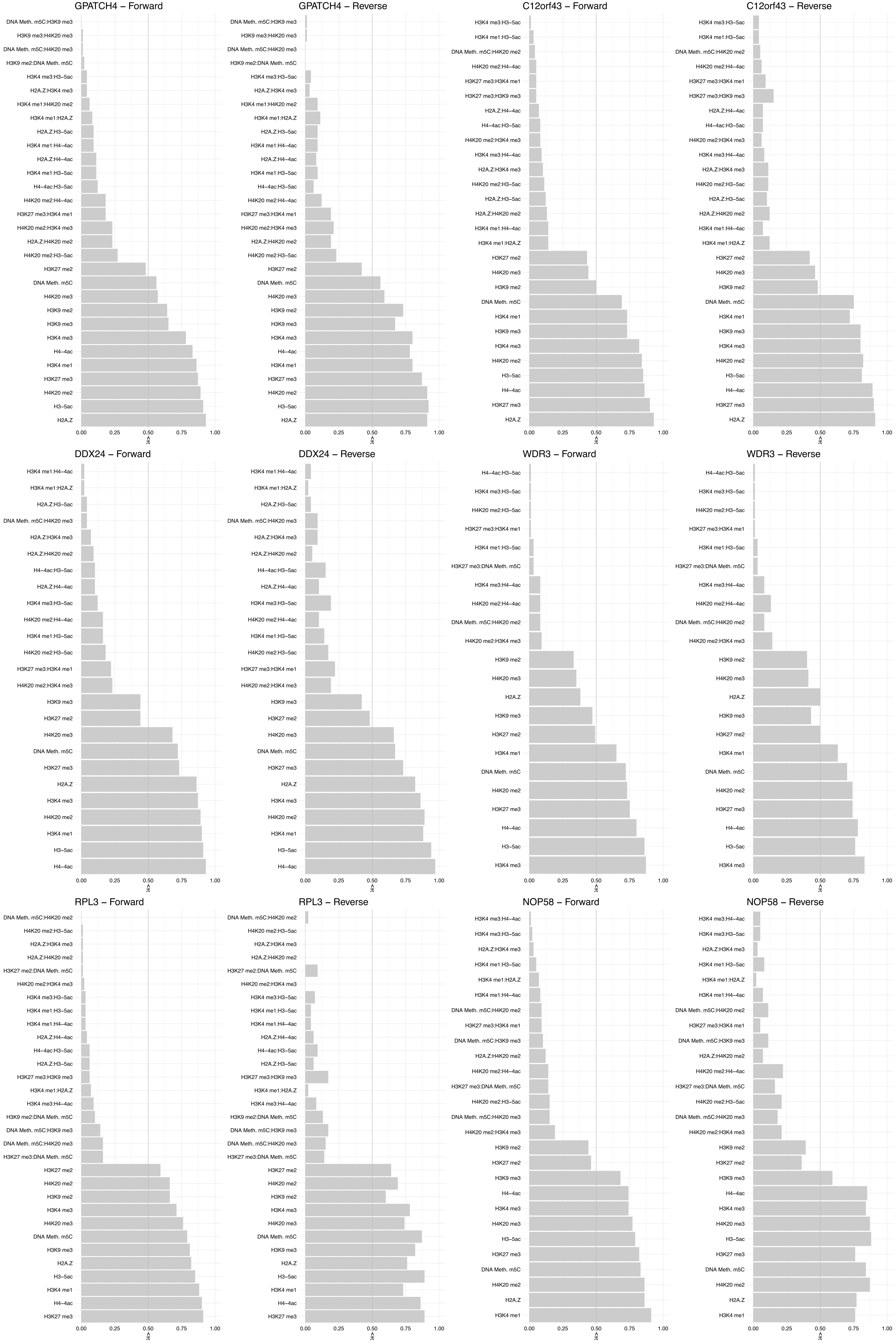
GPATCH4 – Forward
DNA Meth. m5C:H3K9 me3
H3K9 me3:H4K20 me3
DNA Meth. m5C:H4K20 me3
H3K9 me2:DNA Meth. m5C
H3K4 me3:H3–5ac
H2A.Z:H3K4 me3
H3K4 me1:H4K20 me2
H3K4 me1:H2A.Z
H2A.Z:H3–5ac
H3K4 me1:H4–4ac
H2A.Z:H4–4ac
H3K4 me1:H3–5ac
H4–4ac:H3–5ac
H4K20 me2:H4–4ac
H3K27 me3:H3K4 me1
H4K20 me2:H3K4 me3
H2A.Z:H4K20 me2
H4K20 me2:H3–5ac
H3K27 me2
DNA Meth. m5C
H4K20 me3
H3K9 me2
H3K9 me3
H3K4 me3
H4–4ac
H3K4 me1
H3K27 me3
H4K20 me2
H3–5ac
H2A.Z
0.00 0.25 0.50 0.75 1.00
π̂
GPATCH4 – Reverse
C12orf43 – Forward
H3K4 me3:H3–5ac
H3K4 me1:H3–5ac
DNA Meth. m5C:H4K20 me2
H4K20 me2:H4–4ac
H3K27 me3:H3K4 me1
H3K27 me3:H3K9 me3
H2A.Z:H4–4ac
H4–4ac:H3–5ac
H4K20 me2:H3K4 me3
H3K4 me3:H4–4ac
H2A.Z:H3K4 me3
H4K20 me2:H3–5ac
H2A.Z:H3–5ac
H2A.Z:H4K20 me2
H3K4 me1:H4–4ac
H3K4 me1:H2A.Z
H3K27 me2
H4K20 me3
H3K9 me2
DNA Meth. m5C
H3K4 me1
H3K9 me3
H3K4 me3
H4K20 me2
H3–5ac
H4–4ac
H3K27 me3
H2A.Z
C12orf43 – Reverse
DDX24 – Forward
H3K4 me1:H4–4ac
H3K4 me1:H2A.Z
H2A.Z:H3–5ac
DNA Meth. m5C:H4K20 me3
H2A.Z:H3K4 me3
H2A.Z:H4K20 me2
H4–4ac:H3–5ac
H2A.Z:H4–4ac
H3K4 me3:H3–5ac
H4K20 me2:H4–4ac
H3K4 me1:H3–5ac
H4K20 me2:H3–5ac
H3K27 me3:H3K4 me1
H4K20 me2:H3K4 me3
H3K9 me3
H3K27 me2
H4K20 me3
DNA Meth. m5C
H3K27 me3
H2A.Z
H3K4 me3
H4K20 me2
H3K4 me1
H3–5ac
H4–4ac
DDX24 – Reverse
WDR3 – Forward
H4–4ac:H3–5ac
H3K4 me3:H3–5ac
H4K20 me2:H3–5ac
H3K27 me3:H3K4 me1
H3K4 me1:H3–5ac
H3K27 me3:DNA Meth. m5C
H3K4 me3:H4–4ac
H4K20 me2:H4–4ac
DNA Meth. m5C:H4K20 me2
H4K20 me2:H3K4 me3
H3K9 me2
H4K20 me3
H2A.Z
H3K9 me3
H3K27 me2
H3K4 me1
DNA Meth. m5C
H4K20 me2
H3K27 me3
H4–4ac
H3–5ac
H3K4 me3
WDR3 – Reverse
RPL3 – Forward
DNA Meth. m5C:H4K20 me2
H4K20 me2:H3–5ac
H2A.Z:H3K4 me3
H2A.Z:H4K20 me2
H3K27 me2:DNA Meth. m5C
H4K20 me2:H3K4 me3
H3K4 me3:H3–5ac
H3K4 me1:H3–5ac
H3K4 me1:H4–4ac
H2A.Z:H4–4ac
H4–4ac:H3–5ac
H2A.Z:H3–5ac
H3K27 me3:H3K9 me3
H3K4 me1:H2A.Z
H3K4 me3:H4–4ac
H3K9 me2:DNA Meth. m5C
DNA Meth. m5C:H3K9 me3
DNA Meth. m5C:H4K20 me3
H3K27 me3:DNA Meth. m5C
H3K27 me2
H4K20 me2
H3K9 me2
H3K4 me3
H4K20 me3
DNA Meth. m5C
H3K9 me3
H2A.Z
H3–5ac
H3K4 me1
H4–4ac
H3K27 me3
RPL3 – Reverse
NOP58 – Forward
H3K4 me3:H4–4ac
H3K4 me3:H3–5ac
H2A.Z:H3K4 me3
H3K4 me1:H3–5ac
H3K4 me1:H2A.Z
H3K4 me1:H4–4ac
DNA Meth. m5C:H4K20 me2
H3K27 me3:H3K4 me1
DNA Meth. m5C:H3K9 me3
H2A.Z:H4K20 me2
H4K20 me2:H4–4ac
H3K27 me3:DNA Meth. m5C
H4K20 me2:H3–5ac
DNA Meth. m5C:H4K20 me3
H4K20 me2:H3K4 me3
H3K9 me2
H3K27 me2
H3K9 me3
H4–4ac
H3K4 me3
H4K20 me3
H3–5ac
H3K27 me3
DNA Meth. m5C
H4K20 me2
H2A.Z
H3K4 me1
NOP58 – Reverse

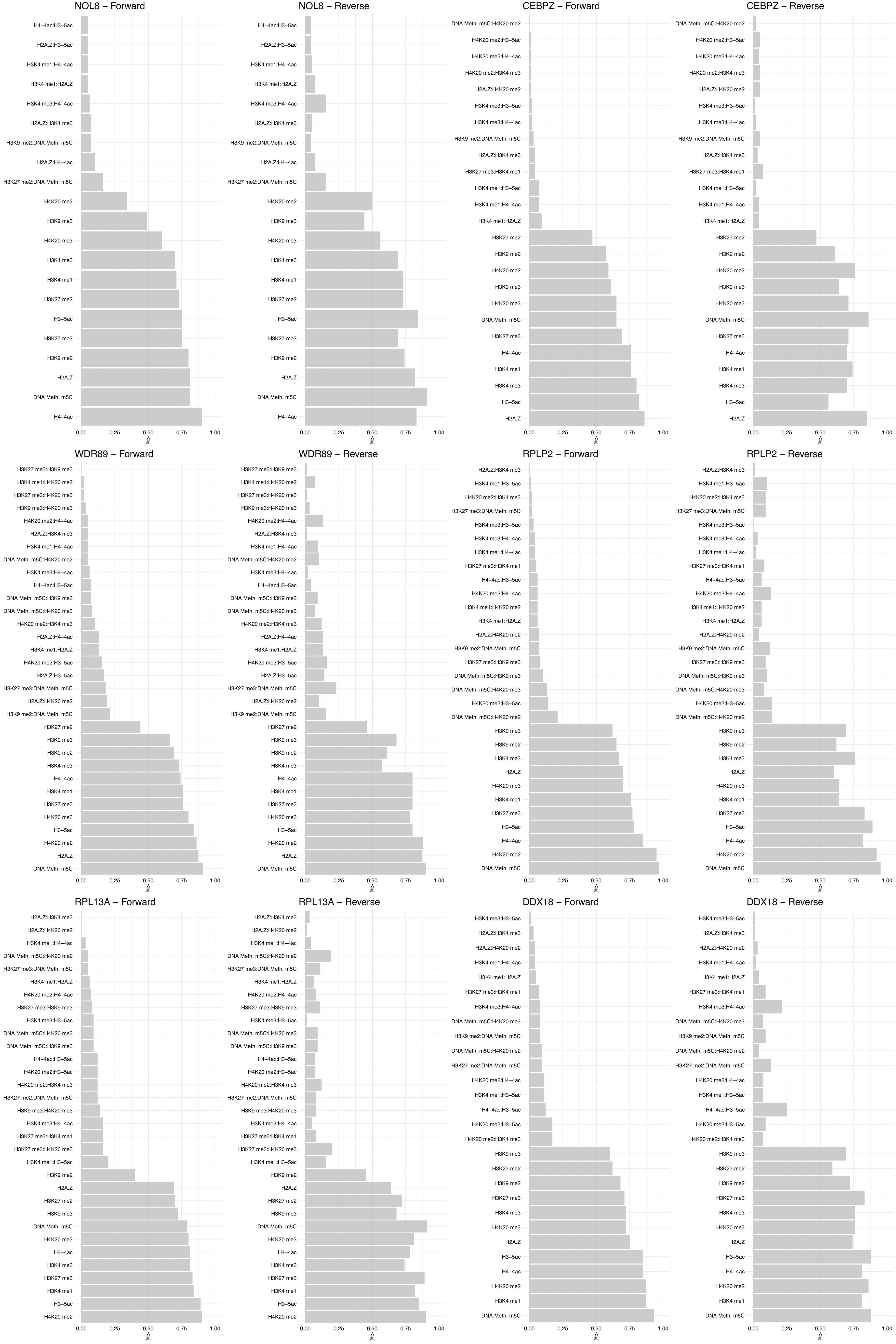
NOL8 – Forward
H4−4ac:H3−5ac
H2A.Z:H3−5ac
H3K4 me1:H4−4ac
H3K4 me1:H2A.Z
H3K4 me3:H4−4ac
H2A.Z:H3K4 me3
H3K9 me2:DNA Meth. m5C
H2A.Z:H4−4ac
H3K27 me2:DNA Meth. m5C
H4K20 me2
H3K9 me3
H4K20 me3
H3K4 me3
H3K4 me1
H3K27 me2
H3−5ac
H3K27 me3
H3K9 me2
H2A.Z
DNA Meth. m5C
H4−4ac
0.00 0.25 0.50 0.75 1.00
$\hat{\pi}$
NOL8 – Reverse
H4−4ac:H3−5ac
H2A.Z:H3−5ac
H3K4 me1:H4−4ac
H3K4 me1:H2A.Z
H3K4 me3:H4−4ac
H2A.Z:H3K4 me3
H3K9 me2:DNA Meth. m5C
H2A.Z:H4−4ac
H3K27 me2:DNA Meth. m5C
H4K20 me2
H3K9 me3
H4K20 me3
H3K4 me3
H3K4 me1
H3K27 me2
H3−5ac
H3K27 me3
H3K9 me2
H2A.Z
DNA Meth. m5C
H4−4ac
0.00 0.25 0.50 0.75 1.00
$\hat{\pi}$
CEBPZ – Forward
DNA Meth. m5C:H4K20 me2
H4K20 me2:H3−5ac
H4K20 me2:H4−4ac
H4K20 me2:H3K4 me3
H2A.Z:H4K20 me2
H3K4 me3:H3−5ac
H3K4 me3:H4−4ac
H3K9 me2:DNA Meth. m5C
H2A.Z:H3K4 me3
H3K27 me3:H3K4 me1
H3K4 me1:H3−5ac
H3K4 me1:H4−4ac
H3K4 me1:H2A.Z
H3K27 me2
H3K9 me2
H4K20 me2
H3K9 me3
H4K20 me3
DNA Meth. m5C
H3K27 me3
H4−4ac
H3K4 me1
H3K4 me3
H3−5ac
H2A.Z
0.00 0.25 0.50 0.75 1.00
$\hat{\pi}$
CEBPZ – Reverse
DNA Meth. m5C:H4K20 me2
H4K20 me2:H3−5ac
H4K20 me2:H4−4ac
H4K20 me2:H3K4 me3
H2A.Z:H4K20 me2
H3K4 me3:H3−5ac
H3K4 me3:H4−4ac
H3K9 me2:DNA Meth. m5C
H2A.Z:H3K4 me3
H3K27 me3:H3K4 me1
H3K4 me1:H3−5ac
H3K4 me1:H4−4ac
H3K4 me1:H2A.Z
H3K27 me2
H3K9 me2
H4K20 me2
H3K9 me3
H4K20 me3
DNA Meth. m5C
H3K27 me3
H4−4ac
H3K4 me1
H3K4 me3
H3−5ac
H2A.Z
0.00 0.25 0.50 0.75 1.00
$\hat{\pi}$
WDR89 – Forward
H3K27 me3:H3K9 me3
H3K4 me1:H4K20 me2
H3K27 me3:H4K20 me3
H3K9 me3:H4K20 me3
H4K20 me2:H4−4ac
H2A.Z:H3K4 me3
H3K4 me1:H4−4ac
DNA Meth. m5C:H4K20 me2
H3K4 me3:H4−4ac
H4−4ac:H3−5ac
DNA Meth. m5C:H3K9 me3
DNA Meth. m5C:H4K20 me3
H4K20 me2:H3K4 me3
H2A.Z:H4−4ac
H3K4 me1:H2A.Z
H4K20 me2:H3−5ac
H2A.Z:H3−5ac
H3K27 me3:DNA Meth. m5C
H2A.Z:H4K20 me2
H3K9 me2:DNA Meth. m5C
H3K27 me2
H3K9 me3
H3K9 me2
H3K4 me3
H4−4ac
H3K4 me1
H3K27 me3
H4K20 me3
H3−5ac
H4K20 me2
H2A.Z
DNA Meth. m5C
0.00 0.25 0.50 0.75 1.00
$\hat{\pi}$
WDR89 – Reverse
H3K27 me3:H3K9 me3
H3K4 me1:H4K20 me2
H3K27 me3:H4K20 me3
H3K9 me3:H4K20 me3
H4K20 me2:H4−4ac
H2A.Z:H3K4 me3
H3K4 me1:H4−4ac
DNA Meth. m5C:H4K20 me2
H3K4 me3:H4−4ac
H4−4ac:H3−5ac
DNA Meth. m5C:H3K9 me3
DNA Meth. m5C:H4K20 me3
H4K20 me2:H3K4 me3
H2A.Z:H4−4ac
H3K4 me1:H2A.Z
H4K20 me2:H3−5ac
H2A.Z:H3−5ac
H3K27 me3:DNA Meth. m5C
H2A.Z:H4K20 me2
H3K9 me2:DNA Meth. m5C
H3K27 me2
H3K9 me3
H3K9 me2
H3K4 me3
H4−4ac
H3K4 me1
H3K27 me3
H4K20 me3
H3−5ac
H4K20 me2
H2A.Z
DNA Meth. m5C
0.00 0.25 0.50 0.75 1.00
$\hat{\pi}$
RPLP2 – Forward
H2A.Z:H3K4 me3
H3K4 me1:H3−5ac
H4K20 me2:H3K4 me3
H3K27 me3:DNA Meth. m5C
H3K4 me3:H3−5ac
H3K4 me3:H4−4ac
H3K4 me1:H4−4ac
H3K27 me3:H3K4 me1
H4−4ac:H3−5ac
H4K20 me2:H4−4ac
H3K4 me1:H4K20 me2
H3K4 me1:H2A.Z
H2A.Z:H4K20 me2
H3K9 me2:DNA Meth. m5C
H3K27 me3:H3K9 me3
DNA Meth. m5C:H3K9 me3
DNA Meth. m5C:H4K20 me3
H4K20 me2:H3−5ac
DNA Meth. m5C:H4K20 me2
H3K9 me3
H3K9 me2
H3K4 me3
H2A.Z
H4K20 me3
H3K4 me1
H3K27 me3
H3−5ac
H4−4ac
H4K20 me2
DNA Meth. m5C
0.00 0.25 0.50 0.75 1.00
$\hat{\pi}$
RPLP2 – Reverse
H2A.Z:H3K4 me3
H3K4 me1:H3−5ac
H4K20 me2:H3K4 me3
H3K27 me3:DNA Meth. m5C
H3K4 me3:H3−5ac
H3K4 me3:H4−4ac
H3K4 me1:H4−4ac
H3K27 me3:H3K4 me1
H4−4ac:H3−5ac
H4K20 me2:H4−4ac
H3K4 me1:H4K20 me2
H3K4 me1:H2A.Z
H2A.Z:H4K20 me2
H3K9 me2:DNA Meth. m5C
H3K27 me3:H3K9 me3
DNA Meth. m5C:H3K9 me3
DNA Meth. m5C:H4K20 me3
H4K20 me2:H3−5ac
DNA Meth. m5C:H4K20 me2
H3K9 me3
H3K9 me2
H3K4 me3
H2A.Z
H4K20 me3
H3K4 me1
H3K27 me3
H3−5ac
H4−4ac
H4K20 me2
DNA Meth. m5C
0.00 0.25 0.50 0.75 1.00
$\hat{\pi}$
RPL13A – Forward
H2A.Z:H3K4 me3
H2A.Z:H4K20 me2
H3K4 me1:H4−4ac
DNA Meth. m5C:H4K20 me2
H3K27 me3:DNA Meth. m5C
H3K4 me1:H2A.Z
H4K20 me2:H4−4ac
H3K27 me3:H3K9 me3
H3K4 me3:H3−5ac
DNA Meth. m5C:H4K20 me3
DNA Meth. m5C:H3K9 me3
H4−4ac:H3−5ac
H4K20 me2:H3−5ac
H4K20 me2:H3K4 me3
H3K27 me2:DNA Meth. m5C
H3K9 me3:H4K20 me3
H3K4 me3:H4−4ac
H3K27 me3:H3K4 me1
H3K27 me3:H4K20 me3
H3K4 me1:H3−5ac
H3K9 me2
H2A.Z
H3K27 me2
H3K9 me3
DNA Meth. m5C
H4K20 me3
H4−4ac
H3K4 me3
H3K27 me3
H3K4 me1
H3−5ac
H4K20 me2
0.00 0.25 0.50 0.75 1.00
$\hat{\pi}$
RPL13A – Reverse
H2A.Z:H3K4 me3
H2A.Z:H4K20 me2
H3K4 me1:H4−4ac
DNA Meth. m5C:H4K20 me2
H3K27 me3:DNA Meth. m5C
H3K4 me1:H2A.Z
H4K20 me2:H4−4ac
H3K27 me3:H3K9 me3
H3K4 me3:H3−5ac
DNA Meth. m5C:H4K20 me3
DNA Meth. m5C:H3K9 me3
H4−4ac:H3−5ac
H4K20 me2:H3−5ac
H4K20 me2:H3K4 me3
H3K27 me2:DNA Meth. m5C
H3K9 me3:H4K20 me3
H3K4 me3:H4−4ac
H3K27 me3:H3K4 me1
H3K27 me3:H4K20 me3
H3K4 me1:H3−5ac
H3K9 me2
H2A.Z
H3K27 me2
H3K9 me3
DNA Meth. m5C
H4K20 me3
H4−4ac
H3K4 me3
H3K27 me3
H3K4 me1
H3−5ac
H4K20 me2
0.00 0.25 0.50 0.75 1.00
$\hat{\pi}$
DDX18 – Forward
H3K4 me3:H3−5ac
H2A.Z:H3K4 me3
H2A.Z:H4K20 me2
H3K4 me1:H4−4ac
H3K4 me1:H2A.Z
H3K27 me3:H3K4 me1
H3K4 me3:H4−4ac
DNA Meth. m5C:H4K20 me3
H3K9 me2:DNA Meth. m5C
DNA Meth. m5C:H4K20 me2
H3K27 me2:DNA Meth. m5C
H4K20 me2:H4−4ac
H3K4 me1:H3−5ac
H4−4ac:H3−5ac
H4K20 me2:H3−5ac
H4K20 me2:H3K4 me3
H3K9 me3
H3K27 me2
H3K9 me2
H3K27 me3
H3K4 me3
H4K20 me3
H2A.Z
H3−5ac
H4−4ac
H4K20 me2
H3K4 me1
DNA Meth. m5C
0.00 0.25 0.50 0.75 1.00
$\hat{\pi}$
DDX18 – Reverse
H3K4 me3:H3−5ac
H2A.Z:H3K4 me3
H2A.Z:H4K20 me2
H3K4 me1:H4−4ac
H3K4 me1:H2A.Z
H3K27 me3:H3K4 me1
H3K4 me3:H4−4ac
DNA Meth. m5C:H4K20 me3
H3K9 me2:DNA Meth. m5C
DNA Meth. m5C:H4K20 me2
H3K27 me2:DNA Meth. m5C
H4K20 me2:H4−4ac
H3K4 me1:H3−5ac
H4−4ac:H3−5ac
H4K20 me2:H3−5ac
H4K20 me2:H3K4 me3
H3K9 me3
H3K27 me2
H3K9 me2
H3K27 me3
H3K4 me3
H4K20 me3
H2A.Z
H3−5ac
H4−4ac
H4K20 me2
H3K4 me1
DNA Meth. m5C
0.00 0.25 0.50 0.75 1.00
$\hat{\pi}$

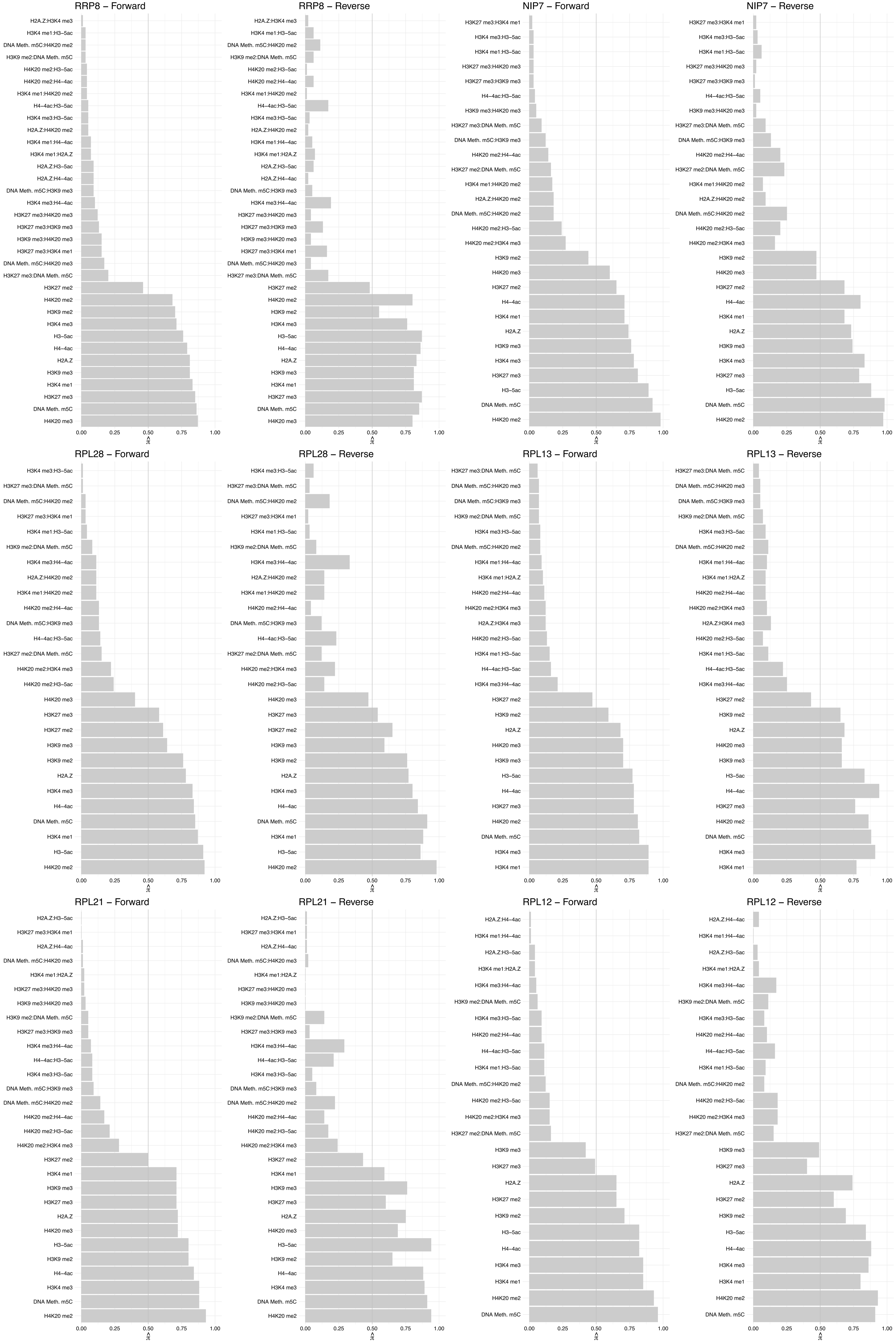
RRP8 – Forward
H2A.Z:H3K4 me3
H3K4 me1:H3–5ac
DNA Meth. m5C:H4K20 me2
H3K9 me2:DNA Meth. m5C
H4K20 me2:H3–5ac
H4K20 me2:H4–4ac
H3K4 me1:H4K20 me2
H4–4ac:H3–5ac
H3K4 me3:H3–5ac
H2A.Z:H4K20 me2
H3K4 me1:H4–4ac
H3K4 me1:H2A.Z
H2A.Z:H3–5ac
H2A.Z:H4–4ac
DNA Meth. m5C:H3K9 me3
H3K4 me3:H4–4ac
H3K27 me3:H4K20 me3
H3K27 me3:H3K9 me3
H3K9 me3:H4K20 me3
H3K27 me3:H3K4 me1
DNA Meth. m5C:H4K20 me3
H3K27 me3:DNA Meth. m5C
H3K27 me2
H4K20 me2
H3K9 me2
H3K4 me3
H3–5ac
H4–4ac
H2A.Z
H3K9 me3
H3K4 me1
H3K27 me3
DNA Meth. m5C
H4K20 me3
0.00
0.25
0.50
0.75
1.00
π̂
RRP8 – Reverse
NIP7 – Forward
H3K27 me3:H3K4 me1
H3K4 me3:H3–5ac
H3K4 me1:H3–5ac
H3K27 me3:H4K20 me3
H3K27 me3:H3K9 me3
H4–4ac:H3–5ac
H3K9 me3:H4K20 me3
H3K27 me3:DNA Meth. m5C
DNA Meth. m5C:H3K9 me3
H4K20 me2:H4–4ac
H3K27 me2:DNA Meth. m5C
H3K4 me1:H4K20 me2
H2A.Z:H4K20 me2
DNA Meth. m5C:H4K20 me2
H4K20 me2:H3–5ac
H4K20 me2:H3K4 me3
H3K9 me2
H4K20 me3
H3K27 me2
H4–4ac
H3K4 me1
H2A.Z
H3K9 me3
H3K4 me3
H3K27 me3
H3–5ac
DNA Meth. m5C
H4K20 me2
NIP7 – Reverse
RPL28 – Forward
H3K4 me3:H3–5ac
H3K27 me3:DNA Meth. m5C
DNA Meth. m5C:H4K20 me2
H3K27 me3:H3K4 me1
H3K4 me1:H3–5ac
H3K9 me2:DNA Meth. m5C
H3K4 me3:H4–4ac
H2A.Z:H4K20 me2
H3K4 me1:H4K20 me2
H4K20 me2:H4–4ac
DNA Meth. m5C:H3K9 me3
H4–4ac:H3–5ac
H3K27 me2:DNA Meth. m5C
H4K20 me2:H3K4 me3
H4K20 me2:H3–5ac
H4K20 me3
H3K27 me3
H3K27 me2
H3K9 me3
H3K9 me2
H2A.Z
H3K4 me3
H4–4ac
DNA Meth. m5C
H3K4 me1
H3–5ac
H4K20 me2
RPL28 – Reverse
RPL13 – Forward
H3K27 me3:DNA Meth. m5C
DNA Meth. m5C:H4K20 me3
DNA Meth. m5C:H3K9 me3
H3K9 me2:DNA Meth. m5C
H3K4 me3:H3–5ac
DNA Meth. m5C:H4K20 me2
H3K4 me1:H4–4ac
H3K4 me1:H2A.Z
H4K20 me2:H4–4ac
H4K20 me2:H3K4 me3
H2A.Z:H3K4 me3
H4K20 me2:H3–5ac
H3K4 me1:H3–5ac
H4–4ac:H3–5ac
H3K4 me3:H4–4ac
H3K27 me2
H3K9 me2
H2A.Z
H4K20 me3
H3K9 me3
H3–5ac
H4–4ac
H3K27 me3
H4K20 me2
DNA Meth. m5C
H3K4 me3
H3K4 me1
RPL13 – Reverse
RPL21 – Forward
H2A.Z:H3–5ac
H3K27 me3:H3K4 me1
H2A.Z:H4–4ac
DNA Meth. m5C:H4K20 me3
H3K4 me1:H2A.Z
H3K27 me3:H4K20 me3
H3K9 me3:H4K20 me3
H3K9 me2:DNA Meth. m5C
H3K27 me3:H3K9 me3
H3K4 me3:H4–4ac
H4–4ac:H3–5ac
H3K4 me3:H3–5ac
DNA Meth. m5C:H3K9 me3
DNA Meth. m5C:H4K20 me2
H4K20 me2:H4–4ac
H4K20 me2:H3–5ac
H4K20 me2:H3K4 me3
H3K27 me2
H3K4 me1
H3K9 me3
H3K27 me3
H2A.Z
H4K20 me3
H3–5ac
H3K9 me2
H4–4ac
H3K4 me3
DNA Meth. m5C
H4K20 me2
RPL21 – Reverse
RPL12 – Forward
H2A.Z:H4–4ac
H3K4 me1:H4–4ac
H2A.Z:H3–5ac
H3K4 me1:H2A.Z
H3K4 me3:H4–4ac
H3K9 me2:DNA Meth. m5C
H3K4 me3:H3–5ac
H4K20 me2:H4–4ac
H4–4ac:H3–5ac
H3K4 me1:H3–5ac
DNA Meth. m5C:H4K20 me2
H4K20 me2:H3–5ac
H4K20 me2:H3K4 me3
H3K27 me2:DNA Meth. m5C
H3K9 me3
H3K27 me3
H2A.Z
H3K27 me2
H3K9 me2
H3–5ac
H4–4ac
H3K4 me3
H3K4 me1
H4K20 me2
DNA Meth. m5C
RPL12 – Reverse

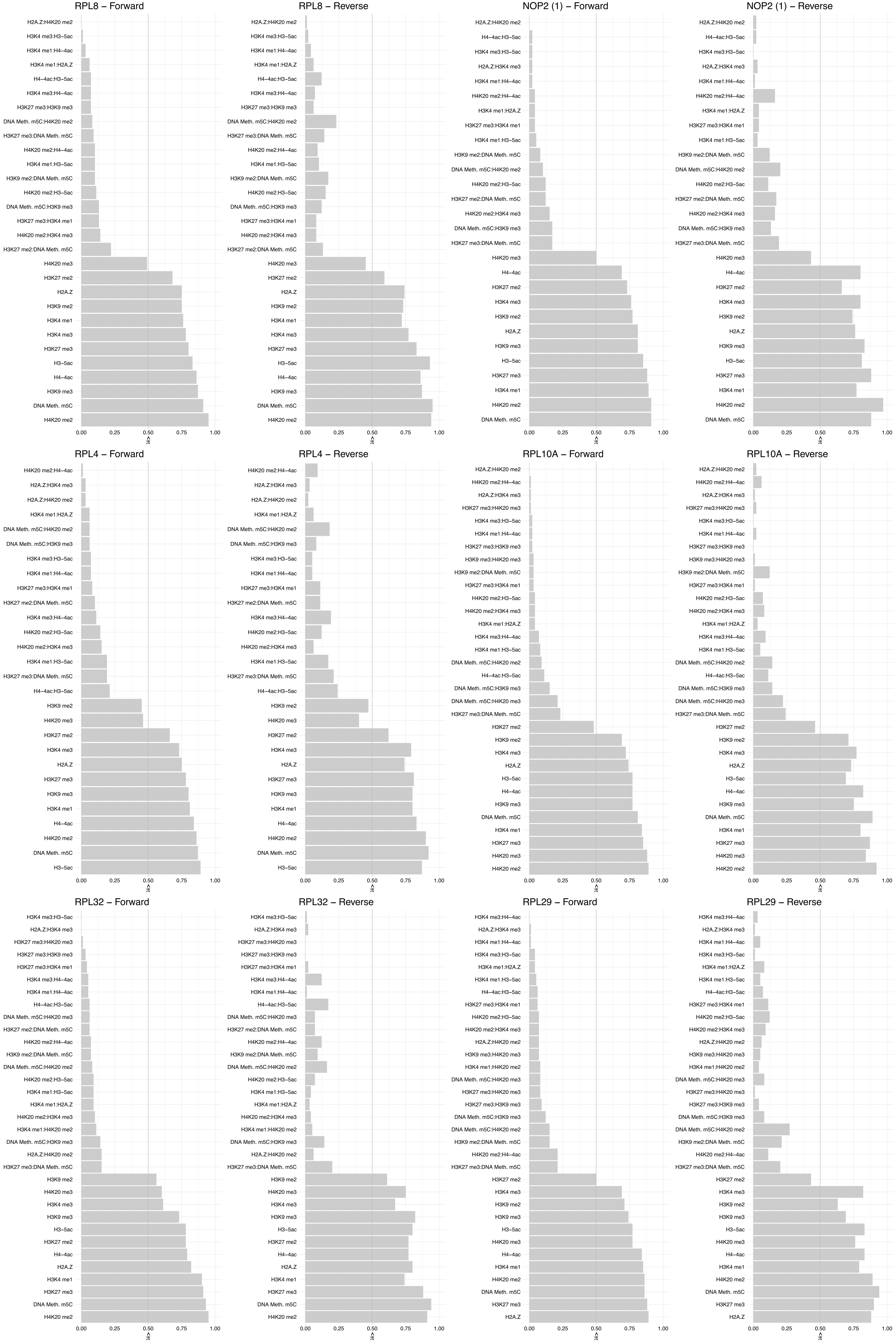
RPL8 – Forward
RPL8 – Reverse
NOP2 (1) – Forward
NOP2 (1) – Reverse
RPL4 – Forward
RPL4 – Reverse
RPL10A – Forward
RPL10A – Reverse
RPL32 – Forward
RPL32 – Reverse
RPL29 – Forward
RPL29 – Reverse
0.00 0.25 0.50 0.75 1.00
π̂

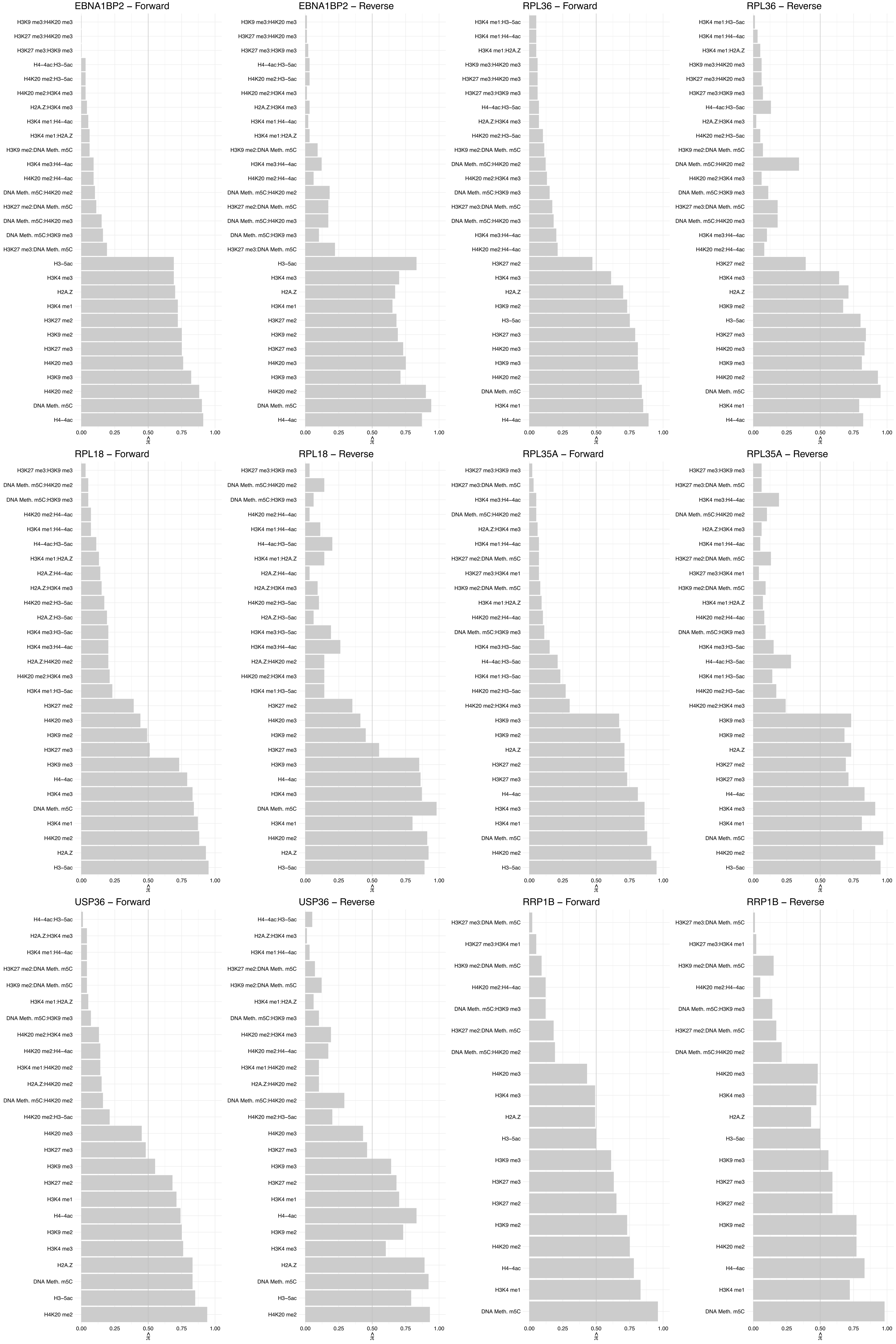
EBNA1BP2 – Forward
H3K9 me3:H4K20 me3
H3K27 me3:H4K20 me3
H3K27 me3:H3K9 me3
H4−4ac:H3−5ac
H4K20 me2:H3−5ac
H4K20 me2:H3K4 me3
H2A.Z:H3K4 me3
H3K4 me1:H4−4ac
H3K4 me1:H2A.Z
H3K9 me2:DNA Meth. m5C
H3K4 me3:H4−4ac
H4K20 me2:H4−4ac
DNA Meth. m5C:H4K20 me2
H3K27 me2:DNA Meth. m5C
DNA Meth. m5C:H4K20 me3
DNA Meth. m5C:H3K9 me3
H3K27 me3:DNA Meth. m5C
H3−5ac
H3K4 me3
H2A.Z
H3K4 me1
H3K27 me2
H3K9 me2
H3K27 me3
H4K20 me3
H3K9 me3
H4K20 me2
DNA Meth. m5C
H4−4ac
0.00
0.25
0.50
0.75
1.00
π̂
EBNA1BP2 – Reverse
H3K9 me3:H4K20 me3
H3K27 me3:H4K20 me3
H3K27 me3:H3K9 me3
H4−4ac:H3−5ac
H4K20 me2:H3−5ac
H4K20 me2:H3K4 me3
H2A.Z:H3K4 me3
H3K4 me1:H4−4ac
H3K4 me1:H2A.Z
H3K9 me2:DNA Meth. m5C
H3K4 me3:H4−4ac
H4K20 me2:H4−4ac
DNA Meth. m5C:H4K20 me2
H3K27 me2:DNA Meth. m5C
DNA Meth. m5C:H4K20 me3
DNA Meth. m5C:H3K9 me3
H3K27 me3:DNA Meth. m5C
H3−5ac
H3K4 me3
H2A.Z
H3K4 me1
H3K27 me2
H3K9 me2
H3K27 me3
H4K20 me3
H3K9 me3
H4K20 me2
DNA Meth. m5C
H4−4ac
0.00
0.25
0.50
0.75
1.00
π̂
RPL36 – Forward
H3K4 me1:H3−5ac
H3K4 me1:H4−4ac
H3K4 me1:H2A.Z
H3K9 me3:H4K20 me3
H3K27 me3:H4K20 me3
H3K27 me3:H3K9 me3
H4−4ac:H3−5ac
H2A.Z:H3K4 me3
H4K20 me2:H3−5ac
H3K9 me2:DNA Meth. m5C
DNA Meth. m5C:H4K20 me2
H4K20 me2:H3K4 me3
DNA Meth. m5C:H3K9 me3
H3K27 me3:DNA Meth. m5C
DNA Meth. m5C:H4K20 me3
H3K4 me3:H4−4ac
H4K20 me2:H4−4ac
H3K27 me2
H3K4 me3
H2A.Z
H3K9 me2
H3−5ac
H3K27 me3
H4K20 me3
H3K9 me3
H4K20 me2
DNA Meth. m5C
H3K4 me1
H4−4ac
0.00
0.25
0.50
0.75
1.00
π̂
RPL36 – Reverse
H3K4 me1:H3−5ac
H3K4 me1:H4−4ac
H3K4 me1:H2A.Z
H3K9 me3:H4K20 me3
H3K27 me3:H4K20 me3
H3K27 me3:H3K9 me3
H4−4ac:H3−5ac
H2A.Z:H3K4 me3
H4K20 me2:H3−5ac
H3K9 me2:DNA Meth. m5C
DNA Meth. m5C:H4K20 me2
H4K20 me2:H3K4 me3
DNA Meth. m5C:H3K9 me3
H3K27 me3:DNA Meth. m5C
DNA Meth. m5C:H4K20 me3
H3K4 me3:H4−4ac
H4K20 me2:H4−4ac
H3K27 me2
H3K4 me3
H2A.Z
H3K9 me2
H3−5ac
H3K27 me3
H4K20 me3
H3K9 me3
H4K20 me2
DNA Meth. m5C
H3K4 me1
H4−4ac
0.00
0.25
0.50
0.75
1.00
π̂
RPL18 – Forward
H3K27 me3:H3K9 me3
DNA Meth. m5C:H4K20 me2
DNA Meth. m5C:H3K9 me3
H4K20 me2:H4−4ac
H3K4 me1:H4−4ac
H4−4ac:H3−5ac
H3K4 me1:H2A.Z
H2A.Z:H4−4ac
H2A.Z:H3K4 me3
H4K20 me2:H3−5ac
H2A.Z:H3−5ac
H3K4 me3:H3−5ac
H3K4 me3:H4−4ac
H2A.Z:H4K20 me2
H4K20 me2:H3K4 me3
H3K4 me1:H3−5ac
H3K27 me2
H4K20 me3
H3K9 me2
H3K27 me3
H3K9 me3
H4−4ac
H3K4 me3
DNA Meth. m5C
H3K4 me1
H4K20 me2
H2A.Z
H3−5ac
0.00
0.25
0.50
0.75
1.00
π̂
RPL18 – Reverse
H3K27 me3:H3K9 me3
DNA Meth. m5C:H4K20 me2
DNA Meth. m5C:H3K9 me3
H4K20 me2:H4−4ac
H3K4 me1:H4−4ac
H4−4ac:H3−5ac
H3K4 me1:H2A.Z
H2A.Z:H4−4ac
H2A.Z:H3K4 me3
H4K20 me2:H3−5ac
H2A.Z:H3−5ac
H3K4 me3:H3−5ac
H3K4 me3:H4−4ac
H2A.Z:H4K20 me2
H4K20 me2:H3K4 me3
H3K4 me1:H3−5ac
H3K27 me2
H4K20 me3
H3K9 me2
H3K27 me3
H3K9 me3
H4−4ac
H3K4 me3
DNA Meth. m5C
H3K4 me1
H4K20 me2
H2A.Z
H3−5ac
0.00
0.25
0.50
0.75
1.00
π̂
RPL35A – Forward
H3K27 me3:H3K9 me3
H3K27 me3:DNA Meth. m5C
H3K4 me3:H4−4ac
DNA Meth. m5C:H4K20 me2
H2A.Z:H3K4 me3
H3K4 me1:H4−4ac
H3K27 me2:DNA Meth. m5C
H3K27 me3:H3K4 me1
H3K9 me2:DNA Meth. m5C
H3K4 me1:H2A.Z
H4K20 me2:H4−4ac
DNA Meth. m5C:H3K9 me3
H3K4 me3:H3−5ac
H4−4ac:H3−5ac
H3K4 me1:H3−5ac
H4K20 me2:H3−5ac
H4K20 me2:H3K4 me3
H3K9 me3
H3K9 me2
H2A.Z
H3K27 me2
H3K27 me3
H4−4ac
H3K4 me3
H3K4 me1
DNA Meth. m5C
H4K20 me2
H3−5ac
0.00
0.25
0.50
0.75
1.00
π̂
RPL35A – Reverse
H3K27 me3:H3K9 me3
H3K27 me3:DNA Meth. m5C
H3K4 me3:H4−4ac
DNA Meth. m5C:H4K20 me2
H2A.Z:H3K4 me3
H3K4 me1:H4−4ac
H3K27 me2:DNA Meth. m5C
H3K27 me3:H3K4 me1
H3K9 me2:DNA Meth. m5C
H3K4 me1:H2A.Z
H4K20 me2:H4−4ac
DNA Meth. m5C:H3K9 me3
H3K4 me3:H3−5ac
H4−4ac:H3−5ac
H3K4 me1:H3−5ac
H4K20 me2:H3−5ac
H4K20 me2:H3K4 me3
H3K9 me3
H3K9 me2
H2A.Z
H3K27 me2
H3K27 me3
H4−4ac
H3K4 me3
H3K4 me1
DNA Meth. m5C
H4K20 me2
H3−5ac
0.00
0.25
0.50
0.75
1.00
π̂
USP36 – Forward
H4−4ac:H3−5ac
H2A.Z:H3K4 me3
H3K4 me1:H4−4ac
H3K27 me2:DNA Meth. m5C
H3K9 me2:DNA Meth. m5C
H3K4 me1:H2A.Z
DNA Meth. m5C:H3K9 me3
H4K20 me2:H3K4 me3
H4K20 me2:H4−4ac
H3K4 me1:H4K20 me2
H2A.Z:H4K20 me2
DNA Meth. m5C:H4K20 me2
H4K20 me2:H3−5ac
H4K20 me3
H3K27 me3
H3K9 me3
H3K27 me2
H3K4 me1
H4−4ac
H3K9 me2
H3K4 me3
H2A.Z
DNA Meth. m5C
H3−5ac
H4K20 me2
0.00
0.25
0.50
0.75
1.00
π̂
USP36 – Reverse
H4−4ac:H3−5ac
H2A.Z:H3K4 me3
H3K4 me1:H4−4ac
H3K27 me2:DNA Meth. m5C
H3K9 me2:DNA Meth. m5C
H3K4 me1:H2A.Z
DNA Meth. m5C:H3K9 me3
H4K20 me2:H3K4 me3
H4K20 me2:H4−4ac
H3K4 me1:H4K20 me2
H2A.Z:H4K20 me2
DNA Meth. m5C:H4K20 me2
H4K20 me2:H3−5ac
H4K20 me3
H3K27 me3
H3K9 me3
H3K27 me2
H3K4 me1
H4−4ac
H3K9 me2
H3K4 me3
H2A.Z
DNA Meth. m5C
H3−5ac
H4K20 me2
0.00
0.25
0.50
0.75
1.00
π̂
RRP1B – Forward
H3K27 me3:DNA Meth. m5C
H3K27 me3:H3K4 me1
H3K9 me2:DNA Meth. m5C
H4K20 me2:H4−4ac
DNA Meth. m5C:H3K9 me3
H3K27 me2:DNA Meth. m5C
DNA Meth. m5C:H4K20 me2
H4K20 me3
H3K4 me3
H2A.Z
H3−5ac
H3K9 me3
H3K27 me3
H3K27 me2
H3K9 me2
H4K20 me2
H4−4ac
H3K4 me1
DNA Meth. m5C
0.00
0.25
0.50
0.75
1.00
π̂
RRP1B – Reverse
H3K27 me3:DNA Meth. m5C
H3K27 me3:H3K4 me1
H3K9 me2:DNA Meth. m5C
H4K20 me2:H4−4ac
DNA Meth. m5C:H3K9 me3
H3K27 me2:DNA Meth. m5C
DNA Meth. m5C:H4K20 me2
H4K20 me3
H3K4 me3
H2A.Z
H3−5ac
H3K9 me3
H3K27 me3
H3K27 me2
H3K9 me2
H4K20 me2
H4−4ac
H3K4 me1
DNA Meth. m5C
0.00
0.25
0.50
0.75
1.00
π̂

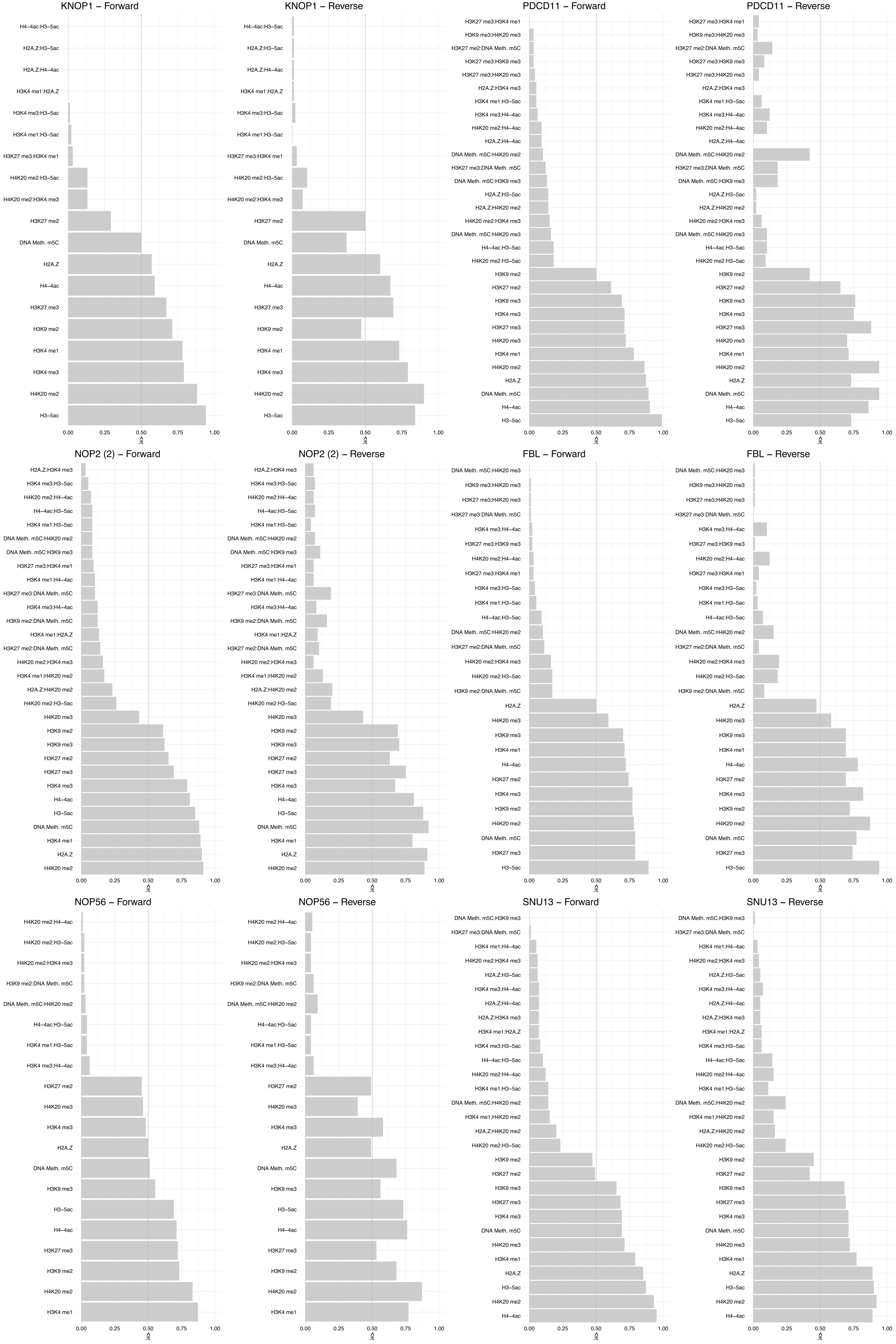
KNOP1 − Forward
H4−4ac:H3−5ac
H2A.Z:H3−5ac
H2A.Z:H4−4ac
H3K4 me1:H2A.Z
H3K4 me3:H3−5ac
H3K4 me1:H3−5ac
H3K27 me3:H3K4 me1
H4K20 me2:H3−5ac
H4K20 me2:H3K4 me3
H3K27 me2
DNA Meth. m5C
H2A.Z
H4−4ac
H3K27 me3
H3K9 me2
H3K4 me1
H3K4 me3
H4K20 me2
H3−5ac
0.00 0.25 0.50 0.75 1.00
π̂
KNOP1 − Reverse
H4−4ac:H3−5ac
H2A.Z:H3−5ac
H2A.Z:H4−4ac
H3K4 me1:H2A.Z
H3K4 me3:H3−5ac
H3K4 me1:H3−5ac
H3K27 me3:H3K4 me1
H4K20 me2:H3−5ac
H4K20 me2:H3K4 me3
H3K27 me2
DNA Meth. m5C
H2A.Z
H4−4ac
H3K27 me3
H3K9 me2
H3K4 me1
H3K4 me3
H4K20 me2
H3−5ac
0.00 0.25 0.50 0.75 1.00
π̂
PDCD11 − Forward
H3K27 me3:H3K4 me1
H3K9 me3:H4K20 me3
H3K27 me2:DNA Meth. m5C
H3K27 me3:H3K9 me3
H3K27 me3:H4K20 me3
H2A.Z:H3K4 me3
H3K4 me1:H3−5ac
H3K4 me3:H4−4ac
H4K20 me2:H4−4ac
H2A.Z:H4−4ac
DNA Meth. m5C:H4K20 me2
H3K27 me3:DNA Meth. m5C
DNA Meth. m5C:H3K9 me3
H2A.Z:H3−5ac
H2A.Z:H4K20 me2
H4K20 me2:H3K4 me3
DNA Meth. m5C:H4K20 me3
H4−4ac:H3−5ac
H4K20 me2:H3−5ac
H3K9 me2
H3K27 me2
H3K9 me3
H3K4 me3
H3K27 me3
H4K20 me3
H3K4 me1
H4K20 me2
H2A.Z
DNA Meth. m5C
H4−4ac
H3−5ac
0.00 0.25 0.50 0.75 1.00
π̂
PDCD11 − Reverse
H3K27 me3:H3K4 me1
H3K9 me3:H4K20 me3
H3K27 me2:DNA Meth. m5C
H3K27 me3:H3K9 me3
H3K27 me3:H4K20 me3
H2A.Z:H3K4 me3
H3K4 me1:H3−5ac
H3K4 me3:H4−4ac
H4K20 me2:H4−4ac
H2A.Z:H4−4ac
DNA Meth. m5C:H4K20 me2
H3K27 me3:DNA Meth. m5C
DNA Meth. m5C:H3K9 me3
H2A.Z:H3−5ac
H2A.Z:H4K20 me2
H4K20 me2:H3K4 me3
DNA Meth. m5C:H4K20 me3
H4−4ac:H3−5ac
H4K20 me2:H3−5ac
H3K9 me2
H3K27 me2
H3K9 me3
H3K4 me3
H3K27 me3
H4K20 me3
H3K4 me1
H4K20 me2
H2A.Z
DNA Meth. m5C
H4−4ac
H3−5ac
0.00 0.25 0.50 0.75 1.00
π̂
NOP2 (2) − Forward
H2A.Z:H3K4 me3
H3K4 me3:H3−5ac
H4K20 me2:H4−4ac
H4−4ac:H3−5ac
H3K4 me1:H3−5ac
DNA Meth. m5C:H4K20 me2
DNA Meth. m5C:H3K9 me3
H3K27 me3:H3K4 me1
H3K4 me1:H4−4ac
H3K27 me3:DNA Meth. m5C
H3K4 me3:H4−4ac
H3K9 me2:DNA Meth. m5C
H3K4 me1:H2A.Z
H3K27 me2:DNA Meth. m5C
H4K20 me2:H3K4 me3
H3K4 me1:H4K20 me2
H2A.Z:H4K20 me2
H4K20 me2:H3−5ac
H4K20 me3
H3K9 me2
H3K9 me3
H3K27 me2
H3K27 me3
H3K4 me3
H4−4ac
H3−5ac
DNA Meth. m5C
H3K4 me1
H2A.Z
H4K20 me2
0.00 0.25 0.50 0.75 1.00
π̂
NOP2 (2) − Reverse
H2A.Z:H3K4 me3
H3K4 me3:H3−5ac
H4K20 me2:H4−4ac
H4−4ac:H3−5ac
H3K4 me1:H3−5ac
DNA Meth. m5C:H4K20 me2
DNA Meth. m5C:H3K9 me3
H3K27 me3:H3K4 me1
H3K4 me1:H4−4ac
H3K27 me3:DNA Meth. m5C
H3K4 me3:H4−4ac
H3K9 me2:DNA Meth. m5C
H3K4 me1:H2A.Z
H3K27 me2:DNA Meth. m5C
H4K20 me2:H3K4 me3
H3K4 me1:H4K20 me2
H2A.Z:H4K20 me2
H4K20 me2:H3−5ac
H4K20 me3
H3K9 me2
H3K9 me3
H3K27 me2
H3K27 me3
H3K4 me3
H4−4ac
H3−5ac
DNA Meth. m5C
H3K4 me1
H2A.Z
H4K20 me2
0.00 0.25 0.50 0.75 1.00
π̂
FBL − Forward
DNA Meth. m5C:H4K20 me3
H3K9 me3:H4K20 me3
H3K27 me3:H4K20 me3
H3K27 me3:DNA Meth. m5C
H3K4 me3:H4−4ac
H3K27 me3:H3K9 me3
H4K20 me2:H4−4ac
H3K27 me3:H3K4 me1
H3K4 me3:H3−5ac
H3K4 me1:H3−5ac
H4−4ac:H3−5ac
DNA Meth. m5C:H4K20 me2
H3K27 me2:DNA Meth. m5C
H4K20 me2:H3K4 me3
H4K20 me2:H3−5ac
H3K9 me2:DNA Meth. m5C
H2A.Z
H4K20 me3
H3K9 me3
H3K4 me1
H4−4ac
H3K27 me2
H3K4 me3
H3K9 me2
H4K20 me2
DNA Meth. m5C
H3K27 me3
H3−5ac
0.00 0.25 0.50 0.75 1.00
π̂
FBL − Reverse
DNA Meth. m5C:H4K20 me3
H3K9 me3:H4K20 me3
H3K27 me3:H4K20 me3
H3K27 me3:DNA Meth. m5C
H3K4 me3:H4−4ac
H3K27 me3:H3K9 me3
H4K20 me2:H4−4ac
H3K27 me3:H3K4 me1
H3K4 me3:H3−5ac
H3K4 me1:H3−5ac
H4−4ac:H3−5ac
DNA Meth. m5C:H4K20 me2
H3K27 me2:DNA Meth. m5C
H4K20 me2:H3K4 me3
H4K20 me2:H3−5ac
H3K9 me2:DNA Meth. m5C
H2A.Z
H4K20 me3
H3K9 me3
H3K4 me1
H4−4ac
H3K27 me2
H3K4 me3
H3K9 me2
H4K20 me2
DNA Meth. m5C
H3K27 me3
H3−5ac
0.00 0.25 0.50 0.75 1.00
π̂
NOP56 − Forward
H4K20 me2:H4−4ac
H4K20 me2:H3−5ac
H4K20 me2:H3K4 me3
H3K9 me2:DNA Meth. m5C
DNA Meth. m5C:H4K20 me2
H4−4ac:H3−5ac
H3K4 me1:H3−5ac
H3K4 me3:H4−4ac
H3K27 me2
H4K20 me3
H3K4 me3
H2A.Z
DNA Meth. m5C
H3K9 me3
H3−5ac
H4−4ac
H3K27 me3
H3K9 me2
H4K20 me2
H3K4 me1
0.00 0.25 0.50 0.75 1.00
π̂
NOP56 − Reverse
H4K20 me2:H4−4ac
H4K20 me2:H3−5ac
H4K20 me2:H3K4 me3
H3K9 me2:DNA Meth. m5C
DNA Meth. m5C:H4K20 me2
H4−4ac:H3−5ac
H3K4 me1:H3−5ac
H3K4 me3:H4−4ac
H3K27 me2
H4K20 me3
H3K4 me3
H2A.Z
DNA Meth. m5C
H3K9 me3
H3−5ac
H4−4ac
H3K27 me3
H3K9 me2
H4K20 me2
H3K4 me1
0.00 0.25 0.50 0.75 1.00
π̂
SNU13 − Forward
DNA Meth. m5C:H3K9 me3
H3K27 me3:DNA Meth. m5C
H3K4 me1:H4−4ac
H4K20 me2:H3K4 me3
H2A.Z:H3−5ac
H3K4 me3:H4−4ac
H2A.Z:H4−4ac
H2A.Z:H3K4 me3
H3K4 me1:H2A.Z
H3K4 me3:H3−5ac
H4−4ac:H3−5ac
H4K20 me2:H4−4ac
H3K4 me1:H3−5ac
DNA Meth. m5C:H4K20 me2
H3K4 me1:H4K20 me2
H2A.Z:H4K20 me2
H4K20 me2:H3−5ac
H3K9 me2
H3K27 me2
H3K9 me3
H3K27 me3
H3K4 me3
DNA Meth. m5C
H4K20 me3
H3K4 me1
H2A.Z
H3−5ac
H4K20 me2
H4−4ac
0.00 0.25 0.50 0.75 1.00
π̂
SNU13 − Reverse
DNA Meth. m5C:H3K9 me3
H3K27 me3:DNA Meth. m5C
H3K4 me1:H4−4ac
H4K20 me2:H3K4 me3
H2A.Z:H3−5ac
H3K4 me3:H4−4ac
H2A.Z:H4−4ac
H2A.Z:H3K4 me3
H3K4 me1:H2A.Z
H3K4 me3:H3−5ac
H4−4ac:H3−5ac
H4K20 me2:H4−4ac
H3K4 me1:H3−5ac
DNA Meth. m5C:H4K20 me2
H3K4 me1:H4K20 me2
H2A.Z:H4K20 me2
H4K20 me2:H3−5ac
H3K9 me2
H3K27 me2
H3K9 me3
H3K27 me3
H3K4 me3
DNA Meth. m5C
H4K20 me3
H3K4 me1
H2A.Z
H3−5ac
H4K20 me2
H4−4ac
0.00 0.25 0.50 0.75 1.00
π̂

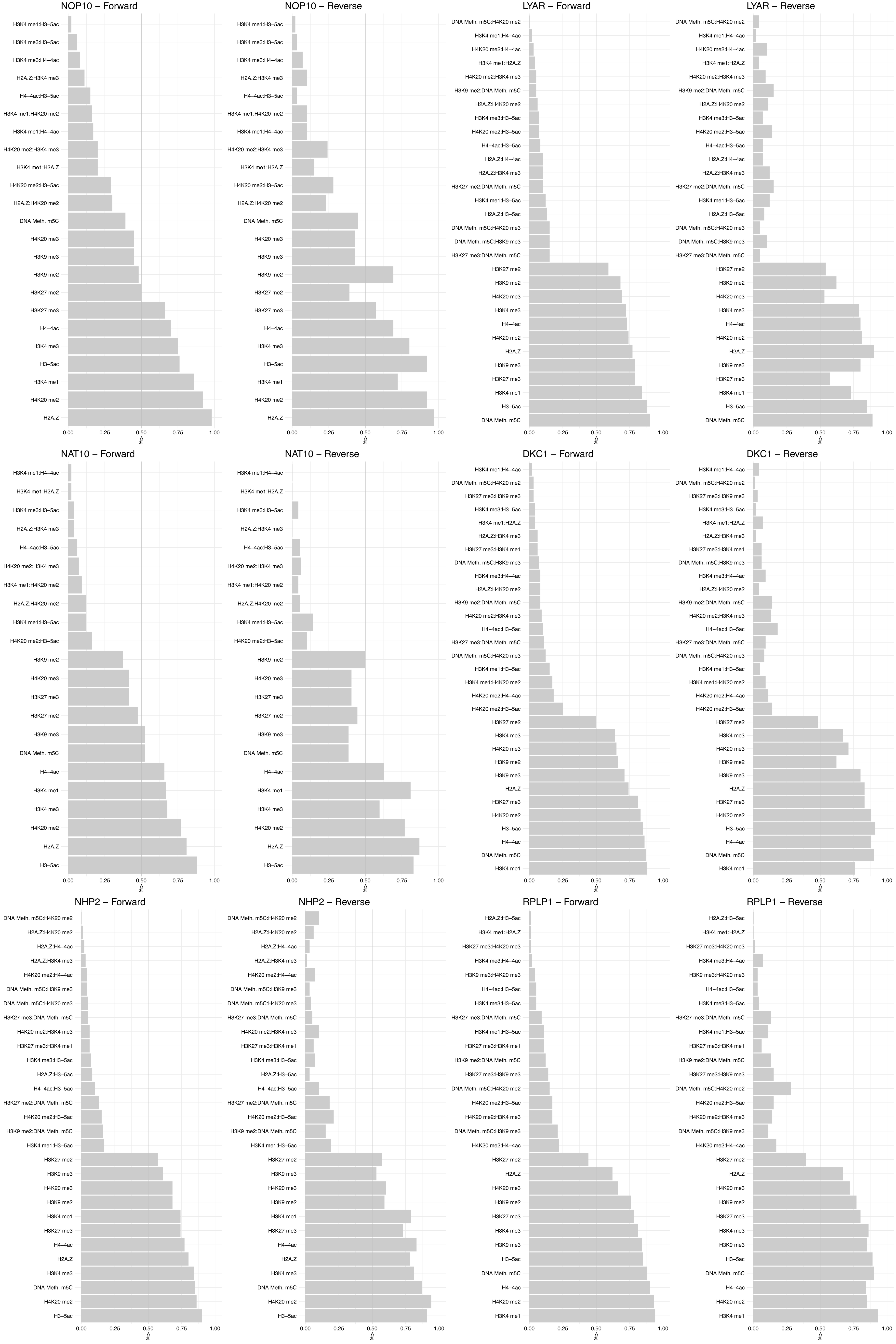
NOP10 – Forward
H3K4 me1:H3-5ac
H3K4 me3:H3-5ac
H3K4 me3:H4-4ac
H2A.Z:H3K4 me3
H4-4ac:H3-5ac
H3K4 me1:H4K20 me2
H3K4 me1:H4-4ac
H4K20 me2:H3K4 me3
H3K4 me1:H2A.Z
H4K20 me2:H3-5ac
H2A.Z:H4K20 me2
DNA Meth. m5C
H4K20 me3
H3K9 me3
H3K9 me2
H3K27 me2
H3K27 me3
H4-4ac
H3K4 me3
H3-5ac
H3K4 me1
H4K20 me2
H2A.Z
0.00 0.25 0.50 0.75 1.00
π̂
NOP10 – Reverse
H3K4 me1:H3-5ac
H3K4 me3:H3-5ac
H3K4 me3:H4-4ac
H2A.Z:H3K4 me3
H4-4ac:H3-5ac
H3K4 me1:H4K20 me2
H3K4 me1:H4-4ac
H4K20 me2:H3K4 me3
H3K4 me1:H2A.Z
H4K20 me2:H3-5ac
H2A.Z:H4K20 me2
DNA Meth. m5C
H4K20 me3
H3K9 me3
H3K9 me2
H3K27 me2
H3K27 me3
H4-4ac
H3K4 me3
H3-5ac
H3K4 me1
H4K20 me2
H2A.Z
0.00 0.25 0.50 0.75 1.00
π̂
LYAR – Forward
DNA Meth. m5C:H4K20 me2
H3K4 me1:H4-4ac
H4K20 me2:H4-4ac
H3K4 me1:H2A.Z
H4K20 me2:H3K4 me3
H3K9 me2:DNA Meth. m5C
H2A.Z:H4K20 me2
H3K4 me3:H3-5ac
H4K20 me2:H3-5ac
H4-4ac:H3-5ac
H2A.Z:H4-4ac
H2A.Z:H3K4 me3
H3K27 me2:DNA Meth. m5C
H3K4 me1:H3-5ac
H2A.Z:H3-5ac
DNA Meth. m5C:H4K20 me3
DNA Meth. m5C:H3K9 me3
H3K27 me3:DNA Meth. m5C
H3K27 me2
H3K9 me2
H4K20 me3
H3K4 me3
H4-4ac
H4K20 me2
H2A.Z
H3K9 me3
H3K27 me3
H3K4 me1
H3-5ac
DNA Meth. m5C
0.00 0.25 0.50 0.75 1.00
π̂
LYAR – Reverse
DNA Meth. m5C:H4K20 me2
H3K4 me1:H4-4ac
H4K20 me2:H4-4ac
H3K4 me1:H2A.Z
H4K20 me2:H3K4 me3
H3K9 me2:DNA Meth. m5C
H2A.Z:H4K20 me2
H3K4 me3:H3-5ac
H4K20 me2:H3-5ac
H4-4ac:H3-5ac
H2A.Z:H4-4ac
H2A.Z:H3K4 me3
H3K27 me2:DNA Meth. m5C
H3K4 me1:H3-5ac
H2A.Z:H3-5ac
DNA Meth. m5C:H4K20 me3
DNA Meth. m5C:H3K9 me3
H3K27 me3:DNA Meth. m5C
H3K27 me2
H3K9 me2
H4K20 me3
H3K4 me3
H4-4ac
H4K20 me2
H2A.Z
H3K9 me3
H3K27 me3
H3K4 me1
H3-5ac
DNA Meth. m5C
0.00 0.25 0.50 0.75 1.00
π̂
NAT10 – Forward
H3K4 me1:H4-4ac
H3K4 me1:H2A.Z
H3K4 me3:H3-5ac
H2A.Z:H3K4 me3
H4-4ac:H3-5ac
H4K20 me2:H3K4 me3
H3K4 me1:H4K20 me2
H2A.Z:H4K20 me2
H3K4 me1:H3-5ac
H4K20 me2:H3-5ac
H3K9 me2
H4K20 me3
H3K27 me3
H3K27 me2
H3K9 me3
DNA Meth. m5C
H4-4ac
H3K4 me1
H3K4 me3
H4K20 me2
H2A.Z
H3-5ac
0.00 0.25 0.50 0.75 1.00
π̂
NAT10 – Reverse
H3K4 me1:H4-4ac
H3K4 me1:H2A.Z
H3K4 me3:H3-5ac
H2A.Z:H3K4 me3
H4-4ac:H3-5ac
H4K20 me2:H3K4 me3
H3K4 me1:H4K20 me2
H2A.Z:H4K20 me2
H3K4 me1:H3-5ac
H4K20 me2:H3-5ac
H3K9 me2
H4K20 me3
H3K27 me3
H3K27 me2
H3K9 me3
DNA Meth. m5C
H4-4ac
H3K4 me1
H3K4 me3
H4K20 me2
H2A.Z
H3-5ac
0.00 0.25 0.50 0.75 1.00
π̂
DKC1 – Forward
H3K4 me1:H4-4ac
DNA Meth. m5C:H4K20 me2
H3K27 me3:H3K9 me3
H3K4 me3:H3-5ac
H3K4 me1:H2A.Z
H2A.Z:H3K4 me3
H3K27 me3:H3K4 me1
DNA Meth. m5C:H3K9 me3
H3K4 me3:H4-4ac
H2A.Z:H4K20 me2
H3K9 me2:DNA Meth. m5C
H4K20 me2:H3K4 me3
H4-4ac:H3-5ac
H3K27 me3:DNA Meth. m5C
DNA Meth. m5C:H4K20 me3
H3K4 me1:H3-5ac
H3K4 me1:H4K20 me2
H4K20 me2:H4-4ac
H4K20 me2:H3-5ac
H3K27 me2
H3K4 me3
H4K20 me3
H3K9 me2
H3K9 me3
H2A.Z
H3K27 me3
H4K20 me2
H3-5ac
H4-4ac
DNA Meth. m5C
H3K4 me1
0.00 0.25 0.50 0.75 1.00
π̂
DKC1 – Reverse
H3K4 me1:H4-4ac
DNA Meth. m5C:H4K20 me2
H3K27 me3:H3K9 me3
H3K4 me3:H3-5ac
H3K4 me1:H2A.Z
H2A.Z:H3K4 me3
H3K27 me3:H3K4 me1
DNA Meth. m5C:H3K9 me3
H3K4 me3:H4-4ac
H2A.Z:H4K20 me2
H3K9 me2:DNA Meth. m5C
H4K20 me2:H3K4 me3
H4-4ac:H3-5ac
H3K27 me3:DNA Meth. m5C
DNA Meth. m5C:H4K20 me3
H3K4 me1:H3-5ac
H3K4 me1:H4K20 me2
H4K20 me2:H4-4ac
H4K20 me2:H3-5ac
H3K27 me2
H3K4 me3
H4K20 me3
H3K9 me2
H3K9 me3
H2A.Z
H3K27 me3
H4K20 me2
H3-5ac
H4-4ac
DNA Meth. m5C
H3K4 me1
0.00 0.25 0.50 0.75 1.00
π̂
NHP2 – Forward
DNA Meth. m5C:H4K20 me2
H2A.Z:H4K20 me2
H2A.Z:H4-4ac
H2A.Z:H3K4 me3
H4K20 me2:H4-4ac
DNA Meth. m5C:H3K9 me3
DNA Meth. m5C:H4K20 me3
H3K27 me3:DNA Meth. m5C
H4K20 me2:H3K4 me3
H3K27 me3:H3K4 me1
H3K4 me3:H3-5ac
H2A.Z:H3-5ac
H4-4ac:H3-5ac
H3K27 me2:DNA Meth. m5C
H4K20 me2:H3-5ac
H3K9 me2:DNA Meth. m5C
H3K4 me1:H3-5ac
H3K27 me2
H3K9 me3
H4K20 me3
H3K9 me2
H3K4 me1
H3K27 me3
H4-4ac
H2A.Z
H3K4 me3
DNA Meth. m5C
H4K20 me2
H3-5ac
0.00 0.25 0.50 0.75 1.00
π̂
NHP2 – Reverse
DNA Meth. m5C:H4K20 me2
H2A.Z:H4K20 me2
H2A.Z:H4-4ac
H2A.Z:H3K4 me3
H4K20 me2:H4-4ac
DNA Meth. m5C:H3K9 me3
DNA Meth. m5C:H4K20 me3
H3K27 me3:DNA Meth. m5C
H4K20 me2:H3K4 me3
H3K27 me3:H3K4 me1
H3K4 me3:H3-5ac
H2A.Z:H3-5ac
H4-4ac:H3-5ac
H3K27 me2:DNA Meth. m5C
H4K20 me2:H3-5ac
H3K9 me2:DNA Meth. m5C
H3K4 me1:H3-5ac
H3K27 me2
H3K9 me3
H4K20 me3
H3K9 me2
H3K4 me1
H3K27 me3
H4-4ac
H2A.Z
H3K4 me3
DNA Meth. m5C
H4K20 me2
H3-5ac
0.00 0.25 0.50 0.75 1.00
π̂
RPLP1 – Forward
H2A.Z:H3-5ac
H3K4 me1:H2A.Z
H3K27 me3:H4K20 me3
H3K4 me3:H4-4ac
H3K9 me3:H4K20 me3
H4-4ac:H3-5ac
H3K4 me3:H3-5ac
H3K27 me3:DNA Meth. m5C
H3K4 me1:H3-5ac
H3K27 me3:H3K4 me1
H3K9 me2:DNA Meth. m5C
H3K27 me3:H3K9 me3
DNA Meth. m5C:H4K20 me2
H4K20 me2:H3-5ac
H4K20 me2:H3K4 me3
DNA Meth. m5C:H3K9 me3
H4K20 me2:H4-4ac
H3K27 me2
H2A.Z
H4K20 me3
H3K9 me2
H3K27 me3
H3K4 me3
H3K9 me3
H3-5ac
DNA Meth. m5C
H4-4ac
H4K20 me2
H3K4 me1
0.00 0.25 0.50 0.75 1.00
π̂
RPLP1 – Reverse
H2A.Z:H3-5ac
H3K4 me1:H2A.Z
H3K27 me3:H4K20 me3
H3K4 me3:H4-4ac
H3K9 me3:H4K20 me3
H4-4ac:H3-5ac
H3K4 me3:H3-5ac
H3K27 me3:DNA Meth. m5C
H3K4 me1:H3-5ac
H3K27 me3:H3K4 me1
H3K9 me2:DNA Meth. m5C
H3K27 me3:H3K9 me3
DNA Meth. m5C:H4K20 me2
H4K20 me2:H3-5ac
H4K20 me2:H3K4 me3
DNA Meth. m5C:H3K9 me3
H4K20 me2:H4-4ac
H3K27 me2
H2A.Z
H4K20 me3
H3K9 me2
H3K27 me3
H3K4 me3
H3K9 me3
H3-5ac
DNA Meth. m5C
H4-4ac
H4K20 me2
H3K4 me1
0.00 0.25 0.50 0.75 1.00
π̂

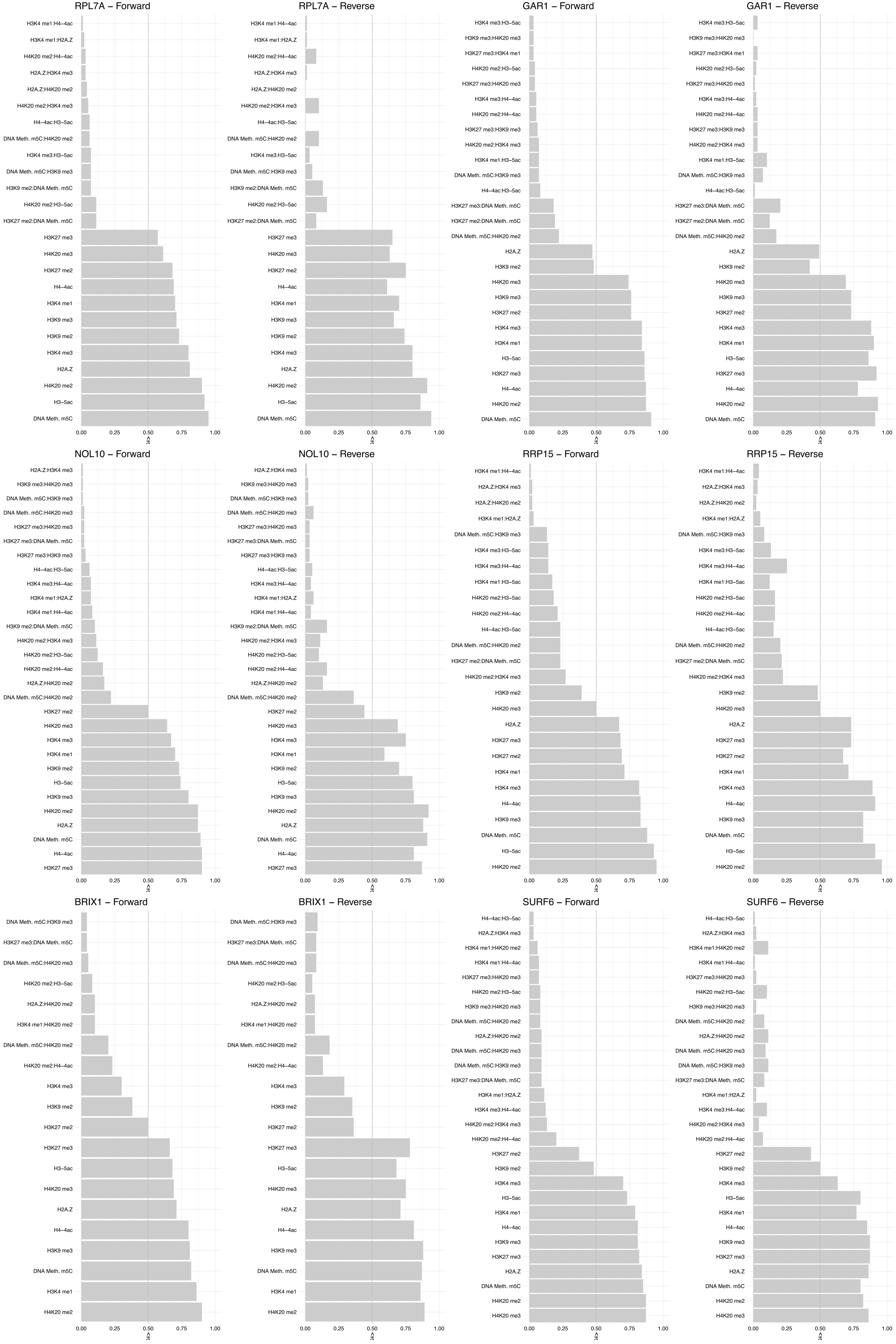
RPL7A – Forward
H3K4 me1:H4–4ac
H3K4 me1:H2A.Z
H4K20 me2:H4–4ac
H2A.Z:H3K4 me3
H2A.Z:H4K20 me2
H4K20 me2:H3K4 me3
H4–4ac:H3–5ac
DNA Meth. m5C:H4K20 me2
H3K4 me3:H3–5ac
DNA Meth. m5C:H3K9 me3
H3K9 me2:DNA Meth. m5C
H4K20 me2:H3–5ac
H3K27 me2:DNA Meth. m5C
H3K27 me3
H4K20 me3
H3K27 me2
H4–4ac
H3K4 me1
H3K9 me3
H3K9 me2
H3K4 me3
H2A.Z
H4K20 me2
H3–5ac
DNA Meth. m5C
0.00
0.25
0.50
0.75
1.00
π̂
RPL7A – Reverse
GAR1 – Forward
H3K4 me3:H3–5ac
H3K9 me3:H4K20 me3
H3K27 me3:H3K4 me1
H4K20 me2:H3–5ac
H3K27 me3:H4K20 me3
H3K4 me3:H4–4ac
H4K20 me2:H4–4ac
H3K27 me3:H3K9 me3
H4K20 me2:H3K4 me3
H3K4 me1:H3–5ac
DNA Meth. m5C:H3K9 me3
H4–4ac:H3–5ac
H3K27 me3:DNA Meth. m5C
H3K27 me2:DNA Meth. m5C
DNA Meth. m5C:H4K20 me2
H2A.Z
H3K9 me2
H4K20 me3
H3K9 me3
H3K27 me2
H3K4 me3
H3K4 me1
H3–5ac
H3K27 me3
H4–4ac
H4K20 me2
DNA Meth. m5C
GAR1 – Reverse
NOL10 – Forward
H2A.Z:H3K4 me3
H3K9 me3:H4K20 me3
DNA Meth. m5C:H3K9 me3
DNA Meth. m5C:H4K20 me3
H3K27 me3:H4K20 me3
H3K27 me3:DNA Meth. m5C
H3K27 me3:H3K9 me3
H4–4ac:H3–5ac
H3K4 me3:H4–4ac
H3K4 me1:H2A.Z
H3K4 me1:H4–4ac
H3K9 me2:DNA Meth. m5C
H4K20 me2:H3K4 me3
H4K20 me2:H3–5ac
H4K20 me2:H4–4ac
H2A.Z:H4K20 me2
DNA Meth. m5C:H4K20 me2
H3K27 me2
H4K20 me3
H3K4 me3
H3K4 me1
H3K9 me2
H3–5ac
H3K9 me3
H4K20 me2
H2A.Z
DNA Meth. m5C
H4–4ac
H3K27 me3
NOL10 – Reverse
RRP15 – Forward
H3K4 me1:H4–4ac
H2A.Z:H3K4 me3
H2A.Z:H4K20 me2
H3K4 me1:H2A.Z
DNA Meth. m5C:H3K9 me3
H3K4 me3:H3–5ac
H3K4 me3:H4–4ac
H3K4 me1:H3–5ac
H4K20 me2:H3–5ac
H4K20 me2:H4–4ac
H4–4ac:H3–5ac
DNA Meth. m5C:H4K20 me2
H3K27 me2:DNA Meth. m5C
H4K20 me2:H3K4 me3
H3K9 me2
H4K20 me3
H2A.Z
H3K27 me3
H3K27 me2
H3K4 me1
H3K4 me3
H4–4ac
H3K9 me3
DNA Meth. m5C
H3–5ac
H4K20 me2
RRP15 – Reverse
BRIX1 – Forward
DNA Meth. m5C:H3K9 me3
H3K27 me3:DNA Meth. m5C
DNA Meth. m5C:H4K20 me3
H4K20 me2:H3–5ac
H2A.Z:H4K20 me2
H3K4 me1:H4K20 me2
DNA Meth. m5C:H4K20 me2
H4K20 me2:H4–4ac
H3K4 me3
H3K9 me2
H3K27 me2
H3K27 me3
H3–5ac
H4K20 me3
H2A.Z
H4–4ac
H3K9 me3
DNA Meth. m5C
H3K4 me1
H4K20 me2
BRIX1 – Reverse
SURF6 – Forward
H4–4ac:H3–5ac
H2A.Z:H3K4 me3
H3K4 me1:H4K20 me2
H3K4 me1:H4–4ac
H3K27 me3:H4K20 me3
H4K20 me2:H3–5ac
H3K9 me3:H4K20 me3
DNA Meth. m5C:H4K20 me2
H2A.Z:H4K20 me2
DNA Meth. m5C:H4K20 me3
DNA Meth. m5C:H3K9 me3
H3K27 me3:DNA Meth. m5C
H3K4 me1:H2A.Z
H3K4 me3:H4–4ac
H4K20 me2:H3K4 me3
H4K20 me2:H4–4ac
H3K27 me2
H3K9 me2
H3K4 me3
H3–5ac
H3K4 me1
H4–4ac
H3K9 me3
H3K27 me3
H2A.Z
DNA Meth. m5C
H4K20 me2
H4K20 me3
SURF6 – Reverse

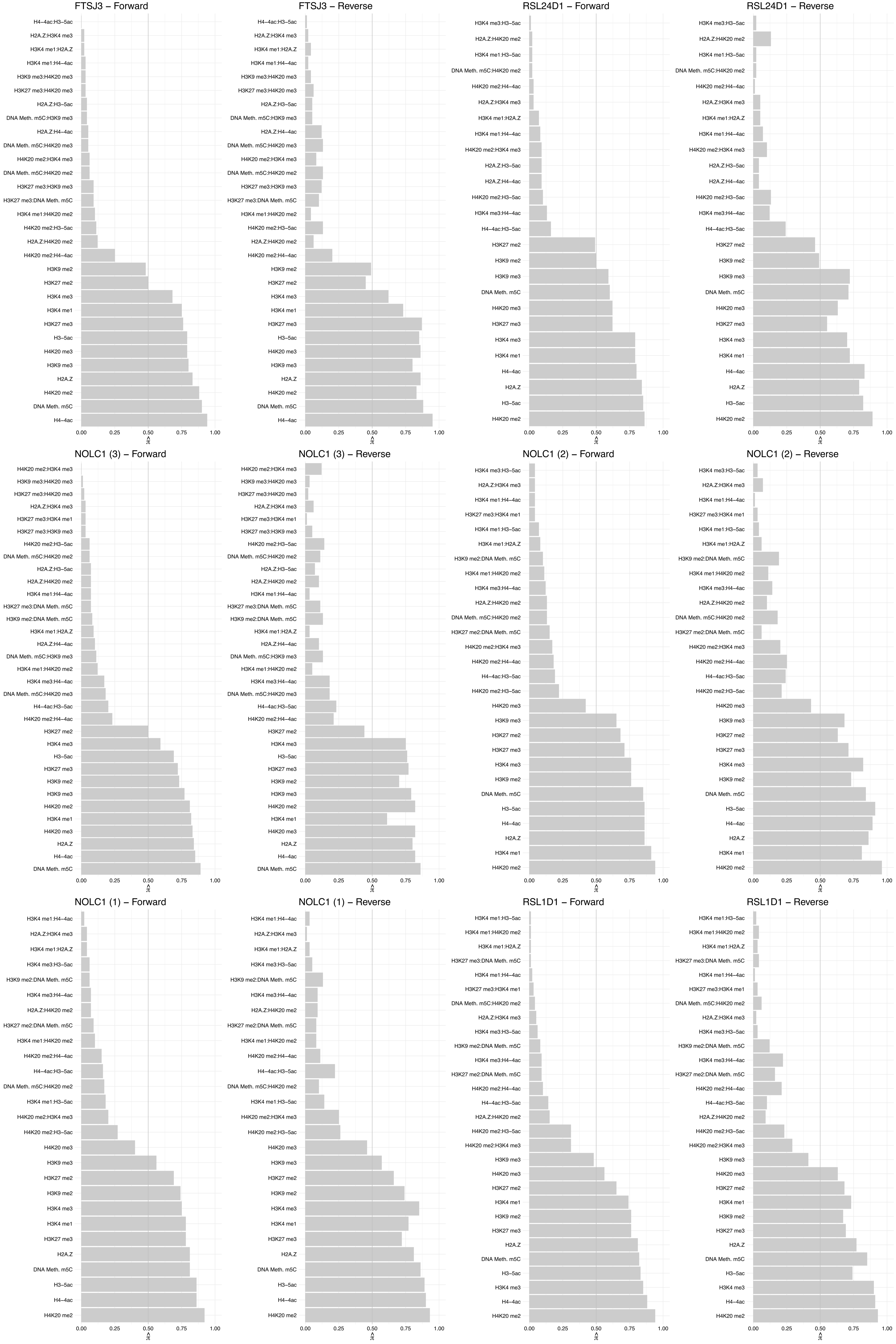
FTSJ3 − Forward
H4−4ac:H3−5ac
H2A.Z:H3K4 me3
H3K4 me1:H2A.Z
H3K4 me1:H4−4ac
H3K9 me3:H4K20 me3
H3K27 me3:H4K20 me3
H2A.Z:H3−5ac
DNA Meth. m5C:H3K9 me3
H2A.Z:H4−4ac
DNA Meth. m5C:H4K20 me3
H4K20 me2:H3K4 me3
DNA Meth. m5C:H4K20 me2
H3K27 me3:H3K9 me3
H3K27 me3:DNA Meth. m5C
H3K4 me1:H4K20 me2
H4K20 me2:H3−5ac
H2A.Z:H4K20 me2
H4K20 me2:H4−4ac
H3K9 me2
H3K27 me2
H3K4 me3
H3K4 me1
H3K27 me3
H3−5ac
H4K20 me3
H3K9 me3
H2A.Z
H4K20 me2
DNA Meth. m5C
H4−4ac
0.00 0.25 0.50 0.75 1.00
π̂
FTSJ3 − Reverse
H4−4ac:H3−5ac
H2A.Z:H3K4 me3
H3K4 me1:H2A.Z
H3K4 me1:H4−4ac
H3K9 me3:H4K20 me3
H3K27 me3:H4K20 me3
H2A.Z:H3−5ac
DNA Meth. m5C:H3K9 me3
H2A.Z:H4−4ac
DNA Meth. m5C:H4K20 me3
H4K20 me2:H3K4 me3
DNA Meth. m5C:H4K20 me2
H3K27 me3:H3K9 me3
H3K27 me3:DNA Meth. m5C
H3K4 me1:H4K20 me2
H4K20 me2:H3−5ac
H2A.Z:H4K20 me2
H4K20 me2:H4−4ac
H3K9 me2
H3K27 me2
H3K4 me3
H3K4 me1
H3K27 me3
H3−5ac
H4K20 me3
H3K9 me3
H2A.Z
H4K20 me2
DNA Meth. m5C
H4−4ac
0.00 0.25 0.50 0.75 1.00
π̂
RSL24D1 − Forward
H3K4 me3:H3−5ac
H2A.Z:H4K20 me2
H3K4 me1:H3−5ac
DNA Meth. m5C:H4K20 me2
H4K20 me2:H4−4ac
H2A.Z:H3K4 me3
H3K4 me1:H2A.Z
H3K4 me1:H4−4ac
H4K20 me2:H3K4 me3
H2A.Z:H3−5ac
H2A.Z:H4−4ac
H4K20 me2:H3−5ac
H3K4 me3:H4−4ac
H4−4ac:H3−5ac
H3K27 me2
H3K9 me2
H3K9 me3
DNA Meth. m5C
H4K20 me3
H3K27 me3
H3K4 me3
H3K4 me1
H4−4ac
H2A.Z
H3−5ac
H4K20 me2
0.00 0.25 0.50 0.75 1.00
π̂
RSL24D1 − Reverse
H3K4 me3:H3−5ac
H2A.Z:H4K20 me2
H3K4 me1:H3−5ac
DNA Meth. m5C:H4K20 me2
H4K20 me2:H4−4ac
H2A.Z:H3K4 me3
H3K4 me1:H2A.Z
H3K4 me1:H4−4ac
H4K20 me2:H3K4 me3
H2A.Z:H3−5ac
H2A.Z:H4−4ac
H4K20 me2:H3−5ac
H3K4 me3:H4−4ac
H4−4ac:H3−5ac
H3K27 me2
H3K9 me2
H3K9 me3
DNA Meth. m5C
H4K20 me3
H3K27 me3
H3K4 me3
H3K4 me1
H4−4ac
H2A.Z
H3−5ac
H4K20 me2
0.00 0.25 0.50 0.75 1.00
π̂
NOLC1 (3) − Forward
H4K20 me2:H3K4 me3
H3K9 me3:H4K20 me3
H3K27 me3:H4K20 me3
H2A.Z:H3K4 me3
H3K27 me3:H3K4 me1
H3K27 me3:H3K9 me3
H4K20 me2:H3−5ac
DNA Meth. m5C:H4K20 me2
H2A.Z:H3−5ac
H2A.Z:H4K20 me2
H3K4 me1:H4−4ac
H3K27 me3:DNA Meth. m5C
H3K9 me2:DNA Meth. m5C
H3K4 me1:H2A.Z
H2A.Z:H4−4ac
DNA Meth. m5C:H3K9 me3
H3K4 me1:H4K20 me2
H3K4 me3:H4−4ac
DNA Meth. m5C:H4K20 me3
H4−4ac:H3−5ac
H4K20 me2:H4−4ac
H3K27 me2
H3K4 me3
H3−5ac
H3K27 me3
H3K9 me2
H3K9 me3
H4K20 me2
H3K4 me1
H4K20 me3
H2A.Z
H4−4ac
DNA Meth. m5C
0.00 0.25 0.50 0.75 1.00
π̂
NOLC1 (3) − Reverse
H4K20 me2:H3K4 me3
H3K9 me3:H4K20 me3
H3K27 me3:H4K20 me3
H2A.Z:H3K4 me3
H3K27 me3:H3K4 me1
H3K27 me3:H3K9 me3
H4K20 me2:H3−5ac
DNA Meth. m5C:H4K20 me2
H2A.Z:H3−5ac
H2A.Z:H4K20 me2
H3K4 me1:H4−4ac
H3K27 me3:DNA Meth. m5C
H3K9 me2:DNA Meth. m5C
H3K4 me1:H2A.Z
H2A.Z:H4−4ac
DNA Meth. m5C:H3K9 me3
H3K4 me1:H4K20 me2
H3K4 me3:H4−4ac
DNA Meth. m5C:H4K20 me3
H4−4ac:H3−5ac
H4K20 me2:H4−4ac
H3K27 me2
H3K4 me3
H3−5ac
H3K27 me3
H3K9 me2
H3K9 me3
H4K20 me2
H3K4 me1
H4K20 me3
H2A.Z
H4−4ac
DNA Meth. m5C
0.00 0.25 0.50 0.75 1.00
π̂
NOLC1 (2) − Forward
H3K4 me3:H3−5ac
H2A.Z:H3K4 me3
H3K4 me1:H4−4ac
H3K27 me3:H3K4 me1
H3K4 me1:H3−5ac
H3K4 me1:H2A.Z
H3K9 me2:DNA Meth. m5C
H3K4 me1:H4K20 me2
H3K4 me3:H4−4ac
H2A.Z:H4K20 me2
DNA Meth. m5C:H4K20 me2
H3K27 me2:DNA Meth. m5C
H4K20 me2:H3K4 me3
H4K20 me2:H4−4ac
H4−4ac:H3−5ac
H4K20 me2:H3−5ac
H4K20 me3
H3K9 me3
H3K27 me2
H3K27 me3
H3K4 me3
H3K9 me2
DNA Meth. m5C
H3−5ac
H4−4ac
H2A.Z
H3K4 me1
H4K20 me2
0.00 0.25 0.50 0.75 1.00
π̂
NOLC1 (2) − Reverse
H3K4 me3:H3−5ac
H2A.Z:H3K4 me3
H3K4 me1:H4−4ac
H3K27 me3:H3K4 me1
H3K4 me1:H3−5ac
H3K4 me1:H2A.Z
H3K9 me2:DNA Meth. m5C
H3K4 me1:H4K20 me2
H3K4 me3:H4−4ac
H2A.Z:H4K20 me2
DNA Meth. m5C:H4K20 me2
H3K27 me2:DNA Meth. m5C
H4K20 me2:H3K4 me3
H4K20 me2:H4−4ac
H4−4ac:H3−5ac
H4K20 me2:H3−5ac
H4K20 me3
H3K9 me3
H3K27 me2
H3K27 me3
H3K4 me3
H3K9 me2
DNA Meth. m5C
H3−5ac
H4−4ac
H2A.Z
H3K4 me1
H4K20 me2
0.00 0.25 0.50 0.75 1.00
π̂
NOLC1 (1) − Forward
H3K4 me1:H4−4ac
H2A.Z:H3K4 me3
H3K4 me1:H2A.Z
H3K4 me3:H3−5ac
H3K9 me2:DNA Meth. m5C
H3K4 me3:H4−4ac
H2A.Z:H4K20 me2
H3K27 me2:DNA Meth. m5C
H3K4 me1:H4K20 me2
H4K20 me2:H4−4ac
H4−4ac:H3−5ac
DNA Meth. m5C:H4K20 me2
H3K4 me1:H3−5ac
H4K20 me2:H3K4 me3
H4K20 me2:H3−5ac
H4K20 me3
H3K9 me3
H3K27 me2
H3K9 me2
H3K4 me3
H3K4 me1
H3K27 me3
H2A.Z
DNA Meth. m5C
H3−5ac
H4−4ac
H4K20 me2
0.00 0.25 0.50 0.75 1.00
π̂
NOLC1 (1) − Reverse
H3K4 me1:H4−4ac
H2A.Z:H3K4 me3
H3K4 me1:H2A.Z
H3K4 me3:H3−5ac
H3K9 me2:DNA Meth. m5C
H3K4 me3:H4−4ac
H2A.Z:H4K20 me2
H3K27 me2:DNA Meth. m5C
H3K4 me1:H4K20 me2
H4K20 me2:H4−4ac
H4−4ac:H3−5ac
DNA Meth. m5C:H4K20 me2
H3K4 me1:H3−5ac
H4K20 me2:H3K4 me3
H4K20 me2:H3−5ac
H4K20 me3
H3K9 me3
H3K27 me2
H3K9 me2
H3K4 me3
H3K4 me1
H3K27 me3
H2A.Z
DNA Meth. m5C
H3−5ac
H4−4ac
H4K20 me2
0.00 0.25 0.50 0.75 1.00
π̂
RSL1D1 − Forward
H3K4 me1:H3−5ac
H3K4 me1:H4K20 me2
H3K4 me1:H2A.Z
H3K27 me3:DNA Meth. m5C
H3K4 me1:H4−4ac
H3K27 me3:H3K4 me1
DNA Meth. m5C:H4K20 me2
H2A.Z:H3K4 me3
H3K4 me3:H3−5ac
H3K9 me2:DNA Meth. m5C
H3K4 me3:H4−4ac
H3K27 me2:DNA Meth. m5C
H4K20 me2:H4−4ac
H4−4ac:H3−5ac
H2A.Z:H4K20 me2
H4K20 me2:H3−5ac
H4K20 me2:H3K4 me3
H3K9 me3
H4K20 me3
H3K27 me2
H3K4 me1
H3K9 me2
H3K27 me3
H2A.Z
DNA Meth. m5C
H3−5ac
H3K4 me3
H4−4ac
H4K20 me2
0.00 0.25 0.50 0.75 1.00
π̂
RSL1D1 − Reverse
H3K4 me1:H3−5ac
H3K4 me1:H4K20 me2
H3K4 me1:H2A.Z
H3K27 me3:DNA Meth. m5C
H3K4 me1:H4−4ac
H3K27 me3:H3K4 me1
DNA Meth. m5C:H4K20 me2
H2A.Z:H3K4 me3
H3K4 me3:H3−5ac
H3K9 me2:DNA Meth. m5C
H3K4 me3:H4−4ac
H3K27 me2:DNA Meth. m5C
H4K20 me2:H4−4ac
H4−4ac:H3−5ac
H2A.Z:H4K20 me2
H4K20 me2:H3−5ac
H4K20 me2:H3K4 me3
H3K9 me3
H4K20 me3
H3K27 me2
H3K4 me1
H3K9 me2
H3K27 me3
H2A.Z
DNA Meth. m5C
H3−5ac
H3K4 me3
H4−4ac
H4K20 me2
0.00 0.25 0.50 0.75 1.00
π̂

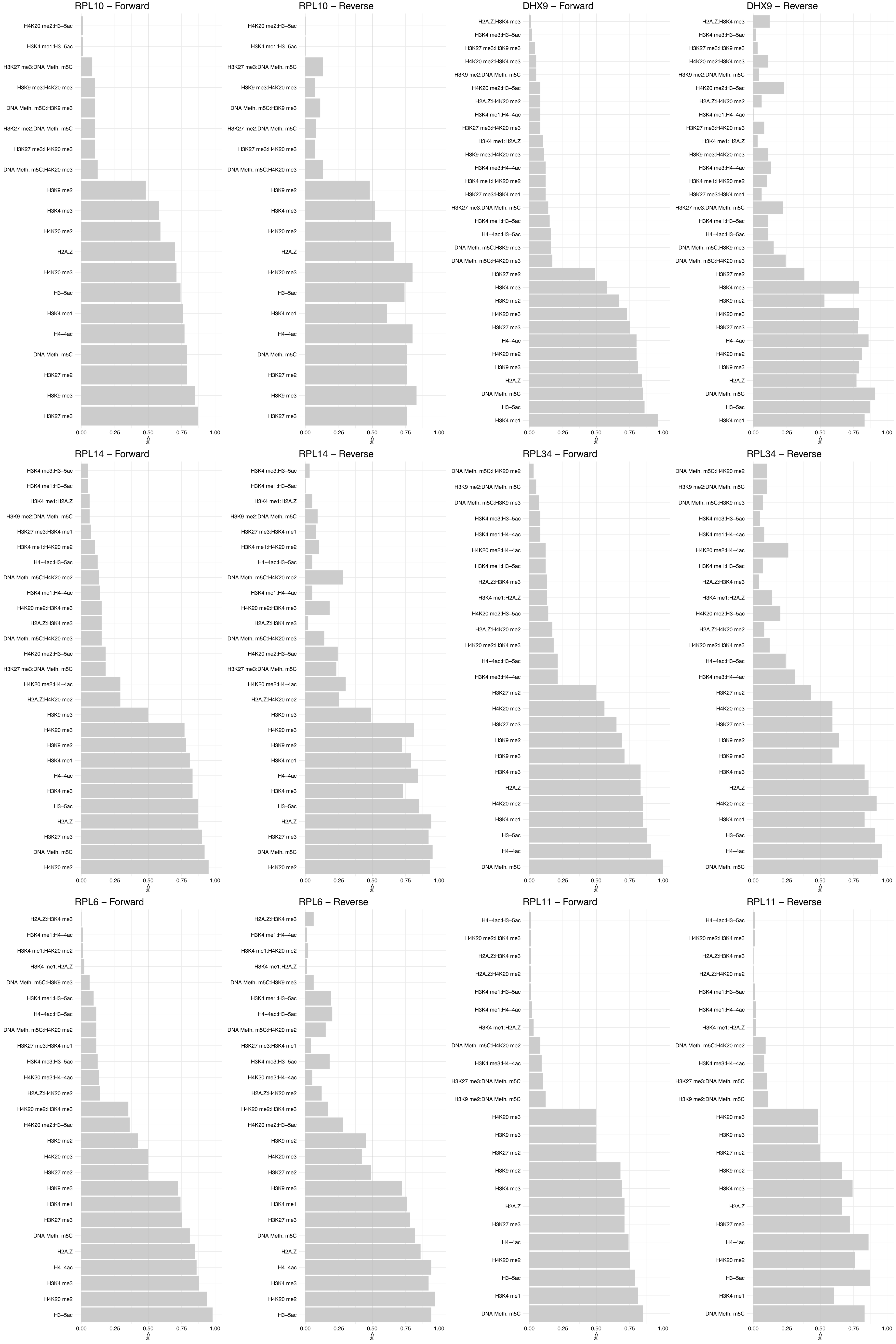
RPL10 – Forward
H4K20 me2:H3–5ac
H3K4 me1:H3–5ac
H3K27 me3:DNA Meth. m5C
H3K9 me3:H4K20 me3
DNA Meth. m5C:H3K9 me3
H3K27 me2:DNA Meth. m5C
H3K27 me3:H4K20 me3
DNA Meth. m5C:H4K20 me3
H3K9 me2
H3K4 me3
H4K20 me2
H2A.Z
H4K20 me3
H3–5ac
H3K4 me1
H4–4ac
DNA Meth. m5C
H3K27 me2
H3K9 me3
H3K27 me3
0.00
0.25
0.50
0.75
1.00
π̂
RPL10 – Reverse
DHX9 – Forward
H2A.Z:H3K4 me3
H3K4 me3:H3–5ac
H3K27 me3:H3K9 me3
H4K20 me2:H3K4 me3
H3K9 me2:DNA Meth. m5C
H4K20 me2:H3–5ac
H2A.Z:H4K20 me2
H3K4 me1:H4–4ac
H3K27 me3:H4K20 me3
H3K4 me1:H2A.Z
H3K9 me3:H4K20 me3
H3K4 me3:H4–4ac
H3K4 me1:H4K20 me2
H3K27 me3:H3K4 me1
H3K27 me3:DNA Meth. m5C
H3K4 me1:H3–5ac
H4–4ac:H3–5ac
DNA Meth. m5C:H3K9 me3
DNA Meth. m5C:H4K20 me3
H3K27 me2
H3K4 me3
H3K9 me2
H4K20 me3
H3K27 me3
H4–4ac
H4K20 me2
H3K9 me3
H2A.Z
DNA Meth. m5C
H3–5ac
H3K4 me1
DHX9 – Reverse
RPL14 – Forward
H3K4 me3:H3–5ac
H3K4 me1:H3–5ac
H3K4 me1:H2A.Z
H3K9 me2:DNA Meth. m5C
H3K27 me3:H3K4 me1
H3K4 me1:H4K20 me2
H4–4ac:H3–5ac
DNA Meth. m5C:H4K20 me2
H3K4 me1:H4–4ac
H4K20 me2:H3K4 me3
H2A.Z:H3K4 me3
DNA Meth. m5C:H4K20 me3
H4K20 me2:H3–5ac
H3K27 me3:DNA Meth. m5C
H4K20 me2:H4–4ac
H2A.Z:H4K20 me2
H3K9 me3
H4K20 me3
H3K9 me2
H3K4 me1
H4–4ac
H3K4 me3
H3–5ac
H2A.Z
H3K27 me3
DNA Meth. m5C
H4K20 me2
RPL14 – Reverse
RPL34 – Forward
DNA Meth. m5C:H4K20 me2
H3K9 me2:DNA Meth. m5C
DNA Meth. m5C:H3K9 me3
H3K4 me3:H3–5ac
H3K4 me1:H4–4ac
H4K20 me2:H4–4ac
H3K4 me1:H3–5ac
H2A.Z:H3K4 me3
H3K4 me1:H2A.Z
H4K20 me2:H3–5ac
H2A.Z:H4K20 me2
H4K20 me2:H3K4 me3
H4–4ac:H3–5ac
H3K4 me3:H4–4ac
H3K27 me2
H4K20 me3
H3K27 me3
H3K9 me2
H3K9 me3
H3K4 me3
H2A.Z
H4K20 me2
H3K4 me1
H3–5ac
H4–4ac
DNA Meth. m5C
RPL34 – Reverse
RPL6 – Forward
H2A.Z:H3K4 me3
H3K4 me1:H4–4ac
H3K4 me1:H4K20 me2
H3K4 me1:H2A.Z
DNA Meth. m5C:H3K9 me3
H3K4 me1:H3–5ac
H4–4ac:H3–5ac
DNA Meth. m5C:H4K20 me2
H3K27 me3:H3K4 me1
H3K4 me3:H3–5ac
H4K20 me2:H4–4ac
H2A.Z:H4K20 me2
H4K20 me2:H3K4 me3
H4K20 me2:H3–5ac
H3K9 me2
H4K20 me3
H3K27 me2
H3K9 me3
H3K4 me1
H3K27 me3
DNA Meth. m5C
H2A.Z
H4–4ac
H3K4 me3
H4K20 me2
H3–5ac
RPL6 – Reverse
RPL11 – Forward
H4–4ac:H3–5ac
H4K20 me2:H3K4 me3
H2A.Z:H3K4 me3
H2A.Z:H4K20 me2
H3K4 me1:H3–5ac
H3K4 me1:H4–4ac
H3K4 me1:H2A.Z
DNA Meth. m5C:H4K20 me2
H3K4 me3:H4–4ac
H3K27 me3:DNA Meth. m5C
H3K9 me2:DNA Meth. m5C
H4K20 me3
H3K9 me3
H3K27 me2
H3K9 me2
H3K4 me3
H2A.Z
H3K27 me3
H4–4ac
H4K20 me2
H3–5ac
H3K4 me1
DNA Meth. m5C
RPL11 – Reverse

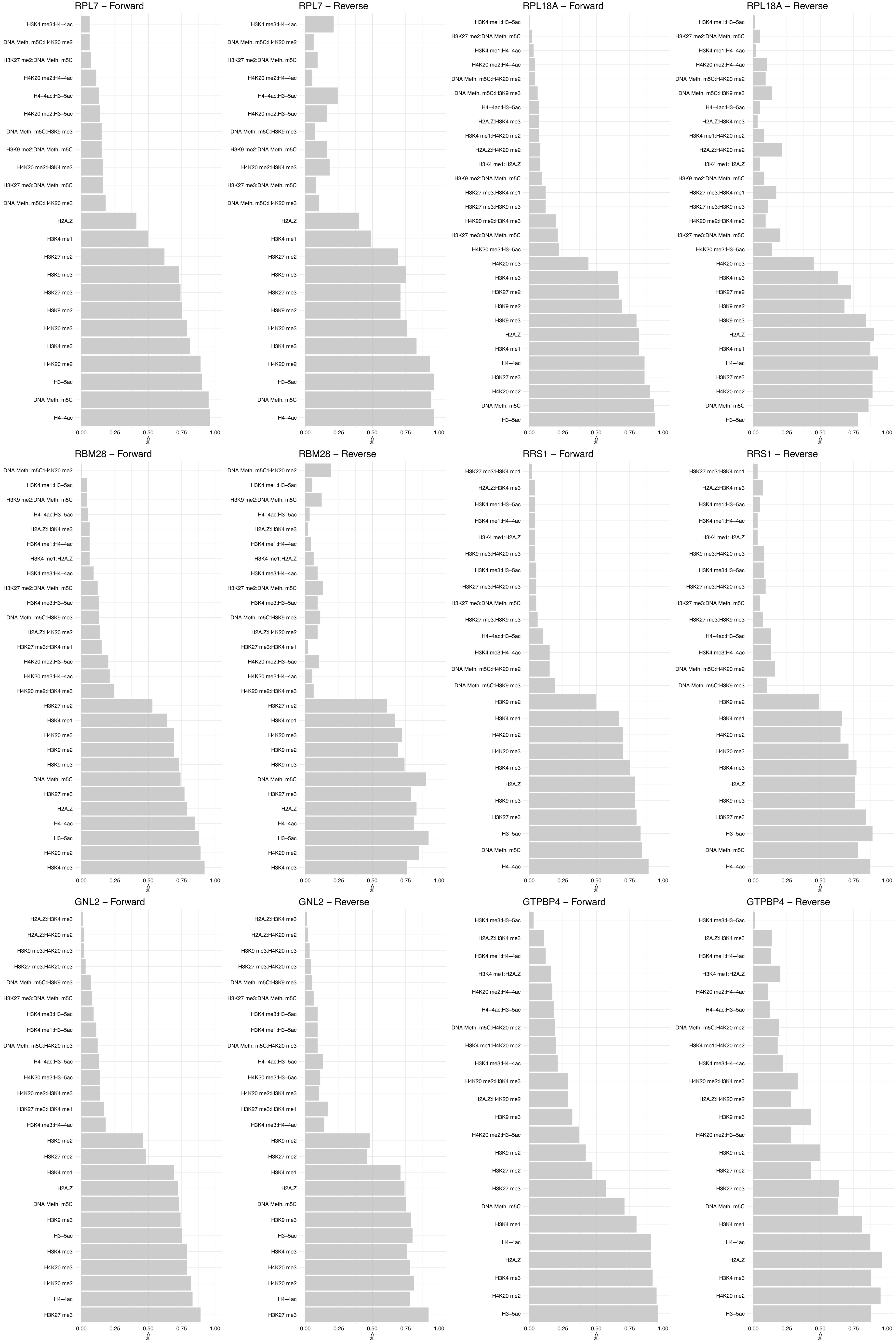
RPL7 – Forward
H3K4 me3:H4–4ac
DNA Meth. m5C:H4K20 me2
H3K27 me2:DNA Meth. m5C
H4K20 me2:H4–4ac
H4–4ac:H3–5ac
H4K20 me2:H3–5ac
DNA Meth. m5C:H3K9 me3
H3K9 me2:DNA Meth. m5C
H4K20 me2:H3K4 me3
H3K27 me3:DNA Meth. m5C
DNA Meth. m5C:H4K20 me3
H2A.Z
H3K4 me1
H3K27 me2
H3K9 me3
H3K27 me3
H3K9 me2
H4K20 me3
H3K4 me3
H4K20 me2
H3–5ac
DNA Meth. m5C
H4–4ac
0.00
0.25
0.50
0.75
1.00
π̂
RPL7 – Reverse
H3K4 me3:H4–4ac
DNA Meth. m5C:H4K20 me2
H3K27 me2:DNA Meth. m5C
H4K20 me2:H4–4ac
H4–4ac:H3–5ac
H4K20 me2:H3–5ac
DNA Meth. m5C:H3K9 me3
H3K9 me2:DNA Meth. m5C
H4K20 me2:H3K4 me3
H3K27 me3:DNA Meth. m5C
DNA Meth. m5C:H4K20 me3
H2A.Z
H3K4 me1
H3K27 me2
H3K9 me3
H3K27 me3
H3K9 me2
H4K20 me3
H3K4 me3
H4K20 me2
H3–5ac
DNA Meth. m5C
H4–4ac
0.00
0.25
0.50
0.75
1.00
π̂
RPL18A – Forward
H3K4 me1:H3–5ac
H3K27 me2:DNA Meth. m5C
H3K4 me1:H4–4ac
H4K20 me2:H4–4ac
DNA Meth. m5C:H4K20 me2
DNA Meth. m5C:H3K9 me3
H4–4ac:H3–5ac
H2A.Z:H3K4 me3
H3K4 me1:H4K20 me2
H2A.Z:H4K20 me2
H3K4 me1:H2A.Z
H3K9 me2:DNA Meth. m5C
H3K27 me3:H3K4 me1
H3K27 me3:H3K9 me3
H4K20 me2:H3K4 me3
H3K27 me3:DNA Meth. m5C
H4K20 me2:H3–5ac
H4K20 me3
H3K4 me3
H3K27 me2
H3K9 me2
H3K9 me3
H2A.Z
H3K4 me1
H4–4ac
H3K27 me3
H4K20 me2
DNA Meth. m5C
H3–5ac
0.00
0.25
0.50
0.75
1.00
π̂
RPL18A – Reverse
H3K4 me1:H3–5ac
H3K27 me2:DNA Meth. m5C
H3K4 me1:H4–4ac
H4K20 me2:H4–4ac
DNA Meth. m5C:H4K20 me2
DNA Meth. m5C:H3K9 me3
H4–4ac:H3–5ac
H2A.Z:H3K4 me3
H3K4 me1:H4K20 me2
H2A.Z:H4K20 me2
H3K4 me1:H2A.Z
H3K9 me2:DNA Meth. m5C
H3K27 me3:H3K4 me1
H3K27 me3:H3K9 me3
H4K20 me2:H3K4 me3
H3K27 me3:DNA Meth. m5C
H4K20 me2:H3–5ac
H4K20 me3
H3K4 me3
H3K27 me2
H3K9 me2
H3K9 me3
H2A.Z
H3K4 me1
H4–4ac
H3K27 me3
H4K20 me2
DNA Meth. m5C
H3–5ac
0.00
0.25
0.50
0.75
1.00
π̂
RBM28 – Forward
DNA Meth. m5C:H4K20 me2
H3K4 me1:H3–5ac
H3K9 me2:DNA Meth. m5C
H4–4ac:H3–5ac
H2A.Z:H3K4 me3
H3K4 me1:H4–4ac
H3K4 me1:H2A.Z
H3K4 me3:H4–4ac
H3K27 me2:DNA Meth. m5C
H3K4 me3:H3–5ac
DNA Meth. m5C:H3K9 me3
H2A.Z:H4K20 me2
H3K27 me3:H3K4 me1
H4K20 me2:H3–5ac
H4K20 me2:H4–4ac
H4K20 me2:H3K4 me3
H3K27 me2
H3K4 me1
H4K20 me3
H3K9 me2
H3K9 me3
DNA Meth. m5C
H3K27 me3
H2A.Z
H4–4ac
H3–5ac
H4K20 me2
H3K4 me3
0.00
0.25
0.50
0.75
1.00
π̂
RBM28 – Reverse
DNA Meth. m5C:H4K20 me2
H3K4 me1:H3–5ac
H3K9 me2:DNA Meth. m5C
H4–4ac:H3–5ac
H2A.Z:H3K4 me3
H3K4 me1:H4–4ac
H3K4 me1:H2A.Z
H3K4 me3:H4–4ac
H3K27 me2:DNA Meth. m5C
H3K4 me3:H3–5ac
DNA Meth. m5C:H3K9 me3
H2A.Z:H4K20 me2
H3K27 me3:H3K4 me1
H4K20 me2:H3–5ac
H4K20 me2:H4–4ac
H4K20 me2:H3K4 me3
H3K27 me2
H3K4 me1
H4K20 me3
H3K9 me2
H3K9 me3
DNA Meth. m5C
H3K27 me3
H2A.Z
H4–4ac
H3–5ac
H4K20 me2
H3K4 me3
0.00
0.25
0.50
0.75
1.00
π̂
RRS1 – Forward
H3K27 me3:H3K4 me1
H2A.Z:H3K4 me3
H3K4 me1:H3–5ac
H3K4 me1:H4–4ac
H3K4 me1:H2A.Z
H3K9 me3:H4K20 me3
H3K4 me3:H3–5ac
H3K27 me3:H4K20 me3
H3K27 me3:DNA Meth. m5C
H3K27 me3:H3K9 me3
H4–4ac:H3–5ac
H3K4 me3:H4–4ac
DNA Meth. m5C:H4K20 me2
DNA Meth. m5C:H3K9 me3
H3K9 me2
H3K4 me1
H4K20 me2
H4K20 me3
H3K4 me3
H2A.Z
H3K9 me3
H3K27 me3
H3–5ac
DNA Meth. m5C
H4–4ac
0.00
0.25
0.50
0.75
1.00
π̂
RRS1 – Reverse
H3K27 me3:H3K4 me1
H2A.Z:H3K4 me3
H3K4 me1:H3–5ac
H3K4 me1:H4–4ac
H3K4 me1:H2A.Z
H3K9 me3:H4K20 me3
H3K4 me3:H3–5ac
H3K27 me3:H4K20 me3
H3K27 me3:DNA Meth. m5C
H3K27 me3:H3K9 me3
H4–4ac:H3–5ac
H3K4 me3:H4–4ac
DNA Meth. m5C:H4K20 me2
DNA Meth. m5C:H3K9 me3
H3K9 me2
H3K4 me1
H4K20 me2
H4K20 me3
H3K4 me3
H2A.Z
H3K9 me3
H3K27 me3
H3–5ac
DNA Meth. m5C
H4–4ac
0.00
0.25
0.50
0.75
1.00
π̂
GNL2 – Forward
H2A.Z:H3K4 me3
H2A.Z:H4K20 me2
H3K9 me3:H4K20 me3
H3K27 me3:H4K20 me3
DNA Meth. m5C:H3K9 me3
H3K27 me3:DNA Meth. m5C
H3K4 me3:H3–5ac
H3K4 me1:H3–5ac
DNA Meth. m5C:H4K20 me3
H4–4ac:H3–5ac
H4K20 me2:H3–5ac
H4K20 me2:H3K4 me3
H3K27 me3:H3K4 me1
H3K4 me3:H4–4ac
H3K9 me2
H3K27 me2
H3K4 me1
H2A.Z
DNA Meth. m5C
H3K9 me3
H3–5ac
H3K4 me3
H4K20 me3
H4K20 me2
H4–4ac
H3K27 me3
0.00
0.25
0.50
0.75
1.00
π̂
GNL2 – Reverse
H2A.Z:H3K4 me3
H2A.Z:H4K20 me2
H3K9 me3:H4K20 me3
H3K27 me3:H4K20 me3
DNA Meth. m5C:H3K9 me3
H3K27 me3:DNA Meth. m5C
H3K4 me3:H3–5ac
H3K4 me1:H3–5ac
DNA Meth. m5C:H4K20 me3
H4–4ac:H3–5ac
H4K20 me2:H3–5ac
H4K20 me2:H3K4 me3
H3K27 me3:H3K4 me1
H3K4 me3:H4–4ac
H3K9 me2
H3K27 me2
H3K4 me1
H2A.Z
DNA Meth. m5C
H3K9 me3
H3–5ac
H3K4 me3
H4K20 me3
H4K20 me2
H4–4ac
H3K27 me3
0.00
0.25
0.50
0.75
1.00
π̂
GTPBP4 – Forward
H3K4 me3:H3–5ac
H2A.Z:H3K4 me3
H3K4 me1:H4–4ac
H3K4 me1:H2A.Z
H4K20 me2:H4–4ac
H4–4ac:H3–5ac
DNA Meth. m5C:H4K20 me2
H3K4 me1:H4K20 me2
H3K4 me3:H4–4ac
H4K20 me2:H3K4 me3
H2A.Z:H4K20 me2
H3K9 me3
H4K20 me2:H3–5ac
H3K9 me2
H3K27 me2
H3K27 me3
DNA Meth. m5C
H3K4 me1
H4–4ac
H2A.Z
H3K4 me3
H4K20 me2
H3–5ac
0.00
0.25
0.50
0.75
1.00
π̂
GTPBP4 – Reverse
H3K4 me3:H3–5ac
H2A.Z:H3K4 me3
H3K4 me1:H4–4ac
H3K4 me1:H2A.Z
H4K20 me2:H4–4ac
H4–4ac:H3–5ac
DNA Meth. m5C:H4K20 me2
H3K4 me1:H4K20 me2
H3K4 me3:H4–4ac
H4K20 me2:H3K4 me3
H2A.Z:H4K20 me2
H3K9 me3
H4K20 me2:H3–5ac
H3K9 me2
H3K27 me2
H3K27 me3
DNA Meth. m5C
H3K4 me1
H4–4ac
H2A.Z
H3K4 me3
H4K20 me2
H3–5ac
0.00
0.25
0.50
0.75
1.00
π̂

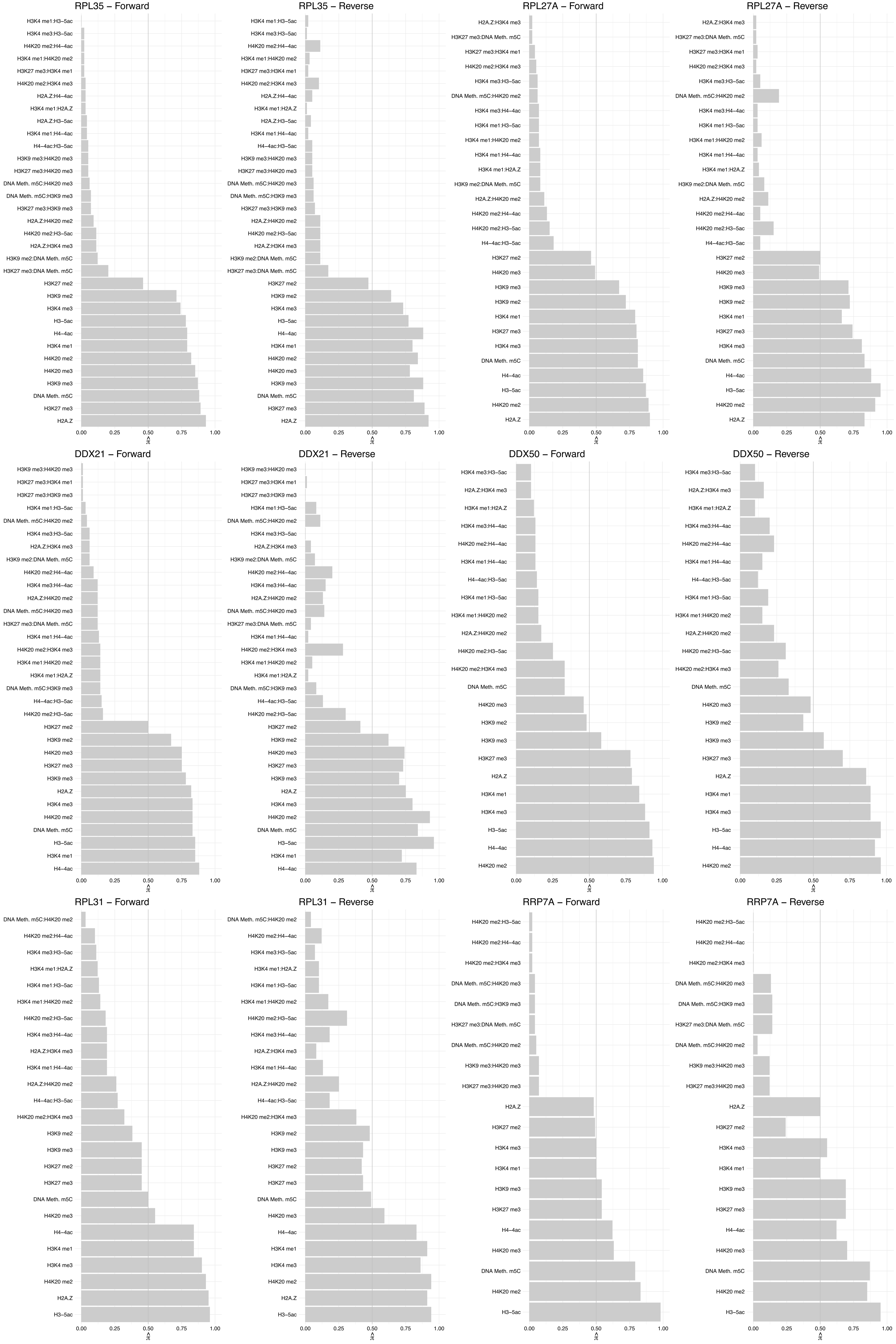
RPL35 – Forward
H3K4 me1:H3−5ac
H3K4 me3:H3−5ac
H4K20 me2:H4−4ac
H3K4 me1:H4K20 me2
H3K27 me3:H3K4 me1
H4K20 me2:H3K4 me3
H2A.Z:H4−4ac
H3K4 me1:H2A.Z
H2A.Z:H3−5ac
H3K4 me1:H4−4ac
H4−4ac:H3−5ac
H3K9 me3:H4K20 me3
H3K27 me3:H4K20 me3
DNA Meth. m5C:H4K20 me3
DNA Meth. m5C:H3K9 me3
H3K27 me3:H3K9 me3
H2A.Z:H4K20 me2
H4K20 me2:H3−5ac
H2A.Z:H3K4 me3
H3K9 me2:DNA Meth. m5C
H3K27 me3:DNA Meth. m5C
H3K27 me2
H3K9 me2
H3K4 me3
H3−5ac
H4−4ac
H3K4 me1
H4K20 me2
H4K20 me3
H3K9 me3
DNA Meth. m5C
H3K27 me3
H2A.Z
0.00 0.25 0.50 0.75 1.00
π̂
RPL35 – Reverse
H3K4 me1:H3−5ac
H3K4 me3:H3−5ac
H4K20 me2:H4−4ac
H3K4 me1:H4K20 me2
H3K27 me3:H3K4 me1
H4K20 me2:H3K4 me3
H2A.Z:H4−4ac
H3K4 me1:H2A.Z
H2A.Z:H3−5ac
H3K4 me1:H4−4ac
H4−4ac:H3−5ac
H3K9 me3:H4K20 me3
H3K27 me3:H4K20 me3
DNA Meth. m5C:H4K20 me3
DNA Meth. m5C:H3K9 me3
H3K27 me3:H3K9 me3
H2A.Z:H4K20 me2
H4K20 me2:H3−5ac
H2A.Z:H3K4 me3
H3K9 me2:DNA Meth. m5C
H3K27 me3:DNA Meth. m5C
H3K27 me2
H3K9 me2
H3K4 me3
H3−5ac
H4−4ac
H3K4 me1
H4K20 me2
H4K20 me3
H3K9 me3
DNA Meth. m5C
H3K27 me3
H2A.Z
0.00 0.25 0.50 0.75 1.00
π̂
RPL27A – Forward
H2A.Z:H3K4 me3
H3K27 me3:DNA Meth. m5C
H3K27 me3:H3K4 me1
H4K20 me2:H3K4 me3
H3K4 me3:H3−5ac
DNA Meth. m5C:H4K20 me2
H3K4 me3:H4−4ac
H3K4 me1:H3−5ac
H3K4 me1:H4K20 me2
H3K4 me1:H4−4ac
H3K4 me1:H2A.Z
H3K9 me2:DNA Meth. m5C
H2A.Z:H4K20 me2
H4K20 me2:H4−4ac
H4K20 me2:H3−5ac
H4−4ac:H3−5ac
H3K27 me2
H4K20 me3
H3K9 me3
H3K9 me2
H3K4 me1
H3K27 me3
H3K4 me3
DNA Meth. m5C
H4−4ac
H3−5ac
H4K20 me2
H2A.Z
0.00 0.25 0.50 0.75 1.00
π̂
RPL27A – Reverse
H2A.Z:H3K4 me3
H3K27 me3:DNA Meth. m5C
H3K27 me3:H3K4 me1
H4K20 me2:H3K4 me3
H3K4 me3:H3−5ac
DNA Meth. m5C:H4K20 me2
H3K4 me3:H4−4ac
H3K4 me1:H3−5ac
H3K4 me1:H4K20 me2
H3K4 me1:H4−4ac
H3K4 me1:H2A.Z
H3K9 me2:DNA Meth. m5C
H2A.Z:H4K20 me2
H4K20 me2:H4−4ac
H4K20 me2:H3−5ac
H4−4ac:H3−5ac
H3K27 me2
H4K20 me3
H3K9 me3
H3K9 me2
H3K4 me1
H3K27 me3
H3K4 me3
DNA Meth. m5C
H4−4ac
H3−5ac
H4K20 me2
H2A.Z
0.00 0.25 0.50 0.75 1.00
π̂
DDX21 – Forward
H3K9 me3:H4K20 me3
H3K27 me3:H3K4 me1
H3K27 me3:H3K9 me3
H3K4 me1:H3−5ac
DNA Meth. m5C:H4K20 me2
H3K4 me3:H3−5ac
H2A.Z:H3K4 me3
H3K9 me2:DNA Meth. m5C
H4K20 me2:H4−4ac
H3K4 me3:H4−4ac
H2A.Z:H4K20 me2
DNA Meth. m5C:H4K20 me3
H3K27 me3:DNA Meth. m5C
H3K4 me1:H4−4ac
H4K20 me2:H3K4 me3
H3K4 me1:H4K20 me2
H3K4 me1:H2A.Z
DNA Meth. m5C:H3K9 me3
H4−4ac:H3−5ac
H4K20 me2:H3−5ac
H3K27 me2
H3K9 me2
H4K20 me3
H3K27 me3
H3K9 me3
H2A.Z
H3K4 me3
H4K20 me2
DNA Meth. m5C
H3−5ac
H3K4 me1
H4−4ac
0.00 0.25 0.50 0.75 1.00
π̂
DDX21 – Reverse
H3K9 me3:H4K20 me3
H3K27 me3:H3K4 me1
H3K27 me3:H3K9 me3
H3K4 me1:H3−5ac
DNA Meth. m5C:H4K20 me2
H3K4 me3:H3−5ac
H2A.Z:H3K4 me3
H3K9 me2:DNA Meth. m5C
H4K20 me2:H4−4ac
H3K4 me3:H4−4ac
H2A.Z:H4K20 me2
DNA Meth. m5C:H4K20 me3
H3K27 me3:DNA Meth. m5C
H3K4 me1:H4−4ac
H4K20 me2:H3K4 me3
H3K4 me1:H4K20 me2
H3K4 me1:H2A.Z
DNA Meth. m5C:H3K9 me3
H4−4ac:H3−5ac
H4K20 me2:H3−5ac
H3K27 me2
H3K9 me2
H4K20 me3
H3K27 me3
H3K9 me3
H2A.Z
H3K4 me3
H4K20 me2
DNA Meth. m5C
H3−5ac
H3K4 me1
H4−4ac
0.00 0.25 0.50 0.75 1.00
π̂
DDX50 – Forward
H3K4 me3:H3−5ac
H2A.Z:H3K4 me3
H3K4 me1:H2A.Z
H3K4 me3:H4−4ac
H4K20 me2:H4−4ac
H3K4 me1:H4−4ac
H4−4ac:H3−5ac
H3K4 me1:H3−5ac
H3K4 me1:H4K20 me2
H2A.Z:H4K20 me2
H4K20 me2:H3−5ac
H4K20 me2:H3K4 me3
DNA Meth. m5C
H4K20 me3
H3K9 me2
H3K9 me3
H3K27 me3
H2A.Z
H3K4 me1
H3K4 me3
H3−5ac
H4−4ac
H4K20 me2
0.00 0.25 0.50 0.75 1.00
DDX50 – Reverse
H3K4 me3:H3−5ac
H2A.Z:H3K4 me3
H3K4 me1:H2A.Z
H3K4 me3:H4−4ac
H4K20 me2:H4−4ac
H3K4 me1:H4−4ac
H4−4ac:H3−5ac
H3K4 me1:H3−5ac
H3K4 me1:H4K20 me2
H2A.Z:H4K20 me2
H4K20 me2:H3−5ac
H4K20 me2:H3K4 me3
DNA Meth. m5C
H4K20 me3
H3K9 me2
H3K9 me3
H3K27 me3
H2A.Z
H3K4 me1
H3K4 me3
H3−5ac
H4−4ac
H4K20 me2
0.00 0.25 0.50 0.75 1.00
RPL31 – Forward
DNA Meth. m5C:H4K20 me2
H4K20 me2:H4−4ac
H3K4 me3:H3−5ac
H3K4 me1:H2A.Z
H3K4 me1:H3−5ac
H3K4 me1:H4K20 me2
H4K20 me2:H3−5ac
H3K4 me3:H4−4ac
H2A.Z:H3K4 me3
H3K4 me1:H4−4ac
H2A.Z:H4K20 me2
H4−4ac:H3−5ac
H4K20 me2:H3K4 me3
H3K9 me2
H3K9 me3
H3K27 me2
H3K27 me3
DNA Meth. m5C
H4K20 me3
H4−4ac
H3K4 me1
H3K4 me3
H4K20 me2
H2A.Z
H3−5ac
0.00 0.25 0.50 0.75 1.00
π̂
RPL31 – Reverse
DNA Meth. m5C:H4K20 me2
H4K20 me2:H4−4ac
H3K4 me3:H3−5ac
H3K4 me1:H2A.Z
H3K4 me1:H3−5ac
H3K4 me1:H4K20 me2
H4K20 me2:H3−5ac
H3K4 me3:H4−4ac
H2A.Z:H3K4 me3
H3K4 me1:H4−4ac
H2A.Z:H4K20 me2
H4−4ac:H3−5ac
H4K20 me2:H3K4 me3
H3K9 me2
H3K9 me3
H3K27 me2
H3K27 me3
DNA Meth. m5C
H4K20 me3
H4−4ac
H3K4 me1
H3K4 me3
H4K20 me2
H2A.Z
H3−5ac
0.00 0.25 0.50 0.75 1.00
π̂
RRP7A – Forward
H4K20 me2:H3−5ac
H4K20 me2:H4−4ac
H4K20 me2:H3K4 me3
DNA Meth. m5C:H4K20 me3
DNA Meth. m5C:H3K9 me3
H3K27 me3:DNA Meth. m5C
DNA Meth. m5C:H4K20 me2
H3K9 me3:H4K20 me3
H3K27 me3:H4K20 me3
H2A.Z
H3K27 me2
H3K4 me3
H3K4 me1
H3K9 me3
H3K27 me3
H4−4ac
H4K20 me3
DNA Meth. m5C
H4K20 me2
H3−5ac
0.00 0.25 0.50 0.75 1.00
π̂
RRP7A – Reverse
H4K20 me2:H3−5ac
H4K20 me2:H4−4ac
H4K20 me2:H3K4 me3
DNA Meth. m5C:H4K20 me3
DNA Meth. m5C:H3K9 me3
H3K27 me3:DNA Meth. m5C
DNA Meth. m5C:H4K20 me2
H3K9 me3:H4K20 me3
H3K27 me3:H4K20 me3
H2A.Z
H3K27 me2
H3K4 me3
H3K4 me1
H3K9 me3
H3K27 me3
H4−4ac
H4K20 me3
DNA Meth. m5C
H4K20 me2
H3−5ac
0.00 0.25 0.50 0.75 1.00
π̂

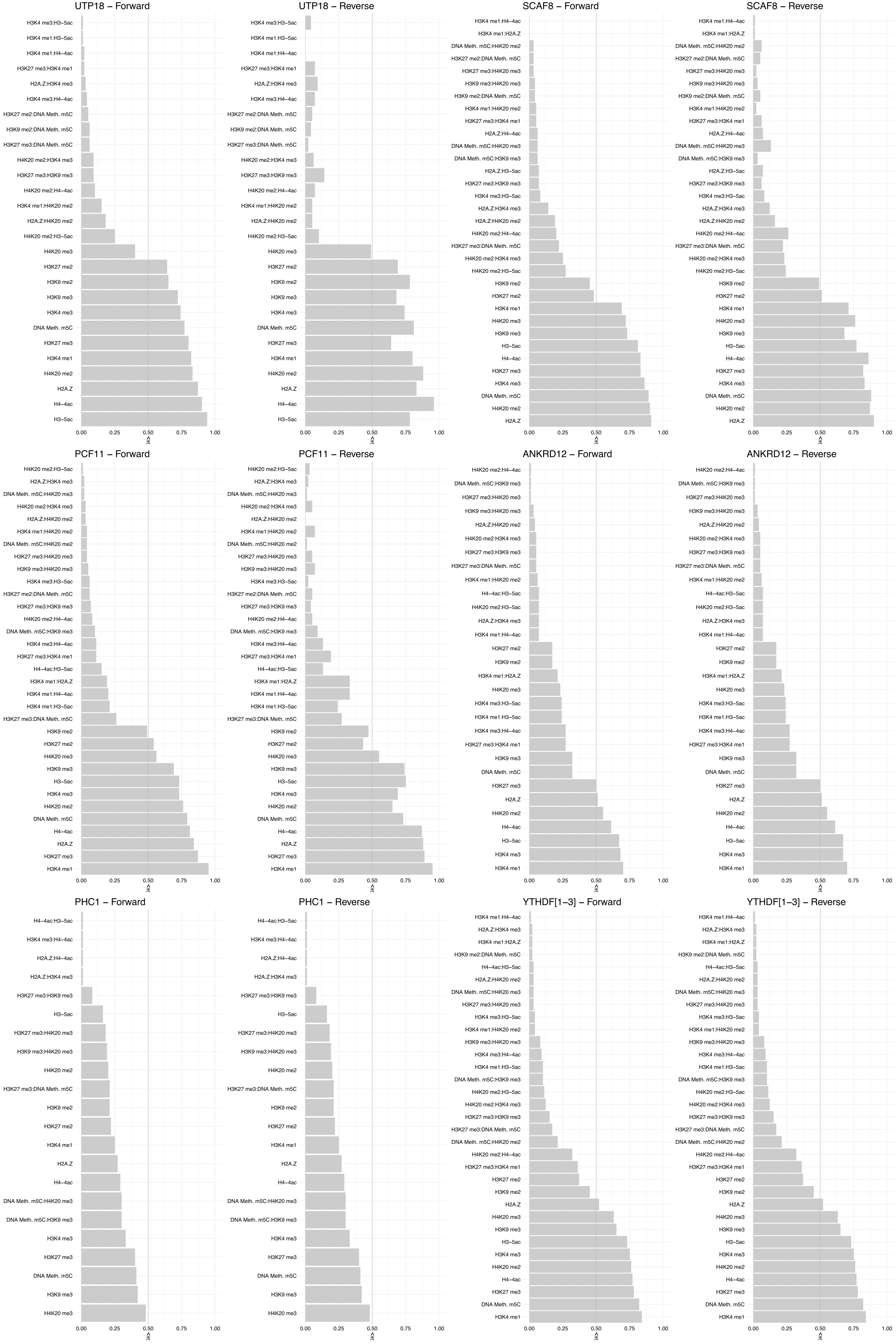
UTP18 – Forward
H3K4 me3:H3−5ac
H3K4 me1:H3−5ac
H3K4 me1:H4−4ac
H3K27 me3:H3K4 me1
H2A.Z:H3K4 me3
H3K4 me3:H4−4ac
H3K27 me2:DNA Meth. m5C
H3K9 me2:DNA Meth. m5C
H3K27 me3:DNA Meth. m5C
H4K20 me2:H3K4 me3
H3K27 me3:H3K9 me3
H4K20 me2:H4−4ac
H3K4 me1:H4K20 me2
H2A.Z:H4K20 me2
H4K20 me2:H3−5ac
H4K20 me3
H3K27 me2
H3K9 me2
H3K9 me3
H3K4 me3
DNA Meth. m5C
H3K27 me3
H3K4 me1
H4K20 me2
H2A.Z
H4−4ac
H3−5ac
0.00 0.25 0.50 0.75 1.00
π̂
UTP18 – Reverse
H3K4 me3:H3−5ac
H3K4 me1:H3−5ac
H3K4 me1:H4−4ac
H3K27 me3:H3K4 me1
H2A.Z:H3K4 me3
H3K4 me3:H4−4ac
H3K27 me2:DNA Meth. m5C
H3K9 me2:DNA Meth. m5C
H3K27 me3:DNA Meth. m5C
H4K20 me2:H3K4 me3
H3K27 me3:H3K9 me3
H4K20 me2:H4−4ac
H3K4 me1:H4K20 me2
H2A.Z:H4K20 me2
H4K20 me2:H3−5ac
H4K20 me3
H3K27 me2
H3K9 me2
H3K9 me3
H3K4 me3
DNA Meth. m5C
H3K27 me3
H3K4 me1
H4K20 me2
H2A.Z
H4−4ac
H3−5ac
0.00 0.25 0.50 0.75 1.00
π̂
SCAF8 – Forward
H3K4 me1:H4−4ac
H3K4 me1:H2A.Z
DNA Meth. m5C:H4K20 me2
H3K27 me2:DNA Meth. m5C
H3K27 me3:H4K20 me3
H3K9 me3:H4K20 me3
H3K9 me2:DNA Meth. m5C
H3K4 me1:H4K20 me2
H3K27 me3:H3K4 me1
H2A.Z:H4−4ac
DNA Meth. m5C:H4K20 me3
DNA Meth. m5C:H3K9 me3
H2A.Z:H3−5ac
H3K27 me3:H3K9 me3
H3K4 me3:H3−5ac
H2A.Z:H3K4 me3
H2A.Z:H4K20 me2
H4K20 me2:H4−4ac
H3K27 me3:DNA Meth. m5C
H4K20 me2:H3K4 me3
H4K20 me2:H3−5ac
H3K9 me2
H3K27 me2
H3K4 me1
H4K20 me3
H3K9 me3
H3−5ac
H4−4ac
H3K27 me3
H3K4 me3
DNA Meth. m5C
H4K20 me2
H2A.Z
0.00 0.25 0.50 0.75 1.00
π̂
SCAF8 – Reverse
H3K4 me1:H4−4ac
H3K4 me1:H2A.Z
DNA Meth. m5C:H4K20 me2
H3K27 me2:DNA Meth. m5C
H3K27 me3:H4K20 me3
H3K9 me3:H4K20 me3
H3K9 me2:DNA Meth. m5C
H3K4 me1:H4K20 me2
H3K27 me3:H3K4 me1
H2A.Z:H4−4ac
DNA Meth. m5C:H4K20 me3
DNA Meth. m5C:H3K9 me3
H2A.Z:H3−5ac
H3K27 me3:H3K9 me3
H3K4 me3:H3−5ac
H2A.Z:H3K4 me3
H2A.Z:H4K20 me2
H4K20 me2:H4−4ac
H3K27 me3:DNA Meth. m5C
H4K20 me2:H3K4 me3
H4K20 me2:H3−5ac
H3K9 me2
H3K27 me2
H3K4 me1
H4K20 me3
H3K9 me3
H3−5ac
H4−4ac
H3K27 me3
H3K4 me3
DNA Meth. m5C
H4K20 me2
H2A.Z
0.00 0.25 0.50 0.75 1.00
π̂
PCF11 – Forward
H4K20 me2:H3−5ac
H2A.Z:H3K4 me3
DNA Meth. m5C:H4K20 me3
H4K20 me2:H3K4 me3
H2A.Z:H4K20 me2
H3K4 me1:H4K20 me2
DNA Meth. m5C:H4K20 me2
H3K27 me3:H4K20 me3
H3K9 me3:H4K20 me3
H3K4 me3:H3−5ac
H3K27 me2:DNA Meth. m5C
H3K27 me3:H3K9 me3
H4K20 me2:H4−4ac
DNA Meth. m5C:H3K9 me3
H3K4 me3:H4−4ac
H3K27 me3:H3K4 me1
H4−4ac:H3−5ac
H3K4 me1:H2A.Z
H3K4 me1:H4−4ac
H3K4 me1:H3−5ac
H3K27 me3:DNA Meth. m5C
H3K9 me2
H3K27 me2
H4K20 me3
H3K9 me3
H3−5ac
H3K4 me3
H4K20 me2
DNA Meth. m5C
H4−4ac
H2A.Z
H3K27 me3
H3K4 me1
0.00 0.25 0.50 0.75 1.00
π̂
PCF11 – Reverse
H4K20 me2:H3−5ac
H2A.Z:H3K4 me3
DNA Meth. m5C:H4K20 me3
H4K20 me2:H3K4 me3
H2A.Z:H4K20 me2
H3K4 me1:H4K20 me2
DNA Meth. m5C:H4K20 me2
H3K27 me3:H4K20 me3
H3K9 me3:H4K20 me3
H3K4 me3:H3−5ac
H3K27 me2:DNA Meth. m5C
H3K27 me3:H3K9 me3
H4K20 me2:H4−4ac
DNA Meth. m5C:H3K9 me3
H3K4 me3:H4−4ac
H3K27 me3:H3K4 me1
H4−4ac:H3−5ac
H3K4 me1:H2A.Z
H3K4 me1:H4−4ac
H3K4 me1:H3−5ac
H3K27 me3:DNA Meth. m5C
H3K9 me2
H3K27 me2
H4K20 me3
H3K9 me3
H3−5ac
H3K4 me3
H4K20 me2
DNA Meth. m5C
H4−4ac
H2A.Z
H3K27 me3
H3K4 me1
0.00 0.25 0.50 0.75 1.00
π̂
ANKRD12 – Forward
H4K20 me2:H4−4ac
DNA Meth. m5C:H3K9 me3
H3K27 me3:H4K20 me3
H3K9 me3:H4K20 me3
H2A.Z:H4K20 me2
H4K20 me2:H3K4 me3
H3K27 me3:H3K9 me3
H3K27 me3:DNA Meth. m5C
H3K4 me1:H4K20 me2
H4−4ac:H3−5ac
H4K20 me2:H3−5ac
H2A.Z:H3K4 me3
H3K4 me1:H4−4ac
H3K27 me2
H3K9 me2
H3K4 me1:H2A.Z
H4K20 me3
H3K4 me3:H3−5ac
H3K4 me1:H3−5ac
H3K4 me3:H4−4ac
H3K27 me3:H3K4 me1
H3K9 me3
DNA Meth. m5C
H3K27 me3
H2A.Z
H4K20 me2
H4−4ac
H3−5ac
H3K4 me3
H3K4 me1
0.00 0.25 0.50 0.75 1.00
π̂
ANKRD12 – Reverse
H4K20 me2:H4−4ac
DNA Meth. m5C:H3K9 me3
H3K27 me3:H4K20 me3
H3K9 me3:H4K20 me3
H2A.Z:H4K20 me2
H4K20 me2:H3K4 me3
H3K27 me3:H3K9 me3
H3K27 me3:DNA Meth. m5C
H3K4 me1:H4K20 me2
H4−4ac:H3−5ac
H4K20 me2:H3−5ac
H2A.Z:H3K4 me3
H3K4 me1:H4−4ac
H3K27 me2
H3K9 me2
H3K4 me1:H2A.Z
H4K20 me3
H3K4 me3:H3−5ac
H3K4 me1:H3−5ac
H3K4 me3:H4−4ac
H3K27 me3:H3K4 me1
H3K9 me3
DNA Meth. m5C
H3K27 me3
H2A.Z
H4K20 me2
H4−4ac
H3−5ac
H3K4 me3
H3K4 me1
0.00 0.25 0.50 0.75 1.00
π̂
PHC1 – Forward
H4−4ac:H3−5ac
H3K4 me3:H4−4ac
H2A.Z:H4−4ac
H2A.Z:H3K4 me3
H3K27 me3:H3K9 me3
H3−5ac
H3K27 me3:H4K20 me3
H3K9 me3:H4K20 me3
H4K20 me2
H3K27 me3:DNA Meth. m5C
H3K9 me2
H3K27 me2
H3K4 me1
H2A.Z
H4−4ac
DNA Meth. m5C:H4K20 me3
DNA Meth. m5C:H3K9 me3
H3K4 me3
H3K27 me3
DNA Meth. m5C
H3K9 me3
H4K20 me3
0.00 0.25 0.50 0.75 1.00
π̂
PHC1 – Reverse
H4−4ac:H3−5ac
H3K4 me3:H4−4ac
H2A.Z:H4−4ac
H2A.Z:H3K4 me3
H3K27 me3:H3K9 me3
H3−5ac
H3K27 me3:H4K20 me3
H3K9 me3:H4K20 me3
H4K20 me2
H3K27 me3:DNA Meth. m5C
H3K9 me2
H3K27 me2
H3K4 me1
H2A.Z
H4−4ac
DNA Meth. m5C:H4K20 me3
DNA Meth. m5C:H3K9 me3
H3K4 me3
H3K27 me3
DNA Meth. m5C
H3K9 me3
H4K20 me3
0.00 0.25 0.50 0.75 1.00
π̂
YTHDF[1−3] – Forward
H3K4 me1:H4−4ac
H2A.Z:H3K4 me3
H3K4 me1:H2A.Z
H3K9 me2:DNA Meth. m5C
H4−4ac:H3−5ac
H2A.Z:H4K20 me2
DNA Meth. m5C:H4K20 me3
H3K27 me3:H4K20 me3
H3K4 me3:H3−5ac
H3K4 me1:H4K20 me2
H3K9 me3:H4K20 me3
H3K4 me3:H4−4ac
H3K4 me1:H3−5ac
DNA Meth. m5C:H3K9 me3
H4K20 me2:H3−5ac
H4K20 me2:H3K4 me3
H3K27 me3:H3K9 me3
H3K27 me3:DNA Meth. m5C
DNA Meth. m5C:H4K20 me2
H4K20 me2:H4−4ac
H3K27 me3:H3K4 me1
H3K27 me2
H3K9 me2
H2A.Z
H4K20 me3
H3K9 me3
H3−5ac
H3K4 me3
H4K20 me2
H4−4ac
H3K27 me3
DNA Meth. m5C
H3K4 me1
0.00 0.25 0.50 0.75 1.00
π̂
YTHDF[1−3] – Reverse
H3K4 me1:H4−4ac
H2A.Z:H3K4 me3
H3K4 me1:H2A.Z
H3K9 me2:DNA Meth. m5C
H4−4ac:H3−5ac
H2A.Z:H4K20 me2
DNA Meth. m5C:H4K20 me3
H3K27 me3:H4K20 me3
H3K4 me3:H3−5ac
H3K4 me1:H4K20 me2
H3K9 me3:H4K20 me3
H3K4 me3:H4−4ac
H3K4 me1:H3−5ac
DNA Meth. m5C:H3K9 me3
H4K20 me2:H3−5ac
H4K20 me2:H3K4 me3
H3K27 me3:H3K9 me3
H3K27 me3:DNA Meth. m5C
DNA Meth. m5C:H4K20 me2
H4K20 me2:H4−4ac
H3K27 me3:H3K4 me1
H3K27 me2
H3K9 me2
H2A.Z
H4K20 me3
H3K9 me3
H3−5ac
H3K4 me3
H4K20 me2
H4−4ac
H3K27 me3
DNA Meth. m5C
H3K4 me1
0.00 0.25 0.50 0.75 1.00
π̂

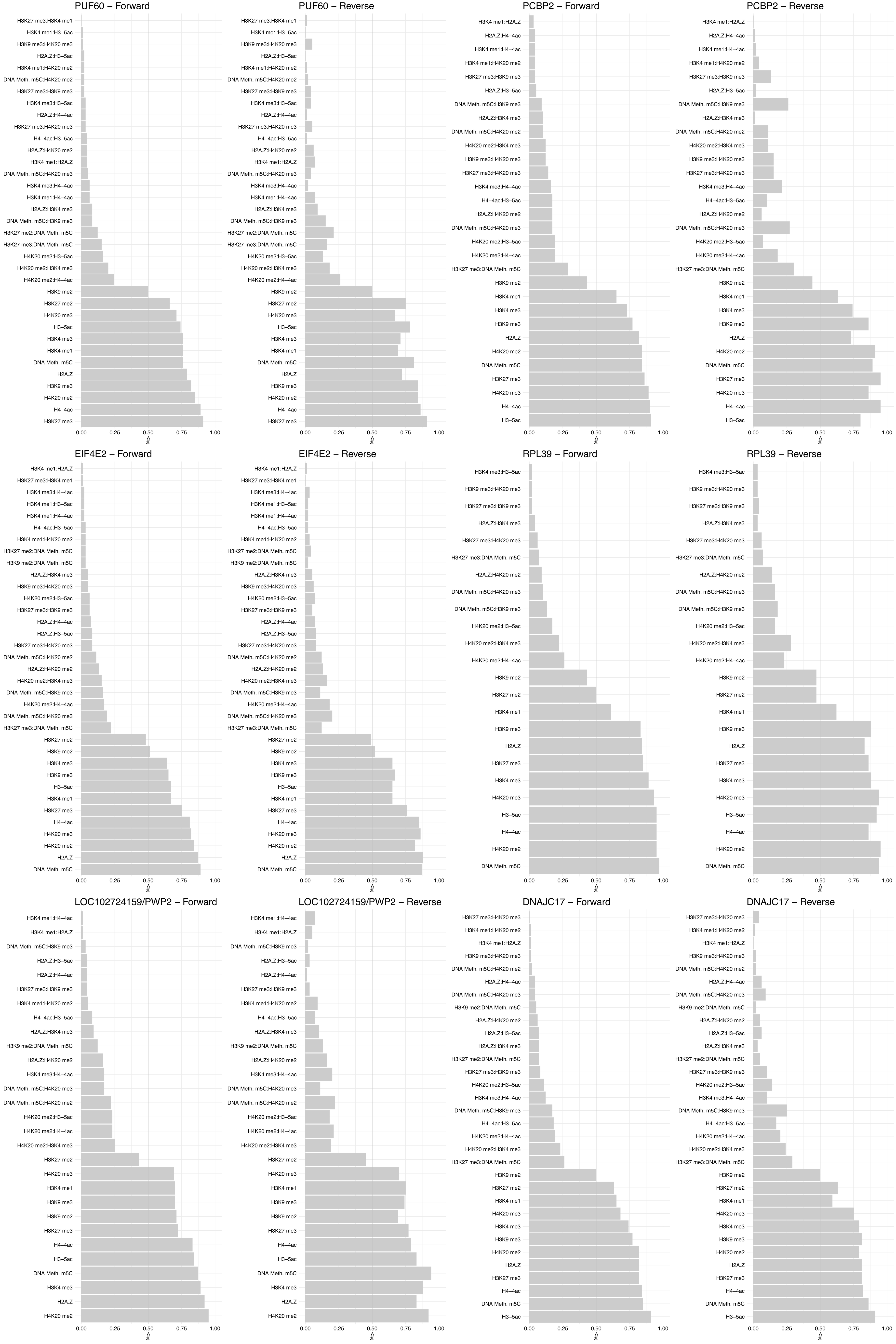
PUF60 – Forward
PUF60 – Reverse
PCBP2 – Forward
PCBP2 – Reverse
EIF4E2 – Forward
EIF4E2 – Reverse
RPL39 – Forward
RPL39 – Reverse
LOC102724159/PWP2 – Forward
LOC102724159/PWP2 – Reverse
DNAJC17 – Forward
DNAJC17 – Reverse
0.00
0.25
0.50
0.75
1.00
$\hat{\pi}$

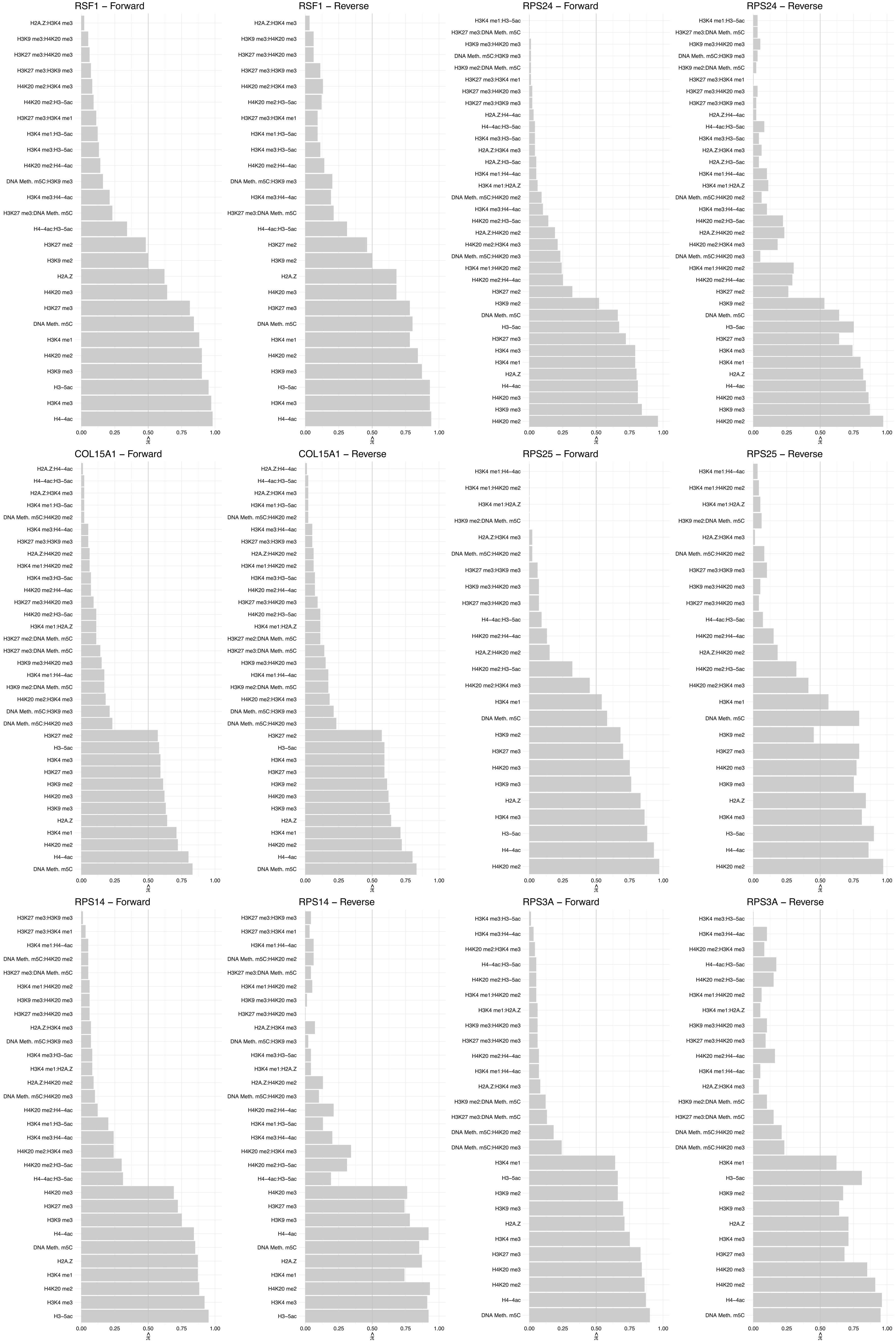
RSF1 – Forward
H2A.Z:H3K4 me3
H3K9 me3:H4K20 me3
H3K27 me3:H4K20 me3
H3K27 me3:H3K9 me3
H4K20 me2:H3K4 me3
H4K20 me2:H3−5ac
H3K27 me3:H3K4 me1
H3K4 me1:H3−5ac
H3K4 me3:H3−5ac
H4K20 me2:H4−4ac
DNA Meth. m5C:H3K9 me3
H3K4 me3:H4−4ac
H3K27 me3:DNA Meth. m5C
H4−4ac:H3−5ac
H3K27 me2
H3K9 me2
H2A.Z
H4K20 me3
H3K27 me3
DNA Meth. m5C
H3K4 me1
H4K20 me2
H3K9 me3
H3−5ac
H3K4 me3
H4−4ac
0.00 0.25 0.50 0.75 1.00
π̂
RSF1 – Reverse
H2A.Z:H3K4 me3
H3K9 me3:H4K20 me3
H3K27 me3:H4K20 me3
H3K27 me3:H3K9 me3
H4K20 me2:H3K4 me3
H4K20 me2:H3−5ac
H3K27 me3:H3K4 me1
H3K4 me1:H3−5ac
H3K4 me3:H3−5ac
H4K20 me2:H4−4ac
DNA Meth. m5C:H3K9 me3
H3K4 me3:H4−4ac
H3K27 me3:DNA Meth. m5C
H4−4ac:H3−5ac
H3K27 me2
H3K9 me2
H2A.Z
H4K20 me3
H3K27 me3
DNA Meth. m5C
H3K4 me1
H4K20 me2
H3K9 me3
H3−5ac
H3K4 me3
H4−4ac
0.00 0.25 0.50 0.75 1.00
π̂
RPS24 – Forward
H3K4 me1:H3−5ac
H3K27 me3:DNA Meth. m5C
H3K9 me3:H4K20 me3
DNA Meth. m5C:H3K9 me3
H3K9 me2:DNA Meth. m5C
H3K27 me3:H3K4 me1
H3K27 me3:H4K20 me3
H3K27 me3:H3K9 me3
H2A.Z:H4−4ac
H4−4ac:H3−5ac
H3K4 me3:H3−5ac
H2A.Z:H3K4 me3
H2A.Z:H3−5ac
H3K4 me1:H4−4ac
H3K4 me1:H2A.Z
DNA Meth. m5C:H4K20 me2
H3K4 me3:H4−4ac
H4K20 me2:H3−5ac
H2A.Z:H4K20 me2
H4K20 me2:H3K4 me3
DNA Meth. m5C:H4K20 me3
H3K4 me1:H4K20 me2
H4K20 me2:H4−4ac
H3K27 me2
H3K9 me2
DNA Meth. m5C
H3−5ac
H3K27 me3
H3K4 me3
H3K4 me1
H2A.Z
H4−4ac
H4K20 me3
H3K9 me3
H4K20 me2
0.00 0.25 0.50 0.75 1.00
π̂
RPS24 – Reverse
H3K4 me1:H3−5ac
H3K27 me3:DNA Meth. m5C
H3K9 me3:H4K20 me3
DNA Meth. m5C:H3K9 me3
H3K9 me2:DNA Meth. m5C
H3K27 me3:H3K4 me1
H3K27 me3:H4K20 me3
H3K27 me3:H3K9 me3
H2A.Z:H4−4ac
H4−4ac:H3−5ac
H3K4 me3:H3−5ac
H2A.Z:H3K4 me3
H2A.Z:H3−5ac
H3K4 me1:H4−4ac
H3K4 me1:H2A.Z
DNA Meth. m5C:H4K20 me2
H3K4 me3:H4−4ac
H4K20 me2:H3−5ac
H2A.Z:H4K20 me2
H4K20 me2:H3K4 me3
DNA Meth. m5C:H4K20 me3
H3K4 me1:H4K20 me2
H4K20 me2:H4−4ac
H3K27 me2
H3K9 me2
DNA Meth. m5C
H3−5ac
H3K27 me3
H3K4 me3
H3K4 me1
H2A.Z
H4−4ac
H4K20 me3
H3K9 me3
H4K20 me2
0.00 0.25 0.50 0.75 1.00
π̂
COL15A1 – Forward
H2A.Z:H4−4ac
H4−4ac:H3−5ac
H2A.Z:H3K4 me3
H3K4 me1:H3−5ac
DNA Meth. m5C:H4K20 me2
H3K4 me3:H4−4ac
H3K27 me3:H3K9 me3
H2A.Z:H4K20 me2
H3K4 me1:H4K20 me2
H3K4 me3:H3−5ac
H4K20 me2:H4−4ac
H3K27 me3:H4K20 me3
H4K20 me2:H3−5ac
H3K4 me1:H2A.Z
H3K27 me2:DNA Meth. m5C
H3K27 me3:DNA Meth. m5C
H3K9 me3:H4K20 me3
H3K4 me1:H4−4ac
H3K9 me2:DNA Meth. m5C
H4K20 me2:H3K4 me3
DNA Meth. m5C:H3K9 me3
DNA Meth. m5C:H4K20 me3
H3K27 me2
H3−5ac
H3K4 me3
H3K27 me3
H3K9 me2
H4K20 me3
H3K9 me3
H2A.Z
H3K4 me1
H4K20 me2
H4−4ac
DNA Meth. m5C
0.00 0.25 0.50 0.75 1.00
π̂
COL15A1 – Reverse
H2A.Z:H4−4ac
H4−4ac:H3−5ac
H2A.Z:H3K4 me3
H3K4 me1:H3−5ac
DNA Meth. m5C:H4K20 me2
H3K4 me3:H4−4ac
H3K27 me3:H3K9 me3
H2A.Z:H4K20 me2
H3K4 me1:H4K20 me2
H3K4 me3:H3−5ac
H4K20 me2:H4−4ac
H3K27 me3:H4K20 me3
H4K20 me2:H3−5ac
H3K4 me1:H2A.Z
H3K27 me2:DNA Meth. m5C
H3K27 me3:DNA Meth. m5C
H3K9 me3:H4K20 me3
H3K4 me1:H4−4ac
H3K9 me2:DNA Meth. m5C
H4K20 me2:H3K4 me3
DNA Meth. m5C:H3K9 me3
DNA Meth. m5C:H4K20 me3
H3K27 me2
H3−5ac
H3K4 me3
H3K27 me3
H3K9 me2
H4K20 me3
H3K9 me3
H2A.Z
H3K4 me1
H4K20 me2
H4−4ac
DNA Meth. m5C
0.00 0.25 0.50 0.75 1.00
π̂
RPS25 – Forward
H3K4 me1:H3−5ac
H3K4 me1:H4K20 me2
H3K4 me1:H2A.Z
H3K9 me2:DNA Meth. m5C
H2A.Z:H3K4 me3
DNA Meth. m5C:H4K20 me2
H3K27 me3:H3K9 me3
H3K9 me3:H4K20 me3
H3K27 me3:H4K20 me3
H4−4ac:H3−5ac
H4K20 me2:H4−4ac
H2A.Z:H4K20 me2
H4K20 me2:H3−5ac
H4K20 me2:H3K4 me3
H3K4 me1
DNA Meth. m5C
H3K9 me2
H3K27 me3
H4K20 me3
H3K9 me3
H2A.Z
H3K4 me3
H3−5ac
H4−4ac
H4K20 me2
0.00 0.25 0.50 0.75 1.00
π̂
RPS25 – Reverse
H3K4 me1:H3−5ac
H3K4 me1:H4K20 me2
H3K4 me1:H2A.Z
H3K9 me2:DNA Meth. m5C
H2A.Z:H3K4 me3
DNA Meth. m5C:H4K20 me2
H3K27 me3:H3K9 me3
H3K9 me3:H4K20 me3
H3K27 me3:H4K20 me3
H4−4ac:H3−5ac
H4K20 me2:H4−4ac
H2A.Z:H4K20 me2
H4K20 me2:H3−5ac
H4K20 me2:H3K4 me3
H3K4 me1
DNA Meth. m5C
H3K9 me2
H3K27 me3
H4K20 me3
H3K9 me3
H2A.Z
H3K4 me3
H3−5ac
H4−4ac
H4K20 me2
0.00 0.25 0.50 0.75 1.00
π̂
RPS14 – Forward
H3K27 me3:H3K9 me3
H3K27 me3:H3K4 me1
H3K4 me1:H4−4ac
DNA Meth. m5C:H4K20 me2
H3K27 me3:DNA Meth. m5C
H3K4 me1:H4K20 me2
H3K9 me3:H4K20 me3
H3K27 me3:H4K20 me3
H2A.Z:H3K4 me3
DNA Meth. m5C:H3K9 me3
H3K4 me3:H3−5ac
H3K4 me1:H2A.Z
H2A.Z:H4K20 me2
DNA Meth. m5C:H4K20 me3
H4K20 me2:H4−4ac
H3K4 me1:H3−5ac
H3K4 me3:H4−4ac
H4K20 me2:H3K4 me3
H4K20 me2:H3−5ac
H4−4ac:H3−5ac
H4K20 me3
H3K27 me3
H3K9 me3
H4−4ac
DNA Meth. m5C
H2A.Z
H3K4 me1
H4K20 me2
H3K4 me3
H3−5ac
0.00 0.25 0.50 0.75 1.00
π̂
RPS14 – Reverse
H3K27 me3:H3K9 me3
H3K27 me3:H3K4 me1
H3K4 me1:H4−4ac
DNA Meth. m5C:H4K20 me2
H3K27 me3:DNA Meth. m5C
H3K4 me1:H4K20 me2
H3K9 me3:H4K20 me3
H3K27 me3:H4K20 me3
H2A.Z:H3K4 me3
DNA Meth. m5C:H3K9 me3
H3K4 me3:H3−5ac
H3K4 me1:H2A.Z
H2A.Z:H4K20 me2
DNA Meth. m5C:H4K20 me3
H4K20 me2:H4−4ac
H3K4 me1:H3−5ac
H3K4 me3:H4−4ac
H4K20 me2:H3K4 me3
H4K20 me2:H3−5ac
H4−4ac:H3−5ac
H4K20 me3
H3K27 me3
H3K9 me3
H4−4ac
DNA Meth. m5C
H2A.Z
H3K4 me1
H4K20 me2
H3K4 me3
H3−5ac
0.00 0.25 0.50 0.75 1.00
π̂
RPS3A – Forward
H3K4 me3:H3−5ac
H3K4 me3:H4−4ac
H4K20 me2:H3K4 me3
H4−4ac:H3−5ac
H4K20 me2:H3−5ac
H3K4 me1:H4K20 me2
H3K4 me1:H2A.Z
H3K9 me3:H4K20 me3
H3K27 me3:H4K20 me3
H4K20 me2:H4−4ac
H3K4 me1:H4−4ac
H2A.Z:H3K4 me3
H3K9 me2:DNA Meth. m5C
H3K27 me3:DNA Meth. m5C
DNA Meth. m5C:H4K20 me2
DNA Meth. m5C:H4K20 me3
H3K4 me1
H3−5ac
H3K9 me2
H3K9 me3
H2A.Z
H3K4 me3
H3K27 me3
H4K20 me3
H4K20 me2
H4−4ac
DNA Meth. m5C
0.00 0.25 0.50 0.75 1.00
π̂
RPS3A – Reverse
H3K4 me3:H3−5ac
H3K4 me3:H4−4ac
H4K20 me2:H3K4 me3
H4−4ac:H3−5ac
H4K20 me2:H3−5ac
H3K4 me1:H4K20 me2
H3K4 me1:H2A.Z
H3K9 me3:H4K20 me3
H3K27 me3:H4K20 me3
H4K20 me2:H4−4ac
H3K4 me1:H4−4ac
H2A.Z:H3K4 me3
H3K9 me2:DNA Meth. m5C
H3K27 me3:DNA Meth. m5C
DNA Meth. m5C:H4K20 me2
DNA Meth. m5C:H4K20 me3
H3K4 me1
H3−5ac
H3K9 me2
H3K9 me3
H2A.Z
H3K4 me3
H3K27 me3
H4K20 me3
H4K20 me2
H4−4ac
DNA Meth. m5C
0.00 0.25 0.50 0.75 1.00
π̂

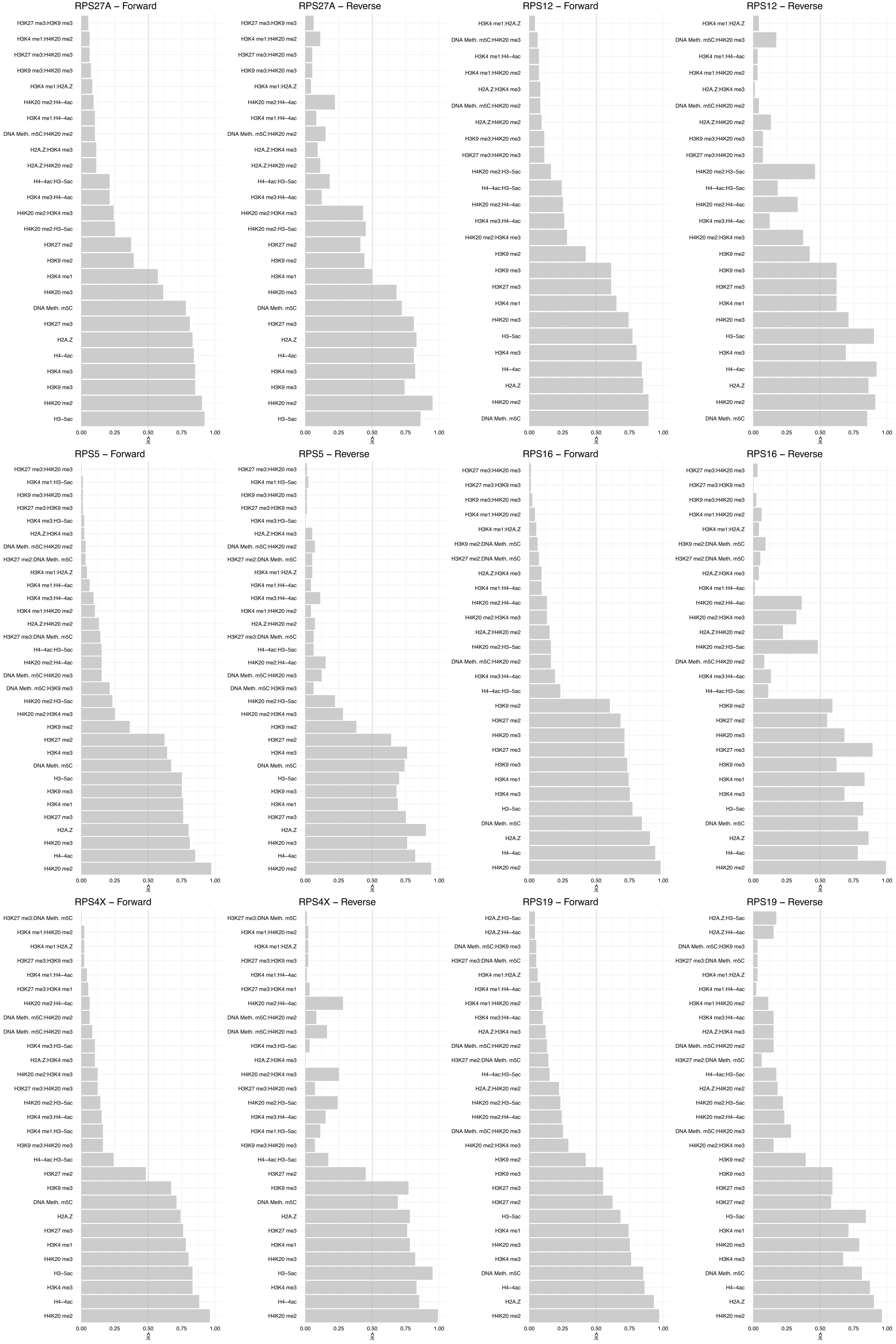

RPS27A – Forward
H3K27 me3:H3K9 me3
H3K4 me1:H4K20 me2
H3K27 me3:H4K20 me3
H3K9 me3:H4K20 me3
H3K4 me1:H2A.Z
H4K20 me2:H4–4ac
H3K4 me1:H4–4ac
DNA Meth. m5C:H4K20 me2
H2A.Z:H3K4 me3
H2A.Z:H4K20 me2
H4–4ac:H3–5ac
H3K4 me3:H4–4ac
H4K20 me2:H3K4 me3
H4K20 me2:H3–5ac
H3K27 me2
H3K9 me2
H3K4 me1
H4K20 me3
DNA Meth. m5C
H3K27 me3
H2A.Z
H4–4ac
H3K4 me3
H3K9 me3
H4K20 me2
H3–5ac
0.00 0.25 0.50 0.75 1.00
π̂
RPS27A – Reverse
RPS12 – Forward
H3K4 me1:H2A.Z
DNA Meth. m5C:H4K20 me3
H3K4 me1:H4–4ac
H3K4 me1:H4K20 me2
H2A.Z:H3K4 me3
DNA Meth. m5C:H4K20 me2
H2A.Z:H4K20 me2
H3K9 me3:H4K20 me3
H3K27 me3:H4K20 me3
H4K20 me2:H3–5ac
H4–4ac:H3–5ac
H4K20 me2:H4–4ac
H3K4 me3:H4–4ac
H4K20 me2:H3K4 me3
H3K9 me2
H3K9 me3
H3K27 me3
H3K4 me1
H4K20 me3
H3–5ac
H3K4 me3
H4–4ac
H2A.Z
H4K20 me2
DNA Meth. m5C
RPS12 – Reverse
RPS5 – Forward
H3K27 me3:H4K20 me3
H3K4 me1:H3–5ac
H3K9 me3:H4K20 me3
H3K27 me3:H3K9 me3
H3K4 me3:H3–5ac
H2A.Z:H3K4 me3
DNA Meth. m5C:H4K20 me2
H3K27 me2:DNA Meth. m5C
H3K4 me1:H2A.Z
H3K4 me1:H4–4ac
H3K4 me3:H4–4ac
H3K4 me1:H4K20 me2
H2A.Z:H4K20 me2
H3K27 me3:DNA Meth. m5C
H4–4ac:H3–5ac
H4K20 me2:H4–4ac
DNA Meth. m5C:H4K20 me3
DNA Meth. m5C:H3K9 me3
H4K20 me2:H3–5ac
H4K20 me2:H3K4 me3
H3K9 me2
H3K27 me2
H3K4 me3
DNA Meth. m5C
H3–5ac
H3K9 me3
H3K4 me1
H3K27 me3
H2A.Z
H4K20 me3
H4–4ac
H4K20 me2
RPS5 – Reverse
RPS16 – Forward
H3K27 me3:H4K20 me3
H3K27 me3:H3K9 me3
H3K9 me3:H4K20 me3
H3K4 me1:H4K20 me2
H3K4 me1:H2A.Z
H3K9 me2:DNA Meth. m5C
H3K27 me2:DNA Meth. m5C
H2A.Z:H3K4 me3
H3K4 me1:H4–4ac
H4K20 me2:H4–4ac
H4K20 me2:H3K4 me3
H2A.Z:H4K20 me2
H4K20 me2:H3–5ac
DNA Meth. m5C:H4K20 me2
H3K4 me3:H4–4ac
H4–4ac:H3–5ac
H3K9 me2
H3K27 me2
H4K20 me3
H3K27 me3
H3K9 me3
H3K4 me1
H3K4 me3
H3–5ac
DNA Meth. m5C
H2A.Z
H4–4ac
H4K20 me2
RPS16 – Reverse
RPS4X – Forward
H3K27 me3:DNA Meth. m5C
H3K4 me1:H4K20 me2
H3K4 me1:H2A.Z
H3K27 me3:H3K9 me3
H3K4 me1:H4–4ac
H3K27 me3:H3K4 me1
H4K20 me2:H4–4ac
DNA Meth. m5C:H4K20 me2
DNA Meth. m5C:H4K20 me3
H3K4 me3:H3–5ac
H2A.Z:H3K4 me3
H4K20 me2:H3K4 me3
H3K27 me3:H4K20 me3
H4K20 me2:H3–5ac
H3K4 me3:H4–4ac
H3K4 me1:H3–5ac
H3K9 me3:H4K20 me3
H4–4ac:H3–5ac
H3K27 me2
H3K9 me3
DNA Meth. m5C
H2A.Z
H3K27 me3
H3K4 me1
H4K20 me3
H3–5ac
H3K4 me3
H4–4ac
H4K20 me2
RPS4X – Reverse
RPS19 – Forward
H2A.Z:H3–5ac
H2A.Z:H4–4ac
DNA Meth. m5C:H3K9 me3
H3K27 me3:DNA Meth. m5C
H3K4 me1:H2A.Z
H3K4 me1:H4–4ac
H3K4 me1:H4K20 me2
H3K4 me3:H4–4ac
H2A.Z:H3K4 me3
DNA Meth. m5C:H4K20 me2
H3K27 me2:DNA Meth. m5C
H4–4ac:H3–5ac
H2A.Z:H4K20 me2
H4K20 me2:H3–5ac
H4K20 me2:H4–4ac
DNA Meth. m5C:H4K20 me3
H4K20 me2:H3K4 me3
H3K9 me2
H3K9 me3
H3K27 me3
H3K27 me2
H3–5ac
H3K4 me1
H4K20 me3
H3K4 me3
DNA Meth. m5C
H4–4ac
H2A.Z
H4K20 me2
RPS19 – Reverse

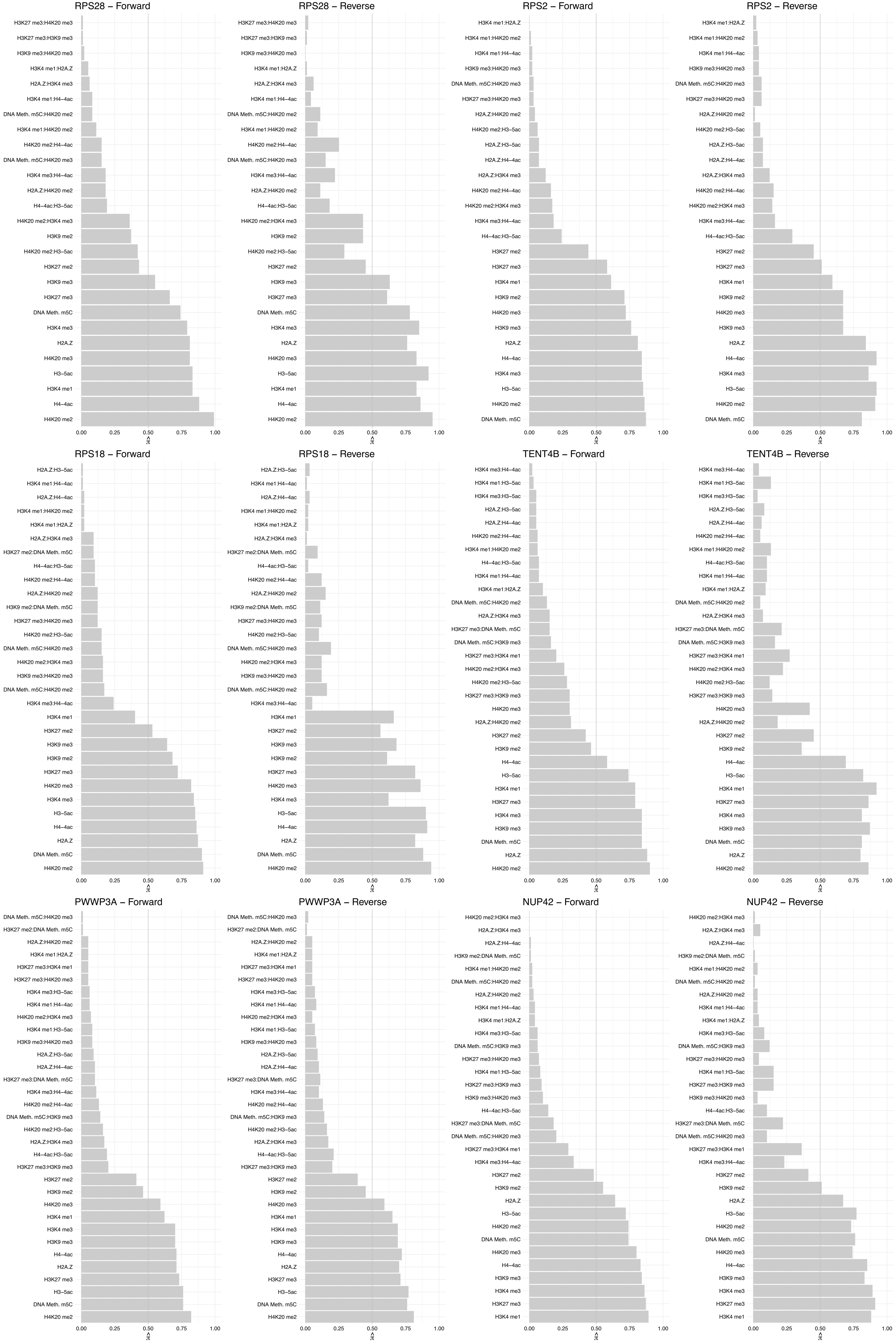
RPS28 − Forward
H3K27 me3:H4K20 me3
H3K27 me3:H3K9 me3
H3K9 me3:H4K20 me3
H3K4 me1:H2A.Z
H2A.Z:H3K4 me3
H3K4 me1:H4−4ac
DNA Meth. m5C:H4K20 me2
H3K4 me1:H4K20 me2
H4K20 me2:H4−4ac
DNA Meth. m5C:H4K20 me3
H3K4 me3:H4−4ac
H2A.Z:H4K20 me2
H4−4ac:H3−5ac
H4K20 me2:H3K4 me3
H3K9 me2
H4K20 me2:H3−5ac
H3K27 me2
H3K9 me3
H3K27 me3
DNA Meth. m5C
H3K4 me3
H2A.Z
H4K20 me3
H3−5ac
H3K4 me1
H4−4ac
H4K20 me2
0.00
0.25
0.50
0.75
1.00
$\hat{\pi}$
RPS28 − Reverse
H3K27 me3:H4K20 me3
H3K27 me3:H3K9 me3
H3K9 me3:H4K20 me3
H3K4 me1:H2A.Z
H2A.Z:H3K4 me3
H3K4 me1:H4−4ac
DNA Meth. m5C:H4K20 me2
H3K4 me1:H4K20 me2
H4K20 me2:H4−4ac
DNA Meth. m5C:H4K20 me3
H3K4 me3:H4−4ac
H2A.Z:H4K20 me2
H4−4ac:H3−5ac
H4K20 me2:H3K4 me3
H3K9 me2
H4K20 me2:H3−5ac
H3K27 me2
H3K9 me3
H3K27 me3
DNA Meth. m5C
H3K4 me3
H2A.Z
H4K20 me3
H3−5ac
H3K4 me1
H4−4ac
H4K20 me2
0.00
0.25
0.50
0.75
1.00
$\hat{\pi}$
RPS2 − Forward
H3K4 me1:H2A.Z
H3K4 me1:H4K20 me2
H3K4 me1:H4−4ac
H3K9 me3:H4K20 me3
DNA Meth. m5C:H4K20 me3
H3K27 me3:H4K20 me3
H2A.Z:H4K20 me2
H4K20 me2:H3−5ac
H2A.Z:H3−5ac
H2A.Z:H4−4ac
H2A.Z:H3K4 me3
H4K20 me2:H4−4ac
H4K20 me2:H3K4 me3
H3K4 me3:H4−4ac
H4−4ac:H3−5ac
H3K27 me2
H3K27 me3
H3K4 me1
H3K9 me2
H4K20 me3
H3K9 me3
H2A.Z
H4−4ac
H3K4 me3
H3−5ac
H4K20 me2
DNA Meth. m5C
0.00
0.25
0.50
0.75
1.00
$\hat{\pi}$
RPS2 − Reverse
H3K4 me1:H2A.Z
H3K4 me1:H4K20 me2
H3K4 me1:H4−4ac
H3K9 me3:H4K20 me3
DNA Meth. m5C:H4K20 me3
H3K27 me3:H4K20 me3
H2A.Z:H4K20 me2
H4K20 me2:H3−5ac
H2A.Z:H3−5ac
H2A.Z:H4−4ac
H2A.Z:H3K4 me3
H4K20 me2:H4−4ac
H4K20 me2:H3K4 me3
H3K4 me3:H4−4ac
H4−4ac:H3−5ac
H3K27 me2
H3K27 me3
H3K4 me1
H3K9 me2
H4K20 me3
H3K9 me3
H2A.Z
H4−4ac
H3K4 me3
H3−5ac
H4K20 me2
DNA Meth. m5C
0.00
0.25
0.50
0.75
1.00
$\hat{\pi}$
RPS18 − Forward
H2A.Z:H3−5ac
H3K4 me1:H4−4ac
H2A.Z:H4−4ac
H3K4 me1:H4K20 me2
H3K4 me1:H2A.Z
H2A.Z:H3K4 me3
H3K27 me2:DNA Meth. m5C
H4−4ac:H3−5ac
H4K20 me2:H4−4ac
H2A.Z:H4K20 me2
H3K9 me2:DNA Meth. m5C
H3K27 me3:H4K20 me3
H4K20 me2:H3−5ac
DNA Meth. m5C:H4K20 me3
H4K20 me2:H3K4 me3
H3K9 me3:H4K20 me3
DNA Meth. m5C:H4K20 me2
H3K4 me3:H4−4ac
H3K4 me1
H3K27 me2
H3K9 me3
H3K9 me2
H3K27 me3
H4K20 me3
H3K4 me3
H3−5ac
H4−4ac
H2A.Z
DNA Meth. m5C
H4K20 me2
0.00
0.25
0.50
0.75
1.00
$\hat{\pi}$
RPS18 − Reverse
H2A.Z:H3−5ac
H3K4 me1:H4−4ac
H2A.Z:H4−4ac
H3K4 me1:H4K20 me2
H3K4 me1:H2A.Z
H2A.Z:H3K4 me3
H3K27 me2:DNA Meth. m5C
H4−4ac:H3−5ac
H4K20 me2:H4−4ac
H2A.Z:H4K20 me2
H3K9 me2:DNA Meth. m5C
H3K27 me3:H4K20 me3
H4K20 me2:H3−5ac
DNA Meth. m5C:H4K20 me3
H4K20 me2:H3K4 me3
H3K9 me3:H4K20 me3
DNA Meth. m5C:H4K20 me2
H3K4 me3:H4−4ac
H3K4 me1
H3K27 me2
H3K9 me3
H3K9 me2
H3K27 me3
H4K20 me3
H3K4 me3
H3−5ac
H4−4ac
H2A.Z
DNA Meth. m5C
H4K20 me2
0.00
0.25
0.50
0.75
1.00
$\hat{\pi}$
TENT4B − Forward
H3K4 me3:H4−4ac
H3K4 me1:H3−5ac
H3K4 me3:H3−5ac
H2A.Z:H3−5ac
H2A.Z:H4−4ac
H4K20 me2:H4−4ac
H3K4 me1:H4K20 me2
H4−4ac:H3−5ac
H3K4 me1:H4−4ac
H3K4 me1:H2A.Z
DNA Meth. m5C:H4K20 me2
H2A.Z:H3K4 me3
H3K27 me3:DNA Meth. m5C
DNA Meth. m5C:H3K9 me3
H3K27 me3:H3K4 me1
H4K20 me2:H3K4 me3
H4K20 me2:H3−5ac
H3K27 me3:H3K9 me3
H4K20 me3
H2A.Z:H4K20 me2
H3K27 me2
H3K9 me2
H4−4ac
H3−5ac
H3K4 me1
H3K27 me3
H3K4 me3
H3K9 me3
DNA Meth. m5C
H2A.Z
H4K20 me2
0.00
0.25
0.50
0.75
1.00
$\hat{\pi}$
TENT4B − Reverse
H3K4 me3:H4−4ac
H3K4 me1:H3−5ac
H3K4 me3:H3−5ac
H2A.Z:H3−5ac
H2A.Z:H4−4ac
H4K20 me2:H4−4ac
H3K4 me1:H4K20 me2
H4−4ac:H3−5ac
H3K4 me1:H4−4ac
H3K4 me1:H2A.Z
DNA Meth. m5C:H4K20 me2
H2A.Z:H3K4 me3
H3K27 me3:DNA Meth. m5C
DNA Meth. m5C:H3K9 me3
H3K27 me3:H3K4 me1
H4K20 me2:H3K4 me3
H4K20 me2:H3−5ac
H3K27 me3:H3K9 me3
H4K20 me3
H2A.Z:H4K20 me2
H3K27 me2
H3K9 me2
H4−4ac
H3−5ac
H3K4 me1
H3K27 me3
H3K4 me3
H3K9 me3
DNA Meth. m5C
H2A.Z
H4K20 me2
0.00
0.25
0.50
0.75
1.00
$\hat{\pi}$
PWWP3A − Forward
DNA Meth. m5C:H4K20 me3
H3K27 me2:DNA Meth. m5C
H2A.Z:H4K20 me2
H3K4 me1:H2A.Z
H3K27 me3:H3K4 me1
H3K27 me3:H4K20 me3
H3K4 me3:H3−5ac
H3K4 me1:H4−4ac
H4K20 me2:H3K4 me3
H3K4 me1:H3−5ac
H3K9 me3:H4K20 me3
H2A.Z:H3−5ac
H2A.Z:H4−4ac
H3K27 me3:DNA Meth. m5C
H3K4 me3:H4−4ac
H4K20 me2:H4−4ac
DNA Meth. m5C:H3K9 me3
H4K20 me2:H3−5ac
H2A.Z:H3K4 me3
H4−4ac:H3−5ac
H3K27 me3:H3K9 me3
H3K27 me2
H3K9 me2
H4K20 me3
H3K4 me1
H3K4 me3
H3K9 me3
H4−4ac
H2A.Z
H3K27 me3
H3−5ac
DNA Meth. m5C
H4K20 me2
0.00
0.25
0.50
0.75
1.00
$\hat{\pi}$
PWWP3A − Reverse
DNA Meth. m5C:H4K20 me3
H3K27 me2:DNA Meth. m5C
H2A.Z:H4K20 me2
H3K4 me1:H2A.Z
H3K27 me3:H3K4 me1
H3K27 me3:H4K20 me3
H3K4 me3:H3−5ac
H3K4 me1:H4−4ac
H4K20 me2:H3K4 me3
H3K4 me1:H3−5ac
H3K9 me3:H4K20 me3
H2A.Z:H3−5ac
H2A.Z:H4−4ac
H3K27 me3:DNA Meth. m5C
H3K4 me3:H4−4ac
H4K20 me2:H4−4ac
DNA Meth. m5C:H3K9 me3
H4K20 me2:H3−5ac
H2A.Z:H3K4 me3
H4−4ac:H3−5ac
H3K27 me3:H3K9 me3
H3K27 me2
H3K9 me2
H4K20 me3
H3K4 me1
H3K4 me3
H3K9 me3
H4−4ac
H2A.Z
H3K27 me3
H3−5ac
DNA Meth. m5C
H4K20 me2
0.00
0.25
0.50
0.75
1.00
$\hat{\pi}$
NUP42 − Forward
H4K20 me2:H3K4 me3
H2A.Z:H3K4 me3
H2A.Z:H4−4ac
H3K9 me2:DNA Meth. m5C
H3K4 me1:H4K20 me2
DNA Meth. m5C:H4K20 me2
H2A.Z:H4K20 me2
H3K4 me1:H4−4ac
H3K4 me1:H2A.Z
H3K4 me3:H3−5ac
DNA Meth. m5C:H3K9 me3
H3K27 me3:H4K20 me3
H3K4 me1:H3−5ac
H3K27 me3:H3K9 me3
H3K9 me3:H4K20 me3
H4−4ac:H3−5ac
H3K27 me3:DNA Meth. m5C
DNA Meth. m5C:H4K20 me3
H3K27 me3:H3K4 me1
H3K4 me3:H4−4ac
H3K27 me2
H3K9 me2
H2A.Z
H3−5ac
H4K20 me2
DNA Meth. m5C
H4K20 me3
H4−4ac
H3K9 me3
H3K4 me3
H3K27 me3
H3K4 me1
0.00
0.25
0.50
0.75
1.00
$\hat{\pi}$
NUP42 − Reverse
H4K20 me2:H3K4 me3
H2A.Z:H3K4 me3
H2A.Z:H4−4ac
H3K9 me2:DNA Meth. m5C
H3K4 me1:H4K20 me2
DNA Meth. m5C:H4K20 me2
H2A.Z:H4K20 me2
H3K4 me1:H4−4ac
H3K4 me1:H2A.Z
H3K4 me3:H3−5ac
DNA Meth. m5C:H3K9 me3
H3K27 me3:H4K20 me3
H3K4 me1:H3−5ac
H3K27 me3:H3K9 me3
H3K9 me3:H4K20 me3
H4−4ac:H3−5ac
H3K27 me3:DNA Meth. m5C
DNA Meth. m5C:H4K20 me3
H3K27 me3:H3K4 me1
H3K4 me3:H4−4ac
H3K27 me2
H3K9 me2
H2A.Z
H3−5ac
H4K20 me2
DNA Meth. m5C
H4K20 me3
H4−4ac
H3K9 me3
H3K4 me3
H3K27 me3
H3K4 me1
0.00
0.25
0.50
0.75
1.00
$\hat{\pi}$

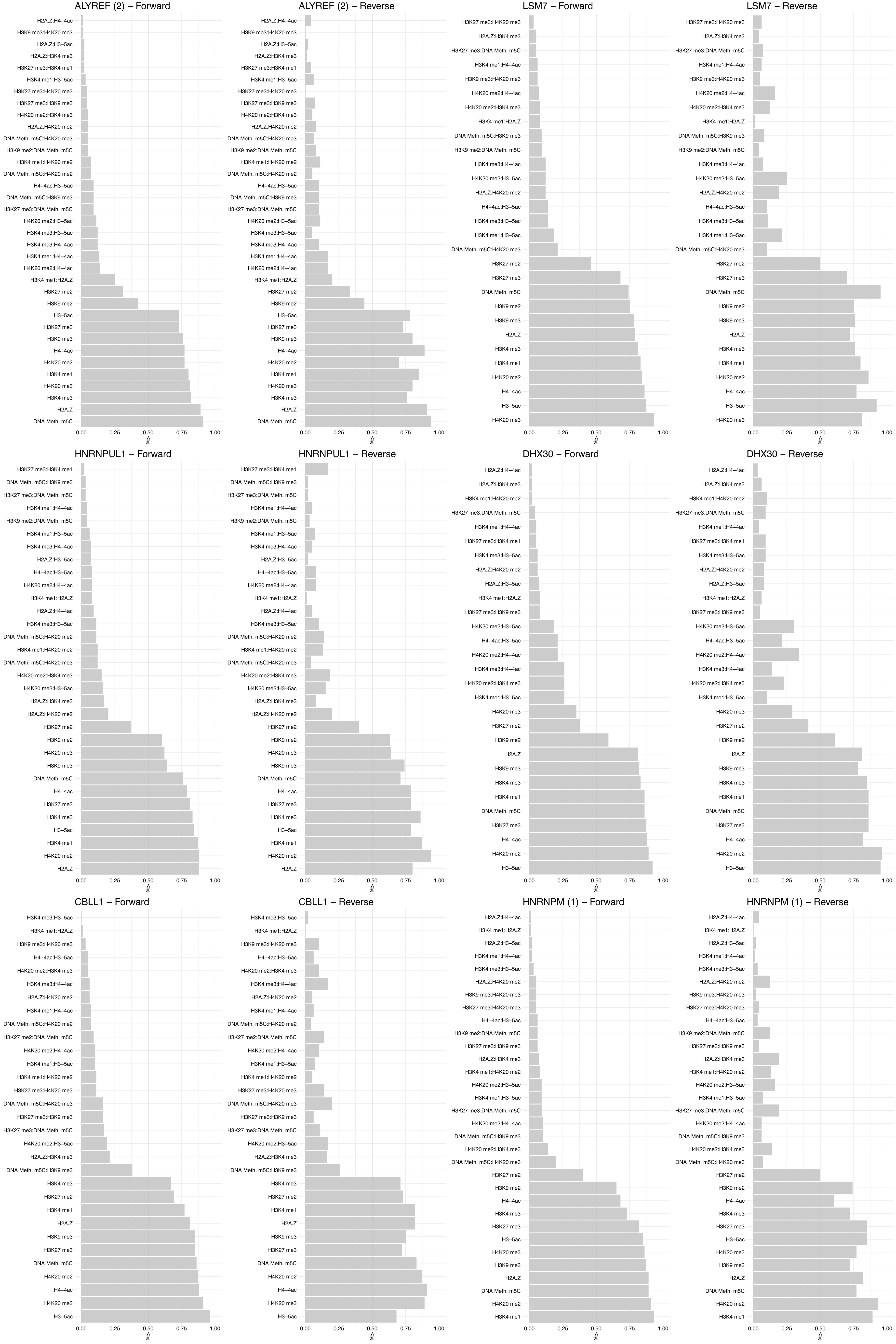
ALYREF (2) − Forward
ALYREF (2) − Reverse
H2A.Z:H4−4ac
H3K9 me3:H4K20 me3
H2A.Z:H3−5ac
H2A.Z:H3K4 me3
H3K27 me3:H3K4 me1
H3K4 me1:H3−5ac
H3K27 me3:H4K20 me3
H3K27 me3:H3K9 me3
H4K20 me2:H3K4 me3
H2A.Z:H4K20 me2
DNA Meth. m5C:H4K20 me3
H3K9 me2:DNA Meth. m5C
H3K4 me1:H4K20 me2
DNA Meth. m5C:H4K20 me2
H4−4ac:H3−5ac
DNA Meth. m5C:H3K9 me3
H3K27 me3:DNA Meth. m5C
H4K20 me2:H3−5ac
H3K4 me3:H3−5ac
H3K4 me3:H4−4ac
H3K4 me1:H4−4ac
H4K20 me2:H4−4ac
H3K4 me1:H2A.Z
H3K27 me2
H3K9 me2
H3−5ac
H3K27 me3
H3K9 me3
H4−4ac
H4K20 me2
H3K4 me1
H4K20 me3
H3K4 me3
H2A.Z
DNA Meth. m5C
LSM7 − Forward
LSM7 − Reverse
H3K27 me3:H4K20 me3
H2A.Z:H3K4 me3
H3K27 me3:DNA Meth. m5C
H3K4 me1:H4−4ac
H3K9 me3:H4K20 me3
H4K20 me2:H4−4ac
H4K20 me2:H3K4 me3
H3K4 me1:H2A.Z
DNA Meth. m5C:H3K9 me3
H3K9 me2:DNA Meth. m5C
H3K4 me3:H4−4ac
H4K20 me2:H3−5ac
H2A.Z:H4K20 me2
H4−4ac:H3−5ac
H3K4 me3:H3−5ac
H3K4 me1:H3−5ac
DNA Meth. m5C:H4K20 me3
H3K27 me2
H3K27 me3
DNA Meth. m5C
H3K9 me2
H3K9 me3
H2A.Z
H3K4 me3
H3K4 me1
H4K20 me2
H4−4ac
H3−5ac
H4K20 me3
HNRNPUL1 − Forward
HNRNPUL1 − Reverse
H3K27 me3:H3K4 me1
DNA Meth. m5C:H3K9 me3
H3K27 me3:DNA Meth. m5C
H3K4 me1:H4−4ac
H3K9 me2:DNA Meth. m5C
H3K4 me1:H3−5ac
H3K4 me3:H4−4ac
H2A.Z:H3−5ac
H4−4ac:H3−5ac
H4K20 me2:H4−4ac
H3K4 me1:H2A.Z
H2A.Z:H4−4ac
H3K4 me3:H3−5ac
DNA Meth. m5C:H4K20 me2
H3K4 me1:H4K20 me2
DNA Meth. m5C:H4K20 me3
H4K20 me2:H3K4 me3
H4K20 me2:H3−5ac
H2A.Z:H3K4 me3
H2A.Z:H4K20 me2
H3K27 me2
H3K9 me2
H4K20 me3
H3K9 me3
DNA Meth. m5C
H4−4ac
H3K27 me3
H3K4 me3
H3−5ac
H3K4 me1
H4K20 me2
H2A.Z
DHX30 − Forward
DHX30 − Reverse
H2A.Z:H4−4ac
H2A.Z:H3K4 me3
H3K4 me1:H4K20 me2
H3K27 me3:DNA Meth. m5C
H3K4 me1:H4−4ac
H3K27 me3:H3K4 me1
H3K4 me3:H3−5ac
H2A.Z:H4K20 me2
H2A.Z:H3−5ac
H3K4 me1:H2A.Z
H3K27 me3:H3K9 me3
H4K20 me2:H3−5ac
H4−4ac:H3−5ac
H4K20 me2:H4−4ac
H3K4 me3:H4−4ac
H4K20 me2:H3K4 me3
H3K4 me1:H3−5ac
H4K20 me3
H3K27 me2
H3K9 me2
H2A.Z
H3K9 me3
H3K4 me3
H3K4 me1
DNA Meth. m5C
H3K27 me3
H4−4ac
H4K20 me2
H3−5ac
CBLL1 − Forward
CBLL1 − Reverse
H3K4 me3:H3−5ac
H3K4 me1:H2A.Z
H3K9 me3:H4K20 me3
H4−4ac:H3−5ac
H4K20 me2:H3K4 me3
H3K4 me3:H4−4ac
H2A.Z:H4K20 me2
H3K4 me1:H4−4ac
DNA Meth. m5C:H4K20 me2
H3K27 me2:DNA Meth. m5C
H4K20 me2:H4−4ac
H3K4 me1:H3−5ac
H3K4 me1:H4K20 me2
H3K27 me3:H4K20 me3
DNA Meth. m5C:H4K20 me3
H3K27 me3:H3K9 me3
H3K27 me3:DNA Meth. m5C
H4K20 me2:H3−5ac
H2A.Z:H3K4 me3
DNA Meth. m5C:H3K9 me3
H3K4 me3
H3K27 me2
H3K4 me1
H2A.Z
H3K9 me3
H3K27 me3
DNA Meth. m5C
H4K20 me2
H4−4ac
H4K20 me3
H3−5ac
HNRNPM (1) − Forward
HNRNPM (1) − Reverse
H2A.Z:H4−4ac
H3K4 me1:H2A.Z
H2A.Z:H3−5ac
H3K4 me1:H4−4ac
H3K4 me3:H3−5ac
H2A.Z:H4K20 me2
H3K9 me3:H4K20 me3
H3K27 me3:H4K20 me3
H4−4ac:H3−5ac
H3K9 me2:DNA Meth. m5C
H3K27 me3:H3K9 me3
H2A.Z:H3K4 me3
H3K4 me1:H4K20 me2
H4K20 me2:H3−5ac
H3K4 me1:H3−5ac
H3K27 me3:DNA Meth. m5C
H4K20 me2:H4−4ac
DNA Meth. m5C:H3K9 me3
H4K20 me2:H3K4 me3
DNA Meth. m5C:H4K20 me3
H3K27 me2
H3K9 me2
H4−4ac
H3K4 me3
H3K27 me3
H3−5ac
H4K20 me3
H3K9 me3
H2A.Z
DNA Meth. m5C
H4K20 me2
H3K4 me1
0.00 0.25 0.50 0.75 1.00
π̂

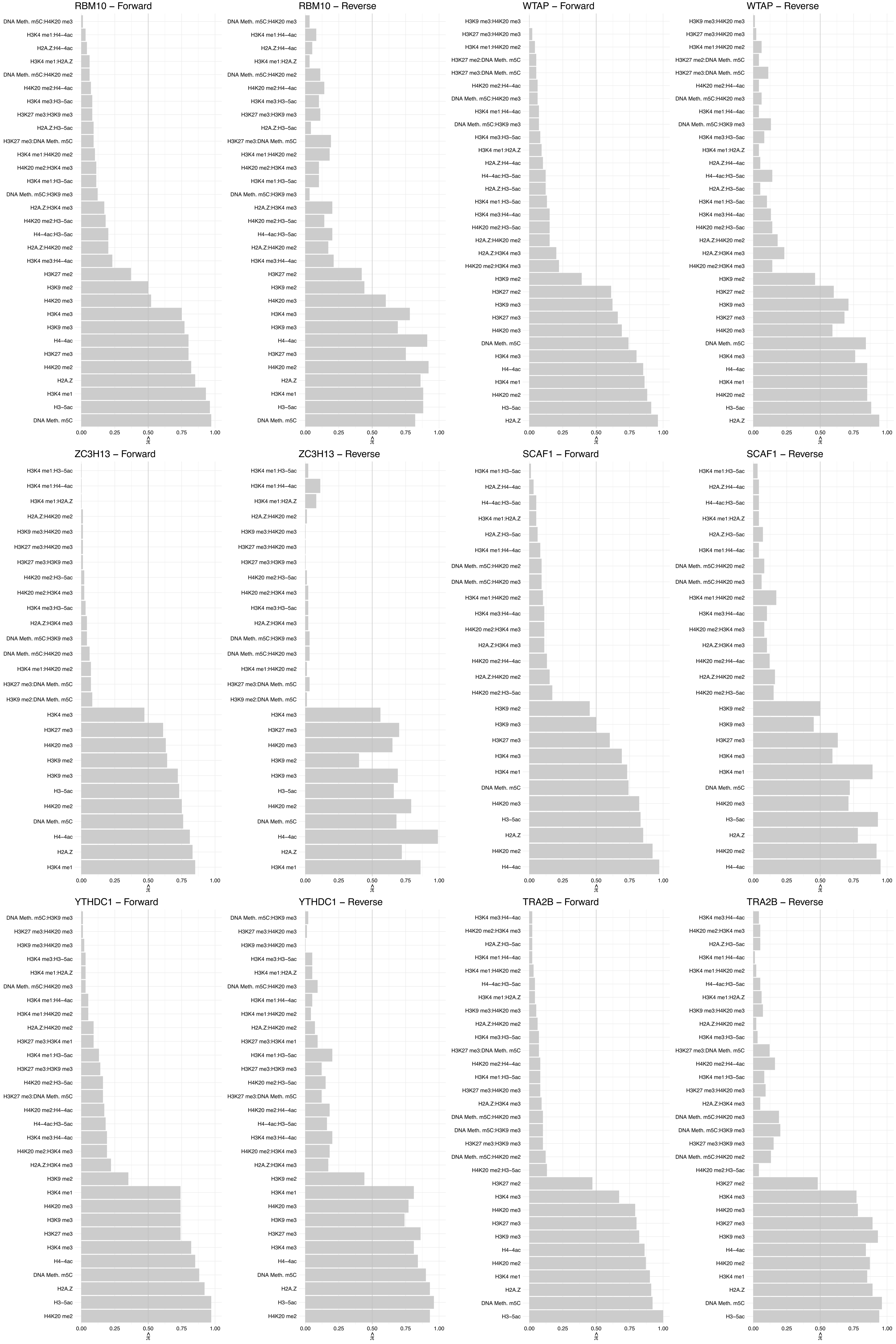
RBM10 – Forward
DNA Meth. m5C:H4K20 me3
H3K4 me1:H4−4ac
H2A.Z:H4−4ac
H3K4 me1:H2A.Z
DNA Meth. m5C:H4K20 me2
H4K20 me2:H4−4ac
H3K4 me3:H3−5ac
H3K27 me3:H3K9 me3
H2A.Z:H3−5ac
H3K27 me3:DNA Meth. m5C
H3K4 me1:H4K20 me2
H4K20 me2:H3K4 me3
H3K4 me1:H3−5ac
DNA Meth. m5C:H3K9 me3
H2A.Z:H3K4 me3
H4K20 me2:H3−5ac
H4−4ac:H3−5ac
H2A.Z:H4K20 me2
H3K4 me3:H4−4ac
H3K27 me2
H3K9 me2
H4K20 me3
H3K4 me3
H3K9 me3
H4−4ac
H3K27 me3
H4K20 me2
H2A.Z
H3K4 me1
H3−5ac
DNA Meth. m5C
0.00
0.25
0.50
0.75
1.00
π̂
RBM10 – Reverse
DNA Meth. m5C:H4K20 me3
H3K4 me1:H4−4ac
H2A.Z:H4−4ac
H3K4 me1:H2A.Z
DNA Meth. m5C:H4K20 me2
H4K20 me2:H4−4ac
H3K4 me3:H3−5ac
H3K27 me3:H3K9 me3
H2A.Z:H3−5ac
H3K27 me3:DNA Meth. m5C
H3K4 me1:H4K20 me2
H4K20 me2:H3K4 me3
H3K4 me1:H3−5ac
DNA Meth. m5C:H3K9 me3
H2A.Z:H3K4 me3
H4K20 me2:H3−5ac
H4−4ac:H3−5ac
H2A.Z:H4K20 me2
H3K4 me3:H4−4ac
H3K27 me2
H3K9 me2
H4K20 me3
H3K4 me3
H3K9 me3
H4−4ac
H3K27 me3
H4K20 me2
H2A.Z
H3K4 me1
H3−5ac
DNA Meth. m5C
0.00
0.25
0.50
0.75
1.00
π̂
WTAP – Forward
H3K9 me3:H4K20 me3
H3K27 me3:H4K20 me3
H3K4 me1:H4K20 me2
H3K27 me2:DNA Meth. m5C
H3K27 me3:DNA Meth. m5C
H4K20 me2:H4−4ac
DNA Meth. m5C:H4K20 me3
H3K4 me1:H4−4ac
DNA Meth. m5C:H3K9 me3
H3K4 me3:H3−5ac
H3K4 me1:H2A.Z
H2A.Z:H4−4ac
H4−4ac:H3−5ac
H2A.Z:H3−5ac
H3K4 me1:H3−5ac
H3K4 me3:H4−4ac
H4K20 me2:H3−5ac
H2A.Z:H4K20 me2
H2A.Z:H3K4 me3
H4K20 me2:H3K4 me3
H3K9 me2
H3K27 me2
H3K9 me3
H3K27 me3
H4K20 me3
DNA Meth. m5C
H3K4 me3
H4−4ac
H3K4 me1
H4K20 me2
H3−5ac
H2A.Z
0.00
0.25
0.50
0.75
1.00
π̂
WTAP – Reverse
H3K9 me3:H4K20 me3
H3K27 me3:H4K20 me3
H3K4 me1:H4K20 me2
H3K27 me2:DNA Meth. m5C
H3K27 me3:DNA Meth. m5C
H4K20 me2:H4−4ac
DNA Meth. m5C:H4K20 me3
H3K4 me1:H4−4ac
DNA Meth. m5C:H3K9 me3
H3K4 me3:H3−5ac
H3K4 me1:H2A.Z
H2A.Z:H4−4ac
H4−4ac:H3−5ac
H2A.Z:H3−5ac
H3K4 me1:H3−5ac
H3K4 me3:H4−4ac
H4K20 me2:H3−5ac
H2A.Z:H4K20 me2
H2A.Z:H3K4 me3
H4K20 me2:H3K4 me3
H3K9 me2
H3K27 me2
H3K9 me3
H3K27 me3
H4K20 me3
DNA Meth. m5C
H3K4 me3
H4−4ac
H3K4 me1
H4K20 me2
H3−5ac
H2A.Z
0.00
0.25
0.50
0.75
1.00
π̂
ZC3H13 – Forward
H3K4 me1:H3−5ac
H3K4 me1:H4−4ac
H3K4 me1:H2A.Z
H2A.Z:H4K20 me2
H3K9 me3:H4K20 me3
H3K27 me3:H4K20 me3
H3K27 me3:H3K9 me3
H4K20 me2:H3−5ac
H4K20 me2:H3K4 me3
H3K4 me3:H3−5ac
H2A.Z:H3K4 me3
DNA Meth. m5C:H3K9 me3
DNA Meth. m5C:H4K20 me3
H3K4 me1:H4K20 me2
H3K27 me3:DNA Meth. m5C
H3K9 me2:DNA Meth. m5C
H3K4 me3
H3K27 me3
H4K20 me3
H3K9 me2
H3K9 me3
H3−5ac
H4K20 me2
DNA Meth. m5C
H4−4ac
H2A.Z
H3K4 me1
0.00
0.25
0.50
0.75
1.00
π̂
ZC3H13 – Reverse
H3K4 me1:H3−5ac
H3K4 me1:H4−4ac
H3K4 me1:H2A.Z
H2A.Z:H4K20 me2
H3K9 me3:H4K20 me3
H3K27 me3:H4K20 me3
H3K27 me3:H3K9 me3
H4K20 me2:H3−5ac
H4K20 me2:H3K4 me3
H3K4 me3:H3−5ac
H2A.Z:H3K4 me3
DNA Meth. m5C:H3K9 me3
DNA Meth. m5C:H4K20 me3
H3K4 me1:H4K20 me2
H3K27 me3:DNA Meth. m5C
H3K9 me2:DNA Meth. m5C
H3K4 me3
H3K27 me3
H4K20 me3
H3K9 me2
H3K9 me3
H3−5ac
H4K20 me2
DNA Meth. m5C
H4−4ac
H2A.Z
H3K4 me1
0.00
0.25
0.50
0.75
1.00
π̂
SCAF1 – Forward
H3K4 me1:H3−5ac
H2A.Z:H4−4ac
H4−4ac:H3−5ac
H3K4 me1:H2A.Z
H2A.Z:H3−5ac
H3K4 me1:H4−4ac
DNA Meth. m5C:H4K20 me2
DNA Meth. m5C:H4K20 me3
H3K4 me1:H4K20 me2
H3K4 me3:H4−4ac
H4K20 me2:H3K4 me3
H2A.Z:H3K4 me3
H4K20 me2:H4−4ac
H2A.Z:H4K20 me2
H4K20 me2:H3−5ac
H3K9 me2
H3K9 me3
H3K27 me3
H3K4 me3
H3K4 me1
DNA Meth. m5C
H4K20 me3
H3−5ac
H2A.Z
H4K20 me2
H4−4ac
0.00
0.25
0.50
0.75
1.00
π̂
SCAF1 – Reverse
H3K4 me1:H3−5ac
H2A.Z:H4−4ac
H4−4ac:H3−5ac
H3K4 me1:H2A.Z
H2A.Z:H3−5ac
H3K4 me1:H4−4ac
DNA Meth. m5C:H4K20 me2
DNA Meth. m5C:H4K20 me3
H3K4 me1:H4K20 me2
H3K4 me3:H4−4ac
H4K20 me2:H3K4 me3
H2A.Z:H3K4 me3
H4K20 me2:H4−4ac
H2A.Z:H4K20 me2
H4K20 me2:H3−5ac
H3K9 me2
H3K9 me3
H3K27 me3
H3K4 me3
H3K4 me1
DNA Meth. m5C
H4K20 me3
H3−5ac
H2A.Z
H4K20 me2
H4−4ac
0.00
0.25
0.50
0.75
1.00
π̂
YTHDC1 – Forward
DNA Meth. m5C:H3K9 me3
H3K27 me3:H4K20 me3
H3K9 me3:H4K20 me3
H3K4 me3:H3−5ac
H3K4 me1:H2A.Z
DNA Meth. m5C:H4K20 me3
H3K4 me1:H4−4ac
H3K4 me1:H4K20 me2
H2A.Z:H4K20 me2
H3K27 me3:H3K4 me1
H3K4 me1:H3−5ac
H3K27 me3:H3K9 me3
H4K20 me2:H3−5ac
H3K27 me3:DNA Meth. m5C
H4K20 me2:H4−4ac
H4−4ac:H3−5ac
H3K4 me3:H4−4ac
H4K20 me2:H3K4 me3
H2A.Z:H3K4 me3
H3K9 me2
H3K4 me1
H4K20 me3
H3K9 me3
H3K27 me3
H3K4 me3
H4−4ac
DNA Meth. m5C
H2A.Z
H3−5ac
H4K20 me2
0.00
0.25
0.50
0.75
1.00
π̂
YTHDC1 – Reverse
DNA Meth. m5C:H3K9 me3
H3K27 me3:H4K20 me3
H3K9 me3:H4K20 me3
H3K4 me3:H3−5ac
H3K4 me1:H2A.Z
DNA Meth. m5C:H4K20 me3
H3K4 me1:H4−4ac
H3K4 me1:H4K20 me2
H2A.Z:H4K20 me2
H3K27 me3:H3K4 me1
H3K4 me1:H3−5ac
H3K27 me3:H3K9 me3
H4K20 me2:H3−5ac
H3K27 me3:DNA Meth. m5C
H4K20 me2:H4−4ac
H4−4ac:H3−5ac
H3K4 me3:H4−4ac
H4K20 me2:H3K4 me3
H2A.Z:H3K4 me3
H3K9 me2
H3K4 me1
H4K20 me3
H3K9 me3
H3K27 me3
H3K4 me3
H4−4ac
DNA Meth. m5C
H2A.Z
H3−5ac
H4K20 me2
0.00
0.25
0.50
0.75
1.00
π̂
TRA2B – Forward
H3K4 me3:H4−4ac
H4K20 me2:H3K4 me3
H2A.Z:H3−5ac
H3K4 me1:H4−4ac
H3K4 me1:H4K20 me2
H4−4ac:H3−5ac
H3K4 me1:H2A.Z
H3K9 me3:H4K20 me3
H2A.Z:H4K20 me2
H3K4 me3:H3−5ac
H3K27 me3:DNA Meth. m5C
H4K20 me2:H4−4ac
H3K4 me1:H3−5ac
H3K27 me3:H4K20 me3
H2A.Z:H3K4 me3
DNA Meth. m5C:H4K20 me3
DNA Meth. m5C:H3K9 me3
H3K27 me3:H3K9 me3
DNA Meth. m5C:H4K20 me2
H4K20 me2:H3−5ac
H3K27 me2
H3K4 me3
H4K20 me3
H3K27 me3
H3K9 me3
H4−4ac
H4K20 me2
H3K4 me1
H2A.Z
DNA Meth. m5C
H3−5ac
0.00
0.25
0.50
0.75
1.00
π̂
TRA2B – Reverse
H3K4 me3:H4−4ac
H4K20 me2:H3K4 me3
H2A.Z:H3−5ac
H3K4 me1:H4−4ac
H3K4 me1:H4K20 me2
H4−4ac:H3−5ac
H3K4 me1:H2A.Z
H3K9 me3:H4K20 me3
H2A.Z:H4K20 me2
H3K4 me3:H3−5ac
H3K27 me3:DNA Meth. m5C
H4K20 me2:H4−4ac
H3K4 me1:H3−5ac
H3K27 me3:H4K20 me3
H2A.Z:H3K4 me3
DNA Meth. m5C:H4K20 me3
DNA Meth. m5C:H3K9 me3
H3K27 me3:H3K9 me3
DNA Meth. m5C:H4K20 me2
H4K20 me2:H3−5ac
H3K27 me2
H3K4 me3
H4K20 me3
H3K27 me3
H3K9 me3
H4−4ac
H4K20 me2
H3K4 me1
H2A.Z
DNA Meth. m5C
H3−5ac
0.00
0.25
0.50
0.75
1.00
π̂

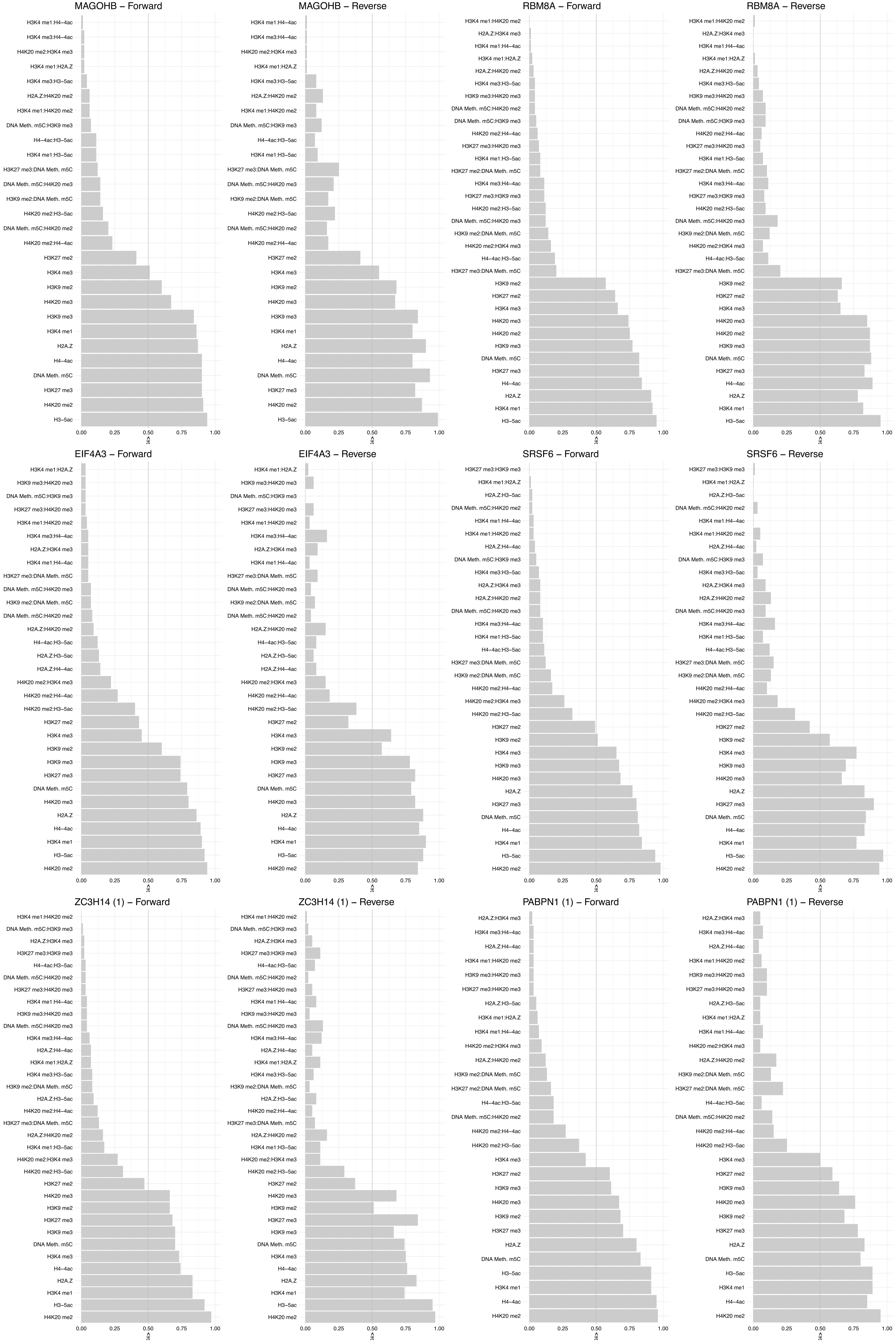
MAGOHB − Forward
H3K4 me1:H4−4ac
H3K4 me3:H4−4ac
H4K20 me2:H3K4 me3
H3K4 me1:H2A.Z
H3K4 me3:H3−5ac
H2A.Z:H4K20 me2
H3K4 me1:H4K20 me2
DNA Meth. m5C:H3K9 me3
H4−4ac:H3−5ac
H3K4 me1:H3−5ac
H3K27 me3:DNA Meth. m5C
DNA Meth. m5C:H4K20 me3
H3K9 me2:DNA Meth. m5C
H4K20 me2:H3−5ac
DNA Meth. m5C:H4K20 me2
H4K20 me2:H4−4ac
H3K27 me2
H3K4 me3
H3K9 me2
H4K20 me3
H3K9 me3
H3K4 me1
H2A.Z
H4−4ac
DNA Meth. m5C
H3K27 me3
H4K20 me2
H3−5ac
0.00 0.25 0.50 0.75 1.00
π̂
MAGOHB − Reverse
H3K4 me1:H4−4ac
H3K4 me3:H4−4ac
H4K20 me2:H3K4 me3
H3K4 me1:H2A.Z
H3K4 me3:H3−5ac
H2A.Z:H4K20 me2
H3K4 me1:H4K20 me2
DNA Meth. m5C:H3K9 me3
H4−4ac:H3−5ac
H3K4 me1:H3−5ac
H3K27 me3:DNA Meth. m5C
DNA Meth. m5C:H4K20 me3
H3K9 me2:DNA Meth. m5C
H4K20 me2:H3−5ac
DNA Meth. m5C:H4K20 me2
H4K20 me2:H4−4ac
H3K27 me2
H3K4 me3
H3K9 me2
H4K20 me3
H3K9 me3
H3K4 me1
H2A.Z
H4−4ac
DNA Meth. m5C
H3K27 me3
H4K20 me2
H3−5ac
0.00 0.25 0.50 0.75 1.00
π̂
RBM8A − Forward
H3K4 me1:H4K20 me2
H2A.Z:H3K4 me3
H3K4 me1:H4−4ac
H3K4 me1:H2A.Z
H2A.Z:H4K20 me2
H3K4 me3:H3−5ac
H3K9 me3:H4K20 me3
DNA Meth. m5C:H4K20 me2
DNA Meth. m5C:H3K9 me3
H4K20 me2:H4−4ac
H3K27 me3:H4K20 me3
H3K4 me1:H3−5ac
H3K27 me2:DNA Meth. m5C
H3K4 me3:H4−4ac
H3K27 me3:H3K9 me3
H4K20 me2:H3−5ac
DNA Meth. m5C:H4K20 me3
H3K9 me2:DNA Meth. m5C
H4K20 me2:H3K4 me3
H4−4ac:H3−5ac
H3K27 me3:DNA Meth. m5C
H3K9 me2
H3K27 me2
H3K4 me3
H4K20 me3
H4K20 me2
H3K9 me3
DNA Meth. m5C
H3K27 me3
H4−4ac
H2A.Z
H3K4 me1
H3−5ac
0.00 0.25 0.50 0.75 1.00
π̂
RBM8A − Reverse
H3K4 me1:H4K20 me2
H2A.Z:H3K4 me3
H3K4 me1:H4−4ac
H3K4 me1:H2A.Z
H2A.Z:H4K20 me2
H3K4 me3:H3−5ac
H3K9 me3:H4K20 me3
DNA Meth. m5C:H4K20 me2
DNA Meth. m5C:H3K9 me3
H4K20 me2:H4−4ac
H3K27 me3:H4K20 me3
H3K4 me1:H3−5ac
H3K27 me2:DNA Meth. m5C
H3K4 me3:H4−4ac
H3K27 me3:H3K9 me3
H4K20 me2:H3−5ac
DNA Meth. m5C:H4K20 me3
H3K9 me2:DNA Meth. m5C
H4K20 me2:H3K4 me3
H4−4ac:H3−5ac
H3K27 me3:DNA Meth. m5C
H3K9 me2
H3K27 me2
H3K4 me3
H4K20 me3
H4K20 me2
H3K9 me3
DNA Meth. m5C
H3K27 me3
H4−4ac
H2A.Z
H3K4 me1
H3−5ac
0.00 0.25 0.50 0.75 1.00
π̂
EIF4A3 − Forward
H3K4 me1:H2A.Z
H3K9 me3:H4K20 me3
DNA Meth. m5C:H3K9 me3
H3K27 me3:H4K20 me3
H3K4 me1:H4K20 me2
H3K4 me3:H4−4ac
H2A.Z:H3K4 me3
H3K4 me1:H4−4ac
H3K27 me3:DNA Meth. m5C
DNA Meth. m5C:H4K20 me3
H3K9 me2:DNA Meth. m5C
DNA Meth. m5C:H4K20 me2
H2A.Z:H4K20 me2
H4−4ac:H3−5ac
H2A.Z:H3−5ac
H2A.Z:H4−4ac
H4K20 me2:H3K4 me3
H4K20 me2:H4−4ac
H4K20 me2:H3−5ac
H3K27 me2
H3K4 me3
H3K9 me2
H3K9 me3
H3K27 me3
DNA Meth. m5C
H4K20 me3
H2A.Z
H4−4ac
H3K4 me1
H3−5ac
H4K20 me2
0.00 0.25 0.50 0.75 1.00
π̂
EIF4A3 − Reverse
H3K4 me1:H2A.Z
H3K9 me3:H4K20 me3
DNA Meth. m5C:H3K9 me3
H3K27 me3:H4K20 me3
H3K4 me1:H4K20 me2
H3K4 me3:H4−4ac
H2A.Z:H3K4 me3
H3K4 me1:H4−4ac
H3K27 me3:DNA Meth. m5C
DNA Meth. m5C:H4K20 me3
H3K9 me2:DNA Meth. m5C
DNA Meth. m5C:H4K20 me2
H2A.Z:H4K20 me2
H4−4ac:H3−5ac
H2A.Z:H3−5ac
H2A.Z:H4−4ac
H4K20 me2:H3K4 me3
H4K20 me2:H4−4ac
H4K20 me2:H3−5ac
H3K27 me2
H3K4 me3
H3K9 me2
H3K9 me3
H3K27 me3
DNA Meth. m5C
H4K20 me3
H2A.Z
H4−4ac
H3K4 me1
H3−5ac
H4K20 me2
0.00 0.25 0.50 0.75 1.00
π̂
SRSF6 − Forward
H3K27 me3:H3K9 me3
H3K4 me1:H2A.Z
H2A.Z:H3−5ac
DNA Meth. m5C:H4K20 me2
H3K4 me1:H4−4ac
H3K4 me1:H4K20 me2
H2A.Z:H4−4ac
DNA Meth. m5C:H3K9 me3
H3K4 me3:H3−5ac
H2A.Z:H3K4 me3
H2A.Z:H4K20 me2
DNA Meth. m5C:H4K20 me3
H3K4 me3:H4−4ac
H3K4 me1:H3−5ac
H4−4ac:H3−5ac
H3K27 me3:DNA Meth. m5C
H3K9 me2:DNA Meth. m5C
H4K20 me2:H4−4ac
H4K20 me2:H3K4 me3
H4K20 me2:H3−5ac
H3K27 me2
H3K9 me2
H3K4 me3
H3K9 me3
H4K20 me3
H2A.Z
H3K27 me3
DNA Meth. m5C
H4−4ac
H3K4 me1
H3−5ac
H4K20 me2
0.00 0.25 0.50 0.75 1.00
π̂
SRSF6 − Reverse
H3K27 me3:H3K9 me3
H3K4 me1:H2A.Z
H2A.Z:H3−5ac
DNA Meth. m5C:H4K20 me2
H3K4 me1:H4−4ac
H3K4 me1:H4K20 me2
H2A.Z:H4−4ac
DNA Meth. m5C:H3K9 me3
H3K4 me3:H3−5ac
H2A.Z:H3K4 me3
H2A.Z:H4K20 me2
DNA Meth. m5C:H4K20 me3
H3K4 me3:H4−4ac
H3K4 me1:H3−5ac
H4−4ac:H3−5ac
H3K27 me3:DNA Meth. m5C
H3K9 me2:DNA Meth. m5C
H4K20 me2:H4−4ac
H4K20 me2:H3K4 me3
H4K20 me2:H3−5ac
H3K27 me2
H3K9 me2
H3K4 me3
H3K9 me3
H4K20 me3
H2A.Z
H3K27 me3
DNA Meth. m5C
H4−4ac
H3K4 me1
H3−5ac
H4K20 me2
0.00 0.25 0.50 0.75 1.00
π̂
ZC3H14 (1) − Forward
H3K4 me1:H4K20 me2
DNA Meth. m5C:H3K9 me3
H2A.Z:H3K4 me3
H3K27 me3:H3K9 me3
H4−4ac:H3−5ac
DNA Meth. m5C:H4K20 me2
H3K27 me3:H4K20 me3
H3K4 me1:H4−4ac
H3K9 me3:H4K20 me3
DNA Meth. m5C:H4K20 me3
H3K4 me3:H4−4ac
H2A.Z:H4−4ac
H3K4 me1:H2A.Z
H3K4 me3:H3−5ac
H3K9 me2:DNA Meth. m5C
H2A.Z:H3−5ac
H4K20 me2:H4−4ac
H3K27 me3:DNA Meth. m5C
H2A.Z:H4K20 me2
H3K4 me1:H3−5ac
H4K20 me2:H3K4 me3
H4K20 me2:H3−5ac
H3K27 me2
H4K20 me3
H3K9 me2
H3K27 me3
H3K9 me3
DNA Meth. m5C
H3K4 me3
H4−4ac
H2A.Z
H3K4 me1
H3−5ac
H4K20 me2
0.00 0.25 0.50 0.75 1.00
π̂
ZC3H14 (1) − Reverse
H3K4 me1:H4K20 me2
DNA Meth. m5C:H3K9 me3
H2A.Z:H3K4 me3
H3K27 me3:H3K9 me3
H4−4ac:H3−5ac
DNA Meth. m5C:H4K20 me2
H3K27 me3:H4K20 me3
H3K4 me1:H4−4ac
H3K9 me3:H4K20 me3
DNA Meth. m5C:H4K20 me3
H3K4 me3:H4−4ac
H2A.Z:H4−4ac
H3K4 me1:H2A.Z
H3K4 me3:H3−5ac
H3K9 me2:DNA Meth. m5C
H2A.Z:H3−5ac
H4K20 me2:H4−4ac
H3K27 me3:DNA Meth. m5C
H2A.Z:H4K20 me2
H3K4 me1:H3−5ac
H4K20 me2:H3K4 me3
H4K20 me2:H3−5ac
H3K27 me2
H4K20 me3
H3K9 me2
H3K27 me3
H3K9 me3
DNA Meth. m5C
H3K4 me3
H4−4ac
H2A.Z
H3K4 me1
H3−5ac
H4K20 me2
0.00 0.25 0.50 0.75 1.00
π̂
PABPN1 (1) − Forward
H2A.Z:H3K4 me3
H3K4 me3:H4−4ac
H2A.Z:H4−4ac
H3K4 me1:H4K20 me2
H3K9 me3:H4K20 me3
H3K27 me3:H4K20 me3
H2A.Z:H3−5ac
H3K4 me1:H2A.Z
H3K4 me1:H4−4ac
H4K20 me2:H3K4 me3
H2A.Z:H4K20 me2
H3K9 me2:DNA Meth. m5C
H3K27 me2:DNA Meth. m5C
H4−4ac:H3−5ac
DNA Meth. m5C:H4K20 me2
H4K20 me2:H4−4ac
H4K20 me2:H3−5ac
H3K4 me3
H3K27 me2
H3K9 me3
H4K20 me3
H3K9 me2
H3K27 me3
H2A.Z
DNA Meth. m5C
H3−5ac
H3K4 me1
H4−4ac
H4K20 me2
0.00 0.25 0.50 0.75 1.00
π̂
PABPN1 (1) − Reverse
H2A.Z:H3K4 me3
H3K4 me3:H4−4ac
H2A.Z:H4−4ac
H3K4 me1:H4K20 me2
H3K9 me3:H4K20 me3
H3K27 me3:H4K20 me3
H2A.Z:H3−5ac
H3K4 me1:H2A.Z
H3K4 me1:H4−4ac
H4K20 me2:H3K4 me3
H2A.Z:H4K20 me2
H3K9 me2:DNA Meth. m5C
H3K27 me2:DNA Meth. m5C
H4−4ac:H3−5ac
DNA Meth. m5C:H4K20 me2
H4K20 me2:H4−4ac
H4K20 me2:H3−5ac
H3K4 me3
H3K27 me2
H3K9 me3
H4K20 me3
H3K9 me2
H3K27 me3
H2A.Z
DNA Meth. m5C
H3−5ac
H3K4 me1
H4−4ac
H4K20 me2
0.00 0.25 0.50 0.75 1.00
π̂

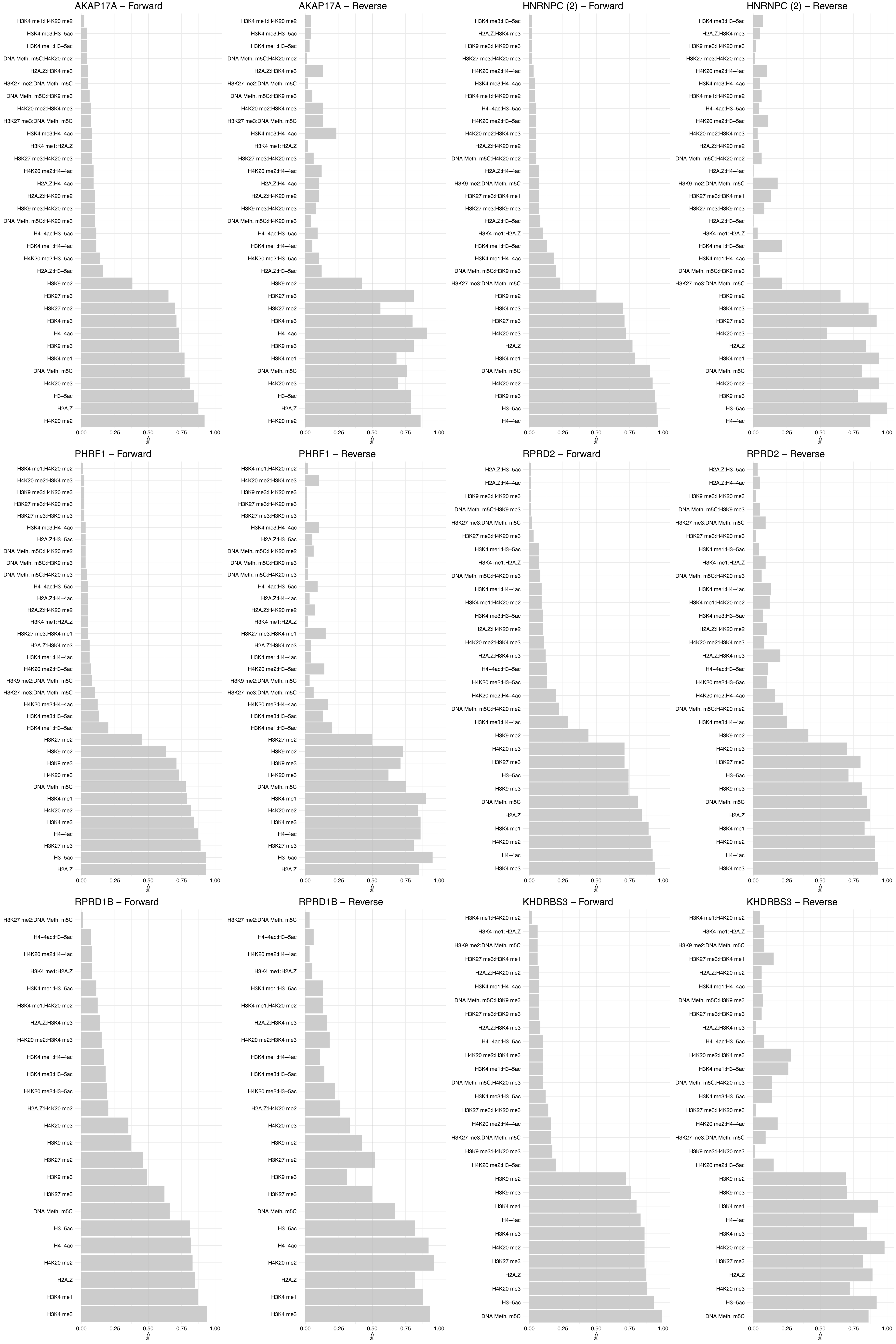

AKAP17A – Forward
H3K4 me1:H4K20 me2
H3K4 me3:H3–5ac
H3K4 me1:H3–5ac
DNA Meth. m5C:H4K20 me2
H2A.Z:H3K4 me3
H3K27 me2:DNA Meth. m5C
DNA Meth. m5C:H3K9 me3
H4K20 me2:H3K4 me3
H3K27 me3:DNA Meth. m5C
H3K4 me3:H4–4ac
H3K4 me1:H2A.Z
H3K27 me3:H4K20 me3
H4K20 me2:H4–4ac
H2A.Z:H4–4ac
H2A.Z:H4K20 me2
H3K9 me3:H4K20 me3
DNA Meth. m5C:H4K20 me3
H4–4ac:H3–5ac
H3K4 me1:H4–4ac
H4K20 me2:H3–5ac
H2A.Z:H3–5ac
H3K9 me2
H3K27 me3
H3K27 me2
H3K4 me3
H4–4ac
H3K9 me3
H3K4 me1
DNA Meth. m5C
H4K20 me3
H3–5ac
H2A.Z
H4K20 me2
0.00
0.25
0.50
0.75
1.00
$\hat{\pi}$
AKAP17A – Reverse
H3K4 me1:H4K20 me2
H3K4 me3:H3–5ac
H3K4 me1:H3–5ac
DNA Meth. m5C:H4K20 me2
H2A.Z:H3K4 me3
H3K27 me2:DNA Meth. m5C
DNA Meth. m5C:H3K9 me3
H4K20 me2:H3K4 me3
H3K27 me3:DNA Meth. m5C
H3K4 me3:H4–4ac
H3K4 me1:H2A.Z
H3K27 me3:H4K20 me3
H4K20 me2:H4–4ac
H2A.Z:H4–4ac
H2A.Z:H4K20 me2
H3K9 me3:H4K20 me3
DNA Meth. m5C:H4K20 me3
H4–4ac:H3–5ac
H3K4 me1:H4–4ac
H4K20 me2:H3–5ac
H2A.Z:H3–5ac
H3K9 me2
H3K27 me3
H3K27 me2
H3K4 me3
H4–4ac
H3K9 me3
H3K4 me1
DNA Meth. m5C
H4K20 me3
H3–5ac
H2A.Z
H4K20 me2
0.00
0.25
0.50
0.75
1.00
$\hat{\pi}$
HNRNPC (2) – Forward
H3K4 me3:H3–5ac
H2A.Z:H3K4 me3
H3K9 me3:H4K20 me3
H3K27 me3:H4K20 me3
H4K20 me2:H4–4ac
H3K4 me3:H4–4ac
H3K4 me1:H4K20 me2
H4–4ac:H3–5ac
H4K20 me2:H3–5ac
H4K20 me2:H3K4 me3
H2A.Z:H4K20 me2
DNA Meth. m5C:H4K20 me2
H2A.Z:H4–4ac
H3K9 me2:DNA Meth. m5C
H3K27 me3:H3K4 me1
H3K27 me3:H3K9 me3
H2A.Z:H3–5ac
H3K4 me1:H2A.Z
H3K4 me1:H3–5ac
H3K4 me1:H4–4ac
DNA Meth. m5C:H3K9 me3
H3K27 me3:DNA Meth. m5C
H3K9 me2
H3K4 me3
H3K27 me3
H4K20 me3
H2A.Z
H3K4 me1
DNA Meth. m5C
H4K20 me2
H3K9 me3
H3–5ac
H4–4ac
0.00
0.25
0.50
0.75
1.00
$\hat{\pi}$
HNRNPC (2) – Reverse
H3K4 me3:H3–5ac
H2A.Z:H3K4 me3
H3K9 me3:H4K20 me3
H3K27 me3:H4K20 me3
H4K20 me2:H4–4ac
H3K4 me3:H4–4ac
H3K4 me1:H4K20 me2
H4–4ac:H3–5ac
H4K20 me2:H3–5ac
H4K20 me2:H3K4 me3
H2A.Z:H4K20 me2
DNA Meth. m5C:H4K20 me2
H2A.Z:H4–4ac
H3K9 me2:DNA Meth. m5C
H3K27 me3:H3K4 me1
H3K27 me3:H3K9 me3
H2A.Z:H3–5ac
H3K4 me1:H2A.Z
H3K4 me1:H3–5ac
H3K4 me1:H4–4ac
DNA Meth. m5C:H3K9 me3
H3K27 me3:DNA Meth. m5C
H3K9 me2
H3K4 me3
H3K27 me3
H4K20 me3
H2A.Z
H3K4 me1
DNA Meth. m5C
H4K20 me2
H3K9 me3
H3–5ac
H4–4ac
0.00
0.25
0.50
0.75
1.00
$\hat{\pi}$
PHRF1 – Forward
H3K4 me1:H4K20 me2
H4K20 me2:H3K4 me3
H3K9 me3:H4K20 me3
H3K27 me3:H4K20 me3
H3K27 me3:H3K9 me3
H3K4 me3:H4–4ac
H2A.Z:H3–5ac
DNA Meth. m5C:H4K20 me2
DNA Meth. m5C:H3K9 me3
DNA Meth. m5C:H4K20 me3
H4–4ac:H3–5ac
H2A.Z:H4–4ac
H2A.Z:H4K20 me2
H3K4 me1:H2A.Z
H3K27 me3:H3K4 me1
H2A.Z:H3K4 me3
H3K4 me1:H4–4ac
H4K20 me2:H3–5ac
H3K9 me2:DNA Meth. m5C
H3K27 me3:DNA Meth. m5C
H4K20 me2:H4–4ac
H3K4 me3:H3–5ac
H3K4 me1:H3–5ac
H3K27 me2
H3K9 me2
H3K9 me3
H4K20 me3
DNA Meth. m5C
H3K4 me1
H4K20 me2
H3K4 me3
H4–4ac
H3K27 me3
H3–5ac
H2A.Z
0.00
0.25
0.50
0.75
1.00
$\hat{\pi}$
PHRF1 – Reverse
H3K4 me1:H4K20 me2
H4K20 me2:H3K4 me3
H3K9 me3:H4K20 me3
H3K27 me3:H4K20 me3
H3K27 me3:H3K9 me3
H3K4 me3:H4–4ac
H2A.Z:H3–5ac
DNA Meth. m5C:H4K20 me2
DNA Meth. m5C:H3K9 me3
DNA Meth. m5C:H4K20 me3
H4–4ac:H3–5ac
H2A.Z:H4–4ac
H2A.Z:H4K20 me2
H3K4 me1:H2A.Z
H3K27 me3:H3K4 me1
H2A.Z:H3K4 me3
H3K4 me1:H4–4ac
H4K20 me2:H3–5ac
H3K9 me2:DNA Meth. m5C
H3K27 me3:DNA Meth. m5C
H4K20 me2:H4–4ac
H3K4 me3:H3–5ac
H3K4 me1:H3–5ac
H3K27 me2
H3K9 me2
H3K9 me3
H4K20 me3
DNA Meth. m5C
H3K4 me1
H4K20 me2
H3K4 me3
H4–4ac
H3K27 me3
H3–5ac
H2A.Z
0.00
0.25
0.50
0.75
1.00
$\hat{\pi}$
RPRD2 – Forward
H2A.Z:H3–5ac
H2A.Z:H4–4ac
H3K9 me3:H4K20 me3
DNA Meth. m5C:H3K9 me3
H3K27 me3:DNA Meth. m5C
H3K27 me3:H4K20 me3
H3K4 me1:H3–5ac
H3K4 me1:H2A.Z
DNA Meth. m5C:H4K20 me3
H3K4 me1:H4–4ac
H3K4 me1:H4K20 me2
H3K4 me3:H3–5ac
H2A.Z:H4K20 me2
H4K20 me2:H3K4 me3
H2A.Z:H3K4 me3
H4–4ac:H3–5ac
H4K20 me2:H3–5ac
H4K20 me2:H4–4ac
DNA Meth. m5C:H4K20 me2
H3K4 me3:H4–4ac
H3K9 me2
H4K20 me3
H3K27 me3
H3–5ac
H3K9 me3
DNA Meth. m5C
H2A.Z
H3K4 me1
H4K20 me2
H4–4ac
H3K4 me3
0.00
0.25
0.50
0.75
1.00
$\hat{\pi}$
RPRD2 – Reverse
H2A.Z:H3–5ac
H2A.Z:H4–4ac
H3K9 me3:H4K20 me3
DNA Meth. m5C:H3K9 me3
H3K27 me3:DNA Meth. m5C
H3K27 me3:H4K20 me3
H3K4 me1:H3–5ac
H3K4 me1:H2A.Z
DNA Meth. m5C:H4K20 me3
H3K4 me1:H4–4ac
H3K4 me1:H4K20 me2
H3K4 me3:H3–5ac
H2A.Z:H4K20 me2
H4K20 me2:H3K4 me3
H2A.Z:H3K4 me3
H4–4ac:H3–5ac
H4K20 me2:H3–5ac
H4K20 me2:H4–4ac
DNA Meth. m5C:H4K20 me2
H3K4 me3:H4–4ac
H3K9 me2
H4K20 me3
H3K27 me3
H3–5ac
H3K9 me3
DNA Meth. m5C
H2A.Z
H3K4 me1
H4K20 me2
H4–4ac
H3K4 me3
0.00
0.25
0.50
0.75
1.00
$\hat{\pi}$
RPRD1B – Forward
H3K27 me2:DNA Meth. m5C
H4–4ac:H3–5ac
H4K20 me2:H4–4ac
H3K4 me1:H2A.Z
H3K4 me1:H3–5ac
H3K4 me1:H4K20 me2
H2A.Z:H3K4 me3
H4K20 me2:H3K4 me3
H3K4 me1:H4–4ac
H3K4 me3:H3–5ac
H4K20 me2:H3–5ac
H2A.Z:H4K20 me2
H4K20 me3
H3K9 me2
H3K27 me2
H3K9 me3
H3K27 me3
DNA Meth. m5C
H3–5ac
H4–4ac
H4K20 me2
H2A.Z
H3K4 me1
H3K4 me3
0.00
0.25
0.50
0.75
1.00
$\hat{\pi}$
RPRD1B – Reverse
H3K27 me2:DNA Meth. m5C
H4–4ac:H3–5ac
H4K20 me2:H4–4ac
H3K4 me1:H2A.Z
H3K4 me1:H3–5ac
H3K4 me1:H4K20 me2
H2A.Z:H3K4 me3
H4K20 me2:H3K4 me3
H3K4 me1:H4–4ac
H3K4 me3:H3–5ac
H4K20 me2:H3–5ac
H2A.Z:H4K20 me2
H4K20 me3
H3K9 me2
H3K27 me2
H3K9 me3
H3K27 me3
DNA Meth. m5C
H3–5ac
H4–4ac
H4K20 me2
H2A.Z
H3K4 me1
H3K4 me3
0.00
0.25
0.50
0.75
1.00
$\hat{\pi}$
KHDRBS3 – Forward
H3K4 me1:H4K20 me2
H3K4 me1:H2A.Z
H3K9 me2:DNA Meth. m5C
H3K27 me3:H3K4 me1
H2A.Z:H4K20 me2
H3K4 me1:H4–4ac
DNA Meth. m5C:H3K9 me3
H3K27 me3:H3K9 me3
H2A.Z:H3K4 me3
H4–4ac:H3–5ac
H4K20 me2:H3K4 me3
H3K4 me1:H3–5ac
DNA Meth. m5C:H4K20 me3
H3K4 me3:H3–5ac
H3K27 me3:H4K20 me3
H4K20 me2:H4–4ac
H3K27 me3:DNA Meth. m5C
H3K9 me3:H4K20 me3
H4K20 me2:H3–5ac
H3K9 me2
H3K9 me3
H3K4 me1
H4–4ac
H3K4 me3
H4K20 me2
H3K27 me3
H2A.Z
H4K20 me3
H3–5ac
DNA Meth. m5C
0.00
0.25
0.50
0.75
1.00
$\hat{\pi}$
KHDRBS3 – Reverse
H3K4 me1:H4K20 me2
H3K4 me1:H2A.Z
H3K9 me2:DNA Meth. m5C
H3K27 me3:H3K4 me1
H2A.Z:H4K20 me2
H3K4 me1:H4–4ac
DNA Meth. m5C:H3K9 me3
H3K27 me3:H3K9 me3
H2A.Z:H3K4 me3
H4–4ac:H3–5ac
H4K20 me2:H3K4 me3
H3K4 me1:H3–5ac
DNA Meth. m5C:H4K20 me3
H3K4 me3:H3–5ac
H3K27 me3:H4K20 me3
H4K20 me2:H4–4ac
H3K27 me3:DNA Meth. m5C
H3K9 me3:H4K20 me3
H4K20 me2:H3–5ac
H3K9 me2
H3K9 me3
H3K4 me1
H4–4ac
H3K4 me3
H4K20 me2
H3K27 me3
H2A.Z
H4K20 me3
H3–5ac
DNA Meth. m5C
0.00
0.25
0.50
0.75
1.00
$\hat{\pi}$

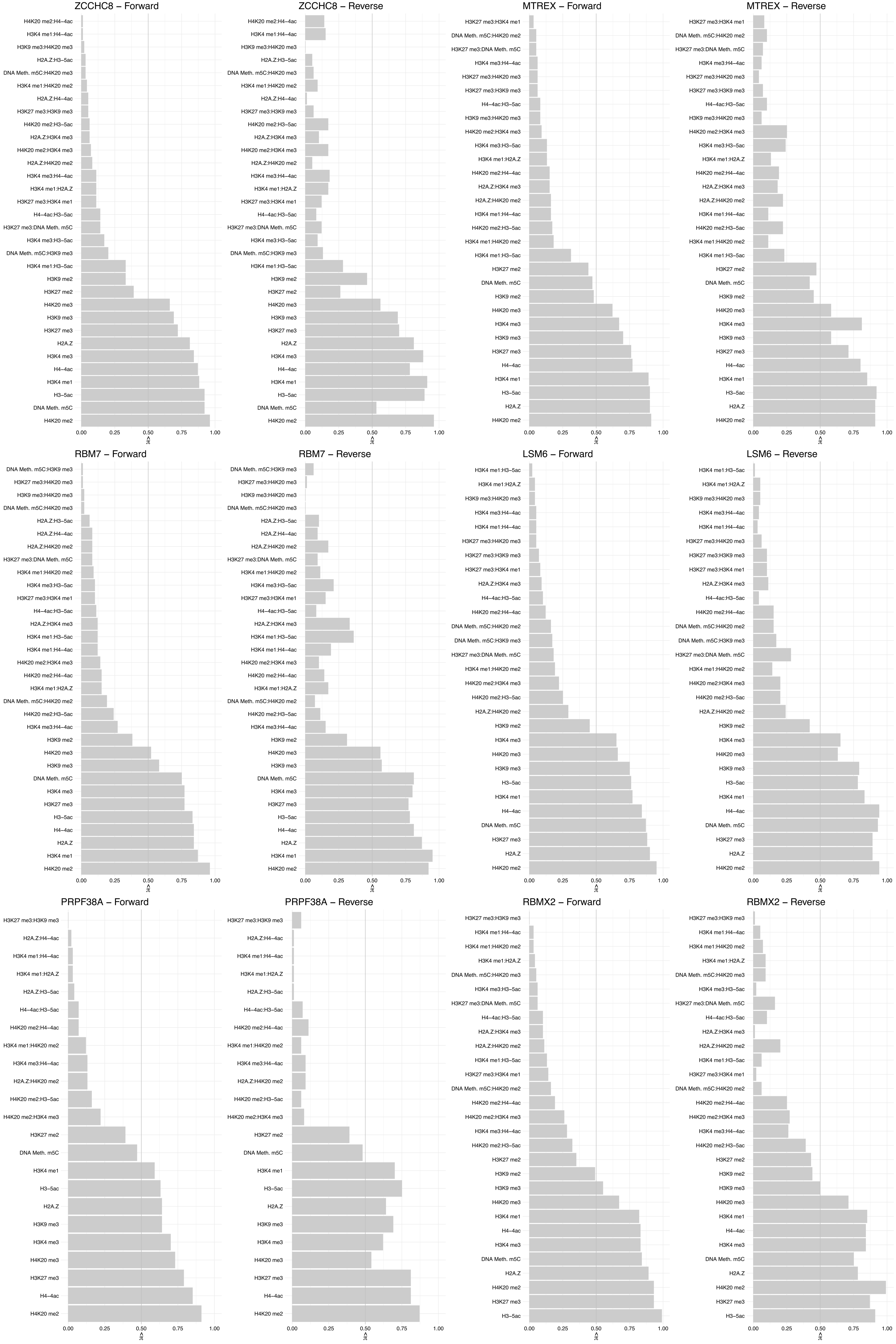

ZCCHC8 – Forward
H4K20 me2:H4–4ac
H3K4 me1:H4–4ac
H3K9 me3:H4K20 me3
H2A.Z:H3–5ac
DNA Meth. m5C:H4K20 me3
H3K4 me1:H4K20 me2
H2A.Z:H4–4ac
H3K27 me3:H3K9 me3
H4K20 me2:H3–5ac
H2A.Z:H3K4 me3
H4K20 me2:H3K4 me3
H2A.Z:H4K20 me2
H3K4 me3:H4–4ac
H3K4 me1:H2A.Z
H3K27 me3:H3K4 me1
H4–4ac:H3–5ac
H3K27 me3:DNA Meth. m5C
H3K4 me3:H3–5ac
DNA Meth. m5C:H3K9 me3
H3K4 me1:H3–5ac
H3K9 me2
H3K27 me2
H4K20 me3
H3K9 me3
H3K27 me3
H2A.Z
H3K4 me3
H4–4ac
H3K4 me1
H3–5ac
DNA Meth. m5C
H4K20 me2
0.00
0.25
0.50
0.75
1.00
π̂
ZCCHC8 – Reverse
H4K20 me2:H4–4ac
H3K4 me1:H4–4ac
H3K9 me3:H4K20 me3
H2A.Z:H3–5ac
DNA Meth. m5C:H4K20 me3
H3K4 me1:H4K20 me2
H2A.Z:H4–4ac
H3K27 me3:H3K9 me3
H4K20 me2:H3–5ac
H2A.Z:H3K4 me3
H4K20 me2:H3K4 me3
H2A.Z:H4K20 me2
H3K4 me3:H4–4ac
H3K4 me1:H2A.Z
H3K27 me3:H3K4 me1
H4–4ac:H3–5ac
H3K27 me3:DNA Meth. m5C
H3K4 me3:H3–5ac
DNA Meth. m5C:H3K9 me3
H3K4 me1:H3–5ac
H3K9 me2
H3K27 me2
H4K20 me3
H3K9 me3
H3K27 me3
H2A.Z
H3K4 me3
H4–4ac
H3K4 me1
H3–5ac
DNA Meth. m5C
H4K20 me2
0.00
0.25
0.50
0.75
1.00
π̂
MTREX – Forward
H3K27 me3:H3K4 me1
DNA Meth. m5C:H4K20 me2
H3K27 me3:DNA Meth. m5C
H3K4 me3:H4–4ac
H3K27 me3:H4K20 me3
H3K27 me3:H3K9 me3
H4–4ac:H3–5ac
H3K9 me3:H4K20 me3
H4K20 me2:H3K4 me3
H3K4 me3:H3–5ac
H3K4 me1:H2A.Z
H4K20 me2:H4–4ac
H2A.Z:H3K4 me3
H2A.Z:H4K20 me2
H3K4 me1:H4–4ac
H4K20 me2:H3–5ac
H3K4 me1:H4K20 me2
H3K4 me1:H3–5ac
H3K27 me2
DNA Meth. m5C
H3K9 me2
H4K20 me3
H3K4 me3
H3K9 me3
H3K27 me3
H4–4ac
H3K4 me1
H3–5ac
H2A.Z
H4K20 me2
0.00
0.25
0.50
0.75
1.00
π̂
MTREX – Reverse
H3K27 me3:H3K4 me1
DNA Meth. m5C:H4K20 me2
H3K27 me3:DNA Meth. m5C
H3K4 me3:H4–4ac
H3K27 me3:H4K20 me3
H3K27 me3:H3K9 me3
H4–4ac:H3–5ac
H3K9 me3:H4K20 me3
H4K20 me2:H3K4 me3
H3K4 me3:H3–5ac
H3K4 me1:H2A.Z
H4K20 me2:H4–4ac
H2A.Z:H3K4 me3
H2A.Z:H4K20 me2
H3K4 me1:H4–4ac
H4K20 me2:H3–5ac
H3K4 me1:H4K20 me2
H3K4 me1:H3–5ac
H3K27 me2
DNA Meth. m5C
H3K9 me2
H4K20 me3
H3K4 me3
H3K9 me3
H3K27 me3
H4–4ac
H3K4 me1
H3–5ac
H2A.Z
H4K20 me2
0.00
0.25
0.50
0.75
1.00
π̂
RBM7 – Forward
DNA Meth. m5C:H3K9 me3
H3K27 me3:H4K20 me3
H3K9 me3:H4K20 me3
DNA Meth. m5C:H4K20 me3
H2A.Z:H3–5ac
H2A.Z:H4–4ac
H2A.Z:H4K20 me2
H3K27 me3:DNA Meth. m5C
H3K4 me1:H4K20 me2
H3K4 me3:H3–5ac
H3K27 me3:H3K4 me1
H4–4ac:H3–5ac
H2A.Z:H3K4 me3
H3K4 me1:H3–5ac
H3K4 me1:H4–4ac
H4K20 me2:H3K4 me3
H4K20 me2:H4–4ac
H3K4 me1:H2A.Z
DNA Meth. m5C:H4K20 me2
H4K20 me2:H3–5ac
H3K4 me3:H4–4ac
H3K9 me2
H4K20 me3
H3K9 me3
DNA Meth. m5C
H3K4 me3
H3K27 me3
H3–5ac
H4–4ac
H2A.Z
H3K4 me1
H4K20 me2
0.00
0.25
0.50
0.75
1.00
π̂
RBM7 – Reverse
DNA Meth. m5C:H3K9 me3
H3K27 me3:H4K20 me3
H3K9 me3:H4K20 me3
DNA Meth. m5C:H4K20 me3
H2A.Z:H3–5ac
H2A.Z:H4–4ac
H2A.Z:H4K20 me2
H3K27 me3:DNA Meth. m5C
H3K4 me1:H4K20 me2
H3K4 me3:H3–5ac
H3K27 me3:H3K4 me1
H4–4ac:H3–5ac
H2A.Z:H3K4 me3
H3K4 me1:H3–5ac
H3K4 me1:H4–4ac
H4K20 me2:H3K4 me3
H4K20 me2:H4–4ac
H3K4 me1:H2A.Z
DNA Meth. m5C:H4K20 me2
H4K20 me2:H3–5ac
H3K4 me3:H4–4ac
H3K9 me2
H4K20 me3
H3K9 me3
DNA Meth. m5C
H3K4 me3
H3K27 me3
H3–5ac
H4–4ac
H2A.Z
H3K4 me1
H4K20 me2
0.00
0.25
0.50
0.75
1.00
π̂
LSM6 – Forward
H3K4 me1:H3–5ac
H3K4 me1:H2A.Z
H3K9 me3:H4K20 me3
H3K4 me3:H4–4ac
H3K4 me1:H4–4ac
H3K27 me3:H4K20 me3
H3K27 me3:H3K9 me3
H3K27 me3:H3K4 me1
H2A.Z:H3K4 me3
H4–4ac:H3–5ac
H4K20 me2:H4–4ac
DNA Meth. m5C:H4K20 me2
DNA Meth. m5C:H3K9 me3
H3K27 me3:DNA Meth. m5C
H3K4 me1:H4K20 me2
H4K20 me2:H3K4 me3
H4K20 me2:H3–5ac
H2A.Z:H4K20 me2
H3K9 me2
H3K4 me3
H4K20 me3
H3K9 me3
H3–5ac
H3K4 me1
H4–4ac
DNA Meth. m5C
H3K27 me3
H2A.Z
H4K20 me2
0.00
0.25
0.50
0.75
1.00
π̂
LSM6 – Reverse
H3K4 me1:H3–5ac
H3K4 me1:H2A.Z
H3K9 me3:H4K20 me3
H3K4 me3:H4–4ac
H3K4 me1:H4–4ac
H3K27 me3:H4K20 me3
H3K27 me3:H3K9 me3
H3K27 me3:H3K4 me1
H2A.Z:H3K4 me3
H4–4ac:H3–5ac
H4K20 me2:H4–4ac
DNA Meth. m5C:H4K20 me2
DNA Meth. m5C:H3K9 me3
H3K27 me3:DNA Meth. m5C
H3K4 me1:H4K20 me2
H4K20 me2:H3K4 me3
H4K20 me2:H3–5ac
H2A.Z:H4K20 me2
H3K9 me2
H3K4 me3
H4K20 me3
H3K9 me3
H3–5ac
H3K4 me1
H4–4ac
DNA Meth. m5C
H3K27 me3
H2A.Z
H4K20 me2
0.00
0.25
0.50
0.75
1.00
π̂
PRPF38A – Forward
H3K27 me3:H3K9 me3
H2A.Z:H4–4ac
H3K4 me1:H4–4ac
H3K4 me1:H2A.Z
H2A.Z:H3–5ac
H4–4ac:H3–5ac
H4K20 me2:H4–4ac
H3K4 me1:H4K20 me2
H3K4 me3:H4–4ac
H2A.Z:H4K20 me2
H4K20 me2:H3–5ac
H4K20 me2:H3K4 me3
H3K27 me2
DNA Meth. m5C
H3K4 me1
H3–5ac
H2A.Z
H3K9 me3
H3K4 me3
H4K20 me3
H3K27 me3
H4–4ac
H4K20 me2
0.00
0.25
0.50
0.75
1.00
π̂
PRPF38A – Reverse
H3K27 me3:H3K9 me3
H2A.Z:H4–4ac
H3K4 me1:H4–4ac
H3K4 me1:H2A.Z
H2A.Z:H3–5ac
H4–4ac:H3–5ac
H4K20 me2:H4–4ac
H3K4 me1:H4K20 me2
H3K4 me3:H4–4ac
H2A.Z:H4K20 me2
H4K20 me2:H3–5ac
H4K20 me2:H3K4 me3
H3K27 me2
DNA Meth. m5C
H3K4 me1
H3–5ac
H2A.Z
H3K9 me3
H3K4 me3
H4K20 me3
H3K27 me3
H4–4ac
H4K20 me2
0.00
0.25
0.50
0.75
1.00
π̂
RBMX2 – Forward
H3K27 me3:H3K9 me3
H3K4 me1:H4–4ac
H3K4 me1:H4K20 me2
H3K4 me1:H2A.Z
DNA Meth. m5C:H4K20 me3
H3K4 me3:H3–5ac
H3K27 me3:DNA Meth. m5C
H4–4ac:H3–5ac
H2A.Z:H3K4 me3
H2A.Z:H4K20 me2
H3K4 me1:H3–5ac
H3K27 me3:H3K4 me1
DNA Meth. m5C:H4K20 me2
H4K20 me2:H4–4ac
H4K20 me2:H3K4 me3
H3K4 me3:H4–4ac
H4K20 me2:H3–5ac
H3K27 me2
H3K9 me2
H3K9 me3
H4K20 me3
H3K4 me1
H4–4ac
H3K4 me3
DNA Meth. m5C
H2A.Z
H4K20 me2
H3K27 me3
H3–5ac
0.00
0.25
0.50
0.75
1.00
π̂
RBMX2 – Reverse
H3K27 me3:H3K9 me3
H3K4 me1:H4–4ac
H3K4 me1:H4K20 me2
H3K4 me1:H2A.Z
DNA Meth. m5C:H4K20 me3
H3K4 me3:H3–5ac
H3K27 me3:DNA Meth. m5C
H4–4ac:H3–5ac
H2A.Z:H3K4 me3
H2A.Z:H4K20 me2
H3K4 me1:H3–5ac
H3K27 me3:H3K4 me1
DNA Meth. m5C:H4K20 me2
H4K20 me2:H4–4ac
H4K20 me2:H3K4 me3
H3K4 me3:H4–4ac
H4K20 me2:H3–5ac
H3K27 me2
H3K9 me2
H3K9 me3
H4K20 me3
H3K4 me1
H4–4ac
H3K4 me3
DNA Meth. m5C
H2A.Z
H4K20 me2
H3K27 me3
H3–5ac
0.00
0.25
0.50
0.75
1.00
π̂

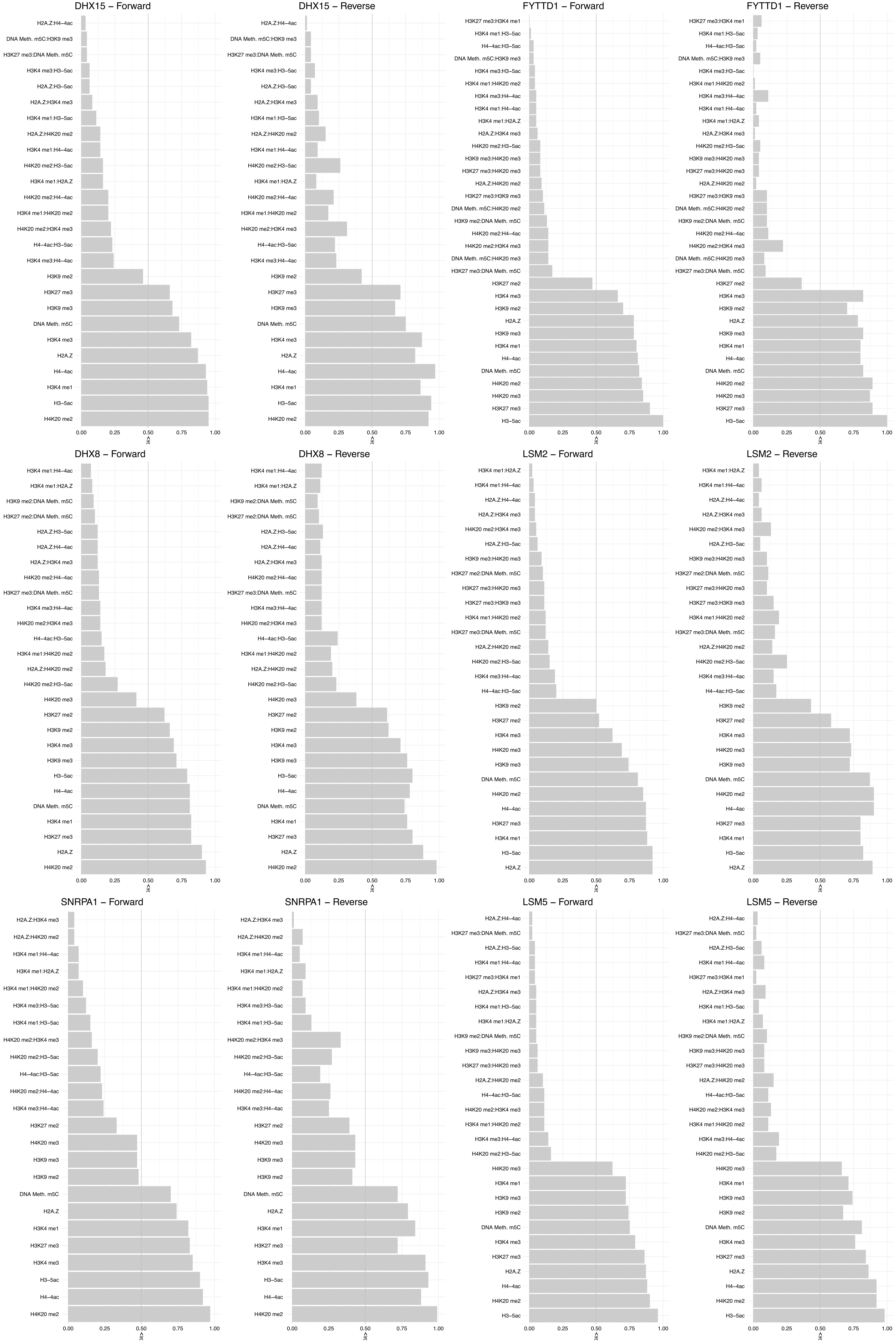
DHX15 – Forward
H2A.Z:H4−4ac
DNA Meth. m5C:H3K9 me3
H3K27 me3:DNA Meth. m5C
H3K4 me3:H3−5ac
H2A.Z:H3−5ac
H2A.Z:H3K4 me3
H3K4 me1:H3−5ac
H2A.Z:H4K20 me2
H3K4 me1:H4−4ac
H4K20 me2:H3−5ac
H3K4 me1:H2A.Z
H4K20 me2:H4−4ac
H3K4 me1:H4K20 me2
H4K20 me2:H3K4 me3
H4−4ac:H3−5ac
H3K4 me3:H4−4ac
H3K9 me2
H3K27 me3
H3K9 me3
DNA Meth. m5C
H3K4 me3
H2A.Z
H4−4ac
H3K4 me1
H3−5ac
H4K20 me2
0.00 0.25 0.50 0.75 1.00
π̂
DHX15 – Reverse
H2A.Z:H4−4ac
DNA Meth. m5C:H3K9 me3
H3K27 me3:DNA Meth. m5C
H3K4 me3:H3−5ac
H2A.Z:H3−5ac
H2A.Z:H3K4 me3
H3K4 me1:H3−5ac
H2A.Z:H4K20 me2
H3K4 me1:H4−4ac
H4K20 me2:H3−5ac
H3K4 me1:H2A.Z
H4K20 me2:H4−4ac
H3K4 me1:H4K20 me2
H4K20 me2:H3K4 me3
H4−4ac:H3−5ac
H3K4 me3:H4−4ac
H3K9 me2
H3K27 me3
H3K9 me3
DNA Meth. m5C
H3K4 me3
H2A.Z
H4−4ac
H3K4 me1
H3−5ac
H4K20 me2
0.00 0.25 0.50 0.75 1.00
π̂
FYTTD1 – Forward
H3K27 me3:H3K4 me1
H3K4 me1:H3−5ac
H4−4ac:H3−5ac
DNA Meth. m5C:H3K9 me3
H3K4 me3:H3−5ac
H3K4 me1:H4K20 me2
H3K4 me3:H4−4ac
H3K4 me1:H4−4ac
H3K4 me1:H2A.Z
H2A.Z:H3K4 me3
H4K20 me2:H3−5ac
H3K9 me3:H4K20 me3
H3K27 me3:H4K20 me3
H2A.Z:H4K20 me2
H3K27 me3:H3K9 me3
DNA Meth. m5C:H4K20 me2
H3K9 me2:DNA Meth. m5C
H4K20 me2:H4−4ac
H4K20 me2:H3K4 me3
DNA Meth. m5C:H4K20 me3
H3K27 me3:DNA Meth. m5C
H3K27 me2
H3K4 me3
H3K9 me2
H2A.Z
H3K9 me3
H3K4 me1
H4−4ac
DNA Meth. m5C
H4K20 me2
H4K20 me3
H3K27 me3
H3−5ac
0.00 0.25 0.50 0.75 1.00
π̂
FYTTD1 – Reverse
H3K27 me3:H3K4 me1
H3K4 me1:H3−5ac
H4−4ac:H3−5ac
DNA Meth. m5C:H3K9 me3
H3K4 me3:H3−5ac
H3K4 me1:H4K20 me2
H3K4 me3:H4−4ac
H3K4 me1:H4−4ac
H3K4 me1:H2A.Z
H2A.Z:H3K4 me3
H4K20 me2:H3−5ac
H3K9 me3:H4K20 me3
H3K27 me3:H4K20 me3
H2A.Z:H4K20 me2
H3K27 me3:H3K9 me3
DNA Meth. m5C:H4K20 me2
H3K9 me2:DNA Meth. m5C
H4K20 me2:H4−4ac
H4K20 me2:H3K4 me3
DNA Meth. m5C:H4K20 me3
H3K27 me3:DNA Meth. m5C
H3K27 me2
H3K4 me3
H3K9 me2
H2A.Z
H3K9 me3
H3K4 me1
H4−4ac
DNA Meth. m5C
H4K20 me2
H4K20 me3
H3K27 me3
H3−5ac
0.00 0.25 0.50 0.75 1.00
π̂
DHX8 – Forward
H3K4 me1:H4−4ac
H3K4 me1:H2A.Z
H3K9 me2:DNA Meth. m5C
H3K27 me2:DNA Meth. m5C
H2A.Z:H3−5ac
H2A.Z:H4−4ac
H2A.Z:H3K4 me3
H4K20 me2:H4−4ac
H3K27 me3:DNA Meth. m5C
H3K4 me3:H4−4ac
H4K20 me2:H3K4 me3
H4−4ac:H3−5ac
H3K4 me1:H4K20 me2
H2A.Z:H4K20 me2
H4K20 me2:H3−5ac
H4K20 me3
H3K27 me2
H3K9 me2
H3K4 me3
H3K9 me3
H3−5ac
H4−4ac
DNA Meth. m5C
H3K4 me1
H3K27 me3
H2A.Z
H4K20 me2
0.00 0.25 0.50 0.75 1.00
π̂
DHX8 – Reverse
H3K4 me1:H4−4ac
H3K4 me1:H2A.Z
H3K9 me2:DNA Meth. m5C
H3K27 me2:DNA Meth. m5C
H2A.Z:H3−5ac
H2A.Z:H4−4ac
H2A.Z:H3K4 me3
H4K20 me2:H4−4ac
H3K27 me3:DNA Meth. m5C
H3K4 me3:H4−4ac
H4K20 me2:H3K4 me3
H4−4ac:H3−5ac
H3K4 me1:H4K20 me2
H2A.Z:H4K20 me2
H4K20 me2:H3−5ac
H4K20 me3
H3K27 me2
H3K9 me2
H3K4 me3
H3K9 me3
H3−5ac
H4−4ac
DNA Meth. m5C
H3K4 me1
H3K27 me3
H2A.Z
H4K20 me2
0.00 0.25 0.50 0.75 1.00
π̂
LSM2 – Forward
H3K4 me1:H2A.Z
H3K4 me1:H4−4ac
H2A.Z:H4−4ac
H2A.Z:H3K4 me3
H4K20 me2:H3K4 me3
H2A.Z:H3−5ac
H3K9 me3:H4K20 me3
H3K27 me2:DNA Meth. m5C
H3K27 me3:H4K20 me3
H3K27 me3:H3K9 me3
H3K4 me1:H4K20 me2
H3K27 me3:DNA Meth. m5C
H2A.Z:H4K20 me2
H4K20 me2:H3−5ac
H3K4 me3:H4−4ac
H4−4ac:H3−5ac
H3K9 me2
H3K27 me2
H3K4 me3
H4K20 me3
H3K9 me3
DNA Meth. m5C
H4K20 me2
H4−4ac
H3K27 me3
H3K4 me1
H3−5ac
H2A.Z
0.00 0.25 0.50 0.75 1.00
π̂
LSM2 – Reverse
H3K4 me1:H2A.Z
H3K4 me1:H4−4ac
H2A.Z:H4−4ac
H2A.Z:H3K4 me3
H4K20 me2:H3K4 me3
H2A.Z:H3−5ac
H3K9 me3:H4K20 me3
H3K27 me2:DNA Meth. m5C
H3K27 me3:H4K20 me3
H3K27 me3:H3K9 me3
H3K4 me1:H4K20 me2
H3K27 me3:DNA Meth. m5C
H2A.Z:H4K20 me2
H4K20 me2:H3−5ac
H3K4 me3:H4−4ac
H4−4ac:H3−5ac
H3K9 me2
H3K27 me2
H3K4 me3
H4K20 me3
H3K9 me3
DNA Meth. m5C
H4K20 me2
H4−4ac
H3K27 me3
H3K4 me1
H3−5ac
H2A.Z
0.00 0.25 0.50 0.75 1.00
π̂
SNRPA1 – Forward
H2A.Z:H3K4 me3
H2A.Z:H4K20 me2
H3K4 me1:H4−4ac
H3K4 me1:H2A.Z
H3K4 me1:H4K20 me2
H3K4 me3:H3−5ac
H3K4 me1:H3−5ac
H4K20 me2:H3K4 me3
H4K20 me2:H3−5ac
H4−4ac:H3−5ac
H4K20 me2:H4−4ac
H3K4 me3:H4−4ac
H3K27 me2
H4K20 me3
H3K9 me3
H3K9 me2
DNA Meth. m5C
H2A.Z
H3K4 me1
H3K27 me3
H3K4 me3
H3−5ac
H4−4ac
H4K20 me2
0.00 0.25 0.50 0.75 1.00
π̂
SNRPA1 – Reverse
H2A.Z:H3K4 me3
H2A.Z:H4K20 me2
H3K4 me1:H4−4ac
H3K4 me1:H2A.Z
H3K4 me1:H4K20 me2
H3K4 me3:H3−5ac
H3K4 me1:H3−5ac
H4K20 me2:H3K4 me3
H4K20 me2:H3−5ac
H4−4ac:H3−5ac
H4K20 me2:H4−4ac
H3K4 me3:H4−4ac
H3K27 me2
H4K20 me3
H3K9 me3
H3K9 me2
DNA Meth. m5C
H2A.Z
H3K4 me1
H3K27 me3
H3K4 me3
H3−5ac
H4−4ac
H4K20 me2
0.00 0.25 0.50 0.75 1.00
π̂
LSM5 – Forward
H2A.Z:H4−4ac
H3K27 me3:DNA Meth. m5C
H2A.Z:H3−5ac
H3K4 me1:H4−4ac
H3K27 me3:H3K4 me1
H2A.Z:H3K4 me3
H3K4 me1:H3−5ac
H3K4 me1:H2A.Z
H3K9 me2:DNA Meth. m5C
H3K9 me3:H4K20 me3
H3K27 me3:H4K20 me3
H2A.Z:H4K20 me2
H4−4ac:H3−5ac
H4K20 me2:H3K4 me3
H3K4 me1:H4K20 me2
H3K4 me3:H4−4ac
H4K20 me2:H3−5ac
H4K20 me3
H3K4 me1
H3K9 me3
H3K9 me2
DNA Meth. m5C
H3K4 me3
H3K27 me3
H2A.Z
H4−4ac
H4K20 me2
H3−5ac
0.00 0.25 0.50 0.75 1.00
π̂
LSM5 – Reverse
H2A.Z:H4−4ac
H3K27 me3:DNA Meth. m5C
H2A.Z:H3−5ac
H3K4 me1:H4−4ac
H3K27 me3:H3K4 me1
H2A.Z:H3K4 me3
H3K4 me1:H3−5ac
H3K4 me1:H2A.Z
H3K9 me2:DNA Meth. m5C
H3K9 me3:H4K20 me3
H3K27 me3:H4K20 me3
H2A.Z:H4K20 me2
H4−4ac:H3−5ac
H4K20 me2:H3K4 me3
H3K4 me1:H4K20 me2
H3K4 me3:H4−4ac
H4K20 me2:H3−5ac
H4K20 me3
H3K4 me1
H3K9 me3
H3K9 me2
DNA Meth. m5C
H3K4 me3
H3K27 me3
H2A.Z
H4−4ac
H4K20 me2
H3−5ac
0.00 0.25 0.50 0.75 1.00
π̂

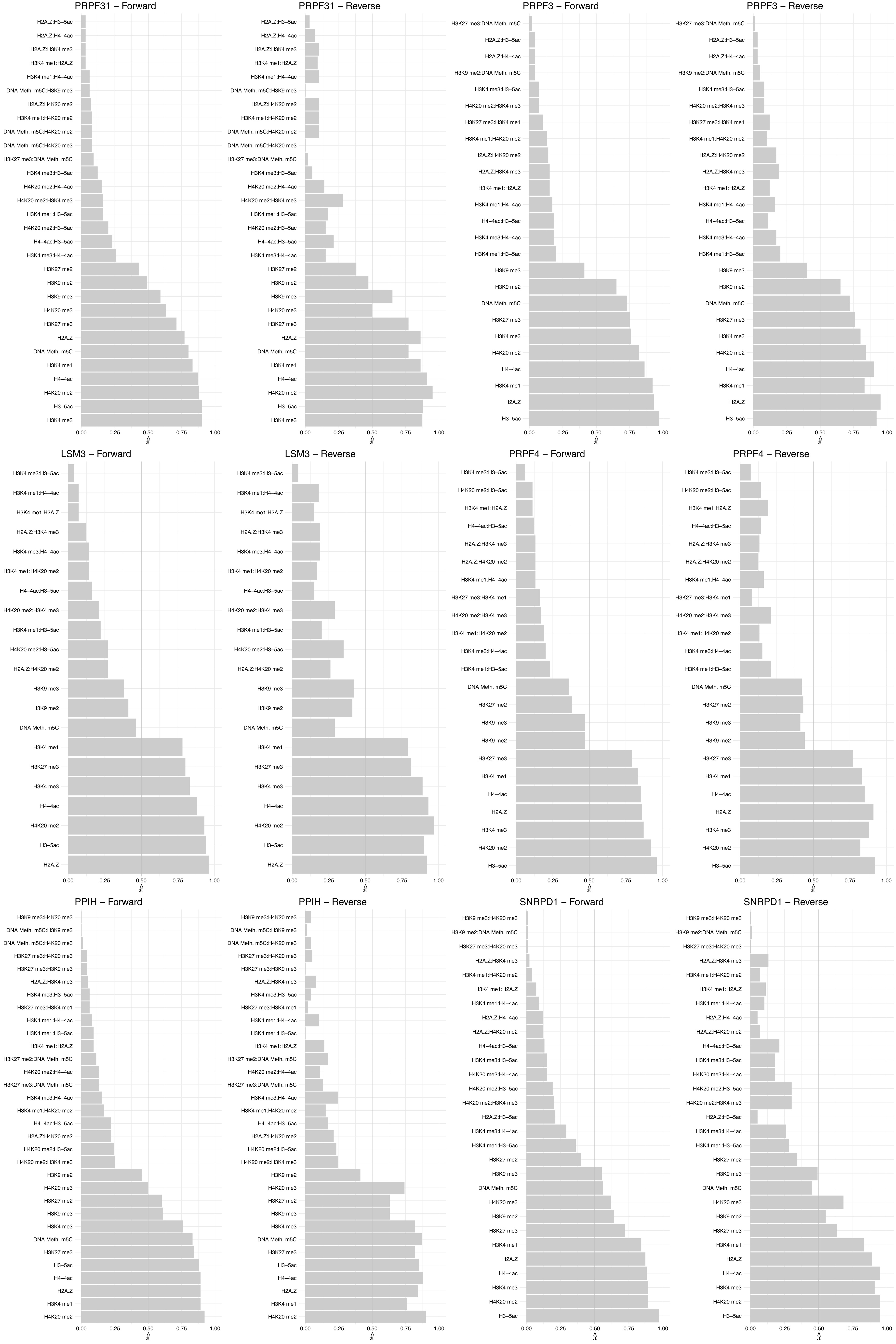
PRPF31 – Forward
H2A.Z:H3–5ac
H2A.Z:H4–4ac
H2A.Z:H3K4 me3
H3K4 me1:H2A.Z
H3K4 me1:H4–4ac
DNA Meth. m5C:H3K9 me3
H2A.Z:H4K20 me2
H3K4 me1:H4K20 me2
DNA Meth. m5C:H4K20 me2
DNA Meth. m5C:H4K20 me3
H3K27 me3:DNA Meth. m5C
H3K4 me3:H3–5ac
H4K20 me2:H4–4ac
H4K20 me2:H3K4 me3
H3K4 me1:H3–5ac
H4K20 me2:H3–5ac
H4–4ac:H3–5ac
H3K4 me3:H4–4ac
H3K27 me2
H3K9 me2
H3K9 me3
H4K20 me3
H3K27 me3
H2A.Z
DNA Meth. m5C
H3K4 me1
H4–4ac
H4K20 me2
H3–5ac
H3K4 me3
0.00
0.25
0.50
0.75
1.00
π̂
PRPF31 – Reverse
H2A.Z:H3–5ac
H2A.Z:H4–4ac
H2A.Z:H3K4 me3
H3K4 me1:H2A.Z
H3K4 me1:H4–4ac
DNA Meth. m5C:H3K9 me3
H2A.Z:H4K20 me2
H3K4 me1:H4K20 me2
DNA Meth. m5C:H4K20 me2
DNA Meth. m5C:H4K20 me3
H3K27 me3:DNA Meth. m5C
H3K4 me3:H3–5ac
H4K20 me2:H4–4ac
H4K20 me2:H3K4 me3
H3K4 me1:H3–5ac
H4K20 me2:H3–5ac
H4–4ac:H3–5ac
H3K4 me3:H4–4ac
H3K27 me2
H3K9 me2
H3K9 me3
H4K20 me3
H3K27 me3
H2A.Z
DNA Meth. m5C
H3K4 me1
H4–4ac
H4K20 me2
H3–5ac
H3K4 me3
0.00
0.25
0.50
0.75
1.00
π̂
PRPF3 – Forward
H3K27 me3:DNA Meth. m5C
H2A.Z:H3–5ac
H2A.Z:H4–4ac
H3K9 me2:DNA Meth. m5C
H3K4 me3:H3–5ac
H4K20 me2:H3K4 me3
H3K27 me3:H3K4 me1
H3K4 me1:H4K20 me2
H2A.Z:H4K20 me2
H2A.Z:H3K4 me3
H3K4 me1:H2A.Z
H3K4 me1:H4–4ac
H4–4ac:H3–5ac
H3K4 me3:H4–4ac
H3K4 me1:H3–5ac
H3K9 me3
H3K9 me2
DNA Meth. m5C
H3K27 me3
H3K4 me3
H4K20 me2
H4–4ac
H3K4 me1
H2A.Z
H3–5ac
0.00
0.25
0.50
0.75
1.00
π̂
PRPF3 – Reverse
H3K27 me3:DNA Meth. m5C
H2A.Z:H3–5ac
H2A.Z:H4–4ac
H3K9 me2:DNA Meth. m5C
H3K4 me3:H3–5ac
H4K20 me2:H3K4 me3
H3K27 me3:H3K4 me1
H3K4 me1:H4K20 me2
H2A.Z:H4K20 me2
H2A.Z:H3K4 me3
H3K4 me1:H2A.Z
H3K4 me1:H4–4ac
H4–4ac:H3–5ac
H3K4 me3:H4–4ac
H3K4 me1:H3–5ac
H3K9 me3
H3K9 me2
DNA Meth. m5C
H3K27 me3
H3K4 me3
H4K20 me2
H4–4ac
H3K4 me1
H2A.Z
H3–5ac
0.00
0.25
0.50
0.75
1.00
π̂
LSM3 – Forward
H3K4 me3:H3–5ac
H3K4 me1:H4–4ac
H3K4 me1:H2A.Z
H2A.Z:H3K4 me3
H3K4 me3:H4–4ac
H3K4 me1:H4K20 me2
H4–4ac:H3–5ac
H4K20 me2:H3K4 me3
H3K4 me1:H3–5ac
H4K20 me2:H3–5ac
H2A.Z:H4K20 me2
H3K9 me3
H3K9 me2
DNA Meth. m5C
H3K4 me1
H3K27 me3
H3K4 me3
H4–4ac
H4K20 me2
H3–5ac
H2A.Z
0.00
0.25
0.50
0.75
1.00
π̂
LSM3 – Reverse
H3K4 me3:H3–5ac
H3K4 me1:H4–4ac
H3K4 me1:H2A.Z
H2A.Z:H3K4 me3
H3K4 me3:H4–4ac
H3K4 me1:H4K20 me2
H4–4ac:H3–5ac
H4K20 me2:H3K4 me3
H3K4 me1:H3–5ac
H4K20 me2:H3–5ac
H2A.Z:H4K20 me2
H3K9 me3
H3K9 me2
DNA Meth. m5C
H3K4 me1
H3K27 me3
H3K4 me3
H4–4ac
H4K20 me2
H3–5ac
H2A.Z
0.00
0.25
0.50
0.75
1.00
π̂
PRPF4 – Forward
H3K4 me3:H3–5ac
H4K20 me2:H3–5ac
H3K4 me1:H2A.Z
H4–4ac:H3–5ac
H2A.Z:H3K4 me3
H2A.Z:H4K20 me2
H3K4 me1:H4–4ac
H3K27 me3:H3K4 me1
H4K20 me2:H3K4 me3
H3K4 me1:H4K20 me2
H3K4 me3:H4–4ac
H3K4 me1:H3–5ac
DNA Meth. m5C
H3K27 me2
H3K9 me3
H3K9 me2
H3K27 me3
H3K4 me1
H4–4ac
H2A.Z
H3K4 me3
H4K20 me2
H3–5ac
0.00
0.25
0.50
0.75
1.00
π̂
PRPF4 – Reverse
H3K4 me3:H3–5ac
H4K20 me2:H3–5ac
H3K4 me1:H2A.Z
H4–4ac:H3–5ac
H2A.Z:H3K4 me3
H2A.Z:H4K20 me2
H3K4 me1:H4–4ac
H3K27 me3:H3K4 me1
H4K20 me2:H3K4 me3
H3K4 me1:H4K20 me2
H3K4 me3:H4–4ac
H3K4 me1:H3–5ac
DNA Meth. m5C
H3K27 me2
H3K9 me3
H3K9 me2
H3K27 me3
H3K4 me1
H4–4ac
H2A.Z
H3K4 me3
H4K20 me2
H3–5ac
0.00
0.25
0.50
0.75
1.00
π̂
PPIH – Forward
H3K9 me3:H4K20 me3
DNA Meth. m5C:H3K9 me3
DNA Meth. m5C:H4K20 me3
H3K27 me3:H4K20 me3
H3K27 me3:H3K9 me3
H2A.Z:H3K4 me3
H3K4 me3:H3–5ac
H3K27 me3:H3K4 me1
H3K4 me1:H4–4ac
H3K4 me1:H3–5ac
H3K4 me1:H2A.Z
H3K27 me2:DNA Meth. m5C
H4K20 me2:H4–4ac
H3K27 me3:DNA Meth. m5C
H3K4 me3:H4–4ac
H3K4 me1:H4K20 me2
H4–4ac:H3–5ac
H2A.Z:H4K20 me2
H4K20 me2:H3–5ac
H4K20 me2:H3K4 me3
H3K9 me2
H4K20 me3
H3K27 me2
H3K9 me3
H3K4 me3
DNA Meth. m5C
H3K27 me3
H3–5ac
H4–4ac
H2A.Z
H3K4 me1
H4K20 me2
0.00
0.25
0.50
0.75
1.00
π̂
PPIH – Reverse
H3K9 me3:H4K20 me3
DNA Meth. m5C:H3K9 me3
DNA Meth. m5C:H4K20 me3
H3K27 me3:H4K20 me3
H3K27 me3:H3K9 me3
H2A.Z:H3K4 me3
H3K4 me3:H3–5ac
H3K27 me3:H3K4 me1
H3K4 me1:H4–4ac
H3K4 me1:H3–5ac
H3K4 me1:H2A.Z
H3K27 me2:DNA Meth. m5C
H4K20 me2:H4–4ac
H3K27 me3:DNA Meth. m5C
H3K4 me3:H4–4ac
H3K4 me1:H4K20 me2
H4–4ac:H3–5ac
H2A.Z:H4K20 me2
H4K20 me2:H3–5ac
H4K20 me2:H3K4 me3
H3K9 me2
H4K20 me3
H3K27 me2
H3K9 me3
H3K4 me3
DNA Meth. m5C
H3K27 me3
H3–5ac
H4–4ac
H2A.Z
H3K4 me1
H4K20 me2
0.00
0.25
0.50
0.75
1.00
π̂
SNRPD1 – Forward
H3K9 me3:H4K20 me3
H3K9 me2:DNA Meth. m5C
H3K27 me3:H4K20 me3
H2A.Z:H3K4 me3
H3K4 me1:H4K20 me2
H3K4 me1:H2A.Z
H3K4 me1:H4–4ac
H2A.Z:H4–4ac
H2A.Z:H4K20 me2
H4–4ac:H3–5ac
H3K4 me3:H3–5ac
H4K20 me2:H4–4ac
H4K20 me2:H3–5ac
H4K20 me2:H3K4 me3
H2A.Z:H3–5ac
H3K4 me3:H4–4ac
H3K4 me1:H3–5ac
H3K27 me2
H3K9 me3
DNA Meth. m5C
H4K20 me3
H3K9 me2
H3K27 me3
H3K4 me1
H2A.Z
H4–4ac
H3K4 me3
H4K20 me2
H3–5ac
0.00
0.25
0.50
0.75
1.00
π̂
SNRPD1 – Reverse
H3K9 me3:H4K20 me3
H3K9 me2:DNA Meth. m5C
H3K27 me3:H4K20 me3
H2A.Z:H3K4 me3
H3K4 me1:H4K20 me2
H3K4 me1:H2A.Z
H3K4 me1:H4–4ac
H2A.Z:H4–4ac
H2A.Z:H4K20 me2
H4–4ac:H3–5ac
H3K4 me3:H3–5ac
H4K20 me2:H4–4ac
H4K20 me2:H3–5ac
H4K20 me2:H3K4 me3
H2A.Z:H3–5ac
H3K4 me3:H4–4ac
H3K4 me1:H3–5ac
H3K27 me2
H3K9 me3
DNA Meth. m5C
H4K20 me3
H3K9 me2
H3K27 me3
H3K4 me1
H2A.Z
H4–4ac
H3K4 me3
H4K20 me2
H3–5ac
0.00
0.25
0.50
0.75
1.00
π̂

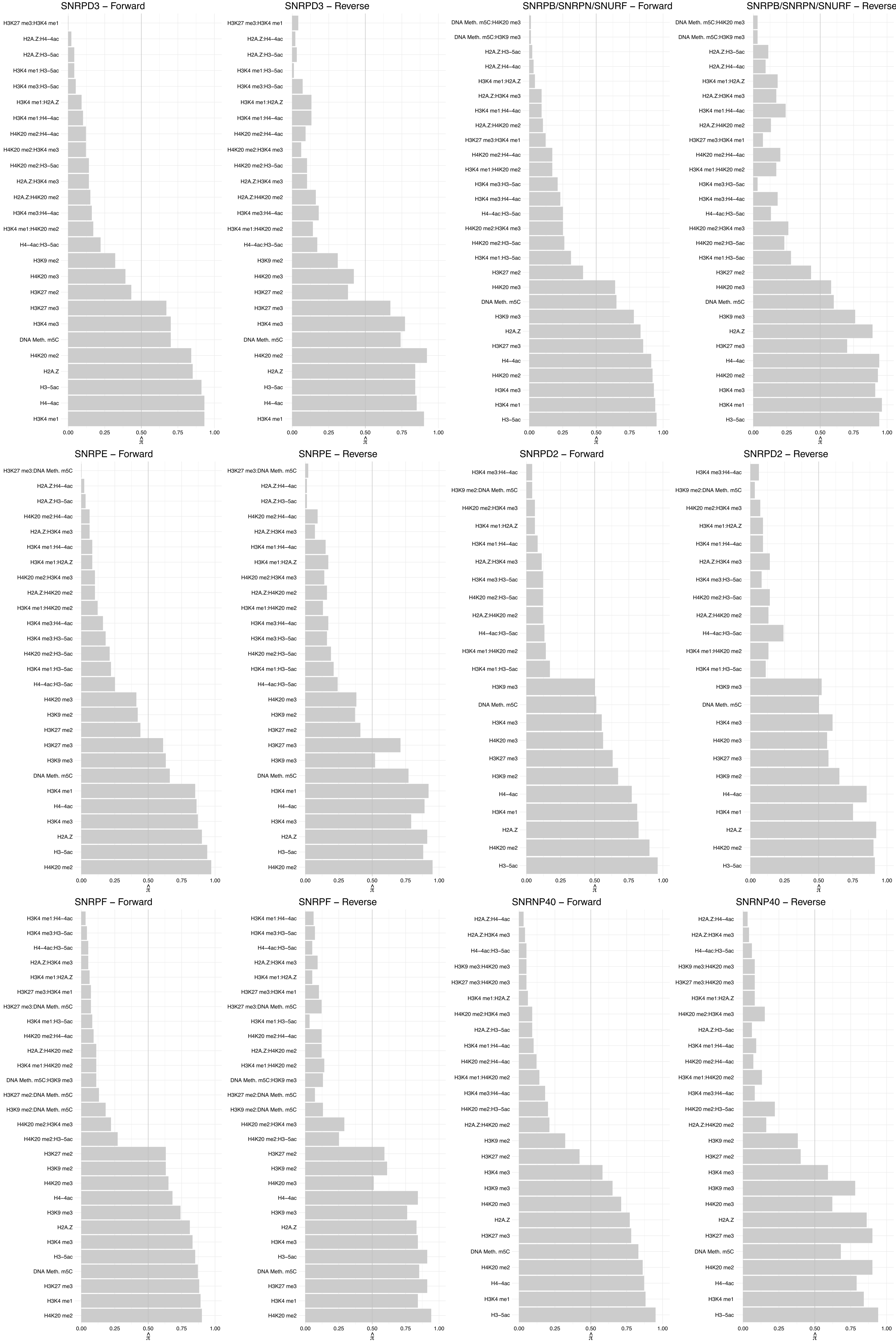
SNRPD3 – Forward
H3K27 me3:H3K4 me1
H2A.Z:H4−4ac
H2A.Z:H3−5ac
H3K4 me1:H3−5ac
H3K4 me3:H3−5ac
H3K4 me1:H2A.Z
H3K4 me1:H4−4ac
H4K20 me2:H4−4ac
H4K20 me2:H3K4 me3
H4K20 me2:H3−5ac
H2A.Z:H3K4 me3
H2A.Z:H4K20 me2
H3K4 me3:H4−4ac
H3K4 me1:H4K20 me2
H4−4ac:H3−5ac
H3K9 me2
H4K20 me3
H3K27 me2
H3K27 me3
H3K4 me3
DNA Meth. m5C
H4K20 me2
H2A.Z
H3−5ac
H4−4ac
H3K4 me1
0.00 0.25 0.50 0.75 1.00
π̂
SNRPD3 – Reverse
H3K27 me3:H3K4 me1
H2A.Z:H4−4ac
H2A.Z:H3−5ac
H3K4 me1:H3−5ac
H3K4 me3:H3−5ac
H3K4 me1:H2A.Z
H3K4 me1:H4−4ac
H4K20 me2:H4−4ac
H4K20 me2:H3K4 me3
H4K20 me2:H3−5ac
H2A.Z:H3K4 me3
H2A.Z:H4K20 me2
H3K4 me3:H4−4ac
H3K4 me1:H4K20 me2
H4−4ac:H3−5ac
H3K9 me2
H4K20 me3
H3K27 me2
H3K27 me3
H3K4 me3
DNA Meth. m5C
H4K20 me2
H2A.Z
H3−5ac
H4−4ac
H3K4 me1
0.00 0.25 0.50 0.75 1.00
π̂
SNRPB/SNRPN/SNURF – Forward
DNA Meth. m5C:H4K20 me3
DNA Meth. m5C:H3K9 me3
H2A.Z:H3−5ac
H2A.Z:H4−4ac
H3K4 me1:H2A.Z
H2A.Z:H3K4 me3
H3K4 me1:H4−4ac
H2A.Z:H4K20 me2
H3K27 me3:H3K4 me1
H4K20 me2:H4−4ac
H3K4 me1:H4K20 me2
H3K4 me3:H3−5ac
H3K4 me3:H4−4ac
H4−4ac:H3−5ac
H4K20 me2:H3K4 me3
H4K20 me2:H3−5ac
H3K4 me1:H3−5ac
H3K27 me2
H4K20 me3
DNA Meth. m5C
H3K9 me3
H2A.Z
H3K27 me3
H4−4ac
H4K20 me2
H3K4 me3
H3K4 me1
H3−5ac
0.00 0.25 0.50 0.75 1.00
π̂
SNRPB/SNRPN/SNURF – Reverse
DNA Meth. m5C:H4K20 me3
DNA Meth. m5C:H3K9 me3
H2A.Z:H3−5ac
H2A.Z:H4−4ac
H3K4 me1:H2A.Z
H2A.Z:H3K4 me3
H3K4 me1:H4−4ac
H2A.Z:H4K20 me2
H3K27 me3:H3K4 me1
H4K20 me2:H4−4ac
H3K4 me1:H4K20 me2
H3K4 me3:H3−5ac
H3K4 me3:H4−4ac
H4−4ac:H3−5ac
H4K20 me2:H3K4 me3
H4K20 me2:H3−5ac
H3K4 me1:H3−5ac
H3K27 me2
H4K20 me3
DNA Meth. m5C
H3K9 me3
H2A.Z
H3K27 me3
H4−4ac
H4K20 me2
H3K4 me3
H3K4 me1
H3−5ac
0.00 0.25 0.50 0.75 1.00
π̂
SNRPE – Forward
H3K27 me3:DNA Meth. m5C
H2A.Z:H4−4ac
H2A.Z:H3−5ac
H4K20 me2:H4−4ac
H2A.Z:H3K4 me3
H3K4 me1:H4−4ac
H3K4 me1:H2A.Z
H4K20 me2:H3K4 me3
H2A.Z:H4K20 me2
H3K4 me1:H4K20 me2
H3K4 me3:H4−4ac
H3K4 me3:H3−5ac
H4K20 me2:H3−5ac
H3K4 me1:H3−5ac
H4−4ac:H3−5ac
H4K20 me3
H3K9 me2
H3K27 me2
H3K27 me3
H3K9 me3
DNA Meth. m5C
H3K4 me1
H4−4ac
H3K4 me3
H2A.Z
H3−5ac
H4K20 me2
0.00 0.25 0.50 0.75 1.00
π̂
SNRPE – Reverse
H3K27 me3:DNA Meth. m5C
H2A.Z:H4−4ac
H2A.Z:H3−5ac
H4K20 me2:H4−4ac
H2A.Z:H3K4 me3
H3K4 me1:H4−4ac
H3K4 me1:H2A.Z
H4K20 me2:H3K4 me3
H2A.Z:H4K20 me2
H3K4 me1:H4K20 me2
H3K4 me3:H4−4ac
H3K4 me3:H3−5ac
H4K20 me2:H3−5ac
H3K4 me1:H3−5ac
H4−4ac:H3−5ac
H4K20 me3
H3K9 me2
H3K27 me2
H3K27 me3
H3K9 me3
DNA Meth. m5C
H3K4 me1
H4−4ac
H3K4 me3
H2A.Z
H3−5ac
H4K20 me2
0.00 0.25 0.50 0.75 1.00
π̂
SNRPD2 – Forward
H3K4 me3:H4−4ac
H3K9 me2:DNA Meth. m5C
H4K20 me2:H3K4 me3
H3K4 me1:H2A.Z
H3K4 me1:H4−4ac
H2A.Z:H3K4 me3
H3K4 me3:H3−5ac
H4K20 me2:H3−5ac
H2A.Z:H4K20 me2
H4−4ac:H3−5ac
H3K4 me1:H4K20 me2
H3K4 me1:H3−5ac
H3K9 me3
DNA Meth. m5C
H3K4 me3
H4K20 me3
H3K27 me3
H3K9 me2
H4−4ac
H3K4 me1
H2A.Z
H4K20 me2
H3−5ac
0.00 0.25 0.50 0.75 1.00
π̂
SNRPD2 – Reverse
H3K4 me3:H4−4ac
H3K9 me2:DNA Meth. m5C
H4K20 me2:H3K4 me3
H3K4 me1:H2A.Z
H3K4 me1:H4−4ac
H2A.Z:H3K4 me3
H3K4 me3:H3−5ac
H4K20 me2:H3−5ac
H2A.Z:H4K20 me2
H4−4ac:H3−5ac
H3K4 me1:H4K20 me2
H3K4 me1:H3−5ac
H3K9 me3
DNA Meth. m5C
H3K4 me3
H4K20 me3
H3K27 me3
H3K9 me2
H4−4ac
H3K4 me1
H2A.Z
H4K20 me2
H3−5ac
0.00 0.25 0.50 0.75 1.00
π̂
SNRPF – Forward
H3K4 me1:H4−4ac
H3K4 me3:H3−5ac
H4−4ac:H3−5ac
H2A.Z:H3K4 me3
H3K4 me1:H2A.Z
H3K27 me3:H3K4 me1
H3K27 me3:DNA Meth. m5C
H3K4 me1:H3−5ac
H4K20 me2:H4−4ac
H2A.Z:H4K20 me2
H3K4 me1:H4K20 me2
DNA Meth. m5C:H3K9 me3
H3K27 me2:DNA Meth. m5C
H3K9 me2:DNA Meth. m5C
H4K20 me2:H3K4 me3
H4K20 me2:H3−5ac
H3K27 me2
H3K9 me2
H4K20 me3
H4−4ac
H3K9 me3
H2A.Z
H3K4 me3
H3−5ac
DNA Meth. m5C
H3K27 me3
H3K4 me1
H4K20 me2
0.00 0.25 0.50 0.75 1.00
π̂
SNRPF – Reverse
H3K4 me1:H4−4ac
H3K4 me3:H3−5ac
H4−4ac:H3−5ac
H2A.Z:H3K4 me3
H3K4 me1:H2A.Z
H3K27 me3:H3K4 me1
H3K27 me3:DNA Meth. m5C
H3K4 me1:H3−5ac
H4K20 me2:H4−4ac
H2A.Z:H4K20 me2
H3K4 me1:H4K20 me2
DNA Meth. m5C:H3K9 me3
H3K27 me2:DNA Meth. m5C
H3K9 me2:DNA Meth. m5C
H4K20 me2:H3K4 me3
H4K20 me2:H3−5ac
H3K27 me2
H3K9 me2
H4K20 me3
H4−4ac
H3K9 me3
H2A.Z
H3K4 me3
H3−5ac
DNA Meth. m5C
H3K27 me3
H3K4 me1
H4K20 me2
0.00 0.25 0.50 0.75 1.00
π̂
SNRNP40 – Forward
H2A.Z:H4−4ac
H2A.Z:H3K4 me3
H4−4ac:H3−5ac
H3K9 me3:H4K20 me3
H3K27 me3:H4K20 me3
H3K4 me1:H2A.Z
H4K20 me2:H3K4 me3
H2A.Z:H3−5ac
H3K4 me1:H4−4ac
H4K20 me2:H4−4ac
H3K4 me1:H4K20 me2
H3K4 me3:H4−4ac
H4K20 me2:H3−5ac
H2A.Z:H4K20 me2
H3K9 me2
H3K27 me2
H3K4 me3
H3K9 me3
H4K20 me3
H2A.Z
H3K27 me3
DNA Meth. m5C
H4K20 me2
H4−4ac
H3K4 me1
H3−5ac
0.00 0.25 0.50 0.75 1.00
π̂
SNRNP40 – Reverse
H2A.Z:H4−4ac
H2A.Z:H3K4 me3
H4−4ac:H3−5ac
H3K9 me3:H4K20 me3
H3K27 me3:H4K20 me3
H3K4 me1:H2A.Z
H4K20 me2:H3K4 me3
H2A.Z:H3−5ac
H3K4 me1:H4−4ac
H4K20 me2:H4−4ac
H3K4 me1:H4K20 me2
H3K4 me3:H4−4ac
H4K20 me2:H3−5ac
H2A.Z:H4K20 me2
H3K9 me2
H3K27 me2
H3K4 me3
H3K9 me3
H4K20 me3
H2A.Z
H3K27 me3
DNA Meth. m5C
H4K20 me2
H4−4ac
H3K4 me1
H3−5ac
0.00 0.25 0.50 0.75 1.00
π̂

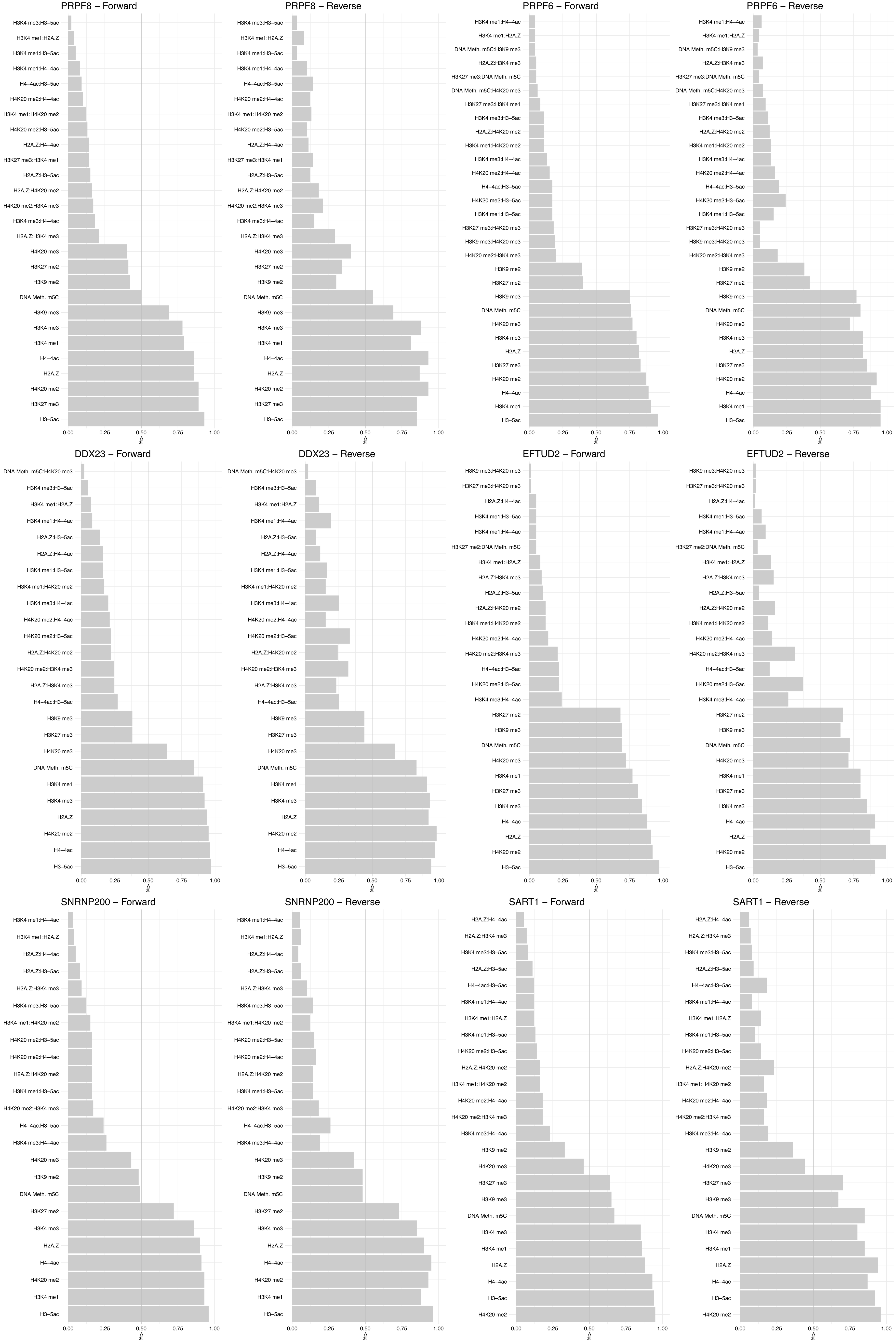
PRPF8 – Forward
H3K4 me3:H3−5ac
H3K4 me1:H2A.Z
H3K4 me1:H3−5ac
H3K4 me1:H4−4ac
H4−4ac:H3−5ac
H4K20 me2:H4−4ac
H3K4 me1:H4K20 me2
H4K20 me2:H3−5ac
H2A.Z:H4−4ac
H3K27 me3:H3K4 me1
H2A.Z:H3−5ac
H2A.Z:H4K20 me2
H4K20 me2:H3K4 me3
H3K4 me3:H4−4ac
H2A.Z:H3K4 me3
H4K20 me3
H3K27 me2
H3K9 me2
DNA Meth. m5C
H3K9 me3
H3K4 me3
H3K4 me1
H4−4ac
H2A.Z
H4K20 me2
H3K27 me3
H3−5ac
0.00 0.25 0.50 0.75 1.00
$\hat{\pi}$
PRPF8 – Reverse
H3K4 me3:H3−5ac
H3K4 me1:H2A.Z
H3K4 me1:H3−5ac
H3K4 me1:H4−4ac
H4−4ac:H3−5ac
H4K20 me2:H4−4ac
H3K4 me1:H4K20 me2
H4K20 me2:H3−5ac
H2A.Z:H4−4ac
H3K27 me3:H3K4 me1
H2A.Z:H3−5ac
H2A.Z:H4K20 me2
H4K20 me2:H3K4 me3
H3K4 me3:H4−4ac
H2A.Z:H3K4 me3
H4K20 me3
H3K27 me2
H3K9 me2
DNA Meth. m5C
H3K9 me3
H3K4 me3
H3K4 me1
H4−4ac
H2A.Z
H4K20 me2
H3K27 me3
H3−5ac
0.00 0.25 0.50 0.75 1.00
$\hat{\pi}$
PRPF6 – Forward
H3K4 me1:H4−4ac
H3K4 me1:H2A.Z
DNA Meth. m5C:H3K9 me3
H2A.Z:H3K4 me3
H3K27 me3:DNA Meth. m5C
DNA Meth. m5C:H4K20 me3
H3K27 me3:H3K4 me1
H3K4 me3:H3−5ac
H2A.Z:H4K20 me2
H3K4 me1:H4K20 me2
H3K4 me3:H4−4ac
H4K20 me2:H4−4ac
H4−4ac:H3−5ac
H4K20 me2:H3−5ac
H3K4 me1:H3−5ac
H3K27 me3:H4K20 me3
H3K9 me3:H4K20 me3
H4K20 me2:H3K4 me3
H3K9 me2
H3K27 me2
H3K9 me3
DNA Meth. m5C
H4K20 me3
H3K4 me3
H2A.Z
H3K27 me3
H4K20 me2
H4−4ac
H3K4 me1
H3−5ac
0.00 0.25 0.50 0.75 1.00
$\hat{\pi}$
PRPF6 – Reverse
H3K4 me1:H4−4ac
H3K4 me1:H2A.Z
DNA Meth. m5C:H3K9 me3
H2A.Z:H3K4 me3
H3K27 me3:DNA Meth. m5C
DNA Meth. m5C:H4K20 me3
H3K27 me3:H3K4 me1
H3K4 me3:H3−5ac
H2A.Z:H4K20 me2
H3K4 me1:H4K20 me2
H3K4 me3:H4−4ac
H4K20 me2:H4−4ac
H4−4ac:H3−5ac
H4K20 me2:H3−5ac
H3K4 me1:H3−5ac
H3K27 me3:H4K20 me3
H3K9 me3:H4K20 me3
H4K20 me2:H3K4 me3
H3K9 me2
H3K27 me2
H3K9 me3
DNA Meth. m5C
H4K20 me3
H3K4 me3
H2A.Z
H3K27 me3
H4K20 me2
H4−4ac
H3K4 me1
H3−5ac
0.00 0.25 0.50 0.75 1.00
$\hat{\pi}$
DDX23 – Forward
DNA Meth. m5C:H4K20 me3
H3K4 me3:H3−5ac
H3K4 me1:H2A.Z
H3K4 me1:H4−4ac
H2A.Z:H3−5ac
H2A.Z:H4−4ac
H3K4 me1:H3−5ac
H3K4 me1:H4K20 me2
H3K4 me3:H4−4ac
H4K20 me2:H4−4ac
H4K20 me2:H3−5ac
H2A.Z:H4K20 me2
H4K20 me2:H3K4 me3
H2A.Z:H3K4 me3
H4−4ac:H3−5ac
H3K9 me3
H3K27 me3
H4K20 me3
DNA Meth. m5C
H3K4 me1
H3K4 me3
H2A.Z
H4K20 me2
H4−4ac
H3−5ac
0.00 0.25 0.50 0.75 1.00
$\hat{\pi}$
DDX23 – Reverse
DNA Meth. m5C:H4K20 me3
H3K4 me3:H3−5ac
H3K4 me1:H2A.Z
H3K4 me1:H4−4ac
H2A.Z:H3−5ac
H2A.Z:H4−4ac
H3K4 me1:H3−5ac
H3K4 me1:H4K20 me2
H3K4 me3:H4−4ac
H4K20 me2:H4−4ac
H4K20 me2:H3−5ac
H2A.Z:H4K20 me2
H4K20 me2:H3K4 me3
H2A.Z:H3K4 me3
H4−4ac:H3−5ac
H3K9 me3
H3K27 me3
H4K20 me3
DNA Meth. m5C
H3K4 me1
H3K4 me3
H2A.Z
H4K20 me2
H4−4ac
H3−5ac
0.00 0.25 0.50 0.75 1.00
$\hat{\pi}$
EFTUD2 – Forward
H3K9 me3:H4K20 me3
H3K27 me3:H4K20 me3
H2A.Z:H4−4ac
H3K4 me1:H3−5ac
H3K4 me1:H4−4ac
H3K27 me2:DNA Meth. m5C
H3K4 me1:H2A.Z
H2A.Z:H3K4 me3
H2A.Z:H3−5ac
H2A.Z:H4K20 me2
H3K4 me1:H4K20 me2
H4K20 me2:H4−4ac
H4K20 me2:H3K4 me3
H4−4ac:H3−5ac
H4K20 me2:H3−5ac
H3K4 me3:H4−4ac
H3K27 me2
H3K9 me3
DNA Meth. m5C
H4K20 me3
H3K4 me1
H3K27 me3
H3K4 me3
H4−4ac
H2A.Z
H4K20 me2
H3−5ac
0.00 0.25 0.50 0.75 1.00
$\hat{\pi}$
EFTUD2 – Reverse
H3K9 me3:H4K20 me3
H3K27 me3:H4K20 me3
H2A.Z:H4−4ac
H3K4 me1:H3−5ac
H3K4 me1:H4−4ac
H3K27 me2:DNA Meth. m5C
H3K4 me1:H2A.Z
H2A.Z:H3K4 me3
H2A.Z:H3−5ac
H2A.Z:H4K20 me2
H3K4 me1:H4K20 me2
H4K20 me2:H4−4ac
H4K20 me2:H3K4 me3
H4−4ac:H3−5ac
H4K20 me2:H3−5ac
H3K4 me3:H4−4ac
H3K27 me2
H3K9 me3
DNA Meth. m5C
H4K20 me3
H3K4 me1
H3K27 me3
H3K4 me3
H4−4ac
H2A.Z
H4K20 me2
H3−5ac
0.00 0.25 0.50 0.75 1.00
$\hat{\pi}$
SNRNP200 – Forward
H3K4 me1:H4−4ac
H3K4 me1:H2A.Z
H2A.Z:H4−4ac
H2A.Z:H3−5ac
H2A.Z:H3K4 me3
H3K4 me3:H3−5ac
H3K4 me1:H4K20 me2
H4K20 me2:H3−5ac
H4K20 me2:H4−4ac
H2A.Z:H4K20 me2
H3K4 me1:H3−5ac
H4K20 me2:H3K4 me3
H4−4ac:H3−5ac
H3K4 me3:H4−4ac
H4K20 me3
H3K9 me2
DNA Meth. m5C
H3K27 me2
H3K4 me3
H2A.Z
H4−4ac
H4K20 me2
H3K4 me1
H3−5ac
0.00 0.25 0.50 0.75 1.00
$\hat{\pi}$
SNRNP200 – Reverse
H3K4 me1:H4−4ac
H3K4 me1:H2A.Z
H2A.Z:H4−4ac
H2A.Z:H3−5ac
H2A.Z:H3K4 me3
H3K4 me3:H3−5ac
H3K4 me1:H4K20 me2
H4K20 me2:H3−5ac
H4K20 me2:H4−4ac
H2A.Z:H4K20 me2
H3K4 me1:H3−5ac
H4K20 me2:H3K4 me3
H4−4ac:H3−5ac
H3K4 me3:H4−4ac
H4K20 me3
H3K9 me2
DNA Meth. m5C
H3K27 me2
H3K4 me3
H2A.Z
H4−4ac
H4K20 me2
H3K4 me1
H3−5ac
0.00 0.25 0.50 0.75 1.00
$\hat{\pi}$
SART1 – Forward
H2A.Z:H4−4ac
H2A.Z:H3K4 me3
H3K4 me3:H3−5ac
H2A.Z:H3−5ac
H4−4ac:H3−5ac
H3K4 me1:H4−4ac
H3K4 me1:H2A.Z
H3K4 me1:H3−5ac
H4K20 me2:H3−5ac
H2A.Z:H4K20 me2
H3K4 me1:H4K20 me2
H4K20 me2:H4−4ac
H4K20 me2:H3K4 me3
H3K4 me3:H4−4ac
H3K9 me2
H4K20 me3
H3K27 me3
H3K9 me3
DNA Meth. m5C
H3K4 me3
H3K4 me1
H2A.Z
H4−4ac
H3−5ac
H4K20 me2
0.00 0.25 0.50 0.75 1.00
$\hat{\pi}$
SART1 – Reverse
H2A.Z:H4−4ac
H2A.Z:H3K4 me3
H3K4 me3:H3−5ac
H2A.Z:H3−5ac
H4−4ac:H3−5ac
H3K4 me1:H4−4ac
H3K4 me1:H2A.Z
H3K4 me1:H3−5ac
H4K20 me2:H3−5ac
H2A.Z:H4K20 me2
H3K4 me1:H4K20 me2
H4K20 me2:H4−4ac
H4K20 me2:H3K4 me3
H3K4 me3:H4−4ac
H3K9 me2
H4K20 me3
H3K27 me3
H3K9 me3
DNA Meth. m5C
H3K4 me3
H3K4 me1
H2A.Z
H4−4ac
H3−5ac
H4K20 me2
0.00 0.25 0.50 0.75 1.00
$\hat{\pi}$

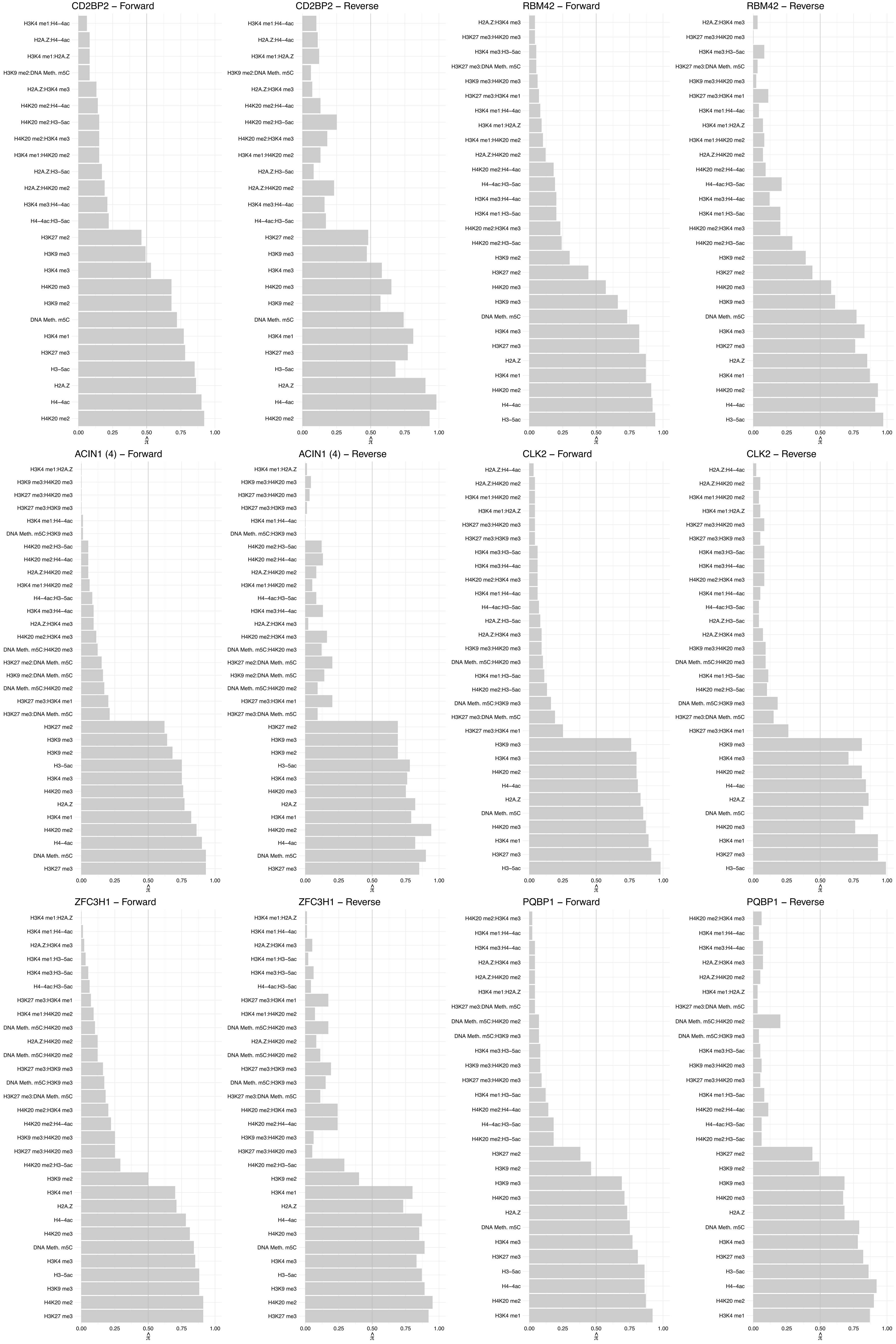
CD2BP2 – Forward
H3K4 me1:H4−4ac
H2A.Z:H4−4ac
H3K4 me1:H2A.Z
H3K9 me2:DNA Meth. m5C
H2A.Z:H3K4 me3
H4K20 me2:H4−4ac
H4K20 me2:H3−5ac
H4K20 me2:H3K4 me3
H3K4 me1:H4K20 me2
H2A.Z:H3−5ac
H2A.Z:H4K20 me2
H3K4 me3:H4−4ac
H4−4ac:H3−5ac
H3K27 me2
H3K9 me3
H3K4 me3
H4K20 me3
H3K9 me2
DNA Meth. m5C
H3K4 me1
H3K27 me3
H3−5ac
H2A.Z
H4−4ac
H4K20 me2
0.00 0.25 0.50 0.75 1.00
π̂
CD2BP2 – Reverse
H3K4 me1:H4−4ac
H2A.Z:H4−4ac
H3K4 me1:H2A.Z
H3K9 me2:DNA Meth. m5C
H2A.Z:H3K4 me3
H4K20 me2:H4−4ac
H4K20 me2:H3−5ac
H4K20 me2:H3K4 me3
H3K4 me1:H4K20 me2
H2A.Z:H3−5ac
H2A.Z:H4K20 me2
H3K4 me3:H4−4ac
H4−4ac:H3−5ac
H3K27 me2
H3K9 me3
H3K4 me3
H4K20 me3
H3K9 me2
DNA Meth. m5C
H3K4 me1
H3K27 me3
H3−5ac
H2A.Z
H4−4ac
H4K20 me2
0.00 0.25 0.50 0.75 1.00
π̂
RBM42 – Forward
H2A.Z:H3K4 me3
H3K27 me3:H4K20 me3
H3K4 me3:H3−5ac
H3K27 me3:DNA Meth. m5C
H3K9 me3:H4K20 me3
H3K27 me3:H3K4 me1
H3K4 me1:H4−4ac
H3K4 me1:H2A.Z
H3K4 me1:H4K20 me2
H2A.Z:H4K20 me2
H4K20 me2:H4−4ac
H4−4ac:H3−5ac
H3K4 me3:H4−4ac
H3K4 me1:H3−5ac
H4K20 me2:H3K4 me3
H4K20 me2:H3−5ac
H3K9 me2
H3K27 me2
H4K20 me3
H3K9 me3
DNA Meth. m5C
H3K4 me3
H3K27 me3
H2A.Z
H3K4 me1
H4K20 me2
H4−4ac
H3−5ac
0.00 0.25 0.50 0.75 1.00
π̂
RBM42 – Reverse
H2A.Z:H3K4 me3
H3K27 me3:H4K20 me3
H3K4 me3:H3−5ac
H3K27 me3:DNA Meth. m5C
H3K9 me3:H4K20 me3
H3K27 me3:H3K4 me1
H3K4 me1:H4−4ac
H3K4 me1:H2A.Z
H3K4 me1:H4K20 me2
H2A.Z:H4K20 me2
H4K20 me2:H4−4ac
H4−4ac:H3−5ac
H3K4 me3:H4−4ac
H3K4 me1:H3−5ac
H4K20 me2:H3K4 me3
H4K20 me2:H3−5ac
H3K9 me2
H3K27 me2
H4K20 me3
H3K9 me3
DNA Meth. m5C
H3K4 me3
H3K27 me3
H2A.Z
H3K4 me1
H4K20 me2
H4−4ac
H3−5ac
0.00 0.25 0.50 0.75 1.00
π̂
ACIN1 (4) – Forward
H3K4 me1:H2A.Z
H3K9 me3:H4K20 me3
H3K27 me3:H4K20 me3
H3K27 me3:H3K9 me3
H3K4 me1:H4−4ac
DNA Meth. m5C:H3K9 me3
H4K20 me2:H3−5ac
H4K20 me2:H4−4ac
H2A.Z:H4K20 me2
H3K4 me1:H4K20 me2
H4−4ac:H3−5ac
H3K4 me3:H4−4ac
H2A.Z:H3K4 me3
H4K20 me2:H3K4 me3
DNA Meth. m5C:H4K20 me3
H3K27 me2:DNA Meth. m5C
H3K9 me2:DNA Meth. m5C
DNA Meth. m5C:H4K20 me2
H3K27 me3:H3K4 me1
H3K27 me3:DNA Meth. m5C
H3K27 me2
H3K9 me3
H3K9 me2
H3−5ac
H3K4 me3
H4K20 me3
H2A.Z
H3K4 me1
H4K20 me2
H4−4ac
DNA Meth. m5C
H3K27 me3
0.00 0.25 0.50 0.75 1.00
π̂
ACIN1 (4) – Reverse
H3K4 me1:H2A.Z
H3K9 me3:H4K20 me3
H3K27 me3:H4K20 me3
H3K27 me3:H3K9 me3
H3K4 me1:H4−4ac
DNA Meth. m5C:H3K9 me3
H4K20 me2:H3−5ac
H4K20 me2:H4−4ac
H2A.Z:H4K20 me2
H3K4 me1:H4K20 me2
H4−4ac:H3−5ac
H3K4 me3:H4−4ac
H2A.Z:H3K4 me3
H4K20 me2:H3K4 me3
DNA Meth. m5C:H4K20 me3
H3K27 me2:DNA Meth. m5C
H3K9 me2:DNA Meth. m5C
DNA Meth. m5C:H4K20 me2
H3K27 me3:H3K4 me1
H3K27 me3:DNA Meth. m5C
H3K27 me2
H3K9 me3
H3K9 me2
H3−5ac
H3K4 me3
H4K20 me3
H2A.Z
H3K4 me1
H4K20 me2
H4−4ac
DNA Meth. m5C
H3K27 me3
0.00 0.25 0.50 0.75 1.00
π̂
CLK2 – Forward
H2A.Z:H4−4ac
H2A.Z:H4K20 me2
H3K4 me1:H4K20 me2
H3K4 me1:H2A.Z
H3K27 me3:H4K20 me3
H3K27 me3:H3K9 me3
H3K4 me3:H3−5ac
H3K4 me3:H4−4ac
H4K20 me2:H3K4 me3
H3K4 me1:H4−4ac
H4−4ac:H3−5ac
H2A.Z:H3−5ac
H2A.Z:H3K4 me3
H3K9 me3:H4K20 me3
DNA Meth. m5C:H4K20 me3
H3K4 me1:H3−5ac
H4K20 me2:H3−5ac
DNA Meth. m5C:H3K9 me3
H3K27 me3:DNA Meth. m5C
H3K27 me3:H3K4 me1
H3K9 me3
H3K4 me3
H4K20 me2
H4−4ac
H2A.Z
DNA Meth. m5C
H4K20 me3
H3K4 me1
H3K27 me3
H3−5ac
0.00 0.25 0.50 0.75 1.00
π̂
CLK2 – Reverse
H2A.Z:H4−4ac
H2A.Z:H4K20 me2
H3K4 me1:H4K20 me2
H3K4 me1:H2A.Z
H3K27 me3:H4K20 me3
H3K27 me3:H3K9 me3
H3K4 me3:H3−5ac
H3K4 me3:H4−4ac
H4K20 me2:H3K4 me3
H3K4 me1:H4−4ac
H4−4ac:H3−5ac
H2A.Z:H3−5ac
H2A.Z:H3K4 me3
H3K9 me3:H4K20 me3
DNA Meth. m5C:H4K20 me3
H3K4 me1:H3−5ac
H4K20 me2:H3−5ac
DNA Meth. m5C:H3K9 me3
H3K27 me3:DNA Meth. m5C
H3K27 me3:H3K4 me1
H3K9 me3
H3K4 me3
H4K20 me2
H4−4ac
H2A.Z
DNA Meth. m5C
H4K20 me3
H3K4 me1
H3K27 me3
H3−5ac
0.00 0.25 0.50 0.75 1.00
π̂
ZFC3H1 – Forward
H3K4 me1:H2A.Z
H3K4 me1:H4−4ac
H2A.Z:H3K4 me3
H3K4 me1:H3−5ac
H3K4 me3:H3−5ac
H4−4ac:H3−5ac
H3K27 me3:H3K4 me1
H3K4 me1:H4K20 me2
DNA Meth. m5C:H4K20 me3
H2A.Z:H4K20 me2
DNA Meth. m5C:H4K20 me2
H3K27 me3:H3K9 me3
DNA Meth. m5C:H3K9 me3
H3K27 me3:DNA Meth. m5C
H4K20 me2:H3K4 me3
H4K20 me2:H4−4ac
H3K9 me3:H4K20 me3
H3K27 me3:H4K20 me3
H4K20 me2:H3−5ac
H3K9 me2
H3K4 me1
H2A.Z
H4−4ac
H4K20 me3
DNA Meth. m5C
H3K4 me3
H3−5ac
H3K9 me3
H4K20 me2
H3K27 me3
0.00 0.25 0.50 0.75 1.00
π̂
ZFC3H1 – Reverse
H3K4 me1:H2A.Z
H3K4 me1:H4−4ac
H2A.Z:H3K4 me3
H3K4 me1:H3−5ac
H3K4 me3:H3−5ac
H4−4ac:H3−5ac
H3K27 me3:H3K4 me1
H3K4 me1:H4K20 me2
DNA Meth. m5C:H4K20 me3
H2A.Z:H4K20 me2
DNA Meth. m5C:H4K20 me2
H3K27 me3:H3K9 me3
DNA Meth. m5C:H3K9 me3
H3K27 me3:DNA Meth. m5C
H4K20 me2:H3K4 me3
H4K20 me2:H4−4ac
H3K9 me3:H4K20 me3
H3K27 me3:H4K20 me3
H4K20 me2:H3−5ac
H3K9 me2
H3K4 me1
H2A.Z
H4−4ac
H4K20 me3
DNA Meth. m5C
H3K4 me3
H3−5ac
H3K9 me3
H4K20 me2
H3K27 me3
0.00 0.25 0.50 0.75 1.00
π̂
PQBP1 – Forward
H4K20 me2:H3K4 me3
H3K4 me1:H4−4ac
H3K4 me3:H4−4ac
H2A.Z:H3K4 me3
H2A.Z:H4K20 me2
H3K4 me1:H2A.Z
H3K27 me3:DNA Meth. m5C
DNA Meth. m5C:H4K20 me2
DNA Meth. m5C:H3K9 me3
H3K4 me3:H3−5ac
H3K9 me3:H4K20 me3
H3K27 me3:H4K20 me3
H3K4 me1:H3−5ac
H4K20 me2:H4−4ac
H4−4ac:H3−5ac
H4K20 me2:H3−5ac
H3K27 me2
H3K9 me2
H3K9 me3
H4K20 me3
H2A.Z
DNA Meth. m5C
H3K4 me3
H3K27 me3
H3−5ac
H4−4ac
H4K20 me2
H3K4 me1
0.00 0.25 0.50 0.75 1.00
π̂
PQBP1 – Reverse
H4K20 me2:H3K4 me3
H3K4 me1:H4−4ac
H3K4 me3:H4−4ac
H2A.Z:H3K4 me3
H2A.Z:H4K20 me2
H3K4 me1:H2A.Z
H3K27 me3:DNA Meth. m5C
DNA Meth. m5C:H4K20 me2
DNA Meth. m5C:H3K9 me3
H3K4 me3:H3−5ac
H3K9 me3:H4K20 me3
H3K27 me3:H4K20 me3
H3K4 me1:H3−5ac
H4K20 me2:H4−4ac
H4−4ac:H3−5ac
H4K20 me2:H3−5ac
H3K27 me2
H3K9 me2
H3K9 me3
H4K20 me3
H2A.Z
DNA Meth. m5C
H3K4 me3
H3K27 me3
H3−5ac
H4−4ac
H4K20 me2
H3K4 me1
0.00 0.25 0.50 0.75 1.00
π̂

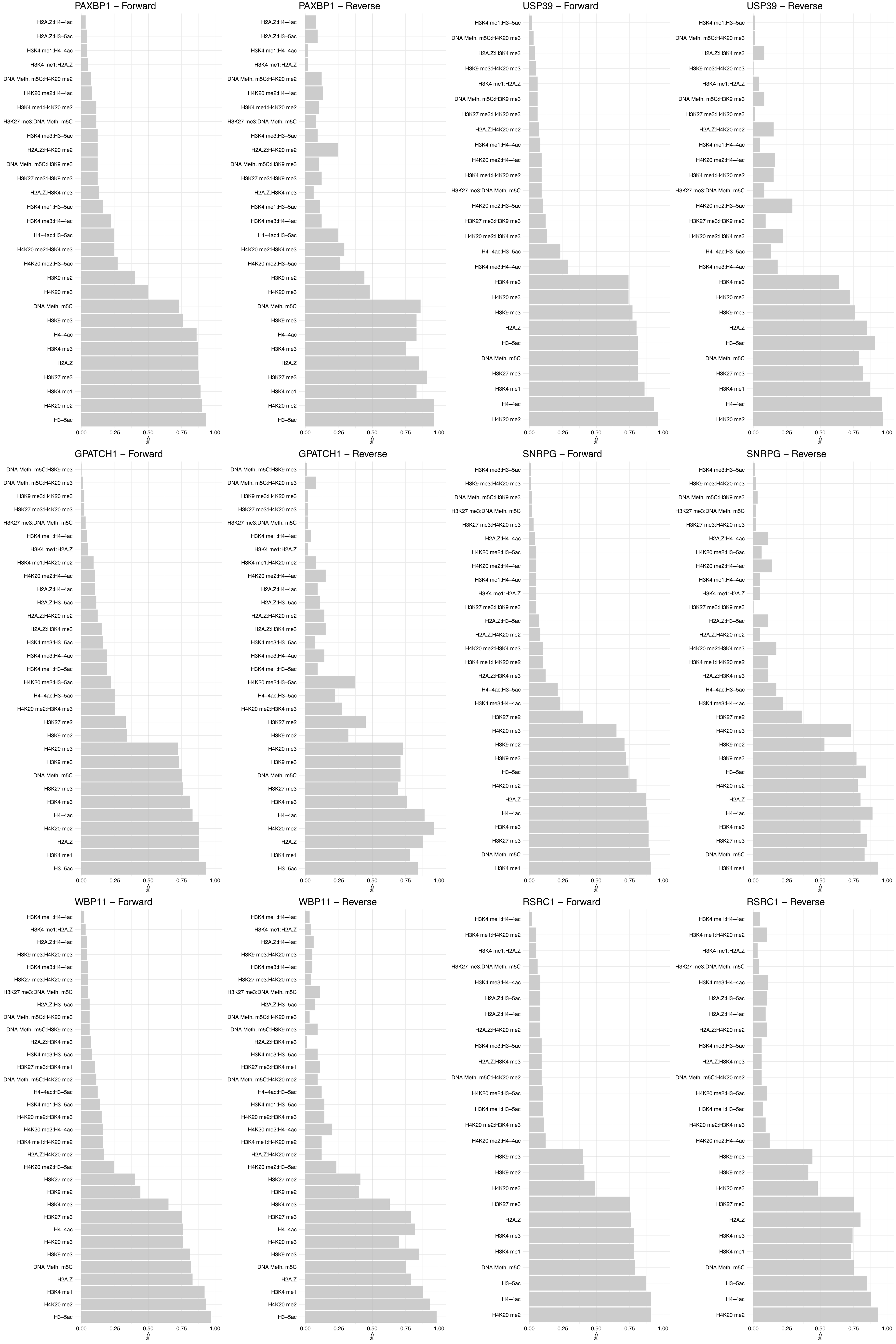
PAXBP1 – Forward
H2A.Z:H4–4ac
H2A.Z:H3–5ac
H3K4 me1:H4–4ac
H3K4 me1:H2A.Z
DNA Meth. m5C:H4K20 me2
H4K20 me2:H4–4ac
H3K4 me1:H4K20 me2
H3K27 me3:DNA Meth. m5C
H3K4 me3:H3–5ac
H2A.Z:H4K20 me2
DNA Meth. m5C:H3K9 me3
H3K27 me3:H3K9 me3
H2A.Z:H3K4 me3
H3K4 me1:H3–5ac
H3K4 me3:H4–4ac
H4–4ac:H3–5ac
H4K20 me2:H3K4 me3
H4K20 me2:H3–5ac
H3K9 me2
H4K20 me3
DNA Meth. m5C
H3K9 me3
H4–4ac
H3K4 me3
H2A.Z
H3K27 me3
H3K4 me1
H4K20 me2
H3–5ac
0.00 0.25 0.50 0.75 1.00
π̂
PAXBP1 – Reverse
H2A.Z:H4–4ac
H2A.Z:H3–5ac
H3K4 me1:H4–4ac
H3K4 me1:H2A.Z
DNA Meth. m5C:H4K20 me2
H4K20 me2:H4–4ac
H3K4 me1:H4K20 me2
H3K27 me3:DNA Meth. m5C
H3K4 me3:H3–5ac
H2A.Z:H4K20 me2
DNA Meth. m5C:H3K9 me3
H3K27 me3:H3K9 me3
H2A.Z:H3K4 me3
H3K4 me1:H3–5ac
H3K4 me3:H4–4ac
H4–4ac:H3–5ac
H4K20 me2:H3K4 me3
H4K20 me2:H3–5ac
H3K9 me2
H4K20 me3
DNA Meth. m5C
H3K9 me3
H4–4ac
H3K4 me3
H2A.Z
H3K27 me3
H3K4 me1
H4K20 me2
H3–5ac
0.00 0.25 0.50 0.75 1.00
π̂
USP39 – Forward
H3K4 me1:H3–5ac
DNA Meth. m5C:H4K20 me3
H2A.Z:H3K4 me3
H3K9 me3:H4K20 me3
H3K4 me1:H2A.Z
DNA Meth. m5C:H3K9 me3
H3K27 me3:H4K20 me3
H2A.Z:H4K20 me2
H3K4 me1:H4–4ac
H4K20 me2:H4–4ac
H3K4 me1:H4K20 me2
H3K27 me3:DNA Meth. m5C
H4K20 me2:H3–5ac
H3K27 me3:H3K9 me3
H4K20 me2:H3K4 me3
H4–4ac:H3–5ac
H3K4 me3:H4–4ac
H3K4 me3
H4K20 me3
H3K9 me3
H2A.Z
H3–5ac
DNA Meth. m5C
H3K27 me3
H3K4 me1
H4–4ac
H4K20 me2
0.00 0.25 0.50 0.75 1.00
π̂
USP39 – Reverse
H3K4 me1:H3–5ac
DNA Meth. m5C:H4K20 me3
H2A.Z:H3K4 me3
H3K9 me3:H4K20 me3
H3K4 me1:H2A.Z
DNA Meth. m5C:H3K9 me3
H3K27 me3:H4K20 me3
H2A.Z:H4K20 me2
H3K4 me1:H4–4ac
H4K20 me2:H4–4ac
H3K4 me1:H4K20 me2
H3K27 me3:DNA Meth. m5C
H4K20 me2:H3–5ac
H3K27 me3:H3K9 me3
H4K20 me2:H3K4 me3
H4–4ac:H3–5ac
H3K4 me3:H4–4ac
H3K4 me3
H4K20 me3
H3K9 me3
H2A.Z
H3–5ac
DNA Meth. m5C
H3K27 me3
H3K4 me1
H4–4ac
H4K20 me2
0.00 0.25 0.50 0.75 1.00
π̂
GPATCH1 – Forward
DNA Meth. m5C:H3K9 me3
DNA Meth. m5C:H4K20 me3
H3K9 me3:H4K20 me3
H3K27 me3:H4K20 me3
H3K27 me3:DNA Meth. m5C
H3K4 me1:H4–4ac
H3K4 me1:H2A.Z
H3K4 me1:H4K20 me2
H4K20 me2:H4–4ac
H2A.Z:H4–4ac
H2A.Z:H3–5ac
H2A.Z:H4K20 me2
H2A.Z:H3K4 me3
H3K4 me3:H3–5ac
H3K4 me3:H4–4ac
H3K4 me1:H3–5ac
H4K20 me2:H3–5ac
H4–4ac:H3–5ac
H4K20 me2:H3K4 me3
H3K27 me2
H3K9 me2
H4K20 me3
H3K9 me3
DNA Meth. m5C
H3K27 me3
H3K4 me3
H4–4ac
H4K20 me2
H2A.Z
H3K4 me1
H3–5ac
0.00 0.25 0.50 0.75 1.00
π̂
GPATCH1 – Reverse
DNA Meth. m5C:H3K9 me3
DNA Meth. m5C:H4K20 me3
H3K9 me3:H4K20 me3
H3K27 me3:H4K20 me3
H3K27 me3:DNA Meth. m5C
H3K4 me1:H4–4ac
H3K4 me1:H2A.Z
H3K4 me1:H4K20 me2
H4K20 me2:H4–4ac
H2A.Z:H4–4ac
H2A.Z:H3–5ac
H2A.Z:H4K20 me2
H2A.Z:H3K4 me3
H3K4 me3:H3–5ac
H3K4 me3:H4–4ac
H3K4 me1:H3–5ac
H4K20 me2:H3–5ac
H4–4ac:H3–5ac
H4K20 me2:H3K4 me3
H3K27 me2
H3K9 me2
H4K20 me3
H3K9 me3
DNA Meth. m5C
H3K27 me3
H3K4 me3
H4–4ac
H4K20 me2
H2A.Z
H3K4 me1
H3–5ac
0.00 0.25 0.50 0.75 1.00
π̂
SNRPG – Forward
H3K4 me3:H3–5ac
H3K9 me3:H4K20 me3
DNA Meth. m5C:H3K9 me3
H3K27 me3:DNA Meth. m5C
H3K27 me3:H4K20 me3
H2A.Z:H4–4ac
H4K20 me2:H3–5ac
H4K20 me2:H4–4ac
H3K4 me1:H4–4ac
H3K4 me1:H2A.Z
H3K27 me3:H3K9 me3
H2A.Z:H3–5ac
H2A.Z:H4K20 me2
H4K20 me2:H3K4 me3
H3K4 me1:H4K20 me2
H2A.Z:H3K4 me3
H4–4ac:H3–5ac
H3K4 me3:H4–4ac
H3K27 me2
H4K20 me3
H3K9 me2
H3K9 me3
H3–5ac
H4K20 me2
H2A.Z
H4–4ac
H3K4 me3
H3K27 me3
DNA Meth. m5C
H3K4 me1
0.00 0.25 0.50 0.75 1.00
π̂
SNRPG – Reverse
H3K4 me3:H3–5ac
H3K9 me3:H4K20 me3
DNA Meth. m5C:H3K9 me3
H3K27 me3:DNA Meth. m5C
H3K27 me3:H4K20 me3
H2A.Z:H4–4ac
H4K20 me2:H3–5ac
H4K20 me2:H4–4ac
H3K4 me1:H4–4ac
H3K4 me1:H2A.Z
H3K27 me3:H3K9 me3
H2A.Z:H3–5ac
H2A.Z:H4K20 me2
H4K20 me2:H3K4 me3
H3K4 me1:H4K20 me2
H2A.Z:H3K4 me3
H4–4ac:H3–5ac
H3K4 me3:H4–4ac
H3K27 me2
H4K20 me3
H3K9 me2
H3K9 me3
H3–5ac
H4K20 me2
H2A.Z
H4–4ac
H3K4 me3
H3K27 me3
DNA Meth. m5C
H3K4 me1
0.00 0.25 0.50 0.75 1.00
π̂
WBP11 – Forward
H3K4 me1:H4–4ac
H3K4 me1:H2A.Z
H2A.Z:H4–4ac
H3K9 me3:H4K20 me3
H3K4 me3:H4–4ac
H3K27 me3:H4K20 me3
H3K27 me3:DNA Meth. m5C
H2A.Z:H3–5ac
DNA Meth. m5C:H4K20 me3
DNA Meth. m5C:H3K9 me3
H2A.Z:H3K4 me3
H3K4 me3:H3–5ac
H3K27 me3:H3K4 me1
DNA Meth. m5C:H4K20 me2
H4–4ac:H3–5ac
H3K4 me1:H3–5ac
H4K20 me2:H3K4 me3
H4K20 me2:H4–4ac
H3K4 me1:H4K20 me2
H2A.Z:H4K20 me2
H4K20 me2:H3–5ac
H3K27 me2
H3K9 me2
H3K4 me3
H3K27 me3
H4–4ac
H4K20 me3
H3K9 me3
DNA Meth. m5C
H2A.Z
H3K4 me1
H4K20 me2
H3–5ac
0.00 0.25 0.50 0.75 1.00
π̂
WBP11 – Reverse
H3K4 me1:H4–4ac
H3K4 me1:H2A.Z
H2A.Z:H4–4ac
H3K9 me3:H4K20 me3
H3K4 me3:H4–4ac
H3K27 me3:H4K20 me3
H3K27 me3:DNA Meth. m5C
H2A.Z:H3–5ac
DNA Meth. m5C:H4K20 me3
DNA Meth. m5C:H3K9 me3
H2A.Z:H3K4 me3
H3K4 me3:H3–5ac
H3K27 me3:H3K4 me1
DNA Meth. m5C:H4K20 me2
H4–4ac:H3–5ac
H3K4 me1:H3–5ac
H4K20 me2:H3K4 me3
H4K20 me2:H4–4ac
H3K4 me1:H4K20 me2
H2A.Z:H4K20 me2
H4K20 me2:H3–5ac
H3K27 me2
H3K9 me2
H3K4 me3
H3K27 me3
H4–4ac
H4K20 me3
H3K9 me3
DNA Meth. m5C
H2A.Z
H3K4 me1
H4K20 me2
H3–5ac
0.00 0.25 0.50 0.75 1.00
π̂
RSRC1 – Forward
H3K4 me1:H4–4ac
H3K4 me1:H4K20 me2
H3K4 me1:H2A.Z
H3K27 me3:DNA Meth. m5C
H3K4 me3:H4–4ac
H2A.Z:H3–5ac
H2A.Z:H4–4ac
H2A.Z:H4K20 me2
H3K4 me3:H3–5ac
H2A.Z:H3K4 me3
DNA Meth. m5C:H4K20 me2
H4K20 me2:H3–5ac
H3K4 me1:H3–5ac
H4K20 me2:H3K4 me3
H4K20 me2:H4–4ac
H3K9 me3
H3K9 me2
H4K20 me3
H3K27 me3
H2A.Z
H3K4 me3
H3K4 me1
DNA Meth. m5C
H3–5ac
H4–4ac
H4K20 me2
0.00 0.25 0.50 0.75 1.00
π̂
RSRC1 – Reverse
H3K4 me1:H4–4ac
H3K4 me1:H4K20 me2
H3K4 me1:H2A.Z
H3K27 me3:DNA Meth. m5C
H3K4 me3:H4–4ac
H2A.Z:H3–5ac
H2A.Z:H4–4ac
H2A.Z:H4K20 me2
H3K4 me3:H3–5ac
H2A.Z:H3K4 me3
DNA Meth. m5C:H4K20 me2
H4K20 me2:H3–5ac
H3K4 me1:H3–5ac
H4K20 me2:H3K4 me3
H4K20 me2:H4–4ac
H3K9 me3
H3K9 me2
H4K20 me3
H3K27 me3
H2A.Z
H3K4 me3
H3K4 me1
DNA Meth. m5C
H3–5ac
H4–4ac
H4K20 me2
0.00 0.25 0.50 0.75 1.00
π̂

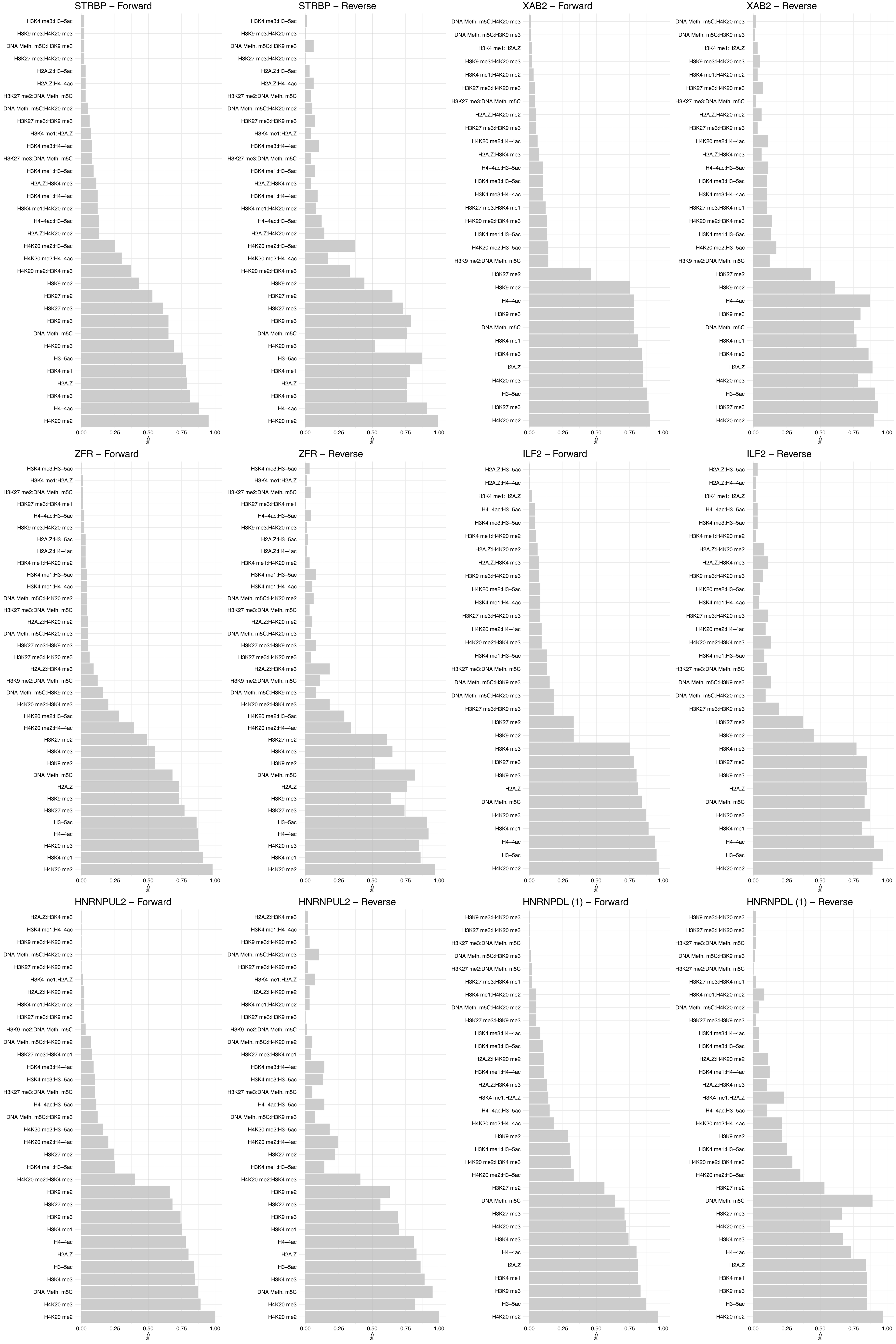
STRBP – Forward
STRBP – Reverse
H3K4 me3:H3–5ac
H3K9 me3:H4K20 me3
DNA Meth. m5C:H3K9 me3
H3K27 me3:H4K20 me3
H2A.Z:H3–5ac
H2A.Z:H4–4ac
H3K27 me2:DNA Meth. m5C
DNA Meth. m5C:H4K20 me2
H3K27 me3:H3K9 me3
H3K4 me1:H2A.Z
H3K4 me3:H4–4ac
H3K27 me3:DNA Meth. m5C
H3K4 me1:H3–5ac
H2A.Z:H3K4 me3
H3K4 me1:H4–4ac
H3K4 me1:H4K20 me2
H4–4ac:H3–5ac
H2A.Z:H4K20 me2
H4K20 me2:H3–5ac
H4K20 me2:H4–4ac
H4K20 me2:H3K4 me3
H3K9 me2
H3K27 me2
H3K27 me3
H3K9 me3
DNA Meth. m5C
H4K20 me3
H3–5ac
H3K4 me1
H2A.Z
H3K4 me3
H4–4ac
H4K20 me2
XAB2 – Forward
XAB2 – Reverse
DNA Meth. m5C:H4K20 me3
DNA Meth. m5C:H3K9 me3
H3K4 me1:H2A.Z
H3K9 me3:H4K20 me3
H3K4 me1:H4K20 me2
H3K27 me3:H4K20 me3
H3K27 me3:DNA Meth. m5C
H2A.Z:H4K20 me2
H3K27 me3:H3K9 me3
H4K20 me2:H4–4ac
H2A.Z:H3K4 me3
H4–4ac:H3–5ac
H3K4 me3:H3–5ac
H3K4 me3:H4–4ac
H3K27 me3:H3K4 me1
H4K20 me2:H3K4 me3
H3K4 me1:H3–5ac
H4K20 me2:H3–5ac
H3K9 me2:DNA Meth. m5C
H3K27 me2
H3K9 me2
H4–4ac
H3K9 me3
DNA Meth. m5C
H3K4 me1
H3K4 me3
H2A.Z
H4K20 me3
H3–5ac
H3K27 me3
H4K20 me2
ZFR – Forward
ZFR – Reverse
H3K4 me3:H3–5ac
H3K4 me1:H2A.Z
H3K27 me2:DNA Meth. m5C
H3K27 me3:H3K4 me1
H4–4ac:H3–5ac
H3K9 me3:H4K20 me3
H2A.Z:H3–5ac
H2A.Z:H4–4ac
H3K4 me1:H4K20 me2
H3K4 me1:H3–5ac
H3K4 me1:H4–4ac
DNA Meth. m5C:H4K20 me2
H3K27 me3:DNA Meth. m5C
H2A.Z:H4K20 me2
DNA Meth. m5C:H4K20 me3
H3K27 me3:H3K9 me3
H3K27 me3:H4K20 me3
H2A.Z:H3K4 me3
H3K9 me2:DNA Meth. m5C
DNA Meth. m5C:H3K9 me3
H4K20 me2:H3K4 me3
H4K20 me2:H3–5ac
H4K20 me2:H4–4ac
H3K27 me2
H3K4 me3
H3K9 me2
DNA Meth. m5C
H2A.Z
H3K9 me3
H3K27 me3
H3–5ac
H4–4ac
H4K20 me3
H3K4 me1
H4K20 me2
ILF2 – Forward
ILF2 – Reverse
H2A.Z:H3–5ac
H2A.Z:H4–4ac
H3K4 me1:H2A.Z
H4–4ac:H3–5ac
H3K4 me3:H3–5ac
H3K4 me1:H4K20 me2
H2A.Z:H4K20 me2
H2A.Z:H3K4 me3
H3K9 me3:H4K20 me3
H4K20 me2:H3–5ac
H3K4 me1:H4–4ac
H3K27 me3:H4K20 me3
H4K20 me2:H4–4ac
H4K20 me2:H3K4 me3
H3K4 me1:H3–5ac
H3K27 me3:DNA Meth. m5C
DNA Meth. m5C:H3K9 me3
DNA Meth. m5C:H4K20 me3
H3K27 me3:H3K9 me3
H3K27 me2
H3K9 me2
H3K4 me3
H3K27 me3
H3K9 me3
H2A.Z
DNA Meth. m5C
H4K20 me3
H3K4 me1
H4–4ac
H3–5ac
H4K20 me2
HNRNPUL2 – Forward
HNRNPUL2 – Reverse
H2A.Z:H3K4 me3
H3K4 me1:H4–4ac
H3K9 me3:H4K20 me3
DNA Meth. m5C:H4K20 me3
H3K27 me3:H4K20 me3
H3K4 me1:H2A.Z
H2A.Z:H4K20 me2
H3K4 me1:H4K20 me2
H3K27 me3:H3K9 me3
H3K9 me2:DNA Meth. m5C
DNA Meth. m5C:H4K20 me2
H3K27 me3:H3K4 me1
H3K4 me3:H4–4ac
H3K4 me3:H3–5ac
H3K27 me3:DNA Meth. m5C
H4–4ac:H3–5ac
DNA Meth. m5C:H3K9 me3
H4K20 me2:H3–5ac
H4K20 me2:H4–4ac
H3K27 me2
H3K4 me1:H3–5ac
H4K20 me2:H3K4 me3
H3K9 me2
H3K27 me3
H3K9 me3
H3K4 me1
H4–4ac
H2A.Z
H3–5ac
H3K4 me3
DNA Meth. m5C
H4K20 me3
H4K20 me2
HNRNPDL (1) – Forward
HNRNPDL (1) – Reverse
H3K9 me3:H4K20 me3
H3K27 me3:H4K20 me3
H3K27 me3:DNA Meth. m5C
DNA Meth. m5C:H3K9 me3
H3K27 me2:DNA Meth. m5C
H3K27 me3:H3K4 me1
H3K4 me1:H4K20 me2
DNA Meth. m5C:H4K20 me2
H3K27 me3:H3K9 me3
H3K4 me3:H4–4ac
H3K4 me3:H3–5ac
H2A.Z:H4K20 me2
H3K4 me1:H4–4ac
H2A.Z:H3K4 me3
H3K4 me1:H2A.Z
H4–4ac:H3–5ac
H4K20 me2:H4–4ac
H3K9 me2
H3K4 me1:H3–5ac
H4K20 me2:H3K4 me3
H4K20 me2:H3–5ac
H3K27 me2
DNA Meth. m5C
H3K27 me3
H4K20 me3
H3K4 me3
H4–4ac
H2A.Z
H3K4 me1
H3K9 me3
H3–5ac
H4K20 me2
0.00
0.25
0.50
0.75
1.00
$\hat{\pi}$

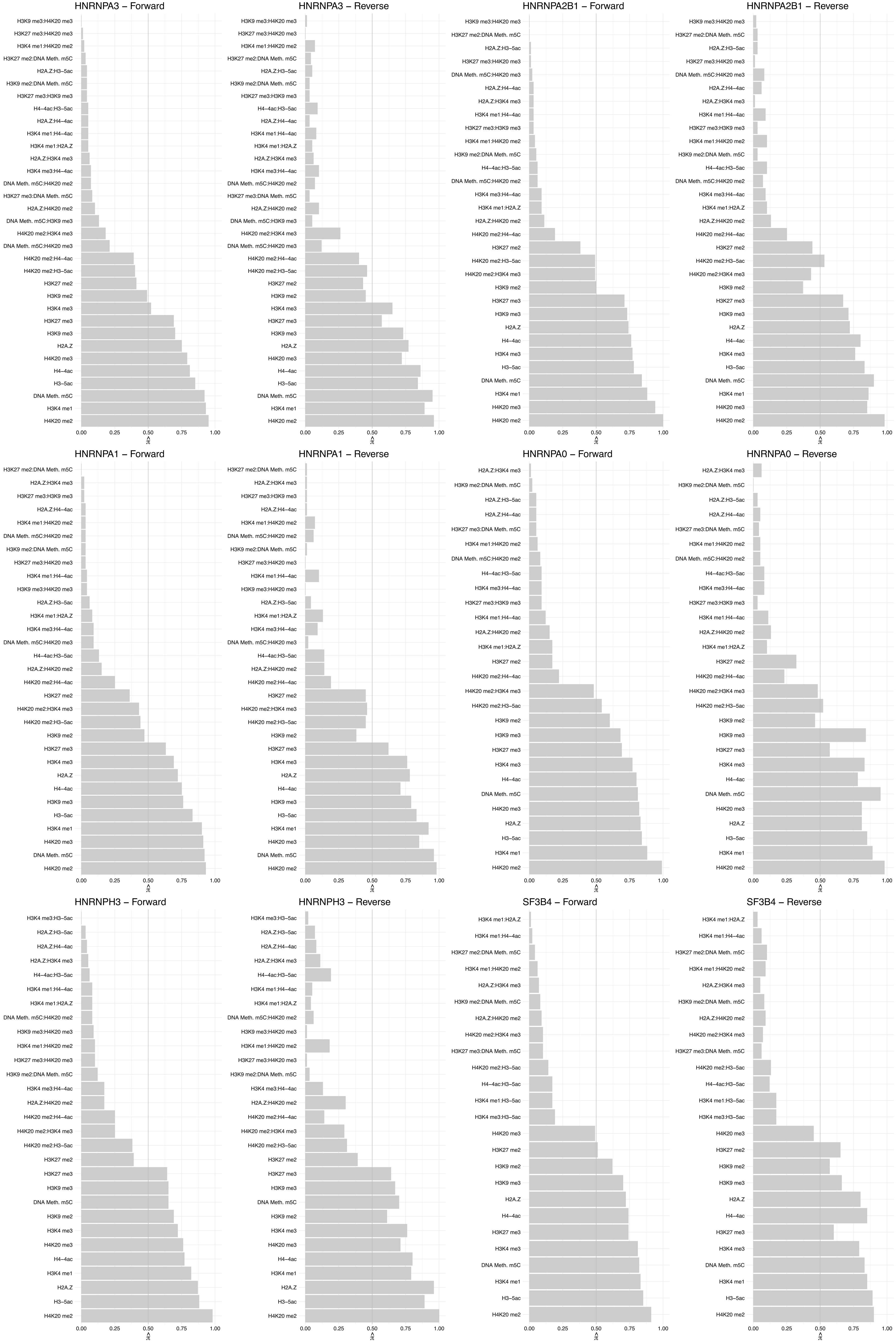
HNRNPA3 − Forward
H3K9 me3:H4K20 me3
H3K27 me3:H4K20 me3
H3K4 me1:H4K20 me2
H3K27 me2:DNA Meth. m5C
H2A.Z:H3−5ac
H3K9 me2:DNA Meth. m5C
H3K27 me3:H3K9 me3
H4−4ac:H3−5ac
H2A.Z:H4−4ac
H3K4 me1:H4−4ac
H3K4 me1:H2A.Z
H2A.Z:H3K4 me3
H3K4 me3:H4−4ac
DNA Meth. m5C:H4K20 me2
H3K27 me3:DNA Meth. m5C
H2A.Z:H4K20 me2
DNA Meth. m5C:H3K9 me3
H4K20 me2:H3K4 me3
DNA Meth. m5C:H4K20 me3
H4K20 me2:H4−4ac
H4K20 me2:H3−5ac
H3K27 me2
H3K9 me2
H3K4 me3
H3K27 me3
H3K9 me3
H2A.Z
H4K20 me3
H4−4ac
H3−5ac
DNA Meth. m5C
H3K4 me1
H4K20 me2
0.00 0.25 0.50 0.75 1.00
$\hat{\pi}$
HNRNPA3 − Reverse
H3K9 me3:H4K20 me3
H3K27 me3:H4K20 me3
H3K4 me1:H4K20 me2
H3K27 me2:DNA Meth. m5C
H2A.Z:H3−5ac
H3K9 me2:DNA Meth. m5C
H3K27 me3:H3K9 me3
H4−4ac:H3−5ac
H2A.Z:H4−4ac
H3K4 me1:H4−4ac
H3K4 me1:H2A.Z
H2A.Z:H3K4 me3
H3K4 me3:H4−4ac
DNA Meth. m5C:H4K20 me2
H3K27 me3:DNA Meth. m5C
H2A.Z:H4K20 me2
DNA Meth. m5C:H3K9 me3
H4K20 me2:H3K4 me3
DNA Meth. m5C:H4K20 me3
H4K20 me2:H4−4ac
H4K20 me2:H3−5ac
H3K27 me2
H3K9 me2
H3K4 me3
H3K27 me3
H3K9 me3
H2A.Z
H4K20 me3
H4−4ac
H3−5ac
DNA Meth. m5C
H3K4 me1
H4K20 me2
0.00 0.25 0.50 0.75 1.00
$\hat{\pi}$
HNRNPA2B1 − Forward
H3K9 me3:H4K20 me3
H3K27 me2:DNA Meth. m5C
H2A.Z:H3−5ac
H3K27 me3:H4K20 me3
DNA Meth. m5C:H4K20 me3
H2A.Z:H4−4ac
H2A.Z:H3K4 me3
H3K4 me1:H4−4ac
H3K27 me3:H3K9 me3
H3K4 me1:H4K20 me2
H3K9 me2:DNA Meth. m5C
H4−4ac:H3−5ac
DNA Meth. m5C:H4K20 me2
H3K4 me3:H4−4ac
H3K4 me1:H2A.Z
H2A.Z:H4K20 me2
H4K20 me2:H4−4ac
H3K27 me2
H4K20 me2:H3−5ac
H4K20 me2:H3K4 me3
H3K9 me2
H3K27 me3
H3K9 me3
H2A.Z
H4−4ac
H3K4 me3
H3−5ac
DNA Meth. m5C
H3K4 me1
H4K20 me3
H4K20 me2
0.00 0.25 0.50 0.75 1.00
$\hat{\pi}$
HNRNPA2B1 − Reverse
H3K9 me3:H4K20 me3
H3K27 me2:DNA Meth. m5C
H2A.Z:H3−5ac
H3K27 me3:H4K20 me3
DNA Meth. m5C:H4K20 me3
H2A.Z:H4−4ac
H2A.Z:H3K4 me3
H3K4 me1:H4−4ac
H3K27 me3:H3K9 me3
H3K4 me1:H4K20 me2
H3K9 me2:DNA Meth. m5C
H4−4ac:H3−5ac
DNA Meth. m5C:H4K20 me2
H3K4 me3:H4−4ac
H3K4 me1:H2A.Z
H2A.Z:H4K20 me2
H4K20 me2:H4−4ac
H3K27 me2
H4K20 me2:H3−5ac
H4K20 me2:H3K4 me3
H3K9 me2
H3K27 me3
H3K9 me3
H2A.Z
H4−4ac
H3K4 me3
H3−5ac
DNA Meth. m5C
H3K4 me1
H4K20 me3
H4K20 me2
0.00 0.25 0.50 0.75 1.00
$\hat{\pi}$
HNRNPA1 − Forward
H3K27 me2:DNA Meth. m5C
H2A.Z:H3K4 me3
H3K27 me3:H3K9 me3
H2A.Z:H4−4ac
H3K4 me1:H4K20 me2
DNA Meth. m5C:H4K20 me2
H3K9 me2:DNA Meth. m5C
H3K27 me3:H4K20 me3
H3K4 me1:H4−4ac
H3K9 me3:H4K20 me3
H2A.Z:H3−5ac
H3K4 me1:H2A.Z
H3K4 me3:H4−4ac
DNA Meth. m5C:H4K20 me3
H4−4ac:H3−5ac
H2A.Z:H4K20 me2
H4K20 me2:H4−4ac
H3K27 me2
H4K20 me2:H3K4 me3
H4K20 me2:H3−5ac
H3K9 me2
H3K27 me3
H3K4 me3
H2A.Z
H4−4ac
H3K9 me3
H3−5ac
H3K4 me1
H4K20 me3
DNA Meth. m5C
H4K20 me2
0.00 0.25 0.50 0.75 1.00
$\hat{\pi}$
HNRNPA1 − Reverse
H3K27 me2:DNA Meth. m5C
H2A.Z:H3K4 me3
H3K27 me3:H3K9 me3
H2A.Z:H4−4ac
H3K4 me1:H4K20 me2
DNA Meth. m5C:H4K20 me2
H3K9 me2:DNA Meth. m5C
H3K27 me3:H4K20 me3
H3K4 me1:H4−4ac
H3K9 me3:H4K20 me3
H2A.Z:H3−5ac
H3K4 me1:H2A.Z
H3K4 me3:H4−4ac
DNA Meth. m5C:H4K20 me3
H4−4ac:H3−5ac
H2A.Z:H4K20 me2
H4K20 me2:H4−4ac
H3K27 me2
H4K20 me2:H3K4 me3
H4K20 me2:H3−5ac
H3K9 me2
H3K27 me3
H3K4 me3
H2A.Z
H4−4ac
H3K9 me3
H3−5ac
H3K4 me1
H4K20 me3
DNA Meth. m5C
H4K20 me2
0.00 0.25 0.50 0.75 1.00
$\hat{\pi}$
HNRNPA0 − Forward
H2A.Z:H3K4 me3
H3K9 me2:DNA Meth. m5C
H2A.Z:H3−5ac
H2A.Z:H4−4ac
H3K27 me3:DNA Meth. m5C
H3K4 me1:H4K20 me2
DNA Meth. m5C:H4K20 me2
H4−4ac:H3−5ac
H3K4 me3:H4−4ac
H3K27 me3:H3K9 me3
H3K4 me1:H4−4ac
H2A.Z:H4K20 me2
H3K4 me1:H2A.Z
H3K27 me2
H4K20 me2:H4−4ac
H4K20 me2:H3K4 me3
H4K20 me2:H3−5ac
H3K9 me2
H3K9 me3
H3K27 me3
H3K4 me3
H4−4ac
DNA Meth. m5C
H4K20 me3
H2A.Z
H3−5ac
H3K4 me1
H4K20 me2
0.00 0.25 0.50 0.75 1.00
$\hat{\pi}$
HNRNPA0 − Reverse
H2A.Z:H3K4 me3
H3K9 me2:DNA Meth. m5C
H2A.Z:H3−5ac
H2A.Z:H4−4ac
H3K27 me3:DNA Meth. m5C
H3K4 me1:H4K20 me2
DNA Meth. m5C:H4K20 me2
H4−4ac:H3−5ac
H3K4 me3:H4−4ac
H3K27 me3:H3K9 me3
H3K4 me1:H4−4ac
H2A.Z:H4K20 me2
H3K4 me1:H2A.Z
H3K27 me2
H4K20 me2:H4−4ac
H4K20 me2:H3K4 me3
H4K20 me2:H3−5ac
H3K9 me2
H3K9 me3
H3K27 me3
H3K4 me3
H4−4ac
DNA Meth. m5C
H4K20 me3
H2A.Z
H3−5ac
H3K4 me1
H4K20 me2
0.00 0.25 0.50 0.75 1.00
$\hat{\pi}$
HNRNPH3 − Forward
H3K4 me3:H3−5ac
H2A.Z:H3−5ac
H2A.Z:H4−4ac
H2A.Z:H3K4 me3
H4−4ac:H3−5ac
H3K4 me1:H4−4ac
H3K4 me1:H2A.Z
DNA Meth. m5C:H4K20 me2
H3K9 me3:H4K20 me3
H3K4 me1:H4K20 me2
H3K27 me3:H4K20 me3
H3K9 me2:DNA Meth. m5C
H3K4 me3:H4−4ac
H2A.Z:H4K20 me2
H4K20 me2:H4−4ac
H4K20 me2:H3K4 me3
H4K20 me2:H3−5ac
H3K27 me2
H3K27 me3
H3K9 me3
DNA Meth. m5C
H3K9 me2
H3K4 me3
H4K20 me3
H4−4ac
H3K4 me1
H2A.Z
H3−5ac
H4K20 me2
0.00 0.25 0.50 0.75 1.00
$\hat{\pi}$
HNRNPH3 − Reverse
H3K4 me3:H3−5ac
H2A.Z:H3−5ac
H2A.Z:H4−4ac
H2A.Z:H3K4 me3
H4−4ac:H3−5ac
H3K4 me1:H4−4ac
H3K4 me1:H2A.Z
DNA Meth. m5C:H4K20 me2
H3K9 me3:H4K20 me3
H3K4 me1:H4K20 me2
H3K27 me3:H4K20 me3
H3K9 me2:DNA Meth. m5C
H3K4 me3:H4−4ac
H2A.Z:H4K20 me2
H4K20 me2:H4−4ac
H4K20 me2:H3K4 me3
H4K20 me2:H3−5ac
H3K27 me2
H3K27 me3
H3K9 me3
DNA Meth. m5C
H3K9 me2
H3K4 me3
H4K20 me3
H4−4ac
H3K4 me1
H2A.Z
H3−5ac
H4K20 me2
0.00 0.25 0.50 0.75 1.00
$\hat{\pi}$
SF3B4 − Forward
H3K4 me1:H2A.Z
H3K4 me1:H4−4ac
H3K27 me2:DNA Meth. m5C
H3K4 me1:H4K20 me2
H2A.Z:H3K4 me3
H3K9 me2:DNA Meth. m5C
H2A.Z:H4K20 me2
H4K20 me2:H3K4 me3
H3K27 me3:DNA Meth. m5C
H4K20 me2:H3−5ac
H4−4ac:H3−5ac
H3K4 me1:H3−5ac
H3K4 me3:H3−5ac
H4K20 me3
H3K27 me2
H3K9 me2
H3K9 me3
H2A.Z
H4−4ac
H3K27 me3
H3K4 me3
DNA Meth. m5C
H3K4 me1
H3−5ac
H4K20 me2
0.00 0.25 0.50 0.75 1.00
$\hat{\pi}$
SF3B4 − Reverse
H3K4 me1:H2A.Z
H3K4 me1:H4−4ac
H3K27 me2:DNA Meth. m5C
H3K4 me1:H4K20 me2
H2A.Z:H3K4 me3
H3K9 me2:DNA Meth. m5C
H2A.Z:H4K20 me2
H4K20 me2:H3K4 me3
H3K27 me3:DNA Meth. m5C
H4K20 me2:H3−5ac
H4−4ac:H3−5ac
H3K4 me1:H3−5ac
H3K4 me3:H3−5ac
H4K20 me3
H3K27 me2
H3K9 me2
H3K9 me3
H2A.Z
H4−4ac
H3K27 me3
H3K4 me3
DNA Meth. m5C
H3K4 me1
H3−5ac
H4K20 me2
0.00 0.25 0.50 0.75 1.00
$\hat{\pi}$

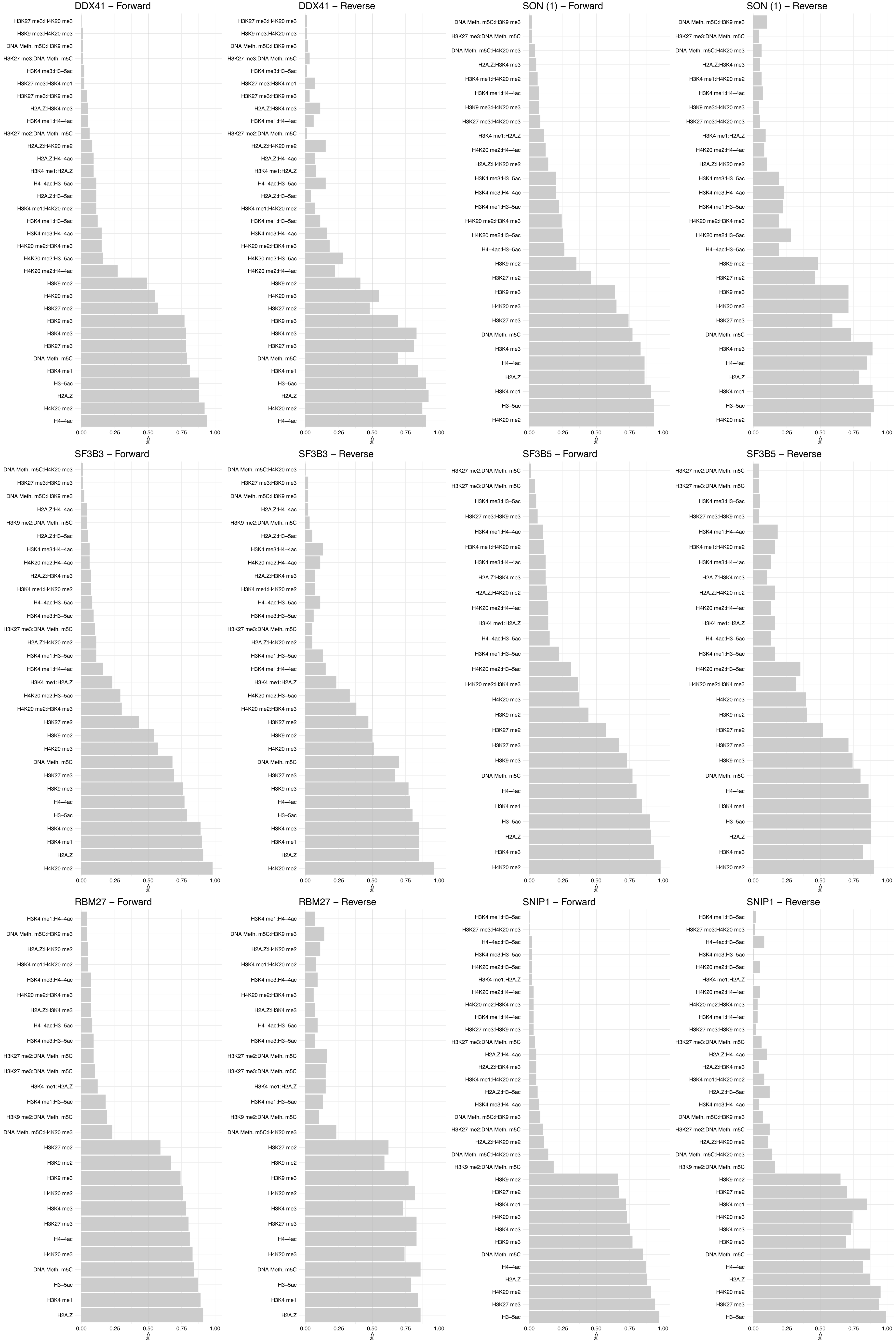
DDX41 – Forward
H3K27 me3:H4K20 me3
H3K9 me3:H4K20 me3
DNA Meth. m5C:H3K9 me3
H3K27 me3:DNA Meth. m5C
H3K4 me3:H3−5ac
H3K27 me3:H3K4 me1
H3K27 me3:H3K9 me3
H2A.Z:H3K4 me3
H3K4 me1:H4−4ac
H3K27 me2:DNA Meth. m5C
H2A.Z:H4K20 me2
H2A.Z:H4−4ac
H3K4 me1:H2A.Z
H4−4ac:H3−5ac
H2A.Z:H3−5ac
H3K4 me1:H4K20 me2
H3K4 me1:H3−5ac
H3K4 me3:H4−4ac
H4K20 me2:H3K4 me3
H4K20 me2:H3−5ac
H4K20 me2:H4−4ac
H3K9 me2
H4K20 me3
H3K27 me2
H3K9 me3
H3K4 me3
H3K27 me3
DNA Meth. m5C
H3K4 me1
H3−5ac
H2A.Z
H4K20 me2
H4−4ac
0.00 0.25 0.50 0.75 1.00
π̂
DDX41 – Reverse
H3K27 me3:H4K20 me3
H3K9 me3:H4K20 me3
DNA Meth. m5C:H3K9 me3
H3K27 me3:DNA Meth. m5C
H3K4 me3:H3−5ac
H3K27 me3:H3K4 me1
H3K27 me3:H3K9 me3
H2A.Z:H3K4 me3
H3K4 me1:H4−4ac
H3K27 me2:DNA Meth. m5C
H2A.Z:H4K20 me2
H2A.Z:H4−4ac
H3K4 me1:H2A.Z
H4−4ac:H3−5ac
H2A.Z:H3−5ac
H3K4 me1:H4K20 me2
H3K4 me1:H3−5ac
H3K4 me3:H4−4ac
H4K20 me2:H3K4 me3
H4K20 me2:H3−5ac
H4K20 me2:H4−4ac
H3K9 me2
H4K20 me3
H3K27 me2
H3K9 me3
H3K4 me3
H3K27 me3
DNA Meth. m5C
H3K4 me1
H3−5ac
H2A.Z
H4K20 me2
H4−4ac
0.00 0.25 0.50 0.75 1.00
π̂
SON (1) – Forward
DNA Meth. m5C:H3K9 me3
H3K27 me3:DNA Meth. m5C
DNA Meth. m5C:H4K20 me3
H2A.Z:H3K4 me3
H3K4 me1:H4K20 me2
H3K4 me1:H4−4ac
H3K9 me3:H4K20 me3
H3K27 me3:H4K20 me3
H3K4 me1:H2A.Z
H4K20 me2:H4−4ac
H2A.Z:H4K20 me2
H3K4 me3:H3−5ac
H3K4 me3:H4−4ac
H3K4 me1:H3−5ac
H4K20 me2:H3K4 me3
H4K20 me2:H3−5ac
H4−4ac:H3−5ac
H3K9 me2
H3K27 me2
H3K9 me3
H4K20 me3
H3K27 me3
DNA Meth. m5C
H3K4 me3
H4−4ac
H2A.Z
H3K4 me1
H3−5ac
H4K20 me2
0.00 0.25 0.50 0.75 1.00
π̂
SON (1) – Reverse
DNA Meth. m5C:H3K9 me3
H3K27 me3:DNA Meth. m5C
DNA Meth. m5C:H4K20 me3
H2A.Z:H3K4 me3
H3K4 me1:H4K20 me2
H3K4 me1:H4−4ac
H3K9 me3:H4K20 me3
H3K27 me3:H4K20 me3
H3K4 me1:H2A.Z
H4K20 me2:H4−4ac
H2A.Z:H4K20 me2
H3K4 me3:H3−5ac
H3K4 me3:H4−4ac
H3K4 me1:H3−5ac
H4K20 me2:H3K4 me3
H4K20 me2:H3−5ac
H4−4ac:H3−5ac
H3K9 me2
H3K27 me2
H3K9 me3
H4K20 me3
H3K27 me3
DNA Meth. m5C
H3K4 me3
H4−4ac
H2A.Z
H3K4 me1
H3−5ac
H4K20 me2
0.00 0.25 0.50 0.75 1.00
π̂
SF3B3 – Forward
DNA Meth. m5C:H4K20 me3
H3K27 me3:H3K9 me3
DNA Meth. m5C:H3K9 me3
H2A.Z:H4−4ac
H3K9 me2:DNA Meth. m5C
H2A.Z:H3−5ac
H3K4 me3:H4−4ac
H4K20 me2:H4−4ac
H2A.Z:H3K4 me3
H3K4 me1:H4K20 me2
H4−4ac:H3−5ac
H3K4 me3:H3−5ac
H3K27 me3:DNA Meth. m5C
H2A.Z:H4K20 me2
H3K4 me1:H3−5ac
H3K4 me1:H4−4ac
H3K4 me1:H2A.Z
H4K20 me2:H3−5ac
H4K20 me2:H3K4 me3
H3K27 me2
H3K9 me2
H4K20 me3
DNA Meth. m5C
H3K27 me3
H3K9 me3
H4−4ac
H3−5ac
H3K4 me3
H3K4 me1
H2A.Z
H4K20 me2
0.00 0.25 0.50 0.75 1.00
π̂
SF3B3 – Reverse
DNA Meth. m5C:H4K20 me3
H3K27 me3:H3K9 me3
DNA Meth. m5C:H3K9 me3
H2A.Z:H4−4ac
H3K9 me2:DNA Meth. m5C
H2A.Z:H3−5ac
H3K4 me3:H4−4ac
H4K20 me2:H4−4ac
H2A.Z:H3K4 me3
H3K4 me1:H4K20 me2
H4−4ac:H3−5ac
H3K4 me3:H3−5ac
H3K27 me3:DNA Meth. m5C
H2A.Z:H4K20 me2
H3K4 me1:H3−5ac
H3K4 me1:H4−4ac
H3K4 me1:H2A.Z
H4K20 me2:H3−5ac
H4K20 me2:H3K4 me3
H3K27 me2
H3K9 me2
H4K20 me3
DNA Meth. m5C
H3K27 me3
H3K9 me3
H4−4ac
H3−5ac
H3K4 me3
H3K4 me1
H2A.Z
H4K20 me2
0.00 0.25 0.50 0.75 1.00
π̂
SF3B5 – Forward
H3K27 me2:DNA Meth. m5C
H3K27 me3:DNA Meth. m5C
H3K4 me3:H3−5ac
H3K27 me3:H3K9 me3
H3K4 me1:H4−4ac
H3K4 me1:H4K20 me2
H3K4 me3:H4−4ac
H2A.Z:H3K4 me3
H2A.Z:H4K20 me2
H4K20 me2:H4−4ac
H3K4 me1:H2A.Z
H4−4ac:H3−5ac
H3K4 me1:H3−5ac
H4K20 me2:H3−5ac
H4K20 me2:H3K4 me3
H4K20 me3
H3K9 me2
H3K27 me2
H3K27 me3
H3K9 me3
DNA Meth. m5C
H4−4ac
H3K4 me1
H3−5ac
H2A.Z
H3K4 me3
H4K20 me2
0.00 0.25 0.50 0.75 1.00
π̂
SF3B5 – Reverse
H3K27 me2:DNA Meth. m5C
H3K27 me3:DNA Meth. m5C
H3K4 me3:H3−5ac
H3K27 me3:H3K9 me3
H3K4 me1:H4−4ac
H3K4 me1:H4K20 me2
H3K4 me3:H4−4ac
H2A.Z:H3K4 me3
H2A.Z:H4K20 me2
H4K20 me2:H4−4ac
H3K4 me1:H2A.Z
H4−4ac:H3−5ac
H3K4 me1:H3−5ac
H4K20 me2:H3−5ac
H4K20 me2:H3K4 me3
H4K20 me3
H3K9 me2
H3K27 me2
H3K27 me3
H3K9 me3
DNA Meth. m5C
H4−4ac
H3K4 me1
H3−5ac
H2A.Z
H3K4 me3
H4K20 me2
0.00 0.25 0.50 0.75 1.00
π̂
RBM27 – Forward
H3K4 me1:H4−4ac
DNA Meth. m5C:H3K9 me3
H2A.Z:H4K20 me2
H3K4 me1:H4K20 me2
H3K4 me3:H4−4ac
H4K20 me2:H3K4 me3
H2A.Z:H3K4 me3
H4−4ac:H3−5ac
H3K4 me3:H3−5ac
H3K27 me2:DNA Meth. m5C
H3K27 me3:DNA Meth. m5C
H3K4 me1:H2A.Z
H3K4 me1:H3−5ac
H3K9 me2:DNA Meth. m5C
DNA Meth. m5C:H4K20 me3
H3K27 me2
H3K9 me2
H3K9 me3
H4K20 me2
H3K4 me3
H3K27 me3
H4−4ac
H4K20 me3
DNA Meth. m5C
H3−5ac
H3K4 me1
H2A.Z
0.00 0.25 0.50 0.75 1.00
π̂
RBM27 – Reverse
H3K4 me1:H4−4ac
DNA Meth. m5C:H3K9 me3
H2A.Z:H4K20 me2
H3K4 me1:H4K20 me2
H3K4 me3:H4−4ac
H4K20 me2:H3K4 me3
H2A.Z:H3K4 me3
H4−4ac:H3−5ac
H3K4 me3:H3−5ac
H3K27 me2:DNA Meth. m5C
H3K27 me3:DNA Meth. m5C
H3K4 me1:H2A.Z
H3K4 me1:H3−5ac
H3K9 me2:DNA Meth. m5C
DNA Meth. m5C:H4K20 me3
H3K27 me2
H3K9 me2
H3K9 me3
H4K20 me2
H3K4 me3
H3K27 me3
H4−4ac
H4K20 me3
DNA Meth. m5C
H3−5ac
H3K4 me1
H2A.Z
0.00 0.25 0.50 0.75 1.00
π̂
SNIP1 – Forward
H3K4 me1:H3−5ac
H3K27 me3:H4K20 me3
H4−4ac:H3−5ac
H3K4 me3:H3−5ac
H4K20 me2:H3−5ac
H3K4 me1:H2A.Z
H4K20 me2:H4−4ac
H4K20 me2:H3K4 me3
H3K4 me1:H4−4ac
H3K27 me3:H3K9 me3
H3K27 me3:DNA Meth. m5C
H2A.Z:H4−4ac
H2A.Z:H3K4 me3
H3K4 me1:H4K20 me2
H2A.Z:H3−5ac
H3K4 me3:H4−4ac
DNA Meth. m5C:H3K9 me3
H3K27 me2:DNA Meth. m5C
H2A.Z:H4K20 me2
DNA Meth. m5C:H4K20 me3
H3K9 me2:DNA Meth. m5C
H3K9 me2
H3K27 me2
H3K4 me1
H4K20 me3
H3K4 me3
H3K9 me3
DNA Meth. m5C
H4−4ac
H2A.Z
H4K20 me2
H3K27 me3
H3−5ac
0.00 0.25 0.50 0.75 1.00
π̂
SNIP1 – Reverse
H3K4 me1:H3−5ac
H3K27 me3:H4K20 me3
H4−4ac:H3−5ac
H3K4 me3:H3−5ac
H4K20 me2:H3−5ac
H3K4 me1:H2A.Z
H4K20 me2:H4−4ac
H4K20 me2:H3K4 me3
H3K4 me1:H4−4ac
H3K27 me3:H3K9 me3
H3K27 me3:DNA Meth. m5C
H2A.Z:H4−4ac
H2A.Z:H3K4 me3
H3K4 me1:H4K20 me2
H2A.Z:H3−5ac
H3K4 me3:H4−4ac
DNA Meth. m5C:H3K9 me3
H3K27 me2:DNA Meth. m5C
H2A.Z:H4K20 me2
DNA Meth. m5C:H4K20 me3
H3K9 me2:DNA Meth. m5C
H3K9 me2
H3K27 me2
H3K4 me1
H4K20 me3
H3K4 me3
H3K9 me3
DNA Meth. m5C
H4−4ac
H2A.Z
H4K20 me2
H3K27 me3
H3−5ac
0.00 0.25 0.50 0.75 1.00
π̂

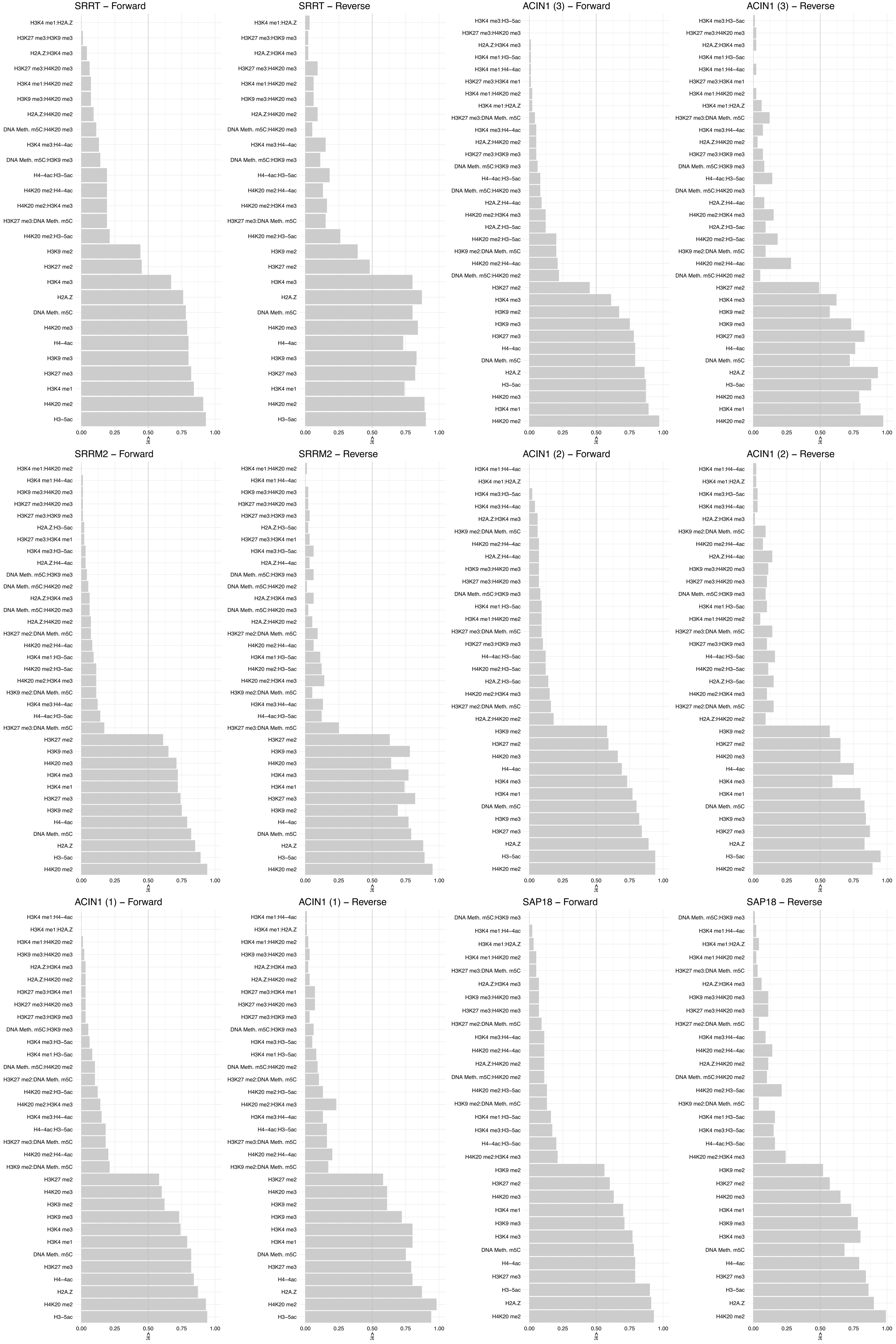
SRRT − Forward
H3K4 me1:H2A.Z
H3K27 me3:H3K9 me3
H2A.Z:H3K4 me3
H3K27 me3:H4K20 me3
H3K4 me1:H4K20 me2
H3K9 me3:H4K20 me3
H2A.Z:H4K20 me2
DNA Meth. m5C:H4K20 me3
H3K4 me3:H4−4ac
DNA Meth. m5C:H3K9 me3
H4−4ac:H3−5ac
H4K20 me2:H4−4ac
H4K20 me2:H3K4 me3
H3K27 me3:DNA Meth. m5C
H4K20 me2:H3−5ac
H3K9 me2
H3K27 me2
H3K4 me3
H2A.Z
DNA Meth. m5C
H4K20 me3
H4−4ac
H3K9 me3
H3K27 me3
H3K4 me1
H4K20 me2
H3−5ac
0.00 0.25 0.50 0.75 1.00
π̂
SRRT − Reverse
H3K4 me1:H2A.Z
H3K27 me3:H3K9 me3
H2A.Z:H3K4 me3
H3K27 me3:H4K20 me3
H3K4 me1:H4K20 me2
H3K9 me3:H4K20 me3
H2A.Z:H4K20 me2
DNA Meth. m5C:H4K20 me3
H3K4 me3:H4−4ac
DNA Meth. m5C:H3K9 me3
H4−4ac:H3−5ac
H4K20 me2:H4−4ac
H4K20 me2:H3K4 me3
H3K27 me3:DNA Meth. m5C
H4K20 me2:H3−5ac
H3K9 me2
H3K27 me2
H3K4 me3
H2A.Z
DNA Meth. m5C
H4K20 me3
H4−4ac
H3K9 me3
H3K27 me3
H3K4 me1
H4K20 me2
H3−5ac
0.00 0.25 0.50 0.75 1.00
π̂
ACIN1 (3) − Forward
H3K4 me3:H3−5ac
H3K27 me3:H4K20 me3
H2A.Z:H3K4 me3
H3K4 me1:H3−5ac
H3K4 me1:H4−4ac
H3K27 me3:H3K4 me1
H3K4 me1:H4K20 me2
H3K4 me1:H2A.Z
H3K27 me3:DNA Meth. m5C
H3K4 me3:H4−4ac
H2A.Z:H4K20 me2
H3K27 me3:H3K9 me3
DNA Meth. m5C:H3K9 me3
H4−4ac:H3−5ac
DNA Meth. m5C:H4K20 me3
H2A.Z:H4−4ac
H4K20 me2:H3K4 me3
H2A.Z:H3−5ac
H4K20 me2:H3−5ac
H3K9 me2:DNA Meth. m5C
H4K20 me2:H4−4ac
DNA Meth. m5C:H4K20 me2
H3K27 me2
H3K4 me3
H3K9 me2
H3K9 me3
H3K27 me3
H4−4ac
DNA Meth. m5C
H2A.Z
H3−5ac
H4K20 me3
H3K4 me1
H4K20 me2
0.00 0.25 0.50 0.75 1.00
π̂
ACIN1 (3) − Reverse
H3K4 me3:H3−5ac
H3K27 me3:H4K20 me3
H2A.Z:H3K4 me3
H3K4 me1:H3−5ac
H3K4 me1:H4−4ac
H3K27 me3:H3K4 me1
H3K4 me1:H4K20 me2
H3K4 me1:H2A.Z
H3K27 me3:DNA Meth. m5C
H3K4 me3:H4−4ac
H2A.Z:H4K20 me2
H3K27 me3:H3K9 me3
DNA Meth. m5C:H3K9 me3
H4−4ac:H3−5ac
DNA Meth. m5C:H4K20 me3
H2A.Z:H4−4ac
H4K20 me2:H3K4 me3
H2A.Z:H3−5ac
H4K20 me2:H3−5ac
H3K9 me2:DNA Meth. m5C
H4K20 me2:H4−4ac
DNA Meth. m5C:H4K20 me2
H3K27 me2
H3K4 me3
H3K9 me2
H3K9 me3
H3K27 me3
H4−4ac
DNA Meth. m5C
H2A.Z
H3−5ac
H4K20 me3
H3K4 me1
H4K20 me2
0.00 0.25 0.50 0.75 1.00
π̂
SRRM2 − Forward
H3K4 me1:H4K20 me2
H3K4 me1:H4−4ac
H3K9 me3:H4K20 me3
H3K27 me3:H4K20 me3
H3K27 me3:H3K9 me3
H2A.Z:H3−5ac
H3K27 me3:H3K4 me1
H3K4 me3:H3−5ac
H2A.Z:H4−4ac
DNA Meth. m5C:H3K9 me3
DNA Meth. m5C:H4K20 me2
H2A.Z:H3K4 me3
DNA Meth. m5C:H4K20 me3
H2A.Z:H4K20 me2
H3K27 me2:DNA Meth. m5C
H4K20 me2:H4−4ac
H3K4 me1:H3−5ac
H4K20 me2:H3−5ac
H4K20 me2:H3K4 me3
H3K9 me2:DNA Meth. m5C
H3K4 me3:H4−4ac
H4−4ac:H3−5ac
H3K27 me3:DNA Meth. m5C
H3K27 me2
H3K9 me3
H4K20 me3
H3K4 me3
H3K4 me1
H3K27 me3
H3K9 me2
H4−4ac
DNA Meth. m5C
H2A.Z
H3−5ac
H4K20 me2
0.00 0.25 0.50 0.75 1.00
π̂
SRRM2 − Reverse
H3K4 me1:H4K20 me2
H3K4 me1:H4−4ac
H3K9 me3:H4K20 me3
H3K27 me3:H4K20 me3
H3K27 me3:H3K9 me3
H2A.Z:H3−5ac
H3K27 me3:H3K4 me1
H3K4 me3:H3−5ac
H2A.Z:H4−4ac
DNA Meth. m5C:H3K9 me3
DNA Meth. m5C:H4K20 me2
H2A.Z:H3K4 me3
DNA Meth. m5C:H4K20 me3
H2A.Z:H4K20 me2
H3K27 me2:DNA Meth. m5C
H4K20 me2:H4−4ac
H3K4 me1:H3−5ac
H4K20 me2:H3−5ac
H4K20 me2:H3K4 me3
H3K9 me2:DNA Meth. m5C
H3K4 me3:H4−4ac
H4−4ac:H3−5ac
H3K27 me3:DNA Meth. m5C
H3K27 me2
H3K9 me3
H4K20 me3
H3K4 me3
H3K4 me1
H3K27 me3
H3K9 me2
H4−4ac
DNA Meth. m5C
H2A.Z
H3−5ac
H4K20 me2
0.00 0.25 0.50 0.75 1.00
π̂
ACIN1 (2) − Forward
H3K4 me1:H4−4ac
H3K4 me1:H2A.Z
H3K4 me3:H3−5ac
H3K4 me3:H4−4ac
H2A.Z:H3K4 me3
H3K9 me2:DNA Meth. m5C
H4K20 me2:H4−4ac
H2A.Z:H4−4ac
H3K9 me3:H4K20 me3
H3K27 me3:H4K20 me3
DNA Meth. m5C:H3K9 me3
H3K4 me1:H3−5ac
H3K4 me1:H4K20 me2
H3K27 me3:DNA Meth. m5C
H3K27 me3:H3K9 me3
H4−4ac:H3−5ac
H4K20 me2:H3−5ac
H2A.Z:H3−5ac
H4K20 me2:H3K4 me3
H3K27 me2:DNA Meth. m5C
H2A.Z:H4K20 me2
H3K9 me2
H3K27 me2
H4K20 me3
H4−4ac
H3K4 me3
H3K4 me1
DNA Meth. m5C
H3K9 me3
H3K27 me3
H2A.Z
H3−5ac
H4K20 me2
0.00 0.25 0.50 0.75 1.00
π̂
ACIN1 (2) − Reverse
H3K4 me1:H4−4ac
H3K4 me1:H2A.Z
H3K4 me3:H3−5ac
H3K4 me3:H4−4ac
H2A.Z:H3K4 me3
H3K9 me2:DNA Meth. m5C
H4K20 me2:H4−4ac
H2A.Z:H4−4ac
H3K9 me3:H4K20 me3
H3K27 me3:H4K20 me3
DNA Meth. m5C:H3K9 me3
H3K4 me1:H3−5ac
H3K4 me1:H4K20 me2
H3K27 me3:DNA Meth. m5C
H3K27 me3:H3K9 me3
H4−4ac:H3−5ac
H4K20 me2:H3−5ac
H2A.Z:H3−5ac
H4K20 me2:H3K4 me3
H3K27 me2:DNA Meth. m5C
H2A.Z:H4K20 me2
H3K9 me2
H3K27 me2
H4K20 me3
H4−4ac
H3K4 me3
H3K4 me1
DNA Meth. m5C
H3K9 me3
H3K27 me3
H2A.Z
H3−5ac
H4K20 me2
0.00 0.25 0.50 0.75 1.00
π̂
ACIN1 (1) − Forward
H3K4 me1:H4−4ac
H3K4 me1:H2A.Z
H3K4 me1:H4K20 me2
H3K9 me3:H4K20 me3
H2A.Z:H3K4 me3
H2A.Z:H4K20 me2
H3K27 me3:H3K4 me1
H3K27 me3:H4K20 me3
H3K27 me3:H3K9 me3
DNA Meth. m5C:H3K9 me3
H3K4 me3:H3−5ac
H3K4 me1:H3−5ac
DNA Meth. m5C:H4K20 me2
H3K27 me2:DNA Meth. m5C
H4K20 me2:H3−5ac
H4K20 me2:H3K4 me3
H3K4 me3:H4−4ac
H4−4ac:H3−5ac
H3K27 me3:DNA Meth. m5C
H4K20 me2:H4−4ac
H3K9 me2:DNA Meth. m5C
H3K27 me2
H4K20 me3
H3K9 me2
H3K9 me3
H3K4 me3
H3K4 me1
DNA Meth. m5C
H3K27 me3
H4−4ac
H2A.Z
H4K20 me2
H3−5ac
0.00 0.25 0.50 0.75 1.00
π̂
ACIN1 (1) − Reverse
H3K4 me1:H4−4ac
H3K4 me1:H2A.Z
H3K4 me1:H4K20 me2
H3K9 me3:H4K20 me3
H2A.Z:H3K4 me3
H2A.Z:H4K20 me2
H3K27 me3:H3K4 me1
H3K27 me3:H4K20 me3
H3K27 me3:H3K9 me3
DNA Meth. m5C:H3K9 me3
H3K4 me3:H3−5ac
H3K4 me1:H3−5ac
DNA Meth. m5C:H4K20 me2
H3K27 me2:DNA Meth. m5C
H4K20 me2:H3−5ac
H4K20 me2:H3K4 me3
H3K4 me3:H4−4ac
H4−4ac:H3−5ac
H3K27 me3:DNA Meth. m5C
H4K20 me2:H4−4ac
H3K9 me2:DNA Meth. m5C
H3K27 me2
H4K20 me3
H3K9 me2
H3K9 me3
H3K4 me3
H3K4 me1
DNA Meth. m5C
H3K27 me3
H4−4ac
H2A.Z
H4K20 me2
H3−5ac
0.00 0.25 0.50 0.75 1.00
π̂
SAP18 − Forward
DNA Meth. m5C:H3K9 me3
H3K4 me1:H4−4ac
H3K4 me1:H2A.Z
H3K4 me1:H4K20 me2
H3K27 me3:DNA Meth. m5C
H2A.Z:H3K4 me3
H3K9 me3:H4K20 me3
H3K27 me3:H4K20 me3
H3K27 me2:DNA Meth. m5C
H3K4 me3:H4−4ac
H4K20 me2:H4−4ac
H2A.Z:H4K20 me2
DNA Meth. m5C:H4K20 me2
H4K20 me2:H3−5ac
H3K9 me2:DNA Meth. m5C
H3K4 me1:H3−5ac
H3K4 me3:H3−5ac
H4−4ac:H3−5ac
H4K20 me2:H3K4 me3
H3K9 me2
H3K27 me2
H4K20 me3
H3K4 me1
H3K9 me3
H3K4 me3
DNA Meth. m5C
H4−4ac
H3K27 me3
H3−5ac
H2A.Z
H4K20 me2
0.00 0.25 0.50 0.75 1.00
π̂
SAP18 − Reverse
DNA Meth. m5C:H3K9 me3
H3K4 me1:H4−4ac
H3K4 me1:H2A.Z
H3K4 me1:H4K20 me2
H3K27 me3:DNA Meth. m5C
H2A.Z:H3K4 me3
H3K9 me3:H4K20 me3
H3K27 me3:H4K20 me3
H3K27 me2:DNA Meth. m5C
H3K4 me3:H4−4ac
H4K20 me2:H4−4ac
H2A.Z:H4K20 me2
DNA Meth. m5C:H4K20 me2
H4K20 me2:H3−5ac
H3K9 me2:DNA Meth. m5C
H3K4 me1:H3−5ac
H3K4 me3:H3−5ac
H4−4ac:H3−5ac
H4K20 me2:H3K4 me3
H3K9 me2
H3K27 me2
H4K20 me3
H3K4 me1
H3K9 me3
H3K4 me3
DNA Meth. m5C
H4−4ac
H3K27 me3
H3−5ac
H2A.Z
H4K20 me2
0.00 0.25 0.50 0.75 1.00
π̂

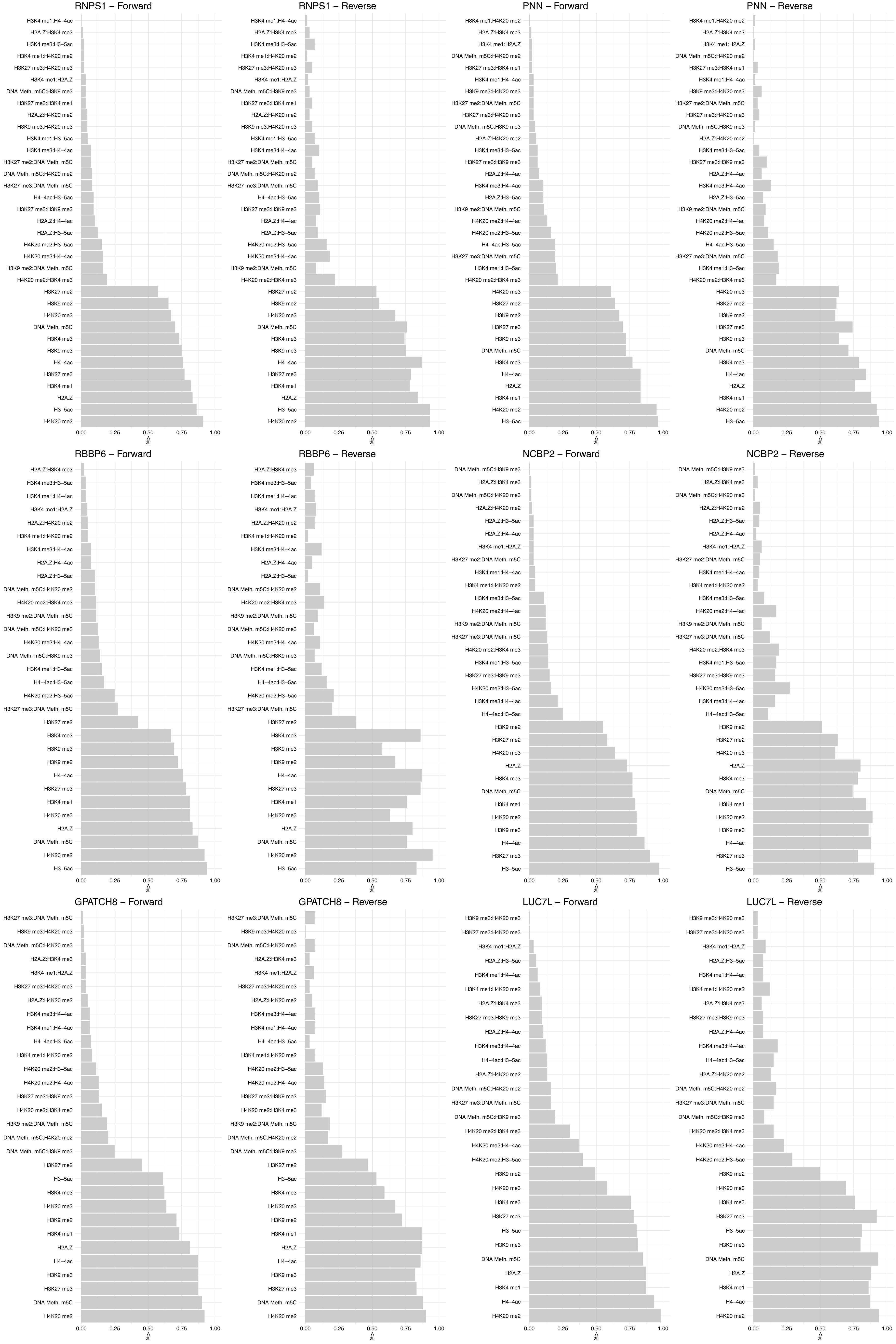

RNPS1 – Forward
H3K4 me1:H4−4ac
H2A.Z:H3K4 me3
H3K4 me3:H3−5ac
H3K4 me1:H4K20 me2
H3K27 me3:H4K20 me3
H3K4 me1:H2A.Z
DNA Meth. m5C:H3K9 me3
H3K27 me3:H3K4 me1
H2A.Z:H4K20 me2
H3K9 me3:H4K20 me3
H3K4 me1:H3−5ac
H3K4 me3:H4−4ac
H3K27 me2:DNA Meth. m5C
DNA Meth. m5C:H4K20 me2
H3K27 me3:DNA Meth. m5C
H4−4ac:H3−5ac
H3K27 me3:H3K9 me3
H2A.Z:H4−4ac
H2A.Z:H3−5ac
H4K20 me2:H3−5ac
H4K20 me2:H4−4ac
H3K9 me2:DNA Meth. m5C
H4K20 me2:H3K4 me3
H3K27 me2
H3K9 me2
H4K20 me3
DNA Meth. m5C
H3K4 me3
H3K9 me3
H4−4ac
H3K27 me3
H3K4 me1
H2A.Z
H3−5ac
H4K20 me2
0.00 0.25 0.50 0.75 1.00
π̂
RNPS1 – Reverse
H3K4 me1:H4−4ac
H2A.Z:H3K4 me3
H3K4 me3:H3−5ac
H3K4 me1:H4K20 me2
H3K27 me3:H4K20 me3
H3K4 me1:H2A.Z
DNA Meth. m5C:H3K9 me3
H3K27 me3:H3K4 me1
H2A.Z:H4K20 me2
H3K9 me3:H4K20 me3
H3K4 me1:H3−5ac
H3K4 me3:H4−4ac
H3K27 me2:DNA Meth. m5C
DNA Meth. m5C:H4K20 me2
H3K27 me3:DNA Meth. m5C
H4−4ac:H3−5ac
H3K27 me3:H3K9 me3
H2A.Z:H4−4ac
H2A.Z:H3−5ac
H4K20 me2:H3−5ac
H4K20 me2:H4−4ac
H3K9 me2:DNA Meth. m5C
H4K20 me2:H3K4 me3
H3K27 me2
H3K9 me2
H4K20 me3
DNA Meth. m5C
H3K4 me3
H3K9 me3
H4−4ac
H3K27 me3
H3K4 me1
H2A.Z
H3−5ac
H4K20 me2
0.00 0.25 0.50 0.75 1.00
π̂
PNN – Forward
H3K4 me1:H4K20 me2
H2A.Z:H3K4 me3
H3K4 me1:H2A.Z
DNA Meth. m5C:H4K20 me2
H3K27 me3:H3K4 me1
H3K4 me1:H4−4ac
H3K9 me3:H4K20 me3
H3K27 me2:DNA Meth. m5C
H3K27 me3:H4K20 me3
DNA Meth. m5C:H3K9 me3
H2A.Z:H4K20 me2
H3K4 me3:H3−5ac
H3K27 me3:H3K9 me3
H2A.Z:H4−4ac
H3K4 me3:H4−4ac
H2A.Z:H3−5ac
H3K9 me2:DNA Meth. m5C
H4K20 me2:H4−4ac
H4K20 me2:H3−5ac
H4−4ac:H3−5ac
H3K27 me3:DNA Meth. m5C
H3K4 me1:H3−5ac
H4K20 me2:H3K4 me3
H4K20 me3
H3K27 me2
H3K9 me2
H3K27 me3
H3K9 me3
DNA Meth. m5C
H3K4 me3
H4−4ac
H2A.Z
H3K4 me1
H4K20 me2
H3−5ac
0.00 0.25 0.50 0.75 1.00
π̂
PNN – Reverse
H3K4 me1:H4K20 me2
H2A.Z:H3K4 me3
H3K4 me1:H2A.Z
DNA Meth. m5C:H4K20 me2
H3K27 me3:H3K4 me1
H3K4 me1:H4−4ac
H3K9 me3:H4K20 me3
H3K27 me2:DNA Meth. m5C
H3K27 me3:H4K20 me3
DNA Meth. m5C:H3K9 me3
H2A.Z:H4K20 me2
H3K4 me3:H3−5ac
H3K27 me3:H3K9 me3
H2A.Z:H4−4ac
H3K4 me3:H4−4ac
H2A.Z:H3−5ac
H3K9 me2:DNA Meth. m5C
H4K20 me2:H4−4ac
H4K20 me2:H3−5ac
H4−4ac:H3−5ac
H3K27 me3:DNA Meth. m5C
H3K4 me1:H3−5ac
H4K20 me2:H3K4 me3
H4K20 me3
H3K27 me2
H3K9 me2
H3K27 me3
H3K9 me3
DNA Meth. m5C
H3K4 me3
H4−4ac
H2A.Z
H3K4 me1
H4K20 me2
H3−5ac
0.00 0.25 0.50 0.75 1.00
π̂
RBBP6 – Forward
H2A.Z:H3K4 me3
H3K4 me3:H3−5ac
H3K4 me1:H4−4ac
H3K4 me1:H2A.Z
H2A.Z:H4K20 me2
H3K4 me1:H4K20 me2
H3K4 me3:H4−4ac
H2A.Z:H4−4ac
H2A.Z:H3−5ac
DNA Meth. m5C:H4K20 me2
H4K20 me2:H3K4 me3
H3K9 me2:DNA Meth. m5C
DNA Meth. m5C:H4K20 me3
H4K20 me2:H4−4ac
DNA Meth. m5C:H3K9 me3
H3K4 me1:H3−5ac
H4−4ac:H3−5ac
H4K20 me2:H3−5ac
H3K27 me3:DNA Meth. m5C
H3K27 me2
H3K4 me3
H3K9 me3
H3K9 me2
H4−4ac
H3K27 me3
H3K4 me1
H4K20 me3
H2A.Z
DNA Meth. m5C
H4K20 me2
H3−5ac
0.00 0.25 0.50 0.75 1.00
π̂
RBBP6 – Reverse
H2A.Z:H3K4 me3
H3K4 me3:H3−5ac
H3K4 me1:H4−4ac
H3K4 me1:H2A.Z
H2A.Z:H4K20 me2
H3K4 me1:H4K20 me2
H3K4 me3:H4−4ac
H2A.Z:H4−4ac
H2A.Z:H3−5ac
DNA Meth. m5C:H4K20 me2
H4K20 me2:H3K4 me3
H3K9 me2:DNA Meth. m5C
DNA Meth. m5C:H4K20 me3
H4K20 me2:H4−4ac
DNA Meth. m5C:H3K9 me3
H3K4 me1:H3−5ac
H4−4ac:H3−5ac
H4K20 me2:H3−5ac
H3K27 me3:DNA Meth. m5C
H3K27 me2
H3K4 me3
H3K9 me3
H3K9 me2
H4−4ac
H3K27 me3
H3K4 me1
H4K20 me3
H2A.Z
DNA Meth. m5C
H4K20 me2
H3−5ac
0.00 0.25 0.50 0.75 1.00
π̂
NCBP2 – Forward
DNA Meth. m5C:H3K9 me3
H2A.Z:H3K4 me3
DNA Meth. m5C:H4K20 me3
H2A.Z:H4K20 me2
H2A.Z:H3−5ac
H2A.Z:H4−4ac
H3K4 me1:H2A.Z
H3K27 me2:DNA Meth. m5C
H3K4 me1:H4−4ac
H3K4 me1:H4K20 me2
H3K4 me3:H3−5ac
H4K20 me2:H4−4ac
H3K9 me2:DNA Meth. m5C
H3K27 me3:DNA Meth. m5C
H4K20 me2:H3K4 me3
H3K4 me1:H3−5ac
H3K27 me3:H3K9 me3
H4K20 me2:H3−5ac
H3K4 me3:H4−4ac
H4−4ac:H3−5ac
H3K9 me2
H3K27 me2
H4K20 me3
H2A.Z
H3K4 me3
DNA Meth. m5C
H3K4 me1
H4K20 me2
H3K9 me3
H4−4ac
H3K27 me3
H3−5ac
0.00 0.25 0.50 0.75 1.00
π̂
NCBP2 – Reverse
DNA Meth. m5C:H3K9 me3
H2A.Z:H3K4 me3
DNA Meth. m5C:H4K20 me3
H2A.Z:H4K20 me2
H2A.Z:H3−5ac
H2A.Z:H4−4ac
H3K4 me1:H2A.Z
H3K27 me2:DNA Meth. m5C
H3K4 me1:H4−4ac
H3K4 me1:H4K20 me2
H3K4 me3:H3−5ac
H4K20 me2:H4−4ac
H3K9 me2:DNA Meth. m5C
H3K27 me3:DNA Meth. m5C
H4K20 me2:H3K4 me3
H3K4 me1:H3−5ac
H3K27 me3:H3K9 me3
H4K20 me2:H3−5ac
H3K4 me3:H4−4ac
H4−4ac:H3−5ac
H3K9 me2
H3K27 me2
H4K20 me3
H2A.Z
H3K4 me3
DNA Meth. m5C
H3K4 me1
H4K20 me2
H3K9 me3
H4−4ac
H3K27 me3
H3−5ac
0.00 0.25 0.50 0.75 1.00
π̂
GPATCH8 – Forward
H3K27 me3:DNA Meth. m5C
H3K9 me3:H4K20 me3
DNA Meth. m5C:H4K20 me3
H2A.Z:H3K4 me3
H3K4 me1:H2A.Z
H3K27 me3:H4K20 me3
H2A.Z:H4K20 me2
H3K4 me3:H4−4ac
H3K4 me1:H4−4ac
H4−4ac:H3−5ac
H3K4 me1:H4K20 me2
H4K20 me2:H3−5ac
H4K20 me2:H4−4ac
H3K27 me3:H3K9 me3
H4K20 me2:H3K4 me3
H3K9 me2:DNA Meth. m5C
DNA Meth. m5C:H4K20 me2
DNA Meth. m5C:H3K9 me3
H3K27 me2
H3−5ac
H3K4 me3
H4K20 me3
H3K9 me2
H3K4 me1
H2A.Z
H4−4ac
H3K9 me3
H3K27 me3
DNA Meth. m5C
H4K20 me2
0.00 0.25 0.50 0.75 1.00
π̂
GPATCH8 – Reverse
H3K27 me3:DNA Meth. m5C
H3K9 me3:H4K20 me3
DNA Meth. m5C:H4K20 me3
H2A.Z:H3K4 me3
H3K4 me1:H2A.Z
H3K27 me3:H4K20 me3
H2A.Z:H4K20 me2
H3K4 me3:H4−4ac
H3K4 me1:H4−4ac
H4−4ac:H3−5ac
H3K4 me1:H4K20 me2
H4K20 me2:H3−5ac
H4K20 me2:H4−4ac
H3K27 me3:H3K9 me3
H4K20 me2:H3K4 me3
H3K9 me2:DNA Meth. m5C
DNA Meth. m5C:H4K20 me2
DNA Meth. m5C:H3K9 me3
H3K27 me2
H3−5ac
H3K4 me3
H4K20 me3
H3K9 me2
H3K4 me1
H2A.Z
H4−4ac
H3K9 me3
H3K27 me3
DNA Meth. m5C
H4K20 me2
0.00 0.25 0.50 0.75 1.00
π̂
LUC7L – Forward
H3K9 me3:H4K20 me3
H3K27 me3:H4K20 me3
H3K4 me1:H2A.Z
H2A.Z:H3−5ac
H3K4 me1:H4−4ac
H3K4 me1:H4K20 me2
H2A.Z:H3K4 me3
H3K27 me3:H3K9 me3
H2A.Z:H4−4ac
H3K4 me3:H4−4ac
H4−4ac:H3−5ac
H2A.Z:H4K20 me2
DNA Meth. m5C:H4K20 me2
H3K27 me3:DNA Meth. m5C
DNA Meth. m5C:H3K9 me3
H4K20 me2:H3K4 me3
H4K20 me2:H4−4ac
H4K20 me2:H3−5ac
H3K9 me2
H4K20 me3
H3K4 me3
H3K27 me3
H3−5ac
H3K9 me3
DNA Meth. m5C
H2A.Z
H3K4 me1
H4−4ac
H4K20 me2
0.00 0.25 0.50 0.75 1.00
π̂
LUC7L – Reverse
H3K9 me3:H4K20 me3
H3K27 me3:H4K20 me3
H3K4 me1:H2A.Z
H2A.Z:H3−5ac
H3K4 me1:H4−4ac
H3K4 me1:H4K20 me2
H2A.Z:H3K4 me3
H3K27 me3:H3K9 me3
H2A.Z:H4−4ac
H3K4 me3:H4−4ac
H4−4ac:H3−5ac
H2A.Z:H4K20 me2
DNA Meth. m5C:H4K20 me2
H3K27 me3:DNA Meth. m5C
DNA Meth. m5C:H3K9 me3
H4K20 me2:H3K4 me3
H4K20 me2:H4−4ac
H4K20 me2:H3−5ac
H3K9 me2
H4K20 me3
H3K4 me3
H3K27 me3
H3−5ac
H3K9 me3
DNA Meth. m5C
H2A.Z
H3K4 me1
H4−4ac
H4K20 me2
0.00 0.25 0.50 0.75 1.00
π̂

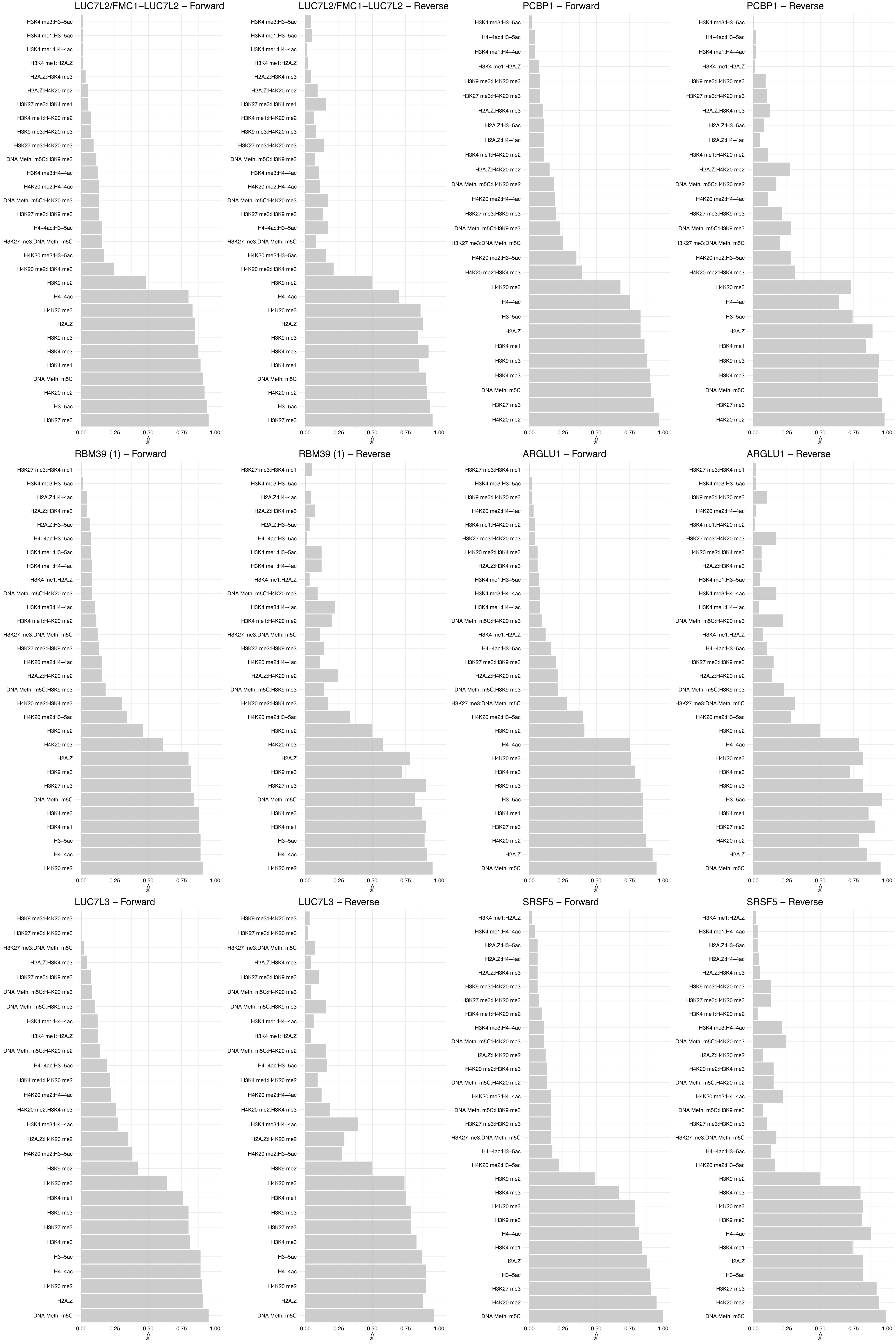
LUC7L2/FMC1−LUC7L2 − Forward
H3K4 me3:H3−5ac
H3K4 me1:H3−5ac
H3K4 me1:H4−4ac
H3K4 me1:H2A.Z
H2A.Z:H3K4 me3
H2A.Z:H4K20 me2
H3K27 me3:H3K4 me1
H3K4 me1:H4K20 me2
H3K9 me3:H4K20 me3
H3K27 me3:H4K20 me3
DNA Meth. m5C:H3K9 me3
H3K4 me3:H4−4ac
H4K20 me2:H4−4ac
DNA Meth. m5C:H4K20 me3
H3K27 me3:H3K9 me3
H4−4ac:H3−5ac
H3K27 me3:DNA Meth. m5C
H4K20 me2:H3−5ac
H4K20 me2:H3K4 me3
H3K9 me2
H4−4ac
H4K20 me3
H2A.Z
H3K9 me3
H3K4 me3
H3K4 me1
DNA Meth. m5C
H4K20 me2
H3−5ac
H3K27 me3
0.00 0.25 0.50 0.75 1.00
π̂
LUC7L2/FMC1−LUC7L2 − Reverse
0.00 0.25 0.50 0.75 1.00
π̂
PCBP1 − Forward
H3K4 me3:H3−5ac
H4−4ac:H3−5ac
H3K4 me1:H4−4ac
H3K4 me1:H2A.Z
H3K9 me3:H4K20 me3
H3K27 me3:H4K20 me3
H2A.Z:H3K4 me3
H2A.Z:H3−5ac
H2A.Z:H4−4ac
H3K4 me1:H4K20 me2
H2A.Z:H4K20 me2
DNA Meth. m5C:H4K20 me2
H4K20 me2:H4−4ac
H3K27 me3:H3K9 me3
DNA Meth. m5C:H3K9 me3
H3K27 me3:DNA Meth. m5C
H4K20 me2:H3−5ac
H4K20 me2:H3K4 me3
H4K20 me3
H4−4ac
H3−5ac
H2A.Z
H3K4 me1
H3K9 me3
H3K4 me3
DNA Meth. m5C
H3K27 me3
H4K20 me2
0.00 0.25 0.50 0.75 1.00
π̂
PCBP1 − Reverse
0.00 0.25 0.50 0.75 1.00
π̂
RBM39 (1) − Forward
H3K27 me3:H3K4 me1
H3K4 me3:H3−5ac
H2A.Z:H4−4ac
H2A.Z:H3K4 me3
H2A.Z:H3−5ac
H4−4ac:H3−5ac
H3K4 me1:H3−5ac
H3K4 me1:H4−4ac
H3K4 me1:H2A.Z
DNA Meth. m5C:H4K20 me3
H3K4 me3:H4−4ac
H3K4 me1:H4K20 me2
H3K27 me3:DNA Meth. m5C
H3K27 me3:H3K9 me3
H4K20 me2:H4−4ac
H2A.Z:H4K20 me2
DNA Meth. m5C:H3K9 me3
H4K20 me2:H3K4 me3
H4K20 me2:H3−5ac
H3K9 me2
H4K20 me3
H2A.Z
H3K9 me3
H3K27 me3
DNA Meth. m5C
H3K4 me3
H3K4 me1
H3−5ac
H4−4ac
H4K20 me2
0.00 0.25 0.50 0.75 1.00
π̂
RBM39 (1) − Reverse
0.00 0.25 0.50 0.75 1.00
π̂
ARGLU1 − Forward
H3K27 me3:H3K4 me1
H3K4 me3:H3−5ac
H3K9 me3:H4K20 me3
H4K20 me2:H4−4ac
H3K4 me1:H4K20 me2
H3K27 me3:H4K20 me3
H4K20 me2:H3K4 me3
H2A.Z:H3K4 me3
H3K4 me1:H3−5ac
H3K4 me3:H4−4ac
H3K4 me1:H4−4ac
DNA Meth. m5C:H4K20 me3
H3K4 me1:H2A.Z
H4−4ac:H3−5ac
H3K27 me3:H3K9 me3
H2A.Z:H4K20 me2
DNA Meth. m5C:H3K9 me3
H3K27 me3:DNA Meth. m5C
H4K20 me2:H3−5ac
H3K9 me2
H4−4ac
H4K20 me3
H3K4 me3
H3K9 me3
H3−5ac
H3K4 me1
H3K27 me3
H4K20 me2
H2A.Z
DNA Meth. m5C
0.00 0.25 0.50 0.75 1.00
π̂
ARGLU1 − Reverse
0.00 0.25 0.50 0.75 1.00
π̂
LUC7L3 − Forward
H3K9 me3:H4K20 me3
H3K27 me3:H4K20 me3
H3K27 me3:DNA Meth. m5C
H2A.Z:H3K4 me3
H3K27 me3:H3K9 me3
DNA Meth. m5C:H4K20 me3
DNA Meth. m5C:H3K9 me3
H3K4 me1:H4−4ac
H3K4 me1:H2A.Z
DNA Meth. m5C:H4K20 me2
H4−4ac:H3−5ac
H3K4 me1:H4K20 me2
H4K20 me2:H4−4ac
H4K20 me2:H3K4 me3
H3K4 me3:H4−4ac
H2A.Z:H4K20 me2
H4K20 me2:H3−5ac
H3K9 me2
H4K20 me3
H3K4 me1
H3K9 me3
H3K27 me3
H3K4 me3
H3−5ac
H4−4ac
H4K20 me2
H2A.Z
DNA Meth. m5C
0.00 0.25 0.50 0.75 1.00
π̂
LUC7L3 − Reverse
0.00 0.25 0.50 0.75 1.00
π̂
SRSF5 − Forward
H3K4 me1:H2A.Z
H3K4 me1:H4−4ac
H2A.Z:H3−5ac
H2A.Z:H4−4ac
H2A.Z:H3K4 me3
H3K9 me3:H4K20 me3
H3K27 me3:H4K20 me3
H3K4 me1:H4K20 me2
H3K4 me3:H4−4ac
DNA Meth. m5C:H4K20 me3
H2A.Z:H4K20 me2
H4K20 me2:H3K4 me3
DNA Meth. m5C:H4K20 me2
H4K20 me2:H4−4ac
DNA Meth. m5C:H3K9 me3
H3K27 me3:H3K9 me3
H3K27 me3:DNA Meth. m5C
H4−4ac:H3−5ac
H4K20 me2:H3−5ac
H3K9 me2
H3K4 me3
H4K20 me3
H3K9 me3
H4−4ac
H3K4 me1
H2A.Z
H3−5ac
H3K27 me3
H4K20 me2
DNA Meth. m5C
0.00 0.25 0.50 0.75 1.00
π̂
SRSF5 − Reverse
0.00 0.25 0.50 0.75 1.00
π̂

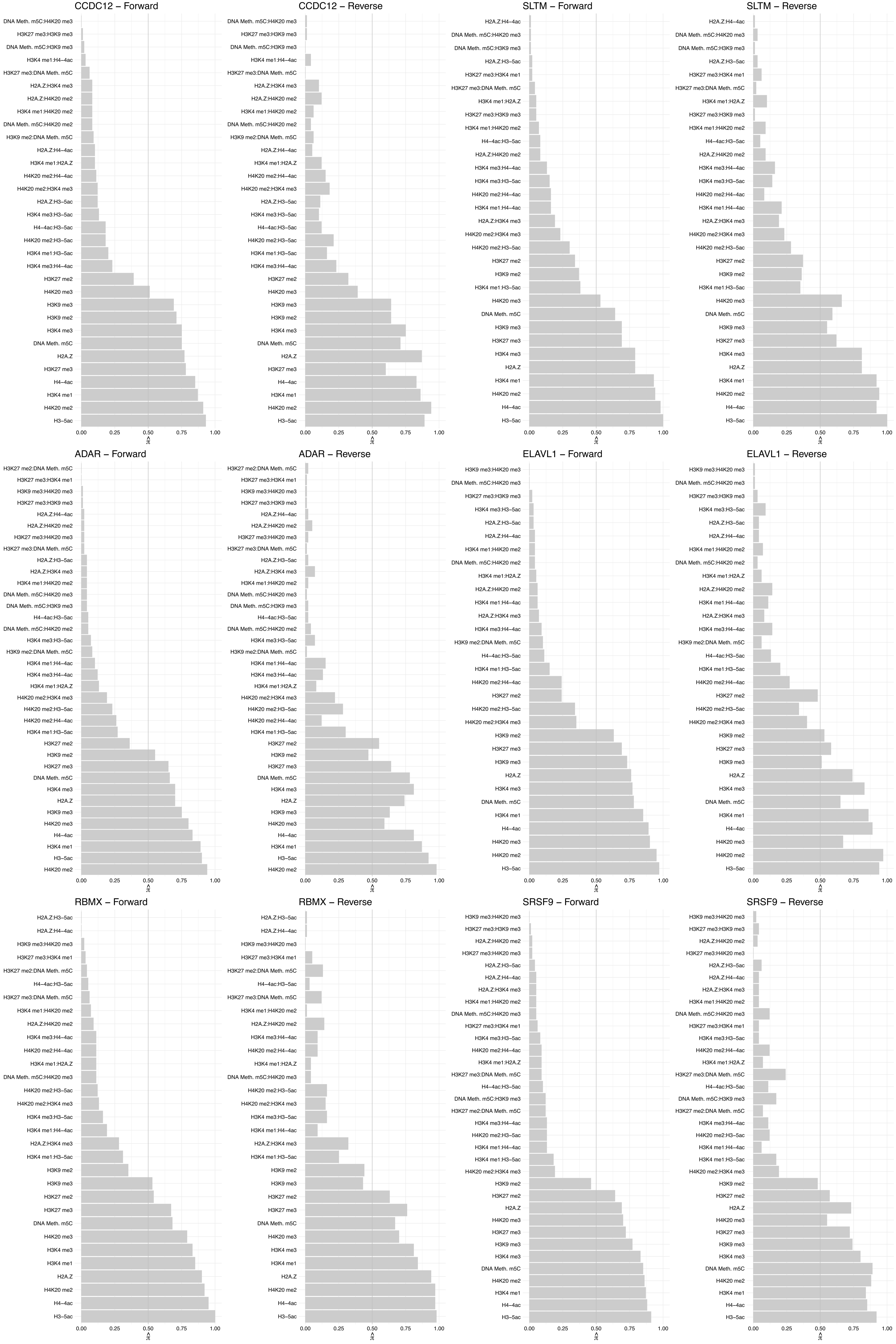

CCDC12 – Forward
DNA Meth. m5C:H4K20 me3
H3K27 me3:H3K9 me3
DNA Meth. m5C:H3K9 me3
H3K4 me1:H4–4ac
H3K27 me3:DNA Meth. m5C
H2A.Z:H3K4 me3
H2A.Z:H4K20 me2
H3K4 me1:H4K20 me2
DNA Meth. m5C:H4K20 me2
H3K9 me2:DNA Meth. m5C
H2A.Z:H4–4ac
H3K4 me1:H2A.Z
H4K20 me2:H4–4ac
H4K20 me2:H3K4 me3
H2A.Z:H3–5ac
H3K4 me3:H3–5ac
H4–4ac:H3–5ac
H4K20 me2:H3–5ac
H3K4 me1:H3–5ac
H3K4 me3:H4–4ac
H3K27 me2
H4K20 me3
H3K9 me3
H3K9 me2
H3K4 me3
DNA Meth. m5C
H2A.Z
H3K27 me3
H4–4ac
H3K4 me1
H4K20 me2
H3–5ac
0.00 0.25 0.50 0.75 1.00
$\hat{\pi}$
CCDC12 – Reverse
DNA Meth. m5C:H4K20 me3
H3K27 me3:H3K9 me3
DNA Meth. m5C:H3K9 me3
H3K4 me1:H4–4ac
H3K27 me3:DNA Meth. m5C
H2A.Z:H3K4 me3
H2A.Z:H4K20 me2
H3K4 me1:H4K20 me2
DNA Meth. m5C:H4K20 me2
H3K9 me2:DNA Meth. m5C
H2A.Z:H4–4ac
H3K4 me1:H2A.Z
H4K20 me2:H4–4ac
H4K20 me2:H3K4 me3
H2A.Z:H3–5ac
H3K4 me3:H3–5ac
H4–4ac:H3–5ac
H4K20 me2:H3–5ac
H3K4 me1:H3–5ac
H3K4 me3:H4–4ac
H3K27 me2
H4K20 me3
H3K9 me3
H3K9 me2
H3K4 me3
DNA Meth. m5C
H2A.Z
H3K27 me3
H4–4ac
H3K4 me1
H4K20 me2
H3–5ac
0.00 0.25 0.50 0.75 1.00
$\hat{\pi}$
SLTM – Forward
H2A.Z:H4–4ac
DNA Meth. m5C:H4K20 me3
DNA Meth. m5C:H3K9 me3
H2A.Z:H3–5ac
H3K27 me3:H3K4 me1
H3K27 me3:DNA Meth. m5C
H3K4 me1:H2A.Z
H3K27 me3:H3K9 me3
H3K4 me1:H4K20 me2
H4–4ac:H3–5ac
H2A.Z:H4K20 me2
H3K4 me3:H4–4ac
H3K4 me3:H3–5ac
H4K20 me2:H4–4ac
H3K4 me1:H4–4ac
H2A.Z:H3K4 me3
H4K20 me2:H3K4 me3
H4K20 me2:H3–5ac
H3K27 me2
H3K9 me2
H3K4 me1:H3–5ac
H4K20 me3
DNA Meth. m5C
H3K9 me3
H3K27 me3
H3K4 me3
H2A.Z
H3K4 me1
H4K20 me2
H4–4ac
H3–5ac
0.00 0.25 0.50 0.75 1.00
$\hat{\pi}$
SLTM – Reverse
H2A.Z:H4–4ac
DNA Meth. m5C:H4K20 me3
DNA Meth. m5C:H3K9 me3
H2A.Z:H3–5ac
H3K27 me3:H3K4 me1
H3K27 me3:DNA Meth. m5C
H3K4 me1:H2A.Z
H3K27 me3:H3K9 me3
H3K4 me1:H4K20 me2
H4–4ac:H3–5ac
H2A.Z:H4K20 me2
H3K4 me3:H4–4ac
H3K4 me3:H3–5ac
H4K20 me2:H4–4ac
H3K4 me1:H4–4ac
H2A.Z:H3K4 me3
H4K20 me2:H3K4 me3
H4K20 me2:H3–5ac
H3K27 me2
H3K9 me2
H3K4 me1:H3–5ac
H4K20 me3
DNA Meth. m5C
H3K9 me3
H3K27 me3
H3K4 me3
H2A.Z
H3K4 me1
H4K20 me2
H4–4ac
H3–5ac
0.00 0.25 0.50 0.75 1.00
$\hat{\pi}$
ADAR – Forward
H3K27 me2:DNA Meth. m5C
H3K27 me3:H3K4 me1
H3K9 me3:H4K20 me3
H3K27 me3:H3K9 me3
H2A.Z:H4–4ac
H2A.Z:H4K20 me2
H3K27 me3:H4K20 me3
H3K27 me3:DNA Meth. m5C
H2A.Z:H3–5ac
H2A.Z:H3K4 me3
H3K4 me1:H4K20 me2
DNA Meth. m5C:H4K20 me3
DNA Meth. m5C:H3K9 me3
H4–4ac:H3–5ac
DNA Meth. m5C:H4K20 me2
H3K4 me3:H3–5ac
H3K9 me2:DNA Meth. m5C
H3K4 me1:H4–4ac
H3K4 me3:H4–4ac
H3K4 me1:H2A.Z
H4K20 me2:H3K4 me3
H4K20 me2:H3–5ac
H4K20 me2:H4–4ac
H3K4 me1:H3–5ac
H3K27 me2
H3K9 me2
H3K27 me3
DNA Meth. m5C
H3K4 me3
H2A.Z
H3K9 me3
H4K20 me3
H4–4ac
H3K4 me1
H3–5ac
H4K20 me2
0.00 0.25 0.50 0.75 1.00
$\hat{\pi}$
ADAR – Reverse
H3K27 me2:DNA Meth. m5C
H3K27 me3:H3K4 me1
H3K9 me3:H4K20 me3
H3K27 me3:H3K9 me3
H2A.Z:H4–4ac
H2A.Z:H4K20 me2
H3K27 me3:H4K20 me3
H3K27 me3:DNA Meth. m5C
H2A.Z:H3–5ac
H2A.Z:H3K4 me3
H3K4 me1:H4K20 me2
DNA Meth. m5C:H4K20 me3
DNA Meth. m5C:H3K9 me3
H4–4ac:H3–5ac
DNA Meth. m5C:H4K20 me2
H3K4 me3:H3–5ac
H3K9 me2:DNA Meth. m5C
H3K4 me1:H4–4ac
H3K4 me3:H4–4ac
H3K4 me1:H2A.Z
H4K20 me2:H3K4 me3
H4K20 me2:H3–5ac
H4K20 me2:H4–4ac
H3K4 me1:H3–5ac
H3K27 me2
H3K9 me2
H3K27 me3
DNA Meth. m5C
H3K4 me3
H2A.Z
H3K9 me3
H4K20 me3
H4–4ac
H3K4 me1
H3–5ac
H4K20 me2
0.00 0.25 0.50 0.75 1.00
$\hat{\pi}$
ELAVL1 – Forward
H3K9 me3:H4K20 me3
DNA Meth. m5C:H4K20 me3
H3K27 me3:H3K9 me3
H3K4 me3:H3–5ac
H2A.Z:H3–5ac
H2A.Z:H4–4ac
H3K4 me1:H4K20 me2
DNA Meth. m5C:H4K20 me2
H3K4 me1:H2A.Z
H2A.Z:H4K20 me2
H3K4 me1:H4–4ac
H2A.Z:H3K4 me3
H3K4 me3:H4–4ac
H3K9 me2:DNA Meth. m5C
H4–4ac:H3–5ac
H3K4 me1:H3–5ac
H4K20 me2:H4–4ac
H3K27 me2
H4K20 me2:H3–5ac
H4K20 me2:H3K4 me3
H3K9 me2
H3K27 me3
H3K9 me3
H2A.Z
H3K4 me3
DNA Meth. m5C
H3K4 me1
H4–4ac
H4K20 me3
H4K20 me2
H3–5ac
0.00 0.25 0.50 0.75 1.00
$\hat{\pi}$
ELAVL1 – Reverse
H3K9 me3:H4K20 me3
DNA Meth. m5C:H4K20 me3
H3K27 me3:H3K9 me3
H3K4 me3:H3–5ac
H2A.Z:H3–5ac
H2A.Z:H4–4ac
H3K4 me1:H4K20 me2
DNA Meth. m5C:H4K20 me2
H3K4 me1:H2A.Z
H2A.Z:H4K20 me2
H3K4 me1:H4–4ac
H2A.Z:H3K4 me3
H3K4 me3:H4–4ac
H3K9 me2:DNA Meth. m5C
H4–4ac:H3–5ac
H3K4 me1:H3–5ac
H4K20 me2:H4–4ac
H3K27 me2
H4K20 me2:H3–5ac
H4K20 me2:H3K4 me3
H3K9 me2
H3K27 me3
H3K9 me3
H2A.Z
H3K4 me3
DNA Meth. m5C
H3K4 me1
H4–4ac
H4K20 me3
H4K20 me2
H3–5ac
0.00 0.25 0.50 0.75 1.00
$\hat{\pi}$
RBMX – Forward
H2A.Z:H3–5ac
H2A.Z:H4–4ac
H3K9 me3:H4K20 me3
H3K27 me3:H3K4 me1
H3K27 me2:DNA Meth. m5C
H4–4ac:H3–5ac
H3K27 me3:DNA Meth. m5C
H3K4 me1:H4K20 me2
H2A.Z:H4K20 me2
H3K4 me3:H4–4ac
H4K20 me2:H4–4ac
H3K4 me1:H2A.Z
DNA Meth. m5C:H4K20 me3
H4K20 me2:H3–5ac
H4K20 me2:H3K4 me3
H3K4 me3:H3–5ac
H3K4 me1:H4–4ac
H2A.Z:H3K4 me3
H3K4 me1:H3–5ac
H3K9 me2
H3K9 me3
H3K27 me2
H3K27 me3
DNA Meth. m5C
H4K20 me3
H3K4 me3
H3K4 me1
H2A.Z
H4K20 me2
H4–4ac
H3–5ac
0.00 0.25 0.50 0.75 1.00
$\hat{\pi}$
RBMX – Reverse
H2A.Z:H3–5ac
H2A.Z:H4–4ac
H3K9 me3:H4K20 me3
H3K27 me3:H3K4 me1
H3K27 me2:DNA Meth. m5C
H4–4ac:H3–5ac
H3K27 me3:DNA Meth. m5C
H3K4 me1:H4K20 me2
H2A.Z:H4K20 me2
H3K4 me3:H4–4ac
H4K20 me2:H4–4ac
H3K4 me1:H2A.Z
DNA Meth. m5C:H4K20 me3
H4K20 me2:H3–5ac
H4K20 me2:H3K4 me3
H3K4 me3:H3–5ac
H3K4 me1:H4–4ac
H2A.Z:H3K4 me3
H3K4 me1:H3–5ac
H3K9 me2
H3K9 me3
H3K27 me2
H3K27 me3
DNA Meth. m5C
H4K20 me3
H3K4 me3
H3K4 me1
H2A.Z
H4K20 me2
H4–4ac
H3–5ac
0.00 0.25 0.50 0.75 1.00
$\hat{\pi}$
SRSF9 – Forward
H3K9 me3:H4K20 me3
H3K27 me3:H3K9 me3
H2A.Z:H4K20 me2
H3K27 me3:H4K20 me3
H2A.Z:H3–5ac
H2A.Z:H4–4ac
H2A.Z:H3K4 me3
H3K4 me1:H4K20 me2
DNA Meth. m5C:H4K20 me3
H3K27 me3:H3K4 me1
H3K4 me3:H3–5ac
H4K20 me2:H4–4ac
H3K4 me1:H2A.Z
H3K27 me3:DNA Meth. m5C
H4–4ac:H3–5ac
DNA Meth. m5C:H3K9 me3
H3K27 me2:DNA Meth. m5C
H3K4 me3:H4–4ac
H4K20 me2:H3–5ac
H3K4 me1:H4–4ac
H3K4 me1:H3–5ac
H4K20 me2:H3K4 me3
H3K9 me2
H3K27 me2
H2A.Z
H4K20 me3
H3K27 me3
H3K9 me3
H3K4 me3
DNA Meth. m5C
H4K20 me2
H3K4 me1
H4–4ac
H3–5ac
0.00 0.25 0.50 0.75 1.00
$\hat{\pi}$
SRSF9 – Reverse
H3K9 me3:H4K20 me3
H3K27 me3:H3K9 me3
H2A.Z:H4K20 me2
H3K27 me3:H4K20 me3
H2A.Z:H3–5ac
H2A.Z:H4–4ac
H2A.Z:H3K4 me3
H3K4 me1:H4K20 me2
DNA Meth. m5C:H4K20 me3
H3K27 me3:H3K4 me1
H3K4 me3:H3–5ac
H4K20 me2:H4–4ac
H3K4 me1:H2A.Z
H3K27 me3:DNA Meth. m5C
H4–4ac:H3–5ac
DNA Meth. m5C:H3K9 me3
H3K27 me2:DNA Meth. m5C
H3K4 me3:H4–4ac
H4K20 me2:H3–5ac
H3K4 me1:H4–4ac
H3K4 me1:H3–5ac
H4K20 me2:H3K4 me3
H3K9 me2
H3K27 me2
H2A.Z
H4K20 me3
H3K27 me3
H3K9 me3
H3K4 me3
DNA Meth. m5C
H4K20 me2
H3K4 me1
H4–4ac
H3–5ac
0.00 0.25 0.50 0.75 1.00
$\hat{\pi}$

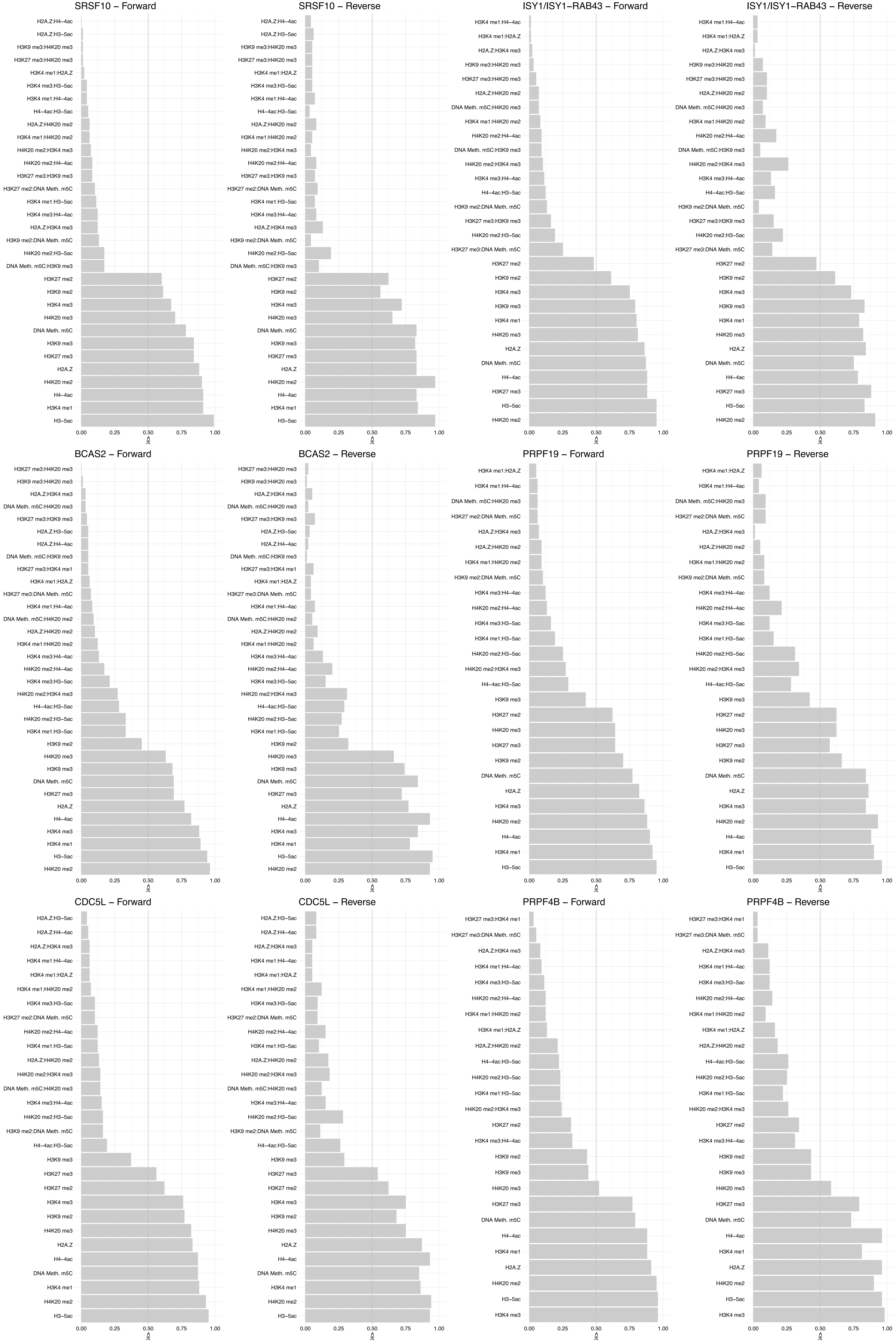
SRSF10 − Forward
H2A.Z:H4−4ac
H2A.Z:H3−5ac
H3K9 me3:H4K20 me3
H3K27 me3:H4K20 me3
H3K4 me1:H2A.Z
H3K4 me3:H3−5ac
H3K4 me1:H4−4ac
H4−4ac:H3−5ac
H2A.Z:H4K20 me2
H3K4 me1:H4K20 me2
H4K20 me2:H3K4 me3
H4K20 me2:H4−4ac
H3K27 me3:H3K9 me3
H3K27 me2:DNA Meth. m5C
H3K4 me1:H3−5ac
H3K4 me3:H4−4ac
H2A.Z:H3K4 me3
H3K9 me2:DNA Meth. m5C
H4K20 me2:H3−5ac
DNA Meth. m5C:H3K9 me3
H3K27 me2
H3K9 me2
H3K4 me3
H4K20 me3
DNA Meth. m5C
H3K9 me3
H3K27 me3
H2A.Z
H4K20 me2
H4−4ac
H3K4 me1
H3−5ac
0.00 0.25 0.50 0.75 1.00
π̂
SRSF10 − Reverse
H2A.Z:H4−4ac
H2A.Z:H3−5ac
H3K9 me3:H4K20 me3
H3K27 me3:H4K20 me3
H3K4 me1:H2A.Z
H3K4 me3:H3−5ac
H3K4 me1:H4−4ac
H4−4ac:H3−5ac
H2A.Z:H4K20 me2
H3K4 me1:H4K20 me2
H4K20 me2:H3K4 me3
H4K20 me2:H4−4ac
H3K27 me3:H3K9 me3
H3K27 me2:DNA Meth. m5C
H3K4 me1:H3−5ac
H3K4 me3:H4−4ac
H2A.Z:H3K4 me3
H3K9 me2:DNA Meth. m5C
H4K20 me2:H3−5ac
DNA Meth. m5C:H3K9 me3
H3K27 me2
H3K9 me2
H3K4 me3
H4K20 me3
DNA Meth. m5C
H3K9 me3
H3K27 me3
H2A.Z
H4K20 me2
H4−4ac
H3K4 me1
H3−5ac
0.00 0.25 0.50 0.75 1.00
π̂
ISY1/ISY1−RAB43 − Forward
H3K4 me1:H4−4ac
H3K4 me1:H2A.Z
H2A.Z:H3K4 me3
H3K9 me3:H4K20 me3
H3K27 me3:H4K20 me3
H2A.Z:H4K20 me2
DNA Meth. m5C:H4K20 me3
H3K4 me1:H4K20 me2
H4K20 me2:H4−4ac
DNA Meth. m5C:H3K9 me3
H4K20 me2:H3K4 me3
H3K4 me3:H4−4ac
H4−4ac:H3−5ac
H3K9 me2:DNA Meth. m5C
H3K27 me3:H3K9 me3
H4K20 me2:H3−5ac
H3K27 me3:DNA Meth. m5C
H3K27 me2
H3K9 me2
H3K4 me3
H3K9 me3
H3K4 me1
H4K20 me3
H2A.Z
DNA Meth. m5C
H4−4ac
H3K27 me3
H3−5ac
H4K20 me2
0.00 0.25 0.50 0.75 1.00
π̂
ISY1/ISY1−RAB43 − Reverse
H3K4 me1:H4−4ac
H3K4 me1:H2A.Z
H2A.Z:H3K4 me3
H3K9 me3:H4K20 me3
H3K27 me3:H4K20 me3
H2A.Z:H4K20 me2
DNA Meth. m5C:H4K20 me3
H3K4 me1:H4K20 me2
H4K20 me2:H4−4ac
DNA Meth. m5C:H3K9 me3
H4K20 me2:H3K4 me3
H3K4 me3:H4−4ac
H4−4ac:H3−5ac
H3K9 me2:DNA Meth. m5C
H3K27 me3:H3K9 me3
H4K20 me2:H3−5ac
H3K27 me3:DNA Meth. m5C
H3K27 me2
H3K9 me2
H3K4 me3
H3K9 me3
H3K4 me1
H4K20 me3
H2A.Z
DNA Meth. m5C
H4−4ac
H3K27 me3
H3−5ac
H4K20 me2
0.00 0.25 0.50 0.75 1.00
π̂
BCAS2 − Forward
H3K27 me3:H4K20 me3
H3K9 me3:H4K20 me3
H2A.Z:H3K4 me3
DNA Meth. m5C:H4K20 me3
H3K27 me3:H3K9 me3
H2A.Z:H3−5ac
H2A.Z:H4−4ac
DNA Meth. m5C:H3K9 me3
H3K27 me3:H3K4 me1
H3K4 me1:H2A.Z
H3K27 me3:DNA Meth. m5C
H3K4 me1:H4−4ac
DNA Meth. m5C:H4K20 me2
H2A.Z:H4K20 me2
H3K4 me1:H4K20 me2
H3K4 me3:H4−4ac
H4K20 me2:H4−4ac
H3K4 me3:H3−5ac
H4K20 me2:H3K4 me3
H4−4ac:H3−5ac
H4K20 me2:H3−5ac
H3K4 me1:H3−5ac
H3K9 me2
H4K20 me3
H3K9 me3
DNA Meth. m5C
H3K27 me3
H2A.Z
H4−4ac
H3K4 me3
H3K4 me1
H3−5ac
H4K20 me2
0.00 0.25 0.50 0.75 1.00
π̂
BCAS2 − Reverse
H3K27 me3:H4K20 me3
H3K9 me3:H4K20 me3
H2A.Z:H3K4 me3
DNA Meth. m5C:H4K20 me3
H3K27 me3:H3K9 me3
H2A.Z:H3−5ac
H2A.Z:H4−4ac
DNA Meth. m5C:H3K9 me3
H3K27 me3:H3K4 me1
H3K4 me1:H2A.Z
H3K27 me3:DNA Meth. m5C
H3K4 me1:H4−4ac
DNA Meth. m5C:H4K20 me2
H2A.Z:H4K20 me2
H3K4 me1:H4K20 me2
H3K4 me3:H4−4ac
H4K20 me2:H4−4ac
H3K4 me3:H3−5ac
H4K20 me2:H3K4 me3
H4−4ac:H3−5ac
H4K20 me2:H3−5ac
H3K4 me1:H3−5ac
H3K9 me2
H4K20 me3
H3K9 me3
DNA Meth. m5C
H3K27 me3
H2A.Z
H4−4ac
H3K4 me3
H3K4 me1
H3−5ac
H4K20 me2
0.00 0.25 0.50 0.75 1.00
π̂
PRPF19 − Forward
H3K4 me1:H2A.Z
H3K4 me1:H4−4ac
DNA Meth. m5C:H4K20 me3
H3K27 me2:DNA Meth. m5C
H2A.Z:H3K4 me3
H2A.Z:H4K20 me2
H3K4 me1:H4K20 me2
H3K9 me2:DNA Meth. m5C
H3K4 me3:H4−4ac
H4K20 me2:H4−4ac
H3K4 me3:H3−5ac
H3K4 me1:H3−5ac
H4K20 me2:H3−5ac
H4K20 me2:H3K4 me3
H4−4ac:H3−5ac
H3K9 me3
H3K27 me2
H4K20 me3
H3K27 me3
H3K9 me2
DNA Meth. m5C
H2A.Z
H3K4 me3
H4K20 me2
H4−4ac
H3K4 me1
H3−5ac
0.00 0.25 0.50 0.75 1.00
π̂
PRPF19 − Reverse
H3K4 me1:H2A.Z
H3K4 me1:H4−4ac
DNA Meth. m5C:H4K20 me3
H3K27 me2:DNA Meth. m5C
H2A.Z:H3K4 me3
H2A.Z:H4K20 me2
H3K4 me1:H4K20 me2
H3K9 me2:DNA Meth. m5C
H3K4 me3:H4−4ac
H4K20 me2:H4−4ac
H3K4 me3:H3−5ac
H3K4 me1:H3−5ac
H4K20 me2:H3−5ac
H4K20 me2:H3K4 me3
H4−4ac:H3−5ac
H3K9 me3
H3K27 me2
H4K20 me3
H3K27 me3
H3K9 me2
DNA Meth. m5C
H2A.Z
H3K4 me3
H4K20 me2
H4−4ac
H3K4 me1
H3−5ac
0.00 0.25 0.50 0.75 1.00
π̂
CDC5L − Forward
H2A.Z:H3−5ac
H2A.Z:H4−4ac
H2A.Z:H3K4 me3
H3K4 me1:H4−4ac
H3K4 me1:H2A.Z
H3K4 me1:H4K20 me2
H3K4 me3:H3−5ac
H3K27 me2:DNA Meth. m5C
H4K20 me2:H4−4ac
H3K4 me1:H3−5ac
H2A.Z:H4K20 me2
H4K20 me2:H3K4 me3
DNA Meth. m5C:H4K20 me3
H3K4 me3:H4−4ac
H4K20 me2:H3−5ac
H3K9 me2:DNA Meth. m5C
H4−4ac:H3−5ac
H3K9 me3
H3K27 me3
H3K27 me2
H3K4 me3
H3K9 me2
H4K20 me3
H2A.Z
H4−4ac
DNA Meth. m5C
H3K4 me1
H4K20 me2
H3−5ac
0.00 0.25 0.50 0.75 1.00
π̂
CDC5L − Reverse
H2A.Z:H3−5ac
H2A.Z:H4−4ac
H2A.Z:H3K4 me3
H3K4 me1:H4−4ac
H3K4 me1:H2A.Z
H3K4 me1:H4K20 me2
H3K4 me3:H3−5ac
H3K27 me2:DNA Meth. m5C
H4K20 me2:H4−4ac
H3K4 me1:H3−5ac
H2A.Z:H4K20 me2
H4K20 me2:H3K4 me3
DNA Meth. m5C:H4K20 me3
H3K4 me3:H4−4ac
H4K20 me2:H3−5ac
H3K9 me2:DNA Meth. m5C
H4−4ac:H3−5ac
H3K9 me3
H3K27 me3
H3K27 me2
H3K4 me3
H3K9 me2
H4K20 me3
H2A.Z
H4−4ac
DNA Meth. m5C
H3K4 me1
H4K20 me2
H3−5ac
0.00 0.25 0.50 0.75 1.00
π̂
PRPF4B − Forward
H3K27 me3:H3K4 me1
H3K27 me3:DNA Meth. m5C
H2A.Z:H3K4 me3
H3K4 me1:H4−4ac
H3K4 me3:H3−5ac
H4K20 me2:H4−4ac
H3K4 me1:H4K20 me2
H3K4 me1:H2A.Z
H2A.Z:H4K20 me2
H4−4ac:H3−5ac
H4K20 me2:H3−5ac
H3K4 me1:H3−5ac
H4K20 me2:H3K4 me3
H3K27 me2
H3K4 me3:H4−4ac
H3K9 me2
H3K9 me3
H4K20 me3
H3K27 me3
DNA Meth. m5C
H4−4ac
H3K4 me1
H2A.Z
H4K20 me2
H3−5ac
H3K4 me3
0.00 0.25 0.50 0.75 1.00
π̂
PRPF4B − Reverse
H3K27 me3:H3K4 me1
H3K27 me3:DNA Meth. m5C
H2A.Z:H3K4 me3
H3K4 me1:H4−4ac
H3K4 me3:H3−5ac
H4K20 me2:H4−4ac
H3K4 me1:H4K20 me2
H3K4 me1:H2A.Z
H2A.Z:H4K20 me2
H4−4ac:H3−5ac
H4K20 me2:H3−5ac
H3K4 me1:H3−5ac
H4K20 me2:H3K4 me3
H3K27 me2
H3K4 me3:H4−4ac
H3K9 me2
H3K9 me3
H4K20 me3
H3K27 me3
DNA Meth. m5C
H4−4ac
H3K4 me1
H2A.Z
H4K20 me2
H3−5ac
H3K4 me3
0.00 0.25 0.50 0.75 1.00
π̂

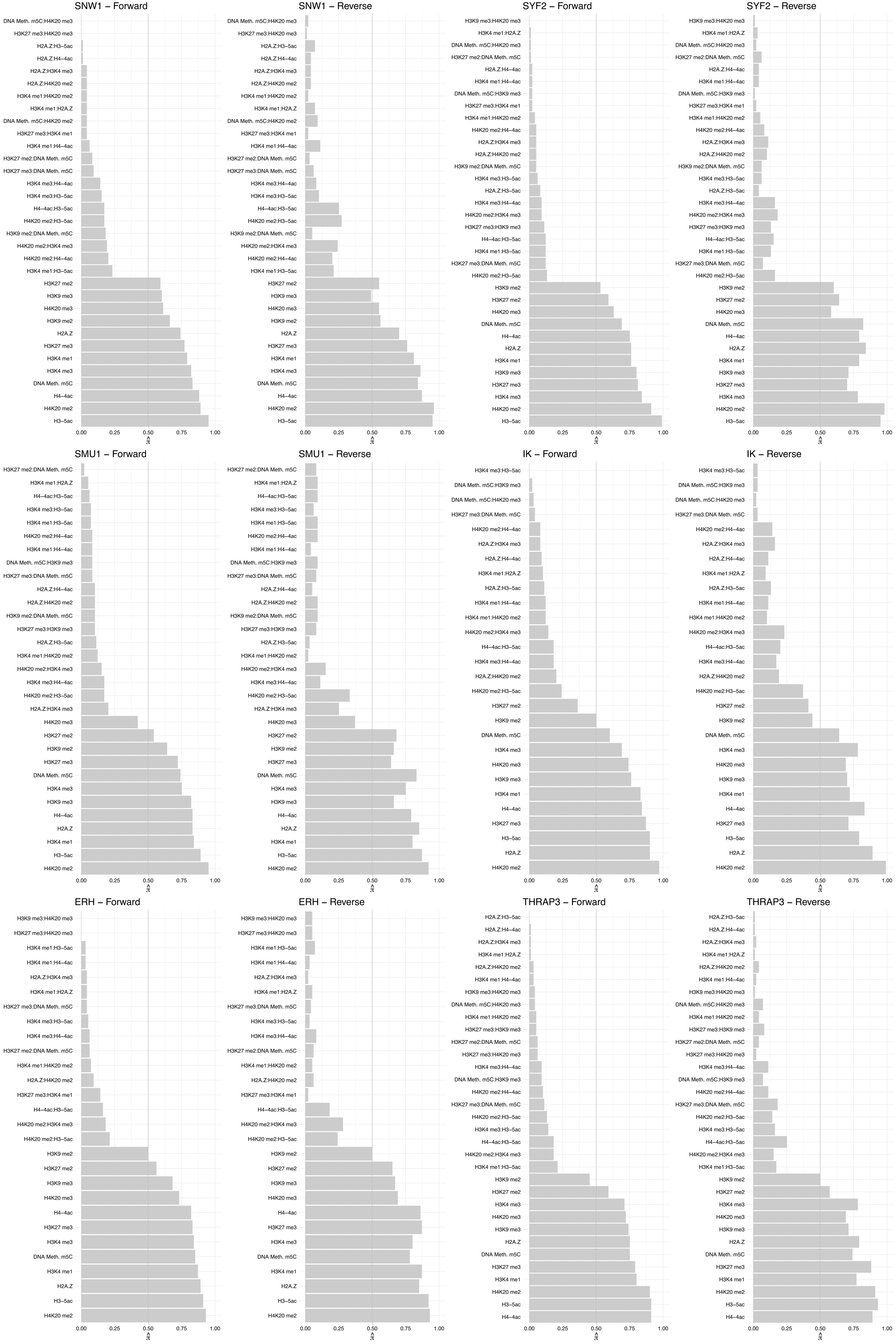
SNW1 – Forward
DNA Meth. m5C:H4K20 me3
H3K27 me3:H4K20 me3
H2A.Z:H3−5ac
H2A.Z:H4−4ac
H2A.Z:H3K4 me3
H2A.Z:H4K20 me2
H3K4 me1:H4K20 me2
H3K4 me1:H2A.Z
DNA Meth. m5C:H4K20 me2
H3K27 me3:H3K4 me1
H3K4 me1:H4−4ac
H3K27 me2:DNA Meth. m5C
H3K27 me3:DNA Meth. m5C
H3K4 me3:H4−4ac
H3K4 me3:H3−5ac
H4−4ac:H3−5ac
H4K20 me2:H3−5ac
H3K9 me2:DNA Meth. m5C
H4K20 me2:H3K4 me3
H4K20 me2:H4−4ac
H3K4 me1:H3−5ac
H3K27 me2
H3K9 me3
H4K20 me3
H3K9 me2
H2A.Z
H3K27 me3
H3K4 me1
H3K4 me3
DNA Meth. m5C
H4−4ac
H4K20 me2
H3−5ac
0.00 0.25 0.50 0.75 1.00
π̂
SNW1 – Reverse
DNA Meth. m5C:H4K20 me3
H3K27 me3:H4K20 me3
H2A.Z:H3−5ac
H2A.Z:H4−4ac
H2A.Z:H3K4 me3
H2A.Z:H4K20 me2
H3K4 me1:H4K20 me2
H3K4 me1:H2A.Z
DNA Meth. m5C:H4K20 me2
H3K27 me3:H3K4 me1
H3K4 me1:H4−4ac
H3K27 me2:DNA Meth. m5C
H3K27 me3:DNA Meth. m5C
H3K4 me3:H4−4ac
H3K4 me3:H3−5ac
H4−4ac:H3−5ac
H4K20 me2:H3−5ac
H3K9 me2:DNA Meth. m5C
H4K20 me2:H3K4 me3
H4K20 me2:H4−4ac
H3K4 me1:H3−5ac
H3K27 me2
H3K9 me3
H4K20 me3
H3K9 me2
H2A.Z
H3K27 me3
H3K4 me1
H3K4 me3
DNA Meth. m5C
H4−4ac
H4K20 me2
H3−5ac
0.00 0.25 0.50 0.75 1.00
π̂
SYF2 – Forward
H3K9 me3:H4K20 me3
H3K4 me1:H2A.Z
DNA Meth. m5C:H4K20 me3
H3K27 me2:DNA Meth. m5C
H2A.Z:H4−4ac
H3K4 me1:H4−4ac
DNA Meth. m5C:H3K9 me3
H3K27 me3:H3K4 me1
H3K4 me1:H4K20 me2
H4K20 me2:H4−4ac
H2A.Z:H3K4 me3
H2A.Z:H4K20 me2
H3K9 me2:DNA Meth. m5C
H3K4 me3:H3−5ac
H2A.Z:H3−5ac
H3K4 me3:H4−4ac
H4K20 me2:H3K4 me3
H3K27 me3:H3K9 me3
H4−4ac:H3−5ac
H3K4 me1:H3−5ac
H3K27 me3:DNA Meth. m5C
H4K20 me2:H3−5ac
H3K9 me2
H3K27 me2
H4K20 me3
DNA Meth. m5C
H4−4ac
H2A.Z
H3K4 me1
H3K9 me3
H3K27 me3
H3K4 me3
H4K20 me2
H3−5ac
0.00 0.25 0.50 0.75 1.00
π̂
SYF2 – Reverse
H3K9 me3:H4K20 me3
H3K4 me1:H2A.Z
DNA Meth. m5C:H4K20 me3
H3K27 me2:DNA Meth. m5C
H2A.Z:H4−4ac
H3K4 me1:H4−4ac
DNA Meth. m5C:H3K9 me3
H3K27 me3:H3K4 me1
H3K4 me1:H4K20 me2
H4K20 me2:H4−4ac
H2A.Z:H3K4 me3
H2A.Z:H4K20 me2
H3K9 me2:DNA Meth. m5C
H3K4 me3:H3−5ac
H2A.Z:H3−5ac
H3K4 me3:H4−4ac
H4K20 me2:H3K4 me3
H3K27 me3:H3K9 me3
H4−4ac:H3−5ac
H3K4 me1:H3−5ac
H3K27 me3:DNA Meth. m5C
H4K20 me2:H3−5ac
H3K9 me2
H3K27 me2
H4K20 me3
DNA Meth. m5C
H4−4ac
H2A.Z
H3K4 me1
H3K9 me3
H3K27 me3
H3K4 me3
H4K20 me2
H3−5ac
0.00 0.25 0.50 0.75 1.00
π̂
SMU1 – Forward
H3K27 me2:DNA Meth. m5C
H3K4 me1:H2A.Z
H4−4ac:H3−5ac
H3K4 me3:H3−5ac
H3K4 me1:H3−5ac
H4K20 me2:H4−4ac
H3K4 me1:H4−4ac
DNA Meth. m5C:H3K9 me3
H3K27 me3:DNA Meth. m5C
H2A.Z:H4−4ac
H2A.Z:H4K20 me2
H3K9 me2:DNA Meth. m5C
H3K27 me3:H3K9 me3
H2A.Z:H3−5ac
H3K4 me1:H4K20 me2
H4K20 me2:H3K4 me3
H3K4 me3:H4−4ac
H4K20 me2:H3−5ac
H2A.Z:H3K4 me3
H4K20 me3
H3K27 me2
H3K9 me2
H3K27 me3
DNA Meth. m5C
H3K4 me3
H3K9 me3
H4−4ac
H2A.Z
H3K4 me1
H3−5ac
H4K20 me2
0.00 0.25 0.50 0.75 1.00
π̂
SMU1 – Reverse
H3K27 me2:DNA Meth. m5C
H3K4 me1:H2A.Z
H4−4ac:H3−5ac
H3K4 me3:H3−5ac
H3K4 me1:H3−5ac
H4K20 me2:H4−4ac
H3K4 me1:H4−4ac
DNA Meth. m5C:H3K9 me3
H3K27 me3:DNA Meth. m5C
H2A.Z:H4−4ac
H2A.Z:H4K20 me2
H3K9 me2:DNA Meth. m5C
H3K27 me3:H3K9 me3
H2A.Z:H3−5ac
H3K4 me1:H4K20 me2
H4K20 me2:H3K4 me3
H3K4 me3:H4−4ac
H4K20 me2:H3−5ac
H2A.Z:H3K4 me3
H4K20 me3
H3K27 me2
H3K9 me2
H3K27 me3
DNA Meth. m5C
H3K4 me3
H3K9 me3
H4−4ac
H2A.Z
H3K4 me1
H3−5ac
H4K20 me2
0.00 0.25 0.50 0.75 1.00
π̂
IK – Forward
H3K4 me3:H3−5ac
DNA Meth. m5C:H3K9 me3
DNA Meth. m5C:H4K20 me3
H3K27 me3:DNA Meth. m5C
H4K20 me2:H4−4ac
H2A.Z:H3K4 me3
H2A.Z:H4−4ac
H3K4 me1:H2A.Z
H2A.Z:H3−5ac
H3K4 me1:H4−4ac
H3K4 me1:H4K20 me2
H4K20 me2:H3K4 me3
H4−4ac:H3−5ac
H3K4 me3:H4−4ac
H2A.Z:H4K20 me2
H4K20 me2:H3−5ac
H3K27 me2
H3K9 me2
DNA Meth. m5C
H3K4 me3
H4K20 me3
H3K9 me3
H3K4 me1
H4−4ac
H3K27 me3
H3−5ac
H2A.Z
H4K20 me2
0.00 0.25 0.50 0.75 1.00
π̂
IK – Reverse
H3K4 me3:H3−5ac
DNA Meth. m5C:H3K9 me3
DNA Meth. m5C:H4K20 me3
H3K27 me3:DNA Meth. m5C
H4K20 me2:H4−4ac
H2A.Z:H3K4 me3
H2A.Z:H4−4ac
H3K4 me1:H2A.Z
H2A.Z:H3−5ac
H3K4 me1:H4−4ac
H3K4 me1:H4K20 me2
H4K20 me2:H3K4 me3
H4−4ac:H3−5ac
H3K4 me3:H4−4ac
H2A.Z:H4K20 me2
H4K20 me2:H3−5ac
H3K27 me2
H3K9 me2
DNA Meth. m5C
H3K4 me3
H4K20 me3
H3K9 me3
H3K4 me1
H4−4ac
H3K27 me3
H3−5ac
H2A.Z
H4K20 me2
0.00 0.25 0.50 0.75 1.00
π̂
ERH – Forward
H3K9 me3:H4K20 me3
H3K27 me3:H4K20 me3
H3K4 me1:H3−5ac
H3K4 me1:H4−4ac
H2A.Z:H3K4 me3
H3K4 me1:H2A.Z
H3K27 me3:DNA Meth. m5C
H3K4 me3:H3−5ac
H3K4 me3:H4−4ac
H3K27 me2:DNA Meth. m5C
H3K4 me1:H4K20 me2
H2A.Z:H4K20 me2
H3K27 me3:H3K4 me1
H4−4ac:H3−5ac
H4K20 me2:H3K4 me3
H4K20 me2:H3−5ac
H3K9 me2
H3K27 me2
H3K9 me3
H4K20 me3
H4−4ac
H3K27 me3
H3K4 me3
DNA Meth. m5C
H3K4 me1
H2A.Z
H3−5ac
H4K20 me2
0.00 0.25 0.50 0.75 1.00
π̂
ERH – Reverse
H3K9 me3:H4K20 me3
H3K27 me3:H4K20 me3
H3K4 me1:H3−5ac
H3K4 me1:H4−4ac
H2A.Z:H3K4 me3
H3K4 me1:H2A.Z
H3K27 me3:DNA Meth. m5C
H3K4 me3:H3−5ac
H3K4 me3:H4−4ac
H3K27 me2:DNA Meth. m5C
H3K4 me1:H4K20 me2
H2A.Z:H4K20 me2
H3K27 me3:H3K4 me1
H4−4ac:H3−5ac
H4K20 me2:H3K4 me3
H4K20 me2:H3−5ac
H3K9 me2
H3K27 me2
H3K9 me3
H4K20 me3
H4−4ac
H3K27 me3
H3K4 me3
DNA Meth. m5C
H3K4 me1
H2A.Z
H3−5ac
H4K20 me2
0.00 0.25 0.50 0.75 1.00
π̂
THRAP3 – Forward
H2A.Z:H3−5ac
H2A.Z:H4−4ac
H2A.Z:H3K4 me3
H3K4 me1:H2A.Z
H2A.Z:H4K20 me2
H3K4 me1:H4−4ac
H3K9 me3:H4K20 me3
DNA Meth. m5C:H4K20 me3
H3K4 me1:H4K20 me2
H3K27 me3:H3K9 me3
H3K27 me2:DNA Meth. m5C
H3K27 me3:H4K20 me3
H3K4 me3:H4−4ac
DNA Meth. m5C:H3K9 me3
H4K20 me2:H4−4ac
H3K27 me3:DNA Meth. m5C
H4K20 me2:H3−5ac
H3K4 me3:H3−5ac
H4−4ac:H3−5ac
H4K20 me2:H3K4 me3
H3K4 me1:H3−5ac
H3K9 me2
H3K27 me2
H3K4 me3
H4K20 me3
H3K9 me3
H2A.Z
DNA Meth. m5C
H3K27 me3
H3K4 me1
H4K20 me2
H3−5ac
H4−4ac
0.00 0.25 0.50 0.75 1.00
π̂
THRAP3 – Reverse
H2A.Z:H3−5ac
H2A.Z:H4−4ac
H2A.Z:H3K4 me3
H3K4 me1:H2A.Z
H2A.Z:H4K20 me2
H3K4 me1:H4−4ac
H3K9 me3:H4K20 me3
DNA Meth. m5C:H4K20 me3
H3K4 me1:H4K20 me2
H3K27 me3:H3K9 me3
H3K27 me2:DNA Meth. m5C
H3K27 me3:H4K20 me3
H3K4 me3:H4−4ac
DNA Meth. m5C:H3K9 me3
H4K20 me2:H4−4ac
H3K27 me3:DNA Meth. m5C
H4K20 me2:H3−5ac
H3K4 me3:H3−5ac
H4−4ac:H3−5ac
H4K20 me2:H3K4 me3
H3K4 me1:H3−5ac
H3K9 me2
H3K27 me2
H3K4 me3
H4K20 me3
H3K9 me3
H2A.Z
DNA Meth. m5C
H3K27 me3
H3K4 me1
H4K20 me2
H3−5ac
H4−4ac
0.00 0.25 0.50 0.75 1.00
π̂

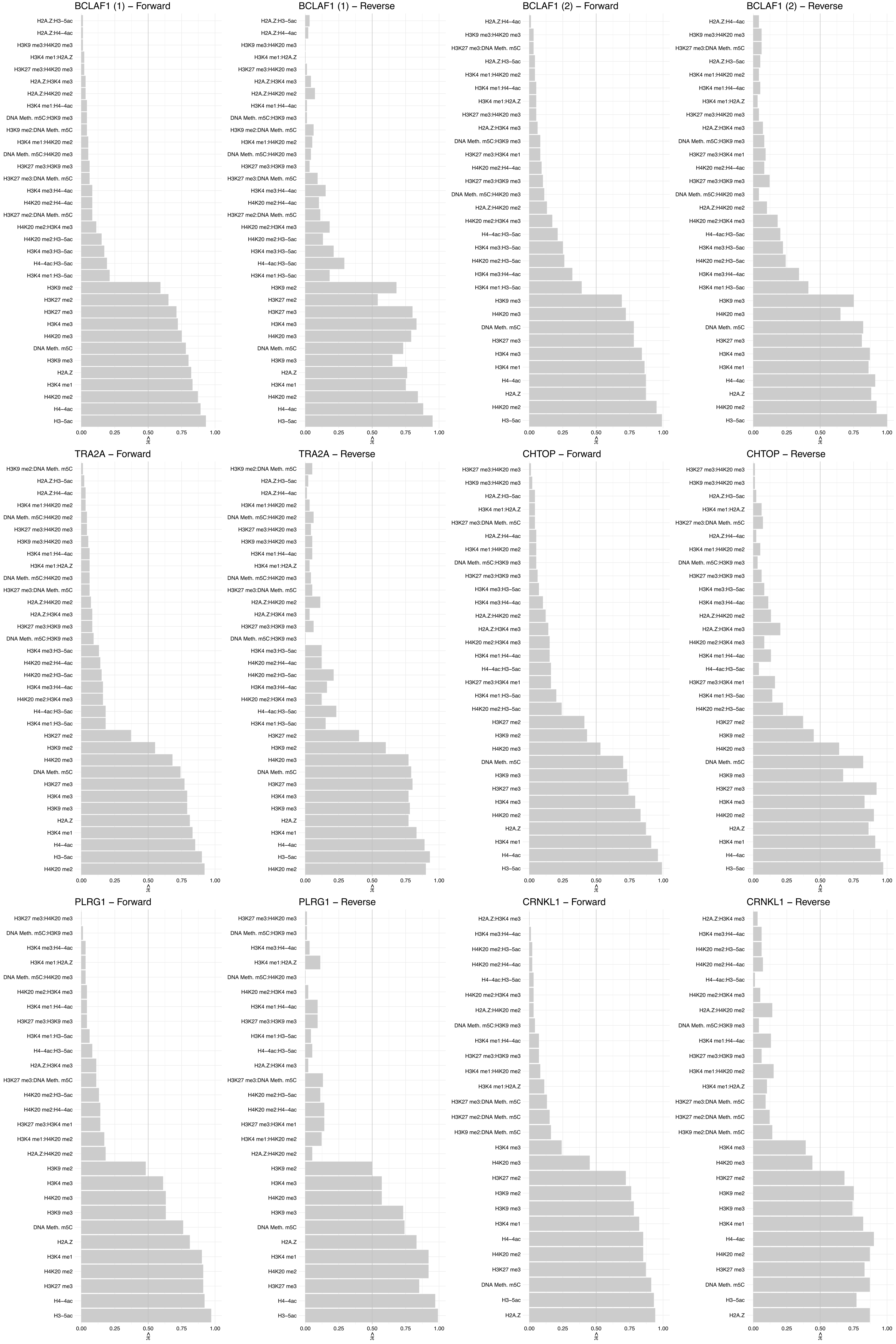
BCLAF1 (1) – Forward
H2A.Z:H3−5ac
H2A.Z:H4−4ac
H3K9 me3:H4K20 me3
H3K4 me1:H2A.Z
H3K27 me3:H4K20 me3
H2A.Z:H3K4 me3
H2A.Z:H4K20 me2
H3K4 me1:H4−4ac
DNA Meth. m5C:H3K9 me3
H3K9 me2:DNA Meth. m5C
H3K4 me1:H4K20 me2
DNA Meth. m5C:H4K20 me3
H3K27 me3:H3K9 me3
H3K27 me3:DNA Meth. m5C
H3K4 me3:H4−4ac
H4K20 me2:H4−4ac
H3K27 me2:DNA Meth. m5C
H4K20 me2:H3K4 me3
H4K20 me2:H3−5ac
H3K4 me3:H3−5ac
H4−4ac:H3−5ac
H3K4 me1:H3−5ac
H3K9 me2
H3K27 me2
H3K27 me3
H3K4 me3
H4K20 me3
DNA Meth. m5C
H3K9 me3
H2A.Z
H3K4 me1
H4K20 me2
H4−4ac
H3−5ac
0.00
0.25
0.50
0.75
1.00
π̂
BCLAF1 (1) – Reverse
H2A.Z:H3−5ac
H2A.Z:H4−4ac
H3K9 me3:H4K20 me3
H3K4 me1:H2A.Z
H3K27 me3:H4K20 me3
H2A.Z:H3K4 me3
H2A.Z:H4K20 me2
H3K4 me1:H4−4ac
DNA Meth. m5C:H3K9 me3
H3K9 me2:DNA Meth. m5C
H3K4 me1:H4K20 me2
DNA Meth. m5C:H4K20 me3
H3K27 me3:H3K9 me3
H3K27 me3:DNA Meth. m5C
H3K4 me3:H4−4ac
H4K20 me2:H4−4ac
H3K27 me2:DNA Meth. m5C
H4K20 me2:H3K4 me3
H4K20 me2:H3−5ac
H3K4 me3:H3−5ac
H4−4ac:H3−5ac
H3K4 me1:H3−5ac
H3K9 me2
H3K27 me2
H3K27 me3
H3K4 me3
H4K20 me3
DNA Meth. m5C
H3K9 me3
H2A.Z
H3K4 me1
H4K20 me2
H4−4ac
H3−5ac
0.00
0.25
0.50
0.75
1.00
π̂
BCLAF1 (2) – Forward
H2A.Z:H4−4ac
H3K9 me3:H4K20 me3
H3K27 me3:DNA Meth. m5C
H2A.Z:H3−5ac
H3K4 me1:H4K20 me2
H3K4 me1:H4−4ac
H3K4 me1:H2A.Z
H3K27 me3:H4K20 me3
H2A.Z:H3K4 me3
DNA Meth. m5C:H3K9 me3
H3K27 me3:H3K4 me1
H4K20 me2:H4−4ac
H3K27 me3:H3K9 me3
DNA Meth. m5C:H4K20 me3
H2A.Z:H4K20 me2
H4K20 me2:H3K4 me3
H4−4ac:H3−5ac
H3K4 me3:H3−5ac
H4K20 me2:H3−5ac
H3K4 me3:H4−4ac
H3K4 me1:H3−5ac
H3K9 me3
H4K20 me3
DNA Meth. m5C
H3K27 me3
H3K4 me3
H3K4 me1
H4−4ac
H2A.Z
H4K20 me2
H3−5ac
0.00
0.25
0.50
0.75
1.00
π̂
BCLAF1 (2) – Reverse
H2A.Z:H4−4ac
H3K9 me3:H4K20 me3
H3K27 me3:DNA Meth. m5C
H2A.Z:H3−5ac
H3K4 me1:H4K20 me2
H3K4 me1:H4−4ac
H3K4 me1:H2A.Z
H3K27 me3:H4K20 me3
H2A.Z:H3K4 me3
DNA Meth. m5C:H3K9 me3
H3K27 me3:H3K4 me1
H4K20 me2:H4−4ac
H3K27 me3:H3K9 me3
DNA Meth. m5C:H4K20 me3
H2A.Z:H4K20 me2
H4K20 me2:H3K4 me3
H4−4ac:H3−5ac
H3K4 me3:H3−5ac
H4K20 me2:H3−5ac
H3K4 me3:H4−4ac
H3K4 me1:H3−5ac
H3K9 me3
H4K20 me3
DNA Meth. m5C
H3K27 me3
H3K4 me3
H3K4 me1
H4−4ac
H2A.Z
H4K20 me2
H3−5ac
0.00
0.25
0.50
0.75
1.00
π̂
TRA2A – Forward
H3K9 me2:DNA Meth. m5C
H2A.Z:H3−5ac
H2A.Z:H4−4ac
H3K4 me1:H4K20 me2
DNA Meth. m5C:H4K20 me2
H3K27 me3:H4K20 me3
H3K9 me3:H4K20 me3
H3K4 me1:H4−4ac
H3K4 me1:H2A.Z
DNA Meth. m5C:H4K20 me3
H3K27 me3:DNA Meth. m5C
H2A.Z:H4K20 me2
H2A.Z:H3K4 me3
H3K27 me3:H3K9 me3
DNA Meth. m5C:H3K9 me3
H3K4 me3:H3−5ac
H4K20 me2:H4−4ac
H4K20 me2:H3−5ac
H3K4 me3:H4−4ac
H4K20 me2:H3K4 me3
H4−4ac:H3−5ac
H3K4 me1:H3−5ac
H3K27 me2
H3K9 me2
H4K20 me3
DNA Meth. m5C
H3K27 me3
H3K4 me3
H3K9 me3
H2A.Z
H3K4 me1
H4−4ac
H3−5ac
H4K20 me2
0.00
0.25
0.50
0.75
1.00
π̂
TRA2A – Reverse
H3K9 me2:DNA Meth. m5C
H2A.Z:H3−5ac
H2A.Z:H4−4ac
H3K4 me1:H4K20 me2
DNA Meth. m5C:H4K20 me2
H3K27 me3:H4K20 me3
H3K9 me3:H4K20 me3
H3K4 me1:H4−4ac
H3K4 me1:H2A.Z
DNA Meth. m5C:H4K20 me3
H3K27 me3:DNA Meth. m5C
H2A.Z:H4K20 me2
H2A.Z:H3K4 me3
H3K27 me3:H3K9 me3
DNA Meth. m5C:H3K9 me3
H3K4 me3:H3−5ac
H4K20 me2:H4−4ac
H4K20 me2:H3−5ac
H3K4 me3:H4−4ac
H4K20 me2:H3K4 me3
H4−4ac:H3−5ac
H3K4 me1:H3−5ac
H3K27 me2
H3K9 me2
H4K20 me3
DNA Meth. m5C
H3K27 me3
H3K4 me3
H3K9 me3
H2A.Z
H3K4 me1
H4−4ac
H3−5ac
H4K20 me2
0.00
0.25
0.50
0.75
1.00
π̂
CHTOP – Forward
H3K27 me3:H4K20 me3
H3K9 me3:H4K20 me3
H2A.Z:H3−5ac
H3K4 me1:H2A.Z
H3K27 me3:DNA Meth. m5C
H2A.Z:H4−4ac
H3K4 me1:H4K20 me2
DNA Meth. m5C:H3K9 me3
H3K27 me3:H3K9 me3
H3K4 me3:H3−5ac
H3K4 me3:H4−4ac
H2A.Z:H4K20 me2
H2A.Z:H3K4 me3
H4K20 me2:H3K4 me3
H3K4 me1:H4−4ac
H4−4ac:H3−5ac
H3K27 me3:H3K4 me1
H3K4 me1:H3−5ac
H4K20 me2:H3−5ac
H3K27 me2
H3K9 me2
H4K20 me3
DNA Meth. m5C
H3K9 me3
H3K27 me3
H3K4 me3
H4K20 me2
H2A.Z
H3K4 me1
H4−4ac
H3−5ac
0.00
0.25
0.50
0.75
1.00
π̂
CHTOP – Reverse
H3K27 me3:H4K20 me3
H3K9 me3:H4K20 me3
H2A.Z:H3−5ac
H3K4 me1:H2A.Z
H3K27 me3:DNA Meth. m5C
H2A.Z:H4−4ac
H3K4 me1:H4K20 me2
DNA Meth. m5C:H3K9 me3
H3K27 me3:H3K9 me3
H3K4 me3:H3−5ac
H3K4 me3:H4−4ac
H2A.Z:H4K20 me2
H2A.Z:H3K4 me3
H4K20 me2:H3K4 me3
H3K4 me1:H4−4ac
H4−4ac:H3−5ac
H3K27 me3:H3K4 me1
H3K4 me1:H3−5ac
H4K20 me2:H3−5ac
H3K27 me2
H3K9 me2
H4K20 me3
DNA Meth. m5C
H3K9 me3
H3K27 me3
H3K4 me3
H4K20 me2
H2A.Z
H3K4 me1
H4−4ac
H3−5ac
0.00
0.25
0.50
0.75
1.00
π̂
PLRG1 – Forward
H3K27 me3:H4K20 me3
DNA Meth. m5C:H3K9 me3
H3K4 me3:H4−4ac
H3K4 me1:H2A.Z
DNA Meth. m5C:H4K20 me3
H4K20 me2:H3K4 me3
H3K4 me1:H4−4ac
H3K27 me3:H3K9 me3
H3K4 me1:H3−5ac
H4−4ac:H3−5ac
H2A.Z:H3K4 me3
H3K27 me3:DNA Meth. m5C
H4K20 me2:H3−5ac
H4K20 me2:H4−4ac
H3K27 me3:H3K4 me1
H3K4 me1:H4K20 me2
H2A.Z:H4K20 me2
H3K9 me2
H3K4 me3
H4K20 me3
H3K9 me3
DNA Meth. m5C
H2A.Z
H3K4 me1
H4K20 me2
H3K27 me3
H4−4ac
H3−5ac
0.00
0.25
0.50
0.75
1.00
π̂
PLRG1 – Reverse
H3K27 me3:H4K20 me3
DNA Meth. m5C:H3K9 me3
H3K4 me3:H4−4ac
H3K4 me1:H2A.Z
DNA Meth. m5C:H4K20 me3
H4K20 me2:H3K4 me3
H3K4 me1:H4−4ac
H3K27 me3:H3K9 me3
H3K4 me1:H3−5ac
H4−4ac:H3−5ac
H2A.Z:H3K4 me3
H3K27 me3:DNA Meth. m5C
H4K20 me2:H3−5ac
H4K20 me2:H4−4ac
H3K27 me3:H3K4 me1
H3K4 me1:H4K20 me2
H2A.Z:H4K20 me2
H3K9 me2
H3K4 me3
H4K20 me3
H3K9 me3
DNA Meth. m5C
H2A.Z
H3K4 me1
H4K20 me2
H3K27 me3
H4−4ac
H3−5ac
0.00
0.25
0.50
0.75
1.00
π̂
CRNKL1 – Forward
H2A.Z:H3K4 me3
H3K4 me3:H4−4ac
H4K20 me2:H3−5ac
H4K20 me2:H4−4ac
H4−4ac:H3−5ac
H4K20 me2:H3K4 me3
H2A.Z:H4K20 me2
DNA Meth. m5C:H3K9 me3
H3K4 me1:H4−4ac
H3K27 me3:H3K9 me3
H3K4 me1:H4K20 me2
H3K4 me1:H2A.Z
H3K27 me3:DNA Meth. m5C
H3K27 me2:DNA Meth. m5C
H3K9 me2:DNA Meth. m5C
H3K4 me3
H4K20 me3
H3K27 me2
H3K9 me2
H3K9 me3
H3K4 me1
H4−4ac
H4K20 me2
H3K27 me3
DNA Meth. m5C
H3−5ac
H2A.Z
0.00
0.25
0.50
0.75
1.00
π̂
CRNKL1 – Reverse
H2A.Z:H3K4 me3
H3K4 me3:H4−4ac
H4K20 me2:H3−5ac
H4K20 me2:H4−4ac
H4−4ac:H3−5ac
H4K20 me2:H3K4 me3
H2A.Z:H4K20 me2
DNA Meth. m5C:H3K9 me3
H3K4 me1:H4−4ac
H3K27 me3:H3K9 me3
H3K4 me1:H4K20 me2
H3K4 me1:H2A.Z
H3K27 me3:DNA Meth. m5C
H3K27 me2:DNA Meth. m5C
H3K9 me2:DNA Meth. m5C
H3K4 me3
H4K20 me3
H3K27 me2
H3K9 me2
H3K9 me3
H3K4 me1
H4−4ac
H4K20 me2
H3K27 me3
DNA Meth. m5C
H3−5ac
H2A.Z
0.00
0.25
0.50
0.75
1.00
π̂

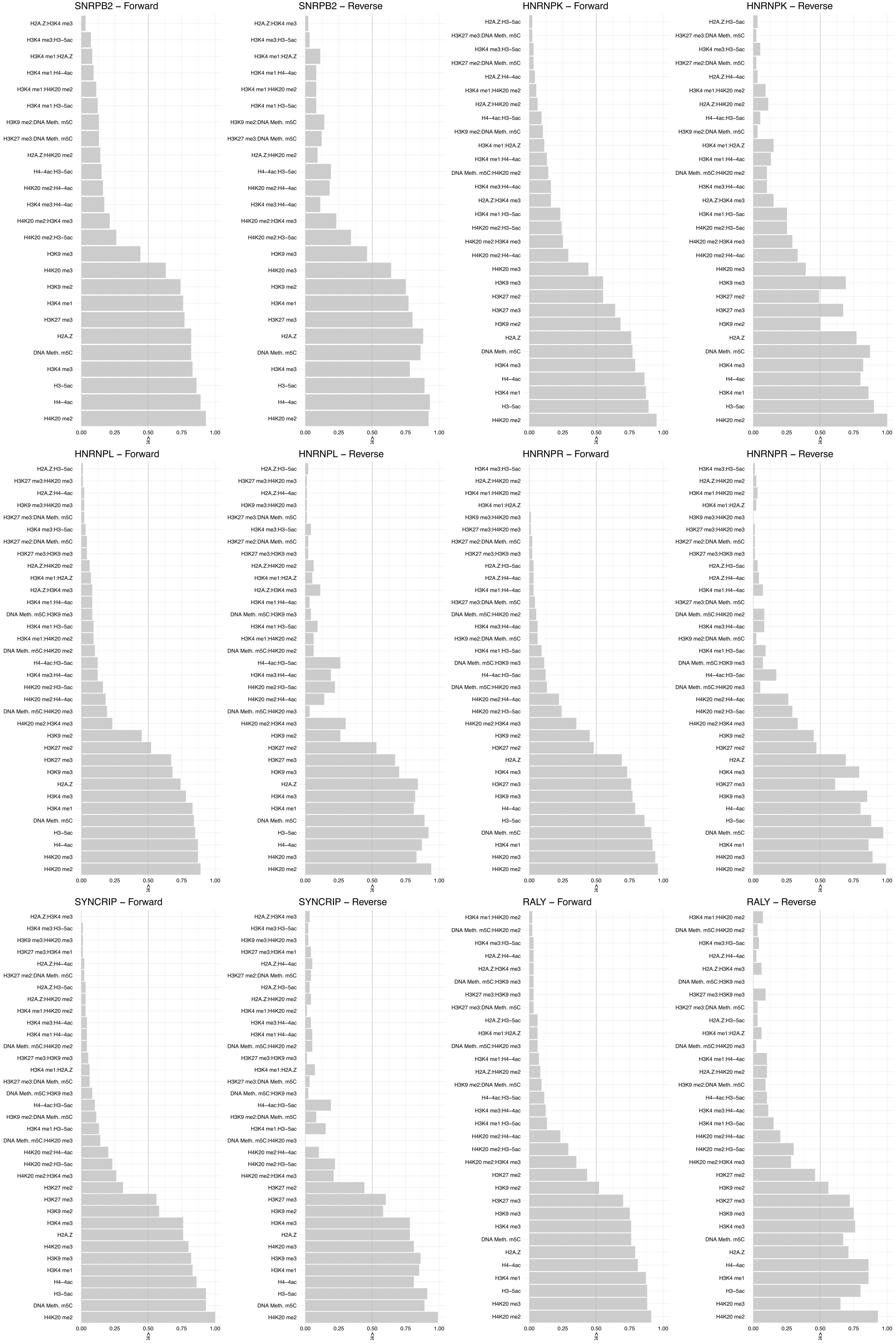
SNRPB2 – Forward
SNRPB2 – Reverse
HNRNPK – Forward
HNRNPK – Reverse
HNRNPL – Forward
HNRNPL – Reverse
HNRNPR – Forward
HNRNPR – Reverse
SYNCRIP – Forward
SYNCRIP – Reverse
RALY – Forward
RALY – Reverse
H4K20 me2
H4–4ac
H3–5ac
H3K4 me3
DNA Meth. m5C
H2A.Z
H3K27 me3
H3K4 me1
H3K9 me2
H4K20 me3
H3K9 me3
H3K27 me2
0.00
0.25
0.50
0.75
1.00

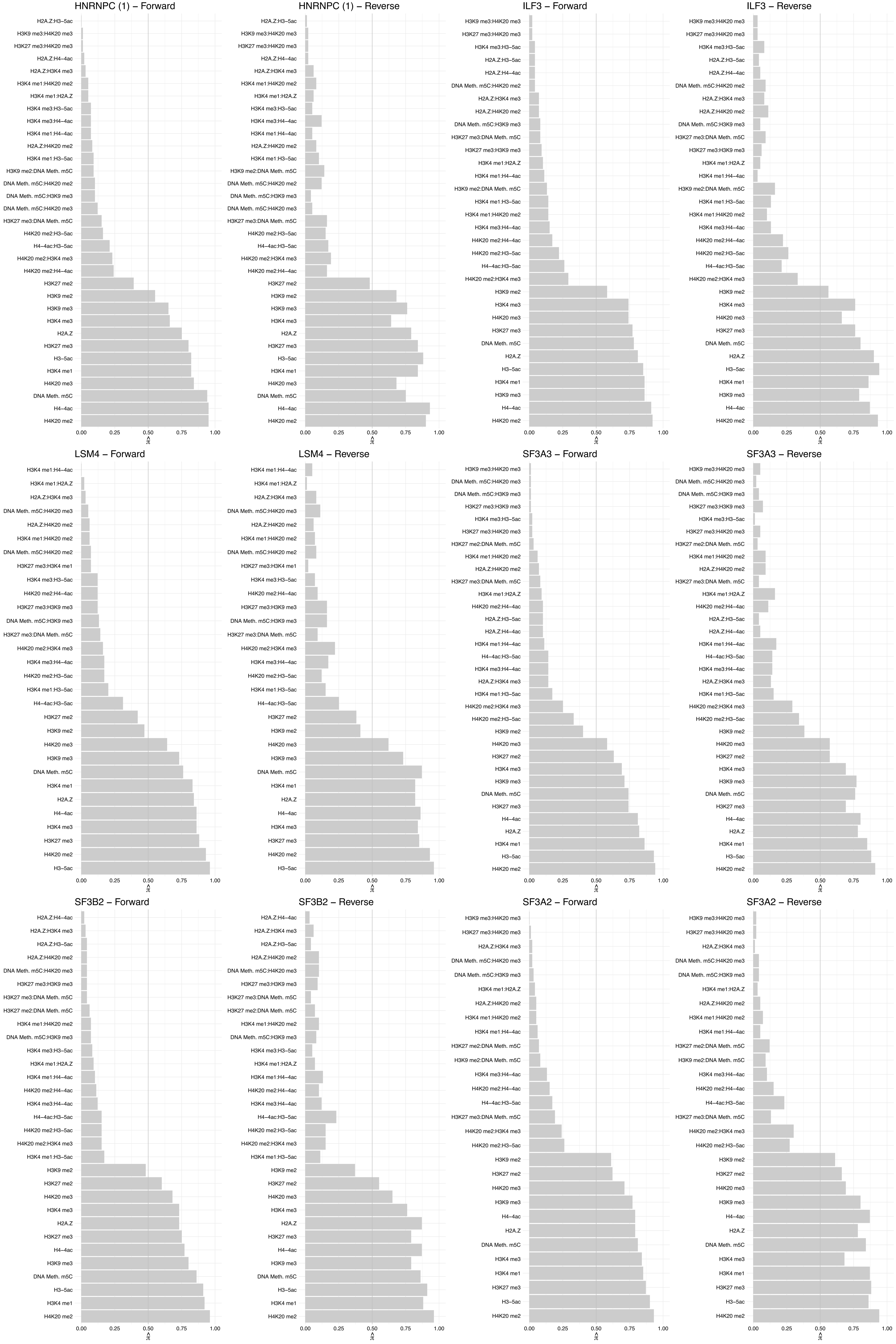

HNRNPC (1) – Forward
H2A.Z:H3−5ac
H3K9 me3:H4K20 me3
H3K27 me3:H4K20 me3
H2A.Z:H4−4ac
H2A.Z:H3K4 me3
H3K4 me1:H4K20 me2
H3K4 me1:H2A.Z
H3K4 me3:H3−5ac
H3K4 me3:H4−4ac
H3K4 me1:H4−4ac
H2A.Z:H4K20 me2
H3K4 me1:H3−5ac
H3K9 me2:DNA Meth. m5C
DNA Meth. m5C:H4K20 me2
DNA Meth. m5C:H3K9 me3
DNA Meth. m5C:H4K20 me3
H3K27 me3:DNA Meth. m5C
H4K20 me2:H3−5ac
H4−4ac:H3−5ac
H4K20 me2:H3K4 me3
H4K20 me2:H4−4ac
H3K27 me2
H3K9 me2
H3K9 me3
H3K4 me3
H2A.Z
H3K27 me3
H3−5ac
H3K4 me1
H4K20 me3
DNA Meth. m5C
H4−4ac
H4K20 me2
0.00 0.25 0.50 0.75 1.00
π̂
HNRNPC (1) – Reverse
H2A.Z:H3−5ac
H3K9 me3:H4K20 me3
H3K27 me3:H4K20 me3
H2A.Z:H4−4ac
H2A.Z:H3K4 me3
H3K4 me1:H4K20 me2
H3K4 me1:H2A.Z
H3K4 me3:H3−5ac
H3K4 me3:H4−4ac
H3K4 me1:H4−4ac
H2A.Z:H4K20 me2
H3K4 me1:H3−5ac
H3K9 me2:DNA Meth. m5C
DNA Meth. m5C:H4K20 me2
DNA Meth. m5C:H3K9 me3
DNA Meth. m5C:H4K20 me3
H3K27 me3:DNA Meth. m5C
H4K20 me2:H3−5ac
H4−4ac:H3−5ac
H4K20 me2:H3K4 me3
H4K20 me2:H4−4ac
H3K27 me2
H3K9 me2
H3K9 me3
H3K4 me3
H2A.Z
H3K27 me3
H3−5ac
H3K4 me1
H4K20 me3
DNA Meth. m5C
H4−4ac
H4K20 me2
0.00 0.25 0.50 0.75 1.00
π̂
ILF3 – Forward
H3K9 me3:H4K20 me3
H3K27 me3:H4K20 me3
H3K4 me3:H3−5ac
H2A.Z:H3−5ac
H2A.Z:H4−4ac
DNA Meth. m5C:H4K20 me2
H2A.Z:H3K4 me3
H2A.Z:H4K20 me2
DNA Meth. m5C:H3K9 me3
H3K27 me3:DNA Meth. m5C
H3K27 me3:H3K9 me3
H3K4 me1:H2A.Z
H3K4 me1:H4−4ac
H3K9 me2:DNA Meth. m5C
H3K4 me1:H3−5ac
H3K4 me1:H4K20 me2
H3K4 me3:H4−4ac
H4K20 me2:H4−4ac
H4K20 me2:H3−5ac
H4−4ac:H3−5ac
H4K20 me2:H3K4 me3
H3K9 me2
H3K4 me3
H4K20 me3
H3K27 me3
DNA Meth. m5C
H2A.Z
H3−5ac
H3K4 me1
H3K9 me3
H4−4ac
H4K20 me2
0.00 0.25 0.50 0.75 1.00
π̂
ILF3 – Reverse
H3K9 me3:H4K20 me3
H3K27 me3:H4K20 me3
H3K4 me3:H3−5ac
H2A.Z:H3−5ac
H2A.Z:H4−4ac
DNA Meth. m5C:H4K20 me2
H2A.Z:H3K4 me3
H2A.Z:H4K20 me2
DNA Meth. m5C:H3K9 me3
H3K27 me3:DNA Meth. m5C
H3K27 me3:H3K9 me3
H3K4 me1:H2A.Z
H3K4 me1:H4−4ac
H3K9 me2:DNA Meth. m5C
H3K4 me1:H3−5ac
H3K4 me1:H4K20 me2
H3K4 me3:H4−4ac
H4K20 me2:H4−4ac
H4K20 me2:H3−5ac
H4−4ac:H3−5ac
H4K20 me2:H3K4 me3
H3K9 me2
H3K4 me3
H4K20 me3
H3K27 me3
DNA Meth. m5C
H2A.Z
H3−5ac
H3K4 me1
H3K9 me3
H4−4ac
H4K20 me2
0.00 0.25 0.50 0.75 1.00
π̂
LSM4 – Forward
H3K4 me1:H4−4ac
H3K4 me1:H2A.Z
H2A.Z:H3K4 me3
DNA Meth. m5C:H4K20 me3
H2A.Z:H4K20 me2
H3K4 me1:H4K20 me2
DNA Meth. m5C:H4K20 me2
H3K27 me3:H3K4 me1
H3K4 me3:H3−5ac
H4K20 me2:H4−4ac
H3K27 me3:H3K9 me3
DNA Meth. m5C:H3K9 me3
H3K27 me3:DNA Meth. m5C
H4K20 me2:H3K4 me3
H3K4 me3:H4−4ac
H4K20 me2:H3−5ac
H3K4 me1:H3−5ac
H4−4ac:H3−5ac
H3K27 me2
H3K9 me2
H4K20 me3
H3K9 me3
DNA Meth. m5C
H3K4 me1
H2A.Z
H4−4ac
H3K4 me3
H3K27 me3
H4K20 me2
H3−5ac
0.00 0.25 0.50 0.75 1.00
π̂
LSM4 – Reverse
H3K4 me1:H4−4ac
H3K4 me1:H2A.Z
H2A.Z:H3K4 me3
DNA Meth. m5C:H4K20 me3
H2A.Z:H4K20 me2
H3K4 me1:H4K20 me2
DNA Meth. m5C:H4K20 me2
H3K27 me3:H3K4 me1
H3K4 me3:H3−5ac
H4K20 me2:H4−4ac
H3K27 me3:H3K9 me3
DNA Meth. m5C:H3K9 me3
H3K27 me3:DNA Meth. m5C
H4K20 me2:H3K4 me3
H3K4 me3:H4−4ac
H4K20 me2:H3−5ac
H3K4 me1:H3−5ac
H4−4ac:H3−5ac
H3K27 me2
H3K9 me2
H4K20 me3
H3K9 me3
DNA Meth. m5C
H3K4 me1
H2A.Z
H4−4ac
H3K4 me3
H3K27 me3
H4K20 me2
H3−5ac
0.00 0.25 0.50 0.75 1.00
π̂
SF3A3 – Forward
H3K9 me3:H4K20 me3
DNA Meth. m5C:H4K20 me3
DNA Meth. m5C:H3K9 me3
H3K27 me3:H3K9 me3
H3K4 me3:H3−5ac
H3K27 me3:H4K20 me3
H3K27 me2:DNA Meth. m5C
H3K4 me1:H4K20 me2
H2A.Z:H4K20 me2
H3K27 me3:DNA Meth. m5C
H3K4 me1:H2A.Z
H4K20 me2:H4−4ac
H2A.Z:H3−5ac
H2A.Z:H4−4ac
H3K4 me1:H4−4ac
H4−4ac:H3−5ac
H3K4 me3:H4−4ac
H2A.Z:H3K4 me3
H3K4 me1:H3−5ac
H4K20 me2:H3K4 me3
H4K20 me2:H3−5ac
H3K9 me2
H4K20 me3
H3K27 me2
H3K4 me3
H3K9 me3
DNA Meth. m5C
H3K27 me3
H4−4ac
H2A.Z
H3K4 me1
H3−5ac
H4K20 me2
0.00 0.25 0.50 0.75 1.00
π̂
SF3A3 – Reverse
H3K9 me3:H4K20 me3
DNA Meth. m5C:H4K20 me3
DNA Meth. m5C:H3K9 me3
H3K27 me3:H3K9 me3
H3K4 me3:H3−5ac
H3K27 me3:H4K20 me3
H3K27 me2:DNA Meth. m5C
H3K4 me1:H4K20 me2
H2A.Z:H4K20 me2
H3K27 me3:DNA Meth. m5C
H3K4 me1:H2A.Z
H4K20 me2:H4−4ac
H2A.Z:H3−5ac
H2A.Z:H4−4ac
H3K4 me1:H4−4ac
H4−4ac:H3−5ac
H3K4 me3:H4−4ac
H2A.Z:H3K4 me3
H3K4 me1:H3−5ac
H4K20 me2:H3K4 me3
H4K20 me2:H3−5ac
H3K9 me2
H4K20 me3
H3K27 me2
H3K4 me3
H3K9 me3
DNA Meth. m5C
H3K27 me3
H4−4ac
H2A.Z
H3K4 me1
H3−5ac
H4K20 me2
0.00 0.25 0.50 0.75 1.00
π̂
SF3B2 – Forward
H2A.Z:H4−4ac
H2A.Z:H3K4 me3
H2A.Z:H3−5ac
H2A.Z:H4K20 me2
DNA Meth. m5C:H4K20 me3
H3K27 me3:H3K9 me3
H3K27 me3:DNA Meth. m5C
H3K27 me2:DNA Meth. m5C
H3K4 me1:H4K20 me2
DNA Meth. m5C:H3K9 me3
H3K4 me3:H3−5ac
H3K4 me1:H2A.Z
H3K4 me1:H4−4ac
H4K20 me2:H4−4ac
H3K4 me3:H4−4ac
H4−4ac:H3−5ac
H4K20 me2:H3−5ac
H4K20 me2:H3K4 me3
H3K4 me1:H3−5ac
H3K9 me2
H3K27 me2
H4K20 me3
H3K4 me3
H2A.Z
H3K27 me3
H4−4ac
H3K9 me3
DNA Meth. m5C
H3−5ac
H3K4 me1
H4K20 me2
0.00 0.25 0.50 0.75 1.00
π̂
SF3B2 – Reverse
H2A.Z:H4−4ac
H2A.Z:H3K4 me3
H2A.Z:H3−5ac
H2A.Z:H4K20 me2
DNA Meth. m5C:H4K20 me3
H3K27 me3:H3K9 me3
H3K27 me3:DNA Meth. m5C
H3K27 me2:DNA Meth. m5C
H3K4 me1:H4K20 me2
DNA Meth. m5C:H3K9 me3
H3K4 me3:H3−5ac
H3K4 me1:H2A.Z
H3K4 me1:H4−4ac
H4K20 me2:H4−4ac
H3K4 me3:H4−4ac
H4−4ac:H3−5ac
H4K20 me2:H3−5ac
H4K20 me2:H3K4 me3
H3K4 me1:H3−5ac
H3K9 me2
H3K27 me2
H4K20 me3
H3K4 me3
H2A.Z
H3K27 me3
H4−4ac
H3K9 me3
DNA Meth. m5C
H3−5ac
H3K4 me1
H4K20 me2
0.00 0.25 0.50 0.75 1.00
π̂
SF3A2 – Forward
H3K9 me3:H4K20 me3
H3K27 me3:H4K20 me3
H2A.Z:H3K4 me3
DNA Meth. m5C:H4K20 me3
DNA Meth. m5C:H3K9 me3
H3K4 me1:H2A.Z
H2A.Z:H4K20 me2
H3K4 me1:H4K20 me2
H3K4 me1:H4−4ac
H3K27 me2:DNA Meth. m5C
H3K9 me2:DNA Meth. m5C
H3K4 me3:H4−4ac
H4K20 me2:H4−4ac
H4−4ac:H3−5ac
H3K27 me3:DNA Meth. m5C
H4K20 me2:H3K4 me3
H4K20 me2:H3−5ac
H3K9 me2
H3K27 me2
H4K20 me3
H3K9 me3
H4−4ac
H2A.Z
DNA Meth. m5C
H3K4 me3
H3K4 me1
H3K27 me3
H3−5ac
H4K20 me2
0.00 0.25 0.50 0.75 1.00
π̂
SF3A2 – Reverse
H3K9 me3:H4K20 me3
H3K27 me3:H4K20 me3
H2A.Z:H3K4 me3
DNA Meth. m5C:H4K20 me3
DNA Meth. m5C:H3K9 me3
H3K4 me1:H2A.Z
H2A.Z:H4K20 me2
H3K4 me1:H4K20 me2
H3K4 me1:H4−4ac
H3K27 me2:DNA Meth. m5C
H3K9 me2:DNA Meth. m5C
H3K4 me3:H4−4ac
H4K20 me2:H4−4ac
H4−4ac:H3−5ac
H3K27 me3:DNA Meth. m5C
H4K20 me2:H3K4 me3
H4K20 me2:H3−5ac
H3K9 me2
H3K27 me2
H4K20 me3
H3K9 me3
H4−4ac
H2A.Z
DNA Meth. m5C
H3K4 me3
H3K4 me1
H3K27 me3
H3−5ac
H4K20 me2
0.00 0.25 0.50 0.75 1.00
π̂

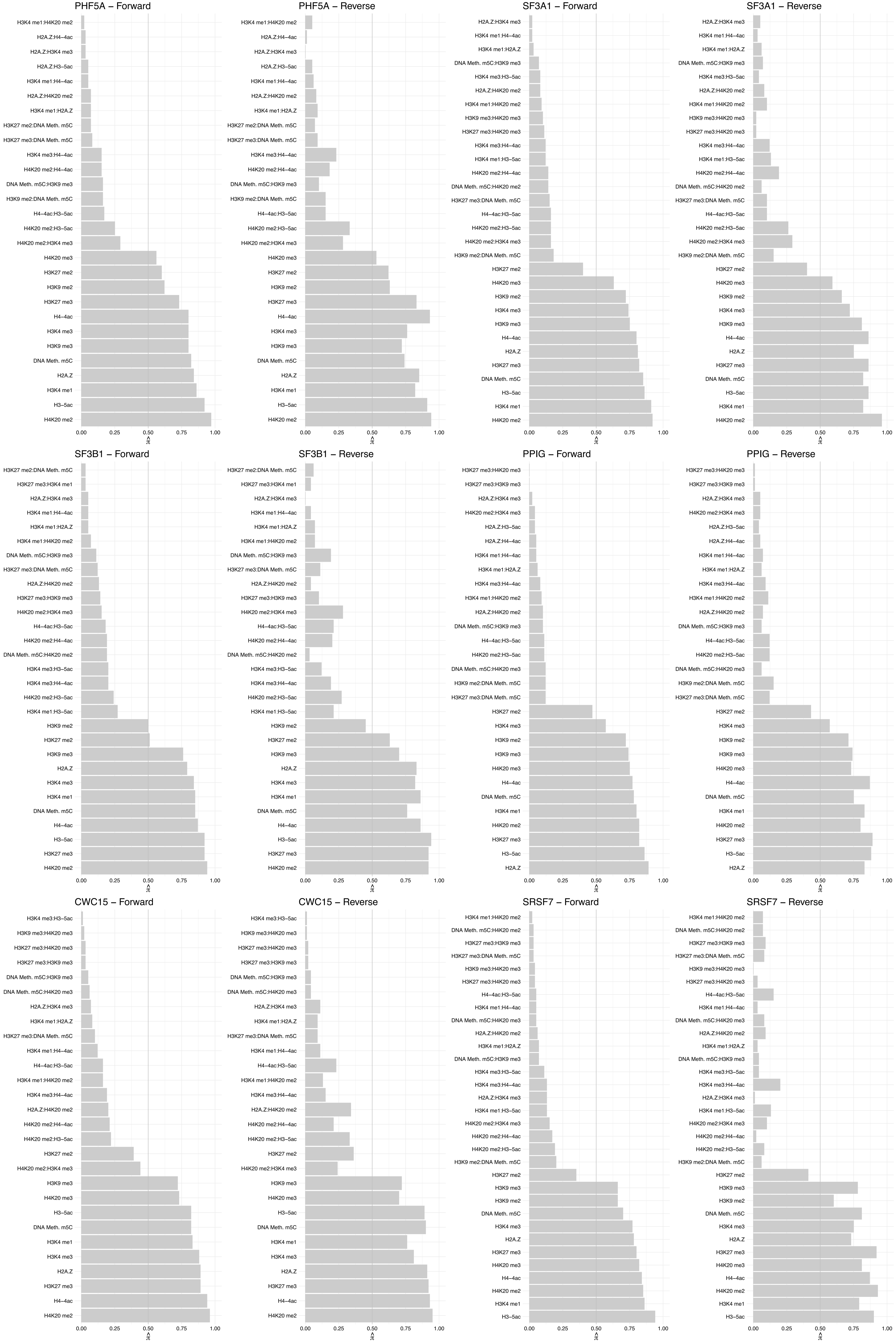
PHF5A – Forward
H3K4 me1:H4K20 me2
H2A.Z:H4–4ac
H2A.Z:H3K4 me3
H2A.Z:H3–5ac
H3K4 me1:H4–4ac
H2A.Z:H4K20 me2
H3K4 me1:H2A.Z
H3K27 me2:DNA Meth. m5C
H3K27 me3:DNA Meth. m5C
H3K4 me3:H4–4ac
H4K20 me2:H4–4ac
DNA Meth. m5C:H3K9 me3
H3K9 me2:DNA Meth. m5C
H4–4ac:H3–5ac
H4K20 me2:H3–5ac
H4K20 me2:H3K4 me3
H4K20 me3
H3K27 me2
H3K9 me2
H3K27 me3
H4–4ac
H3K4 me3
H3K9 me3
DNA Meth. m5C
H2A.Z
H3K4 me1
H3–5ac
H4K20 me2
0.00 0.25 0.50 0.75 1.00
π̂
PHF5A – Reverse
H3K4 me1:H4K20 me2
H2A.Z:H4–4ac
H2A.Z:H3K4 me3
H2A.Z:H3–5ac
H3K4 me1:H4–4ac
H2A.Z:H4K20 me2
H3K4 me1:H2A.Z
H3K27 me2:DNA Meth. m5C
H3K27 me3:DNA Meth. m5C
H3K4 me3:H4–4ac
H4K20 me2:H4–4ac
DNA Meth. m5C:H3K9 me3
H3K9 me2:DNA Meth. m5C
H4–4ac:H3–5ac
H4K20 me2:H3–5ac
H4K20 me2:H3K4 me3
H4K20 me3
H3K27 me2
H3K9 me2
H3K27 me3
H4–4ac
H3K4 me3
H3K9 me3
DNA Meth. m5C
H2A.Z
H3K4 me1
H3–5ac
H4K20 me2
0.00 0.25 0.50 0.75 1.00
π̂
SF3A1 – Forward
H2A.Z:H3K4 me3
H3K4 me1:H4–4ac
H3K4 me1:H2A.Z
DNA Meth. m5C:H3K9 me3
H3K4 me3:H3–5ac
H2A.Z:H4K20 me2
H3K4 me1:H4K20 me2
H3K9 me3:H4K20 me3
H3K27 me3:H4K20 me3
H3K4 me3:H4–4ac
H3K4 me1:H3–5ac
H4K20 me2:H4–4ac
DNA Meth. m5C:H4K20 me2
H3K27 me3:DNA Meth. m5C
H4–4ac:H3–5ac
H4K20 me2:H3–5ac
H4K20 me2:H3K4 me3
H3K9 me2:DNA Meth. m5C
H3K27 me2
H4K20 me3
H3K9 me2
H3K4 me3
H3K9 me3
H4–4ac
H2A.Z
H3K27 me3
DNA Meth. m5C
H3–5ac
H3K4 me1
H4K20 me2
0.00 0.25 0.50 0.75 1.00
π̂
SF3A1 – Reverse
H2A.Z:H3K4 me3
H3K4 me1:H4–4ac
H3K4 me1:H2A.Z
DNA Meth. m5C:H3K9 me3
H3K4 me3:H3–5ac
H2A.Z:H4K20 me2
H3K4 me1:H4K20 me2
H3K9 me3:H4K20 me3
H3K27 me3:H4K20 me3
H3K4 me3:H4–4ac
H3K4 me1:H3–5ac
H4K20 me2:H4–4ac
DNA Meth. m5C:H4K20 me2
H3K27 me3:DNA Meth. m5C
H4–4ac:H3–5ac
H4K20 me2:H3–5ac
H4K20 me2:H3K4 me3
H3K9 me2:DNA Meth. m5C
H3K27 me2
H4K20 me3
H3K9 me2
H3K4 me3
H3K9 me3
H4–4ac
H2A.Z
H3K27 me3
DNA Meth. m5C
H3–5ac
H3K4 me1
H4K20 me2
0.00 0.25 0.50 0.75 1.00
π̂
SF3B1 – Forward
H3K27 me2:DNA Meth. m5C
H3K27 me3:H3K4 me1
H2A.Z:H3K4 me3
H3K4 me1:H4–4ac
H3K4 me1:H2A.Z
H3K4 me1:H4K20 me2
DNA Meth. m5C:H3K9 me3
H3K27 me3:DNA Meth. m5C
H2A.Z:H4K20 me2
H3K27 me3:H3K9 me3
H4K20 me2:H3K4 me3
H4–4ac:H3–5ac
H4K20 me2:H4–4ac
DNA Meth. m5C:H4K20 me2
H3K4 me3:H3–5ac
H3K4 me3:H4–4ac
H4K20 me2:H3–5ac
H3K4 me1:H3–5ac
H3K9 me2
H3K27 me2
H3K9 me3
H2A.Z
H3K4 me3
H3K4 me1
DNA Meth. m5C
H4–4ac
H3–5ac
H3K27 me3
H4K20 me2
0.00 0.25 0.50 0.75 1.00
π̂
SF3B1 – Reverse
H3K27 me2:DNA Meth. m5C
H3K27 me3:H3K4 me1
H2A.Z:H3K4 me3
H3K4 me1:H4–4ac
H3K4 me1:H2A.Z
H3K4 me1:H4K20 me2
DNA Meth. m5C:H3K9 me3
H3K27 me3:DNA Meth. m5C
H2A.Z:H4K20 me2
H3K27 me3:H3K9 me3
H4K20 me2:H3K4 me3
H4–4ac:H3–5ac
H4K20 me2:H4–4ac
DNA Meth. m5C:H4K20 me2
H3K4 me3:H3–5ac
H3K4 me3:H4–4ac
H4K20 me2:H3–5ac
H3K4 me1:H3–5ac
H3K9 me2
H3K27 me2
H3K9 me3
H2A.Z
H3K4 me3
H3K4 me1
DNA Meth. m5C
H4–4ac
H3–5ac
H3K27 me3
H4K20 me2
0.00 0.25 0.50 0.75 1.00
π̂
PPIG – Forward
H3K27 me3:H4K20 me3
H3K27 me3:H3K9 me3
H2A.Z:H3K4 me3
H4K20 me2:H3K4 me3
H2A.Z:H3–5ac
H2A.Z:H4–4ac
H3K4 me1:H4–4ac
H3K4 me1:H2A.Z
H3K4 me3:H4–4ac
H3K4 me1:H4K20 me2
H2A.Z:H4K20 me2
DNA Meth. m5C:H3K9 me3
H4–4ac:H3–5ac
H4K20 me2:H3–5ac
DNA Meth. m5C:H4K20 me3
H3K9 me2:DNA Meth. m5C
H3K27 me3:DNA Meth. m5C
H3K27 me2
H3K4 me3
H3K9 me2
H3K9 me3
H4K20 me3
H4–4ac
DNA Meth. m5C
H3K4 me1
H4K20 me2
H3K27 me3
H3–5ac
H2A.Z
0.00 0.25 0.50 0.75 1.00
π̂
PPIG – Reverse
H3K27 me3:H4K20 me3
H3K27 me3:H3K9 me3
H2A.Z:H3K4 me3
H4K20 me2:H3K4 me3
H2A.Z:H3–5ac
H2A.Z:H4–4ac
H3K4 me1:H4–4ac
H3K4 me1:H2A.Z
H3K4 me3:H4–4ac
H3K4 me1:H4K20 me2
H2A.Z:H4K20 me2
DNA Meth. m5C:H3K9 me3
H4–4ac:H3–5ac
H4K20 me2:H3–5ac
DNA Meth. m5C:H4K20 me3
H3K9 me2:DNA Meth. m5C
H3K27 me3:DNA Meth. m5C
H3K27 me2
H3K4 me3
H3K9 me2
H3K9 me3
H4K20 me3
H4–4ac
DNA Meth. m5C
H3K4 me1
H4K20 me2
H3K27 me3
H3–5ac
H2A.Z
0.00 0.25 0.50 0.75 1.00
π̂
CWC15 – Forward
H3K4 me3:H3–5ac
H3K9 me3:H4K20 me3
H3K27 me3:H4K20 me3
H3K27 me3:H3K9 me3
DNA Meth. m5C:H3K9 me3
DNA Meth. m5C:H4K20 me3
H2A.Z:H3K4 me3
H3K4 me1:H2A.Z
H3K27 me3:DNA Meth. m5C
H3K4 me1:H4–4ac
H4–4ac:H3–5ac
H3K4 me1:H4K20 me2
H3K4 me3:H4–4ac
H2A.Z:H4K20 me2
H4K20 me2:H4–4ac
H4K20 me2:H3–5ac
H3K27 me2
H4K20 me2:H3K4 me3
H3K9 me3
H4K20 me3
H3–5ac
DNA Meth. m5C
H3K4 me1
H3K4 me3
H2A.Z
H3K27 me3
H4–4ac
H4K20 me2
0.00 0.25 0.50 0.75 1.00
π̂
CWC15 – Reverse
H3K4 me3:H3–5ac
H3K9 me3:H4K20 me3
H3K27 me3:H4K20 me3
H3K27 me3:H3K9 me3
DNA Meth. m5C:H3K9 me3
DNA Meth. m5C:H4K20 me3
H2A.Z:H3K4 me3
H3K4 me1:H2A.Z
H3K27 me3:DNA Meth. m5C
H3K4 me1:H4–4ac
H4–4ac:H3–5ac
H3K4 me1:H4K20 me2
H3K4 me3:H4–4ac
H2A.Z:H4K20 me2
H4K20 me2:H4–4ac
H4K20 me2:H3–5ac
H3K27 me2
H4K20 me2:H3K4 me3
H3K9 me3
H4K20 me3
H3–5ac
DNA Meth. m5C
H3K4 me1
H3K4 me3
H2A.Z
H3K27 me3
H4–4ac
H4K20 me2
0.00 0.25 0.50 0.75 1.00
π̂
SRSF7 – Forward
H3K4 me1:H4K20 me2
DNA Meth. m5C:H4K20 me2
H3K27 me3:H3K9 me3
H3K27 me3:DNA Meth. m5C
H3K9 me3:H4K20 me3
H3K27 me3:H4K20 me3
H4–4ac:H3–5ac
H3K4 me1:H4–4ac
DNA Meth. m5C:H4K20 me3
H2A.Z:H4K20 me2
H3K4 me1:H2A.Z
DNA Meth. m5C:H3K9 me3
H3K4 me3:H3–5ac
H3K4 me3:H4–4ac
H2A.Z:H3K4 me3
H3K4 me1:H3–5ac
H4K20 me2:H3K4 me3
H4K20 me2:H4–4ac
H4K20 me2:H3–5ac
H3K9 me2:DNA Meth. m5C
H3K27 me2
H3K9 me3
H3K9 me2
DNA Meth. m5C
H3K4 me3
H2A.Z
H3K27 me3
H4K20 me3
H4–4ac
H4K20 me2
H3K4 me1
H3–5ac
0.00 0.25 0.50 0.75 1.00
π̂
SRSF7 – Reverse
H3K4 me1:H4K20 me2
DNA Meth. m5C:H4K20 me2
H3K27 me3:H3K9 me3
H3K27 me3:DNA Meth. m5C
H3K9 me3:H4K20 me3
H3K27 me3:H4K20 me3
H4–4ac:H3–5ac
H3K4 me1:H4–4ac
DNA Meth. m5C:H4K20 me3
H2A.Z:H4K20 me2
H3K4 me1:H2A.Z
DNA Meth. m5C:H3K9 me3
H3K4 me3:H3–5ac
H3K4 me3:H4–4ac
H2A.Z:H3K4 me3
H3K4 me1:H3–5ac
H4K20 me2:H3K4 me3
H4K20 me2:H4–4ac
H4K20 me2:H3–5ac
H3K9 me2:DNA Meth. m5C
H3K27 me2
H3K9 me3
H3K9 me2
DNA Meth. m5C
H3K4 me3
H2A.Z
H3K27 me3
H4K20 me3
H4–4ac
H4K20 me2
H3K4 me1
H3–5ac
0.00 0.25 0.50 0.75 1.00
π̂

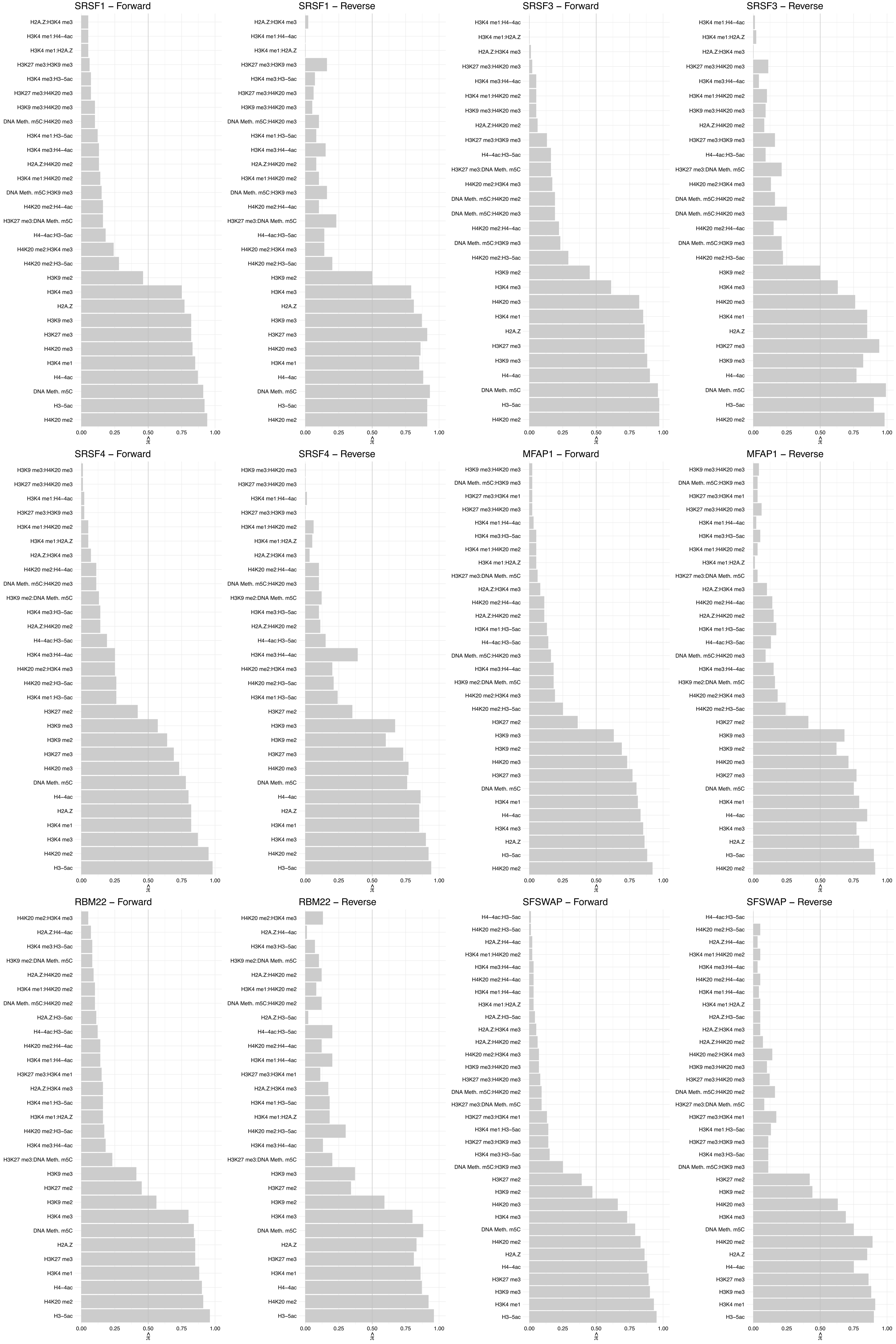
SRSF1 – Forward
H2A.Z:H3K4 me3
H3K4 me1:H4–4ac
H3K4 me1:H2A.Z
H3K27 me3:H3K9 me3
H3K4 me3:H3–5ac
H3K27 me3:H4K20 me3
H3K9 me3:H4K20 me3
DNA Meth. m5C:H4K20 me3
H3K4 me1:H3–5ac
H3K4 me3:H4–4ac
H2A.Z:H4K20 me2
H3K4 me1:H4K20 me2
DNA Meth. m5C:H3K9 me3
H4K20 me2:H4–4ac
H3K27 me3:DNA Meth. m5C
H4–4ac:H3–5ac
H4K20 me2:H3K4 me3
H4K20 me2:H3–5ac
H3K9 me2
H3K4 me3
H2A.Z
H3K9 me3
H3K27 me3
H4K20 me3
H3K4 me1
H4–4ac
DNA Meth. m5C
H3–5ac
H4K20 me2
0.00 0.25 0.50 0.75 1.00
π̂
SRSF1 – Reverse
H2A.Z:H3K4 me3
H3K4 me1:H4–4ac
H3K4 me1:H2A.Z
H3K27 me3:H3K9 me3
H3K4 me3:H3–5ac
H3K27 me3:H4K20 me3
H3K9 me3:H4K20 me3
DNA Meth. m5C:H4K20 me3
H3K4 me1:H3–5ac
H3K4 me3:H4–4ac
H2A.Z:H4K20 me2
H3K4 me1:H4K20 me2
DNA Meth. m5C:H3K9 me3
H4K20 me2:H4–4ac
H3K27 me3:DNA Meth. m5C
H4–4ac:H3–5ac
H4K20 me2:H3K4 me3
H4K20 me2:H3–5ac
H3K9 me2
H3K4 me3
H2A.Z
H3K9 me3
H3K27 me3
H4K20 me3
H3K4 me1
H4–4ac
DNA Meth. m5C
H3–5ac
H4K20 me2
0.00 0.25 0.50 0.75 1.00
π̂
SRSF3 – Forward
H3K4 me1:H4–4ac
H3K4 me1:H2A.Z
H2A.Z:H3K4 me3
H3K27 me3:H4K20 me3
H3K4 me3:H4–4ac
H3K4 me1:H4K20 me2
H3K9 me3:H4K20 me3
H2A.Z:H4K20 me2
H3K27 me3:H3K9 me3
H4–4ac:H3–5ac
H3K27 me3:DNA Meth. m5C
H4K20 me2:H3K4 me3
DNA Meth. m5C:H4K20 me2
DNA Meth. m5C:H4K20 me3
H4K20 me2:H4–4ac
DNA Meth. m5C:H3K9 me3
H4K20 me2:H3–5ac
H3K9 me2
H3K4 me3
H4K20 me3
H3K4 me1
H2A.Z
H3K27 me3
H3K9 me3
H4–4ac
DNA Meth. m5C
H3–5ac
H4K20 me2
0.00 0.25 0.50 0.75 1.00
π̂
SRSF3 – Reverse
H3K4 me1:H4–4ac
H3K4 me1:H2A.Z
H2A.Z:H3K4 me3
H3K27 me3:H4K20 me3
H3K4 me3:H4–4ac
H3K4 me1:H4K20 me2
H3K9 me3:H4K20 me3
H2A.Z:H4K20 me2
H3K27 me3:H3K9 me3
H4–4ac:H3–5ac
H3K27 me3:DNA Meth. m5C
H4K20 me2:H3K4 me3
DNA Meth. m5C:H4K20 me2
DNA Meth. m5C:H4K20 me3
H4K20 me2:H4–4ac
DNA Meth. m5C:H3K9 me3
H4K20 me2:H3–5ac
H3K9 me2
H3K4 me3
H4K20 me3
H3K4 me1
H2A.Z
H3K27 me3
H3K9 me3
H4–4ac
DNA Meth. m5C
H3–5ac
H4K20 me2
0.00 0.25 0.50 0.75 1.00
π̂
SRSF4 – Forward
H3K9 me3:H4K20 me3
H3K27 me3:H4K20 me3
H3K4 me1:H4–4ac
H3K27 me3:H3K9 me3
H3K4 me1:H4K20 me2
H3K4 me1:H2A.Z
H2A.Z:H3K4 me3
H4K20 me2:H4–4ac
DNA Meth. m5C:H4K20 me3
H3K9 me2:DNA Meth. m5C
H3K4 me3:H3–5ac
H2A.Z:H4K20 me2
H4–4ac:H3–5ac
H3K4 me3:H4–4ac
H4K20 me2:H3K4 me3
H4K20 me2:H3–5ac
H3K4 me1:H3–5ac
H3K27 me2
H3K9 me3
H3K9 me2
H3K27 me3
H4K20 me3
DNA Meth. m5C
H4–4ac
H2A.Z
H3K4 me1
H3K4 me3
H4K20 me2
H3–5ac
0.00 0.25 0.50 0.75 1.00
π̂
SRSF4 – Reverse
H3K9 me3:H4K20 me3
H3K27 me3:H4K20 me3
H3K4 me1:H4–4ac
H3K27 me3:H3K9 me3
H3K4 me1:H4K20 me2
H3K4 me1:H2A.Z
H2A.Z:H3K4 me3
H4K20 me2:H4–4ac
DNA Meth. m5C:H4K20 me3
H3K9 me2:DNA Meth. m5C
H3K4 me3:H3–5ac
H2A.Z:H4K20 me2
H4–4ac:H3–5ac
H3K4 me3:H4–4ac
H4K20 me2:H3K4 me3
H4K20 me2:H3–5ac
H3K4 me1:H3–5ac
H3K27 me2
H3K9 me3
H3K9 me2
H3K27 me3
H4K20 me3
DNA Meth. m5C
H4–4ac
H2A.Z
H3K4 me1
H3K4 me3
H4K20 me2
H3–5ac
0.00 0.25 0.50 0.75 1.00
π̂
MFAP1 – Forward
H3K9 me3:H4K20 me3
DNA Meth. m5C:H3K9 me3
H3K27 me3:H3K4 me1
H3K27 me3:H4K20 me3
H3K4 me1:H4–4ac
H3K4 me3:H3–5ac
H3K4 me1:H4K20 me2
H3K4 me1:H2A.Z
H3K27 me3:DNA Meth. m5C
H2A.Z:H3K4 me3
H4K20 me2:H4–4ac
H2A.Z:H4K20 me2
H3K4 me1:H3–5ac
H4–4ac:H3–5ac
DNA Meth. m5C:H4K20 me3
H3K4 me3:H4–4ac
H3K9 me2:DNA Meth. m5C
H4K20 me2:H3K4 me3
H4K20 me2:H3–5ac
H3K27 me2
H3K9 me3
H3K9 me2
H4K20 me3
H3K27 me3
DNA Meth. m5C
H3K4 me1
H4–4ac
H3K4 me3
H2A.Z
H3–5ac
H4K20 me2
0.00 0.25 0.50 0.75 1.00
π̂
MFAP1 – Reverse
H3K9 me3:H4K20 me3
DNA Meth. m5C:H3K9 me3
H3K27 me3:H3K4 me1
H3K27 me3:H4K20 me3
H3K4 me1:H4–4ac
H3K4 me3:H3–5ac
H3K4 me1:H4K20 me2
H3K4 me1:H2A.Z
H3K27 me3:DNA Meth. m5C
H2A.Z:H3K4 me3
H4K20 me2:H4–4ac
H2A.Z:H4K20 me2
H3K4 me1:H3–5ac
H4–4ac:H3–5ac
DNA Meth. m5C:H4K20 me3
H3K4 me3:H4–4ac
H3K9 me2:DNA Meth. m5C
H4K20 me2:H3K4 me3
H4K20 me2:H3–5ac
H3K27 me2
H3K9 me3
H3K9 me2
H4K20 me3
H3K27 me3
DNA Meth. m5C
H3K4 me1
H4–4ac
H3K4 me3
H2A.Z
H3–5ac
H4K20 me2
0.00 0.25 0.50 0.75 1.00
π̂
RBM22 – Forward
H4K20 me2:H3K4 me3
H2A.Z:H4–4ac
H3K4 me3:H3–5ac
H3K9 me2:DNA Meth. m5C
H2A.Z:H4K20 me2
H3K4 me1:H4K20 me2
DNA Meth. m5C:H4K20 me2
H2A.Z:H3–5ac
H4–4ac:H3–5ac
H4K20 me2:H4–4ac
H3K4 me1:H4–4ac
H3K27 me3:H3K4 me1
H2A.Z:H3K4 me3
H3K4 me1:H3–5ac
H3K4 me1:H2A.Z
H4K20 me2:H3–5ac
H3K4 me3:H4–4ac
H3K27 me3:DNA Meth. m5C
H3K9 me3
H3K27 me2
H3K9 me2
H3K4 me3
DNA Meth. m5C
H2A.Z
H3K27 me3
H3K4 me1
H4–4ac
H4K20 me2
H3–5ac
0.00 0.25 0.50 0.75 1.00
π̂
RBM22 – Reverse
H4K20 me2:H3K4 me3
H2A.Z:H4–4ac
H3K4 me3:H3–5ac
H3K9 me2:DNA Meth. m5C
H2A.Z:H4K20 me2
H3K4 me1:H4K20 me2
DNA Meth. m5C:H4K20 me2
H2A.Z:H3–5ac
H4–4ac:H3–5ac
H4K20 me2:H4–4ac
H3K4 me1:H4–4ac
H3K27 me3:H3K4 me1
H2A.Z:H3K4 me3
H3K4 me1:H3–5ac
H3K4 me1:H2A.Z
H4K20 me2:H3–5ac
H3K4 me3:H4–4ac
H3K27 me3:DNA Meth. m5C
H3K9 me3
H3K27 me2
H3K9 me2
H3K4 me3
DNA Meth. m5C
H2A.Z
H3K27 me3
H3K4 me1
H4–4ac
H4K20 me2
H3–5ac
0.00 0.25 0.50 0.75 1.00
π̂
SFSWAP – Forward
H4–4ac:H3–5ac
H4K20 me2:H3–5ac
H2A.Z:H4–4ac
H3K4 me1:H4K20 me2
H3K4 me3:H4–4ac
H4K20 me2:H4–4ac
H3K4 me1:H4–4ac
H3K4 me1:H2A.Z
H2A.Z:H3–5ac
H2A.Z:H3K4 me3
H2A.Z:H4K20 me2
H4K20 me2:H3K4 me3
H3K9 me3:H4K20 me3
H3K27 me3:H4K20 me3
DNA Meth. m5C:H4K20 me2
H3K27 me3:DNA Meth. m5C
H3K27 me3:H3K4 me1
H3K4 me1:H3–5ac
H3K27 me3:H3K9 me3
H3K4 me3:H3–5ac
DNA Meth. m5C:H3K9 me3
H3K27 me2
H3K9 me2
H4K20 me3
H3K4 me3
DNA Meth. m5C
H4K20 me2
H2A.Z
H4–4ac
H3K27 me3
H3K9 me3
H3K4 me1
H3–5ac
0.00 0.25 0.50 0.75 1.00
π̂
SFSWAP – Reverse
H4–4ac:H3–5ac
H4K20 me2:H3–5ac
H2A.Z:H4–4ac
H3K4 me1:H4K20 me2
H3K4 me3:H4–4ac
H4K20 me2:H4–4ac
H3K4 me1:H4–4ac
H3K4 me1:H2A.Z
H2A.Z:H3–5ac
H2A.Z:H3K4 me3
H2A.Z:H4K20 me2
H4K20 me2:H3K4 me3
H3K9 me3:H4K20 me3
H3K27 me3:H4K20 me3
DNA Meth. m5C:H4K20 me2
H3K27 me3:DNA Meth. m5C
H3K27 me3:H3K4 me1
H3K4 me1:H3–5ac
H3K27 me3:H3K9 me3
H3K4 me3:H3–5ac
DNA Meth. m5C:H3K9 me3
H3K27 me2
H3K9 me2
H4K20 me3
H3K4 me3
DNA Meth. m5C
H4K20 me2
H2A.Z
H4–4ac
H3K27 me3
H3K9 me3
H3K4 me1
H3–5ac
0.00 0.25 0.50 0.75 1.00
π̂

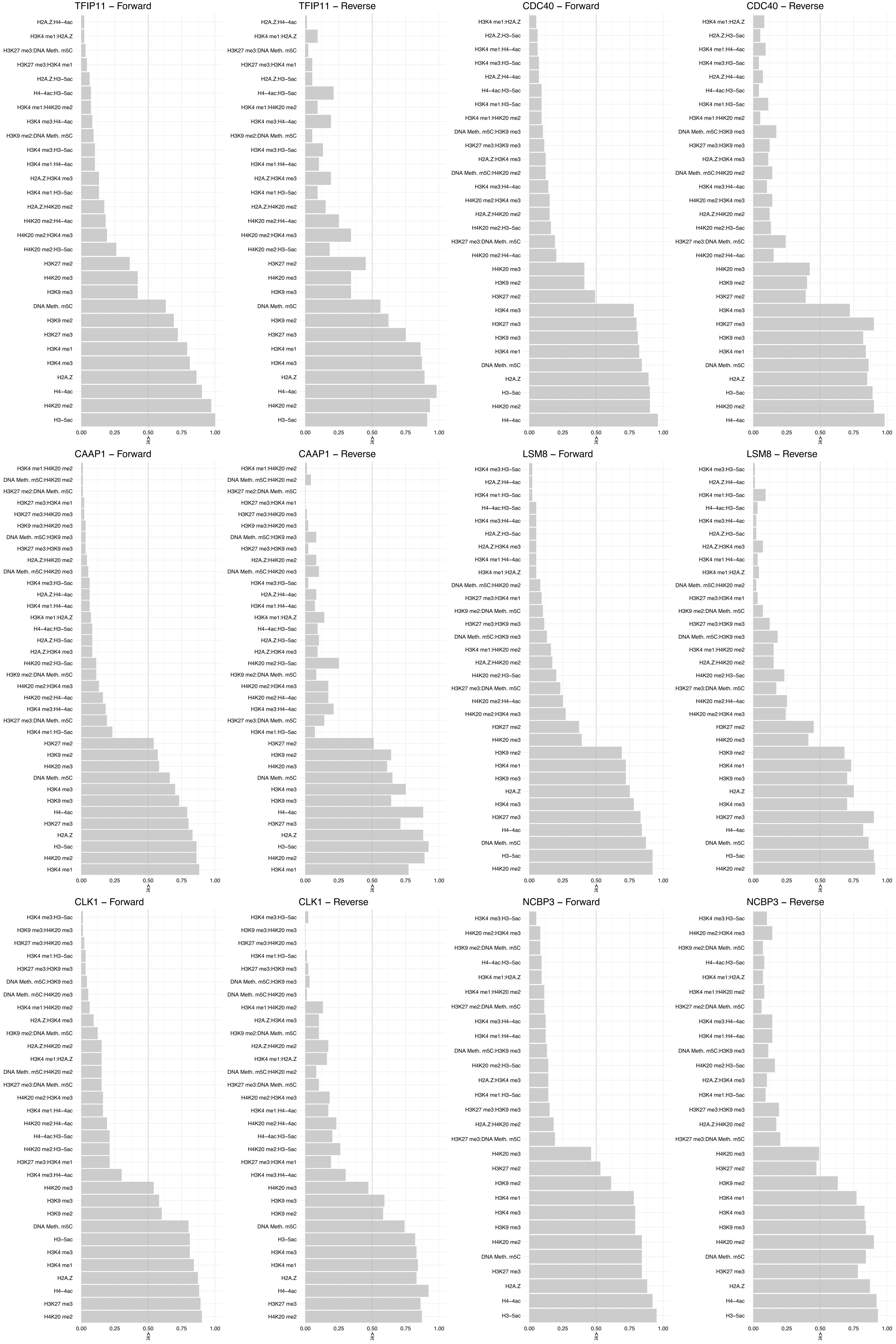
TFIP11 – Forward
H2A.Z:H4–4ac
H3K4 me1:H2A.Z
H3K27 me3:DNA Meth. m5C
H3K27 me3:H3K4 me1
H2A.Z:H3–5ac
H4–4ac:H3–5ac
H3K4 me1:H4K20 me2
H3K4 me3:H4–4ac
H3K9 me2:DNA Meth. m5C
H3K4 me3:H3–5ac
H3K4 me1:H4–4ac
H2A.Z:H3K4 me3
H3K4 me1:H3–5ac
H2A.Z:H4K20 me2
H4K20 me2:H4–4ac
H4K20 me2:H3K4 me3
H4K20 me2:H3–5ac
H3K27 me2
H4K20 me3
H3K9 me3
DNA Meth. m5C
H3K9 me2
H3K27 me3
H3K4 me1
H3K4 me3
H2A.Z
H4–4ac
H4K20 me2
H3–5ac
0.00 0.25 0.50 0.75 1.00
$\hat{\pi}$
TFIP11 – Reverse
H2A.Z:H4–4ac
H3K4 me1:H2A.Z
H3K27 me3:DNA Meth. m5C
H3K27 me3:H3K4 me1
H2A.Z:H3–5ac
H4–4ac:H3–5ac
H3K4 me1:H4K20 me2
H3K4 me3:H4–4ac
H3K9 me2:DNA Meth. m5C
H3K4 me3:H3–5ac
H3K4 me1:H4–4ac
H2A.Z:H3K4 me3
H3K4 me1:H3–5ac
H2A.Z:H4K20 me2
H4K20 me2:H4–4ac
H4K20 me2:H3K4 me3
H4K20 me2:H3–5ac
H3K27 me2
H4K20 me3
H3K9 me3
DNA Meth. m5C
H3K9 me2
H3K27 me3
H3K4 me1
H3K4 me3
H2A.Z
H4–4ac
H4K20 me2
H3–5ac
0.00 0.25 0.50 0.75 1.00
$\hat{\pi}$
CDC40 – Forward
H3K4 me1:H2A.Z
H2A.Z:H3–5ac
H3K4 me1:H4–4ac
H3K4 me3:H3–5ac
H2A.Z:H4–4ac
H4–4ac:H3–5ac
H3K4 me1:H3–5ac
H3K4 me1:H4K20 me2
DNA Meth. m5C:H3K9 me3
H3K27 me3:H3K9 me3
H2A.Z:H3K4 me3
DNA Meth. m5C:H4K20 me2
H3K4 me3:H4–4ac
H4K20 me2:H3K4 me3
H2A.Z:H4K20 me2
H4K20 me2:H3–5ac
H3K27 me3:DNA Meth. m5C
H4K20 me2:H4–4ac
H4K20 me3
H3K9 me2
H3K27 me2
H3K4 me3
H3K27 me3
H3K9 me3
H3K4 me1
DNA Meth. m5C
H2A.Z
H3–5ac
H4K20 me2
H4–4ac
0.00 0.25 0.50 0.75 1.00
$\hat{\pi}$
CDC40 – Reverse
H3K4 me1:H2A.Z
H2A.Z:H3–5ac
H3K4 me1:H4–4ac
H3K4 me3:H3–5ac
H2A.Z:H4–4ac
H4–4ac:H3–5ac
H3K4 me1:H3–5ac
H3K4 me1:H4K20 me2
DNA Meth. m5C:H3K9 me3
H3K27 me3:H3K9 me3
H2A.Z:H3K4 me3
DNA Meth. m5C:H4K20 me2
H3K4 me3:H4–4ac
H4K20 me2:H3K4 me3
H2A.Z:H4K20 me2
H4K20 me2:H3–5ac
H3K27 me3:DNA Meth. m5C
H4K20 me2:H4–4ac
H4K20 me3
H3K9 me2
H3K27 me2
H3K4 me3
H3K27 me3
H3K9 me3
H3K4 me1
DNA Meth. m5C
H2A.Z
H3–5ac
H4K20 me2
H4–4ac
0.00 0.25 0.50 0.75 1.00
$\hat{\pi}$
CAAP1 – Forward
H3K4 me1:H4K20 me2
DNA Meth. m5C:H4K20 me2
H3K27 me2:DNA Meth. m5C
H3K27 me3:H3K4 me1
H3K27 me3:H4K20 me3
H3K9 me3:H4K20 me3
DNA Meth. m5C:H3K9 me3
H3K27 me3:H3K9 me3
H2A.Z:H4K20 me2
DNA Meth. m5C:H4K20 me3
H3K4 me3:H3–5ac
H2A.Z:H4–4ac
H3K4 me1:H4–4ac
H3K4 me1:H2A.Z
H4–4ac:H3–5ac
H2A.Z:H3–5ac
H2A.Z:H3K4 me3
H4K20 me2:H3–5ac
H3K9 me2:DNA Meth. m5C
H4K20 me2:H3K4 me3
H4K20 me2:H4–4ac
H3K4 me3:H4–4ac
H3K27 me3:DNA Meth. m5C
H3K4 me1:H3–5ac
H3K27 me2
H3K9 me2
H4K20 me3
DNA Meth. m5C
H3K4 me3
H3K9 me3
H4–4ac
H3K27 me3
H2A.Z
H3–5ac
H4K20 me2
H3K4 me1
0.00 0.25 0.50 0.75 1.00
$\hat{\pi}$
CAAP1 – Reverse
H3K4 me1:H4K20 me2
DNA Meth. m5C:H4K20 me2
H3K27 me2:DNA Meth. m5C
H3K27 me3:H3K4 me1
H3K27 me3:H4K20 me3
H3K9 me3:H4K20 me3
DNA Meth. m5C:H3K9 me3
H3K27 me3:H3K9 me3
H2A.Z:H4K20 me2
DNA Meth. m5C:H4K20 me3
H3K4 me3:H3–5ac
H2A.Z:H4–4ac
H3K4 me1:H4–4ac
H3K4 me1:H2A.Z
H4–4ac:H3–5ac
H2A.Z:H3–5ac
H2A.Z:H3K4 me3
H4K20 me2:H3–5ac
H3K9 me2:DNA Meth. m5C
H4K20 me2:H3K4 me3
H4K20 me2:H4–4ac
H3K4 me3:H4–4ac
H3K27 me3:DNA Meth. m5C
H3K4 me1:H3–5ac
H3K27 me2
H3K9 me2
H4K20 me3
DNA Meth. m5C
H3K4 me3
H3K9 me3
H4–4ac
H3K27 me3
H2A.Z
H3–5ac
H4K20 me2
H3K4 me1
0.00 0.25 0.50 0.75 1.00
$\hat{\pi}$
LSM8 – Forward
H3K4 me3:H3–5ac
H2A.Z:H4–4ac
H3K4 me1:H3–5ac
H4–4ac:H3–5ac
H3K4 me3:H4–4ac
H2A.Z:H3–5ac
H2A.Z:H3K4 me3
H3K4 me1:H4–4ac
H3K4 me1:H2A.Z
DNA Meth. m5C:H4K20 me2
H3K27 me3:H3K4 me1
H3K9 me2:DNA Meth. m5C
H3K27 me3:H3K9 me3
DNA Meth. m5C:H3K9 me3
H3K4 me1:H4K20 me2
H2A.Z:H4K20 me2
H4K20 me2:H3–5ac
H3K27 me3:DNA Meth. m5C
H4K20 me2:H4–4ac
H4K20 me2:H3K4 me3
H3K27 me2
H4K20 me3
H3K9 me2
H3K4 me1
H3K9 me3
H2A.Z
H3K4 me3
H3K27 me3
H4–4ac
DNA Meth. m5C
H3–5ac
H4K20 me2
0.00 0.25 0.50 0.75 1.00
$\hat{\pi}$
LSM8 – Reverse
H3K4 me3:H3–5ac
H2A.Z:H4–4ac
H3K4 me1:H3–5ac
H4–4ac:H3–5ac
H3K4 me3:H4–4ac
H2A.Z:H3–5ac
H2A.Z:H3K4 me3
H3K4 me1:H4–4ac
H3K4 me1:H2A.Z
DNA Meth. m5C:H4K20 me2
H3K27 me3:H3K4 me1
H3K9 me2:DNA Meth. m5C
H3K27 me3:H3K9 me3
DNA Meth. m5C:H3K9 me3
H3K4 me1:H4K20 me2
H2A.Z:H4K20 me2
H4K20 me2:H3–5ac
H3K27 me3:DNA Meth. m5C
H4K20 me2:H4–4ac
H4K20 me2:H3K4 me3
H3K27 me2
H4K20 me3
H3K9 me2
H3K4 me1
H3K9 me3
H2A.Z
H3K4 me3
H3K27 me3
H4–4ac
DNA Meth. m5C
H3–5ac
H4K20 me2
0.00 0.25 0.50 0.75 1.00
$\hat{\pi}$
CLK1 – Forward
H3K4 me3:H3–5ac
H3K9 me3:H4K20 me3
H3K27 me3:H4K20 me3
H3K4 me1:H3–5ac
H3K27 me3:H3K9 me3
DNA Meth. m5C:H3K9 me3
DNA Meth. m5C:H4K20 me3
H3K4 me1:H4K20 me2
H2A.Z:H3K4 me3
H3K9 me2:DNA Meth. m5C
H2A.Z:H4K20 me2
H3K4 me1:H2A.Z
DNA Meth. m5C:H4K20 me2
H3K27 me3:DNA Meth. m5C
H4K20 me2:H3K4 me3
H3K4 me1:H4–4ac
H4K20 me2:H4–4ac
H4–4ac:H3–5ac
H4K20 me2:H3–5ac
H3K27 me3:H3K4 me1
H3K4 me3:H4–4ac
H4K20 me3
H3K9 me3
H3K9 me2
DNA Meth. m5C
H3–5ac
H3K4 me3
H3K4 me1
H2A.Z
H4–4ac
H3K27 me3
H4K20 me2
0.00 0.25 0.50 0.75 1.00
$\hat{\pi}$
CLK1 – Reverse
H3K4 me3:H3–5ac
H3K9 me3:H4K20 me3
H3K27 me3:H4K20 me3
H3K4 me1:H3–5ac
H3K27 me3:H3K9 me3
DNA Meth. m5C:H3K9 me3
DNA Meth. m5C:H4K20 me3
H3K4 me1:H4K20 me2
H2A.Z:H3K4 me3
H3K9 me2:DNA Meth. m5C
H2A.Z:H4K20 me2
H3K4 me1:H2A.Z
DNA Meth. m5C:H4K20 me2
H3K27 me3:DNA Meth. m5C
H4K20 me2:H3K4 me3
H3K4 me1:H4–4ac
H4K20 me2:H4–4ac
H4–4ac:H3–5ac
H4K20 me2:H3–5ac
H3K27 me3:H3K4 me1
H3K4 me3:H4–4ac
H4K20 me3
H3K9 me3
H3K9 me2
DNA Meth. m5C
H3–5ac
H3K4 me3
H3K4 me1
H2A.Z
H4–4ac
H3K27 me3
H4K20 me2
0.00 0.25 0.50 0.75 1.00
$\hat{\pi}$
NCBP3 – Forward
H3K4 me3:H3–5ac
H4K20 me2:H3K4 me3
H3K9 me2:DNA Meth. m5C
H4–4ac:H3–5ac
H3K4 me1:H2A.Z
H3K4 me1:H4K20 me2
H3K27 me2:DNA Meth. m5C
H3K4 me3:H4–4ac
H3K4 me1:H4–4ac
DNA Meth. m5C:H3K9 me3
H4K20 me2:H3–5ac
H2A.Z:H3K4 me3
H3K4 me1:H3–5ac
H3K27 me3:H3K9 me3
H2A.Z:H4K20 me2
H3K27 me3:DNA Meth. m5C
H4K20 me3
H3K27 me2
H3K9 me2
H3K4 me1
H3K4 me3
H3K9 me3
H4K20 me2
DNA Meth. m5C
H3K27 me3
H2A.Z
H4–4ac
H3–5ac
0.00 0.25 0.50 0.75 1.00
$\hat{\pi}$
NCBP3 – Reverse
H3K4 me3:H3–5ac
H4K20 me2:H3K4 me3
H3K9 me2:DNA Meth. m5C
H4–4ac:H3–5ac
H3K4 me1:H2A.Z
H3K4 me1:H4K20 me2
H3K27 me2:DNA Meth. m5C
H3K4 me3:H4–4ac
H3K4 me1:H4–4ac
DNA Meth. m5C:H3K9 me3
H4K20 me2:H3–5ac
H2A.Z:H3K4 me3
H3K4 me1:H3–5ac
H3K27 me3:H3K9 me3
H2A.Z:H4K20 me2
H3K27 me3:DNA Meth. m5C
H4K20 me3
H3K27 me2
H3K9 me2
H3K4 me1
H3K4 me3
H3K9 me3
H4K20 me2
DNA Meth. m5C
H3K27 me3
H2A.Z
H4–4ac
H3–5ac
0.00 0.25 0.50 0.75 1.00
$\hat{\pi}$

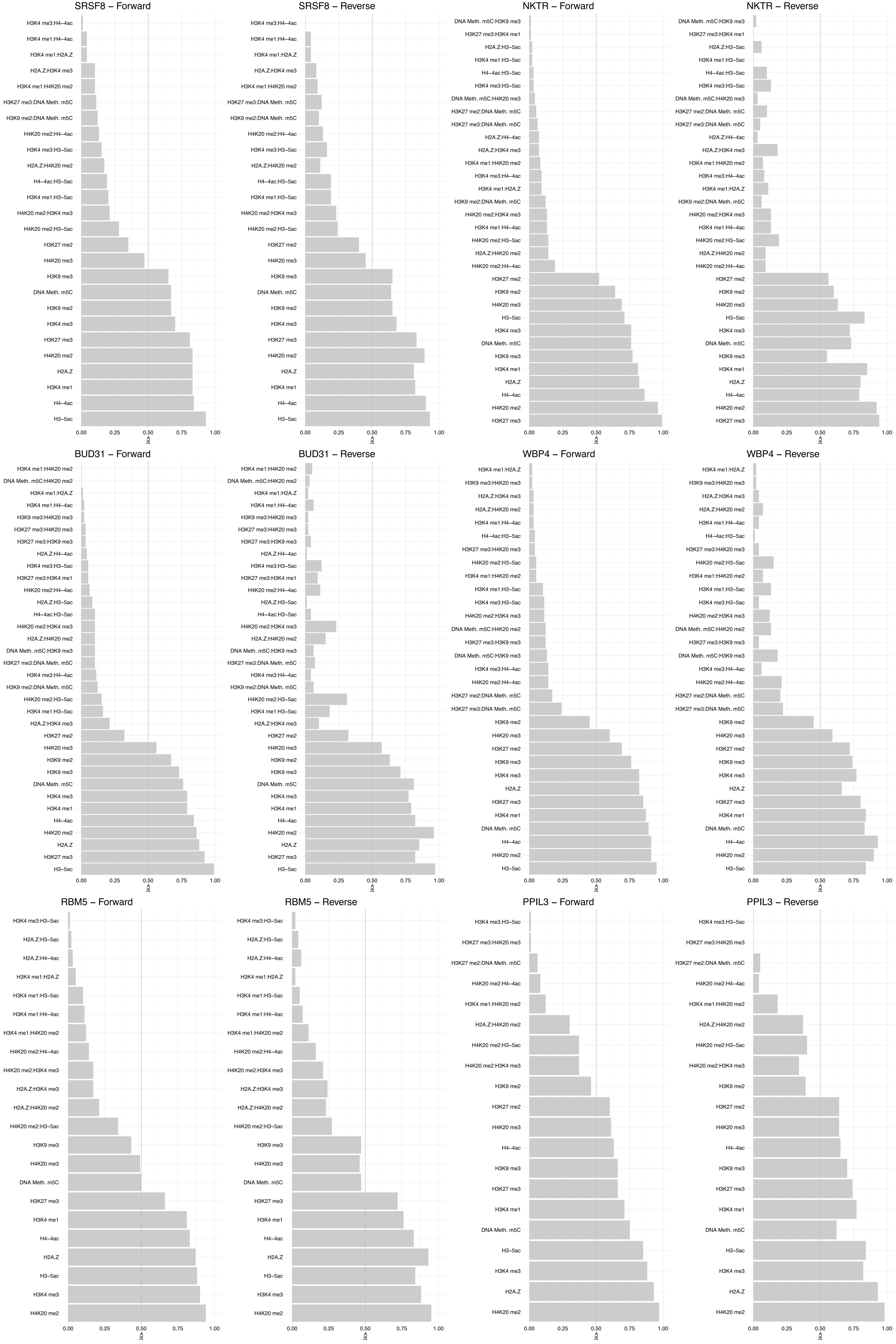
SRSF8 − Forward
H3K4 me3:H4−4ac
H3K4 me1:H4−4ac
H3K4 me1:H2A.Z
H2A.Z:H3K4 me3
H3K4 me1:H4K20 me2
H3K27 me3:DNA Meth. m5C
H3K9 me2:DNA Meth. m5C
H4K20 me2:H4−4ac
H3K4 me3:H3−5ac
H2A.Z:H4K20 me2
H4−4ac:H3−5ac
H3K4 me1:H3−5ac
H4K20 me2:H3K4 me3
H4K20 me2:H3−5ac
H3K27 me2
H4K20 me3
H3K9 me3
DNA Meth. m5C
H3K9 me2
H3K4 me3
H3K27 me3
H4K20 me2
H2A.Z
H3K4 me1
H4−4ac
H3−5ac
0.00 0.25 0.50 0.75 1.00
π̂
SRSF8 − Reverse
H3K4 me3:H4−4ac
H3K4 me1:H4−4ac
H3K4 me1:H2A.Z
H2A.Z:H3K4 me3
H3K4 me1:H4K20 me2
H3K27 me3:DNA Meth. m5C
H3K9 me2:DNA Meth. m5C
H4K20 me2:H4−4ac
H3K4 me3:H3−5ac
H2A.Z:H4K20 me2
H4−4ac:H3−5ac
H3K4 me1:H3−5ac
H4K20 me2:H3K4 me3
H4K20 me2:H3−5ac
H3K27 me2
H4K20 me3
H3K9 me3
DNA Meth. m5C
H3K9 me2
H3K4 me3
H3K27 me3
H4K20 me2
H2A.Z
H3K4 me1
H4−4ac
H3−5ac
0.00 0.25 0.50 0.75 1.00
π̂
NKTR − Forward
DNA Meth. m5C:H3K9 me3
H3K27 me3:H3K4 me1
H2A.Z:H3−5ac
H3K4 me1:H3−5ac
H4−4ac:H3−5ac
H3K4 me3:H3−5ac
DNA Meth. m5C:H4K20 me3
H3K27 me2:DNA Meth. m5C
H3K27 me3:DNA Meth. m5C
H2A.Z:H4−4ac
H2A.Z:H3K4 me3
H3K4 me1:H4K20 me2
H3K4 me3:H4−4ac
H3K4 me1:H2A.Z
H3K9 me2:DNA Meth. m5C
H4K20 me2:H3K4 me3
H3K4 me1:H4−4ac
H4K20 me2:H3−5ac
H2A.Z:H4K20 me2
H4K20 me2:H4−4ac
H3K27 me2
H3K9 me2
H4K20 me3
H3−5ac
H3K4 me3
DNA Meth. m5C
H3K9 me3
H3K4 me1
H2A.Z
H4−4ac
H4K20 me2
H3K27 me3
0.00 0.25 0.50 0.75 1.00
π̂
NKTR − Reverse
DNA Meth. m5C:H3K9 me3
H3K27 me3:H3K4 me1
H2A.Z:H3−5ac
H3K4 me1:H3−5ac
H4−4ac:H3−5ac
H3K4 me3:H3−5ac
DNA Meth. m5C:H4K20 me3
H3K27 me2:DNA Meth. m5C
H3K27 me3:DNA Meth. m5C
H2A.Z:H4−4ac
H2A.Z:H3K4 me3
H3K4 me1:H4K20 me2
H3K4 me3:H4−4ac
H3K4 me1:H2A.Z
H3K9 me2:DNA Meth. m5C
H4K20 me2:H3K4 me3
H3K4 me1:H4−4ac
H4K20 me2:H3−5ac
H2A.Z:H4K20 me2
H4K20 me2:H4−4ac
H3K27 me2
H3K9 me2
H4K20 me3
H3−5ac
H3K4 me3
DNA Meth. m5C
H3K9 me3
H3K4 me1
H2A.Z
H4−4ac
H4K20 me2
H3K27 me3
0.00 0.25 0.50 0.75 1.00
π̂
BUD31 − Forward
H3K4 me1:H4K20 me2
DNA Meth. m5C:H4K20 me2
H3K4 me1:H2A.Z
H3K4 me1:H4−4ac
H3K9 me3:H4K20 me3
H3K27 me3:H4K20 me3
H3K27 me3:H3K9 me3
H2A.Z:H4−4ac
H3K4 me3:H3−5ac
H3K27 me3:H3K4 me1
H4K20 me2:H4−4ac
H2A.Z:H3−5ac
H4−4ac:H3−5ac
H4K20 me2:H3K4 me3
H2A.Z:H4K20 me2
DNA Meth. m5C:H3K9 me3
H3K27 me3:DNA Meth. m5C
H3K4 me3:H4−4ac
H3K9 me2:DNA Meth. m5C
H4K20 me2:H3−5ac
H3K4 me1:H3−5ac
H2A.Z:H3K4 me3
H3K27 me2
H4K20 me3
H3K9 me2
H3K9 me3
DNA Meth. m5C
H3K4 me3
H3K4 me1
H4−4ac
H4K20 me2
H2A.Z
H3K27 me3
H3−5ac
0.00 0.25 0.50 0.75 1.00
π̂
BUD31 − Reverse
H3K4 me1:H4K20 me2
DNA Meth. m5C:H4K20 me2
H3K4 me1:H2A.Z
H3K4 me1:H4−4ac
H3K9 me3:H4K20 me3
H3K27 me3:H4K20 me3
H3K27 me3:H3K9 me3
H2A.Z:H4−4ac
H3K4 me3:H3−5ac
H3K27 me3:H3K4 me1
H4K20 me2:H4−4ac
H2A.Z:H3−5ac
H4−4ac:H3−5ac
H4K20 me2:H3K4 me3
H2A.Z:H4K20 me2
DNA Meth. m5C:H3K9 me3
H3K27 me3:DNA Meth. m5C
H3K4 me3:H4−4ac
H3K9 me2:DNA Meth. m5C
H4K20 me2:H3−5ac
H3K4 me1:H3−5ac
H2A.Z:H3K4 me3
H3K27 me2
H4K20 me3
H3K9 me2
H3K9 me3
DNA Meth. m5C
H3K4 me3
H3K4 me1
H4−4ac
H4K20 me2
H2A.Z
H3K27 me3
H3−5ac
0.00 0.25 0.50 0.75 1.00
π̂
WBP4 − Forward
H3K4 me1:H2A.Z
H3K9 me3:H4K20 me3
H2A.Z:H3K4 me3
H2A.Z:H4K20 me2
H3K4 me1:H4−4ac
H4−4ac:H3−5ac
H3K27 me3:H4K20 me3
H4K20 me2:H3−5ac
H3K4 me1:H4K20 me2
H3K4 me1:H3−5ac
H3K4 me3:H3−5ac
H4K20 me2:H3K4 me3
DNA Meth. m5C:H4K20 me2
H3K27 me3:H3K9 me3
DNA Meth. m5C:H3K9 me3
H3K4 me3:H4−4ac
H4K20 me2:H4−4ac
H3K27 me2:DNA Meth. m5C
H3K27 me3:DNA Meth. m5C
H3K9 me2
H4K20 me3
H3K27 me2
H3K9 me3
H3K4 me3
H2A.Z
H3K27 me3
H3K4 me1
DNA Meth. m5C
H4−4ac
H4K20 me2
H3−5ac
0.00 0.25 0.50 0.75 1.00
π̂
WBP4 − Reverse
H3K4 me1:H2A.Z
H3K9 me3:H4K20 me3
H2A.Z:H3K4 me3
H2A.Z:H4K20 me2
H3K4 me1:H4−4ac
H4−4ac:H3−5ac
H3K27 me3:H4K20 me3
H4K20 me2:H3−5ac
H3K4 me1:H4K20 me2
H3K4 me1:H3−5ac
H3K4 me3:H3−5ac
H4K20 me2:H3K4 me3
DNA Meth. m5C:H4K20 me2
H3K27 me3:H3K9 me3
DNA Meth. m5C:H3K9 me3
H3K4 me3:H4−4ac
H4K20 me2:H4−4ac
H3K27 me2:DNA Meth. m5C
H3K27 me3:DNA Meth. m5C
H3K9 me2
H4K20 me3
H3K27 me2
H3K9 me3
H3K4 me3
H2A.Z
H3K27 me3
H3K4 me1
DNA Meth. m5C
H4−4ac
H4K20 me2
H3−5ac
0.00 0.25 0.50 0.75 1.00
π̂
RBM5 − Forward
H3K4 me3:H3−5ac
H2A.Z:H3−5ac
H2A.Z:H4−4ac
H3K4 me1:H2A.Z
H3K4 me1:H3−5ac
H3K4 me1:H4−4ac
H3K4 me1:H4K20 me2
H4K20 me2:H4−4ac
H4K20 me2:H3K4 me3
H2A.Z:H3K4 me3
H2A.Z:H4K20 me2
H4K20 me2:H3−5ac
H3K9 me3
H4K20 me3
DNA Meth. m5C
H3K27 me3
H3K4 me1
H4−4ac
H2A.Z
H3−5ac
H3K4 me3
H4K20 me2
0.00 0.25 0.50 0.75 1.00
π̂
RBM5 − Reverse
H3K4 me3:H3−5ac
H2A.Z:H3−5ac
H2A.Z:H4−4ac
H3K4 me1:H2A.Z
H3K4 me1:H3−5ac
H3K4 me1:H4−4ac
H3K4 me1:H4K20 me2
H4K20 me2:H4−4ac
H4K20 me2:H3K4 me3
H2A.Z:H3K4 me3
H2A.Z:H4K20 me2
H4K20 me2:H3−5ac
H3K9 me3
H4K20 me3
DNA Meth. m5C
H3K27 me3
H3K4 me1
H4−4ac
H2A.Z
H3−5ac
H3K4 me3
H4K20 me2
0.00 0.25 0.50 0.75 1.00
π̂
PPIL3 − Forward
H3K4 me3:H3−5ac
H3K27 me3:H4K20 me3
H3K27 me2:DNA Meth. m5C
H4K20 me2:H4−4ac
H3K4 me1:H4K20 me2
H2A.Z:H4K20 me2
H4K20 me2:H3−5ac
H4K20 me2:H3K4 me3
H3K9 me2
H3K27 me2
H4K20 me3
H4−4ac
H3K9 me3
H3K27 me3
H3K4 me1
DNA Meth. m5C
H3−5ac
H3K4 me3
H2A.Z
H4K20 me2
0.00 0.25 0.50 0.75 1.00
π̂
PPIL3 − Reverse
H3K4 me3:H3−5ac
H3K27 me3:H4K20 me3
H3K27 me2:DNA Meth. m5C
H4K20 me2:H4−4ac
H3K4 me1:H4K20 me2
H2A.Z:H4K20 me2
H4K20 me2:H3−5ac
H4K20 me2:H3K4 me3
H3K9 me2
H3K27 me2
H4K20 me3
H4−4ac
H3K9 me3
H3K27 me3
H3K4 me1
DNA Meth. m5C
H3−5ac
H3K4 me3
H2A.Z
H4K20 me2
0.00 0.25 0.50 0.75 1.00
π̂

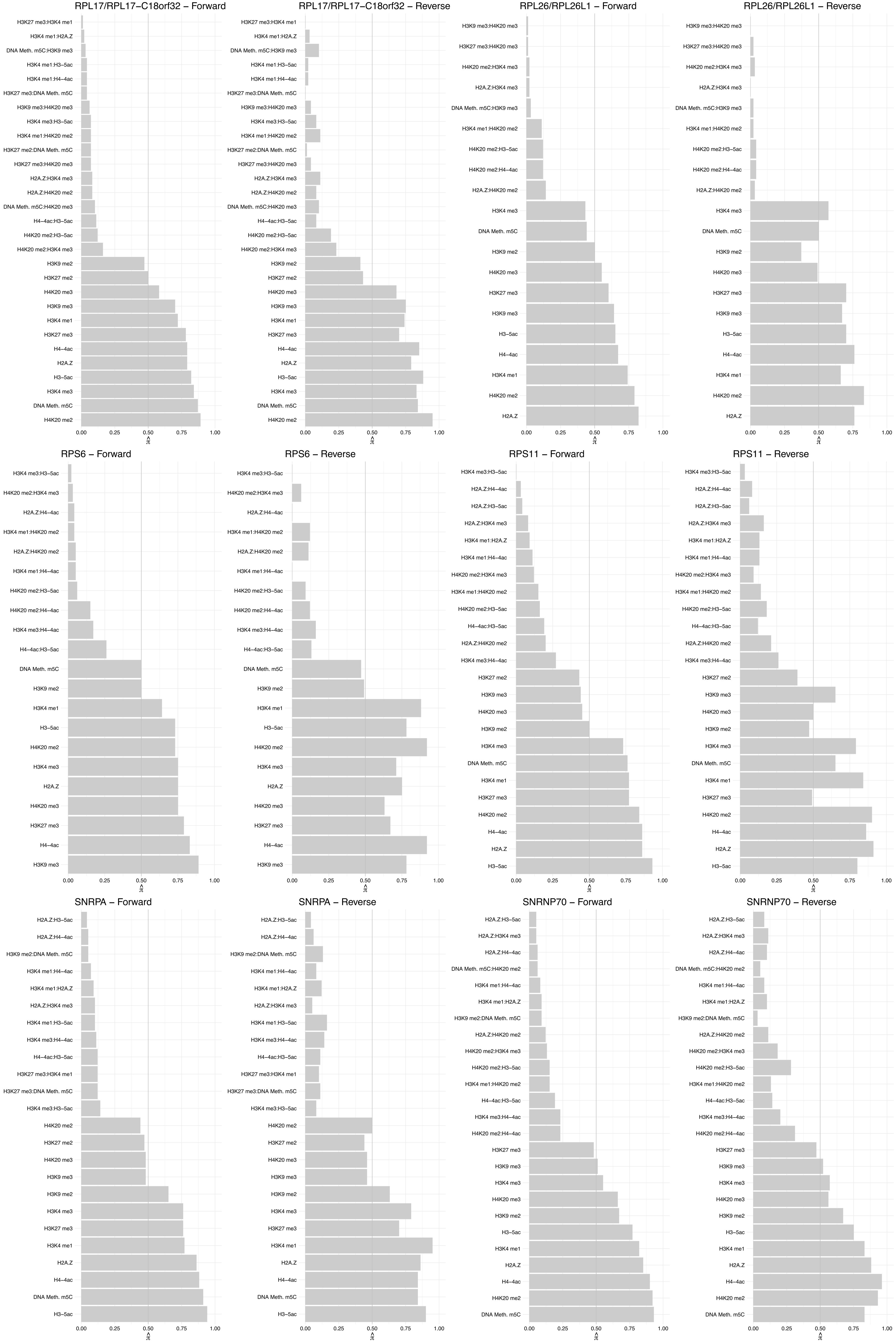
RPL17/RPL17−C18orf32 – Forward
H3K27 me3:H3K4 me1
H3K4 me1:H2A.Z
DNA Meth. m5C:H3K9 me3
H3K4 me1:H3−5ac
H3K4 me1:H4−4ac
H3K27 me3:DNA Meth. m5C
H3K9 me3:H4K20 me3
H3K4 me3:H3−5ac
H3K4 me1:H4K20 me2
H3K27 me2:DNA Meth. m5C
H3K27 me3:H4K20 me3
H2A.Z:H3K4 me3
H2A.Z:H4K20 me2
DNA Meth. m5C:H4K20 me3
H4−4ac:H3−5ac
H4K20 me2:H3−5ac
H4K20 me2:H3K4 me3
H3K9 me2
H3K27 me2
H4K20 me3
H3K9 me3
H3K4 me1
H3K27 me3
H4−4ac
H2A.Z
H3−5ac
H3K4 me3
DNA Meth. m5C
H4K20 me2
0.00 0.25 0.50 0.75 1.00
π̂
RPL17/RPL17−C18orf32 – Reverse
H3K27 me3:H3K4 me1
H3K4 me1:H2A.Z
DNA Meth. m5C:H3K9 me3
H3K4 me1:H3−5ac
H3K4 me1:H4−4ac
H3K27 me3:DNA Meth. m5C
H3K9 me3:H4K20 me3
H3K4 me3:H3−5ac
H3K4 me1:H4K20 me2
H3K27 me2:DNA Meth. m5C
H3K27 me3:H4K20 me3
H2A.Z:H3K4 me3
H2A.Z:H4K20 me2
DNA Meth. m5C:H4K20 me3
H4−4ac:H3−5ac
H4K20 me2:H3−5ac
H4K20 me2:H3K4 me3
H3K9 me2
H3K27 me2
H4K20 me3
H3K9 me3
H3K4 me1
H3K27 me3
H4−4ac
H2A.Z
H3−5ac
H3K4 me3
DNA Meth. m5C
H4K20 me2
0.00 0.25 0.50 0.75 1.00
π̂
RPL26/RPL26L1 – Forward
H3K9 me3:H4K20 me3
H3K27 me3:H4K20 me3
H4K20 me2:H3K4 me3
H2A.Z:H3K4 me3
DNA Meth. m5C:H3K9 me3
H3K4 me1:H4K20 me2
H4K20 me2:H3−5ac
H4K20 me2:H4−4ac
H2A.Z:H4K20 me2
H3K4 me3
DNA Meth. m5C
H3K9 me2
H4K20 me3
H3K27 me3
H3K9 me3
H3−5ac
H4−4ac
H3K4 me1
H4K20 me2
H2A.Z
0.00 0.25 0.50 0.75 1.00
π̂
RPL26/RPL26L1 – Reverse
H3K9 me3:H4K20 me3
H3K27 me3:H4K20 me3
H4K20 me2:H3K4 me3
H2A.Z:H3K4 me3
DNA Meth. m5C:H3K9 me3
H3K4 me1:H4K20 me2
H4K20 me2:H3−5ac
H4K20 me2:H4−4ac
H2A.Z:H4K20 me2
H3K4 me3
DNA Meth. m5C
H3K9 me2
H4K20 me3
H3K27 me3
H3K9 me3
H3−5ac
H4−4ac
H3K4 me1
H4K20 me2
H2A.Z
0.00 0.25 0.50 0.75 1.00
π̂
RPS6 – Forward
H3K4 me3:H3−5ac
H4K20 me2:H3K4 me3
H2A.Z:H4−4ac
H3K4 me1:H4K20 me2
H2A.Z:H4K20 me2
H3K4 me1:H4−4ac
H4K20 me2:H3−5ac
H4K20 me2:H4−4ac
H3K4 me3:H4−4ac
H4−4ac:H3−5ac
DNA Meth. m5C
H3K9 me2
H3K4 me1
H3−5ac
H4K20 me2
H3K4 me3
H2A.Z
H4K20 me3
H3K27 me3
H4−4ac
H3K9 me3
0.00 0.25 0.50 0.75 1.00
π̂
RPS6 – Reverse
H3K4 me3:H3−5ac
H4K20 me2:H3K4 me3
H2A.Z:H4−4ac
H3K4 me1:H4K20 me2
H2A.Z:H4K20 me2
H3K4 me1:H4−4ac
H4K20 me2:H3−5ac
H4K20 me2:H4−4ac
H3K4 me3:H4−4ac
H4−4ac:H3−5ac
DNA Meth. m5C
H3K9 me2
H3K4 me1
H3−5ac
H4K20 me2
H3K4 me3
H2A.Z
H4K20 me3
H3K27 me3
H4−4ac
H3K9 me3
0.00 0.25 0.50 0.75 1.00
π̂
RPS11 – Forward
H3K4 me3:H3−5ac
H2A.Z:H4−4ac
H2A.Z:H3−5ac
H2A.Z:H3K4 me3
H3K4 me1:H2A.Z
H3K4 me1:H4−4ac
H4K20 me2:H3K4 me3
H3K4 me1:H4K20 me2
H4K20 me2:H3−5ac
H4−4ac:H3−5ac
H2A.Z:H4K20 me2
H3K4 me3:H4−4ac
H3K27 me2
H3K9 me3
H4K20 me3
H3K9 me2
H3K4 me3
DNA Meth. m5C
H3K4 me1
H3K27 me3
H4K20 me2
H4−4ac
H2A.Z
H3−5ac
0.00 0.25 0.50 0.75 1.00
π̂
RPS11 – Reverse
H3K4 me3:H3−5ac
H2A.Z:H4−4ac
H2A.Z:H3−5ac
H2A.Z:H3K4 me3
H3K4 me1:H2A.Z
H3K4 me1:H4−4ac
H4K20 me2:H3K4 me3
H3K4 me1:H4K20 me2
H4K20 me2:H3−5ac
H4−4ac:H3−5ac
H2A.Z:H4K20 me2
H3K4 me3:H4−4ac
H3K27 me2
H3K9 me3
H4K20 me3
H3K9 me2
H3K4 me3
DNA Meth. m5C
H3K4 me1
H3K27 me3
H4K20 me2
H4−4ac
H2A.Z
H3−5ac
0.00 0.25 0.50 0.75 1.00
π̂
SNRPA – Forward
H2A.Z:H3−5ac
H2A.Z:H4−4ac
H3K9 me2:DNA Meth. m5C
H3K4 me1:H4−4ac
H3K4 me1:H2A.Z
H2A.Z:H3K4 me3
H3K4 me1:H3−5ac
H3K4 me3:H4−4ac
H4−4ac:H3−5ac
H3K27 me3:H3K4 me1
H3K27 me3:DNA Meth. m5C
H3K4 me3:H3−5ac
H4K20 me2
H3K27 me2
H4K20 me3
H3K9 me3
H3K9 me2
H3K4 me3
H3K27 me3
H3K4 me1
H2A.Z
H4−4ac
DNA Meth. m5C
H3−5ac
0.00 0.25 0.50 0.75 1.00
π̂
SNRPA – Reverse
H2A.Z:H3−5ac
H2A.Z:H4−4ac
H3K9 me2:DNA Meth. m5C
H3K4 me1:H4−4ac
H3K4 me1:H2A.Z
H2A.Z:H3K4 me3
H3K4 me1:H3−5ac
H3K4 me3:H4−4ac
H4−4ac:H3−5ac
H3K27 me3:H3K4 me1
H3K27 me3:DNA Meth. m5C
H3K4 me3:H3−5ac
H4K20 me2
H3K27 me2
H4K20 me3
H3K9 me3
H3K9 me2
H3K4 me3
H3K27 me3
H3K4 me1
H2A.Z
H4−4ac
DNA Meth. m5C
H3−5ac
0.00 0.25 0.50 0.75 1.00
π̂
SNRNP70 – Forward
H2A.Z:H3−5ac
H2A.Z:H3K4 me3
H2A.Z:H4−4ac
DNA Meth. m5C:H4K20 me2
H3K4 me1:H4−4ac
H3K4 me1:H2A.Z
H3K9 me2:DNA Meth. m5C
H2A.Z:H4K20 me2
H4K20 me2:H3K4 me3
H4K20 me2:H3−5ac
H3K4 me1:H4K20 me2
H4−4ac:H3−5ac
H3K4 me3:H4−4ac
H4K20 me2:H4−4ac
H3K27 me3
H3K9 me3
H3K4 me3
H4K20 me3
H3K9 me2
H3−5ac
H3K4 me1
H2A.Z
H4−4ac
H4K20 me2
DNA Meth. m5C
0.00 0.25 0.50 0.75 1.00
π̂
SNRNP70 – Reverse
H2A.Z:H3−5ac
H2A.Z:H3K4 me3
H2A.Z:H4−4ac
DNA Meth. m5C:H4K20 me2
H3K4 me1:H4−4ac
H3K4 me1:H2A.Z
H3K9 me2:DNA Meth. m5C
H2A.Z:H4K20 me2
H4K20 me2:H3K4 me3
H4K20 me2:H3−5ac
H3K4 me1:H4K20 me2
H4−4ac:H3−5ac
H3K4 me3:H4−4ac
H4K20 me2:H4−4ac
H3K27 me3
H3K9 me3
H3K4 me3
H4K20 me3
H3K9 me2
H3−5ac
H3K4 me1
H2A.Z
H4−4ac
H4K20 me2
DNA Meth. m5C
0.00 0.25 0.50 0.75 1.00
π̂

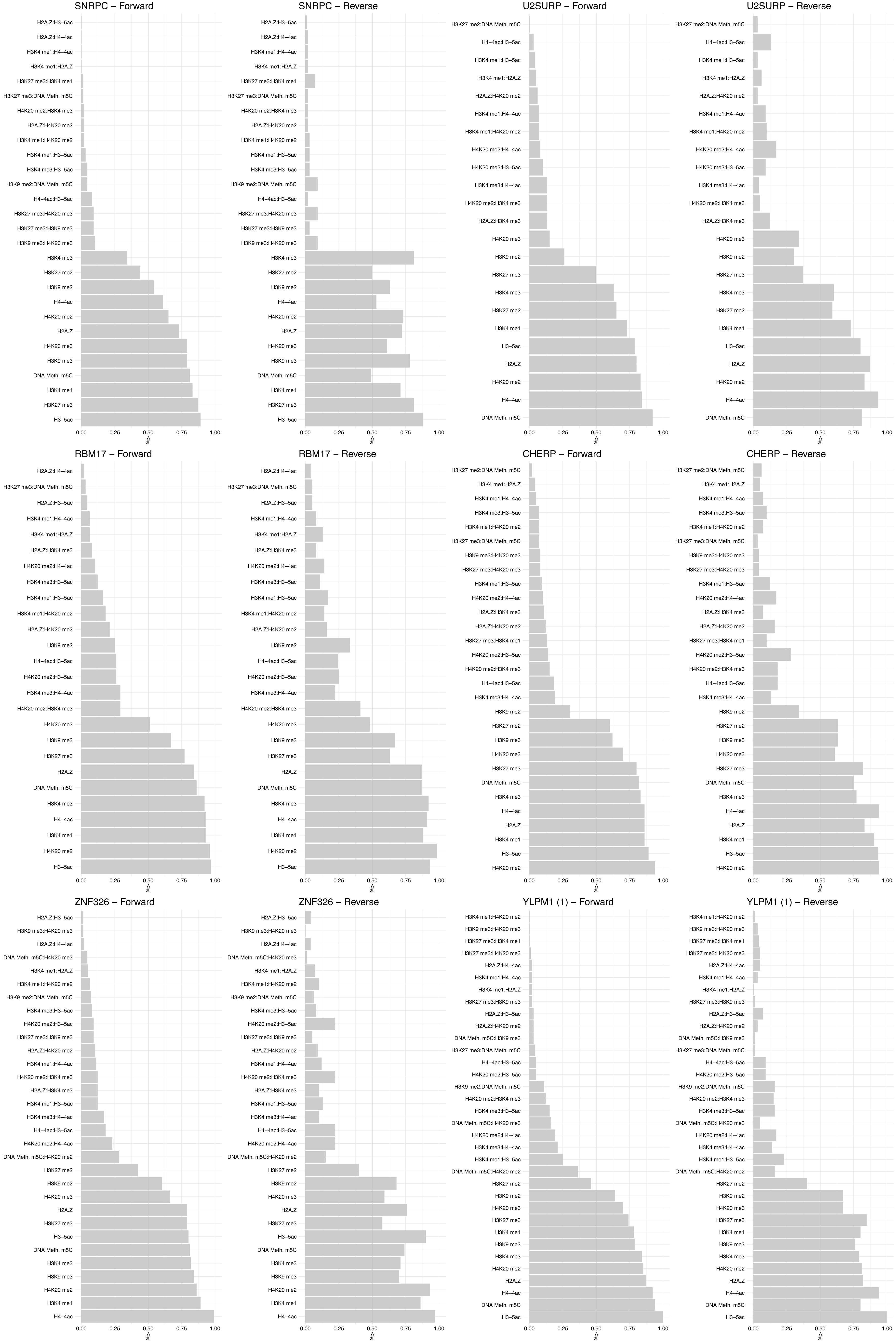
SNRPC – Forward
H2A.Z:H3–5ac
H2A.Z:H4–4ac
H3K4 me1:H4–4ac
H3K4 me1:H2A.Z
H3K27 me3:H3K4 me1
H3K27 me3:DNA Meth. m5C
H4K20 me2:H3K4 me3
H2A.Z:H4K20 me2
H3K4 me1:H4K20 me2
H3K4 me1:H3–5ac
H3K4 me3:H3–5ac
H3K9 me2:DNA Meth. m5C
H4–4ac:H3–5ac
H3K27 me3:H4K20 me3
H3K27 me3:H3K9 me3
H3K9 me3:H4K20 me3
H3K4 me3
H3K27 me2
H3K9 me2
H4–4ac
H4K20 me2
H2A.Z
H4K20 me3
H3K9 me3
DNA Meth. m5C
H3K4 me1
H3K27 me3
H3–5ac
0.00 0.25 0.50 0.75 1.00
$\hat{\pi}$
SNRPC – Reverse
H2A.Z:H3–5ac
H2A.Z:H4–4ac
H3K4 me1:H4–4ac
H3K4 me1:H2A.Z
H3K27 me3:H3K4 me1
H3K27 me3:DNA Meth. m5C
H4K20 me2:H3K4 me3
H2A.Z:H4K20 me2
H3K4 me1:H4K20 me2
H3K4 me1:H3–5ac
H3K4 me3:H3–5ac
H3K9 me2:DNA Meth. m5C
H4–4ac:H3–5ac
H3K27 me3:H4K20 me3
H3K27 me3:H3K9 me3
H3K9 me3:H4K20 me3
H3K4 me3
H3K27 me2
H3K9 me2
H4–4ac
H4K20 me2
H2A.Z
H4K20 me3
H3K9 me3
DNA Meth. m5C
H3K4 me1
H3K27 me3
H3–5ac
0.00 0.25 0.50 0.75 1.00
$\hat{\pi}$
U2SURP – Forward
H3K27 me2:DNA Meth. m5C
H4–4ac:H3–5ac
H3K4 me1:H3–5ac
H3K4 me1:H2A.Z
H2A.Z:H4K20 me2
H3K4 me1:H4–4ac
H3K4 me1:H4K20 me2
H4K20 me2:H4–4ac
H4K20 me2:H3–5ac
H3K4 me3:H4–4ac
H4K20 me2:H3K4 me3
H2A.Z:H3K4 me3
H4K20 me3
H3K9 me2
H3K27 me3
H3K4 me3
H3K27 me2
H3K4 me1
H3–5ac
H2A.Z
H4K20 me2
H4–4ac
DNA Meth. m5C
0.00 0.25 0.50 0.75 1.00
$\hat{\pi}$
U2SURP – Reverse
H3K27 me2:DNA Meth. m5C
H4–4ac:H3–5ac
H3K4 me1:H3–5ac
H3K4 me1:H2A.Z
H2A.Z:H4K20 me2
H3K4 me1:H4–4ac
H3K4 me1:H4K20 me2
H4K20 me2:H4–4ac
H4K20 me2:H3–5ac
H3K4 me3:H4–4ac
H4K20 me2:H3K4 me3
H2A.Z:H3K4 me3
H4K20 me3
H3K9 me2
H3K27 me3
H3K4 me3
H3K27 me2
H3K4 me1
H3–5ac
H2A.Z
H4K20 me2
H4–4ac
DNA Meth. m5C
0.00 0.25 0.50 0.75 1.00
$\hat{\pi}$
RBM17 – Forward
H2A.Z:H4–4ac
H3K27 me3:DNA Meth. m5C
H2A.Z:H3–5ac
H3K4 me1:H4–4ac
H3K4 me1:H2A.Z
H2A.Z:H3K4 me3
H4K20 me2:H4–4ac
H3K4 me3:H3–5ac
H3K4 me1:H3–5ac
H3K4 me1:H4K20 me2
H2A.Z:H4K20 me2
H3K9 me2
H4–4ac:H3–5ac
H4K20 me2:H3–5ac
H3K4 me3:H4–4ac
H4K20 me2:H3K4 me3
H4K20 me3
H3K9 me3
H3K27 me3
H2A.Z
DNA Meth. m5C
H3K4 me3
H4–4ac
H3K4 me1
H4K20 me2
H3–5ac
0.00 0.25 0.50 0.75 1.00
$\hat{\pi}$
RBM17 – Reverse
H2A.Z:H4–4ac
H3K27 me3:DNA Meth. m5C
H2A.Z:H3–5ac
H3K4 me1:H4–4ac
H3K4 me1:H2A.Z
H2A.Z:H3K4 me3
H4K20 me2:H4–4ac
H3K4 me3:H3–5ac
H3K4 me1:H3–5ac
H3K4 me1:H4K20 me2
H2A.Z:H4K20 me2
H3K9 me2
H4–4ac:H3–5ac
H4K20 me2:H3–5ac
H3K4 me3:H4–4ac
H4K20 me2:H3K4 me3
H4K20 me3
H3K9 me3
H3K27 me3
H2A.Z
DNA Meth. m5C
H3K4 me3
H4–4ac
H3K4 me1
H4K20 me2
H3–5ac
0.00 0.25 0.50 0.75 1.00
$\hat{\pi}$
CHERP – Forward
H3K27 me2:DNA Meth. m5C
H3K4 me1:H2A.Z
H3K4 me1:H4–4ac
H3K4 me3:H3–5ac
H3K4 me1:H4K20 me2
H3K27 me3:DNA Meth. m5C
H3K9 me3:H4K20 me3
H3K27 me3:H4K20 me3
H3K4 me1:H3–5ac
H4K20 me2:H4–4ac
H2A.Z:H3K4 me3
H2A.Z:H4K20 me2
H3K27 me3:H3K4 me1
H4K20 me2:H3–5ac
H4K20 me2:H3K4 me3
H4–4ac:H3–5ac
H3K4 me3:H4–4ac
H3K9 me2
H3K27 me2
H3K9 me3
H4K20 me3
H3K27 me3
DNA Meth. m5C
H3K4 me3
H4–4ac
H2A.Z
H3K4 me1
H3–5ac
H4K20 me2
0.00 0.25 0.50 0.75 1.00
$\hat{\pi}$
CHERP – Reverse
H3K27 me2:DNA Meth. m5C
H3K4 me1:H2A.Z
H3K4 me1:H4–4ac
H3K4 me3:H3–5ac
H3K4 me1:H4K20 me2
H3K27 me3:DNA Meth. m5C
H3K9 me3:H4K20 me3
H3K27 me3:H4K20 me3
H3K4 me1:H3–5ac
H4K20 me2:H4–4ac
H2A.Z:H3K4 me3
H2A.Z:H4K20 me2
H3K27 me3:H3K4 me1
H4K20 me2:H3–5ac
H4K20 me2:H3K4 me3
H4–4ac:H3–5ac
H3K4 me3:H4–4ac
H3K9 me2
H3K27 me2
H3K9 me3
H4K20 me3
H3K27 me3
DNA Meth. m5C
H3K4 me3
H4–4ac
H2A.Z
H3K4 me1
H3–5ac
H4K20 me2
0.00 0.25 0.50 0.75 1.00
$\hat{\pi}$
ZNF326 – Forward
H2A.Z:H3–5ac
H3K9 me3:H4K20 me3
H2A.Z:H4–4ac
DNA Meth. m5C:H4K20 me3
H3K4 me1:H2A.Z
H3K4 me1:H4K20 me2
H3K9 me2:DNA Meth. m5C
H3K4 me3:H3–5ac
H4K20 me2:H3–5ac
H3K27 me3:H3K9 me3
H2A.Z:H4K20 me2
H3K4 me1:H4–4ac
H4K20 me2:H3K4 me3
H2A.Z:H3K4 me3
H3K4 me1:H3–5ac
H3K4 me3:H4–4ac
H4–4ac:H3–5ac
H4K20 me2:H4–4ac
DNA Meth. m5C:H4K20 me2
H3K27 me2
H3K9 me2
H4K20 me3
H2A.Z
H3K27 me3
H3–5ac
DNA Meth. m5C
H3K4 me3
H3K9 me3
H4K20 me2
H3K4 me1
H4–4ac
0.00 0.25 0.50 0.75 1.00
$\hat{\pi}$
ZNF326 – Reverse
H2A.Z:H3–5ac
H3K9 me3:H4K20 me3
H2A.Z:H4–4ac
DNA Meth. m5C:H4K20 me3
H3K4 me1:H2A.Z
H3K4 me1:H4K20 me2
H3K9 me2:DNA Meth. m5C
H3K4 me3:H3–5ac
H4K20 me2:H3–5ac
H3K27 me3:H3K9 me3
H2A.Z:H4K20 me2
H3K4 me1:H4–4ac
H4K20 me2:H3K4 me3
H2A.Z:H3K4 me3
H3K4 me1:H3–5ac
H3K4 me3:H4–4ac
H4–4ac:H3–5ac
H4K20 me2:H4–4ac
DNA Meth. m5C:H4K20 me2
H3K27 me2
H3K9 me2
H4K20 me3
H2A.Z
H3K27 me3
H3–5ac
DNA Meth. m5C
H3K4 me3
H3K9 me3
H4K20 me2
H3K4 me1
H4–4ac
0.00 0.25 0.50 0.75 1.00
$\hat{\pi}$
YLPM1 (1) – Forward
H3K4 me1:H4K20 me2
H3K9 me3:H4K20 me3
H3K27 me3:H3K4 me1
H3K27 me3:H4K20 me3
H2A.Z:H4–4ac
H3K4 me1:H4–4ac
H3K4 me1:H2A.Z
H3K27 me3:H3K9 me3
H2A.Z:H3–5ac
H2A.Z:H4K20 me2
DNA Meth. m5C:H3K9 me3
H3K27 me3:DNA Meth. m5C
H4–4ac:H3–5ac
H4K20 me2:H3–5ac
H3K9 me2:DNA Meth. m5C
H4K20 me2:H3K4 me3
H3K4 me3:H3–5ac
DNA Meth. m5C:H4K20 me3
H4K20 me2:H4–4ac
H3K4 me3:H4–4ac
H3K4 me1:H3–5ac
DNA Meth. m5C:H4K20 me2
H3K27 me2
H3K9 me2
H4K20 me3
H3K27 me3
H3K4 me1
H3K9 me3
H3K4 me3
H4K20 me2
H2A.Z
H4–4ac
DNA Meth. m5C
H3–5ac
0.00 0.25 0.50 0.75 1.00
$\hat{\pi}$
YLPM1 (1) – Reverse
H3K4 me1:H4K20 me2
H3K9 me3:H4K20 me3
H3K27 me3:H3K4 me1
H3K27 me3:H4K20 me3
H2A.Z:H4–4ac
H3K4 me1:H4–4ac
H3K4 me1:H2A.Z
H3K27 me3:H3K9 me3
H2A.Z:H3–5ac
H2A.Z:H4K20 me2
DNA Meth. m5C:H3K9 me3
H3K27 me3:DNA Meth. m5C
H4–4ac:H3–5ac
H4K20 me2:H3–5ac
H3K9 me2:DNA Meth. m5C
H4K20 me2:H3K4 me3
H3K4 me3:H3–5ac
DNA Meth. m5C:H4K20 me3
H4K20 me2:H4–4ac
H3K4 me3:H4–4ac
H3K4 me1:H3–5ac
DNA Meth. m5C:H4K20 me2
H3K27 me2
H3K9 me2
H4K20 me3
H3K27 me3
H3K4 me1
H3K9 me3
H3K4 me3
H4K20 me2
H2A.Z
H4–4ac
DNA Meth. m5C
H3–5ac
0.00 0.25 0.50 0.75 1.00
$\hat{\pi}$

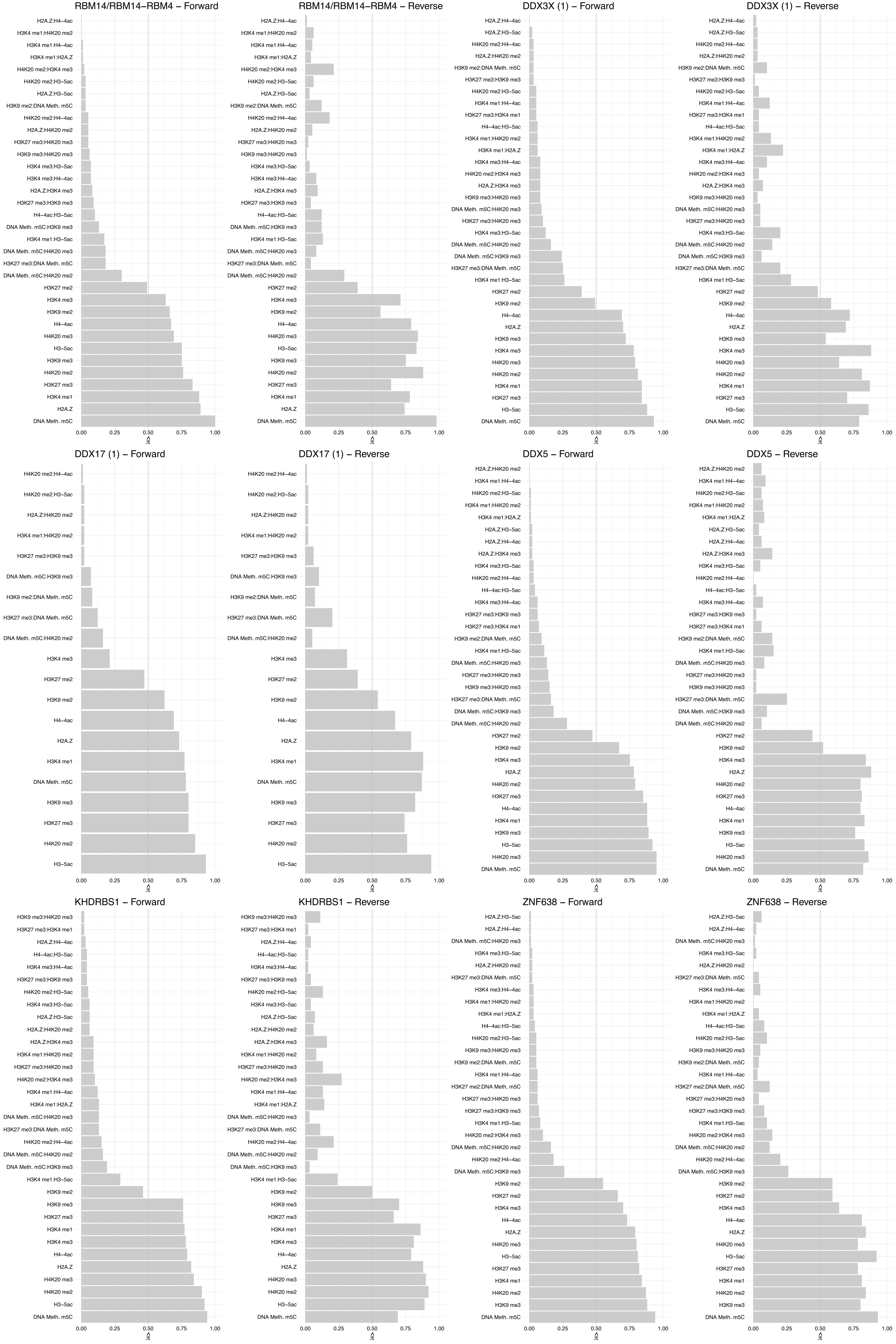

RBM14/RBM14−RBM4 − Forward
RBM14/RBM14−RBM4 − Reverse
DDX3X (1) − Forward
DDX3X (1) − Reverse
DDX17 (1) − Forward
DDX17 (1) − Reverse
DDX5 − Forward
DDX5 − Reverse
KHDRBS1 − Forward
KHDRBS1 − Reverse
ZNF638 − Forward
ZNF638 − Reverse
0.00
0.25
0.50
0.75
1.00
$\hat{\pi}$

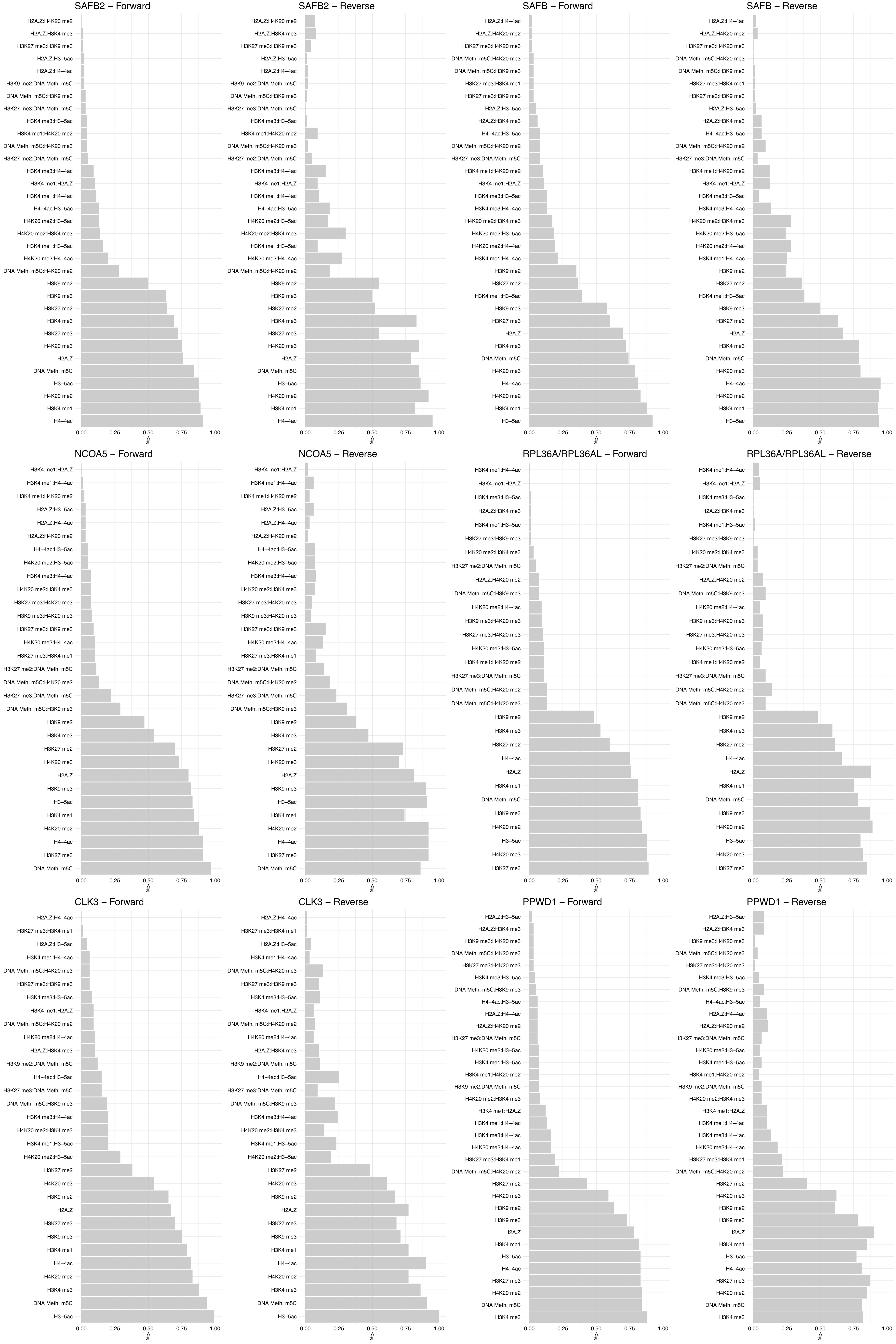

SAFB2 – Forward
SAFB2 – Reverse
SAFB – Forward
SAFB – Reverse
NCOA5 – Forward
NCOA5 – Reverse
RPL36A/RPL36AL – Forward
RPL36A/RPL36AL – Reverse
CLK3 – Forward
CLK3 – Reverse
PPWD1 – Forward
PPWD1 – Reverse
0.00
0.25
0.50
0.75
1.00
$\hat{\pi}$

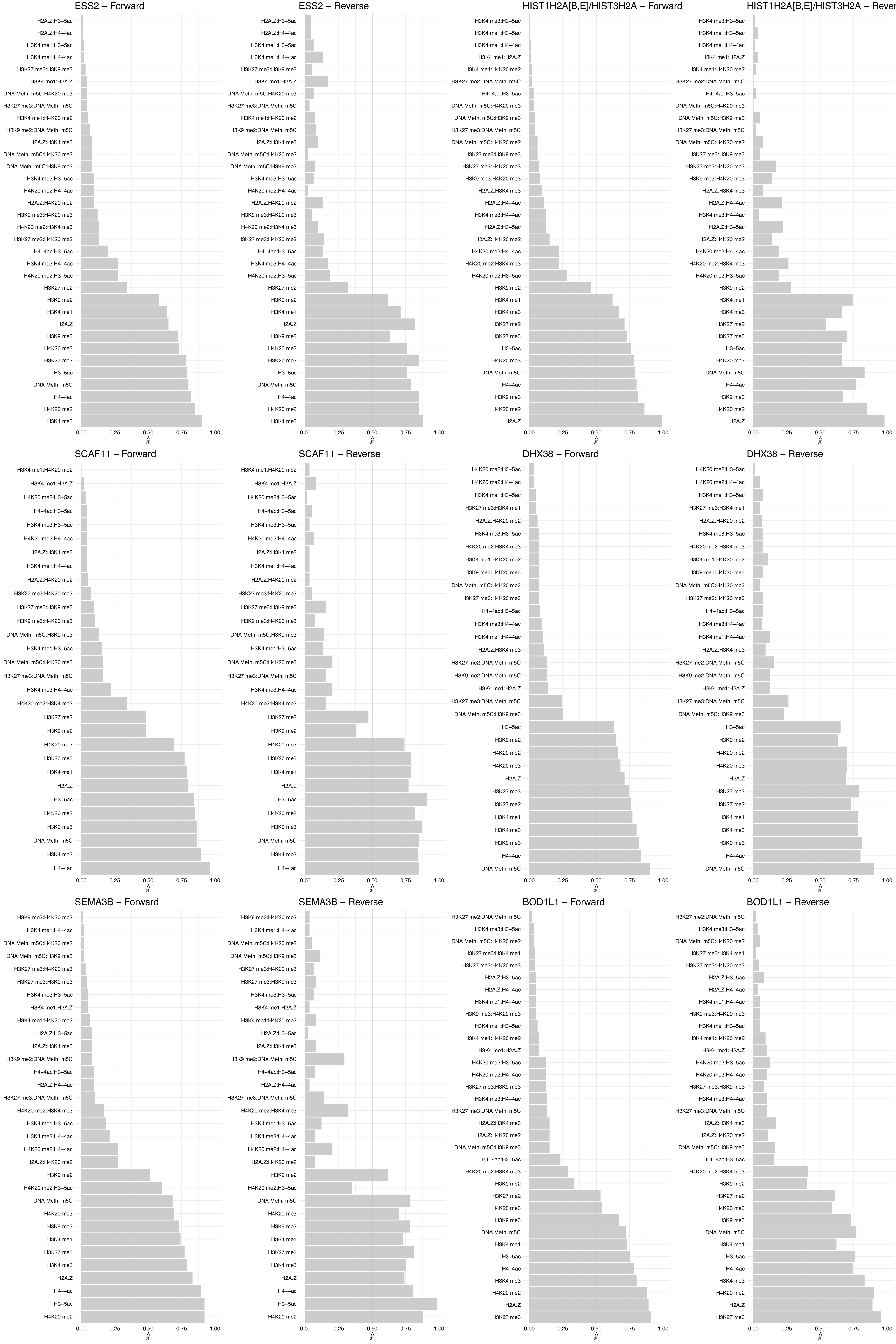
ESS2 – Forward
ESS2 – Reverse
HIST1H2A[B,E]/HIST3H2A – Forward
HIST1H2A[B,E]/HIST3H2A – Rever
SCAF11 – Forward
SCAF11 – Reverse
DHX38 – Forward
DHX38 – Reverse
SEMA3B – Forward
SEMA3B – Reverse
BOD1L1 – Forward
BOD1L1 – Reverse
0.00
0.25
0.50
0.75
1.00
$\hat{\pi}$

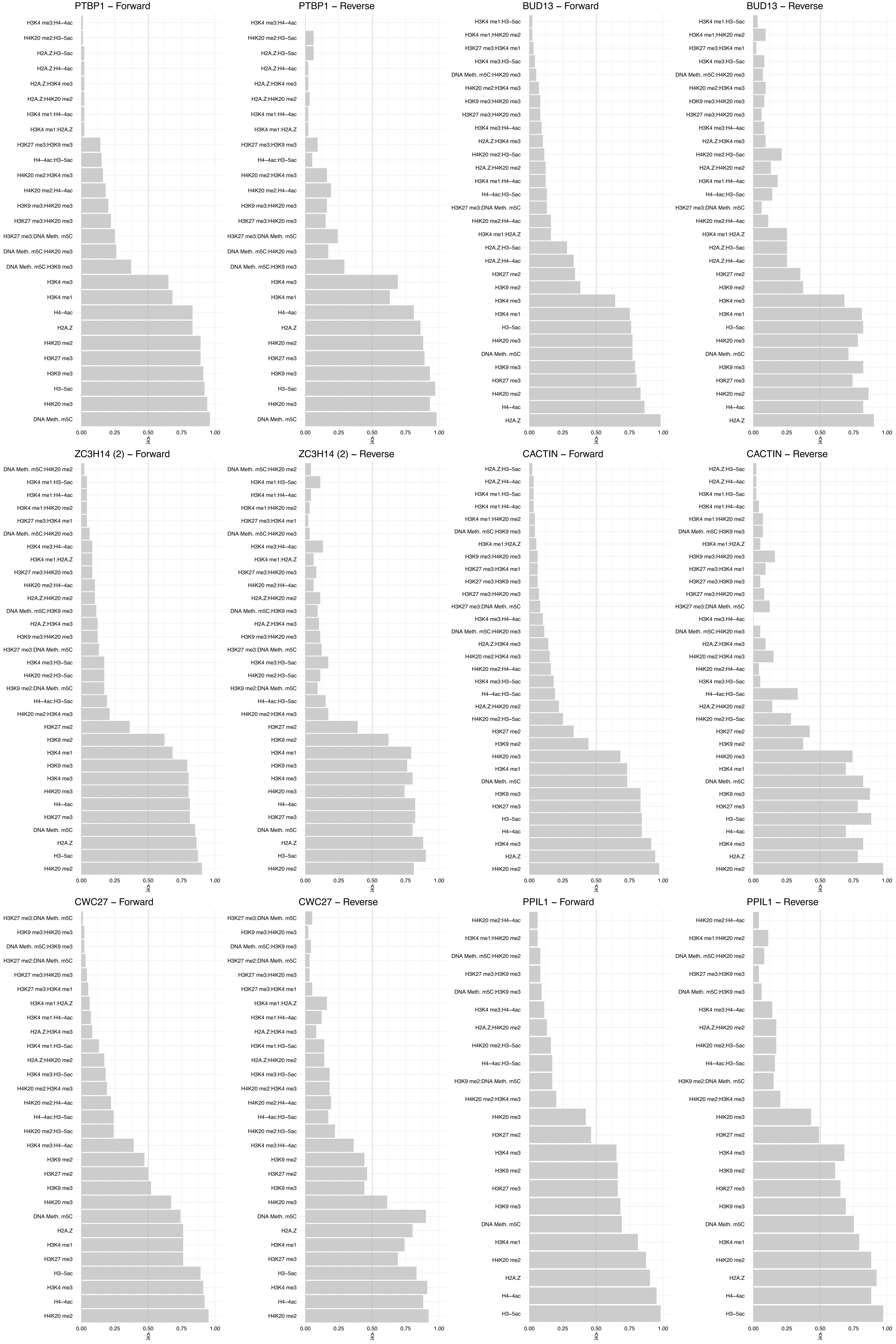
PTBP1 – Forward
H3K4 me3:H4−4ac
H4K20 me2:H3−5ac
H2A.Z:H3−5ac
H2A.Z:H4−4ac
H2A.Z:H3K4 me3
H2A.Z:H4K20 me2
H3K4 me1:H4−4ac
H3K4 me1:H2A.Z
H3K27 me3:H3K9 me3
H4−4ac:H3−5ac
H4K20 me2:H3K4 me3
H4K20 me2:H4−4ac
H3K9 me3:H4K20 me3
H3K27 me3:H4K20 me3
H3K27 me3:DNA Meth. m5C
DNA Meth. m5C:H4K20 me3
DNA Meth. m5C:H3K9 me3
H3K4 me3
H3K4 me1
H4−4ac
H2A.Z
H4K20 me2
H3K27 me3
H3K9 me3
H3−5ac
H4K20 me3
DNA Meth. m5C
0.00
0.25
0.50
0.75
1.00
π̂
PTBP1 – Reverse
H3K4 me3:H4−4ac
H4K20 me2:H3−5ac
H2A.Z:H3−5ac
H2A.Z:H4−4ac
H2A.Z:H3K4 me3
H2A.Z:H4K20 me2
H3K4 me1:H4−4ac
H3K4 me1:H2A.Z
H3K27 me3:H3K9 me3
H4−4ac:H3−5ac
H4K20 me2:H3K4 me3
H4K20 me2:H4−4ac
H3K9 me3:H4K20 me3
H3K27 me3:H4K20 me3
H3K27 me3:DNA Meth. m5C
DNA Meth. m5C:H4K20 me3
DNA Meth. m5C:H3K9 me3
H3K4 me3
H3K4 me1
H4−4ac
H2A.Z
H4K20 me2
H3K27 me3
H3K9 me3
H3−5ac
H4K20 me3
DNA Meth. m5C
0.00
0.25
0.50
0.75
1.00
π̂
BUD13 – Forward
H3K4 me1:H3−5ac
H3K4 me1:H4K20 me2
H3K27 me3:H3K4 me1
H3K4 me3:H3−5ac
DNA Meth. m5C:H4K20 me3
H4K20 me2:H3K4 me3
H3K9 me3:H4K20 me3
H3K27 me3:H4K20 me3
H3K4 me3:H4−4ac
H2A.Z:H3K4 me3
H4K20 me2:H3−5ac
H2A.Z:H4K20 me2
H3K4 me1:H4−4ac
H4−4ac:H3−5ac
H3K27 me3:DNA Meth. m5C
H4K20 me2:H4−4ac
H3K4 me1:H2A.Z
H2A.Z:H3−5ac
H2A.Z:H4−4ac
H3K27 me2
H3K9 me2
H3K4 me3
H3K4 me1
H3−5ac
H4K20 me3
DNA Meth. m5C
H3K9 me3
H3K27 me3
H4K20 me2
H4−4ac
H2A.Z
0.00
0.25
0.50
0.75
1.00
π̂
BUD13 – Reverse
H3K4 me1:H3−5ac
H3K4 me1:H4K20 me2
H3K27 me3:H3K4 me1
H3K4 me3:H3−5ac
DNA Meth. m5C:H4K20 me3
H4K20 me2:H3K4 me3
H3K9 me3:H4K20 me3
H3K27 me3:H4K20 me3
H3K4 me3:H4−4ac
H2A.Z:H3K4 me3
H4K20 me2:H3−5ac
H2A.Z:H4K20 me2
H3K4 me1:H4−4ac
H4−4ac:H3−5ac
H3K27 me3:DNA Meth. m5C
H4K20 me2:H4−4ac
H3K4 me1:H2A.Z
H2A.Z:H3−5ac
H2A.Z:H4−4ac
H3K27 me2
H3K9 me2
H3K4 me3
H3K4 me1
H3−5ac
H4K20 me3
DNA Meth. m5C
H3K9 me3
H3K27 me3
H4K20 me2
H4−4ac
H2A.Z
0.00
0.25
0.50
0.75
1.00
π̂
ZC3H14 (2) – Forward
DNA Meth. m5C:H4K20 me2
H3K4 me1:H3−5ac
H3K4 me1:H4−4ac
H3K4 me1:H4K20 me2
H3K27 me3:H3K4 me1
DNA Meth. m5C:H4K20 me3
H3K4 me3:H4−4ac
H3K4 me1:H2A.Z
H3K27 me3:H4K20 me3
H4K20 me2:H4−4ac
H2A.Z:H4K20 me2
DNA Meth. m5C:H3K9 me3
H2A.Z:H3K4 me3
H3K9 me3:H4K20 me3
H3K27 me3:DNA Meth. m5C
H3K4 me3:H3−5ac
H4K20 me2:H3−5ac
H3K9 me2:DNA Meth. m5C
H4−4ac:H3−5ac
H4K20 me2:H3K4 me3
H3K27 me2
H3K9 me2
H3K4 me1
H3K9 me3
H3K4 me3
H4K20 me3
H4−4ac
H3K27 me3
DNA Meth. m5C
H2A.Z
H3−5ac
H4K20 me2
0.00
0.25
0.50
0.75
1.00
π̂
ZC3H14 (2) – Reverse
DNA Meth. m5C:H4K20 me2
H3K4 me1:H3−5ac
H3K4 me1:H4−4ac
H3K4 me1:H4K20 me2
H3K27 me3:H3K4 me1
DNA Meth. m5C:H4K20 me3
H3K4 me3:H4−4ac
H3K4 me1:H2A.Z
H3K27 me3:H4K20 me3
H4K20 me2:H4−4ac
H2A.Z:H4K20 me2
DNA Meth. m5C:H3K9 me3
H2A.Z:H3K4 me3
H3K9 me3:H4K20 me3
H3K27 me3:DNA Meth. m5C
H3K4 me3:H3−5ac
H4K20 me2:H3−5ac
H3K9 me2:DNA Meth. m5C
H4−4ac:H3−5ac
H4K20 me2:H3K4 me3
H3K27 me2
H3K9 me2
H3K4 me1
H3K9 me3
H3K4 me3
H4K20 me3
H4−4ac
H3K27 me3
DNA Meth. m5C
H2A.Z
H3−5ac
H4K20 me2
0.00
0.25
0.50
0.75
1.00
π̂
CACTIN – Forward
H2A.Z:H3−5ac
H2A.Z:H4−4ac
H3K4 me1:H3−5ac
H3K4 me1:H4−4ac
H3K4 me1:H4K20 me2
DNA Meth. m5C:H3K9 me3
H3K4 me1:H2A.Z
H3K9 me3:H4K20 me3
H3K27 me3:H3K4 me1
H3K27 me3:H3K9 me3
H3K27 me3:H4K20 me3
H3K27 me3:DNA Meth. m5C
H3K4 me3:H4−4ac
DNA Meth. m5C:H4K20 me3
H2A.Z:H3K4 me3
H4K20 me2:H3K4 me3
H4K20 me2:H4−4ac
H3K4 me3:H3−5ac
H4−4ac:H3−5ac
H2A.Z:H4K20 me2
H4K20 me2:H3−5ac
H3K27 me2
H3K9 me2
H4K20 me3
H3K4 me1
DNA Meth. m5C
H3K9 me3
H3K27 me3
H3−5ac
H4−4ac
H3K4 me3
H2A.Z
H4K20 me2
0.00
0.25
0.50
0.75
1.00
π̂
CACTIN – Reverse
H2A.Z:H3−5ac
H2A.Z:H4−4ac
H3K4 me1:H3−5ac
H3K4 me1:H4−4ac
H3K4 me1:H4K20 me2
DNA Meth. m5C:H3K9 me3
H3K4 me1:H2A.Z
H3K9 me3:H4K20 me3
H3K27 me3:H3K4 me1
H3K27 me3:H3K9 me3
H3K27 me3:H4K20 me3
H3K27 me3:DNA Meth. m5C
H3K4 me3:H4−4ac
DNA Meth. m5C:H4K20 me3
H2A.Z:H3K4 me3
H4K20 me2:H3K4 me3
H4K20 me2:H4−4ac
H3K4 me3:H3−5ac
H4−4ac:H3−5ac
H2A.Z:H4K20 me2
H4K20 me2:H3−5ac
H3K27 me2
H3K9 me2
H4K20 me3
H3K4 me1
DNA Meth. m5C
H3K9 me3
H3K27 me3
H3−5ac
H4−4ac
H3K4 me3
H2A.Z
H4K20 me2
0.00
0.25
0.50
0.75
1.00
π̂
CWC27 – Forward
H3K27 me3:DNA Meth. m5C
H3K9 me3:H4K20 me3
DNA Meth. m5C:H3K9 me3
H3K27 me2:DNA Meth. m5C
H3K27 me3:H4K20 me3
H3K27 me3:H3K4 me1
H3K4 me1:H2A.Z
H3K4 me1:H4−4ac
H2A.Z:H3K4 me3
H3K4 me1:H3−5ac
H2A.Z:H4K20 me2
H3K4 me3:H3−5ac
H4K20 me2:H3K4 me3
H4K20 me2:H4−4ac
H4−4ac:H3−5ac
H4K20 me2:H3−5ac
H3K4 me3:H4−4ac
H3K9 me2
H3K27 me2
H3K9 me3
H4K20 me3
DNA Meth. m5C
H2A.Z
H3K4 me1
H3K27 me3
H3−5ac
H3K4 me3
H4−4ac
H4K20 me2
0.00
0.25
0.50
0.75
1.00
π̂
CWC27 – Reverse
H3K27 me3:DNA Meth. m5C
H3K9 me3:H4K20 me3
DNA Meth. m5C:H3K9 me3
H3K27 me2:DNA Meth. m5C
H3K27 me3:H4K20 me3
H3K27 me3:H3K4 me1
H3K4 me1:H2A.Z
H3K4 me1:H4−4ac
H2A.Z:H3K4 me3
H3K4 me1:H3−5ac
H2A.Z:H4K20 me2
H3K4 me3:H3−5ac
H4K20 me2:H3K4 me3
H4K20 me2:H4−4ac
H4−4ac:H3−5ac
H4K20 me2:H3−5ac
H3K4 me3:H4−4ac
H3K9 me2
H3K27 me2
H3K9 me3
H4K20 me3
DNA Meth. m5C
H2A.Z
H3K4 me1
H3K27 me3
H3−5ac
H3K4 me3
H4−4ac
H4K20 me2
0.00
0.25
0.50
0.75
1.00
π̂
PPIL1 – Forward
H4K20 me2:H4−4ac
H3K4 me1:H4K20 me2
DNA Meth. m5C:H4K20 me2
H3K27 me3:H3K9 me3
DNA Meth. m5C:H3K9 me3
H3K4 me3:H4−4ac
H2A.Z:H4K20 me2
H4K20 me2:H3−5ac
H4−4ac:H3−5ac
H3K9 me2:DNA Meth. m5C
H4K20 me2:H3K4 me3
H4K20 me3
H3K27 me2
H3K4 me3
H3K9 me2
H3K27 me3
H3K9 me3
DNA Meth. m5C
H3K4 me1
H4K20 me2
H2A.Z
H4−4ac
H3−5ac
0.00
0.25
0.50
0.75
1.00
π̂
PPIL1 – Reverse
H4K20 me2:H4−4ac
H3K4 me1:H4K20 me2
DNA Meth. m5C:H4K20 me2
H3K27 me3:H3K9 me3
DNA Meth. m5C:H3K9 me3
H3K4 me3:H4−4ac
H2A.Z:H4K20 me2
H4K20 me2:H3−5ac
H4−4ac:H3−5ac
H3K9 me2:DNA Meth. m5C
H4K20 me2:H3K4 me3
H4K20 me3
H3K27 me2
H3K4 me3
H3K9 me2
H3K27 me3
H3K9 me3
DNA Meth. m5C
H3K4 me1
H4K20 me2
H2A.Z
H4−4ac
H3−5ac
0.00
0.25
0.50
0.75
1.00
π̂

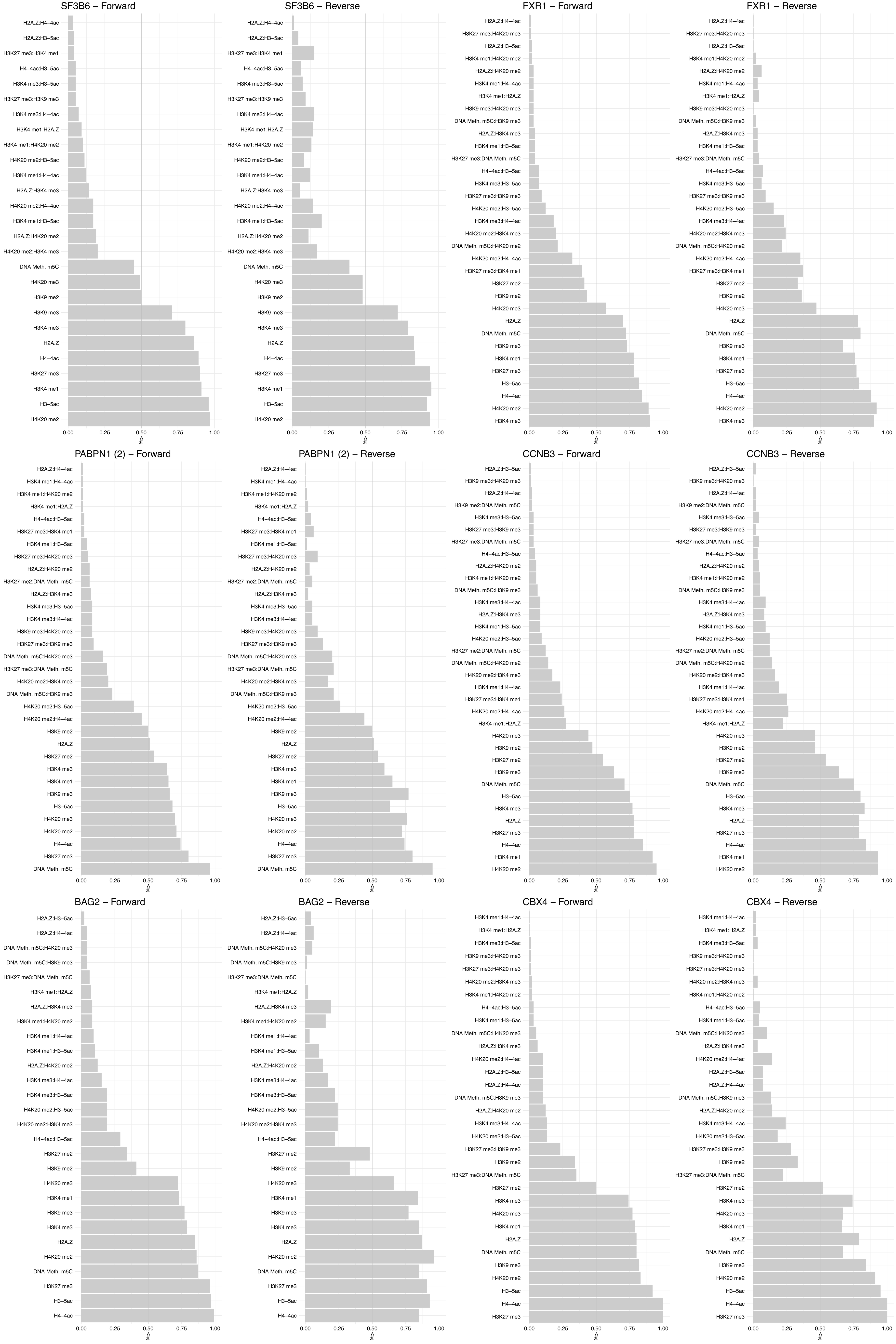
SF3B6 – Forward
H2A.Z:H4–4ac
H2A.Z:H3–5ac
H3K27 me3:H3K4 me1
H4–4ac:H3–5ac
H3K4 me3:H3–5ac
H3K27 me3:H3K9 me3
H3K4 me3:H4–4ac
H3K4 me1:H2A.Z
H3K4 me1:H4K20 me2
H4K20 me2:H3–5ac
H3K4 me1:H4–4ac
H2A.Z:H3K4 me3
H4K20 me2:H4–4ac
H3K4 me1:H3–5ac
H2A.Z:H4K20 me2
H4K20 me2:H3K4 me3
DNA Meth. m5C
H4K20 me3
H3K9 me2
H3K9 me3
H3K4 me3
H2A.Z
H4–4ac
H3K27 me3
H3K4 me1
H3–5ac
H4K20 me2
0.00
0.25
0.50
0.75
1.00
$\hat{\pi}$
SF3B6 – Reverse
H2A.Z:H4–4ac
H2A.Z:H3–5ac
H3K27 me3:H3K4 me1
H4–4ac:H3–5ac
H3K4 me3:H3–5ac
H3K27 me3:H3K9 me3
H3K4 me3:H4–4ac
H3K4 me1:H2A.Z
H3K4 me1:H4K20 me2
H4K20 me2:H3–5ac
H3K4 me1:H4–4ac
H2A.Z:H3K4 me3
H4K20 me2:H4–4ac
H3K4 me1:H3–5ac
H2A.Z:H4K20 me2
H4K20 me2:H3K4 me3
DNA Meth. m5C
H4K20 me3
H3K9 me2
H3K9 me3
H3K4 me3
H2A.Z
H4–4ac
H3K27 me3
H3K4 me1
H3–5ac
H4K20 me2
0.00
0.25
0.50
0.75
1.00
$\hat{\pi}$
FXR1 – Forward
H2A.Z:H4–4ac
H3K27 me3:H4K20 me3
H2A.Z:H3–5ac
H3K4 me1:H4K20 me2
H2A.Z:H4K20 me2
H3K4 me1:H4–4ac
H3K4 me1:H2A.Z
H3K9 me3:H4K20 me3
DNA Meth. m5C:H3K9 me3
H2A.Z:H3K4 me3
H3K4 me1:H3–5ac
H3K27 me3:DNA Meth. m5C
H4–4ac:H3–5ac
H3K4 me3:H3–5ac
H3K27 me3:H3K9 me3
H4K20 me2:H3–5ac
H3K4 me3:H4–4ac
H4K20 me2:H3K4 me3
DNA Meth. m5C:H4K20 me2
H4K20 me2:H4–4ac
H3K27 me3:H3K4 me1
H3K27 me2
H3K9 me2
H4K20 me3
H2A.Z
DNA Meth. m5C
H3K9 me3
H3K4 me1
H3K27 me3
H3–5ac
H4–4ac
H4K20 me2
H3K4 me3
0.00
0.25
0.50
0.75
1.00
$\hat{\pi}$
FXR1 – Reverse
H2A.Z:H4–4ac
H3K27 me3:H4K20 me3
H2A.Z:H3–5ac
H3K4 me1:H4K20 me2
H2A.Z:H4K20 me2
H3K4 me1:H4–4ac
H3K4 me1:H2A.Z
H3K9 me3:H4K20 me3
DNA Meth. m5C:H3K9 me3
H2A.Z:H3K4 me3
H3K4 me1:H3–5ac
H3K27 me3:DNA Meth. m5C
H4–4ac:H3–5ac
H3K4 me3:H3–5ac
H3K27 me3:H3K9 me3
H4K20 me2:H3–5ac
H3K4 me3:H4–4ac
H4K20 me2:H3K4 me3
DNA Meth. m5C:H4K20 me2
H4K20 me2:H4–4ac
H3K27 me3:H3K4 me1
H3K27 me2
H3K9 me2
H4K20 me3
H2A.Z
DNA Meth. m5C
H3K9 me3
H3K4 me1
H3K27 me3
H3–5ac
H4–4ac
H4K20 me2
H3K4 me3
0.00
0.25
0.50
0.75
1.00
$\hat{\pi}$
PABPN1 (2) – Forward
H2A.Z:H4–4ac
H3K4 me1:H4–4ac
H3K4 me1:H4K20 me2
H3K4 me1:H2A.Z
H4–4ac:H3–5ac
H3K27 me3:H3K4 me1
H3K4 me1:H3–5ac
H3K27 me3:H4K20 me3
H2A.Z:H4K20 me2
H3K27 me2:DNA Meth. m5C
H2A.Z:H3K4 me3
H3K4 me3:H3–5ac
H3K4 me3:H4–4ac
H3K9 me3:H4K20 me3
H3K27 me3:H3K9 me3
DNA Meth. m5C:H4K20 me3
H3K27 me3:DNA Meth. m5C
H4K20 me2:H3K4 me3
DNA Meth. m5C:H3K9 me3
H4K20 me2:H3–5ac
H4K20 me2:H4–4ac
H3K9 me2
H2A.Z
H3K27 me2
H3K4 me3
H3K4 me1
H3K9 me3
H3–5ac
H4K20 me3
H4K20 me2
H4–4ac
H3K27 me3
DNA Meth. m5C
0.00
0.25
0.50
0.75
1.00
$\hat{\pi}$
PABPN1 (2) – Reverse
H2A.Z:H4–4ac
H3K4 me1:H4–4ac
H3K4 me1:H4K20 me2
H3K4 me1:H2A.Z
H4–4ac:H3–5ac
H3K27 me3:H3K4 me1
H3K4 me1:H3–5ac
H3K27 me3:H4K20 me3
H2A.Z:H4K20 me2
H3K27 me2:DNA Meth. m5C
H2A.Z:H3K4 me3
H3K4 me3:H3–5ac
H3K4 me3:H4–4ac
H3K9 me3:H4K20 me3
H3K27 me3:H3K9 me3
DNA Meth. m5C:H4K20 me3
H3K27 me3:DNA Meth. m5C
H4K20 me2:H3K4 me3
DNA Meth. m5C:H3K9 me3
H4K20 me2:H3–5ac
H4K20 me2:H4–4ac
H3K9 me2
H2A.Z
H3K27 me2
H3K4 me3
H3K4 me1
H3K9 me3
H3–5ac
H4K20 me3
H4K20 me2
H4–4ac
H3K27 me3
DNA Meth. m5C
0.00
0.25
0.50
0.75
1.00
$\hat{\pi}$
CCNB3 – Forward
H2A.Z:H3–5ac
H3K9 me3:H4K20 me3
H2A.Z:H4–4ac
H3K9 me2:DNA Meth. m5C
H3K4 me3:H3–5ac
H3K27 me3:H3K9 me3
H3K27 me3:DNA Meth. m5C
H4–4ac:H3–5ac
H2A.Z:H4K20 me2
H3K4 me1:H4K20 me2
DNA Meth. m5C:H3K9 me3
H3K4 me3:H4–4ac
H2A.Z:H3K4 me3
H3K4 me1:H3–5ac
H4K20 me2:H3–5ac
H3K27 me2:DNA Meth. m5C
DNA Meth. m5C:H4K20 me2
H4K20 me2:H3K4 me3
H3K4 me1:H4–4ac
H3K27 me3:H3K4 me1
H4K20 me2:H4–4ac
H3K4 me1:H2A.Z
H4K20 me3
H3K9 me2
H3K27 me2
H3K9 me3
DNA Meth. m5C
H3–5ac
H3K4 me3
H2A.Z
H3K27 me3
H4–4ac
H3K4 me1
H4K20 me2
0.00
0.25
0.50
0.75
1.00
$\hat{\pi}$
CCNB3 – Reverse
H2A.Z:H3–5ac
H3K9 me3:H4K20 me3
H2A.Z:H4–4ac
H3K9 me2:DNA Meth. m5C
H3K4 me3:H3–5ac
H3K27 me3:H3K9 me3
H3K27 me3:DNA Meth. m5C
H4–4ac:H3–5ac
H2A.Z:H4K20 me2
H3K4 me1:H4K20 me2
DNA Meth. m5C:H3K9 me3
H3K4 me3:H4–4ac
H2A.Z:H3K4 me3
H3K4 me1:H3–5ac
H4K20 me2:H3–5ac
H3K27 me2:DNA Meth. m5C
DNA Meth. m5C:H4K20 me2
H4K20 me2:H3K4 me3
H3K4 me1:H4–4ac
H3K27 me3:H3K4 me1
H4K20 me2:H4–4ac
H3K4 me1:H2A.Z
H4K20 me3
H3K9 me2
H3K27 me2
H3K9 me3
DNA Meth. m5C
H3–5ac
H3K4 me3
H2A.Z
H3K27 me3
H4–4ac
H3K4 me1
H4K20 me2
0.00
0.25
0.50
0.75
1.00
$\hat{\pi}$
BAG2 – Forward
H2A.Z:H3–5ac
H2A.Z:H4–4ac
DNA Meth. m5C:H4K20 me3
DNA Meth. m5C:H3K9 me3
H3K27 me3:DNA Meth. m5C
H3K4 me1:H2A.Z
H2A.Z:H3K4 me3
H3K4 me1:H4K20 me2
H3K4 me1:H4–4ac
H3K4 me1:H3–5ac
H2A.Z:H4K20 me2
H3K4 me3:H4–4ac
H3K4 me3:H3–5ac
H4K20 me2:H3–5ac
H4K20 me2:H3K4 me3
H4–4ac:H3–5ac
H3K27 me2
H3K9 me2
H4K20 me3
H3K4 me1
H3K9 me3
H3K4 me3
H2A.Z
H4K20 me2
DNA Meth. m5C
H3K27 me3
H3–5ac
H4–4ac
0.00
0.25
0.50
0.75
1.00
$\hat{\pi}$
BAG2 – Reverse
H2A.Z:H3–5ac
H2A.Z:H4–4ac
DNA Meth. m5C:H4K20 me3
DNA Meth. m5C:H3K9 me3
H3K27 me3:DNA Meth. m5C
H3K4 me1:H2A.Z
H2A.Z:H3K4 me3
H3K4 me1:H4K20 me2
H3K4 me1:H4–4ac
H3K4 me1:H3–5ac
H2A.Z:H4K20 me2
H3K4 me3:H4–4ac
H3K4 me3:H3–5ac
H4K20 me2:H3–5ac
H4K20 me2:H3K4 me3
H4–4ac:H3–5ac
H3K27 me2
H3K9 me2
H4K20 me3
H3K4 me1
H3K9 me3
H3K4 me3
H2A.Z
H4K20 me2
DNA Meth. m5C
H3K27 me3
H3–5ac
H4–4ac
0.00
0.25
0.50
0.75
1.00
$\hat{\pi}$
CBX4 – Forward
H3K4 me1:H4–4ac
H3K4 me1:H2A.Z
H3K4 me3:H3–5ac
H3K9 me3:H4K20 me3
H3K27 me3:H4K20 me3
H4K20 me2:H3K4 me3
H3K4 me1:H4K20 me2
H4–4ac:H3–5ac
H3K4 me1:H3–5ac
DNA Meth. m5C:H4K20 me3
H2A.Z:H3K4 me3
H4K20 me2:H4–4ac
H2A.Z:H3–5ac
H2A.Z:H4–4ac
DNA Meth. m5C:H3K9 me3
H2A.Z:H4K20 me2
H3K4 me3:H4–4ac
H4K20 me2:H3–5ac
H3K27 me3:H3K9 me3
H3K9 me2
H3K27 me3:DNA Meth. m5C
H3K27 me2
H3K4 me3
H4K20 me3
H3K4 me1
H2A.Z
DNA Meth. m5C
H3K9 me3
H4K20 me2
H3–5ac
H4–4ac
H3K27 me3
0.00
0.25
0.50
0.75
1.00
$\hat{\pi}$
CBX4 – Reverse
H3K4 me1:H4–4ac
H3K4 me1:H2A.Z
H3K4 me3:H3–5ac
H3K9 me3:H4K20 me3
H3K27 me3:H4K20 me3
H4K20 me2:H3K4 me3
H3K4 me1:H4K20 me2
H4–4ac:H3–5ac
H3K4 me1:H3–5ac
DNA Meth. m5C:H4K20 me3
H2A.Z:H3K4 me3
H4K20 me2:H4–4ac
H2A.Z:H3–5ac
H2A.Z:H4–4ac
DNA Meth. m5C:H3K9 me3
H2A.Z:H4K20 me2
H3K4 me3:H4–4ac
H4K20 me2:H3–5ac
H3K27 me3:H3K9 me3
H3K9 me2
H3K27 me3:DNA Meth. m5C
H3K27 me2
H3K4 me3
H4K20 me3
H3K4 me1
H2A.Z
DNA Meth. m5C
H3K9 me3
H4K20 me2
H3–5ac
H4–4ac
H3K27 me3
0.00
0.25
0.50
0.75
1.00
$\hat{\pi}$

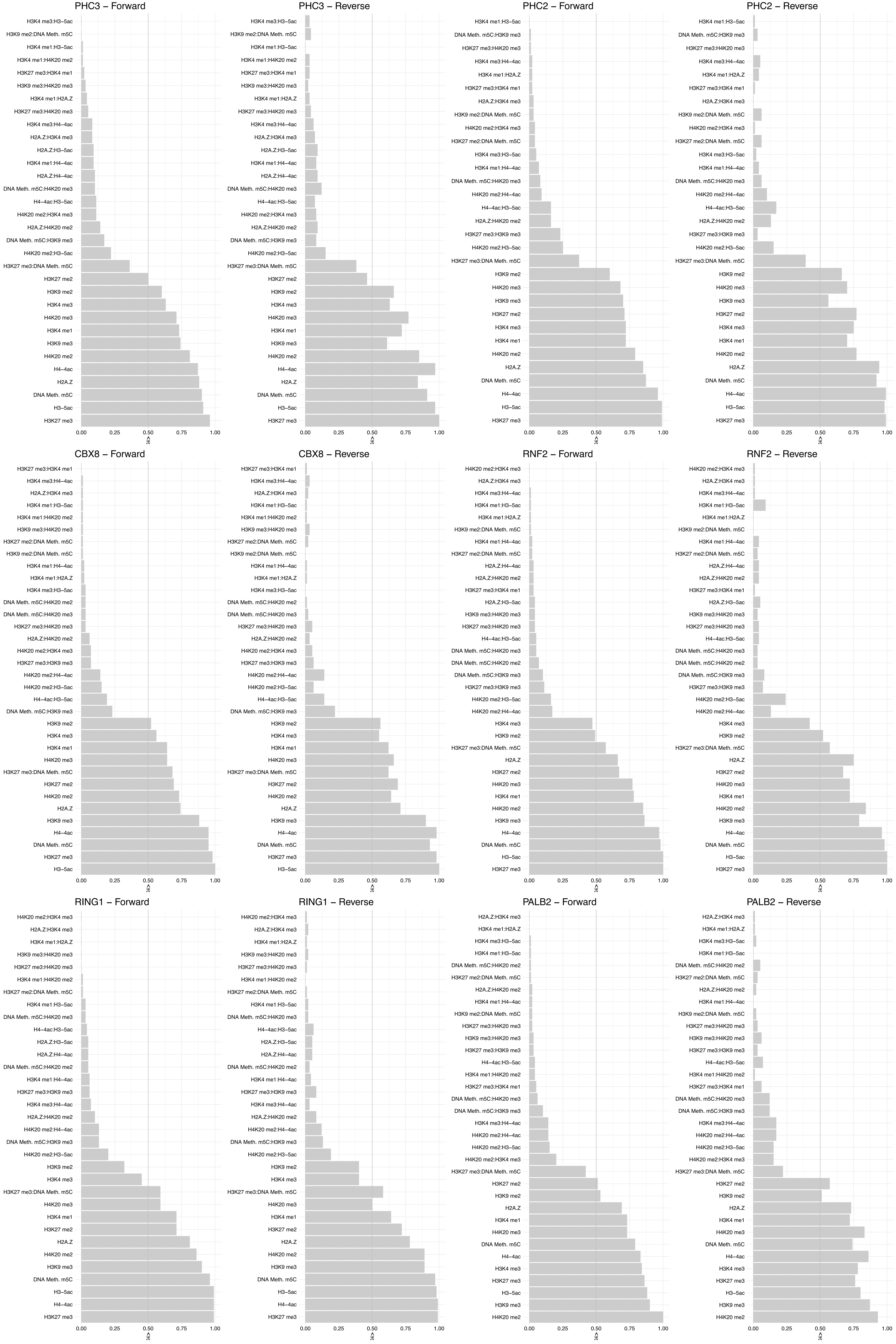
PHC3 – Forward
H3K4 me3:H3−5ac
H3K9 me2:DNA Meth. m5C
H3K4 me1:H3−5ac
H3K4 me1:H4K20 me2
H3K27 me3:H3K4 me1
H3K9 me3:H4K20 me3
H3K4 me1:H2A.Z
H3K27 me3:H4K20 me3
H3K4 me3:H4−4ac
H2A.Z:H3K4 me3
H2A.Z:H3−5ac
H3K4 me1:H4−4ac
H2A.Z:H4−4ac
DNA Meth. m5C:H4K20 me3
H4−4ac:H3−5ac
H4K20 me2:H3K4 me3
H2A.Z:H4K20 me2
DNA Meth. m5C:H3K9 me3
H4K20 me2:H3−5ac
H3K27 me3:DNA Meth. m5C
H3K27 me2
H3K9 me2
H3K4 me3
H4K20 me3
H3K4 me1
H3K9 me3
H4K20 me2
H4−4ac
H2A.Z
DNA Meth. m5C
H3−5ac
H3K27 me3
0.00 0.25 0.50 0.75 1.00
π̂
PHC3 – Reverse
H3K4 me3:H3−5ac
H3K9 me2:DNA Meth. m5C
H3K4 me1:H3−5ac
H3K4 me1:H4K20 me2
H3K27 me3:H3K4 me1
H3K9 me3:H4K20 me3
H3K4 me1:H2A.Z
H3K27 me3:H4K20 me3
H3K4 me3:H4−4ac
H2A.Z:H3K4 me3
H2A.Z:H3−5ac
H3K4 me1:H4−4ac
H2A.Z:H4−4ac
DNA Meth. m5C:H4K20 me3
H4−4ac:H3−5ac
H4K20 me2:H3K4 me3
H2A.Z:H4K20 me2
DNA Meth. m5C:H3K9 me3
H4K20 me2:H3−5ac
H3K27 me3:DNA Meth. m5C
H3K27 me2
H3K9 me2
H3K4 me3
H4K20 me3
H3K4 me1
H3K9 me3
H4K20 me2
H4−4ac
H2A.Z
DNA Meth. m5C
H3−5ac
H3K27 me3
0.00 0.25 0.50 0.75 1.00
π̂
PHC2 – Forward
H3K4 me3:H3−5ac
DNA Meth. m5C:H3K9 me3
H3K27 me3:H4K20 me3
H3K4 me3:H4−4ac
H3K4 me1:H2A.Z
H3K27 me3:H3K4 me1
H2A.Z:H3K4 me3
H3K9 me2:DNA Meth. m5C
H4K20 me2:H3K4 me3
H3K27 me2:DNA Meth. m5C
H3K4 me3:H3−5ac
H3K4 me1:H4−4ac
DNA Meth. m5C:H4K20 me3
H4K20 me2:H4−4ac
H4−4ac:H3−5ac
H2A.Z:H4K20 me2
H3K27 me3:H3K9 me3
H4K20 me2:H3−5ac
H3K27 me3:DNA Meth. m5C
H3K9 me2
H4K20 me3
H3K9 me3
H3K27 me2
H3K4 me3
H3K4 me1
H4K20 me2
H2A.Z
DNA Meth. m5C
H4−4ac
H3−5ac
H3K27 me3
0.00 0.25 0.50 0.75 1.00
π̂
PHC2 – Reverse
H3K4 me3:H3−5ac
DNA Meth. m5C:H3K9 me3
H3K27 me3:H4K20 me3
H3K4 me3:H4−4ac
H3K4 me1:H2A.Z
H3K27 me3:H3K4 me1
H2A.Z:H3K4 me3
H3K9 me2:DNA Meth. m5C
H4K20 me2:H3K4 me3
H3K27 me2:DNA Meth. m5C
H3K4 me3:H3−5ac
H3K4 me1:H4−4ac
DNA Meth. m5C:H4K20 me3
H4K20 me2:H4−4ac
H4−4ac:H3−5ac
H2A.Z:H4K20 me2
H3K27 me3:H3K9 me3
H4K20 me2:H3−5ac
H3K27 me3:DNA Meth. m5C
H3K9 me2
H4K20 me3
H3K9 me3
H3K27 me2
H3K4 me3
H3K4 me1
H4K20 me2
H2A.Z
DNA Meth. m5C
H4−4ac
H3−5ac
H3K27 me3
0.00 0.25 0.50 0.75 1.00
π̂
CBX8 – Forward
H3K27 me3:H3K4 me1
H3K4 me3:H4−4ac
H2A.Z:H3K4 me3
H3K4 me1:H3−5ac
H3K4 me1:H4K20 me2
H3K9 me3:H4K20 me3
H3K27 me2:DNA Meth. m5C
H3K9 me2:DNA Meth. m5C
H3K4 me1:H4−4ac
H3K4 me1:H2A.Z
H3K4 me3:H3−5ac
DNA Meth. m5C:H4K20 me2
DNA Meth. m5C:H4K20 me3
H3K27 me3:H4K20 me3
H2A.Z:H4K20 me2
H4K20 me2:H3K4 me3
H3K27 me3:H3K9 me3
H4K20 me2:H4−4ac
H4K20 me2:H3−5ac
H4−4ac:H3−5ac
DNA Meth. m5C:H3K9 me3
H3K9 me2
H3K4 me3
H3K4 me1
H4K20 me3
H3K27 me3:DNA Meth. m5C
H3K27 me2
H4K20 me2
H2A.Z
H3K9 me3
H4−4ac
DNA Meth. m5C
H3K27 me3
H3−5ac
0.00 0.25 0.50 0.75 1.00
π̂
CBX8 – Reverse
H3K27 me3:H3K4 me1
H3K4 me3:H4−4ac
H2A.Z:H3K4 me3
H3K4 me1:H3−5ac
H3K4 me1:H4K20 me2
H3K9 me3:H4K20 me3
H3K27 me2:DNA Meth. m5C
H3K9 me2:DNA Meth. m5C
H3K4 me1:H4−4ac
H3K4 me1:H2A.Z
H3K4 me3:H3−5ac
DNA Meth. m5C:H4K20 me2
DNA Meth. m5C:H4K20 me3
H3K27 me3:H4K20 me3
H2A.Z:H4K20 me2
H4K20 me2:H3K4 me3
H3K27 me3:H3K9 me3
H4K20 me2:H4−4ac
H4K20 me2:H3−5ac
H4−4ac:H3−5ac
DNA Meth. m5C:H3K9 me3
H3K9 me2
H3K4 me3
H3K4 me1
H4K20 me3
H3K27 me3:DNA Meth. m5C
H3K27 me2
H4K20 me2
H2A.Z
H3K9 me3
H4−4ac
DNA Meth. m5C
H3K27 me3
H3−5ac
0.00 0.25 0.50 0.75 1.00
π̂
RNF2 – Forward
H4K20 me2:H3K4 me3
H2A.Z:H3K4 me3
H3K4 me3:H4−4ac
H3K4 me1:H3−5ac
H3K4 me1:H2A.Z
H3K9 me2:DNA Meth. m5C
H3K4 me1:H4−4ac
H3K27 me2:DNA Meth. m5C
H2A.Z:H4−4ac
H2A.Z:H4K20 me2
H3K27 me3:H3K4 me1
H2A.Z:H3−5ac
H3K9 me3:H4K20 me3
H3K27 me3:H4K20 me3
H4−4ac:H3−5ac
DNA Meth. m5C:H4K20 me3
DNA Meth. m5C:H4K20 me2
DNA Meth. m5C:H3K9 me3
H3K27 me3:H3K9 me3
H4K20 me2:H3−5ac
H4K20 me2:H4−4ac
H3K4 me3
H3K9 me2
H3K27 me3:DNA Meth. m5C
H2A.Z
H3K27 me2
H4K20 me3
H3K4 me1
H4K20 me2
H3K9 me3
H4−4ac
DNA Meth. m5C
H3−5ac
H3K27 me3
0.00 0.25 0.50 0.75 1.00
π̂
RNF2 – Reverse
H4K20 me2:H3K4 me3
H2A.Z:H3K4 me3
H3K4 me3:H4−4ac
H3K4 me1:H3−5ac
H3K4 me1:H2A.Z
H3K9 me2:DNA Meth. m5C
H3K4 me1:H4−4ac
H3K27 me2:DNA Meth. m5C
H2A.Z:H4−4ac
H2A.Z:H4K20 me2
H3K27 me3:H3K4 me1
H2A.Z:H3−5ac
H3K9 me3:H4K20 me3
H3K27 me3:H4K20 me3
H4−4ac:H3−5ac
DNA Meth. m5C:H4K20 me3
DNA Meth. m5C:H4K20 me2
DNA Meth. m5C:H3K9 me3
H3K27 me3:H3K9 me3
H4K20 me2:H3−5ac
H4K20 me2:H4−4ac
H3K4 me3
H3K9 me2
H3K27 me3:DNA Meth. m5C
H2A.Z
H3K27 me2
H4K20 me3
H3K4 me1
H4K20 me2
H3K9 me3
H4−4ac
DNA Meth. m5C
H3−5ac
H3K27 me3
0.00 0.25 0.50 0.75 1.00
π̂
RING1 – Forward
H4K20 me2:H3K4 me3
H2A.Z:H3K4 me3
H3K4 me1:H2A.Z
H3K9 me3:H4K20 me3
H3K27 me3:H4K20 me3
H3K4 me1:H4K20 me2
H3K27 me2:DNA Meth. m5C
H3K4 me1:H3−5ac
DNA Meth. m5C:H4K20 me3
H4−4ac:H3−5ac
H2A.Z:H3−5ac
H2A.Z:H4−4ac
DNA Meth. m5C:H4K20 me2
H3K4 me1:H4−4ac
H3K27 me3:H3K9 me3
H3K4 me3:H4−4ac
H2A.Z:H4K20 me2
H4K20 me2:H4−4ac
DNA Meth. m5C:H3K9 me3
H4K20 me2:H3−5ac
H3K9 me2
H3K4 me3
H3K27 me3:DNA Meth. m5C
H4K20 me3
H3K4 me1
H3K27 me2
H2A.Z
H4K20 me2
H3K9 me3
DNA Meth. m5C
H3−5ac
H4−4ac
H3K27 me3
0.00 0.25 0.50 0.75 1.00
π̂
RING1 – Reverse
H4K20 me2:H3K4 me3
H2A.Z:H3K4 me3
H3K4 me1:H2A.Z
H3K9 me3:H4K20 me3
H3K27 me3:H4K20 me3
H3K4 me1:H4K20 me2
H3K27 me2:DNA Meth. m5C
H3K4 me1:H3−5ac
DNA Meth. m5C:H4K20 me3
H4−4ac:H3−5ac
H2A.Z:H3−5ac
H2A.Z:H4−4ac
DNA Meth. m5C:H4K20 me2
H3K4 me1:H4−4ac
H3K27 me3:H3K9 me3
H3K4 me3:H4−4ac
H2A.Z:H4K20 me2
H4K20 me2:H4−4ac
DNA Meth. m5C:H3K9 me3
H4K20 me2:H3−5ac
H3K9 me2
H3K4 me3
H3K27 me3:DNA Meth. m5C
H4K20 me3
H3K4 me1
H3K27 me2
H2A.Z
H4K20 me2
H3K9 me3
DNA Meth. m5C
H3−5ac
H4−4ac
H3K27 me3
0.00 0.25 0.50 0.75 1.00
π̂
PALB2 – Forward
H2A.Z:H3K4 me3
H3K4 me1:H2A.Z
H3K4 me3:H3−5ac
H3K4 me1:H3−5ac
DNA Meth. m5C:H4K20 me2
H3K27 me2:DNA Meth. m5C
H2A.Z:H4K20 me2
H3K4 me1:H4−4ac
H3K9 me2:DNA Meth. m5C
H3K27 me3:H4K20 me3
H3K9 me3:H4K20 me3
H3K27 me3:H3K9 me3
H4−4ac:H3−5ac
H3K4 me1:H4K20 me2
H3K27 me3:H3K4 me1
DNA Meth. m5C:H4K20 me3
DNA Meth. m5C:H3K9 me3
H3K4 me3:H4−4ac
H4K20 me2:H4−4ac
H4K20 me2:H3−5ac
H4K20 me2:H3K4 me3
H3K27 me3:DNA Meth. m5C
H3K27 me2
H3K9 me2
H2A.Z
H3K4 me1
H4K20 me3
DNA Meth. m5C
H4−4ac
H3K4 me3
H3K27 me3
H3−5ac
H3K9 me3
H4K20 me2
0.00 0.25 0.50 0.75 1.00
π̂
PALB2 – Reverse
H2A.Z:H3K4 me3
H3K4 me1:H2A.Z
H3K4 me3:H3−5ac
H3K4 me1:H3−5ac
DNA Meth. m5C:H4K20 me2
H3K27 me2:DNA Meth. m5C
H2A.Z:H4K20 me2
H3K4 me1:H4−4ac
H3K9 me2:DNA Meth. m5C
H3K27 me3:H4K20 me3
H3K9 me3:H4K20 me3
H3K27 me3:H3K9 me3
H4−4ac:H3−5ac
H3K4 me1:H4K20 me2
H3K27 me3:H3K4 me1
DNA Meth. m5C:H4K20 me3
DNA Meth. m5C:H3K9 me3
H3K4 me3:H4−4ac
H4K20 me2:H4−4ac
H4K20 me2:H3−5ac
H4K20 me2:H3K4 me3
H3K27 me3:DNA Meth. m5C
H3K27 me2
H3K9 me2
H2A.Z
H3K4 me1
H4K20 me3
DNA Meth. m5C
H4−4ac
H3K4 me3
H3K27 me3
H3−5ac
H3K9 me3
H4K20 me2
0.00 0.25 0.50 0.75 1.00
π̂

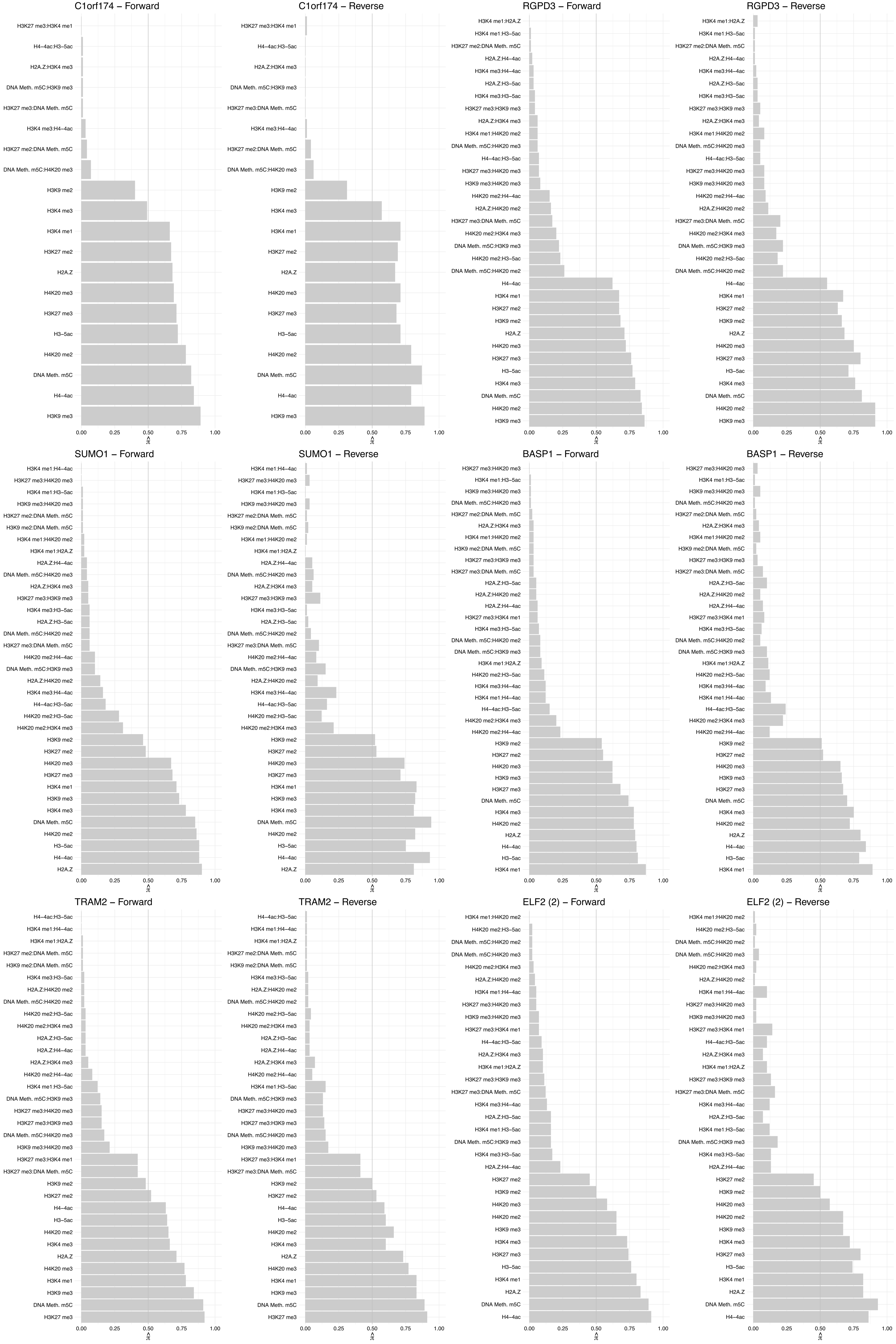
C1orf174 – Forward
H3K27 me3:H3K4 me1
H4−4ac:H3−5ac
H2A.Z:H3K4 me3
DNA Meth. m5C:H3K9 me3
H3K27 me3:DNA Meth. m5C
H3K4 me3:H4−4ac
H3K27 me2:DNA Meth. m5C
DNA Meth. m5C:H4K20 me3
H3K9 me2
H3K4 me3
H3K4 me1
H3K27 me2
H2A.Z
H4K20 me3
H3K27 me3
H3−5ac
H4K20 me2
DNA Meth. m5C
H4−4ac
H3K9 me3
0.00 0.25 0.50 0.75 1.00
π̂
C1orf174 – Reverse
H3K27 me3:H3K4 me1
H4−4ac:H3−5ac
H2A.Z:H3K4 me3
DNA Meth. m5C:H3K9 me3
H3K27 me3:DNA Meth. m5C
H3K4 me3:H4−4ac
H3K27 me2:DNA Meth. m5C
DNA Meth. m5C:H4K20 me3
H3K9 me2
H3K4 me3
H3K4 me1
H3K27 me2
H2A.Z
H4K20 me3
H3K27 me3
H3−5ac
H4K20 me2
DNA Meth. m5C
H4−4ac
H3K9 me3
0.00 0.25 0.50 0.75 1.00
π̂
RGPD3 – Forward
H3K4 me1:H2A.Z
H3K4 me1:H3−5ac
H3K27 me2:DNA Meth. m5C
H2A.Z:H4−4ac
H3K4 me3:H4−4ac
H2A.Z:H3−5ac
H3K4 me3:H3−5ac
H3K27 me3:H3K9 me3
H2A.Z:H3K4 me3
H3K4 me1:H4K20 me2
DNA Meth. m5C:H4K20 me3
H4−4ac:H3−5ac
H3K27 me3:H4K20 me3
H3K9 me3:H4K20 me3
H4K20 me2:H4−4ac
H2A.Z:H4K20 me2
H3K27 me3:DNA Meth. m5C
H4K20 me2:H3K4 me3
DNA Meth. m5C:H3K9 me3
H4K20 me2:H3−5ac
DNA Meth. m5C:H4K20 me2
H4−4ac
H3K4 me1
H3K27 me2
H3K9 me2
H2A.Z
H4K20 me3
H3K27 me3
H3−5ac
H3K4 me3
DNA Meth. m5C
H4K20 me2
H3K9 me3
0.00 0.25 0.50 0.75 1.00
π̂
RGPD3 – Reverse
H3K4 me1:H2A.Z
H3K4 me1:H3−5ac
H3K27 me2:DNA Meth. m5C
H2A.Z:H4−4ac
H3K4 me3:H4−4ac
H2A.Z:H3−5ac
H3K4 me3:H3−5ac
H3K27 me3:H3K9 me3
H2A.Z:H3K4 me3
H3K4 me1:H4K20 me2
DNA Meth. m5C:H4K20 me3
H4−4ac:H3−5ac
H3K27 me3:H4K20 me3
H3K9 me3:H4K20 me3
H4K20 me2:H4−4ac
H2A.Z:H4K20 me2
H3K27 me3:DNA Meth. m5C
H4K20 me2:H3K4 me3
DNA Meth. m5C:H3K9 me3
H4K20 me2:H3−5ac
DNA Meth. m5C:H4K20 me2
H4−4ac
H3K4 me1
H3K27 me2
H3K9 me2
H2A.Z
H4K20 me3
H3K27 me3
H3−5ac
H3K4 me3
DNA Meth. m5C
H4K20 me2
H3K9 me3
0.00 0.25 0.50 0.75 1.00
π̂
SUMO1 – Forward
H3K4 me1:H4−4ac
H3K27 me3:H4K20 me3
H3K4 me1:H3−5ac
H3K9 me3:H4K20 me3
H3K27 me2:DNA Meth. m5C
H3K9 me2:DNA Meth. m5C
H3K4 me1:H4K20 me2
H3K4 me1:H2A.Z
H2A.Z:H4−4ac
DNA Meth. m5C:H4K20 me3
H2A.Z:H3K4 me3
H3K27 me3:H3K9 me3
H3K4 me3:H3−5ac
H2A.Z:H3−5ac
DNA Meth. m5C:H4K20 me2
H3K27 me3:DNA Meth. m5C
H4K20 me2:H4−4ac
DNA Meth. m5C:H3K9 me3
H2A.Z:H4K20 me2
H3K4 me3:H4−4ac
H4−4ac:H3−5ac
H4K20 me2:H3−5ac
H4K20 me2:H3K4 me3
H3K9 me2
H3K27 me2
H4K20 me3
H3K27 me3
H3K4 me1
H3K9 me3
H3K4 me3
DNA Meth. m5C
H4K20 me2
H3−5ac
H4−4ac
H2A.Z
0.00 0.25 0.50 0.75 1.00
π̂
SUMO1 – Reverse
H3K4 me1:H4−4ac
H3K27 me3:H4K20 me3
H3K4 me1:H3−5ac
H3K9 me3:H4K20 me3
H3K27 me2:DNA Meth. m5C
H3K9 me2:DNA Meth. m5C
H3K4 me1:H4K20 me2
H3K4 me1:H2A.Z
H2A.Z:H4−4ac
DNA Meth. m5C:H4K20 me3
H2A.Z:H3K4 me3
H3K27 me3:H3K9 me3
H3K4 me3:H3−5ac
H2A.Z:H3−5ac
DNA Meth. m5C:H4K20 me2
H3K27 me3:DNA Meth. m5C
H4K20 me2:H4−4ac
DNA Meth. m5C:H3K9 me3
H2A.Z:H4K20 me2
H3K4 me3:H4−4ac
H4−4ac:H3−5ac
H4K20 me2:H3−5ac
H4K20 me2:H3K4 me3
H3K9 me2
H3K27 me2
H4K20 me3
H3K27 me3
H3K4 me1
H3K9 me3
H3K4 me3
DNA Meth. m5C
H4K20 me2
H3−5ac
H4−4ac
H2A.Z
0.00 0.25 0.50 0.75 1.00
π̂
BASP1 – Forward
H3K27 me3:H4K20 me3
H3K4 me1:H3−5ac
H3K9 me3:H4K20 me3
DNA Meth. m5C:H4K20 me3
H3K27 me2:DNA Meth. m5C
H2A.Z:H3K4 me3
H3K4 me1:H4K20 me2
H3K9 me2:DNA Meth. m5C
H3K27 me3:H3K9 me3
H3K27 me3:DNA Meth. m5C
H2A.Z:H3−5ac
H2A.Z:H4K20 me2
H2A.Z:H4−4ac
H3K27 me3:H3K4 me1
H3K4 me3:H3−5ac
DNA Meth. m5C:H4K20 me2
DNA Meth. m5C:H3K9 me3
H3K4 me1:H2A.Z
H4K20 me2:H3−5ac
H3K4 me3:H4−4ac
H3K4 me1:H4−4ac
H4−4ac:H3−5ac
H4K20 me2:H3K4 me3
H4K20 me2:H4−4ac
H3K9 me2
H3K27 me2
H4K20 me3
H3K9 me3
H3K27 me3
DNA Meth. m5C
H3K4 me3
H4K20 me2
H2A.Z
H4−4ac
H3−5ac
H3K4 me1
0.00 0.25 0.50 0.75 1.00
π̂
BASP1 – Reverse
H3K27 me3:H4K20 me3
H3K4 me1:H3−5ac
H3K9 me3:H4K20 me3
DNA Meth. m5C:H4K20 me3
H3K27 me2:DNA Meth. m5C
H2A.Z:H3K4 me3
H3K4 me1:H4K20 me2
H3K9 me2:DNA Meth. m5C
H3K27 me3:H3K9 me3
H3K27 me3:DNA Meth. m5C
H2A.Z:H3−5ac
H2A.Z:H4K20 me2
H2A.Z:H4−4ac
H3K27 me3:H3K4 me1
H3K4 me3:H3−5ac
DNA Meth. m5C:H4K20 me2
DNA Meth. m5C:H3K9 me3
H3K4 me1:H2A.Z
H4K20 me2:H3−5ac
H3K4 me3:H4−4ac
H3K4 me1:H4−4ac
H4−4ac:H3−5ac
H4K20 me2:H3K4 me3
H4K20 me2:H4−4ac
H3K9 me2
H3K27 me2
H4K20 me3
H3K9 me3
H3K27 me3
DNA Meth. m5C
H3K4 me3
H4K20 me2
H2A.Z
H4−4ac
H3−5ac
H3K4 me1
0.00 0.25 0.50 0.75 1.00
π̂
TRAM2 – Forward
H4−4ac:H3−5ac
H3K4 me1:H4−4ac
H3K4 me1:H2A.Z
H3K27 me2:DNA Meth. m5C
H3K9 me2:DNA Meth. m5C
H3K4 me3:H3−5ac
H2A.Z:H4K20 me2
DNA Meth. m5C:H4K20 me2
H4K20 me2:H3−5ac
H4K20 me2:H3K4 me3
H2A.Z:H3−5ac
H2A.Z:H4−4ac
H2A.Z:H3K4 me3
H4K20 me2:H4−4ac
H3K4 me1:H3−5ac
DNA Meth. m5C:H3K9 me3
H3K27 me3:H4K20 me3
H3K27 me3:H3K9 me3
DNA Meth. m5C:H4K20 me3
H3K9 me3:H4K20 me3
H3K27 me3:H3K4 me1
H3K27 me3:DNA Meth. m5C
H3K9 me2
H3K27 me2
H4−4ac
H3−5ac
H4K20 me2
H3K4 me3
H2A.Z
H4K20 me3
H3K4 me1
H3K9 me3
DNA Meth. m5C
H3K27 me3
0.00 0.25 0.50 0.75 1.00
π̂
TRAM2 – Reverse
H4−4ac:H3−5ac
H3K4 me1:H4−4ac
H3K4 me1:H2A.Z
H3K27 me2:DNA Meth. m5C
H3K9 me2:DNA Meth. m5C
H3K4 me3:H3−5ac
H2A.Z:H4K20 me2
DNA Meth. m5C:H4K20 me2
H4K20 me2:H3−5ac
H4K20 me2:H3K4 me3
H2A.Z:H3−5ac
H2A.Z:H4−4ac
H2A.Z:H3K4 me3
H4K20 me2:H4−4ac
H3K4 me1:H3−5ac
DNA Meth. m5C:H3K9 me3
H3K27 me3:H4K20 me3
H3K27 me3:H3K9 me3
DNA Meth. m5C:H4K20 me3
H3K9 me3:H4K20 me3
H3K27 me3:H3K4 me1
H3K27 me3:DNA Meth. m5C
H3K9 me2
H3K27 me2
H4−4ac
H3−5ac
H4K20 me2
H3K4 me3
H2A.Z
H4K20 me3
H3K4 me1
H3K9 me3
DNA Meth. m5C
H3K27 me3
0.00 0.25 0.50 0.75 1.00
π̂
ELF2 (2) – Forward
H3K4 me1:H4K20 me2
H4K20 me2:H3−5ac
DNA Meth. m5C:H4K20 me2
DNA Meth. m5C:H4K20 me3
H4K20 me2:H3K4 me3
H2A.Z:H4K20 me2
H3K4 me1:H4−4ac
H3K27 me3:H4K20 me3
H3K9 me3:H4K20 me3
H3K27 me3:H3K4 me1
H4−4ac:H3−5ac
H2A.Z:H3K4 me3
H3K4 me1:H2A.Z
H3K27 me3:H3K9 me3
H3K27 me3:DNA Meth. m5C
H3K4 me3:H4−4ac
H2A.Z:H3−5ac
H3K4 me1:H3−5ac
DNA Meth. m5C:H3K9 me3
H3K4 me3:H3−5ac
H2A.Z:H4−4ac
H3K27 me2
H3K9 me2
H4K20 me3
H4K20 me2
H3K9 me3
H3K4 me3
H3K27 me3
H3−5ac
H3K4 me1
H2A.Z
DNA Meth. m5C
H4−4ac
0.00 0.25 0.50 0.75 1.00
π̂
ELF2 (2) – Reverse
H3K4 me1:H4K20 me2
H4K20 me2:H3−5ac
DNA Meth. m5C:H4K20 me2
DNA Meth. m5C:H4K20 me3
H4K20 me2:H3K4 me3
H2A.Z:H4K20 me2
H3K4 me1:H4−4ac
H3K27 me3:H4K20 me3
H3K9 me3:H4K20 me3
H3K27 me3:H3K4 me1
H4−4ac:H3−5ac
H2A.Z:H3K4 me3
H3K4 me1:H2A.Z
H3K27 me3:H3K9 me3
H3K27 me3:DNA Meth. m5C
H3K4 me3:H4−4ac
H2A.Z:H3−5ac
H3K4 me1:H3−5ac
DNA Meth. m5C:H3K9 me3
H3K4 me3:H3−5ac
H2A.Z:H4−4ac
H3K27 me2
H3K9 me2
H4K20 me3
H4K20 me2
H3K9 me3
H3K4 me3
H3K27 me3
H3−5ac
H3K4 me1
H2A.Z
DNA Meth. m5C
H4−4ac
0.00 0.25 0.50 0.75 1.00
π̂

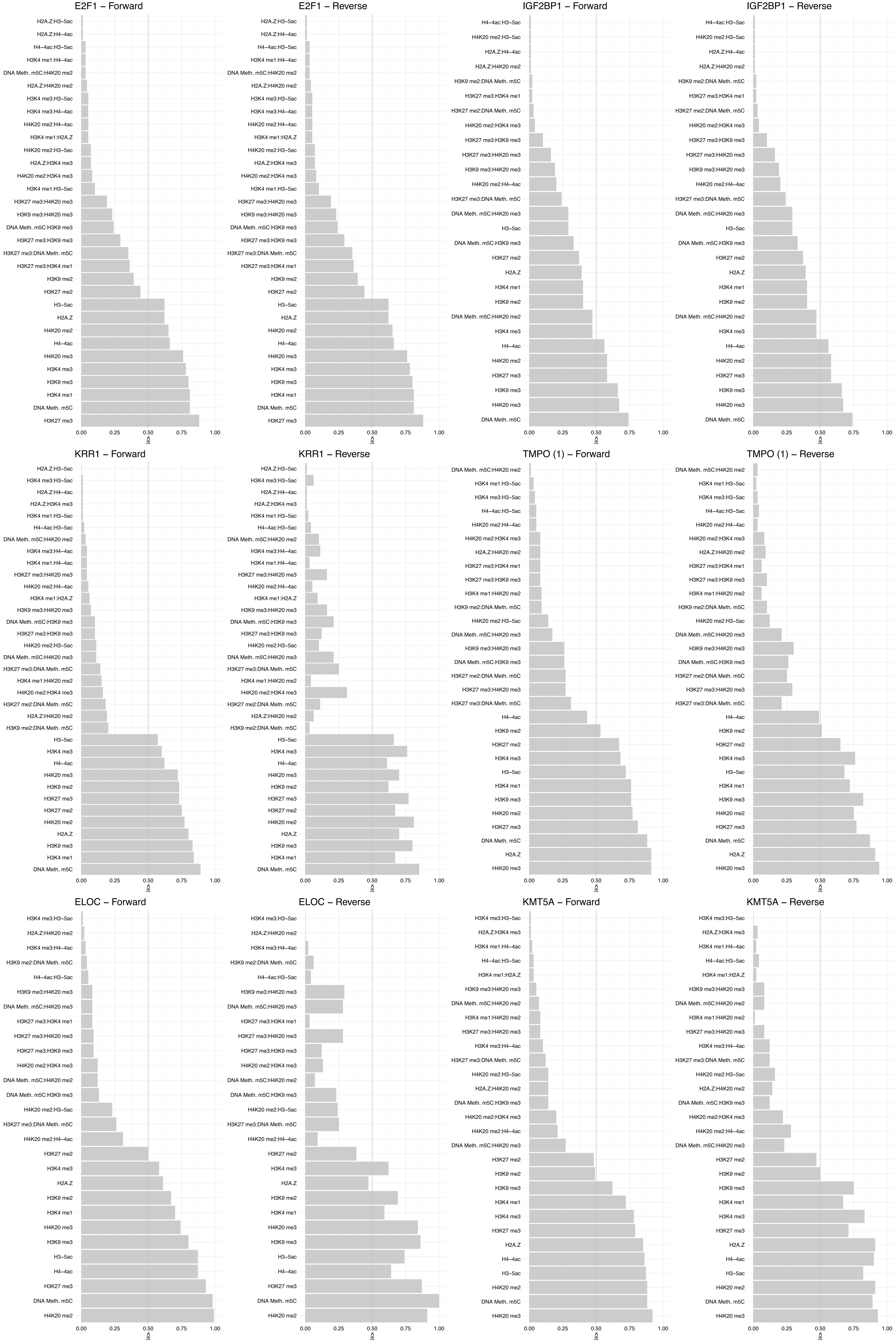
E2F1 – Forward
H2A.Z:H3–5ac
H2A.Z:H4–4ac
H4–4ac:H3–5ac
H3K4 me1:H4–4ac
DNA Meth. m5C:H4K20 me2
H2A.Z:H4K20 me2
H3K4 me3:H3–5ac
H3K4 me3:H4–4ac
H4K20 me2:H4–4ac
H3K4 me1:H2A.Z
H4K20 me2:H3–5ac
H2A.Z:H3K4 me3
H4K20 me2:H3K4 me3
H3K4 me1:H3–5ac
H3K27 me3:H4K20 me3
H3K9 me3:H4K20 me3
DNA Meth. m5C:H3K9 me3
H3K27 me3:H3K9 me3
H3K27 me3:DNA Meth. m5C
H3K27 me3:H3K4 me1
H3K9 me2
H3K27 me2
H3–5ac
H2A.Z
H4K20 me2
H4–4ac
H4K20 me3
H3K4 me3
H3K9 me3
H3K4 me1
DNA Meth. m5C
H3K27 me3
0.00
0.25
0.50
0.75
1.00
π̂
E2F1 – Reverse
IGF2BP1 – Forward
H4–4ac:H3–5ac
H4K20 me2:H3–5ac
H2A.Z:H4–4ac
H2A.Z:H4K20 me2
H3K9 me2:DNA Meth. m5C
H3K27 me3:H3K4 me1
H3K27 me2:DNA Meth. m5C
H4K20 me2:H3K4 me3
H3K27 me3:H3K9 me3
H3K27 me3:H4K20 me3
H3K9 me3:H4K20 me3
H4K20 me2:H4–4ac
H3K27 me3:DNA Meth. m5C
DNA Meth. m5C:H4K20 me3
H3–5ac
DNA Meth. m5C:H3K9 me3
H3K27 me2
H2A.Z
H3K4 me1
H3K9 me2
DNA Meth. m5C:H4K20 me2
H3K4 me3
H4–4ac
H4K20 me2
H3K27 me3
H3K9 me3
H4K20 me3
DNA Meth. m5C
IGF2BP1 – Reverse
KRR1 – Forward
H2A.Z:H3–5ac
H3K4 me3:H3–5ac
H2A.Z:H4–4ac
H2A.Z:H3K4 me3
H3K4 me1:H3–5ac
H4–4ac:H3–5ac
DNA Meth. m5C:H4K20 me2
H3K4 me3:H4–4ac
H3K4 me1:H4–4ac
H3K27 me3:H4K20 me3
H4K20 me2:H4–4ac
H3K4 me1:H2A.Z
H3K9 me3:H4K20 me3
DNA Meth. m5C:H3K9 me3
H3K27 me3:H3K9 me3
H4K20 me2:H3–5ac
DNA Meth. m5C:H4K20 me3
H3K27 me3:DNA Meth. m5C
H3K4 me1:H4K20 me2
H4K20 me2:H3K4 me3
H3K27 me2:DNA Meth. m5C
H2A.Z:H4K20 me2
H3K9 me2:DNA Meth. m5C
H3–5ac
H3K4 me3
H4–4ac
H4K20 me3
H3K9 me2
H3K27 me3
H3K27 me2
H4K20 me2
H2A.Z
H3K9 me3
H3K4 me1
DNA Meth. m5C
KRR1 – Reverse
TMPO (1) – Forward
DNA Meth. m5C:H4K20 me2
H3K4 me1:H3–5ac
H3K4 me3:H3–5ac
H4–4ac:H3–5ac
H4K20 me2:H4–4ac
H4K20 me2:H3K4 me3
H2A.Z:H4K20 me2
H3K27 me3:H3K4 me1
H3K27 me3:H3K9 me3
H3K4 me1:H4K20 me2
H3K9 me2:DNA Meth. m5C
H4K20 me2:H3–5ac
DNA Meth. m5C:H4K20 me3
H3K9 me3:H4K20 me3
DNA Meth. m5C:H3K9 me3
H3K27 me2:DNA Meth. m5C
H3K27 me3:H4K20 me3
H3K27 me3:DNA Meth. m5C
H4–4ac
H3K9 me2
H3K27 me2
H3K4 me3
H3–5ac
H3K4 me1
H3K9 me3
H4K20 me2
H3K27 me3
DNA Meth. m5C
H2A.Z
H4K20 me3
TMPO (1) – Reverse
ELOC – Forward
H3K4 me3:H3–5ac
H2A.Z:H4K20 me2
H3K4 me3:H4–4ac
H3K9 me2:DNA Meth. m5C
H4–4ac:H3–5ac
H3K9 me3:H4K20 me3
DNA Meth. m5C:H4K20 me3
H3K27 me3:H3K4 me1
H3K27 me3:H4K20 me3
H3K27 me3:H3K9 me3
H4K20 me2:H3K4 me3
DNA Meth. m5C:H4K20 me2
DNA Meth. m5C:H3K9 me3
H4K20 me2:H3–5ac
H3K27 me3:DNA Meth. m5C
H4K20 me2:H4–4ac
H3K27 me2
H3K4 me3
H2A.Z
H3K9 me2
H3K4 me1
H4K20 me3
H3K9 me3
H3–5ac
H4–4ac
H3K27 me3
DNA Meth. m5C
H4K20 me2
ELOC – Reverse
KMT5A – Forward
H3K4 me3:H3–5ac
H2A.Z:H3K4 me3
H3K4 me1:H4–4ac
H4–4ac:H3–5ac
H3K4 me1:H2A.Z
H3K9 me3:H4K20 me3
DNA Meth. m5C:H4K20 me2
H3K4 me1:H4K20 me2
H3K27 me3:H4K20 me3
H3K4 me3:H4–4ac
H3K27 me3:DNA Meth. m5C
H4K20 me2:H3–5ac
H2A.Z:H4K20 me2
DNA Meth. m5C:H3K9 me3
H4K20 me2:H3K4 me3
H4K20 me2:H4–4ac
DNA Meth. m5C:H4K20 me3
H3K27 me2
H3K9 me2
H3K9 me3
H3K4 me1
H3K4 me3
H3K27 me3
H2A.Z
H4–4ac
H3–5ac
H4K20 me2
DNA Meth. m5C
H4K20 me3
KMT5A – Reverse

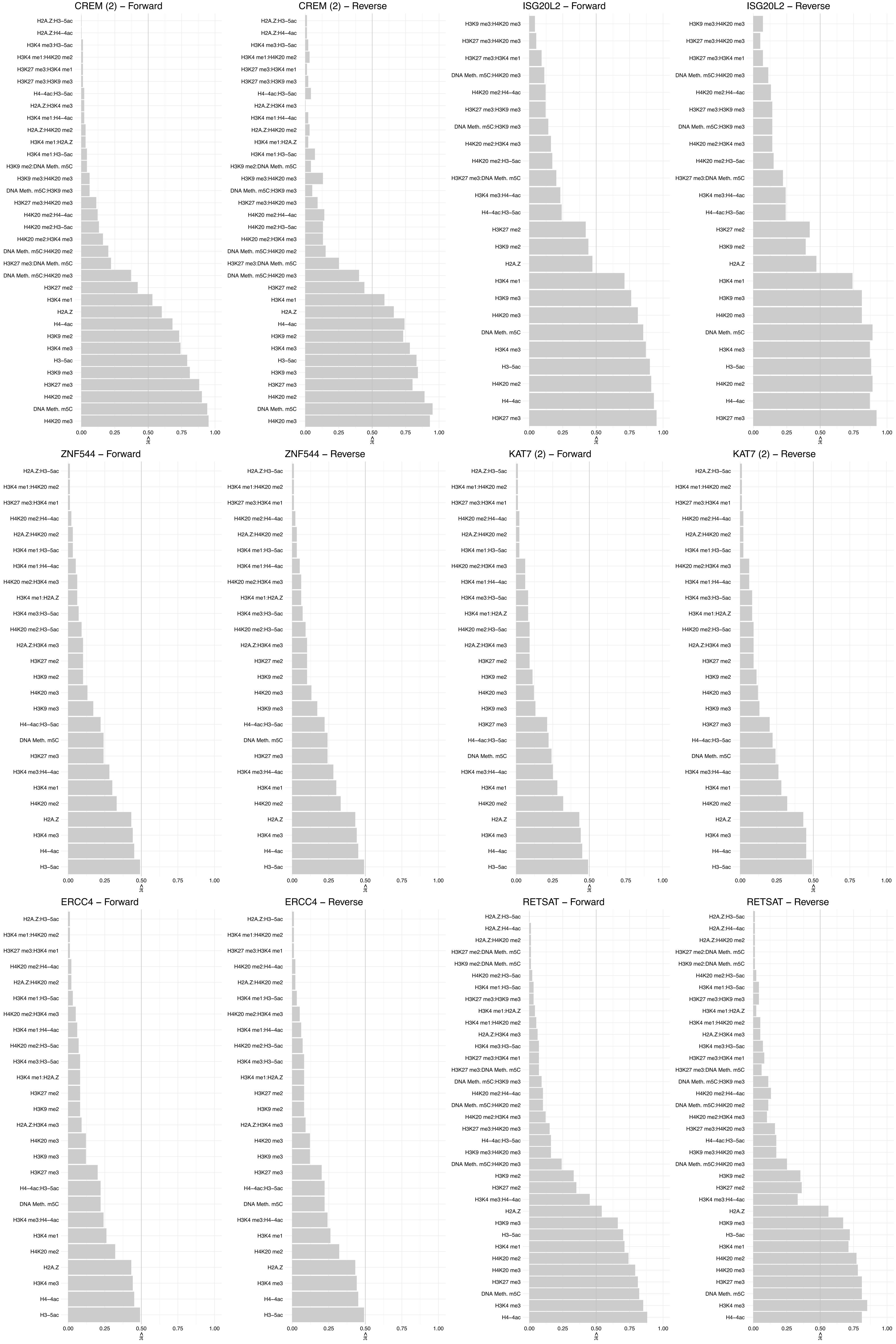
CREM (2) – Forward
CREM (2) – Reverse
ISG20L2 – Forward
ISG20L2 – Reverse
ZNF544 – Forward
ZNF544 – Reverse
KAT7 (2) – Forward
KAT7 (2) – Reverse
ERCC4 – Forward
ERCC4 – Reverse
RETSAT – Forward
RETSAT – Reverse
0.00 0.25 0.50 0.75 1.00
π̂

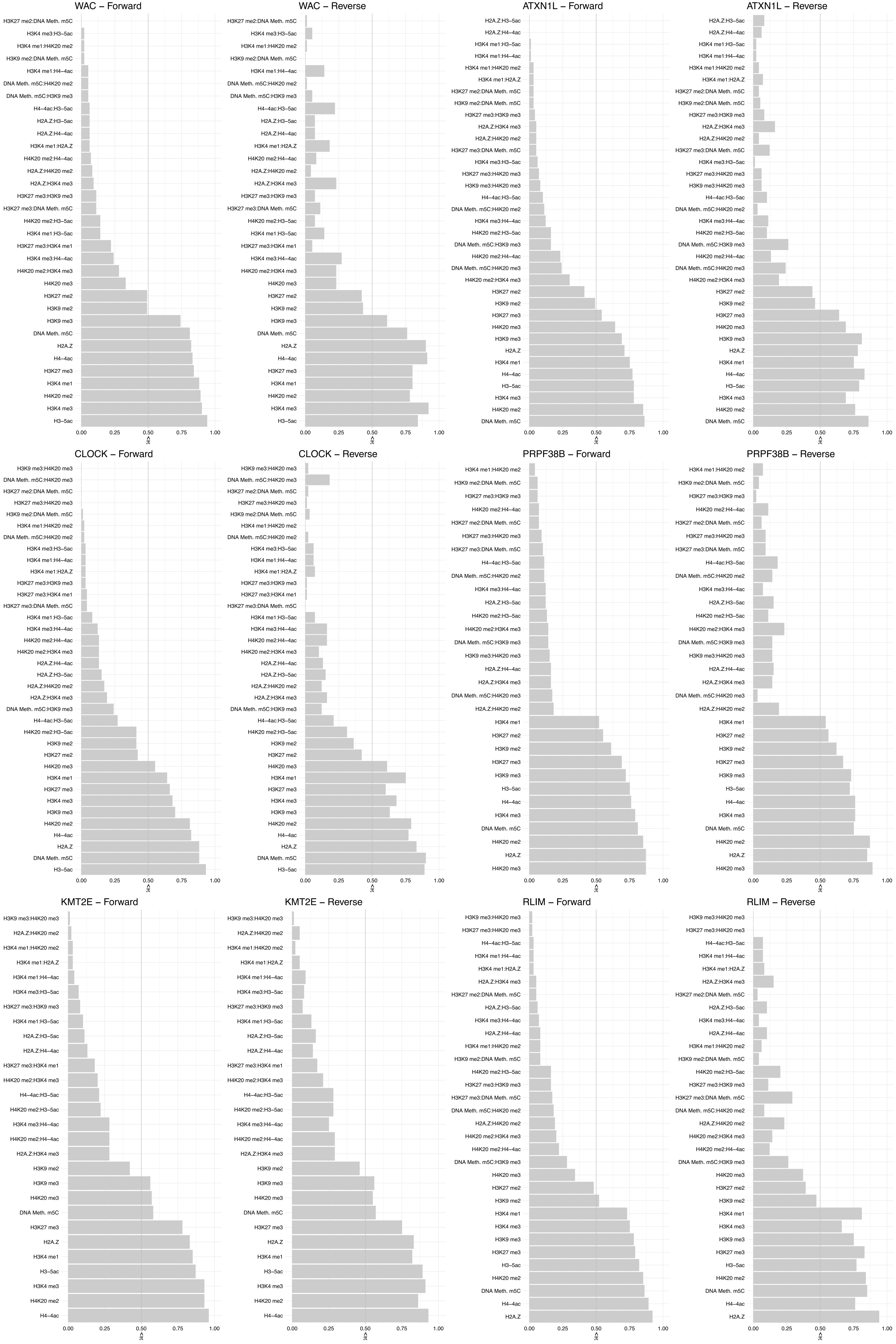
WAC – Forward
H3K27 me2:DNA Meth. m5C
H3K4 me3:H3−5ac
H3K4 me1:H4K20 me2
H3K9 me2:DNA Meth. m5C
H3K4 me1:H4−4ac
DNA Meth. m5C:H4K20 me2
DNA Meth. m5C:H3K9 me3
H4−4ac:H3−5ac
H2A.Z:H3−5ac
H2A.Z:H4−4ac
H3K4 me1:H2A.Z
H4K20 me2:H4−4ac
H2A.Z:H4K20 me2
H2A.Z:H3K4 me3
H3K27 me3:H3K9 me3
H3K27 me3:DNA Meth. m5C
H4K20 me2:H3−5ac
H3K4 me1:H3−5ac
H3K27 me3:H3K4 me1
H3K4 me3:H4−4ac
H4K20 me2:H3K4 me3
H4K20 me3
H3K27 me2
H3K9 me2
H3K9 me3
DNA Meth. m5C
H2A.Z
H4−4ac
H3K27 me3
H3K4 me1
H4K20 me2
H3K4 me3
H3−5ac
0.00 0.25 0.50 0.75 1.00
π̂
WAC – Reverse
H3K27 me2:DNA Meth. m5C
H3K4 me3:H3−5ac
H3K4 me1:H4K20 me2
H3K9 me2:DNA Meth. m5C
H3K4 me1:H4−4ac
DNA Meth. m5C:H4K20 me2
DNA Meth. m5C:H3K9 me3
H4−4ac:H3−5ac
H2A.Z:H3−5ac
H2A.Z:H4−4ac
H3K4 me1:H2A.Z
H4K20 me2:H4−4ac
H2A.Z:H4K20 me2
H2A.Z:H3K4 me3
H3K27 me3:H3K9 me3
H3K27 me3:DNA Meth. m5C
H4K20 me2:H3−5ac
H3K4 me1:H3−5ac
H3K27 me3:H3K4 me1
H3K4 me3:H4−4ac
H4K20 me2:H3K4 me3
H4K20 me3
H3K27 me2
H3K9 me2
H3K9 me3
DNA Meth. m5C
H2A.Z
H4−4ac
H3K27 me3
H3K4 me1
H4K20 me2
H3K4 me3
H3−5ac
0.00 0.25 0.50 0.75 1.00
π̂
ATXN1L – Forward
H2A.Z:H3−5ac
H2A.Z:H4−4ac
H3K4 me1:H3−5ac
H3K4 me1:H4−4ac
H3K4 me1:H4K20 me2
H3K4 me1:H2A.Z
H3K27 me2:DNA Meth. m5C
H3K9 me2:DNA Meth. m5C
H3K27 me3:H3K9 me3
H2A.Z:H3K4 me3
H2A.Z:H4K20 me2
H3K27 me3:DNA Meth. m5C
H3K4 me3:H3−5ac
H3K27 me3:H4K20 me3
H3K9 me3:H4K20 me3
H4−4ac:H3−5ac
DNA Meth. m5C:H4K20 me2
H3K4 me3:H4−4ac
H4K20 me2:H3−5ac
DNA Meth. m5C:H3K9 me3
H4K20 me2:H4−4ac
DNA Meth. m5C:H4K20 me3
H4K20 me2:H3K4 me3
H3K27 me2
H3K9 me2
H3K27 me3
H4K20 me3
H3K9 me3
H2A.Z
H3K4 me1
H4−4ac
H3−5ac
H3K4 me3
H4K20 me2
DNA Meth. m5C
0.00 0.25 0.50 0.75 1.00
π̂
ATXN1L – Reverse
H2A.Z:H3−5ac
H2A.Z:H4−4ac
H3K4 me1:H3−5ac
H3K4 me1:H4−4ac
H3K4 me1:H4K20 me2
H3K4 me1:H2A.Z
H3K27 me2:DNA Meth. m5C
H3K9 me2:DNA Meth. m5C
H3K27 me3:H3K9 me3
H2A.Z:H3K4 me3
H2A.Z:H4K20 me2
H3K27 me3:DNA Meth. m5C
H3K4 me3:H3−5ac
H3K27 me3:H4K20 me3
H3K9 me3:H4K20 me3
H4−4ac:H3−5ac
DNA Meth. m5C:H4K20 me2
H3K4 me3:H4−4ac
H4K20 me2:H3−5ac
DNA Meth. m5C:H3K9 me3
H4K20 me2:H4−4ac
DNA Meth. m5C:H4K20 me3
H4K20 me2:H3K4 me3
H3K27 me2
H3K9 me2
H3K27 me3
H4K20 me3
H3K9 me3
H2A.Z
H3K4 me1
H4−4ac
H3−5ac
H3K4 me3
H4K20 me2
DNA Meth. m5C
0.00 0.25 0.50 0.75 1.00
π̂
CLOCK – Forward
H3K9 me3:H4K20 me3
DNA Meth. m5C:H4K20 me3
H3K27 me2:DNA Meth. m5C
H3K27 me3:H4K20 me3
H3K9 me2:DNA Meth. m5C
H3K4 me1:H4K20 me2
DNA Meth. m5C:H4K20 me2
H3K4 me3:H3−5ac
H3K4 me1:H4−4ac
H3K4 me1:H2A.Z
H3K27 me3:H3K9 me3
H3K27 me3:H3K4 me1
H3K27 me3:DNA Meth. m5C
H3K4 me1:H3−5ac
H3K4 me3:H4−4ac
H4K20 me2:H4−4ac
H4K20 me2:H3K4 me3
H2A.Z:H4−4ac
H2A.Z:H3−5ac
H2A.Z:H4K20 me2
H2A.Z:H3K4 me3
DNA Meth. m5C:H3K9 me3
H4−4ac:H3−5ac
H4K20 me2:H3−5ac
H3K9 me2
H3K27 me2
H4K20 me3
H3K4 me1
H3K27 me3
H3K4 me3
H3K9 me3
H4K20 me2
H4−4ac
H2A.Z
DNA Meth. m5C
H3−5ac
0.00 0.25 0.50 0.75 1.00
π̂
CLOCK – Reverse
H3K9 me3:H4K20 me3
DNA Meth. m5C:H4K20 me3
H3K27 me2:DNA Meth. m5C
H3K27 me3:H4K20 me3
H3K9 me2:DNA Meth. m5C
H3K4 me1:H4K20 me2
DNA Meth. m5C:H4K20 me2
H3K4 me3:H3−5ac
H3K4 me1:H4−4ac
H3K4 me1:H2A.Z
H3K27 me3:H3K9 me3
H3K27 me3:H3K4 me1
H3K27 me3:DNA Meth. m5C
H3K4 me1:H3−5ac
H3K4 me3:H4−4ac
H4K20 me2:H4−4ac
H4K20 me2:H3K4 me3
H2A.Z:H4−4ac
H2A.Z:H3−5ac
H2A.Z:H4K20 me2
H2A.Z:H3K4 me3
DNA Meth. m5C:H3K9 me3
H4−4ac:H3−5ac
H4K20 me2:H3−5ac
H3K9 me2
H3K27 me2
H4K20 me3
H3K4 me1
H3K27 me3
H3K4 me3
H3K9 me3
H4K20 me2
H4−4ac
H2A.Z
DNA Meth. m5C
H3−5ac
0.00 0.25 0.50 0.75 1.00
π̂
PRPF38B – Forward
H3K4 me1:H4K20 me2
H3K9 me2:DNA Meth. m5C
H3K27 me3:H3K9 me3
H4K20 me2:H4−4ac
H3K27 me2:DNA Meth. m5C
H3K27 me3:H4K20 me3
H3K27 me3:DNA Meth. m5C
H4−4ac:H3−5ac
DNA Meth. m5C:H4K20 me2
H3K4 me3:H4−4ac
H2A.Z:H3−5ac
H4K20 me2:H3−5ac
H4K20 me2:H3K4 me3
DNA Meth. m5C:H3K9 me3
H3K9 me3:H4K20 me3
H2A.Z:H4−4ac
H2A.Z:H3K4 me3
DNA Meth. m5C:H4K20 me3
H2A.Z:H4K20 me2
H3K4 me1
H3K27 me2
H3K9 me2
H3K27 me3
H3K9 me3
H3−5ac
H4−4ac
H3K4 me3
DNA Meth. m5C
H4K20 me2
H2A.Z
H4K20 me3
0.00 0.25 0.50 0.75 1.00
π̂
PRPF38B – Reverse
H3K4 me1:H4K20 me2
H3K9 me2:DNA Meth. m5C
H3K27 me3:H3K9 me3
H4K20 me2:H4−4ac
H3K27 me2:DNA Meth. m5C
H3K27 me3:H4K20 me3
H3K27 me3:DNA Meth. m5C
H4−4ac:H3−5ac
DNA Meth. m5C:H4K20 me2
H3K4 me3:H4−4ac
H2A.Z:H3−5ac
H4K20 me2:H3−5ac
H4K20 me2:H3K4 me3
DNA Meth. m5C:H3K9 me3
H3K9 me3:H4K20 me3
H2A.Z:H4−4ac
H2A.Z:H3K4 me3
DNA Meth. m5C:H4K20 me3
H2A.Z:H4K20 me2
H3K4 me1
H3K27 me2
H3K9 me2
H3K27 me3
H3K9 me3
H3−5ac
H4−4ac
H3K4 me3
DNA Meth. m5C
H4K20 me2
H2A.Z
H4K20 me3
0.00 0.25 0.50 0.75 1.00
π̂
KMT2E – Forward
H3K9 me3:H4K20 me3
H2A.Z:H4K20 me2
H3K4 me1:H4K20 me2
H3K4 me1:H2A.Z
H3K4 me1:H4−4ac
H3K4 me3:H3−5ac
H3K27 me3:H3K9 me3
H3K4 me1:H3−5ac
H2A.Z:H3−5ac
H2A.Z:H4−4ac
H3K27 me3:H3K4 me1
H4K20 me2:H3K4 me3
H4−4ac:H3−5ac
H4K20 me2:H3−5ac
H3K4 me3:H4−4ac
H4K20 me2:H4−4ac
H2A.Z:H3K4 me3
H3K9 me2
H3K9 me3
H4K20 me3
DNA Meth. m5C
H3K27 me3
H2A.Z
H3K4 me1
H3−5ac
H3K4 me3
H4K20 me2
H4−4ac
0.00 0.25 0.50 0.75 1.00
π̂
KMT2E – Reverse
H3K9 me3:H4K20 me3
H2A.Z:H4K20 me2
H3K4 me1:H4K20 me2
H3K4 me1:H2A.Z
H3K4 me1:H4−4ac
H3K4 me3:H3−5ac
H3K27 me3:H3K9 me3
H3K4 me1:H3−5ac
H2A.Z:H3−5ac
H2A.Z:H4−4ac
H3K27 me3:H3K4 me1
H4K20 me2:H3K4 me3
H4−4ac:H3−5ac
H4K20 me2:H3−5ac
H3K4 me3:H4−4ac
H4K20 me2:H4−4ac
H2A.Z:H3K4 me3
H3K9 me2
H3K9 me3
H4K20 me3
DNA Meth. m5C
H3K27 me3
H2A.Z
H3K4 me1
H3−5ac
H3K4 me3
H4K20 me2
H4−4ac
0.00 0.25 0.50 0.75 1.00
π̂
RLIM – Forward
H3K9 me3:H4K20 me3
H3K27 me3:H4K20 me3
H4−4ac:H3−5ac
H3K4 me1:H4−4ac
H3K4 me1:H2A.Z
H2A.Z:H3K4 me3
H3K27 me2:DNA Meth. m5C
H2A.Z:H3−5ac
H3K4 me3:H4−4ac
H2A.Z:H4−4ac
H3K4 me1:H4K20 me2
H3K9 me2:DNA Meth. m5C
H4K20 me2:H3−5ac
H3K27 me3:H3K9 me3
H3K27 me3:DNA Meth. m5C
DNA Meth. m5C:H4K20 me2
H2A.Z:H4K20 me2
H4K20 me2:H3K4 me3
H4K20 me2:H4−4ac
DNA Meth. m5C:H3K9 me3
H4K20 me3
H3K27 me2
H3K9 me2
H3K4 me1
H3K4 me3
H3K9 me3
H3K27 me3
H3−5ac
H4K20 me2
DNA Meth. m5C
H4−4ac
H2A.Z
0.00 0.25 0.50 0.75 1.00
π̂
RLIM – Reverse
H3K9 me3:H4K20 me3
H3K27 me3:H4K20 me3
H4−4ac:H3−5ac
H3K4 me1:H4−4ac
H3K4 me1:H2A.Z
H2A.Z:H3K4 me3
H3K27 me2:DNA Meth. m5C
H2A.Z:H3−5ac
H3K4 me3:H4−4ac
H2A.Z:H4−4ac
H3K4 me1:H4K20 me2
H3K9 me2:DNA Meth. m5C
H4K20 me2:H3−5ac
H3K27 me3:H3K9 me3
H3K27 me3:DNA Meth. m5C
DNA Meth. m5C:H4K20 me2
H2A.Z:H4K20 me2
H4K20 me2:H3K4 me3
H4K20 me2:H4−4ac
DNA Meth. m5C:H3K9 me3
H4K20 me3
H3K27 me2
H3K9 me2
H3K4 me1
H3K4 me3
H3K9 me3
H3K27 me3
H3−5ac
H4K20 me2
DNA Meth. m5C
H4−4ac
H2A.Z
0.00 0.25 0.50 0.75 1.00
π̂

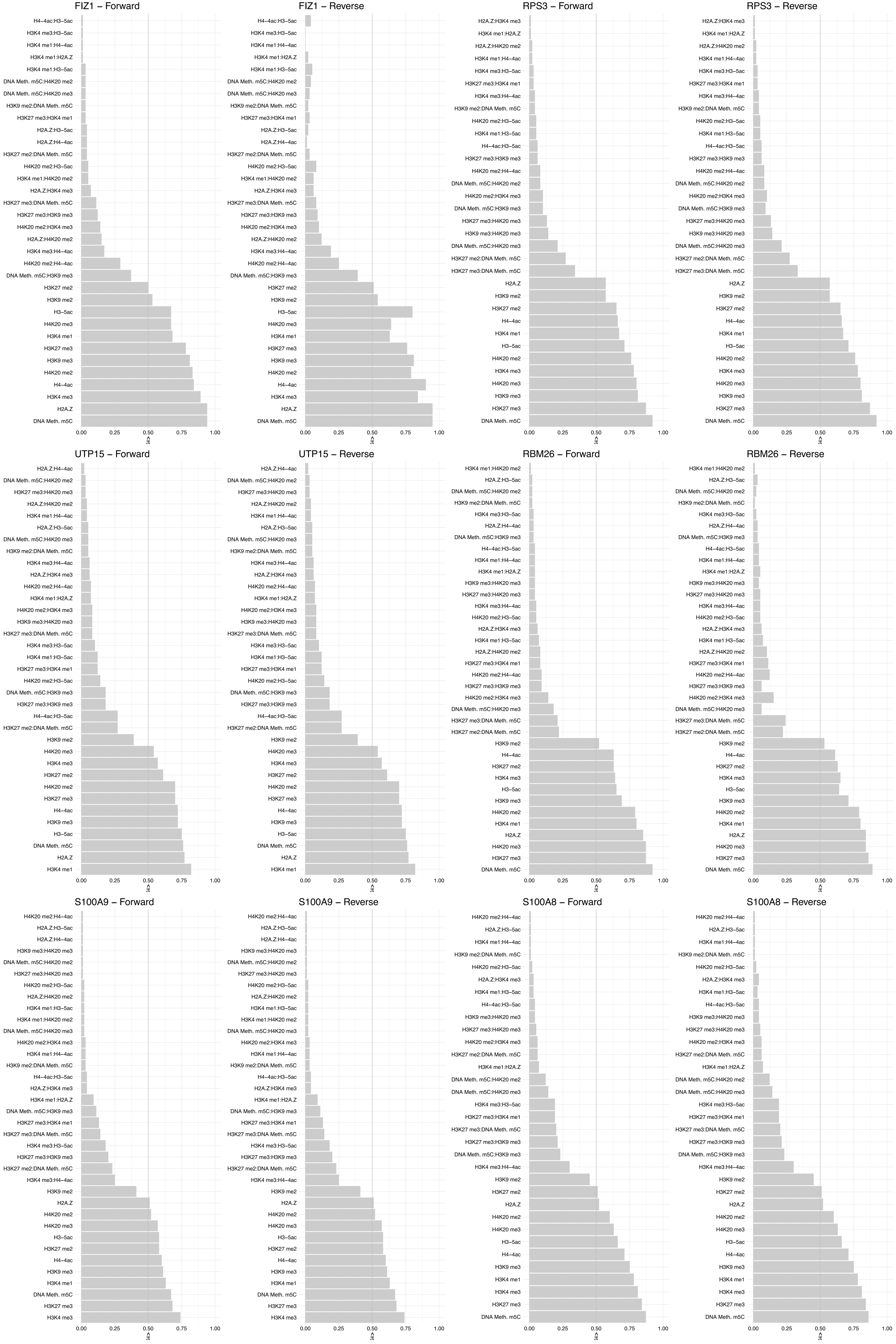
FIZ1 − Forward
H4−4ac:H3−5ac
H3K4 me3:H3−5ac
H3K4 me1:H4−4ac
H3K4 me1:H2A.Z
H3K4 me1:H3−5ac
DNA Meth. m5C:H4K20 me2
DNA Meth. m5C:H4K20 me3
H3K9 me2:DNA Meth. m5C
H3K27 me3:H3K4 me1
H2A.Z:H3−5ac
H2A.Z:H4−4ac
H3K27 me2:DNA Meth. m5C
H4K20 me2:H3−5ac
H3K4 me1:H4K20 me2
H2A.Z:H3K4 me3
H3K27 me3:DNA Meth. m5C
H3K27 me3:H3K9 me3
H4K20 me2:H3K4 me3
H2A.Z:H4K20 me2
H3K4 me3:H4−4ac
H4K20 me2:H4−4ac
DNA Meth. m5C:H3K9 me3
H3K27 me2
H3K9 me2
H3−5ac
H4K20 me3
H3K4 me1
H3K27 me3
H3K9 me3
H4K20 me2
H4−4ac
H3K4 me3
H2A.Z
DNA Meth. m5C
0.00
0.25
0.50
0.75
1.00
π̂
FIZ1 − Reverse
H4−4ac:H3−5ac
H3K4 me3:H3−5ac
H3K4 me1:H4−4ac
H3K4 me1:H2A.Z
H3K4 me1:H3−5ac
DNA Meth. m5C:H4K20 me2
DNA Meth. m5C:H4K20 me3
H3K9 me2:DNA Meth. m5C
H3K27 me3:H3K4 me1
H2A.Z:H3−5ac
H2A.Z:H4−4ac
H3K27 me2:DNA Meth. m5C
H4K20 me2:H3−5ac
H3K4 me1:H4K20 me2
H2A.Z:H3K4 me3
H3K27 me3:DNA Meth. m5C
H3K27 me3:H3K9 me3
H4K20 me2:H3K4 me3
H2A.Z:H4K20 me2
H3K4 me3:H4−4ac
H4K20 me2:H4−4ac
DNA Meth. m5C:H3K9 me3
H3K27 me2
H3K9 me2
H3−5ac
H4K20 me3
H3K4 me1
H3K27 me3
H3K9 me3
H4K20 me2
H4−4ac
H3K4 me3
H2A.Z
DNA Meth. m5C
0.00
0.25
0.50
0.75
1.00
π̂
RPS3 − Forward
H2A.Z:H3K4 me3
H3K4 me1:H2A.Z
H2A.Z:H4K20 me2
H3K4 me1:H4−4ac
H3K4 me3:H3−5ac
H3K27 me3:H3K4 me1
H3K4 me3:H4−4ac
H3K9 me2:DNA Meth. m5C
H4K20 me2:H3−5ac
H3K4 me1:H3−5ac
H4−4ac:H3−5ac
H3K27 me3:H3K9 me3
H4K20 me2:H4−4ac
DNA Meth. m5C:H4K20 me2
H4K20 me2:H3K4 me3
DNA Meth. m5C:H3K9 me3
H3K27 me3:H4K20 me3
H3K9 me3:H4K20 me3
DNA Meth. m5C:H4K20 me3
H3K27 me2:DNA Meth. m5C
H3K27 me3:DNA Meth. m5C
H2A.Z
H3K9 me2
H3K27 me2
H4−4ac
H3K4 me1
H3−5ac
H4K20 me2
H3K4 me3
H4K20 me3
H3K9 me3
H3K27 me3
DNA Meth. m5C
0.00
0.25
0.50
0.75
1.00
π̂
RPS3 − Reverse
H2A.Z:H3K4 me3
H3K4 me1:H2A.Z
H2A.Z:H4K20 me2
H3K4 me1:H4−4ac
H3K4 me3:H3−5ac
H3K27 me3:H3K4 me1
H3K4 me3:H4−4ac
H3K9 me2:DNA Meth. m5C
H4K20 me2:H3−5ac
H3K4 me1:H3−5ac
H4−4ac:H3−5ac
H3K27 me3:H3K9 me3
H4K20 me2:H4−4ac
DNA Meth. m5C:H4K20 me2
H4K20 me2:H3K4 me3
DNA Meth. m5C:H3K9 me3
H3K27 me3:H4K20 me3
H3K9 me3:H4K20 me3
DNA Meth. m5C:H4K20 me3
H3K27 me2:DNA Meth. m5C
H3K27 me3:DNA Meth. m5C
H2A.Z
H3K9 me2
H3K27 me2
H4−4ac
H3K4 me1
H3−5ac
H4K20 me2
H3K4 me3
H4K20 me3
H3K9 me3
H3K27 me3
DNA Meth. m5C
0.00
0.25
0.50
0.75
1.00
π̂
UTP15 − Forward
H2A.Z:H4−4ac
DNA Meth. m5C:H4K20 me2
H3K27 me3:H4K20 me3
H2A.Z:H4K20 me2
H3K4 me1:H4−4ac
H2A.Z:H3−5ac
DNA Meth. m5C:H4K20 me3
H3K9 me2:DNA Meth. m5C
H3K4 me3:H4−4ac
H2A.Z:H3K4 me3
H4K20 me2:H4−4ac
H3K4 me1:H2A.Z
H4K20 me2:H3K4 me3
H3K9 me3:H4K20 me3
H3K27 me3:DNA Meth. m5C
H3K4 me3:H3−5ac
H3K4 me1:H3−5ac
H3K27 me3:H3K4 me1
H4K20 me2:H3−5ac
DNA Meth. m5C:H3K9 me3
H3K27 me3:H3K9 me3
H4−4ac:H3−5ac
H3K27 me2:DNA Meth. m5C
H3K9 me2
H4K20 me3
H3K4 me3
H3K27 me2
H4K20 me2
H3K27 me3
H4−4ac
H3K9 me3
H3−5ac
DNA Meth. m5C
H2A.Z
H3K4 me1
0.00
0.25
0.50
0.75
1.00
π̂
UTP15 − Reverse
H2A.Z:H4−4ac
DNA Meth. m5C:H4K20 me2
H3K27 me3:H4K20 me3
H2A.Z:H4K20 me2
H3K4 me1:H4−4ac
H2A.Z:H3−5ac
DNA Meth. m5C:H4K20 me3
H3K9 me2:DNA Meth. m5C
H3K4 me3:H4−4ac
H2A.Z:H3K4 me3
H4K20 me2:H4−4ac
H3K4 me1:H2A.Z
H4K20 me2:H3K4 me3
H3K9 me3:H4K20 me3
H3K27 me3:DNA Meth. m5C
H3K4 me3:H3−5ac
H3K4 me1:H3−5ac
H3K27 me3:H3K4 me1
H4K20 me2:H3−5ac
DNA Meth. m5C:H3K9 me3
H3K27 me3:H3K9 me3
H4−4ac:H3−5ac
H3K27 me2:DNA Meth. m5C
H3K9 me2
H4K20 me3
H3K4 me3
H3K27 me2
H4K20 me2
H3K27 me3
H4−4ac
H3K9 me3
H3−5ac
DNA Meth. m5C
H2A.Z
H3K4 me1
0.00
0.25
0.50
0.75
1.00
π̂
RBM26 − Forward
H3K4 me1:H4K20 me2
H2A.Z:H3−5ac
DNA Meth. m5C:H4K20 me2
H3K9 me2:DNA Meth. m5C
H3K4 me3:H3−5ac
H2A.Z:H4−4ac
DNA Meth. m5C:H3K9 me3
H4−4ac:H3−5ac
H3K4 me1:H4−4ac
H3K4 me1:H2A.Z
H3K9 me3:H4K20 me3
H3K27 me3:H4K20 me3
H3K4 me3:H4−4ac
H4K20 me2:H3−5ac
H2A.Z:H3K4 me3
H3K4 me1:H3−5ac
H2A.Z:H4K20 me2
H3K27 me3:H3K4 me1
H4K20 me2:H4−4ac
H3K27 me3:H3K9 me3
H4K20 me2:H3K4 me3
DNA Meth. m5C:H4K20 me3
H3K27 me3:DNA Meth. m5C
H3K27 me2:DNA Meth. m5C
H3K9 me2
H4−4ac
H3K27 me2
H3K4 me3
H3−5ac
H3K9 me3
H4K20 me2
H3K4 me1
H2A.Z
H4K20 me3
H3K27 me3
DNA Meth. m5C
0.00
0.25
0.50
0.75
1.00
π̂
RBM26 − Reverse
H3K4 me1:H4K20 me2
H2A.Z:H3−5ac
DNA Meth. m5C:H4K20 me2
H3K9 me2:DNA Meth. m5C
H3K4 me3:H3−5ac
H2A.Z:H4−4ac
DNA Meth. m5C:H3K9 me3
H4−4ac:H3−5ac
H3K4 me1:H4−4ac
H3K4 me1:H2A.Z
H3K9 me3:H4K20 me3
H3K27 me3:H4K20 me3
H3K4 me3:H4−4ac
H4K20 me2:H3−5ac
H2A.Z:H3K4 me3
H3K4 me1:H3−5ac
H2A.Z:H4K20 me2
H3K27 me3:H3K4 me1
H4K20 me2:H4−4ac
H3K27 me3:H3K9 me3
H4K20 me2:H3K4 me3
DNA Meth. m5C:H4K20 me3
H3K27 me3:DNA Meth. m5C
H3K27 me2:DNA Meth. m5C
H3K9 me2
H4−4ac
H3K27 me2
H3K4 me3
H3−5ac
H3K9 me3
H4K20 me2
H3K4 me1
H2A.Z
H4K20 me3
H3K27 me3
DNA Meth. m5C
0.00
0.25
0.50
0.75
1.00
π̂
S100A9 − Forward
H4K20 me2:H4−4ac
H2A.Z:H3−5ac
H2A.Z:H4−4ac
H3K9 me3:H4K20 me3
DNA Meth. m5C:H4K20 me2
H3K27 me3:H4K20 me3
H4K20 me2:H3−5ac
H2A.Z:H4K20 me2
H3K4 me1:H3−5ac
H3K4 me1:H4K20 me2
DNA Meth. m5C:H4K20 me3
H4K20 me2:H3K4 me3
H3K4 me1:H4−4ac
H3K9 me2:DNA Meth. m5C
H4−4ac:H3−5ac
H2A.Z:H3K4 me3
H3K4 me1:H2A.Z
DNA Meth. m5C:H3K9 me3
H3K27 me3:H3K4 me1
H3K27 me3:DNA Meth. m5C
H3K4 me3:H3−5ac
H3K27 me3:H3K9 me3
H3K27 me2:DNA Meth. m5C
H3K4 me3:H4−4ac
H3K9 me2
H2A.Z
H4K20 me2
H4K20 me3
H3−5ac
H3K27 me2
H4−4ac
H3K9 me3
H3K4 me1
DNA Meth. m5C
H3K27 me3
H3K4 me3
0.00
0.25
0.50
0.75
1.00
π̂
S100A9 − Reverse
H4K20 me2:H4−4ac
H2A.Z:H3−5ac
H2A.Z:H4−4ac
H3K9 me3:H4K20 me3
DNA Meth. m5C:H4K20 me2
H3K27 me3:H4K20 me3
H4K20 me2:H3−5ac
H2A.Z:H4K20 me2
H3K4 me1:H3−5ac
H3K4 me1:H4K20 me2
DNA Meth. m5C:H4K20 me3
H4K20 me2:H3K4 me3
H3K4 me1:H4−4ac
H3K9 me2:DNA Meth. m5C
H4−4ac:H3−5ac
H2A.Z:H3K4 me3
H3K4 me1:H2A.Z
DNA Meth. m5C:H3K9 me3
H3K27 me3:H3K4 me1
H3K27 me3:DNA Meth. m5C
H3K4 me3:H3−5ac
H3K27 me3:H3K9 me3
H3K27 me2:DNA Meth. m5C
H3K4 me3:H4−4ac
H3K9 me2
H2A.Z
H4K20 me2
H4K20 me3
H3−5ac
H3K27 me2
H4−4ac
H3K9 me3
H3K4 me1
DNA Meth. m5C
H3K27 me3
H3K4 me3
0.00
0.25
0.50
0.75
1.00
π̂
S100A8 − Forward
H4K20 me2:H4−4ac
H2A.Z:H3−5ac
H3K4 me1:H4−4ac
H3K9 me2:DNA Meth. m5C
H4K20 me2:H3−5ac
H2A.Z:H3K4 me3
H3K4 me1:H3−5ac
H4−4ac:H3−5ac
H3K9 me3:H4K20 me3
H3K27 me3:H4K20 me3
H4K20 me2:H3K4 me3
H3K27 me2:DNA Meth. m5C
H3K4 me1:H2A.Z
DNA Meth. m5C:H4K20 me2
DNA Meth. m5C:H4K20 me3
H3K4 me3:H3−5ac
H3K27 me3:H3K4 me1
H3K27 me3:DNA Meth. m5C
H3K27 me3:H3K9 me3
DNA Meth. m5C:H3K9 me3
H3K4 me3:H4−4ac
H3K9 me2
H3K27 me2
H2A.Z
H4K20 me2
H4K20 me3
H3−5ac
H4−4ac
H3K9 me3
H3K4 me1
H3K4 me3
H3K27 me3
DNA Meth. m5C
0.00
0.25
0.50
0.75
1.00
π̂
S100A8 − Reverse
H4K20 me2:H4−4ac
H2A.Z:H3−5ac
H3K4 me1:H4−4ac
H3K9 me2:DNA Meth. m5C
H4K20 me2:H3−5ac
H2A.Z:H3K4 me3
H3K4 me1:H3−5ac
H4−4ac:H3−5ac
H3K9 me3:H4K20 me3
H3K27 me3:H4K20 me3
H4K20 me2:H3K4 me3
H3K27 me2:DNA Meth. m5C
H3K4 me1:H2A.Z
DNA Meth. m5C:H4K20 me2
DNA Meth. m5C:H4K20 me3
H3K4 me3:H3−5ac
H3K27 me3:H3K4 me1
H3K27 me3:DNA Meth. m5C
H3K27 me3:H3K9 me3
DNA Meth. m5C:H3K9 me3
H3K4 me3:H4−4ac
H3K9 me2
H3K27 me2
H2A.Z
H4K20 me2
H4K20 me3
H3−5ac
H4−4ac
H3K9 me3
H3K4 me1
H3K4 me3
H3K27 me3
DNA Meth. m5C
0.00
0.25
0.50
0.75
1.00
π̂

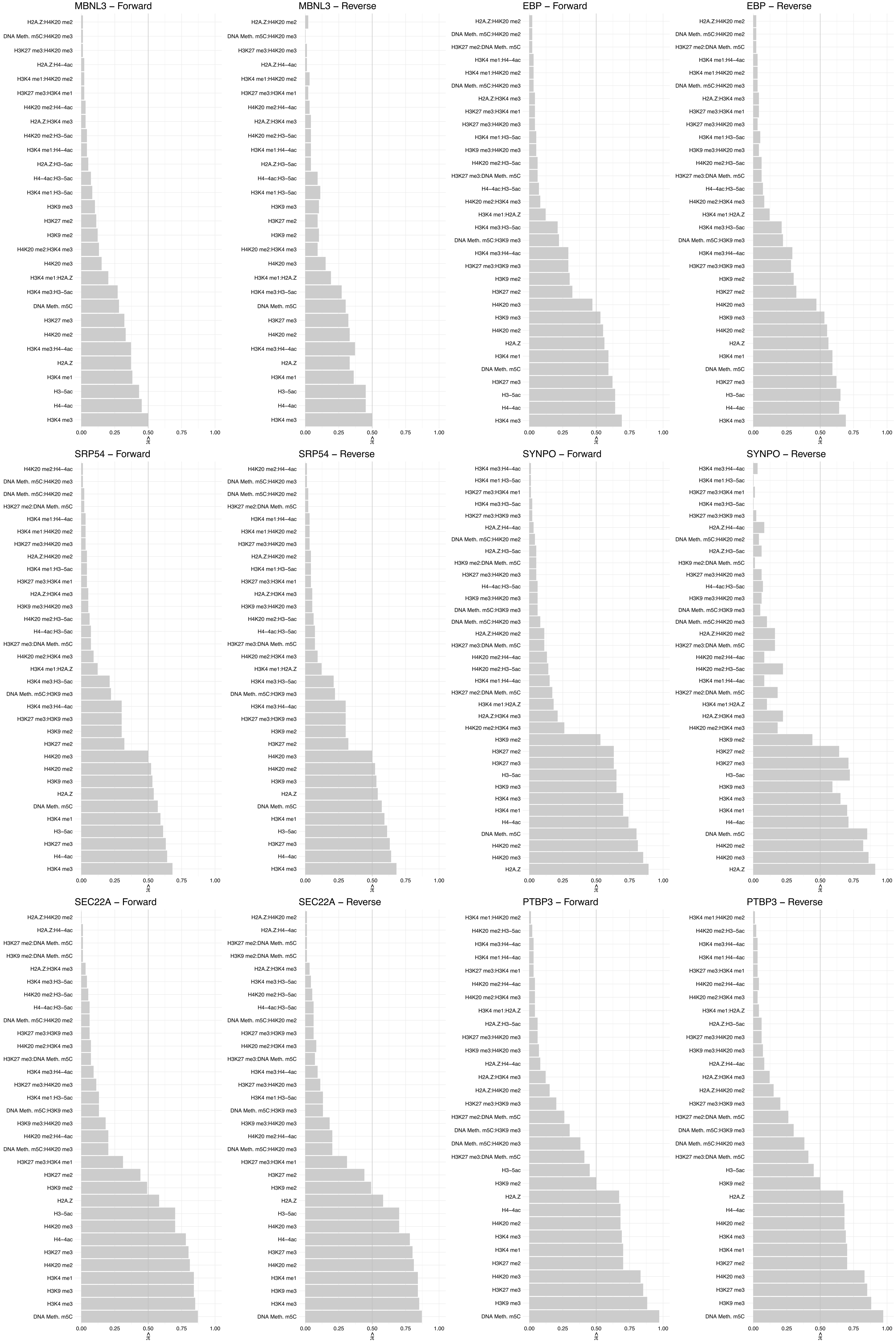
MBNL3 − Forward
H2A.Z:H4K20 me2
DNA Meth. m5C:H4K20 me3
H3K27 me3:H4K20 me3
H2A.Z:H4−4ac
H3K4 me1:H4K20 me2
H3K27 me3:H3K4 me1
H4K20 me2:H4−4ac
H2A.Z:H3K4 me3
H4K20 me2:H3−5ac
H3K4 me1:H4−4ac
H2A.Z:H3−5ac
H4−4ac:H3−5ac
H3K4 me1:H3−5ac
H3K9 me3
H3K27 me2
H3K9 me2
H4K20 me2:H3K4 me3
H4K20 me3
H3K4 me1:H2A.Z
H3K4 me3:H3−5ac
DNA Meth. m5C
H3K27 me3
H4K20 me2
H3K4 me3:H4−4ac
H2A.Z
H3K4 me1
H3−5ac
H4−4ac
H3K4 me3
0.00
0.25
0.50
0.75
1.00
π̂
MBNL3 − Reverse
H2A.Z:H4K20 me2
DNA Meth. m5C:H4K20 me3
H3K27 me3:H4K20 me3
H2A.Z:H4−4ac
H3K4 me1:H4K20 me2
H3K27 me3:H3K4 me1
H4K20 me2:H4−4ac
H2A.Z:H3K4 me3
H4K20 me2:H3−5ac
H3K4 me1:H4−4ac
H2A.Z:H3−5ac
H4−4ac:H3−5ac
H3K4 me1:H3−5ac
H3K9 me3
H3K27 me2
H3K9 me2
H4K20 me2:H3K4 me3
H4K20 me3
H3K4 me1:H2A.Z
H3K4 me3:H3−5ac
DNA Meth. m5C
H3K27 me3
H4K20 me2
H3K4 me3:H4−4ac
H2A.Z
H3K4 me1
H3−5ac
H4−4ac
H3K4 me3
0.00
0.25
0.50
0.75
1.00
π̂
EBP − Forward
H2A.Z:H4K20 me2
DNA Meth. m5C:H4K20 me2
H3K27 me2:DNA Meth. m5C
H3K4 me1:H4−4ac
H3K4 me1:H4K20 me2
DNA Meth. m5C:H4K20 me3
H2A.Z:H3K4 me3
H3K27 me3:H3K4 me1
H3K27 me3:H4K20 me3
H3K4 me1:H3−5ac
H3K9 me3:H4K20 me3
H4K20 me2:H3−5ac
H3K27 me3:DNA Meth. m5C
H4−4ac:H3−5ac
H4K20 me2:H3K4 me3
H3K4 me1:H2A.Z
H3K4 me3:H3−5ac
DNA Meth. m5C:H3K9 me3
H3K4 me3:H4−4ac
H3K27 me3:H3K9 me3
H3K9 me2
H3K27 me2
H4K20 me3
H3K9 me3
H4K20 me2
H2A.Z
H3K4 me1
DNA Meth. m5C
H3K27 me3
H3−5ac
H4−4ac
H3K4 me3
0.00
0.25
0.50
0.75
1.00
π̂
EBP − Reverse
H2A.Z:H4K20 me2
DNA Meth. m5C:H4K20 me2
H3K27 me2:DNA Meth. m5C
H3K4 me1:H4−4ac
H3K4 me1:H4K20 me2
DNA Meth. m5C:H4K20 me3
H2A.Z:H3K4 me3
H3K27 me3:H3K4 me1
H3K27 me3:H4K20 me3
H3K4 me1:H3−5ac
H3K9 me3:H4K20 me3
H4K20 me2:H3−5ac
H3K27 me3:DNA Meth. m5C
H4−4ac:H3−5ac
H4K20 me2:H3K4 me3
H3K4 me1:H2A.Z
H3K4 me3:H3−5ac
DNA Meth. m5C:H3K9 me3
H3K4 me3:H4−4ac
H3K27 me3:H3K9 me3
H3K9 me2
H3K27 me2
H4K20 me3
H3K9 me3
H4K20 me2
H2A.Z
H3K4 me1
DNA Meth. m5C
H3K27 me3
H3−5ac
H4−4ac
H3K4 me3
0.00
0.25
0.50
0.75
1.00
π̂
SRP54 − Forward
H4K20 me2:H4−4ac
DNA Meth. m5C:H4K20 me3
DNA Meth. m5C:H4K20 me2
H3K27 me2:DNA Meth. m5C
H3K4 me1:H4−4ac
H3K4 me1:H4K20 me2
H3K27 me3:H4K20 me3
H2A.Z:H4K20 me2
H3K4 me1:H3−5ac
H3K27 me3:H3K4 me1
H2A.Z:H3K4 me3
H3K9 me3:H4K20 me3
H4K20 me2:H3−5ac
H4−4ac:H3−5ac
H3K27 me3:DNA Meth. m5C
H4K20 me2:H3K4 me3
H3K4 me1:H2A.Z
H3K4 me3:H3−5ac
DNA Meth. m5C:H3K9 me3
H3K4 me3:H4−4ac
H3K27 me3:H3K9 me3
H3K9 me2
H3K27 me2
H4K20 me3
H4K20 me2
H3K9 me3
H2A.Z
DNA Meth. m5C
H3K4 me1
H3−5ac
H3K27 me3
H4−4ac
H3K4 me3
0.00
0.25
0.50
0.75
1.00
π̂
SRP54 − Reverse
H4K20 me2:H4−4ac
DNA Meth. m5C:H4K20 me3
DNA Meth. m5C:H4K20 me2
H3K27 me2:DNA Meth. m5C
H3K4 me1:H4−4ac
H3K4 me1:H4K20 me2
H3K27 me3:H4K20 me3
H2A.Z:H4K20 me2
H3K4 me1:H3−5ac
H3K27 me3:H3K4 me1
H2A.Z:H3K4 me3
H3K9 me3:H4K20 me3
H4K20 me2:H3−5ac
H4−4ac:H3−5ac
H3K27 me3:DNA Meth. m5C
H4K20 me2:H3K4 me3
H3K4 me1:H2A.Z
H3K4 me3:H3−5ac
DNA Meth. m5C:H3K9 me3
H3K4 me3:H4−4ac
H3K27 me3:H3K9 me3
H3K9 me2
H3K27 me2
H4K20 me3
H4K20 me2
H3K9 me3
H2A.Z
DNA Meth. m5C
H3K4 me1
H3−5ac
H3K27 me3
H4−4ac
H3K4 me3
0.00
0.25
0.50
0.75
1.00
π̂
SYNPO − Forward
H3K4 me3:H4−4ac
H3K4 me1:H3−5ac
H3K27 me3:H3K4 me1
H3K4 me3:H3−5ac
H3K27 me3:H3K9 me3
H2A.Z:H4−4ac
DNA Meth. m5C:H4K20 me2
H2A.Z:H3−5ac
H3K9 me2:DNA Meth. m5C
H3K27 me3:H4K20 me3
H4−4ac:H3−5ac
H3K9 me3:H4K20 me3
DNA Meth. m5C:H3K9 me3
DNA Meth. m5C:H4K20 me3
H2A.Z:H4K20 me2
H3K27 me3:DNA Meth. m5C
H4K20 me2:H4−4ac
H4K20 me2:H3−5ac
H3K4 me1:H4−4ac
H3K27 me2:DNA Meth. m5C
H3K4 me1:H2A.Z
H2A.Z:H3K4 me3
H4K20 me2:H3K4 me3
H3K9 me2
H3K27 me2
H3K27 me3
H3−5ac
H3K9 me3
H3K4 me3
H3K4 me1
H4−4ac
DNA Meth. m5C
H4K20 me2
H4K20 me3
H2A.Z
0.00
0.25
0.50
0.75
1.00
π̂
SYNPO − Reverse
H3K4 me3:H4−4ac
H3K4 me1:H3−5ac
H3K27 me3:H3K4 me1
H3K4 me3:H3−5ac
H3K27 me3:H3K9 me3
H2A.Z:H4−4ac
DNA Meth. m5C:H4K20 me2
H2A.Z:H3−5ac
H3K9 me2:DNA Meth. m5C
H3K27 me3:H4K20 me3
H4−4ac:H3−5ac
H3K9 me3:H4K20 me3
DNA Meth. m5C:H3K9 me3
DNA Meth. m5C:H4K20 me3
H2A.Z:H4K20 me2
H3K27 me3:DNA Meth. m5C
H4K20 me2:H4−4ac
H4K20 me2:H3−5ac
H3K4 me1:H4−4ac
H3K27 me2:DNA Meth. m5C
H3K4 me1:H2A.Z
H2A.Z:H3K4 me3
H4K20 me2:H3K4 me3
H3K9 me2
H3K27 me2
H3K27 me3
H3−5ac
H3K9 me3
H3K4 me3
H3K4 me1
H4−4ac
DNA Meth. m5C
H4K20 me2
H4K20 me3
H2A.Z
0.00
0.25
0.50
0.75
1.00
π̂
SEC22A − Forward
H2A.Z:H4K20 me2
H2A.Z:H4−4ac
H3K27 me2:DNA Meth. m5C
H3K9 me2:DNA Meth. m5C
H2A.Z:H3K4 me3
H3K4 me3:H3−5ac
H4K20 me2:H3−5ac
H4−4ac:H3−5ac
DNA Meth. m5C:H4K20 me2
H3K27 me3:H3K9 me3
H4K20 me2:H3K4 me3
H3K27 me3:DNA Meth. m5C
H3K4 me3:H4−4ac
H3K27 me3:H4K20 me3
H3K4 me1:H3−5ac
DNA Meth. m5C:H3K9 me3
H3K9 me3:H4K20 me3
H4K20 me2:H4−4ac
DNA Meth. m5C:H4K20 me3
H3K27 me3:H3K4 me1
H3K27 me2
H3K9 me2
H2A.Z
H3−5ac
H4K20 me3
H4−4ac
H3K27 me3
H4K20 me2
H3K4 me1
H3K9 me3
H3K4 me3
DNA Meth. m5C
0.00
0.25
0.50
0.75
1.00
π̂
SEC22A − Reverse
H2A.Z:H4K20 me2
H2A.Z:H4−4ac
H3K27 me2:DNA Meth. m5C
H3K9 me2:DNA Meth. m5C
H2A.Z:H3K4 me3
H3K4 me3:H3−5ac
H4K20 me2:H3−5ac
H4−4ac:H3−5ac
DNA Meth. m5C:H4K20 me2
H3K27 me3:H3K9 me3
H4K20 me2:H3K4 me3
H3K27 me3:DNA Meth. m5C
H3K4 me3:H4−4ac
H3K27 me3:H4K20 me3
H3K4 me1:H3−5ac
DNA Meth. m5C:H3K9 me3
H3K9 me3:H4K20 me3
H4K20 me2:H4−4ac
DNA Meth. m5C:H4K20 me3
H3K27 me3:H3K4 me1
H3K27 me2
H3K9 me2
H2A.Z
H3−5ac
H4K20 me3
H4−4ac
H3K27 me3
H4K20 me2
H3K4 me1
H3K9 me3
H3K4 me3
DNA Meth. m5C
0.00
0.25
0.50
0.75
1.00
π̂
PTBP3 − Forward
H3K4 me1:H4K20 me2
H4K20 me2:H3−5ac
H3K4 me3:H4−4ac
H3K4 me1:H4−4ac
H3K27 me3:H3K4 me1
H4K20 me2:H4−4ac
H4K20 me2:H3K4 me3
H3K4 me1:H2A.Z
H2A.Z:H3−5ac
H3K27 me3:H4K20 me3
H3K9 me3:H4K20 me3
H2A.Z:H4−4ac
H2A.Z:H3K4 me3
H2A.Z:H4K20 me2
H3K27 me3:H3K9 me3
H3K27 me2:DNA Meth. m5C
DNA Meth. m5C:H3K9 me3
DNA Meth. m5C:H4K20 me3
H3K27 me3:DNA Meth. m5C
H3−5ac
H3K9 me2
H2A.Z
H4−4ac
H4K20 me2
H3K4 me3
H3K4 me1
H3K27 me2
H4K20 me3
H3K27 me3
H3K9 me3
DNA Meth. m5C
0.00
0.25
0.50
0.75
1.00
π̂
PTBP3 − Reverse
H3K4 me1:H4K20 me2
H4K20 me2:H3−5ac
H3K4 me3:H4−4ac
H3K4 me1:H4−4ac
H3K27 me3:H3K4 me1
H4K20 me2:H4−4ac
H4K20 me2:H3K4 me3
H3K4 me1:H2A.Z
H2A.Z:H3−5ac
H3K27 me3:H4K20 me3
H3K9 me3:H4K20 me3
H2A.Z:H4−4ac
H2A.Z:H3K4 me3
H2A.Z:H4K20 me2
H3K27 me3:H3K9 me3
H3K27 me2:DNA Meth. m5C
DNA Meth. m5C:H3K9 me3
DNA Meth. m5C:H4K20 me3
H3K27 me3:DNA Meth. m5C
H3−5ac
H3K9 me2
H2A.Z
H4−4ac
H4K20 me2
H3K4 me3
H3K4 me1
H3K27 me2
H4K20 me3
H3K27 me3
H3K9 me3
DNA Meth. m5C
0.00
0.25
0.50
0.75
1.00
π̂

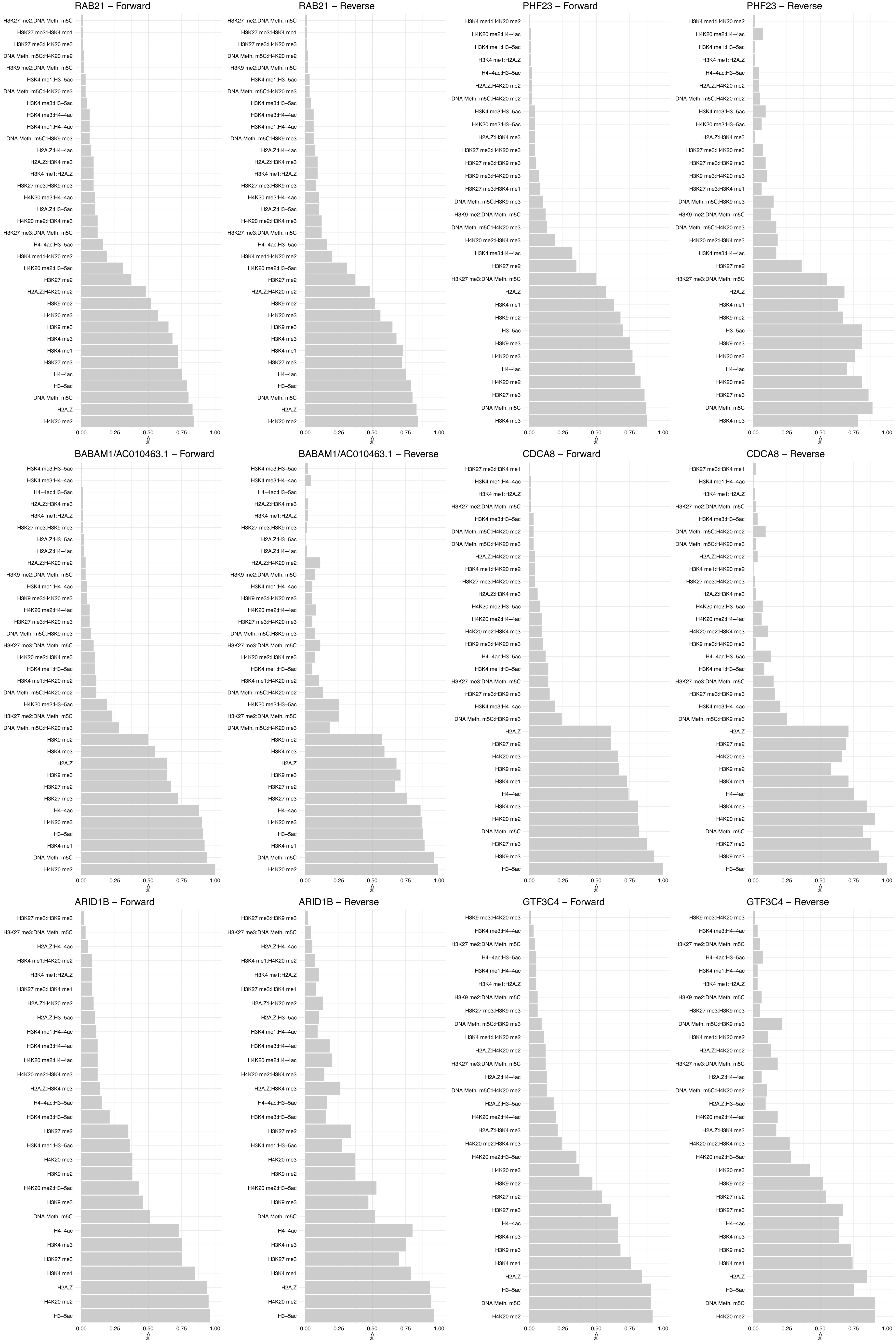
RAB21 – Forward
RAB21 – Reverse
PHF23 – Forward
PHF23 – Reverse
BABAM1/AC010463.1 – Forward
BABAM1/AC010463.1 – Reverse
CDCA8 – Forward
CDCA8 – Reverse
ARID1B – Forward
ARID1B – Reverse
GTF3C4 – Forward
GTF3C4 – Reverse

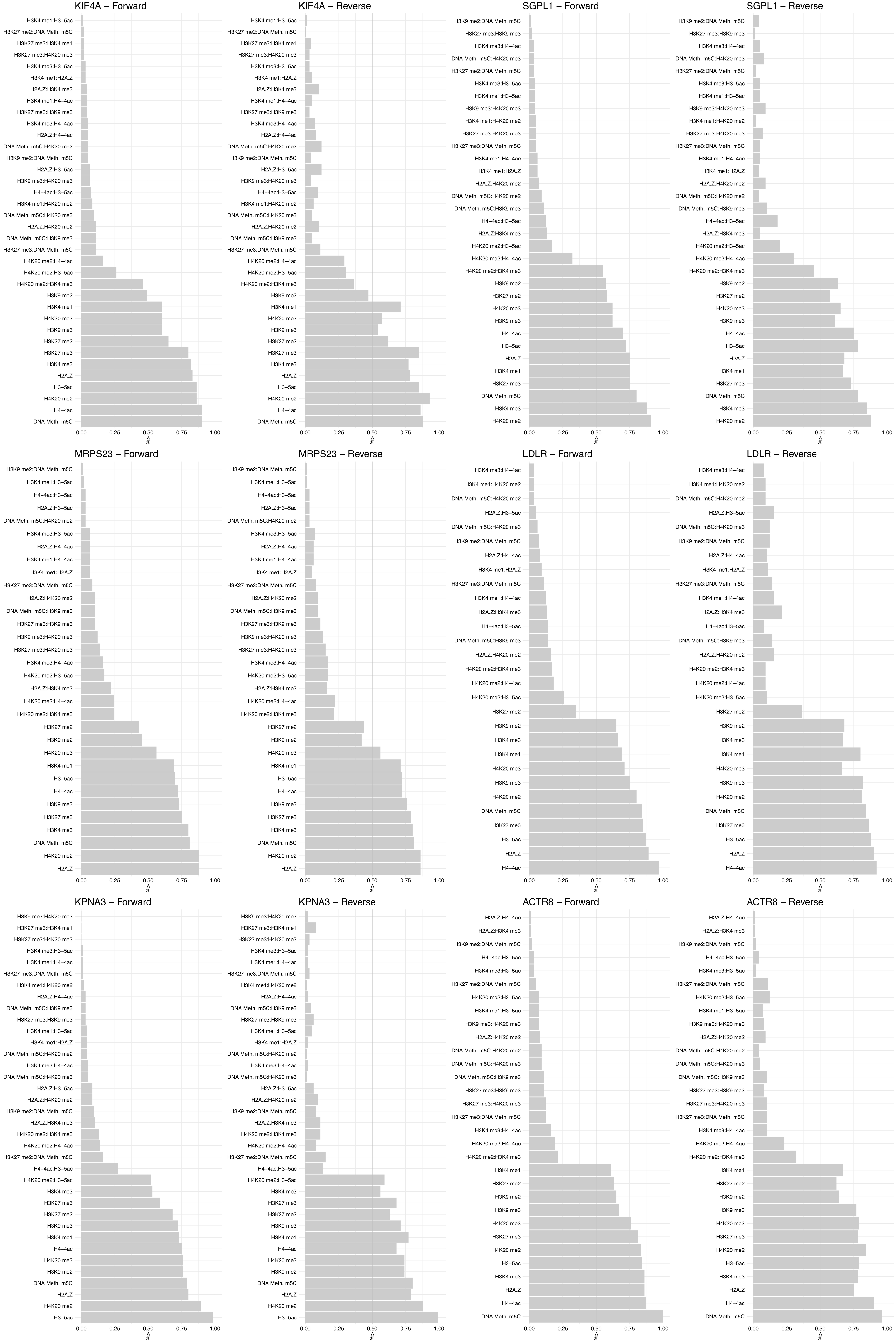
KIF4A – Forward
H3K4 me1:H3–5ac
H3K27 me2:DNA Meth. m5C
H3K27 me3:H3K4 me1
H3K27 me3:H4K20 me3
H3K4 me3:H3–5ac
H3K4 me1:H2A.Z
H2A.Z:H3K4 me3
H3K4 me1:H4–4ac
H3K27 me3:H3K9 me3
H3K4 me3:H4–4ac
H2A.Z:H4–4ac
DNA Meth. m5C:H4K20 me2
H3K9 me2:DNA Meth. m5C
H2A.Z:H3–5ac
H3K9 me3:H4K20 me3
H4–4ac:H3–5ac
H3K4 me1:H4K20 me2
DNA Meth. m5C:H4K20 me3
H2A.Z:H4K20 me2
DNA Meth. m5C:H3K9 me3
H3K27 me3:DNA Meth. m5C
H4K20 me2:H4–4ac
H4K20 me2:H3–5ac
H4K20 me2:H3K4 me3
H3K9 me2
H3K4 me1
H4K20 me3
H3K9 me3
H3K27 me2
H3K27 me3
H3K4 me3
H2A.Z
H3–5ac
H4K20 me2
H4–4ac
DNA Meth. m5C
0.00 0.25 0.50 0.75 1.00
π̂
KIF4A – Reverse
H3K4 me1:H3–5ac
H3K27 me2:DNA Meth. m5C
H3K27 me3:H3K4 me1
H3K27 me3:H4K20 me3
H3K4 me3:H3–5ac
H3K4 me1:H2A.Z
H2A.Z:H3K4 me3
H3K4 me1:H4–4ac
H3K27 me3:H3K9 me3
H3K4 me3:H4–4ac
H2A.Z:H4–4ac
DNA Meth. m5C:H4K20 me2
H3K9 me2:DNA Meth. m5C
H2A.Z:H3–5ac
H3K9 me3:H4K20 me3
H4–4ac:H3–5ac
H3K4 me1:H4K20 me2
DNA Meth. m5C:H4K20 me3
H2A.Z:H4K20 me2
DNA Meth. m5C:H3K9 me3
H3K27 me3:DNA Meth. m5C
H4K20 me2:H4–4ac
H4K20 me2:H3–5ac
H4K20 me2:H3K4 me3
H3K9 me2
H3K4 me1
H4K20 me3
H3K9 me3
H3K27 me2
H3K27 me3
H3K4 me3
H2A.Z
H3–5ac
H4K20 me2
H4–4ac
DNA Meth. m5C
0.00 0.25 0.50 0.75 1.00
π̂
SGPL1 – Forward
H3K9 me2:DNA Meth. m5C
H3K27 me3:H3K9 me3
H3K4 me3:H4–4ac
DNA Meth. m5C:H4K20 me3
H3K27 me2:DNA Meth. m5C
H3K4 me3:H3–5ac
H3K4 me1:H3–5ac
H3K9 me3:H4K20 me3
H3K4 me1:H4K20 me2
H3K27 me3:H4K20 me3
H3K27 me3:DNA Meth. m5C
H3K4 me1:H4–4ac
H3K4 me1:H2A.Z
H2A.Z:H4K20 me2
DNA Meth. m5C:H4K20 me2
DNA Meth. m5C:H3K9 me3
H4–4ac:H3–5ac
H2A.Z:H3K4 me3
H4K20 me2:H3–5ac
H4K20 me2:H4–4ac
H4K20 me2:H3K4 me3
H3K9 me2
H3K27 me2
H4K20 me3
H3K9 me3
H4–4ac
H3–5ac
H2A.Z
H3K4 me1
H3K27 me3
DNA Meth. m5C
H3K4 me3
H4K20 me2
0.00 0.25 0.50 0.75 1.00
π̂
SGPL1 – Reverse
H3K9 me2:DNA Meth. m5C
H3K27 me3:H3K9 me3
H3K4 me3:H4–4ac
DNA Meth. m5C:H4K20 me3
H3K27 me2:DNA Meth. m5C
H3K4 me3:H3–5ac
H3K4 me1:H3–5ac
H3K9 me3:H4K20 me3
H3K4 me1:H4K20 me2
H3K27 me3:H4K20 me3
H3K27 me3:DNA Meth. m5C
H3K4 me1:H4–4ac
H3K4 me1:H2A.Z
H2A.Z:H4K20 me2
DNA Meth. m5C:H4K20 me2
DNA Meth. m5C:H3K9 me3
H4–4ac:H3–5ac
H2A.Z:H3K4 me3
H4K20 me2:H3–5ac
H4K20 me2:H4–4ac
H4K20 me2:H3K4 me3
H3K9 me2
H3K27 me2
H4K20 me3
H3K9 me3
H4–4ac
H3–5ac
H2A.Z
H3K4 me1
H3K27 me3
DNA Meth. m5C
H3K4 me3
H4K20 me2
0.00 0.25 0.50 0.75 1.00
π̂
MRPS23 – Forward
H3K9 me2:DNA Meth. m5C
H3K4 me1:H3–5ac
H4–4ac:H3–5ac
H2A.Z:H3–5ac
DNA Meth. m5C:H4K20 me2
H3K4 me3:H3–5ac
H2A.Z:H4–4ac
H3K4 me1:H4–4ac
H3K4 me1:H2A.Z
H3K27 me3:DNA Meth. m5C
H2A.Z:H4K20 me2
DNA Meth. m5C:H3K9 me3
H3K27 me3:H3K9 me3
H3K9 me3:H4K20 me3
H3K27 me3:H4K20 me3
H3K4 me3:H4–4ac
H4K20 me2:H3–5ac
H2A.Z:H3K4 me3
H4K20 me2:H4–4ac
H4K20 me2:H3K4 me3
H3K27 me2
H3K9 me2
H4K20 me3
H3K4 me1
H3–5ac
H4–4ac
H3K9 me3
H3K27 me3
H3K4 me3
DNA Meth. m5C
H4K20 me2
H2A.Z
0.00 0.25 0.50 0.75 1.00
π̂
MRPS23 – Reverse
H3K9 me2:DNA Meth. m5C
H3K4 me1:H3–5ac
H4–4ac:H3–5ac
H2A.Z:H3–5ac
DNA Meth. m5C:H4K20 me2
H3K4 me3:H3–5ac
H2A.Z:H4–4ac
H3K4 me1:H4–4ac
H3K4 me1:H2A.Z
H3K27 me3:DNA Meth. m5C
H2A.Z:H4K20 me2
DNA Meth. m5C:H3K9 me3
H3K27 me3:H3K9 me3
H3K9 me3:H4K20 me3
H3K27 me3:H4K20 me3
H3K4 me3:H4–4ac
H4K20 me2:H3–5ac
H2A.Z:H3K4 me3
H4K20 me2:H4–4ac
H4K20 me2:H3K4 me3
H3K27 me2
H3K9 me2
H4K20 me3
H3K4 me1
H3–5ac
H4–4ac
H3K9 me3
H3K27 me3
H3K4 me3
DNA Meth. m5C
H4K20 me2
H2A.Z
0.00 0.25 0.50 0.75 1.00
π̂
LDLR – Forward
H3K4 me3:H4–4ac
H3K4 me1:H4K20 me2
DNA Meth. m5C:H4K20 me2
H2A.Z:H3–5ac
DNA Meth. m5C:H4K20 me3
H3K9 me2:DNA Meth. m5C
H2A.Z:H4–4ac
H3K4 me1:H2A.Z
H3K27 me3:DNA Meth. m5C
H3K4 me1:H4–4ac
H2A.Z:H3K4 me3
H4–4ac:H3–5ac
DNA Meth. m5C:H3K9 me3
H2A.Z:H4K20 me2
H4K20 me2:H3K4 me3
H4K20 me2:H4–4ac
H4K20 me2:H3–5ac
H3K27 me2
H3K9 me2
H3K4 me3
H3K4 me1
H4K20 me3
H3K9 me3
H4K20 me2
DNA Meth. m5C
H3K27 me3
H3–5ac
H2A.Z
H4–4ac
0.00 0.25 0.50 0.75 1.00
π̂
LDLR – Reverse
H3K4 me3:H4–4ac
H3K4 me1:H4K20 me2
DNA Meth. m5C:H4K20 me2
H2A.Z:H3–5ac
DNA Meth. m5C:H4K20 me3
H3K9 me2:DNA Meth. m5C
H2A.Z:H4–4ac
H3K4 me1:H2A.Z
H3K27 me3:DNA Meth. m5C
H3K4 me1:H4–4ac
H2A.Z:H3K4 me3
H4–4ac:H3–5ac
DNA Meth. m5C:H3K9 me3
H2A.Z:H4K20 me2
H4K20 me2:H3K4 me3
H4K20 me2:H4–4ac
H4K20 me2:H3–5ac
H3K27 me2
H3K9 me2
H3K4 me3
H3K4 me1
H4K20 me3
H3K9 me3
H4K20 me2
DNA Meth. m5C
H3K27 me3
H3–5ac
H2A.Z
H4–4ac
0.00 0.25 0.50 0.75 1.00
π̂
KPNA3 – Forward
H3K9 me3:H4K20 me3
H3K27 me3:H3K4 me1
H3K27 me3:H4K20 me3
H3K4 me3:H3–5ac
H3K4 me1:H4–4ac
H3K27 me3:DNA Meth. m5C
H3K4 me1:H4K20 me2
H2A.Z:H4–4ac
DNA Meth. m5C:H3K9 me3
H3K27 me3:H3K9 me3
H3K4 me1:H3–5ac
H3K4 me1:H2A.Z
DNA Meth. m5C:H4K20 me2
H3K4 me3:H4–4ac
DNA Meth. m5C:H4K20 me3
H2A.Z:H3–5ac
H2A.Z:H4K20 me2
H3K9 me2:DNA Meth. m5C
H2A.Z:H3K4 me3
H4K20 me2:H3K4 me3
H4K20 me2:H4–4ac
H3K27 me2:DNA Meth. m5C
H4–4ac:H3–5ac
H4K20 me2:H3–5ac
H3K4 me3
H3K27 me3
H3K27 me2
H3K9 me3
H3K4 me1
H4–4ac
H4K20 me3
H3K9 me2
DNA Meth. m5C
H2A.Z
H4K20 me2
H3–5ac
0.00 0.25 0.50 0.75 1.00
π̂
KPNA3 – Reverse
H3K9 me3:H4K20 me3
H3K27 me3:H3K4 me1
H3K27 me3:H4K20 me3
H3K4 me3:H3–5ac
H3K4 me1:H4–4ac
H3K27 me3:DNA Meth. m5C
H3K4 me1:H4K20 me2
H2A.Z:H4–4ac
DNA Meth. m5C:H3K9 me3
H3K27 me3:H3K9 me3
H3K4 me1:H3–5ac
H3K4 me1:H2A.Z
DNA Meth. m5C:H4K20 me2
H3K4 me3:H4–4ac
DNA Meth. m5C:H4K20 me3
H2A.Z:H3–5ac
H2A.Z:H4K20 me2
H3K9 me2:DNA Meth. m5C
H2A.Z:H3K4 me3
H4K20 me2:H3K4 me3
H4K20 me2:H4–4ac
H3K27 me2:DNA Meth. m5C
H4–4ac:H3–5ac
H4K20 me2:H3–5ac
H3K4 me3
H3K27 me3
H3K27 me2
H3K9 me3
H3K4 me1
H4–4ac
H4K20 me3
H3K9 me2
DNA Meth. m5C
H2A.Z
H4K20 me2
H3–5ac
0.00 0.25 0.50 0.75 1.00
π̂
ACTR8 – Forward
H2A.Z:H4–4ac
H2A.Z:H3K4 me3
H3K9 me2:DNA Meth. m5C
H4–4ac:H3–5ac
H3K4 me3:H3–5ac
H3K27 me2:DNA Meth. m5C
H4K20 me2:H3–5ac
H3K4 me1:H3–5ac
H3K9 me3:H4K20 me3
H2A.Z:H4K20 me2
DNA Meth. m5C:H4K20 me2
DNA Meth. m5C:H4K20 me3
DNA Meth. m5C:H3K9 me3
H3K27 me3:H3K9 me3
H3K27 me3:H4K20 me3
H3K27 me3:DNA Meth. m5C
H3K4 me3:H4–4ac
H4K20 me2:H4–4ac
H4K20 me2:H3K4 me3
H3K4 me1
H3K27 me2
H3K9 me2
H3K9 me3
H4K20 me3
H3K27 me3
H4K20 me2
H3–5ac
H3K4 me3
H2A.Z
H4–4ac
DNA Meth. m5C
0.00 0.25 0.50 0.75 1.00
π̂
ACTR8 – Reverse
H2A.Z:H4–4ac
H2A.Z:H3K4 me3
H3K9 me2:DNA Meth. m5C
H4–4ac:H3–5ac
H3K4 me3:H3–5ac
H3K27 me2:DNA Meth. m5C
H4K20 me2:H3–5ac
H3K4 me1:H3–5ac
H3K9 me3:H4K20 me3
H2A.Z:H4K20 me2
DNA Meth. m5C:H4K20 me2
DNA Meth. m5C:H4K20 me3
DNA Meth. m5C:H3K9 me3
H3K27 me3:H3K9 me3
H3K27 me3:H4K20 me3
H3K27 me3:DNA Meth. m5C
H3K4 me3:H4–4ac
H4K20 me2:H4–4ac
H4K20 me2:H3K4 me3
H3K4 me1
H3K27 me2
H3K9 me2
H3K9 me3
H4K20 me3
H3K27 me3
H4K20 me2
H3–5ac
H3K4 me3
H2A.Z
H4–4ac
DNA Meth. m5C
0.00 0.25 0.50 0.75 1.00
π̂

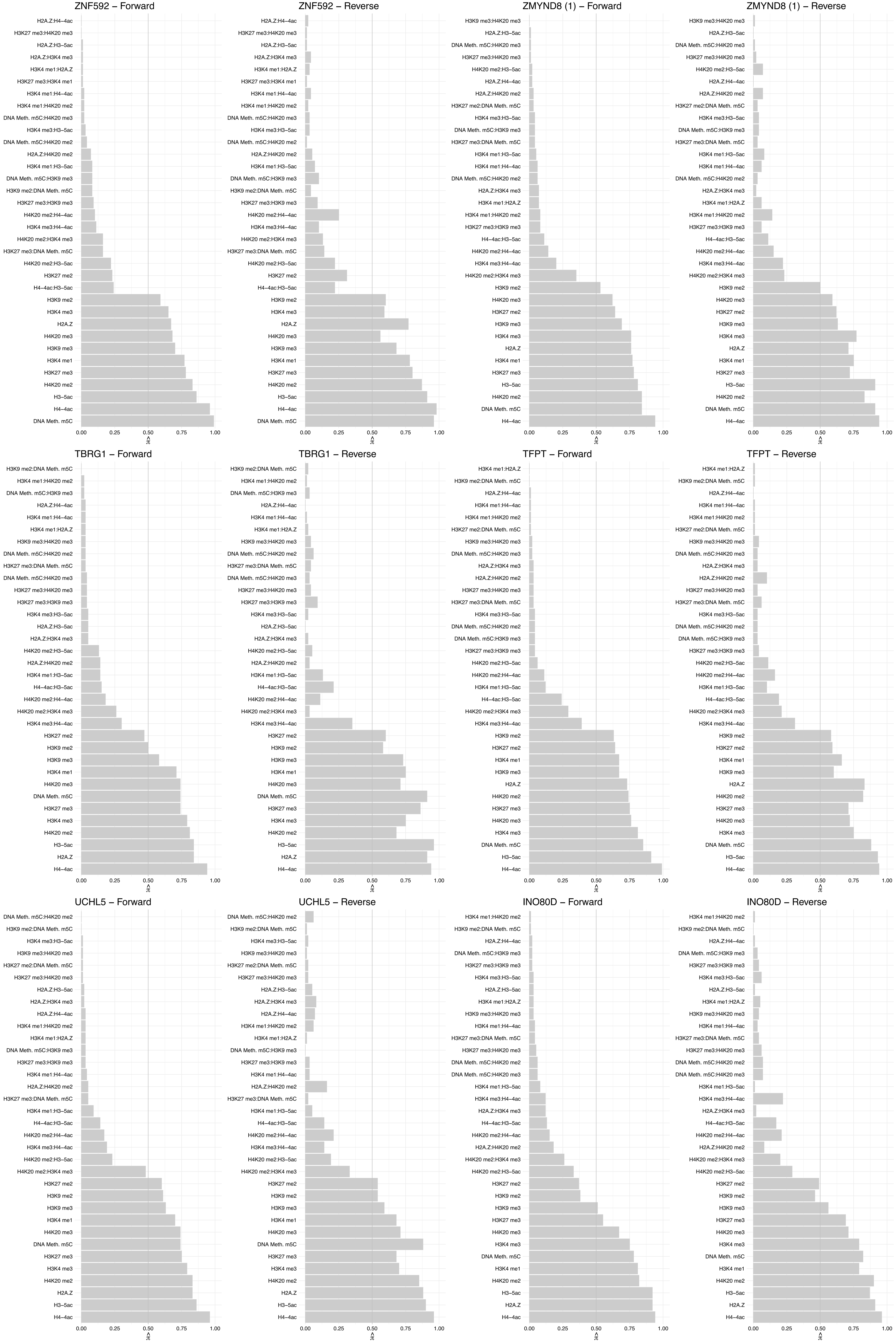
ZNF592 – Forward
H2A.Z:H4–4ac
H3K27 me3:H4K20 me3
H2A.Z:H3–5ac
H2A.Z:H3K4 me3
H3K4 me1:H2A.Z
H3K27 me3:H3K4 me1
H3K4 me1:H4–4ac
H3K4 me1:H4K20 me2
DNA Meth. m5C:H4K20 me3
H3K4 me3:H3–5ac
DNA Meth. m5C:H4K20 me2
H2A.Z:H4K20 me2
H3K4 me1:H3–5ac
DNA Meth. m5C:H3K9 me3
H3K9 me2:DNA Meth. m5C
H3K27 me3:H3K9 me3
H4K20 me2:H4–4ac
H3K4 me3:H4–4ac
H4K20 me2:H3K4 me3
H3K27 me3:DNA Meth. m5C
H4K20 me2:H3–5ac
H3K27 me2
H4–4ac:H3–5ac
H3K9 me2
H3K4 me3
H2A.Z
H4K20 me3
H3K9 me3
H3K4 me1
H3K27 me3
H4K20 me2
H3–5ac
H4–4ac
DNA Meth. m5C
0.00 0.25 0.50 0.75 1.00
π̂
ZNF592 – Reverse
H2A.Z:H4–4ac
H3K27 me3:H4K20 me3
H2A.Z:H3–5ac
H2A.Z:H3K4 me3
H3K4 me1:H2A.Z
H3K27 me3:H3K4 me1
H3K4 me1:H4–4ac
H3K4 me1:H4K20 me2
DNA Meth. m5C:H4K20 me3
H3K4 me3:H3–5ac
DNA Meth. m5C:H4K20 me2
H2A.Z:H4K20 me2
H3K4 me1:H3–5ac
DNA Meth. m5C:H3K9 me3
H3K9 me2:DNA Meth. m5C
H3K27 me3:H3K9 me3
H4K20 me2:H4–4ac
H3K4 me3:H4–4ac
H4K20 me2:H3K4 me3
H3K27 me3:DNA Meth. m5C
H4K20 me2:H3–5ac
H3K27 me2
H4–4ac:H3–5ac
H3K9 me2
H3K4 me3
H2A.Z
H4K20 me3
H3K9 me3
H3K4 me1
H3K27 me3
H4K20 me2
H3–5ac
H4–4ac
DNA Meth. m5C
0.00 0.25 0.50 0.75 1.00
π̂
ZMYND8 (1) – Forward
H3K9 me3:H4K20 me3
H2A.Z:H3–5ac
DNA Meth. m5C:H4K20 me3
H3K27 me3:H4K20 me3
H4K20 me2:H3–5ac
H2A.Z:H4–4ac
H2A.Z:H4K20 me2
H3K27 me2:DNA Meth. m5C
H3K4 me3:H3–5ac
DNA Meth. m5C:H3K9 me3
H3K27 me3:DNA Meth. m5C
H3K4 me1:H3–5ac
H3K4 me1:H4–4ac
DNA Meth. m5C:H4K20 me2
H2A.Z:H3K4 me3
H3K4 me1:H2A.Z
H3K4 me1:H4K20 me2
H3K27 me3:H3K9 me3
H4–4ac:H3–5ac
H4K20 me2:H4–4ac
H3K4 me3:H4–4ac
H4K20 me2:H3K4 me3
H3K9 me2
H4K20 me3
H3K27 me2
H3K9 me3
H3K4 me3
H2A.Z
H3K4 me1
H3K27 me3
H3–5ac
H4K20 me2
DNA Meth. m5C
H4–4ac
0.00 0.25 0.50 0.75 1.00
π̂
ZMYND8 (1) – Reverse
H3K9 me3:H4K20 me3
H2A.Z:H3–5ac
DNA Meth. m5C:H4K20 me3
H3K27 me3:H4K20 me3
H4K20 me2:H3–5ac
H2A.Z:H4–4ac
H2A.Z:H4K20 me2
H3K27 me2:DNA Meth. m5C
H3K4 me3:H3–5ac
DNA Meth. m5C:H3K9 me3
H3K27 me3:DNA Meth. m5C
H3K4 me1:H3–5ac
H3K4 me1:H4–4ac
DNA Meth. m5C:H4K20 me2
H2A.Z:H3K4 me3
H3K4 me1:H2A.Z
H3K4 me1:H4K20 me2
H3K27 me3:H3K9 me3
H4–4ac:H3–5ac
H4K20 me2:H4–4ac
H3K4 me3:H4–4ac
H4K20 me2:H3K4 me3
H3K9 me2
H4K20 me3
H3K27 me2
H3K9 me3
H3K4 me3
H2A.Z
H3K4 me1
H3K27 me3
H3–5ac
H4K20 me2
DNA Meth. m5C
H4–4ac
0.00 0.25 0.50 0.75 1.00
π̂
TBRG1 – Forward
H3K9 me2:DNA Meth. m5C
H3K4 me1:H4K20 me2
DNA Meth. m5C:H3K9 me3
H2A.Z:H4–4ac
H3K4 me1:H4–4ac
H3K4 me1:H2A.Z
H3K9 me3:H4K20 me3
DNA Meth. m5C:H4K20 me2
H3K27 me3:DNA Meth. m5C
DNA Meth. m5C:H4K20 me3
H3K27 me3:H4K20 me3
H3K27 me3:H3K9 me3
H3K4 me3:H3–5ac
H2A.Z:H3–5ac
H2A.Z:H3K4 me3
H4K20 me2:H3–5ac
H2A.Z:H4K20 me2
H3K4 me1:H3–5ac
H4–4ac:H3–5ac
H4K20 me2:H4–4ac
H4K20 me2:H3K4 me3
H3K4 me3:H4–4ac
H3K27 me2
H3K9 me2
H3K9 me3
H3K4 me1
H4K20 me3
DNA Meth. m5C
H3K27 me3
H3K4 me3
H4K20 me2
H3–5ac
H2A.Z
H4–4ac
0.00 0.25 0.50 0.75 1.00
π̂
TBRG1 – Reverse
H3K9 me2:DNA Meth. m5C
H3K4 me1:H4K20 me2
DNA Meth. m5C:H3K9 me3
H2A.Z:H4–4ac
H3K4 me1:H4–4ac
H3K4 me1:H2A.Z
H3K9 me3:H4K20 me3
DNA Meth. m5C:H4K20 me2
H3K27 me3:DNA Meth. m5C
DNA Meth. m5C:H4K20 me3
H3K27 me3:H4K20 me3
H3K27 me3:H3K9 me3
H3K4 me3:H3–5ac
H2A.Z:H3–5ac
H2A.Z:H3K4 me3
H4K20 me2:H3–5ac
H2A.Z:H4K20 me2
H3K4 me1:H3–5ac
H4–4ac:H3–5ac
H4K20 me2:H4–4ac
H4K20 me2:H3K4 me3
H3K4 me3:H4–4ac
H3K27 me2
H3K9 me2
H3K9 me3
H3K4 me1
H4K20 me3
DNA Meth. m5C
H3K27 me3
H3K4 me3
H4K20 me2
H3–5ac
H2A.Z
H4–4ac
0.00 0.25 0.50 0.75 1.00
π̂
TFPT – Forward
H3K4 me1:H2A.Z
H3K9 me2:DNA Meth. m5C
H2A.Z:H4–4ac
H3K4 me1:H4–4ac
H3K4 me1:H4K20 me2
H3K27 me2:DNA Meth. m5C
H3K9 me3:H4K20 me3
DNA Meth. m5C:H4K20 me3
H2A.Z:H3K4 me3
H2A.Z:H4K20 me2
H3K27 me3:H4K20 me3
H3K27 me3:DNA Meth. m5C
H3K4 me3:H3–5ac
DNA Meth. m5C:H4K20 me2
DNA Meth. m5C:H3K9 me3
H3K27 me3:H3K9 me3
H4K20 me2:H3–5ac
H4K20 me2:H4–4ac
H3K4 me1:H3–5ac
H4–4ac:H3–5ac
H4K20 me2:H3K4 me3
H3K4 me3:H4–4ac
H3K9 me2
H3K27 me2
H3K4 me1
H3K9 me3
H2A.Z
H4K20 me2
H3K27 me3
H4K20 me3
H3K4 me3
DNA Meth. m5C
H3–5ac
H4–4ac
0.00 0.25 0.50 0.75 1.00
π̂
TFPT – Reverse
H3K4 me1:H2A.Z
H3K9 me2:DNA Meth. m5C
H2A.Z:H4–4ac
H3K4 me1:H4–4ac
H3K4 me1:H4K20 me2
H3K27 me2:DNA Meth. m5C
H3K9 me3:H4K20 me3
DNA Meth. m5C:H4K20 me3
H2A.Z:H3K4 me3
H2A.Z:H4K20 me2
H3K27 me3:H4K20 me3
H3K27 me3:DNA Meth. m5C
H3K4 me3:H3–5ac
DNA Meth. m5C:H4K20 me2
DNA Meth. m5C:H3K9 me3
H3K27 me3:H3K9 me3
H4K20 me2:H3–5ac
H4K20 me2:H4–4ac
H3K4 me1:H3–5ac
H4–4ac:H3–5ac
H4K20 me2:H3K4 me3
H3K4 me3:H4–4ac
H3K9 me2
H3K27 me2
H3K4 me1
H3K9 me3
H2A.Z
H4K20 me2
H3K27 me3
H4K20 me3
H3K4 me3
DNA Meth. m5C
H3–5ac
H4–4ac
0.00 0.25 0.50 0.75 1.00
π̂
UCHL5 – Forward
DNA Meth. m5C:H4K20 me2
H3K9 me2:DNA Meth. m5C
H3K4 me3:H3–5ac
H3K9 me3:H4K20 me3
H3K27 me2:DNA Meth. m5C
H3K27 me3:H4K20 me3
H2A.Z:H3–5ac
H2A.Z:H3K4 me3
H2A.Z:H4–4ac
H3K4 me1:H4K20 me2
H3K4 me1:H2A.Z
DNA Meth. m5C:H3K9 me3
H3K27 me3:H3K9 me3
H3K4 me1:H4–4ac
H2A.Z:H4K20 me2
H3K27 me3:DNA Meth. m5C
H3K4 me1:H3–5ac
H4–4ac:H3–5ac
H4K20 me2:H4–4ac
H3K4 me3:H4–4ac
H4K20 me2:H3–5ac
H4K20 me2:H3K4 me3
H3K27 me2
H3K9 me2
H3K9 me3
H3K4 me1
H4K20 me3
DNA Meth. m5C
H3K27 me3
H3K4 me3
H4K20 me2
H2A.Z
H3–5ac
H4–4ac
0.00 0.25 0.50 0.75 1.00
π̂
UCHL5 – Reverse
DNA Meth. m5C:H4K20 me2
H3K9 me2:DNA Meth. m5C
H3K4 me3:H3–5ac
H3K9 me3:H4K20 me3
H3K27 me2:DNA Meth. m5C
H3K27 me3:H4K20 me3
H2A.Z:H3–5ac
H2A.Z:H3K4 me3
H2A.Z:H4–4ac
H3K4 me1:H4K20 me2
H3K4 me1:H2A.Z
DNA Meth. m5C:H3K9 me3
H3K27 me3:H3K9 me3
H3K4 me1:H4–4ac
H2A.Z:H4K20 me2
H3K27 me3:DNA Meth. m5C
H3K4 me1:H3–5ac
H4–4ac:H3–5ac
H4K20 me2:H4–4ac
H3K4 me3:H4–4ac
H4K20 me2:H3–5ac
H4K20 me2:H3K4 me3
H3K27 me2
H3K9 me2
H3K9 me3
H3K4 me1
H4K20 me3
DNA Meth. m5C
H3K27 me3
H3K4 me3
H4K20 me2
H2A.Z
H3–5ac
H4–4ac
0.00 0.25 0.50 0.75 1.00
π̂
INO80D – Forward
H3K4 me1:H4K20 me2
H3K9 me2:DNA Meth. m5C
H2A.Z:H4–4ac
DNA Meth. m5C:H3K9 me3
H3K27 me3:H3K9 me3
H3K4 me3:H3–5ac
H2A.Z:H3–5ac
H3K4 me1:H2A.Z
H3K9 me3:H4K20 me3
H3K4 me1:H4–4ac
H3K27 me3:DNA Meth. m5C
H3K27 me3:H4K20 me3
DNA Meth. m5C:H4K20 me2
DNA Meth. m5C:H4K20 me3
H3K4 me1:H3–5ac
H3K4 me3:H4–4ac
H2A.Z:H3K4 me3
H4–4ac:H3–5ac
H4K20 me2:H4–4ac
H2A.Z:H4K20 me2
H4K20 me2:H3K4 me3
H4K20 me2:H3–5ac
H3K27 me2
H3K9 me2
H3K9 me3
H3K27 me3
H4K20 me3
H3K4 me3
DNA Meth. m5C
H3K4 me1
H4K20 me2
H3–5ac
H2A.Z
H4–4ac
0.00 0.25 0.50 0.75 1.00
π̂
INO80D – Reverse
H3K4 me1:H4K20 me2
H3K9 me2:DNA Meth. m5C
H2A.Z:H4–4ac
DNA Meth. m5C:H3K9 me3
H3K27 me3:H3K9 me3
H3K4 me3:H3–5ac
H2A.Z:H3–5ac
H3K4 me1:H2A.Z
H3K9 me3:H4K20 me3
H3K4 me1:H4–4ac
H3K27 me3:DNA Meth. m5C
H3K27 me3:H4K20 me3
DNA Meth. m5C:H4K20 me2
DNA Meth. m5C:H4K20 me3
H3K4 me1:H3–5ac
H3K4 me3:H4–4ac
H2A.Z:H3K4 me3
H4–4ac:H3–5ac
H4K20 me2:H4–4ac
H2A.Z:H4K20 me2
H4K20 me2:H3K4 me3
H4K20 me2:H3–5ac
H3K27 me2
H3K9 me2
H3K9 me3
H3K27 me3
H4K20 me3
H3K4 me3
DNA Meth. m5C
H3K4 me1
H4K20 me2
H3–5ac
H2A.Z
H4–4ac
0.00 0.25 0.50 0.75 1.00
π̂

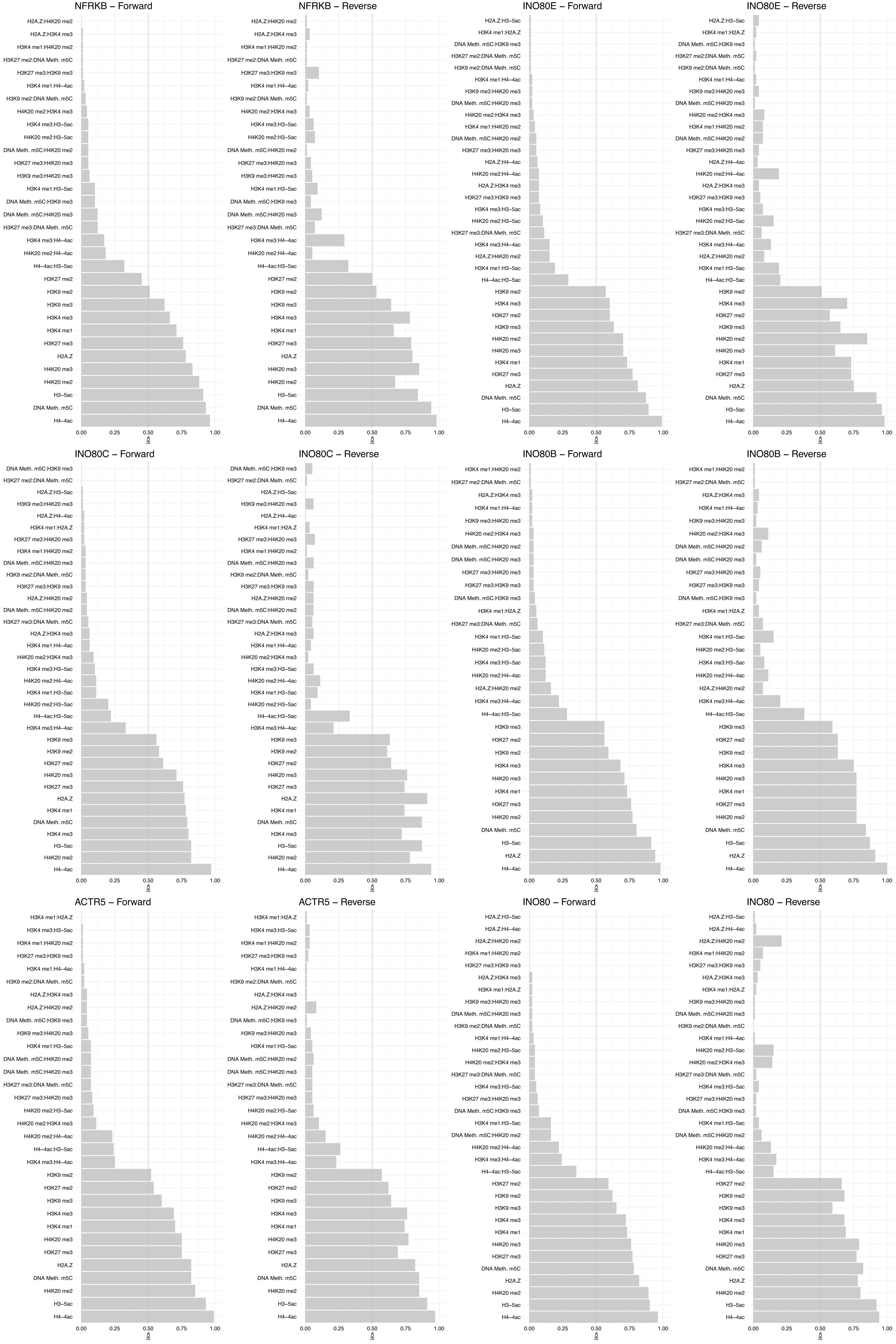
NFRKB – Forward
NFRKB – Reverse
INO80E – Forward
INO80E – Reverse
INO80C – Forward
INO80C – Reverse
INO80B – Forward
INO80B – Reverse
ACTR5 – Forward
ACTR5 – Reverse
INO80 – Forward
INO80 – Reverse
0.00
0.25
0.50
0.75
1.00

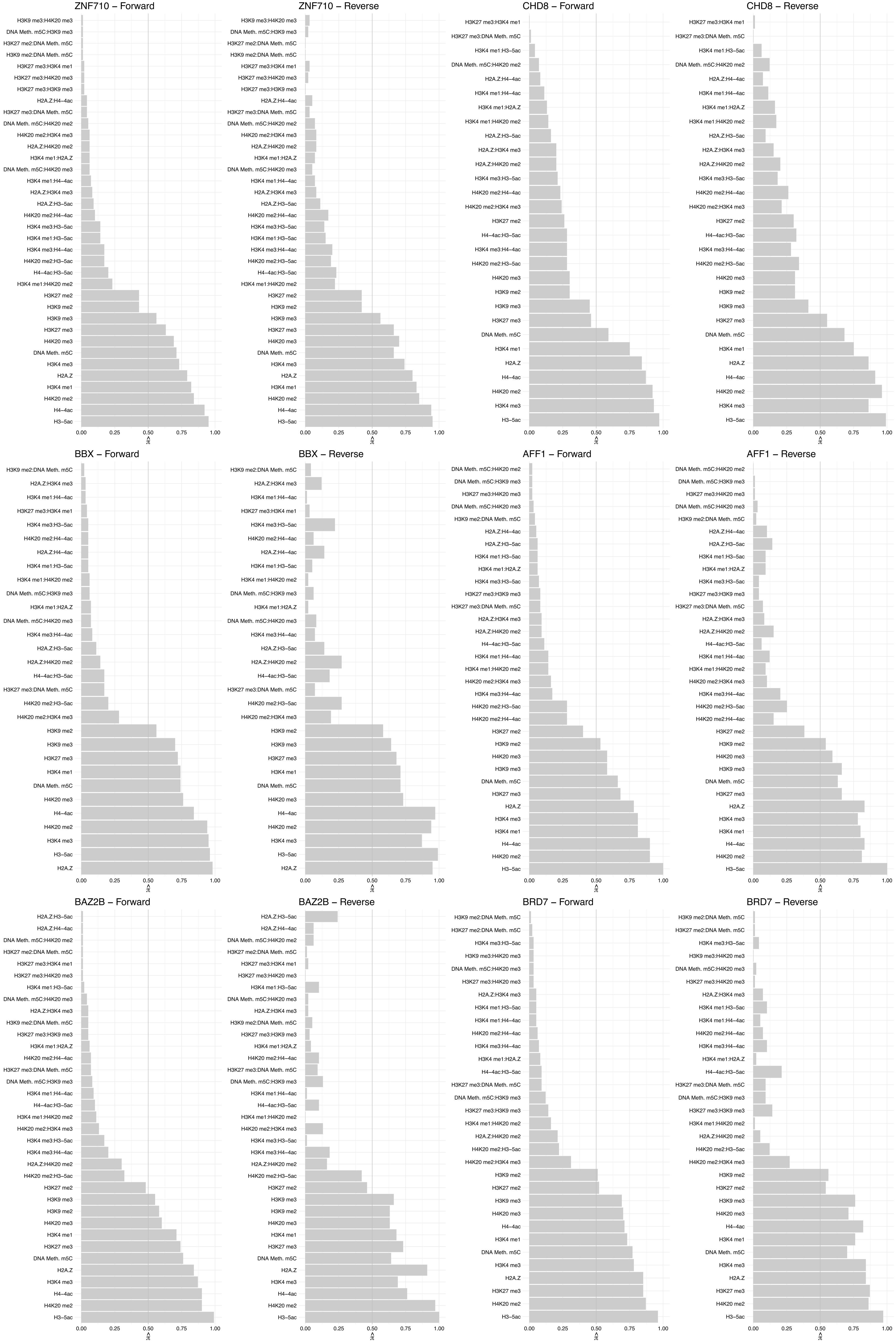
ZNF710 − Forward
H3K9 me3:H4K20 me3
DNA Meth. m5C:H3K9 me3
H3K27 me2:DNA Meth. m5C
H3K9 me2:DNA Meth. m5C
H3K27 me3:H3K4 me1
H3K27 me3:H4K20 me3
H3K27 me3:H3K9 me3
H2A.Z:H4−4ac
H3K27 me3:DNA Meth. m5C
DNA Meth. m5C:H4K20 me2
H4K20 me2:H3K4 me3
H2A.Z:H4K20 me2
H3K4 me1:H2A.Z
DNA Meth. m5C:H4K20 me3
H3K4 me1:H4−4ac
H2A.Z:H3K4 me3
H2A.Z:H3−5ac
H4K20 me2:H4−4ac
H3K4 me3:H3−5ac
H3K4 me1:H3−5ac
H3K4 me3:H4−4ac
H4K20 me2:H3−5ac
H4−4ac:H3−5ac
H3K4 me1:H4K20 me2
H3K27 me2
H3K9 me2
H3K9 me3
H3K27 me3
H4K20 me3
DNA Meth. m5C
H3K4 me3
H2A.Z
H3K4 me1
H4K20 me2
H4−4ac
H3−5ac
0.00 0.25 0.50 0.75 1.00
$\hat{\pi}$
ZNF710 − Reverse
H3K9 me3:H4K20 me3
DNA Meth. m5C:H3K9 me3
H3K27 me2:DNA Meth. m5C
H3K9 me2:DNA Meth. m5C
H3K27 me3:H3K4 me1
H3K27 me3:H4K20 me3
H3K27 me3:H3K9 me3
H2A.Z:H4−4ac
H3K27 me3:DNA Meth. m5C
DNA Meth. m5C:H4K20 me2
H4K20 me2:H3K4 me3
H2A.Z:H4K20 me2
H3K4 me1:H2A.Z
DNA Meth. m5C:H4K20 me3
H3K4 me1:H4−4ac
H2A.Z:H3K4 me3
H2A.Z:H3−5ac
H4K20 me2:H4−4ac
H3K4 me3:H3−5ac
H3K4 me1:H3−5ac
H3K4 me3:H4−4ac
H4K20 me2:H3−5ac
H4−4ac:H3−5ac
H3K4 me1:H4K20 me2
H3K27 me2
H3K9 me2
H3K9 me3
H3K27 me3
H4K20 me3
DNA Meth. m5C
H3K4 me3
H2A.Z
H3K4 me1
H4K20 me2
H4−4ac
H3−5ac
0.00 0.25 0.50 0.75 1.00
$\hat{\pi}$
CHD8 − Forward
H3K27 me3:H3K4 me1
H3K27 me3:DNA Meth. m5C
H3K4 me1:H3−5ac
DNA Meth. m5C:H4K20 me2
H2A.Z:H4−4ac
H3K4 me1:H4−4ac
H3K4 me1:H2A.Z
H3K4 me1:H4K20 me2
H2A.Z:H3−5ac
H2A.Z:H3K4 me3
H2A.Z:H4K20 me2
H3K4 me3:H3−5ac
H4K20 me2:H4−4ac
H4K20 me2:H3K4 me3
H3K27 me2
H4−4ac:H3−5ac
H3K4 me3:H4−4ac
H4K20 me2:H3−5ac
H4K20 me3
H3K9 me2
H3K9 me3
H3K27 me3
DNA Meth. m5C
H3K4 me1
H2A.Z
H4−4ac
H4K20 me2
H3K4 me3
H3−5ac
0.00 0.25 0.50 0.75 1.00
$\hat{\pi}$
CHD8 − Reverse
H3K27 me3:H3K4 me1
H3K27 me3:DNA Meth. m5C
H3K4 me1:H3−5ac
DNA Meth. m5C:H4K20 me2
H2A.Z:H4−4ac
H3K4 me1:H4−4ac
H3K4 me1:H2A.Z
H3K4 me1:H4K20 me2
H2A.Z:H3−5ac
H2A.Z:H3K4 me3
H2A.Z:H4K20 me2
H3K4 me3:H3−5ac
H4K20 me2:H4−4ac
H4K20 me2:H3K4 me3
H3K27 me2
H4−4ac:H3−5ac
H3K4 me3:H4−4ac
H4K20 me2:H3−5ac
H4K20 me3
H3K9 me2
H3K9 me3
H3K27 me3
DNA Meth. m5C
H3K4 me1
H2A.Z
H4−4ac
H4K20 me2
H3K4 me3
H3−5ac
0.00 0.25 0.50 0.75 1.00
$\hat{\pi}$
BBX − Forward
H3K9 me2:DNA Meth. m5C
H2A.Z:H3K4 me3
H3K4 me1:H4−4ac
H3K27 me3:H3K4 me1
H3K4 me3:H3−5ac
H4K20 me2:H4−4ac
H2A.Z:H4−4ac
H3K4 me1:H3−5ac
H3K4 me1:H4K20 me2
DNA Meth. m5C:H3K9 me3
H3K4 me1:H2A.Z
DNA Meth. m5C:H4K20 me3
H3K4 me3:H4−4ac
H2A.Z:H3−5ac
H2A.Z:H4K20 me2
H4−4ac:H3−5ac
H3K27 me3:DNA Meth. m5C
H4K20 me2:H3−5ac
H4K20 me2:H3K4 me3
H3K9 me2
H3K9 me3
H3K27 me3
H3K4 me1
DNA Meth. m5C
H4K20 me3
H4−4ac
H4K20 me2
H3K4 me3
H3−5ac
H2A.Z
0.00 0.25 0.50 0.75 1.00
$\hat{\pi}$
BBX − Reverse
H3K9 me2:DNA Meth. m5C
H2A.Z:H3K4 me3
H3K4 me1:H4−4ac
H3K27 me3:H3K4 me1
H3K4 me3:H3−5ac
H4K20 me2:H4−4ac
H2A.Z:H4−4ac
H3K4 me1:H3−5ac
H3K4 me1:H4K20 me2
DNA Meth. m5C:H3K9 me3
H3K4 me1:H2A.Z
DNA Meth. m5C:H4K20 me3
H3K4 me3:H4−4ac
H2A.Z:H3−5ac
H2A.Z:H4K20 me2
H4−4ac:H3−5ac
H3K27 me3:DNA Meth. m5C
H4K20 me2:H3−5ac
H4K20 me2:H3K4 me3
H3K9 me2
H3K9 me3
H3K27 me3
H3K4 me1
DNA Meth. m5C
H4K20 me3
H4−4ac
H4K20 me2
H3K4 me3
H3−5ac
H2A.Z
0.00 0.25 0.50 0.75 1.00
$\hat{\pi}$
AFF1 − Forward
DNA Meth. m5C:H4K20 me2
DNA Meth. m5C:H3K9 me3
H3K27 me3:H4K20 me3
DNA Meth. m5C:H4K20 me3
H3K9 me2:DNA Meth. m5C
H2A.Z:H4−4ac
H2A.Z:H3−5ac
H3K4 me1:H3−5ac
H3K4 me1:H2A.Z
H3K4 me3:H3−5ac
H3K27 me3:H3K9 me3
H3K27 me3:DNA Meth. m5C
H2A.Z:H3K4 me3
H2A.Z:H4K20 me2
H4−4ac:H3−5ac
H3K4 me1:H4−4ac
H3K4 me1:H4K20 me2
H4K20 me2:H3K4 me3
H3K4 me3:H4−4ac
H4K20 me2:H3−5ac
H4K20 me2:H4−4ac
H3K27 me2
H3K9 me2
H4K20 me3
H3K9 me3
DNA Meth. m5C
H3K27 me3
H2A.Z
H3K4 me3
H3K4 me1
H4−4ac
H4K20 me2
H3−5ac
0.00 0.25 0.50 0.75 1.00
$\hat{\pi}$
AFF1 − Reverse
DNA Meth. m5C:H4K20 me2
DNA Meth. m5C:H3K9 me3
H3K27 me3:H4K20 me3
DNA Meth. m5C:H4K20 me3
H3K9 me2:DNA Meth. m5C
H2A.Z:H4−4ac
H2A.Z:H3−5ac
H3K4 me1:H3−5ac
H3K4 me1:H2A.Z
H3K4 me3:H3−5ac
H3K27 me3:H3K9 me3
H3K27 me3:DNA Meth. m5C
H2A.Z:H3K4 me3
H2A.Z:H4K20 me2
H4−4ac:H3−5ac
H3K4 me1:H4−4ac
H3K4 me1:H4K20 me2
H4K20 me2:H3K4 me3
H3K4 me3:H4−4ac
H4K20 me2:H3−5ac
H4K20 me2:H4−4ac
H3K27 me2
H3K9 me2
H4K20 me3
H3K9 me3
DNA Meth. m5C
H3K27 me3
H2A.Z
H3K4 me3
H3K4 me1
H4−4ac
H4K20 me2
H3−5ac
0.00 0.25 0.50 0.75 1.00
$\hat{\pi}$
BAZ2B − Forward
H2A.Z:H3−5ac
H2A.Z:H4−4ac
DNA Meth. m5C:H4K20 me2
H3K27 me2:DNA Meth. m5C
H3K27 me3:H3K4 me1
H3K27 me3:H4K20 me3
H3K4 me1:H3−5ac
DNA Meth. m5C:H4K20 me3
H2A.Z:H3K4 me3
H3K9 me2:DNA Meth. m5C
H3K27 me3:H3K9 me3
H3K4 me1:H2A.Z
H4K20 me2:H4−4ac
H3K27 me3:DNA Meth. m5C
DNA Meth. m5C:H3K9 me3
H3K4 me1:H4−4ac
H4−4ac:H3−5ac
H3K4 me1:H4K20 me2
H4K20 me2:H3K4 me3
H3K4 me3:H3−5ac
H3K4 me3:H4−4ac
H2A.Z:H4K20 me2
H4K20 me2:H3−5ac
H3K27 me2
H3K9 me3
H3K9 me2
H4K20 me3
H3K4 me1
H3K27 me3
DNA Meth. m5C
H2A.Z
H3K4 me3
H4−4ac
H4K20 me2
H3−5ac
0.00 0.25 0.50 0.75 1.00
$\hat{\pi}$
BAZ2B − Reverse
H2A.Z:H3−5ac
H2A.Z:H4−4ac
DNA Meth. m5C:H4K20 me2
H3K27 me2:DNA Meth. m5C
H3K27 me3:H3K4 me1
H3K27 me3:H4K20 me3
H3K4 me1:H3−5ac
DNA Meth. m5C:H4K20 me3
H2A.Z:H3K4 me3
H3K9 me2:DNA Meth. m5C
H3K27 me3:H3K9 me3
H3K4 me1:H2A.Z
H4K20 me2:H4−4ac
H3K27 me3:DNA Meth. m5C
DNA Meth. m5C:H3K9 me3
H3K4 me1:H4−4ac
H4−4ac:H3−5ac
H3K4 me1:H4K20 me2
H4K20 me2:H3K4 me3
H3K4 me3:H3−5ac
H3K4 me3:H4−4ac
H2A.Z:H4K20 me2
H4K20 me2:H3−5ac
H3K27 me2
H3K9 me3
H3K9 me2
H4K20 me3
H3K4 me1
H3K27 me3
DNA Meth. m5C
H2A.Z
H3K4 me3
H4−4ac
H4K20 me2
H3−5ac
0.00 0.25 0.50 0.75 1.00
$\hat{\pi}$
BRD7 − Forward
H3K9 me2:DNA Meth. m5C
H3K27 me2:DNA Meth. m5C
H3K4 me3:H3−5ac
H3K9 me3:H4K20 me3
DNA Meth. m5C:H4K20 me3
H3K27 me3:H4K20 me3
H2A.Z:H3K4 me3
H3K4 me1:H3−5ac
H3K4 me1:H4−4ac
H4K20 me2:H4−4ac
H3K4 me3:H4−4ac
H3K4 me1:H2A.Z
H4−4ac:H3−5ac
H3K27 me3:DNA Meth. m5C
DNA Meth. m5C:H3K9 me3
H3K27 me3:H3K9 me3
H3K4 me1:H4K20 me2
H2A.Z:H4K20 me2
H4K20 me2:H3−5ac
H4K20 me2:H3K4 me3
H3K9 me2
H3K27 me2
H3K9 me3
H4K20 me3
H4−4ac
H3K4 me1
DNA Meth. m5C
H3K4 me3
H2A.Z
H3K27 me3
H4K20 me2
H3−5ac
0.00 0.25 0.50 0.75 1.00
$\hat{\pi}$
BRD7 − Reverse
H3K9 me2:DNA Meth. m5C
H3K27 me2:DNA Meth. m5C
H3K4 me3:H3−5ac
H3K9 me3:H4K20 me3
DNA Meth. m5C:H4K20 me3
H3K27 me3:H4K20 me3
H2A.Z:H3K4 me3
H3K4 me1:H3−5ac
H3K4 me1:H4−4ac
H4K20 me2:H4−4ac
H3K4 me3:H4−4ac
H3K4 me1:H2A.Z
H4−4ac:H3−5ac
H3K27 me3:DNA Meth. m5C
DNA Meth. m5C:H3K9 me3
H3K27 me3:H3K9 me3
H3K4 me1:H4K20 me2
H2A.Z:H4K20 me2
H4K20 me2:H3−5ac
H4K20 me2:H3K4 me3
H3K9 me2
H3K27 me2
H3K9 me3
H4K20 me3
H4−4ac
H3K4 me1
DNA Meth. m5C
H3K4 me3
H2A.Z
H3K27 me3
H4K20 me2
H3−5ac
0.00 0.25 0.50 0.75 1.00
$\hat{\pi}$

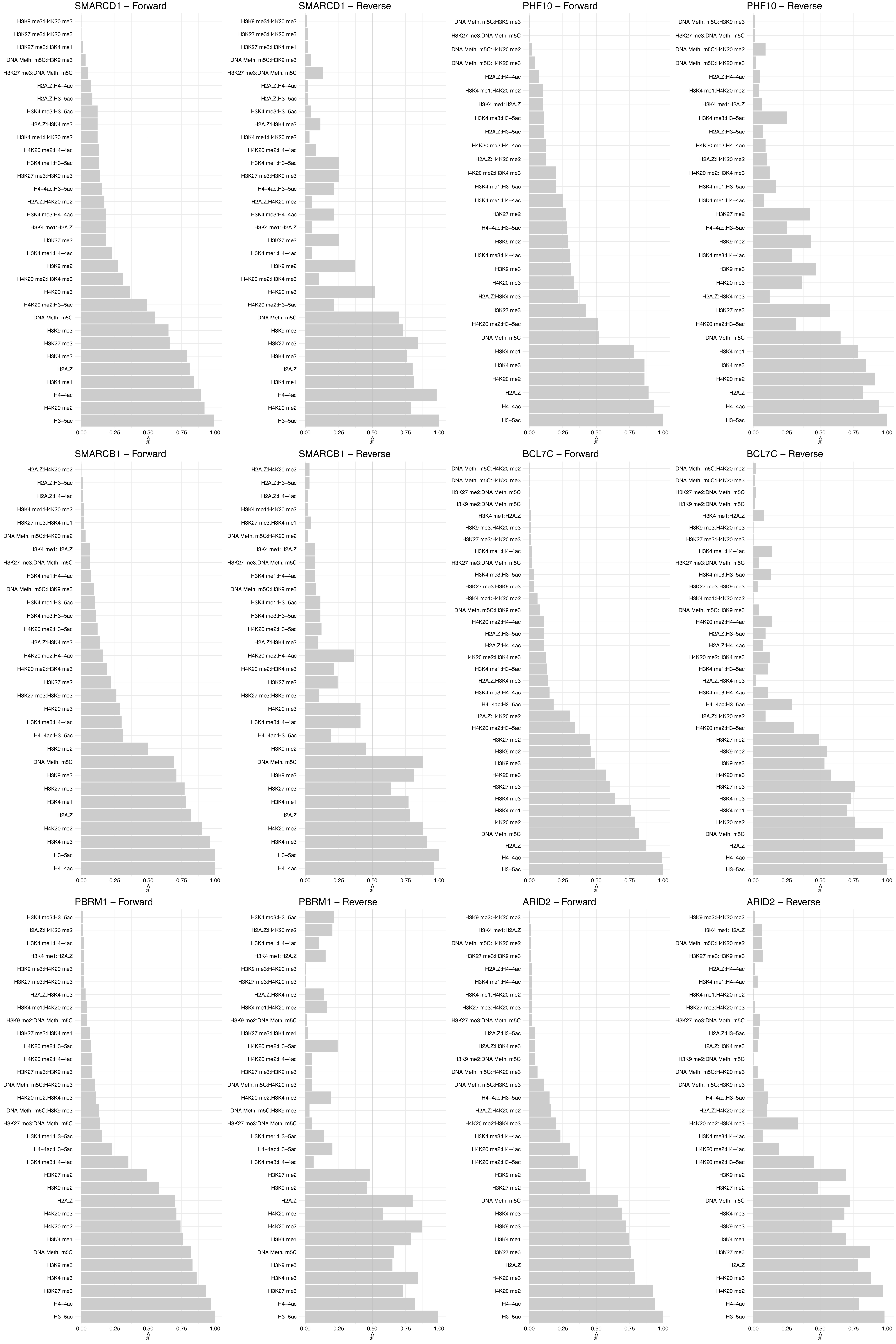
SMARCD1 – Forward
SMARCD1 – Reverse
PHF10 – Forward
PHF10 – Reverse
SMARCB1 – Forward
SMARCB1 – Reverse
BCL7C – Forward
BCL7C – Reverse
PBRM1 – Forward
PBRM1 – Reverse
ARID2 – Forward
ARID2 – Reverse
0.00 0.25 0.50 0.75 1.00
$\hat{\pi}$

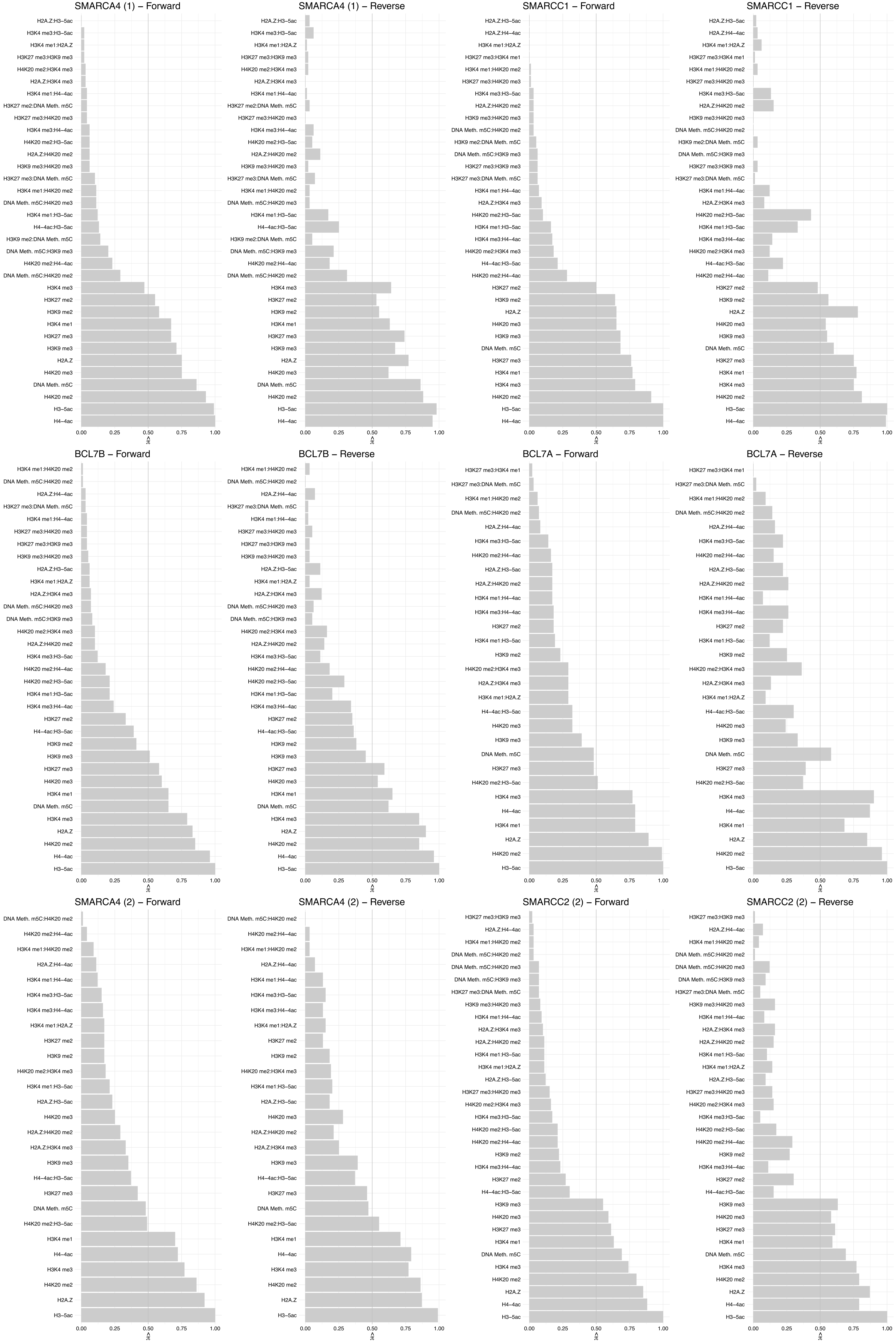
SMARCA4 (1) – Forward
H2A.Z:H3−5ac
H3K4 me3:H3−5ac
H3K4 me1:H2A.Z
H3K27 me3:H3K9 me3
H4K20 me2:H3K4 me3
H2A.Z:H3K4 me3
H3K4 me1:H4−4ac
H3K27 me2:DNA Meth. m5C
H3K27 me3:H4K20 me3
H3K4 me3:H4−4ac
H4K20 me2:H3−5ac
H2A.Z:H4K20 me2
H3K9 me3:H4K20 me3
H3K27 me3:DNA Meth. m5C
H3K4 me1:H4K20 me2
DNA Meth. m5C:H4K20 me3
H3K4 me1:H3−5ac
H4−4ac:H3−5ac
H3K9 me2:DNA Meth. m5C
DNA Meth. m5C:H3K9 me3
H4K20 me2:H4−4ac
DNA Meth. m5C:H4K20 me2
H3K4 me3
H3K27 me2
H3K9 me2
H3K4 me1
H3K27 me3
H3K9 me3
H2A.Z
H4K20 me3
DNA Meth. m5C
H4K20 me2
H3−5ac
H4−4ac
0.00 0.25 0.50 0.75 1.00
π̂
SMARCA4 (1) – Reverse
H2A.Z:H3−5ac
H3K4 me3:H3−5ac
H3K4 me1:H2A.Z
H3K27 me3:H3K9 me3
H4K20 me2:H3K4 me3
H2A.Z:H3K4 me3
H3K4 me1:H4−4ac
H3K27 me2:DNA Meth. m5C
H3K27 me3:H4K20 me3
H3K4 me3:H4−4ac
H4K20 me2:H3−5ac
H2A.Z:H4K20 me2
H3K9 me3:H4K20 me3
H3K27 me3:DNA Meth. m5C
H3K4 me1:H4K20 me2
DNA Meth. m5C:H4K20 me3
H3K4 me1:H3−5ac
H4−4ac:H3−5ac
H3K9 me2:DNA Meth. m5C
DNA Meth. m5C:H3K9 me3
H4K20 me2:H4−4ac
DNA Meth. m5C:H4K20 me2
H3K4 me3
H3K27 me2
H3K9 me2
H3K4 me1
H3K27 me3
H3K9 me3
H2A.Z
H4K20 me3
DNA Meth. m5C
H4K20 me2
H3−5ac
H4−4ac
0.00 0.25 0.50 0.75 1.00
π̂
SMARCC1 – Forward
H2A.Z:H3−5ac
H2A.Z:H4−4ac
H3K4 me1:H2A.Z
H3K27 me3:H3K4 me1
H3K4 me1:H4K20 me2
H3K27 me3:H4K20 me3
H3K4 me3:H3−5ac
H2A.Z:H4K20 me2
H3K9 me3:H4K20 me3
DNA Meth. m5C:H4K20 me2
H3K9 me2:DNA Meth. m5C
DNA Meth. m5C:H3K9 me3
H3K27 me3:H3K9 me3
H3K27 me3:DNA Meth. m5C
H3K4 me1:H4−4ac
H2A.Z:H3K4 me3
H4K20 me2:H3−5ac
H3K4 me1:H3−5ac
H3K4 me3:H4−4ac
H4K20 me2:H3K4 me3
H4−4ac:H3−5ac
H4K20 me2:H4−4ac
H3K27 me2
H3K9 me2
H2A.Z
H4K20 me3
H3K9 me3
DNA Meth. m5C
H3K27 me3
H3K4 me1
H3K4 me3
H4K20 me2
H3−5ac
H4−4ac
0.00 0.25 0.50 0.75 1.00
π̂
SMARCC1 – Reverse
H2A.Z:H3−5ac
H2A.Z:H4−4ac
H3K4 me1:H2A.Z
H3K27 me3:H3K4 me1
H3K4 me1:H4K20 me2
H3K27 me3:H4K20 me3
H3K4 me3:H3−5ac
H2A.Z:H4K20 me2
H3K9 me3:H4K20 me3
DNA Meth. m5C:H4K20 me2
H3K9 me2:DNA Meth. m5C
DNA Meth. m5C:H3K9 me3
H3K27 me3:H3K9 me3
H3K27 me3:DNA Meth. m5C
H3K4 me1:H4−4ac
H2A.Z:H3K4 me3
H4K20 me2:H3−5ac
H3K4 me1:H3−5ac
H3K4 me3:H4−4ac
H4K20 me2:H3K4 me3
H4−4ac:H3−5ac
H4K20 me2:H4−4ac
H3K27 me2
H3K9 me2
H2A.Z
H4K20 me3
H3K9 me3
DNA Meth. m5C
H3K27 me3
H3K4 me1
H3K4 me3
H4K20 me2
H3−5ac
H4−4ac
0.00 0.25 0.50 0.75 1.00
π̂
BCL7B – Forward
H3K4 me1:H4K20 me2
DNA Meth. m5C:H4K20 me2
H2A.Z:H4−4ac
H3K27 me3:DNA Meth. m5C
H3K4 me1:H4−4ac
H3K27 me3:H4K20 me3
H3K27 me3:H3K9 me3
H3K9 me3:H4K20 me3
H2A.Z:H3−5ac
H3K4 me1:H2A.Z
H2A.Z:H3K4 me3
DNA Meth. m5C:H4K20 me3
DNA Meth. m5C:H3K9 me3
H4K20 me2:H3K4 me3
H2A.Z:H4K20 me2
H3K4 me3:H3−5ac
H4K20 me2:H4−4ac
H4K20 me2:H3−5ac
H3K4 me1:H3−5ac
H3K4 me3:H4−4ac
H3K27 me2
H4−4ac:H3−5ac
H3K9 me2
H3K9 me3
H3K27 me3
H4K20 me3
H3K4 me1
DNA Meth. m5C
H3K4 me3
H2A.Z
H4K20 me2
H4−4ac
H3−5ac
0.00 0.25 0.50 0.75 1.00
π̂
BCL7B – Reverse
H3K4 me1:H4K20 me2
DNA Meth. m5C:H4K20 me2
H2A.Z:H4−4ac
H3K27 me3:DNA Meth. m5C
H3K4 me1:H4−4ac
H3K27 me3:H4K20 me3
H3K27 me3:H3K9 me3
H3K9 me3:H4K20 me3
H2A.Z:H3−5ac
H3K4 me1:H2A.Z
H2A.Z:H3K4 me3
DNA Meth. m5C:H4K20 me3
DNA Meth. m5C:H3K9 me3
H4K20 me2:H3K4 me3
H2A.Z:H4K20 me2
H3K4 me3:H3−5ac
H4K20 me2:H4−4ac
H4K20 me2:H3−5ac
H3K4 me1:H3−5ac
H3K4 me3:H4−4ac
H3K27 me2
H4−4ac:H3−5ac
H3K9 me2
H3K9 me3
H3K27 me3
H4K20 me3
H3K4 me1
DNA Meth. m5C
H3K4 me3
H2A.Z
H4K20 me2
H4−4ac
H3−5ac
0.00 0.25 0.50 0.75 1.00
π̂
BCL7A – Forward
H3K27 me3:H3K4 me1
H3K27 me3:DNA Meth. m5C
H3K4 me1:H4K20 me2
DNA Meth. m5C:H4K20 me2
H2A.Z:H4−4ac
H3K4 me3:H3−5ac
H4K20 me2:H4−4ac
H2A.Z:H3−5ac
H2A.Z:H4K20 me2
H3K4 me1:H4−4ac
H3K4 me3:H4−4ac
H3K27 me2
H3K4 me1:H3−5ac
H3K9 me2
H4K20 me2:H3K4 me3
H2A.Z:H3K4 me3
H3K4 me1:H2A.Z
H4−4ac:H3−5ac
H4K20 me3
H3K9 me3
DNA Meth. m5C
H3K27 me3
H4K20 me2:H3−5ac
H3K4 me3
H4−4ac
H3K4 me1
H2A.Z
H4K20 me2
H3−5ac
0.00 0.25 0.50 0.75 1.00
π̂
BCL7A – Reverse
H3K27 me3:H3K4 me1
H3K27 me3:DNA Meth. m5C
H3K4 me1:H4K20 me2
DNA Meth. m5C:H4K20 me2
H2A.Z:H4−4ac
H3K4 me3:H3−5ac
H4K20 me2:H4−4ac
H2A.Z:H3−5ac
H2A.Z:H4K20 me2
H3K4 me1:H4−4ac
H3K4 me3:H4−4ac
H3K27 me2
H3K4 me1:H3−5ac
H3K9 me2
H4K20 me2:H3K4 me3
H2A.Z:H3K4 me3
H3K4 me1:H2A.Z
H4−4ac:H3−5ac
H4K20 me3
H3K9 me3
DNA Meth. m5C
H3K27 me3
H4K20 me2:H3−5ac
H3K4 me3
H4−4ac
H3K4 me1
H2A.Z
H4K20 me2
H3−5ac
0.00 0.25 0.50 0.75 1.00
π̂
SMARCA4 (2) – Forward
DNA Meth. m5C:H4K20 me2
H4K20 me2:H4−4ac
H3K4 me1:H4K20 me2
H2A.Z:H4−4ac
H3K4 me1:H4−4ac
H3K4 me3:H3−5ac
H3K4 me3:H4−4ac
H3K4 me1:H2A.Z
H3K27 me2
H3K9 me2
H4K20 me2:H3K4 me3
H3K4 me1:H3−5ac
H2A.Z:H3−5ac
H4K20 me3
H2A.Z:H4K20 me2
H2A.Z:H3K4 me3
H3K9 me3
H4−4ac:H3−5ac
H3K27 me3
DNA Meth. m5C
H4K20 me2:H3−5ac
H3K4 me1
H4−4ac
H3K4 me3
H4K20 me2
H2A.Z
H3−5ac
0.00 0.25 0.50 0.75 1.00
π̂
SMARCA4 (2) – Reverse
DNA Meth. m5C:H4K20 me2
H4K20 me2:H4−4ac
H3K4 me1:H4K20 me2
H2A.Z:H4−4ac
H3K4 me1:H4−4ac
H3K4 me3:H3−5ac
H3K4 me3:H4−4ac
H3K4 me1:H2A.Z
H3K27 me2
H3K9 me2
H4K20 me2:H3K4 me3
H3K4 me1:H3−5ac
H2A.Z:H3−5ac
H4K20 me3
H2A.Z:H4K20 me2
H2A.Z:H3K4 me3
H3K9 me3
H4−4ac:H3−5ac
H3K27 me3
DNA Meth. m5C
H4K20 me2:H3−5ac
H3K4 me1
H4−4ac
H3K4 me3
H4K20 me2
H2A.Z
H3−5ac
0.00 0.25 0.50 0.75 1.00
π̂
SMARCC2 (2) – Forward
H3K27 me3:H3K9 me3
H2A.Z:H4−4ac
H3K4 me1:H4K20 me2
DNA Meth. m5C:H4K20 me2
DNA Meth. m5C:H4K20 me3
DNA Meth. m5C:H3K9 me3
H3K27 me3:DNA Meth. m5C
H3K9 me3:H4K20 me3
H3K4 me1:H4−4ac
H2A.Z:H3K4 me3
H2A.Z:H4K20 me2
H3K4 me1:H3−5ac
H3K4 me1:H2A.Z
H2A.Z:H3−5ac
H3K27 me3:H4K20 me3
H4K20 me2:H3K4 me3
H3K4 me3:H3−5ac
H4K20 me2:H3−5ac
H4K20 me2:H4−4ac
H3K9 me2
H3K4 me3:H4−4ac
H3K27 me2
H4−4ac:H3−5ac
H3K9 me3
H4K20 me3
H3K27 me3
H3K4 me1
DNA Meth. m5C
H3K4 me3
H4K20 me2
H2A.Z
H4−4ac
H3−5ac
0.00 0.25 0.50 0.75 1.00
π̂
SMARCC2 (2) – Reverse
H3K27 me3:H3K9 me3
H2A.Z:H4−4ac
H3K4 me1:H4K20 me2
DNA Meth. m5C:H4K20 me2
DNA Meth. m5C:H4K20 me3
DNA Meth. m5C:H3K9 me3
H3K27 me3:DNA Meth. m5C
H3K9 me3:H4K20 me3
H3K4 me1:H4−4ac
H2A.Z:H3K4 me3
H2A.Z:H4K20 me2
H3K4 me1:H3−5ac
H3K4 me1:H2A.Z
H2A.Z:H3−5ac
H3K27 me3:H4K20 me3
H4K20 me2:H3K4 me3
H3K4 me3:H3−5ac
H4K20 me2:H3−5ac
H4K20 me2:H4−4ac
H3K9 me2
H3K4 me3:H4−4ac
H3K27 me2
H4−4ac:H3−5ac
H3K9 me3
H4K20 me3
H3K27 me3
H3K4 me1
DNA Meth. m5C
H3K4 me3
H4K20 me2
H2A.Z
H4−4ac
H3−5ac
0.00 0.25 0.50 0.75 1.00
π̂

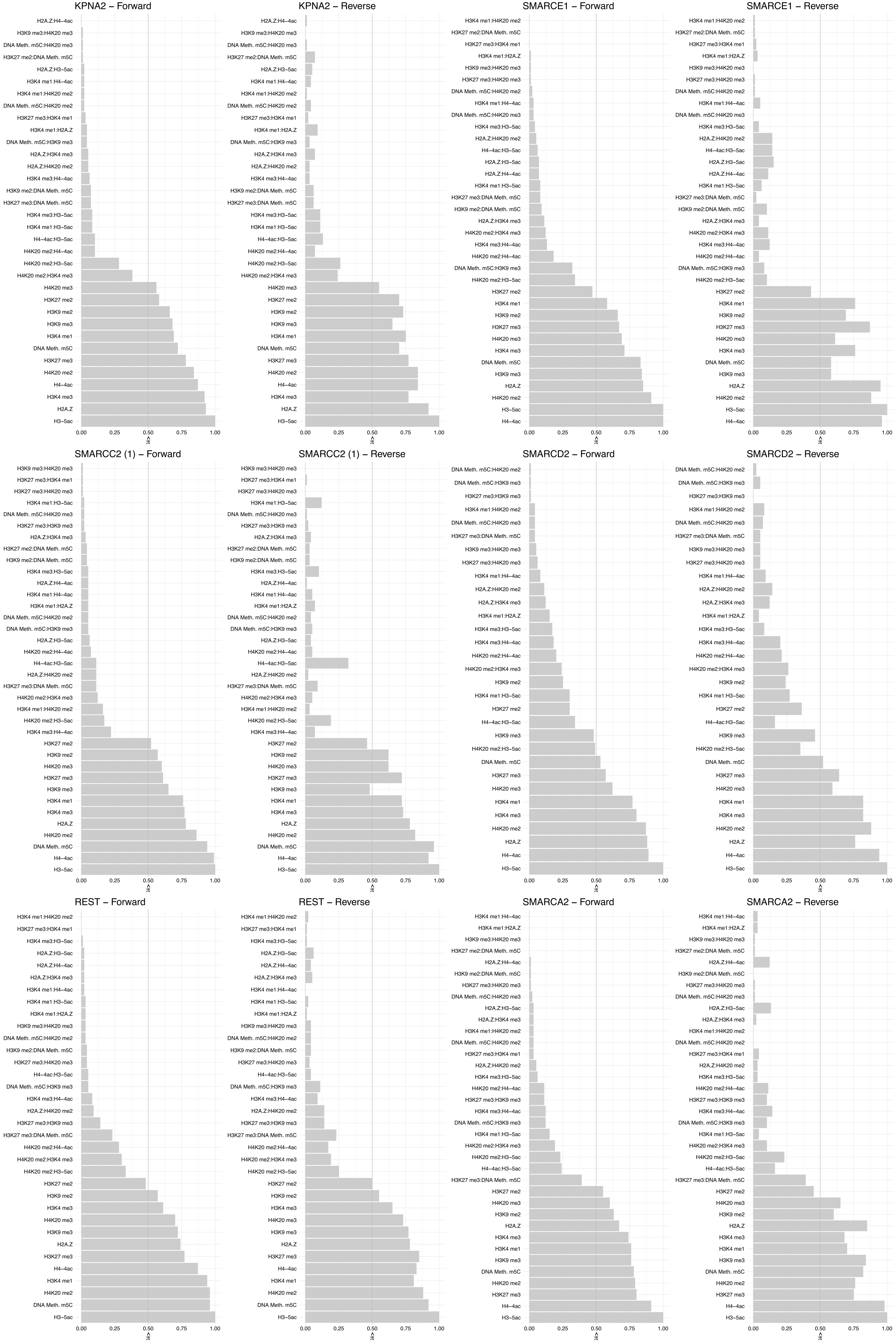

KPNA2 – Forward
H2A.Z:H4–4ac
H3K9 me3:H4K20 me3
DNA Meth. m5C:H4K20 me3
H3K27 me2:DNA Meth. m5C
H2A.Z:H3–5ac
H3K4 me1:H4–4ac
H3K4 me1:H4K20 me2
DNA Meth. m5C:H4K20 me2
H3K27 me3:H3K4 me1
H3K4 me1:H2A.Z
DNA Meth. m5C:H3K9 me3
H2A.Z:H3K4 me3
H2A.Z:H4K20 me2
H3K4 me3:H4–4ac
H3K9 me2:DNA Meth. m5C
H3K27 me3:DNA Meth. m5C
H3K4 me3:H3–5ac
H3K4 me1:H3–5ac
H4–4ac:H3–5ac
H4K20 me2:H4–4ac
H4K20 me2:H3–5ac
H4K20 me2:H3K4 me3
H4K20 me3
H3K27 me2
H3K9 me2
H3K9 me3
H3K4 me1
DNA Meth. m5C
H3K27 me3
H4K20 me2
H4–4ac
H3K4 me3
H2A.Z
H3–5ac
0.00 0.25 0.50 0.75 1.00
π̂
KPNA2 – Reverse
H2A.Z:H4–4ac
H3K9 me3:H4K20 me3
DNA Meth. m5C:H4K20 me3
H3K27 me2:DNA Meth. m5C
H2A.Z:H3–5ac
H3K4 me1:H4–4ac
H3K4 me1:H4K20 me2
DNA Meth. m5C:H4K20 me2
H3K27 me3:H3K4 me1
H3K4 me1:H2A.Z
DNA Meth. m5C:H3K9 me3
H2A.Z:H3K4 me3
H2A.Z:H4K20 me2
H3K4 me3:H4–4ac
H3K9 me2:DNA Meth. m5C
H3K27 me3:DNA Meth. m5C
H3K4 me3:H3–5ac
H3K4 me1:H3–5ac
H4–4ac:H3–5ac
H4K20 me2:H4–4ac
H4K20 me2:H3–5ac
H4K20 me2:H3K4 me3
H4K20 me3
H3K27 me2
H3K9 me2
H3K9 me3
H3K4 me1
DNA Meth. m5C
H3K27 me3
H4K20 me2
H4–4ac
H3K4 me3
H2A.Z
H3–5ac
0.00 0.25 0.50 0.75 1.00
π̂
SMARCE1 – Forward
H3K4 me1:H4K20 me2
H3K27 me2:DNA Meth. m5C
H3K27 me3:H3K4 me1
H3K4 me1:H2A.Z
H3K9 me3:H4K20 me3
H3K27 me3:H4K20 me3
DNA Meth. m5C:H4K20 me2
H3K4 me1:H4–4ac
DNA Meth. m5C:H4K20 me3
H3K4 me3:H3–5ac
H2A.Z:H4K20 me2
H4–4ac:H3–5ac
H2A.Z:H3–5ac
H2A.Z:H4–4ac
H3K4 me1:H3–5ac
H3K27 me3:DNA Meth. m5C
H3K9 me2:DNA Meth. m5C
H2A.Z:H3K4 me3
H4K20 me2:H3K4 me3
H3K4 me3:H4–4ac
H4K20 me2:H4–4ac
DNA Meth. m5C:H3K9 me3
H4K20 me2:H3–5ac
H3K27 me2
H3K4 me1
H3K9 me2
H3K27 me3
H4K20 me3
H3K4 me3
DNA Meth. m5C
H3K9 me3
H2A.Z
H4K20 me2
H3–5ac
H4–4ac
0.00 0.25 0.50 0.75 1.00
π̂
SMARCE1 – Reverse
H3K4 me1:H4K20 me2
H3K27 me2:DNA Meth. m5C
H3K27 me3:H3K4 me1
H3K4 me1:H2A.Z
H3K9 me3:H4K20 me3
H3K27 me3:H4K20 me3
DNA Meth. m5C:H4K20 me2
H3K4 me1:H4–4ac
DNA Meth. m5C:H4K20 me3
H3K4 me3:H3–5ac
H2A.Z:H4K20 me2
H4–4ac:H3–5ac
H2A.Z:H3–5ac
H2A.Z:H4–4ac
H3K4 me1:H3–5ac
H3K27 me3:DNA Meth. m5C
H3K9 me2:DNA Meth. m5C
H2A.Z:H3K4 me3
H4K20 me2:H3K4 me3
H3K4 me3:H4–4ac
H4K20 me2:H4–4ac
DNA Meth. m5C:H3K9 me3
H4K20 me2:H3–5ac
H3K27 me2
H3K4 me1
H3K9 me2
H3K27 me3
H4K20 me3
H3K4 me3
DNA Meth. m5C
H3K9 me3
H2A.Z
H4K20 me2
H3–5ac
H4–4ac
0.00 0.25 0.50 0.75 1.00
π̂
SMARCC2 (1) – Forward
H3K9 me3:H4K20 me3
H3K27 me3:H3K4 me1
H3K27 me3:H4K20 me3
H3K4 me1:H3–5ac
DNA Meth. m5C:H4K20 me3
H3K27 me3:H3K9 me3
H2A.Z:H3K4 me3
H3K27 me2:DNA Meth. m5C
H3K9 me2:DNA Meth. m5C
H3K4 me3:H3–5ac
H2A.Z:H4–4ac
H3K4 me1:H4–4ac
H3K4 me1:H2A.Z
DNA Meth. m5C:H4K20 me2
DNA Meth. m5C:H3K9 me3
H2A.Z:H3–5ac
H4K20 me2:H4–4ac
H4–4ac:H3–5ac
H2A.Z:H4K20 me2
H3K27 me3:DNA Meth. m5C
H4K20 me2:H3K4 me3
H3K4 me1:H4K20 me2
H4K20 me2:H3–5ac
H3K4 me3:H4–4ac
H3K27 me2
H3K9 me2
H4K20 me3
H3K27 me3
H3K9 me3
H3K4 me1
H3K4 me3
H2A.Z
H4K20 me2
DNA Meth. m5C
H4–4ac
H3–5ac
0.00 0.25 0.50 0.75 1.00
π̂
SMARCC2 (1) – Reverse
H3K9 me3:H4K20 me3
H3K27 me3:H3K4 me1
H3K27 me3:H4K20 me3
H3K4 me1:H3–5ac
DNA Meth. m5C:H4K20 me3
H3K27 me3:H3K9 me3
H2A.Z:H3K4 me3
H3K27 me2:DNA Meth. m5C
H3K9 me2:DNA Meth. m5C
H3K4 me3:H3–5ac
H2A.Z:H4–4ac
H3K4 me1:H4–4ac
H3K4 me1:H2A.Z
DNA Meth. m5C:H4K20 me2
DNA Meth. m5C:H3K9 me3
H2A.Z:H3–5ac
H4K20 me2:H4–4ac
H4–4ac:H3–5ac
H2A.Z:H4K20 me2
H3K27 me3:DNA Meth. m5C
H4K20 me2:H3K4 me3
H3K4 me1:H4K20 me2
H4K20 me2:H3–5ac
H3K4 me3:H4–4ac
H3K27 me2
H3K9 me2
H4K20 me3
H3K27 me3
H3K9 me3
H3K4 me1
H3K4 me3
H2A.Z
H4K20 me2
DNA Meth. m5C
H4–4ac
H3–5ac
0.00 0.25 0.50 0.75 1.00
π̂
SMARCD2 – Forward
DNA Meth. m5C:H4K20 me2
DNA Meth. m5C:H3K9 me3
H3K27 me3:H3K9 me3
H3K4 me1:H4K20 me2
DNA Meth. m5C:H4K20 me3
H3K27 me3:DNA Meth. m5C
H3K9 me3:H4K20 me3
H3K27 me3:H4K20 me3
H3K4 me1:H4–4ac
H2A.Z:H4K20 me2
H2A.Z:H3K4 me3
H3K4 me1:H2A.Z
H3K4 me3:H3–5ac
H3K4 me3:H4–4ac
H4K20 me2:H4–4ac
H4K20 me2:H3K4 me3
H3K9 me2
H3K4 me1:H3–5ac
H3K27 me2
H4–4ac:H3–5ac
H3K9 me3
H4K20 me2:H3–5ac
DNA Meth. m5C
H3K27 me3
H4K20 me3
H3K4 me1
H3K4 me3
H4K20 me2
H2A.Z
H4–4ac
H3–5ac
0.00 0.25 0.50 0.75 1.00
π̂
SMARCD2 – Reverse
DNA Meth. m5C:H4K20 me2
DNA Meth. m5C:H3K9 me3
H3K27 me3:H3K9 me3
H3K4 me1:H4K20 me2
DNA Meth. m5C:H4K20 me3
H3K27 me3:DNA Meth. m5C
H3K9 me3:H4K20 me3
H3K27 me3:H4K20 me3
H3K4 me1:H4–4ac
H2A.Z:H4K20 me2
H2A.Z:H3K4 me3
H3K4 me1:H2A.Z
H3K4 me3:H3–5ac
H3K4 me3:H4–4ac
H4K20 me2:H4–4ac
H4K20 me2:H3K4 me3
H3K9 me2
H3K4 me1:H3–5ac
H3K27 me2
H4–4ac:H3–5ac
H3K9 me3
H4K20 me2:H3–5ac
DNA Meth. m5C
H3K27 me3
H4K20 me3
H3K4 me1
H3K4 me3
H4K20 me2
H2A.Z
H4–4ac
H3–5ac
0.00 0.25 0.50 0.75 1.00
π̂
REST – Forward
H3K4 me1:H4K20 me2
H3K27 me3:H3K4 me1
H3K4 me3:H3–5ac
H2A.Z:H3–5ac
H2A.Z:H4–4ac
H2A.Z:H3K4 me3
H3K4 me1:H4–4ac
H3K4 me1:H3–5ac
H3K4 me1:H2A.Z
H3K9 me3:H4K20 me3
DNA Meth. m5C:H4K20 me2
H3K9 me2:DNA Meth. m5C
H3K27 me3:H4K20 me3
H4–4ac:H3–5ac
DNA Meth. m5C:H3K9 me3
H3K4 me3:H4–4ac
H2A.Z:H4K20 me2
H3K27 me3:H3K9 me3
H3K27 me3:DNA Meth. m5C
H4K20 me2:H4–4ac
H4K20 me2:H3K4 me3
H4K20 me2:H3–5ac
H3K27 me2
H3K9 me2
H3K4 me3
H4K20 me3
H3K9 me3
H2A.Z
H3K27 me3
H4–4ac
H3K4 me1
H4K20 me2
DNA Meth. m5C
H3–5ac
0.00 0.25 0.50 0.75 1.00
π̂
REST – Reverse
H3K4 me1:H4K20 me2
H3K27 me3:H3K4 me1
H3K4 me3:H3–5ac
H2A.Z:H3–5ac
H2A.Z:H4–4ac
H2A.Z:H3K4 me3
H3K4 me1:H4–4ac
H3K4 me1:H3–5ac
H3K4 me1:H2A.Z
H3K9 me3:H4K20 me3
DNA Meth. m5C:H4K20 me2
H3K9 me2:DNA Meth. m5C
H3K27 me3:H4K20 me3
H4–4ac:H3–5ac
DNA Meth. m5C:H3K9 me3
H3K4 me3:H4–4ac
H2A.Z:H4K20 me2
H3K27 me3:H3K9 me3
H3K27 me3:DNA Meth. m5C
H4K20 me2:H4–4ac
H4K20 me2:H3K4 me3
H4K20 me2:H3–5ac
H3K27 me2
H3K9 me2
H3K4 me3
H4K20 me3
H3K9 me3
H2A.Z
H3K27 me3
H4–4ac
H3K4 me1
H4K20 me2
DNA Meth. m5C
H3–5ac
0.00 0.25 0.50 0.75 1.00
π̂
SMARCA2 – Forward
H3K4 me1:H4–4ac
H3K4 me1:H2A.Z
H3K9 me3:H4K20 me3
H3K27 me2:DNA Meth. m5C
H2A.Z:H4–4ac
H3K9 me2:DNA Meth. m5C
H3K27 me3:H4K20 me3
DNA Meth. m5C:H4K20 me3
H2A.Z:H3–5ac
H2A.Z:H3K4 me3
H3K4 me1:H4K20 me2
DNA Meth. m5C:H4K20 me2
H3K27 me3:H3K4 me1
H2A.Z:H4K20 me2
H3K4 me3:H3–5ac
H4K20 me2:H4–4ac
H3K27 me3:H3K9 me3
H3K4 me3:H4–4ac
DNA Meth. m5C:H3K9 me3
H3K4 me1:H3–5ac
H4K20 me2:H3K4 me3
H4K20 me2:H3–5ac
H4–4ac:H3–5ac
H3K27 me3:DNA Meth. m5C
H3K27 me2
H4K20 me3
H3K9 me2
H2A.Z
H3K4 me3
H3K4 me1
H3K9 me3
DNA Meth. m5C
H4K20 me2
H3K27 me3
H4–4ac
H3–5ac
0.00 0.25 0.50 0.75 1.00
π̂
SMARCA2 – Reverse
H3K4 me1:H4–4ac
H3K4 me1:H2A.Z
H3K9 me3:H4K20 me3
H3K27 me2:DNA Meth. m5C
H2A.Z:H4–4ac
H3K9 me2:DNA Meth. m5C
H3K27 me3:H4K20 me3
DNA Meth. m5C:H4K20 me3
H2A.Z:H3–5ac
H2A.Z:H3K4 me3
H3K4 me1:H4K20 me2
DNA Meth. m5C:H4K20 me2
H3K27 me3:H3K4 me1
H2A.Z:H4K20 me2
H3K4 me3:H3–5ac
H4K20 me2:H4–4ac
H3K27 me3:H3K9 me3
H3K4 me3:H4–4ac
DNA Meth. m5C:H3K9 me3
H3K4 me1:H3–5ac
H4K20 me2:H3K4 me3
H4K20 me2:H3–5ac
H4–4ac:H3–5ac
H3K27 me3:DNA Meth. m5C
H3K27 me2
H4K20 me3
H3K9 me2
H2A.Z
H3K4 me3
H3K4 me1
H3K9 me3
DNA Meth. m5C
H4K20 me2
H3K27 me3
H4–4ac
H3–5ac
0.00 0.25 0.50 0.75 1.00
π̂

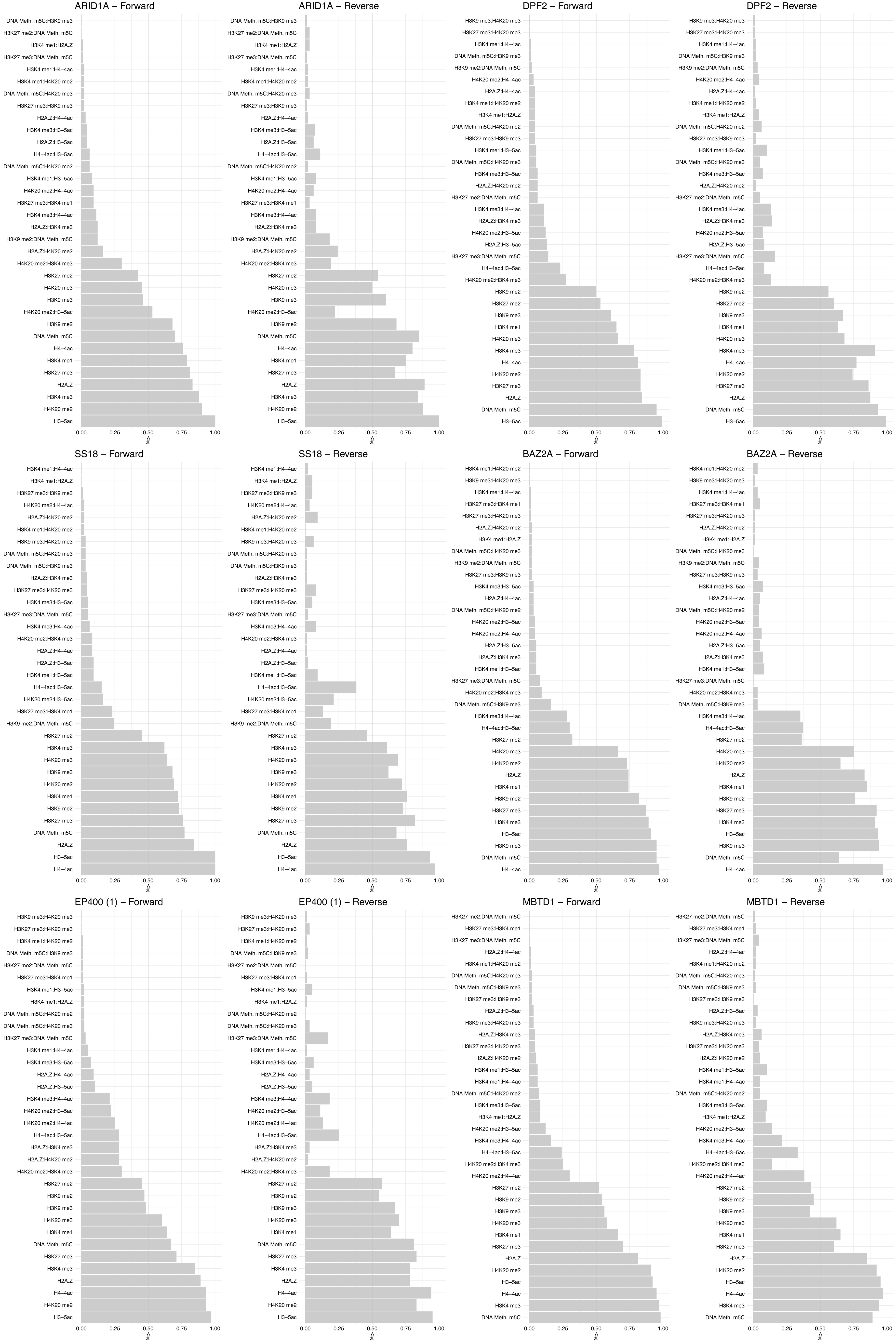
ARID1A – Forward
ARID1A – Reverse
DPF2 – Forward
DPF2 – Reverse
SS18 – Forward
SS18 – Reverse
BAZ2A – Forward
BAZ2A – Reverse
EP400 (1) – Forward
EP400 (1) – Reverse
MBTD1 – Forward
MBTD1 – Reverse
0.00 0.25 0.50 0.75 1.00
$\hat{\pi}$

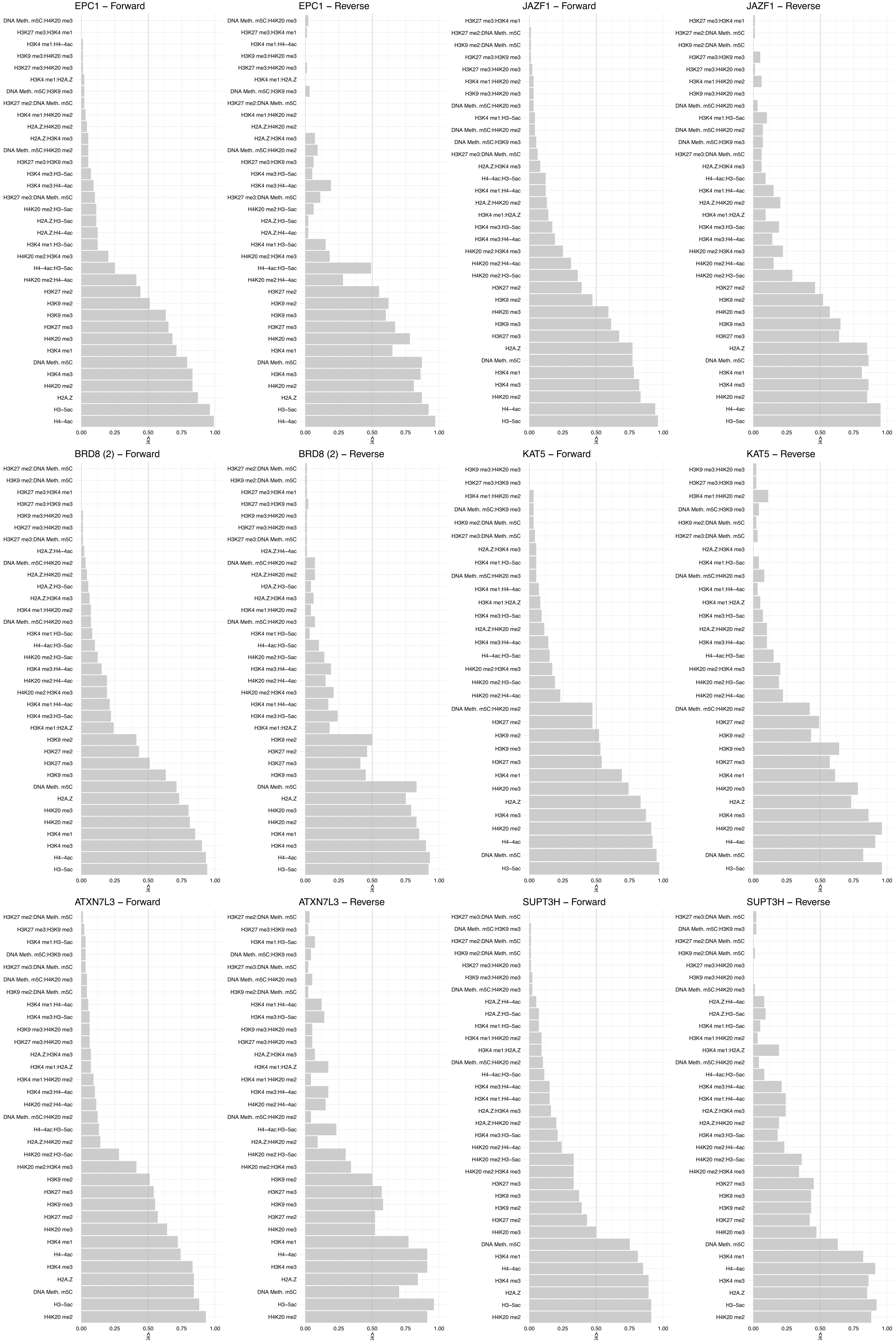

EPC1 – Forward
EPC1 – Reverse
JAZF1 – Forward
JAZF1 – Reverse
BRD8 (2) – Forward
BRD8 (2) – Reverse
KAT5 – Forward
KAT5 – Reverse
ATXN7L3 – Forward
ATXN7L3 – Reverse
SUPT3H – Forward
SUPT3H – Reverse
0.00 0.25 0.50 0.75 1.00
$\hat{\pi}$

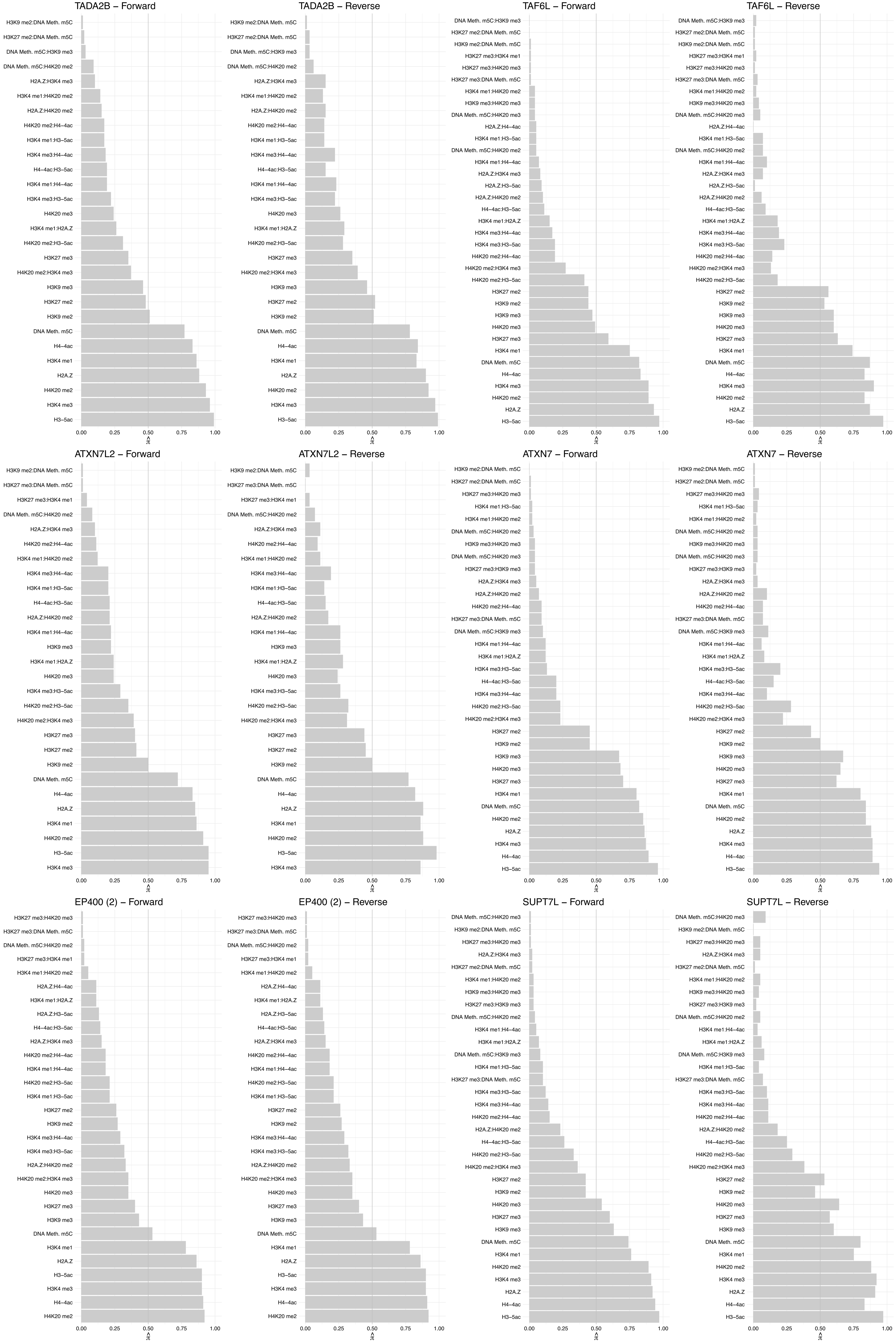
TADA2B – Forward
H3K9 me2:DNA Meth. m5C
H3K27 me2:DNA Meth. m5C
DNA Meth. m5C:H3K9 me3
DNA Meth. m5C:H4K20 me2
H2A.Z:H3K4 me3
H3K4 me1:H4K20 me2
H2A.Z:H4K20 me2
H4K20 me2:H4−4ac
H3K4 me1:H3−5ac
H3K4 me3:H4−4ac
H4−4ac:H3−5ac
H3K4 me1:H4−4ac
H3K4 me3:H3−5ac
H4K20 me3
H3K4 me1:H2A.Z
H4K20 me2:H3−5ac
H3K27 me3
H4K20 me2:H3K4 me3
H3K9 me3
H3K27 me2
H3K9 me2
DNA Meth. m5C
H4−4ac
H3K4 me1
H2A.Z
H4K20 me2
H3K4 me3
H3−5ac
0.00 0.25 0.50 0.75 1.00
π̂
TADA2B – Reverse
H3K9 me2:DNA Meth. m5C
H3K27 me2:DNA Meth. m5C
DNA Meth. m5C:H3K9 me3
DNA Meth. m5C:H4K20 me2
H2A.Z:H3K4 me3
H3K4 me1:H4K20 me2
H2A.Z:H4K20 me2
H4K20 me2:H4−4ac
H3K4 me1:H3−5ac
H3K4 me3:H4−4ac
H4−4ac:H3−5ac
H3K4 me1:H4−4ac
H3K4 me3:H3−5ac
H4K20 me3
H3K4 me1:H2A.Z
H4K20 me2:H3−5ac
H3K27 me3
H4K20 me2:H3K4 me3
H3K9 me3
H3K27 me2
H3K9 me2
DNA Meth. m5C
H4−4ac
H3K4 me1
H2A.Z
H4K20 me2
H3K4 me3
H3−5ac
0.00 0.25 0.50 0.75 1.00
π̂
TAF6L – Forward
DNA Meth. m5C:H3K9 me3
H3K27 me2:DNA Meth. m5C
H3K9 me2:DNA Meth. m5C
H3K27 me3:H3K4 me1
H3K27 me3:H4K20 me3
H3K27 me3:DNA Meth. m5C
H3K4 me1:H4K20 me2
H3K9 me3:H4K20 me3
DNA Meth. m5C:H4K20 me3
H2A.Z:H4−4ac
H3K4 me1:H3−5ac
DNA Meth. m5C:H4K20 me2
H3K4 me1:H4−4ac
H2A.Z:H3K4 me3
H2A.Z:H3−5ac
H2A.Z:H4K20 me2
H4−4ac:H3−5ac
H3K4 me1:H2A.Z
H3K4 me3:H4−4ac
H3K4 me3:H3−5ac
H4K20 me2:H4−4ac
H4K20 me2:H3K4 me3
H4K20 me2:H3−5ac
H3K27 me2
H3K9 me2
H3K9 me3
H4K20 me3
H3K27 me3
H3K4 me1
DNA Meth. m5C
H4−4ac
H3K4 me3
H4K20 me2
H2A.Z
H3−5ac
0.00 0.25 0.50 0.75 1.00
π̂
TAF6L – Reverse
DNA Meth. m5C:H3K9 me3
H3K27 me2:DNA Meth. m5C
H3K9 me2:DNA Meth. m5C
H3K27 me3:H3K4 me1
H3K27 me3:H4K20 me3
H3K27 me3:DNA Meth. m5C
H3K4 me1:H4K20 me2
H3K9 me3:H4K20 me3
DNA Meth. m5C:H4K20 me3
H2A.Z:H4−4ac
H3K4 me1:H3−5ac
DNA Meth. m5C:H4K20 me2
H3K4 me1:H4−4ac
H2A.Z:H3K4 me3
H2A.Z:H3−5ac
H2A.Z:H4K20 me2
H4−4ac:H3−5ac
H3K4 me1:H2A.Z
H3K4 me3:H4−4ac
H3K4 me3:H3−5ac
H4K20 me2:H4−4ac
H4K20 me2:H3K4 me3
H4K20 me2:H3−5ac
H3K27 me2
H3K9 me2
H3K9 me3
H4K20 me3
H3K27 me3
H3K4 me1
DNA Meth. m5C
H4−4ac
H3K4 me3
H4K20 me2
H2A.Z
H3−5ac
0.00 0.25 0.50 0.75 1.00
π̂
ATXN7L2 – Forward
H3K9 me2:DNA Meth. m5C
H3K27 me3:DNA Meth. m5C
H3K27 me3:H3K4 me1
DNA Meth. m5C:H4K20 me2
H2A.Z:H3K4 me3
H4K20 me2:H4−4ac
H3K4 me1:H4K20 me2
H3K4 me3:H4−4ac
H3K4 me1:H3−5ac
H4−4ac:H3−5ac
H2A.Z:H4K20 me2
H3K4 me1:H4−4ac
H3K9 me3
H3K4 me1:H2A.Z
H4K20 me3
H3K4 me3:H3−5ac
H4K20 me2:H3−5ac
H4K20 me2:H3K4 me3
H3K27 me3
H3K27 me2
H3K9 me2
DNA Meth. m5C
H4−4ac
H2A.Z
H3K4 me1
H4K20 me2
H3−5ac
H3K4 me3
0.00 0.25 0.50 0.75 1.00
π̂
ATXN7L2 – Reverse
H3K9 me2:DNA Meth. m5C
H3K27 me3:DNA Meth. m5C
H3K27 me3:H3K4 me1
DNA Meth. m5C:H4K20 me2
H2A.Z:H3K4 me3
H4K20 me2:H4−4ac
H3K4 me1:H4K20 me2
H3K4 me3:H4−4ac
H3K4 me1:H3−5ac
H4−4ac:H3−5ac
H2A.Z:H4K20 me2
H3K4 me1:H4−4ac
H3K9 me3
H3K4 me1:H2A.Z
H4K20 me3
H3K4 me3:H3−5ac
H4K20 me2:H3−5ac
H4K20 me2:H3K4 me3
H3K27 me3
H3K27 me2
H3K9 me2
DNA Meth. m5C
H4−4ac
H2A.Z
H3K4 me1
H4K20 me2
H3−5ac
H3K4 me3
0.00 0.25 0.50 0.75 1.00
π̂
ATXN7 – Forward
H3K9 me2:DNA Meth. m5C
H3K27 me2:DNA Meth. m5C
H3K27 me3:H4K20 me3
H3K4 me1:H3−5ac
H3K4 me1:H4K20 me2
DNA Meth. m5C:H4K20 me2
H3K9 me3:H4K20 me3
DNA Meth. m5C:H4K20 me3
H3K27 me3:H3K9 me3
H2A.Z:H3K4 me3
H2A.Z:H4K20 me2
H4K20 me2:H4−4ac
H3K27 me3:DNA Meth. m5C
DNA Meth. m5C:H3K9 me3
H3K4 me1:H4−4ac
H3K4 me1:H2A.Z
H3K4 me3:H3−5ac
H4−4ac:H3−5ac
H3K4 me3:H4−4ac
H4K20 me2:H3−5ac
H4K20 me2:H3K4 me3
H3K27 me2
H3K9 me2
H3K9 me3
H4K20 me3
H3K27 me3
H3K4 me1
DNA Meth. m5C
H4K20 me2
H2A.Z
H3K4 me3
H4−4ac
H3−5ac
0.00 0.25 0.50 0.75 1.00
π̂
ATXN7 – Reverse
H3K9 me2:DNA Meth. m5C
H3K27 me2:DNA Meth. m5C
H3K27 me3:H4K20 me3
H3K4 me1:H3−5ac
H3K4 me1:H4K20 me2
DNA Meth. m5C:H4K20 me2
H3K9 me3:H4K20 me3
DNA Meth. m5C:H4K20 me3
H3K27 me3:H3K9 me3
H2A.Z:H3K4 me3
H2A.Z:H4K20 me2
H4K20 me2:H4−4ac
H3K27 me3:DNA Meth. m5C
DNA Meth. m5C:H3K9 me3
H3K4 me1:H4−4ac
H3K4 me1:H2A.Z
H3K4 me3:H3−5ac
H4−4ac:H3−5ac
H3K4 me3:H4−4ac
H4K20 me2:H3−5ac
H4K20 me2:H3K4 me3
H3K27 me2
H3K9 me2
H3K9 me3
H4K20 me3
H3K27 me3
H3K4 me1
DNA Meth. m5C
H4K20 me2
H2A.Z
H3K4 me3
H4−4ac
H3−5ac
0.00 0.25 0.50 0.75 1.00
π̂
EP400 (2) – Forward
H3K27 me3:H4K20 me3
H3K27 me3:DNA Meth. m5C
DNA Meth. m5C:H4K20 me2
H3K27 me3:H3K4 me1
H3K4 me1:H4K20 me2
H2A.Z:H4−4ac
H3K4 me1:H2A.Z
H2A.Z:H3−5ac
H4−4ac:H3−5ac
H2A.Z:H3K4 me3
H4K20 me2:H4−4ac
H3K4 me1:H4−4ac
H4K20 me2:H3−5ac
H3K4 me1:H3−5ac
H3K27 me2
H3K9 me2
H3K4 me3:H4−4ac
H3K4 me3:H3−5ac
H2A.Z:H4K20 me2
H4K20 me2:H3K4 me3
H4K20 me3
H3K27 me3
H3K9 me3
DNA Meth. m5C
H3K4 me1
H2A.Z
H3−5ac
H3K4 me3
H4−4ac
H4K20 me2
0.00 0.25 0.50 0.75 1.00
π̂
EP400 (2) – Reverse
H3K27 me3:H4K20 me3
H3K27 me3:DNA Meth. m5C
DNA Meth. m5C:H4K20 me2
H3K27 me3:H3K4 me1
H3K4 me1:H4K20 me2
H2A.Z:H4−4ac
H3K4 me1:H2A.Z
H2A.Z:H3−5ac
H4−4ac:H3−5ac
H2A.Z:H3K4 me3
H4K20 me2:H4−4ac
H3K4 me1:H4−4ac
H4K20 me2:H3−5ac
H3K4 me1:H3−5ac
H3K27 me2
H3K9 me2
H3K4 me3:H4−4ac
H3K4 me3:H3−5ac
H2A.Z:H4K20 me2
H4K20 me2:H3K4 me3
H4K20 me3
H3K27 me3
H3K9 me3
DNA Meth. m5C
H3K4 me1
H2A.Z
H3−5ac
H3K4 me3
H4−4ac
H4K20 me2
0.00 0.25 0.50 0.75 1.00
π̂
SUPT7L – Forward
DNA Meth. m5C:H4K20 me3
H3K9 me2:DNA Meth. m5C
H3K27 me3:H4K20 me3
H2A.Z:H3K4 me3
H3K27 me2:DNA Meth. m5C
H3K4 me1:H4K20 me2
H3K9 me3:H4K20 me3
H3K27 me3:H3K9 me3
DNA Meth. m5C:H4K20 me2
H3K4 me1:H4−4ac
H3K4 me1:H2A.Z
DNA Meth. m5C:H3K9 me3
H3K4 me1:H3−5ac
H3K27 me3:DNA Meth. m5C
H3K4 me3:H3−5ac
H3K4 me3:H4−4ac
H4K20 me2:H4−4ac
H2A.Z:H4K20 me2
H4−4ac:H3−5ac
H4K20 me2:H3−5ac
H4K20 me2:H3K4 me3
H3K27 me2
H3K9 me2
H4K20 me3
H3K27 me3
H3K9 me3
DNA Meth. m5C
H3K4 me1
H4K20 me2
H3K4 me3
H2A.Z
H4−4ac
H3−5ac
0.00 0.25 0.50 0.75 1.00
π̂
SUPT7L – Reverse
DNA Meth. m5C:H4K20 me3
H3K9 me2:DNA Meth. m5C
H3K27 me3:H4K20 me3
H2A.Z:H3K4 me3
H3K27 me2:DNA Meth. m5C
H3K4 me1:H4K20 me2
H3K9 me3:H4K20 me3
H3K27 me3:H3K9 me3
DNA Meth. m5C:H4K20 me2
H3K4 me1:H4−4ac
H3K4 me1:H2A.Z
DNA Meth. m5C:H3K9 me3
H3K4 me1:H3−5ac
H3K27 me3:DNA Meth. m5C
H3K4 me3:H3−5ac
H3K4 me3:H4−4ac
H4K20 me2:H4−4ac
H2A.Z:H4K20 me2
H4−4ac:H3−5ac
H4K20 me2:H3−5ac
H4K20 me2:H3K4 me3
H3K27 me2
H3K9 me2
H4K20 me3
H3K27 me3
H3K9 me3
DNA Meth. m5C
H3K4 me1
H4K20 me2
H3K4 me3
H2A.Z
H4−4ac
H3−5ac
0.00 0.25 0.50 0.75 1.00
π̂

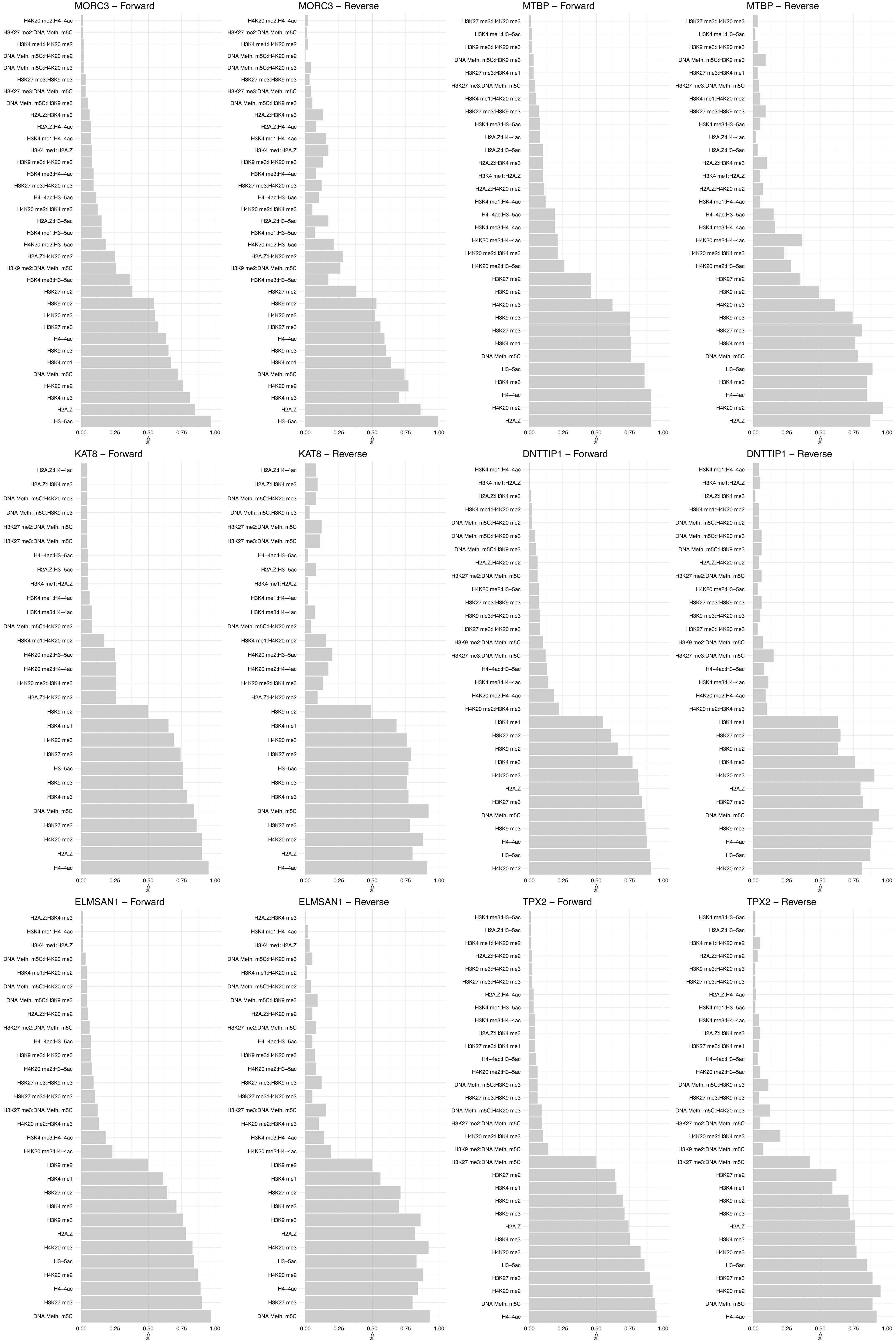

MORC3 – Forward
H4K20 me2:H4–4ac
H3K27 me2:DNA Meth. m5C
H3K4 me1:H4K20 me2
DNA Meth. m5C:H4K20 me2
DNA Meth. m5C:H4K20 me3
H3K27 me3:H3K9 me3
H3K27 me3:DNA Meth. m5C
DNA Meth. m5C:H3K9 me3
H2A.Z:H3K4 me3
H2A.Z:H4–4ac
H3K4 me1:H4–4ac
H3K4 me1:H2A.Z
H3K9 me3:H4K20 me3
H3K4 me3:H4–4ac
H3K27 me3:H4K20 me3
H4–4ac:H3–5ac
H4K20 me2:H3K4 me3
H2A.Z:H3–5ac
H3K4 me1:H3–5ac
H4K20 me2:H3–5ac
H2A.Z:H4K20 me2
H3K9 me2:DNA Meth. m5C
H3K4 me3:H3–5ac
H3K27 me2
H3K9 me2
H4K20 me3
H3K27 me3
H4–4ac
H3K9 me3
H3K4 me1
DNA Meth. m5C
H4K20 me2
H3K4 me3
H2A.Z
H3–5ac
0.00 0.25 0.50 0.75 1.00
π̂
MORC3 – Reverse
H4K20 me2:H4–4ac
H3K27 me2:DNA Meth. m5C
H3K4 me1:H4K20 me2
DNA Meth. m5C:H4K20 me2
DNA Meth. m5C:H4K20 me3
H3K27 me3:H3K9 me3
H3K27 me3:DNA Meth. m5C
DNA Meth. m5C:H3K9 me3
H2A.Z:H3K4 me3
H2A.Z:H4–4ac
H3K4 me1:H4–4ac
H3K4 me1:H2A.Z
H3K9 me3:H4K20 me3
H3K4 me3:H4–4ac
H3K27 me3:H4K20 me3
H4–4ac:H3–5ac
H4K20 me2:H3K4 me3
H2A.Z:H3–5ac
H3K4 me1:H3–5ac
H4K20 me2:H3–5ac
H2A.Z:H4K20 me2
H3K9 me2:DNA Meth. m5C
H3K4 me3:H3–5ac
H3K27 me2
H3K9 me2
H4K20 me3
H3K27 me3
H4–4ac
H3K9 me3
H3K4 me1
DNA Meth. m5C
H4K20 me2
H3K4 me3
H2A.Z
H3–5ac
0.00 0.25 0.50 0.75 1.00
π̂
MTBP – Forward
H3K27 me3:H4K20 me3
H3K4 me1:H3–5ac
H3K9 me3:H4K20 me3
DNA Meth. m5C:H3K9 me3
H3K27 me3:H3K4 me1
H3K27 me3:DNA Meth. m5C
H3K4 me1:H4K20 me2
H3K27 me3:H3K9 me3
H3K4 me3:H3–5ac
H2A.Z:H4–4ac
H2A.Z:H3–5ac
H2A.Z:H3K4 me3
H3K4 me1:H2A.Z
H2A.Z:H4K20 me2
H3K4 me1:H4–4ac
H4–4ac:H3–5ac
H3K4 me3:H4–4ac
H4K20 me2:H4–4ac
H4K20 me2:H3K4 me3
H4K20 me2:H3–5ac
H3K27 me2
H3K9 me2
H4K20 me3
H3K9 me3
H3K27 me3
H3K4 me1
DNA Meth. m5C
H3–5ac
H3K4 me3
H4–4ac
H4K20 me2
H2A.Z
0.00 0.25 0.50 0.75 1.00
π̂
MTBP – Reverse
H3K27 me3:H4K20 me3
H3K4 me1:H3–5ac
H3K9 me3:H4K20 me3
DNA Meth. m5C:H3K9 me3
H3K27 me3:H3K4 me1
H3K27 me3:DNA Meth. m5C
H3K4 me1:H4K20 me2
H3K27 me3:H3K9 me3
H3K4 me3:H3–5ac
H2A.Z:H4–4ac
H2A.Z:H3–5ac
H2A.Z:H3K4 me3
H3K4 me1:H2A.Z
H2A.Z:H4K20 me2
H3K4 me1:H4–4ac
H4–4ac:H3–5ac
H3K4 me3:H4–4ac
H4K20 me2:H4–4ac
H4K20 me2:H3K4 me3
H4K20 me2:H3–5ac
H3K27 me2
H3K9 me2
H4K20 me3
H3K9 me3
H3K27 me3
H3K4 me1
DNA Meth. m5C
H3–5ac
H3K4 me3
H4–4ac
H4K20 me2
H2A.Z
0.00 0.25 0.50 0.75 1.00
π̂
KAT8 – Forward
H2A.Z:H4–4ac
H2A.Z:H3K4 me3
DNA Meth. m5C:H4K20 me3
DNA Meth. m5C:H3K9 me3
H3K27 me2:DNA Meth. m5C
H3K27 me3:DNA Meth. m5C
H4–4ac:H3–5ac
H2A.Z:H3–5ac
H3K4 me1:H2A.Z
H3K4 me1:H4–4ac
H3K4 me3:H4–4ac
DNA Meth. m5C:H4K20 me2
H3K4 me1:H4K20 me2
H4K20 me2:H3–5ac
H4K20 me2:H4–4ac
H4K20 me2:H3K4 me3
H2A.Z:H4K20 me2
H3K9 me2
H3K4 me1
H4K20 me3
H3K27 me2
H3–5ac
H3K9 me3
H3K4 me3
DNA Meth. m5C
H3K27 me3
H4K20 me2
H2A.Z
H4–4ac
0.00 0.25 0.50 0.75 1.00
π̂
KAT8 – Reverse
H2A.Z:H4–4ac
H2A.Z:H3K4 me3
DNA Meth. m5C:H4K20 me3
DNA Meth. m5C:H3K9 me3
H3K27 me2:DNA Meth. m5C
H3K27 me3:DNA Meth. m5C
H4–4ac:H3–5ac
H2A.Z:H3–5ac
H3K4 me1:H2A.Z
H3K4 me1:H4–4ac
H3K4 me3:H4–4ac
DNA Meth. m5C:H4K20 me2
H3K4 me1:H4K20 me2
H4K20 me2:H3–5ac
H4K20 me2:H4–4ac
H4K20 me2:H3K4 me3
H2A.Z:H4K20 me2
H3K9 me2
H3K4 me1
H4K20 me3
H3K27 me2
H3–5ac
H3K9 me3
H3K4 me3
DNA Meth. m5C
H3K27 me3
H4K20 me2
H2A.Z
H4–4ac
0.00 0.25 0.50 0.75 1.00
π̂
DNTTIP1 – Forward
H3K4 me1:H4–4ac
H3K4 me1:H2A.Z
H2A.Z:H3K4 me3
H3K4 me1:H4K20 me2
DNA Meth. m5C:H4K20 me2
DNA Meth. m5C:H4K20 me3
DNA Meth. m5C:H3K9 me3
H2A.Z:H4K20 me2
H3K27 me2:DNA Meth. m5C
H4K20 me2:H3–5ac
H3K27 me3:H3K9 me3
H3K9 me3:H4K20 me3
H3K27 me3:H4K20 me3
H3K9 me2:DNA Meth. m5C
H3K27 me3:DNA Meth. m5C
H4–4ac:H3–5ac
H3K4 me3:H4–4ac
H4K20 me2:H4–4ac
H4K20 me2:H3K4 me3
H3K4 me1
H3K27 me2
H3K9 me2
H3K4 me3
H4K20 me3
H2A.Z
H3K27 me3
DNA Meth. m5C
H3K9 me3
H4–4ac
H3–5ac
H4K20 me2
0.00 0.25 0.50 0.75 1.00
π̂
DNTTIP1 – Reverse
H3K4 me1:H4–4ac
H3K4 me1:H2A.Z
H2A.Z:H3K4 me3
H3K4 me1:H4K20 me2
DNA Meth. m5C:H4K20 me2
DNA Meth. m5C:H4K20 me3
DNA Meth. m5C:H3K9 me3
H2A.Z:H4K20 me2
H3K27 me2:DNA Meth. m5C
H4K20 me2:H3–5ac
H3K27 me3:H3K9 me3
H3K9 me3:H4K20 me3
H3K27 me3:H4K20 me3
H3K9 me2:DNA Meth. m5C
H3K27 me3:DNA Meth. m5C
H4–4ac:H3–5ac
H3K4 me3:H4–4ac
H4K20 me2:H4–4ac
H4K20 me2:H3K4 me3
H3K4 me1
H3K27 me2
H3K9 me2
H3K4 me3
H4K20 me3
H2A.Z
H3K27 me3
DNA Meth. m5C
H3K9 me3
H4–4ac
H3–5ac
H4K20 me2
0.00 0.25 0.50 0.75 1.00
π̂
ELMSAN1 – Forward
H2A.Z:H3K4 me3
H3K4 me1:H4–4ac
H3K4 me1:H2A.Z
DNA Meth. m5C:H4K20 me3
H3K4 me1:H4K20 me2
DNA Meth. m5C:H4K20 me2
DNA Meth. m5C:H3K9 me3
H2A.Z:H4K20 me2
H3K27 me2:DNA Meth. m5C
H4–4ac:H3–5ac
H3K9 me3:H4K20 me3
H4K20 me2:H3–5ac
H3K27 me3:H3K9 me3
H3K27 me3:H4K20 me3
H3K27 me3:DNA Meth. m5C
H4K20 me2:H3K4 me3
H3K4 me3:H4–4ac
H4K20 me2:H4–4ac
H3K9 me2
H3K4 me1
H3K27 me2
H3K4 me3
H3K9 me3
H2A.Z
H4K20 me3
H3–5ac
H4K20 me2
H4–4ac
H3K27 me3
DNA Meth. m5C
0.00 0.25 0.50 0.75 1.00
π̂
ELMSAN1 – Reverse
H2A.Z:H3K4 me3
H3K4 me1:H4–4ac
H3K4 me1:H2A.Z
DNA Meth. m5C:H4K20 me3
H3K4 me1:H4K20 me2
DNA Meth. m5C:H4K20 me2
DNA Meth. m5C:H3K9 me3
H2A.Z:H4K20 me2
H3K27 me2:DNA Meth. m5C
H4–4ac:H3–5ac
H3K9 me3:H4K20 me3
H4K20 me2:H3–5ac
H3K27 me3:H3K9 me3
H3K27 me3:H4K20 me3
H3K27 me3:DNA Meth. m5C
H4K20 me2:H3K4 me3
H3K4 me3:H4–4ac
H4K20 me2:H4–4ac
H3K9 me2
H3K4 me1
H3K27 me2
H3K4 me3
H3K9 me3
H2A.Z
H4K20 me3
H3–5ac
H4K20 me2
H4–4ac
H3K27 me3
DNA Meth. m5C
0.00 0.25 0.50 0.75 1.00
π̂
TPX2 – Forward
H3K4 me3:H3–5ac
H2A.Z:H3–5ac
H3K4 me1:H4K20 me2
H2A.Z:H4K20 me2
H3K9 me3:H4K20 me3
H3K27 me3:H4K20 me3
H2A.Z:H4–4ac
H3K4 me1:H3–5ac
H3K4 me3:H4–4ac
H2A.Z:H3K4 me3
H3K27 me3:H3K4 me1
H4–4ac:H3–5ac
H4K20 me2:H3–5ac
DNA Meth. m5C:H3K9 me3
H3K27 me3:H3K9 me3
DNA Meth. m5C:H4K20 me3
H3K27 me2:DNA Meth. m5C
H4K20 me2:H3K4 me3
H3K9 me2:DNA Meth. m5C
H3K27 me3:DNA Meth. m5C
H3K27 me2
H3K4 me1
H3K9 me2
H3K9 me3
H2A.Z
H3K4 me3
H4K20 me3
H3–5ac
H3K27 me3
H4K20 me2
DNA Meth. m5C
H4–4ac
0.00 0.25 0.50 0.75 1.00
π̂
TPX2 – Reverse
H3K4 me3:H3–5ac
H2A.Z:H3–5ac
H3K4 me1:H4K20 me2
H2A.Z:H4K20 me2
H3K9 me3:H4K20 me3
H3K27 me3:H4K20 me3
H2A.Z:H4–4ac
H3K4 me1:H3–5ac
H3K4 me3:H4–4ac
H2A.Z:H3K4 me3
H3K27 me3:H3K4 me1
H4–4ac:H3–5ac
H4K20 me2:H3–5ac
DNA Meth. m5C:H3K9 me3
H3K27 me3:H3K9 me3
DNA Meth. m5C:H4K20 me3
H3K27 me2:DNA Meth. m5C
H4K20 me2:H3K4 me3
H3K9 me2:DNA Meth. m5C
H3K27 me3:DNA Meth. m5C
H3K27 me2
H3K4 me1
H3K9 me2
H3K9 me3
H2A.Z
H3K4 me3
H4K20 me3
H3–5ac
H3K27 me3
H4K20 me2
DNA Meth. m5C
H4–4ac
0.00 0.25 0.50 0.75 1.00
π̂

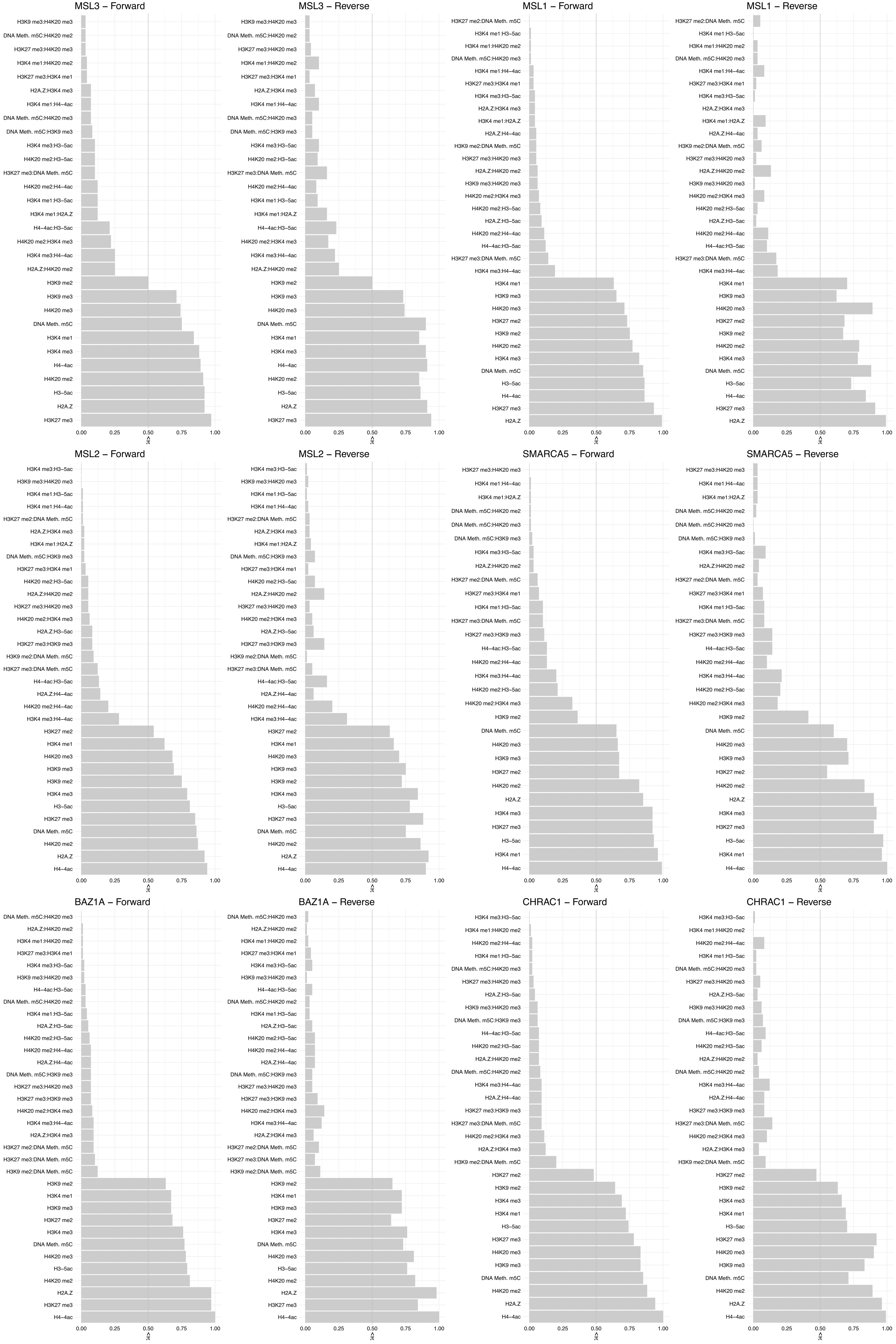
MSL3 – Forward
H3K9 me3:H4K20 me3
DNA Meth. m5C:H4K20 me2
H3K27 me3:H4K20 me3
H3K4 me1:H4K20 me2
H3K27 me3:H3K4 me1
H2A.Z:H3K4 me3
H3K4 me1:H4–4ac
DNA Meth. m5C:H4K20 me3
DNA Meth. m5C:H3K9 me3
H3K4 me3:H3–5ac
H4K20 me2:H3–5ac
H3K27 me3:DNA Meth. m5C
H4K20 me2:H4–4ac
H3K4 me1:H3–5ac
H3K4 me1:H2A.Z
H4–4ac:H3–5ac
H4K20 me2:H3K4 me3
H3K4 me3:H4–4ac
H2A.Z:H4K20 me2
H3K9 me2
H3K9 me3
H4K20 me3
DNA Meth. m5C
H3K4 me1
H3K4 me3
H4–4ac
H4K20 me2
H3–5ac
H2A.Z
H3K27 me3
0.00 0.25 0.50 0.75 1.00
π̂
MSL3 – Reverse
H3K9 me3:H4K20 me3
DNA Meth. m5C:H4K20 me2
H3K27 me3:H4K20 me3
H3K4 me1:H4K20 me2
H3K27 me3:H3K4 me1
H2A.Z:H3K4 me3
H3K4 me1:H4–4ac
DNA Meth. m5C:H4K20 me3
DNA Meth. m5C:H3K9 me3
H3K4 me3:H3–5ac
H4K20 me2:H3–5ac
H3K27 me3:DNA Meth. m5C
H4K20 me2:H4–4ac
H3K4 me1:H3–5ac
H3K4 me1:H2A.Z
H4–4ac:H3–5ac
H4K20 me2:H3K4 me3
H3K4 me3:H4–4ac
H2A.Z:H4K20 me2
H3K9 me2
H3K9 me3
H4K20 me3
DNA Meth. m5C
H3K4 me1
H3K4 me3
H4–4ac
H4K20 me2
H3–5ac
H2A.Z
H3K27 me3
0.00 0.25 0.50 0.75 1.00
π̂
MSL1 – Forward
H3K27 me2:DNA Meth. m5C
H3K4 me1:H3–5ac
H3K4 me1:H4K20 me2
DNA Meth. m5C:H4K20 me3
H3K4 me1:H4–4ac
H3K27 me3:H3K4 me1
H3K4 me3:H3–5ac
H2A.Z:H3K4 me3
H3K4 me1:H2A.Z
H2A.Z:H4–4ac
H3K9 me2:DNA Meth. m5C
H3K27 me3:H4K20 me3
H2A.Z:H4K20 me2
H3K9 me3:H4K20 me3
H4K20 me2:H3K4 me3
H4K20 me2:H3–5ac
H2A.Z:H3–5ac
H4K20 me2:H4–4ac
H4–4ac:H3–5ac
H3K27 me3:DNA Meth. m5C
H3K4 me3:H4–4ac
H3K4 me1
H3K9 me3
H4K20 me3
H3K27 me2
H3K9 me2
H4K20 me2
H3K4 me3
DNA Meth. m5C
H3–5ac
H4–4ac
H3K27 me3
H2A.Z
0.00 0.25 0.50 0.75 1.00
π̂
MSL1 – Reverse
H3K27 me2:DNA Meth. m5C
H3K4 me1:H3–5ac
H3K4 me1:H4K20 me2
DNA Meth. m5C:H4K20 me3
H3K4 me1:H4–4ac
H3K27 me3:H3K4 me1
H3K4 me3:H3–5ac
H2A.Z:H3K4 me3
H3K4 me1:H2A.Z
H2A.Z:H4–4ac
H3K9 me2:DNA Meth. m5C
H3K27 me3:H4K20 me3
H2A.Z:H4K20 me2
H3K9 me3:H4K20 me3
H4K20 me2:H3K4 me3
H4K20 me2:H3–5ac
H2A.Z:H3–5ac
H4K20 me2:H4–4ac
H4–4ac:H3–5ac
H3K27 me3:DNA Meth. m5C
H3K4 me3:H4–4ac
H3K4 me1
H3K9 me3
H4K20 me3
H3K27 me2
H3K9 me2
H4K20 me2
H3K4 me3
DNA Meth. m5C
H3–5ac
H4–4ac
H3K27 me3
H2A.Z
0.00 0.25 0.50 0.75 1.00
π̂
MSL2 – Forward
H3K4 me3:H3–5ac
H3K9 me3:H4K20 me3
H3K4 me1:H3–5ac
H3K4 me1:H4–4ac
H3K27 me2:DNA Meth. m5C
H2A.Z:H3K4 me3
H3K4 me1:H2A.Z
DNA Meth. m5C:H3K9 me3
H3K27 me3:H3K4 me1
H4K20 me2:H3–5ac
H2A.Z:H4K20 me2
H3K27 me3:H4K20 me3
H4K20 me2:H3K4 me3
H2A.Z:H3–5ac
H3K27 me3:H3K9 me3
H3K9 me2:DNA Meth. m5C
H3K27 me3:DNA Meth. m5C
H4–4ac:H3–5ac
H2A.Z:H4–4ac
H4K20 me2:H4–4ac
H3K4 me3:H4–4ac
H3K27 me2
H3K4 me1
H4K20 me3
H3K9 me3
H3K9 me2
H3K4 me3
H3–5ac
H3K27 me3
DNA Meth. m5C
H4K20 me2
H2A.Z
H4–4ac
0.00 0.25 0.50 0.75 1.00
π̂
MSL2 – Reverse
H3K4 me3:H3–5ac
H3K9 me3:H4K20 me3
H3K4 me1:H3–5ac
H3K4 me1:H4–4ac
H3K27 me2:DNA Meth. m5C
H2A.Z:H3K4 me3
H3K4 me1:H2A.Z
DNA Meth. m5C:H3K9 me3
H3K27 me3:H3K4 me1
H4K20 me2:H3–5ac
H2A.Z:H4K20 me2
H3K27 me3:H4K20 me3
H4K20 me2:H3K4 me3
H2A.Z:H3–5ac
H3K27 me3:H3K9 me3
H3K9 me2:DNA Meth. m5C
H3K27 me3:DNA Meth. m5C
H4–4ac:H3–5ac
H2A.Z:H4–4ac
H4K20 me2:H4–4ac
H3K4 me3:H4–4ac
H3K27 me2
H3K4 me1
H4K20 me3
H3K9 me3
H3K9 me2
H3K4 me3
H3–5ac
H3K27 me3
DNA Meth. m5C
H4K20 me2
H2A.Z
H4–4ac
0.00 0.25 0.50 0.75 1.00
π̂
SMARCA5 – Forward
H3K27 me3:H4K20 me3
H3K4 me1:H4–4ac
H3K4 me1:H2A.Z
DNA Meth. m5C:H4K20 me2
DNA Meth. m5C:H4K20 me3
DNA Meth. m5C:H3K9 me3
H3K4 me3:H3–5ac
H2A.Z:H4K20 me2
H3K27 me2:DNA Meth. m5C
H3K27 me3:H3K4 me1
H3K4 me1:H3–5ac
H3K27 me3:DNA Meth. m5C
H3K27 me3:H3K9 me3
H4–4ac:H3–5ac
H4K20 me2:H4–4ac
H3K4 me3:H4–4ac
H4K20 me2:H3–5ac
H4K20 me2:H3K4 me3
H3K9 me2
DNA Meth. m5C
H4K20 me3
H3K9 me3
H3K27 me2
H4K20 me2
H2A.Z
H3K4 me3
H3K27 me3
H3–5ac
H3K4 me1
H4–4ac
0.00 0.25 0.50 0.75 1.00
π̂
SMARCA5 – Reverse
H3K27 me3:H4K20 me3
H3K4 me1:H4–4ac
H3K4 me1:H2A.Z
DNA Meth. m5C:H4K20 me2
DNA Meth. m5C:H4K20 me3
DNA Meth. m5C:H3K9 me3
H3K4 me3:H3–5ac
H2A.Z:H4K20 me2
H3K27 me2:DNA Meth. m5C
H3K27 me3:H3K4 me1
H3K4 me1:H3–5ac
H3K27 me3:DNA Meth. m5C
H3K27 me3:H3K9 me3
H4–4ac:H3–5ac
H4K20 me2:H4–4ac
H3K4 me3:H4–4ac
H4K20 me2:H3–5ac
H4K20 me2:H3K4 me3
H3K9 me2
DNA Meth. m5C
H4K20 me3
H3K9 me3
H3K27 me2
H4K20 me2
H2A.Z
H3K4 me3
H3K27 me3
H3–5ac
H3K4 me1
H4–4ac
0.00 0.25 0.50 0.75 1.00
π̂
BAZ1A – Forward
DNA Meth. m5C:H4K20 me3
H2A.Z:H4K20 me2
H3K4 me1:H4K20 me2
H3K27 me3:H3K4 me1
H3K4 me3:H3–5ac
H3K9 me3:H4K20 me3
H4–4ac:H3–5ac
DNA Meth. m5C:H4K20 me2
H3K4 me1:H3–5ac
H2A.Z:H3–5ac
H4K20 me2:H3–5ac
H4K20 me2:H4–4ac
H2A.Z:H4–4ac
DNA Meth. m5C:H3K9 me3
H3K27 me3:H4K20 me3
H3K27 me3:H3K9 me3
H4K20 me2:H3K4 me3
H3K4 me3:H4–4ac
H2A.Z:H3K4 me3
H3K27 me2:DNA Meth. m5C
H3K27 me3:DNA Meth. m5C
H3K9 me2:DNA Meth. m5C
H3K9 me2
H3K4 me1
H3K9 me3
H3K27 me2
H3K4 me3
DNA Meth. m5C
H4K20 me3
H3–5ac
H4K20 me2
H2A.Z
H3K27 me3
H4–4ac
0.00 0.25 0.50 0.75 1.00
π̂
BAZ1A – Reverse
DNA Meth. m5C:H4K20 me3
H2A.Z:H4K20 me2
H3K4 me1:H4K20 me2
H3K27 me3:H3K4 me1
H3K4 me3:H3–5ac
H3K9 me3:H4K20 me3
H4–4ac:H3–5ac
DNA Meth. m5C:H4K20 me2
H3K4 me1:H3–5ac
H2A.Z:H3–5ac
H4K20 me2:H3–5ac
H4K20 me2:H4–4ac
H2A.Z:H4–4ac
DNA Meth. m5C:H3K9 me3
H3K27 me3:H4K20 me3
H3K27 me3:H3K9 me3
H4K20 me2:H3K4 me3
H3K4 me3:H4–4ac
H2A.Z:H3K4 me3
H3K27 me2:DNA Meth. m5C
H3K27 me3:DNA Meth. m5C
H3K9 me2:DNA Meth. m5C
H3K9 me2
H3K4 me1
H3K9 me3
H3K27 me2
H3K4 me3
DNA Meth. m5C
H4K20 me3
H3–5ac
H4K20 me2
H2A.Z
H3K27 me3
H4–4ac
0.00 0.25 0.50 0.75 1.00
π̂
CHRAC1 – Forward
H3K4 me3:H3–5ac
H3K4 me1:H4K20 me2
H4K20 me2:H4–4ac
H3K4 me1:H3–5ac
DNA Meth. m5C:H4K20 me3
H3K27 me3:H4K20 me3
H2A.Z:H3–5ac
H3K9 me3:H4K20 me3
DNA Meth. m5C:H3K9 me3
H4–4ac:H3–5ac
H4K20 me2:H3–5ac
H2A.Z:H4K20 me2
DNA Meth. m5C:H4K20 me2
H3K4 me3:H4–4ac
H2A.Z:H4–4ac
H3K27 me3:H3K9 me3
H3K27 me3:DNA Meth. m5C
H4K20 me2:H3K4 me3
H2A.Z:H3K4 me3
H3K9 me2:DNA Meth. m5C
H3K27 me2
H3K9 me2
H3K4 me3
H3K4 me1
H3–5ac
H3K27 me3
H4K20 me3
H3K9 me3
DNA Meth. m5C
H4K20 me2
H2A.Z
H4–4ac
0.00 0.25 0.50 0.75 1.00
π̂
CHRAC1 – Reverse
H3K4 me3:H3–5ac
H3K4 me1:H4K20 me2
H4K20 me2:H4–4ac
H3K4 me1:H3–5ac
DNA Meth. m5C:H4K20 me3
H3K27 me3:H4K20 me3
H2A.Z:H3–5ac
H3K9 me3:H4K20 me3
DNA Meth. m5C:H3K9 me3
H4–4ac:H3–5ac
H4K20 me2:H3–5ac
H2A.Z:H4K20 me2
DNA Meth. m5C:H4K20 me2
H3K4 me3:H4–4ac
H2A.Z:H4–4ac
H3K27 me3:H3K9 me3
H3K27 me3:DNA Meth. m5C
H4K20 me2:H3K4 me3
H2A.Z:H3K4 me3
H3K9 me2:DNA Meth. m5C
H3K27 me2
H3K9 me2
H3K4 me3
H3K4 me1
H3–5ac
H3K27 me3
H4K20 me3
H3K9 me3
DNA Meth. m5C
H4K20 me2
H2A.Z
H4–4ac
0.00 0.25 0.50 0.75 1.00
π̂

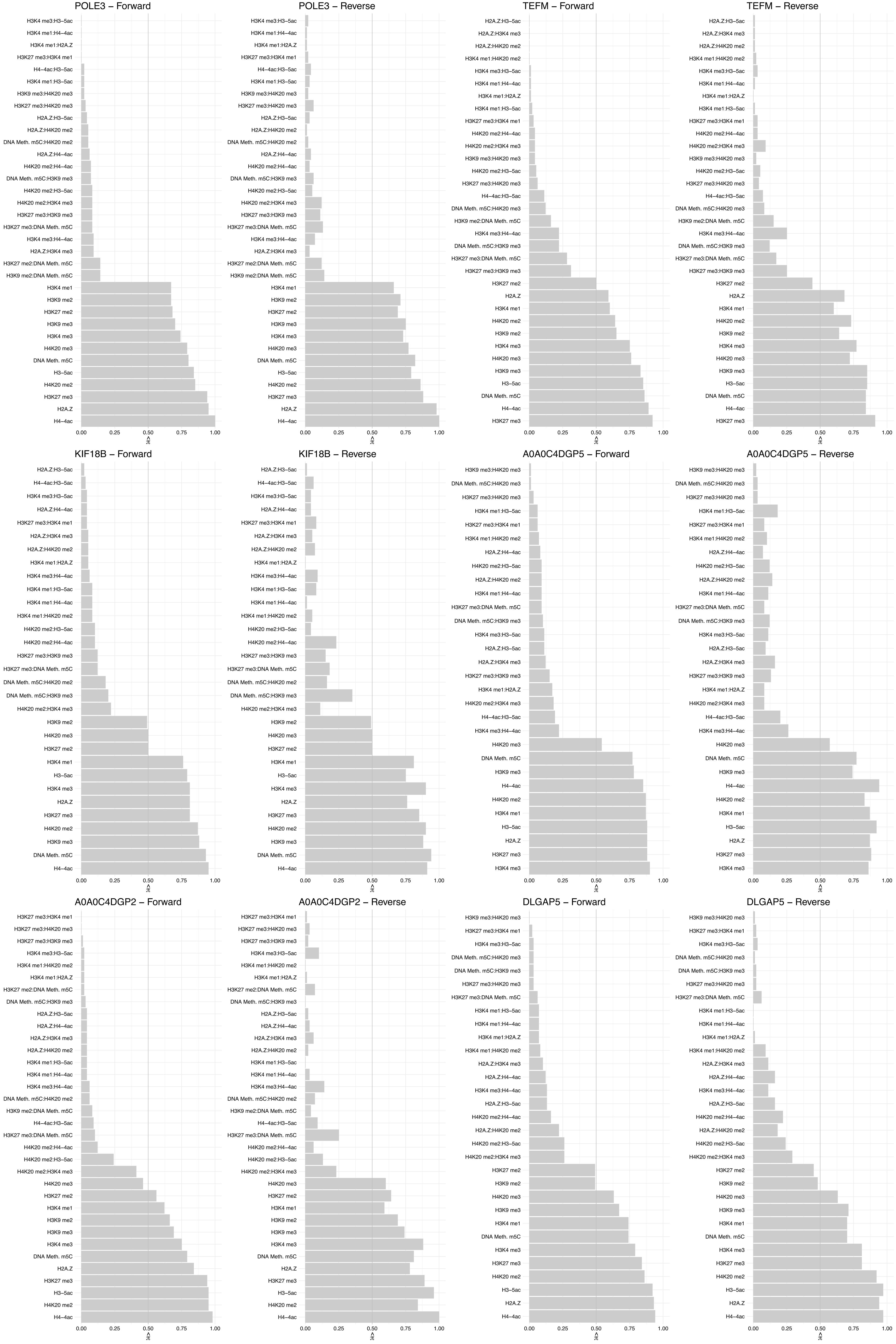
POLE3 – Forward
H3K4 me3:H3−5ac
H3K4 me1:H4−4ac
H3K4 me1:H2A.Z
H3K27 me3:H3K4 me1
H4−4ac:H3−5ac
H3K4 me1:H3−5ac
H3K9 me3:H4K20 me3
H3K27 me3:H4K20 me3
H2A.Z:H3−5ac
H2A.Z:H4K20 me2
DNA Meth. m5C:H4K20 me2
H2A.Z:H4−4ac
H4K20 me2:H4−4ac
DNA Meth. m5C:H3K9 me3
H4K20 me2:H3−5ac
H4K20 me2:H3K4 me3
H3K27 me3:H3K9 me3
H3K27 me3:DNA Meth. m5C
H3K4 me3:H4−4ac
H2A.Z:H3K4 me3
H3K27 me2:DNA Meth. m5C
H3K9 me2:DNA Meth. m5C
H3K4 me1
H3K9 me2
H3K27 me2
H3K9 me3
H3K4 me3
H4K20 me3
DNA Meth. m5C
H3−5ac
H4K20 me2
H3K27 me3
H2A.Z
H4−4ac
0.00 0.25 0.50 0.75 1.00
π̂
POLE3 – Reverse
H3K4 me3:H3−5ac
H3K4 me1:H4−4ac
H3K4 me1:H2A.Z
H3K27 me3:H3K4 me1
H4−4ac:H3−5ac
H3K4 me1:H3−5ac
H3K9 me3:H4K20 me3
H3K27 me3:H4K20 me3
H2A.Z:H3−5ac
H2A.Z:H4K20 me2
DNA Meth. m5C:H4K20 me2
H2A.Z:H4−4ac
H4K20 me2:H4−4ac
DNA Meth. m5C:H3K9 me3
H4K20 me2:H3−5ac
H4K20 me2:H3K4 me3
H3K27 me3:H3K9 me3
H3K27 me3:DNA Meth. m5C
H3K4 me3:H4−4ac
H2A.Z:H3K4 me3
H3K27 me2:DNA Meth. m5C
H3K9 me2:DNA Meth. m5C
H3K4 me1
H3K9 me2
H3K27 me2
H3K9 me3
H3K4 me3
H4K20 me3
DNA Meth. m5C
H3−5ac
H4K20 me2
H3K27 me3
H2A.Z
H4−4ac
0.00 0.25 0.50 0.75 1.00
π̂
TEFM – Forward
H2A.Z:H3−5ac
H2A.Z:H3K4 me3
H2A.Z:H4K20 me2
H3K4 me1:H4K20 me2
H3K4 me3:H3−5ac
H3K4 me1:H4−4ac
H3K4 me1:H2A.Z
H3K4 me1:H3−5ac
H3K27 me3:H3K4 me1
H4K20 me2:H4−4ac
H4K20 me2:H3K4 me3
H3K9 me3:H4K20 me3
H4K20 me2:H3−5ac
H3K27 me3:H4K20 me3
H4−4ac:H3−5ac
DNA Meth. m5C:H4K20 me3
H3K9 me2:DNA Meth. m5C
H3K4 me3:H4−4ac
DNA Meth. m5C:H3K9 me3
H3K27 me3:DNA Meth. m5C
H3K27 me3:H3K9 me3
H3K27 me2
H2A.Z
H3K4 me1
H4K20 me2
H3K9 me2
H3K4 me3
H4K20 me3
H3K9 me3
H3−5ac
DNA Meth. m5C
H4−4ac
H3K27 me3
0.00 0.25 0.50 0.75 1.00
π̂
TEFM – Reverse
H2A.Z:H3−5ac
H2A.Z:H3K4 me3
H2A.Z:H4K20 me2
H3K4 me1:H4K20 me2
H3K4 me3:H3−5ac
H3K4 me1:H4−4ac
H3K4 me1:H2A.Z
H3K4 me1:H3−5ac
H3K27 me3:H3K4 me1
H4K20 me2:H4−4ac
H4K20 me2:H3K4 me3
H3K9 me3:H4K20 me3
H4K20 me2:H3−5ac
H3K27 me3:H4K20 me3
H4−4ac:H3−5ac
DNA Meth. m5C:H4K20 me3
H3K9 me2:DNA Meth. m5C
H3K4 me3:H4−4ac
DNA Meth. m5C:H3K9 me3
H3K27 me3:DNA Meth. m5C
H3K27 me3:H3K9 me3
H3K27 me2
H2A.Z
H3K4 me1
H4K20 me2
H3K9 me2
H3K4 me3
H4K20 me3
H3K9 me3
H3−5ac
DNA Meth. m5C
H4−4ac
H3K27 me3
0.00 0.25 0.50 0.75 1.00
π̂
KIF18B – Forward
H2A.Z:H3−5ac
H4−4ac:H3−5ac
H3K4 me3:H3−5ac
H2A.Z:H4−4ac
H3K27 me3:H3K4 me1
H2A.Z:H3K4 me3
H2A.Z:H4K20 me2
H3K4 me1:H2A.Z
H3K4 me3:H4−4ac
H3K4 me1:H3−5ac
H3K4 me1:H4−4ac
H3K4 me1:H4K20 me2
H4K20 me2:H3−5ac
H4K20 me2:H4−4ac
H3K27 me3:H3K9 me3
H3K27 me3:DNA Meth. m5C
DNA Meth. m5C:H4K20 me2
DNA Meth. m5C:H3K9 me3
H4K20 me2:H3K4 me3
H3K9 me2
H4K20 me3
H3K27 me2
H3K4 me1
H3−5ac
H3K4 me3
H2A.Z
H3K27 me3
H4K20 me2
H3K9 me3
DNA Meth. m5C
H4−4ac
0.00 0.25 0.50 0.75 1.00
π̂
KIF18B – Reverse
H2A.Z:H3−5ac
H4−4ac:H3−5ac
H3K4 me3:H3−5ac
H2A.Z:H4−4ac
H3K27 me3:H3K4 me1
H2A.Z:H3K4 me3
H2A.Z:H4K20 me2
H3K4 me1:H2A.Z
H3K4 me3:H4−4ac
H3K4 me1:H3−5ac
H3K4 me1:H4−4ac
H3K4 me1:H4K20 me2
H4K20 me2:H3−5ac
H4K20 me2:H4−4ac
H3K27 me3:H3K9 me3
H3K27 me3:DNA Meth. m5C
DNA Meth. m5C:H4K20 me2
DNA Meth. m5C:H3K9 me3
H4K20 me2:H3K4 me3
H3K9 me2
H4K20 me3
H3K27 me2
H3K4 me1
H3−5ac
H3K4 me3
H2A.Z
H3K27 me3
H4K20 me2
H3K9 me3
DNA Meth. m5C
H4−4ac
0.00 0.25 0.50 0.75 1.00
π̂
A0A0C4DGP5 – Forward
H3K9 me3:H4K20 me3
DNA Meth. m5C:H4K20 me3
H3K27 me3:H4K20 me3
H3K4 me1:H3−5ac
H3K27 me3:H3K4 me1
H3K4 me1:H4K20 me2
H2A.Z:H4−4ac
H4K20 me2:H3−5ac
H2A.Z:H4K20 me2
H3K4 me1:H4−4ac
H3K27 me3:DNA Meth. m5C
DNA Meth. m5C:H3K9 me3
H3K4 me3:H3−5ac
H2A.Z:H3−5ac
H2A.Z:H3K4 me3
H3K27 me3:H3K9 me3
H3K4 me1:H2A.Z
H4K20 me2:H3K4 me3
H4−4ac:H3−5ac
H3K4 me3:H4−4ac
H4K20 me3
DNA Meth. m5C
H3K9 me3
H4−4ac
H4K20 me2
H3K4 me1
H3−5ac
H2A.Z
H3K27 me3
H3K4 me3
0.00 0.25 0.50 0.75 1.00
π̂
A0A0C4DGP5 – Reverse
H3K9 me3:H4K20 me3
DNA Meth. m5C:H4K20 me3
H3K27 me3:H4K20 me3
H3K4 me1:H3−5ac
H3K27 me3:H3K4 me1
H3K4 me1:H4K20 me2
H2A.Z:H4−4ac
H4K20 me2:H3−5ac
H2A.Z:H4K20 me2
H3K4 me1:H4−4ac
H3K27 me3:DNA Meth. m5C
DNA Meth. m5C:H3K9 me3
H3K4 me3:H3−5ac
H2A.Z:H3−5ac
H2A.Z:H3K4 me3
H3K27 me3:H3K9 me3
H3K4 me1:H2A.Z
H4K20 me2:H3K4 me3
H4−4ac:H3−5ac
H3K4 me3:H4−4ac
H4K20 me3
DNA Meth. m5C
H3K9 me3
H4−4ac
H4K20 me2
H3K4 me1
H3−5ac
H2A.Z
H3K27 me3
H3K4 me3
0.00 0.25 0.50 0.75 1.00
π̂
A0A0C4DGP2 – Forward
H3K27 me3:H3K4 me1
H3K27 me3:H4K20 me3
H3K27 me3:H3K9 me3
H3K4 me3:H3−5ac
H3K4 me1:H4K20 me2
H3K4 me1:H2A.Z
H3K27 me2:DNA Meth. m5C
DNA Meth. m5C:H3K9 me3
H2A.Z:H3−5ac
H2A.Z:H4−4ac
H2A.Z:H3K4 me3
H2A.Z:H4K20 me2
H3K4 me1:H3−5ac
H3K4 me1:H4−4ac
H3K4 me3:H4−4ac
DNA Meth. m5C:H4K20 me2
H3K9 me2:DNA Meth. m5C
H4−4ac:H3−5ac
H3K27 me3:DNA Meth. m5C
H4K20 me2:H4−4ac
H4K20 me2:H3−5ac
H4K20 me2:H3K4 me3
H4K20 me3
H3K27 me2
H3K4 me1
H3K9 me2
H3K9 me3
H3K4 me3
DNA Meth. m5C
H2A.Z
H3K27 me3
H3−5ac
H4K20 me2
H4−4ac
0.00 0.25 0.50 0.75 1.00
π̂
A0A0C4DGP2 – Reverse
H3K27 me3:H3K4 me1
H3K27 me3:H4K20 me3
H3K27 me3:H3K9 me3
H3K4 me3:H3−5ac
H3K4 me1:H4K20 me2
H3K4 me1:H2A.Z
H3K27 me2:DNA Meth. m5C
DNA Meth. m5C:H3K9 me3
H2A.Z:H3−5ac
H2A.Z:H4−4ac
H2A.Z:H3K4 me3
H2A.Z:H4K20 me2
H3K4 me1:H3−5ac
H3K4 me1:H4−4ac
H3K4 me3:H4−4ac
DNA Meth. m5C:H4K20 me2
H3K9 me2:DNA Meth. m5C
H4−4ac:H3−5ac
H3K27 me3:DNA Meth. m5C
H4K20 me2:H4−4ac
H4K20 me2:H3−5ac
H4K20 me2:H3K4 me3
H4K20 me3
H3K27 me2
H3K4 me1
H3K9 me2
H3K9 me3
H3K4 me3
DNA Meth. m5C
H2A.Z
H3K27 me3
H3−5ac
H4K20 me2
H4−4ac
0.00 0.25 0.50 0.75 1.00
π̂
DLGAP5 – Forward
H3K9 me3:H4K20 me3
H3K27 me3:H3K4 me1
H3K4 me3:H3−5ac
DNA Meth. m5C:H4K20 me3
DNA Meth. m5C:H3K9 me3
H3K27 me3:H4K20 me3
H3K27 me3:DNA Meth. m5C
H3K4 me1:H3−5ac
H3K4 me1:H4−4ac
H3K4 me1:H2A.Z
H3K4 me1:H4K20 me2
H2A.Z:H3K4 me3
H2A.Z:H4−4ac
H3K4 me3:H4−4ac
H2A.Z:H3−5ac
H4K20 me2:H4−4ac
H2A.Z:H4K20 me2
H4K20 me2:H3−5ac
H4K20 me2:H3K4 me3
H3K27 me2
H3K9 me2
H4K20 me3
H3K9 me3
H3K4 me1
DNA Meth. m5C
H3K4 me3
H3K27 me3
H4K20 me2
H3−5ac
H2A.Z
H4−4ac
0.00 0.25 0.50 0.75 1.00
π̂
DLGAP5 – Reverse
H3K9 me3:H4K20 me3
H3K27 me3:H3K4 me1
H3K4 me3:H3−5ac
DNA Meth. m5C:H4K20 me3
DNA Meth. m5C:H3K9 me3
H3K27 me3:H4K20 me3
H3K27 me3:DNA Meth. m5C
H3K4 me1:H3−5ac
H3K4 me1:H4−4ac
H3K4 me1:H2A.Z
H3K4 me1:H4K20 me2
H2A.Z:H3K4 me3
H2A.Z:H4−4ac
H3K4 me3:H4−4ac
H2A.Z:H3−5ac
H4K20 me2:H4−4ac
H2A.Z:H4K20 me2
H4K20 me2:H3−5ac
H4K20 me2:H3K4 me3
H3K27 me2
H3K9 me2
H4K20 me3
H3K9 me3
H3K4 me1
DNA Meth. m5C
H3K4 me3
H3K27 me3
H4K20 me2
H3−5ac
H2A.Z
H4−4ac
0.00 0.25 0.50 0.75 1.00
π̂

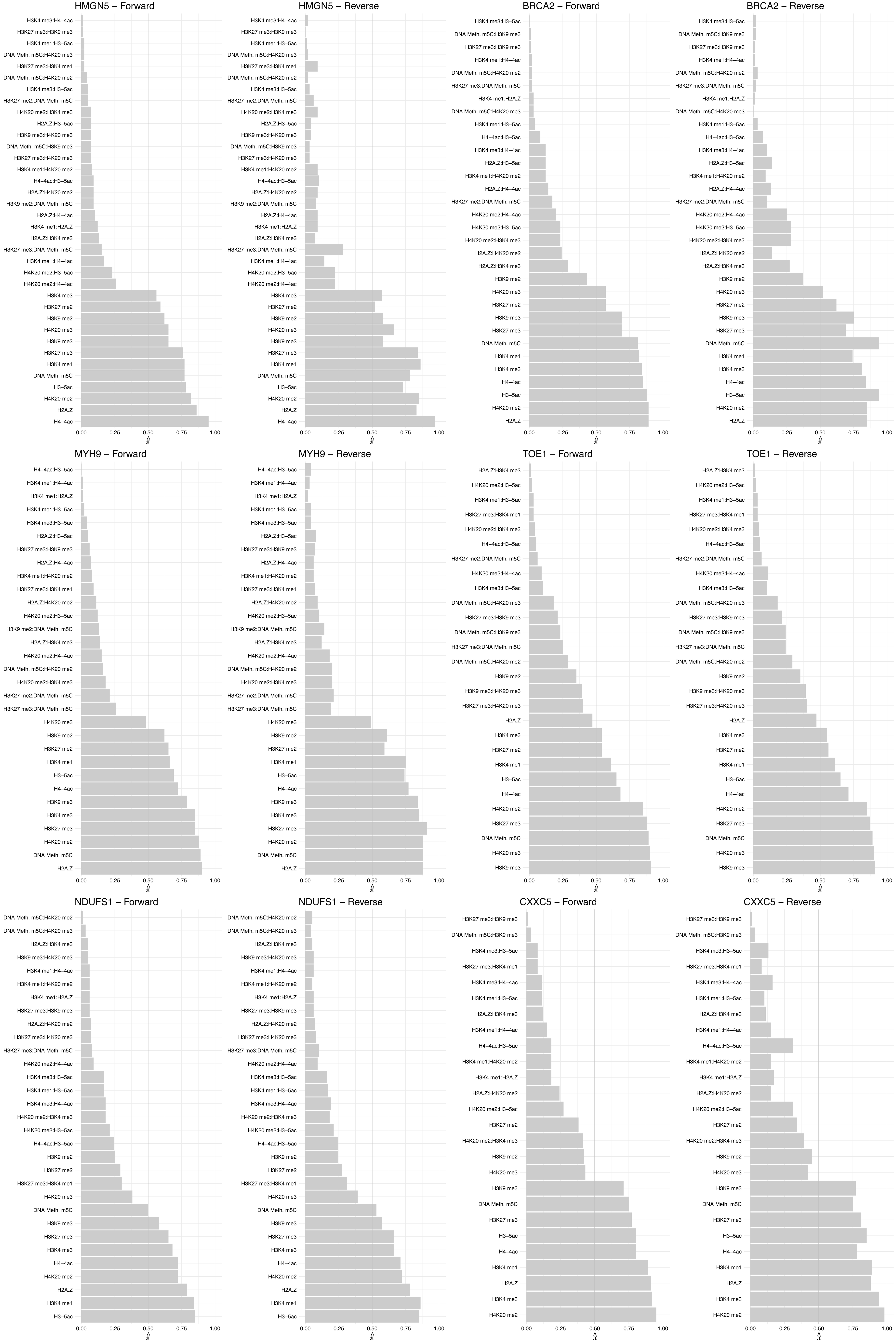
HMGN5 – Forward
H3K4 me3:H4–4ac
H3K27 me3:H3K9 me3
H3K4 me1:H3–5ac
DNA Meth. m5C:H4K20 me3
H3K27 me3:H3K4 me1
DNA Meth. m5C:H4K20 me2
H3K4 me3:H3–5ac
H3K27 me2:DNA Meth. m5C
H4K20 me2:H3K4 me3
H2A.Z:H3–5ac
H3K9 me3:H4K20 me3
DNA Meth. m5C:H3K9 me3
H3K27 me3:H4K20 me3
H3K4 me1:H4K20 me2
H4–4ac:H3–5ac
H2A.Z:H4K20 me2
H3K9 me2:DNA Meth. m5C
H2A.Z:H4–4ac
H3K4 me1:H2A.Z
H2A.Z:H3K4 me3
H3K27 me3:DNA Meth. m5C
H3K4 me1:H4–4ac
H4K20 me2:H3–5ac
H4K20 me2:H4–4ac
H3K4 me3
H3K27 me2
H3K9 me2
H4K20 me3
H3K9 me3
H3K27 me3
H3K4 me1
DNA Meth. m5C
H3–5ac
H4K20 me2
H2A.Z
H4–4ac
0.00 0.25 0.50 0.75 1.00
π̂
HMGN5 – Reverse
H3K4 me3:H4–4ac
H3K27 me3:H3K9 me3
H3K4 me1:H3–5ac
DNA Meth. m5C:H4K20 me3
H3K27 me3:H3K4 me1
DNA Meth. m5C:H4K20 me2
H3K4 me3:H3–5ac
H3K27 me2:DNA Meth. m5C
H4K20 me2:H3K4 me3
H2A.Z:H3–5ac
H3K9 me3:H4K20 me3
DNA Meth. m5C:H3K9 me3
H3K27 me3:H4K20 me3
H3K4 me1:H4K20 me2
H4–4ac:H3–5ac
H2A.Z:H4K20 me2
H3K9 me2:DNA Meth. m5C
H2A.Z:H4–4ac
H3K4 me1:H2A.Z
H2A.Z:H3K4 me3
H3K27 me3:DNA Meth. m5C
H3K4 me1:H4–4ac
H4K20 me2:H3–5ac
H4K20 me2:H4–4ac
H3K4 me3
H3K27 me2
H3K9 me2
H4K20 me3
H3K9 me3
H3K27 me3
H3K4 me1
DNA Meth. m5C
H3–5ac
H4K20 me2
H2A.Z
H4–4ac
0.00 0.25 0.50 0.75 1.00
π̂
BRCA2 – Forward
H3K4 me3:H3–5ac
DNA Meth. m5C:H3K9 me3
H3K27 me3:H3K9 me3
H3K4 me1:H4–4ac
DNA Meth. m5C:H4K20 me2
H3K27 me3:DNA Meth. m5C
H3K4 me1:H2A.Z
DNA Meth. m5C:H4K20 me3
H3K4 me1:H3–5ac
H4–4ac:H3–5ac
H3K4 me3:H4–4ac
H2A.Z:H3–5ac
H3K4 me1:H4K20 me2
H2A.Z:H4–4ac
H3K27 me2:DNA Meth. m5C
H4K20 me2:H4–4ac
H4K20 me2:H3–5ac
H4K20 me2:H3K4 me3
H2A.Z:H4K20 me2
H2A.Z:H3K4 me3
H3K9 me2
H4K20 me3
H3K27 me2
H3K9 me3
H3K27 me3
DNA Meth. m5C
H3K4 me1
H3K4 me3
H4–4ac
H3–5ac
H4K20 me2
H2A.Z
0.00 0.25 0.50 0.75 1.00
π̂
BRCA2 – Reverse
H3K4 me3:H3–5ac
DNA Meth. m5C:H3K9 me3
H3K27 me3:H3K9 me3
H3K4 me1:H4–4ac
DNA Meth. m5C:H4K20 me2
H3K27 me3:DNA Meth. m5C
H3K4 me1:H2A.Z
DNA Meth. m5C:H4K20 me3
H3K4 me1:H3–5ac
H4–4ac:H3–5ac
H3K4 me3:H4–4ac
H2A.Z:H3–5ac
H3K4 me1:H4K20 me2
H2A.Z:H4–4ac
H3K27 me2:DNA Meth. m5C
H4K20 me2:H4–4ac
H4K20 me2:H3–5ac
H4K20 me2:H3K4 me3
H2A.Z:H4K20 me2
H2A.Z:H3K4 me3
H3K9 me2
H4K20 me3
H3K27 me2
H3K9 me3
H3K27 me3
DNA Meth. m5C
H3K4 me1
H3K4 me3
H4–4ac
H3–5ac
H4K20 me2
H2A.Z
0.00 0.25 0.50 0.75 1.00
π̂
MYH9 – Forward
H4–4ac:H3–5ac
H3K4 me1:H4–4ac
H3K4 me1:H2A.Z
H3K4 me1:H3–5ac
H3K4 me3:H3–5ac
H2A.Z:H3–5ac
H3K27 me3:H3K9 me3
H2A.Z:H4–4ac
H3K4 me1:H4K20 me2
H3K27 me3:H3K4 me1
H2A.Z:H4K20 me2
H4K20 me2:H3–5ac
H3K9 me2:DNA Meth. m5C
H2A.Z:H3K4 me3
H4K20 me2:H4–4ac
DNA Meth. m5C:H4K20 me2
H4K20 me2:H3K4 me3
H3K27 me2:DNA Meth. m5C
H3K27 me3:DNA Meth. m5C
H4K20 me3
H3K9 me2
H3K27 me2
H3K4 me1
H3–5ac
H4–4ac
H3K9 me3
H3K4 me3
H3K27 me3
H4K20 me2
DNA Meth. m5C
H2A.Z
0.00 0.25 0.50 0.75 1.00
π̂
MYH9 – Reverse
H4–4ac:H3–5ac
H3K4 me1:H4–4ac
H3K4 me1:H2A.Z
H3K4 me1:H3–5ac
H3K4 me3:H3–5ac
H2A.Z:H3–5ac
H3K27 me3:H3K9 me3
H2A.Z:H4–4ac
H3K4 me1:H4K20 me2
H3K27 me3:H3K4 me1
H2A.Z:H4K20 me2
H4K20 me2:H3–5ac
H3K9 me2:DNA Meth. m5C
H2A.Z:H3K4 me3
H4K20 me2:H4–4ac
DNA Meth. m5C:H4K20 me2
H4K20 me2:H3K4 me3
H3K27 me2:DNA Meth. m5C
H3K27 me3:DNA Meth. m5C
H4K20 me3
H3K9 me2
H3K27 me2
H3K4 me1
H3–5ac
H4–4ac
H3K9 me3
H3K4 me3
H3K27 me3
H4K20 me2
DNA Meth. m5C
H2A.Z
0.00 0.25 0.50 0.75 1.00
π̂
TOE1 – Forward
H2A.Z:H3K4 me3
H4K20 me2:H3–5ac
H3K4 me1:H3–5ac
H3K27 me3:H3K4 me1
H4K20 me2:H3K4 me3
H4–4ac:H3–5ac
H3K27 me2:DNA Meth. m5C
H4K20 me2:H4–4ac
H3K4 me3:H3–5ac
DNA Meth. m5C:H4K20 me3
H3K27 me3:H3K9 me3
DNA Meth. m5C:H3K9 me3
H3K27 me3:DNA Meth. m5C
DNA Meth. m5C:H4K20 me2
H3K9 me2
H3K9 me3:H4K20 me3
H3K27 me3:H4K20 me3
H2A.Z
H3K4 me3
H3K27 me2
H3K4 me1
H3–5ac
H4–4ac
H4K20 me2
H3K27 me3
DNA Meth. m5C
H4K20 me3
H3K9 me3
0.00 0.25 0.50 0.75 1.00
π̂
TOE1 – Reverse
H2A.Z:H3K4 me3
H4K20 me2:H3–5ac
H3K4 me1:H3–5ac
H3K27 me3:H3K4 me1
H4K20 me2:H3K4 me3
H4–4ac:H3–5ac
H3K27 me2:DNA Meth. m5C
H4K20 me2:H4–4ac
H3K4 me3:H3–5ac
DNA Meth. m5C:H4K20 me3
H3K27 me3:H3K9 me3
DNA Meth. m5C:H3K9 me3
H3K27 me3:DNA Meth. m5C
DNA Meth. m5C:H4K20 me2
H3K9 me2
H3K9 me3:H4K20 me3
H3K27 me3:H4K20 me3
H2A.Z
H3K4 me3
H3K27 me2
H3K4 me1
H3–5ac
H4–4ac
H4K20 me2
H3K27 me3
DNA Meth. m5C
H4K20 me3
H3K9 me3
0.00 0.25 0.50 0.75 1.00
π̂
NDUFS1 – Forward
DNA Meth. m5C:H4K20 me2
DNA Meth. m5C:H4K20 me3
H2A.Z:H3K4 me3
H3K9 me3:H4K20 me3
H3K4 me1:H4–4ac
H3K4 me1:H4K20 me2
H3K4 me1:H2A.Z
H3K27 me3:H3K9 me3
H2A.Z:H4K20 me2
H3K27 me3:H4K20 me3
H3K27 me3:DNA Meth. m5C
H4K20 me2:H4–4ac
H3K4 me3:H3–5ac
H3K4 me1:H3–5ac
H3K4 me3:H4–4ac
H4K20 me2:H3K4 me3
H4K20 me2:H3–5ac
H4–4ac:H3–5ac
H3K9 me2
H3K27 me2
H3K27 me3:H3K4 me1
H4K20 me3
DNA Meth. m5C
H3K9 me3
H3K27 me3
H3K4 me3
H4–4ac
H4K20 me2
H2A.Z
H3K4 me1
H3–5ac
0.00 0.25 0.50 0.75 1.00
π̂
NDUFS1 – Reverse
DNA Meth. m5C:H4K20 me2
DNA Meth. m5C:H4K20 me3
H2A.Z:H3K4 me3
H3K9 me3:H4K20 me3
H3K4 me1:H4–4ac
H3K4 me1:H4K20 me2
H3K4 me1:H2A.Z
H3K27 me3:H3K9 me3
H2A.Z:H4K20 me2
H3K27 me3:H4K20 me3
H3K27 me3:DNA Meth. m5C
H4K20 me2:H4–4ac
H3K4 me3:H3–5ac
H3K4 me1:H3–5ac
H3K4 me3:H4–4ac
H4K20 me2:H3K4 me3
H4K20 me2:H3–5ac
H4–4ac:H3–5ac
H3K9 me2
H3K27 me2
H3K27 me3:H3K4 me1
H4K20 me3
DNA Meth. m5C
H3K9 me3
H3K27 me3
H3K4 me3
H4–4ac
H4K20 me2
H2A.Z
H3K4 me1
H3–5ac
0.00 0.25 0.50 0.75 1.00
π̂
CXXC5 – Forward
H3K27 me3:H3K9 me3
DNA Meth. m5C:H3K9 me3
H3K4 me3:H3–5ac
H3K27 me3:H3K4 me1
H3K4 me3:H4–4ac
H3K4 me1:H3–5ac
H2A.Z:H3K4 me3
H3K4 me1:H4–4ac
H4–4ac:H3–5ac
H3K4 me1:H4K20 me2
H3K4 me1:H2A.Z
H2A.Z:H4K20 me2
H4K20 me2:H3–5ac
H3K27 me2
H4K20 me2:H3K4 me3
H3K9 me2
H4K20 me3
H3K9 me3
DNA Meth. m5C
H3K27 me3
H3–5ac
H4–4ac
H3K4 me1
H2A.Z
H3K4 me3
H4K20 me2
0.00 0.25 0.50 0.75 1.00
π̂
CXXC5 – Reverse
H3K27 me3:H3K9 me3
DNA Meth. m5C:H3K9 me3
H3K4 me3:H3–5ac
H3K27 me3:H3K4 me1
H3K4 me3:H4–4ac
H3K4 me1:H3–5ac
H2A.Z:H3K4 me3
H3K4 me1:H4–4ac
H4–4ac:H3–5ac
H3K4 me1:H4K20 me2
H3K4 me1:H2A.Z
H2A.Z:H4K20 me2
H4K20 me2:H3–5ac
H3K27 me2
H4K20 me2:H3K4 me3
H3K9 me2
H4K20 me3
H3K9 me3
DNA Meth. m5C
H3K27 me3
H3–5ac
H4–4ac
H3K4 me1
H2A.Z
H3K4 me3
H4K20 me2
0.00 0.25 0.50 0.75 1.00
π̂

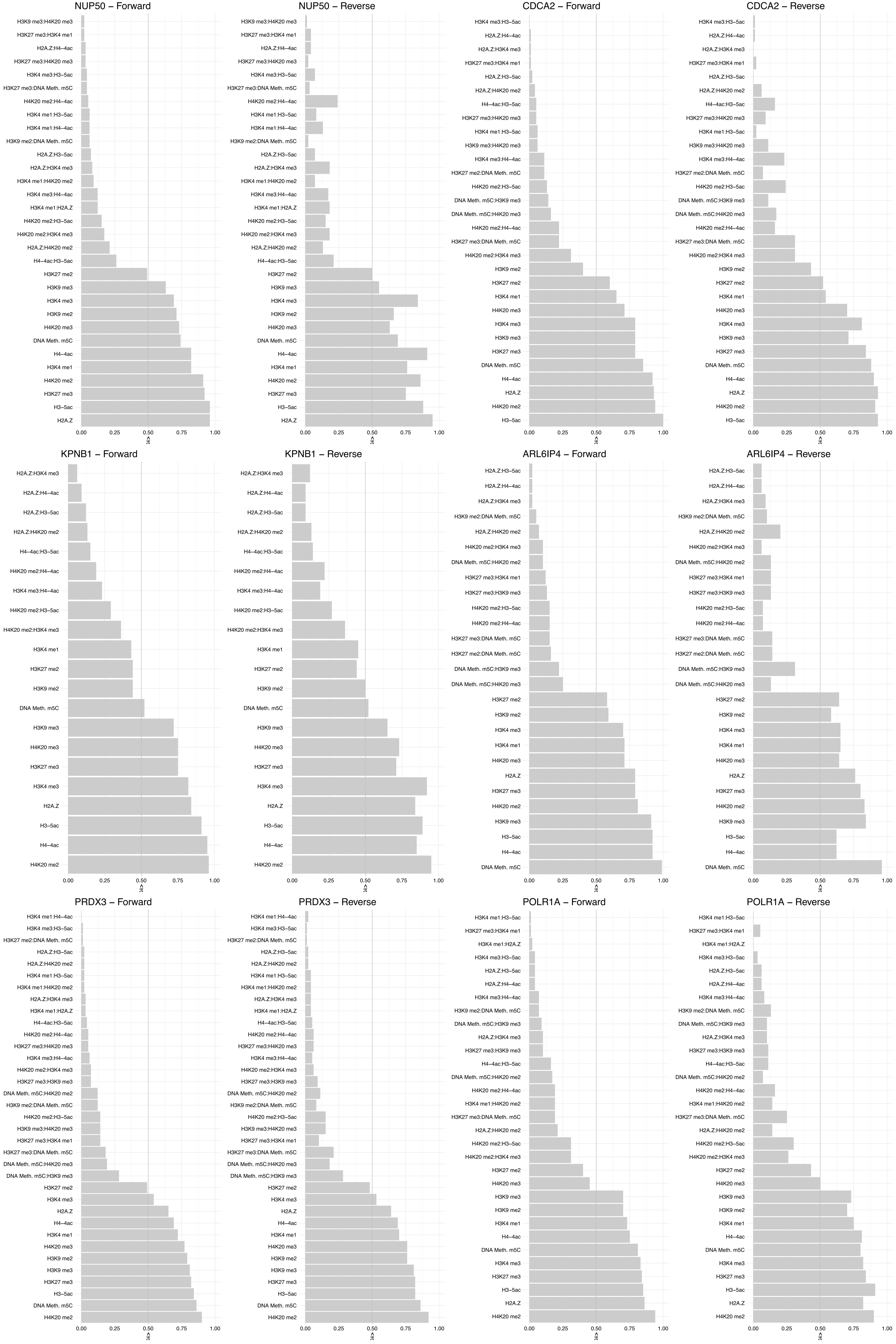
NUP50 – Forward
H3K9 me3:H4K20 me3
H3K27 me3:H3K4 me1
H2A.Z:H4−4ac
H3K27 me3:H4K20 me3
H3K4 me3:H3−5ac
H3K27 me3:DNA Meth. m5C
H4K20 me2:H4−4ac
H3K4 me1:H3−5ac
H3K4 me1:H4−4ac
H3K9 me2:DNA Meth. m5C
H2A.Z:H3−5ac
H2A.Z:H3K4 me3
H3K4 me1:H4K20 me2
H3K4 me3:H4−4ac
H3K4 me1:H2A.Z
H4K20 me2:H3−5ac
H4K20 me2:H3K4 me3
H2A.Z:H4K20 me2
H4−4ac:H3−5ac
H3K27 me2
H3K9 me3
H3K4 me3
H3K9 me2
H4K20 me3
DNA Meth. m5C
H4−4ac
H3K4 me1
H4K20 me2
H3K27 me3
H3−5ac
H2A.Z
0.00 0.25 0.50 0.75 1.00
π̂
NUP50 – Reverse
H3K9 me3:H4K20 me3
H3K27 me3:H3K4 me1
H2A.Z:H4−4ac
H3K27 me3:H4K20 me3
H3K4 me3:H3−5ac
H3K27 me3:DNA Meth. m5C
H4K20 me2:H4−4ac
H3K4 me1:H3−5ac
H3K4 me1:H4−4ac
H3K9 me2:DNA Meth. m5C
H2A.Z:H3−5ac
H2A.Z:H3K4 me3
H3K4 me1:H4K20 me2
H3K4 me3:H4−4ac
H3K4 me1:H2A.Z
H4K20 me2:H3−5ac
H4K20 me2:H3K4 me3
H2A.Z:H4K20 me2
H4−4ac:H3−5ac
H3K27 me2
H3K9 me3
H3K4 me3
H3K9 me2
H4K20 me3
DNA Meth. m5C
H4−4ac
H3K4 me1
H4K20 me2
H3K27 me3
H3−5ac
H2A.Z
0.00 0.25 0.50 0.75 1.00
π̂
CDCA2 – Forward
H3K4 me3:H3−5ac
H2A.Z:H4−4ac
H2A.Z:H3K4 me3
H3K27 me3:H3K4 me1
H2A.Z:H3−5ac
H2A.Z:H4K20 me2
H4−4ac:H3−5ac
H3K27 me3:H4K20 me3
H3K4 me1:H3−5ac
H3K9 me3:H4K20 me3
H3K4 me3:H4−4ac
H3K27 me2:DNA Meth. m5C
H4K20 me2:H3−5ac
DNA Meth. m5C:H3K9 me3
DNA Meth. m5C:H4K20 me3
H4K20 me2:H4−4ac
H3K27 me3:DNA Meth. m5C
H4K20 me2:H3K4 me3
H3K9 me2
H3K27 me2
H3K4 me1
H4K20 me3
H3K4 me3
H3K9 me3
H3K27 me3
DNA Meth. m5C
H4−4ac
H2A.Z
H4K20 me2
H3−5ac
0.00 0.25 0.50 0.75 1.00
π̂
CDCA2 – Reverse
H3K4 me3:H3−5ac
H2A.Z:H4−4ac
H2A.Z:H3K4 me3
H3K27 me3:H3K4 me1
H2A.Z:H3−5ac
H2A.Z:H4K20 me2
H4−4ac:H3−5ac
H3K27 me3:H4K20 me3
H3K4 me1:H3−5ac
H3K9 me3:H4K20 me3
H3K4 me3:H4−4ac
H3K27 me2:DNA Meth. m5C
H4K20 me2:H3−5ac
DNA Meth. m5C:H3K9 me3
DNA Meth. m5C:H4K20 me3
H4K20 me2:H4−4ac
H3K27 me3:DNA Meth. m5C
H4K20 me2:H3K4 me3
H3K9 me2
H3K27 me2
H3K4 me1
H4K20 me3
H3K4 me3
H3K9 me3
H3K27 me3
DNA Meth. m5C
H4−4ac
H2A.Z
H4K20 me2
H3−5ac
0.00 0.25 0.50 0.75 1.00
π̂
KPNB1 – Forward
H2A.Z:H3K4 me3
H2A.Z:H4−4ac
H2A.Z:H3−5ac
H2A.Z:H4K20 me2
H4−4ac:H3−5ac
H4K20 me2:H4−4ac
H3K4 me3:H4−4ac
H4K20 me2:H3−5ac
H4K20 me2:H3K4 me3
H3K4 me1
H3K27 me2
H3K9 me2
DNA Meth. m5C
H3K9 me3
H4K20 me3
H3K27 me3
H3K4 me3
H2A.Z
H3−5ac
H4−4ac
H4K20 me2
0.00 0.25 0.50 0.75 1.00
π̂
KPNB1 – Reverse
H2A.Z:H3K4 me3
H2A.Z:H4−4ac
H2A.Z:H3−5ac
H2A.Z:H4K20 me2
H4−4ac:H3−5ac
H4K20 me2:H4−4ac
H3K4 me3:H4−4ac
H4K20 me2:H3−5ac
H4K20 me2:H3K4 me3
H3K4 me1
H3K27 me2
H3K9 me2
DNA Meth. m5C
H3K9 me3
H4K20 me3
H3K27 me3
H3K4 me3
H2A.Z
H3−5ac
H4−4ac
H4K20 me2
0.00 0.25 0.50 0.75 1.00
π̂
ARL6IP4 – Forward
H2A.Z:H3−5ac
H2A.Z:H4−4ac
H2A.Z:H3K4 me3
H3K9 me2:DNA Meth. m5C
H2A.Z:H4K20 me2
H4K20 me2:H3K4 me3
DNA Meth. m5C:H4K20 me2
H3K27 me3:H3K4 me1
H3K27 me3:H3K9 me3
H4K20 me2:H3−5ac
H4K20 me2:H4−4ac
H3K27 me3:DNA Meth. m5C
H3K27 me2:DNA Meth. m5C
DNA Meth. m5C:H3K9 me3
DNA Meth. m5C:H4K20 me3
H3K27 me2
H3K9 me2
H3K4 me3
H3K4 me1
H4K20 me3
H2A.Z
H3K27 me3
H4K20 me2
H3K9 me3
H3−5ac
H4−4ac
DNA Meth. m5C
0.00 0.25 0.50 0.75 1.00
π̂
ARL6IP4 – Reverse
H2A.Z:H3−5ac
H2A.Z:H4−4ac
H2A.Z:H3K4 me3
H3K9 me2:DNA Meth. m5C
H2A.Z:H4K20 me2
H4K20 me2:H3K4 me3
DNA Meth. m5C:H4K20 me2
H3K27 me3:H3K4 me1
H3K27 me3:H3K9 me3
H4K20 me2:H3−5ac
H4K20 me2:H4−4ac
H3K27 me3:DNA Meth. m5C
H3K27 me2:DNA Meth. m5C
DNA Meth. m5C:H3K9 me3
DNA Meth. m5C:H4K20 me3
H3K27 me2
H3K9 me2
H3K4 me3
H3K4 me1
H4K20 me3
H2A.Z
H3K27 me3
H4K20 me2
H3K9 me3
H3−5ac
H4−4ac
DNA Meth. m5C
0.00 0.25 0.50 0.75 1.00
π̂
PRDX3 – Forward
H3K4 me1:H4−4ac
H3K4 me3:H3−5ac
H3K27 me2:DNA Meth. m5C
H2A.Z:H3−5ac
H2A.Z:H4K20 me2
H3K4 me1:H3−5ac
H3K4 me1:H4K20 me2
H2A.Z:H3K4 me3
H3K4 me1:H2A.Z
H4−4ac:H3−5ac
H4K20 me2:H4−4ac
H3K27 me3:H4K20 me3
H3K4 me3:H4−4ac
H4K20 me2:H3K4 me3
H3K27 me3:H3K9 me3
DNA Meth. m5C:H4K20 me2
H3K9 me2:DNA Meth. m5C
H4K20 me2:H3−5ac
H3K9 me3:H4K20 me3
H3K27 me3:H3K4 me1
H3K27 me3:DNA Meth. m5C
DNA Meth. m5C:H4K20 me3
DNA Meth. m5C:H3K9 me3
H3K27 me2
H3K4 me3
H2A.Z
H4−4ac
H3K4 me1
H4K20 me3
H3K9 me2
H3K9 me3
H3K27 me3
H3−5ac
DNA Meth. m5C
H4K20 me2
0.00 0.25 0.50 0.75 1.00
π̂
PRDX3 – Reverse
H3K4 me1:H4−4ac
H3K4 me3:H3−5ac
H3K27 me2:DNA Meth. m5C
H2A.Z:H3−5ac
H2A.Z:H4K20 me2
H3K4 me1:H3−5ac
H3K4 me1:H4K20 me2
H2A.Z:H3K4 me3
H3K4 me1:H2A.Z
H4−4ac:H3−5ac
H4K20 me2:H4−4ac
H3K27 me3:H4K20 me3
H3K4 me3:H4−4ac
H4K20 me2:H3K4 me3
H3K27 me3:H3K9 me3
DNA Meth. m5C:H4K20 me2
H3K9 me2:DNA Meth. m5C
H4K20 me2:H3−5ac
H3K9 me3:H4K20 me3
H3K27 me3:H3K4 me1
H3K27 me3:DNA Meth. m5C
DNA Meth. m5C:H4K20 me3
DNA Meth. m5C:H3K9 me3
H3K27 me2
H3K4 me3
H2A.Z
H4−4ac
H3K4 me1
H4K20 me3
H3K9 me2
H3K9 me3
H3K27 me3
H3−5ac
DNA Meth. m5C
H4K20 me2
0.00 0.25 0.50 0.75 1.00
π̂
POLR1A – Forward
H3K4 me1:H3−5ac
H3K27 me3:H3K4 me1
H3K4 me1:H2A.Z
H3K4 me3:H3−5ac
H2A.Z:H3−5ac
H2A.Z:H4−4ac
H3K4 me3:H4−4ac
H3K9 me2:DNA Meth. m5C
DNA Meth. m5C:H3K9 me3
H2A.Z:H3K4 me3
H3K27 me3:H3K9 me3
H4−4ac:H3−5ac
DNA Meth. m5C:H4K20 me2
H4K20 me2:H4−4ac
H3K4 me1:H4K20 me2
H3K27 me3:DNA Meth. m5C
H2A.Z:H4K20 me2
H4K20 me2:H3−5ac
H4K20 me2:H3K4 me3
H3K27 me2
H4K20 me3
H3K9 me3
H3K9 me2
H3K4 me1
H4−4ac
DNA Meth. m5C
H3K4 me3
H3K27 me3
H3−5ac
H2A.Z
H4K20 me2
0.00 0.25 0.50 0.75 1.00
π̂
POLR1A – Reverse
H3K4 me1:H3−5ac
H3K27 me3:H3K4 me1
H3K4 me1:H2A.Z
H3K4 me3:H3−5ac
H2A.Z:H3−5ac
H2A.Z:H4−4ac
H3K4 me3:H4−4ac
H3K9 me2:DNA Meth. m5C
DNA Meth. m5C:H3K9 me3
H2A.Z:H3K4 me3
H3K27 me3:H3K9 me3
H4−4ac:H3−5ac
DNA Meth. m5C:H4K20 me2
H4K20 me2:H4−4ac
H3K4 me1:H4K20 me2
H3K27 me3:DNA Meth. m5C
H2A.Z:H4K20 me2
H4K20 me2:H3−5ac
H4K20 me2:H3K4 me3
H3K27 me2
H4K20 me3
H3K9 me3
H3K9 me2
H3K4 me1
H4−4ac
DNA Meth. m5C
H3K4 me3
H3K27 me3
H3−5ac
H2A.Z
H4K20 me2
0.00 0.25 0.50 0.75 1.00
π̂

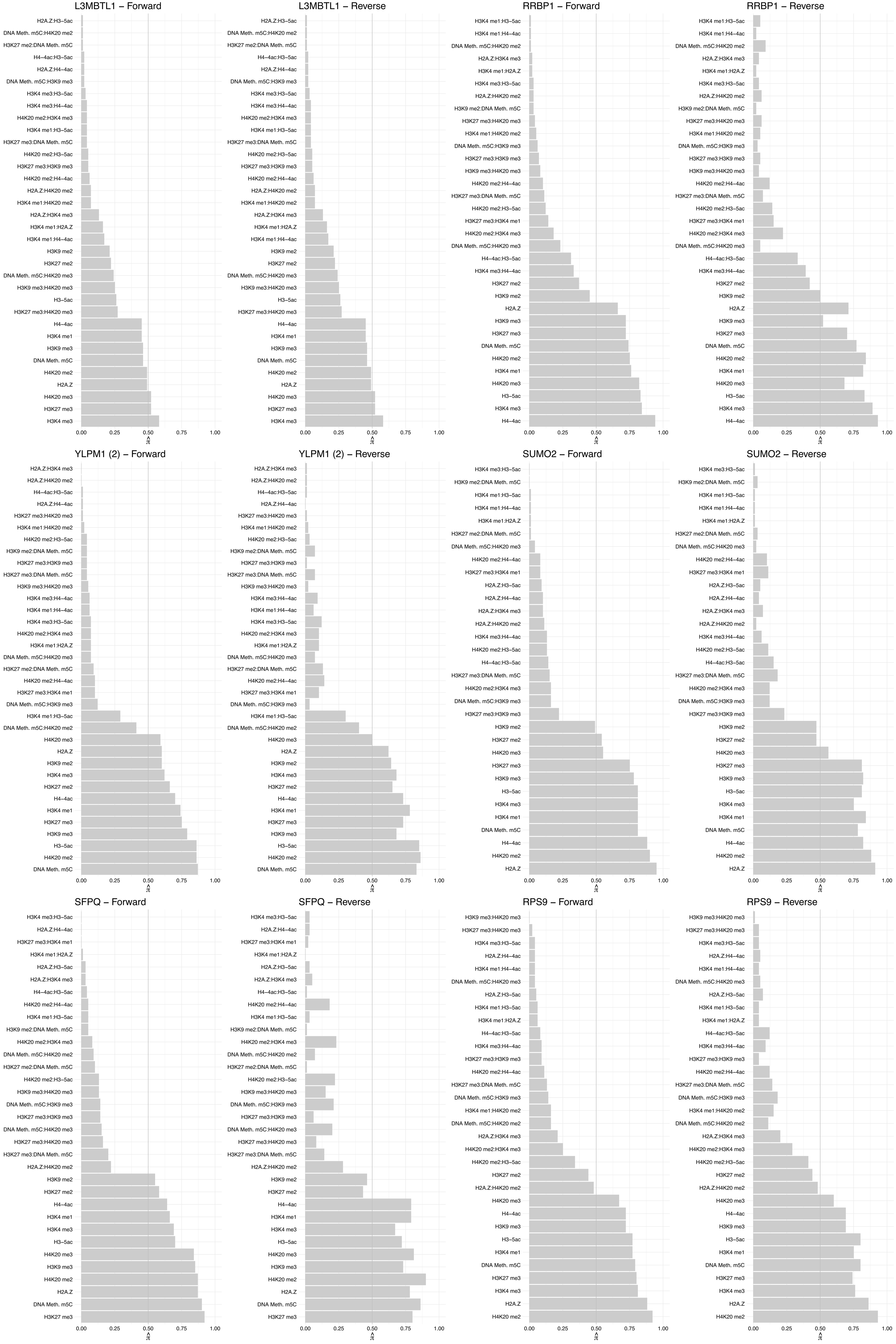
L3MBTL1 − Forward
H2A.Z:H3−5ac
DNA Meth. m5C:H4K20 me2
H3K27 me2:DNA Meth. m5C
H4−4ac:H3−5ac
H2A.Z:H4−4ac
DNA Meth. m5C:H3K9 me3
H3K4 me3:H3−5ac
H3K4 me3:H4−4ac
H4K20 me2:H3K4 me3
H3K4 me1:H3−5ac
H3K27 me3:DNA Meth. m5C
H4K20 me2:H3−5ac
H3K27 me3:H3K9 me3
H4K20 me2:H4−4ac
H2A.Z:H4K20 me2
H3K4 me1:H4K20 me2
H2A.Z:H3K4 me3
H3K4 me1:H2A.Z
H3K4 me1:H4−4ac
H3K9 me2
H3K27 me2
DNA Meth. m5C:H4K20 me3
H3K9 me3:H4K20 me3
H3−5ac
H3K27 me3:H4K20 me3
H4−4ac
H3K4 me1
H3K9 me3
DNA Meth. m5C
H4K20 me2
H2A.Z
H4K20 me3
H3K27 me3
H3K4 me3
0.00 0.25 0.50 0.75 1.00
$\hat{\pi}$
L3MBTL1 − Reverse
H2A.Z:H3−5ac
DNA Meth. m5C:H4K20 me2
H3K27 me2:DNA Meth. m5C
H4−4ac:H3−5ac
H2A.Z:H4−4ac
DNA Meth. m5C:H3K9 me3
H3K4 me3:H3−5ac
H3K4 me3:H4−4ac
H4K20 me2:H3K4 me3
H3K4 me1:H3−5ac
H3K27 me3:DNA Meth. m5C
H4K20 me2:H3−5ac
H3K27 me3:H3K9 me3
H4K20 me2:H4−4ac
H2A.Z:H4K20 me2
H3K4 me1:H4K20 me2
H2A.Z:H3K4 me3
H3K4 me1:H2A.Z
H3K4 me1:H4−4ac
H3K9 me2
H3K27 me2
DNA Meth. m5C:H4K20 me3
H3K9 me3:H4K20 me3
H3−5ac
H3K27 me3:H4K20 me3
H4−4ac
H3K4 me1
H3K9 me3
DNA Meth. m5C
H4K20 me2
H2A.Z
H4K20 me3
H3K27 me3
H3K4 me3
0.00 0.25 0.50 0.75 1.00
$\hat{\pi}$
RRBP1 − Forward
H3K4 me1:H3−5ac
H3K4 me1:H4−4ac
DNA Meth. m5C:H4K20 me2
H2A.Z:H3K4 me3
H3K4 me1:H2A.Z
H3K4 me3:H3−5ac
H2A.Z:H4K20 me2
H3K9 me2:DNA Meth. m5C
H3K27 me3:H4K20 me3
H3K4 me1:H4K20 me2
DNA Meth. m5C:H3K9 me3
H3K27 me3:H3K9 me3
H3K9 me3:H4K20 me3
H4K20 me2:H4−4ac
H3K27 me3:DNA Meth. m5C
H4K20 me2:H3−5ac
H3K27 me3:H3K4 me1
H4K20 me2:H3K4 me3
DNA Meth. m5C:H4K20 me3
H4−4ac:H3−5ac
H3K4 me3:H4−4ac
H3K27 me2
H3K9 me2
H2A.Z
H3K9 me3
H3K27 me3
DNA Meth. m5C
H4K20 me2
H3K4 me1
H4K20 me3
H3−5ac
H3K4 me3
H4−4ac
0.00 0.25 0.50 0.75 1.00
$\hat{\pi}$
RRBP1 − Reverse
H3K4 me1:H3−5ac
H3K4 me1:H4−4ac
DNA Meth. m5C:H4K20 me2
H2A.Z:H3K4 me3
H3K4 me1:H2A.Z
H3K4 me3:H3−5ac
H2A.Z:H4K20 me2
H3K9 me2:DNA Meth. m5C
H3K27 me3:H4K20 me3
H3K4 me1:H4K20 me2
DNA Meth. m5C:H3K9 me3
H3K27 me3:H3K9 me3
H3K9 me3:H4K20 me3
H4K20 me2:H4−4ac
H3K27 me3:DNA Meth. m5C
H4K20 me2:H3−5ac
H3K27 me3:H3K4 me1
H4K20 me2:H3K4 me3
DNA Meth. m5C:H4K20 me3
H4−4ac:H3−5ac
H3K4 me3:H4−4ac
H3K27 me2
H3K9 me2
H2A.Z
H3K9 me3
H3K27 me3
DNA Meth. m5C
H4K20 me2
H3K4 me1
H4K20 me3
H3−5ac
H3K4 me3
H4−4ac
0.00 0.25 0.50 0.75 1.00
$\hat{\pi}$
YLPM1 (2) − Forward
H2A.Z:H3K4 me3
H2A.Z:H4K20 me2
H4−4ac:H3−5ac
H2A.Z:H4−4ac
H3K27 me3:H4K20 me3
H3K4 me1:H4K20 me2
H4K20 me2:H3−5ac
H3K9 me2:DNA Meth. m5C
H3K27 me3:H3K9 me3
H3K27 me3:DNA Meth. m5C
H3K9 me3:H4K20 me3
H3K4 me3:H4−4ac
H3K4 me1:H4−4ac
H3K4 me3:H3−5ac
H4K20 me2:H3K4 me3
H3K4 me1:H2A.Z
DNA Meth. m5C:H4K20 me3
H3K27 me2:DNA Meth. m5C
H4K20 me2:H4−4ac
H3K27 me3:H3K4 me1
DNA Meth. m5C:H3K9 me3
H3K4 me1:H3−5ac
DNA Meth. m5C:H4K20 me2
H4K20 me3
H2A.Z
H3K9 me2
H3K4 me3
H3K27 me2
H4−4ac
H3K4 me1
H3K27 me3
H3K9 me3
H3−5ac
H4K20 me2
DNA Meth. m5C
0.00 0.25 0.50 0.75 1.00
$\hat{\pi}$
YLPM1 (2) − Reverse
H2A.Z:H3K4 me3
H2A.Z:H4K20 me2
H4−4ac:H3−5ac
H2A.Z:H4−4ac
H3K27 me3:H4K20 me3
H3K4 me1:H4K20 me2
H4K20 me2:H3−5ac
H3K9 me2:DNA Meth. m5C
H3K27 me3:H3K9 me3
H3K27 me3:DNA Meth. m5C
H3K9 me3:H4K20 me3
H3K4 me3:H4−4ac
H3K4 me1:H4−4ac
H3K4 me3:H3−5ac
H4K20 me2:H3K4 me3
H3K4 me1:H2A.Z
DNA Meth. m5C:H4K20 me3
H3K27 me2:DNA Meth. m5C
H4K20 me2:H4−4ac
H3K27 me3:H3K4 me1
DNA Meth. m5C:H3K9 me3
H3K4 me1:H3−5ac
DNA Meth. m5C:H4K20 me2
H4K20 me3
H2A.Z
H3K9 me2
H3K4 me3
H3K27 me2
H4−4ac
H3K4 me1
H3K27 me3
H3K9 me3
H3−5ac
H4K20 me2
DNA Meth. m5C
0.00 0.25 0.50 0.75 1.00
$\hat{\pi}$
SUMO2 − Forward
H3K4 me3:H3−5ac
H3K9 me2:DNA Meth. m5C
H3K4 me1:H3−5ac
H3K4 me1:H4−4ac
H3K4 me1:H2A.Z
H3K27 me2:DNA Meth. m5C
DNA Meth. m5C:H4K20 me3
H4K20 me2:H4−4ac
H3K27 me3:H3K4 me1
H2A.Z:H3−5ac
H2A.Z:H4−4ac
H2A.Z:H3K4 me3
H2A.Z:H4K20 me2
H3K4 me3:H4−4ac
H4K20 me2:H3−5ac
H4−4ac:H3−5ac
H3K27 me3:DNA Meth. m5C
H4K20 me2:H3K4 me3
DNA Meth. m5C:H3K9 me3
H3K27 me3:H3K9 me3
H3K9 me2
H3K27 me2
H4K20 me3
H3K27 me3
H3K9 me3
H3−5ac
H3K4 me3
H3K4 me1
DNA Meth. m5C
H4−4ac
H4K20 me2
H2A.Z
0.00 0.25 0.50 0.75 1.00
$\hat{\pi}$
SUMO2 − Reverse
H3K4 me3:H3−5ac
H3K9 me2:DNA Meth. m5C
H3K4 me1:H3−5ac
H3K4 me1:H4−4ac
H3K4 me1:H2A.Z
H3K27 me2:DNA Meth. m5C
DNA Meth. m5C:H4K20 me3
H4K20 me2:H4−4ac
H3K27 me3:H3K4 me1
H2A.Z:H3−5ac
H2A.Z:H4−4ac
H2A.Z:H3K4 me3
H2A.Z:H4K20 me2
H3K4 me3:H4−4ac
H4K20 me2:H3−5ac
H4−4ac:H3−5ac
H3K27 me3:DNA Meth. m5C
H4K20 me2:H3K4 me3
DNA Meth. m5C:H3K9 me3
H3K27 me3:H3K9 me3
H3K9 me2
H3K27 me2
H4K20 me3
H3K27 me3
H3K9 me3
H3−5ac
H3K4 me3
H3K4 me1
DNA Meth. m5C
H4−4ac
H4K20 me2
H2A.Z
0.00 0.25 0.50 0.75 1.00
$\hat{\pi}$
SFPQ − Forward
H3K4 me3:H3−5ac
H2A.Z:H4−4ac
H3K27 me3:H3K4 me1
H3K4 me1:H2A.Z
H2A.Z:H3−5ac
H2A.Z:H3K4 me3
H4−4ac:H3−5ac
H4K20 me2:H4−4ac
H3K4 me1:H3−5ac
H3K9 me2:DNA Meth. m5C
H4K20 me2:H3K4 me3
DNA Meth. m5C:H4K20 me2
H3K27 me2:DNA Meth. m5C
H4K20 me2:H3−5ac
H3K9 me3:H4K20 me3
DNA Meth. m5C:H3K9 me3
H3K27 me3:H3K9 me3
DNA Meth. m5C:H4K20 me3
H3K27 me3:H4K20 me3
H3K27 me3:DNA Meth. m5C
H2A.Z:H4K20 me2
H3K9 me2
H3K27 me2
H4−4ac
H3K4 me1
H3K4 me3
H3−5ac
H4K20 me3
H3K9 me3
H4K20 me2
H2A.Z
DNA Meth. m5C
H3K27 me3
0.00 0.25 0.50 0.75 1.00
$\hat{\pi}$
SFPQ − Reverse
H3K4 me3:H3−5ac
H2A.Z:H4−4ac
H3K27 me3:H3K4 me1
H3K4 me1:H2A.Z
H2A.Z:H3−5ac
H2A.Z:H3K4 me3
H4−4ac:H3−5ac
H4K20 me2:H4−4ac
H3K4 me1:H3−5ac
H3K9 me2:DNA Meth. m5C
H4K20 me2:H3K4 me3
DNA Meth. m5C:H4K20 me2
H3K27 me2:DNA Meth. m5C
H4K20 me2:H3−5ac
H3K9 me3:H4K20 me3
DNA Meth. m5C:H3K9 me3
H3K27 me3:H3K9 me3
DNA Meth. m5C:H4K20 me3
H3K27 me3:H4K20 me3
H3K27 me3:DNA Meth. m5C
H2A.Z:H4K20 me2
H3K9 me2
H3K27 me2
H4−4ac
H3K4 me1
H3K4 me3
H3−5ac
H4K20 me3
H3K9 me3
H4K20 me2
H2A.Z
DNA Meth. m5C
H3K27 me3
0.00 0.25 0.50 0.75 1.00
$\hat{\pi}$
RPS9 − Forward
H3K9 me3:H4K20 me3
H3K27 me3:H4K20 me3
H3K4 me3:H3−5ac
H2A.Z:H4−4ac
H3K4 me1:H4−4ac
DNA Meth. m5C:H4K20 me3
H2A.Z:H3−5ac
H3K4 me1:H3−5ac
H3K4 me1:H2A.Z
H4−4ac:H3−5ac
H3K4 me3:H4−4ac
H3K27 me3:H3K9 me3
H4K20 me2:H4−4ac
H3K27 me3:DNA Meth. m5C
DNA Meth. m5C:H3K9 me3
H3K4 me1:H4K20 me2
DNA Meth. m5C:H4K20 me2
H2A.Z:H3K4 me3
H4K20 me2:H3K4 me3
H4K20 me2:H3−5ac
H3K27 me2
H2A.Z:H4K20 me2
H4K20 me3
H4−4ac
H3K9 me3
H3−5ac
H3K4 me1
DNA Meth. m5C
H3K27 me3
H3K4 me3
H2A.Z
H4K20 me2
0.00 0.25 0.50 0.75 1.00
$\hat{\pi}$
RPS9 − Reverse
H3K9 me3:H4K20 me3
H3K27 me3:H4K20 me3
H3K4 me3:H3−5ac
H2A.Z:H4−4ac
H3K4 me1:H4−4ac
DNA Meth. m5C:H4K20 me3
H2A.Z:H3−5ac
H3K4 me1:H3−5ac
H3K4 me1:H2A.Z
H4−4ac:H3−5ac
H3K4 me3:H4−4ac
H3K27 me3:H3K9 me3
H4K20 me2:H4−4ac
H3K27 me3:DNA Meth. m5C
DNA Meth. m5C:H3K9 me3
H3K4 me1:H4K20 me2
DNA Meth. m5C:H4K20 me2
H2A.Z:H3K4 me3
H4K20 me2:H3K4 me3
H4K20 me2:H3−5ac
H3K27 me2
H2A.Z:H4K20 me2
H4K20 me3
H4−4ac
H3K9 me3
H3−5ac
H3K4 me1
DNA Meth. m5C
H3K27 me3
H3K4 me3
H2A.Z
H4K20 me2
0.00 0.25 0.50 0.75 1.00
$\hat{\pi}$

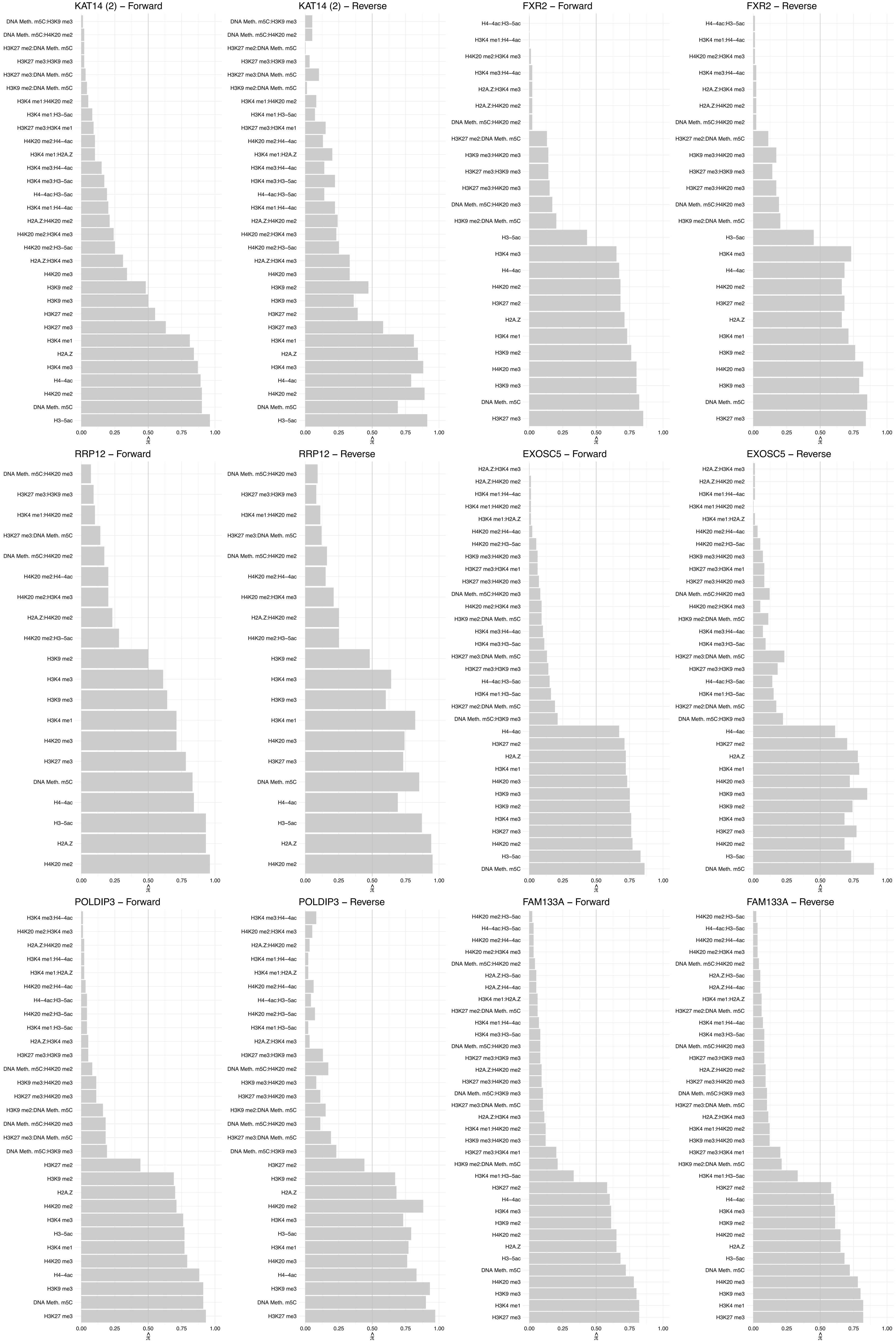
KAT14 (2) – Forward
KAT14 (2) – Reverse
DNA Meth. m5C:H3K9 me3
DNA Meth. m5C:H4K20 me2
H3K27 me2:DNA Meth. m5C
H3K27 me3:H3K9 me3
H3K27 me3:DNA Meth. m5C
H3K9 me2:DNA Meth. m5C
H3K4 me1:H4K20 me2
H3K4 me1:H3−5ac
H3K27 me3:H3K4 me1
H4K20 me2:H4−4ac
H3K4 me1:H2A.Z
H3K4 me3:H4−4ac
H3K4 me3:H3−5ac
H4−4ac:H3−5ac
H3K4 me1:H4−4ac
H2A.Z:H4K20 me2
H4K20 me2:H3K4 me3
H4K20 me2:H3−5ac
H2A.Z:H3K4 me3
H4K20 me3
H3K9 me2
H3K9 me3
H3K27 me2
H3K27 me3
H3K4 me1
H2A.Z
H3K4 me3
H4−4ac
H4K20 me2
DNA Meth. m5C
H3−5ac
FXR2 – Forward
FXR2 – Reverse
H4−4ac:H3−5ac
H3K4 me1:H4−4ac
H4K20 me2:H3K4 me3
H3K4 me3:H4−4ac
H2A.Z:H3K4 me3
H2A.Z:H4K20 me2
DNA Meth. m5C:H4K20 me2
H3K27 me2:DNA Meth. m5C
H3K9 me3:H4K20 me3
H3K27 me3:H3K9 me3
H3K27 me3:H4K20 me3
DNA Meth. m5C:H4K20 me3
H3K9 me2:DNA Meth. m5C
H3−5ac
H3K4 me3
H4−4ac
H4K20 me2
H3K27 me2
H2A.Z
H3K4 me1
H3K9 me2
H4K20 me3
H3K9 me3
DNA Meth. m5C
H3K27 me3
RRP12 – Forward
RRP12 – Reverse
DNA Meth. m5C:H4K20 me3
H3K27 me3:H3K9 me3
H3K4 me1:H4K20 me2
H3K27 me3:DNA Meth. m5C
DNA Meth. m5C:H4K20 me2
H4K20 me2:H4−4ac
H4K20 me2:H3K4 me3
H2A.Z:H4K20 me2
H4K20 me2:H3−5ac
H3K9 me2
H3K4 me3
H3K9 me3
H3K4 me1
H4K20 me3
H3K27 me3
DNA Meth. m5C
H4−4ac
H3−5ac
H2A.Z
H4K20 me2
EXOSC5 – Forward
EXOSC5 – Reverse
H2A.Z:H3K4 me3
H2A.Z:H4K20 me2
H3K4 me1:H4−4ac
H3K4 me1:H4K20 me2
H3K4 me1:H2A.Z
H4K20 me2:H4−4ac
H4K20 me2:H3−5ac
H3K9 me3:H4K20 me3
H3K27 me3:H3K4 me1
H3K27 me3:H4K20 me3
DNA Meth. m5C:H4K20 me3
H4K20 me2:H3K4 me3
H3K9 me2:DNA Meth. m5C
H3K4 me3:H4−4ac
H3K4 me3:H3−5ac
H3K27 me3:DNA Meth. m5C
H3K27 me3:H3K9 me3
H4−4ac:H3−5ac
H3K4 me1:H3−5ac
H3K27 me2:DNA Meth. m5C
DNA Meth. m5C:H3K9 me3
H4−4ac
H3K27 me2
H2A.Z
H3K4 me1
H4K20 me3
H3K9 me3
H3K9 me2
H3K4 me3
H3K27 me3
H4K20 me2
H3−5ac
DNA Meth. m5C
POLDIP3 – Forward
POLDIP3 – Reverse
H3K4 me3:H4−4ac
H4K20 me2:H3K4 me3
H2A.Z:H4K20 me2
H3K4 me1:H4−4ac
H3K4 me1:H2A.Z
H4K20 me2:H4−4ac
H4−4ac:H3−5ac
H4K20 me2:H3−5ac
H3K4 me1:H3−5ac
H2A.Z:H3K4 me3
H3K27 me3:H3K9 me3
DNA Meth. m5C:H4K20 me2
H3K9 me3:H4K20 me3
H3K27 me3:H4K20 me3
H3K9 me2:DNA Meth. m5C
DNA Meth. m5C:H4K20 me3
H3K27 me3:DNA Meth. m5C
DNA Meth. m5C:H3K9 me3
H3K27 me2
H3K9 me2
H2A.Z
H4K20 me2
H3K4 me3
H3−5ac
H3K4 me1
H4K20 me3
H4−4ac
H3K9 me3
DNA Meth. m5C
H3K27 me3
FAM133A – Forward
FAM133A – Reverse
H4K20 me2:H3−5ac
H4−4ac:H3−5ac
H4K20 me2:H4−4ac
H4K20 me2:H3K4 me3
DNA Meth. m5C:H4K20 me2
H2A.Z:H3−5ac
H2A.Z:H4−4ac
H3K4 me1:H2A.Z
H3K27 me2:DNA Meth. m5C
H3K4 me1:H4−4ac
H3K4 me3:H3−5ac
DNA Meth. m5C:H4K20 me3
H3K27 me3:H3K9 me3
H2A.Z:H4K20 me2
H3K27 me3:H4K20 me3
DNA Meth. m5C:H3K9 me3
H3K27 me3:DNA Meth. m5C
H2A.Z:H3K4 me3
H3K4 me1:H4K20 me2
H3K9 me3:H4K20 me3
H3K27 me3:H3K4 me1
H3K9 me2:DNA Meth. m5C
H3K4 me1:H3−5ac
H3K27 me2
H4−4ac
H3K4 me3
H3K9 me2
H4K20 me2
H2A.Z
H3−5ac
DNA Meth. m5C
H4K20 me3
H3K9 me3
H3K4 me1
H3K27 me3
0.00
0.25
0.50
0.75
1.00
π̂

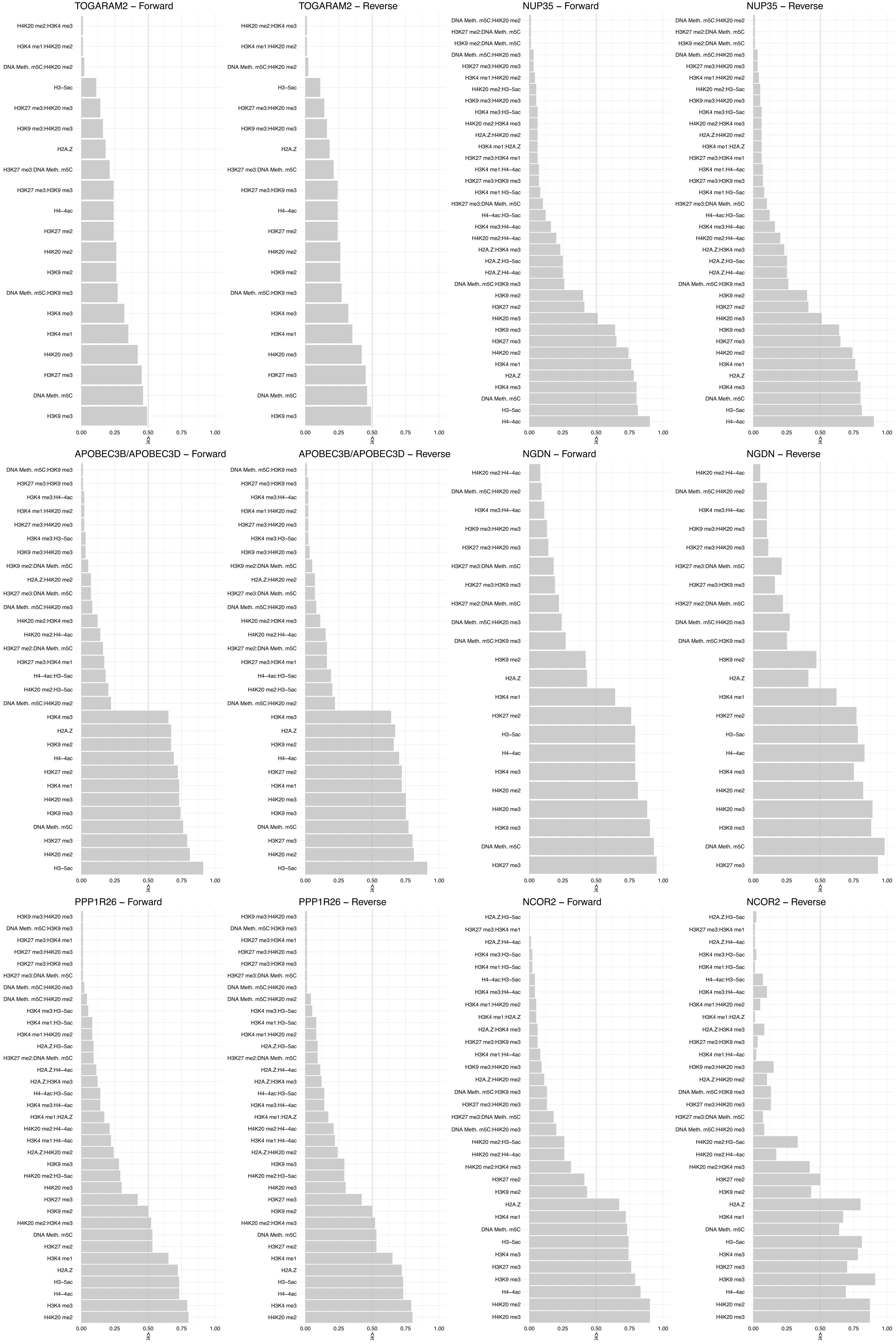
TOGARAM2 – Forward
H4K20 me2:H3K4 me3
H3K4 me1:H4K20 me2
DNA Meth. m5C:H4K20 me2
H3–5ac
H3K27 me3:H4K20 me3
H3K9 me3:H4K20 me3
H2A.Z
H3K27 me3:DNA Meth. m5C
H3K27 me3:H3K9 me3
H4–4ac
H3K27 me2
H4K20 me2
H3K9 me2
DNA Meth. m5C:H3K9 me3
H3K4 me3
H3K4 me1
H4K20 me3
H3K27 me3
DNA Meth. m5C
H3K9 me3
0.00 0.25 0.50 0.75 1.00
π̂
TOGARAM2 – Reverse
H4K20 me2:H3K4 me3
H3K4 me1:H4K20 me2
DNA Meth. m5C:H4K20 me2
H3–5ac
H3K27 me3:H4K20 me3
H3K9 me3:H4K20 me3
H2A.Z
H3K27 me3:DNA Meth. m5C
H3K27 me3:H3K9 me3
H4–4ac
H3K27 me2
H4K20 me2
H3K9 me2
DNA Meth. m5C:H3K9 me3
H3K4 me3
H3K4 me1
H4K20 me3
H3K27 me3
DNA Meth. m5C
H3K9 me3
0.00 0.25 0.50 0.75 1.00
π̂
NUP35 – Forward
DNA Meth. m5C:H4K20 me2
H3K27 me2:DNA Meth. m5C
H3K9 me2:DNA Meth. m5C
DNA Meth. m5C:H4K20 me3
H3K27 me3:H4K20 me3
H3K4 me1:H4K20 me2
H4K20 me2:H3–5ac
H3K9 me3:H4K20 me3
H3K4 me3:H3–5ac
H4K20 me2:H3K4 me3
H2A.Z:H4K20 me2
H3K4 me1:H2A.Z
H3K27 me3:H3K4 me1
H3K4 me1:H4–4ac
H3K27 me3:H3K9 me3
H3K4 me1:H3–5ac
H3K27 me3:DNA Meth. m5C
H4–4ac:H3–5ac
H3K4 me3:H4–4ac
H4K20 me2:H4–4ac
H2A.Z:H3K4 me3
H2A.Z:H3–5ac
H2A.Z:H4–4ac
DNA Meth. m5C:H3K9 me3
H3K9 me2
H3K27 me2
H4K20 me3
H3K9 me3
H3K27 me3
H4K20 me2
H3K4 me1
H2A.Z
H3K4 me3
DNA Meth. m5C
H3–5ac
H4–4ac
0.00 0.25 0.50 0.75 1.00
π̂
NUP35 – Reverse
DNA Meth. m5C:H4K20 me2
H3K27 me2:DNA Meth. m5C
H3K9 me2:DNA Meth. m5C
DNA Meth. m5C:H4K20 me3
H3K27 me3:H4K20 me3
H3K4 me1:H4K20 me2
H4K20 me2:H3–5ac
H3K9 me3:H4K20 me3
H3K4 me3:H3–5ac
H4K20 me2:H3K4 me3
H2A.Z:H4K20 me2
H3K4 me1:H2A.Z
H3K27 me3:H3K4 me1
H3K4 me1:H4–4ac
H3K27 me3:H3K9 me3
H3K4 me1:H3–5ac
H3K27 me3:DNA Meth. m5C
H4–4ac:H3–5ac
H3K4 me3:H4–4ac
H4K20 me2:H4–4ac
H2A.Z:H3K4 me3
H2A.Z:H3–5ac
H2A.Z:H4–4ac
DNA Meth. m5C:H3K9 me3
H3K9 me2
H3K27 me2
H4K20 me3
H3K9 me3
H3K27 me3
H4K20 me2
H3K4 me1
H2A.Z
H3K4 me3
DNA Meth. m5C
H3–5ac
H4–4ac
0.00 0.25 0.50 0.75 1.00
π̂
APOBEC3B/APOBEC3D – Forward
DNA Meth. m5C:H3K9 me3
H3K27 me3:H3K9 me3
H3K4 me3:H4–4ac
H3K4 me1:H4K20 me2
H3K27 me3:H4K20 me3
H3K4 me3:H3–5ac
H3K9 me3:H4K20 me3
H3K9 me2:DNA Meth. m5C
H2A.Z:H4K20 me2
H3K27 me3:DNA Meth. m5C
DNA Meth. m5C:H4K20 me3
H4K20 me2:H3K4 me3
H4K20 me2:H4–4ac
H3K27 me2:DNA Meth. m5C
H3K27 me3:H3K4 me1
H4–4ac:H3–5ac
H4K20 me2:H3–5ac
DNA Meth. m5C:H4K20 me2
H3K4 me3
H2A.Z
H3K9 me2
H4–4ac
H3K27 me2
H3K4 me1
H4K20 me3
H3K9 me3
DNA Meth. m5C
H3K27 me3
H4K20 me2
H3–5ac
0.00 0.25 0.50 0.75 1.00
π̂
APOBEC3B/APOBEC3D – Reverse
DNA Meth. m5C:H3K9 me3
H3K27 me3:H3K9 me3
H3K4 me3:H4–4ac
H3K4 me1:H4K20 me2
H3K27 me3:H4K20 me3
H3K4 me3:H3–5ac
H3K9 me3:H4K20 me3
H3K9 me2:DNA Meth. m5C
H2A.Z:H4K20 me2
H3K27 me3:DNA Meth. m5C
DNA Meth. m5C:H4K20 me3
H4K20 me2:H3K4 me3
H4K20 me2:H4–4ac
H3K27 me2:DNA Meth. m5C
H3K27 me3:H3K4 me1
H4–4ac:H3–5ac
H4K20 me2:H3–5ac
DNA Meth. m5C:H4K20 me2
H3K4 me3
H2A.Z
H3K9 me2
H4–4ac
H3K27 me2
H3K4 me1
H4K20 me3
H3K9 me3
DNA Meth. m5C
H3K27 me3
H4K20 me2
H3–5ac
0.00 0.25 0.50 0.75 1.00
π̂
NGDN – Forward
H4K20 me2:H4–4ac
DNA Meth. m5C:H4K20 me2
H3K4 me3:H4–4ac
H3K9 me3:H4K20 me3
H3K27 me3:H4K20 me3
H3K27 me3:DNA Meth. m5C
H3K27 me3:H3K9 me3
H3K27 me2:DNA Meth. m5C
DNA Meth. m5C:H4K20 me3
DNA Meth. m5C:H3K9 me3
H3K9 me2
H2A.Z
H3K4 me1
H3K27 me2
H3–5ac
H4–4ac
H3K4 me3
H4K20 me2
H4K20 me3
H3K9 me3
DNA Meth. m5C
H3K27 me3
0.00 0.25 0.50 0.75 1.00
π̂
NGDN – Reverse
H4K20 me2:H4–4ac
DNA Meth. m5C:H4K20 me2
H3K4 me3:H4–4ac
H3K9 me3:H4K20 me3
H3K27 me3:H4K20 me3
H3K27 me3:DNA Meth. m5C
H3K27 me3:H3K9 me3
H3K27 me2:DNA Meth. m5C
DNA Meth. m5C:H4K20 me3
DNA Meth. m5C:H3K9 me3
H3K9 me2
H2A.Z
H3K4 me1
H3K27 me2
H3–5ac
H4–4ac
H3K4 me3
H4K20 me2
H4K20 me3
H3K9 me3
DNA Meth. m5C
H3K27 me3
0.00 0.25 0.50 0.75 1.00
π̂
PPP1R26 – Forward
H3K9 me3:H4K20 me3
DNA Meth. m5C:H3K9 me3
H3K27 me3:H3K4 me1
H3K27 me3:H4K20 me3
H3K27 me3:H3K9 me3
H3K27 me3:DNA Meth. m5C
DNA Meth. m5C:H4K20 me3
DNA Meth. m5C:H4K20 me2
H3K4 me3:H3–5ac
H3K4 me1:H3–5ac
H3K4 me1:H4K20 me2
H2A.Z:H3–5ac
H3K27 me2:DNA Meth. m5C
H2A.Z:H4–4ac
H2A.Z:H3K4 me3
H4–4ac:H3–5ac
H3K4 me3:H4–4ac
H3K4 me1:H2A.Z
H4K20 me2:H4–4ac
H3K4 me1:H4–4ac
H2A.Z:H4K20 me2
H3K9 me3
H4K20 me2:H3–5ac
H4K20 me3
H3K27 me3
H3K9 me2
H4K20 me2:H3K4 me3
DNA Meth. m5C
H3K27 me2
H3K4 me1
H2A.Z
H3–5ac
H4–4ac
H3K4 me3
H4K20 me2
0.00 0.25 0.50 0.75 1.00
π̂
PPP1R26 – Reverse
H3K9 me3:H4K20 me3
DNA Meth. m5C:H3K9 me3
H3K27 me3:H3K4 me1
H3K27 me3:H4K20 me3
H3K27 me3:H3K9 me3
H3K27 me3:DNA Meth. m5C
DNA Meth. m5C:H4K20 me3
DNA Meth. m5C:H4K20 me2
H3K4 me3:H3–5ac
H3K4 me1:H3–5ac
H3K4 me1:H4K20 me2
H2A.Z:H3–5ac
H3K27 me2:DNA Meth. m5C
H2A.Z:H4–4ac
H2A.Z:H3K4 me3
H4–4ac:H3–5ac
H3K4 me3:H4–4ac
H3K4 me1:H2A.Z
H4K20 me2:H4–4ac
H3K4 me1:H4–4ac
H2A.Z:H4K20 me2
H3K9 me3
H4K20 me2:H3–5ac
H4K20 me3
H3K27 me3
H3K9 me2
H4K20 me2:H3K4 me3
DNA Meth. m5C
H3K27 me2
H3K4 me1
H2A.Z
H3–5ac
H4–4ac
H3K4 me3
H4K20 me2
0.00 0.25 0.50 0.75 1.00
π̂
NCOR2 – Forward
H2A.Z:H3–5ac
H3K27 me3:H3K4 me1
H2A.Z:H4–4ac
H3K4 me3:H3–5ac
H3K4 me1:H3–5ac
H4–4ac:H3–5ac
H3K4 me3:H4–4ac
H3K4 me1:H4K20 me2
H3K4 me1:H2A.Z
H2A.Z:H3K4 me3
H3K27 me3:H3K9 me3
H3K4 me1:H4–4ac
H3K9 me3:H4K20 me3
H2A.Z:H4K20 me2
DNA Meth. m5C:H3K9 me3
H3K27 me3:H4K20 me3
H3K27 me3:DNA Meth. m5C
DNA Meth. m5C:H4K20 me3
H4K20 me2:H3–5ac
H4K20 me2:H4–4ac
H4K20 me2:H3K4 me3
H3K27 me2
H3K9 me2
H2A.Z
H3K4 me1
DNA Meth. m5C
H3–5ac
H3K4 me3
H3K27 me3
H3K9 me3
H4–4ac
H4K20 me2
H4K20 me3
0.00 0.25 0.50 0.75 1.00
π̂
NCOR2 – Reverse
H2A.Z:H3–5ac
H3K27 me3:H3K4 me1
H2A.Z:H4–4ac
H3K4 me3:H3–5ac
H3K4 me1:H3–5ac
H4–4ac:H3–5ac
H3K4 me3:H4–4ac
H3K4 me1:H4K20 me2
H3K4 me1:H2A.Z
H2A.Z:H3K4 me3
H3K27 me3:H3K9 me3
H3K4 me1:H4–4ac
H3K9 me3:H4K20 me3
H2A.Z:H4K20 me2
DNA Meth. m5C:H3K9 me3
H3K27 me3:H4K20 me3
H3K27 me3:DNA Meth. m5C
DNA Meth. m5C:H4K20 me3
H4K20 me2:H3–5ac
H4K20 me2:H4–4ac
H4K20 me2:H3K4 me3
H3K27 me2
H3K9 me2
H2A.Z
H3K4 me1
DNA Meth. m5C
H3–5ac
H3K4 me3
H3K27 me3
H3K9 me3
H4–4ac
H4K20 me2
H4K20 me3
0.00 0.25 0.50 0.75 1.00
π̂

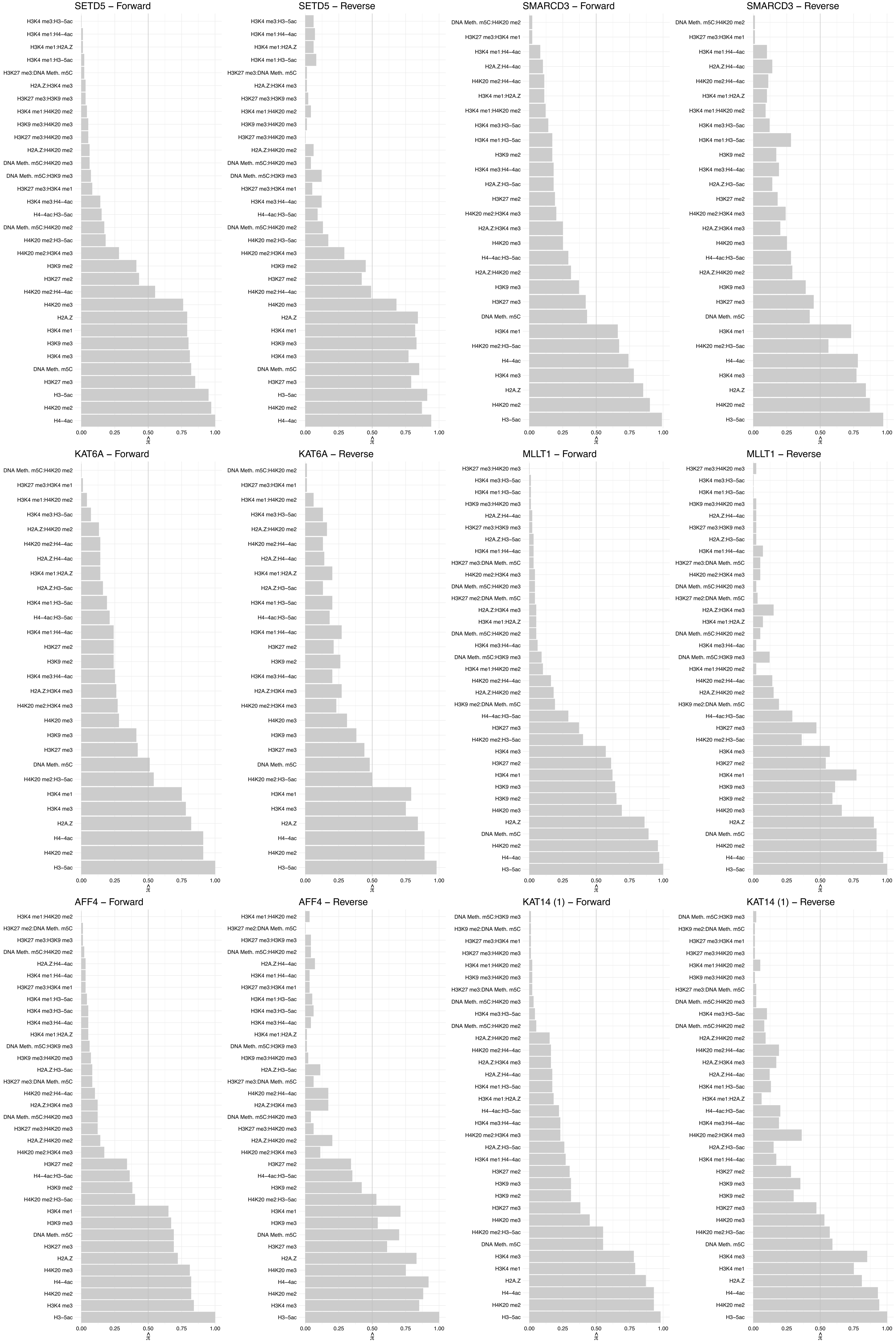
SETD5 – Forward
H3K4 me3:H3−5ac
H3K4 me1:H4−4ac
H3K4 me1:H2A.Z
H3K4 me1:H3−5ac
H3K27 me3:DNA Meth. m5C
H2A.Z:H3K4 me3
H3K27 me3:H3K9 me3
H3K4 me1:H4K20 me2
H3K9 me3:H4K20 me3
H3K27 me3:H4K20 me3
H2A.Z:H4K20 me2
DNA Meth. m5C:H4K20 me3
DNA Meth. m5C:H3K9 me3
H3K27 me3:H3K4 me1
H3K4 me3:H4−4ac
H4−4ac:H3−5ac
DNA Meth. m5C:H4K20 me2
H4K20 me2:H3−5ac
H4K20 me2:H3K4 me3
H3K9 me2
H3K27 me2
H4K20 me2:H4−4ac
H4K20 me3
H2A.Z
H3K4 me1
H3K9 me3
H3K4 me3
DNA Meth. m5C
H3K27 me3
H3−5ac
H4K20 me2
H4−4ac
0.00
0.25
0.50
0.75
1.00
π̂
SETD5 – Reverse
H3K4 me3:H3−5ac
H3K4 me1:H4−4ac
H3K4 me1:H2A.Z
H3K4 me1:H3−5ac
H3K27 me3:DNA Meth. m5C
H2A.Z:H3K4 me3
H3K27 me3:H3K9 me3
H3K4 me1:H4K20 me2
H3K9 me3:H4K20 me3
H3K27 me3:H4K20 me3
H2A.Z:H4K20 me2
DNA Meth. m5C:H4K20 me3
DNA Meth. m5C:H3K9 me3
H3K27 me3:H3K4 me1
H3K4 me3:H4−4ac
H4−4ac:H3−5ac
DNA Meth. m5C:H4K20 me2
H4K20 me2:H3−5ac
H4K20 me2:H3K4 me3
H3K9 me2
H3K27 me2
H4K20 me2:H4−4ac
H4K20 me3
H2A.Z
H3K4 me1
H3K9 me3
H3K4 me3
DNA Meth. m5C
H3K27 me3
H3−5ac
H4K20 me2
H4−4ac
0.00
0.25
0.50
0.75
1.00
π̂
SMARCD3 – Forward
DNA Meth. m5C:H4K20 me2
H3K27 me3:H3K4 me1
H3K4 me1:H4−4ac
H2A.Z:H4−4ac
H4K20 me2:H4−4ac
H3K4 me1:H2A.Z
H3K4 me1:H4K20 me2
H3K4 me3:H3−5ac
H3K4 me1:H3−5ac
H3K9 me2
H3K4 me3:H4−4ac
H2A.Z:H3−5ac
H3K27 me2
H4K20 me2:H3K4 me3
H2A.Z:H3K4 me3
H4K20 me3
H4−4ac:H3−5ac
H2A.Z:H4K20 me2
H3K9 me3
H3K27 me3
DNA Meth. m5C
H3K4 me1
H4K20 me2:H3−5ac
H4−4ac
H3K4 me3
H2A.Z
H4K20 me2
H3−5ac
0.00
0.25
0.50
0.75
1.00
π̂
SMARCD3 – Reverse
DNA Meth. m5C:H4K20 me2
H3K27 me3:H3K4 me1
H3K4 me1:H4−4ac
H2A.Z:H4−4ac
H4K20 me2:H4−4ac
H3K4 me1:H2A.Z
H3K4 me1:H4K20 me2
H3K4 me3:H3−5ac
H3K4 me1:H3−5ac
H3K9 me2
H3K4 me3:H4−4ac
H2A.Z:H3−5ac
H3K27 me2
H4K20 me2:H3K4 me3
H2A.Z:H3K4 me3
H4K20 me3
H4−4ac:H3−5ac
H2A.Z:H4K20 me2
H3K9 me3
H3K27 me3
DNA Meth. m5C
H3K4 me1
H4K20 me2:H3−5ac
H4−4ac
H3K4 me3
H2A.Z
H4K20 me2
H3−5ac
0.00
0.25
0.50
0.75
1.00
π̂
KAT6A – Forward
DNA Meth. m5C:H4K20 me2
H3K27 me3:H3K4 me1
H3K4 me1:H4K20 me2
H3K4 me3:H3−5ac
H2A.Z:H4K20 me2
H4K20 me2:H4−4ac
H2A.Z:H4−4ac
H3K4 me1:H2A.Z
H2A.Z:H3−5ac
H3K4 me1:H3−5ac
H4−4ac:H3−5ac
H3K4 me1:H4−4ac
H3K27 me2
H3K9 me2
H3K4 me3:H4−4ac
H2A.Z:H3K4 me3
H4K20 me2:H3K4 me3
H4K20 me3
H3K9 me3
H3K27 me3
DNA Meth. m5C
H4K20 me2:H3−5ac
H3K4 me1
H3K4 me3
H2A.Z
H4−4ac
H4K20 me2
H3−5ac
0.00
0.25
0.50
0.75
1.00
π̂
KAT6A – Reverse
DNA Meth. m5C:H4K20 me2
H3K27 me3:H3K4 me1
H3K4 me1:H4K20 me2
H3K4 me3:H3−5ac
H2A.Z:H4K20 me2
H4K20 me2:H4−4ac
H2A.Z:H4−4ac
H3K4 me1:H2A.Z
H2A.Z:H3−5ac
H3K4 me1:H3−5ac
H4−4ac:H3−5ac
H3K4 me1:H4−4ac
H3K27 me2
H3K9 me2
H3K4 me3:H4−4ac
H2A.Z:H3K4 me3
H4K20 me2:H3K4 me3
H4K20 me3
H3K9 me3
H3K27 me3
DNA Meth. m5C
H4K20 me2:H3−5ac
H3K4 me1
H3K4 me3
H2A.Z
H4−4ac
H4K20 me2
H3−5ac
0.00
0.25
0.50
0.75
1.00
π̂
MLLT1 – Forward
H3K27 me3:H4K20 me3
H3K4 me3:H3−5ac
H3K4 me1:H3−5ac
H3K9 me3:H4K20 me3
H2A.Z:H4−4ac
H3K27 me3:H3K9 me3
H2A.Z:H3−5ac
H3K4 me1:H4−4ac
H3K27 me3:DNA Meth. m5C
H4K20 me2:H3K4 me3
DNA Meth. m5C:H4K20 me3
H3K27 me2:DNA Meth. m5C
H2A.Z:H3K4 me3
H3K4 me1:H2A.Z
DNA Meth. m5C:H4K20 me2
H3K4 me3:H4−4ac
DNA Meth. m5C:H3K9 me3
H3K4 me1:H4K20 me2
H4K20 me2:H4−4ac
H2A.Z:H4K20 me2
H3K9 me2:DNA Meth. m5C
H4−4ac:H3−5ac
H3K27 me3
H4K20 me2:H3−5ac
H3K4 me3
H3K27 me2
H3K4 me1
H3K9 me3
H3K9 me2
H4K20 me3
H2A.Z
DNA Meth. m5C
H4K20 me2
H4−4ac
H3−5ac
0.00
0.25
0.50
0.75
1.00
π̂
MLLT1 – Reverse
H3K27 me3:H4K20 me3
H3K4 me3:H3−5ac
H3K4 me1:H3−5ac
H3K9 me3:H4K20 me3
H2A.Z:H4−4ac
H3K27 me3:H3K9 me3
H2A.Z:H3−5ac
H3K4 me1:H4−4ac
H3K27 me3:DNA Meth. m5C
H4K20 me2:H3K4 me3
DNA Meth. m5C:H4K20 me3
H3K27 me2:DNA Meth. m5C
H2A.Z:H3K4 me3
H3K4 me1:H2A.Z
DNA Meth. m5C:H4K20 me2
H3K4 me3:H4−4ac
DNA Meth. m5C:H3K9 me3
H3K4 me1:H4K20 me2
H4K20 me2:H4−4ac
H2A.Z:H4K20 me2
H3K9 me2:DNA Meth. m5C
H4−4ac:H3−5ac
H3K27 me3
H4K20 me2:H3−5ac
H3K4 me3
H3K27 me2
H3K4 me1
H3K9 me3
H3K9 me2
H4K20 me3
H2A.Z
DNA Meth. m5C
H4K20 me2
H4−4ac
H3−5ac
0.00
0.25
0.50
0.75
1.00
π̂
AFF4 – Forward
H3K4 me1:H4K20 me2
H3K27 me2:DNA Meth. m5C
H3K27 me3:H3K9 me3
DNA Meth. m5C:H4K20 me2
H2A.Z:H4−4ac
H3K4 me1:H4−4ac
H3K27 me3:H3K4 me1
H3K4 me1:H3−5ac
H3K4 me3:H3−5ac
H3K4 me3:H4−4ac
H3K4 me1:H2A.Z
DNA Meth. m5C:H3K9 me3
H3K9 me3:H4K20 me3
H2A.Z:H3−5ac
H3K27 me3:DNA Meth. m5C
H4K20 me2:H4−4ac
H2A.Z:H3K4 me3
DNA Meth. m5C:H4K20 me3
H3K27 me3:H4K20 me3
H2A.Z:H4K20 me2
H4K20 me2:H3K4 me3
H3K27 me2
H4−4ac:H3−5ac
H3K9 me2
H4K20 me2:H3−5ac
H3K4 me1
H3K9 me3
DNA Meth. m5C
H3K27 me3
H2A.Z
H4K20 me3
H4−4ac
H4K20 me2
H3K4 me3
H3−5ac
0.00
0.25
0.50
0.75
1.00
π̂
AFF4 – Reverse
H3K4 me1:H4K20 me2
H3K27 me2:DNA Meth. m5C
H3K27 me3:H3K9 me3
DNA Meth. m5C:H4K20 me2
H2A.Z:H4−4ac
H3K4 me1:H4−4ac
H3K27 me3:H3K4 me1
H3K4 me1:H3−5ac
H3K4 me3:H3−5ac
H3K4 me3:H4−4ac
H3K4 me1:H2A.Z
DNA Meth. m5C:H3K9 me3
H3K9 me3:H4K20 me3
H2A.Z:H3−5ac
H3K27 me3:DNA Meth. m5C
H4K20 me2:H4−4ac
H2A.Z:H3K4 me3
DNA Meth. m5C:H4K20 me3
H3K27 me3:H4K20 me3
H2A.Z:H4K20 me2
H4K20 me2:H3K4 me3
H3K27 me2
H4−4ac:H3−5ac
H3K9 me2
H4K20 me2:H3−5ac
H3K4 me1
H3K9 me3
DNA Meth. m5C
H3K27 me3
H2A.Z
H4K20 me3
H4−4ac
H4K20 me2
H3K4 me3
H3−5ac
0.00
0.25
0.50
0.75
1.00
π̂
KAT14 (1) – Forward
DNA Meth. m5C:H3K9 me3
H3K9 me2:DNA Meth. m5C
H3K27 me3:H3K4 me1
H3K27 me3:H4K20 me3
H3K4 me1:H4K20 me2
H3K9 me3:H4K20 me3
H3K27 me3:DNA Meth. m5C
DNA Meth. m5C:H4K20 me3
H3K4 me3:H3−5ac
DNA Meth. m5C:H4K20 me2
H2A.Z:H4K20 me2
H4K20 me2:H4−4ac
H2A.Z:H3K4 me3
H2A.Z:H4−4ac
H3K4 me1:H3−5ac
H3K4 me1:H2A.Z
H4−4ac:H3−5ac
H3K4 me3:H4−4ac
H4K20 me2:H3K4 me3
H2A.Z:H3−5ac
H3K4 me1:H4−4ac
H3K27 me2
H3K9 me3
H3K9 me2
H3K27 me3
H4K20 me3
H4K20 me2:H3−5ac
DNA Meth. m5C
H3K4 me3
H3K4 me1
H2A.Z
H4−4ac
H4K20 me2
H3−5ac
0.00
0.25
0.50
0.75
1.00
π̂
KAT14 (1) – Reverse
DNA Meth. m5C:H3K9 me3
H3K9 me2:DNA Meth. m5C
H3K27 me3:H3K4 me1
H3K27 me3:H4K20 me3
H3K4 me1:H4K20 me2
H3K9 me3:H4K20 me3
H3K27 me3:DNA Meth. m5C
DNA Meth. m5C:H4K20 me3
H3K4 me3:H3−5ac
DNA Meth. m5C:H4K20 me2
H2A.Z:H4K20 me2
H4K20 me2:H4−4ac
H2A.Z:H3K4 me3
H2A.Z:H4−4ac
H3K4 me1:H3−5ac
H3K4 me1:H2A.Z
H4−4ac:H3−5ac
H3K4 me3:H4−4ac
H4K20 me2:H3K4 me3
H2A.Z:H3−5ac
H3K4 me1:H4−4ac
H3K27 me2
H3K9 me3
H3K9 me2
H3K27 me3
H4K20 me3
H4K20 me2:H3−5ac
DNA Meth. m5C
H3K4 me3
H3K4 me1
H2A.Z
H4−4ac
H4K20 me2
H3−5ac
0.00
0.25
0.50
0.75
1.00
π̂

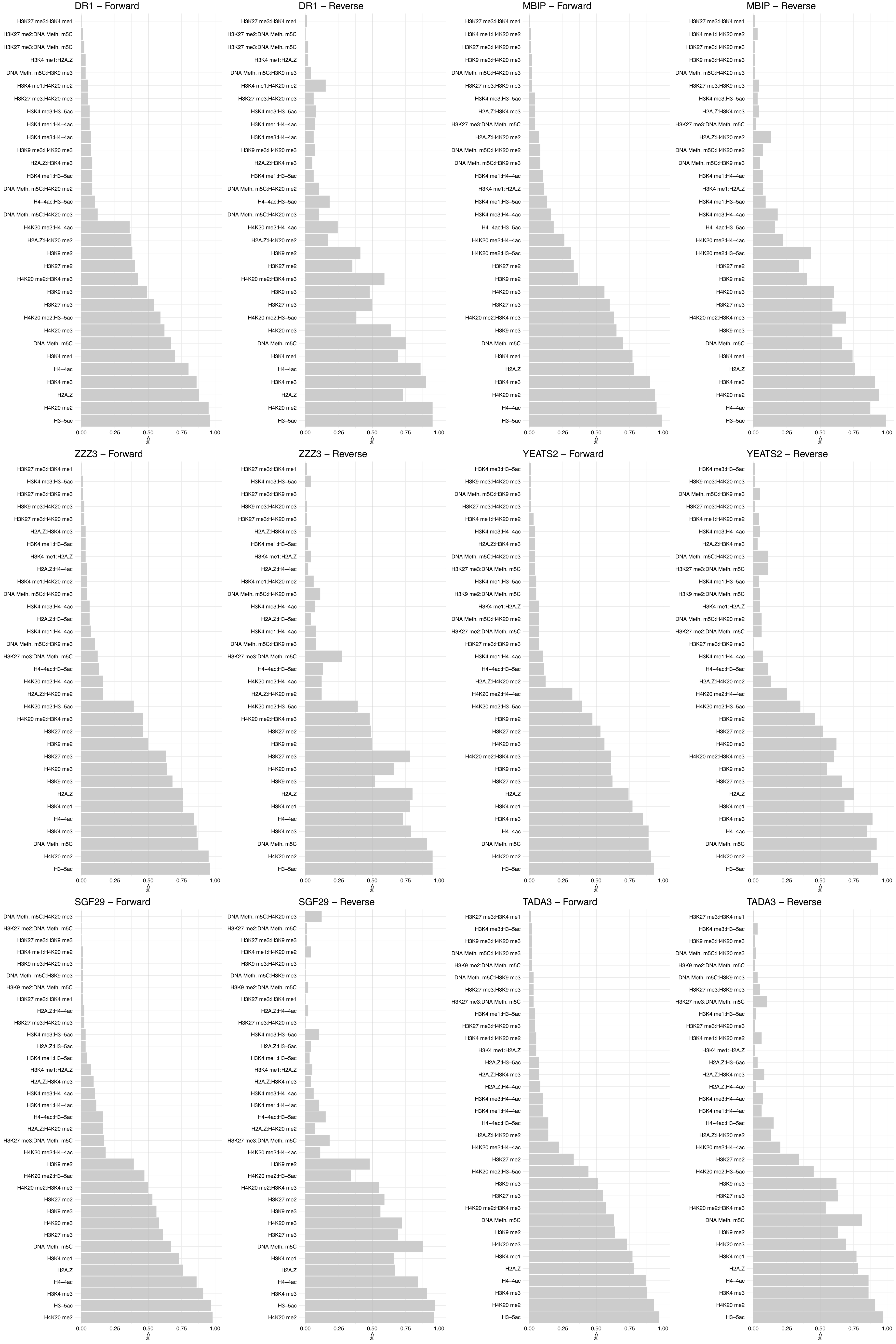
DR1 – Forward
H3K27 me3:H3K4 me1
H3K27 me2:DNA Meth. m5C
H3K27 me3:DNA Meth. m5C
H3K4 me1:H2A.Z
DNA Meth. m5C:H3K9 me3
H3K4 me1:H4K20 me2
H3K27 me3:H4K20 me3
H3K4 me3:H3−5ac
H3K4 me1:H4−4ac
H3K4 me3:H4−4ac
H3K9 me3:H4K20 me3
H2A.Z:H3K4 me3
H3K4 me1:H3−5ac
DNA Meth. m5C:H4K20 me2
H4−4ac:H3−5ac
DNA Meth. m5C:H4K20 me3
H4K20 me2:H4−4ac
H2A.Z:H4K20 me2
H3K9 me2
H3K27 me2
H4K20 me2:H3K4 me3
H3K9 me3
H3K27 me3
H4K20 me2:H3−5ac
H4K20 me3
DNA Meth. m5C
H3K4 me1
H4−4ac
H3K4 me3
H2A.Z
H4K20 me2
H3−5ac
0.00 0.25 0.50 0.75 1.00
π̂
DR1 – Reverse
H3K27 me3:H3K4 me1
H3K27 me2:DNA Meth. m5C
H3K27 me3:DNA Meth. m5C
H3K4 me1:H2A.Z
DNA Meth. m5C:H3K9 me3
H3K4 me1:H4K20 me2
H3K27 me3:H4K20 me3
H3K4 me3:H3−5ac
H3K4 me1:H4−4ac
H3K4 me3:H4−4ac
H3K9 me3:H4K20 me3
H2A.Z:H3K4 me3
H3K4 me1:H3−5ac
DNA Meth. m5C:H4K20 me2
H4−4ac:H3−5ac
DNA Meth. m5C:H4K20 me3
H4K20 me2:H4−4ac
H2A.Z:H4K20 me2
H3K9 me2
H3K27 me2
H4K20 me2:H3K4 me3
H3K9 me3
H3K27 me3
H4K20 me2:H3−5ac
H4K20 me3
DNA Meth. m5C
H3K4 me1
H4−4ac
H3K4 me3
H2A.Z
H4K20 me2
H3−5ac
0.00 0.25 0.50 0.75 1.00
π̂
MBIP – Forward
H3K27 me3:H3K4 me1
H3K4 me1:H4K20 me2
H3K27 me3:H4K20 me3
H3K9 me3:H4K20 me3
DNA Meth. m5C:H4K20 me3
H3K27 me3:H3K9 me3
H3K4 me3:H3−5ac
H2A.Z:H3K4 me3
H3K27 me3:DNA Meth. m5C
H2A.Z:H4K20 me2
DNA Meth. m5C:H4K20 me2
DNA Meth. m5C:H3K9 me3
H3K4 me1:H4−4ac
H3K4 me1:H2A.Z
H3K4 me1:H3−5ac
H3K4 me3:H4−4ac
H4−4ac:H3−5ac
H4K20 me2:H4−4ac
H4K20 me2:H3−5ac
H3K27 me2
H3K9 me2
H4K20 me3
H3K27 me3
H4K20 me2:H3K4 me3
H3K9 me3
DNA Meth. m5C
H3K4 me1
H2A.Z
H3K4 me3
H4K20 me2
H4−4ac
H3−5ac
0.00 0.25 0.50 0.75 1.00
π̂
MBIP – Reverse
H3K27 me3:H3K4 me1
H3K4 me1:H4K20 me2
H3K27 me3:H4K20 me3
H3K9 me3:H4K20 me3
DNA Meth. m5C:H4K20 me3
H3K27 me3:H3K9 me3
H3K4 me3:H3−5ac
H2A.Z:H3K4 me3
H3K27 me3:DNA Meth. m5C
H2A.Z:H4K20 me2
DNA Meth. m5C:H4K20 me2
DNA Meth. m5C:H3K9 me3
H3K4 me1:H4−4ac
H3K4 me1:H2A.Z
H3K4 me1:H3−5ac
H3K4 me3:H4−4ac
H4−4ac:H3−5ac
H4K20 me2:H4−4ac
H4K20 me2:H3−5ac
H3K27 me2
H3K9 me2
H4K20 me3
H3K27 me3
H4K20 me2:H3K4 me3
H3K9 me3
DNA Meth. m5C
H3K4 me1
H2A.Z
H3K4 me3
H4K20 me2
H4−4ac
H3−5ac
0.00 0.25 0.50 0.75 1.00
π̂
ZZZ3 – Forward
H3K27 me3:H3K4 me1
H3K4 me3:H3−5ac
H3K27 me3:H3K9 me3
H3K9 me3:H4K20 me3
H3K27 me3:H4K20 me3
H2A.Z:H3K4 me3
H3K4 me1:H3−5ac
H3K4 me1:H2A.Z
H2A.Z:H4−4ac
H3K4 me1:H4K20 me2
DNA Meth. m5C:H4K20 me3
H3K4 me3:H4−4ac
H2A.Z:H3−5ac
H3K4 me1:H4−4ac
DNA Meth. m5C:H3K9 me3
H3K27 me3:DNA Meth. m5C
H4−4ac:H3−5ac
H4K20 me2:H4−4ac
H2A.Z:H4K20 me2
H4K20 me2:H3−5ac
H4K20 me2:H3K4 me3
H3K27 me2
H3K9 me2
H3K27 me3
H4K20 me3
H3K9 me3
H2A.Z
H3K4 me1
H4−4ac
H3K4 me3
DNA Meth. m5C
H4K20 me2
H3−5ac
0.00 0.25 0.50 0.75 1.00
π̂
ZZZ3 – Reverse
H3K27 me3:H3K4 me1
H3K4 me3:H3−5ac
H3K27 me3:H3K9 me3
H3K9 me3:H4K20 me3
H3K27 me3:H4K20 me3
H2A.Z:H3K4 me3
H3K4 me1:H3−5ac
H3K4 me1:H2A.Z
H2A.Z:H4−4ac
H3K4 me1:H4K20 me2
DNA Meth. m5C:H4K20 me3
H3K4 me3:H4−4ac
H2A.Z:H3−5ac
H3K4 me1:H4−4ac
DNA Meth. m5C:H3K9 me3
H3K27 me3:DNA Meth. m5C
H4−4ac:H3−5ac
H4K20 me2:H4−4ac
H2A.Z:H4K20 me2
H4K20 me2:H3−5ac
H4K20 me2:H3K4 me3
H3K27 me2
H3K9 me2
H3K27 me3
H4K20 me3
H3K9 me3
H2A.Z
H3K4 me1
H4−4ac
H3K4 me3
DNA Meth. m5C
H4K20 me2
H3−5ac
0.00 0.25 0.50 0.75 1.00
π̂
YEATS2 – Forward
H3K4 me3:H3−5ac
H3K9 me3:H4K20 me3
DNA Meth. m5C:H3K9 me3
H3K27 me3:H4K20 me3
H3K4 me1:H4K20 me2
H3K4 me3:H4−4ac
H2A.Z:H3K4 me3
DNA Meth. m5C:H4K20 me3
H3K27 me3:DNA Meth. m5C
H3K4 me1:H3−5ac
H3K9 me2:DNA Meth. m5C
H3K4 me1:H2A.Z
DNA Meth. m5C:H4K20 me2
H3K27 me2:DNA Meth. m5C
H3K27 me3:H3K9 me3
H3K4 me1:H4−4ac
H4−4ac:H3−5ac
H2A.Z:H4K20 me2
H4K20 me2:H4−4ac
H4K20 me2:H3−5ac
H3K9 me2
H3K27 me2
H4K20 me3
H4K20 me2:H3K4 me3
H3K9 me3
H3K27 me3
H2A.Z
H3K4 me1
H3K4 me3
H4−4ac
DNA Meth. m5C
H4K20 me2
H3−5ac
0.00 0.25 0.50 0.75 1.00
π̂
YEATS2 – Reverse
H3K4 me3:H3−5ac
H3K9 me3:H4K20 me3
DNA Meth. m5C:H3K9 me3
H3K27 me3:H4K20 me3
H3K4 me1:H4K20 me2
H3K4 me3:H4−4ac
H2A.Z:H3K4 me3
DNA Meth. m5C:H4K20 me3
H3K27 me3:DNA Meth. m5C
H3K4 me1:H3−5ac
H3K9 me2:DNA Meth. m5C
H3K4 me1:H2A.Z
DNA Meth. m5C:H4K20 me2
H3K27 me2:DNA Meth. m5C
H3K27 me3:H3K9 me3
H3K4 me1:H4−4ac
H4−4ac:H3−5ac
H2A.Z:H4K20 me2
H4K20 me2:H4−4ac
H4K20 me2:H3−5ac
H3K9 me2
H3K27 me2
H4K20 me3
H4K20 me2:H3K4 me3
H3K9 me3
H3K27 me3
H2A.Z
H3K4 me1
H3K4 me3
H4−4ac
DNA Meth. m5C
H4K20 me2
H3−5ac
0.00 0.25 0.50 0.75 1.00
π̂
SGF29 – Forward
DNA Meth. m5C:H4K20 me3
H3K27 me2:DNA Meth. m5C
H3K27 me3:H3K9 me3
H3K4 me1:H4K20 me2
H3K9 me3:H4K20 me3
DNA Meth. m5C:H3K9 me3
H3K9 me2:DNA Meth. m5C
H3K27 me3:H3K4 me1
H2A.Z:H4−4ac
H3K27 me3:H4K20 me3
H3K4 me3:H3−5ac
H2A.Z:H3−5ac
H3K4 me1:H3−5ac
H3K4 me1:H2A.Z
H2A.Z:H3K4 me3
H3K4 me3:H4−4ac
H3K4 me1:H4−4ac
H4−4ac:H3−5ac
H2A.Z:H4K20 me2
H3K27 me3:DNA Meth. m5C
H4K20 me2:H4−4ac
H3K9 me2
H4K20 me2:H3−5ac
H4K20 me2:H3K4 me3
H3K27 me2
H3K9 me3
H4K20 me3
H3K27 me3
DNA Meth. m5C
H3K4 me1
H2A.Z
H4−4ac
H3K4 me3
H3−5ac
H4K20 me2
0.00 0.25 0.50 0.75 1.00
π̂
SGF29 – Reverse
DNA Meth. m5C:H4K20 me3
H3K27 me2:DNA Meth. m5C
H3K27 me3:H3K9 me3
H3K4 me1:H4K20 me2
H3K9 me3:H4K20 me3
DNA Meth. m5C:H3K9 me3
H3K9 me2:DNA Meth. m5C
H3K27 me3:H3K4 me1
H2A.Z:H4−4ac
H3K27 me3:H4K20 me3
H3K4 me3:H3−5ac
H2A.Z:H3−5ac
H3K4 me1:H3−5ac
H3K4 me1:H2A.Z
H2A.Z:H3K4 me3
H3K4 me3:H4−4ac
H3K4 me1:H4−4ac
H4−4ac:H3−5ac
H2A.Z:H4K20 me2
H3K27 me3:DNA Meth. m5C
H4K20 me2:H4−4ac
H3K9 me2
H4K20 me2:H3−5ac
H4K20 me2:H3K4 me3
H3K27 me2
H3K9 me3
H4K20 me3
H3K27 me3
DNA Meth. m5C
H3K4 me1
H2A.Z
H4−4ac
H3K4 me3
H3−5ac
H4K20 me2
0.00 0.25 0.50 0.75 1.00
π̂
TADA3 – Forward
H3K27 me3:H3K4 me1
H3K4 me3:H3−5ac
H3K9 me3:H4K20 me3
DNA Meth. m5C:H4K20 me3
H3K9 me2:DNA Meth. m5C
DNA Meth. m5C:H3K9 me3
H3K27 me3:H3K9 me3
H3K27 me3:DNA Meth. m5C
H3K4 me1:H3−5ac
H3K27 me3:H4K20 me3
H3K4 me1:H4K20 me2
H3K4 me1:H2A.Z
H2A.Z:H3−5ac
H2A.Z:H3K4 me3
H2A.Z:H4−4ac
H3K4 me3:H4−4ac
H3K4 me1:H4−4ac
H4−4ac:H3−5ac
H2A.Z:H4K20 me2
H4K20 me2:H4−4ac
H3K27 me2
H4K20 me2:H3−5ac
H3K9 me3
H3K27 me3
H4K20 me2:H3K4 me3
DNA Meth. m5C
H3K9 me2
H4K20 me3
H3K4 me1
H2A.Z
H4−4ac
H3K4 me3
H4K20 me2
H3−5ac
0.00 0.25 0.50 0.75 1.00
π̂
TADA3 – Reverse
H3K27 me3:H3K4 me1
H3K4 me3:H3−5ac
H3K9 me3:H4K20 me3
DNA Meth. m5C:H4K20 me3
H3K9 me2:DNA Meth. m5C
DNA Meth. m5C:H3K9 me3
H3K27 me3:H3K9 me3
H3K27 me3:DNA Meth. m5C
H3K4 me1:H3−5ac
H3K27 me3:H4K20 me3
H3K4 me1:H4K20 me2
H3K4 me1:H2A.Z
H2A.Z:H3−5ac
H2A.Z:H3K4 me3
H2A.Z:H4−4ac
H3K4 me3:H4−4ac
H3K4 me1:H4−4ac
H4−4ac:H3−5ac
H2A.Z:H4K20 me2
H4K20 me2:H4−4ac
H3K27 me2
H4K20 me2:H3−5ac
H3K9 me3
H3K27 me3
H4K20 me2:H3K4 me3
DNA Meth. m5C
H3K9 me2
H4K20 me3
H3K4 me1
H2A.Z
H4−4ac
H3K4 me3
H4K20 me2
H3−5ac
0.00 0.25 0.50 0.75 1.00
π̂

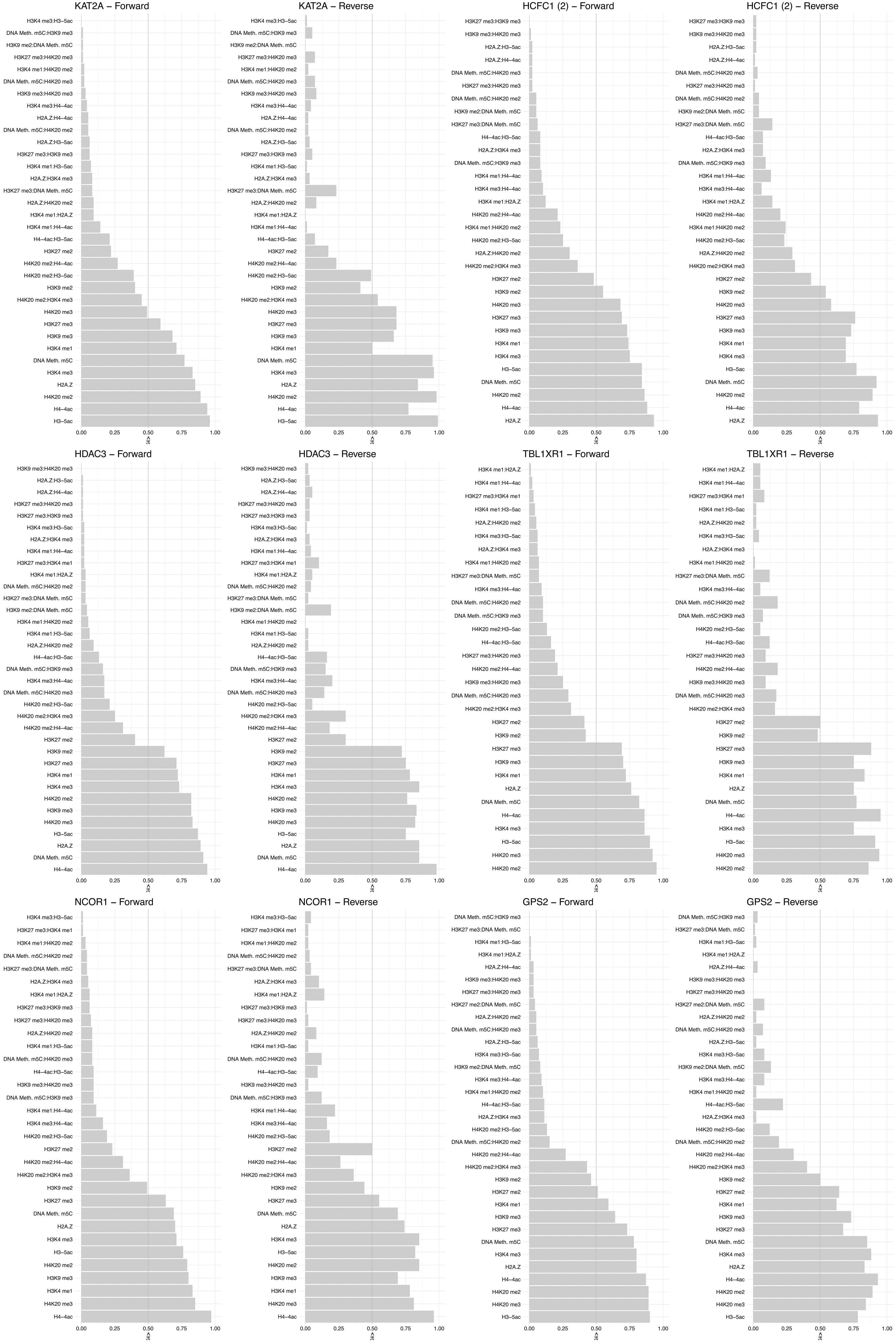
KAT2A – Forward
KAT2A – Reverse
HCFC1 (2) – Forward
HCFC1 (2) – Reverse
HDAC3 – Forward
HDAC3 – Reverse
TBL1XR1 – Forward
TBL1XR1 – Reverse
NCOR1 – Forward
NCOR1 – Reverse
GPS2 – Forward
GPS2 – Reverse

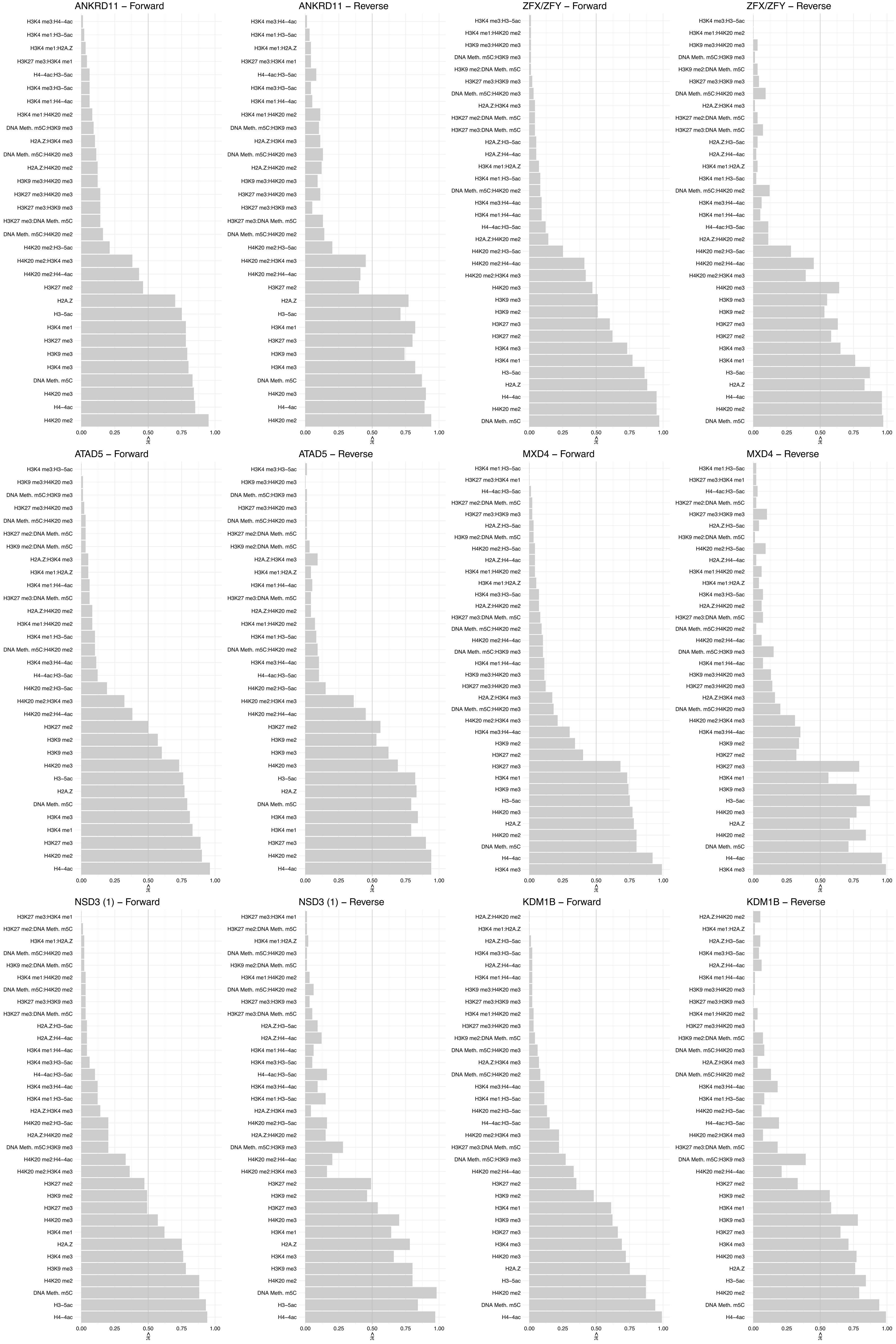
ANKRD11 – Forward
ANKRD11 – Reverse
ZFX/ZFY – Forward
ZFX/ZFY – Reverse
ATAD5 – Forward
ATAD5 – Reverse
MXD4 – Forward
MXD4 – Reverse
NSD3 (1) – Forward
NSD3 (1) – Reverse
KDM1B – Forward
KDM1B – Reverse
0.00
0.25
0.50
0.75
1.00
π̂

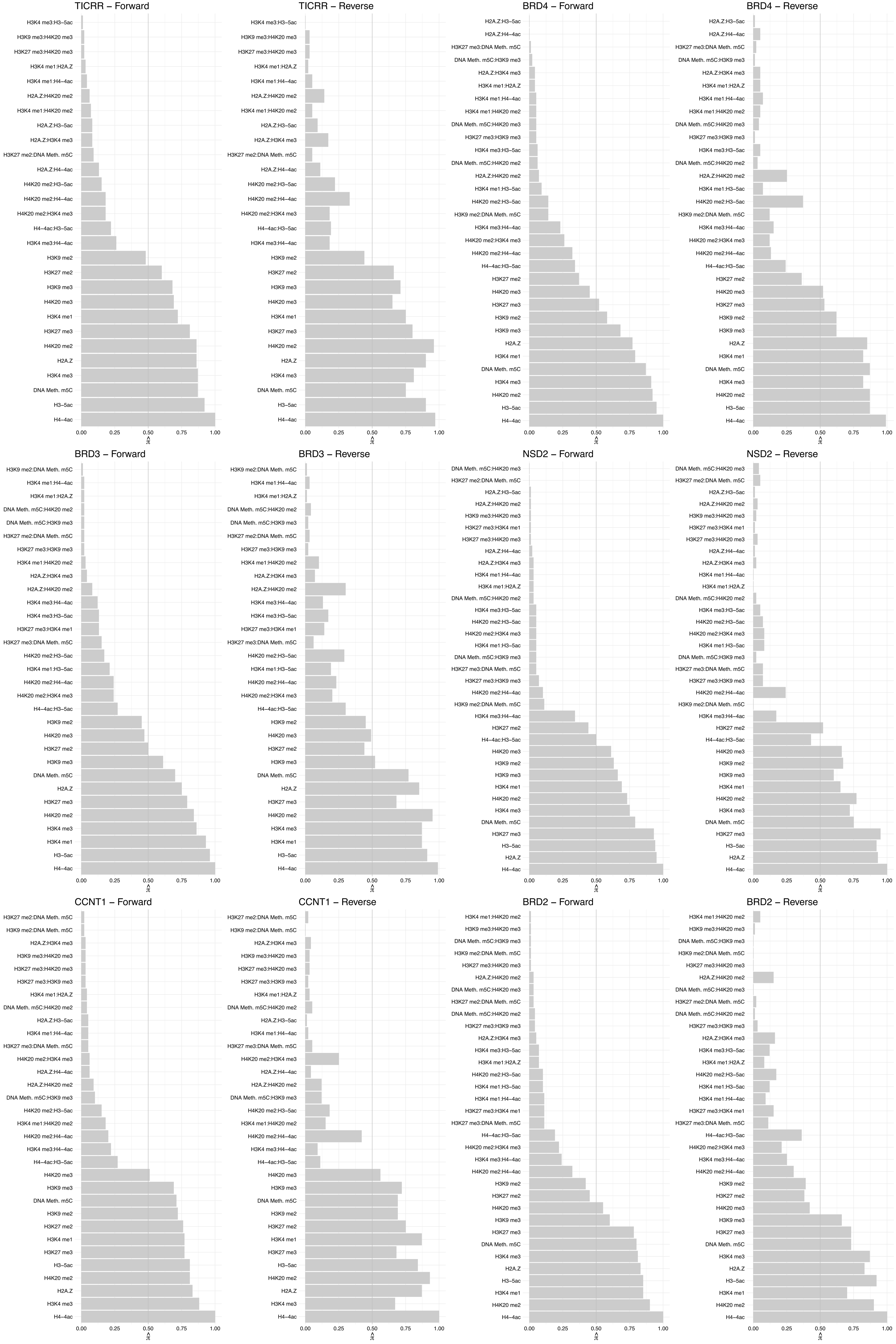
TICRR – Forward
H3K4 me3:H3−5ac
H3K9 me3:H4K20 me3
H3K27 me3:H4K20 me3
H3K4 me1:H2A.Z
H3K4 me1:H4−4ac
H2A.Z:H4K20 me2
H3K4 me1:H4K20 me2
H2A.Z:H3−5ac
H2A.Z:H3K4 me3
H3K27 me2:DNA Meth. m5C
H2A.Z:H4−4ac
H4K20 me2:H3−5ac
H4K20 me2:H4−4ac
H4K20 me2:H3K4 me3
H4−4ac:H3−5ac
H3K4 me3:H4−4ac
H3K9 me2
H3K27 me2
H3K9 me3
H4K20 me3
H3K4 me1
H3K27 me3
H4K20 me2
H2A.Z
H3K4 me3
DNA Meth. m5C
H3−5ac
H4−4ac
0.00 0.25 0.50 0.75 1.00
$\hat{\pi}$
TICRR – Reverse
H3K4 me3:H3−5ac
H3K9 me3:H4K20 me3
H3K27 me3:H4K20 me3
H3K4 me1:H2A.Z
H3K4 me1:H4−4ac
H2A.Z:H4K20 me2
H3K4 me1:H4K20 me2
H2A.Z:H3−5ac
H2A.Z:H3K4 me3
H3K27 me2:DNA Meth. m5C
H2A.Z:H4−4ac
H4K20 me2:H3−5ac
H4K20 me2:H4−4ac
H4K20 me2:H3K4 me3
H4−4ac:H3−5ac
H3K4 me3:H4−4ac
H3K9 me2
H3K27 me2
H3K9 me3
H4K20 me3
H3K4 me1
H3K27 me3
H4K20 me2
H2A.Z
H3K4 me3
DNA Meth. m5C
H3−5ac
H4−4ac
0.00 0.25 0.50 0.75 1.00
$\hat{\pi}$
BRD4 – Forward
H2A.Z:H3−5ac
H2A.Z:H4−4ac
H3K27 me3:DNA Meth. m5C
DNA Meth. m5C:H3K9 me3
H2A.Z:H3K4 me3
H3K4 me1:H2A.Z
H3K4 me1:H4−4ac
H3K4 me1:H4K20 me2
DNA Meth. m5C:H4K20 me3
H3K27 me3:H3K9 me3
H3K4 me3:H3−5ac
DNA Meth. m5C:H4K20 me2
H2A.Z:H4K20 me2
H3K4 me1:H3−5ac
H4K20 me2:H3−5ac
H3K9 me2:DNA Meth. m5C
H3K4 me3:H4−4ac
H4K20 me2:H3K4 me3
H4K20 me2:H4−4ac
H4−4ac:H3−5ac
H3K27 me2
H4K20 me3
H3K27 me3
H3K9 me2
H3K9 me3
H2A.Z
H3K4 me1
DNA Meth. m5C
H3K4 me3
H4K20 me2
H3−5ac
H4−4ac
0.00 0.25 0.50 0.75 1.00
$\hat{\pi}$
BRD4 – Reverse
H2A.Z:H3−5ac
H2A.Z:H4−4ac
H3K27 me3:DNA Meth. m5C
DNA Meth. m5C:H3K9 me3
H2A.Z:H3K4 me3
H3K4 me1:H2A.Z
H3K4 me1:H4−4ac
H3K4 me1:H4K20 me2
DNA Meth. m5C:H4K20 me3
H3K27 me3:H3K9 me3
H3K4 me3:H3−5ac
DNA Meth. m5C:H4K20 me2
H2A.Z:H4K20 me2
H3K4 me1:H3−5ac
H4K20 me2:H3−5ac
H3K9 me2:DNA Meth. m5C
H3K4 me3:H4−4ac
H4K20 me2:H3K4 me3
H4K20 me2:H4−4ac
H4−4ac:H3−5ac
H3K27 me2
H4K20 me3
H3K27 me3
H3K9 me2
H3K9 me3
H2A.Z
H3K4 me1
DNA Meth. m5C
H3K4 me3
H4K20 me2
H3−5ac
H4−4ac
0.00 0.25 0.50 0.75 1.00
$\hat{\pi}$
BRD3 – Forward
H3K9 me2:DNA Meth. m5C
H3K4 me1:H4−4ac
H3K4 me1:H2A.Z
DNA Meth. m5C:H4K20 me2
DNA Meth. m5C:H3K9 me3
H3K27 me2:DNA Meth. m5C
H3K27 me3:H3K9 me3
H3K4 me1:H4K20 me2
H2A.Z:H3K4 me3
H2A.Z:H4K20 me2
H3K4 me3:H4−4ac
H3K4 me3:H3−5ac
H3K27 me3:H3K4 me1
H3K27 me3:DNA Meth. m5C
H4K20 me2:H3−5ac
H3K4 me1:H3−5ac
H4K20 me2:H4−4ac
H4K20 me2:H3K4 me3
H4−4ac:H3−5ac
H3K9 me2
H4K20 me3
H3K27 me2
H3K9 me3
DNA Meth. m5C
H2A.Z
H3K27 me3
H4K20 me2
H3K4 me3
H3K4 me1
H3−5ac
H4−4ac
0.00 0.25 0.50 0.75 1.00
$\hat{\pi}$
BRD3 – Reverse
H3K9 me2:DNA Meth. m5C
H3K4 me1:H4−4ac
H3K4 me1:H2A.Z
DNA Meth. m5C:H4K20 me2
DNA Meth. m5C:H3K9 me3
H3K27 me2:DNA Meth. m5C
H3K27 me3:H3K9 me3
H3K4 me1:H4K20 me2
H2A.Z:H3K4 me3
H2A.Z:H4K20 me2
H3K4 me3:H4−4ac
H3K4 me3:H3−5ac
H3K27 me3:H3K4 me1
H3K27 me3:DNA Meth. m5C
H4K20 me2:H3−5ac
H3K4 me1:H3−5ac
H4K20 me2:H4−4ac
H4K20 me2:H3K4 me3
H4−4ac:H3−5ac
H3K9 me2
H4K20 me3
H3K27 me2
H3K9 me3
DNA Meth. m5C
H2A.Z
H3K27 me3
H4K20 me2
H3K4 me3
H3K4 me1
H3−5ac
H4−4ac
0.00 0.25 0.50 0.75 1.00
$\hat{\pi}$
NSD2 – Forward
DNA Meth. m5C:H4K20 me3
H3K27 me2:DNA Meth. m5C
H2A.Z:H3−5ac
H2A.Z:H4K20 me2
H3K9 me3:H4K20 me3
H3K27 me3:H3K4 me1
H3K27 me3:H4K20 me3
H2A.Z:H4−4ac
H2A.Z:H3K4 me3
H3K4 me1:H4−4ac
H3K4 me1:H2A.Z
DNA Meth. m5C:H4K20 me2
H3K4 me3:H3−5ac
H4K20 me2:H3−5ac
H4K20 me2:H3K4 me3
H3K4 me1:H3−5ac
DNA Meth. m5C:H3K9 me3
H3K27 me3:DNA Meth. m5C
H3K27 me3:H3K9 me3
H4K20 me2:H4−4ac
H3K9 me2:DNA Meth. m5C
H3K4 me3:H4−4ac
H3K27 me2
H4−4ac:H3−5ac
H4K20 me3
H3K9 me2
H3K9 me3
H3K4 me1
H4K20 me2
H3K4 me3
DNA Meth. m5C
H3K27 me3
H3−5ac
H2A.Z
H4−4ac
0.00 0.25 0.50 0.75 1.00
$\hat{\pi}$
NSD2 – Reverse
DNA Meth. m5C:H4K20 me3
H3K27 me2:DNA Meth. m5C
H2A.Z:H3−5ac
H2A.Z:H4K20 me2
H3K9 me3:H4K20 me3
H3K27 me3:H3K4 me1
H3K27 me3:H4K20 me3
H2A.Z:H4−4ac
H2A.Z:H3K4 me3
H3K4 me1:H4−4ac
H3K4 me1:H2A.Z
DNA Meth. m5C:H4K20 me2
H3K4 me3:H3−5ac
H4K20 me2:H3−5ac
H4K20 me2:H3K4 me3
H3K4 me1:H3−5ac
DNA Meth. m5C:H3K9 me3
H3K27 me3:DNA Meth. m5C
H3K27 me3:H3K9 me3
H4K20 me2:H4−4ac
H3K9 me2:DNA Meth. m5C
H3K4 me3:H4−4ac
H3K27 me2
H4−4ac:H3−5ac
H4K20 me3
H3K9 me2
H3K9 me3
H3K4 me1
H4K20 me2
H3K4 me3
DNA Meth. m5C
H3K27 me3
H3−5ac
H2A.Z
H4−4ac
0.00 0.25 0.50 0.75 1.00
$\hat{\pi}$
CCNT1 – Forward
H3K27 me2:DNA Meth. m5C
H3K9 me2:DNA Meth. m5C
H2A.Z:H3K4 me3
H3K9 me3:H4K20 me3
H3K27 me3:H4K20 me3
H3K27 me3:H3K9 me3
H3K4 me1:H2A.Z
DNA Meth. m5C:H4K20 me2
H2A.Z:H3−5ac
H3K4 me1:H4−4ac
H3K27 me3:DNA Meth. m5C
H4K20 me2:H3K4 me3
H2A.Z:H4−4ac
H2A.Z:H4K20 me2
DNA Meth. m5C:H3K9 me3
H4K20 me2:H3−5ac
H3K4 me1:H4K20 me2
H4K20 me2:H4−4ac
H3K4 me3:H4−4ac
H4−4ac:H3−5ac
H4K20 me3
H3K9 me3
DNA Meth. m5C
H3K9 me2
H3K27 me2
H3K4 me1
H3K27 me3
H3−5ac
H4K20 me2
H2A.Z
H3K4 me3
H4−4ac
0.00 0.25 0.50 0.75 1.00
$\hat{\pi}$
CCNT1 – Reverse
H3K27 me2:DNA Meth. m5C
H3K9 me2:DNA Meth. m5C
H2A.Z:H3K4 me3
H3K9 me3:H4K20 me3
H3K27 me3:H4K20 me3
H3K27 me3:H3K9 me3
H3K4 me1:H2A.Z
DNA Meth. m5C:H4K20 me2
H2A.Z:H3−5ac
H3K4 me1:H4−4ac
H3K27 me3:DNA Meth. m5C
H4K20 me2:H3K4 me3
H2A.Z:H4−4ac
H2A.Z:H4K20 me2
DNA Meth. m5C:H3K9 me3
H4K20 me2:H3−5ac
H3K4 me1:H4K20 me2
H4K20 me2:H4−4ac
H3K4 me3:H4−4ac
H4−4ac:H3−5ac
H4K20 me3
H3K9 me3
DNA Meth. m5C
H3K9 me2
H3K27 me2
H3K4 me1
H3K27 me3
H3−5ac
H4K20 me2
H2A.Z
H3K4 me3
H4−4ac
0.00 0.25 0.50 0.75 1.00
$\hat{\pi}$
BRD2 – Forward
H3K4 me1:H4K20 me2
H3K9 me3:H4K20 me3
DNA Meth. m5C:H3K9 me3
H3K9 me2:DNA Meth. m5C
H3K27 me3:H4K20 me3
H2A.Z:H4K20 me2
DNA Meth. m5C:H4K20 me3
H3K27 me2:DNA Meth. m5C
DNA Meth. m5C:H4K20 me2
H3K27 me3:H3K9 me3
H2A.Z:H3K4 me3
H3K4 me3:H3−5ac
H3K4 me1:H2A.Z
H4K20 me2:H3−5ac
H3K4 me1:H3−5ac
H3K4 me1:H4−4ac
H3K27 me3:H3K4 me1
H3K27 me3:DNA Meth. m5C
H4−4ac:H3−5ac
H4K20 me2:H3K4 me3
H3K4 me3:H4−4ac
H4K20 me2:H4−4ac
H3K9 me2
H3K27 me2
H4K20 me3
H3K9 me3
H3K27 me3
DNA Meth. m5C
H3K4 me3
H2A.Z
H3−5ac
H3K4 me1
H4K20 me2
H4−4ac
0.00 0.25 0.50 0.75 1.00
$\hat{\pi}$
BRD2 – Reverse
H3K4 me1:H4K20 me2
H3K9 me3:H4K20 me3
DNA Meth. m5C:H3K9 me3
H3K9 me2:DNA Meth. m5C
H3K27 me3:H4K20 me3
H2A.Z:H4K20 me2
DNA Meth. m5C:H4K20 me3
H3K27 me2:DNA Meth. m5C
DNA Meth. m5C:H4K20 me2
H3K27 me3:H3K9 me3
H2A.Z:H3K4 me3
H3K4 me3:H3−5ac
H3K4 me1:H2A.Z
H4K20 me2:H3−5ac
H3K4 me1:H3−5ac
H3K4 me1:H4−4ac
H3K27 me3:H3K4 me1
H3K27 me3:DNA Meth. m5C
H4−4ac:H3−5ac
H4K20 me2:H3K4 me3
H3K4 me3:H4−4ac
H4K20 me2:H4−4ac
H3K9 me2
H3K27 me2
H4K20 me3
H3K9 me3
H3K27 me3
DNA Meth. m5C
H3K4 me3
H2A.Z
H3−5ac
H3K4 me1
H4K20 me2
H4−4ac
0.00 0.25 0.50 0.75 1.00
$\hat{\pi}$

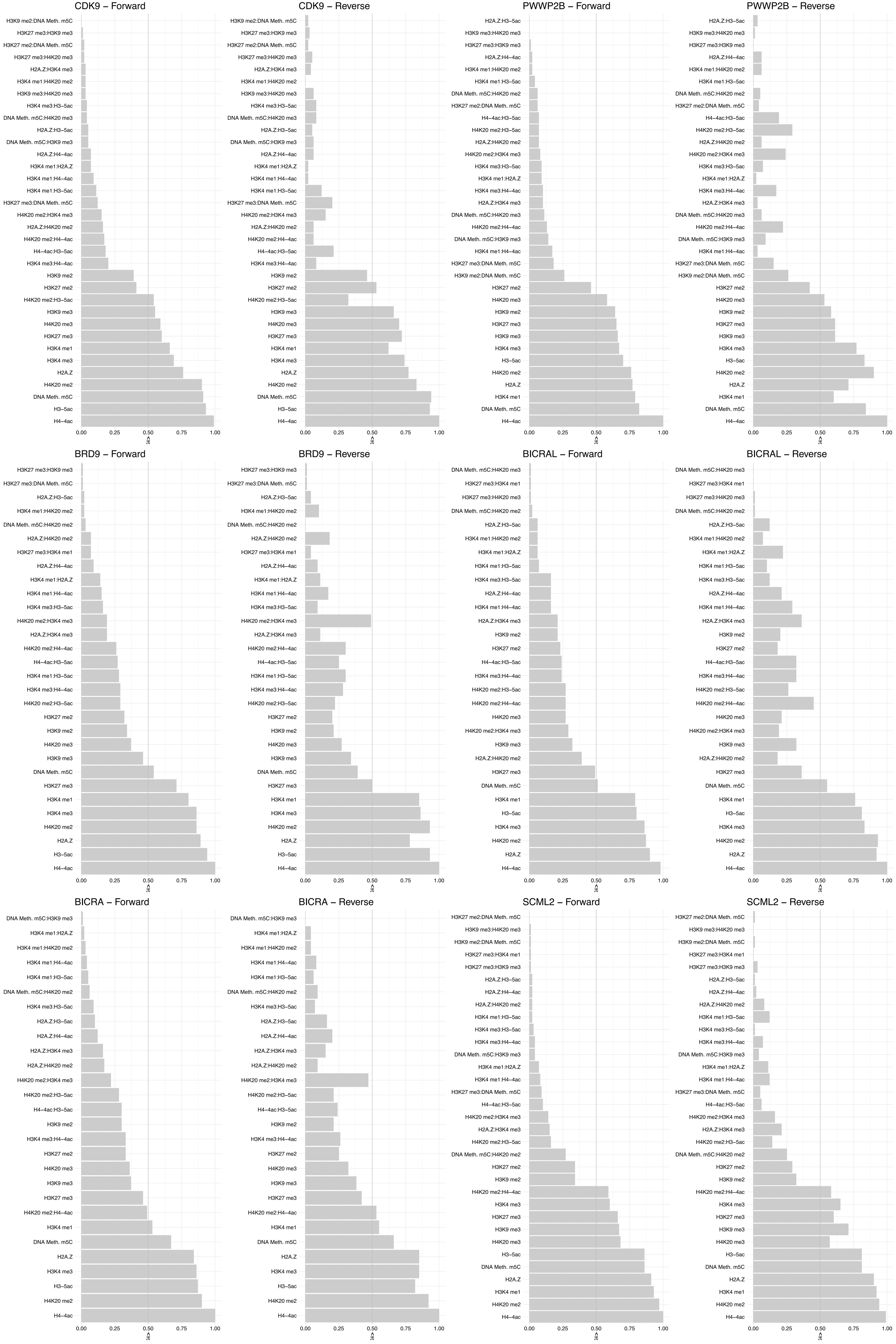
CDK9 − Forward
H3K9 me2:DNA Meth. m5C
H3K27 me3:H3K9 me3
H3K27 me2:DNA Meth. m5C
H3K27 me3:H4K20 me3
H2A.Z:H3K4 me3
H3K4 me1:H4K20 me2
H3K9 me3:H4K20 me3
H3K4 me3:H3−5ac
DNA Meth. m5C:H4K20 me3
H2A.Z:H3−5ac
DNA Meth. m5C:H3K9 me3
H2A.Z:H4−4ac
H3K4 me1:H2A.Z
H3K4 me1:H4−4ac
H3K4 me1:H3−5ac
H3K27 me3:DNA Meth. m5C
H4K20 me2:H3K4 me3
H2A.Z:H4K20 me2
H4K20 me2:H4−4ac
H4−4ac:H3−5ac
H3K4 me3:H4−4ac
H3K9 me2
H3K27 me2
H4K20 me2:H3−5ac
H3K9 me3
H4K20 me3
H3K27 me3
H3K4 me1
H3K4 me3
H2A.Z
H4K20 me2
DNA Meth. m5C
H3−5ac
H4−4ac
0.00 0.25 0.50 0.75 1.00
$\hat{\pi}$
CDK9 − Reverse
H3K9 me2:DNA Meth. m5C
H3K27 me3:H3K9 me3
H3K27 me2:DNA Meth. m5C
H3K27 me3:H4K20 me3
H2A.Z:H3K4 me3
H3K4 me1:H4K20 me2
H3K9 me3:H4K20 me3
H3K4 me3:H3−5ac
DNA Meth. m5C:H4K20 me3
H2A.Z:H3−5ac
DNA Meth. m5C:H3K9 me3
H2A.Z:H4−4ac
H3K4 me1:H2A.Z
H3K4 me1:H4−4ac
H3K4 me1:H3−5ac
H3K27 me3:DNA Meth. m5C
H4K20 me2:H3K4 me3
H2A.Z:H4K20 me2
H4K20 me2:H4−4ac
H4−4ac:H3−5ac
H3K4 me3:H4−4ac
H3K9 me2
H3K27 me2
H4K20 me2:H3−5ac
H3K9 me3
H4K20 me3
H3K27 me3
H3K4 me1
H3K4 me3
H2A.Z
H4K20 me2
DNA Meth. m5C
H3−5ac
H4−4ac
0.00 0.25 0.50 0.75 1.00
$\hat{\pi}$
PWWP2B − Forward
H2A.Z:H3−5ac
H3K9 me3:H4K20 me3
H3K27 me3:H3K9 me3
H2A.Z:H4−4ac
H3K4 me1:H4K20 me2
H3K4 me1:H3−5ac
DNA Meth. m5C:H4K20 me2
H3K27 me2:DNA Meth. m5C
H4−4ac:H3−5ac
H4K20 me2:H3−5ac
H2A.Z:H4K20 me2
H4K20 me2:H3K4 me3
H3K4 me3:H3−5ac
H3K4 me1:H2A.Z
H3K4 me3:H4−4ac
H2A.Z:H3K4 me3
DNA Meth. m5C:H4K20 me3
H4K20 me2:H4−4ac
DNA Meth. m5C:H3K9 me3
H3K4 me1:H4−4ac
H3K27 me3:DNA Meth. m5C
H3K9 me2:DNA Meth. m5C
H3K27 me2
H4K20 me3
H3K9 me2
H3K27 me3
H3K9 me3
H3K4 me3
H3−5ac
H4K20 me2
H2A.Z
H3K4 me1
DNA Meth. m5C
H4−4ac
0.00 0.25 0.50 0.75 1.00
$\hat{\pi}$
PWWP2B − Reverse
H2A.Z:H3−5ac
H3K9 me3:H4K20 me3
H3K27 me3:H3K9 me3
H2A.Z:H4−4ac
H3K4 me1:H4K20 me2
H3K4 me1:H3−5ac
DNA Meth. m5C:H4K20 me2
H3K27 me2:DNA Meth. m5C
H4−4ac:H3−5ac
H4K20 me2:H3−5ac
H2A.Z:H4K20 me2
H4K20 me2:H3K4 me3
H3K4 me3:H3−5ac
H3K4 me1:H2A.Z
H3K4 me3:H4−4ac
H2A.Z:H3K4 me3
DNA Meth. m5C:H4K20 me3
H4K20 me2:H4−4ac
DNA Meth. m5C:H3K9 me3
H3K4 me1:H4−4ac
H3K27 me3:DNA Meth. m5C
H3K9 me2:DNA Meth. m5C
H3K27 me2
H4K20 me3
H3K9 me2
H3K27 me3
H3K9 me3
H3K4 me3
H3−5ac
H4K20 me2
H2A.Z
H3K4 me1
DNA Meth. m5C
H4−4ac
0.00 0.25 0.50 0.75 1.00
$\hat{\pi}$
BRD9 − Forward
H3K27 me3:H3K9 me3
H3K27 me3:DNA Meth. m5C
H2A.Z:H3−5ac
H3K4 me1:H4K20 me2
DNA Meth. m5C:H4K20 me2
H2A.Z:H4K20 me2
H3K27 me3:H3K4 me1
H2A.Z:H4−4ac
H3K4 me1:H2A.Z
H3K4 me1:H4−4ac
H3K4 me3:H3−5ac
H4K20 me2:H3K4 me3
H2A.Z:H3K4 me3
H4K20 me2:H4−4ac
H4−4ac:H3−5ac
H3K4 me1:H3−5ac
H3K4 me3:H4−4ac
H4K20 me2:H3−5ac
H3K27 me2
H3K9 me2
H4K20 me3
H3K9 me3
DNA Meth. m5C
H3K27 me3
H3K4 me1
H3K4 me3
H4K20 me2
H2A.Z
H3−5ac
H4−4ac
0.00 0.25 0.50 0.75 1.00
$\hat{\pi}$
BRD9 − Reverse
H3K27 me3:H3K9 me3
H3K27 me3:DNA Meth. m5C
H2A.Z:H3−5ac
H3K4 me1:H4K20 me2
DNA Meth. m5C:H4K20 me2
H2A.Z:H4K20 me2
H3K27 me3:H3K4 me1
H2A.Z:H4−4ac
H3K4 me1:H2A.Z
H3K4 me1:H4−4ac
H3K4 me3:H3−5ac
H4K20 me2:H3K4 me3
H2A.Z:H3K4 me3
H4K20 me2:H4−4ac
H4−4ac:H3−5ac
H3K4 me1:H3−5ac
H3K4 me3:H4−4ac
H4K20 me2:H3−5ac
H3K27 me2
H3K9 me2
H4K20 me3
H3K9 me3
DNA Meth. m5C
H3K27 me3
H3K4 me1
H3K4 me3
H4K20 me2
H2A.Z
H3−5ac
H4−4ac
0.00 0.25 0.50 0.75 1.00
$\hat{\pi}$
BICRAL − Forward
DNA Meth. m5C:H4K20 me3
H3K27 me3:H3K4 me1
H3K27 me3:H4K20 me3
DNA Meth. m5C:H4K20 me2
H2A.Z:H3−5ac
H3K4 me1:H4K20 me2
H3K4 me1:H2A.Z
H3K4 me1:H3−5ac
H3K4 me3:H3−5ac
H2A.Z:H4−4ac
H3K4 me1:H4−4ac
H2A.Z:H3K4 me3
H3K9 me2
H3K27 me2
H4−4ac:H3−5ac
H3K4 me3:H4−4ac
H4K20 me2:H3−5ac
H4K20 me2:H4−4ac
H4K20 me3
H4K20 me2:H3K4 me3
H3K9 me3
H2A.Z:H4K20 me2
H3K27 me3
DNA Meth. m5C
H3K4 me1
H3−5ac
H3K4 me3
H4K20 me2
H2A.Z
H4−4ac
0.00 0.25 0.50 0.75 1.00
$\hat{\pi}$
BICRAL − Reverse
DNA Meth. m5C:H4K20 me3
H3K27 me3:H3K4 me1
H3K27 me3:H4K20 me3
DNA Meth. m5C:H4K20 me2
H2A.Z:H3−5ac
H3K4 me1:H4K20 me2
H3K4 me1:H2A.Z
H3K4 me1:H3−5ac
H3K4 me3:H3−5ac
H2A.Z:H4−4ac
H3K4 me1:H4−4ac
H2A.Z:H3K4 me3
H3K9 me2
H3K27 me2
H4−4ac:H3−5ac
H3K4 me3:H4−4ac
H4K20 me2:H3−5ac
H4K20 me2:H4−4ac
H4K20 me3
H4K20 me2:H3K4 me3
H3K9 me3
H2A.Z:H4K20 me2
H3K27 me3
DNA Meth. m5C
H3K4 me1
H3−5ac
H3K4 me3
H4K20 me2
H2A.Z
H4−4ac
0.00 0.25 0.50 0.75 1.00
$\hat{\pi}$
BICRA − Forward
DNA Meth. m5C:H3K9 me3
H3K4 me1:H2A.Z
H3K4 me1:H4K20 me2
H3K4 me1:H4−4ac
H3K4 me1:H3−5ac
DNA Meth. m5C:H4K20 me2
H3K4 me3:H3−5ac
H2A.Z:H3−5ac
H2A.Z:H4−4ac
H2A.Z:H3K4 me3
H2A.Z:H4K20 me2
H4K20 me2:H3K4 me3
H4K20 me2:H3−5ac
H4−4ac:H3−5ac
H3K9 me2
H3K4 me3:H4−4ac
H3K27 me2
H4K20 me3
H3K9 me3
H3K27 me3
H4K20 me2:H4−4ac
H3K4 me1
DNA Meth. m5C
H2A.Z
H3K4 me3
H3−5ac
H4K20 me2
H4−4ac
0.00 0.25 0.50 0.75 1.00
$\hat{\pi}$
BICRA − Reverse
DNA Meth. m5C:H3K9 me3
H3K4 me1:H2A.Z
H3K4 me1:H4K20 me2
H3K4 me1:H4−4ac
H3K4 me1:H3−5ac
DNA Meth. m5C:H4K20 me2
H3K4 me3:H3−5ac
H2A.Z:H3−5ac
H2A.Z:H4−4ac
H2A.Z:H3K4 me3
H2A.Z:H4K20 me2
H4K20 me2:H3K4 me3
H4K20 me2:H3−5ac
H4−4ac:H3−5ac
H3K9 me2
H3K4 me3:H4−4ac
H3K27 me2
H4K20 me3
H3K9 me3
H3K27 me3
H4K20 me2:H4−4ac
H3K4 me1
DNA Meth. m5C
H2A.Z
H3K4 me3
H3−5ac
H4K20 me2
H4−4ac
0.00 0.25 0.50 0.75 1.00
$\hat{\pi}$
SCML2 − Forward
H3K27 me2:DNA Meth. m5C
H3K9 me3:H4K20 me3
H3K9 me2:DNA Meth. m5C
H3K27 me3:H3K4 me1
H3K27 me3:H3K9 me3
H2A.Z:H3−5ac
H2A.Z:H4−4ac
H2A.Z:H4K20 me2
H3K4 me1:H3−5ac
H3K4 me3:H3−5ac
H3K4 me3:H4−4ac
DNA Meth. m5C:H3K9 me3
H3K4 me1:H2A.Z
H3K4 me1:H4−4ac
H3K27 me3:DNA Meth. m5C
H4−4ac:H3−5ac
H4K20 me2:H3K4 me3
H2A.Z:H3K4 me3
H4K20 me2:H3−5ac
DNA Meth. m5C:H4K20 me2
H3K27 me2
H3K9 me2
H4K20 me2:H4−4ac
H3K4 me3
H3K27 me3
H3K9 me3
H4K20 me3
H3−5ac
DNA Meth. m5C
H2A.Z
H3K4 me1
H4K20 me2
H4−4ac
0.00 0.25 0.50 0.75 1.00
$\hat{\pi}$
SCML2 − Reverse
H3K27 me2:DNA Meth. m5C
H3K9 me3:H4K20 me3
H3K9 me2:DNA Meth. m5C
H3K27 me3:H3K4 me1
H3K27 me3:H3K9 me3
H2A.Z:H3−5ac
H2A.Z:H4−4ac
H2A.Z:H4K20 me2
H3K4 me1:H3−5ac
H3K4 me3:H3−5ac
H3K4 me3:H4−4ac
DNA Meth. m5C:H3K9 me3
H3K4 me1:H2A.Z
H3K4 me1:H4−4ac
H3K27 me3:DNA Meth. m5C
H4−4ac:H3−5ac
H4K20 me2:H3K4 me3
H2A.Z:H3K4 me3
H4K20 me2:H3−5ac
DNA Meth. m5C:H4K20 me2
H3K27 me2
H3K9 me2
H4K20 me2:H4−4ac
H3K4 me3
H3K27 me3
H3K9 me3
H4K20 me3
H3−5ac
DNA Meth. m5C
H2A.Z
H3K4 me1
H4K20 me2
H4−4ac
0.00 0.25 0.50 0.75 1.00
$\hat{\pi}$

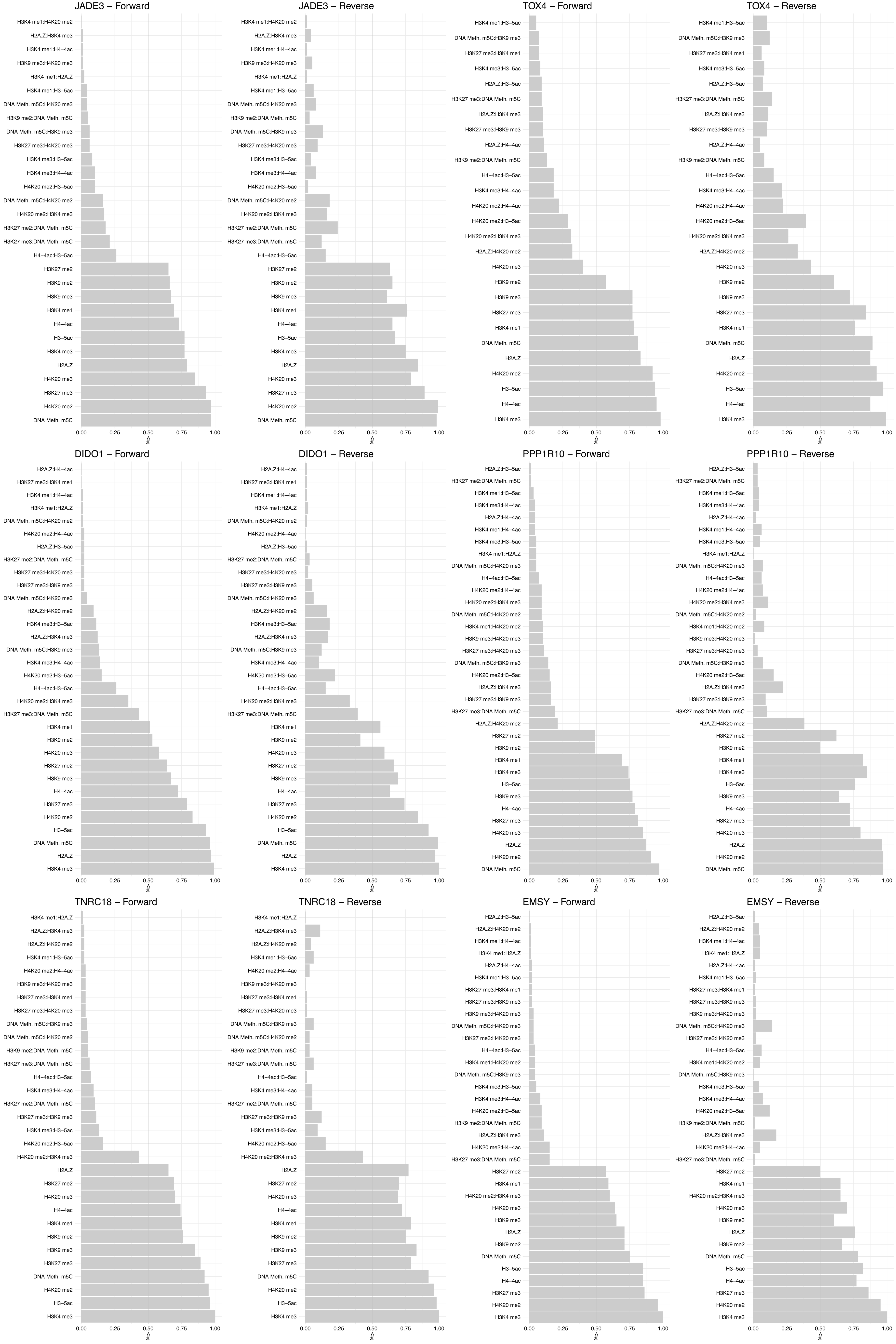
JADE3 – Forward
H3K4 me1:H4K20 me2
H2A.Z:H3K4 me3
H3K4 me1:H4–4ac
H3K9 me3:H4K20 me3
H3K4 me1:H2A.Z
H3K4 me1:H3–5ac
DNA Meth. m5C:H4K20 me3
H3K9 me2:DNA Meth. m5C
DNA Meth. m5C:H3K9 me3
H3K27 me3:H4K20 me3
H3K4 me3:H3–5ac
H3K4 me3:H4–4ac
H4K20 me2:H3–5ac
DNA Meth. m5C:H4K20 me2
H4K20 me2:H3K4 me3
H3K27 me2:DNA Meth. m5C
H3K27 me3:DNA Meth. m5C
H4–4ac:H3–5ac
H3K27 me2
H3K9 me2
H3K9 me3
H3K4 me1
H4–4ac
H3–5ac
H3K4 me3
H2A.Z
H4K20 me3
H3K27 me3
H4K20 me2
DNA Meth. m5C
0.00 0.25 0.50 0.75 1.00
π̂
JADE3 – Reverse
H3K4 me1:H4K20 me2
H2A.Z:H3K4 me3
H3K4 me1:H4–4ac
H3K9 me3:H4K20 me3
H3K4 me1:H2A.Z
H3K4 me1:H3–5ac
DNA Meth. m5C:H4K20 me3
H3K9 me2:DNA Meth. m5C
DNA Meth. m5C:H3K9 me3
H3K27 me3:H4K20 me3
H3K4 me3:H3–5ac
H3K4 me3:H4–4ac
H4K20 me2:H3–5ac
DNA Meth. m5C:H4K20 me2
H4K20 me2:H3K4 me3
H3K27 me2:DNA Meth. m5C
H3K27 me3:DNA Meth. m5C
H4–4ac:H3–5ac
H3K27 me2
H3K9 me2
H3K9 me3
H3K4 me1
H4–4ac
H3–5ac
H3K4 me3
H2A.Z
H4K20 me3
H3K27 me3
H4K20 me2
DNA Meth. m5C
0.00 0.25 0.50 0.75 1.00
π̂
TOX4 – Forward
H3K4 me1:H3–5ac
DNA Meth. m5C:H3K9 me3
H3K27 me3:H3K4 me1
H3K4 me3:H3–5ac
H2A.Z:H3–5ac
H3K27 me3:DNA Meth. m5C
H2A.Z:H3K4 me3
H3K27 me3:H3K9 me3
H2A.Z:H4–4ac
H3K9 me2:DNA Meth. m5C
H4–4ac:H3–5ac
H3K4 me3:H4–4ac
H4K20 me2:H4–4ac
H4K20 me2:H3–5ac
H4K20 me2:H3K4 me3
H2A.Z:H4K20 me2
H4K20 me3
H3K9 me2
H3K9 me3
H3K27 me3
H3K4 me1
DNA Meth. m5C
H2A.Z
H4K20 me2
H3–5ac
H4–4ac
H3K4 me3
0.00 0.25 0.50 0.75 1.00
π̂
TOX4 – Reverse
H3K4 me1:H3–5ac
DNA Meth. m5C:H3K9 me3
H3K27 me3:H3K4 me1
H3K4 me3:H3–5ac
H2A.Z:H3–5ac
H3K27 me3:DNA Meth. m5C
H2A.Z:H3K4 me3
H3K27 me3:H3K9 me3
H2A.Z:H4–4ac
H3K9 me2:DNA Meth. m5C
H4–4ac:H3–5ac
H3K4 me3:H4–4ac
H4K20 me2:H4–4ac
H4K20 me2:H3–5ac
H4K20 me2:H3K4 me3
H2A.Z:H4K20 me2
H4K20 me3
H3K9 me2
H3K9 me3
H3K27 me3
H3K4 me1
DNA Meth. m5C
H2A.Z
H4K20 me2
H3–5ac
H4–4ac
H3K4 me3
0.00 0.25 0.50 0.75 1.00
π̂
DIDO1 – Forward
H2A.Z:H4–4ac
H3K27 me3:H3K4 me1
H3K4 me1:H4–4ac
H3K4 me1:H2A.Z
DNA Meth. m5C:H4K20 me2
H4K20 me2:H4–4ac
H2A.Z:H3–5ac
H3K27 me2:DNA Meth. m5C
H3K27 me3:H4K20 me3
H3K27 me3:H3K9 me3
DNA Meth. m5C:H4K20 me3
H2A.Z:H4K20 me2
H3K4 me3:H3–5ac
H2A.Z:H3K4 me3
DNA Meth. m5C:H3K9 me3
H3K4 me3:H4–4ac
H4K20 me2:H3–5ac
H4–4ac:H3–5ac
H4K20 me2:H3K4 me3
H3K27 me3:DNA Meth. m5C
H3K4 me1
H3K9 me2
H4K20 me3
H3K27 me2
H3K9 me3
H4–4ac
H3K27 me3
H4K20 me2
H3–5ac
DNA Meth. m5C
H2A.Z
H3K4 me3
0.00 0.25 0.50 0.75 1.00
π̂
DIDO1 – Reverse
H2A.Z:H4–4ac
H3K27 me3:H3K4 me1
H3K4 me1:H4–4ac
H3K4 me1:H2A.Z
DNA Meth. m5C:H4K20 me2
H4K20 me2:H4–4ac
H2A.Z:H3–5ac
H3K27 me2:DNA Meth. m5C
H3K27 me3:H4K20 me3
H3K27 me3:H3K9 me3
DNA Meth. m5C:H4K20 me3
H2A.Z:H4K20 me2
H3K4 me3:H3–5ac
H2A.Z:H3K4 me3
DNA Meth. m5C:H3K9 me3
H3K4 me3:H4–4ac
H4K20 me2:H3–5ac
H4–4ac:H3–5ac
H4K20 me2:H3K4 me3
H3K27 me3:DNA Meth. m5C
H3K4 me1
H3K9 me2
H4K20 me3
H3K27 me2
H3K9 me3
H4–4ac
H3K27 me3
H4K20 me2
H3–5ac
DNA Meth. m5C
H2A.Z
H3K4 me3
0.00 0.25 0.50 0.75 1.00
π̂
PPP1R10 – Forward
H2A.Z:H3–5ac
H3K27 me2:DNA Meth. m5C
H3K4 me1:H3–5ac
H3K4 me3:H4–4ac
H2A.Z:H4–4ac
H3K4 me1:H4–4ac
H3K4 me3:H3–5ac
H3K4 me1:H2A.Z
DNA Meth. m5C:H4K20 me3
H4–4ac:H3–5ac
H4K20 me2:H4–4ac
H4K20 me2:H3K4 me3
DNA Meth. m5C:H4K20 me2
H3K4 me1:H4K20 me2
H3K9 me3:H4K20 me3
H3K27 me3:H4K20 me3
DNA Meth. m5C:H3K9 me3
H4K20 me2:H3–5ac
H2A.Z:H3K4 me3
H3K27 me3:H3K9 me3
H3K27 me3:DNA Meth. m5C
H2A.Z:H4K20 me2
H3K27 me2
H3K9 me2
H3K4 me1
H3K4 me3
H3–5ac
H3K9 me3
H4–4ac
H3K27 me3
H4K20 me3
H2A.Z
H4K20 me2
DNA Meth. m5C
0.00 0.25 0.50 0.75 1.00
π̂
PPP1R10 – Reverse
H2A.Z:H3–5ac
H3K27 me2:DNA Meth. m5C
H3K4 me1:H3–5ac
H3K4 me3:H4–4ac
H2A.Z:H4–4ac
H3K4 me1:H4–4ac
H3K4 me3:H3–5ac
H3K4 me1:H2A.Z
DNA Meth. m5C:H4K20 me3
H4–4ac:H3–5ac
H4K20 me2:H4–4ac
H4K20 me2:H3K4 me3
DNA Meth. m5C:H4K20 me2
H3K4 me1:H4K20 me2
H3K9 me3:H4K20 me3
H3K27 me3:H4K20 me3
DNA Meth. m5C:H3K9 me3
H4K20 me2:H3–5ac
H2A.Z:H3K4 me3
H3K27 me3:H3K9 me3
H3K27 me3:DNA Meth. m5C
H2A.Z:H4K20 me2
H3K27 me2
H3K9 me2
H3K4 me1
H3K4 me3
H3–5ac
H3K9 me3
H4–4ac
H3K27 me3
H4K20 me3
H2A.Z
H4K20 me2
DNA Meth. m5C
0.00 0.25 0.50 0.75 1.00
π̂
TNRC18 – Forward
H3K4 me1:H2A.Z
H2A.Z:H3K4 me3
H2A.Z:H4K20 me2
H3K4 me1:H3–5ac
H4K20 me2:H4–4ac
H3K9 me3:H4K20 me3
H3K27 me3:H3K4 me1
H3K27 me3:H4K20 me3
DNA Meth. m5C:H3K9 me3
DNA Meth. m5C:H4K20 me2
H3K9 me2:DNA Meth. m5C
H3K27 me3:DNA Meth. m5C
H4–4ac:H3–5ac
H3K4 me3:H4–4ac
H3K27 me2:DNA Meth. m5C
H3K27 me3:H3K9 me3
H3K4 me3:H3–5ac
H4K20 me2:H3–5ac
H4K20 me2:H3K4 me3
H2A.Z
H3K27 me2
H4K20 me3
H4–4ac
H3K4 me1
H3K9 me2
H3K9 me3
H3K27 me3
DNA Meth. m5C
H4K20 me2
H3–5ac
H3K4 me3
0.00 0.25 0.50 0.75 1.00
π̂
TNRC18 – Reverse
H3K4 me1:H2A.Z
H2A.Z:H3K4 me3
H2A.Z:H4K20 me2
H3K4 me1:H3–5ac
H4K20 me2:H4–4ac
H3K9 me3:H4K20 me3
H3K27 me3:H3K4 me1
H3K27 me3:H4K20 me3
DNA Meth. m5C:H3K9 me3
DNA Meth. m5C:H4K20 me2
H3K9 me2:DNA Meth. m5C
H3K27 me3:DNA Meth. m5C
H4–4ac:H3–5ac
H3K4 me3:H4–4ac
H3K27 me2:DNA Meth. m5C
H3K27 me3:H3K9 me3
H3K4 me3:H3–5ac
H4K20 me2:H3–5ac
H4K20 me2:H3K4 me3
H2A.Z
H3K27 me2
H4K20 me3
H4–4ac
H3K4 me1
H3K9 me2
H3K9 me3
H3K27 me3
DNA Meth. m5C
H4K20 me2
H3–5ac
H3K4 me3
0.00 0.25 0.50 0.75 1.00
π̂
EMSY – Forward
H2A.Z:H3–5ac
H2A.Z:H4K20 me2
H3K4 me1:H4–4ac
H3K4 me1:H2A.Z
H2A.Z:H4–4ac
H3K4 me1:H3–5ac
H3K27 me3:H3K4 me1
H3K27 me3:H3K9 me3
H3K9 me3:H4K20 me3
DNA Meth. m5C:H4K20 me3
H3K27 me3:H4K20 me3
H4–4ac:H3–5ac
H3K4 me1:H4K20 me2
DNA Meth. m5C:H3K9 me3
H3K4 me3:H3–5ac
H3K4 me3:H4–4ac
H4K20 me2:H3–5ac
H3K9 me2:DNA Meth. m5C
H2A.Z:H3K4 me3
H4K20 me2:H4–4ac
H3K27 me3:DNA Meth. m5C
H3K27 me2
H3K4 me1
H4K20 me2:H3K4 me3
H4K20 me3
H3K9 me3
H2A.Z
H3K9 me2
DNA Meth. m5C
H3–5ac
H4–4ac
H3K27 me3
H4K20 me2
H3K4 me3
0.00 0.25 0.50 0.75 1.00
π̂
EMSY – Reverse
H2A.Z:H3–5ac
H2A.Z:H4K20 me2
H3K4 me1:H4–4ac
H3K4 me1:H2A.Z
H2A.Z:H4–4ac
H3K4 me1:H3–5ac
H3K27 me3:H3K4 me1
H3K27 me3:H3K9 me3
H3K9 me3:H4K20 me3
DNA Meth. m5C:H4K20 me3
H3K27 me3:H4K20 me3
H4–4ac:H3–5ac
H3K4 me1:H4K20 me2
DNA Meth. m5C:H3K9 me3
H3K4 me3:H3–5ac
H3K4 me3:H4–4ac
H4K20 me2:H3–5ac
H3K9 me2:DNA Meth. m5C
H2A.Z:H3K4 me3
H4K20 me2:H4–4ac
H3K27 me3:DNA Meth. m5C
H3K27 me2
H3K4 me1
H4K20 me2:H3K4 me3
H4K20 me3
H3K9 me3
H2A.Z
H3K9 me2
DNA Meth. m5C
H3–5ac
H4–4ac
H3K27 me3
H4K20 me2
H3K4 me3
0.00 0.25 0.50 0.75 1.00
π̂

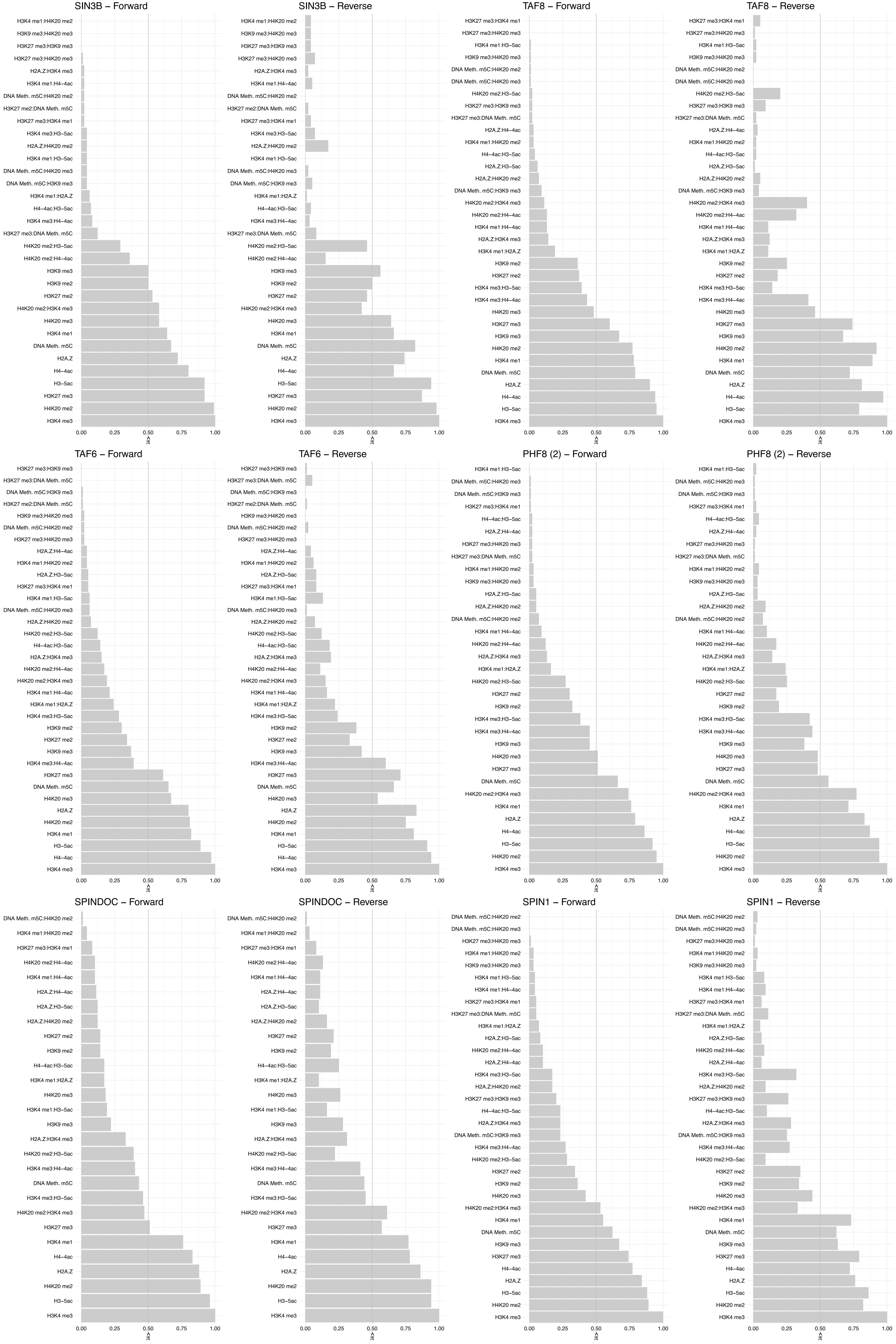
SIN3B – Forward
H3K4 me1:H4K20 me2
H3K9 me3:H4K20 me3
H3K27 me3:H3K9 me3
H3K27 me3:H4K20 me3
H2A.Z:H3K4 me3
H3K4 me1:H4−4ac
DNA Meth. m5C:H4K20 me2
H3K27 me2:DNA Meth. m5C
H3K27 me3:H3K4 me1
H3K4 me3:H3−5ac
H2A.Z:H4K20 me2
H3K4 me1:H3−5ac
DNA Meth. m5C:H4K20 me3
DNA Meth. m5C:H3K9 me3
H3K4 me1:H2A.Z
H4−4ac:H3−5ac
H3K4 me3:H4−4ac
H3K27 me3:DNA Meth. m5C
H4K20 me2:H3−5ac
H4K20 me2:H4−4ac
H3K9 me3
H3K9 me2
H3K27 me2
H4K20 me2:H3K4 me3
H4K20 me3
H3K4 me1
DNA Meth. m5C
H2A.Z
H4−4ac
H3−5ac
H3K27 me3
H4K20 me2
H3K4 me3
0.00
0.25
0.50
0.75
1.00
$\hat{\pi}$
SIN3B – Reverse
H3K4 me1:H4K20 me2
H3K9 me3:H4K20 me3
H3K27 me3:H3K9 me3
H3K27 me3:H4K20 me3
H2A.Z:H3K4 me3
H3K4 me1:H4−4ac
DNA Meth. m5C:H4K20 me2
H3K27 me2:DNA Meth. m5C
H3K27 me3:H3K4 me1
H3K4 me3:H3−5ac
H2A.Z:H4K20 me2
H3K4 me1:H3−5ac
DNA Meth. m5C:H4K20 me3
DNA Meth. m5C:H3K9 me3
H3K4 me1:H2A.Z
H4−4ac:H3−5ac
H3K4 me3:H4−4ac
H3K27 me3:DNA Meth. m5C
H4K20 me2:H3−5ac
H4K20 me2:H4−4ac
H3K9 me3
H3K9 me2
H3K27 me2
H4K20 me2:H3K4 me3
H4K20 me3
H3K4 me1
DNA Meth. m5C
H2A.Z
H4−4ac
H3−5ac
H3K27 me3
H4K20 me2
H3K4 me3
0.00
0.25
0.50
0.75
1.00
$\hat{\pi}$
TAF8 – Forward
H3K27 me3:H3K4 me1
H3K27 me3:H4K20 me3
H3K4 me1:H3−5ac
H3K9 me3:H4K20 me3
DNA Meth. m5C:H4K20 me2
DNA Meth. m5C:H4K20 me3
H4K20 me2:H3−5ac
H3K27 me3:H3K9 me3
H3K27 me3:DNA Meth. m5C
H2A.Z:H4−4ac
H3K4 me1:H4K20 me2
H4−4ac:H3−5ac
H2A.Z:H3−5ac
H2A.Z:H4K20 me2
DNA Meth. m5C:H3K9 me3
H4K20 me2:H3K4 me3
H4K20 me2:H4−4ac
H3K4 me1:H4−4ac
H2A.Z:H3K4 me3
H3K4 me1:H2A.Z
H3K9 me2
H3K27 me2
H3K4 me3:H3−5ac
H3K4 me3:H4−4ac
H4K20 me3
H3K27 me3
H3K9 me3
H4K20 me2
H3K4 me1
DNA Meth. m5C
H2A.Z
H4−4ac
H3−5ac
H3K4 me3
0.00
0.25
0.50
0.75
1.00
$\hat{\pi}$
TAF8 – Reverse
H3K27 me3:H3K4 me1
H3K27 me3:H4K20 me3
H3K4 me1:H3−5ac
H3K9 me3:H4K20 me3
DNA Meth. m5C:H4K20 me2
DNA Meth. m5C:H4K20 me3
H4K20 me2:H3−5ac
H3K27 me3:H3K9 me3
H3K27 me3:DNA Meth. m5C
H2A.Z:H4−4ac
H3K4 me1:H4K20 me2
H4−4ac:H3−5ac
H2A.Z:H3−5ac
H2A.Z:H4K20 me2
DNA Meth. m5C:H3K9 me3
H4K20 me2:H3K4 me3
H4K20 me2:H4−4ac
H3K4 me1:H4−4ac
H2A.Z:H3K4 me3
H3K4 me1:H2A.Z
H3K9 me2
H3K27 me2
H3K4 me3:H3−5ac
H3K4 me3:H4−4ac
H4K20 me3
H3K27 me3
H3K9 me3
H4K20 me2
H3K4 me1
DNA Meth. m5C
H2A.Z
H4−4ac
H3−5ac
H3K4 me3
0.00
0.25
0.50
0.75
1.00
$\hat{\pi}$
TAF6 – Forward
H3K27 me3:H3K9 me3
H3K27 me3:DNA Meth. m5C
DNA Meth. m5C:H3K9 me3
H3K27 me2:DNA Meth. m5C
H3K9 me3:H4K20 me3
DNA Meth. m5C:H4K20 me2
H3K27 me3:H4K20 me3
H2A.Z:H4−4ac
H3K4 me1:H4K20 me2
H2A.Z:H3−5ac
H3K27 me3:H3K4 me1
H3K4 me1:H3−5ac
DNA Meth. m5C:H4K20 me3
H2A.Z:H4K20 me2
H4K20 me2:H3−5ac
H4−4ac:H3−5ac
H2A.Z:H3K4 me3
H4K20 me2:H4−4ac
H4K20 me2:H3K4 me3
H3K4 me1:H4−4ac
H3K4 me1:H2A.Z
H3K4 me3:H3−5ac
H3K9 me2
H3K27 me2
H3K9 me3
H3K4 me3:H4−4ac
H3K27 me3
DNA Meth. m5C
H4K20 me3
H2A.Z
H4K20 me2
H3K4 me1
H3−5ac
H4−4ac
H3K4 me3
0.00
0.25
0.50
0.75
1.00
$\hat{\pi}$
TAF6 – Reverse
H3K27 me3:H3K9 me3
H3K27 me3:DNA Meth. m5C
DNA Meth. m5C:H3K9 me3
H3K27 me2:DNA Meth. m5C
H3K9 me3:H4K20 me3
DNA Meth. m5C:H4K20 me2
H3K27 me3:H4K20 me3
H2A.Z:H4−4ac
H3K4 me1:H4K20 me2
H2A.Z:H3−5ac
H3K27 me3:H3K4 me1
H3K4 me1:H3−5ac
DNA Meth. m5C:H4K20 me3
H2A.Z:H4K20 me2
H4K20 me2:H3−5ac
H4−4ac:H3−5ac
H2A.Z:H3K4 me3
H4K20 me2:H4−4ac
H4K20 me2:H3K4 me3
H3K4 me1:H4−4ac
H3K4 me1:H2A.Z
H3K4 me3:H3−5ac
H3K9 me2
H3K27 me2
H3K9 me3
H3K4 me3:H4−4ac
H3K27 me3
DNA Meth. m5C
H4K20 me3
H2A.Z
H4K20 me2
H3K4 me1
H3−5ac
H4−4ac
H3K4 me3
0.00
0.25
0.50
0.75
1.00
$\hat{\pi}$
PHF8 (2) – Forward
H3K4 me1:H3−5ac
DNA Meth. m5C:H4K20 me3
DNA Meth. m5C:H3K9 me3
H3K27 me3:H3K4 me1
H4−4ac:H3−5ac
H2A.Z:H4−4ac
H3K27 me3:H4K20 me3
H3K27 me3:DNA Meth. m5C
H3K4 me1:H4K20 me2
H3K9 me3:H4K20 me3
H2A.Z:H3−5ac
H2A.Z:H4K20 me2
DNA Meth. m5C:H4K20 me2
H3K4 me1:H4−4ac
H4K20 me2:H4−4ac
H2A.Z:H3K4 me3
H3K4 me1:H2A.Z
H4K20 me2:H3−5ac
H3K27 me2
H3K9 me2
H3K4 me3:H3−5ac
H3K4 me3:H4−4ac
H3K9 me3
H4K20 me3
H3K27 me3
DNA Meth. m5C
H4K20 me2:H3K4 me3
H3K4 me1
H2A.Z
H4−4ac
H3−5ac
H4K20 me2
H3K4 me3
0.00
0.25
0.50
0.75
1.00
$\hat{\pi}$
PHF8 (2) – Reverse
H3K4 me1:H3−5ac
DNA Meth. m5C:H4K20 me3
DNA Meth. m5C:H3K9 me3
H3K27 me3:H3K4 me1
H4−4ac:H3−5ac
H2A.Z:H4−4ac
H3K27 me3:H4K20 me3
H3K27 me3:DNA Meth. m5C
H3K4 me1:H4K20 me2
H3K9 me3:H4K20 me3
H2A.Z:H3−5ac
H2A.Z:H4K20 me2
DNA Meth. m5C:H4K20 me2
H3K4 me1:H4−4ac
H4K20 me2:H4−4ac
H2A.Z:H3K4 me3
H3K4 me1:H2A.Z
H4K20 me2:H3−5ac
H3K27 me2
H3K9 me2
H3K4 me3:H3−5ac
H3K4 me3:H4−4ac
H3K9 me3
H4K20 me3
H3K27 me3
DNA Meth. m5C
H4K20 me2:H3K4 me3
H3K4 me1
H2A.Z
H4−4ac
H3−5ac
H4K20 me2
H3K4 me3
0.00
0.25
0.50
0.75
1.00
$\hat{\pi}$
SPINDOC – Forward
DNA Meth. m5C:H4K20 me2
H3K4 me1:H4K20 me2
H3K27 me3:H3K4 me1
H4K20 me2:H4−4ac
H3K4 me1:H4−4ac
H2A.Z:H4−4ac
H2A.Z:H3−5ac
H2A.Z:H4K20 me2
H3K27 me2
H3K9 me2
H4−4ac:H3−5ac
H3K4 me1:H2A.Z
H4K20 me3
H3K4 me1:H3−5ac
H3K9 me3
H2A.Z:H3K4 me3
H4K20 me2:H3−5ac
H3K4 me3:H4−4ac
DNA Meth. m5C
H3K4 me3:H3−5ac
H4K20 me2:H3K4 me3
H3K27 me3
H3K4 me1
H4−4ac
H2A.Z
H4K20 me2
H3−5ac
H3K4 me3
0.00
0.25
0.50
0.75
1.00
$\hat{\pi}$
SPINDOC – Reverse
DNA Meth. m5C:H4K20 me2
H3K4 me1:H4K20 me2
H3K27 me3:H3K4 me1
H4K20 me2:H4−4ac
H3K4 me1:H4−4ac
H2A.Z:H4−4ac
H2A.Z:H3−5ac
H2A.Z:H4K20 me2
H3K27 me2
H3K9 me2
H4−4ac:H3−5ac
H3K4 me1:H2A.Z
H4K20 me3
H3K4 me1:H3−5ac
H3K9 me3
H2A.Z:H3K4 me3
H4K20 me2:H3−5ac
H3K4 me3:H4−4ac
DNA Meth. m5C
H3K4 me3:H3−5ac
H4K20 me2:H3K4 me3
H3K27 me3
H3K4 me1
H4−4ac
H2A.Z
H4K20 me2
H3−5ac
H3K4 me3
0.00
0.25
0.50
0.75
1.00
$\hat{\pi}$
SPIN1 – Forward
DNA Meth. m5C:H4K20 me2
DNA Meth. m5C:H4K20 me3
H3K27 me3:H4K20 me3
H3K4 me1:H4K20 me2
H3K9 me3:H4K20 me3
H3K4 me1:H3−5ac
H3K4 me1:H4−4ac
H3K27 me3:H3K4 me1
H3K27 me3:DNA Meth. m5C
H3K4 me1:H2A.Z
H2A.Z:H3−5ac
H4K20 me2:H4−4ac
H2A.Z:H4−4ac
H3K4 me3:H3−5ac
H2A.Z:H4K20 me2
H3K27 me3:H3K9 me3
H4−4ac:H3−5ac
H2A.Z:H3K4 me3
DNA Meth. m5C:H3K9 me3
H3K4 me3:H4−4ac
H4K20 me2:H3−5ac
H3K27 me2
H3K9 me2
H4K20 me3
H4K20 me2:H3K4 me3
H3K4 me1
DNA Meth. m5C
H3K9 me3
H3K27 me3
H4−4ac
H2A.Z
H3−5ac
H4K20 me2
H3K4 me3
0.00
0.25
0.50
0.75
1.00
$\hat{\pi}$
SPIN1 – Reverse
DNA Meth. m5C:H4K20 me2
DNA Meth. m5C:H4K20 me3
H3K27 me3:H4K20 me3
H3K4 me1:H4K20 me2
H3K9 me3:H4K20 me3
H3K4 me1:H3−5ac
H3K4 me1:H4−4ac
H3K27 me3:H3K4 me1
H3K27 me3:DNA Meth. m5C
H3K4 me1:H2A.Z
H2A.Z:H3−5ac
H4K20 me2:H4−4ac
H2A.Z:H4−4ac
H3K4 me3:H3−5ac
H2A.Z:H4K20 me2
H3K27 me3:H3K9 me3
H4−4ac:H3−5ac
H2A.Z:H3K4 me3
DNA Meth. m5C:H3K9 me3
H3K4 me3:H4−4ac
H4K20 me2:H3−5ac
H3K27 me2
H3K9 me2
H4K20 me3
H4K20 me2:H3K4 me3
H3K4 me1
DNA Meth. m5C
H3K9 me3
H3K27 me3
H4−4ac
H2A.Z
H3−5ac
H4K20 me2
H3K4 me3
0.00
0.25
0.50
0.75
1.00
$\hat{\pi}$

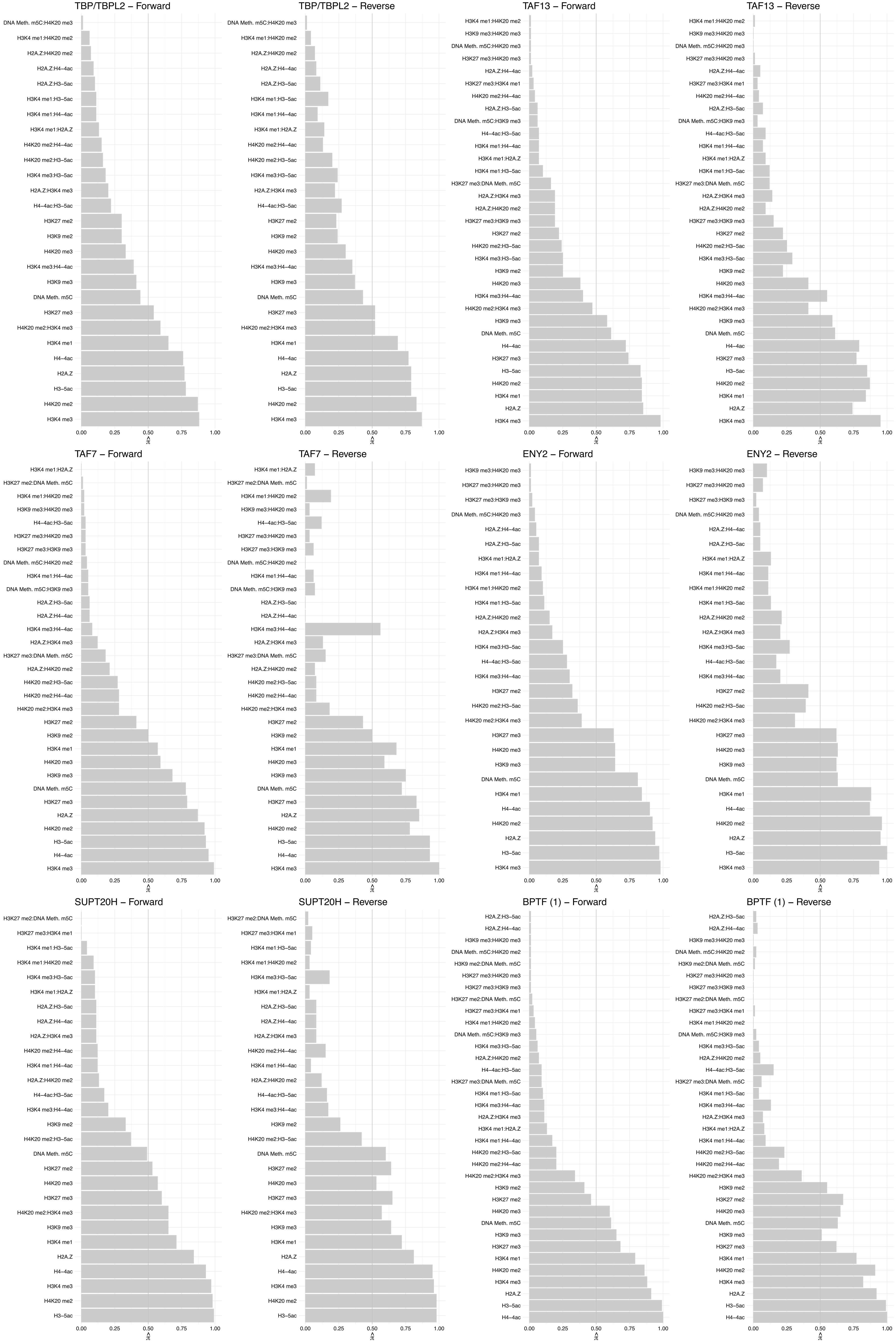

TBP/TBPL2 – Forward
TBP/TBPL2 – Reverse
DNA Meth. m5C:H4K20 me3
H3K4 me1:H4K20 me2
H2A.Z:H4K20 me2
H2A.Z:H4–4ac
H2A.Z:H3–5ac
H3K4 me1:H3–5ac
H3K4 me1:H4–4ac
H3K4 me1:H2A.Z
H4K20 me2:H4–4ac
H4K20 me2:H3–5ac
H3K4 me3:H3–5ac
H2A.Z:H3K4 me3
H4–4ac:H3–5ac
H3K27 me2
H3K9 me2
H4K20 me3
H3K4 me3:H4–4ac
H3K9 me3
DNA Meth. m5C
H3K27 me3
H4K20 me2:H3K4 me3
H3K4 me1
H4–4ac
H2A.Z
H3–5ac
H4K20 me2
H3K4 me3
TAF13 – Forward
TAF13 – Reverse
H3K4 me1:H4K20 me2
H3K9 me3:H4K20 me3
DNA Meth. m5C:H4K20 me3
H3K27 me3:H4K20 me3
H2A.Z:H4–4ac
H3K27 me3:H3K4 me1
H4K20 me2:H4–4ac
H2A.Z:H3–5ac
DNA Meth. m5C:H3K9 me3
H4–4ac:H3–5ac
H3K4 me1:H4–4ac
H3K4 me1:H2A.Z
H3K4 me1:H3–5ac
H3K27 me3:DNA Meth. m5C
H2A.Z:H3K4 me3
H2A.Z:H4K20 me2
H3K27 me3:H3K9 me3
H3K27 me2
H4K20 me2:H3–5ac
H3K4 me3:H3–5ac
H3K9 me2
H4K20 me3
H3K4 me3:H4–4ac
H4K20 me2:H3K4 me3
H3K9 me3
DNA Meth. m5C
H4–4ac
H3K27 me3
H3–5ac
H4K20 me2
H3K4 me1
H2A.Z
H3K4 me3
TAF7 – Forward
TAF7 – Reverse
H3K4 me1:H2A.Z
H3K27 me2:DNA Meth. m5C
H3K4 me1:H4K20 me2
H3K9 me3:H4K20 me3
H4–4ac:H3–5ac
H3K27 me3:H4K20 me3
H3K27 me3:H3K9 me3
DNA Meth. m5C:H4K20 me2
H3K4 me1:H4–4ac
DNA Meth. m5C:H3K9 me3
H2A.Z:H3–5ac
H2A.Z:H4–4ac
H3K4 me3:H4–4ac
H2A.Z:H3K4 me3
H3K27 me3:DNA Meth. m5C
H2A.Z:H4K20 me2
H4K20 me2:H3–5ac
H4K20 me2:H4–4ac
H4K20 me2:H3K4 me3
H3K27 me2
H3K9 me2
H3K4 me1
H4K20 me3
H3K9 me3
DNA Meth. m5C
H3K27 me3
H2A.Z
H4K20 me2
H3–5ac
H4–4ac
H3K4 me3
ENY2 – Forward
ENY2 – Reverse
H3K9 me3:H4K20 me3
H3K27 me3:H4K20 me3
H3K27 me3:H3K9 me3
DNA Meth. m5C:H4K20 me3
H2A.Z:H4–4ac
H2A.Z:H3–5ac
H3K4 me1:H2A.Z
H3K4 me1:H4–4ac
H3K4 me1:H4K20 me2
H3K4 me1:H3–5ac
H2A.Z:H4K20 me2
H2A.Z:H3K4 me3
H3K4 me3:H3–5ac
H4–4ac:H3–5ac
H3K4 me3:H4–4ac
H3K27 me2
H4K20 me2:H3–5ac
H4K20 me2:H3K4 me3
H3K27 me3
H4K20 me3
H3K9 me3
DNA Meth. m5C
H3K4 me1
H4–4ac
H4K20 me2
H2A.Z
H3–5ac
H3K4 me3
SUPT20H – Forward
SUPT20H – Reverse
H3K27 me2:DNA Meth. m5C
H3K27 me3:H3K4 me1
H3K4 me1:H3–5ac
H3K4 me1:H4K20 me2
H3K4 me3:H3–5ac
H3K4 me1:H2A.Z
H2A.Z:H3–5ac
H2A.Z:H4–4ac
H2A.Z:H3K4 me3
H4K20 me2:H4–4ac
H3K4 me1:H4–4ac
H2A.Z:H4K20 me2
H4–4ac:H3–5ac
H3K4 me3:H4–4ac
H3K9 me2
H4K20 me2:H3–5ac
DNA Meth. m5C
H3K27 me2
H4K20 me3
H3K27 me3
H4K20 me2:H3K4 me3
H3K9 me3
H3K4 me1
H2A.Z
H4–4ac
H3K4 me3
H4K20 me2
H3–5ac
BPTF (1) – Forward
BPTF (1) – Reverse
H2A.Z:H3–5ac
H2A.Z:H4–4ac
H3K9 me3:H4K20 me3
DNA Meth. m5C:H4K20 me2
H3K9 me2:DNA Meth. m5C
H3K27 me3:H4K20 me3
H3K27 me3:H3K9 me3
H3K27 me2:DNA Meth. m5C
H3K27 me3:H3K4 me1
H3K4 me1:H4K20 me2
DNA Meth. m5C:H3K9 me3
H3K4 me3:H3–5ac
H2A.Z:H4K20 me2
H4–4ac:H3–5ac
H3K27 me3:DNA Meth. m5C
H3K4 me1:H3–5ac
H3K4 me3:H4–4ac
H2A.Z:H3K4 me3
H3K4 me1:H2A.Z
H3K4 me1:H4–4ac
H4K20 me2:H3–5ac
H4K20 me2:H4–4ac
H4K20 me2:H3K4 me3
H3K9 me2
H3K27 me2
H4K20 me3
DNA Meth. m5C
H3K9 me3
H3K27 me3
H3K4 me1
H4K20 me2
H3K4 me3
H2A.Z
H3–5ac
H4–4ac
0.00 0.25 0.50 0.75 1.00
π̂

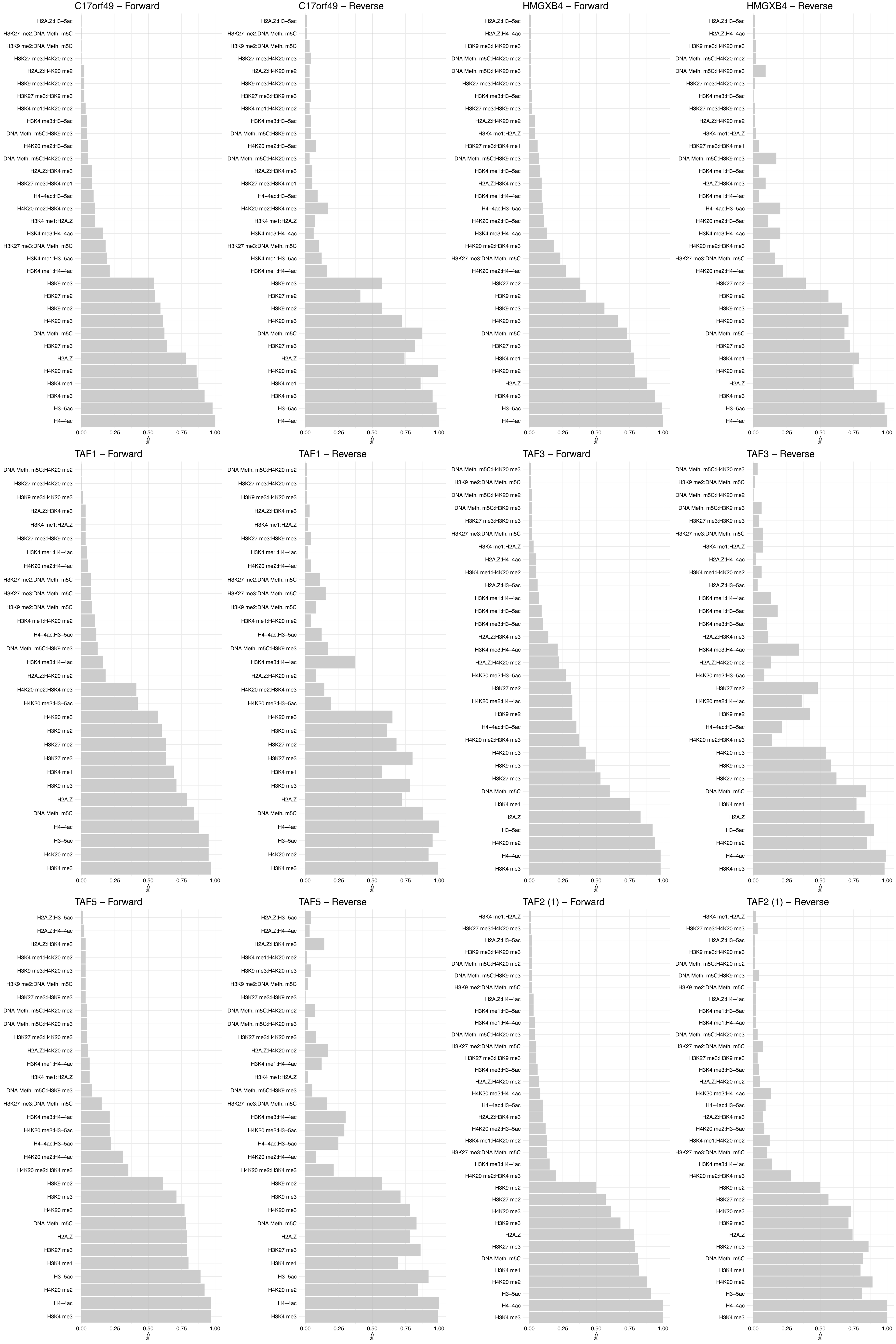
C17orf49 – Forward
H2A.Z:H3–5ac
H3K27 me2:DNA Meth. m5C
H3K9 me2:DNA Meth. m5C
H3K27 me3:H4K20 me3
H2A.Z:H4K20 me2
H3K9 me3:H4K20 me3
H3K27 me3:H3K9 me3
H3K4 me1:H4K20 me2
H3K4 me3:H3–5ac
DNA Meth. m5C:H3K9 me3
H4K20 me2:H3–5ac
DNA Meth. m5C:H4K20 me3
H2A.Z:H3K4 me3
H3K27 me3:H3K4 me1
H4–4ac:H3–5ac
H4K20 me2:H3K4 me3
H3K4 me1:H2A.Z
H3K4 me3:H4–4ac
H3K27 me3:DNA Meth. m5C
H3K4 me1:H3–5ac
H3K4 me1:H4–4ac
H3K9 me3
H3K27 me2
H3K9 me2
H4K20 me3
DNA Meth. m5C
H3K27 me3
H2A.Z
H4K20 me2
H3K4 me1
H3K4 me3
H3–5ac
H4–4ac
0.00
0.25
0.50
0.75
1.00
$\hat{\pi}$
C17orf49 – Reverse
H2A.Z:H3–5ac
H3K27 me2:DNA Meth. m5C
H3K9 me2:DNA Meth. m5C
H3K27 me3:H4K20 me3
H2A.Z:H4K20 me2
H3K9 me3:H4K20 me3
H3K27 me3:H3K9 me3
H3K4 me1:H4K20 me2
H3K4 me3:H3–5ac
DNA Meth. m5C:H3K9 me3
H4K20 me2:H3–5ac
DNA Meth. m5C:H4K20 me3
H2A.Z:H3K4 me3
H3K27 me3:H3K4 me1
H4–4ac:H3–5ac
H4K20 me2:H3K4 me3
H3K4 me1:H2A.Z
H3K4 me3:H4–4ac
H3K27 me3:DNA Meth. m5C
H3K4 me1:H3–5ac
H3K4 me1:H4–4ac
H3K9 me3
H3K27 me2
H3K9 me2
H4K20 me3
DNA Meth. m5C
H3K27 me3
H2A.Z
H4K20 me2
H3K4 me1
H3K4 me3
H3–5ac
H4–4ac
0.00
0.25
0.50
0.75
1.00
$\hat{\pi}$
HMGXB4 – Forward
H2A.Z:H3–5ac
H2A.Z:H4–4ac
H3K9 me3:H4K20 me3
DNA Meth. m5C:H4K20 me2
DNA Meth. m5C:H4K20 me3
H3K27 me3:H4K20 me3
H3K4 me3:H3–5ac
H3K27 me3:H3K9 me3
H2A.Z:H4K20 me2
H3K4 me1:H2A.Z
H3K27 me3:H3K4 me1
DNA Meth. m5C:H3K9 me3
H3K4 me1:H3–5ac
H2A.Z:H3K4 me3
H3K4 me1:H4–4ac
H4–4ac:H3–5ac
H4K20 me2:H3–5ac
H3K4 me3:H4–4ac
H4K20 me2:H3K4 me3
H3K27 me3:DNA Meth. m5C
H4K20 me2:H4–4ac
H3K27 me2
H3K9 me2
H3K9 me3
H4K20 me3
DNA Meth. m5C
H3K27 me3
H3K4 me1
H4K20 me2
H2A.Z
H3K4 me3
H3–5ac
H4–4ac
0.00
0.25
0.50
0.75
1.00
$\hat{\pi}$
HMGXB4 – Reverse
H2A.Z:H3–5ac
H2A.Z:H4–4ac
H3K9 me3:H4K20 me3
DNA Meth. m5C:H4K20 me2
DNA Meth. m5C:H4K20 me3
H3K27 me3:H4K20 me3
H3K4 me3:H3–5ac
H3K27 me3:H3K9 me3
H2A.Z:H4K20 me2
H3K4 me1:H2A.Z
H3K27 me3:H3K4 me1
DNA Meth. m5C:H3K9 me3
H3K4 me1:H3–5ac
H2A.Z:H3K4 me3
H3K4 me1:H4–4ac
H4–4ac:H3–5ac
H4K20 me2:H3–5ac
H3K4 me3:H4–4ac
H4K20 me2:H3K4 me3
H3K27 me3:DNA Meth. m5C
H4K20 me2:H4–4ac
H3K27 me2
H3K9 me2
H3K9 me3
H4K20 me3
DNA Meth. m5C
H3K27 me3
H3K4 me1
H4K20 me2
H2A.Z
H3K4 me3
H3–5ac
H4–4ac
0.00
0.25
0.50
0.75
1.00
$\hat{\pi}$
TAF1 – Forward
DNA Meth. m5C:H4K20 me2
H3K27 me3:H4K20 me3
H3K9 me3:H4K20 me3
H2A.Z:H3K4 me3
H3K4 me1:H2A.Z
H3K27 me3:H3K9 me3
H3K4 me1:H4–4ac
H4K20 me2:H4–4ac
H3K27 me2:DNA Meth. m5C
H3K27 me3:DNA Meth. m5C
H3K9 me2:DNA Meth. m5C
H3K4 me1:H4K20 me2
H4–4ac:H3–5ac
DNA Meth. m5C:H3K9 me3
H3K4 me3:H4–4ac
H2A.Z:H4K20 me2
H4K20 me2:H3K4 me3
H4K20 me2:H3–5ac
H4K20 me3
H3K9 me2
H3K27 me2
H3K27 me3
H3K4 me1
H3K9 me3
H2A.Z
DNA Meth. m5C
H4–4ac
H3–5ac
H4K20 me2
H3K4 me3
0.00
0.25
0.50
0.75
1.00
$\hat{\pi}$
TAF1 – Reverse
DNA Meth. m5C:H4K20 me2
H3K27 me3:H4K20 me3
H3K9 me3:H4K20 me3
H2A.Z:H3K4 me3
H3K4 me1:H2A.Z
H3K27 me3:H3K9 me3
H3K4 me1:H4–4ac
H4K20 me2:H4–4ac
H3K27 me2:DNA Meth. m5C
H3K27 me3:DNA Meth. m5C
H3K9 me2:DNA Meth. m5C
H3K4 me1:H4K20 me2
H4–4ac:H3–5ac
DNA Meth. m5C:H3K9 me3
H3K4 me3:H4–4ac
H2A.Z:H4K20 me2
H4K20 me2:H3K4 me3
H4K20 me2:H3–5ac
H4K20 me3
H3K9 me2
H3K27 me2
H3K27 me3
H3K4 me1
H3K9 me3
H2A.Z
DNA Meth. m5C
H4–4ac
H3–5ac
H4K20 me2
H3K4 me3
0.00
0.25
0.50
0.75
1.00
$\hat{\pi}$
TAF3 – Forward
DNA Meth. m5C:H4K20 me3
H3K9 me2:DNA Meth. m5C
DNA Meth. m5C:H4K20 me2
DNA Meth. m5C:H3K9 me3
H3K27 me3:H3K9 me3
H3K27 me3:DNA Meth. m5C
H3K4 me1:H2A.Z
H2A.Z:H4–4ac
H3K4 me1:H4K20 me2
H2A.Z:H3–5ac
H3K4 me1:H4–4ac
H3K4 me1:H3–5ac
H3K4 me3:H3–5ac
H2A.Z:H3K4 me3
H3K4 me3:H4–4ac
H2A.Z:H4K20 me2
H4K20 me2:H3–5ac
H3K27 me2
H4K20 me2:H4–4ac
H3K9 me2
H4–4ac:H3–5ac
H4K20 me2:H3K4 me3
H4K20 me3
H3K9 me3
H3K27 me3
DNA Meth. m5C
H3K4 me1
H2A.Z
H3–5ac
H4K20 me2
H4–4ac
H3K4 me3
0.00
0.25
0.50
0.75
1.00
$\hat{\pi}$
TAF3 – Reverse
DNA Meth. m5C:H4K20 me3
H3K9 me2:DNA Meth. m5C
DNA Meth. m5C:H4K20 me2
DNA Meth. m5C:H3K9 me3
H3K27 me3:H3K9 me3
H3K27 me3:DNA Meth. m5C
H3K4 me1:H2A.Z
H2A.Z:H4–4ac
H3K4 me1:H4K20 me2
H2A.Z:H3–5ac
H3K4 me1:H4–4ac
H3K4 me1:H3–5ac
H3K4 me3:H3–5ac
H2A.Z:H3K4 me3
H3K4 me3:H4–4ac
H2A.Z:H4K20 me2
H4K20 me2:H3–5ac
H3K27 me2
H4K20 me2:H4–4ac
H3K9 me2
H4–4ac:H3–5ac
H4K20 me2:H3K4 me3
H4K20 me3
H3K9 me3
H3K27 me3
DNA Meth. m5C
H3K4 me1
H2A.Z
H3–5ac
H4K20 me2
H4–4ac
H3K4 me3
0.00
0.25
0.50
0.75
1.00
$\hat{\pi}$
TAF5 – Forward
H2A.Z:H3–5ac
H2A.Z:H4–4ac
H2A.Z:H3K4 me3
H3K4 me1:H4K20 me2
H3K9 me3:H4K20 me3
H3K9 me2:DNA Meth. m5C
H3K27 me3:H3K9 me3
DNA Meth. m5C:H4K20 me2
DNA Meth. m5C:H4K20 me3
H3K27 me3:H4K20 me3
H2A.Z:H4K20 me2
H3K4 me1:H4–4ac
H3K4 me1:H2A.Z
DNA Meth. m5C:H3K9 me3
H3K27 me3:DNA Meth. m5C
H3K4 me3:H4–4ac
H4K20 me2:H3–5ac
H4–4ac:H3–5ac
H4K20 me2:H4–4ac
H4K20 me2:H3K4 me3
H3K9 me2
H3K9 me3
H4K20 me3
DNA Meth. m5C
H2A.Z
H3K27 me3
H3K4 me1
H3–5ac
H4K20 me2
H4–4ac
H3K4 me3
0.00
0.25
0.50
0.75
1.00
$\hat{\pi}$
TAF5 – Reverse
H2A.Z:H3–5ac
H2A.Z:H4–4ac
H2A.Z:H3K4 me3
H3K4 me1:H4K20 me2
H3K9 me3:H4K20 me3
H3K9 me2:DNA Meth. m5C
H3K27 me3:H3K9 me3
DNA Meth. m5C:H4K20 me2
DNA Meth. m5C:H4K20 me3
H3K27 me3:H4K20 me3
H2A.Z:H4K20 me2
H3K4 me1:H4–4ac
H3K4 me1:H2A.Z
DNA Meth. m5C:H3K9 me3
H3K27 me3:DNA Meth. m5C
H3K4 me3:H4–4ac
H4K20 me2:H3–5ac
H4–4ac:H3–5ac
H4K20 me2:H4–4ac
H4K20 me2:H3K4 me3
H3K9 me2
H3K9 me3
H4K20 me3
DNA Meth. m5C
H2A.Z
H3K27 me3
H3K4 me1
H3–5ac
H4K20 me2
H4–4ac
H3K4 me3
0.00
0.25
0.50
0.75
1.00
$\hat{\pi}$
TAF2 (1) – Forward
H3K4 me1:H2A.Z
H3K27 me3:H4K20 me3
H2A.Z:H3–5ac
H3K9 me3:H4K20 me3
DNA Meth. m5C:H4K20 me2
DNA Meth. m5C:H3K9 me3
H3K9 me2:DNA Meth. m5C
H2A.Z:H4–4ac
H3K4 me1:H3–5ac
H3K4 me1:H4–4ac
DNA Meth. m5C:H4K20 me3
H3K27 me2:DNA Meth. m5C
H3K27 me3:H3K9 me3
H3K4 me3:H3–5ac
H2A.Z:H4K20 me2
H4K20 me2:H4–4ac
H4–4ac:H3–5ac
H2A.Z:H3K4 me3
H4K20 me2:H3–5ac
H3K4 me1:H4K20 me2
H3K27 me3:DNA Meth. m5C
H3K4 me3:H4–4ac
H4K20 me2:H3K4 me3
H3K9 me2
H3K27 me2
H4K20 me3
H3K9 me3
H2A.Z
H3K27 me3
DNA Meth. m5C
H3K4 me1
H4K20 me2
H3–5ac
H4–4ac
H3K4 me3
0.00
0.25
0.50
0.75
1.00
$\hat{\pi}$
TAF2 (1) – Reverse
H3K4 me1:H2A.Z
H3K27 me3:H4K20 me3
H2A.Z:H3–5ac
H3K9 me3:H4K20 me3
DNA Meth. m5C:H4K20 me2
DNA Meth. m5C:H3K9 me3
H3K9 me2:DNA Meth. m5C
H2A.Z:H4–4ac
H3K4 me1:H3–5ac
H3K4 me1:H4–4ac
DNA Meth. m5C:H4K20 me3
H3K27 me2:DNA Meth. m5C
H3K27 me3:H3K9 me3
H3K4 me3:H3–5ac
H2A.Z:H4K20 me2
H4K20 me2:H4–4ac
H4–4ac:H3–5ac
H2A.Z:H3K4 me3
H4K20 me2:H3–5ac
H3K4 me1:H4K20 me2
H3K27 me3:DNA Meth. m5C
H3K4 me3:H4–4ac
H4K20 me2:H3K4 me3
H3K9 me2
H3K27 me2
H4K20 me3
H3K9 me3
H2A.Z
H3K27 me3
DNA Meth. m5C
H3K4 me1
H4K20 me2
H3–5ac
H4–4ac
H3K4 me3
0.00
0.25
0.50
0.75
1.00
$\hat{\pi}$

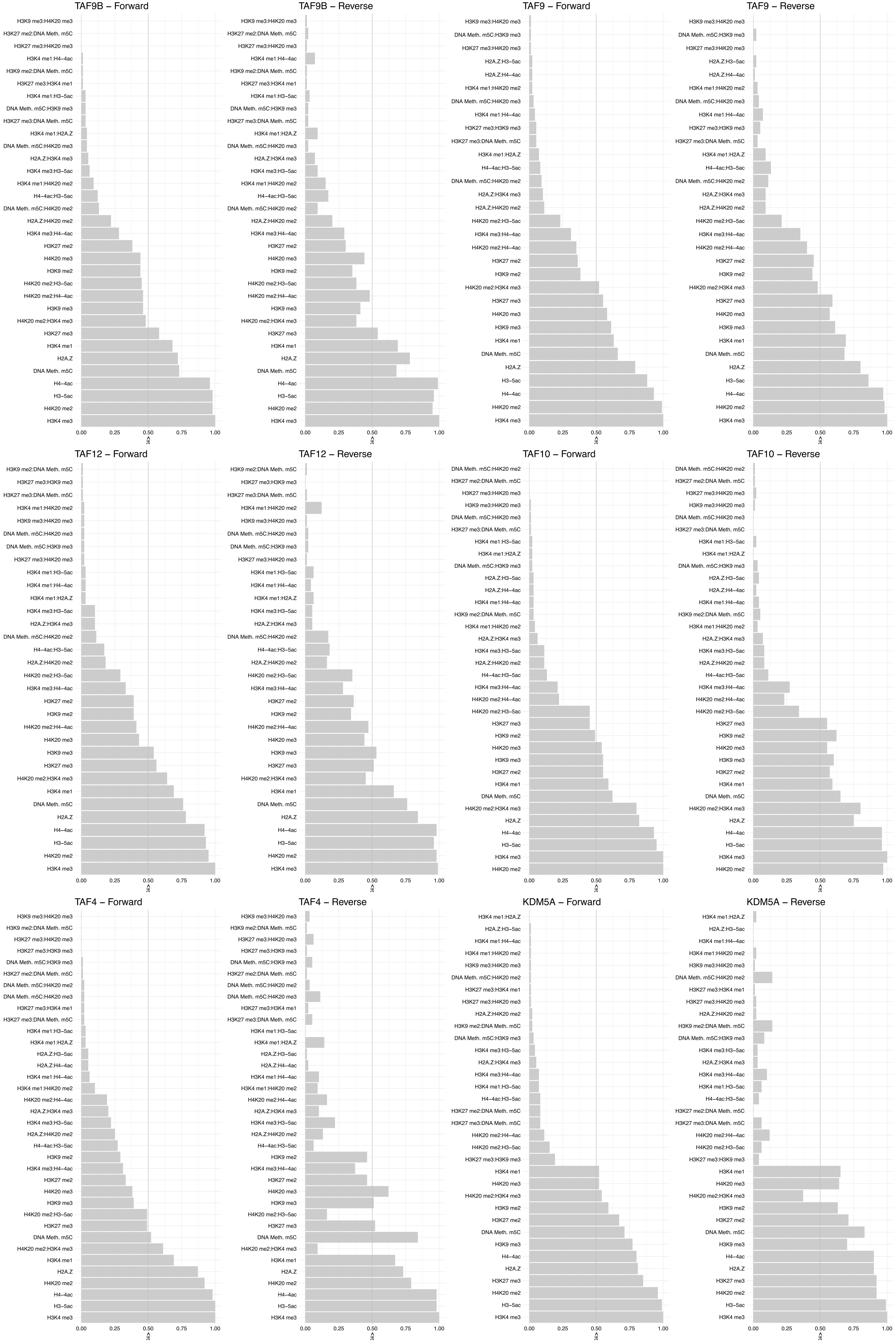
TAF9B – Forward
H3K9 me3:H4K20 me3
H3K27 me2:DNA Meth. m5C
H3K27 me3:H4K20 me3
H3K4 me1:H4−4ac
H3K9 me2:DNA Meth. m5C
H3K27 me3:H3K4 me1
H3K4 me1:H3−5ac
DNA Meth. m5C:H3K9 me3
H3K27 me3:DNA Meth. m5C
H3K4 me1:H2A.Z
DNA Meth. m5C:H4K20 me3
H2A.Z:H3K4 me3
H3K4 me3:H3−5ac
H3K4 me1:H4K20 me2
H4−4ac:H3−5ac
DNA Meth. m5C:H4K20 me2
H2A.Z:H4K20 me2
H3K4 me3:H4−4ac
H3K27 me2
H4K20 me3
H3K9 me2
H4K20 me2:H3−5ac
H4K20 me2:H4−4ac
H3K9 me3
H4K20 me2:H3K4 me3
H3K27 me3
H3K4 me1
H2A.Z
DNA Meth. m5C
H4−4ac
H3−5ac
H4K20 me2
H3K4 me3
0.00 0.25 0.50 0.75 1.00
$\hat{\pi}$
TAF9B – Reverse
H3K9 me3:H4K20 me3
H3K27 me2:DNA Meth. m5C
H3K27 me3:H4K20 me3
H3K4 me1:H4−4ac
H3K9 me2:DNA Meth. m5C
H3K27 me3:H3K4 me1
H3K4 me1:H3−5ac
DNA Meth. m5C:H3K9 me3
H3K27 me3:DNA Meth. m5C
H3K4 me1:H2A.Z
DNA Meth. m5C:H4K20 me3
H2A.Z:H3K4 me3
H3K4 me3:H3−5ac
H3K4 me1:H4K20 me2
H4−4ac:H3−5ac
DNA Meth. m5C:H4K20 me2
H2A.Z:H4K20 me2
H3K4 me3:H4−4ac
H3K27 me2
H4K20 me3
H3K9 me2
H4K20 me2:H3−5ac
H4K20 me2:H4−4ac
H3K9 me3
H4K20 me2:H3K4 me3
H3K27 me3
H3K4 me1
H2A.Z
DNA Meth. m5C
H4−4ac
H3−5ac
H4K20 me2
H3K4 me3
0.00 0.25 0.50 0.75 1.00
$\hat{\pi}$
TAF9 – Forward
H3K9 me3:H4K20 me3
DNA Meth. m5C:H3K9 me3
H3K27 me3:H4K20 me3
H2A.Z:H3−5ac
H2A.Z:H4−4ac
H3K4 me1:H4K20 me2
DNA Meth. m5C:H4K20 me3
H3K4 me1:H4−4ac
H3K27 me3:H3K9 me3
H3K27 me3:DNA Meth. m5C
H3K4 me1:H2A.Z
H4−4ac:H3−5ac
DNA Meth. m5C:H4K20 me2
H2A.Z:H3K4 me3
H2A.Z:H4K20 me2
H4K20 me2:H3−5ac
H3K4 me3:H4−4ac
H4K20 me2:H4−4ac
H3K27 me2
H3K9 me2
H4K20 me2:H3K4 me3
H3K27 me3
H4K20 me3
H3K9 me3
H3K4 me1
DNA Meth. m5C
H2A.Z
H3−5ac
H4−4ac
H4K20 me2
H3K4 me3
0.00 0.25 0.50 0.75 1.00
$\hat{\pi}$
TAF9 – Reverse
H3K9 me3:H4K20 me3
DNA Meth. m5C:H3K9 me3
H3K27 me3:H4K20 me3
H2A.Z:H3−5ac
H2A.Z:H4−4ac
H3K4 me1:H4K20 me2
DNA Meth. m5C:H4K20 me3
H3K4 me1:H4−4ac
H3K27 me3:H3K9 me3
H3K27 me3:DNA Meth. m5C
H3K4 me1:H2A.Z
H4−4ac:H3−5ac
DNA Meth. m5C:H4K20 me2
H2A.Z:H3K4 me3
H2A.Z:H4K20 me2
H4K20 me2:H3−5ac
H3K4 me3:H4−4ac
H4K20 me2:H4−4ac
H3K27 me2
H3K9 me2
H4K20 me2:H3K4 me3
H3K27 me3
H4K20 me3
H3K9 me3
H3K4 me1
DNA Meth. m5C
H2A.Z
H3−5ac
H4−4ac
H4K20 me2
H3K4 me3
0.00 0.25 0.50 0.75 1.00
$\hat{\pi}$
TAF12 – Forward
H3K9 me2:DNA Meth. m5C
H3K27 me3:H3K9 me3
H3K27 me3:DNA Meth. m5C
H3K4 me1:H4K20 me2
H3K9 me3:H4K20 me3
DNA Meth. m5C:H4K20 me3
DNA Meth. m5C:H3K9 me3
H3K27 me3:H4K20 me3
H3K4 me1:H3−5ac
H3K4 me1:H4−4ac
H3K4 me1:H2A.Z
H3K4 me3:H3−5ac
H2A.Z:H3K4 me3
DNA Meth. m5C:H4K20 me2
H4−4ac:H3−5ac
H2A.Z:H4K20 me2
H4K20 me2:H3−5ac
H3K4 me3:H4−4ac
H3K27 me2
H3K9 me2
H4K20 me2:H4−4ac
H4K20 me3
H3K9 me3
H3K27 me3
H4K20 me2:H3K4 me3
H3K4 me1
DNA Meth. m5C
H2A.Z
H4−4ac
H3−5ac
H4K20 me2
H3K4 me3
0.00 0.25 0.50 0.75 1.00
$\hat{\pi}$
TAF12 – Reverse
H3K9 me2:DNA Meth. m5C
H3K27 me3:H3K9 me3
H3K27 me3:DNA Meth. m5C
H3K4 me1:H4K20 me2
H3K9 me3:H4K20 me3
DNA Meth. m5C:H4K20 me3
DNA Meth. m5C:H3K9 me3
H3K27 me3:H4K20 me3
H3K4 me1:H3−5ac
H3K4 me1:H4−4ac
H3K4 me1:H2A.Z
H3K4 me3:H3−5ac
H2A.Z:H3K4 me3
DNA Meth. m5C:H4K20 me2
H4−4ac:H3−5ac
H2A.Z:H4K20 me2
H4K20 me2:H3−5ac
H3K4 me3:H4−4ac
H3K27 me2
H3K9 me2
H4K20 me2:H4−4ac
H4K20 me3
H3K9 me3
H3K27 me3
H4K20 me2:H3K4 me3
H3K4 me1
DNA Meth. m5C
H2A.Z
H4−4ac
H3−5ac
H4K20 me2
H3K4 me3
0.00 0.25 0.50 0.75 1.00
$\hat{\pi}$
TAF10 – Forward
DNA Meth. m5C:H4K20 me2
H3K27 me2:DNA Meth. m5C
H3K27 me3:H4K20 me3
H3K9 me3:H4K20 me3
DNA Meth. m5C:H4K20 me3
H3K27 me3:DNA Meth. m5C
H3K4 me1:H3−5ac
H3K4 me1:H2A.Z
DNA Meth. m5C:H3K9 me3
H2A.Z:H3−5ac
H2A.Z:H4−4ac
H3K4 me1:H4−4ac
H3K9 me2:DNA Meth. m5C
H3K4 me1:H4K20 me2
H2A.Z:H3K4 me3
H3K4 me3:H3−5ac
H2A.Z:H4K20 me2
H4−4ac:H3−5ac
H3K4 me3:H4−4ac
H4K20 me2:H4−4ac
H4K20 me2:H3−5ac
H3K27 me3
H3K9 me2
H4K20 me3
H3K9 me3
H3K27 me2
H3K4 me1
DNA Meth. m5C
H4K20 me2:H3K4 me3
H2A.Z
H4−4ac
H3−5ac
H3K4 me3
H4K20 me2
0.00 0.25 0.50 0.75 1.00
$\hat{\pi}$
TAF10 – Reverse
DNA Meth. m5C:H4K20 me2
H3K27 me2:DNA Meth. m5C
H3K27 me3:H4K20 me3
H3K9 me3:H4K20 me3
DNA Meth. m5C:H4K20 me3
H3K27 me3:DNA Meth. m5C
H3K4 me1:H3−5ac
H3K4 me1:H2A.Z
DNA Meth. m5C:H3K9 me3
H2A.Z:H3−5ac
H2A.Z:H4−4ac
H3K4 me1:H4−4ac
H3K9 me2:DNA Meth. m5C
H3K4 me1:H4K20 me2
H2A.Z:H3K4 me3
H3K4 me3:H3−5ac
H2A.Z:H4K20 me2
H4−4ac:H3−5ac
H3K4 me3:H4−4ac
H4K20 me2:H4−4ac
H4K20 me2:H3−5ac
H3K27 me3
H3K9 me2
H4K20 me3
H3K9 me3
H3K27 me2
H3K4 me1
DNA Meth. m5C
H4K20 me2:H3K4 me3
H2A.Z
H4−4ac
H3−5ac
H3K4 me3
H4K20 me2
0.00 0.25 0.50 0.75 1.00
$\hat{\pi}$
TAF4 – Forward
H3K9 me3:H4K20 me3
H3K9 me2:DNA Meth. m5C
H3K27 me3:H4K20 me3
H3K27 me3:H3K9 me3
DNA Meth. m5C:H3K9 me3
H3K27 me2:DNA Meth. m5C
DNA Meth. m5C:H4K20 me2
DNA Meth. m5C:H4K20 me3
H3K27 me3:H3K4 me1
H3K27 me3:DNA Meth. m5C
H3K4 me1:H3−5ac
H3K4 me1:H2A.Z
H2A.Z:H3−5ac
H2A.Z:H4−4ac
H3K4 me1:H4−4ac
H3K4 me1:H4K20 me2
H4K20 me2:H4−4ac
H2A.Z:H3K4 me3
H3K4 me3:H3−5ac
H2A.Z:H4K20 me2
H4−4ac:H3−5ac
H3K9 me2
H3K4 me3:H4−4ac
H3K27 me2
H4K20 me3
H3K9 me3
H4K20 me2:H3−5ac
H3K27 me3
DNA Meth. m5C
H4K20 me2:H3K4 me3
H3K4 me1
H2A.Z
H4K20 me2
H4−4ac
H3−5ac
H3K4 me3
0.00 0.25 0.50 0.75 1.00
$\hat{\pi}$
TAF4 – Reverse
H3K9 me3:H4K20 me3
H3K9 me2:DNA Meth. m5C
H3K27 me3:H4K20 me3
H3K27 me3:H3K9 me3
DNA Meth. m5C:H3K9 me3
H3K27 me2:DNA Meth. m5C
DNA Meth. m5C:H4K20 me2
DNA Meth. m5C:H4K20 me3
H3K27 me3:H3K4 me1
H3K27 me3:DNA Meth. m5C
H3K4 me1:H3−5ac
H3K4 me1:H2A.Z
H2A.Z:H3−5ac
H2A.Z:H4−4ac
H3K4 me1:H4−4ac
H3K4 me1:H4K20 me2
H4K20 me2:H4−4ac
H2A.Z:H3K4 me3
H3K4 me3:H3−5ac
H2A.Z:H4K20 me2
H4−4ac:H3−5ac
H3K9 me2
H3K4 me3:H4−4ac
H3K27 me2
H4K20 me3
H3K9 me3
H4K20 me2:H3−5ac
H3K27 me3
DNA Meth. m5C
H4K20 me2:H3K4 me3
H3K4 me1
H2A.Z
H4K20 me2
H4−4ac
H3−5ac
H3K4 me3
0.00 0.25 0.50 0.75 1.00
$\hat{\pi}$
KDM5A – Forward
H3K4 me1:H2A.Z
H2A.Z:H3−5ac
H3K4 me1:H4−4ac
H3K4 me1:H4K20 me2
H3K9 me3:H4K20 me3
DNA Meth. m5C:H4K20 me2
H3K27 me3:H3K4 me1
H3K27 me3:H4K20 me3
H2A.Z:H4K20 me2
H3K9 me2:DNA Meth. m5C
DNA Meth. m5C:H3K9 me3
H3K4 me3:H3−5ac
H2A.Z:H3K4 me3
H3K4 me3:H4−4ac
H3K4 me1:H3−5ac
H4−4ac:H3−5ac
H3K27 me2:DNA Meth. m5C
H3K27 me3:DNA Meth. m5C
H4K20 me2:H4−4ac
H4K20 me2:H3−5ac
H3K27 me3:H3K9 me3
H3K4 me1
H4K20 me3
H4K20 me2:H3K4 me3
H3K9 me2
H3K27 me2
DNA Meth. m5C
H3K9 me3
H4−4ac
H2A.Z
H3K27 me3
H4K20 me2
H3−5ac
H3K4 me3
0.00 0.25 0.50 0.75 1.00
$\hat{\pi}$
KDM5A – Reverse
H3K4 me1:H2A.Z
H2A.Z:H3−5ac
H3K4 me1:H4−4ac
H3K4 me1:H4K20 me2
H3K9 me3:H4K20 me3
DNA Meth. m5C:H4K20 me2
H3K27 me3:H3K4 me1
H3K27 me3:H4K20 me3
H2A.Z:H4K20 me2
H3K9 me2:DNA Meth. m5C
DNA Meth. m5C:H3K9 me3
H3K4 me3:H3−5ac
H2A.Z:H3K4 me3
H3K4 me3:H4−4ac
H3K4 me1:H3−5ac
H4−4ac:H3−5ac
H3K27 me2:DNA Meth. m5C
H3K27 me3:DNA Meth. m5C
H4K20 me2:H4−4ac
H4K20 me2:H3−5ac
H3K27 me3:H3K9 me3
H3K4 me1
H4K20 me3
H4K20 me2:H3K4 me3
H3K9 me2
H3K27 me2
DNA Meth. m5C
H3K9 me3
H4−4ac
H2A.Z
H3K27 me3
H4K20 me2
H3−5ac
H3K4 me3
0.00 0.25 0.50 0.75 1.00
$\hat{\pi}$

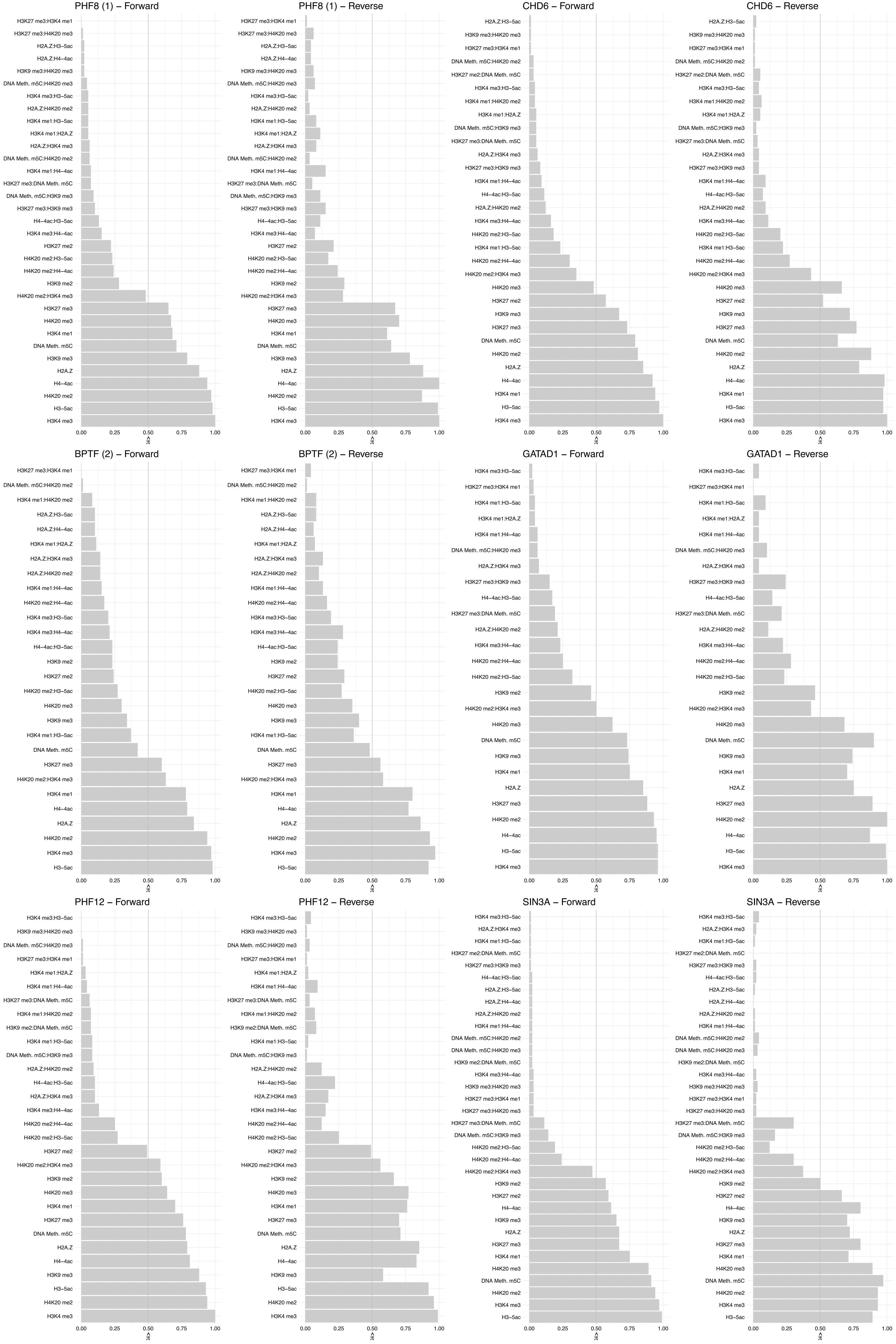
PHF8 (1) – Forward
H3K27 me3:H3K4 me1
H3K27 me3:H4K20 me3
H2A.Z:H3−5ac
H2A.Z:H4−4ac
H3K9 me3:H4K20 me3
DNA Meth. m5C:H4K20 me3
H3K4 me3:H3−5ac
H2A.Z:H4K20 me2
H3K4 me1:H3−5ac
H3K4 me1:H2A.Z
H2A.Z:H3K4 me3
DNA Meth. m5C:H4K20 me2
H3K4 me1:H4−4ac
H3K27 me3:DNA Meth. m5C
DNA Meth. m5C:H3K9 me3
H3K27 me3:H3K9 me3
H4−4ac:H3−5ac
H3K4 me3:H4−4ac
H3K27 me2
H4K20 me2:H3−5ac
H4K20 me2:H4−4ac
H3K9 me2
H4K20 me2:H3K4 me3
H3K27 me3
H4K20 me3
H3K4 me1
DNA Meth. m5C
H3K9 me3
H2A.Z
H4−4ac
H4K20 me2
H3−5ac
H3K4 me3
0.00
0.25
0.50
0.75
1.00
π̂
PHF8 (1) – Reverse
H3K27 me3:H3K4 me1
H3K27 me3:H4K20 me3
H2A.Z:H3−5ac
H2A.Z:H4−4ac
H3K9 me3:H4K20 me3
DNA Meth. m5C:H4K20 me3
H3K4 me3:H3−5ac
H2A.Z:H4K20 me2
H3K4 me1:H3−5ac
H3K4 me1:H2A.Z
H2A.Z:H3K4 me3
DNA Meth. m5C:H4K20 me2
H3K4 me1:H4−4ac
H3K27 me3:DNA Meth. m5C
DNA Meth. m5C:H3K9 me3
H3K27 me3:H3K9 me3
H4−4ac:H3−5ac
H3K4 me3:H4−4ac
H3K27 me2
H4K20 me2:H3−5ac
H4K20 me2:H4−4ac
H3K9 me2
H4K20 me2:H3K4 me3
H3K27 me3
H4K20 me3
H3K4 me1
DNA Meth. m5C
H3K9 me3
H2A.Z
H4−4ac
H4K20 me2
H3−5ac
H3K4 me3
0.00
0.25
0.50
0.75
1.00
π̂
CHD6 – Forward
H2A.Z:H3−5ac
H3K9 me3:H4K20 me3
H3K27 me3:H3K4 me1
DNA Meth. m5C:H4K20 me2
H3K27 me2:DNA Meth. m5C
H3K4 me3:H3−5ac
H3K4 me1:H4K20 me2
H3K4 me1:H2A.Z
DNA Meth. m5C:H3K9 me3
H3K27 me3:DNA Meth. m5C
H2A.Z:H3K4 me3
H3K27 me3:H3K9 me3
H3K4 me1:H4−4ac
H4−4ac:H3−5ac
H2A.Z:H4K20 me2
H3K4 me3:H4−4ac
H4K20 me2:H3−5ac
H3K4 me1:H3−5ac
H4K20 me2:H4−4ac
H4K20 me2:H3K4 me3
H4K20 me3
H3K27 me2
H3K9 me3
H3K27 me3
DNA Meth. m5C
H4K20 me2
H2A.Z
H4−4ac
H3K4 me1
H3−5ac
H3K4 me3
0.00
0.25
0.50
0.75
1.00
π̂
CHD6 – Reverse
H2A.Z:H3−5ac
H3K9 me3:H4K20 me3
H3K27 me3:H3K4 me1
DNA Meth. m5C:H4K20 me2
H3K27 me2:DNA Meth. m5C
H3K4 me3:H3−5ac
H3K4 me1:H4K20 me2
H3K4 me1:H2A.Z
DNA Meth. m5C:H3K9 me3
H3K27 me3:DNA Meth. m5C
H2A.Z:H3K4 me3
H3K27 me3:H3K9 me3
H3K4 me1:H4−4ac
H4−4ac:H3−5ac
H2A.Z:H4K20 me2
H3K4 me3:H4−4ac
H4K20 me2:H3−5ac
H3K4 me1:H3−5ac
H4K20 me2:H4−4ac
H4K20 me2:H3K4 me3
H4K20 me3
H3K27 me2
H3K9 me3
H3K27 me3
DNA Meth. m5C
H4K20 me2
H2A.Z
H4−4ac
H3K4 me1
H3−5ac
H3K4 me3
0.00
0.25
0.50
0.75
1.00
π̂
BPTF (2) – Forward
H3K27 me3:H3K4 me1
DNA Meth. m5C:H4K20 me2
H3K4 me1:H4K20 me2
H2A.Z:H3−5ac
H2A.Z:H4−4ac
H3K4 me1:H2A.Z
H2A.Z:H3K4 me3
H2A.Z:H4K20 me2
H3K4 me1:H4−4ac
H4K20 me2:H4−4ac
H3K4 me3:H3−5ac
H3K4 me3:H4−4ac
H4−4ac:H3−5ac
H3K9 me2
H3K27 me2
H4K20 me2:H3−5ac
H4K20 me3
H3K9 me3
H3K4 me1:H3−5ac
DNA Meth. m5C
H3K27 me3
H4K20 me2:H3K4 me3
H3K4 me1
H4−4ac
H2A.Z
H4K20 me2
H3K4 me3
H3−5ac
0.00
0.25
0.50
0.75
1.00
π̂
BPTF (2) – Reverse
H3K27 me3:H3K4 me1
DNA Meth. m5C:H4K20 me2
H3K4 me1:H4K20 me2
H2A.Z:H3−5ac
H2A.Z:H4−4ac
H3K4 me1:H2A.Z
H2A.Z:H3K4 me3
H2A.Z:H4K20 me2
H3K4 me1:H4−4ac
H4K20 me2:H4−4ac
H3K4 me3:H3−5ac
H3K4 me3:H4−4ac
H4−4ac:H3−5ac
H3K9 me2
H3K27 me2
H4K20 me2:H3−5ac
H4K20 me3
H3K9 me3
H3K4 me1:H3−5ac
DNA Meth. m5C
H3K27 me3
H4K20 me2:H3K4 me3
H3K4 me1
H4−4ac
H2A.Z
H4K20 me2
H3K4 me3
H3−5ac
0.00
0.25
0.50
0.75
1.00
π̂
GATAD1 – Forward
H3K4 me3:H3−5ac
H3K27 me3:H3K4 me1
H3K4 me1:H3−5ac
H3K4 me1:H2A.Z
H3K4 me1:H4−4ac
DNA Meth. m5C:H4K20 me3
H2A.Z:H3K4 me3
H3K27 me3:H3K9 me3
H4−4ac:H3−5ac
H3K27 me3:DNA Meth. m5C
H2A.Z:H4K20 me2
H3K4 me3:H4−4ac
H4K20 me2:H4−4ac
H4K20 me2:H3−5ac
H3K9 me2
H4K20 me2:H3K4 me3
H4K20 me3
DNA Meth. m5C
H3K9 me3
H3K4 me1
H2A.Z
H3K27 me3
H4K20 me2
H4−4ac
H3−5ac
H3K4 me3
0.00
0.25
0.50
0.75
1.00
π̂
GATAD1 – Reverse
H3K4 me3:H3−5ac
H3K27 me3:H3K4 me1
H3K4 me1:H3−5ac
H3K4 me1:H2A.Z
H3K4 me1:H4−4ac
DNA Meth. m5C:H4K20 me3
H2A.Z:H3K4 me3
H3K27 me3:H3K9 me3
H4−4ac:H3−5ac
H3K27 me3:DNA Meth. m5C
H2A.Z:H4K20 me2
H3K4 me3:H4−4ac
H4K20 me2:H4−4ac
H4K20 me2:H3−5ac
H3K9 me2
H4K20 me2:H3K4 me3
H4K20 me3
DNA Meth. m5C
H3K9 me3
H3K4 me1
H2A.Z
H3K27 me3
H4K20 me2
H4−4ac
H3−5ac
H3K4 me3
0.00
0.25
0.50
0.75
1.00
π̂
PHF12 – Forward
H3K4 me3:H3−5ac
H3K9 me3:H4K20 me3
DNA Meth. m5C:H4K20 me3
H3K27 me3:H3K4 me1
H3K4 me1:H2A.Z
H3K4 me1:H4−4ac
H3K27 me3:DNA Meth. m5C
H3K4 me1:H4K20 me2
H3K9 me2:DNA Meth. m5C
H3K4 me1:H3−5ac
DNA Meth. m5C:H3K9 me3
H2A.Z:H4K20 me2
H4−4ac:H3−5ac
H2A.Z:H3K4 me3
H3K4 me3:H4−4ac
H4K20 me2:H4−4ac
H4K20 me2:H3−5ac
H3K27 me2
H4K20 me2:H3K4 me3
H3K9 me2
H4K20 me3
H3K4 me1
H3K27 me3
DNA Meth. m5C
H2A.Z
H4−4ac
H3K9 me3
H3−5ac
H4K20 me2
H3K4 me3
0.00
0.25
0.50
0.75
1.00
π̂
PHF12 – Reverse
H3K4 me3:H3−5ac
H3K9 me3:H4K20 me3
DNA Meth. m5C:H4K20 me3
H3K27 me3:H3K4 me1
H3K4 me1:H2A.Z
H3K4 me1:H4−4ac
H3K27 me3:DNA Meth. m5C
H3K4 me1:H4K20 me2
H3K9 me2:DNA Meth. m5C
H3K4 me1:H3−5ac
DNA Meth. m5C:H3K9 me3
H2A.Z:H4K20 me2
H4−4ac:H3−5ac
H2A.Z:H3K4 me3
H3K4 me3:H4−4ac
H4K20 me2:H4−4ac
H4K20 me2:H3−5ac
H3K27 me2
H4K20 me2:H3K4 me3
H3K9 me2
H4K20 me3
H3K4 me1
H3K27 me3
DNA Meth. m5C
H2A.Z
H4−4ac
H3K9 me3
H3−5ac
H4K20 me2
H3K4 me3
0.00
0.25
0.50
0.75
1.00
π̂
SIN3A – Forward
H3K4 me3:H3−5ac
H2A.Z:H3K4 me3
H3K4 me1:H3−5ac
H3K27 me2:DNA Meth. m5C
H3K27 me3:H3K9 me3
H4−4ac:H3−5ac
H2A.Z:H3−5ac
H2A.Z:H4−4ac
H2A.Z:H4K20 me2
H3K4 me1:H4−4ac
DNA Meth. m5C:H4K20 me2
DNA Meth. m5C:H4K20 me3
H3K9 me2:DNA Meth. m5C
H3K4 me3:H4−4ac
H3K9 me3:H4K20 me3
H3K27 me3:H3K4 me1
H3K27 me3:H4K20 me3
H3K27 me3:DNA Meth. m5C
DNA Meth. m5C:H3K9 me3
H4K20 me2:H3−5ac
H4K20 me2:H4−4ac
H4K20 me2:H3K4 me3
H3K9 me2
H3K27 me2
H4−4ac
H3K9 me3
H2A.Z
H3K27 me3
H3K4 me1
H4K20 me3
DNA Meth. m5C
H4K20 me2
H3K4 me3
H3−5ac
0.00
0.25
0.50
0.75
1.00
π̂
SIN3A – Reverse
H3K4 me3:H3−5ac
H2A.Z:H3K4 me3
H3K4 me1:H3−5ac
H3K27 me2:DNA Meth. m5C
H3K27 me3:H3K9 me3
H4−4ac:H3−5ac
H2A.Z:H3−5ac
H2A.Z:H4−4ac
H2A.Z:H4K20 me2
H3K4 me1:H4−4ac
DNA Meth. m5C:H4K20 me2
DNA Meth. m5C:H4K20 me3
H3K9 me2:DNA Meth. m5C
H3K4 me3:H4−4ac
H3K9 me3:H4K20 me3
H3K27 me3:H3K4 me1
H3K27 me3:H4K20 me3
H3K27 me3:DNA Meth. m5C
DNA Meth. m5C:H3K9 me3
H4K20 me2:H3−5ac
H4K20 me2:H4−4ac
H4K20 me2:H3K4 me3
H3K9 me2
H3K27 me2
H4−4ac
H3K9 me3
H2A.Z
H3K27 me3
H3K4 me1
H4K20 me3
DNA Meth. m5C
H4K20 me2
H3K4 me3
H3−5ac
0.00
0.25
0.50
0.75
1.00
π̂

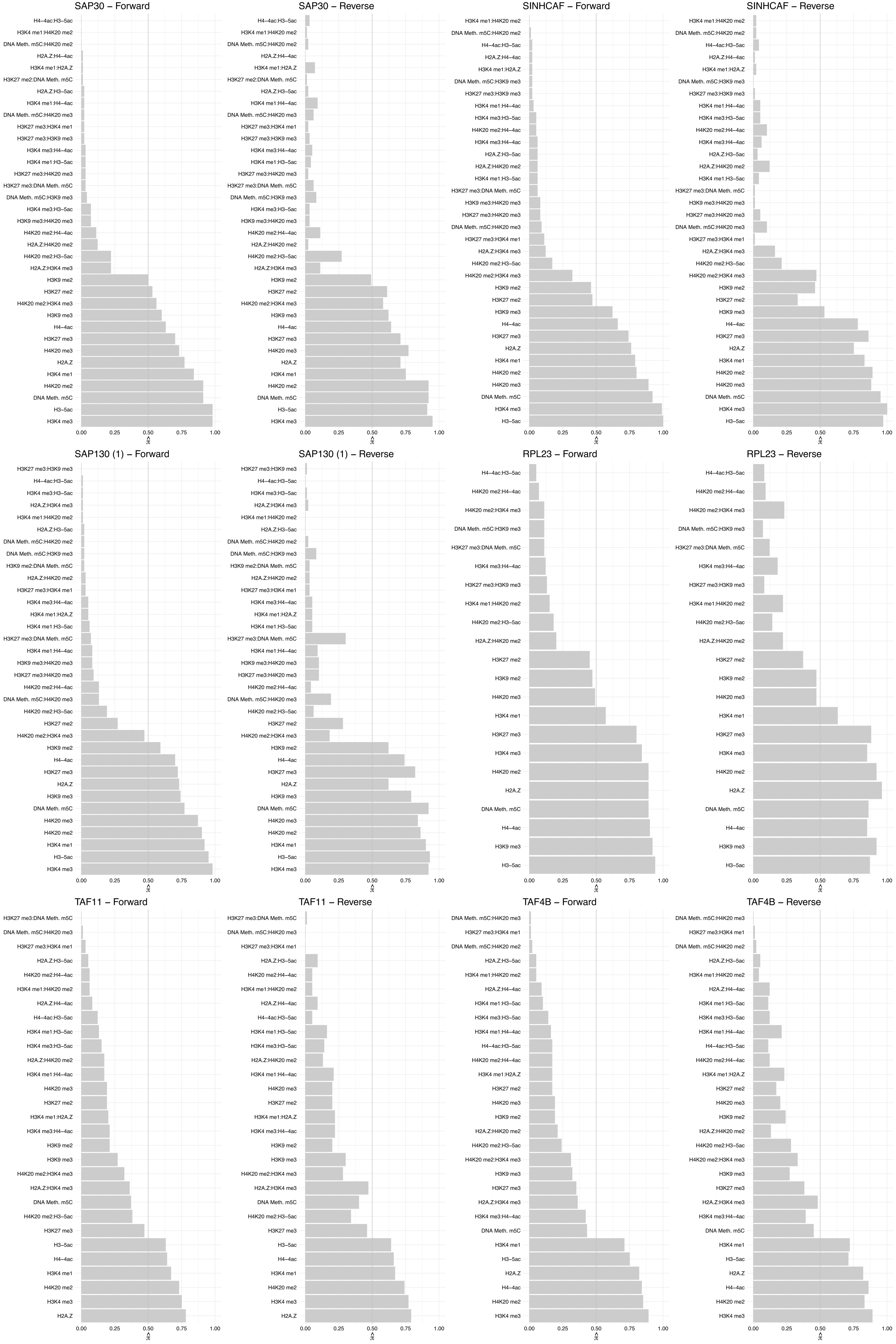
SAP30 − Forward
H4−4ac:H3−5ac
H3K4 me1:H4K20 me2
DNA Meth. m5C:H4K20 me2
H2A.Z:H4−4ac
H3K4 me1:H2A.Z
H3K27 me2:DNA Meth. m5C
H2A.Z:H3−5ac
H3K4 me1:H4−4ac
DNA Meth. m5C:H4K20 me3
H3K27 me3:H3K4 me1
H3K27 me3:H3K9 me3
H3K4 me3:H4−4ac
H3K4 me1:H3−5ac
H3K27 me3:H4K20 me3
H3K27 me3:DNA Meth. m5C
DNA Meth. m5C:H3K9 me3
H3K4 me3:H3−5ac
H3K9 me3:H4K20 me3
H4K20 me2:H4−4ac
H2A.Z:H4K20 me2
H4K20 me2:H3−5ac
H2A.Z:H3K4 me3
H3K9 me2
H3K27 me2
H4K20 me2:H3K4 me3
H3K9 me3
H4−4ac
H3K27 me3
H4K20 me3
H2A.Z
H3K4 me1
H4K20 me2
DNA Meth. m5C
H3−5ac
H3K4 me3
0.00 0.25 0.50 0.75 1.00
π̂
SAP30 − Reverse
H4−4ac:H3−5ac
H3K4 me1:H4K20 me2
DNA Meth. m5C:H4K20 me2
H2A.Z:H4−4ac
H3K4 me1:H2A.Z
H3K27 me2:DNA Meth. m5C
H2A.Z:H3−5ac
H3K4 me1:H4−4ac
DNA Meth. m5C:H4K20 me3
H3K27 me3:H3K4 me1
H3K27 me3:H3K9 me3
H3K4 me3:H4−4ac
H3K4 me1:H3−5ac
H3K27 me3:H4K20 me3
H3K27 me3:DNA Meth. m5C
DNA Meth. m5C:H3K9 me3
H3K4 me3:H3−5ac
H3K9 me3:H4K20 me3
H4K20 me2:H4−4ac
H2A.Z:H4K20 me2
H4K20 me2:H3−5ac
H2A.Z:H3K4 me3
H3K9 me2
H3K27 me2
H4K20 me2:H3K4 me3
H3K9 me3
H4−4ac
H3K27 me3
H4K20 me3
H2A.Z
H3K4 me1
H4K20 me2
DNA Meth. m5C
H3−5ac
H3K4 me3
0.00 0.25 0.50 0.75 1.00
π̂
SINHCAF − Forward
H3K4 me1:H4K20 me2
DNA Meth. m5C:H4K20 me2
H4−4ac:H3−5ac
H2A.Z:H4−4ac
H3K4 me1:H2A.Z
DNA Meth. m5C:H3K9 me3
H3K27 me3:H3K9 me3
H3K4 me1:H4−4ac
H3K4 me3:H3−5ac
H4K20 me2:H4−4ac
H3K4 me3:H4−4ac
H2A.Z:H3−5ac
H2A.Z:H4K20 me2
H3K4 me1:H3−5ac
H3K27 me3:DNA Meth. m5C
H3K9 me3:H4K20 me3
H3K27 me3:H4K20 me3
DNA Meth. m5C:H4K20 me3
H3K27 me3:H3K4 me1
H2A.Z:H3K4 me3
H4K20 me2:H3−5ac
H4K20 me2:H3K4 me3
H3K9 me2
H3K27 me2
H3K9 me3
H4−4ac
H3K27 me3
H2A.Z
H3K4 me1
H4K20 me2
H4K20 me3
DNA Meth. m5C
H3K4 me3
H3−5ac
0.00 0.25 0.50 0.75 1.00
π̂
SINHCAF − Reverse
H3K4 me1:H4K20 me2
DNA Meth. m5C:H4K20 me2
H4−4ac:H3−5ac
H2A.Z:H4−4ac
H3K4 me1:H2A.Z
DNA Meth. m5C:H3K9 me3
H3K27 me3:H3K9 me3
H3K4 me1:H4−4ac
H3K4 me3:H3−5ac
H4K20 me2:H4−4ac
H3K4 me3:H4−4ac
H2A.Z:H3−5ac
H2A.Z:H4K20 me2
H3K4 me1:H3−5ac
H3K27 me3:DNA Meth. m5C
H3K9 me3:H4K20 me3
H3K27 me3:H4K20 me3
DNA Meth. m5C:H4K20 me3
H3K27 me3:H3K4 me1
H2A.Z:H3K4 me3
H4K20 me2:H3−5ac
H4K20 me2:H3K4 me3
H3K9 me2
H3K27 me2
H3K9 me3
H4−4ac
H3K27 me3
H2A.Z
H3K4 me1
H4K20 me2
H4K20 me3
DNA Meth. m5C
H3K4 me3
H3−5ac
0.00 0.25 0.50 0.75 1.00
π̂
SAP130 (1) − Forward
H3K27 me3:H3K9 me3
H4−4ac:H3−5ac
H3K4 me3:H3−5ac
H2A.Z:H3K4 me3
H3K4 me1:H4K20 me2
H2A.Z:H3−5ac
DNA Meth. m5C:H4K20 me2
DNA Meth. m5C:H3K9 me3
H3K9 me2:DNA Meth. m5C
H2A.Z:H4K20 me2
H3K27 me3:H3K4 me1
H3K4 me3:H4−4ac
H3K4 me1:H2A.Z
H3K4 me1:H3−5ac
H3K27 me3:DNA Meth. m5C
H3K4 me1:H4−4ac
H3K9 me3:H4K20 me3
H3K27 me3:H4K20 me3
H4K20 me2:H4−4ac
DNA Meth. m5C:H4K20 me3
H4K20 me2:H3−5ac
H3K27 me2
H4K20 me2:H3K4 me3
H3K9 me2
H4−4ac
H3K27 me3
H2A.Z
H3K9 me3
DNA Meth. m5C
H4K20 me3
H4K20 me2
H3K4 me1
H3−5ac
H3K4 me3
0.00 0.25 0.50 0.75 1.00
π̂
SAP130 (1) − Reverse
H3K27 me3:H3K9 me3
H4−4ac:H3−5ac
H3K4 me3:H3−5ac
H2A.Z:H3K4 me3
H3K4 me1:H4K20 me2
H2A.Z:H3−5ac
DNA Meth. m5C:H4K20 me2
DNA Meth. m5C:H3K9 me3
H3K9 me2:DNA Meth. m5C
H2A.Z:H4K20 me2
H3K27 me3:H3K4 me1
H3K4 me3:H4−4ac
H3K4 me1:H2A.Z
H3K4 me1:H3−5ac
H3K27 me3:DNA Meth. m5C
H3K4 me1:H4−4ac
H3K9 me3:H4K20 me3
H3K27 me3:H4K20 me3
H4K20 me2:H4−4ac
DNA Meth. m5C:H4K20 me3
H4K20 me2:H3−5ac
H3K27 me2
H4K20 me2:H3K4 me3
H3K9 me2
H4−4ac
H3K27 me3
H2A.Z
H3K9 me3
DNA Meth. m5C
H4K20 me3
H4K20 me2
H3K4 me1
H3−5ac
H3K4 me3
0.00 0.25 0.50 0.75 1.00
π̂
RPL23 − Forward
H4−4ac:H3−5ac
H4K20 me2:H4−4ac
H4K20 me2:H3K4 me3
DNA Meth. m5C:H3K9 me3
H3K27 me3:DNA Meth. m5C
H3K4 me3:H4−4ac
H3K27 me3:H3K9 me3
H3K4 me1:H4K20 me2
H4K20 me2:H3−5ac
H2A.Z:H4K20 me2
H3K27 me2
H3K9 me2
H4K20 me3
H3K4 me1
H3K27 me3
H3K4 me3
H4K20 me2
H2A.Z
DNA Meth. m5C
H4−4ac
H3K9 me3
H3−5ac
0.00 0.25 0.50 0.75 1.00
π̂
RPL23 − Reverse
H4−4ac:H3−5ac
H4K20 me2:H4−4ac
H4K20 me2:H3K4 me3
DNA Meth. m5C:H3K9 me3
H3K27 me3:DNA Meth. m5C
H3K4 me3:H4−4ac
H3K27 me3:H3K9 me3
H3K4 me1:H4K20 me2
H4K20 me2:H3−5ac
H2A.Z:H4K20 me2
H3K27 me2
H3K9 me2
H4K20 me3
H3K4 me1
H3K27 me3
H3K4 me3
H4K20 me2
H2A.Z
DNA Meth. m5C
H4−4ac
H3K9 me3
H3−5ac
0.00 0.25 0.50 0.75 1.00
π̂
TAF11 − Forward
H3K27 me3:DNA Meth. m5C
DNA Meth. m5C:H4K20 me3
H3K27 me3:H3K4 me1
H2A.Z:H3−5ac
H4K20 me2:H4−4ac
H3K4 me1:H4K20 me2
H2A.Z:H4−4ac
H4−4ac:H3−5ac
H3K4 me1:H3−5ac
H3K4 me3:H3−5ac
H2A.Z:H4K20 me2
H3K4 me1:H4−4ac
H4K20 me3
H3K27 me2
H3K4 me1:H2A.Z
H3K4 me3:H4−4ac
H3K9 me2
H3K9 me3
H4K20 me2:H3K4 me3
H2A.Z:H3K4 me3
DNA Meth. m5C
H4K20 me2:H3−5ac
H3K27 me3
H3−5ac
H4−4ac
H3K4 me1
H4K20 me2
H3K4 me3
H2A.Z
0.00 0.25 0.50 0.75 1.00
π̂
TAF11 − Reverse
H3K27 me3:DNA Meth. m5C
DNA Meth. m5C:H4K20 me3
H3K27 me3:H3K4 me1
H2A.Z:H3−5ac
H4K20 me2:H4−4ac
H3K4 me1:H4K20 me2
H2A.Z:H4−4ac
H4−4ac:H3−5ac
H3K4 me1:H3−5ac
H3K4 me3:H3−5ac
H2A.Z:H4K20 me2
H3K4 me1:H4−4ac
H4K20 me3
H3K27 me2
H3K4 me1:H2A.Z
H3K4 me3:H4−4ac
H3K9 me2
H3K9 me3
H4K20 me2:H3K4 me3
H2A.Z:H3K4 me3
DNA Meth. m5C
H4K20 me2:H3−5ac
H3K27 me3
H3−5ac
H4−4ac
H3K4 me1
H4K20 me2
H3K4 me3
H2A.Z
0.00 0.25 0.50 0.75 1.00
π̂
TAF4B − Forward
DNA Meth. m5C:H4K20 me3
H3K27 me3:H3K4 me1
DNA Meth. m5C:H4K20 me2
H2A.Z:H3−5ac
H3K4 me1:H4K20 me2
H2A.Z:H4−4ac
H3K4 me1:H3−5ac
H3K4 me3:H3−5ac
H3K4 me1:H4−4ac
H4−4ac:H3−5ac
H4K20 me2:H4−4ac
H3K4 me1:H2A.Z
H3K27 me2
H4K20 me3
H3K9 me2
H2A.Z:H4K20 me2
H4K20 me2:H3−5ac
H4K20 me2:H3K4 me3
H3K9 me3
H3K27 me3
H2A.Z:H3K4 me3
H3K4 me3:H4−4ac
DNA Meth. m5C
H3K4 me1
H3−5ac
H2A.Z
H4−4ac
H4K20 me2
H3K4 me3
0.00 0.25 0.50 0.75 1.00
π̂
TAF4B − Reverse
DNA Meth. m5C:H4K20 me3
H3K27 me3:H3K4 me1
DNA Meth. m5C:H4K20 me2
H2A.Z:H3−5ac
H3K4 me1:H4K20 me2
H2A.Z:H4−4ac
H3K4 me1:H3−5ac
H3K4 me3:H3−5ac
H3K4 me1:H4−4ac
H4−4ac:H3−5ac
H4K20 me2:H4−4ac
H3K4 me1:H2A.Z
H3K27 me2
H4K20 me3
H3K9 me2
H2A.Z:H4K20 me2
H4K20 me2:H3−5ac
H4K20 me2:H3K4 me3
H3K9 me3
H3K27 me3
H2A.Z:H3K4 me3
H3K4 me3:H4−4ac
DNA Meth. m5C
H3K4 me1
H3−5ac
H2A.Z
H4−4ac
H4K20 me2
H3K4 me3
0.00 0.25 0.50 0.75 1.00
π̂

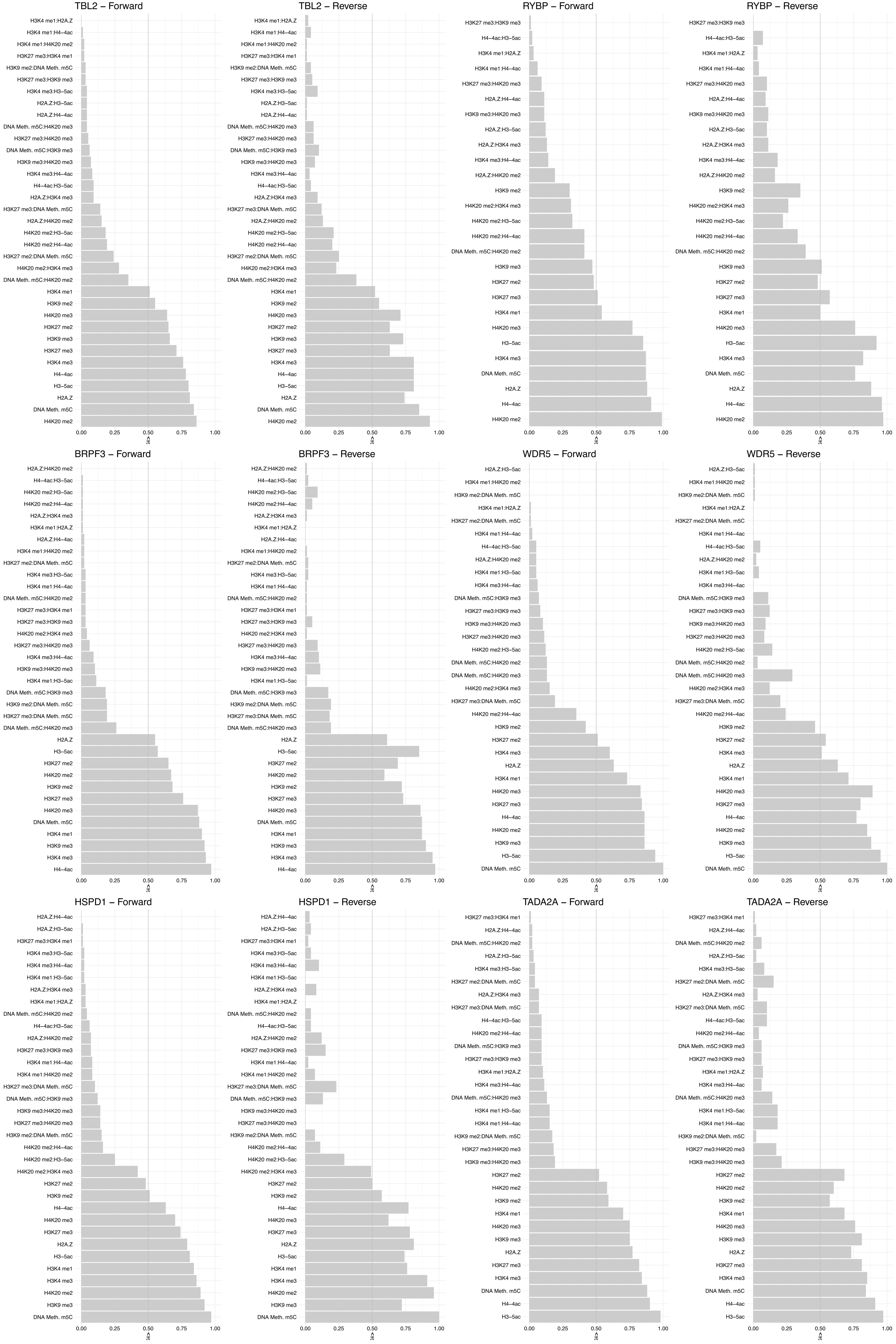
TBL2 − Forward
H3K4 me1:H2A.Z
H3K4 me1:H4−4ac
H3K4 me1:H4K20 me2
H3K27 me3:H3K4 me1
H3K9 me2:DNA Meth. m5C
H3K27 me3:H3K9 me3
H3K4 me3:H3−5ac
H2A.Z:H3−5ac
H2A.Z:H4−4ac
DNA Meth. m5C:H4K20 me3
H3K27 me3:H4K20 me3
DNA Meth. m5C:H3K9 me3
H3K9 me3:H4K20 me3
H3K4 me3:H4−4ac
H4−4ac:H3−5ac
H2A.Z:H3K4 me3
H3K27 me3:DNA Meth. m5C
H2A.Z:H4K20 me2
H4K20 me2:H3−5ac
H4K20 me2:H4−4ac
H3K27 me2:DNA Meth. m5C
H4K20 me2:H3K4 me3
DNA Meth. m5C:H4K20 me2
H3K4 me1
H3K9 me2
H4K20 me3
H3K27 me2
H3K9 me3
H3K27 me3
H3K4 me3
H4−4ac
H3−5ac
H2A.Z
DNA Meth. m5C
H4K20 me2
0.00 0.25 0.50 0.75 1.00
π̂
TBL2 − Reverse
H3K4 me1:H2A.Z
H3K4 me1:H4−4ac
H3K4 me1:H4K20 me2
H3K27 me3:H3K4 me1
H3K9 me2:DNA Meth. m5C
H3K27 me3:H3K9 me3
H3K4 me3:H3−5ac
H2A.Z:H3−5ac
H2A.Z:H4−4ac
DNA Meth. m5C:H4K20 me3
H3K27 me3:H4K20 me3
DNA Meth. m5C:H3K9 me3
H3K9 me3:H4K20 me3
H3K4 me3:H4−4ac
H4−4ac:H3−5ac
H2A.Z:H3K4 me3
H3K27 me3:DNA Meth. m5C
H2A.Z:H4K20 me2
H4K20 me2:H3−5ac
H4K20 me2:H4−4ac
H3K27 me2:DNA Meth. m5C
H4K20 me2:H3K4 me3
DNA Meth. m5C:H4K20 me2
H3K4 me1
H3K9 me2
H4K20 me3
H3K27 me2
H3K9 me3
H3K27 me3
H3K4 me3
H4−4ac
H3−5ac
H2A.Z
DNA Meth. m5C
H4K20 me2
0.00 0.25 0.50 0.75 1.00
π̂
RYBP − Forward
H3K27 me3:H3K9 me3
H4−4ac:H3−5ac
H3K4 me1:H2A.Z
H3K4 me1:H4−4ac
H3K27 me3:H4K20 me3
H2A.Z:H4−4ac
H3K9 me3:H4K20 me3
H2A.Z:H3−5ac
H2A.Z:H3K4 me3
H3K4 me3:H4−4ac
H2A.Z:H4K20 me2
H3K9 me2
H4K20 me2:H3K4 me3
H4K20 me2:H3−5ac
H4K20 me2:H4−4ac
DNA Meth. m5C:H4K20 me2
H3K9 me3
H3K27 me2
H3K27 me3
H3K4 me1
H4K20 me3
H3−5ac
H3K4 me3
DNA Meth. m5C
H2A.Z
H4−4ac
H4K20 me2
0.00 0.25 0.50 0.75 1.00
π̂
RYBP − Reverse
H3K27 me3:H3K9 me3
H4−4ac:H3−5ac
H3K4 me1:H2A.Z
H3K4 me1:H4−4ac
H3K27 me3:H4K20 me3
H2A.Z:H4−4ac
H3K9 me3:H4K20 me3
H2A.Z:H3−5ac
H2A.Z:H3K4 me3
H3K4 me3:H4−4ac
H2A.Z:H4K20 me2
H3K9 me2
H4K20 me2:H3K4 me3
H4K20 me2:H3−5ac
H4K20 me2:H4−4ac
DNA Meth. m5C:H4K20 me2
H3K9 me3
H3K27 me2
H3K27 me3
H3K4 me1
H4K20 me3
H3−5ac
H3K4 me3
DNA Meth. m5C
H2A.Z
H4−4ac
H4K20 me2
0.00 0.25 0.50 0.75 1.00
π̂
BRPF3 − Forward
H2A.Z:H4K20 me2
H4−4ac:H3−5ac
H4K20 me2:H3−5ac
H4K20 me2:H4−4ac
H2A.Z:H3K4 me3
H3K4 me1:H2A.Z
H2A.Z:H4−4ac
H3K4 me1:H4K20 me2
H3K27 me2:DNA Meth. m5C
H3K4 me3:H3−5ac
H3K4 me1:H4−4ac
DNA Meth. m5C:H4K20 me2
H3K27 me3:H3K4 me1
H3K27 me3:H3K9 me3
H4K20 me2:H3K4 me3
H3K27 me3:H4K20 me3
H3K4 me3:H4−4ac
H3K9 me3:H4K20 me3
H3K4 me1:H3−5ac
DNA Meth. m5C:H3K9 me3
H3K9 me2:DNA Meth. m5C
H3K27 me3:DNA Meth. m5C
DNA Meth. m5C:H4K20 me3
H2A.Z
H3−5ac
H3K27 me2
H4K20 me2
H3K9 me2
H3K27 me3
H4K20 me3
DNA Meth. m5C
H3K4 me1
H3K9 me3
H3K4 me3
H4−4ac
0.00 0.25 0.50 0.75 1.00
π̂
BRPF3 − Reverse
H2A.Z:H4K20 me2
H4−4ac:H3−5ac
H4K20 me2:H3−5ac
H4K20 me2:H4−4ac
H2A.Z:H3K4 me3
H3K4 me1:H2A.Z
H2A.Z:H4−4ac
H3K4 me1:H4K20 me2
H3K27 me2:DNA Meth. m5C
H3K4 me3:H3−5ac
H3K4 me1:H4−4ac
DNA Meth. m5C:H4K20 me2
H3K27 me3:H3K4 me1
H3K27 me3:H3K9 me3
H4K20 me2:H3K4 me3
H3K27 me3:H4K20 me3
H3K4 me3:H4−4ac
H3K9 me3:H4K20 me3
H3K4 me1:H3−5ac
DNA Meth. m5C:H3K9 me3
H3K9 me2:DNA Meth. m5C
H3K27 me3:DNA Meth. m5C
DNA Meth. m5C:H4K20 me3
H2A.Z
H3−5ac
H3K27 me2
H4K20 me2
H3K9 me2
H3K27 me3
H4K20 me3
DNA Meth. m5C
H3K4 me1
H3K9 me3
H3K4 me3
H4−4ac
0.00 0.25 0.50 0.75 1.00
π̂
WDR5 − Forward
H2A.Z:H3−5ac
H3K4 me1:H4K20 me2
H3K9 me2:DNA Meth. m5C
H3K4 me1:H2A.Z
H3K27 me2:DNA Meth. m5C
H3K4 me1:H4−4ac
H4−4ac:H3−5ac
H2A.Z:H4K20 me2
H3K4 me1:H3−5ac
H3K4 me3:H4−4ac
DNA Meth. m5C:H3K9 me3
H3K27 me3:H3K9 me3
H3K9 me3:H4K20 me3
H3K27 me3:H4K20 me3
H4K20 me2:H3−5ac
DNA Meth. m5C:H4K20 me2
DNA Meth. m5C:H4K20 me3
H4K20 me2:H3K4 me3
H3K27 me3:DNA Meth. m5C
H4K20 me2:H4−4ac
H3K9 me2
H3K27 me2
H3K4 me3
H2A.Z
H3K4 me1
H4K20 me3
H3K27 me3
H4−4ac
H4K20 me2
H3K9 me3
H3−5ac
DNA Meth. m5C
0.00 0.25 0.50 0.75 1.00
π̂
WDR5 − Reverse
H2A.Z:H3−5ac
H3K4 me1:H4K20 me2
H3K9 me2:DNA Meth. m5C
H3K4 me1:H2A.Z
H3K27 me2:DNA Meth. m5C
H3K4 me1:H4−4ac
H4−4ac:H3−5ac
H2A.Z:H4K20 me2
H3K4 me1:H3−5ac
H3K4 me3:H4−4ac
DNA Meth. m5C:H3K9 me3
H3K27 me3:H3K9 me3
H3K9 me3:H4K20 me3
H3K27 me3:H4K20 me3
H4K20 me2:H3−5ac
DNA Meth. m5C:H4K20 me2
DNA Meth. m5C:H4K20 me3
H4K20 me2:H3K4 me3
H3K27 me3:DNA Meth. m5C
H4K20 me2:H4−4ac
H3K9 me2
H3K27 me2
H3K4 me3
H2A.Z
H3K4 me1
H4K20 me3
H3K27 me3
H4−4ac
H4K20 me2
H3K9 me3
H3−5ac
DNA Meth. m5C
0.00 0.25 0.50 0.75 1.00
π̂
HSPD1 − Forward
H2A.Z:H4−4ac
H2A.Z:H3−5ac
H3K27 me3:H3K4 me1
H3K4 me3:H3−5ac
H3K4 me3:H4−4ac
H3K4 me1:H3−5ac
H2A.Z:H3K4 me3
H3K4 me1:H2A.Z
DNA Meth. m5C:H4K20 me2
H4−4ac:H3−5ac
H2A.Z:H4K20 me2
H3K27 me3:H3K9 me3
H3K4 me1:H4−4ac
H3K4 me1:H4K20 me2
H3K27 me3:DNA Meth. m5C
DNA Meth. m5C:H3K9 me3
H3K9 me3:H4K20 me3
H3K27 me3:H4K20 me3
H3K9 me2:DNA Meth. m5C
H4K20 me2:H4−4ac
H4K20 me2:H3−5ac
H4K20 me2:H3K4 me3
H3K27 me2
H3K9 me2
H4−4ac
H4K20 me3
H3K27 me3
H2A.Z
H3−5ac
H3K4 me1
H3K4 me3
H4K20 me2
H3K9 me3
DNA Meth. m5C
0.00 0.25 0.50 0.75 1.00
π̂
HSPD1 − Reverse
H2A.Z:H4−4ac
H2A.Z:H3−5ac
H3K27 me3:H3K4 me1
H3K4 me3:H3−5ac
H3K4 me3:H4−4ac
H3K4 me1:H3−5ac
H2A.Z:H3K4 me3
H3K4 me1:H2A.Z
DNA Meth. m5C:H4K20 me2
H4−4ac:H3−5ac
H2A.Z:H4K20 me2
H3K27 me3:H3K9 me3
H3K4 me1:H4−4ac
H3K4 me1:H4K20 me2
H3K27 me3:DNA Meth. m5C
DNA Meth. m5C:H3K9 me3
H3K9 me3:H4K20 me3
H3K27 me3:H4K20 me3
H3K9 me2:DNA Meth. m5C
H4K20 me2:H4−4ac
H4K20 me2:H3−5ac
H4K20 me2:H3K4 me3
H3K27 me2
H3K9 me2
H4−4ac
H4K20 me3
H3K27 me3
H2A.Z
H3−5ac
H3K4 me1
H3K4 me3
H4K20 me2
H3K9 me3
DNA Meth. m5C
0.00 0.25 0.50 0.75 1.00
π̂
TADA2A − Forward
H3K27 me3:H3K4 me1
H2A.Z:H4−4ac
DNA Meth. m5C:H4K20 me2
H2A.Z:H3−5ac
H3K4 me3:H3−5ac
H3K27 me2:DNA Meth. m5C
H2A.Z:H3K4 me3
H3K27 me3:DNA Meth. m5C
H4−4ac:H3−5ac
H4K20 me2:H4−4ac
DNA Meth. m5C:H3K9 me3
H3K27 me3:H3K9 me3
H3K4 me1:H2A.Z
H3K4 me3:H4−4ac
DNA Meth. m5C:H4K20 me3
H3K4 me1:H3−5ac
H3K4 me1:H4−4ac
H3K9 me2:DNA Meth. m5C
H3K27 me3:H4K20 me3
H3K9 me3:H4K20 me3
H3K27 me2
H4K20 me2
H3K9 me2
H3K4 me1
H4K20 me3
H3K9 me3
H2A.Z
H3K27 me3
H3K4 me3
DNA Meth. m5C
H4−4ac
H3−5ac
0.00 0.25 0.50 0.75 1.00
π̂
TADA2A − Reverse
H3K27 me3:H3K4 me1
H2A.Z:H4−4ac
DNA Meth. m5C:H4K20 me2
H2A.Z:H3−5ac
H3K4 me3:H3−5ac
H3K27 me2:DNA Meth. m5C
H2A.Z:H3K4 me3
H3K27 me3:DNA Meth. m5C
H4−4ac:H3−5ac
H4K20 me2:H4−4ac
DNA Meth. m5C:H3K9 me3
H3K27 me3:H3K9 me3
H3K4 me1:H2A.Z
H3K4 me3:H4−4ac
DNA Meth. m5C:H4K20 me3
H3K4 me1:H3−5ac
H3K4 me1:H4−4ac
H3K9 me2:DNA Meth. m5C
H3K27 me3:H4K20 me3
H3K9 me3:H4K20 me3
H3K27 me2
H4K20 me2
H3K9 me2
H3K4 me1
H4K20 me3
H3K9 me3
H2A.Z
H3K27 me3
H3K4 me3
DNA Meth. m5C
H4−4ac
H3−5ac
0.00 0.25 0.50 0.75 1.00
π̂

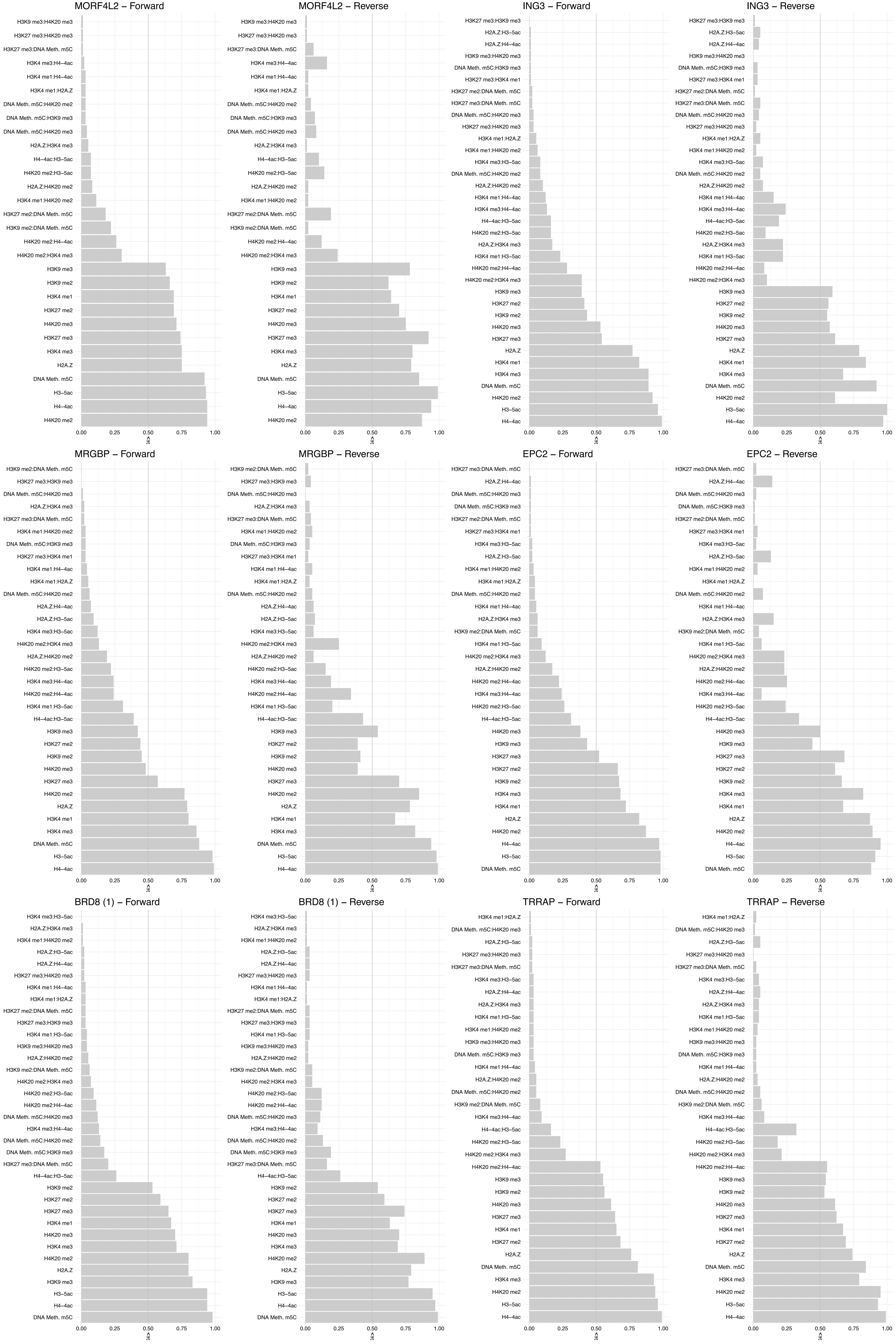
MORF4L2 – Forward
MORF4L2 – Reverse
ING3 – Forward
ING3 – Reverse
MRGBP – Forward
MRGBP – Reverse
EPC2 – Forward
EPC2 – Reverse
BRD8 (1) – Forward
BRD8 (1) – Reverse
TRRAP – Forward
TRRAP – Reverse
0.00
0.25
0.50
0.75
1.00
$\hat{\pi}$

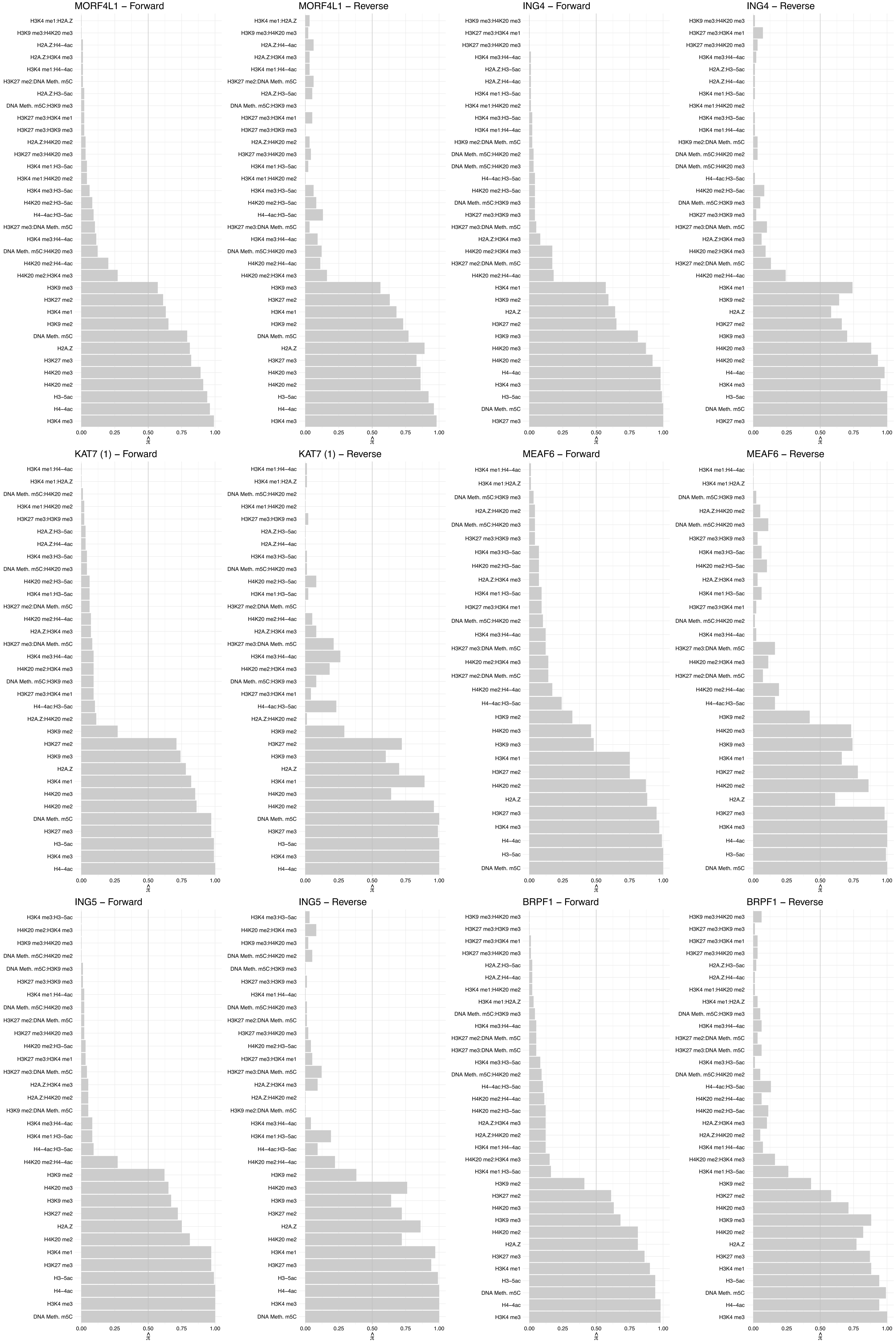
MORF4L1 – Forward
MORF4L1 – Reverse
ING4 – Forward
ING4 – Reverse
KAT7 (1) – Forward
KAT7 (1) – Reverse
MEAF6 – Forward
MEAF6 – Reverse
ING5 – Forward
ING5 – Reverse
BRPF1 – Forward
BRPF1 – Reverse
0.00 0.25 0.50 0.75 1.00
$\hat{\pi}$
H3K4 me3
H4–4ac
H3–5ac
H4K20 me2
H4K20 me3
H3K27 me3
H2A.Z
DNA Meth. m5C
H3K9 me2
H3K4 me1
H3K27 me2
H3K9 me3
H4K20 me2:H3K4 me3
H4K20 me2:H4–4ac
DNA Meth. m5C:H4K20 me3
H3K4 me3:H4–4ac
H3K27 me3:DNA Meth. m5C
H4–4ac:H3–5ac
H4K20 me2:H3–5ac
H3K4 me3:H3–5ac
H3K4 me1:H4K20 me2
H3K4 me1:H3–5ac
H3K27 me3:H4K20 me3
H2A.Z:H4K20 me2
H3K27 me3:H3K9 me3
H3K27 me3:H3K4 me1
DNA Meth. m5C:H3K9 me3
H2A.Z:H3–5ac
H3K27 me2:DNA Meth. m5C
H3K4 me1:H4–4ac
H2A.Z:H3K4 me3
H2A.Z:H4–4ac
H3K9 me3:H4K20 me3
H3K4 me1:H2A.Z
H3K9 me2:DNA Meth. m5C
DNA Meth. m5C:H4K20 me2

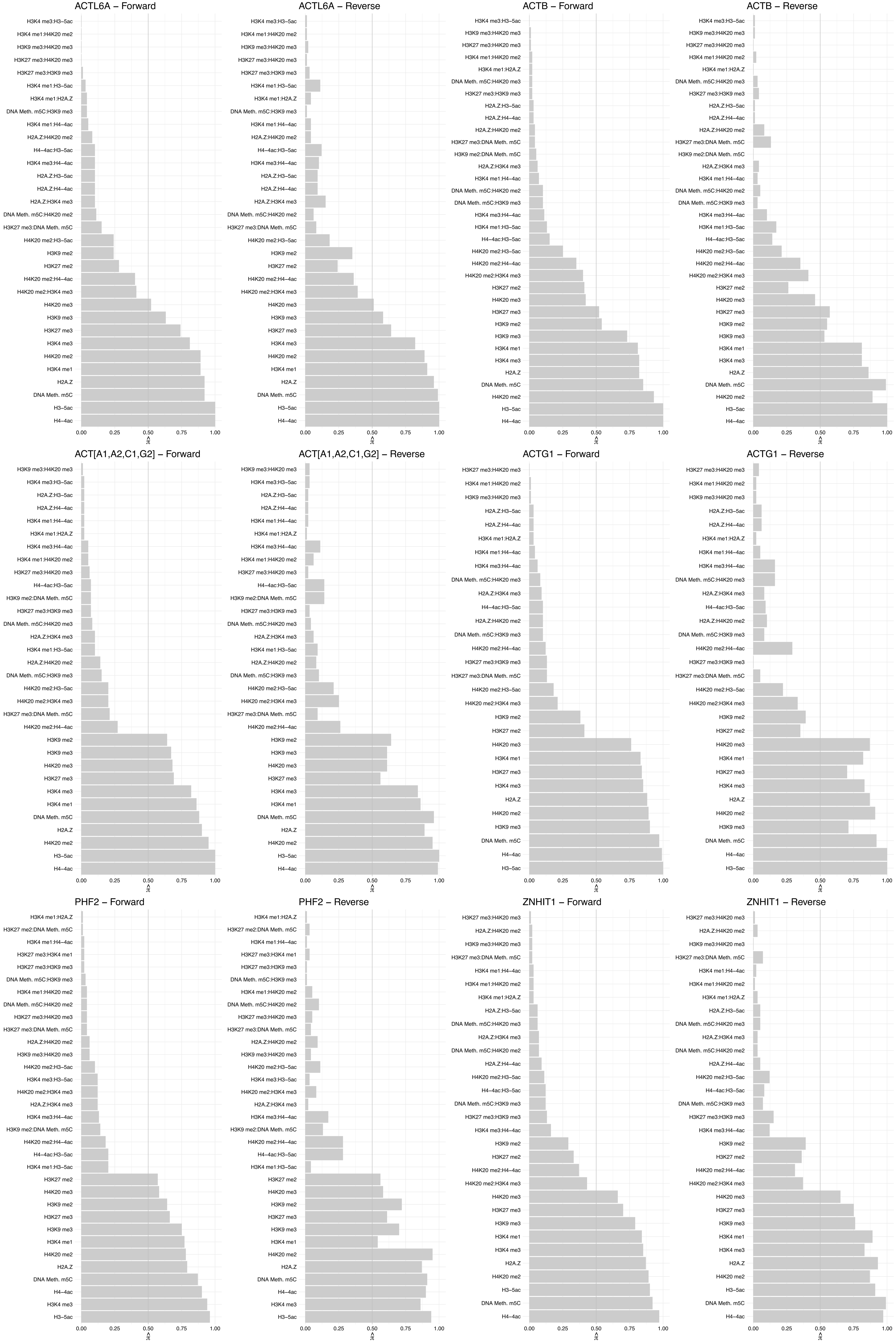
ACTL6A – Forward
ACTL6A – Reverse
ACTB – Forward
ACTB – Reverse
ACT[A1,A2,C1,G2] – Forward
ACT[A1,A2,C1,G2] – Reverse
ACTG1 – Forward
ACTG1 – Reverse
PHF2 – Forward
PHF2 – Reverse
ZNHIT1 – Forward
ZNHIT1 – Reverse
0.00
0.25
0.50
0.75
1.00
$\hat{\pi}$

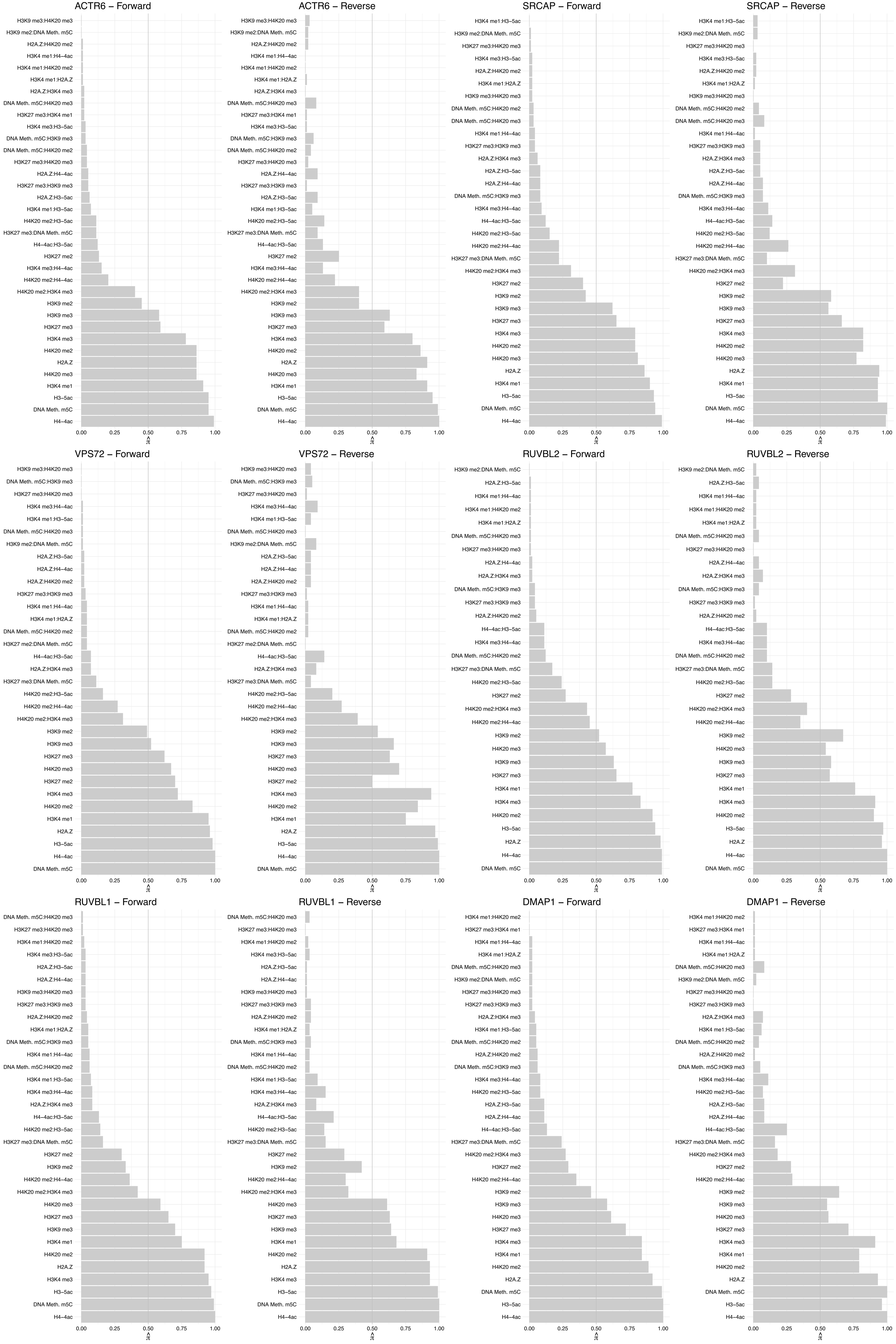
ACTR6 − Forward
H3K9 me3:H4K20 me3
H3K9 me2:DNA Meth. m5C
H2A.Z:H4K20 me2
H3K4 me1:H4−4ac
H3K4 me1:H4K20 me2
H3K4 me1:H2A.Z
H2A.Z:H3K4 me3
DNA Meth. m5C:H4K20 me3
H3K27 me3:H3K4 me1
H3K4 me3:H3−5ac
DNA Meth. m5C:H3K9 me3
DNA Meth. m5C:H4K20 me2
H3K27 me3:H4K20 me3
H2A.Z:H4−4ac
H3K27 me3:H3K9 me3
H2A.Z:H3−5ac
H3K4 me1:H3−5ac
H4K20 me2:H3−5ac
H3K27 me3:DNA Meth. m5C
H4−4ac:H3−5ac
H3K27 me2
H3K4 me3:H4−4ac
H4K20 me2:H4−4ac
H4K20 me2:H3K4 me3
H3K9 me2
H3K9 me3
H3K27 me3
H3K4 me3
H4K20 me2
H2A.Z
H4K20 me3
H3K4 me1
H3−5ac
DNA Meth. m5C
H4−4ac
0.00 0.25 0.50 0.75 1.00
π̂
ACTR6 − Reverse
H3K9 me3:H4K20 me3
H3K9 me2:DNA Meth. m5C
H2A.Z:H4K20 me2
H3K4 me1:H4−4ac
H3K4 me1:H4K20 me2
H3K4 me1:H2A.Z
H2A.Z:H3K4 me3
DNA Meth. m5C:H4K20 me3
H3K27 me3:H3K4 me1
H3K4 me3:H3−5ac
DNA Meth. m5C:H3K9 me3
DNA Meth. m5C:H4K20 me2
H3K27 me3:H4K20 me3
H2A.Z:H4−4ac
H3K27 me3:H3K9 me3
H2A.Z:H3−5ac
H3K4 me1:H3−5ac
H4K20 me2:H3−5ac
H3K27 me3:DNA Meth. m5C
H4−4ac:H3−5ac
H3K27 me2
H3K4 me3:H4−4ac
H4K20 me2:H4−4ac
H4K20 me2:H3K4 me3
H3K9 me2
H3K9 me3
H3K27 me3
H3K4 me3
H4K20 me2
H2A.Z
H4K20 me3
H3K4 me1
H3−5ac
DNA Meth. m5C
H4−4ac
0.00 0.25 0.50 0.75 1.00
π̂
SRCAP − Forward
H3K4 me1:H3−5ac
H3K9 me2:DNA Meth. m5C
H3K27 me3:H4K20 me3
H3K4 me3:H3−5ac
H2A.Z:H4K20 me2
H3K4 me1:H2A.Z
H3K9 me3:H4K20 me3
DNA Meth. m5C:H4K20 me2
DNA Meth. m5C:H4K20 me3
H3K4 me1:H4−4ac
H3K27 me3:H3K9 me3
H2A.Z:H3K4 me3
H2A.Z:H3−5ac
H2A.Z:H4−4ac
DNA Meth. m5C:H3K9 me3
H3K4 me3:H4−4ac
H4−4ac:H3−5ac
H4K20 me2:H3−5ac
H4K20 me2:H4−4ac
H3K27 me3:DNA Meth. m5C
H4K20 me2:H3K4 me3
H3K27 me2
H3K9 me2
H3K9 me3
H3K27 me3
H3K4 me3
H4K20 me2
H4K20 me3
H2A.Z
H3K4 me1
H3−5ac
DNA Meth. m5C
H4−4ac
0.00 0.25 0.50 0.75 1.00
π̂
SRCAP − Reverse
H3K4 me1:H3−5ac
H3K9 me2:DNA Meth. m5C
H3K27 me3:H4K20 me3
H3K4 me3:H3−5ac
H2A.Z:H4K20 me2
H3K4 me1:H2A.Z
H3K9 me3:H4K20 me3
DNA Meth. m5C:H4K20 me2
DNA Meth. m5C:H4K20 me3
H3K4 me1:H4−4ac
H3K27 me3:H3K9 me3
H2A.Z:H3K4 me3
H2A.Z:H3−5ac
H2A.Z:H4−4ac
DNA Meth. m5C:H3K9 me3
H3K4 me3:H4−4ac
H4−4ac:H3−5ac
H4K20 me2:H3−5ac
H4K20 me2:H4−4ac
H3K27 me3:DNA Meth. m5C
H4K20 me2:H3K4 me3
H3K27 me2
H3K9 me2
H3K9 me3
H3K27 me3
H3K4 me3
H4K20 me2
H4K20 me3
H2A.Z
H3K4 me1
H3−5ac
DNA Meth. m5C
H4−4ac
0.00 0.25 0.50 0.75 1.00
π̂
VPS72 − Forward
H3K9 me3:H4K20 me3
DNA Meth. m5C:H3K9 me3
H3K27 me3:H4K20 me3
H3K4 me3:H4−4ac
H3K4 me1:H3−5ac
DNA Meth. m5C:H4K20 me3
H3K9 me2:DNA Meth. m5C
H2A.Z:H3−5ac
H2A.Z:H4−4ac
H2A.Z:H4K20 me2
H3K27 me3:H3K9 me3
H3K4 me1:H4−4ac
H3K4 me1:H2A.Z
DNA Meth. m5C:H4K20 me2
H3K27 me2:DNA Meth. m5C
H4−4ac:H3−5ac
H2A.Z:H3K4 me3
H3K27 me3:DNA Meth. m5C
H4K20 me2:H3−5ac
H4K20 me2:H4−4ac
H4K20 me2:H3K4 me3
H3K9 me2
H3K9 me3
H3K27 me3
H4K20 me3
H3K27 me2
H3K4 me3
H4K20 me2
H3K4 me1
H2A.Z
H3−5ac
H4−4ac
DNA Meth. m5C
0.00 0.25 0.50 0.75 1.00
π̂
VPS72 − Reverse
H3K9 me3:H4K20 me3
DNA Meth. m5C:H3K9 me3
H3K27 me3:H4K20 me3
H3K4 me3:H4−4ac
H3K4 me1:H3−5ac
DNA Meth. m5C:H4K20 me3
H3K9 me2:DNA Meth. m5C
H2A.Z:H3−5ac
H2A.Z:H4−4ac
H2A.Z:H4K20 me2
H3K27 me3:H3K9 me3
H3K4 me1:H4−4ac
H3K4 me1:H2A.Z
DNA Meth. m5C:H4K20 me2
H3K27 me2:DNA Meth. m5C
H4−4ac:H3−5ac
H2A.Z:H3K4 me3
H3K27 me3:DNA Meth. m5C
H4K20 me2:H3−5ac
H4K20 me2:H4−4ac
H4K20 me2:H3K4 me3
H3K9 me2
H3K9 me3
H3K27 me3
H4K20 me3
H3K27 me2
H3K4 me3
H4K20 me2
H3K4 me1
H2A.Z
H3−5ac
H4−4ac
DNA Meth. m5C
0.00 0.25 0.50 0.75 1.00
π̂
RUVBL2 − Forward
H3K9 me2:DNA Meth. m5C
H2A.Z:H3−5ac
H3K4 me1:H4−4ac
H3K4 me1:H4K20 me2
H3K4 me1:H2A.Z
DNA Meth. m5C:H4K20 me3
H3K27 me3:H4K20 me3
H2A.Z:H4−4ac
H2A.Z:H3K4 me3
DNA Meth. m5C:H3K9 me3
H3K27 me3:H3K9 me3
H2A.Z:H4K20 me2
H4−4ac:H3−5ac
H3K4 me3:H4−4ac
DNA Meth. m5C:H4K20 me2
H3K27 me3:DNA Meth. m5C
H4K20 me2:H3−5ac
H3K27 me2
H4K20 me2:H3K4 me3
H4K20 me2:H4−4ac
H3K9 me2
H4K20 me3
H3K9 me3
H3K27 me3
H3K4 me1
H3K4 me3
H4K20 me2
H3−5ac
H2A.Z
H4−4ac
DNA Meth. m5C
0.00 0.25 0.50 0.75 1.00
π̂
RUVBL2 − Reverse
H3K9 me2:DNA Meth. m5C
H2A.Z:H3−5ac
H3K4 me1:H4−4ac
H3K4 me1:H4K20 me2
H3K4 me1:H2A.Z
DNA Meth. m5C:H4K20 me3
H3K27 me3:H4K20 me3
H2A.Z:H4−4ac
H2A.Z:H3K4 me3
DNA Meth. m5C:H3K9 me3
H3K27 me3:H3K9 me3
H2A.Z:H4K20 me2
H4−4ac:H3−5ac
H3K4 me3:H4−4ac
DNA Meth. m5C:H4K20 me2
H3K27 me3:DNA Meth. m5C
H4K20 me2:H3−5ac
H3K27 me2
H4K20 me2:H3K4 me3
H4K20 me2:H4−4ac
H3K9 me2
H4K20 me3
H3K9 me3
H3K27 me3
H3K4 me1
H3K4 me3
H4K20 me2
H3−5ac
H2A.Z
H4−4ac
DNA Meth. m5C
0.00 0.25 0.50 0.75 1.00
π̂
RUVBL1 − Forward
DNA Meth. m5C:H4K20 me3
H3K27 me3:H4K20 me3
H3K4 me1:H4K20 me2
H3K4 me3:H3−5ac
H2A.Z:H3−5ac
H2A.Z:H4−4ac
H3K9 me3:H4K20 me3
H3K27 me3:H3K9 me3
H2A.Z:H4K20 me2
H3K4 me1:H2A.Z
DNA Meth. m5C:H3K9 me3
H3K4 me1:H4−4ac
DNA Meth. m5C:H4K20 me2
H3K4 me1:H3−5ac
H3K4 me3:H4−4ac
H2A.Z:H3K4 me3
H4−4ac:H3−5ac
H4K20 me2:H3−5ac
H3K27 me3:DNA Meth. m5C
H3K27 me2
H3K9 me2
H4K20 me2:H4−4ac
H4K20 me2:H3K4 me3
H4K20 me3
H3K27 me3
H3K9 me3
H3K4 me1
H4K20 me2
H2A.Z
H3K4 me3
H3−5ac
DNA Meth. m5C
H4−4ac
0.00 0.25 0.50 0.75 1.00
π̂
RUVBL1 − Reverse
DNA Meth. m5C:H4K20 me3
H3K27 me3:H4K20 me3
H3K4 me1:H4K20 me2
H3K4 me3:H3−5ac
H2A.Z:H3−5ac
H2A.Z:H4−4ac
H3K9 me3:H4K20 me3
H3K27 me3:H3K9 me3
H2A.Z:H4K20 me2
H3K4 me1:H2A.Z
DNA Meth. m5C:H3K9 me3
H3K4 me1:H4−4ac
DNA Meth. m5C:H4K20 me2
H3K4 me1:H3−5ac
H3K4 me3:H4−4ac
H2A.Z:H3K4 me3
H4−4ac:H3−5ac
H4K20 me2:H3−5ac
H3K27 me3:DNA Meth. m5C
H3K27 me2
H3K9 me2
H4K20 me2:H4−4ac
H4K20 me2:H3K4 me3
H4K20 me3
H3K27 me3
H3K9 me3
H3K4 me1
H4K20 me2
H2A.Z
H3K4 me3
H3−5ac
DNA Meth. m5C
H4−4ac
0.00 0.25 0.50 0.75 1.00
π̂
DMAP1 − Forward
H3K4 me1:H4K20 me2
H3K27 me3:H3K4 me1
H3K4 me1:H4−4ac
H3K4 me1:H2A.Z
DNA Meth. m5C:H4K20 me3
H3K9 me2:DNA Meth. m5C
H3K27 me3:H4K20 me3
H3K27 me3:H3K9 me3
H2A.Z:H3K4 me3
H3K4 me1:H3−5ac
DNA Meth. m5C:H4K20 me2
H2A.Z:H4K20 me2
DNA Meth. m5C:H3K9 me3
H3K4 me3:H4−4ac
H4K20 me2:H3−5ac
H2A.Z:H3−5ac
H2A.Z:H4−4ac
H4−4ac:H3−5ac
H3K27 me3:DNA Meth. m5C
H4K20 me2:H3K4 me3
H3K27 me2
H4K20 me2:H4−4ac
H3K9 me2
H3K9 me3
H4K20 me3
H3K27 me3
H3K4 me3
H3K4 me1
H4K20 me2
H2A.Z
DNA Meth. m5C
H3−5ac
H4−4ac
0.00 0.25 0.50 0.75 1.00
π̂
DMAP1 − Reverse
H3K4 me1:H4K20 me2
H3K27 me3:H3K4 me1
H3K4 me1:H4−4ac
H3K4 me1:H2A.Z
DNA Meth. m5C:H4K20 me3
H3K9 me2:DNA Meth. m5C
H3K27 me3:H4K20 me3
H3K27 me3:H3K9 me3
H2A.Z:H3K4 me3
H3K4 me1:H3−5ac
DNA Meth. m5C:H4K20 me2
H2A.Z:H4K20 me2
DNA Meth. m5C:H3K9 me3
H3K4 me3:H4−4ac
H4K20 me2:H3−5ac
H2A.Z:H3−5ac
H2A.Z:H4−4ac
H4−4ac:H3−5ac
H3K27 me3:DNA Meth. m5C
H4K20 me2:H3K4 me3
H3K27 me2
H4K20 me2:H4−4ac
H3K9 me2
H3K9 me3
H4K20 me3
H3K27 me3
H3K4 me3
H3K4 me1
H4K20 me2
H2A.Z
DNA Meth. m5C
H3−5ac
H4−4ac
0.00 0.25 0.50 0.75 1.00
π̂

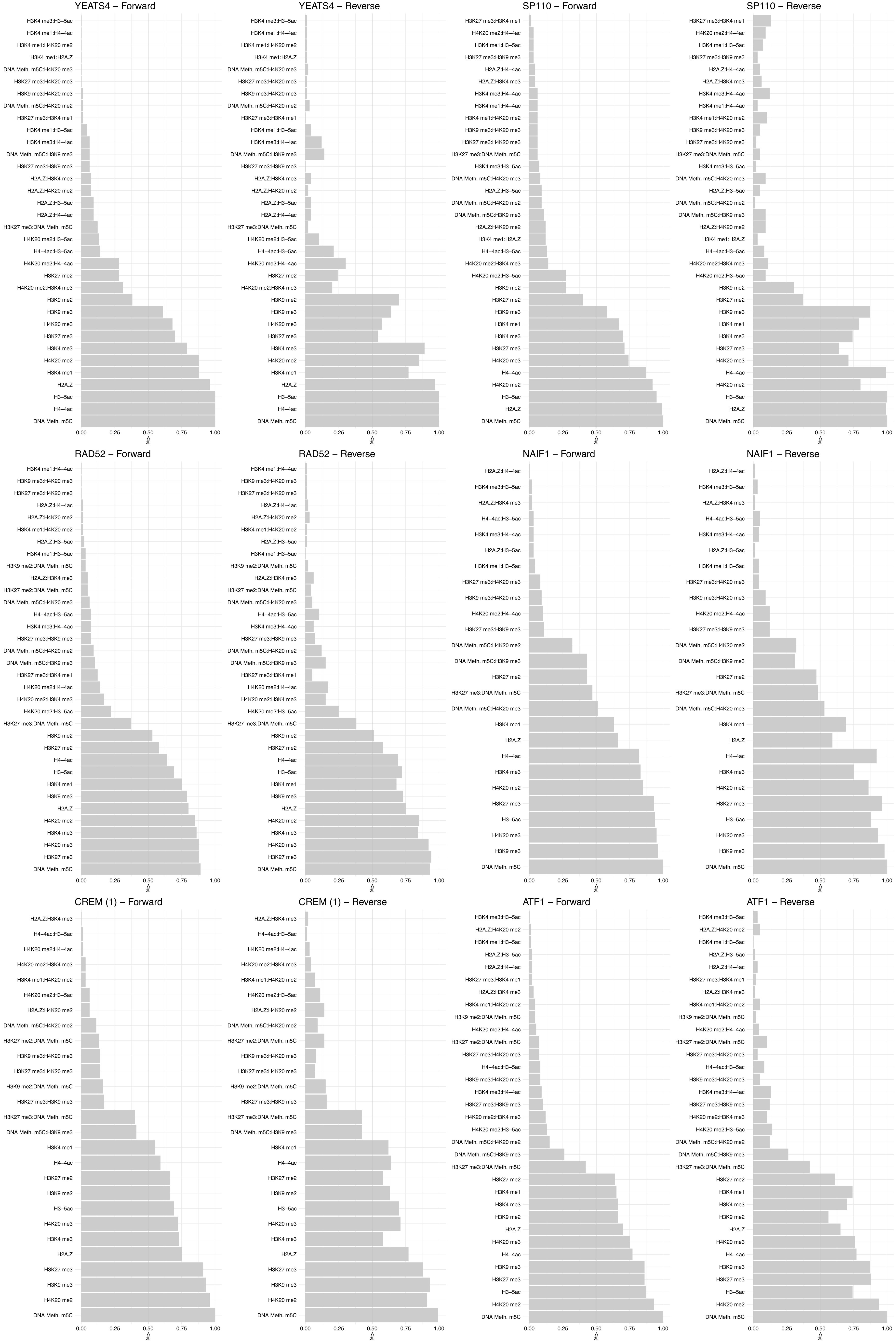

YEATS4 – Forward
H3K4 me3:H3–5ac
H3K4 me1:H4–4ac
H3K4 me1:H4K20 me2
H3K4 me1:H2A.Z
DNA Meth. m5C:H4K20 me3
H3K27 me3:H4K20 me3
H3K9 me3:H4K20 me3
DNA Meth. m5C:H4K20 me2
H3K27 me3:H3K4 me1
H3K4 me1:H3–5ac
H3K4 me3:H4–4ac
DNA Meth. m5C:H3K9 me3
H3K27 me3:H3K9 me3
H2A.Z:H3K4 me3
H2A.Z:H4K20 me2
H2A.Z:H3–5ac
H2A.Z:H4–4ac
H3K27 me3:DNA Meth. m5C
H4K20 me2:H3–5ac
H4–4ac:H3–5ac
H4K20 me2:H4–4ac
H3K27 me2
H4K20 me2:H3K4 me3
H3K9 me2
H3K9 me3
H4K20 me3
H3K27 me3
H3K4 me3
H4K20 me2
H3K4 me1
H2A.Z
H3–5ac
H4–4ac
DNA Meth. m5C
0.00
0.25
0.50
0.75
1.00
$\hat{\pi}$
YEATS4 – Reverse
SP110 – Forward
H3K27 me3:H3K4 me1
H4K20 me2:H4–4ac
H3K4 me1:H3–5ac
H3K27 me3:H3K9 me3
H2A.Z:H4–4ac
H2A.Z:H3K4 me3
H3K4 me3:H4–4ac
H3K4 me1:H4–4ac
H3K4 me1:H4K20 me2
H3K9 me3:H4K20 me3
H3K27 me3:H4K20 me3
H3K27 me3:DNA Meth. m5C
H3K4 me3:H3–5ac
DNA Meth. m5C:H4K20 me3
H2A.Z:H3–5ac
DNA Meth. m5C:H4K20 me2
DNA Meth. m5C:H3K9 me3
H2A.Z:H4K20 me2
H3K4 me1:H2A.Z
H4–4ac:H3–5ac
H4K20 me2:H3K4 me3
H4K20 me2:H3–5ac
H3K9 me2
H3K27 me2
H3K9 me3
H3K4 me1
H3K4 me3
H3K27 me3
H4K20 me3
H4–4ac
H4K20 me2
H3–5ac
H2A.Z
DNA Meth. m5C
SP110 – Reverse
RAD52 – Forward
H3K4 me1:H4–4ac
H3K9 me3:H4K20 me3
H3K27 me3:H4K20 me3
H2A.Z:H4–4ac
H2A.Z:H4K20 me2
H3K4 me1:H4K20 me2
H2A.Z:H3–5ac
H3K4 me1:H3–5ac
H3K9 me2:DNA Meth. m5C
H2A.Z:H3K4 me3
H3K27 me2:DNA Meth. m5C
DNA Meth. m5C:H4K20 me3
H4–4ac:H3–5ac
H3K4 me3:H4–4ac
H3K27 me3:H3K9 me3
DNA Meth. m5C:H4K20 me2
DNA Meth. m5C:H3K9 me3
H3K27 me3:H3K4 me1
H4K20 me2:H4–4ac
H4K20 me2:H3K4 me3
H4K20 me2:H3–5ac
H3K27 me3:DNA Meth. m5C
H3K9 me2
H3K27 me2
H4–4ac
H3–5ac
H3K4 me1
H3K9 me3
H2A.Z
H4K20 me2
H3K4 me3
H4K20 me3
H3K27 me3
DNA Meth. m5C
RAD52 – Reverse
NAIF1 – Forward
H2A.Z:H4–4ac
H3K4 me3:H3–5ac
H2A.Z:H3K4 me3
H4–4ac:H3–5ac
H3K4 me3:H4–4ac
H2A.Z:H3–5ac
H3K4 me1:H3–5ac
H3K27 me3:H4K20 me3
H3K9 me3:H4K20 me3
H4K20 me2:H4–4ac
H3K27 me3:H3K9 me3
DNA Meth. m5C:H4K20 me2
DNA Meth. m5C:H3K9 me3
H3K27 me2
H3K27 me3:DNA Meth. m5C
DNA Meth. m5C:H4K20 me3
H3K4 me1
H2A.Z
H4–4ac
H3K4 me3
H4K20 me2
H3K27 me3
H3–5ac
H4K20 me3
H3K9 me3
DNA Meth. m5C
NAIF1 – Reverse
CREM (1) – Forward
H2A.Z:H3K4 me3
H4–4ac:H3–5ac
H4K20 me2:H4–4ac
H4K20 me2:H3K4 me3
H3K4 me1:H4K20 me2
H4K20 me2:H3–5ac
H2A.Z:H4K20 me2
DNA Meth. m5C:H4K20 me2
H3K27 me2:DNA Meth. m5C
H3K9 me3:H4K20 me3
H3K27 me3:H4K20 me3
H3K9 me2:DNA Meth. m5C
H3K27 me3:H3K9 me3
H3K27 me3:DNA Meth. m5C
DNA Meth. m5C:H3K9 me3
H3K4 me1
H4–4ac
H3K27 me2
H3K9 me2
H3–5ac
H4K20 me3
H3K4 me3
H2A.Z
H3K27 me3
H3K9 me3
H4K20 me2
DNA Meth. m5C
CREM (1) – Reverse
ATF1 – Forward
H3K4 me3:H3–5ac
H2A.Z:H4K20 me2
H3K4 me1:H3–5ac
H2A.Z:H3–5ac
H2A.Z:H4–4ac
H3K27 me3:H3K4 me1
H2A.Z:H3K4 me3
H3K4 me1:H4K20 me2
H3K9 me2:DNA Meth. m5C
H4K20 me2:H4–4ac
H3K27 me2:DNA Meth. m5C
H3K27 me3:H4K20 me3
H4–4ac:H3–5ac
H3K9 me3:H4K20 me3
H3K4 me3:H4–4ac
H3K27 me3:H3K9 me3
H4K20 me2:H3K4 me3
H4K20 me2:H3–5ac
DNA Meth. m5C:H4K20 me2
DNA Meth. m5C:H3K9 me3
H3K27 me3:DNA Meth. m5C
H3K27 me2
H3K4 me1
H3K4 me3
H3K9 me2
H2A.Z
H4K20 me3
H4–4ac
H3K9 me3
H3K27 me3
H3–5ac
H4K20 me2
DNA Meth. m5C
ATF1 – Reverse

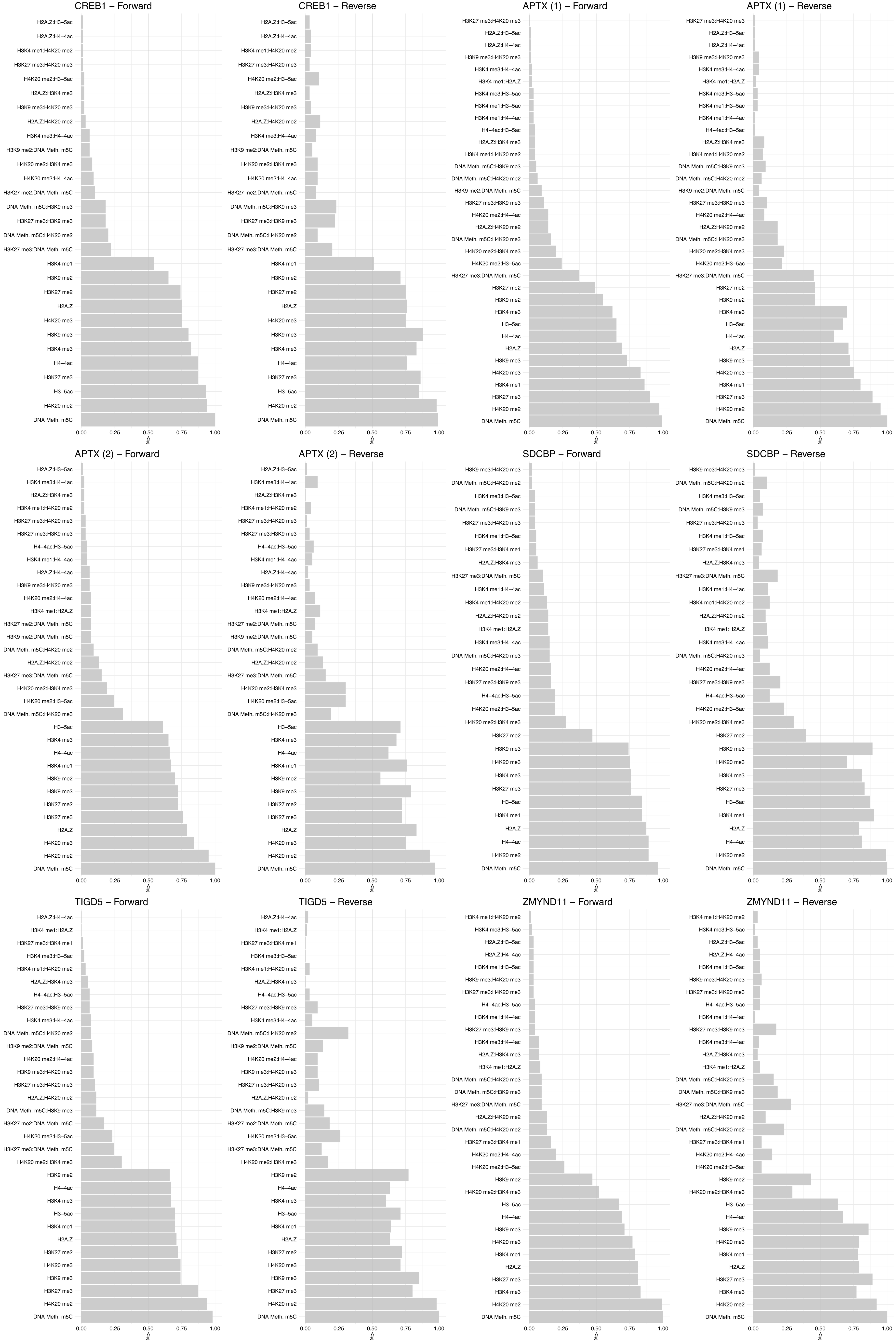

CREB1 – Forward
CREB1 – Reverse
H2A.Z:H3–5ac
H2A.Z:H4–4ac
H3K4 me1:H4K20 me2
H3K27 me3:H4K20 me3
H4K20 me2:H3–5ac
H2A.Z:H3K4 me3
H3K9 me3:H4K20 me3
H2A.Z:H4K20 me2
H3K4 me3:H4–4ac
H3K9 me2:DNA Meth. m5C
H4K20 me2:H3K4 me3
H4K20 me2:H4–4ac
H3K27 me2:DNA Meth. m5C
DNA Meth. m5C:H3K9 me3
H3K27 me3:H3K9 me3
DNA Meth. m5C:H4K20 me2
H3K27 me3:DNA Meth. m5C
H3K4 me1
H3K9 me2
H3K27 me2
H2A.Z
H4K20 me3
H3K9 me3
H3K4 me3
H4–4ac
H3K27 me3
H3–5ac
H4K20 me2
DNA Meth. m5C
APTX (1) – Forward
APTX (1) – Reverse
H3K27 me3:H4K20 me3
H2A.Z:H3–5ac
H2A.Z:H4–4ac
H3K9 me3:H4K20 me3
H3K4 me3:H4–4ac
H3K4 me1:H2A.Z
H3K4 me3:H3–5ac
H3K4 me1:H3–5ac
H3K4 me1:H4–4ac
H4–4ac:H3–5ac
H2A.Z:H3K4 me3
H3K4 me1:H4K20 me2
DNA Meth. m5C:H3K9 me3
DNA Meth. m5C:H4K20 me2
H3K9 me2:DNA Meth. m5C
H3K27 me3:H3K9 me3
H4K20 me2:H4–4ac
H2A.Z:H4K20 me2
DNA Meth. m5C:H4K20 me3
H4K20 me2:H3K4 me3
H4K20 me2:H3–5ac
H3K27 me3:DNA Meth. m5C
H3K27 me2
H3K9 me2
H3K4 me3
H3–5ac
H4–4ac
H2A.Z
H3K9 me3
H4K20 me3
H3K4 me1
H3K27 me3
H4K20 me2
DNA Meth. m5C
APTX (2) – Forward
APTX (2) – Reverse
H2A.Z:H3–5ac
H3K4 me3:H4–4ac
H2A.Z:H3K4 me3
H3K4 me1:H4K20 me2
H3K27 me3:H4K20 me3
H3K27 me3:H3K9 me3
H4–4ac:H3–5ac
H3K4 me1:H4–4ac
H2A.Z:H4–4ac
H3K9 me3:H4K20 me3
H4K20 me2:H4–4ac
H3K4 me1:H2A.Z
H3K27 me2:DNA Meth. m5C
H3K9 me2:DNA Meth. m5C
DNA Meth. m5C:H4K20 me2
H2A.Z:H4K20 me2
H3K27 me3:DNA Meth. m5C
H4K20 me2:H3K4 me3
H4K20 me2:H3–5ac
DNA Meth. m5C:H4K20 me3
H3–5ac
H3K4 me3
H4–4ac
H3K4 me1
H3K9 me2
H3K9 me3
H3K27 me2
H3K27 me3
H2A.Z
H4K20 me3
H4K20 me2
DNA Meth. m5C
SDCBP – Forward
SDCBP – Reverse
H3K9 me3:H4K20 me3
DNA Meth. m5C:H4K20 me2
H3K4 me3:H3–5ac
DNA Meth. m5C:H3K9 me3
H3K27 me3:H4K20 me3
H3K4 me1:H3–5ac
H3K27 me3:H3K4 me1
H2A.Z:H3K4 me3
H3K27 me3:DNA Meth. m5C
H3K4 me1:H4–4ac
H3K4 me1:H4K20 me2
H2A.Z:H4K20 me2
H3K4 me1:H2A.Z
H3K4 me3:H4–4ac
DNA Meth. m5C:H4K20 me3
H4K20 me2:H4–4ac
H3K27 me3:H3K9 me3
H4–4ac:H3–5ac
H4K20 me2:H3–5ac
H4K20 me2:H3K4 me3
H3K27 me2
H3K9 me3
H4K20 me3
H3K4 me3
H3K27 me3
H3–5ac
H3K4 me1
H2A.Z
H4–4ac
H4K20 me2
DNA Meth. m5C
TIGD5 – Forward
TIGD5 – Reverse
H2A.Z:H4–4ac
H3K4 me1:H2A.Z
H3K27 me3:H3K4 me1
H3K4 me3:H3–5ac
H3K4 me1:H4K20 me2
H2A.Z:H3K4 me3
H4–4ac:H3–5ac
H3K27 me3:H3K9 me3
H3K4 me3:H4–4ac
DNA Meth. m5C:H4K20 me2
H3K9 me2:DNA Meth. m5C
H4K20 me2:H4–4ac
H3K9 me3:H4K20 me3
H3K27 me3:H4K20 me3
H2A.Z:H4K20 me2
DNA Meth. m5C:H3K9 me3
H3K27 me2:DNA Meth. m5C
H4K20 me2:H3–5ac
H3K27 me3:DNA Meth. m5C
H4K20 me2:H3K4 me3
H3K9 me2
H4–4ac
H3K4 me3
H3–5ac
H3K4 me1
H2A.Z
H3K27 me2
H4K20 me3
H3K9 me3
H3K27 me3
H4K20 me2
DNA Meth. m5C
ZMYND11 – Forward
ZMYND11 – Reverse
H3K4 me1:H4K20 me2
H3K4 me3:H3–5ac
H2A.Z:H3–5ac
H2A.Z:H4–4ac
H3K4 me1:H3–5ac
H3K9 me3:H4K20 me3
H3K27 me3:H4K20 me3
H4–4ac:H3–5ac
H3K4 me1:H4–4ac
H3K27 me3:H3K9 me3
H3K4 me3:H4–4ac
H2A.Z:H3K4 me3
H3K4 me1:H2A.Z
DNA Meth. m5C:H4K20 me3
DNA Meth. m5C:H3K9 me3
H3K27 me3:DNA Meth. m5C
H2A.Z:H4K20 me2
DNA Meth. m5C:H4K20 me2
H3K27 me3:H3K4 me1
H4K20 me2:H4–4ac
H4K20 me2:H3–5ac
H3K9 me2
H4K20 me2:H3K4 me3
H3–5ac
H4–4ac
H3K9 me3
H4K20 me3
H3K4 me1
H2A.Z
H3K27 me3
H3K4 me3
H4K20 me2
DNA Meth. m5C
0.00
0.25
0.50
0.75
1.00
$\hat{\pi}$

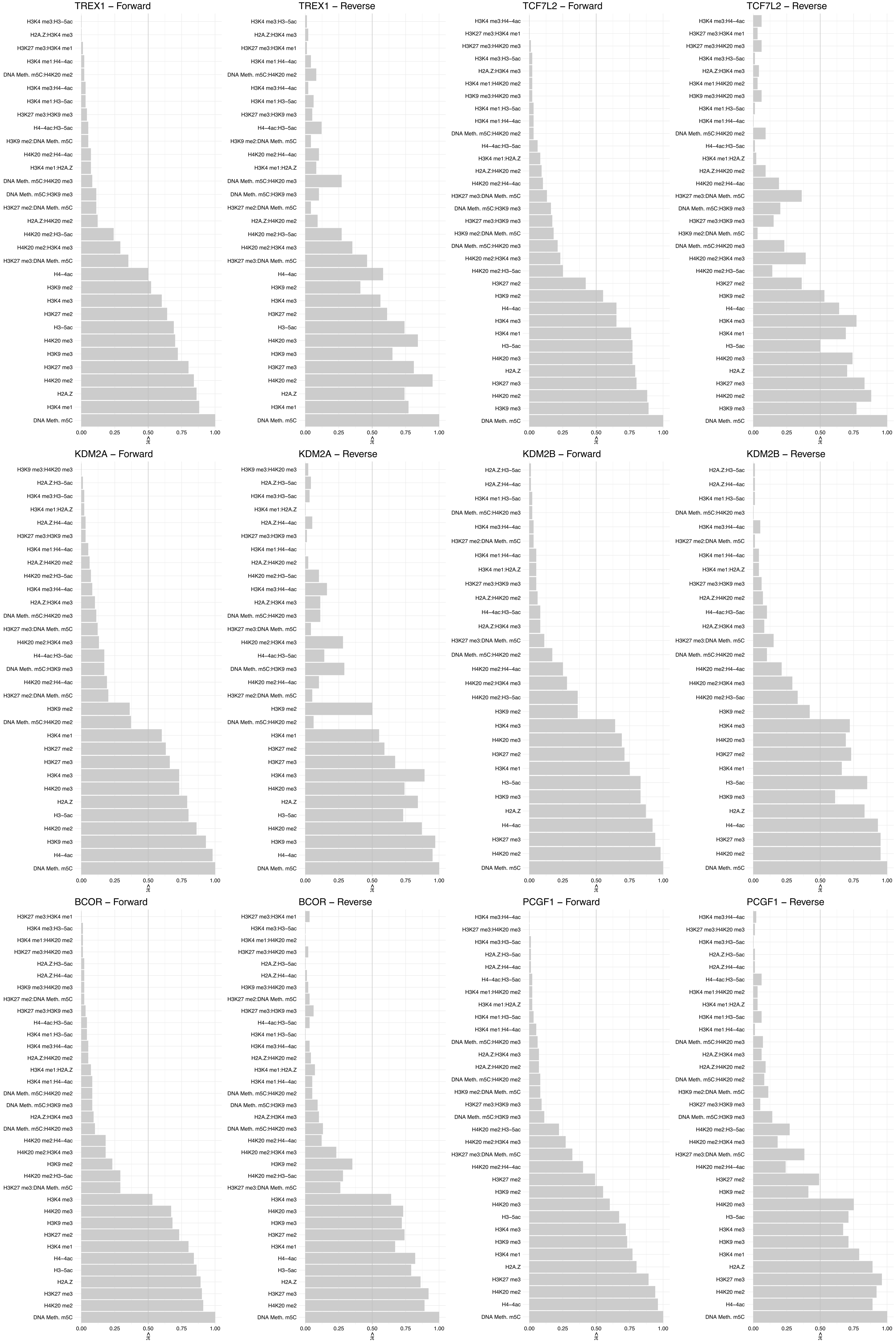
TREX1 – Forward
TREX1 – Reverse
TCF7L2 – Forward
TCF7L2 – Reverse
KDM2A – Forward
KDM2A – Reverse
KDM2B – Forward
KDM2B – Reverse
BCOR – Forward
BCOR – Reverse
PCGF1 – Forward
PCGF1 – Reverse
0.00 0.25 0.50 0.75 1.00
$\hat{\pi}$

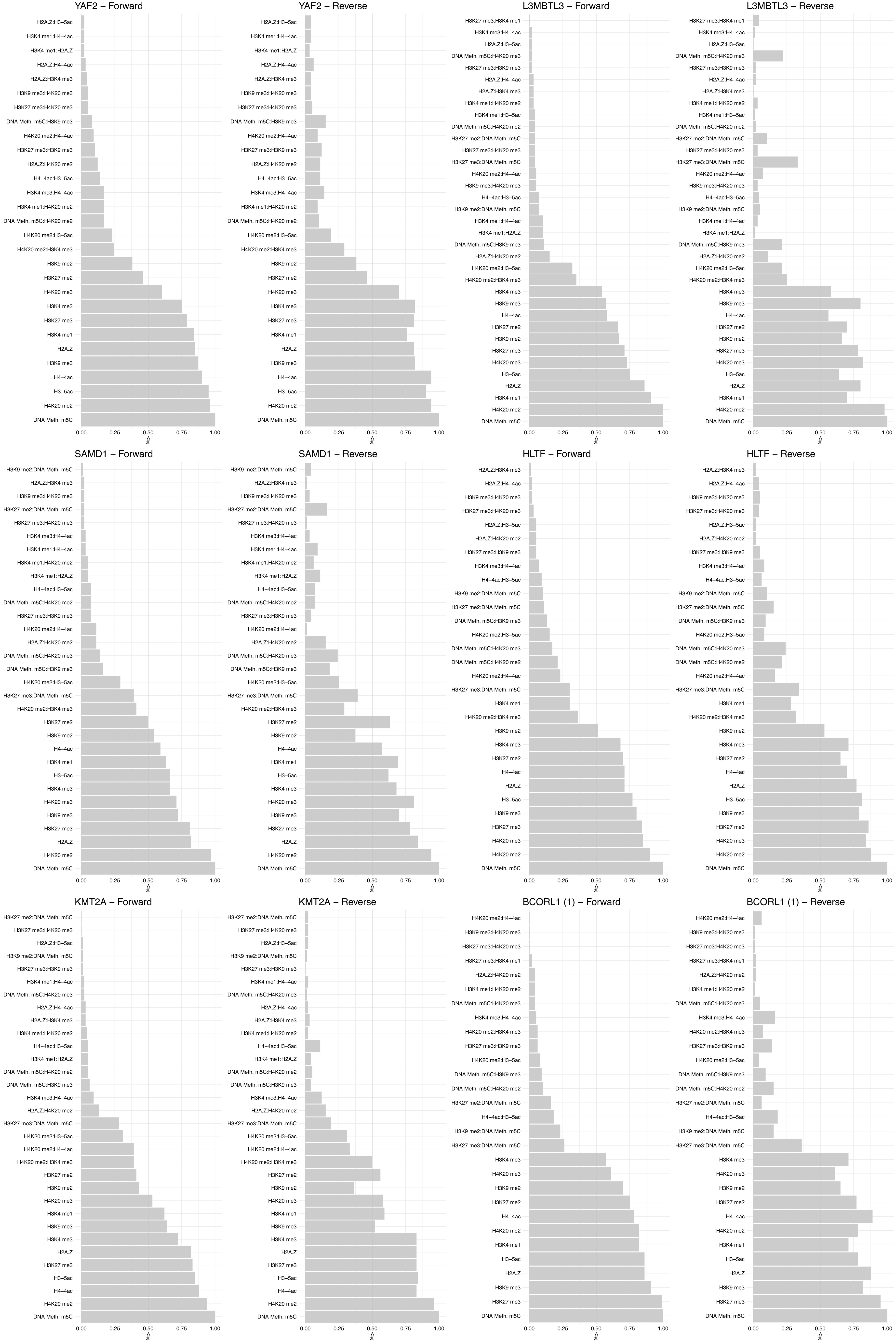

YAF2 – Forward
H2A.Z:H3–5ac
H3K4 me1:H4–4ac
H3K4 me1:H2A.Z
H2A.Z:H4–4ac
H2A.Z:H3K4 me3
H3K9 me3:H4K20 me3
H3K27 me3:H4K20 me3
DNA Meth. m5C:H3K9 me3
H4K20 me2:H4–4ac
H3K27 me3:H3K9 me3
H2A.Z:H4K20 me2
H4–4ac:H3–5ac
H3K4 me3:H4–4ac
H3K4 me1:H4K20 me2
DNA Meth. m5C:H4K20 me2
H4K20 me2:H3–5ac
H4K20 me2:H3K4 me3
H3K9 me2
H3K27 me2
H4K20 me3
H3K4 me3
H3K27 me3
H3K4 me1
H2A.Z
H3K9 me3
H4–4ac
H3–5ac
H4K20 me2
DNA Meth. m5C
0.00
0.25
0.50
0.75
1.00
π̂
YAF2 – Reverse
L3MBTL3 – Forward
H3K27 me3:H3K4 me1
H3K4 me3:H4–4ac
H2A.Z:H3–5ac
DNA Meth. m5C:H4K20 me3
H3K27 me3:H3K9 me3
H2A.Z:H4–4ac
H2A.Z:H3K4 me3
H3K4 me1:H4K20 me2
H3K4 me1:H3–5ac
DNA Meth. m5C:H4K20 me2
H3K27 me2:DNA Meth. m5C
H3K27 me3:H4K20 me3
H3K27 me3:DNA Meth. m5C
H4K20 me2:H4–4ac
H3K9 me3:H4K20 me3
H4–4ac:H3–5ac
H3K9 me2:DNA Meth. m5C
H3K4 me1:H4–4ac
H3K4 me1:H2A.Z
DNA Meth. m5C:H3K9 me3
H2A.Z:H4K20 me2
H4K20 me2:H3–5ac
H4K20 me2:H3K4 me3
H3K4 me3
H3K9 me3
H4–4ac
H3K27 me2
H3K9 me2
H3K27 me3
H4K20 me3
H3–5ac
H2A.Z
H3K4 me1
H4K20 me2
DNA Meth. m5C
L3MBTL3 – Reverse
SAMD1 – Forward
H3K9 me2:DNA Meth. m5C
H2A.Z:H3K4 me3
H3K9 me3:H4K20 me3
H3K27 me2:DNA Meth. m5C
H3K27 me3:H4K20 me3
H3K4 me3:H4–4ac
H3K4 me1:H4–4ac
H3K4 me1:H4K20 me2
H3K4 me1:H2A.Z
H4–4ac:H3–5ac
DNA Meth. m5C:H4K20 me2
H3K27 me3:H3K9 me3
H4K20 me2:H4–4ac
H2A.Z:H4K20 me2
DNA Meth. m5C:H4K20 me3
DNA Meth. m5C:H3K9 me3
H4K20 me2:H3–5ac
H3K27 me3:DNA Meth. m5C
H4K20 me2:H3K4 me3
H3K27 me2
H3K9 me2
H4–4ac
H3K4 me1
H3–5ac
H3K4 me3
H4K20 me3
H3K9 me3
H3K27 me3
H2A.Z
H4K20 me2
DNA Meth. m5C
SAMD1 – Reverse
HLTF – Forward
H2A.Z:H3K4 me3
H2A.Z:H4–4ac
H3K9 me3:H4K20 me3
H3K27 me3:H4K20 me3
H2A.Z:H3–5ac
H2A.Z:H4K20 me2
H3K27 me3:H3K9 me3
H3K4 me3:H4–4ac
H4–4ac:H3–5ac
H3K9 me2:DNA Meth. m5C
H3K27 me2:DNA Meth. m5C
DNA Meth. m5C:H3K9 me3
H4K20 me2:H3–5ac
DNA Meth. m5C:H4K20 me3
DNA Meth. m5C:H4K20 me2
H4K20 me2:H4–4ac
H3K27 me3:DNA Meth. m5C
H3K4 me1
H4K20 me2:H3K4 me3
H3K9 me2
H3K4 me3
H3K27 me2
H4–4ac
H2A.Z
H3–5ac
H3K9 me3
H3K27 me3
H4K20 me3
H4K20 me2
DNA Meth. m5C
HLTF – Reverse
KMT2A – Forward
H3K27 me2:DNA Meth. m5C
H3K27 me3:H4K20 me3
H2A.Z:H3–5ac
H3K9 me2:DNA Meth. m5C
H3K27 me3:H3K9 me3
H3K4 me1:H4–4ac
DNA Meth. m5C:H4K20 me3
H2A.Z:H4–4ac
H2A.Z:H3K4 me3
H3K4 me1:H4K20 me2
H4–4ac:H3–5ac
H3K4 me1:H2A.Z
DNA Meth. m5C:H4K20 me2
DNA Meth. m5C:H3K9 me3
H3K4 me3:H4–4ac
H2A.Z:H4K20 me2
H3K27 me3:DNA Meth. m5C
H4K20 me2:H3–5ac
H4K20 me2:H4–4ac
H4K20 me2:H3K4 me3
H3K27 me2
H3K9 me2
H4K20 me3
H3K4 me1
H3K9 me3
H3K4 me3
H2A.Z
H3K27 me3
H3–5ac
H4–4ac
H4K20 me2
DNA Meth. m5C
KMT2A – Reverse
BCORL1 (1) – Forward
H4K20 me2:H4–4ac
H3K9 me3:H4K20 me3
H3K27 me3:H4K20 me3
H3K27 me3:H3K4 me1
H2A.Z:H4K20 me2
H3K4 me1:H4K20 me2
DNA Meth. m5C:H4K20 me3
H3K4 me3:H4–4ac
H4K20 me2:H3K4 me3
H3K27 me3:H3K9 me3
H4K20 me2:H3–5ac
DNA Meth. m5C:H3K9 me3
DNA Meth. m5C:H4K20 me2
H3K27 me2:DNA Meth. m5C
H4–4ac:H3–5ac
H3K9 me2:DNA Meth. m5C
H3K27 me3:DNA Meth. m5C
H3K4 me3
H4K20 me3
H3K9 me2
H3K27 me2
H4–4ac
H4K20 me2
H3K4 me1
H3–5ac
H2A.Z
H3K9 me3
H3K27 me3
DNA Meth. m5C
BCORL1 (1) – Reverse

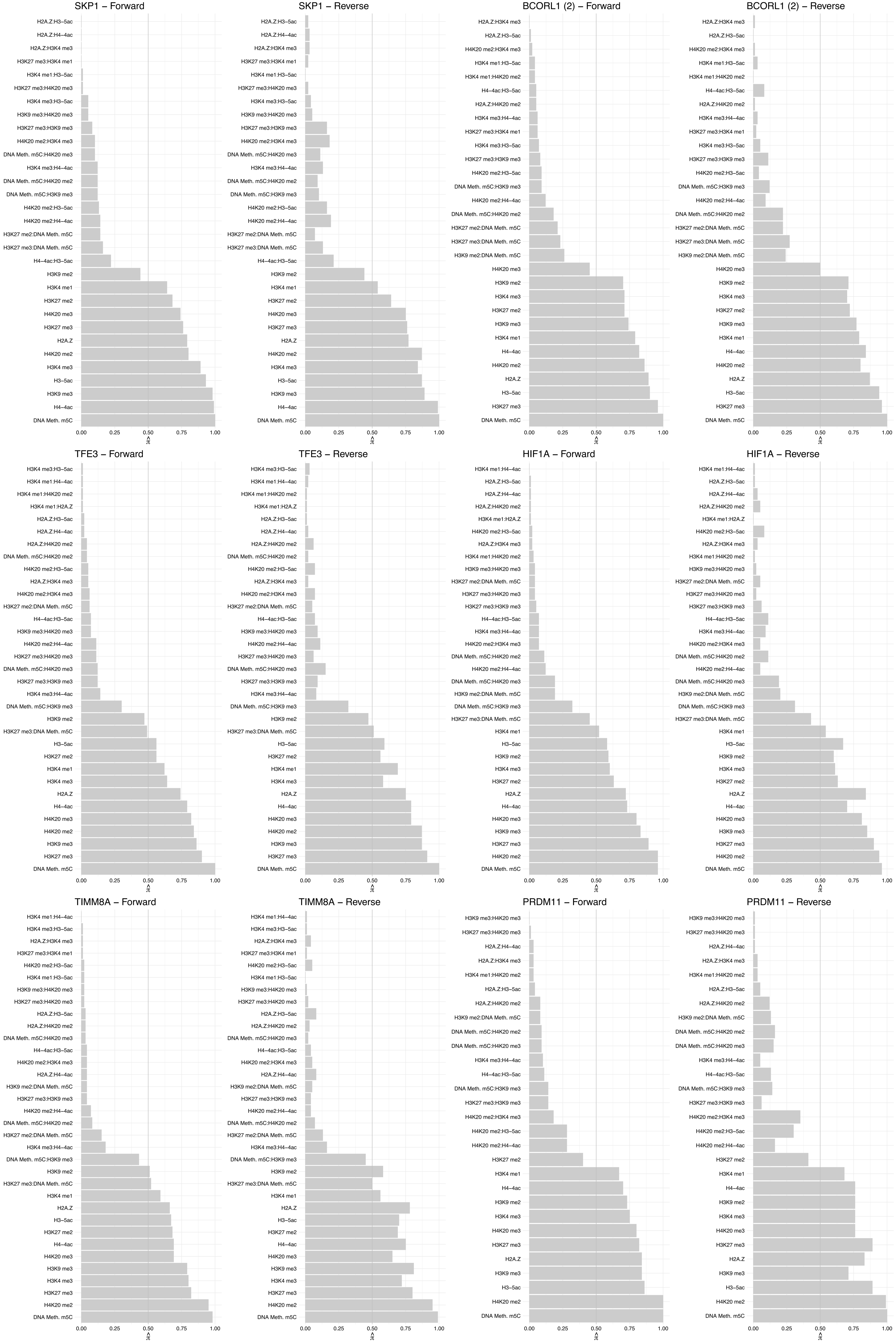

SKP1 – Forward
SKP1 – Reverse
BCORL1 (2) – Forward
BCORL1 (2) – Reverse
TFE3 – Forward
TFE3 – Reverse
HIF1A – Forward
HIF1A – Reverse
TIMM8A – Forward
TIMM8A – Reverse
PRDM11 – Forward
PRDM11 – Reverse
0.00
0.25
0.50
0.75
1.00
$\hat{\pi}$

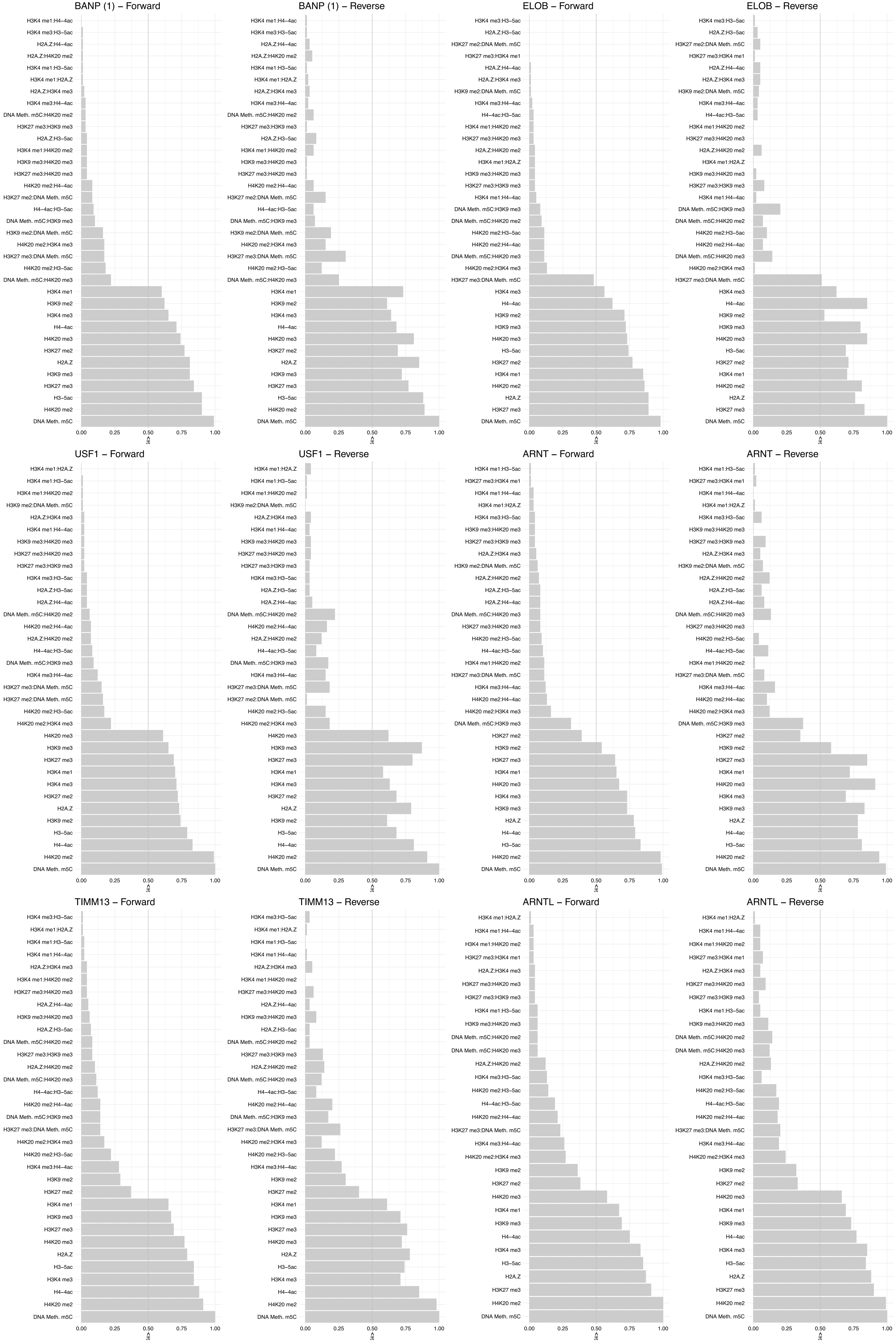
BANP (1) – Forward
H3K4 me1:H4–4ac
H3K4 me3:H3–5ac
H2A.Z:H4–4ac
H2A.Z:H4K20 me2
H3K4 me1:H3–5ac
H3K4 me1:H2A.Z
H2A.Z:H3K4 me3
H3K4 me3:H4–4ac
DNA Meth. m5C:H4K20 me2
H3K27 me3:H3K9 me3
H2A.Z:H3–5ac
H3K4 me1:H4K20 me2
H3K9 me3:H4K20 me3
H3K27 me3:H4K20 me3
H4K20 me2:H4–4ac
H3K27 me2:DNA Meth. m5C
H4–4ac:H3–5ac
DNA Meth. m5C:H3K9 me3
H3K9 me2:DNA Meth. m5C
H4K20 me2:H3K4 me3
H3K27 me3:DNA Meth. m5C
H4K20 me2:H3–5ac
DNA Meth. m5C:H4K20 me3
H3K4 me1
H3K9 me2
H3K4 me3
H4–4ac
H4K20 me3
H3K27 me2
H2A.Z
H3K9 me3
H3K27 me3
H3–5ac
H4K20 me2
DNA Meth. m5C
0.00 0.25 0.50 0.75 1.00
π̂
BANP (1) – Reverse
H3K4 me1:H4–4ac
H3K4 me3:H3–5ac
H2A.Z:H4–4ac
H2A.Z:H4K20 me2
H3K4 me1:H3–5ac
H3K4 me1:H2A.Z
H2A.Z:H3K4 me3
H3K4 me3:H4–4ac
DNA Meth. m5C:H4K20 me2
H3K27 me3:H3K9 me3
H2A.Z:H3–5ac
H3K4 me1:H4K20 me2
H3K9 me3:H4K20 me3
H3K27 me3:H4K20 me3
H4K20 me2:H4–4ac
H3K27 me2:DNA Meth. m5C
H4–4ac:H3–5ac
DNA Meth. m5C:H3K9 me3
H3K9 me2:DNA Meth. m5C
H4K20 me2:H3K4 me3
H3K27 me3:DNA Meth. m5C
H4K20 me2:H3–5ac
DNA Meth. m5C:H4K20 me3
H3K4 me1
H3K9 me2
H3K4 me3
H4–4ac
H4K20 me3
H3K27 me2
H2A.Z
H3K9 me3
H3K27 me3
H3–5ac
H4K20 me2
DNA Meth. m5C
0.00 0.25 0.50 0.75 1.00
π̂
ELOB – Forward
H3K4 me3:H3–5ac
H2A.Z:H3–5ac
H3K27 me2:DNA Meth. m5C
H3K27 me3:H3K4 me1
H2A.Z:H4–4ac
H2A.Z:H3K4 me3
H3K9 me2:DNA Meth. m5C
H3K4 me3:H4–4ac
H4–4ac:H3–5ac
H3K4 me1:H4K20 me2
H3K27 me3:H4K20 me3
H2A.Z:H4K20 me2
H3K4 me1:H2A.Z
H3K9 me3:H4K20 me3
H3K27 me3:H3K9 me3
H3K4 me1:H4–4ac
DNA Meth. m5C:H3K9 me3
DNA Meth. m5C:H4K20 me2
H4K20 me2:H3–5ac
H4K20 me2:H4–4ac
DNA Meth. m5C:H4K20 me3
H4K20 me2:H3K4 me3
H3K27 me3:DNA Meth. m5C
H3K4 me3
H4–4ac
H3K9 me2
H3K9 me3
H4K20 me3
H3–5ac
H3K27 me2
H3K4 me1
H4K20 me2
H2A.Z
H3K27 me3
DNA Meth. m5C
0.00 0.25 0.50 0.75 1.00
π̂
ELOB – Reverse
H3K4 me3:H3–5ac
H2A.Z:H3–5ac
H3K27 me2:DNA Meth. m5C
H3K27 me3:H3K4 me1
H2A.Z:H4–4ac
H2A.Z:H3K4 me3
H3K9 me2:DNA Meth. m5C
H3K4 me3:H4–4ac
H4–4ac:H3–5ac
H3K4 me1:H4K20 me2
H3K27 me3:H4K20 me3
H2A.Z:H4K20 me2
H3K4 me1:H2A.Z
H3K9 me3:H4K20 me3
H3K27 me3:H3K9 me3
H3K4 me1:H4–4ac
DNA Meth. m5C:H3K9 me3
DNA Meth. m5C:H4K20 me2
H4K20 me2:H3–5ac
H4K20 me2:H4–4ac
DNA Meth. m5C:H4K20 me3
H4K20 me2:H3K4 me3
H3K27 me3:DNA Meth. m5C
H3K4 me3
H4–4ac
H3K9 me2
H3K9 me3
H4K20 me3
H3–5ac
H3K27 me2
H3K4 me1
H4K20 me2
H2A.Z
H3K27 me3
DNA Meth. m5C
0.00 0.25 0.50 0.75 1.00
π̂
USF1 – Forward
H3K4 me1:H2A.Z
H3K4 me1:H3–5ac
H3K4 me1:H4K20 me2
H3K9 me2:DNA Meth. m5C
H2A.Z:H3K4 me3
H3K4 me1:H4–4ac
H3K9 me3:H4K20 me3
H3K27 me3:H4K20 me3
H3K27 me3:H3K9 me3
H3K4 me3:H3–5ac
H2A.Z:H3–5ac
H2A.Z:H4–4ac
DNA Meth. m5C:H4K20 me2
H4K20 me2:H4–4ac
H2A.Z:H4K20 me2
H4–4ac:H3–5ac
DNA Meth. m5C:H3K9 me3
H3K4 me3:H4–4ac
H3K27 me3:DNA Meth. m5C
H3K27 me2:DNA Meth. m5C
H4K20 me2:H3–5ac
H4K20 me2:H3K4 me3
H4K20 me3
H3K9 me3
H3K27 me3
H3K4 me1
H3K4 me3
H3K27 me2
H2A.Z
H3K9 me2
H3–5ac
H4–4ac
H4K20 me2
DNA Meth. m5C
0.00 0.25 0.50 0.75 1.00
π̂
USF1 – Reverse
H3K4 me1:H2A.Z
H3K4 me1:H3–5ac
H3K4 me1:H4K20 me2
H3K9 me2:DNA Meth. m5C
H2A.Z:H3K4 me3
H3K4 me1:H4–4ac
H3K9 me3:H4K20 me3
H3K27 me3:H4K20 me3
H3K27 me3:H3K9 me3
H3K4 me3:H3–5ac
H2A.Z:H3–5ac
H2A.Z:H4–4ac
DNA Meth. m5C:H4K20 me2
H4K20 me2:H4–4ac
H2A.Z:H4K20 me2
H4–4ac:H3–5ac
DNA Meth. m5C:H3K9 me3
H3K4 me3:H4–4ac
H3K27 me3:DNA Meth. m5C
H3K27 me2:DNA Meth. m5C
H4K20 me2:H3–5ac
H4K20 me2:H3K4 me3
H4K20 me3
H3K9 me3
H3K27 me3
H3K4 me1
H3K4 me3
H3K27 me2
H2A.Z
H3K9 me2
H3–5ac
H4–4ac
H4K20 me2
DNA Meth. m5C
0.00 0.25 0.50 0.75 1.00
π̂
ARNT – Forward
H3K4 me1:H3–5ac
H3K27 me3:H3K4 me1
H3K4 me1:H4–4ac
H3K4 me1:H2A.Z
H3K4 me3:H3–5ac
H3K9 me3:H4K20 me3
H3K27 me3:H3K9 me3
H2A.Z:H3K4 me3
H3K9 me2:DNA Meth. m5C
H2A.Z:H4K20 me2
H2A.Z:H3–5ac
H2A.Z:H4–4ac
DNA Meth. m5C:H4K20 me3
H3K27 me3:H4K20 me3
H4K20 me2:H3–5ac
H4–4ac:H3–5ac
H3K4 me1:H4K20 me2
H3K27 me3:DNA Meth. m5C
H3K4 me3:H4–4ac
H4K20 me2:H4–4ac
H4K20 me2:H3K4 me3
DNA Meth. m5C:H3K9 me3
H3K27 me2
H3K9 me2
H3K27 me3
H3K4 me1
H4K20 me3
H3K4 me3
H3K9 me3
H2A.Z
H4–4ac
H3–5ac
H4K20 me2
DNA Meth. m5C
0.00 0.25 0.50 0.75 1.00
π̂
ARNT – Reverse
H3K4 me1:H3–5ac
H3K27 me3:H3K4 me1
H3K4 me1:H4–4ac
H3K4 me1:H2A.Z
H3K4 me3:H3–5ac
H3K9 me3:H4K20 me3
H3K27 me3:H3K9 me3
H2A.Z:H3K4 me3
H3K9 me2:DNA Meth. m5C
H2A.Z:H4K20 me2
H2A.Z:H3–5ac
H2A.Z:H4–4ac
DNA Meth. m5C:H4K20 me3
H3K27 me3:H4K20 me3
H4K20 me2:H3–5ac
H4–4ac:H3–5ac
H3K4 me1:H4K20 me2
H3K27 me3:DNA Meth. m5C
H3K4 me3:H4–4ac
H4K20 me2:H4–4ac
H4K20 me2:H3K4 me3
DNA Meth. m5C:H3K9 me3
H3K27 me2
H3K9 me2
H3K27 me3
H3K4 me1
H4K20 me3
H3K4 me3
H3K9 me3
H2A.Z
H4–4ac
H3–5ac
H4K20 me2
DNA Meth. m5C
0.00 0.25 0.50 0.75 1.00
π̂
TIMM13 – Forward
H3K4 me3:H3–5ac
H3K4 me1:H2A.Z
H3K4 me1:H3–5ac
H3K4 me1:H4–4ac
H2A.Z:H3K4 me3
H3K4 me1:H4K20 me2
H3K27 me3:H4K20 me3
H2A.Z:H4–4ac
H3K9 me3:H4K20 me3
H2A.Z:H3–5ac
DNA Meth. m5C:H4K20 me2
H3K27 me3:H3K9 me3
H2A.Z:H4K20 me2
DNA Meth. m5C:H4K20 me3
H4–4ac:H3–5ac
H4K20 me2:H4–4ac
DNA Meth. m5C:H3K9 me3
H3K27 me3:DNA Meth. m5C
H4K20 me2:H3K4 me3
H4K20 me2:H3–5ac
H3K4 me3:H4–4ac
H3K9 me2
H3K27 me2
H3K4 me1
H3K9 me3
H3K27 me3
H4K20 me3
H2A.Z
H3–5ac
H3K4 me3
H4–4ac
H4K20 me2
DNA Meth. m5C
0.00 0.25 0.50 0.75 1.00
π̂
TIMM13 – Reverse
H3K4 me3:H3–5ac
H3K4 me1:H2A.Z
H3K4 me1:H3–5ac
H3K4 me1:H4–4ac
H2A.Z:H3K4 me3
H3K4 me1:H4K20 me2
H3K27 me3:H4K20 me3
H2A.Z:H4–4ac
H3K9 me3:H4K20 me3
H2A.Z:H3–5ac
DNA Meth. m5C:H4K20 me2
H3K27 me3:H3K9 me3
H2A.Z:H4K20 me2
DNA Meth. m5C:H4K20 me3
H4–4ac:H3–5ac
H4K20 me2:H4–4ac
DNA Meth. m5C:H3K9 me3
H3K27 me3:DNA Meth. m5C
H4K20 me2:H3K4 me3
H4K20 me2:H3–5ac
H3K4 me3:H4–4ac
H3K9 me2
H3K27 me2
H3K4 me1
H3K9 me3
H3K27 me3
H4K20 me3
H2A.Z
H3–5ac
H3K4 me3
H4–4ac
H4K20 me2
DNA Meth. m5C
0.00 0.25 0.50 0.75 1.00
π̂
ARNTL – Forward
H3K4 me1:H2A.Z
H3K4 me1:H4–4ac
H3K4 me1:H4K20 me2
H3K27 me3:H3K4 me1
H2A.Z:H3K4 me3
H3K27 me3:H4K20 me3
H3K27 me3:H3K9 me3
H3K4 me1:H3–5ac
H3K9 me3:H4K20 me3
DNA Meth. m5C:H4K20 me2
DNA Meth. m5C:H4K20 me3
H2A.Z:H4K20 me2
H3K4 me3:H3–5ac
H4K20 me2:H3–5ac
H4–4ac:H3–5ac
H4K20 me2:H4–4ac
H3K27 me3:DNA Meth. m5C
H3K4 me3:H4–4ac
H4K20 me2:H3K4 me3
H3K9 me2
H3K27 me2
H4K20 me3
H3K4 me1
H3K9 me3
H4–4ac
H3K4 me3
H3–5ac
H2A.Z
H3K27 me3
H4K20 me2
DNA Meth. m5C
0.00 0.25 0.50 0.75 1.00
π̂
ARNTL – Reverse
H3K4 me1:H2A.Z
H3K4 me1:H4–4ac
H3K4 me1:H4K20 me2
H3K27 me3:H3K4 me1
H2A.Z:H3K4 me3
H3K27 me3:H4K20 me3
H3K27 me3:H3K9 me3
H3K4 me1:H3–5ac
H3K9 me3:H4K20 me3
DNA Meth. m5C:H4K20 me2
DNA Meth. m5C:H4K20 me3
H2A.Z:H4K20 me2
H3K4 me3:H3–5ac
H4K20 me2:H3–5ac
H4–4ac:H3–5ac
H4K20 me2:H4–4ac
H3K27 me3:DNA Meth. m5C
H3K4 me3:H4–4ac
H4K20 me2:H3K4 me3
H3K9 me2
H3K27 me2
H4K20 me3
H3K4 me1
H3K9 me3
H4–4ac
H3K4 me3
H3–5ac
H2A.Z
H3K27 me3
H4K20 me2
DNA Meth. m5C
0.00 0.25 0.50 0.75 1.00
π̂

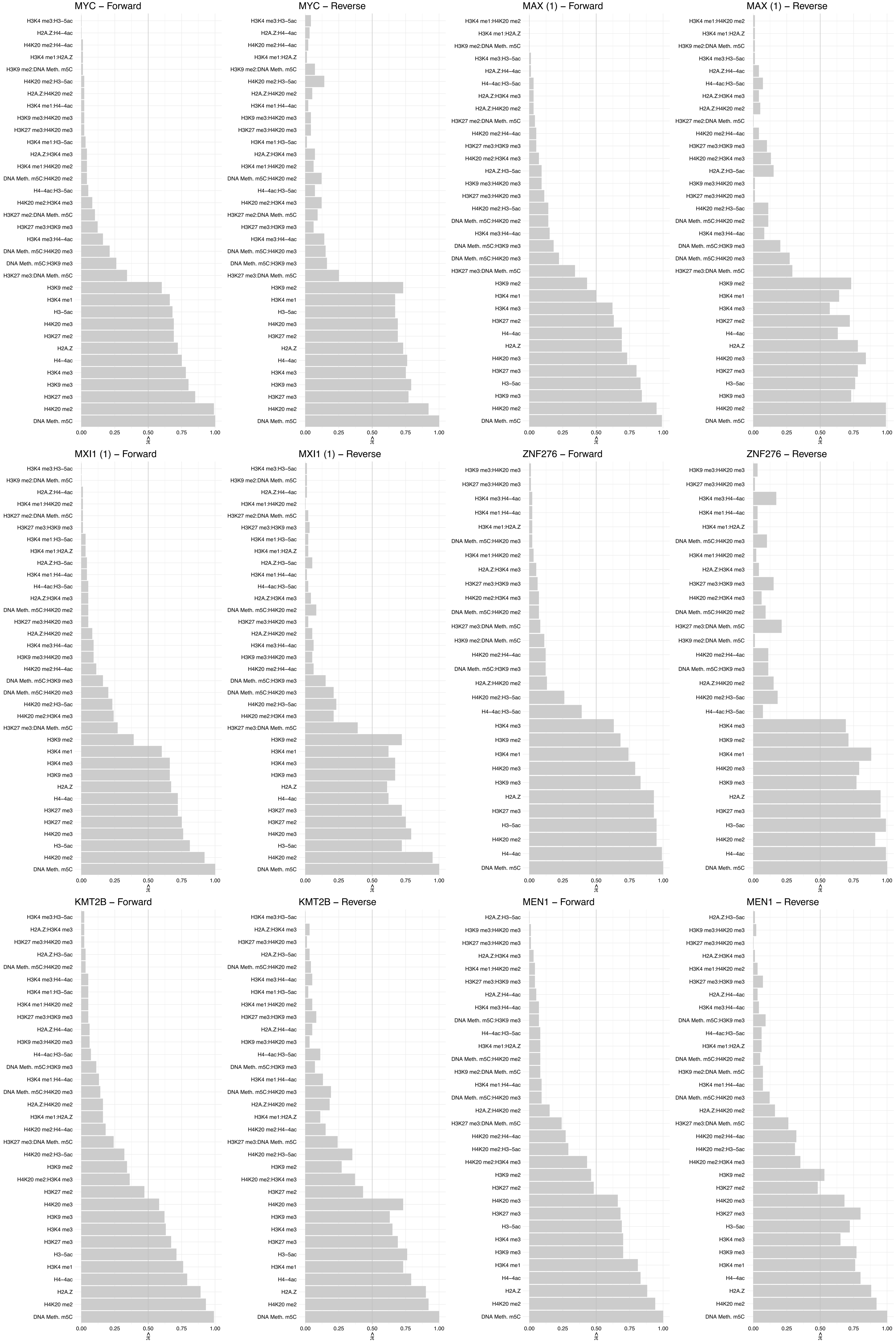
MYC – Forward
H3K4 me3:H3−5ac
H2A.Z:H4−4ac
H4K20 me2:H4−4ac
H3K4 me1:H2A.Z
H3K9 me2:DNA Meth. m5C
H4K20 me2:H3−5ac
H2A.Z:H4K20 me2
H3K4 me1:H4−4ac
H3K9 me3:H4K20 me3
H3K27 me3:H4K20 me3
H3K4 me1:H3−5ac
H2A.Z:H3K4 me3
H3K4 me1:H4K20 me2
DNA Meth. m5C:H4K20 me2
H4−4ac:H3−5ac
H4K20 me2:H3K4 me3
H3K27 me2:DNA Meth. m5C
H3K27 me3:H3K9 me3
H3K4 me3:H4−4ac
DNA Meth. m5C:H4K20 me3
DNA Meth. m5C:H3K9 me3
H3K27 me3:DNA Meth. m5C
H3K9 me2
H3K4 me1
H3−5ac
H4K20 me3
H3K27 me2
H2A.Z
H4−4ac
H3K4 me3
H3K9 me3
H3K27 me3
H4K20 me2
DNA Meth. m5C
0.00 0.25 0.50 0.75 1.00
$\hat{\pi}$
MYC – Reverse
H3K4 me3:H3−5ac
H2A.Z:H4−4ac
H4K20 me2:H4−4ac
H3K4 me1:H2A.Z
H3K9 me2:DNA Meth. m5C
H4K20 me2:H3−5ac
H2A.Z:H4K20 me2
H3K4 me1:H4−4ac
H3K9 me3:H4K20 me3
H3K27 me3:H4K20 me3
H3K4 me1:H3−5ac
H2A.Z:H3K4 me3
H3K4 me1:H4K20 me2
DNA Meth. m5C:H4K20 me2
H4−4ac:H3−5ac
H4K20 me2:H3K4 me3
H3K27 me2:DNA Meth. m5C
H3K27 me3:H3K9 me3
H3K4 me3:H4−4ac
DNA Meth. m5C:H4K20 me3
DNA Meth. m5C:H3K9 me3
H3K27 me3:DNA Meth. m5C
H3K9 me2
H3K4 me1
H3−5ac
H4K20 me3
H3K27 me2
H2A.Z
H4−4ac
H3K4 me3
H3K9 me3
H3K27 me3
H4K20 me2
DNA Meth. m5C
0.00 0.25 0.50 0.75 1.00
$\hat{\pi}$
MAX (1) – Forward
H3K4 me1:H4K20 me2
H3K4 me1:H2A.Z
H3K9 me2:DNA Meth. m5C
H3K4 me3:H3−5ac
H2A.Z:H4−4ac
H4−4ac:H3−5ac
H2A.Z:H3K4 me3
H2A.Z:H4K20 me2
H3K27 me2:DNA Meth. m5C
H4K20 me2:H4−4ac
H3K27 me3:H3K9 me3
H4K20 me2:H3K4 me3
H2A.Z:H3−5ac
H3K9 me3:H4K20 me3
H3K27 me3:H4K20 me3
H4K20 me2:H3−5ac
DNA Meth. m5C:H4K20 me2
H3K4 me3:H4−4ac
DNA Meth. m5C:H3K9 me3
DNA Meth. m5C:H4K20 me3
H3K27 me3:DNA Meth. m5C
H3K9 me2
H3K4 me1
H3K4 me3
H3K27 me2
H4−4ac
H2A.Z
H4K20 me3
H3K27 me3
H3−5ac
H3K9 me3
H4K20 me2
DNA Meth. m5C
0.00 0.25 0.50 0.75 1.00
$\hat{\pi}$
MAX (1) – Reverse
H3K4 me1:H4K20 me2
H3K4 me1:H2A.Z
H3K9 me2:DNA Meth. m5C
H3K4 me3:H3−5ac
H2A.Z:H4−4ac
H4−4ac:H3−5ac
H2A.Z:H3K4 me3
H2A.Z:H4K20 me2
H3K27 me2:DNA Meth. m5C
H4K20 me2:H4−4ac
H3K27 me3:H3K9 me3
H4K20 me2:H3K4 me3
H2A.Z:H3−5ac
H3K9 me3:H4K20 me3
H3K27 me3:H4K20 me3
H4K20 me2:H3−5ac
DNA Meth. m5C:H4K20 me2
H3K4 me3:H4−4ac
DNA Meth. m5C:H3K9 me3
DNA Meth. m5C:H4K20 me3
H3K27 me3:DNA Meth. m5C
H3K9 me2
H3K4 me1
H3K4 me3
H3K27 me2
H4−4ac
H2A.Z
H4K20 me3
H3K27 me3
H3−5ac
H3K9 me3
H4K20 me2
DNA Meth. m5C
0.00 0.25 0.50 0.75 1.00
$\hat{\pi}$
MXI1 (1) – Forward
H3K4 me3:H3−5ac
H3K9 me2:DNA Meth. m5C
H2A.Z:H4−4ac
H3K4 me1:H4K20 me2
H3K27 me2:DNA Meth. m5C
H3K27 me3:H3K9 me3
H3K4 me1:H3−5ac
H3K4 me1:H2A.Z
H2A.Z:H3−5ac
H3K4 me1:H4−4ac
H4−4ac:H3−5ac
H2A.Z:H3K4 me3
DNA Meth. m5C:H4K20 me2
H3K27 me3:H4K20 me3
H2A.Z:H4K20 me2
H3K4 me3:H4−4ac
H3K9 me3:H4K20 me3
H4K20 me2:H4−4ac
DNA Meth. m5C:H3K9 me3
DNA Meth. m5C:H4K20 me3
H4K20 me2:H3−5ac
H4K20 me2:H3K4 me3
H3K27 me3:DNA Meth. m5C
H3K9 me2
H3K4 me1
H3K4 me3
H3K9 me3
H2A.Z
H4−4ac
H3K27 me3
H3K27 me2
H4K20 me3
H3−5ac
H4K20 me2
DNA Meth. m5C
0.00 0.25 0.50 0.75 1.00
$\hat{\pi}$
MXI1 (1) – Reverse
H3K4 me3:H3−5ac
H3K9 me2:DNA Meth. m5C
H2A.Z:H4−4ac
H3K4 me1:H4K20 me2
H3K27 me2:DNA Meth. m5C
H3K27 me3:H3K9 me3
H3K4 me1:H3−5ac
H3K4 me1:H2A.Z
H2A.Z:H3−5ac
H3K4 me1:H4−4ac
H4−4ac:H3−5ac
H2A.Z:H3K4 me3
DNA Meth. m5C:H4K20 me2
H3K27 me3:H4K20 me3
H2A.Z:H4K20 me2
H3K4 me3:H4−4ac
H3K9 me3:H4K20 me3
H4K20 me2:H4−4ac
DNA Meth. m5C:H3K9 me3
DNA Meth. m5C:H4K20 me3
H4K20 me2:H3−5ac
H4K20 me2:H3K4 me3
H3K27 me3:DNA Meth. m5C
H3K9 me2
H3K4 me1
H3K4 me3
H3K9 me3
H2A.Z
H4−4ac
H3K27 me3
H3K27 me2
H4K20 me3
H3−5ac
H4K20 me2
DNA Meth. m5C
0.00 0.25 0.50 0.75 1.00
$\hat{\pi}$
ZNF276 – Forward
H3K9 me3:H4K20 me3
H3K27 me3:H4K20 me3
H3K4 me3:H4−4ac
H3K4 me1:H4−4ac
H3K4 me1:H2A.Z
DNA Meth. m5C:H4K20 me3
H3K4 me1:H4K20 me2
H2A.Z:H3K4 me3
H3K27 me3:H3K9 me3
H4K20 me2:H3K4 me3
DNA Meth. m5C:H4K20 me2
H3K27 me3:DNA Meth. m5C
H3K9 me2:DNA Meth. m5C
H4K20 me2:H4−4ac
DNA Meth. m5C:H3K9 me3
H2A.Z:H4K20 me2
H4K20 me2:H3−5ac
H4−4ac:H3−5ac
H3K4 me3
H3K9 me2
H3K4 me1
H4K20 me3
H3K9 me3
H2A.Z
H3K27 me3
H3−5ac
H4K20 me2
H4−4ac
DNA Meth. m5C
0.00 0.25 0.50 0.75 1.00
$\hat{\pi}$
ZNF276 – Reverse
H3K9 me3:H4K20 me3
H3K27 me3:H4K20 me3
H3K4 me3:H4−4ac
H3K4 me1:H4−4ac
H3K4 me1:H2A.Z
DNA Meth. m5C:H4K20 me3
H3K4 me1:H4K20 me2
H2A.Z:H3K4 me3
H3K27 me3:H3K9 me3
H4K20 me2:H3K4 me3
DNA Meth. m5C:H4K20 me2
H3K27 me3:DNA Meth. m5C
H3K9 me2:DNA Meth. m5C
H4K20 me2:H4−4ac
DNA Meth. m5C:H3K9 me3
H2A.Z:H4K20 me2
H4K20 me2:H3−5ac
H4−4ac:H3−5ac
H3K4 me3
H3K9 me2
H3K4 me1
H4K20 me3
H3K9 me3
H2A.Z
H3K27 me3
H3−5ac
H4K20 me2
H4−4ac
DNA Meth. m5C
0.00 0.25 0.50 0.75 1.00
$\hat{\pi}$
KMT2B – Forward
H3K4 me3:H3−5ac
H2A.Z:H3K4 me3
H3K27 me3:H4K20 me3
H2A.Z:H3−5ac
DNA Meth. m5C:H4K20 me2
H3K4 me3:H4−4ac
H3K4 me1:H3−5ac
H3K4 me1:H4K20 me2
H3K27 me3:H3K9 me3
H2A.Z:H4−4ac
H3K9 me3:H4K20 me3
H4−4ac:H3−5ac
DNA Meth. m5C:H3K9 me3
H3K4 me1:H4−4ac
DNA Meth. m5C:H4K20 me3
H2A.Z:H4K20 me2
H3K4 me1:H2A.Z
H4K20 me2:H4−4ac
H3K27 me3:DNA Meth. m5C
H4K20 me2:H3−5ac
H3K9 me2
H4K20 me2:H3K4 me3
H3K27 me2
H4K20 me3
H3K9 me3
H3K4 me3
H3K27 me3
H3−5ac
H3K4 me1
H4−4ac
H2A.Z
H4K20 me2
DNA Meth. m5C
0.00 0.25 0.50 0.75 1.00
$\hat{\pi}$
KMT2B – Reverse
H3K4 me3:H3−5ac
H2A.Z:H3K4 me3
H3K27 me3:H4K20 me3
H2A.Z:H3−5ac
DNA Meth. m5C:H4K20 me2
H3K4 me3:H4−4ac
H3K4 me1:H3−5ac
H3K4 me1:H4K20 me2
H3K27 me3:H3K9 me3
H2A.Z:H4−4ac
H3K9 me3:H4K20 me3
H4−4ac:H3−5ac
DNA Meth. m5C:H3K9 me3
H3K4 me1:H4−4ac
DNA Meth. m5C:H4K20 me3
H2A.Z:H4K20 me2
H3K4 me1:H2A.Z
H4K20 me2:H4−4ac
H3K27 me3:DNA Meth. m5C
H4K20 me2:H3−5ac
H3K9 me2
H4K20 me2:H3K4 me3
H3K27 me2
H4K20 me3
H3K9 me3
H3K4 me3
H3K27 me3
H3−5ac
H3K4 me1
H4−4ac
H2A.Z
H4K20 me2
DNA Meth. m5C
0.00 0.25 0.50 0.75 1.00
$\hat{\pi}$
MEN1 – Forward
H2A.Z:H3−5ac
H3K9 me3:H4K20 me3
H3K27 me3:H4K20 me3
H2A.Z:H3K4 me3
H3K4 me1:H4K20 me2
H3K27 me3:H3K9 me3
H2A.Z:H4−4ac
H3K4 me3:H4−4ac
DNA Meth. m5C:H3K9 me3
H4−4ac:H3−5ac
H3K4 me1:H2A.Z
DNA Meth. m5C:H4K20 me2
H3K9 me2:DNA Meth. m5C
H3K4 me1:H4−4ac
DNA Meth. m5C:H4K20 me3
H2A.Z:H4K20 me2
H3K27 me3:DNA Meth. m5C
H4K20 me2:H4−4ac
H4K20 me2:H3−5ac
H4K20 me2:H3K4 me3
H3K9 me2
H3K27 me2
H4K20 me3
H3K27 me3
H3−5ac
H3K4 me3
H3K9 me3
H3K4 me1
H4−4ac
H2A.Z
H4K20 me2
DNA Meth. m5C
0.00 0.25 0.50 0.75 1.00
$\hat{\pi}$
MEN1 – Reverse
H2A.Z:H3−5ac
H3K9 me3:H4K20 me3
H3K27 me3:H4K20 me3
H2A.Z:H3K4 me3
H3K4 me1:H4K20 me2
H3K27 me3:H3K9 me3
H2A.Z:H4−4ac
H3K4 me3:H4−4ac
DNA Meth. m5C:H3K9 me3
H4−4ac:H3−5ac
H3K4 me1:H2A.Z
DNA Meth. m5C:H4K20 me2
H3K9 me2:DNA Meth. m5C
H3K4 me1:H4−4ac
DNA Meth. m5C:H4K20 me3
H2A.Z:H4K20 me2
H3K27 me3:DNA Meth. m5C
H4K20 me2:H4−4ac
H4K20 me2:H3−5ac
H4K20 me2:H3K4 me3
H3K9 me2
H3K27 me2
H4K20 me3
H3K27 me3
H3−5ac
H3K4 me3
H3K9 me3
H3K4 me1
H4−4ac
H2A.Z
H4K20 me2
DNA Meth. m5C
0.00 0.25 0.50 0.75 1.00
$\hat{\pi}$

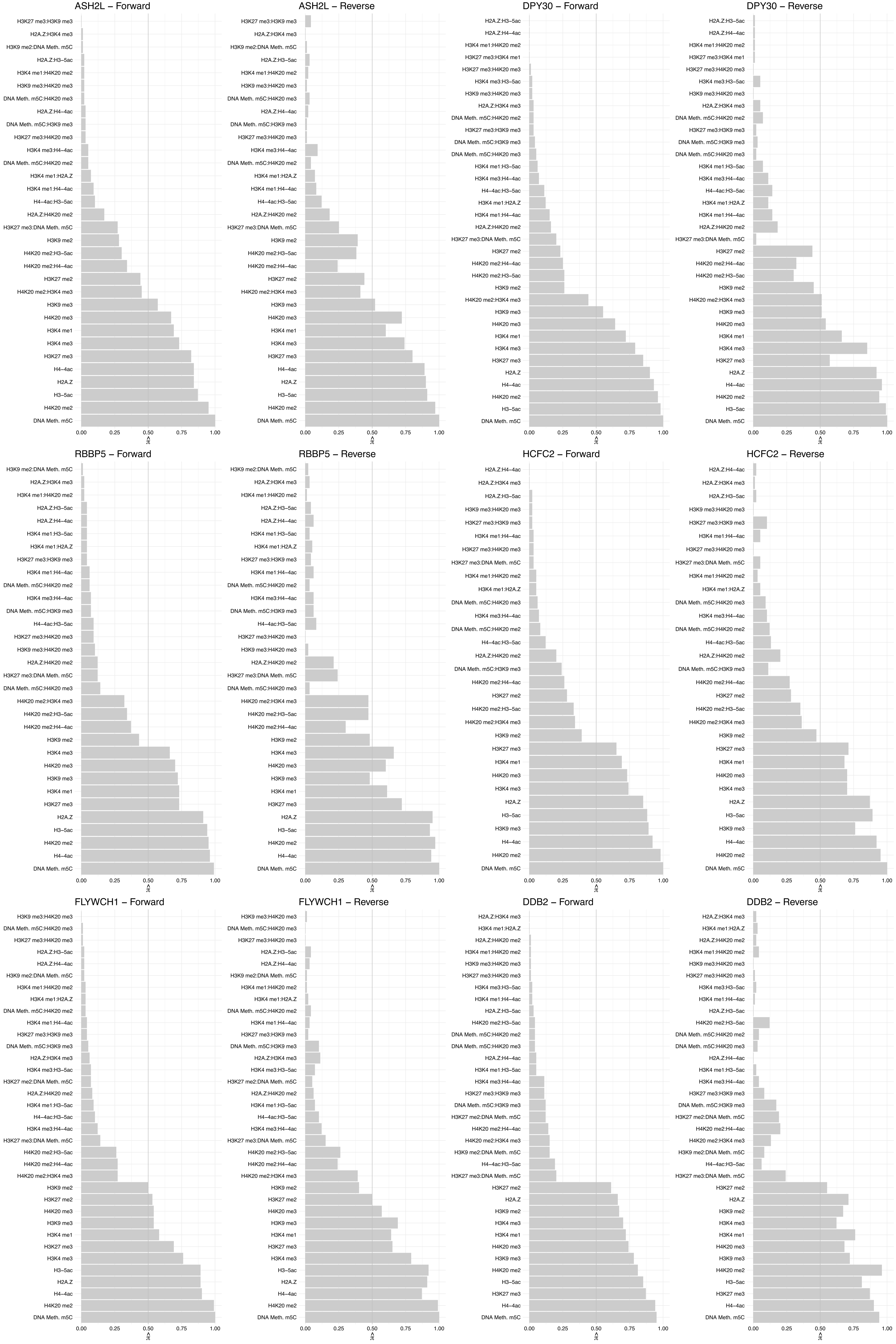
ASH2L – Forward
H3K27 me3:H3K9 me3
H2A.Z:H3K4 me3
H3K9 me2:DNA Meth. m5C
H2A.Z:H3–5ac
H3K4 me1:H4K20 me2
H3K9 me3:H4K20 me3
DNA Meth. m5C:H4K20 me3
H2A.Z:H4–4ac
DNA Meth. m5C:H3K9 me3
H3K27 me3:H4K20 me3
H3K4 me3:H4–4ac
DNA Meth. m5C:H4K20 me2
H3K4 me1:H2A.Z
H3K4 me1:H4–4ac
H4–4ac:H3–5ac
H2A.Z:H4K20 me2
H3K27 me3:DNA Meth. m5C
H3K9 me2
H4K20 me2:H3–5ac
H4K20 me2:H4–4ac
H3K27 me2
H4K20 me2:H3K4 me3
H3K9 me3
H4K20 me3
H3K4 me1
H3K4 me3
H3K27 me3
H4–4ac
H2A.Z
H3–5ac
H4K20 me2
DNA Meth. m5C
0.00 0.25 0.50 0.75 1.00
π̂
ASH2L – Reverse
DPY30 – Forward
H2A.Z:H3–5ac
H2A.Z:H4–4ac
H3K4 me1:H4K20 me2
H3K27 me3:H3K4 me1
H3K27 me3:H4K20 me3
H3K4 me3:H3–5ac
H3K9 me3:H4K20 me3
H2A.Z:H3K4 me3
DNA Meth. m5C:H4K20 me2
H3K27 me3:H3K9 me3
DNA Meth. m5C:H3K9 me3
DNA Meth. m5C:H4K20 me3
H3K4 me1:H3–5ac
H3K4 me3:H4–4ac
H4–4ac:H3–5ac
H3K4 me1:H2A.Z
H3K4 me1:H4–4ac
H2A.Z:H4K20 me2
H3K27 me3:DNA Meth. m5C
H3K27 me2
H4K20 me2:H4–4ac
H4K20 me2:H3–5ac
H3K9 me2
H4K20 me2:H3K4 me3
H3K9 me3
H4K20 me3
H3K4 me1
H3K4 me3
H3K27 me3
H2A.Z
H4–4ac
H4K20 me2
H3–5ac
DNA Meth. m5C
DPY30 – Reverse
RBBP5 – Forward
H3K9 me2:DNA Meth. m5C
H2A.Z:H3K4 me3
H3K4 me1:H4K20 me2
H2A.Z:H3–5ac
H2A.Z:H4–4ac
H3K4 me1:H3–5ac
H3K4 me1:H2A.Z
H3K27 me3:H3K9 me3
H3K4 me1:H4–4ac
DNA Meth. m5C:H4K20 me2
H3K4 me3:H4–4ac
DNA Meth. m5C:H3K9 me3
H4–4ac:H3–5ac
H3K27 me3:H4K20 me3
H3K9 me3:H4K20 me3
H2A.Z:H4K20 me2
H3K27 me3:DNA Meth. m5C
DNA Meth. m5C:H4K20 me3
H4K20 me2:H3K4 me3
H4K20 me2:H3–5ac
H4K20 me2:H4–4ac
H3K9 me2
H3K4 me3
H4K20 me3
H3K9 me3
H3K4 me1
H3K27 me3
H2A.Z
H3–5ac
H4K20 me2
H4–4ac
DNA Meth. m5C
RBBP5 – Reverse
HCFC2 – Forward
H2A.Z:H4–4ac
H2A.Z:H3K4 me3
H2A.Z:H3–5ac
H3K9 me3:H4K20 me3
H3K27 me3:H3K9 me3
H3K4 me1:H4–4ac
H3K27 me3:H4K20 me3
H3K27 me3:DNA Meth. m5C
H3K4 me1:H4K20 me2
H3K4 me1:H2A.Z
DNA Meth. m5C:H4K20 me3
H3K4 me3:H4–4ac
DNA Meth. m5C:H4K20 me2
H4–4ac:H3–5ac
H2A.Z:H4K20 me2
DNA Meth. m5C:H3K9 me3
H4K20 me2:H4–4ac
H3K27 me2
H4K20 me2:H3–5ac
H4K20 me2:H3K4 me3
H3K9 me2
H3K27 me3
H3K4 me1
H4K20 me3
H3K4 me3
H2A.Z
H3–5ac
H3K9 me3
H4–4ac
H4K20 me2
DNA Meth. m5C
HCFC2 – Reverse
FLYWCH1 – Forward
H3K9 me3:H4K20 me3
DNA Meth. m5C:H4K20 me3
H3K27 me3:H4K20 me3
H2A.Z:H3–5ac
H2A.Z:H4–4ac
H3K9 me2:DNA Meth. m5C
H3K4 me1:H4K20 me2
H3K4 me1:H2A.Z
DNA Meth. m5C:H4K20 me2
H3K4 me1:H4–4ac
H3K27 me3:H3K9 me3
DNA Meth. m5C:H3K9 me3
H2A.Z:H3K4 me3
H3K4 me3:H3–5ac
H3K27 me2:DNA Meth. m5C
H2A.Z:H4K20 me2
H3K4 me1:H3–5ac
H4–4ac:H3–5ac
H3K4 me3:H4–4ac
H3K27 me3:DNA Meth. m5C
H4K20 me2:H3–5ac
H4K20 me2:H4–4ac
H4K20 me2:H3K4 me3
H3K9 me2
H3K27 me2
H4K20 me3
H3K9 me3
H3K4 me1
H3K27 me3
H3K4 me3
H3–5ac
H2A.Z
H4–4ac
H4K20 me2
DNA Meth. m5C
FLYWCH1 – Reverse
DDB2 – Forward
H2A.Z:H3K4 me3
H3K4 me1:H2A.Z
H2A.Z:H4K20 me2
H3K4 me1:H4K20 me2
H3K9 me3:H4K20 me3
H3K27 me3:H4K20 me3
H3K4 me3:H3–5ac
H3K4 me1:H4–4ac
H2A.Z:H3–5ac
H4K20 me2:H3–5ac
DNA Meth. m5C:H4K20 me2
DNA Meth. m5C:H4K20 me3
H2A.Z:H4–4ac
H3K4 me1:H3–5ac
H3K4 me3:H4–4ac
H3K27 me3:H3K9 me3
DNA Meth. m5C:H3K9 me3
H3K27 me2:DNA Meth. m5C
H4K20 me2:H4–4ac
H4K20 me2:H3K4 me3
H3K9 me2:DNA Meth. m5C
H4–4ac:H3–5ac
H3K27 me3:DNA Meth. m5C
H3K27 me2
H2A.Z
H3K9 me2
H3K4 me3
H3K4 me1
H4K20 me3
H3K9 me3
H4K20 me2
H3–5ac
H3K27 me3
H4–4ac
DNA Meth. m5C
DDB2 – Reverse

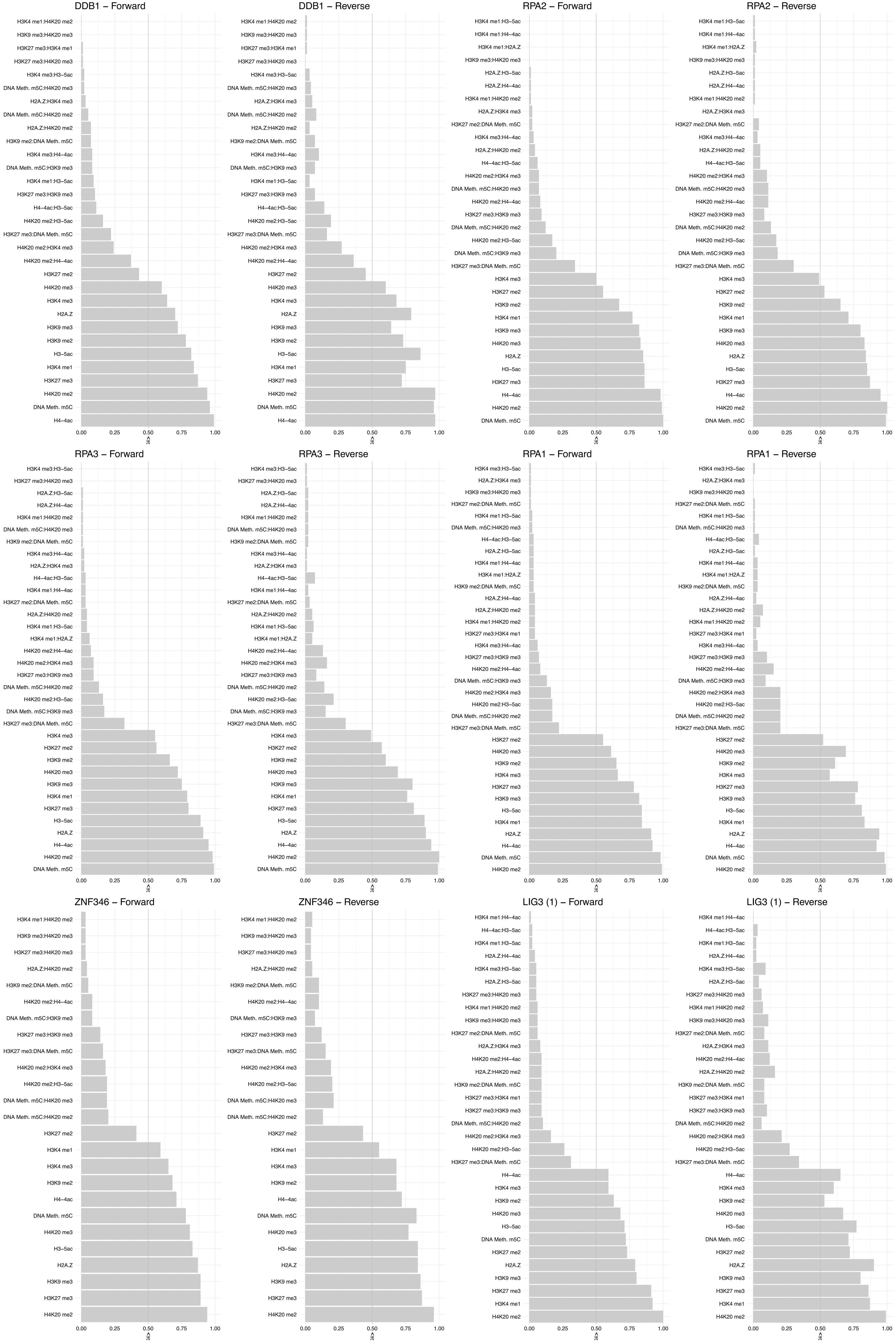
DDB1 – Forward
H3K4 me1:H4K20 me2
H3K9 me3:H4K20 me3
H3K27 me3:H3K4 me1
H3K27 me3:H4K20 me3
H3K4 me3:H3−5ac
DNA Meth. m5C:H4K20 me3
H2A.Z:H3K4 me3
DNA Meth. m5C:H4K20 me2
H2A.Z:H4K20 me2
H3K9 me2:DNA Meth. m5C
H3K4 me3:H4−4ac
DNA Meth. m5C:H3K9 me3
H3K4 me1:H3−5ac
H3K27 me3:H3K9 me3
H4−4ac:H3−5ac
H4K20 me2:H3−5ac
H3K27 me3:DNA Meth. m5C
H4K20 me2:H3K4 me3
H4K20 me2:H4−4ac
H3K27 me2
H4K20 me3
H3K4 me3
H2A.Z
H3K9 me3
H3K9 me2
H3−5ac
H3K4 me1
H3K27 me3
H4K20 me2
DNA Meth. m5C
H4−4ac
0.00 0.25 0.50 0.75 1.00
$\hat{\pi}$
DDB1 – Reverse
H3K4 me1:H4K20 me2
H3K9 me3:H4K20 me3
H3K27 me3:H3K4 me1
H3K27 me3:H4K20 me3
H3K4 me3:H3−5ac
DNA Meth. m5C:H4K20 me3
H2A.Z:H3K4 me3
DNA Meth. m5C:H4K20 me2
H2A.Z:H4K20 me2
H3K9 me2:DNA Meth. m5C
H3K4 me3:H4−4ac
DNA Meth. m5C:H3K9 me3
H3K4 me1:H3−5ac
H3K27 me3:H3K9 me3
H4−4ac:H3−5ac
H4K20 me2:H3−5ac
H3K27 me3:DNA Meth. m5C
H4K20 me2:H3K4 me3
H4K20 me2:H4−4ac
H3K27 me2
H4K20 me3
H3K4 me3
H2A.Z
H3K9 me3
H3K9 me2
H3−5ac
H3K4 me1
H3K27 me3
H4K20 me2
DNA Meth. m5C
H4−4ac
0.00 0.25 0.50 0.75 1.00
$\hat{\pi}$
RPA2 – Forward
H3K4 me1:H3−5ac
H3K4 me1:H4−4ac
H3K4 me1:H2A.Z
H3K9 me3:H4K20 me3
H2A.Z:H3−5ac
H2A.Z:H4−4ac
H3K4 me1:H4K20 me2
H2A.Z:H3K4 me3
H3K27 me2:DNA Meth. m5C
H3K4 me3:H4−4ac
H2A.Z:H4K20 me2
H4−4ac:H3−5ac
H4K20 me2:H3K4 me3
DNA Meth. m5C:H4K20 me3
H4K20 me2:H4−4ac
H3K27 me3:H3K9 me3
DNA Meth. m5C:H4K20 me2
H4K20 me2:H3−5ac
DNA Meth. m5C:H3K9 me3
H3K27 me3:DNA Meth. m5C
H3K4 me3
H3K27 me2
H3K9 me2
H3K4 me1
H3K9 me3
H4K20 me3
H2A.Z
H3−5ac
H3K27 me3
H4−4ac
H4K20 me2
DNA Meth. m5C
0.00 0.25 0.50 0.75 1.00
$\hat{\pi}$
RPA2 – Reverse
H3K4 me1:H3−5ac
H3K4 me1:H4−4ac
H3K4 me1:H2A.Z
H3K9 me3:H4K20 me3
H2A.Z:H3−5ac
H2A.Z:H4−4ac
H3K4 me1:H4K20 me2
H2A.Z:H3K4 me3
H3K27 me2:DNA Meth. m5C
H3K4 me3:H4−4ac
H2A.Z:H4K20 me2
H4−4ac:H3−5ac
H4K20 me2:H3K4 me3
DNA Meth. m5C:H4K20 me3
H4K20 me2:H4−4ac
H3K27 me3:H3K9 me3
DNA Meth. m5C:H4K20 me2
H4K20 me2:H3−5ac
DNA Meth. m5C:H3K9 me3
H3K27 me3:DNA Meth. m5C
H3K4 me3
H3K27 me2
H3K9 me2
H3K4 me1
H3K9 me3
H4K20 me3
H2A.Z
H3−5ac
H3K27 me3
H4−4ac
H4K20 me2
DNA Meth. m5C
0.00 0.25 0.50 0.75 1.00
$\hat{\pi}$
RPA3 – Forward
H3K4 me3:H3−5ac
H3K27 me3:H4K20 me3
H2A.Z:H3−5ac
H2A.Z:H4−4ac
H3K4 me1:H4K20 me2
DNA Meth. m5C:H4K20 me3
H3K9 me2:DNA Meth. m5C
H3K4 me3:H4−4ac
H2A.Z:H3K4 me3
H4−4ac:H3−5ac
H3K4 me1:H4−4ac
H3K27 me2:DNA Meth. m5C
H2A.Z:H4K20 me2
H3K4 me1:H3−5ac
H3K4 me1:H2A.Z
H4K20 me2:H4−4ac
H4K20 me2:H3K4 me3
H3K27 me3:H3K9 me3
DNA Meth. m5C:H4K20 me2
H4K20 me2:H3−5ac
DNA Meth. m5C:H3K9 me3
H3K27 me3:DNA Meth. m5C
H3K4 me3
H3K27 me2
H3K9 me2
H4K20 me3
H3K9 me3
H3K4 me1
H3K27 me3
H3−5ac
H2A.Z
H4−4ac
H4K20 me2
DNA Meth. m5C
0.00 0.25 0.50 0.75 1.00
$\hat{\pi}$
RPA3 – Reverse
H3K4 me3:H3−5ac
H3K27 me3:H4K20 me3
H2A.Z:H3−5ac
H2A.Z:H4−4ac
H3K4 me1:H4K20 me2
DNA Meth. m5C:H4K20 me3
H3K9 me2:DNA Meth. m5C
H3K4 me3:H4−4ac
H2A.Z:H3K4 me3
H4−4ac:H3−5ac
H3K4 me1:H4−4ac
H3K27 me2:DNA Meth. m5C
H2A.Z:H4K20 me2
H3K4 me1:H3−5ac
H3K4 me1:H2A.Z
H4K20 me2:H4−4ac
H4K20 me2:H3K4 me3
H3K27 me3:H3K9 me3
DNA Meth. m5C:H4K20 me2
H4K20 me2:H3−5ac
DNA Meth. m5C:H3K9 me3
H3K27 me3:DNA Meth. m5C
H3K4 me3
H3K27 me2
H3K9 me2
H4K20 me3
H3K9 me3
H3K4 me1
H3K27 me3
H3−5ac
H2A.Z
H4−4ac
H4K20 me2
DNA Meth. m5C
0.00 0.25 0.50 0.75 1.00
$\hat{\pi}$
RPA1 – Forward
H3K4 me3:H3−5ac
H2A.Z:H3K4 me3
H3K9 me3:H4K20 me3
H3K27 me2:DNA Meth. m5C
H3K4 me1:H3−5ac
DNA Meth. m5C:H4K20 me3
H4−4ac:H3−5ac
H2A.Z:H3−5ac
H3K4 me1:H4−4ac
H3K4 me1:H2A.Z
H3K9 me2:DNA Meth. m5C
H2A.Z:H4−4ac
H2A.Z:H4K20 me2
H3K4 me1:H4K20 me2
H3K27 me3:H3K4 me1
H3K4 me3:H4−4ac
H3K27 me3:H3K9 me3
H4K20 me2:H4−4ac
DNA Meth. m5C:H3K9 me3
H4K20 me2:H3K4 me3
H4K20 me2:H3−5ac
DNA Meth. m5C:H4K20 me2
H3K27 me3:DNA Meth. m5C
H3K27 me2
H4K20 me3
H3K9 me2
H3K4 me3
H3K27 me3
H3K9 me3
H3−5ac
H3K4 me1
H2A.Z
H4−4ac
DNA Meth. m5C
H4K20 me2
0.00 0.25 0.50 0.75 1.00
$\hat{\pi}$
RPA1 – Reverse
H3K4 me3:H3−5ac
H2A.Z:H3K4 me3
H3K9 me3:H4K20 me3
H3K27 me2:DNA Meth. m5C
H3K4 me1:H3−5ac
DNA Meth. m5C:H4K20 me3
H4−4ac:H3−5ac
H2A.Z:H3−5ac
H3K4 me1:H4−4ac
H3K4 me1:H2A.Z
H3K9 me2:DNA Meth. m5C
H2A.Z:H4−4ac
H2A.Z:H4K20 me2
H3K4 me1:H4K20 me2
H3K27 me3:H3K4 me1
H3K4 me3:H4−4ac
H3K27 me3:H3K9 me3
H4K20 me2:H4−4ac
DNA Meth. m5C:H3K9 me3
H4K20 me2:H3K4 me3
H4K20 me2:H3−5ac
DNA Meth. m5C:H4K20 me2
H3K27 me3:DNA Meth. m5C
H3K27 me2
H4K20 me3
H3K9 me2
H3K4 me3
H3K27 me3
H3K9 me3
H3−5ac
H3K4 me1
H2A.Z
H4−4ac
DNA Meth. m5C
H4K20 me2
0.00 0.25 0.50 0.75 1.00
$\hat{\pi}$
ZNF346 – Forward
H3K4 me1:H4K20 me2
H3K9 me3:H4K20 me3
H3K27 me3:H4K20 me3
H2A.Z:H4K20 me2
H3K9 me2:DNA Meth. m5C
H4K20 me2:H4−4ac
DNA Meth. m5C:H3K9 me3
H3K27 me3:H3K9 me3
H3K27 me3:DNA Meth. m5C
H4K20 me2:H3K4 me3
H4K20 me2:H3−5ac
DNA Meth. m5C:H4K20 me3
DNA Meth. m5C:H4K20 me2
H3K27 me2
H3K4 me1
H3K4 me3
H3K9 me2
H4−4ac
DNA Meth. m5C
H4K20 me3
H3−5ac
H2A.Z
H3K9 me3
H3K27 me3
H4K20 me2
0.00 0.25 0.50 0.75 1.00
$\hat{\pi}$
ZNF346 – Reverse
H3K4 me1:H4K20 me2
H3K9 me3:H4K20 me3
H3K27 me3:H4K20 me3
H2A.Z:H4K20 me2
H3K9 me2:DNA Meth. m5C
H4K20 me2:H4−4ac
DNA Meth. m5C:H3K9 me3
H3K27 me3:H3K9 me3
H3K27 me3:DNA Meth. m5C
H4K20 me2:H3K4 me3
H4K20 me2:H3−5ac
DNA Meth. m5C:H4K20 me3
DNA Meth. m5C:H4K20 me2
H3K27 me2
H3K4 me1
H3K4 me3
H3K9 me2
H4−4ac
DNA Meth. m5C
H4K20 me3
H3−5ac
H2A.Z
H3K9 me3
H3K27 me3
H4K20 me2
0.00 0.25 0.50 0.75 1.00
$\hat{\pi}$
LIG3 (1) – Forward
H3K4 me1:H4−4ac
H4−4ac:H3−5ac
H3K4 me1:H3−5ac
H2A.Z:H4−4ac
H3K4 me3:H3−5ac
H2A.Z:H3−5ac
H3K27 me3:H4K20 me3
H3K4 me1:H4K20 me2
H3K9 me3:H4K20 me3
H3K27 me2:DNA Meth. m5C
H2A.Z:H3K4 me3
H4K20 me2:H4−4ac
H2A.Z:H4K20 me2
H3K9 me2:DNA Meth. m5C
H3K27 me3:H3K4 me1
H3K27 me3:H3K9 me3
DNA Meth. m5C:H4K20 me2
H4K20 me2:H3K4 me3
H4K20 me2:H3−5ac
H3K27 me3:DNA Meth. m5C
H4−4ac
H3K4 me3
H3K9 me2
H4K20 me3
H3−5ac
DNA Meth. m5C
H3K27 me2
H2A.Z
H3K9 me3
H3K27 me3
H3K4 me1
H4K20 me2
0.00 0.25 0.50 0.75 1.00
$\hat{\pi}$
LIG3 (1) – Reverse
H3K4 me1:H4−4ac
H4−4ac:H3−5ac
H3K4 me1:H3−5ac
H2A.Z:H4−4ac
H3K4 me3:H3−5ac
H2A.Z:H3−5ac
H3K27 me3:H4K20 me3
H3K4 me1:H4K20 me2
H3K9 me3:H4K20 me3
H3K27 me2:DNA Meth. m5C
H2A.Z:H3K4 me3
H4K20 me2:H4−4ac
H2A.Z:H4K20 me2
H3K9 me2:DNA Meth. m5C
H3K27 me3:H3K4 me1
H3K27 me3:H3K9 me3
DNA Meth. m5C:H4K20 me2
H4K20 me2:H3K4 me3
H4K20 me2:H3−5ac
H3K27 me3:DNA Meth. m5C
H4−4ac
H3K4 me3
H3K9 me2
H4K20 me3
H3−5ac
DNA Meth. m5C
H3K27 me2
H2A.Z
H3K9 me3
H3K27 me3
H3K4 me1
H4K20 me2
0.00 0.25 0.50 0.75 1.00
$\hat{\pi}$

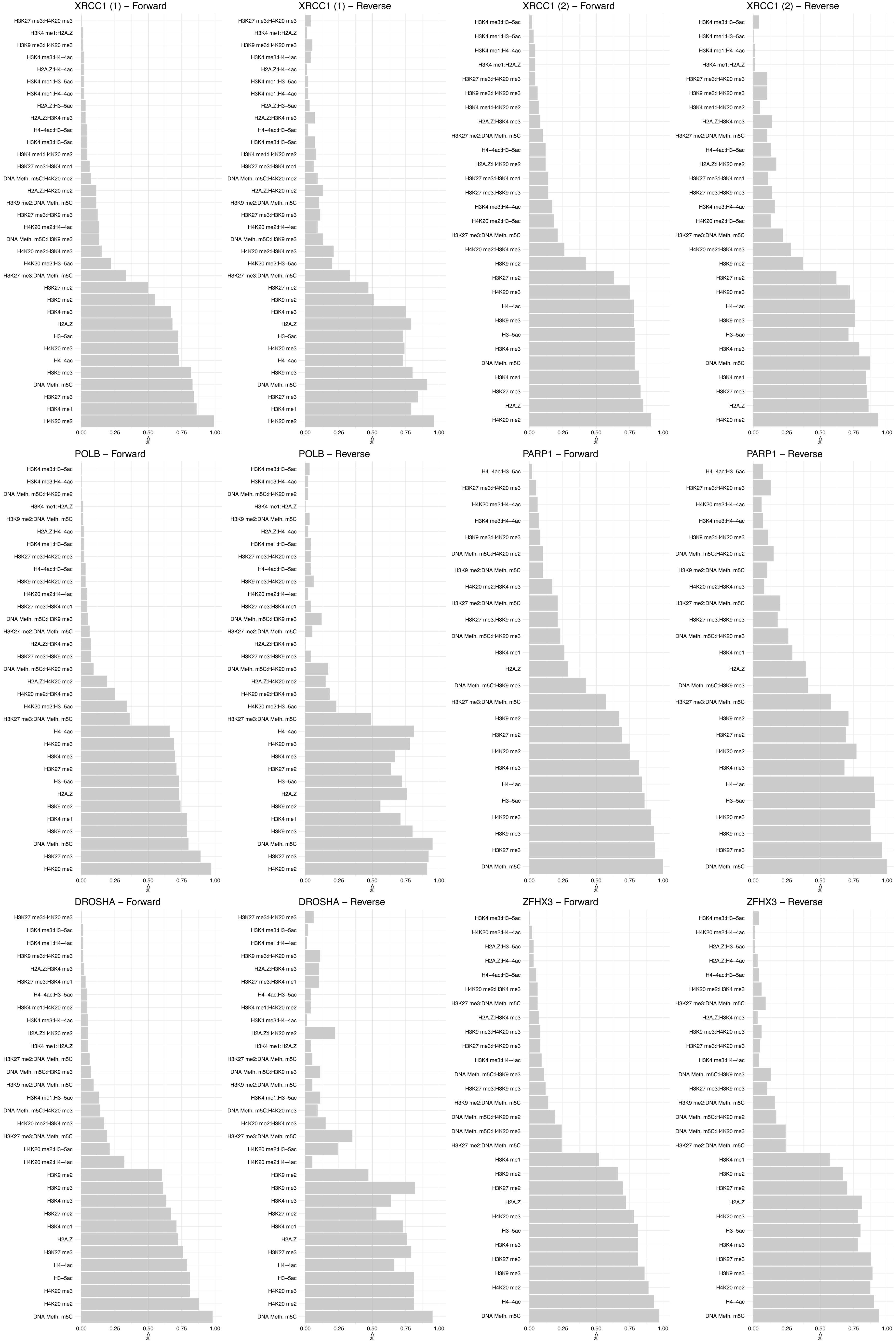
XRCC1 (1) – Forward
H3K27 me3:H4K20 me3
H3K4 me1:H2A.Z
H3K9 me3:H4K20 me3
H3K4 me3:H4–4ac
H2A.Z:H4–4ac
H3K4 me1:H3–5ac
H3K4 me1:H4–4ac
H2A.Z:H3–5ac
H2A.Z:H3K4 me3
H4–4ac:H3–5ac
H3K4 me3:H3–5ac
H3K4 me1:H4K20 me2
H3K27 me3:H3K4 me1
DNA Meth. m5C:H4K20 me2
H2A.Z:H4K20 me2
H3K9 me2:DNA Meth. m5C
H3K27 me3:H3K9 me3
H4K20 me2:H4–4ac
DNA Meth. m5C:H3K9 me3
H4K20 me2:H3K4 me3
H4K20 me2:H3–5ac
H3K27 me3:DNA Meth. m5C
H3K27 me2
H3K9 me2
H3K4 me3
H2A.Z
H3–5ac
H4K20 me3
H4–4ac
H3K9 me3
DNA Meth. m5C
H3K27 me3
H3K4 me1
H4K20 me2
0.00
0.25
0.50
0.75
1.00
π̂
XRCC1 (1) – Reverse
H3K27 me3:H4K20 me3
H3K4 me1:H2A.Z
H3K9 me3:H4K20 me3
H3K4 me3:H4–4ac
H2A.Z:H4–4ac
H3K4 me1:H3–5ac
H3K4 me1:H4–4ac
H2A.Z:H3–5ac
H2A.Z:H3K4 me3
H4–4ac:H3–5ac
H3K4 me3:H3–5ac
H3K4 me1:H4K20 me2
H3K27 me3:H3K4 me1
DNA Meth. m5C:H4K20 me2
H2A.Z:H4K20 me2
H3K9 me2:DNA Meth. m5C
H3K27 me3:H3K9 me3
H4K20 me2:H4–4ac
DNA Meth. m5C:H3K9 me3
H4K20 me2:H3K4 me3
H4K20 me2:H3–5ac
H3K27 me3:DNA Meth. m5C
H3K27 me2
H3K9 me2
H3K4 me3
H2A.Z
H3–5ac
H4K20 me3
H4–4ac
H3K9 me3
DNA Meth. m5C
H3K27 me3
H3K4 me1
H4K20 me2
0.00
0.25
0.50
0.75
1.00
π̂
XRCC1 (2) – Forward
H3K4 me3:H3–5ac
H3K4 me1:H3–5ac
H3K4 me1:H4–4ac
H3K4 me1:H2A.Z
H3K27 me3:H4K20 me3
H3K9 me3:H4K20 me3
H3K4 me1:H4K20 me2
H2A.Z:H3K4 me3
H3K27 me2:DNA Meth. m5C
H4–4ac:H3–5ac
H2A.Z:H4K20 me2
H3K27 me3:H3K4 me1
H3K27 me3:H3K9 me3
H3K4 me3:H4–4ac
H4K20 me2:H3–5ac
H3K27 me3:DNA Meth. m5C
H4K20 me2:H3K4 me3
H3K9 me2
H3K27 me2
H4K20 me3
H4–4ac
H3K9 me3
H3–5ac
H3K4 me3
DNA Meth. m5C
H3K4 me1
H3K27 me3
H2A.Z
H4K20 me2
0.00
0.25
0.50
0.75
1.00
π̂
XRCC1 (2) – Reverse
H3K4 me3:H3–5ac
H3K4 me1:H3–5ac
H3K4 me1:H4–4ac
H3K4 me1:H2A.Z
H3K27 me3:H4K20 me3
H3K9 me3:H4K20 me3
H3K4 me1:H4K20 me2
H2A.Z:H3K4 me3
H3K27 me2:DNA Meth. m5C
H4–4ac:H3–5ac
H2A.Z:H4K20 me2
H3K27 me3:H3K4 me1
H3K27 me3:H3K9 me3
H3K4 me3:H4–4ac
H4K20 me2:H3–5ac
H3K27 me3:DNA Meth. m5C
H4K20 me2:H3K4 me3
H3K9 me2
H3K27 me2
H4K20 me3
H4–4ac
H3K9 me3
H3–5ac
H3K4 me3
DNA Meth. m5C
H3K4 me1
H3K27 me3
H2A.Z
H4K20 me2
0.00
0.25
0.50
0.75
1.00
π̂
POLB – Forward
H3K4 me3:H3–5ac
H3K4 me3:H4–4ac
DNA Meth. m5C:H4K20 me2
H3K4 me1:H2A.Z
H3K9 me2:DNA Meth. m5C
H2A.Z:H4–4ac
H3K4 me1:H3–5ac
H3K27 me3:H4K20 me3
H4–4ac:H3–5ac
H3K9 me3:H4K20 me3
H4K20 me2:H4–4ac
H3K27 me3:H3K4 me1
DNA Meth. m5C:H3K9 me3
H3K27 me2:DNA Meth. m5C
H2A.Z:H3K4 me3
H3K27 me3:H3K9 me3
DNA Meth. m5C:H4K20 me3
H2A.Z:H4K20 me2
H4K20 me2:H3K4 me3
H4K20 me2:H3–5ac
H3K27 me3:DNA Meth. m5C
H4–4ac
H4K20 me3
H3K4 me3
H3K27 me2
H3–5ac
H2A.Z
H3K9 me2
H3K4 me1
H3K9 me3
DNA Meth. m5C
H3K27 me3
H4K20 me2
0.00
0.25
0.50
0.75
1.00
π̂
POLB – Reverse
H3K4 me3:H3–5ac
H3K4 me3:H4–4ac
DNA Meth. m5C:H4K20 me2
H3K4 me1:H2A.Z
H3K9 me2:DNA Meth. m5C
H2A.Z:H4–4ac
H3K4 me1:H3–5ac
H3K27 me3:H4K20 me3
H4–4ac:H3–5ac
H3K9 me3:H4K20 me3
H4K20 me2:H4–4ac
H3K27 me3:H3K4 me1
DNA Meth. m5C:H3K9 me3
H3K27 me2:DNA Meth. m5C
H2A.Z:H3K4 me3
H3K27 me3:H3K9 me3
DNA Meth. m5C:H4K20 me3
H2A.Z:H4K20 me2
H4K20 me2:H3K4 me3
H4K20 me2:H3–5ac
H3K27 me3:DNA Meth. m5C
H4–4ac
H4K20 me3
H3K4 me3
H3K27 me2
H3–5ac
H2A.Z
H3K9 me2
H3K4 me1
H3K9 me3
DNA Meth. m5C
H3K27 me3
H4K20 me2
0.00
0.25
0.50
0.75
1.00
π̂
PARP1 – Forward
H4–4ac:H3–5ac
H3K27 me3:H4K20 me3
H4K20 me2:H4–4ac
H3K4 me3:H4–4ac
H3K9 me3:H4K20 me3
DNA Meth. m5C:H4K20 me2
H3K9 me2:DNA Meth. m5C
H4K20 me2:H3K4 me3
H3K27 me2:DNA Meth. m5C
H3K27 me3:H3K9 me3
DNA Meth. m5C:H4K20 me3
H3K4 me1
H2A.Z
DNA Meth. m5C:H3K9 me3
H3K27 me3:DNA Meth. m5C
H3K9 me2
H3K27 me2
H4K20 me2
H3K4 me3
H4–4ac
H3–5ac
H4K20 me3
H3K9 me3
H3K27 me3
DNA Meth. m5C
0.00
0.25
0.50
0.75
1.00
π̂
PARP1 – Reverse
H4–4ac:H3–5ac
H3K27 me3:H4K20 me3
H4K20 me2:H4–4ac
H3K4 me3:H4–4ac
H3K9 me3:H4K20 me3
DNA Meth. m5C:H4K20 me2
H3K9 me2:DNA Meth. m5C
H4K20 me2:H3K4 me3
H3K27 me2:DNA Meth. m5C
H3K27 me3:H3K9 me3
DNA Meth. m5C:H4K20 me3
H3K4 me1
H2A.Z
DNA Meth. m5C:H3K9 me3
H3K27 me3:DNA Meth. m5C
H3K9 me2
H3K27 me2
H4K20 me2
H3K4 me3
H4–4ac
H3–5ac
H4K20 me3
H3K9 me3
H3K27 me3
DNA Meth. m5C
0.00
0.25
0.50
0.75
1.00
π̂
DROSHA – Forward
H3K27 me3:H4K20 me3
H3K4 me3:H3–5ac
H3K4 me1:H4–4ac
H3K9 me3:H4K20 me3
H2A.Z:H3K4 me3
H3K27 me3:H3K4 me1
H4–4ac:H3–5ac
H3K4 me1:H4K20 me2
H3K4 me3:H4–4ac
H2A.Z:H4K20 me2
H3K4 me1:H2A.Z
H3K27 me2:DNA Meth. m5C
DNA Meth. m5C:H3K9 me3
H3K9 me2:DNA Meth. m5C
H3K4 me1:H3–5ac
DNA Meth. m5C:H4K20 me3
H4K20 me2:H3K4 me3
H3K27 me3:DNA Meth. m5C
H4K20 me2:H3–5ac
H4K20 me2:H4–4ac
H3K9 me2
H3K9 me3
H3K4 me3
H3K27 me2
H3K4 me1
H2A.Z
H3K27 me3
H4–4ac
H3–5ac
H4K20 me3
H4K20 me2
DNA Meth. m5C
0.00
0.25
0.50
0.75
1.00
π̂
DROSHA – Reverse
H3K27 me3:H4K20 me3
H3K4 me3:H3–5ac
H3K4 me1:H4–4ac
H3K9 me3:H4K20 me3
H2A.Z:H3K4 me3
H3K27 me3:H3K4 me1
H4–4ac:H3–5ac
H3K4 me1:H4K20 me2
H3K4 me3:H4–4ac
H2A.Z:H4K20 me2
H3K4 me1:H2A.Z
H3K27 me2:DNA Meth. m5C
DNA Meth. m5C:H3K9 me3
H3K9 me2:DNA Meth. m5C
H3K4 me1:H3–5ac
DNA Meth. m5C:H4K20 me3
H4K20 me2:H3K4 me3
H3K27 me3:DNA Meth. m5C
H4K20 me2:H3–5ac
H4K20 me2:H4–4ac
H3K9 me2
H3K9 me3
H3K4 me3
H3K27 me2
H3K4 me1
H2A.Z
H3K27 me3
H4–4ac
H3–5ac
H4K20 me3
H4K20 me2
DNA Meth. m5C
0.00
0.25
0.50
0.75
1.00
π̂
ZFHX3 – Forward
H3K4 me3:H3–5ac
H4K20 me2:H4–4ac
H2A.Z:H3–5ac
H2A.Z:H4–4ac
H4–4ac:H3–5ac
H4K20 me2:H3K4 me3
H3K27 me3:DNA Meth. m5C
H2A.Z:H3K4 me3
H3K9 me3:H4K20 me3
H3K27 me3:H4K20 me3
H3K4 me3:H4–4ac
DNA Meth. m5C:H3K9 me3
H3K27 me3:H3K9 me3
H3K9 me2:DNA Meth. m5C
DNA Meth. m5C:H4K20 me2
DNA Meth. m5C:H4K20 me3
H3K27 me2:DNA Meth. m5C
H3K4 me1
H3K9 me2
H3K27 me2
H2A.Z
H4K20 me3
H3–5ac
H3K4 me3
H3K27 me3
H3K9 me3
H4K20 me2
H4–4ac
DNA Meth. m5C
0.00
0.25
0.50
0.75
1.00
π̂
ZFHX3 – Reverse
H3K4 me3:H3–5ac
H4K20 me2:H4–4ac
H2A.Z:H3–5ac
H2A.Z:H4–4ac
H4–4ac:H3–5ac
H4K20 me2:H3K4 me3
H3K27 me3:DNA Meth. m5C
H2A.Z:H3K4 me3
H3K9 me3:H4K20 me3
H3K27 me3:H4K20 me3
H3K4 me3:H4–4ac
DNA Meth. m5C:H3K9 me3
H3K27 me3:H3K9 me3
H3K9 me2:DNA Meth. m5C
DNA Meth. m5C:H4K20 me2
DNA Meth. m5C:H4K20 me3
H3K27 me2:DNA Meth. m5C
H3K4 me1
H3K9 me2
H3K27 me2
H2A.Z
H4K20 me3
H3–5ac
H3K4 me3
H3K27 me3
H3K9 me3
H4K20 me2
H4–4ac
DNA Meth. m5C
0.00
0.25
0.50
0.75
1.00
π̂

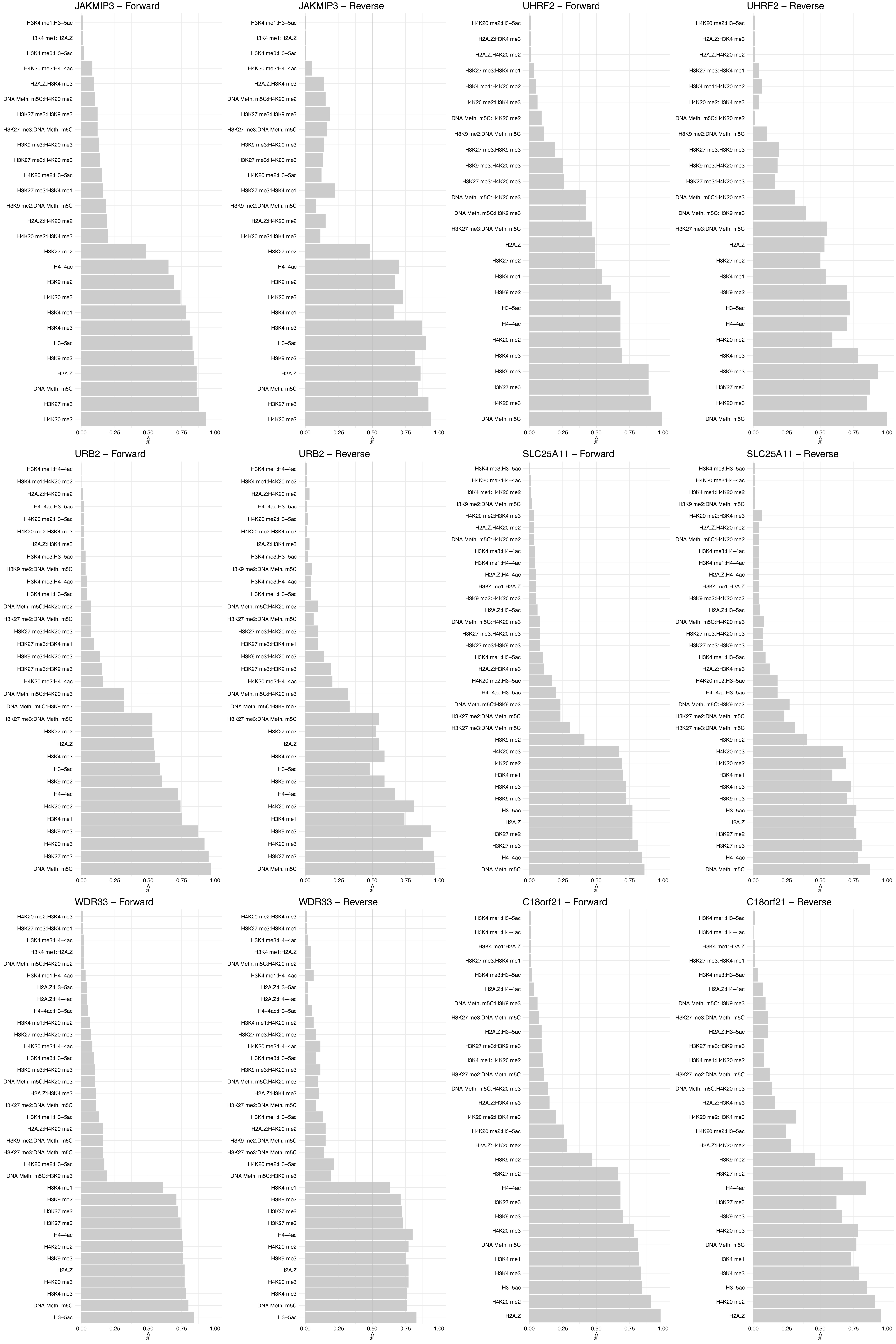
JAKMIP3 – Forward
H3K4 me1:H3−5ac
H3K4 me1:H2A.Z
H3K4 me3:H3−5ac
H4K20 me2:H4−4ac
H2A.Z:H3K4 me3
DNA Meth. m5C:H4K20 me2
H3K27 me3:H3K9 me3
H3K27 me3:DNA Meth. m5C
H3K9 me3:H4K20 me3
H3K27 me3:H4K20 me3
H4K20 me2:H3−5ac
H3K27 me3:H3K4 me1
H3K9 me2:DNA Meth. m5C
H2A.Z:H4K20 me2
H4K20 me2:H3K4 me3
H3K27 me2
H4−4ac
H3K9 me2
H4K20 me3
H3K4 me1
H3K4 me3
H3−5ac
H3K9 me3
H2A.Z
DNA Meth. m5C
H3K27 me3
H4K20 me2
0.00
0.25
0.50
0.75
1.00
π̂
JAKMIP3 – Reverse
H3K4 me1:H3−5ac
H3K4 me1:H2A.Z
H3K4 me3:H3−5ac
H4K20 me2:H4−4ac
H2A.Z:H3K4 me3
DNA Meth. m5C:H4K20 me2
H3K27 me3:H3K9 me3
H3K27 me3:DNA Meth. m5C
H3K9 me3:H4K20 me3
H3K27 me3:H4K20 me3
H4K20 me2:H3−5ac
H3K27 me3:H3K4 me1
H3K9 me2:DNA Meth. m5C
H2A.Z:H4K20 me2
H4K20 me2:H3K4 me3
H3K27 me2
H4−4ac
H3K9 me2
H4K20 me3
H3K4 me1
H3K4 me3
H3−5ac
H3K9 me3
H2A.Z
DNA Meth. m5C
H3K27 me3
H4K20 me2
0.00
0.25
0.50
0.75
1.00
π̂
UHRF2 – Forward
H4K20 me2:H3−5ac
H2A.Z:H3K4 me3
H2A.Z:H4K20 me2
H3K27 me3:H3K4 me1
H3K4 me1:H4K20 me2
H4K20 me2:H3K4 me3
DNA Meth. m5C:H4K20 me2
H3K9 me2:DNA Meth. m5C
H3K27 me3:H3K9 me3
H3K9 me3:H4K20 me3
H3K27 me3:H4K20 me3
DNA Meth. m5C:H4K20 me3
DNA Meth. m5C:H3K9 me3
H3K27 me3:DNA Meth. m5C
H2A.Z
H3K27 me2
H3K4 me1
H3K9 me2
H3−5ac
H4−4ac
H4K20 me2
H3K4 me3
H3K9 me3
H3K27 me3
H4K20 me3
DNA Meth. m5C
0.00
0.25
0.50
0.75
1.00
π̂
UHRF2 – Reverse
H4K20 me2:H3−5ac
H2A.Z:H3K4 me3
H2A.Z:H4K20 me2
H3K27 me3:H3K4 me1
H3K4 me1:H4K20 me2
H4K20 me2:H3K4 me3
DNA Meth. m5C:H4K20 me2
H3K9 me2:DNA Meth. m5C
H3K27 me3:H3K9 me3
H3K9 me3:H4K20 me3
H3K27 me3:H4K20 me3
DNA Meth. m5C:H4K20 me3
DNA Meth. m5C:H3K9 me3
H3K27 me3:DNA Meth. m5C
H2A.Z
H3K27 me2
H3K4 me1
H3K9 me2
H3−5ac
H4−4ac
H4K20 me2
H3K4 me3
H3K9 me3
H3K27 me3
H4K20 me3
DNA Meth. m5C
0.00
0.25
0.50
0.75
1.00
π̂
URB2 – Forward
H3K4 me1:H4−4ac
H3K4 me1:H4K20 me2
H2A.Z:H4K20 me2
H4−4ac:H3−5ac
H4K20 me2:H3−5ac
H4K20 me2:H3K4 me3
H2A.Z:H3K4 me3
H3K4 me3:H3−5ac
H3K9 me2:DNA Meth. m5C
H3K4 me3:H4−4ac
H3K4 me1:H3−5ac
DNA Meth. m5C:H4K20 me2
H3K27 me2:DNA Meth. m5C
H3K27 me3:H4K20 me3
H3K27 me3:H3K4 me1
H3K9 me3:H4K20 me3
H3K27 me3:H3K9 me3
H4K20 me2:H4−4ac
DNA Meth. m5C:H4K20 me3
DNA Meth. m5C:H3K9 me3
H3K27 me3:DNA Meth. m5C
H3K27 me2
H2A.Z
H3K4 me3
H3−5ac
H3K9 me2
H4−4ac
H4K20 me2
H3K4 me1
H3K9 me3
H4K20 me3
H3K27 me3
DNA Meth. m5C
0.00
0.25
0.50
0.75
1.00
π̂
URB2 – Reverse
H3K4 me1:H4−4ac
H3K4 me1:H4K20 me2
H2A.Z:H4K20 me2
H4−4ac:H3−5ac
H4K20 me2:H3−5ac
H4K20 me2:H3K4 me3
H2A.Z:H3K4 me3
H3K4 me3:H3−5ac
H3K9 me2:DNA Meth. m5C
H3K4 me3:H4−4ac
H3K4 me1:H3−5ac
DNA Meth. m5C:H4K20 me2
H3K27 me2:DNA Meth. m5C
H3K27 me3:H4K20 me3
H3K27 me3:H3K4 me1
H3K9 me3:H4K20 me3
H3K27 me3:H3K9 me3
H4K20 me2:H4−4ac
DNA Meth. m5C:H4K20 me3
DNA Meth. m5C:H3K9 me3
H3K27 me3:DNA Meth. m5C
H3K27 me2
H2A.Z
H3K4 me3
H3−5ac
H3K9 me2
H4−4ac
H4K20 me2
H3K4 me1
H3K9 me3
H4K20 me3
H3K27 me3
DNA Meth. m5C
0.00
0.25
0.50
0.75
1.00
π̂
SLC25A11 – Forward
H3K4 me3:H3−5ac
H4K20 me2:H4−4ac
H3K4 me1:H4K20 me2
H3K9 me2:DNA Meth. m5C
H4K20 me2:H3K4 me3
H2A.Z:H4K20 me2
DNA Meth. m5C:H4K20 me2
H3K4 me3:H4−4ac
H3K4 me1:H4−4ac
H2A.Z:H4−4ac
H3K4 me1:H2A.Z
H3K9 me3:H4K20 me3
H2A.Z:H3−5ac
DNA Meth. m5C:H4K20 me3
H3K27 me3:H4K20 me3
H3K27 me3:H3K9 me3
H3K4 me1:H3−5ac
H2A.Z:H3K4 me3
H4K20 me2:H3−5ac
H4−4ac:H3−5ac
DNA Meth. m5C:H3K9 me3
H3K27 me2:DNA Meth. m5C
H3K27 me3:DNA Meth. m5C
H3K9 me2
H4K20 me3
H4K20 me2
H3K4 me1
H3K4 me3
H3K9 me3
H3−5ac
H2A.Z
H3K27 me2
H3K27 me3
H4−4ac
DNA Meth. m5C
0.00
0.25
0.50
0.75
1.00
π̂
SLC25A11 – Reverse
H3K4 me3:H3−5ac
H4K20 me2:H4−4ac
H3K4 me1:H4K20 me2
H3K9 me2:DNA Meth. m5C
H4K20 me2:H3K4 me3
H2A.Z:H4K20 me2
DNA Meth. m5C:H4K20 me2
H3K4 me3:H4−4ac
H3K4 me1:H4−4ac
H2A.Z:H4−4ac
H3K4 me1:H2A.Z
H3K9 me3:H4K20 me3
H2A.Z:H3−5ac
DNA Meth. m5C:H4K20 me3
H3K27 me3:H4K20 me3
H3K27 me3:H3K9 me3
H3K4 me1:H3−5ac
H2A.Z:H3K4 me3
H4K20 me2:H3−5ac
H4−4ac:H3−5ac
DNA Meth. m5C:H3K9 me3
H3K27 me2:DNA Meth. m5C
H3K27 me3:DNA Meth. m5C
H3K9 me2
H4K20 me3
H4K20 me2
H3K4 me1
H3K4 me3
H3K9 me3
H3−5ac
H2A.Z
H3K27 me2
H3K27 me3
H4−4ac
DNA Meth. m5C
0.00
0.25
0.50
0.75
1.00
π̂
WDR33 – Forward
H4K20 me2:H3K4 me3
H3K27 me3:H3K4 me1
H3K4 me3:H4−4ac
H3K4 me1:H2A.Z
DNA Meth. m5C:H4K20 me2
H3K4 me1:H4−4ac
H2A.Z:H3−5ac
H2A.Z:H4−4ac
H4−4ac:H3−5ac
H3K4 me1:H4K20 me2
H3K27 me3:H4K20 me3
H4K20 me2:H4−4ac
H3K4 me3:H3−5ac
H3K9 me3:H4K20 me3
DNA Meth. m5C:H4K20 me3
H2A.Z:H3K4 me3
H3K27 me2:DNA Meth. m5C
H3K4 me1:H3−5ac
H2A.Z:H4K20 me2
H3K9 me2:DNA Meth. m5C
H3K27 me3:DNA Meth. m5C
H4K20 me2:H3−5ac
DNA Meth. m5C:H3K9 me3
H3K4 me1
H3K9 me2
H3K27 me2
H3K27 me3
H4−4ac
H4K20 me2
H3K9 me3
H2A.Z
H4K20 me3
H3K4 me3
DNA Meth. m5C
H3−5ac
0.00
0.25
0.50
0.75
1.00
π̂
WDR33 – Reverse
H4K20 me2:H3K4 me3
H3K27 me3:H3K4 me1
H3K4 me3:H4−4ac
H3K4 me1:H2A.Z
DNA Meth. m5C:H4K20 me2
H3K4 me1:H4−4ac
H2A.Z:H3−5ac
H2A.Z:H4−4ac
H4−4ac:H3−5ac
H3K4 me1:H4K20 me2
H3K27 me3:H4K20 me3
H4K20 me2:H4−4ac
H3K4 me3:H3−5ac
H3K9 me3:H4K20 me3
DNA Meth. m5C:H4K20 me3
H2A.Z:H3K4 me3
H3K27 me2:DNA Meth. m5C
H3K4 me1:H3−5ac
H2A.Z:H4K20 me2
H3K9 me2:DNA Meth. m5C
H3K27 me3:DNA Meth. m5C
H4K20 me2:H3−5ac
DNA Meth. m5C:H3K9 me3
H3K4 me1
H3K9 me2
H3K27 me2
H3K27 me3
H4−4ac
H4K20 me2
H3K9 me3
H2A.Z
H4K20 me3
H3K4 me3
DNA Meth. m5C
H3−5ac
0.00
0.25
0.50
0.75
1.00
π̂
C18orf21 – Forward
H3K4 me1:H3−5ac
H3K4 me1:H4−4ac
H3K4 me1:H2A.Z
H3K27 me3:H3K4 me1
H3K4 me3:H3−5ac
H2A.Z:H4−4ac
DNA Meth. m5C:H3K9 me3
H3K27 me3:DNA Meth. m5C
H2A.Z:H3−5ac
H3K27 me3:H3K9 me3
H3K4 me1:H4K20 me2
H3K27 me2:DNA Meth. m5C
DNA Meth. m5C:H4K20 me3
H2A.Z:H3K4 me3
H4K20 me2:H3K4 me3
H4K20 me2:H3−5ac
H2A.Z:H4K20 me2
H3K9 me2
H3K27 me2
H4−4ac
H3K27 me3
H3K9 me3
H4K20 me3
DNA Meth. m5C
H3K4 me1
H3K4 me3
H3−5ac
H4K20 me2
H2A.Z
0.00
0.25
0.50
0.75
1.00
π̂
C18orf21 – Reverse
H3K4 me1:H3−5ac
H3K4 me1:H4−4ac
H3K4 me1:H2A.Z
H3K27 me3:H3K4 me1
H3K4 me3:H3−5ac
H2A.Z:H4−4ac
DNA Meth. m5C:H3K9 me3
H3K27 me3:DNA Meth. m5C
H2A.Z:H3−5ac
H3K27 me3:H3K9 me3
H3K4 me1:H4K20 me2
H3K27 me2:DNA Meth. m5C
DNA Meth. m5C:H4K20 me3
H2A.Z:H3K4 me3
H4K20 me2:H3K4 me3
H4K20 me2:H3−5ac
H2A.Z:H4K20 me2
H3K9 me2
H3K27 me2
H4−4ac
H3K27 me3
H3K9 me3
H4K20 me3
DNA Meth. m5C
H3K4 me1
H3K4 me3
H3−5ac
H4K20 me2
H2A.Z
0.00
0.25
0.50
0.75
1.00
π̂

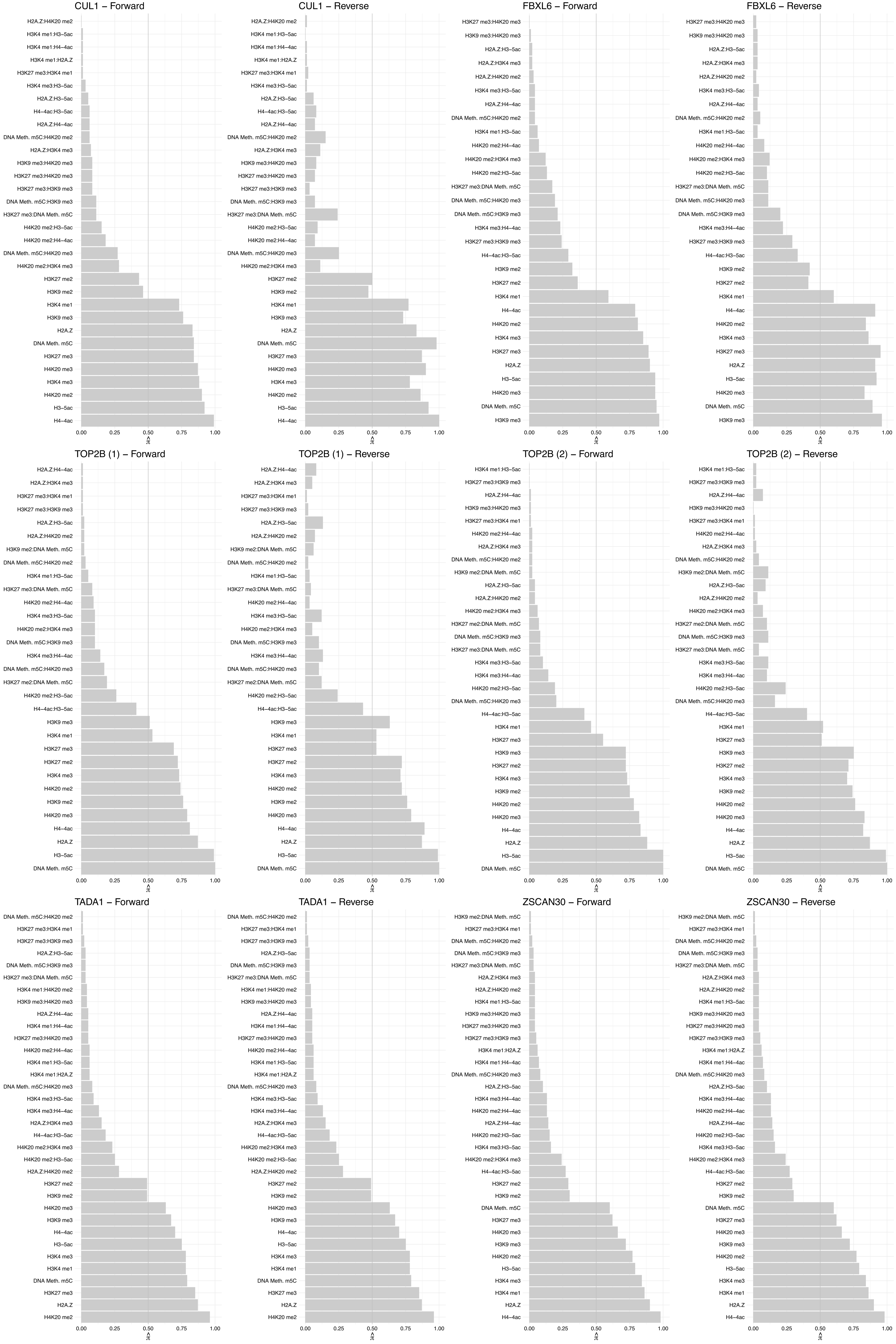
CUL1 – Forward
H2A.Z:H4K20 me2
H3K4 me1:H3–5ac
H3K4 me1:H4–4ac
H3K4 me1:H2A.Z
H3K27 me3:H3K4 me1
H3K4 me3:H3–5ac
H2A.Z:H3–5ac
H4–4ac:H3–5ac
H2A.Z:H4–4ac
DNA Meth. m5C:H4K20 me2
H2A.Z:H3K4 me3
H3K9 me3:H4K20 me3
H3K27 me3:H4K20 me3
H3K27 me3:H3K9 me3
DNA Meth. m5C:H3K9 me3
H3K27 me3:DNA Meth. m5C
H4K20 me2:H3–5ac
H4K20 me2:H4–4ac
DNA Meth. m5C:H4K20 me3
H4K20 me2:H3K4 me3
H3K27 me2
H3K9 me2
H3K4 me1
H3K9 me3
H2A.Z
DNA Meth. m5C
H3K27 me3
H4K20 me3
H3K4 me3
H4K20 me2
H3–5ac
H4–4ac
0.00 0.25 0.50 0.75 1.00
$\hat{\pi}$
CUL1 – Reverse
H2A.Z:H4K20 me2
H3K4 me1:H3–5ac
H3K4 me1:H4–4ac
H3K4 me1:H2A.Z
H3K27 me3:H3K4 me1
H3K4 me3:H3–5ac
H2A.Z:H3–5ac
H4–4ac:H3–5ac
H2A.Z:H4–4ac
DNA Meth. m5C:H4K20 me2
H2A.Z:H3K4 me3
H3K9 me3:H4K20 me3
H3K27 me3:H4K20 me3
H3K27 me3:H3K9 me3
DNA Meth. m5C:H3K9 me3
H3K27 me3:DNA Meth. m5C
H4K20 me2:H3–5ac
H4K20 me2:H4–4ac
DNA Meth. m5C:H4K20 me3
H4K20 me2:H3K4 me3
H3K27 me2
H3K9 me2
H3K4 me1
H3K9 me3
H2A.Z
DNA Meth. m5C
H3K27 me3
H4K20 me3
H3K4 me3
H4K20 me2
H3–5ac
H4–4ac
0.00 0.25 0.50 0.75 1.00
$\hat{\pi}$
FBXL6 – Forward
H3K27 me3:H4K20 me3
H3K9 me3:H4K20 me3
H2A.Z:H3–5ac
H2A.Z:H3K4 me3
H2A.Z:H4K20 me2
H3K4 me3:H3–5ac
H2A.Z:H4–4ac
DNA Meth. m5C:H4K20 me2
H3K4 me1:H3–5ac
H4K20 me2:H4–4ac
H4K20 me2:H3K4 me3
H4K20 me2:H3–5ac
H3K27 me3:DNA Meth. m5C
DNA Meth. m5C:H4K20 me3
DNA Meth. m5C:H3K9 me3
H3K4 me3:H4–4ac
H3K27 me3:H3K9 me3
H4–4ac:H3–5ac
H3K9 me2
H3K27 me2
H3K4 me1
H4–4ac
H4K20 me2
H3K4 me3
H3K27 me3
H2A.Z
H3–5ac
H4K20 me3
DNA Meth. m5C
H3K9 me3
0.00 0.25 0.50 0.75 1.00
$\hat{\pi}$
FBXL6 – Reverse
H3K27 me3:H4K20 me3
H3K9 me3:H4K20 me3
H2A.Z:H3–5ac
H2A.Z:H3K4 me3
H2A.Z:H4K20 me2
H3K4 me3:H3–5ac
H2A.Z:H4–4ac
DNA Meth. m5C:H4K20 me2
H3K4 me1:H3–5ac
H4K20 me2:H4–4ac
H4K20 me2:H3K4 me3
H4K20 me2:H3–5ac
H3K27 me3:DNA Meth. m5C
DNA Meth. m5C:H4K20 me3
DNA Meth. m5C:H3K9 me3
H3K4 me3:H4–4ac
H3K27 me3:H3K9 me3
H4–4ac:H3–5ac
H3K9 me2
H3K27 me2
H3K4 me1
H4–4ac
H4K20 me2
H3K4 me3
H3K27 me3
H2A.Z
H3–5ac
H4K20 me3
DNA Meth. m5C
H3K9 me3
0.00 0.25 0.50 0.75 1.00
$\hat{\pi}$
TOP2B (1) – Forward
H2A.Z:H4–4ac
H2A.Z:H3K4 me3
H3K27 me3:H3K4 me1
H3K27 me3:H3K9 me3
H2A.Z:H3–5ac
H2A.Z:H4K20 me2
H3K9 me2:DNA Meth. m5C
DNA Meth. m5C:H4K20 me2
H3K4 me1:H3–5ac
H3K27 me3:DNA Meth. m5C
H4K20 me2:H4–4ac
H3K4 me3:H3–5ac
H4K20 me2:H3K4 me3
DNA Meth. m5C:H3K9 me3
H3K4 me3:H4–4ac
DNA Meth. m5C:H4K20 me3
H3K27 me2:DNA Meth. m5C
H4K20 me2:H3–5ac
H4–4ac:H3–5ac
H3K9 me3
H3K4 me1
H3K27 me3
H3K27 me2
H3K4 me3
H4K20 me2
H3K9 me2
H4K20 me3
H4–4ac
H2A.Z
H3–5ac
DNA Meth. m5C
0.00 0.25 0.50 0.75 1.00
$\hat{\pi}$
TOP2B (1) – Reverse
H2A.Z:H4–4ac
H2A.Z:H3K4 me3
H3K27 me3:H3K4 me1
H3K27 me3:H3K9 me3
H2A.Z:H3–5ac
H2A.Z:H4K20 me2
H3K9 me2:DNA Meth. m5C
DNA Meth. m5C:H4K20 me2
H3K4 me1:H3–5ac
H3K27 me3:DNA Meth. m5C
H4K20 me2:H4–4ac
H3K4 me3:H3–5ac
H4K20 me2:H3K4 me3
DNA Meth. m5C:H3K9 me3
H3K4 me3:H4–4ac
DNA Meth. m5C:H4K20 me3
H3K27 me2:DNA Meth. m5C
H4K20 me2:H3–5ac
H4–4ac:H3–5ac
H3K9 me3
H3K4 me1
H3K27 me3
H3K27 me2
H3K4 me3
H4K20 me2
H3K9 me2
H4K20 me3
H4–4ac
H2A.Z
H3–5ac
DNA Meth. m5C
0.00 0.25 0.50 0.75 1.00
$\hat{\pi}$
TOP2B (2) – Forward
H3K4 me1:H3–5ac
H3K27 me3:H3K9 me3
H2A.Z:H4–4ac
H3K9 me3:H4K20 me3
H3K27 me3:H3K4 me1
H4K20 me2:H4–4ac
H2A.Z:H3K4 me3
DNA Meth. m5C:H4K20 me2
H3K9 me2:DNA Meth. m5C
H2A.Z:H3–5ac
H2A.Z:H4K20 me2
H4K20 me2:H3K4 me3
H3K27 me2:DNA Meth. m5C
DNA Meth. m5C:H3K9 me3
H3K27 me3:DNA Meth. m5C
H3K4 me3:H3–5ac
H3K4 me3:H4–4ac
H4K20 me2:H3–5ac
DNA Meth. m5C:H4K20 me3
H4–4ac:H3–5ac
H3K4 me1
H3K27 me3
H3K9 me3
H3K27 me2
H3K4 me3
H3K9 me2
H4K20 me2
H4K20 me3
H4–4ac
H2A.Z
H3–5ac
DNA Meth. m5C
0.00 0.25 0.50 0.75 1.00
$\hat{\pi}$
TOP2B (2) – Reverse
H3K4 me1:H3–5ac
H3K27 me3:H3K9 me3
H2A.Z:H4–4ac
H3K9 me3:H4K20 me3
H3K27 me3:H3K4 me1
H4K20 me2:H4–4ac
H2A.Z:H3K4 me3
DNA Meth. m5C:H4K20 me2
H3K9 me2:DNA Meth. m5C
H2A.Z:H3–5ac
H2A.Z:H4K20 me2
H4K20 me2:H3K4 me3
H3K27 me2:DNA Meth. m5C
DNA Meth. m5C:H3K9 me3
H3K27 me3:DNA Meth. m5C
H3K4 me3:H3–5ac
H3K4 me3:H4–4ac
H4K20 me2:H3–5ac
DNA Meth. m5C:H4K20 me3
H4–4ac:H3–5ac
H3K4 me1
H3K27 me3
H3K9 me3
H3K27 me2
H3K4 me3
H3K9 me2
H4K20 me2
H4K20 me3
H4–4ac
H2A.Z
H3–5ac
DNA Meth. m5C
0.00 0.25 0.50 0.75 1.00
$\hat{\pi}$
TADA1 – Forward
DNA Meth. m5C:H4K20 me2
H3K27 me3:H3K4 me1
H3K27 me3:H3K9 me3
H2A.Z:H3–5ac
DNA Meth. m5C:H3K9 me3
H3K27 me3:DNA Meth. m5C
H3K4 me1:H4K20 me2
H3K9 me3:H4K20 me3
H2A.Z:H4–4ac
H3K4 me1:H4–4ac
H3K27 me3:H4K20 me3
H4K20 me2:H4–4ac
H3K4 me1:H3–5ac
H3K4 me1:H2A.Z
DNA Meth. m5C:H4K20 me3
H3K4 me3:H3–5ac
H3K4 me3:H4–4ac
H2A.Z:H3K4 me3
H4–4ac:H3–5ac
H4K20 me2:H3K4 me3
H4K20 me2:H3–5ac
H2A.Z:H4K20 me2
H3K27 me2
H3K9 me2
H4K20 me3
H3K9 me3
H4–4ac
H3–5ac
H3K4 me3
H3K4 me1
DNA Meth. m5C
H3K27 me3
H2A.Z
H4K20 me2
0.00 0.25 0.50 0.75 1.00
$\hat{\pi}$
TADA1 – Reverse
DNA Meth. m5C:H4K20 me2
H3K27 me3:H3K4 me1
H3K27 me3:H3K9 me3
H2A.Z:H3–5ac
DNA Meth. m5C:H3K9 me3
H3K27 me3:DNA Meth. m5C
H3K4 me1:H4K20 me2
H3K9 me3:H4K20 me3
H2A.Z:H4–4ac
H3K4 me1:H4–4ac
H3K27 me3:H4K20 me3
H4K20 me2:H4–4ac
H3K4 me1:H3–5ac
H3K4 me1:H2A.Z
DNA Meth. m5C:H4K20 me3
H3K4 me3:H3–5ac
H3K4 me3:H4–4ac
H2A.Z:H3K4 me3
H4–4ac:H3–5ac
H4K20 me2:H3K4 me3
H4K20 me2:H3–5ac
H2A.Z:H4K20 me2
H3K27 me2
H3K9 me2
H4K20 me3
H3K9 me3
H4–4ac
H3–5ac
H3K4 me3
H3K4 me1
DNA Meth. m5C
H3K27 me3
H2A.Z
H4K20 me2
0.00 0.25 0.50 0.75 1.00
$\hat{\pi}$
ZSCAN30 – Forward
H3K9 me2:DNA Meth. m5C
H3K27 me3:H3K4 me1
DNA Meth. m5C:H4K20 me2
DNA Meth. m5C:H3K9 me3
H3K27 me3:DNA Meth. m5C
H2A.Z:H3K4 me3
H2A.Z:H4K20 me2
H3K4 me1:H3–5ac
H3K9 me3:H4K20 me3
H3K27 me3:H4K20 me3
H3K27 me3:H3K9 me3
H3K4 me1:H2A.Z
H3K4 me1:H4–4ac
DNA Meth. m5C:H4K20 me3
H2A.Z:H3–5ac
H3K4 me3:H4–4ac
H4K20 me2:H4–4ac
H2A.Z:H4–4ac
H4K20 me2:H3–5ac
H3K4 me3:H3–5ac
H4K20 me2:H3K4 me3
H4–4ac:H3–5ac
H3K27 me2
H3K9 me2
DNA Meth. m5C
H3K27 me3
H4K20 me3
H3K9 me3
H4K20 me2
H3–5ac
H3K4 me3
H3K4 me1
H2A.Z
H4–4ac
0.00 0.25 0.50 0.75 1.00
$\hat{\pi}$
ZSCAN30 – Reverse
H3K9 me2:DNA Meth. m5C
H3K27 me3:H3K4 me1
DNA Meth. m5C:H4K20 me2
DNA Meth. m5C:H3K9 me3
H3K27 me3:DNA Meth. m5C
H2A.Z:H3K4 me3
H2A.Z:H4K20 me2
H3K4 me1:H3–5ac
H3K9 me3:H4K20 me3
H3K27 me3:H4K20 me3
H3K27 me3:H3K9 me3
H3K4 me1:H2A.Z
H3K4 me1:H4–4ac
DNA Meth. m5C:H4K20 me3
H2A.Z:H3–5ac
H3K4 me3:H4–4ac
H4K20 me2:H4–4ac
H2A.Z:H4–4ac
H4K20 me2:H3–5ac
H3K4 me3:H3–5ac
H4K20 me2:H3K4 me3
H4–4ac:H3–5ac
H3K27 me2
H3K9 me2
DNA Meth. m5C
H3K27 me3
H4K20 me3
H3K9 me3
H4K20 me2
H3–5ac
H3K4 me3
H3K4 me1
H2A.Z
H4–4ac
0.00 0.25 0.50 0.75 1.00
$\hat{\pi}$

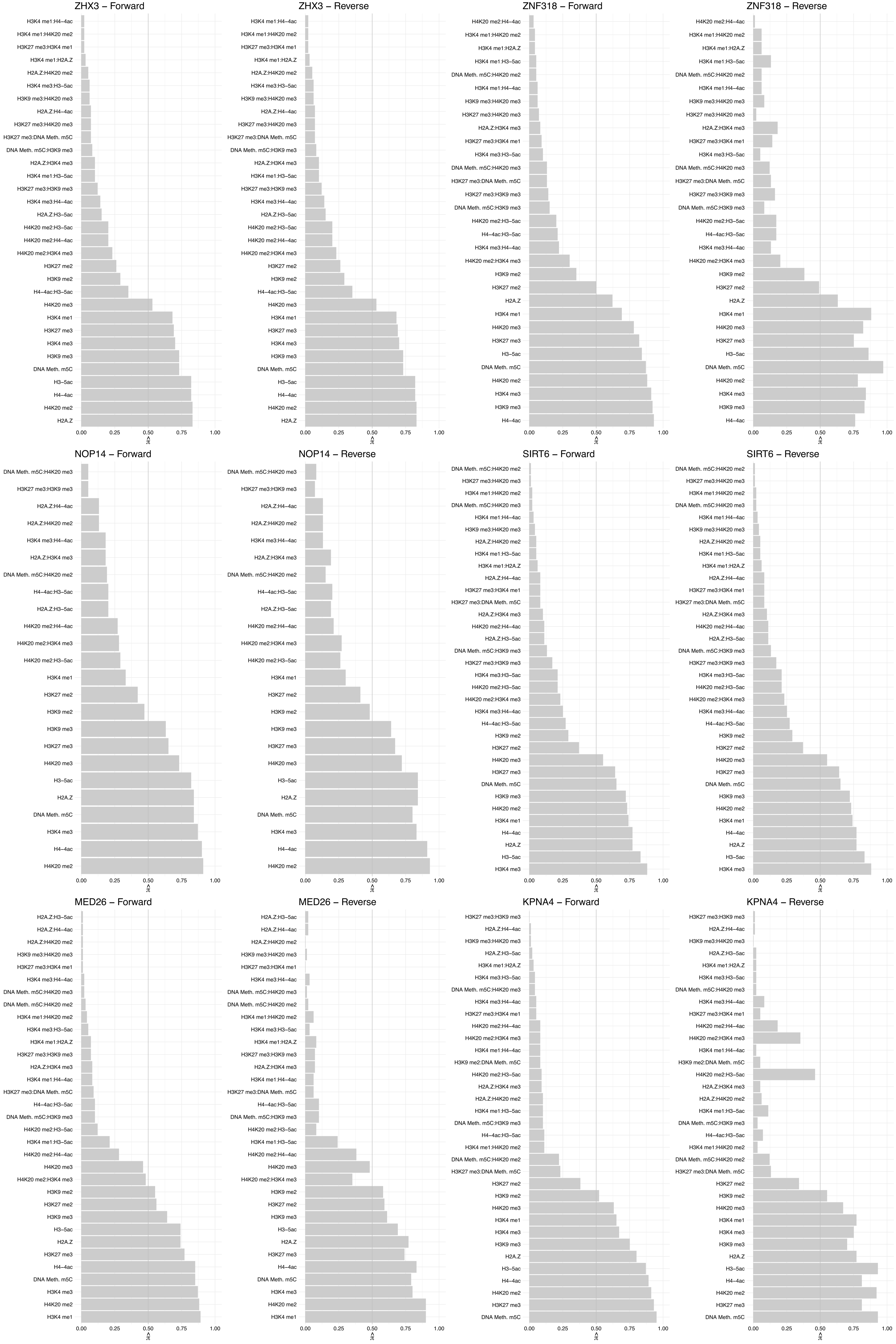
ZHX3 − Forward
H3K4 me1:H4−4ac
H3K4 me1:H4K20 me2
H3K27 me3:H3K4 me1
H3K4 me1:H2A.Z
H2A.Z:H4K20 me2
H3K4 me3:H3−5ac
H3K9 me3:H4K20 me3
H2A.Z:H4−4ac
H3K27 me3:H4K20 me3
H3K27 me3:DNA Meth. m5C
DNA Meth. m5C:H3K9 me3
H2A.Z:H3K4 me3
H3K4 me1:H3−5ac
H3K27 me3:H3K9 me3
H3K4 me3:H4−4ac
H2A.Z:H3−5ac
H4K20 me2:H3−5ac
H4K20 me2:H4−4ac
H4K20 me2:H3K4 me3
H3K27 me2
H3K9 me2
H4−4ac:H3−5ac
H4K20 me3
H3K4 me1
H3K27 me3
H3K4 me3
H3K9 me3
DNA Meth. m5C
H3−5ac
H4−4ac
H4K20 me2
H2A.Z
0.00 0.25 0.50 0.75 1.00
π̂
ZHX3 − Reverse
H3K4 me1:H4−4ac
H3K4 me1:H4K20 me2
H3K27 me3:H3K4 me1
H3K4 me1:H2A.Z
H2A.Z:H4K20 me2
H3K4 me3:H3−5ac
H3K9 me3:H4K20 me3
H2A.Z:H4−4ac
H3K27 me3:H4K20 me3
H3K27 me3:DNA Meth. m5C
DNA Meth. m5C:H3K9 me3
H2A.Z:H3K4 me3
H3K4 me1:H3−5ac
H3K27 me3:H3K9 me3
H3K4 me3:H4−4ac
H2A.Z:H3−5ac
H4K20 me2:H3−5ac
H4K20 me2:H4−4ac
H4K20 me2:H3K4 me3
H3K27 me2
H3K9 me2
H4−4ac:H3−5ac
H4K20 me3
H3K4 me1
H3K27 me3
H3K4 me3
H3K9 me3
DNA Meth. m5C
H3−5ac
H4−4ac
H4K20 me2
H2A.Z
0.00 0.25 0.50 0.75 1.00
π̂
ZNF318 − Forward
H4K20 me2:H4−4ac
H3K4 me1:H4K20 me2
H3K4 me1:H2A.Z
H3K4 me1:H3−5ac
DNA Meth. m5C:H4K20 me2
H3K4 me1:H4−4ac
H3K9 me3:H4K20 me3
H3K27 me3:H4K20 me3
H2A.Z:H3K4 me3
H3K27 me3:H3K4 me1
H3K4 me3:H3−5ac
DNA Meth. m5C:H4K20 me3
H3K27 me3:DNA Meth. m5C
H3K27 me3:H3K9 me3
DNA Meth. m5C:H3K9 me3
H4K20 me2:H3−5ac
H4−4ac:H3−5ac
H3K4 me3:H4−4ac
H4K20 me2:H3K4 me3
H3K9 me2
H3K27 me2
H2A.Z
H3K4 me1
H4K20 me3
H3K27 me3
H3−5ac
DNA Meth. m5C
H4K20 me2
H3K4 me3
H3K9 me3
H4−4ac
0.00 0.25 0.50 0.75 1.00
π̂
ZNF318 − Reverse
H4K20 me2:H4−4ac
H3K4 me1:H4K20 me2
H3K4 me1:H2A.Z
H3K4 me1:H3−5ac
DNA Meth. m5C:H4K20 me2
H3K4 me1:H4−4ac
H3K9 me3:H4K20 me3
H3K27 me3:H4K20 me3
H2A.Z:H3K4 me3
H3K27 me3:H3K4 me1
H3K4 me3:H3−5ac
DNA Meth. m5C:H4K20 me3
H3K27 me3:DNA Meth. m5C
H3K27 me3:H3K9 me3
DNA Meth. m5C:H3K9 me3
H4K20 me2:H3−5ac
H4−4ac:H3−5ac
H3K4 me3:H4−4ac
H4K20 me2:H3K4 me3
H3K9 me2
H3K27 me2
H2A.Z
H3K4 me1
H4K20 me3
H3K27 me3
H3−5ac
DNA Meth. m5C
H4K20 me2
H3K4 me3
H3K9 me3
H4−4ac
0.00 0.25 0.50 0.75 1.00
π̂
NOP14 − Forward
DNA Meth. m5C:H4K20 me3
H3K27 me3:H3K9 me3
H2A.Z:H4−4ac
H2A.Z:H4K20 me2
H3K4 me3:H4−4ac
H2A.Z:H3K4 me3
DNA Meth. m5C:H4K20 me2
H4−4ac:H3−5ac
H2A.Z:H3−5ac
H4K20 me2:H4−4ac
H4K20 me2:H3K4 me3
H4K20 me2:H3−5ac
H3K4 me1
H3K27 me2
H3K9 me2
H3K9 me3
H3K27 me3
H4K20 me3
H3−5ac
H2A.Z
DNA Meth. m5C
H3K4 me3
H4−4ac
H4K20 me2
0.00 0.25 0.50 0.75 1.00
π̂
NOP14 − Reverse
DNA Meth. m5C:H4K20 me3
H3K27 me3:H3K9 me3
H2A.Z:H4−4ac
H2A.Z:H4K20 me2
H3K4 me3:H4−4ac
H2A.Z:H3K4 me3
DNA Meth. m5C:H4K20 me2
H4−4ac:H3−5ac
H2A.Z:H3−5ac
H4K20 me2:H4−4ac
H4K20 me2:H3K4 me3
H4K20 me2:H3−5ac
H3K4 me1
H3K27 me2
H3K9 me2
H3K9 me3
H3K27 me3
H4K20 me3
H3−5ac
H2A.Z
DNA Meth. m5C
H3K4 me3
H4−4ac
H4K20 me2
0.00 0.25 0.50 0.75 1.00
π̂
SIRT6 − Forward
DNA Meth. m5C:H4K20 me2
H3K27 me3:H4K20 me3
H3K4 me1:H4K20 me2
DNA Meth. m5C:H4K20 me3
H3K4 me1:H4−4ac
H3K9 me3:H4K20 me3
H2A.Z:H4K20 me2
H3K4 me1:H3−5ac
H3K4 me1:H2A.Z
H2A.Z:H4−4ac
H3K27 me3:H3K4 me1
H3K27 me3:DNA Meth. m5C
H2A.Z:H3K4 me3
H4K20 me2:H4−4ac
H2A.Z:H3−5ac
DNA Meth. m5C:H3K9 me3
H3K27 me3:H3K9 me3
H3K4 me3:H3−5ac
H4K20 me2:H3−5ac
H4K20 me2:H3K4 me3
H3K4 me3:H4−4ac
H4−4ac:H3−5ac
H3K9 me2
H3K27 me2
H4K20 me3
H3K27 me3
DNA Meth. m5C
H3K9 me3
H4K20 me2
H3K4 me1
H4−4ac
H2A.Z
H3−5ac
H3K4 me3
0.00 0.25 0.50 0.75 1.00
π̂
SIRT6 − Reverse
DNA Meth. m5C:H4K20 me2
H3K27 me3:H4K20 me3
H3K4 me1:H4K20 me2
DNA Meth. m5C:H4K20 me3
H3K4 me1:H4−4ac
H3K9 me3:H4K20 me3
H2A.Z:H4K20 me2
H3K4 me1:H3−5ac
H3K4 me1:H2A.Z
H2A.Z:H4−4ac
H3K27 me3:H3K4 me1
H3K27 me3:DNA Meth. m5C
H2A.Z:H3K4 me3
H4K20 me2:H4−4ac
H2A.Z:H3−5ac
DNA Meth. m5C:H3K9 me3
H3K27 me3:H3K9 me3
H3K4 me3:H3−5ac
H4K20 me2:H3−5ac
H4K20 me2:H3K4 me3
H3K4 me3:H4−4ac
H4−4ac:H3−5ac
H3K9 me2
H3K27 me2
H4K20 me3
H3K27 me3
DNA Meth. m5C
H3K9 me3
H4K20 me2
H3K4 me1
H4−4ac
H2A.Z
H3−5ac
H3K4 me3
0.00 0.25 0.50 0.75 1.00
π̂
MED26 − Forward
H2A.Z:H3−5ac
H2A.Z:H4−4ac
H2A.Z:H4K20 me2
H3K9 me3:H4K20 me3
H3K27 me3:H3K4 me1
H3K4 me3:H4−4ac
DNA Meth. m5C:H4K20 me3
DNA Meth. m5C:H4K20 me2
H3K4 me1:H4K20 me2
H3K4 me3:H3−5ac
H3K4 me1:H2A.Z
H3K27 me3:H3K9 me3
H2A.Z:H3K4 me3
H3K4 me1:H4−4ac
H3K27 me3:DNA Meth. m5C
H4−4ac:H3−5ac
DNA Meth. m5C:H3K9 me3
H4K20 me2:H3−5ac
H3K4 me1:H3−5ac
H4K20 me2:H4−4ac
H4K20 me3
H4K20 me2:H3K4 me3
H3K9 me2
H3K27 me2
H3K9 me3
H3−5ac
H2A.Z
H3K27 me3
H4−4ac
DNA Meth. m5C
H3K4 me3
H4K20 me2
H3K4 me1
0.00 0.25 0.50 0.75 1.00
π̂
MED26 − Reverse
H2A.Z:H3−5ac
H2A.Z:H4−4ac
H2A.Z:H4K20 me2
H3K9 me3:H4K20 me3
H3K27 me3:H3K4 me1
H3K4 me3:H4−4ac
DNA Meth. m5C:H4K20 me3
DNA Meth. m5C:H4K20 me2
H3K4 me1:H4K20 me2
H3K4 me3:H3−5ac
H3K4 me1:H2A.Z
H3K27 me3:H3K9 me3
H2A.Z:H3K4 me3
H3K4 me1:H4−4ac
H3K27 me3:DNA Meth. m5C
H4−4ac:H3−5ac
DNA Meth. m5C:H3K9 me3
H4K20 me2:H3−5ac
H3K4 me1:H3−5ac
H4K20 me2:H4−4ac
H4K20 me3
H4K20 me2:H3K4 me3
H3K9 me2
H3K27 me2
H3K9 me3
H3−5ac
H2A.Z
H3K27 me3
H4−4ac
DNA Meth. m5C
H3K4 me3
H4K20 me2
H3K4 me1
0.00 0.25 0.50 0.75 1.00
π̂
KPNA4 − Forward
H3K27 me3:H3K9 me3
H2A.Z:H4−4ac
H3K9 me3:H4K20 me3
H2A.Z:H3−5ac
H3K4 me1:H2A.Z
H3K4 me3:H3−5ac
DNA Meth. m5C:H4K20 me3
H3K4 me3:H4−4ac
H3K27 me3:H3K4 me1
H4K20 me2:H4−4ac
H4K20 me2:H3K4 me3
H3K4 me1:H4−4ac
H3K9 me2:DNA Meth. m5C
H4K20 me2:H3−5ac
H2A.Z:H3K4 me3
H2A.Z:H4K20 me2
H3K4 me1:H3−5ac
DNA Meth. m5C:H3K9 me3
H4−4ac:H3−5ac
H3K4 me1:H4K20 me2
DNA Meth. m5C:H4K20 me2
H3K27 me3:DNA Meth. m5C
H3K27 me2
H3K9 me2
H4K20 me3
H3K4 me1
H3K4 me3
H3K9 me3
H2A.Z
H3−5ac
H4−4ac
H4K20 me2
H3K27 me3
DNA Meth. m5C
0.00 0.25 0.50 0.75 1.00
π̂
KPNA4 − Reverse
H3K27 me3:H3K9 me3
H2A.Z:H4−4ac
H3K9 me3:H4K20 me3
H2A.Z:H3−5ac
H3K4 me1:H2A.Z
H3K4 me3:H3−5ac
DNA Meth. m5C:H4K20 me3
H3K4 me3:H4−4ac
H3K27 me3:H3K4 me1
H4K20 me2:H4−4ac
H4K20 me2:H3K4 me3
H3K4 me1:H4−4ac
H3K9 me2:DNA Meth. m5C
H4K20 me2:H3−5ac
H2A.Z:H3K4 me3
H2A.Z:H4K20 me2
H3K4 me1:H3−5ac
DNA Meth. m5C:H3K9 me3
H4−4ac:H3−5ac
H3K4 me1:H4K20 me2
DNA Meth. m5C:H4K20 me2
H3K27 me3:DNA Meth. m5C
H3K27 me2
H3K9 me2
H4K20 me3
H3K4 me1
H3K4 me3
H3K9 me3
H2A.Z
H3−5ac
H4−4ac
H4K20 me2
H3K27 me3
DNA Meth. m5C
0.00 0.25 0.50 0.75 1.00
π̂

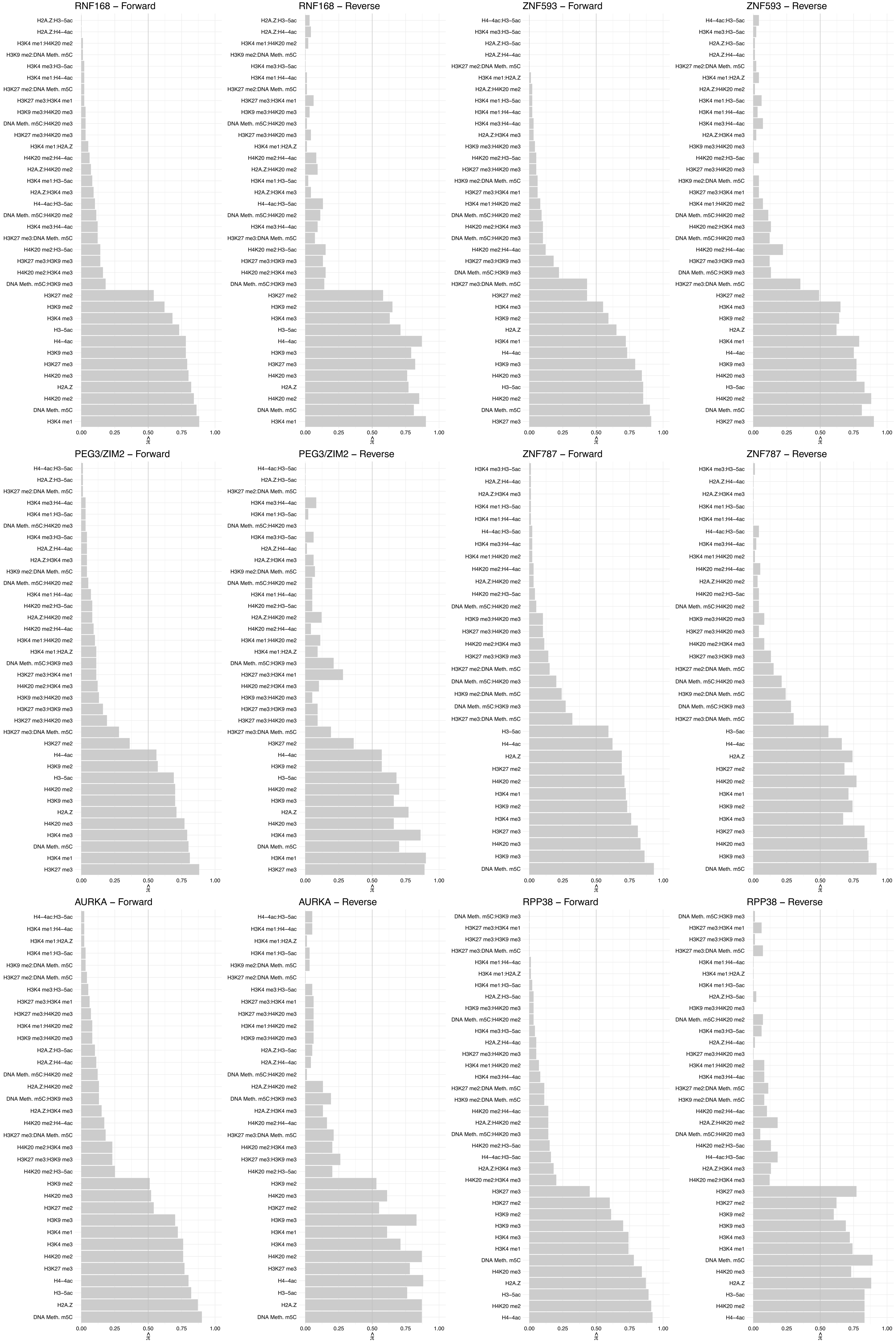
RNF168 – Forward
H2A.Z:H3-5ac
H2A.Z:H4-4ac
H3K4 me1:H4K20 me2
H3K9 me2:DNA Meth. m5C
H3K4 me3:H3-5ac
H3K4 me1:H4-4ac
H3K27 me2:DNA Meth. m5C
H3K27 me3:H3K4 me1
H3K9 me3:H4K20 me3
DNA Meth. m5C:H4K20 me3
H3K27 me3:H4K20 me3
H3K4 me1:H2A.Z
H4K20 me2:H4-4ac
H2A.Z:H4K20 me2
H3K4 me1:H3-5ac
H2A.Z:H3K4 me3
H4-4ac:H3-5ac
DNA Meth. m5C:H4K20 me2
H3K4 me3:H4-4ac
H3K27 me3:DNA Meth. m5C
H4K20 me2:H3-5ac
H3K27 me3:H3K9 me3
H4K20 me2:H3K4 me3
DNA Meth. m5C:H3K9 me3
H3K27 me2
H3K9 me2
H3K4 me3
H3-5ac
H4-4ac
H3K9 me3
H3K27 me3
H4K20 me3
H2A.Z
H4K20 me2
DNA Meth. m5C
H3K4 me1
0.00 0.25 0.50 0.75 1.00
π̂
RNF168 – Reverse
H2A.Z:H3-5ac
H2A.Z:H4-4ac
H3K4 me1:H4K20 me2
H3K9 me2:DNA Meth. m5C
H3K4 me3:H3-5ac
H3K4 me1:H4-4ac
H3K27 me2:DNA Meth. m5C
H3K27 me3:H3K4 me1
H3K9 me3:H4K20 me3
DNA Meth. m5C:H4K20 me3
H3K27 me3:H4K20 me3
H3K4 me1:H2A.Z
H4K20 me2:H4-4ac
H2A.Z:H4K20 me2
H3K4 me1:H3-5ac
H2A.Z:H3K4 me3
H4-4ac:H3-5ac
DNA Meth. m5C:H4K20 me2
H3K4 me3:H4-4ac
H3K27 me3:DNA Meth. m5C
H4K20 me2:H3-5ac
H3K27 me3:H3K9 me3
H4K20 me2:H3K4 me3
DNA Meth. m5C:H3K9 me3
H3K27 me2
H3K9 me2
H3K4 me3
H3-5ac
H4-4ac
H3K9 me3
H3K27 me3
H4K20 me3
H2A.Z
H4K20 me2
DNA Meth. m5C
H3K4 me1
0.00 0.25 0.50 0.75 1.00
π̂
ZNF593 – Forward
H4-4ac:H3-5ac
H3K4 me3:H3-5ac
H2A.Z:H3-5ac
H2A.Z:H4-4ac
H3K27 me2:DNA Meth. m5C
H3K4 me1:H2A.Z
H2A.Z:H4K20 me2
H3K4 me1:H3-5ac
H3K4 me1:H4-4ac
H3K4 me3:H4-4ac
H2A.Z:H3K4 me3
H3K9 me3:H4K20 me3
H4K20 me2:H3-5ac
H3K27 me3:H4K20 me3
H3K9 me2:DNA Meth. m5C
H3K27 me3:H3K4 me1
H3K4 me1:H4K20 me2
DNA Meth. m5C:H4K20 me2
H4K20 me2:H3K4 me3
DNA Meth. m5C:H4K20 me3
H4K20 me2:H4-4ac
H3K27 me3:H3K9 me3
DNA Meth. m5C:H3K9 me3
H3K27 me3:DNA Meth. m5C
H3K27 me2
H3K4 me3
H3K9 me2
H2A.Z
H3K4 me1
H4-4ac
H3K9 me3
H4K20 me3
H3-5ac
H4K20 me2
DNA Meth. m5C
H3K27 me3
0.00 0.25 0.50 0.75 1.00
π̂
ZNF593 – Reverse
H4-4ac:H3-5ac
H3K4 me3:H3-5ac
H2A.Z:H3-5ac
H2A.Z:H4-4ac
H3K27 me2:DNA Meth. m5C
H3K4 me1:H2A.Z
H2A.Z:H4K20 me2
H3K4 me1:H3-5ac
H3K4 me1:H4-4ac
H3K4 me3:H4-4ac
H2A.Z:H3K4 me3
H3K9 me3:H4K20 me3
H4K20 me2:H3-5ac
H3K27 me3:H4K20 me3
H3K9 me2:DNA Meth. m5C
H3K27 me3:H3K4 me1
H3K4 me1:H4K20 me2
DNA Meth. m5C:H4K20 me2
H4K20 me2:H3K4 me3
DNA Meth. m5C:H4K20 me3
H4K20 me2:H4-4ac
H3K27 me3:H3K9 me3
DNA Meth. m5C:H3K9 me3
H3K27 me3:DNA Meth. m5C
H3K27 me2
H3K4 me3
H3K9 me2
H2A.Z
H3K4 me1
H4-4ac
H3K9 me3
H4K20 me3
H3-5ac
H4K20 me2
DNA Meth. m5C
H3K27 me3
0.00 0.25 0.50 0.75 1.00
π̂
PEG3/ZIM2 – Forward
H4-4ac:H3-5ac
H2A.Z:H3-5ac
H3K27 me2:DNA Meth. m5C
H3K4 me3:H4-4ac
H3K4 me1:H3-5ac
DNA Meth. m5C:H4K20 me3
H3K4 me3:H3-5ac
H2A.Z:H4-4ac
H2A.Z:H3K4 me3
H3K9 me2:DNA Meth. m5C
DNA Meth. m5C:H4K20 me2
H3K4 me1:H4-4ac
H4K20 me2:H3-5ac
H2A.Z:H4K20 me2
H4K20 me2:H4-4ac
H3K4 me1:H4K20 me2
H3K4 me1:H2A.Z
DNA Meth. m5C:H3K9 me3
H3K27 me3:H3K4 me1
H4K20 me2:H3K4 me3
H3K9 me3:H4K20 me3
H3K27 me3:H3K9 me3
H3K27 me3:H4K20 me3
H3K27 me3:DNA Meth. m5C
H3K27 me2
H4-4ac
H3K9 me2
H3-5ac
H4K20 me2
H3K9 me3
H2A.Z
H4K20 me3
H3K4 me3
DNA Meth. m5C
H3K4 me1
H3K27 me3
0.00 0.25 0.50 0.75 1.00
π̂
PEG3/ZIM2 – Reverse
H4-4ac:H3-5ac
H2A.Z:H3-5ac
H3K27 me2:DNA Meth. m5C
H3K4 me3:H4-4ac
H3K4 me1:H3-5ac
DNA Meth. m5C:H4K20 me3
H3K4 me3:H3-5ac
H2A.Z:H4-4ac
H2A.Z:H3K4 me3
H3K9 me2:DNA Meth. m5C
DNA Meth. m5C:H4K20 me2
H3K4 me1:H4-4ac
H4K20 me2:H3-5ac
H2A.Z:H4K20 me2
H4K20 me2:H4-4ac
H3K4 me1:H4K20 me2
H3K4 me1:H2A.Z
DNA Meth. m5C:H3K9 me3
H3K27 me3:H3K4 me1
H4K20 me2:H3K4 me3
H3K9 me3:H4K20 me3
H3K27 me3:H3K9 me3
H3K27 me3:H4K20 me3
H3K27 me3:DNA Meth. m5C
H3K27 me2
H4-4ac
H3K9 me2
H3-5ac
H4K20 me2
H3K9 me3
H2A.Z
H4K20 me3
H3K4 me3
DNA Meth. m5C
H3K4 me1
H3K27 me3
0.00 0.25 0.50 0.75 1.00
π̂
ZNF787 – Forward
H3K4 me3:H3-5ac
H2A.Z:H4-4ac
H2A.Z:H3K4 me3
H3K4 me1:H3-5ac
H3K4 me1:H4-4ac
H4-4ac:H3-5ac
H3K4 me3:H4-4ac
H3K4 me1:H4K20 me2
H4K20 me2:H4-4ac
H2A.Z:H4K20 me2
H4K20 me2:H3-5ac
DNA Meth. m5C:H4K20 me2
H3K9 me3:H4K20 me3
H3K27 me3:H4K20 me3
H4K20 me2:H3K4 me3
H3K27 me3:H3K9 me3
H3K27 me2:DNA Meth. m5C
DNA Meth. m5C:H4K20 me3
H3K9 me2:DNA Meth. m5C
DNA Meth. m5C:H3K9 me3
H3K27 me3:DNA Meth. m5C
H3-5ac
H4-4ac
H2A.Z
H3K27 me2
H4K20 me2
H3K4 me1
H3K9 me2
H3K4 me3
H3K27 me3
H4K20 me3
H3K9 me3
DNA Meth. m5C
0.00 0.25 0.50 0.75 1.00
π̂
ZNF787 – Reverse
H3K4 me3:H3-5ac
H2A.Z:H4-4ac
H2A.Z:H3K4 me3
H3K4 me1:H3-5ac
H3K4 me1:H4-4ac
H4-4ac:H3-5ac
H3K4 me3:H4-4ac
H3K4 me1:H4K20 me2
H4K20 me2:H4-4ac
H2A.Z:H4K20 me2
H4K20 me2:H3-5ac
DNA Meth. m5C:H4K20 me2
H3K9 me3:H4K20 me3
H3K27 me3:H4K20 me3
H4K20 me2:H3K4 me3
H3K27 me3:H3K9 me3
H3K27 me2:DNA Meth. m5C
DNA Meth. m5C:H4K20 me3
H3K9 me2:DNA Meth. m5C
DNA Meth. m5C:H3K9 me3
H3K27 me3:DNA Meth. m5C
H3-5ac
H4-4ac
H2A.Z
H3K27 me2
H4K20 me2
H3K4 me1
H3K9 me2
H3K4 me3
H3K27 me3
H4K20 me3
H3K9 me3
DNA Meth. m5C
0.00 0.25 0.50 0.75 1.00
π̂
AURKA – Forward
H4-4ac:H3-5ac
H3K4 me1:H4-4ac
H3K4 me1:H2A.Z
H3K4 me1:H3-5ac
H3K9 me2:DNA Meth. m5C
H3K27 me2:DNA Meth. m5C
H3K4 me3:H3-5ac
H3K27 me3:H3K4 me1
H3K27 me3:H4K20 me3
H3K4 me1:H4K20 me2
H3K9 me3:H4K20 me3
H2A.Z:H3-5ac
H2A.Z:H4-4ac
DNA Meth. m5C:H4K20 me2
H2A.Z:H4K20 me2
DNA Meth. m5C:H3K9 me3
H2A.Z:H3K4 me3
H4K20 me2:H4-4ac
H3K27 me3:DNA Meth. m5C
H4K20 me2:H3K4 me3
H3K27 me3:H3K9 me3
H4K20 me2:H3-5ac
H3K9 me2
H4K20 me3
H3K27 me2
H3K9 me3
H3K4 me1
H3K4 me3
H4K20 me2
H3K27 me3
H4-4ac
H3-5ac
H2A.Z
DNA Meth. m5C
0.00 0.25 0.50 0.75 1.00
π̂
AURKA – Reverse
H4-4ac:H3-5ac
H3K4 me1:H4-4ac
H3K4 me1:H2A.Z
H3K4 me1:H3-5ac
H3K9 me2:DNA Meth. m5C
H3K27 me2:DNA Meth. m5C
H3K4 me3:H3-5ac
H3K27 me3:H3K4 me1
H3K27 me3:H4K20 me3
H3K4 me1:H4K20 me2
H3K9 me3:H4K20 me3
H2A.Z:H3-5ac
H2A.Z:H4-4ac
DNA Meth. m5C:H4K20 me2
H2A.Z:H4K20 me2
DNA Meth. m5C:H3K9 me3
H2A.Z:H3K4 me3
H4K20 me2:H4-4ac
H3K27 me3:DNA Meth. m5C
H4K20 me2:H3K4 me3
H3K27 me3:H3K9 me3
H4K20 me2:H3-5ac
H3K9 me2
H4K20 me3
H3K27 me2
H3K9 me3
H3K4 me1
H3K4 me3
H4K20 me2
H3K27 me3
H4-4ac
H3-5ac
H2A.Z
DNA Meth. m5C
0.00 0.25 0.50 0.75 1.00
π̂
RPP38 – Forward
DNA Meth. m5C:H3K9 me3
H3K27 me3:H3K4 me1
H3K27 me3:H3K9 me3
H3K27 me3:DNA Meth. m5C
H3K4 me1:H4-4ac
H3K4 me1:H2A.Z
H3K4 me1:H3-5ac
H2A.Z:H3-5ac
H3K9 me3:H4K20 me3
DNA Meth. m5C:H4K20 me2
H3K4 me3:H3-5ac
H2A.Z:H4-4ac
H3K27 me3:H4K20 me3
H3K4 me1:H4K20 me2
H3K4 me3:H4-4ac
H3K27 me2:DNA Meth. m5C
H3K9 me2:DNA Meth. m5C
H4K20 me2:H4-4ac
H2A.Z:H4K20 me2
DNA Meth. m5C:H4K20 me3
H4K20 me2:H3-5ac
H4-4ac:H3-5ac
H2A.Z:H3K4 me3
H4K20 me2:H3K4 me3
H3K27 me3
H3K27 me2
H3K9 me2
H3K9 me3
H3K4 me3
H3K4 me1
DNA Meth. m5C
H4K20 me3
H2A.Z
H3-5ac
H4K20 me2
H4-4ac
0.00 0.25 0.50 0.75 1.00
π̂
RPP38 – Reverse
DNA Meth. m5C:H3K9 me3
H3K27 me3:H3K4 me1
H3K27 me3:H3K9 me3
H3K27 me3:DNA Meth. m5C
H3K4 me1:H4-4ac
H3K4 me1:H2A.Z
H3K4 me1:H3-5ac
H2A.Z:H3-5ac
H3K9 me3:H4K20 me3
DNA Meth. m5C:H4K20 me2
H3K4 me3:H3-5ac
H2A.Z:H4-4ac
H3K27 me3:H4K20 me3
H3K4 me1:H4K20 me2
H3K4 me3:H4-4ac
H3K27 me2:DNA Meth. m5C
H3K9 me2:DNA Meth. m5C
H4K20 me2:H4-4ac
H2A.Z:H4K20 me2
DNA Meth. m5C:H4K20 me3
H4K20 me2:H3-5ac
H4-4ac:H3-5ac
H2A.Z:H3K4 me3
H4K20 me2:H3K4 me3
H3K27 me3
H3K27 me2
H3K9 me2
H3K9 me3
H3K4 me3
H3K4 me1
DNA Meth. m5C
H4K20 me3
H2A.Z
H3-5ac
H4K20 me2
H4-4ac
0.00 0.25 0.50 0.75 1.00
π̂

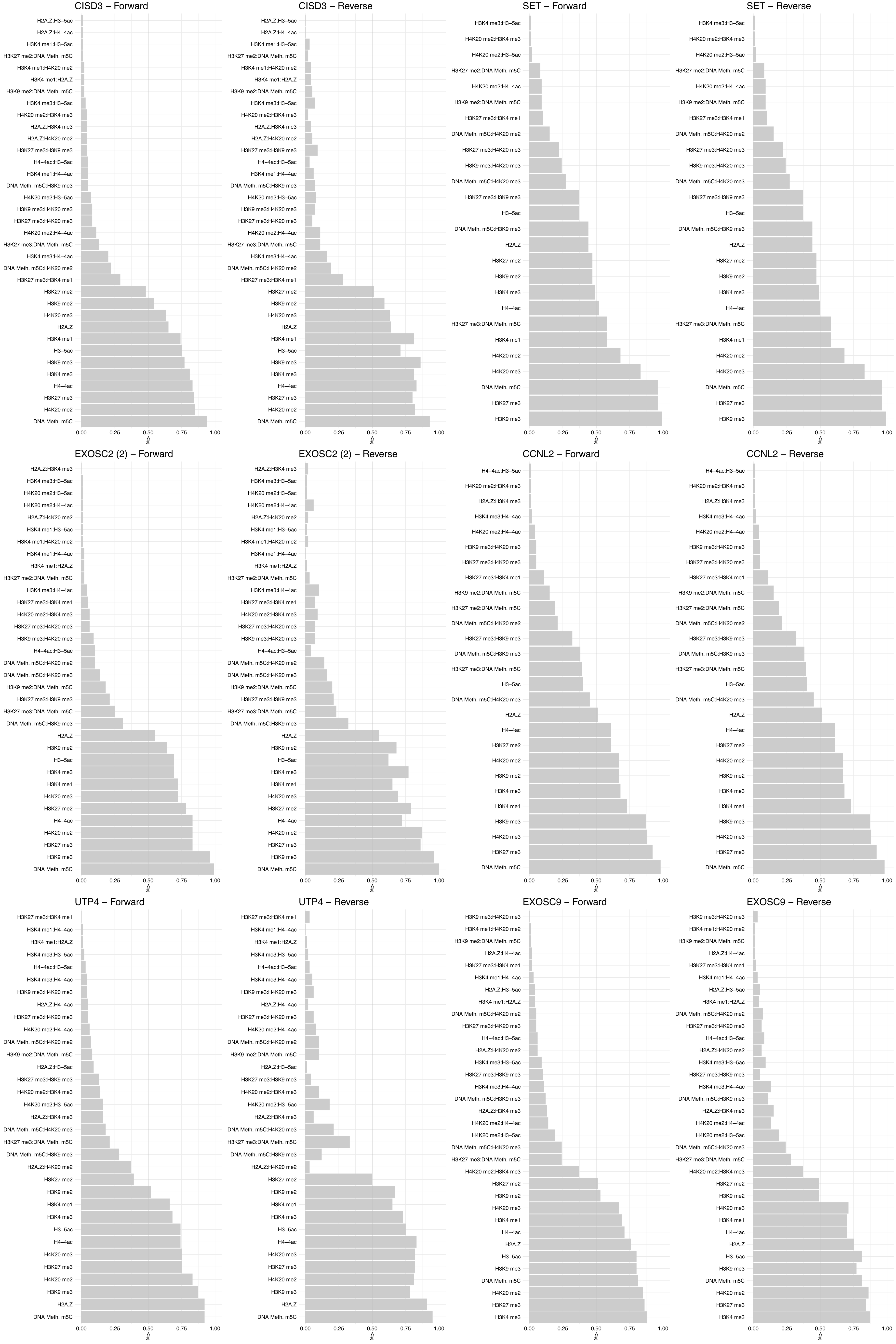
CISD3 – Forward
H2A.Z:H3−5ac
H2A.Z:H4−4ac
H3K4 me1:H3−5ac
H3K27 me2:DNA Meth. m5C
H3K4 me1:H4K20 me2
H3K4 me1:H2A.Z
H3K9 me2:DNA Meth. m5C
H3K4 me3:H3−5ac
H4K20 me2:H3K4 me3
H2A.Z:H3K4 me3
H2A.Z:H4K20 me2
H3K27 me3:H3K9 me3
H4−4ac:H3−5ac
H3K4 me1:H4−4ac
DNA Meth. m5C:H3K9 me3
H4K20 me2:H3−5ac
H3K9 me3:H4K20 me3
H3K27 me3:H4K20 me3
H4K20 me2:H4−4ac
H3K27 me3:DNA Meth. m5C
H3K4 me3:H4−4ac
DNA Meth. m5C:H4K20 me2
H3K27 me3:H3K4 me1
H3K27 me2
H3K9 me2
H4K20 me3
H2A.Z
H3K4 me1
H3−5ac
H3K9 me3
H3K4 me3
H4−4ac
H3K27 me3
H4K20 me2
DNA Meth. m5C
0.00 0.25 0.50 0.75 1.00
π̂
CISD3 – Reverse
H2A.Z:H3−5ac
H2A.Z:H4−4ac
H3K4 me1:H3−5ac
H3K27 me2:DNA Meth. m5C
H3K4 me1:H4K20 me2
H3K4 me1:H2A.Z
H3K9 me2:DNA Meth. m5C
H3K4 me3:H3−5ac
H4K20 me2:H3K4 me3
H2A.Z:H3K4 me3
H2A.Z:H4K20 me2
H3K27 me3:H3K9 me3
H4−4ac:H3−5ac
H3K4 me1:H4−4ac
DNA Meth. m5C:H3K9 me3
H4K20 me2:H3−5ac
H3K9 me3:H4K20 me3
H3K27 me3:H4K20 me3
H4K20 me2:H4−4ac
H3K27 me3:DNA Meth. m5C
H3K4 me3:H4−4ac
DNA Meth. m5C:H4K20 me2
H3K27 me3:H3K4 me1
H3K27 me2
H3K9 me2
H4K20 me3
H2A.Z
H3K4 me1
H3−5ac
H3K9 me3
H3K4 me3
H4−4ac
H3K27 me3
H4K20 me2
DNA Meth. m5C
0.00 0.25 0.50 0.75 1.00
π̂
SET – Forward
H3K4 me3:H3−5ac
H4K20 me2:H3K4 me3
H4K20 me2:H3−5ac
H3K27 me2:DNA Meth. m5C
H4K20 me2:H4−4ac
H3K9 me2:DNA Meth. m5C
H3K27 me3:H3K4 me1
DNA Meth. m5C:H4K20 me2
H3K27 me3:H4K20 me3
H3K9 me3:H4K20 me3
DNA Meth. m5C:H4K20 me3
H3K27 me3:H3K9 me3
H3−5ac
DNA Meth. m5C:H3K9 me3
H2A.Z
H3K27 me2
H3K9 me2
H3K4 me3
H4−4ac
H3K27 me3:DNA Meth. m5C
H3K4 me1
H4K20 me2
H4K20 me3
DNA Meth. m5C
H3K27 me3
H3K9 me3
0.00 0.25 0.50 0.75 1.00
π̂
SET – Reverse
H3K4 me3:H3−5ac
H4K20 me2:H3K4 me3
H4K20 me2:H3−5ac
H3K27 me2:DNA Meth. m5C
H4K20 me2:H4−4ac
H3K9 me2:DNA Meth. m5C
H3K27 me3:H3K4 me1
DNA Meth. m5C:H4K20 me2
H3K27 me3:H4K20 me3
H3K9 me3:H4K20 me3
DNA Meth. m5C:H4K20 me3
H3K27 me3:H3K9 me3
H3−5ac
DNA Meth. m5C:H3K9 me3
H2A.Z
H3K27 me2
H3K9 me2
H3K4 me3
H4−4ac
H3K27 me3:DNA Meth. m5C
H3K4 me1
H4K20 me2
H4K20 me3
DNA Meth. m5C
H3K27 me3
H3K9 me3
0.00 0.25 0.50 0.75 1.00
π̂
EXOSC2 (2) – Forward
H2A.Z:H3K4 me3
H3K4 me3:H3−5ac
H4K20 me2:H3−5ac
H4K20 me2:H4−4ac
H2A.Z:H4K20 me2
H3K4 me1:H3−5ac
H3K4 me1:H4K20 me2
H3K4 me1:H4−4ac
H3K4 me1:H2A.Z
H3K27 me2:DNA Meth. m5C
H3K4 me3:H4−4ac
H3K27 me3:H3K4 me1
H4K20 me2:H3K4 me3
H3K27 me3:H4K20 me3
H3K9 me3:H4K20 me3
H4−4ac:H3−5ac
DNA Meth. m5C:H4K20 me2
DNA Meth. m5C:H4K20 me3
H3K9 me2:DNA Meth. m5C
H3K27 me3:H3K9 me3
H3K27 me3:DNA Meth. m5C
DNA Meth. m5C:H3K9 me3
H2A.Z
H3K9 me2
H3−5ac
H3K4 me3
H3K4 me1
H4K20 me3
H3K27 me2
H4−4ac
H4K20 me2
H3K27 me3
H3K9 me3
DNA Meth. m5C
0.00 0.25 0.50 0.75 1.00
π̂
EXOSC2 (2) – Reverse
H2A.Z:H3K4 me3
H3K4 me3:H3−5ac
H4K20 me2:H3−5ac
H4K20 me2:H4−4ac
H2A.Z:H4K20 me2
H3K4 me1:H3−5ac
H3K4 me1:H4K20 me2
H3K4 me1:H4−4ac
H3K4 me1:H2A.Z
H3K27 me2:DNA Meth. m5C
H3K4 me3:H4−4ac
H3K27 me3:H3K4 me1
H4K20 me2:H3K4 me3
H3K27 me3:H4K20 me3
H3K9 me3:H4K20 me3
H4−4ac:H3−5ac
DNA Meth. m5C:H4K20 me2
DNA Meth. m5C:H4K20 me3
H3K9 me2:DNA Meth. m5C
H3K27 me3:H3K9 me3
H3K27 me3:DNA Meth. m5C
DNA Meth. m5C:H3K9 me3
H2A.Z
H3K9 me2
H3−5ac
H3K4 me3
H3K4 me1
H4K20 me3
H3K27 me2
H4−4ac
H4K20 me2
H3K27 me3
H3K9 me3
DNA Meth. m5C
0.00 0.25 0.50 0.75 1.00
π̂
CCNL2 – Forward
H4−4ac:H3−5ac
H4K20 me2:H3K4 me3
H2A.Z:H3K4 me3
H3K4 me3:H4−4ac
H4K20 me2:H4−4ac
H3K9 me3:H4K20 me3
H3K27 me3:H4K20 me3
H3K27 me3:H3K4 me1
H3K9 me2:DNA Meth. m5C
H3K27 me2:DNA Meth. m5C
DNA Meth. m5C:H4K20 me2
H3K27 me3:H3K9 me3
DNA Meth. m5C:H3K9 me3
H3K27 me3:DNA Meth. m5C
H3−5ac
DNA Meth. m5C:H4K20 me3
H2A.Z
H4−4ac
H3K27 me2
H4K20 me2
H3K9 me2
H3K4 me3
H3K4 me1
H3K9 me3
H4K20 me3
H3K27 me3
DNA Meth. m5C
0.00 0.25 0.50 0.75 1.00
π̂
CCNL2 – Reverse
H4−4ac:H3−5ac
H4K20 me2:H3K4 me3
H2A.Z:H3K4 me3
H3K4 me3:H4−4ac
H4K20 me2:H4−4ac
H3K9 me3:H4K20 me3
H3K27 me3:H4K20 me3
H3K27 me3:H3K4 me1
H3K9 me2:DNA Meth. m5C
H3K27 me2:DNA Meth. m5C
DNA Meth. m5C:H4K20 me2
H3K27 me3:H3K9 me3
DNA Meth. m5C:H3K9 me3
H3K27 me3:DNA Meth. m5C
H3−5ac
DNA Meth. m5C:H4K20 me3
H2A.Z
H4−4ac
H3K27 me2
H4K20 me2
H3K9 me2
H3K4 me3
H3K4 me1
H3K9 me3
H4K20 me3
H3K27 me3
DNA Meth. m5C
0.00 0.25 0.50 0.75 1.00
π̂
UTP4 – Forward
H3K27 me3:H3K4 me1
H3K4 me1:H4−4ac
H3K4 me1:H2A.Z
H3K4 me3:H3−5ac
H4−4ac:H3−5ac
H3K4 me3:H4−4ac
H3K9 me3:H4K20 me3
H2A.Z:H4−4ac
H3K27 me3:H4K20 me3
H4K20 me2:H4−4ac
DNA Meth. m5C:H4K20 me2
H3K9 me2:DNA Meth. m5C
H2A.Z:H3−5ac
H3K27 me3:H3K9 me3
H4K20 me2:H3K4 me3
H4K20 me2:H3−5ac
H2A.Z:H3K4 me3
DNA Meth. m5C:H4K20 me3
H3K27 me3:DNA Meth. m5C
DNA Meth. m5C:H3K9 me3
H2A.Z:H4K20 me2
H3K27 me2
H3K9 me2
H3K4 me1
H3K4 me3
H3−5ac
H4−4ac
H4K20 me3
H3K27 me3
H4K20 me2
H3K9 me3
H2A.Z
DNA Meth. m5C
0.00 0.25 0.50 0.75 1.00
π̂
UTP4 – Reverse
H3K27 me3:H3K4 me1
H3K4 me1:H4−4ac
H3K4 me1:H2A.Z
H3K4 me3:H3−5ac
H4−4ac:H3−5ac
H3K4 me3:H4−4ac
H3K9 me3:H4K20 me3
H2A.Z:H4−4ac
H3K27 me3:H4K20 me3
H4K20 me2:H4−4ac
DNA Meth. m5C:H4K20 me2
H3K9 me2:DNA Meth. m5C
H2A.Z:H3−5ac
H3K27 me3:H3K9 me3
H4K20 me2:H3K4 me3
H4K20 me2:H3−5ac
H2A.Z:H3K4 me3
DNA Meth. m5C:H4K20 me3
H3K27 me3:DNA Meth. m5C
DNA Meth. m5C:H3K9 me3
H2A.Z:H4K20 me2
H3K27 me2
H3K9 me2
H3K4 me1
H3K4 me3
H3−5ac
H4−4ac
H4K20 me3
H3K27 me3
H4K20 me2
H3K9 me3
H2A.Z
DNA Meth. m5C
0.00 0.25 0.50 0.75 1.00
π̂
EXOSC9 – Forward
H3K9 me3:H4K20 me3
H3K4 me1:H4K20 me2
H3K9 me2:DNA Meth. m5C
H2A.Z:H4−4ac
H3K27 me3:H3K4 me1
H3K4 me1:H4−4ac
H2A.Z:H3−5ac
H3K4 me1:H2A.Z
DNA Meth. m5C:H4K20 me2
H3K27 me3:H4K20 me3
H4−4ac:H3−5ac
H2A.Z:H4K20 me2
H3K4 me3:H3−5ac
H3K27 me3:H3K9 me3
H3K4 me3:H4−4ac
DNA Meth. m5C:H3K9 me3
H2A.Z:H3K4 me3
H4K20 me2:H4−4ac
H4K20 me2:H3−5ac
DNA Meth. m5C:H4K20 me3
H3K27 me3:DNA Meth. m5C
H4K20 me2:H3K4 me3
H3K27 me2
H3K9 me2
H4K20 me3
H3K4 me1
H4−4ac
H2A.Z
H3−5ac
H3K9 me3
DNA Meth. m5C
H4K20 me2
H3K27 me3
H3K4 me3
0.00 0.25 0.50 0.75 1.00
π̂
EXOSC9 – Reverse
H3K9 me3:H4K20 me3
H3K4 me1:H4K20 me2
H3K9 me2:DNA Meth. m5C
H2A.Z:H4−4ac
H3K27 me3:H3K4 me1
H3K4 me1:H4−4ac
H2A.Z:H3−5ac
H3K4 me1:H2A.Z
DNA Meth. m5C:H4K20 me2
H3K27 me3:H4K20 me3
H4−4ac:H3−5ac
H2A.Z:H4K20 me2
H3K4 me3:H3−5ac
H3K27 me3:H3K9 me3
H3K4 me3:H4−4ac
DNA Meth. m5C:H3K9 me3
H2A.Z:H3K4 me3
H4K20 me2:H4−4ac
H4K20 me2:H3−5ac
DNA Meth. m5C:H4K20 me3
H3K27 me3:DNA Meth. m5C
H4K20 me2:H3K4 me3
H3K27 me2
H3K9 me2
H4K20 me3
H3K4 me1
H4−4ac
H2A.Z
H3−5ac
H3K9 me3
DNA Meth. m5C
H4K20 me2
H3K27 me3
H3K4 me3
0.00 0.25 0.50 0.75 1.00
π̂

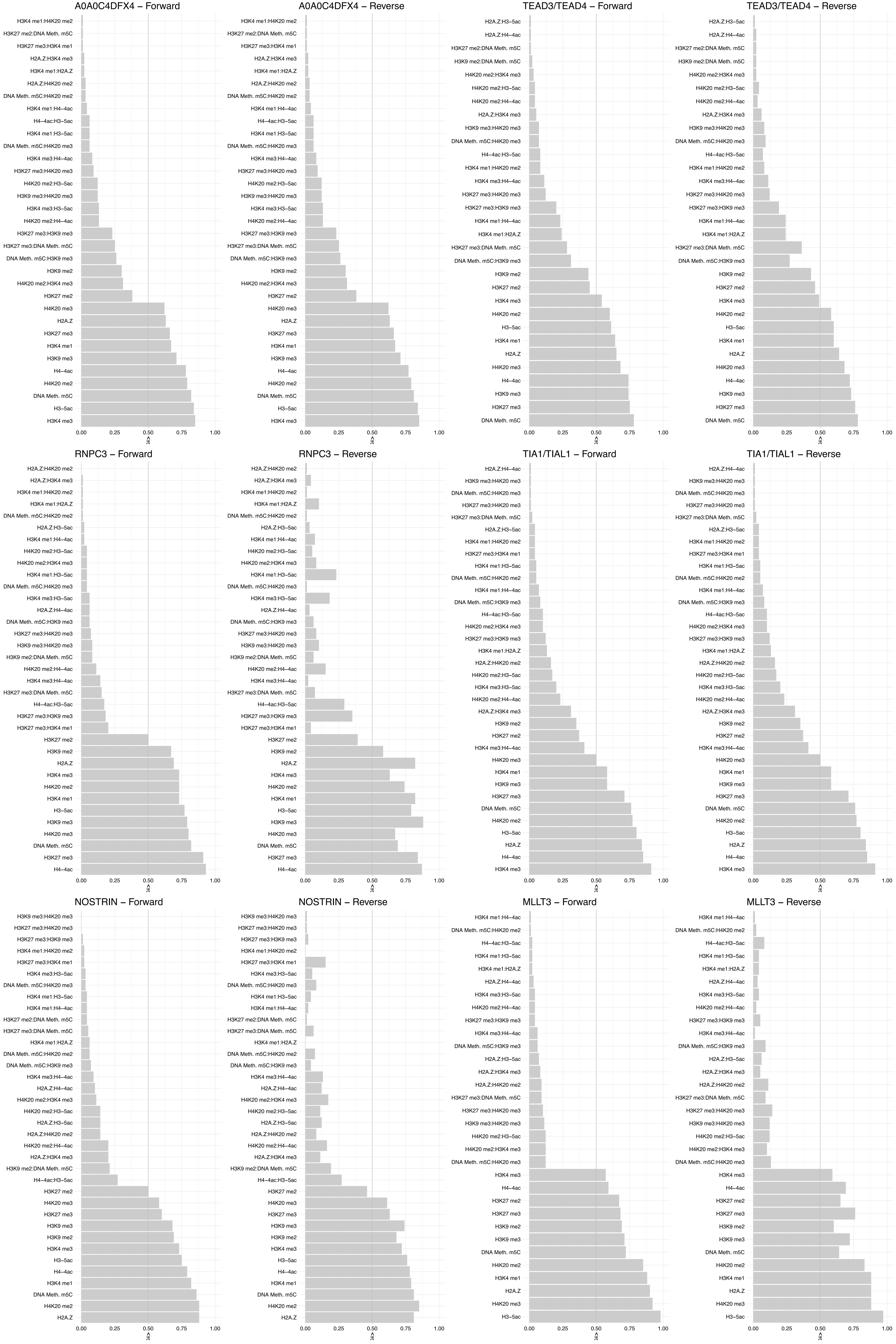
A0A0C4DFX4 – Forward
H3K4 me1:H4K20 me2
H3K27 me2:DNA Meth. m5C
H3K27 me3:H3K4 me1
H2A.Z:H3K4 me3
H3K4 me1:H2A.Z
H2A.Z:H4K20 me2
DNA Meth. m5C:H4K20 me2
H3K4 me1:H4–4ac
H4–4ac:H3–5ac
H3K4 me1:H3–5ac
DNA Meth. m5C:H4K20 me3
H3K4 me3:H4–4ac
H3K27 me3:H4K20 me3
H4K20 me2:H3–5ac
H3K9 me3:H4K20 me3
H3K4 me3:H3–5ac
H4K20 me2:H4–4ac
H3K27 me3:H3K9 me3
H3K27 me3:DNA Meth. m5C
DNA Meth. m5C:H3K9 me3
H3K9 me2
H4K20 me2:H3K4 me3
H3K27 me2
H4K20 me3
H2A.Z
H3K27 me3
H3K4 me1
H3K9 me3
H4–4ac
H4K20 me2
DNA Meth. m5C
H3–5ac
H3K4 me3
0.00 0.25 0.50 0.75 1.00
$\hat{\pi}$
A0A0C4DFX4 – Reverse
H3K4 me1:H4K20 me2
H3K27 me2:DNA Meth. m5C
H3K27 me3:H3K4 me1
H2A.Z:H3K4 me3
H3K4 me1:H2A.Z
H2A.Z:H4K20 me2
DNA Meth. m5C:H4K20 me2
H3K4 me1:H4–4ac
H4–4ac:H3–5ac
H3K4 me1:H3–5ac
DNA Meth. m5C:H4K20 me3
H3K4 me3:H4–4ac
H3K27 me3:H4K20 me3
H4K20 me2:H3–5ac
H3K9 me3:H4K20 me3
H3K4 me3:H3–5ac
H4K20 me2:H4–4ac
H3K27 me3:H3K9 me3
H3K27 me3:DNA Meth. m5C
DNA Meth. m5C:H3K9 me3
H3K9 me2
H4K20 me2:H3K4 me3
H3K27 me2
H4K20 me3
H2A.Z
H3K27 me3
H3K4 me1
H3K9 me3
H4–4ac
H4K20 me2
DNA Meth. m5C
H3–5ac
H3K4 me3
0.00 0.25 0.50 0.75 1.00
$\hat{\pi}$
TEAD3/TEAD4 – Forward
H2A.Z:H3–5ac
H2A.Z:H4–4ac
H3K27 me2:DNA Meth. m5C
H3K9 me2:DNA Meth. m5C
H4K20 me2:H3K4 me3
H4K20 me2:H3–5ac
H4K20 me2:H4–4ac
H2A.Z:H3K4 me3
H3K9 me3:H4K20 me3
DNA Meth. m5C:H4K20 me3
H4–4ac:H3–5ac
H3K4 me1:H4K20 me2
H3K4 me3:H4–4ac
H3K27 me3:H4K20 me3
H3K27 me3:H3K9 me3
H3K4 me1:H4–4ac
H3K4 me1:H2A.Z
H3K27 me3:DNA Meth. m5C
DNA Meth. m5C:H3K9 me3
H3K9 me2
H3K27 me2
H3K4 me3
H4K20 me2
H3–5ac
H3K4 me1
H2A.Z
H4K20 me3
H4–4ac
H3K9 me3
H3K27 me3
DNA Meth. m5C
0.00 0.25 0.50 0.75 1.00
$\hat{\pi}$
TEAD3/TEAD4 – Reverse
H2A.Z:H3–5ac
H2A.Z:H4–4ac
H3K27 me2:DNA Meth. m5C
H3K9 me2:DNA Meth. m5C
H4K20 me2:H3K4 me3
H4K20 me2:H3–5ac
H4K20 me2:H4–4ac
H2A.Z:H3K4 me3
H3K9 me3:H4K20 me3
DNA Meth. m5C:H4K20 me3
H4–4ac:H3–5ac
H3K4 me1:H4K20 me2
H3K4 me3:H4–4ac
H3K27 me3:H4K20 me3
H3K27 me3:H3K9 me3
H3K4 me1:H4–4ac
H3K4 me1:H2A.Z
H3K27 me3:DNA Meth. m5C
DNA Meth. m5C:H3K9 me3
H3K9 me2
H3K27 me2
H3K4 me3
H4K20 me2
H3–5ac
H3K4 me1
H2A.Z
H4K20 me3
H4–4ac
H3K9 me3
H3K27 me3
DNA Meth. m5C
0.00 0.25 0.50 0.75 1.00
$\hat{\pi}$
RNPC3 – Forward
H2A.Z:H4K20 me2
H2A.Z:H3K4 me3
H3K4 me1:H4K20 me2
H3K4 me1:H2A.Z
DNA Meth. m5C:H4K20 me2
H2A.Z:H3–5ac
H3K4 me1:H4–4ac
H4K20 me2:H3–5ac
H4K20 me2:H3K4 me3
H3K4 me1:H3–5ac
DNA Meth. m5C:H4K20 me3
H3K4 me3:H3–5ac
H2A.Z:H4–4ac
DNA Meth. m5C:H3K9 me3
H3K27 me3:H4K20 me3
H3K9 me3:H4K20 me3
H3K9 me2:DNA Meth. m5C
H4K20 me2:H4–4ac
H3K4 me3:H4–4ac
H3K27 me3:DNA Meth. m5C
H4–4ac:H3–5ac
H3K27 me3:H3K9 me3
H3K27 me3:H3K4 me1
H3K27 me2
H3K9 me2
H2A.Z
H3K4 me3
H4K20 me2
H3K4 me1
H3–5ac
H3K9 me3
H4K20 me3
DNA Meth. m5C
H3K27 me3
H4–4ac
0.00 0.25 0.50 0.75 1.00
$\hat{\pi}$
RNPC3 – Reverse
H2A.Z:H4K20 me2
H2A.Z:H3K4 me3
H3K4 me1:H4K20 me2
H3K4 me1:H2A.Z
DNA Meth. m5C:H4K20 me2
H2A.Z:H3–5ac
H3K4 me1:H4–4ac
H4K20 me2:H3–5ac
H4K20 me2:H3K4 me3
H3K4 me1:H3–5ac
DNA Meth. m5C:H4K20 me3
H3K4 me3:H3–5ac
H2A.Z:H4–4ac
DNA Meth. m5C:H3K9 me3
H3K27 me3:H4K20 me3
H3K9 me3:H4K20 me3
H3K9 me2:DNA Meth. m5C
H4K20 me2:H4–4ac
H3K4 me3:H4–4ac
H3K27 me3:DNA Meth. m5C
H4–4ac:H3–5ac
H3K27 me3:H3K9 me3
H3K27 me3:H3K4 me1
H3K27 me2
H3K9 me2
H2A.Z
H3K4 me3
H4K20 me2
H3K4 me1
H3–5ac
H3K9 me3
H4K20 me3
DNA Meth. m5C
H3K27 me3
H4–4ac
0.00 0.25 0.50 0.75 1.00
$\hat{\pi}$
TIA1/TIAL1 – Forward
H2A.Z:H4–4ac
H3K9 me3:H4K20 me3
DNA Meth. m5C:H4K20 me3
H3K27 me3:H4K20 me3
H3K27 me3:DNA Meth. m5C
H2A.Z:H3–5ac
H3K4 me1:H4K20 me2
H3K27 me3:H3K4 me1
H3K4 me1:H3–5ac
DNA Meth. m5C:H4K20 me2
H3K4 me1:H4–4ac
DNA Meth. m5C:H3K9 me3
H4–4ac:H3–5ac
H4K20 me2:H3K4 me3
H3K27 me3:H3K9 me3
H3K4 me1:H2A.Z
H2A.Z:H4K20 me2
H4K20 me2:H3–5ac
H3K4 me3:H3–5ac
H4K20 me2:H4–4ac
H2A.Z:H3K4 me3
H3K9 me2
H3K27 me2
H3K4 me3:H4–4ac
H4K20 me3
H3K4 me1
H3K9 me3
H3K27 me3
DNA Meth. m5C
H4K20 me2
H3–5ac
H2A.Z
H4–4ac
H3K4 me3
0.00 0.25 0.50 0.75 1.00
$\hat{\pi}$
TIA1/TIAL1 – Reverse
H2A.Z:H4–4ac
H3K9 me3:H4K20 me3
DNA Meth. m5C:H4K20 me3
H3K27 me3:H4K20 me3
H3K27 me3:DNA Meth. m5C
H2A.Z:H3–5ac
H3K4 me1:H4K20 me2
H3K27 me3:H3K4 me1
H3K4 me1:H3–5ac
DNA Meth. m5C:H4K20 me2
H3K4 me1:H4–4ac
DNA Meth. m5C:H3K9 me3
H4–4ac:H3–5ac
H4K20 me2:H3K4 me3
H3K27 me3:H3K9 me3
H3K4 me1:H2A.Z
H2A.Z:H4K20 me2
H4K20 me2:H3–5ac
H3K4 me3:H3–5ac
H4K20 me2:H4–4ac
H2A.Z:H3K4 me3
H3K9 me2
H3K27 me2
H3K4 me3:H4–4ac
H4K20 me3
H3K4 me1
H3K9 me3
H3K27 me3
DNA Meth. m5C
H4K20 me2
H3–5ac
H2A.Z
H4–4ac
H3K4 me3
0.00 0.25 0.50 0.75 1.00
$\hat{\pi}$
NOSTRIN – Forward
H3K9 me3:H4K20 me3
H3K27 me3:H4K20 me3
H3K27 me3:H3K9 me3
H3K4 me1:H4K20 me2
H3K27 me3:H3K4 me1
H3K4 me3:H3–5ac
DNA Meth. m5C:H4K20 me3
H3K4 me1:H3–5ac
H3K4 me1:H4–4ac
H3K27 me2:DNA Meth. m5C
H3K27 me3:DNA Meth. m5C
H3K4 me1:H2A.Z
DNA Meth. m5C:H4K20 me2
DNA Meth. m5C:H3K9 me3
H3K4 me3:H4–4ac
H2A.Z:H4–4ac
H4K20 me2:H3K4 me3
H4K20 me2:H3–5ac
H2A.Z:H3–5ac
H2A.Z:H4K20 me2
H4K20 me2:H4–4ac
H2A.Z:H3K4 me3
H3K9 me2:DNA Meth. m5C
H4–4ac:H3–5ac
H3K27 me2
H4K20 me3
H3K27 me3
H3K9 me3
H3K9 me2
H3K4 me3
H3–5ac
H4–4ac
H3K4 me1
DNA Meth. m5C
H4K20 me2
H2A.Z
0.00 0.25 0.50 0.75 1.00
$\hat{\pi}$
NOSTRIN – Reverse
H3K9 me3:H4K20 me3
H3K27 me3:H4K20 me3
H3K27 me3:H3K9 me3
H3K4 me1:H4K20 me2
H3K27 me3:H3K4 me1
H3K4 me3:H3–5ac
DNA Meth. m5C:H4K20 me3
H3K4 me1:H3–5ac
H3K4 me1:H4–4ac
H3K27 me2:DNA Meth. m5C
H3K27 me3:DNA Meth. m5C
H3K4 me1:H2A.Z
DNA Meth. m5C:H4K20 me2
DNA Meth. m5C:H3K9 me3
H3K4 me3:H4–4ac
H2A.Z:H4–4ac
H4K20 me2:H3K4 me3
H4K20 me2:H3–5ac
H2A.Z:H3–5ac
H2A.Z:H4K20 me2
H4K20 me2:H4–4ac
H2A.Z:H3K4 me3
H3K9 me2:DNA Meth. m5C
H4–4ac:H3–5ac
H3K27 me2
H4K20 me3
H3K27 me3
H3K9 me3
H3K9 me2
H3K4 me3
H3–5ac
H4–4ac
H3K4 me1
DNA Meth. m5C
H4K20 me2
H2A.Z
0.00 0.25 0.50 0.75 1.00
$\hat{\pi}$
MLLT3 – Forward
H3K4 me1:H4–4ac
DNA Meth. m5C:H4K20 me2
H4–4ac:H3–5ac
H3K4 me1:H3–5ac
H3K4 me1:H2A.Z
H2A.Z:H4–4ac
H3K4 me3:H3–5ac
H4K20 me2:H4–4ac
H3K27 me3:H3K9 me3
H3K4 me3:H4–4ac
DNA Meth. m5C:H3K9 me3
H2A.Z:H3–5ac
H2A.Z:H3K4 me3
H2A.Z:H4K20 me2
H3K27 me3:DNA Meth. m5C
H3K27 me3:H4K20 me3
H3K9 me3:H4K20 me3
H4K20 me2:H3–5ac
H4K20 me2:H3K4 me3
DNA Meth. m5C:H4K20 me3
H3K4 me3
H4–4ac
H3K27 me2
H3K27 me3
H3K9 me2
H3K9 me3
DNA Meth. m5C
H4K20 me2
H3K4 me1
H2A.Z
H4K20 me3
H3–5ac
0.00 0.25 0.50 0.75 1.00
$\hat{\pi}$
MLLT3 – Reverse
H3K4 me1:H4–4ac
DNA Meth. m5C:H4K20 me2
H4–4ac:H3–5ac
H3K4 me1:H3–5ac
H3K4 me1:H2A.Z
H2A.Z:H4–4ac
H3K4 me3:H3–5ac
H4K20 me2:H4–4ac
H3K27 me3:H3K9 me3
H3K4 me3:H4–4ac
DNA Meth. m5C:H3K9 me3
H2A.Z:H3–5ac
H2A.Z:H3K4 me3
H2A.Z:H4K20 me2
H3K27 me3:DNA Meth. m5C
H3K27 me3:H4K20 me3
H3K9 me3:H4K20 me3
H4K20 me2:H3–5ac
H4K20 me2:H3K4 me3
DNA Meth. m5C:H4K20 me3
H3K4 me3
H4–4ac
H3K27 me2
H3K27 me3
H3K9 me2
H3K9 me3
DNA Meth. m5C
H4K20 me2
H3K4 me1
H2A.Z
H4K20 me3
H3–5ac
0.00 0.25 0.50 0.75 1.00
$\hat{\pi}$

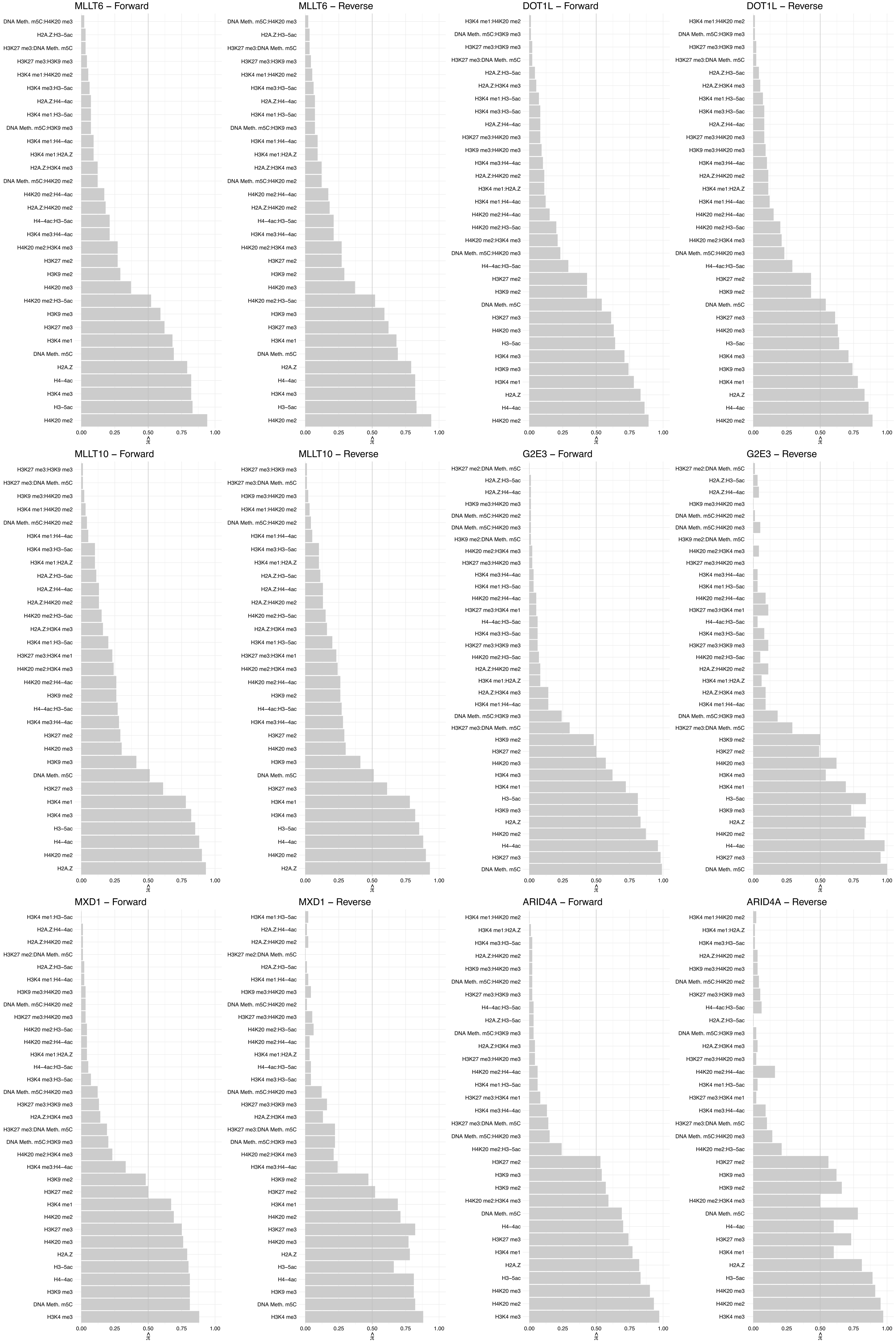
MLLT6 – Forward
DNA Meth. m5C:H4K20 me3
H2A.Z:H3–5ac
H3K27 me3:DNA Meth. m5C
H3K27 me3:H3K9 me3
H3K4 me1:H4K20 me2
H3K4 me3:H3–5ac
H2A.Z:H4–4ac
H3K4 me1:H3–5ac
DNA Meth. m5C:H3K9 me3
H3K4 me1:H4–4ac
H3K4 me1:H2A.Z
H2A.Z:H3K4 me3
DNA Meth. m5C:H4K20 me2
H4K20 me2:H4–4ac
H2A.Z:H4K20 me2
H4–4ac:H3–5ac
H3K4 me3:H4–4ac
H4K20 me2:H3K4 me3
H3K27 me2
H3K9 me2
H4K20 me3
H4K20 me2:H3–5ac
H3K9 me3
H3K27 me3
H3K4 me1
DNA Meth. m5C
H2A.Z
H4–4ac
H3K4 me3
H3–5ac
H4K20 me2
0.00 0.25 0.50 0.75 1.00
π̂
MLLT6 – Reverse
DNA Meth. m5C:H4K20 me3
H2A.Z:H3–5ac
H3K27 me3:DNA Meth. m5C
H3K27 me3:H3K9 me3
H3K4 me1:H4K20 me2
H3K4 me3:H3–5ac
H2A.Z:H4–4ac
H3K4 me1:H3–5ac
DNA Meth. m5C:H3K9 me3
H3K4 me1:H4–4ac
H3K4 me1:H2A.Z
H2A.Z:H3K4 me3
DNA Meth. m5C:H4K20 me2
H4K20 me2:H4–4ac
H2A.Z:H4K20 me2
H4–4ac:H3–5ac
H3K4 me3:H4–4ac
H4K20 me2:H3K4 me3
H3K27 me2
H3K9 me2
H4K20 me3
H4K20 me2:H3–5ac
H3K9 me3
H3K27 me3
H3K4 me1
DNA Meth. m5C
H2A.Z
H4–4ac
H3K4 me3
H3–5ac
H4K20 me2
0.00 0.25 0.50 0.75 1.00
π̂
DOT1L – Forward
H3K4 me1:H4K20 me2
DNA Meth. m5C:H3K9 me3
H3K27 me3:H3K9 me3
H3K27 me3:DNA Meth. m5C
H2A.Z:H3–5ac
H2A.Z:H3K4 me3
H3K4 me1:H3–5ac
H3K4 me3:H3–5ac
H2A.Z:H4–4ac
H3K27 me3:H4K20 me3
H3K9 me3:H4K20 me3
H3K4 me3:H4–4ac
H2A.Z:H4K20 me2
H3K4 me1:H2A.Z
H3K4 me1:H4–4ac
H4K20 me2:H4–4ac
H4K20 me2:H3–5ac
H4K20 me2:H3K4 me3
DNA Meth. m5C:H4K20 me3
H4–4ac:H3–5ac
H3K27 me2
H3K9 me2
DNA Meth. m5C
H3K27 me3
H4K20 me3
H3–5ac
H3K4 me3
H3K9 me3
H3K4 me1
H2A.Z
H4–4ac
H4K20 me2
0.00 0.25 0.50 0.75 1.00
π̂
DOT1L – Reverse
H3K4 me1:H4K20 me2
DNA Meth. m5C:H3K9 me3
H3K27 me3:H3K9 me3
H3K27 me3:DNA Meth. m5C
H2A.Z:H3–5ac
H2A.Z:H3K4 me3
H3K4 me1:H3–5ac
H3K4 me3:H3–5ac
H2A.Z:H4–4ac
H3K27 me3:H4K20 me3
H3K9 me3:H4K20 me3
H3K4 me3:H4–4ac
H2A.Z:H4K20 me2
H3K4 me1:H2A.Z
H3K4 me1:H4–4ac
H4K20 me2:H4–4ac
H4K20 me2:H3–5ac
H4K20 me2:H3K4 me3
DNA Meth. m5C:H4K20 me3
H4–4ac:H3–5ac
H3K27 me2
H3K9 me2
DNA Meth. m5C
H3K27 me3
H4K20 me3
H3–5ac
H3K4 me3
H3K9 me3
H3K4 me1
H2A.Z
H4–4ac
H4K20 me2
0.00 0.25 0.50 0.75 1.00
π̂
MLLT10 – Forward
H3K27 me3:H3K9 me3
H3K27 me3:DNA Meth. m5C
H3K9 me3:H4K20 me3
H3K4 me1:H4K20 me2
DNA Meth. m5C:H4K20 me2
H3K4 me1:H4–4ac
H3K4 me3:H3–5ac
H3K4 me1:H2A.Z
H2A.Z:H3–5ac
H2A.Z:H4–4ac
H2A.Z:H4K20 me2
H4K20 me2:H3–5ac
H2A.Z:H3K4 me3
H3K4 me1:H3–5ac
H3K27 me3:H3K4 me1
H4K20 me2:H3K4 me3
H4K20 me2:H4–4ac
H3K9 me2
H4–4ac:H3–5ac
H3K4 me3:H4–4ac
H3K27 me2
H4K20 me3
H3K9 me3
DNA Meth. m5C
H3K27 me3
H3K4 me1
H3K4 me3
H3–5ac
H4–4ac
H4K20 me2
H2A.Z
0.00 0.25 0.50 0.75 1.00
π̂
MLLT10 – Reverse
H3K27 me3:H3K9 me3
H3K27 me3:DNA Meth. m5C
H3K9 me3:H4K20 me3
H3K4 me1:H4K20 me2
DNA Meth. m5C:H4K20 me2
H3K4 me1:H4–4ac
H3K4 me3:H3–5ac
H3K4 me1:H2A.Z
H2A.Z:H3–5ac
H2A.Z:H4–4ac
H2A.Z:H4K20 me2
H4K20 me2:H3–5ac
H2A.Z:H3K4 me3
H3K4 me1:H3–5ac
H3K27 me3:H3K4 me1
H4K20 me2:H3K4 me3
H4K20 me2:H4–4ac
H3K9 me2
H4–4ac:H3–5ac
H3K4 me3:H4–4ac
H3K27 me2
H4K20 me3
H3K9 me3
DNA Meth. m5C
H3K27 me3
H3K4 me1
H3K4 me3
H3–5ac
H4–4ac
H4K20 me2
H2A.Z
0.00 0.25 0.50 0.75 1.00
π̂
G2E3 – Forward
H3K27 me2:DNA Meth. m5C
H2A.Z:H3–5ac
H2A.Z:H4–4ac
H3K9 me3:H4K20 me3
DNA Meth. m5C:H4K20 me2
DNA Meth. m5C:H4K20 me3
H3K9 me2:DNA Meth. m5C
H4K20 me2:H3K4 me3
H3K27 me3:H4K20 me3
H3K4 me3:H4–4ac
H3K4 me1:H3–5ac
H4K20 me2:H4–4ac
H3K27 me3:H3K4 me1
H4–4ac:H3–5ac
H3K4 me3:H3–5ac
H3K27 me3:H3K9 me3
H4K20 me2:H3–5ac
H2A.Z:H4K20 me2
H3K4 me1:H2A.Z
H2A.Z:H3K4 me3
H3K4 me1:H4–4ac
DNA Meth. m5C:H3K9 me3
H3K27 me3:DNA Meth. m5C
H3K9 me2
H3K27 me2
H4K20 me3
H3K4 me3
H3K4 me1
H3–5ac
H3K9 me3
H2A.Z
H4K20 me2
H4–4ac
H3K27 me3
DNA Meth. m5C
0.00 0.25 0.50 0.75 1.00
π̂
G2E3 – Reverse
H3K27 me2:DNA Meth. m5C
H2A.Z:H3–5ac
H2A.Z:H4–4ac
H3K9 me3:H4K20 me3
DNA Meth. m5C:H4K20 me2
DNA Meth. m5C:H4K20 me3
H3K9 me2:DNA Meth. m5C
H4K20 me2:H3K4 me3
H3K27 me3:H4K20 me3
H3K4 me3:H4–4ac
H3K4 me1:H3–5ac
H4K20 me2:H4–4ac
H3K27 me3:H3K4 me1
H4–4ac:H3–5ac
H3K4 me3:H3–5ac
H3K27 me3:H3K9 me3
H4K20 me2:H3–5ac
H2A.Z:H4K20 me2
H3K4 me1:H2A.Z
H2A.Z:H3K4 me3
H3K4 me1:H4–4ac
DNA Meth. m5C:H3K9 me3
H3K27 me3:DNA Meth. m5C
H3K9 me2
H3K27 me2
H4K20 me3
H3K4 me3
H3K4 me1
H3–5ac
H3K9 me3
H2A.Z
H4K20 me2
H4–4ac
H3K27 me3
DNA Meth. m5C
0.00 0.25 0.50 0.75 1.00
π̂
MXD1 – Forward
H3K4 me1:H3–5ac
H2A.Z:H4–4ac
H2A.Z:H4K20 me2
H3K27 me2:DNA Meth. m5C
H2A.Z:H3–5ac
H3K4 me1:H4–4ac
H3K9 me3:H4K20 me3
DNA Meth. m5C:H4K20 me2
H3K27 me3:H4K20 me3
H4K20 me2:H3–5ac
H4K20 me2:H4–4ac
H3K4 me1:H2A.Z
H4–4ac:H3–5ac
H3K4 me3:H3–5ac
DNA Meth. m5C:H4K20 me3
H3K27 me3:H3K9 me3
H2A.Z:H3K4 me3
H3K27 me3:DNA Meth. m5C
DNA Meth. m5C:H3K9 me3
H4K20 me2:H3K4 me3
H3K4 me3:H4–4ac
H3K9 me2
H3K27 me2
H3K4 me1
H4K20 me2
H3K27 me3
H4K20 me3
H2A.Z
H3–5ac
H4–4ac
H3K9 me3
DNA Meth. m5C
H3K4 me3
0.00 0.25 0.50 0.75 1.00
π̂
MXD1 – Reverse
H3K4 me1:H3–5ac
H2A.Z:H4–4ac
H2A.Z:H4K20 me2
H3K27 me2:DNA Meth. m5C
H2A.Z:H3–5ac
H3K4 me1:H4–4ac
H3K9 me3:H4K20 me3
DNA Meth. m5C:H4K20 me2
H3K27 me3:H4K20 me3
H4K20 me2:H3–5ac
H4K20 me2:H4–4ac
H3K4 me1:H2A.Z
H4–4ac:H3–5ac
H3K4 me3:H3–5ac
DNA Meth. m5C:H4K20 me3
H3K27 me3:H3K9 me3
H2A.Z:H3K4 me3
H3K27 me3:DNA Meth. m5C
DNA Meth. m5C:H3K9 me3
H4K20 me2:H3K4 me3
H3K4 me3:H4–4ac
H3K9 me2
H3K27 me2
H3K4 me1
H4K20 me2
H3K27 me3
H4K20 me3
H2A.Z
H3–5ac
H4–4ac
H3K9 me3
DNA Meth. m5C
H3K4 me3
0.00 0.25 0.50 0.75 1.00
π̂
ARID4A – Forward
H3K4 me1:H4K20 me2
H3K4 me1:H2A.Z
H3K4 me3:H3–5ac
H2A.Z:H4K20 me2
H3K9 me3:H4K20 me3
DNA Meth. m5C:H4K20 me2
H3K27 me3:H3K9 me3
H4–4ac:H3–5ac
H2A.Z:H3–5ac
DNA Meth. m5C:H3K9 me3
H2A.Z:H3K4 me3
H3K27 me3:H4K20 me3
H4K20 me2:H4–4ac
H3K4 me1:H3–5ac
H3K27 me3:H3K4 me1
H3K4 me3:H4–4ac
H3K27 me3:DNA Meth. m5C
DNA Meth. m5C:H4K20 me3
H4K20 me2:H3–5ac
H3K27 me2
H3K9 me3
H3K9 me2
H4K20 me2:H3K4 me3
DNA Meth. m5C
H4–4ac
H3K27 me3
H3K4 me1
H2A.Z
H3–5ac
H4K20 me3
H4K20 me2
H3K4 me3
0.00 0.25 0.50 0.75 1.00
π̂
ARID4A – Reverse
H3K4 me1:H4K20 me2
H3K4 me1:H2A.Z
H3K4 me3:H3–5ac
H2A.Z:H4K20 me2
H3K9 me3:H4K20 me3
DNA Meth. m5C:H4K20 me2
H3K27 me3:H3K9 me3
H4–4ac:H3–5ac
H2A.Z:H3–5ac
DNA Meth. m5C:H3K9 me3
H2A.Z:H3K4 me3
H3K27 me3:H4K20 me3
H4K20 me2:H4–4ac
H3K4 me1:H3–5ac
H3K27 me3:H3K4 me1
H3K4 me3:H4–4ac
H3K27 me3:DNA Meth. m5C
DNA Meth. m5C:H4K20 me3
H4K20 me2:H3–5ac
H3K27 me2
H3K9 me3
H3K9 me2
H4K20 me2:H3K4 me3
DNA Meth. m5C
H4–4ac
H3K27 me3
H3K4 me1
H2A.Z
H3–5ac
H4K20 me3
H4K20 me2
H3K4 me3
0.00 0.25 0.50 0.75 1.00
π̂

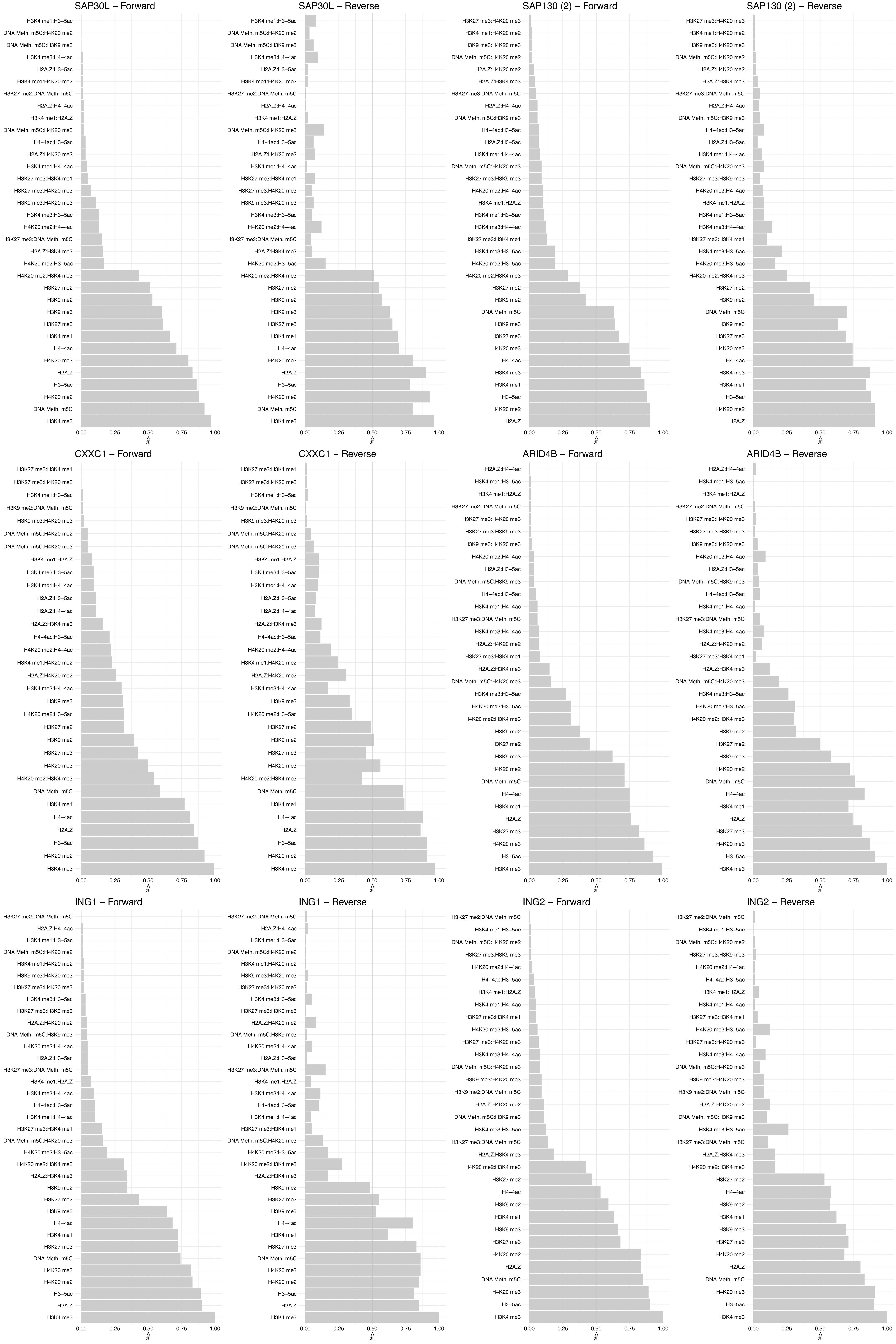
SAP30L − Forward
H3K4 me1:H3−5ac
DNA Meth. m5C:H4K20 me2
DNA Meth. m5C:H3K9 me3
H3K4 me3:H4−4ac
H2A.Z:H3−5ac
H3K4 me1:H4K20 me2
H3K27 me2:DNA Meth. m5C
H2A.Z:H4−4ac
H3K4 me1:H2A.Z
DNA Meth. m5C:H4K20 me3
H4−4ac:H3−5ac
H2A.Z:H4K20 me2
H3K4 me1:H4−4ac
H3K27 me3:H3K4 me1
H3K27 me3:H4K20 me3
H3K9 me3:H4K20 me3
H3K4 me3:H3−5ac
H4K20 me2:H4−4ac
H3K27 me3:DNA Meth. m5C
H2A.Z:H3K4 me3
H4K20 me2:H3−5ac
H4K20 me2:H3K4 me3
H3K27 me2
H3K9 me2
H3K9 me3
H3K27 me3
H3K4 me1
H4−4ac
H4K20 me3
H2A.Z
H3−5ac
H4K20 me2
DNA Meth. m5C
H3K4 me3
0.00 0.25 0.50 0.75 1.00
π̂
SAP30L − Reverse
H3K4 me1:H3−5ac
DNA Meth. m5C:H4K20 me2
DNA Meth. m5C:H3K9 me3
H3K4 me3:H4−4ac
H2A.Z:H3−5ac
H3K4 me1:H4K20 me2
H3K27 me2:DNA Meth. m5C
H2A.Z:H4−4ac
H3K4 me1:H2A.Z
DNA Meth. m5C:H4K20 me3
H4−4ac:H3−5ac
H2A.Z:H4K20 me2
H3K4 me1:H4−4ac
H3K27 me3:H3K4 me1
H3K27 me3:H4K20 me3
H3K9 me3:H4K20 me3
H3K4 me3:H3−5ac
H4K20 me2:H4−4ac
H3K27 me3:DNA Meth. m5C
H2A.Z:H3K4 me3
H4K20 me2:H3−5ac
H4K20 me2:H3K4 me3
H3K27 me2
H3K9 me2
H3K9 me3
H3K27 me3
H3K4 me1
H4−4ac
H4K20 me3
H2A.Z
H3−5ac
H4K20 me2
DNA Meth. m5C
H3K4 me3
0.00 0.25 0.50 0.75 1.00
π̂
SAP130 (2) − Forward
H3K27 me3:H4K20 me3
H3K4 me1:H4K20 me2
H3K9 me3:H4K20 me3
DNA Meth. m5C:H4K20 me2
H2A.Z:H4K20 me2
H2A.Z:H3K4 me3
H3K27 me3:DNA Meth. m5C
H2A.Z:H4−4ac
DNA Meth. m5C:H3K9 me3
H4−4ac:H3−5ac
H2A.Z:H3−5ac
H3K4 me1:H4−4ac
DNA Meth. m5C:H4K20 me3
H3K27 me3:H3K9 me3
H4K20 me2:H4−4ac
H3K4 me1:H2A.Z
H3K4 me1:H3−5ac
H3K4 me3:H4−4ac
H3K27 me3:H3K4 me1
H3K4 me3:H3−5ac
H4K20 me2:H3−5ac
H4K20 me2:H3K4 me3
H3K27 me2
H3K9 me2
DNA Meth. m5C
H3K9 me3
H3K27 me3
H4K20 me3
H4−4ac
H3K4 me3
H3K4 me1
H3−5ac
H4K20 me2
H2A.Z
0.00 0.25 0.50 0.75 1.00
π̂
SAP130 (2) − Reverse
H3K27 me3:H4K20 me3
H3K4 me1:H4K20 me2
H3K9 me3:H4K20 me3
DNA Meth. m5C:H4K20 me2
H2A.Z:H4K20 me2
H2A.Z:H3K4 me3
H3K27 me3:DNA Meth. m5C
H2A.Z:H4−4ac
DNA Meth. m5C:H3K9 me3
H4−4ac:H3−5ac
H2A.Z:H3−5ac
H3K4 me1:H4−4ac
DNA Meth. m5C:H4K20 me3
H3K27 me3:H3K9 me3
H4K20 me2:H4−4ac
H3K4 me1:H2A.Z
H3K4 me1:H3−5ac
H3K4 me3:H4−4ac
H3K27 me3:H3K4 me1
H3K4 me3:H3−5ac
H4K20 me2:H3−5ac
H4K20 me2:H3K4 me3
H3K27 me2
H3K9 me2
DNA Meth. m5C
H3K9 me3
H3K27 me3
H4K20 me3
H4−4ac
H3K4 me3
H3K4 me1
H3−5ac
H4K20 me2
H2A.Z
0.00 0.25 0.50 0.75 1.00
π̂
CXXC1 − Forward
H3K27 me3:H3K4 me1
H3K27 me3:H4K20 me3
H3K4 me1:H3−5ac
H3K9 me2:DNA Meth. m5C
H3K9 me3:H4K20 me3
DNA Meth. m5C:H4K20 me2
DNA Meth. m5C:H4K20 me3
H3K4 me1:H2A.Z
H3K4 me3:H3−5ac
H3K4 me1:H4−4ac
H2A.Z:H3−5ac
H2A.Z:H4−4ac
H2A.Z:H3K4 me3
H4−4ac:H3−5ac
H4K20 me2:H4−4ac
H3K4 me1:H4K20 me2
H2A.Z:H4K20 me2
H3K4 me3:H4−4ac
H3K9 me3
H4K20 me2:H3−5ac
H3K27 me2
H3K9 me2
H3K27 me3
H4K20 me3
H4K20 me2:H3K4 me3
DNA Meth. m5C
H3K4 me1
H4−4ac
H2A.Z
H3−5ac
H4K20 me2
H3K4 me3
0.00 0.25 0.50 0.75 1.00
π̂
CXXC1 − Reverse
H3K27 me3:H3K4 me1
H3K27 me3:H4K20 me3
H3K4 me1:H3−5ac
H3K9 me2:DNA Meth. m5C
H3K9 me3:H4K20 me3
DNA Meth. m5C:H4K20 me2
DNA Meth. m5C:H4K20 me3
H3K4 me1:H2A.Z
H3K4 me3:H3−5ac
H3K4 me1:H4−4ac
H2A.Z:H3−5ac
H2A.Z:H4−4ac
H2A.Z:H3K4 me3
H4−4ac:H3−5ac
H4K20 me2:H4−4ac
H3K4 me1:H4K20 me2
H2A.Z:H4K20 me2
H3K4 me3:H4−4ac
H3K9 me3
H4K20 me2:H3−5ac
H3K27 me2
H3K9 me2
H3K27 me3
H4K20 me3
H4K20 me2:H3K4 me3
DNA Meth. m5C
H3K4 me1
H4−4ac
H2A.Z
H3−5ac
H4K20 me2
H3K4 me3
0.00 0.25 0.50 0.75 1.00
π̂
ARID4B − Forward
H2A.Z:H4−4ac
H3K4 me1:H3−5ac
H3K4 me1:H2A.Z
H3K27 me2:DNA Meth. m5C
H3K27 me3:H4K20 me3
H3K27 me3:H3K9 me3
H3K9 me3:H4K20 me3
H4K20 me2:H4−4ac
H2A.Z:H3−5ac
DNA Meth. m5C:H3K9 me3
H4−4ac:H3−5ac
H3K4 me1:H4−4ac
H3K27 me3:DNA Meth. m5C
H3K4 me3:H4−4ac
H2A.Z:H4K20 me2
H3K27 me3:H3K4 me1
H2A.Z:H3K4 me3
DNA Meth. m5C:H4K20 me3
H3K4 me3:H3−5ac
H4K20 me2:H3−5ac
H4K20 me2:H3K4 me3
H3K9 me2
H3K27 me2
H3K9 me3
H4K20 me2
DNA Meth. m5C
H4−4ac
H3K4 me1
H2A.Z
H3K27 me3
H4K20 me3
H3−5ac
H3K4 me3
0.00 0.25 0.50 0.75 1.00
π̂
ARID4B − Reverse
H2A.Z:H4−4ac
H3K4 me1:H3−5ac
H3K4 me1:H2A.Z
H3K27 me2:DNA Meth. m5C
H3K27 me3:H4K20 me3
H3K27 me3:H3K9 me3
H3K9 me3:H4K20 me3
H4K20 me2:H4−4ac
H2A.Z:H3−5ac
DNA Meth. m5C:H3K9 me3
H4−4ac:H3−5ac
H3K4 me1:H4−4ac
H3K27 me3:DNA Meth. m5C
H3K4 me3:H4−4ac
H2A.Z:H4K20 me2
H3K27 me3:H3K4 me1
H2A.Z:H3K4 me3
DNA Meth. m5C:H4K20 me3
H3K4 me3:H3−5ac
H4K20 me2:H3−5ac
H4K20 me2:H3K4 me3
H3K9 me2
H3K27 me2
H3K9 me3
H4K20 me2
DNA Meth. m5C
H4−4ac
H3K4 me1
H2A.Z
H3K27 me3
H4K20 me3
H3−5ac
H3K4 me3
0.00 0.25 0.50 0.75 1.00
π̂
ING1 − Forward
H3K27 me2:DNA Meth. m5C
H2A.Z:H4−4ac
H3K4 me1:H3−5ac
DNA Meth. m5C:H4K20 me2
H3K4 me1:H4K20 me2
H3K9 me3:H4K20 me3
H3K27 me3:H4K20 me3
H3K4 me3:H3−5ac
H3K27 me3:H3K9 me3
H2A.Z:H4K20 me2
DNA Meth. m5C:H3K9 me3
H4K20 me2:H4−4ac
H2A.Z:H3−5ac
H3K27 me3:DNA Meth. m5C
H3K4 me1:H2A.Z
H3K4 me3:H4−4ac
H4−4ac:H3−5ac
H3K4 me1:H4−4ac
H3K27 me3:H3K4 me1
DNA Meth. m5C:H4K20 me3
H4K20 me2:H3−5ac
H4K20 me2:H3K4 me3
H2A.Z:H3K4 me3
H3K9 me2
H3K27 me2
H3K9 me3
H4−4ac
H3K4 me1
H3K27 me3
DNA Meth. m5C
H4K20 me3
H4K20 me2
H3−5ac
H2A.Z
H3K4 me3
0.00 0.25 0.50 0.75 1.00
π̂
ING1 − Reverse
H3K27 me2:DNA Meth. m5C
H2A.Z:H4−4ac
H3K4 me1:H3−5ac
DNA Meth. m5C:H4K20 me2
H3K4 me1:H4K20 me2
H3K9 me3:H4K20 me3
H3K27 me3:H4K20 me3
H3K4 me3:H3−5ac
H3K27 me3:H3K9 me3
H2A.Z:H4K20 me2
DNA Meth. m5C:H3K9 me3
H4K20 me2:H4−4ac
H2A.Z:H3−5ac
H3K27 me3:DNA Meth. m5C
H3K4 me1:H2A.Z
H3K4 me3:H4−4ac
H4−4ac:H3−5ac
H3K4 me1:H4−4ac
H3K27 me3:H3K4 me1
DNA Meth. m5C:H4K20 me3
H4K20 me2:H3−5ac
H4K20 me2:H3K4 me3
H2A.Z:H3K4 me3
H3K9 me2
H3K27 me2
H3K9 me3
H4−4ac
H3K4 me1
H3K27 me3
DNA Meth. m5C
H4K20 me3
H4K20 me2
H3−5ac
H2A.Z
H3K4 me3
0.00 0.25 0.50 0.75 1.00
π̂
ING2 − Forward
H3K27 me2:DNA Meth. m5C
H3K4 me1:H3−5ac
DNA Meth. m5C:H4K20 me2
H3K27 me3:H3K9 me3
H4K20 me2:H4−4ac
H4−4ac:H3−5ac
H3K4 me1:H2A.Z
H3K4 me1:H4−4ac
H3K27 me3:H3K4 me1
H4K20 me2:H3−5ac
H3K27 me3:H4K20 me3
H3K4 me3:H4−4ac
DNA Meth. m5C:H4K20 me3
H3K9 me3:H4K20 me3
H3K9 me2:DNA Meth. m5C
H2A.Z:H4K20 me2
DNA Meth. m5C:H3K9 me3
H3K4 me3:H3−5ac
H3K27 me3:DNA Meth. m5C
H2A.Z:H3K4 me3
H4K20 me2:H3K4 me3
H3K27 me2
H4−4ac
H3K9 me2
H3K4 me1
H3K9 me3
H3K27 me3
H4K20 me2
H2A.Z
DNA Meth. m5C
H4K20 me3
H3−5ac
H3K4 me3
0.00 0.25 0.50 0.75 1.00
π̂
ING2 − Reverse
H3K27 me2:DNA Meth. m5C
H3K4 me1:H3−5ac
DNA Meth. m5C:H4K20 me2
H3K27 me3:H3K9 me3
H4K20 me2:H4−4ac
H4−4ac:H3−5ac
H3K4 me1:H2A.Z
H3K4 me1:H4−4ac
H3K27 me3:H3K4 me1
H4K20 me2:H3−5ac
H3K27 me3:H4K20 me3
H3K4 me3:H4−4ac
DNA Meth. m5C:H4K20 me3
H3K9 me3:H4K20 me3
H3K9 me2:DNA Meth. m5C
H2A.Z:H4K20 me2
DNA Meth. m5C:H3K9 me3
H3K4 me3:H3−5ac
H3K27 me3:DNA Meth. m5C
H2A.Z:H3K4 me3
H4K20 me2:H3K4 me3
H3K27 me2
H4−4ac
H3K9 me2
H3K4 me1
H3K9 me3
H3K27 me3
H4K20 me2
H2A.Z
DNA Meth. m5C
H4K20 me3
H3−5ac
H3K4 me3
0.00 0.25 0.50 0.75 1.00
π̂

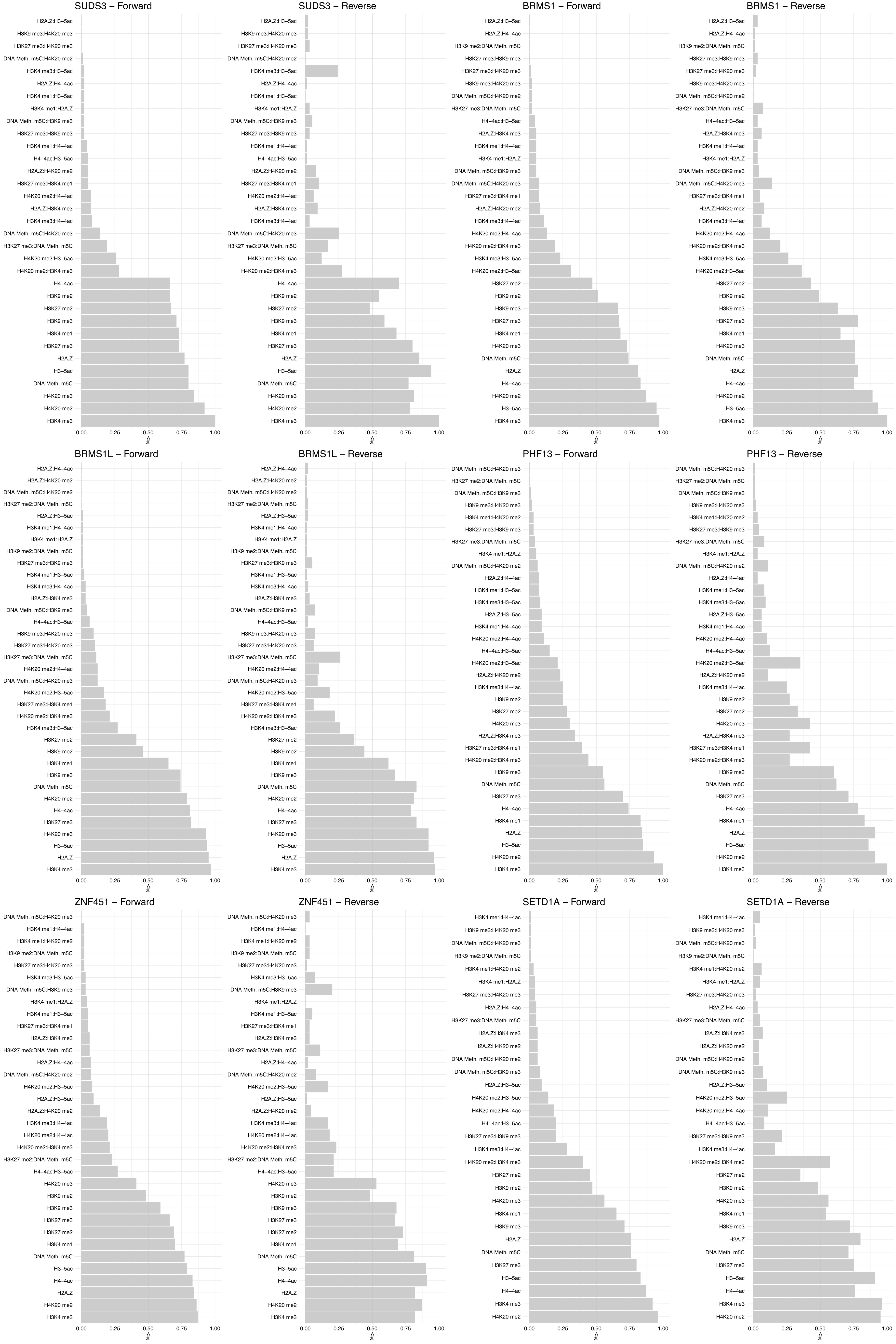
SUDS3 – Forward
H2A.Z:H3–5ac
H3K9 me3:H4K20 me3
H3K27 me3:H4K20 me3
DNA Meth. m5C:H4K20 me2
H3K4 me3:H3–5ac
H2A.Z:H4–4ac
H3K4 me1:H3–5ac
H3K4 me1:H2A.Z
DNA Meth. m5C:H3K9 me3
H3K27 me3:H3K9 me3
H3K4 me1:H4–4ac
H4–4ac:H3–5ac
H2A.Z:H4K20 me2
H3K27 me3:H3K4 me1
H4K20 me2:H4–4ac
H2A.Z:H3K4 me3
H3K4 me3:H4–4ac
DNA Meth. m5C:H4K20 me3
H3K27 me3:DNA Meth. m5C
H4K20 me2:H3–5ac
H4K20 me2:H3K4 me3
H4–4ac
H3K9 me2
H3K27 me2
H3K9 me3
H3K4 me1
H3K27 me3
H2A.Z
H3–5ac
DNA Meth. m5C
H4K20 me3
H4K20 me2
H3K4 me3
0.00 0.25 0.50 0.75 1.00
$\hat{\pi}$
SUDS3 – Reverse
H2A.Z:H3–5ac
H3K9 me3:H4K20 me3
H3K27 me3:H4K20 me3
DNA Meth. m5C:H4K20 me2
H3K4 me3:H3–5ac
H2A.Z:H4–4ac
H3K4 me1:H3–5ac
H3K4 me1:H2A.Z
DNA Meth. m5C:H3K9 me3
H3K27 me3:H3K9 me3
H3K4 me1:H4–4ac
H4–4ac:H3–5ac
H2A.Z:H4K20 me2
H3K27 me3:H3K4 me1
H4K20 me2:H4–4ac
H2A.Z:H3K4 me3
H3K4 me3:H4–4ac
DNA Meth. m5C:H4K20 me3
H3K27 me3:DNA Meth. m5C
H4K20 me2:H3–5ac
H4K20 me2:H3K4 me3
H4–4ac
H3K9 me2
H3K27 me2
H3K9 me3
H3K4 me1
H3K27 me3
H2A.Z
H3–5ac
DNA Meth. m5C
H4K20 me3
H4K20 me2
H3K4 me3
0.00 0.25 0.50 0.75 1.00
$\hat{\pi}$
BRMS1 – Forward
H2A.Z:H3–5ac
H2A.Z:H4–4ac
H3K9 me2:DNA Meth. m5C
H3K27 me3:H3K9 me3
H3K27 me3:H4K20 me3
H3K9 me3:H4K20 me3
DNA Meth. m5C:H4K20 me2
H3K27 me3:DNA Meth. m5C
H4–4ac:H3–5ac
H2A.Z:H3K4 me3
H3K4 me1:H4–4ac
H3K4 me1:H2A.Z
DNA Meth. m5C:H3K9 me3
DNA Meth. m5C:H4K20 me3
H3K27 me3:H3K4 me1
H2A.Z:H4K20 me2
H3K4 me3:H4–4ac
H4K20 me2:H4–4ac
H4K20 me2:H3K4 me3
H3K4 me3:H3–5ac
H4K20 me2:H3–5ac
H3K27 me2
H3K9 me2
H3K9 me3
H3K27 me3
H3K4 me1
H4K20 me3
DNA Meth. m5C
H2A.Z
H4–4ac
H4K20 me2
H3–5ac
H3K4 me3
0.00 0.25 0.50 0.75 1.00
$\hat{\pi}$
BRMS1 – Reverse
H2A.Z:H3–5ac
H2A.Z:H4–4ac
H3K9 me2:DNA Meth. m5C
H3K27 me3:H3K9 me3
H3K27 me3:H4K20 me3
H3K9 me3:H4K20 me3
DNA Meth. m5C:H4K20 me2
H3K27 me3:DNA Meth. m5C
H4–4ac:H3–5ac
H2A.Z:H3K4 me3
H3K4 me1:H4–4ac
H3K4 me1:H2A.Z
DNA Meth. m5C:H3K9 me3
DNA Meth. m5C:H4K20 me3
H3K27 me3:H3K4 me1
H2A.Z:H4K20 me2
H3K4 me3:H4–4ac
H4K20 me2:H4–4ac
H4K20 me2:H3K4 me3
H3K4 me3:H3–5ac
H4K20 me2:H3–5ac
H3K27 me2
H3K9 me2
H3K9 me3
H3K27 me3
H3K4 me1
H4K20 me3
DNA Meth. m5C
H2A.Z
H4–4ac
H4K20 me2
H3–5ac
H3K4 me3
0.00 0.25 0.50 0.75 1.00
$\hat{\pi}$
BRMS1L – Forward
H2A.Z:H4–4ac
H2A.Z:H4K20 me2
DNA Meth. m5C:H4K20 me2
H3K27 me2:DNA Meth. m5C
H2A.Z:H3–5ac
H3K4 me1:H4–4ac
H3K4 me1:H2A.Z
H3K9 me2:DNA Meth. m5C
H3K27 me3:H3K9 me3
H3K4 me1:H3–5ac
H3K4 me3:H4–4ac
H2A.Z:H3K4 me3
DNA Meth. m5C:H3K9 me3
H4–4ac:H3–5ac
H3K9 me3:H4K20 me3
H3K27 me3:H4K20 me3
H3K27 me3:DNA Meth. m5C
H4K20 me2:H4–4ac
DNA Meth. m5C:H4K20 me3
H4K20 me2:H3–5ac
H3K27 me3:H3K4 me1
H4K20 me2:H3K4 me3
H3K4 me3:H3–5ac
H3K27 me2
H3K9 me2
H3K4 me1
H3K9 me3
DNA Meth. m5C
H4K20 me2
H4–4ac
H3K27 me3
H4K20 me3
H3–5ac
H2A.Z
H3K4 me3
0.00 0.25 0.50 0.75 1.00
$\hat{\pi}$
BRMS1L – Reverse
H2A.Z:H4–4ac
H2A.Z:H4K20 me2
DNA Meth. m5C:H4K20 me2
H3K27 me2:DNA Meth. m5C
H2A.Z:H3–5ac
H3K4 me1:H4–4ac
H3K4 me1:H2A.Z
H3K9 me2:DNA Meth. m5C
H3K27 me3:H3K9 me3
H3K4 me1:H3–5ac
H3K4 me3:H4–4ac
H2A.Z:H3K4 me3
DNA Meth. m5C:H3K9 me3
H4–4ac:H3–5ac
H3K9 me3:H4K20 me3
H3K27 me3:H4K20 me3
H3K27 me3:DNA Meth. m5C
H4K20 me2:H4–4ac
DNA Meth. m5C:H4K20 me3
H4K20 me2:H3–5ac
H3K27 me3:H3K4 me1
H4K20 me2:H3K4 me3
H3K4 me3:H3–5ac
H3K27 me2
H3K9 me2
H3K4 me1
H3K9 me3
DNA Meth. m5C
H4K20 me2
H4–4ac
H3K27 me3
H4K20 me3
H3–5ac
H2A.Z
H3K4 me3
0.00 0.25 0.50 0.75 1.00
$\hat{\pi}$
PHF13 – Forward
DNA Meth. m5C:H4K20 me3
H3K27 me2:DNA Meth. m5C
DNA Meth. m5C:H3K9 me3
H3K9 me3:H4K20 me3
H3K4 me1:H4K20 me2
H3K27 me3:H3K9 me3
H3K27 me3:DNA Meth. m5C
H3K4 me1:H2A.Z
DNA Meth. m5C:H4K20 me2
H2A.Z:H4–4ac
H3K4 me1:H3–5ac
H3K4 me3:H3–5ac
H2A.Z:H3–5ac
H3K4 me1:H4–4ac
H4K20 me2:H4–4ac
H4–4ac:H3–5ac
H4K20 me2:H3–5ac
H2A.Z:H4K20 me2
H3K4 me3:H4–4ac
H3K9 me2
H3K27 me2
H4K20 me3
H2A.Z:H3K4 me3
H3K27 me3:H3K4 me1
H4K20 me2:H3K4 me3
H3K9 me3
DNA Meth. m5C
H3K27 me3
H4–4ac
H3K4 me1
H2A.Z
H3–5ac
H4K20 me2
H3K4 me3
0.00 0.25 0.50 0.75 1.00
$\hat{\pi}$
PHF13 – Reverse
DNA Meth. m5C:H4K20 me3
H3K27 me2:DNA Meth. m5C
DNA Meth. m5C:H3K9 me3
H3K9 me3:H4K20 me3
H3K4 me1:H4K20 me2
H3K27 me3:H3K9 me3
H3K27 me3:DNA Meth. m5C
H3K4 me1:H2A.Z
DNA Meth. m5C:H4K20 me2
H2A.Z:H4–4ac
H3K4 me1:H3–5ac
H3K4 me3:H3–5ac
H2A.Z:H3–5ac
H3K4 me1:H4–4ac
H4K20 me2:H4–4ac
H4–4ac:H3–5ac
H4K20 me2:H3–5ac
H2A.Z:H4K20 me2
H3K4 me3:H4–4ac
H3K9 me2
H3K27 me2
H4K20 me3
H2A.Z:H3K4 me3
H3K27 me3:H3K4 me1
H4K20 me2:H3K4 me3
H3K9 me3
DNA Meth. m5C
H3K27 me3
H4–4ac
H3K4 me1
H2A.Z
H3–5ac
H4K20 me2
H3K4 me3
0.00 0.25 0.50 0.75 1.00
$\hat{\pi}$
ZNF451 – Forward
DNA Meth. m5C:H4K20 me3
H3K4 me1:H4–4ac
H3K4 me1:H4K20 me2
H3K9 me2:DNA Meth. m5C
H3K27 me3:H4K20 me3
H3K4 me3:H3–5ac
DNA Meth. m5C:H3K9 me3
H3K4 me1:H2A.Z
H3K4 me1:H3–5ac
H3K27 me3:H3K4 me1
H2A.Z:H3K4 me3
H3K27 me3:DNA Meth. m5C
H2A.Z:H4–4ac
DNA Meth. m5C:H4K20 me2
H4K20 me2:H3–5ac
H2A.Z:H3–5ac
H2A.Z:H4K20 me2
H3K4 me3:H4–4ac
H4K20 me2:H4–4ac
H4K20 me2:H3K4 me3
H3K27 me2:DNA Meth. m5C
H4–4ac:H3–5ac
H4K20 me3
H3K9 me2
H3K9 me3
H3K27 me3
H3K27 me2
H3K4 me1
DNA Meth. m5C
H3–5ac
H4–4ac
H2A.Z
H4K20 me2
H3K4 me3
0.00 0.25 0.50 0.75 1.00
$\hat{\pi}$
ZNF451 – Reverse
DNA Meth. m5C:H4K20 me3
H3K4 me1:H4–4ac
H3K4 me1:H4K20 me2
H3K9 me2:DNA Meth. m5C
H3K27 me3:H4K20 me3
H3K4 me3:H3–5ac
DNA Meth. m5C:H3K9 me3
H3K4 me1:H2A.Z
H3K4 me1:H3–5ac
H3K27 me3:H3K4 me1
H2A.Z:H3K4 me3
H3K27 me3:DNA Meth. m5C
H2A.Z:H4–4ac
DNA Meth. m5C:H4K20 me2
H4K20 me2:H3–5ac
H2A.Z:H3–5ac
H2A.Z:H4K20 me2
H3K4 me3:H4–4ac
H4K20 me2:H4–4ac
H4K20 me2:H3K4 me3
H3K27 me2:DNA Meth. m5C
H4–4ac:H3–5ac
H4K20 me3
H3K9 me2
H3K9 me3
H3K27 me3
H3K27 me2
H3K4 me1
DNA Meth. m5C
H3–5ac
H4–4ac
H2A.Z
H4K20 me2
H3K4 me3
0.00 0.25 0.50 0.75 1.00
$\hat{\pi}$
SETD1A – Forward
H3K4 me1:H4–4ac
H3K9 me3:H4K20 me3
DNA Meth. m5C:H4K20 me3
H3K9 me2:DNA Meth. m5C
H3K4 me1:H4K20 me2
H3K4 me1:H2A.Z
H3K27 me3:H4K20 me3
H2A.Z:H4–4ac
H3K27 me3:DNA Meth. m5C
H2A.Z:H3K4 me3
H2A.Z:H4K20 me2
DNA Meth. m5C:H4K20 me2
DNA Meth. m5C:H3K9 me3
H2A.Z:H3–5ac
H4K20 me2:H3–5ac
H4K20 me2:H4–4ac
H4–4ac:H3–5ac
H3K27 me3:H3K9 me3
H3K4 me3:H4–4ac
H4K20 me2:H3K4 me3
H3K27 me2
H3K9 me2
H4K20 me3
H3K4 me1
H3K9 me3
H2A.Z
DNA Meth. m5C
H3K27 me3
H3–5ac
H4–4ac
H3K4 me3
H4K20 me2
0.00 0.25 0.50 0.75 1.00
$\hat{\pi}$
SETD1A – Reverse
H3K4 me1:H4–4ac
H3K9 me3:H4K20 me3
DNA Meth. m5C:H4K20 me3
H3K9 me2:DNA Meth. m5C
H3K4 me1:H4K20 me2
H3K4 me1:H2A.Z
H3K27 me3:H4K20 me3
H2A.Z:H4–4ac
H3K27 me3:DNA Meth. m5C
H2A.Z:H3K4 me3
H2A.Z:H4K20 me2
DNA Meth. m5C:H4K20 me2
DNA Meth. m5C:H3K9 me3
H2A.Z:H3–5ac
H4K20 me2:H3–5ac
H4K20 me2:H4–4ac
H4–4ac:H3–5ac
H3K27 me3:H3K9 me3
H3K4 me3:H4–4ac
H4K20 me2:H3K4 me3
H3K27 me2
H3K9 me2
H4K20 me3
H3K4 me1
H3K9 me3
H2A.Z
DNA Meth. m5C
H3K27 me3
H3–5ac
H4–4ac
H3K4 me3
H4K20 me2
0.00 0.25 0.50 0.75 1.00
$\hat{\pi}$

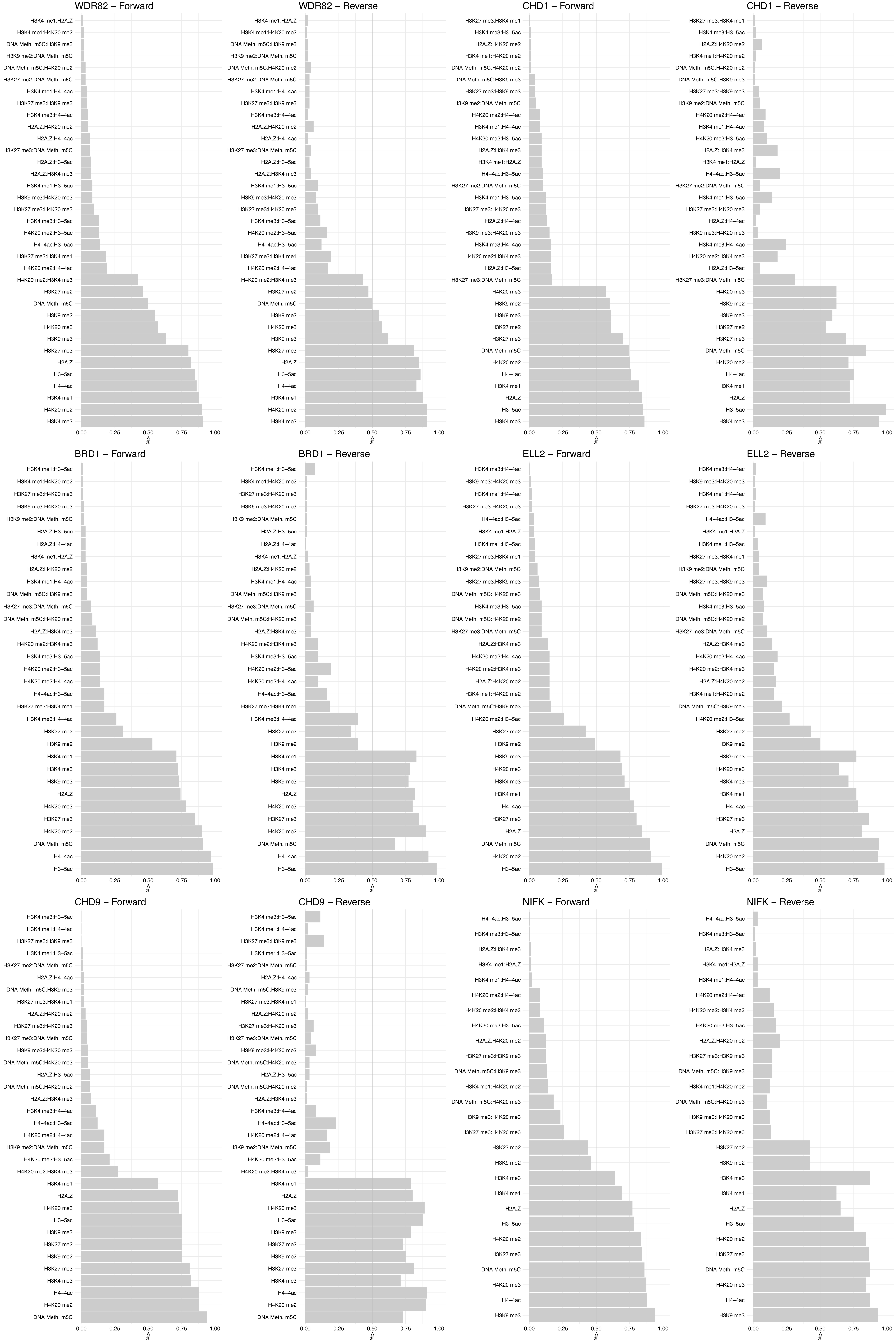
WDR82 − Forward
H3K4 me1:H2A.Z
H3K4 me1:H4K20 me2
DNA Meth. m5C:H3K9 me3
H3K9 me2:DNA Meth. m5C
DNA Meth. m5C:H4K20 me2
H3K27 me2:DNA Meth. m5C
H3K4 me1:H4−4ac
H3K27 me3:H3K9 me3
H3K4 me3:H4−4ac
H2A.Z:H4K20 me2
H2A.Z:H4−4ac
H3K27 me3:DNA Meth. m5C
H2A.Z:H3−5ac
H2A.Z:H3K4 me3
H3K4 me1:H3−5ac
H3K9 me3:H4K20 me3
H3K27 me3:H4K20 me3
H3K4 me3:H3−5ac
H4K20 me2:H3−5ac
H4−4ac:H3−5ac
H3K27 me3:H3K4 me1
H4K20 me2:H4−4ac
H4K20 me2:H3K4 me3
H3K27 me2
DNA Meth. m5C
H3K9 me2
H4K20 me3
H3K9 me3
H3K27 me3
H2A.Z
H3−5ac
H4−4ac
H3K4 me1
H4K20 me2
H3K4 me3
0.00 0.25 0.50 0.75 1.00
π̂
WDR82 − Reverse
H3K4 me1:H2A.Z
H3K4 me1:H4K20 me2
DNA Meth. m5C:H3K9 me3
H3K9 me2:DNA Meth. m5C
DNA Meth. m5C:H4K20 me2
H3K27 me2:DNA Meth. m5C
H3K4 me1:H4−4ac
H3K27 me3:H3K9 me3
H3K4 me3:H4−4ac
H2A.Z:H4K20 me2
H2A.Z:H4−4ac
H3K27 me3:DNA Meth. m5C
H2A.Z:H3−5ac
H2A.Z:H3K4 me3
H3K4 me1:H3−5ac
H3K9 me3:H4K20 me3
H3K27 me3:H4K20 me3
H3K4 me3:H3−5ac
H4K20 me2:H3−5ac
H4−4ac:H3−5ac
H3K27 me3:H3K4 me1
H4K20 me2:H4−4ac
H4K20 me2:H3K4 me3
H3K27 me2
DNA Meth. m5C
H3K9 me2
H4K20 me3
H3K9 me3
H3K27 me3
H2A.Z
H3−5ac
H4−4ac
H3K4 me1
H4K20 me2
H3K4 me3
0.00 0.25 0.50 0.75 1.00
π̂
CHD1 − Forward
H3K27 me3:H3K4 me1
H3K4 me3:H3−5ac
H2A.Z:H4K20 me2
H3K4 me1:H4K20 me2
DNA Meth. m5C:H4K20 me2
DNA Meth. m5C:H3K9 me3
H3K27 me3:H3K9 me3
H3K9 me2:DNA Meth. m5C
H4K20 me2:H4−4ac
H3K4 me1:H4−4ac
H4K20 me2:H3−5ac
H2A.Z:H3K4 me3
H3K4 me1:H2A.Z
H4−4ac:H3−5ac
H3K27 me2:DNA Meth. m5C
H3K4 me1:H3−5ac
H3K27 me3:H4K20 me3
H2A.Z:H4−4ac
H3K9 me3:H4K20 me3
H3K4 me3:H4−4ac
H4K20 me2:H3K4 me3
H2A.Z:H3−5ac
H3K27 me3:DNA Meth. m5C
H4K20 me3
H3K9 me2
H3K9 me3
H3K27 me2
H3K27 me3
DNA Meth. m5C
H4K20 me2
H4−4ac
H3K4 me1
H2A.Z
H3−5ac
H3K4 me3
0.00 0.25 0.50 0.75 1.00
π̂
CHD1 − Reverse
H3K27 me3:H3K4 me1
H3K4 me3:H3−5ac
H2A.Z:H4K20 me2
H3K4 me1:H4K20 me2
DNA Meth. m5C:H4K20 me2
DNA Meth. m5C:H3K9 me3
H3K27 me3:H3K9 me3
H3K9 me2:DNA Meth. m5C
H4K20 me2:H4−4ac
H3K4 me1:H4−4ac
H4K20 me2:H3−5ac
H2A.Z:H3K4 me3
H3K4 me1:H2A.Z
H4−4ac:H3−5ac
H3K27 me2:DNA Meth. m5C
H3K4 me1:H3−5ac
H3K27 me3:H4K20 me3
H2A.Z:H4−4ac
H3K9 me3:H4K20 me3
H3K4 me3:H4−4ac
H4K20 me2:H3K4 me3
H2A.Z:H3−5ac
H3K27 me3:DNA Meth. m5C
H4K20 me3
H3K9 me2
H3K9 me3
H3K27 me2
H3K27 me3
DNA Meth. m5C
H4K20 me2
H4−4ac
H3K4 me1
H2A.Z
H3−5ac
H3K4 me3
0.00 0.25 0.50 0.75 1.00
π̂
BRD1 − Forward
H3K4 me1:H3−5ac
H3K4 me1:H4K20 me2
H3K27 me3:H4K20 me3
H3K9 me3:H4K20 me3
H3K9 me2:DNA Meth. m5C
H2A.Z:H3−5ac
H2A.Z:H4−4ac
H3K4 me1:H2A.Z
H2A.Z:H4K20 me2
H3K4 me1:H4−4ac
DNA Meth. m5C:H3K9 me3
H3K27 me3:DNA Meth. m5C
DNA Meth. m5C:H4K20 me2
H2A.Z:H3K4 me3
H4K20 me2:H3K4 me3
H3K4 me3:H3−5ac
H4K20 me2:H3−5ac
H4K20 me2:H4−4ac
H4−4ac:H3−5ac
H3K27 me3:H3K4 me1
H3K4 me3:H4−4ac
H3K27 me2
H3K9 me2
H3K4 me1
H3K4 me3
H3K9 me3
H2A.Z
H4K20 me3
H3K27 me3
H4K20 me2
DNA Meth. m5C
H4−4ac
H3−5ac
0.00 0.25 0.50 0.75 1.00
π̂
BRD1 − Reverse
H3K4 me1:H3−5ac
H3K4 me1:H4K20 me2
H3K27 me3:H4K20 me3
H3K9 me3:H4K20 me3
H3K9 me2:DNA Meth. m5C
H2A.Z:H3−5ac
H2A.Z:H4−4ac
H3K4 me1:H2A.Z
H2A.Z:H4K20 me2
H3K4 me1:H4−4ac
DNA Meth. m5C:H3K9 me3
H3K27 me3:DNA Meth. m5C
DNA Meth. m5C:H4K20 me2
H2A.Z:H3K4 me3
H4K20 me2:H3K4 me3
H3K4 me3:H3−5ac
H4K20 me2:H3−5ac
H4K20 me2:H4−4ac
H4−4ac:H3−5ac
H3K27 me3:H3K4 me1
H3K4 me3:H4−4ac
H3K27 me2
H3K9 me2
H3K4 me1
H3K4 me3
H3K9 me3
H2A.Z
H4K20 me3
H3K27 me3
H4K20 me2
DNA Meth. m5C
H4−4ac
H3−5ac
0.00 0.25 0.50 0.75 1.00
π̂
ELL2 − Forward
H3K4 me3:H4−4ac
H3K9 me3:H4K20 me3
H3K4 me1:H4−4ac
H3K27 me3:H4K20 me3
H4−4ac:H3−5ac
H3K4 me1:H2A.Z
H3K4 me1:H3−5ac
H3K27 me3:H3K4 me1
H3K9 me2:DNA Meth. m5C
H3K27 me3:H3K9 me3
DNA Meth. m5C:H4K20 me3
H3K4 me3:H3−5ac
DNA Meth. m5C:H4K20 me2
H3K27 me3:DNA Meth. m5C
H2A.Z:H3K4 me3
H4K20 me2:H4−4ac
H4K20 me2:H3K4 me3
H2A.Z:H4K20 me2
H3K4 me1:H4K20 me2
DNA Meth. m5C:H3K9 me3
H4K20 me2:H3−5ac
H3K27 me2
H3K9 me2
H3K9 me3
H4K20 me3
H3K4 me3
H3K4 me1
H4−4ac
H3K27 me3
H2A.Z
DNA Meth. m5C
H4K20 me2
H3−5ac
0.00 0.25 0.50 0.75 1.00
π̂
ELL2 − Reverse
H3K4 me3:H4−4ac
H3K9 me3:H4K20 me3
H3K4 me1:H4−4ac
H3K27 me3:H4K20 me3
H4−4ac:H3−5ac
H3K4 me1:H2A.Z
H3K4 me1:H3−5ac
H3K27 me3:H3K4 me1
H3K9 me2:DNA Meth. m5C
H3K27 me3:H3K9 me3
DNA Meth. m5C:H4K20 me3
H3K4 me3:H3−5ac
DNA Meth. m5C:H4K20 me2
H3K27 me3:DNA Meth. m5C
H2A.Z:H3K4 me3
H4K20 me2:H4−4ac
H4K20 me2:H3K4 me3
H2A.Z:H4K20 me2
H3K4 me1:H4K20 me2
DNA Meth. m5C:H3K9 me3
H4K20 me2:H3−5ac
H3K27 me2
H3K9 me2
H3K9 me3
H4K20 me3
H3K4 me3
H3K4 me1
H4−4ac
H3K27 me3
H2A.Z
DNA Meth. m5C
H4K20 me2
H3−5ac
0.00 0.25 0.50 0.75 1.00
π̂
CHD9 − Forward
H3K4 me3:H3−5ac
H3K4 me1:H4−4ac
H3K27 me3:H3K9 me3
H3K4 me1:H3−5ac
H3K27 me2:DNA Meth. m5C
H2A.Z:H4−4ac
DNA Meth. m5C:H3K9 me3
H3K27 me3:H3K4 me1
H2A.Z:H4K20 me2
H3K27 me3:H4K20 me3
H3K27 me3:DNA Meth. m5C
H3K9 me3:H4K20 me3
DNA Meth. m5C:H4K20 me3
H2A.Z:H3−5ac
DNA Meth. m5C:H4K20 me2
H2A.Z:H3K4 me3
H3K4 me3:H4−4ac
H4−4ac:H3−5ac
H4K20 me2:H4−4ac
H3K9 me2:DNA Meth. m5C
H4K20 me2:H3−5ac
H4K20 me2:H3K4 me3
H3K4 me1
H2A.Z
H4K20 me3
H3−5ac
H3K9 me3
H3K27 me2
H3K9 me2
H3K27 me3
H3K4 me3
H4−4ac
H4K20 me2
DNA Meth. m5C
0.00 0.25 0.50 0.75 1.00
π̂
CHD9 − Reverse
H3K4 me3:H3−5ac
H3K4 me1:H4−4ac
H3K27 me3:H3K9 me3
H3K4 me1:H3−5ac
H3K27 me2:DNA Meth. m5C
H2A.Z:H4−4ac
DNA Meth. m5C:H3K9 me3
H3K27 me3:H3K4 me1
H2A.Z:H4K20 me2
H3K27 me3:H4K20 me3
H3K27 me3:DNA Meth. m5C
H3K9 me3:H4K20 me3
DNA Meth. m5C:H4K20 me3
H2A.Z:H3−5ac
DNA Meth. m5C:H4K20 me2
H2A.Z:H3K4 me3
H3K4 me3:H4−4ac
H4−4ac:H3−5ac
H4K20 me2:H4−4ac
H3K9 me2:DNA Meth. m5C
H4K20 me2:H3−5ac
H4K20 me2:H3K4 me3
H3K4 me1
H2A.Z
H4K20 me3
H3−5ac
H3K9 me3
H3K27 me2
H3K9 me2
H3K27 me3
H3K4 me3
H4−4ac
H4K20 me2
DNA Meth. m5C
0.00 0.25 0.50 0.75 1.00
π̂
NIFK − Forward
H4−4ac:H3−5ac
H3K4 me3:H3−5ac
H2A.Z:H3K4 me3
H3K4 me1:H2A.Z
H3K4 me1:H4−4ac
H4K20 me2:H4−4ac
H4K20 me2:H3K4 me3
H4K20 me2:H3−5ac
H2A.Z:H4K20 me2
H3K27 me3:H3K9 me3
DNA Meth. m5C:H3K9 me3
H3K4 me1:H4K20 me2
DNA Meth. m5C:H4K20 me3
H3K9 me3:H4K20 me3
H3K27 me3:H4K20 me3
H3K27 me2
H3K9 me2
H3K4 me3
H3K4 me1
H2A.Z
H3−5ac
H4K20 me2
H3K27 me3
DNA Meth. m5C
H4K20 me3
H4−4ac
H3K9 me3
0.00 0.25 0.50 0.75 1.00
π̂
NIFK − Reverse
H4−4ac:H3−5ac
H3K4 me3:H3−5ac
H2A.Z:H3K4 me3
H3K4 me1:H2A.Z
H3K4 me1:H4−4ac
H4K20 me2:H4−4ac
H4K20 me2:H3K4 me3
H4K20 me2:H3−5ac
H2A.Z:H4K20 me2
H3K27 me3:H3K9 me3
DNA Meth. m5C:H3K9 me3
H3K4 me1:H4K20 me2
DNA Meth. m5C:H4K20 me3
H3K9 me3:H4K20 me3
H3K27 me3:H4K20 me3
H3K27 me2
H3K9 me2
H3K4 me3
H3K4 me1
H2A.Z
H3−5ac
H4K20 me2
H3K27 me3
DNA Meth. m5C
H4K20 me3
H4−4ac
H3K9 me3
0.00 0.25 0.50 0.75 1.00
π̂

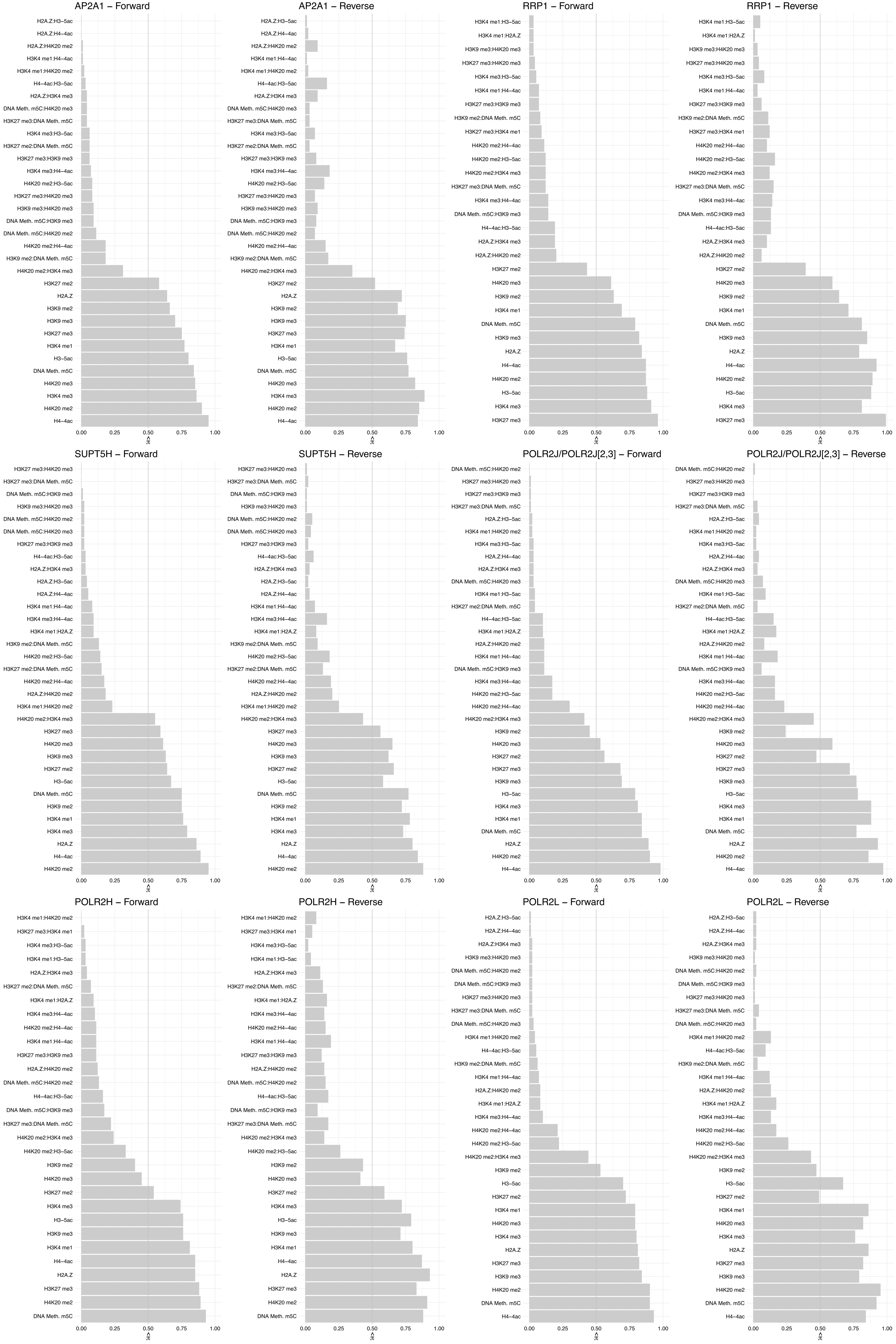

AP2A1 − Forward
H2A.Z:H3−5ac
H2A.Z:H4−4ac
H2A.Z:H4K20 me2
H3K4 me1:H4−4ac
H3K4 me1:H4K20 me2
H4−4ac:H3−5ac
H2A.Z:H3K4 me3
DNA Meth. m5C:H4K20 me3
H3K27 me3:DNA Meth. m5C
H3K4 me3:H3−5ac
H3K27 me2:DNA Meth. m5C
H3K27 me3:H3K9 me3
H3K4 me3:H4−4ac
H4K20 me2:H3−5ac
H3K27 me3:H4K20 me3
H3K9 me3:H4K20 me3
DNA Meth. m5C:H3K9 me3
DNA Meth. m5C:H4K20 me2
H4K20 me2:H4−4ac
H3K9 me2:DNA Meth. m5C
H4K20 me2:H3K4 me3
H3K27 me2
H2A.Z
H3K9 me2
H3K9 me3
H3K27 me3
H3K4 me1
H3−5ac
DNA Meth. m5C
H4K20 me3
H3K4 me3
H4K20 me2
H4−4ac
0.00 0.25 0.50 0.75 1.00
$\hat{\pi}$
AP2A1 − Reverse
H2A.Z:H3−5ac
H2A.Z:H4−4ac
H2A.Z:H4K20 me2
H3K4 me1:H4−4ac
H3K4 me1:H4K20 me2
H4−4ac:H3−5ac
H2A.Z:H3K4 me3
DNA Meth. m5C:H4K20 me3
H3K27 me3:DNA Meth. m5C
H3K4 me3:H3−5ac
H3K27 me2:DNA Meth. m5C
H3K27 me3:H3K9 me3
H3K4 me3:H4−4ac
H4K20 me2:H3−5ac
H3K27 me3:H4K20 me3
H3K9 me3:H4K20 me3
DNA Meth. m5C:H3K9 me3
DNA Meth. m5C:H4K20 me2
H4K20 me2:H4−4ac
H3K9 me2:DNA Meth. m5C
H4K20 me2:H3K4 me3
H3K27 me2
H2A.Z
H3K9 me2
H3K9 me3
H3K27 me3
H3K4 me1
H3−5ac
DNA Meth. m5C
H4K20 me3
H3K4 me3
H4K20 me2
H4−4ac
0.00 0.25 0.50 0.75 1.00
$\hat{\pi}$
RRP1 − Forward
H3K4 me1:H3−5ac
H3K4 me1:H2A.Z
H3K9 me3:H4K20 me3
H3K27 me3:H4K20 me3
H3K4 me3:H3−5ac
H3K4 me1:H4−4ac
H3K27 me3:H3K9 me3
H3K9 me2:DNA Meth. m5C
H3K27 me3:H3K4 me1
H4K20 me2:H4−4ac
H4K20 me2:H3−5ac
H4K20 me2:H3K4 me3
H3K27 me3:DNA Meth. m5C
H3K4 me3:H4−4ac
DNA Meth. m5C:H3K9 me3
H4−4ac:H3−5ac
H2A.Z:H3K4 me3
H2A.Z:H4K20 me2
H3K27 me2
H4K20 me3
H3K9 me2
H3K4 me1
DNA Meth. m5C
H3K9 me3
H2A.Z
H4−4ac
H4K20 me2
H3−5ac
H3K4 me3
H3K27 me3
0.00 0.25 0.50 0.75 1.00
$\hat{\pi}$
RRP1 − Reverse
H3K4 me1:H3−5ac
H3K4 me1:H2A.Z
H3K9 me3:H4K20 me3
H3K27 me3:H4K20 me3
H3K4 me3:H3−5ac
H3K4 me1:H4−4ac
H3K27 me3:H3K9 me3
H3K9 me2:DNA Meth. m5C
H3K27 me3:H3K4 me1
H4K20 me2:H4−4ac
H4K20 me2:H3−5ac
H4K20 me2:H3K4 me3
H3K27 me3:DNA Meth. m5C
H3K4 me3:H4−4ac
DNA Meth. m5C:H3K9 me3
H4−4ac:H3−5ac
H2A.Z:H3K4 me3
H2A.Z:H4K20 me2
H3K27 me2
H4K20 me3
H3K9 me2
H3K4 me1
DNA Meth. m5C
H3K9 me3
H2A.Z
H4−4ac
H4K20 me2
H3−5ac
H3K4 me3
H3K27 me3
0.00 0.25 0.50 0.75 1.00
$\hat{\pi}$
SUPT5H − Forward
H3K27 me3:H4K20 me3
H3K27 me3:DNA Meth. m5C
DNA Meth. m5C:H3K9 me3
H3K9 me3:H4K20 me3
DNA Meth. m5C:H4K20 me2
DNA Meth. m5C:H4K20 me3
H3K27 me3:H3K9 me3
H4−4ac:H3−5ac
H2A.Z:H3K4 me3
H2A.Z:H3−5ac
H2A.Z:H4−4ac
H3K4 me1:H4−4ac
H3K4 me3:H4−4ac
H3K4 me1:H2A.Z
H3K9 me2:DNA Meth. m5C
H4K20 me2:H3−5ac
H3K27 me2:DNA Meth. m5C
H4K20 me2:H4−4ac
H2A.Z:H4K20 me2
H3K4 me1:H4K20 me2
H4K20 me2:H3K4 me3
H3K27 me3
H4K20 me3
H3K9 me3
H3K27 me2
H3−5ac
DNA Meth. m5C
H3K9 me2
H3K4 me1
H3K4 me3
H2A.Z
H4−4ac
H4K20 me2
0.00 0.25 0.50 0.75 1.00
$\hat{\pi}$
SUPT5H − Reverse
H3K27 me3:H4K20 me3
H3K27 me3:DNA Meth. m5C
DNA Meth. m5C:H3K9 me3
H3K9 me3:H4K20 me3
DNA Meth. m5C:H4K20 me2
DNA Meth. m5C:H4K20 me3
H3K27 me3:H3K9 me3
H4−4ac:H3−5ac
H2A.Z:H3K4 me3
H2A.Z:H3−5ac
H2A.Z:H4−4ac
H3K4 me1:H4−4ac
H3K4 me3:H4−4ac
H3K4 me1:H2A.Z
H3K9 me2:DNA Meth. m5C
H4K20 me2:H3−5ac
H3K27 me2:DNA Meth. m5C
H4K20 me2:H4−4ac
H2A.Z:H4K20 me2
H3K4 me1:H4K20 me2
H4K20 me2:H3K4 me3
H3K27 me3
H4K20 me3
H3K9 me3
H3K27 me2
H3−5ac
DNA Meth. m5C
H3K9 me2
H3K4 me1
H3K4 me3
H2A.Z
H4−4ac
H4K20 me2
0.00 0.25 0.50 0.75 1.00
$\hat{\pi}$
POLR2J/POLR2J[2,3] − Forward
DNA Meth. m5C:H4K20 me2
H3K27 me3:H4K20 me3
H3K27 me3:H3K9 me3
H3K27 me3:DNA Meth. m5C
H2A.Z:H3−5ac
H3K4 me1:H4K20 me2
H3K4 me3:H3−5ac
H2A.Z:H4−4ac
H2A.Z:H3K4 me3
DNA Meth. m5C:H4K20 me3
H3K4 me1:H3−5ac
H3K27 me2:DNA Meth. m5C
H4−4ac:H3−5ac
H3K4 me1:H2A.Z
H2A.Z:H4K20 me2
H3K4 me1:H4−4ac
DNA Meth. m5C:H3K9 me3
H3K4 me3:H4−4ac
H4K20 me2:H3−5ac
H4K20 me2:H4−4ac
H4K20 me2:H3K4 me3
H3K9 me2
H4K20 me3
H3K27 me2
H3K27 me3
H3K9 me3
H3−5ac
H3K4 me3
H3K4 me1
DNA Meth. m5C
H2A.Z
H4K20 me2
H4−4ac
0.00 0.25 0.50 0.75 1.00
$\hat{\pi}$
POLR2J/POLR2J[2,3] − Reverse
DNA Meth. m5C:H4K20 me2
H3K27 me3:H4K20 me3
H3K27 me3:H3K9 me3
H3K27 me3:DNA Meth. m5C
H2A.Z:H3−5ac
H3K4 me1:H4K20 me2
H3K4 me3:H3−5ac
H2A.Z:H4−4ac
H2A.Z:H3K4 me3
DNA Meth. m5C:H4K20 me3
H3K4 me1:H3−5ac
H3K27 me2:DNA Meth. m5C
H4−4ac:H3−5ac
H3K4 me1:H2A.Z
H2A.Z:H4K20 me2
H3K4 me1:H4−4ac
DNA Meth. m5C:H3K9 me3
H3K4 me3:H4−4ac
H4K20 me2:H3−5ac
H4K20 me2:H4−4ac
H4K20 me2:H3K4 me3
H3K9 me2
H4K20 me3
H3K27 me2
H3K27 me3
H3K9 me3
H3−5ac
H3K4 me3
H3K4 me1
DNA Meth. m5C
H2A.Z
H4K20 me2
H4−4ac
0.00 0.25 0.50 0.75 1.00
$\hat{\pi}$
POLR2H − Forward
H3K4 me1:H4K20 me2
H3K27 me3:H3K4 me1
H3K4 me3:H3−5ac
H3K4 me1:H3−5ac
H2A.Z:H3K4 me3
H3K27 me2:DNA Meth. m5C
H3K4 me1:H2A.Z
H3K4 me3:H4−4ac
H4K20 me2:H4−4ac
H3K4 me1:H4−4ac
H3K27 me3:H3K9 me3
H2A.Z:H4K20 me2
DNA Meth. m5C:H4K20 me2
H4−4ac:H3−5ac
DNA Meth. m5C:H3K9 me3
H3K27 me3:DNA Meth. m5C
H4K20 me2:H3K4 me3
H4K20 me2:H3−5ac
H3K9 me2
H4K20 me3
H3K27 me2
H3K4 me3
H3−5ac
H3K9 me3
H3K4 me1
H4−4ac
H2A.Z
H3K27 me3
H4K20 me2
DNA Meth. m5C
0.00 0.25 0.50 0.75 1.00
$\hat{\pi}$
POLR2H − Reverse
H3K4 me1:H4K20 me2
H3K27 me3:H3K4 me1
H3K4 me3:H3−5ac
H3K4 me1:H3−5ac
H2A.Z:H3K4 me3
H3K27 me2:DNA Meth. m5C
H3K4 me1:H2A.Z
H3K4 me3:H4−4ac
H4K20 me2:H4−4ac
H3K4 me1:H4−4ac
H3K27 me3:H3K9 me3
H2A.Z:H4K20 me2
DNA Meth. m5C:H4K20 me2
H4−4ac:H3−5ac
DNA Meth. m5C:H3K9 me3
H3K27 me3:DNA Meth. m5C
H4K20 me2:H3K4 me3
H4K20 me2:H3−5ac
H3K9 me2
H4K20 me3
H3K27 me2
H3K4 me3
H3−5ac
H3K9 me3
H3K4 me1
H4−4ac
H2A.Z
H3K27 me3
H4K20 me2
DNA Meth. m5C
0.00 0.25 0.50 0.75 1.00
$\hat{\pi}$
POLR2L − Forward
H2A.Z:H3−5ac
H2A.Z:H4−4ac
H2A.Z:H3K4 me3
H3K9 me3:H4K20 me3
DNA Meth. m5C:H4K20 me2
DNA Meth. m5C:H3K9 me3
H3K27 me3:H4K20 me3
H3K27 me3:DNA Meth. m5C
DNA Meth. m5C:H4K20 me3
H3K4 me1:H4K20 me2
H4−4ac:H3−5ac
H3K9 me2:DNA Meth. m5C
H3K4 me1:H4−4ac
H2A.Z:H4K20 me2
H3K4 me1:H2A.Z
H3K4 me3:H4−4ac
H4K20 me2:H4−4ac
H4K20 me2:H3−5ac
H4K20 me2:H3K4 me3
H3K9 me2
H3−5ac
H3K27 me2
H3K4 me1
H4K20 me3
H3K4 me3
H2A.Z
H3K27 me3
H3K9 me3
H4K20 me2
DNA Meth. m5C
H4−4ac
0.00 0.25 0.50 0.75 1.00
$\hat{\pi}$
POLR2L − Reverse
H2A.Z:H3−5ac
H2A.Z:H4−4ac
H2A.Z:H3K4 me3
H3K9 me3:H4K20 me3
DNA Meth. m5C:H4K20 me2
DNA Meth. m5C:H3K9 me3
H3K27 me3:H4K20 me3
H3K27 me3:DNA Meth. m5C
DNA Meth. m5C:H4K20 me3
H3K4 me1:H4K20 me2
H4−4ac:H3−5ac
H3K9 me2:DNA Meth. m5C
H3K4 me1:H4−4ac
H2A.Z:H4K20 me2
H3K4 me1:H2A.Z
H3K4 me3:H4−4ac
H4K20 me2:H4−4ac
H4K20 me2:H3−5ac
H4K20 me2:H3K4 me3
H3K9 me2
H3−5ac
H3K27 me2
H3K4 me1
H4K20 me3
H3K4 me3
H2A.Z
H3K27 me3
H3K9 me3
H4K20 me2
DNA Meth. m5C
H4−4ac
0.00 0.25 0.50 0.75 1.00
$\hat{\pi}$

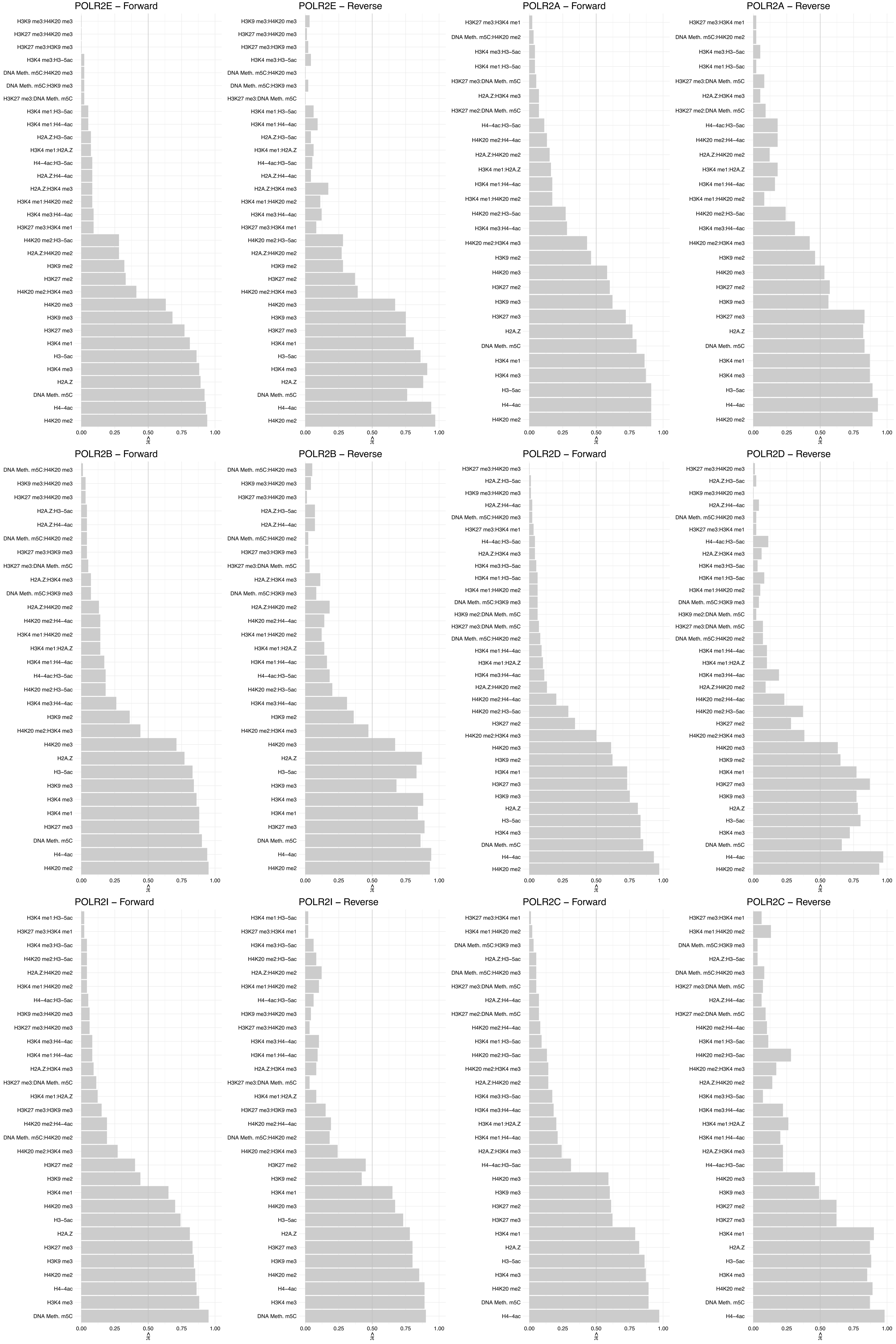

POLR2E – Forward
POLR2E – Reverse
H3K9 me3:H4K20 me3
H3K27 me3:H4K20 me3
H3K27 me3:H3K9 me3
H3K4 me3:H3−5ac
DNA Meth. m5C:H4K20 me3
DNA Meth. m5C:H3K9 me3
H3K27 me3:DNA Meth. m5C
H3K4 me1:H3−5ac
H3K4 me1:H4−4ac
H2A.Z:H3−5ac
H3K4 me1:H2A.Z
H4−4ac:H3−5ac
H2A.Z:H4−4ac
H2A.Z:H3K4 me3
H3K4 me1:H4K20 me2
H3K4 me3:H4−4ac
H3K27 me3:H3K4 me1
H4K20 me2:H3−5ac
H2A.Z:H4K20 me2
H3K9 me2
H3K27 me2
H4K20 me2:H3K4 me3
H4K20 me3
H3K9 me3
H3K27 me3
H3K4 me1
H3−5ac
H3K4 me3
H2A.Z
DNA Meth. m5C
H4−4ac
H4K20 me2
POLR2A – Forward
POLR2A – Reverse
H3K27 me3:H3K4 me1
DNA Meth. m5C:H4K20 me2
H3K4 me3:H3−5ac
H3K4 me1:H3−5ac
H3K27 me3:DNA Meth. m5C
H2A.Z:H3K4 me3
H3K27 me2:DNA Meth. m5C
H4−4ac:H3−5ac
H4K20 me2:H4−4ac
H2A.Z:H4K20 me2
H3K4 me1:H2A.Z
H3K4 me1:H4−4ac
H3K4 me1:H4K20 me2
H4K20 me2:H3−5ac
H3K4 me3:H4−4ac
H4K20 me2:H3K4 me3
H3K9 me2
H4K20 me3
H3K27 me2
H3K9 me3
H3K27 me3
H2A.Z
DNA Meth. m5C
H3K4 me1
H3K4 me3
H3−5ac
H4−4ac
H4K20 me2
POLR2B – Forward
POLR2B – Reverse
DNA Meth. m5C:H4K20 me3
H3K9 me3:H4K20 me3
H3K27 me3:H4K20 me3
H2A.Z:H3−5ac
H2A.Z:H4−4ac
DNA Meth. m5C:H4K20 me2
H3K27 me3:H3K9 me3
H3K27 me3:DNA Meth. m5C
H2A.Z:H3K4 me3
DNA Meth. m5C:H3K9 me3
H2A.Z:H4K20 me2
H4K20 me2:H4−4ac
H3K4 me1:H4K20 me2
H3K4 me1:H2A.Z
H3K4 me1:H4−4ac
H4−4ac:H3−5ac
H4K20 me2:H3−5ac
H3K4 me3:H4−4ac
H3K9 me2
H4K20 me2:H3K4 me3
H4K20 me3
H2A.Z
H3−5ac
H3K9 me3
H3K4 me3
H3K4 me1
H3K27 me3
DNA Meth. m5C
H4−4ac
H4K20 me2
POLR2D – Forward
POLR2D – Reverse
H3K27 me3:H4K20 me3
H2A.Z:H3−5ac
H3K9 me3:H4K20 me3
H2A.Z:H4−4ac
DNA Meth. m5C:H4K20 me3
H3K27 me3:H3K4 me1
H4−4ac:H3−5ac
H2A.Z:H3K4 me3
H3K4 me3:H3−5ac
H3K4 me1:H3−5ac
H3K4 me1:H4K20 me2
DNA Meth. m5C:H3K9 me3
H3K9 me2:DNA Meth. m5C
H3K27 me3:DNA Meth. m5C
DNA Meth. m5C:H4K20 me2
H3K4 me1:H4−4ac
H3K4 me1:H2A.Z
H3K4 me3:H4−4ac
H2A.Z:H4K20 me2
H4K20 me2:H4−4ac
H4K20 me2:H3−5ac
H3K27 me2
H4K20 me2:H3K4 me3
H4K20 me3
H3K9 me2
H3K4 me1
H3K27 me3
H3K9 me3
H2A.Z
H3−5ac
H3K4 me3
DNA Meth. m5C
H4−4ac
H4K20 me2
POLR2I – Forward
POLR2I – Reverse
H3K4 me1:H3−5ac
H3K27 me3:H3K4 me1
H3K4 me3:H3−5ac
H4K20 me2:H3−5ac
H2A.Z:H4K20 me2
H3K4 me1:H4K20 me2
H4−4ac:H3−5ac
H3K9 me3:H4K20 me3
H3K27 me3:H4K20 me3
H3K4 me3:H4−4ac
H3K4 me1:H4−4ac
H2A.Z:H3K4 me3
H3K27 me3:DNA Meth. m5C
H3K4 me1:H2A.Z
H3K27 me3:H3K9 me3
H4K20 me2:H4−4ac
DNA Meth. m5C:H4K20 me2
H4K20 me2:H3K4 me3
H3K27 me2
H3K9 me2
H3K4 me1
H4K20 me3
H3−5ac
H2A.Z
H3K27 me3
H3K9 me3
H4K20 me2
H4−4ac
H3K4 me3
DNA Meth. m5C
POLR2C – Forward
POLR2C – Reverse
H3K27 me3:H3K4 me1
H3K4 me1:H4K20 me2
DNA Meth. m5C:H3K9 me3
H2A.Z:H3−5ac
DNA Meth. m5C:H4K20 me3
H3K27 me3:DNA Meth. m5C
H2A.Z:H4−4ac
H3K27 me2:DNA Meth. m5C
H4K20 me2:H4−4ac
H3K4 me1:H3−5ac
H4K20 me2:H3−5ac
H4K20 me2:H3K4 me3
H2A.Z:H4K20 me2
H3K4 me3:H3−5ac
H3K4 me3:H4−4ac
H3K4 me1:H2A.Z
H3K4 me1:H4−4ac
H2A.Z:H3K4 me3
H4−4ac:H3−5ac
H4K20 me3
H3K9 me3
H3K27 me2
H3K27 me3
H3K4 me1
H2A.Z
H3−5ac
H3K4 me3
H4K20 me2
DNA Meth. m5C
H4−4ac
0.00
0.25
0.50
0.75
1.00
$\hat{\pi}$

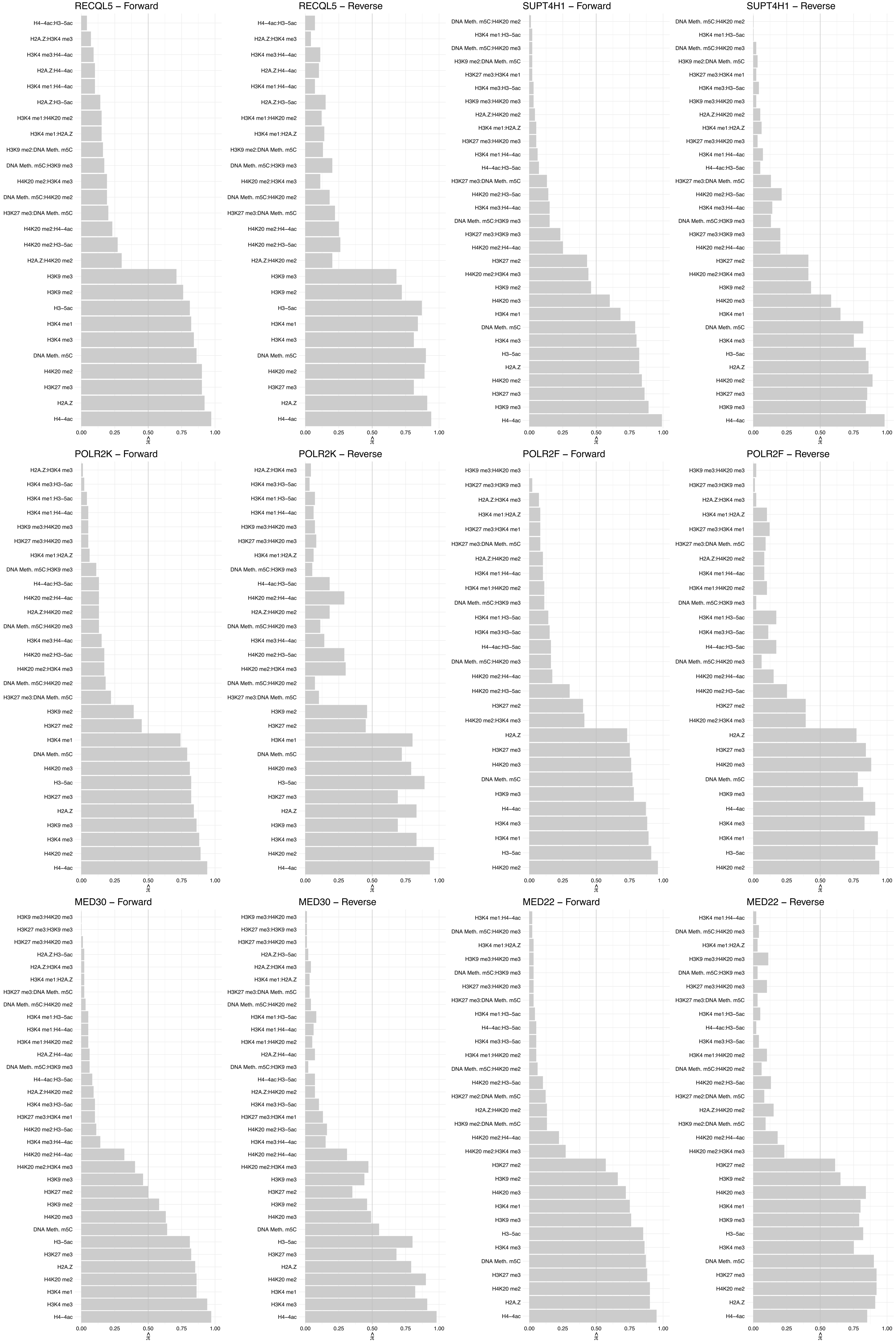
RECQL5 – Forward
H4–4ac:H3–5ac
H2A.Z:H3K4 me3
H3K4 me3:H4–4ac
H2A.Z:H4–4ac
H3K4 me1:H4–4ac
H2A.Z:H3–5ac
H3K4 me1:H4K20 me2
H3K4 me1:H2A.Z
H3K9 me2:DNA Meth. m5C
DNA Meth. m5C:H3K9 me3
H4K20 me2:H3K4 me3
DNA Meth. m5C:H4K20 me2
H3K27 me3:DNA Meth. m5C
H4K20 me2:H4–4ac
H4K20 me2:H3–5ac
H2A.Z:H4K20 me2
H3K9 me3
H3K9 me2
H3–5ac
H3K4 me1
H3K4 me3
DNA Meth. m5C
H4K20 me2
H3K27 me3
H2A.Z
H4–4ac
0.00
0.25
0.50
0.75
1.00
π̂
RECQL5 – Reverse
SUPT4H1 – Forward
DNA Meth. m5C:H4K20 me2
H3K4 me1:H3–5ac
DNA Meth. m5C:H4K20 me3
H3K9 me2:DNA Meth. m5C
H3K27 me3:H3K4 me1
H3K4 me3:H3–5ac
H3K9 me3:H4K20 me3
H2A.Z:H4K20 me2
H3K4 me1:H2A.Z
H3K27 me3:H4K20 me3
H3K4 me1:H4–4ac
H4–4ac:H3–5ac
H3K27 me3:DNA Meth. m5C
H4K20 me2:H3–5ac
H3K4 me3:H4–4ac
DNA Meth. m5C:H3K9 me3
H3K27 me3:H3K9 me3
H4K20 me2:H4–4ac
H3K27 me2
H4K20 me2:H3K4 me3
H3K9 me2
H4K20 me3
H3K4 me1
DNA Meth. m5C
H3K4 me3
H3–5ac
H2A.Z
H4K20 me2
H3K27 me3
H3K9 me3
H4–4ac
SUPT4H1 – Reverse
POLR2K – Forward
H2A.Z:H3K4 me3
H3K4 me3:H3–5ac
H3K4 me1:H3–5ac
H3K4 me1:H4–4ac
H3K9 me3:H4K20 me3
H3K27 me3:H4K20 me3
H3K4 me1:H2A.Z
DNA Meth. m5C:H3K9 me3
H4–4ac:H3–5ac
H4K20 me2:H4–4ac
H2A.Z:H4K20 me2
DNA Meth. m5C:H4K20 me3
H3K4 me3:H4–4ac
H4K20 me2:H3–5ac
H4K20 me2:H3K4 me3
DNA Meth. m5C:H4K20 me2
H3K27 me3:DNA Meth. m5C
H3K9 me2
H3K27 me2
H3K4 me1
DNA Meth. m5C
H4K20 me3
H3–5ac
H3K27 me3
H2A.Z
H3K9 me3
H3K4 me3
H4K20 me2
H4–4ac
POLR2K – Reverse
POLR2F – Forward
H3K9 me3:H4K20 me3
H3K27 me3:H3K9 me3
H2A.Z:H3K4 me3
H3K4 me1:H2A.Z
H3K27 me3:H3K4 me1
H3K27 me3:DNA Meth. m5C
H2A.Z:H4K20 me2
H3K4 me1:H4–4ac
H3K4 me1:H4K20 me2
DNA Meth. m5C:H3K9 me3
H3K4 me1:H3–5ac
H3K4 me3:H3–5ac
H4–4ac:H3–5ac
DNA Meth. m5C:H4K20 me3
H4K20 me2:H4–4ac
H4K20 me2:H3–5ac
H3K27 me2
H4K20 me2:H3K4 me3
H2A.Z
H3K27 me3
H4K20 me3
DNA Meth. m5C
H3K9 me3
H4–4ac
H3K4 me3
H3K4 me1
H3–5ac
H4K20 me2
POLR2F – Reverse
MED30 – Forward
H3K9 me3:H4K20 me3
H3K27 me3:H3K9 me3
H3K27 me3:H4K20 me3
H2A.Z:H3–5ac
H2A.Z:H3K4 me3
H3K4 me1:H2A.Z
H3K27 me3:DNA Meth. m5C
DNA Meth. m5C:H4K20 me2
H3K4 me1:H3–5ac
H3K4 me1:H4–4ac
H3K4 me1:H4K20 me2
H2A.Z:H4–4ac
DNA Meth. m5C:H3K9 me3
H4–4ac:H3–5ac
H2A.Z:H4K20 me2
H3K4 me3:H3–5ac
H3K27 me3:H3K4 me1
H4K20 me2:H3–5ac
H3K4 me3:H4–4ac
H4K20 me2:H4–4ac
H4K20 me2:H3K4 me3
H3K9 me3
H3K27 me2
H3K9 me2
H4K20 me3
DNA Meth. m5C
H3–5ac
H3K27 me3
H2A.Z
H4K20 me2
H3K4 me1
H3K4 me3
H4–4ac
MED30 – Reverse
MED22 – Forward
H3K4 me1:H4–4ac
DNA Meth. m5C:H4K20 me3
H3K4 me1:H2A.Z
H3K9 me3:H4K20 me3
DNA Meth. m5C:H3K9 me3
H3K27 me3:H4K20 me3
H3K27 me3:DNA Meth. m5C
H3K4 me1:H3–5ac
H4–4ac:H3–5ac
H3K4 me3:H3–5ac
H3K4 me1:H4K20 me2
DNA Meth. m5C:H4K20 me2
H4K20 me2:H3–5ac
H3K27 me2:DNA Meth. m5C
H2A.Z:H4K20 me2
H3K9 me2:DNA Meth. m5C
H4K20 me2:H4–4ac
H4K20 me2:H3K4 me3
H3K27 me2
H3K9 me2
H4K20 me3
H3K4 me1
H3K9 me3
H3–5ac
H3K4 me3
DNA Meth. m5C
H3K27 me3
H4K20 me2
H2A.Z
H4–4ac
MED22 – Reverse

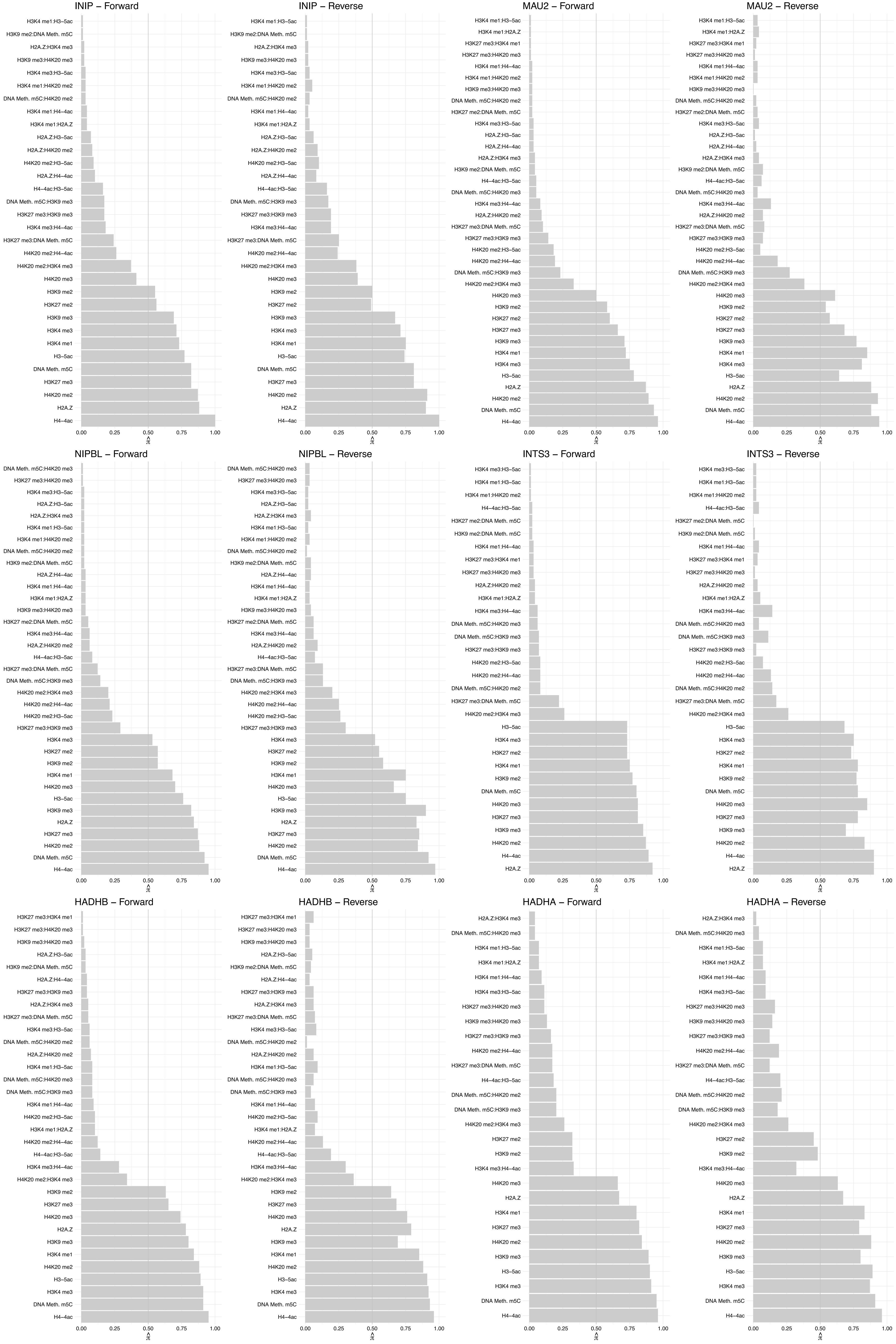
INIP – Forward
H3K4 me1:H3−5ac
H3K9 me2:DNA Meth. m5C
H2A.Z:H3K4 me3
H3K9 me3:H4K20 me3
H3K4 me3:H3−5ac
H3K4 me1:H4K20 me2
DNA Meth. m5C:H4K20 me2
H3K4 me1:H4−4ac
H3K4 me1:H2A.Z
H2A.Z:H3−5ac
H2A.Z:H4K20 me2
H4K20 me2:H3−5ac
H2A.Z:H4−4ac
H4−4ac:H3−5ac
DNA Meth. m5C:H3K9 me3
H3K27 me3:H3K9 me3
H3K4 me3:H4−4ac
H3K27 me3:DNA Meth. m5C
H4K20 me2:H4−4ac
H4K20 me2:H3K4 me3
H4K20 me3
H3K9 me2
H3K27 me2
H3K9 me3
H3K4 me3
H3K4 me1
H3−5ac
DNA Meth. m5C
H3K27 me3
H4K20 me2
H2A.Z
H4−4ac
0.00
0.25
0.50
0.75
1.00
π̂
INIP – Reverse
H3K4 me1:H3−5ac
H3K9 me2:DNA Meth. m5C
H2A.Z:H3K4 me3
H3K9 me3:H4K20 me3
H3K4 me3:H3−5ac
H3K4 me1:H4K20 me2
DNA Meth. m5C:H4K20 me2
H3K4 me1:H4−4ac
H3K4 me1:H2A.Z
H2A.Z:H3−5ac
H2A.Z:H4K20 me2
H4K20 me2:H3−5ac
H2A.Z:H4−4ac
H4−4ac:H3−5ac
DNA Meth. m5C:H3K9 me3
H3K27 me3:H3K9 me3
H3K4 me3:H4−4ac
H3K27 me3:DNA Meth. m5C
H4K20 me2:H4−4ac
H4K20 me2:H3K4 me3
H4K20 me3
H3K9 me2
H3K27 me2
H3K9 me3
H3K4 me3
H3K4 me1
H3−5ac
DNA Meth. m5C
H3K27 me3
H4K20 me2
H2A.Z
H4−4ac
MAU2 – Forward
H3K4 me1:H3−5ac
H3K4 me1:H2A.Z
H3K27 me3:H3K4 me1
H3K27 me3:H4K20 me3
H3K4 me1:H4−4ac
H3K4 me1:H4K20 me2
H3K9 me3:H4K20 me3
DNA Meth. m5C:H4K20 me2
H3K27 me2:DNA Meth. m5C
H3K4 me3:H3−5ac
H2A.Z:H3−5ac
H2A.Z:H4−4ac
H2A.Z:H3K4 me3
H3K9 me2:DNA Meth. m5C
H4−4ac:H3−5ac
DNA Meth. m5C:H4K20 me3
H3K4 me3:H4−4ac
H2A.Z:H4K20 me2
H3K27 me3:DNA Meth. m5C
H3K27 me3:H3K9 me3
H4K20 me2:H3−5ac
H4K20 me2:H4−4ac
DNA Meth. m5C:H3K9 me3
H4K20 me2:H3K4 me3
H4K20 me3
H3K9 me2
H3K27 me2
H3K27 me3
H3K9 me3
H3K4 me1
H3K4 me3
H3−5ac
H2A.Z
H4K20 me2
DNA Meth. m5C
H4−4ac
MAU2 – Reverse
H3K4 me1:H3−5ac
H3K4 me1:H2A.Z
H3K27 me3:H3K4 me1
H3K27 me3:H4K20 me3
H3K4 me1:H4−4ac
H3K4 me1:H4K20 me2
H3K9 me3:H4K20 me3
DNA Meth. m5C:H4K20 me2
H3K27 me2:DNA Meth. m5C
H3K4 me3:H3−5ac
H2A.Z:H3−5ac
H2A.Z:H4−4ac
H2A.Z:H3K4 me3
H3K9 me2:DNA Meth. m5C
H4−4ac:H3−5ac
DNA Meth. m5C:H4K20 me3
H3K4 me3:H4−4ac
H2A.Z:H4K20 me2
H3K27 me3:DNA Meth. m5C
H3K27 me3:H3K9 me3
H4K20 me2:H3−5ac
H4K20 me2:H4−4ac
DNA Meth. m5C:H3K9 me3
H4K20 me2:H3K4 me3
H4K20 me3
H3K9 me2
H3K27 me2
H3K27 me3
H3K9 me3
H3K4 me1
H3K4 me3
H3−5ac
H2A.Z
H4K20 me2
DNA Meth. m5C
H4−4ac
NIPBL – Forward
DNA Meth. m5C:H4K20 me3
H3K27 me3:H4K20 me3
H3K4 me3:H3−5ac
H2A.Z:H3−5ac
H2A.Z:H3K4 me3
H3K4 me1:H3−5ac
H3K4 me1:H4K20 me2
DNA Meth. m5C:H4K20 me2
H3K9 me2:DNA Meth. m5C
H2A.Z:H4−4ac
H3K4 me1:H4−4ac
H3K4 me1:H2A.Z
H3K9 me3:H4K20 me3
H3K27 me2:DNA Meth. m5C
H3K4 me3:H4−4ac
H2A.Z:H4K20 me2
H4−4ac:H3−5ac
H3K27 me3:DNA Meth. m5C
DNA Meth. m5C:H3K9 me3
H4K20 me2:H3K4 me3
H4K20 me2:H4−4ac
H4K20 me2:H3−5ac
H3K27 me3:H3K9 me3
H3K4 me3
H3K27 me2
H3K9 me2
H3K4 me1
H4K20 me3
H3−5ac
H3K9 me3
H2A.Z
H3K27 me3
H4K20 me2
DNA Meth. m5C
H4−4ac
NIPBL – Reverse
DNA Meth. m5C:H4K20 me3
H3K27 me3:H4K20 me3
H3K4 me3:H3−5ac
H2A.Z:H3−5ac
H2A.Z:H3K4 me3
H3K4 me1:H3−5ac
H3K4 me1:H4K20 me2
DNA Meth. m5C:H4K20 me2
H3K9 me2:DNA Meth. m5C
H2A.Z:H4−4ac
H3K4 me1:H4−4ac
H3K4 me1:H2A.Z
H3K9 me3:H4K20 me3
H3K27 me2:DNA Meth. m5C
H3K4 me3:H4−4ac
H2A.Z:H4K20 me2
H4−4ac:H3−5ac
H3K27 me3:DNA Meth. m5C
DNA Meth. m5C:H3K9 me3
H4K20 me2:H3K4 me3
H4K20 me2:H4−4ac
H4K20 me2:H3−5ac
H3K27 me3:H3K9 me3
H3K4 me3
H3K27 me2
H3K9 me2
H3K4 me1
H4K20 me3
H3−5ac
H3K9 me3
H2A.Z
H3K27 me3
H4K20 me2
DNA Meth. m5C
H4−4ac
INTS3 – Forward
H3K4 me3:H3−5ac
H3K4 me1:H3−5ac
H3K4 me1:H4K20 me2
H4−4ac:H3−5ac
H3K27 me2:DNA Meth. m5C
H3K9 me2:DNA Meth. m5C
H3K4 me1:H4−4ac
H3K27 me3:H3K4 me1
H3K27 me3:H4K20 me3
H2A.Z:H4K20 me2
H3K4 me1:H2A.Z
H3K4 me3:H4−4ac
DNA Meth. m5C:H4K20 me3
DNA Meth. m5C:H3K9 me3
H3K27 me3:H3K9 me3
H4K20 me2:H3−5ac
H4K20 me2:H4−4ac
DNA Meth. m5C:H4K20 me2
H3K27 me3:DNA Meth. m5C
H4K20 me2:H3K4 me3
H3−5ac
H3K4 me3
H3K27 me2
H3K4 me1
H3K9 me2
DNA Meth. m5C
H4K20 me3
H3K27 me3
H3K9 me3
H4K20 me2
H4−4ac
H2A.Z
INTS3 – Reverse
H3K4 me3:H3−5ac
H3K4 me1:H3−5ac
H3K4 me1:H4K20 me2
H4−4ac:H3−5ac
H3K27 me2:DNA Meth. m5C
H3K9 me2:DNA Meth. m5C
H3K4 me1:H4−4ac
H3K27 me3:H3K4 me1
H3K27 me3:H4K20 me3
H2A.Z:H4K20 me2
H3K4 me1:H2A.Z
H3K4 me3:H4−4ac
DNA Meth. m5C:H4K20 me3
DNA Meth. m5C:H3K9 me3
H3K27 me3:H3K9 me3
H4K20 me2:H3−5ac
H4K20 me2:H4−4ac
DNA Meth. m5C:H4K20 me2
H3K27 me3:DNA Meth. m5C
H4K20 me2:H3K4 me3
H3−5ac
H3K4 me3
H3K27 me2
H3K4 me1
H3K9 me2
DNA Meth. m5C
H4K20 me3
H3K27 me3
H3K9 me3
H4K20 me2
H4−4ac
H2A.Z
HADHB – Forward
H3K27 me3:H3K4 me1
H3K27 me3:H4K20 me3
H3K9 me3:H4K20 me3
H2A.Z:H3−5ac
H3K9 me2:DNA Meth. m5C
H2A.Z:H4−4ac
H3K27 me3:H3K9 me3
H2A.Z:H3K4 me3
H3K27 me3:DNA Meth. m5C
H3K4 me3:H3−5ac
DNA Meth. m5C:H4K20 me2
H2A.Z:H4K20 me2
H3K4 me1:H3−5ac
DNA Meth. m5C:H4K20 me3
DNA Meth. m5C:H3K9 me3
H3K4 me1:H4−4ac
H4K20 me2:H3−5ac
H3K4 me1:H2A.Z
H4K20 me2:H4−4ac
H4−4ac:H3−5ac
H3K4 me3:H4−4ac
H4K20 me2:H3K4 me3
H3K9 me2
H3K27 me3
H4K20 me3
H2A.Z
H3K9 me3
H3K4 me1
H4K20 me2
H3−5ac
H3K4 me3
DNA Meth. m5C
H4−4ac
HADHB – Reverse
H3K27 me3:H3K4 me1
H3K27 me3:H4K20 me3
H3K9 me3:H4K20 me3
H2A.Z:H3−5ac
H3K9 me2:DNA Meth. m5C
H2A.Z:H4−4ac
H3K27 me3:H3K9 me3
H2A.Z:H3K4 me3
H3K27 me3:DNA Meth. m5C
H3K4 me3:H3−5ac
DNA Meth. m5C:H4K20 me2
H2A.Z:H4K20 me2
H3K4 me1:H3−5ac
DNA Meth. m5C:H4K20 me3
DNA Meth. m5C:H3K9 me3
H3K4 me1:H4−4ac
H4K20 me2:H3−5ac
H3K4 me1:H2A.Z
H4K20 me2:H4−4ac
H4−4ac:H3−5ac
H3K4 me3:H4−4ac
H4K20 me2:H3K4 me3
H3K9 me2
H3K27 me3
H4K20 me3
H2A.Z
H3K9 me3
H3K4 me1
H4K20 me2
H3−5ac
H3K4 me3
DNA Meth. m5C
H4−4ac
HADHA – Forward
H2A.Z:H3K4 me3
DNA Meth. m5C:H4K20 me3
H3K4 me1:H3−5ac
H3K4 me1:H2A.Z
H3K4 me1:H4−4ac
H3K4 me3:H3−5ac
H3K27 me3:H4K20 me3
H3K9 me3:H4K20 me3
H3K27 me3:H3K9 me3
H4K20 me2:H4−4ac
H3K27 me3:DNA Meth. m5C
H4−4ac:H3−5ac
DNA Meth. m5C:H4K20 me2
DNA Meth. m5C:H3K9 me3
H4K20 me2:H3K4 me3
H3K27 me2
H3K9 me2
H3K4 me3:H4−4ac
H4K20 me3
H2A.Z
H3K4 me1
H3K27 me3
H4K20 me2
H3K9 me3
H3−5ac
H3K4 me3
DNA Meth. m5C
H4−4ac
HADHA – Reverse
H2A.Z:H3K4 me3
DNA Meth. m5C:H4K20 me3
H3K4 me1:H3−5ac
H3K4 me1:H2A.Z
H3K4 me1:H4−4ac
H3K4 me3:H3−5ac
H3K27 me3:H4K20 me3
H3K9 me3:H4K20 me3
H3K27 me3:H3K9 me3
H4K20 me2:H4−4ac
H3K27 me3:DNA Meth. m5C
H4−4ac:H3−5ac
DNA Meth. m5C:H4K20 me2
DNA Meth. m5C:H3K9 me3
H4K20 me2:H3K4 me3
H3K27 me2
H3K9 me2
H3K4 me3:H4−4ac
H4K20 me3
H2A.Z
H3K4 me1
H3K27 me3
H4K20 me2
H3K9 me3
H3−5ac
H3K4 me3
DNA Meth. m5C
H4−4ac

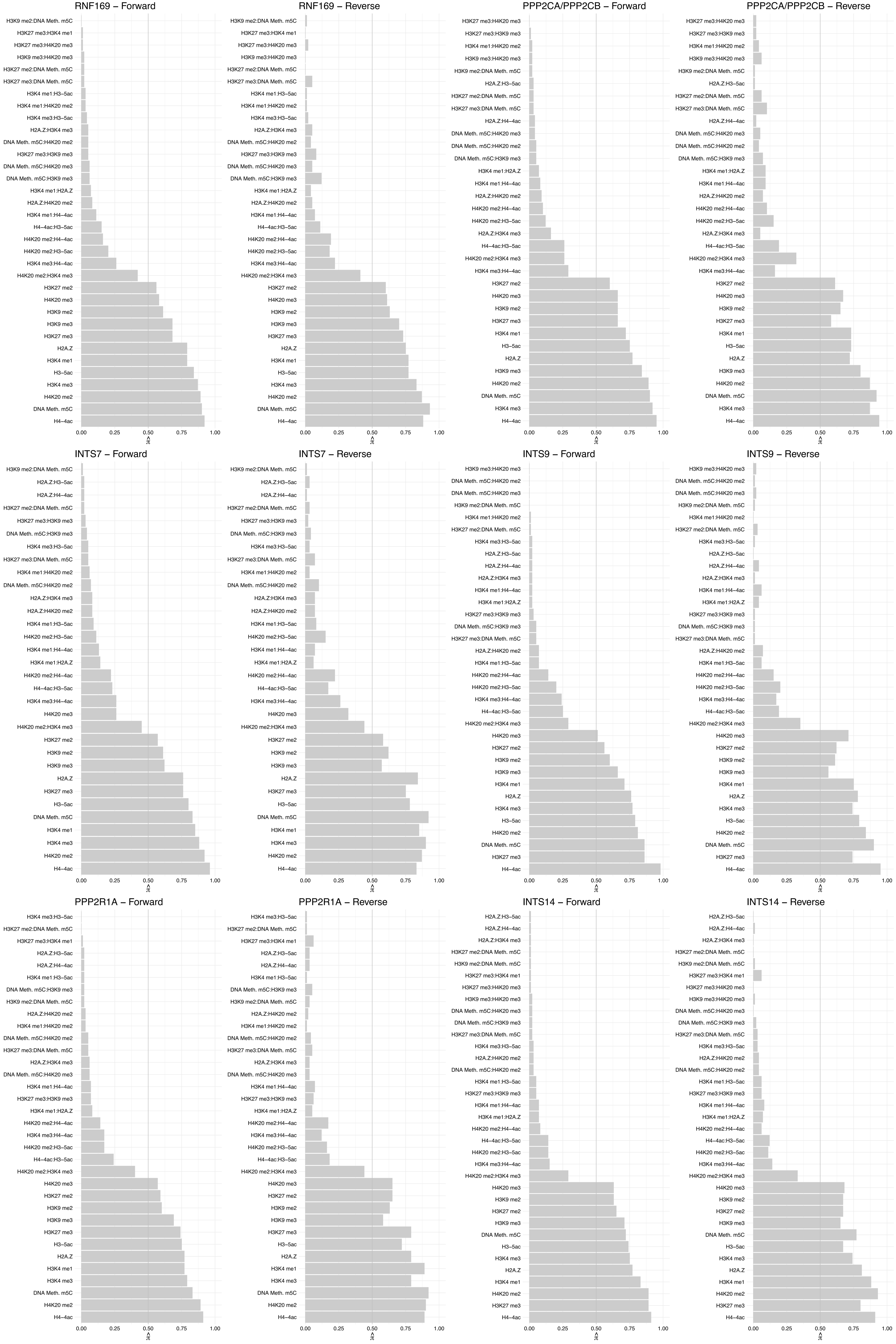
RNF169 – Forward
RNF169 – Reverse
PPP2CA/PPP2CB – Forward
PPP2CA/PPP2CB – Reverse
INTS7 – Forward
INTS7 – Reverse
INTS9 – Forward
INTS9 – Reverse
PPP2R1A – Forward
PPP2R1A – Reverse
INTS14 – Forward
INTS14 – Reverse

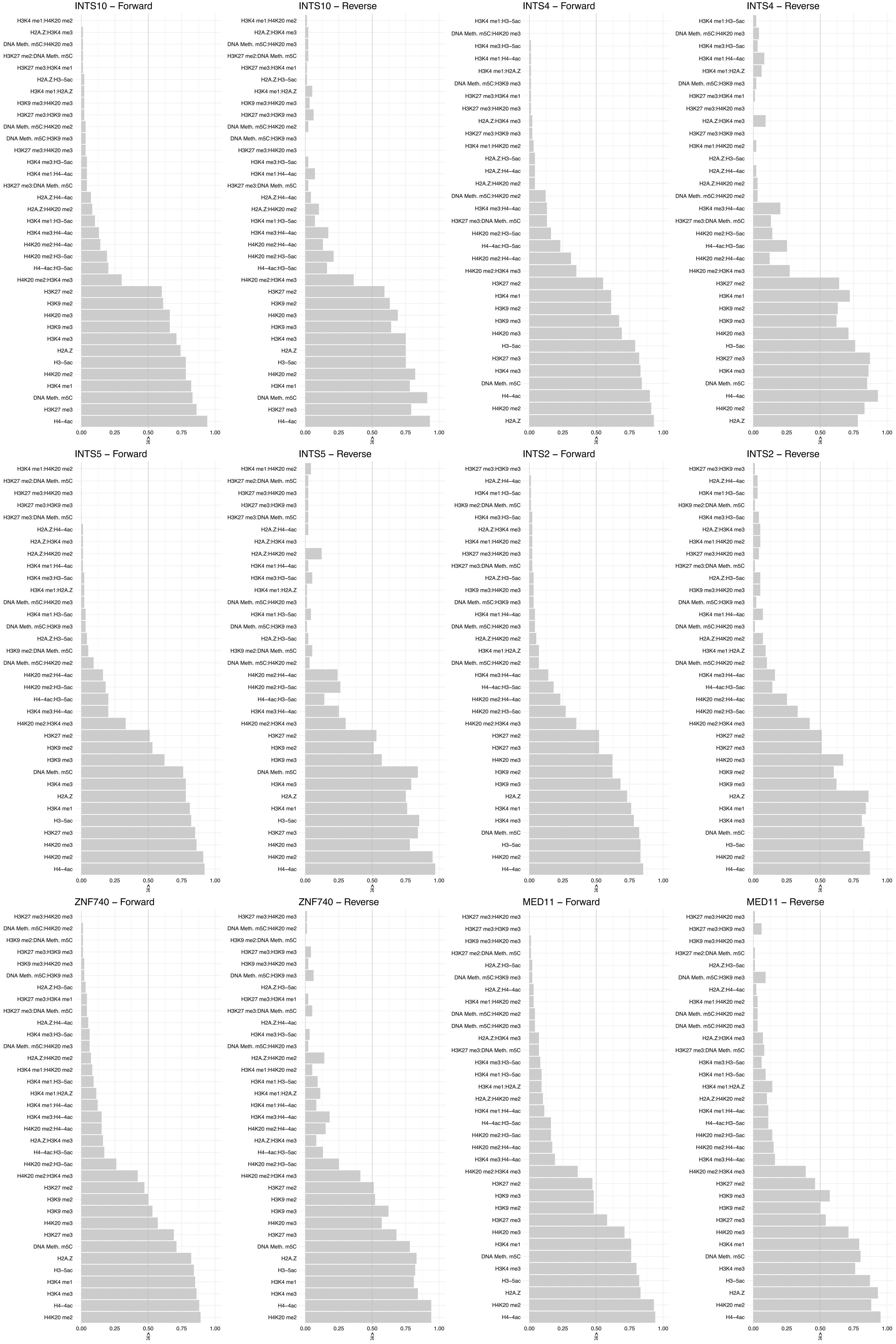
INTS10 – Forward
H3K4 me1:H4K20 me2
H2A.Z:H3K4 me3
DNA Meth. m5C:H4K20 me3
H3K27 me2:DNA Meth. m5C
H3K27 me3:H3K4 me1
H2A.Z:H3−5ac
H3K4 me1:H2A.Z
H3K9 me3:H4K20 me3
H3K27 me3:H3K9 me3
DNA Meth. m5C:H4K20 me2
DNA Meth. m5C:H3K9 me3
H3K27 me3:H4K20 me3
H3K4 me3:H3−5ac
H3K4 me1:H4−4ac
H3K27 me3:DNA Meth. m5C
H2A.Z:H4−4ac
H2A.Z:H4K20 me2
H3K4 me1:H3−5ac
H3K4 me3:H4−4ac
H4K20 me2:H4−4ac
H4K20 me2:H3−5ac
H4−4ac:H3−5ac
H4K20 me2:H3K4 me3
H3K27 me2
H3K9 me2
H4K20 me3
H3K9 me3
H3K4 me3
H2A.Z
H3−5ac
H4K20 me2
H3K4 me1
DNA Meth. m5C
H3K27 me3
H4−4ac
0.00 0.25 0.50 0.75 1.00
$\hat{\pi}$
INTS10 – Reverse
H3K4 me1:H4K20 me2
H2A.Z:H3K4 me3
DNA Meth. m5C:H4K20 me3
H3K27 me2:DNA Meth. m5C
H3K27 me3:H3K4 me1
H2A.Z:H3−5ac
H3K4 me1:H2A.Z
H3K9 me3:H4K20 me3
H3K27 me3:H3K9 me3
DNA Meth. m5C:H4K20 me2
DNA Meth. m5C:H3K9 me3
H3K27 me3:H4K20 me3
H3K4 me3:H3−5ac
H3K4 me1:H4−4ac
H3K27 me3:DNA Meth. m5C
H2A.Z:H4−4ac
H2A.Z:H4K20 me2
H3K4 me1:H3−5ac
H3K4 me3:H4−4ac
H4K20 me2:H4−4ac
H4K20 me2:H3−5ac
H4−4ac:H3−5ac
H4K20 me2:H3K4 me3
H3K27 me2
H3K9 me2
H4K20 me3
H3K9 me3
H3K4 me3
H2A.Z
H3−5ac
H4K20 me2
H3K4 me1
DNA Meth. m5C
H3K27 me3
H4−4ac
0.00 0.25 0.50 0.75 1.00
$\hat{\pi}$
INTS4 – Forward
H3K4 me1:H3−5ac
DNA Meth. m5C:H4K20 me3
H3K4 me3:H3−5ac
H3K4 me1:H4−4ac
H3K4 me1:H2A.Z
DNA Meth. m5C:H3K9 me3
H3K27 me3:H3K4 me1
H3K27 me3:H4K20 me3
H2A.Z:H3K4 me3
H3K27 me3:H3K9 me3
H3K4 me1:H4K20 me2
H2A.Z:H3−5ac
H2A.Z:H4−4ac
H2A.Z:H4K20 me2
DNA Meth. m5C:H4K20 me2
H3K4 me3:H4−4ac
H3K27 me3:DNA Meth. m5C
H4K20 me2:H3−5ac
H4−4ac:H3−5ac
H4K20 me2:H4−4ac
H4K20 me2:H3K4 me3
H3K27 me2
H3K4 me1
H3K9 me2
H3K9 me3
H4K20 me3
H3−5ac
H3K27 me3
H3K4 me3
DNA Meth. m5C
H4−4ac
H4K20 me2
H2A.Z
0.00 0.25 0.50 0.75 1.00
$\hat{\pi}$
INTS4 – Reverse
H3K4 me1:H3−5ac
DNA Meth. m5C:H4K20 me3
H3K4 me3:H3−5ac
H3K4 me1:H4−4ac
H3K4 me1:H2A.Z
DNA Meth. m5C:H3K9 me3
H3K27 me3:H3K4 me1
H3K27 me3:H4K20 me3
H2A.Z:H3K4 me3
H3K27 me3:H3K9 me3
H3K4 me1:H4K20 me2
H2A.Z:H3−5ac
H2A.Z:H4−4ac
H2A.Z:H4K20 me2
DNA Meth. m5C:H4K20 me2
H3K4 me3:H4−4ac
H3K27 me3:DNA Meth. m5C
H4K20 me2:H3−5ac
H4−4ac:H3−5ac
H4K20 me2:H4−4ac
H4K20 me2:H3K4 me3
H3K27 me2
H3K4 me1
H3K9 me2
H3K9 me3
H4K20 me3
H3−5ac
H3K27 me3
H3K4 me3
DNA Meth. m5C
H4−4ac
H4K20 me2
H2A.Z
0.00 0.25 0.50 0.75 1.00
$\hat{\pi}$
INTS5 – Forward
H3K4 me1:H4K20 me2
H3K27 me2:DNA Meth. m5C
H3K27 me3:H4K20 me3
H3K27 me3:H3K9 me3
H3K27 me3:DNA Meth. m5C
H2A.Z:H4−4ac
H2A.Z:H3K4 me3
H2A.Z:H4K20 me2
H3K4 me1:H4−4ac
H3K4 me3:H3−5ac
H3K4 me1:H2A.Z
DNA Meth. m5C:H4K20 me3
H3K4 me1:H3−5ac
DNA Meth. m5C:H3K9 me3
H2A.Z:H3−5ac
H3K9 me2:DNA Meth. m5C
DNA Meth. m5C:H4K20 me2
H4K20 me2:H4−4ac
H4K20 me2:H3−5ac
H4−4ac:H3−5ac
H3K4 me3:H4−4ac
H4K20 me2:H3K4 me3
H3K27 me2
H3K9 me2
H3K9 me3
DNA Meth. m5C
H3K4 me3
H2A.Z
H3K4 me1
H3−5ac
H3K27 me3
H4K20 me3
H4K20 me2
H4−4ac
0.00 0.25 0.50 0.75 1.00
$\hat{\pi}$
INTS5 – Reverse
H3K4 me1:H4K20 me2
H3K27 me2:DNA Meth. m5C
H3K27 me3:H4K20 me3
H3K27 me3:H3K9 me3
H3K27 me3:DNA Meth. m5C
H2A.Z:H4−4ac
H2A.Z:H3K4 me3
H2A.Z:H4K20 me2
H3K4 me1:H4−4ac
H3K4 me3:H3−5ac
H3K4 me1:H2A.Z
DNA Meth. m5C:H4K20 me3
H3K4 me1:H3−5ac
DNA Meth. m5C:H3K9 me3
H2A.Z:H3−5ac
H3K9 me2:DNA Meth. m5C
DNA Meth. m5C:H4K20 me2
H4K20 me2:H4−4ac
H4K20 me2:H3−5ac
H4−4ac:H3−5ac
H3K4 me3:H4−4ac
H4K20 me2:H3K4 me3
H3K27 me2
H3K9 me2
H3K9 me3
DNA Meth. m5C
H3K4 me3
H2A.Z
H3K4 me1
H3−5ac
H3K27 me3
H4K20 me3
H4K20 me2
H4−4ac
0.00 0.25 0.50 0.75 1.00
$\hat{\pi}$
INTS2 – Forward
H3K27 me3:H3K9 me3
H2A.Z:H4−4ac
H3K4 me1:H3−5ac
H3K9 me2:DNA Meth. m5C
H3K4 me3:H3−5ac
H2A.Z:H3K4 me3
H3K4 me1:H4K20 me2
H3K27 me3:H4K20 me3
H3K27 me3:DNA Meth. m5C
H2A.Z:H3−5ac
H3K9 me3:H4K20 me3
DNA Meth. m5C:H3K9 me3
H3K4 me1:H4−4ac
DNA Meth. m5C:H4K20 me3
H2A.Z:H4K20 me2
H3K4 me1:H2A.Z
DNA Meth. m5C:H4K20 me2
H3K4 me3:H4−4ac
H4−4ac:H3−5ac
H4K20 me2:H4−4ac
H4K20 me2:H3−5ac
H4K20 me2:H3K4 me3
H3K27 me2
H3K27 me3
H4K20 me3
H3K9 me2
H3K9 me3
H2A.Z
H3K4 me1
H3K4 me3
DNA Meth. m5C
H3−5ac
H4K20 me2
H4−4ac
0.00 0.25 0.50 0.75 1.00
$\hat{\pi}$
INTS2 – Reverse
H3K27 me3:H3K9 me3
H2A.Z:H4−4ac
H3K4 me1:H3−5ac
H3K9 me2:DNA Meth. m5C
H3K4 me3:H3−5ac
H2A.Z:H3K4 me3
H3K4 me1:H4K20 me2
H3K27 me3:H4K20 me3
H3K27 me3:DNA Meth. m5C
H2A.Z:H3−5ac
H3K9 me3:H4K20 me3
DNA Meth. m5C:H3K9 me3
H3K4 me1:H4−4ac
DNA Meth. m5C:H4K20 me3
H2A.Z:H4K20 me2
H3K4 me1:H2A.Z
DNA Meth. m5C:H4K20 me2
H3K4 me3:H4−4ac
H4−4ac:H3−5ac
H4K20 me2:H4−4ac
H4K20 me2:H3−5ac
H4K20 me2:H3K4 me3
H3K27 me2
H3K27 me3
H4K20 me3
H3K9 me2
H3K9 me3
H2A.Z
H3K4 me1
H3K4 me3
DNA Meth. m5C
H3−5ac
H4K20 me2
H4−4ac
0.00 0.25 0.50 0.75 1.00
$\hat{\pi}$
ZNF740 – Forward
H3K27 me3:H4K20 me3
DNA Meth. m5C:H4K20 me2
H3K9 me2:DNA Meth. m5C
H3K27 me3:H3K9 me3
H3K9 me3:H4K20 me3
DNA Meth. m5C:H3K9 me3
H2A.Z:H3−5ac
H3K27 me3:H3K4 me1
H3K27 me3:DNA Meth. m5C
H2A.Z:H4−4ac
H3K4 me3:H3−5ac
DNA Meth. m5C:H4K20 me3
H2A.Z:H4K20 me2
H3K4 me1:H4K20 me2
H3K4 me1:H3−5ac
H3K4 me1:H2A.Z
H3K4 me1:H4−4ac
H3K4 me3:H4−4ac
H4K20 me2:H4−4ac
H2A.Z:H3K4 me3
H4−4ac:H3−5ac
H4K20 me2:H3−5ac
H4K20 me2:H3K4 me3
H3K27 me2
H3K9 me2
H3K9 me3
H4K20 me3
H3K27 me3
DNA Meth. m5C
H2A.Z
H3−5ac
H3K4 me1
H3K4 me3
H4−4ac
H4K20 me2
0.00 0.25 0.50 0.75 1.00
$\hat{\pi}$
ZNF740 – Reverse
H3K27 me3:H4K20 me3
DNA Meth. m5C:H4K20 me2
H3K9 me2:DNA Meth. m5C
H3K27 me3:H3K9 me3
H3K9 me3:H4K20 me3
DNA Meth. m5C:H3K9 me3
H2A.Z:H3−5ac
H3K27 me3:H3K4 me1
H3K27 me3:DNA Meth. m5C
H2A.Z:H4−4ac
H3K4 me3:H3−5ac
DNA Meth. m5C:H4K20 me3
H2A.Z:H4K20 me2
H3K4 me1:H4K20 me2
H3K4 me1:H3−5ac
H3K4 me1:H2A.Z
H3K4 me1:H4−4ac
H3K4 me3:H4−4ac
H4K20 me2:H4−4ac
H2A.Z:H3K4 me3
H4−4ac:H3−5ac
H4K20 me2:H3−5ac
H4K20 me2:H3K4 me3
H3K27 me2
H3K9 me2
H3K9 me3
H4K20 me3
H3K27 me3
DNA Meth. m5C
H2A.Z
H3−5ac
H3K4 me1
H3K4 me3
H4−4ac
H4K20 me2
0.00 0.25 0.50 0.75 1.00
$\hat{\pi}$
MED11 – Forward
H3K27 me3:H4K20 me3
H3K27 me3:H3K9 me3
H3K9 me3:H4K20 me3
H3K27 me2:DNA Meth. m5C
H2A.Z:H3−5ac
DNA Meth. m5C:H3K9 me3
H2A.Z:H4−4ac
H3K4 me1:H4K20 me2
DNA Meth. m5C:H4K20 me2
DNA Meth. m5C:H4K20 me3
H2A.Z:H3K4 me3
H3K27 me3:DNA Meth. m5C
H3K4 me3:H3−5ac
H3K4 me1:H3−5ac
H3K4 me1:H2A.Z
H2A.Z:H4K20 me2
H3K4 me1:H4−4ac
H4−4ac:H3−5ac
H4K20 me2:H3−5ac
H4K20 me2:H4−4ac
H3K4 me3:H4−4ac
H4K20 me2:H3K4 me3
H3K27 me2
H3K9 me3
H3K9 me2
H3K27 me3
H4K20 me3
H3K4 me1
DNA Meth. m5C
H3K4 me3
H3−5ac
H2A.Z
H4K20 me2
H4−4ac
0.00 0.25 0.50 0.75 1.00
$\hat{\pi}$
MED11 – Reverse
H3K27 me3:H4K20 me3
H3K27 me3:H3K9 me3
H3K9 me3:H4K20 me3
H3K27 me2:DNA Meth. m5C
H2A.Z:H3−5ac
DNA Meth. m5C:H3K9 me3
H2A.Z:H4−4ac
H3K4 me1:H4K20 me2
DNA Meth. m5C:H4K20 me2
DNA Meth. m5C:H4K20 me3
H2A.Z:H3K4 me3
H3K27 me3:DNA Meth. m5C
H3K4 me3:H3−5ac
H3K4 me1:H3−5ac
H3K4 me1:H2A.Z
H2A.Z:H4K20 me2
H3K4 me1:H4−4ac
H4−4ac:H3−5ac
H4K20 me2:H3−5ac
H4K20 me2:H4−4ac
H3K4 me3:H4−4ac
H4K20 me2:H3K4 me3
H3K27 me2
H3K9 me3
H3K9 me2
H3K27 me3
H4K20 me3
H3K4 me1
DNA Meth. m5C
H3K4 me3
H3−5ac
H2A.Z
H4K20 me2
H4−4ac
0.00 0.25 0.50 0.75 1.00
$\hat{\pi}$

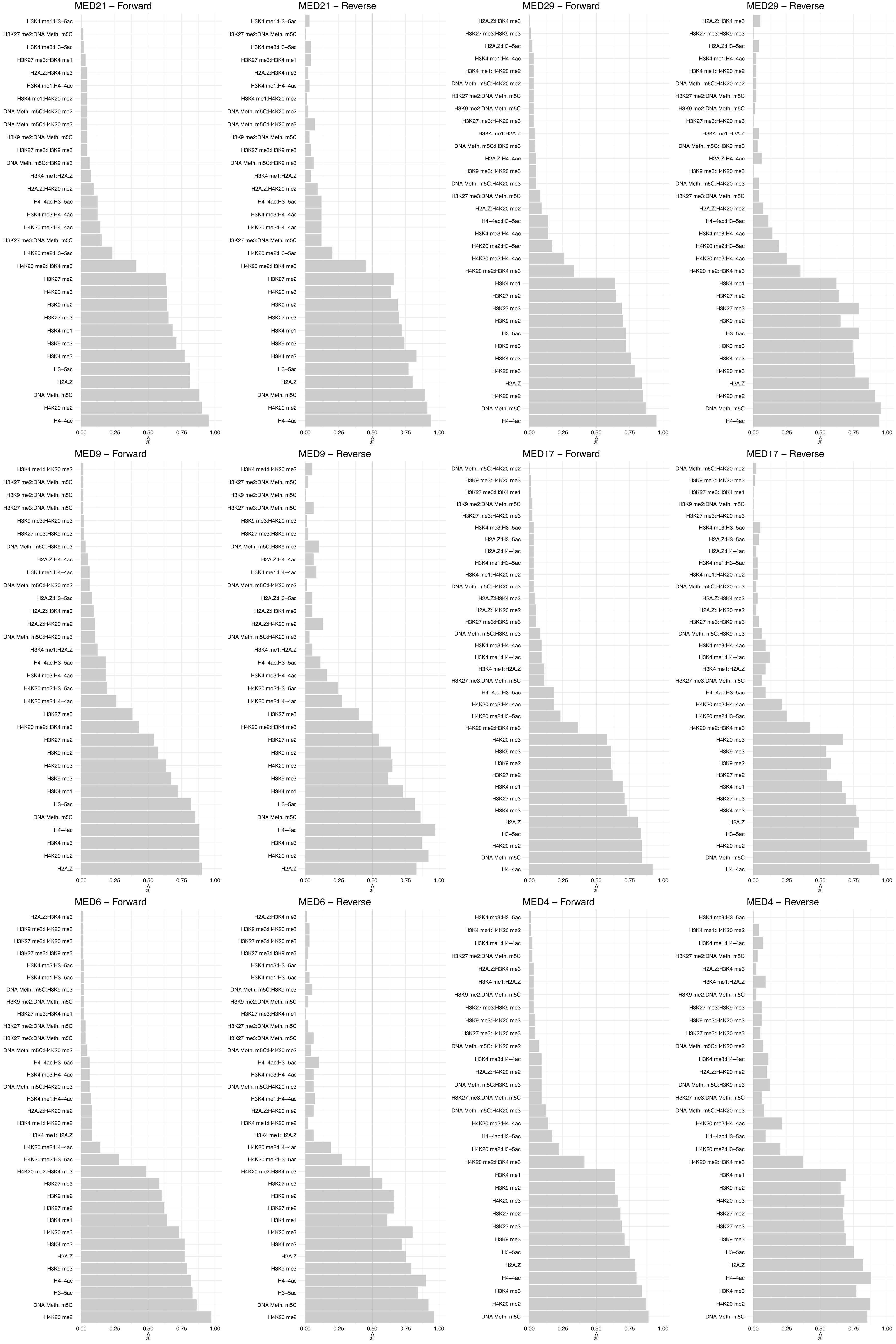
MED21 – Forward
MED21 – Reverse
H3K4 me1:H3−5ac
H3K27 me2:DNA Meth. m5C
H3K4 me3:H3−5ac
H3K27 me3:H3K4 me1
H2A.Z:H3K4 me3
H3K4 me1:H4−4ac
H3K4 me1:H4K20 me2
DNA Meth. m5C:H4K20 me2
DNA Meth. m5C:H4K20 me3
H3K9 me2:DNA Meth. m5C
H3K27 me3:H3K9 me3
DNA Meth. m5C:H3K9 me3
H3K4 me1:H2A.Z
H2A.Z:H4K20 me2
H4−4ac:H3−5ac
H3K4 me3:H4−4ac
H4K20 me2:H4−4ac
H3K27 me3:DNA Meth. m5C
H4K20 me2:H3−5ac
H4K20 me2:H3K4 me3
H3K27 me2
H4K20 me3
H3K9 me2
H3K27 me3
H3K4 me1
H3K9 me3
H3K4 me3
H3−5ac
H2A.Z
DNA Meth. m5C
H4K20 me2
H4−4ac
0.00
0.25
0.50
0.75
1.00
π̂
MED29 – Forward
MED29 – Reverse
H2A.Z:H3K4 me3
H3K27 me3:H3K9 me3
H2A.Z:H3−5ac
H3K4 me1:H4−4ac
H3K4 me1:H4K20 me2
DNA Meth. m5C:H4K20 me2
H3K27 me2:DNA Meth. m5C
H3K9 me2:DNA Meth. m5C
H3K27 me3:H4K20 me3
H3K4 me1:H2A.Z
DNA Meth. m5C:H3K9 me3
H2A.Z:H4−4ac
H3K9 me3:H4K20 me3
DNA Meth. m5C:H4K20 me3
H3K27 me3:DNA Meth. m5C
H2A.Z:H4K20 me2
H4−4ac:H3−5ac
H3K4 me3:H4−4ac
H4K20 me2:H3−5ac
H4K20 me2:H4−4ac
H4K20 me2:H3K4 me3
H3K4 me1
H3K27 me2
H3K27 me3
H3K9 me2
H3−5ac
H3K9 me3
H3K4 me3
H4K20 me3
H2A.Z
H4K20 me2
DNA Meth. m5C
H4−4ac
MED9 – Forward
MED9 – Reverse
H3K4 me1:H4K20 me2
H3K27 me2:DNA Meth. m5C
H3K9 me2:DNA Meth. m5C
H3K27 me3:DNA Meth. m5C
H3K9 me3:H4K20 me3
H3K27 me3:H3K9 me3
DNA Meth. m5C:H3K9 me3
H2A.Z:H4−4ac
H3K4 me1:H4−4ac
DNA Meth. m5C:H4K20 me2
H2A.Z:H3−5ac
H2A.Z:H3K4 me3
H2A.Z:H4K20 me2
DNA Meth. m5C:H4K20 me3
H3K4 me1:H2A.Z
H4−4ac:H3−5ac
H3K4 me3:H4−4ac
H4K20 me2:H3−5ac
H4K20 me2:H4−4ac
H3K27 me3
H4K20 me2:H3K4 me3
H3K27 me2
H3K9 me2
H4K20 me3
H3K9 me3
H3K4 me1
H3−5ac
DNA Meth. m5C
H4−4ac
H3K4 me3
H4K20 me2
H2A.Z
MED17 – Forward
MED17 – Reverse
DNA Meth. m5C:H4K20 me2
H3K9 me3:H4K20 me3
H3K27 me3:H3K4 me1
H3K9 me2:DNA Meth. m5C
H3K27 me3:H4K20 me3
H3K4 me3:H3−5ac
H2A.Z:H3−5ac
H2A.Z:H4−4ac
H3K4 me1:H3−5ac
H3K4 me1:H4K20 me2
DNA Meth. m5C:H4K20 me3
H2A.Z:H3K4 me3
H2A.Z:H4K20 me2
H3K27 me3:H3K9 me3
DNA Meth. m5C:H3K9 me3
H3K4 me3:H4−4ac
H3K4 me1:H4−4ac
H3K4 me1:H2A.Z
H3K27 me3:DNA Meth. m5C
H4−4ac:H3−5ac
H4K20 me2:H4−4ac
H4K20 me2:H3−5ac
H4K20 me2:H3K4 me3
H4K20 me3
H3K9 me3
H3K9 me2
H3K27 me2
H3K4 me1
H3K27 me3
H3K4 me3
H2A.Z
H3−5ac
H4K20 me2
DNA Meth. m5C
H4−4ac
MED6 – Forward
MED6 – Reverse
H2A.Z:H3K4 me3
H3K9 me3:H4K20 me3
H3K27 me3:H4K20 me3
H3K27 me3:H3K9 me3
H3K4 me3:H3−5ac
H3K4 me1:H3−5ac
DNA Meth. m5C:H3K9 me3
H3K9 me2:DNA Meth. m5C
H3K27 me3:H3K4 me1
H3K27 me2:DNA Meth. m5C
H3K27 me3:DNA Meth. m5C
DNA Meth. m5C:H4K20 me2
H4−4ac:H3−5ac
H3K4 me3:H4−4ac
DNA Meth. m5C:H4K20 me3
H3K4 me1:H4−4ac
H2A.Z:H4K20 me2
H3K4 me1:H4K20 me2
H3K4 me1:H2A.Z
H4K20 me2:H4−4ac
H4K20 me2:H3−5ac
H4K20 me2:H3K4 me3
H3K27 me3
H3K9 me2
H3K27 me2
H3K4 me1
H4K20 me3
H3K4 me3
H2A.Z
H3K9 me3
H4−4ac
H3−5ac
DNA Meth. m5C
H4K20 me2
MED4 – Forward
MED4 – Reverse
H3K4 me3:H3−5ac
H3K4 me1:H4K20 me2
H3K4 me1:H4−4ac
H3K27 me2:DNA Meth. m5C
H2A.Z:H3K4 me3
H3K4 me1:H2A.Z
H3K9 me2:DNA Meth. m5C
H3K27 me3:H3K9 me3
H3K9 me3:H4K20 me3
H3K27 me3:H4K20 me3
DNA Meth. m5C:H4K20 me2
H3K4 me3:H4−4ac
H2A.Z:H4K20 me2
DNA Meth. m5C:H3K9 me3
H3K27 me3:DNA Meth. m5C
DNA Meth. m5C:H4K20 me3
H4K20 me2:H4−4ac
H4−4ac:H3−5ac
H4K20 me2:H3−5ac
H4K20 me2:H3K4 me3
H3K4 me1
H3K9 me2
H4K20 me3
H3K27 me2
H3K27 me3
H3K9 me3
H3−5ac
H2A.Z
H4−4ac
H3K4 me3
H4K20 me2
DNA Meth. m5C

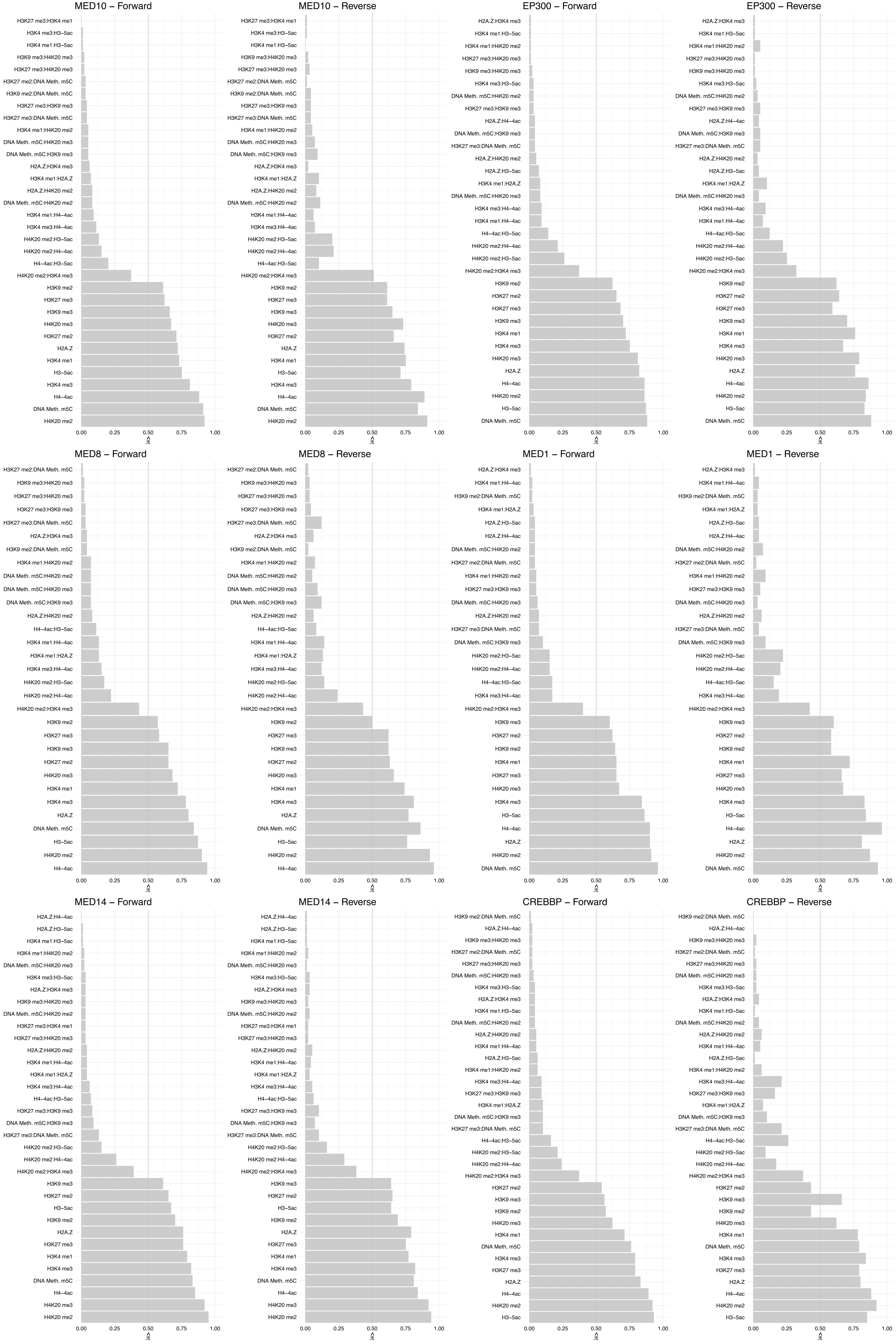
MED10 – Forward
H3K27 me3:H3K4 me1
H3K4 me3:H3–5ac
H3K4 me1:H3–5ac
H3K9 me3:H4K20 me3
H3K27 me3:H4K20 me3
H3K27 me2:DNA Meth. m5C
H3K9 me2:DNA Meth. m5C
H3K27 me3:H3K9 me3
H3K27 me3:DNA Meth. m5C
H3K4 me1:H4K20 me2
DNA Meth. m5C:H4K20 me3
DNA Meth. m5C:H3K9 me3
H2A.Z:H3K4 me3
H3K4 me1:H2A.Z
H2A.Z:H4K20 me2
DNA Meth. m5C:H4K20 me2
H3K4 me1:H4–4ac
H3K4 me3:H4–4ac
H4K20 me2:H3–5ac
H4K20 me2:H4–4ac
H4–4ac:H3–5ac
H4K20 me2:H3K4 me3
H3K9 me2
H3K27 me3
H3K9 me3
H4K20 me3
H3K27 me2
H2A.Z
H3K4 me1
H3–5ac
H3K4 me3
H4–4ac
DNA Meth. m5C
H4K20 me2
0.00 0.25 0.50 0.75 1.00
π̂
MED10 – Reverse
H3K27 me3:H3K4 me1
H3K4 me3:H3–5ac
H3K4 me1:H3–5ac
H3K9 me3:H4K20 me3
H3K27 me3:H4K20 me3
H3K27 me2:DNA Meth. m5C
H3K9 me2:DNA Meth. m5C
H3K27 me3:H3K9 me3
H3K27 me3:DNA Meth. m5C
H3K4 me1:H4K20 me2
DNA Meth. m5C:H4K20 me3
DNA Meth. m5C:H3K9 me3
H2A.Z:H3K4 me3
H3K4 me1:H2A.Z
H2A.Z:H4K20 me2
DNA Meth. m5C:H4K20 me2
H3K4 me1:H4–4ac
H3K4 me3:H4–4ac
H4K20 me2:H3–5ac
H4K20 me2:H4–4ac
H4–4ac:H3–5ac
H4K20 me2:H3K4 me3
H3K9 me2
H3K27 me3
H3K9 me3
H4K20 me3
H3K27 me2
H2A.Z
H3K4 me1
H3–5ac
H3K4 me3
H4–4ac
DNA Meth. m5C
H4K20 me2
0.00 0.25 0.50 0.75 1.00
π̂
EP300 – Forward
H2A.Z:H3K4 me3
H3K4 me1:H3–5ac
H3K4 me1:H4K20 me2
H3K27 me3:H4K20 me3
H3K9 me3:H4K20 me3
H3K4 me3:H3–5ac
DNA Meth. m5C:H4K20 me2
H3K27 me3:H3K9 me3
H2A.Z:H4–4ac
DNA Meth. m5C:H3K9 me3
H3K27 me3:DNA Meth. m5C
H2A.Z:H4K20 me2
H2A.Z:H3–5ac
H3K4 me1:H2A.Z
DNA Meth. m5C:H4K20 me3
H3K4 me3:H4–4ac
H3K4 me1:H4–4ac
H4–4ac:H3–5ac
H4K20 me2:H4–4ac
H4K20 me2:H3–5ac
H4K20 me2:H3K4 me3
H3K9 me2
H3K27 me2
H3K27 me3
H3K9 me3
H3K4 me1
H3K4 me3
H4K20 me3
H2A.Z
H4–4ac
H4K20 me2
H3–5ac
DNA Meth. m5C
0.00 0.25 0.50 0.75 1.00
π̂
EP300 – Reverse
H2A.Z:H3K4 me3
H3K4 me1:H3–5ac
H3K4 me1:H4K20 me2
H3K27 me3:H4K20 me3
H3K9 me3:H4K20 me3
H3K4 me3:H3–5ac
DNA Meth. m5C:H4K20 me2
H3K27 me3:H3K9 me3
H2A.Z:H4–4ac
DNA Meth. m5C:H3K9 me3
H3K27 me3:DNA Meth. m5C
H2A.Z:H4K20 me2
H2A.Z:H3–5ac
H3K4 me1:H2A.Z
DNA Meth. m5C:H4K20 me3
H3K4 me3:H4–4ac
H3K4 me1:H4–4ac
H4–4ac:H3–5ac
H4K20 me2:H4–4ac
H4K20 me2:H3–5ac
H4K20 me2:H3K4 me3
H3K9 me2
H3K27 me2
H3K27 me3
H3K9 me3
H3K4 me1
H3K4 me3
H4K20 me3
H2A.Z
H4–4ac
H4K20 me2
H3–5ac
DNA Meth. m5C
0.00 0.25 0.50 0.75 1.00
π̂
MED8 – Forward
H3K27 me2:DNA Meth. m5C
H3K9 me3:H4K20 me3
H3K27 me3:H4K20 me3
H3K27 me3:H3K9 me3
H3K27 me3:DNA Meth. m5C
H2A.Z:H3K4 me3
H3K9 me2:DNA Meth. m5C
H3K4 me1:H4K20 me2
DNA Meth. m5C:H4K20 me2
DNA Meth. m5C:H4K20 me3
DNA Meth. m5C:H3K9 me3
H2A.Z:H4K20 me2
H4–4ac:H3–5ac
H3K4 me1:H4–4ac
H3K4 me1:H2A.Z
H3K4 me3:H4–4ac
H4K20 me2:H3–5ac
H4K20 me2:H4–4ac
H4K20 me2:H3K4 me3
H3K9 me2
H3K27 me3
H3K9 me3
H3K27 me2
H4K20 me3
H3K4 me1
H3K4 me3
H2A.Z
DNA Meth. m5C
H3–5ac
H4K20 me2
H4–4ac
0.00 0.25 0.50 0.75 1.00
π̂
MED8 – Reverse
H3K27 me2:DNA Meth. m5C
H3K9 me3:H4K20 me3
H3K27 me3:H4K20 me3
H3K27 me3:H3K9 me3
H3K27 me3:DNA Meth. m5C
H2A.Z:H3K4 me3
H3K9 me2:DNA Meth. m5C
H3K4 me1:H4K20 me2
DNA Meth. m5C:H4K20 me2
DNA Meth. m5C:H4K20 me3
DNA Meth. m5C:H3K9 me3
H2A.Z:H4K20 me2
H4–4ac:H3–5ac
H3K4 me1:H4–4ac
H3K4 me1:H2A.Z
H3K4 me3:H4–4ac
H4K20 me2:H3–5ac
H4K20 me2:H4–4ac
H4K20 me2:H3K4 me3
H3K9 me2
H3K27 me3
H3K9 me3
H3K27 me2
H4K20 me3
H3K4 me1
H3K4 me3
H2A.Z
DNA Meth. m5C
H3–5ac
H4K20 me2
H4–4ac
0.00 0.25 0.50 0.75 1.00
π̂
MED1 – Forward
H2A.Z:H3K4 me3
H3K4 me1:H4–4ac
H3K9 me2:DNA Meth. m5C
H3K4 me1:H2A.Z
H2A.Z:H3–5ac
H2A.Z:H4–4ac
DNA Meth. m5C:H4K20 me2
H3K27 me2:DNA Meth. m5C
H3K4 me1:H4K20 me2
H3K27 me3:H3K9 me3
DNA Meth. m5C:H4K20 me3
H2A.Z:H4K20 me2
H3K27 me3:DNA Meth. m5C
DNA Meth. m5C:H3K9 me3
H4K20 me2:H3–5ac
H4K20 me2:H4–4ac
H4–4ac:H3–5ac
H3K4 me3:H4–4ac
H4K20 me2:H3K4 me3
H3K9 me3
H3K27 me2
H3K9 me2
H3K4 me1
H3K27 me3
H4K20 me3
H3K4 me3
H3–5ac
H4–4ac
H2A.Z
H4K20 me2
DNA Meth. m5C
0.00 0.25 0.50 0.75 1.00
π̂
MED1 – Reverse
H2A.Z:H3K4 me3
H3K4 me1:H4–4ac
H3K9 me2:DNA Meth. m5C
H3K4 me1:H2A.Z
H2A.Z:H3–5ac
H2A.Z:H4–4ac
DNA Meth. m5C:H4K20 me2
H3K27 me2:DNA Meth. m5C
H3K4 me1:H4K20 me2
H3K27 me3:H3K9 me3
DNA Meth. m5C:H4K20 me3
H2A.Z:H4K20 me2
H3K27 me3:DNA Meth. m5C
DNA Meth. m5C:H3K9 me3
H4K20 me2:H3–5ac
H4K20 me2:H4–4ac
H4–4ac:H3–5ac
H3K4 me3:H4–4ac
H4K20 me2:H3K4 me3
H3K9 me3
H3K27 me2
H3K9 me2
H3K4 me1
H3K27 me3
H4K20 me3
H3K4 me3
H3–5ac
H4–4ac
H2A.Z
H4K20 me2
DNA Meth. m5C
0.00 0.25 0.50 0.75 1.00
π̂
MED14 – Forward
H2A.Z:H4–4ac
H2A.Z:H3–5ac
H3K4 me1:H3–5ac
H3K4 me1:H4K20 me2
DNA Meth. m5C:H4K20 me3
H3K4 me3:H3–5ac
H2A.Z:H3K4 me3
H3K9 me3:H4K20 me3
DNA Meth. m5C:H4K20 me2
H3K27 me3:H3K4 me1
H3K27 me3:H4K20 me3
H2A.Z:H4K20 me2
H3K4 me1:H4–4ac
H3K4 me1:H2A.Z
H3K4 me3:H4–4ac
H4–4ac:H3–5ac
H3K27 me3:H3K9 me3
DNA Meth. m5C:H3K9 me3
H3K27 me3:DNA Meth. m5C
H4K20 me2:H3–5ac
H4K20 me2:H4–4ac
H4K20 me2:H3K4 me3
H3K9 me3
H3K27 me2
H3–5ac
H3K9 me2
H2A.Z
H3K27 me3
H3K4 me1
H3K4 me3
DNA Meth. m5C
H4–4ac
H4K20 me3
H4K20 me2
0.00 0.25 0.50 0.75 1.00
π̂
MED14 – Reverse
H2A.Z:H4–4ac
H2A.Z:H3–5ac
H3K4 me1:H3–5ac
H3K4 me1:H4K20 me2
DNA Meth. m5C:H4K20 me3
H3K4 me3:H3–5ac
H2A.Z:H3K4 me3
H3K9 me3:H4K20 me3
DNA Meth. m5C:H4K20 me2
H3K27 me3:H3K4 me1
H3K27 me3:H4K20 me3
H2A.Z:H4K20 me2
H3K4 me1:H4–4ac
H3K4 me1:H2A.Z
H3K4 me3:H4–4ac
H4–4ac:H3–5ac
H3K27 me3:H3K9 me3
DNA Meth. m5C:H3K9 me3
H3K27 me3:DNA Meth. m5C
H4K20 me2:H3–5ac
H4K20 me2:H4–4ac
H4K20 me2:H3K4 me3
H3K9 me3
H3K27 me2
H3–5ac
H3K9 me2
H2A.Z
H3K27 me3
H3K4 me1
H3K4 me3
DNA Meth. m5C
H4–4ac
H4K20 me3
H4K20 me2
0.00 0.25 0.50 0.75 1.00
π̂
CREBBP – Forward
H3K9 me2:DNA Meth. m5C
H2A.Z:H4–4ac
H3K9 me3:H4K20 me3
H3K27 me2:DNA Meth. m5C
H3K27 me3:H4K20 me3
DNA Meth. m5C:H4K20 me3
H3K4 me3:H3–5ac
H2A.Z:H3K4 me3
H3K4 me1:H3–5ac
DNA Meth. m5C:H4K20 me2
H2A.Z:H4K20 me2
H3K4 me1:H4–4ac
H2A.Z:H3–5ac
H3K4 me1:H4K20 me2
H3K4 me3:H4–4ac
H3K27 me3:H3K9 me3
H3K4 me1:H2A.Z
DNA Meth. m5C:H3K9 me3
H3K27 me3:DNA Meth. m5C
H4–4ac:H3–5ac
H4K20 me2:H3–5ac
H4K20 me2:H4–4ac
H4K20 me2:H3K4 me3
H3K27 me2
H3K9 me3
H3K9 me2
H4K20 me3
H3K4 me1
DNA Meth. m5C
H3K4 me3
H3K27 me3
H2A.Z
H4–4ac
H4K20 me2
H3–5ac
0.00 0.25 0.50 0.75 1.00
π̂
CREBBP – Reverse
H3K9 me2:DNA Meth. m5C
H2A.Z:H4–4ac
H3K9 me3:H4K20 me3
H3K27 me2:DNA Meth. m5C
H3K27 me3:H4K20 me3
DNA Meth. m5C:H4K20 me3
H3K4 me3:H3–5ac
H2A.Z:H3K4 me3
H3K4 me1:H3–5ac
DNA Meth. m5C:H4K20 me2
H2A.Z:H4K20 me2
H3K4 me1:H4–4ac
H2A.Z:H3–5ac
H3K4 me1:H4K20 me2
H3K4 me3:H4–4ac
H3K27 me3:H3K9 me3
H3K4 me1:H2A.Z
DNA Meth. m5C:H3K9 me3
H3K27 me3:DNA Meth. m5C
H4–4ac:H3–5ac
H4K20 me2:H3–5ac
H4K20 me2:H4–4ac
H4K20 me2:H3K4 me3
H3K27 me2
H3K9 me3
H3K9 me2
H4K20 me3
H3K4 me1
DNA Meth. m5C
H3K4 me3
H3K27 me3
H2A.Z
H4–4ac
H4K20 me2
H3–5ac
0.00 0.25 0.50 0.75 1.00
π̂

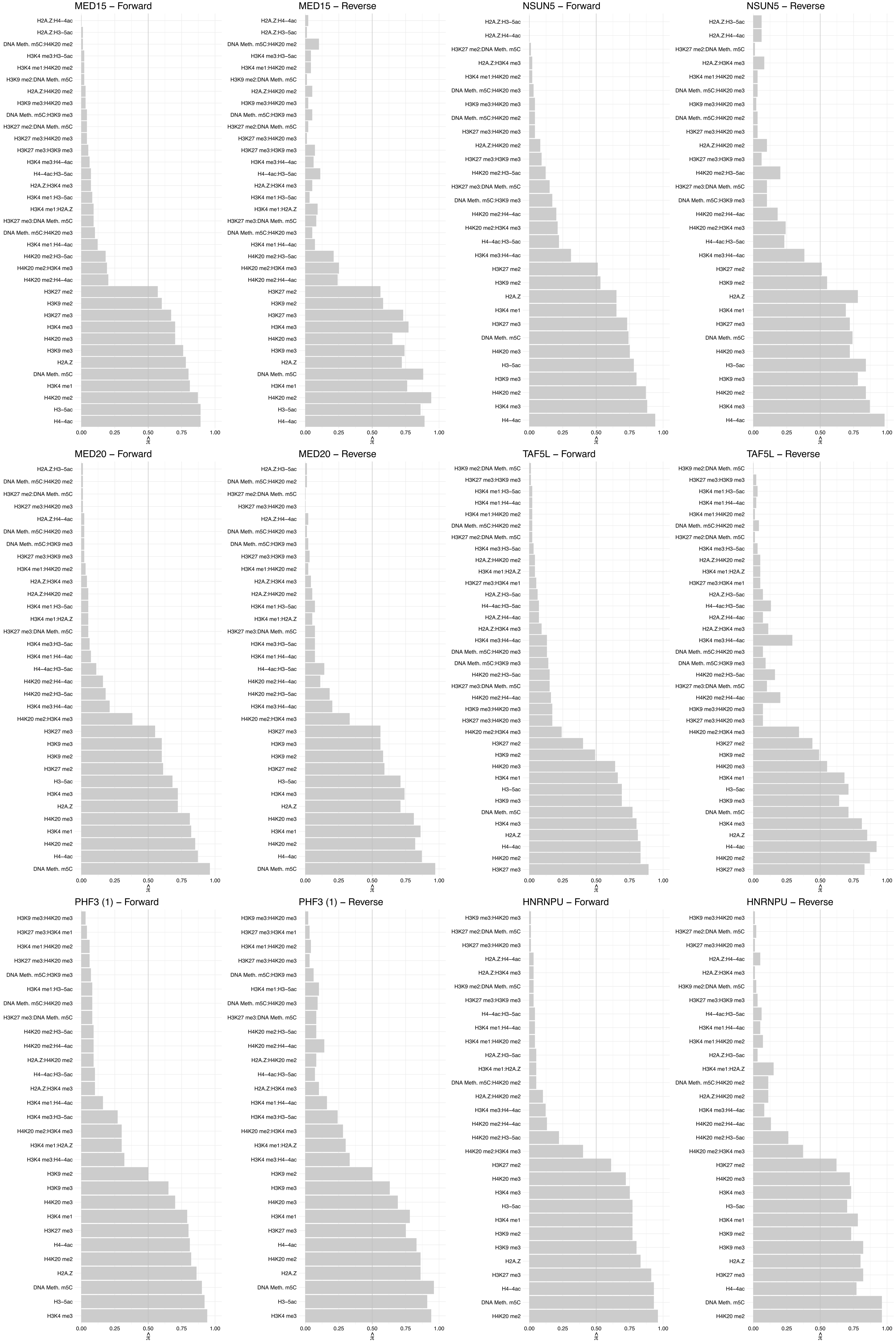
MED15 – Forward
MED15 – Reverse
NSUN5 – Forward
NSUN5 – Reverse
MED20 – Forward
MED20 – Reverse
TAF5L – Forward
TAF5L – Reverse
PHF3 (1) – Forward
PHF3 (1) – Reverse
HNRNPU – Forward
HNRNPU – Reverse
0.00
0.25
0.50
0.75
1.00
$\hat{\pi}$

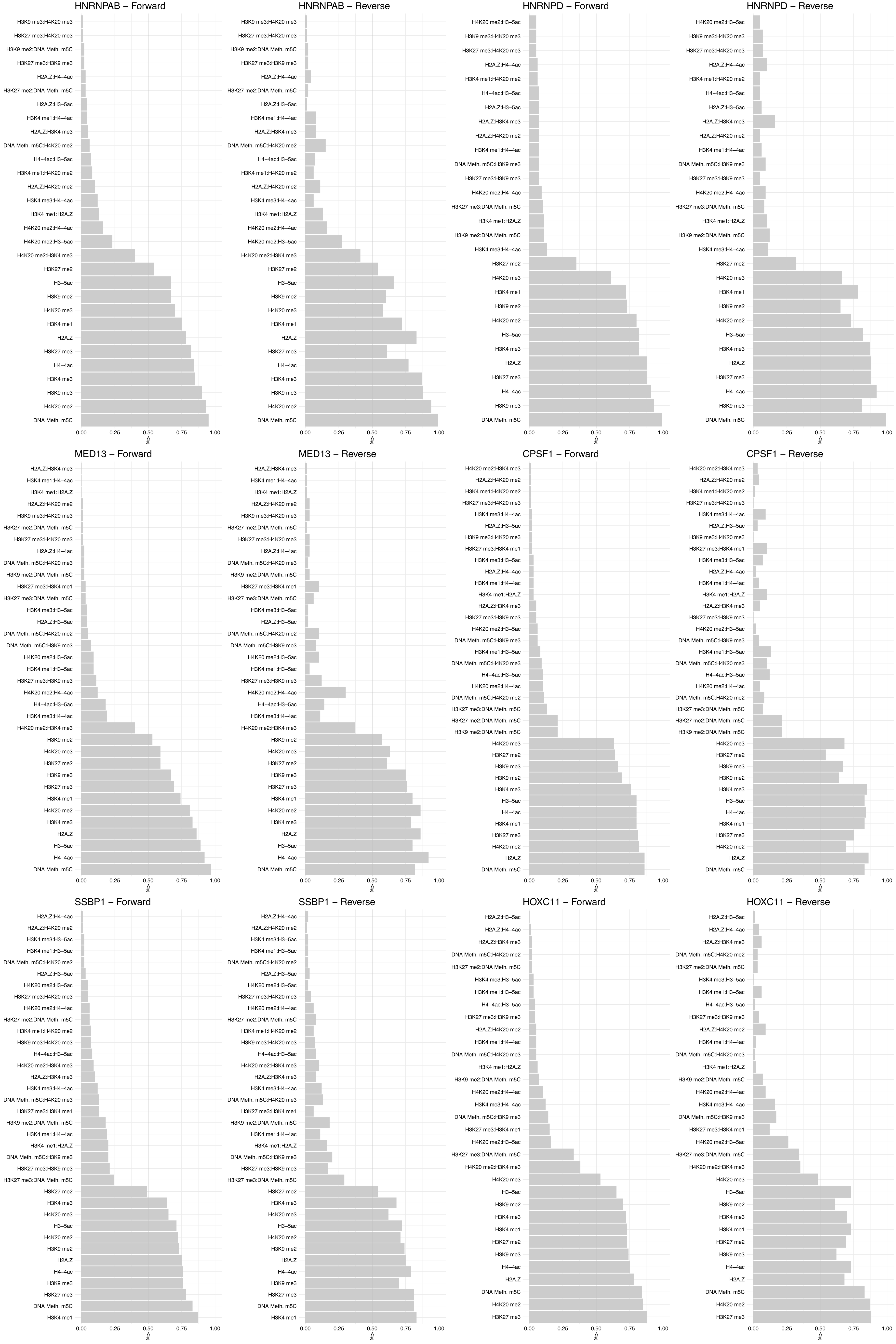
HNRNPAB – Forward
H3K9 me3:H4K20 me3
H3K27 me3:H4K20 me3
H3K9 me2:DNA Meth. m5C
H3K27 me3:H3K9 me3
H2A.Z:H4–4ac
H3K27 me2:DNA Meth. m5C
H2A.Z:H3–5ac
H3K4 me1:H4–4ac
H2A.Z:H3K4 me3
DNA Meth. m5C:H4K20 me2
H4–4ac:H3–5ac
H3K4 me1:H4K20 me2
H2A.Z:H4K20 me2
H3K4 me3:H4–4ac
H3K4 me1:H2A.Z
H4K20 me2:H4–4ac
H4K20 me2:H3–5ac
H4K20 me2:H3K4 me3
H3K27 me2
H3–5ac
H3K9 me2
H4K20 me3
H3K4 me1
H2A.Z
H3K27 me3
H4–4ac
H3K4 me3
H3K9 me3
H4K20 me2
DNA Meth. m5C
0.00 0.25 0.50 0.75 1.00
π̂
HNRNPAB – Reverse
H3K9 me3:H4K20 me3
H3K27 me3:H4K20 me3
H3K9 me2:DNA Meth. m5C
H3K27 me3:H3K9 me3
H2A.Z:H4–4ac
H3K27 me2:DNA Meth. m5C
H2A.Z:H3–5ac
H3K4 me1:H4–4ac
H2A.Z:H3K4 me3
DNA Meth. m5C:H4K20 me2
H4–4ac:H3–5ac
H3K4 me1:H4K20 me2
H2A.Z:H4K20 me2
H3K4 me3:H4–4ac
H3K4 me1:H2A.Z
H4K20 me2:H4–4ac
H4K20 me2:H3–5ac
H4K20 me2:H3K4 me3
H3K27 me2
H3–5ac
H3K9 me2
H4K20 me3
H3K4 me1
H2A.Z
H3K27 me3
H4–4ac
H3K4 me3
H3K9 me3
H4K20 me2
DNA Meth. m5C
0.00 0.25 0.50 0.75 1.00
π̂
HNRNPD – Forward
H4K20 me2:H3–5ac
H3K9 me3:H4K20 me3
H3K27 me3:H4K20 me3
H2A.Z:H4–4ac
H3K4 me1:H4K20 me2
H4–4ac:H3–5ac
H2A.Z:H3–5ac
H2A.Z:H3K4 me3
H2A.Z:H4K20 me2
H3K4 me1:H4–4ac
DNA Meth. m5C:H3K9 me3
H3K27 me3:H3K9 me3
H4K20 me2:H4–4ac
H3K27 me3:DNA Meth. m5C
H3K4 me1:H2A.Z
H3K9 me2:DNA Meth. m5C
H3K4 me3:H4–4ac
H3K27 me2
H4K20 me3
H3K4 me1
H3K9 me2
H4K20 me2
H3–5ac
H3K4 me3
H2A.Z
H3K27 me3
H4–4ac
H3K9 me3
DNA Meth. m5C
0.00 0.25 0.50 0.75 1.00
π̂
HNRNPD – Reverse
H4K20 me2:H3–5ac
H3K9 me3:H4K20 me3
H3K27 me3:H4K20 me3
H2A.Z:H4–4ac
H3K4 me1:H4K20 me2
H4–4ac:H3–5ac
H2A.Z:H3–5ac
H2A.Z:H3K4 me3
H2A.Z:H4K20 me2
H3K4 me1:H4–4ac
DNA Meth. m5C:H3K9 me3
H3K27 me3:H3K9 me3
H4K20 me2:H4–4ac
H3K27 me3:DNA Meth. m5C
H3K4 me1:H2A.Z
H3K9 me2:DNA Meth. m5C
H3K4 me3:H4–4ac
H3K27 me2
H4K20 me3
H3K4 me1
H3K9 me2
H4K20 me2
H3–5ac
H3K4 me3
H2A.Z
H3K27 me3
H4–4ac
H3K9 me3
DNA Meth. m5C
0.00 0.25 0.50 0.75 1.00
π̂
MED13 – Forward
H2A.Z:H3K4 me3
H3K4 me1:H4–4ac
H3K4 me1:H2A.Z
H2A.Z:H4K20 me2
H3K9 me3:H4K20 me3
H3K27 me2:DNA Meth. m5C
H3K27 me3:H4K20 me3
H2A.Z:H4–4ac
DNA Meth. m5C:H4K20 me3
H3K9 me2:DNA Meth. m5C
H3K27 me3:H3K4 me1
H3K27 me3:DNA Meth. m5C
H3K4 me3:H3–5ac
H2A.Z:H3–5ac
DNA Meth. m5C:H4K20 me2
DNA Meth. m5C:H3K9 me3
H4K20 me2:H3–5ac
H3K4 me1:H3–5ac
H3K27 me3:H3K9 me3
H4K20 me2:H4–4ac
H4–4ac:H3–5ac
H3K4 me3:H4–4ac
H4K20 me2:H3K4 me3
H3K9 me2
H4K20 me3
H3K27 me2
H3K9 me3
H3K27 me3
H3K4 me1
H4K20 me2
H3K4 me3
H2A.Z
H3–5ac
H4–4ac
DNA Meth. m5C
0.00 0.25 0.50 0.75 1.00
π̂
MED13 – Reverse
H2A.Z:H3K4 me3
H3K4 me1:H4–4ac
H3K4 me1:H2A.Z
H2A.Z:H4K20 me2
H3K9 me3:H4K20 me3
H3K27 me2:DNA Meth. m5C
H3K27 me3:H4K20 me3
H2A.Z:H4–4ac
DNA Meth. m5C:H4K20 me3
H3K9 me2:DNA Meth. m5C
H3K27 me3:H3K4 me1
H3K27 me3:DNA Meth. m5C
H3K4 me3:H3–5ac
H2A.Z:H3–5ac
DNA Meth. m5C:H4K20 me2
DNA Meth. m5C:H3K9 me3
H4K20 me2:H3–5ac
H3K4 me1:H3–5ac
H3K27 me3:H3K9 me3
H4K20 me2:H4–4ac
H4–4ac:H3–5ac
H3K4 me3:H4–4ac
H4K20 me2:H3K4 me3
H3K9 me2
H4K20 me3
H3K27 me2
H3K9 me3
H3K27 me3
H3K4 me1
H4K20 me2
H3K4 me3
H2A.Z
H3–5ac
H4–4ac
DNA Meth. m5C
0.00 0.25 0.50 0.75 1.00
π̂
CPSF1 – Forward
H4K20 me2:H3K4 me3
H2A.Z:H4K20 me2
H3K4 me1:H4K20 me2
H3K27 me3:H4K20 me3
H3K4 me3:H4–4ac
H2A.Z:H3–5ac
H3K9 me3:H4K20 me3
H3K27 me3:H3K4 me1
H3K4 me3:H3–5ac
H2A.Z:H4–4ac
H3K4 me1:H4–4ac
H3K4 me1:H2A.Z
H2A.Z:H3K4 me3
H3K27 me3:H3K9 me3
H4K20 me2:H3–5ac
DNA Meth. m5C:H3K9 me3
H3K4 me1:H3–5ac
DNA Meth. m5C:H4K20 me3
H4–4ac:H3–5ac
H4K20 me2:H4–4ac
DNA Meth. m5C:H4K20 me2
H3K27 me3:DNA Meth. m5C
H3K27 me2:DNA Meth. m5C
H3K9 me2:DNA Meth. m5C
H4K20 me3
H3K27 me2
H3K9 me3
H3K9 me2
H3K4 me3
H3–5ac
H4–4ac
H3K4 me1
H3K27 me3
H4K20 me2
H2A.Z
DNA Meth. m5C
0.00 0.25 0.50 0.75 1.00
π̂
CPSF1 – Reverse
H4K20 me2:H3K4 me3
H2A.Z:H4K20 me2
H3K4 me1:H4K20 me2
H3K27 me3:H4K20 me3
H3K4 me3:H4–4ac
H2A.Z:H3–5ac
H3K9 me3:H4K20 me3
H3K27 me3:H3K4 me1
H3K4 me3:H3–5ac
H2A.Z:H4–4ac
H3K4 me1:H4–4ac
H3K4 me1:H2A.Z
H2A.Z:H3K4 me3
H3K27 me3:H3K9 me3
H4K20 me2:H3–5ac
DNA Meth. m5C:H3K9 me3
H3K4 me1:H3–5ac
DNA Meth. m5C:H4K20 me3
H4–4ac:H3–5ac
H4K20 me2:H4–4ac
DNA Meth. m5C:H4K20 me2
H3K27 me3:DNA Meth. m5C
H3K27 me2:DNA Meth. m5C
H3K9 me2:DNA Meth. m5C
H4K20 me3
H3K27 me2
H3K9 me3
H3K9 me2
H3K4 me3
H3–5ac
H4–4ac
H3K4 me1
H3K27 me3
H4K20 me2
H2A.Z
DNA Meth. m5C
0.00 0.25 0.50 0.75 1.00
π̂
SSBP1 – Forward
H2A.Z:H4–4ac
H2A.Z:H4K20 me2
H3K4 me3:H3–5ac
H3K4 me1:H3–5ac
DNA Meth. m5C:H4K20 me2
H2A.Z:H3–5ac
H4K20 me2:H3–5ac
H3K27 me3:H4K20 me3
H4K20 me2:H4–4ac
H3K27 me2:DNA Meth. m5C
H3K4 me1:H4K20 me2
H3K9 me3:H4K20 me3
H4–4ac:H3–5ac
H4K20 me2:H3K4 me3
H2A.Z:H3K4 me3
H3K4 me3:H4–4ac
DNA Meth. m5C:H4K20 me3
H3K27 me3:H3K4 me1
H3K9 me2:DNA Meth. m5C
H3K4 me1:H4–4ac
H3K4 me1:H2A.Z
DNA Meth. m5C:H3K9 me3
H3K27 me3:H3K9 me3
H3K27 me3:DNA Meth. m5C
H3K27 me2
H3K4 me3
H4K20 me3
H3–5ac
H4K20 me2
H3K9 me2
H2A.Z
H4–4ac
H3K9 me3
H3K27 me3
DNA Meth. m5C
H3K4 me1
0.00 0.25 0.50 0.75 1.00
π̂
SSBP1 – Reverse
H2A.Z:H4–4ac
H2A.Z:H4K20 me2
H3K4 me3:H3–5ac
H3K4 me1:H3–5ac
DNA Meth. m5C:H4K20 me2
H2A.Z:H3–5ac
H4K20 me2:H3–5ac
H3K27 me3:H4K20 me3
H4K20 me2:H4–4ac
H3K27 me2:DNA Meth. m5C
H3K4 me1:H4K20 me2
H3K9 me3:H4K20 me3
H4–4ac:H3–5ac
H4K20 me2:H3K4 me3
H2A.Z:H3K4 me3
H3K4 me3:H4–4ac
DNA Meth. m5C:H4K20 me3
H3K27 me3:H3K4 me1
H3K9 me2:DNA Meth. m5C
H3K4 me1:H4–4ac
H3K4 me1:H2A.Z
DNA Meth. m5C:H3K9 me3
H3K27 me3:H3K9 me3
H3K27 me3:DNA Meth. m5C
H3K27 me2
H3K4 me3
H4K20 me3
H3–5ac
H4K20 me2
H3K9 me2
H2A.Z
H4–4ac
H3K9 me3
H3K27 me3
DNA Meth. m5C
H3K4 me1
0.00 0.25 0.50 0.75 1.00
π̂
HOXC11 – Forward
H2A.Z:H3–5ac
H2A.Z:H4–4ac
H2A.Z:H3K4 me3
DNA Meth. m5C:H4K20 me2
H3K27 me2:DNA Meth. m5C
H3K4 me3:H3–5ac
H3K4 me1:H3–5ac
H4–4ac:H3–5ac
H3K27 me3:H3K9 me3
H2A.Z:H4K20 me2
H3K4 me1:H4–4ac
DNA Meth. m5C:H4K20 me3
H3K4 me1:H2A.Z
H3K9 me2:DNA Meth. m5C
H4K20 me2:H4–4ac
H3K4 me3:H4–4ac
DNA Meth. m5C:H3K9 me3
H3K27 me3:H3K4 me1
H4K20 me2:H3–5ac
H3K27 me3:DNA Meth. m5C
H4K20 me2:H3K4 me3
H4K20 me3
H3–5ac
H3K9 me2
H3K4 me3
H3K4 me1
H3K27 me2
H3K9 me3
H4–4ac
H2A.Z
DNA Meth. m5C
H4K20 me2
H3K27 me3
0.00 0.25 0.50 0.75 1.00
π̂
HOXC11 – Reverse
H2A.Z:H3–5ac
H2A.Z:H4–4ac
H2A.Z:H3K4 me3
DNA Meth. m5C:H4K20 me2
H3K27 me2:DNA Meth. m5C
H3K4 me3:H3–5ac
H3K4 me1:H3–5ac
H4–4ac:H3–5ac
H3K27 me3:H3K9 me3
H2A.Z:H4K20 me2
H3K4 me1:H4–4ac
DNA Meth. m5C:H4K20 me3
H3K4 me1:H2A.Z
H3K9 me2:DNA Meth. m5C
H4K20 me2:H4–4ac
H3K4 me3:H4–4ac
DNA Meth. m5C:H3K9 me3
H3K27 me3:H3K4 me1
H4K20 me2:H3–5ac
H3K27 me3:DNA Meth. m5C
H4K20 me2:H3K4 me3
H4K20 me3
H3–5ac
H3K9 me2
H3K4 me3
H3K4 me1
H3K27 me2
H3K9 me3
H4–4ac
H2A.Z
DNA Meth. m5C
H4K20 me2
H3K27 me3
0.00 0.25 0.50 0.75 1.00
π̂

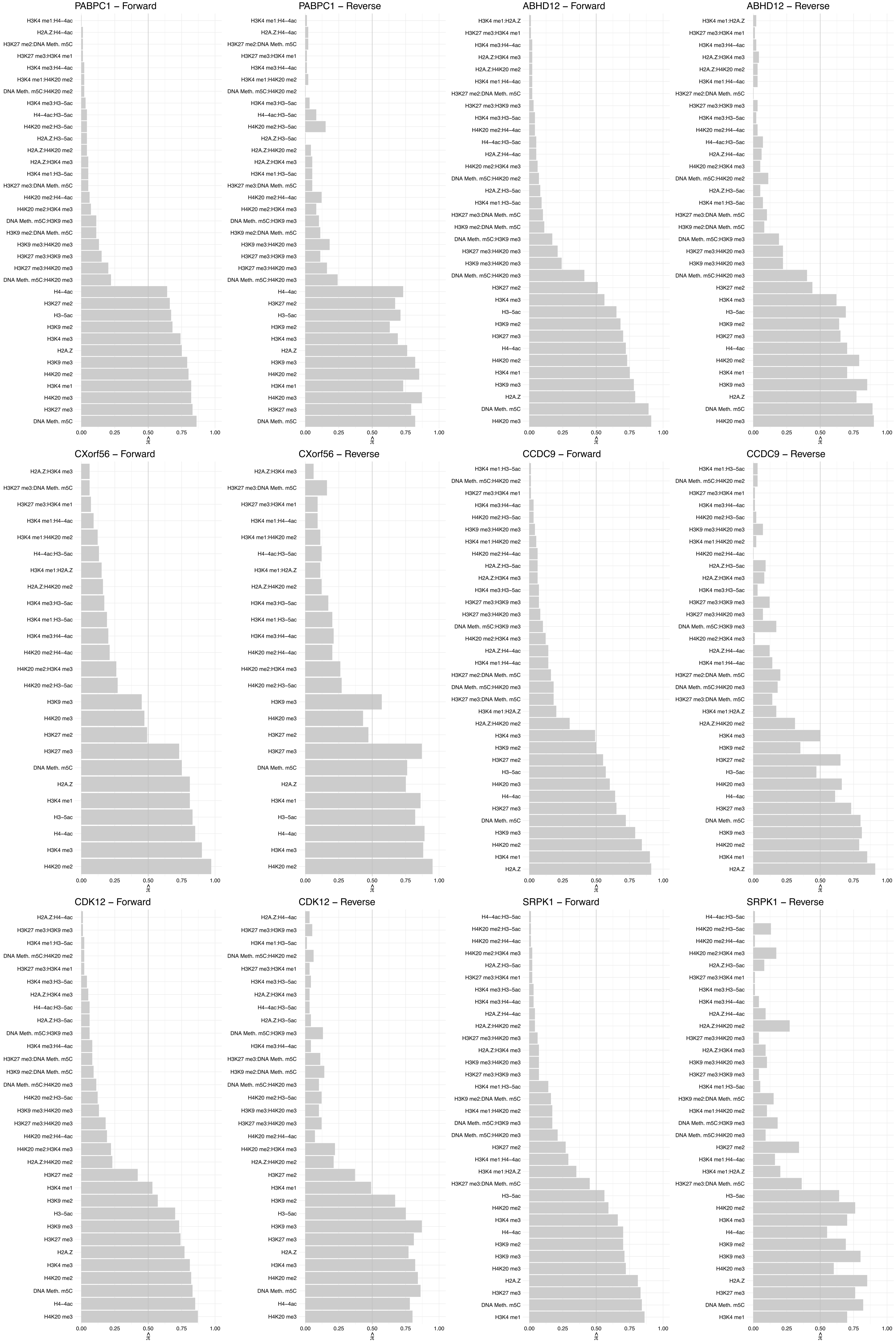
PABPC1 – Forward
H3K4 me1:H4−4ac
H2A.Z:H4−4ac
H3K27 me2:DNA Meth. m5C
H3K27 me3:H3K4 me1
H3K4 me3:H4−4ac
H3K4 me1:H4K20 me2
DNA Meth. m5C:H4K20 me2
H3K4 me3:H3−5ac
H4−4ac:H3−5ac
H4K20 me2:H3−5ac
H2A.Z:H3−5ac
H2A.Z:H4K20 me2
H2A.Z:H3K4 me3
H3K4 me1:H3−5ac
H3K27 me3:DNA Meth. m5C
H4K20 me2:H4−4ac
H4K20 me2:H3K4 me3
DNA Meth. m5C:H3K9 me3
H3K9 me2:DNA Meth. m5C
H3K9 me3:H4K20 me3
H3K27 me3:H3K9 me3
H3K27 me3:H4K20 me3
DNA Meth. m5C:H4K20 me3
H4−4ac
H3K27 me2
H3−5ac
H3K9 me2
H3K4 me3
H2A.Z
H3K9 me3
H4K20 me2
H3K4 me1
H4K20 me3
H3K27 me3
DNA Meth. m5C
0.00 0.25 0.50 0.75 1.00
π̂
PABPC1 – Reverse
H3K4 me1:H4−4ac
H2A.Z:H4−4ac
H3K27 me2:DNA Meth. m5C
H3K27 me3:H3K4 me1
H3K4 me3:H4−4ac
H3K4 me1:H4K20 me2
DNA Meth. m5C:H4K20 me2
H3K4 me3:H3−5ac
H4−4ac:H3−5ac
H4K20 me2:H3−5ac
H2A.Z:H3−5ac
H2A.Z:H4K20 me2
H2A.Z:H3K4 me3
H3K4 me1:H3−5ac
H3K27 me3:DNA Meth. m5C
H4K20 me2:H4−4ac
H4K20 me2:H3K4 me3
DNA Meth. m5C:H3K9 me3
H3K9 me2:DNA Meth. m5C
H3K9 me3:H4K20 me3
H3K27 me3:H3K9 me3
H3K27 me3:H4K20 me3
DNA Meth. m5C:H4K20 me3
H4−4ac
H3K27 me2
H3−5ac
H3K9 me2
H3K4 me3
H2A.Z
H3K9 me3
H4K20 me2
H3K4 me1
H4K20 me3
H3K27 me3
DNA Meth. m5C
0.00 0.25 0.50 0.75 1.00
π̂
ABHD12 – Forward
H3K4 me1:H2A.Z
H3K27 me3:H3K4 me1
H3K4 me3:H4−4ac
H2A.Z:H3K4 me3
H2A.Z:H4K20 me2
H3K4 me1:H4−4ac
H3K27 me2:DNA Meth. m5C
H3K27 me3:H3K9 me3
H3K4 me3:H3−5ac
H4K20 me2:H4−4ac
H4−4ac:H3−5ac
H2A.Z:H4−4ac
H4K20 me2:H3K4 me3
DNA Meth. m5C:H4K20 me2
H2A.Z:H3−5ac
H3K4 me1:H3−5ac
H3K27 me3:DNA Meth. m5C
H3K9 me2:DNA Meth. m5C
DNA Meth. m5C:H3K9 me3
H3K27 me3:H4K20 me3
H3K9 me3:H4K20 me3
DNA Meth. m5C:H4K20 me3
H3K27 me2
H3K4 me3
H3−5ac
H3K9 me2
H3K27 me3
H4−4ac
H4K20 me2
H3K4 me1
H3K9 me3
H2A.Z
DNA Meth. m5C
H4K20 me3
0.00 0.25 0.50 0.75 1.00
π̂
ABHD12 – Reverse
H3K4 me1:H2A.Z
H3K27 me3:H3K4 me1
H3K4 me3:H4−4ac
H2A.Z:H3K4 me3
H2A.Z:H4K20 me2
H3K4 me1:H4−4ac
H3K27 me2:DNA Meth. m5C
H3K27 me3:H3K9 me3
H3K4 me3:H3−5ac
H4K20 me2:H4−4ac
H4−4ac:H3−5ac
H2A.Z:H4−4ac
H4K20 me2:H3K4 me3
DNA Meth. m5C:H4K20 me2
H2A.Z:H3−5ac
H3K4 me1:H3−5ac
H3K27 me3:DNA Meth. m5C
H3K9 me2:DNA Meth. m5C
DNA Meth. m5C:H3K9 me3
H3K27 me3:H4K20 me3
H3K9 me3:H4K20 me3
DNA Meth. m5C:H4K20 me3
H3K27 me2
H3K4 me3
H3−5ac
H3K9 me2
H3K27 me3
H4−4ac
H4K20 me2
H3K4 me1
H3K9 me3
H2A.Z
DNA Meth. m5C
H4K20 me3
0.00 0.25 0.50 0.75 1.00
π̂
CXorf56 – Forward
H2A.Z:H3K4 me3
H3K27 me3:DNA Meth. m5C
H3K27 me3:H3K4 me1
H3K4 me1:H4−4ac
H3K4 me1:H4K20 me2
H4−4ac:H3−5ac
H3K4 me1:H2A.Z
H2A.Z:H4K20 me2
H3K4 me3:H3−5ac
H3K4 me1:H3−5ac
H3K4 me3:H4−4ac
H4K20 me2:H4−4ac
H4K20 me2:H3K4 me3
H4K20 me2:H3−5ac
H3K9 me3
H4K20 me3
H3K27 me2
H3K27 me3
DNA Meth. m5C
H2A.Z
H3K4 me1
H3−5ac
H4−4ac
H3K4 me3
H4K20 me2
0.00 0.25 0.50 0.75 1.00
π̂
CXorf56 – Reverse
H2A.Z:H3K4 me3
H3K27 me3:DNA Meth. m5C
H3K27 me3:H3K4 me1
H3K4 me1:H4−4ac
H3K4 me1:H4K20 me2
H4−4ac:H3−5ac
H3K4 me1:H2A.Z
H2A.Z:H4K20 me2
H3K4 me3:H3−5ac
H3K4 me1:H3−5ac
H3K4 me3:H4−4ac
H4K20 me2:H4−4ac
H4K20 me2:H3K4 me3
H4K20 me2:H3−5ac
H3K9 me3
H4K20 me3
H3K27 me2
H3K27 me3
DNA Meth. m5C
H2A.Z
H3K4 me1
H3−5ac
H4−4ac
H3K4 me3
H4K20 me2
0.00 0.25 0.50 0.75 1.00
π̂
CCDC9 – Forward
H3K4 me1:H3−5ac
DNA Meth. m5C:H4K20 me2
H3K27 me3:H3K4 me1
H3K4 me3:H4−4ac
H4K20 me2:H3−5ac
H3K9 me3:H4K20 me3
H3K4 me1:H4K20 me2
H4K20 me2:H4−4ac
H2A.Z:H3−5ac
H2A.Z:H3K4 me3
H3K4 me3:H3−5ac
H3K27 me3:H3K9 me3
H3K27 me3:H4K20 me3
DNA Meth. m5C:H3K9 me3
H4K20 me2:H3K4 me3
H2A.Z:H4−4ac
H3K4 me1:H4−4ac
H3K27 me2:DNA Meth. m5C
DNA Meth. m5C:H4K20 me3
H3K27 me3:DNA Meth. m5C
H3K4 me1:H2A.Z
H2A.Z:H4K20 me2
H3K4 me3
H3K9 me2
H3K27 me2
H3−5ac
H4K20 me3
H4−4ac
H3K27 me3
DNA Meth. m5C
H3K9 me3
H4K20 me2
H3K4 me1
H2A.Z
0.00 0.25 0.50 0.75 1.00
π̂
CCDC9 – Reverse
H3K4 me1:H3−5ac
DNA Meth. m5C:H4K20 me2
H3K27 me3:H3K4 me1
H3K4 me3:H4−4ac
H4K20 me2:H3−5ac
H3K9 me3:H4K20 me3
H3K4 me1:H4K20 me2
H4K20 me2:H4−4ac
H2A.Z:H3−5ac
H2A.Z:H3K4 me3
H3K4 me3:H3−5ac
H3K27 me3:H3K9 me3
H3K27 me3:H4K20 me3
DNA Meth. m5C:H3K9 me3
H4K20 me2:H3K4 me3
H2A.Z:H4−4ac
H3K4 me1:H4−4ac
H3K27 me2:DNA Meth. m5C
DNA Meth. m5C:H4K20 me3
H3K27 me3:DNA Meth. m5C
H3K4 me1:H2A.Z
H2A.Z:H4K20 me2
H3K4 me3
H3K9 me2
H3K27 me2
H3−5ac
H4K20 me3
H4−4ac
H3K27 me3
DNA Meth. m5C
H3K9 me3
H4K20 me2
H3K4 me1
H2A.Z
0.00 0.25 0.50 0.75 1.00
π̂
CDK12 – Forward
H2A.Z:H4−4ac
H3K27 me3:H3K9 me3
H3K4 me1:H3−5ac
DNA Meth. m5C:H4K20 me2
H3K27 me3:H3K4 me1
H3K4 me3:H3−5ac
H2A.Z:H3K4 me3
H4−4ac:H3−5ac
H2A.Z:H3−5ac
DNA Meth. m5C:H3K9 me3
H3K4 me3:H4−4ac
H3K27 me3:DNA Meth. m5C
H3K9 me2:DNA Meth. m5C
DNA Meth. m5C:H4K20 me3
H4K20 me2:H3−5ac
H3K9 me3:H4K20 me3
H3K27 me3:H4K20 me3
H4K20 me2:H4−4ac
H4K20 me2:H3K4 me3
H2A.Z:H4K20 me2
H3K27 me2
H3K4 me1
H3K9 me2
H3−5ac
H3K9 me3
H3K27 me3
H2A.Z
H3K4 me3
H4K20 me2
DNA Meth. m5C
H4−4ac
H4K20 me3
0.00 0.25 0.50 0.75 1.00
π̂
CDK12 – Reverse
H2A.Z:H4−4ac
H3K27 me3:H3K9 me3
H3K4 me1:H3−5ac
DNA Meth. m5C:H4K20 me2
H3K27 me3:H3K4 me1
H3K4 me3:H3−5ac
H2A.Z:H3K4 me3
H4−4ac:H3−5ac
H2A.Z:H3−5ac
DNA Meth. m5C:H3K9 me3
H3K4 me3:H4−4ac
H3K27 me3:DNA Meth. m5C
H3K9 me2:DNA Meth. m5C
DNA Meth. m5C:H4K20 me3
H4K20 me2:H3−5ac
H3K9 me3:H4K20 me3
H3K27 me3:H4K20 me3
H4K20 me2:H4−4ac
H4K20 me2:H3K4 me3
H2A.Z:H4K20 me2
H3K27 me2
H3K4 me1
H3K9 me2
H3−5ac
H3K9 me3
H3K27 me3
H2A.Z
H3K4 me3
H4K20 me2
DNA Meth. m5C
H4−4ac
H4K20 me3
0.00 0.25 0.50 0.75 1.00
π̂
SRPK1 – Forward
H4−4ac:H3−5ac
H4K20 me2:H3−5ac
H4K20 me2:H4−4ac
H4K20 me2:H3K4 me3
H2A.Z:H3−5ac
H3K27 me3:H3K4 me1
H3K4 me3:H3−5ac
H3K4 me3:H4−4ac
H2A.Z:H4−4ac
H2A.Z:H4K20 me2
H3K27 me3:H4K20 me3
H2A.Z:H3K4 me3
H3K9 me3:H4K20 me3
H3K27 me3:H3K9 me3
H3K4 me1:H3−5ac
H3K9 me2:DNA Meth. m5C
H3K4 me1:H4K20 me2
DNA Meth. m5C:H3K9 me3
DNA Meth. m5C:H4K20 me3
H3K27 me2
H3K4 me1:H4−4ac
H3K4 me1:H2A.Z
H3K27 me3:DNA Meth. m5C
H3−5ac
H4K20 me2
H3K4 me3
H4−4ac
H3K9 me2
H3K9 me3
H4K20 me3
H2A.Z
H3K27 me3
DNA Meth. m5C
H3K4 me1
0.00 0.25 0.50 0.75 1.00
π̂
SRPK1 – Reverse
H4−4ac:H3−5ac
H4K20 me2:H3−5ac
H4K20 me2:H4−4ac
H4K20 me2:H3K4 me3
H2A.Z:H3−5ac
H3K27 me3:H3K4 me1
H3K4 me3:H3−5ac
H3K4 me3:H4−4ac
H2A.Z:H4−4ac
H2A.Z:H4K20 me2
H3K27 me3:H4K20 me3
H2A.Z:H3K4 me3
H3K9 me3:H4K20 me3
H3K27 me3:H3K9 me3
H3K4 me1:H3−5ac
H3K9 me2:DNA Meth. m5C
H3K4 me1:H4K20 me2
DNA Meth. m5C:H3K9 me3
DNA Meth. m5C:H4K20 me3
H3K27 me2
H3K4 me1:H4−4ac
H3K4 me1:H2A.Z
H3K27 me3:DNA Meth. m5C
H3−5ac
H4K20 me2
H3K4 me3
H4−4ac
H3K9 me2
H3K9 me3
H4K20 me3
H2A.Z
H3K27 me3
DNA Meth. m5C
H3K4 me1
0.00 0.25 0.50 0.75 1.00
π̂

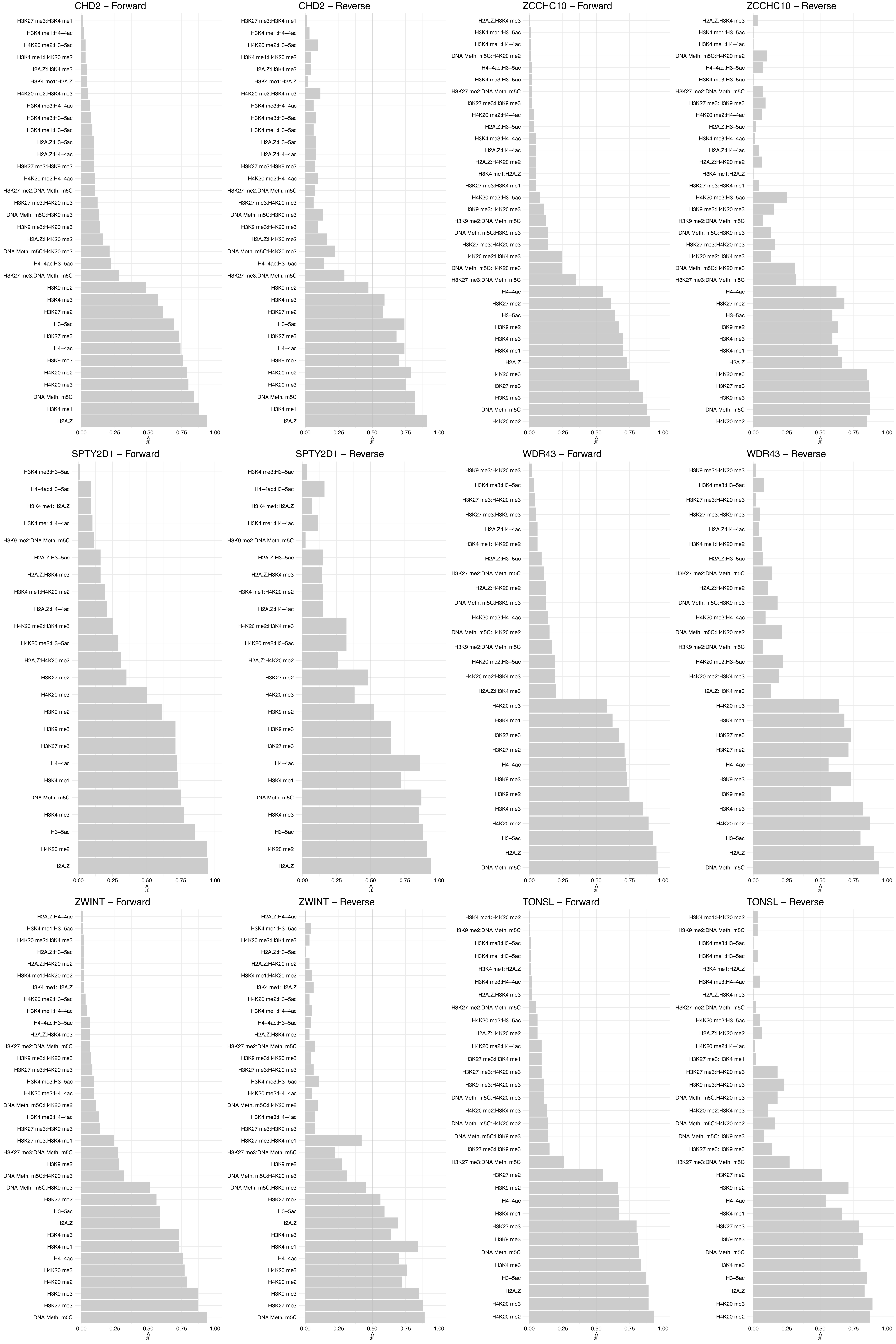
CHD2 – Forward
H3K27 me3:H3K4 me1
H3K4 me1:H4−4ac
H4K20 me2:H3−5ac
H3K4 me1:H4K20 me2
H2A.Z:H3K4 me3
H3K4 me1:H2A.Z
H4K20 me2:H3K4 me3
H3K4 me3:H4−4ac
H3K4 me3:H3−5ac
H3K4 me1:H3−5ac
H2A.Z:H3−5ac
H2A.Z:H4−4ac
H3K27 me3:H3K9 me3
H4K20 me2:H4−4ac
H3K27 me2:DNA Meth. m5C
H3K27 me3:H4K20 me3
DNA Meth. m5C:H3K9 me3
H3K9 me3:H4K20 me3
H2A.Z:H4K20 me2
DNA Meth. m5C:H4K20 me3
H4−4ac:H3−5ac
H3K27 me3:DNA Meth. m5C
H3K9 me2
H3K4 me3
H3K27 me2
H3−5ac
H3K27 me3
H4−4ac
H3K9 me3
H4K20 me2
H4K20 me3
DNA Meth. m5C
H3K4 me1
H2A.Z
0.00 0.25 0.50 0.75 1.00
$\hat{\pi}$
CHD2 – Reverse
H3K27 me3:H3K4 me1
H3K4 me1:H4−4ac
H4K20 me2:H3−5ac
H3K4 me1:H4K20 me2
H2A.Z:H3K4 me3
H3K4 me1:H2A.Z
H4K20 me2:H3K4 me3
H3K4 me3:H4−4ac
H3K4 me3:H3−5ac
H3K4 me1:H3−5ac
H2A.Z:H3−5ac
H2A.Z:H4−4ac
H3K27 me3:H3K9 me3
H4K20 me2:H4−4ac
H3K27 me2:DNA Meth. m5C
H3K27 me3:H4K20 me3
DNA Meth. m5C:H3K9 me3
H3K9 me3:H4K20 me3
H2A.Z:H4K20 me2
DNA Meth. m5C:H4K20 me3
H4−4ac:H3−5ac
H3K27 me3:DNA Meth. m5C
H3K9 me2
H3K4 me3
H3K27 me2
H3−5ac
H3K27 me3
H4−4ac
H3K9 me3
H4K20 me2
H4K20 me3
DNA Meth. m5C
H3K4 me1
H2A.Z
0.00 0.25 0.50 0.75 1.00
$\hat{\pi}$
ZCCHC10 – Forward
H2A.Z:H3K4 me3
H3K4 me1:H3−5ac
H3K4 me1:H4−4ac
DNA Meth. m5C:H4K20 me2
H4−4ac:H3−5ac
H3K4 me3:H3−5ac
H3K27 me2:DNA Meth. m5C
H3K27 me3:H3K9 me3
H4K20 me2:H4−4ac
H2A.Z:H3−5ac
H3K4 me3:H4−4ac
H2A.Z:H4−4ac
H2A.Z:H4K20 me2
H3K4 me1:H2A.Z
H3K27 me3:H3K4 me1
H4K20 me2:H3−5ac
H3K9 me3:H4K20 me3
H3K9 me2:DNA Meth. m5C
DNA Meth. m5C:H3K9 me3
H3K27 me3:H4K20 me3
H4K20 me2:H3K4 me3
DNA Meth. m5C:H4K20 me3
H3K27 me3:DNA Meth. m5C
H4−4ac
H3K27 me2
H3−5ac
H3K9 me2
H3K4 me3
H3K4 me1
H2A.Z
H4K20 me3
H3K27 me3
H3K9 me3
DNA Meth. m5C
H4K20 me2
0.00 0.25 0.50 0.75 1.00
$\hat{\pi}$
ZCCHC10 – Reverse
H2A.Z:H3K4 me3
H3K4 me1:H3−5ac
H3K4 me1:H4−4ac
DNA Meth. m5C:H4K20 me2
H4−4ac:H3−5ac
H3K4 me3:H3−5ac
H3K27 me2:DNA Meth. m5C
H3K27 me3:H3K9 me3
H4K20 me2:H4−4ac
H2A.Z:H3−5ac
H3K4 me3:H4−4ac
H2A.Z:H4−4ac
H2A.Z:H4K20 me2
H3K4 me1:H2A.Z
H3K27 me3:H3K4 me1
H4K20 me2:H3−5ac
H3K9 me3:H4K20 me3
H3K9 me2:DNA Meth. m5C
DNA Meth. m5C:H3K9 me3
H3K27 me3:H4K20 me3
H4K20 me2:H3K4 me3
DNA Meth. m5C:H4K20 me3
H3K27 me3:DNA Meth. m5C
H4−4ac
H3K27 me2
H3−5ac
H3K9 me2
H3K4 me3
H3K4 me1
H2A.Z
H4K20 me3
H3K27 me3
H3K9 me3
DNA Meth. m5C
H4K20 me2
0.00 0.25 0.50 0.75 1.00
$\hat{\pi}$
SPTY2D1 – Forward
H3K4 me3:H3−5ac
H4−4ac:H3−5ac
H3K4 me1:H2A.Z
H3K4 me1:H4−4ac
H3K9 me2:DNA Meth. m5C
H2A.Z:H3−5ac
H2A.Z:H3K4 me3
H3K4 me1:H4K20 me2
H2A.Z:H4−4ac
H4K20 me2:H3K4 me3
H4K20 me2:H3−5ac
H2A.Z:H4K20 me2
H3K27 me2
H4K20 me3
H3K9 me2
H3K9 me3
H3K27 me3
H4−4ac
H3K4 me1
DNA Meth. m5C
H3K4 me3
H3−5ac
H4K20 me2
H2A.Z
0.00 0.25 0.50 0.75 1.00
$\hat{\pi}$
SPTY2D1 – Reverse
H3K4 me3:H3−5ac
H4−4ac:H3−5ac
H3K4 me1:H2A.Z
H3K4 me1:H4−4ac
H3K9 me2:DNA Meth. m5C
H2A.Z:H3−5ac
H2A.Z:H3K4 me3
H3K4 me1:H4K20 me2
H2A.Z:H4−4ac
H4K20 me2:H3K4 me3
H4K20 me2:H3−5ac
H2A.Z:H4K20 me2
H3K27 me2
H4K20 me3
H3K9 me2
H3K9 me3
H3K27 me3
H4−4ac
H3K4 me1
DNA Meth. m5C
H3K4 me3
H3−5ac
H4K20 me2
H2A.Z
0.00 0.25 0.50 0.75 1.00
$\hat{\pi}$
WDR43 – Forward
H3K9 me3:H4K20 me3
H3K4 me3:H3−5ac
H3K27 me3:H4K20 me3
H3K27 me3:H3K9 me3
H2A.Z:H4−4ac
H3K4 me1:H4K20 me2
H2A.Z:H3−5ac
H3K27 me2:DNA Meth. m5C
H2A.Z:H4K20 me2
DNA Meth. m5C:H3K9 me3
H4K20 me2:H4−4ac
DNA Meth. m5C:H4K20 me2
H3K9 me2:DNA Meth. m5C
H4K20 me2:H3−5ac
H4K20 me2:H3K4 me3
H2A.Z:H3K4 me3
H4K20 me3
H3K4 me1
H3K27 me3
H3K27 me2
H4−4ac
H3K9 me3
H3K9 me2
H3K4 me3
H4K20 me2
H3−5ac
H2A.Z
DNA Meth. m5C
0.00 0.25 0.50 0.75 1.00
$\hat{\pi}$
WDR43 – Reverse
H3K9 me3:H4K20 me3
H3K4 me3:H3−5ac
H3K27 me3:H4K20 me3
H3K27 me3:H3K9 me3
H2A.Z:H4−4ac
H3K4 me1:H4K20 me2
H2A.Z:H3−5ac
H3K27 me2:DNA Meth. m5C
H2A.Z:H4K20 me2
DNA Meth. m5C:H3K9 me3
H4K20 me2:H4−4ac
DNA Meth. m5C:H4K20 me2
H3K9 me2:DNA Meth. m5C
H4K20 me2:H3−5ac
H4K20 me2:H3K4 me3
H2A.Z:H3K4 me3
H4K20 me3
H3K4 me1
H3K27 me3
H3K27 me2
H4−4ac
H3K9 me3
H3K9 me2
H3K4 me3
H4K20 me2
H3−5ac
H2A.Z
DNA Meth. m5C
0.00 0.25 0.50 0.75 1.00
$\hat{\pi}$
ZWINT – Forward
H2A.Z:H4−4ac
H3K4 me1:H3−5ac
H4K20 me2:H3K4 me3
H2A.Z:H3−5ac
H2A.Z:H4K20 me2
H3K4 me1:H4K20 me2
H3K4 me1:H2A.Z
H4K20 me2:H3−5ac
H3K4 me1:H4−4ac
H4−4ac:H3−5ac
H2A.Z:H3K4 me3
H3K27 me2:DNA Meth. m5C
H3K9 me3:H4K20 me3
H3K27 me3:H4K20 me3
H3K4 me3:H3−5ac
H4K20 me2:H4−4ac
DNA Meth. m5C:H4K20 me2
H3K4 me3:H4−4ac
H3K27 me3:H3K9 me3
H3K27 me3:H3K4 me1
H3K27 me3:DNA Meth. m5C
H3K9 me2
DNA Meth. m5C:H4K20 me3
DNA Meth. m5C:H3K9 me3
H3K27 me2
H3−5ac
H2A.Z
H3K4 me3
H3K4 me1
H4−4ac
H4K20 me3
H4K20 me2
H3K9 me3
H3K27 me3
DNA Meth. m5C
0.00 0.25 0.50 0.75 1.00
$\hat{\pi}$
ZWINT – Reverse
H2A.Z:H4−4ac
H3K4 me1:H3−5ac
H4K20 me2:H3K4 me3
H2A.Z:H3−5ac
H2A.Z:H4K20 me2
H3K4 me1:H4K20 me2
H3K4 me1:H2A.Z
H4K20 me2:H3−5ac
H3K4 me1:H4−4ac
H4−4ac:H3−5ac
H2A.Z:H3K4 me3
H3K27 me2:DNA Meth. m5C
H3K9 me3:H4K20 me3
H3K27 me3:H4K20 me3
H3K4 me3:H3−5ac
H4K20 me2:H4−4ac
DNA Meth. m5C:H4K20 me2
H3K4 me3:H4−4ac
H3K27 me3:H3K9 me3
H3K27 me3:H3K4 me1
H3K27 me3:DNA Meth. m5C
H3K9 me2
DNA Meth. m5C:H4K20 me3
DNA Meth. m5C:H3K9 me3
H3K27 me2
H3−5ac
H2A.Z
H3K4 me3
H3K4 me1
H4−4ac
H4K20 me3
H4K20 me2
H3K9 me3
H3K27 me3
DNA Meth. m5C
0.00 0.25 0.50 0.75 1.00
$\hat{\pi}$
TONSL – Forward
H3K4 me1:H4K20 me2
H3K9 me2:DNA Meth. m5C
H3K4 me3:H3−5ac
H3K4 me1:H3−5ac
H3K4 me1:H2A.Z
H3K4 me3:H4−4ac
H2A.Z:H3K4 me3
H3K27 me2:DNA Meth. m5C
H4K20 me2:H3−5ac
H2A.Z:H4K20 me2
H4K20 me2:H4−4ac
H3K27 me3:H3K4 me1
H3K27 me3:H4K20 me3
H3K9 me3:H4K20 me3
DNA Meth. m5C:H4K20 me3
H4K20 me2:H3K4 me3
DNA Meth. m5C:H4K20 me2
DNA Meth. m5C:H3K9 me3
H3K27 me3:H3K9 me3
H3K27 me3:DNA Meth. m5C
H3K27 me2
H3K9 me2
H4−4ac
H3K4 me1
H3K27 me3
H3K9 me3
DNA Meth. m5C
H3K4 me3
H3−5ac
H2A.Z
H4K20 me3
H4K20 me2
0.00 0.25 0.50 0.75 1.00
$\hat{\pi}$
TONSL – Reverse
H3K4 me1:H4K20 me2
H3K9 me2:DNA Meth. m5C
H3K4 me3:H3−5ac
H3K4 me1:H3−5ac
H3K4 me1:H2A.Z
H3K4 me3:H4−4ac
H2A.Z:H3K4 me3
H3K27 me2:DNA Meth. m5C
H4K20 me2:H3−5ac
H2A.Z:H4K20 me2
H4K20 me2:H4−4ac
H3K27 me3:H3K4 me1
H3K27 me3:H4K20 me3
H3K9 me3:H4K20 me3
DNA Meth. m5C:H4K20 me3
H4K20 me2:H3K4 me3
DNA Meth. m5C:H4K20 me2
DNA Meth. m5C:H3K9 me3
H3K27 me3:H3K9 me3
H3K27 me3:DNA Meth. m5C
H3K27 me2
H3K9 me2
H4−4ac
H3K4 me1
H3K27 me3
H3K9 me3
DNA Meth. m5C
H3K4 me3
H3−5ac
H2A.Z
H4K20 me3
H4K20 me2
0.00 0.25 0.50 0.75 1.00
$\hat{\pi}$

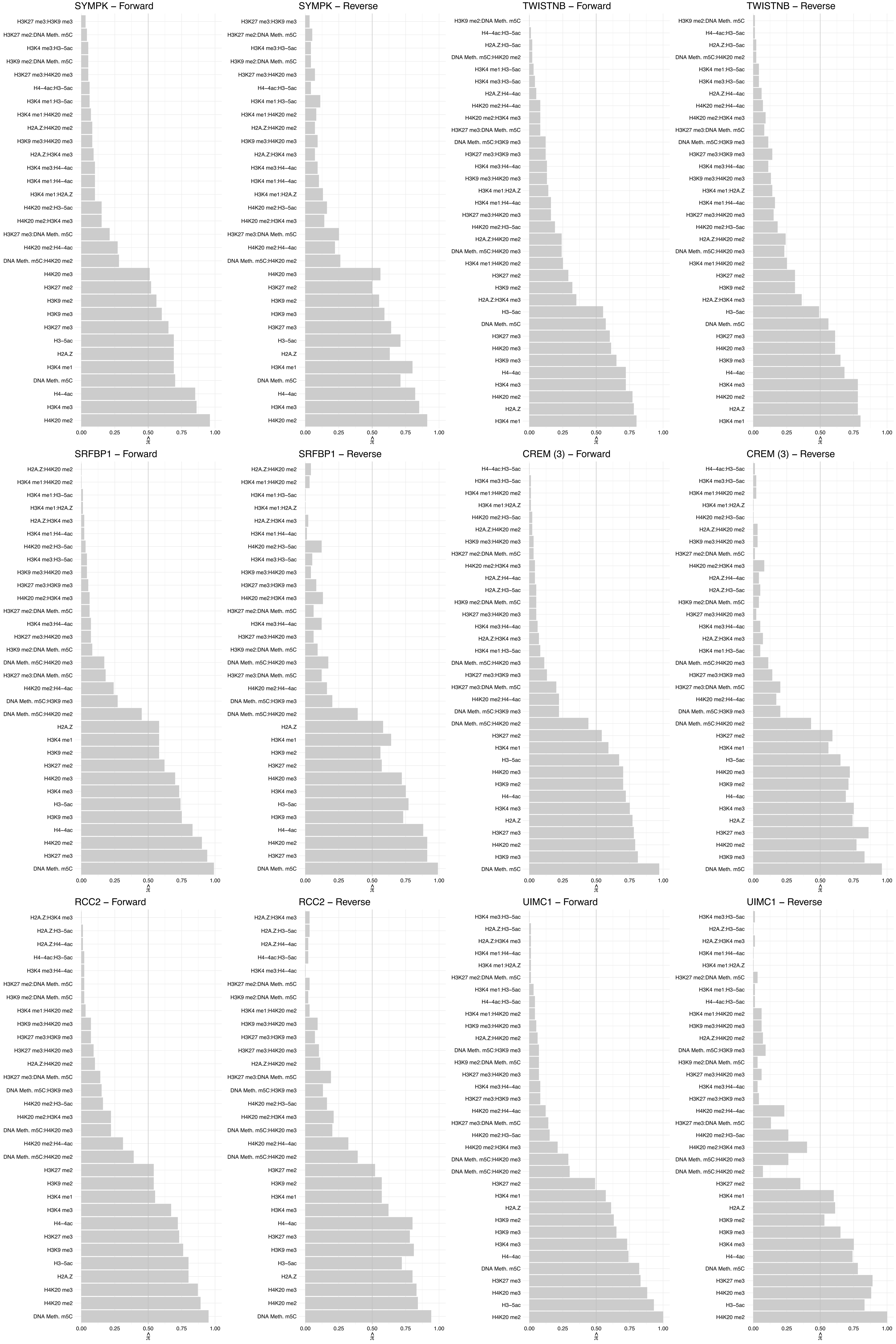
SYMPK – Forward
H3K27 me3:H3K9 me3
H3K27 me2:DNA Meth. m5C
H3K4 me3:H3–5ac
H3K9 me2:DNA Meth. m5C
H3K27 me3:H4K20 me3
H4–4ac:H3–5ac
H3K4 me1:H3–5ac
H3K4 me1:H4K20 me2
H2A.Z:H4K20 me2
H3K9 me3:H4K20 me3
H2A.Z:H3K4 me3
H3K4 me3:H4–4ac
H3K4 me1:H4–4ac
H3K4 me1:H2A.Z
H4K20 me2:H3–5ac
H4K20 me2:H3K4 me3
H3K27 me3:DNA Meth. m5C
H4K20 me2:H4–4ac
DNA Meth. m5C:H4K20 me2
H4K20 me3
H3K27 me2
H3K9 me2
H3K9 me3
H3K27 me3
H3–5ac
H2A.Z
H3K4 me1
DNA Meth. m5C
H4–4ac
H3K4 me3
H4K20 me2
0.00 0.25 0.50 0.75 1.00
π̂
SYMPK – Reverse
H3K27 me3:H3K9 me3
H3K27 me2:DNA Meth. m5C
H3K4 me3:H3–5ac
H3K9 me2:DNA Meth. m5C
H3K27 me3:H4K20 me3
H4–4ac:H3–5ac
H3K4 me1:H3–5ac
H3K4 me1:H4K20 me2
H2A.Z:H4K20 me2
H3K9 me3:H4K20 me3
H2A.Z:H3K4 me3
H3K4 me3:H4–4ac
H3K4 me1:H4–4ac
H3K4 me1:H2A.Z
H4K20 me2:H3–5ac
H4K20 me2:H3K4 me3
H3K27 me3:DNA Meth. m5C
H4K20 me2:H4–4ac
DNA Meth. m5C:H4K20 me2
H4K20 me3
H3K27 me2
H3K9 me2
H3K9 me3
H3K27 me3
H3–5ac
H2A.Z
H3K4 me1
DNA Meth. m5C
H4–4ac
H3K4 me3
H4K20 me2
0.00 0.25 0.50 0.75 1.00
π̂
TWISTNB – Forward
H3K9 me2:DNA Meth. m5C
H4–4ac:H3–5ac
H2A.Z:H3–5ac
DNA Meth. m5C:H4K20 me2
H3K4 me1:H3–5ac
H3K4 me3:H3–5ac
H2A.Z:H4–4ac
H4K20 me2:H4–4ac
H4K20 me2:H3K4 me3
H3K27 me3:DNA Meth. m5C
DNA Meth. m5C:H3K9 me3
H3K27 me3:H3K9 me3
H3K4 me3:H4–4ac
H3K9 me3:H4K20 me3
H3K4 me1:H2A.Z
H3K4 me1:H4–4ac
H3K27 me3:H4K20 me3
H4K20 me2:H3–5ac
H2A.Z:H4K20 me2
DNA Meth. m5C:H4K20 me3
H3K4 me1:H4K20 me2
H3K27 me2
H3K9 me2
H2A.Z:H3K4 me3
H3–5ac
DNA Meth. m5C
H3K27 me3
H4K20 me3
H3K9 me3
H4–4ac
H3K4 me3
H4K20 me2
H2A.Z
H3K4 me1
0.00 0.25 0.50 0.75 1.00
π̂
TWISTNB – Reverse
H3K9 me2:DNA Meth. m5C
H4–4ac:H3–5ac
H2A.Z:H3–5ac
DNA Meth. m5C:H4K20 me2
H3K4 me1:H3–5ac
H3K4 me3:H3–5ac
H2A.Z:H4–4ac
H4K20 me2:H4–4ac
H4K20 me2:H3K4 me3
H3K27 me3:DNA Meth. m5C
DNA Meth. m5C:H3K9 me3
H3K27 me3:H3K9 me3
H3K4 me3:H4–4ac
H3K9 me3:H4K20 me3
H3K4 me1:H2A.Z
H3K4 me1:H4–4ac
H3K27 me3:H4K20 me3
H4K20 me2:H3–5ac
H2A.Z:H4K20 me2
DNA Meth. m5C:H4K20 me3
H3K4 me1:H4K20 me2
H3K27 me2
H3K9 me2
H2A.Z:H3K4 me3
H3–5ac
DNA Meth. m5C
H3K27 me3
H4K20 me3
H3K9 me3
H4–4ac
H3K4 me3
H4K20 me2
H2A.Z
H3K4 me1
0.00 0.25 0.50 0.75 1.00
π̂
SRFBP1 – Forward
H2A.Z:H4K20 me2
H3K4 me1:H4K20 me2
H3K4 me1:H3–5ac
H3K4 me1:H2A.Z
H2A.Z:H3K4 me3
H3K4 me1:H4–4ac
H4K20 me2:H3–5ac
H3K4 me3:H3–5ac
H3K9 me3:H4K20 me3
H3K27 me3:H3K9 me3
H4K20 me2:H3K4 me3
H3K27 me2:DNA Meth. m5C
H3K4 me3:H4–4ac
H3K27 me3:H4K20 me3
H3K9 me2:DNA Meth. m5C
DNA Meth. m5C:H4K20 me3
H3K27 me3:DNA Meth. m5C
H4K20 me2:H4–4ac
DNA Meth. m5C:H3K9 me3
DNA Meth. m5C:H4K20 me2
H2A.Z
H3K4 me1
H3K9 me2
H3K27 me2
H4K20 me3
H3K4 me3
H3–5ac
H3K9 me3
H4–4ac
H4K20 me2
H3K27 me3
DNA Meth. m5C
0.00 0.25 0.50 0.75 1.00
π̂
SRFBP1 – Reverse
H2A.Z:H4K20 me2
H3K4 me1:H4K20 me2
H3K4 me1:H3–5ac
H3K4 me1:H2A.Z
H2A.Z:H3K4 me3
H3K4 me1:H4–4ac
H4K20 me2:H3–5ac
H3K4 me3:H3–5ac
H3K9 me3:H4K20 me3
H3K27 me3:H3K9 me3
H4K20 me2:H3K4 me3
H3K27 me2:DNA Meth. m5C
H3K4 me3:H4–4ac
H3K27 me3:H4K20 me3
H3K9 me2:DNA Meth. m5C
DNA Meth. m5C:H4K20 me3
H3K27 me3:DNA Meth. m5C
H4K20 me2:H4–4ac
DNA Meth. m5C:H3K9 me3
DNA Meth. m5C:H4K20 me2
H2A.Z
H3K4 me1
H3K9 me2
H3K27 me2
H4K20 me3
H3K4 me3
H3–5ac
H3K9 me3
H4–4ac
H4K20 me2
H3K27 me3
DNA Meth. m5C
0.00 0.25 0.50 0.75 1.00
π̂
CREM (3) – Forward
H4–4ac:H3–5ac
H3K4 me3:H3–5ac
H3K4 me1:H4K20 me2
H3K4 me1:H2A.Z
H4K20 me2:H3–5ac
H2A.Z:H4K20 me2
H3K9 me3:H4K20 me3
H3K27 me2:DNA Meth. m5C
H4K20 me2:H3K4 me3
H2A.Z:H4–4ac
H2A.Z:H3–5ac
H3K9 me2:DNA Meth. m5C
H3K27 me3:H4K20 me3
H3K4 me3:H4–4ac
H2A.Z:H3K4 me3
H3K4 me1:H3–5ac
DNA Meth. m5C:H4K20 me3
H3K27 me3:H3K9 me3
H3K27 me3:DNA Meth. m5C
H4K20 me2:H4–4ac
DNA Meth. m5C:H3K9 me3
DNA Meth. m5C:H4K20 me2
H3K27 me2
H3K4 me1
H3–5ac
H4K20 me3
H3K9 me2
H4–4ac
H3K4 me3
H2A.Z
H3K27 me3
H4K20 me2
H3K9 me3
DNA Meth. m5C
0.00 0.25 0.50 0.75 1.00
π̂
CREM (3) – Reverse
H4–4ac:H3–5ac
H3K4 me3:H3–5ac
H3K4 me1:H4K20 me2
H3K4 me1:H2A.Z
H4K20 me2:H3–5ac
H2A.Z:H4K20 me2
H3K9 me3:H4K20 me3
H3K27 me2:DNA Meth. m5C
H4K20 me2:H3K4 me3
H2A.Z:H4–4ac
H2A.Z:H3–5ac
H3K9 me2:DNA Meth. m5C
H3K27 me3:H4K20 me3
H3K4 me3:H4–4ac
H2A.Z:H3K4 me3
H3K4 me1:H3–5ac
DNA Meth. m5C:H4K20 me3
H3K27 me3:H3K9 me3
H3K27 me3:DNA Meth. m5C
H4K20 me2:H4–4ac
DNA Meth. m5C:H3K9 me3
DNA Meth. m5C:H4K20 me2
H3K27 me2
H3K4 me1
H3–5ac
H4K20 me3
H3K9 me2
H4–4ac
H3K4 me3
H2A.Z
H3K27 me3
H4K20 me2
H3K9 me3
DNA Meth. m5C
0.00 0.25 0.50 0.75 1.00
π̂
RCC2 – Forward
H2A.Z:H3K4 me3
H2A.Z:H3–5ac
H2A.Z:H4–4ac
H4–4ac:H3–5ac
H3K4 me3:H4–4ac
H3K27 me2:DNA Meth. m5C
H3K9 me2:DNA Meth. m5C
H3K4 me1:H4K20 me2
H3K9 me3:H4K20 me3
H3K27 me3:H3K9 me3
H3K27 me3:H4K20 me3
H2A.Z:H4K20 me2
H3K27 me3:DNA Meth. m5C
DNA Meth. m5C:H3K9 me3
H4K20 me2:H3–5ac
H4K20 me2:H3K4 me3
DNA Meth. m5C:H4K20 me3
H4K20 me2:H4–4ac
DNA Meth. m5C:H4K20 me2
H3K27 me2
H3K9 me2
H3K4 me1
H3K4 me3
H4–4ac
H3K27 me3
H3K9 me3
H3–5ac
H2A.Z
H4K20 me3
H4K20 me2
DNA Meth. m5C
0.00 0.25 0.50 0.75 1.00
π̂
RCC2 – Reverse
H2A.Z:H3K4 me3
H2A.Z:H3–5ac
H2A.Z:H4–4ac
H4–4ac:H3–5ac
H3K4 me3:H4–4ac
H3K27 me2:DNA Meth. m5C
H3K9 me2:DNA Meth. m5C
H3K4 me1:H4K20 me2
H3K9 me3:H4K20 me3
H3K27 me3:H3K9 me3
H3K27 me3:H4K20 me3
H2A.Z:H4K20 me2
H3K27 me3:DNA Meth. m5C
DNA Meth. m5C:H3K9 me3
H4K20 me2:H3–5ac
H4K20 me2:H3K4 me3
DNA Meth. m5C:H4K20 me3
H4K20 me2:H4–4ac
DNA Meth. m5C:H4K20 me2
H3K27 me2
H3K9 me2
H3K4 me1
H3K4 me3
H4–4ac
H3K27 me3
H3K9 me3
H3–5ac
H2A.Z
H4K20 me3
H4K20 me2
DNA Meth. m5C
0.00 0.25 0.50 0.75 1.00
π̂
UIMC1 – Forward
H3K4 me3:H3–5ac
H2A.Z:H3–5ac
H2A.Z:H3K4 me3
H3K4 me1:H4–4ac
H3K4 me1:H2A.Z
H3K27 me2:DNA Meth. m5C
H3K4 me1:H3–5ac
H4–4ac:H3–5ac
H3K4 me1:H4K20 me2
H3K9 me3:H4K20 me3
H2A.Z:H4K20 me2
DNA Meth. m5C:H3K9 me3
H3K9 me2:DNA Meth. m5C
H3K27 me3:H4K20 me3
H3K4 me3:H4–4ac
H3K27 me3:H3K9 me3
H4K20 me2:H4–4ac
H3K27 me3:DNA Meth. m5C
H4K20 me2:H3–5ac
H4K20 me2:H3K4 me3
DNA Meth. m5C:H4K20 me3
DNA Meth. m5C:H4K20 me2
H3K27 me2
H3K4 me1
H2A.Z
H3K9 me2
H3K9 me3
H3K4 me3
H4–4ac
DNA Meth. m5C
H3K27 me3
H4K20 me3
H3–5ac
H4K20 me2
0.00 0.25 0.50 0.75 1.00
π̂
UIMC1 – Reverse
H3K4 me3:H3–5ac
H2A.Z:H3–5ac
H2A.Z:H3K4 me3
H3K4 me1:H4–4ac
H3K4 me1:H2A.Z
H3K27 me2:DNA Meth. m5C
H3K4 me1:H3–5ac
H4–4ac:H3–5ac
H3K4 me1:H4K20 me2
H3K9 me3:H4K20 me3
H2A.Z:H4K20 me2
DNA Meth. m5C:H3K9 me3
H3K9 me2:DNA Meth. m5C
H3K27 me3:H4K20 me3
H3K4 me3:H4–4ac
H3K27 me3:H3K9 me3
H4K20 me2:H4–4ac
H3K27 me3:DNA Meth. m5C
H4K20 me2:H3–5ac
H4K20 me2:H3K4 me3
DNA Meth. m5C:H4K20 me3
DNA Meth. m5C:H4K20 me2
H3K27 me2
H3K4 me1
H2A.Z
H3K9 me2
H3K9 me3
H3K4 me3
H4–4ac
DNA Meth. m5C
H3K27 me3
H4K20 me3
H3–5ac
H4K20 me2
0.00 0.25 0.50 0.75 1.00
π̂

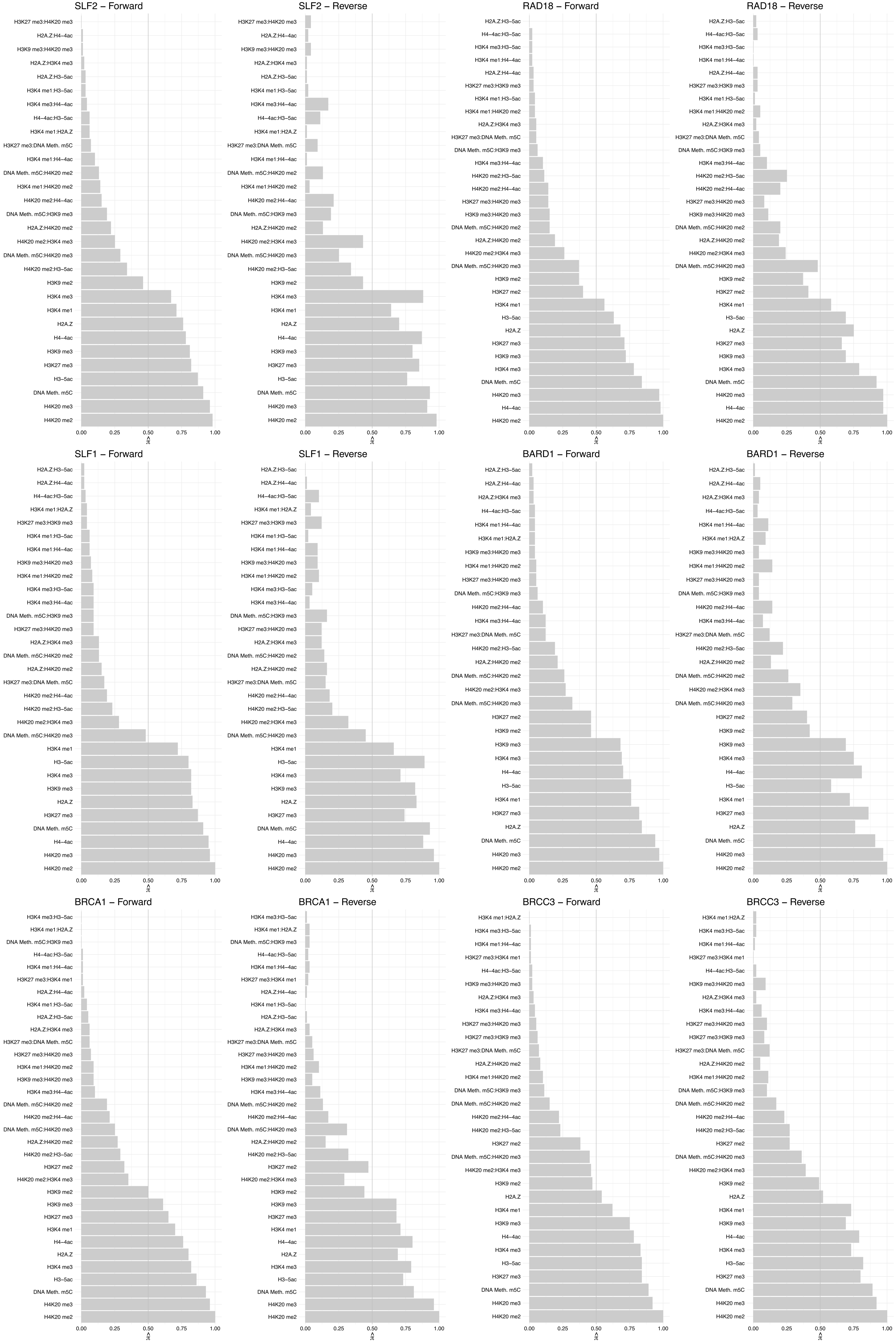
SLF2 – Forward
H3K27 me3:H4K20 me3
H2A.Z:H4–4ac
H3K9 me3:H4K20 me3
H2A.Z:H3K4 me3
H2A.Z:H3–5ac
H3K4 me1:H3–5ac
H3K4 me3:H4–4ac
H4–4ac:H3–5ac
H3K4 me1:H2A.Z
H3K27 me3:DNA Meth. m5C
H3K4 me1:H4–4ac
DNA Meth. m5C:H4K20 me2
H3K4 me1:H4K20 me2
H4K20 me2:H4–4ac
DNA Meth. m5C:H3K9 me3
H2A.Z:H4K20 me2
H4K20 me2:H3K4 me3
DNA Meth. m5C:H4K20 me3
H4K20 me2:H3–5ac
H3K9 me2
H3K4 me3
H3K4 me1
H2A.Z
H4–4ac
H3K9 me3
H3K27 me3
H3–5ac
DNA Meth. m5C
H4K20 me3
H4K20 me2
0.00
0.25
0.50
0.75
1.00
$\hat{\pi}$
SLF2 – Reverse
H3K27 me3:H4K20 me3
H2A.Z:H4–4ac
H3K9 me3:H4K20 me3
H2A.Z:H3K4 me3
H2A.Z:H3–5ac
H3K4 me1:H3–5ac
H3K4 me3:H4–4ac
H4–4ac:H3–5ac
H3K4 me1:H2A.Z
H3K27 me3:DNA Meth. m5C
H3K4 me1:H4–4ac
DNA Meth. m5C:H4K20 me2
H3K4 me1:H4K20 me2
H4K20 me2:H4–4ac
DNA Meth. m5C:H3K9 me3
H2A.Z:H4K20 me2
H4K20 me2:H3K4 me3
DNA Meth. m5C:H4K20 me3
H4K20 me2:H3–5ac
H3K9 me2
H3K4 me3
H3K4 me1
H2A.Z
H4–4ac
H3K9 me3
H3K27 me3
H3–5ac
DNA Meth. m5C
H4K20 me3
H4K20 me2
0.00
0.25
0.50
0.75
1.00
$\hat{\pi}$
RAD18 – Forward
H2A.Z:H3–5ac
H4–4ac:H3–5ac
H3K4 me3:H3–5ac
H3K4 me1:H4–4ac
H2A.Z:H4–4ac
H3K27 me3:H3K9 me3
H3K4 me1:H3–5ac
H3K4 me1:H4K20 me2
H2A.Z:H3K4 me3
H3K27 me3:DNA Meth. m5C
DNA Meth. m5C:H3K9 me3
H3K4 me3:H4–4ac
H4K20 me2:H3–5ac
H4K20 me2:H4–4ac
H3K27 me3:H4K20 me3
H3K9 me3:H4K20 me3
DNA Meth. m5C:H4K20 me2
H2A.Z:H4K20 me2
H4K20 me2:H3K4 me3
DNA Meth. m5C:H4K20 me3
H3K9 me2
H3K27 me2
H3K4 me1
H3–5ac
H2A.Z
H3K27 me3
H3K9 me3
H3K4 me3
DNA Meth. m5C
H4K20 me3
H4–4ac
H4K20 me2
0.00
0.25
0.50
0.75
1.00
$\hat{\pi}$
RAD18 – Reverse
H2A.Z:H3–5ac
H4–4ac:H3–5ac
H3K4 me3:H3–5ac
H3K4 me1:H4–4ac
H2A.Z:H4–4ac
H3K27 me3:H3K9 me3
H3K4 me1:H3–5ac
H3K4 me1:H4K20 me2
H2A.Z:H3K4 me3
H3K27 me3:DNA Meth. m5C
DNA Meth. m5C:H3K9 me3
H3K4 me3:H4–4ac
H4K20 me2:H3–5ac
H4K20 me2:H4–4ac
H3K27 me3:H4K20 me3
H3K9 me3:H4K20 me3
DNA Meth. m5C:H4K20 me2
H2A.Z:H4K20 me2
H4K20 me2:H3K4 me3
DNA Meth. m5C:H4K20 me3
H3K9 me2
H3K27 me2
H3K4 me1
H3–5ac
H2A.Z
H3K27 me3
H3K9 me3
H3K4 me3
DNA Meth. m5C
H4K20 me3
H4–4ac
H4K20 me2
0.00
0.25
0.50
0.75
1.00
$\hat{\pi}$
SLF1 – Forward
H2A.Z:H3–5ac
H2A.Z:H4–4ac
H4–4ac:H3–5ac
H3K4 me1:H2A.Z
H3K27 me3:H3K9 me3
H3K4 me1:H3–5ac
H3K4 me1:H4–4ac
H3K9 me3:H4K20 me3
H3K4 me1:H4K20 me2
H3K4 me3:H3–5ac
H3K4 me3:H4–4ac
DNA Meth. m5C:H3K9 me3
H3K27 me3:H4K20 me3
H2A.Z:H3K4 me3
DNA Meth. m5C:H4K20 me2
H2A.Z:H4K20 me2
H3K27 me3:DNA Meth. m5C
H4K20 me2:H4–4ac
H4K20 me2:H3–5ac
H4K20 me2:H3K4 me3
DNA Meth. m5C:H4K20 me3
H3K4 me1
H3–5ac
H3K4 me3
H3K9 me3
H2A.Z
H3K27 me3
DNA Meth. m5C
H4–4ac
H4K20 me3
H4K20 me2
0.00
0.25
0.50
0.75
1.00
$\hat{\pi}$
SLF1 – Reverse
H2A.Z:H3–5ac
H2A.Z:H4–4ac
H4–4ac:H3–5ac
H3K4 me1:H2A.Z
H3K27 me3:H3K9 me3
H3K4 me1:H3–5ac
H3K4 me1:H4–4ac
H3K9 me3:H4K20 me3
H3K4 me1:H4K20 me2
H3K4 me3:H3–5ac
H3K4 me3:H4–4ac
DNA Meth. m5C:H3K9 me3
H3K27 me3:H4K20 me3
H2A.Z:H3K4 me3
DNA Meth. m5C:H4K20 me2
H2A.Z:H4K20 me2
H3K27 me3:DNA Meth. m5C
H4K20 me2:H4–4ac
H4K20 me2:H3–5ac
H4K20 me2:H3K4 me3
DNA Meth. m5C:H4K20 me3
H3K4 me1
H3–5ac
H3K4 me3
H3K9 me3
H2A.Z
H3K27 me3
DNA Meth. m5C
H4–4ac
H4K20 me3
H4K20 me2
0.00
0.25
0.50
0.75
1.00
$\hat{\pi}$
BARD1 – Forward
H2A.Z:H3–5ac
H2A.Z:H4–4ac
H2A.Z:H3K4 me3
H4–4ac:H3–5ac
H3K4 me1:H4–4ac
H3K4 me1:H2A.Z
H3K9 me3:H4K20 me3
H3K4 me1:H4K20 me2
H3K27 me3:H4K20 me3
DNA Meth. m5C:H3K9 me3
H4K20 me2:H4–4ac
H3K4 me3:H4–4ac
H3K27 me3:DNA Meth. m5C
H4K20 me2:H3–5ac
H2A.Z:H4K20 me2
DNA Meth. m5C:H4K20 me2
H4K20 me2:H3K4 me3
DNA Meth. m5C:H4K20 me3
H3K27 me2
H3K9 me2
H3K9 me3
H3K4 me3
H4–4ac
H3–5ac
H3K4 me1
H3K27 me3
H2A.Z
DNA Meth. m5C
H4K20 me3
H4K20 me2
0.00
0.25
0.50
0.75
1.00
$\hat{\pi}$
BARD1 – Reverse
H2A.Z:H3–5ac
H2A.Z:H4–4ac
H2A.Z:H3K4 me3
H4–4ac:H3–5ac
H3K4 me1:H4–4ac
H3K4 me1:H2A.Z
H3K9 me3:H4K20 me3
H3K4 me1:H4K20 me2
H3K27 me3:H4K20 me3
DNA Meth. m5C:H3K9 me3
H4K20 me2:H4–4ac
H3K4 me3:H4–4ac
H3K27 me3:DNA Meth. m5C
H4K20 me2:H3–5ac
H2A.Z:H4K20 me2
DNA Meth. m5C:H4K20 me2
H4K20 me2:H3K4 me3
DNA Meth. m5C:H4K20 me3
H3K27 me2
H3K9 me2
H3K9 me3
H3K4 me3
H4–4ac
H3–5ac
H3K4 me1
H3K27 me3
H2A.Z
DNA Meth. m5C
H4K20 me3
H4K20 me2
0.00
0.25
0.50
0.75
1.00
$\hat{\pi}$
BRCA1 – Forward
H3K4 me3:H3–5ac
H3K4 me1:H2A.Z
DNA Meth. m5C:H3K9 me3
H4–4ac:H3–5ac
H3K4 me1:H4–4ac
H3K27 me3:H3K4 me1
H2A.Z:H4–4ac
H3K4 me1:H3–5ac
H2A.Z:H3–5ac
H2A.Z:H3K4 me3
H3K27 me3:DNA Meth. m5C
H3K27 me3:H4K20 me3
H3K4 me1:H4K20 me2
H3K9 me3:H4K20 me3
H3K4 me3:H4–4ac
DNA Meth. m5C:H4K20 me2
H4K20 me2:H4–4ac
DNA Meth. m5C:H4K20 me3
H2A.Z:H4K20 me2
H4K20 me2:H3–5ac
H3K27 me2
H4K20 me2:H3K4 me3
H3K9 me2
H3K9 me3
H3K27 me3
H3K4 me1
H4–4ac
H2A.Z
H3K4 me3
H3–5ac
DNA Meth. m5C
H4K20 me3
H4K20 me2
0.00
0.25
0.50
0.75
1.00
$\hat{\pi}$
BRCA1 – Reverse
H3K4 me3:H3–5ac
H3K4 me1:H2A.Z
DNA Meth. m5C:H3K9 me3
H4–4ac:H3–5ac
H3K4 me1:H4–4ac
H3K27 me3:H3K4 me1
H2A.Z:H4–4ac
H3K4 me1:H3–5ac
H2A.Z:H3–5ac
H2A.Z:H3K4 me3
H3K27 me3:DNA Meth. m5C
H3K27 me3:H4K20 me3
H3K4 me1:H4K20 me2
H3K9 me3:H4K20 me3
H3K4 me3:H4–4ac
DNA Meth. m5C:H4K20 me2
H4K20 me2:H4–4ac
DNA Meth. m5C:H4K20 me3
H2A.Z:H4K20 me2
H4K20 me2:H3–5ac
H3K27 me2
H4K20 me2:H3K4 me3
H3K9 me2
H3K9 me3
H3K27 me3
H3K4 me1
H4–4ac
H2A.Z
H3K4 me3
H3–5ac
DNA Meth. m5C
H4K20 me3
H4K20 me2
0.00
0.25
0.50
0.75
1.00
$\hat{\pi}$
BRCC3 – Forward
H3K4 me1:H2A.Z
H3K4 me3:H3–5ac
H3K4 me1:H4–4ac
H3K27 me3:H3K4 me1
H4–4ac:H3–5ac
H3K9 me3:H4K20 me3
H2A.Z:H3K4 me3
H3K4 me3:H4–4ac
H3K27 me3:H4K20 me3
H3K27 me3:H3K9 me3
H3K27 me3:DNA Meth. m5C
H2A.Z:H4K20 me2
H3K4 me1:H4K20 me2
DNA Meth. m5C:H3K9 me3
DNA Meth. m5C:H4K20 me2
H4K20 me2:H4–4ac
H4K20 me2:H3–5ac
H3K27 me2
DNA Meth. m5C:H4K20 me3
H4K20 me2:H3K4 me3
H3K9 me2
H2A.Z
H3K4 me1
H3K9 me3
H4–4ac
H3K4 me3
H3–5ac
H3K27 me3
DNA Meth. m5C
H4K20 me3
H4K20 me2
0.00
0.25
0.50
0.75
1.00
$\hat{\pi}$
BRCC3 – Reverse
H3K4 me1:H2A.Z
H3K4 me3:H3–5ac
H3K4 me1:H4–4ac
H3K27 me3:H3K4 me1
H4–4ac:H3–5ac
H3K9 me3:H4K20 me3
H2A.Z:H3K4 me3
H3K4 me3:H4–4ac
H3K27 me3:H4K20 me3
H3K27 me3:H3K9 me3
H3K27 me3:DNA Meth. m5C
H2A.Z:H4K20 me2
H3K4 me1:H4K20 me2
DNA Meth. m5C:H3K9 me3
DNA Meth. m5C:H4K20 me2
H4K20 me2:H4–4ac
H4K20 me2:H3–5ac
H3K27 me2
DNA Meth. m5C:H4K20 me3
H4K20 me2:H3K4 me3
H3K9 me2
H2A.Z
H3K4 me1
H3K9 me3
H4–4ac
H3K4 me3
H3–5ac
H3K27 me3
DNA Meth. m5C
H4K20 me3
H4K20 me2
0.00
0.25
0.50
0.75
1.00
$\hat{\pi}$

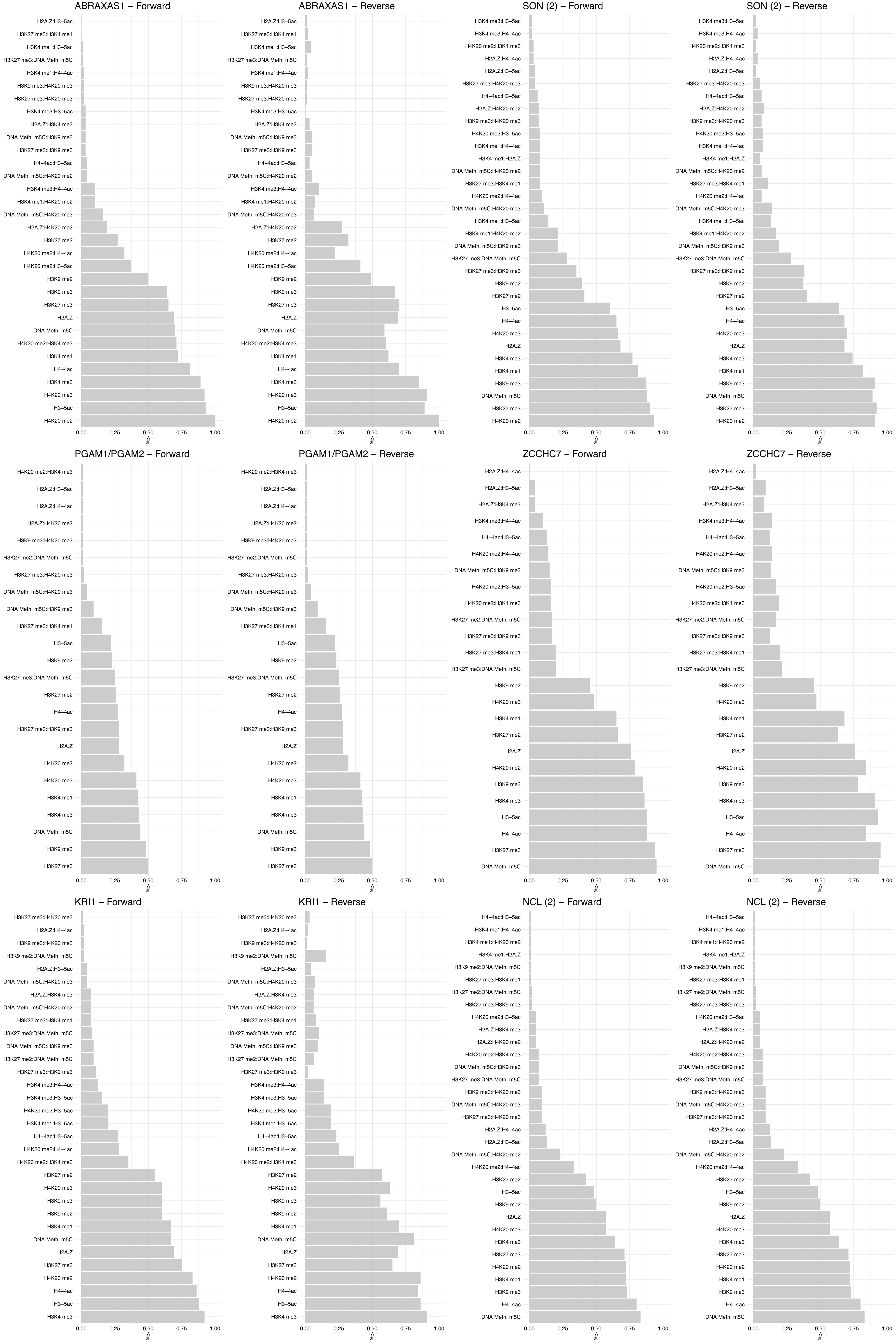

ABRAXAS1 – Forward
H2A.Z:H3–5ac
H3K27 me3:H3K4 me1
H3K4 me1:H3–5ac
H3K27 me3:DNA Meth. m5C
H3K4 me1:H4–4ac
H3K9 me3:H4K20 me3
H3K27 me3:H4K20 me3
H3K4 me3:H3–5ac
H2A.Z:H3K4 me3
DNA Meth. m5C:H3K9 me3
H3K27 me3:H3K9 me3
H4–4ac:H3–5ac
DNA Meth. m5C:H4K20 me2
H3K4 me3:H4–4ac
H3K4 me1:H4K20 me2
DNA Meth. m5C:H4K20 me3
H2A.Z:H4K20 me2
H3K27 me2
H4K20 me2:H4–4ac
H4K20 me2:H3–5ac
H3K9 me2
H3K9 me3
H3K27 me3
H2A.Z
DNA Meth. m5C
H4K20 me2:H3K4 me3
H3K4 me1
H4–4ac
H3K4 me3
H4K20 me3
H3–5ac
H4K20 me2
0.00 0.25 0.50 0.75 1.00
$\hat{\pi}$
ABRAXAS1 – Reverse
H2A.Z:H3–5ac
H3K27 me3:H3K4 me1
H3K4 me1:H3–5ac
H3K27 me3:DNA Meth. m5C
H3K4 me1:H4–4ac
H3K9 me3:H4K20 me3
H3K27 me3:H4K20 me3
H3K4 me3:H3–5ac
H2A.Z:H3K4 me3
DNA Meth. m5C:H3K9 me3
H3K27 me3:H3K9 me3
H4–4ac:H3–5ac
DNA Meth. m5C:H4K20 me2
H3K4 me3:H4–4ac
H3K4 me1:H4K20 me2
DNA Meth. m5C:H4K20 me3
H2A.Z:H4K20 me2
H3K27 me2
H4K20 me2:H4–4ac
H4K20 me2:H3–5ac
H3K9 me2
H3K9 me3
H3K27 me3
H2A.Z
DNA Meth. m5C
H4K20 me2:H3K4 me3
H3K4 me1
H4–4ac
H3K4 me3
H4K20 me3
H3–5ac
H4K20 me2
0.00 0.25 0.50 0.75 1.00
$\hat{\pi}$
SON (2) – Forward
H3K4 me3:H3–5ac
H3K4 me3:H4–4ac
H4K20 me2:H3K4 me3
H2A.Z:H4–4ac
H2A.Z:H3–5ac
H3K27 me3:H4K20 me3
H4–4ac:H3–5ac
H2A.Z:H4K20 me2
H3K9 me3:H4K20 me3
H4K20 me2:H3–5ac
H3K4 me1:H4–4ac
H3K4 me1:H2A.Z
DNA Meth. m5C:H4K20 me2
H3K27 me3:H3K4 me1
H4K20 me2:H4–4ac
DNA Meth. m5C:H4K20 me3
H3K4 me1:H3–5ac
H3K4 me1:H4K20 me2
DNA Meth. m5C:H3K9 me3
H3K27 me3:DNA Meth. m5C
H3K27 me3:H3K9 me3
H3K9 me2
H3K27 me2
H3–5ac
H4–4ac
H4K20 me3
H2A.Z
H3K4 me3
H3K4 me1
H3K9 me3
DNA Meth. m5C
H3K27 me3
H4K20 me2
0.00 0.25 0.50 0.75 1.00
$\hat{\pi}$
SON (2) – Reverse
H3K4 me3:H3–5ac
H3K4 me3:H4–4ac
H4K20 me2:H3K4 me3
H2A.Z:H4–4ac
H2A.Z:H3–5ac
H3K27 me3:H4K20 me3
H4–4ac:H3–5ac
H2A.Z:H4K20 me2
H3K9 me3:H4K20 me3
H4K20 me2:H3–5ac
H3K4 me1:H4–4ac
H3K4 me1:H2A.Z
DNA Meth. m5C:H4K20 me2
H3K27 me3:H3K4 me1
H4K20 me2:H4–4ac
DNA Meth. m5C:H4K20 me3
H3K4 me1:H3–5ac
H3K4 me1:H4K20 me2
DNA Meth. m5C:H3K9 me3
H3K27 me3:DNA Meth. m5C
H3K27 me3:H3K9 me3
H3K9 me2
H3K27 me2
H3–5ac
H4–4ac
H4K20 me3
H2A.Z
H3K4 me3
H3K4 me1
H3K9 me3
DNA Meth. m5C
H3K27 me3
H4K20 me2
0.00 0.25 0.50 0.75 1.00
$\hat{\pi}$
PGAM1/PGAM2 – Forward
H4K20 me2:H3K4 me3
H2A.Z:H3–5ac
H2A.Z:H4–4ac
H2A.Z:H4K20 me2
H3K9 me3:H4K20 me3
H3K27 me2:DNA Meth. m5C
H3K27 me3:H4K20 me3
DNA Meth. m5C:H4K20 me3
DNA Meth. m5C:H3K9 me3
H3K27 me3:H3K4 me1
H3–5ac
H3K9 me2
H3K27 me3:DNA Meth. m5C
H3K27 me2
H4–4ac
H3K27 me3:H3K9 me3
H2A.Z
H4K20 me2
H4K20 me3
H3K4 me1
H3K4 me3
DNA Meth. m5C
H3K9 me3
H3K27 me3
0.00 0.25 0.50 0.75 1.00
$\hat{\pi}$
PGAM1/PGAM2 – Reverse
H4K20 me2:H3K4 me3
H2A.Z:H3–5ac
H2A.Z:H4–4ac
H2A.Z:H4K20 me2
H3K9 me3:H4K20 me3
H3K27 me2:DNA Meth. m5C
H3K27 me3:H4K20 me3
DNA Meth. m5C:H4K20 me3
DNA Meth. m5C:H3K9 me3
H3K27 me3:H3K4 me1
H3–5ac
H3K9 me2
H3K27 me3:DNA Meth. m5C
H3K27 me2
H4–4ac
H3K27 me3:H3K9 me3
H2A.Z
H4K20 me2
H4K20 me3
H3K4 me1
H3K4 me3
DNA Meth. m5C
H3K9 me3
H3K27 me3
0.00 0.25 0.50 0.75 1.00
$\hat{\pi}$
ZCCHC7 – Forward
H2A.Z:H4–4ac
H2A.Z:H3–5ac
H2A.Z:H3K4 me3
H3K4 me3:H4–4ac
H4–4ac:H3–5ac
H4K20 me2:H4–4ac
DNA Meth. m5C:H3K9 me3
H4K20 me2:H3–5ac
H4K20 me2:H3K4 me3
H3K27 me2:DNA Meth. m5C
H3K27 me3:H3K9 me3
H3K27 me3:H3K4 me1
H3K27 me3:DNA Meth. m5C
H3K9 me2
H4K20 me3
H3K4 me1
H3K27 me2
H2A.Z
H4K20 me2
H3K9 me3
H3K4 me3
H3–5ac
H4–4ac
H3K27 me3
DNA Meth. m5C
0.00 0.25 0.50 0.75 1.00
$\hat{\pi}$
ZCCHC7 – Reverse
H2A.Z:H4–4ac
H2A.Z:H3–5ac
H2A.Z:H3K4 me3
H3K4 me3:H4–4ac
H4–4ac:H3–5ac
H4K20 me2:H4–4ac
DNA Meth. m5C:H3K9 me3
H4K20 me2:H3–5ac
H4K20 me2:H3K4 me3
H3K27 me2:DNA Meth. m5C
H3K27 me3:H3K9 me3
H3K27 me3:H3K4 me1
H3K27 me3:DNA Meth. m5C
H3K9 me2
H4K20 me3
H3K4 me1
H3K27 me2
H2A.Z
H4K20 me2
H3K9 me3
H3K4 me3
H3–5ac
H4–4ac
H3K27 me3
DNA Meth. m5C
0.00 0.25 0.50 0.75 1.00
$\hat{\pi}$
KRI1 – Forward
H3K27 me3:H4K20 me3
H2A.Z:H4–4ac
H3K9 me3:H4K20 me3
H3K9 me2:DNA Meth. m5C
H2A.Z:H3–5ac
DNA Meth. m5C:H4K20 me3
H2A.Z:H3K4 me3
DNA Meth. m5C:H4K20 me2
H3K27 me3:H3K4 me1
H3K27 me3:DNA Meth. m5C
DNA Meth. m5C:H3K9 me3
H3K27 me2:DNA Meth. m5C
H3K27 me3:H3K9 me3
H3K4 me3:H4–4ac
H3K4 me3:H3–5ac
H4K20 me2:H3–5ac
H3K4 me1:H3–5ac
H4–4ac:H3–5ac
H4K20 me2:H4–4ac
H4K20 me2:H3K4 me3
H3K27 me2
H4K20 me3
H3K9 me3
H3K9 me2
H3K4 me1
DNA Meth. m5C
H2A.Z
H3K27 me3
H4K20 me2
H4–4ac
H3–5ac
H3K4 me3
0.00 0.25 0.50 0.75 1.00
$\hat{\pi}$
KRI1 – Reverse
H3K27 me3:H4K20 me3
H2A.Z:H4–4ac
H3K9 me3:H4K20 me3
H3K9 me2:DNA Meth. m5C
H2A.Z:H3–5ac
DNA Meth. m5C:H4K20 me3
H2A.Z:H3K4 me3
DNA Meth. m5C:H4K20 me2
H3K27 me3:H3K4 me1
H3K27 me3:DNA Meth. m5C
DNA Meth. m5C:H3K9 me3
H3K27 me2:DNA Meth. m5C
H3K27 me3:H3K9 me3
H3K4 me3:H4–4ac
H3K4 me3:H3–5ac
H4K20 me2:H3–5ac
H3K4 me1:H3–5ac
H4–4ac:H3–5ac
H4K20 me2:H4–4ac
H4K20 me2:H3K4 me3
H3K27 me2
H4K20 me3
H3K9 me3
H3K9 me2
H3K4 me1
DNA Meth. m5C
H2A.Z
H3K27 me3
H4K20 me2
H4–4ac
H3–5ac
H3K4 me3
0.00 0.25 0.50 0.75 1.00
$\hat{\pi}$
NCL (2) – Forward
H4–4ac:H3–5ac
H3K4 me1:H4–4ac
H3K4 me1:H4K20 me2
H3K4 me1:H2A.Z
H3K9 me2:DNA Meth. m5C
H3K27 me3:H3K4 me1
H3K27 me2:DNA Meth. m5C
H3K27 me3:H3K9 me3
H4K20 me2:H3–5ac
H2A.Z:H3K4 me3
H2A.Z:H4K20 me2
H4K20 me2:H3K4 me3
DNA Meth. m5C:H3K9 me3
H3K27 me3:DNA Meth. m5C
H3K9 me3:H4K20 me3
DNA Meth. m5C:H4K20 me3
H3K27 me3:H4K20 me3
H2A.Z:H4–4ac
H2A.Z:H3–5ac
DNA Meth. m5C:H4K20 me2
H4K20 me2:H4–4ac
H3K27 me2
H3–5ac
H3K9 me2
H2A.Z
H4K20 me3
H3K4 me3
H3K27 me3
H4K20 me2
H3K4 me1
H3K9 me3
H4–4ac
DNA Meth. m5C
0.00 0.25 0.50 0.75 1.00
$\hat{\pi}$
NCL (2) – Reverse
H4–4ac:H3–5ac
H3K4 me1:H4–4ac
H3K4 me1:H4K20 me2
H3K4 me1:H2A.Z
H3K9 me2:DNA Meth. m5C
H3K27 me3:H3K4 me1
H3K27 me2:DNA Meth. m5C
H3K27 me3:H3K9 me3
H4K20 me2:H3–5ac
H2A.Z:H3K4 me3
H2A.Z:H4K20 me2
H4K20 me2:H3K4 me3
DNA Meth. m5C:H3K9 me3
H3K27 me3:DNA Meth. m5C
H3K9 me3:H4K20 me3
DNA Meth. m5C:H4K20 me3
H3K27 me3:H4K20 me3
H2A.Z:H4–4ac
H2A.Z:H3–5ac
DNA Meth. m5C:H4K20 me2
H4K20 me2:H4–4ac
H3K27 me2
H3–5ac
H3K9 me2
H2A.Z
H4K20 me3
H3K4 me3
H3K27 me3
H4K20 me2
H3K4 me1
H3K9 me3
H4–4ac
DNA Meth. m5C
0.00 0.25 0.50 0.75 1.00
$\hat{\pi}$

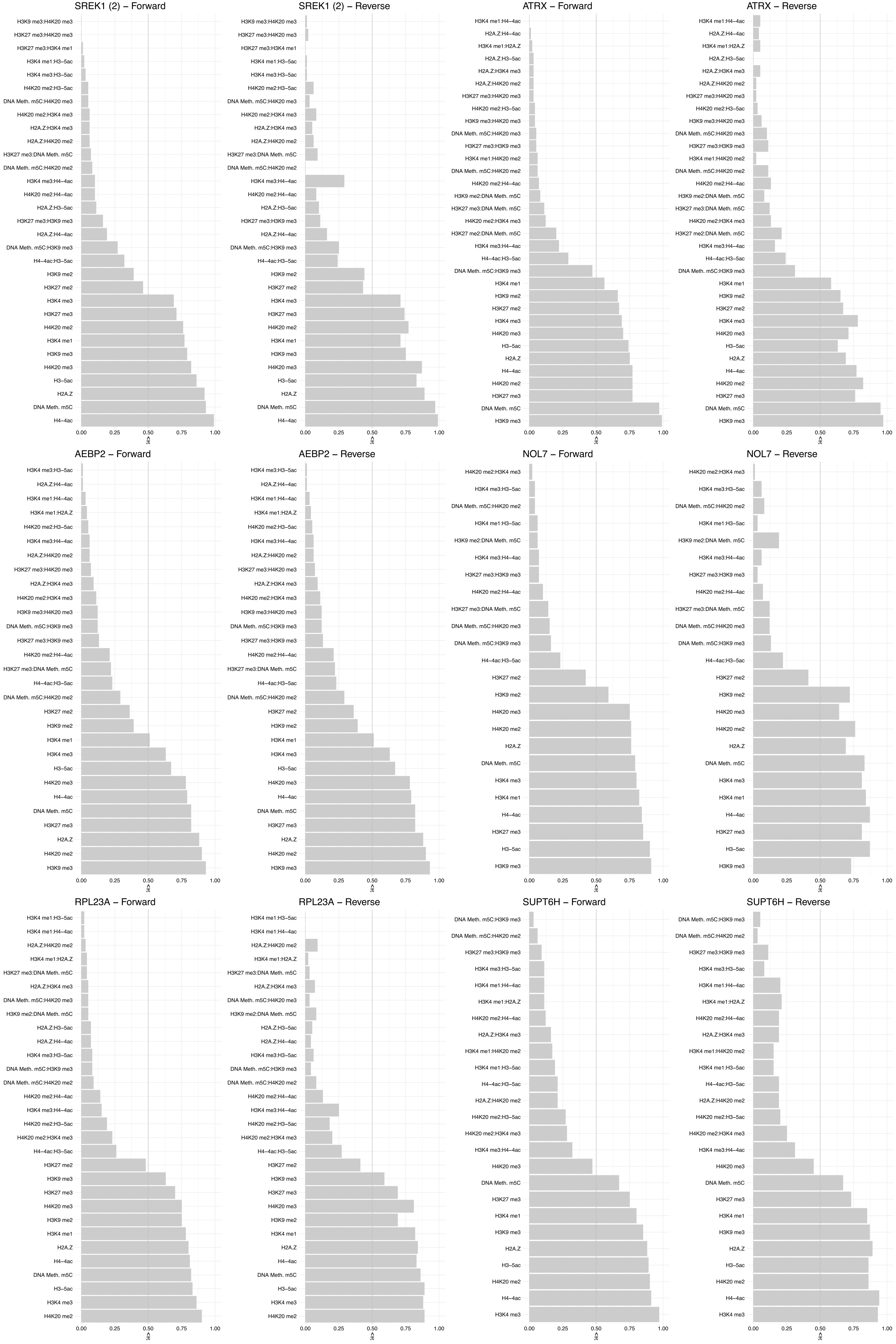
SREK1 (2) – Forward
H3K9 me3:H4K20 me3
H3K27 me3:H4K20 me3
H3K27 me3:H3K4 me1
H3K4 me1:H3−5ac
H3K4 me3:H3−5ac
H4K20 me2:H3−5ac
DNA Meth. m5C:H4K20 me3
H4K20 me2:H3K4 me3
H2A.Z:H3K4 me3
H2A.Z:H4K20 me2
H3K27 me3:DNA Meth. m5C
DNA Meth. m5C:H4K20 me2
H3K4 me3:H4−4ac
H4K20 me2:H4−4ac
H2A.Z:H3−5ac
H3K27 me3:H3K9 me3
H2A.Z:H4−4ac
DNA Meth. m5C:H3K9 me3
H4−4ac:H3−5ac
H3K9 me2
H3K27 me2
H3K4 me3
H3K27 me3
H4K20 me2
H3K4 me1
H3K9 me3
H4K20 me3
H3−5ac
H2A.Z
DNA Meth. m5C
H4−4ac
0.00
0.25
0.50
0.75
1.00
$\hat{\pi}$
SREK1 (2) – Reverse
H3K9 me3:H4K20 me3
H3K27 me3:H4K20 me3
H3K27 me3:H3K4 me1
H3K4 me1:H3−5ac
H3K4 me3:H3−5ac
H4K20 me2:H3−5ac
DNA Meth. m5C:H4K20 me3
H4K20 me2:H3K4 me3
H2A.Z:H3K4 me3
H2A.Z:H4K20 me2
H3K27 me3:DNA Meth. m5C
DNA Meth. m5C:H4K20 me2
H3K4 me3:H4−4ac
H4K20 me2:H4−4ac
H2A.Z:H3−5ac
H3K27 me3:H3K9 me3
H2A.Z:H4−4ac
DNA Meth. m5C:H3K9 me3
H4−4ac:H3−5ac
H3K9 me2
H3K27 me2
H3K4 me3
H3K27 me3
H4K20 me2
H3K4 me1
H3K9 me3
H4K20 me3
H3−5ac
H2A.Z
DNA Meth. m5C
H4−4ac
0.00
0.25
0.50
0.75
1.00
$\hat{\pi}$
ATRX – Forward
H3K4 me1:H4−4ac
H2A.Z:H4−4ac
H3K4 me1:H2A.Z
H2A.Z:H3−5ac
H2A.Z:H3K4 me3
H2A.Z:H4K20 me2
H3K27 me3:H4K20 me3
H4K20 me2:H3−5ac
H3K9 me3:H4K20 me3
DNA Meth. m5C:H4K20 me3
H3K27 me3:H3K9 me3
H3K4 me1:H4K20 me2
DNA Meth. m5C:H4K20 me2
H4K20 me2:H4−4ac
H3K9 me2:DNA Meth. m5C
H3K27 me3:DNA Meth. m5C
H4K20 me2:H3K4 me3
H3K27 me2:DNA Meth. m5C
H3K4 me3:H4−4ac
H4−4ac:H3−5ac
DNA Meth. m5C:H3K9 me3
H3K4 me1
H3K9 me2
H3K27 me2
H3K4 me3
H4K20 me3
H3−5ac
H2A.Z
H4−4ac
H4K20 me2
H3K27 me3
DNA Meth. m5C
H3K9 me3
0.00
0.25
0.50
0.75
1.00
$\hat{\pi}$
ATRX – Reverse
H3K4 me1:H4−4ac
H2A.Z:H4−4ac
H3K4 me1:H2A.Z
H2A.Z:H3−5ac
H2A.Z:H3K4 me3
H2A.Z:H4K20 me2
H3K27 me3:H4K20 me3
H4K20 me2:H3−5ac
H3K9 me3:H4K20 me3
DNA Meth. m5C:H4K20 me3
H3K27 me3:H3K9 me3
H3K4 me1:H4K20 me2
DNA Meth. m5C:H4K20 me2
H4K20 me2:H4−4ac
H3K9 me2:DNA Meth. m5C
H3K27 me3:DNA Meth. m5C
H4K20 me2:H3K4 me3
H3K27 me2:DNA Meth. m5C
H3K4 me3:H4−4ac
H4−4ac:H3−5ac
DNA Meth. m5C:H3K9 me3
H3K4 me1
H3K9 me2
H3K27 me2
H3K4 me3
H4K20 me3
H3−5ac
H2A.Z
H4−4ac
H4K20 me2
H3K27 me3
DNA Meth. m5C
H3K9 me3
0.00
0.25
0.50
0.75
1.00
$\hat{\pi}$
AEBP2 – Forward
H3K4 me3:H3−5ac
H2A.Z:H4−4ac
H3K4 me1:H4−4ac
H3K4 me1:H2A.Z
H4K20 me2:H3−5ac
H3K4 me3:H4−4ac
H2A.Z:H4K20 me2
H3K27 me3:H4K20 me3
H2A.Z:H3K4 me3
H4K20 me2:H3K4 me3
H3K9 me3:H4K20 me3
DNA Meth. m5C:H3K9 me3
H3K27 me3:H3K9 me3
H4K20 me2:H4−4ac
H3K27 me3:DNA Meth. m5C
H4−4ac:H3−5ac
DNA Meth. m5C:H4K20 me2
H3K27 me2
H3K9 me2
H3K4 me1
H3K4 me3
H3−5ac
H4K20 me3
H4−4ac
DNA Meth. m5C
H3K27 me3
H2A.Z
H4K20 me2
H3K9 me3
0.00
0.25
0.50
0.75
1.00
$\hat{\pi}$
AEBP2 – Reverse
H3K4 me3:H3−5ac
H2A.Z:H4−4ac
H3K4 me1:H4−4ac
H3K4 me1:H2A.Z
H4K20 me2:H3−5ac
H3K4 me3:H4−4ac
H2A.Z:H4K20 me2
H3K27 me3:H4K20 me3
H2A.Z:H3K4 me3
H4K20 me2:H3K4 me3
H3K9 me3:H4K20 me3
DNA Meth. m5C:H3K9 me3
H3K27 me3:H3K9 me3
H4K20 me2:H4−4ac
H3K27 me3:DNA Meth. m5C
H4−4ac:H3−5ac
DNA Meth. m5C:H4K20 me2
H3K27 me2
H3K9 me2
H3K4 me1
H3K4 me3
H3−5ac
H4K20 me3
H4−4ac
DNA Meth. m5C
H3K27 me3
H2A.Z
H4K20 me2
H3K9 me3
0.00
0.25
0.50
0.75
1.00
$\hat{\pi}$
NOL7 – Forward
H4K20 me2:H3K4 me3
H3K4 me3:H3−5ac
DNA Meth. m5C:H4K20 me2
H3K4 me1:H3−5ac
H3K9 me2:DNA Meth. m5C
H3K4 me3:H4−4ac
H3K27 me3:H3K9 me3
H4K20 me2:H4−4ac
H3K27 me3:DNA Meth. m5C
DNA Meth. m5C:H4K20 me3
DNA Meth. m5C:H3K9 me3
H4−4ac:H3−5ac
H3K27 me2
H3K9 me2
H4K20 me3
H4K20 me2
H2A.Z
DNA Meth. m5C
H3K4 me3
H3K4 me1
H4−4ac
H3K27 me3
H3−5ac
H3K9 me3
0.00
0.25
0.50
0.75
1.00
$\hat{\pi}$
NOL7 – Reverse
H4K20 me2:H3K4 me3
H3K4 me3:H3−5ac
DNA Meth. m5C:H4K20 me2
H3K4 me1:H3−5ac
H3K9 me2:DNA Meth. m5C
H3K4 me3:H4−4ac
H3K27 me3:H3K9 me3
H4K20 me2:H4−4ac
H3K27 me3:DNA Meth. m5C
DNA Meth. m5C:H4K20 me3
DNA Meth. m5C:H3K9 me3
H4−4ac:H3−5ac
H3K27 me2
H3K9 me2
H4K20 me3
H4K20 me2
H2A.Z
DNA Meth. m5C
H3K4 me3
H3K4 me1
H4−4ac
H3K27 me3
H3−5ac
H3K9 me3
0.00
0.25
0.50
0.75
1.00
$\hat{\pi}$
RPL23A – Forward
H3K4 me1:H3−5ac
H3K4 me1:H4−4ac
H2A.Z:H4K20 me2
H3K4 me1:H2A.Z
H3K27 me3:DNA Meth. m5C
H2A.Z:H3K4 me3
DNA Meth. m5C:H4K20 me3
H3K9 me2:DNA Meth. m5C
H2A.Z:H3−5ac
H2A.Z:H4−4ac
H3K4 me3:H3−5ac
DNA Meth. m5C:H3K9 me3
DNA Meth. m5C:H4K20 me2
H4K20 me2:H4−4ac
H3K4 me3:H4−4ac
H4K20 me2:H3−5ac
H4K20 me2:H3K4 me3
H4−4ac:H3−5ac
H3K27 me2
H3K9 me3
H3K27 me3
H4K20 me3
H3K9 me2
H3K4 me1
H2A.Z
H4−4ac
DNA Meth. m5C
H3−5ac
H3K4 me3
H4K20 me2
0.00
0.25
0.50
0.75
1.00
$\hat{\pi}$
RPL23A – Reverse
H3K4 me1:H3−5ac
H3K4 me1:H4−4ac
H2A.Z:H4K20 me2
H3K4 me1:H2A.Z
H3K27 me3:DNA Meth. m5C
H2A.Z:H3K4 me3
DNA Meth. m5C:H4K20 me3
H3K9 me2:DNA Meth. m5C
H2A.Z:H3−5ac
H2A.Z:H4−4ac
H3K4 me3:H3−5ac
DNA Meth. m5C:H3K9 me3
DNA Meth. m5C:H4K20 me2
H4K20 me2:H4−4ac
H3K4 me3:H4−4ac
H4K20 me2:H3−5ac
H4K20 me2:H3K4 me3
H4−4ac:H3−5ac
H3K27 me2
H3K9 me3
H3K27 me3
H4K20 me3
H3K9 me2
H3K4 me1
H2A.Z
H4−4ac
DNA Meth. m5C
H3−5ac
H3K4 me3
H4K20 me2
0.00
0.25
0.50
0.75
1.00
$\hat{\pi}$
SUPT6H – Forward
DNA Meth. m5C:H3K9 me3
DNA Meth. m5C:H4K20 me2
H3K27 me3:H3K9 me3
H3K4 me3:H3−5ac
H3K4 me1:H4−4ac
H3K4 me1:H2A.Z
H4K20 me2:H4−4ac
H2A.Z:H3K4 me3
H3K4 me1:H4K20 me2
H3K4 me1:H3−5ac
H4−4ac:H3−5ac
H2A.Z:H4K20 me2
H4K20 me2:H3−5ac
H4K20 me2:H3K4 me3
H3K4 me3:H4−4ac
H4K20 me3
DNA Meth. m5C
H3K27 me3
H3K4 me1
H3K9 me3
H2A.Z
H3−5ac
H4K20 me2
H4−4ac
H3K4 me3
0.00
0.25
0.50
0.75
1.00
$\hat{\pi}$
SUPT6H – Reverse
DNA Meth. m5C:H3K9 me3
DNA Meth. m5C:H4K20 me2
H3K27 me3:H3K9 me3
H3K4 me3:H3−5ac
H3K4 me1:H4−4ac
H3K4 me1:H2A.Z
H4K20 me2:H4−4ac
H2A.Z:H3K4 me3
H3K4 me1:H4K20 me2
H3K4 me1:H3−5ac
H4−4ac:H3−5ac
H2A.Z:H4K20 me2
H4K20 me2:H3−5ac
H4K20 me2:H3K4 me3
H3K4 me3:H4−4ac
H4K20 me3
DNA Meth. m5C
H3K27 me3
H3K4 me1
H3K9 me3
H2A.Z
H3−5ac
H4K20 me2
H4−4ac
H3K4 me3
0.00
0.25
0.50
0.75
1.00
$\hat{\pi}$

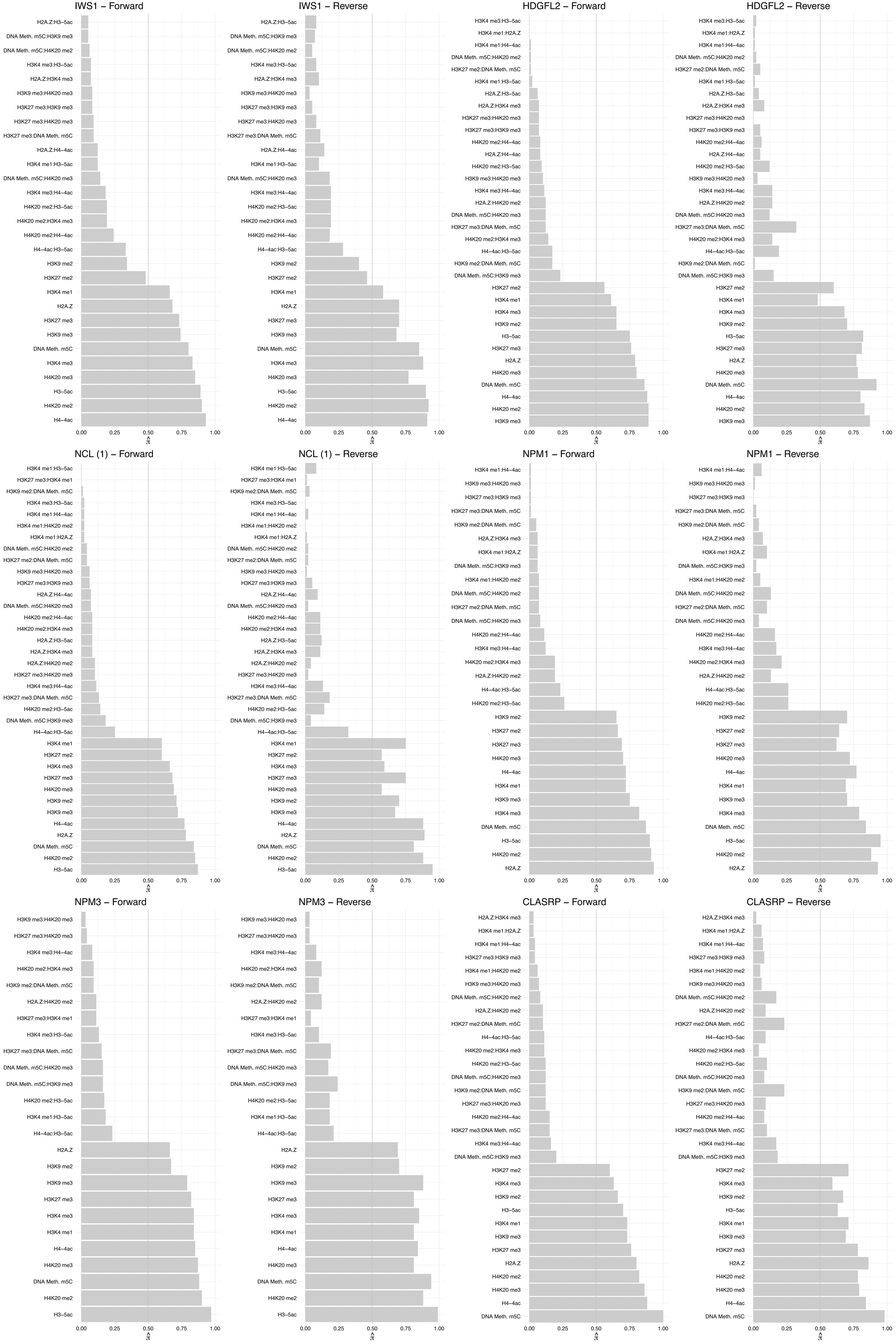
IWS1 – Forward
H2A.Z:H3−5ac
DNA Meth. m5C:H3K9 me3
DNA Meth. m5C:H4K20 me2
H3K4 me3:H3−5ac
H2A.Z:H3K4 me3
H3K9 me3:H4K20 me3
H3K27 me3:H3K9 me3
H3K27 me3:H4K20 me3
H3K27 me3:DNA Meth. m5C
H2A.Z:H4−4ac
H3K4 me1:H3−5ac
DNA Meth. m5C:H4K20 me3
H3K4 me3:H4−4ac
H4K20 me2:H3−5ac
H4K20 me2:H3K4 me3
H4K20 me2:H4−4ac
H4−4ac:H3−5ac
H3K9 me2
H3K27 me2
H3K4 me1
H2A.Z
H3K27 me3
H3K9 me3
DNA Meth. m5C
H3K4 me3
H4K20 me3
H3−5ac
H4K20 me2
H4−4ac
0.00 0.25 0.50 0.75 1.00
$\hat{\pi}$
IWS1 – Reverse
H2A.Z:H3−5ac
DNA Meth. m5C:H3K9 me3
DNA Meth. m5C:H4K20 me2
H3K4 me3:H3−5ac
H2A.Z:H3K4 me3
H3K9 me3:H4K20 me3
H3K27 me3:H3K9 me3
H3K27 me3:H4K20 me3
H3K27 me3:DNA Meth. m5C
H2A.Z:H4−4ac
H3K4 me1:H3−5ac
DNA Meth. m5C:H4K20 me3
H3K4 me3:H4−4ac
H4K20 me2:H3−5ac
H4K20 me2:H3K4 me3
H4K20 me2:H4−4ac
H4−4ac:H3−5ac
H3K9 me2
H3K27 me2
H3K4 me1
H2A.Z
H3K27 me3
H3K9 me3
DNA Meth. m5C
H3K4 me3
H4K20 me3
H3−5ac
H4K20 me2
H4−4ac
0.00 0.25 0.50 0.75 1.00
$\hat{\pi}$
HDGFL2 – Forward
H3K4 me3:H3−5ac
H3K4 me1:H2A.Z
H3K4 me1:H4−4ac
DNA Meth. m5C:H4K20 me2
H3K27 me2:DNA Meth. m5C
H3K4 me1:H3−5ac
H2A.Z:H3−5ac
H2A.Z:H3K4 me3
H3K27 me3:H4K20 me3
H3K27 me3:H3K9 me3
H4K20 me2:H4−4ac
H2A.Z:H4−4ac
H4K20 me2:H3−5ac
H3K9 me3:H4K20 me3
H3K4 me3:H4−4ac
H2A.Z:H4K20 me2
DNA Meth. m5C:H4K20 me3
H3K27 me3:DNA Meth. m5C
H4K20 me2:H3K4 me3
H4−4ac:H3−5ac
H3K9 me2:DNA Meth. m5C
DNA Meth. m5C:H3K9 me3
H3K27 me2
H3K4 me1
H3K4 me3
H3K9 me2
H3−5ac
H3K27 me3
H2A.Z
H4K20 me3
DNA Meth. m5C
H4−4ac
H4K20 me2
H3K9 me3
0.00 0.25 0.50 0.75 1.00
$\hat{\pi}$
HDGFL2 – Reverse
H3K4 me3:H3−5ac
H3K4 me1:H2A.Z
H3K4 me1:H4−4ac
DNA Meth. m5C:H4K20 me2
H3K27 me2:DNA Meth. m5C
H3K4 me1:H3−5ac
H2A.Z:H3−5ac
H2A.Z:H3K4 me3
H3K27 me3:H4K20 me3
H3K27 me3:H3K9 me3
H4K20 me2:H4−4ac
H2A.Z:H4−4ac
H4K20 me2:H3−5ac
H3K9 me3:H4K20 me3
H3K4 me3:H4−4ac
H2A.Z:H4K20 me2
DNA Meth. m5C:H4K20 me3
H3K27 me3:DNA Meth. m5C
H4K20 me2:H3K4 me3
H4−4ac:H3−5ac
H3K9 me2:DNA Meth. m5C
DNA Meth. m5C:H3K9 me3
H3K27 me2
H3K4 me1
H3K4 me3
H3K9 me2
H3−5ac
H3K27 me3
H2A.Z
H4K20 me3
DNA Meth. m5C
H4−4ac
H4K20 me2
H3K9 me3
0.00 0.25 0.50 0.75 1.00
$\hat{\pi}$
NCL (1) – Forward
H3K4 me1:H3−5ac
H3K27 me3:H3K4 me1
H3K9 me2:DNA Meth. m5C
H3K4 me3:H3−5ac
H3K4 me1:H4−4ac
H3K4 me1:H4K20 me2
H3K4 me1:H2A.Z
DNA Meth. m5C:H4K20 me2
H3K27 me2:DNA Meth. m5C
H3K9 me3:H4K20 me3
H3K27 me3:H3K9 me3
H2A.Z:H4−4ac
DNA Meth. m5C:H4K20 me3
H4K20 me2:H4−4ac
H4K20 me2:H3K4 me3
H2A.Z:H3−5ac
H2A.Z:H3K4 me3
H2A.Z:H4K20 me2
H3K27 me3:H4K20 me3
H3K4 me3:H4−4ac
H3K27 me3:DNA Meth. m5C
H4K20 me2:H3−5ac
DNA Meth. m5C:H3K9 me3
H4−4ac:H3−5ac
H3K4 me1
H3K27 me2
H3K4 me3
H3K27 me3
H4K20 me3
H3K9 me2
H3K9 me3
H4−4ac
H2A.Z
DNA Meth. m5C
H4K20 me2
H3−5ac
0.00 0.25 0.50 0.75 1.00
$\hat{\pi}$
NCL (1) – Reverse
H3K4 me1:H3−5ac
H3K27 me3:H3K4 me1
H3K9 me2:DNA Meth. m5C
H3K4 me3:H3−5ac
H3K4 me1:H4−4ac
H3K4 me1:H4K20 me2
H3K4 me1:H2A.Z
DNA Meth. m5C:H4K20 me2
H3K27 me2:DNA Meth. m5C
H3K9 me3:H4K20 me3
H3K27 me3:H3K9 me3
H2A.Z:H4−4ac
DNA Meth. m5C:H4K20 me3
H4K20 me2:H4−4ac
H4K20 me2:H3K4 me3
H2A.Z:H3−5ac
H2A.Z:H3K4 me3
H2A.Z:H4K20 me2
H3K27 me3:H4K20 me3
H3K4 me3:H4−4ac
H3K27 me3:DNA Meth. m5C
H4K20 me2:H3−5ac
DNA Meth. m5C:H3K9 me3
H4−4ac:H3−5ac
H3K4 me1
H3K27 me2
H3K4 me3
H3K27 me3
H4K20 me3
H3K9 me2
H3K9 me3
H4−4ac
H2A.Z
DNA Meth. m5C
H4K20 me2
H3−5ac
0.00 0.25 0.50 0.75 1.00
$\hat{\pi}$
NPM1 – Forward
H3K4 me1:H4−4ac
H3K9 me3:H4K20 me3
H3K27 me3:H3K9 me3
H3K27 me3:DNA Meth. m5C
H3K9 me2:DNA Meth. m5C
H2A.Z:H3K4 me3
H3K4 me1:H2A.Z
DNA Meth. m5C:H3K9 me3
H3K4 me1:H4K20 me2
DNA Meth. m5C:H4K20 me2
H3K27 me2:DNA Meth. m5C
DNA Meth. m5C:H4K20 me3
H4K20 me2:H4−4ac
H3K4 me3:H4−4ac
H4K20 me2:H3K4 me3
H2A.Z:H4K20 me2
H4−4ac:H3−5ac
H4K20 me2:H3−5ac
H3K9 me2
H3K27 me2
H3K27 me3
H4K20 me3
H4−4ac
H3K4 me1
H3K9 me3
H3K4 me3
DNA Meth. m5C
H3−5ac
H4K20 me2
H2A.Z
0.00 0.25 0.50 0.75 1.00
$\hat{\pi}$
NPM1 – Reverse
H3K4 me1:H4−4ac
H3K9 me3:H4K20 me3
H3K27 me3:H3K9 me3
H3K27 me3:DNA Meth. m5C
H3K9 me2:DNA Meth. m5C
H2A.Z:H3K4 me3
H3K4 me1:H2A.Z
DNA Meth. m5C:H3K9 me3
H3K4 me1:H4K20 me2
DNA Meth. m5C:H4K20 me2
H3K27 me2:DNA Meth. m5C
DNA Meth. m5C:H4K20 me3
H4K20 me2:H4−4ac
H3K4 me3:H4−4ac
H4K20 me2:H3K4 me3
H2A.Z:H4K20 me2
H4−4ac:H3−5ac
H4K20 me2:H3−5ac
H3K9 me2
H3K27 me2
H3K27 me3
H4K20 me3
H4−4ac
H3K4 me1
H3K9 me3
H3K4 me3
DNA Meth. m5C
H3−5ac
H4K20 me2
H2A.Z
0.00 0.25 0.50 0.75 1.00
$\hat{\pi}$
NPM3 – Forward
H3K9 me3:H4K20 me3
H3K27 me3:H4K20 me3
H3K4 me3:H4−4ac
H4K20 me2:H3K4 me3
H3K9 me2:DNA Meth. m5C
H2A.Z:H4K20 me2
H3K27 me3:H3K4 me1
H3K4 me3:H3−5ac
H3K27 me3:DNA Meth. m5C
DNA Meth. m5C:H4K20 me3
DNA Meth. m5C:H3K9 me3
H4K20 me2:H3−5ac
H3K4 me1:H3−5ac
H4−4ac:H3−5ac
H2A.Z
H3K9 me2
H3K9 me3
H3K27 me3
H3K4 me3
H3K4 me1
H4−4ac
H4K20 me3
DNA Meth. m5C
H4K20 me2
H3−5ac
0.00 0.25 0.50 0.75 1.00
$\hat{\pi}$
NPM3 – Reverse
H3K9 me3:H4K20 me3
H3K27 me3:H4K20 me3
H3K4 me3:H4−4ac
H4K20 me2:H3K4 me3
H3K9 me2:DNA Meth. m5C
H2A.Z:H4K20 me2
H3K27 me3:H3K4 me1
H3K4 me3:H3−5ac
H3K27 me3:DNA Meth. m5C
DNA Meth. m5C:H4K20 me3
DNA Meth. m5C:H3K9 me3
H4K20 me2:H3−5ac
H3K4 me1:H3−5ac
H4−4ac:H3−5ac
H2A.Z
H3K9 me2
H3K9 me3
H3K27 me3
H3K4 me3
H3K4 me1
H4−4ac
H4K20 me3
DNA Meth. m5C
H4K20 me2
H3−5ac
0.00 0.25 0.50 0.75 1.00
$\hat{\pi}$
CLASRP – Forward
H2A.Z:H3K4 me3
H3K4 me1:H2A.Z
H3K4 me1:H4−4ac
H3K27 me3:H3K9 me3
H3K4 me1:H4K20 me2
H3K9 me3:H4K20 me3
DNA Meth. m5C:H4K20 me2
H2A.Z:H4K20 me2
H3K27 me2:DNA Meth. m5C
H4−4ac:H3−5ac
H4K20 me2:H3K4 me3
H4K20 me2:H3−5ac
DNA Meth. m5C:H4K20 me3
H3K9 me2:DNA Meth. m5C
H3K27 me3:H4K20 me3
H4K20 me2:H4−4ac
H3K27 me3:DNA Meth. m5C
H3K4 me3:H4−4ac
DNA Meth. m5C:H3K9 me3
H3K27 me2
H3K4 me3
H3K9 me2
H3−5ac
H3K4 me1
H3K9 me3
H3K27 me3
H2A.Z
H4K20 me2
H4K20 me3
H4−4ac
DNA Meth. m5C
0.00 0.25 0.50 0.75 1.00
$\hat{\pi}$
CLASRP – Reverse
H2A.Z:H3K4 me3
H3K4 me1:H2A.Z
H3K4 me1:H4−4ac
H3K27 me3:H3K9 me3
H3K4 me1:H4K20 me2
H3K9 me3:H4K20 me3
DNA Meth. m5C:H4K20 me2
H2A.Z:H4K20 me2
H3K27 me2:DNA Meth. m5C
H4−4ac:H3−5ac
H4K20 me2:H3K4 me3
H4K20 me2:H3−5ac
DNA Meth. m5C:H4K20 me3
H3K9 me2:DNA Meth. m5C
H3K27 me3:H4K20 me3
H4K20 me2:H4−4ac
H3K27 me3:DNA Meth. m5C
H3K4 me3:H4−4ac
DNA Meth. m5C:H3K9 me3
H3K27 me2
H3K4 me3
H3K9 me2
H3−5ac
H3K4 me1
H3K9 me3
H3K27 me3
H2A.Z
H4K20 me2
H4K20 me3
H4−4ac
DNA Meth. m5C
0.00 0.25 0.50 0.75 1.00
$\hat{\pi}$

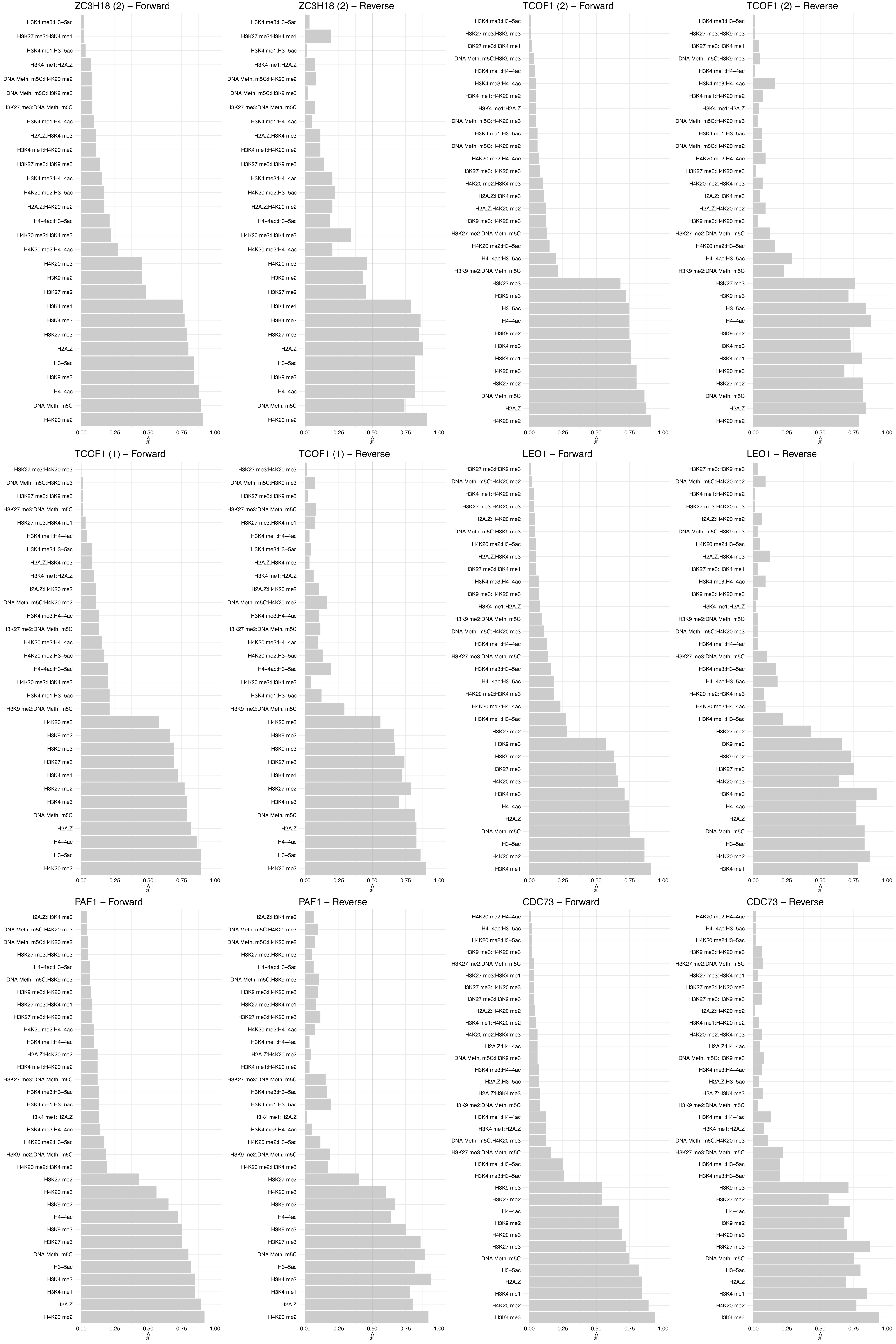
ZC3H18 (2) – Forward
H3K4 me3:H3–5ac
H3K27 me3:H3K4 me1
H3K4 me1:H3–5ac
H3K4 me1:H2A.Z
DNA Meth. m5C:H4K20 me2
DNA Meth. m5C:H3K9 me3
H3K27 me3:DNA Meth. m5C
H3K4 me1:H4–4ac
H2A.Z:H3K4 me3
H3K4 me1:H4K20 me2
H3K27 me3:H3K9 me3
H3K4 me3:H4–4ac
H4K20 me2:H3–5ac
H2A.Z:H4K20 me2
H4–4ac:H3–5ac
H4K20 me2:H3K4 me3
H4K20 me2:H4–4ac
H4K20 me3
H3K9 me2
H3K27 me2
H3K4 me1
H3K4 me3
H3K27 me3
H2A.Z
H3–5ac
H3K9 me3
H4–4ac
DNA Meth. m5C
H4K20 me2
0.00 0.25 0.50 0.75 1.00
$\hat{\pi}$
ZC3H18 (2) – Reverse
H3K4 me3:H3–5ac
H3K27 me3:H3K4 me1
H3K4 me1:H3–5ac
H3K4 me1:H2A.Z
DNA Meth. m5C:H4K20 me2
DNA Meth. m5C:H3K9 me3
H3K27 me3:DNA Meth. m5C
H3K4 me1:H4–4ac
H2A.Z:H3K4 me3
H3K4 me1:H4K20 me2
H3K27 me3:H3K9 me3
H3K4 me3:H4–4ac
H4K20 me2:H3–5ac
H2A.Z:H4K20 me2
H4–4ac:H3–5ac
H4K20 me2:H3K4 me3
H4K20 me2:H4–4ac
H4K20 me3
H3K9 me2
H3K27 me2
H3K4 me1
H3K4 me3
H3K27 me3
H2A.Z
H3–5ac
H3K9 me3
H4–4ac
DNA Meth. m5C
H4K20 me2
0.00 0.25 0.50 0.75 1.00
$\hat{\pi}$
TCOF1 (2) – Forward
H3K4 me3:H3–5ac
H3K27 me3:H3K9 me3
H3K27 me3:H3K4 me1
DNA Meth. m5C:H3K9 me3
H3K4 me1:H4–4ac
H3K4 me3:H4–4ac
H3K4 me1:H4K20 me2
H3K4 me1:H2A.Z
DNA Meth. m5C:H4K20 me3
H3K4 me1:H3–5ac
DNA Meth. m5C:H4K20 me2
H4K20 me2:H4–4ac
H3K27 me3:H4K20 me3
H4K20 me2:H3K4 me3
H2A.Z:H3K4 me3
H2A.Z:H4K20 me2
H3K9 me3:H4K20 me3
H3K27 me2:DNA Meth. m5C
H4K20 me2:H3–5ac
H4–4ac:H3–5ac
H3K9 me2:DNA Meth. m5C
H3K27 me3
H3K9 me3
H3–5ac
H4–4ac
H3K9 me2
H3K4 me3
H3K4 me1
H4K20 me3
H3K27 me2
DNA Meth. m5C
H2A.Z
H4K20 me2
0.00 0.25 0.50 0.75 1.00
$\hat{\pi}$
TCOF1 (2) – Reverse
H3K4 me3:H3–5ac
H3K27 me3:H3K9 me3
H3K27 me3:H3K4 me1
DNA Meth. m5C:H3K9 me3
H3K4 me1:H4–4ac
H3K4 me3:H4–4ac
H3K4 me1:H4K20 me2
H3K4 me1:H2A.Z
DNA Meth. m5C:H4K20 me3
H3K4 me1:H3–5ac
DNA Meth. m5C:H4K20 me2
H4K20 me2:H4–4ac
H3K27 me3:H4K20 me3
H4K20 me2:H3K4 me3
H2A.Z:H3K4 me3
H2A.Z:H4K20 me2
H3K9 me3:H4K20 me3
H3K27 me2:DNA Meth. m5C
H4K20 me2:H3–5ac
H4–4ac:H3–5ac
H3K9 me2:DNA Meth. m5C
H3K27 me3
H3K9 me3
H3–5ac
H4–4ac
H3K9 me2
H3K4 me3
H3K4 me1
H4K20 me3
H3K27 me2
DNA Meth. m5C
H2A.Z
H4K20 me2
0.00 0.25 0.50 0.75 1.00
$\hat{\pi}$
TCOF1 (1) – Forward
H3K27 me3:H4K20 me3
DNA Meth. m5C:H3K9 me3
H3K27 me3:H3K9 me3
H3K27 me3:DNA Meth. m5C
H3K27 me3:H3K4 me1
H3K4 me1:H4–4ac
H3K4 me3:H3–5ac
H2A.Z:H3K4 me3
H3K4 me1:H2A.Z
H2A.Z:H4K20 me2
DNA Meth. m5C:H4K20 me2
H3K4 me3:H4–4ac
H3K27 me2:DNA Meth. m5C
H4K20 me2:H4–4ac
H4K20 me2:H3–5ac
H4–4ac:H3–5ac
H4K20 me2:H3K4 me3
H3K4 me1:H3–5ac
H3K9 me2:DNA Meth. m5C
H4K20 me3
H3K9 me2
H3K9 me3
H3K27 me3
H3K4 me1
H3K27 me2
H3K4 me3
DNA Meth. m5C
H2A.Z
H4–4ac
H3–5ac
H4K20 me2
0.00 0.25 0.50 0.75 1.00
$\hat{\pi}$
TCOF1 (1) – Reverse
H3K27 me3:H4K20 me3
DNA Meth. m5C:H3K9 me3
H3K27 me3:H3K9 me3
H3K27 me3:DNA Meth. m5C
H3K27 me3:H3K4 me1
H3K4 me1:H4–4ac
H3K4 me3:H3–5ac
H2A.Z:H3K4 me3
H3K4 me1:H2A.Z
H2A.Z:H4K20 me2
DNA Meth. m5C:H4K20 me2
H3K4 me3:H4–4ac
H3K27 me2:DNA Meth. m5C
H4K20 me2:H4–4ac
H4K20 me2:H3–5ac
H4–4ac:H3–5ac
H4K20 me2:H3K4 me3
H3K4 me1:H3–5ac
H3K9 me2:DNA Meth. m5C
H4K20 me3
H3K9 me2
H3K9 me3
H3K27 me3
H3K4 me1
H3K27 me2
H3K4 me3
DNA Meth. m5C
H2A.Z
H4–4ac
H3–5ac
H4K20 me2
0.00 0.25 0.50 0.75 1.00
$\hat{\pi}$
LEO1 – Forward
H3K27 me3:H3K9 me3
DNA Meth. m5C:H4K20 me2
H3K4 me1:H4K20 me2
H3K27 me3:H4K20 me3
H2A.Z:H4K20 me2
DNA Meth. m5C:H3K9 me3
H4K20 me2:H3–5ac
H2A.Z:H3K4 me3
H3K27 me3:H3K4 me1
H3K4 me3:H4–4ac
H3K9 me3:H4K20 me3
H3K4 me1:H2A.Z
H3K9 me2:DNA Meth. m5C
DNA Meth. m5C:H4K20 me3
H3K4 me1:H4–4ac
H3K27 me3:DNA Meth. m5C
H3K4 me3:H3–5ac
H4–4ac:H3–5ac
H4K20 me2:H3K4 me3
H4K20 me2:H4–4ac
H3K4 me1:H3–5ac
H3K27 me2
H3K9 me3
H3K9 me2
H3K27 me3
H4K20 me3
H3K4 me3
H4–4ac
H2A.Z
DNA Meth. m5C
H3–5ac
H4K20 me2
H3K4 me1
0.00 0.25 0.50 0.75 1.00
$\hat{\pi}$
LEO1 – Reverse
H3K27 me3:H3K9 me3
DNA Meth. m5C:H4K20 me2
H3K4 me1:H4K20 me2
H3K27 me3:H4K20 me3
H2A.Z:H4K20 me2
DNA Meth. m5C:H3K9 me3
H4K20 me2:H3–5ac
H2A.Z:H3K4 me3
H3K27 me3:H3K4 me1
H3K4 me3:H4–4ac
H3K9 me3:H4K20 me3
H3K4 me1:H2A.Z
H3K9 me2:DNA Meth. m5C
DNA Meth. m5C:H4K20 me3
H3K4 me1:H4–4ac
H3K27 me3:DNA Meth. m5C
H3K4 me3:H3–5ac
H4–4ac:H3–5ac
H4K20 me2:H3K4 me3
H4K20 me2:H4–4ac
H3K4 me1:H3–5ac
H3K27 me2
H3K9 me3
H3K9 me2
H3K27 me3
H4K20 me3
H3K4 me3
H4–4ac
H2A.Z
DNA Meth. m5C
H3–5ac
H4K20 me2
H3K4 me1
0.00 0.25 0.50 0.75 1.00
$\hat{\pi}$
PAF1 – Forward
H2A.Z:H3K4 me3
DNA Meth. m5C:H4K20 me3
DNA Meth. m5C:H4K20 me2
H3K27 me3:H3K9 me3
H4–4ac:H3–5ac
DNA Meth. m5C:H3K9 me3
H3K9 me3:H4K20 me3
H3K27 me3:H3K4 me1
H3K27 me3:H4K20 me3
H4K20 me2:H4–4ac
H3K4 me1:H4–4ac
H2A.Z:H4K20 me2
H3K4 me1:H4K20 me2
H3K27 me3:DNA Meth. m5C
H3K4 me3:H3–5ac
H3K4 me1:H3–5ac
H3K4 me1:H2A.Z
H3K4 me3:H4–4ac
H4K20 me2:H3–5ac
H3K9 me2:DNA Meth. m5C
H4K20 me2:H3K4 me3
H3K27 me2
H4K20 me3
H3K9 me2
H4–4ac
H3K9 me3
H3K27 me3
DNA Meth. m5C
H3–5ac
H3K4 me3
H3K4 me1
H2A.Z
H4K20 me2
0.00 0.25 0.50 0.75 1.00
$\hat{\pi}$
PAF1 – Reverse
H2A.Z:H3K4 me3
DNA Meth. m5C:H4K20 me3
DNA Meth. m5C:H4K20 me2
H3K27 me3:H3K9 me3
H4–4ac:H3–5ac
DNA Meth. m5C:H3K9 me3
H3K9 me3:H4K20 me3
H3K27 me3:H3K4 me1
H3K27 me3:H4K20 me3
H4K20 me2:H4–4ac
H3K4 me1:H4–4ac
H2A.Z:H4K20 me2
H3K4 me1:H4K20 me2
H3K27 me3:DNA Meth. m5C
H3K4 me3:H3–5ac
H3K4 me1:H3–5ac
H3K4 me1:H2A.Z
H3K4 me3:H4–4ac
H4K20 me2:H3–5ac
H3K9 me2:DNA Meth. m5C
H4K20 me2:H3K4 me3
H3K27 me2
H4K20 me3
H3K9 me2
H4–4ac
H3K9 me3
H3K27 me3
DNA Meth. m5C
H3–5ac
H3K4 me3
H3K4 me1
H2A.Z
H4K20 me2
0.00 0.25 0.50 0.75 1.00
$\hat{\pi}$
CDC73 – Forward
H4K20 me2:H4–4ac
H4–4ac:H3–5ac
H4K20 me2:H3–5ac
H3K9 me3:H4K20 me3
H3K27 me2:DNA Meth. m5C
H3K27 me3:H3K4 me1
H3K27 me3:H4K20 me3
H3K27 me3:H3K9 me3
H2A.Z:H4K20 me2
H3K4 me1:H4K20 me2
H4K20 me2:H3K4 me3
H2A.Z:H4–4ac
DNA Meth. m5C:H3K9 me3
H3K4 me3:H4–4ac
H2A.Z:H3–5ac
H2A.Z:H3K4 me3
H3K9 me2:DNA Meth. m5C
H3K4 me1:H4–4ac
H3K4 me1:H2A.Z
DNA Meth. m5C:H4K20 me3
H3K27 me3:DNA Meth. m5C
H3K4 me1:H3–5ac
H3K4 me3:H3–5ac
H3K9 me3
H3K27 me2
H4–4ac
H3K9 me2
H4K20 me3
H3K27 me3
DNA Meth. m5C
H3–5ac
H2A.Z
H3K4 me1
H4K20 me2
H3K4 me3
0.00 0.25 0.50 0.75 1.00
$\hat{\pi}$
CDC73 – Reverse
H4K20 me2:H4–4ac
H4–4ac:H3–5ac
H4K20 me2:H3–5ac
H3K9 me3:H4K20 me3
H3K27 me2:DNA Meth. m5C
H3K27 me3:H3K4 me1
H3K27 me3:H4K20 me3
H3K27 me3:H3K9 me3
H2A.Z:H4K20 me2
H3K4 me1:H4K20 me2
H4K20 me2:H3K4 me3
H2A.Z:H4–4ac
DNA Meth. m5C:H3K9 me3
H3K4 me3:H4–4ac
H2A.Z:H3–5ac
H2A.Z:H3K4 me3
H3K9 me2:DNA Meth. m5C
H3K4 me1:H4–4ac
H3K4 me1:H2A.Z
DNA Meth. m5C:H4K20 me3
H3K27 me3:DNA Meth. m5C
H3K4 me1:H3–5ac
H3K4 me3:H3–5ac
H3K9 me3
H3K27 me2
H4–4ac
H3K9 me2
H4K20 me3
H3K27 me3
DNA Meth. m5C
H3–5ac
H2A.Z
H3K4 me1
H4K20 me2
H3K4 me3
0.00 0.25 0.50 0.75 1.00
$\hat{\pi}$

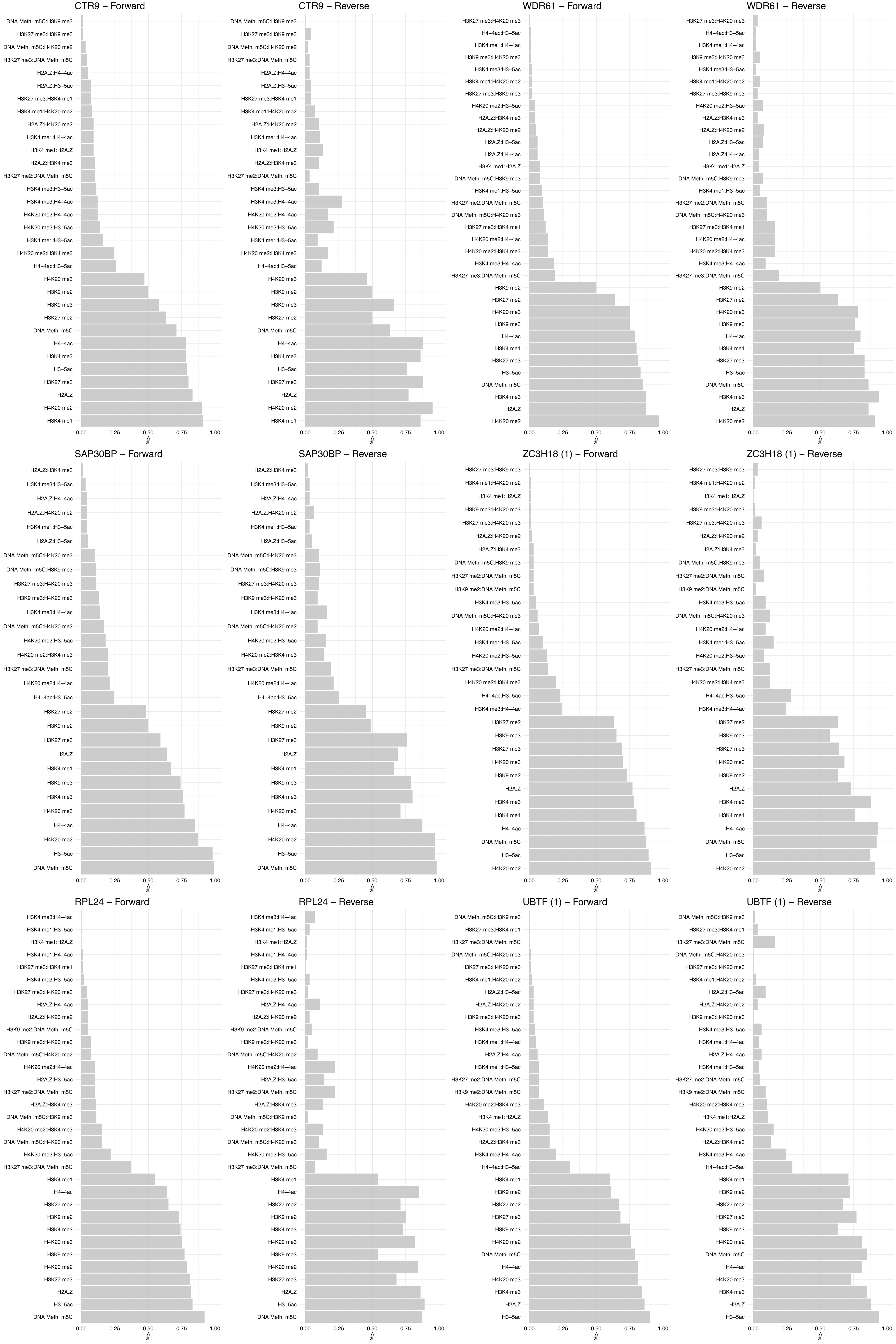
CTR9 – Forward
DNA Meth. m5C:H3K9 me3
H3K27 me3:H3K9 me3
DNA Meth. m5C:H4K20 me2
H3K27 me3:DNA Meth. m5C
H2A.Z:H4−4ac
H2A.Z:H3−5ac
H3K27 me3:H3K4 me1
H3K4 me1:H4K20 me2
H2A.Z:H4K20 me2
H3K4 me1:H4−4ac
H3K4 me1:H2A.Z
H2A.Z:H3K4 me3
H3K27 me2:DNA Meth. m5C
H3K4 me3:H3−5ac
H3K4 me3:H4−4ac
H4K20 me2:H4−4ac
H4K20 me2:H3−5ac
H3K4 me1:H3−5ac
H4K20 me2:H3K4 me3
H4−4ac:H3−5ac
H4K20 me3
H3K9 me2
H3K9 me3
H3K27 me2
DNA Meth. m5C
H4−4ac
H3K4 me3
H3−5ac
H3K27 me3
H2A.Z
H4K20 me2
H3K4 me1
0.00
0.25
0.50
0.75
1.00
π̂
CTR9 – Reverse
DNA Meth. m5C:H3K9 me3
H3K27 me3:H3K9 me3
DNA Meth. m5C:H4K20 me2
H3K27 me3:DNA Meth. m5C
H2A.Z:H4−4ac
H2A.Z:H3−5ac
H3K27 me3:H3K4 me1
H3K4 me1:H4K20 me2
H2A.Z:H4K20 me2
H3K4 me1:H4−4ac
H3K4 me1:H2A.Z
H2A.Z:H3K4 me3
H3K27 me2:DNA Meth. m5C
H3K4 me3:H3−5ac
H3K4 me3:H4−4ac
H4K20 me2:H4−4ac
H4K20 me2:H3−5ac
H3K4 me1:H3−5ac
H4K20 me2:H3K4 me3
H4−4ac:H3−5ac
H4K20 me3
H3K9 me2
H3K9 me3
H3K27 me2
DNA Meth. m5C
H4−4ac
H3K4 me3
H3−5ac
H3K27 me3
H2A.Z
H4K20 me2
H3K4 me1
0.00
0.25
0.50
0.75
1.00
π̂
WDR61 – Forward
H3K27 me3:H4K20 me3
H4−4ac:H3−5ac
H3K4 me1:H4−4ac
H3K9 me3:H4K20 me3
H3K4 me3:H3−5ac
H3K4 me1:H4K20 me2
H3K27 me3:H3K9 me3
H4K20 me2:H3−5ac
H2A.Z:H3K4 me3
H2A.Z:H4K20 me2
H2A.Z:H3−5ac
H2A.Z:H4−4ac
H3K4 me1:H2A.Z
DNA Meth. m5C:H3K9 me3
H3K4 me1:H3−5ac
H3K27 me2:DNA Meth. m5C
DNA Meth. m5C:H4K20 me3
H3K27 me3:H3K4 me1
H4K20 me2:H4−4ac
H4K20 me2:H3K4 me3
H3K4 me3:H4−4ac
H3K27 me3:DNA Meth. m5C
H3K9 me2
H3K27 me2
H4K20 me3
H3K9 me3
H4−4ac
H3K4 me1
H3K27 me3
H3−5ac
DNA Meth. m5C
H3K4 me3
H2A.Z
H4K20 me2
0.00
0.25
0.50
0.75
1.00
π̂
WDR61 – Reverse
H3K27 me3:H4K20 me3
H4−4ac:H3−5ac
H3K4 me1:H4−4ac
H3K9 me3:H4K20 me3
H3K4 me3:H3−5ac
H3K4 me1:H4K20 me2
H3K27 me3:H3K9 me3
H4K20 me2:H3−5ac
H2A.Z:H3K4 me3
H2A.Z:H4K20 me2
H2A.Z:H3−5ac
H2A.Z:H4−4ac
H3K4 me1:H2A.Z
DNA Meth. m5C:H3K9 me3
H3K4 me1:H3−5ac
H3K27 me2:DNA Meth. m5C
DNA Meth. m5C:H4K20 me3
H3K27 me3:H3K4 me1
H4K20 me2:H4−4ac
H4K20 me2:H3K4 me3
H3K4 me3:H4−4ac
H3K27 me3:DNA Meth. m5C
H3K9 me2
H3K27 me2
H4K20 me3
H3K9 me3
H4−4ac
H3K4 me1
H3K27 me3
H3−5ac
DNA Meth. m5C
H3K4 me3
H2A.Z
H4K20 me2
0.00
0.25
0.50
0.75
1.00
π̂
SAP30BP – Forward
H2A.Z:H3K4 me3
H3K4 me3:H3−5ac
H2A.Z:H4−4ac
H2A.Z:H4K20 me2
H3K4 me1:H3−5ac
H2A.Z:H3−5ac
DNA Meth. m5C:H4K20 me3
DNA Meth. m5C:H3K9 me3
H3K27 me3:H4K20 me3
H3K9 me3:H4K20 me3
H3K4 me3:H4−4ac
DNA Meth. m5C:H4K20 me2
H4K20 me2:H3−5ac
H4K20 me2:H3K4 me3
H3K27 me3:DNA Meth. m5C
H4K20 me2:H4−4ac
H4−4ac:H3−5ac
H3K27 me2
H3K9 me2
H3K27 me3
H2A.Z
H3K4 me1
H3K9 me3
H3K4 me3
H4K20 me3
H4−4ac
H4K20 me2
H3−5ac
DNA Meth. m5C
0.00
0.25
0.50
0.75
1.00
π̂
SAP30BP – Reverse
H2A.Z:H3K4 me3
H3K4 me3:H3−5ac
H2A.Z:H4−4ac
H2A.Z:H4K20 me2
H3K4 me1:H3−5ac
H2A.Z:H3−5ac
DNA Meth. m5C:H4K20 me3
DNA Meth. m5C:H3K9 me3
H3K27 me3:H4K20 me3
H3K9 me3:H4K20 me3
H3K4 me3:H4−4ac
DNA Meth. m5C:H4K20 me2
H4K20 me2:H3−5ac
H4K20 me2:H3K4 me3
H3K27 me3:DNA Meth. m5C
H4K20 me2:H4−4ac
H4−4ac:H3−5ac
H3K27 me2
H3K9 me2
H3K27 me3
H2A.Z
H3K4 me1
H3K9 me3
H3K4 me3
H4K20 me3
H4−4ac
H4K20 me2
H3−5ac
DNA Meth. m5C
0.00
0.25
0.50
0.75
1.00
π̂
ZC3H18 (1) – Forward
H3K27 me3:H3K9 me3
H3K4 me1:H4K20 me2
H3K4 me1:H2A.Z
H3K9 me3:H4K20 me3
H3K27 me3:H4K20 me3
H2A.Z:H4K20 me2
H2A.Z:H3K4 me3
DNA Meth. m5C:H3K9 me3
H3K27 me2:DNA Meth. m5C
H3K9 me2:DNA Meth. m5C
H3K4 me3:H3−5ac
DNA Meth. m5C:H4K20 me3
H4K20 me2:H4−4ac
H3K4 me1:H3−5ac
H4K20 me2:H3−5ac
H3K27 me3:DNA Meth. m5C
H4K20 me2:H3K4 me3
H4−4ac:H3−5ac
H3K4 me3:H4−4ac
H3K27 me2
H3K9 me3
H3K27 me3
H4K20 me3
H3K9 me2
H2A.Z
H3K4 me3
H3K4 me1
H4−4ac
DNA Meth. m5C
H3−5ac
H4K20 me2
0.00
0.25
0.50
0.75
1.00
π̂
ZC3H18 (1) – Reverse
H3K27 me3:H3K9 me3
H3K4 me1:H4K20 me2
H3K4 me1:H2A.Z
H3K9 me3:H4K20 me3
H3K27 me3:H4K20 me3
H2A.Z:H4K20 me2
H2A.Z:H3K4 me3
DNA Meth. m5C:H3K9 me3
H3K27 me2:DNA Meth. m5C
H3K9 me2:DNA Meth. m5C
H3K4 me3:H3−5ac
DNA Meth. m5C:H4K20 me3
H4K20 me2:H4−4ac
H3K4 me1:H3−5ac
H4K20 me2:H3−5ac
H3K27 me3:DNA Meth. m5C
H4K20 me2:H3K4 me3
H4−4ac:H3−5ac
H3K4 me3:H4−4ac
H3K27 me2
H3K9 me3
H3K27 me3
H4K20 me3
H3K9 me2
H2A.Z
H3K4 me3
H3K4 me1
H4−4ac
DNA Meth. m5C
H3−5ac
H4K20 me2
0.00
0.25
0.50
0.75
1.00
π̂
RPL24 – Forward
H3K4 me3:H4−4ac
H3K4 me1:H3−5ac
H3K4 me1:H2A.Z
H3K4 me1:H4−4ac
H3K27 me3:H3K4 me1
H3K4 me3:H3−5ac
H3K27 me3:H4K20 me3
H2A.Z:H4−4ac
H2A.Z:H4K20 me2
H3K9 me2:DNA Meth. m5C
H3K9 me3:H4K20 me3
DNA Meth. m5C:H4K20 me2
H4K20 me2:H4−4ac
H2A.Z:H3−5ac
H3K27 me2:DNA Meth. m5C
H2A.Z:H3K4 me3
DNA Meth. m5C:H3K9 me3
H4K20 me2:H3K4 me3
DNA Meth. m5C:H4K20 me3
H4K20 me2:H3−5ac
H3K27 me3:DNA Meth. m5C
H3K4 me1
H4−4ac
H3K27 me2
H3K9 me2
H3K4 me3
H4K20 me3
H3K9 me3
H4K20 me2
H3K27 me3
H2A.Z
H3−5ac
DNA Meth. m5C
0.00
0.25
0.50
0.75
1.00
π̂
RPL24 – Reverse
H3K4 me3:H4−4ac
H3K4 me1:H3−5ac
H3K4 me1:H2A.Z
H3K4 me1:H4−4ac
H3K27 me3:H3K4 me1
H3K4 me3:H3−5ac
H3K27 me3:H4K20 me3
H2A.Z:H4−4ac
H2A.Z:H4K20 me2
H3K9 me2:DNA Meth. m5C
H3K9 me3:H4K20 me3
DNA Meth. m5C:H4K20 me2
H4K20 me2:H4−4ac
H2A.Z:H3−5ac
H3K27 me2:DNA Meth. m5C
H2A.Z:H3K4 me3
DNA Meth. m5C:H3K9 me3
H4K20 me2:H3K4 me3
DNA Meth. m5C:H4K20 me3
H4K20 me2:H3−5ac
H3K27 me3:DNA Meth. m5C
H3K4 me1
H4−4ac
H3K27 me2
H3K9 me2
H3K4 me3
H4K20 me3
H3K9 me3
H4K20 me2
H3K27 me3
H2A.Z
H3−5ac
DNA Meth. m5C
0.00
0.25
0.50
0.75
1.00
π̂
UBTF (1) – Forward
DNA Meth. m5C:H3K9 me3
H3K27 me3:H3K4 me1
H3K27 me3:DNA Meth. m5C
DNA Meth. m5C:H4K20 me3
H3K27 me3:H4K20 me3
H3K4 me1:H4K20 me2
H2A.Z:H3−5ac
H2A.Z:H4K20 me2
H3K9 me3:H4K20 me3
H3K4 me3:H3−5ac
H3K4 me1:H4−4ac
H2A.Z:H4−4ac
H3K4 me1:H3−5ac
H3K27 me2:DNA Meth. m5C
H3K9 me2:DNA Meth. m5C
H4K20 me2:H3K4 me3
H3K4 me1:H2A.Z
H4K20 me2:H3−5ac
H2A.Z:H3K4 me3
H3K4 me3:H4−4ac
H4−4ac:H3−5ac
H3K4 me1
H3K9 me2
H3K27 me2
H3K27 me3
H3K9 me3
H4K20 me2
DNA Meth. m5C
H4−4ac
H4K20 me3
H3K4 me3
H2A.Z
H3−5ac
0.00
0.25
0.50
0.75
1.00
π̂
UBTF (1) – Reverse
DNA Meth. m5C:H3K9 me3
H3K27 me3:H3K4 me1
H3K27 me3:DNA Meth. m5C
DNA Meth. m5C:H4K20 me3
H3K27 me3:H4K20 me3
H3K4 me1:H4K20 me2
H2A.Z:H3−5ac
H2A.Z:H4K20 me2
H3K9 me3:H4K20 me3
H3K4 me3:H3−5ac
H3K4 me1:H4−4ac
H2A.Z:H4−4ac
H3K4 me1:H3−5ac
H3K27 me2:DNA Meth. m5C
H3K9 me2:DNA Meth. m5C
H4K20 me2:H3K4 me3
H3K4 me1:H2A.Z
H4K20 me2:H3−5ac
H2A.Z:H3K4 me3
H3K4 me3:H4−4ac
H4−4ac:H3−5ac
H3K4 me1
H3K9 me2
H3K27 me2
H3K27 me3
H3K9 me3
H4K20 me2
DNA Meth. m5C
H4−4ac
H4K20 me3
H3K4 me3
H2A.Z
H3−5ac
0.00
0.25
0.50
0.75
1.00
π̂

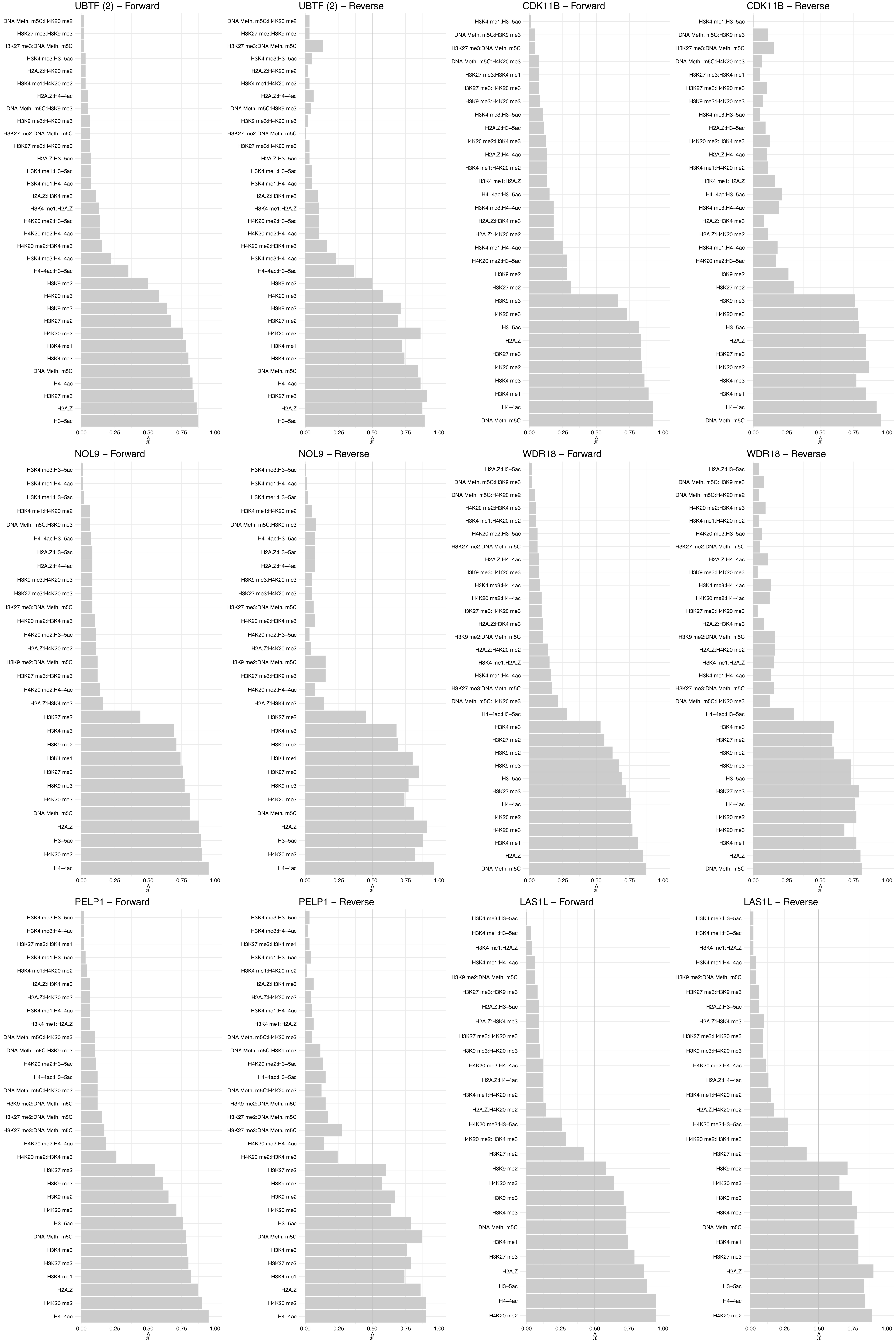
UBTF (2) – Forward
DNA Meth. m5C:H4K20 me2
H3K27 me3:H3K9 me3
H3K27 me3:DNA Meth. m5C
H3K4 me3:H3–5ac
H2A.Z:H4K20 me2
H3K4 me1:H4K20 me2
H2A.Z:H4–4ac
DNA Meth. m5C:H3K9 me3
H3K9 me3:H4K20 me3
H3K27 me2:DNA Meth. m5C
H3K27 me3:H4K20 me3
H2A.Z:H3–5ac
H3K4 me1:H3–5ac
H3K4 me1:H4–4ac
H2A.Z:H3K4 me3
H3K4 me1:H2A.Z
H4K20 me2:H3–5ac
H4K20 me2:H4–4ac
H4K20 me2:H3K4 me3
H3K4 me3:H4–4ac
H4–4ac:H3–5ac
H3K9 me2
H4K20 me3
H3K9 me3
H3K27 me2
H4K20 me2
H3K4 me1
H3K4 me3
DNA Meth. m5C
H4–4ac
H3K27 me3
H2A.Z
H3–5ac
0.00 0.25 0.50 0.75 1.00
π̂
UBTF (2) – Reverse
DNA Meth. m5C:H4K20 me2
H3K27 me3:H3K9 me3
H3K27 me3:DNA Meth. m5C
H3K4 me3:H3–5ac
H2A.Z:H4K20 me2
H3K4 me1:H4K20 me2
H2A.Z:H4–4ac
DNA Meth. m5C:H3K9 me3
H3K9 me3:H4K20 me3
H3K27 me2:DNA Meth. m5C
H3K27 me3:H4K20 me3
H2A.Z:H3–5ac
H3K4 me1:H3–5ac
H3K4 me1:H4–4ac
H2A.Z:H3K4 me3
H3K4 me1:H2A.Z
H4K20 me2:H3–5ac
H4K20 me2:H4–4ac
H4K20 me2:H3K4 me3
H3K4 me3:H4–4ac
H4–4ac:H3–5ac
H3K9 me2
H4K20 me3
H3K9 me3
H3K27 me2
H4K20 me2
H3K4 me1
H3K4 me3
DNA Meth. m5C
H4–4ac
H3K27 me3
H2A.Z
H3–5ac
0.00 0.25 0.50 0.75 1.00
π̂
CDK11B – Forward
H3K4 me1:H3–5ac
DNA Meth. m5C:H3K9 me3
H3K27 me3:DNA Meth. m5C
DNA Meth. m5C:H4K20 me3
H3K27 me3:H3K4 me1
H3K27 me3:H4K20 me3
H3K9 me3:H4K20 me3
H3K4 me3:H3–5ac
H2A.Z:H3–5ac
H4K20 me2:H3K4 me3
H2A.Z:H4–4ac
H3K4 me1:H4K20 me2
H3K4 me1:H2A.Z
H4–4ac:H3–5ac
H3K4 me3:H4–4ac
H2A.Z:H3K4 me3
H2A.Z:H4K20 me2
H3K4 me1:H4–4ac
H4K20 me2:H3–5ac
H3K9 me2
H3K27 me2
H3K9 me3
H4K20 me3
H3–5ac
H2A.Z
H3K27 me3
H4K20 me2
H3K4 me3
H3K4 me1
H4–4ac
DNA Meth. m5C
0.00 0.25 0.50 0.75 1.00
π̂
CDK11B – Reverse
H3K4 me1:H3–5ac
DNA Meth. m5C:H3K9 me3
H3K27 me3:DNA Meth. m5C
DNA Meth. m5C:H4K20 me3
H3K27 me3:H3K4 me1
H3K27 me3:H4K20 me3
H3K9 me3:H4K20 me3
H3K4 me3:H3–5ac
H2A.Z:H3–5ac
H4K20 me2:H3K4 me3
H2A.Z:H4–4ac
H3K4 me1:H4K20 me2
H3K4 me1:H2A.Z
H4–4ac:H3–5ac
H3K4 me3:H4–4ac
H2A.Z:H3K4 me3
H2A.Z:H4K20 me2
H3K4 me1:H4–4ac
H4K20 me2:H3–5ac
H3K9 me2
H3K27 me2
H3K9 me3
H4K20 me3
H3–5ac
H2A.Z
H3K27 me3
H4K20 me2
H3K4 me3
H3K4 me1
H4–4ac
DNA Meth. m5C
0.00 0.25 0.50 0.75 1.00
π̂
NOL9 – Forward
H3K4 me3:H3–5ac
H3K4 me1:H4–4ac
H3K4 me1:H3–5ac
H3K4 me1:H4K20 me2
DNA Meth. m5C:H3K9 me3
H4–4ac:H3–5ac
H2A.Z:H3–5ac
H2A.Z:H4–4ac
H3K9 me3:H4K20 me3
H3K27 me3:H4K20 me3
H3K27 me3:DNA Meth. m5C
H4K20 me2:H3K4 me3
H4K20 me2:H3–5ac
H2A.Z:H4K20 me2
H3K9 me2:DNA Meth. m5C
H3K27 me3:H3K9 me3
H4K20 me2:H4–4ac
H2A.Z:H3K4 me3
H3K27 me2
H3K4 me3
H3K9 me2
H3K4 me1
H3K27 me3
H3K9 me3
H4K20 me3
DNA Meth. m5C
H2A.Z
H3–5ac
H4K20 me2
H4–4ac
0.00 0.25 0.50 0.75 1.00
π̂
NOL9 – Reverse
H3K4 me3:H3–5ac
H3K4 me1:H4–4ac
H3K4 me1:H3–5ac
H3K4 me1:H4K20 me2
DNA Meth. m5C:H3K9 me3
H4–4ac:H3–5ac
H2A.Z:H3–5ac
H2A.Z:H4–4ac
H3K9 me3:H4K20 me3
H3K27 me3:H4K20 me3
H3K27 me3:DNA Meth. m5C
H4K20 me2:H3K4 me3
H4K20 me2:H3–5ac
H2A.Z:H4K20 me2
H3K9 me2:DNA Meth. m5C
H3K27 me3:H3K9 me3
H4K20 me2:H4–4ac
H2A.Z:H3K4 me3
H3K27 me2
H3K4 me3
H3K9 me2
H3K4 me1
H3K27 me3
H3K9 me3
H4K20 me3
DNA Meth. m5C
H2A.Z
H3–5ac
H4K20 me2
H4–4ac
0.00 0.25 0.50 0.75 1.00
π̂
WDR18 – Forward
H2A.Z:H3–5ac
DNA Meth. m5C:H3K9 me3
DNA Meth. m5C:H4K20 me2
H4K20 me2:H3K4 me3
H3K4 me1:H4K20 me2
H4K20 me2:H3–5ac
H3K27 me2:DNA Meth. m5C
H2A.Z:H4–4ac
H3K9 me3:H4K20 me3
H3K4 me3:H4–4ac
H4K20 me2:H4–4ac
H3K27 me3:H4K20 me3
H2A.Z:H3K4 me3
H3K9 me2:DNA Meth. m5C
H2A.Z:H4K20 me2
H3K4 me1:H2A.Z
H3K4 me1:H4–4ac
H3K27 me3:DNA Meth. m5C
DNA Meth. m5C:H4K20 me3
H4–4ac:H3–5ac
H3K4 me3
H3K27 me2
H3K9 me2
H3K9 me3
H3–5ac
H3K27 me3
H4–4ac
H4K20 me2
H4K20 me3
H3K4 me1
H2A.Z
DNA Meth. m5C
0.00 0.25 0.50 0.75 1.00
π̂
WDR18 – Reverse
H2A.Z:H3–5ac
DNA Meth. m5C:H3K9 me3
DNA Meth. m5C:H4K20 me2
H4K20 me2:H3K4 me3
H3K4 me1:H4K20 me2
H4K20 me2:H3–5ac
H3K27 me2:DNA Meth. m5C
H2A.Z:H4–4ac
H3K9 me3:H4K20 me3
H3K4 me3:H4–4ac
H4K20 me2:H4–4ac
H3K27 me3:H4K20 me3
H2A.Z:H3K4 me3
H3K9 me2:DNA Meth. m5C
H2A.Z:H4K20 me2
H3K4 me1:H2A.Z
H3K4 me1:H4–4ac
H3K27 me3:DNA Meth. m5C
DNA Meth. m5C:H4K20 me3
H4–4ac:H3–5ac
H3K4 me3
H3K27 me2
H3K9 me2
H3K9 me3
H3–5ac
H3K27 me3
H4–4ac
H4K20 me2
H4K20 me3
H3K4 me1
H2A.Z
DNA Meth. m5C
0.00 0.25 0.50 0.75 1.00
π̂
PELP1 – Forward
H3K4 me3:H3–5ac
H3K4 me3:H4–4ac
H3K27 me3:H3K4 me1
H3K4 me1:H3–5ac
H3K4 me1:H4K20 me2
H2A.Z:H3K4 me3
H2A.Z:H4K20 me2
H3K4 me1:H4–4ac
H3K4 me1:H2A.Z
DNA Meth. m5C:H4K20 me3
DNA Meth. m5C:H3K9 me3
H4K20 me2:H3–5ac
H4–4ac:H3–5ac
DNA Meth. m5C:H4K20 me2
H3K9 me2:DNA Meth. m5C
H3K27 me2:DNA Meth. m5C
H3K27 me3:DNA Meth. m5C
H4K20 me2:H4–4ac
H4K20 me2:H3K4 me3
H3K27 me2
H3K9 me3
H3K9 me2
H4K20 me3
H3–5ac
DNA Meth. m5C
H3K4 me3
H3K27 me3
H3K4 me1
H2A.Z
H4K20 me2
H4–4ac
0.00 0.25 0.50 0.75 1.00
π̂
PELP1 – Reverse
H3K4 me3:H3–5ac
H3K4 me3:H4–4ac
H3K27 me3:H3K4 me1
H3K4 me1:H3–5ac
H3K4 me1:H4K20 me2
H2A.Z:H3K4 me3
H2A.Z:H4K20 me2
H3K4 me1:H4–4ac
H3K4 me1:H2A.Z
DNA Meth. m5C:H4K20 me3
DNA Meth. m5C:H3K9 me3
H4K20 me2:H3–5ac
H4–4ac:H3–5ac
DNA Meth. m5C:H4K20 me2
H3K9 me2:DNA Meth. m5C
H3K27 me2:DNA Meth. m5C
H3K27 me3:DNA Meth. m5C
H4K20 me2:H4–4ac
H4K20 me2:H3K4 me3
H3K27 me2
H3K9 me3
H3K9 me2
H4K20 me3
H3–5ac
DNA Meth. m5C
H3K4 me3
H3K27 me3
H3K4 me1
H2A.Z
H4K20 me2
H4–4ac
0.00 0.25 0.50 0.75 1.00
π̂
LAS1L – Forward
H3K4 me3:H3–5ac
H3K4 me1:H3–5ac
H3K4 me1:H2A.Z
H3K4 me1:H4–4ac
H3K9 me2:DNA Meth. m5C
H3K27 me3:H3K9 me3
H2A.Z:H3–5ac
H2A.Z:H3K4 me3
H3K27 me3:H4K20 me3
H3K9 me3:H4K20 me3
H4K20 me2:H4–4ac
H2A.Z:H4–4ac
H3K4 me1:H4K20 me2
H2A.Z:H4K20 me2
H4K20 me2:H3–5ac
H4K20 me2:H3K4 me3
H3K27 me2
H3K9 me2
H4K20 me3
H3K9 me3
H3K4 me3
DNA Meth. m5C
H3K4 me1
H3K27 me3
H2A.Z
H3–5ac
H4–4ac
H4K20 me2
0.00 0.25 0.50 0.75 1.00
π̂
LAS1L – Reverse
H3K4 me3:H3–5ac
H3K4 me1:H3–5ac
H3K4 me1:H2A.Z
H3K4 me1:H4–4ac
H3K9 me2:DNA Meth. m5C
H3K27 me3:H3K9 me3
H2A.Z:H3–5ac
H2A.Z:H3K4 me3
H3K27 me3:H4K20 me3
H3K9 me3:H4K20 me3
H4K20 me2:H4–4ac
H2A.Z:H4–4ac
H3K4 me1:H4K20 me2
H2A.Z:H4K20 me2
H4K20 me2:H3–5ac
H4K20 me2:H3K4 me3
H3K27 me2
H3K9 me2
H4K20 me3
H3K9 me3
H3K4 me3
DNA Meth. m5C
H3K4 me1
H3K27 me3
H2A.Z
H3–5ac
H4–4ac
H4K20 me2
0.00 0.25 0.50 0.75 1.00
π̂

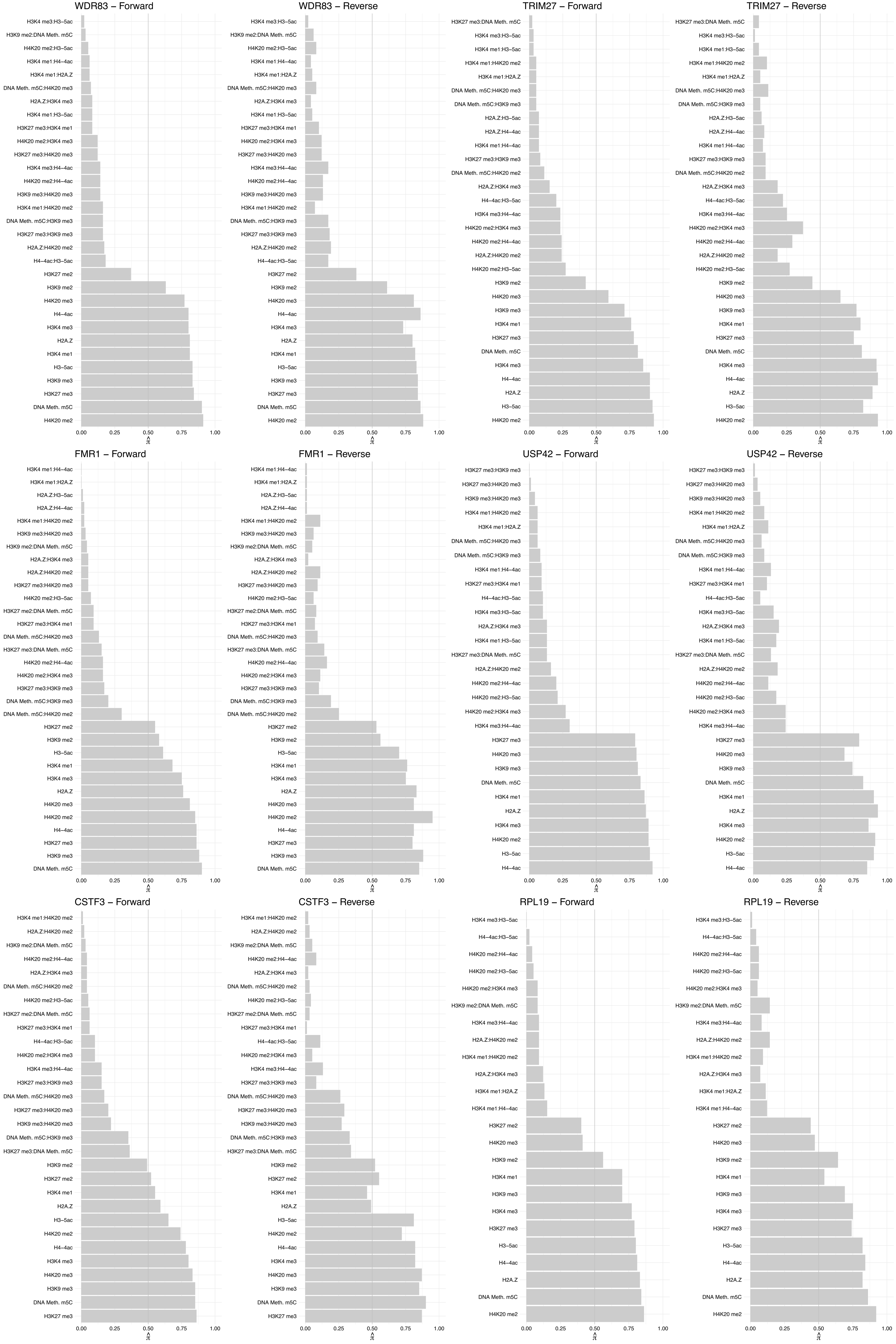
WDR83 – Forward
H3K4 me3:H3−5ac
H3K9 me2:DNA Meth. m5C
H4K20 me2:H3−5ac
H3K4 me1:H4−4ac
H3K4 me1:H2A.Z
DNA Meth. m5C:H4K20 me3
H2A.Z:H3K4 me3
H3K4 me1:H3−5ac
H3K27 me3:H3K4 me1
H4K20 me2:H3K4 me3
H3K27 me3:H4K20 me3
H3K4 me3:H4−4ac
H4K20 me2:H4−4ac
H3K9 me3:H4K20 me3
H3K4 me1:H4K20 me2
DNA Meth. m5C:H3K9 me3
H3K27 me3:H3K9 me3
H2A.Z:H4K20 me2
H4−4ac:H3−5ac
H3K27 me2
H3K9 me2
H4K20 me3
H4−4ac
H3K4 me3
H2A.Z
H3K4 me1
H3−5ac
H3K9 me3
H3K27 me3
DNA Meth. m5C
H4K20 me2
0.00 0.25 0.50 0.75 1.00
π̂
WDR83 – Reverse
H3K4 me3:H3−5ac
H3K9 me2:DNA Meth. m5C
H4K20 me2:H3−5ac
H3K4 me1:H4−4ac
H3K4 me1:H2A.Z
DNA Meth. m5C:H4K20 me3
H2A.Z:H3K4 me3
H3K4 me1:H3−5ac
H3K27 me3:H3K4 me1
H4K20 me2:H3K4 me3
H3K27 me3:H4K20 me3
H3K4 me3:H4−4ac
H4K20 me2:H4−4ac
H3K9 me3:H4K20 me3
H3K4 me1:H4K20 me2
DNA Meth. m5C:H3K9 me3
H3K27 me3:H3K9 me3
H2A.Z:H4K20 me2
H4−4ac:H3−5ac
H3K27 me2
H3K9 me2
H4K20 me3
H4−4ac
H3K4 me3
H2A.Z
H3K4 me1
H3−5ac
H3K9 me3
H3K27 me3
DNA Meth. m5C
H4K20 me2
0.00 0.25 0.50 0.75 1.00
π̂
TRIM27 – Forward
H3K27 me3:DNA Meth. m5C
H3K4 me3:H3−5ac
H3K4 me1:H3−5ac
H3K4 me1:H4K20 me2
H3K4 me1:H2A.Z
DNA Meth. m5C:H4K20 me3
DNA Meth. m5C:H3K9 me3
H2A.Z:H3−5ac
H2A.Z:H4−4ac
H3K4 me1:H4−4ac
H3K27 me3:H3K9 me3
DNA Meth. m5C:H4K20 me2
H2A.Z:H3K4 me3
H4−4ac:H3−5ac
H3K4 me3:H4−4ac
H4K20 me2:H3K4 me3
H4K20 me2:H4−4ac
H2A.Z:H4K20 me2
H4K20 me2:H3−5ac
H3K9 me2
H4K20 me3
H3K9 me3
H3K4 me1
H3K27 me3
DNA Meth. m5C
H3K4 me3
H4−4ac
H2A.Z
H3−5ac
H4K20 me2
0.00 0.25 0.50 0.75 1.00
π̂
TRIM27 – Reverse
H3K27 me3:DNA Meth. m5C
H3K4 me3:H3−5ac
H3K4 me1:H3−5ac
H3K4 me1:H4K20 me2
H3K4 me1:H2A.Z
DNA Meth. m5C:H4K20 me3
DNA Meth. m5C:H3K9 me3
H2A.Z:H3−5ac
H2A.Z:H4−4ac
H3K4 me1:H4−4ac
H3K27 me3:H3K9 me3
DNA Meth. m5C:H4K20 me2
H2A.Z:H3K4 me3
H4−4ac:H3−5ac
H3K4 me3:H4−4ac
H4K20 me2:H3K4 me3
H4K20 me2:H4−4ac
H2A.Z:H4K20 me2
H4K20 me2:H3−5ac
H3K9 me2
H4K20 me3
H3K9 me3
H3K4 me1
H3K27 me3
DNA Meth. m5C
H3K4 me3
H4−4ac
H2A.Z
H3−5ac
H4K20 me2
0.00 0.25 0.50 0.75 1.00
π̂
FMR1 – Forward
H3K4 me1:H4−4ac
H3K4 me1:H2A.Z
H2A.Z:H3−5ac
H2A.Z:H4−4ac
H3K4 me1:H4K20 me2
H3K9 me3:H4K20 me3
H3K9 me2:DNA Meth. m5C
H2A.Z:H3K4 me3
H2A.Z:H4K20 me2
H3K27 me3:H4K20 me3
H4K20 me2:H3−5ac
H3K27 me2:DNA Meth. m5C
H3K27 me3:H3K4 me1
DNA Meth. m5C:H4K20 me3
H3K27 me3:DNA Meth. m5C
H4K20 me2:H4−4ac
H4K20 me2:H3K4 me3
H3K27 me3:H3K9 me3
DNA Meth. m5C:H3K9 me3
DNA Meth. m5C:H4K20 me2
H3K27 me2
H3K9 me2
H3−5ac
H3K4 me1
H3K4 me3
H2A.Z
H4K20 me3
H4K20 me2
H4−4ac
H3K27 me3
H3K9 me3
DNA Meth. m5C
0.00 0.25 0.50 0.75 1.00
π̂
FMR1 – Reverse
H3K4 me1:H4−4ac
H3K4 me1:H2A.Z
H2A.Z:H3−5ac
H2A.Z:H4−4ac
H3K4 me1:H4K20 me2
H3K9 me3:H4K20 me3
H3K9 me2:DNA Meth. m5C
H2A.Z:H3K4 me3
H2A.Z:H4K20 me2
H3K27 me3:H4K20 me3
H4K20 me2:H3−5ac
H3K27 me2:DNA Meth. m5C
H3K27 me3:H3K4 me1
DNA Meth. m5C:H4K20 me3
H3K27 me3:DNA Meth. m5C
H4K20 me2:H4−4ac
H4K20 me2:H3K4 me3
H3K27 me3:H3K9 me3
DNA Meth. m5C:H3K9 me3
DNA Meth. m5C:H4K20 me2
H3K27 me2
H3K9 me2
H3−5ac
H3K4 me1
H3K4 me3
H2A.Z
H4K20 me3
H4K20 me2
H4−4ac
H3K27 me3
H3K9 me3
DNA Meth. m5C
0.00 0.25 0.50 0.75 1.00
π̂
USP42 – Forward
H3K27 me3:H3K9 me3
H3K27 me3:H4K20 me3
H3K9 me3:H4K20 me3
H3K4 me1:H4K20 me2
H3K4 me1:H2A.Z
DNA Meth. m5C:H4K20 me3
DNA Meth. m5C:H3K9 me3
H3K4 me1:H4−4ac
H3K27 me3:H3K4 me1
H4−4ac:H3−5ac
H3K4 me3:H3−5ac
H2A.Z:H3K4 me3
H3K4 me1:H3−5ac
H3K27 me3:DNA Meth. m5C
H2A.Z:H4K20 me2
H4K20 me2:H4−4ac
H4K20 me2:H3−5ac
H4K20 me2:H3K4 me3
H3K4 me3:H4−4ac
H3K27 me3
H4K20 me3
H3K9 me3
DNA Meth. m5C
H3K4 me1
H2A.Z
H3K4 me3
H4K20 me2
H3−5ac
H4−4ac
0.00 0.25 0.50 0.75 1.00
π̂
USP42 – Reverse
H3K27 me3:H3K9 me3
H3K27 me3:H4K20 me3
H3K9 me3:H4K20 me3
H3K4 me1:H4K20 me2
H3K4 me1:H2A.Z
DNA Meth. m5C:H4K20 me3
DNA Meth. m5C:H3K9 me3
H3K4 me1:H4−4ac
H3K27 me3:H3K4 me1
H4−4ac:H3−5ac
H3K4 me3:H3−5ac
H2A.Z:H3K4 me3
H3K4 me1:H3−5ac
H3K27 me3:DNA Meth. m5C
H2A.Z:H4K20 me2
H4K20 me2:H4−4ac
H4K20 me2:H3−5ac
H4K20 me2:H3K4 me3
H3K4 me3:H4−4ac
H3K27 me3
H4K20 me3
H3K9 me3
DNA Meth. m5C
H3K4 me1
H2A.Z
H3K4 me3
H4K20 me2
H3−5ac
H4−4ac
0.00 0.25 0.50 0.75 1.00
π̂
CSTF3 – Forward
H3K4 me1:H4K20 me2
H2A.Z:H4K20 me2
H3K9 me2:DNA Meth. m5C
H4K20 me2:H4−4ac
H2A.Z:H3K4 me3
DNA Meth. m5C:H4K20 me2
H4K20 me2:H3−5ac
H3K27 me2:DNA Meth. m5C
H3K27 me3:H3K4 me1
H4−4ac:H3−5ac
H4K20 me2:H3K4 me3
H3K4 me3:H4−4ac
H3K27 me3:H3K9 me3
DNA Meth. m5C:H4K20 me3
H3K27 me3:H4K20 me3
H3K9 me3:H4K20 me3
DNA Meth. m5C:H3K9 me3
H3K27 me3:DNA Meth. m5C
H3K9 me2
H3K27 me2
H3K4 me1
H2A.Z
H3−5ac
H4K20 me2
H4−4ac
H3K4 me3
H4K20 me3
H3K9 me3
DNA Meth. m5C
H3K27 me3
0.00 0.25 0.50 0.75 1.00
π̂
CSTF3 – Reverse
H3K4 me1:H4K20 me2
H2A.Z:H4K20 me2
H3K9 me2:DNA Meth. m5C
H4K20 me2:H4−4ac
H2A.Z:H3K4 me3
DNA Meth. m5C:H4K20 me2
H4K20 me2:H3−5ac
H3K27 me2:DNA Meth. m5C
H3K27 me3:H3K4 me1
H4−4ac:H3−5ac
H4K20 me2:H3K4 me3
H3K4 me3:H4−4ac
H3K27 me3:H3K9 me3
DNA Meth. m5C:H4K20 me3
H3K27 me3:H4K20 me3
H3K9 me3:H4K20 me3
DNA Meth. m5C:H3K9 me3
H3K27 me3:DNA Meth. m5C
H3K9 me2
H3K27 me2
H3K4 me1
H2A.Z
H3−5ac
H4K20 me2
H4−4ac
H3K4 me3
H4K20 me3
H3K9 me3
DNA Meth. m5C
H3K27 me3
0.00 0.25 0.50 0.75 1.00
π̂
RPL19 – Forward
H3K4 me3:H3−5ac
H4−4ac:H3−5ac
H4K20 me2:H4−4ac
H4K20 me2:H3−5ac
H4K20 me2:H3K4 me3
H3K9 me2:DNA Meth. m5C
H3K4 me3:H4−4ac
H2A.Z:H4K20 me2
H3K4 me1:H4K20 me2
H2A.Z:H3K4 me3
H3K4 me1:H2A.Z
H3K4 me1:H4−4ac
H3K27 me2
H4K20 me3
H3K9 me2
H3K4 me1
H3K9 me3
H3K4 me3
H3K27 me3
H3−5ac
H4−4ac
H2A.Z
DNA Meth. m5C
H4K20 me2
0.00 0.25 0.50 0.75 1.00
π̂
RPL19 – Reverse
H3K4 me3:H3−5ac
H4−4ac:H3−5ac
H4K20 me2:H4−4ac
H4K20 me2:H3−5ac
H4K20 me2:H3K4 me3
H3K9 me2:DNA Meth. m5C
H3K4 me3:H4−4ac
H2A.Z:H4K20 me2
H3K4 me1:H4K20 me2
H2A.Z:H3K4 me3
H3K4 me1:H2A.Z
H3K4 me1:H4−4ac
H3K27 me2
H4K20 me3
H3K9 me2
H3K4 me1
H3K9 me3
H3K4 me3
H3K27 me3
H3−5ac
H4−4ac
H2A.Z
DNA Meth. m5C
H4K20 me2
0.00 0.25 0.50 0.75 1.00
π̂

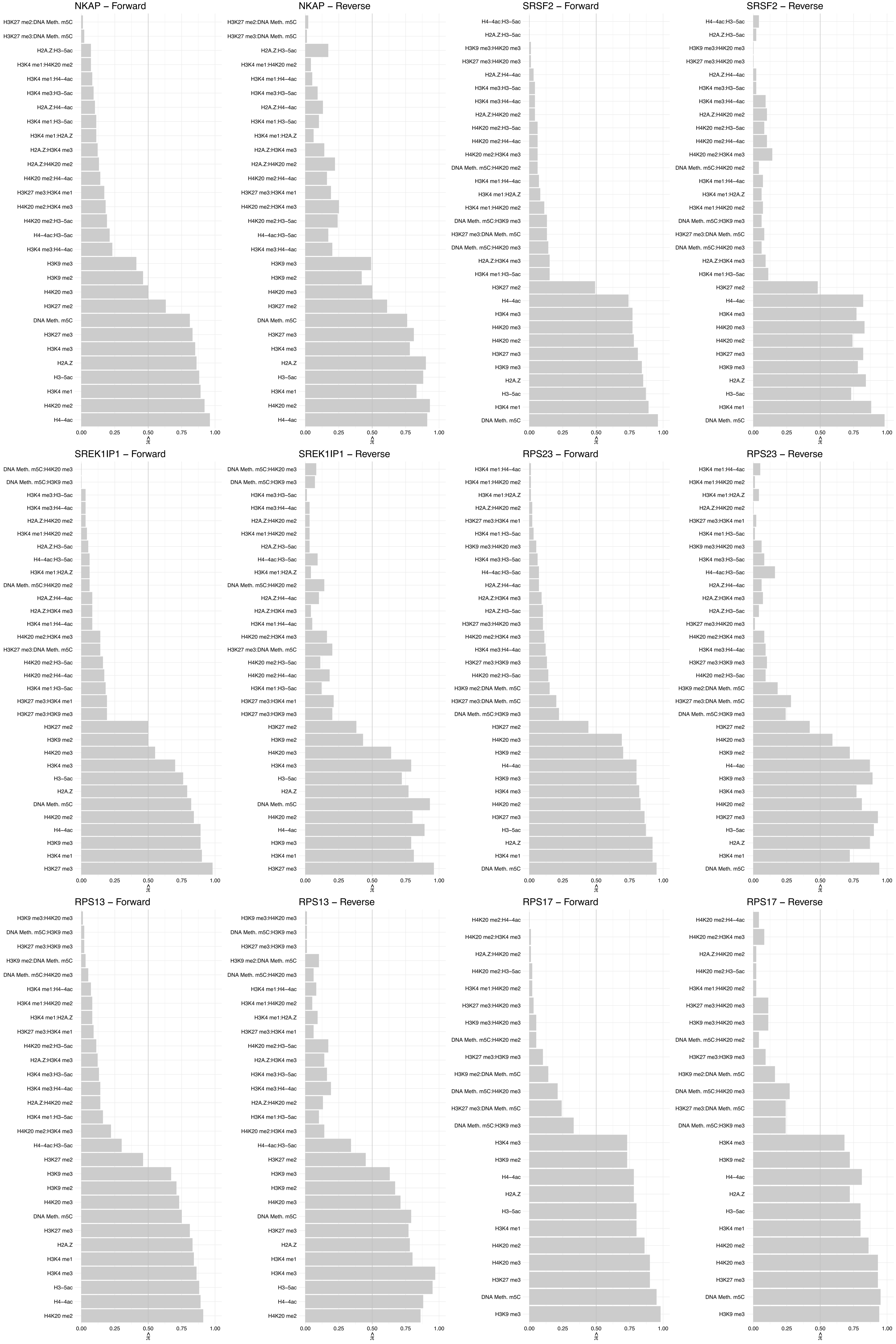
NKAP − Forward
H3K27 me2:DNA Meth. m5C
H3K27 me3:DNA Meth. m5C
H2A.Z:H3−5ac
H3K4 me1:H4K20 me2
H3K4 me1:H4−4ac
H3K4 me3:H3−5ac
H2A.Z:H4−4ac
H3K4 me1:H3−5ac
H3K4 me1:H2A.Z
H2A.Z:H3K4 me3
H2A.Z:H4K20 me2
H4K20 me2:H4−4ac
H3K27 me3:H3K4 me1
H4K20 me2:H3K4 me3
H4K20 me2:H3−5ac
H4−4ac:H3−5ac
H3K4 me3:H4−4ac
H3K9 me3
H3K9 me2
H4K20 me3
H3K27 me2
DNA Meth. m5C
H3K27 me3
H3K4 me3
H2A.Z
H3−5ac
H3K4 me1
H4K20 me2
H4−4ac
0.00 0.25 0.50 0.75 1.00
π̂
NKAP − Reverse
H3K27 me2:DNA Meth. m5C
H3K27 me3:DNA Meth. m5C
H2A.Z:H3−5ac
H3K4 me1:H4K20 me2
H3K4 me1:H4−4ac
H3K4 me3:H3−5ac
H2A.Z:H4−4ac
H3K4 me1:H3−5ac
H3K4 me1:H2A.Z
H2A.Z:H3K4 me3
H2A.Z:H4K20 me2
H4K20 me2:H4−4ac
H3K27 me3:H3K4 me1
H4K20 me2:H3K4 me3
H4K20 me2:H3−5ac
H4−4ac:H3−5ac
H3K4 me3:H4−4ac
H3K9 me3
H3K9 me2
H4K20 me3
H3K27 me2
DNA Meth. m5C
H3K27 me3
H3K4 me3
H2A.Z
H3−5ac
H3K4 me1
H4K20 me2
H4−4ac
0.00 0.25 0.50 0.75 1.00
π̂
SRSF2 − Forward
H4−4ac:H3−5ac
H2A.Z:H3−5ac
H3K9 me3:H4K20 me3
H3K27 me3:H4K20 me3
H2A.Z:H4−4ac
H3K4 me3:H3−5ac
H3K4 me3:H4−4ac
H2A.Z:H4K20 me2
H4K20 me2:H3−5ac
H4K20 me2:H4−4ac
H4K20 me2:H3K4 me3
DNA Meth. m5C:H4K20 me2
H3K4 me1:H4−4ac
H3K4 me1:H2A.Z
H3K4 me1:H4K20 me2
DNA Meth. m5C:H3K9 me3
H3K27 me3:DNA Meth. m5C
DNA Meth. m5C:H4K20 me3
H2A.Z:H3K4 me3
H3K4 me1:H3−5ac
H3K27 me2
H4−4ac
H3K4 me3
H4K20 me3
H4K20 me2
H3K27 me3
H3K9 me3
H2A.Z
H3−5ac
H3K4 me1
DNA Meth. m5C
0.00 0.25 0.50 0.75 1.00
π̂
SRSF2 − Reverse
H4−4ac:H3−5ac
H2A.Z:H3−5ac
H3K9 me3:H4K20 me3
H3K27 me3:H4K20 me3
H2A.Z:H4−4ac
H3K4 me3:H3−5ac
H3K4 me3:H4−4ac
H2A.Z:H4K20 me2
H4K20 me2:H3−5ac
H4K20 me2:H4−4ac
H4K20 me2:H3K4 me3
DNA Meth. m5C:H4K20 me2
H3K4 me1:H4−4ac
H3K4 me1:H2A.Z
H3K4 me1:H4K20 me2
DNA Meth. m5C:H3K9 me3
H3K27 me3:DNA Meth. m5C
DNA Meth. m5C:H4K20 me3
H2A.Z:H3K4 me3
H3K4 me1:H3−5ac
H3K27 me2
H4−4ac
H3K4 me3
H4K20 me3
H4K20 me2
H3K27 me3
H3K9 me3
H2A.Z
H3−5ac
H3K4 me1
DNA Meth. m5C
0.00 0.25 0.50 0.75 1.00
π̂
SREK1IP1 − Forward
DNA Meth. m5C:H4K20 me3
DNA Meth. m5C:H3K9 me3
H3K4 me3:H3−5ac
H3K4 me3:H4−4ac
H2A.Z:H4K20 me2
H3K4 me1:H4K20 me2
H2A.Z:H3−5ac
H4−4ac:H3−5ac
H3K4 me1:H2A.Z
DNA Meth. m5C:H4K20 me2
H2A.Z:H4−4ac
H2A.Z:H3K4 me3
H3K4 me1:H4−4ac
H4K20 me2:H3K4 me3
H3K27 me3:DNA Meth. m5C
H4K20 me2:H3−5ac
H4K20 me2:H4−4ac
H3K4 me1:H3−5ac
H3K27 me3:H3K4 me1
H3K27 me3:H3K9 me3
H3K27 me2
H3K9 me2
H4K20 me3
H3K4 me3
H3−5ac
H2A.Z
DNA Meth. m5C
H4K20 me2
H4−4ac
H3K9 me3
H3K4 me1
H3K27 me3
0.00 0.25 0.50 0.75 1.00
π̂
SREK1IP1 − Reverse
DNA Meth. m5C:H4K20 me3
DNA Meth. m5C:H3K9 me3
H3K4 me3:H3−5ac
H3K4 me3:H4−4ac
H2A.Z:H4K20 me2
H3K4 me1:H4K20 me2
H2A.Z:H3−5ac
H4−4ac:H3−5ac
H3K4 me1:H2A.Z
DNA Meth. m5C:H4K20 me2
H2A.Z:H4−4ac
H2A.Z:H3K4 me3
H3K4 me1:H4−4ac
H4K20 me2:H3K4 me3
H3K27 me3:DNA Meth. m5C
H4K20 me2:H3−5ac
H4K20 me2:H4−4ac
H3K4 me1:H3−5ac
H3K27 me3:H3K4 me1
H3K27 me3:H3K9 me3
H3K27 me2
H3K9 me2
H4K20 me3
H3K4 me3
H3−5ac
H2A.Z
DNA Meth. m5C
H4K20 me2
H4−4ac
H3K9 me3
H3K4 me1
H3K27 me3
0.00 0.25 0.50 0.75 1.00
π̂
RPS23 − Forward
H3K4 me1:H4−4ac
H3K4 me1:H4K20 me2
H3K4 me1:H2A.Z
H2A.Z:H4K20 me2
H3K27 me3:H3K4 me1
H3K4 me1:H3−5ac
H3K9 me3:H4K20 me3
H3K4 me3:H3−5ac
H4−4ac:H3−5ac
H2A.Z:H4−4ac
H2A.Z:H3K4 me3
H2A.Z:H3−5ac
H3K27 me3:H4K20 me3
H4K20 me2:H3K4 me3
H3K4 me3:H4−4ac
H3K27 me3:H3K9 me3
H4K20 me2:H3−5ac
H3K9 me2:DNA Meth. m5C
H3K27 me3:DNA Meth. m5C
DNA Meth. m5C:H3K9 me3
H3K27 me2
H4K20 me3
H3K9 me2
H4−4ac
H3K9 me3
H3K4 me3
H4K20 me2
H3K27 me3
H3−5ac
H2A.Z
H3K4 me1
DNA Meth. m5C
0.00 0.25 0.50 0.75 1.00
π̂
RPS23 − Reverse
H3K4 me1:H4−4ac
H3K4 me1:H4K20 me2
H3K4 me1:H2A.Z
H2A.Z:H4K20 me2
H3K27 me3:H3K4 me1
H3K4 me1:H3−5ac
H3K9 me3:H4K20 me3
H3K4 me3:H3−5ac
H4−4ac:H3−5ac
H2A.Z:H4−4ac
H2A.Z:H3K4 me3
H2A.Z:H3−5ac
H3K27 me3:H4K20 me3
H4K20 me2:H3K4 me3
H3K4 me3:H4−4ac
H3K27 me3:H3K9 me3
H4K20 me2:H3−5ac
H3K9 me2:DNA Meth. m5C
H3K27 me3:DNA Meth. m5C
DNA Meth. m5C:H3K9 me3
H3K27 me2
H4K20 me3
H3K9 me2
H4−4ac
H3K9 me3
H3K4 me3
H4K20 me2
H3K27 me3
H3−5ac
H2A.Z
H3K4 me1
DNA Meth. m5C
0.00 0.25 0.50 0.75 1.00
π̂
RPS13 − Forward
H3K9 me3:H4K20 me3
DNA Meth. m5C:H3K9 me3
H3K27 me3:H3K9 me3
H3K9 me2:DNA Meth. m5C
DNA Meth. m5C:H4K20 me3
H3K4 me1:H4−4ac
H3K4 me1:H4K20 me2
H3K4 me1:H2A.Z
H3K27 me3:H3K4 me1
H4K20 me2:H3−5ac
H2A.Z:H3K4 me3
H3K4 me3:H3−5ac
H3K4 me3:H4−4ac
H2A.Z:H4K20 me2
H3K4 me1:H3−5ac
H4K20 me2:H3K4 me3
H4−4ac:H3−5ac
H3K27 me2
H3K9 me3
H3K9 me2
H4K20 me3
DNA Meth. m5C
H3K27 me3
H2A.Z
H3K4 me1
H3K4 me3
H3−5ac
H4−4ac
H4K20 me2
0.00 0.25 0.50 0.75 1.00
π̂
RPS13 − Reverse
H3K9 me3:H4K20 me3
DNA Meth. m5C:H3K9 me3
H3K27 me3:H3K9 me3
H3K9 me2:DNA Meth. m5C
DNA Meth. m5C:H4K20 me3
H3K4 me1:H4−4ac
H3K4 me1:H4K20 me2
H3K4 me1:H2A.Z
H3K27 me3:H3K4 me1
H4K20 me2:H3−5ac
H2A.Z:H3K4 me3
H3K4 me3:H3−5ac
H3K4 me3:H4−4ac
H2A.Z:H4K20 me2
H3K4 me1:H3−5ac
H4K20 me2:H3K4 me3
H4−4ac:H3−5ac
H3K27 me2
H3K9 me3
H3K9 me2
H4K20 me3
DNA Meth. m5C
H3K27 me3
H2A.Z
H3K4 me1
H3K4 me3
H3−5ac
H4−4ac
H4K20 me2
0.00 0.25 0.50 0.75 1.00
π̂
RPS17 − Forward
H4K20 me2:H4−4ac
H4K20 me2:H3K4 me3
H2A.Z:H4K20 me2
H4K20 me2:H3−5ac
H3K4 me1:H4K20 me2
H3K27 me3:H4K20 me3
H3K9 me3:H4K20 me3
DNA Meth. m5C:H4K20 me2
H3K27 me3:H3K9 me3
H3K9 me2:DNA Meth. m5C
DNA Meth. m5C:H4K20 me3
H3K27 me3:DNA Meth. m5C
DNA Meth. m5C:H3K9 me3
H3K4 me3
H3K9 me2
H4−4ac
H2A.Z
H3−5ac
H3K4 me1
H4K20 me2
H4K20 me3
H3K27 me3
DNA Meth. m5C
H3K9 me3
0.00 0.25 0.50 0.75 1.00
π̂
RPS17 − Reverse
H4K20 me2:H4−4ac
H4K20 me2:H3K4 me3
H2A.Z:H4K20 me2
H4K20 me2:H3−5ac
H3K4 me1:H4K20 me2
H3K27 me3:H4K20 me3
H3K9 me3:H4K20 me3
DNA Meth. m5C:H4K20 me2
H3K27 me3:H3K9 me3
H3K9 me2:DNA Meth. m5C
DNA Meth. m5C:H4K20 me3
H3K27 me3:DNA Meth. m5C
DNA Meth. m5C:H3K9 me3
H3K4 me3
H3K9 me2
H4−4ac
H2A.Z
H3−5ac
H3K4 me1
H4K20 me2
H4K20 me3
H3K27 me3
DNA Meth. m5C
H3K9 me3
0.00 0.25 0.50 0.75 1.00
π̂

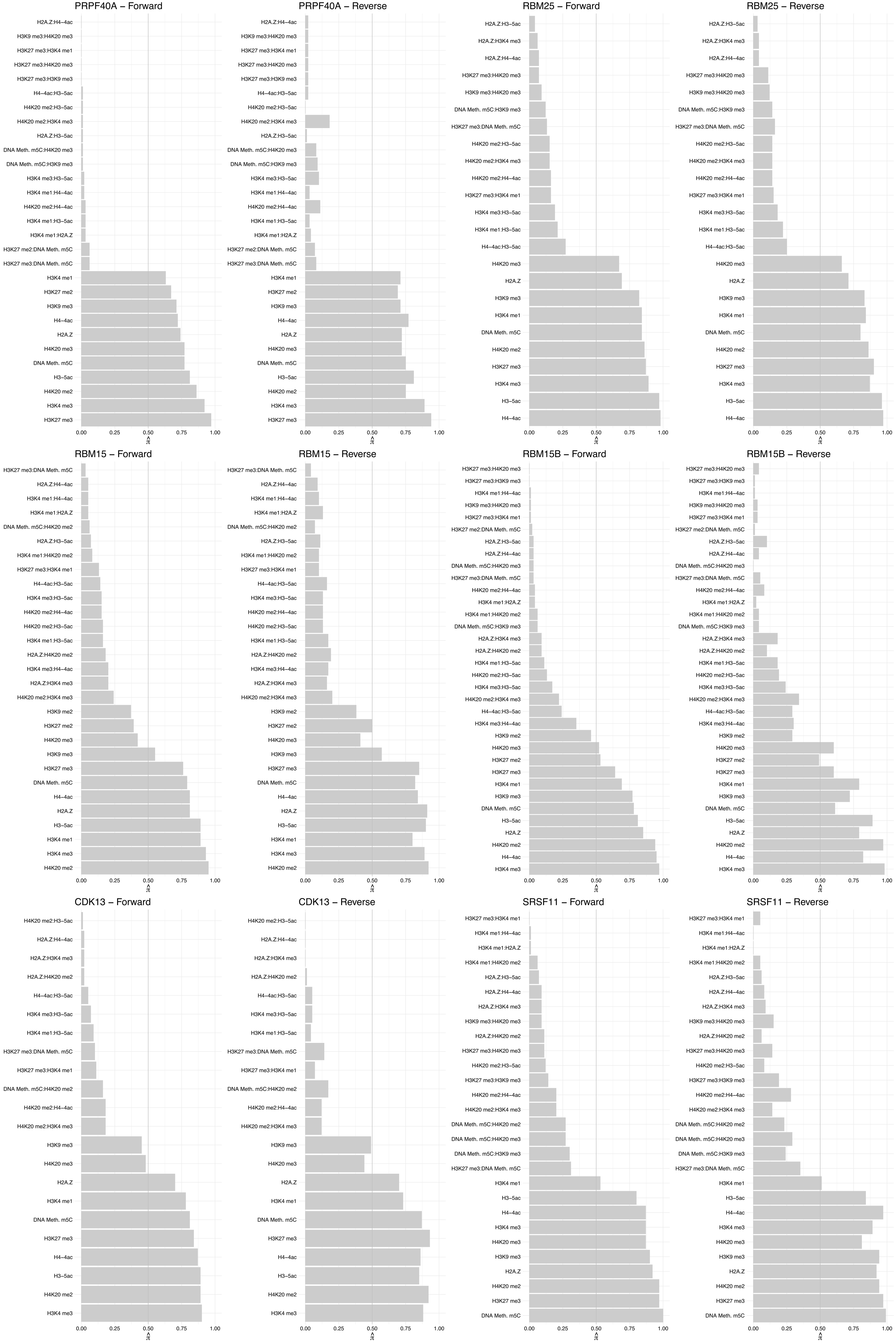
PRPF40A – Forward
H2A.Z:H4−4ac
H3K9 me3:H4K20 me3
H3K27 me3:H3K4 me1
H3K27 me3:H4K20 me3
H3K27 me3:H3K9 me3
H4−4ac:H3−5ac
H4K20 me2:H3−5ac
H4K20 me2:H3K4 me3
H2A.Z:H3−5ac
DNA Meth. m5C:H4K20 me3
DNA Meth. m5C:H3K9 me3
H3K4 me3:H3−5ac
H3K4 me1:H4−4ac
H4K20 me2:H4−4ac
H3K4 me1:H3−5ac
H3K4 me1:H2A.Z
H3K27 me2:DNA Meth. m5C
H3K27 me3:DNA Meth. m5C
H3K4 me1
H3K27 me2
H3K9 me3
H4−4ac
H2A.Z
H4K20 me3
DNA Meth. m5C
H3−5ac
H4K20 me2
H3K4 me3
H3K27 me3
0.00 0.25 0.50 0.75 1.00
$\hat{\pi}$
PRPF40A – Reverse
H2A.Z:H4−4ac
H3K9 me3:H4K20 me3
H3K27 me3:H3K4 me1
H3K27 me3:H4K20 me3
H3K27 me3:H3K9 me3
H4−4ac:H3−5ac
H4K20 me2:H3−5ac
H4K20 me2:H3K4 me3
H2A.Z:H3−5ac
DNA Meth. m5C:H4K20 me3
DNA Meth. m5C:H3K9 me3
H3K4 me3:H3−5ac
H3K4 me1:H4−4ac
H4K20 me2:H4−4ac
H3K4 me1:H3−5ac
H3K4 me1:H2A.Z
H3K27 me2:DNA Meth. m5C
H3K27 me3:DNA Meth. m5C
H3K4 me1
H3K27 me2
H3K9 me3
H4−4ac
H2A.Z
H4K20 me3
DNA Meth. m5C
H3−5ac
H4K20 me2
H3K4 me3
H3K27 me3
0.00 0.25 0.50 0.75 1.00
$\hat{\pi}$
RBM25 – Forward
H2A.Z:H3−5ac
H2A.Z:H3K4 me3
H2A.Z:H4−4ac
H3K27 me3:H4K20 me3
H3K9 me3:H4K20 me3
DNA Meth. m5C:H3K9 me3
H3K27 me3:DNA Meth. m5C
H4K20 me2:H3−5ac
H4K20 me2:H3K4 me3
H4K20 me2:H4−4ac
H3K27 me3:H3K4 me1
H3K4 me3:H3−5ac
H3K4 me1:H3−5ac
H4−4ac:H3−5ac
H4K20 me3
H2A.Z
H3K9 me3
H3K4 me1
DNA Meth. m5C
H4K20 me2
H3K27 me3
H3K4 me3
H3−5ac
H4−4ac
0.00 0.25 0.50 0.75 1.00
$\hat{\pi}$
RBM25 – Reverse
H2A.Z:H3−5ac
H2A.Z:H3K4 me3
H2A.Z:H4−4ac
H3K27 me3:H4K20 me3
H3K9 me3:H4K20 me3
DNA Meth. m5C:H3K9 me3
H3K27 me3:DNA Meth. m5C
H4K20 me2:H3−5ac
H4K20 me2:H3K4 me3
H4K20 me2:H4−4ac
H3K27 me3:H3K4 me1
H3K4 me3:H3−5ac
H3K4 me1:H3−5ac
H4−4ac:H3−5ac
H4K20 me3
H2A.Z
H3K9 me3
H3K4 me1
DNA Meth. m5C
H4K20 me2
H3K27 me3
H3K4 me3
H3−5ac
H4−4ac
0.00 0.25 0.50 0.75 1.00
$\hat{\pi}$
RBM15 – Forward
H3K27 me3:DNA Meth. m5C
H2A.Z:H4−4ac
H3K4 me1:H4−4ac
H3K4 me1:H2A.Z
DNA Meth. m5C:H4K20 me2
H2A.Z:H3−5ac
H3K4 me1:H4K20 me2
H3K27 me3:H3K4 me1
H4−4ac:H3−5ac
H3K4 me3:H3−5ac
H4K20 me2:H4−4ac
H4K20 me2:H3−5ac
H3K4 me1:H3−5ac
H2A.Z:H4K20 me2
H3K4 me3:H4−4ac
H2A.Z:H3K4 me3
H4K20 me2:H3K4 me3
H3K9 me2
H3K27 me2
H4K20 me3
H3K9 me3
H3K27 me3
DNA Meth. m5C
H4−4ac
H2A.Z
H3−5ac
H3K4 me1
H3K4 me3
H4K20 me2
0.00 0.25 0.50 0.75 1.00
$\hat{\pi}$
RBM15 – Reverse
H3K27 me3:DNA Meth. m5C
H2A.Z:H4−4ac
H3K4 me1:H4−4ac
H3K4 me1:H2A.Z
DNA Meth. m5C:H4K20 me2
H2A.Z:H3−5ac
H3K4 me1:H4K20 me2
H3K27 me3:H3K4 me1
H4−4ac:H3−5ac
H3K4 me3:H3−5ac
H4K20 me2:H4−4ac
H4K20 me2:H3−5ac
H3K4 me1:H3−5ac
H2A.Z:H4K20 me2
H3K4 me3:H4−4ac
H2A.Z:H3K4 me3
H4K20 me2:H3K4 me3
H3K9 me2
H3K27 me2
H4K20 me3
H3K9 me3
H3K27 me3
DNA Meth. m5C
H4−4ac
H2A.Z
H3−5ac
H3K4 me1
H3K4 me3
H4K20 me2
0.00 0.25 0.50 0.75 1.00
$\hat{\pi}$
RBM15B – Forward
H3K27 me3:H4K20 me3
H3K27 me3:H3K9 me3
H3K4 me1:H4−4ac
H3K9 me3:H4K20 me3
H3K27 me3:H3K4 me1
H3K27 me2:DNA Meth. m5C
H2A.Z:H3−5ac
H2A.Z:H4−4ac
DNA Meth. m5C:H4K20 me3
H3K27 me3:DNA Meth. m5C
H4K20 me2:H4−4ac
H3K4 me1:H2A.Z
H3K4 me1:H4K20 me2
DNA Meth. m5C:H3K9 me3
H2A.Z:H3K4 me3
H2A.Z:H4K20 me2
H3K4 me1:H3−5ac
H4K20 me2:H3−5ac
H3K4 me3:H3−5ac
H4K20 me2:H3K4 me3
H4−4ac:H3−5ac
H3K4 me3:H4−4ac
H3K9 me2
H4K20 me3
H3K27 me2
H3K27 me3
H3K4 me1
H3K9 me3
DNA Meth. m5C
H3−5ac
H2A.Z
H4K20 me2
H4−4ac
H3K4 me3
0.00 0.25 0.50 0.75 1.00
$\hat{\pi}$
RBM15B – Reverse
H3K27 me3:H4K20 me3
H3K27 me3:H3K9 me3
H3K4 me1:H4−4ac
H3K9 me3:H4K20 me3
H3K27 me3:H3K4 me1
H3K27 me2:DNA Meth. m5C
H2A.Z:H3−5ac
H2A.Z:H4−4ac
DNA Meth. m5C:H4K20 me3
H3K27 me3:DNA Meth. m5C
H4K20 me2:H4−4ac
H3K4 me1:H2A.Z
H3K4 me1:H4K20 me2
DNA Meth. m5C:H3K9 me3
H2A.Z:H3K4 me3
H2A.Z:H4K20 me2
H3K4 me1:H3−5ac
H4K20 me2:H3−5ac
H3K4 me3:H3−5ac
H4K20 me2:H3K4 me3
H4−4ac:H3−5ac
H3K4 me3:H4−4ac
H3K9 me2
H4K20 me3
H3K27 me2
H3K27 me3
H3K4 me1
H3K9 me3
DNA Meth. m5C
H3−5ac
H2A.Z
H4K20 me2
H4−4ac
H3K4 me3
0.00 0.25 0.50 0.75 1.00
$\hat{\pi}$
CDK13 – Forward
H4K20 me2:H3−5ac
H2A.Z:H4−4ac
H2A.Z:H3K4 me3
H2A.Z:H4K20 me2
H4−4ac:H3−5ac
H3K4 me3:H3−5ac
H3K4 me1:H3−5ac
H3K27 me3:DNA Meth. m5C
H3K27 me3:H3K4 me1
DNA Meth. m5C:H4K20 me2
H4K20 me2:H4−4ac
H4K20 me2:H3K4 me3
H3K9 me3
H4K20 me3
H2A.Z
H3K4 me1
DNA Meth. m5C
H3K27 me3
H4−4ac
H3−5ac
H4K20 me2
H3K4 me3
0.00 0.25 0.50 0.75 1.00
$\hat{\pi}$
CDK13 – Reverse
H4K20 me2:H3−5ac
H2A.Z:H4−4ac
H2A.Z:H3K4 me3
H2A.Z:H4K20 me2
H4−4ac:H3−5ac
H3K4 me3:H3−5ac
H3K4 me1:H3−5ac
H3K27 me3:DNA Meth. m5C
H3K27 me3:H3K4 me1
DNA Meth. m5C:H4K20 me2
H4K20 me2:H4−4ac
H4K20 me2:H3K4 me3
H3K9 me3
H4K20 me3
H2A.Z
H3K4 me1
DNA Meth. m5C
H3K27 me3
H4−4ac
H3−5ac
H4K20 me2
H3K4 me3
0.00 0.25 0.50 0.75 1.00
$\hat{\pi}$
SRSF11 – Forward
H3K27 me3:H3K4 me1
H3K4 me1:H4−4ac
H3K4 me1:H2A.Z
H3K4 me1:H4K20 me2
H2A.Z:H3−5ac
H2A.Z:H4−4ac
H2A.Z:H3K4 me3
H3K9 me3:H4K20 me3
H2A.Z:H4K20 me2
H3K27 me3:H4K20 me3
H4K20 me2:H3−5ac
H3K27 me3:H3K9 me3
H4K20 me2:H4−4ac
H4K20 me2:H3K4 me3
DNA Meth. m5C:H4K20 me2
DNA Meth. m5C:H4K20 me3
DNA Meth. m5C:H3K9 me3
H3K27 me3:DNA Meth. m5C
H3K4 me1
H3−5ac
H4−4ac
H3K4 me3
H4K20 me3
H3K9 me3
H2A.Z
H4K20 me2
H3K27 me3
DNA Meth. m5C
0.00 0.25 0.50 0.75 1.00
$\hat{\pi}$
SRSF11 – Reverse
H3K27 me3:H3K4 me1
H3K4 me1:H4−4ac
H3K4 me1:H2A.Z
H3K4 me1:H4K20 me2
H2A.Z:H3−5ac
H2A.Z:H4−4ac
H2A.Z:H3K4 me3
H3K9 me3:H4K20 me3
H2A.Z:H4K20 me2
H3K27 me3:H4K20 me3
H4K20 me2:H3−5ac
H3K27 me3:H3K9 me3
H4K20 me2:H4−4ac
H4K20 me2:H3K4 me3
DNA Meth. m5C:H4K20 me2
DNA Meth. m5C:H4K20 me3
DNA Meth. m5C:H3K9 me3
H3K27 me3:DNA Meth. m5C
H3K4 me1
H3−5ac
H4−4ac
H3K4 me3
H4K20 me3
H3K9 me3
H2A.Z
H4K20 me2
H3K27 me3
DNA Meth. m5C
0.00 0.25 0.50 0.75 1.00
$\hat{\pi}$

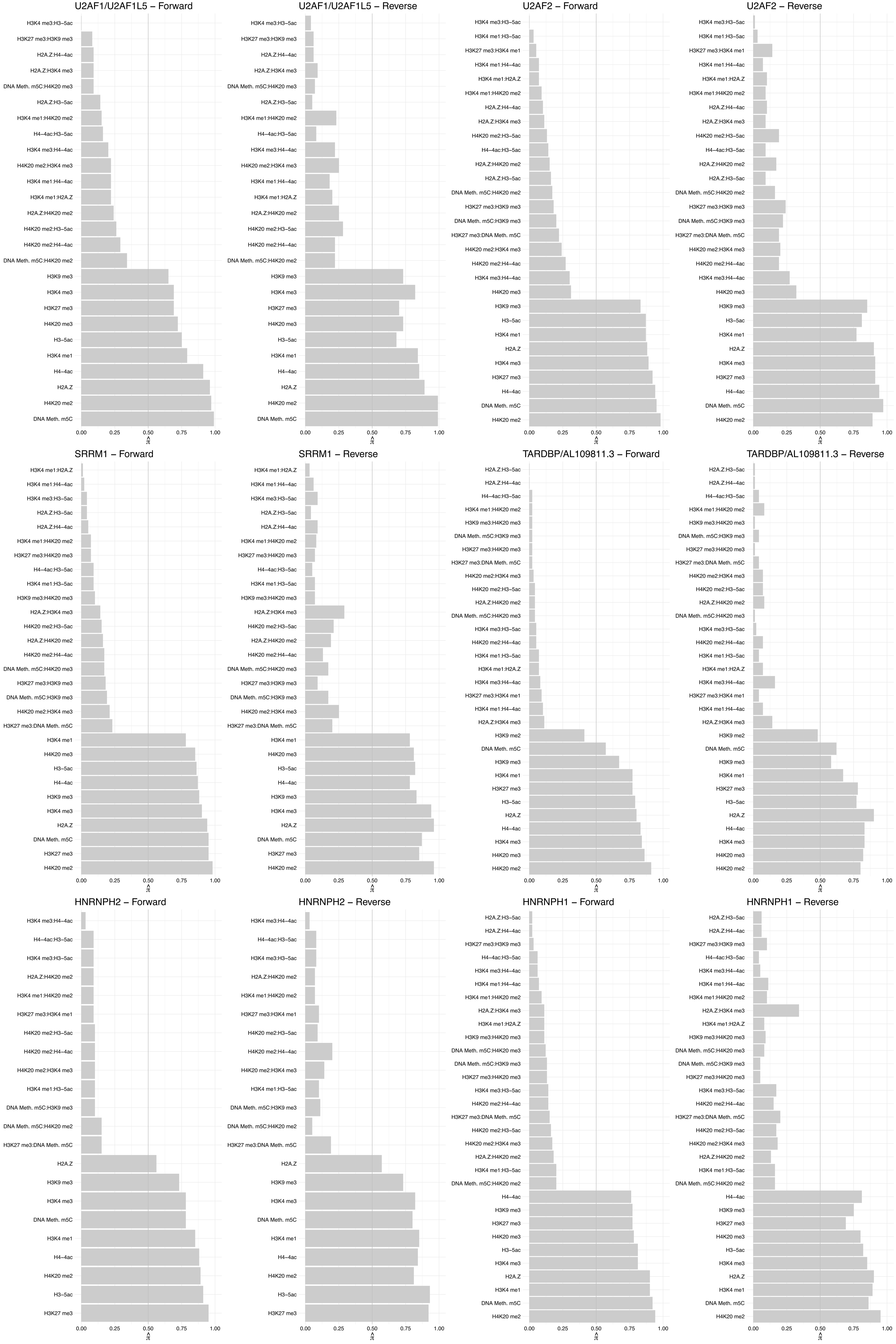

U2AF1/U2AF1L5 – Forward
H3K4 me3:H3−5ac
H3K27 me3:H3K9 me3
H2A.Z:H4−4ac
H2A.Z:H3K4 me3
DNA Meth. m5C:H4K20 me3
H2A.Z:H3−5ac
H3K4 me1:H4K20 me2
H4−4ac:H3−5ac
H3K4 me3:H4−4ac
H4K20 me2:H3K4 me3
H3K4 me1:H4−4ac
H3K4 me1:H2A.Z
H2A.Z:H4K20 me2
H4K20 me2:H3−5ac
H4K20 me2:H4−4ac
DNA Meth. m5C:H4K20 me2
H3K9 me3
H3K4 me3
H3K27 me3
H4K20 me3
H3−5ac
H3K4 me1
H4−4ac
H2A.Z
H4K20 me2
DNA Meth. m5C
0.00 0.25 0.50 0.75 1.00
π̂
U2AF1/U2AF1L5 – Reverse
H3K4 me3:H3−5ac
H3K27 me3:H3K9 me3
H2A.Z:H4−4ac
H2A.Z:H3K4 me3
DNA Meth. m5C:H4K20 me3
H2A.Z:H3−5ac
H3K4 me1:H4K20 me2
H4−4ac:H3−5ac
H3K4 me3:H4−4ac
H4K20 me2:H3K4 me3
H3K4 me1:H4−4ac
H3K4 me1:H2A.Z
H2A.Z:H4K20 me2
H4K20 me2:H3−5ac
H4K20 me2:H4−4ac
DNA Meth. m5C:H4K20 me2
H3K9 me3
H3K4 me3
H3K27 me3
H4K20 me3
H3−5ac
H3K4 me1
H4−4ac
H2A.Z
H4K20 me2
DNA Meth. m5C
0.00 0.25 0.50 0.75 1.00
π̂
U2AF2 – Forward
H3K4 me3:H3−5ac
H3K4 me1:H3−5ac
H3K27 me3:H3K4 me1
H3K4 me1:H4−4ac
H3K4 me1:H2A.Z
H3K4 me1:H4K20 me2
H2A.Z:H4−4ac
H2A.Z:H3K4 me3
H4K20 me2:H3−5ac
H4−4ac:H3−5ac
H2A.Z:H4K20 me2
H2A.Z:H3−5ac
DNA Meth. m5C:H4K20 me2
H3K27 me3:H3K9 me3
DNA Meth. m5C:H3K9 me3
H3K27 me3:DNA Meth. m5C
H4K20 me2:H3K4 me3
H4K20 me2:H4−4ac
H3K4 me3:H4−4ac
H4K20 me3
H3K9 me3
H3−5ac
H3K4 me1
H2A.Z
H3K4 me3
H3K27 me3
H4−4ac
DNA Meth. m5C
H4K20 me2
0.00 0.25 0.50 0.75 1.00
π̂
U2AF2 – Reverse
H3K4 me3:H3−5ac
H3K4 me1:H3−5ac
H3K27 me3:H3K4 me1
H3K4 me1:H4−4ac
H3K4 me1:H2A.Z
H3K4 me1:H4K20 me2
H2A.Z:H4−4ac
H2A.Z:H3K4 me3
H4K20 me2:H3−5ac
H4−4ac:H3−5ac
H2A.Z:H4K20 me2
H2A.Z:H3−5ac
DNA Meth. m5C:H4K20 me2
H3K27 me3:H3K9 me3
DNA Meth. m5C:H3K9 me3
H3K27 me3:DNA Meth. m5C
H4K20 me2:H3K4 me3
H4K20 me2:H4−4ac
H3K4 me3:H4−4ac
H4K20 me3
H3K9 me3
H3−5ac
H3K4 me1
H2A.Z
H3K4 me3
H3K27 me3
H4−4ac
DNA Meth. m5C
H4K20 me2
0.00 0.25 0.50 0.75 1.00
π̂
SRRM1 – Forward
H3K4 me1:H2A.Z
H3K4 me1:H4−4ac
H3K4 me3:H3−5ac
H2A.Z:H3−5ac
H2A.Z:H4−4ac
H3K4 me1:H4K20 me2
H3K27 me3:H4K20 me3
H4−4ac:H3−5ac
H3K4 me1:H3−5ac
H3K9 me3:H4K20 me3
H2A.Z:H3K4 me3
H4K20 me2:H3−5ac
H2A.Z:H4K20 me2
H4K20 me2:H4−4ac
DNA Meth. m5C:H4K20 me3
H3K27 me3:H3K9 me3
DNA Meth. m5C:H3K9 me3
H4K20 me2:H3K4 me3
H3K27 me3:DNA Meth. m5C
H3K4 me1
H4K20 me3
H3−5ac
H4−4ac
H3K9 me3
H3K4 me3
H2A.Z
DNA Meth. m5C
H3K27 me3
H4K20 me2
0.00 0.25 0.50 0.75 1.00
π̂
SRRM1 – Reverse
H3K4 me1:H2A.Z
H3K4 me1:H4−4ac
H3K4 me3:H3−5ac
H2A.Z:H3−5ac
H2A.Z:H4−4ac
H3K4 me1:H4K20 me2
H3K27 me3:H4K20 me3
H4−4ac:H3−5ac
H3K4 me1:H3−5ac
H3K9 me3:H4K20 me3
H2A.Z:H3K4 me3
H4K20 me2:H3−5ac
H2A.Z:H4K20 me2
H4K20 me2:H4−4ac
DNA Meth. m5C:H4K20 me3
H3K27 me3:H3K9 me3
DNA Meth. m5C:H3K9 me3
H4K20 me2:H3K4 me3
H3K27 me3:DNA Meth. m5C
H3K4 me1
H4K20 me3
H3−5ac
H4−4ac
H3K9 me3
H3K4 me3
H2A.Z
DNA Meth. m5C
H3K27 me3
H4K20 me2
0.00 0.25 0.50 0.75 1.00
π̂
TARDBP/AL109811.3 – Forward
H2A.Z:H3−5ac
H2A.Z:H4−4ac
H4−4ac:H3−5ac
H3K4 me1:H4K20 me2
H3K9 me3:H4K20 me3
DNA Meth. m5C:H3K9 me3
H3K27 me3:H4K20 me3
H3K27 me3:DNA Meth. m5C
H4K20 me2:H3K4 me3
H4K20 me2:H3−5ac
H2A.Z:H4K20 me2
DNA Meth. m5C:H4K20 me3
H3K4 me3:H3−5ac
H4K20 me2:H4−4ac
H3K4 me1:H3−5ac
H3K4 me1:H2A.Z
H3K4 me3:H4−4ac
H3K27 me3:H3K4 me1
H3K4 me1:H4−4ac
H2A.Z:H3K4 me3
H3K9 me2
DNA Meth. m5C
H3K9 me3
H3K4 me1
H3K27 me3
H3−5ac
H2A.Z
H4−4ac
H3K4 me3
H4K20 me3
H4K20 me2
0.00 0.25 0.50 0.75 1.00
π̂
TARDBP/AL109811.3 – Reverse
H2A.Z:H3−5ac
H2A.Z:H4−4ac
H4−4ac:H3−5ac
H3K4 me1:H4K20 me2
H3K9 me3:H4K20 me3
DNA Meth. m5C:H3K9 me3
H3K27 me3:H4K20 me3
H3K27 me3:DNA Meth. m5C
H4K20 me2:H3K4 me3
H4K20 me2:H3−5ac
H2A.Z:H4K20 me2
DNA Meth. m5C:H4K20 me3
H3K4 me3:H3−5ac
H4K20 me2:H4−4ac
H3K4 me1:H3−5ac
H3K4 me1:H2A.Z
H3K4 me3:H4−4ac
H3K27 me3:H3K4 me1
H3K4 me1:H4−4ac
H2A.Z:H3K4 me3
H3K9 me2
DNA Meth. m5C
H3K9 me3
H3K4 me1
H3K27 me3
H3−5ac
H2A.Z
H4−4ac
H3K4 me3
H4K20 me3
H4K20 me2
0.00 0.25 0.50 0.75 1.00
π̂
HNRNPH2 – Forward
H3K4 me3:H4−4ac
H4−4ac:H3−5ac
H3K4 me3:H3−5ac
H2A.Z:H4K20 me2
H3K4 me1:H4K20 me2
H3K27 me3:H3K4 me1
H4K20 me2:H3−5ac
H4K20 me2:H4−4ac
H4K20 me2:H3K4 me3
H3K4 me1:H3−5ac
DNA Meth. m5C:H3K9 me3
DNA Meth. m5C:H4K20 me2
H3K27 me3:DNA Meth. m5C
H2A.Z
H3K9 me3
H3K4 me3
DNA Meth. m5C
H3K4 me1
H4−4ac
H4K20 me2
H3−5ac
H3K27 me3
0.00 0.25 0.50 0.75 1.00
π̂
HNRNPH2 – Reverse
H3K4 me3:H4−4ac
H4−4ac:H3−5ac
H3K4 me3:H3−5ac
H2A.Z:H4K20 me2
H3K4 me1:H4K20 me2
H3K27 me3:H3K4 me1
H4K20 me2:H3−5ac
H4K20 me2:H4−4ac
H4K20 me2:H3K4 me3
H3K4 me1:H3−5ac
DNA Meth. m5C:H3K9 me3
DNA Meth. m5C:H4K20 me2
H3K27 me3:DNA Meth. m5C
H2A.Z
H3K9 me3
H3K4 me3
DNA Meth. m5C
H3K4 me1
H4−4ac
H4K20 me2
H3−5ac
H3K27 me3
0.00 0.25 0.50 0.75 1.00
π̂
HNRNPH1 – Forward
H2A.Z:H3−5ac
H2A.Z:H4−4ac
H3K27 me3:H3K9 me3
H4−4ac:H3−5ac
H3K4 me3:H4−4ac
H3K4 me1:H4−4ac
H3K4 me1:H4K20 me2
H2A.Z:H3K4 me3
H3K4 me1:H2A.Z
H3K9 me3:H4K20 me3
DNA Meth. m5C:H4K20 me3
DNA Meth. m5C:H3K9 me3
H3K27 me3:H4K20 me3
H3K4 me3:H3−5ac
H4K20 me2:H4−4ac
H3K27 me3:DNA Meth. m5C
H4K20 me2:H3−5ac
H4K20 me2:H3K4 me3
H2A.Z:H4K20 me2
H3K4 me1:H3−5ac
DNA Meth. m5C:H4K20 me2
H4−4ac
H3K9 me3
H3K27 me3
H4K20 me3
H3−5ac
H3K4 me3
H2A.Z
H3K4 me1
DNA Meth. m5C
H4K20 me2
0.00 0.25 0.50 0.75 1.00
π̂
HNRNPH1 – Reverse
H2A.Z:H3−5ac
H2A.Z:H4−4ac
H3K27 me3:H3K9 me3
H4−4ac:H3−5ac
H3K4 me3:H4−4ac
H3K4 me1:H4−4ac
H3K4 me1:H4K20 me2
H2A.Z:H3K4 me3
H3K4 me1:H2A.Z
H3K9 me3:H4K20 me3
DNA Meth. m5C:H4K20 me3
DNA Meth. m5C:H3K9 me3
H3K27 me3:H4K20 me3
H3K4 me3:H3−5ac
H4K20 me2:H4−4ac
H3K27 me3:DNA Meth. m5C
H4K20 me2:H3−5ac
H4K20 me2:H3K4 me3
H2A.Z:H4K20 me2
H3K4 me1:H3−5ac
DNA Meth. m5C:H4K20 me2
H4−4ac
H3K9 me3
H3K27 me3
H4K20 me3
H3−5ac
H3K4 me3
H2A.Z
H3K4 me1
DNA Meth. m5C
H4K20 me2
0.00 0.25 0.50 0.75 1.00
π̂

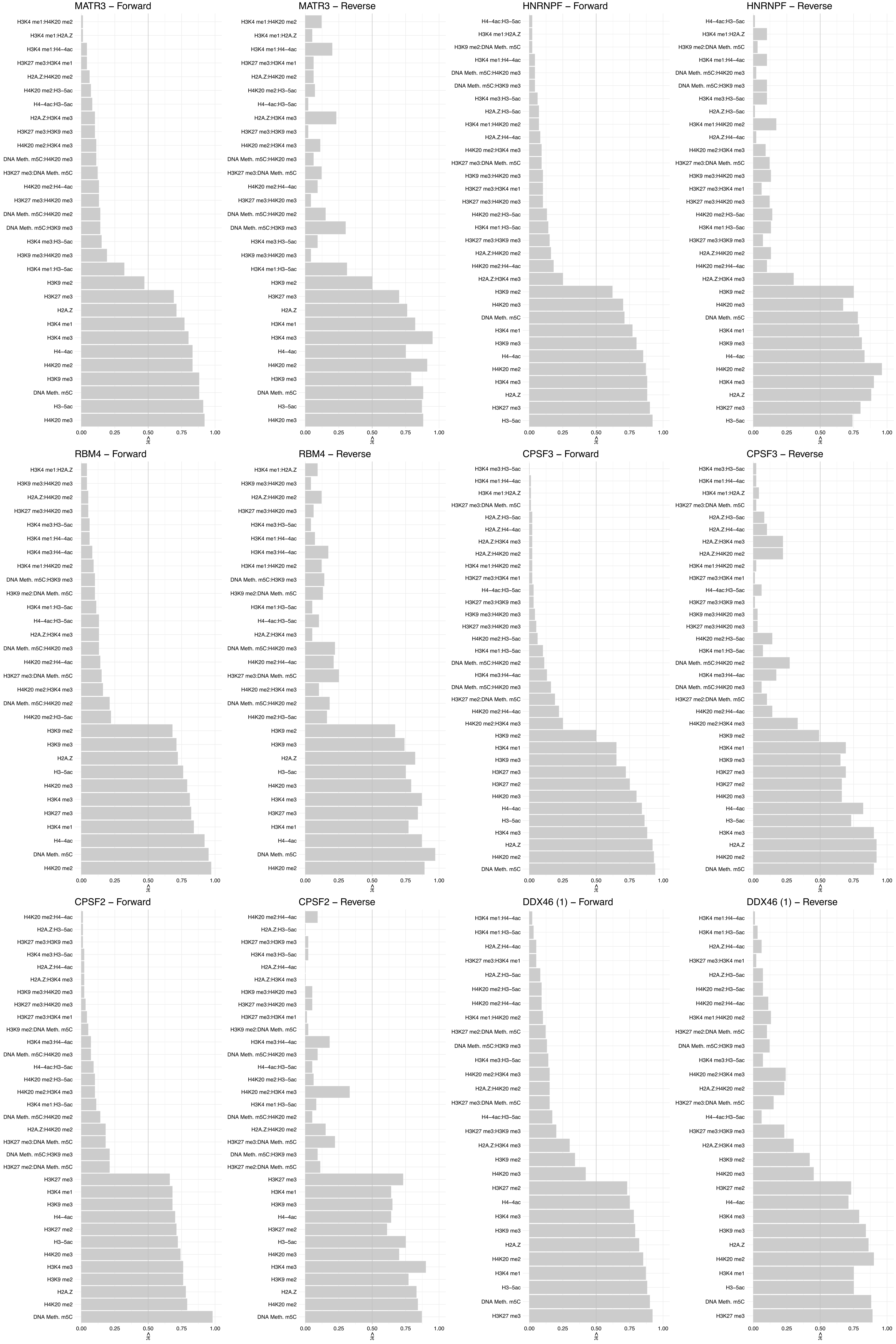
MATR3 – Forward
H3K4 me1:H4K20 me2
H3K4 me1:H2A.Z
H3K4 me1:H4–4ac
H3K27 me3:H3K4 me1
H2A.Z:H4K20 me2
H4K20 me2:H3–5ac
H4–4ac:H3–5ac
H2A.Z:H3K4 me3
H3K27 me3:H3K9 me3
H4K20 me2:H3K4 me3
DNA Meth. m5C:H4K20 me3
H3K27 me3:DNA Meth. m5C
H4K20 me2:H4–4ac
H3K27 me3:H4K20 me3
DNA Meth. m5C:H4K20 me2
DNA Meth. m5C:H3K9 me3
H3K4 me3:H3–5ac
H3K9 me3:H4K20 me3
H3K4 me1:H3–5ac
H3K9 me2
H3K27 me3
H2A.Z
H3K4 me1
H3K4 me3
H4–4ac
H4K20 me2
H3K9 me3
DNA Meth. m5C
H3–5ac
H4K20 me3
0.00 0.25 0.50 0.75 1.00
π̂
MATR3 – Reverse
H3K4 me1:H4K20 me2
H3K4 me1:H2A.Z
H3K4 me1:H4–4ac
H3K27 me3:H3K4 me1
H2A.Z:H4K20 me2
H4K20 me2:H3–5ac
H4–4ac:H3–5ac
H2A.Z:H3K4 me3
H3K27 me3:H3K9 me3
H4K20 me2:H3K4 me3
DNA Meth. m5C:H4K20 me3
H3K27 me3:DNA Meth. m5C
H4K20 me2:H4–4ac
H3K27 me3:H4K20 me3
DNA Meth. m5C:H4K20 me2
DNA Meth. m5C:H3K9 me3
H3K4 me3:H3–5ac
H3K9 me3:H4K20 me3
H3K4 me1:H3–5ac
H3K9 me2
H3K27 me3
H2A.Z
H3K4 me1
H3K4 me3
H4–4ac
H4K20 me2
H3K9 me3
DNA Meth. m5C
H3–5ac
H4K20 me3
0.00 0.25 0.50 0.75 1.00
π̂
HNRNPF – Forward
H4–4ac:H3–5ac
H3K4 me1:H2A.Z
H3K9 me2:DNA Meth. m5C
H3K4 me1:H4–4ac
DNA Meth. m5C:H4K20 me3
DNA Meth. m5C:H3K9 me3
H3K4 me3:H3–5ac
H2A.Z:H3–5ac
H3K4 me1:H4K20 me2
H2A.Z:H4–4ac
H4K20 me2:H3K4 me3
H3K27 me3:DNA Meth. m5C
H3K9 me3:H4K20 me3
H3K27 me3:H3K4 me1
H3K27 me3:H4K20 me3
H4K20 me2:H3–5ac
H3K4 me1:H3–5ac
H3K27 me3:H3K9 me3
H2A.Z:H4K20 me2
H4K20 me2:H4–4ac
H2A.Z:H3K4 me3
H3K9 me2
H4K20 me3
DNA Meth. m5C
H3K4 me1
H3K9 me3
H4–4ac
H4K20 me2
H3K4 me3
H2A.Z
H3K27 me3
H3–5ac
0.00 0.25 0.50 0.75 1.00
π̂
HNRNPF – Reverse
H4–4ac:H3–5ac
H3K4 me1:H2A.Z
H3K9 me2:DNA Meth. m5C
H3K4 me1:H4–4ac
DNA Meth. m5C:H4K20 me3
DNA Meth. m5C:H3K9 me3
H3K4 me3:H3–5ac
H2A.Z:H3–5ac
H3K4 me1:H4K20 me2
H2A.Z:H4–4ac
H4K20 me2:H3K4 me3
H3K27 me3:DNA Meth. m5C
H3K9 me3:H4K20 me3
H3K27 me3:H3K4 me1
H3K27 me3:H4K20 me3
H4K20 me2:H3–5ac
H3K4 me1:H3–5ac
H3K27 me3:H3K9 me3
H2A.Z:H4K20 me2
H4K20 me2:H4–4ac
H2A.Z:H3K4 me3
H3K9 me2
H4K20 me3
DNA Meth. m5C
H3K4 me1
H3K9 me3
H4–4ac
H4K20 me2
H3K4 me3
H2A.Z
H3K27 me3
H3–5ac
0.00 0.25 0.50 0.75 1.00
π̂
RBM4 – Forward
H3K4 me1:H2A.Z
H3K9 me3:H4K20 me3
H2A.Z:H4K20 me2
H3K27 me3:H4K20 me3
H3K4 me3:H3–5ac
H3K4 me1:H4–4ac
H3K4 me3:H4–4ac
H3K4 me1:H4K20 me2
DNA Meth. m5C:H3K9 me3
H3K9 me2:DNA Meth. m5C
H3K4 me1:H3–5ac
H4–4ac:H3–5ac
H2A.Z:H3K4 me3
DNA Meth. m5C:H4K20 me3
H4K20 me2:H4–4ac
H3K27 me3:DNA Meth. m5C
H4K20 me2:H3K4 me3
DNA Meth. m5C:H4K20 me2
H4K20 me2:H3–5ac
H3K9 me2
H3K9 me3
H2A.Z
H3–5ac
H4K20 me3
H3K4 me3
H3K27 me3
H3K4 me1
H4–4ac
DNA Meth. m5C
H4K20 me2
0.00 0.25 0.50 0.75 1.00
π̂
RBM4 – Reverse
H3K4 me1:H2A.Z
H3K9 me3:H4K20 me3
H2A.Z:H4K20 me2
H3K27 me3:H4K20 me3
H3K4 me3:H3–5ac
H3K4 me1:H4–4ac
H3K4 me3:H4–4ac
H3K4 me1:H4K20 me2
DNA Meth. m5C:H3K9 me3
H3K9 me2:DNA Meth. m5C
H3K4 me1:H3–5ac
H4–4ac:H3–5ac
H2A.Z:H3K4 me3
DNA Meth. m5C:H4K20 me3
H4K20 me2:H4–4ac
H3K27 me3:DNA Meth. m5C
H4K20 me2:H3K4 me3
DNA Meth. m5C:H4K20 me2
H4K20 me2:H3–5ac
H3K9 me2
H3K9 me3
H2A.Z
H3–5ac
H4K20 me3
H3K4 me3
H3K27 me3
H3K4 me1
H4–4ac
DNA Meth. m5C
H4K20 me2
0.00 0.25 0.50 0.75 1.00
π̂
CPSF3 – Forward
H3K4 me3:H3–5ac
H3K4 me1:H4–4ac
H3K4 me1:H2A.Z
H3K27 me3:DNA Meth. m5C
H2A.Z:H3–5ac
H2A.Z:H4–4ac
H2A.Z:H3K4 me3
H2A.Z:H4K20 me2
H3K4 me1:H4K20 me2
H3K27 me3:H3K4 me1
H4–4ac:H3–5ac
H3K27 me3:H3K9 me3
H3K9 me3:H4K20 me3
H3K27 me3:H4K20 me3
H4K20 me2:H3–5ac
H3K4 me1:H3–5ac
DNA Meth. m5C:H4K20 me2
H3K4 me3:H4–4ac
DNA Meth. m5C:H4K20 me3
H3K27 me2:DNA Meth. m5C
H4K20 me2:H4–4ac
H4K20 me2:H3K4 me3
H3K9 me2
H3K4 me1
H3K9 me3
H3K27 me3
H3K27 me2
H4K20 me3
H4–4ac
H3–5ac
H3K4 me3
H2A.Z
H4K20 me2
DNA Meth. m5C
0.00 0.25 0.50 0.75 1.00
π̂
CPSF3 – Reverse
H3K4 me3:H3–5ac
H3K4 me1:H4–4ac
H3K4 me1:H2A.Z
H3K27 me3:DNA Meth. m5C
H2A.Z:H3–5ac
H2A.Z:H4–4ac
H2A.Z:H3K4 me3
H2A.Z:H4K20 me2
H3K4 me1:H4K20 me2
H3K27 me3:H3K4 me1
H4–4ac:H3–5ac
H3K27 me3:H3K9 me3
H3K9 me3:H4K20 me3
H3K27 me3:H4K20 me3
H4K20 me2:H3–5ac
H3K4 me1:H3–5ac
DNA Meth. m5C:H4K20 me2
H3K4 me3:H4–4ac
DNA Meth. m5C:H4K20 me3
H3K27 me2:DNA Meth. m5C
H4K20 me2:H4–4ac
H4K20 me2:H3K4 me3
H3K9 me2
H3K4 me1
H3K9 me3
H3K27 me3
H3K27 me2
H4K20 me3
H4–4ac
H3–5ac
H3K4 me3
H2A.Z
H4K20 me2
DNA Meth. m5C
0.00 0.25 0.50 0.75 1.00
π̂
CPSF2 – Forward
H4K20 me2:H4–4ac
H2A.Z:H3–5ac
H3K27 me3:H3K9 me3
H3K4 me3:H3–5ac
H2A.Z:H4–4ac
H2A.Z:H3K4 me3
H3K9 me3:H4K20 me3
H3K27 me3:H4K20 me3
H3K27 me3:H3K4 me1
H3K9 me2:DNA Meth. m5C
H3K4 me3:H4–4ac
DNA Meth. m5C:H4K20 me3
H4–4ac:H3–5ac
H4K20 me2:H3–5ac
H4K20 me2:H3K4 me3
H3K4 me1:H3–5ac
DNA Meth. m5C:H4K20 me2
H2A.Z:H4K20 me2
H3K27 me3:DNA Meth. m5C
DNA Meth. m5C:H3K9 me3
H3K27 me2:DNA Meth. m5C
H3K27 me3
H3K4 me1
H3K9 me3
H4–4ac
H3K27 me2
H3–5ac
H4K20 me3
H3K4 me3
H3K9 me2
H2A.Z
H4K20 me2
DNA Meth. m5C
0.00 0.25 0.50 0.75 1.00
π̂
CPSF2 – Reverse
H4K20 me2:H4–4ac
H2A.Z:H3–5ac
H3K27 me3:H3K9 me3
H3K4 me3:H3–5ac
H2A.Z:H4–4ac
H2A.Z:H3K4 me3
H3K9 me3:H4K20 me3
H3K27 me3:H4K20 me3
H3K27 me3:H3K4 me1
H3K9 me2:DNA Meth. m5C
H3K4 me3:H4–4ac
DNA Meth. m5C:H4K20 me3
H4–4ac:H3–5ac
H4K20 me2:H3–5ac
H4K20 me2:H3K4 me3
H3K4 me1:H3–5ac
DNA Meth. m5C:H4K20 me2
H2A.Z:H4K20 me2
H3K27 me3:DNA Meth. m5C
DNA Meth. m5C:H3K9 me3
H3K27 me2:DNA Meth. m5C
H3K27 me3
H3K4 me1
H3K9 me3
H4–4ac
H3K27 me2
H3–5ac
H4K20 me3
H3K4 me3
H3K9 me2
H2A.Z
H4K20 me2
DNA Meth. m5C
0.00 0.25 0.50 0.75 1.00
π̂
DDX46 (1) – Forward
H3K4 me1:H4–4ac
H3K4 me1:H3–5ac
H2A.Z:H4–4ac
H3K27 me3:H3K4 me1
H2A.Z:H3–5ac
H4K20 me2:H3–5ac
H4K20 me2:H4–4ac
H3K4 me1:H4K20 me2
H3K27 me2:DNA Meth. m5C
DNA Meth. m5C:H3K9 me3
H3K4 me3:H3–5ac
H4K20 me2:H3K4 me3
H2A.Z:H4K20 me2
H3K27 me3:DNA Meth. m5C
H4–4ac:H3–5ac
H3K27 me3:H3K9 me3
H2A.Z:H3K4 me3
H3K9 me2
H4K20 me3
H3K27 me2
H4–4ac
H3K4 me3
H3K9 me3
H2A.Z
H4K20 me2
H3K4 me1
H3–5ac
DNA Meth. m5C
H3K27 me3
0.00 0.25 0.50 0.75 1.00
π̂
DDX46 (1) – Reverse
H3K4 me1:H4–4ac
H3K4 me1:H3–5ac
H2A.Z:H4–4ac
H3K27 me3:H3K4 me1
H2A.Z:H3–5ac
H4K20 me2:H3–5ac
H4K20 me2:H4–4ac
H3K4 me1:H4K20 me2
H3K27 me2:DNA Meth. m5C
DNA Meth. m5C:H3K9 me3
H3K4 me3:H3–5ac
H4K20 me2:H3K4 me3
H2A.Z:H4K20 me2
H3K27 me3:DNA Meth. m5C
H4–4ac:H3–5ac
H3K27 me3:H3K9 me3
H2A.Z:H3K4 me3
H3K9 me2
H4K20 me3
H3K27 me2
H4–4ac
H3K4 me3
H3K9 me3
H2A.Z
H4K20 me2
H3K4 me1
H3–5ac
DNA Meth. m5C
H3K27 me3
0.00 0.25 0.50 0.75 1.00
π̂

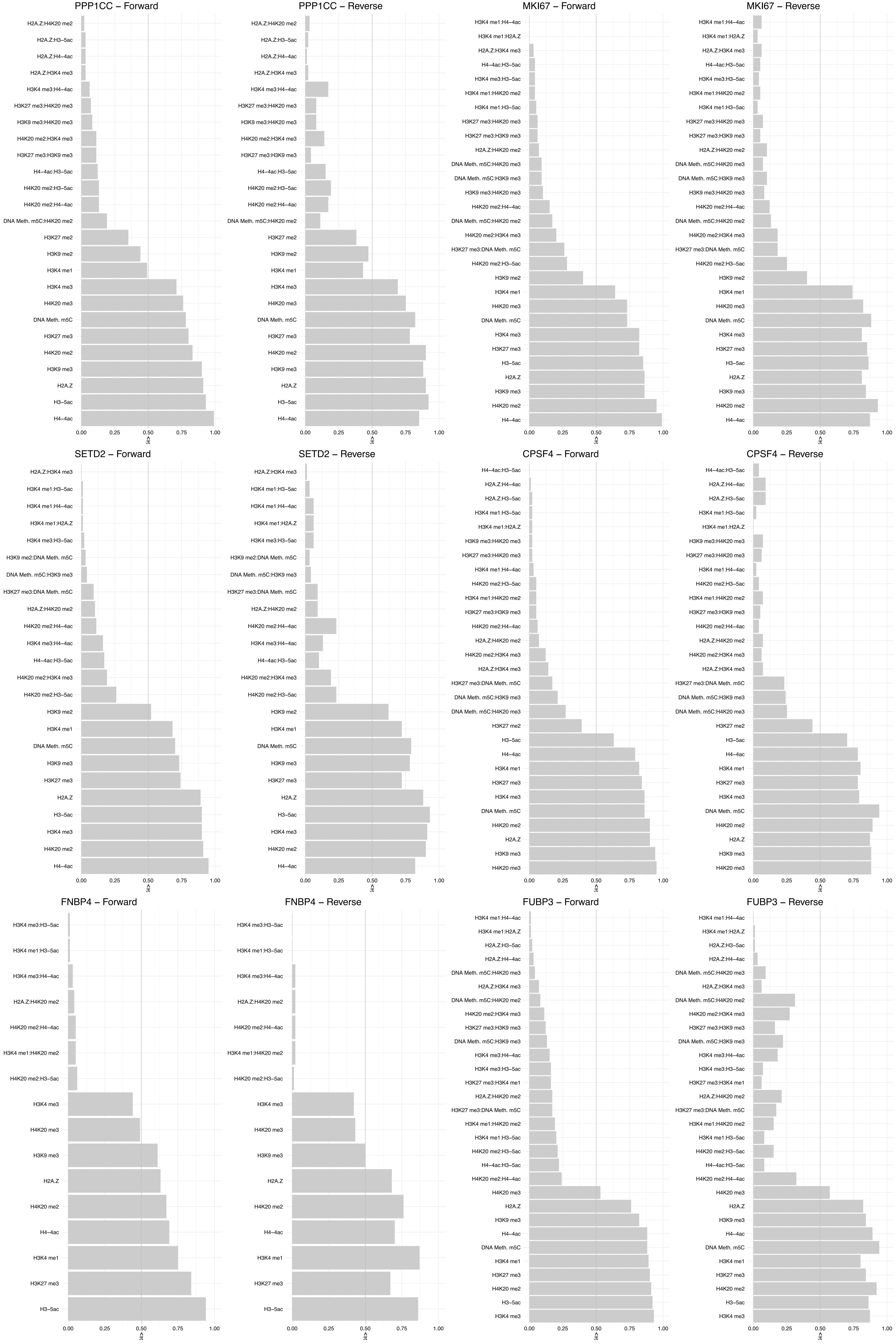

PPP1CC – Forward
H2A.Z:H4K20 me2
H2A.Z:H3–5ac
H2A.Z:H4–4ac
H2A.Z:H3K4 me3
H3K4 me3:H4–4ac
H3K27 me3:H4K20 me3
H3K9 me3:H4K20 me3
H4K20 me2:H3K4 me3
H3K27 me3:H3K9 me3
H4–4ac:H3–5ac
H4K20 me2:H3–5ac
H4K20 me2:H4–4ac
DNA Meth. m5C:H4K20 me2
H3K27 me2
H3K9 me2
H3K4 me1
H3K4 me3
H4K20 me3
DNA Meth. m5C
H3K27 me3
H4K20 me2
H3K9 me3
H2A.Z
H3–5ac
H4–4ac
0.00 0.25 0.50 0.75 1.00
π̂
PPP1CC – Reverse
H2A.Z:H4K20 me2
H2A.Z:H3–5ac
H2A.Z:H4–4ac
H2A.Z:H3K4 me3
H3K4 me3:H4–4ac
H3K27 me3:H4K20 me3
H3K9 me3:H4K20 me3
H4K20 me2:H3K4 me3
H3K27 me3:H3K9 me3
H4–4ac:H3–5ac
H4K20 me2:H3–5ac
H4K20 me2:H4–4ac
DNA Meth. m5C:H4K20 me2
H3K27 me2
H3K9 me2
H3K4 me1
H3K4 me3
H4K20 me3
DNA Meth. m5C
H3K27 me3
H4K20 me2
H3K9 me3
H2A.Z
H3–5ac
H4–4ac
0.00 0.25 0.50 0.75 1.00
π̂
MKI67 – Forward
H3K4 me1:H4–4ac
H3K4 me1:H2A.Z
H2A.Z:H3K4 me3
H4–4ac:H3–5ac
H3K4 me3:H3–5ac
H3K4 me1:H4K20 me2
H3K4 me1:H3–5ac
H3K27 me3:H4K20 me3
H3K27 me3:H3K9 me3
H2A.Z:H4K20 me2
DNA Meth. m5C:H4K20 me3
DNA Meth. m5C:H3K9 me3
H3K9 me3:H4K20 me3
H4K20 me2:H4–4ac
DNA Meth. m5C:H4K20 me2
H4K20 me2:H3K4 me3
H3K27 me3:DNA Meth. m5C
H4K20 me2:H3–5ac
H3K9 me2
H3K4 me1
H4K20 me3
DNA Meth. m5C
H3K4 me3
H3K27 me3
H3–5ac
H2A.Z
H3K9 me3
H4K20 me2
H4–4ac
0.00 0.25 0.50 0.75 1.00
π̂
MKI67 – Reverse
H3K4 me1:H4–4ac
H3K4 me1:H2A.Z
H2A.Z:H3K4 me3
H4–4ac:H3–5ac
H3K4 me3:H3–5ac
H3K4 me1:H4K20 me2
H3K4 me1:H3–5ac
H3K27 me3:H4K20 me3
H3K27 me3:H3K9 me3
H2A.Z:H4K20 me2
DNA Meth. m5C:H4K20 me3
DNA Meth. m5C:H3K9 me3
H3K9 me3:H4K20 me3
H4K20 me2:H4–4ac
DNA Meth. m5C:H4K20 me2
H4K20 me2:H3K4 me3
H3K27 me3:DNA Meth. m5C
H4K20 me2:H3–5ac
H3K9 me2
H3K4 me1
H4K20 me3
DNA Meth. m5C
H3K4 me3
H3K27 me3
H3–5ac
H2A.Z
H3K9 me3
H4K20 me2
H4–4ac
0.00 0.25 0.50 0.75 1.00
π̂
SETD2 – Forward
H2A.Z:H3K4 me3
H3K4 me1:H3–5ac
H3K4 me1:H4–4ac
H3K4 me1:H2A.Z
H3K4 me3:H3–5ac
H3K9 me2:DNA Meth. m5C
DNA Meth. m5C:H3K9 me3
H3K27 me3:DNA Meth. m5C
H2A.Z:H4K20 me2
H4K20 me2:H4–4ac
H3K4 me3:H4–4ac
H4–4ac:H3–5ac
H4K20 me2:H3K4 me3
H4K20 me2:H3–5ac
H3K9 me2
H3K4 me1
DNA Meth. m5C
H3K9 me3
H3K27 me3
H2A.Z
H3–5ac
H3K4 me3
H4K20 me2
H4–4ac
0.00 0.25 0.50 0.75 1.00
π̂
SETD2 – Reverse
H2A.Z:H3K4 me3
H3K4 me1:H3–5ac
H3K4 me1:H4–4ac
H3K4 me1:H2A.Z
H3K4 me3:H3–5ac
H3K9 me2:DNA Meth. m5C
DNA Meth. m5C:H3K9 me3
H3K27 me3:DNA Meth. m5C
H2A.Z:H4K20 me2
H4K20 me2:H4–4ac
H3K4 me3:H4–4ac
H4–4ac:H3–5ac
H4K20 me2:H3K4 me3
H4K20 me2:H3–5ac
H3K9 me2
H3K4 me1
DNA Meth. m5C
H3K9 me3
H3K27 me3
H2A.Z
H3–5ac
H3K4 me3
H4K20 me2
H4–4ac
0.00 0.25 0.50 0.75 1.00
π̂
CPSF4 – Forward
H4–4ac:H3–5ac
H2A.Z:H4–4ac
H2A.Z:H3–5ac
H3K4 me1:H3–5ac
H3K4 me1:H2A.Z
H3K9 me3:H4K20 me3
H3K27 me3:H4K20 me3
H3K4 me1:H4–4ac
H4K20 me2:H3–5ac
H3K4 me1:H4K20 me2
H3K27 me3:H3K9 me3
H4K20 me2:H4–4ac
H2A.Z:H4K20 me2
H4K20 me2:H3K4 me3
H2A.Z:H3K4 me3
H3K27 me3:DNA Meth. m5C
DNA Meth. m5C:H3K9 me3
DNA Meth. m5C:H4K20 me3
H3K27 me2
H3–5ac
H4–4ac
H3K4 me1
H3K27 me3
H3K4 me3
DNA Meth. m5C
H4K20 me2
H2A.Z
H3K9 me3
H4K20 me3
0.00 0.25 0.50 0.75 1.00
π̂
CPSF4 – Reverse
H4–4ac:H3–5ac
H2A.Z:H4–4ac
H2A.Z:H3–5ac
H3K4 me1:H3–5ac
H3K4 me1:H2A.Z
H3K9 me3:H4K20 me3
H3K27 me3:H4K20 me3
H3K4 me1:H4–4ac
H4K20 me2:H3–5ac
H3K4 me1:H4K20 me2
H3K27 me3:H3K9 me3
H4K20 me2:H4–4ac
H2A.Z:H4K20 me2
H4K20 me2:H3K4 me3
H2A.Z:H3K4 me3
H3K27 me3:DNA Meth. m5C
DNA Meth. m5C:H3K9 me3
DNA Meth. m5C:H4K20 me3
H3K27 me2
H3–5ac
H4–4ac
H3K4 me1
H3K27 me3
H3K4 me3
DNA Meth. m5C
H4K20 me2
H2A.Z
H3K9 me3
H4K20 me3
0.00 0.25 0.50 0.75 1.00
π̂
FNBP4 – Forward
H3K4 me3:H3–5ac
H3K4 me1:H3–5ac
H3K4 me3:H4–4ac
H2A.Z:H4K20 me2
H4K20 me2:H4–4ac
H3K4 me1:H4K20 me2
H4K20 me2:H3–5ac
H3K4 me3
H4K20 me3
H3K9 me3
H2A.Z
H4K20 me2
H4–4ac
H3K4 me1
H3K27 me3
H3–5ac
0.00 0.25 0.50 0.75 1.00
π̂
FNBP4 – Reverse
H3K4 me3:H3–5ac
H3K4 me1:H3–5ac
H3K4 me3:H4–4ac
H2A.Z:H4K20 me2
H4K20 me2:H4–4ac
H3K4 me1:H4K20 me2
H4K20 me2:H3–5ac
H3K4 me3
H4K20 me3
H3K9 me3
H2A.Z
H4K20 me2
H4–4ac
H3K4 me1
H3K27 me3
H3–5ac
0.00 0.25 0.50 0.75 1.00
π̂
FUBP3 – Forward
H3K4 me1:H4–4ac
H3K4 me1:H2A.Z
H2A.Z:H3–5ac
H2A.Z:H4–4ac
DNA Meth. m5C:H4K20 me3
H2A.Z:H3K4 me3
DNA Meth. m5C:H4K20 me2
H4K20 me2:H3K4 me3
H3K27 me3:H3K9 me3
DNA Meth. m5C:H3K9 me3
H3K4 me3:H4–4ac
H3K4 me3:H3–5ac
H3K27 me3:H3K4 me1
H2A.Z:H4K20 me2
H3K27 me3:DNA Meth. m5C
H3K4 me1:H4K20 me2
H3K4 me1:H3–5ac
H4K20 me2:H3–5ac
H4–4ac:H3–5ac
H4K20 me2:H4–4ac
H4K20 me3
H2A.Z
H3K9 me3
H4–4ac
DNA Meth. m5C
H3K4 me1
H3K27 me3
H4K20 me2
H3–5ac
H3K4 me3
0.00 0.25 0.50 0.75 1.00
π̂
FUBP3 – Reverse
H3K4 me1:H4–4ac
H3K4 me1:H2A.Z
H2A.Z:H3–5ac
H2A.Z:H4–4ac
DNA Meth. m5C:H4K20 me3
H2A.Z:H3K4 me3
DNA Meth. m5C:H4K20 me2
H4K20 me2:H3K4 me3
H3K27 me3:H3K9 me3
DNA Meth. m5C:H3K9 me3
H3K4 me3:H4–4ac
H3K4 me3:H3–5ac
H3K27 me3:H3K4 me1
H2A.Z:H4K20 me2
H3K27 me3:DNA Meth. m5C
H3K4 me1:H4K20 me2
H3K4 me1:H3–5ac
H4K20 me2:H3–5ac
H4–4ac:H3–5ac
H4K20 me2:H4–4ac
H4K20 me3
H2A.Z
H3K9 me3
H4–4ac
DNA Meth. m5C
H3K4 me1
H3K27 me3
H4K20 me2
H3–5ac
H3K4 me3
0.00 0.25 0.50 0.75 1.00
π̂

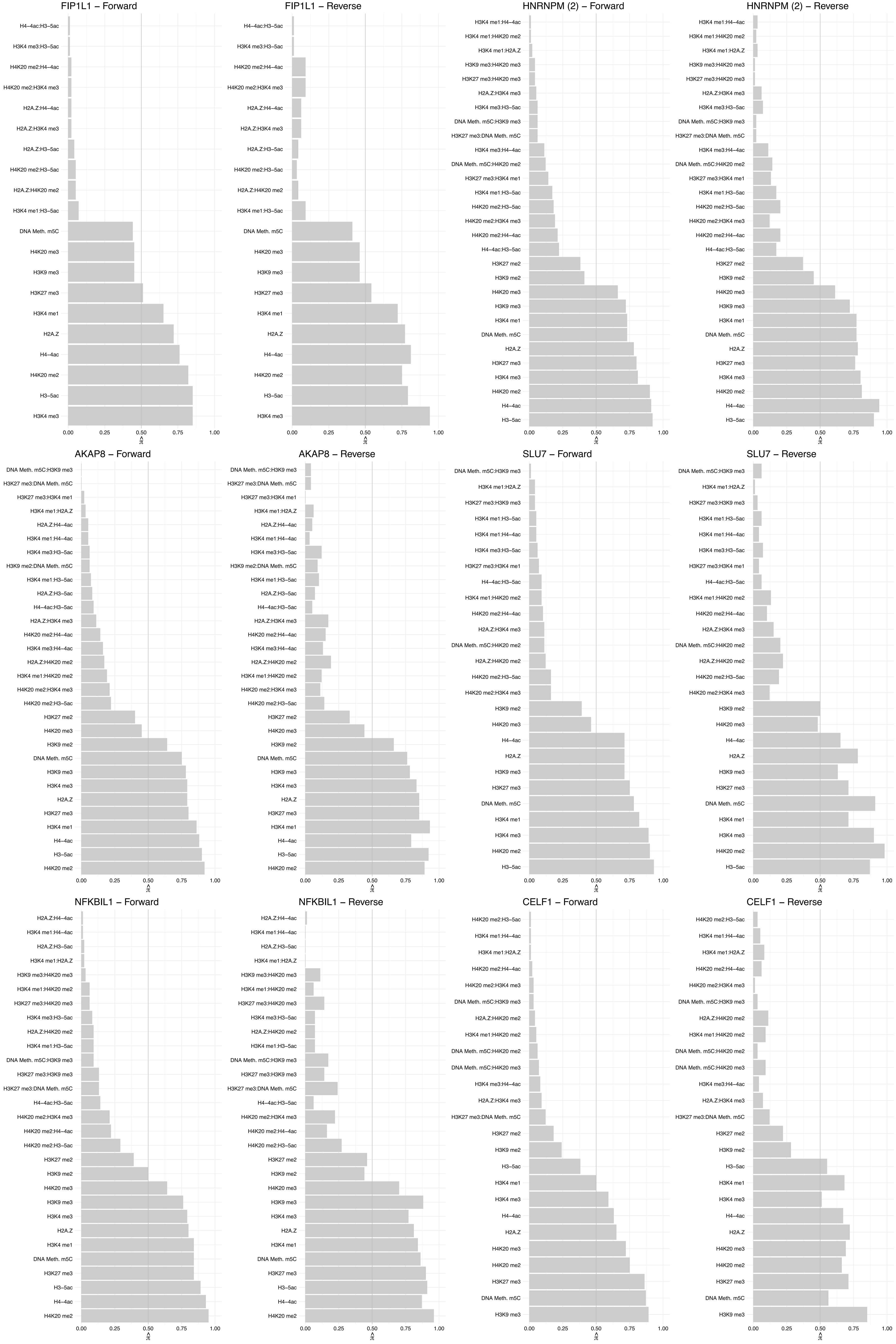
FIP1L1 – Forward
H4−4ac:H3−5ac
H3K4 me3:H3−5ac
H4K20 me2:H4−4ac
H4K20 me2:H3K4 me3
H2A.Z:H4−4ac
H2A.Z:H3K4 me3
H2A.Z:H3−5ac
H4K20 me2:H3−5ac
H2A.Z:H4K20 me2
H3K4 me1:H3−5ac
DNA Meth. m5C
H4K20 me3
H3K9 me3
H3K27 me3
H3K4 me1
H2A.Z
H4−4ac
H4K20 me2
H3−5ac
H3K4 me3
0.00
0.25
0.50
0.75
1.00
π̂
FIP1L1 – Reverse
H4−4ac:H3−5ac
H3K4 me3:H3−5ac
H4K20 me2:H4−4ac
H4K20 me2:H3K4 me3
H2A.Z:H4−4ac
H2A.Z:H3K4 me3
H2A.Z:H3−5ac
H4K20 me2:H3−5ac
H2A.Z:H4K20 me2
H3K4 me1:H3−5ac
DNA Meth. m5C
H4K20 me3
H3K9 me3
H3K27 me3
H3K4 me1
H2A.Z
H4−4ac
H4K20 me2
H3−5ac
H3K4 me3
0.00
0.25
0.50
0.75
1.00
π̂
HNRNPM (2) – Forward
H3K4 me1:H4−4ac
H3K4 me1:H4K20 me2
H3K4 me1:H2A.Z
H3K9 me3:H4K20 me3
H3K27 me3:H4K20 me3
H2A.Z:H3K4 me3
H3K4 me3:H3−5ac
DNA Meth. m5C:H3K9 me3
H3K27 me3:DNA Meth. m5C
H3K4 me3:H4−4ac
DNA Meth. m5C:H4K20 me2
H3K27 me3:H3K4 me1
H3K4 me1:H3−5ac
H4K20 me2:H3−5ac
H4K20 me2:H3K4 me3
H4K20 me2:H4−4ac
H4−4ac:H3−5ac
H3K27 me2
H3K9 me2
H4K20 me3
H3K9 me3
H3K4 me1
DNA Meth. m5C
H2A.Z
H3K27 me3
H3K4 me3
H4K20 me2
H4−4ac
H3−5ac
0.00
0.25
0.50
0.75
1.00
π̂
HNRNPM (2) – Reverse
H3K4 me1:H4−4ac
H3K4 me1:H4K20 me2
H3K4 me1:H2A.Z
H3K9 me3:H4K20 me3
H3K27 me3:H4K20 me3
H2A.Z:H3K4 me3
H3K4 me3:H3−5ac
DNA Meth. m5C:H3K9 me3
H3K27 me3:DNA Meth. m5C
H3K4 me3:H4−4ac
DNA Meth. m5C:H4K20 me2
H3K27 me3:H3K4 me1
H3K4 me1:H3−5ac
H4K20 me2:H3−5ac
H4K20 me2:H3K4 me3
H4K20 me2:H4−4ac
H4−4ac:H3−5ac
H3K27 me2
H3K9 me2
H4K20 me3
H3K9 me3
H3K4 me1
DNA Meth. m5C
H2A.Z
H3K27 me3
H3K4 me3
H4K20 me2
H4−4ac
H3−5ac
0.00
0.25
0.50
0.75
1.00
π̂
AKAP8 – Forward
DNA Meth. m5C:H3K9 me3
H3K27 me3:DNA Meth. m5C
H3K27 me3:H3K4 me1
H3K4 me1:H2A.Z
H2A.Z:H4−4ac
H3K4 me1:H4−4ac
H3K4 me3:H3−5ac
H3K9 me2:DNA Meth. m5C
H3K4 me1:H3−5ac
H2A.Z:H3−5ac
H4−4ac:H3−5ac
H2A.Z:H3K4 me3
H4K20 me2:H4−4ac
H3K4 me3:H4−4ac
H2A.Z:H4K20 me2
H3K4 me1:H4K20 me2
H4K20 me2:H3K4 me3
H4K20 me2:H3−5ac
H3K27 me2
H4K20 me3
H3K9 me2
DNA Meth. m5C
H3K9 me3
H3K4 me3
H2A.Z
H3K27 me3
H3K4 me1
H4−4ac
H3−5ac
H4K20 me2
0.00
0.25
0.50
0.75
1.00
π̂
AKAP8 – Reverse
DNA Meth. m5C:H3K9 me3
H3K27 me3:DNA Meth. m5C
H3K27 me3:H3K4 me1
H3K4 me1:H2A.Z
H2A.Z:H4−4ac
H3K4 me1:H4−4ac
H3K4 me3:H3−5ac
H3K9 me2:DNA Meth. m5C
H3K4 me1:H3−5ac
H2A.Z:H3−5ac
H4−4ac:H3−5ac
H2A.Z:H3K4 me3
H4K20 me2:H4−4ac
H3K4 me3:H4−4ac
H2A.Z:H4K20 me2
H3K4 me1:H4K20 me2
H4K20 me2:H3K4 me3
H4K20 me2:H3−5ac
H3K27 me2
H4K20 me3
H3K9 me2
DNA Meth. m5C
H3K9 me3
H3K4 me3
H2A.Z
H3K27 me3
H3K4 me1
H4−4ac
H3−5ac
H4K20 me2
0.00
0.25
0.50
0.75
1.00
π̂
SLU7 – Forward
DNA Meth. m5C:H3K9 me3
H3K4 me1:H2A.Z
H3K27 me3:H3K9 me3
H3K4 me1:H3−5ac
H3K4 me1:H4−4ac
H3K4 me3:H3−5ac
H3K27 me3:H3K4 me1
H4−4ac:H3−5ac
H3K4 me1:H4K20 me2
H4K20 me2:H4−4ac
H2A.Z:H3K4 me3
DNA Meth. m5C:H4K20 me2
H2A.Z:H4K20 me2
H4K20 me2:H3−5ac
H4K20 me2:H3K4 me3
H3K9 me2
H4K20 me3
H4−4ac
H2A.Z
H3K9 me3
H3K27 me3
DNA Meth. m5C
H3K4 me1
H3K4 me3
H4K20 me2
H3−5ac
0.00
0.25
0.50
0.75
1.00
π̂
SLU7 – Reverse
DNA Meth. m5C:H3K9 me3
H3K4 me1:H2A.Z
H3K27 me3:H3K9 me3
H3K4 me1:H3−5ac
H3K4 me1:H4−4ac
H3K4 me3:H3−5ac
H3K27 me3:H3K4 me1
H4−4ac:H3−5ac
H3K4 me1:H4K20 me2
H4K20 me2:H4−4ac
H2A.Z:H3K4 me3
DNA Meth. m5C:H4K20 me2
H2A.Z:H4K20 me2
H4K20 me2:H3−5ac
H4K20 me2:H3K4 me3
H3K9 me2
H4K20 me3
H4−4ac
H2A.Z
H3K9 me3
H3K27 me3
DNA Meth. m5C
H3K4 me1
H3K4 me3
H4K20 me2
H3−5ac
0.00
0.25
0.50
0.75
1.00
π̂
NFKBIL1 – Forward
H2A.Z:H4−4ac
H3K4 me1:H4−4ac
H2A.Z:H3−5ac
H3K4 me1:H2A.Z
H3K9 me3:H4K20 me3
H3K4 me1:H4K20 me2
H3K27 me3:H4K20 me3
H3K4 me3:H3−5ac
H2A.Z:H4K20 me2
H3K4 me1:H3−5ac
DNA Meth. m5C:H3K9 me3
H3K27 me3:H3K9 me3
H3K27 me3:DNA Meth. m5C
H4−4ac:H3−5ac
H4K20 me2:H3K4 me3
H4K20 me2:H4−4ac
H4K20 me2:H3−5ac
H3K27 me2
H3K9 me2
H4K20 me3
H3K9 me3
H3K4 me3
H2A.Z
H3K4 me1
DNA Meth. m5C
H3K27 me3
H3−5ac
H4−4ac
H4K20 me2
0.00
0.25
0.50
0.75
1.00
π̂
NFKBIL1 – Reverse
H2A.Z:H4−4ac
H3K4 me1:H4−4ac
H2A.Z:H3−5ac
H3K4 me1:H2A.Z
H3K9 me3:H4K20 me3
H3K4 me1:H4K20 me2
H3K27 me3:H4K20 me3
H3K4 me3:H3−5ac
H2A.Z:H4K20 me2
H3K4 me1:H3−5ac
DNA Meth. m5C:H3K9 me3
H3K27 me3:H3K9 me3
H3K27 me3:DNA Meth. m5C
H4−4ac:H3−5ac
H4K20 me2:H3K4 me3
H4K20 me2:H4−4ac
H4K20 me2:H3−5ac
H3K27 me2
H3K9 me2
H4K20 me3
H3K9 me3
H3K4 me3
H2A.Z
H3K4 me1
DNA Meth. m5C
H3K27 me3
H3−5ac
H4−4ac
H4K20 me2
0.00
0.25
0.50
0.75
1.00
π̂
CELF1 – Forward
H4K20 me2:H3−5ac
H3K4 me1:H4−4ac
H3K4 me1:H2A.Z
H4K20 me2:H4−4ac
H4K20 me2:H3K4 me3
DNA Meth. m5C:H3K9 me3
H2A.Z:H4K20 me2
H3K4 me1:H4K20 me2
DNA Meth. m5C:H4K20 me2
DNA Meth. m5C:H4K20 me3
H3K4 me3:H4−4ac
H2A.Z:H3K4 me3
H3K27 me3:DNA Meth. m5C
H3K27 me2
H3K9 me2
H3−5ac
H3K4 me1
H3K4 me3
H4−4ac
H2A.Z
H4K20 me3
H4K20 me2
H3K27 me3
DNA Meth. m5C
H3K9 me3
0.00
0.25
0.50
0.75
1.00
π̂
CELF1 – Reverse
H4K20 me2:H3−5ac
H3K4 me1:H4−4ac
H3K4 me1:H2A.Z
H4K20 me2:H4−4ac
H4K20 me2:H3K4 me3
DNA Meth. m5C:H3K9 me3
H2A.Z:H4K20 me2
H3K4 me1:H4K20 me2
DNA Meth. m5C:H4K20 me2
DNA Meth. m5C:H4K20 me3
H3K4 me3:H4−4ac
H2A.Z:H3K4 me3
H3K27 me3:DNA Meth. m5C
H3K27 me2
H3K9 me2
H3−5ac
H3K4 me1
H3K4 me3
H4−4ac
H2A.Z
H4K20 me3
H4K20 me2
H3K27 me3
DNA Meth. m5C
H3K9 me3
0.00
0.25
0.50
0.75
1.00
π̂

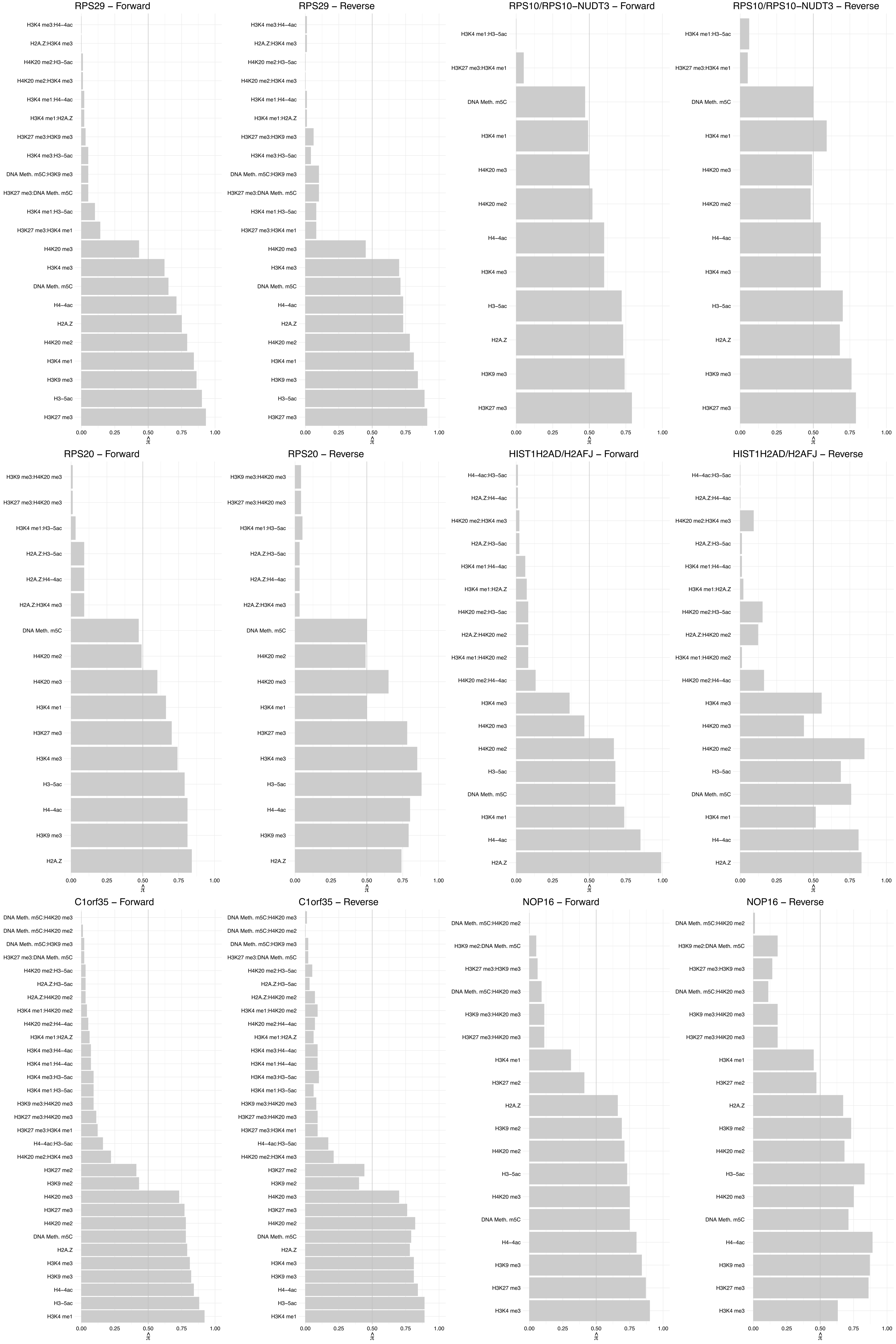
RPS29 – Forward
H3K4 me3:H4–4ac
H2A.Z:H3K4 me3
H4K20 me2:H3–5ac
H4K20 me2:H3K4 me3
H3K4 me1:H4–4ac
H3K4 me1:H2A.Z
H3K27 me3:H3K9 me3
H3K4 me3:H3–5ac
DNA Meth. m5C:H3K9 me3
H3K27 me3:DNA Meth. m5C
H3K4 me1:H3–5ac
H3K27 me3:H3K4 me1
H4K20 me3
H3K4 me3
DNA Meth. m5C
H4–4ac
H2A.Z
H4K20 me2
H3K4 me1
H3K9 me3
H3–5ac
H3K27 me3
0.00 0.25 0.50 0.75 1.00
π^
RPS29 – Reverse
H3K4 me3:H4–4ac
H2A.Z:H3K4 me3
H4K20 me2:H3–5ac
H4K20 me2:H3K4 me3
H3K4 me1:H4–4ac
H3K4 me1:H2A.Z
H3K27 me3:H3K9 me3
H3K4 me3:H3–5ac
DNA Meth. m5C:H3K9 me3
H3K27 me3:DNA Meth. m5C
H3K4 me1:H3–5ac
H3K27 me3:H3K4 me1
H4K20 me3
H3K4 me3
DNA Meth. m5C
H4–4ac
H2A.Z
H4K20 me2
H3K4 me1
H3K9 me3
H3–5ac
H3K27 me3
0.00 0.25 0.50 0.75 1.00
π^
RPS10/RPS10–NUDT3 – Forward
H3K4 me1:H3–5ac
H3K27 me3:H3K4 me1
DNA Meth. m5C
H3K4 me1
H4K20 me3
H4K20 me2
H4–4ac
H3K4 me3
H3–5ac
H2A.Z
H3K9 me3
H3K27 me3
0.00 0.25 0.50 0.75 1.00
π^
RPS10/RPS10–NUDT3 – Reverse
H3K4 me1:H3–5ac
H3K27 me3:H3K4 me1
DNA Meth. m5C
H3K4 me1
H4K20 me3
H4K20 me2
H4–4ac
H3K4 me3
H3–5ac
H2A.Z
H3K9 me3
H3K27 me3
0.00 0.25 0.50 0.75 1.00
π^
RPS20 – Forward
H3K9 me3:H4K20 me3
H3K27 me3:H4K20 me3
H3K4 me1:H3–5ac
H2A.Z:H3–5ac
H2A.Z:H4–4ac
H2A.Z:H3K4 me3
DNA Meth. m5C
H4K20 me2
H4K20 me3
H3K4 me1
H3K27 me3
H3K4 me3
H3–5ac
H4–4ac
H3K9 me3
H2A.Z
0.00 0.25 0.50 0.75 1.00
π^
RPS20 – Reverse
H3K9 me3:H4K20 me3
H3K27 me3:H4K20 me3
H3K4 me1:H3–5ac
H2A.Z:H3–5ac
H2A.Z:H4–4ac
H2A.Z:H3K4 me3
DNA Meth. m5C
H4K20 me2
H4K20 me3
H3K4 me1
H3K27 me3
H3K4 me3
H3–5ac
H4–4ac
H3K9 me3
H2A.Z
0.00 0.25 0.50 0.75 1.00
π^
HIST1H2AD/H2AFJ – Forward
H4–4ac:H3–5ac
H2A.Z:H4–4ac
H4K20 me2:H3K4 me3
H2A.Z:H3–5ac
H3K4 me1:H4–4ac
H3K4 me1:H2A.Z
H4K20 me2:H3–5ac
H2A.Z:H4K20 me2
H3K4 me1:H4K20 me2
H4K20 me2:H4–4ac
H3K4 me3
H4K20 me3
H4K20 me2
H3–5ac
DNA Meth. m5C
H3K4 me1
H4–4ac
H2A.Z
0.00 0.25 0.50 0.75 1.00
π^
HIST1H2AD/H2AFJ – Reverse
H4–4ac:H3–5ac
H2A.Z:H4–4ac
H4K20 me2:H3K4 me3
H2A.Z:H3–5ac
H3K4 me1:H4–4ac
H3K4 me1:H2A.Z
H4K20 me2:H3–5ac
H2A.Z:H4K20 me2
H3K4 me1:H4K20 me2
H4K20 me2:H4–4ac
H3K4 me3
H4K20 me3
H4K20 me2
H3–5ac
DNA Meth. m5C
H3K4 me1
H4–4ac
H2A.Z
0.00 0.25 0.50 0.75 1.00
π^
C1orf35 – Forward
DNA Meth. m5C:H4K20 me3
DNA Meth. m5C:H4K20 me2
DNA Meth. m5C:H3K9 me3
H3K27 me3:DNA Meth. m5C
H4K20 me2:H3–5ac
H2A.Z:H3–5ac
H2A.Z:H4K20 me2
H3K4 me1:H4K20 me2
H4K20 me2:H4–4ac
H3K4 me1:H2A.Z
H3K4 me3:H4–4ac
H3K4 me1:H4–4ac
H3K4 me3:H3–5ac
H3K4 me1:H3–5ac
H3K9 me3:H4K20 me3
H3K27 me3:H4K20 me3
H3K27 me3:H3K4 me1
H4–4ac:H3–5ac
H4K20 me2:H3K4 me3
H3K27 me2
H3K9 me2
H4K20 me3
H3K27 me3
H4K20 me2
DNA Meth. m5C
H2A.Z
H3K4 me3
H3K9 me3
H4–4ac
H3–5ac
H3K4 me1
0.00 0.25 0.50 0.75 1.00
π^
C1orf35 – Reverse
DNA Meth. m5C:H4K20 me3
DNA Meth. m5C:H4K20 me2
DNA Meth. m5C:H3K9 me3
H3K27 me3:DNA Meth. m5C
H4K20 me2:H3–5ac
H2A.Z:H3–5ac
H2A.Z:H4K20 me2
H3K4 me1:H4K20 me2
H4K20 me2:H4–4ac
H3K4 me1:H2A.Z
H3K4 me3:H4–4ac
H3K4 me1:H4–4ac
H3K4 me3:H3–5ac
H3K4 me1:H3–5ac
H3K9 me3:H4K20 me3
H3K27 me3:H4K20 me3
H3K27 me3:H3K4 me1
H4–4ac:H3–5ac
H4K20 me2:H3K4 me3
H3K27 me2
H3K9 me2
H4K20 me3
H3K27 me3
H4K20 me2
DNA Meth. m5C
H2A.Z
H3K4 me3
H3K9 me3
H4–4ac
H3–5ac
H3K4 me1
0.00 0.25 0.50 0.75 1.00
π^
NOP16 – Forward
DNA Meth. m5C:H4K20 me2
H3K9 me2:DNA Meth. m5C
H3K27 me3:H3K9 me3
DNA Meth. m5C:H4K20 me3
H3K9 me3:H4K20 me3
H3K27 me3:H4K20 me3
H3K4 me1
H3K27 me2
H2A.Z
H3K9 me2
H4K20 me2
H3–5ac
H4K20 me3
DNA Meth. m5C
H4–4ac
H3K9 me3
H3K27 me3
H3K4 me3
0.00 0.25 0.50 0.75 1.00
π^
NOP16 – Reverse
DNA Meth. m5C:H4K20 me2
H3K9 me2:DNA Meth. m5C
H3K27 me3:H3K9 me3
DNA Meth. m5C:H4K20 me3
H3K9 me3:H4K20 me3
H3K27 me3:H4K20 me3
H3K4 me1
H3K27 me2
H2A.Z
H3K9 me2
H4K20 me2
H3–5ac
H4K20 me3
DNA Meth. m5C
H4–4ac
H3K9 me3
H3K27 me3
H3K4 me3
0.00 0.25 0.50 0.75 1.00
π^

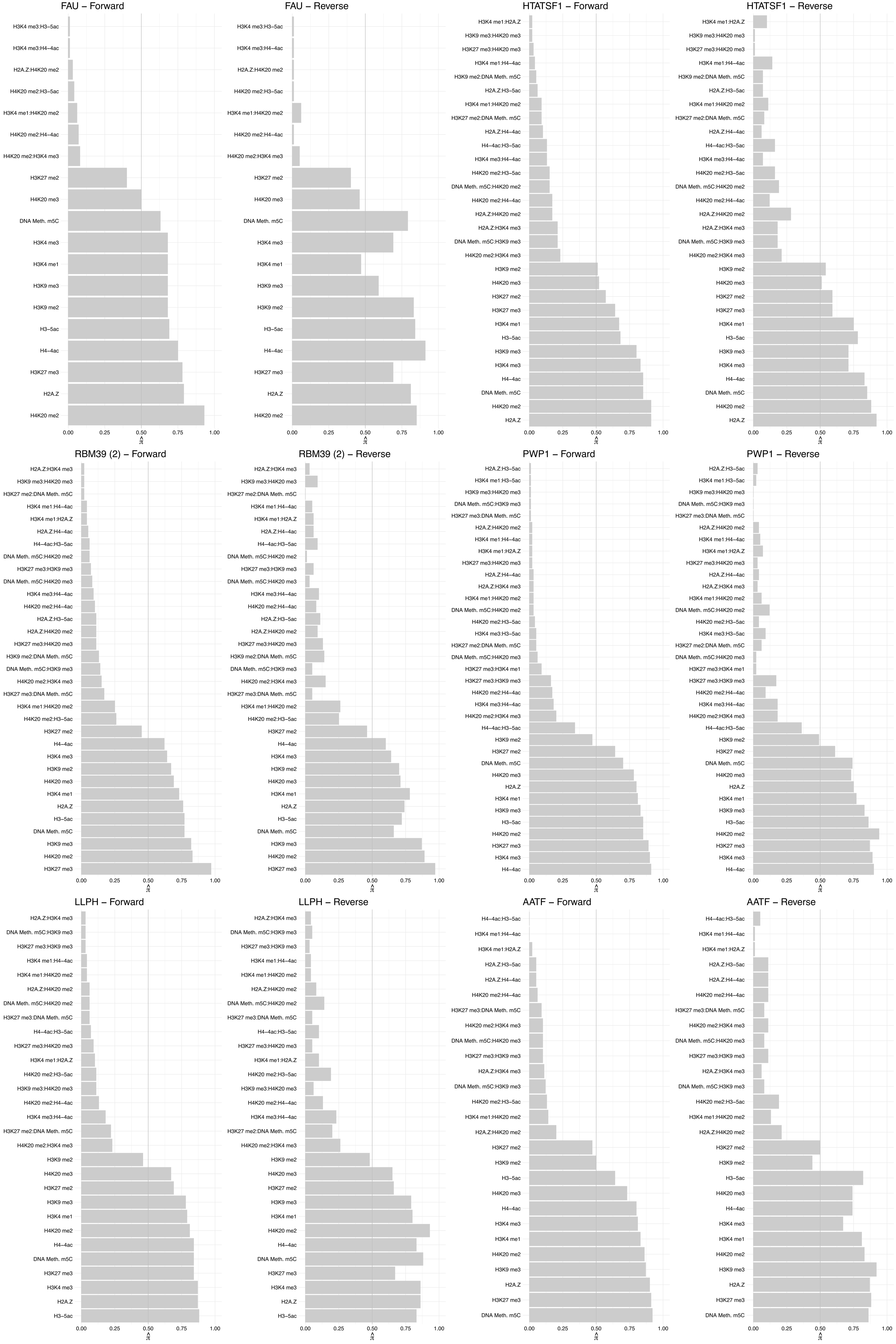

FAU – Forward
H3K4 me3:H3–5ac
H3K4 me3:H4–4ac
H2A.Z:H4K20 me2
H4K20 me2:H3–5ac
H3K4 me1:H4K20 me2
H4K20 me2:H4–4ac
H4K20 me2:H3K4 me3
H3K27 me2
H4K20 me3
DNA Meth. m5C
H3K4 me3
H3K4 me1
H3K9 me3
H3K9 me2
H3–5ac
H4–4ac
H3K27 me3
H2A.Z
H4K20 me2
0.00
0.25
0.50
0.75
1.00
$\hat{\pi}$
FAU – Reverse
H3K4 me3:H3–5ac
H3K4 me3:H4–4ac
H2A.Z:H4K20 me2
H4K20 me2:H3–5ac
H3K4 me1:H4K20 me2
H4K20 me2:H4–4ac
H4K20 me2:H3K4 me3
H3K27 me2
H4K20 me3
DNA Meth. m5C
H3K4 me3
H3K4 me1
H3K9 me3
H3K9 me2
H3–5ac
H4–4ac
H3K27 me3
H2A.Z
H4K20 me2
0.00
0.25
0.50
0.75
1.00
$\hat{\pi}$
HTATSF1 – Forward
H3K4 me1:H2A.Z
H3K9 me3:H4K20 me3
H3K27 me3:H4K20 me3
H3K4 me1:H4–4ac
H3K9 me2:DNA Meth. m5C
H2A.Z:H3–5ac
H3K4 me1:H4K20 me2
H3K27 me2:DNA Meth. m5C
H2A.Z:H4–4ac
H4–4ac:H3–5ac
H3K4 me3:H4–4ac
H4K20 me2:H3–5ac
DNA Meth. m5C:H4K20 me2
H4K20 me2:H4–4ac
H2A.Z:H4K20 me2
H2A.Z:H3K4 me3
DNA Meth. m5C:H3K9 me3
H4K20 me2:H3K4 me3
H3K9 me2
H4K20 me3
H3K27 me2
H3K27 me3
H3K4 me1
H3–5ac
H3K9 me3
H3K4 me3
H4–4ac
DNA Meth. m5C
H4K20 me2
H2A.Z
0.00
0.25
0.50
0.75
1.00
$\hat{\pi}$
HTATSF1 – Reverse
H3K4 me1:H2A.Z
H3K9 me3:H4K20 me3
H3K27 me3:H4K20 me3
H3K4 me1:H4–4ac
H3K9 me2:DNA Meth. m5C
H2A.Z:H3–5ac
H3K4 me1:H4K20 me2
H3K27 me2:DNA Meth. m5C
H2A.Z:H4–4ac
H4–4ac:H3–5ac
H3K4 me3:H4–4ac
H4K20 me2:H3–5ac
DNA Meth. m5C:H4K20 me2
H4K20 me2:H4–4ac
H2A.Z:H4K20 me2
H2A.Z:H3K4 me3
DNA Meth. m5C:H3K9 me3
H4K20 me2:H3K4 me3
H3K9 me2
H4K20 me3
H3K27 me2
H3K27 me3
H3K4 me1
H3–5ac
H3K9 me3
H3K4 me3
H4–4ac
DNA Meth. m5C
H4K20 me2
H2A.Z
0.00
0.25
0.50
0.75
1.00
$\hat{\pi}$
RBM39 (2) – Forward
H2A.Z:H3K4 me3
H3K9 me3:H4K20 me3
H3K27 me2:DNA Meth. m5C
H3K4 me1:H4–4ac
H3K4 me1:H2A.Z
H2A.Z:H4–4ac
H4–4ac:H3–5ac
DNA Meth. m5C:H4K20 me2
H3K27 me3:H3K9 me3
DNA Meth. m5C:H4K20 me3
H3K4 me3:H4–4ac
H4K20 me2:H4–4ac
H2A.Z:H3–5ac
H2A.Z:H4K20 me2
H3K27 me3:H4K20 me3
H3K9 me2:DNA Meth. m5C
DNA Meth. m5C:H3K9 me3
H4K20 me2:H3K4 me3
H3K27 me3:DNA Meth. m5C
H3K4 me1:H4K20 me2
H4K20 me2:H3–5ac
H3K27 me2
H4–4ac
H3K4 me3
H3K9 me2
H4K20 me3
H3K4 me1
H2A.Z
H3–5ac
DNA Meth. m5C
H3K9 me3
H4K20 me2
H3K27 me3
0.00
0.25
0.50
0.75
1.00
$\hat{\pi}$
RBM39 (2) – Reverse
H2A.Z:H3K4 me3
H3K9 me3:H4K20 me3
H3K27 me2:DNA Meth. m5C
H3K4 me1:H4–4ac
H3K4 me1:H2A.Z
H2A.Z:H4–4ac
H4–4ac:H3–5ac
DNA Meth. m5C:H4K20 me2
H3K27 me3:H3K9 me3
DNA Meth. m5C:H4K20 me3
H3K4 me3:H4–4ac
H4K20 me2:H4–4ac
H2A.Z:H3–5ac
H2A.Z:H4K20 me2
H3K27 me3:H4K20 me3
H3K9 me2:DNA Meth. m5C
DNA Meth. m5C:H3K9 me3
H4K20 me2:H3K4 me3
H3K27 me3:DNA Meth. m5C
H3K4 me1:H4K20 me2
H4K20 me2:H3–5ac
H3K27 me2
H4–4ac
H3K4 me3
H3K9 me2
H4K20 me3
H3K4 me1
H2A.Z
H3–5ac
DNA Meth. m5C
H3K9 me3
H4K20 me2
H3K27 me3
0.00
0.25
0.50
0.75
1.00
$\hat{\pi}$
PWP1 – Forward
H2A.Z:H3–5ac
H3K4 me1:H3–5ac
H3K9 me3:H4K20 me3
DNA Meth. m5C:H3K9 me3
H3K27 me3:DNA Meth. m5C
H2A.Z:H4K20 me2
H3K4 me1:H4–4ac
H3K4 me1:H2A.Z
H3K27 me3:H4K20 me3
H2A.Z:H4–4ac
H2A.Z:H3K4 me3
H3K4 me1:H4K20 me2
DNA Meth. m5C:H4K20 me2
H4K20 me2:H3–5ac
H3K4 me3:H3–5ac
H3K27 me2:DNA Meth. m5C
DNA Meth. m5C:H4K20 me3
H3K27 me3:H3K4 me1
H3K27 me3:H3K9 me3
H4K20 me2:H4–4ac
H3K4 me3:H4–4ac
H4K20 me2:H3K4 me3
H4–4ac:H3–5ac
H3K9 me2
H3K27 me2
DNA Meth. m5C
H4K20 me3
H2A.Z
H3K4 me1
H3K9 me3
H3–5ac
H4K20 me2
H3K27 me3
H3K4 me3
H4–4ac
0.00
0.25
0.50
0.75
1.00
$\hat{\pi}$
PWP1 – Reverse
H2A.Z:H3–5ac
H3K4 me1:H3–5ac
H3K9 me3:H4K20 me3
DNA Meth. m5C:H3K9 me3
H3K27 me3:DNA Meth. m5C
H2A.Z:H4K20 me2
H3K4 me1:H4–4ac
H3K4 me1:H2A.Z
H3K27 me3:H4K20 me3
H2A.Z:H4–4ac
H2A.Z:H3K4 me3
H3K4 me1:H4K20 me2
DNA Meth. m5C:H4K20 me2
H4K20 me2:H3–5ac
H3K4 me3:H3–5ac
H3K27 me2:DNA Meth. m5C
DNA Meth. m5C:H4K20 me3
H3K27 me3:H3K4 me1
H3K27 me3:H3K9 me3
H4K20 me2:H4–4ac
H3K4 me3:H4–4ac
H4K20 me2:H3K4 me3
H4–4ac:H3–5ac
H3K9 me2
H3K27 me2
DNA Meth. m5C
H4K20 me3
H2A.Z
H3K4 me1
H3K9 me3
H3–5ac
H4K20 me2
H3K27 me3
H3K4 me3
H4–4ac
0.00
0.25
0.50
0.75
1.00
$\hat{\pi}$
LLPH – Forward
H2A.Z:H3K4 me3
DNA Meth. m5C:H3K9 me3
H3K27 me3:H3K9 me3
H3K4 me1:H4–4ac
H3K4 me1:H4K20 me2
H2A.Z:H4K20 me2
DNA Meth. m5C:H4K20 me2
H3K27 me3:DNA Meth. m5C
H4–4ac:H3–5ac
H3K27 me3:H4K20 me3
H3K4 me1:H2A.Z
H4K20 me2:H3–5ac
H3K9 me3:H4K20 me3
H4K20 me2:H4–4ac
H3K4 me3:H4–4ac
H3K27 me2:DNA Meth. m5C
H4K20 me2:H3K4 me3
H3K9 me2
H4K20 me3
H3K27 me2
H3K9 me3
H3K4 me1
H4K20 me2
H4–4ac
DNA Meth. m5C
H3K27 me3
H3K4 me3
H2A.Z
H3–5ac
0.00
0.25
0.50
0.75
1.00
$\hat{\pi}$
LLPH – Reverse
H2A.Z:H3K4 me3
DNA Meth. m5C:H3K9 me3
H3K27 me3:H3K9 me3
H3K4 me1:H4–4ac
H3K4 me1:H4K20 me2
H2A.Z:H4K20 me2
DNA Meth. m5C:H4K20 me2
H3K27 me3:DNA Meth. m5C
H4–4ac:H3–5ac
H3K27 me3:H4K20 me3
H3K4 me1:H2A.Z
H4K20 me2:H3–5ac
H3K9 me3:H4K20 me3
H4K20 me2:H4–4ac
H3K4 me3:H4–4ac
H3K27 me2:DNA Meth. m5C
H4K20 me2:H3K4 me3
H3K9 me2
H4K20 me3
H3K27 me2
H3K9 me3
H3K4 me1
H4K20 me2
H4–4ac
DNA Meth. m5C
H3K27 me3
H3K4 me3
H2A.Z
H3–5ac
0.00
0.25
0.50
0.75
1.00
$\hat{\pi}$
AATF – Forward
H4–4ac:H3–5ac
H3K4 me1:H4–4ac
H3K4 me1:H2A.Z
H2A.Z:H3–5ac
H2A.Z:H4–4ac
H4K20 me2:H4–4ac
H3K27 me3:DNA Meth. m5C
H4K20 me2:H3K4 me3
DNA Meth. m5C:H4K20 me3
H3K27 me3:H3K9 me3
H2A.Z:H3K4 me3
DNA Meth. m5C:H3K9 me3
H4K20 me2:H3–5ac
H3K4 me1:H4K20 me2
H2A.Z:H4K20 me2
H3K27 me2
H3K9 me2
H3–5ac
H4K20 me3
H4–4ac
H3K4 me3
H3K4 me1
H4K20 me2
H3K9 me3
H2A.Z
H3K27 me3
DNA Meth. m5C
0.00
0.25
0.50
0.75
1.00
$\hat{\pi}$
AATF – Reverse
H4–4ac:H3–5ac
H3K4 me1:H4–4ac
H3K4 me1:H2A.Z
H2A.Z:H3–5ac
H2A.Z:H4–4ac
H4K20 me2:H4–4ac
H3K27 me3:DNA Meth. m5C
H4K20 me2:H3K4 me3
DNA Meth. m5C:H4K20 me3
H3K27 me3:H3K9 me3
H2A.Z:H3K4 me3
DNA Meth. m5C:H3K9 me3
H4K20 me2:H3–5ac
H3K4 me1:H4K20 me2
H2A.Z:H4K20 me2
H3K27 me2
H3K9 me2
H3–5ac
H4K20 me3
H4–4ac
H3K4 me3
H3K4 me1
H4K20 me2
H3K9 me3
H2A.Z
H3K27 me3
DNA Meth. m5C
0.00
0.25
0.50
0.75
1.00
$\hat{\pi}$

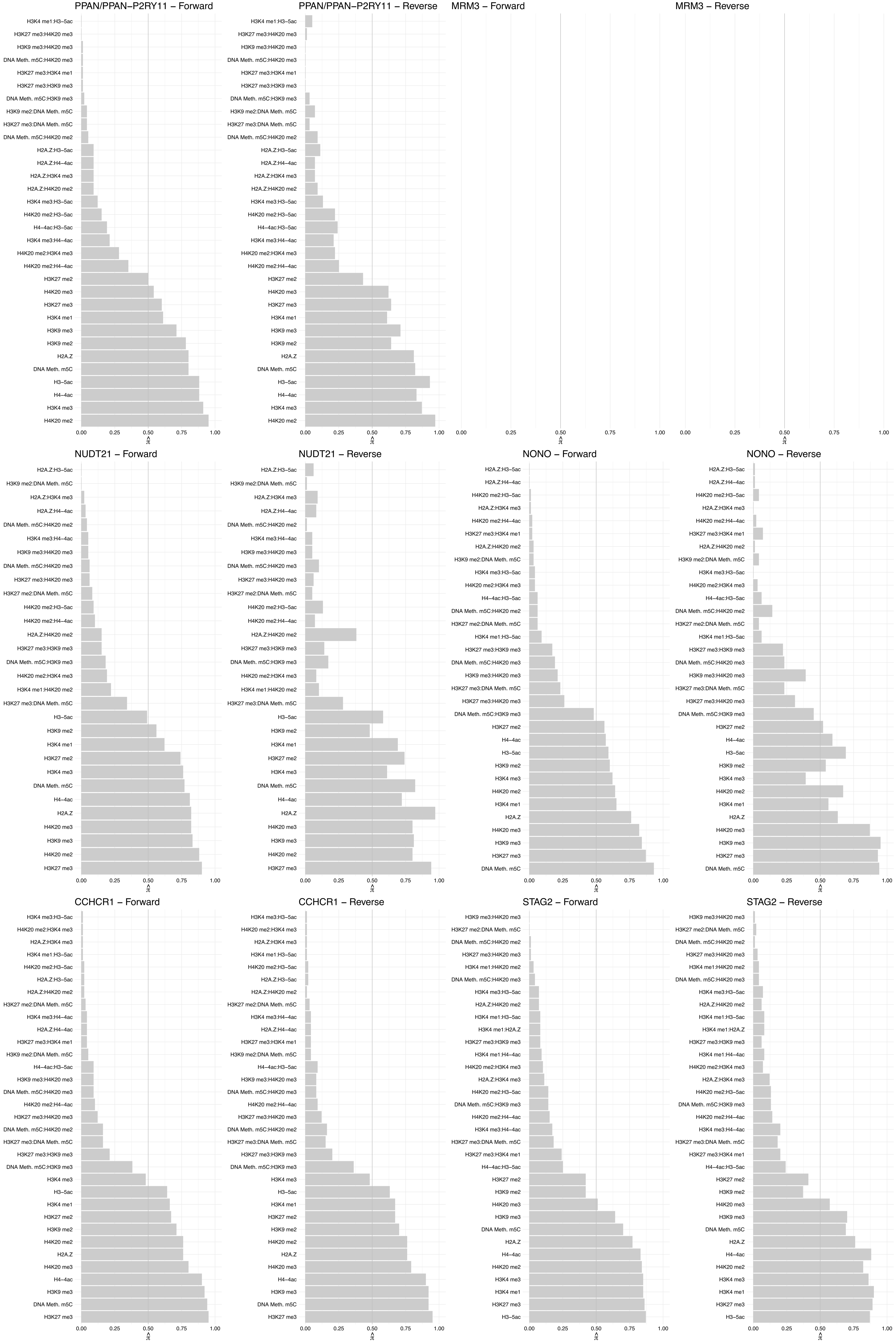
PPAN/PPAN−P2RY11 − Forward
H3K4 me1:H3−5ac
H3K27 me3:H4K20 me3
H3K9 me3:H4K20 me3
DNA Meth. m5C:H4K20 me3
H3K27 me3:H3K4 me1
H3K27 me3:H3K9 me3
DNA Meth. m5C:H3K9 me3
H3K9 me2:DNA Meth. m5C
H3K27 me3:DNA Meth. m5C
DNA Meth. m5C:H4K20 me2
H2A.Z:H3−5ac
H2A.Z:H4−4ac
H2A.Z:H3K4 me3
H2A.Z:H4K20 me2
H3K4 me3:H3−5ac
H4K20 me2:H3−5ac
H4−4ac:H3−5ac
H3K4 me3:H4−4ac
H4K20 me2:H3K4 me3
H4K20 me2:H4−4ac
H3K27 me2
H4K20 me3
H3K27 me3
H3K4 me1
H3K9 me3
H3K9 me2
H2A.Z
DNA Meth. m5C
H3−5ac
H4−4ac
H3K4 me3
H4K20 me2
0.00 0.25 0.50 0.75 1.00
π̂
PPAN/PPAN−P2RY11 − Reverse
MRM3 − Forward
0.00 0.25 0.50 0.75 1.00
π̂
MRM3 − Reverse
0.00 0.25 0.50 0.75 1.00
π̂
NUDT21 − Forward
H2A.Z:H3−5ac
H3K9 me2:DNA Meth. m5C
H2A.Z:H3K4 me3
H2A.Z:H4−4ac
DNA Meth. m5C:H4K20 me2
H3K4 me3:H4−4ac
H3K9 me3:H4K20 me3
DNA Meth. m5C:H4K20 me3
H3K27 me3:H4K20 me3
H3K27 me2:DNA Meth. m5C
H4K20 me2:H3−5ac
H4K20 me2:H4−4ac
H2A.Z:H4K20 me2
H3K27 me3:H3K9 me3
DNA Meth. m5C:H3K9 me3
H4K20 me2:H3K4 me3
H3K4 me1:H4K20 me2
H3K27 me3:DNA Meth. m5C
H3−5ac
H3K9 me2
H3K4 me1
H3K27 me2
H3K4 me3
DNA Meth. m5C
H4−4ac
H2A.Z
H4K20 me3
H3K9 me3
H4K20 me2
H3K27 me3
NUDT21 − Reverse
NONO − Forward
H2A.Z:H3−5ac
H2A.Z:H4−4ac
H4K20 me2:H3−5ac
H2A.Z:H3K4 me3
H4K20 me2:H4−4ac
H3K27 me3:H3K4 me1
H2A.Z:H4K20 me2
H3K9 me2:DNA Meth. m5C
H3K4 me3:H3−5ac
H4K20 me2:H3K4 me3
H4−4ac:H3−5ac
DNA Meth. m5C:H4K20 me2
H3K27 me2:DNA Meth. m5C
H3K4 me1:H3−5ac
H3K27 me3:H3K9 me3
DNA Meth. m5C:H4K20 me3
H3K9 me3:H4K20 me3
H3K27 me3:DNA Meth. m5C
H3K27 me3:H4K20 me3
DNA Meth. m5C:H3K9 me3
H3K27 me2
H4−4ac
H3−5ac
H3K9 me2
H3K4 me3
H4K20 me2
H3K4 me1
H2A.Z
H4K20 me3
H3K9 me3
H3K27 me3
DNA Meth. m5C
NONO − Reverse
CCHCR1 − Forward
H3K4 me3:H3−5ac
H4K20 me2:H3K4 me3
H2A.Z:H3K4 me3
H3K4 me1:H3−5ac
H4K20 me2:H3−5ac
H2A.Z:H3−5ac
H2A.Z:H4K20 me2
H3K27 me2:DNA Meth. m5C
H3K4 me3:H4−4ac
H2A.Z:H4−4ac
H3K27 me3:H3K4 me1
H3K9 me2:DNA Meth. m5C
H4−4ac:H3−5ac
H3K9 me3:H4K20 me3
DNA Meth. m5C:H4K20 me3
H4K20 me2:H4−4ac
H3K27 me3:H4K20 me3
DNA Meth. m5C:H4K20 me2
H3K27 me3:DNA Meth. m5C
H3K27 me3:H3K9 me3
DNA Meth. m5C:H3K9 me3
H3K4 me3
H3−5ac
H3K4 me1
H3K27 me2
H3K9 me2
H4K20 me2
H2A.Z
H4K20 me3
H4−4ac
H3K9 me3
DNA Meth. m5C
H3K27 me3
CCHCR1 − Reverse
STAG2 − Forward
H3K9 me3:H4K20 me3
H3K27 me2:DNA Meth. m5C
DNA Meth. m5C:H4K20 me2
H3K27 me3:H4K20 me3
H3K4 me1:H4K20 me2
DNA Meth. m5C:H4K20 me3
H3K4 me3:H3−5ac
H2A.Z:H4K20 me2
H3K4 me1:H3−5ac
H3K4 me1:H2A.Z
H3K27 me3:H3K9 me3
H3K4 me1:H4−4ac
H4K20 me2:H3K4 me3
H2A.Z:H3K4 me3
H4K20 me2:H3−5ac
DNA Meth. m5C:H3K9 me3
H4K20 me2:H4−4ac
H3K4 me3:H4−4ac
H3K27 me3:DNA Meth. m5C
H3K27 me3:H3K4 me1
H4−4ac:H3−5ac
H3K27 me2
H3K9 me2
H4K20 me3
H3K9 me3
DNA Meth. m5C
H2A.Z
H4−4ac
H4K20 me2
H3K4 me3
H3K4 me1
H3K27 me3
H3−5ac
STAG2 − Reverse

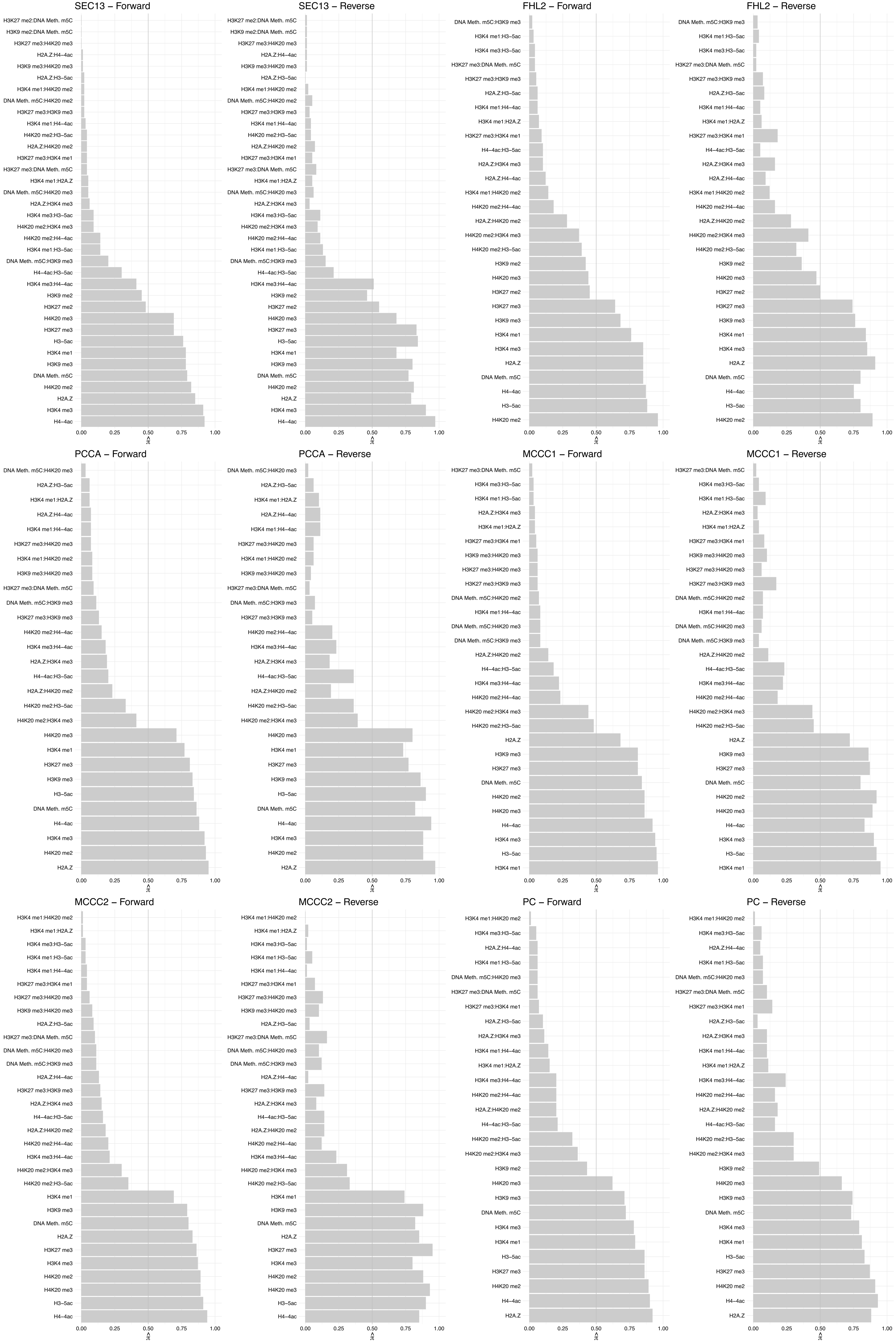

SEC13 – Forward
SEC13 – Reverse
FHL2 – Forward
FHL2 – Reverse
PCCA – Forward
PCCA – Reverse
MCCC1 – Forward
MCCC1 – Reverse
MCCC2 – Forward
MCCC2 – Reverse
PC – Forward
PC – Reverse
0.00 0.25 0.50 0.75 1.00
$\hat{\pi}$

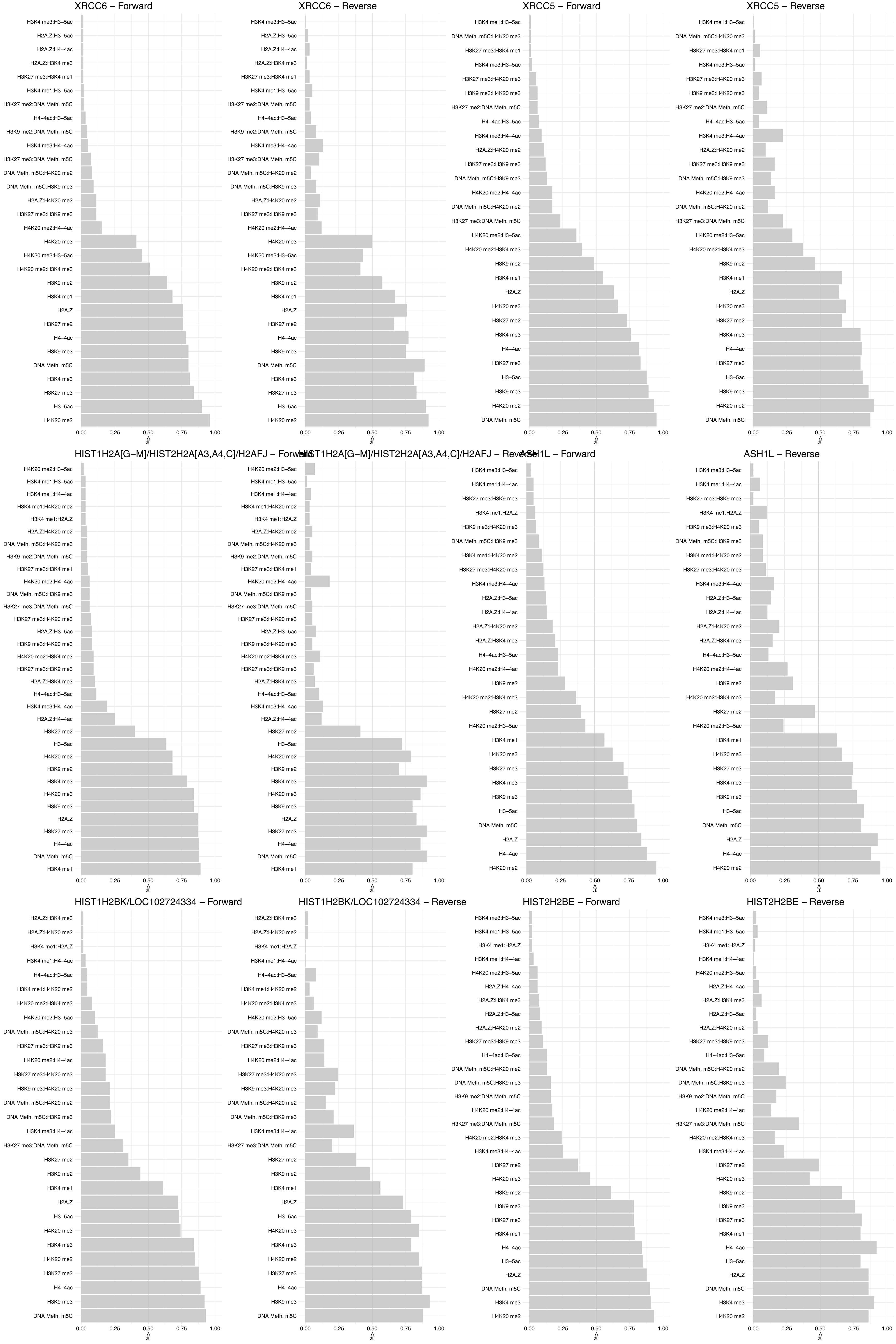
XRCC6 – Forward
XRCC6 – Reverse
XRCC5 – Forward
XRCC5 – Reverse
HIST1H2A[G–M]/HIST2H2A[A3,A4,C]/H2AFJ – Forward
HIST1H2A[G–M]/HIST2H2A[A3,A4,C]/H2AFJ – Reverse
ASH1L – Forward
ASH1L – Reverse
HIST1H2BK/LOC102724334 – Forward
HIST1H2BK/LOC102724334 – Reverse
HIST2H2BE – Forward
HIST2H2BE – Reverse
0.00 0.25 0.50 0.75 1.00
$\hat{\pi}$

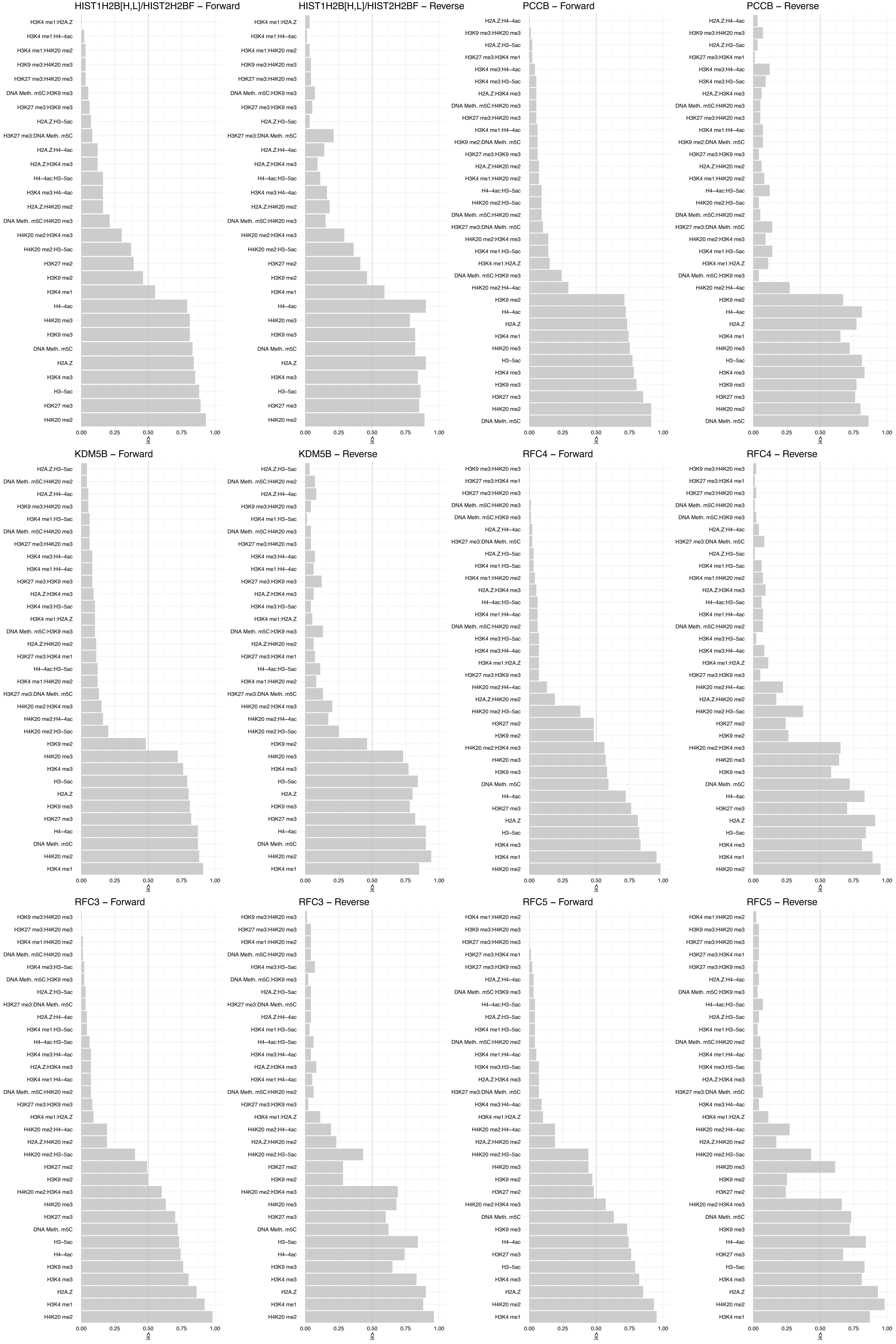

HIST1H2B[H,L]/HIST2H2BF – Forward
H3K4 me1:H2A.Z
H3K4 me1:H4−4ac
H3K4 me1:H4K20 me2
H3K9 me3:H4K20 me3
H3K27 me3:H4K20 me3
DNA Meth. m5C:H3K9 me3
H3K27 me3:H3K9 me3
H2A.Z:H3−5ac
H3K27 me3:DNA Meth. m5C
H2A.Z:H4−4ac
H2A.Z:H3K4 me3
H4−4ac:H3−5ac
H3K4 me3:H4−4ac
H2A.Z:H4K20 me2
DNA Meth. m5C:H4K20 me3
H4K20 me2:H3K4 me3
H4K20 me2:H3−5ac
H3K27 me2
H3K9 me2
H3K4 me1
H4−4ac
H4K20 me3
H3K9 me3
DNA Meth. m5C
H2A.Z
H3K4 me3
H3−5ac
H3K27 me3
H4K20 me2
0.00 0.25 0.50 0.75 1.00
π̂
HIST1H2B[H,L]/HIST2H2BF – Reverse
H3K4 me1:H2A.Z
H3K4 me1:H4−4ac
H3K4 me1:H4K20 me2
H3K9 me3:H4K20 me3
H3K27 me3:H4K20 me3
DNA Meth. m5C:H3K9 me3
H3K27 me3:H3K9 me3
H2A.Z:H3−5ac
H3K27 me3:DNA Meth. m5C
H2A.Z:H4−4ac
H2A.Z:H3K4 me3
H4−4ac:H3−5ac
H3K4 me3:H4−4ac
H2A.Z:H4K20 me2
DNA Meth. m5C:H4K20 me3
H4K20 me2:H3K4 me3
H4K20 me2:H3−5ac
H3K27 me2
H3K9 me2
H3K4 me1
H4−4ac
H4K20 me3
H3K9 me3
DNA Meth. m5C
H2A.Z
H3K4 me3
H3−5ac
H3K27 me3
H4K20 me2
0.00 0.25 0.50 0.75 1.00
π̂
PCCB – Forward
H2A.Z:H4−4ac
H3K9 me3:H4K20 me3
H2A.Z:H3−5ac
H3K27 me3:H3K4 me1
H3K4 me3:H4−4ac
H3K4 me3:H3−5ac
H2A.Z:H3K4 me3
DNA Meth. m5C:H4K20 me3
H3K27 me3:H4K20 me3
H3K4 me1:H4−4ac
H3K9 me2:DNA Meth. m5C
H3K27 me3:H3K9 me3
H2A.Z:H4K20 me2
H3K4 me1:H4K20 me2
H4−4ac:H3−5ac
H4K20 me2:H3−5ac
DNA Meth. m5C:H4K20 me2
H3K27 me3:DNA Meth. m5C
H4K20 me2:H3K4 me3
H3K4 me1:H3−5ac
H3K4 me1:H2A.Z
DNA Meth. m5C:H3K9 me3
H4K20 me2:H4−4ac
H3K9 me2
H4−4ac
H2A.Z
H3K4 me1
H4K20 me3
H3−5ac
H3K4 me3
H3K9 me3
H3K27 me3
H4K20 me2
DNA Meth. m5C
0.00 0.25 0.50 0.75 1.00
π̂
PCCB – Reverse
H2A.Z:H4−4ac
H3K9 me3:H4K20 me3
H2A.Z:H3−5ac
H3K27 me3:H3K4 me1
H3K4 me3:H4−4ac
H3K4 me3:H3−5ac
H2A.Z:H3K4 me3
DNA Meth. m5C:H4K20 me3
H3K27 me3:H4K20 me3
H3K4 me1:H4−4ac
H3K9 me2:DNA Meth. m5C
H3K27 me3:H3K9 me3
H2A.Z:H4K20 me2
H3K4 me1:H4K20 me2
H4−4ac:H3−5ac
H4K20 me2:H3−5ac
DNA Meth. m5C:H4K20 me2
H3K27 me3:DNA Meth. m5C
H4K20 me2:H3K4 me3
H3K4 me1:H3−5ac
H3K4 me1:H2A.Z
DNA Meth. m5C:H3K9 me3
H4K20 me2:H4−4ac
H3K9 me2
H4−4ac
H2A.Z
H3K4 me1
H4K20 me3
H3−5ac
H3K4 me3
H3K9 me3
H3K27 me3
H4K20 me2
DNA Meth. m5C
0.00 0.25 0.50 0.75 1.00
π̂
KDM5B – Forward
H2A.Z:H3−5ac
DNA Meth. m5C:H4K20 me2
H2A.Z:H4−4ac
H3K9 me3:H4K20 me3
H3K4 me1:H3−5ac
DNA Meth. m5C:H4K20 me3
H3K27 me3:H4K20 me3
H3K4 me3:H4−4ac
H3K4 me1:H4−4ac
H3K27 me3:H3K9 me3
H2A.Z:H3K4 me3
H3K4 me3:H3−5ac
H3K4 me1:H2A.Z
DNA Meth. m5C:H3K9 me3
H2A.Z:H4K20 me2
H3K27 me3:H3K4 me1
H4−4ac:H3−5ac
H3K4 me1:H4K20 me2
H3K27 me3:DNA Meth. m5C
H4K20 me2:H3K4 me3
H4K20 me2:H4−4ac
H4K20 me2:H3−5ac
H3K9 me2
H4K20 me3
H3K4 me3
H3−5ac
H2A.Z
H3K9 me3
H3K27 me3
H4−4ac
DNA Meth. m5C
H4K20 me2
H3K4 me1
0.00 0.25 0.50 0.75 1.00
π̂
KDM5B – Reverse
H2A.Z:H3−5ac
DNA Meth. m5C:H4K20 me2
H2A.Z:H4−4ac
H3K9 me3:H4K20 me3
H3K4 me1:H3−5ac
DNA Meth. m5C:H4K20 me3
H3K27 me3:H4K20 me3
H3K4 me3:H4−4ac
H3K4 me1:H4−4ac
H3K27 me3:H3K9 me3
H2A.Z:H3K4 me3
H3K4 me3:H3−5ac
H3K4 me1:H2A.Z
DNA Meth. m5C:H3K9 me3
H2A.Z:H4K20 me2
H3K27 me3:H3K4 me1
H4−4ac:H3−5ac
H3K4 me1:H4K20 me2
H3K27 me3:DNA Meth. m5C
H4K20 me2:H3K4 me3
H4K20 me2:H4−4ac
H4K20 me2:H3−5ac
H3K9 me2
H4K20 me3
H3K4 me3
H3−5ac
H2A.Z
H3K9 me3
H3K27 me3
H4−4ac
DNA Meth. m5C
H4K20 me2
H3K4 me1
0.00 0.25 0.50 0.75 1.00
π̂
RFC4 – Forward
H3K9 me3:H4K20 me3
H3K27 me3:H3K4 me1
H3K27 me3:H4K20 me3
DNA Meth. m5C:H4K20 me3
DNA Meth. m5C:H3K9 me3
H2A.Z:H4−4ac
H3K27 me3:DNA Meth. m5C
H2A.Z:H3−5ac
H3K4 me1:H3−5ac
H3K4 me1:H4K20 me2
H2A.Z:H3K4 me3
H4−4ac:H3−5ac
H3K4 me1:H4−4ac
DNA Meth. m5C:H4K20 me2
H3K4 me3:H3−5ac
H3K4 me3:H4−4ac
H3K4 me1:H2A.Z
H3K27 me3:H3K9 me3
H4K20 me2:H4−4ac
H2A.Z:H4K20 me2
H4K20 me2:H3−5ac
H3K27 me2
H3K9 me2
H4K20 me2:H3K4 me3
H4K20 me3
H3K9 me3
DNA Meth. m5C
H4−4ac
H3K27 me3
H2A.Z
H3−5ac
H3K4 me3
H3K4 me1
H4K20 me2
0.00 0.25 0.50 0.75 1.00
π̂
RFC4 – Reverse
H3K9 me3:H4K20 me3
H3K27 me3:H3K4 me1
H3K27 me3:H4K20 me3
DNA Meth. m5C:H4K20 me3
DNA Meth. m5C:H3K9 me3
H2A.Z:H4−4ac
H3K27 me3:DNA Meth. m5C
H2A.Z:H3−5ac
H3K4 me1:H3−5ac
H3K4 me1:H4K20 me2
H2A.Z:H3K4 me3
H4−4ac:H3−5ac
H3K4 me1:H4−4ac
DNA Meth. m5C:H4K20 me2
H3K4 me3:H3−5ac
H3K4 me3:H4−4ac
H3K4 me1:H2A.Z
H3K27 me3:H3K9 me3
H4K20 me2:H4−4ac
H2A.Z:H4K20 me2
H4K20 me2:H3−5ac
H3K27 me2
H3K9 me2
H4K20 me2:H3K4 me3
H4K20 me3
H3K9 me3
DNA Meth. m5C
H4−4ac
H3K27 me3
H2A.Z
H3−5ac
H3K4 me3
H3K4 me1
H4K20 me2
0.00 0.25 0.50 0.75 1.00
π̂
RFC3 – Forward
H3K9 me3:H4K20 me3
H3K27 me3:H4K20 me3
H3K4 me1:H4K20 me2
DNA Meth. m5C:H4K20 me3
H3K4 me3:H3−5ac
DNA Meth. m5C:H3K9 me3
H2A.Z:H3−5ac
H3K27 me3:DNA Meth. m5C
H2A.Z:H4−4ac
H3K4 me1:H3−5ac
H4−4ac:H3−5ac
H3K4 me3:H4−4ac
H2A.Z:H3K4 me3
H3K4 me1:H4−4ac
DNA Meth. m5C:H4K20 me2
H3K27 me3:H3K9 me3
H3K4 me1:H2A.Z
H4K20 me2:H4−4ac
H2A.Z:H4K20 me2
H4K20 me2:H3−5ac
H3K27 me2
H3K9 me2
H4K20 me2:H3K4 me3
H4K20 me3
H3K27 me3
DNA Meth. m5C
H3−5ac
H4−4ac
H3K9 me3
H3K4 me3
H2A.Z
H3K4 me1
H4K20 me2
0.00 0.25 0.50 0.75 1.00
π̂
RFC3 – Reverse
H3K9 me3:H4K20 me3
H3K27 me3:H4K20 me3
H3K4 me1:H4K20 me2
DNA Meth. m5C:H4K20 me3
H3K4 me3:H3−5ac
DNA Meth. m5C:H3K9 me3
H2A.Z:H3−5ac
H3K27 me3:DNA Meth. m5C
H2A.Z:H4−4ac
H3K4 me1:H3−5ac
H4−4ac:H3−5ac
H3K4 me3:H4−4ac
H2A.Z:H3K4 me3
H3K4 me1:H4−4ac
DNA Meth. m5C:H4K20 me2
H3K27 me3:H3K9 me3
H3K4 me1:H2A.Z
H4K20 me2:H4−4ac
H2A.Z:H4K20 me2
H4K20 me2:H3−5ac
H3K27 me2
H3K9 me2
H4K20 me2:H3K4 me3
H4K20 me3
H3K27 me3
DNA Meth. m5C
H3−5ac
H4−4ac
H3K9 me3
H3K4 me3
H2A.Z
H3K4 me1
H4K20 me2
0.00 0.25 0.50 0.75 1.00
π̂
RFC5 – Forward
H3K4 me1:H4K20 me2
H3K9 me3:H4K20 me3
H3K27 me3:H4K20 me3
H3K27 me3:H3K4 me1
H3K27 me3:H3K9 me3
H2A.Z:H4−4ac
DNA Meth. m5C:H3K9 me3
H4−4ac:H3−5ac
H2A.Z:H3−5ac
H3K4 me1:H3−5ac
DNA Meth. m5C:H4K20 me2
H3K4 me1:H4−4ac
H3K4 me3:H3−5ac
H2A.Z:H3K4 me3
H3K27 me3:DNA Meth. m5C
H3K4 me3:H4−4ac
H3K4 me1:H2A.Z
H4K20 me2:H4−4ac
H2A.Z:H4K20 me2
H4K20 me2:H3−5ac
H4K20 me3
H3K9 me2
H3K27 me2
H4K20 me2:H3K4 me3
DNA Meth. m5C
H3K9 me3
H4−4ac
H3K27 me3
H3−5ac
H3K4 me3
H2A.Z
H4K20 me2
H3K4 me1
0.00 0.25 0.50 0.75 1.00
π̂
RFC5 – Reverse
H3K4 me1:H4K20 me2
H3K9 me3:H4K20 me3
H3K27 me3:H4K20 me3
H3K27 me3:H3K4 me1
H3K27 me3:H3K9 me3
H2A.Z:H4−4ac
DNA Meth. m5C:H3K9 me3
H4−4ac:H3−5ac
H2A.Z:H3−5ac
H3K4 me1:H3−5ac
DNA Meth. m5C:H4K20 me2
H3K4 me1:H4−4ac
H3K4 me3:H3−5ac
H2A.Z:H3K4 me3
H3K27 me3:DNA Meth. m5C
H3K4 me3:H4−4ac
H3K4 me1:H2A.Z
H4K20 me2:H4−4ac
H2A.Z:H4K20 me2
H4K20 me2:H3−5ac
H4K20 me3
H3K9 me2
H3K27 me2
H4K20 me2:H3K4 me3
DNA Meth. m5C
H3K9 me3
H4−4ac
H3K27 me3
H3−5ac
H3K4 me3
H2A.Z
H4K20 me2
H3K4 me1
0.00 0.25 0.50 0.75 1.00
π̂

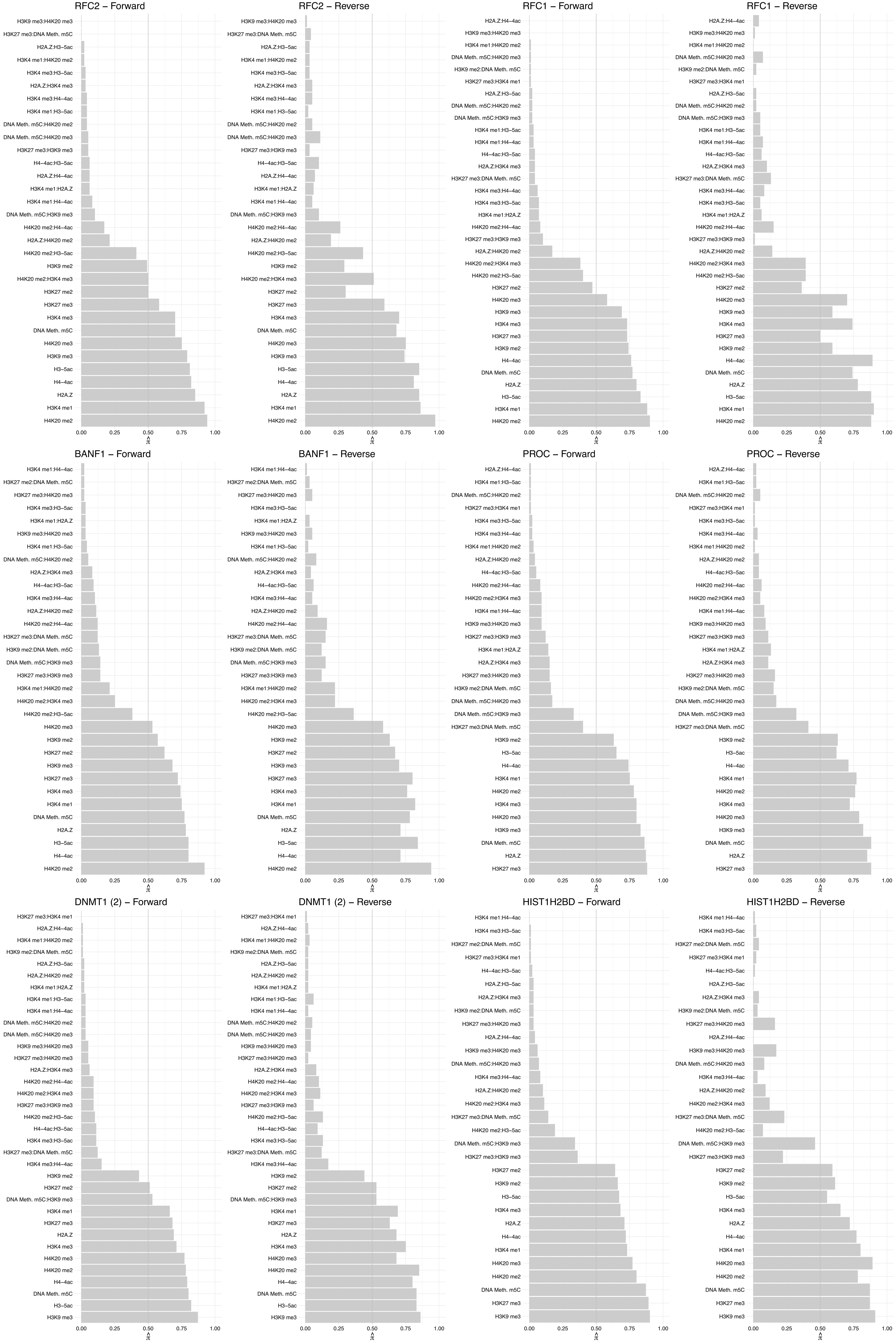
RFC2 – Forward
H3K9 me3:H4K20 me3
H3K27 me3:DNA Meth. m5C
H2A.Z:H3−5ac
H3K4 me1:H4K20 me2
H3K4 me3:H3−5ac
H2A.Z:H3K4 me3
H3K4 me3:H4−4ac
H3K4 me1:H3−5ac
DNA Meth. m5C:H4K20 me2
DNA Meth. m5C:H4K20 me3
H3K27 me3:H3K9 me3
H4−4ac:H3−5ac
H2A.Z:H4−4ac
H3K4 me1:H2A.Z
H3K4 me1:H4−4ac
DNA Meth. m5C:H3K9 me3
H4K20 me2:H4−4ac
H2A.Z:H4K20 me2
H4K20 me2:H3−5ac
H3K9 me2
H4K20 me2:H3K4 me3
H3K27 me2
H3K27 me3
H3K4 me3
DNA Meth. m5C
H4K20 me3
H3K9 me3
H3−5ac
H4−4ac
H2A.Z
H3K4 me1
H4K20 me2
0.00 0.25 0.50 0.75 1.00
π̂
RFC2 – Reverse
H3K9 me3:H4K20 me3
H3K27 me3:DNA Meth. m5C
H2A.Z:H3−5ac
H3K4 me1:H4K20 me2
H3K4 me3:H3−5ac
H2A.Z:H3K4 me3
H3K4 me3:H4−4ac
H3K4 me1:H3−5ac
DNA Meth. m5C:H4K20 me2
DNA Meth. m5C:H4K20 me3
H3K27 me3:H3K9 me3
H4−4ac:H3−5ac
H2A.Z:H4−4ac
H3K4 me1:H2A.Z
H3K4 me1:H4−4ac
DNA Meth. m5C:H3K9 me3
H4K20 me2:H4−4ac
H2A.Z:H4K20 me2
H4K20 me2:H3−5ac
H3K9 me2
H4K20 me2:H3K4 me3
H3K27 me2
H3K27 me3
H3K4 me3
DNA Meth. m5C
H4K20 me3
H3K9 me3
H3−5ac
H4−4ac
H2A.Z
H3K4 me1
H4K20 me2
0.00 0.25 0.50 0.75 1.00
π̂
RFC1 – Forward
H2A.Z:H4−4ac
H3K9 me3:H4K20 me3
H3K4 me1:H4K20 me2
DNA Meth. m5C:H4K20 me3
H3K9 me2:DNA Meth. m5C
H3K27 me3:H3K4 me1
H2A.Z:H3−5ac
DNA Meth. m5C:H4K20 me2
DNA Meth. m5C:H3K9 me3
H3K4 me1:H3−5ac
H3K4 me1:H4−4ac
H4−4ac:H3−5ac
H2A.Z:H3K4 me3
H3K27 me3:DNA Meth. m5C
H3K4 me3:H4−4ac
H3K4 me3:H3−5ac
H3K4 me1:H2A.Z
H4K20 me2:H4−4ac
H3K27 me3:H3K9 me3
H2A.Z:H4K20 me2
H4K20 me2:H3K4 me3
H4K20 me2:H3−5ac
H3K27 me2
H4K20 me3
H3K9 me3
H3K4 me3
H3K27 me3
H3K9 me2
H4−4ac
DNA Meth. m5C
H2A.Z
H3−5ac
H3K4 me1
H4K20 me2
0.00 0.25 0.50 0.75 1.00
π̂
RFC1 – Reverse
H2A.Z:H4−4ac
H3K9 me3:H4K20 me3
H3K4 me1:H4K20 me2
DNA Meth. m5C:H4K20 me3
H3K9 me2:DNA Meth. m5C
H3K27 me3:H3K4 me1
H2A.Z:H3−5ac
DNA Meth. m5C:H4K20 me2
DNA Meth. m5C:H3K9 me3
H3K4 me1:H3−5ac
H3K4 me1:H4−4ac
H4−4ac:H3−5ac
H2A.Z:H3K4 me3
H3K27 me3:DNA Meth. m5C
H3K4 me3:H4−4ac
H3K4 me3:H3−5ac
H3K4 me1:H2A.Z
H4K20 me2:H4−4ac
H3K27 me3:H3K9 me3
H2A.Z:H4K20 me2
H4K20 me2:H3K4 me3
H4K20 me2:H3−5ac
H3K27 me2
H4K20 me3
H3K9 me3
H3K4 me3
H3K27 me3
H3K9 me2
H4−4ac
DNA Meth. m5C
H2A.Z
H3−5ac
H3K4 me1
H4K20 me2
0.00 0.25 0.50 0.75 1.00
π̂
BANF1 – Forward
H3K4 me1:H4−4ac
H3K27 me2:DNA Meth. m5C
H3K27 me3:H4K20 me3
H3K4 me3:H3−5ac
H3K4 me1:H2A.Z
H3K9 me3:H4K20 me3
H3K4 me1:H3−5ac
DNA Meth. m5C:H4K20 me2
H2A.Z:H3K4 me3
H4−4ac:H3−5ac
H3K4 me3:H4−4ac
H2A.Z:H4K20 me2
H4K20 me2:H4−4ac
H3K27 me3:DNA Meth. m5C
H3K9 me2:DNA Meth. m5C
DNA Meth. m5C:H3K9 me3
H3K27 me3:H3K9 me3
H3K4 me1:H4K20 me2
H4K20 me2:H3K4 me3
H4K20 me2:H3−5ac
H4K20 me3
H3K9 me2
H3K27 me2
H3K9 me3
H3K27 me3
H3K4 me3
H3K4 me1
DNA Meth. m5C
H2A.Z
H3−5ac
H4−4ac
H4K20 me2
0.00 0.25 0.50 0.75 1.00
π̂
BANF1 – Reverse
H3K4 me1:H4−4ac
H3K27 me2:DNA Meth. m5C
H3K27 me3:H4K20 me3
H3K4 me3:H3−5ac
H3K4 me1:H2A.Z
H3K9 me3:H4K20 me3
H3K4 me1:H3−5ac
DNA Meth. m5C:H4K20 me2
H2A.Z:H3K4 me3
H4−4ac:H3−5ac
H3K4 me3:H4−4ac
H2A.Z:H4K20 me2
H4K20 me2:H4−4ac
H3K27 me3:DNA Meth. m5C
H3K9 me2:DNA Meth. m5C
DNA Meth. m5C:H3K9 me3
H3K27 me3:H3K9 me3
H3K4 me1:H4K20 me2
H4K20 me2:H3K4 me3
H4K20 me2:H3−5ac
H4K20 me3
H3K9 me2
H3K27 me2
H3K9 me3
H3K27 me3
H3K4 me3
H3K4 me1
DNA Meth. m5C
H2A.Z
H3−5ac
H4−4ac
H4K20 me2
0.00 0.25 0.50 0.75 1.00
π̂
PROC – Forward
H2A.Z:H4−4ac
H3K4 me1:H3−5ac
DNA Meth. m5C:H4K20 me2
H3K27 me3:H3K4 me1
H3K4 me3:H3−5ac
H3K4 me3:H4−4ac
H3K4 me1:H4K20 me2
H2A.Z:H4K20 me2
H4−4ac:H3−5ac
H4K20 me2:H4−4ac
H4K20 me2:H3K4 me3
H3K4 me1:H4−4ac
H3K9 me3:H4K20 me3
H3K27 me3:H3K9 me3
H3K4 me1:H2A.Z
H2A.Z:H3K4 me3
H3K27 me3:H4K20 me3
H3K9 me2:DNA Meth. m5C
DNA Meth. m5C:H4K20 me3
DNA Meth. m5C:H3K9 me3
H3K27 me3:DNA Meth. m5C
H3K9 me2
H3−5ac
H4−4ac
H3K4 me1
H4K20 me2
H3K4 me3
H4K20 me3
H3K9 me3
DNA Meth. m5C
H2A.Z
H3K27 me3
0.00 0.25 0.50 0.75 1.00
π̂
PROC – Reverse
H2A.Z:H4−4ac
H3K4 me1:H3−5ac
DNA Meth. m5C:H4K20 me2
H3K27 me3:H3K4 me1
H3K4 me3:H3−5ac
H3K4 me3:H4−4ac
H3K4 me1:H4K20 me2
H2A.Z:H4K20 me2
H4−4ac:H3−5ac
H4K20 me2:H4−4ac
H4K20 me2:H3K4 me3
H3K4 me1:H4−4ac
H3K9 me3:H4K20 me3
H3K27 me3:H3K9 me3
H3K4 me1:H2A.Z
H2A.Z:H3K4 me3
H3K27 me3:H4K20 me3
H3K9 me2:DNA Meth. m5C
DNA Meth. m5C:H4K20 me3
DNA Meth. m5C:H3K9 me3
H3K27 me3:DNA Meth. m5C
H3K9 me2
H3−5ac
H4−4ac
H3K4 me1
H4K20 me2
H3K4 me3
H4K20 me3
H3K9 me3
DNA Meth. m5C
H2A.Z
H3K27 me3
0.00 0.25 0.50 0.75 1.00
π̂
DNMT1 (2) – Forward
H3K27 me3:H3K4 me1
H2A.Z:H4−4ac
H3K4 me1:H4K20 me2
H3K9 me2:DNA Meth. m5C
H2A.Z:H3−5ac
H2A.Z:H4K20 me2
H3K4 me1:H2A.Z
H3K4 me1:H3−5ac
H3K4 me1:H4−4ac
DNA Meth. m5C:H4K20 me2
DNA Meth. m5C:H4K20 me3
H3K9 me3:H4K20 me3
H3K27 me3:H4K20 me3
H2A.Z:H3K4 me3
H4K20 me2:H4−4ac
H4K20 me2:H3K4 me3
H3K27 me3:H3K9 me3
H4K20 me2:H3−5ac
H4−4ac:H3−5ac
H3K4 me3:H3−5ac
H3K27 me3:DNA Meth. m5C
H3K4 me3:H4−4ac
H3K9 me2
H3K27 me2
DNA Meth. m5C:H3K9 me3
H3K4 me1
H3K27 me3
H2A.Z
H3K4 me3
H4K20 me3
H4K20 me2
H4−4ac
DNA Meth. m5C
H3−5ac
H3K9 me3
0.00 0.25 0.50 0.75 1.00
π̂
DNMT1 (2) – Reverse
H3K27 me3:H3K4 me1
H2A.Z:H4−4ac
H3K4 me1:H4K20 me2
H3K9 me2:DNA Meth. m5C
H2A.Z:H3−5ac
H2A.Z:H4K20 me2
H3K4 me1:H2A.Z
H3K4 me1:H3−5ac
H3K4 me1:H4−4ac
DNA Meth. m5C:H4K20 me2
DNA Meth. m5C:H4K20 me3
H3K9 me3:H4K20 me3
H3K27 me3:H4K20 me3
H2A.Z:H3K4 me3
H4K20 me2:H4−4ac
H4K20 me2:H3K4 me3
H3K27 me3:H3K9 me3
H4K20 me2:H3−5ac
H4−4ac:H3−5ac
H3K4 me3:H3−5ac
H3K27 me3:DNA Meth. m5C
H3K4 me3:H4−4ac
H3K9 me2
H3K27 me2
DNA Meth. m5C:H3K9 me3
H3K4 me1
H3K27 me3
H2A.Z
H3K4 me3
H4K20 me3
H4K20 me2
H4−4ac
DNA Meth. m5C
H3−5ac
H3K9 me3
0.00 0.25 0.50 0.75 1.00
π̂
HIST1H2BD – Forward
H3K4 me1:H4−4ac
H3K4 me3:H3−5ac
H3K27 me2:DNA Meth. m5C
H3K27 me3:H3K4 me1
H4−4ac:H3−5ac
H2A.Z:H3−5ac
H2A.Z:H3K4 me3
H3K9 me2:DNA Meth. m5C
H3K27 me3:H4K20 me3
H2A.Z:H4−4ac
H3K9 me3:H4K20 me3
DNA Meth. m5C:H4K20 me3
H3K4 me3:H4−4ac
H2A.Z:H4K20 me2
H4K20 me2:H3K4 me3
H3K27 me3:DNA Meth. m5C
H4K20 me2:H3−5ac
DNA Meth. m5C:H3K9 me3
H3K27 me3:H3K9 me3
H3K27 me2
H3K9 me2
H3−5ac
H3K4 me3
H2A.Z
H4−4ac
H3K4 me1
H4K20 me3
H4K20 me2
DNA Meth. m5C
H3K27 me3
H3K9 me3
0.00 0.25 0.50 0.75 1.00
π̂
HIST1H2BD – Reverse
H3K4 me1:H4−4ac
H3K4 me3:H3−5ac
H3K27 me2:DNA Meth. m5C
H3K27 me3:H3K4 me1
H4−4ac:H3−5ac
H2A.Z:H3−5ac
H2A.Z:H3K4 me3
H3K9 me2:DNA Meth. m5C
H3K27 me3:H4K20 me3
H2A.Z:H4−4ac
H3K9 me3:H4K20 me3
DNA Meth. m5C:H4K20 me3
H3K4 me3:H4−4ac
H2A.Z:H4K20 me2
H4K20 me2:H3K4 me3
H3K27 me3:DNA Meth. m5C
H4K20 me2:H3−5ac
DNA Meth. m5C:H3K9 me3
H3K27 me3:H3K9 me3
H3K27 me2
H3K9 me2
H3−5ac
H3K4 me3
H2A.Z
H4−4ac
H3K4 me1
H4K20 me3
H4K20 me2
DNA Meth. m5C
H3K27 me3
H3K9 me3
0.00 0.25 0.50 0.75 1.00
π̂

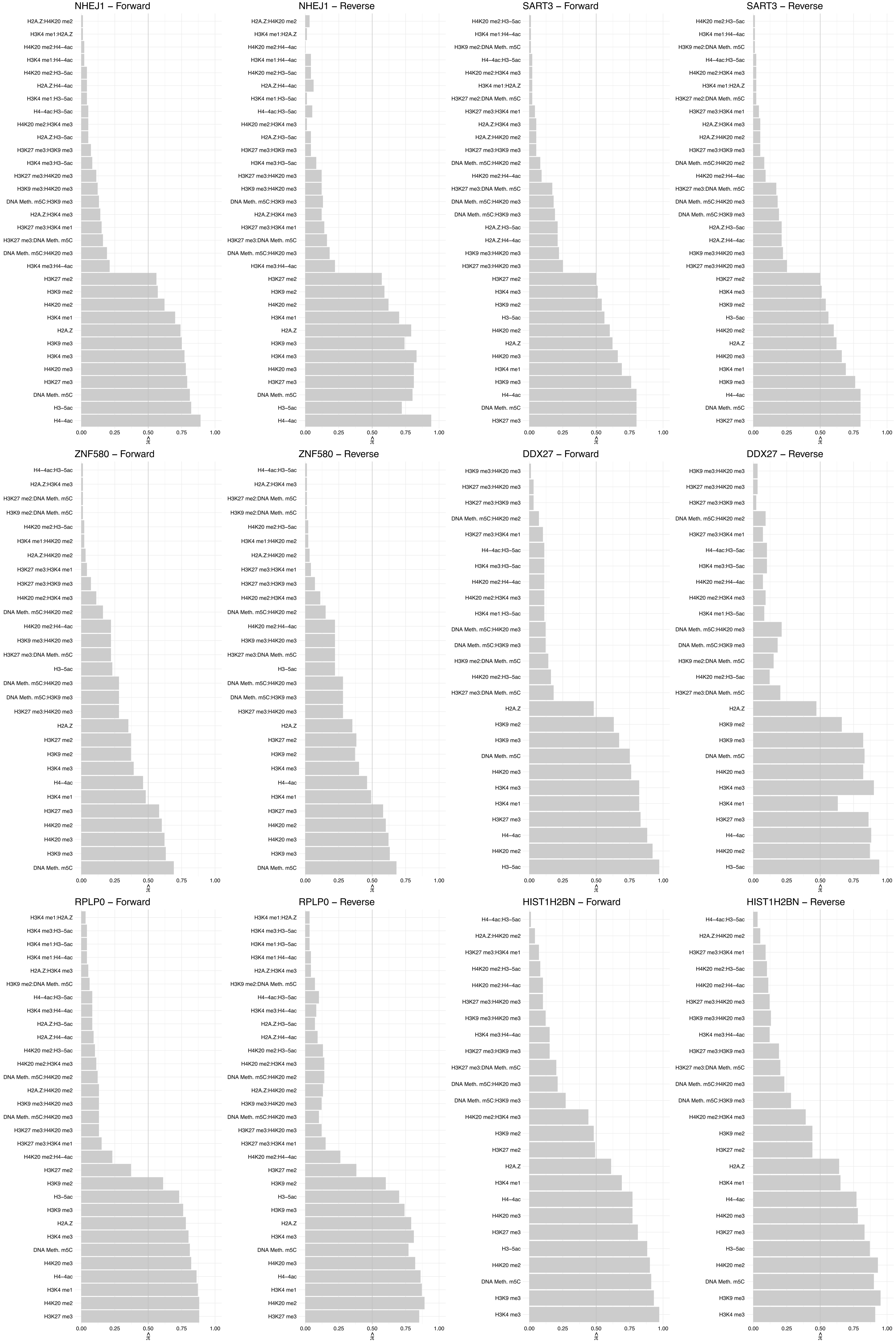
NHEJ1 – Forward
H2A.Z:H4K20 me2
H3K4 me1:H2A.Z
H4K20 me2:H4–4ac
H3K4 me1:H4–4ac
H4K20 me2:H3–5ac
H2A.Z:H4–4ac
H3K4 me1:H3–5ac
H4–4ac:H3–5ac
H4K20 me2:H3K4 me3
H2A.Z:H3–5ac
H3K27 me3:H3K9 me3
H3K4 me3:H3–5ac
H3K27 me3:H4K20 me3
H3K9 me3:H4K20 me3
DNA Meth. m5C:H3K9 me3
H2A.Z:H3K4 me3
H3K27 me3:H3K4 me1
H3K27 me3:DNA Meth. m5C
DNA Meth. m5C:H4K20 me3
H3K4 me3:H4–4ac
H3K27 me2
H3K9 me2
H4K20 me2
H3K4 me1
H2A.Z
H3K9 me3
H3K4 me3
H4K20 me3
H3K27 me3
DNA Meth. m5C
H3–5ac
H4–4ac
0.00 0.25 0.50 0.75 1.00
$\hat{\pi}$
NHEJ1 – Reverse
H2A.Z:H4K20 me2
H3K4 me1:H2A.Z
H4K20 me2:H4–4ac
H3K4 me1:H4–4ac
H4K20 me2:H3–5ac
H2A.Z:H4–4ac
H3K4 me1:H3–5ac
H4–4ac:H3–5ac
H4K20 me2:H3K4 me3
H2A.Z:H3–5ac
H3K27 me3:H3K9 me3
H3K4 me3:H3–5ac
H3K27 me3:H4K20 me3
H3K9 me3:H4K20 me3
DNA Meth. m5C:H3K9 me3
H2A.Z:H3K4 me3
H3K27 me3:H3K4 me1
H3K27 me3:DNA Meth. m5C
DNA Meth. m5C:H4K20 me3
H3K4 me3:H4–4ac
H3K27 me2
H3K9 me2
H4K20 me2
H3K4 me1
H2A.Z
H3K9 me3
H3K4 me3
H4K20 me3
H3K27 me3
DNA Meth. m5C
H3–5ac
H4–4ac
0.00 0.25 0.50 0.75 1.00
$\hat{\pi}$
SART3 – Forward
H4K20 me2:H3–5ac
H3K4 me1:H4–4ac
H3K9 me2:DNA Meth. m5C
H4–4ac:H3–5ac
H4K20 me2:H3K4 me3
H3K4 me1:H2A.Z
H3K27 me2:DNA Meth. m5C
H3K27 me3:H3K4 me1
H2A.Z:H3K4 me3
H2A.Z:H4K20 me2
H3K27 me3:H3K9 me3
DNA Meth. m5C:H4K20 me2
H4K20 me2:H4–4ac
H3K27 me3:DNA Meth. m5C
DNA Meth. m5C:H4K20 me3
DNA Meth. m5C:H3K9 me3
H2A.Z:H3–5ac
H2A.Z:H4–4ac
H3K9 me3:H4K20 me3
H3K27 me3:H4K20 me3
H3K27 me2
H3K4 me3
H3K9 me2
H3–5ac
H4K20 me2
H2A.Z
H4K20 me3
H3K4 me1
H3K9 me3
H4–4ac
DNA Meth. m5C
H3K27 me3
0.00 0.25 0.50 0.75 1.00
$\hat{\pi}$
SART3 – Reverse
H4K20 me2:H3–5ac
H3K4 me1:H4–4ac
H3K9 me2:DNA Meth. m5C
H4–4ac:H3–5ac
H4K20 me2:H3K4 me3
H3K4 me1:H2A.Z
H3K27 me2:DNA Meth. m5C
H3K27 me3:H3K4 me1
H2A.Z:H3K4 me3
H2A.Z:H4K20 me2
H3K27 me3:H3K9 me3
DNA Meth. m5C:H4K20 me2
H4K20 me2:H4–4ac
H3K27 me3:DNA Meth. m5C
DNA Meth. m5C:H4K20 me3
DNA Meth. m5C:H3K9 me3
H2A.Z:H3–5ac
H2A.Z:H4–4ac
H3K9 me3:H4K20 me3
H3K27 me3:H4K20 me3
H3K27 me2
H3K4 me3
H3K9 me2
H3–5ac
H4K20 me2
H2A.Z
H4K20 me3
H3K4 me1
H3K9 me3
H4–4ac
DNA Meth. m5C
H3K27 me3
0.00 0.25 0.50 0.75 1.00
$\hat{\pi}$
ZNF580 – Forward
H4–4ac:H3–5ac
H2A.Z:H3K4 me3
H3K27 me2:DNA Meth. m5C
H3K9 me2:DNA Meth. m5C
H4K20 me2:H3–5ac
H3K4 me1:H4K20 me2
H2A.Z:H4K20 me2
H3K27 me3:H3K4 me1
H3K27 me3:H3K9 me3
H4K20 me2:H3K4 me3
DNA Meth. m5C:H4K20 me2
H4K20 me2:H4–4ac
H3K9 me3:H4K20 me3
H3K27 me3:DNA Meth. m5C
H3–5ac
DNA Meth. m5C:H4K20 me3
DNA Meth. m5C:H3K9 me3
H3K27 me3:H4K20 me3
H2A.Z
H3K27 me2
H3K9 me2
H3K4 me3
H4–4ac
H3K4 me1
H3K27 me3
H4K20 me2
H4K20 me3
H3K9 me3
DNA Meth. m5C
0.00 0.25 0.50 0.75 1.00
$\hat{\pi}$
ZNF580 – Reverse
H4–4ac:H3–5ac
H2A.Z:H3K4 me3
H3K27 me2:DNA Meth. m5C
H3K9 me2:DNA Meth. m5C
H4K20 me2:H3–5ac
H3K4 me1:H4K20 me2
H2A.Z:H4K20 me2
H3K27 me3:H3K4 me1
H3K27 me3:H3K9 me3
H4K20 me2:H3K4 me3
DNA Meth. m5C:H4K20 me2
H4K20 me2:H4–4ac
H3K9 me3:H4K20 me3
H3K27 me3:DNA Meth. m5C
H3–5ac
DNA Meth. m5C:H4K20 me3
DNA Meth. m5C:H3K9 me3
H3K27 me3:H4K20 me3
H2A.Z
H3K27 me2
H3K9 me2
H3K4 me3
H4–4ac
H3K4 me1
H3K27 me3
H4K20 me2
H4K20 me3
H3K9 me3
DNA Meth. m5C
0.00 0.25 0.50 0.75 1.00
$\hat{\pi}$
DDX27 – Forward
H3K9 me3:H4K20 me3
H3K27 me3:H4K20 me3
H3K27 me3:H3K9 me3
DNA Meth. m5C:H4K20 me2
H3K27 me3:H3K4 me1
H4–4ac:H3–5ac
H3K4 me3:H3–5ac
H4K20 me2:H4–4ac
H4K20 me2:H3K4 me3
H3K4 me1:H3–5ac
DNA Meth. m5C:H4K20 me3
DNA Meth. m5C:H3K9 me3
H3K9 me2:DNA Meth. m5C
H4K20 me2:H3–5ac
H3K27 me3:DNA Meth. m5C
H2A.Z
H3K9 me2
H3K9 me3
DNA Meth. m5C
H4K20 me3
H3K4 me3
H3K4 me1
H3K27 me3
H4–4ac
H4K20 me2
H3–5ac
0.00 0.25 0.50 0.75 1.00
$\hat{\pi}$
DDX27 – Reverse
H3K9 me3:H4K20 me3
H3K27 me3:H4K20 me3
H3K27 me3:H3K9 me3
DNA Meth. m5C:H4K20 me2
H3K27 me3:H3K4 me1
H4–4ac:H3–5ac
H3K4 me3:H3–5ac
H4K20 me2:H4–4ac
H4K20 me2:H3K4 me3
H3K4 me1:H3–5ac
DNA Meth. m5C:H4K20 me3
DNA Meth. m5C:H3K9 me3
H3K9 me2:DNA Meth. m5C
H4K20 me2:H3–5ac
H3K27 me3:DNA Meth. m5C
H2A.Z
H3K9 me2
H3K9 me3
DNA Meth. m5C
H4K20 me3
H3K4 me3
H3K4 me1
H3K27 me3
H4–4ac
H4K20 me2
H3–5ac
0.00 0.25 0.50 0.75 1.00
$\hat{\pi}$
RPLP0 – Forward
H3K4 me1:H2A.Z
H3K4 me3:H3–5ac
H3K4 me1:H3–5ac
H3K4 me1:H4–4ac
H2A.Z:H3K4 me3
H3K9 me2:DNA Meth. m5C
H4–4ac:H3–5ac
H3K4 me3:H4–4ac
H2A.Z:H3–5ac
H2A.Z:H4–4ac
H4K20 me2:H3–5ac
H4K20 me2:H3K4 me3
DNA Meth. m5C:H4K20 me2
H2A.Z:H4K20 me2
H3K9 me3:H4K20 me3
DNA Meth. m5C:H4K20 me3
H3K27 me3:H4K20 me3
H3K27 me3:H3K4 me1
H4K20 me2:H4–4ac
H3K27 me2
H3K9 me2
H3–5ac
H3K9 me3
H2A.Z
H3K4 me3
DNA Meth. m5C
H4K20 me3
H4–4ac
H3K4 me1
H4K20 me2
H3K27 me3
0.00 0.25 0.50 0.75 1.00
$\hat{\pi}$
RPLP0 – Reverse
H3K4 me1:H2A.Z
H3K4 me3:H3–5ac
H3K4 me1:H3–5ac
H3K4 me1:H4–4ac
H2A.Z:H3K4 me3
H3K9 me2:DNA Meth. m5C
H4–4ac:H3–5ac
H3K4 me3:H4–4ac
H2A.Z:H3–5ac
H2A.Z:H4–4ac
H4K20 me2:H3–5ac
H4K20 me2:H3K4 me3
DNA Meth. m5C:H4K20 me2
H2A.Z:H4K20 me2
H3K9 me3:H4K20 me3
DNA Meth. m5C:H4K20 me3
H3K27 me3:H4K20 me3
H3K27 me3:H3K4 me1
H4K20 me2:H4–4ac
H3K27 me2
H3K9 me2
H3–5ac
H3K9 me3
H2A.Z
H3K4 me3
DNA Meth. m5C
H4K20 me3
H4–4ac
H3K4 me1
H4K20 me2
H3K27 me3
0.00 0.25 0.50 0.75 1.00
$\hat{\pi}$
HIST1H2BN – Forward
H4–4ac:H3–5ac
H2A.Z:H4K20 me2
H3K27 me3:H3K4 me1
H4K20 me2:H3–5ac
H4K20 me2:H4–4ac
H3K27 me3:H4K20 me3
H3K9 me3:H4K20 me3
H3K4 me3:H4–4ac
H3K27 me3:H3K9 me3
H3K27 me3:DNA Meth. m5C
DNA Meth. m5C:H4K20 me3
DNA Meth. m5C:H3K9 me3
H4K20 me2:H3K4 me3
H3K9 me2
H3K27 me2
H2A.Z
H3K4 me1
H4–4ac
H4K20 me3
H3K27 me3
H3–5ac
H4K20 me2
DNA Meth. m5C
H3K9 me3
H3K4 me3
0.00 0.25 0.50 0.75 1.00
$\hat{\pi}$
HIST1H2BN – Reverse
H4–4ac:H3–5ac
H2A.Z:H4K20 me2
H3K27 me3:H3K4 me1
H4K20 me2:H3–5ac
H4K20 me2:H4–4ac
H3K27 me3:H4K20 me3
H3K9 me3:H4K20 me3
H3K4 me3:H4–4ac
H3K27 me3:H3K9 me3
H3K27 me3:DNA Meth. m5C
DNA Meth. m5C:H4K20 me3
DNA Meth. m5C:H3K9 me3
H4K20 me2:H3K4 me3
H3K9 me2
H3K27 me2
H2A.Z
H3K4 me1
H4–4ac
H4K20 me3
H3K27 me3
H3–5ac
H4K20 me2
DNA Meth. m5C
H3K9 me3
H3K4 me3
0.00 0.25 0.50 0.75 1.00
$\hat{\pi}$

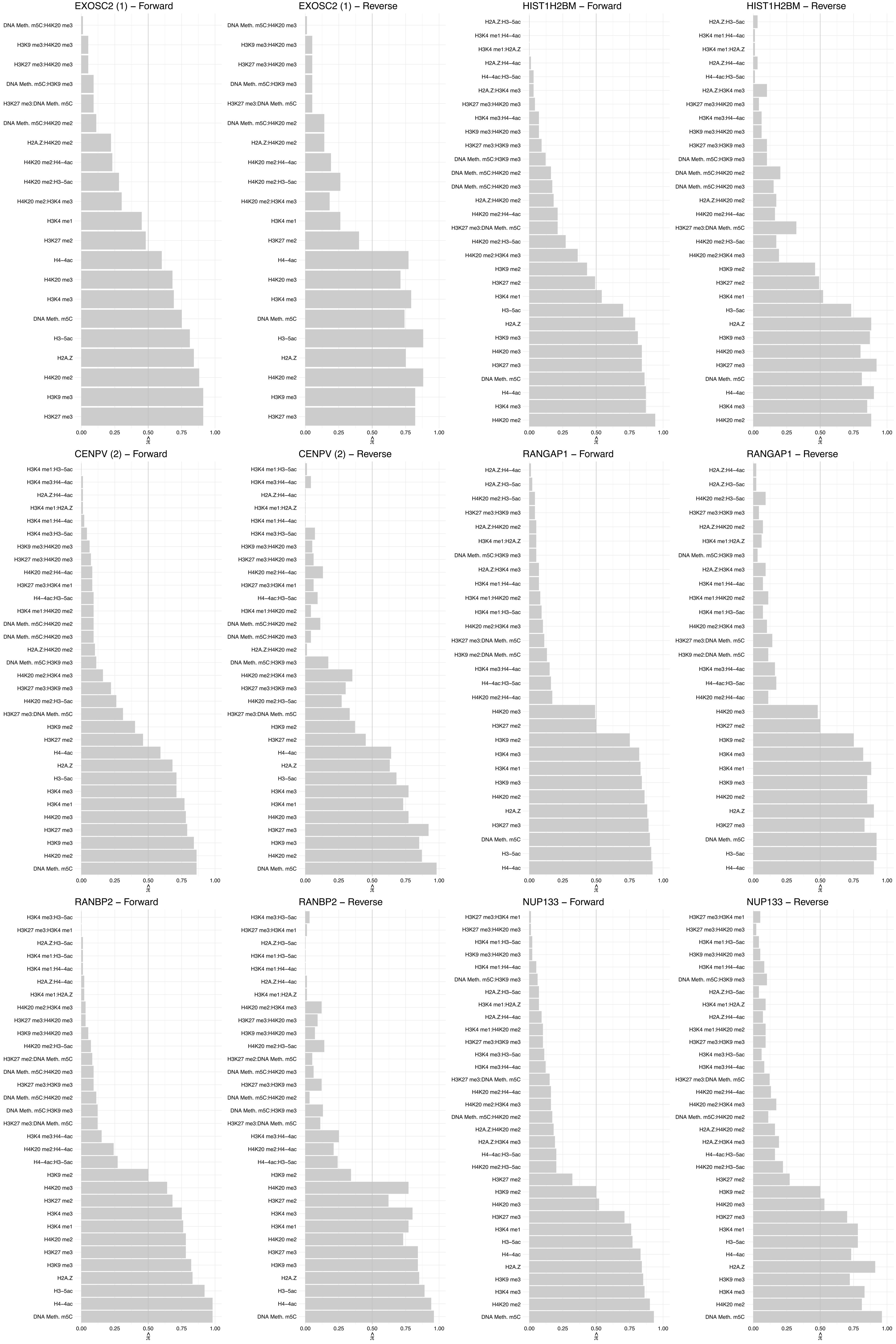
EXOSC2 (1) – Forward
EXOSC2 (1) – Reverse
HIST1H2BM – Forward
HIST1H2BM – Reverse
CENPV (2) – Forward
CENPV (2) – Reverse
RANGAP1 – Forward
RANGAP1 – Reverse
RANBP2 – Forward
RANBP2 – Reverse
NUP133 – Forward
NUP133 – Reverse
0.00 0.25 0.50 0.75 1.00
$\hat{\pi}$

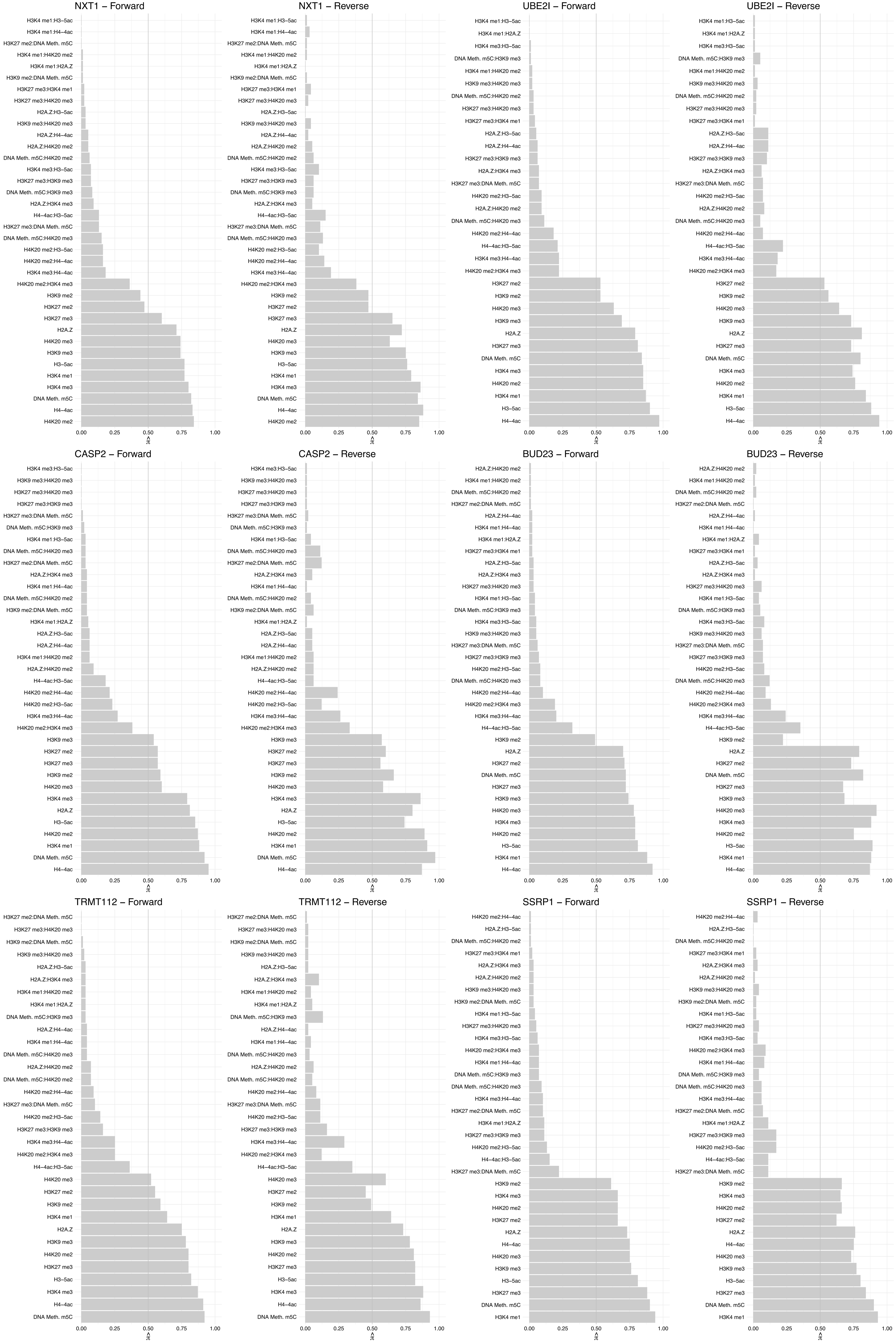

NXT1 – Forward
NXT1 – Reverse
H3K4 me1:H3−5ac
H3K4 me1:H4−4ac
H3K27 me2:DNA Meth. m5C
H3K4 me1:H4K20 me2
H3K4 me1:H2A.Z
H3K9 me2:DNA Meth. m5C
H3K27 me3:H3K4 me1
H3K27 me3:H4K20 me3
H2A.Z:H3−5ac
H3K9 me3:H4K20 me3
H2A.Z:H4−4ac
H2A.Z:H4K20 me2
DNA Meth. m5C:H4K20 me2
H3K4 me3:H3−5ac
H3K27 me3:H3K9 me3
DNA Meth. m5C:H3K9 me3
H2A.Z:H3K4 me3
H4−4ac:H3−5ac
H3K27 me3:DNA Meth. m5C
DNA Meth. m5C:H4K20 me3
H4K20 me2:H3−5ac
H4K20 me2:H4−4ac
H3K4 me3:H4−4ac
H4K20 me2:H3K4 me3
H3K9 me2
H3K27 me2
H3K27 me3
H2A.Z
H4K20 me3
H3K9 me3
H3−5ac
H3K4 me1
H3K4 me3
DNA Meth. m5C
H4−4ac
H4K20 me2
0.00
0.25
0.50
0.75
1.00
$\hat{\pi}$
UBE2I – Forward
UBE2I – Reverse
H3K4 me1:H3−5ac
H3K4 me1:H2A.Z
H3K4 me3:H3−5ac
DNA Meth. m5C:H3K9 me3
H3K4 me1:H4K20 me2
H3K9 me3:H4K20 me3
DNA Meth. m5C:H4K20 me2
H3K27 me3:H4K20 me3
H3K27 me3:H3K4 me1
H2A.Z:H3−5ac
H2A.Z:H4−4ac
H3K27 me3:H3K9 me3
H2A.Z:H3K4 me3
H3K27 me3:DNA Meth. m5C
H4K20 me2:H3−5ac
H2A.Z:H4K20 me2
DNA Meth. m5C:H4K20 me3
H4K20 me2:H4−4ac
H4−4ac:H3−5ac
H3K4 me3:H4−4ac
H4K20 me2:H3K4 me3
H3K27 me2
H3K9 me2
H4K20 me3
H3K9 me3
H2A.Z
H3K27 me3
DNA Meth. m5C
H3K4 me3
H4K20 me2
H3K4 me1
H3−5ac
H4−4ac
CASP2 – Forward
CASP2 – Reverse
H3K4 me3:H3−5ac
H3K9 me3:H4K20 me3
H3K27 me3:H4K20 me3
H3K27 me3:H3K9 me3
H3K27 me3:DNA Meth. m5C
DNA Meth. m5C:H3K9 me3
H3K4 me1:H3−5ac
DNA Meth. m5C:H4K20 me3
H3K27 me2:DNA Meth. m5C
H2A.Z:H3K4 me3
H3K4 me1:H4−4ac
DNA Meth. m5C:H4K20 me2
H3K9 me2:DNA Meth. m5C
H3K4 me1:H2A.Z
H2A.Z:H3−5ac
H2A.Z:H4−4ac
H3K4 me1:H4K20 me2
H2A.Z:H4K20 me2
H4−4ac:H3−5ac
H4K20 me2:H4−4ac
H4K20 me2:H3−5ac
H3K4 me3:H4−4ac
H4K20 me2:H3K4 me3
H3K9 me3
H3K27 me2
H3K27 me3
H3K9 me2
H4K20 me3
H3K4 me3
H2A.Z
H3−5ac
H4K20 me2
H3K4 me1
DNA Meth. m5C
H4−4ac
BUD23 – Forward
BUD23 – Reverse
H2A.Z:H4K20 me2
H3K4 me1:H4K20 me2
DNA Meth. m5C:H4K20 me2
H3K27 me2:DNA Meth. m5C
H2A.Z:H4−4ac
H3K4 me1:H4−4ac
H3K4 me1:H2A.Z
H3K27 me3:H3K4 me1
H2A.Z:H3−5ac
H2A.Z:H3K4 me3
H3K27 me3:H4K20 me3
H3K4 me1:H3−5ac
DNA Meth. m5C:H3K9 me3
H3K4 me3:H3−5ac
H3K9 me3:H4K20 me3
H3K27 me3:DNA Meth. m5C
H3K27 me3:H3K9 me3
H4K20 me2:H3−5ac
DNA Meth. m5C:H4K20 me3
H4K20 me2:H4−4ac
H4K20 me2:H3K4 me3
H3K4 me3:H4−4ac
H4−4ac:H3−5ac
H3K9 me2
H2A.Z
H3K27 me2
DNA Meth. m5C
H3K27 me3
H3K9 me3
H4K20 me3
H3K4 me3
H4K20 me2
H3−5ac
H3K4 me1
H4−4ac
TRMT112 – Forward
TRMT112 – Reverse
H3K27 me2:DNA Meth. m5C
H3K27 me3:H4K20 me3
H3K9 me2:DNA Meth. m5C
H3K9 me3:H4K20 me3
H2A.Z:H3−5ac
H2A.Z:H3K4 me3
H3K4 me1:H4K20 me2
H3K4 me1:H2A.Z
DNA Meth. m5C:H3K9 me3
H2A.Z:H4−4ac
H3K4 me1:H4−4ac
DNA Meth. m5C:H4K20 me3
H2A.Z:H4K20 me2
DNA Meth. m5C:H4K20 me2
H4K20 me2:H4−4ac
H3K27 me3:DNA Meth. m5C
H4K20 me2:H3−5ac
H3K27 me3:H3K9 me3
H3K4 me3:H4−4ac
H4K20 me2:H3K4 me3
H4−4ac:H3−5ac
H4K20 me3
H3K27 me2
H3K9 me2
H3K4 me1
H2A.Z
H3K9 me3
H4K20 me2
H3K27 me3
H3−5ac
H3K4 me3
H4−4ac
DNA Meth. m5C
SSRP1 – Forward
SSRP1 – Reverse
H4K20 me2:H4−4ac
H2A.Z:H3−5ac
DNA Meth. m5C:H4K20 me2
H3K27 me3:H3K4 me1
H2A.Z:H3K4 me3
H2A.Z:H4K20 me2
H3K9 me3:H4K20 me3
H3K9 me2:DNA Meth. m5C
H3K4 me1:H3−5ac
H3K27 me3:H4K20 me3
H3K4 me3:H3−5ac
H4K20 me2:H3K4 me3
H3K4 me1:H4−4ac
DNA Meth. m5C:H3K9 me3
DNA Meth. m5C:H4K20 me3
H3K4 me3:H4−4ac
H3K27 me2:DNA Meth. m5C
H3K4 me1:H2A.Z
H3K27 me3:H3K9 me3
H4K20 me2:H3−5ac
H4−4ac:H3−5ac
H3K27 me3:DNA Meth. m5C
H3K9 me2
H3K4 me3
H4K20 me2
H3K27 me2
H2A.Z
H4−4ac
H4K20 me3
H3K9 me3
H3−5ac
H3K27 me3
DNA Meth. m5C
H3K4 me1

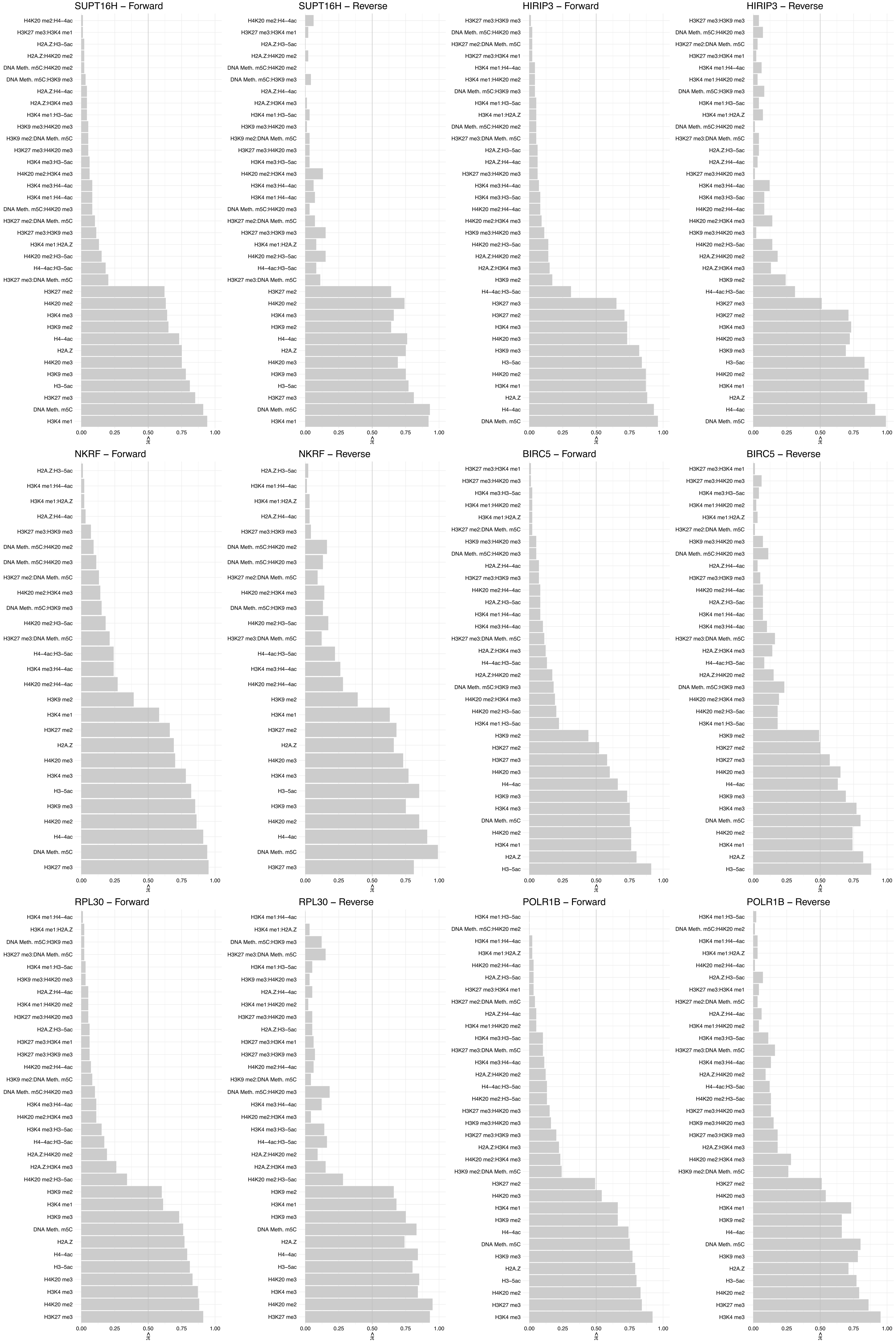
SUPT16H – Forward
H4K20 me2:H4−4ac
H3K27 me3:H3K4 me1
H2A.Z:H3−5ac
H2A.Z:H4K20 me2
DNA Meth. m5C:H4K20 me2
DNA Meth. m5C:H3K9 me3
H2A.Z:H4−4ac
H2A.Z:H3K4 me3
H3K4 me1:H3−5ac
H3K9 me3:H4K20 me3
H3K9 me2:DNA Meth. m5C
H3K27 me3:H4K20 me3
H3K4 me3:H3−5ac
H4K20 me2:H3K4 me3
H3K4 me3:H4−4ac
H3K4 me1:H4−4ac
DNA Meth. m5C:H4K20 me3
H3K27 me2:DNA Meth. m5C
H3K27 me3:H3K9 me3
H3K4 me1:H2A.Z
H4K20 me2:H3−5ac
H4−4ac:H3−5ac
H3K27 me3:DNA Meth. m5C
H3K27 me2
H4K20 me2
H3K4 me3
H3K9 me2
H4−4ac
H2A.Z
H4K20 me3
H3K9 me3
H3−5ac
H3K27 me3
DNA Meth. m5C
H3K4 me1
0.00
0.25
0.50
0.75
1.00
$\hat{\pi}$
SUPT16H – Reverse
H4K20 me2:H4−4ac
H3K27 me3:H3K4 me1
H2A.Z:H3−5ac
H2A.Z:H4K20 me2
DNA Meth. m5C:H4K20 me2
DNA Meth. m5C:H3K9 me3
H2A.Z:H4−4ac
H2A.Z:H3K4 me3
H3K4 me1:H3−5ac
H3K9 me3:H4K20 me3
H3K9 me2:DNA Meth. m5C
H3K27 me3:H4K20 me3
H3K4 me3:H3−5ac
H4K20 me2:H3K4 me3
H3K4 me3:H4−4ac
H3K4 me1:H4−4ac
DNA Meth. m5C:H4K20 me3
H3K27 me2:DNA Meth. m5C
H3K27 me3:H3K9 me3
H3K4 me1:H2A.Z
H4K20 me2:H3−5ac
H4−4ac:H3−5ac
H3K27 me3:DNA Meth. m5C
H3K27 me2
H4K20 me2
H3K4 me3
H3K9 me2
H4−4ac
H2A.Z
H4K20 me3
H3K9 me3
H3−5ac
H3K27 me3
DNA Meth. m5C
H3K4 me1
0.00
0.25
0.50
0.75
1.00
$\hat{\pi}$
HIRIP3 – Forward
H3K27 me3:H3K9 me3
DNA Meth. m5C:H4K20 me3
H3K27 me2:DNA Meth. m5C
H3K27 me3:H3K4 me1
H3K4 me1:H4−4ac
H3K4 me1:H4K20 me2
DNA Meth. m5C:H3K9 me3
H3K4 me1:H3−5ac
H3K4 me1:H2A.Z
DNA Meth. m5C:H4K20 me2
H3K27 me3:DNA Meth. m5C
H2A.Z:H3−5ac
H2A.Z:H4−4ac
H3K27 me3:H4K20 me3
H3K4 me3:H4−4ac
H3K4 me3:H3−5ac
H4K20 me2:H4−4ac
H4K20 me2:H3K4 me3
H3K9 me3:H4K20 me3
H4K20 me2:H3−5ac
H2A.Z:H4K20 me2
H2A.Z:H3K4 me3
H3K9 me2
H4−4ac:H3−5ac
H3K27 me3
H3K27 me2
H3K4 me3
H4K20 me3
H3K9 me3
H3−5ac
H4K20 me2
H3K4 me1
H2A.Z
H4−4ac
DNA Meth. m5C
0.00
0.25
0.50
0.75
1.00
$\hat{\pi}$
HIRIP3 – Reverse
H3K27 me3:H3K9 me3
DNA Meth. m5C:H4K20 me3
H3K27 me2:DNA Meth. m5C
H3K27 me3:H3K4 me1
H3K4 me1:H4−4ac
H3K4 me1:H4K20 me2
DNA Meth. m5C:H3K9 me3
H3K4 me1:H3−5ac
H3K4 me1:H2A.Z
DNA Meth. m5C:H4K20 me2
H3K27 me3:DNA Meth. m5C
H2A.Z:H3−5ac
H2A.Z:H4−4ac
H3K27 me3:H4K20 me3
H3K4 me3:H4−4ac
H3K4 me3:H3−5ac
H4K20 me2:H4−4ac
H4K20 me2:H3K4 me3
H3K9 me3:H4K20 me3
H4K20 me2:H3−5ac
H2A.Z:H4K20 me2
H2A.Z:H3K4 me3
H3K9 me2
H4−4ac:H3−5ac
H3K27 me3
H3K27 me2
H3K4 me3
H4K20 me3
H3K9 me3
H3−5ac
H4K20 me2
H3K4 me1
H2A.Z
H4−4ac
DNA Meth. m5C
0.00
0.25
0.50
0.75
1.00
$\hat{\pi}$
NKRF – Forward
H2A.Z:H3−5ac
H3K4 me1:H4−4ac
H3K4 me1:H2A.Z
H2A.Z:H4−4ac
H3K27 me3:H3K9 me3
DNA Meth. m5C:H4K20 me2
DNA Meth. m5C:H4K20 me3
H3K27 me2:DNA Meth. m5C
H4K20 me2:H3K4 me3
DNA Meth. m5C:H3K9 me3
H4K20 me2:H3−5ac
H3K27 me3:DNA Meth. m5C
H4−4ac:H3−5ac
H3K4 me3:H4−4ac
H4K20 me2:H4−4ac
H3K9 me2
H3K4 me1
H3K27 me2
H2A.Z
H4K20 me3
H3K4 me3
H3−5ac
H3K9 me3
H4K20 me2
H4−4ac
DNA Meth. m5C
H3K27 me3
0.00
0.25
0.50
0.75
1.00
$\hat{\pi}$
NKRF – Reverse
H2A.Z:H3−5ac
H3K4 me1:H4−4ac
H3K4 me1:H2A.Z
H2A.Z:H4−4ac
H3K27 me3:H3K9 me3
DNA Meth. m5C:H4K20 me2
DNA Meth. m5C:H4K20 me3
H3K27 me2:DNA Meth. m5C
H4K20 me2:H3K4 me3
DNA Meth. m5C:H3K9 me3
H4K20 me2:H3−5ac
H3K27 me3:DNA Meth. m5C
H4−4ac:H3−5ac
H3K4 me3:H4−4ac
H4K20 me2:H4−4ac
H3K9 me2
H3K4 me1
H3K27 me2
H2A.Z
H4K20 me3
H3K4 me3
H3−5ac
H3K9 me3
H4K20 me2
H4−4ac
DNA Meth. m5C
H3K27 me3
0.00
0.25
0.50
0.75
1.00
$\hat{\pi}$
BIRC5 – Forward
H3K27 me3:H3K4 me1
H3K27 me3:H4K20 me3
H3K4 me3:H3−5ac
H3K4 me1:H4K20 me2
H3K4 me1:H2A.Z
H3K27 me2:DNA Meth. m5C
H3K9 me3:H4K20 me3
DNA Meth. m5C:H4K20 me3
H2A.Z:H4−4ac
H3K27 me3:H3K9 me3
H4K20 me2:H4−4ac
H2A.Z:H3−5ac
H3K4 me1:H4−4ac
H3K4 me3:H4−4ac
H3K27 me3:DNA Meth. m5C
H2A.Z:H3K4 me3
H4−4ac:H3−5ac
H2A.Z:H4K20 me2
DNA Meth. m5C:H3K9 me3
H4K20 me2:H3K4 me3
H4K20 me2:H3−5ac
H3K4 me1:H3−5ac
H3K9 me2
H3K27 me2
H3K27 me3
H4K20 me3
H4−4ac
H3K9 me3
H3K4 me3
DNA Meth. m5C
H4K20 me2
H3K4 me1
H2A.Z
H3−5ac
0.00
0.25
0.50
0.75
1.00
$\hat{\pi}$
BIRC5 – Reverse
H3K27 me3:H3K4 me1
H3K27 me3:H4K20 me3
H3K4 me3:H3−5ac
H3K4 me1:H4K20 me2
H3K4 me1:H2A.Z
H3K27 me2:DNA Meth. m5C
H3K9 me3:H4K20 me3
DNA Meth. m5C:H4K20 me3
H2A.Z:H4−4ac
H3K27 me3:H3K9 me3
H4K20 me2:H4−4ac
H2A.Z:H3−5ac
H3K4 me1:H4−4ac
H3K4 me3:H4−4ac
H3K27 me3:DNA Meth. m5C
H2A.Z:H3K4 me3
H4−4ac:H3−5ac
H2A.Z:H4K20 me2
DNA Meth. m5C:H3K9 me3
H4K20 me2:H3K4 me3
H4K20 me2:H3−5ac
H3K4 me1:H3−5ac
H3K9 me2
H3K27 me2
H3K27 me3
H4K20 me3
H4−4ac
H3K9 me3
H3K4 me3
DNA Meth. m5C
H4K20 me2
H3K4 me1
H2A.Z
H3−5ac
0.00
0.25
0.50
0.75
1.00
$\hat{\pi}$
RPL30 – Forward
H3K4 me1:H4−4ac
H3K4 me1:H2A.Z
DNA Meth. m5C:H3K9 me3
H3K27 me3:DNA Meth. m5C
H3K4 me1:H3−5ac
H3K9 me3:H4K20 me3
H2A.Z:H4−4ac
H3K4 me1:H4K20 me2
H3K27 me3:H4K20 me3
H2A.Z:H3−5ac
H3K27 me3:H3K4 me1
H3K27 me3:H3K9 me3
H4K20 me2:H4−4ac
H3K9 me2:DNA Meth. m5C
DNA Meth. m5C:H4K20 me3
H3K4 me3:H4−4ac
H4K20 me2:H3K4 me3
H3K4 me3:H3−5ac
H4−4ac:H3−5ac
H2A.Z:H4K20 me2
H2A.Z:H3K4 me3
H4K20 me2:H3−5ac
H3K9 me2
H3K4 me1
H3K9 me3
DNA Meth. m5C
H2A.Z
H4−4ac
H3−5ac
H4K20 me3
H3K4 me3
H4K20 me2
H3K27 me3
0.00
0.25
0.50
0.75
1.00
$\hat{\pi}$
RPL30 – Reverse
H3K4 me1:H4−4ac
H3K4 me1:H2A.Z
DNA Meth. m5C:H3K9 me3
H3K27 me3:DNA Meth. m5C
H3K4 me1:H3−5ac
H3K9 me3:H4K20 me3
H2A.Z:H4−4ac
H3K4 me1:H4K20 me2
H3K27 me3:H4K20 me3
H2A.Z:H3−5ac
H3K27 me3:H3K4 me1
H3K27 me3:H3K9 me3
H4K20 me2:H4−4ac
H3K9 me2:DNA Meth. m5C
DNA Meth. m5C:H4K20 me3
H3K4 me3:H4−4ac
H4K20 me2:H3K4 me3
H3K4 me3:H3−5ac
H4−4ac:H3−5ac
H2A.Z:H4K20 me2
H2A.Z:H3K4 me3
H4K20 me2:H3−5ac
H3K9 me2
H3K4 me1
H3K9 me3
DNA Meth. m5C
H2A.Z
H4−4ac
H3−5ac
H4K20 me3
H3K4 me3
H4K20 me2
H3K27 me3
0.00
0.25
0.50
0.75
1.00
$\hat{\pi}$
POLR1B – Forward
H3K4 me1:H3−5ac
DNA Meth. m5C:H4K20 me2
H3K4 me1:H4−4ac
H3K4 me1:H2A.Z
H4K20 me2:H4−4ac
H2A.Z:H3−5ac
H3K27 me3:H3K4 me1
H3K27 me2:DNA Meth. m5C
H2A.Z:H4−4ac
H3K4 me1:H4K20 me2
H3K4 me3:H3−5ac
H3K27 me3:DNA Meth. m5C
H3K4 me3:H4−4ac
H2A.Z:H4K20 me2
H4−4ac:H3−5ac
H4K20 me2:H3−5ac
H3K27 me3:H4K20 me3
H3K9 me3:H4K20 me3
H3K27 me3:H3K9 me3
H2A.Z:H3K4 me3
H4K20 me2:H3K4 me3
H3K9 me2:DNA Meth. m5C
H3K27 me2
H4K20 me3
H3K4 me1
H3K9 me2
H4−4ac
DNA Meth. m5C
H3K9 me3
H2A.Z
H3−5ac
H4K20 me2
H3K27 me3
H3K4 me3
0.00
0.25
0.50
0.75
1.00
$\hat{\pi}$
POLR1B – Reverse
H3K4 me1:H3−5ac
DNA Meth. m5C:H4K20 me2
H3K4 me1:H4−4ac
H3K4 me1:H2A.Z
H4K20 me2:H4−4ac
H2A.Z:H3−5ac
H3K27 me3:H3K4 me1
H3K27 me2:DNA Meth. m5C
H2A.Z:H4−4ac
H3K4 me1:H4K20 me2
H3K4 me3:H3−5ac
H3K27 me3:DNA Meth. m5C
H3K4 me3:H4−4ac
H2A.Z:H4K20 me2
H4−4ac:H3−5ac
H4K20 me2:H3−5ac
H3K27 me3:H4K20 me3
H3K9 me3:H4K20 me3
H3K27 me3:H3K9 me3
H2A.Z:H3K4 me3
H4K20 me2:H3K4 me3
H3K9 me2:DNA Meth. m5C
H3K27 me2
H4K20 me3
H3K4 me1
H3K9 me2
H4−4ac
DNA Meth. m5C
H3K9 me3
H2A.Z
H3−5ac
H4K20 me2
H3K27 me3
H3K4 me3
0.00
0.25
0.50
0.75
1.00
$\hat{\pi}$

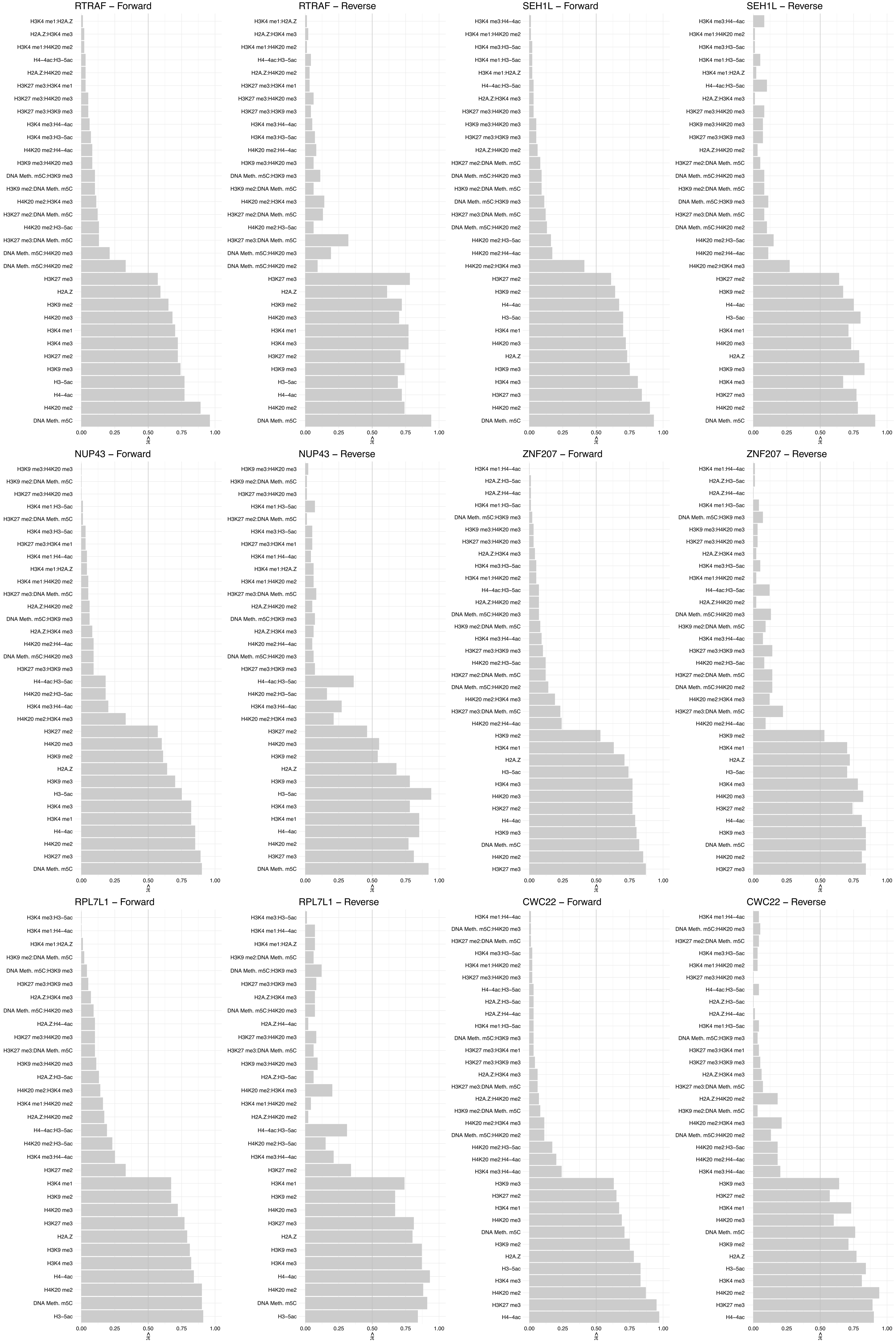
RTRAF – Forward
H3K4 me1:H2A.Z
H2A.Z:H3K4 me3
H3K4 me1:H4K20 me2
H4−4ac:H3−5ac
H2A.Z:H4K20 me2
H3K27 me3:H3K4 me1
H3K27 me3:H4K20 me3
H3K27 me3:H3K9 me3
H3K4 me3:H4−4ac
H3K4 me3:H3−5ac
H4K20 me2:H4−4ac
H3K9 me3:H4K20 me3
DNA Meth. m5C:H3K9 me3
H3K9 me2:DNA Meth. m5C
H4K20 me2:H3K4 me3
H3K27 me2:DNA Meth. m5C
H4K20 me2:H3−5ac
H3K27 me3:DNA Meth. m5C
DNA Meth. m5C:H4K20 me3
DNA Meth. m5C:H4K20 me2
H3K27 me3
H2A.Z
H3K9 me2
H4K20 me3
H3K4 me1
H3K4 me3
H3K27 me2
H3K9 me3
H3−5ac
H4−4ac
H4K20 me2
DNA Meth. m5C
0.00 0.25 0.50 0.75 1.00
π̂
RTRAF – Reverse
H3K4 me1:H2A.Z
H2A.Z:H3K4 me3
H3K4 me1:H4K20 me2
H4−4ac:H3−5ac
H2A.Z:H4K20 me2
H3K27 me3:H3K4 me1
H3K27 me3:H4K20 me3
H3K27 me3:H3K9 me3
H3K4 me3:H4−4ac
H3K4 me3:H3−5ac
H4K20 me2:H4−4ac
H3K9 me3:H4K20 me3
DNA Meth. m5C:H3K9 me3
H3K9 me2:DNA Meth. m5C
H4K20 me2:H3K4 me3
H3K27 me2:DNA Meth. m5C
H4K20 me2:H3−5ac
H3K27 me3:DNA Meth. m5C
DNA Meth. m5C:H4K20 me3
DNA Meth. m5C:H4K20 me2
H3K27 me3
H2A.Z
H3K9 me2
H4K20 me3
H3K4 me1
H3K4 me3
H3K27 me2
H3K9 me3
H3−5ac
H4−4ac
H4K20 me2
DNA Meth. m5C
0.00 0.25 0.50 0.75 1.00
π̂
SEH1L – Forward
H3K4 me3:H4−4ac
H3K4 me1:H4K20 me2
H3K4 me3:H3−5ac
H3K4 me1:H3−5ac
H3K4 me1:H2A.Z
H4−4ac:H3−5ac
H2A.Z:H3K4 me3
H3K27 me3:H4K20 me3
H3K9 me3:H4K20 me3
H3K27 me3:H3K9 me3
H2A.Z:H4K20 me2
H3K27 me2:DNA Meth. m5C
DNA Meth. m5C:H4K20 me3
H3K9 me2:DNA Meth. m5C
DNA Meth. m5C:H3K9 me3
H3K27 me3:DNA Meth. m5C
DNA Meth. m5C:H4K20 me2
H4K20 me2:H3−5ac
H4K20 me2:H4−4ac
H4K20 me2:H3K4 me3
H3K27 me2
H3K9 me2
H4−4ac
H3−5ac
H3K4 me1
H4K20 me3
H2A.Z
H3K9 me3
H3K4 me3
H3K27 me3
H4K20 me2
DNA Meth. m5C
0.00 0.25 0.50 0.75 1.00
π̂
SEH1L – Reverse
H3K4 me3:H4−4ac
H3K4 me1:H4K20 me2
H3K4 me3:H3−5ac
H3K4 me1:H3−5ac
H3K4 me1:H2A.Z
H4−4ac:H3−5ac
H2A.Z:H3K4 me3
H3K27 me3:H4K20 me3
H3K9 me3:H4K20 me3
H3K27 me3:H3K9 me3
H2A.Z:H4K20 me2
H3K27 me2:DNA Meth. m5C
DNA Meth. m5C:H4K20 me3
H3K9 me2:DNA Meth. m5C
DNA Meth. m5C:H3K9 me3
H3K27 me3:DNA Meth. m5C
DNA Meth. m5C:H4K20 me2
H4K20 me2:H3−5ac
H4K20 me2:H4−4ac
H4K20 me2:H3K4 me3
H3K27 me2
H3K9 me2
H4−4ac
H3−5ac
H3K4 me1
H4K20 me3
H2A.Z
H3K9 me3
H3K4 me3
H3K27 me3
H4K20 me2
DNA Meth. m5C
0.00 0.25 0.50 0.75 1.00
π̂
NUP43 – Forward
H3K9 me3:H4K20 me3
H3K9 me2:DNA Meth. m5C
H3K27 me3:H4K20 me3
H3K4 me1:H3−5ac
H3K27 me2:DNA Meth. m5C
H3K4 me3:H3−5ac
H3K27 me3:H3K4 me1
H3K4 me1:H4−4ac
H3K4 me1:H2A.Z
H3K4 me1:H4K20 me2
H3K27 me3:DNA Meth. m5C
H2A.Z:H4K20 me2
DNA Meth. m5C:H3K9 me3
H2A.Z:H3K4 me3
H4K20 me2:H4−4ac
DNA Meth. m5C:H4K20 me3
H3K27 me3:H3K9 me3
H4−4ac:H3−5ac
H4K20 me2:H3−5ac
H3K4 me3:H4−4ac
H4K20 me2:H3K4 me3
H3K27 me2
H4K20 me3
H3K9 me2
H2A.Z
H3K9 me3
H3−5ac
H3K4 me3
H3K4 me1
H4−4ac
H4K20 me2
H3K27 me3
DNA Meth. m5C
0.00 0.25 0.50 0.75 1.00
π̂
NUP43 – Reverse
H3K9 me3:H4K20 me3
H3K9 me2:DNA Meth. m5C
H3K27 me3:H4K20 me3
H3K4 me1:H3−5ac
H3K27 me2:DNA Meth. m5C
H3K4 me3:H3−5ac
H3K27 me3:H3K4 me1
H3K4 me1:H4−4ac
H3K4 me1:H2A.Z
H3K4 me1:H4K20 me2
H3K27 me3:DNA Meth. m5C
H2A.Z:H4K20 me2
DNA Meth. m5C:H3K9 me3
H2A.Z:H3K4 me3
H4K20 me2:H4−4ac
DNA Meth. m5C:H4K20 me3
H3K27 me3:H3K9 me3
H4−4ac:H3−5ac
H4K20 me2:H3−5ac
H3K4 me3:H4−4ac
H4K20 me2:H3K4 me3
H3K27 me2
H4K20 me3
H3K9 me2
H2A.Z
H3K9 me3
H3−5ac
H3K4 me3
H3K4 me1
H4−4ac
H4K20 me2
H3K27 me3
DNA Meth. m5C
0.00 0.25 0.50 0.75 1.00
π̂
ZNF207 – Forward
H3K4 me1:H4−4ac
H2A.Z:H3−5ac
H2A.Z:H4−4ac
H3K4 me1:H3−5ac
DNA Meth. m5C:H3K9 me3
H3K9 me3:H4K20 me3
H3K27 me3:H4K20 me3
H2A.Z:H3K4 me3
H3K4 me3:H3−5ac
H3K4 me1:H4K20 me2
H4−4ac:H3−5ac
H2A.Z:H4K20 me2
DNA Meth. m5C:H4K20 me3
H3K9 me2:DNA Meth. m5C
H3K4 me3:H4−4ac
H3K27 me3:H3K9 me3
H4K20 me2:H3−5ac
H3K27 me2:DNA Meth. m5C
DNA Meth. m5C:H4K20 me2
H4K20 me2:H3K4 me3
H3K27 me3:DNA Meth. m5C
H4K20 me2:H4−4ac
H3K9 me2
H3K4 me1
H2A.Z
H3−5ac
H3K4 me3
H4K20 me3
H3K27 me2
H4−4ac
H3K9 me3
DNA Meth. m5C
H4K20 me2
H3K27 me3
0.00 0.25 0.50 0.75 1.00
π̂
ZNF207 – Reverse
H3K4 me1:H4−4ac
H2A.Z:H3−5ac
H2A.Z:H4−4ac
H3K4 me1:H3−5ac
DNA Meth. m5C:H3K9 me3
H3K9 me3:H4K20 me3
H3K27 me3:H4K20 me3
H2A.Z:H3K4 me3
H3K4 me3:H3−5ac
H3K4 me1:H4K20 me2
H4−4ac:H3−5ac
H2A.Z:H4K20 me2
DNA Meth. m5C:H4K20 me3
H3K9 me2:DNA Meth. m5C
H3K4 me3:H4−4ac
H3K27 me3:H3K9 me3
H4K20 me2:H3−5ac
H3K27 me2:DNA Meth. m5C
DNA Meth. m5C:H4K20 me2
H4K20 me2:H3K4 me3
H3K27 me3:DNA Meth. m5C
H4K20 me2:H4−4ac
H3K9 me2
H3K4 me1
H2A.Z
H3−5ac
H3K4 me3
H4K20 me3
H3K27 me2
H4−4ac
H3K9 me3
DNA Meth. m5C
H4K20 me2
H3K27 me3
0.00 0.25 0.50 0.75 1.00
π̂
RPL7L1 – Forward
H3K4 me3:H3−5ac
H3K4 me1:H4−4ac
H3K4 me1:H2A.Z
H3K9 me2:DNA Meth. m5C
DNA Meth. m5C:H3K9 me3
H3K27 me3:H3K9 me3
H2A.Z:H3K4 me3
DNA Meth. m5C:H4K20 me3
H2A.Z:H4−4ac
H3K27 me3:H4K20 me3
H3K27 me3:DNA Meth. m5C
H3K9 me3:H4K20 me3
H2A.Z:H3−5ac
H4K20 me2:H3K4 me3
H3K4 me1:H4K20 me2
H2A.Z:H4K20 me2
H4−4ac:H3−5ac
H4K20 me2:H3−5ac
H3K4 me3:H4−4ac
H3K27 me2
H3K4 me1
H3K9 me2
H4K20 me3
H3K27 me3
H2A.Z
H3K9 me3
H3K4 me3
H4−4ac
H4K20 me2
DNA Meth. m5C
H3−5ac
0.00 0.25 0.50 0.75 1.00
π̂
RPL7L1 – Reverse
H3K4 me3:H3−5ac
H3K4 me1:H4−4ac
H3K4 me1:H2A.Z
H3K9 me2:DNA Meth. m5C
DNA Meth. m5C:H3K9 me3
H3K27 me3:H3K9 me3
H2A.Z:H3K4 me3
DNA Meth. m5C:H4K20 me3
H2A.Z:H4−4ac
H3K27 me3:H4K20 me3
H3K27 me3:DNA Meth. m5C
H3K9 me3:H4K20 me3
H2A.Z:H3−5ac
H4K20 me2:H3K4 me3
H3K4 me1:H4K20 me2
H2A.Z:H4K20 me2
H4−4ac:H3−5ac
H4K20 me2:H3−5ac
H3K4 me3:H4−4ac
H3K27 me2
H3K4 me1
H3K9 me2
H4K20 me3
H3K27 me3
H2A.Z
H3K9 me3
H3K4 me3
H4−4ac
H4K20 me2
DNA Meth. m5C
H3−5ac
0.00 0.25 0.50 0.75 1.00
π̂
CWC22 – Forward
H3K4 me1:H4−4ac
DNA Meth. m5C:H4K20 me3
H3K27 me2:DNA Meth. m5C
H3K4 me3:H3−5ac
H3K4 me1:H4K20 me2
H3K27 me3:H4K20 me3
H4−4ac:H3−5ac
H2A.Z:H3−5ac
H2A.Z:H4−4ac
H3K4 me1:H3−5ac
DNA Meth. m5C:H3K9 me3
H3K27 me3:H3K4 me1
H3K27 me3:H3K9 me3
H2A.Z:H3K4 me3
H3K27 me3:DNA Meth. m5C
H2A.Z:H4K20 me2
H3K9 me2:DNA Meth. m5C
H4K20 me2:H3K4 me3
DNA Meth. m5C:H4K20 me2
H4K20 me2:H3−5ac
H4K20 me2:H4−4ac
H3K4 me3:H4−4ac
H3K9 me3
H3K27 me2
H3K4 me1
H4K20 me3
DNA Meth. m5C
H3K9 me2
H2A.Z
H3−5ac
H3K4 me3
H4K20 me2
H3K27 me3
H4−4ac
0.00 0.25 0.50 0.75 1.00
π̂
CWC22 – Reverse
H3K4 me1:H4−4ac
DNA Meth. m5C:H4K20 me3
H3K27 me2:DNA Meth. m5C
H3K4 me3:H3−5ac
H3K4 me1:H4K20 me2
H3K27 me3:H4K20 me3
H4−4ac:H3−5ac
H2A.Z:H3−5ac
H2A.Z:H4−4ac
H3K4 me1:H3−5ac
DNA Meth. m5C:H3K9 me3
H3K27 me3:H3K4 me1
H3K27 me3:H3K9 me3
H2A.Z:H3K4 me3
H3K27 me3:DNA Meth. m5C
H2A.Z:H4K20 me2
H3K9 me2:DNA Meth. m5C
H4K20 me2:H3K4 me3
DNA Meth. m5C:H4K20 me2
H4K20 me2:H3−5ac
H4K20 me2:H4−4ac
H3K4 me3:H4−4ac
H3K9 me3
H3K27 me2
H3K4 me1
H4K20 me3
DNA Meth. m5C
H3K9 me2
H2A.Z
H3−5ac
H3K4 me3
H4K20 me2
H3K27 me3
H4−4ac
0.00 0.25 0.50 0.75 1.00
π̂

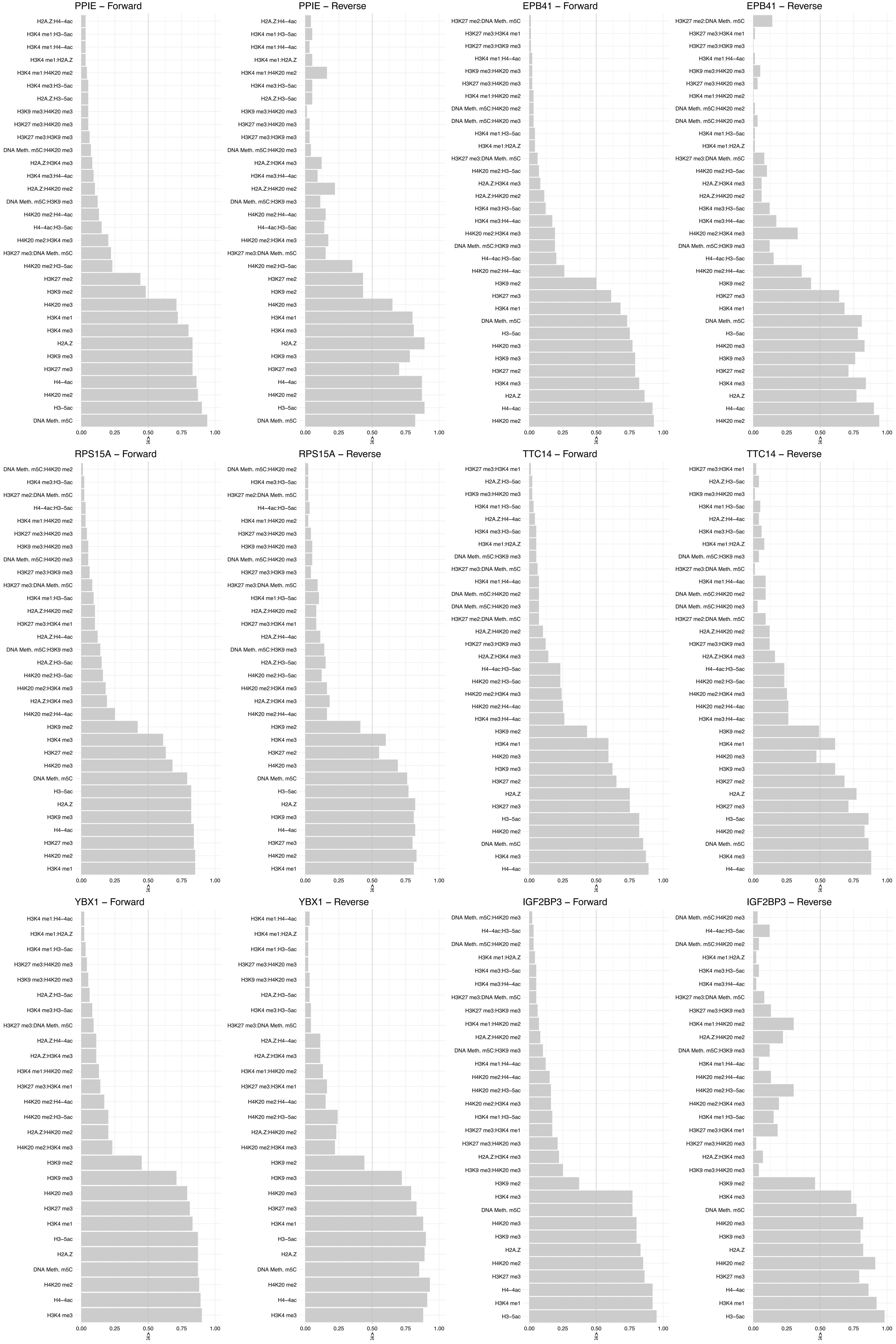
PPIE – Forward
H2A.Z:H4−4ac
H3K4 me1:H3−5ac
H3K4 me1:H4−4ac
H3K4 me1:H2A.Z
H3K4 me1:H4K20 me2
H3K4 me3:H3−5ac
H2A.Z:H3−5ac
H3K9 me3:H4K20 me3
H3K27 me3:H4K20 me3
H3K27 me3:H3K9 me3
DNA Meth. m5C:H4K20 me3
H2A.Z:H3K4 me3
H3K4 me3:H4−4ac
H2A.Z:H4K20 me2
DNA Meth. m5C:H3K9 me3
H4K20 me2:H4−4ac
H4−4ac:H3−5ac
H4K20 me2:H3K4 me3
H3K27 me3:DNA Meth. m5C
H4K20 me2:H3−5ac
H3K27 me2
H3K9 me2
H4K20 me3
H3K4 me1
H3K4 me3
H2A.Z
H3K9 me3
H3K27 me3
H4−4ac
H4K20 me2
H3−5ac
DNA Meth. m5C
0.00 0.25 0.50 0.75 1.00
$\hat{\pi}$
PPIE – Reverse
H2A.Z:H4−4ac
H3K4 me1:H3−5ac
H3K4 me1:H4−4ac
H3K4 me1:H2A.Z
H3K4 me1:H4K20 me2
H3K4 me3:H3−5ac
H2A.Z:H3−5ac
H3K9 me3:H4K20 me3
H3K27 me3:H4K20 me3
H3K27 me3:H3K9 me3
DNA Meth. m5C:H4K20 me3
H2A.Z:H3K4 me3
H3K4 me3:H4−4ac
H2A.Z:H4K20 me2
DNA Meth. m5C:H3K9 me3
H4K20 me2:H4−4ac
H4−4ac:H3−5ac
H4K20 me2:H3K4 me3
H3K27 me3:DNA Meth. m5C
H4K20 me2:H3−5ac
H3K27 me2
H3K9 me2
H4K20 me3
H3K4 me1
H3K4 me3
H2A.Z
H3K9 me3
H3K27 me3
H4−4ac
H4K20 me2
H3−5ac
DNA Meth. m5C
0.00 0.25 0.50 0.75 1.00
$\hat{\pi}$
EPB41 – Forward
H3K27 me2:DNA Meth. m5C
H3K27 me3:H3K4 me1
H3K27 me3:H3K9 me3
H3K4 me1:H4−4ac
H3K9 me3:H4K20 me3
H3K27 me3:H4K20 me3
H3K4 me1:H4K20 me2
DNA Meth. m5C:H4K20 me2
DNA Meth. m5C:H4K20 me3
H3K4 me1:H3−5ac
H3K4 me1:H2A.Z
H3K27 me3:DNA Meth. m5C
H4K20 me2:H3−5ac
H2A.Z:H3K4 me3
H2A.Z:H4K20 me2
H3K4 me3:H3−5ac
H3K4 me3:H4−4ac
H4K20 me2:H3K4 me3
DNA Meth. m5C:H3K9 me3
H4−4ac:H3−5ac
H4K20 me2:H4−4ac
H3K9 me2
H3K27 me3
H3K4 me1
DNA Meth. m5C
H3−5ac
H4K20 me3
H3K9 me3
H3K27 me2
H3K4 me3
H2A.Z
H4−4ac
H4K20 me2
0.00 0.25 0.50 0.75 1.00
$\hat{\pi}$
EPB41 – Reverse
H3K27 me2:DNA Meth. m5C
H3K27 me3:H3K4 me1
H3K27 me3:H3K9 me3
H3K4 me1:H4−4ac
H3K9 me3:H4K20 me3
H3K27 me3:H4K20 me3
H3K4 me1:H4K20 me2
DNA Meth. m5C:H4K20 me2
DNA Meth. m5C:H4K20 me3
H3K4 me1:H3−5ac
H3K4 me1:H2A.Z
H3K27 me3:DNA Meth. m5C
H4K20 me2:H3−5ac
H2A.Z:H3K4 me3
H2A.Z:H4K20 me2
H3K4 me3:H3−5ac
H3K4 me3:H4−4ac
H4K20 me2:H3K4 me3
DNA Meth. m5C:H3K9 me3
H4−4ac:H3−5ac
H4K20 me2:H4−4ac
H3K9 me2
H3K27 me3
H3K4 me1
DNA Meth. m5C
H3−5ac
H4K20 me3
H3K9 me3
H3K27 me2
H3K4 me3
H2A.Z
H4−4ac
H4K20 me2
0.00 0.25 0.50 0.75 1.00
$\hat{\pi}$
RPS15A – Forward
DNA Meth. m5C:H4K20 me2
H3K4 me3:H3−5ac
H3K27 me2:DNA Meth. m5C
H4−4ac:H3−5ac
H3K4 me1:H4K20 me2
H3K27 me3:H4K20 me3
H3K9 me3:H4K20 me3
DNA Meth. m5C:H4K20 me3
H3K27 me3:H3K9 me3
H3K27 me3:DNA Meth. m5C
H3K4 me1:H3−5ac
H2A.Z:H4K20 me2
H3K27 me3:H3K4 me1
H2A.Z:H4−4ac
DNA Meth. m5C:H3K9 me3
H2A.Z:H3−5ac
H4K20 me2:H3−5ac
H4K20 me2:H3K4 me3
H2A.Z:H3K4 me3
H4K20 me2:H4−4ac
H3K9 me2
H3K4 me3
H3K27 me2
H4K20 me3
DNA Meth. m5C
H3−5ac
H2A.Z
H3K9 me3
H4−4ac
H3K27 me3
H4K20 me2
H3K4 me1
0.00 0.25 0.50 0.75 1.00
$\hat{\pi}$
RPS15A – Reverse
DNA Meth. m5C:H4K20 me2
H3K4 me3:H3−5ac
H3K27 me2:DNA Meth. m5C
H4−4ac:H3−5ac
H3K4 me1:H4K20 me2
H3K27 me3:H4K20 me3
H3K9 me3:H4K20 me3
DNA Meth. m5C:H4K20 me3
H3K27 me3:H3K9 me3
H3K27 me3:DNA Meth. m5C
H3K4 me1:H3−5ac
H2A.Z:H4K20 me2
H3K27 me3:H3K4 me1
H2A.Z:H4−4ac
DNA Meth. m5C:H3K9 me3
H2A.Z:H3−5ac
H4K20 me2:H3−5ac
H4K20 me2:H3K4 me3
H2A.Z:H3K4 me3
H4K20 me2:H4−4ac
H3K9 me2
H3K4 me3
H3K27 me2
H4K20 me3
DNA Meth. m5C
H3−5ac
H2A.Z
H3K9 me3
H4−4ac
H3K27 me3
H4K20 me2
H3K4 me1
0.00 0.25 0.50 0.75 1.00
$\hat{\pi}$
TTC14 – Forward
H3K27 me3:H3K4 me1
H2A.Z:H3−5ac
H3K9 me3:H4K20 me3
H3K4 me1:H3−5ac
H2A.Z:H4−4ac
H3K4 me3:H3−5ac
H3K4 me1:H2A.Z
DNA Meth. m5C:H3K9 me3
H3K27 me3:DNA Meth. m5C
H3K4 me1:H4−4ac
DNA Meth. m5C:H4K20 me2
DNA Meth. m5C:H4K20 me3
H3K27 me2:DNA Meth. m5C
H2A.Z:H4K20 me2
H3K27 me3:H3K9 me3
H2A.Z:H3K4 me3
H4−4ac:H3−5ac
H4K20 me2:H3−5ac
H4K20 me2:H3K4 me3
H4K20 me2:H4−4ac
H3K4 me3:H4−4ac
H3K9 me2
H3K4 me1
H4K20 me3
H3K9 me3
H3K27 me2
H2A.Z
H3K27 me3
H3−5ac
H4K20 me2
DNA Meth. m5C
H3K4 me3
H4−4ac
0.00 0.25 0.50 0.75 1.00
$\hat{\pi}$
TTC14 – Reverse
H3K27 me3:H3K4 me1
H2A.Z:H3−5ac
H3K9 me3:H4K20 me3
H3K4 me1:H3−5ac
H2A.Z:H4−4ac
H3K4 me3:H3−5ac
H3K4 me1:H2A.Z
DNA Meth. m5C:H3K9 me3
H3K27 me3:DNA Meth. m5C
H3K4 me1:H4−4ac
DNA Meth. m5C:H4K20 me2
DNA Meth. m5C:H4K20 me3
H3K27 me2:DNA Meth. m5C
H2A.Z:H4K20 me2
H3K27 me3:H3K9 me3
H2A.Z:H3K4 me3
H4−4ac:H3−5ac
H4K20 me2:H3−5ac
H4K20 me2:H3K4 me3
H4K20 me2:H4−4ac
H3K4 me3:H4−4ac
H3K9 me2
H3K4 me1
H4K20 me3
H3K9 me3
H3K27 me2
H2A.Z
H3K27 me3
H3−5ac
H4K20 me2
DNA Meth. m5C
H3K4 me3
H4−4ac
0.00 0.25 0.50 0.75 1.00
$\hat{\pi}$
YBX1 – Forward
H3K4 me1:H4−4ac
H3K4 me1:H2A.Z
H3K4 me1:H3−5ac
H3K27 me3:H4K20 me3
H3K9 me3:H4K20 me3
H2A.Z:H3−5ac
H3K4 me3:H3−5ac
H3K27 me3:DNA Meth. m5C
H2A.Z:H4−4ac
H2A.Z:H3K4 me3
H3K4 me1:H4K20 me2
H3K27 me3:H3K4 me1
H4K20 me2:H4−4ac
H4K20 me2:H3−5ac
H2A.Z:H4K20 me2
H4K20 me2:H3K4 me3
H3K9 me2
H3K9 me3
H4K20 me3
H3K27 me3
H3K4 me1
H3−5ac
H2A.Z
DNA Meth. m5C
H4K20 me2
H4−4ac
H3K4 me3
0.00 0.25 0.50 0.75 1.00
$\hat{\pi}$
YBX1 – Reverse
H3K4 me1:H4−4ac
H3K4 me1:H2A.Z
H3K4 me1:H3−5ac
H3K27 me3:H4K20 me3
H3K9 me3:H4K20 me3
H2A.Z:H3−5ac
H3K4 me3:H3−5ac
H3K27 me3:DNA Meth. m5C
H2A.Z:H4−4ac
H2A.Z:H3K4 me3
H3K4 me1:H4K20 me2
H3K27 me3:H3K4 me1
H4K20 me2:H4−4ac
H4K20 me2:H3−5ac
H2A.Z:H4K20 me2
H4K20 me2:H3K4 me3
H3K9 me2
H3K9 me3
H4K20 me3
H3K27 me3
H3K4 me1
H3−5ac
H2A.Z
DNA Meth. m5C
H4K20 me2
H4−4ac
H3K4 me3
0.00 0.25 0.50 0.75 1.00
$\hat{\pi}$
IGF2BP3 – Forward
DNA Meth. m5C:H4K20 me3
H4−4ac:H3−5ac
DNA Meth. m5C:H4K20 me2
H3K4 me1:H2A.Z
H3K4 me3:H3−5ac
H3K4 me3:H4−4ac
H3K27 me3:DNA Meth. m5C
H3K27 me3:H3K9 me3
H3K4 me1:H4K20 me2
H2A.Z:H4K20 me2
DNA Meth. m5C:H3K9 me3
H3K4 me1:H4−4ac
H4K20 me2:H4−4ac
H4K20 me2:H3−5ac
H4K20 me2:H3K4 me3
H3K4 me1:H3−5ac
H3K27 me3:H3K4 me1
H3K27 me3:H4K20 me3
H2A.Z:H3K4 me3
H3K9 me3:H4K20 me3
H3K9 me2
H3K4 me3
DNA Meth. m5C
H4K20 me3
H3K9 me3
H2A.Z
H4K20 me2
H3K27 me3
H4−4ac
H3K4 me1
H3−5ac
0.00 0.25 0.50 0.75 1.00
$\hat{\pi}$
IGF2BP3 – Reverse
DNA Meth. m5C:H4K20 me3
H4−4ac:H3−5ac
DNA Meth. m5C:H4K20 me2
H3K4 me1:H2A.Z
H3K4 me3:H3−5ac
H3K4 me3:H4−4ac
H3K27 me3:DNA Meth. m5C
H3K27 me3:H3K9 me3
H3K4 me1:H4K20 me2
H2A.Z:H4K20 me2
DNA Meth. m5C:H3K9 me3
H3K4 me1:H4−4ac
H4K20 me2:H4−4ac
H4K20 me2:H3−5ac
H4K20 me2:H3K4 me3
H3K4 me1:H3−5ac
H3K27 me3:H3K4 me1
H3K27 me3:H4K20 me3
H2A.Z:H3K4 me3
H3K9 me3:H4K20 me3
H3K9 me2
H3K4 me3
DNA Meth. m5C
H4K20 me3
H3K9 me3
H2A.Z
H4K20 me2
H3K27 me3
H4−4ac
H3K4 me1
H3−5ac
0.00 0.25 0.50 0.75 1.00
$\hat{\pi}$

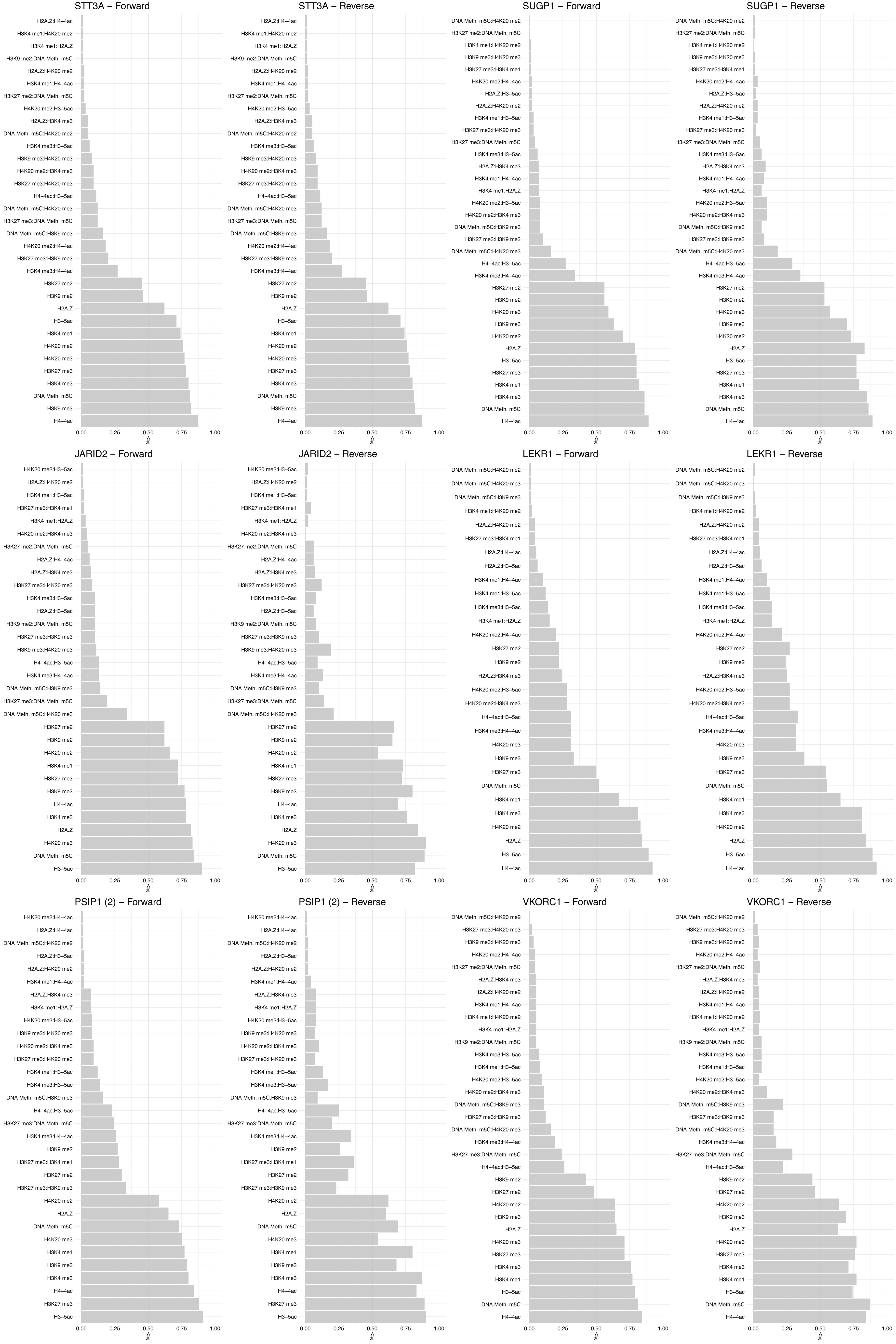
STT3A – Forward
H2A.Z:H4−4ac
H3K4 me1:H4K20 me2
H3K4 me1:H2A.Z
H3K9 me2:DNA Meth. m5C
H2A.Z:H4K20 me2
H3K4 me1:H4−4ac
H3K27 me2:DNA Meth. m5C
H4K20 me2:H3−5ac
H2A.Z:H3K4 me3
DNA Meth. m5C:H4K20 me2
H3K4 me3:H3−5ac
H3K9 me3:H4K20 me3
H4K20 me2:H3K4 me3
H3K27 me3:H4K20 me3
H4−4ac:H3−5ac
DNA Meth. m5C:H4K20 me3
H3K27 me3:DNA Meth. m5C
DNA Meth. m5C:H3K9 me3
H4K20 me2:H4−4ac
H3K27 me3:H3K9 me3
H3K4 me3:H4−4ac
H3K27 me2
H3K9 me2
H2A.Z
H3−5ac
H3K4 me1
H4K20 me2
H4K20 me3
H3K27 me3
H3K4 me3
DNA Meth. m5C
H3K9 me3
H4−4ac
0.00
0.25
0.50
0.75
1.00
π̂
STT3A – Reverse
H2A.Z:H4−4ac
H3K4 me1:H4K20 me2
H3K4 me1:H2A.Z
H3K9 me2:DNA Meth. m5C
H2A.Z:H4K20 me2
H3K4 me1:H4−4ac
H3K27 me2:DNA Meth. m5C
H4K20 me2:H3−5ac
H2A.Z:H3K4 me3
DNA Meth. m5C:H4K20 me2
H3K4 me3:H3−5ac
H3K9 me3:H4K20 me3
H4K20 me2:H3K4 me3
H3K27 me3:H4K20 me3
H4−4ac:H3−5ac
DNA Meth. m5C:H4K20 me3
H3K27 me3:DNA Meth. m5C
DNA Meth. m5C:H3K9 me3
H4K20 me2:H4−4ac
H3K27 me3:H3K9 me3
H3K4 me3:H4−4ac
H3K27 me2
H3K9 me2
H2A.Z
H3−5ac
H3K4 me1
H4K20 me2
H4K20 me3
H3K27 me3
H3K4 me3
DNA Meth. m5C
H3K9 me3
H4−4ac
0.00
0.25
0.50
0.75
1.00
π̂
SUGP1 – Forward
DNA Meth. m5C:H4K20 me2
H3K27 me2:DNA Meth. m5C
H3K4 me1:H4K20 me2
H3K9 me3:H4K20 me3
H3K27 me3:H3K4 me1
H4K20 me2:H4−4ac
H2A.Z:H3−5ac
H2A.Z:H4K20 me2
H3K4 me1:H3−5ac
H3K27 me3:H4K20 me3
H3K27 me3:DNA Meth. m5C
H3K4 me3:H3−5ac
H2A.Z:H3K4 me3
H3K4 me1:H4−4ac
H3K4 me1:H2A.Z
H4K20 me2:H3−5ac
H4K20 me2:H3K4 me3
DNA Meth. m5C:H3K9 me3
H3K27 me3:H3K9 me3
DNA Meth. m5C:H4K20 me3
H4−4ac:H3−5ac
H3K4 me3:H4−4ac
H3K27 me2
H3K9 me2
H4K20 me3
H3K9 me3
H4K20 me2
H2A.Z
H3−5ac
H3K27 me3
H3K4 me1
H3K4 me3
DNA Meth. m5C
H4−4ac
0.00
0.25
0.50
0.75
1.00
π̂
SUGP1 – Reverse
DNA Meth. m5C:H4K20 me2
H3K27 me2:DNA Meth. m5C
H3K4 me1:H4K20 me2
H3K9 me3:H4K20 me3
H3K27 me3:H3K4 me1
H4K20 me2:H4−4ac
H2A.Z:H3−5ac
H2A.Z:H4K20 me2
H3K4 me1:H3−5ac
H3K27 me3:H4K20 me3
H3K27 me3:DNA Meth. m5C
H3K4 me3:H3−5ac
H2A.Z:H3K4 me3
H3K4 me1:H4−4ac
H3K4 me1:H2A.Z
H4K20 me2:H3−5ac
H4K20 me2:H3K4 me3
DNA Meth. m5C:H3K9 me3
H3K27 me3:H3K9 me3
DNA Meth. m5C:H4K20 me3
H4−4ac:H3−5ac
H3K4 me3:H4−4ac
H3K27 me2
H3K9 me2
H4K20 me3
H3K9 me3
H4K20 me2
H2A.Z
H3−5ac
H3K27 me3
H3K4 me1
H3K4 me3
DNA Meth. m5C
H4−4ac
0.00
0.25
0.50
0.75
1.00
π̂
JARID2 – Forward
H4K20 me2:H3−5ac
H2A.Z:H4K20 me2
H3K4 me1:H3−5ac
H3K27 me3:H3K4 me1
H3K4 me1:H2A.Z
H4K20 me2:H3K4 me3
H3K27 me2:DNA Meth. m5C
H2A.Z:H4−4ac
H2A.Z:H3K4 me3
H3K27 me3:H4K20 me3
H3K4 me3:H3−5ac
H2A.Z:H3−5ac
H3K9 me2:DNA Meth. m5C
H3K27 me3:H3K9 me3
H3K9 me3:H4K20 me3
H4−4ac:H3−5ac
H3K4 me3:H4−4ac
DNA Meth. m5C:H3K9 me3
H3K27 me3:DNA Meth. m5C
DNA Meth. m5C:H4K20 me3
H3K27 me2
H3K9 me2
H4K20 me2
H3K4 me1
H3K27 me3
H3K9 me3
H4−4ac
H3K4 me3
H2A.Z
H4K20 me3
DNA Meth. m5C
H3−5ac
0.00
0.25
0.50
0.75
1.00
π̂
JARID2 – Reverse
H4K20 me2:H3−5ac
H2A.Z:H4K20 me2
H3K4 me1:H3−5ac
H3K27 me3:H3K4 me1
H3K4 me1:H2A.Z
H4K20 me2:H3K4 me3
H3K27 me2:DNA Meth. m5C
H2A.Z:H4−4ac
H2A.Z:H3K4 me3
H3K27 me3:H4K20 me3
H3K4 me3:H3−5ac
H2A.Z:H3−5ac
H3K9 me2:DNA Meth. m5C
H3K27 me3:H3K9 me3
H3K9 me3:H4K20 me3
H4−4ac:H3−5ac
H3K4 me3:H4−4ac
DNA Meth. m5C:H3K9 me3
H3K27 me3:DNA Meth. m5C
DNA Meth. m5C:H4K20 me3
H3K27 me2
H3K9 me2
H4K20 me2
H3K4 me1
H3K27 me3
H3K9 me3
H4−4ac
H3K4 me3
H2A.Z
H4K20 me3
DNA Meth. m5C
H3−5ac
0.00
0.25
0.50
0.75
1.00
π̂
LEKR1 – Forward
DNA Meth. m5C:H4K20 me2
DNA Meth. m5C:H4K20 me3
DNA Meth. m5C:H3K9 me3
H3K4 me1:H4K20 me2
H2A.Z:H4K20 me2
H3K27 me3:H3K4 me1
H2A.Z:H4−4ac
H2A.Z:H3−5ac
H3K4 me1:H4−4ac
H3K4 me1:H3−5ac
H3K4 me3:H3−5ac
H3K4 me1:H2A.Z
H4K20 me2:H4−4ac
H3K27 me2
H3K9 me2
H2A.Z:H3K4 me3
H4K20 me2:H3−5ac
H4K20 me2:H3K4 me3
H4−4ac:H3−5ac
H3K4 me3:H4−4ac
H4K20 me3
H3K9 me3
H3K27 me3
DNA Meth. m5C
H3K4 me1
H3K4 me3
H4K20 me2
H2A.Z
H3−5ac
H4−4ac
0.00
0.25
0.50
0.75
1.00
π̂
LEKR1 – Reverse
DNA Meth. m5C:H4K20 me2
DNA Meth. m5C:H4K20 me3
DNA Meth. m5C:H3K9 me3
H3K4 me1:H4K20 me2
H2A.Z:H4K20 me2
H3K27 me3:H3K4 me1
H2A.Z:H4−4ac
H2A.Z:H3−5ac
H3K4 me1:H4−4ac
H3K4 me1:H3−5ac
H3K4 me3:H3−5ac
H3K4 me1:H2A.Z
H4K20 me2:H4−4ac
H3K27 me2
H3K9 me2
H2A.Z:H3K4 me3
H4K20 me2:H3−5ac
H4K20 me2:H3K4 me3
H4−4ac:H3−5ac
H3K4 me3:H4−4ac
H4K20 me3
H3K9 me3
H3K27 me3
DNA Meth. m5C
H3K4 me1
H3K4 me3
H4K20 me2
H2A.Z
H3−5ac
H4−4ac
0.00
0.25
0.50
0.75
1.00
π̂
PSIP1 (2) – Forward
H4K20 me2:H4−4ac
H2A.Z:H4−4ac
DNA Meth. m5C:H4K20 me2
H2A.Z:H3−5ac
H2A.Z:H4K20 me2
H3K4 me1:H4−4ac
H2A.Z:H3K4 me3
H3K4 me1:H2A.Z
H4K20 me2:H3−5ac
H3K9 me3:H4K20 me3
H4K20 me2:H3K4 me3
H3K27 me3:H4K20 me3
H3K4 me1:H3−5ac
H3K4 me3:H3−5ac
DNA Meth. m5C:H3K9 me3
H4−4ac:H3−5ac
H3K27 me3:DNA Meth. m5C
H3K4 me3:H4−4ac
H3K9 me2
H3K27 me3:H3K4 me1
H3K27 me2
H3K27 me3:H3K9 me3
H4K20 me2
H2A.Z
DNA Meth. m5C
H4K20 me3
H3K4 me1
H3K9 me3
H3K4 me3
H4−4ac
H3K27 me3
H3−5ac
0.00
0.25
0.50
0.75
1.00
π̂
PSIP1 (2) – Reverse
H4K20 me2:H4−4ac
H2A.Z:H4−4ac
DNA Meth. m5C:H4K20 me2
H2A.Z:H3−5ac
H2A.Z:H4K20 me2
H3K4 me1:H4−4ac
H2A.Z:H3K4 me3
H3K4 me1:H2A.Z
H4K20 me2:H3−5ac
H3K9 me3:H4K20 me3
H4K20 me2:H3K4 me3
H3K27 me3:H4K20 me3
H3K4 me1:H3−5ac
H3K4 me3:H3−5ac
DNA Meth. m5C:H3K9 me3
H4−4ac:H3−5ac
H3K27 me3:DNA Meth. m5C
H3K4 me3:H4−4ac
H3K9 me2
H3K27 me3:H3K4 me1
H3K27 me2
H3K27 me3:H3K9 me3
H4K20 me2
H2A.Z
DNA Meth. m5C
H4K20 me3
H3K4 me1
H3K9 me3
H3K4 me3
H4−4ac
H3K27 me3
H3−5ac
0.00
0.25
0.50
0.75
1.00
π̂
VKORC1 – Forward
DNA Meth. m5C:H4K20 me2
H3K27 me3:H4K20 me3
H3K9 me3:H4K20 me3
H4K20 me2:H4−4ac
H3K27 me2:DNA Meth. m5C
H2A.Z:H3K4 me3
H2A.Z:H4K20 me2
H3K4 me1:H4−4ac
H3K4 me1:H4K20 me2
H3K4 me1:H2A.Z
H3K9 me2:DNA Meth. m5C
H3K4 me3:H3−5ac
H3K4 me1:H3−5ac
H4K20 me2:H3−5ac
H4K20 me2:H3K4 me3
DNA Meth. m5C:H3K9 me3
H3K27 me3:H3K9 me3
DNA Meth. m5C:H4K20 me3
H3K4 me3:H4−4ac
H3K27 me3:DNA Meth. m5C
H4−4ac:H3−5ac
H3K9 me2
H3K27 me2
H4K20 me2
H3K9 me3
H2A.Z
H4K20 me3
H3K27 me3
H3K4 me3
H3K4 me1
H3−5ac
DNA Meth. m5C
H4−4ac
0.00
0.25
0.50
0.75
1.00
π̂
VKORC1 – Reverse
DNA Meth. m5C:H4K20 me2
H3K27 me3:H4K20 me3
H3K9 me3:H4K20 me3
H4K20 me2:H4−4ac
H3K27 me2:DNA Meth. m5C
H2A.Z:H3K4 me3
H2A.Z:H4K20 me2
H3K4 me1:H4−4ac
H3K4 me1:H4K20 me2
H3K4 me1:H2A.Z
H3K9 me2:DNA Meth. m5C
H3K4 me3:H3−5ac
H3K4 me1:H3−5ac
H4K20 me2:H3−5ac
H4K20 me2:H3K4 me3
DNA Meth. m5C:H3K9 me3
H3K27 me3:H3K9 me3
DNA Meth. m5C:H4K20 me3
H3K4 me3:H4−4ac
H3K27 me3:DNA Meth. m5C
H4−4ac:H3−5ac
H3K9 me2
H3K27 me2
H4K20 me2
H3K9 me3
H2A.Z
H4K20 me3
H3K27 me3
H3K4 me3
H3K4 me1
H3−5ac
DNA Meth. m5C
H4−4ac
0.00
0.25
0.50
0.75
1.00
π̂

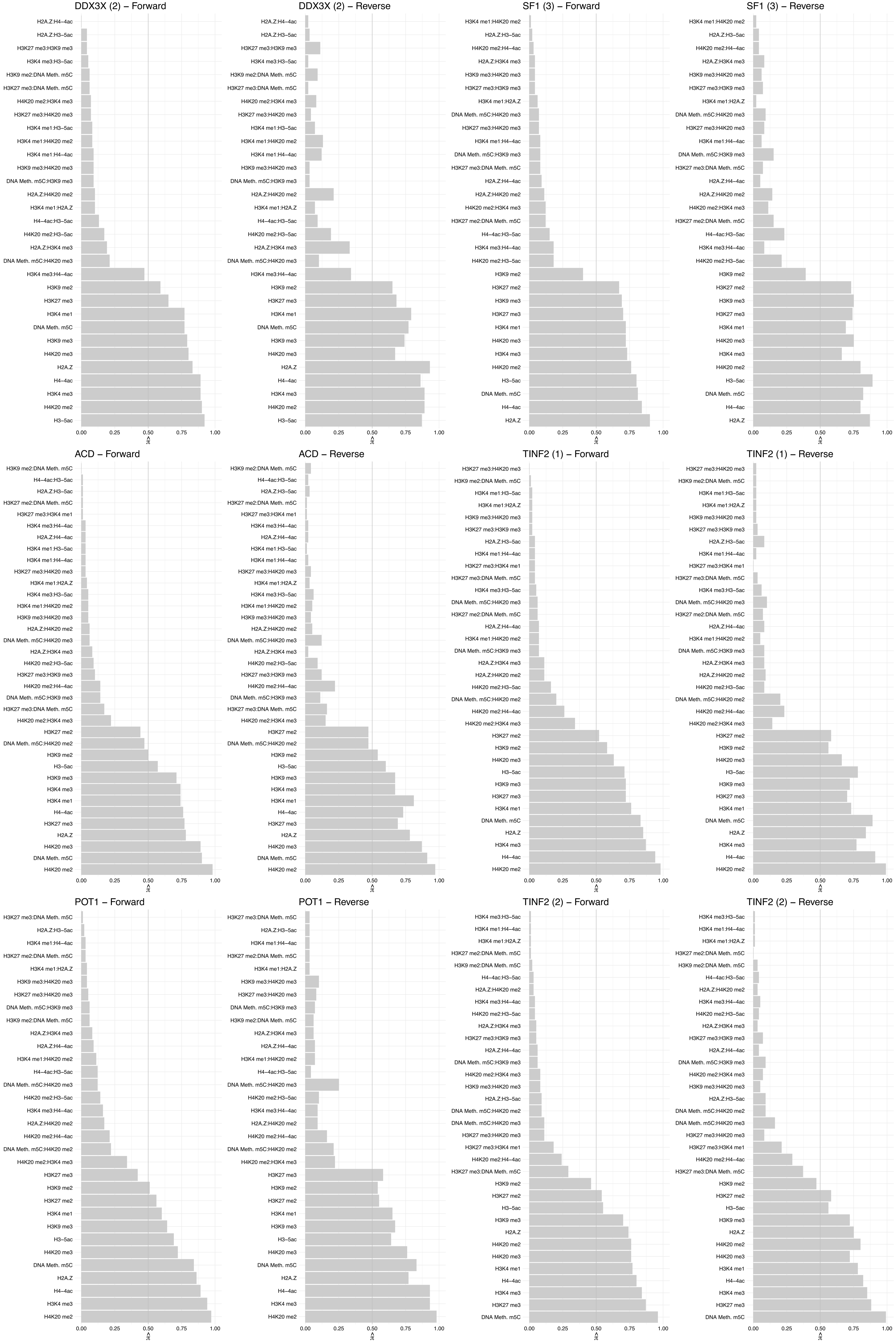
DDX3X (2) – Forward
H2A.Z:H4–4ac
H2A.Z:H3–5ac
H3K27 me3:H3K9 me3
H3K4 me3:H3–5ac
H3K9 me2:DNA Meth. m5C
H3K27 me3:DNA Meth. m5C
H4K20 me2:H3K4 me3
H3K27 me3:H4K20 me3
H3K4 me1:H3–5ac
H3K4 me1:H4K20 me2
H3K4 me1:H4–4ac
H3K9 me3:H4K20 me3
DNA Meth. m5C:H3K9 me3
H2A.Z:H4K20 me2
H3K4 me1:H2A.Z
H4–4ac:H3–5ac
H4K20 me2:H3–5ac
H2A.Z:H3K4 me3
DNA Meth. m5C:H4K20 me3
H3K4 me3:H4–4ac
H3K9 me2
H3K27 me3
H3K4 me1
DNA Meth. m5C
H3K9 me3
H4K20 me3
H2A.Z
H4–4ac
H3K4 me3
H4K20 me2
H3–5ac
0.00
0.25
0.50
0.75
1.00
$\hat{\pi}$
DDX3X (2) – Reverse
H2A.Z:H4–4ac
H2A.Z:H3–5ac
H3K27 me3:H3K9 me3
H3K4 me3:H3–5ac
H3K9 me2:DNA Meth. m5C
H3K27 me3:DNA Meth. m5C
H4K20 me2:H3K4 me3
H3K27 me3:H4K20 me3
H3K4 me1:H3–5ac
H3K4 me1:H4K20 me2
H3K4 me1:H4–4ac
H3K9 me3:H4K20 me3
DNA Meth. m5C:H3K9 me3
H2A.Z:H4K20 me2
H3K4 me1:H2A.Z
H4–4ac:H3–5ac
H4K20 me2:H3–5ac
H2A.Z:H3K4 me3
DNA Meth. m5C:H4K20 me3
H3K4 me3:H4–4ac
H3K9 me2
H3K27 me3
H3K4 me1
DNA Meth. m5C
H3K9 me3
H4K20 me3
H2A.Z
H4–4ac
H3K4 me3
H4K20 me2
H3–5ac
0.00
0.25
0.50
0.75
1.00
$\hat{\pi}$
SF1 (3) – Forward
H3K4 me1:H4K20 me2
H2A.Z:H3–5ac
H4K20 me2:H4–4ac
H2A.Z:H3K4 me3
H3K9 me3:H4K20 me3
H3K27 me3:H3K9 me3
H3K4 me1:H2A.Z
DNA Meth. m5C:H4K20 me3
H3K27 me3:H4K20 me3
H3K4 me1:H4–4ac
DNA Meth. m5C:H3K9 me3
H3K27 me3:DNA Meth. m5C
H2A.Z:H4–4ac
H2A.Z:H4K20 me2
H4K20 me2:H3K4 me3
H3K27 me2:DNA Meth. m5C
H4–4ac:H3–5ac
H3K4 me3:H4–4ac
H4K20 me2:H3–5ac
H3K9 me2
H3K27 me2
H3K9 me3
H3K27 me3
H3K4 me1
H4K20 me3
H3K4 me3
H4K20 me2
H3–5ac
DNA Meth. m5C
H4–4ac
H2A.Z
0.00
0.25
0.50
0.75
1.00
$\hat{\pi}$
SF1 (3) – Reverse
H3K4 me1:H4K20 me2
H2A.Z:H3–5ac
H4K20 me2:H4–4ac
H2A.Z:H3K4 me3
H3K9 me3:H4K20 me3
H3K27 me3:H3K9 me3
H3K4 me1:H2A.Z
DNA Meth. m5C:H4K20 me3
H3K27 me3:H4K20 me3
H3K4 me1:H4–4ac
DNA Meth. m5C:H3K9 me3
H3K27 me3:DNA Meth. m5C
H2A.Z:H4–4ac
H2A.Z:H4K20 me2
H4K20 me2:H3K4 me3
H3K27 me2:DNA Meth. m5C
H4–4ac:H3–5ac
H3K4 me3:H4–4ac
H4K20 me2:H3–5ac
H3K9 me2
H3K27 me2
H3K9 me3
H3K27 me3
H3K4 me1
H4K20 me3
H3K4 me3
H4K20 me2
H3–5ac
DNA Meth. m5C
H4–4ac
H2A.Z
0.00
0.25
0.50
0.75
1.00
$\hat{\pi}$
ACD – Forward
H3K9 me2:DNA Meth. m5C
H4–4ac:H3–5ac
H2A.Z:H3–5ac
H3K27 me2:DNA Meth. m5C
H3K27 me3:H3K4 me1
H3K4 me3:H4–4ac
H2A.Z:H4–4ac
H3K4 me1:H3–5ac
H3K4 me1:H4–4ac
H3K27 me3:H4K20 me3
H3K4 me1:H2A.Z
H3K4 me3:H3–5ac
H3K4 me1:H4K20 me2
H3K9 me3:H4K20 me3
H2A.Z:H4K20 me2
DNA Meth. m5C:H4K20 me3
H2A.Z:H3K4 me3
H4K20 me2:H3–5ac
H3K27 me3:H3K9 me3
H4K20 me2:H4–4ac
DNA Meth. m5C:H3K9 me3
H3K27 me3:DNA Meth. m5C
H4K20 me2:H3K4 me3
H3K27 me2
DNA Meth. m5C:H4K20 me2
H3K9 me2
H3–5ac
H3K9 me3
H3K4 me3
H3K4 me1
H4–4ac
H3K27 me3
H2A.Z
H4K20 me3
DNA Meth. m5C
H4K20 me2
0.00
0.25
0.50
0.75
1.00
$\hat{\pi}$
ACD – Reverse
H3K9 me2:DNA Meth. m5C
H4–4ac:H3–5ac
H2A.Z:H3–5ac
H3K27 me2:DNA Meth. m5C
H3K27 me3:H3K4 me1
H3K4 me3:H4–4ac
H2A.Z:H4–4ac
H3K4 me1:H3–5ac
H3K4 me1:H4–4ac
H3K27 me3:H4K20 me3
H3K4 me1:H2A.Z
H3K4 me3:H3–5ac
H3K4 me1:H4K20 me2
H3K9 me3:H4K20 me3
H2A.Z:H4K20 me2
DNA Meth. m5C:H4K20 me3
H2A.Z:H3K4 me3
H4K20 me2:H3–5ac
H3K27 me3:H3K9 me3
H4K20 me2:H4–4ac
DNA Meth. m5C:H3K9 me3
H3K27 me3:DNA Meth. m5C
H4K20 me2:H3K4 me3
H3K27 me2
DNA Meth. m5C:H4K20 me2
H3K9 me2
H3–5ac
H3K9 me3
H3K4 me3
H3K4 me1
H4–4ac
H3K27 me3
H2A.Z
H4K20 me3
DNA Meth. m5C
H4K20 me2
0.00
0.25
0.50
0.75
1.00
$\hat{\pi}$
TINF2 (1) – Forward
H3K27 me3:H4K20 me3
H3K9 me2:DNA Meth. m5C
H3K4 me1:H3–5ac
H3K4 me1:H2A.Z
H3K9 me3:H4K20 me3
H3K27 me3:H3K9 me3
H2A.Z:H3–5ac
H3K4 me1:H4–4ac
H3K27 me3:H3K4 me1
H3K27 me3:DNA Meth. m5C
H3K4 me3:H3–5ac
DNA Meth. m5C:H4K20 me3
H3K27 me2:DNA Meth. m5C
H2A.Z:H4–4ac
H3K4 me1:H4K20 me2
DNA Meth. m5C:H3K9 me3
H2A.Z:H3K4 me3
H2A.Z:H4K20 me2
H4K20 me2:H3–5ac
DNA Meth. m5C:H4K20 me2
H4K20 me2:H4–4ac
H4K20 me2:H3K4 me3
H3K27 me2
H3K9 me2
H4K20 me3
H3–5ac
H3K9 me3
H3K27 me3
H3K4 me1
DNA Meth. m5C
H2A.Z
H3K4 me3
H4–4ac
H4K20 me2
0.00
0.25
0.50
0.75
1.00
$\hat{\pi}$
TINF2 (1) – Reverse
H3K27 me3:H4K20 me3
H3K9 me2:DNA Meth. m5C
H3K4 me1:H3–5ac
H3K4 me1:H2A.Z
H3K9 me3:H4K20 me3
H3K27 me3:H3K9 me3
H2A.Z:H3–5ac
H3K4 me1:H4–4ac
H3K27 me3:H3K4 me1
H3K27 me3:DNA Meth. m5C
H3K4 me3:H3–5ac
DNA Meth. m5C:H4K20 me3
H3K27 me2:DNA Meth. m5C
H2A.Z:H4–4ac
H3K4 me1:H4K20 me2
DNA Meth. m5C:H3K9 me3
H2A.Z:H3K4 me3
H2A.Z:H4K20 me2
H4K20 me2:H3–5ac
DNA Meth. m5C:H4K20 me2
H4K20 me2:H4–4ac
H4K20 me2:H3K4 me3
H3K27 me2
H3K9 me2
H4K20 me3
H3–5ac
H3K9 me3
H3K27 me3
H3K4 me1
DNA Meth. m5C
H2A.Z
H3K4 me3
H4–4ac
H4K20 me2
0.00
0.25
0.50
0.75
1.00
$\hat{\pi}$
POT1 – Forward
H3K27 me3:DNA Meth. m5C
H2A.Z:H3–5ac
H3K4 me1:H4–4ac
H3K27 me2:DNA Meth. m5C
H3K4 me1:H2A.Z
H3K9 me3:H4K20 me3
H3K27 me3:H4K20 me3
DNA Meth. m5C:H3K9 me3
H3K9 me2:DNA Meth. m5C
H2A.Z:H3K4 me3
H2A.Z:H4–4ac
H3K4 me1:H4K20 me2
H4–4ac:H3–5ac
DNA Meth. m5C:H4K20 me3
H4K20 me2:H3–5ac
H3K4 me3:H4–4ac
H2A.Z:H4K20 me2
H4K20 me2:H4–4ac
DNA Meth. m5C:H4K20 me2
H4K20 me2:H3K4 me3
H3K27 me3
H3K9 me2
H3K27 me2
H3K4 me1
H3K9 me3
H3–5ac
H4K20 me3
DNA Meth. m5C
H2A.Z
H4–4ac
H3K4 me3
H4K20 me2
0.00
0.25
0.50
0.75
1.00
$\hat{\pi}$
POT1 – Reverse
H3K27 me3:DNA Meth. m5C
H2A.Z:H3–5ac
H3K4 me1:H4–4ac
H3K27 me2:DNA Meth. m5C
H3K4 me1:H2A.Z
H3K9 me3:H4K20 me3
H3K27 me3:H4K20 me3
DNA Meth. m5C:H3K9 me3
H3K9 me2:DNA Meth. m5C
H2A.Z:H3K4 me3
H2A.Z:H4–4ac
H3K4 me1:H4K20 me2
H4–4ac:H3–5ac
DNA Meth. m5C:H4K20 me3
H4K20 me2:H3–5ac
H3K4 me3:H4–4ac
H2A.Z:H4K20 me2
H4K20 me2:H4–4ac
DNA Meth. m5C:H4K20 me2
H4K20 me2:H3K4 me3
H3K27 me3
H3K9 me2
H3K27 me2
H3K4 me1
H3K9 me3
H3–5ac
H4K20 me3
DNA Meth. m5C
H2A.Z
H4–4ac
H3K4 me3
H4K20 me2
0.00
0.25
0.50
0.75
1.00
$\hat{\pi}$
TINF2 (2) – Forward
H3K4 me3:H3–5ac
H3K4 me1:H4–4ac
H3K4 me1:H2A.Z
H3K27 me2:DNA Meth. m5C
H3K9 me2:DNA Meth. m5C
H4–4ac:H3–5ac
H2A.Z:H4K20 me2
H3K4 me3:H4–4ac
H4K20 me2:H3–5ac
H2A.Z:H3K4 me3
H3K27 me3:H3K9 me3
H2A.Z:H4–4ac
DNA Meth. m5C:H3K9 me3
H4K20 me2:H3K4 me3
H3K9 me3:H4K20 me3
H2A.Z:H3–5ac
DNA Meth. m5C:H4K20 me2
DNA Meth. m5C:H4K20 me3
H3K27 me3:H4K20 me3
H3K27 me3:H3K4 me1
H4K20 me2:H4–4ac
H3K27 me3:DNA Meth. m5C
H3K9 me2
H3K27 me2
H3–5ac
H3K9 me3
H2A.Z
H4K20 me2
H4K20 me3
H3K4 me1
H4–4ac
H3K4 me3
H3K27 me3
DNA Meth. m5C
0.00
0.25
0.50
0.75
1.00
$\hat{\pi}$
TINF2 (2) – Reverse
H3K4 me3:H3–5ac
H3K4 me1:H4–4ac
H3K4 me1:H2A.Z
H3K27 me2:DNA Meth. m5C
H3K9 me2:DNA Meth. m5C
H4–4ac:H3–5ac
H2A.Z:H4K20 me2
H3K4 me3:H4–4ac
H4K20 me2:H3–5ac
H2A.Z:H3K4 me3
H3K27 me3:H3K9 me3
H2A.Z:H4–4ac
DNA Meth. m5C:H3K9 me3
H4K20 me2:H3K4 me3
H3K9 me3:H4K20 me3
H2A.Z:H3–5ac
DNA Meth. m5C:H4K20 me2
DNA Meth. m5C:H4K20 me3
H3K27 me3:H4K20 me3
H3K27 me3:H3K4 me1
H4K20 me2:H4–4ac
H3K27 me3:DNA Meth. m5C
H3K9 me2
H3K27 me2
H3–5ac
H3K9 me3
H2A.Z
H4K20 me2
H4K20 me3
H3K4 me1
H4–4ac
H3K4 me3
H3K27 me3
DNA Meth. m5C
0.00
0.25
0.50
0.75
1.00
$\hat{\pi}$

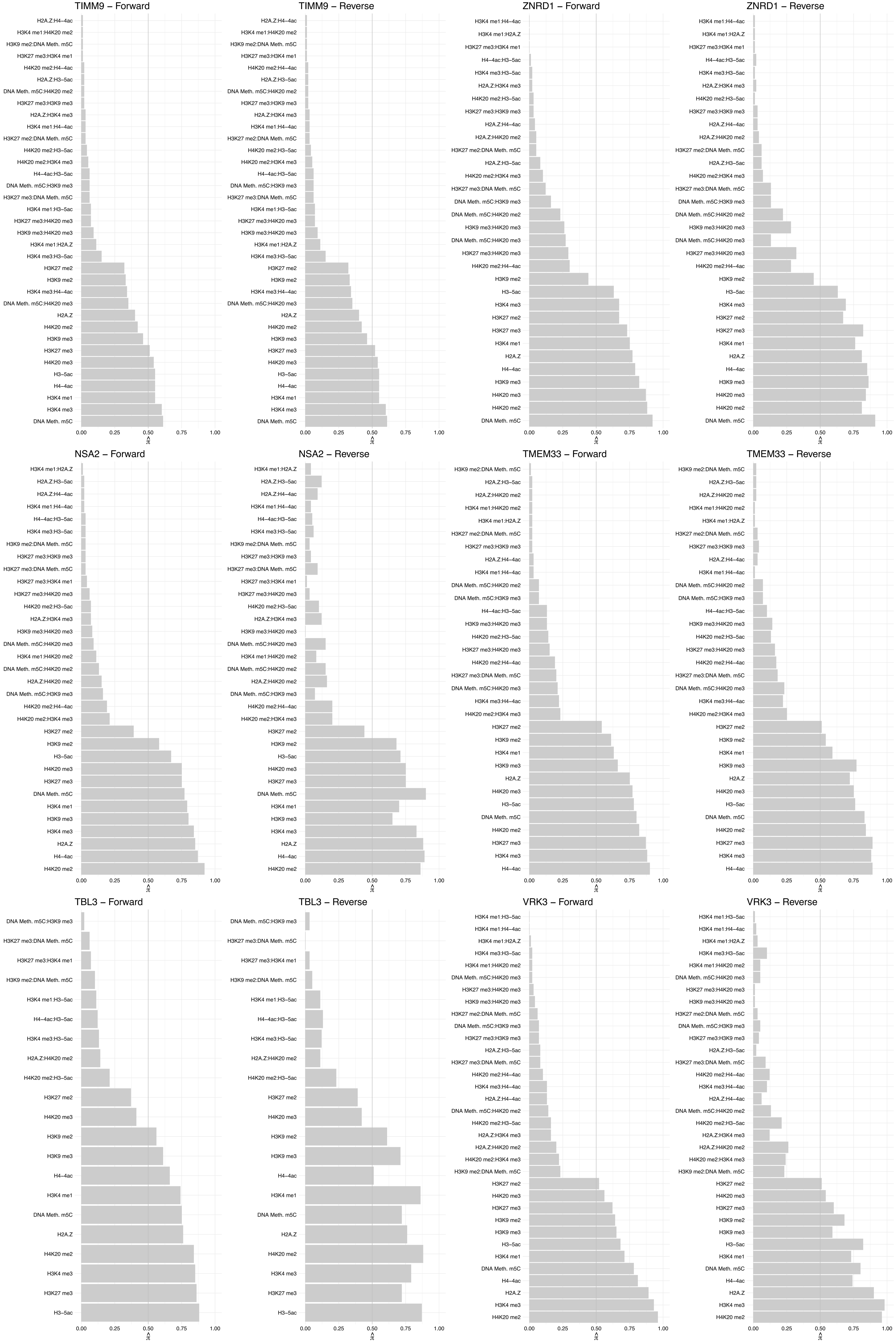

TIMM9 – Forward
H2A.Z:H4−4ac
H3K4 me1:H4K20 me2
H3K9 me2:DNA Meth. m5C
H3K27 me3:H3K4 me1
H4K20 me2:H4−4ac
H2A.Z:H3−5ac
DNA Meth. m5C:H4K20 me2
H3K27 me3:H3K9 me3
H2A.Z:H3K4 me3
H3K4 me1:H4−4ac
H3K27 me2:DNA Meth. m5C
H4K20 me2:H3−5ac
H4K20 me2:H3K4 me3
H4−4ac:H3−5ac
DNA Meth. m5C:H3K9 me3
H3K27 me3:DNA Meth. m5C
H3K4 me1:H3−5ac
H3K27 me3:H4K20 me3
H3K9 me3:H4K20 me3
H3K4 me1:H2A.Z
H3K4 me3:H3−5ac
H3K27 me2
H3K9 me2
H3K4 me3:H4−4ac
DNA Meth. m5C:H4K20 me3
H2A.Z
H4K20 me2
H3K9 me3
H3K27 me3
H4K20 me3
H3−5ac
H4−4ac
H3K4 me1
H3K4 me3
DNA Meth. m5C
0.00 0.25 0.50 0.75 1.00
π̂
TIMM9 – Reverse
H2A.Z:H4−4ac
H3K4 me1:H4K20 me2
H3K9 me2:DNA Meth. m5C
H3K27 me3:H3K4 me1
H4K20 me2:H4−4ac
H2A.Z:H3−5ac
DNA Meth. m5C:H4K20 me2
H3K27 me3:H3K9 me3
H2A.Z:H3K4 me3
H3K4 me1:H4−4ac
H3K27 me2:DNA Meth. m5C
H4K20 me2:H3−5ac
H4K20 me2:H3K4 me3
H4−4ac:H3−5ac
DNA Meth. m5C:H3K9 me3
H3K27 me3:DNA Meth. m5C
H3K4 me1:H3−5ac
H3K27 me3:H4K20 me3
H3K9 me3:H4K20 me3
H3K4 me1:H2A.Z
H3K4 me3:H3−5ac
H3K27 me2
H3K9 me2
H3K4 me3:H4−4ac
DNA Meth. m5C:H4K20 me3
H2A.Z
H4K20 me2
H3K9 me3
H3K27 me3
H4K20 me3
H3−5ac
H4−4ac
H3K4 me1
H3K4 me3
DNA Meth. m5C
0.00 0.25 0.50 0.75 1.00
π̂
ZNRD1 – Forward
H3K4 me1:H4−4ac
H3K4 me1:H2A.Z
H3K27 me3:H3K4 me1
H4−4ac:H3−5ac
H3K4 me3:H3−5ac
H2A.Z:H3K4 me3
H4K20 me2:H3−5ac
H3K27 me3:H3K9 me3
H2A.Z:H4−4ac
H2A.Z:H4K20 me2
H3K27 me2:DNA Meth. m5C
H2A.Z:H3−5ac
H4K20 me2:H3K4 me3
H3K27 me3:DNA Meth. m5C
DNA Meth. m5C:H3K9 me3
DNA Meth. m5C:H4K20 me2
H3K9 me3:H4K20 me3
DNA Meth. m5C:H4K20 me3
H3K27 me3:H4K20 me3
H4K20 me2:H4−4ac
H3K9 me2
H3−5ac
H3K4 me3
H3K27 me2
H3K27 me3
H3K4 me1
H2A.Z
H4−4ac
H3K9 me3
H4K20 me3
H4K20 me2
DNA Meth. m5C
0.00 0.25 0.50 0.75 1.00
π̂
ZNRD1 – Reverse
H3K4 me1:H4−4ac
H3K4 me1:H2A.Z
H3K27 me3:H3K4 me1
H4−4ac:H3−5ac
H3K4 me3:H3−5ac
H2A.Z:H3K4 me3
H4K20 me2:H3−5ac
H3K27 me3:H3K9 me3
H2A.Z:H4−4ac
H2A.Z:H4K20 me2
H3K27 me2:DNA Meth. m5C
H2A.Z:H3−5ac
H4K20 me2:H3K4 me3
H3K27 me3:DNA Meth. m5C
DNA Meth. m5C:H3K9 me3
DNA Meth. m5C:H4K20 me2
H3K9 me3:H4K20 me3
DNA Meth. m5C:H4K20 me3
H3K27 me3:H4K20 me3
H4K20 me2:H4−4ac
H3K9 me2
H3−5ac
H3K4 me3
H3K27 me2
H3K27 me3
H3K4 me1
H2A.Z
H4−4ac
H3K9 me3
H4K20 me3
H4K20 me2
DNA Meth. m5C
0.00 0.25 0.50 0.75 1.00
π̂
NSA2 – Forward
H3K4 me1:H2A.Z
H2A.Z:H3−5ac
H2A.Z:H4−4ac
H3K4 me1:H4−4ac
H4−4ac:H3−5ac
H3K4 me3:H3−5ac
H3K9 me2:DNA Meth. m5C
H3K27 me3:H3K9 me3
H3K27 me3:DNA Meth. m5C
H3K27 me3:H3K4 me1
H3K27 me3:H4K20 me3
H4K20 me2:H3−5ac
H2A.Z:H3K4 me3
H3K9 me3:H4K20 me3
DNA Meth. m5C:H4K20 me3
H3K4 me1:H4K20 me2
DNA Meth. m5C:H4K20 me2
H2A.Z:H4K20 me2
DNA Meth. m5C:H3K9 me3
H4K20 me2:H4−4ac
H4K20 me2:H3K4 me3
H3K27 me2
H3K9 me2
H3−5ac
H4K20 me3
H3K27 me3
DNA Meth. m5C
H3K4 me1
H3K9 me3
H3K4 me3
H2A.Z
H4−4ac
H4K20 me2
0.00 0.25 0.50 0.75 1.00
π̂
NSA2 – Reverse
H3K4 me1:H2A.Z
H2A.Z:H3−5ac
H2A.Z:H4−4ac
H3K4 me1:H4−4ac
H4−4ac:H3−5ac
H3K4 me3:H3−5ac
H3K9 me2:DNA Meth. m5C
H3K27 me3:H3K9 me3
H3K27 me3:DNA Meth. m5C
H3K27 me3:H3K4 me1
H3K27 me3:H4K20 me3
H4K20 me2:H3−5ac
H2A.Z:H3K4 me3
H3K9 me3:H4K20 me3
DNA Meth. m5C:H4K20 me3
H3K4 me1:H4K20 me2
DNA Meth. m5C:H4K20 me2
H2A.Z:H4K20 me2
DNA Meth. m5C:H3K9 me3
H4K20 me2:H4−4ac
H4K20 me2:H3K4 me3
H3K27 me2
H3K9 me2
H3−5ac
H4K20 me3
H3K27 me3
DNA Meth. m5C
H3K4 me1
H3K9 me3
H3K4 me3
H2A.Z
H4−4ac
H4K20 me2
0.00 0.25 0.50 0.75 1.00
π̂
TMEM33 – Forward
H3K9 me2:DNA Meth. m5C
H2A.Z:H3−5ac
H2A.Z:H4K20 me2
H3K4 me1:H4K20 me2
H3K4 me1:H2A.Z
H3K27 me2:DNA Meth. m5C
H3K27 me3:H3K9 me3
H2A.Z:H4−4ac
H3K4 me1:H4−4ac
DNA Meth. m5C:H4K20 me2
DNA Meth. m5C:H3K9 me3
H4−4ac:H3−5ac
H3K9 me3:H4K20 me3
H4K20 me2:H3−5ac
H3K27 me3:H4K20 me3
H4K20 me2:H4−4ac
H3K27 me3:DNA Meth. m5C
DNA Meth. m5C:H4K20 me3
H3K4 me3:H4−4ac
H4K20 me2:H3K4 me3
H3K27 me2
H3K9 me2
H3K4 me1
H3K9 me3
H2A.Z
H4K20 me3
H3−5ac
DNA Meth. m5C
H4K20 me2
H3K27 me3
H3K4 me3
H4−4ac
0.00 0.25 0.50 0.75 1.00
π̂
TMEM33 – Reverse
H3K9 me2:DNA Meth. m5C
H2A.Z:H3−5ac
H2A.Z:H4K20 me2
H3K4 me1:H4K20 me2
H3K4 me1:H2A.Z
H3K27 me2:DNA Meth. m5C
H3K27 me3:H3K9 me3
H2A.Z:H4−4ac
H3K4 me1:H4−4ac
DNA Meth. m5C:H4K20 me2
DNA Meth. m5C:H3K9 me3
H4−4ac:H3−5ac
H3K9 me3:H4K20 me3
H4K20 me2:H3−5ac
H3K27 me3:H4K20 me3
H4K20 me2:H4−4ac
H3K27 me3:DNA Meth. m5C
DNA Meth. m5C:H4K20 me3
H3K4 me3:H4−4ac
H4K20 me2:H3K4 me3
H3K27 me2
H3K9 me2
H3K4 me1
H3K9 me3
H2A.Z
H4K20 me3
H3−5ac
DNA Meth. m5C
H4K20 me2
H3K27 me3
H3K4 me3
H4−4ac
0.00 0.25 0.50 0.75 1.00
π̂
TBL3 – Forward
DNA Meth. m5C:H3K9 me3
H3K27 me3:DNA Meth. m5C
H3K27 me3:H3K4 me1
H3K9 me2:DNA Meth. m5C
H3K4 me1:H3−5ac
H4−4ac:H3−5ac
H3K4 me3:H3−5ac
H2A.Z:H4K20 me2
H4K20 me2:H3−5ac
H3K27 me2
H4K20 me3
H3K9 me2
H3K9 me3
H4−4ac
H3K4 me1
DNA Meth. m5C
H2A.Z
H4K20 me2
H3K4 me3
H3K27 me3
H3−5ac
0.00 0.25 0.50 0.75 1.00
π̂
TBL3 – Reverse
DNA Meth. m5C:H3K9 me3
H3K27 me3:DNA Meth. m5C
H3K27 me3:H3K4 me1
H3K9 me2:DNA Meth. m5C
H3K4 me1:H3−5ac
H4−4ac:H3−5ac
H3K4 me3:H3−5ac
H2A.Z:H4K20 me2
H4K20 me2:H3−5ac
H3K27 me2
H4K20 me3
H3K9 me2
H3K9 me3
H4−4ac
H3K4 me1
DNA Meth. m5C
H2A.Z
H4K20 me2
H3K4 me3
H3K27 me3
H3−5ac
0.00 0.25 0.50 0.75 1.00
π̂
VRK3 – Forward
H3K4 me1:H3−5ac
H3K4 me1:H4−4ac
H3K4 me1:H2A.Z
H3K4 me3:H3−5ac
H3K4 me1:H4K20 me2
DNA Meth. m5C:H4K20 me3
H3K27 me3:H4K20 me3
H3K9 me3:H4K20 me3
H3K27 me2:DNA Meth. m5C
DNA Meth. m5C:H3K9 me3
H3K27 me3:H3K9 me3
H2A.Z:H3−5ac
H3K27 me3:DNA Meth. m5C
H4K20 me2:H4−4ac
H3K4 me3:H4−4ac
H2A.Z:H4−4ac
DNA Meth. m5C:H4K20 me2
H4K20 me2:H3−5ac
H2A.Z:H3K4 me3
H2A.Z:H4K20 me2
H4K20 me2:H3K4 me3
H3K9 me2:DNA Meth. m5C
H3K27 me2
H4K20 me3
H3K27 me3
H3K9 me2
H3K9 me3
H3−5ac
H3K4 me1
DNA Meth. m5C
H4−4ac
H2A.Z
H3K4 me3
H4K20 me2
0.00 0.25 0.50 0.75 1.00
π̂
VRK3 – Reverse
H3K4 me1:H3−5ac
H3K4 me1:H4−4ac
H3K4 me1:H2A.Z
H3K4 me3:H3−5ac
H3K4 me1:H4K20 me2
DNA Meth. m5C:H4K20 me3
H3K27 me3:H4K20 me3
H3K9 me3:H4K20 me3
H3K27 me2:DNA Meth. m5C
DNA Meth. m5C:H3K9 me3
H3K27 me3:H3K9 me3
H2A.Z:H3−5ac
H3K27 me3:DNA Meth. m5C
H4K20 me2:H4−4ac
H3K4 me3:H4−4ac
H2A.Z:H4−4ac
DNA Meth. m5C:H4K20 me2
H4K20 me2:H3−5ac
H2A.Z:H3K4 me3
H2A.Z:H4K20 me2
H4K20 me2:H3K4 me3
H3K9 me2:DNA Meth. m5C
H3K27 me2
H4K20 me3
H3K27 me3
H3K9 me2
H3K9 me3
H3−5ac
H3K4 me1
DNA Meth. m5C
H4−4ac
H2A.Z
H3K4 me3
H4K20 me2
0.00 0.25 0.50 0.75 1.00
π̂

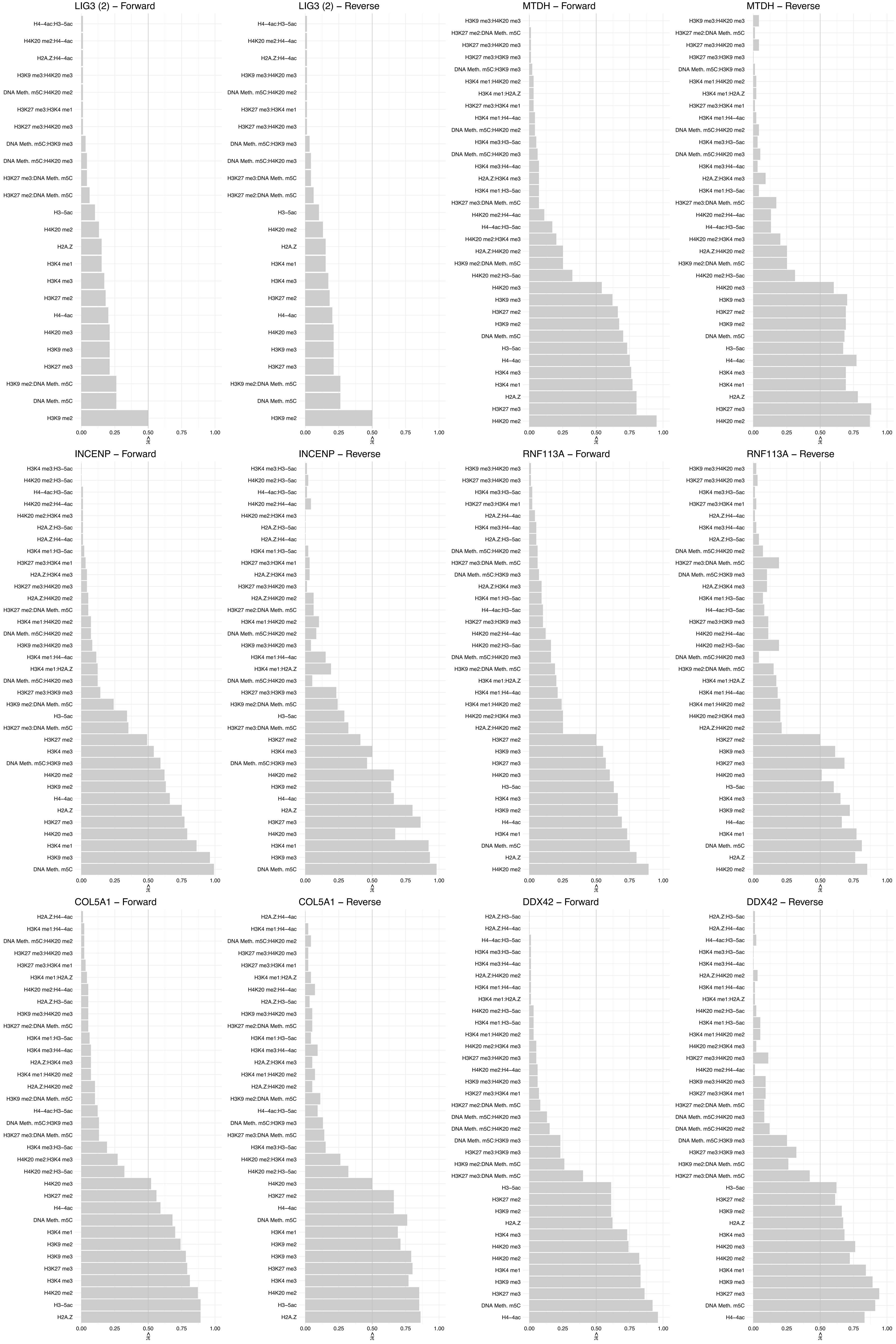
LIG3 (2) – Forward
H4−4ac:H3−5ac
H4K20 me2:H4−4ac
H2A.Z:H4−4ac
H3K9 me3:H4K20 me3
DNA Meth. m5C:H4K20 me2
H3K27 me3:H3K4 me1
H3K27 me3:H4K20 me3
DNA Meth. m5C:H3K9 me3
DNA Meth. m5C:H4K20 me3
H3K27 me3:DNA Meth. m5C
H3K27 me2:DNA Meth. m5C
H3−5ac
H4K20 me2
H2A.Z
H3K4 me1
H3K4 me3
H3K27 me2
H4−4ac
H4K20 me3
H3K9 me3
H3K27 me3
H3K9 me2:DNA Meth. m5C
DNA Meth. m5C
H3K9 me2
0.00 0.25 0.50 0.75 1.00
$\hat{\pi}$
LIG3 (2) – Reverse
MTDH – Forward
H3K9 me3:H4K20 me3
H3K27 me2:DNA Meth. m5C
H3K27 me3:H4K20 me3
H3K27 me3:H3K9 me3
DNA Meth. m5C:H3K9 me3
H3K4 me1:H4K20 me2
H3K4 me1:H2A.Z
H3K27 me3:H3K4 me1
H3K4 me1:H4−4ac
DNA Meth. m5C:H4K20 me2
H3K4 me3:H3−5ac
DNA Meth. m5C:H4K20 me3
H3K4 me3:H4−4ac
H2A.Z:H3K4 me3
H3K4 me1:H3−5ac
H3K27 me3:DNA Meth. m5C
H4K20 me2:H4−4ac
H4−4ac:H3−5ac
H4K20 me2:H3K4 me3
H2A.Z:H4K20 me2
H3K9 me2:DNA Meth. m5C
H4K20 me2:H3−5ac
H4K20 me3
H3K9 me3
H3K27 me2
H3K9 me2
DNA Meth. m5C
H3−5ac
H4−4ac
H3K4 me3
H3K4 me1
H2A.Z
H3K27 me3
H4K20 me2
MTDH – Reverse
INCENP – Forward
H3K4 me3:H3−5ac
H4K20 me2:H3−5ac
H4−4ac:H3−5ac
H4K20 me2:H4−4ac
H4K20 me2:H3K4 me3
H2A.Z:H3−5ac
H2A.Z:H4−4ac
H3K4 me1:H3−5ac
H3K27 me3:H3K4 me1
H2A.Z:H3K4 me3
H3K27 me3:H4K20 me3
H2A.Z:H4K20 me2
H3K27 me2:DNA Meth. m5C
H3K4 me1:H4K20 me2
DNA Meth. m5C:H4K20 me2
H3K9 me3:H4K20 me3
H3K4 me1:H4−4ac
H3K4 me1:H2A.Z
DNA Meth. m5C:H4K20 me3
H3K27 me3:H3K9 me3
H3K9 me2:DNA Meth. m5C
H3−5ac
H3K27 me3:DNA Meth. m5C
H3K27 me2
H3K4 me3
DNA Meth. m5C:H3K9 me3
H4K20 me2
H3K9 me2
H4−4ac
H2A.Z
H3K27 me3
H4K20 me3
H3K4 me1
H3K9 me3
DNA Meth. m5C
INCENP – Reverse
RNF113A – Forward
H3K9 me3:H4K20 me3
H3K27 me3:H4K20 me3
H3K4 me3:H3−5ac
H3K27 me3:H3K4 me1
H2A.Z:H4−4ac
H3K4 me3:H4−4ac
H2A.Z:H3−5ac
DNA Meth. m5C:H4K20 me2
H3K27 me3:DNA Meth. m5C
DNA Meth. m5C:H3K9 me3
H2A.Z:H3K4 me3
H3K4 me1:H3−5ac
H4−4ac:H3−5ac
H3K27 me3:H3K9 me3
H4K20 me2:H4−4ac
H4K20 me2:H3−5ac
DNA Meth. m5C:H4K20 me3
H3K9 me2:DNA Meth. m5C
H3K4 me1:H2A.Z
H3K4 me1:H4−4ac
H3K4 me1:H4K20 me2
H4K20 me2:H3K4 me3
H2A.Z:H4K20 me2
H3K27 me2
H3K9 me3
H3K27 me3
H4K20 me3
H3−5ac
H3K4 me3
H3K9 me2
H4−4ac
H3K4 me1
DNA Meth. m5C
H2A.Z
H4K20 me2
RNF113A – Reverse
COL5A1 – Forward
H2A.Z:H4−4ac
H3K4 me1:H4−4ac
DNA Meth. m5C:H4K20 me2
H3K27 me3:H4K20 me3
H3K27 me3:H3K4 me1
H3K4 me1:H2A.Z
H4K20 me2:H4−4ac
H2A.Z:H3−5ac
H3K9 me3:H4K20 me3
H3K27 me2:DNA Meth. m5C
H3K4 me1:H3−5ac
H3K4 me3:H4−4ac
H2A.Z:H3K4 me3
H3K4 me1:H4K20 me2
H2A.Z:H4K20 me2
H3K9 me2:DNA Meth. m5C
H4−4ac:H3−5ac
DNA Meth. m5C:H3K9 me3
H3K27 me3:DNA Meth. m5C
H3K4 me3:H3−5ac
H4K20 me2:H3K4 me3
H4K20 me2:H3−5ac
H4K20 me3
H3K27 me2
H4−4ac
DNA Meth. m5C
H3K4 me1
H3K9 me2
H3K9 me3
H3K27 me3
H3K4 me3
H4K20 me2
H3−5ac
H2A.Z
COL5A1 – Reverse
DDX42 – Forward
H2A.Z:H3−5ac
H2A.Z:H4−4ac
H4−4ac:H3−5ac
H3K4 me3:H3−5ac
H3K4 me3:H4−4ac
H2A.Z:H4K20 me2
H3K4 me1:H4−4ac
H3K4 me1:H2A.Z
H4K20 me2:H3−5ac
H3K4 me1:H3−5ac
H3K4 me1:H4K20 me2
H4K20 me2:H3K4 me3
H3K27 me3:H4K20 me3
H4K20 me2:H4−4ac
H3K9 me3:H4K20 me3
H3K27 me3:H3K4 me1
H3K27 me2:DNA Meth. m5C
DNA Meth. m5C:H4K20 me3
DNA Meth. m5C:H4K20 me2
DNA Meth. m5C:H3K9 me3
H3K27 me3:H3K9 me3
H3K9 me2:DNA Meth. m5C
H3K27 me3:DNA Meth. m5C
H3−5ac
H3K27 me2
H3K9 me2
H2A.Z
H3K4 me3
H4K20 me3
H4K20 me2
H3K4 me1
H3K9 me3
H3K27 me3
DNA Meth. m5C
H4−4ac
DDX42 – Reverse

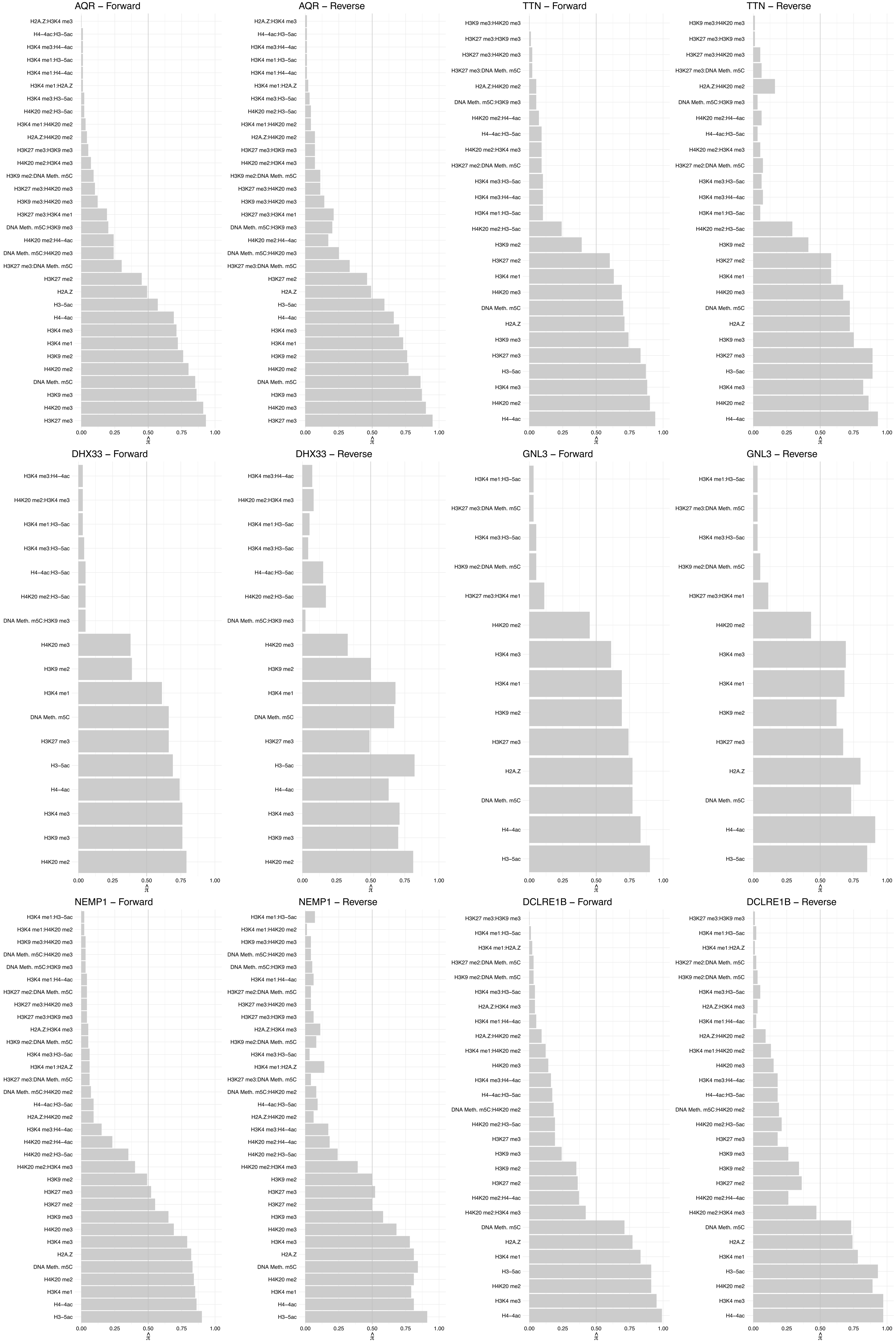
AQR – Forward
H2A.Z:H3K4 me3
H4–4ac:H3–5ac
H3K4 me3:H4–4ac
H3K4 me1:H3–5ac
H3K4 me1:H4–4ac
H3K4 me1:H2A.Z
H3K4 me3:H3–5ac
H4K20 me2:H3–5ac
H3K4 me1:H4K20 me2
H2A.Z:H4K20 me2
H3K27 me3:H3K9 me3
H4K20 me2:H3K4 me3
H3K9 me2:DNA Meth. m5C
H3K27 me3:H4K20 me3
H3K9 me3:H4K20 me3
H3K27 me3:H3K4 me1
DNA Meth. m5C:H3K9 me3
H4K20 me2:H4–4ac
DNA Meth. m5C:H4K20 me3
H3K27 me3:DNA Meth. m5C
H3K27 me2
H2A.Z
H3–5ac
H4–4ac
H3K4 me3
H3K4 me1
H3K9 me2
H4K20 me2
DNA Meth. m5C
H3K9 me3
H4K20 me3
H3K27 me3
0.00
0.25
0.50
0.75
1.00
π̂
AQR – Reverse
TTN – Forward
H3K9 me3:H4K20 me3
H3K27 me3:H3K9 me3
H3K27 me3:H4K20 me3
H3K27 me3:DNA Meth. m5C
H2A.Z:H4K20 me2
DNA Meth. m5C:H3K9 me3
H4K20 me2:H4–4ac
H4–4ac:H3–5ac
H4K20 me2:H3K4 me3
H3K27 me2:DNA Meth. m5C
H3K4 me3:H3–5ac
H3K4 me3:H4–4ac
H3K4 me1:H3–5ac
H4K20 me2:H3–5ac
H3K9 me2
H3K27 me2
H3K4 me1
H4K20 me3
DNA Meth. m5C
H2A.Z
H3K9 me3
H3K27 me3
H3–5ac
H3K4 me3
H4K20 me2
H4–4ac
TTN – Reverse
DHX33 – Forward
H3K4 me3:H4–4ac
H4K20 me2:H3K4 me3
H3K4 me1:H3–5ac
H3K4 me3:H3–5ac
H4–4ac:H3–5ac
H4K20 me2:H3–5ac
DNA Meth. m5C:H3K9 me3
H4K20 me3
H3K9 me2
H3K4 me1
DNA Meth. m5C
H3K27 me3
H3–5ac
H4–4ac
H3K4 me3
H3K9 me3
H4K20 me2
DHX33 – Reverse
GNL3 – Forward
H3K4 me1:H3–5ac
H3K27 me3:DNA Meth. m5C
H3K4 me3:H3–5ac
H3K9 me2:DNA Meth. m5C
H3K27 me3:H3K4 me1
H4K20 me2
H3K4 me3
H3K4 me1
H3K9 me2
H3K27 me3
H2A.Z
DNA Meth. m5C
H4–4ac
H3–5ac
GNL3 – Reverse
NEMP1 – Forward
H3K4 me1:H3–5ac
H3K4 me1:H4K20 me2
H3K9 me3:H4K20 me3
DNA Meth. m5C:H4K20 me3
DNA Meth. m5C:H3K9 me3
H3K4 me1:H4–4ac
H3K27 me2:DNA Meth. m5C
H3K27 me3:H4K20 me3
H3K27 me3:H3K9 me3
H2A.Z:H3K4 me3
H3K9 me2:DNA Meth. m5C
H3K4 me3:H3–5ac
H3K4 me1:H2A.Z
H3K27 me3:DNA Meth. m5C
DNA Meth. m5C:H4K20 me2
H4–4ac:H3–5ac
H2A.Z:H4K20 me2
H3K4 me3:H4–4ac
H4K20 me2:H4–4ac
H4K20 me2:H3–5ac
H4K20 me2:H3K4 me3
H3K9 me2
H3K27 me3
H3K27 me2
H3K9 me3
H4K20 me3
H3K4 me3
H2A.Z
DNA Meth. m5C
H4K20 me2
H3K4 me1
H4–4ac
H3–5ac
NEMP1 – Reverse
DCLRE1B – Forward
H3K27 me3:H3K9 me3
H3K4 me1:H3–5ac
H3K4 me1:H2A.Z
H3K27 me2:DNA Meth. m5C
H3K9 me2:DNA Meth. m5C
H3K4 me3:H3–5ac
H2A.Z:H3K4 me3
H3K4 me1:H4–4ac
H2A.Z:H4K20 me2
H3K4 me1:H4K20 me2
H4K20 me3
H3K4 me3:H4–4ac
H4–4ac:H3–5ac
DNA Meth. m5C:H4K20 me2
H4K20 me2:H3–5ac
H3K27 me3
H3K9 me3
H3K9 me2
H3K27 me2
H4K20 me2:H4–4ac
H4K20 me2:H3K4 me3
DNA Meth. m5C
H2A.Z
H3K4 me1
H3–5ac
H4K20 me2
H3K4 me3
H4–4ac
DCLRE1B – Reverse

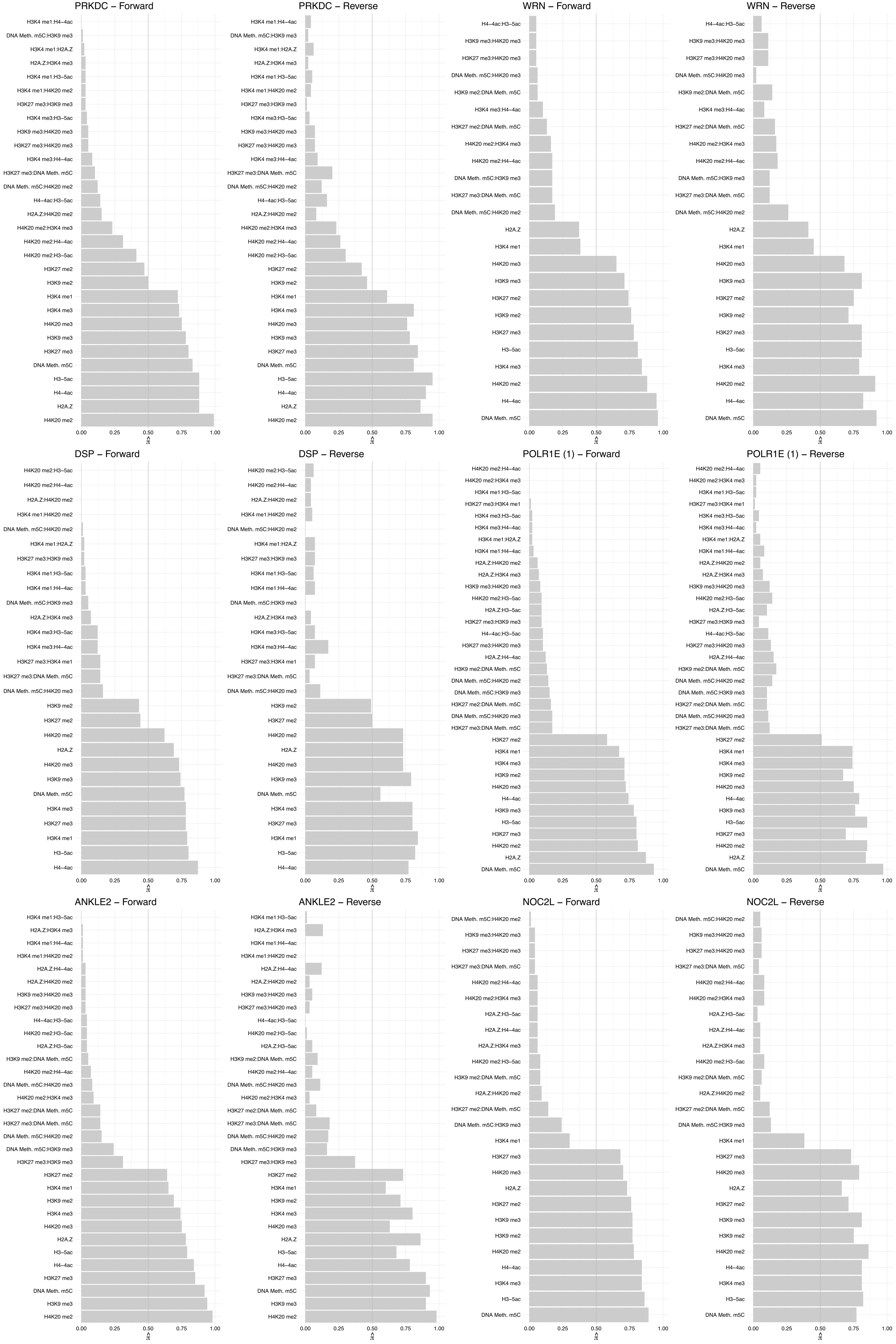

PRKDC – Forward
H3K4 me1:H4–4ac
DNA Meth. m5C:H3K9 me3
H3K4 me1:H2A.Z
H2A.Z:H3K4 me3
H3K4 me1:H3–5ac
H3K4 me1:H4K20 me2
H3K27 me3:H3K9 me3
H3K4 me3:H3–5ac
H3K9 me3:H4K20 me3
H3K27 me3:H4K20 me3
H3K4 me3:H4–4ac
H3K27 me3:DNA Meth. m5C
DNA Meth. m5C:H4K20 me2
H4–4ac:H3–5ac
H2A.Z:H4K20 me2
H4K20 me2:H3K4 me3
H4K20 me2:H4–4ac
H4K20 me2:H3–5ac
H3K27 me2
H3K9 me2
H3K4 me1
H3K4 me3
H4K20 me3
H3K9 me3
H3K27 me3
DNA Meth. m5C
H3–5ac
H4–4ac
H2A.Z
H4K20 me2
0.00
0.25
0.50
0.75
1.00
$\hat{\pi}$
PRKDC – Reverse
WRN – Forward
H4–4ac:H3–5ac
H3K9 me3:H4K20 me3
H3K27 me3:H4K20 me3
DNA Meth. m5C:H4K20 me3
H3K9 me2:DNA Meth. m5C
H3K4 me3:H4–4ac
H3K27 me2:DNA Meth. m5C
H4K20 me2:H3K4 me3
H4K20 me2:H4–4ac
DNA Meth. m5C:H3K9 me3
H3K27 me3:DNA Meth. m5C
DNA Meth. m5C:H4K20 me2
H2A.Z
H3K4 me1
H4K20 me3
H3K9 me3
H3K27 me2
H3K9 me2
H3K27 me3
H3–5ac
H3K4 me3
H4K20 me2
H4–4ac
DNA Meth. m5C
WRN – Reverse
DSP – Forward
H4K20 me2:H3–5ac
H4K20 me2:H4–4ac
H2A.Z:H4K20 me2
H3K4 me1:H4K20 me2
DNA Meth. m5C:H4K20 me2
H3K4 me1:H2A.Z
H3K27 me3:H3K9 me3
H3K4 me1:H3–5ac
H3K4 me1:H4–4ac
DNA Meth. m5C:H3K9 me3
H2A.Z:H3K4 me3
H3K4 me3:H3–5ac
H3K4 me3:H4–4ac
H3K27 me3:H3K4 me1
H3K27 me3:DNA Meth. m5C
DNA Meth. m5C:H4K20 me3
H3K9 me2
H3K27 me2
H4K20 me2
H2A.Z
H4K20 me3
H3K9 me3
DNA Meth. m5C
H3K4 me3
H3K27 me3
H3K4 me1
H3–5ac
H4–4ac
DSP – Reverse
POLR1E (1) – Forward
H4K20 me2:H4–4ac
H4K20 me2:H3K4 me3
H3K4 me1:H3–5ac
H3K27 me3:H3K4 me1
H3K4 me3:H3–5ac
H3K4 me3:H4–4ac
H3K4 me1:H2A.Z
H3K4 me1:H4–4ac
H2A.Z:H4K20 me2
H2A.Z:H3K4 me3
H3K9 me3:H4K20 me3
H4K20 me2:H3–5ac
H2A.Z:H3–5ac
H3K27 me3:H3K9 me3
H4–4ac:H3–5ac
H3K27 me3:H4K20 me3
H2A.Z:H4–4ac
H3K9 me2:DNA Meth. m5C
DNA Meth. m5C:H4K20 me2
DNA Meth. m5C:H3K9 me3
H3K27 me2:DNA Meth. m5C
DNA Meth. m5C:H4K20 me3
H3K27 me3:DNA Meth. m5C
H3K27 me2
H3K4 me1
H3K4 me3
H3K9 me2
H4K20 me3
H4–4ac
H3K9 me3
H3–5ac
H3K27 me3
H4K20 me2
H2A.Z
DNA Meth. m5C
POLR1E (1) – Reverse
ANKLE2 – Forward
H3K4 me1:H3–5ac
H2A.Z:H3K4 me3
H3K4 me1:H4–4ac
H3K4 me1:H4K20 me2
H2A.Z:H4–4ac
H2A.Z:H4K20 me2
H3K9 me3:H4K20 me3
H3K27 me3:H4K20 me3
H4–4ac:H3–5ac
H4K20 me2:H3–5ac
H2A.Z:H3–5ac
H3K9 me2:DNA Meth. m5C
H4K20 me2:H4–4ac
DNA Meth. m5C:H4K20 me3
H4K20 me2:H3K4 me3
H3K27 me2:DNA Meth. m5C
H3K27 me3:DNA Meth. m5C
DNA Meth. m5C:H4K20 me2
DNA Meth. m5C:H3K9 me3
H3K27 me3:H3K9 me3
H3K27 me2
H3K4 me1
H3K9 me2
H3K4 me3
H4K20 me3
H2A.Z
H3–5ac
H4–4ac
H3K27 me3
DNA Meth. m5C
H3K9 me3
H4K20 me2
ANKLE2 – Reverse
NOC2L – Forward
DNA Meth. m5C:H4K20 me2
H3K9 me3:H4K20 me3
H3K27 me3:H4K20 me3
H3K27 me3:DNA Meth. m5C
H4K20 me2:H4–4ac
H4K20 me2:H3K4 me3
H2A.Z:H3–5ac
H2A.Z:H4–4ac
H2A.Z:H3K4 me3
H4K20 me2:H3–5ac
H3K9 me2:DNA Meth. m5C
H2A.Z:H4K20 me2
H3K27 me2:DNA Meth. m5C
DNA Meth. m5C:H3K9 me3
H3K4 me1
H3K27 me3
H4K20 me3
H2A.Z
H3K27 me2
H3K9 me3
H3K9 me2
H4K20 me2
H4–4ac
H3K4 me3
H3–5ac
DNA Meth. m5C
NOC2L – Reverse

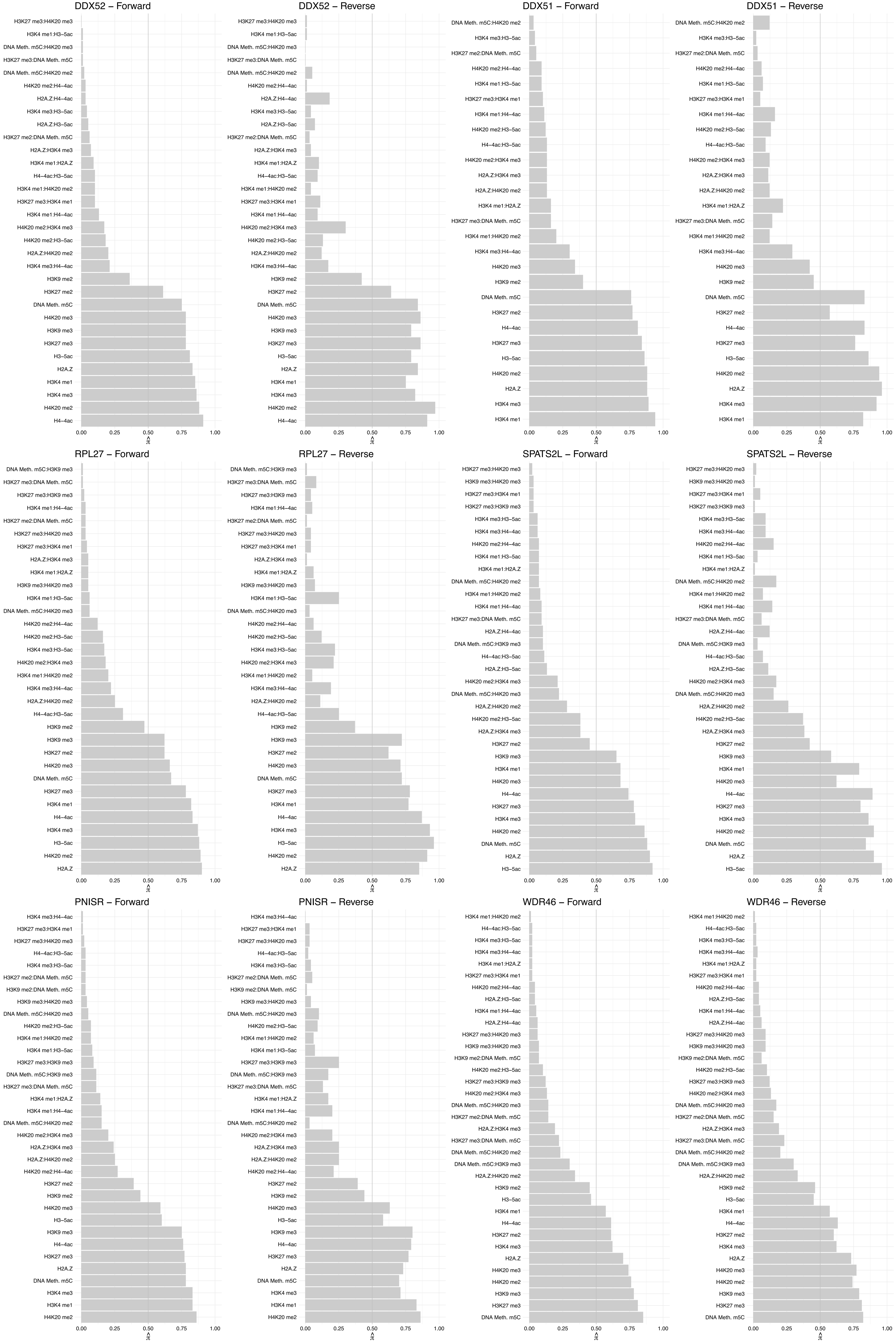
DDX52 − Forward
H3K27 me3:H4K20 me3
H3K4 me1:H3−5ac
DNA Meth. m5C:H4K20 me3
H3K27 me3:DNA Meth. m5C
DNA Meth. m5C:H4K20 me2
H4K20 me2:H4−4ac
H2A.Z:H4−4ac
H3K4 me3:H3−5ac
H2A.Z:H3−5ac
H3K27 me2:DNA Meth. m5C
H2A.Z:H3K4 me3
H3K4 me1:H2A.Z
H4−4ac:H3−5ac
H3K4 me1:H4K20 me2
H3K27 me3:H3K4 me1
H3K4 me1:H4−4ac
H4K20 me2:H3K4 me3
H4K20 me2:H3−5ac
H2A.Z:H4K20 me2
H3K4 me3:H4−4ac
H3K9 me2
H3K27 me2
DNA Meth. m5C
H4K20 me3
H3K9 me3
H3K27 me3
H3−5ac
H2A.Z
H3K4 me1
H3K4 me3
H4K20 me2
H4−4ac
0.00 0.25 0.50 0.75 1.00
π̂
DDX52 − Reverse
H3K27 me3:H4K20 me3
H3K4 me1:H3−5ac
DNA Meth. m5C:H4K20 me3
H3K27 me3:DNA Meth. m5C
DNA Meth. m5C:H4K20 me2
H4K20 me2:H4−4ac
H2A.Z:H4−4ac
H3K4 me3:H3−5ac
H2A.Z:H3−5ac
H3K27 me2:DNA Meth. m5C
H2A.Z:H3K4 me3
H3K4 me1:H2A.Z
H4−4ac:H3−5ac
H3K4 me1:H4K20 me2
H3K27 me3:H3K4 me1
H3K4 me1:H4−4ac
H4K20 me2:H3K4 me3
H4K20 me2:H3−5ac
H2A.Z:H4K20 me2
H3K4 me3:H4−4ac
H3K9 me2
H3K27 me2
DNA Meth. m5C
H4K20 me3
H3K9 me3
H3K27 me3
H3−5ac
H2A.Z
H3K4 me1
H3K4 me3
H4K20 me2
H4−4ac
0.00 0.25 0.50 0.75 1.00
π̂
DDX51 − Forward
DNA Meth. m5C:H4K20 me2
H3K4 me3:H3−5ac
H3K27 me2:DNA Meth. m5C
H4K20 me2:H4−4ac
H3K4 me1:H3−5ac
H3K27 me3:H3K4 me1
H3K4 me1:H4−4ac
H4K20 me2:H3−5ac
H4−4ac:H3−5ac
H4K20 me2:H3K4 me3
H2A.Z:H3K4 me3
H2A.Z:H4K20 me2
H3K4 me1:H2A.Z
H3K27 me3:DNA Meth. m5C
H3K4 me1:H4K20 me2
H3K4 me3:H4−4ac
H4K20 me3
H3K9 me2
DNA Meth. m5C
H3K27 me2
H4−4ac
H3K27 me3
H3−5ac
H4K20 me2
H2A.Z
H3K4 me3
H3K4 me1
0.00 0.25 0.50 0.75 1.00
π̂
DDX51 − Reverse
DNA Meth. m5C:H4K20 me2
H3K4 me3:H3−5ac
H3K27 me2:DNA Meth. m5C
H4K20 me2:H4−4ac
H3K4 me1:H3−5ac
H3K27 me3:H3K4 me1
H3K4 me1:H4−4ac
H4K20 me2:H3−5ac
H4−4ac:H3−5ac
H4K20 me2:H3K4 me3
H2A.Z:H3K4 me3
H2A.Z:H4K20 me2
H3K4 me1:H2A.Z
H3K27 me3:DNA Meth. m5C
H3K4 me1:H4K20 me2
H3K4 me3:H4−4ac
H4K20 me3
H3K9 me2
DNA Meth. m5C
H3K27 me2
H4−4ac
H3K27 me3
H3−5ac
H4K20 me2
H2A.Z
H3K4 me3
H3K4 me1
0.00 0.25 0.50 0.75 1.00
π̂
RPL27 − Forward
DNA Meth. m5C:H3K9 me3
H3K27 me3:DNA Meth. m5C
H3K27 me3:H3K9 me3
H3K4 me1:H4−4ac
H3K27 me2:DNA Meth. m5C
H3K27 me3:H4K20 me3
H3K27 me3:H3K4 me1
H2A.Z:H3K4 me3
H3K4 me1:H2A.Z
H3K9 me3:H4K20 me3
H3K4 me1:H3−5ac
DNA Meth. m5C:H4K20 me3
H4K20 me2:H4−4ac
H4K20 me2:H3−5ac
H3K4 me3:H3−5ac
H4K20 me2:H3K4 me3
H3K4 me1:H4K20 me2
H3K4 me3:H4−4ac
H2A.Z:H4K20 me2
H4−4ac:H3−5ac
H3K9 me2
H3K9 me3
H3K27 me2
H4K20 me3
DNA Meth. m5C
H3K27 me3
H3K4 me1
H4−4ac
H3K4 me3
H3−5ac
H4K20 me2
H2A.Z
0.00 0.25 0.50 0.75 1.00
π̂
RPL27 − Reverse
DNA Meth. m5C:H3K9 me3
H3K27 me3:DNA Meth. m5C
H3K27 me3:H3K9 me3
H3K4 me1:H4−4ac
H3K27 me2:DNA Meth. m5C
H3K27 me3:H4K20 me3
H3K27 me3:H3K4 me1
H2A.Z:H3K4 me3
H3K4 me1:H2A.Z
H3K9 me3:H4K20 me3
H3K4 me1:H3−5ac
DNA Meth. m5C:H4K20 me3
H4K20 me2:H4−4ac
H4K20 me2:H3−5ac
H3K4 me3:H3−5ac
H4K20 me2:H3K4 me3
H3K4 me1:H4K20 me2
H3K4 me3:H4−4ac
H2A.Z:H4K20 me2
H4−4ac:H3−5ac
H3K9 me2
H3K9 me3
H3K27 me2
H4K20 me3
DNA Meth. m5C
H3K27 me3
H3K4 me1
H4−4ac
H3K4 me3
H3−5ac
H4K20 me2
H2A.Z
0.00 0.25 0.50 0.75 1.00
π̂
SPATS2L − Forward
H3K27 me3:H4K20 me3
H3K9 me3:H4K20 me3
H3K27 me3:H3K4 me1
H3K27 me3:H3K9 me3
H3K4 me3:H3−5ac
H3K4 me3:H4−4ac
H4K20 me2:H4−4ac
H3K4 me1:H3−5ac
H3K4 me1:H2A.Z
DNA Meth. m5C:H4K20 me2
H3K4 me1:H4K20 me2
H3K4 me1:H4−4ac
H3K27 me3:DNA Meth. m5C
H2A.Z:H4−4ac
DNA Meth. m5C:H3K9 me3
H4−4ac:H3−5ac
H2A.Z:H3−5ac
H4K20 me2:H3K4 me3
DNA Meth. m5C:H4K20 me3
H2A.Z:H4K20 me2
H4K20 me2:H3−5ac
H2A.Z:H3K4 me3
H3K27 me2
H3K9 me3
H3K4 me1
H4K20 me3
H4−4ac
H3K27 me3
H3K4 me3
H4K20 me2
DNA Meth. m5C
H2A.Z
H3−5ac
0.00 0.25 0.50 0.75 1.00
π̂
SPATS2L − Reverse
H3K27 me3:H4K20 me3
H3K9 me3:H4K20 me3
H3K27 me3:H3K4 me1
H3K27 me3:H3K9 me3
H3K4 me3:H3−5ac
H3K4 me3:H4−4ac
H4K20 me2:H4−4ac
H3K4 me1:H3−5ac
H3K4 me1:H2A.Z
DNA Meth. m5C:H4K20 me2
H3K4 me1:H4K20 me2
H3K4 me1:H4−4ac
H3K27 me3:DNA Meth. m5C
H2A.Z:H4−4ac
DNA Meth. m5C:H3K9 me3
H4−4ac:H3−5ac
H2A.Z:H3−5ac
H4K20 me2:H3K4 me3
DNA Meth. m5C:H4K20 me3
H2A.Z:H4K20 me2
H4K20 me2:H3−5ac
H2A.Z:H3K4 me3
H3K27 me2
H3K9 me3
H3K4 me1
H4K20 me3
H4−4ac
H3K27 me3
H3K4 me3
H4K20 me2
DNA Meth. m5C
H2A.Z
H3−5ac
0.00 0.25 0.50 0.75 1.00
π̂
PNISR − Forward
H3K4 me3:H4−4ac
H3K27 me3:H3K4 me1
H3K27 me3:H4K20 me3
H4−4ac:H3−5ac
H3K4 me3:H3−5ac
H3K27 me2:DNA Meth. m5C
H3K9 me2:DNA Meth. m5C
H3K9 me3:H4K20 me3
DNA Meth. m5C:H4K20 me3
H4K20 me2:H3−5ac
H3K4 me1:H4K20 me2
H3K4 me1:H3−5ac
H3K27 me3:H3K9 me3
DNA Meth. m5C:H3K9 me3
H3K27 me3:DNA Meth. m5C
H3K4 me1:H2A.Z
H3K4 me1:H4−4ac
DNA Meth. m5C:H4K20 me2
H4K20 me2:H3K4 me3
H2A.Z:H3K4 me3
H2A.Z:H4K20 me2
H4K20 me2:H4−4ac
H3K27 me2
H3K9 me2
H4K20 me3
H3−5ac
H3K9 me3
H4−4ac
H3K27 me3
H2A.Z
DNA Meth. m5C
H3K4 me3
H3K4 me1
H4K20 me2
0.00 0.25 0.50 0.75 1.00
π̂
PNISR − Reverse
H3K4 me3:H4−4ac
H3K27 me3:H3K4 me1
H3K27 me3:H4K20 me3
H4−4ac:H3−5ac
H3K4 me3:H3−5ac
H3K27 me2:DNA Meth. m5C
H3K9 me2:DNA Meth. m5C
H3K9 me3:H4K20 me3
DNA Meth. m5C:H4K20 me3
H4K20 me2:H3−5ac
H3K4 me1:H4K20 me2
H3K4 me1:H3−5ac
H3K27 me3:H3K9 me3
DNA Meth. m5C:H3K9 me3
H3K27 me3:DNA Meth. m5C
H3K4 me1:H2A.Z
H3K4 me1:H4−4ac
DNA Meth. m5C:H4K20 me2
H4K20 me2:H3K4 me3
H2A.Z:H3K4 me3
H2A.Z:H4K20 me2
H4K20 me2:H4−4ac
H3K27 me2
H3K9 me2
H4K20 me3
H3−5ac
H3K9 me3
H4−4ac
H3K27 me3
H2A.Z
DNA Meth. m5C
H3K4 me3
H3K4 me1
H4K20 me2
0.00 0.25 0.50 0.75 1.00
π̂
WDR46 − Forward
H3K4 me1:H4K20 me2
H4−4ac:H3−5ac
H3K4 me3:H3−5ac
H3K4 me3:H4−4ac
H3K4 me1:H2A.Z
H3K27 me3:H3K4 me1
H4K20 me2:H4−4ac
H2A.Z:H3−5ac
H3K4 me1:H4−4ac
H2A.Z:H4−4ac
H3K27 me3:H4K20 me3
H3K9 me3:H4K20 me3
H3K9 me2:DNA Meth. m5C
H4K20 me2:H3−5ac
H3K27 me3:H3K9 me3
H4K20 me2:H3K4 me3
DNA Meth. m5C:H4K20 me3
H3K27 me2:DNA Meth. m5C
H2A.Z:H3K4 me3
H3K27 me3:DNA Meth. m5C
DNA Meth. m5C:H4K20 me2
DNA Meth. m5C:H3K9 me3
H2A.Z:H4K20 me2
H3K9 me2
H3−5ac
H3K4 me1
H4−4ac
H3K27 me2
H3K4 me3
H2A.Z
H4K20 me3
H4K20 me2
H3K9 me3
H3K27 me3
DNA Meth. m5C
0.00 0.25 0.50 0.75 1.00
π̂
WDR46 − Reverse
H3K4 me1:H4K20 me2
H4−4ac:H3−5ac
H3K4 me3:H3−5ac
H3K4 me3:H4−4ac
H3K4 me1:H2A.Z
H3K27 me3:H3K4 me1
H4K20 me2:H4−4ac
H2A.Z:H3−5ac
H3K4 me1:H4−4ac
H2A.Z:H4−4ac
H3K27 me3:H4K20 me3
H3K9 me3:H4K20 me3
H3K9 me2:DNA Meth. m5C
H4K20 me2:H3−5ac
H3K27 me3:H3K9 me3
H4K20 me2:H3K4 me3
DNA Meth. m5C:H4K20 me3
H3K27 me2:DNA Meth. m5C
H2A.Z:H3K4 me3
H3K27 me3:DNA Meth. m5C
DNA Meth. m5C:H4K20 me2
DNA Meth. m5C:H3K9 me3
H2A.Z:H4K20 me2
H3K9 me2
H3−5ac
H3K4 me1
H4−4ac
H3K27 me2
H3K4 me3
H2A.Z
H4K20 me3
H4K20 me2
H3K9 me3
H3K27 me3
DNA Meth. m5C
0.00 0.25 0.50 0.75 1.00
π̂

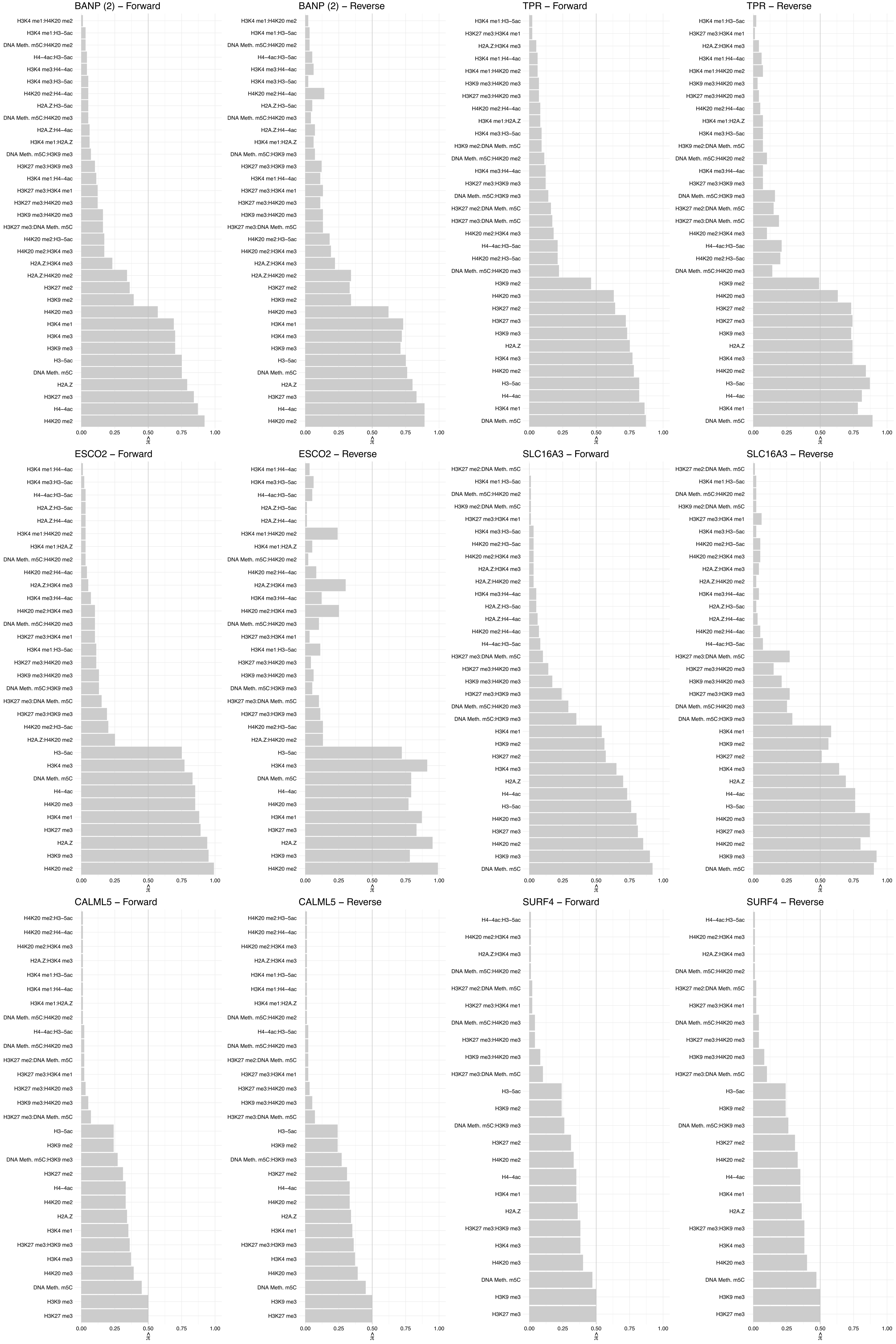
BANP (2) – Forward
H3K4 me1:H4K20 me2
H3K4 me1:H3–5ac
DNA Meth. m5C:H4K20 me2
H4–4ac:H3–5ac
H3K4 me3:H4–4ac
H3K4 me3:H3–5ac
H4K20 me2:H4–4ac
H2A.Z:H3–5ac
DNA Meth. m5C:H4K20 me3
H2A.Z:H4–4ac
H3K4 me1:H2A.Z
DNA Meth. m5C:H3K9 me3
H3K27 me3:H3K9 me3
H3K4 me1:H4–4ac
H3K27 me3:H3K4 me1
H3K27 me3:H4K20 me3
H3K9 me3:H4K20 me3
H3K27 me3:DNA Meth. m5C
H4K20 me2:H3–5ac
H4K20 me2:H3K4 me3
H2A.Z:H3K4 me3
H2A.Z:H4K20 me2
H3K27 me2
H3K9 me2
H4K20 me3
H3K4 me1
H3K4 me3
H3K9 me3
H3–5ac
DNA Meth. m5C
H2A.Z
H3K27 me3
H4–4ac
H4K20 me2
0.00 0.25 0.50 0.75 1.00
$\hat{\pi}$
BANP (2) – Reverse
H3K4 me1:H4K20 me2
H3K4 me1:H3–5ac
DNA Meth. m5C:H4K20 me2
H4–4ac:H3–5ac
H3K4 me3:H4–4ac
H3K4 me3:H3–5ac
H4K20 me2:H4–4ac
H2A.Z:H3–5ac
DNA Meth. m5C:H4K20 me3
H2A.Z:H4–4ac
H3K4 me1:H2A.Z
DNA Meth. m5C:H3K9 me3
H3K27 me3:H3K9 me3
H3K4 me1:H4–4ac
H3K27 me3:H3K4 me1
H3K27 me3:H4K20 me3
H3K9 me3:H4K20 me3
H3K27 me3:DNA Meth. m5C
H4K20 me2:H3–5ac
H4K20 me2:H3K4 me3
H2A.Z:H3K4 me3
H2A.Z:H4K20 me2
H3K27 me2
H3K9 me2
H4K20 me3
H3K4 me1
H3K4 me3
H3K9 me3
H3–5ac
DNA Meth. m5C
H2A.Z
H3K27 me3
H4–4ac
H4K20 me2
0.00 0.25 0.50 0.75 1.00
$\hat{\pi}$
TPR – Forward
H3K4 me1:H3–5ac
H3K27 me3:H3K4 me1
H2A.Z:H3K4 me3
H3K4 me1:H4–4ac
H3K4 me1:H4K20 me2
H3K9 me3:H4K20 me3
H3K27 me3:H4K20 me3
H4K20 me2:H4–4ac
H3K4 me1:H2A.Z
H3K4 me3:H3–5ac
H3K9 me2:DNA Meth. m5C
DNA Meth. m5C:H4K20 me2
H3K4 me3:H4–4ac
H3K27 me3:H3K9 me3
DNA Meth. m5C:H3K9 me3
H3K27 me2:DNA Meth. m5C
H3K27 me3:DNA Meth. m5C
H4K20 me2:H3K4 me3
H4–4ac:H3–5ac
H4K20 me2:H3–5ac
DNA Meth. m5C:H4K20 me3
H3K9 me2
H4K20 me3
H3K27 me2
H3K27 me3
H3K9 me3
H2A.Z
H3K4 me3
H4K20 me2
H3–5ac
H4–4ac
H3K4 me1
DNA Meth. m5C
0.00 0.25 0.50 0.75 1.00
$\hat{\pi}$
TPR – Reverse
H3K4 me1:H3–5ac
H3K27 me3:H3K4 me1
H2A.Z:H3K4 me3
H3K4 me1:H4–4ac
H3K4 me1:H4K20 me2
H3K9 me3:H4K20 me3
H3K27 me3:H4K20 me3
H4K20 me2:H4–4ac
H3K4 me1:H2A.Z
H3K4 me3:H3–5ac
H3K9 me2:DNA Meth. m5C
DNA Meth. m5C:H4K20 me2
H3K4 me3:H4–4ac
H3K27 me3:H3K9 me3
DNA Meth. m5C:H3K9 me3
H3K27 me2:DNA Meth. m5C
H3K27 me3:DNA Meth. m5C
H4K20 me2:H3K4 me3
H4–4ac:H3–5ac
H4K20 me2:H3–5ac
DNA Meth. m5C:H4K20 me3
H3K9 me2
H4K20 me3
H3K27 me2
H3K27 me3
H3K9 me3
H2A.Z
H3K4 me3
H4K20 me2
H3–5ac
H4–4ac
H3K4 me1
DNA Meth. m5C
0.00 0.25 0.50 0.75 1.00
$\hat{\pi}$
ESCO2 – Forward
H3K4 me1:H4–4ac
H3K4 me3:H3–5ac
H4–4ac:H3–5ac
H2A.Z:H3–5ac
H2A.Z:H4–4ac
H3K4 me1:H4K20 me2
H3K4 me1:H2A.Z
DNA Meth. m5C:H4K20 me2
H4K20 me2:H4–4ac
H2A.Z:H3K4 me3
H3K4 me3:H4–4ac
H4K20 me2:H3K4 me3
DNA Meth. m5C:H4K20 me3
H3K27 me3:H3K4 me1
H3K4 me1:H3–5ac
H3K27 me3:H4K20 me3
H3K9 me3:H4K20 me3
DNA Meth. m5C:H3K9 me3
H3K27 me3:DNA Meth. m5C
H3K27 me3:H3K9 me3
H4K20 me2:H3–5ac
H2A.Z:H4K20 me2
H3–5ac
H3K4 me3
DNA Meth. m5C
H4–4ac
H4K20 me3
H3K4 me1
H3K27 me3
H2A.Z
H3K9 me3
H4K20 me2
0.00 0.25 0.50 0.75 1.00
$\hat{\pi}$
ESCO2 – Reverse
H3K4 me1:H4–4ac
H3K4 me3:H3–5ac
H4–4ac:H3–5ac
H2A.Z:H3–5ac
H2A.Z:H4–4ac
H3K4 me1:H4K20 me2
H3K4 me1:H2A.Z
DNA Meth. m5C:H4K20 me2
H4K20 me2:H4–4ac
H2A.Z:H3K4 me3
H3K4 me3:H4–4ac
H4K20 me2:H3K4 me3
DNA Meth. m5C:H4K20 me3
H3K27 me3:H3K4 me1
H3K4 me1:H3–5ac
H3K27 me3:H4K20 me3
H3K9 me3:H4K20 me3
DNA Meth. m5C:H3K9 me3
H3K27 me3:DNA Meth. m5C
H3K27 me3:H3K9 me3
H4K20 me2:H3–5ac
H2A.Z:H4K20 me2
H3–5ac
H3K4 me3
DNA Meth. m5C
H4–4ac
H4K20 me3
H3K4 me1
H3K27 me3
H2A.Z
H3K9 me3
H4K20 me2
0.00 0.25 0.50 0.75 1.00
$\hat{\pi}$
SLC16A3 – Forward
H3K27 me2:DNA Meth. m5C
H3K4 me1:H3–5ac
DNA Meth. m5C:H4K20 me2
H3K9 me2:DNA Meth. m5C
H3K27 me3:H3K4 me1
H3K4 me3:H3–5ac
H4K20 me2:H3–5ac
H4K20 me2:H3K4 me3
H2A.Z:H3K4 me3
H2A.Z:H4K20 me2
H3K4 me3:H4–4ac
H2A.Z:H3–5ac
H2A.Z:H4–4ac
H4K20 me2:H4–4ac
H4–4ac:H3–5ac
H3K27 me3:DNA Meth. m5C
H3K27 me3:H4K20 me3
H3K9 me3:H4K20 me3
H3K27 me3:H3K9 me3
DNA Meth. m5C:H4K20 me3
DNA Meth. m5C:H3K9 me3
H3K4 me1
H3K9 me2
H3K27 me2
H3K4 me3
H2A.Z
H4–4ac
H3–5ac
H4K20 me3
H3K27 me3
H4K20 me2
H3K9 me3
DNA Meth. m5C
0.00 0.25 0.50 0.75 1.00
$\hat{\pi}$
SLC16A3 – Reverse
H3K27 me2:DNA Meth. m5C
H3K4 me1:H3–5ac
DNA Meth. m5C:H4K20 me2
H3K9 me2:DNA Meth. m5C
H3K27 me3:H3K4 me1
H3K4 me3:H3–5ac
H4K20 me2:H3–5ac
H4K20 me2:H3K4 me3
H2A.Z:H3K4 me3
H2A.Z:H4K20 me2
H3K4 me3:H4–4ac
H2A.Z:H3–5ac
H2A.Z:H4–4ac
H4K20 me2:H4–4ac
H4–4ac:H3–5ac
H3K27 me3:DNA Meth. m5C
H3K27 me3:H4K20 me3
H3K9 me3:H4K20 me3
H3K27 me3:H3K9 me3
DNA Meth. m5C:H4K20 me3
DNA Meth. m5C:H3K9 me3
H3K4 me1
H3K9 me2
H3K27 me2
H3K4 me3
H2A.Z
H4–4ac
H3–5ac
H4K20 me3
H3K27 me3
H4K20 me2
H3K9 me3
DNA Meth. m5C
0.00 0.25 0.50 0.75 1.00
$\hat{\pi}$
CALML5 – Forward
H4K20 me2:H3–5ac
H4K20 me2:H4–4ac
H4K20 me2:H3K4 me3
H2A.Z:H3K4 me3
H3K4 me1:H3–5ac
H3K4 me1:H4–4ac
H3K4 me1:H2A.Z
DNA Meth. m5C:H4K20 me2
H4–4ac:H3–5ac
DNA Meth. m5C:H4K20 me3
H3K27 me2:DNA Meth. m5C
H3K27 me3:H3K4 me1
H3K27 me3:H4K20 me3
H3K9 me3:H4K20 me3
H3K27 me3:DNA Meth. m5C
H3–5ac
H3K9 me2
DNA Meth. m5C:H3K9 me3
H3K27 me2
H4–4ac
H4K20 me2
H2A.Z
H3K4 me1
H3K27 me3:H3K9 me3
H3K4 me3
H4K20 me3
DNA Meth. m5C
H3K9 me3
H3K27 me3
0.00 0.25 0.50 0.75 1.00
$\hat{\pi}$
CALML5 – Reverse
H4K20 me2:H3–5ac
H4K20 me2:H4–4ac
H4K20 me2:H3K4 me3
H2A.Z:H3K4 me3
H3K4 me1:H3–5ac
H3K4 me1:H4–4ac
H3K4 me1:H2A.Z
DNA Meth. m5C:H4K20 me2
H4–4ac:H3–5ac
DNA Meth. m5C:H4K20 me3
H3K27 me2:DNA Meth. m5C
H3K27 me3:H3K4 me1
H3K27 me3:H4K20 me3
H3K9 me3:H4K20 me3
H3K27 me3:DNA Meth. m5C
H3–5ac
H3K9 me2
DNA Meth. m5C:H3K9 me3
H3K27 me2
H4–4ac
H4K20 me2
H2A.Z
H3K4 me1
H3K27 me3:H3K9 me3
H3K4 me3
H4K20 me3
DNA Meth. m5C
H3K9 me3
H3K27 me3
0.00 0.25 0.50 0.75 1.00
$\hat{\pi}$
SURF4 – Forward
H4–4ac:H3–5ac
H4K20 me2:H3K4 me3
H2A.Z:H3K4 me3
DNA Meth. m5C:H4K20 me2
H3K27 me2:DNA Meth. m5C
H3K27 me3:H3K4 me1
DNA Meth. m5C:H4K20 me3
H3K27 me3:H4K20 me3
H3K9 me3:H4K20 me3
H3K27 me3:DNA Meth. m5C
H3–5ac
H3K9 me2
DNA Meth. m5C:H3K9 me3
H3K27 me2
H4K20 me2
H4–4ac
H3K4 me1
H2A.Z
H3K27 me3:H3K9 me3
H3K4 me3
H4K20 me3
DNA Meth. m5C
H3K9 me3
H3K27 me3
0.00 0.25 0.50 0.75 1.00
$\hat{\pi}$
SURF4 – Reverse
H4–4ac:H3–5ac
H4K20 me2:H3K4 me3
H2A.Z:H3K4 me3
DNA Meth. m5C:H4K20 me2
H3K27 me2:DNA Meth. m5C
H3K27 me3:H3K4 me1
DNA Meth. m5C:H4K20 me3
H3K27 me3:H4K20 me3
H3K9 me3:H4K20 me3
H3K27 me3:DNA Meth. m5C
H3–5ac
H3K9 me2
DNA Meth. m5C:H3K9 me3
H3K27 me2
H4K20 me2
H4–4ac
H3K4 me1
H2A.Z
H3K27 me3:H3K9 me3
H3K4 me3
H4K20 me3
DNA Meth. m5C
H3K9 me3
H3K27 me3
0.00 0.25 0.50 0.75 1.00
$\hat{\pi}$

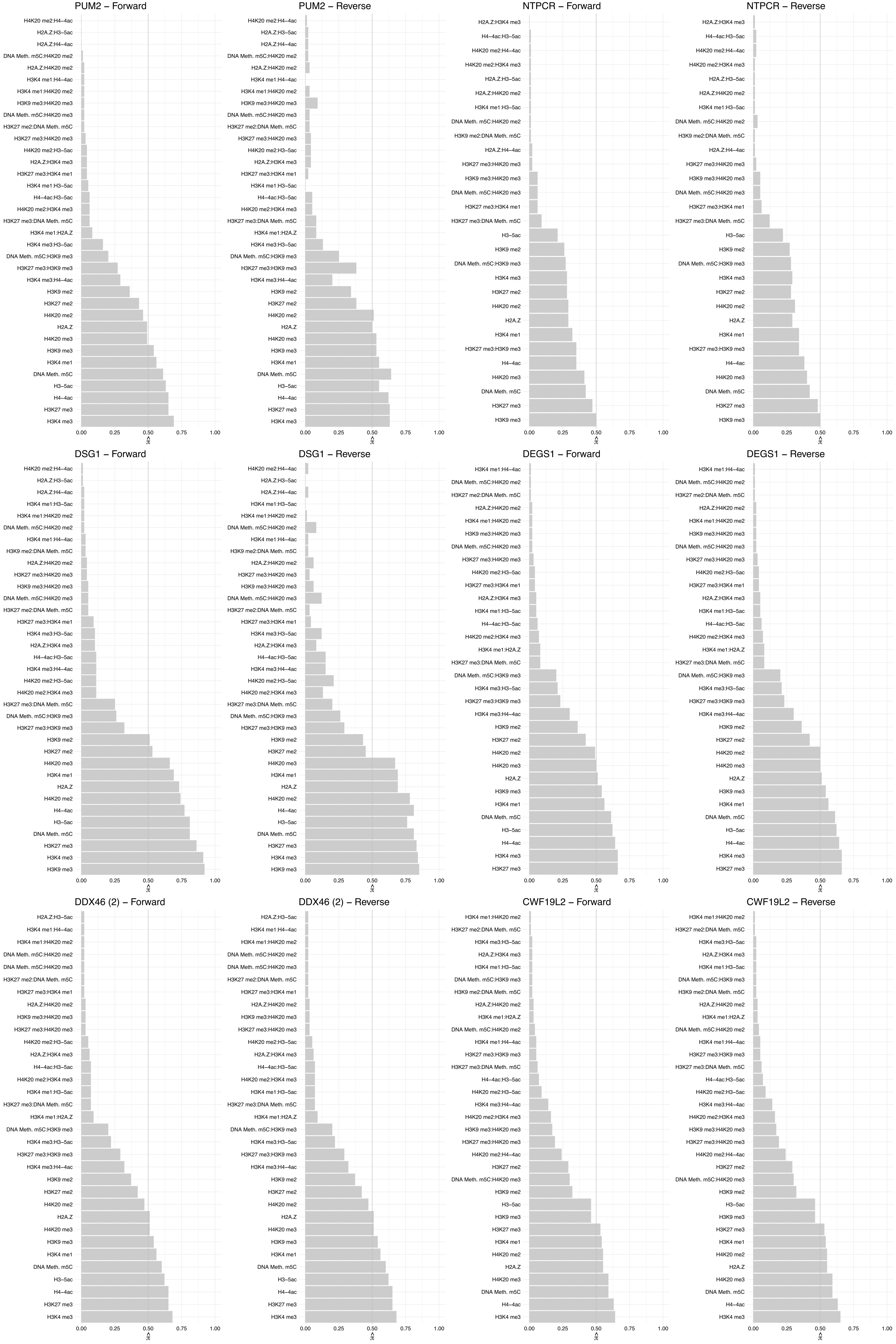
PUM2 – Forward
PUM2 – Reverse
NTPCR – Forward
NTPCR – Reverse
DSG1 – Forward
DSG1 – Reverse
DEGS1 – Forward
DEGS1 – Reverse
DDX46 (2) – Forward
DDX46 (2) – Reverse
CWF19L2 – Forward
CWF19L2 – Reverse
0.00 0.25 0.50 0.75 1.00
$\hat{\pi}$

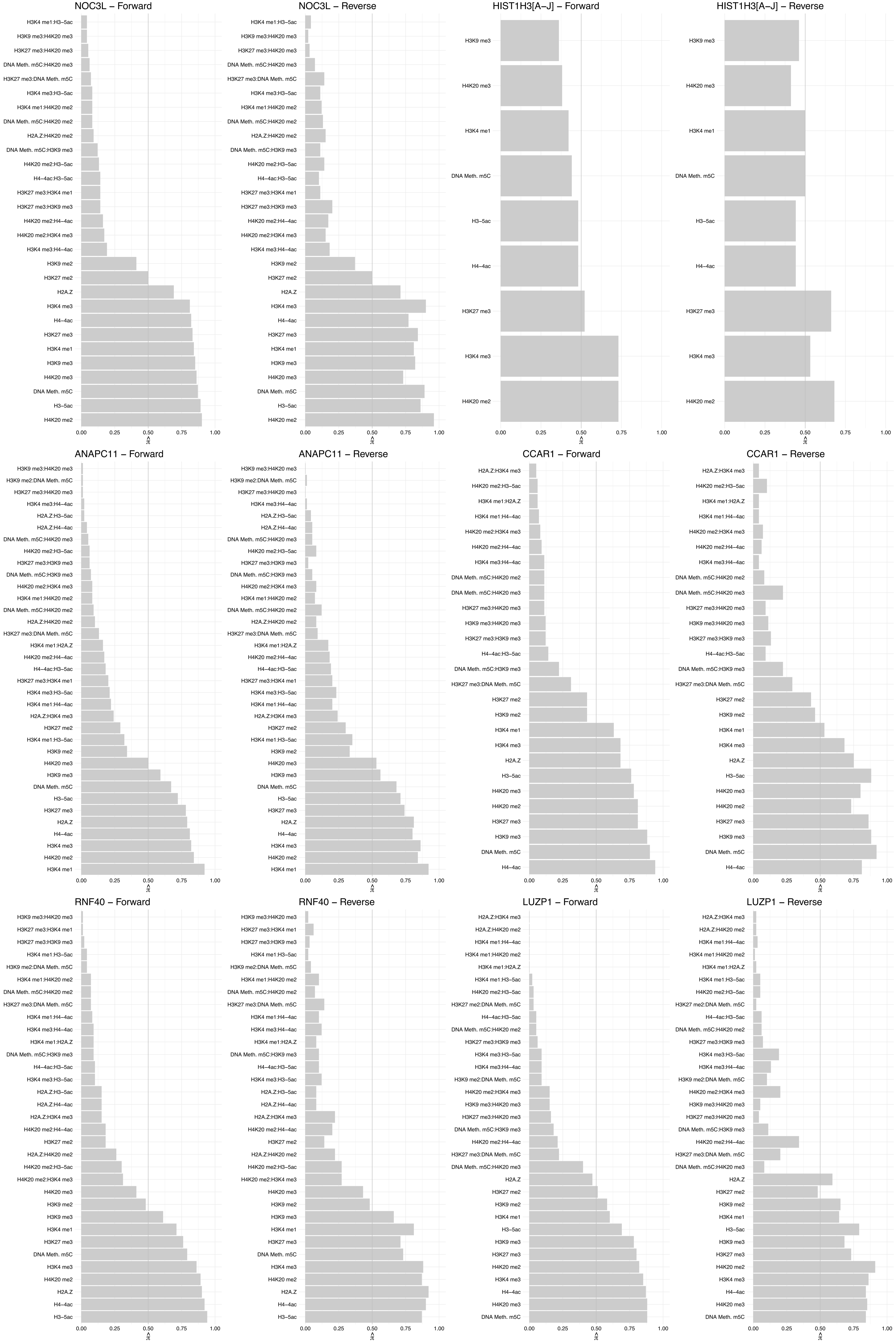
NOC3L – Forward
H3K4 me1:H3–5ac
H3K9 me3:H4K20 me3
H3K27 me3:H4K20 me3
DNA Meth. m5C:H4K20 me3
H3K27 me3:DNA Meth. m5C
H3K4 me3:H3–5ac
H3K4 me1:H4K20 me2
DNA Meth. m5C:H4K20 me2
H2A.Z:H4K20 me2
DNA Meth. m5C:H3K9 me3
H4K20 me2:H3–5ac
H4–4ac:H3–5ac
H3K27 me3:H3K4 me1
H3K27 me3:H3K9 me3
H4K20 me2:H4–4ac
H4K20 me2:H3K4 me3
H3K4 me3:H4–4ac
H3K9 me2
H3K27 me2
H2A.Z
H3K4 me3
H4–4ac
H3K27 me3
H3K4 me1
H3K9 me3
H4K20 me3
DNA Meth. m5C
H3–5ac
H4K20 me2
0.00 0.25 0.50 0.75 1.00
π̂
NOC3L – Reverse
H3K4 me1:H3–5ac
H3K9 me3:H4K20 me3
H3K27 me3:H4K20 me3
DNA Meth. m5C:H4K20 me3
H3K27 me3:DNA Meth. m5C
H3K4 me3:H3–5ac
H3K4 me1:H4K20 me2
DNA Meth. m5C:H4K20 me2
H2A.Z:H4K20 me2
DNA Meth. m5C:H3K9 me3
H4K20 me2:H3–5ac
H4–4ac:H3–5ac
H3K27 me3:H3K4 me1
H3K27 me3:H3K9 me3
H4K20 me2:H4–4ac
H4K20 me2:H3K4 me3
H3K4 me3:H4–4ac
H3K9 me2
H3K27 me2
H2A.Z
H3K4 me3
H4–4ac
H3K27 me3
H3K4 me1
H3K9 me3
H4K20 me3
DNA Meth. m5C
H3–5ac
H4K20 me2
0.00 0.25 0.50 0.75 1.00
π̂
HIST1H3[A–J] – Forward
H3K9 me3
H4K20 me3
H3K4 me1
DNA Meth. m5C
H3–5ac
H4–4ac
H3K27 me3
H3K4 me3
H4K20 me2
0.00 0.25 0.50 0.75 1.00
π̂
HIST1H3[A–J] – Reverse
H3K9 me3
H4K20 me3
H3K4 me1
DNA Meth. m5C
H3–5ac
H4–4ac
H3K27 me3
H3K4 me3
H4K20 me2
0.00 0.25 0.50 0.75 1.00
π̂
ANAPC11 – Forward
H3K9 me3:H4K20 me3
H3K9 me2:DNA Meth. m5C
H3K27 me3:H4K20 me3
H3K4 me3:H4–4ac
H2A.Z:H3–5ac
H2A.Z:H4–4ac
DNA Meth. m5C:H4K20 me3
H4K20 me2:H3–5ac
H3K27 me3:H3K9 me3
DNA Meth. m5C:H3K9 me3
H4K20 me2:H3K4 me3
H3K4 me1:H4K20 me2
DNA Meth. m5C:H4K20 me2
H2A.Z:H4K20 me2
H3K27 me3:DNA Meth. m5C
H3K4 me1:H2A.Z
H4K20 me2:H4–4ac
H4–4ac:H3–5ac
H3K27 me3:H3K4 me1
H3K4 me3:H3–5ac
H3K4 me1:H4–4ac
H2A.Z:H3K4 me3
H3K27 me2
H3K4 me1:H3–5ac
H3K9 me2
H4K20 me3
H3K9 me3
DNA Meth. m5C
H3–5ac
H3K27 me3
H2A.Z
H4–4ac
H3K4 me3
H4K20 me2
H3K4 me1
0.00 0.25 0.50 0.75 1.00
π̂
ANAPC11 – Reverse
H3K9 me3:H4K20 me3
H3K9 me2:DNA Meth. m5C
H3K27 me3:H4K20 me3
H3K4 me3:H4–4ac
H2A.Z:H3–5ac
H2A.Z:H4–4ac
DNA Meth. m5C:H4K20 me3
H4K20 me2:H3–5ac
H3K27 me3:H3K9 me3
DNA Meth. m5C:H3K9 me3
H4K20 me2:H3K4 me3
H3K4 me1:H4K20 me2
DNA Meth. m5C:H4K20 me2
H2A.Z:H4K20 me2
H3K27 me3:DNA Meth. m5C
H3K4 me1:H2A.Z
H4K20 me2:H4–4ac
H4–4ac:H3–5ac
H3K27 me3:H3K4 me1
H3K4 me3:H3–5ac
H3K4 me1:H4–4ac
H2A.Z:H3K4 me3
H3K27 me2
H3K4 me1:H3–5ac
H3K9 me2
H4K20 me3
H3K9 me3
DNA Meth. m5C
H3–5ac
H3K27 me3
H2A.Z
H4–4ac
H3K4 me3
H4K20 me2
H3K4 me1
0.00 0.25 0.50 0.75 1.00
π̂
CCAR1 – Forward
H2A.Z:H3K4 me3
H4K20 me2:H3–5ac
H3K4 me1:H2A.Z
H3K4 me1:H4–4ac
H4K20 me2:H3K4 me3
H4K20 me2:H4–4ac
H3K4 me3:H4–4ac
DNA Meth. m5C:H4K20 me2
DNA Meth. m5C:H4K20 me3
H3K27 me3:H4K20 me3
H3K9 me3:H4K20 me3
H3K27 me3:H3K9 me3
H4–4ac:H3–5ac
DNA Meth. m5C:H3K9 me3
H3K27 me3:DNA Meth. m5C
H3K27 me2
H3K9 me2
H3K4 me1
H3K4 me3
H2A.Z
H3–5ac
H4K20 me3
H4K20 me2
H3K27 me3
H3K9 me3
DNA Meth. m5C
H4–4ac
0.00 0.25 0.50 0.75 1.00
π̂
CCAR1 – Reverse
H2A.Z:H3K4 me3
H4K20 me2:H3–5ac
H3K4 me1:H2A.Z
H3K4 me1:H4–4ac
H4K20 me2:H3K4 me3
H4K20 me2:H4–4ac
H3K4 me3:H4–4ac
DNA Meth. m5C:H4K20 me2
DNA Meth. m5C:H4K20 me3
H3K27 me3:H4K20 me3
H3K9 me3:H4K20 me3
H3K27 me3:H3K9 me3
H4–4ac:H3–5ac
DNA Meth. m5C:H3K9 me3
H3K27 me3:DNA Meth. m5C
H3K27 me2
H3K9 me2
H3K4 me1
H3K4 me3
H2A.Z
H3–5ac
H4K20 me3
H4K20 me2
H3K27 me3
H3K9 me3
DNA Meth. m5C
H4–4ac
0.00 0.25 0.50 0.75 1.00
π̂
RNF40 – Forward
H3K9 me3:H4K20 me3
H3K27 me3:H3K4 me1
H3K27 me3:H3K9 me3
H3K4 me1:H3–5ac
H3K9 me2:DNA Meth. m5C
H3K4 me1:H4K20 me2
DNA Meth. m5C:H4K20 me2
H3K27 me3:DNA Meth. m5C
H3K4 me1:H4–4ac
H3K4 me3:H4–4ac
H3K4 me1:H2A.Z
DNA Meth. m5C:H3K9 me3
H4–4ac:H3–5ac
H3K4 me3:H3–5ac
H2A.Z:H3–5ac
H2A.Z:H4–4ac
H2A.Z:H3K4 me3
H4K20 me2:H4–4ac
H3K27 me2
H2A.Z:H4K20 me2
H4K20 me2:H3–5ac
H4K20 me2:H3K4 me3
H4K20 me3
H3K9 me2
H3K9 me3
H3K4 me1
H3K27 me3
DNA Meth. m5C
H3K4 me3
H4K20 me2
H2A.Z
H4–4ac
H3–5ac
0.00 0.25 0.50 0.75 1.00
π̂
RNF40 – Reverse
H3K9 me3:H4K20 me3
H3K27 me3:H3K4 me1
H3K27 me3:H3K9 me3
H3K4 me1:H3–5ac
H3K9 me2:DNA Meth. m5C
H3K4 me1:H4K20 me2
DNA Meth. m5C:H4K20 me2
H3K27 me3:DNA Meth. m5C
H3K4 me1:H4–4ac
H3K4 me3:H4–4ac
H3K4 me1:H2A.Z
DNA Meth. m5C:H3K9 me3
H4–4ac:H3–5ac
H3K4 me3:H3–5ac
H2A.Z:H3–5ac
H2A.Z:H4–4ac
H2A.Z:H3K4 me3
H4K20 me2:H4–4ac
H3K27 me2
H2A.Z:H4K20 me2
H4K20 me2:H3–5ac
H4K20 me2:H3K4 me3
H4K20 me3
H3K9 me2
H3K9 me3
H3K4 me1
H3K27 me3
DNA Meth. m5C
H3K4 me3
H4K20 me2
H2A.Z
H4–4ac
H3–5ac
0.00 0.25 0.50 0.75 1.00
π̂
LUZP1 – Forward
H2A.Z:H3K4 me3
H2A.Z:H4K20 me2
H3K4 me1:H4–4ac
H3K4 me1:H4K20 me2
H3K4 me1:H2A.Z
H3K4 me1:H3–5ac
H4K20 me2:H3–5ac
H3K27 me2:DNA Meth. m5C
H4–4ac:H3–5ac
DNA Meth. m5C:H4K20 me2
H3K27 me3:H3K9 me3
H3K4 me3:H3–5ac
H3K4 me3:H4–4ac
H3K9 me2:DNA Meth. m5C
H4K20 me2:H3K4 me3
H3K9 me3:H4K20 me3
H3K27 me3:H4K20 me3
DNA Meth. m5C:H3K9 me3
H4K20 me2:H4–4ac
H3K27 me3:DNA Meth. m5C
DNA Meth. m5C:H4K20 me3
H2A.Z
H3K27 me2
H3K9 me2
H3K4 me1
H3–5ac
H3K9 me3
H3K27 me3
H4K20 me2
H3K4 me3
H4–4ac
H4K20 me3
DNA Meth. m5C
0.00 0.25 0.50 0.75 1.00
π̂
LUZP1 – Reverse
H2A.Z:H3K4 me3
H2A.Z:H4K20 me2
H3K4 me1:H4–4ac
H3K4 me1:H4K20 me2
H3K4 me1:H2A.Z
H3K4 me1:H3–5ac
H4K20 me2:H3–5ac
H3K27 me2:DNA Meth. m5C
H4–4ac:H3–5ac
DNA Meth. m5C:H4K20 me2
H3K27 me3:H3K9 me3
H3K4 me3:H3–5ac
H3K4 me3:H4–4ac
H3K9 me2:DNA Meth. m5C
H4K20 me2:H3K4 me3
H3K9 me3:H4K20 me3
H3K27 me3:H4K20 me3
DNA Meth. m5C:H3K9 me3
H4K20 me2:H4–4ac
H3K27 me3:DNA Meth. m5C
DNA Meth. m5C:H4K20 me3
H2A.Z
H3K27 me2
H3K9 me2
H3K4 me1
H3–5ac
H3K9 me3
H3K27 me3
H4K20 me2
H3K4 me3
H4–4ac
H4K20 me3
DNA Meth. m5C
0.00 0.25 0.50 0.75 1.00
π̂

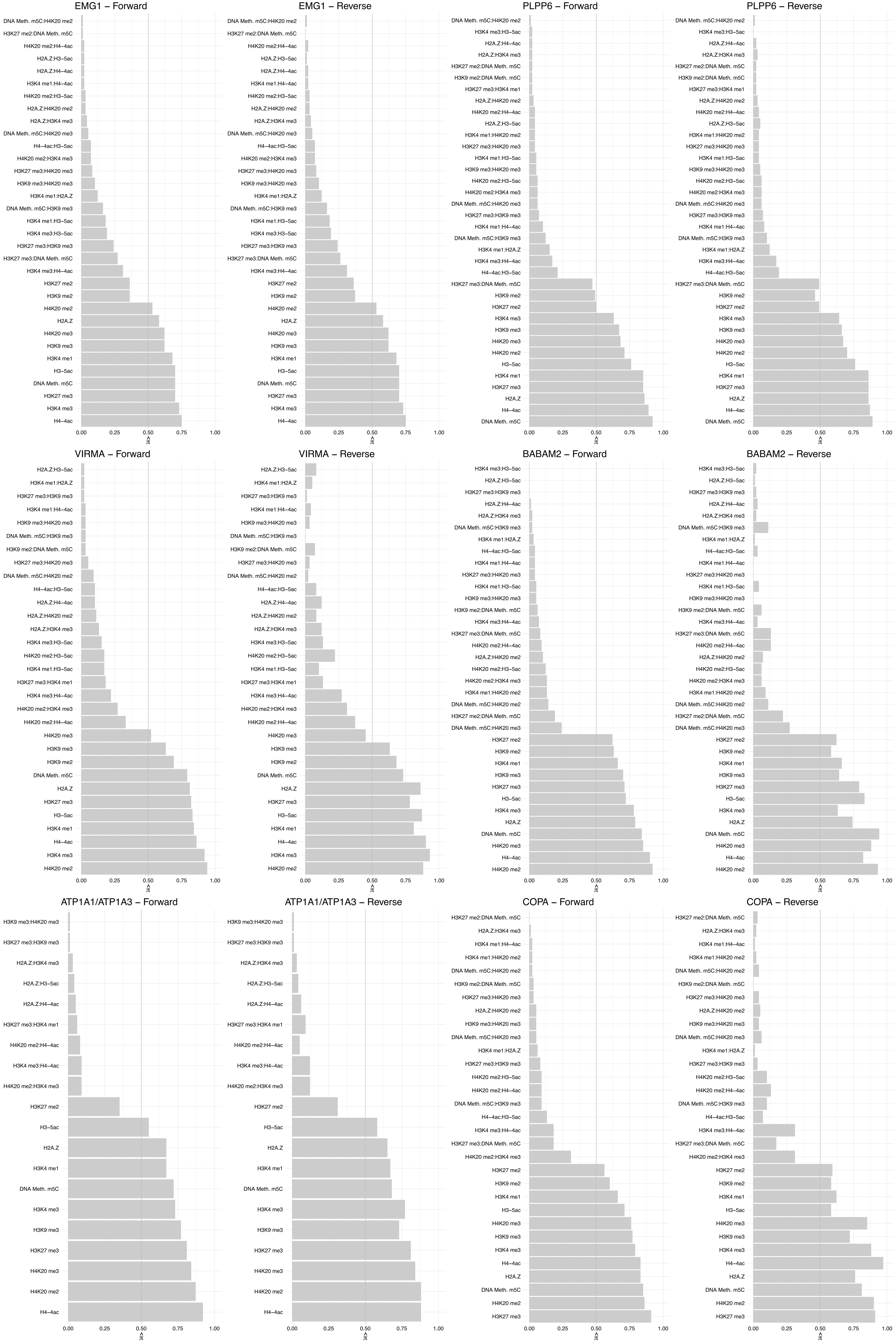
EMG1 – Forward
DNA Meth. m5C:H4K20 me2
H3K27 me2:DNA Meth. m5C
H4K20 me2:H4–4ac
H2A.Z:H3–5ac
H2A.Z:H4–4ac
H3K4 me1:H4–4ac
H4K20 me2:H3–5ac
H2A.Z:H4K20 me2
H2A.Z:H3K4 me3
DNA Meth. m5C:H4K20 me3
H4–4ac:H3–5ac
H4K20 me2:H3K4 me3
H3K27 me3:H4K20 me3
H3K9 me3:H4K20 me3
H3K4 me1:H2A.Z
DNA Meth. m5C:H3K9 me3
H3K4 me1:H3–5ac
H3K4 me3:H3–5ac
H3K27 me3:H3K9 me3
H3K27 me3:DNA Meth. m5C
H3K4 me3:H4–4ac
H3K27 me2
H3K9 me2
H4K20 me2
H2A.Z
H4K20 me3
H3K9 me3
H3K4 me1
H3–5ac
DNA Meth. m5C
H3K27 me3
H3K4 me3
H4–4ac
0.00
0.25
0.50
0.75
1.00
π̂
EMG1 – Reverse
DNA Meth. m5C:H4K20 me2
H3K27 me2:DNA Meth. m5C
H4K20 me2:H4–4ac
H2A.Z:H3–5ac
H2A.Z:H4–4ac
H3K4 me1:H4–4ac
H4K20 me2:H3–5ac
H2A.Z:H4K20 me2
H2A.Z:H3K4 me3
DNA Meth. m5C:H4K20 me3
H4–4ac:H3–5ac
H4K20 me2:H3K4 me3
H3K27 me3:H4K20 me3
H3K9 me3:H4K20 me3
H3K4 me1:H2A.Z
DNA Meth. m5C:H3K9 me3
H3K4 me1:H3–5ac
H3K4 me3:H3–5ac
H3K27 me3:H3K9 me3
H3K27 me3:DNA Meth. m5C
H3K4 me3:H4–4ac
H3K27 me2
H3K9 me2
H4K20 me2
H2A.Z
H4K20 me3
H3K9 me3
H3K4 me1
H3–5ac
DNA Meth. m5C
H3K27 me3
H3K4 me3
H4–4ac
0.00
0.25
0.50
0.75
1.00
π̂
PLPP6 – Forward
DNA Meth. m5C:H4K20 me2
H3K4 me3:H3–5ac
H2A.Z:H4–4ac
H2A.Z:H3K4 me3
H3K27 me2:DNA Meth. m5C
H3K9 me2:DNA Meth. m5C
H3K27 me3:H3K4 me1
H2A.Z:H4K20 me2
H4K20 me2:H4–4ac
H2A.Z:H3–5ac
H3K4 me1:H4K20 me2
H3K27 me3:H4K20 me3
H3K4 me1:H3–5ac
H3K9 me3:H4K20 me3
H4K20 me2:H3–5ac
H4K20 me2:H3K4 me3
DNA Meth. m5C:H4K20 me3
H3K27 me3:H3K9 me3
H3K4 me1:H4–4ac
DNA Meth. m5C:H3K9 me3
H3K4 me1:H2A.Z
H3K4 me3:H4–4ac
H4–4ac:H3–5ac
H3K27 me3:DNA Meth. m5C
H3K9 me2
H3K27 me2
H3K4 me3
H3K9 me3
H4K20 me3
H4K20 me2
H3–5ac
H3K4 me1
H3K27 me3
H2A.Z
H4–4ac
DNA Meth. m5C
0.00
0.25
0.50
0.75
1.00
π̂
PLPP6 – Reverse
DNA Meth. m5C:H4K20 me2
H3K4 me3:H3–5ac
H2A.Z:H4–4ac
H2A.Z:H3K4 me3
H3K27 me2:DNA Meth. m5C
H3K9 me2:DNA Meth. m5C
H3K27 me3:H3K4 me1
H2A.Z:H4K20 me2
H4K20 me2:H4–4ac
H2A.Z:H3–5ac
H3K4 me1:H4K20 me2
H3K27 me3:H4K20 me3
H3K4 me1:H3–5ac
H3K9 me3:H4K20 me3
H4K20 me2:H3–5ac
H4K20 me2:H3K4 me3
DNA Meth. m5C:H4K20 me3
H3K27 me3:H3K9 me3
H3K4 me1:H4–4ac
DNA Meth. m5C:H3K9 me3
H3K4 me1:H2A.Z
H3K4 me3:H4–4ac
H4–4ac:H3–5ac
H3K27 me3:DNA Meth. m5C
H3K9 me2
H3K27 me2
H3K4 me3
H3K9 me3
H4K20 me3
H4K20 me2
H3–5ac
H3K4 me1
H3K27 me3
H2A.Z
H4–4ac
DNA Meth. m5C
0.00
0.25
0.50
0.75
1.00
π̂
VIRMA – Forward
H2A.Z:H3–5ac
H3K4 me1:H2A.Z
H3K27 me3:H3K9 me3
H3K4 me1:H4–4ac
H3K9 me3:H4K20 me3
DNA Meth. m5C:H3K9 me3
H3K9 me2:DNA Meth. m5C
H3K27 me3:H4K20 me3
DNA Meth. m5C:H4K20 me2
H4–4ac:H3–5ac
H2A.Z:H4–4ac
H2A.Z:H4K20 me2
H2A.Z:H3K4 me3
H3K4 me3:H3–5ac
H4K20 me2:H3–5ac
H3K4 me1:H3–5ac
H3K27 me3:H3K4 me1
H3K4 me3:H4–4ac
H4K20 me2:H3K4 me3
H4K20 me2:H4–4ac
H4K20 me3
H3K9 me3
H3K9 me2
DNA Meth. m5C
H2A.Z
H3K27 me3
H3–5ac
H3K4 me1
H4–4ac
H3K4 me3
H4K20 me2
0.00
0.25
0.50
0.75
1.00
π̂
VIRMA – Reverse
H2A.Z:H3–5ac
H3K4 me1:H2A.Z
H3K27 me3:H3K9 me3
H3K4 me1:H4–4ac
H3K9 me3:H4K20 me3
DNA Meth. m5C:H3K9 me3
H3K9 me2:DNA Meth. m5C
H3K27 me3:H4K20 me3
DNA Meth. m5C:H4K20 me2
H4–4ac:H3–5ac
H2A.Z:H4–4ac
H2A.Z:H4K20 me2
H2A.Z:H3K4 me3
H3K4 me3:H3–5ac
H4K20 me2:H3–5ac
H3K4 me1:H3–5ac
H3K27 me3:H3K4 me1
H3K4 me3:H4–4ac
H4K20 me2:H3K4 me3
H4K20 me2:H4–4ac
H4K20 me3
H3K9 me3
H3K9 me2
DNA Meth. m5C
H2A.Z
H3K27 me3
H3–5ac
H3K4 me1
H4–4ac
H3K4 me3
H4K20 me2
0.00
0.25
0.50
0.75
1.00
π̂
BABAM2 – Forward
H3K4 me3:H3–5ac
H2A.Z:H3–5ac
H3K27 me3:H3K9 me3
H2A.Z:H4–4ac
H2A.Z:H3K4 me3
DNA Meth. m5C:H3K9 me3
H3K4 me1:H2A.Z
H4–4ac:H3–5ac
H3K4 me1:H4–4ac
H3K27 me3:H4K20 me3
H3K4 me1:H3–5ac
H3K9 me3:H4K20 me3
H3K9 me2:DNA Meth. m5C
H3K4 me3:H4–4ac
H3K27 me3:DNA Meth. m5C
H4K20 me2:H4–4ac
H2A.Z:H4K20 me2
H4K20 me2:H3–5ac
H4K20 me2:H3K4 me3
H3K4 me1:H4K20 me2
DNA Meth. m5C:H4K20 me2
H3K27 me2:DNA Meth. m5C
DNA Meth. m5C:H4K20 me3
H3K27 me2
H3K9 me2
H3K4 me1
H3K9 me3
H3K27 me3
H3–5ac
H3K4 me3
H2A.Z
DNA Meth. m5C
H4K20 me3
H4–4ac
H4K20 me2
0.00
0.25
0.50
0.75
1.00
π̂
BABAM2 – Reverse
H3K4 me3:H3–5ac
H2A.Z:H3–5ac
H3K27 me3:H3K9 me3
H2A.Z:H4–4ac
H2A.Z:H3K4 me3
DNA Meth. m5C:H3K9 me3
H3K4 me1:H2A.Z
H4–4ac:H3–5ac
H3K4 me1:H4–4ac
H3K27 me3:H4K20 me3
H3K4 me1:H3–5ac
H3K9 me3:H4K20 me3
H3K9 me2:DNA Meth. m5C
H3K4 me3:H4–4ac
H3K27 me3:DNA Meth. m5C
H4K20 me2:H4–4ac
H2A.Z:H4K20 me2
H4K20 me2:H3–5ac
H4K20 me2:H3K4 me3
H3K4 me1:H4K20 me2
DNA Meth. m5C:H4K20 me2
H3K27 me2:DNA Meth. m5C
DNA Meth. m5C:H4K20 me3
H3K27 me2
H3K9 me2
H3K4 me1
H3K9 me3
H3K27 me3
H3–5ac
H3K4 me3
H2A.Z
DNA Meth. m5C
H4K20 me3
H4–4ac
H4K20 me2
0.00
0.25
0.50
0.75
1.00
π̂
ATP1A1/ATP1A3 – Forward
H3K9 me3:H4K20 me3
H3K27 me3:H3K9 me3
H2A.Z:H3K4 me3
H2A.Z:H3–5ac
H2A.Z:H4–4ac
H3K27 me3:H3K4 me1
H4K20 me2:H4–4ac
H3K4 me3:H4–4ac
H4K20 me2:H3K4 me3
H3K27 me2
H3–5ac
H2A.Z
H3K4 me1
DNA Meth. m5C
H3K4 me3
H3K9 me3
H3K27 me3
H4K20 me3
H4K20 me2
H4–4ac
0.00
0.25
0.50
0.75
1.00
π̂
ATP1A1/ATP1A3 – Reverse
H3K9 me3:H4K20 me3
H3K27 me3:H3K9 me3
H2A.Z:H3K4 me3
H2A.Z:H3–5ac
H2A.Z:H4–4ac
H3K27 me3:H3K4 me1
H4K20 me2:H4–4ac
H3K4 me3:H4–4ac
H4K20 me2:H3K4 me3
H3K27 me2
H3–5ac
H2A.Z
H3K4 me1
DNA Meth. m5C
H3K4 me3
H3K9 me3
H3K27 me3
H4K20 me3
H4K20 me2
H4–4ac
0.00
0.25
0.50
0.75
1.00
π̂
COPA – Forward
H3K27 me2:DNA Meth. m5C
H2A.Z:H3K4 me3
H3K4 me1:H4–4ac
H3K4 me1:H4K20 me2
DNA Meth. m5C:H4K20 me2
H3K9 me2:DNA Meth. m5C
H3K27 me3:H4K20 me3
H2A.Z:H4K20 me2
H3K9 me3:H4K20 me3
DNA Meth. m5C:H4K20 me3
H3K4 me1:H2A.Z
H3K27 me3:H3K9 me3
H4K20 me2:H3–5ac
H4K20 me2:H4–4ac
DNA Meth. m5C:H3K9 me3
H4–4ac:H3–5ac
H3K4 me3:H4–4ac
H3K27 me3:DNA Meth. m5C
H4K20 me2:H3K4 me3
H3K27 me2
H3K9 me2
H3K4 me1
H3–5ac
H4K20 me3
H3K9 me3
H3K4 me3
H4–4ac
H2A.Z
DNA Meth. m5C
H4K20 me2
H3K27 me3
0.00
0.25
0.50
0.75
1.00
π̂
COPA – Reverse
H3K27 me2:DNA Meth. m5C
H2A.Z:H3K4 me3
H3K4 me1:H4–4ac
H3K4 me1:H4K20 me2
DNA Meth. m5C:H4K20 me2
H3K9 me2:DNA Meth. m5C
H3K27 me3:H4K20 me3
H2A.Z:H4K20 me2
H3K9 me3:H4K20 me3
DNA Meth. m5C:H4K20 me3
H3K4 me1:H2A.Z
H3K27 me3:H3K9 me3
H4K20 me2:H3–5ac
H4K20 me2:H4–4ac
DNA Meth. m5C:H3K9 me3
H4–4ac:H3–5ac
H3K4 me3:H4–4ac
H3K27 me3:DNA Meth. m5C
H4K20 me2:H3K4 me3
H3K27 me2
H3K9 me2
H3K4 me1
H3–5ac
H4K20 me3
H3K9 me3
H3K4 me3
H4–4ac
H2A.Z
DNA Meth. m5C
H4K20 me2
H3K27 me3
0.00
0.25
0.50
0.75
1.00
π̂

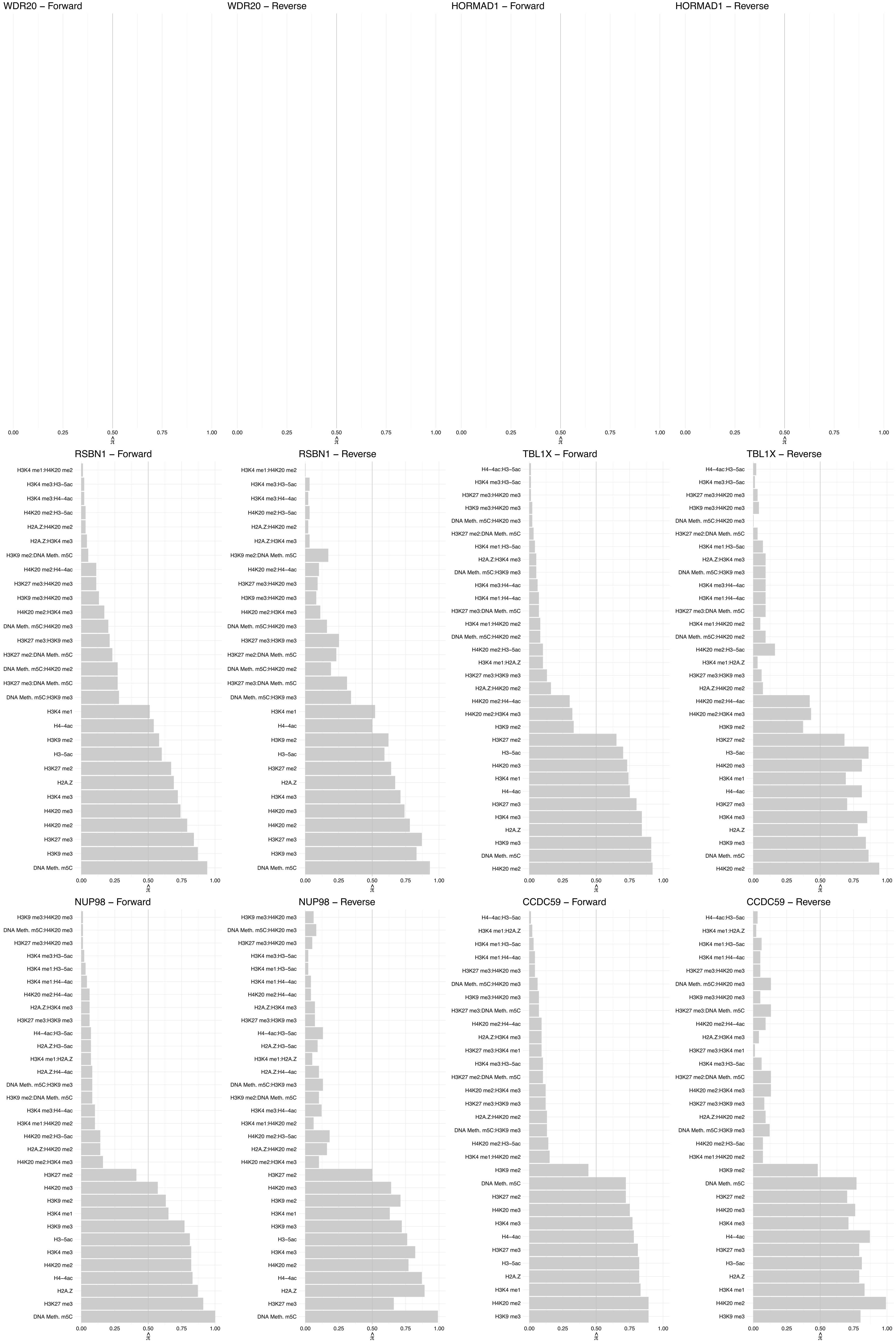

WDR20 – Forward
WDR20 – Reverse
HORMAD1 – Forward
HORMAD1 – Reverse
0.00
0.25
0.50
0.75
1.00
$\hat{\pi}$
RSBN1 – Forward
RSBN1 – Reverse
H3K4 me1:H4K20 me2
H3K4 me3:H3–5ac
H3K4 me3:H4–4ac
H4K20 me2:H3–5ac
H2A.Z:H4K20 me2
H2A.Z:H3K4 me3
H3K9 me2:DNA Meth. m5C
H4K20 me2:H4–4ac
H3K27 me3:H4K20 me3
H3K9 me3:H4K20 me3
H4K20 me2:H3K4 me3
DNA Meth. m5C:H4K20 me3
H3K27 me3:H3K9 me3
H3K27 me2:DNA Meth. m5C
DNA Meth. m5C:H4K20 me2
H3K27 me3:DNA Meth. m5C
DNA Meth. m5C:H3K9 me3
H3K4 me1
H4–4ac
H3K9 me2
H3–5ac
H3K27 me2
H2A.Z
H3K4 me3
H4K20 me3
H4K20 me2
H3K27 me3
H3K9 me3
DNA Meth. m5C
TBL1X – Forward
TBL1X – Reverse
H4–4ac:H3–5ac
H3K4 me3:H3–5ac
H3K27 me3:H4K20 me3
H3K9 me3:H4K20 me3
DNA Meth. m5C:H4K20 me3
H3K27 me2:DNA Meth. m5C
H3K4 me1:H3–5ac
H2A.Z:H3K4 me3
DNA Meth. m5C:H3K9 me3
H3K4 me3:H4–4ac
H3K4 me1:H4–4ac
H3K27 me3:DNA Meth. m5C
H3K4 me1:H4K20 me2
DNA Meth. m5C:H4K20 me2
H4K20 me2:H3–5ac
H3K4 me1:H2A.Z
H3K27 me3:H3K9 me3
H2A.Z:H4K20 me2
H4K20 me2:H4–4ac
H4K20 me2:H3K4 me3
H3K9 me2
H3K27 me2
H3–5ac
H4K20 me3
H3K4 me1
H4–4ac
H3K27 me3
H3K4 me3
H2A.Z
H3K9 me3
DNA Meth. m5C
H4K20 me2
NUP98 – Forward
NUP98 – Reverse
H3K9 me3:H4K20 me3
DNA Meth. m5C:H4K20 me3
H3K27 me3:H4K20 me3
H3K4 me3:H3–5ac
H3K4 me1:H3–5ac
H3K4 me1:H4–4ac
H4K20 me2:H4–4ac
H2A.Z:H3K4 me3
H3K27 me3:H3K9 me3
H4–4ac:H3–5ac
H2A.Z:H3–5ac
H3K4 me1:H2A.Z
H2A.Z:H4–4ac
DNA Meth. m5C:H3K9 me3
H3K9 me2:DNA Meth. m5C
H3K4 me3:H4–4ac
H3K4 me1:H4K20 me2
H4K20 me2:H3–5ac
H2A.Z:H4K20 me2
H4K20 me2:H3K4 me3
H3K27 me2
H4K20 me3
H3K9 me2
H3K4 me1
H3K9 me3
H3–5ac
H3K4 me3
H4K20 me2
H4–4ac
H2A.Z
H3K27 me3
DNA Meth. m5C
CCDC59 – Forward
CCDC59 – Reverse
H4–4ac:H3–5ac
H3K4 me1:H2A.Z
H3K4 me1:H3–5ac
H3K4 me1:H4–4ac
H3K27 me3:H4K20 me3
DNA Meth. m5C:H4K20 me3
H3K9 me3:H4K20 me3
H3K27 me3:DNA Meth. m5C
H4K20 me2:H4–4ac
H2A.Z:H3K4 me3
H3K27 me3:H3K4 me1
H3K4 me3:H3–5ac
H3K27 me2:DNA Meth. m5C
H4K20 me2:H3K4 me3
H3K27 me3:H3K9 me3
H2A.Z:H4K20 me2
DNA Meth. m5C:H3K9 me3
H4K20 me2:H3–5ac
H3K4 me1:H4K20 me2
H3K9 me2
DNA Meth. m5C
H3K27 me2
H4K20 me3
H3K4 me3
H4–4ac
H3K27 me3
H3–5ac
H2A.Z
H3K4 me1
H4K20 me2
H3K9 me3

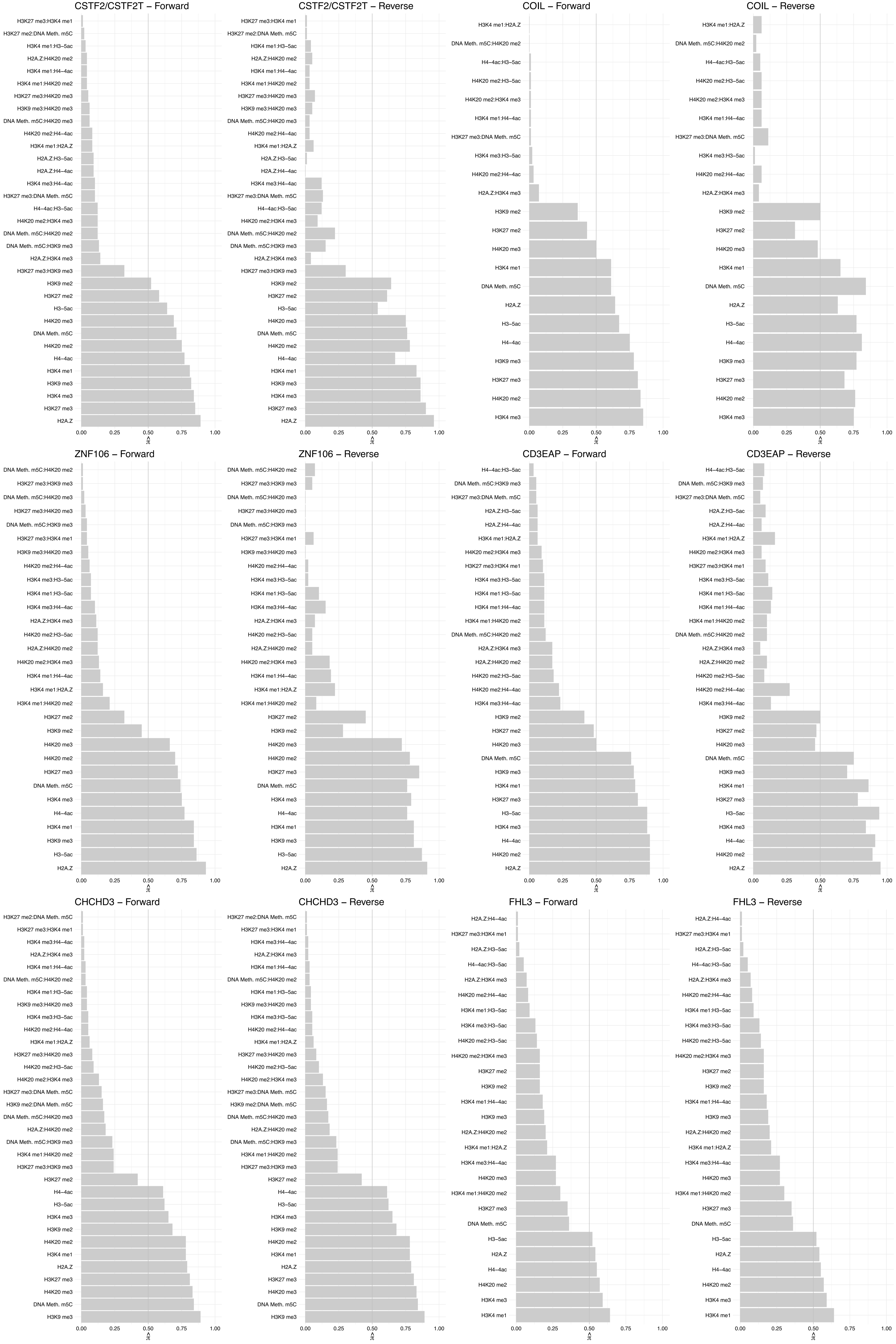
CSTF2/CSTF2T – Forward
H3K27 me3:H3K4 me1
H3K27 me2:DNA Meth. m5C
H3K4 me1:H3–5ac
H2A.Z:H4K20 me2
H3K4 me1:H4–4ac
H3K4 me1:H4K20 me2
H3K27 me3:H4K20 me3
H3K9 me3:H4K20 me3
DNA Meth. m5C:H4K20 me3
H4K20 me2:H4–4ac
H3K4 me1:H2A.Z
H2A.Z:H3–5ac
H2A.Z:H4–4ac
H3K4 me3:H4–4ac
H3K27 me3:DNA Meth. m5C
H4–4ac:H3–5ac
H4K20 me2:H3K4 me3
DNA Meth. m5C:H4K20 me2
DNA Meth. m5C:H3K9 me3
H2A.Z:H3K4 me3
H3K27 me3:H3K9 me3
H3K9 me2
H3K27 me2
H3–5ac
H4K20 me3
DNA Meth. m5C
H4K20 me2
H4–4ac
H3K4 me1
H3K9 me3
H3K4 me3
H3K27 me3
H2A.Z
0.00
0.25
0.50
0.75
1.00
$\hat{\pi}$
CSTF2/CSTF2T – Reverse
H3K27 me3:H3K4 me1
H3K27 me2:DNA Meth. m5C
H3K4 me1:H3–5ac
H2A.Z:H4K20 me2
H3K4 me1:H4–4ac
H3K4 me1:H4K20 me2
H3K27 me3:H4K20 me3
H3K9 me3:H4K20 me3
DNA Meth. m5C:H4K20 me3
H4K20 me2:H4–4ac
H3K4 me1:H2A.Z
H2A.Z:H3–5ac
H2A.Z:H4–4ac
H3K4 me3:H4–4ac
H3K27 me3:DNA Meth. m5C
H4–4ac:H3–5ac
H4K20 me2:H3K4 me3
DNA Meth. m5C:H4K20 me2
DNA Meth. m5C:H3K9 me3
H2A.Z:H3K4 me3
H3K27 me3:H3K9 me3
H3K9 me2
H3K27 me2
H3–5ac
H4K20 me3
DNA Meth. m5C
H4K20 me2
H4–4ac
H3K4 me1
H3K9 me3
H3K4 me3
H3K27 me3
H2A.Z
0.00
0.25
0.50
0.75
1.00
$\hat{\pi}$
COIL – Forward
H3K4 me1:H2A.Z
DNA Meth. m5C:H4K20 me2
H4–4ac:H3–5ac
H4K20 me2:H3–5ac
H4K20 me2:H3K4 me3
H3K4 me1:H4–4ac
H3K27 me3:DNA Meth. m5C
H3K4 me3:H3–5ac
H4K20 me2:H4–4ac
H2A.Z:H3K4 me3
H3K9 me2
H3K27 me2
H4K20 me3
H3K4 me1
DNA Meth. m5C
H2A.Z
H3–5ac
H4–4ac
H3K9 me3
H3K27 me3
H4K20 me2
H3K4 me3
0.00
0.25
0.50
0.75
1.00
$\hat{\pi}$
COIL – Reverse
H3K4 me1:H2A.Z
DNA Meth. m5C:H4K20 me2
H4–4ac:H3–5ac
H4K20 me2:H3–5ac
H4K20 me2:H3K4 me3
H3K4 me1:H4–4ac
H3K27 me3:DNA Meth. m5C
H3K4 me3:H3–5ac
H4K20 me2:H4–4ac
H2A.Z:H3K4 me3
H3K9 me2
H3K27 me2
H4K20 me3
H3K4 me1
DNA Meth. m5C
H2A.Z
H3–5ac
H4–4ac
H3K9 me3
H3K27 me3
H4K20 me2
H3K4 me3
0.00
0.25
0.50
0.75
1.00
$\hat{\pi}$
ZNF106 – Forward
DNA Meth. m5C:H4K20 me2
H3K27 me3:H3K9 me3
DNA Meth. m5C:H4K20 me3
H3K27 me3:H4K20 me3
DNA Meth. m5C:H3K9 me3
H3K27 me3:H3K4 me1
H3K9 me3:H4K20 me3
H4K20 me2:H4–4ac
H3K4 me3:H3–5ac
H3K4 me1:H3–5ac
H3K4 me3:H4–4ac
H2A.Z:H3K4 me3
H4K20 me2:H3–5ac
H2A.Z:H4K20 me2
H4K20 me2:H3K4 me3
H3K4 me1:H4–4ac
H3K4 me1:H2A.Z
H3K4 me1:H4K20 me2
H3K27 me2
H3K9 me2
H4K20 me3
H4K20 me2
H3K27 me3
DNA Meth. m5C
H3K4 me3
H4–4ac
H3K4 me1
H3K9 me3
H3–5ac
H2A.Z
0.00
0.25
0.50
0.75
1.00
$\hat{\pi}$
ZNF106 – Reverse
DNA Meth. m5C:H4K20 me2
H3K27 me3:H3K9 me3
DNA Meth. m5C:H4K20 me3
H3K27 me3:H4K20 me3
DNA Meth. m5C:H3K9 me3
H3K27 me3:H3K4 me1
H3K9 me3:H4K20 me3
H4K20 me2:H4–4ac
H3K4 me3:H3–5ac
H3K4 me1:H3–5ac
H3K4 me3:H4–4ac
H2A.Z:H3K4 me3
H4K20 me2:H3–5ac
H2A.Z:H4K20 me2
H4K20 me2:H3K4 me3
H3K4 me1:H4–4ac
H3K4 me1:H2A.Z
H3K4 me1:H4K20 me2
H3K27 me2
H3K9 me2
H4K20 me3
H4K20 me2
H3K27 me3
DNA Meth. m5C
H3K4 me3
H4–4ac
H3K4 me1
H3K9 me3
H3–5ac
H2A.Z
0.00
0.25
0.50
0.75
1.00
$\hat{\pi}$
CD3EAP – Forward
H4–4ac:H3–5ac
DNA Meth. m5C:H3K9 me3
H3K27 me3:DNA Meth. m5C
H2A.Z:H3–5ac
H2A.Z:H4–4ac
H3K4 me1:H2A.Z
H4K20 me2:H3K4 me3
H3K27 me3:H3K4 me1
H3K4 me3:H3–5ac
H3K4 me1:H3–5ac
H3K4 me1:H4–4ac
H3K4 me1:H4K20 me2
DNA Meth. m5C:H4K20 me2
H2A.Z:H3K4 me3
H2A.Z:H4K20 me2
H4K20 me2:H3–5ac
H4K20 me2:H4–4ac
H3K4 me3:H4–4ac
H3K9 me2
H3K27 me2
H4K20 me3
DNA Meth. m5C
H3K9 me3
H3K4 me1
H3K27 me3
H3–5ac
H3K4 me3
H4–4ac
H4K20 me2
H2A.Z
0.00
0.25
0.50
0.75
1.00
$\hat{\pi}$
CD3EAP – Reverse
H4–4ac:H3–5ac
DNA Meth. m5C:H3K9 me3
H3K27 me3:DNA Meth. m5C
H2A.Z:H3–5ac
H2A.Z:H4–4ac
H3K4 me1:H2A.Z
H4K20 me2:H3K4 me3
H3K27 me3:H3K4 me1
H3K4 me3:H3–5ac
H3K4 me1:H3–5ac
H3K4 me1:H4–4ac
H3K4 me1:H4K20 me2
DNA Meth. m5C:H4K20 me2
H2A.Z:H3K4 me3
H2A.Z:H4K20 me2
H4K20 me2:H3–5ac
H4K20 me2:H4–4ac
H3K4 me3:H4–4ac
H3K9 me2
H3K27 me2
H4K20 me3
DNA Meth. m5C
H3K9 me3
H3K4 me1
H3K27 me3
H3–5ac
H3K4 me3
H4–4ac
H4K20 me2
H2A.Z
0.00
0.25
0.50
0.75
1.00
$\hat{\pi}$
CHCHD3 – Forward
H3K27 me2:DNA Meth. m5C
H3K27 me3:H3K4 me1
H3K4 me3:H4–4ac
H2A.Z:H3K4 me3
H3K4 me1:H4–4ac
DNA Meth. m5C:H4K20 me2
H3K4 me1:H3–5ac
H3K9 me3:H4K20 me3
H3K4 me3:H3–5ac
H4K20 me2:H4–4ac
H3K4 me1:H2A.Z
H3K27 me3:H4K20 me3
H4K20 me2:H3–5ac
H4K20 me2:H3K4 me3
H3K27 me3:DNA Meth. m5C
H3K9 me2:DNA Meth. m5C
DNA Meth. m5C:H4K20 me3
H2A.Z:H4K20 me2
DNA Meth. m5C:H3K9 me3
H3K4 me1:H4K20 me2
H3K27 me3:H3K9 me3
H3K27 me2
H4–4ac
H3–5ac
H3K4 me3
H3K9 me2
H4K20 me2
H3K4 me1
H2A.Z
H3K27 me3
H4K20 me3
DNA Meth. m5C
H3K9 me3
0.00
0.25
0.50
0.75
1.00
$\hat{\pi}$
CHCHD3 – Reverse
H3K27 me2:DNA Meth. m5C
H3K27 me3:H3K4 me1
H3K4 me3:H4–4ac
H2A.Z:H3K4 me3
H3K4 me1:H4–4ac
DNA Meth. m5C:H4K20 me2
H3K4 me1:H3–5ac
H3K9 me3:H4K20 me3
H3K4 me3:H3–5ac
H4K20 me2:H4–4ac
H3K4 me1:H2A.Z
H3K27 me3:H4K20 me3
H4K20 me2:H3–5ac
H4K20 me2:H3K4 me3
H3K27 me3:DNA Meth. m5C
H3K9 me2:DNA Meth. m5C
DNA Meth. m5C:H4K20 me3
H2A.Z:H4K20 me2
DNA Meth. m5C:H3K9 me3
H3K4 me1:H4K20 me2
H3K27 me3:H3K9 me3
H3K27 me2
H4–4ac
H3–5ac
H3K4 me3
H3K9 me2
H4K20 me2
H3K4 me1
H2A.Z
H3K27 me3
H4K20 me3
DNA Meth. m5C
H3K9 me3
0.00
0.25
0.50
0.75
1.00
$\hat{\pi}$
FHL3 – Forward
H2A.Z:H4–4ac
H3K27 me3:H3K4 me1
H2A.Z:H3–5ac
H4–4ac:H3–5ac
H2A.Z:H3K4 me3
H4K20 me2:H4–4ac
H3K4 me1:H3–5ac
H3K4 me3:H3–5ac
H4K20 me2:H3–5ac
H4K20 me2:H3K4 me3
H3K27 me2
H3K9 me2
H3K4 me1:H4–4ac
H3K9 me3
H2A.Z:H4K20 me2
H3K4 me1:H2A.Z
H3K4 me3:H4–4ac
H4K20 me3
H3K4 me1:H4K20 me2
H3K27 me3
DNA Meth. m5C
H3–5ac
H2A.Z
H4–4ac
H4K20 me2
H3K4 me3
H3K4 me1
0.00
0.25
0.50
0.75
1.00
$\hat{\pi}$
FHL3 – Reverse
H2A.Z:H4–4ac
H3K27 me3:H3K4 me1
H2A.Z:H3–5ac
H4–4ac:H3–5ac
H2A.Z:H3K4 me3
H4K20 me2:H4–4ac
H3K4 me1:H3–5ac
H3K4 me3:H3–5ac
H4K20 me2:H3–5ac
H4K20 me2:H3K4 me3
H3K27 me2
H3K9 me2
H3K4 me1:H4–4ac
H3K9 me3
H2A.Z:H4K20 me2
H3K4 me1:H2A.Z
H3K4 me3:H4–4ac
H4K20 me3
H3K4 me1:H4K20 me2
H3K27 me3
DNA Meth. m5C
H3–5ac
H2A.Z
H4–4ac
H4K20 me2
H3K4 me3
H3K4 me1
0.00
0.25
0.50
0.75
1.00
$\hat{\pi}$

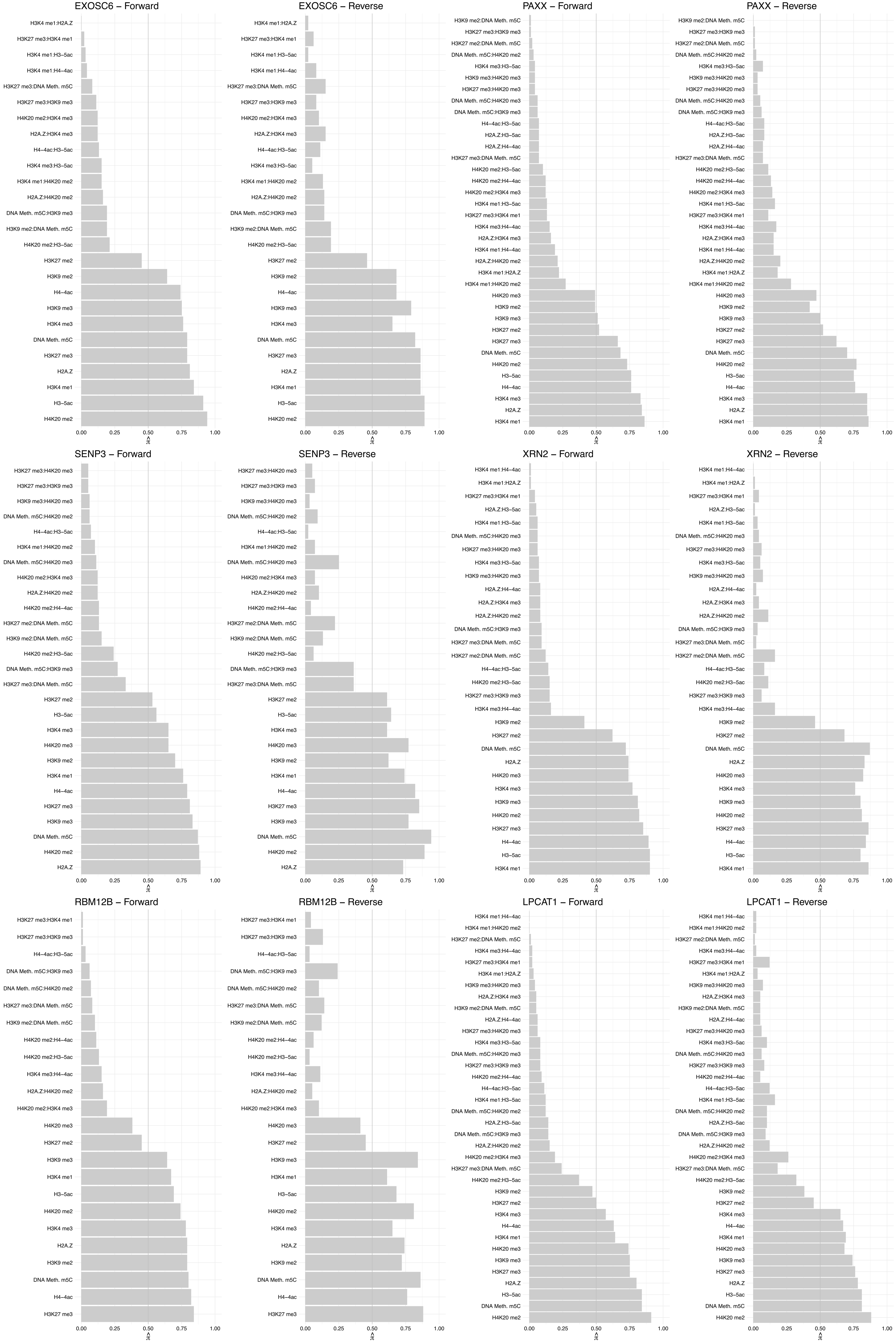

EXOSC6 – Forward
H3K4 me1:H2A.Z
H3K27 me3:H3K4 me1
H3K4 me1:H3–5ac
H3K4 me1:H4–4ac
H3K27 me3:DNA Meth. m5C
H3K27 me3:H3K9 me3
H4K20 me2:H3K4 me3
H2A.Z:H3K4 me3
H4–4ac:H3–5ac
H3K4 me3:H3–5ac
H3K4 me1:H4K20 me2
H2A.Z:H4K20 me2
DNA Meth. m5C:H3K9 me3
H3K9 me2:DNA Meth. m5C
H4K20 me2:H3–5ac
H3K27 me2
H3K9 me2
H4–4ac
H3K9 me3
H3K4 me3
DNA Meth. m5C
H3K27 me3
H2A.Z
H3K4 me1
H3–5ac
H4K20 me2
0.00
0.25
0.50
0.75
1.00
π̂
EXOSC6 – Reverse
H3K4 me1:H2A.Z
H3K27 me3:H3K4 me1
H3K4 me1:H3–5ac
H3K4 me1:H4–4ac
H3K27 me3:DNA Meth. m5C
H3K27 me3:H3K9 me3
H4K20 me2:H3K4 me3
H2A.Z:H3K4 me3
H4–4ac:H3–5ac
H3K4 me3:H3–5ac
H3K4 me1:H4K20 me2
H2A.Z:H4K20 me2
DNA Meth. m5C:H3K9 me3
H3K9 me2:DNA Meth. m5C
H4K20 me2:H3–5ac
H3K27 me2
H3K9 me2
H4–4ac
H3K9 me3
H3K4 me3
DNA Meth. m5C
H3K27 me3
H2A.Z
H3K4 me1
H3–5ac
H4K20 me2
0.00
0.25
0.50
0.75
1.00
π̂
PAXX – Forward
H3K9 me2:DNA Meth. m5C
H3K27 me3:H3K9 me3
H3K27 me2:DNA Meth. m5C
DNA Meth. m5C:H4K20 me2
H3K4 me3:H3–5ac
H3K9 me3:H4K20 me3
H3K27 me3:H4K20 me3
DNA Meth. m5C:H4K20 me3
DNA Meth. m5C:H3K9 me3
H4–4ac:H3–5ac
H2A.Z:H3–5ac
H2A.Z:H4–4ac
H3K27 me3:DNA Meth. m5C
H4K20 me2:H3–5ac
H4K20 me2:H4–4ac
H4K20 me2:H3K4 me3
H3K4 me1:H3–5ac
H3K27 me3:H3K4 me1
H3K4 me3:H4–4ac
H2A.Z:H3K4 me3
H3K4 me1:H4–4ac
H2A.Z:H4K20 me2
H3K4 me1:H2A.Z
H3K4 me1:H4K20 me2
H4K20 me3
H3K9 me2
H3K9 me3
H3K27 me2
H3K27 me3
DNA Meth. m5C
H4K20 me2
H3–5ac
H4–4ac
H3K4 me3
H2A.Z
H3K4 me1
0.00
0.25
0.50
0.75
1.00
π̂
PAXX – Reverse
H3K9 me2:DNA Meth. m5C
H3K27 me3:H3K9 me3
H3K27 me2:DNA Meth. m5C
DNA Meth. m5C:H4K20 me2
H3K4 me3:H3–5ac
H3K9 me3:H4K20 me3
H3K27 me3:H4K20 me3
DNA Meth. m5C:H4K20 me3
DNA Meth. m5C:H3K9 me3
H4–4ac:H3–5ac
H2A.Z:H3–5ac
H2A.Z:H4–4ac
H3K27 me3:DNA Meth. m5C
H4K20 me2:H3–5ac
H4K20 me2:H4–4ac
H4K20 me2:H3K4 me3
H3K4 me1:H3–5ac
H3K27 me3:H3K4 me1
H3K4 me3:H4–4ac
H2A.Z:H3K4 me3
H3K4 me1:H4–4ac
H2A.Z:H4K20 me2
H3K4 me1:H2A.Z
H3K4 me1:H4K20 me2
H4K20 me3
H3K9 me2
H3K9 me3
H3K27 me2
H3K27 me3
DNA Meth. m5C
H4K20 me2
H3–5ac
H4–4ac
H3K4 me3
H2A.Z
H3K4 me1
0.00
0.25
0.50
0.75
1.00
π̂
SENP3 – Forward
H3K27 me3:H4K20 me3
H3K27 me3:H3K9 me3
H3K9 me3:H4K20 me3
DNA Meth. m5C:H4K20 me2
H4–4ac:H3–5ac
H3K4 me1:H4K20 me2
DNA Meth. m5C:H4K20 me3
H4K20 me2:H3K4 me3
H2A.Z:H4K20 me2
H4K20 me2:H4–4ac
H3K27 me2:DNA Meth. m5C
H3K9 me2:DNA Meth. m5C
H4K20 me2:H3–5ac
DNA Meth. m5C:H3K9 me3
H3K27 me3:DNA Meth. m5C
H3K27 me2
H3–5ac
H3K4 me3
H4K20 me3
H3K9 me2
H3K4 me1
H4–4ac
H3K27 me3
H3K9 me3
DNA Meth. m5C
H4K20 me2
H2A.Z
0.00
0.25
0.50
0.75
1.00
π̂
SENP3 – Reverse
H3K27 me3:H4K20 me3
H3K27 me3:H3K9 me3
H3K9 me3:H4K20 me3
DNA Meth. m5C:H4K20 me2
H4–4ac:H3–5ac
H3K4 me1:H4K20 me2
DNA Meth. m5C:H4K20 me3
H4K20 me2:H3K4 me3
H2A.Z:H4K20 me2
H4K20 me2:H4–4ac
H3K27 me2:DNA Meth. m5C
H3K9 me2:DNA Meth. m5C
H4K20 me2:H3–5ac
DNA Meth. m5C:H3K9 me3
H3K27 me3:DNA Meth. m5C
H3K27 me2
H3–5ac
H3K4 me3
H4K20 me3
H3K9 me2
H3K4 me1
H4–4ac
H3K27 me3
H3K9 me3
DNA Meth. m5C
H4K20 me2
H2A.Z
0.00
0.25
0.50
0.75
1.00
π̂
XRN2 – Forward
H3K4 me1:H4–4ac
H3K4 me1:H2A.Z
H3K27 me3:H3K4 me1
H2A.Z:H3–5ac
H3K4 me1:H3–5ac
DNA Meth. m5C:H4K20 me3
H3K27 me3:H4K20 me3
H3K4 me3:H3–5ac
H3K9 me3:H4K20 me3
H2A.Z:H4–4ac
H2A.Z:H3K4 me3
H2A.Z:H4K20 me2
DNA Meth. m5C:H3K9 me3
H3K27 me3:DNA Meth. m5C
H3K27 me2:DNA Meth. m5C
H4–4ac:H3–5ac
H4K20 me2:H3–5ac
H3K27 me3:H3K9 me3
H3K4 me3:H4–4ac
H3K9 me2
H3K27 me2
DNA Meth. m5C
H2A.Z
H4K20 me3
H3K4 me3
H3K9 me3
H4K20 me2
H3K27 me3
H4–4ac
H3–5ac
H3K4 me1
0.00
0.25
0.50
0.75
1.00
π̂
XRN2 – Reverse
H3K4 me1:H4–4ac
H3K4 me1:H2A.Z
H3K27 me3:H3K4 me1
H2A.Z:H3–5ac
H3K4 me1:H3–5ac
DNA Meth. m5C:H4K20 me3
H3K27 me3:H4K20 me3
H3K4 me3:H3–5ac
H3K9 me3:H4K20 me3
H2A.Z:H4–4ac
H2A.Z:H3K4 me3
H2A.Z:H4K20 me2
DNA Meth. m5C:H3K9 me3
H3K27 me3:DNA Meth. m5C
H3K27 me2:DNA Meth. m5C
H4–4ac:H3–5ac
H4K20 me2:H3–5ac
H3K27 me3:H3K9 me3
H3K4 me3:H4–4ac
H3K9 me2
H3K27 me2
DNA Meth. m5C
H2A.Z
H4K20 me3
H3K4 me3
H3K9 me3
H4K20 me2
H3K27 me3
H4–4ac
H3–5ac
H3K4 me1
0.00
0.25
0.50
0.75
1.00
π̂
RBM12B – Forward
H3K27 me3:H3K4 me1
H3K27 me3:H3K9 me3
H4–4ac:H3–5ac
DNA Meth. m5C:H3K9 me3
DNA Meth. m5C:H4K20 me2
H3K27 me3:DNA Meth. m5C
H3K9 me2:DNA Meth. m5C
H4K20 me2:H4–4ac
H4K20 me2:H3–5ac
H3K4 me3:H4–4ac
H2A.Z:H4K20 me2
H4K20 me2:H3K4 me3
H4K20 me3
H3K27 me2
H3K9 me3
H3K4 me1
H3–5ac
H4K20 me2
H3K4 me3
H2A.Z
H3K9 me2
DNA Meth. m5C
H4–4ac
H3K27 me3
0.00
0.25
0.50
0.75
1.00
π̂
RBM12B – Reverse
H3K27 me3:H3K4 me1
H3K27 me3:H3K9 me3
H4–4ac:H3–5ac
DNA Meth. m5C:H3K9 me3
DNA Meth. m5C:H4K20 me2
H3K27 me3:DNA Meth. m5C
H3K9 me2:DNA Meth. m5C
H4K20 me2:H4–4ac
H4K20 me2:H3–5ac
H3K4 me3:H4–4ac
H2A.Z:H4K20 me2
H4K20 me2:H3K4 me3
H4K20 me3
H3K27 me2
H3K9 me3
H3K4 me1
H3–5ac
H4K20 me2
H3K4 me3
H2A.Z
H3K9 me2
DNA Meth. m5C
H4–4ac
H3K27 me3
0.00
0.25
0.50
0.75
1.00
π̂
LPCAT1 – Forward
H3K4 me1:H4–4ac
H3K4 me1:H4K20 me2
H3K27 me2:DNA Meth. m5C
H3K4 me3:H4–4ac
H3K27 me3:H3K4 me1
H3K4 me1:H2A.Z
H3K9 me3:H4K20 me3
H2A.Z:H3K4 me3
H3K9 me2:DNA Meth. m5C
H2A.Z:H4–4ac
H3K27 me3:H4K20 me3
H3K4 me3:H3–5ac
DNA Meth. m5C:H4K20 me3
H3K27 me3:H3K9 me3
H4K20 me2:H4–4ac
H4–4ac:H3–5ac
H3K4 me1:H3–5ac
DNA Meth. m5C:H4K20 me2
H2A.Z:H3–5ac
DNA Meth. m5C:H3K9 me3
H2A.Z:H4K20 me2
H4K20 me2:H3K4 me3
H3K27 me3:DNA Meth. m5C
H4K20 me2:H3–5ac
H3K9 me2
H3K27 me2
H3K4 me3
H4–4ac
H3K4 me1
H4K20 me3
H3K9 me3
H3K27 me3
H2A.Z
H3–5ac
DNA Meth. m5C
H4K20 me2
0.00
0.25
0.50
0.75
1.00
π̂
LPCAT1 – Reverse
H3K4 me1:H4–4ac
H3K4 me1:H4K20 me2
H3K27 me2:DNA Meth. m5C
H3K4 me3:H4–4ac
H3K27 me3:H3K4 me1
H3K4 me1:H2A.Z
H3K9 me3:H4K20 me3
H2A.Z:H3K4 me3
H3K9 me2:DNA Meth. m5C
H2A.Z:H4–4ac
H3K27 me3:H4K20 me3
H3K4 me3:H3–5ac
DNA Meth. m5C:H4K20 me3
H3K27 me3:H3K9 me3
H4K20 me2:H4–4ac
H4–4ac:H3–5ac
H3K4 me1:H3–5ac
DNA Meth. m5C:H4K20 me2
H2A.Z:H3–5ac
DNA Meth. m5C:H3K9 me3
H2A.Z:H4K20 me2
H4K20 me2:H3K4 me3
H3K27 me3:DNA Meth. m5C
H4K20 me2:H3–5ac
H3K9 me2
H3K27 me2
H3K4 me3
H4–4ac
H3K4 me1
H4K20 me3
H3K9 me3
H3K27 me3
H2A.Z
H3–5ac
DNA Meth. m5C
H4K20 me2
0.00
0.25
0.50
0.75
1.00
π̂

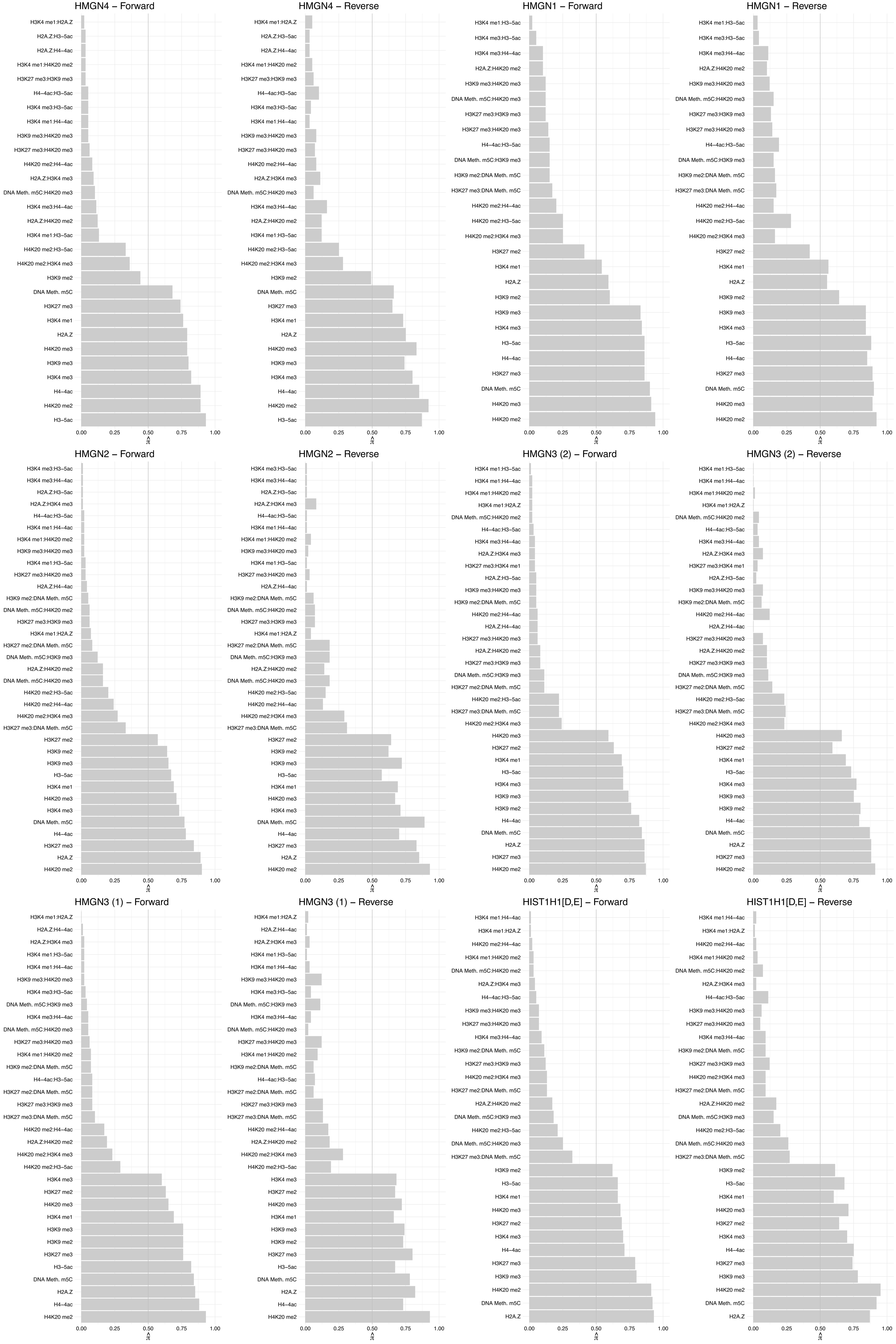
HMGN4 – Forward
H3K4 me1:H2A.Z
H2A.Z:H3–5ac
H2A.Z:H4–4ac
H3K4 me1:H4K20 me2
H3K27 me3:H3K9 me3
H4–4ac:H3–5ac
H3K4 me3:H3–5ac
H3K4 me1:H4–4ac
H3K9 me3:H4K20 me3
H3K27 me3:H4K20 me3
H4K20 me2:H4–4ac
H2A.Z:H3K4 me3
DNA Meth. m5C:H4K20 me3
H3K4 me3:H4–4ac
H2A.Z:H4K20 me2
H3K4 me1:H3–5ac
H4K20 me2:H3–5ac
H4K20 me2:H3K4 me3
H3K9 me2
DNA Meth. m5C
H3K27 me3
H3K4 me1
H2A.Z
H4K20 me3
H3K9 me3
H3K4 me3
H4–4ac
H4K20 me2
H3–5ac
0.00
0.25
0.50
0.75
1.00
$\hat{\pi}$
HMGN4 – Reverse
H3K4 me1:H2A.Z
H2A.Z:H3–5ac
H2A.Z:H4–4ac
H3K4 me1:H4K20 me2
H3K27 me3:H3K9 me3
H4–4ac:H3–5ac
H3K4 me3:H3–5ac
H3K4 me1:H4–4ac
H3K9 me3:H4K20 me3
H3K27 me3:H4K20 me3
H4K20 me2:H4–4ac
H2A.Z:H3K4 me3
DNA Meth. m5C:H4K20 me3
H3K4 me3:H4–4ac
H2A.Z:H4K20 me2
H3K4 me1:H3–5ac
H4K20 me2:H3–5ac
H4K20 me2:H3K4 me3
H3K9 me2
DNA Meth. m5C
H3K27 me3
H3K4 me1
H2A.Z
H4K20 me3
H3K9 me3
H3K4 me3
H4–4ac
H4K20 me2
H3–5ac
0.00
0.25
0.50
0.75
1.00
$\hat{\pi}$
HMGN1 – Forward
H3K4 me1:H3–5ac
H3K4 me3:H3–5ac
H3K4 me3:H4–4ac
H2A.Z:H4K20 me2
H3K9 me3:H4K20 me3
DNA Meth. m5C:H4K20 me3
H3K27 me3:H3K9 me3
H3K27 me3:H4K20 me3
H4–4ac:H3–5ac
DNA Meth. m5C:H3K9 me3
H3K9 me2:DNA Meth. m5C
H3K27 me3:DNA Meth. m5C
H4K20 me2:H4–4ac
H4K20 me2:H3–5ac
H4K20 me2:H3K4 me3
H3K27 me2
H3K4 me1
H2A.Z
H3K9 me2
H3K9 me3
H3K4 me3
H3–5ac
H4–4ac
H3K27 me3
DNA Meth. m5C
H4K20 me3
H4K20 me2
0.00
0.25
0.50
0.75
1.00
$\hat{\pi}$
HMGN1 – Reverse
H3K4 me1:H3–5ac
H3K4 me3:H3–5ac
H3K4 me3:H4–4ac
H2A.Z:H4K20 me2
H3K9 me3:H4K20 me3
DNA Meth. m5C:H4K20 me3
H3K27 me3:H3K9 me3
H3K27 me3:H4K20 me3
H4–4ac:H3–5ac
DNA Meth. m5C:H3K9 me3
H3K9 me2:DNA Meth. m5C
H3K27 me3:DNA Meth. m5C
H4K20 me2:H4–4ac
H4K20 me2:H3–5ac
H4K20 me2:H3K4 me3
H3K27 me2
H3K4 me1
H2A.Z
H3K9 me2
H3K9 me3
H3K4 me3
H3–5ac
H4–4ac
H3K27 me3
DNA Meth. m5C
H4K20 me3
H4K20 me2
0.00
0.25
0.50
0.75
1.00
$\hat{\pi}$
HMGN2 – Forward
H3K4 me3:H3–5ac
H3K4 me3:H4–4ac
H2A.Z:H3–5ac
H2A.Z:H3K4 me3
H4–4ac:H3–5ac
H3K4 me1:H4–4ac
H3K4 me1:H4K20 me2
H3K9 me3:H4K20 me3
H3K4 me1:H3–5ac
H3K27 me3:H4K20 me3
H2A.Z:H4–4ac
H3K9 me2:DNA Meth. m5C
DNA Meth. m5C:H4K20 me2
H3K27 me3:H3K9 me3
H3K4 me1:H2A.Z
H3K27 me2:DNA Meth. m5C
DNA Meth. m5C:H3K9 me3
H2A.Z:H4K20 me2
DNA Meth. m5C:H4K20 me3
H4K20 me2:H3–5ac
H4K20 me2:H4–4ac
H4K20 me2:H3K4 me3
H3K27 me3:DNA Meth. m5C
H3K27 me2
H3K9 me2
H3K9 me3
H3–5ac
H3K4 me1
H4K20 me3
H3K4 me3
DNA Meth. m5C
H4–4ac
H3K27 me3
H2A.Z
H4K20 me2
0.00
0.25
0.50
0.75
1.00
$\hat{\pi}$
HMGN2 – Reverse
H3K4 me3:H3–5ac
H3K4 me3:H4–4ac
H2A.Z:H3–5ac
H2A.Z:H3K4 me3
H4–4ac:H3–5ac
H3K4 me1:H4–4ac
H3K4 me1:H4K20 me2
H3K9 me3:H4K20 me3
H3K4 me1:H3–5ac
H3K27 me3:H4K20 me3
H2A.Z:H4–4ac
H3K9 me2:DNA Meth. m5C
DNA Meth. m5C:H4K20 me2
H3K27 me3:H3K9 me3
H3K4 me1:H2A.Z
H3K27 me2:DNA Meth. m5C
DNA Meth. m5C:H3K9 me3
H2A.Z:H4K20 me2
DNA Meth. m5C:H4K20 me3
H4K20 me2:H3–5ac
H4K20 me2:H4–4ac
H4K20 me2:H3K4 me3
H3K27 me3:DNA Meth. m5C
H3K27 me2
H3K9 me2
H3K9 me3
H3–5ac
H3K4 me1
H4K20 me3
H3K4 me3
DNA Meth. m5C
H4–4ac
H3K27 me3
H2A.Z
H4K20 me2
0.00
0.25
0.50
0.75
1.00
$\hat{\pi}$
HMGN3 (2) – Forward
H3K4 me1:H3–5ac
H3K4 me1:H4–4ac
H3K4 me1:H4K20 me2
H3K4 me1:H2A.Z
DNA Meth. m5C:H4K20 me2
H4–4ac:H3–5ac
H3K4 me3:H4–4ac
H2A.Z:H3K4 me3
H3K27 me3:H3K4 me1
H2A.Z:H3–5ac
H3K9 me3:H4K20 me3
H3K9 me2:DNA Meth. m5C
H4K20 me2:H4–4ac
H2A.Z:H4–4ac
H3K27 me3:H4K20 me3
H2A.Z:H4K20 me2
H3K27 me3:H3K9 me3
DNA Meth. m5C:H3K9 me3
H3K27 me2:DNA Meth. m5C
H4K20 me2:H3–5ac
H3K27 me3:DNA Meth. m5C
H4K20 me2:H3K4 me3
H4K20 me3
H3K27 me2
H3K4 me1
H3–5ac
H3K4 me3
H3K9 me3
H3K9 me2
H4–4ac
DNA Meth. m5C
H2A.Z
H3K27 me3
H4K20 me2
0.00
0.25
0.50
0.75
1.00
$\hat{\pi}$
HMGN3 (2) – Reverse
H3K4 me1:H3–5ac
H3K4 me1:H4–4ac
H3K4 me1:H4K20 me2
H3K4 me1:H2A.Z
DNA Meth. m5C:H4K20 me2
H4–4ac:H3–5ac
H3K4 me3:H4–4ac
H2A.Z:H3K4 me3
H3K27 me3:H3K4 me1
H2A.Z:H3–5ac
H3K9 me3:H4K20 me3
H3K9 me2:DNA Meth. m5C
H4K20 me2:H4–4ac
H2A.Z:H4–4ac
H3K27 me3:H4K20 me3
H2A.Z:H4K20 me2
H3K27 me3:H3K9 me3
DNA Meth. m5C:H3K9 me3
H3K27 me2:DNA Meth. m5C
H4K20 me2:H3–5ac
H3K27 me3:DNA Meth. m5C
H4K20 me2:H3K4 me3
H4K20 me3
H3K27 me2
H3K4 me1
H3–5ac
H3K4 me3
H3K9 me3
H3K9 me2
H4–4ac
DNA Meth. m5C
H2A.Z
H3K27 me3
H4K20 me2
0.00
0.25
0.50
0.75
1.00
$\hat{\pi}$
HMGN3 (1) – Forward
H3K4 me1:H2A.Z
H2A.Z:H4–4ac
H2A.Z:H3K4 me3
H3K4 me1:H3–5ac
H3K4 me1:H4–4ac
H3K9 me3:H4K20 me3
H3K4 me3:H3–5ac
DNA Meth. m5C:H3K9 me3
H3K4 me3:H4–4ac
DNA Meth. m5C:H4K20 me3
H3K27 me3:H4K20 me3
H3K4 me1:H4K20 me2
H3K9 me2:DNA Meth. m5C
H4–4ac:H3–5ac
H3K27 me2:DNA Meth. m5C
H3K27 me3:H3K9 me3
H3K27 me3:DNA Meth. m5C
H4K20 me2:H4–4ac
H2A.Z:H4K20 me2
H4K20 me2:H3K4 me3
H4K20 me2:H3–5ac
H3K4 me3
H3K27 me2
H4K20 me3
H3K4 me1
H3K9 me3
H3K9 me2
H3K27 me3
H3–5ac
DNA Meth. m5C
H2A.Z
H4–4ac
H4K20 me2
0.00
0.25
0.50
0.75
1.00
$\hat{\pi}$
HMGN3 (1) – Reverse
H3K4 me1:H2A.Z
H2A.Z:H4–4ac
H2A.Z:H3K4 me3
H3K4 me1:H3–5ac
H3K4 me1:H4–4ac
H3K9 me3:H4K20 me3
H3K4 me3:H3–5ac
DNA Meth. m5C:H3K9 me3
H3K4 me3:H4–4ac
DNA Meth. m5C:H4K20 me3
H3K27 me3:H4K20 me3
H3K4 me1:H4K20 me2
H3K9 me2:DNA Meth. m5C
H4–4ac:H3–5ac
H3K27 me2:DNA Meth. m5C
H3K27 me3:H3K9 me3
H3K27 me3:DNA Meth. m5C
H4K20 me2:H4–4ac
H2A.Z:H4K20 me2
H4K20 me2:H3K4 me3
H4K20 me2:H3–5ac
H3K4 me3
H3K27 me2
H4K20 me3
H3K4 me1
H3K9 me3
H3K9 me2
H3K27 me3
H3–5ac
DNA Meth. m5C
H2A.Z
H4–4ac
H4K20 me2
0.00
0.25
0.50
0.75
1.00
$\hat{\pi}$
HIST1H1[D,E] – Forward
H3K4 me1:H4–4ac
H3K4 me1:H2A.Z
H4K20 me2:H4–4ac
H3K4 me1:H4K20 me2
DNA Meth. m5C:H4K20 me2
H2A.Z:H3K4 me3
H4–4ac:H3–5ac
H3K9 me3:H4K20 me3
H3K27 me3:H4K20 me3
H3K4 me3:H4–4ac
H3K9 me2:DNA Meth. m5C
H3K27 me3:H3K9 me3
H4K20 me2:H3K4 me3
H3K27 me2:DNA Meth. m5C
H2A.Z:H4K20 me2
DNA Meth. m5C:H3K9 me3
H4K20 me2:H3–5ac
DNA Meth. m5C:H4K20 me3
H3K27 me3:DNA Meth. m5C
H3K9 me2
H3–5ac
H3K4 me1
H4K20 me3
H3K27 me2
H3K4 me3
H4–4ac
H3K27 me3
H3K9 me3
H4K20 me2
DNA Meth. m5C
H2A.Z
0.00
0.25
0.50
0.75
1.00
$\hat{\pi}$
HIST1H1[D,E] – Reverse
H3K4 me1:H4–4ac
H3K4 me1:H2A.Z
H4K20 me2:H4–4ac
H3K4 me1:H4K20 me2
DNA Meth. m5C:H4K20 me2
H2A.Z:H3K4 me3
H4–4ac:H3–5ac
H3K9 me3:H4K20 me3
H3K27 me3:H4K20 me3
H3K4 me3:H4–4ac
H3K9 me2:DNA Meth. m5C
H3K27 me3:H3K9 me3
H4K20 me2:H3K4 me3
H3K27 me2:DNA Meth. m5C
H2A.Z:H4K20 me2
DNA Meth. m5C:H3K9 me3
H4K20 me2:H3–5ac
DNA Meth. m5C:H4K20 me3
H3K27 me3:DNA Meth. m5C
H3K9 me2
H3–5ac
H3K4 me1
H4K20 me3
H3K27 me2
H3K4 me3
H4–4ac
H3K27 me3
H3K9 me3
H4K20 me2
DNA Meth. m5C
H2A.Z
0.00
0.25
0.50
0.75
1.00
$\hat{\pi}$

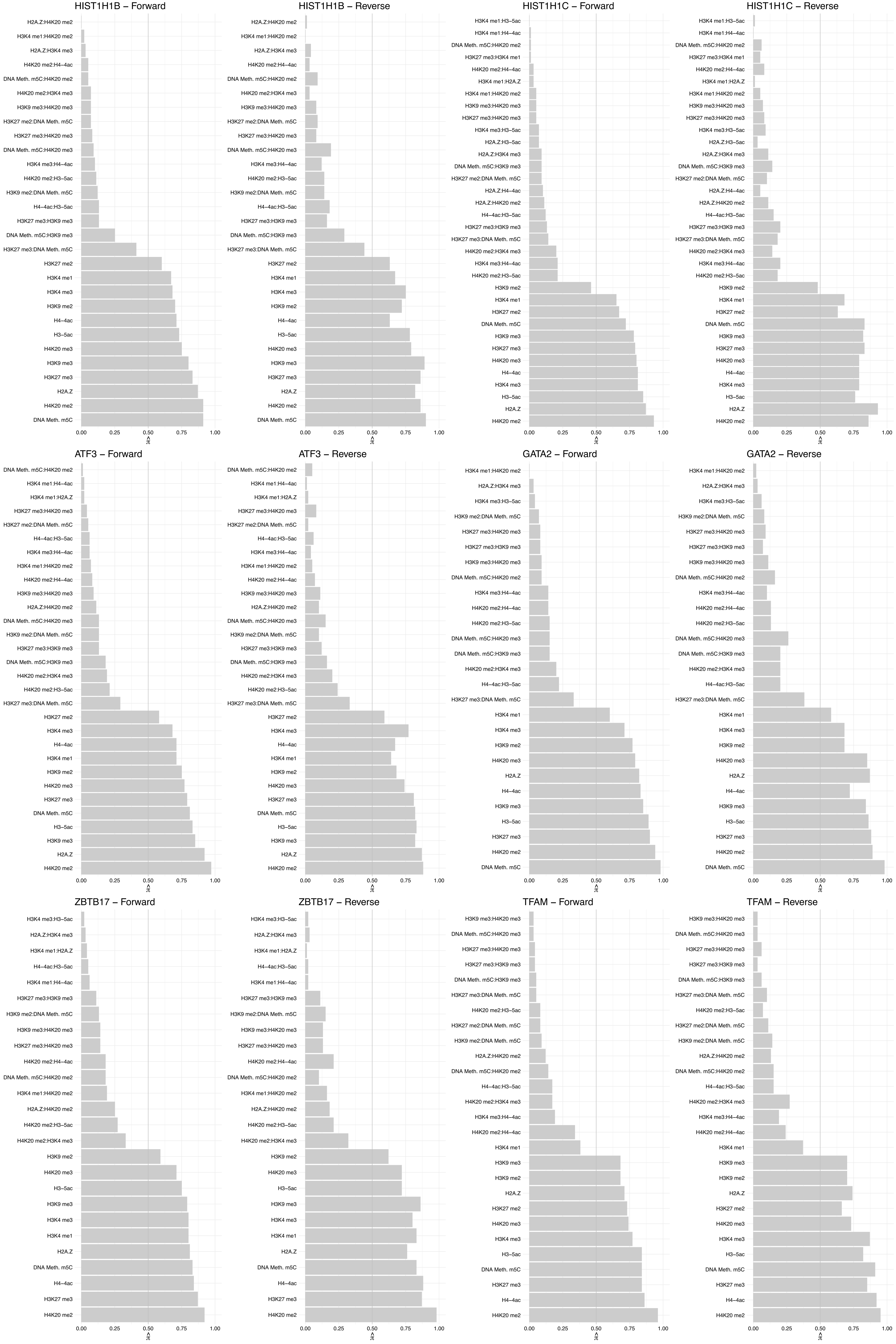
HIST1H1B – Forward
H2A.Z:H4K20 me2
H3K4 me1:H4K20 me2
H2A.Z:H3K4 me3
H4K20 me2:H4−4ac
DNA Meth. m5C:H4K20 me2
H4K20 me2:H3K4 me3
H3K9 me3:H4K20 me3
H3K27 me2:DNA Meth. m5C
H3K27 me3:H4K20 me3
DNA Meth. m5C:H4K20 me3
H3K4 me3:H4−4ac
H4K20 me2:H3−5ac
H3K9 me2:DNA Meth. m5C
H4−4ac:H3−5ac
H3K27 me3:H3K9 me3
DNA Meth. m5C:H3K9 me3
H3K27 me3:DNA Meth. m5C
H3K27 me2
H3K4 me1
H3K4 me3
H3K9 me2
H4−4ac
H3−5ac
H4K20 me3
H3K9 me3
H3K27 me3
H2A.Z
H4K20 me2
DNA Meth. m5C
0.00 0.25 0.50 0.75 1.00
$\hat{\pi}$
HIST1H1B – Reverse
H2A.Z:H4K20 me2
H3K4 me1:H4K20 me2
H2A.Z:H3K4 me3
H4K20 me2:H4−4ac
DNA Meth. m5C:H4K20 me2
H4K20 me2:H3K4 me3
H3K9 me3:H4K20 me3
H3K27 me2:DNA Meth. m5C
H3K27 me3:H4K20 me3
DNA Meth. m5C:H4K20 me3
H3K4 me3:H4−4ac
H4K20 me2:H3−5ac
H3K9 me2:DNA Meth. m5C
H4−4ac:H3−5ac
H3K27 me3:H3K9 me3
DNA Meth. m5C:H3K9 me3
H3K27 me3:DNA Meth. m5C
H3K27 me2
H3K4 me1
H3K4 me3
H3K9 me2
H4−4ac
H3−5ac
H4K20 me3
H3K9 me3
H3K27 me3
H2A.Z
H4K20 me2
DNA Meth. m5C
0.00 0.25 0.50 0.75 1.00
$\hat{\pi}$
HIST1H1C – Forward
H3K4 me1:H3−5ac
H3K4 me1:H4−4ac
DNA Meth. m5C:H4K20 me2
H3K27 me3:H3K4 me1
H4K20 me2:H4−4ac
H3K4 me1:H2A.Z
H3K4 me1:H4K20 me2
H3K9 me3:H4K20 me3
H3K27 me3:H4K20 me3
H3K4 me3:H3−5ac
H2A.Z:H3−5ac
H2A.Z:H3K4 me3
DNA Meth. m5C:H3K9 me3
H3K27 me2:DNA Meth. m5C
H2A.Z:H4−4ac
H2A.Z:H4K20 me2
H4−4ac:H3−5ac
H3K27 me3:H3K9 me3
H3K27 me3:DNA Meth. m5C
H4K20 me2:H3K4 me3
H3K4 me3:H4−4ac
H4K20 me2:H3−5ac
H3K9 me2
H3K4 me1
H3K27 me2
DNA Meth. m5C
H3K9 me3
H3K27 me3
H4K20 me3
H4−4ac
H3K4 me3
H3−5ac
H2A.Z
H4K20 me2
0.00 0.25 0.50 0.75 1.00
$\hat{\pi}$
HIST1H1C – Reverse
H3K4 me1:H3−5ac
H3K4 me1:H4−4ac
DNA Meth. m5C:H4K20 me2
H3K27 me3:H3K4 me1
H4K20 me2:H4−4ac
H3K4 me1:H2A.Z
H3K4 me1:H4K20 me2
H3K9 me3:H4K20 me3
H3K27 me3:H4K20 me3
H3K4 me3:H3−5ac
H2A.Z:H3−5ac
H2A.Z:H3K4 me3
DNA Meth. m5C:H3K9 me3
H3K27 me2:DNA Meth. m5C
H2A.Z:H4−4ac
H2A.Z:H4K20 me2
H4−4ac:H3−5ac
H3K27 me3:H3K9 me3
H3K27 me3:DNA Meth. m5C
H4K20 me2:H3K4 me3
H3K4 me3:H4−4ac
H4K20 me2:H3−5ac
H3K9 me2
H3K4 me1
H3K27 me2
DNA Meth. m5C
H3K9 me3
H3K27 me3
H4K20 me3
H4−4ac
H3K4 me3
H3−5ac
H2A.Z
H4K20 me2
0.00 0.25 0.50 0.75 1.00
$\hat{\pi}$
ATF3 – Forward
DNA Meth. m5C:H4K20 me2
H3K4 me1:H4−4ac
H3K4 me1:H2A.Z
H3K27 me3:H4K20 me3
H3K27 me2:DNA Meth. m5C
H4−4ac:H3−5ac
H3K4 me3:H4−4ac
H3K4 me1:H4K20 me2
H4K20 me2:H4−4ac
H3K9 me3:H4K20 me3
H2A.Z:H4K20 me2
DNA Meth. m5C:H4K20 me3
H3K9 me2:DNA Meth. m5C
H3K27 me3:H3K9 me3
DNA Meth. m5C:H3K9 me3
H4K20 me2:H3K4 me3
H4K20 me2:H3−5ac
H3K27 me3:DNA Meth. m5C
H3K27 me2
H3K4 me3
H4−4ac
H3K4 me1
H3K9 me2
H4K20 me3
H3K27 me3
DNA Meth. m5C
H3−5ac
H3K9 me3
H2A.Z
H4K20 me2
0.00 0.25 0.50 0.75 1.00
$\hat{\pi}$
ATF3 – Reverse
DNA Meth. m5C:H4K20 me2
H3K4 me1:H4−4ac
H3K4 me1:H2A.Z
H3K27 me3:H4K20 me3
H3K27 me2:DNA Meth. m5C
H4−4ac:H3−5ac
H3K4 me3:H4−4ac
H3K4 me1:H4K20 me2
H4K20 me2:H4−4ac
H3K9 me3:H4K20 me3
H2A.Z:H4K20 me2
DNA Meth. m5C:H4K20 me3
H3K9 me2:DNA Meth. m5C
H3K27 me3:H3K9 me3
DNA Meth. m5C:H3K9 me3
H4K20 me2:H3K4 me3
H4K20 me2:H3−5ac
H3K27 me3:DNA Meth. m5C
H3K27 me2
H3K4 me3
H4−4ac
H3K4 me1
H3K9 me2
H4K20 me3
H3K27 me3
DNA Meth. m5C
H3−5ac
H3K9 me3
H2A.Z
H4K20 me2
0.00 0.25 0.50 0.75 1.00
$\hat{\pi}$
GATA2 – Forward
H3K4 me1:H4K20 me2
H2A.Z:H3K4 me3
H3K4 me3:H3−5ac
H3K9 me2:DNA Meth. m5C
H3K27 me3:H4K20 me3
H3K27 me3:H3K9 me3
H3K9 me3:H4K20 me3
DNA Meth. m5C:H4K20 me2
H3K4 me3:H4−4ac
H4K20 me2:H4−4ac
H4K20 me2:H3−5ac
DNA Meth. m5C:H4K20 me3
DNA Meth. m5C:H3K9 me3
H4K20 me2:H3K4 me3
H4−4ac:H3−5ac
H3K27 me3:DNA Meth. m5C
H3K4 me1
H3K4 me3
H3K9 me2
H4K20 me3
H2A.Z
H4−4ac
H3K9 me3
H3−5ac
H3K27 me3
H4K20 me2
DNA Meth. m5C
0.00 0.25 0.50 0.75 1.00
$\hat{\pi}$
GATA2 – Reverse
H3K4 me1:H4K20 me2
H2A.Z:H3K4 me3
H3K4 me3:H3−5ac
H3K9 me2:DNA Meth. m5C
H3K27 me3:H4K20 me3
H3K27 me3:H3K9 me3
H3K9 me3:H4K20 me3
DNA Meth. m5C:H4K20 me2
H3K4 me3:H4−4ac
H4K20 me2:H4−4ac
H4K20 me2:H3−5ac
DNA Meth. m5C:H4K20 me3
DNA Meth. m5C:H3K9 me3
H4K20 me2:H3K4 me3
H4−4ac:H3−5ac
H3K27 me3:DNA Meth. m5C
H3K4 me1
H3K4 me3
H3K9 me2
H4K20 me3
H2A.Z
H4−4ac
H3K9 me3
H3−5ac
H3K27 me3
H4K20 me2
DNA Meth. m5C
0.00 0.25 0.50 0.75 1.00
$\hat{\pi}$
ZBTB17 – Forward
H3K4 me3:H3−5ac
H2A.Z:H3K4 me3
H3K4 me1:H2A.Z
H4−4ac:H3−5ac
H3K4 me1:H4−4ac
H3K27 me3:H3K9 me3
H3K9 me2:DNA Meth. m5C
H3K9 me3:H4K20 me3
H3K27 me3:H4K20 me3
H4K20 me2:H4−4ac
DNA Meth. m5C:H4K20 me2
H3K4 me1:H4K20 me2
H2A.Z:H4K20 me2
H4K20 me2:H3−5ac
H4K20 me2:H3K4 me3
H3K9 me2
H4K20 me3
H3−5ac
H3K9 me3
H3K4 me3
H3K4 me1
H2A.Z
DNA Meth. m5C
H4−4ac
H3K27 me3
H4K20 me2
0.00 0.25 0.50 0.75 1.00
$\hat{\pi}$
ZBTB17 – Reverse
H3K4 me3:H3−5ac
H2A.Z:H3K4 me3
H3K4 me1:H2A.Z
H4−4ac:H3−5ac
H3K4 me1:H4−4ac
H3K27 me3:H3K9 me3
H3K9 me2:DNA Meth. m5C
H3K9 me3:H4K20 me3
H3K27 me3:H4K20 me3
H4K20 me2:H4−4ac
DNA Meth. m5C:H4K20 me2
H3K4 me1:H4K20 me2
H2A.Z:H4K20 me2
H4K20 me2:H3−5ac
H4K20 me2:H3K4 me3
H3K9 me2
H4K20 me3
H3−5ac
H3K9 me3
H3K4 me3
H3K4 me1
H2A.Z
DNA Meth. m5C
H4−4ac
H3K27 me3
H4K20 me2
0.00 0.25 0.50 0.75 1.00
$\hat{\pi}$
TFAM – Forward
H3K9 me3:H4K20 me3
DNA Meth. m5C:H4K20 me3
H3K27 me3:H4K20 me3
H3K27 me3:H3K9 me3
DNA Meth. m5C:H3K9 me3
H3K27 me3:DNA Meth. m5C
H4K20 me2:H3−5ac
H3K27 me2:DNA Meth. m5C
H3K9 me2:DNA Meth. m5C
H2A.Z:H4K20 me2
DNA Meth. m5C:H4K20 me2
H4−4ac:H3−5ac
H4K20 me2:H3K4 me3
H3K4 me3:H4−4ac
H4K20 me2:H4−4ac
H3K4 me1
H3K9 me3
H3K9 me2
H2A.Z
H3K27 me2
H4K20 me3
H3K4 me3
H3−5ac
DNA Meth. m5C
H3K27 me3
H4−4ac
H4K20 me2
0.00 0.25 0.50 0.75 1.00
$\hat{\pi}$
TFAM – Reverse
H3K9 me3:H4K20 me3
DNA Meth. m5C:H4K20 me3
H3K27 me3:H4K20 me3
H3K27 me3:H3K9 me3
DNA Meth. m5C:H3K9 me3
H3K27 me3:DNA Meth. m5C
H4K20 me2:H3−5ac
H3K27 me2:DNA Meth. m5C
H3K9 me2:DNA Meth. m5C
H2A.Z:H4K20 me2
DNA Meth. m5C:H4K20 me2
H4−4ac:H3−5ac
H4K20 me2:H3K4 me3
H3K4 me3:H4−4ac
H4K20 me2:H4−4ac
H3K4 me1
H3K9 me3
H3K9 me2
H2A.Z
H3K27 me2
H4K20 me3
H3K4 me3
H3−5ac
DNA Meth. m5C
H3K27 me3
H4−4ac
H4K20 me2
0.00 0.25 0.50 0.75 1.00
$\hat{\pi}$

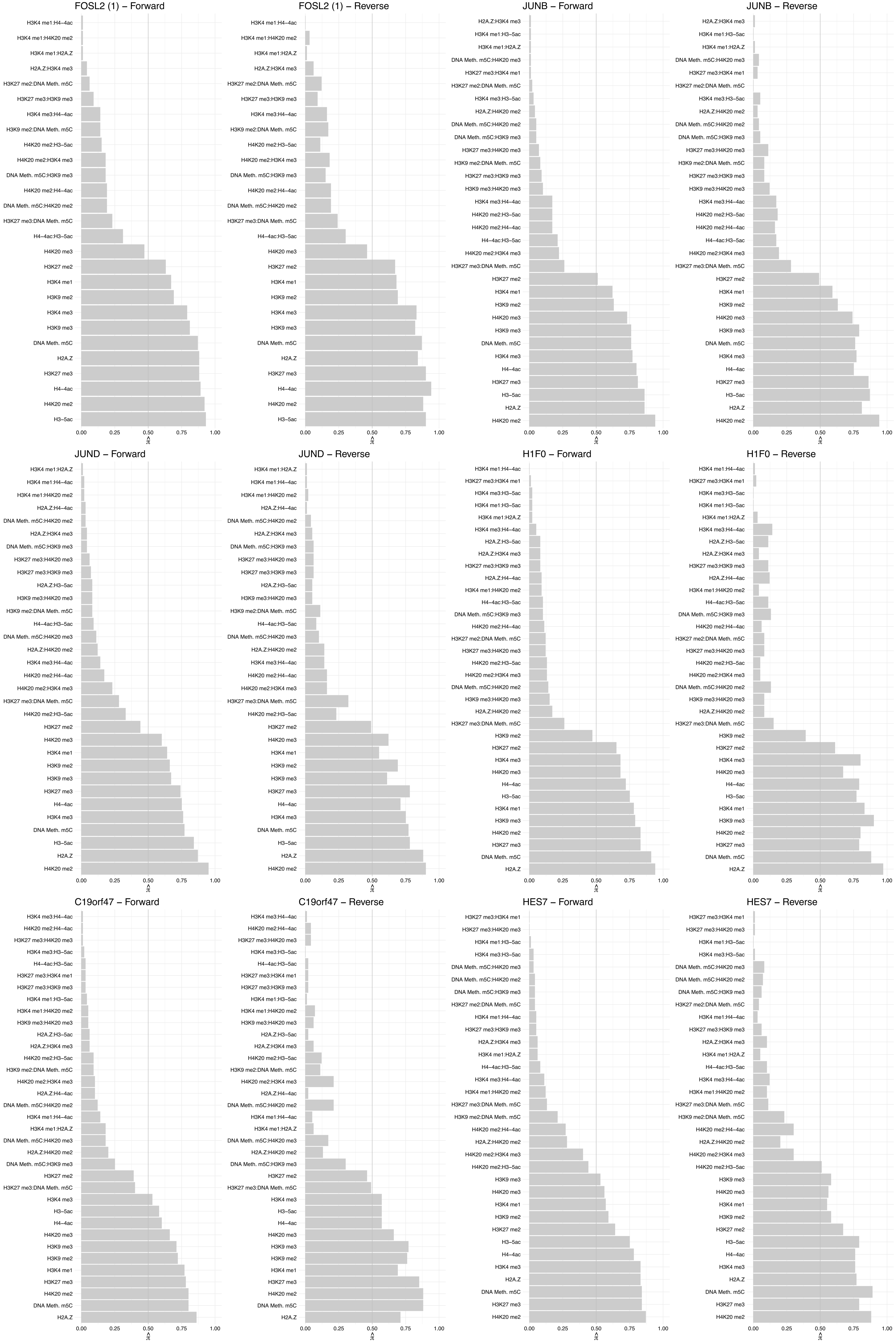

FOSL2 (1) – Forward
H3K4 me1:H4−4ac
H3K4 me1:H4K20 me2
H3K4 me1:H2A.Z
H2A.Z:H3K4 me3
H3K27 me2:DNA Meth. m5C
H3K27 me3:H3K9 me3
H3K4 me3:H4−4ac
H3K9 me2:DNA Meth. m5C
H4K20 me2:H3−5ac
H4K20 me2:H3K4 me3
DNA Meth. m5C:H3K9 me3
H4K20 me2:H4−4ac
DNA Meth. m5C:H4K20 me2
H3K27 me3:DNA Meth. m5C
H4−4ac:H3−5ac
H4K20 me3
H3K27 me2
H3K4 me1
H3K9 me2
H3K4 me3
H3K9 me3
DNA Meth. m5C
H2A.Z
H3K27 me3
H4−4ac
H4K20 me2
H3−5ac
0.00 0.25 0.50 0.75 1.00
π̂
FOSL2 (1) – Reverse
H3K4 me1:H4−4ac
H3K4 me1:H4K20 me2
H3K4 me1:H2A.Z
H2A.Z:H3K4 me3
H3K27 me2:DNA Meth. m5C
H3K27 me3:H3K9 me3
H3K4 me3:H4−4ac
H3K9 me2:DNA Meth. m5C
H4K20 me2:H3−5ac
H4K20 me2:H3K4 me3
DNA Meth. m5C:H3K9 me3
H4K20 me2:H4−4ac
DNA Meth. m5C:H4K20 me2
H3K27 me3:DNA Meth. m5C
H4−4ac:H3−5ac
H4K20 me3
H3K27 me2
H3K4 me1
H3K9 me2
H3K4 me3
H3K9 me3
DNA Meth. m5C
H2A.Z
H3K27 me3
H4−4ac
H4K20 me2
H3−5ac
0.00 0.25 0.50 0.75 1.00
π̂
JUNB – Forward
H2A.Z:H3K4 me3
H3K4 me1:H3−5ac
H3K4 me1:H2A.Z
DNA Meth. m5C:H4K20 me3
H3K27 me3:H3K4 me1
H3K27 me2:DNA Meth. m5C
H3K4 me3:H3−5ac
H2A.Z:H4K20 me2
DNA Meth. m5C:H4K20 me2
DNA Meth. m5C:H3K9 me3
H3K27 me3:H4K20 me3
H3K9 me2:DNA Meth. m5C
H3K27 me3:H3K9 me3
H3K9 me3:H4K20 me3
H3K4 me3:H4−4ac
H4K20 me2:H3−5ac
H4K20 me2:H4−4ac
H4−4ac:H3−5ac
H4K20 me2:H3K4 me3
H3K27 me3:DNA Meth. m5C
H3K27 me2
H3K4 me1
H3K9 me2
H4K20 me3
H3K9 me3
DNA Meth. m5C
H3K4 me3
H4−4ac
H3K27 me3
H3−5ac
H2A.Z
H4K20 me2
0.00 0.25 0.50 0.75 1.00
π̂
JUNB – Reverse
H2A.Z:H3K4 me3
H3K4 me1:H3−5ac
H3K4 me1:H2A.Z
DNA Meth. m5C:H4K20 me3
H3K27 me3:H3K4 me1
H3K27 me2:DNA Meth. m5C
H3K4 me3:H3−5ac
H2A.Z:H4K20 me2
DNA Meth. m5C:H4K20 me2
DNA Meth. m5C:H3K9 me3
H3K27 me3:H4K20 me3
H3K9 me2:DNA Meth. m5C
H3K27 me3:H3K9 me3
H3K9 me3:H4K20 me3
H3K4 me3:H4−4ac
H4K20 me2:H3−5ac
H4K20 me2:H4−4ac
H4−4ac:H3−5ac
H4K20 me2:H3K4 me3
H3K27 me3:DNA Meth. m5C
H3K27 me2
H3K4 me1
H3K9 me2
H4K20 me3
H3K9 me3
DNA Meth. m5C
H3K4 me3
H4−4ac
H3K27 me3
H3−5ac
H2A.Z
H4K20 me2
0.00 0.25 0.50 0.75 1.00
π̂
JUND – Forward
H3K4 me1:H2A.Z
H3K4 me1:H4−4ac
H3K4 me1:H4K20 me2
H2A.Z:H4−4ac
DNA Meth. m5C:H4K20 me2
H2A.Z:H3K4 me3
DNA Meth. m5C:H3K9 me3
H3K27 me3:H4K20 me3
H3K27 me3:H3K9 me3
H2A.Z:H3−5ac
H3K9 me3:H4K20 me3
H3K9 me2:DNA Meth. m5C
H4−4ac:H3−5ac
DNA Meth. m5C:H4K20 me3
H2A.Z:H4K20 me2
H3K4 me3:H4−4ac
H4K20 me2:H4−4ac
H4K20 me2:H3K4 me3
H3K27 me3:DNA Meth. m5C
H4K20 me2:H3−5ac
H3K27 me2
H4K20 me3
H3K4 me1
H3K9 me2
H3K9 me3
H3K27 me3
H4−4ac
H3K4 me3
DNA Meth. m5C
H3−5ac
H2A.Z
H4K20 me2
0.00 0.25 0.50 0.75 1.00
π̂
JUND – Reverse
H3K4 me1:H2A.Z
H3K4 me1:H4−4ac
H3K4 me1:H4K20 me2
H2A.Z:H4−4ac
DNA Meth. m5C:H4K20 me2
H2A.Z:H3K4 me3
DNA Meth. m5C:H3K9 me3
H3K27 me3:H4K20 me3
H3K27 me3:H3K9 me3
H2A.Z:H3−5ac
H3K9 me3:H4K20 me3
H3K9 me2:DNA Meth. m5C
H4−4ac:H3−5ac
DNA Meth. m5C:H4K20 me3
H2A.Z:H4K20 me2
H3K4 me3:H4−4ac
H4K20 me2:H4−4ac
H4K20 me2:H3K4 me3
H3K27 me3:DNA Meth. m5C
H4K20 me2:H3−5ac
H3K27 me2
H4K20 me3
H3K4 me1
H3K9 me2
H3K9 me3
H3K27 me3
H4−4ac
H3K4 me3
DNA Meth. m5C
H3−5ac
H2A.Z
H4K20 me2
0.00 0.25 0.50 0.75 1.00
π̂
H1F0 – Forward
H3K4 me1:H4−4ac
H3K27 me3:H3K4 me1
H3K4 me3:H3−5ac
H3K4 me1:H3−5ac
H3K4 me1:H2A.Z
H3K4 me3:H4−4ac
H2A.Z:H3−5ac
H2A.Z:H3K4 me3
H3K27 me3:H3K9 me3
H2A.Z:H4−4ac
H3K4 me1:H4K20 me2
H4−4ac:H3−5ac
DNA Meth. m5C:H3K9 me3
H4K20 me2:H4−4ac
H3K27 me2:DNA Meth. m5C
H3K27 me3:H4K20 me3
H4K20 me2:H3−5ac
H4K20 me2:H3K4 me3
DNA Meth. m5C:H4K20 me2
H3K9 me3:H4K20 me3
H2A.Z:H4K20 me2
H3K27 me3:DNA Meth. m5C
H3K9 me2
H3K27 me2
H3K4 me3
H4K20 me3
H4−4ac
H3−5ac
H3K4 me1
H3K9 me3
H4K20 me2
H3K27 me3
DNA Meth. m5C
H2A.Z
0.00 0.25 0.50 0.75 1.00
π̂
H1F0 – Reverse
H3K4 me1:H4−4ac
H3K27 me3:H3K4 me1
H3K4 me3:H3−5ac
H3K4 me1:H3−5ac
H3K4 me1:H2A.Z
H3K4 me3:H4−4ac
H2A.Z:H3−5ac
H2A.Z:H3K4 me3
H3K27 me3:H3K9 me3
H2A.Z:H4−4ac
H3K4 me1:H4K20 me2
H4−4ac:H3−5ac
DNA Meth. m5C:H3K9 me3
H4K20 me2:H4−4ac
H3K27 me2:DNA Meth. m5C
H3K27 me3:H4K20 me3
H4K20 me2:H3−5ac
H4K20 me2:H3K4 me3
DNA Meth. m5C:H4K20 me2
H3K9 me3:H4K20 me3
H2A.Z:H4K20 me2
H3K27 me3:DNA Meth. m5C
H3K9 me2
H3K27 me2
H3K4 me3
H4K20 me3
H4−4ac
H3−5ac
H3K4 me1
H3K9 me3
H4K20 me2
H3K27 me3
DNA Meth. m5C
H2A.Z
0.00 0.25 0.50 0.75 1.00
π̂
C19orf47 – Forward
H3K4 me3:H4−4ac
H4K20 me2:H4−4ac
H3K27 me3:H4K20 me3
H3K4 me3:H3−5ac
H4−4ac:H3−5ac
H3K27 me3:H3K4 me1
H3K27 me3:H3K9 me3
H3K4 me1:H3−5ac
H3K4 me1:H4K20 me2
H3K9 me3:H4K20 me3
H2A.Z:H3−5ac
H2A.Z:H3K4 me3
H4K20 me2:H3−5ac
H3K9 me2:DNA Meth. m5C
H4K20 me2:H3K4 me3
H2A.Z:H4−4ac
DNA Meth. m5C:H4K20 me2
H3K4 me1:H4−4ac
H3K4 me1:H2A.Z
DNA Meth. m5C:H4K20 me3
H2A.Z:H4K20 me2
DNA Meth. m5C:H3K9 me3
H3K27 me2
H3K27 me3:DNA Meth. m5C
H3K4 me3
H3−5ac
H4−4ac
H4K20 me3
H3K9 me3
H3K9 me2
H3K4 me1
H3K27 me3
H4K20 me2
DNA Meth. m5C
H2A.Z
0.00 0.25 0.50 0.75 1.00
π̂
C19orf47 – Reverse
H3K4 me3:H4−4ac
H4K20 me2:H4−4ac
H3K27 me3:H4K20 me3
H3K4 me3:H3−5ac
H4−4ac:H3−5ac
H3K27 me3:H3K4 me1
H3K27 me3:H3K9 me3
H3K4 me1:H3−5ac
H3K4 me1:H4K20 me2
H3K9 me3:H4K20 me3
H2A.Z:H3−5ac
H2A.Z:H3K4 me3
H4K20 me2:H3−5ac
H3K9 me2:DNA Meth. m5C
H4K20 me2:H3K4 me3
H2A.Z:H4−4ac
DNA Meth. m5C:H4K20 me2
H3K4 me1:H4−4ac
H3K4 me1:H2A.Z
DNA Meth. m5C:H4K20 me3
H2A.Z:H4K20 me2
DNA Meth. m5C:H3K9 me3
H3K27 me2
H3K27 me3:DNA Meth. m5C
H3K4 me3
H3−5ac
H4−4ac
H4K20 me3
H3K9 me3
H3K9 me2
H3K4 me1
H3K27 me3
H4K20 me2
DNA Meth. m5C
H2A.Z
0.00 0.25 0.50 0.75 1.00
π̂
HES7 – Forward
H3K27 me3:H3K4 me1
H3K27 me3:H4K20 me3
H3K4 me1:H3−5ac
H3K4 me3:H3−5ac
DNA Meth. m5C:H4K20 me3
DNA Meth. m5C:H4K20 me2
DNA Meth. m5C:H3K9 me3
H3K27 me2:DNA Meth. m5C
H3K4 me1:H4−4ac
H3K27 me3:H3K9 me3
H2A.Z:H3K4 me3
H3K4 me1:H2A.Z
H4−4ac:H3−5ac
H3K4 me3:H4−4ac
H3K4 me1:H4K20 me2
H3K27 me3:DNA Meth. m5C
H3K9 me2:DNA Meth. m5C
H4K20 me2:H4−4ac
H2A.Z:H4K20 me2
H4K20 me2:H3K4 me3
H4K20 me2:H3−5ac
H3K9 me3
H4K20 me3
H3K4 me1
H3K9 me2
H3K27 me2
H3−5ac
H4−4ac
H3K4 me3
H2A.Z
DNA Meth. m5C
H3K27 me3
H4K20 me2
0.00 0.25 0.50 0.75 1.00
π̂
HES7 – Reverse
H3K27 me3:H3K4 me1
H3K27 me3:H4K20 me3
H3K4 me1:H3−5ac
H3K4 me3:H3−5ac
DNA Meth. m5C:H4K20 me3
DNA Meth. m5C:H4K20 me2
DNA Meth. m5C:H3K9 me3
H3K27 me2:DNA Meth. m5C
H3K4 me1:H4−4ac
H3K27 me3:H3K9 me3
H2A.Z:H3K4 me3
H3K4 me1:H2A.Z
H4−4ac:H3−5ac
H3K4 me3:H4−4ac
H3K4 me1:H4K20 me2
H3K27 me3:DNA Meth. m5C
H3K9 me2:DNA Meth. m5C
H4K20 me2:H4−4ac
H2A.Z:H4K20 me2
H4K20 me2:H3K4 me3
H4K20 me2:H3−5ac
H3K9 me3
H4K20 me3
H3K4 me1
H3K9 me2
H3K27 me2
H3−5ac
H4−4ac
H3K4 me3
H2A.Z
DNA Meth. m5C
H3K27 me3
H4K20 me2
0.00 0.25 0.50 0.75 1.00
π̂

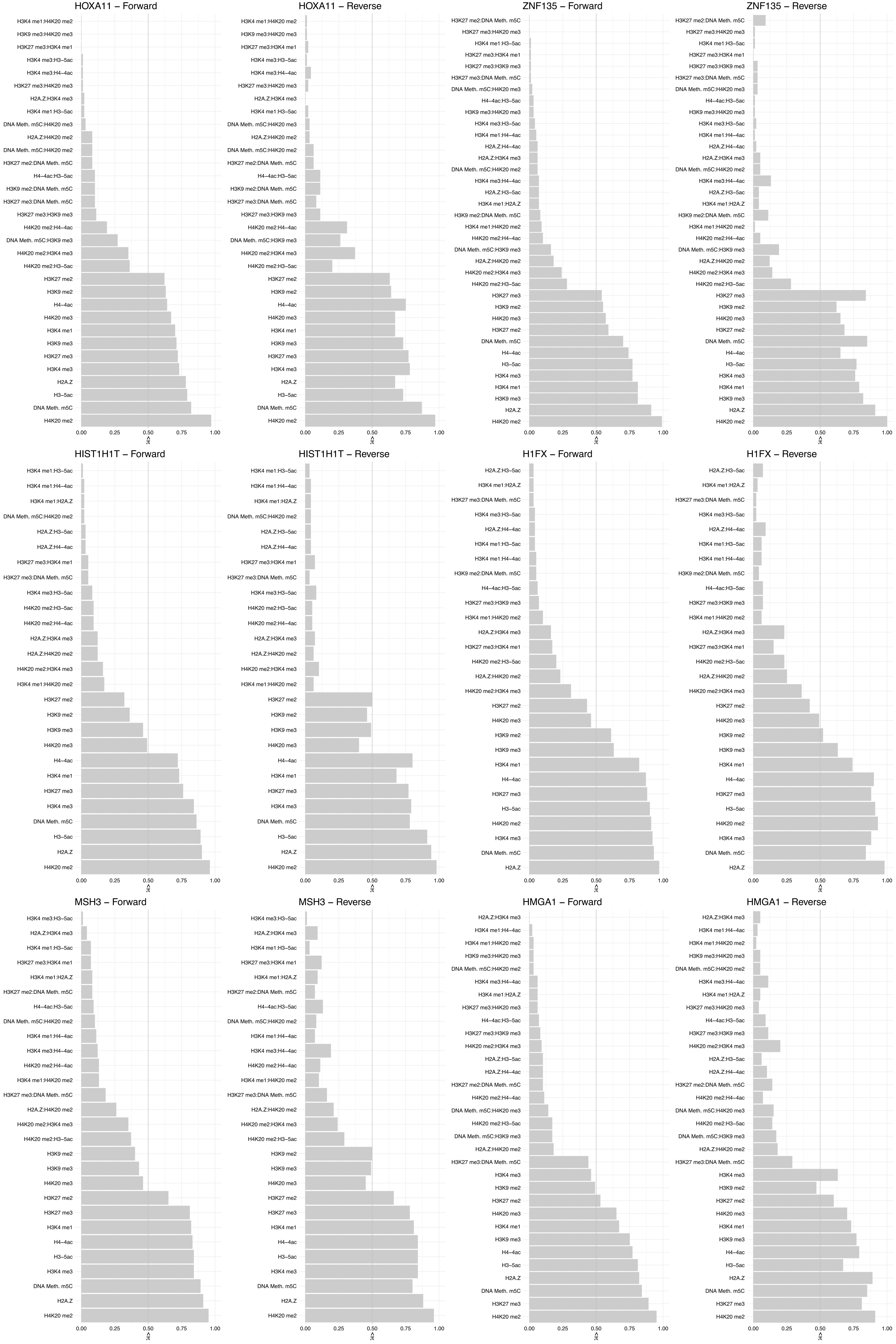
HOXA11 – Forward
H3K4 me1:H4K20 me2
H3K9 me3:H4K20 me3
H3K27 me3:H3K4 me1
H3K4 me3:H3–5ac
H3K4 me3:H4–4ac
H3K27 me3:H4K20 me3
H2A.Z:H3K4 me3
H3K4 me1:H3–5ac
DNA Meth. m5C:H4K20 me3
H2A.Z:H4K20 me2
DNA Meth. m5C:H4K20 me2
H3K27 me2:DNA Meth. m5C
H4–4ac:H3–5ac
H3K9 me2:DNA Meth. m5C
H3K27 me3:DNA Meth. m5C
H3K27 me3:H3K9 me3
H4K20 me2:H4–4ac
DNA Meth. m5C:H3K9 me3
H4K20 me2:H3K4 me3
H4K20 me2:H3–5ac
H3K27 me2
H3K9 me2
H4–4ac
H4K20 me3
H3K4 me1
H3K9 me3
H3K27 me3
H3K4 me3
H2A.Z
H3–5ac
DNA Meth. m5C
H4K20 me2
0.00 0.25 0.50 0.75 1.00
$\hat{\pi}$
HOXA11 – Reverse
H3K4 me1:H4K20 me2
H3K9 me3:H4K20 me3
H3K27 me3:H3K4 me1
H3K4 me3:H3–5ac
H3K4 me3:H4–4ac
H3K27 me3:H4K20 me3
H2A.Z:H3K4 me3
H3K4 me1:H3–5ac
DNA Meth. m5C:H4K20 me3
H2A.Z:H4K20 me2
DNA Meth. m5C:H4K20 me2
H3K27 me2:DNA Meth. m5C
H4–4ac:H3–5ac
H3K9 me2:DNA Meth. m5C
H3K27 me3:DNA Meth. m5C
H3K27 me3:H3K9 me3
H4K20 me2:H4–4ac
DNA Meth. m5C:H3K9 me3
H4K20 me2:H3K4 me3
H4K20 me2:H3–5ac
H3K27 me2
H3K9 me2
H4–4ac
H4K20 me3
H3K4 me1
H3K9 me3
H3K27 me3
H3K4 me3
H2A.Z
H3–5ac
DNA Meth. m5C
H4K20 me2
0.00 0.25 0.50 0.75 1.00
$\hat{\pi}$
ZNF135 – Forward
H3K27 me2:DNA Meth. m5C
H3K27 me3:H4K20 me3
H3K4 me1:H3–5ac
H3K27 me3:H3K4 me1
H3K27 me3:H3K9 me3
H3K27 me3:DNA Meth. m5C
DNA Meth. m5C:H4K20 me3
H4–4ac:H3–5ac
H3K9 me3:H4K20 me3
H3K4 me3:H3–5ac
H3K4 me1:H4–4ac
H2A.Z:H4–4ac
H2A.Z:H3K4 me3
DNA Meth. m5C:H4K20 me2
H3K4 me3:H4–4ac
H2A.Z:H3–5ac
H3K4 me1:H2A.Z
H3K9 me2:DNA Meth. m5C
H3K4 me1:H4K20 me2
H4K20 me2:H4–4ac
DNA Meth. m5C:H3K9 me3
H2A.Z:H4K20 me2
H4K20 me2:H3K4 me3
H4K20 me2:H3–5ac
H3K27 me3
H3K9 me2
H4K20 me3
H3K27 me2
DNA Meth. m5C
H4–4ac
H3–5ac
H3K4 me3
H3K4 me1
H3K9 me3
H2A.Z
H4K20 me2
0.00 0.25 0.50 0.75 1.00
$\hat{\pi}$
ZNF135 – Reverse
H3K27 me2:DNA Meth. m5C
H3K27 me3:H4K20 me3
H3K4 me1:H3–5ac
H3K27 me3:H3K4 me1
H3K27 me3:H3K9 me3
H3K27 me3:DNA Meth. m5C
DNA Meth. m5C:H4K20 me3
H4–4ac:H3–5ac
H3K9 me3:H4K20 me3
H3K4 me3:H3–5ac
H3K4 me1:H4–4ac
H2A.Z:H4–4ac
H2A.Z:H3K4 me3
DNA Meth. m5C:H4K20 me2
H3K4 me3:H4–4ac
H2A.Z:H3–5ac
H3K4 me1:H2A.Z
H3K9 me2:DNA Meth. m5C
H3K4 me1:H4K20 me2
H4K20 me2:H4–4ac
DNA Meth. m5C:H3K9 me3
H2A.Z:H4K20 me2
H4K20 me2:H3K4 me3
H4K20 me2:H3–5ac
H3K27 me3
H3K9 me2
H4K20 me3
H3K27 me2
DNA Meth. m5C
H4–4ac
H3–5ac
H3K4 me3
H3K4 me1
H3K9 me3
H2A.Z
H4K20 me2
0.00 0.25 0.50 0.75 1.00
$\hat{\pi}$
HIST1H1T – Forward
H3K4 me1:H3–5ac
H3K4 me1:H4–4ac
H3K4 me1:H2A.Z
DNA Meth. m5C:H4K20 me2
H2A.Z:H3–5ac
H2A.Z:H4–4ac
H3K27 me3:H3K4 me1
H3K27 me3:DNA Meth. m5C
H3K4 me3:H3–5ac
H4K20 me2:H3–5ac
H4K20 me2:H4–4ac
H2A.Z:H3K4 me3
H2A.Z:H4K20 me2
H4K20 me2:H3K4 me3
H3K4 me1:H4K20 me2
H3K27 me2
H3K9 me2
H3K9 me3
H4K20 me3
H4–4ac
H3K4 me1
H3K27 me3
H3K4 me3
DNA Meth. m5C
H3–5ac
H2A.Z
H4K20 me2
0.00 0.25 0.50 0.75 1.00
$\hat{\pi}$
HIST1H1T – Reverse
H3K4 me1:H3–5ac
H3K4 me1:H4–4ac
H3K4 me1:H2A.Z
DNA Meth. m5C:H4K20 me2
H2A.Z:H3–5ac
H2A.Z:H4–4ac
H3K27 me3:H3K4 me1
H3K27 me3:DNA Meth. m5C
H3K4 me3:H3–5ac
H4K20 me2:H3–5ac
H4K20 me2:H4–4ac
H2A.Z:H3K4 me3
H2A.Z:H4K20 me2
H4K20 me2:H3K4 me3
H3K4 me1:H4K20 me2
H3K27 me2
H3K9 me2
H3K9 me3
H4K20 me3
H4–4ac
H3K4 me1
H3K27 me3
H3K4 me3
DNA Meth. m5C
H3–5ac
H2A.Z
H4K20 me2
0.00 0.25 0.50 0.75 1.00
$\hat{\pi}$
H1FX – Forward
H2A.Z:H3–5ac
H3K4 me1:H2A.Z
H3K27 me3:DNA Meth. m5C
H3K4 me3:H3–5ac
H2A.Z:H4–4ac
H3K4 me1:H3–5ac
H3K4 me1:H4–4ac
H3K9 me2:DNA Meth. m5C
H4–4ac:H3–5ac
H3K27 me3:H3K9 me3
H3K4 me1:H4K20 me2
H2A.Z:H3K4 me3
H3K27 me3:H3K4 me1
H4K20 me2:H3–5ac
H2A.Z:H4K20 me2
H4K20 me2:H3K4 me3
H3K27 me2
H4K20 me3
H3K9 me2
H3K9 me3
H3K4 me1
H4–4ac
H3K27 me3
H3–5ac
H4K20 me2
H3K4 me3
DNA Meth. m5C
H2A.Z
0.00 0.25 0.50 0.75 1.00
$\hat{\pi}$
H1FX – Reverse
H2A.Z:H3–5ac
H3K4 me1:H2A.Z
H3K27 me3:DNA Meth. m5C
H3K4 me3:H3–5ac
H2A.Z:H4–4ac
H3K4 me1:H3–5ac
H3K4 me1:H4–4ac
H3K9 me2:DNA Meth. m5C
H4–4ac:H3–5ac
H3K27 me3:H3K9 me3
H3K4 me1:H4K20 me2
H2A.Z:H3K4 me3
H3K27 me3:H3K4 me1
H4K20 me2:H3–5ac
H2A.Z:H4K20 me2
H4K20 me2:H3K4 me3
H3K27 me2
H4K20 me3
H3K9 me2
H3K9 me3
H3K4 me1
H4–4ac
H3K27 me3
H3–5ac
H4K20 me2
H3K4 me3
DNA Meth. m5C
H2A.Z
0.00 0.25 0.50 0.75 1.00
$\hat{\pi}$
MSH3 – Forward
H3K4 me3:H3–5ac
H2A.Z:H3K4 me3
H3K4 me1:H3–5ac
H3K27 me3:H3K4 me1
H3K4 me1:H2A.Z
H3K27 me2:DNA Meth. m5C
H4–4ac:H3–5ac
DNA Meth. m5C:H4K20 me2
H3K4 me1:H4–4ac
H3K4 me3:H4–4ac
H4K20 me2:H4–4ac
H3K4 me1:H4K20 me2
H3K27 me3:DNA Meth. m5C
H2A.Z:H4K20 me2
H4K20 me2:H3K4 me3
H4K20 me2:H3–5ac
H3K9 me2
H3K9 me3
H4K20 me3
H3K27 me2
H3K27 me3
H3K4 me1
H4–4ac
H3–5ac
H3K4 me3
DNA Meth. m5C
H2A.Z
H4K20 me2
0.00 0.25 0.50 0.75 1.00
$\hat{\pi}$
MSH3 – Reverse
H3K4 me3:H3–5ac
H2A.Z:H3K4 me3
H3K4 me1:H3–5ac
H3K27 me3:H3K4 me1
H3K4 me1:H2A.Z
H3K27 me2:DNA Meth. m5C
H4–4ac:H3–5ac
DNA Meth. m5C:H4K20 me2
H3K4 me1:H4–4ac
H3K4 me3:H4–4ac
H4K20 me2:H4–4ac
H3K4 me1:H4K20 me2
H3K27 me3:DNA Meth. m5C
H2A.Z:H4K20 me2
H4K20 me2:H3K4 me3
H4K20 me2:H3–5ac
H3K9 me2
H3K9 me3
H4K20 me3
H3K27 me2
H3K27 me3
H3K4 me1
H4–4ac
H3–5ac
H3K4 me3
DNA Meth. m5C
H2A.Z
H4K20 me2
0.00 0.25 0.50 0.75 1.00
$\hat{\pi}$
HMGA1 – Forward
H2A.Z:H3K4 me3
H3K4 me1:H4–4ac
H3K4 me1:H4K20 me2
H3K9 me3:H4K20 me3
DNA Meth. m5C:H4K20 me2
H3K4 me3:H4–4ac
H3K4 me1:H2A.Z
H3K27 me3:H4K20 me3
H4–4ac:H3–5ac
H3K27 me3:H3K9 me3
H4K20 me2:H3K4 me3
H2A.Z:H3–5ac
H2A.Z:H4–4ac
H3K27 me2:DNA Meth. m5C
H4K20 me2:H4–4ac
DNA Meth. m5C:H4K20 me3
H4K20 me2:H3–5ac
DNA Meth. m5C:H3K9 me3
H2A.Z:H4K20 me2
H3K27 me3:DNA Meth. m5C
H3K4 me3
H3K9 me2
H3K27 me2
H4K20 me3
H3K4 me1
H3K9 me3
H4–4ac
H3–5ac
H2A.Z
DNA Meth. m5C
H3K27 me3
H4K20 me2
0.00 0.25 0.50 0.75 1.00
$\hat{\pi}$
HMGA1 – Reverse
H2A.Z:H3K4 me3
H3K4 me1:H4–4ac
H3K4 me1:H4K20 me2
H3K9 me3:H4K20 me3
DNA Meth. m5C:H4K20 me2
H3K4 me3:H4–4ac
H3K4 me1:H2A.Z
H3K27 me3:H4K20 me3
H4–4ac:H3–5ac
H3K27 me3:H3K9 me3
H4K20 me2:H3K4 me3
H2A.Z:H3–5ac
H2A.Z:H4–4ac
H3K27 me2:DNA Meth. m5C
H4K20 me2:H4–4ac
DNA Meth. m5C:H4K20 me3
H4K20 me2:H3–5ac
DNA Meth. m5C:H3K9 me3
H2A.Z:H4K20 me2
H3K27 me3:DNA Meth. m5C
H3K4 me3
H3K9 me2
H3K27 me2
H4K20 me3
H3K4 me1
H3K9 me3
H4–4ac
H3–5ac
H2A.Z
DNA Meth. m5C
H3K27 me3
H4K20 me2
0.00 0.25 0.50 0.75 1.00
$\hat{\pi}$

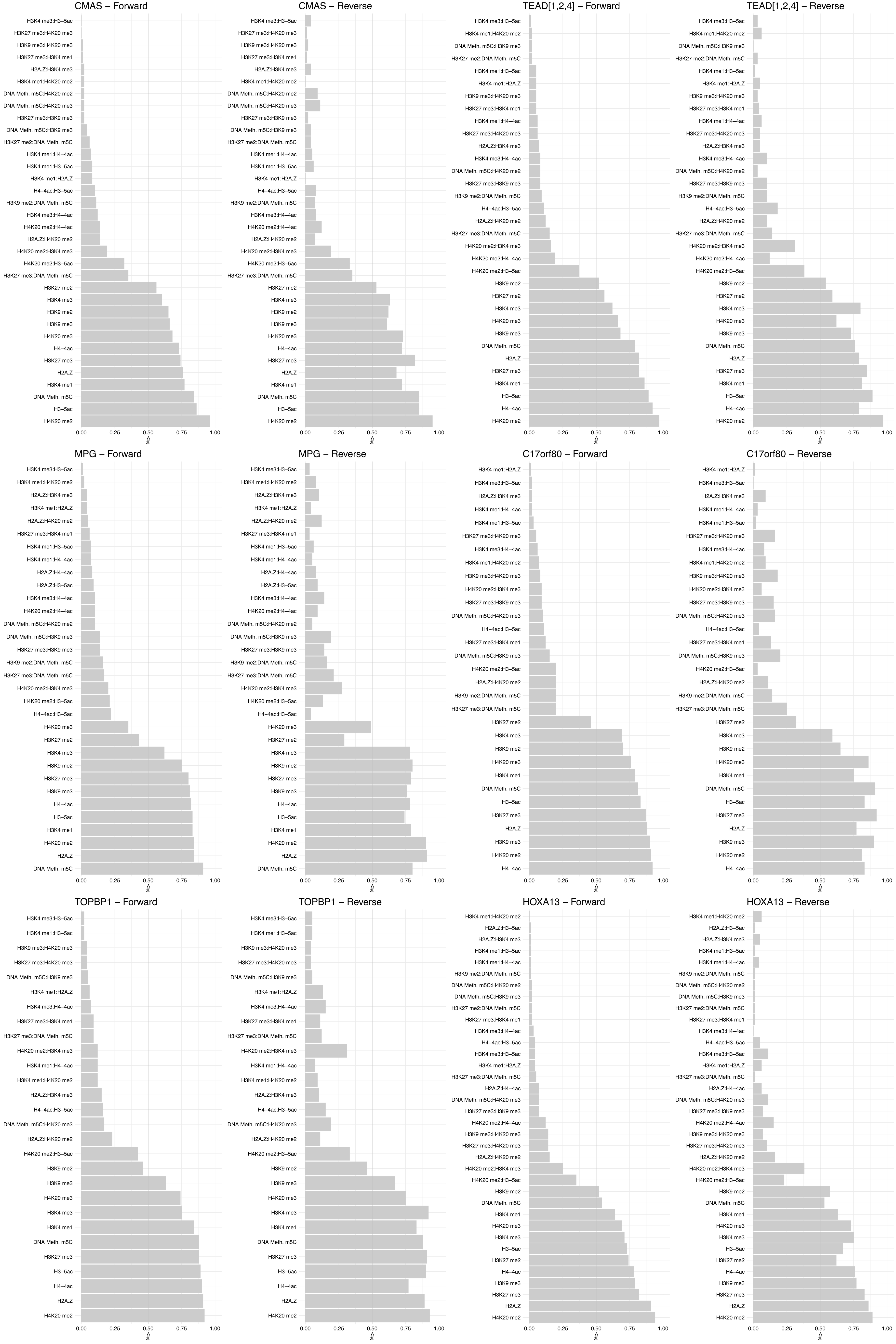
CMAS – Forward
H3K4 me3:H3−5ac
H3K27 me3:H4K20 me3
H3K9 me3:H4K20 me3
H3K27 me3:H3K4 me1
H2A.Z:H3K4 me3
H3K4 me1:H4K20 me2
DNA Meth. m5C:H4K20 me2
DNA Meth. m5C:H4K20 me3
H3K27 me3:H3K9 me3
DNA Meth. m5C:H3K9 me3
H3K27 me2:DNA Meth. m5C
H3K4 me1:H4−4ac
H3K4 me1:H3−5ac
H3K4 me1:H2A.Z
H4−4ac:H3−5ac
H3K9 me2:DNA Meth. m5C
H3K4 me3:H4−4ac
H4K20 me2:H4−4ac
H2A.Z:H4K20 me2
H4K20 me2:H3K4 me3
H4K20 me2:H3−5ac
H3K27 me3:DNA Meth. m5C
H3K27 me2
H3K4 me3
H3K9 me2
H3K9 me3
H4K20 me3
H4−4ac
H3K27 me3
H2A.Z
H3K4 me1
DNA Meth. m5C
H3−5ac
H4K20 me2
0.00
0.25
0.50
0.75
1.00
π̂
CMAS – Reverse
H3K4 me3:H3−5ac
H3K27 me3:H4K20 me3
H3K9 me3:H4K20 me3
H3K27 me3:H3K4 me1
H2A.Z:H3K4 me3
H3K4 me1:H4K20 me2
DNA Meth. m5C:H4K20 me2
DNA Meth. m5C:H4K20 me3
H3K27 me3:H3K9 me3
DNA Meth. m5C:H3K9 me3
H3K27 me2:DNA Meth. m5C
H3K4 me1:H4−4ac
H3K4 me1:H3−5ac
H3K4 me1:H2A.Z
H4−4ac:H3−5ac
H3K9 me2:DNA Meth. m5C
H3K4 me3:H4−4ac
H4K20 me2:H4−4ac
H2A.Z:H4K20 me2
H4K20 me2:H3K4 me3
H4K20 me2:H3−5ac
H3K27 me3:DNA Meth. m5C
H3K27 me2
H3K4 me3
H3K9 me2
H3K9 me3
H4K20 me3
H4−4ac
H3K27 me3
H2A.Z
H3K4 me1
DNA Meth. m5C
H3−5ac
H4K20 me2
0.00
0.25
0.50
0.75
1.00
π̂
TEAD[1,2,4] – Forward
H3K4 me3:H3−5ac
H3K4 me1:H4K20 me2
DNA Meth. m5C:H3K9 me3
H3K27 me2:DNA Meth. m5C
H3K4 me1:H3−5ac
H3K4 me1:H2A.Z
H3K9 me3:H4K20 me3
H3K27 me3:H3K4 me1
H3K4 me1:H4−4ac
H3K27 me3:H4K20 me3
H2A.Z:H3K4 me3
H3K4 me3:H4−4ac
DNA Meth. m5C:H4K20 me2
H3K27 me3:H3K9 me3
H3K9 me2:DNA Meth. m5C
H4−4ac:H3−5ac
H2A.Z:H4K20 me2
H3K27 me3:DNA Meth. m5C
H4K20 me2:H3K4 me3
H4K20 me2:H4−4ac
H4K20 me2:H3−5ac
H3K9 me2
H3K27 me2
H3K4 me3
H4K20 me3
H3K9 me3
DNA Meth. m5C
H2A.Z
H3K27 me3
H3K4 me1
H3−5ac
H4−4ac
H4K20 me2
0.00
0.25
0.50
0.75
1.00
π̂
TEAD[1,2,4] – Reverse
H3K4 me3:H3−5ac
H3K4 me1:H4K20 me2
DNA Meth. m5C:H3K9 me3
H3K27 me2:DNA Meth. m5C
H3K4 me1:H3−5ac
H3K4 me1:H2A.Z
H3K9 me3:H4K20 me3
H3K27 me3:H3K4 me1
H3K4 me1:H4−4ac
H3K27 me3:H4K20 me3
H2A.Z:H3K4 me3
H3K4 me3:H4−4ac
DNA Meth. m5C:H4K20 me2
H3K27 me3:H3K9 me3
H3K9 me2:DNA Meth. m5C
H4−4ac:H3−5ac
H2A.Z:H4K20 me2
H3K27 me3:DNA Meth. m5C
H4K20 me2:H3K4 me3
H4K20 me2:H4−4ac
H4K20 me2:H3−5ac
H3K9 me2
H3K27 me2
H3K4 me3
H4K20 me3
H3K9 me3
DNA Meth. m5C
H2A.Z
H3K27 me3
H3K4 me1
H3−5ac
H4−4ac
H4K20 me2
0.00
0.25
0.50
0.75
1.00
π̂
MPG – Forward
H3K4 me3:H3−5ac
H3K4 me1:H4K20 me2
H2A.Z:H3K4 me3
H3K4 me1:H2A.Z
H2A.Z:H4K20 me2
H3K27 me3:H3K4 me1
H3K4 me1:H3−5ac
H3K4 me1:H4−4ac
H2A.Z:H4−4ac
H2A.Z:H3−5ac
H3K4 me3:H4−4ac
H4K20 me2:H4−4ac
DNA Meth. m5C:H4K20 me2
DNA Meth. m5C:H3K9 me3
H3K27 me3:H3K9 me3
H3K9 me2:DNA Meth. m5C
H3K27 me3:DNA Meth. m5C
H4K20 me2:H3K4 me3
H4K20 me2:H3−5ac
H4−4ac:H3−5ac
H4K20 me3
H3K27 me2
H3K4 me3
H3K9 me2
H3K27 me3
H3K9 me3
H4−4ac
H3−5ac
H3K4 me1
H4K20 me2
H2A.Z
DNA Meth. m5C
0.00
0.25
0.50
0.75
1.00
π̂
MPG – Reverse
H3K4 me3:H3−5ac
H3K4 me1:H4K20 me2
H2A.Z:H3K4 me3
H3K4 me1:H2A.Z
H2A.Z:H4K20 me2
H3K27 me3:H3K4 me1
H3K4 me1:H3−5ac
H3K4 me1:H4−4ac
H2A.Z:H4−4ac
H2A.Z:H3−5ac
H3K4 me3:H4−4ac
H4K20 me2:H4−4ac
DNA Meth. m5C:H4K20 me2
DNA Meth. m5C:H3K9 me3
H3K27 me3:H3K9 me3
H3K9 me2:DNA Meth. m5C
H3K27 me3:DNA Meth. m5C
H4K20 me2:H3K4 me3
H4K20 me2:H3−5ac
H4−4ac:H3−5ac
H4K20 me3
H3K27 me2
H3K4 me3
H3K9 me2
H3K27 me3
H3K9 me3
H4−4ac
H3−5ac
H3K4 me1
H4K20 me2
H2A.Z
DNA Meth. m5C
0.00
0.25
0.50
0.75
1.00
π̂
C17orf80 – Forward
H3K4 me1:H2A.Z
H3K4 me3:H3−5ac
H2A.Z:H3K4 me3
H3K4 me1:H4−4ac
H3K4 me1:H3−5ac
H3K27 me3:H4K20 me3
H3K4 me3:H4−4ac
H3K4 me1:H4K20 me2
H3K9 me3:H4K20 me3
H4K20 me2:H3K4 me3
H3K27 me3:H3K9 me3
DNA Meth. m5C:H4K20 me3
H4−4ac:H3−5ac
H3K27 me3:H3K4 me1
DNA Meth. m5C:H3K9 me3
H4K20 me2:H3−5ac
H2A.Z:H4K20 me2
H3K9 me2:DNA Meth. m5C
H3K27 me3:DNA Meth. m5C
H3K27 me2
H3K4 me3
H3K9 me2
H4K20 me3
H3K4 me1
DNA Meth. m5C
H3−5ac
H3K27 me3
H2A.Z
H3K9 me3
H4K20 me2
H4−4ac
0.00
0.25
0.50
0.75
1.00
π̂
C17orf80 – Reverse
H3K4 me1:H2A.Z
H3K4 me3:H3−5ac
H2A.Z:H3K4 me3
H3K4 me1:H4−4ac
H3K4 me1:H3−5ac
H3K27 me3:H4K20 me3
H3K4 me3:H4−4ac
H3K4 me1:H4K20 me2
H3K9 me3:H4K20 me3
H4K20 me2:H3K4 me3
H3K27 me3:H3K9 me3
DNA Meth. m5C:H4K20 me3
H4−4ac:H3−5ac
H3K27 me3:H3K4 me1
DNA Meth. m5C:H3K9 me3
H4K20 me2:H3−5ac
H2A.Z:H4K20 me2
H3K9 me2:DNA Meth. m5C
H3K27 me3:DNA Meth. m5C
H3K27 me2
H3K4 me3
H3K9 me2
H4K20 me3
H3K4 me1
DNA Meth. m5C
H3−5ac
H3K27 me3
H2A.Z
H3K9 me3
H4K20 me2
H4−4ac
0.00
0.25
0.50
0.75
1.00
π̂
TOPBP1 – Forward
H3K4 me3:H3−5ac
H3K4 me1:H3−5ac
H3K9 me3:H4K20 me3
H3K27 me3:H4K20 me3
DNA Meth. m5C:H3K9 me3
H3K4 me1:H2A.Z
H3K4 me3:H4−4ac
H3K27 me3:H3K4 me1
H3K27 me3:DNA Meth. m5C
H4K20 me2:H3K4 me3
H3K4 me1:H4−4ac
H3K4 me1:H4K20 me2
H2A.Z:H3K4 me3
H4−4ac:H3−5ac
DNA Meth. m5C:H4K20 me3
H2A.Z:H4K20 me2
H4K20 me2:H3−5ac
H3K9 me2
H3K9 me3
H4K20 me3
H3K4 me3
H3K4 me1
DNA Meth. m5C
H3K27 me3
H3−5ac
H4−4ac
H2A.Z
H4K20 me2
0.00
0.25
0.50
0.75
1.00
π̂
TOPBP1 – Reverse
H3K4 me3:H3−5ac
H3K4 me1:H3−5ac
H3K9 me3:H4K20 me3
H3K27 me3:H4K20 me3
DNA Meth. m5C:H3K9 me3
H3K4 me1:H2A.Z
H3K4 me3:H4−4ac
H3K27 me3:H3K4 me1
H3K27 me3:DNA Meth. m5C
H4K20 me2:H3K4 me3
H3K4 me1:H4−4ac
H3K4 me1:H4K20 me2
H2A.Z:H3K4 me3
H4−4ac:H3−5ac
DNA Meth. m5C:H4K20 me3
H2A.Z:H4K20 me2
H4K20 me2:H3−5ac
H3K9 me2
H3K9 me3
H4K20 me3
H3K4 me3
H3K4 me1
DNA Meth. m5C
H3K27 me3
H3−5ac
H4−4ac
H2A.Z
H4K20 me2
0.00
0.25
0.50
0.75
1.00
π̂
HOXA13 – Forward
H3K4 me1:H4K20 me2
H2A.Z:H3−5ac
H2A.Z:H3K4 me3
H3K4 me1:H3−5ac
H3K4 me1:H4−4ac
H3K9 me2:DNA Meth. m5C
DNA Meth. m5C:H4K20 me2
DNA Meth. m5C:H3K9 me3
H3K27 me2:DNA Meth. m5C
H3K27 me3:H3K4 me1
H3K4 me3:H4−4ac
H4−4ac:H3−5ac
H3K4 me3:H3−5ac
H3K4 me1:H2A.Z
H3K27 me3:DNA Meth. m5C
H2A.Z:H4−4ac
DNA Meth. m5C:H4K20 me3
H3K27 me3:H3K9 me3
H4K20 me2:H4−4ac
H3K9 me3:H4K20 me3
H3K27 me3:H4K20 me3
H2A.Z:H4K20 me2
H4K20 me2:H3K4 me3
H4K20 me2:H3−5ac
H3K9 me2
DNA Meth. m5C
H3K4 me1
H4K20 me3
H3K4 me3
H3−5ac
H3K27 me2
H4−4ac
H3K9 me3
H3K27 me3
H2A.Z
H4K20 me2
0.00
0.25
0.50
0.75
1.00
π̂
HOXA13 – Reverse
H3K4 me1:H4K20 me2
H2A.Z:H3−5ac
H2A.Z:H3K4 me3
H3K4 me1:H3−5ac
H3K4 me1:H4−4ac
H3K9 me2:DNA Meth. m5C
DNA Meth. m5C:H4K20 me2
DNA Meth. m5C:H3K9 me3
H3K27 me2:DNA Meth. m5C
H3K27 me3:H3K4 me1
H3K4 me3:H4−4ac
H4−4ac:H3−5ac
H3K4 me3:H3−5ac
H3K4 me1:H2A.Z
H3K27 me3:DNA Meth. m5C
H2A.Z:H4−4ac
DNA Meth. m5C:H4K20 me3
H3K27 me3:H3K9 me3
H4K20 me2:H4−4ac
H3K9 me3:H4K20 me3
H3K27 me3:H4K20 me3
H2A.Z:H4K20 me2
H4K20 me2:H3K4 me3
H4K20 me2:H3−5ac
H3K9 me2
DNA Meth. m5C
H3K4 me1
H4K20 me3
H3K4 me3
H3−5ac
H3K27 me2
H4−4ac
H3K9 me3
H3K27 me3
H2A.Z
H4K20 me2
0.00
0.25
0.50
0.75
1.00
π̂

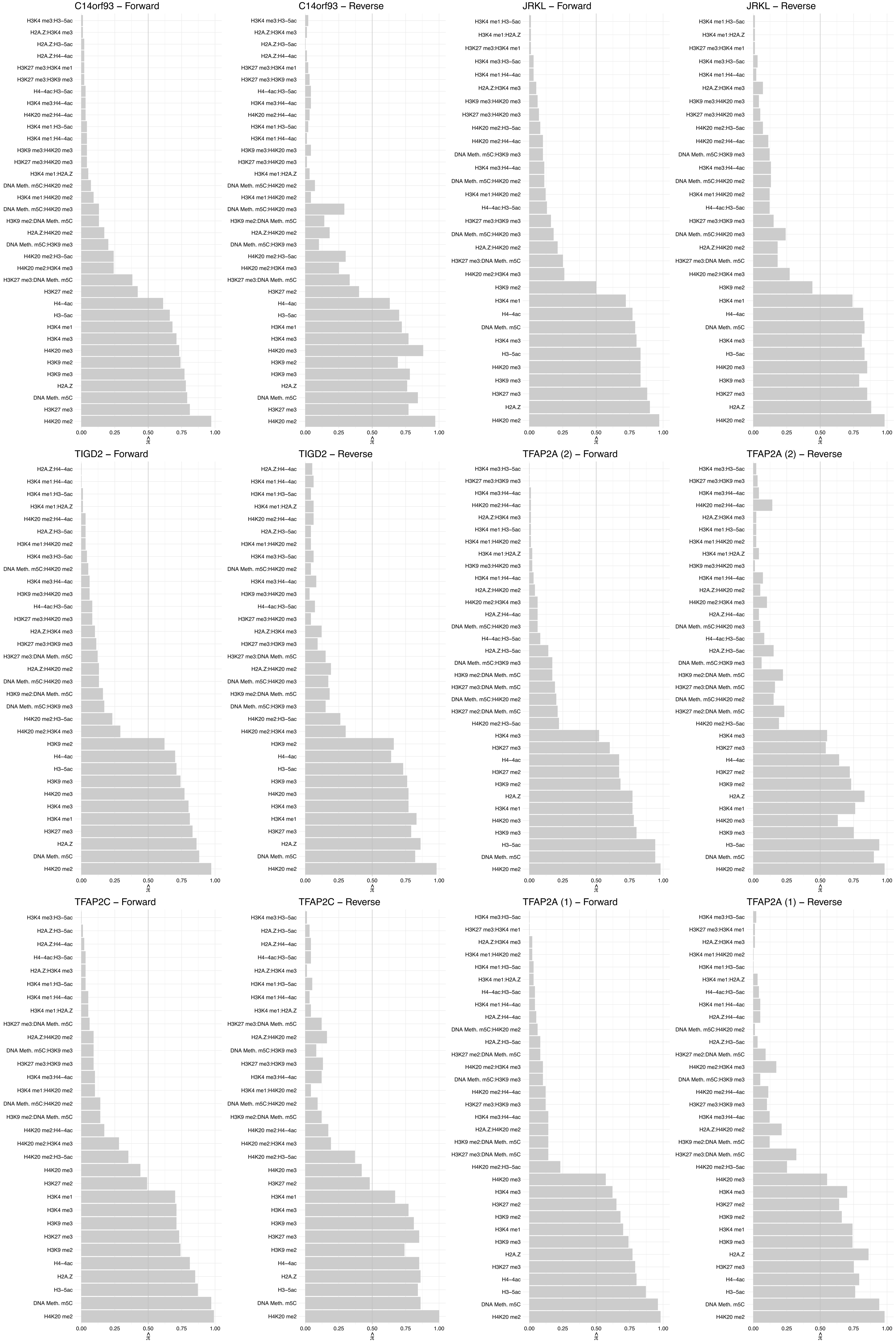

C14orf93 – Forward
H3K4 me3:H3–5ac
H2A.Z:H3K4 me3
H2A.Z:H3–5ac
H2A.Z:H4–4ac
H3K27 me3:H3K4 me1
H3K27 me3:H3K9 me3
H4–4ac:H3–5ac
H3K4 me3:H4–4ac
H4K20 me2:H4–4ac
H3K4 me1:H3–5ac
H3K4 me1:H4–4ac
H3K9 me3:H4K20 me3
H3K27 me3:H4K20 me3
H3K4 me1:H2A.Z
DNA Meth. m5C:H4K20 me2
H3K4 me1:H4K20 me2
DNA Meth. m5C:H4K20 me3
H3K9 me2:DNA Meth. m5C
H2A.Z:H4K20 me2
DNA Meth. m5C:H3K9 me3
H4K20 me2:H3–5ac
H4K20 me2:H3K4 me3
H3K27 me3:DNA Meth. m5C
H3K27 me2
H4–4ac
H3–5ac
H3K4 me1
H3K4 me3
H4K20 me3
H3K9 me2
H3K9 me3
H2A.Z
DNA Meth. m5C
H3K27 me3
H4K20 me2
0.00 0.25 0.50 0.75 1.00
π̂
C14orf93 – Reverse
H3K4 me3:H3–5ac
H2A.Z:H3K4 me3
H2A.Z:H3–5ac
H2A.Z:H4–4ac
H3K27 me3:H3K4 me1
H3K27 me3:H3K9 me3
H4–4ac:H3–5ac
H3K4 me3:H4–4ac
H4K20 me2:H4–4ac
H3K4 me1:H3–5ac
H3K4 me1:H4–4ac
H3K9 me3:H4K20 me3
H3K27 me3:H4K20 me3
H3K4 me1:H2A.Z
DNA Meth. m5C:H4K20 me2
H3K4 me1:H4K20 me2
DNA Meth. m5C:H4K20 me3
H3K9 me2:DNA Meth. m5C
H2A.Z:H4K20 me2
DNA Meth. m5C:H3K9 me3
H4K20 me2:H3–5ac
H4K20 me2:H3K4 me3
H3K27 me3:DNA Meth. m5C
H3K27 me2
H4–4ac
H3–5ac
H3K4 me1
H3K4 me3
H4K20 me3
H3K9 me2
H3K9 me3
H2A.Z
DNA Meth. m5C
H3K27 me3
H4K20 me2
0.00 0.25 0.50 0.75 1.00
π̂
JRKL – Forward
H3K4 me1:H3–5ac
H3K4 me1:H2A.Z
H3K27 me3:H3K4 me1
H3K4 me3:H3–5ac
H3K4 me1:H4–4ac
H2A.Z:H3K4 me3
H3K9 me3:H4K20 me3
H3K27 me3:H4K20 me3
H4K20 me2:H3–5ac
H4K20 me2:H4–4ac
DNA Meth. m5C:H3K9 me3
H3K4 me3:H4–4ac
DNA Meth. m5C:H4K20 me2
H3K4 me1:H4K20 me2
H4–4ac:H3–5ac
H3K27 me3:H3K9 me3
DNA Meth. m5C:H4K20 me3
H2A.Z:H4K20 me2
H3K27 me3:DNA Meth. m5C
H4K20 me2:H3K4 me3
H3K9 me2
H3K4 me1
H4–4ac
DNA Meth. m5C
H3K4 me3
H3–5ac
H4K20 me3
H3K9 me3
H3K27 me3
H2A.Z
H4K20 me2
0.00 0.25 0.50 0.75 1.00
π̂
JRKL – Reverse
H3K4 me1:H3–5ac
H3K4 me1:H2A.Z
H3K27 me3:H3K4 me1
H3K4 me3:H3–5ac
H3K4 me1:H4–4ac
H2A.Z:H3K4 me3
H3K9 me3:H4K20 me3
H3K27 me3:H4K20 me3
H4K20 me2:H3–5ac
H4K20 me2:H4–4ac
DNA Meth. m5C:H3K9 me3
H3K4 me3:H4–4ac
DNA Meth. m5C:H4K20 me2
H3K4 me1:H4K20 me2
H4–4ac:H3–5ac
H3K27 me3:H3K9 me3
DNA Meth. m5C:H4K20 me3
H2A.Z:H4K20 me2
H3K27 me3:DNA Meth. m5C
H4K20 me2:H3K4 me3
H3K9 me2
H3K4 me1
H4–4ac
DNA Meth. m5C
H3K4 me3
H3–5ac
H4K20 me3
H3K9 me3
H3K27 me3
H2A.Z
H4K20 me2
0.00 0.25 0.50 0.75 1.00
π̂
TIGD2 – Forward
H2A.Z:H4–4ac
H3K4 me1:H4–4ac
H3K4 me1:H3–5ac
H3K4 me1:H2A.Z
H4K20 me2:H4–4ac
H2A.Z:H3–5ac
H3K4 me1:H4K20 me2
H3K4 me3:H3–5ac
DNA Meth. m5C:H4K20 me2
H3K4 me3:H4–4ac
H3K9 me3:H4K20 me3
H4–4ac:H3–5ac
H3K27 me3:H4K20 me3
H2A.Z:H3K4 me3
H3K27 me3:H3K9 me3
H3K27 me3:DNA Meth. m5C
H2A.Z:H4K20 me2
DNA Meth. m5C:H4K20 me3
H3K9 me2:DNA Meth. m5C
DNA Meth. m5C:H3K9 me3
H4K20 me2:H3–5ac
H4K20 me2:H3K4 me3
H3K9 me2
H4–4ac
H3–5ac
H3K9 me3
H4K20 me3
H3K4 me3
H3K4 me1
H3K27 me3
H2A.Z
DNA Meth. m5C
H4K20 me2
0.00 0.25 0.50 0.75 1.00
π̂
TIGD2 – Reverse
H2A.Z:H4–4ac
H3K4 me1:H4–4ac
H3K4 me1:H3–5ac
H3K4 me1:H2A.Z
H4K20 me2:H4–4ac
H2A.Z:H3–5ac
H3K4 me1:H4K20 me2
H3K4 me3:H3–5ac
DNA Meth. m5C:H4K20 me2
H3K4 me3:H4–4ac
H3K9 me3:H4K20 me3
H4–4ac:H3–5ac
H3K27 me3:H4K20 me3
H2A.Z:H3K4 me3
H3K27 me3:H3K9 me3
H3K27 me3:DNA Meth. m5C
H2A.Z:H4K20 me2
DNA Meth. m5C:H4K20 me3
H3K9 me2:DNA Meth. m5C
DNA Meth. m5C:H3K9 me3
H4K20 me2:H3–5ac
H4K20 me2:H3K4 me3
H3K9 me2
H4–4ac
H3–5ac
H3K9 me3
H4K20 me3
H3K4 me3
H3K4 me1
H3K27 me3
H2A.Z
DNA Meth. m5C
H4K20 me2
0.00 0.25 0.50 0.75 1.00
π̂
TFAP2A (2) – Forward
H3K4 me3:H3–5ac
H3K27 me3:H3K9 me3
H3K4 me3:H4–4ac
H4K20 me2:H4–4ac
H2A.Z:H3K4 me3
H3K4 me1:H3–5ac
H3K4 me1:H4K20 me2
H3K4 me1:H2A.Z
H3K9 me3:H4K20 me3
H3K4 me1:H4–4ac
H2A.Z:H4K20 me2
H4K20 me2:H3K4 me3
H2A.Z:H4–4ac
DNA Meth. m5C:H4K20 me3
H4–4ac:H3–5ac
H2A.Z:H3–5ac
DNA Meth. m5C:H3K9 me3
H3K9 me2:DNA Meth. m5C
H3K27 me3:DNA Meth. m5C
DNA Meth. m5C:H4K20 me2
H3K27 me2:DNA Meth. m5C
H4K20 me2:H3–5ac
H3K4 me3
H3K27 me3
H4–4ac
H3K27 me2
H3K9 me2
H2A.Z
H3K4 me1
H4K20 me3
H3K9 me3
H3–5ac
DNA Meth. m5C
H4K20 me2
0.00 0.25 0.50 0.75 1.00
π̂
TFAP2A (2) – Reverse
H3K4 me3:H3–5ac
H3K27 me3:H3K9 me3
H3K4 me3:H4–4ac
H4K20 me2:H4–4ac
H2A.Z:H3K4 me3
H3K4 me1:H3–5ac
H3K4 me1:H4K20 me2
H3K4 me1:H2A.Z
H3K9 me3:H4K20 me3
H3K4 me1:H4–4ac
H2A.Z:H4K20 me2
H4K20 me2:H3K4 me3
H2A.Z:H4–4ac
DNA Meth. m5C:H4K20 me3
H4–4ac:H3–5ac
H2A.Z:H3–5ac
DNA Meth. m5C:H3K9 me3
H3K9 me2:DNA Meth. m5C
H3K27 me3:DNA Meth. m5C
DNA Meth. m5C:H4K20 me2
H3K27 me2:DNA Meth. m5C
H4K20 me2:H3–5ac
H3K4 me3
H3K27 me3
H4–4ac
H3K27 me2
H3K9 me2
H2A.Z
H3K4 me1
H4K20 me3
H3K9 me3
H3–5ac
DNA Meth. m5C
H4K20 me2
0.00 0.25 0.50 0.75 1.00
π̂
TFAP2C – Forward
H3K4 me3:H3–5ac
H2A.Z:H3–5ac
H2A.Z:H4–4ac
H4–4ac:H3–5ac
H2A.Z:H3K4 me3
H3K4 me1:H3–5ac
H3K4 me1:H4–4ac
H3K4 me1:H2A.Z
H3K27 me3:DNA Meth. m5C
H2A.Z:H4K20 me2
DNA Meth. m5C:H3K9 me3
H3K27 me3:H3K9 me3
H3K4 me3:H4–4ac
H3K4 me1:H4K20 me2
DNA Meth. m5C:H4K20 me2
H3K9 me2:DNA Meth. m5C
H4K20 me2:H4–4ac
H4K20 me2:H3K4 me3
H4K20 me2:H3–5ac
H4K20 me3
H3K27 me2
H3K4 me1
H3K4 me3
H3K9 me3
H3K27 me3
H3K9 me2
H4–4ac
H2A.Z
H3–5ac
DNA Meth. m5C
H4K20 me2
0.00 0.25 0.50 0.75 1.00
π̂
TFAP2C – Reverse
H3K4 me3:H3–5ac
H2A.Z:H3–5ac
H2A.Z:H4–4ac
H4–4ac:H3–5ac
H2A.Z:H3K4 me3
H3K4 me1:H3–5ac
H3K4 me1:H4–4ac
H3K4 me1:H2A.Z
H3K27 me3:DNA Meth. m5C
H2A.Z:H4K20 me2
DNA Meth. m5C:H3K9 me3
H3K27 me3:H3K9 me3
H3K4 me3:H4–4ac
H3K4 me1:H4K20 me2
DNA Meth. m5C:H4K20 me2
H3K9 me2:DNA Meth. m5C
H4K20 me2:H4–4ac
H4K20 me2:H3K4 me3
H4K20 me2:H3–5ac
H4K20 me3
H3K27 me2
H3K4 me1
H3K4 me3
H3K9 me3
H3K27 me3
H3K9 me2
H4–4ac
H2A.Z
H3–5ac
DNA Meth. m5C
H4K20 me2
0.00 0.25 0.50 0.75 1.00
π̂
TFAP2A (1) – Forward
H3K4 me3:H3–5ac
H3K27 me3:H3K4 me1
H2A.Z:H3K4 me3
H3K4 me1:H4K20 me2
H3K4 me1:H3–5ac
H3K4 me1:H2A.Z
H4–4ac:H3–5ac
H3K4 me1:H4–4ac
H2A.Z:H4–4ac
DNA Meth. m5C:H4K20 me2
H2A.Z:H3–5ac
H3K27 me2:DNA Meth. m5C
H4K20 me2:H3K4 me3
DNA Meth. m5C:H3K9 me3
H4K20 me2:H4–4ac
H3K27 me3:H3K9 me3
H3K4 me3:H4–4ac
H2A.Z:H4K20 me2
H3K9 me2:DNA Meth. m5C
H3K27 me3:DNA Meth. m5C
H4K20 me2:H3–5ac
H4K20 me3
H3K4 me3
H3K27 me2
H3K9 me2
H3K4 me1
H3K9 me3
H2A.Z
H3K27 me3
H4–4ac
H3–5ac
DNA Meth. m5C
H4K20 me2
0.00 0.25 0.50 0.75 1.00
π̂
TFAP2A (1) – Reverse
H3K4 me3:H3–5ac
H3K27 me3:H3K4 me1
H2A.Z:H3K4 me3
H3K4 me1:H4K20 me2
H3K4 me1:H3–5ac
H3K4 me1:H2A.Z
H4–4ac:H3–5ac
H3K4 me1:H4–4ac
H2A.Z:H4–4ac
DNA Meth. m5C:H4K20 me2
H2A.Z:H3–5ac
H3K27 me2:DNA Meth. m5C
H4K20 me2:H3K4 me3
DNA Meth. m5C:H3K9 me3
H4K20 me2:H4–4ac
H3K27 me3:H3K9 me3
H3K4 me3:H4–4ac
H2A.Z:H4K20 me2
H3K9 me2:DNA Meth. m5C
H3K27 me3:DNA Meth. m5C
H4K20 me2:H3–5ac
H4K20 me3
H3K4 me3
H3K27 me2
H3K9 me2
H3K4 me1
H3K9 me3
H2A.Z
H3K27 me3
H4–4ac
H3–5ac
DNA Meth. m5C
H4K20 me2
0.00 0.25 0.50 0.75 1.00
π̂

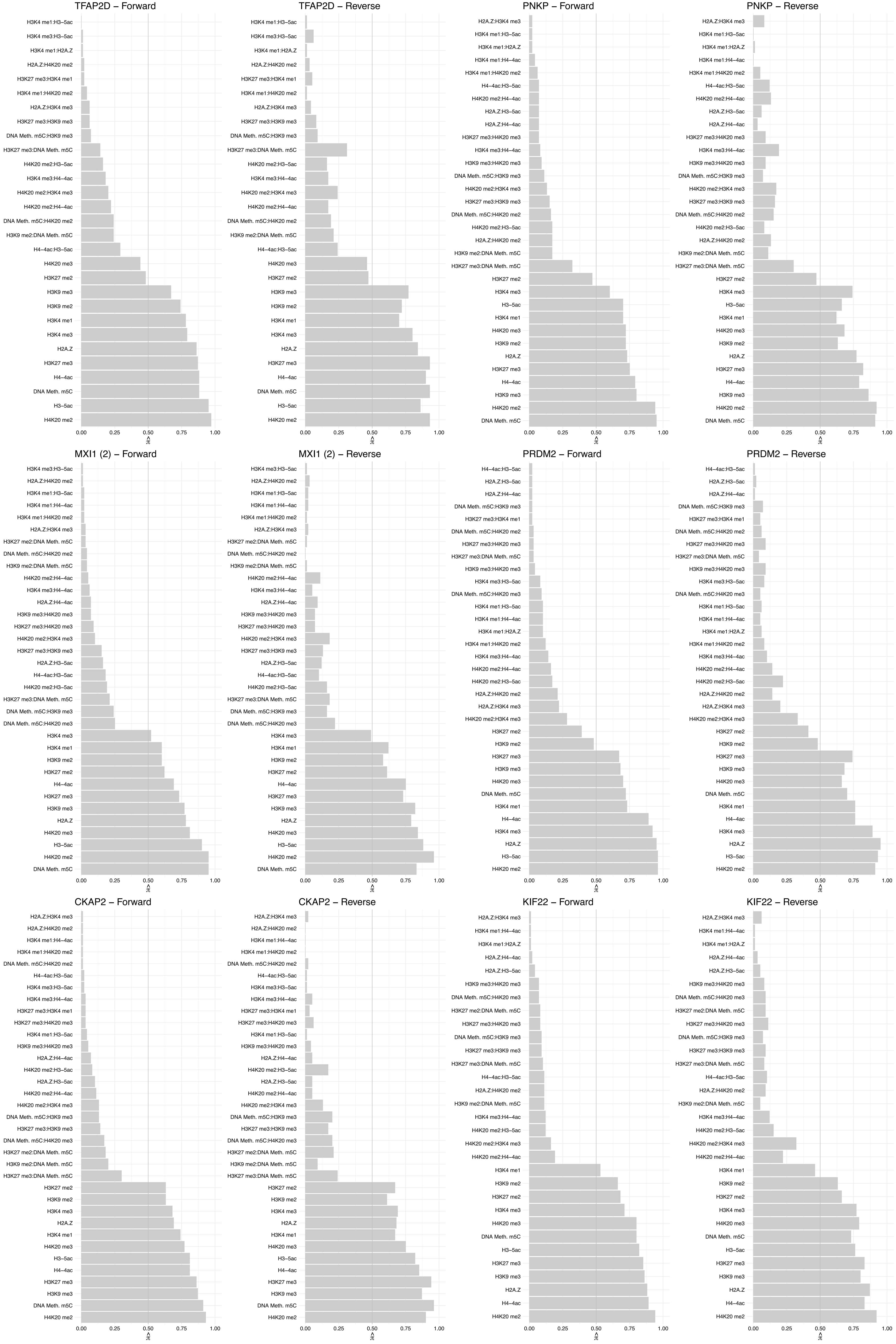

TFAP2D – Forward
TFAP2D – Reverse
PNKP – Forward
PNKP – Reverse
MXI1 (2) – Forward
MXI1 (2) – Reverse
PRDM2 – Forward
PRDM2 – Reverse
CKAP2 – Forward
CKAP2 – Reverse
KIF22 – Forward
KIF22 – Reverse
0.00
0.25
0.50
0.75
1.00
$\hat{\pi}$

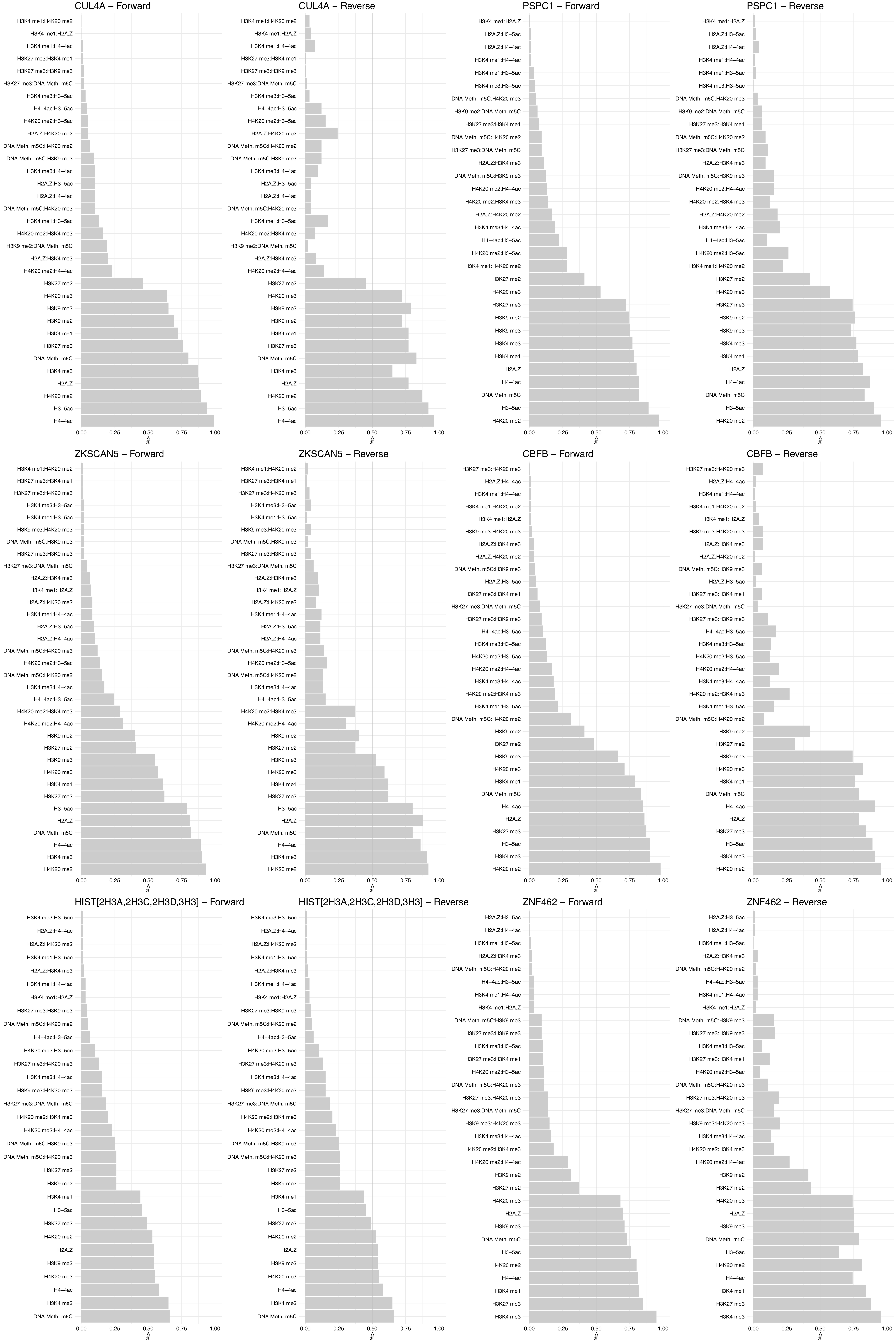
CUL4A – Forward
H3K4 me1:H4K20 me2
H3K4 me1:H2A.Z
H3K4 me1:H4–4ac
H3K27 me3:H3K4 me1
H3K27 me3:H3K9 me3
H3K27 me3:DNA Meth. m5C
H3K4 me3:H3–5ac
H4–4ac:H3–5ac
H4K20 me2:H3–5ac
H2A.Z:H4K20 me2
DNA Meth. m5C:H4K20 me2
DNA Meth. m5C:H3K9 me3
H3K4 me3:H4–4ac
H2A.Z:H3–5ac
H2A.Z:H4–4ac
DNA Meth. m5C:H4K20 me3
H3K4 me1:H3–5ac
H4K20 me2:H3K4 me3
H3K9 me2:DNA Meth. m5C
H2A.Z:H3K4 me3
H4K20 me2:H4–4ac
H3K27 me2
H4K20 me3
H3K9 me3
H3K9 me2
H3K4 me1
H3K27 me3
DNA Meth. m5C
H3K4 me3
H2A.Z
H4K20 me2
H3–5ac
H4–4ac
0.00 0.25 0.50 0.75 1.00
π̂
CUL4A – Reverse
PSPC1 – Forward
H3K4 me1:H2A.Z
H2A.Z:H3–5ac
H2A.Z:H4–4ac
H3K4 me1:H4–4ac
H3K4 me1:H3–5ac
H3K4 me3:H3–5ac
DNA Meth. m5C:H4K20 me3
H3K9 me2:DNA Meth. m5C
H3K27 me3:H3K4 me1
DNA Meth. m5C:H4K20 me2
H3K27 me3:DNA Meth. m5C
H2A.Z:H3K4 me3
DNA Meth. m5C:H3K9 me3
H4K20 me2:H4–4ac
H4K20 me2:H3K4 me3
H2A.Z:H4K20 me2
H3K4 me3:H4–4ac
H4–4ac:H3–5ac
H4K20 me2:H3–5ac
H3K4 me1:H4K20 me2
H3K27 me2
H4K20 me3
H3K27 me3
H3K9 me2
H3K9 me3
H3K4 me3
H3K4 me1
H2A.Z
H4–4ac
DNA Meth. m5C
H3–5ac
H4K20 me2
PSPC1 – Reverse
ZKSCAN5 – Forward
H3K4 me1:H4K20 me2
H3K27 me3:H3K4 me1
H3K27 me3:H4K20 me3
H3K4 me3:H3–5ac
H3K4 me1:H3–5ac
H3K9 me3:H4K20 me3
DNA Meth. m5C:H3K9 me3
H3K27 me3:H3K9 me3
H3K27 me3:DNA Meth. m5C
H2A.Z:H3K4 me3
H3K4 me1:H2A.Z
H2A.Z:H4K20 me2
H3K4 me1:H4–4ac
H2A.Z:H3–5ac
H2A.Z:H4–4ac
DNA Meth. m5C:H4K20 me3
H4K20 me2:H3–5ac
DNA Meth. m5C:H4K20 me2
H3K4 me3:H4–4ac
H4–4ac:H3–5ac
H4K20 me2:H3K4 me3
H4K20 me2:H4–4ac
H3K9 me2
H3K27 me2
H3K9 me3
H4K20 me3
H3K4 me1
H3K27 me3
H3–5ac
H2A.Z
DNA Meth. m5C
H4–4ac
H3K4 me3
H4K20 me2
ZKSCAN5 – Reverse
CBFB – Forward
H3K27 me3:H4K20 me3
H2A.Z:H4–4ac
H3K4 me1:H4–4ac
H3K4 me1:H4K20 me2
H3K4 me1:H2A.Z
H3K9 me3:H4K20 me3
H2A.Z:H3K4 me3
H2A.Z:H4K20 me2
DNA Meth. m5C:H3K9 me3
H2A.Z:H3–5ac
H3K27 me3:H3K4 me1
H3K27 me3:DNA Meth. m5C
H3K27 me3:H3K9 me3
H4–4ac:H3–5ac
H3K4 me3:H3–5ac
H4K20 me2:H3–5ac
H4K20 me2:H4–4ac
H3K4 me3:H4–4ac
H4K20 me2:H3K4 me3
H3K4 me1:H3–5ac
DNA Meth. m5C:H4K20 me2
H3K9 me2
H3K27 me2
H3K9 me3
H4K20 me3
H3K4 me1
DNA Meth. m5C
H4–4ac
H2A.Z
H3K27 me3
H3–5ac
H3K4 me3
H4K20 me2
CBFB – Reverse
HIST[2H3A,2H3C,2H3D,3H3] – Forward
H3K4 me3:H3–5ac
H2A.Z:H4–4ac
H2A.Z:H4K20 me2
H3K4 me1:H3–5ac
H2A.Z:H3K4 me3
H3K4 me1:H4–4ac
H3K4 me1:H2A.Z
H3K27 me3:H3K9 me3
DNA Meth. m5C:H4K20 me2
H4–4ac:H3–5ac
H4K20 me2:H3–5ac
H3K27 me3:H4K20 me3
H3K4 me3:H4–4ac
H3K9 me3:H4K20 me3
H3K27 me3:DNA Meth. m5C
H4K20 me2:H3K4 me3
H4K20 me2:H4–4ac
DNA Meth. m5C:H3K9 me3
DNA Meth. m5C:H4K20 me3
H3K27 me2
H3K9 me2
H3K4 me1
H3–5ac
H3K27 me3
H4K20 me2
H2A.Z
H3K9 me3
H4K20 me3
H4–4ac
H3K4 me3
DNA Meth. m5C
HIST[2H3A,2H3C,2H3D,3H3] – Reverse
ZNF462 – Forward
H2A.Z:H3–5ac
H2A.Z:H4–4ac
H3K4 me1:H3–5ac
H2A.Z:H3K4 me3
DNA Meth. m5C:H4K20 me2
H4–4ac:H3–5ac
H3K4 me1:H4–4ac
H3K4 me1:H2A.Z
DNA Meth. m5C:H3K9 me3
H3K27 me3:H3K9 me3
H3K4 me3:H3–5ac
H3K27 me3:H3K4 me1
H4K20 me2:H3–5ac
DNA Meth. m5C:H4K20 me3
H3K27 me3:H4K20 me3
H3K27 me3:DNA Meth. m5C
H3K9 me3:H4K20 me3
H3K4 me3:H4–4ac
H4K20 me2:H3K4 me3
H4K20 me2:H4–4ac
H3K9 me2
H3K27 me2
H4K20 me3
H2A.Z
H3K9 me3
DNA Meth. m5C
H3–5ac
H4K20 me2
H4–4ac
H3K4 me1
H3K27 me3
H3K4 me3
ZNF462 – Reverse

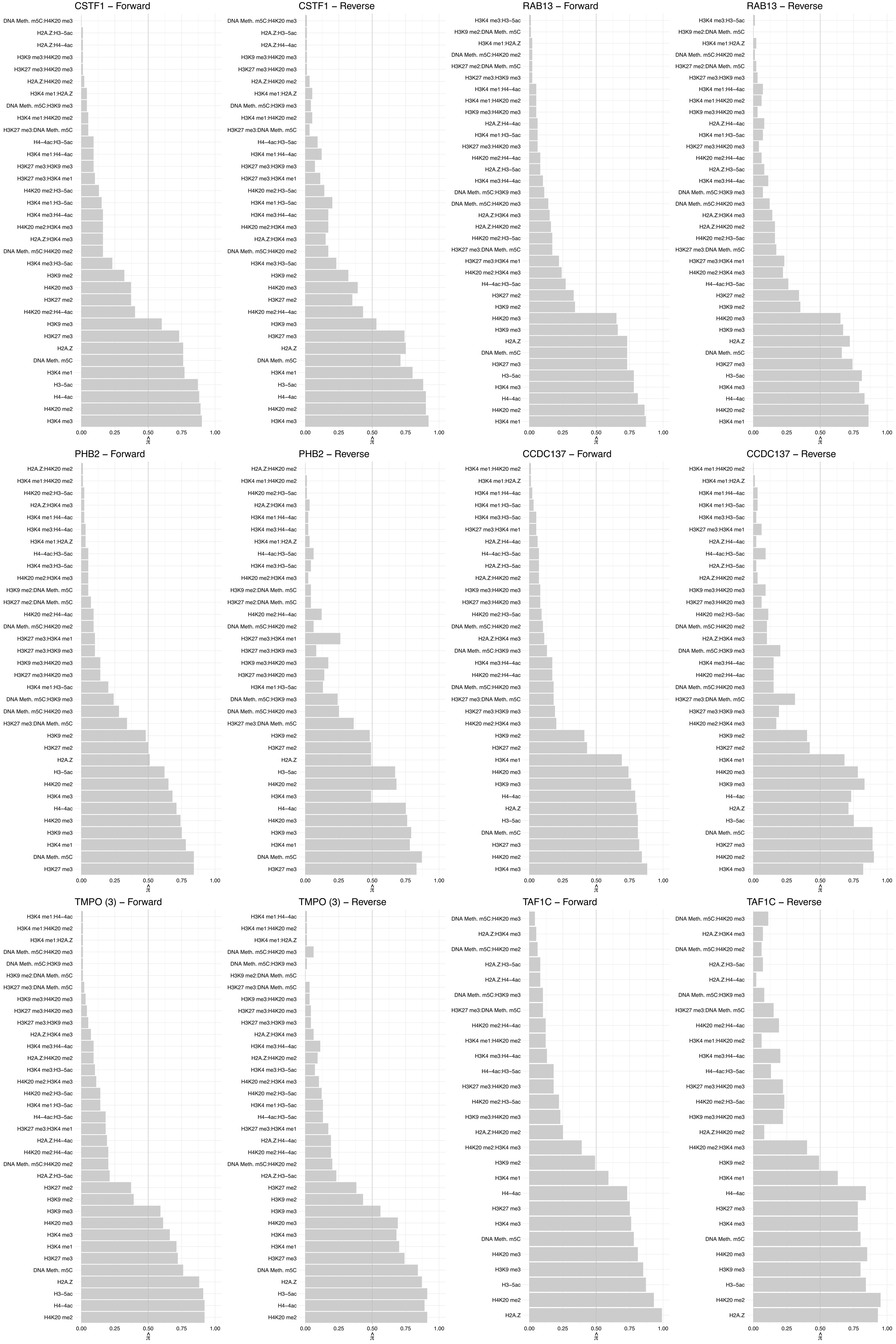
CSTF1 – Forward
DNA Meth. m5C:H4K20 me3
H2A.Z:H3−5ac
H2A.Z:H4−4ac
H3K9 me3:H4K20 me3
H3K27 me3:H4K20 me3
H2A.Z:H4K20 me2
H3K4 me1:H2A.Z
DNA Meth. m5C:H3K9 me3
H3K4 me1:H4K20 me2
H3K27 me3:DNA Meth. m5C
H4−4ac:H3−5ac
H3K4 me1:H4−4ac
H3K27 me3:H3K9 me3
H3K27 me3:H3K4 me1
H4K20 me2:H3−5ac
H3K4 me1:H3−5ac
H3K4 me3:H4−4ac
H4K20 me2:H3K4 me3
H2A.Z:H3K4 me3
DNA Meth. m5C:H4K20 me2
H3K4 me3:H3−5ac
H3K9 me2
H4K20 me3
H3K27 me2
H4K20 me2:H4−4ac
H3K9 me3
H3K27 me3
H2A.Z
DNA Meth. m5C
H3K4 me1
H3−5ac
H4−4ac
H4K20 me2
H3K4 me3
0.00 0.25 0.50 0.75 1.00
π̂
CSTF1 – Reverse
DNA Meth. m5C:H4K20 me3
H2A.Z:H3−5ac
H2A.Z:H4−4ac
H3K9 me3:H4K20 me3
H3K27 me3:H4K20 me3
H2A.Z:H4K20 me2
H3K4 me1:H2A.Z
DNA Meth. m5C:H3K9 me3
H3K4 me1:H4K20 me2
H3K27 me3:DNA Meth. m5C
H4−4ac:H3−5ac
H3K4 me1:H4−4ac
H3K27 me3:H3K9 me3
H3K27 me3:H3K4 me1
H4K20 me2:H3−5ac
H3K4 me1:H3−5ac
H3K4 me3:H4−4ac
H4K20 me2:H3K4 me3
H2A.Z:H3K4 me3
DNA Meth. m5C:H4K20 me2
H3K4 me3:H3−5ac
H3K9 me2
H4K20 me3
H3K27 me2
H4K20 me2:H4−4ac
H3K9 me3
H3K27 me3
H2A.Z
DNA Meth. m5C
H3K4 me1
H3−5ac
H4−4ac
H4K20 me2
H3K4 me3
0.00 0.25 0.50 0.75 1.00
π̂
RAB13 – Forward
H3K4 me3:H3−5ac
H3K9 me2:DNA Meth. m5C
H3K4 me1:H2A.Z
DNA Meth. m5C:H4K20 me2
H3K27 me2:DNA Meth. m5C
H3K27 me3:H3K9 me3
H3K4 me1:H4−4ac
H3K4 me1:H4K20 me2
H3K9 me3:H4K20 me3
H2A.Z:H4−4ac
H3K4 me1:H3−5ac
H3K27 me3:H4K20 me3
H4K20 me2:H4−4ac
H2A.Z:H3−5ac
H3K4 me3:H4−4ac
DNA Meth. m5C:H3K9 me3
DNA Meth. m5C:H4K20 me3
H2A.Z:H3K4 me3
H2A.Z:H4K20 me2
H4K20 me2:H3−5ac
H3K27 me3:DNA Meth. m5C
H3K27 me3:H3K4 me1
H4K20 me2:H3K4 me3
H4−4ac:H3−5ac
H3K27 me2
H3K9 me2
H4K20 me3
H3K9 me3
H2A.Z
DNA Meth. m5C
H3K27 me3
H3−5ac
H3K4 me3
H4−4ac
H4K20 me2
H3K4 me1
0.00 0.25 0.50 0.75 1.00
π̂
RAB13 – Reverse
H3K4 me3:H3−5ac
H3K9 me2:DNA Meth. m5C
H3K4 me1:H2A.Z
DNA Meth. m5C:H4K20 me2
H3K27 me2:DNA Meth. m5C
H3K27 me3:H3K9 me3
H3K4 me1:H4−4ac
H3K4 me1:H4K20 me2
H3K9 me3:H4K20 me3
H2A.Z:H4−4ac
H3K4 me1:H3−5ac
H3K27 me3:H4K20 me3
H4K20 me2:H4−4ac
H2A.Z:H3−5ac
H3K4 me3:H4−4ac
DNA Meth. m5C:H3K9 me3
DNA Meth. m5C:H4K20 me3
H2A.Z:H3K4 me3
H2A.Z:H4K20 me2
H4K20 me2:H3−5ac
H3K27 me3:DNA Meth. m5C
H3K27 me3:H3K4 me1
H4K20 me2:H3K4 me3
H4−4ac:H3−5ac
H3K27 me2
H3K9 me2
H4K20 me3
H3K9 me3
H2A.Z
DNA Meth. m5C
H3K27 me3
H3−5ac
H3K4 me3
H4−4ac
H4K20 me2
H3K4 me1
0.00 0.25 0.50 0.75 1.00
π̂
PHB2 – Forward
H2A.Z:H4K20 me2
H3K4 me1:H4K20 me2
H4K20 me2:H3−5ac
H2A.Z:H3K4 me3
H3K4 me1:H4−4ac
H3K4 me3:H4−4ac
H3K4 me1:H2A.Z
H4−4ac:H3−5ac
H3K4 me3:H3−5ac
H4K20 me2:H3K4 me3
H3K9 me2:DNA Meth. m5C
H3K27 me2:DNA Meth. m5C
H4K20 me2:H4−4ac
DNA Meth. m5C:H4K20 me2
H3K27 me3:H3K4 me1
H3K27 me3:H3K9 me3
H3K9 me3:H4K20 me3
H3K27 me3:H4K20 me3
H3K4 me1:H3−5ac
DNA Meth. m5C:H3K9 me3
DNA Meth. m5C:H4K20 me3
H3K27 me3:DNA Meth. m5C
H3K9 me2
H3K27 me2
H2A.Z
H3−5ac
H4K20 me2
H3K4 me3
H4−4ac
H4K20 me3
H3K9 me3
H3K4 me1
DNA Meth. m5C
H3K27 me3
0.00 0.25 0.50 0.75 1.00
π̂
PHB2 – Reverse
H2A.Z:H4K20 me2
H3K4 me1:H4K20 me2
H4K20 me2:H3−5ac
H2A.Z:H3K4 me3
H3K4 me1:H4−4ac
H3K4 me3:H4−4ac
H3K4 me1:H2A.Z
H4−4ac:H3−5ac
H3K4 me3:H3−5ac
H4K20 me2:H3K4 me3
H3K9 me2:DNA Meth. m5C
H3K27 me2:DNA Meth. m5C
H4K20 me2:H4−4ac
DNA Meth. m5C:H4K20 me2
H3K27 me3:H3K4 me1
H3K27 me3:H3K9 me3
H3K9 me3:H4K20 me3
H3K27 me3:H4K20 me3
H3K4 me1:H3−5ac
DNA Meth. m5C:H3K9 me3
DNA Meth. m5C:H4K20 me3
H3K27 me3:DNA Meth. m5C
H3K9 me2
H3K27 me2
H2A.Z
H3−5ac
H4K20 me2
H3K4 me3
H4−4ac
H4K20 me3
H3K9 me3
H3K4 me1
DNA Meth. m5C
H3K27 me3
0.00 0.25 0.50 0.75 1.00
π̂
CCDC137 – Forward
H3K4 me1:H4K20 me2
H3K4 me1:H2A.Z
H3K4 me1:H4−4ac
H3K4 me1:H3−5ac
H3K4 me3:H3−5ac
H3K27 me3:H3K4 me1
H2A.Z:H4−4ac
H4−4ac:H3−5ac
H2A.Z:H3−5ac
H2A.Z:H4K20 me2
H3K9 me3:H4K20 me3
H3K27 me3:H4K20 me3
H4K20 me2:H3−5ac
DNA Meth. m5C:H4K20 me2
H2A.Z:H3K4 me3
DNA Meth. m5C:H3K9 me3
H3K4 me3:H4−4ac
H4K20 me2:H4−4ac
DNA Meth. m5C:H4K20 me3
H3K27 me3:DNA Meth. m5C
H3K27 me3:H3K9 me3
H4K20 me2:H3K4 me3
H3K9 me2
H3K27 me2
H3K4 me1
H4K20 me3
H3K9 me3
H4−4ac
H2A.Z
H3−5ac
DNA Meth. m5C
H3K27 me3
H4K20 me2
H3K4 me3
0.00 0.25 0.50 0.75 1.00
π̂
CCDC137 – Reverse
H3K4 me1:H4K20 me2
H3K4 me1:H2A.Z
H3K4 me1:H4−4ac
H3K4 me1:H3−5ac
H3K4 me3:H3−5ac
H3K27 me3:H3K4 me1
H2A.Z:H4−4ac
H4−4ac:H3−5ac
H2A.Z:H3−5ac
H2A.Z:H4K20 me2
H3K9 me3:H4K20 me3
H3K27 me3:H4K20 me3
H4K20 me2:H3−5ac
DNA Meth. m5C:H4K20 me2
H2A.Z:H3K4 me3
DNA Meth. m5C:H3K9 me3
H3K4 me3:H4−4ac
H4K20 me2:H4−4ac
DNA Meth. m5C:H4K20 me3
H3K27 me3:DNA Meth. m5C
H3K27 me3:H3K9 me3
H4K20 me2:H3K4 me3
H3K9 me2
H3K27 me2
H3K4 me1
H4K20 me3
H3K9 me3
H4−4ac
H2A.Z
H3−5ac
DNA Meth. m5C
H3K27 me3
H4K20 me2
H3K4 me3
0.00 0.25 0.50 0.75 1.00
π̂
TMPO (3) – Forward
H3K4 me1:H4−4ac
H3K4 me1:H4K20 me2
H3K4 me1:H2A.Z
DNA Meth. m5C:H4K20 me3
DNA Meth. m5C:H3K9 me3
H3K9 me2:DNA Meth. m5C
H3K27 me3:DNA Meth. m5C
H3K9 me3:H4K20 me3
H3K27 me3:H4K20 me3
H3K27 me3:H3K9 me3
H2A.Z:H3K4 me3
H3K4 me3:H4−4ac
H2A.Z:H4K20 me2
H3K4 me3:H3−5ac
H4K20 me2:H3K4 me3
H4K20 me2:H3−5ac
H3K4 me1:H3−5ac
H4−4ac:H3−5ac
H3K27 me3:H3K4 me1
H2A.Z:H4−4ac
H4K20 me2:H4−4ac
DNA Meth. m5C:H4K20 me2
H2A.Z:H3−5ac
H3K27 me2
H3K9 me2
H3K9 me3
H4K20 me3
H3K4 me3
H3K4 me1
H3K27 me3
DNA Meth. m5C
H2A.Z
H3−5ac
H4−4ac
H4K20 me2
0.00 0.25 0.50 0.75 1.00
π̂
TMPO (3) – Reverse
H3K4 me1:H4−4ac
H3K4 me1:H4K20 me2
H3K4 me1:H2A.Z
DNA Meth. m5C:H4K20 me3
DNA Meth. m5C:H3K9 me3
H3K9 me2:DNA Meth. m5C
H3K27 me3:DNA Meth. m5C
H3K9 me3:H4K20 me3
H3K27 me3:H4K20 me3
H3K27 me3:H3K9 me3
H2A.Z:H3K4 me3
H3K4 me3:H4−4ac
H2A.Z:H4K20 me2
H3K4 me3:H3−5ac
H4K20 me2:H3K4 me3
H4K20 me2:H3−5ac
H3K4 me1:H3−5ac
H4−4ac:H3−5ac
H3K27 me3:H3K4 me1
H2A.Z:H4−4ac
H4K20 me2:H4−4ac
DNA Meth. m5C:H4K20 me2
H2A.Z:H3−5ac
H3K27 me2
H3K9 me2
H3K9 me3
H4K20 me3
H3K4 me3
H3K4 me1
H3K27 me3
DNA Meth. m5C
H2A.Z
H3−5ac
H4−4ac
H4K20 me2
0.00 0.25 0.50 0.75 1.00
π̂
TAF1C – Forward
DNA Meth. m5C:H4K20 me3
H2A.Z:H3K4 me3
DNA Meth. m5C:H4K20 me2
H2A.Z:H3−5ac
H2A.Z:H4−4ac
DNA Meth. m5C:H3K9 me3
H3K27 me3:DNA Meth. m5C
H4K20 me2:H4−4ac
H3K4 me1:H4K20 me2
H3K4 me3:H4−4ac
H4−4ac:H3−5ac
H3K27 me3:H4K20 me3
H4K20 me2:H3−5ac
H3K9 me3:H4K20 me3
H2A.Z:H4K20 me2
H4K20 me2:H3K4 me3
H3K9 me2
H3K4 me1
H4−4ac
H3K27 me3
H3K4 me3
DNA Meth. m5C
H4K20 me3
H3K9 me3
H3−5ac
H4K20 me2
H2A.Z
0.00 0.25 0.50 0.75 1.00
π̂
TAF1C – Reverse
DNA Meth. m5C:H4K20 me3
H2A.Z:H3K4 me3
DNA Meth. m5C:H4K20 me2
H2A.Z:H3−5ac
H2A.Z:H4−4ac
DNA Meth. m5C:H3K9 me3
H3K27 me3:DNA Meth. m5C
H4K20 me2:H4−4ac
H3K4 me1:H4K20 me2
H3K4 me3:H4−4ac
H4−4ac:H3−5ac
H3K27 me3:H4K20 me3
H4K20 me2:H3−5ac
H3K9 me3:H4K20 me3
H2A.Z:H4K20 me2
H4K20 me2:H3K4 me3
H3K9 me2
H3K4 me1
H4−4ac
H3K27 me3
H3K4 me3
DNA Meth. m5C
H4K20 me3
H3K9 me3
H3−5ac
H4K20 me2
H2A.Z
0.00 0.25 0.50 0.75 1.00
π̂

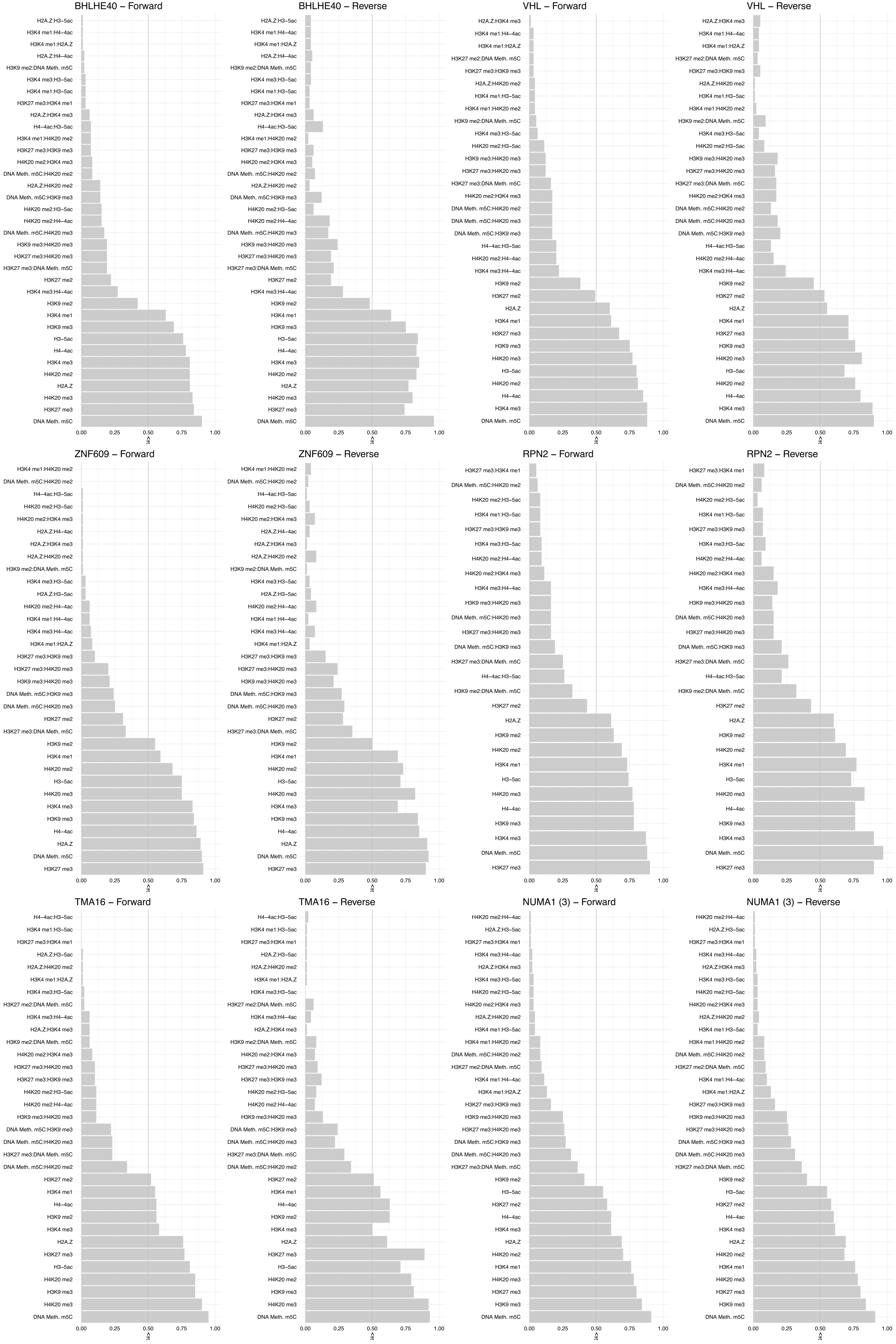
BHLHE40 – Forward
H2A.Z:H3–5ac
H3K4 me1:H4–4ac
H3K4 me1:H2A.Z
H2A.Z:H4–4ac
H3K9 me2:DNA Meth. m5C
H3K4 me3:H3–5ac
H3K4 me1:H3–5ac
H3K27 me3:H3K4 me1
H2A.Z:H3K4 me3
H4–4ac:H3–5ac
H3K4 me1:H4K20 me2
H3K27 me3:H3K9 me3
H4K20 me2:H3K4 me3
DNA Meth. m5C:H4K20 me2
H2A.Z:H4K20 me2
DNA Meth. m5C:H3K9 me3
H4K20 me2:H3–5ac
H4K20 me2:H4–4ac
DNA Meth. m5C:H4K20 me3
H3K9 me3:H4K20 me3
H3K27 me3:H4K20 me3
H3K27 me3:DNA Meth. m5C
H3K27 me2
H3K4 me3:H4–4ac
H3K9 me2
H3K4 me1
H3K9 me3
H3–5ac
H4–4ac
H3K4 me3
H4K20 me2
H2A.Z
H4K20 me3
H3K27 me3
DNA Meth. m5C
0.00 0.25 0.50 0.75 1.00
π̂
BHLHE40 – Reverse
VHL – Forward
VHL – Reverse
ZNF609 – Forward
ZNF609 – Reverse
RPN2 – Forward
RPN2 – Reverse
TMA16 – Forward
TMA16 – Reverse
NUMA1 (3) – Forward
NUMA1 (3) – Reverse

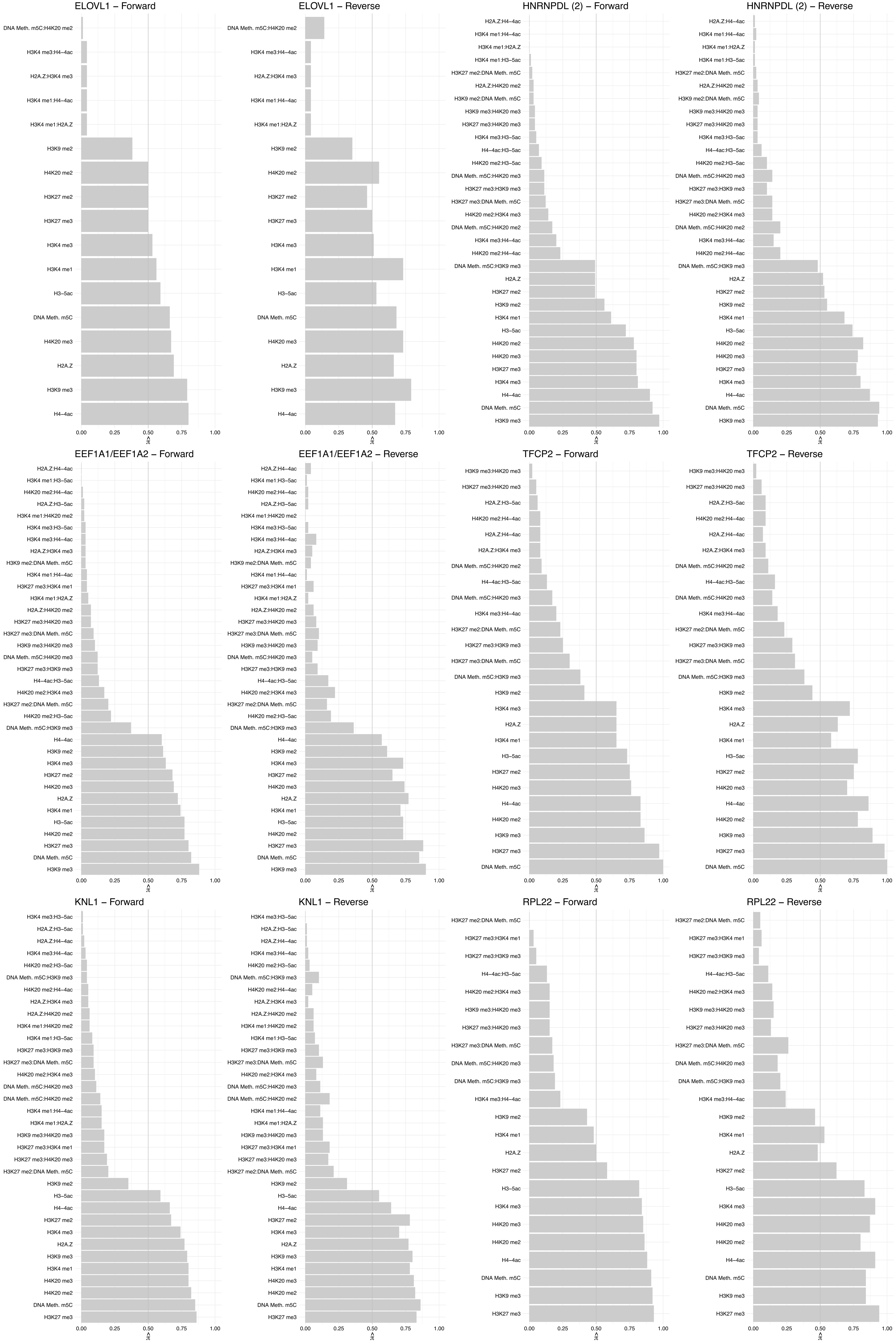

ELOVL1 – Forward
DNA Meth. m5C:H4K20 me2
H3K4 me3:H4–4ac
H2A.Z:H3K4 me3
H3K4 me1:H4–4ac
H3K4 me1:H2A.Z
H3K9 me2
H4K20 me2
H3K27 me2
H3K27 me3
H3K4 me3
H3K4 me1
H3–5ac
DNA Meth. m5C
H4K20 me3
H2A.Z
H3K9 me3
H4–4ac
ELOVL1 – Reverse
HNRNPDL (2) – Forward
H2A.Z:H4–4ac
H3K4 me1:H4–4ac
H3K4 me1:H2A.Z
H3K4 me1:H3–5ac
H3K27 me2:DNA Meth. m5C
H2A.Z:H4K20 me2
H3K9 me2:DNA Meth. m5C
H3K9 me3:H4K20 me3
H3K27 me3:H4K20 me3
H3K4 me3:H3–5ac
H4–4ac:H3–5ac
H4K20 me2:H3–5ac
DNA Meth. m5C:H4K20 me3
H3K27 me3:H3K9 me3
H3K27 me3:DNA Meth. m5C
H4K20 me2:H3K4 me3
DNA Meth. m5C:H4K20 me2
H3K4 me3:H4–4ac
H4K20 me2:H4–4ac
DNA Meth. m5C:H3K9 me3
H2A.Z
H3K27 me2
H3K9 me2
H3K4 me1
H3–5ac
H4K20 me2
H4K20 me3
H3K27 me3
H3K4 me3
H4–4ac
DNA Meth. m5C
H3K9 me3
HNRNPDL (2) – Reverse
EEF1A1/EEF1A2 – Forward
H2A.Z:H4–4ac
H3K4 me1:H3–5ac
H4K20 me2:H4–4ac
H2A.Z:H3–5ac
H3K4 me1:H4K20 me2
H3K4 me3:H3–5ac
H3K4 me3:H4–4ac
H2A.Z:H3K4 me3
H3K9 me2:DNA Meth. m5C
H3K4 me1:H4–4ac
H3K27 me3:H3K4 me1
H3K4 me1:H2A.Z
H2A.Z:H4K20 me2
H3K27 me3:H4K20 me3
H3K27 me3:DNA Meth. m5C
H3K9 me3:H4K20 me3
DNA Meth. m5C:H4K20 me3
H3K27 me3:H3K9 me3
H4–4ac:H3–5ac
H4K20 me2:H3K4 me3
H3K27 me2:DNA Meth. m5C
H4K20 me2:H3–5ac
DNA Meth. m5C:H3K9 me3
H4–4ac
H3K9 me2
H3K4 me3
H3K27 me2
H4K20 me3
H2A.Z
H3K4 me1
H3–5ac
H4K20 me2
H3K27 me3
DNA Meth. m5C
H3K9 me3
EEF1A1/EEF1A2 – Reverse
TFCP2 – Forward
H3K9 me3:H4K20 me3
H3K27 me3:H4K20 me3
H2A.Z:H3–5ac
H4K20 me2:H4–4ac
H2A.Z:H4–4ac
H2A.Z:H3K4 me3
DNA Meth. m5C:H4K20 me2
H4–4ac:H3–5ac
DNA Meth. m5C:H4K20 me3
H3K4 me3:H4–4ac
H3K27 me2:DNA Meth. m5C
H3K27 me3:H3K9 me3
H3K27 me3:DNA Meth. m5C
DNA Meth. m5C:H3K9 me3
H3K9 me2
H3K4 me3
H2A.Z
H3K4 me1
H3–5ac
H3K27 me2
H4K20 me3
H4–4ac
H4K20 me2
H3K9 me3
H3K27 me3
DNA Meth. m5C
TFCP2 – Reverse
KNL1 – Forward
H3K4 me3:H3–5ac
H2A.Z:H3–5ac
H2A.Z:H4–4ac
H3K4 me3:H4–4ac
H4K20 me2:H3–5ac
DNA Meth. m5C:H3K9 me3
H4K20 me2:H4–4ac
H2A.Z:H3K4 me3
H2A.Z:H4K20 me2
H3K4 me1:H4K20 me2
H3K4 me1:H3–5ac
H3K27 me3:H3K9 me3
H3K27 me3:DNA Meth. m5C
H4K20 me2:H3K4 me3
DNA Meth. m5C:H4K20 me3
DNA Meth. m5C:H4K20 me2
H3K4 me1:H4–4ac
H3K4 me1:H2A.Z
H3K9 me3:H4K20 me3
H3K27 me3:H3K4 me1
H3K27 me3:H4K20 me3
H3K27 me2:DNA Meth. m5C
H3K9 me2
H3–5ac
H4–4ac
H3K27 me2
H3K4 me3
H2A.Z
H3K9 me3
H3K4 me1
H4K20 me3
H4K20 me2
DNA Meth. m5C
H3K27 me3
KNL1 – Reverse
RPL22 – Forward
H3K27 me2:DNA Meth. m5C
H3K27 me3:H3K4 me1
H3K27 me3:H3K9 me3
H4–4ac:H3–5ac
H4K20 me2:H3K4 me3
H3K9 me3:H4K20 me3
H3K27 me3:H4K20 me3
H3K27 me3:DNA Meth. m5C
DNA Meth. m5C:H4K20 me3
DNA Meth. m5C:H3K9 me3
H3K4 me3:H4–4ac
H3K9 me2
H3K4 me1
H2A.Z
H3K27 me2
H3–5ac
H3K4 me3
H4K20 me3
H4K20 me2
H4–4ac
DNA Meth. m5C
H3K9 me3
H3K27 me3
RPL22 – Reverse
0.00
0.25
0.50
0.75
1.00
π̂

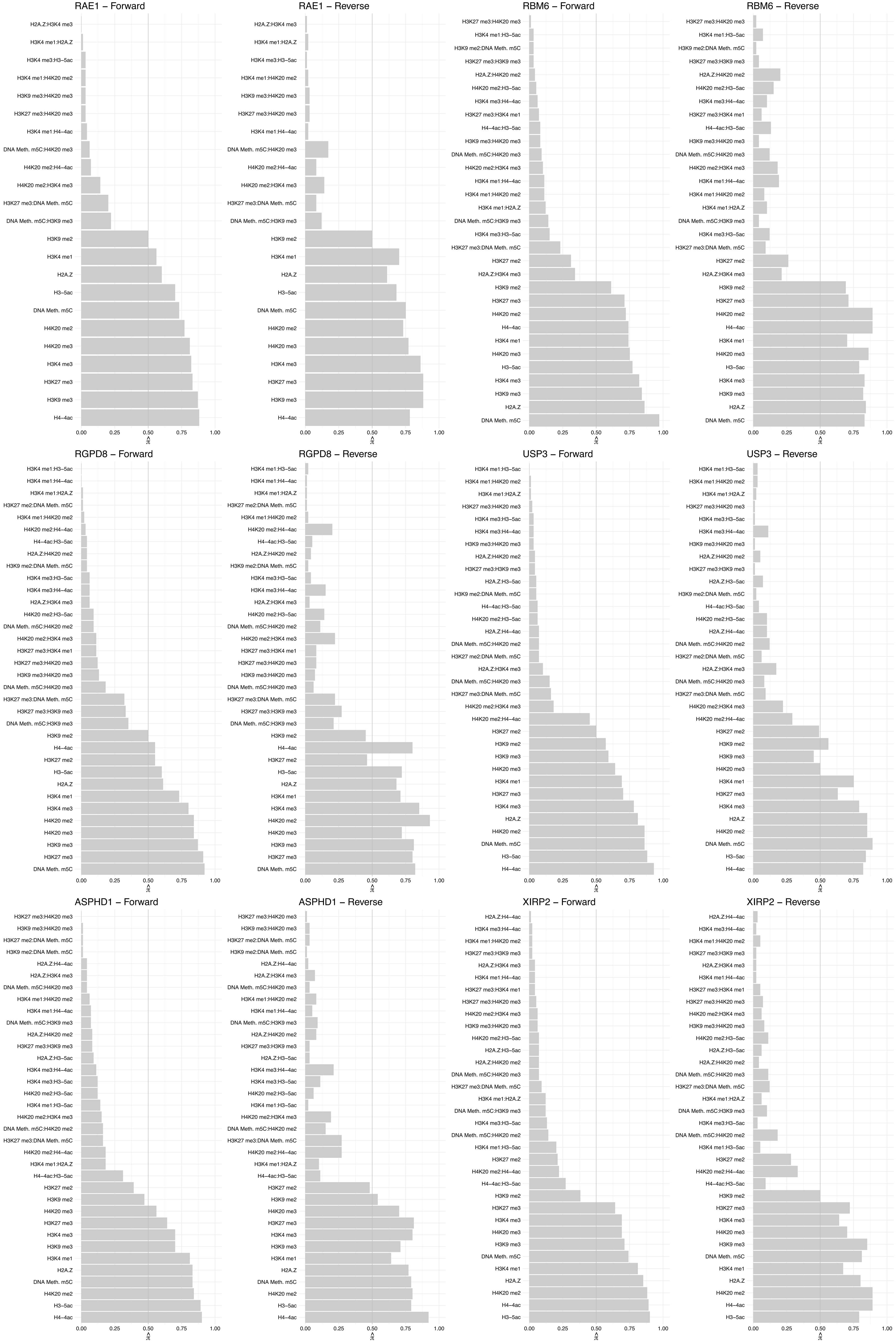
RAE1 – Forward
H2A.Z:H3K4 me3
H3K4 me1:H2A.Z
H3K4 me3:H3−5ac
H3K4 me1:H4K20 me2
H3K9 me3:H4K20 me3
H3K27 me3:H4K20 me3
H3K4 me1:H4−4ac
DNA Meth. m5C:H4K20 me3
H4K20 me2:H4−4ac
H4K20 me2:H3K4 me3
H3K27 me3:DNA Meth. m5C
DNA Meth. m5C:H3K9 me3
H3K9 me2
H3K4 me1
H2A.Z
H3−5ac
DNA Meth. m5C
H4K20 me2
H4K20 me3
H3K4 me3
H3K27 me3
H3K9 me3
H4−4ac
0.00 0.25 0.50 0.75 1.00
π̂
RAE1 – Reverse
H2A.Z:H3K4 me3
H3K4 me1:H2A.Z
H3K4 me3:H3−5ac
H3K4 me1:H4K20 me2
H3K9 me3:H4K20 me3
H3K27 me3:H4K20 me3
H3K4 me1:H4−4ac
DNA Meth. m5C:H4K20 me3
H4K20 me2:H4−4ac
H4K20 me2:H3K4 me3
H3K27 me3:DNA Meth. m5C
DNA Meth. m5C:H3K9 me3
H3K9 me2
H3K4 me1
H2A.Z
H3−5ac
DNA Meth. m5C
H4K20 me2
H4K20 me3
H3K4 me3
H3K27 me3
H3K9 me3
H4−4ac
0.00 0.25 0.50 0.75 1.00
π̂
RBM6 – Forward
H3K27 me3:H4K20 me3
H3K4 me1:H3−5ac
H3K9 me2:DNA Meth. m5C
H3K27 me3:H3K9 me3
H2A.Z:H4K20 me2
H4K20 me2:H3−5ac
H3K4 me3:H4−4ac
H3K27 me3:H3K4 me1
H4−4ac:H3−5ac
H3K9 me3:H4K20 me3
DNA Meth. m5C:H4K20 me3
H4K20 me2:H3K4 me3
H3K4 me1:H4−4ac
H3K4 me1:H4K20 me2
H3K4 me1:H2A.Z
DNA Meth. m5C:H3K9 me3
H3K4 me3:H3−5ac
H3K27 me3:DNA Meth. m5C
H3K27 me2
H2A.Z:H3K4 me3
H3K9 me2
H3K27 me3
H4K20 me2
H4−4ac
H3K4 me1
H4K20 me3
H3−5ac
H3K4 me3
H3K9 me3
H2A.Z
DNA Meth. m5C
0.00 0.25 0.50 0.75 1.00
π̂
RBM6 – Reverse
H3K27 me3:H4K20 me3
H3K4 me1:H3−5ac
H3K9 me2:DNA Meth. m5C
H3K27 me3:H3K9 me3
H2A.Z:H4K20 me2
H4K20 me2:H3−5ac
H3K4 me3:H4−4ac
H3K27 me3:H3K4 me1
H4−4ac:H3−5ac
H3K9 me3:H4K20 me3
DNA Meth. m5C:H4K20 me3
H4K20 me2:H3K4 me3
H3K4 me1:H4−4ac
H3K4 me1:H4K20 me2
H3K4 me1:H2A.Z
DNA Meth. m5C:H3K9 me3
H3K4 me3:H3−5ac
H3K27 me3:DNA Meth. m5C
H3K27 me2
H2A.Z:H3K4 me3
H3K9 me2
H3K27 me3
H4K20 me2
H4−4ac
H3K4 me1
H4K20 me3
H3−5ac
H3K4 me3
H3K9 me3
H2A.Z
DNA Meth. m5C
0.00 0.25 0.50 0.75 1.00
π̂
RGPD8 – Forward
H3K4 me1:H3−5ac
H3K4 me1:H4−4ac
H3K4 me1:H2A.Z
H3K27 me2:DNA Meth. m5C
H3K4 me1:H4K20 me2
H4K20 me2:H4−4ac
H4−4ac:H3−5ac
H2A.Z:H4K20 me2
H3K9 me2:DNA Meth. m5C
H3K4 me3:H3−5ac
H3K4 me3:H4−4ac
H2A.Z:H3K4 me3
H4K20 me2:H3−5ac
DNA Meth. m5C:H4K20 me2
H4K20 me2:H3K4 me3
H3K27 me3:H3K4 me1
H3K27 me3:H4K20 me3
H3K9 me3:H4K20 me3
DNA Meth. m5C:H4K20 me3
H3K27 me3:DNA Meth. m5C
H3K27 me3:H3K9 me3
DNA Meth. m5C:H3K9 me3
H3K9 me2
H4−4ac
H3K27 me2
H3−5ac
H2A.Z
H3K4 me1
H3K4 me3
H4K20 me2
H4K20 me3
H3K9 me3
H3K27 me3
DNA Meth. m5C
0.00 0.25 0.50 0.75 1.00
π̂
RGPD8 – Reverse
H3K4 me1:H3−5ac
H3K4 me1:H4−4ac
H3K4 me1:H2A.Z
H3K27 me2:DNA Meth. m5C
H3K4 me1:H4K20 me2
H4K20 me2:H4−4ac
H4−4ac:H3−5ac
H2A.Z:H4K20 me2
H3K9 me2:DNA Meth. m5C
H3K4 me3:H3−5ac
H3K4 me3:H4−4ac
H2A.Z:H3K4 me3
H4K20 me2:H3−5ac
DNA Meth. m5C:H4K20 me2
H4K20 me2:H3K4 me3
H3K27 me3:H3K4 me1
H3K27 me3:H4K20 me3
H3K9 me3:H4K20 me3
DNA Meth. m5C:H4K20 me3
H3K27 me3:DNA Meth. m5C
H3K27 me3:H3K9 me3
DNA Meth. m5C:H3K9 me3
H3K9 me2
H4−4ac
H3K27 me2
H3−5ac
H2A.Z
H3K4 me1
H3K4 me3
H4K20 me2
H4K20 me3
H3K9 me3
H3K27 me3
DNA Meth. m5C
0.00 0.25 0.50 0.75 1.00
π̂
USP3 – Forward
H3K4 me1:H3−5ac
H3K4 me1:H4K20 me2
H3K4 me1:H2A.Z
H3K27 me3:H4K20 me3
H3K4 me3:H3−5ac
H3K4 me3:H4−4ac
H3K9 me3:H4K20 me3
H2A.Z:H4K20 me2
H3K27 me3:H3K9 me3
H2A.Z:H3−5ac
H3K9 me2:DNA Meth. m5C
H4−4ac:H3−5ac
H4K20 me2:H3−5ac
H2A.Z:H4−4ac
DNA Meth. m5C:H4K20 me2
H3K27 me2:DNA Meth. m5C
H2A.Z:H3K4 me3
DNA Meth. m5C:H4K20 me3
H3K27 me3:DNA Meth. m5C
H4K20 me2:H3K4 me3
H4K20 me2:H4−4ac
H3K27 me2
H3K9 me2
H3K9 me3
H4K20 me3
H3K4 me1
H3K27 me3
H3K4 me3
H2A.Z
H4K20 me2
DNA Meth. m5C
H3−5ac
H4−4ac
0.00 0.25 0.50 0.75 1.00
π̂
USP3 – Reverse
H3K4 me1:H3−5ac
H3K4 me1:H4K20 me2
H3K4 me1:H2A.Z
H3K27 me3:H4K20 me3
H3K4 me3:H3−5ac
H3K4 me3:H4−4ac
H3K9 me3:H4K20 me3
H2A.Z:H4K20 me2
H3K27 me3:H3K9 me3
H2A.Z:H3−5ac
H3K9 me2:DNA Meth. m5C
H4−4ac:H3−5ac
H4K20 me2:H3−5ac
H2A.Z:H4−4ac
DNA Meth. m5C:H4K20 me2
H3K27 me2:DNA Meth. m5C
H2A.Z:H3K4 me3
DNA Meth. m5C:H4K20 me3
H3K27 me3:DNA Meth. m5C
H4K20 me2:H3K4 me3
H4K20 me2:H4−4ac
H3K27 me2
H3K9 me2
H3K9 me3
H4K20 me3
H3K4 me1
H3K27 me3
H3K4 me3
H2A.Z
H4K20 me2
DNA Meth. m5C
H3−5ac
H4−4ac
0.00 0.25 0.50 0.75 1.00
π̂
ASPHD1 – Forward
H3K27 me3:H4K20 me3
H3K9 me3:H4K20 me3
H3K27 me2:DNA Meth. m5C
H3K9 me2:DNA Meth. m5C
H2A.Z:H4−4ac
H2A.Z:H3K4 me3
DNA Meth. m5C:H4K20 me3
H3K4 me1:H4K20 me2
H3K4 me1:H4−4ac
DNA Meth. m5C:H3K9 me3
H2A.Z:H4K20 me2
H3K27 me3:H3K9 me3
H2A.Z:H3−5ac
H3K4 me3:H4−4ac
H3K4 me3:H3−5ac
H4K20 me2:H3−5ac
H3K4 me1:H3−5ac
H4K20 me2:H3K4 me3
DNA Meth. m5C:H4K20 me2
H3K27 me3:DNA Meth. m5C
H4K20 me2:H4−4ac
H3K4 me1:H2A.Z
H4−4ac:H3−5ac
H3K27 me2
H3K9 me2
H4K20 me3
H3K27 me3
H3K4 me3
H3K9 me3
H3K4 me1
H2A.Z
DNA Meth. m5C
H4K20 me2
H3−5ac
H4−4ac
0.00 0.25 0.50 0.75 1.00
π̂
ASPHD1 – Reverse
H3K27 me3:H4K20 me3
H3K9 me3:H4K20 me3
H3K27 me2:DNA Meth. m5C
H3K9 me2:DNA Meth. m5C
H2A.Z:H4−4ac
H2A.Z:H3K4 me3
DNA Meth. m5C:H4K20 me3
H3K4 me1:H4K20 me2
H3K4 me1:H4−4ac
DNA Meth. m5C:H3K9 me3
H2A.Z:H4K20 me2
H3K27 me3:H3K9 me3
H2A.Z:H3−5ac
H3K4 me3:H4−4ac
H3K4 me3:H3−5ac
H4K20 me2:H3−5ac
H3K4 me1:H3−5ac
H4K20 me2:H3K4 me3
DNA Meth. m5C:H4K20 me2
H3K27 me3:DNA Meth. m5C
H4K20 me2:H4−4ac
H3K4 me1:H2A.Z
H4−4ac:H3−5ac
H3K27 me2
H3K9 me2
H4K20 me3
H3K27 me3
H3K4 me3
H3K9 me3
H3K4 me1
H2A.Z
DNA Meth. m5C
H4K20 me2
H3−5ac
H4−4ac
0.00 0.25 0.50 0.75 1.00
π̂
XIRP2 – Forward
H2A.Z:H4−4ac
H3K4 me3:H4−4ac
H3K4 me1:H4K20 me2
H3K27 me3:H3K9 me3
H2A.Z:H3K4 me3
H3K4 me1:H4−4ac
H3K27 me3:H3K4 me1
H3K27 me3:H4K20 me3
H4K20 me2:H3K4 me3
H3K9 me3:H4K20 me3
H4K20 me2:H3−5ac
H2A.Z:H3−5ac
H2A.Z:H4K20 me2
DNA Meth. m5C:H4K20 me3
H3K27 me3:DNA Meth. m5C
H3K4 me1:H2A.Z
DNA Meth. m5C:H3K9 me3
H3K4 me3:H3−5ac
DNA Meth. m5C:H4K20 me2
H3K4 me1:H3−5ac
H3K27 me2
H4K20 me2:H4−4ac
H4−4ac:H3−5ac
H3K9 me2
H3K27 me3
H3K4 me3
H4K20 me3
H3K9 me3
DNA Meth. m5C
H3K4 me1
H2A.Z
H4K20 me2
H4−4ac
H3−5ac
0.00 0.25 0.50 0.75 1.00
π̂
XIRP2 – Reverse
H2A.Z:H4−4ac
H3K4 me3:H4−4ac
H3K4 me1:H4K20 me2
H3K27 me3:H3K9 me3
H2A.Z:H3K4 me3
H3K4 me1:H4−4ac
H3K27 me3:H3K4 me1
H3K27 me3:H4K20 me3
H4K20 me2:H3K4 me3
H3K9 me3:H4K20 me3
H4K20 me2:H3−5ac
H2A.Z:H3−5ac
H2A.Z:H4K20 me2
DNA Meth. m5C:H4K20 me3
H3K27 me3:DNA Meth. m5C
H3K4 me1:H2A.Z
DNA Meth. m5C:H3K9 me3
H3K4 me3:H3−5ac
DNA Meth. m5C:H4K20 me2
H3K4 me1:H3−5ac
H3K27 me2
H4K20 me2:H4−4ac
H4−4ac:H3−5ac
H3K9 me2
H3K27 me3
H3K4 me3
H4K20 me3
H3K9 me3
DNA Meth. m5C
H3K4 me1
H2A.Z
H4K20 me2
H4−4ac
H3−5ac
0.00 0.25 0.50 0.75 1.00
π̂

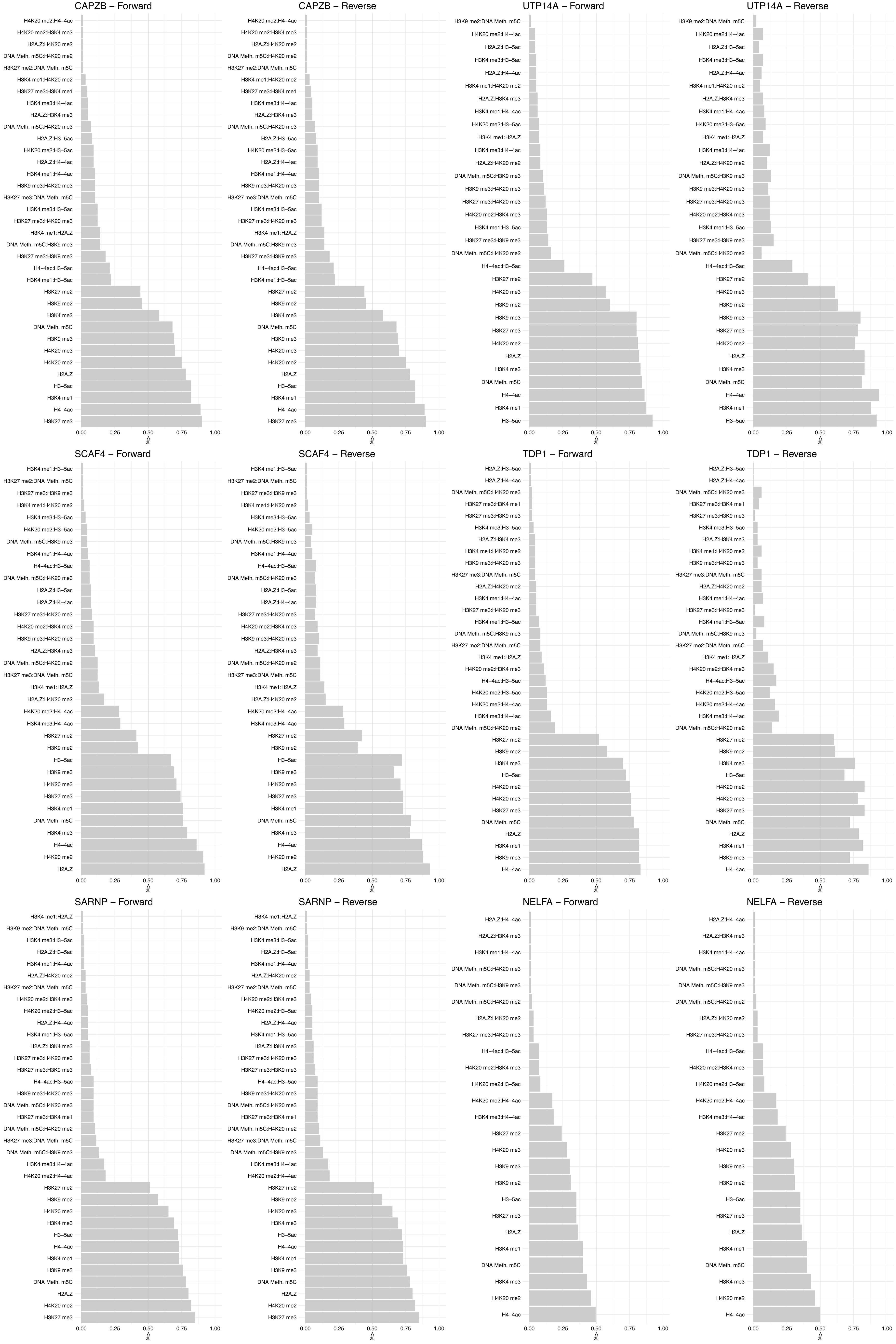
CAPZB – Forward
H4K20 me2:H4−4ac
H4K20 me2:H3K4 me3
H2A.Z:H4K20 me2
DNA Meth. m5C:H4K20 me2
H3K27 me2:DNA Meth. m5C
H3K4 me1:H4K20 me2
H3K27 me3:H3K4 me1
H3K4 me3:H4−4ac
H2A.Z:H3K4 me3
DNA Meth. m5C:H4K20 me3
H2A.Z:H3−5ac
H4K20 me2:H3−5ac
H2A.Z:H4−4ac
H3K4 me1:H4−4ac
H3K9 me3:H4K20 me3
H3K27 me3:DNA Meth. m5C
H3K4 me3:H3−5ac
H3K27 me3:H4K20 me3
H3K4 me1:H2A.Z
DNA Meth. m5C:H3K9 me3
H3K27 me3:H3K9 me3
H4−4ac:H3−5ac
H3K4 me1:H3−5ac
H3K27 me2
H3K9 me2
H3K4 me3
DNA Meth. m5C
H3K9 me3
H4K20 me3
H4K20 me2
H2A.Z
H3−5ac
H3K4 me1
H4−4ac
H3K27 me3
0.00 0.25 0.50 0.75 1.00
$\hat{\pi}$
CAPZB – Reverse
H4K20 me2:H4−4ac
H4K20 me2:H3K4 me3
H2A.Z:H4K20 me2
DNA Meth. m5C:H4K20 me2
H3K27 me2:DNA Meth. m5C
H3K4 me1:H4K20 me2
H3K27 me3:H3K4 me1
H3K4 me3:H4−4ac
H2A.Z:H3K4 me3
DNA Meth. m5C:H4K20 me3
H2A.Z:H3−5ac
H4K20 me2:H3−5ac
H2A.Z:H4−4ac
H3K4 me1:H4−4ac
H3K9 me3:H4K20 me3
H3K27 me3:DNA Meth. m5C
H3K4 me3:H3−5ac
H3K27 me3:H4K20 me3
H3K4 me1:H2A.Z
DNA Meth. m5C:H3K9 me3
H3K27 me3:H3K9 me3
H4−4ac:H3−5ac
H3K4 me1:H3−5ac
H3K27 me2
H3K9 me2
H3K4 me3
DNA Meth. m5C
H3K9 me3
H4K20 me3
H4K20 me2
H2A.Z
H3−5ac
H3K4 me1
H4−4ac
H3K27 me3
0.00 0.25 0.50 0.75 1.00
$\hat{\pi}$
UTP14A – Forward
H3K9 me2:DNA Meth. m5C
H4K20 me2:H4−4ac
H2A.Z:H3−5ac
H3K4 me3:H3−5ac
H2A.Z:H4−4ac
H3K4 me1:H4K20 me2
H2A.Z:H3K4 me3
H3K4 me1:H4−4ac
H4K20 me2:H3−5ac
H3K4 me1:H2A.Z
H3K4 me3:H4−4ac
H2A.Z:H4K20 me2
DNA Meth. m5C:H3K9 me3
H3K9 me3:H4K20 me3
H3K27 me3:H4K20 me3
H4K20 me2:H3K4 me3
H3K4 me1:H3−5ac
H3K27 me3:H3K9 me3
DNA Meth. m5C:H4K20 me2
H4−4ac:H3−5ac
H3K27 me2
H4K20 me3
H3K9 me2
H3K9 me3
H3K27 me3
H4K20 me2
H2A.Z
H3K4 me3
DNA Meth. m5C
H4−4ac
H3K4 me1
H3−5ac
0.00 0.25 0.50 0.75 1.00
$\hat{\pi}$
UTP14A – Reverse
H3K9 me2:DNA Meth. m5C
H4K20 me2:H4−4ac
H2A.Z:H3−5ac
H3K4 me3:H3−5ac
H2A.Z:H4−4ac
H3K4 me1:H4K20 me2
H2A.Z:H3K4 me3
H3K4 me1:H4−4ac
H4K20 me2:H3−5ac
H3K4 me1:H2A.Z
H3K4 me3:H4−4ac
H2A.Z:H4K20 me2
DNA Meth. m5C:H3K9 me3
H3K9 me3:H4K20 me3
H3K27 me3:H4K20 me3
H4K20 me2:H3K4 me3
H3K4 me1:H3−5ac
H3K27 me3:H3K9 me3
DNA Meth. m5C:H4K20 me2
H4−4ac:H3−5ac
H3K27 me2
H4K20 me3
H3K9 me2
H3K9 me3
H3K27 me3
H4K20 me2
H2A.Z
H3K4 me3
DNA Meth. m5C
H4−4ac
H3K4 me1
H3−5ac
0.00 0.25 0.50 0.75 1.00
$\hat{\pi}$
SCAF4 – Forward
H3K4 me1:H3−5ac
H3K27 me2:DNA Meth. m5C
H3K27 me3:H3K9 me3
H3K4 me1:H4K20 me2
H3K4 me3:H3−5ac
H4K20 me2:H3−5ac
DNA Meth. m5C:H3K9 me3
H3K4 me1:H4−4ac
H4−4ac:H3−5ac
DNA Meth. m5C:H4K20 me3
H2A.Z:H3−5ac
H2A.Z:H4−4ac
H3K27 me3:H4K20 me3
H4K20 me2:H3K4 me3
H3K9 me3:H4K20 me3
H2A.Z:H3K4 me3
DNA Meth. m5C:H4K20 me2
H3K27 me3:DNA Meth. m5C
H3K4 me1:H2A.Z
H2A.Z:H4K20 me2
H4K20 me2:H4−4ac
H3K4 me3:H4−4ac
H3K27 me2
H3K9 me2
H3−5ac
H3K9 me3
H4K20 me3
H3K27 me3
H3K4 me1
DNA Meth. m5C
H3K4 me3
H4−4ac
H4K20 me2
H2A.Z
0.00 0.25 0.50 0.75 1.00
$\hat{\pi}$
SCAF4 – Reverse
H3K4 me1:H3−5ac
H3K27 me2:DNA Meth. m5C
H3K27 me3:H3K9 me3
H3K4 me1:H4K20 me2
H3K4 me3:H3−5ac
H4K20 me2:H3−5ac
DNA Meth. m5C:H3K9 me3
H3K4 me1:H4−4ac
H4−4ac:H3−5ac
DNA Meth. m5C:H4K20 me3
H2A.Z:H3−5ac
H2A.Z:H4−4ac
H3K27 me3:H4K20 me3
H4K20 me2:H3K4 me3
H3K9 me3:H4K20 me3
H2A.Z:H3K4 me3
DNA Meth. m5C:H4K20 me2
H3K27 me3:DNA Meth. m5C
H3K4 me1:H2A.Z
H2A.Z:H4K20 me2
H4K20 me2:H4−4ac
H3K4 me3:H4−4ac
H3K27 me2
H3K9 me2
H3−5ac
H3K9 me3
H4K20 me3
H3K27 me3
H3K4 me1
DNA Meth. m5C
H3K4 me3
H4−4ac
H4K20 me2
H2A.Z
0.00 0.25 0.50 0.75 1.00
$\hat{\pi}$
TDP1 – Forward
H2A.Z:H3−5ac
H2A.Z:H4−4ac
DNA Meth. m5C:H4K20 me3
H3K27 me3:H3K4 me1
H3K27 me3:H3K9 me3
H3K4 me3:H3−5ac
H2A.Z:H3K4 me3
H3K4 me1:H4K20 me2
H3K9 me3:H4K20 me3
H3K27 me3:DNA Meth. m5C
H2A.Z:H4K20 me2
H3K4 me1:H4−4ac
H3K27 me3:H4K20 me3
H3K4 me1:H3−5ac
DNA Meth. m5C:H3K9 me3
H3K27 me2:DNA Meth. m5C
H3K4 me1:H2A.Z
H4K20 me2:H3K4 me3
H4−4ac:H3−5ac
H4K20 me2:H3−5ac
H4K20 me2:H4−4ac
H3K4 me3:H4−4ac
DNA Meth. m5C:H4K20 me2
H3K27 me2
H3K9 me2
H3K4 me3
H3−5ac
H4K20 me2
H4K20 me3
H3K27 me3
DNA Meth. m5C
H2A.Z
H3K4 me1
H3K9 me3
H4−4ac
0.00 0.25 0.50 0.75 1.00
$\hat{\pi}$
TDP1 – Reverse
H2A.Z:H3−5ac
H2A.Z:H4−4ac
DNA Meth. m5C:H4K20 me3
H3K27 me3:H3K4 me1
H3K27 me3:H3K9 me3
H3K4 me3:H3−5ac
H2A.Z:H3K4 me3
H3K4 me1:H4K20 me2
H3K9 me3:H4K20 me3
H3K27 me3:DNA Meth. m5C
H2A.Z:H4K20 me2
H3K4 me1:H4−4ac
H3K27 me3:H4K20 me3
H3K4 me1:H3−5ac
DNA Meth. m5C:H3K9 me3
H3K27 me2:DNA Meth. m5C
H3K4 me1:H2A.Z
H4K20 me2:H3K4 me3
H4−4ac:H3−5ac
H4K20 me2:H3−5ac
H4K20 me2:H4−4ac
H3K4 me3:H4−4ac
DNA Meth. m5C:H4K20 me2
H3K27 me2
H3K9 me2
H3K4 me3
H3−5ac
H4K20 me2
H4K20 me3
H3K27 me3
DNA Meth. m5C
H2A.Z
H3K4 me1
H3K9 me3
H4−4ac
0.00 0.25 0.50 0.75 1.00
$\hat{\pi}$
SARNP – Forward
H3K4 me1:H2A.Z
H3K9 me2:DNA Meth. m5C
H3K4 me3:H3−5ac
H2A.Z:H3−5ac
H3K4 me1:H4−4ac
H2A.Z:H4K20 me2
H3K27 me2:DNA Meth. m5C
H4K20 me2:H3K4 me3
H4K20 me2:H3−5ac
H2A.Z:H4−4ac
H3K4 me1:H3−5ac
H2A.Z:H3K4 me3
H3K27 me3:H4K20 me3
H3K27 me3:H3K9 me3
H4−4ac:H3−5ac
H3K9 me3:H4K20 me3
DNA Meth. m5C:H4K20 me3
H3K27 me3:H3K4 me1
DNA Meth. m5C:H4K20 me2
H3K27 me3:DNA Meth. m5C
DNA Meth. m5C:H3K9 me3
H3K4 me3:H4−4ac
H4K20 me2:H4−4ac
H3K27 me2
H3K9 me2
H4K20 me3
H3K4 me3
H3−5ac
H4−4ac
H3K4 me1
H3K9 me3
DNA Meth. m5C
H2A.Z
H4K20 me2
H3K27 me3
0.00 0.25 0.50 0.75 1.00
$\hat{\pi}$
SARNP – Reverse
H3K4 me1:H2A.Z
H3K9 me2:DNA Meth. m5C
H3K4 me3:H3−5ac
H2A.Z:H3−5ac
H3K4 me1:H4−4ac
H2A.Z:H4K20 me2
H3K27 me2:DNA Meth. m5C
H4K20 me2:H3K4 me3
H4K20 me2:H3−5ac
H2A.Z:H4−4ac
H3K4 me1:H3−5ac
H2A.Z:H3K4 me3
H3K27 me3:H4K20 me3
H3K27 me3:H3K9 me3
H4−4ac:H3−5ac
H3K9 me3:H4K20 me3
DNA Meth. m5C:H4K20 me3
H3K27 me3:H3K4 me1
DNA Meth. m5C:H4K20 me2
H3K27 me3:DNA Meth. m5C
DNA Meth. m5C:H3K9 me3
H3K4 me3:H4−4ac
H4K20 me2:H4−4ac
H3K27 me2
H3K9 me2
H4K20 me3
H3K4 me3
H3−5ac
H4−4ac
H3K4 me1
H3K9 me3
DNA Meth. m5C
H2A.Z
H4K20 me2
H3K27 me3
0.00 0.25 0.50 0.75 1.00
$\hat{\pi}$
NELFA – Forward
H2A.Z:H4−4ac
H2A.Z:H3K4 me3
H3K4 me1:H4−4ac
DNA Meth. m5C:H4K20 me3
DNA Meth. m5C:H3K9 me3
DNA Meth. m5C:H4K20 me2
H2A.Z:H4K20 me2
H3K27 me3:H4K20 me3
H4−4ac:H3−5ac
H4K20 me2:H3K4 me3
H4K20 me2:H3−5ac
H4K20 me2:H4−4ac
H3K4 me3:H4−4ac
H3K27 me2
H4K20 me3
H3K9 me3
H3K9 me2
H3−5ac
H3K27 me3
H2A.Z
H3K4 me1
DNA Meth. m5C
H3K4 me3
H4K20 me2
H4−4ac
0.00 0.25 0.50 0.75 1.00
$\hat{\pi}$
NELFA – Reverse
H2A.Z:H4−4ac
H2A.Z:H3K4 me3
H3K4 me1:H4−4ac
DNA Meth. m5C:H4K20 me3
DNA Meth. m5C:H3K9 me3
DNA Meth. m5C:H4K20 me2
H2A.Z:H4K20 me2
H3K27 me3:H4K20 me3
H4−4ac:H3−5ac
H4K20 me2:H3K4 me3
H4K20 me2:H3−5ac
H4K20 me2:H4−4ac
H3K4 me3:H4−4ac
H3K27 me2
H4K20 me3
H3K9 me3
H3K9 me2
H3−5ac
H3K27 me3
H2A.Z
H3K4 me1
DNA Meth. m5C
H3K4 me3
H4K20 me2
H4−4ac
0.00 0.25 0.50 0.75 1.00
$\hat{\pi}$

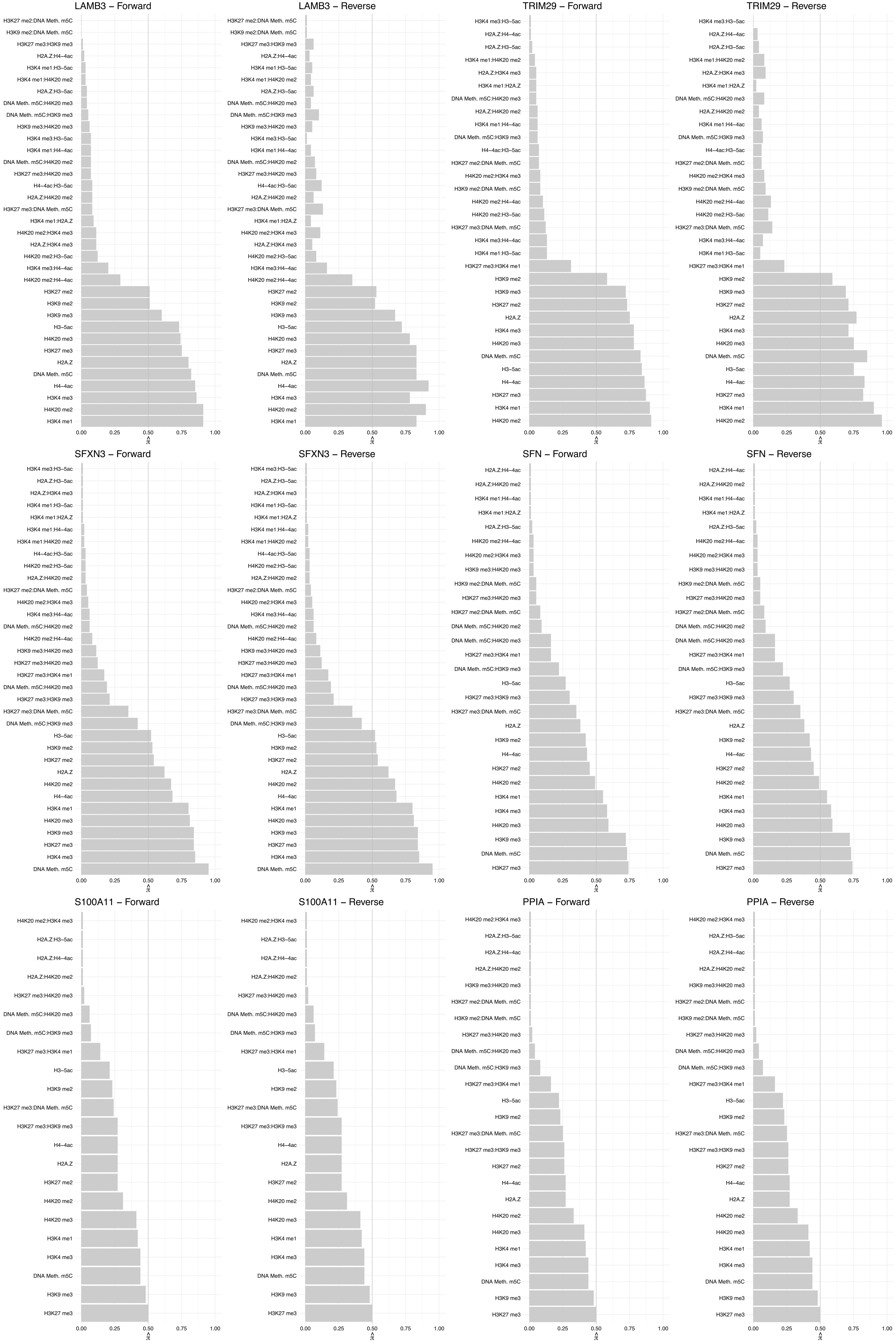

LAMB3 – Forward
H3K27 me2:DNA Meth. m5C
H3K9 me2:DNA Meth. m5C
H3K27 me3:H3K9 me3
H2A.Z:H4−4ac
H3K4 me1:H3−5ac
H3K4 me1:H4K20 me2
H2A.Z:H3−5ac
DNA Meth. m5C:H4K20 me3
DNA Meth. m5C:H3K9 me3
H3K9 me3:H4K20 me3
H3K4 me3:H3−5ac
H3K4 me1:H4−4ac
DNA Meth. m5C:H4K20 me2
H3K27 me3:H4K20 me3
H4−4ac:H3−5ac
H2A.Z:H4K20 me2
H3K27 me3:DNA Meth. m5C
H3K4 me1:H2A.Z
H4K20 me2:H3K4 me3
H2A.Z:H3K4 me3
H4K20 me2:H3−5ac
H3K4 me3:H4−4ac
H4K20 me2:H4−4ac
H3K27 me2
H3K9 me2
H3K9 me3
H3−5ac
H4K20 me3
H3K27 me3
H2A.Z
DNA Meth. m5C
H4−4ac
H3K4 me3
H4K20 me2
H3K4 me1
0.00 0.25 0.50 0.75 1.00
π̂
LAMB3 – Reverse
H3K27 me2:DNA Meth. m5C
H3K9 me2:DNA Meth. m5C
H3K27 me3:H3K9 me3
H2A.Z:H4−4ac
H3K4 me1:H3−5ac
H3K4 me1:H4K20 me2
H2A.Z:H3−5ac
DNA Meth. m5C:H4K20 me3
DNA Meth. m5C:H3K9 me3
H3K9 me3:H4K20 me3
H3K4 me3:H3−5ac
H3K4 me1:H4−4ac
DNA Meth. m5C:H4K20 me2
H3K27 me3:H4K20 me3
H4−4ac:H3−5ac
H2A.Z:H4K20 me2
H3K27 me3:DNA Meth. m5C
H3K4 me1:H2A.Z
H4K20 me2:H3K4 me3
H2A.Z:H3K4 me3
H4K20 me2:H3−5ac
H3K4 me3:H4−4ac
H4K20 me2:H4−4ac
H3K27 me2
H3K9 me2
H3K9 me3
H3−5ac
H4K20 me3
H3K27 me3
H2A.Z
DNA Meth. m5C
H4−4ac
H3K4 me3
H4K20 me2
H3K4 me1
0.00 0.25 0.50 0.75 1.00
π̂
TRIM29 – Forward
H3K4 me3:H3−5ac
H2A.Z:H4−4ac
H2A.Z:H3−5ac
H3K4 me1:H4K20 me2
H2A.Z:H3K4 me3
H3K4 me1:H2A.Z
DNA Meth. m5C:H4K20 me3
H2A.Z:H4K20 me2
H3K4 me1:H4−4ac
DNA Meth. m5C:H3K9 me3
H4−4ac:H3−5ac
H3K27 me2:DNA Meth. m5C
H4K20 me2:H3K4 me3
H3K9 me2:DNA Meth. m5C
H4K20 me2:H4−4ac
H4K20 me2:H3−5ac
H3K27 me3:DNA Meth. m5C
H3K4 me3:H4−4ac
H3K4 me1:H3−5ac
H3K27 me3:H3K4 me1
H3K9 me2
H3K9 me3
H3K27 me2
H2A.Z
H3K4 me3
H4K20 me3
DNA Meth. m5C
H3−5ac
H4−4ac
H3K27 me3
H3K4 me1
H4K20 me2
0.00 0.25 0.50 0.75 1.00
π̂
TRIM29 – Reverse
H3K4 me3:H3−5ac
H2A.Z:H4−4ac
H2A.Z:H3−5ac
H3K4 me1:H4K20 me2
H2A.Z:H3K4 me3
H3K4 me1:H2A.Z
DNA Meth. m5C:H4K20 me3
H2A.Z:H4K20 me2
H3K4 me1:H4−4ac
DNA Meth. m5C:H3K9 me3
H4−4ac:H3−5ac
H3K27 me2:DNA Meth. m5C
H4K20 me2:H3K4 me3
H3K9 me2:DNA Meth. m5C
H4K20 me2:H4−4ac
H4K20 me2:H3−5ac
H3K27 me3:DNA Meth. m5C
H3K4 me3:H4−4ac
H3K4 me1:H3−5ac
H3K27 me3:H3K4 me1
H3K9 me2
H3K9 me3
H3K27 me2
H2A.Z
H3K4 me3
H4K20 me3
DNA Meth. m5C
H3−5ac
H4−4ac
H3K27 me3
H3K4 me1
H4K20 me2
0.00 0.25 0.50 0.75 1.00
π̂
SFXN3 – Forward
H3K4 me3:H3−5ac
H2A.Z:H3−5ac
H2A.Z:H3K4 me3
H3K4 me1:H3−5ac
H3K4 me1:H2A.Z
H3K4 me1:H4−4ac
H3K4 me1:H4K20 me2
H4−4ac:H3−5ac
H4K20 me2:H3−5ac
H2A.Z:H4K20 me2
H3K27 me2:DNA Meth. m5C
H4K20 me2:H3K4 me3
H3K4 me3:H4−4ac
DNA Meth. m5C:H4K20 me2
H4K20 me2:H4−4ac
H3K9 me3:H4K20 me3
H3K27 me3:H4K20 me3
H3K27 me3:H3K4 me1
DNA Meth. m5C:H4K20 me3
H3K27 me3:H3K9 me3
H3K27 me3:DNA Meth. m5C
DNA Meth. m5C:H3K9 me3
H3−5ac
H3K9 me2
H3K27 me2
H2A.Z
H4K20 me2
H4−4ac
H3K4 me1
H4K20 me3
H3K9 me3
H3K27 me3
H3K4 me3
DNA Meth. m5C
0.00 0.25 0.50 0.75 1.00
π̂
SFXN3 – Reverse
H3K4 me3:H3−5ac
H2A.Z:H3−5ac
H2A.Z:H3K4 me3
H3K4 me1:H3−5ac
H3K4 me1:H2A.Z
H3K4 me1:H4−4ac
H3K4 me1:H4K20 me2
H4−4ac:H3−5ac
H4K20 me2:H3−5ac
H2A.Z:H4K20 me2
H3K27 me2:DNA Meth. m5C
H4K20 me2:H3K4 me3
H3K4 me3:H4−4ac
DNA Meth. m5C:H4K20 me2
H4K20 me2:H4−4ac
H3K9 me3:H4K20 me3
H3K27 me3:H4K20 me3
H3K27 me3:H3K4 me1
DNA Meth. m5C:H4K20 me3
H3K27 me3:H3K9 me3
H3K27 me3:DNA Meth. m5C
DNA Meth. m5C:H3K9 me3
H3−5ac
H3K9 me2
H3K27 me2
H2A.Z
H4K20 me2
H4−4ac
H3K4 me1
H4K20 me3
H3K9 me3
H3K27 me3
H3K4 me3
DNA Meth. m5C
0.00 0.25 0.50 0.75 1.00
π̂
SFN – Forward
H2A.Z:H4−4ac
H2A.Z:H4K20 me2
H3K4 me1:H4−4ac
H3K4 me1:H2A.Z
H2A.Z:H3−5ac
H4K20 me2:H4−4ac
H4K20 me2:H3K4 me3
H3K9 me3:H4K20 me3
H3K9 me2:DNA Meth. m5C
H3K27 me3:H4K20 me3
H3K27 me2:DNA Meth. m5C
DNA Meth. m5C:H4K20 me2
DNA Meth. m5C:H4K20 me3
H3K27 me3:H3K4 me1
DNA Meth. m5C:H3K9 me3
H3−5ac
H3K27 me3:H3K9 me3
H3K27 me3:DNA Meth. m5C
H2A.Z
H3K9 me2
H4−4ac
H3K27 me2
H4K20 me2
H3K4 me1
H3K4 me3
H4K20 me3
H3K9 me3
DNA Meth. m5C
H3K27 me3
0.00 0.25 0.50 0.75 1.00
π̂
SFN – Reverse
H2A.Z:H4−4ac
H2A.Z:H4K20 me2
H3K4 me1:H4−4ac
H3K4 me1:H2A.Z
H2A.Z:H3−5ac
H4K20 me2:H4−4ac
H4K20 me2:H3K4 me3
H3K9 me3:H4K20 me3
H3K9 me2:DNA Meth. m5C
H3K27 me3:H4K20 me3
H3K27 me2:DNA Meth. m5C
DNA Meth. m5C:H4K20 me2
DNA Meth. m5C:H4K20 me3
H3K27 me3:H3K4 me1
DNA Meth. m5C:H3K9 me3
H3−5ac
H3K27 me3:H3K9 me3
H3K27 me3:DNA Meth. m5C
H2A.Z
H3K9 me2
H4−4ac
H3K27 me2
H4K20 me2
H3K4 me1
H3K4 me3
H4K20 me3
H3K9 me3
DNA Meth. m5C
H3K27 me3
0.00 0.25 0.50 0.75 1.00
π̂
S100A11 – Forward
H4K20 me2:H3K4 me3
H2A.Z:H3−5ac
H2A.Z:H4−4ac
H2A.Z:H4K20 me2
H3K27 me3:H4K20 me3
DNA Meth. m5C:H4K20 me3
DNA Meth. m5C:H3K9 me3
H3K27 me3:H3K4 me1
H3−5ac
H3K9 me2
H3K27 me3:DNA Meth. m5C
H3K27 me3:H3K9 me3
H4−4ac
H2A.Z
H3K27 me2
H4K20 me2
H4K20 me3
H3K4 me1
H3K4 me3
DNA Meth. m5C
H3K9 me3
H3K27 me3
0.00 0.25 0.50 0.75 1.00
π̂
S100A11 – Reverse
H4K20 me2:H3K4 me3
H2A.Z:H3−5ac
H2A.Z:H4−4ac
H2A.Z:H4K20 me2
H3K27 me3:H4K20 me3
DNA Meth. m5C:H4K20 me3
DNA Meth. m5C:H3K9 me3
H3K27 me3:H3K4 me1
H3−5ac
H3K9 me2
H3K27 me3:DNA Meth. m5C
H3K27 me3:H3K9 me3
H4−4ac
H2A.Z
H3K27 me2
H4K20 me2
H4K20 me3
H3K4 me1
H3K4 me3
DNA Meth. m5C
H3K9 me3
H3K27 me3
0.00 0.25 0.50 0.75 1.00
π̂
PPIA – Forward
H4K20 me2:H3K4 me3
H2A.Z:H3−5ac
H2A.Z:H4−4ac
H2A.Z:H4K20 me2
H3K9 me3:H4K20 me3
H3K27 me2:DNA Meth. m5C
H3K9 me2:DNA Meth. m5C
H3K27 me3:H4K20 me3
DNA Meth. m5C:H4K20 me3
DNA Meth. m5C:H3K9 me3
H3K27 me3:H3K4 me1
H3−5ac
H3K9 me2
H3K27 me3:DNA Meth. m5C
H3K27 me3:H3K9 me3
H3K27 me2
H4−4ac
H2A.Z
H4K20 me2
H4K20 me3
H3K4 me1
H3K4 me3
DNA Meth. m5C
H3K9 me3
H3K27 me3
0.00 0.25 0.50 0.75 1.00
π̂
PPIA – Reverse
H4K20 me2:H3K4 me3
H2A.Z:H3−5ac
H2A.Z:H4−4ac
H2A.Z:H4K20 me2
H3K9 me3:H4K20 me3
H3K27 me2:DNA Meth. m5C
H3K9 me2:DNA Meth. m5C
H3K27 me3:H4K20 me3
DNA Meth. m5C:H4K20 me3
DNA Meth. m5C:H3K9 me3
H3K27 me3:H3K4 me1
H3−5ac
H3K9 me2
H3K27 me3:DNA Meth. m5C
H3K27 me3:H3K9 me3
H3K27 me2
H4−4ac
H2A.Z
H4K20 me2
H4K20 me3
H3K4 me1
H3K4 me3
DNA Meth. m5C
H3K9 me3
H3K27 me3
0.00 0.25 0.50 0.75 1.00
π̂

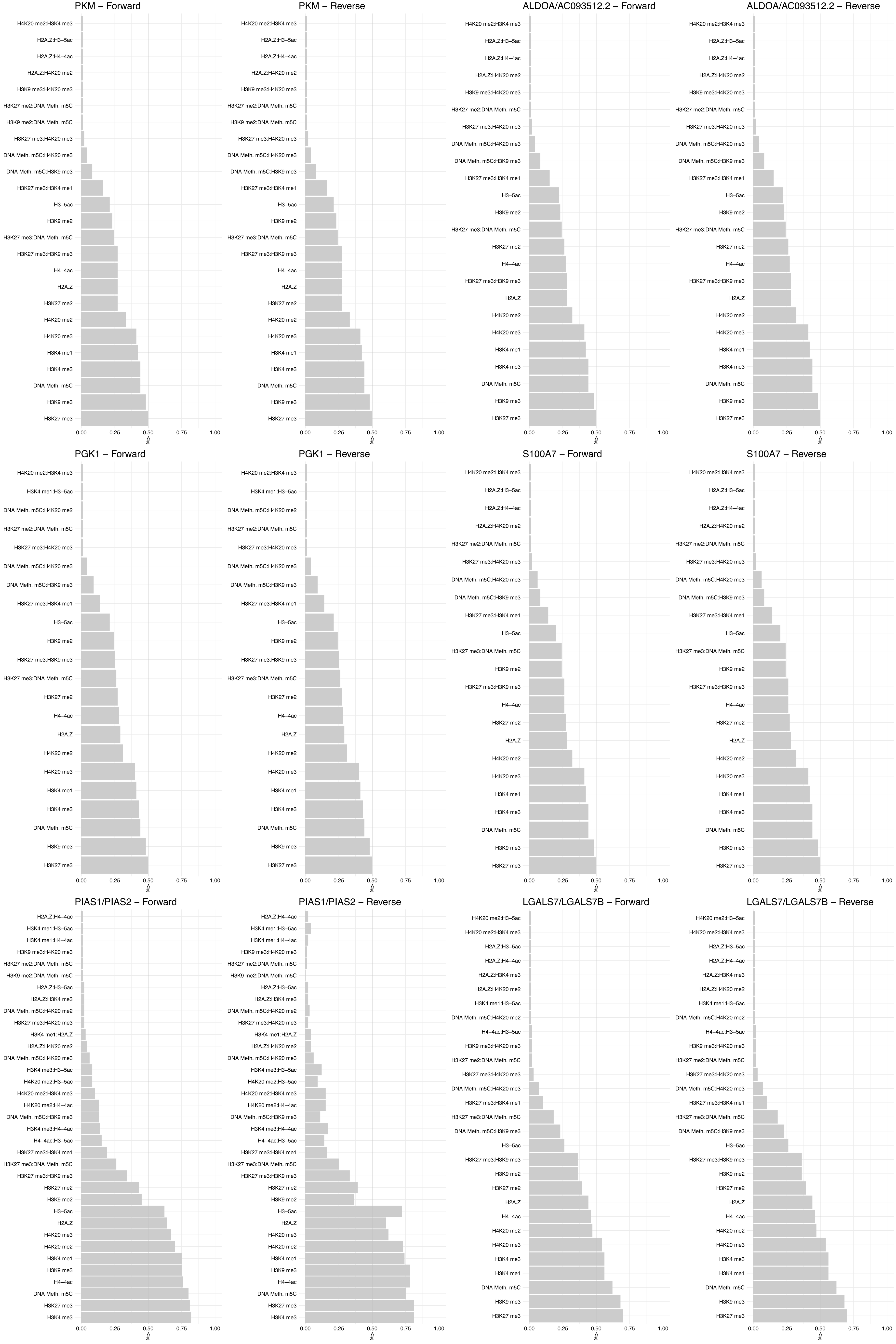
PKM – Forward
PKM – Reverse
ALDOA/AC093512.2 – Forward
ALDOA/AC093512.2 – Reverse
PGK1 – Forward
PGK1 – Reverse
S100A7 – Forward
S100A7 – Reverse
PIAS1/PIAS2 – Forward
PIAS1/PIAS2 – Reverse
LGALS7/LGALS7B – Forward
LGALS7/LGALS7B – Reverse
H4K20 me2:H3K4 me3
H2A.Z:H3–5ac
H2A.Z:H4–4ac
H2A.Z:H4K20 me2
H3K9 me3:H4K20 me3
H3K27 me2:DNA Meth. m5C
H3K9 me2:DNA Meth. m5C
H3K27 me3:H4K20 me3
DNA Meth. m5C:H4K20 me3
DNA Meth. m5C:H3K9 me3
H3K27 me3:H3K4 me1
H3–5ac
H3K9 me2
H3K27 me3:DNA Meth. m5C
H3K27 me3:H3K9 me3
H4–4ac
H2A.Z
H3K27 me2
H4K20 me2
H4K20 me3
H3K4 me1
H3K4 me3
DNA Meth. m5C
H3K9 me3
H3K27 me3
H3K4 me1:H3–5ac
DNA Meth. m5C:H4K20 me2
H3K4 me1:H4–4ac
H2A.Z:H3K4 me3
H3K4 me1:H2A.Z
H3K4 me3:H3–5ac
H4K20 me2:H3–5ac
H4K20 me2:H4–4ac
H3K4 me3:H4–4ac
H4–4ac:H3–5ac
0.00
0.25
0.50
0.75
1.00
$\hat{\pi}$

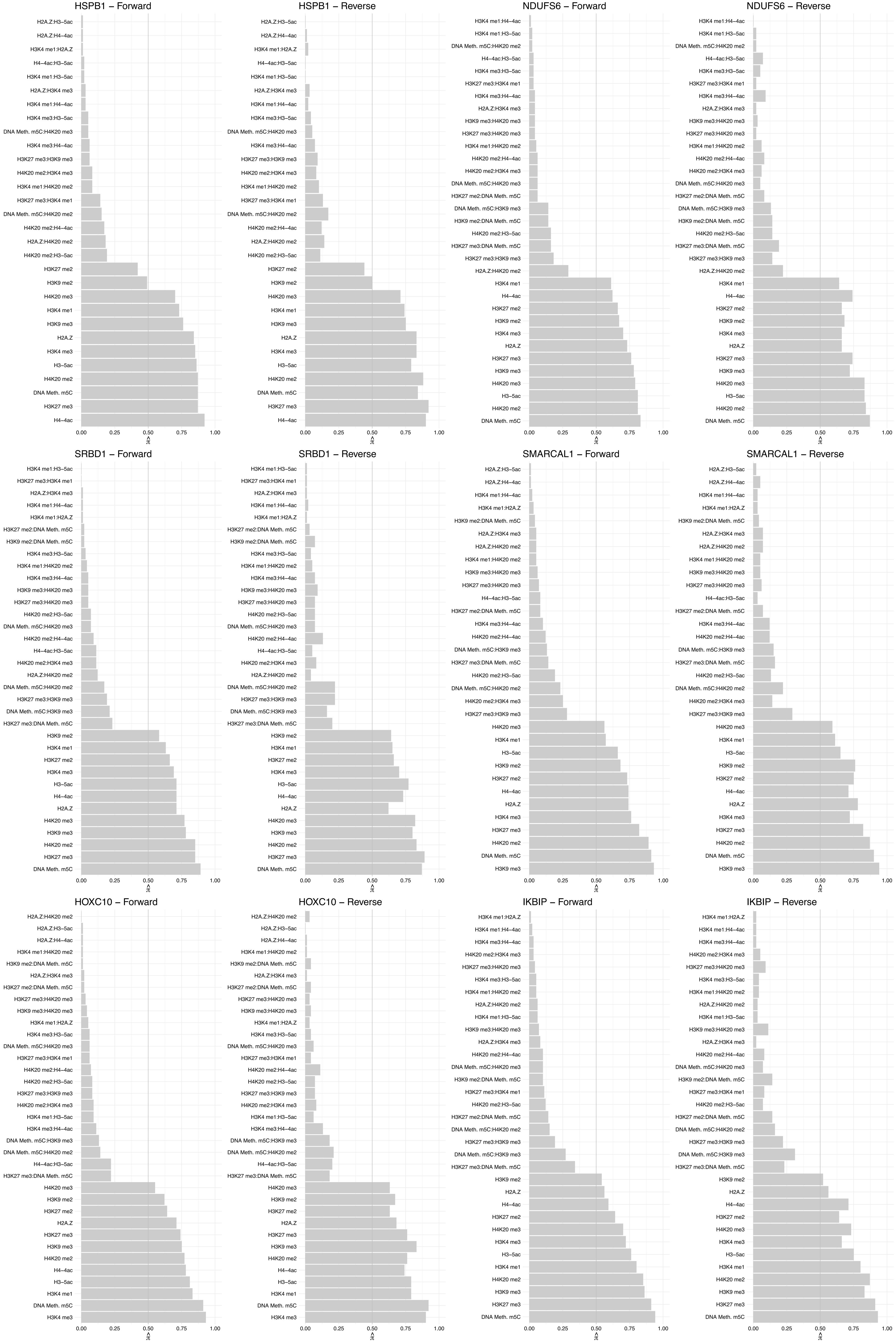
HSPB1 – Forward
HSPB1 – Reverse
H2A.Z:H3−5ac
H2A.Z:H4−4ac
H3K4 me1:H2A.Z
H4−4ac:H3−5ac
H3K4 me1:H3−5ac
H2A.Z:H3K4 me3
H3K4 me1:H4−4ac
H3K4 me3:H3−5ac
DNA Meth. m5C:H4K20 me3
H3K4 me3:H4−4ac
H3K27 me3:H3K9 me3
H4K20 me2:H3K4 me3
H3K4 me1:H4K20 me2
H3K27 me3:H3K4 me1
DNA Meth. m5C:H4K20 me2
H4K20 me2:H4−4ac
H2A.Z:H4K20 me2
H4K20 me2:H3−5ac
H3K27 me2
H3K9 me2
H4K20 me3
H3K4 me1
H3K9 me3
H2A.Z
H3K4 me3
H3−5ac
H4K20 me2
DNA Meth. m5C
H3K27 me3
H4−4ac
NDUFS6 – Forward
NDUFS6 – Reverse
H3K4 me1:H4−4ac
H3K4 me1:H3−5ac
DNA Meth. m5C:H4K20 me2
H4−4ac:H3−5ac
H3K4 me3:H3−5ac
H3K27 me3:H3K4 me1
H3K4 me3:H4−4ac
H2A.Z:H3K4 me3
H3K9 me3:H4K20 me3
H3K27 me3:H4K20 me3
H3K4 me1:H4K20 me2
H4K20 me2:H4−4ac
H4K20 me2:H3K4 me3
DNA Meth. m5C:H4K20 me3
H3K27 me2:DNA Meth. m5C
DNA Meth. m5C:H3K9 me3
H3K9 me2:DNA Meth. m5C
H4K20 me2:H3−5ac
H3K27 me3:DNA Meth. m5C
H3K27 me3:H3K9 me3
H2A.Z:H4K20 me2
H3K4 me1
H4−4ac
H3K27 me2
H3K9 me2
H3K4 me3
H2A.Z
H3K27 me3
H3K9 me3
H4K20 me3
H3−5ac
H4K20 me2
DNA Meth. m5C
SRBD1 – Forward
SRBD1 – Reverse
H3K4 me1:H3−5ac
H3K27 me3:H3K4 me1
H2A.Z:H3K4 me3
H3K4 me1:H4−4ac
H3K4 me1:H2A.Z
H3K27 me2:DNA Meth. m5C
H3K9 me2:DNA Meth. m5C
H3K4 me3:H3−5ac
H3K4 me1:H4K20 me2
H3K4 me3:H4−4ac
H3K9 me3:H4K20 me3
H3K27 me3:H4K20 me3
H4K20 me2:H3−5ac
DNA Meth. m5C:H4K20 me3
H4K20 me2:H4−4ac
H4−4ac:H3−5ac
H4K20 me2:H3K4 me3
H2A.Z:H4K20 me2
DNA Meth. m5C:H4K20 me2
H3K27 me3:H3K9 me3
DNA Meth. m5C:H3K9 me3
H3K27 me3:DNA Meth. m5C
H3K9 me2
H3K4 me1
H3K27 me2
H3K4 me3
H3−5ac
H4−4ac
H2A.Z
H4K20 me3
H3K9 me3
H4K20 me2
H3K27 me3
DNA Meth. m5C
SMARCAL1 – Forward
SMARCAL1 – Reverse
H2A.Z:H3−5ac
H2A.Z:H4−4ac
H3K4 me1:H4−4ac
H3K4 me1:H2A.Z
H3K9 me2:DNA Meth. m5C
H2A.Z:H3K4 me3
H2A.Z:H4K20 me2
H3K4 me1:H4K20 me2
H3K9 me3:H4K20 me3
H3K27 me3:H4K20 me3
H4−4ac:H3−5ac
H3K27 me2:DNA Meth. m5C
H3K4 me3:H4−4ac
H4K20 me2:H4−4ac
DNA Meth. m5C:H3K9 me3
H3K27 me3:DNA Meth. m5C
H4K20 me2:H3−5ac
DNA Meth. m5C:H4K20 me2
H4K20 me2:H3K4 me3
H3K27 me3:H3K9 me3
H4K20 me3
H3K4 me1
H3−5ac
H3K9 me2
H3K27 me2
H4−4ac
H2A.Z
H3K4 me3
H3K27 me3
H4K20 me2
DNA Meth. m5C
H3K9 me3
HOXC10 – Forward
HOXC10 – Reverse
H2A.Z:H4K20 me2
H2A.Z:H3−5ac
H2A.Z:H4−4ac
H3K4 me1:H4K20 me2
H3K9 me2:DNA Meth. m5C
H2A.Z:H3K4 me3
H3K27 me2:DNA Meth. m5C
H3K27 me3:H4K20 me3
H3K9 me3:H4K20 me3
H3K4 me1:H2A.Z
H3K4 me3:H3−5ac
DNA Meth. m5C:H4K20 me3
H3K27 me3:H3K4 me1
H4K20 me2:H4−4ac
H4K20 me2:H3−5ac
H3K27 me3:H3K9 me3
H4K20 me2:H3K4 me3
H3K4 me1:H3−5ac
H3K4 me3:H4−4ac
DNA Meth. m5C:H3K9 me3
DNA Meth. m5C:H4K20 me2
H4−4ac:H3−5ac
H3K27 me3:DNA Meth. m5C
H4K20 me3
H3K9 me2
H3K27 me2
H2A.Z
H3K27 me3
H3K9 me3
H4K20 me2
H4−4ac
H3−5ac
H3K4 me1
DNA Meth. m5C
H3K4 me3
IKBIP – Forward
IKBIP – Reverse
H3K4 me1:H2A.Z
H3K4 me1:H4−4ac
H3K4 me3:H4−4ac
H4K20 me2:H3K4 me3
H3K27 me3:H4K20 me3
H3K4 me3:H3−5ac
H3K4 me1:H4K20 me2
H2A.Z:H4K20 me2
H3K4 me1:H3−5ac
H3K9 me3:H4K20 me3
H2A.Z:H3K4 me3
H4K20 me2:H4−4ac
DNA Meth. m5C:H4K20 me3
H3K9 me2:DNA Meth. m5C
H3K27 me3:H3K4 me1
H4K20 me2:H3−5ac
H3K27 me2:DNA Meth. m5C
DNA Meth. m5C:H4K20 me2
H3K27 me3:H3K9 me3
DNA Meth. m5C:H3K9 me3
H3K27 me3:DNA Meth. m5C
H3K9 me2
H2A.Z
H4−4ac
H3K27 me2
H4K20 me3
H3K4 me3
H3−5ac
H3K4 me1
H4K20 me2
H3K9 me3
H3K27 me3
DNA Meth. m5C
0.00 0.25 0.50 0.75 1.00
π̂

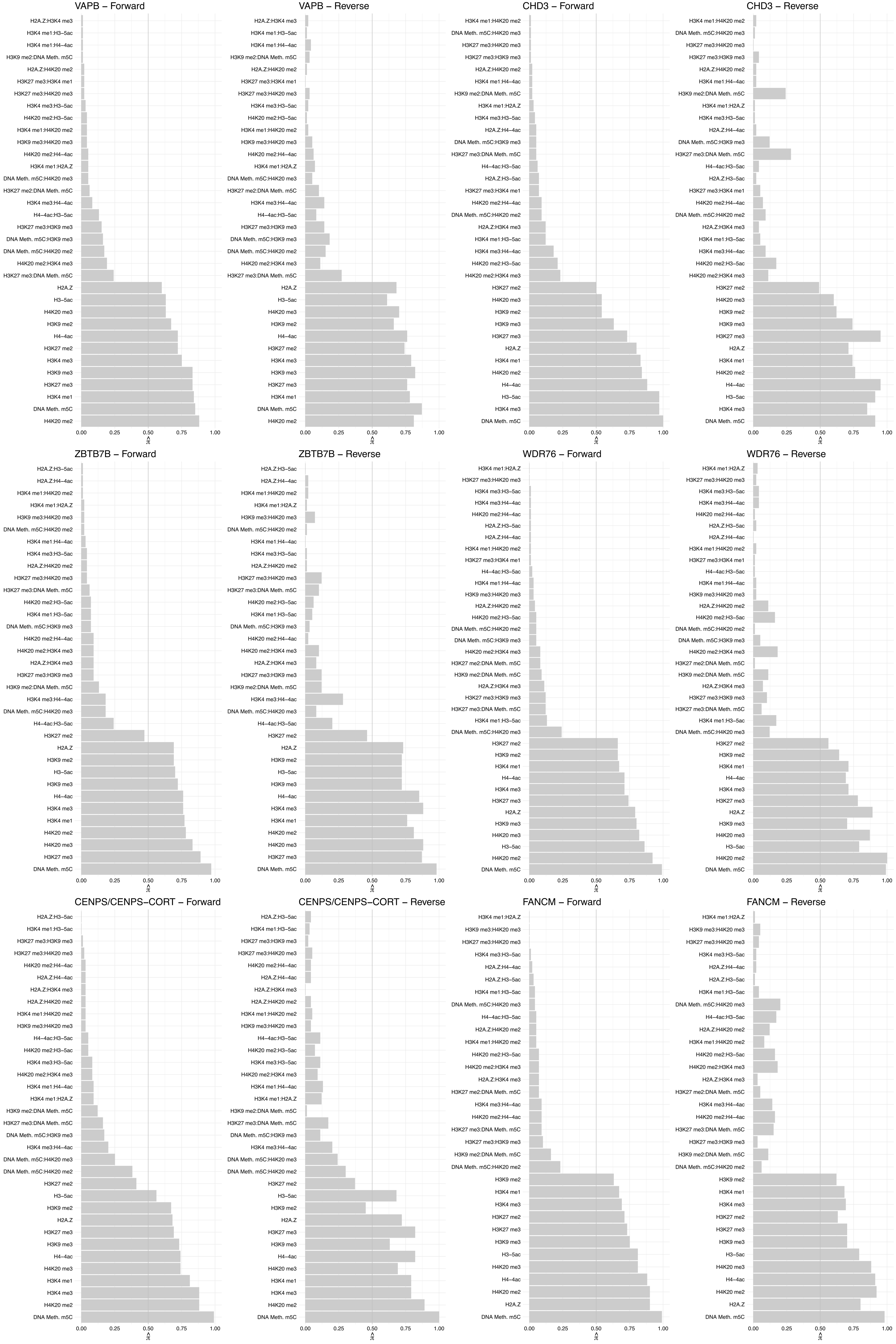
VAPB – Forward
VAPB – Reverse
CHD3 – Forward
CHD3 – Reverse
ZBTB7B – Forward
ZBTB7B – Reverse
WDR76 – Forward
WDR76 – Reverse
CENPS/CENPS–CORT – Forward
CENPS/CENPS–CORT – Reverse
FANCM – Forward
FANCM – Reverse
0.00
0.25
0.50
0.75
1.00
$\hat{\pi}$

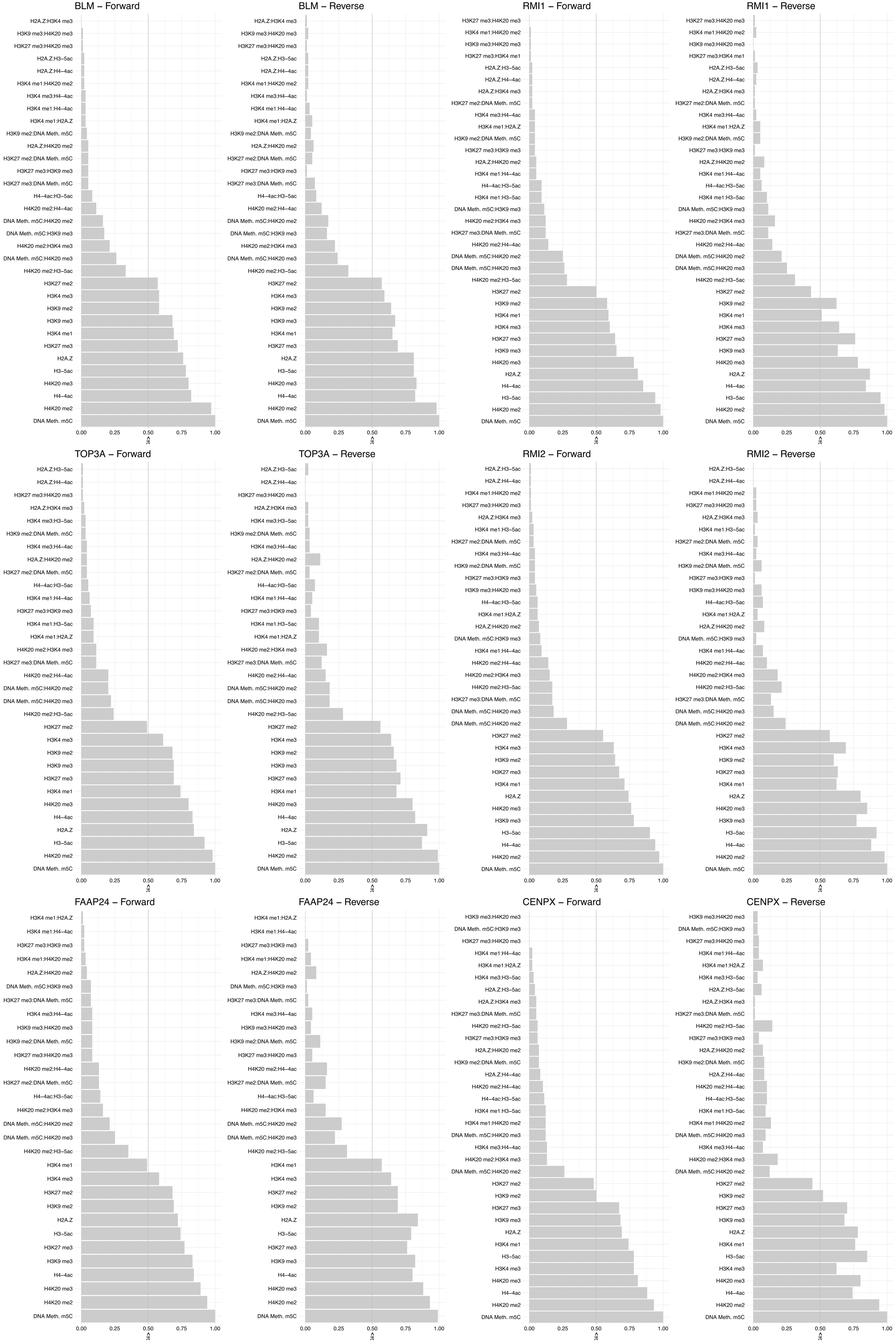

BLM – Forward
BLM – Reverse
H2A.Z:H3K4 me3
H3K9 me3:H4K20 me3
H3K27 me3:H4K20 me3
H2A.Z:H3−5ac
H2A.Z:H4−4ac
H3K4 me1:H4K20 me2
H3K4 me3:H4−4ac
H3K4 me1:H4−4ac
H3K4 me1:H2A.Z
H3K9 me2:DNA Meth. m5C
H2A.Z:H4K20 me2
H3K27 me2:DNA Meth. m5C
H3K27 me3:H3K9 me3
H3K27 me3:DNA Meth. m5C
H4−4ac:H3−5ac
H4K20 me2:H4−4ac
DNA Meth. m5C:H4K20 me2
DNA Meth. m5C:H3K9 me3
H4K20 me2:H3K4 me3
DNA Meth. m5C:H4K20 me3
H4K20 me2:H3−5ac
H3K27 me2
H3K4 me3
H3K9 me2
H3K9 me3
H3K4 me1
H3K27 me3
H2A.Z
H3−5ac
H4K20 me3
H4−4ac
H4K20 me2
DNA Meth. m5C
0.00
0.25
0.50
0.75
1.00
π̂
RMI1 – Forward
RMI1 – Reverse
H3K27 me3:H4K20 me3
H3K4 me1:H4K20 me2
H3K9 me3:H4K20 me3
H3K27 me3:H3K4 me1
H2A.Z:H3−5ac
H2A.Z:H4−4ac
H2A.Z:H3K4 me3
H3K27 me2:DNA Meth. m5C
H3K4 me3:H4−4ac
H3K4 me1:H2A.Z
H3K9 me2:DNA Meth. m5C
H3K27 me3:H3K9 me3
H2A.Z:H4K20 me2
H3K4 me1:H4−4ac
H4−4ac:H3−5ac
H3K4 me1:H3−5ac
DNA Meth. m5C:H3K9 me3
H4K20 me2:H3K4 me3
H3K27 me3:DNA Meth. m5C
H4K20 me2:H4−4ac
DNA Meth. m5C:H4K20 me2
DNA Meth. m5C:H4K20 me3
H4K20 me2:H3−5ac
H3K27 me2
H3K9 me2
H3K4 me1
H3K4 me3
H3K27 me3
H3K9 me3
H4K20 me3
H2A.Z
H4−4ac
H3−5ac
H4K20 me2
DNA Meth. m5C
TOP3A – Forward
TOP3A – Reverse
H2A.Z:H3−5ac
H2A.Z:H4−4ac
H3K27 me3:H4K20 me3
H2A.Z:H3K4 me3
H3K4 me3:H3−5ac
H3K9 me2:DNA Meth. m5C
H3K4 me3:H4−4ac
H2A.Z:H4K20 me2
H3K27 me2:DNA Meth. m5C
H4−4ac:H3−5ac
H3K4 me1:H4−4ac
H3K27 me3:H3K9 me3
H3K4 me1:H3−5ac
H3K4 me1:H2A.Z
H4K20 me2:H3K4 me3
H3K27 me3:DNA Meth. m5C
H4K20 me2:H4−4ac
DNA Meth. m5C:H4K20 me2
DNA Meth. m5C:H4K20 me3
H4K20 me2:H3−5ac
H3K27 me2
H3K4 me3
H3K9 me2
H3K9 me3
H3K27 me3
H3K4 me1
H4K20 me3
H4−4ac
H2A.Z
H3−5ac
H4K20 me2
DNA Meth. m5C
RMI2 – Forward
RMI2 – Reverse
H2A.Z:H3−5ac
H2A.Z:H4−4ac
H3K4 me1:H4K20 me2
H3K27 me3:H4K20 me3
H2A.Z:H3K4 me3
H3K4 me1:H3−5ac
H3K27 me2:DNA Meth. m5C
H3K4 me3:H4−4ac
H3K9 me2:DNA Meth. m5C
H3K27 me3:H3K9 me3
H3K9 me3:H4K20 me3
H4−4ac:H3−5ac
H3K4 me1:H2A.Z
H2A.Z:H4K20 me2
DNA Meth. m5C:H3K9 me3
H3K4 me1:H4−4ac
H4K20 me2:H4−4ac
H4K20 me2:H3K4 me3
H4K20 me2:H3−5ac
H3K27 me3:DNA Meth. m5C
DNA Meth. m5C:H4K20 me3
DNA Meth. m5C:H4K20 me2
H3K27 me2
H3K4 me3
H3K9 me2
H3K27 me3
H3K4 me1
H2A.Z
H4K20 me3
H3K9 me3
H3−5ac
H4−4ac
H4K20 me2
DNA Meth. m5C
FAAP24 – Forward
FAAP24 – Reverse
H3K4 me1:H2A.Z
H3K4 me1:H4−4ac
H3K27 me3:H3K9 me3
H3K4 me1:H4K20 me2
H2A.Z:H4K20 me2
DNA Meth. m5C:H3K9 me3
H3K27 me3:DNA Meth. m5C
H3K4 me3:H4−4ac
H3K9 me3:H4K20 me3
H3K9 me2:DNA Meth. m5C
H3K27 me3:H4K20 me3
H4K20 me2:H4−4ac
H3K27 me2:DNA Meth. m5C
H4−4ac:H3−5ac
H4K20 me2:H3K4 me3
DNA Meth. m5C:H4K20 me2
DNA Meth. m5C:H4K20 me3
H4K20 me2:H3−5ac
H3K4 me1
H3K4 me3
H3K27 me2
H3K9 me2
H2A.Z
H3−5ac
H3K27 me3
H3K9 me3
H4−4ac
H4K20 me3
H4K20 me2
DNA Meth. m5C
CENPX – Forward
CENPX – Reverse
H3K9 me3:H4K20 me3
DNA Meth. m5C:H3K9 me3
H3K27 me3:H4K20 me3
H3K4 me1:H4−4ac
H3K4 me1:H2A.Z
H3K4 me3:H3−5ac
H2A.Z:H3−5ac
H2A.Z:H3K4 me3
H3K27 me3:DNA Meth. m5C
H4K20 me2:H3−5ac
H3K27 me3:H3K9 me3
H2A.Z:H4K20 me2
H3K9 me2:DNA Meth. m5C
H2A.Z:H4−4ac
H4K20 me2:H4−4ac
H4−4ac:H3−5ac
H3K4 me1:H3−5ac
H3K4 me1:H4K20 me2
DNA Meth. m5C:H4K20 me3
H3K4 me3:H4−4ac
H4K20 me2:H3K4 me3
DNA Meth. m5C:H4K20 me2
H3K27 me2
H3K9 me2
H3K27 me3
H3K9 me3
H2A.Z
H3K4 me1
H3−5ac
H3K4 me3
H4K20 me3
H4−4ac
H4K20 me2
DNA Meth. m5C

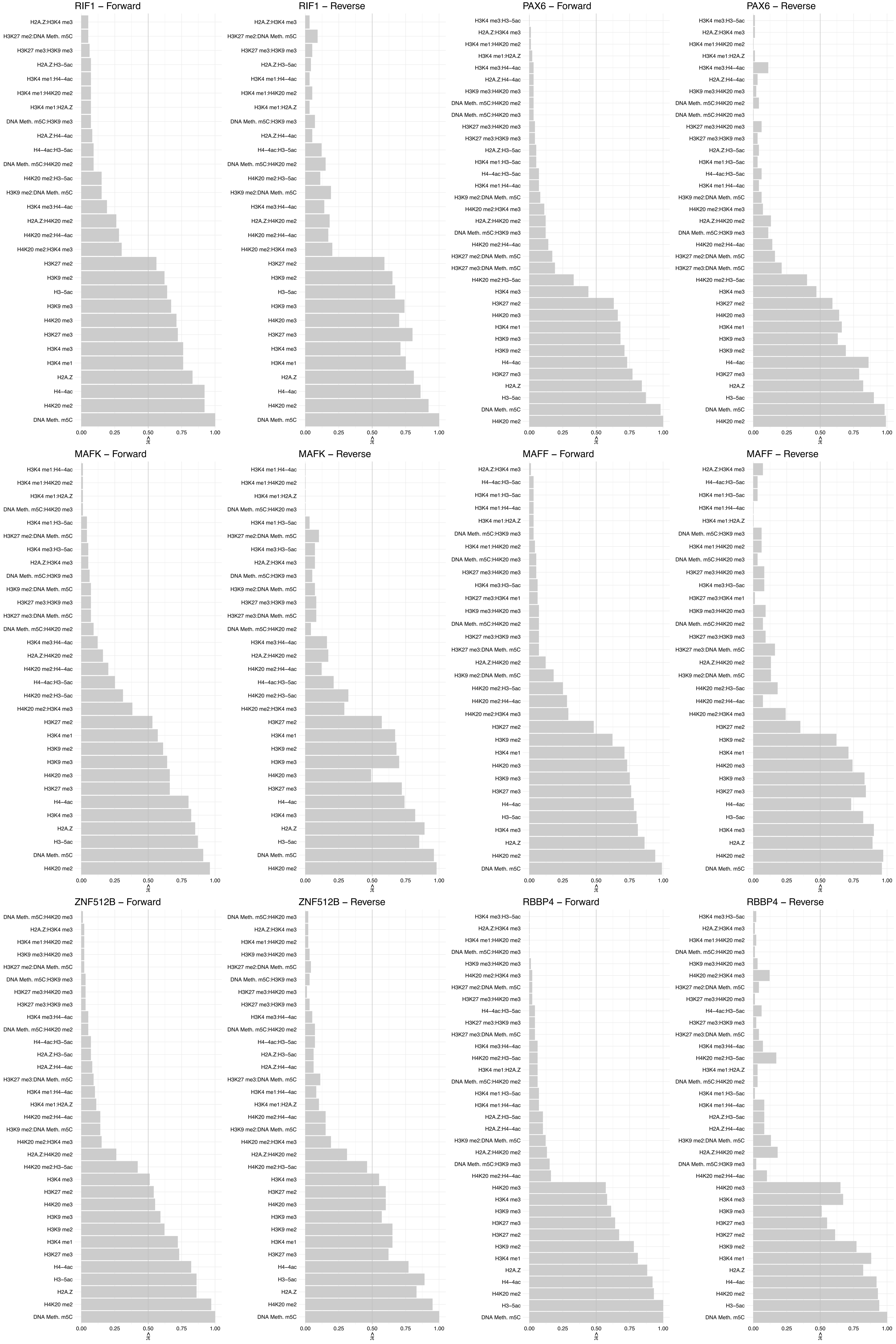
RIF1 – Forward
H2A.Z:H3K4 me3
H3K27 me2:DNA Meth. m5C
H3K27 me3:H3K9 me3
H2A.Z:H3−5ac
H3K4 me1:H4−4ac
H3K4 me1:H4K20 me2
H3K4 me1:H2A.Z
DNA Meth. m5C:H3K9 me3
H2A.Z:H4−4ac
H4−4ac:H3−5ac
DNA Meth. m5C:H4K20 me2
H4K20 me2:H3−5ac
H3K9 me2:DNA Meth. m5C
H3K4 me3:H4−4ac
H2A.Z:H4K20 me2
H4K20 me2:H4−4ac
H4K20 me2:H3K4 me3
H3K27 me2
H3K9 me2
H3−5ac
H3K9 me3
H4K20 me3
H3K27 me3
H3K4 me3
H3K4 me1
H2A.Z
H4−4ac
H4K20 me2
DNA Meth. m5C
0.00
0.25
0.50
0.75
1.00
π̂
RIF1 – Reverse
H2A.Z:H3K4 me3
H3K27 me2:DNA Meth. m5C
H3K27 me3:H3K9 me3
H2A.Z:H3−5ac
H3K4 me1:H4−4ac
H3K4 me1:H4K20 me2
H3K4 me1:H2A.Z
DNA Meth. m5C:H3K9 me3
H2A.Z:H4−4ac
H4−4ac:H3−5ac
DNA Meth. m5C:H4K20 me2
H4K20 me2:H3−5ac
H3K9 me2:DNA Meth. m5C
H3K4 me3:H4−4ac
H2A.Z:H4K20 me2
H4K20 me2:H4−4ac
H4K20 me2:H3K4 me3
H3K27 me2
H3K9 me2
H3−5ac
H3K9 me3
H4K20 me3
H3K27 me3
H3K4 me3
H3K4 me1
H2A.Z
H4−4ac
H4K20 me2
DNA Meth. m5C
0.00
0.25
0.50
0.75
1.00
π̂
PAX6 – Forward
H3K4 me3:H3−5ac
H2A.Z:H3K4 me3
H3K4 me1:H4K20 me2
H3K4 me1:H2A.Z
H3K4 me3:H4−4ac
H2A.Z:H4−4ac
H3K9 me3:H4K20 me3
DNA Meth. m5C:H4K20 me2
DNA Meth. m5C:H4K20 me3
H3K27 me3:H4K20 me3
H3K27 me3:H3K9 me3
H2A.Z:H3−5ac
H3K4 me1:H3−5ac
H4−4ac:H3−5ac
H3K4 me1:H4−4ac
H3K9 me2:DNA Meth. m5C
H4K20 me2:H3K4 me3
H2A.Z:H4K20 me2
DNA Meth. m5C:H3K9 me3
H4K20 me2:H4−4ac
H3K27 me2:DNA Meth. m5C
H3K27 me3:DNA Meth. m5C
H4K20 me2:H3−5ac
H3K4 me3
H3K27 me2
H4K20 me3
H3K4 me1
H3K9 me3
H3K9 me2
H4−4ac
H3K27 me3
H2A.Z
H3−5ac
DNA Meth. m5C
H4K20 me2
0.00
0.25
0.50
0.75
1.00
π̂
PAX6 – Reverse
H3K4 me3:H3−5ac
H2A.Z:H3K4 me3
H3K4 me1:H4K20 me2
H3K4 me1:H2A.Z
H3K4 me3:H4−4ac
H2A.Z:H4−4ac
H3K9 me3:H4K20 me3
DNA Meth. m5C:H4K20 me2
DNA Meth. m5C:H4K20 me3
H3K27 me3:H4K20 me3
H3K27 me3:H3K9 me3
H2A.Z:H3−5ac
H3K4 me1:H3−5ac
H4−4ac:H3−5ac
H3K4 me1:H4−4ac
H3K9 me2:DNA Meth. m5C
H4K20 me2:H3K4 me3
H2A.Z:H4K20 me2
DNA Meth. m5C:H3K9 me3
H4K20 me2:H4−4ac
H3K27 me2:DNA Meth. m5C
H3K27 me3:DNA Meth. m5C
H4K20 me2:H3−5ac
H3K4 me3
H3K27 me2
H4K20 me3
H3K4 me1
H3K9 me3
H3K9 me2
H4−4ac
H3K27 me3
H2A.Z
H3−5ac
DNA Meth. m5C
H4K20 me2
0.00
0.25
0.50
0.75
1.00
π̂
MAFK – Forward
H3K4 me1:H4−4ac
H3K4 me1:H4K20 me2
H3K4 me1:H2A.Z
DNA Meth. m5C:H4K20 me3
H3K4 me1:H3−5ac
H3K27 me2:DNA Meth. m5C
H3K4 me3:H3−5ac
H2A.Z:H3K4 me3
DNA Meth. m5C:H3K9 me3
H3K9 me2:DNA Meth. m5C
H3K27 me3:H3K9 me3
H3K27 me3:DNA Meth. m5C
DNA Meth. m5C:H4K20 me2
H3K4 me3:H4−4ac
H2A.Z:H4K20 me2
H4K20 me2:H4−4ac
H4−4ac:H3−5ac
H4K20 me2:H3−5ac
H4K20 me2:H3K4 me3
H3K27 me2
H3K4 me1
H3K9 me2
H3K9 me3
H4K20 me3
H3K27 me3
H4−4ac
H3K4 me3
H2A.Z
H3−5ac
DNA Meth. m5C
H4K20 me2
0.00
0.25
0.50
0.75
1.00
π̂
MAFK – Reverse
H3K4 me1:H4−4ac
H3K4 me1:H4K20 me2
H3K4 me1:H2A.Z
DNA Meth. m5C:H4K20 me3
H3K4 me1:H3−5ac
H3K27 me2:DNA Meth. m5C
H3K4 me3:H3−5ac
H2A.Z:H3K4 me3
DNA Meth. m5C:H3K9 me3
H3K9 me2:DNA Meth. m5C
H3K27 me3:H3K9 me3
H3K27 me3:DNA Meth. m5C
DNA Meth. m5C:H4K20 me2
H3K4 me3:H4−4ac
H2A.Z:H4K20 me2
H4K20 me2:H4−4ac
H4−4ac:H3−5ac
H4K20 me2:H3−5ac
H4K20 me2:H3K4 me3
H3K27 me2
H3K4 me1
H3K9 me2
H3K9 me3
H4K20 me3
H3K27 me3
H4−4ac
H3K4 me3
H2A.Z
H3−5ac
DNA Meth. m5C
H4K20 me2
0.00
0.25
0.50
0.75
1.00
π̂
MAFF – Forward
H2A.Z:H3K4 me3
H4−4ac:H3−5ac
H3K4 me1:H3−5ac
H3K4 me1:H4−4ac
H3K4 me1:H2A.Z
DNA Meth. m5C:H3K9 me3
H3K4 me1:H4K20 me2
DNA Meth. m5C:H4K20 me3
H3K27 me3:H4K20 me3
H3K4 me3:H3−5ac
H3K27 me3:H3K4 me1
H3K9 me3:H4K20 me3
DNA Meth. m5C:H4K20 me2
H3K27 me3:H3K9 me3
H3K27 me3:DNA Meth. m5C
H2A.Z:H4K20 me2
H3K9 me2:DNA Meth. m5C
H4K20 me2:H3−5ac
H4K20 me2:H4−4ac
H4K20 me2:H3K4 me3
H3K27 me2
H3K9 me2
H3K4 me1
H4K20 me3
H3K9 me3
H3K27 me3
H4−4ac
H3−5ac
H3K4 me3
H2A.Z
H4K20 me2
DNA Meth. m5C
0.00
0.25
0.50
0.75
1.00
π̂
MAFF – Reverse
H2A.Z:H3K4 me3
H4−4ac:H3−5ac
H3K4 me1:H3−5ac
H3K4 me1:H4−4ac
H3K4 me1:H2A.Z
DNA Meth. m5C:H3K9 me3
H3K4 me1:H4K20 me2
DNA Meth. m5C:H4K20 me3
H3K27 me3:H4K20 me3
H3K4 me3:H3−5ac
H3K27 me3:H3K4 me1
H3K9 me3:H4K20 me3
DNA Meth. m5C:H4K20 me2
H3K27 me3:H3K9 me3
H3K27 me3:DNA Meth. m5C
H2A.Z:H4K20 me2
H3K9 me2:DNA Meth. m5C
H4K20 me2:H3−5ac
H4K20 me2:H4−4ac
H4K20 me2:H3K4 me3
H3K27 me2
H3K9 me2
H3K4 me1
H4K20 me3
H3K9 me3
H3K27 me3
H4−4ac
H3−5ac
H3K4 me3
H2A.Z
H4K20 me2
DNA Meth. m5C
0.00
0.25
0.50
0.75
1.00
π̂
ZNF512B – Forward
DNA Meth. m5C:H4K20 me3
H2A.Z:H3K4 me3
H3K4 me1:H4K20 me2
H3K9 me3:H4K20 me3
H3K27 me2:DNA Meth. m5C
DNA Meth. m5C:H3K9 me3
H3K27 me3:H4K20 me3
H3K27 me3:H3K9 me3
H3K4 me3:H4−4ac
DNA Meth. m5C:H4K20 me2
H4−4ac:H3−5ac
H2A.Z:H3−5ac
H2A.Z:H4−4ac
H3K27 me3:DNA Meth. m5C
H3K4 me1:H4−4ac
H3K4 me1:H2A.Z
H4K20 me2:H4−4ac
H3K9 me2:DNA Meth. m5C
H4K20 me2:H3K4 me3
H2A.Z:H4K20 me2
H4K20 me2:H3−5ac
H3K4 me3
H3K27 me2
H4K20 me3
H3K9 me3
H3K9 me2
H3K4 me1
H3K27 me3
H4−4ac
H3−5ac
H2A.Z
H4K20 me2
DNA Meth. m5C
0.00
0.25
0.50
0.75
1.00
π̂
ZNF512B – Reverse
DNA Meth. m5C:H4K20 me3
H2A.Z:H3K4 me3
H3K4 me1:H4K20 me2
H3K9 me3:H4K20 me3
H3K27 me2:DNA Meth. m5C
DNA Meth. m5C:H3K9 me3
H3K27 me3:H4K20 me3
H3K27 me3:H3K9 me3
H3K4 me3:H4−4ac
DNA Meth. m5C:H4K20 me2
H4−4ac:H3−5ac
H2A.Z:H3−5ac
H2A.Z:H4−4ac
H3K27 me3:DNA Meth. m5C
H3K4 me1:H4−4ac
H3K4 me1:H2A.Z
H4K20 me2:H4−4ac
H3K9 me2:DNA Meth. m5C
H4K20 me2:H3K4 me3
H2A.Z:H4K20 me2
H4K20 me2:H3−5ac
H3K4 me3
H3K27 me2
H4K20 me3
H3K9 me3
H3K9 me2
H3K4 me1
H3K27 me3
H4−4ac
H3−5ac
H2A.Z
H4K20 me2
DNA Meth. m5C
0.00
0.25
0.50
0.75
1.00
π̂
RBBP4 – Forward
H3K4 me3:H3−5ac
H2A.Z:H3K4 me3
H3K4 me1:H4K20 me2
DNA Meth. m5C:H4K20 me3
H3K9 me3:H4K20 me3
H4K20 me2:H3K4 me3
H3K27 me2:DNA Meth. m5C
H3K27 me3:H4K20 me3
H4−4ac:H3−5ac
H3K27 me3:H3K9 me3
H3K27 me3:DNA Meth. m5C
H3K4 me3:H4−4ac
H4K20 me2:H3−5ac
H3K4 me1:H2A.Z
DNA Meth. m5C:H4K20 me2
H3K4 me1:H3−5ac
H3K4 me1:H4−4ac
H2A.Z:H3−5ac
H2A.Z:H4−4ac
H3K9 me2:DNA Meth. m5C
H2A.Z:H4K20 me2
DNA Meth. m5C:H3K9 me3
H4K20 me2:H4−4ac
H4K20 me3
H3K4 me3
H3K9 me3
H3K27 me3
H3K27 me2
H3K9 me2
H3K4 me1
H2A.Z
H4−4ac
H4K20 me2
H3−5ac
DNA Meth. m5C
0.00
0.25
0.50
0.75
1.00
π̂
RBBP4 – Reverse
H3K4 me3:H3−5ac
H2A.Z:H3K4 me3
H3K4 me1:H4K20 me2
DNA Meth. m5C:H4K20 me3
H3K9 me3:H4K20 me3
H4K20 me2:H3K4 me3
H3K27 me2:DNA Meth. m5C
H3K27 me3:H4K20 me3
H4−4ac:H3−5ac
H3K27 me3:H3K9 me3
H3K27 me3:DNA Meth. m5C
H3K4 me3:H4−4ac
H4K20 me2:H3−5ac
H3K4 me1:H2A.Z
DNA Meth. m5C:H4K20 me2
H3K4 me1:H3−5ac
H3K4 me1:H4−4ac
H2A.Z:H3−5ac
H2A.Z:H4−4ac
H3K9 me2:DNA Meth. m5C
H2A.Z:H4K20 me2
DNA Meth. m5C:H3K9 me3
H4K20 me2:H4−4ac
H4K20 me3
H3K4 me3
H3K9 me3
H3K27 me3
H3K27 me2
H3K9 me2
H3K4 me1
H2A.Z
H4−4ac
H4K20 me2
H3−5ac
DNA Meth. m5C
0.00
0.25
0.50
0.75
1.00
π̂

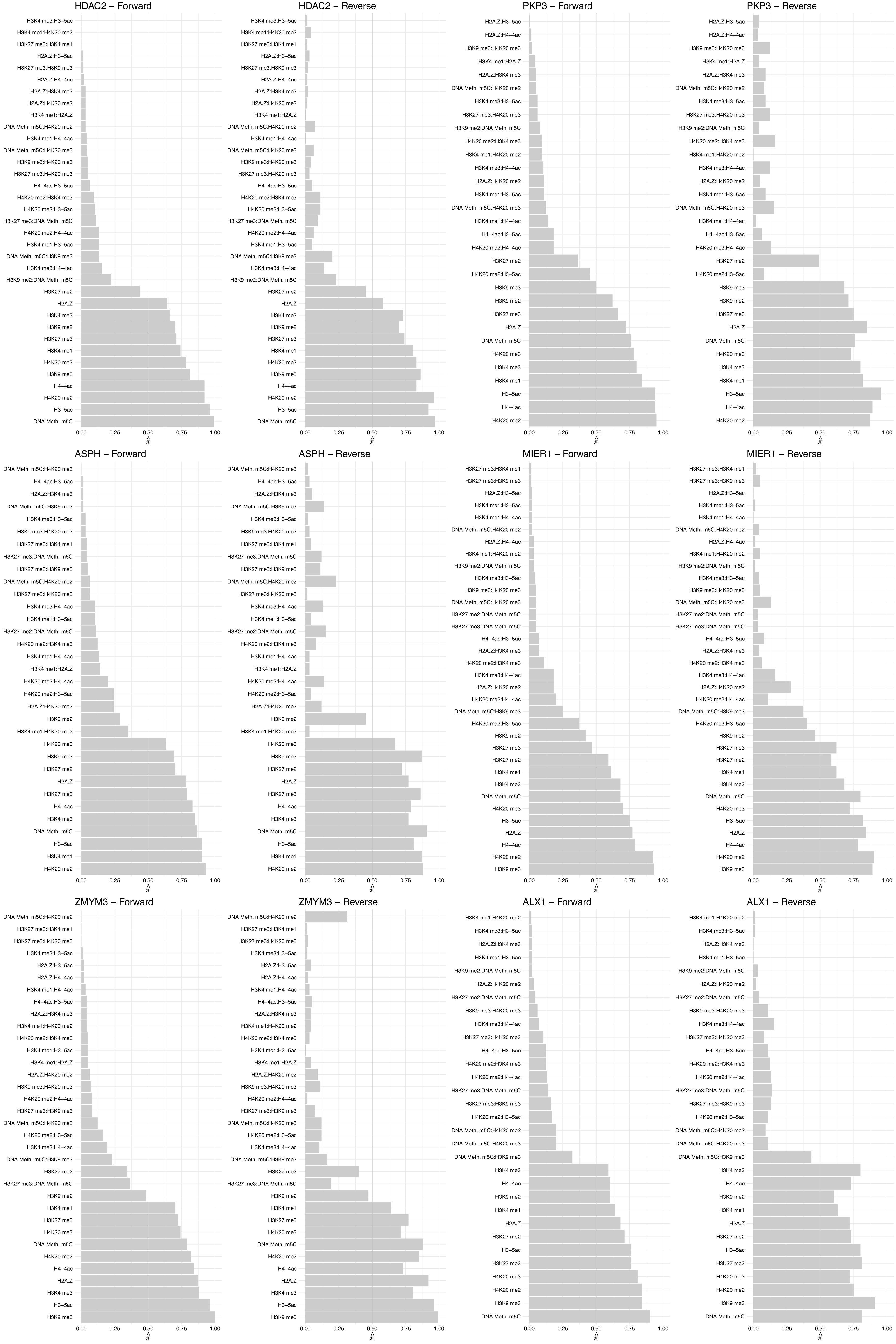
HDAC2 – Forward
HDAC2 – Reverse
H3K4 me3:H3–5ac
H3K4 me1:H4K20 me2
H3K27 me3:H3K4 me1
H2A.Z:H3–5ac
H3K27 me3:H3K9 me3
H2A.Z:H4–4ac
H2A.Z:H3K4 me3
H2A.Z:H4K20 me2
H3K4 me1:H2A.Z
DNA Meth. m5C:H4K20 me2
H3K4 me1:H4–4ac
DNA Meth. m5C:H4K20 me3
H3K9 me3:H4K20 me3
H3K27 me3:H4K20 me3
H4–4ac:H3–5ac
H4K20 me2:H3K4 me3
H4K20 me2:H3–5ac
H3K27 me3:DNA Meth. m5C
H4K20 me2:H4–4ac
H3K4 me1:H3–5ac
DNA Meth. m5C:H3K9 me3
H3K4 me3:H4–4ac
H3K9 me2:DNA Meth. m5C
H3K27 me2
H2A.Z
H3K4 me3
H3K9 me2
H3K27 me3
H3K4 me1
H4K20 me3
H3K9 me3
H4–4ac
H4K20 me2
H3–5ac
DNA Meth. m5C
PKP3 – Forward
PKP3 – Reverse
H2A.Z:H3–5ac
H2A.Z:H4–4ac
H3K9 me3:H4K20 me3
H3K4 me1:H2A.Z
H2A.Z:H3K4 me3
DNA Meth. m5C:H4K20 me2
H3K4 me3:H3–5ac
H3K27 me3:H4K20 me3
H3K9 me2:DNA Meth. m5C
H4K20 me2:H3K4 me3
H3K4 me1:H4K20 me2
H3K4 me3:H4–4ac
H2A.Z:H4K20 me2
H3K4 me1:H3–5ac
DNA Meth. m5C:H4K20 me3
H3K4 me1:H4–4ac
H4–4ac:H3–5ac
H4K20 me2:H4–4ac
H3K27 me2
H4K20 me2:H3–5ac
H3K9 me3
H3K9 me2
H3K27 me3
H2A.Z
DNA Meth. m5C
H4K20 me3
H3K4 me3
H3K4 me1
H3–5ac
H4–4ac
H4K20 me2
ASPH – Forward
ASPH – Reverse
DNA Meth. m5C:H4K20 me3
H4–4ac:H3–5ac
H2A.Z:H3K4 me3
DNA Meth. m5C:H3K9 me3
H3K4 me3:H3–5ac
H3K9 me3:H4K20 me3
H3K27 me3:H3K4 me1
H3K27 me3:DNA Meth. m5C
H3K27 me3:H3K9 me3
DNA Meth. m5C:H4K20 me2
H3K27 me3:H4K20 me3
H3K4 me3:H4–4ac
H3K4 me1:H3–5ac
H3K27 me2:DNA Meth. m5C
H4K20 me2:H3K4 me3
H3K4 me1:H4–4ac
H3K4 me1:H2A.Z
H4K20 me2:H4–4ac
H4K20 me2:H3–5ac
H2A.Z:H4K20 me2
H3K9 me2
H3K4 me1:H4K20 me2
H4K20 me3
H3K9 me3
H3K27 me2
H2A.Z
H3K27 me3
H4–4ac
H3K4 me3
DNA Meth. m5C
H3–5ac
H3K4 me1
H4K20 me2
MIER1 – Forward
MIER1 – Reverse
H3K27 me3:H3K4 me1
H3K27 me3:H3K9 me3
H2A.Z:H3–5ac
H3K4 me1:H3–5ac
H3K4 me1:H4–4ac
DNA Meth. m5C:H4K20 me2
H2A.Z:H4–4ac
H3K4 me1:H4K20 me2
H3K9 me2:DNA Meth. m5C
H3K4 me3:H3–5ac
H3K9 me3:H4K20 me3
DNA Meth. m5C:H4K20 me3
H3K27 me2:DNA Meth. m5C
H3K27 me3:DNA Meth. m5C
H4–4ac:H3–5ac
H2A.Z:H3K4 me3
H4K20 me2:H3K4 me3
H3K4 me3:H4–4ac
H2A.Z:H4K20 me2
H4K20 me2:H4–4ac
DNA Meth. m5C:H3K9 me3
H4K20 me2:H3–5ac
H3K9 me2
H3K27 me3
H3K27 me2
H3K4 me1
H3K4 me3
DNA Meth. m5C
H4K20 me3
H3–5ac
H2A.Z
H4–4ac
H4K20 me2
H3K9 me3
ZMYM3 – Forward
ZMYM3 – Reverse
DNA Meth. m5C:H4K20 me2
H3K27 me3:H3K4 me1
H3K27 me3:H4K20 me3
H3K4 me3:H3–5ac
H2A.Z:H3–5ac
H2A.Z:H4–4ac
H3K4 me1:H4–4ac
H4–4ac:H3–5ac
H2A.Z:H3K4 me3
H3K4 me1:H4K20 me2
H4K20 me2:H3K4 me3
H3K4 me1:H3–5ac
H3K4 me1:H2A.Z
H2A.Z:H4K20 me2
H3K9 me3:H4K20 me3
H4K20 me2:H4–4ac
H3K27 me3:H3K9 me3
DNA Meth. m5C:H4K20 me3
H4K20 me2:H3–5ac
H3K4 me3:H4–4ac
DNA Meth. m5C:H3K9 me3
H3K27 me2
H3K27 me3:DNA Meth. m5C
H3K9 me2
H3K4 me1
H3K27 me3
H4K20 me3
DNA Meth. m5C
H4K20 me2
H4–4ac
H2A.Z
H3K4 me3
H3–5ac
H3K9 me3
ALX1 – Forward
ALX1 – Reverse
H3K4 me1:H4K20 me2
H3K4 me3:H3–5ac
H2A.Z:H3K4 me3
H3K4 me1:H3–5ac
H3K9 me2:DNA Meth. m5C
H2A.Z:H4K20 me2
H3K27 me2:DNA Meth. m5C
H3K9 me3:H4K20 me3
H3K4 me3:H4–4ac
H3K27 me3:H4K20 me3
H4–4ac:H3–5ac
H4K20 me2:H3K4 me3
H4K20 me2:H4–4ac
H3K27 me3:DNA Meth. m5C
H3K27 me3:H3K9 me3
H4K20 me2:H3–5ac
DNA Meth. m5C:H4K20 me2
DNA Meth. m5C:H4K20 me3
DNA Meth. m5C:H3K9 me3
H3K4 me3
H4–4ac
H3K9 me2
H3K4 me1
H2A.Z
H3K27 me2
H3–5ac
H3K27 me3
H4K20 me3
H4K20 me2
H3K9 me3
DNA Meth. m5C
0.00
0.25
0.50
0.75
1.00
$\hat{\pi}$

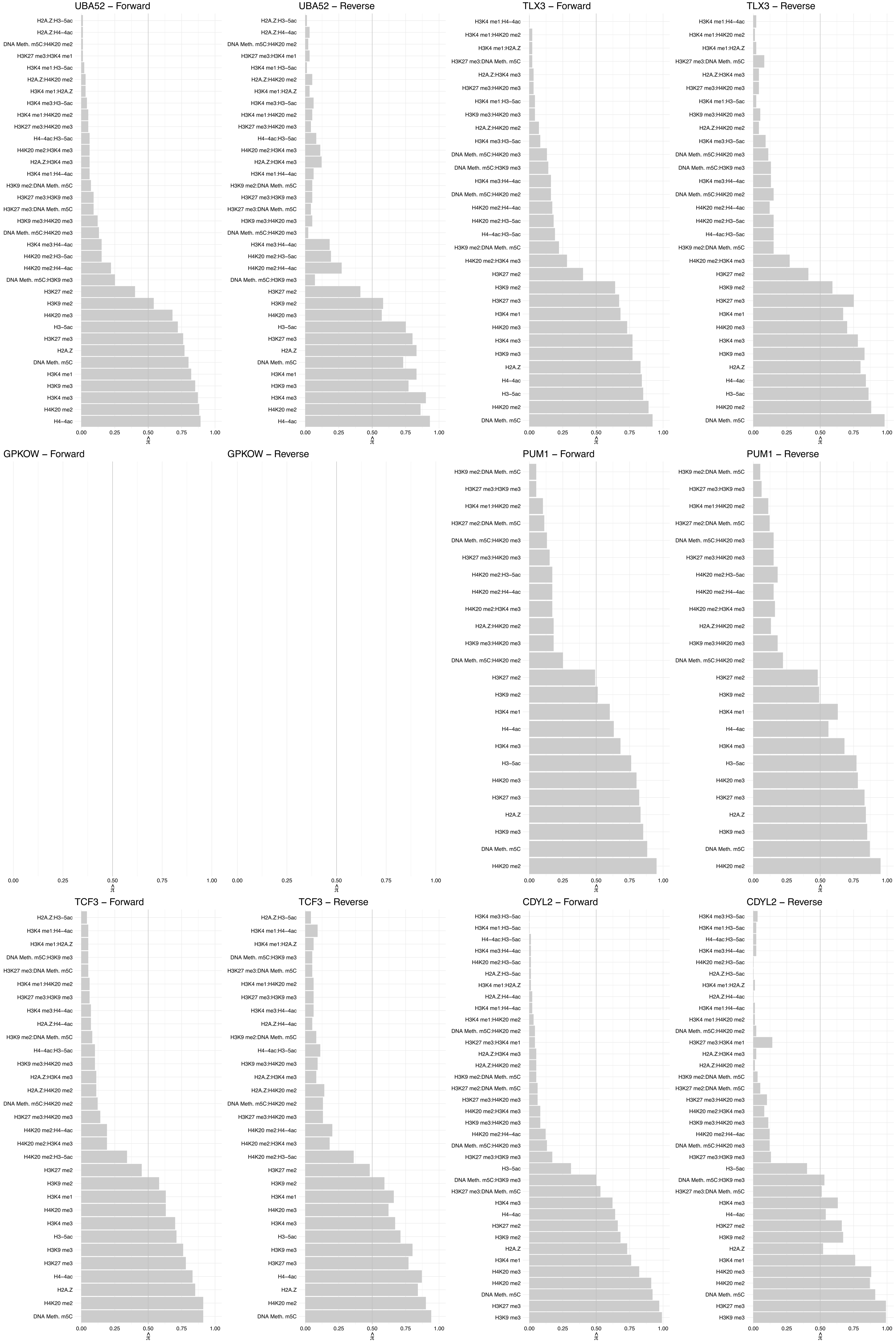

UBA52 – Forward
H2A.Z:H3–5ac
H2A.Z:H4–4ac
DNA Meth. m5C:H4K20 me2
H3K27 me3:H3K4 me1
H3K4 me1:H3–5ac
H2A.Z:H4K20 me2
H3K4 me1:H2A.Z
H3K4 me3:H3–5ac
H3K4 me1:H4K20 me2
H3K27 me3:H4K20 me3
H4–4ac:H3–5ac
H4K20 me2:H3K4 me3
H2A.Z:H3K4 me3
H3K4 me1:H4–4ac
H3K9 me2:DNA Meth. m5C
H3K27 me3:H3K9 me3
H3K27 me3:DNA Meth. m5C
H3K9 me3:H4K20 me3
DNA Meth. m5C:H4K20 me3
H3K4 me3:H4–4ac
H4K20 me2:H3–5ac
H4K20 me2:H4–4ac
DNA Meth. m5C:H3K9 me3
H3K27 me2
H3K9 me2
H4K20 me3
H3–5ac
H3K27 me3
H2A.Z
DNA Meth. m5C
H3K4 me1
H3K9 me3
H3K4 me3
H4K20 me2
H4–4ac
0.00 0.25 0.50 0.75 1.00
π̂
UBA52 – Reverse
TLX3 – Forward
TLX3 – Reverse
GPKOW – Forward
GPKOW – Reverse
PUM1 – Forward
PUM1 – Reverse
TCF3 – Forward
TCF3 – Reverse
CDYL2 – Forward
CDYL2 – Reverse

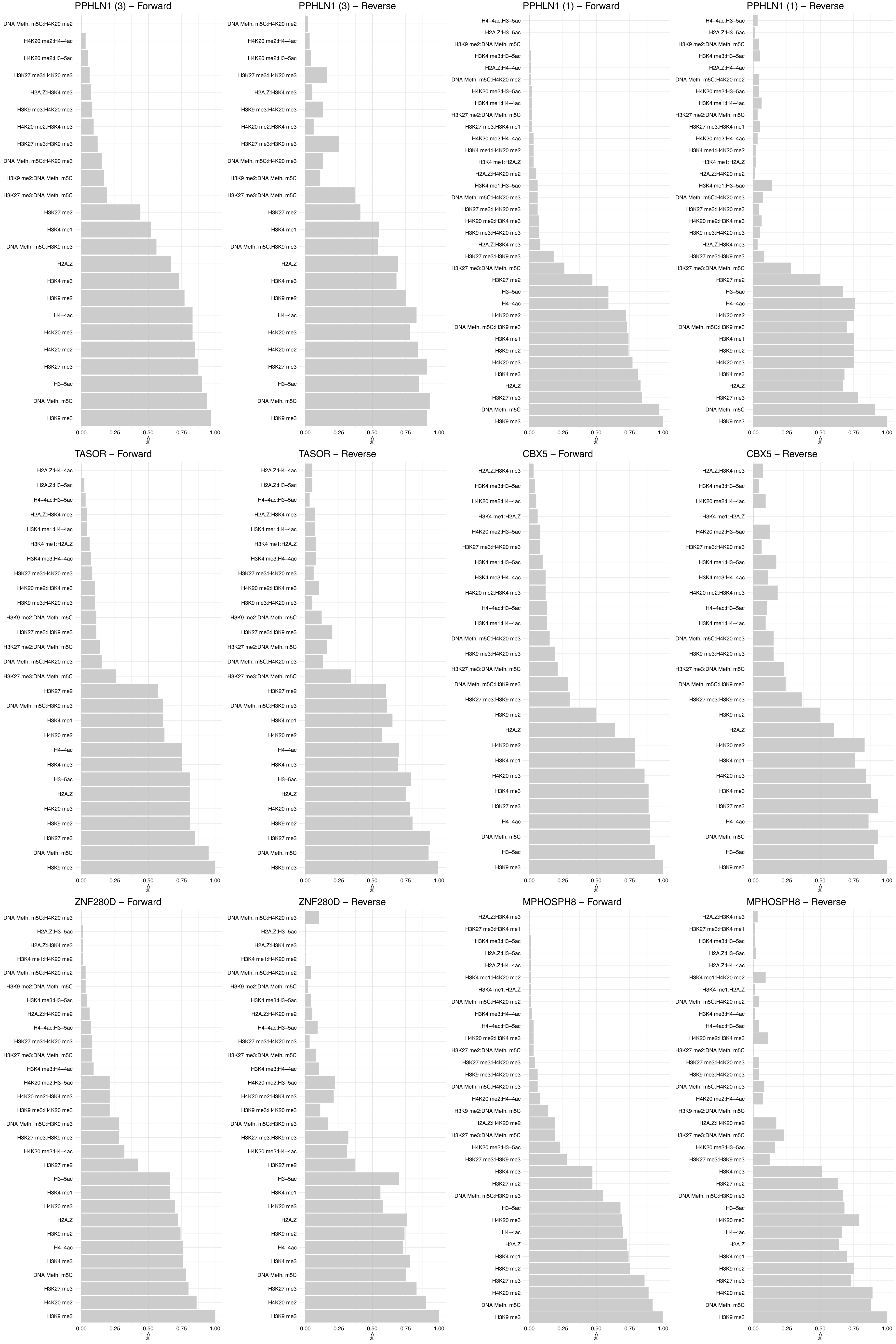
PPHLN1 (3) – Forward
PPHLN1 (3) – Reverse
PPHLN1 (1) – Forward
PPHLN1 (1) – Reverse
TASOR – Forward
TASOR – Reverse
CBX5 – Forward
CBX5 – Reverse
ZNF280D – Forward
ZNF280D – Reverse
MPHOSPH8 – Forward
MPHOSPH8 – Reverse
0.00 0.25 0.50 0.75 1.00
$\hat{\pi}$

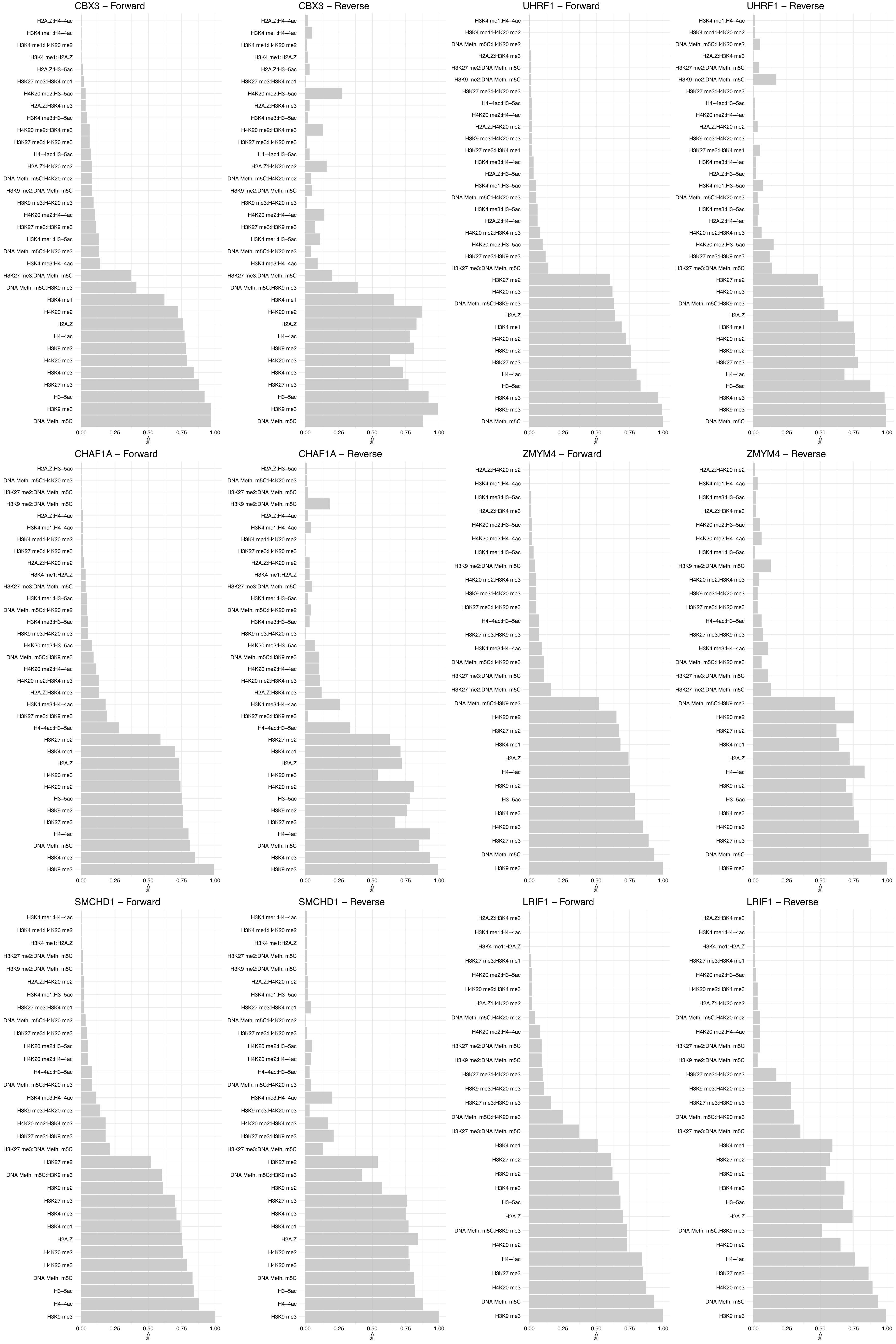
CBX3 − Forward
CBX3 − Reverse
H2A.Z:H4−4ac
H3K4 me1:H4−4ac
H3K4 me1:H4K20 me2
H3K4 me1:H2A.Z
H2A.Z:H3−5ac
H3K27 me3:H3K4 me1
H4K20 me2:H3−5ac
H2A.Z:H3K4 me3
H3K4 me3:H3−5ac
H4K20 me2:H3K4 me3
H3K27 me3:H4K20 me3
H4−4ac:H3−5ac
H2A.Z:H4K20 me2
DNA Meth. m5C:H4K20 me2
H3K9 me2:DNA Meth. m5C
H3K9 me3:H4K20 me3
H4K20 me2:H4−4ac
H3K27 me3:H3K9 me3
H3K4 me1:H3−5ac
DNA Meth. m5C:H4K20 me3
H3K4 me3:H4−4ac
H3K27 me3:DNA Meth. m5C
DNA Meth. m5C:H3K9 me3
H3K4 me1
H4K20 me2
H2A.Z
H4−4ac
H3K9 me2
H4K20 me3
H3K4 me3
H3K27 me3
H3−5ac
H3K9 me3
DNA Meth. m5C
UHRF1 − Forward
UHRF1 − Reverse
H3K4 me1:H4−4ac
H3K4 me1:H4K20 me2
DNA Meth. m5C:H4K20 me2
H2A.Z:H3K4 me3
H3K27 me2:DNA Meth. m5C
H3K9 me2:DNA Meth. m5C
H3K27 me3:H4K20 me3
H4−4ac:H3−5ac
H4K20 me2:H4−4ac
H2A.Z:H4K20 me2
H3K9 me3:H4K20 me3
H3K27 me3:H3K4 me1
H3K4 me3:H4−4ac
H2A.Z:H3−5ac
H3K4 me1:H3−5ac
DNA Meth. m5C:H4K20 me3
H3K4 me3:H3−5ac
H2A.Z:H4−4ac
H4K20 me2:H3K4 me3
H4K20 me2:H3−5ac
H3K27 me3:H3K9 me3
H3K27 me3:DNA Meth. m5C
H3K27 me2
H4K20 me3
DNA Meth. m5C:H3K9 me3
H2A.Z
H3K4 me1
H4K20 me2
H3K9 me2
H3K27 me3
H4−4ac
H3−5ac
H3K4 me3
H3K9 me3
DNA Meth. m5C
CHAF1A − Forward
CHAF1A − Reverse
H2A.Z:H3−5ac
DNA Meth. m5C:H4K20 me3
H3K27 me2:DNA Meth. m5C
H3K9 me2:DNA Meth. m5C
H2A.Z:H4−4ac
H3K4 me1:H4−4ac
H3K4 me1:H4K20 me2
H3K27 me3:H4K20 me3
H2A.Z:H4K20 me2
H3K4 me1:H2A.Z
H3K27 me3:DNA Meth. m5C
H3K4 me1:H3−5ac
DNA Meth. m5C:H4K20 me2
H3K4 me3:H3−5ac
H3K9 me3:H4K20 me3
H4K20 me2:H3−5ac
DNA Meth. m5C:H3K9 me3
H4K20 me2:H4−4ac
H4K20 me2:H3K4 me3
H2A.Z:H3K4 me3
H3K4 me3:H4−4ac
H3K27 me3:H3K9 me3
H4−4ac:H3−5ac
H3K27 me2
H3K4 me1
H2A.Z
H4K20 me3
H4K20 me2
H3−5ac
H3K9 me2
H3K27 me3
H4−4ac
DNA Meth. m5C
H3K4 me3
H3K9 me3
ZMYM4 − Forward
ZMYM4 − Reverse
H2A.Z:H4K20 me2
H3K4 me1:H4−4ac
H3K4 me3:H3−5ac
H2A.Z:H3K4 me3
H4K20 me2:H3−5ac
H4K20 me2:H4−4ac
H3K4 me1:H3−5ac
H3K9 me2:DNA Meth. m5C
H4K20 me2:H3K4 me3
H3K9 me3:H4K20 me3
H3K27 me3:H4K20 me3
H4−4ac:H3−5ac
H3K27 me3:H3K9 me3
H3K4 me3:H4−4ac
DNA Meth. m5C:H4K20 me3
H3K27 me3:DNA Meth. m5C
H3K27 me2:DNA Meth. m5C
DNA Meth. m5C:H3K9 me3
H4K20 me2
H3K27 me2
H3K4 me1
H2A.Z
H4−4ac
H3K9 me2
H3−5ac
H3K4 me3
H4K20 me3
H3K27 me3
DNA Meth. m5C
H3K9 me3
SMCHD1 − Forward
SMCHD1 − Reverse
H3K4 me1:H4−4ac
H3K4 me1:H4K20 me2
H3K4 me1:H2A.Z
H3K27 me2:DNA Meth. m5C
H3K9 me2:DNA Meth. m5C
H2A.Z:H4K20 me2
H3K4 me1:H3−5ac
H3K27 me3:H3K4 me1
DNA Meth. m5C:H4K20 me2
H3K27 me3:H4K20 me3
H4K20 me2:H3−5ac
H4K20 me2:H4−4ac
H4−4ac:H3−5ac
DNA Meth. m5C:H4K20 me3
H3K4 me3:H4−4ac
H3K9 me3:H4K20 me3
H4K20 me2:H3K4 me3
H3K27 me3:H3K9 me3
H3K27 me3:DNA Meth. m5C
H3K27 me2
DNA Meth. m5C:H3K9 me3
H3K9 me2
H3K27 me3
H3K4 me3
H3K4 me1
H2A.Z
H4K20 me2
H4K20 me3
DNA Meth. m5C
H3−5ac
H4−4ac
H3K9 me3
LRIF1 − Forward
LRIF1 − Reverse
H2A.Z:H3K4 me3
H3K4 me1:H4−4ac
H3K4 me1:H2A.Z
H3K27 me3:H3K4 me1
H4K20 me2:H3−5ac
H4K20 me2:H3K4 me3
H2A.Z:H4K20 me2
DNA Meth. m5C:H4K20 me2
H4K20 me2:H4−4ac
H3K27 me2:DNA Meth. m5C
H3K9 me2:DNA Meth. m5C
H3K27 me3:H4K20 me3
H3K9 me3:H4K20 me3
H3K27 me3:H3K9 me3
DNA Meth. m5C:H4K20 me3
H3K27 me3:DNA Meth. m5C
H3K4 me1
H3K27 me2
H3K9 me2
H3K4 me3
H3−5ac
H2A.Z
DNA Meth. m5C:H3K9 me3
H4K20 me2
H4−4ac
H3K27 me3
H4K20 me3
DNA Meth. m5C
H3K9 me3
0.00
0.25
0.50
0.75
1.00
π̂

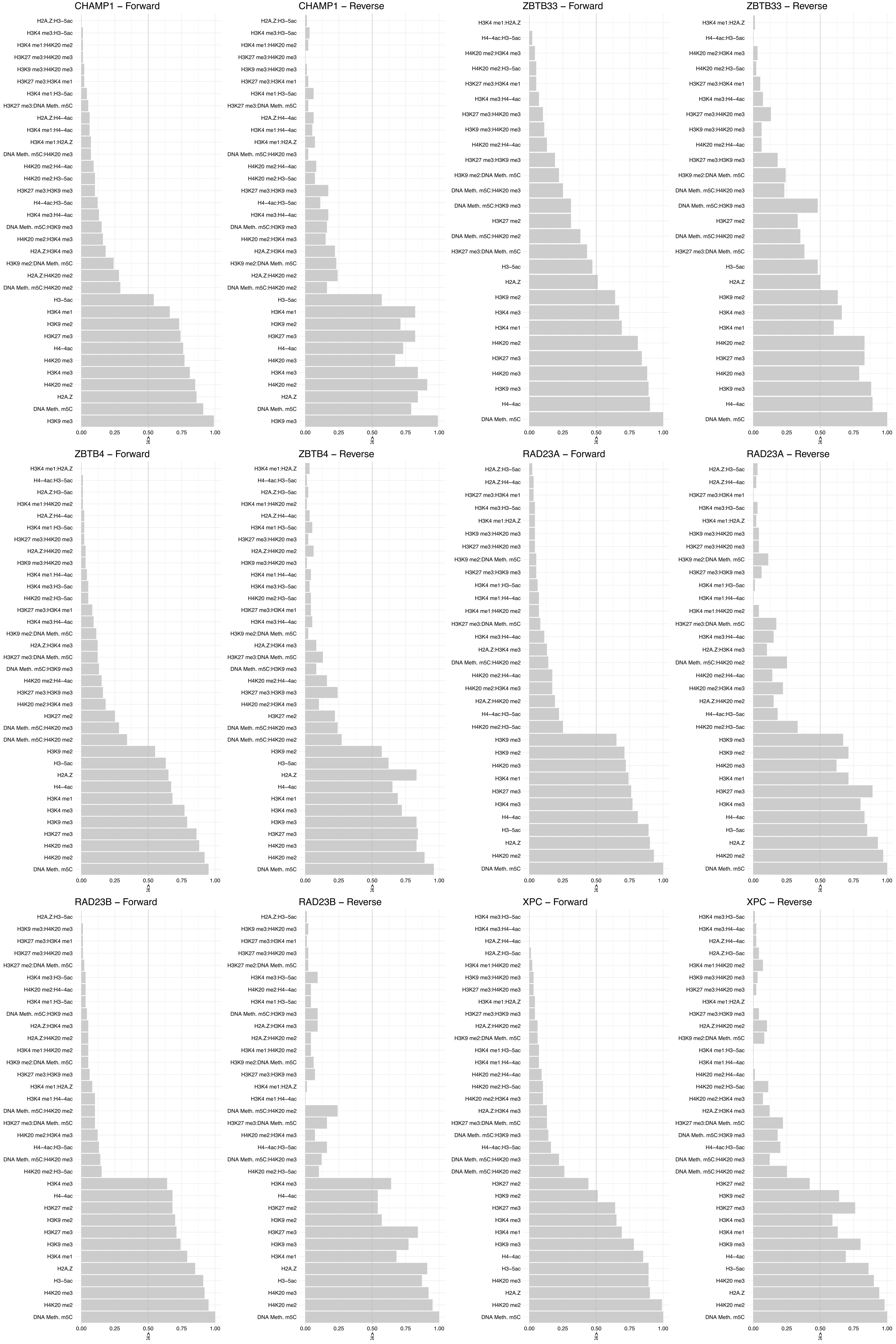
CHAMP1 – Forward
CHAMP1 – Reverse
ZBTB33 – Forward
ZBTB33 – Reverse
ZBTB4 – Forward
ZBTB4 – Reverse
RAD23A – Forward
RAD23A – Reverse
RAD23B – Forward
RAD23B – Reverse
XPC – Forward
XPC – Reverse
0.00
0.25
0.50
0.75
1.00
$\hat{\pi}$

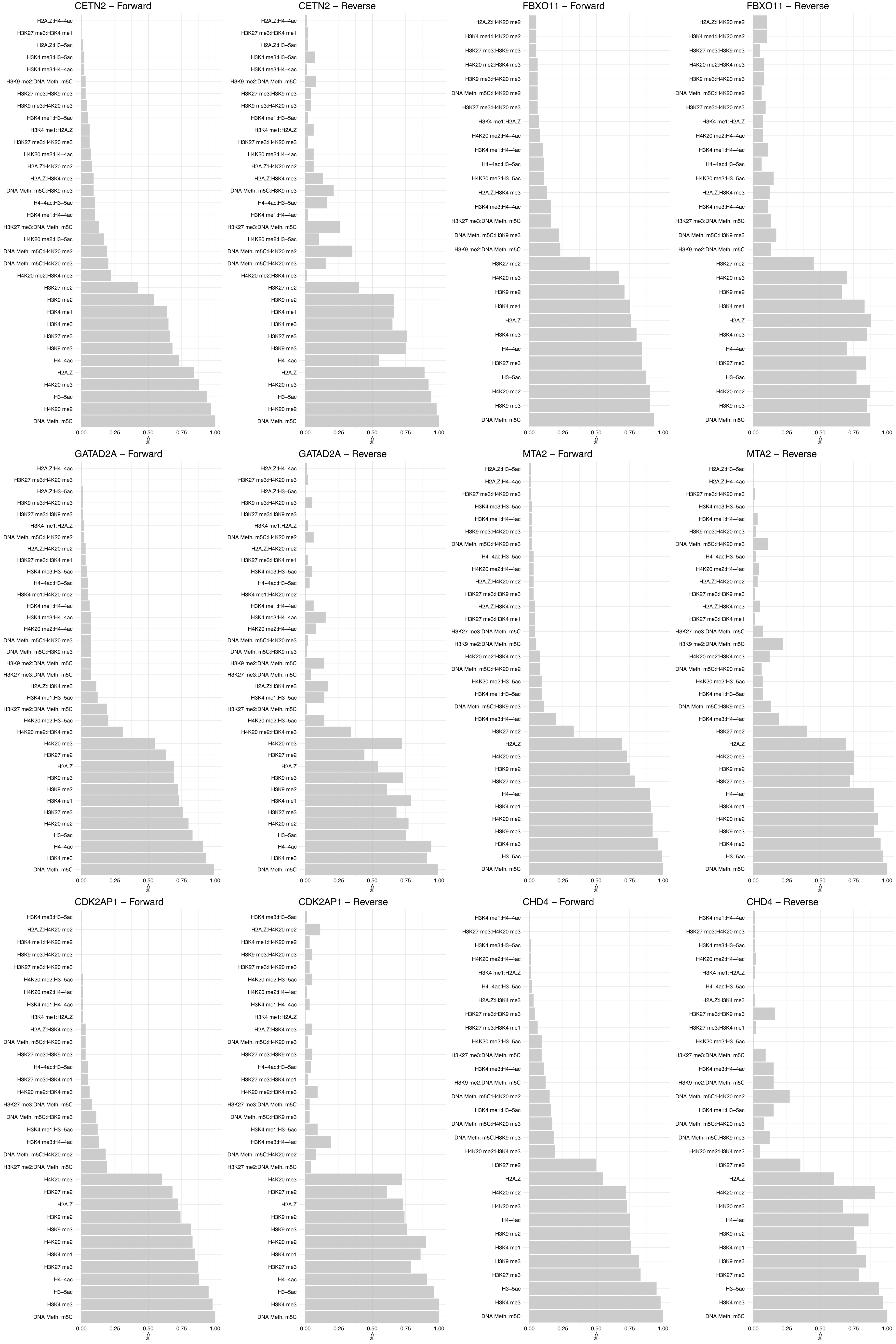
CETN2 – Forward
H2A.Z:H4–4ac
H3K27 me3:H3K4 me1
H2A.Z:H3–5ac
H3K4 me3:H3–5ac
H3K4 me3:H4–4ac
H3K9 me2:DNA Meth. m5C
H3K27 me3:H3K9 me3
H3K9 me3:H4K20 me3
H3K4 me1:H3–5ac
H3K4 me1:H2A.Z
H3K27 me3:H4K20 me3
H4K20 me2:H4–4ac
H2A.Z:H4K20 me2
H2A.Z:H3K4 me3
DNA Meth. m5C:H3K9 me3
H4–4ac:H3–5ac
H3K4 me1:H4–4ac
H3K27 me3:DNA Meth. m5C
H4K20 me2:H3–5ac
DNA Meth. m5C:H4K20 me2
DNA Meth. m5C:H4K20 me3
H4K20 me2:H3K4 me3
H3K27 me2
H3K9 me2
H3K4 me1
H3K4 me3
H3K27 me3
H3K9 me3
H4–4ac
H2A.Z
H4K20 me3
H3–5ac
H4K20 me2
DNA Meth. m5C
0.00 0.25 0.50 0.75 1.00
π̂
CETN2 – Reverse
H2A.Z:H4–4ac
H3K27 me3:H3K4 me1
H2A.Z:H3–5ac
H3K4 me3:H3–5ac
H3K4 me3:H4–4ac
H3K9 me2:DNA Meth. m5C
H3K27 me3:H3K9 me3
H3K9 me3:H4K20 me3
H3K4 me1:H3–5ac
H3K4 me1:H2A.Z
H3K27 me3:H4K20 me3
H4K20 me2:H4–4ac
H2A.Z:H4K20 me2
H2A.Z:H3K4 me3
DNA Meth. m5C:H3K9 me3
H4–4ac:H3–5ac
H3K4 me1:H4–4ac
H3K27 me3:DNA Meth. m5C
H4K20 me2:H3–5ac
DNA Meth. m5C:H4K20 me2
DNA Meth. m5C:H4K20 me3
H4K20 me2:H3K4 me3
H3K27 me2
H3K9 me2
H3K4 me1
H3K4 me3
H3K27 me3
H3K9 me3
H4–4ac
H2A.Z
H4K20 me3
H3–5ac
H4K20 me2
DNA Meth. m5C
0.00 0.25 0.50 0.75 1.00
π̂
FBXO11 – Forward
H2A.Z:H4K20 me2
H3K4 me1:H4K20 me2
H3K27 me3:H3K9 me3
H4K20 me2:H3K4 me3
H3K9 me3:H4K20 me3
DNA Meth. m5C:H4K20 me2
H3K27 me3:H4K20 me3
H3K4 me1:H2A.Z
H4K20 me2:H4–4ac
H3K4 me1:H4–4ac
H4–4ac:H3–5ac
H4K20 me2:H3–5ac
H2A.Z:H3K4 me3
H3K4 me3:H4–4ac
H3K27 me3:DNA Meth. m5C
DNA Meth. m5C:H3K9 me3
H3K9 me2:DNA Meth. m5C
H3K27 me2
H4K20 me3
H3K9 me2
H3K4 me1
H2A.Z
H3K4 me3
H4–4ac
H3K27 me3
H3–5ac
H4K20 me2
H3K9 me3
DNA Meth. m5C
0.00 0.25 0.50 0.75 1.00
π̂
FBXO11 – Reverse
H2A.Z:H4K20 me2
H3K4 me1:H4K20 me2
H3K27 me3:H3K9 me3
H4K20 me2:H3K4 me3
H3K9 me3:H4K20 me3
DNA Meth. m5C:H4K20 me2
H3K27 me3:H4K20 me3
H3K4 me1:H2A.Z
H4K20 me2:H4–4ac
H3K4 me1:H4–4ac
H4–4ac:H3–5ac
H4K20 me2:H3–5ac
H2A.Z:H3K4 me3
H3K4 me3:H4–4ac
H3K27 me3:DNA Meth. m5C
DNA Meth. m5C:H3K9 me3
H3K9 me2:DNA Meth. m5C
H3K27 me2
H4K20 me3
H3K9 me2
H3K4 me1
H2A.Z
H3K4 me3
H4–4ac
H3K27 me3
H3–5ac
H4K20 me2
H3K9 me3
DNA Meth. m5C
0.00 0.25 0.50 0.75 1.00
π̂
GATAD2A – Forward
H2A.Z:H4–4ac
H3K27 me3:H4K20 me3
H2A.Z:H3–5ac
H3K9 me3:H4K20 me3
H3K27 me3:H3K9 me3
H3K4 me1:H2A.Z
DNA Meth. m5C:H4K20 me2
H2A.Z:H4K20 me2
H3K27 me3:H3K4 me1
H3K4 me3:H3–5ac
H4–4ac:H3–5ac
H3K4 me1:H4K20 me2
H3K4 me1:H4–4ac
H3K4 me3:H4–4ac
H4K20 me2:H4–4ac
DNA Meth. m5C:H4K20 me3
DNA Meth. m5C:H3K9 me3
H3K9 me2:DNA Meth. m5C
H3K27 me3:DNA Meth. m5C
H2A.Z:H3K4 me3
H3K4 me1:H3–5ac
H3K27 me2:DNA Meth. m5C
H4K20 me2:H3–5ac
H4K20 me2:H3K4 me3
H4K20 me3
H3K27 me2
H2A.Z
H3K9 me3
H3K9 me2
H3K4 me1
H3K27 me3
H4K20 me2
H3–5ac
H4–4ac
H3K4 me3
DNA Meth. m5C
0.00 0.25 0.50 0.75 1.00
π̂
GATAD2A – Reverse
H2A.Z:H4–4ac
H3K27 me3:H4K20 me3
H2A.Z:H3–5ac
H3K9 me3:H4K20 me3
H3K27 me3:H3K9 me3
H3K4 me1:H2A.Z
DNA Meth. m5C:H4K20 me2
H2A.Z:H4K20 me2
H3K27 me3:H3K4 me1
H3K4 me3:H3–5ac
H4–4ac:H3–5ac
H3K4 me1:H4K20 me2
H3K4 me1:H4–4ac
H3K4 me3:H4–4ac
H4K20 me2:H4–4ac
DNA Meth. m5C:H4K20 me3
DNA Meth. m5C:H3K9 me3
H3K9 me2:DNA Meth. m5C
H3K27 me3:DNA Meth. m5C
H2A.Z:H3K4 me3
H3K4 me1:H3–5ac
H3K27 me2:DNA Meth. m5C
H4K20 me2:H3–5ac
H4K20 me2:H3K4 me3
H4K20 me3
H3K27 me2
H2A.Z
H3K9 me3
H3K9 me2
H3K4 me1
H3K27 me3
H4K20 me2
H3–5ac
H4–4ac
H3K4 me3
DNA Meth. m5C
0.00 0.25 0.50 0.75 1.00
π̂
MTA2 – Forward
H2A.Z:H3–5ac
H2A.Z:H4–4ac
H3K27 me3:H4K20 me3
H3K4 me3:H3–5ac
H3K4 me1:H4–4ac
H3K9 me3:H4K20 me3
DNA Meth. m5C:H4K20 me3
H4–4ac:H3–5ac
H4K20 me2:H4–4ac
H2A.Z:H4K20 me2
H3K27 me3:H3K9 me3
H2A.Z:H3K4 me3
H3K27 me3:H3K4 me1
H3K27 me3:DNA Meth. m5C
H3K9 me2:DNA Meth. m5C
H4K20 me2:H3K4 me3
DNA Meth. m5C:H4K20 me2
H4K20 me2:H3–5ac
H3K4 me1:H3–5ac
DNA Meth. m5C:H3K9 me3
H3K4 me3:H4–4ac
H3K27 me2
H2A.Z
H4K20 me3
H3K9 me2
H3K27 me3
H4–4ac
H3K4 me1
H4K20 me2
H3K9 me3
H3K4 me3
H3–5ac
DNA Meth. m5C
0.00 0.25 0.50 0.75 1.00
π̂
MTA2 – Reverse
H2A.Z:H3–5ac
H2A.Z:H4–4ac
H3K27 me3:H4K20 me3
H3K4 me3:H3–5ac
H3K4 me1:H4–4ac
H3K9 me3:H4K20 me3
DNA Meth. m5C:H4K20 me3
H4–4ac:H3–5ac
H4K20 me2:H4–4ac
H2A.Z:H4K20 me2
H3K27 me3:H3K9 me3
H2A.Z:H3K4 me3
H3K27 me3:H3K4 me1
H3K27 me3:DNA Meth. m5C
H3K9 me2:DNA Meth. m5C
H4K20 me2:H3K4 me3
DNA Meth. m5C:H4K20 me2
H4K20 me2:H3–5ac
H3K4 me1:H3–5ac
DNA Meth. m5C:H3K9 me3
H3K4 me3:H4–4ac
H3K27 me2
H2A.Z
H4K20 me3
H3K9 me2
H3K27 me3
H4–4ac
H3K4 me1
H4K20 me2
H3K9 me3
H3K4 me3
H3–5ac
DNA Meth. m5C
0.00 0.25 0.50 0.75 1.00
π̂
CDK2AP1 – Forward
H3K4 me3:H3–5ac
H2A.Z:H4K20 me2
H3K4 me1:H4K20 me2
H3K9 me3:H4K20 me3
H3K27 me3:H4K20 me3
H4K20 me2:H3–5ac
H4K20 me2:H4–4ac
H3K4 me1:H4–4ac
H3K4 me1:H2A.Z
H2A.Z:H3K4 me3
DNA Meth. m5C:H4K20 me3
H3K27 me3:H3K9 me3
H4–4ac:H3–5ac
H3K27 me3:H3K4 me1
H4K20 me2:H3K4 me3
H3K27 me3:DNA Meth. m5C
DNA Meth. m5C:H3K9 me3
H3K4 me1:H3–5ac
H3K4 me3:H4–4ac
DNA Meth. m5C:H4K20 me2
H3K27 me2:DNA Meth. m5C
H4K20 me3
H3K27 me2
H2A.Z
H3K9 me2
H3K9 me3
H4K20 me2
H3K4 me1
H3K27 me3
H4–4ac
H3–5ac
H3K4 me3
DNA Meth. m5C
0.00 0.25 0.50 0.75 1.00
π̂
CDK2AP1 – Reverse
H3K4 me3:H3–5ac
H2A.Z:H4K20 me2
H3K4 me1:H4K20 me2
H3K9 me3:H4K20 me3
H3K27 me3:H4K20 me3
H4K20 me2:H3–5ac
H4K20 me2:H4–4ac
H3K4 me1:H4–4ac
H3K4 me1:H2A.Z
H2A.Z:H3K4 me3
DNA Meth. m5C:H4K20 me3
H3K27 me3:H3K9 me3
H4–4ac:H3–5ac
H3K27 me3:H3K4 me1
H4K20 me2:H3K4 me3
H3K27 me3:DNA Meth. m5C
DNA Meth. m5C:H3K9 me3
H3K4 me1:H3–5ac
H3K4 me3:H4–4ac
DNA Meth. m5C:H4K20 me2
H3K27 me2:DNA Meth. m5C
H4K20 me3
H3K27 me2
H2A.Z
H3K9 me2
H3K9 me3
H4K20 me2
H3K4 me1
H3K27 me3
H4–4ac
H3–5ac
H3K4 me3
DNA Meth. m5C
0.00 0.25 0.50 0.75 1.00
π̂
CHD4 – Forward
H3K4 me1:H4–4ac
H3K27 me3:H4K20 me3
H3K4 me3:H3–5ac
H4K20 me2:H4–4ac
H3K4 me1:H2A.Z
H4–4ac:H3–5ac
H2A.Z:H3K4 me3
H3K27 me3:H3K9 me3
H3K27 me3:H3K4 me1
H4K20 me2:H3–5ac
H3K27 me3:DNA Meth. m5C
H3K4 me3:H4–4ac
H3K9 me2:DNA Meth. m5C
DNA Meth. m5C:H4K20 me2
H3K4 me1:H3–5ac
DNA Meth. m5C:H4K20 me3
DNA Meth. m5C:H3K9 me3
H4K20 me2:H3K4 me3
H3K27 me2
H2A.Z
H4K20 me2
H4K20 me3
H4–4ac
H3K9 me2
H3K4 me1
H3K9 me3
H3K27 me3
H3–5ac
H3K4 me3
DNA Meth. m5C
0.00 0.25 0.50 0.75 1.00
π̂
CHD4 – Reverse
H3K4 me1:H4–4ac
H3K27 me3:H4K20 me3
H3K4 me3:H3–5ac
H4K20 me2:H4–4ac
H3K4 me1:H2A.Z
H4–4ac:H3–5ac
H2A.Z:H3K4 me3
H3K27 me3:H3K9 me3
H3K27 me3:H3K4 me1
H4K20 me2:H3–5ac
H3K27 me3:DNA Meth. m5C
H3K4 me3:H4–4ac
H3K9 me2:DNA Meth. m5C
DNA Meth. m5C:H4K20 me2
H3K4 me1:H3–5ac
DNA Meth. m5C:H4K20 me3
DNA Meth. m5C:H3K9 me3
H4K20 me2:H3K4 me3
H3K27 me2
H2A.Z
H4K20 me2
H4K20 me3
H4–4ac
H3K9 me2
H3K4 me1
H3K9 me3
H3K27 me3
H3–5ac
H3K4 me3
DNA Meth. m5C
0.00 0.25 0.50 0.75 1.00
π̂

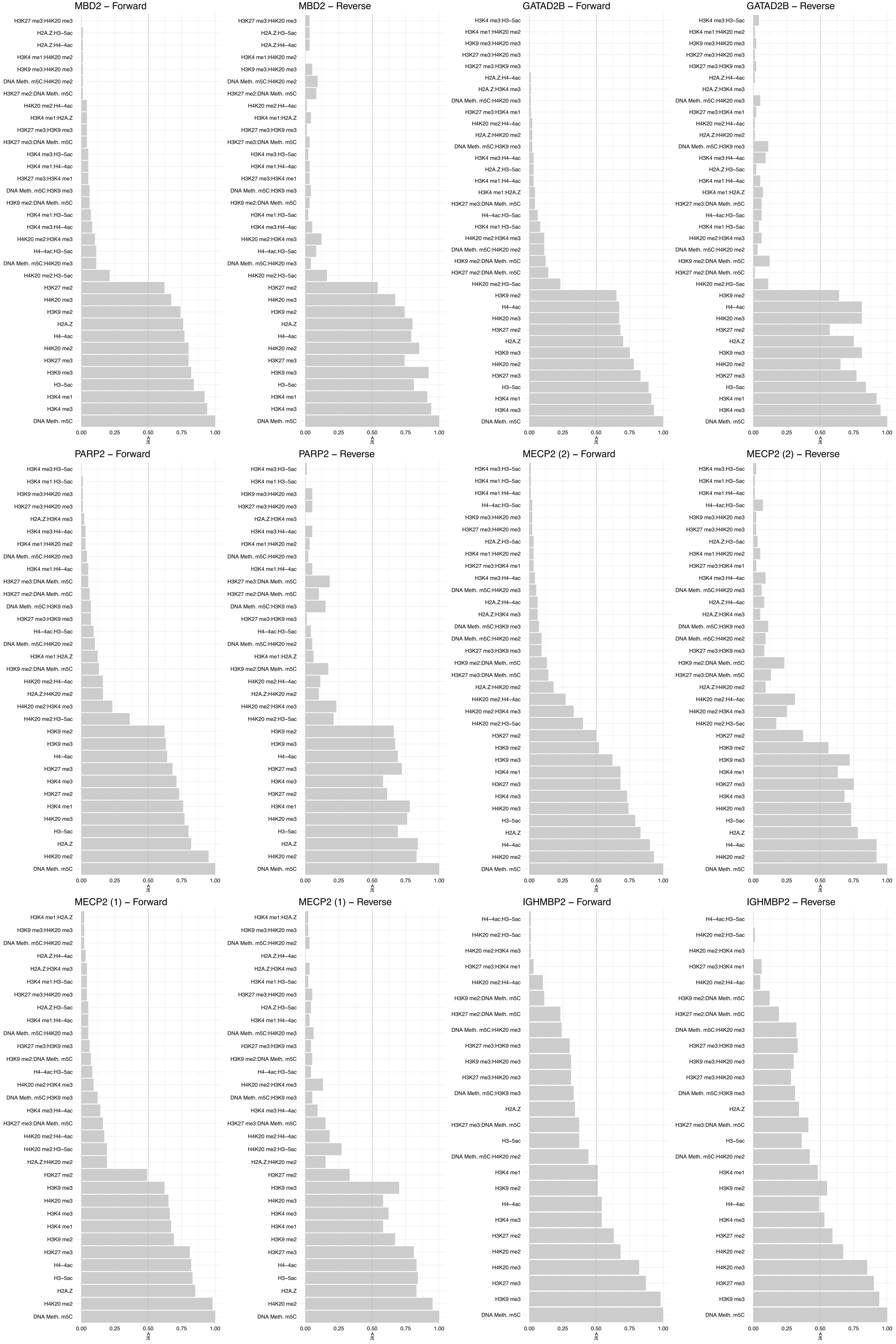
MBD2 – Forward
H3K27 me3:H4K20 me3
H2A.Z:H3−5ac
H2A.Z:H4−4ac
H3K4 me1:H4K20 me2
H3K9 me3:H4K20 me3
DNA Meth. m5C:H4K20 me2
H3K27 me2:DNA Meth. m5C
H4K20 me2:H4−4ac
H3K4 me1:H2A.Z
H3K27 me3:H3K9 me3
H3K27 me3:DNA Meth. m5C
H3K4 me3:H3−5ac
H3K4 me1:H4−4ac
H3K27 me3:H3K4 me1
DNA Meth. m5C:H3K9 me3
H3K9 me2:DNA Meth. m5C
H3K4 me1:H3−5ac
H3K4 me3:H4−4ac
H4K20 me2:H3K4 me3
H4−4ac:H3−5ac
DNA Meth. m5C:H4K20 me3
H4K20 me2:H3−5ac
H3K27 me2
H4K20 me3
H3K9 me2
H2A.Z
H4−4ac
H4K20 me2
H3K27 me3
H3K9 me3
H3−5ac
H3K4 me1
H3K4 me3
DNA Meth. m5C
0.00 0.25 0.50 0.75 1.00
π̂
MBD2 – Reverse
H3K27 me3:H4K20 me3
H2A.Z:H3−5ac
H2A.Z:H4−4ac
H3K4 me1:H4K20 me2
H3K9 me3:H4K20 me3
DNA Meth. m5C:H4K20 me2
H3K27 me2:DNA Meth. m5C
H4K20 me2:H4−4ac
H3K4 me1:H2A.Z
H3K27 me3:H3K9 me3
H3K27 me3:DNA Meth. m5C
H3K4 me3:H3−5ac
H3K4 me1:H4−4ac
H3K27 me3:H3K4 me1
DNA Meth. m5C:H3K9 me3
H3K9 me2:DNA Meth. m5C
H3K4 me1:H3−5ac
H3K4 me3:H4−4ac
H4K20 me2:H3K4 me3
H4−4ac:H3−5ac
DNA Meth. m5C:H4K20 me3
H4K20 me2:H3−5ac
H3K27 me2
H4K20 me3
H3K9 me2
H2A.Z
H4−4ac
H4K20 me2
H3K27 me3
H3K9 me3
H3−5ac
H3K4 me1
H3K4 me3
DNA Meth. m5C
0.00 0.25 0.50 0.75 1.00
π̂
GATAD2B – Forward
H3K4 me3:H3−5ac
H3K4 me1:H4K20 me2
H3K9 me3:H4K20 me3
H3K27 me3:H4K20 me3
H3K27 me3:H3K9 me3
H2A.Z:H4−4ac
H2A.Z:H3K4 me3
DNA Meth. m5C:H4K20 me3
H3K27 me3:H3K4 me1
H4K20 me2:H4−4ac
H2A.Z:H4K20 me2
DNA Meth. m5C:H3K9 me3
H3K4 me3:H4−4ac
H2A.Z:H3−5ac
H3K4 me1:H4−4ac
H3K4 me1:H2A.Z
H3K27 me3:DNA Meth. m5C
H4−4ac:H3−5ac
H3K4 me1:H3−5ac
H4K20 me2:H3K4 me3
DNA Meth. m5C:H4K20 me2
H3K9 me2:DNA Meth. m5C
H3K27 me2:DNA Meth. m5C
H4K20 me2:H3−5ac
H3K9 me2
H4−4ac
H4K20 me3
H3K27 me2
H2A.Z
H3K9 me3
H4K20 me2
H3K27 me3
H3−5ac
H3K4 me1
H3K4 me3
DNA Meth. m5C
0.00 0.25 0.50 0.75 1.00
π̂
GATAD2B – Reverse
H3K4 me3:H3−5ac
H3K4 me1:H4K20 me2
H3K9 me3:H4K20 me3
H3K27 me3:H4K20 me3
H3K27 me3:H3K9 me3
H2A.Z:H4−4ac
H2A.Z:H3K4 me3
DNA Meth. m5C:H4K20 me3
H3K27 me3:H3K4 me1
H4K20 me2:H4−4ac
H2A.Z:H4K20 me2
DNA Meth. m5C:H3K9 me3
H3K4 me3:H4−4ac
H2A.Z:H3−5ac
H3K4 me1:H4−4ac
H3K4 me1:H2A.Z
H3K27 me3:DNA Meth. m5C
H4−4ac:H3−5ac
H3K4 me1:H3−5ac
H4K20 me2:H3K4 me3
DNA Meth. m5C:H4K20 me2
H3K9 me2:DNA Meth. m5C
H3K27 me2:DNA Meth. m5C
H4K20 me2:H3−5ac
H3K9 me2
H4−4ac
H4K20 me3
H3K27 me2
H2A.Z
H3K9 me3
H4K20 me2
H3K27 me3
H3−5ac
H3K4 me1
H3K4 me3
DNA Meth. m5C
0.00 0.25 0.50 0.75 1.00
π̂
PARP2 – Forward
H3K4 me3:H3−5ac
H3K4 me1:H3−5ac
H3K9 me3:H4K20 me3
H3K27 me3:H4K20 me3
H2A.Z:H3K4 me3
H3K4 me3:H4−4ac
H3K4 me1:H4K20 me2
DNA Meth. m5C:H4K20 me3
H3K4 me1:H4−4ac
H3K27 me3:DNA Meth. m5C
H3K27 me2:DNA Meth. m5C
DNA Meth. m5C:H3K9 me3
H3K27 me3:H3K9 me3
H4−4ac:H3−5ac
DNA Meth. m5C:H4K20 me2
H3K4 me1:H2A.Z
H3K9 me2:DNA Meth. m5C
H4K20 me2:H4−4ac
H2A.Z:H4K20 me2
H4K20 me2:H3K4 me3
H4K20 me2:H3−5ac
H3K9 me2
H3K9 me3
H4−4ac
H3K27 me3
H3K4 me3
H3K27 me2
H3K4 me1
H4K20 me3
H3−5ac
H2A.Z
H4K20 me2
DNA Meth. m5C
0.00 0.25 0.50 0.75 1.00
π̂
PARP2 – Reverse
H3K4 me3:H3−5ac
H3K4 me1:H3−5ac
H3K9 me3:H4K20 me3
H3K27 me3:H4K20 me3
H2A.Z:H3K4 me3
H3K4 me3:H4−4ac
H3K4 me1:H4K20 me2
DNA Meth. m5C:H4K20 me3
H3K4 me1:H4−4ac
H3K27 me3:DNA Meth. m5C
H3K27 me2:DNA Meth. m5C
DNA Meth. m5C:H3K9 me3
H3K27 me3:H3K9 me3
H4−4ac:H3−5ac
DNA Meth. m5C:H4K20 me2
H3K4 me1:H2A.Z
H3K9 me2:DNA Meth. m5C
H4K20 me2:H4−4ac
H2A.Z:H4K20 me2
H4K20 me2:H3K4 me3
H4K20 me2:H3−5ac
H3K9 me2
H3K9 me3
H4−4ac
H3K27 me3
H3K4 me3
H3K27 me2
H3K4 me1
H4K20 me3
H3−5ac
H2A.Z
H4K20 me2
DNA Meth. m5C
0.00 0.25 0.50 0.75 1.00
π̂
MECP2 (2) – Forward
H3K4 me3:H3−5ac
H3K4 me1:H3−5ac
H3K4 me1:H4−4ac
H4−4ac:H3−5ac
H3K9 me3:H4K20 me3
H3K27 me3:H4K20 me3
H2A.Z:H3−5ac
H3K4 me1:H4K20 me2
H3K27 me3:H3K4 me1
H3K4 me3:H4−4ac
DNA Meth. m5C:H4K20 me3
H2A.Z:H4−4ac
H2A.Z:H3K4 me3
DNA Meth. m5C:H3K9 me3
DNA Meth. m5C:H4K20 me2
H3K27 me3:H3K9 me3
H3K9 me2:DNA Meth. m5C
H3K27 me3:DNA Meth. m5C
H2A.Z:H4K20 me2
H4K20 me2:H4−4ac
H4K20 me2:H3K4 me3
H4K20 me2:H3−5ac
H3K27 me2
H3K9 me2
H3K9 me3
H3K4 me1
H3K27 me3
H3K4 me3
H4K20 me3
H3−5ac
H2A.Z
H4−4ac
H4K20 me2
DNA Meth. m5C
0.00 0.25 0.50 0.75 1.00
π̂
MECP2 (2) – Reverse
H3K4 me3:H3−5ac
H3K4 me1:H3−5ac
H3K4 me1:H4−4ac
H4−4ac:H3−5ac
H3K9 me3:H4K20 me3
H3K27 me3:H4K20 me3
H2A.Z:H3−5ac
H3K4 me1:H4K20 me2
H3K27 me3:H3K4 me1
H3K4 me3:H4−4ac
DNA Meth. m5C:H4K20 me3
H2A.Z:H4−4ac
H2A.Z:H3K4 me3
DNA Meth. m5C:H3K9 me3
DNA Meth. m5C:H4K20 me2
H3K27 me3:H3K9 me3
H3K9 me2:DNA Meth. m5C
H3K27 me3:DNA Meth. m5C
H2A.Z:H4K20 me2
H4K20 me2:H4−4ac
H4K20 me2:H3K4 me3
H4K20 me2:H3−5ac
H3K27 me2
H3K9 me2
H3K9 me3
H3K4 me1
H3K27 me3
H3K4 me3
H4K20 me3
H3−5ac
H2A.Z
H4−4ac
H4K20 me2
DNA Meth. m5C
0.00 0.25 0.50 0.75 1.00
π̂
MECP2 (1) – Forward
H3K4 me1:H2A.Z
H3K9 me3:H4K20 me3
DNA Meth. m5C:H4K20 me2
H2A.Z:H4−4ac
H2A.Z:H3K4 me3
H3K4 me1:H3−5ac
H3K27 me3:H4K20 me3
H2A.Z:H3−5ac
H3K4 me1:H4−4ac
DNA Meth. m5C:H4K20 me3
H3K27 me3:H3K9 me3
H3K9 me2:DNA Meth. m5C
H4−4ac:H3−5ac
H4K20 me2:H3K4 me3
DNA Meth. m5C:H3K9 me3
H3K4 me3:H4−4ac
H3K27 me3:DNA Meth. m5C
H4K20 me2:H4−4ac
H4K20 me2:H3−5ac
H2A.Z:H4K20 me2
H3K27 me2
H3K9 me3
H4K20 me3
H3K4 me3
H3K4 me1
H3K9 me2
H3K27 me3
H4−4ac
H3−5ac
H2A.Z
H4K20 me2
DNA Meth. m5C
0.00 0.25 0.50 0.75 1.00
π̂
MECP2 (1) – Reverse
H3K4 me1:H2A.Z
H3K9 me3:H4K20 me3
DNA Meth. m5C:H4K20 me2
H2A.Z:H4−4ac
H2A.Z:H3K4 me3
H3K4 me1:H3−5ac
H3K27 me3:H4K20 me3
H2A.Z:H3−5ac
H3K4 me1:H4−4ac
DNA Meth. m5C:H4K20 me3
H3K27 me3:H3K9 me3
H3K9 me2:DNA Meth. m5C
H4−4ac:H3−5ac
H4K20 me2:H3K4 me3
DNA Meth. m5C:H3K9 me3
H3K4 me3:H4−4ac
H3K27 me3:DNA Meth. m5C
H4K20 me2:H4−4ac
H4K20 me2:H3−5ac
H2A.Z:H4K20 me2
H3K27 me2
H3K9 me3
H4K20 me3
H3K4 me3
H3K4 me1
H3K9 me2
H3K27 me3
H4−4ac
H3−5ac
H2A.Z
H4K20 me2
DNA Meth. m5C
0.00 0.25 0.50 0.75 1.00
π̂
IGHMBP2 – Forward
H4−4ac:H3−5ac
H4K20 me2:H3−5ac
H4K20 me2:H3K4 me3
H3K27 me3:H3K4 me1
H4K20 me2:H4−4ac
H3K9 me2:DNA Meth. m5C
H3K27 me2:DNA Meth. m5C
DNA Meth. m5C:H4K20 me3
H3K27 me3:H3K9 me3
H3K9 me3:H4K20 me3
H3K27 me3:H4K20 me3
DNA Meth. m5C:H3K9 me3
H2A.Z
H3K27 me3:DNA Meth. m5C
H3−5ac
DNA Meth. m5C:H4K20 me2
H3K4 me1
H3K9 me2
H4−4ac
H3K4 me3
H3K27 me2
H4K20 me2
H4K20 me3
H3K27 me3
H3K9 me3
DNA Meth. m5C
0.00 0.25 0.50 0.75 1.00
π̂
IGHMBP2 – Reverse
H4−4ac:H3−5ac
H4K20 me2:H3−5ac
H4K20 me2:H3K4 me3
H3K27 me3:H3K4 me1
H4K20 me2:H4−4ac
H3K9 me2:DNA Meth. m5C
H3K27 me2:DNA Meth. m5C
DNA Meth. m5C:H4K20 me3
H3K27 me3:H3K9 me3
H3K9 me3:H4K20 me3
H3K27 me3:H4K20 me3
DNA Meth. m5C:H3K9 me3
H2A.Z
H3K27 me3:DNA Meth. m5C
H3−5ac
DNA Meth. m5C:H4K20 me2
H3K4 me1
H3K9 me2
H4−4ac
H3K4 me3
H3K27 me2
H4K20 me2
H4K20 me3
H3K27 me3
H3K9 me3
DNA Meth. m5C
0.00 0.25 0.50 0.75 1.00
π̂

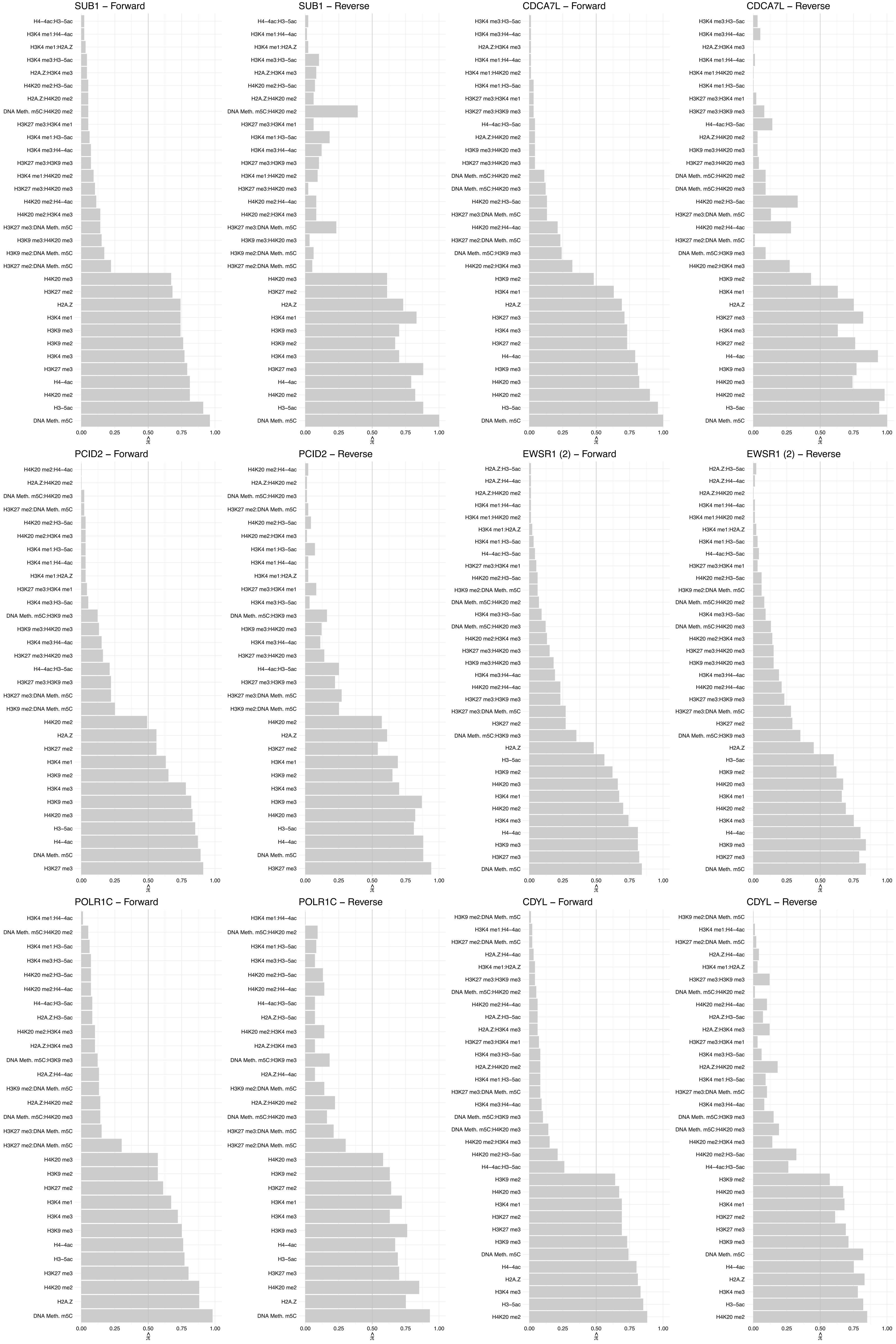

SUB1 – Forward
H4–4ac:H3–5ac
H3K4 me1:H4–4ac
H3K4 me1:H2A.Z
H3K4 me3:H3–5ac
H2A.Z:H3K4 me3
H4K20 me2:H3–5ac
H2A.Z:H4K20 me2
DNA Meth. m5C:H4K20 me2
H3K27 me3:H3K4 me1
H3K4 me1:H3–5ac
H3K4 me3:H4–4ac
H3K27 me3:H3K9 me3
H3K4 me1:H4K20 me2
H3K27 me3:H4K20 me3
H4K20 me2:H4–4ac
H4K20 me2:H3K4 me3
H3K27 me3:DNA Meth. m5C
H3K9 me3:H4K20 me3
H3K9 me2:DNA Meth. m5C
H3K27 me2:DNA Meth. m5C
H4K20 me3
H3K27 me2
H2A.Z
H3K4 me1
H3K9 me3
H3K9 me2
H3K4 me3
H3K27 me3
H4–4ac
H4K20 me2
H3–5ac
DNA Meth. m5C
0.00
0.25
0.50
0.75
1.00
π̂
SUB1 – Reverse
H4–4ac:H3–5ac
H3K4 me1:H4–4ac
H3K4 me1:H2A.Z
H3K4 me3:H3–5ac
H2A.Z:H3K4 me3
H4K20 me2:H3–5ac
H2A.Z:H4K20 me2
DNA Meth. m5C:H4K20 me2
H3K27 me3:H3K4 me1
H3K4 me1:H3–5ac
H3K4 me3:H4–4ac
H3K27 me3:H3K9 me3
H3K4 me1:H4K20 me2
H3K27 me3:H4K20 me3
H4K20 me2:H4–4ac
H4K20 me2:H3K4 me3
H3K27 me3:DNA Meth. m5C
H3K9 me3:H4K20 me3
H3K9 me2:DNA Meth. m5C
H3K27 me2:DNA Meth. m5C
H4K20 me3
H3K27 me2
H2A.Z
H3K4 me1
H3K9 me3
H3K9 me2
H3K4 me3
H3K27 me3
H4–4ac
H4K20 me2
H3–5ac
DNA Meth. m5C
0.00
0.25
0.50
0.75
1.00
π̂
CDCA7L – Forward
H3K4 me3:H3–5ac
H3K4 me3:H4–4ac
H2A.Z:H3K4 me3
H3K4 me1:H4–4ac
H3K4 me1:H4K20 me2
H3K4 me1:H3–5ac
H3K27 me3:H3K4 me1
H3K27 me3:H3K9 me3
H4–4ac:H3–5ac
H2A.Z:H4K20 me2
H3K9 me3:H4K20 me3
H3K27 me3:H4K20 me3
DNA Meth. m5C:H4K20 me2
DNA Meth. m5C:H4K20 me3
H4K20 me2:H3–5ac
H3K27 me3:DNA Meth. m5C
H4K20 me2:H4–4ac
H3K27 me2:DNA Meth. m5C
DNA Meth. m5C:H3K9 me3
H4K20 me2:H3K4 me3
H3K9 me2
H3K4 me1
H2A.Z
H3K27 me3
H3K4 me3
H3K27 me2
H4–4ac
H3K9 me3
H4K20 me3
H4K20 me2
H3–5ac
DNA Meth. m5C
0.00
0.25
0.50
0.75
1.00
π̂
CDCA7L – Reverse
H3K4 me3:H3–5ac
H3K4 me3:H4–4ac
H2A.Z:H3K4 me3
H3K4 me1:H4–4ac
H3K4 me1:H4K20 me2
H3K4 me1:H3–5ac
H3K27 me3:H3K4 me1
H3K27 me3:H3K9 me3
H4–4ac:H3–5ac
H2A.Z:H4K20 me2
H3K9 me3:H4K20 me3
H3K27 me3:H4K20 me3
DNA Meth. m5C:H4K20 me2
DNA Meth. m5C:H4K20 me3
H4K20 me2:H3–5ac
H3K27 me3:DNA Meth. m5C
H4K20 me2:H4–4ac
H3K27 me2:DNA Meth. m5C
DNA Meth. m5C:H3K9 me3
H4K20 me2:H3K4 me3
H3K9 me2
H3K4 me1
H2A.Z
H3K27 me3
H3K4 me3
H3K27 me2
H4–4ac
H3K9 me3
H4K20 me3
H4K20 me2
H3–5ac
DNA Meth. m5C
0.00
0.25
0.50
0.75
1.00
π̂
PCID2 – Forward
H4K20 me2:H4–4ac
H2A.Z:H4K20 me2
DNA Meth. m5C:H4K20 me3
H3K27 me2:DNA Meth. m5C
H4K20 me2:H3–5ac
H4K20 me2:H3K4 me3
H3K4 me1:H3–5ac
H3K4 me1:H4–4ac
H3K4 me1:H2A.Z
H3K27 me3:H3K4 me1
H3K4 me3:H3–5ac
DNA Meth. m5C:H3K9 me3
H3K9 me3:H4K20 me3
H3K4 me3:H4–4ac
H3K27 me3:H4K20 me3
H4–4ac:H3–5ac
H3K27 me3:H3K9 me3
H3K27 me3:DNA Meth. m5C
H3K9 me2:DNA Meth. m5C
H4K20 me2
H2A.Z
H3K27 me2
H3K4 me1
H3K9 me2
H3K4 me3
H3K9 me3
H4K20 me3
H3–5ac
H4–4ac
DNA Meth. m5C
H3K27 me3
0.00
0.25
0.50
0.75
1.00
π̂
PCID2 – Reverse
H4K20 me2:H4–4ac
H2A.Z:H4K20 me2
DNA Meth. m5C:H4K20 me3
H3K27 me2:DNA Meth. m5C
H4K20 me2:H3–5ac
H4K20 me2:H3K4 me3
H3K4 me1:H3–5ac
H3K4 me1:H4–4ac
H3K4 me1:H2A.Z
H3K27 me3:H3K4 me1
H3K4 me3:H3–5ac
DNA Meth. m5C:H3K9 me3
H3K9 me3:H4K20 me3
H3K4 me3:H4–4ac
H3K27 me3:H4K20 me3
H4–4ac:H3–5ac
H3K27 me3:H3K9 me3
H3K27 me3:DNA Meth. m5C
H3K9 me2:DNA Meth. m5C
H4K20 me2
H2A.Z
H3K27 me2
H3K4 me1
H3K9 me2
H3K4 me3
H3K9 me3
H4K20 me3
H3–5ac
H4–4ac
DNA Meth. m5C
H3K27 me3
0.00
0.25
0.50
0.75
1.00
π̂
EWSR1 (2) – Forward
H2A.Z:H3–5ac
H2A.Z:H4–4ac
H2A.Z:H4K20 me2
H3K4 me1:H4–4ac
H3K4 me1:H4K20 me2
H3K4 me1:H2A.Z
H3K4 me1:H3–5ac
H4–4ac:H3–5ac
H3K27 me3:H3K4 me1
H4K20 me2:H3–5ac
H3K9 me2:DNA Meth. m5C
DNA Meth. m5C:H4K20 me2
H3K4 me3:H3–5ac
DNA Meth. m5C:H4K20 me3
H4K20 me2:H3K4 me3
H3K27 me3:H4K20 me3
H3K9 me3:H4K20 me3
H3K4 me3:H4–4ac
H4K20 me2:H4–4ac
H3K27 me3:H3K9 me3
H3K27 me3:DNA Meth. m5C
H3K27 me2
DNA Meth. m5C:H3K9 me3
H2A.Z
H3–5ac
H3K9 me2
H4K20 me3
H3K4 me1
H4K20 me2
H3K4 me3
H4–4ac
H3K9 me3
H3K27 me3
DNA Meth. m5C
0.00
0.25
0.50
0.75
1.00
π̂
EWSR1 (2) – Reverse
H2A.Z:H3–5ac
H2A.Z:H4–4ac
H2A.Z:H4K20 me2
H3K4 me1:H4–4ac
H3K4 me1:H4K20 me2
H3K4 me1:H2A.Z
H3K4 me1:H3–5ac
H4–4ac:H3–5ac
H3K27 me3:H3K4 me1
H4K20 me2:H3–5ac
H3K9 me2:DNA Meth. m5C
DNA Meth. m5C:H4K20 me2
H3K4 me3:H3–5ac
DNA Meth. m5C:H4K20 me3
H4K20 me2:H3K4 me3
H3K27 me3:H4K20 me3
H3K9 me3:H4K20 me3
H3K4 me3:H4–4ac
H4K20 me2:H4–4ac
H3K27 me3:H3K9 me3
H3K27 me3:DNA Meth. m5C
H3K27 me2
DNA Meth. m5C:H3K9 me3
H2A.Z
H3–5ac
H3K9 me2
H4K20 me3
H3K4 me1
H4K20 me2
H3K4 me3
H4–4ac
H3K9 me3
H3K27 me3
DNA Meth. m5C
0.00
0.25
0.50
0.75
1.00
π̂
POLR1C – Forward
H3K4 me1:H4–4ac
DNA Meth. m5C:H4K20 me2
H3K4 me1:H3–5ac
H3K4 me3:H3–5ac
H4K20 me2:H3–5ac
H4K20 me2:H4–4ac
H4–4ac:H3–5ac
H2A.Z:H3–5ac
H4K20 me2:H3K4 me3
H2A.Z:H3K4 me3
DNA Meth. m5C:H3K9 me3
H2A.Z:H4–4ac
H3K9 me2:DNA Meth. m5C
H2A.Z:H4K20 me2
DNA Meth. m5C:H4K20 me3
H3K27 me3:DNA Meth. m5C
H3K27 me2:DNA Meth. m5C
H4K20 me3
H3K9 me2
H3K27 me2
H3K4 me1
H3K4 me3
H3K9 me3
H4–4ac
H3–5ac
H3K27 me3
H4K20 me2
H2A.Z
DNA Meth. m5C
0.00
0.25
0.50
0.75
1.00
π̂
POLR1C – Reverse
H3K4 me1:H4–4ac
DNA Meth. m5C:H4K20 me2
H3K4 me1:H3–5ac
H3K4 me3:H3–5ac
H4K20 me2:H3–5ac
H4K20 me2:H4–4ac
H4–4ac:H3–5ac
H2A.Z:H3–5ac
H4K20 me2:H3K4 me3
H2A.Z:H3K4 me3
DNA Meth. m5C:H3K9 me3
H2A.Z:H4–4ac
H3K9 me2:DNA Meth. m5C
H2A.Z:H4K20 me2
DNA Meth. m5C:H4K20 me3
H3K27 me3:DNA Meth. m5C
H3K27 me2:DNA Meth. m5C
H4K20 me3
H3K9 me2
H3K27 me2
H3K4 me1
H3K4 me3
H3K9 me3
H4–4ac
H3–5ac
H3K27 me3
H4K20 me2
H2A.Z
DNA Meth. m5C
0.00
0.25
0.50
0.75
1.00
π̂
CDYL – Forward
H3K9 me2:DNA Meth. m5C
H3K4 me1:H4–4ac
H3K27 me2:DNA Meth. m5C
H2A.Z:H4–4ac
H3K4 me1:H2A.Z
H3K27 me3:H3K9 me3
DNA Meth. m5C:H4K20 me2
H4K20 me2:H4–4ac
H2A.Z:H3–5ac
H2A.Z:H3K4 me3
H3K27 me3:H3K4 me1
H3K4 me3:H3–5ac
H2A.Z:H4K20 me2
H3K4 me1:H3–5ac
H3K27 me3:DNA Meth. m5C
H3K4 me3:H4–4ac
DNA Meth. m5C:H3K9 me3
DNA Meth. m5C:H4K20 me3
H4K20 me2:H3K4 me3
H4K20 me2:H3–5ac
H4–4ac:H3–5ac
H3K9 me2
H4K20 me3
H3K4 me1
H3K27 me2
H3K27 me3
H3K9 me3
DNA Meth. m5C
H4–4ac
H2A.Z
H3K4 me3
H3–5ac
H4K20 me2
0.00
0.25
0.50
0.75
1.00
π̂
CDYL – Reverse
H3K9 me2:DNA Meth. m5C
H3K4 me1:H4–4ac
H3K27 me2:DNA Meth. m5C
H2A.Z:H4–4ac
H3K4 me1:H2A.Z
H3K27 me3:H3K9 me3
DNA Meth. m5C:H4K20 me2
H4K20 me2:H4–4ac
H2A.Z:H3–5ac
H2A.Z:H3K4 me3
H3K27 me3:H3K4 me1
H3K4 me3:H3–5ac
H2A.Z:H4K20 me2
H3K4 me1:H3–5ac
H3K27 me3:DNA Meth. m5C
H3K4 me3:H4–4ac
DNA Meth. m5C:H3K9 me3
DNA Meth. m5C:H4K20 me3
H4K20 me2:H3K4 me3
H4K20 me2:H3–5ac
H4–4ac:H3–5ac
H3K9 me2
H4K20 me3
H3K4 me1
H3K27 me2
H3K27 me3
H3K9 me3
DNA Meth. m5C
H4–4ac
H2A.Z
H3K4 me3
H3–5ac
H4K20 me2
0.00
0.25
0.50
0.75
1.00
π̂

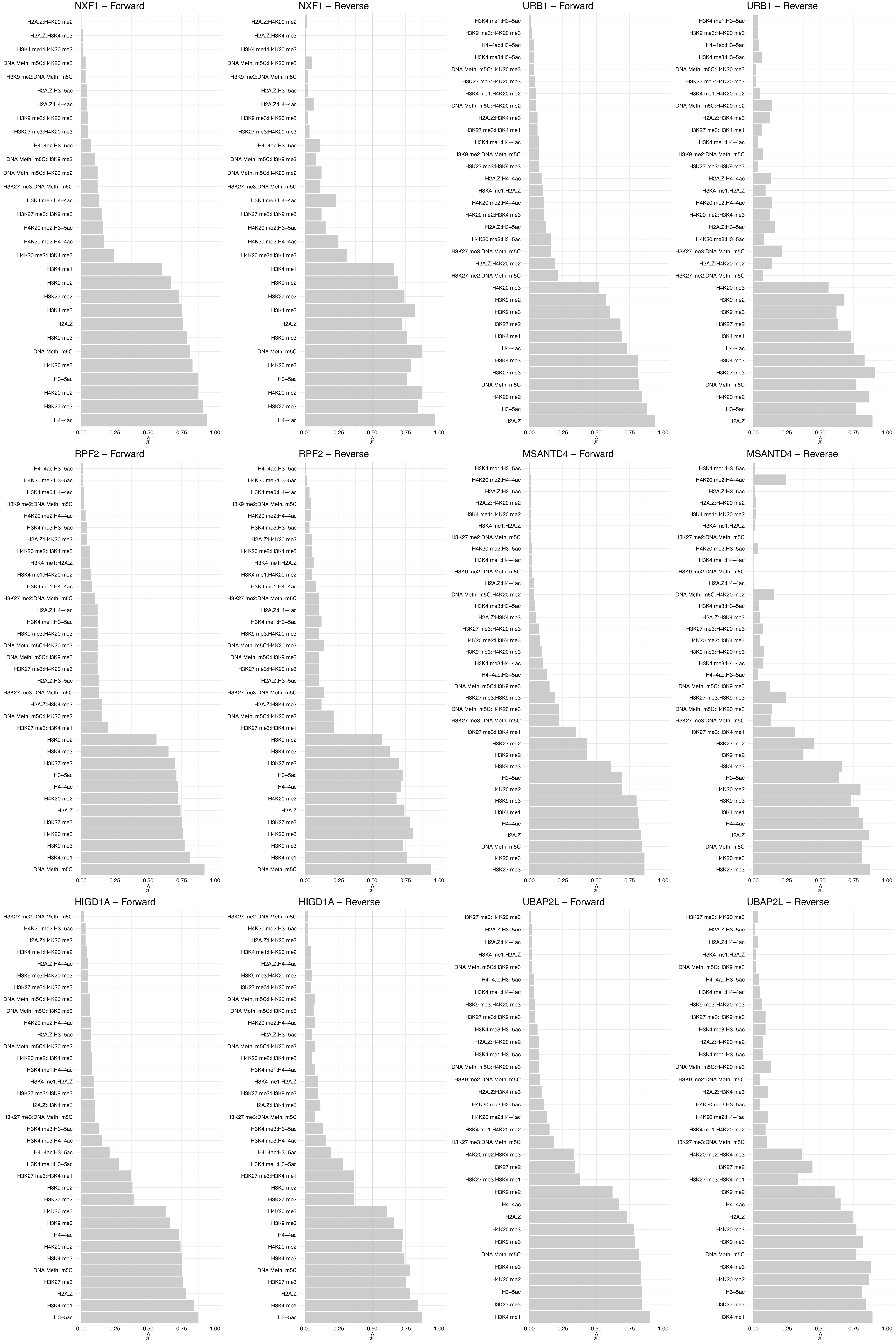
NXF1 – Forward
H2A.Z:H4K20 me2
H2A.Z:H3K4 me3
H3K4 me1:H4K20 me2
DNA Meth. m5C:H4K20 me3
H3K9 me2:DNA Meth. m5C
H2A.Z:H3−5ac
H2A.Z:H4−4ac
H3K9 me3:H4K20 me3
H3K27 me3:H4K20 me3
H4−4ac:H3−5ac
DNA Meth. m5C:H3K9 me3
DNA Meth. m5C:H4K20 me2
H3K27 me3:DNA Meth. m5C
H3K4 me3:H4−4ac
H3K27 me3:H3K9 me3
H4K20 me2:H3−5ac
H4K20 me2:H4−4ac
H4K20 me2:H3K4 me3
H3K4 me1
H3K9 me2
H3K27 me2
H3K4 me3
H2A.Z
H3K9 me3
DNA Meth. m5C
H4K20 me3
H3−5ac
H4K20 me2
H3K27 me3
H4−4ac
0.00 0.25 0.50 0.75 1.00
$\hat{\pi}$
NXF1 – Reverse
H2A.Z:H4K20 me2
H2A.Z:H3K4 me3
H3K4 me1:H4K20 me2
DNA Meth. m5C:H4K20 me3
H3K9 me2:DNA Meth. m5C
H2A.Z:H3−5ac
H2A.Z:H4−4ac
H3K9 me3:H4K20 me3
H3K27 me3:H4K20 me3
H4−4ac:H3−5ac
DNA Meth. m5C:H3K9 me3
DNA Meth. m5C:H4K20 me2
H3K27 me3:DNA Meth. m5C
H3K4 me3:H4−4ac
H3K27 me3:H3K9 me3
H4K20 me2:H3−5ac
H4K20 me2:H4−4ac
H4K20 me2:H3K4 me3
H3K4 me1
H3K9 me2
H3K27 me2
H3K4 me3
H2A.Z
H3K9 me3
DNA Meth. m5C
H4K20 me3
H3−5ac
H4K20 me2
H3K27 me3
H4−4ac
0.00 0.25 0.50 0.75 1.00
$\hat{\pi}$
URB1 – Forward
H3K4 me1:H3−5ac
H3K9 me3:H4K20 me3
H4−4ac:H3−5ac
H3K4 me3:H3−5ac
DNA Meth. m5C:H4K20 me3
H3K27 me3:H4K20 me3
H3K4 me1:H4K20 me2
DNA Meth. m5C:H4K20 me2
H2A.Z:H3K4 me3
H3K27 me3:H3K4 me1
H3K4 me1:H4−4ac
H3K9 me2:DNA Meth. m5C
H3K27 me3:H3K9 me3
H2A.Z:H4−4ac
H3K4 me1:H2A.Z
H4K20 me2:H4−4ac
H4K20 me2:H3K4 me3
H2A.Z:H3−5ac
H4K20 me2:H3−5ac
H3K27 me3:DNA Meth. m5C
H2A.Z:H4K20 me2
H3K27 me2:DNA Meth. m5C
H4K20 me3
H3K9 me2
H3K9 me3
H3K27 me2
H3K4 me1
H4−4ac
H3K4 me3
H3K27 me3
DNA Meth. m5C
H4K20 me2
H3−5ac
H2A.Z
0.00 0.25 0.50 0.75 1.00
$\hat{\pi}$
URB1 – Reverse
H3K4 me1:H3−5ac
H3K9 me3:H4K20 me3
H4−4ac:H3−5ac
H3K4 me3:H3−5ac
DNA Meth. m5C:H4K20 me3
H3K27 me3:H4K20 me3
H3K4 me1:H4K20 me2
DNA Meth. m5C:H4K20 me2
H2A.Z:H3K4 me3
H3K27 me3:H3K4 me1
H3K4 me1:H4−4ac
H3K9 me2:DNA Meth. m5C
H3K27 me3:H3K9 me3
H2A.Z:H4−4ac
H3K4 me1:H2A.Z
H4K20 me2:H4−4ac
H4K20 me2:H3K4 me3
H2A.Z:H3−5ac
H4K20 me2:H3−5ac
H3K27 me3:DNA Meth. m5C
H2A.Z:H4K20 me2
H3K27 me2:DNA Meth. m5C
H4K20 me3
H3K9 me2
H3K9 me3
H3K27 me2
H3K4 me1
H4−4ac
H3K4 me3
H3K27 me3
DNA Meth. m5C
H4K20 me2
H3−5ac
H2A.Z
0.00 0.25 0.50 0.75 1.00
$\hat{\pi}$
RPF2 – Forward
H4−4ac:H3−5ac
H4K20 me2:H3−5ac
H3K4 me3:H4−4ac
H3K9 me2:DNA Meth. m5C
H4K20 me2:H4−4ac
H3K4 me3:H3−5ac
H2A.Z:H4K20 me2
H4K20 me2:H3K4 me3
H3K4 me1:H2A.Z
H3K4 me1:H4K20 me2
H3K4 me1:H4−4ac
H3K27 me2:DNA Meth. m5C
H2A.Z:H4−4ac
H3K4 me1:H3−5ac
H3K9 me3:H4K20 me3
DNA Meth. m5C:H4K20 me3
DNA Meth. m5C:H3K9 me3
H3K27 me3:H4K20 me3
H2A.Z:H3−5ac
H3K27 me3:DNA Meth. m5C
H2A.Z:H3K4 me3
DNA Meth. m5C:H4K20 me2
H3K27 me3:H3K4 me1
H3K9 me2
H3K4 me3
H3K27 me2
H3−5ac
H4−4ac
H4K20 me2
H2A.Z
H3K27 me3
H4K20 me3
H3K9 me3
H3K4 me1
DNA Meth. m5C
0.00 0.25 0.50 0.75 1.00
$\hat{\pi}$
RPF2 – Reverse
H4−4ac:H3−5ac
H4K20 me2:H3−5ac
H3K4 me3:H4−4ac
H3K9 me2:DNA Meth. m5C
H4K20 me2:H4−4ac
H3K4 me3:H3−5ac
H2A.Z:H4K20 me2
H4K20 me2:H3K4 me3
H3K4 me1:H2A.Z
H3K4 me1:H4K20 me2
H3K4 me1:H4−4ac
H3K27 me2:DNA Meth. m5C
H2A.Z:H4−4ac
H3K4 me1:H3−5ac
H3K9 me3:H4K20 me3
DNA Meth. m5C:H4K20 me3
DNA Meth. m5C:H3K9 me3
H3K27 me3:H4K20 me3
H2A.Z:H3−5ac
H3K27 me3:DNA Meth. m5C
H2A.Z:H3K4 me3
DNA Meth. m5C:H4K20 me2
H3K27 me3:H3K4 me1
H3K9 me2
H3K4 me3
H3K27 me2
H3−5ac
H4−4ac
H4K20 me2
H2A.Z
H3K27 me3
H4K20 me3
H3K9 me3
H3K4 me1
DNA Meth. m5C
0.00 0.25 0.50 0.75 1.00
$\hat{\pi}$
MSANTD4 – Forward
H3K4 me1:H3−5ac
H4K20 me2:H4−4ac
H2A.Z:H3−5ac
H2A.Z:H4K20 me2
H3K4 me1:H4K20 me2
H3K4 me1:H2A.Z
H3K27 me2:DNA Meth. m5C
H4K20 me2:H3−5ac
H3K4 me1:H4−4ac
H3K9 me2:DNA Meth. m5C
H2A.Z:H4−4ac
DNA Meth. m5C:H4K20 me2
H3K4 me3:H3−5ac
H2A.Z:H3K4 me3
H3K27 me3:H4K20 me3
H4K20 me2:H3K4 me3
H3K9 me3:H4K20 me3
H3K4 me3:H4−4ac
H4−4ac:H3−5ac
DNA Meth. m5C:H3K9 me3
H3K27 me3:H3K9 me3
DNA Meth. m5C:H4K20 me3
H3K27 me3:DNA Meth. m5C
H3K27 me3:H3K4 me1
H3K27 me2
H3K9 me2
H3K4 me3
H3−5ac
H4K20 me2
H3K9 me3
H3K4 me1
H4−4ac
H2A.Z
DNA Meth. m5C
H4K20 me3
H3K27 me3
0.00 0.25 0.50 0.75 1.00
$\hat{\pi}$
MSANTD4 – Reverse
H3K4 me1:H3−5ac
H4K20 me2:H4−4ac
H2A.Z:H3−5ac
H2A.Z:H4K20 me2
H3K4 me1:H4K20 me2
H3K4 me1:H2A.Z
H3K27 me2:DNA Meth. m5C
H4K20 me2:H3−5ac
H3K4 me1:H4−4ac
H3K9 me2:DNA Meth. m5C
H2A.Z:H4−4ac
DNA Meth. m5C:H4K20 me2
H3K4 me3:H3−5ac
H2A.Z:H3K4 me3
H3K27 me3:H4K20 me3
H4K20 me2:H3K4 me3
H3K9 me3:H4K20 me3
H3K4 me3:H4−4ac
H4−4ac:H3−5ac
DNA Meth. m5C:H3K9 me3
H3K27 me3:H3K9 me3
DNA Meth. m5C:H4K20 me3
H3K27 me3:DNA Meth. m5C
H3K27 me3:H3K4 me1
H3K27 me2
H3K9 me2
H3K4 me3
H3−5ac
H4K20 me2
H3K9 me3
H3K4 me1
H4−4ac
H2A.Z
DNA Meth. m5C
H4K20 me3
H3K27 me3
0.00 0.25 0.50 0.75 1.00
$\hat{\pi}$
HIGD1A – Forward
H3K27 me2:DNA Meth. m5C
H4K20 me2:H3−5ac
H2A.Z:H4K20 me2
H3K4 me1:H4K20 me2
H2A.Z:H4−4ac
H3K9 me3:H4K20 me3
H3K27 me3:H4K20 me3
DNA Meth. m5C:H4K20 me3
DNA Meth. m5C:H3K9 me3
H4K20 me2:H4−4ac
H2A.Z:H3−5ac
DNA Meth. m5C:H4K20 me2
H4K20 me2:H3K4 me3
H3K4 me1:H4−4ac
H3K4 me1:H2A.Z
H3K27 me3:H3K9 me3
H2A.Z:H3K4 me3
H3K27 me3:DNA Meth. m5C
H3K4 me3:H3−5ac
H3K4 me3:H4−4ac
H4−4ac:H3−5ac
H3K4 me1:H3−5ac
H3K27 me3:H3K4 me1
H3K9 me2
H3K27 me2
H4K20 me3
H3K9 me3
H4−4ac
H4K20 me2
H3K4 me3
DNA Meth. m5C
H3K27 me3
H2A.Z
H3K4 me1
H3−5ac
0.00 0.25 0.50 0.75 1.00
$\hat{\pi}$
HIGD1A – Reverse
H3K27 me2:DNA Meth. m5C
H4K20 me2:H3−5ac
H2A.Z:H4K20 me2
H3K4 me1:H4K20 me2
H2A.Z:H4−4ac
H3K9 me3:H4K20 me3
H3K27 me3:H4K20 me3
DNA Meth. m5C:H4K20 me3
DNA Meth. m5C:H3K9 me3
H4K20 me2:H4−4ac
H2A.Z:H3−5ac
DNA Meth. m5C:H4K20 me2
H4K20 me2:H3K4 me3
H3K4 me1:H4−4ac
H3K4 me1:H2A.Z
H3K27 me3:H3K9 me3
H2A.Z:H3K4 me3
H3K27 me3:DNA Meth. m5C
H3K4 me3:H3−5ac
H3K4 me3:H4−4ac
H4−4ac:H3−5ac
H3K4 me1:H3−5ac
H3K27 me3:H3K4 me1
H3K9 me2
H3K27 me2
H4K20 me3
H3K9 me3
H4−4ac
H4K20 me2
H3K4 me3
DNA Meth. m5C
H3K27 me3
H2A.Z
H3K4 me1
H3−5ac
0.00 0.25 0.50 0.75 1.00
$\hat{\pi}$
UBAP2L – Forward
H3K27 me3:H4K20 me3
H2A.Z:H3−5ac
H2A.Z:H4−4ac
H3K4 me1:H2A.Z
DNA Meth. m5C:H3K9 me3
H4−4ac:H3−5ac
H3K4 me1:H4−4ac
H3K9 me3:H4K20 me3
H3K27 me3:H3K9 me3
H3K4 me3:H3−5ac
H2A.Z:H4K20 me2
H3K4 me1:H3−5ac
DNA Meth. m5C:H4K20 me3
H3K9 me2:DNA Meth. m5C
H2A.Z:H3K4 me3
H4K20 me2:H3−5ac
H4K20 me2:H4−4ac
H3K4 me1:H4K20 me2
H3K27 me3:DNA Meth. m5C
H4K20 me2:H3K4 me3
H3K27 me2
H3K27 me3:H3K4 me1
H3K9 me2
H4−4ac
H2A.Z
H4K20 me3
H3K9 me3
DNA Meth. m5C
H3K4 me3
H4K20 me2
H3−5ac
H3K27 me3
H3K4 me1
0.00 0.25 0.50 0.75 1.00
$\hat{\pi}$
UBAP2L – Reverse
H3K27 me3:H4K20 me3
H2A.Z:H3−5ac
H2A.Z:H4−4ac
H3K4 me1:H2A.Z
DNA Meth. m5C:H3K9 me3
H4−4ac:H3−5ac
H3K4 me1:H4−4ac
H3K9 me3:H4K20 me3
H3K27 me3:H3K9 me3
H3K4 me3:H3−5ac
H2A.Z:H4K20 me2
H3K4 me1:H3−5ac
DNA Meth. m5C:H4K20 me3
H3K9 me2:DNA Meth. m5C
H2A.Z:H3K4 me3
H4K20 me2:H3−5ac
H4K20 me2:H4−4ac
H3K4 me1:H4K20 me2
H3K27 me3:DNA Meth. m5C
H4K20 me2:H3K4 me3
H3K27 me2
H3K27 me3:H3K4 me1
H3K9 me2
H4−4ac
H2A.Z
H4K20 me3
H3K9 me3
DNA Meth. m5C
H3K4 me3
H4K20 me2
H3−5ac
H3K27 me3
H3K4 me1
0.00 0.25 0.50 0.75 1.00
$\hat{\pi}$

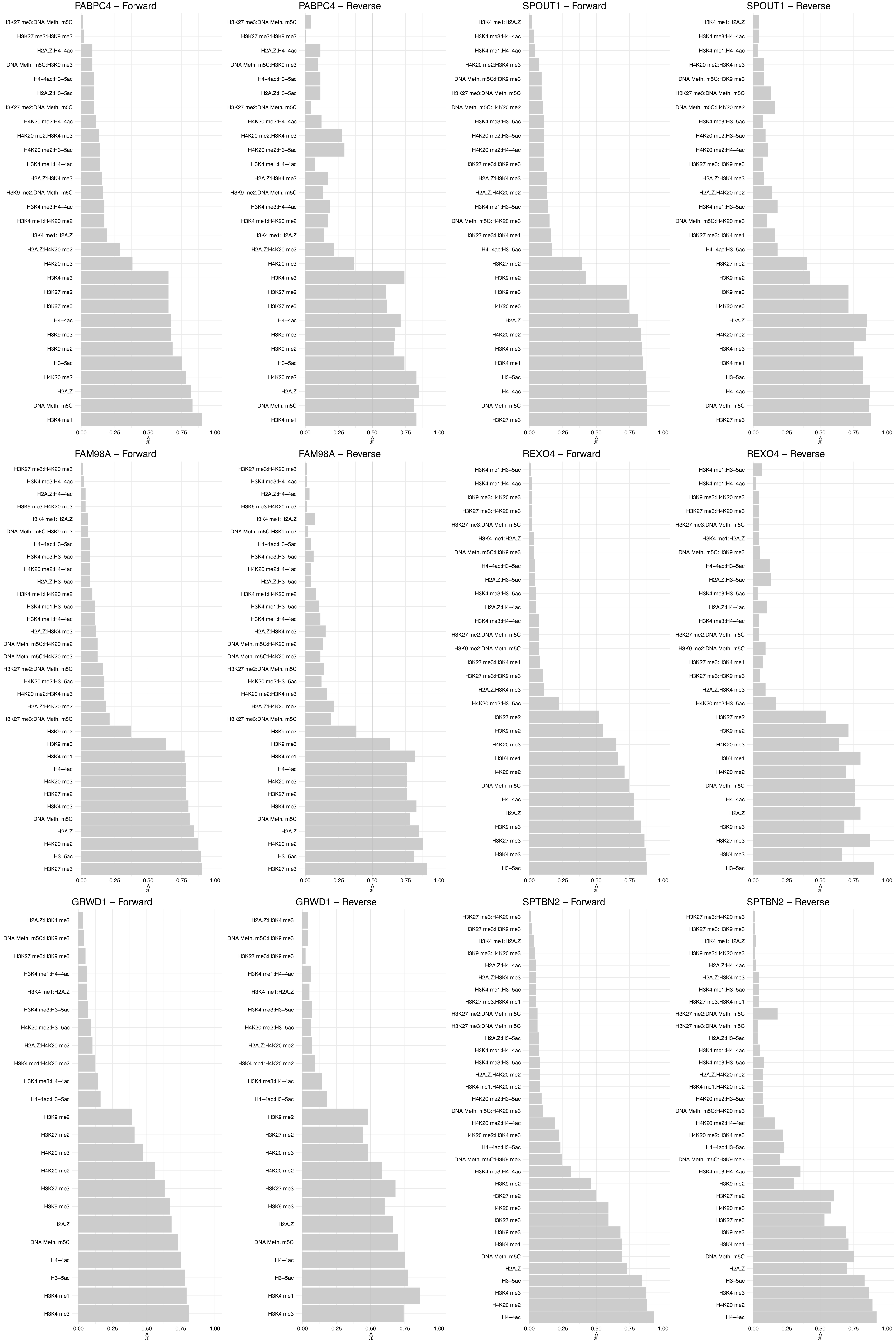
PABPC4 – Forward
H3K27 me3:DNA Meth. m5C
H3K27 me3:H3K9 me3
H2A.Z:H4–4ac
DNA Meth. m5C:H3K9 me3
H4–4ac:H3–5ac
H2A.Z:H3–5ac
H3K27 me2:DNA Meth. m5C
H4K20 me2:H4–4ac
H4K20 me2:H3K4 me3
H4K20 me2:H3–5ac
H3K4 me1:H4–4ac
H2A.Z:H3K4 me3
H3K9 me2:DNA Meth. m5C
H3K4 me3:H4–4ac
H3K4 me1:H4K20 me2
H3K4 me1:H2A.Z
H2A.Z:H4K20 me2
H4K20 me3
H3K4 me3
H3K27 me2
H3K27 me3
H4–4ac
H3K9 me3
H3K9 me2
H3–5ac
H4K20 me2
H2A.Z
DNA Meth. m5C
H3K4 me1
0.00 0.25 0.50 0.75 1.00
$\hat{\pi}$
PABPC4 – Reverse
H3K27 me3:DNA Meth. m5C
H3K27 me3:H3K9 me3
H2A.Z:H4–4ac
DNA Meth. m5C:H3K9 me3
H4–4ac:H3–5ac
H2A.Z:H3–5ac
H3K27 me2:DNA Meth. m5C
H4K20 me2:H4–4ac
H4K20 me2:H3K4 me3
H4K20 me2:H3–5ac
H3K4 me1:H4–4ac
H2A.Z:H3K4 me3
H3K9 me2:DNA Meth. m5C
H3K4 me3:H4–4ac
H3K4 me1:H4K20 me2
H3K4 me1:H2A.Z
H2A.Z:H4K20 me2
H4K20 me3
H3K4 me3
H3K27 me2
H3K27 me3
H4–4ac
H3K9 me3
H3K9 me2
H3–5ac
H4K20 me2
H2A.Z
DNA Meth. m5C
H3K4 me1
0.00 0.25 0.50 0.75 1.00
$\hat{\pi}$
SPOUT1 – Forward
H3K4 me1:H2A.Z
H3K4 me3:H4–4ac
H3K4 me1:H4–4ac
H4K20 me2:H3K4 me3
DNA Meth. m5C:H3K9 me3
H3K27 me3:DNA Meth. m5C
DNA Meth. m5C:H4K20 me2
H3K4 me3:H3–5ac
H4K20 me2:H3–5ac
H4K20 me2:H4–4ac
H3K27 me3:H3K9 me3
H2A.Z:H3K4 me3
H2A.Z:H4K20 me2
H3K4 me1:H3–5ac
DNA Meth. m5C:H4K20 me3
H3K27 me3:H3K4 me1
H4–4ac:H3–5ac
H3K27 me2
H3K9 me2
H3K9 me3
H4K20 me3
H2A.Z
H4K20 me2
H3K4 me3
H3K4 me1
H3–5ac
H4–4ac
DNA Meth. m5C
H3K27 me3
0.00 0.25 0.50 0.75 1.00
$\hat{\pi}$
SPOUT1 – Reverse
H3K4 me1:H2A.Z
H3K4 me3:H4–4ac
H3K4 me1:H4–4ac
H4K20 me2:H3K4 me3
DNA Meth. m5C:H3K9 me3
H3K27 me3:DNA Meth. m5C
DNA Meth. m5C:H4K20 me2
H3K4 me3:H3–5ac
H4K20 me2:H3–5ac
H4K20 me2:H4–4ac
H3K27 me3:H3K9 me3
H2A.Z:H3K4 me3
H2A.Z:H4K20 me2
H3K4 me1:H3–5ac
DNA Meth. m5C:H4K20 me3
H3K27 me3:H3K4 me1
H4–4ac:H3–5ac
H3K27 me2
H3K9 me2
H3K9 me3
H4K20 me3
H2A.Z
H4K20 me2
H3K4 me3
H3K4 me1
H3–5ac
H4–4ac
DNA Meth. m5C
H3K27 me3
0.00 0.25 0.50 0.75 1.00
$\hat{\pi}$
FAM98A – Forward
H3K27 me3:H4K20 me3
H3K4 me3:H4–4ac
H2A.Z:H4–4ac
H3K9 me3:H4K20 me3
H3K4 me1:H2A.Z
DNA Meth. m5C:H3K9 me3
H4–4ac:H3–5ac
H3K4 me3:H3–5ac
H4K20 me2:H4–4ac
H2A.Z:H3–5ac
H3K4 me1:H4K20 me2
H3K4 me1:H3–5ac
H3K4 me1:H4–4ac
H2A.Z:H3K4 me3
DNA Meth. m5C:H4K20 me2
DNA Meth. m5C:H4K20 me3
H3K27 me2:DNA Meth. m5C
H4K20 me2:H3–5ac
H4K20 me2:H3K4 me3
H2A.Z:H4K20 me2
H3K27 me3:DNA Meth. m5C
H3K9 me2
H3K9 me3
H3K4 me1
H4–4ac
H4K20 me3
H3K27 me2
H3K4 me3
DNA Meth. m5C
H2A.Z
H4K20 me2
H3–5ac
H3K27 me3
0.00 0.25 0.50 0.75 1.00
$\hat{\pi}$
FAM98A – Reverse
H3K27 me3:H4K20 me3
H3K4 me3:H4–4ac
H2A.Z:H4–4ac
H3K9 me3:H4K20 me3
H3K4 me1:H2A.Z
DNA Meth. m5C:H3K9 me3
H4–4ac:H3–5ac
H3K4 me3:H3–5ac
H4K20 me2:H4–4ac
H2A.Z:H3–5ac
H3K4 me1:H4K20 me2
H3K4 me1:H3–5ac
H3K4 me1:H4–4ac
H2A.Z:H3K4 me3
DNA Meth. m5C:H4K20 me2
DNA Meth. m5C:H4K20 me3
H3K27 me2:DNA Meth. m5C
H4K20 me2:H3–5ac
H4K20 me2:H3K4 me3
H2A.Z:H4K20 me2
H3K27 me3:DNA Meth. m5C
H3K9 me2
H3K9 me3
H3K4 me1
H4–4ac
H4K20 me3
H3K27 me2
H3K4 me3
DNA Meth. m5C
H2A.Z
H4K20 me2
H3–5ac
H3K27 me3
0.00 0.25 0.50 0.75 1.00
$\hat{\pi}$
REXO4 – Forward
H3K4 me1:H3–5ac
H3K4 me1:H4–4ac
H3K9 me3:H4K20 me3
H3K27 me3:H4K20 me3
H3K27 me3:DNA Meth. m5C
H3K4 me1:H2A.Z
DNA Meth. m5C:H3K9 me3
H4–4ac:H3–5ac
H2A.Z:H3–5ac
H3K4 me3:H3–5ac
H2A.Z:H4–4ac
H3K4 me3:H4–4ac
H3K27 me2:DNA Meth. m5C
H3K9 me2:DNA Meth. m5C
H3K27 me3:H3K4 me1
H3K27 me3:H3K9 me3
H2A.Z:H3K4 me3
H4K20 me2:H3–5ac
H3K27 me2
H3K9 me2
H4K20 me3
H3K4 me1
H4K20 me2
DNA Meth. m5C
H4–4ac
H2A.Z
H3K9 me3
H3K27 me3
H3K4 me3
H3–5ac
0.00 0.25 0.50 0.75 1.00
$\hat{\pi}$
REXO4 – Reverse
H3K4 me1:H3–5ac
H3K4 me1:H4–4ac
H3K9 me3:H4K20 me3
H3K27 me3:H4K20 me3
H3K27 me3:DNA Meth. m5C
H3K4 me1:H2A.Z
DNA Meth. m5C:H3K9 me3
H4–4ac:H3–5ac
H2A.Z:H3–5ac
H3K4 me3:H3–5ac
H2A.Z:H4–4ac
H3K4 me3:H4–4ac
H3K27 me2:DNA Meth. m5C
H3K9 me2:DNA Meth. m5C
H3K27 me3:H3K4 me1
H3K27 me3:H3K9 me3
H2A.Z:H3K4 me3
H4K20 me2:H3–5ac
H3K27 me2
H3K9 me2
H4K20 me3
H3K4 me1
H4K20 me2
DNA Meth. m5C
H4–4ac
H2A.Z
H3K9 me3
H3K27 me3
H3K4 me3
H3–5ac
0.00 0.25 0.50 0.75 1.00
$\hat{\pi}$
GRWD1 – Forward
H2A.Z:H3K4 me3
DNA Meth. m5C:H3K9 me3
H3K27 me3:H3K9 me3
H3K4 me1:H4–4ac
H3K4 me1:H2A.Z
H3K4 me3:H3–5ac
H4K20 me2:H3–5ac
H2A.Z:H4K20 me2
H3K4 me1:H4K20 me2
H3K4 me3:H4–4ac
H4–4ac:H3–5ac
H3K9 me2
H3K27 me2
H4K20 me3
H4K20 me2
H3K27 me3
H3K9 me3
H2A.Z
DNA Meth. m5C
H4–4ac
H3–5ac
H3K4 me1
H3K4 me3
0.00 0.25 0.50 0.75 1.00
$\hat{\pi}$
GRWD1 – Reverse
H2A.Z:H3K4 me3
DNA Meth. m5C:H3K9 me3
H3K27 me3:H3K9 me3
H3K4 me1:H4–4ac
H3K4 me1:H2A.Z
H3K4 me3:H3–5ac
H4K20 me2:H3–5ac
H2A.Z:H4K20 me2
H3K4 me1:H4K20 me2
H3K4 me3:H4–4ac
H4–4ac:H3–5ac
H3K9 me2
H3K27 me2
H4K20 me3
H4K20 me2
H3K27 me3
H3K9 me3
H2A.Z
DNA Meth. m5C
H4–4ac
H3–5ac
H3K4 me1
H3K4 me3
0.00 0.25 0.50 0.75 1.00
$\hat{\pi}$
SPTBN2 – Forward
H3K27 me3:H4K20 me3
H3K27 me3:H3K9 me3
H3K4 me1:H2A.Z
H3K9 me3:H4K20 me3
H2A.Z:H4–4ac
H2A.Z:H3K4 me3
H3K4 me1:H3–5ac
H3K27 me3:H3K4 me1
H3K27 me2:DNA Meth. m5C
H3K27 me3:DNA Meth. m5C
H2A.Z:H3–5ac
H3K4 me1:H4–4ac
H3K4 me3:H3–5ac
H2A.Z:H4K20 me2
H3K4 me1:H4K20 me2
H4K20 me2:H3–5ac
DNA Meth. m5C:H4K20 me3
H4K20 me2:H4–4ac
H4K20 me2:H3K4 me3
H4–4ac:H3–5ac
DNA Meth. m5C:H3K9 me3
H3K4 me3:H4–4ac
H3K9 me2
H3K27 me2
H4K20 me3
H3K27 me3
H3K9 me3
H3K4 me1
DNA Meth. m5C
H2A.Z
H3–5ac
H3K4 me3
H4K20 me2
H4–4ac
0.00 0.25 0.50 0.75 1.00
$\hat{\pi}$
SPTBN2 – Reverse
H3K27 me3:H4K20 me3
H3K27 me3:H3K9 me3
H3K4 me1:H2A.Z
H3K9 me3:H4K20 me3
H2A.Z:H4–4ac
H2A.Z:H3K4 me3
H3K4 me1:H3–5ac
H3K27 me3:H3K4 me1
H3K27 me2:DNA Meth. m5C
H3K27 me3:DNA Meth. m5C
H2A.Z:H3–5ac
H3K4 me1:H4–4ac
H3K4 me3:H3–5ac
H2A.Z:H4K20 me2
H3K4 me1:H4K20 me2
H4K20 me2:H3–5ac
DNA Meth. m5C:H4K20 me3
H4K20 me2:H4–4ac
H4K20 me2:H3K4 me3
H4–4ac:H3–5ac
DNA Meth. m5C:H3K9 me3
H3K4 me3:H4–4ac
H3K9 me2
H3K27 me2
H4K20 me3
H3K27 me3
H3K9 me3
H3K4 me1
DNA Meth. m5C
H2A.Z
H3–5ac
H3K4 me3
H4K20 me2
H4–4ac
0.00 0.25 0.50 0.75 1.00
$\hat{\pi}$

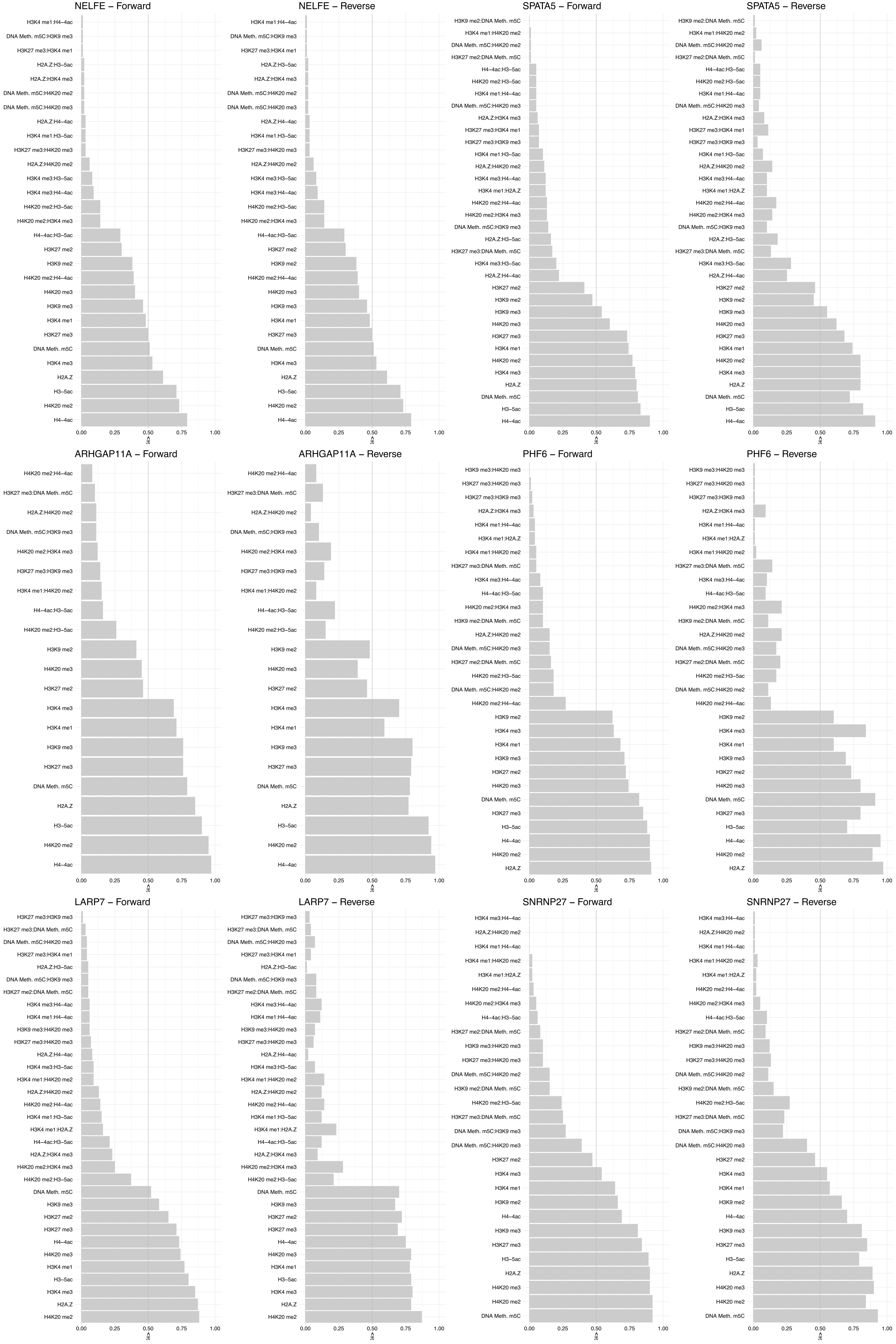

NELFE – Forward
H3K4 me1:H4–4ac
DNA Meth. m5C:H3K9 me3
H3K27 me3:H3K4 me1
H2A.Z:H3–5ac
H2A.Z:H3K4 me3
DNA Meth. m5C:H4K20 me2
DNA Meth. m5C:H4K20 me3
H2A.Z:H4–4ac
H3K4 me1:H3–5ac
H3K27 me3:H4K20 me3
H2A.Z:H4K20 me2
H3K4 me3:H3–5ac
H3K4 me3:H4–4ac
H4K20 me2:H3–5ac
H4K20 me2:H3K4 me3
H4–4ac:H3–5ac
H3K27 me2
H3K9 me2
H4K20 me2:H4–4ac
H4K20 me3
H3K9 me3
H3K4 me1
H3K27 me3
DNA Meth. m5C
H3K4 me3
H2A.Z
H3–5ac
H4K20 me2
H4–4ac
0.00
0.25
0.50
0.75
1.00
π̂
NELFE – Reverse
SPATA5 – Forward
H3K9 me2:DNA Meth. m5C
H3K4 me1:H4K20 me2
DNA Meth. m5C:H4K20 me2
H3K27 me2:DNA Meth. m5C
H4–4ac:H3–5ac
H4K20 me2:H3–5ac
H3K4 me1:H4–4ac
DNA Meth. m5C:H4K20 me3
H2A.Z:H3K4 me3
H3K27 me3:H3K4 me1
H3K27 me3:H3K9 me3
H3K4 me1:H3–5ac
H2A.Z:H4K20 me2
H3K4 me3:H4–4ac
H3K4 me1:H2A.Z
H4K20 me2:H4–4ac
H4K20 me2:H3K4 me3
DNA Meth. m5C:H3K9 me3
H2A.Z:H3–5ac
H3K27 me3:DNA Meth. m5C
H3K4 me3:H3–5ac
H2A.Z:H4–4ac
H3K27 me2
H3K9 me2
H3K9 me3
H4K20 me3
H3K27 me3
H3K4 me1
H4K20 me2
H3K4 me3
H2A.Z
DNA Meth. m5C
H3–5ac
H4–4ac
SPATA5 – Reverse
ARHGAP11A – Forward
H4K20 me2:H4–4ac
H3K27 me3:DNA Meth. m5C
H2A.Z:H4K20 me2
DNA Meth. m5C:H3K9 me3
H4K20 me2:H3K4 me3
H3K27 me3:H3K9 me3
H3K4 me1:H4K20 me2
H4–4ac:H3–5ac
H4K20 me2:H3–5ac
H3K9 me2
H4K20 me3
H3K27 me2
H3K4 me3
H3K4 me1
H3K9 me3
H3K27 me3
DNA Meth. m5C
H2A.Z
H3–5ac
H4K20 me2
H4–4ac
ARHGAP11A – Reverse
PHF6 – Forward
H3K9 me3:H4K20 me3
H3K27 me3:H4K20 me3
H3K27 me3:H3K9 me3
H2A.Z:H3K4 me3
H3K4 me1:H4–4ac
H3K4 me1:H2A.Z
H3K4 me1:H4K20 me2
H3K27 me3:DNA Meth. m5C
H3K4 me3:H4–4ac
H4–4ac:H3–5ac
H4K20 me2:H3K4 me3
H3K9 me2:DNA Meth. m5C
H2A.Z:H4K20 me2
DNA Meth. m5C:H4K20 me3
H3K27 me2:DNA Meth. m5C
H4K20 me2:H3–5ac
DNA Meth. m5C:H4K20 me2
H4K20 me2:H4–4ac
H3K9 me2
H3K4 me3
H3K4 me1
H3K9 me3
H3K27 me2
H4K20 me3
DNA Meth. m5C
H3K27 me3
H3–5ac
H4–4ac
H4K20 me2
H2A.Z
PHF6 – Reverse
LARP7 – Forward
H3K27 me3:H3K9 me3
H3K27 me3:DNA Meth. m5C
DNA Meth. m5C:H4K20 me3
H3K27 me3:H3K4 me1
H2A.Z:H3–5ac
DNA Meth. m5C:H3K9 me3
H3K27 me2:DNA Meth. m5C
H3K4 me3:H4–4ac
H3K4 me1:H4–4ac
H3K9 me3:H4K20 me3
H3K27 me3:H4K20 me3
H2A.Z:H4–4ac
H3K4 me3:H3–5ac
H3K4 me1:H4K20 me2
H2A.Z:H4K20 me2
H4K20 me2:H4–4ac
H3K4 me1:H3–5ac
H3K4 me1:H2A.Z
H4–4ac:H3–5ac
H2A.Z:H3K4 me3
H4K20 me2:H3K4 me3
H4K20 me2:H3–5ac
DNA Meth. m5C
H3K9 me3
H3K27 me2
H3K27 me3
H4–4ac
H4K20 me3
H3K4 me1
H3–5ac
H3K4 me3
H2A.Z
H4K20 me2
LARP7 – Reverse
SNRNP27 – Forward
H3K4 me3:H4–4ac
H2A.Z:H4K20 me2
H3K4 me1:H4–4ac
H3K4 me1:H4K20 me2
H3K4 me1:H2A.Z
H4K20 me2:H4–4ac
H4K20 me2:H3K4 me3
H4–4ac:H3–5ac
H3K27 me2:DNA Meth. m5C
H3K9 me3:H4K20 me3
H3K27 me3:H4K20 me3
DNA Meth. m5C:H4K20 me2
H3K9 me2:DNA Meth. m5C
H4K20 me2:H3–5ac
H3K27 me3:DNA Meth. m5C
DNA Meth. m5C:H3K9 me3
DNA Meth. m5C:H4K20 me3
H3K27 me2
H3K4 me3
H3K4 me1
H3K9 me2
H4–4ac
H3K9 me3
H3K27 me3
H3–5ac
H2A.Z
H4K20 me3
H4K20 me2
DNA Meth. m5C
SNRNP27 – Reverse

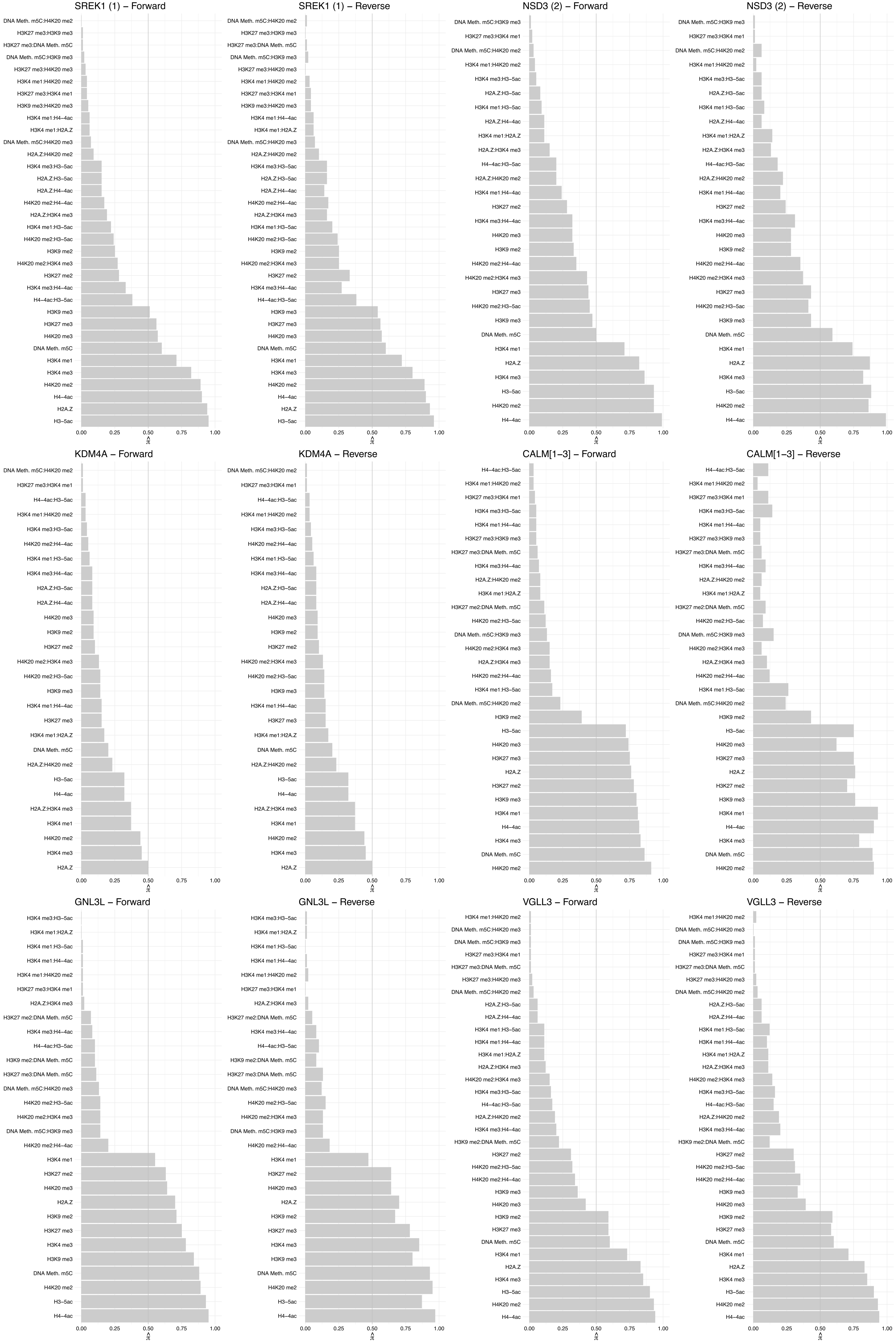
SREK1 (1) – Forward
DNA Meth. m5C:H4K20 me2
H3K27 me3:H3K9 me3
H3K27 me3:DNA Meth. m5C
DNA Meth. m5C:H3K9 me3
H3K27 me3:H4K20 me3
H3K4 me1:H4K20 me2
H3K27 me3:H3K4 me1
H3K9 me3:H4K20 me3
H3K4 me1:H4−4ac
H3K4 me1:H2A.Z
DNA Meth. m5C:H4K20 me3
H2A.Z:H4K20 me2
H3K4 me3:H3−5ac
H2A.Z:H3−5ac
H2A.Z:H4−4ac
H4K20 me2:H4−4ac
H2A.Z:H3K4 me3
H3K4 me1:H3−5ac
H4K20 me2:H3−5ac
H3K9 me2
H4K20 me2:H3K4 me3
H3K27 me2
H3K4 me3:H4−4ac
H4−4ac:H3−5ac
H3K9 me3
H3K27 me3
H4K20 me3
DNA Meth. m5C
H3K4 me1
H3K4 me3
H4K20 me2
H4−4ac
H2A.Z
H3−5ac
0.00 0.25 0.50 0.75 1.00
$\hat{\pi}$
SREK1 (1) – Reverse
DNA Meth. m5C:H4K20 me2
H3K27 me3:H3K9 me3
H3K27 me3:DNA Meth. m5C
DNA Meth. m5C:H3K9 me3
H3K27 me3:H4K20 me3
H3K4 me1:H4K20 me2
H3K27 me3:H3K4 me1
H3K9 me3:H4K20 me3
H3K4 me1:H4−4ac
H3K4 me1:H2A.Z
DNA Meth. m5C:H4K20 me3
H2A.Z:H4K20 me2
H3K4 me3:H3−5ac
H2A.Z:H3−5ac
H2A.Z:H4−4ac
H4K20 me2:H4−4ac
H2A.Z:H3K4 me3
H3K4 me1:H3−5ac
H4K20 me2:H3−5ac
H3K9 me2
H4K20 me2:H3K4 me3
H3K27 me2
H3K4 me3:H4−4ac
H4−4ac:H3−5ac
H3K9 me3
H3K27 me3
H4K20 me3
DNA Meth. m5C
H3K4 me1
H3K4 me3
H4K20 me2
H4−4ac
H2A.Z
H3−5ac
0.00 0.25 0.50 0.75 1.00
$\hat{\pi}$
NSD3 (2) – Forward
DNA Meth. m5C:H3K9 me3
H3K27 me3:H3K4 me1
DNA Meth. m5C:H4K20 me2
H3K4 me1:H4K20 me2
H3K4 me3:H3−5ac
H2A.Z:H3−5ac
H3K4 me1:H3−5ac
H2A.Z:H4−4ac
H3K4 me1:H2A.Z
H2A.Z:H3K4 me3
H4−4ac:H3−5ac
H2A.Z:H4K20 me2
H3K4 me1:H4−4ac
H3K27 me2
H3K4 me3:H4−4ac
H4K20 me3
H3K9 me2
H4K20 me2:H4−4ac
H4K20 me2:H3K4 me3
H3K27 me3
H4K20 me2:H3−5ac
H3K9 me3
DNA Meth. m5C
H3K4 me1
H2A.Z
H3K4 me3
H3−5ac
H4K20 me2
H4−4ac
0.00 0.25 0.50 0.75 1.00
$\hat{\pi}$
NSD3 (2) – Reverse
DNA Meth. m5C:H3K9 me3
H3K27 me3:H3K4 me1
DNA Meth. m5C:H4K20 me2
H3K4 me1:H4K20 me2
H3K4 me3:H3−5ac
H2A.Z:H3−5ac
H3K4 me1:H3−5ac
H2A.Z:H4−4ac
H3K4 me1:H2A.Z
H2A.Z:H3K4 me3
H4−4ac:H3−5ac
H2A.Z:H4K20 me2
H3K4 me1:H4−4ac
H3K27 me2
H3K4 me3:H4−4ac
H4K20 me3
H3K9 me2
H4K20 me2:H4−4ac
H4K20 me2:H3K4 me3
H3K27 me3
H4K20 me2:H3−5ac
H3K9 me3
DNA Meth. m5C
H3K4 me1
H2A.Z
H3K4 me3
H3−5ac
H4K20 me2
H4−4ac
0.00 0.25 0.50 0.75 1.00
$\hat{\pi}$
KDM4A – Forward
DNA Meth. m5C:H4K20 me2
H3K27 me3:H3K4 me1
H4−4ac:H3−5ac
H3K4 me1:H4K20 me2
H3K4 me3:H3−5ac
H4K20 me2:H4−4ac
H3K4 me1:H3−5ac
H3K4 me3:H4−4ac
H2A.Z:H3−5ac
H2A.Z:H4−4ac
H4K20 me3
H3K9 me2
H3K27 me2
H4K20 me2:H3K4 me3
H4K20 me2:H3−5ac
H3K9 me3
H3K4 me1:H4−4ac
H3K27 me3
H3K4 me1:H2A.Z
DNA Meth. m5C
H2A.Z:H4K20 me2
H3−5ac
H4−4ac
H2A.Z:H3K4 me3
H3K4 me1
H4K20 me2
H3K4 me3
H2A.Z
0.00 0.25 0.50 0.75 1.00
$\hat{\pi}$
KDM4A – Reverse
DNA Meth. m5C:H4K20 me2
H3K27 me3:H3K4 me1
H4−4ac:H3−5ac
H3K4 me1:H4K20 me2
H3K4 me3:H3−5ac
H4K20 me2:H4−4ac
H3K4 me1:H3−5ac
H3K4 me3:H4−4ac
H2A.Z:H3−5ac
H2A.Z:H4−4ac
H4K20 me3
H3K9 me2
H3K27 me2
H4K20 me2:H3K4 me3
H4K20 me2:H3−5ac
H3K9 me3
H3K4 me1:H4−4ac
H3K27 me3
H3K4 me1:H2A.Z
DNA Meth. m5C
H2A.Z:H4K20 me2
H3−5ac
H4−4ac
H2A.Z:H3K4 me3
H3K4 me1
H4K20 me2
H3K4 me3
H2A.Z
0.00 0.25 0.50 0.75 1.00
$\hat{\pi}$
CALM[1−3] – Forward
H4−4ac:H3−5ac
H3K4 me1:H4K20 me2
H3K27 me3:H3K4 me1
H3K4 me3:H3−5ac
H3K4 me1:H4−4ac
H3K27 me3:H3K9 me3
H3K27 me3:DNA Meth. m5C
H3K4 me3:H4−4ac
H2A.Z:H4K20 me2
H3K4 me1:H2A.Z
H3K27 me2:DNA Meth. m5C
H4K20 me2:H3−5ac
DNA Meth. m5C:H3K9 me3
H4K20 me2:H3K4 me3
H2A.Z:H3K4 me3
H4K20 me2:H4−4ac
H3K4 me1:H3−5ac
DNA Meth. m5C:H4K20 me2
H3K9 me2
H3−5ac
H4K20 me3
H3K27 me3
H2A.Z
H3K27 me2
H3K9 me3
H3K4 me1
H4−4ac
H3K4 me3
DNA Meth. m5C
H4K20 me2
0.00 0.25 0.50 0.75 1.00
$\hat{\pi}$
CALM[1−3] – Reverse
H4−4ac:H3−5ac
H3K4 me1:H4K20 me2
H3K27 me3:H3K4 me1
H3K4 me3:H3−5ac
H3K4 me1:H4−4ac
H3K27 me3:H3K9 me3
H3K27 me3:DNA Meth. m5C
H3K4 me3:H4−4ac
H2A.Z:H4K20 me2
H3K4 me1:H2A.Z
H3K27 me2:DNA Meth. m5C
H4K20 me2:H3−5ac
DNA Meth. m5C:H3K9 me3
H4K20 me2:H3K4 me3
H2A.Z:H3K4 me3
H4K20 me2:H4−4ac
H3K4 me1:H3−5ac
DNA Meth. m5C:H4K20 me2
H3K9 me2
H3−5ac
H4K20 me3
H3K27 me3
H2A.Z
H3K27 me2
H3K9 me3
H3K4 me1
H4−4ac
H3K4 me3
DNA Meth. m5C
H4K20 me2
0.00 0.25 0.50 0.75 1.00
$\hat{\pi}$
GNL3L – Forward
H3K4 me3:H3−5ac
H3K4 me1:H2A.Z
H3K4 me1:H3−5ac
H3K4 me1:H4−4ac
H3K4 me1:H4K20 me2
H3K27 me3:H3K4 me1
H2A.Z:H3K4 me3
H3K27 me2:DNA Meth. m5C
H3K4 me3:H4−4ac
H4−4ac:H3−5ac
H3K9 me2:DNA Meth. m5C
H3K27 me3:DNA Meth. m5C
DNA Meth. m5C:H4K20 me3
H4K20 me2:H3−5ac
H4K20 me2:H3K4 me3
DNA Meth. m5C:H3K9 me3
H4K20 me2:H4−4ac
H3K4 me1
H3K27 me2
H4K20 me3
H2A.Z
H3K9 me2
H3K27 me3
H3K4 me3
H3K9 me3
DNA Meth. m5C
H4K20 me2
H3−5ac
H4−4ac
0.00 0.25 0.50 0.75 1.00
$\hat{\pi}$
GNL3L – Reverse
H3K4 me3:H3−5ac
H3K4 me1:H2A.Z
H3K4 me1:H3−5ac
H3K4 me1:H4−4ac
H3K4 me1:H4K20 me2
H3K27 me3:H3K4 me1
H2A.Z:H3K4 me3
H3K27 me2:DNA Meth. m5C
H3K4 me3:H4−4ac
H4−4ac:H3−5ac
H3K9 me2:DNA Meth. m5C
H3K27 me3:DNA Meth. m5C
DNA Meth. m5C:H4K20 me3
H4K20 me2:H3−5ac
H4K20 me2:H3K4 me3
DNA Meth. m5C:H3K9 me3
H4K20 me2:H4−4ac
H3K4 me1
H3K27 me2
H4K20 me3
H2A.Z
H3K9 me2
H3K27 me3
H3K4 me3
H3K9 me3
DNA Meth. m5C
H4K20 me2
H3−5ac
H4−4ac
0.00 0.25 0.50 0.75 1.00
$\hat{\pi}$
VGLL3 – Forward
H3K4 me1:H4K20 me2
DNA Meth. m5C:H4K20 me3
DNA Meth. m5C:H3K9 me3
H3K27 me3:H3K4 me1
H3K27 me3:DNA Meth. m5C
H3K27 me3:H4K20 me3
DNA Meth. m5C:H4K20 me2
H2A.Z:H3−5ac
H2A.Z:H4−4ac
H3K4 me1:H3−5ac
H3K4 me1:H4−4ac
H3K4 me1:H2A.Z
H2A.Z:H3K4 me3
H4K20 me2:H3K4 me3
H3K4 me3:H3−5ac
H4−4ac:H3−5ac
H2A.Z:H4K20 me2
H3K4 me3:H4−4ac
H3K9 me2:DNA Meth. m5C
H3K27 me2
H4K20 me2:H3−5ac
H4K20 me2:H4−4ac
H3K9 me3
H4K20 me3
H3K9 me2
H3K27 me3
DNA Meth. m5C
H3K4 me1
H2A.Z
H3K4 me3
H3−5ac
H4K20 me2
H4−4ac
0.00 0.25 0.50 0.75 1.00
$\hat{\pi}$
VGLL3 – Reverse
H3K4 me1:H4K20 me2
DNA Meth. m5C:H4K20 me3
DNA Meth. m5C:H3K9 me3
H3K27 me3:H3K4 me1
H3K27 me3:DNA Meth. m5C
H3K27 me3:H4K20 me3
DNA Meth. m5C:H4K20 me2
H2A.Z:H3−5ac
H2A.Z:H4−4ac
H3K4 me1:H3−5ac
H3K4 me1:H4−4ac
H3K4 me1:H2A.Z
H2A.Z:H3K4 me3
H4K20 me2:H3K4 me3
H3K4 me3:H3−5ac
H4−4ac:H3−5ac
H2A.Z:H4K20 me2
H3K4 me3:H4−4ac
H3K9 me2:DNA Meth. m5C
H3K27 me2
H4K20 me2:H3−5ac
H4K20 me2:H4−4ac
H3K9 me3
H4K20 me3
H3K9 me2
H3K27 me3
DNA Meth. m5C
H3K4 me1
H2A.Z
H3K4 me3
H3−5ac
H4K20 me2
H4−4ac
0.00 0.25 0.50 0.75 1.00
$\hat{\pi}$

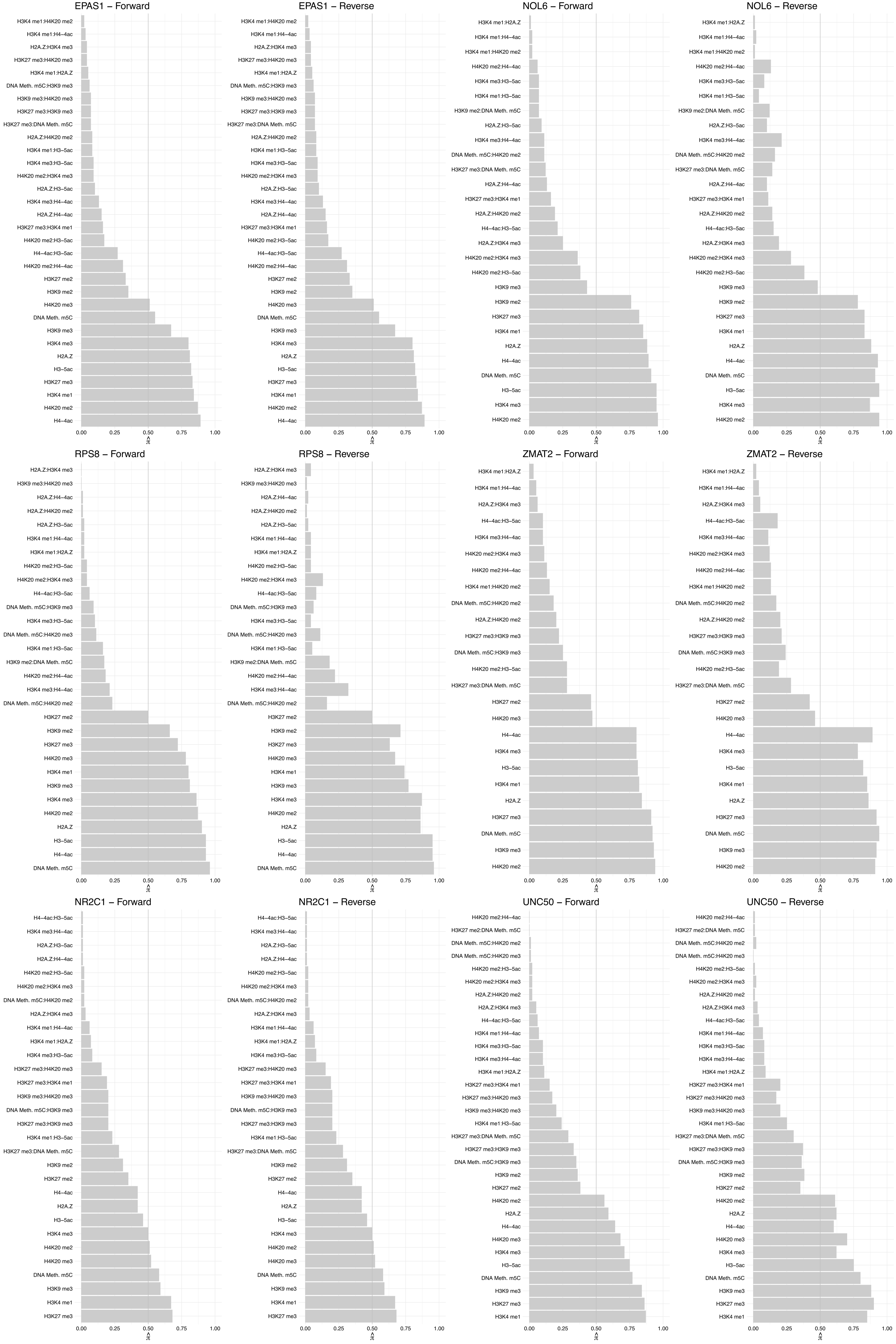
EPAS1 – Forward
H3K4 me1:H4K20 me2
H3K4 me1:H4−4ac
H2A.Z:H3K4 me3
H3K27 me3:H4K20 me3
H3K4 me1:H2A.Z
DNA Meth. m5C:H3K9 me3
H3K9 me3:H4K20 me3
H3K27 me3:H3K9 me3
H3K27 me3:DNA Meth. m5C
H2A.Z:H4K20 me2
H3K4 me1:H3−5ac
H3K4 me3:H3−5ac
H4K20 me2:H3K4 me3
H2A.Z:H3−5ac
H3K4 me3:H4−4ac
H2A.Z:H4−4ac
H3K27 me3:H3K4 me1
H4K20 me2:H3−5ac
H4−4ac:H3−5ac
H4K20 me2:H4−4ac
H3K27 me2
H3K9 me2
H4K20 me3
DNA Meth. m5C
H3K9 me3
H3K4 me3
H2A.Z
H3−5ac
H3K27 me3
H3K4 me1
H4K20 me2
H4−4ac
0.00 0.25 0.50 0.75 1.00
π̂
EPAS1 – Reverse
H3K4 me1:H4K20 me2
H3K4 me1:H4−4ac
H2A.Z:H3K4 me3
H3K27 me3:H4K20 me3
H3K4 me1:H2A.Z
DNA Meth. m5C:H3K9 me3
H3K9 me3:H4K20 me3
H3K27 me3:H3K9 me3
H3K27 me3:DNA Meth. m5C
H2A.Z:H4K20 me2
H3K4 me1:H3−5ac
H3K4 me3:H3−5ac
H4K20 me2:H3K4 me3
H2A.Z:H3−5ac
H3K4 me3:H4−4ac
H2A.Z:H4−4ac
H3K27 me3:H3K4 me1
H4K20 me2:H3−5ac
H4−4ac:H3−5ac
H4K20 me2:H4−4ac
H3K27 me2
H3K9 me2
H4K20 me3
DNA Meth. m5C
H3K9 me3
H3K4 me3
H2A.Z
H3−5ac
H3K27 me3
H3K4 me1
H4K20 me2
H4−4ac
0.00 0.25 0.50 0.75 1.00
π̂
NOL6 – Forward
H3K4 me1:H2A.Z
H3K4 me1:H4−4ac
H3K4 me1:H4K20 me2
H4K20 me2:H4−4ac
H3K4 me3:H3−5ac
H3K4 me1:H3−5ac
H3K9 me2:DNA Meth. m5C
H2A.Z:H3−5ac
H3K4 me3:H4−4ac
DNA Meth. m5C:H4K20 me2
H3K27 me3:DNA Meth. m5C
H2A.Z:H4−4ac
H3K27 me3:H3K4 me1
H2A.Z:H4K20 me2
H4−4ac:H3−5ac
H2A.Z:H3K4 me3
H4K20 me2:H3K4 me3
H4K20 me2:H3−5ac
H3K9 me3
H3K9 me2
H3K27 me3
H3K4 me1
H2A.Z
H4−4ac
DNA Meth. m5C
H3−5ac
H3K4 me3
H4K20 me2
0.00 0.25 0.50 0.75 1.00
π̂
NOL6 – Reverse
H3K4 me1:H2A.Z
H3K4 me1:H4−4ac
H3K4 me1:H4K20 me2
H4K20 me2:H4−4ac
H3K4 me3:H3−5ac
H3K4 me1:H3−5ac
H3K9 me2:DNA Meth. m5C
H2A.Z:H3−5ac
H3K4 me3:H4−4ac
DNA Meth. m5C:H4K20 me2
H3K27 me3:DNA Meth. m5C
H2A.Z:H4−4ac
H3K27 me3:H3K4 me1
H2A.Z:H4K20 me2
H4−4ac:H3−5ac
H2A.Z:H3K4 me3
H4K20 me2:H3K4 me3
H4K20 me2:H3−5ac
H3K9 me3
H3K9 me2
H3K27 me3
H3K4 me1
H2A.Z
H4−4ac
DNA Meth. m5C
H3−5ac
H3K4 me3
H4K20 me2
0.00 0.25 0.50 0.75 1.00
π̂
RPS8 – Forward
H2A.Z:H3K4 me3
H3K9 me3:H4K20 me3
H2A.Z:H4−4ac
H2A.Z:H4K20 me2
H2A.Z:H3−5ac
H3K4 me1:H4−4ac
H3K4 me1:H2A.Z
H4K20 me2:H3−5ac
H4K20 me2:H3K4 me3
H4−4ac:H3−5ac
DNA Meth. m5C:H3K9 me3
H3K4 me3:H3−5ac
DNA Meth. m5C:H4K20 me3
H3K4 me1:H3−5ac
H3K9 me2:DNA Meth. m5C
H4K20 me2:H4−4ac
H3K4 me3:H4−4ac
DNA Meth. m5C:H4K20 me2
H3K27 me2
H3K9 me2
H3K27 me3
H4K20 me3
H3K4 me1
H3K9 me3
H3K4 me3
H4K20 me2
H2A.Z
H3−5ac
H4−4ac
DNA Meth. m5C
0.00 0.25 0.50 0.75 1.00
π̂
RPS8 – Reverse
H2A.Z:H3K4 me3
H3K9 me3:H4K20 me3
H2A.Z:H4−4ac
H2A.Z:H4K20 me2
H2A.Z:H3−5ac
H3K4 me1:H4−4ac
H3K4 me1:H2A.Z
H4K20 me2:H3−5ac
H4K20 me2:H3K4 me3
H4−4ac:H3−5ac
DNA Meth. m5C:H3K9 me3
H3K4 me3:H3−5ac
DNA Meth. m5C:H4K20 me3
H3K4 me1:H3−5ac
H3K9 me2:DNA Meth. m5C
H4K20 me2:H4−4ac
H3K4 me3:H4−4ac
DNA Meth. m5C:H4K20 me2
H3K27 me2
H3K9 me2
H3K27 me3
H4K20 me3
H3K4 me1
H3K9 me3
H3K4 me3
H4K20 me2
H2A.Z
H3−5ac
H4−4ac
DNA Meth. m5C
0.00 0.25 0.50 0.75 1.00
π̂
ZMAT2 – Forward
H3K4 me1:H2A.Z
H3K4 me1:H4−4ac
H2A.Z:H3K4 me3
H4−4ac:H3−5ac
H3K4 me3:H4−4ac
H4K20 me2:H3K4 me3
H4K20 me2:H4−4ac
H3K4 me1:H4K20 me2
DNA Meth. m5C:H4K20 me2
H2A.Z:H4K20 me2
H3K27 me3:H3K9 me3
DNA Meth. m5C:H3K9 me3
H4K20 me2:H3−5ac
H3K27 me3:DNA Meth. m5C
H3K27 me2
H4K20 me3
H4−4ac
H3K4 me3
H3−5ac
H3K4 me1
H2A.Z
H3K27 me3
DNA Meth. m5C
H3K9 me3
H4K20 me2
0.00 0.25 0.50 0.75 1.00
π̂
ZMAT2 – Reverse
H3K4 me1:H2A.Z
H3K4 me1:H4−4ac
H2A.Z:H3K4 me3
H4−4ac:H3−5ac
H3K4 me3:H4−4ac
H4K20 me2:H3K4 me3
H4K20 me2:H4−4ac
H3K4 me1:H4K20 me2
DNA Meth. m5C:H4K20 me2
H2A.Z:H4K20 me2
H3K27 me3:H3K9 me3
DNA Meth. m5C:H3K9 me3
H4K20 me2:H3−5ac
H3K27 me3:DNA Meth. m5C
H3K27 me2
H4K20 me3
H4−4ac
H3K4 me3
H3−5ac
H3K4 me1
H2A.Z
H3K27 me3
DNA Meth. m5C
H3K9 me3
H4K20 me2
0.00 0.25 0.50 0.75 1.00
π̂
NR2C1 – Forward
H4−4ac:H3−5ac
H3K4 me3:H4−4ac
H2A.Z:H3−5ac
H2A.Z:H4−4ac
H4K20 me2:H3−5ac
H4K20 me2:H3K4 me3
DNA Meth. m5C:H4K20 me2
H2A.Z:H3K4 me3
H3K4 me1:H4−4ac
H3K4 me1:H2A.Z
H3K4 me3:H3−5ac
H3K27 me3:H4K20 me3
H3K27 me3:H3K4 me1
H3K9 me3:H4K20 me3
DNA Meth. m5C:H3K9 me3
H3K27 me3:H3K9 me3
H3K4 me1:H3−5ac
H3K27 me3:DNA Meth. m5C
H3K9 me2
H3K27 me2
H4−4ac
H2A.Z
H3−5ac
H3K4 me3
H4K20 me2
H4K20 me3
DNA Meth. m5C
H3K9 me3
H3K4 me1
H3K27 me3
0.00 0.25 0.50 0.75 1.00
π̂
NR2C1 – Reverse
H4−4ac:H3−5ac
H3K4 me3:H4−4ac
H2A.Z:H3−5ac
H2A.Z:H4−4ac
H4K20 me2:H3−5ac
H4K20 me2:H3K4 me3
DNA Meth. m5C:H4K20 me2
H2A.Z:H3K4 me3
H3K4 me1:H4−4ac
H3K4 me1:H2A.Z
H3K4 me3:H3−5ac
H3K27 me3:H4K20 me3
H3K27 me3:H3K4 me1
H3K9 me3:H4K20 me3
DNA Meth. m5C:H3K9 me3
H3K27 me3:H3K9 me3
H3K4 me1:H3−5ac
H3K27 me3:DNA Meth. m5C
H3K9 me2
H3K27 me2
H4−4ac
H2A.Z
H3−5ac
H3K4 me3
H4K20 me2
H4K20 me3
DNA Meth. m5C
H3K9 me3
H3K4 me1
H3K27 me3
0.00 0.25 0.50 0.75 1.00
π̂
UNC50 – Forward
H4K20 me2:H4−4ac
H3K27 me2:DNA Meth. m5C
DNA Meth. m5C:H4K20 me2
DNA Meth. m5C:H4K20 me3
H4K20 me2:H3−5ac
H4K20 me2:H3K4 me3
H2A.Z:H4K20 me2
H2A.Z:H3K4 me3
H4−4ac:H3−5ac
H3K4 me1:H4−4ac
H3K4 me3:H3−5ac
H3K4 me3:H4−4ac
H3K4 me1:H2A.Z
H3K27 me3:H3K4 me1
H3K27 me3:H4K20 me3
H3K9 me3:H4K20 me3
H3K4 me1:H3−5ac
H3K27 me3:DNA Meth. m5C
H3K27 me3:H3K9 me3
DNA Meth. m5C:H3K9 me3
H3K9 me2
H3K27 me2
H4K20 me2
H2A.Z
H4−4ac
H4K20 me3
H3K4 me3
H3−5ac
DNA Meth. m5C
H3K9 me3
H3K27 me3
H3K4 me1
0.00 0.25 0.50 0.75 1.00
π̂
UNC50 – Reverse
H4K20 me2:H4−4ac
H3K27 me2:DNA Meth. m5C
DNA Meth. m5C:H4K20 me2
DNA Meth. m5C:H4K20 me3
H4K20 me2:H3−5ac
H4K20 me2:H3K4 me3
H2A.Z:H4K20 me2
H2A.Z:H3K4 me3
H4−4ac:H3−5ac
H3K4 me1:H4−4ac
H3K4 me3:H3−5ac
H3K4 me3:H4−4ac
H3K4 me1:H2A.Z
H3K27 me3:H3K4 me1
H3K27 me3:H4K20 me3
H3K9 me3:H4K20 me3
H3K4 me1:H3−5ac
H3K27 me3:DNA Meth. m5C
H3K27 me3:H3K9 me3
DNA Meth. m5C:H3K9 me3
H3K9 me2
H3K27 me2
H4K20 me2
H2A.Z
H4−4ac
H4K20 me3
H3K4 me3
H3−5ac
DNA Meth. m5C
H3K9 me3
H3K27 me3
H3K4 me1
0.00 0.25 0.50 0.75 1.00
π̂

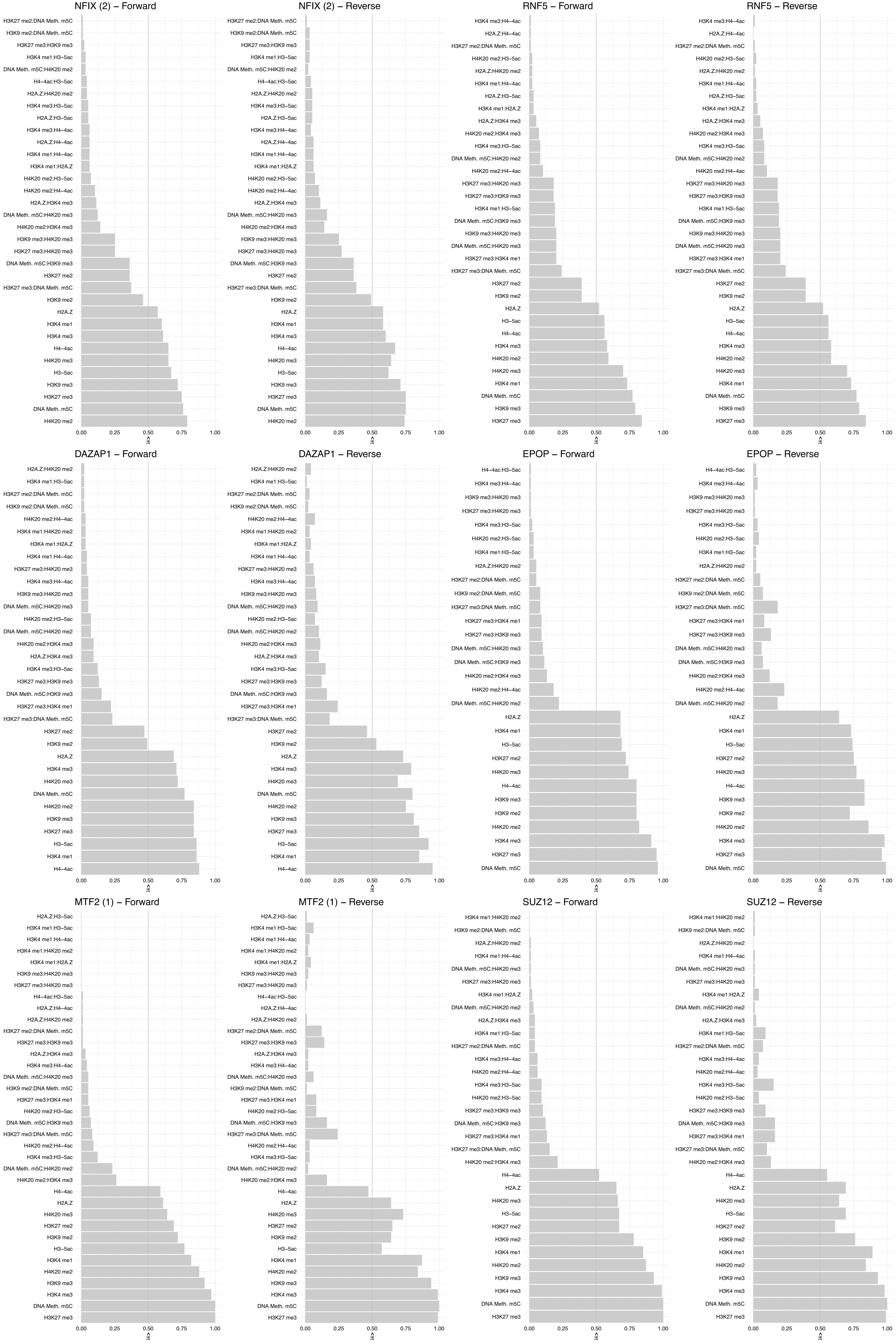
NFIX (2) – Forward
H3K27 me2:DNA Meth. m5C
H3K9 me2:DNA Meth. m5C
H3K27 me3:H3K9 me3
H3K4 me1:H3–5ac
DNA Meth. m5C:H4K20 me2
H4–4ac:H3–5ac
H2A.Z:H4K20 me2
H3K4 me3:H3–5ac
H2A.Z:H3–5ac
H3K4 me3:H4–4ac
H2A.Z:H4–4ac
H3K4 me1:H4–4ac
H3K4 me1:H2A.Z
H4K20 me2:H3–5ac
H4K20 me2:H4–4ac
H2A.Z:H3K4 me3
DNA Meth. m5C:H4K20 me3
H4K20 me2:H3K4 me3
H3K9 me3:H4K20 me3
H3K27 me3:H4K20 me3
DNA Meth. m5C:H3K9 me3
H3K27 me2
H3K27 me3:DNA Meth. m5C
H3K9 me2
H2A.Z
H3K4 me1
H3K4 me3
H4–4ac
H4K20 me3
H3–5ac
H3K9 me3
H3K27 me3
DNA Meth. m5C
H4K20 me2
0.00 0.25 0.50 0.75 1.00
π̂
NFIX (2) – Reverse
H3K27 me2:DNA Meth. m5C
H3K9 me2:DNA Meth. m5C
H3K27 me3:H3K9 me3
H3K4 me1:H3–5ac
DNA Meth. m5C:H4K20 me2
H4–4ac:H3–5ac
H2A.Z:H4K20 me2
H3K4 me3:H3–5ac
H2A.Z:H3–5ac
H3K4 me3:H4–4ac
H2A.Z:H4–4ac
H3K4 me1:H4–4ac
H3K4 me1:H2A.Z
H4K20 me2:H3–5ac
H4K20 me2:H4–4ac
H2A.Z:H3K4 me3
DNA Meth. m5C:H4K20 me3
H4K20 me2:H3K4 me3
H3K9 me3:H4K20 me3
H3K27 me3:H4K20 me3
DNA Meth. m5C:H3K9 me3
H3K27 me2
H3K27 me3:DNA Meth. m5C
H3K9 me2
H2A.Z
H3K4 me1
H3K4 me3
H4–4ac
H4K20 me3
H3–5ac
H3K9 me3
H3K27 me3
DNA Meth. m5C
H4K20 me2
0.00 0.25 0.50 0.75 1.00
π̂
RNF5 – Forward
H3K4 me3:H4–4ac
H2A.Z:H4–4ac
H3K27 me2:DNA Meth. m5C
H4K20 me2:H3–5ac
H2A.Z:H4K20 me2
H3K4 me1:H4–4ac
H2A.Z:H3–5ac
H3K4 me1:H2A.Z
H2A.Z:H3K4 me3
H4K20 me2:H3K4 me3
H3K4 me3:H3–5ac
DNA Meth. m5C:H4K20 me2
H4K20 me2:H4–4ac
H3K27 me3:H4K20 me3
H3K27 me3:H3K9 me3
H3K4 me1:H3–5ac
DNA Meth. m5C:H3K9 me3
H3K9 me3:H4K20 me3
DNA Meth. m5C:H4K20 me3
H3K27 me3:H3K4 me1
H3K27 me3:DNA Meth. m5C
H3K27 me2
H3K9 me2
H2A.Z
H3–5ac
H4–4ac
H3K4 me3
H4K20 me2
H4K20 me3
H3K4 me1
DNA Meth. m5C
H3K9 me3
H3K27 me3
0.00 0.25 0.50 0.75 1.00
π̂
RNF5 – Reverse
H3K4 me3:H4–4ac
H2A.Z:H4–4ac
H3K27 me2:DNA Meth. m5C
H4K20 me2:H3–5ac
H2A.Z:H4K20 me2
H3K4 me1:H4–4ac
H2A.Z:H3–5ac
H3K4 me1:H2A.Z
H2A.Z:H3K4 me3
H4K20 me2:H3K4 me3
H3K4 me3:H3–5ac
DNA Meth. m5C:H4K20 me2
H4K20 me2:H4–4ac
H3K27 me3:H4K20 me3
H3K27 me3:H3K9 me3
H3K4 me1:H3–5ac
DNA Meth. m5C:H3K9 me3
H3K9 me3:H4K20 me3
DNA Meth. m5C:H4K20 me3
H3K27 me3:H3K4 me1
H3K27 me3:DNA Meth. m5C
H3K27 me2
H3K9 me2
H2A.Z
H3–5ac
H4–4ac
H3K4 me3
H4K20 me2
H4K20 me3
H3K4 me1
DNA Meth. m5C
H3K9 me3
H3K27 me3
0.00 0.25 0.50 0.75 1.00
π̂
DAZAP1 – Forward
H2A.Z:H4K20 me2
H3K4 me1:H3–5ac
H3K27 me2:DNA Meth. m5C
H3K9 me2:DNA Meth. m5C
H4K20 me2:H4–4ac
H3K4 me1:H4K20 me2
H3K4 me1:H2A.Z
H3K4 me1:H4–4ac
H3K27 me3:H4K20 me3
H3K4 me3:H4–4ac
H3K9 me3:H4K20 me3
DNA Meth. m5C:H4K20 me3
H4K20 me2:H3–5ac
DNA Meth. m5C:H4K20 me2
H4K20 me2:H3K4 me3
H2A.Z:H3K4 me3
H3K4 me3:H3–5ac
H3K27 me3:H3K9 me3
DNA Meth. m5C:H3K9 me3
H3K27 me3:H3K4 me1
H3K27 me3:DNA Meth. m5C
H3K27 me2
H3K9 me2
H2A.Z
H3K4 me3
H4K20 me3
DNA Meth. m5C
H4K20 me2
H3K9 me3
H3K27 me3
H3–5ac
H3K4 me1
H4–4ac
0.00 0.25 0.50 0.75 1.00
π̂
DAZAP1 – Reverse
H2A.Z:H4K20 me2
H3K4 me1:H3–5ac
H3K27 me2:DNA Meth. m5C
H3K9 me2:DNA Meth. m5C
H4K20 me2:H4–4ac
H3K4 me1:H4K20 me2
H3K4 me1:H2A.Z
H3K4 me1:H4–4ac
H3K27 me3:H4K20 me3
H3K4 me3:H4–4ac
H3K9 me3:H4K20 me3
DNA Meth. m5C:H4K20 me3
H4K20 me2:H3–5ac
DNA Meth. m5C:H4K20 me2
H4K20 me2:H3K4 me3
H2A.Z:H3K4 me3
H3K4 me3:H3–5ac
H3K27 me3:H3K9 me3
DNA Meth. m5C:H3K9 me3
H3K27 me3:H3K4 me1
H3K27 me3:DNA Meth. m5C
H3K27 me2
H3K9 me2
H2A.Z
H3K4 me3
H4K20 me3
DNA Meth. m5C
H4K20 me2
H3K9 me3
H3K27 me3
H3–5ac
H3K4 me1
H4–4ac
0.00 0.25 0.50 0.75 1.00
π̂
EPOP – Forward
H4–4ac:H3–5ac
H3K4 me3:H4–4ac
H3K9 me3:H4K20 me3
H3K27 me3:H4K20 me3
H3K4 me3:H3–5ac
H4K20 me2:H3–5ac
H3K4 me1:H3–5ac
H2A.Z:H4K20 me2
H3K27 me2:DNA Meth. m5C
H3K9 me2:DNA Meth. m5C
H3K27 me3:DNA Meth. m5C
H3K27 me3:H3K4 me1
H3K27 me3:H3K9 me3
DNA Meth. m5C:H4K20 me3
DNA Meth. m5C:H3K9 me3
H4K20 me2:H3K4 me3
H4K20 me2:H4–4ac
DNA Meth. m5C:H4K20 me2
H2A.Z
H3K4 me1
H3–5ac
H3K27 me2
H4K20 me3
H4–4ac
H3K9 me3
H3K9 me2
H4K20 me2
H3K4 me3
H3K27 me3
DNA Meth. m5C
0.00 0.25 0.50 0.75 1.00
π̂
EPOP – Reverse
H4–4ac:H3–5ac
H3K4 me3:H4–4ac
H3K9 me3:H4K20 me3
H3K27 me3:H4K20 me3
H3K4 me3:H3–5ac
H4K20 me2:H3–5ac
H3K4 me1:H3–5ac
H2A.Z:H4K20 me2
H3K27 me2:DNA Meth. m5C
H3K9 me2:DNA Meth. m5C
H3K27 me3:DNA Meth. m5C
H3K27 me3:H3K4 me1
H3K27 me3:H3K9 me3
DNA Meth. m5C:H4K20 me3
DNA Meth. m5C:H3K9 me3
H4K20 me2:H3K4 me3
H4K20 me2:H4–4ac
DNA Meth. m5C:H4K20 me2
H2A.Z
H3K4 me1
H3–5ac
H3K27 me2
H4K20 me3
H4–4ac
H3K9 me3
H3K9 me2
H4K20 me2
H3K4 me3
H3K27 me3
DNA Meth. m5C
0.00 0.25 0.50 0.75 1.00
π̂
MTF2 (1) – Forward
H2A.Z:H3–5ac
H3K4 me1:H3–5ac
H3K4 me1:H4–4ac
H3K4 me1:H4K20 me2
H3K4 me1:H2A.Z
H3K9 me3:H4K20 me3
H3K27 me3:H4K20 me3
H4–4ac:H3–5ac
H2A.Z:H4–4ac
H2A.Z:H4K20 me2
H3K27 me2:DNA Meth. m5C
H3K27 me3:H3K9 me3
H2A.Z:H3K4 me3
H3K4 me3:H4–4ac
DNA Meth. m5C:H4K20 me3
H3K9 me2:DNA Meth. m5C
H3K27 me3:H3K4 me1
H4K20 me2:H3–5ac
DNA Meth. m5C:H3K9 me3
H3K27 me3:DNA Meth. m5C
H4K20 me2:H4–4ac
H3K4 me3:H3–5ac
DNA Meth. m5C:H4K20 me2
H4K20 me2:H3K4 me3
H4–4ac
H2A.Z
H4K20 me3
H3K27 me2
H3K9 me2
H3–5ac
H3K4 me1
H4K20 me2
H3K9 me3
H3K4 me3
DNA Meth. m5C
H3K27 me3
0.00 0.25 0.50 0.75 1.00
π̂
MTF2 (1) – Reverse
H2A.Z:H3–5ac
H3K4 me1:H3–5ac
H3K4 me1:H4–4ac
H3K4 me1:H4K20 me2
H3K4 me1:H2A.Z
H3K9 me3:H4K20 me3
H3K27 me3:H4K20 me3
H4–4ac:H3–5ac
H2A.Z:H4–4ac
H2A.Z:H4K20 me2
H3K27 me2:DNA Meth. m5C
H3K27 me3:H3K9 me3
H2A.Z:H3K4 me3
H3K4 me3:H4–4ac
DNA Meth. m5C:H4K20 me3
H3K9 me2:DNA Meth. m5C
H3K27 me3:H3K4 me1
H4K20 me2:H3–5ac
DNA Meth. m5C:H3K9 me3
H3K27 me3:DNA Meth. m5C
H4K20 me2:H4–4ac
H3K4 me3:H3–5ac
DNA Meth. m5C:H4K20 me2
H4K20 me2:H3K4 me3
H4–4ac
H2A.Z
H4K20 me3
H3K27 me2
H3K9 me2
H3–5ac
H3K4 me1
H4K20 me2
H3K9 me3
H3K4 me3
DNA Meth. m5C
H3K27 me3
0.00 0.25 0.50 0.75 1.00
π̂
SUZ12 – Forward
H3K4 me1:H4K20 me2
H3K9 me2:DNA Meth. m5C
H2A.Z:H4K20 me2
H3K4 me1:H4–4ac
DNA Meth. m5C:H4K20 me3
H3K27 me3:H4K20 me3
H3K4 me1:H2A.Z
DNA Meth. m5C:H4K20 me2
H2A.Z:H3K4 me3
H3K4 me1:H3–5ac
H3K27 me2:DNA Meth. m5C
H3K4 me3:H4–4ac
H4K20 me2:H4–4ac
H3K4 me3:H3–5ac
H4K20 me2:H3–5ac
H3K27 me3:H3K9 me3
DNA Meth. m5C:H3K9 me3
H3K27 me3:H3K4 me1
H3K27 me3:DNA Meth. m5C
H4K20 me2:H3K4 me3
H4–4ac
H2A.Z
H4K20 me3
H3–5ac
H3K27 me2
H3K9 me2
H3K4 me1
H4K20 me2
H3K9 me3
H3K4 me3
DNA Meth. m5C
H3K27 me3
0.00 0.25 0.50 0.75 1.00
π̂
SUZ12 – Reverse
H3K4 me1:H4K20 me2
H3K9 me2:DNA Meth. m5C
H2A.Z:H4K20 me2
H3K4 me1:H4–4ac
DNA Meth. m5C:H4K20 me3
H3K27 me3:H4K20 me3
H3K4 me1:H2A.Z
DNA Meth. m5C:H4K20 me2
H2A.Z:H3K4 me3
H3K4 me1:H3–5ac
H3K27 me2:DNA Meth. m5C
H3K4 me3:H4–4ac
H4K20 me2:H4–4ac
H3K4 me3:H3–5ac
H4K20 me2:H3–5ac
H3K27 me3:H3K9 me3
DNA Meth. m5C:H3K9 me3
H3K27 me3:H3K4 me1
H3K27 me3:DNA Meth. m5C
H4K20 me2:H3K4 me3
H4–4ac
H2A.Z
H4K20 me3
H3–5ac
H3K27 me2
H3K9 me2
H3K4 me1
H4K20 me2
H3K9 me3
H3K4 me3
DNA Meth. m5C
H3K27 me3
0.00 0.25 0.50 0.75 1.00
π̂

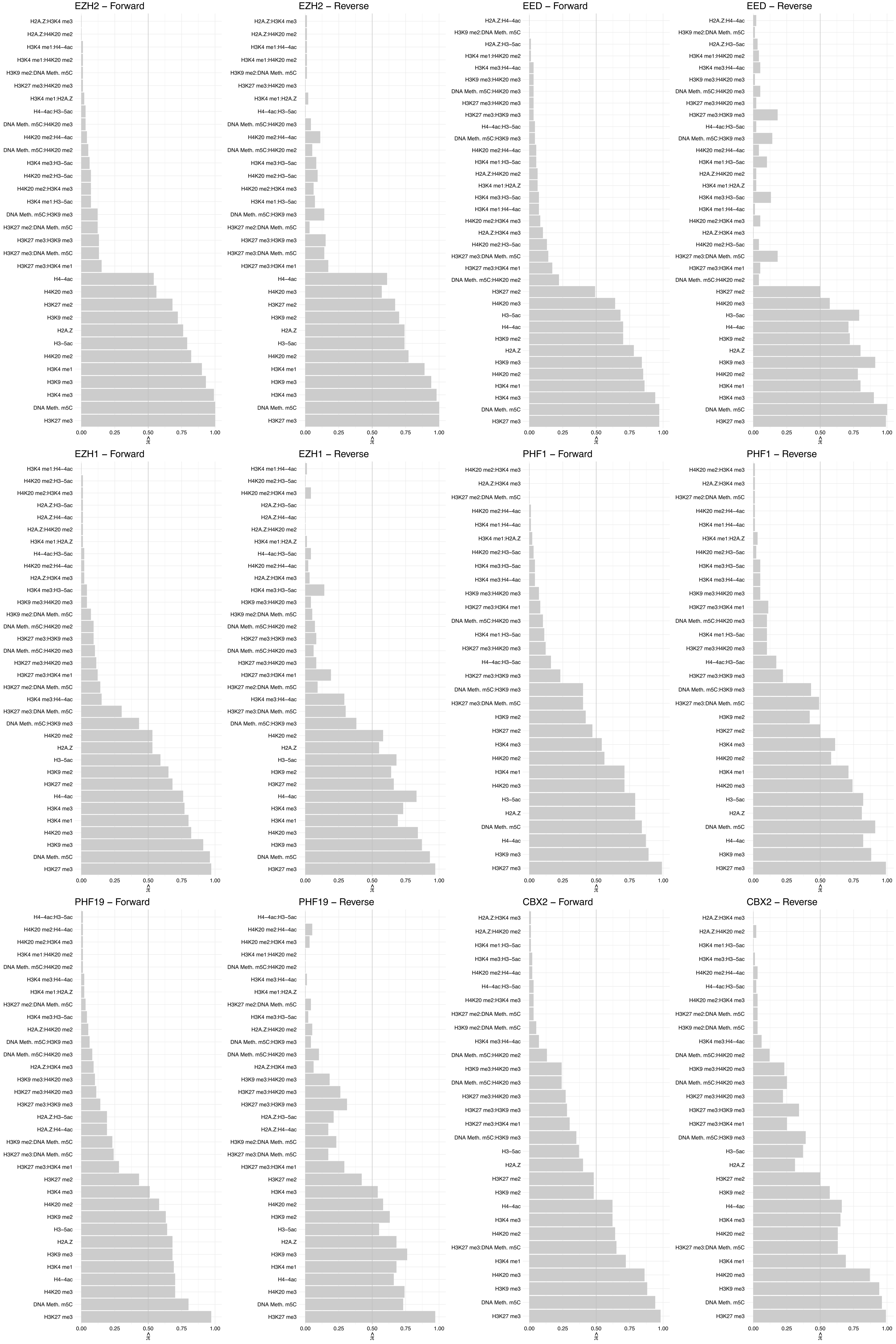
EZH2 − Forward
H2A.Z:H3K4 me3
H2A.Z:H4K20 me2
H3K4 me1:H4−4ac
H3K4 me1:H4K20 me2
H3K9 me2:DNA Meth. m5C
H3K27 me3:H4K20 me3
H3K4 me1:H2A.Z
H4−4ac:H3−5ac
DNA Meth. m5C:H4K20 me3
H4K20 me2:H4−4ac
DNA Meth. m5C:H4K20 me2
H3K4 me3:H3−5ac
H4K20 me2:H3−5ac
H4K20 me2:H3K4 me3
H3K4 me1:H3−5ac
DNA Meth. m5C:H3K9 me3
H3K27 me2:DNA Meth. m5C
H3K27 me3:H3K9 me3
H3K27 me3:DNA Meth. m5C
H3K27 me3:H3K4 me1
H4−4ac
H4K20 me3
H3K27 me2
H3K9 me2
H2A.Z
H3−5ac
H4K20 me2
H3K4 me1
H3K9 me3
H3K4 me3
DNA Meth. m5C
H3K27 me3
0.00 0.25 0.50 0.75 1.00
$\hat{\pi}$
EZH2 − Reverse
H2A.Z:H3K4 me3
H2A.Z:H4K20 me2
H3K4 me1:H4−4ac
H3K4 me1:H4K20 me2
H3K9 me2:DNA Meth. m5C
H3K27 me3:H4K20 me3
H3K4 me1:H2A.Z
H4−4ac:H3−5ac
DNA Meth. m5C:H4K20 me3
H4K20 me2:H4−4ac
DNA Meth. m5C:H4K20 me2
H3K4 me3:H3−5ac
H4K20 me2:H3−5ac
H4K20 me2:H3K4 me3
H3K4 me1:H3−5ac
DNA Meth. m5C:H3K9 me3
H3K27 me2:DNA Meth. m5C
H3K27 me3:H3K9 me3
H3K27 me3:DNA Meth. m5C
H3K27 me3:H3K4 me1
H4−4ac
H4K20 me3
H3K27 me2
H3K9 me2
H2A.Z
H3−5ac
H4K20 me2
H3K4 me1
H3K9 me3
H3K4 me3
DNA Meth. m5C
H3K27 me3
0.00 0.25 0.50 0.75 1.00
$\hat{\pi}$
EED − Forward
H2A.Z:H4−4ac
H3K9 me2:DNA Meth. m5C
H2A.Z:H3−5ac
H3K4 me1:H4K20 me2
H3K4 me3:H4−4ac
H3K9 me3:H4K20 me3
DNA Meth. m5C:H4K20 me3
H3K27 me3:H4K20 me3
H3K27 me3:H3K9 me3
H4−4ac:H3−5ac
DNA Meth. m5C:H3K9 me3
H4K20 me2:H4−4ac
H3K4 me1:H3−5ac
H2A.Z:H4K20 me2
H3K4 me1:H2A.Z
H3K4 me3:H3−5ac
H3K4 me1:H4−4ac
H4K20 me2:H3K4 me3
H2A.Z:H3K4 me3
H4K20 me2:H3−5ac
H3K27 me3:DNA Meth. m5C
H3K27 me3:H3K4 me1
DNA Meth. m5C:H4K20 me2
H3K27 me2
H4K20 me3
H3−5ac
H4−4ac
H3K9 me2
H2A.Z
H3K9 me3
H4K20 me2
H3K4 me1
H3K4 me3
DNA Meth. m5C
H3K27 me3
0.00 0.25 0.50 0.75 1.00
$\hat{\pi}$
EED − Reverse
H2A.Z:H4−4ac
H3K9 me2:DNA Meth. m5C
H2A.Z:H3−5ac
H3K4 me1:H4K20 me2
H3K4 me3:H4−4ac
H3K9 me3:H4K20 me3
DNA Meth. m5C:H4K20 me3
H3K27 me3:H4K20 me3
H3K27 me3:H3K9 me3
H4−4ac:H3−5ac
DNA Meth. m5C:H3K9 me3
H4K20 me2:H4−4ac
H3K4 me1:H3−5ac
H2A.Z:H4K20 me2
H3K4 me1:H2A.Z
H3K4 me3:H3−5ac
H3K4 me1:H4−4ac
H4K20 me2:H3K4 me3
H2A.Z:H3K4 me3
H4K20 me2:H3−5ac
H3K27 me3:DNA Meth. m5C
H3K27 me3:H3K4 me1
DNA Meth. m5C:H4K20 me2
H3K27 me2
H4K20 me3
H3−5ac
H4−4ac
H3K9 me2
H2A.Z
H3K9 me3
H4K20 me2
H3K4 me1
H3K4 me3
DNA Meth. m5C
H3K27 me3
0.00 0.25 0.50 0.75 1.00
$\hat{\pi}$
EZH1 − Forward
H3K4 me1:H4−4ac
H4K20 me2:H3−5ac
H4K20 me2:H3K4 me3
H2A.Z:H3−5ac
H2A.Z:H4−4ac
H2A.Z:H4K20 me2
H3K4 me1:H2A.Z
H4−4ac:H3−5ac
H4K20 me2:H4−4ac
H2A.Z:H3K4 me3
H3K4 me3:H3−5ac
H3K9 me3:H4K20 me3
H3K9 me2:DNA Meth. m5C
DNA Meth. m5C:H4K20 me2
H3K27 me3:H3K9 me3
DNA Meth. m5C:H4K20 me3
H3K27 me3:H4K20 me3
H3K27 me3:H3K4 me1
H3K27 me2:DNA Meth. m5C
H3K4 me3:H4−4ac
H3K27 me3:DNA Meth. m5C
DNA Meth. m5C:H3K9 me3
H4K20 me2
H2A.Z
H3−5ac
H3K9 me2
H3K27 me2
H4−4ac
H3K4 me3
H3K4 me1
H4K20 me3
H3K9 me3
DNA Meth. m5C
H3K27 me3
0.00 0.25 0.50 0.75 1.00
$\hat{\pi}$
EZH1 − Reverse
H3K4 me1:H4−4ac
H4K20 me2:H3−5ac
H4K20 me2:H3K4 me3
H2A.Z:H3−5ac
H2A.Z:H4−4ac
H2A.Z:H4K20 me2
H3K4 me1:H2A.Z
H4−4ac:H3−5ac
H4K20 me2:H4−4ac
H2A.Z:H3K4 me3
H3K4 me3:H3−5ac
H3K9 me3:H4K20 me3
H3K9 me2:DNA Meth. m5C
DNA Meth. m5C:H4K20 me2
H3K27 me3:H3K9 me3
DNA Meth. m5C:H4K20 me3
H3K27 me3:H4K20 me3
H3K27 me3:H3K4 me1
H3K27 me2:DNA Meth. m5C
H3K4 me3:H4−4ac
H3K27 me3:DNA Meth. m5C
DNA Meth. m5C:H3K9 me3
H4K20 me2
H2A.Z
H3−5ac
H3K9 me2
H3K27 me2
H4−4ac
H3K4 me3
H3K4 me1
H4K20 me3
H3K9 me3
DNA Meth. m5C
H3K27 me3
0.00 0.25 0.50 0.75 1.00
$\hat{\pi}$
PHF1 − Forward
H4K20 me2:H3K4 me3
H2A.Z:H3K4 me3
H3K27 me2:DNA Meth. m5C
H4K20 me2:H4−4ac
H3K4 me1:H4−4ac
H3K4 me1:H2A.Z
H4K20 me2:H3−5ac
H3K4 me3:H3−5ac
H3K4 me3:H4−4ac
H3K9 me3:H4K20 me3
H3K27 me3:H3K4 me1
DNA Meth. m5C:H4K20 me3
H3K4 me1:H3−5ac
H3K27 me3:H4K20 me3
H4−4ac:H3−5ac
H3K27 me3:H3K9 me3
DNA Meth. m5C:H3K9 me3
H3K27 me3:DNA Meth. m5C
H3K9 me2
H3K27 me2
H3K4 me3
H4K20 me2
H3K4 me1
H4K20 me3
H3−5ac
H2A.Z
DNA Meth. m5C
H4−4ac
H3K9 me3
H3K27 me3
0.00 0.25 0.50 0.75 1.00
$\hat{\pi}$
PHF1 − Reverse
H4K20 me2:H3K4 me3
H2A.Z:H3K4 me3
H3K27 me2:DNA Meth. m5C
H4K20 me2:H4−4ac
H3K4 me1:H4−4ac
H3K4 me1:H2A.Z
H4K20 me2:H3−5ac
H3K4 me3:H3−5ac
H3K4 me3:H4−4ac
H3K9 me3:H4K20 me3
H3K27 me3:H3K4 me1
DNA Meth. m5C:H4K20 me3
H3K4 me1:H3−5ac
H3K27 me3:H4K20 me3
H4−4ac:H3−5ac
H3K27 me3:H3K9 me3
DNA Meth. m5C:H3K9 me3
H3K27 me3:DNA Meth. m5C
H3K9 me2
H3K27 me2
H3K4 me3
H4K20 me2
H3K4 me1
H4K20 me3
H3−5ac
H2A.Z
DNA Meth. m5C
H4−4ac
H3K9 me3
H3K27 me3
0.00 0.25 0.50 0.75 1.00
$\hat{\pi}$
PHF19 − Forward
H4−4ac:H3−5ac
H4K20 me2:H4−4ac
H4K20 me2:H3K4 me3
H3K4 me1:H4K20 me2
DNA Meth. m5C:H4K20 me2
H3K4 me3:H4−4ac
H3K4 me1:H2A.Z
H3K27 me2:DNA Meth. m5C
H3K4 me3:H3−5ac
H2A.Z:H4K20 me2
DNA Meth. m5C:H3K9 me3
DNA Meth. m5C:H4K20 me3
H2A.Z:H3K4 me3
H3K9 me3:H4K20 me3
H3K27 me3:H4K20 me3
H3K27 me3:H3K9 me3
H2A.Z:H3−5ac
H2A.Z:H4−4ac
H3K9 me2:DNA Meth. m5C
H3K27 me3:DNA Meth. m5C
H3K27 me3:H3K4 me1
H3K27 me2
H3K4 me3
H4K20 me2
H3K9 me2
H3−5ac
H2A.Z
H3K9 me3
H3K4 me1
H4−4ac
H4K20 me3
DNA Meth. m5C
H3K27 me3
0.00 0.25 0.50 0.75 1.00
$\hat{\pi}$
PHF19 − Reverse
H4−4ac:H3−5ac
H4K20 me2:H4−4ac
H4K20 me2:H3K4 me3
H3K4 me1:H4K20 me2
DNA Meth. m5C:H4K20 me2
H3K4 me3:H4−4ac
H3K4 me1:H2A.Z
H3K27 me2:DNA Meth. m5C
H3K4 me3:H3−5ac
H2A.Z:H4K20 me2
DNA Meth. m5C:H3K9 me3
DNA Meth. m5C:H4K20 me3
H2A.Z:H3K4 me3
H3K9 me3:H4K20 me3
H3K27 me3:H4K20 me3
H3K27 me3:H3K9 me3
H2A.Z:H3−5ac
H2A.Z:H4−4ac
H3K9 me2:DNA Meth. m5C
H3K27 me3:DNA Meth. m5C
H3K27 me3:H3K4 me1
H3K27 me2
H3K4 me3
H4K20 me2
H3K9 me2
H3−5ac
H2A.Z
H3K9 me3
H3K4 me1
H4−4ac
H4K20 me3
DNA Meth. m5C
H3K27 me3
0.00 0.25 0.50 0.75 1.00
$\hat{\pi}$
CBX2 − Forward
H2A.Z:H3K4 me3
H2A.Z:H4K20 me2
H3K4 me1:H3−5ac
H3K4 me3:H3−5ac
H4K20 me2:H4−4ac
H4−4ac:H3−5ac
H4K20 me2:H3K4 me3
H3K27 me2:DNA Meth. m5C
H3K9 me2:DNA Meth. m5C
H3K4 me3:H4−4ac
DNA Meth. m5C:H4K20 me2
H3K9 me3:H4K20 me3
DNA Meth. m5C:H4K20 me3
H3K27 me3:H4K20 me3
H3K27 me3:H3K9 me3
H3K27 me3:H3K4 me1
DNA Meth. m5C:H3K9 me3
H3−5ac
H2A.Z
H3K27 me2
H3K9 me2
H4−4ac
H3K4 me3
H4K20 me2
H3K27 me3:DNA Meth. m5C
H3K4 me1
H4K20 me3
H3K9 me3
DNA Meth. m5C
H3K27 me3
0.00 0.25 0.50 0.75 1.00
$\hat{\pi}$
CBX2 − Reverse
H2A.Z:H3K4 me3
H2A.Z:H4K20 me2
H3K4 me1:H3−5ac
H3K4 me3:H3−5ac
H4K20 me2:H4−4ac
H4−4ac:H3−5ac
H4K20 me2:H3K4 me3
H3K27 me2:DNA Meth. m5C
H3K9 me2:DNA Meth. m5C
H3K4 me3:H4−4ac
DNA Meth. m5C:H4K20 me2
H3K9 me3:H4K20 me3
DNA Meth. m5C:H4K20 me3
H3K27 me3:H4K20 me3
H3K27 me3:H3K9 me3
H3K27 me3:H3K4 me1
DNA Meth. m5C:H3K9 me3
H3−5ac
H2A.Z
H3K27 me2
H3K9 me2
H4−4ac
H3K4 me3
H4K20 me2
H3K27 me3:DNA Meth. m5C
H3K4 me1
H4K20 me3
H3K9 me3
DNA Meth. m5C
H3K27 me3
0.00 0.25 0.50 0.75 1.00
$\hat{\pi}$

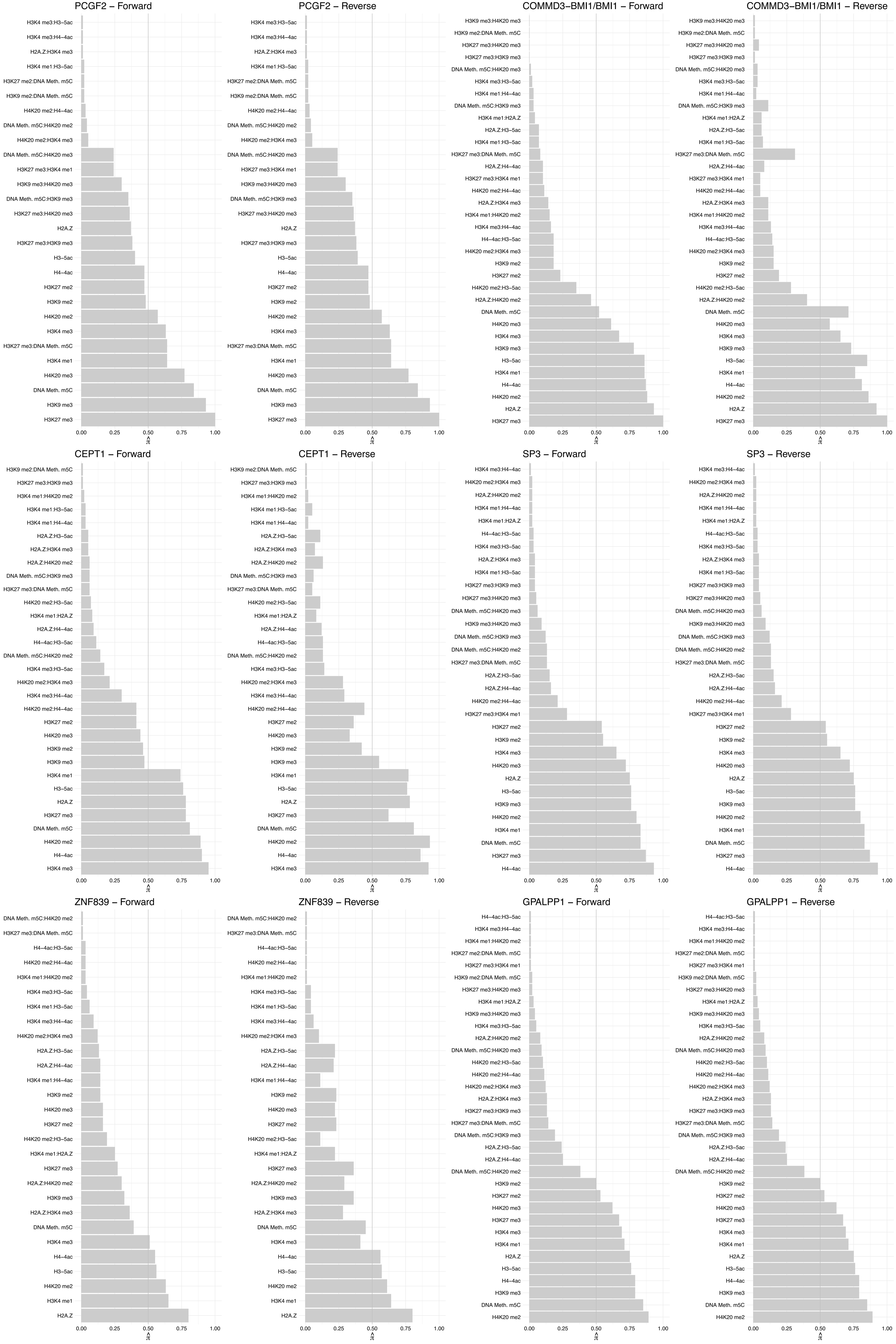

PCGF2 − Forward
PCGF2 − Reverse
H3K4 me3:H3−5ac
H3K4 me3:H4−4ac
H2A.Z:H3K4 me3
H3K4 me1:H3−5ac
H3K27 me2:DNA Meth. m5C
H3K9 me2:DNA Meth. m5C
H4K20 me2:H4−4ac
DNA Meth. m5C:H4K20 me2
H4K20 me2:H3K4 me3
DNA Meth. m5C:H4K20 me3
H3K27 me3:H3K4 me1
H3K9 me3:H4K20 me3
DNA Meth. m5C:H3K9 me3
H3K27 me3:H4K20 me3
H2A.Z
H3K27 me3:H3K9 me3
H3−5ac
H4−4ac
H3K27 me2
H3K9 me2
H4K20 me2
H3K4 me3
H3K27 me3:DNA Meth. m5C
H3K4 me1
H4K20 me3
DNA Meth. m5C
H3K9 me3
H3K27 me3
COMMD3−BMI1/BMI1 − Forward
COMMD3−BMI1/BMI1 − Reverse
H3K9 me3:H4K20 me3
H3K9 me2:DNA Meth. m5C
H3K27 me3:H4K20 me3
H3K27 me3:H3K9 me3
DNA Meth. m5C:H4K20 me3
H3K4 me3:H3−5ac
H3K4 me1:H4−4ac
DNA Meth. m5C:H3K9 me3
H3K4 me1:H2A.Z
H2A.Z:H3−5ac
H3K4 me1:H3−5ac
H3K27 me3:DNA Meth. m5C
H2A.Z:H4−4ac
H3K27 me3:H3K4 me1
H4K20 me2:H4−4ac
H2A.Z:H3K4 me3
H3K4 me1:H4K20 me2
H3K4 me3:H4−4ac
H4−4ac:H3−5ac
H4K20 me2:H3K4 me3
H3K9 me2
H3K27 me2
H4K20 me2:H3−5ac
H2A.Z:H4K20 me2
DNA Meth. m5C
H4K20 me3
H3K4 me3
H3K9 me3
H3−5ac
H3K4 me1
H4−4ac
H4K20 me2
H2A.Z
H3K27 me3
CEPT1 − Forward
CEPT1 − Reverse
H3K9 me2:DNA Meth. m5C
H3K27 me3:H3K9 me3
H3K4 me1:H4K20 me2
H3K4 me1:H3−5ac
H3K4 me1:H4−4ac
H2A.Z:H3−5ac
H2A.Z:H3K4 me3
H2A.Z:H4K20 me2
DNA Meth. m5C:H3K9 me3
H3K27 me3:DNA Meth. m5C
H4K20 me2:H3−5ac
H3K4 me1:H2A.Z
H2A.Z:H4−4ac
H4−4ac:H3−5ac
DNA Meth. m5C:H4K20 me2
H3K4 me3:H3−5ac
H4K20 me2:H3K4 me3
H3K4 me3:H4−4ac
H4K20 me2:H4−4ac
H3K27 me2
H4K20 me3
H3K9 me2
H3K9 me3
H3K4 me1
H3−5ac
H2A.Z
H3K27 me3
DNA Meth. m5C
H4K20 me2
H4−4ac
H3K4 me3
SP3 − Forward
SP3 − Reverse
H3K4 me3:H4−4ac
H4K20 me2:H3K4 me3
H2A.Z:H4K20 me2
H3K4 me1:H4−4ac
H3K4 me1:H2A.Z
H4−4ac:H3−5ac
H3K4 me3:H3−5ac
H2A.Z:H3K4 me3
H3K4 me1:H3−5ac
H3K27 me3:H3K9 me3
H3K27 me3:H4K20 me3
DNA Meth. m5C:H4K20 me3
H3K9 me3:H4K20 me3
DNA Meth. m5C:H3K9 me3
DNA Meth. m5C:H4K20 me2
H3K27 me3:DNA Meth. m5C
H2A.Z:H3−5ac
H2A.Z:H4−4ac
H4K20 me2:H4−4ac
H3K27 me3:H3K4 me1
H3K27 me2
H3K9 me2
H3K4 me3
H4K20 me3
H2A.Z
H3−5ac
H3K9 me3
H4K20 me2
H3K4 me1
DNA Meth. m5C
H3K27 me3
H4−4ac
ZNF839 − Forward
ZNF839 − Reverse
DNA Meth. m5C:H4K20 me2
H3K27 me3:DNA Meth. m5C
H4−4ac:H3−5ac
H4K20 me2:H4−4ac
H3K4 me1:H4K20 me2
H3K4 me3:H3−5ac
H3K4 me1:H3−5ac
H3K4 me3:H4−4ac
H4K20 me2:H3K4 me3
H2A.Z:H3−5ac
H2A.Z:H4−4ac
H3K4 me1:H4−4ac
H3K9 me2
H4K20 me3
H3K27 me2
H4K20 me2:H3−5ac
H3K4 me1:H2A.Z
H3K27 me3
H2A.Z:H4K20 me2
H3K9 me3
H2A.Z:H3K4 me3
DNA Meth. m5C
H3K4 me3
H4−4ac
H3−5ac
H4K20 me2
H3K4 me1
H2A.Z
GPALPP1 − Forward
GPALPP1 − Reverse
H4−4ac:H3−5ac
H3K4 me3:H4−4ac
H3K4 me1:H4K20 me2
H3K27 me2:DNA Meth. m5C
H3K27 me3:H3K4 me1
H3K9 me2:DNA Meth. m5C
H3K27 me3:H4K20 me3
H3K4 me1:H2A.Z
H3K9 me3:H4K20 me3
H3K4 me3:H3−5ac
H2A.Z:H4K20 me2
DNA Meth. m5C:H4K20 me3
H4K20 me2:H3−5ac
H4K20 me2:H4−4ac
H4K20 me2:H3K4 me3
H2A.Z:H3K4 me3
H3K27 me3:H3K9 me3
H3K27 me3:DNA Meth. m5C
DNA Meth. m5C:H3K9 me3
H2A.Z:H3−5ac
H2A.Z:H4−4ac
DNA Meth. m5C:H4K20 me2
H3K9 me2
H3K27 me2
H4K20 me3
H3K27 me3
H3K4 me3
H3K4 me1
H2A.Z
H3−5ac
H4−4ac
H3K9 me3
DNA Meth. m5C
H4K20 me2
0.00 0.25 0.50 0.75 1.00
π̂

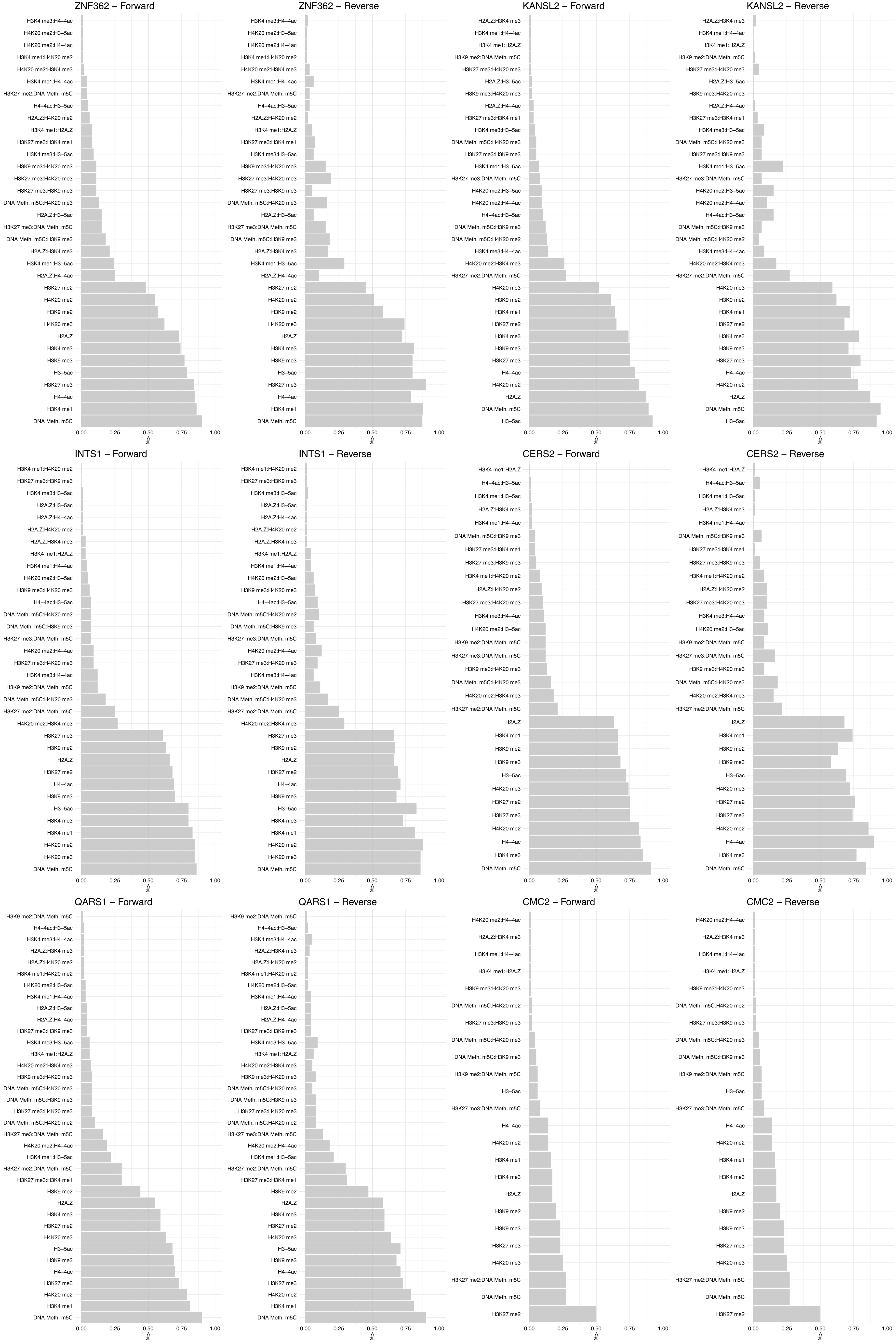
ZNF362 – Forward
ZNF362 – Reverse
H3K4 me3:H4–4ac
H4K20 me2:H3–5ac
H4K20 me2:H4–4ac
H3K4 me1:H4K20 me2
H4K20 me2:H3K4 me3
H3K4 me1:H4–4ac
H3K27 me2:DNA Meth. m5C
H4–4ac:H3–5ac
H2A.Z:H4K20 me2
H3K4 me1:H2A.Z
H3K27 me3:H3K4 me1
H3K4 me3:H3–5ac
H3K9 me3:H4K20 me3
H3K27 me3:H4K20 me3
H3K27 me3:H3K9 me3
DNA Meth. m5C:H4K20 me3
H2A.Z:H3–5ac
H3K27 me3:DNA Meth. m5C
DNA Meth. m5C:H3K9 me3
H2A.Z:H3K4 me3
H3K4 me1:H3–5ac
H2A.Z:H4–4ac
H3K27 me2
H4K20 me2
H3K9 me2
H4K20 me3
H2A.Z
H3K4 me3
H3K9 me3
H3–5ac
H3K27 me3
H4–4ac
H3K4 me1
DNA Meth. m5C
KANSL2 – Forward
KANSL2 – Reverse
H2A.Z:H3K4 me3
H3K4 me1:H4–4ac
H3K4 me1:H2A.Z
H3K9 me2:DNA Meth. m5C
H3K27 me3:H4K20 me3
H2A.Z:H3–5ac
H3K9 me3:H4K20 me3
H2A.Z:H4–4ac
H3K27 me3:H3K4 me1
H3K4 me3:H3–5ac
DNA Meth. m5C:H4K20 me3
H3K27 me3:H3K9 me3
H3K4 me1:H3–5ac
H3K27 me3:DNA Meth. m5C
H4K20 me2:H3–5ac
H4K20 me2:H4–4ac
H4–4ac:H3–5ac
DNA Meth. m5C:H3K9 me3
DNA Meth. m5C:H4K20 me2
H3K4 me3:H4–4ac
H4K20 me2:H3K4 me3
H3K27 me2:DNA Meth. m5C
H4K20 me3
H3K9 me2
H3K4 me1
H3K27 me2
H3K4 me3
H3K9 me3
H3K27 me3
H4–4ac
H4K20 me2
H2A.Z
DNA Meth. m5C
H3–5ac
INTS1 – Forward
INTS1 – Reverse
H3K4 me1:H4K20 me2
H3K27 me3:H3K9 me3
H3K4 me3:H3–5ac
H2A.Z:H3–5ac
H2A.Z:H4–4ac
H2A.Z:H4K20 me2
H2A.Z:H3K4 me3
H3K4 me1:H2A.Z
H3K4 me1:H4–4ac
H4K20 me2:H3–5ac
H3K9 me3:H4K20 me3
H4–4ac:H3–5ac
DNA Meth. m5C:H4K20 me2
DNA Meth. m5C:H3K9 me3
H3K27 me3:DNA Meth. m5C
H4K20 me2:H4–4ac
H3K27 me3:H4K20 me3
H3K4 me3:H4–4ac
H3K9 me2:DNA Meth. m5C
DNA Meth. m5C:H4K20 me3
H3K27 me2:DNA Meth. m5C
H4K20 me2:H3K4 me3
H3K27 me3
H3K9 me2
H2A.Z
H3K27 me2
H4–4ac
H3K9 me3
H3–5ac
H3K4 me3
H3K4 me1
H4K20 me2
H4K20 me3
DNA Meth. m5C
CERS2 – Forward
CERS2 – Reverse
H3K4 me1:H2A.Z
H4–4ac:H3–5ac
H3K4 me1:H3–5ac
H2A.Z:H3K4 me3
H3K4 me1:H4–4ac
DNA Meth. m5C:H3K9 me3
H3K27 me3:H3K4 me1
H3K27 me3:H3K9 me3
H3K4 me1:H4K20 me2
H2A.Z:H4K20 me2
H3K27 me3:H4K20 me3
H3K4 me3:H4–4ac
H4K20 me2:H3–5ac
H3K9 me2:DNA Meth. m5C
H3K27 me3:DNA Meth. m5C
H3K9 me3:H4K20 me3
DNA Meth. m5C:H4K20 me3
H4K20 me2:H3K4 me3
H3K27 me2:DNA Meth. m5C
H2A.Z
H3K4 me1
H3K9 me2
H3K9 me3
H3–5ac
H4K20 me3
H3K27 me2
H3K27 me3
H4K20 me2
H4–4ac
H3K4 me3
DNA Meth. m5C
QARS1 – Forward
QARS1 – Reverse
H3K9 me2:DNA Meth. m5C
H4–4ac:H3–5ac
H3K4 me3:H4–4ac
H2A.Z:H3K4 me3
H2A.Z:H4K20 me2
H3K4 me1:H4K20 me2
H4K20 me2:H3–5ac
H3K4 me1:H4–4ac
H2A.Z:H3–5ac
H2A.Z:H4–4ac
H3K27 me3:H3K9 me3
H3K4 me3:H3–5ac
H3K4 me1:H2A.Z
H4K20 me2:H3K4 me3
H3K9 me3:H4K20 me3
DNA Meth. m5C:H4K20 me3
DNA Meth. m5C:H3K9 me3
H3K27 me3:H4K20 me3
DNA Meth. m5C:H4K20 me2
H3K27 me3:DNA Meth. m5C
H4K20 me2:H4–4ac
H3K4 me1:H3–5ac
H3K27 me2:DNA Meth. m5C
H3K27 me3:H3K4 me1
H3K9 me2
H2A.Z
H3K4 me3
H3K27 me2
H4K20 me3
H3–5ac
H3K9 me3
H4–4ac
H3K27 me3
H4K20 me2
H3K4 me1
DNA Meth. m5C
CMC2 – Forward
CMC2 – Reverse
H4K20 me2:H4–4ac
H2A.Z:H3K4 me3
H3K4 me1:H4–4ac
H3K4 me1:H2A.Z
H3K9 me3:H4K20 me3
DNA Meth. m5C:H4K20 me2
H3K27 me3:H3K9 me3
DNA Meth. m5C:H4K20 me3
DNA Meth. m5C:H3K9 me3
H3K9 me2:DNA Meth. m5C
H3–5ac
H3K27 me3:DNA Meth. m5C
H4–4ac
H4K20 me2
H3K4 me1
H3K4 me3
H2A.Z
H3K9 me2
H3K9 me3
H3K27 me3
H4K20 me3
H3K27 me2:DNA Meth. m5C
DNA Meth. m5C
H3K27 me2
0.00
0.25
0.50
0.75
1.00
π̂

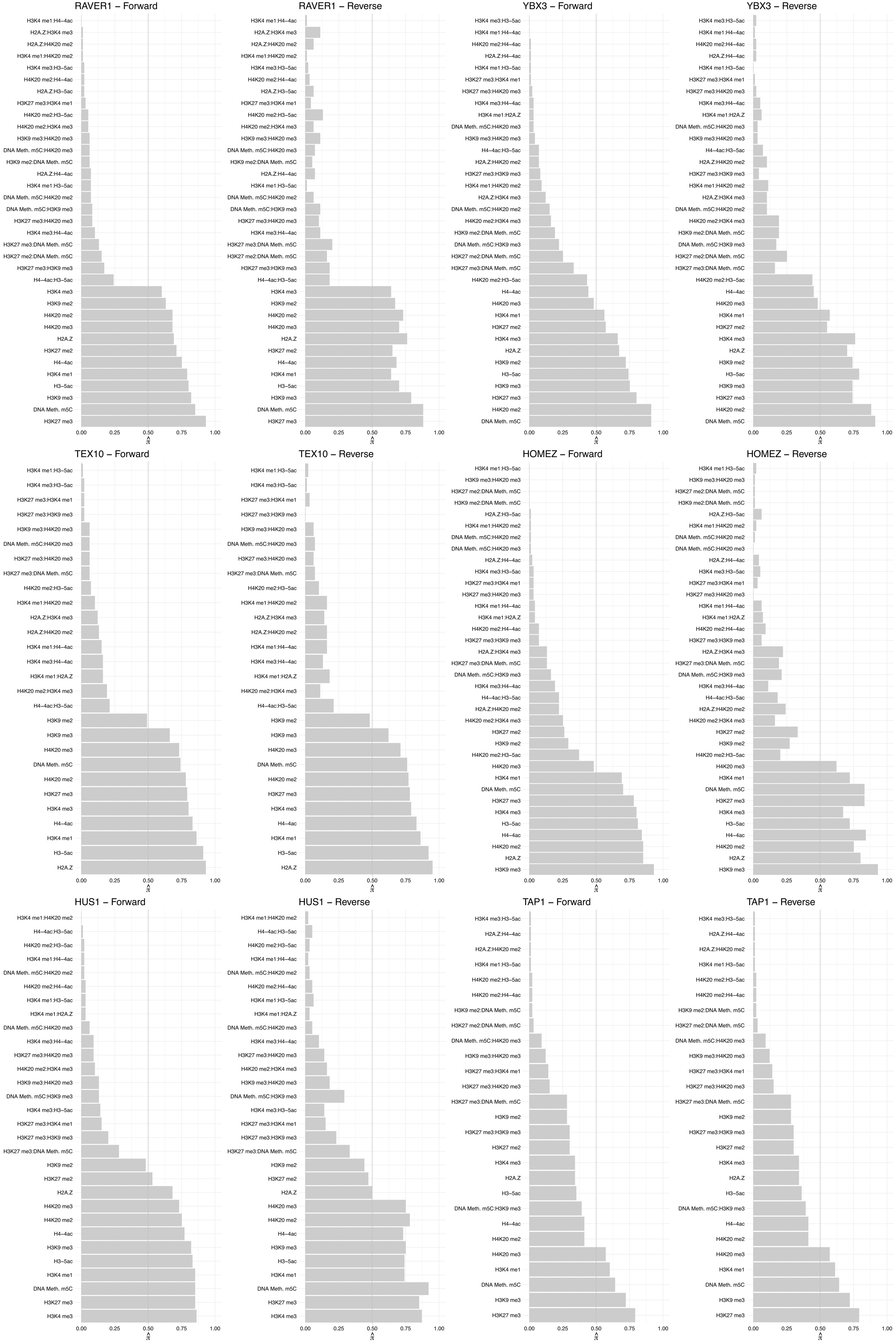
RAVER1 − Forward
H3K4 me1:H4−4ac
H2A.Z:H3K4 me3
H2A.Z:H4K20 me2
H3K4 me1:H4K20 me2
H3K4 me3:H3−5ac
H4K20 me2:H4−4ac
H2A.Z:H3−5ac
H3K27 me3:H3K4 me1
H4K20 me2:H3−5ac
H4K20 me2:H3K4 me3
H3K9 me3:H4K20 me3
DNA Meth. m5C:H4K20 me3
H3K9 me2:DNA Meth. m5C
H2A.Z:H4−4ac
H3K4 me1:H3−5ac
DNA Meth. m5C:H4K20 me2
DNA Meth. m5C:H3K9 me3
H3K27 me3:H4K20 me3
H3K4 me3:H4−4ac
H3K27 me3:DNA Meth. m5C
H3K27 me2:DNA Meth. m5C
H3K27 me3:H3K9 me3
H4−4ac:H3−5ac
H3K4 me3
H3K9 me2
H4K20 me2
H4K20 me3
H2A.Z
H3K27 me2
H4−4ac
H3K4 me1
H3−5ac
H3K9 me3
DNA Meth. m5C
H3K27 me3
0.00
0.25
0.50
0.75
1.00
$\hat{\pi}$
RAVER1 − Reverse
H3K4 me1:H4−4ac
H2A.Z:H3K4 me3
H2A.Z:H4K20 me2
H3K4 me1:H4K20 me2
H3K4 me3:H3−5ac
H4K20 me2:H4−4ac
H2A.Z:H3−5ac
H3K27 me3:H3K4 me1
H4K20 me2:H3−5ac
H4K20 me2:H3K4 me3
H3K9 me3:H4K20 me3
DNA Meth. m5C:H4K20 me3
H3K9 me2:DNA Meth. m5C
H2A.Z:H4−4ac
H3K4 me1:H3−5ac
DNA Meth. m5C:H4K20 me2
DNA Meth. m5C:H3K9 me3
H3K27 me3:H4K20 me3
H3K4 me3:H4−4ac
H3K27 me3:DNA Meth. m5C
H3K27 me2:DNA Meth. m5C
H3K27 me3:H3K9 me3
H4−4ac:H3−5ac
H3K4 me3
H3K9 me2
H4K20 me2
H4K20 me3
H2A.Z
H3K27 me2
H4−4ac
H3K4 me1
H3−5ac
H3K9 me3
DNA Meth. m5C
H3K27 me3
0.00
0.25
0.50
0.75
1.00
$\hat{\pi}$
YBX3 − Forward
H3K4 me3:H3−5ac
H3K4 me1:H4−4ac
H4K20 me2:H4−4ac
H2A.Z:H4−4ac
H3K4 me1:H3−5ac
H3K27 me3:H3K4 me1
H3K27 me3:H4K20 me3
H3K4 me3:H4−4ac
H3K4 me1:H2A.Z
DNA Meth. m5C:H4K20 me3
H3K9 me3:H4K20 me3
H4−4ac:H3−5ac
H2A.Z:H4K20 me2
H3K27 me3:H3K9 me3
H3K4 me1:H4K20 me2
H2A.Z:H3K4 me3
DNA Meth. m5C:H4K20 me2
H4K20 me2:H3K4 me3
H3K9 me2:DNA Meth. m5C
DNA Meth. m5C:H3K9 me3
H3K27 me2:DNA Meth. m5C
H3K27 me3:DNA Meth. m5C
H4K20 me2:H3−5ac
H4−4ac
H4K20 me3
H3K4 me1
H3K27 me2
H3K4 me3
H2A.Z
H3K9 me2
H3−5ac
H3K9 me3
H3K27 me3
H4K20 me2
DNA Meth. m5C
0.00
0.25
0.50
0.75
1.00
$\hat{\pi}$
YBX3 − Reverse
H3K4 me3:H3−5ac
H3K4 me1:H4−4ac
H4K20 me2:H4−4ac
H2A.Z:H4−4ac
H3K4 me1:H3−5ac
H3K27 me3:H3K4 me1
H3K27 me3:H4K20 me3
H3K4 me3:H4−4ac
H3K4 me1:H2A.Z
DNA Meth. m5C:H4K20 me3
H3K9 me3:H4K20 me3
H4−4ac:H3−5ac
H2A.Z:H4K20 me2
H3K27 me3:H3K9 me3
H3K4 me1:H4K20 me2
H2A.Z:H3K4 me3
DNA Meth. m5C:H4K20 me2
H4K20 me2:H3K4 me3
H3K9 me2:DNA Meth. m5C
DNA Meth. m5C:H3K9 me3
H3K27 me2:DNA Meth. m5C
H3K27 me3:DNA Meth. m5C
H4K20 me2:H3−5ac
H4−4ac
H4K20 me3
H3K4 me1
H3K27 me2
H3K4 me3
H2A.Z
H3K9 me2
H3−5ac
H3K9 me3
H3K27 me3
H4K20 me2
DNA Meth. m5C
0.00
0.25
0.50
0.75
1.00
$\hat{\pi}$
TEX10 − Forward
H3K4 me1:H3−5ac
H3K4 me3:H3−5ac
H3K27 me3:H3K4 me1
H3K27 me3:H3K9 me3
H3K9 me3:H4K20 me3
DNA Meth. m5C:H4K20 me3
H3K27 me3:H4K20 me3
H3K27 me3:DNA Meth. m5C
H4K20 me2:H3−5ac
H3K4 me1:H4K20 me2
H2A.Z:H3K4 me3
H2A.Z:H4K20 me2
H3K4 me1:H4−4ac
H3K4 me3:H4−4ac
H3K4 me1:H2A.Z
H4K20 me2:H3K4 me3
H4−4ac:H3−5ac
H3K9 me2
H3K9 me3
H4K20 me3
DNA Meth. m5C
H4K20 me2
H3K27 me3
H3K4 me3
H4−4ac
H3K4 me1
H3−5ac
H2A.Z
0.00
0.25
0.50
0.75
1.00
$\hat{\pi}$
TEX10 − Reverse
H3K4 me1:H3−5ac
H3K4 me3:H3−5ac
H3K27 me3:H3K4 me1
H3K27 me3:H3K9 me3
H3K9 me3:H4K20 me3
DNA Meth. m5C:H4K20 me3
H3K27 me3:H4K20 me3
H3K27 me3:DNA Meth. m5C
H4K20 me2:H3−5ac
H3K4 me1:H4K20 me2
H2A.Z:H3K4 me3
H2A.Z:H4K20 me2
H3K4 me1:H4−4ac
H3K4 me3:H4−4ac
H3K4 me1:H2A.Z
H4K20 me2:H3K4 me3
H4−4ac:H3−5ac
H3K9 me2
H3K9 me3
H4K20 me3
DNA Meth. m5C
H4K20 me2
H3K27 me3
H3K4 me3
H4−4ac
H3K4 me1
H3−5ac
H2A.Z
0.00
0.25
0.50
0.75
1.00
$\hat{\pi}$
HOMEZ − Forward
H3K4 me1:H3−5ac
H3K9 me3:H4K20 me3
H3K27 me2:DNA Meth. m5C
H3K9 me2:DNA Meth. m5C
H2A.Z:H3−5ac
H3K4 me1:H4K20 me2
DNA Meth. m5C:H4K20 me2
DNA Meth. m5C:H4K20 me3
H2A.Z:H4−4ac
H3K4 me3:H3−5ac
H3K27 me3:H3K4 me1
H3K27 me3:H4K20 me3
H3K4 me1:H4−4ac
H3K4 me1:H2A.Z
H4K20 me2:H4−4ac
H3K27 me3:H3K9 me3
H2A.Z:H3K4 me3
H3K27 me3:DNA Meth. m5C
DNA Meth. m5C:H3K9 me3
H3K4 me3:H4−4ac
H4−4ac:H3−5ac
H2A.Z:H4K20 me2
H4K20 me2:H3K4 me3
H3K27 me2
H3K9 me2
H4K20 me2:H3−5ac
H4K20 me3
H3K4 me1
DNA Meth. m5C
H3K27 me3
H3K4 me3
H3−5ac
H4−4ac
H4K20 me2
H2A.Z
H3K9 me3
0.00
0.25
0.50
0.75
1.00
$\hat{\pi}$
HOMEZ − Reverse
H3K4 me1:H3−5ac
H3K9 me3:H4K20 me3
H3K27 me2:DNA Meth. m5C
H3K9 me2:DNA Meth. m5C
H2A.Z:H3−5ac
H3K4 me1:H4K20 me2
DNA Meth. m5C:H4K20 me2
DNA Meth. m5C:H4K20 me3
H2A.Z:H4−4ac
H3K4 me3:H3−5ac
H3K27 me3:H3K4 me1
H3K27 me3:H4K20 me3
H3K4 me1:H4−4ac
H3K4 me1:H2A.Z
H4K20 me2:H4−4ac
H3K27 me3:H3K9 me3
H2A.Z:H3K4 me3
H3K27 me3:DNA Meth. m5C
DNA Meth. m5C:H3K9 me3
H3K4 me3:H4−4ac
H4−4ac:H3−5ac
H2A.Z:H4K20 me2
H4K20 me2:H3K4 me3
H3K27 me2
H3K9 me2
H4K20 me2:H3−5ac
H4K20 me3
H3K4 me1
DNA Meth. m5C
H3K27 me3
H3K4 me3
H3−5ac
H4−4ac
H4K20 me2
H2A.Z
H3K9 me3
0.00
0.25
0.50
0.75
1.00
$\hat{\pi}$
HUS1 − Forward
H3K4 me1:H4K20 me2
H4−4ac:H3−5ac
H4K20 me2:H3−5ac
H3K4 me1:H4−4ac
DNA Meth. m5C:H4K20 me2
H4K20 me2:H4−4ac
H3K4 me1:H3−5ac
H3K4 me1:H2A.Z
DNA Meth. m5C:H4K20 me3
H3K4 me3:H4−4ac
H3K27 me3:H4K20 me3
H4K20 me2:H3K4 me3
H3K9 me3:H4K20 me3
DNA Meth. m5C:H3K9 me3
H3K4 me3:H3−5ac
H3K27 me3:H3K4 me1
H3K27 me3:H3K9 me3
H3K27 me3:DNA Meth. m5C
H3K9 me2
H3K27 me2
H2A.Z
H4K20 me3
H4K20 me2
H4−4ac
H3K9 me3
H3−5ac
H3K4 me1
DNA Meth. m5C
H3K27 me3
H3K4 me3
0.00
0.25
0.50
0.75
1.00
$\hat{\pi}$
HUS1 − Reverse
H3K4 me1:H4K20 me2
H4−4ac:H3−5ac
H4K20 me2:H3−5ac
H3K4 me1:H4−4ac
DNA Meth. m5C:H4K20 me2
H4K20 me2:H4−4ac
H3K4 me1:H3−5ac
H3K4 me1:H2A.Z
DNA Meth. m5C:H4K20 me3
H3K4 me3:H4−4ac
H3K27 me3:H4K20 me3
H4K20 me2:H3K4 me3
H3K9 me3:H4K20 me3
DNA Meth. m5C:H3K9 me3
H3K4 me3:H3−5ac
H3K27 me3:H3K4 me1
H3K27 me3:H3K9 me3
H3K27 me3:DNA Meth. m5C
H3K9 me2
H3K27 me2
H2A.Z
H4K20 me3
H4K20 me2
H4−4ac
H3K9 me3
H3−5ac
H3K4 me1
DNA Meth. m5C
H3K27 me3
H3K4 me3
0.00
0.25
0.50
0.75
1.00
$\hat{\pi}$
TAP1 − Forward
H3K4 me3:H3−5ac
H2A.Z:H4−4ac
H2A.Z:H4K20 me2
H3K4 me1:H3−5ac
H4K20 me2:H3−5ac
H4K20 me2:H4−4ac
H3K9 me2:DNA Meth. m5C
H3K27 me2:DNA Meth. m5C
DNA Meth. m5C:H4K20 me3
H3K9 me3:H4K20 me3
H3K27 me3:H3K4 me1
H3K27 me3:H4K20 me3
H3K27 me3:DNA Meth. m5C
H3K9 me2
H3K27 me3:H3K9 me3
H3K27 me2
H3K4 me3
H2A.Z
H3−5ac
DNA Meth. m5C:H3K9 me3
H4−4ac
H4K20 me2
H4K20 me3
H3K4 me1
DNA Meth. m5C
H3K9 me3
H3K27 me3
0.00
0.25
0.50
0.75
1.00
$\hat{\pi}$
TAP1 − Reverse
H3K4 me3:H3−5ac
H2A.Z:H4−4ac
H2A.Z:H4K20 me2
H3K4 me1:H3−5ac
H4K20 me2:H3−5ac
H4K20 me2:H4−4ac
H3K9 me2:DNA Meth. m5C
H3K27 me2:DNA Meth. m5C
DNA Meth. m5C:H4K20 me3
H3K9 me3:H4K20 me3
H3K27 me3:H3K4 me1
H3K27 me3:H4K20 me3
H3K27 me3:DNA Meth. m5C
H3K9 me2
H3K27 me3:H3K9 me3
H3K27 me2
H3K4 me3
H2A.Z
H3−5ac
DNA Meth. m5C:H3K9 me3
H4−4ac
H4K20 me2
H4K20 me3
H3K4 me1
DNA Meth. m5C
H3K9 me3
H3K27 me3
0.00
0.25
0.50
0.75
1.00
$\hat{\pi}$

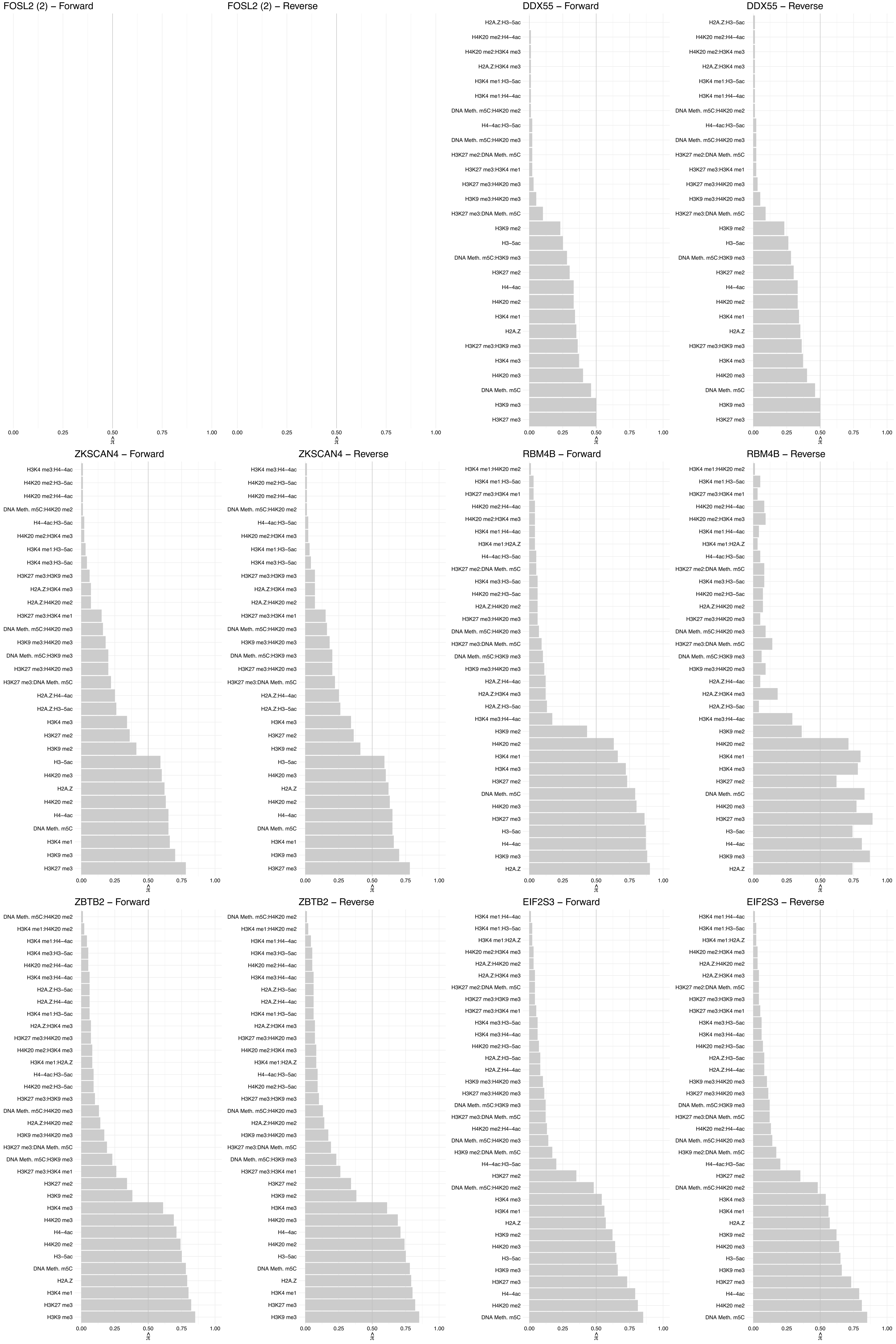
FOSL2 (2) – Forward
0.00
0.25
0.50
0.75
1.00
π̂
FOSL2 (2) – Reverse
0.00
0.25
0.50
0.75
1.00
π̂
DDX55 – Forward
H2A.Z:H3–5ac
H4K20 me2:H4–4ac
H4K20 me2:H3K4 me3
H2A.Z:H3K4 me3
H3K4 me1:H3–5ac
H3K4 me1:H4–4ac
DNA Meth. m5C:H4K20 me2
H4–4ac:H3–5ac
DNA Meth. m5C:H4K20 me3
H3K27 me2:DNA Meth. m5C
H3K27 me3:H3K4 me1
H3K27 me3:H4K20 me3
H3K9 me3:H4K20 me3
H3K27 me3:DNA Meth. m5C
H3K9 me2
H3–5ac
DNA Meth. m5C:H3K9 me3
H3K27 me2
H4–4ac
H4K20 me2
H3K4 me1
H2A.Z
H3K27 me3:H3K9 me3
H3K4 me3
H4K20 me3
DNA Meth. m5C
H3K9 me3
H3K27 me3
0.00
0.25
0.50
0.75
1.00
π̂
DDX55 – Reverse
H2A.Z:H3–5ac
H4K20 me2:H4–4ac
H4K20 me2:H3K4 me3
H2A.Z:H3K4 me3
H3K4 me1:H3–5ac
H3K4 me1:H4–4ac
DNA Meth. m5C:H4K20 me2
H4–4ac:H3–5ac
DNA Meth. m5C:H4K20 me3
H3K27 me2:DNA Meth. m5C
H3K27 me3:H3K4 me1
H3K27 me3:H4K20 me3
H3K9 me3:H4K20 me3
H3K27 me3:DNA Meth. m5C
H3K9 me2
H3–5ac
DNA Meth. m5C:H3K9 me3
H3K27 me2
H4–4ac
H4K20 me2
H3K4 me1
H2A.Z
H3K27 me3:H3K9 me3
H3K4 me3
H4K20 me3
DNA Meth. m5C
H3K9 me3
H3K27 me3
0.00
0.25
0.50
0.75
1.00
π̂
ZKSCAN4 – Forward
H3K4 me3:H4–4ac
H4K20 me2:H3–5ac
H4K20 me2:H4–4ac
DNA Meth. m5C:H4K20 me2
H4–4ac:H3–5ac
H4K20 me2:H3K4 me3
H3K4 me1:H3–5ac
H3K4 me3:H3–5ac
H3K27 me3:H3K9 me3
H2A.Z:H3K4 me3
H2A.Z:H4K20 me2
H3K27 me3:H3K4 me1
DNA Meth. m5C:H4K20 me3
H3K9 me3:H4K20 me3
DNA Meth. m5C:H3K9 me3
H3K27 me3:H4K20 me3
H3K27 me3:DNA Meth. m5C
H2A.Z:H4–4ac
H2A.Z:H3–5ac
H3K4 me3
H3K27 me2
H3K9 me2
H3–5ac
H4K20 me3
H2A.Z
H4K20 me2
H4–4ac
DNA Meth. m5C
H3K4 me1
H3K9 me3
H3K27 me3
0.00
0.25
0.50
0.75
1.00
π̂
ZKSCAN4 – Reverse
H3K4 me3:H4–4ac
H4K20 me2:H3–5ac
H4K20 me2:H4–4ac
DNA Meth. m5C:H4K20 me2
H4–4ac:H3–5ac
H4K20 me2:H3K4 me3
H3K4 me1:H3–5ac
H3K4 me3:H3–5ac
H3K27 me3:H3K9 me3
H2A.Z:H3K4 me3
H2A.Z:H4K20 me2
H3K27 me3:H3K4 me1
DNA Meth. m5C:H4K20 me3
H3K9 me3:H4K20 me3
DNA Meth. m5C:H3K9 me3
H3K27 me3:H4K20 me3
H3K27 me3:DNA Meth. m5C
H2A.Z:H4–4ac
H2A.Z:H3–5ac
H3K4 me3
H3K27 me2
H3K9 me2
H3–5ac
H4K20 me3
H2A.Z
H4K20 me2
H4–4ac
DNA Meth. m5C
H3K4 me1
H3K9 me3
H3K27 me3
0.00
0.25
0.50
0.75
1.00
π̂
RBM4B – Forward
H3K4 me1:H4K20 me2
H3K4 me1:H3–5ac
H3K27 me3:H3K4 me1
H4K20 me2:H4–4ac
H4K20 me2:H3K4 me3
H3K4 me1:H4–4ac
H3K4 me1:H2A.Z
H4–4ac:H3–5ac
H3K27 me2:DNA Meth. m5C
H3K4 me3:H3–5ac
H4K20 me2:H3–5ac
H2A.Z:H4K20 me2
H3K27 me3:H4K20 me3
DNA Meth. m5C:H4K20 me3
H3K27 me3:DNA Meth. m5C
DNA Meth. m5C:H3K9 me3
H3K9 me3:H4K20 me3
H2A.Z:H4–4ac
H2A.Z:H3K4 me3
H2A.Z:H3–5ac
H3K4 me3:H4–4ac
H3K9 me2
H4K20 me2
H3K4 me1
H3K4 me3
H3K27 me2
DNA Meth. m5C
H4K20 me3
H3K27 me3
H3–5ac
H4–4ac
H3K9 me3
H2A.Z
0.00
0.25
0.50
0.75
1.00
π̂
RBM4B – Reverse
H3K4 me1:H4K20 me2
H3K4 me1:H3–5ac
H3K27 me3:H3K4 me1
H4K20 me2:H4–4ac
H4K20 me2:H3K4 me3
H3K4 me1:H4–4ac
H3K4 me1:H2A.Z
H4–4ac:H3–5ac
H3K27 me2:DNA Meth. m5C
H3K4 me3:H3–5ac
H4K20 me2:H3–5ac
H2A.Z:H4K20 me2
H3K27 me3:H4K20 me3
DNA Meth. m5C:H4K20 me3
H3K27 me3:DNA Meth. m5C
DNA Meth. m5C:H3K9 me3
H3K9 me3:H4K20 me3
H2A.Z:H4–4ac
H2A.Z:H3K4 me3
H2A.Z:H3–5ac
H3K4 me3:H4–4ac
H3K9 me2
H4K20 me2
H3K4 me1
H3K4 me3
H3K27 me2
DNA Meth. m5C
H4K20 me3
H3K27 me3
H3–5ac
H4–4ac
H3K9 me3
H2A.Z
0.00
0.25
0.50
0.75
1.00
π̂
ZBTB2 – Forward
DNA Meth. m5C:H4K20 me2
H3K4 me1:H4K20 me2
H3K4 me1:H4–4ac
H3K4 me3:H3–5ac
H4K20 me2:H4–4ac
H3K4 me3:H4–4ac
H2A.Z:H3–5ac
H2A.Z:H4–4ac
H3K4 me1:H3–5ac
H2A.Z:H3K4 me3
H3K27 me3:H4K20 me3
H4K20 me2:H3K4 me3
H3K4 me1:H2A.Z
H4–4ac:H3–5ac
H4K20 me2:H3–5ac
H3K27 me3:H3K9 me3
DNA Meth. m5C:H4K20 me3
H2A.Z:H4K20 me2
H3K9 me3:H4K20 me3
H3K27 me3:DNA Meth. m5C
DNA Meth. m5C:H3K9 me3
H3K27 me3:H3K4 me1
H3K27 me2
H3K9 me2
H3K4 me3
H4K20 me3
H4–4ac
H4K20 me2
H3–5ac
DNA Meth. m5C
H2A.Z
H3K4 me1
H3K27 me3
H3K9 me3
0.00
0.25
0.50
0.75
1.00
π̂
ZBTB2 – Reverse
DNA Meth. m5C:H4K20 me2
H3K4 me1:H4K20 me2
H3K4 me1:H4–4ac
H3K4 me3:H3–5ac
H4K20 me2:H4–4ac
H3K4 me3:H4–4ac
H2A.Z:H3–5ac
H2A.Z:H4–4ac
H3K4 me1:H3–5ac
H2A.Z:H3K4 me3
H3K27 me3:H4K20 me3
H4K20 me2:H3K4 me3
H3K4 me1:H2A.Z
H4–4ac:H3–5ac
H4K20 me2:H3–5ac
H3K27 me3:H3K9 me3
DNA Meth. m5C:H4K20 me3
H2A.Z:H4K20 me2
H3K9 me3:H4K20 me3
H3K27 me3:DNA Meth. m5C
DNA Meth. m5C:H3K9 me3
H3K27 me3:H3K4 me1
H3K27 me2
H3K9 me2
H3K4 me3
H4K20 me3
H4–4ac
H4K20 me2
H3–5ac
DNA Meth. m5C
H2A.Z
H3K4 me1
H3K27 me3
H3K9 me3
0.00
0.25
0.50
0.75
1.00
π̂
EIF2S3 – Forward
H3K4 me1:H4–4ac
H3K4 me1:H3–5ac
H3K4 me1:H2A.Z
H4K20 me2:H3K4 me3
H2A.Z:H4K20 me2
H2A.Z:H3K4 me3
H3K27 me2:DNA Meth. m5C
H3K27 me3:H3K9 me3
H3K27 me3:H3K4 me1
H3K4 me3:H3–5ac
H3K4 me3:H4–4ac
H4K20 me2:H3–5ac
H2A.Z:H3–5ac
H2A.Z:H4–4ac
H3K9 me3:H4K20 me3
H3K27 me3:H4K20 me3
DNA Meth. m5C:H3K9 me3
H3K27 me3:DNA Meth. m5C
H4K20 me2:H4–4ac
DNA Meth. m5C:H4K20 me3
H3K9 me2:DNA Meth. m5C
H4–4ac:H3–5ac
H3K27 me2
DNA Meth. m5C:H4K20 me2
H3K4 me3
H3K4 me1
H2A.Z
H3K9 me2
H4K20 me3
H3–5ac
H3K9 me3
H3K27 me3
H4–4ac
H4K20 me2
DNA Meth. m5C
0.00
0.25
0.50
0.75
1.00
π̂
EIF2S3 – Reverse
H3K4 me1:H4–4ac
H3K4 me1:H3–5ac
H3K4 me1:H2A.Z
H4K20 me2:H3K4 me3
H2A.Z:H4K20 me2
H2A.Z:H3K4 me3
H3K27 me2:DNA Meth. m5C
H3K27 me3:H3K9 me3
H3K27 me3:H3K4 me1
H3K4 me3:H3–5ac
H3K4 me3:H4–4ac
H4K20 me2:H3–5ac
H2A.Z:H3–5ac
H2A.Z:H4–4ac
H3K9 me3:H4K20 me3
H3K27 me3:H4K20 me3
DNA Meth. m5C:H3K9 me3
H3K27 me3:DNA Meth. m5C
H4K20 me2:H4–4ac
DNA Meth. m5C:H4K20 me3
H3K9 me2:DNA Meth. m5C
H4–4ac:H3–5ac
H3K27 me2
DNA Meth. m5C:H4K20 me2
H3K4 me3
H3K4 me1
H2A.Z
H3K9 me2
H4K20 me3
H3–5ac
H3K9 me3
H3K27 me3
H4–4ac
H4K20 me2
DNA Meth. m5C
0.00
0.25
0.50
0.75
1.00
π̂

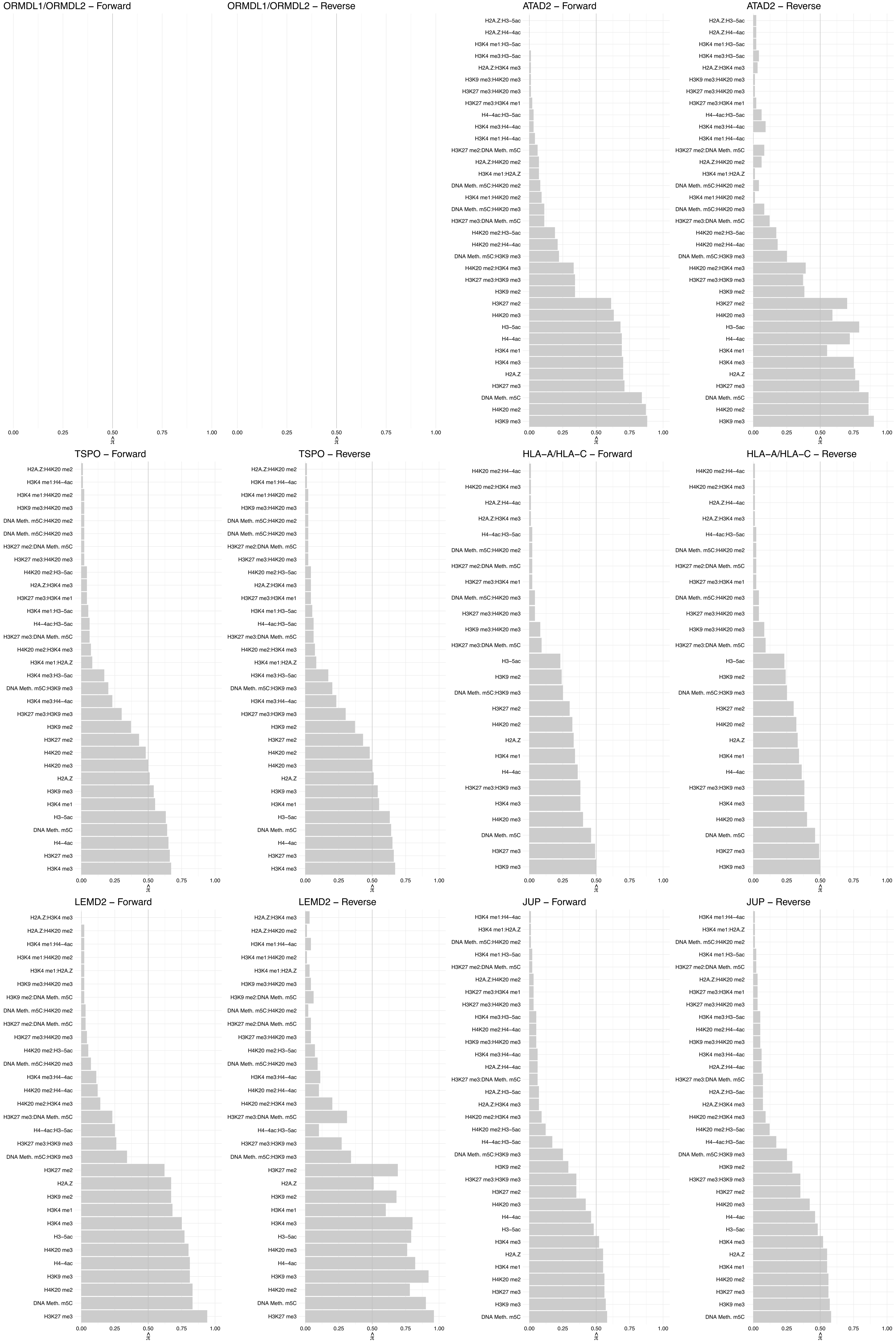
ORMDL1/ORMDL2 − Forward
ORMDL1/ORMDL2 − Reverse
ATAD2 − Forward
ATAD2 − Reverse
H2A.Z:H3−5ac
H2A.Z:H4−4ac
H3K4 me1:H3−5ac
H3K4 me3:H3−5ac
H2A.Z:H3K4 me3
H3K9 me3:H4K20 me3
H3K27 me3:H4K20 me3
H3K27 me3:H3K4 me1
H4−4ac:H3−5ac
H3K4 me3:H4−4ac
H3K4 me1:H4−4ac
H3K27 me2:DNA Meth. m5C
H2A.Z:H4K20 me2
H3K4 me1:H2A.Z
DNA Meth. m5C:H4K20 me2
H3K4 me1:H4K20 me2
DNA Meth. m5C:H4K20 me3
H3K27 me3:DNA Meth. m5C
H4K20 me2:H3−5ac
H4K20 me2:H4−4ac
DNA Meth. m5C:H3K9 me3
H4K20 me2:H3K4 me3
H3K27 me3:H3K9 me3
H3K9 me2
H3K27 me2
H4K20 me3
H3−5ac
H4−4ac
H3K4 me1
H3K4 me3
H2A.Z
H3K27 me3
DNA Meth. m5C
H4K20 me2
H3K9 me3
TSPO − Forward
TSPO − Reverse
H2A.Z:H4K20 me2
H3K4 me1:H4−4ac
H3K4 me1:H4K20 me2
H3K9 me3:H4K20 me3
DNA Meth. m5C:H4K20 me2
DNA Meth. m5C:H4K20 me3
H3K27 me2:DNA Meth. m5C
H3K27 me3:H4K20 me3
H4K20 me2:H3−5ac
H2A.Z:H3K4 me3
H3K27 me3:H3K4 me1
H3K4 me1:H3−5ac
H4−4ac:H3−5ac
H3K27 me3:DNA Meth. m5C
H4K20 me2:H3K4 me3
H3K4 me1:H2A.Z
H3K4 me3:H3−5ac
DNA Meth. m5C:H3K9 me3
H3K4 me3:H4−4ac
H3K27 me3:H3K9 me3
H3K9 me2
H3K27 me2
H4K20 me2
H4K20 me3
H2A.Z
H3K9 me3
H3K4 me1
H3−5ac
DNA Meth. m5C
H4−4ac
H3K27 me3
H3K4 me3
HLA−A/HLA−C − Forward
HLA−A/HLA−C − Reverse
H4K20 me2:H4−4ac
H4K20 me2:H3K4 me3
H2A.Z:H4−4ac
H2A.Z:H3K4 me3
H4−4ac:H3−5ac
DNA Meth. m5C:H4K20 me2
H3K27 me2:DNA Meth. m5C
H3K27 me3:H3K4 me1
DNA Meth. m5C:H4K20 me3
H3K27 me3:H4K20 me3
H3K9 me3:H4K20 me3
H3K27 me3:DNA Meth. m5C
H3−5ac
H3K9 me2
DNA Meth. m5C:H3K9 me3
H3K27 me2
H4K20 me2
H2A.Z
H3K4 me1
H4−4ac
H3K27 me3:H3K9 me3
H3K4 me3
H4K20 me3
DNA Meth. m5C
H3K27 me3
H3K9 me3
LEMD2 − Forward
LEMD2 − Reverse
H2A.Z:H3K4 me3
H2A.Z:H4K20 me2
H3K4 me1:H4−4ac
H3K4 me1:H4K20 me2
H3K4 me1:H2A.Z
H3K9 me3:H4K20 me3
H3K9 me2:DNA Meth. m5C
DNA Meth. m5C:H4K20 me2
H3K27 me2:DNA Meth. m5C
H3K27 me3:H4K20 me3
H4K20 me2:H3−5ac
DNA Meth. m5C:H4K20 me3
H3K4 me3:H4−4ac
H4K20 me2:H4−4ac
H4K20 me2:H3K4 me3
H3K27 me3:DNA Meth. m5C
H4−4ac:H3−5ac
H3K27 me3:H3K9 me3
DNA Meth. m5C:H3K9 me3
H3K27 me2
H2A.Z
H3K9 me2
H3K4 me1
H3K4 me3
H3−5ac
H4K20 me3
H4−4ac
H3K9 me3
H4K20 me2
DNA Meth. m5C
H3K27 me3
JUP − Forward
JUP − Reverse
H3K4 me1:H4−4ac
H3K4 me1:H2A.Z
DNA Meth. m5C:H4K20 me2
H3K4 me1:H3−5ac
H3K27 me2:DNA Meth. m5C
H2A.Z:H4K20 me2
H3K27 me3:H3K4 me1
H3K27 me3:H4K20 me3
H3K4 me3:H3−5ac
H4K20 me2:H4−4ac
H3K9 me3:H4K20 me3
H3K4 me3:H4−4ac
H2A.Z:H4−4ac
H3K27 me3:DNA Meth. m5C
H2A.Z:H3−5ac
H2A.Z:H3K4 me3
H4K20 me2:H3K4 me3
H4K20 me2:H3−5ac
H4−4ac:H3−5ac
DNA Meth. m5C:H3K9 me3
H3K9 me2
H3K27 me3:H3K9 me3
H3K27 me2
H4K20 me3
H4−4ac
H3−5ac
H3K4 me3
H2A.Z
H3K4 me1
H4K20 me2
H3K27 me3
H3K9 me3
DNA Meth. m5C
0.00
0.25
0.50
0.75
1.00
π̂

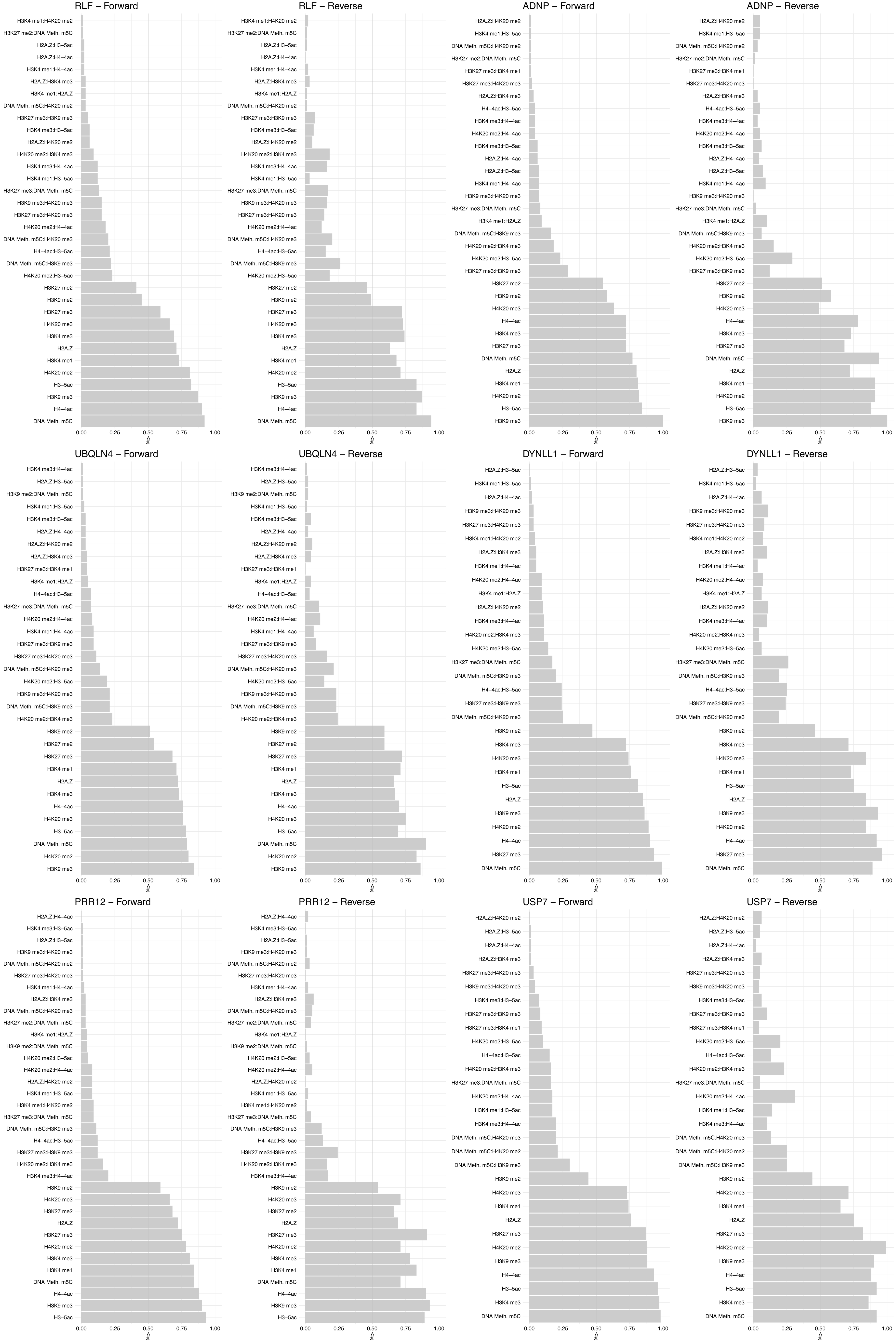

RLF – Forward
H3K4 me1:H4K20 me2
H3K27 me2:DNA Meth. m5C
H2A.Z:H3–5ac
H2A.Z:H4–4ac
H3K4 me1:H4–4ac
H2A.Z:H3K4 me3
H3K4 me1:H2A.Z
DNA Meth. m5C:H4K20 me2
H3K27 me3:H3K9 me3
H3K4 me3:H3–5ac
H2A.Z:H4K20 me2
H4K20 me2:H3K4 me3
H3K4 me3:H4–4ac
H3K4 me1:H3–5ac
H3K27 me3:DNA Meth. m5C
H3K9 me3:H4K20 me3
H3K27 me3:H4K20 me3
H4K20 me2:H4–4ac
DNA Meth. m5C:H4K20 me3
H4–4ac:H3–5ac
DNA Meth. m5C:H3K9 me3
H4K20 me2:H3–5ac
H3K27 me2
H3K9 me2
H3K27 me3
H4K20 me3
H3K4 me3
H2A.Z
H3K4 me1
H4K20 me2
H3–5ac
H3K9 me3
H4–4ac
DNA Meth. m5C
0.00 0.25 0.50 0.75 1.00
π̂
RLF – Reverse
H3K4 me1:H4K20 me2
H3K27 me2:DNA Meth. m5C
H2A.Z:H3–5ac
H2A.Z:H4–4ac
H3K4 me1:H4–4ac
H2A.Z:H3K4 me3
H3K4 me1:H2A.Z
DNA Meth. m5C:H4K20 me2
H3K27 me3:H3K9 me3
H3K4 me3:H3–5ac
H2A.Z:H4K20 me2
H4K20 me2:H3K4 me3
H3K4 me3:H4–4ac
H3K4 me1:H3–5ac
H3K27 me3:DNA Meth. m5C
H3K9 me3:H4K20 me3
H3K27 me3:H4K20 me3
H4K20 me2:H4–4ac
DNA Meth. m5C:H4K20 me3
H4–4ac:H3–5ac
DNA Meth. m5C:H3K9 me3
H4K20 me2:H3–5ac
H3K27 me2
H3K9 me2
H3K27 me3
H4K20 me3
H3K4 me3
H2A.Z
H3K4 me1
H4K20 me2
H3–5ac
H3K9 me3
H4–4ac
DNA Meth. m5C
0.00 0.25 0.50 0.75 1.00
π̂
ADNP – Forward
H2A.Z:H4K20 me2
H3K4 me1:H3–5ac
DNA Meth. m5C:H4K20 me2
H3K27 me2:DNA Meth. m5C
H3K27 me3:H3K4 me1
H3K27 me3:H4K20 me3
H2A.Z:H3K4 me3
H4–4ac:H3–5ac
H3K4 me3:H4–4ac
H4K20 me2:H4–4ac
H3K4 me3:H3–5ac
H2A.Z:H4–4ac
H2A.Z:H3–5ac
H3K4 me1:H4–4ac
H3K9 me3:H4K20 me3
H3K27 me3:DNA Meth. m5C
H3K4 me1:H2A.Z
DNA Meth. m5C:H3K9 me3
H4K20 me2:H3K4 me3
H4K20 me2:H3–5ac
H3K27 me3:H3K9 me3
H3K27 me2
H3K9 me2
H4K20 me3
H4–4ac
H3K4 me3
H3K27 me3
DNA Meth. m5C
H2A.Z
H3K4 me1
H4K20 me2
H3–5ac
H3K9 me3
0.00 0.25 0.50 0.75 1.00
π̂
ADNP – Reverse
H2A.Z:H4K20 me2
H3K4 me1:H3–5ac
DNA Meth. m5C:H4K20 me2
H3K27 me2:DNA Meth. m5C
H3K27 me3:H3K4 me1
H3K27 me3:H4K20 me3
H2A.Z:H3K4 me3
H4–4ac:H3–5ac
H3K4 me3:H4–4ac
H4K20 me2:H4–4ac
H3K4 me3:H3–5ac
H2A.Z:H4–4ac
H2A.Z:H3–5ac
H3K4 me1:H4–4ac
H3K9 me3:H4K20 me3
H3K27 me3:DNA Meth. m5C
H3K4 me1:H2A.Z
DNA Meth. m5C:H3K9 me3
H4K20 me2:H3K4 me3
H4K20 me2:H3–5ac
H3K27 me3:H3K9 me3
H3K27 me2
H3K9 me2
H4K20 me3
H4–4ac
H3K4 me3
H3K27 me3
DNA Meth. m5C
H2A.Z
H3K4 me1
H4K20 me2
H3–5ac
H3K9 me3
0.00 0.25 0.50 0.75 1.00
π̂
UBQLN4 – Forward
H3K4 me3:H4–4ac
H2A.Z:H3–5ac
H3K9 me2:DNA Meth. m5C
H3K4 me1:H3–5ac
H3K4 me3:H3–5ac
H2A.Z:H4–4ac
H2A.Z:H4K20 me2
H2A.Z:H3K4 me3
H3K27 me3:H3K4 me1
H3K4 me1:H2A.Z
H4–4ac:H3–5ac
H3K27 me3:DNA Meth. m5C
H4K20 me2:H4–4ac
H3K4 me1:H4–4ac
H3K27 me3:H3K9 me3
H3K27 me3:H4K20 me3
DNA Meth. m5C:H4K20 me3
H4K20 me2:H3–5ac
H3K9 me3:H4K20 me3
DNA Meth. m5C:H3K9 me3
H4K20 me2:H3K4 me3
H3K9 me2
H3K27 me2
H3K27 me3
H3K4 me1
H2A.Z
H3K4 me3
H4–4ac
H4K20 me3
H3–5ac
DNA Meth. m5C
H4K20 me2
H3K9 me3
0.00 0.25 0.50 0.75 1.00
π̂
UBQLN4 – Reverse
H3K4 me3:H4–4ac
H2A.Z:H3–5ac
H3K9 me2:DNA Meth. m5C
H3K4 me1:H3–5ac
H3K4 me3:H3–5ac
H2A.Z:H4–4ac
H2A.Z:H4K20 me2
H2A.Z:H3K4 me3
H3K27 me3:H3K4 me1
H3K4 me1:H2A.Z
H4–4ac:H3–5ac
H3K27 me3:DNA Meth. m5C
H4K20 me2:H4–4ac
H3K4 me1:H4–4ac
H3K27 me3:H3K9 me3
H3K27 me3:H4K20 me3
DNA Meth. m5C:H4K20 me3
H4K20 me2:H3–5ac
H3K9 me3:H4K20 me3
DNA Meth. m5C:H3K9 me3
H4K20 me2:H3K4 me3
H3K9 me2
H3K27 me2
H3K27 me3
H3K4 me1
H2A.Z
H3K4 me3
H4–4ac
H4K20 me3
H3–5ac
DNA Meth. m5C
H4K20 me2
H3K9 me3
0.00 0.25 0.50 0.75 1.00
π̂
DYNLL1 – Forward
H2A.Z:H3–5ac
H3K4 me1:H3–5ac
H2A.Z:H4–4ac
H3K9 me3:H4K20 me3
H3K27 me3:H4K20 me3
H3K4 me1:H4K20 me2
H2A.Z:H3K4 me3
H3K4 me1:H4–4ac
H4K20 me2:H4–4ac
H3K4 me1:H2A.Z
H2A.Z:H4K20 me2
H3K4 me3:H4–4ac
H4K20 me2:H3K4 me3
H4K20 me2:H3–5ac
H3K27 me3:DNA Meth. m5C
DNA Meth. m5C:H3K9 me3
H4–4ac:H3–5ac
H3K27 me3:H3K9 me3
DNA Meth. m5C:H4K20 me3
H3K9 me2
H3K4 me3
H4K20 me3
H3K4 me1
H3–5ac
H2A.Z
H3K9 me3
H4K20 me2
H4–4ac
H3K27 me3
DNA Meth. m5C
0.00 0.25 0.50 0.75 1.00
π̂
DYNLL1 – Reverse
H2A.Z:H3–5ac
H3K4 me1:H3–5ac
H2A.Z:H4–4ac
H3K9 me3:H4K20 me3
H3K27 me3:H4K20 me3
H3K4 me1:H4K20 me2
H2A.Z:H3K4 me3
H3K4 me1:H4–4ac
H4K20 me2:H4–4ac
H3K4 me1:H2A.Z
H2A.Z:H4K20 me2
H3K4 me3:H4–4ac
H4K20 me2:H3K4 me3
H4K20 me2:H3–5ac
H3K27 me3:DNA Meth. m5C
DNA Meth. m5C:H3K9 me3
H4–4ac:H3–5ac
H3K27 me3:H3K9 me3
DNA Meth. m5C:H4K20 me3
H3K9 me2
H3K4 me3
H4K20 me3
H3K4 me1
H3–5ac
H2A.Z
H3K9 me3
H4K20 me2
H4–4ac
H3K27 me3
DNA Meth. m5C
0.00 0.25 0.50 0.75 1.00
π̂
PRR12 – Forward
H2A.Z:H4–4ac
H3K4 me3:H3–5ac
H2A.Z:H3–5ac
H3K9 me3:H4K20 me3
DNA Meth. m5C:H4K20 me2
H3K27 me3:H4K20 me3
H3K4 me1:H4–4ac
H2A.Z:H3K4 me3
DNA Meth. m5C:H4K20 me3
H3K27 me2:DNA Meth. m5C
H3K4 me1:H2A.Z
H3K9 me2:DNA Meth. m5C
H4K20 me2:H3–5ac
H4K20 me2:H4–4ac
H2A.Z:H4K20 me2
H3K4 me1:H3–5ac
H3K4 me1:H4K20 me2
H3K27 me3:DNA Meth. m5C
DNA Meth. m5C:H3K9 me3
H4–4ac:H3–5ac
H3K27 me3:H3K9 me3
H4K20 me2:H3K4 me3
H3K4 me3:H4–4ac
H3K9 me2
H4K20 me3
H3K27 me2
H2A.Z
H3K27 me3
H4K20 me2
H3K4 me3
H3K4 me1
DNA Meth. m5C
H4–4ac
H3K9 me3
H3–5ac
0.00 0.25 0.50 0.75 1.00
π̂
PRR12 – Reverse
H2A.Z:H4–4ac
H3K4 me3:H3–5ac
H2A.Z:H3–5ac
H3K9 me3:H4K20 me3
DNA Meth. m5C:H4K20 me2
H3K27 me3:H4K20 me3
H3K4 me1:H4–4ac
H2A.Z:H3K4 me3
DNA Meth. m5C:H4K20 me3
H3K27 me2:DNA Meth. m5C
H3K4 me1:H2A.Z
H3K9 me2:DNA Meth. m5C
H4K20 me2:H3–5ac
H4K20 me2:H4–4ac
H2A.Z:H4K20 me2
H3K4 me1:H3–5ac
H3K4 me1:H4K20 me2
H3K27 me3:DNA Meth. m5C
DNA Meth. m5C:H3K9 me3
H4–4ac:H3–5ac
H3K27 me3:H3K9 me3
H4K20 me2:H3K4 me3
H3K4 me3:H4–4ac
H3K9 me2
H4K20 me3
H3K27 me2
H2A.Z
H3K27 me3
H4K20 me2
H3K4 me3
H3K4 me1
DNA Meth. m5C
H4–4ac
H3K9 me3
H3–5ac
0.00 0.25 0.50 0.75 1.00
π̂
USP7 – Forward
H2A.Z:H4K20 me2
H2A.Z:H3–5ac
H2A.Z:H4–4ac
H2A.Z:H3K4 me3
H3K27 me3:H4K20 me3
H3K9 me3:H4K20 me3
H3K4 me3:H3–5ac
H3K27 me3:H3K9 me3
H3K27 me3:H3K4 me1
H4K20 me2:H3–5ac
H4–4ac:H3–5ac
H4K20 me2:H3K4 me3
H3K27 me3:DNA Meth. m5C
H4K20 me2:H4–4ac
H3K4 me1:H3–5ac
H3K4 me3:H4–4ac
DNA Meth. m5C:H4K20 me3
DNA Meth. m5C:H4K20 me2
DNA Meth. m5C:H3K9 me3
H3K9 me2
H4K20 me3
H3K4 me1
H2A.Z
H3K27 me3
H4K20 me2
H3K9 me3
H4–4ac
H3–5ac
H3K4 me3
DNA Meth. m5C
0.00 0.25 0.50 0.75 1.00
π̂
USP7 – Reverse
H2A.Z:H4K20 me2
H2A.Z:H3–5ac
H2A.Z:H4–4ac
H2A.Z:H3K4 me3
H3K27 me3:H4K20 me3
H3K9 me3:H4K20 me3
H3K4 me3:H3–5ac
H3K27 me3:H3K9 me3
H3K27 me3:H3K4 me1
H4K20 me2:H3–5ac
H4–4ac:H3–5ac
H4K20 me2:H3K4 me3
H3K27 me3:DNA Meth. m5C
H4K20 me2:H4–4ac
H3K4 me1:H3–5ac
H3K4 me3:H4–4ac
DNA Meth. m5C:H4K20 me3
DNA Meth. m5C:H4K20 me2
DNA Meth. m5C:H3K9 me3
H3K9 me2
H4K20 me3
H3K4 me1
H2A.Z
H3K27 me3
H4K20 me2
H3K9 me3
H4–4ac
H3–5ac
H3K4 me3
DNA Meth. m5C
0.00 0.25 0.50 0.75 1.00
π̂

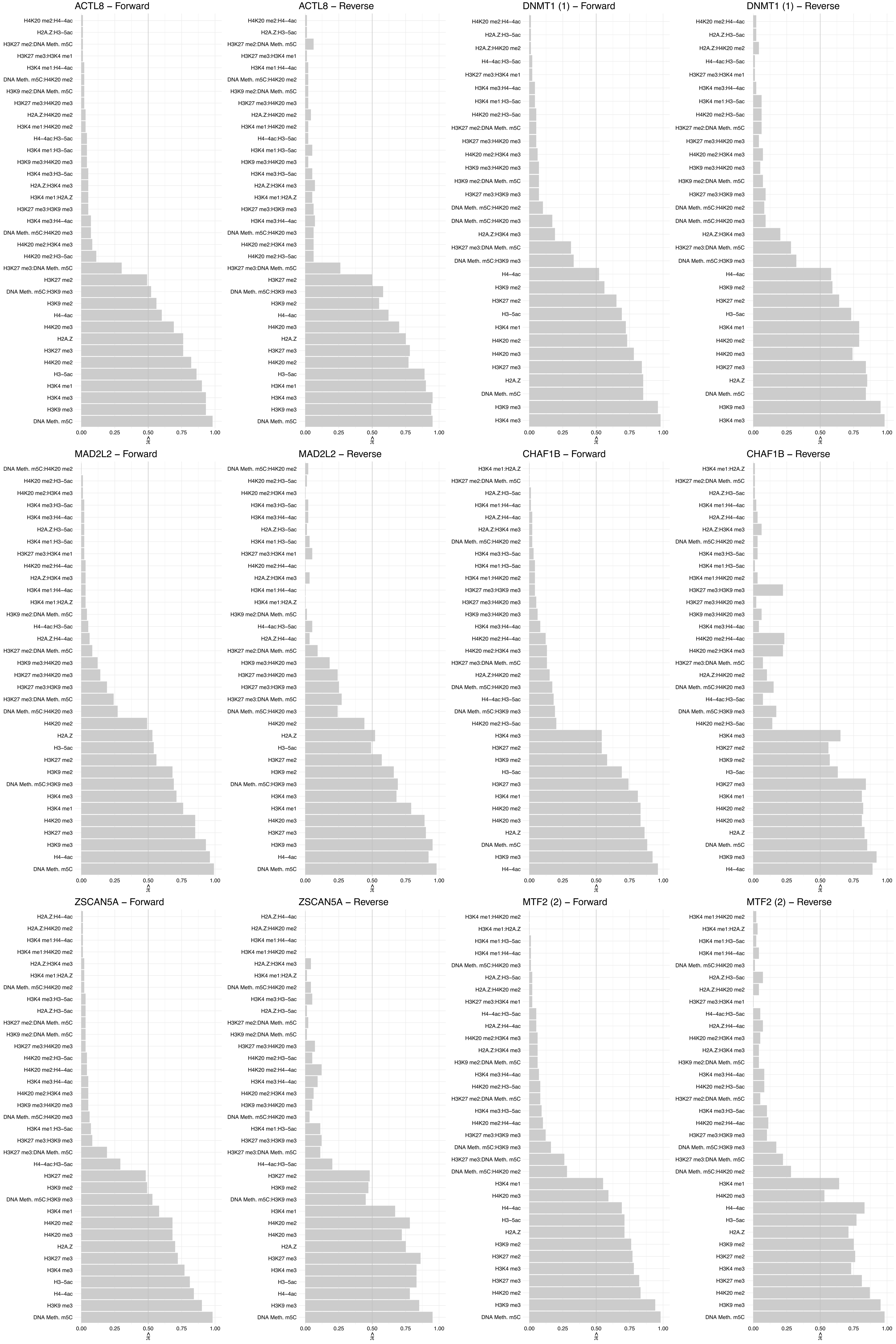
ACTL8 – Forward
H4K20 me2:H4–4ac
H2A.Z:H3–5ac
H3K27 me2:DNA Meth. m5C
H3K27 me3:H3K4 me1
H3K4 me1:H4–4ac
DNA Meth. m5C:H4K20 me2
H3K9 me2:DNA Meth. m5C
H3K27 me3:H4K20 me3
H2A.Z:H4K20 me2
H3K4 me1:H4K20 me2
H4–4ac:H3–5ac
H3K4 me1:H3–5ac
H3K9 me3:H4K20 me3
H3K4 me3:H3–5ac
H2A.Z:H3K4 me3
H3K4 me1:H2A.Z
H3K27 me3:H3K9 me3
H3K4 me3:H4–4ac
DNA Meth. m5C:H4K20 me3
H4K20 me2:H3K4 me3
H4K20 me2:H3–5ac
H3K27 me3:DNA Meth. m5C
H3K27 me2
DNA Meth. m5C:H3K9 me3
H3K9 me2
H4–4ac
H4K20 me3
H2A.Z
H3K27 me3
H4K20 me2
H3–5ac
H3K4 me1
H3K4 me3
H3K9 me3
DNA Meth. m5C
0.00
0.25
0.50
0.75
1.00
π̂
ACTL8 – Reverse
H4K20 me2:H4–4ac
H2A.Z:H3–5ac
H3K27 me2:DNA Meth. m5C
H3K27 me3:H3K4 me1
H3K4 me1:H4–4ac
DNA Meth. m5C:H4K20 me2
H3K9 me2:DNA Meth. m5C
H3K27 me3:H4K20 me3
H2A.Z:H4K20 me2
H3K4 me1:H4K20 me2
H4–4ac:H3–5ac
H3K4 me1:H3–5ac
H3K9 me3:H4K20 me3
H3K4 me3:H3–5ac
H2A.Z:H3K4 me3
H3K4 me1:H2A.Z
H3K27 me3:H3K9 me3
H3K4 me3:H4–4ac
DNA Meth. m5C:H4K20 me3
H4K20 me2:H3K4 me3
H4K20 me2:H3–5ac
H3K27 me3:DNA Meth. m5C
H3K27 me2
DNA Meth. m5C:H3K9 me3
H3K9 me2
H4–4ac
H4K20 me3
H2A.Z
H3K27 me3
H4K20 me2
H3–5ac
H3K4 me1
H3K4 me3
H3K9 me3
DNA Meth. m5C
0.00
0.25
0.50
0.75
1.00
π̂
DNMT1 (1) – Forward
H4K20 me2:H4–4ac
H2A.Z:H3–5ac
H2A.Z:H4K20 me2
H4–4ac:H3–5ac
H3K27 me3:H3K4 me1
H3K4 me3:H4–4ac
H3K4 me1:H3–5ac
H4K20 me2:H3–5ac
H3K27 me2:DNA Meth. m5C
H3K27 me3:H4K20 me3
H4K20 me2:H3K4 me3
H3K9 me3:H4K20 me3
H3K9 me2:DNA Meth. m5C
H3K27 me3:H3K9 me3
DNA Meth. m5C:H4K20 me2
DNA Meth. m5C:H4K20 me3
H2A.Z:H3K4 me3
H3K27 me3:DNA Meth. m5C
DNA Meth. m5C:H3K9 me3
H4–4ac
H3K9 me2
H3K27 me2
H3–5ac
H3K4 me1
H4K20 me2
H4K20 me3
H3K27 me3
H2A.Z
DNA Meth. m5C
H3K9 me3
H3K4 me3
0.00
0.25
0.50
0.75
1.00
π̂
DNMT1 (1) – Reverse
H4K20 me2:H4–4ac
H2A.Z:H3–5ac
H2A.Z:H4K20 me2
H4–4ac:H3–5ac
H3K27 me3:H3K4 me1
H3K4 me3:H4–4ac
H3K4 me1:H3–5ac
H4K20 me2:H3–5ac
H3K27 me2:DNA Meth. m5C
H3K27 me3:H4K20 me3
H4K20 me2:H3K4 me3
H3K9 me3:H4K20 me3
H3K9 me2:DNA Meth. m5C
H3K27 me3:H3K9 me3
DNA Meth. m5C:H4K20 me2
DNA Meth. m5C:H4K20 me3
H2A.Z:H3K4 me3
H3K27 me3:DNA Meth. m5C
DNA Meth. m5C:H3K9 me3
H4–4ac
H3K9 me2
H3K27 me2
H3–5ac
H3K4 me1
H4K20 me2
H4K20 me3
H3K27 me3
H2A.Z
DNA Meth. m5C
H3K9 me3
H3K4 me3
0.00
0.25
0.50
0.75
1.00
π̂
MAD2L2 – Forward
DNA Meth. m5C:H4K20 me2
H4K20 me2:H3–5ac
H4K20 me2:H3K4 me3
H3K4 me3:H3–5ac
H3K4 me3:H4–4ac
H2A.Z:H3–5ac
H3K4 me1:H3–5ac
H3K27 me3:H3K4 me1
H4K20 me2:H4–4ac
H2A.Z:H3K4 me3
H3K4 me1:H4–4ac
H3K4 me1:H2A.Z
H3K9 me2:DNA Meth. m5C
H4–4ac:H3–5ac
H2A.Z:H4–4ac
H3K27 me2:DNA Meth. m5C
H3K9 me3:H4K20 me3
H3K27 me3:H4K20 me3
H3K27 me3:H3K9 me3
H3K27 me3:DNA Meth. m5C
DNA Meth. m5C:H4K20 me3
H4K20 me2
H2A.Z
H3–5ac
H3K27 me2
H3K9 me2
DNA Meth. m5C:H3K9 me3
H3K4 me3
H3K4 me1
H4K20 me3
H3K27 me3
H3K9 me3
H4–4ac
DNA Meth. m5C
0.00
0.25
0.50
0.75
1.00
π̂
MAD2L2 – Reverse
DNA Meth. m5C:H4K20 me2
H4K20 me2:H3–5ac
H4K20 me2:H3K4 me3
H3K4 me3:H3–5ac
H3K4 me3:H4–4ac
H2A.Z:H3–5ac
H3K4 me1:H3–5ac
H3K27 me3:H3K4 me1
H4K20 me2:H4–4ac
H2A.Z:H3K4 me3
H3K4 me1:H4–4ac
H3K4 me1:H2A.Z
H3K9 me2:DNA Meth. m5C
H4–4ac:H3–5ac
H2A.Z:H4–4ac
H3K27 me2:DNA Meth. m5C
H3K9 me3:H4K20 me3
H3K27 me3:H4K20 me3
H3K27 me3:H3K9 me3
H3K27 me3:DNA Meth. m5C
DNA Meth. m5C:H4K20 me3
H4K20 me2
H2A.Z
H3–5ac
H3K27 me2
H3K9 me2
DNA Meth. m5C:H3K9 me3
H3K4 me3
H3K4 me1
H4K20 me3
H3K27 me3
H3K9 me3
H4–4ac
DNA Meth. m5C
0.00
0.25
0.50
0.75
1.00
π̂
CHAF1B – Forward
H3K4 me1:H2A.Z
H3K27 me2:DNA Meth. m5C
H2A.Z:H3–5ac
H3K4 me1:H4–4ac
H2A.Z:H4–4ac
H2A.Z:H3K4 me3
DNA Meth. m5C:H4K20 me2
H3K4 me3:H3–5ac
H3K4 me1:H3–5ac
H3K4 me1:H4K20 me2
H3K27 me3:H3K9 me3
H3K27 me3:H4K20 me3
H3K9 me3:H4K20 me3
H3K4 me3:H4–4ac
H4K20 me2:H4–4ac
H4K20 me2:H3K4 me3
H3K27 me3:DNA Meth. m5C
H2A.Z:H4K20 me2
DNA Meth. m5C:H4K20 me3
H4–4ac:H3–5ac
DNA Meth. m5C:H3K9 me3
H4K20 me2:H3–5ac
H3K4 me3
H3K27 me2
H3K9 me2
H3–5ac
H3K27 me3
H3K4 me1
H4K20 me2
H4K20 me3
H2A.Z
DNA Meth. m5C
H3K9 me3
H4–4ac
0.00
0.25
0.50
0.75
1.00
π̂
CHAF1B – Reverse
H3K4 me1:H2A.Z
H3K27 me2:DNA Meth. m5C
H2A.Z:H3–5ac
H3K4 me1:H4–4ac
H2A.Z:H4–4ac
H2A.Z:H3K4 me3
DNA Meth. m5C:H4K20 me2
H3K4 me3:H3–5ac
H3K4 me1:H3–5ac
H3K4 me1:H4K20 me2
H3K27 me3:H3K9 me3
H3K27 me3:H4K20 me3
H3K9 me3:H4K20 me3
H3K4 me3:H4–4ac
H4K20 me2:H4–4ac
H4K20 me2:H3K4 me3
H3K27 me3:DNA Meth. m5C
H2A.Z:H4K20 me2
DNA Meth. m5C:H4K20 me3
H4–4ac:H3–5ac
DNA Meth. m5C:H3K9 me3
H4K20 me2:H3–5ac
H3K4 me3
H3K27 me2
H3K9 me2
H3–5ac
H3K27 me3
H3K4 me1
H4K20 me2
H4K20 me3
H2A.Z
DNA Meth. m5C
H3K9 me3
H4–4ac
0.00
0.25
0.50
0.75
1.00
π̂
ZSCAN5A – Forward
H2A.Z:H4–4ac
H2A.Z:H4K20 me2
H3K4 me1:H4–4ac
H3K4 me1:H4K20 me2
H2A.Z:H3K4 me3
H3K4 me1:H2A.Z
DNA Meth. m5C:H4K20 me2
H3K4 me3:H3–5ac
H2A.Z:H3–5ac
H3K27 me2:DNA Meth. m5C
H3K9 me2:DNA Meth. m5C
H3K27 me3:H4K20 me3
H4K20 me2:H3–5ac
H4K20 me2:H4–4ac
H3K4 me3:H4–4ac
H4K20 me2:H3K4 me3
H3K9 me3:H4K20 me3
DNA Meth. m5C:H4K20 me3
H3K4 me1:H3–5ac
H3K27 me3:H3K9 me3
H3K27 me3:DNA Meth. m5C
H4–4ac:H3–5ac
H3K27 me2
H3K9 me2
DNA Meth. m5C:H3K9 me3
H3K4 me1
H4K20 me2
H4K20 me3
H2A.Z
H3K27 me3
H3K4 me3
H3–5ac
H4–4ac
H3K9 me3
DNA Meth. m5C
0.00
0.25
0.50
0.75
1.00
π̂
ZSCAN5A – Reverse
H2A.Z:H4–4ac
H2A.Z:H4K20 me2
H3K4 me1:H4–4ac
H3K4 me1:H4K20 me2
H2A.Z:H3K4 me3
H3K4 me1:H2A.Z
DNA Meth. m5C:H4K20 me2
H3K4 me3:H3–5ac
H2A.Z:H3–5ac
H3K27 me2:DNA Meth. m5C
H3K9 me2:DNA Meth. m5C
H3K27 me3:H4K20 me3
H4K20 me2:H3–5ac
H4K20 me2:H4–4ac
H3K4 me3:H4–4ac
H4K20 me2:H3K4 me3
H3K9 me3:H4K20 me3
DNA Meth. m5C:H4K20 me3
H3K4 me1:H3–5ac
H3K27 me3:H3K9 me3
H3K27 me3:DNA Meth. m5C
H4–4ac:H3–5ac
H3K27 me2
H3K9 me2
DNA Meth. m5C:H3K9 me3
H3K4 me1
H4K20 me2
H4K20 me3
H2A.Z
H3K27 me3
H3K4 me3
H3–5ac
H4–4ac
H3K9 me3
DNA Meth. m5C
0.00
0.25
0.50
0.75
1.00
π̂
MTF2 (2) – Forward
H3K4 me1:H4K20 me2
H3K4 me1:H2A.Z
H3K4 me1:H3–5ac
H3K4 me1:H4–4ac
DNA Meth. m5C:H4K20 me3
H2A.Z:H3–5ac
H2A.Z:H4K20 me2
H3K27 me3:H3K4 me1
H4–4ac:H3–5ac
H2A.Z:H4–4ac
H4K20 me2:H3K4 me3
H2A.Z:H3K4 me3
H3K9 me2:DNA Meth. m5C
H3K4 me3:H4–4ac
H4K20 me2:H3–5ac
H3K27 me2:DNA Meth. m5C
H3K4 me3:H3–5ac
H4K20 me2:H4–4ac
H3K27 me3:H3K9 me3
DNA Meth. m5C:H3K9 me3
H3K27 me3:DNA Meth. m5C
DNA Meth. m5C:H4K20 me2
H3K4 me1
H4K20 me3
H4–4ac
H3–5ac
H2A.Z
H3K9 me2
H3K27 me2
H3K4 me3
H3K27 me3
H4K20 me2
H3K9 me3
DNA Meth. m5C
0.00
0.25
0.50
0.75
1.00
π̂
MTF2 (2) – Reverse
H3K4 me1:H4K20 me2
H3K4 me1:H2A.Z
H3K4 me1:H3–5ac
H3K4 me1:H4–4ac
DNA Meth. m5C:H4K20 me3
H2A.Z:H3–5ac
H2A.Z:H4K20 me2
H3K27 me3:H3K4 me1
H4–4ac:H3–5ac
H2A.Z:H4–4ac
H4K20 me2:H3K4 me3
H2A.Z:H3K4 me3
H3K9 me2:DNA Meth. m5C
H3K4 me3:H4–4ac
H4K20 me2:H3–5ac
H3K27 me2:DNA Meth. m5C
H3K4 me3:H3–5ac
H4K20 me2:H4–4ac
H3K27 me3:H3K9 me3
DNA Meth. m5C:H3K9 me3
H3K27 me3:DNA Meth. m5C
DNA Meth. m5C:H4K20 me2
H3K4 me1
H4K20 me3
H4–4ac
H3–5ac
H2A.Z
H3K9 me2
H3K27 me2
H3K4 me3
H3K27 me3
H4K20 me2
H3K9 me3
DNA Meth. m5C
0.00
0.25
0.50
0.75
1.00
π̂

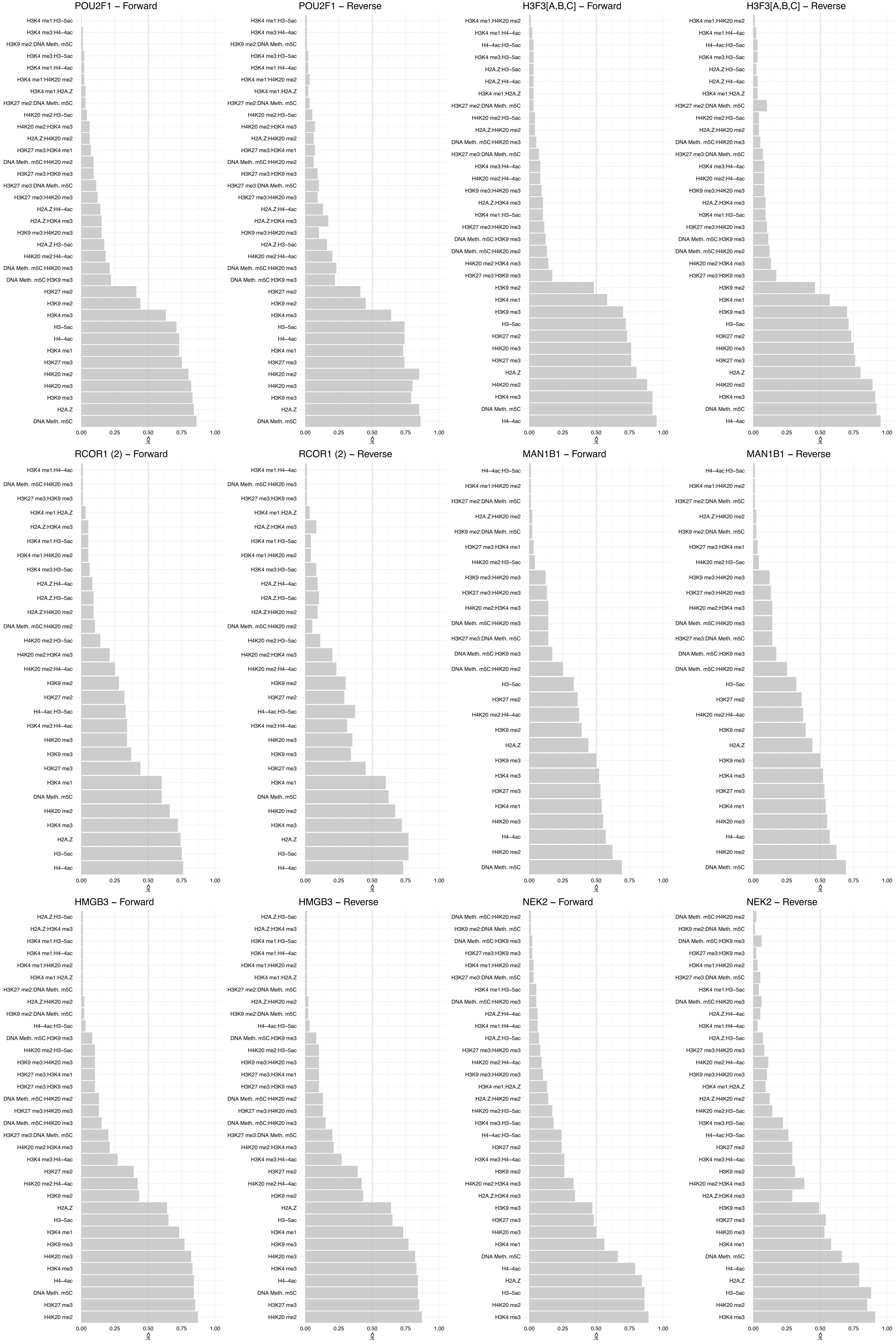
POU2F1 – Forward
H3K4 me1:H3–5ac
H3K4 me3:H4–4ac
H3K9 me2:DNA Meth. m5C
H3K4 me3:H3–5ac
H3K4 me1:H4–4ac
H3K4 me1:H4K20 me2
H3K4 me1:H2A.Z
H3K27 me2:DNA Meth. m5C
H4K20 me2:H3–5ac
H4K20 me2:H3K4 me3
H2A.Z:H4K20 me2
H3K27 me3:H3K4 me1
DNA Meth. m5C:H4K20 me2
H3K27 me3:H3K9 me3
H3K27 me3:DNA Meth. m5C
H3K27 me3:H4K20 me3
H2A.Z:H4–4ac
H2A.Z:H3K4 me3
H3K9 me3:H4K20 me3
H2A.Z:H3–5ac
H4K20 me2:H4–4ac
DNA Meth. m5C:H4K20 me3
DNA Meth. m5C:H3K9 me3
H3K27 me2
H3K9 me2
H3K4 me3
H3–5ac
H4–4ac
H3K4 me1
H3K27 me3
H4K20 me2
H4K20 me3
H3K9 me3
H2A.Z
DNA Meth. m5C
0.00
0.25
0.50
0.75
1.00
$\hat{\pi}$
POU2F1 – Reverse
H3K4 me1:H3–5ac
H3K4 me3:H4–4ac
H3K9 me2:DNA Meth. m5C
H3K4 me3:H3–5ac
H3K4 me1:H4–4ac
H3K4 me1:H4K20 me2
H3K4 me1:H2A.Z
H3K27 me2:DNA Meth. m5C
H4K20 me2:H3–5ac
H4K20 me2:H3K4 me3
H2A.Z:H4K20 me2
H3K27 me3:H3K4 me1
DNA Meth. m5C:H4K20 me2
H3K27 me3:H3K9 me3
H3K27 me3:DNA Meth. m5C
H3K27 me3:H4K20 me3
H2A.Z:H4–4ac
H2A.Z:H3K4 me3
H3K9 me3:H4K20 me3
H2A.Z:H3–5ac
H4K20 me2:H4–4ac
DNA Meth. m5C:H4K20 me3
DNA Meth. m5C:H3K9 me3
H3K27 me2
H3K9 me2
H3K4 me3
H3–5ac
H4–4ac
H3K4 me1
H3K27 me3
H4K20 me2
H4K20 me3
H3K9 me3
H2A.Z
DNA Meth. m5C
0.00
0.25
0.50
0.75
1.00
$\hat{\pi}$
H3F3[A,B,C] – Forward
H3K4 me1:H4K20 me2
H3K4 me1:H4–4ac
H4–4ac:H3–5ac
H3K4 me3:H3–5ac
H2A.Z:H3–5ac
H2A.Z:H4–4ac
H3K4 me1:H2A.Z
H3K27 me2:DNA Meth. m5C
H4K20 me2:H3–5ac
H2A.Z:H4K20 me2
DNA Meth. m5C:H4K20 me3
H3K27 me3:DNA Meth. m5C
H3K4 me3:H4–4ac
H4K20 me2:H4–4ac
H3K9 me3:H4K20 me3
H2A.Z:H3K4 me3
H3K4 me1:H3–5ac
H3K27 me3:H4K20 me3
DNA Meth. m5C:H3K9 me3
DNA Meth. m5C:H4K20 me2
H4K20 me2:H3K4 me3
H3K27 me3:H3K9 me3
H3K9 me2
H3K4 me1
H3K9 me3
H3–5ac
H3K27 me2
H4K20 me3
H3K27 me3
H2A.Z
H4K20 me2
H3K4 me3
DNA Meth. m5C
H4–4ac
0.00
0.25
0.50
0.75
1.00
$\hat{\pi}$
H3F3[A,B,C] – Reverse
H3K4 me1:H4K20 me2
H3K4 me1:H4–4ac
H4–4ac:H3–5ac
H3K4 me3:H3–5ac
H2A.Z:H3–5ac
H2A.Z:H4–4ac
H3K4 me1:H2A.Z
H3K27 me2:DNA Meth. m5C
H4K20 me2:H3–5ac
H2A.Z:H4K20 me2
DNA Meth. m5C:H4K20 me3
H3K27 me3:DNA Meth. m5C
H3K4 me3:H4–4ac
H4K20 me2:H4–4ac
H3K9 me3:H4K20 me3
H2A.Z:H3K4 me3
H3K4 me1:H3–5ac
H3K27 me3:H4K20 me3
DNA Meth. m5C:H3K9 me3
DNA Meth. m5C:H4K20 me2
H4K20 me2:H3K4 me3
H3K27 me3:H3K9 me3
H3K9 me2
H3K4 me1
H3K9 me3
H3–5ac
H3K27 me2
H4K20 me3
H3K27 me3
H2A.Z
H4K20 me2
H3K4 me3
DNA Meth. m5C
H4–4ac
0.00
0.25
0.50
0.75
1.00
$\hat{\pi}$
RCOR1 (2) – Forward
H3K4 me1:H4–4ac
DNA Meth. m5C:H4K20 me3
H3K27 me3:H3K9 me3
H3K4 me1:H2A.Z
H2A.Z:H3K4 me3
H3K4 me1:H3–5ac
H3K4 me1:H4K20 me2
H3K4 me3:H3–5ac
H2A.Z:H4–4ac
H2A.Z:H3–5ac
H2A.Z:H4K20 me2
DNA Meth. m5C:H4K20 me2
H4K20 me2:H3–5ac
H4K20 me2:H3K4 me3
H4K20 me2:H4–4ac
H3K9 me2
H3K27 me2
H4–4ac:H3–5ac
H3K4 me3:H4–4ac
H4K20 me3
H3K9 me3
H3K27 me3
H3K4 me1
DNA Meth. m5C
H4K20 me2
H3K4 me3
H2A.Z
H3–5ac
H4–4ac
0.00
0.25
0.50
0.75
1.00
$\hat{\pi}$
RCOR1 (2) – Reverse
H3K4 me1:H4–4ac
DNA Meth. m5C:H4K20 me3
H3K27 me3:H3K9 me3
H3K4 me1:H2A.Z
H2A.Z:H3K4 me3
H3K4 me1:H3–5ac
H3K4 me1:H4K20 me2
H3K4 me3:H3–5ac
H2A.Z:H4–4ac
H2A.Z:H3–5ac
H2A.Z:H4K20 me2
DNA Meth. m5C:H4K20 me2
H4K20 me2:H3–5ac
H4K20 me2:H3K4 me3
H4K20 me2:H4–4ac
H3K9 me2
H3K27 me2
H4–4ac:H3–5ac
H3K4 me3:H4–4ac
H4K20 me3
H3K9 me3
H3K27 me3
H3K4 me1
DNA Meth. m5C
H4K20 me2
H3K4 me3
H2A.Z
H3–5ac
H4–4ac
0.00
0.25
0.50
0.75
1.00
$\hat{\pi}$
MAN1B1 – Forward
H4–4ac:H3–5ac
H3K4 me1:H4K20 me2
H3K27 me2:DNA Meth. m5C
H2A.Z:H4K20 me2
H3K9 me2:DNA Meth. m5C
H3K27 me3:H3K4 me1
H4K20 me2:H3–5ac
H3K9 me3:H4K20 me3
H3K27 me3:H4K20 me3
H4K20 me2:H3K4 me3
DNA Meth. m5C:H4K20 me3
H3K27 me3:DNA Meth. m5C
DNA Meth. m5C:H3K9 me3
DNA Meth. m5C:H4K20 me2
H3–5ac
H3K27 me2
H4K20 me2:H4–4ac
H3K9 me2
H2A.Z
H3K9 me3
H3K4 me3
H3K27 me3
H3K4 me1
H4K20 me3
H4–4ac
H4K20 me2
DNA Meth. m5C
0.00
0.25
0.50
0.75
1.00
$\hat{\pi}$
MAN1B1 – Reverse
H4–4ac:H3–5ac
H3K4 me1:H4K20 me2
H3K27 me2:DNA Meth. m5C
H2A.Z:H4K20 me2
H3K9 me2:DNA Meth. m5C
H3K27 me3:H3K4 me1
H4K20 me2:H3–5ac
H3K9 me3:H4K20 me3
H3K27 me3:H4K20 me3
H4K20 me2:H3K4 me3
DNA Meth. m5C:H4K20 me3
H3K27 me3:DNA Meth. m5C
DNA Meth. m5C:H3K9 me3
DNA Meth. m5C:H4K20 me2
H3–5ac
H3K27 me2
H4K20 me2:H4–4ac
H3K9 me2
H2A.Z
H3K9 me3
H3K4 me3
H3K27 me3
H3K4 me1
H4K20 me3
H4–4ac
H4K20 me2
DNA Meth. m5C
0.00
0.25
0.50
0.75
1.00
$\hat{\pi}$
HMGB3 – Forward
H2A.Z:H3–5ac
H2A.Z:H3K4 me3
H3K4 me1:H3–5ac
H3K4 me1:H4–4ac
H3K4 me1:H4K20 me2
H3K4 me1:H2A.Z
H3K27 me2:DNA Meth. m5C
H2A.Z:H4K20 me2
H3K9 me2:DNA Meth. m5C
H4–4ac:H3–5ac
DNA Meth. m5C:H3K9 me3
H4K20 me2:H3–5ac
H3K9 me3:H4K20 me3
H3K27 me3:H3K4 me1
H3K27 me3:H3K9 me3
DNA Meth. m5C:H4K20 me2
H3K27 me3:H4K20 me3
DNA Meth. m5C:H4K20 me3
H3K27 me3:DNA Meth. m5C
H4K20 me2:H3K4 me3
H3K4 me3:H4–4ac
H3K27 me2
H4K20 me2:H4–4ac
H3K9 me2
H2A.Z
H3–5ac
H3K4 me1
H3K9 me3
H4K20 me3
H3K4 me3
H4–4ac
DNA Meth. m5C
H3K27 me3
H4K20 me2
0.00
0.25
0.50
0.75
1.00
$\hat{\pi}$
HMGB3 – Reverse
H2A.Z:H3–5ac
H2A.Z:H3K4 me3
H3K4 me1:H3–5ac
H3K4 me1:H4–4ac
H3K4 me1:H4K20 me2
H3K4 me1:H2A.Z
H3K27 me2:DNA Meth. m5C
H2A.Z:H4K20 me2
H3K9 me2:DNA Meth. m5C
H4–4ac:H3–5ac
DNA Meth. m5C:H3K9 me3
H4K20 me2:H3–5ac
H3K9 me3:H4K20 me3
H3K27 me3:H3K4 me1
H3K27 me3:H3K9 me3
DNA Meth. m5C:H4K20 me2
H3K27 me3:H4K20 me3
DNA Meth. m5C:H4K20 me3
H3K27 me3:DNA Meth. m5C
H4K20 me2:H3K4 me3
H3K4 me3:H4–4ac
H3K27 me2
H4K20 me2:H4–4ac
H3K9 me2
H2A.Z
H3–5ac
H3K4 me1
H3K9 me3
H4K20 me3
H3K4 me3
H4–4ac
DNA Meth. m5C
H3K27 me3
H4K20 me2
0.00
0.25
0.50
0.75
1.00
$\hat{\pi}$
NEK2 – Forward
DNA Meth. m5C:H4K20 me2
H3K9 me2:DNA Meth. m5C
DNA Meth. m5C:H3K9 me3
H3K27 me3:H3K9 me3
H3K4 me1:H4K20 me2
H3K27 me3:DNA Meth. m5C
H3K4 me1:H3–5ac
DNA Meth. m5C:H4K20 me3
H2A.Z:H4–4ac
H3K4 me1:H4–4ac
H2A.Z:H3–5ac
H3K27 me3:H4K20 me3
H4K20 me2:H4–4ac
H3K9 me3:H4K20 me3
H3K4 me1:H2A.Z
H2A.Z:H4K20 me2
H4K20 me2:H3–5ac
H3K4 me3:H3–5ac
H4–4ac:H3–5ac
H3K27 me2
H3K4 me3:H4–4ac
H3K9 me2
H4K20 me2:H3K4 me3
H2A.Z:H3K4 me3
H3K9 me3
H3K27 me3
H4K20 me3
H3K4 me1
DNA Meth. m5C
H4–4ac
H2A.Z
H3–5ac
H4K20 me2
H3K4 me3
0.00
0.25
0.50
0.75
1.00
$\hat{\pi}$
NEK2 – Reverse
DNA Meth. m5C:H4K20 me2
H3K9 me2:DNA Meth. m5C
DNA Meth. m5C:H3K9 me3
H3K27 me3:H3K9 me3
H3K4 me1:H4K20 me2
H3K27 me3:DNA Meth. m5C
H3K4 me1:H3–5ac
DNA Meth. m5C:H4K20 me3
H2A.Z:H4–4ac
H3K4 me1:H4–4ac
H2A.Z:H3–5ac
H3K27 me3:H4K20 me3
H4K20 me2:H4–4ac
H3K9 me3:H4K20 me3
H3K4 me1:H2A.Z
H2A.Z:H4K20 me2
H4K20 me2:H3–5ac
H3K4 me3:H3–5ac
H4–4ac:H3–5ac
H3K27 me2
H3K4 me3:H4–4ac
H3K9 me2
H4K20 me2:H3K4 me3
H2A.Z:H3K4 me3
H3K9 me3
H3K27 me3
H4K20 me3
H3K4 me1
DNA Meth. m5C
H4–4ac
H2A.Z
H3–5ac
H4K20 me2
H3K4 me3
0.00
0.25
0.50
0.75
1.00
$\hat{\pi}$

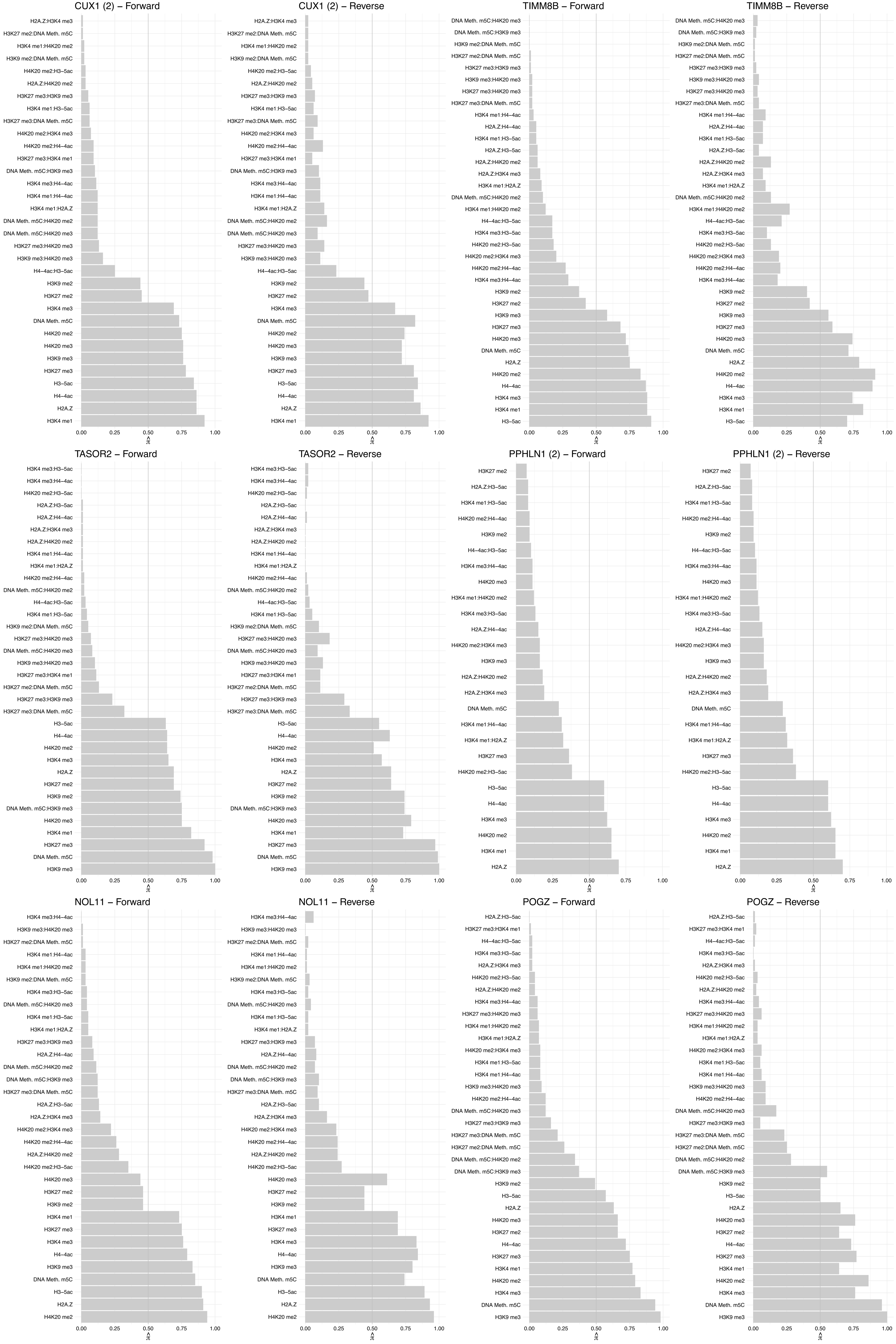
CUX1 (2) – Forward
H2A.Z:H3K4 me3
H3K27 me2:DNA Meth. m5C
H3K4 me1:H4K20 me2
H3K9 me2:DNA Meth. m5C
H4K20 me2:H3−5ac
H2A.Z:H4K20 me2
H3K27 me3:H3K9 me3
H3K4 me1:H3−5ac
H3K27 me3:DNA Meth. m5C
H4K20 me2:H3K4 me3
H4K20 me2:H4−4ac
H3K27 me3:H3K4 me1
DNA Meth. m5C:H3K9 me3
H3K4 me3:H4−4ac
H3K4 me1:H4−4ac
H3K4 me1:H2A.Z
DNA Meth. m5C:H4K20 me2
DNA Meth. m5C:H4K20 me3
H3K27 me3:H4K20 me3
H3K9 me3:H4K20 me3
H4−4ac:H3−5ac
H3K9 me2
H3K27 me2
H3K4 me3
DNA Meth. m5C
H4K20 me2
H4K20 me3
H3K9 me3
H3K27 me3
H3−5ac
H4−4ac
H2A.Z
H3K4 me1
0.00 0.25 0.50 0.75 1.00
π̂
CUX1 (2) – Reverse
H2A.Z:H3K4 me3
H3K27 me2:DNA Meth. m5C
H3K4 me1:H4K20 me2
H3K9 me2:DNA Meth. m5C
H4K20 me2:H3−5ac
H2A.Z:H4K20 me2
H3K27 me3:H3K9 me3
H3K4 me1:H3−5ac
H3K27 me3:DNA Meth. m5C
H4K20 me2:H3K4 me3
H4K20 me2:H4−4ac
H3K27 me3:H3K4 me1
DNA Meth. m5C:H3K9 me3
H3K4 me3:H4−4ac
H3K4 me1:H4−4ac
H3K4 me1:H2A.Z
DNA Meth. m5C:H4K20 me2
DNA Meth. m5C:H4K20 me3
H3K27 me3:H4K20 me3
H3K9 me3:H4K20 me3
H4−4ac:H3−5ac
H3K9 me2
H3K27 me2
H3K4 me3
DNA Meth. m5C
H4K20 me2
H4K20 me3
H3K9 me3
H3K27 me3
H3−5ac
H4−4ac
H2A.Z
H3K4 me1
0.00 0.25 0.50 0.75 1.00
π̂
TIMM8B – Forward
DNA Meth. m5C:H4K20 me3
DNA Meth. m5C:H3K9 me3
H3K9 me2:DNA Meth. m5C
H3K27 me2:DNA Meth. m5C
H3K27 me3:H3K9 me3
H3K9 me3:H4K20 me3
H3K27 me3:H4K20 me3
H3K27 me3:DNA Meth. m5C
H3K4 me1:H4−4ac
H2A.Z:H4−4ac
H3K4 me1:H3−5ac
H2A.Z:H3−5ac
H2A.Z:H4K20 me2
H2A.Z:H3K4 me3
H3K4 me1:H2A.Z
DNA Meth. m5C:H4K20 me2
H3K4 me1:H4K20 me2
H4−4ac:H3−5ac
H3K4 me3:H3−5ac
H4K20 me2:H3−5ac
H4K20 me2:H3K4 me3
H4K20 me2:H4−4ac
H3K4 me3:H4−4ac
H3K9 me2
H3K27 me2
H3K9 me3
H3K27 me3
H4K20 me3
DNA Meth. m5C
H2A.Z
H4K20 me2
H4−4ac
H3K4 me3
H3K4 me1
H3−5ac
0.00 0.25 0.50 0.75 1.00
π̂
TIMM8B – Reverse
DNA Meth. m5C:H4K20 me3
DNA Meth. m5C:H3K9 me3
H3K9 me2:DNA Meth. m5C
H3K27 me2:DNA Meth. m5C
H3K27 me3:H3K9 me3
H3K9 me3:H4K20 me3
H3K27 me3:H4K20 me3
H3K27 me3:DNA Meth. m5C
H3K4 me1:H4−4ac
H2A.Z:H4−4ac
H3K4 me1:H3−5ac
H2A.Z:H3−5ac
H2A.Z:H4K20 me2
H2A.Z:H3K4 me3
H3K4 me1:H2A.Z
DNA Meth. m5C:H4K20 me2
H3K4 me1:H4K20 me2
H4−4ac:H3−5ac
H3K4 me3:H3−5ac
H4K20 me2:H3−5ac
H4K20 me2:H3K4 me3
H4K20 me2:H4−4ac
H3K4 me3:H4−4ac
H3K9 me2
H3K27 me2
H3K9 me3
H3K27 me3
H4K20 me3
DNA Meth. m5C
H2A.Z
H4K20 me2
H4−4ac
H3K4 me3
H3K4 me1
H3−5ac
0.00 0.25 0.50 0.75 1.00
π̂
TASOR2 – Forward
H3K4 me3:H3−5ac
H3K4 me3:H4−4ac
H4K20 me2:H3−5ac
H2A.Z:H3−5ac
H2A.Z:H4−4ac
H2A.Z:H3K4 me3
H2A.Z:H4K20 me2
H3K4 me1:H4−4ac
H3K4 me1:H2A.Z
H4K20 me2:H4−4ac
DNA Meth. m5C:H4K20 me2
H4−4ac:H3−5ac
H3K4 me1:H3−5ac
H3K9 me2:DNA Meth. m5C
H3K27 me3:H4K20 me3
DNA Meth. m5C:H4K20 me3
H3K9 me3:H4K20 me3
H3K27 me3:H3K4 me1
H3K27 me2:DNA Meth. m5C
H3K27 me3:H3K9 me3
H3K27 me3:DNA Meth. m5C
H3−5ac
H4−4ac
H4K20 me2
H3K4 me3
H2A.Z
H3K27 me2
H3K9 me2
DNA Meth. m5C:H3K9 me3
H4K20 me3
H3K4 me1
H3K27 me3
DNA Meth. m5C
H3K9 me3
0.00 0.25 0.50 0.75 1.00
π̂
TASOR2 – Reverse
H3K4 me3:H3−5ac
H3K4 me3:H4−4ac
H4K20 me2:H3−5ac
H2A.Z:H3−5ac
H2A.Z:H4−4ac
H2A.Z:H3K4 me3
H2A.Z:H4K20 me2
H3K4 me1:H4−4ac
H3K4 me1:H2A.Z
H4K20 me2:H4−4ac
DNA Meth. m5C:H4K20 me2
H4−4ac:H3−5ac
H3K4 me1:H3−5ac
H3K9 me2:DNA Meth. m5C
H3K27 me3:H4K20 me3
DNA Meth. m5C:H4K20 me3
H3K9 me3:H4K20 me3
H3K27 me3:H3K4 me1
H3K27 me2:DNA Meth. m5C
H3K27 me3:H3K9 me3
H3K27 me3:DNA Meth. m5C
H3−5ac
H4−4ac
H4K20 me2
H3K4 me3
H2A.Z
H3K27 me2
H3K9 me2
DNA Meth. m5C:H3K9 me3
H4K20 me3
H3K4 me1
H3K27 me3
DNA Meth. m5C
H3K9 me3
0.00 0.25 0.50 0.75 1.00
π̂
PPHLN1 (2) – Forward
H3K27 me2
H2A.Z:H3−5ac
H3K4 me1:H3−5ac
H4K20 me2:H4−4ac
H3K9 me2
H4−4ac:H3−5ac
H3K4 me3:H4−4ac
H4K20 me3
H3K4 me1:H4K20 me2
H3K4 me3:H3−5ac
H2A.Z:H4−4ac
H4K20 me2:H3K4 me3
H3K9 me3
H2A.Z:H4K20 me2
H2A.Z:H3K4 me3
DNA Meth. m5C
H3K4 me1:H4−4ac
H3K4 me1:H2A.Z
H3K27 me3
H4K20 me2:H3−5ac
H3−5ac
H4−4ac
H3K4 me3
H4K20 me2
H3K4 me1
H2A.Z
0.00 0.25 0.50 0.75 1.00
π̂
PPHLN1 (2) – Reverse
H3K27 me2
H2A.Z:H3−5ac
H3K4 me1:H3−5ac
H4K20 me2:H4−4ac
H3K9 me2
H4−4ac:H3−5ac
H3K4 me3:H4−4ac
H4K20 me3
H3K4 me1:H4K20 me2
H3K4 me3:H3−5ac
H2A.Z:H4−4ac
H4K20 me2:H3K4 me3
H3K9 me3
H2A.Z:H4K20 me2
H2A.Z:H3K4 me3
DNA Meth. m5C
H3K4 me1:H4−4ac
H3K4 me1:H2A.Z
H3K27 me3
H4K20 me2:H3−5ac
H3−5ac
H4−4ac
H3K4 me3
H4K20 me2
H3K4 me1
H2A.Z
0.00 0.25 0.50 0.75 1.00
π̂
NOL11 – Forward
H3K4 me3:H4−4ac
H3K9 me3:H4K20 me3
H3K27 me2:DNA Meth. m5C
H3K4 me1:H4−4ac
H3K4 me1:H4K20 me2
H3K9 me2:DNA Meth. m5C
H3K4 me3:H3−5ac
DNA Meth. m5C:H4K20 me3
H3K4 me1:H3−5ac
H3K4 me1:H2A.Z
H3K27 me3:H3K9 me3
H2A.Z:H4−4ac
DNA Meth. m5C:H4K20 me2
DNA Meth. m5C:H3K9 me3
H3K27 me3:DNA Meth. m5C
H2A.Z:H3−5ac
H2A.Z:H3K4 me3
H4K20 me2:H3K4 me3
H4K20 me2:H4−4ac
H2A.Z:H4K20 me2
H4K20 me2:H3−5ac
H4K20 me3
H3K27 me2
H3K9 me2
H3K4 me1
H3K27 me3
H3K4 me3
H4−4ac
H3K9 me3
DNA Meth. m5C
H3−5ac
H2A.Z
H4K20 me2
0.00 0.25 0.50 0.75 1.00
π̂
NOL11 – Reverse
H3K4 me3:H4−4ac
H3K9 me3:H4K20 me3
H3K27 me2:DNA Meth. m5C
H3K4 me1:H4−4ac
H3K4 me1:H4K20 me2
H3K9 me2:DNA Meth. m5C
H3K4 me3:H3−5ac
DNA Meth. m5C:H4K20 me3
H3K4 me1:H3−5ac
H3K4 me1:H2A.Z
H3K27 me3:H3K9 me3
H2A.Z:H4−4ac
DNA Meth. m5C:H4K20 me2
DNA Meth. m5C:H3K9 me3
H3K27 me3:DNA Meth. m5C
H2A.Z:H3−5ac
H2A.Z:H3K4 me3
H4K20 me2:H3K4 me3
H4K20 me2:H4−4ac
H2A.Z:H4K20 me2
H4K20 me2:H3−5ac
H4K20 me3
H3K27 me2
H3K9 me2
H3K4 me1
H3K27 me3
H3K4 me3
H4−4ac
H3K9 me3
DNA Meth. m5C
H3−5ac
H2A.Z
H4K20 me2
0.00 0.25 0.50 0.75 1.00
π̂
POGZ – Forward
H2A.Z:H3−5ac
H3K27 me3:H3K4 me1
H4−4ac:H3−5ac
H3K4 me3:H3−5ac
H2A.Z:H3K4 me3
H4K20 me2:H3−5ac
H2A.Z:H4K20 me2
H3K4 me3:H4−4ac
H3K27 me3:H4K20 me3
H3K4 me1:H4K20 me2
H3K4 me1:H2A.Z
H4K20 me2:H3K4 me3
H3K4 me1:H3−5ac
H3K4 me1:H4−4ac
H3K9 me3:H4K20 me3
H4K20 me2:H4−4ac
DNA Meth. m5C:H4K20 me3
H3K27 me3:H3K9 me3
H3K27 me3:DNA Meth. m5C
H3K27 me2:DNA Meth. m5C
DNA Meth. m5C:H4K20 me2
DNA Meth. m5C:H3K9 me3
H3K9 me2
H3−5ac
H2A.Z
H4K20 me3
H3K27 me2
H4−4ac
H3K27 me3
H3K4 me1
H4K20 me2
H3K4 me3
DNA Meth. m5C
H3K9 me3
0.00 0.25 0.50 0.75 1.00
π̂
POGZ – Reverse
H2A.Z:H3−5ac
H3K27 me3:H3K4 me1
H4−4ac:H3−5ac
H3K4 me3:H3−5ac
H2A.Z:H3K4 me3
H4K20 me2:H3−5ac
H2A.Z:H4K20 me2
H3K4 me3:H4−4ac
H3K27 me3:H4K20 me3
H3K4 me1:H4K20 me2
H3K4 me1:H2A.Z
H4K20 me2:H3K4 me3
H3K4 me1:H3−5ac
H3K4 me1:H4−4ac
H3K9 me3:H4K20 me3
H4K20 me2:H4−4ac
DNA Meth. m5C:H4K20 me3
H3K27 me3:H3K9 me3
H3K27 me3:DNA Meth. m5C
H3K27 me2:DNA Meth. m5C
DNA Meth. m5C:H4K20 me2
DNA Meth. m5C:H3K9 me3
H3K9 me2
H3−5ac
H2A.Z
H4K20 me3
H3K27 me2
H4−4ac
H3K27 me3
H3K4 me1
H4K20 me2
H3K4 me3
DNA Meth. m5C
H3K9 me3
0.00 0.25 0.50 0.75 1.00
π̂

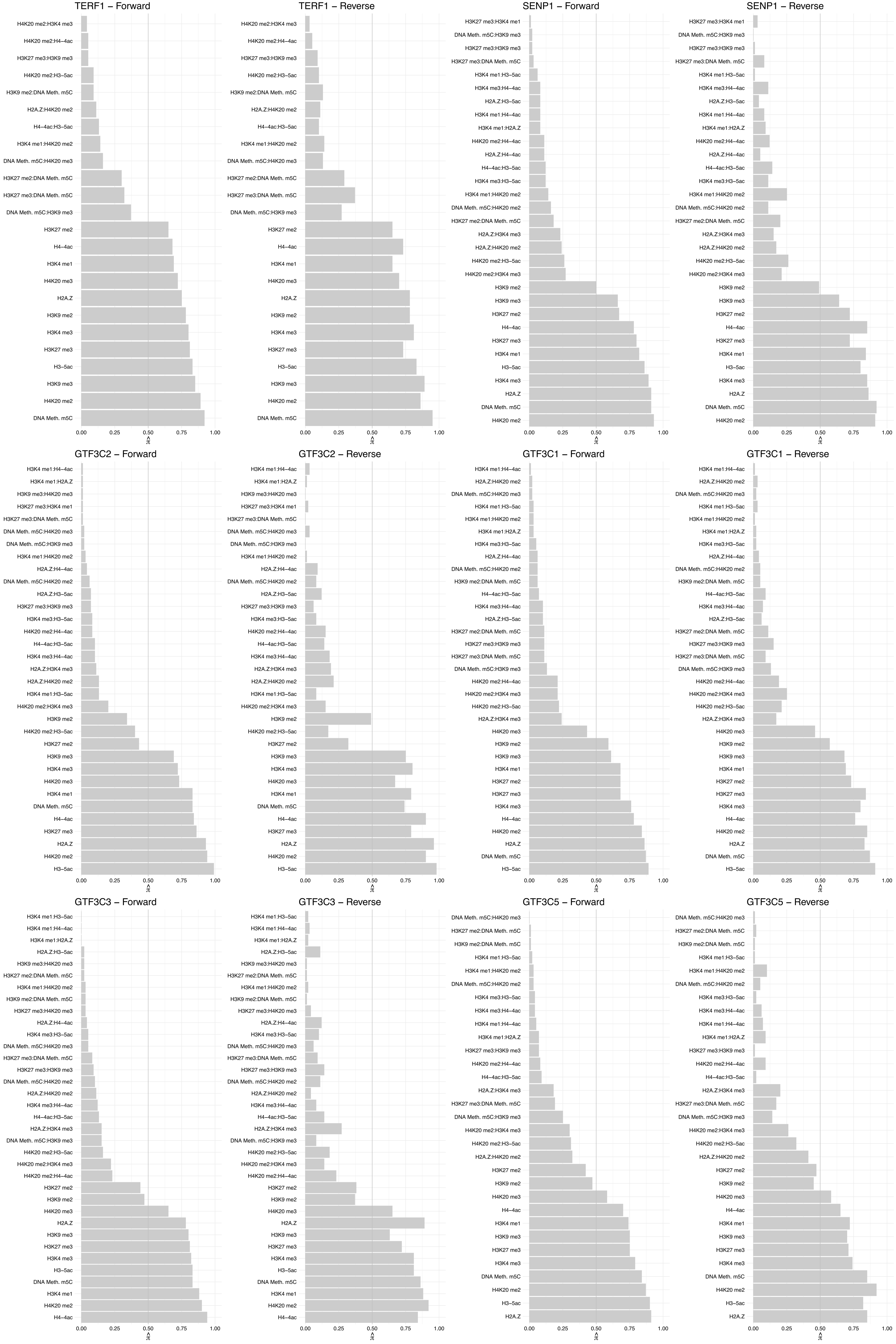
TERF1 − Forward
H4K20 me2:H3K4 me3
H4K20 me2:H4−4ac
H3K27 me3:H3K9 me3
H4K20 me2:H3−5ac
H3K9 me2:DNA Meth. m5C
H2A.Z:H4K20 me2
H4−4ac:H3−5ac
H3K4 me1:H4K20 me2
DNA Meth. m5C:H4K20 me3
H3K27 me2:DNA Meth. m5C
H3K27 me3:DNA Meth. m5C
DNA Meth. m5C:H3K9 me3
H3K27 me2
H4−4ac
H3K4 me1
H4K20 me3
H2A.Z
H3K9 me2
H3K4 me3
H3K27 me3
H3−5ac
H3K9 me3
H4K20 me2
DNA Meth. m5C
0.00 0.25 0.50 0.75 1.00
π̂
TERF1 − Reverse
H4K20 me2:H3K4 me3
H4K20 me2:H4−4ac
H3K27 me3:H3K9 me3
H4K20 me2:H3−5ac
H3K9 me2:DNA Meth. m5C
H2A.Z:H4K20 me2
H4−4ac:H3−5ac
H3K4 me1:H4K20 me2
DNA Meth. m5C:H4K20 me3
H3K27 me2:DNA Meth. m5C
H3K27 me3:DNA Meth. m5C
DNA Meth. m5C:H3K9 me3
H3K27 me2
H4−4ac
H3K4 me1
H4K20 me3
H2A.Z
H3K9 me2
H3K4 me3
H3K27 me3
H3−5ac
H3K9 me3
H4K20 me2
DNA Meth. m5C
0.00 0.25 0.50 0.75 1.00
π̂
SENP1 − Forward
H3K27 me3:H3K4 me1
DNA Meth. m5C:H3K9 me3
H3K27 me3:H3K9 me3
H3K27 me3:DNA Meth. m5C
H3K4 me1:H3−5ac
H3K4 me3:H4−4ac
H2A.Z:H3−5ac
H3K4 me1:H4−4ac
H3K4 me1:H2A.Z
H4K20 me2:H4−4ac
H2A.Z:H4−4ac
H4−4ac:H3−5ac
H3K4 me3:H3−5ac
H3K4 me1:H4K20 me2
DNA Meth. m5C:H4K20 me2
H3K27 me2:DNA Meth. m5C
H2A.Z:H3K4 me3
H2A.Z:H4K20 me2
H4K20 me2:H3−5ac
H4K20 me2:H3K4 me3
H3K9 me2
H3K9 me3
H3K27 me2
H4−4ac
H3K27 me3
H3K4 me1
H3−5ac
H3K4 me3
H2A.Z
DNA Meth. m5C
H4K20 me2
0.00 0.25 0.50 0.75 1.00
π̂
SENP1 − Reverse
H3K27 me3:H3K4 me1
DNA Meth. m5C:H3K9 me3
H3K27 me3:H3K9 me3
H3K27 me3:DNA Meth. m5C
H3K4 me1:H3−5ac
H3K4 me3:H4−4ac
H2A.Z:H3−5ac
H3K4 me1:H4−4ac
H3K4 me1:H2A.Z
H4K20 me2:H4−4ac
H2A.Z:H4−4ac
H4−4ac:H3−5ac
H3K4 me3:H3−5ac
H3K4 me1:H4K20 me2
DNA Meth. m5C:H4K20 me2
H3K27 me2:DNA Meth. m5C
H2A.Z:H3K4 me3
H2A.Z:H4K20 me2
H4K20 me2:H3−5ac
H4K20 me2:H3K4 me3
H3K9 me2
H3K9 me3
H3K27 me2
H4−4ac
H3K27 me3
H3K4 me1
H3−5ac
H3K4 me3
H2A.Z
DNA Meth. m5C
H4K20 me2
0.00 0.25 0.50 0.75 1.00
π̂
GTF3C2 − Forward
H3K4 me1:H4−4ac
H3K4 me1:H2A.Z
H3K9 me3:H4K20 me3
H3K27 me3:H3K4 me1
H3K27 me3:DNA Meth. m5C
DNA Meth. m5C:H4K20 me3
DNA Meth. m5C:H3K9 me3
H3K4 me1:H4K20 me2
H2A.Z:H4−4ac
DNA Meth. m5C:H4K20 me2
H2A.Z:H3−5ac
H3K27 me3:H3K9 me3
H3K4 me3:H3−5ac
H4K20 me2:H4−4ac
H4−4ac:H3−5ac
H3K4 me3:H4−4ac
H2A.Z:H3K4 me3
H2A.Z:H4K20 me2
H3K4 me1:H3−5ac
H4K20 me2:H3K4 me3
H3K9 me2
H4K20 me2:H3−5ac
H3K27 me2
H3K9 me3
H3K4 me3
H4K20 me3
H3K4 me1
DNA Meth. m5C
H4−4ac
H3K27 me3
H2A.Z
H4K20 me2
H3−5ac
0.00 0.25 0.50 0.75 1.00
π̂
GTF3C2 − Reverse
H3K4 me1:H4−4ac
H3K4 me1:H2A.Z
H3K9 me3:H4K20 me3
H3K27 me3:H3K4 me1
H3K27 me3:DNA Meth. m5C
DNA Meth. m5C:H4K20 me3
DNA Meth. m5C:H3K9 me3
H3K4 me1:H4K20 me2
H2A.Z:H4−4ac
DNA Meth. m5C:H4K20 me2
H2A.Z:H3−5ac
H3K27 me3:H3K9 me3
H3K4 me3:H3−5ac
H4K20 me2:H4−4ac
H4−4ac:H3−5ac
H3K4 me3:H4−4ac
H2A.Z:H3K4 me3
H2A.Z:H4K20 me2
H3K4 me1:H3−5ac
H4K20 me2:H3K4 me3
H3K9 me2
H4K20 me2:H3−5ac
H3K27 me2
H3K9 me3
H3K4 me3
H4K20 me3
H3K4 me1
DNA Meth. m5C
H4−4ac
H3K27 me3
H2A.Z
H4K20 me2
H3−5ac
0.00 0.25 0.50 0.75 1.00
π̂
GTF3C1 − Forward
H3K4 me1:H4−4ac
H2A.Z:H4K20 me2
DNA Meth. m5C:H4K20 me3
H3K4 me1:H3−5ac
H3K4 me1:H4K20 me2
H3K4 me1:H2A.Z
H3K4 me3:H3−5ac
H2A.Z:H4−4ac
DNA Meth. m5C:H4K20 me2
H3K9 me2:DNA Meth. m5C
H4−4ac:H3−5ac
H3K4 me3:H4−4ac
H2A.Z:H3−5ac
H3K27 me2:DNA Meth. m5C
H3K27 me3:H3K9 me3
H3K27 me3:DNA Meth. m5C
DNA Meth. m5C:H3K9 me3
H4K20 me2:H4−4ac
H4K20 me2:H3K4 me3
H4K20 me2:H3−5ac
H2A.Z:H3K4 me3
H4K20 me3
H3K9 me2
H3K9 me3
H3K4 me1
H3K27 me2
H3K27 me3
H3K4 me3
H4−4ac
H4K20 me2
H2A.Z
DNA Meth. m5C
H3−5ac
0.00 0.25 0.50 0.75 1.00
π̂
GTF3C1 − Reverse
H3K4 me1:H4−4ac
H2A.Z:H4K20 me2
DNA Meth. m5C:H4K20 me3
H3K4 me1:H3−5ac
H3K4 me1:H4K20 me2
H3K4 me1:H2A.Z
H3K4 me3:H3−5ac
H2A.Z:H4−4ac
DNA Meth. m5C:H4K20 me2
H3K9 me2:DNA Meth. m5C
H4−4ac:H3−5ac
H3K4 me3:H4−4ac
H2A.Z:H3−5ac
H3K27 me2:DNA Meth. m5C
H3K27 me3:H3K9 me3
H3K27 me3:DNA Meth. m5C
DNA Meth. m5C:H3K9 me3
H4K20 me2:H4−4ac
H4K20 me2:H3K4 me3
H4K20 me2:H3−5ac
H2A.Z:H3K4 me3
H4K20 me3
H3K9 me2
H3K9 me3
H3K4 me1
H3K27 me2
H3K27 me3
H3K4 me3
H4−4ac
H4K20 me2
H2A.Z
DNA Meth. m5C
H3−5ac
0.00 0.25 0.50 0.75 1.00
π̂
GTF3C3 − Forward
H3K4 me1:H3−5ac
H3K4 me1:H4−4ac
H3K4 me1:H2A.Z
H2A.Z:H3−5ac
H3K9 me3:H4K20 me3
H3K27 me2:DNA Meth. m5C
H3K4 me1:H4K20 me2
H3K9 me2:DNA Meth. m5C
H3K27 me3:H4K20 me3
H2A.Z:H4−4ac
H3K4 me3:H3−5ac
DNA Meth. m5C:H4K20 me3
H3K27 me3:DNA Meth. m5C
H3K27 me3:H3K9 me3
DNA Meth. m5C:H4K20 me2
H2A.Z:H4K20 me2
H3K4 me3:H4−4ac
H4−4ac:H3−5ac
H2A.Z:H3K4 me3
DNA Meth. m5C:H3K9 me3
H4K20 me2:H3−5ac
H4K20 me2:H3K4 me3
H4K20 me2:H4−4ac
H3K27 me2
H3K9 me2
H4K20 me3
H2A.Z
H3K9 me3
H3K27 me3
H3K4 me3
H3−5ac
DNA Meth. m5C
H3K4 me1
H4K20 me2
H4−4ac
0.00 0.25 0.50 0.75 1.00
π̂
GTF3C3 − Reverse
H3K4 me1:H3−5ac
H3K4 me1:H4−4ac
H3K4 me1:H2A.Z
H2A.Z:H3−5ac
H3K9 me3:H4K20 me3
H3K27 me2:DNA Meth. m5C
H3K4 me1:H4K20 me2
H3K9 me2:DNA Meth. m5C
H3K27 me3:H4K20 me3
H2A.Z:H4−4ac
H3K4 me3:H3−5ac
DNA Meth. m5C:H4K20 me3
H3K27 me3:DNA Meth. m5C
H3K27 me3:H3K9 me3
DNA Meth. m5C:H4K20 me2
H2A.Z:H4K20 me2
H3K4 me3:H4−4ac
H4−4ac:H3−5ac
H2A.Z:H3K4 me3
DNA Meth. m5C:H3K9 me3
H4K20 me2:H3−5ac
H4K20 me2:H3K4 me3
H4K20 me2:H4−4ac
H3K27 me2
H3K9 me2
H4K20 me3
H2A.Z
H3K9 me3
H3K27 me3
H3K4 me3
H3−5ac
DNA Meth. m5C
H3K4 me1
H4K20 me2
H4−4ac
0.00 0.25 0.50 0.75 1.00
π̂
GTF3C5 − Forward
DNA Meth. m5C:H4K20 me3
H3K27 me2:DNA Meth. m5C
H3K9 me2:DNA Meth. m5C
H3K4 me1:H3−5ac
H3K4 me1:H4K20 me2
DNA Meth. m5C:H4K20 me2
H3K4 me3:H3−5ac
H3K4 me3:H4−4ac
H3K4 me1:H4−4ac
H3K4 me1:H2A.Z
H3K27 me3:H3K9 me3
H4K20 me2:H4−4ac
H4−4ac:H3−5ac
H2A.Z:H3K4 me3
H3K27 me3:DNA Meth. m5C
DNA Meth. m5C:H3K9 me3
H4K20 me2:H3K4 me3
H4K20 me2:H3−5ac
H2A.Z:H4K20 me2
H3K27 me2
H3K9 me2
H4K20 me3
H4−4ac
H3K4 me1
H3K9 me3
H3K27 me3
H3K4 me3
DNA Meth. m5C
H4K20 me2
H3−5ac
H2A.Z
0.00 0.25 0.50 0.75 1.00
π̂
GTF3C5 − Reverse
DNA Meth. m5C:H4K20 me3
H3K27 me2:DNA Meth. m5C
H3K9 me2:DNA Meth. m5C
H3K4 me1:H3−5ac
H3K4 me1:H4K20 me2
DNA Meth. m5C:H4K20 me2
H3K4 me3:H3−5ac
H3K4 me3:H4−4ac
H3K4 me1:H4−4ac
H3K4 me1:H2A.Z
H3K27 me3:H3K9 me3
H4K20 me2:H4−4ac
H4−4ac:H3−5ac
H2A.Z:H3K4 me3
H3K27 me3:DNA Meth. m5C
DNA Meth. m5C:H3K9 me3
H4K20 me2:H3K4 me3
H4K20 me2:H3−5ac
H2A.Z:H4K20 me2
H3K27 me2
H3K9 me2
H4K20 me3
H4−4ac
H3K4 me1
H3K9 me3
H3K27 me3
H3K4 me3
DNA Meth. m5C
H4K20 me2
H3−5ac
H2A.Z
0.00 0.25 0.50 0.75 1.00
π̂

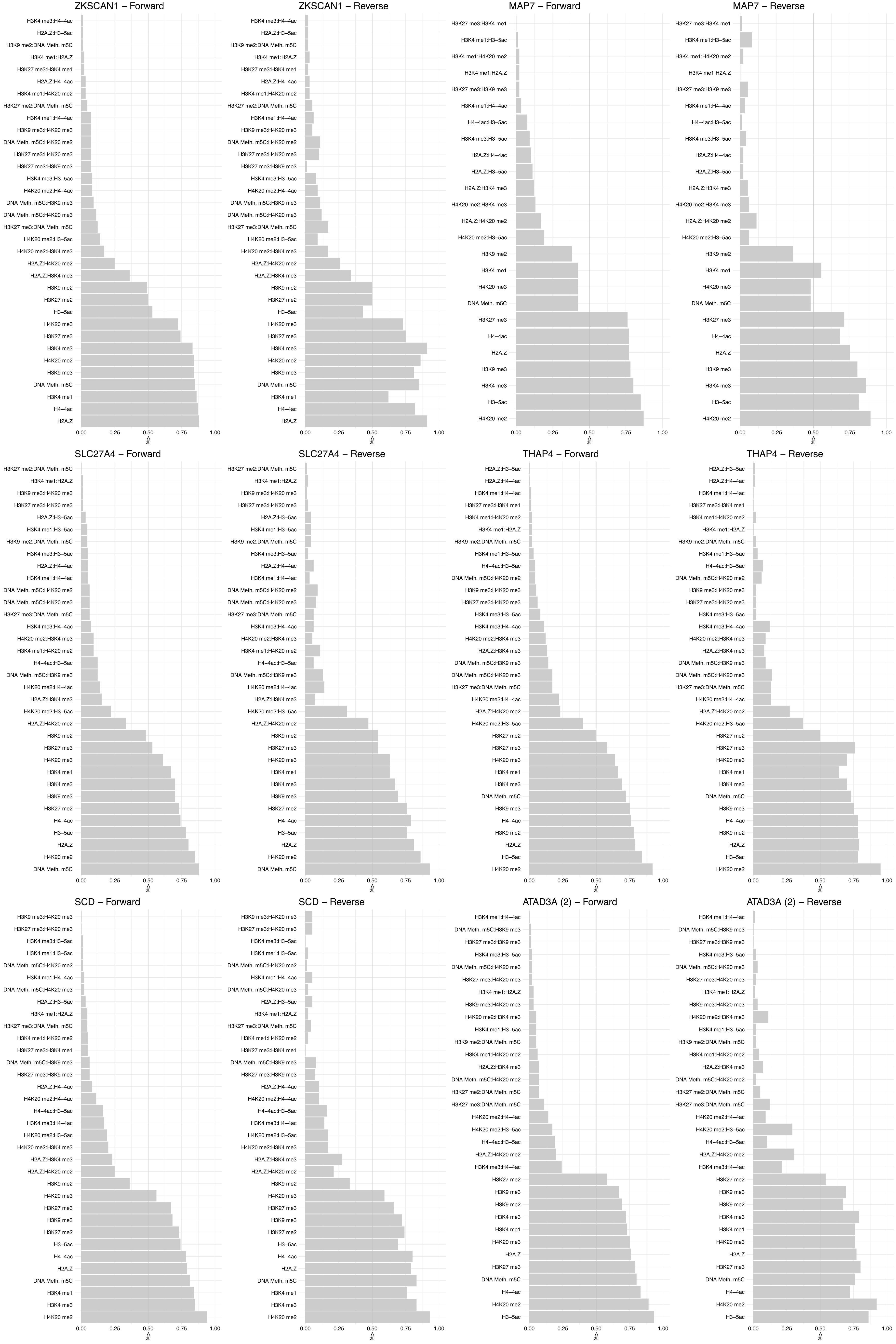

ZKSCAN1 – Forward
H3K4 me3:H4–4ac
H2A.Z:H3–5ac
H3K9 me2:DNA Meth. m5C
H3K4 me1:H2A.Z
H3K27 me3:H3K4 me1
H2A.Z:H4–4ac
H3K4 me1:H4K20 me2
H3K27 me2:DNA Meth. m5C
H3K4 me1:H4–4ac
H3K9 me3:H4K20 me3
DNA Meth. m5C:H4K20 me2
H3K27 me3:H4K20 me3
H3K27 me3:H3K9 me3
H3K4 me3:H3–5ac
H4K20 me2:H4–4ac
DNA Meth. m5C:H3K9 me3
DNA Meth. m5C:H4K20 me3
H3K27 me3:DNA Meth. m5C
H4K20 me2:H3–5ac
H4K20 me2:H3K4 me3
H2A.Z:H4K20 me2
H2A.Z:H3K4 me3
H3K9 me2
H3K27 me2
H3–5ac
H4K20 me3
H3K27 me3
H3K4 me3
H4K20 me2
H3K9 me3
DNA Meth. m5C
H3K4 me1
H4–4ac
H2A.Z
0.00 0.25 0.50 0.75 1.00
$\hat{\pi}$
ZKSCAN1 – Reverse
H3K4 me3:H4–4ac
H2A.Z:H3–5ac
H3K9 me2:DNA Meth. m5C
H3K4 me1:H2A.Z
H3K27 me3:H3K4 me1
H2A.Z:H4–4ac
H3K4 me1:H4K20 me2
H3K27 me2:DNA Meth. m5C
H3K4 me1:H4–4ac
H3K9 me3:H4K20 me3
DNA Meth. m5C:H4K20 me2
H3K27 me3:H4K20 me3
H3K27 me3:H3K9 me3
H3K4 me3:H3–5ac
H4K20 me2:H4–4ac
DNA Meth. m5C:H3K9 me3
DNA Meth. m5C:H4K20 me3
H3K27 me3:DNA Meth. m5C
H4K20 me2:H3–5ac
H4K20 me2:H3K4 me3
H2A.Z:H4K20 me2
H2A.Z:H3K4 me3
H3K9 me2
H3K27 me2
H3–5ac
H4K20 me3
H3K27 me3
H3K4 me3
H4K20 me2
H3K9 me3
DNA Meth. m5C
H3K4 me1
H4–4ac
H2A.Z
0.00 0.25 0.50 0.75 1.00
$\hat{\pi}$
MAP7 – Forward
H3K27 me3:H3K4 me1
H3K4 me1:H3–5ac
H3K4 me1:H4K20 me2
H3K4 me1:H2A.Z
H3K27 me3:H3K9 me3
H3K4 me1:H4–4ac
H4–4ac:H3–5ac
H3K4 me3:H3–5ac
H2A.Z:H4–4ac
H2A.Z:H3–5ac
H2A.Z:H3K4 me3
H4K20 me2:H3K4 me3
H2A.Z:H4K20 me2
H4K20 me2:H3–5ac
H3K9 me2
H3K4 me1
H4K20 me3
DNA Meth. m5C
H3K27 me3
H4–4ac
H2A.Z
H3K9 me3
H3K4 me3
H3–5ac
H4K20 me2
0.00 0.25 0.50 0.75 1.00
$\hat{\pi}$
MAP7 – Reverse
H3K27 me3:H3K4 me1
H3K4 me1:H3–5ac
H3K4 me1:H4K20 me2
H3K4 me1:H2A.Z
H3K27 me3:H3K9 me3
H3K4 me1:H4–4ac
H4–4ac:H3–5ac
H3K4 me3:H3–5ac
H2A.Z:H4–4ac
H2A.Z:H3–5ac
H2A.Z:H3K4 me3
H4K20 me2:H3K4 me3
H2A.Z:H4K20 me2
H4K20 me2:H3–5ac
H3K9 me2
H3K4 me1
H4K20 me3
DNA Meth. m5C
H3K27 me3
H4–4ac
H2A.Z
H3K9 me3
H3K4 me3
H3–5ac
H4K20 me2
0.00 0.25 0.50 0.75 1.00
$\hat{\pi}$
SLC27A4 – Forward
H3K27 me2:DNA Meth. m5C
H3K4 me1:H2A.Z
H3K9 me3:H4K20 me3
H3K27 me3:H4K20 me3
H2A.Z:H3–5ac
H3K4 me1:H3–5ac
H3K9 me2:DNA Meth. m5C
H3K4 me3:H3–5ac
H2A.Z:H4–4ac
H3K4 me1:H4–4ac
DNA Meth. m5C:H4K20 me2
DNA Meth. m5C:H4K20 me3
H3K27 me3:DNA Meth. m5C
H3K4 me3:H4–4ac
H4K20 me2:H3K4 me3
H3K4 me1:H4K20 me2
H4–4ac:H3–5ac
DNA Meth. m5C:H3K9 me3
H4K20 me2:H4–4ac
H2A.Z:H3K4 me3
H4K20 me2:H3–5ac
H2A.Z:H4K20 me2
H3K9 me2
H3K27 me3
H4K20 me3
H3K4 me1
H3K4 me3
H3K9 me3
H3K27 me2
H4–4ac
H3–5ac
H2A.Z
H4K20 me2
DNA Meth. m5C
0.00 0.25 0.50 0.75 1.00
$\hat{\pi}$
SLC27A4 – Reverse
H3K27 me2:DNA Meth. m5C
H3K4 me1:H2A.Z
H3K9 me3:H4K20 me3
H3K27 me3:H4K20 me3
H2A.Z:H3–5ac
H3K4 me1:H3–5ac
H3K9 me2:DNA Meth. m5C
H3K4 me3:H3–5ac
H2A.Z:H4–4ac
H3K4 me1:H4–4ac
DNA Meth. m5C:H4K20 me2
DNA Meth. m5C:H4K20 me3
H3K27 me3:DNA Meth. m5C
H3K4 me3:H4–4ac
H4K20 me2:H3K4 me3
H3K4 me1:H4K20 me2
H4–4ac:H3–5ac
DNA Meth. m5C:H3K9 me3
H4K20 me2:H4–4ac
H2A.Z:H3K4 me3
H4K20 me2:H3–5ac
H2A.Z:H4K20 me2
H3K9 me2
H3K27 me3
H4K20 me3
H3K4 me1
H3K4 me3
H3K9 me3
H3K27 me2
H4–4ac
H3–5ac
H2A.Z
H4K20 me2
DNA Meth. m5C
0.00 0.25 0.50 0.75 1.00
$\hat{\pi}$
THAP4 – Forward
H2A.Z:H3–5ac
H2A.Z:H4–4ac
H3K4 me1:H4–4ac
H3K27 me3:H3K4 me1
H3K4 me1:H4K20 me2
H3K4 me1:H2A.Z
H3K9 me2:DNA Meth. m5C
H3K4 me1:H3–5ac
H4–4ac:H3–5ac
DNA Meth. m5C:H4K20 me2
H3K9 me3:H4K20 me3
H3K27 me3:H4K20 me3
H3K4 me3:H3–5ac
H3K4 me3:H4–4ac
H4K20 me2:H3K4 me3
H2A.Z:H3K4 me3
DNA Meth. m5C:H3K9 me3
DNA Meth. m5C:H4K20 me3
H3K27 me3:DNA Meth. m5C
H4K20 me2:H4–4ac
H2A.Z:H4K20 me2
H4K20 me2:H3–5ac
H3K27 me2
H3K27 me3
H4K20 me3
H3K4 me1
H3K4 me3
DNA Meth. m5C
H3K9 me3
H4–4ac
H3K9 me2
H2A.Z
H3–5ac
H4K20 me2
0.00 0.25 0.50 0.75 1.00
$\hat{\pi}$
THAP4 – Reverse
H2A.Z:H3–5ac
H2A.Z:H4–4ac
H3K4 me1:H4–4ac
H3K27 me3:H3K4 me1
H3K4 me1:H4K20 me2
H3K4 me1:H2A.Z
H3K9 me2:DNA Meth. m5C
H3K4 me1:H3–5ac
H4–4ac:H3–5ac
DNA Meth. m5C:H4K20 me2
H3K9 me3:H4K20 me3
H3K27 me3:H4K20 me3
H3K4 me3:H3–5ac
H3K4 me3:H4–4ac
H4K20 me2:H3K4 me3
H2A.Z:H3K4 me3
DNA Meth. m5C:H3K9 me3
DNA Meth. m5C:H4K20 me3
H3K27 me3:DNA Meth. m5C
H4K20 me2:H4–4ac
H2A.Z:H4K20 me2
H4K20 me2:H3–5ac
H3K27 me2
H3K27 me3
H4K20 me3
H3K4 me1
H3K4 me3
DNA Meth. m5C
H3K9 me3
H4–4ac
H3K9 me2
H2A.Z
H3–5ac
H4K20 me2
0.00 0.25 0.50 0.75 1.00
$\hat{\pi}$
SCD – Forward
H3K9 me3:H4K20 me3
H3K27 me3:H4K20 me3
H3K4 me3:H3–5ac
H3K4 me1:H3–5ac
DNA Meth. m5C:H4K20 me2
H3K4 me1:H4–4ac
DNA Meth. m5C:H4K20 me3
H2A.Z:H3–5ac
H3K4 me1:H2A.Z
H3K27 me3:DNA Meth. m5C
H3K4 me1:H4K20 me2
H3K27 me3:H3K4 me1
DNA Meth. m5C:H3K9 me3
H3K27 me3:H3K9 me3
H2A.Z:H4–4ac
H4K20 me2:H4–4ac
H4–4ac:H3–5ac
H3K4 me3:H4–4ac
H4K20 me2:H3–5ac
H4K20 me2:H3K4 me3
H2A.Z:H3K4 me3
H2A.Z:H4K20 me2
H3K9 me2
H4K20 me3
H3K27 me3
H3K9 me3
H3K27 me2
H3–5ac
H4–4ac
H2A.Z
DNA Meth. m5C
H3K4 me1
H3K4 me3
H4K20 me2
0.00 0.25 0.50 0.75 1.00
$\hat{\pi}$
SCD – Reverse
H3K9 me3:H4K20 me3
H3K27 me3:H4K20 me3
H3K4 me3:H3–5ac
H3K4 me1:H3–5ac
DNA Meth. m5C:H4K20 me2
H3K4 me1:H4–4ac
DNA Meth. m5C:H4K20 me3
H2A.Z:H3–5ac
H3K4 me1:H2A.Z
H3K27 me3:DNA Meth. m5C
H3K4 me1:H4K20 me2
H3K27 me3:H3K4 me1
DNA Meth. m5C:H3K9 me3
H3K27 me3:H3K9 me3
H2A.Z:H4–4ac
H4K20 me2:H4–4ac
H4–4ac:H3–5ac
H3K4 me3:H4–4ac
H4K20 me2:H3–5ac
H4K20 me2:H3K4 me3
H2A.Z:H3K4 me3
H2A.Z:H4K20 me2
H3K9 me2
H4K20 me3
H3K27 me3
H3K9 me3
H3K27 me2
H3–5ac
H4–4ac
H2A.Z
DNA Meth. m5C
H3K4 me1
H3K4 me3
H4K20 me2
0.00 0.25 0.50 0.75 1.00
$\hat{\pi}$
ATAD3A (2) – Forward
H3K4 me1:H4–4ac
DNA Meth. m5C:H3K9 me3
H3K27 me3:H3K9 me3
H3K4 me3:H3–5ac
DNA Meth. m5C:H4K20 me3
H3K27 me3:H4K20 me3
H3K4 me1:H2A.Z
H3K9 me3:H4K20 me3
H4K20 me2:H3K4 me3
H3K4 me1:H3–5ac
H3K9 me2:DNA Meth. m5C
H3K4 me1:H4K20 me2
H2A.Z:H3K4 me3
DNA Meth. m5C:H4K20 me2
H3K27 me2:DNA Meth. m5C
H3K27 me3:DNA Meth. m5C
H4K20 me2:H4–4ac
H4K20 me2:H3–5ac
H4–4ac:H3–5ac
H2A.Z:H4K20 me2
H3K4 me3:H4–4ac
H3K27 me2
H3K9 me3
H3K9 me2
H3K4 me3
H3K4 me1
H4K20 me3
H2A.Z
H3K27 me3
DNA Meth. m5C
H4–4ac
H4K20 me2
H3–5ac
0.00 0.25 0.50 0.75 1.00
$\hat{\pi}$
ATAD3A (2) – Reverse
H3K4 me1:H4–4ac
DNA Meth. m5C:H3K9 me3
H3K27 me3:H3K9 me3
H3K4 me3:H3–5ac
DNA Meth. m5C:H4K20 me3
H3K27 me3:H4K20 me3
H3K4 me1:H2A.Z
H3K9 me3:H4K20 me3
H4K20 me2:H3K4 me3
H3K4 me1:H3–5ac
H3K9 me2:DNA Meth. m5C
H3K4 me1:H4K20 me2
H2A.Z:H3K4 me3
DNA Meth. m5C:H4K20 me2
H3K27 me2:DNA Meth. m5C
H3K27 me3:DNA Meth. m5C
H4K20 me2:H4–4ac
H4K20 me2:H3–5ac
H4–4ac:H3–5ac
H2A.Z:H4K20 me2
H3K4 me3:H4–4ac
H3K27 me2
H3K9 me3
H3K9 me2
H3K4 me3
H3K4 me1
H4K20 me3
H2A.Z
H3K27 me3
DNA Meth. m5C
H4–4ac
H4K20 me2
H3–5ac
0.00 0.25 0.50 0.75 1.00
$\hat{\pi}$

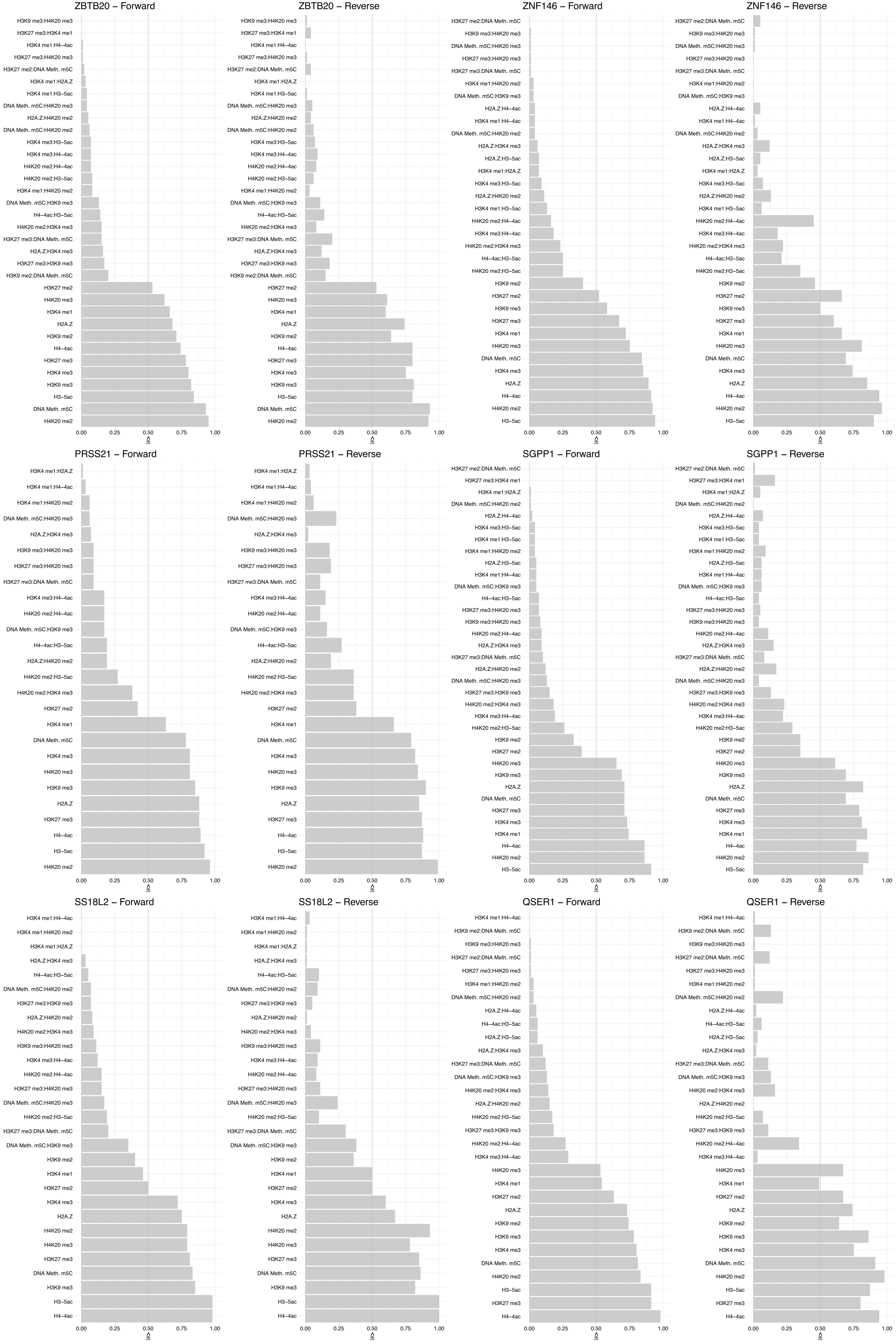
ZBTB20 – Forward
ZBTB20 – Reverse
H3K9 me3:H4K20 me3
H3K27 me3:H3K4 me1
H3K4 me1:H4–4ac
H3K27 me3:H4K20 me3
H3K27 me2:DNA Meth. m5C
H3K4 me1:H2A.Z
H3K4 me1:H3–5ac
DNA Meth. m5C:H4K20 me3
H2A.Z:H4K20 me2
DNA Meth. m5C:H4K20 me2
H3K4 me3:H3–5ac
H3K4 me3:H4–4ac
H4K20 me2:H4–4ac
H4K20 me2:H3–5ac
H3K4 me1:H4K20 me2
DNA Meth. m5C:H3K9 me3
H4–4ac:H3–5ac
H4K20 me2:H3K4 me3
H3K27 me3:DNA Meth. m5C
H2A.Z:H3K4 me3
H3K27 me3:H3K9 me3
H3K9 me2:DNA Meth. m5C
H3K27 me2
H4K20 me3
H3K4 me1
H2A.Z
H3K9 me2
H4–4ac
H3K27 me3
H3K4 me3
H3K9 me3
H3–5ac
DNA Meth. m5C
H4K20 me2
ZNF146 – Forward
ZNF146 – Reverse
H3K27 me2:DNA Meth. m5C
H3K9 me3:H4K20 me3
DNA Meth. m5C:H4K20 me3
H3K27 me3:H4K20 me3
H3K27 me3:DNA Meth. m5C
H3K4 me1:H4K20 me2
DNA Meth. m5C:H3K9 me3
H2A.Z:H4–4ac
H3K4 me1:H4–4ac
DNA Meth. m5C:H4K20 me2
H2A.Z:H3K4 me3
H2A.Z:H3–5ac
H3K4 me1:H2A.Z
H3K4 me3:H3–5ac
H2A.Z:H4K20 me2
H3K4 me1:H3–5ac
H4K20 me2:H4–4ac
H3K4 me3:H4–4ac
H4K20 me2:H3K4 me3
H4–4ac:H3–5ac
H4K20 me2:H3–5ac
H3K9 me2
H3K27 me2
H3K9 me3
H3K27 me3
H3K4 me1
H4K20 me3
DNA Meth. m5C
H3K4 me3
H2A.Z
H4–4ac
H4K20 me2
H3–5ac
PRSS21 – Forward
PRSS21 – Reverse
H3K4 me1:H2A.Z
H3K4 me1:H4–4ac
H3K4 me1:H4K20 me2
DNA Meth. m5C:H4K20 me3
H2A.Z:H3K4 me3
H3K9 me3:H4K20 me3
H3K27 me3:H4K20 me3
H3K27 me3:DNA Meth. m5C
H3K4 me3:H4–4ac
H4K20 me2:H4–4ac
DNA Meth. m5C:H3K9 me3
H4–4ac:H3–5ac
H2A.Z:H4K20 me2
H4K20 me2:H3–5ac
H4K20 me2:H3K4 me3
H3K27 me2
H3K4 me1
DNA Meth. m5C
H3K4 me3
H4K20 me3
H3K9 me3
H2A.Z
H3K27 me3
H4–4ac
H3–5ac
H4K20 me2
SGPP1 – Forward
SGPP1 – Reverse
H3K27 me2:DNA Meth. m5C
H3K27 me3:H3K4 me1
H3K4 me1:H2A.Z
DNA Meth. m5C:H4K20 me2
H2A.Z:H4–4ac
H3K4 me3:H3–5ac
H3K4 me1:H3–5ac
H3K4 me1:H4K20 me2
H2A.Z:H3–5ac
H3K4 me1:H4–4ac
DNA Meth. m5C:H3K9 me3
H4–4ac:H3–5ac
H3K27 me3:H4K20 me3
H3K9 me3:H4K20 me3
H4K20 me2:H4–4ac
H2A.Z:H3K4 me3
H3K27 me3:DNA Meth. m5C
H2A.Z:H4K20 me2
DNA Meth. m5C:H4K20 me3
H3K27 me3:H3K9 me3
H4K20 me2:H3K4 me3
H3K4 me3:H4–4ac
H4K20 me2:H3–5ac
H3K9 me2
H3K27 me2
H4K20 me3
H3K9 me3
H2A.Z
DNA Meth. m5C
H3K27 me3
H3K4 me3
H3K4 me1
H4–4ac
H4K20 me2
H3–5ac
SS18L2 – Forward
SS18L2 – Reverse
H3K4 me1:H4–4ac
H3K4 me1:H4K20 me2
H3K4 me1:H2A.Z
H2A.Z:H3K4 me3
H4–4ac:H3–5ac
DNA Meth. m5C:H4K20 me2
H3K27 me3:H3K9 me3
H2A.Z:H4K20 me2
H4K20 me2:H3K4 me3
H3K9 me3:H4K20 me3
H3K4 me3:H4–4ac
H4K20 me2:H4–4ac
H3K27 me3:H4K20 me3
DNA Meth. m5C:H4K20 me3
H4K20 me2:H3–5ac
H3K27 me3:DNA Meth. m5C
DNA Meth. m5C:H3K9 me3
H3K9 me2
H3K4 me1
H3K27 me2
H3K4 me3
H2A.Z
H4K20 me2
H4K20 me3
H3K27 me3
DNA Meth. m5C
H3K9 me3
H3–5ac
H4–4ac
QSER1 – Forward
QSER1 – Reverse
H3K4 me1:H4–4ac
H3K9 me2:DNA Meth. m5C
H3K9 me3:H4K20 me3
H3K27 me2:DNA Meth. m5C
H3K27 me3:H4K20 me3
H3K4 me1:H4K20 me2
DNA Meth. m5C:H4K20 me2
H2A.Z:H4–4ac
H4–4ac:H3–5ac
H2A.Z:H3–5ac
H2A.Z:H3K4 me3
H3K27 me3:DNA Meth. m5C
DNA Meth. m5C:H3K9 me3
H4K20 me2:H3K4 me3
H2A.Z:H4K20 me2
H4K20 me2:H3–5ac
H3K27 me3:H3K9 me3
H4K20 me2:H4–4ac
H3K4 me3:H4–4ac
H4K20 me3
H3K4 me1
H3K27 me2
H2A.Z
H3K9 me2
H3K9 me3
H3K4 me3
DNA Meth. m5C
H4K20 me2
H3–5ac
H3K27 me3
H4–4ac
0.00
0.25
0.50
0.75
1.00
π̂

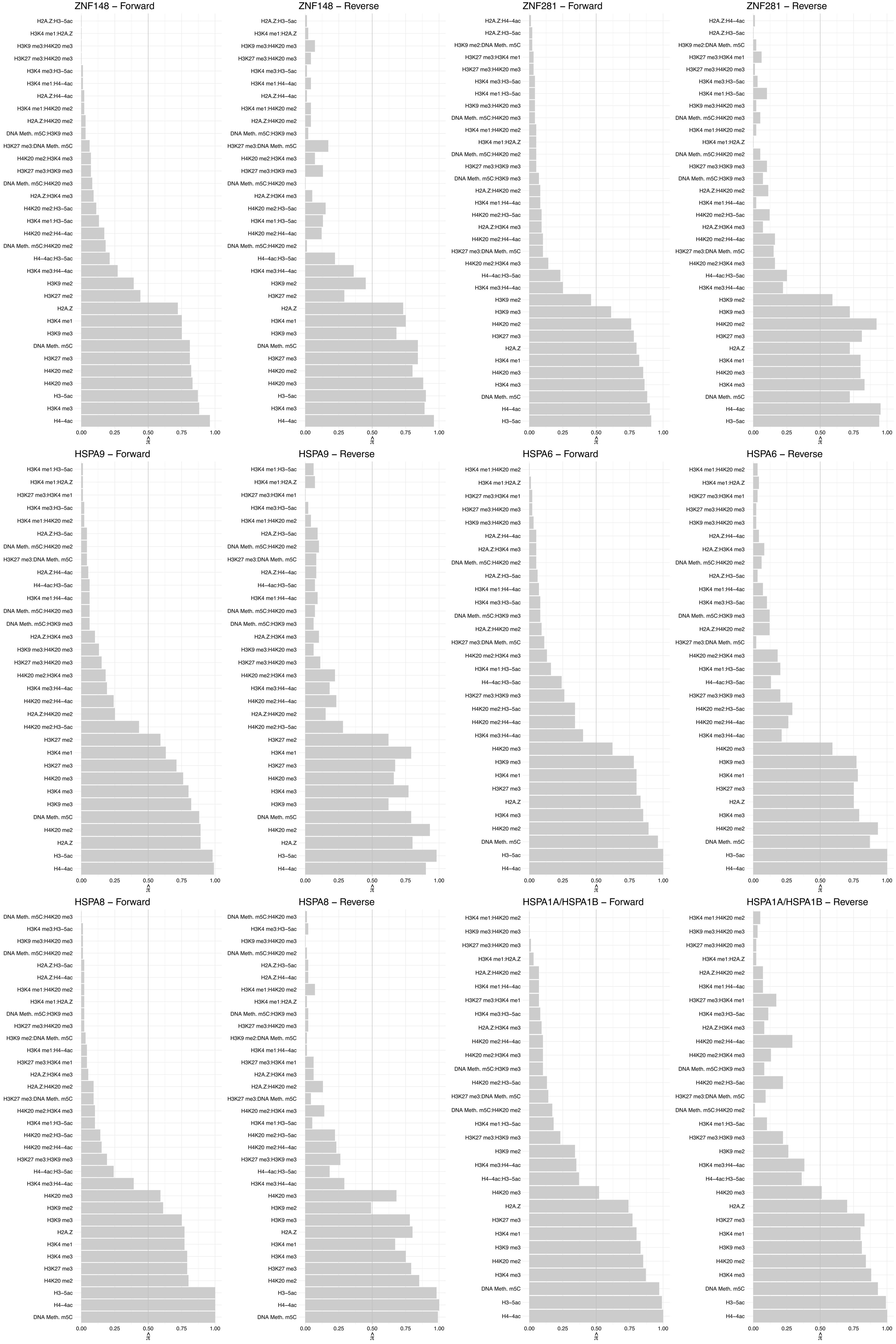
ZNF148 − Forward
H2A.Z:H3−5ac
H3K4 me1:H2A.Z
H3K9 me3:H4K20 me3
H3K27 me3:H4K20 me3
H3K4 me3:H3−5ac
H3K4 me1:H4−4ac
H2A.Z:H4−4ac
H3K4 me1:H4K20 me2
H2A.Z:H4K20 me2
DNA Meth. m5C:H3K9 me3
H3K27 me3:DNA Meth. m5C
H4K20 me2:H3K4 me3
H3K27 me3:H3K9 me3
DNA Meth. m5C:H4K20 me3
H2A.Z:H3K4 me3
H4K20 me2:H3−5ac
H3K4 me1:H3−5ac
H4K20 me2:H4−4ac
DNA Meth. m5C:H4K20 me2
H4−4ac:H3−5ac
H3K4 me3:H4−4ac
H3K9 me2
H3K27 me2
H2A.Z
H3K4 me1
H3K9 me3
DNA Meth. m5C
H3K27 me3
H4K20 me2
H4K20 me3
H3−5ac
H3K4 me3
H4−4ac
0.00 0.25 0.50 0.75 1.00
π̂
ZNF148 − Reverse
H2A.Z:H3−5ac
H3K4 me1:H2A.Z
H3K9 me3:H4K20 me3
H3K27 me3:H4K20 me3
H3K4 me3:H3−5ac
H3K4 me1:H4−4ac
H2A.Z:H4−4ac
H3K4 me1:H4K20 me2
H2A.Z:H4K20 me2
DNA Meth. m5C:H3K9 me3
H3K27 me3:DNA Meth. m5C
H4K20 me2:H3K4 me3
H3K27 me3:H3K9 me3
DNA Meth. m5C:H4K20 me3
H2A.Z:H3K4 me3
H4K20 me2:H3−5ac
H3K4 me1:H3−5ac
H4K20 me2:H4−4ac
DNA Meth. m5C:H4K20 me2
H4−4ac:H3−5ac
H3K4 me3:H4−4ac
H3K9 me2
H3K27 me2
H2A.Z
H3K4 me1
H3K9 me3
DNA Meth. m5C
H3K27 me3
H4K20 me2
H4K20 me3
H3−5ac
H3K4 me3
H4−4ac
0.00 0.25 0.50 0.75 1.00
π̂
ZNF281 − Forward
H2A.Z:H4−4ac
H2A.Z:H3−5ac
H3K9 me2:DNA Meth. m5C
H3K27 me3:H3K4 me1
H3K27 me3:H4K20 me3
H3K4 me3:H3−5ac
H3K4 me1:H3−5ac
H3K9 me3:H4K20 me3
DNA Meth. m5C:H4K20 me3
H3K4 me1:H4K20 me2
H3K4 me1:H2A.Z
DNA Meth. m5C:H4K20 me2
H3K27 me3:H3K9 me3
DNA Meth. m5C:H3K9 me3
H2A.Z:H4K20 me2
H3K4 me1:H4−4ac
H4K20 me2:H3−5ac
H2A.Z:H3K4 me3
H4K20 me2:H4−4ac
H3K27 me3:DNA Meth. m5C
H4K20 me2:H3K4 me3
H4−4ac:H3−5ac
H3K4 me3:H4−4ac
H3K9 me2
H3K9 me3
H4K20 me2
H3K27 me3
H2A.Z
H3K4 me1
H4K20 me3
H3K4 me3
DNA Meth. m5C
H4−4ac
H3−5ac
0.00 0.25 0.50 0.75 1.00
π̂
ZNF281 − Reverse
H2A.Z:H4−4ac
H2A.Z:H3−5ac
H3K9 me2:DNA Meth. m5C
H3K27 me3:H3K4 me1
H3K27 me3:H4K20 me3
H3K4 me3:H3−5ac
H3K4 me1:H3−5ac
H3K9 me3:H4K20 me3
DNA Meth. m5C:H4K20 me3
H3K4 me1:H4K20 me2
H3K4 me1:H2A.Z
DNA Meth. m5C:H4K20 me2
H3K27 me3:H3K9 me3
DNA Meth. m5C:H3K9 me3
H2A.Z:H4K20 me2
H3K4 me1:H4−4ac
H4K20 me2:H3−5ac
H2A.Z:H3K4 me3
H4K20 me2:H4−4ac
H3K27 me3:DNA Meth. m5C
H4K20 me2:H3K4 me3
H4−4ac:H3−5ac
H3K4 me3:H4−4ac
H3K9 me2
H3K9 me3
H4K20 me2
H3K27 me3
H2A.Z
H3K4 me1
H4K20 me3
H3K4 me3
DNA Meth. m5C
H4−4ac
H3−5ac
0.00 0.25 0.50 0.75 1.00
π̂
HSPA9 − Forward
H3K4 me1:H3−5ac
H3K4 me1:H2A.Z
H3K27 me3:H3K4 me1
H3K4 me3:H3−5ac
H3K4 me1:H4K20 me2
H2A.Z:H3−5ac
DNA Meth. m5C:H4K20 me2
H3K27 me3:DNA Meth. m5C
H2A.Z:H4−4ac
H4−4ac:H3−5ac
H3K4 me1:H4−4ac
DNA Meth. m5C:H4K20 me3
DNA Meth. m5C:H3K9 me3
H2A.Z:H3K4 me3
H3K9 me3:H4K20 me3
H3K27 me3:H4K20 me3
H4K20 me2:H3K4 me3
H3K4 me3:H4−4ac
H4K20 me2:H4−4ac
H2A.Z:H4K20 me2
H4K20 me2:H3−5ac
H3K27 me2
H3K4 me1
H3K27 me3
H4K20 me3
H3K4 me3
H3K9 me3
DNA Meth. m5C
H4K20 me2
H2A.Z
H3−5ac
H4−4ac
0.00 0.25 0.50 0.75 1.00
π̂
HSPA9 − Reverse
H3K4 me1:H3−5ac
H3K4 me1:H2A.Z
H3K27 me3:H3K4 me1
H3K4 me3:H3−5ac
H3K4 me1:H4K20 me2
H2A.Z:H3−5ac
DNA Meth. m5C:H4K20 me2
H3K27 me3:DNA Meth. m5C
H2A.Z:H4−4ac
H4−4ac:H3−5ac
H3K4 me1:H4−4ac
DNA Meth. m5C:H4K20 me3
DNA Meth. m5C:H3K9 me3
H2A.Z:H3K4 me3
H3K9 me3:H4K20 me3
H3K27 me3:H4K20 me3
H4K20 me2:H3K4 me3
H3K4 me3:H4−4ac
H4K20 me2:H4−4ac
H2A.Z:H4K20 me2
H4K20 me2:H3−5ac
H3K27 me2
H3K4 me1
H3K27 me3
H4K20 me3
H3K4 me3
H3K9 me3
DNA Meth. m5C
H4K20 me2
H2A.Z
H3−5ac
H4−4ac
0.00 0.25 0.50 0.75 1.00
π̂
HSPA6 − Forward
H3K4 me1:H4K20 me2
H3K4 me1:H2A.Z
H3K27 me3:H3K4 me1
H3K27 me3:H4K20 me3
H3K9 me3:H4K20 me3
H2A.Z:H4−4ac
H2A.Z:H3K4 me3
DNA Meth. m5C:H4K20 me2
H2A.Z:H3−5ac
H3K4 me1:H4−4ac
H3K4 me3:H3−5ac
DNA Meth. m5C:H3K9 me3
H2A.Z:H4K20 me2
H3K27 me3:DNA Meth. m5C
H4K20 me2:H3K4 me3
H3K4 me1:H3−5ac
H4−4ac:H3−5ac
H3K27 me3:H3K9 me3
H4K20 me2:H3−5ac
H4K20 me2:H4−4ac
H3K4 me3:H4−4ac
H4K20 me3
H3K9 me3
H3K4 me1
H3K27 me3
H2A.Z
H3K4 me3
H4K20 me2
DNA Meth. m5C
H3−5ac
H4−4ac
0.00 0.25 0.50 0.75 1.00
π̂
HSPA6 − Reverse
H3K4 me1:H4K20 me2
H3K4 me1:H2A.Z
H3K27 me3:H3K4 me1
H3K27 me3:H4K20 me3
H3K9 me3:H4K20 me3
H2A.Z:H4−4ac
H2A.Z:H3K4 me3
DNA Meth. m5C:H4K20 me2
H2A.Z:H3−5ac
H3K4 me1:H4−4ac
H3K4 me3:H3−5ac
DNA Meth. m5C:H3K9 me3
H2A.Z:H4K20 me2
H3K27 me3:DNA Meth. m5C
H4K20 me2:H3K4 me3
H3K4 me1:H3−5ac
H4−4ac:H3−5ac
H3K27 me3:H3K9 me3
H4K20 me2:H3−5ac
H4K20 me2:H4−4ac
H3K4 me3:H4−4ac
H4K20 me3
H3K9 me3
H3K4 me1
H3K27 me3
H2A.Z
H3K4 me3
H4K20 me2
DNA Meth. m5C
H3−5ac
H4−4ac
0.00 0.25 0.50 0.75 1.00
π̂
HSPA8 − Forward
DNA Meth. m5C:H4K20 me3
H3K4 me3:H3−5ac
H3K9 me3:H4K20 me3
DNA Meth. m5C:H4K20 me2
H2A.Z:H3−5ac
H2A.Z:H4−4ac
H3K4 me1:H4K20 me2
H3K4 me1:H2A.Z
DNA Meth. m5C:H3K9 me3
H3K27 me3:H4K20 me3
H3K9 me2:DNA Meth. m5C
H3K4 me1:H4−4ac
H3K27 me3:H3K4 me1
H2A.Z:H3K4 me3
H2A.Z:H4K20 me2
H3K27 me3:DNA Meth. m5C
H4K20 me2:H3K4 me3
H3K4 me1:H3−5ac
H4K20 me2:H3−5ac
H4K20 me2:H4−4ac
H3K27 me3:H3K9 me3
H4−4ac:H3−5ac
H3K4 me3:H4−4ac
H4K20 me3
H3K9 me2
H3K9 me3
H2A.Z
H3K4 me1
H3K4 me3
H3K27 me3
H4K20 me2
H3−5ac
H4−4ac
DNA Meth. m5C
0.00 0.25 0.50 0.75 1.00
π̂
HSPA8 − Reverse
DNA Meth. m5C:H4K20 me3
H3K4 me3:H3−5ac
H3K9 me3:H4K20 me3
DNA Meth. m5C:H4K20 me2
H2A.Z:H3−5ac
H2A.Z:H4−4ac
H3K4 me1:H4K20 me2
H3K4 me1:H2A.Z
DNA Meth. m5C:H3K9 me3
H3K27 me3:H4K20 me3
H3K9 me2:DNA Meth. m5C
H3K4 me1:H4−4ac
H3K27 me3:H3K4 me1
H2A.Z:H3K4 me3
H2A.Z:H4K20 me2
H3K27 me3:DNA Meth. m5C
H4K20 me2:H3K4 me3
H3K4 me1:H3−5ac
H4K20 me2:H3−5ac
H4K20 me2:H4−4ac
H3K27 me3:H3K9 me3
H4−4ac:H3−5ac
H3K4 me3:H4−4ac
H4K20 me3
H3K9 me2
H3K9 me3
H2A.Z
H3K4 me1
H3K4 me3
H3K27 me3
H4K20 me2
H3−5ac
H4−4ac
DNA Meth. m5C
0.00 0.25 0.50 0.75 1.00
π̂
HSPA1A/HSPA1B − Forward
H3K4 me1:H4K20 me2
H3K9 me3:H4K20 me3
H3K27 me3:H4K20 me3
H3K4 me1:H2A.Z
H2A.Z:H4K20 me2
H3K4 me1:H4−4ac
H3K27 me3:H3K4 me1
H3K4 me3:H3−5ac
H2A.Z:H3K4 me3
H4K20 me2:H4−4ac
H4K20 me2:H3K4 me3
DNA Meth. m5C:H3K9 me3
H4K20 me2:H3−5ac
H3K27 me3:DNA Meth. m5C
DNA Meth. m5C:H4K20 me2
H3K4 me1:H3−5ac
H3K27 me3:H3K9 me3
H3K9 me2
H3K4 me3:H4−4ac
H4−4ac:H3−5ac
H4K20 me3
H2A.Z
H3K27 me3
H3K4 me1
H3K9 me3
H4K20 me2
H3K4 me3
DNA Meth. m5C
H3−5ac
H4−4ac
0.00 0.25 0.50 0.75 1.00
π̂
HSPA1A/HSPA1B − Reverse
H3K4 me1:H4K20 me2
H3K9 me3:H4K20 me3
H3K27 me3:H4K20 me3
H3K4 me1:H2A.Z
H2A.Z:H4K20 me2
H3K4 me1:H4−4ac
H3K27 me3:H3K4 me1
H3K4 me3:H3−5ac
H2A.Z:H3K4 me3
H4K20 me2:H4−4ac
H4K20 me2:H3K4 me3
DNA Meth. m5C:H3K9 me3
H4K20 me2:H3−5ac
H3K27 me3:DNA Meth. m5C
DNA Meth. m5C:H4K20 me2
H3K4 me1:H3−5ac
H3K27 me3:H3K9 me3
H3K9 me2
H3K4 me3:H4−4ac
H4−4ac:H3−5ac
H4K20 me3
H2A.Z
H3K27 me3
H3K4 me1
H3K9 me3
H4K20 me2
H3K4 me3
DNA Meth. m5C
H3−5ac
H4−4ac
0.00 0.25 0.50 0.75 1.00
π̂

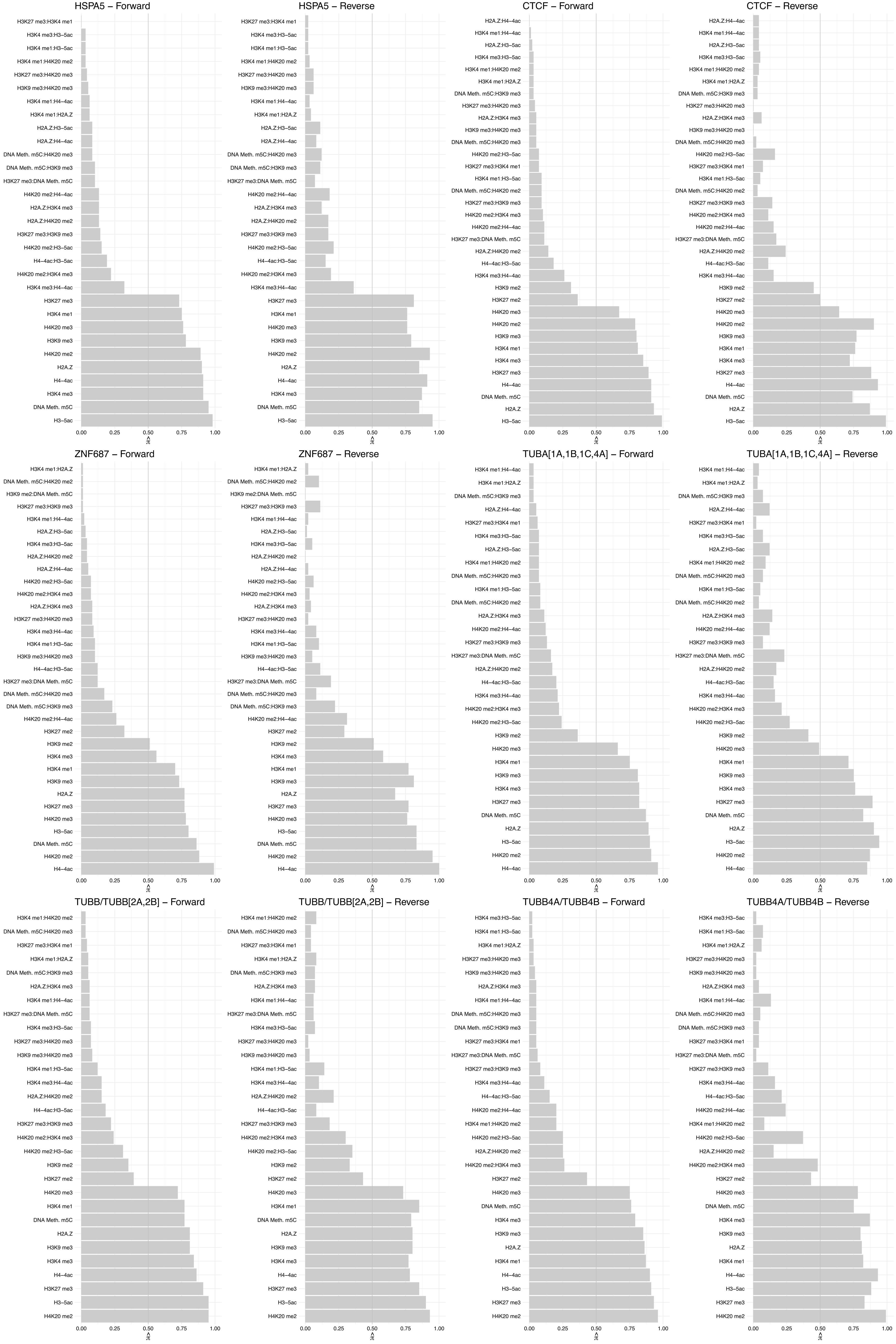
HSPA5 – Forward
H3K27 me3:H3K4 me1
H3K4 me3:H3−5ac
H3K4 me1:H3−5ac
H3K4 me1:H4K20 me2
H3K27 me3:H4K20 me3
H3K9 me3:H4K20 me3
H3K4 me1:H4−4ac
H3K4 me1:H2A.Z
H2A.Z:H3−5ac
H2A.Z:H4−4ac
DNA Meth. m5C:H4K20 me3
DNA Meth. m5C:H3K9 me3
H3K27 me3:DNA Meth. m5C
H4K20 me2:H4−4ac
H2A.Z:H3K4 me3
H2A.Z:H4K20 me2
H3K27 me3:H3K9 me3
H4K20 me2:H3−5ac
H4−4ac:H3−5ac
H4K20 me2:H3K4 me3
H3K4 me3:H4−4ac
H3K27 me3
H3K4 me1
H4K20 me3
H3K9 me3
H4K20 me2
H2A.Z
H4−4ac
H3K4 me3
DNA Meth. m5C
H3−5ac
0.00
0.25
0.50
0.75
1.00
π̂
HSPA5 – Reverse
H3K27 me3:H3K4 me1
H3K4 me3:H3−5ac
H3K4 me1:H3−5ac
H3K4 me1:H4K20 me2
H3K27 me3:H4K20 me3
H3K9 me3:H4K20 me3
H3K4 me1:H4−4ac
H3K4 me1:H2A.Z
H2A.Z:H3−5ac
H2A.Z:H4−4ac
DNA Meth. m5C:H4K20 me3
DNA Meth. m5C:H3K9 me3
H3K27 me3:DNA Meth. m5C
H4K20 me2:H4−4ac
H2A.Z:H3K4 me3
H2A.Z:H4K20 me2
H3K27 me3:H3K9 me3
H4K20 me2:H3−5ac
H4−4ac:H3−5ac
H4K20 me2:H3K4 me3
H3K4 me3:H4−4ac
H3K27 me3
H3K4 me1
H4K20 me3
H3K9 me3
H4K20 me2
H2A.Z
H4−4ac
H3K4 me3
DNA Meth. m5C
H3−5ac
0.00
0.25
0.50
0.75
1.00
π̂
CTCF – Forward
H2A.Z:H4−4ac
H3K4 me1:H4−4ac
H2A.Z:H3−5ac
H3K4 me3:H3−5ac
H3K4 me1:H4K20 me2
H3K4 me1:H2A.Z
DNA Meth. m5C:H3K9 me3
H3K27 me3:H4K20 me3
H2A.Z:H3K4 me3
H3K9 me3:H4K20 me3
DNA Meth. m5C:H4K20 me3
H4K20 me2:H3−5ac
H3K27 me3:H3K4 me1
H3K4 me1:H3−5ac
DNA Meth. m5C:H4K20 me2
H3K27 me3:H3K9 me3
H4K20 me2:H3K4 me3
H4K20 me2:H4−4ac
H3K27 me3:DNA Meth. m5C
H2A.Z:H4K20 me2
H4−4ac:H3−5ac
H3K4 me3:H4−4ac
H3K9 me2
H3K27 me2
H4K20 me3
H4K20 me2
H3K9 me3
H3K4 me1
H3K4 me3
H3K27 me3
H4−4ac
DNA Meth. m5C
H2A.Z
H3−5ac
0.00
0.25
0.50
0.75
1.00
π̂
CTCF – Reverse
H2A.Z:H4−4ac
H3K4 me1:H4−4ac
H2A.Z:H3−5ac
H3K4 me3:H3−5ac
H3K4 me1:H4K20 me2
H3K4 me1:H2A.Z
DNA Meth. m5C:H3K9 me3
H3K27 me3:H4K20 me3
H2A.Z:H3K4 me3
H3K9 me3:H4K20 me3
DNA Meth. m5C:H4K20 me3
H4K20 me2:H3−5ac
H3K27 me3:H3K4 me1
H3K4 me1:H3−5ac
DNA Meth. m5C:H4K20 me2
H3K27 me3:H3K9 me3
H4K20 me2:H3K4 me3
H4K20 me2:H4−4ac
H3K27 me3:DNA Meth. m5C
H2A.Z:H4K20 me2
H4−4ac:H3−5ac
H3K4 me3:H4−4ac
H3K9 me2
H3K27 me2
H4K20 me3
H4K20 me2
H3K9 me3
H3K4 me1
H3K4 me3
H3K27 me3
H4−4ac
DNA Meth. m5C
H2A.Z
H3−5ac
0.00
0.25
0.50
0.75
1.00
π̂
ZNF687 – Forward
H3K4 me1:H2A.Z
DNA Meth. m5C:H4K20 me2
H3K9 me2:DNA Meth. m5C
H3K27 me3:H3K9 me3
H3K4 me1:H4−4ac
H2A.Z:H3−5ac
H3K4 me3:H3−5ac
H2A.Z:H4K20 me2
H2A.Z:H4−4ac
H4K20 me2:H3−5ac
H4K20 me2:H3K4 me3
H2A.Z:H3K4 me3
H3K27 me3:H4K20 me3
H3K4 me3:H4−4ac
H3K4 me1:H3−5ac
H3K9 me3:H4K20 me3
H4−4ac:H3−5ac
H3K27 me3:DNA Meth. m5C
DNA Meth. m5C:H4K20 me3
DNA Meth. m5C:H3K9 me3
H4K20 me2:H4−4ac
H3K27 me2
H3K9 me2
H3K4 me3
H3K4 me1
H3K9 me3
H2A.Z
H3K27 me3
H4K20 me3
H3−5ac
DNA Meth. m5C
H4K20 me2
H4−4ac
0.00
0.25
0.50
0.75
1.00
π̂
ZNF687 – Reverse
H3K4 me1:H2A.Z
DNA Meth. m5C:H4K20 me2
H3K9 me2:DNA Meth. m5C
H3K27 me3:H3K9 me3
H3K4 me1:H4−4ac
H2A.Z:H3−5ac
H3K4 me3:H3−5ac
H2A.Z:H4K20 me2
H2A.Z:H4−4ac
H4K20 me2:H3−5ac
H4K20 me2:H3K4 me3
H2A.Z:H3K4 me3
H3K27 me3:H4K20 me3
H3K4 me3:H4−4ac
H3K4 me1:H3−5ac
H3K9 me3:H4K20 me3
H4−4ac:H3−5ac
H3K27 me3:DNA Meth. m5C
DNA Meth. m5C:H4K20 me3
DNA Meth. m5C:H3K9 me3
H4K20 me2:H4−4ac
H3K27 me2
H3K9 me2
H3K4 me3
H3K4 me1
H3K9 me3
H2A.Z
H3K27 me3
H4K20 me3
H3−5ac
DNA Meth. m5C
H4K20 me2
H4−4ac
0.00
0.25
0.50
0.75
1.00
π̂
TUBA[1A,1B,1C,4A] – Forward
H3K4 me1:H4−4ac
H3K4 me1:H2A.Z
DNA Meth. m5C:H3K9 me3
H2A.Z:H4−4ac
H3K27 me3:H3K4 me1
H3K4 me3:H3−5ac
H2A.Z:H3−5ac
H3K4 me1:H4K20 me2
DNA Meth. m5C:H4K20 me3
H3K4 me1:H3−5ac
DNA Meth. m5C:H4K20 me2
H2A.Z:H3K4 me3
H4K20 me2:H4−4ac
H3K27 me3:H3K9 me3
H3K27 me3:DNA Meth. m5C
H2A.Z:H4K20 me2
H4−4ac:H3−5ac
H3K4 me3:H4−4ac
H4K20 me2:H3K4 me3
H4K20 me2:H3−5ac
H3K9 me2
H4K20 me3
H3K4 me1
H3K9 me3
H3K4 me3
H3K27 me3
DNA Meth. m5C
H2A.Z
H3−5ac
H4K20 me2
H4−4ac
0.00
0.25
0.50
0.75
1.00
π̂
TUBA[1A,1B,1C,4A] – Reverse
H3K4 me1:H4−4ac
H3K4 me1:H2A.Z
DNA Meth. m5C:H3K9 me3
H2A.Z:H4−4ac
H3K27 me3:H3K4 me1
H3K4 me3:H3−5ac
H2A.Z:H3−5ac
H3K4 me1:H4K20 me2
DNA Meth. m5C:H4K20 me3
H3K4 me1:H3−5ac
DNA Meth. m5C:H4K20 me2
H2A.Z:H3K4 me3
H4K20 me2:H4−4ac
H3K27 me3:H3K9 me3
H3K27 me3:DNA Meth. m5C
H2A.Z:H4K20 me2
H4−4ac:H3−5ac
H3K4 me3:H4−4ac
H4K20 me2:H3K4 me3
H4K20 me2:H3−5ac
H3K9 me2
H4K20 me3
H3K4 me1
H3K9 me3
H3K4 me3
H3K27 me3
DNA Meth. m5C
H2A.Z
H3−5ac
H4K20 me2
H4−4ac
0.00
0.25
0.50
0.75
1.00
π̂
TUBB/TUBB[2A,2B] – Forward
H3K4 me1:H4K20 me2
DNA Meth. m5C:H4K20 me3
H3K27 me3:H3K4 me1
H3K4 me1:H2A.Z
DNA Meth. m5C:H3K9 me3
H2A.Z:H3K4 me3
H3K4 me1:H4−4ac
H3K27 me3:DNA Meth. m5C
H3K4 me3:H3−5ac
H3K27 me3:H4K20 me3
H3K9 me3:H4K20 me3
H3K4 me1:H3−5ac
H3K4 me3:H4−4ac
H2A.Z:H4K20 me2
H4−4ac:H3−5ac
H3K27 me3:H3K9 me3
H4K20 me2:H3K4 me3
H4K20 me2:H3−5ac
H3K9 me2
H3K27 me2
H4K20 me3
H3K4 me1
DNA Meth. m5C
H2A.Z
H3K9 me3
H3K4 me3
H4−4ac
H3K27 me3
H3−5ac
H4K20 me2
0.00
0.25
0.50
0.75
1.00
π̂
TUBB/TUBB[2A,2B] – Reverse
H3K4 me1:H4K20 me2
DNA Meth. m5C:H4K20 me3
H3K27 me3:H3K4 me1
H3K4 me1:H2A.Z
DNA Meth. m5C:H3K9 me3
H2A.Z:H3K4 me3
H3K4 me1:H4−4ac
H3K27 me3:DNA Meth. m5C
H3K4 me3:H3−5ac
H3K27 me3:H4K20 me3
H3K9 me3:H4K20 me3
H3K4 me1:H3−5ac
H3K4 me3:H4−4ac
H2A.Z:H4K20 me2
H4−4ac:H3−5ac
H3K27 me3:H3K9 me3
H4K20 me2:H3K4 me3
H4K20 me2:H3−5ac
H3K9 me2
H3K27 me2
H4K20 me3
H3K4 me1
DNA Meth. m5C
H2A.Z
H3K9 me3
H3K4 me3
H4−4ac
H3K27 me3
H3−5ac
H4K20 me2
0.00
0.25
0.50
0.75
1.00
π̂
TUBB4A/TUBB4B – Forward
H3K4 me3:H3−5ac
H3K4 me1:H3−5ac
H3K4 me1:H2A.Z
H3K27 me3:H4K20 me3
H3K9 me3:H4K20 me3
H2A.Z:H3K4 me3
H3K4 me1:H4−4ac
DNA Meth. m5C:H4K20 me3
DNA Meth. m5C:H3K9 me3
H3K27 me3:H3K4 me1
H3K27 me3:DNA Meth. m5C
H3K27 me3:H3K9 me3
H3K4 me3:H4−4ac
H4−4ac:H3−5ac
H4K20 me2:H4−4ac
H3K4 me1:H4K20 me2
H4K20 me2:H3−5ac
H2A.Z:H4K20 me2
H4K20 me2:H3K4 me3
H3K27 me2
H4K20 me3
DNA Meth. m5C
H3K4 me3
H3K9 me3
H2A.Z
H3K4 me1
H4−4ac
H3−5ac
H3K27 me3
H4K20 me2
0.00
0.25
0.50
0.75
1.00
π̂
TUBB4A/TUBB4B – Reverse
H3K4 me3:H3−5ac
H3K4 me1:H3−5ac
H3K4 me1:H2A.Z
H3K27 me3:H4K20 me3
H3K9 me3:H4K20 me3
H2A.Z:H3K4 me3
H3K4 me1:H4−4ac
DNA Meth. m5C:H4K20 me3
DNA Meth. m5C:H3K9 me3
H3K27 me3:H3K4 me1
H3K27 me3:DNA Meth. m5C
H3K27 me3:H3K9 me3
H3K4 me3:H4−4ac
H4−4ac:H3−5ac
H4K20 me2:H4−4ac
H3K4 me1:H4K20 me2
H4K20 me2:H3−5ac
H2A.Z:H4K20 me2
H4K20 me2:H3K4 me3
H3K27 me2
H4K20 me3
DNA Meth. m5C
H3K4 me3
H3K9 me3
H2A.Z
H3K4 me1
H4−4ac
H3−5ac
H3K27 me3
H4K20 me2
0.00
0.25
0.50
0.75
1.00
π̂

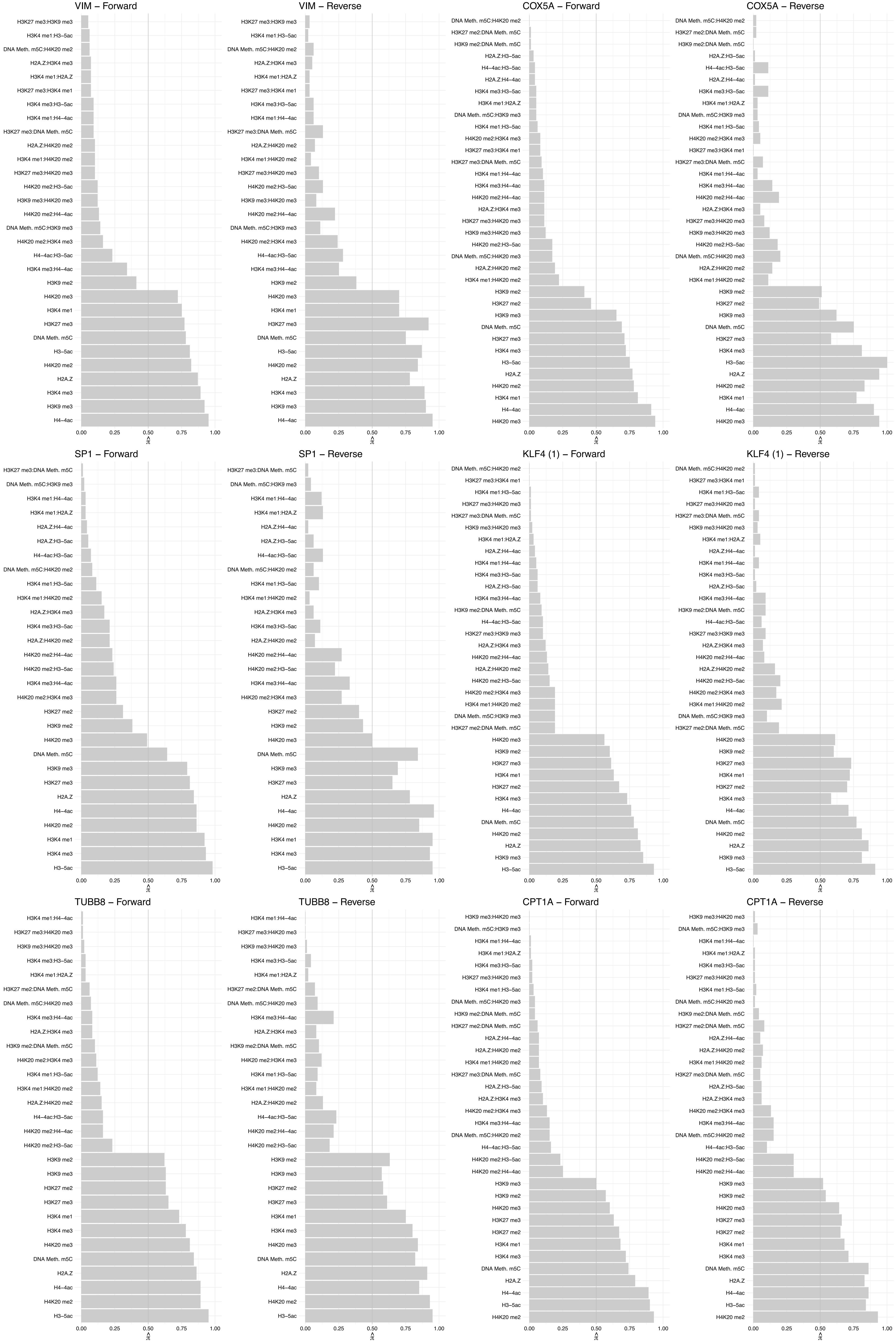
VIM – Forward
H3K27 me3:H3K9 me3
H3K4 me1:H3–5ac
DNA Meth. m5C:H4K20 me2
H2A.Z:H3K4 me3
H3K4 me1:H2A.Z
H3K27 me3:H3K4 me1
H3K4 me3:H3–5ac
H3K4 me1:H4–4ac
H3K27 me3:DNA Meth. m5C
H2A.Z:H4K20 me2
H3K4 me1:H4K20 me2
H3K27 me3:H4K20 me3
H4K20 me2:H3–5ac
H3K9 me3:H4K20 me3
H4K20 me2:H4–4ac
DNA Meth. m5C:H3K9 me3
H4K20 me2:H3K4 me3
H4–4ac:H3–5ac
H3K4 me3:H4–4ac
H3K9 me2
H4K20 me3
H3K4 me1
H3K27 me3
DNA Meth. m5C
H3–5ac
H4K20 me2
H2A.Z
H3K4 me3
H3K9 me3
H4–4ac
VIM – Reverse
H3K27 me3:H3K9 me3
H3K4 me1:H3–5ac
DNA Meth. m5C:H4K20 me2
H2A.Z:H3K4 me3
H3K4 me1:H2A.Z
H3K27 me3:H3K4 me1
H3K4 me3:H3–5ac
H3K4 me1:H4–4ac
H3K27 me3:DNA Meth. m5C
H2A.Z:H4K20 me2
H3K4 me1:H4K20 me2
H3K27 me3:H4K20 me3
H4K20 me2:H3–5ac
H3K9 me3:H4K20 me3
H4K20 me2:H4–4ac
DNA Meth. m5C:H3K9 me3
H4K20 me2:H3K4 me3
H4–4ac:H3–5ac
H3K4 me3:H4–4ac
H3K9 me2
H4K20 me3
H3K4 me1
H3K27 me3
DNA Meth. m5C
H3–5ac
H4K20 me2
H2A.Z
H3K4 me3
H3K9 me3
H4–4ac
COX5A – Forward
DNA Meth. m5C:H4K20 me2
H3K27 me2:DNA Meth. m5C
H3K9 me2:DNA Meth. m5C
H2A.Z:H3–5ac
H4–4ac:H3–5ac
H2A.Z:H4–4ac
H3K4 me3:H3–5ac
H3K4 me1:H2A.Z
DNA Meth. m5C:H3K9 me3
H3K4 me1:H3–5ac
H4K20 me2:H3K4 me3
H3K27 me3:H3K4 me1
H3K27 me3:DNA Meth. m5C
H3K4 me1:H4–4ac
H3K4 me3:H4–4ac
H4K20 me2:H4–4ac
H2A.Z:H3K4 me3
H3K27 me3:H4K20 me3
H3K9 me3:H4K20 me3
H4K20 me2:H3–5ac
DNA Meth. m5C:H4K20 me3
H2A.Z:H4K20 me2
H3K4 me1:H4K20 me2
H3K9 me2
H3K27 me2
H3K9 me3
DNA Meth. m5C
H3K27 me3
H3K4 me3
H3–5ac
H2A.Z
H4K20 me2
H3K4 me1
H4–4ac
H4K20 me3
COX5A – Reverse
DNA Meth. m5C:H4K20 me2
H3K27 me2:DNA Meth. m5C
H3K9 me2:DNA Meth. m5C
H2A.Z:H3–5ac
H4–4ac:H3–5ac
H2A.Z:H4–4ac
H3K4 me3:H3–5ac
H3K4 me1:H2A.Z
DNA Meth. m5C:H3K9 me3
H3K4 me1:H3–5ac
H4K20 me2:H3K4 me3
H3K27 me3:H3K4 me1
H3K27 me3:DNA Meth. m5C
H3K4 me1:H4–4ac
H3K4 me3:H4–4ac
H4K20 me2:H4–4ac
H2A.Z:H3K4 me3
H3K27 me3:H4K20 me3
H3K9 me3:H4K20 me3
H4K20 me2:H3–5ac
DNA Meth. m5C:H4K20 me3
H2A.Z:H4K20 me2
H3K4 me1:H4K20 me2
H3K9 me2
H3K27 me2
H3K9 me3
DNA Meth. m5C
H3K27 me3
H3K4 me3
H3–5ac
H2A.Z
H4K20 me2
H3K4 me1
H4–4ac
H4K20 me3
SP1 – Forward
H3K27 me3:DNA Meth. m5C
DNA Meth. m5C:H3K9 me3
H3K4 me1:H4–4ac
H3K4 me1:H2A.Z
H2A.Z:H4–4ac
H2A.Z:H3–5ac
H4–4ac:H3–5ac
DNA Meth. m5C:H4K20 me2
H3K4 me1:H3–5ac
H3K4 me1:H4K20 me2
H2A.Z:H3K4 me3
H3K4 me3:H3–5ac
H2A.Z:H4K20 me2
H4K20 me2:H4–4ac
H4K20 me2:H3–5ac
H3K4 me3:H4–4ac
H4K20 me2:H3K4 me3
H3K27 me2
H3K9 me2
H4K20 me3
DNA Meth. m5C
H3K9 me3
H3K27 me3
H2A.Z
H4–4ac
H4K20 me2
H3K4 me1
H3K4 me3
H3–5ac
SP1 – Reverse
H3K27 me3:DNA Meth. m5C
DNA Meth. m5C:H3K9 me3
H3K4 me1:H4–4ac
H3K4 me1:H2A.Z
H2A.Z:H4–4ac
H2A.Z:H3–5ac
H4–4ac:H3–5ac
DNA Meth. m5C:H4K20 me2
H3K4 me1:H3–5ac
H3K4 me1:H4K20 me2
H2A.Z:H3K4 me3
H3K4 me3:H3–5ac
H2A.Z:H4K20 me2
H4K20 me2:H4–4ac
H4K20 me2:H3–5ac
H3K4 me3:H4–4ac
H4K20 me2:H3K4 me3
H3K27 me2
H3K9 me2
H4K20 me3
DNA Meth. m5C
H3K9 me3
H3K27 me3
H2A.Z
H4–4ac
H4K20 me2
H3K4 me1
H3K4 me3
H3–5ac
KLF4 (1) – Forward
DNA Meth. m5C:H4K20 me2
H3K27 me3:H3K4 me1
H3K4 me1:H3–5ac
H3K27 me3:H4K20 me3
H3K27 me3:DNA Meth. m5C
H3K9 me3:H4K20 me3
H3K4 me1:H2A.Z
H2A.Z:H4–4ac
H3K4 me1:H4–4ac
H3K4 me3:H3–5ac
H2A.Z:H3–5ac
H3K4 me3:H4–4ac
H3K9 me2:DNA Meth. m5C
H4–4ac:H3–5ac
H3K27 me3:H3K9 me3
H2A.Z:H3K4 me3
H4K20 me2:H4–4ac
H2A.Z:H4K20 me2
H4K20 me2:H3–5ac
H4K20 me2:H3K4 me3
H3K4 me1:H4K20 me2
DNA Meth. m5C:H3K9 me3
H3K27 me2:DNA Meth. m5C
H4K20 me3
H3K9 me2
H3K27 me3
H3K4 me1
H3K27 me2
H3K4 me3
H4–4ac
DNA Meth. m5C
H4K20 me2
H2A.Z
H3K9 me3
H3–5ac
KLF4 (1) – Reverse
DNA Meth. m5C:H4K20 me2
H3K27 me3:H3K4 me1
H3K4 me1:H3–5ac
H3K27 me3:H4K20 me3
H3K27 me3:DNA Meth. m5C
H3K9 me3:H4K20 me3
H3K4 me1:H2A.Z
H2A.Z:H4–4ac
H3K4 me1:H4–4ac
H3K4 me3:H3–5ac
H2A.Z:H3–5ac
H3K4 me3:H4–4ac
H3K9 me2:DNA Meth. m5C
H4–4ac:H3–5ac
H3K27 me3:H3K9 me3
H2A.Z:H3K4 me3
H4K20 me2:H4–4ac
H2A.Z:H4K20 me2
H4K20 me2:H3–5ac
H4K20 me2:H3K4 me3
H3K4 me1:H4K20 me2
DNA Meth. m5C:H3K9 me3
H3K27 me2:DNA Meth. m5C
H4K20 me3
H3K9 me2
H3K27 me3
H3K4 me1
H3K27 me2
H3K4 me3
H4–4ac
DNA Meth. m5C
H4K20 me2
H2A.Z
H3K9 me3
H3–5ac
TUBB8 – Forward
H3K4 me1:H4–4ac
H3K27 me3:H4K20 me3
H3K9 me3:H4K20 me3
H3K4 me3:H3–5ac
H3K4 me1:H2A.Z
H3K27 me2:DNA Meth. m5C
DNA Meth. m5C:H4K20 me3
H3K4 me3:H4–4ac
H2A.Z:H3K4 me3
H3K9 me2:DNA Meth. m5C
H4K20 me2:H3K4 me3
H3K4 me1:H3–5ac
H3K4 me1:H4K20 me2
H2A.Z:H4K20 me2
H4–4ac:H3–5ac
H4K20 me2:H4–4ac
H4K20 me2:H3–5ac
H3K9 me2
H3K9 me3
H3K27 me2
H3K27 me3
H3K4 me1
H3K4 me3
H4K20 me3
DNA Meth. m5C
H2A.Z
H4–4ac
H4K20 me2
H3–5ac
TUBB8 – Reverse
H3K4 me1:H4–4ac
H3K27 me3:H4K20 me3
H3K9 me3:H4K20 me3
H3K4 me3:H3–5ac
H3K4 me1:H2A.Z
H3K27 me2:DNA Meth. m5C
DNA Meth. m5C:H4K20 me3
H3K4 me3:H4–4ac
H2A.Z:H3K4 me3
H3K9 me2:DNA Meth. m5C
H4K20 me2:H3K4 me3
H3K4 me1:H3–5ac
H3K4 me1:H4K20 me2
H2A.Z:H4K20 me2
H4–4ac:H3–5ac
H4K20 me2:H4–4ac
H4K20 me2:H3–5ac
H3K9 me2
H3K9 me3
H3K27 me2
H3K27 me3
H3K4 me1
H3K4 me3
H4K20 me3
DNA Meth. m5C
H2A.Z
H4–4ac
H4K20 me2
H3–5ac
CPT1A – Forward
H3K9 me3:H4K20 me3
DNA Meth. m5C:H3K9 me3
H3K4 me1:H4–4ac
H3K4 me1:H2A.Z
H3K4 me3:H3–5ac
H3K27 me3:H4K20 me3
H3K4 me1:H3–5ac
DNA Meth. m5C:H4K20 me3
H3K9 me2:DNA Meth. m5C
H3K27 me2:DNA Meth. m5C
H2A.Z:H4–4ac
H2A.Z:H4K20 me2
H3K4 me1:H4K20 me2
H3K27 me3:DNA Meth. m5C
H2A.Z:H3–5ac
H2A.Z:H3K4 me3
H4K20 me2:H3K4 me3
H3K4 me3:H4–4ac
DNA Meth. m5C:H4K20 me2
H4–4ac:H3–5ac
H4K20 me2:H3–5ac
H4K20 me2:H4–4ac
H3K9 me3
H3K9 me2
H4K20 me3
H3K27 me3
H3K27 me2
H3K4 me1
H3K4 me3
DNA Meth. m5C
H2A.Z
H4–4ac
H3–5ac
H4K20 me2
CPT1A – Reverse
H3K9 me3:H4K20 me3
DNA Meth. m5C:H3K9 me3
H3K4 me1:H4–4ac
H3K4 me1:H2A.Z
H3K4 me3:H3–5ac
H3K27 me3:H4K20 me3
H3K4 me1:H3–5ac
DNA Meth. m5C:H4K20 me3
H3K9 me2:DNA Meth. m5C
H3K27 me2:DNA Meth. m5C
H2A.Z:H4–4ac
H2A.Z:H4K20 me2
H3K4 me1:H4K20 me2
H3K27 me3:DNA Meth. m5C
H2A.Z:H3–5ac
H2A.Z:H3K4 me3
H4K20 me2:H3K4 me3
H3K4 me3:H4–4ac
DNA Meth. m5C:H4K20 me2
H4–4ac:H3–5ac
H4K20 me2:H3–5ac
H4K20 me2:H4–4ac
H3K9 me3
H3K9 me2
H4K20 me3
H3K27 me3
H3K27 me2
H3K4 me1
H3K4 me3
DNA Meth. m5C
H2A.Z
H4–4ac
H3–5ac
H4K20 me2
0.00
0.25
0.50
0.75
1.00
$\hat{\pi}$

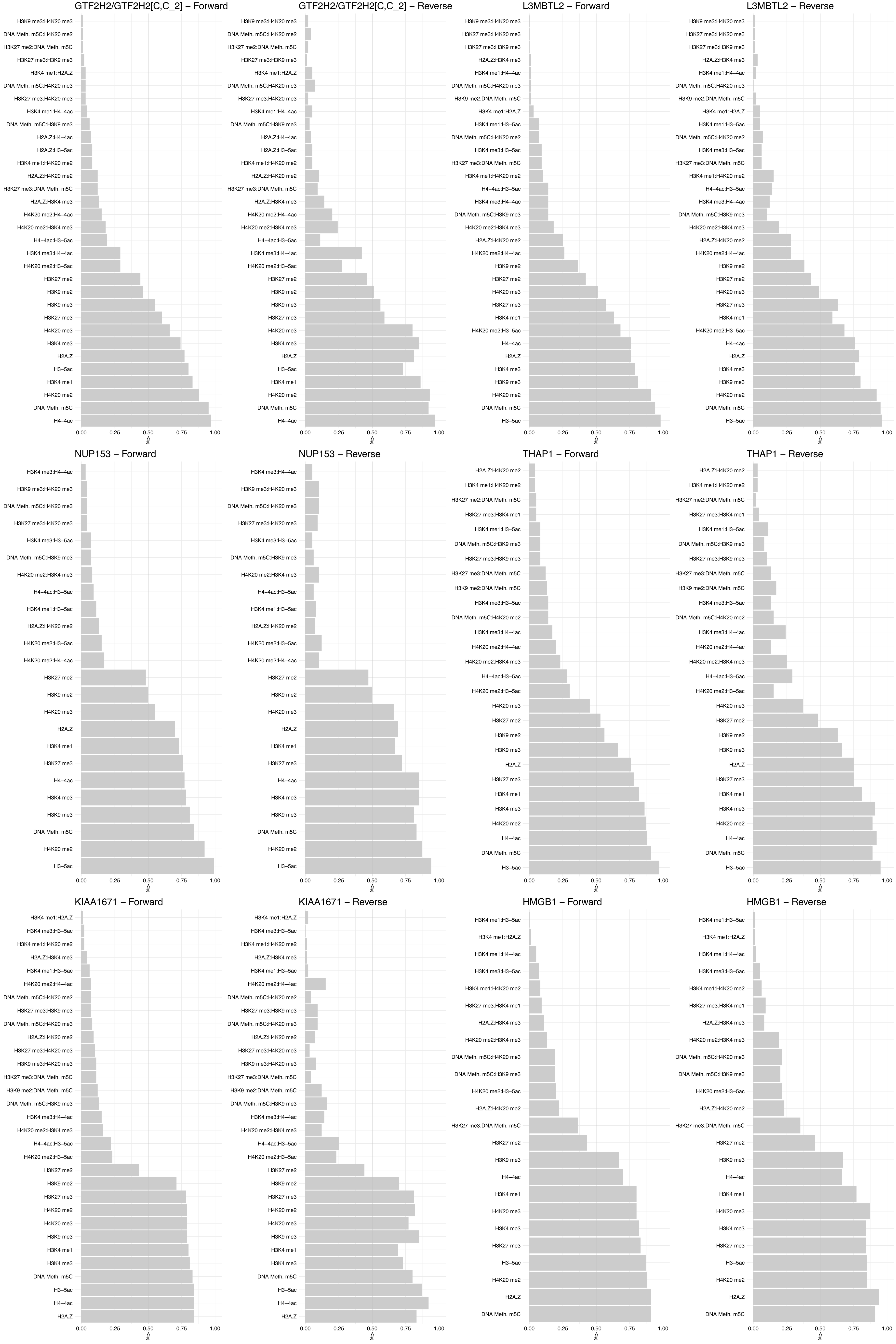
GTF2H2/GTF2H2[C,C_2] – Forward
H3K9 me3:H4K20 me3
DNA Meth. m5C:H4K20 me2
H3K27 me2:DNA Meth. m5C
H3K27 me3:H3K9 me3
H3K4 me1:H2A.Z
DNA Meth. m5C:H4K20 me3
H3K27 me3:H4K20 me3
H3K4 me1:H4−4ac
DNA Meth. m5C:H3K9 me3
H2A.Z:H4−4ac
H2A.Z:H3−5ac
H3K4 me1:H4K20 me2
H2A.Z:H4K20 me2
H3K27 me3:DNA Meth. m5C
H2A.Z:H3K4 me3
H4K20 me2:H4−4ac
H4K20 me2:H3K4 me3
H4−4ac:H3−5ac
H3K4 me3:H4−4ac
H4K20 me2:H3−5ac
H3K27 me2
H3K9 me2
H3K9 me3
H3K27 me3
H4K20 me3
H3K4 me3
H2A.Z
H3−5ac
H3K4 me1
H4K20 me2
DNA Meth. m5C
H4−4ac
0.00 0.25 0.50 0.75 1.00
π̂
GTF2H2/GTF2H2[C,C_2] – Reverse
H3K9 me3:H4K20 me3
DNA Meth. m5C:H4K20 me2
H3K27 me2:DNA Meth. m5C
H3K27 me3:H3K9 me3
H3K4 me1:H2A.Z
DNA Meth. m5C:H4K20 me3
H3K27 me3:H4K20 me3
H3K4 me1:H4−4ac
DNA Meth. m5C:H3K9 me3
H2A.Z:H4−4ac
H2A.Z:H3−5ac
H3K4 me1:H4K20 me2
H2A.Z:H4K20 me2
H3K27 me3:DNA Meth. m5C
H2A.Z:H3K4 me3
H4K20 me2:H4−4ac
H4K20 me2:H3K4 me3
H4−4ac:H3−5ac
H3K4 me3:H4−4ac
H4K20 me2:H3−5ac
H3K27 me2
H3K9 me2
H3K9 me3
H3K27 me3
H4K20 me3
H3K4 me3
H2A.Z
H3−5ac
H3K4 me1
H4K20 me2
DNA Meth. m5C
H4−4ac
0.00 0.25 0.50 0.75 1.00
π̂
L3MBTL2 – Forward
H3K9 me3:H4K20 me3
H3K27 me3:H4K20 me3
H3K27 me3:H3K9 me3
H2A.Z:H3K4 me3
H3K4 me1:H4−4ac
DNA Meth. m5C:H4K20 me3
H3K9 me2:DNA Meth. m5C
H3K4 me1:H2A.Z
H3K4 me1:H3−5ac
DNA Meth. m5C:H4K20 me2
H3K4 me3:H3−5ac
H3K27 me3:DNA Meth. m5C
H3K4 me1:H4K20 me2
H4−4ac:H3−5ac
H3K4 me3:H4−4ac
DNA Meth. m5C:H3K9 me3
H4K20 me2:H3K4 me3
H2A.Z:H4K20 me2
H4K20 me2:H4−4ac
H3K9 me2
H3K27 me2
H4K20 me3
H3K27 me3
H3K4 me1
H4K20 me2:H3−5ac
H4−4ac
H2A.Z
H3K4 me3
H3K9 me3
H4K20 me2
DNA Meth. m5C
H3−5ac
0.00 0.25 0.50 0.75 1.00
π̂
L3MBTL2 – Reverse
H3K9 me3:H4K20 me3
H3K27 me3:H4K20 me3
H3K27 me3:H3K9 me3
H2A.Z:H3K4 me3
H3K4 me1:H4−4ac
DNA Meth. m5C:H4K20 me3
H3K9 me2:DNA Meth. m5C
H3K4 me1:H2A.Z
H3K4 me1:H3−5ac
DNA Meth. m5C:H4K20 me2
H3K4 me3:H3−5ac
H3K27 me3:DNA Meth. m5C
H3K4 me1:H4K20 me2
H4−4ac:H3−5ac
H3K4 me3:H4−4ac
DNA Meth. m5C:H3K9 me3
H4K20 me2:H3K4 me3
H2A.Z:H4K20 me2
H4K20 me2:H4−4ac
H3K9 me2
H3K27 me2
H4K20 me3
H3K27 me3
H3K4 me1
H4K20 me2:H3−5ac
H4−4ac
H2A.Z
H3K4 me3
H3K9 me3
H4K20 me2
DNA Meth. m5C
H3−5ac
0.00 0.25 0.50 0.75 1.00
π̂
NUP153 – Forward
H3K4 me3:H4−4ac
H3K9 me3:H4K20 me3
DNA Meth. m5C:H4K20 me3
H3K27 me3:H4K20 me3
H3K4 me3:H3−5ac
DNA Meth. m5C:H3K9 me3
H4K20 me2:H3K4 me3
H4−4ac:H3−5ac
H3K4 me1:H3−5ac
H2A.Z:H4K20 me2
H4K20 me2:H3−5ac
H4K20 me2:H4−4ac
H3K27 me2
H3K9 me2
H4K20 me3
H2A.Z
H3K4 me1
H3K27 me3
H4−4ac
H3K4 me3
H3K9 me3
DNA Meth. m5C
H4K20 me2
H3−5ac
0.00 0.25 0.50 0.75 1.00
π̂
NUP153 – Reverse
H3K4 me3:H4−4ac
H3K9 me3:H4K20 me3
DNA Meth. m5C:H4K20 me3
H3K27 me3:H4K20 me3
H3K4 me3:H3−5ac
DNA Meth. m5C:H3K9 me3
H4K20 me2:H3K4 me3
H4−4ac:H3−5ac
H3K4 me1:H3−5ac
H2A.Z:H4K20 me2
H4K20 me2:H3−5ac
H4K20 me2:H4−4ac
H3K27 me2
H3K9 me2
H4K20 me3
H2A.Z
H3K4 me1
H3K27 me3
H4−4ac
H3K4 me3
H3K9 me3
DNA Meth. m5C
H4K20 me2
H3−5ac
0.00 0.25 0.50 0.75 1.00
π̂
THAP1 – Forward
H2A.Z:H4K20 me2
H3K4 me1:H4K20 me2
H3K27 me2:DNA Meth. m5C
H3K27 me3:H3K4 me1
H3K4 me1:H3−5ac
DNA Meth. m5C:H3K9 me3
H3K27 me3:H3K9 me3
H3K27 me3:DNA Meth. m5C
H3K9 me2:DNA Meth. m5C
H3K4 me3:H3−5ac
DNA Meth. m5C:H4K20 me2
H3K4 me3:H4−4ac
H4K20 me2:H4−4ac
H4K20 me2:H3K4 me3
H4−4ac:H3−5ac
H4K20 me2:H3−5ac
H4K20 me3
H3K27 me2
H3K9 me2
H3K9 me3
H2A.Z
H3K27 me3
H3K4 me1
H3K4 me3
H4K20 me2
H4−4ac
DNA Meth. m5C
H3−5ac
0.00 0.25 0.50 0.75 1.00
π̂
THAP1 – Reverse
H2A.Z:H4K20 me2
H3K4 me1:H4K20 me2
H3K27 me2:DNA Meth. m5C
H3K27 me3:H3K4 me1
H3K4 me1:H3−5ac
DNA Meth. m5C:H3K9 me3
H3K27 me3:H3K9 me3
H3K27 me3:DNA Meth. m5C
H3K9 me2:DNA Meth. m5C
H3K4 me3:H3−5ac
DNA Meth. m5C:H4K20 me2
H3K4 me3:H4−4ac
H4K20 me2:H4−4ac
H4K20 me2:H3K4 me3
H4−4ac:H3−5ac
H4K20 me2:H3−5ac
H4K20 me3
H3K27 me2
H3K9 me2
H3K9 me3
H2A.Z
H3K27 me3
H3K4 me1
H3K4 me3
H4K20 me2
H4−4ac
DNA Meth. m5C
H3−5ac
0.00 0.25 0.50 0.75 1.00
π̂
KIAA1671 – Forward
H3K4 me1:H2A.Z
H3K4 me3:H3−5ac
H3K4 me1:H4K20 me2
H2A.Z:H3K4 me3
H3K4 me1:H3−5ac
H4K20 me2:H4−4ac
DNA Meth. m5C:H4K20 me2
H3K27 me3:H3K9 me3
DNA Meth. m5C:H4K20 me3
H2A.Z:H4K20 me2
H3K27 me3:H4K20 me3
H3K9 me3:H4K20 me3
H3K27 me3:DNA Meth. m5C
H3K9 me2:DNA Meth. m5C
DNA Meth. m5C:H3K9 me3
H3K4 me3:H4−4ac
H4K20 me2:H3K4 me3
H4−4ac:H3−5ac
H4K20 me2:H3−5ac
H3K27 me2
H3K9 me2
H3K27 me3
H4K20 me2
H4K20 me3
H3K9 me3
H3K4 me1
H3K4 me3
DNA Meth. m5C
H3−5ac
H4−4ac
H2A.Z
0.00 0.25 0.50 0.75 1.00
π̂
KIAA1671 – Reverse
H3K4 me1:H2A.Z
H3K4 me3:H3−5ac
H3K4 me1:H4K20 me2
H2A.Z:H3K4 me3
H3K4 me1:H3−5ac
H4K20 me2:H4−4ac
DNA Meth. m5C:H4K20 me2
H3K27 me3:H3K9 me3
DNA Meth. m5C:H4K20 me3
H2A.Z:H4K20 me2
H3K27 me3:H4K20 me3
H3K9 me3:H4K20 me3
H3K27 me3:DNA Meth. m5C
H3K9 me2:DNA Meth. m5C
DNA Meth. m5C:H3K9 me3
H3K4 me3:H4−4ac
H4K20 me2:H3K4 me3
H4−4ac:H3−5ac
H4K20 me2:H3−5ac
H3K27 me2
H3K9 me2
H3K27 me3
H4K20 me2
H4K20 me3
H3K9 me3
H3K4 me1
H3K4 me3
DNA Meth. m5C
H3−5ac
H4−4ac
H2A.Z
0.00 0.25 0.50 0.75 1.00
π̂
HMGB1 – Forward
H3K4 me1:H3−5ac
H3K4 me1:H2A.Z
H3K4 me1:H4−4ac
H3K4 me3:H3−5ac
H3K4 me1:H4K20 me2
H3K27 me3:H3K4 me1
H2A.Z:H3K4 me3
H4K20 me2:H3K4 me3
DNA Meth. m5C:H4K20 me3
DNA Meth. m5C:H3K9 me3
H4K20 me2:H3−5ac
H2A.Z:H4K20 me2
H3K27 me3:DNA Meth. m5C
H3K27 me2
H3K9 me3
H4−4ac
H3K4 me1
H4K20 me3
H3K4 me3
H3K27 me3
H3−5ac
H4K20 me2
H2A.Z
DNA Meth. m5C
0.00 0.25 0.50 0.75 1.00
π̂
HMGB1 – Reverse
H3K4 me1:H3−5ac
H3K4 me1:H2A.Z
H3K4 me1:H4−4ac
H3K4 me3:H3−5ac
H3K4 me1:H4K20 me2
H3K27 me3:H3K4 me1
H2A.Z:H3K4 me3
H4K20 me2:H3K4 me3
DNA Meth. m5C:H4K20 me3
DNA Meth. m5C:H3K9 me3
H4K20 me2:H3−5ac
H2A.Z:H4K20 me2
H3K27 me3:DNA Meth. m5C
H3K27 me2
H3K9 me3
H4−4ac
H3K4 me1
H4K20 me3
H3K4 me3
H3K27 me3
H3−5ac
H4K20 me2
H2A.Z
DNA Meth. m5C
0.00 0.25 0.50 0.75 1.00
π̂

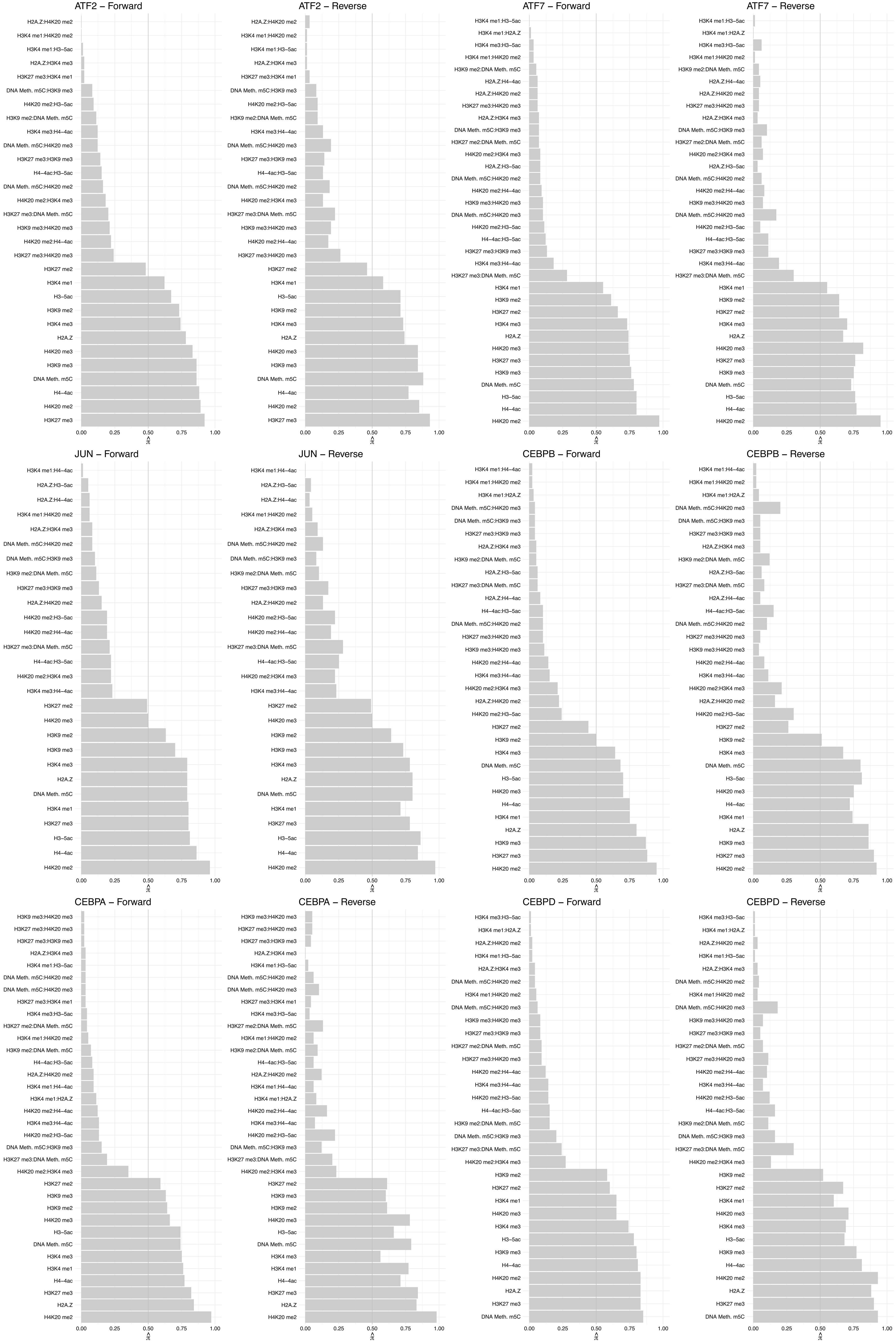
ATF2 – Forward
H2A.Z:H4K20 me2
H3K4 me1:H4K20 me2
H3K4 me1:H3–5ac
H2A.Z:H3K4 me3
H3K27 me3:H3K4 me1
DNA Meth. m5C:H3K9 me3
H4K20 me2:H3–5ac
H3K9 me2:DNA Meth. m5C
H3K4 me3:H4–4ac
DNA Meth. m5C:H4K20 me3
H3K27 me3:H3K9 me3
H4–4ac:H3–5ac
DNA Meth. m5C:H4K20 me2
H4K20 me2:H3K4 me3
H3K27 me3:DNA Meth. m5C
H3K9 me3:H4K20 me3
H4K20 me2:H4–4ac
H3K27 me3:H4K20 me3
H3K27 me2
H3K4 me1
H3–5ac
H3K9 me2
H3K4 me3
H2A.Z
H4K20 me3
H3K9 me3
DNA Meth. m5C
H4–4ac
H4K20 me2
H3K27 me3
0.00
0.25
0.50
0.75
1.00
π̂
ATF2 – Reverse
ATF7 – Forward
H3K4 me1:H3–5ac
H3K4 me1:H2A.Z
H3K4 me3:H3–5ac
H3K4 me1:H4K20 me2
H3K9 me2:DNA Meth. m5C
H2A.Z:H4–4ac
H2A.Z:H4K20 me2
H3K27 me3:H4K20 me3
H2A.Z:H3K4 me3
DNA Meth. m5C:H3K9 me3
H3K27 me2:DNA Meth. m5C
H4K20 me2:H3K4 me3
H2A.Z:H3–5ac
DNA Meth. m5C:H4K20 me2
H4K20 me2:H4–4ac
H3K9 me3:H4K20 me3
DNA Meth. m5C:H4K20 me3
H4K20 me2:H3–5ac
H4–4ac:H3–5ac
H3K27 me3:H3K9 me3
H3K4 me3:H4–4ac
H3K27 me3:DNA Meth. m5C
H3K4 me1
H3K9 me2
H3K27 me2
H3K4 me3
H2A.Z
H4K20 me3
H3K27 me3
H3K9 me3
DNA Meth. m5C
H3–5ac
H4–4ac
H4K20 me2
ATF7 – Reverse
JUN – Forward
H3K4 me1:H4–4ac
H2A.Z:H3–5ac
H2A.Z:H4–4ac
H3K4 me1:H4K20 me2
H2A.Z:H3K4 me3
DNA Meth. m5C:H4K20 me2
DNA Meth. m5C:H3K9 me3
H3K9 me2:DNA Meth. m5C
H3K27 me3:H3K9 me3
H2A.Z:H4K20 me2
H4K20 me2:H3–5ac
H4K20 me2:H4–4ac
H3K27 me3:DNA Meth. m5C
H4–4ac:H3–5ac
H4K20 me2:H3K4 me3
H3K4 me3:H4–4ac
H3K27 me2
H4K20 me3
H3K9 me2
H3K9 me3
H3K4 me3
H2A.Z
DNA Meth. m5C
H3K4 me1
H3K27 me3
H3–5ac
H4–4ac
H4K20 me2
JUN – Reverse
CEBPB – Forward
H3K4 me1:H4–4ac
H3K4 me1:H4K20 me2
H3K4 me1:H2A.Z
DNA Meth. m5C:H4K20 me3
DNA Meth. m5C:H3K9 me3
H3K27 me3:H3K9 me3
H2A.Z:H3K4 me3
H3K9 me2:DNA Meth. m5C
H2A.Z:H3–5ac
H3K27 me3:DNA Meth. m5C
H2A.Z:H4–4ac
H4–4ac:H3–5ac
DNA Meth. m5C:H4K20 me2
H3K27 me3:H4K20 me3
H3K9 me3:H4K20 me3
H4K20 me2:H4–4ac
H3K4 me3:H4–4ac
H4K20 me2:H3K4 me3
H2A.Z:H4K20 me2
H4K20 me2:H3–5ac
H3K27 me2
H3K9 me2
H3K4 me3
DNA Meth. m5C
H3–5ac
H4K20 me3
H4–4ac
H3K4 me1
H2A.Z
H3K9 me3
H3K27 me3
H4K20 me2
CEBPB – Reverse
CEBPA – Forward
H3K9 me3:H4K20 me3
H3K27 me3:H4K20 me3
H3K27 me3:H3K9 me3
H2A.Z:H3K4 me3
H3K4 me1:H3–5ac
DNA Meth. m5C:H4K20 me2
DNA Meth. m5C:H4K20 me3
H3K27 me3:H3K4 me1
H3K4 me3:H3–5ac
H3K27 me2:DNA Meth. m5C
H3K4 me1:H4K20 me2
H3K9 me2:DNA Meth. m5C
H4–4ac:H3–5ac
H2A.Z:H4K20 me2
H3K4 me1:H4–4ac
H3K4 me1:H2A.Z
H4K20 me2:H4–4ac
H3K4 me3:H4–4ac
H4K20 me2:H3–5ac
DNA Meth. m5C:H3K9 me3
H3K27 me3:DNA Meth. m5C
H4K20 me2:H3K4 me3
H3K27 me2
H3K9 me3
H3K9 me2
H4K20 me3
H3–5ac
DNA Meth. m5C
H3K4 me3
H3K4 me1
H4–4ac
H3K27 me3
H2A.Z
H4K20 me2
CEBPA – Reverse
CEBPD – Forward
H3K4 me3:H3–5ac
H3K4 me1:H2A.Z
H2A.Z:H4K20 me2
H3K4 me1:H3–5ac
H2A.Z:H3K4 me3
DNA Meth. m5C:H4K20 me2
H3K4 me1:H4K20 me2
DNA Meth. m5C:H4K20 me3
H3K9 me3:H4K20 me3
H3K27 me3:H3K9 me3
H3K27 me2:DNA Meth. m5C
H3K27 me3:H4K20 me3
H4K20 me2:H4–4ac
H3K4 me3:H4–4ac
H4K20 me2:H3–5ac
H4–4ac:H3–5ac
H3K9 me2:DNA Meth. m5C
DNA Meth. m5C:H3K9 me3
H3K27 me3:DNA Meth. m5C
H4K20 me2:H3K4 me3
H3K9 me2
H3K27 me2
H3K4 me1
H4K20 me3
H3K4 me3
H3–5ac
H3K9 me3
H4–4ac
H4K20 me2
H2A.Z
H3K27 me3
DNA Meth. m5C
CEBPD – Reverse

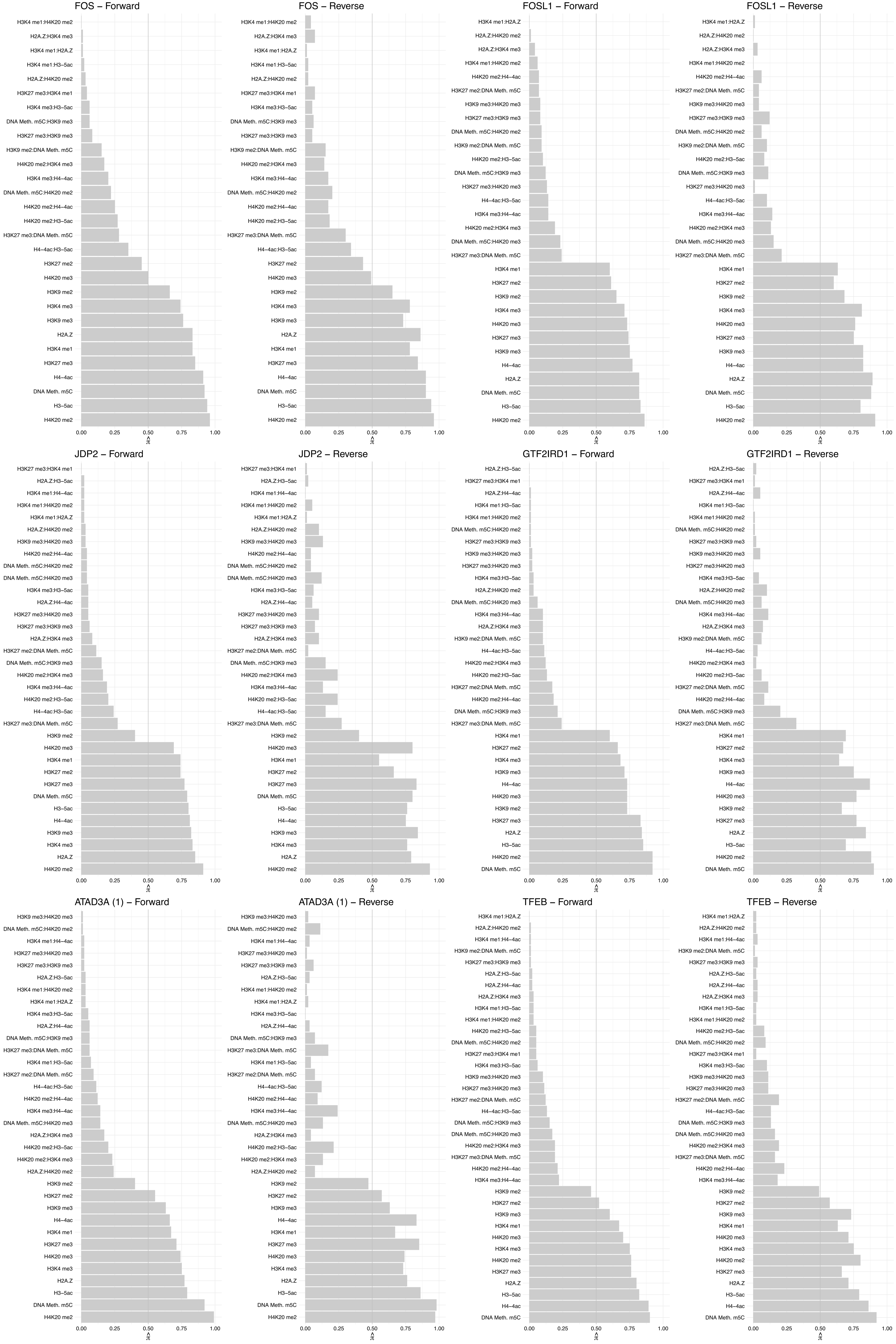
FOS – Forward
H3K4 me1:H4K20 me2
H2A.Z:H3K4 me3
H3K4 me1:H2A.Z
H3K4 me1:H3–5ac
H2A.Z:H4K20 me2
H3K27 me3:H3K4 me1
H3K4 me3:H3–5ac
DNA Meth. m5C:H3K9 me3
H3K27 me3:H3K9 me3
H3K9 me2:DNA Meth. m5C
H4K20 me2:H3K4 me3
H3K4 me3:H4–4ac
DNA Meth. m5C:H4K20 me2
H4K20 me2:H4–4ac
H4K20 me2:H3–5ac
H3K27 me3:DNA Meth. m5C
H4–4ac:H3–5ac
H3K27 me2
H4K20 me3
H3K9 me2
H3K4 me3
H3K9 me3
H2A.Z
H3K4 me1
H3K27 me3
H4–4ac
DNA Meth. m5C
H3–5ac
H4K20 me2
0.00 0.25 0.50 0.75 1.00
π̂
FOS – Reverse
H3K4 me1:H4K20 me2
H2A.Z:H3K4 me3
H3K4 me1:H2A.Z
H3K4 me1:H3–5ac
H2A.Z:H4K20 me2
H3K27 me3:H3K4 me1
H3K4 me3:H3–5ac
DNA Meth. m5C:H3K9 me3
H3K27 me3:H3K9 me3
H3K9 me2:DNA Meth. m5C
H4K20 me2:H3K4 me3
H3K4 me3:H4–4ac
DNA Meth. m5C:H4K20 me2
H4K20 me2:H4–4ac
H4K20 me2:H3–5ac
H3K27 me3:DNA Meth. m5C
H4–4ac:H3–5ac
H3K27 me2
H4K20 me3
H3K9 me2
H3K4 me3
H3K9 me3
H2A.Z
H3K4 me1
H3K27 me3
H4–4ac
DNA Meth. m5C
H3–5ac
H4K20 me2
0.00 0.25 0.50 0.75 1.00
π̂
FOSL1 – Forward
H3K4 me1:H2A.Z
H2A.Z:H4K20 me2
H2A.Z:H3K4 me3
H3K4 me1:H4K20 me2
H4K20 me2:H4–4ac
H3K27 me2:DNA Meth. m5C
H3K9 me3:H4K20 me3
H3K27 me3:H3K9 me3
DNA Meth. m5C:H4K20 me2
H3K9 me2:DNA Meth. m5C
H4K20 me2:H3–5ac
DNA Meth. m5C:H3K9 me3
H3K27 me3:H4K20 me3
H4–4ac:H3–5ac
H3K4 me3:H4–4ac
H4K20 me2:H3K4 me3
DNA Meth. m5C:H4K20 me3
H3K27 me3:DNA Meth. m5C
H3K4 me1
H3K27 me2
H3K9 me2
H3K4 me3
H4K20 me3
H3K27 me3
H3K9 me3
H4–4ac
H2A.Z
DNA Meth. m5C
H3–5ac
H4K20 me2
0.00 0.25 0.50 0.75 1.00
π̂
FOSL1 – Reverse
H3K4 me1:H2A.Z
H2A.Z:H4K20 me2
H2A.Z:H3K4 me3
H3K4 me1:H4K20 me2
H4K20 me2:H4–4ac
H3K27 me2:DNA Meth. m5C
H3K9 me3:H4K20 me3
H3K27 me3:H3K9 me3
DNA Meth. m5C:H4K20 me2
H3K9 me2:DNA Meth. m5C
H4K20 me2:H3–5ac
DNA Meth. m5C:H3K9 me3
H3K27 me3:H4K20 me3
H4–4ac:H3–5ac
H3K4 me3:H4–4ac
H4K20 me2:H3K4 me3
DNA Meth. m5C:H4K20 me3
H3K27 me3:DNA Meth. m5C
H3K4 me1
H3K27 me2
H3K9 me2
H3K4 me3
H4K20 me3
H3K27 me3
H3K9 me3
H4–4ac
H2A.Z
DNA Meth. m5C
H3–5ac
H4K20 me2
0.00 0.25 0.50 0.75 1.00
π̂
JDP2 – Forward
H3K27 me3:H3K4 me1
H2A.Z:H3–5ac
H3K4 me1:H4–4ac
H3K4 me1:H4K20 me2
H3K4 me1:H2A.Z
H2A.Z:H4K20 me2
H3K9 me3:H4K20 me3
H4K20 me2:H4–4ac
DNA Meth. m5C:H4K20 me2
DNA Meth. m5C:H4K20 me3
H3K4 me3:H3–5ac
H2A.Z:H4–4ac
H3K27 me3:H4K20 me3
H3K27 me3:H3K9 me3
H2A.Z:H3K4 me3
H3K27 me2:DNA Meth. m5C
DNA Meth. m5C:H3K9 me3
H4K20 me2:H3K4 me3
H3K4 me3:H4–4ac
H4K20 me2:H3–5ac
H4–4ac:H3–5ac
H3K27 me3:DNA Meth. m5C
H3K9 me2
H4K20 me3
H3K4 me1
H3K27 me2
H3K27 me3
DNA Meth. m5C
H3–5ac
H4–4ac
H3K9 me3
H3K4 me3
H2A.Z
H4K20 me2
0.00 0.25 0.50 0.75 1.00
π̂
JDP2 – Reverse
H3K27 me3:H3K4 me1
H2A.Z:H3–5ac
H3K4 me1:H4–4ac
H3K4 me1:H4K20 me2
H3K4 me1:H2A.Z
H2A.Z:H4K20 me2
H3K9 me3:H4K20 me3
H4K20 me2:H4–4ac
DNA Meth. m5C:H4K20 me2
DNA Meth. m5C:H4K20 me3
H3K4 me3:H3–5ac
H2A.Z:H4–4ac
H3K27 me3:H4K20 me3
H3K27 me3:H3K9 me3
H2A.Z:H3K4 me3
H3K27 me2:DNA Meth. m5C
DNA Meth. m5C:H3K9 me3
H4K20 me2:H3K4 me3
H3K4 me3:H4–4ac
H4K20 me2:H3–5ac
H4–4ac:H3–5ac
H3K27 me3:DNA Meth. m5C
H3K9 me2
H4K20 me3
H3K4 me1
H3K27 me2
H3K27 me3
DNA Meth. m5C
H3–5ac
H4–4ac
H3K9 me3
H3K4 me3
H2A.Z
H4K20 me2
0.00 0.25 0.50 0.75 1.00
π̂
GTF2IRD1 – Forward
H2A.Z:H3–5ac
H3K27 me3:H3K4 me1
H2A.Z:H4–4ac
H3K4 me1:H3–5ac
H3K4 me1:H4K20 me2
DNA Meth. m5C:H4K20 me2
H3K27 me3:H3K9 me3
H3K9 me3:H4K20 me3
H3K27 me3:H4K20 me3
H3K4 me3:H3–5ac
H2A.Z:H4K20 me2
DNA Meth. m5C:H4K20 me3
H3K4 me3:H4–4ac
H2A.Z:H3K4 me3
H3K9 me2:DNA Meth. m5C
H4–4ac:H3–5ac
H4K20 me2:H3K4 me3
H4K20 me2:H3–5ac
H3K27 me2:DNA Meth. m5C
H4K20 me2:H4–4ac
DNA Meth. m5C:H3K9 me3
H3K27 me3:DNA Meth. m5C
H3K4 me1
H3K27 me2
H3K4 me3
H3K9 me3
H4–4ac
H4K20 me3
H3K9 me2
H3K27 me3
H2A.Z
H3–5ac
H4K20 me2
DNA Meth. m5C
0.00 0.25 0.50 0.75 1.00
π̂
GTF2IRD1 – Reverse
H2A.Z:H3–5ac
H3K27 me3:H3K4 me1
H2A.Z:H4–4ac
H3K4 me1:H3–5ac
H3K4 me1:H4K20 me2
DNA Meth. m5C:H4K20 me2
H3K27 me3:H3K9 me3
H3K9 me3:H4K20 me3
H3K27 me3:H4K20 me3
H3K4 me3:H3–5ac
H2A.Z:H4K20 me2
DNA Meth. m5C:H4K20 me3
H3K4 me3:H4–4ac
H2A.Z:H3K4 me3
H3K9 me2:DNA Meth. m5C
H4–4ac:H3–5ac
H4K20 me2:H3K4 me3
H4K20 me2:H3–5ac
H3K27 me2:DNA Meth. m5C
H4K20 me2:H4–4ac
DNA Meth. m5C:H3K9 me3
H3K27 me3:DNA Meth. m5C
H3K4 me1
H3K27 me2
H3K4 me3
H3K9 me3
H4–4ac
H4K20 me3
H3K9 me2
H3K27 me3
H2A.Z
H3–5ac
H4K20 me2
DNA Meth. m5C
0.00 0.25 0.50 0.75 1.00
π̂
ATAD3A (1) – Forward
H3K9 me3:H4K20 me3
DNA Meth. m5C:H4K20 me2
H3K4 me1:H4–4ac
H3K27 me3:H4K20 me3
H3K27 me3:H3K9 me3
H2A.Z:H3–5ac
H3K4 me1:H4K20 me2
H3K4 me1:H2A.Z
H3K4 me3:H3–5ac
H2A.Z:H4–4ac
DNA Meth. m5C:H3K9 me3
H3K27 me3:DNA Meth. m5C
H3K4 me1:H3–5ac
H3K27 me2:DNA Meth. m5C
H4–4ac:H3–5ac
H4K20 me2:H4–4ac
H3K4 me3:H4–4ac
DNA Meth. m5C:H4K20 me3
H2A.Z:H3K4 me3
H4K20 me2:H3–5ac
H4K20 me2:H3K4 me3
H2A.Z:H4K20 me2
H3K9 me2
H3K27 me2
H3K9 me3
H4–4ac
H3K4 me1
H3K27 me3
H4K20 me3
H3K4 me3
H2A.Z
H3–5ac
DNA Meth. m5C
H4K20 me2
0.00 0.25 0.50 0.75 1.00
π̂
ATAD3A (1) – Reverse
H3K9 me3:H4K20 me3
DNA Meth. m5C:H4K20 me2
H3K4 me1:H4–4ac
H3K27 me3:H4K20 me3
H3K27 me3:H3K9 me3
H2A.Z:H3–5ac
H3K4 me1:H4K20 me2
H3K4 me1:H2A.Z
H3K4 me3:H3–5ac
H2A.Z:H4–4ac
DNA Meth. m5C:H3K9 me3
H3K27 me3:DNA Meth. m5C
H3K4 me1:H3–5ac
H3K27 me2:DNA Meth. m5C
H4–4ac:H3–5ac
H4K20 me2:H4–4ac
H3K4 me3:H4–4ac
DNA Meth. m5C:H4K20 me3
H2A.Z:H3K4 me3
H4K20 me2:H3–5ac
H4K20 me2:H3K4 me3
H2A.Z:H4K20 me2
H3K9 me2
H3K27 me2
H3K9 me3
H4–4ac
H3K4 me1
H3K27 me3
H4K20 me3
H3K4 me3
H2A.Z
H3–5ac
DNA Meth. m5C
H4K20 me2
0.00 0.25 0.50 0.75 1.00
π̂
TFEB – Forward
H3K4 me1:H2A.Z
H2A.Z:H4K20 me2
H3K4 me1:H4–4ac
H3K9 me2:DNA Meth. m5C
H3K27 me3:H3K9 me3
H2A.Z:H3–5ac
H2A.Z:H4–4ac
H2A.Z:H3K4 me3
H3K4 me1:H3–5ac
H3K4 me1:H4K20 me2
H4K20 me2:H3–5ac
DNA Meth. m5C:H4K20 me2
H3K27 me3:H3K4 me1
H3K4 me3:H3–5ac
H3K9 me3:H4K20 me3
H3K27 me3:H4K20 me3
H3K27 me2:DNA Meth. m5C
H4–4ac:H3–5ac
DNA Meth. m5C:H3K9 me3
DNA Meth. m5C:H4K20 me3
H4K20 me2:H3K4 me3
H3K27 me3:DNA Meth. m5C
H4K20 me2:H4–4ac
H3K4 me3:H4–4ac
H3K9 me2
H3K27 me2
H3K9 me3
H3K4 me1
H4K20 me3
H3K4 me3
H4K20 me2
H3K27 me3
H2A.Z
H3–5ac
H4–4ac
DNA Meth. m5C
0.00 0.25 0.50 0.75 1.00
π̂
TFEB – Reverse
H3K4 me1:H2A.Z
H2A.Z:H4K20 me2
H3K4 me1:H4–4ac
H3K9 me2:DNA Meth. m5C
H3K27 me3:H3K9 me3
H2A.Z:H3–5ac
H2A.Z:H4–4ac
H2A.Z:H3K4 me3
H3K4 me1:H3–5ac
H3K4 me1:H4K20 me2
H4K20 me2:H3–5ac
DNA Meth. m5C:H4K20 me2
H3K27 me3:H3K4 me1
H3K4 me3:H3–5ac
H3K9 me3:H4K20 me3
H3K27 me3:H4K20 me3
H3K27 me2:DNA Meth. m5C
H4–4ac:H3–5ac
DNA Meth. m5C:H3K9 me3
DNA Meth. m5C:H4K20 me3
H4K20 me2:H3K4 me3
H3K27 me3:DNA Meth. m5C
H4K20 me2:H4–4ac
H3K4 me3:H4–4ac
H3K9 me2
H3K27 me2
H3K9 me3
H3K4 me1
H4K20 me3
H3K4 me3
H4K20 me2
H3K27 me3
H2A.Z
H3–5ac
H4–4ac
DNA Meth. m5C
0.00 0.25 0.50 0.75 1.00
π̂

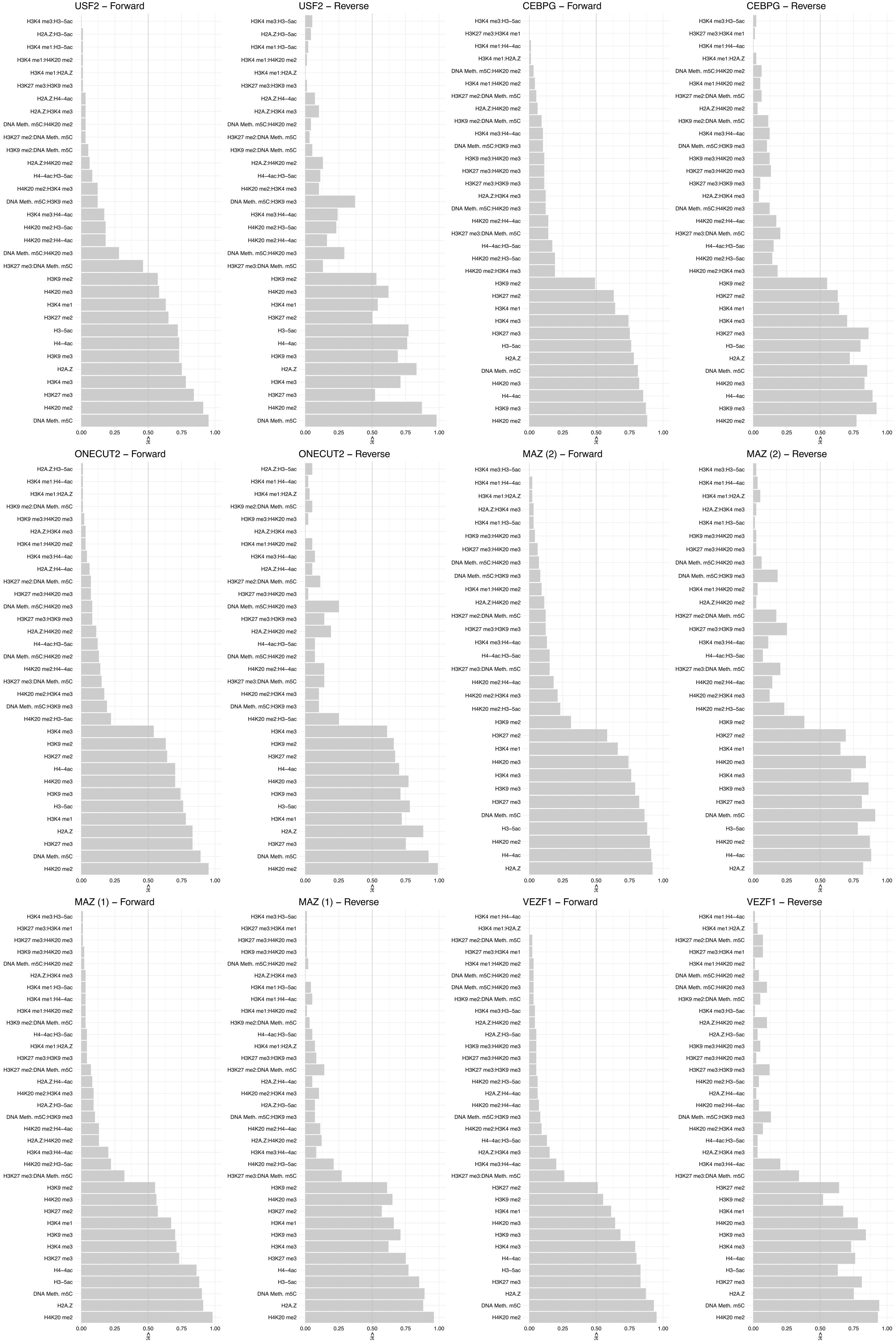
USF2 – Forward
H3K4 me3:H3–5ac
H2A.Z:H3–5ac
H3K4 me1:H3–5ac
H3K4 me1:H4K20 me2
H3K4 me1:H2A.Z
H3K27 me3:H3K9 me3
H2A.Z:H4–4ac
H2A.Z:H3K4 me3
DNA Meth. m5C:H4K20 me2
H3K27 me2:DNA Meth. m5C
H3K9 me2:DNA Meth. m5C
H2A.Z:H4K20 me2
H4–4ac:H3–5ac
H4K20 me2:H3K4 me3
DNA Meth. m5C:H3K9 me3
H3K4 me3:H4–4ac
H4K20 me2:H3–5ac
H4K20 me2:H4–4ac
DNA Meth. m5C:H4K20 me3
H3K27 me3:DNA Meth. m5C
H3K9 me2
H4K20 me3
H3K4 me1
H3K27 me2
H3–5ac
H4–4ac
H3K9 me3
H2A.Z
H3K4 me3
H3K27 me3
H4K20 me2
DNA Meth. m5C
0.00 0.25 0.50 0.75 1.00
π̂
USF2 – Reverse
H3K4 me3:H3–5ac
H2A.Z:H3–5ac
H3K4 me1:H3–5ac
H3K4 me1:H4K20 me2
H3K4 me1:H2A.Z
H3K27 me3:H3K9 me3
H2A.Z:H4–4ac
H2A.Z:H3K4 me3
DNA Meth. m5C:H4K20 me2
H3K27 me2:DNA Meth. m5C
H3K9 me2:DNA Meth. m5C
H2A.Z:H4K20 me2
H4–4ac:H3–5ac
H4K20 me2:H3K4 me3
DNA Meth. m5C:H3K9 me3
H3K4 me3:H4–4ac
H4K20 me2:H3–5ac
H4K20 me2:H4–4ac
DNA Meth. m5C:H4K20 me3
H3K27 me3:DNA Meth. m5C
H3K9 me2
H4K20 me3
H3K4 me1
H3K27 me2
H3–5ac
H4–4ac
H3K9 me3
H2A.Z
H3K4 me3
H3K27 me3
H4K20 me2
DNA Meth. m5C
0.00 0.25 0.50 0.75 1.00
π̂
CEBPG – Forward
H3K4 me3:H3–5ac
H3K27 me3:H3K4 me1
H3K4 me1:H4–4ac
H3K4 me1:H2A.Z
DNA Meth. m5C:H4K20 me2
H3K4 me1:H4K20 me2
H3K27 me2:DNA Meth. m5C
H2A.Z:H4K20 me2
H3K9 me2:DNA Meth. m5C
H3K4 me3:H4–4ac
DNA Meth. m5C:H3K9 me3
H3K9 me3:H4K20 me3
H3K27 me3:H4K20 me3
H3K27 me3:H3K9 me3
H2A.Z:H3K4 me3
DNA Meth. m5C:H4K20 me3
H4K20 me2:H4–4ac
H3K27 me3:DNA Meth. m5C
H4–4ac:H3–5ac
H4K20 me2:H3–5ac
H4K20 me2:H3K4 me3
H3K9 me2
H3K27 me2
H3K4 me1
H3K4 me3
H3K27 me3
H3–5ac
H2A.Z
DNA Meth. m5C
H4K20 me3
H4–4ac
H3K9 me3
H4K20 me2
0.00 0.25 0.50 0.75 1.00
π̂
CEBPG – Reverse
H3K4 me3:H3–5ac
H3K27 me3:H3K4 me1
H3K4 me1:H4–4ac
H3K4 me1:H2A.Z
DNA Meth. m5C:H4K20 me2
H3K4 me1:H4K20 me2
H3K27 me2:DNA Meth. m5C
H2A.Z:H4K20 me2
H3K9 me2:DNA Meth. m5C
H3K4 me3:H4–4ac
DNA Meth. m5C:H3K9 me3
H3K9 me3:H4K20 me3
H3K27 me3:H4K20 me3
H3K27 me3:H3K9 me3
H2A.Z:H3K4 me3
DNA Meth. m5C:H4K20 me3
H4K20 me2:H4–4ac
H3K27 me3:DNA Meth. m5C
H4–4ac:H3–5ac
H4K20 me2:H3–5ac
H4K20 me2:H3K4 me3
H3K9 me2
H3K27 me2
H3K4 me1
H3K4 me3
H3K27 me3
H3–5ac
H2A.Z
DNA Meth. m5C
H4K20 me3
H4–4ac
H3K9 me3
H4K20 me2
0.00 0.25 0.50 0.75 1.00
π̂
ONECUT2 – Forward
H2A.Z:H3–5ac
H3K4 me1:H4–4ac
H3K4 me1:H2A.Z
H3K9 me2:DNA Meth. m5C
H3K9 me3:H4K20 me3
H2A.Z:H3K4 me3
H3K4 me1:H4K20 me2
H3K4 me3:H4–4ac
H2A.Z:H4–4ac
H3K27 me2:DNA Meth. m5C
H3K27 me3:H4K20 me3
DNA Meth. m5C:H4K20 me3
H3K27 me3:H3K9 me3
H2A.Z:H4K20 me2
H4–4ac:H3–5ac
DNA Meth. m5C:H4K20 me2
H4K20 me2:H4–4ac
H3K27 me3:DNA Meth. m5C
H4K20 me2:H3K4 me3
DNA Meth. m5C:H3K9 me3
H4K20 me2:H3–5ac
H3K4 me3
H3K9 me2
H3K27 me2
H4–4ac
H4K20 me3
H3K9 me3
H3–5ac
H3K4 me1
H2A.Z
H3K27 me3
DNA Meth. m5C
H4K20 me2
0.00 0.25 0.50 0.75 1.00
π̂
ONECUT2 – Reverse
H2A.Z:H3–5ac
H3K4 me1:H4–4ac
H3K4 me1:H2A.Z
H3K9 me2:DNA Meth. m5C
H3K9 me3:H4K20 me3
H2A.Z:H3K4 me3
H3K4 me1:H4K20 me2
H3K4 me3:H4–4ac
H2A.Z:H4–4ac
H3K27 me2:DNA Meth. m5C
H3K27 me3:H4K20 me3
DNA Meth. m5C:H4K20 me3
H3K27 me3:H3K9 me3
H2A.Z:H4K20 me2
H4–4ac:H3–5ac
DNA Meth. m5C:H4K20 me2
H4K20 me2:H4–4ac
H3K27 me3:DNA Meth. m5C
H4K20 me2:H3K4 me3
DNA Meth. m5C:H3K9 me3
H4K20 me2:H3–5ac
H3K4 me3
H3K9 me2
H3K27 me2
H4–4ac
H4K20 me3
H3K9 me3
H3–5ac
H3K4 me1
H2A.Z
H3K27 me3
DNA Meth. m5C
H4K20 me2
0.00 0.25 0.50 0.75 1.00
π̂
MAZ (2) – Forward
H3K4 me3:H3–5ac
H3K4 me1:H4–4ac
H3K4 me1:H2A.Z
H2A.Z:H3K4 me3
H3K4 me1:H3–5ac
H3K9 me3:H4K20 me3
H3K27 me3:H4K20 me3
DNA Meth. m5C:H4K20 me3
DNA Meth. m5C:H3K9 me3
H3K4 me1:H4K20 me2
H2A.Z:H4K20 me2
H3K27 me2:DNA Meth. m5C
H3K27 me3:H3K9 me3
H3K4 me3:H4–4ac
H4–4ac:H3–5ac
H3K27 me3:DNA Meth. m5C
H4K20 me2:H4–4ac
H4K20 me2:H3K4 me3
H4K20 me2:H3–5ac
H3K9 me2
H3K27 me2
H3K4 me1
H4K20 me3
H3K4 me3
H3K9 me3
H3K27 me3
DNA Meth. m5C
H3–5ac
H4K20 me2
H4–4ac
H2A.Z
0.00 0.25 0.50 0.75 1.00
π̂
MAZ (2) – Reverse
H3K4 me3:H3–5ac
H3K4 me1:H4–4ac
H3K4 me1:H2A.Z
H2A.Z:H3K4 me3
H3K4 me1:H3–5ac
H3K9 me3:H4K20 me3
H3K27 me3:H4K20 me3
DNA Meth. m5C:H4K20 me3
DNA Meth. m5C:H3K9 me3
H3K4 me1:H4K20 me2
H2A.Z:H4K20 me2
H3K27 me2:DNA Meth. m5C
H3K27 me3:H3K9 me3
H3K4 me3:H4–4ac
H4–4ac:H3–5ac
H3K27 me3:DNA Meth. m5C
H4K20 me2:H4–4ac
H4K20 me2:H3K4 me3
H4K20 me2:H3–5ac
H3K9 me2
H3K27 me2
H3K4 me1
H4K20 me3
H3K4 me3
H3K9 me3
H3K27 me3
DNA Meth. m5C
H3–5ac
H4K20 me2
H4–4ac
H2A.Z
0.00 0.25 0.50 0.75 1.00
π̂
MAZ (1) – Forward
H3K4 me3:H3–5ac
H3K27 me3:H3K4 me1
H3K27 me3:H4K20 me3
H3K9 me3:H4K20 me3
DNA Meth. m5C:H4K20 me2
H2A.Z:H3K4 me3
H3K4 me1:H3–5ac
H3K4 me1:H4–4ac
H3K4 me1:H4K20 me2
H3K9 me2:DNA Meth. m5C
H4–4ac:H3–5ac
H3K4 me1:H2A.Z
H3K27 me3:H3K9 me3
H3K27 me2:DNA Meth. m5C
H2A.Z:H4–4ac
H4K20 me2:H3K4 me3
H2A.Z:H3–5ac
DNA Meth. m5C:H3K9 me3
H4K20 me2:H4–4ac
H2A.Z:H4K20 me2
H3K4 me3:H4–4ac
H4K20 me2:H3–5ac
H3K27 me3:DNA Meth. m5C
H3K9 me2
H4K20 me3
H3K27 me2
H3K4 me1
H3K9 me3
H3K4 me3
H3K27 me3
H4–4ac
H3–5ac
DNA Meth. m5C
H2A.Z
H4K20 me2
0.00 0.25 0.50 0.75 1.00
π̂
MAZ (1) – Reverse
H3K4 me3:H3–5ac
H3K27 me3:H3K4 me1
H3K27 me3:H4K20 me3
H3K9 me3:H4K20 me3
DNA Meth. m5C:H4K20 me2
H2A.Z:H3K4 me3
H3K4 me1:H3–5ac
H3K4 me1:H4–4ac
H3K4 me1:H4K20 me2
H3K9 me2:DNA Meth. m5C
H4–4ac:H3–5ac
H3K4 me1:H2A.Z
H3K27 me3:H3K9 me3
H3K27 me2:DNA Meth. m5C
H2A.Z:H4–4ac
H4K20 me2:H3K4 me3
H2A.Z:H3–5ac
DNA Meth. m5C:H3K9 me3
H4K20 me2:H4–4ac
H2A.Z:H4K20 me2
H3K4 me3:H4–4ac
H4K20 me2:H3–5ac
H3K27 me3:DNA Meth. m5C
H3K9 me2
H4K20 me3
H3K27 me2
H3K4 me1
H3K9 me3
H3K4 me3
H3K27 me3
H4–4ac
H3–5ac
DNA Meth. m5C
H2A.Z
H4K20 me2
0.00 0.25 0.50 0.75 1.00
π̂
VEZF1 – Forward
H3K4 me1:H4–4ac
H3K4 me1:H2A.Z
H3K27 me2:DNA Meth. m5C
H3K27 me3:H3K4 me1
H3K4 me1:H4K20 me2
DNA Meth. m5C:H4K20 me2
DNA Meth. m5C:H4K20 me3
H3K9 me2:DNA Meth. m5C
H3K4 me3:H3–5ac
H2A.Z:H4K20 me2
H2A.Z:H3–5ac
H3K9 me3:H4K20 me3
H3K27 me3:H4K20 me3
H3K27 me3:H3K9 me3
H4K20 me2:H3–5ac
H2A.Z:H4–4ac
H4K20 me2:H4–4ac
DNA Meth. m5C:H3K9 me3
H4K20 me2:H3K4 me3
H4–4ac:H3–5ac
H2A.Z:H3K4 me3
H3K4 me3:H4–4ac
H3K27 me3:DNA Meth. m5C
H3K27 me2
H3K9 me2
H3K4 me1
H4K20 me3
H3K9 me3
H3K4 me3
H4–4ac
H3–5ac
H3K27 me3
H2A.Z
DNA Meth. m5C
H4K20 me2
0.00 0.25 0.50 0.75 1.00
π̂
VEZF1 – Reverse
H3K4 me1:H4–4ac
H3K4 me1:H2A.Z
H3K27 me2:DNA Meth. m5C
H3K27 me3:H3K4 me1
H3K4 me1:H4K20 me2
DNA Meth. m5C:H4K20 me2
DNA Meth. m5C:H4K20 me3
H3K9 me2:DNA Meth. m5C
H3K4 me3:H3–5ac
H2A.Z:H4K20 me2
H2A.Z:H3–5ac
H3K9 me3:H4K20 me3
H3K27 me3:H4K20 me3
H3K27 me3:H3K9 me3
H4K20 me2:H3–5ac
H2A.Z:H4–4ac
H4K20 me2:H4–4ac
DNA Meth. m5C:H3K9 me3
H4K20 me2:H3K4 me3
H4–4ac:H3–5ac
H2A.Z:H3K4 me3
H3K4 me3:H4–4ac
H3K27 me3:DNA Meth. m5C
H3K27 me2
H3K9 me2
H3K4 me1
H4K20 me3
H3K9 me3
H3K4 me3
H4–4ac
H3–5ac
H3K27 me3
H2A.Z
DNA Meth. m5C
H4K20 me2
0.00 0.25 0.50 0.75 1.00
π̂

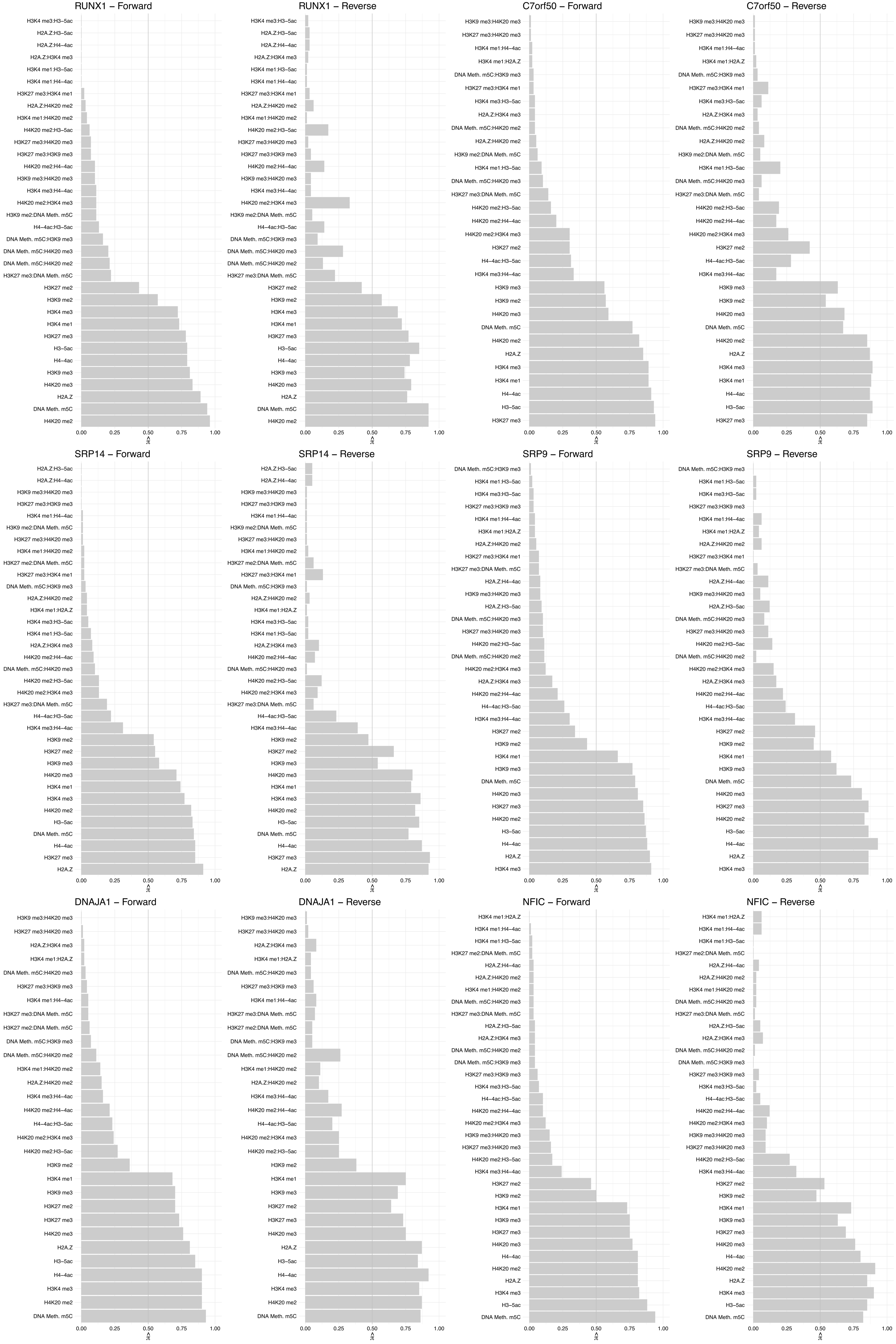
RUNX1 – Forward
H3K4 me3:H3–5ac
H2A.Z:H3–5ac
H2A.Z:H4–4ac
H2A.Z:H3K4 me3
H3K4 me1:H3–5ac
H3K4 me1:H4–4ac
H3K27 me3:H3K4 me1
H2A.Z:H4K20 me2
H3K4 me1:H4K20 me2
H4K20 me2:H3–5ac
H3K27 me3:H4K20 me3
H3K27 me3:H3K9 me3
H4K20 me2:H4–4ac
H3K9 me3:H4K20 me3
H3K4 me3:H4–4ac
H4K20 me2:H3K4 me3
H3K9 me2:DNA Meth. m5C
H4–4ac:H3–5ac
DNA Meth. m5C:H3K9 me3
DNA Meth. m5C:H4K20 me3
DNA Meth. m5C:H4K20 me2
H3K27 me3:DNA Meth. m5C
H3K27 me2
H3K9 me2
H3K4 me3
H3K4 me1
H3K27 me3
H3–5ac
H4–4ac
H3K9 me3
H4K20 me3
H2A.Z
DNA Meth. m5C
H4K20 me2
0.00 0.25 0.50 0.75 1.00
$\hat{\pi}$
RUNX1 – Reverse
H3K4 me3:H3–5ac
H2A.Z:H3–5ac
H2A.Z:H4–4ac
H2A.Z:H3K4 me3
H3K4 me1:H3–5ac
H3K4 me1:H4–4ac
H3K27 me3:H3K4 me1
H2A.Z:H4K20 me2
H3K4 me1:H4K20 me2
H4K20 me2:H3–5ac
H3K27 me3:H4K20 me3
H3K27 me3:H3K9 me3
H4K20 me2:H4–4ac
H3K9 me3:H4K20 me3
H3K4 me3:H4–4ac
H4K20 me2:H3K4 me3
H3K9 me2:DNA Meth. m5C
H4–4ac:H3–5ac
DNA Meth. m5C:H3K9 me3
DNA Meth. m5C:H4K20 me3
DNA Meth. m5C:H4K20 me2
H3K27 me3:DNA Meth. m5C
H3K27 me2
H3K9 me2
H3K4 me3
H3K4 me1
H3K27 me3
H3–5ac
H4–4ac
H3K9 me3
H4K20 me3
H2A.Z
DNA Meth. m5C
H4K20 me2
0.00 0.25 0.50 0.75 1.00
$\hat{\pi}$
C7orf50 – Forward
H3K9 me3:H4K20 me3
H3K27 me3:H4K20 me3
H3K4 me1:H4–4ac
H3K4 me1:H2A.Z
DNA Meth. m5C:H3K9 me3
H3K27 me3:H3K4 me1
H3K4 me3:H3–5ac
H2A.Z:H3K4 me3
DNA Meth. m5C:H4K20 me2
H2A.Z:H4K20 me2
H3K9 me2:DNA Meth. m5C
H3K4 me1:H3–5ac
DNA Meth. m5C:H4K20 me3
H3K27 me3:DNA Meth. m5C
H4K20 me2:H3–5ac
H4K20 me2:H4–4ac
H4K20 me2:H3K4 me3
H3K27 me2
H4–4ac:H3–5ac
H3K4 me3:H4–4ac
H3K9 me3
H3K9 me2
H4K20 me3
DNA Meth. m5C
H4K20 me2
H2A.Z
H3K4 me3
H3K4 me1
H4–4ac
H3–5ac
H3K27 me3
0.00 0.25 0.50 0.75 1.00
$\hat{\pi}$
C7orf50 – Reverse
H3K9 me3:H4K20 me3
H3K27 me3:H4K20 me3
H3K4 me1:H4–4ac
H3K4 me1:H2A.Z
DNA Meth. m5C:H3K9 me3
H3K27 me3:H3K4 me1
H3K4 me3:H3–5ac
H2A.Z:H3K4 me3
DNA Meth. m5C:H4K20 me2
H2A.Z:H4K20 me2
H3K9 me2:DNA Meth. m5C
H3K4 me1:H3–5ac
DNA Meth. m5C:H4K20 me3
H3K27 me3:DNA Meth. m5C
H4K20 me2:H3–5ac
H4K20 me2:H4–4ac
H4K20 me2:H3K4 me3
H3K27 me2
H4–4ac:H3–5ac
H3K4 me3:H4–4ac
H3K9 me3
H3K9 me2
H4K20 me3
DNA Meth. m5C
H4K20 me2
H2A.Z
H3K4 me3
H3K4 me1
H4–4ac
H3–5ac
H3K27 me3
0.00 0.25 0.50 0.75 1.00
$\hat{\pi}$
SRP14 – Forward
H2A.Z:H3–5ac
H2A.Z:H4–4ac
H3K9 me3:H4K20 me3
H3K27 me3:H3K9 me3
H3K4 me1:H4–4ac
H3K9 me2:DNA Meth. m5C
H3K27 me3:H4K20 me3
H3K4 me1:H4K20 me2
H3K27 me2:DNA Meth. m5C
H3K27 me3:H3K4 me1
DNA Meth. m5C:H3K9 me3
H2A.Z:H4K20 me2
H3K4 me1:H2A.Z
H3K4 me3:H3–5ac
H3K4 me1:H3–5ac
H2A.Z:H3K4 me3
H4K20 me2:H4–4ac
DNA Meth. m5C:H4K20 me3
H4K20 me2:H3–5ac
H4K20 me2:H3K4 me3
H3K27 me3:DNA Meth. m5C
H4–4ac:H3–5ac
H3K4 me3:H4–4ac
H3K9 me2
H3K27 me2
H3K9 me3
H4K20 me3
H3K4 me1
H3K4 me3
H4K20 me2
H3–5ac
DNA Meth. m5C
H4–4ac
H3K27 me3
H2A.Z
0.00 0.25 0.50 0.75 1.00
$\hat{\pi}$
SRP14 – Reverse
H2A.Z:H3–5ac
H2A.Z:H4–4ac
H3K9 me3:H4K20 me3
H3K27 me3:H3K9 me3
H3K4 me1:H4–4ac
H3K9 me2:DNA Meth. m5C
H3K27 me3:H4K20 me3
H3K4 me1:H4K20 me2
H3K27 me2:DNA Meth. m5C
H3K27 me3:H3K4 me1
DNA Meth. m5C:H3K9 me3
H2A.Z:H4K20 me2
H3K4 me1:H2A.Z
H3K4 me3:H3–5ac
H3K4 me1:H3–5ac
H2A.Z:H3K4 me3
H4K20 me2:H4–4ac
DNA Meth. m5C:H4K20 me3
H4K20 me2:H3–5ac
H4K20 me2:H3K4 me3
H3K27 me3:DNA Meth. m5C
H4–4ac:H3–5ac
H3K4 me3:H4–4ac
H3K9 me2
H3K27 me2
H3K9 me3
H4K20 me3
H3K4 me1
H3K4 me3
H4K20 me2
H3–5ac
DNA Meth. m5C
H4–4ac
H3K27 me3
H2A.Z
0.00 0.25 0.50 0.75 1.00
$\hat{\pi}$
SRP9 – Forward
DNA Meth. m5C:H3K9 me3
H3K4 me1:H3–5ac
H3K4 me3:H3–5ac
H3K27 me3:H3K9 me3
H3K4 me1:H4–4ac
H3K4 me1:H2A.Z
H2A.Z:H4K20 me2
H3K27 me3:H3K4 me1
H3K27 me3:DNA Meth. m5C
H2A.Z:H4–4ac
H3K9 me3:H4K20 me3
H2A.Z:H3–5ac
DNA Meth. m5C:H4K20 me3
H3K27 me3:H4K20 me3
H4K20 me2:H3–5ac
DNA Meth. m5C:H4K20 me2
H4K20 me2:H3K4 me3
H2A.Z:H3K4 me3
H4K20 me2:H4–4ac
H4–4ac:H3–5ac
H3K4 me3:H4–4ac
H3K27 me2
H3K9 me2
H3K4 me1
H3K9 me3
DNA Meth. m5C
H4K20 me3
H3K27 me3
H4K20 me2
H3–5ac
H4–4ac
H2A.Z
H3K4 me3
0.00 0.25 0.50 0.75 1.00
$\hat{\pi}$
SRP9 – Reverse
DNA Meth. m5C:H3K9 me3
H3K4 me1:H3–5ac
H3K4 me3:H3–5ac
H3K27 me3:H3K9 me3
H3K4 me1:H4–4ac
H3K4 me1:H2A.Z
H2A.Z:H4K20 me2
H3K27 me3:H3K4 me1
H3K27 me3:DNA Meth. m5C
H2A.Z:H4–4ac
H3K9 me3:H4K20 me3
H2A.Z:H3–5ac
DNA Meth. m5C:H4K20 me3
H3K27 me3:H4K20 me3
H4K20 me2:H3–5ac
DNA Meth. m5C:H4K20 me2
H4K20 me2:H3K4 me3
H2A.Z:H3K4 me3
H4K20 me2:H4–4ac
H4–4ac:H3–5ac
H3K4 me3:H4–4ac
H3K27 me2
H3K9 me2
H3K4 me1
H3K9 me3
DNA Meth. m5C
H4K20 me3
H3K27 me3
H4K20 me2
H3–5ac
H4–4ac
H2A.Z
H3K4 me3
0.00 0.25 0.50 0.75 1.00
$\hat{\pi}$
DNAJA1 – Forward
H3K9 me3:H4K20 me3
H3K27 me3:H4K20 me3
H2A.Z:H3K4 me3
H3K4 me1:H2A.Z
DNA Meth. m5C:H4K20 me3
H3K27 me3:H3K9 me3
H3K4 me1:H4–4ac
H3K27 me3:DNA Meth. m5C
H3K27 me2:DNA Meth. m5C
DNA Meth. m5C:H3K9 me3
DNA Meth. m5C:H4K20 me2
H3K4 me1:H4K20 me2
H2A.Z:H4K20 me2
H3K4 me3:H4–4ac
H4K20 me2:H4–4ac
H4–4ac:H3–5ac
H4K20 me2:H3K4 me3
H4K20 me2:H3–5ac
H3K9 me2
H3K4 me1
H3K9 me3
H3K27 me2
H3K27 me3
H4K20 me3
H2A.Z
H3–5ac
H4–4ac
H3K4 me3
H4K20 me2
DNA Meth. m5C
0.00 0.25 0.50 0.75 1.00
$\hat{\pi}$
DNAJA1 – Reverse
H3K9 me3:H4K20 me3
H3K27 me3:H4K20 me3
H2A.Z:H3K4 me3
H3K4 me1:H2A.Z
DNA Meth. m5C:H4K20 me3
H3K27 me3:H3K9 me3
H3K4 me1:H4–4ac
H3K27 me3:DNA Meth. m5C
H3K27 me2:DNA Meth. m5C
DNA Meth. m5C:H3K9 me3
DNA Meth. m5C:H4K20 me2
H3K4 me1:H4K20 me2
H2A.Z:H4K20 me2
H3K4 me3:H4–4ac
H4K20 me2:H4–4ac
H4–4ac:H3–5ac
H4K20 me2:H3K4 me3
H4K20 me2:H3–5ac
H3K9 me2
H3K4 me1
H3K9 me3
H3K27 me2
H3K27 me3
H4K20 me3
H2A.Z
H3–5ac
H4–4ac
H3K4 me3
H4K20 me2
DNA Meth. m5C
0.00 0.25 0.50 0.75 1.00
$\hat{\pi}$
NFIC – Forward
H3K4 me1:H2A.Z
H3K4 me1:H4–4ac
H3K4 me1:H3–5ac
H3K27 me2:DNA Meth. m5C
H2A.Z:H4–4ac
H2A.Z:H4K20 me2
H3K4 me1:H4K20 me2
DNA Meth. m5C:H4K20 me3
H3K27 me3:DNA Meth. m5C
H2A.Z:H3–5ac
H2A.Z:H3K4 me3
DNA Meth. m5C:H4K20 me2
DNA Meth. m5C:H3K9 me3
H3K27 me3:H3K9 me3
H3K4 me3:H3–5ac
H4–4ac:H3–5ac
H4K20 me2:H4–4ac
H4K20 me2:H3K4 me3
H3K9 me3:H4K20 me3
H3K27 me3:H4K20 me3
H4K20 me2:H3–5ac
H3K4 me3:H4–4ac
H3K27 me2
H3K9 me2
H3K4 me1
H3K9 me3
H3K27 me3
H4K20 me3
H4–4ac
H4K20 me2
H2A.Z
H3K4 me3
H3–5ac
DNA Meth. m5C
0.00 0.25 0.50 0.75 1.00
$\hat{\pi}$
NFIC – Reverse
H3K4 me1:H2A.Z
H3K4 me1:H4–4ac
H3K4 me1:H3–5ac
H3K27 me2:DNA Meth. m5C
H2A.Z:H4–4ac
H2A.Z:H4K20 me2
H3K4 me1:H4K20 me2
DNA Meth. m5C:H4K20 me3
H3K27 me3:DNA Meth. m5C
H2A.Z:H3–5ac
H2A.Z:H3K4 me3
DNA Meth. m5C:H4K20 me2
DNA Meth. m5C:H3K9 me3
H3K27 me3:H3K9 me3
H3K4 me3:H3–5ac
H4–4ac:H3–5ac
H4K20 me2:H4–4ac
H4K20 me2:H3K4 me3
H3K9 me3:H4K20 me3
H3K27 me3:H4K20 me3
H4K20 me2:H3–5ac
H3K4 me3:H4–4ac
H3K27 me2
H3K9 me2
H3K4 me1
H3K9 me3
H3K27 me3
H4K20 me3
H4–4ac
H4K20 me2
H2A.Z
H3K4 me3
H3–5ac
DNA Meth. m5C
0.00 0.25 0.50 0.75 1.00
$\hat{\pi}$

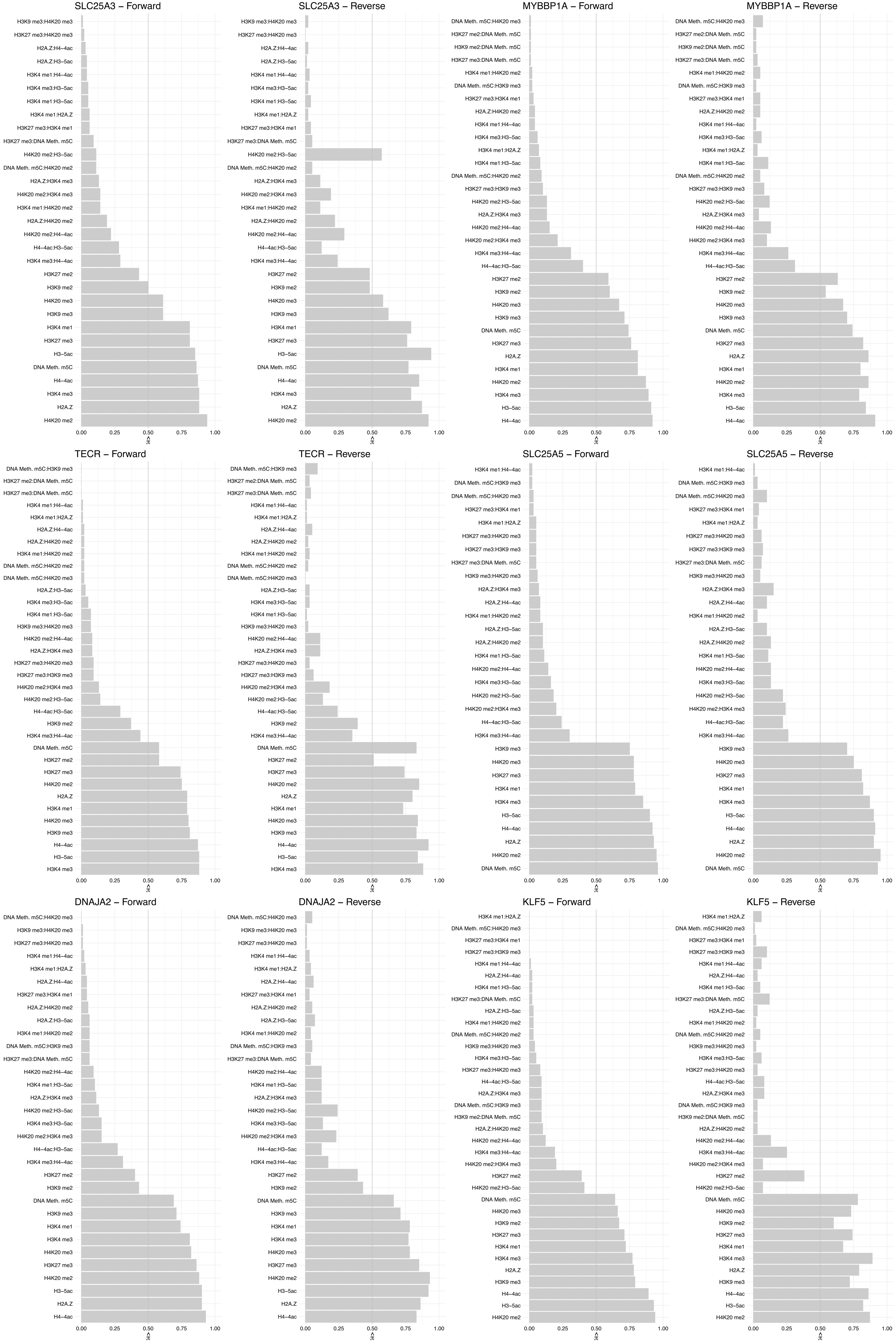
SLC25A3 – Forward
H3K9 me3:H4K20 me3
H3K27 me3:H4K20 me3
H2A.Z:H4−4ac
H2A.Z:H3−5ac
H3K4 me1:H4−4ac
H3K4 me3:H3−5ac
H3K4 me1:H3−5ac
H3K4 me1:H2A.Z
H3K27 me3:H3K4 me1
H3K27 me3:DNA Meth. m5C
H4K20 me2:H3−5ac
DNA Meth. m5C:H4K20 me2
H2A.Z:H3K4 me3
H4K20 me2:H3K4 me3
H3K4 me1:H4K20 me2
H2A.Z:H4K20 me2
H4K20 me2:H4−4ac
H4−4ac:H3−5ac
H3K4 me3:H4−4ac
H3K27 me2
H3K9 me2
H4K20 me3
H3K9 me3
H3K4 me1
H3K27 me3
H3−5ac
DNA Meth. m5C
H4−4ac
H3K4 me3
H2A.Z
H4K20 me2
0.00 0.25 0.50 0.75 1.00
π̂
SLC25A3 – Reverse
H3K9 me3:H4K20 me3
H3K27 me3:H4K20 me3
H2A.Z:H4−4ac
H2A.Z:H3−5ac
H3K4 me1:H4−4ac
H3K4 me3:H3−5ac
H3K4 me1:H3−5ac
H3K4 me1:H2A.Z
H3K27 me3:H3K4 me1
H3K27 me3:DNA Meth. m5C
H4K20 me2:H3−5ac
DNA Meth. m5C:H4K20 me2
H2A.Z:H3K4 me3
H4K20 me2:H3K4 me3
H3K4 me1:H4K20 me2
H2A.Z:H4K20 me2
H4K20 me2:H4−4ac
H4−4ac:H3−5ac
H3K4 me3:H4−4ac
H3K27 me2
H3K9 me2
H4K20 me3
H3K9 me3
H3K4 me1
H3K27 me3
H3−5ac
DNA Meth. m5C
H4−4ac
H3K4 me3
H2A.Z
H4K20 me2
0.00 0.25 0.50 0.75 1.00
π̂
MYBBP1A – Forward
DNA Meth. m5C:H4K20 me3
H3K27 me2:DNA Meth. m5C
H3K9 me2:DNA Meth. m5C
H3K27 me3:DNA Meth. m5C
H3K4 me1:H4K20 me2
DNA Meth. m5C:H3K9 me3
H3K27 me3:H3K4 me1
H2A.Z:H4K20 me2
H3K4 me1:H4−4ac
H3K4 me3:H3−5ac
H3K4 me1:H2A.Z
H3K4 me1:H3−5ac
DNA Meth. m5C:H4K20 me2
H3K27 me3:H3K9 me3
H4K20 me2:H3−5ac
H2A.Z:H3K4 me3
H4K20 me2:H4−4ac
H4K20 me2:H3K4 me3
H3K4 me3:H4−4ac
H4−4ac:H3−5ac
H3K27 me2
H3K9 me2
H4K20 me3
H3K9 me3
DNA Meth. m5C
H3K27 me3
H2A.Z
H3K4 me1
H4K20 me2
H3K4 me3
H3−5ac
H4−4ac
0.00 0.25 0.50 0.75 1.00
π̂
MYBBP1A – Reverse
DNA Meth. m5C:H4K20 me3
H3K27 me2:DNA Meth. m5C
H3K9 me2:DNA Meth. m5C
H3K27 me3:DNA Meth. m5C
H3K4 me1:H4K20 me2
DNA Meth. m5C:H3K9 me3
H3K27 me3:H3K4 me1
H2A.Z:H4K20 me2
H3K4 me1:H4−4ac
H3K4 me3:H3−5ac
H3K4 me1:H2A.Z
H3K4 me1:H3−5ac
DNA Meth. m5C:H4K20 me2
H3K27 me3:H3K9 me3
H4K20 me2:H3−5ac
H2A.Z:H3K4 me3
H4K20 me2:H4−4ac
H4K20 me2:H3K4 me3
H3K4 me3:H4−4ac
H4−4ac:H3−5ac
H3K27 me2
H3K9 me2
H4K20 me3
H3K9 me3
DNA Meth. m5C
H3K27 me3
H2A.Z
H3K4 me1
H4K20 me2
H3K4 me3
H3−5ac
H4−4ac
0.00 0.25 0.50 0.75 1.00
π̂
TECR – Forward
DNA Meth. m5C:H3K9 me3
H3K27 me2:DNA Meth. m5C
H3K27 me3:DNA Meth. m5C
H3K4 me1:H4−4ac
H3K4 me1:H2A.Z
H2A.Z:H4−4ac
H2A.Z:H4K20 me2
H3K4 me1:H4K20 me2
DNA Meth. m5C:H4K20 me2
DNA Meth. m5C:H4K20 me3
H2A.Z:H3−5ac
H3K4 me3:H3−5ac
H3K4 me1:H3−5ac
H3K9 me3:H4K20 me3
H4K20 me2:H4−4ac
H2A.Z:H3K4 me3
H3K27 me3:H4K20 me3
H3K27 me3:H3K9 me3
H4K20 me2:H3K4 me3
H4K20 me2:H3−5ac
H4−4ac:H3−5ac
H3K9 me2
H3K4 me3:H4−4ac
DNA Meth. m5C
H3K27 me2
H3K27 me3
H4K20 me2
H2A.Z
H3K4 me1
H4K20 me3
H3K9 me3
H4−4ac
H3−5ac
H3K4 me3
0.00 0.25 0.50 0.75 1.00
π̂
TECR – Reverse
DNA Meth. m5C:H3K9 me3
H3K27 me2:DNA Meth. m5C
H3K27 me3:DNA Meth. m5C
H3K4 me1:H4−4ac
H3K4 me1:H2A.Z
H2A.Z:H4−4ac
H2A.Z:H4K20 me2
H3K4 me1:H4K20 me2
DNA Meth. m5C:H4K20 me2
DNA Meth. m5C:H4K20 me3
H2A.Z:H3−5ac
H3K4 me3:H3−5ac
H3K4 me1:H3−5ac
H3K9 me3:H4K20 me3
H4K20 me2:H4−4ac
H2A.Z:H3K4 me3
H3K27 me3:H4K20 me3
H3K27 me3:H3K9 me3
H4K20 me2:H3K4 me3
H4K20 me2:H3−5ac
H4−4ac:H3−5ac
H3K9 me2
H3K4 me3:H4−4ac
DNA Meth. m5C
H3K27 me2
H3K27 me3
H4K20 me2
H2A.Z
H3K4 me1
H4K20 me3
H3K9 me3
H4−4ac
H3−5ac
H3K4 me3
0.00 0.25 0.50 0.75 1.00
π̂
SLC25A5 – Forward
H3K4 me1:H4−4ac
DNA Meth. m5C:H3K9 me3
DNA Meth. m5C:H4K20 me3
H3K27 me3:H3K4 me1
H3K4 me1:H2A.Z
H3K27 me3:H4K20 me3
H3K27 me3:H3K9 me3
H3K27 me3:DNA Meth. m5C
H3K9 me3:H4K20 me3
H2A.Z:H3K4 me3
H2A.Z:H4−4ac
H3K4 me1:H4K20 me2
H2A.Z:H3−5ac
H2A.Z:H4K20 me2
H3K4 me1:H3−5ac
H4K20 me2:H4−4ac
H3K4 me3:H3−5ac
H4K20 me2:H3−5ac
H4K20 me2:H3K4 me3
H4−4ac:H3−5ac
H3K4 me3:H4−4ac
H3K9 me3
H4K20 me3
H3K27 me3
H3K4 me1
H3K4 me3
H3−5ac
H4−4ac
H2A.Z
H4K20 me2
DNA Meth. m5C
0.00 0.25 0.50 0.75 1.00
π̂
SLC25A5 – Reverse
H3K4 me1:H4−4ac
DNA Meth. m5C:H3K9 me3
DNA Meth. m5C:H4K20 me3
H3K27 me3:H3K4 me1
H3K4 me1:H2A.Z
H3K27 me3:H4K20 me3
H3K27 me3:H3K9 me3
H3K27 me3:DNA Meth. m5C
H3K9 me3:H4K20 me3
H2A.Z:H3K4 me3
H2A.Z:H4−4ac
H3K4 me1:H4K20 me2
H2A.Z:H3−5ac
H2A.Z:H4K20 me2
H3K4 me1:H3−5ac
H4K20 me2:H4−4ac
H3K4 me3:H3−5ac
H4K20 me2:H3−5ac
H4K20 me2:H3K4 me3
H4−4ac:H3−5ac
H3K4 me3:H4−4ac
H3K9 me3
H4K20 me3
H3K27 me3
H3K4 me1
H3K4 me3
H3−5ac
H4−4ac
H2A.Z
H4K20 me2
DNA Meth. m5C
0.00 0.25 0.50 0.75 1.00
π̂
DNAJA2 – Forward
DNA Meth. m5C:H4K20 me3
H3K9 me3:H4K20 me3
H3K27 me3:H4K20 me3
H3K4 me1:H4−4ac
H3K4 me1:H2A.Z
H2A.Z:H4−4ac
H3K27 me3:H3K4 me1
H2A.Z:H4K20 me2
H2A.Z:H3−5ac
H3K4 me1:H4K20 me2
DNA Meth. m5C:H3K9 me3
H3K27 me3:DNA Meth. m5C
H4K20 me2:H4−4ac
H3K4 me1:H3−5ac
H2A.Z:H3K4 me3
H4K20 me2:H3−5ac
H3K4 me3:H3−5ac
H4K20 me2:H3K4 me3
H4−4ac:H3−5ac
H3K4 me3:H4−4ac
H3K27 me2
H3K9 me2
DNA Meth. m5C
H3K9 me3
H3K4 me1
H3K4 me3
H4K20 me3
H3K27 me3
H4K20 me2
H3−5ac
H2A.Z
H4−4ac
0.00 0.25 0.50 0.75 1.00
π̂
DNAJA2 – Reverse
DNA Meth. m5C:H4K20 me3
H3K9 me3:H4K20 me3
H3K27 me3:H4K20 me3
H3K4 me1:H4−4ac
H3K4 me1:H2A.Z
H2A.Z:H4−4ac
H3K27 me3:H3K4 me1
H2A.Z:H4K20 me2
H2A.Z:H3−5ac
H3K4 me1:H4K20 me2
DNA Meth. m5C:H3K9 me3
H3K27 me3:DNA Meth. m5C
H4K20 me2:H4−4ac
H3K4 me1:H3−5ac
H2A.Z:H3K4 me3
H4K20 me2:H3−5ac
H3K4 me3:H3−5ac
H4K20 me2:H3K4 me3
H4−4ac:H3−5ac
H3K4 me3:H4−4ac
H3K27 me2
H3K9 me2
DNA Meth. m5C
H3K9 me3
H3K4 me1
H3K4 me3
H4K20 me3
H3K27 me3
H4K20 me2
H3−5ac
H2A.Z
H4−4ac
0.00 0.25 0.50 0.75 1.00
π̂
KLF5 – Forward
H3K4 me1:H2A.Z
DNA Meth. m5C:H4K20 me3
H3K27 me3:H3K4 me1
H3K27 me3:H3K9 me3
H3K4 me1:H4−4ac
H2A.Z:H4−4ac
H3K4 me1:H3−5ac
H3K27 me3:DNA Meth. m5C
H2A.Z:H3−5ac
H3K4 me1:H4K20 me2
DNA Meth. m5C:H4K20 me2
H3K9 me3:H4K20 me3
H3K4 me3:H3−5ac
H3K27 me3:H4K20 me3
H4−4ac:H3−5ac
H2A.Z:H3K4 me3
DNA Meth. m5C:H3K9 me3
H3K9 me2:DNA Meth. m5C
H2A.Z:H4K20 me2
H4K20 me2:H4−4ac
H3K4 me3:H4−4ac
H4K20 me2:H3K4 me3
H3K27 me2
H4K20 me2:H3−5ac
DNA Meth. m5C
H4K20 me3
H3K9 me2
H3K27 me3
H3K4 me1
H3K4 me3
H2A.Z
H3K9 me3
H4−4ac
H3−5ac
H4K20 me2
0.00 0.25 0.50 0.75 1.00
π̂
KLF5 – Reverse
H3K4 me1:H2A.Z
DNA Meth. m5C:H4K20 me3
H3K27 me3:H3K4 me1
H3K27 me3:H3K9 me3
H3K4 me1:H4−4ac
H2A.Z:H4−4ac
H3K4 me1:H3−5ac
H3K27 me3:DNA Meth. m5C
H2A.Z:H3−5ac
H3K4 me1:H4K20 me2
DNA Meth. m5C:H4K20 me2
H3K9 me3:H4K20 me3
H3K4 me3:H3−5ac
H3K27 me3:H4K20 me3
H4−4ac:H3−5ac
H2A.Z:H3K4 me3
DNA Meth. m5C:H3K9 me3
H3K9 me2:DNA Meth. m5C
H2A.Z:H4K20 me2
H4K20 me2:H4−4ac
H3K4 me3:H4−4ac
H4K20 me2:H3K4 me3
H3K27 me2
H4K20 me2:H3−5ac
DNA Meth. m5C
H4K20 me3
H3K9 me2
H3K27 me3
H3K4 me1
H3K4 me3
H2A.Z
H3K9 me3
H4−4ac
H3−5ac
H4K20 me2
0.00 0.25 0.50 0.75 1.00
π̂

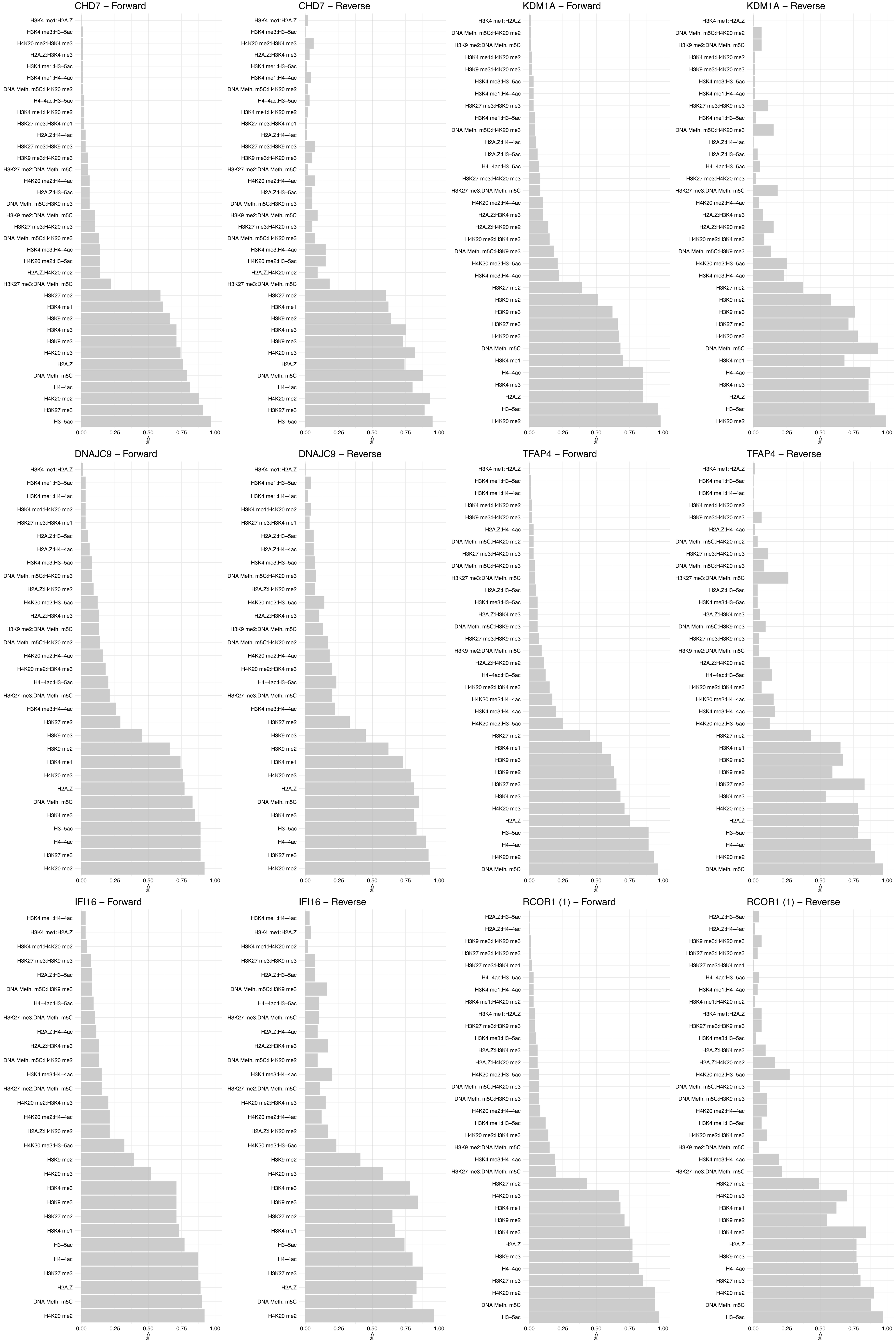
CHD7 − Forward
H3K4 me1:H2A.Z
H3K4 me3:H3−5ac
H4K20 me2:H3K4 me3
H2A.Z:H3K4 me3
H3K4 me1:H3−5ac
H3K4 me1:H4−4ac
DNA Meth. m5C:H4K20 me2
H4−4ac:H3−5ac
H3K4 me1:H4K20 me2
H3K27 me3:H3K4 me1
H2A.Z:H4−4ac
H3K27 me3:H3K9 me3
H3K9 me3:H4K20 me3
H3K27 me2:DNA Meth. m5C
H4K20 me2:H4−4ac
H2A.Z:H3−5ac
DNA Meth. m5C:H3K9 me3
H3K9 me2:DNA Meth. m5C
H3K27 me3:H4K20 me3
DNA Meth. m5C:H4K20 me3
H3K4 me3:H4−4ac
H4K20 me2:H3−5ac
H2A.Z:H4K20 me2
H3K27 me3:DNA Meth. m5C
H3K27 me2
H3K4 me1
H3K9 me2
H3K4 me3
H3K9 me3
H4K20 me3
H2A.Z
DNA Meth. m5C
H4−4ac
H4K20 me2
H3K27 me3
H3−5ac
0.00
0.25
0.50
0.75
1.00
π̂
CHD7 − Reverse
H3K4 me1:H2A.Z
H3K4 me3:H3−5ac
H4K20 me2:H3K4 me3
H2A.Z:H3K4 me3
H3K4 me1:H3−5ac
H3K4 me1:H4−4ac
DNA Meth. m5C:H4K20 me2
H4−4ac:H3−5ac
H3K4 me1:H4K20 me2
H3K27 me3:H3K4 me1
H2A.Z:H4−4ac
H3K27 me3:H3K9 me3
H3K9 me3:H4K20 me3
H3K27 me2:DNA Meth. m5C
H4K20 me2:H4−4ac
H2A.Z:H3−5ac
DNA Meth. m5C:H3K9 me3
H3K9 me2:DNA Meth. m5C
H3K27 me3:H4K20 me3
DNA Meth. m5C:H4K20 me3
H3K4 me3:H4−4ac
H4K20 me2:H3−5ac
H2A.Z:H4K20 me2
H3K27 me3:DNA Meth. m5C
H3K27 me2
H3K4 me1
H3K9 me2
H3K4 me3
H3K9 me3
H4K20 me3
H2A.Z
DNA Meth. m5C
H4−4ac
H4K20 me2
H3K27 me3
H3−5ac
0.00
0.25
0.50
0.75
1.00
π̂
KDM1A − Forward
H3K4 me1:H2A.Z
DNA Meth. m5C:H4K20 me2
H3K9 me2:DNA Meth. m5C
H3K4 me1:H4K20 me2
H3K9 me3:H4K20 me3
H3K4 me3:H3−5ac
H3K4 me1:H4−4ac
H3K27 me3:H3K9 me3
H3K4 me1:H3−5ac
DNA Meth. m5C:H4K20 me3
H2A.Z:H4−4ac
H2A.Z:H3−5ac
H4−4ac:H3−5ac
H3K27 me3:H4K20 me3
H3K27 me3:DNA Meth. m5C
H4K20 me2:H4−4ac
H2A.Z:H3K4 me3
H2A.Z:H4K20 me2
H4K20 me2:H3K4 me3
DNA Meth. m5C:H3K9 me3
H4K20 me2:H3−5ac
H3K4 me3:H4−4ac
H3K27 me2
H3K9 me2
H3K9 me3
H3K27 me3
H4K20 me3
DNA Meth. m5C
H3K4 me1
H4−4ac
H3K4 me3
H2A.Z
H3−5ac
H4K20 me2
0.00
0.25
0.50
0.75
1.00
π̂
KDM1A − Reverse
H3K4 me1:H2A.Z
DNA Meth. m5C:H4K20 me2
H3K9 me2:DNA Meth. m5C
H3K4 me1:H4K20 me2
H3K9 me3:H4K20 me3
H3K4 me3:H3−5ac
H3K4 me1:H4−4ac
H3K27 me3:H3K9 me3
H3K4 me1:H3−5ac
DNA Meth. m5C:H4K20 me3
H2A.Z:H4−4ac
H2A.Z:H3−5ac
H4−4ac:H3−5ac
H3K27 me3:H4K20 me3
H3K27 me3:DNA Meth. m5C
H4K20 me2:H4−4ac
H2A.Z:H3K4 me3
H2A.Z:H4K20 me2
H4K20 me2:H3K4 me3
DNA Meth. m5C:H3K9 me3
H4K20 me2:H3−5ac
H3K4 me3:H4−4ac
H3K27 me2
H3K9 me2
H3K9 me3
H3K27 me3
H4K20 me3
DNA Meth. m5C
H3K4 me1
H4−4ac
H3K4 me3
H2A.Z
H3−5ac
H4K20 me2
0.00
0.25
0.50
0.75
1.00
π̂
DNAJC9 − Forward
H3K4 me1:H2A.Z
H3K4 me1:H3−5ac
H3K4 me1:H4−4ac
H3K4 me1:H4K20 me2
H3K27 me3:H3K4 me1
H2A.Z:H3−5ac
H2A.Z:H4−4ac
H3K4 me3:H3−5ac
DNA Meth. m5C:H4K20 me3
H2A.Z:H4K20 me2
H4K20 me2:H3−5ac
H2A.Z:H3K4 me3
H3K9 me2:DNA Meth. m5C
DNA Meth. m5C:H4K20 me2
H4K20 me2:H4−4ac
H4K20 me2:H3K4 me3
H4−4ac:H3−5ac
H3K27 me3:DNA Meth. m5C
H3K4 me3:H4−4ac
H3K27 me2
H3K9 me3
H3K9 me2
H3K4 me1
H4K20 me3
H2A.Z
DNA Meth. m5C
H3K4 me3
H3−5ac
H4−4ac
H3K27 me3
H4K20 me2
0.00
0.25
0.50
0.75
1.00
π̂
DNAJC9 − Reverse
H3K4 me1:H2A.Z
H3K4 me1:H3−5ac
H3K4 me1:H4−4ac
H3K4 me1:H4K20 me2
H3K27 me3:H3K4 me1
H2A.Z:H3−5ac
H2A.Z:H4−4ac
H3K4 me3:H3−5ac
DNA Meth. m5C:H4K20 me3
H2A.Z:H4K20 me2
H4K20 me2:H3−5ac
H2A.Z:H3K4 me3
H3K9 me2:DNA Meth. m5C
DNA Meth. m5C:H4K20 me2
H4K20 me2:H4−4ac
H4K20 me2:H3K4 me3
H4−4ac:H3−5ac
H3K27 me3:DNA Meth. m5C
H3K4 me3:H4−4ac
H3K27 me2
H3K9 me3
H3K9 me2
H3K4 me1
H4K20 me3
H2A.Z
DNA Meth. m5C
H3K4 me3
H3−5ac
H4−4ac
H3K27 me3
H4K20 me2
0.00
0.25
0.50
0.75
1.00
π̂
TFAP4 − Forward
H3K4 me1:H2A.Z
H3K4 me1:H3−5ac
H3K4 me1:H4−4ac
H3K4 me1:H4K20 me2
H3K9 me3:H4K20 me3
H2A.Z:H4−4ac
DNA Meth. m5C:H4K20 me2
H3K27 me3:H4K20 me3
DNA Meth. m5C:H4K20 me3
H3K27 me3:DNA Meth. m5C
H2A.Z:H3−5ac
H3K4 me3:H3−5ac
H2A.Z:H3K4 me3
DNA Meth. m5C:H3K9 me3
H3K27 me3:H3K9 me3
H3K9 me2:DNA Meth. m5C
H2A.Z:H4K20 me2
H4−4ac:H3−5ac
H4K20 me2:H3K4 me3
H4K20 me2:H4−4ac
H3K4 me3:H4−4ac
H4K20 me2:H3−5ac
H3K27 me2
H3K4 me1
H3K9 me3
H3K9 me2
H3K27 me3
H3K4 me3
H4K20 me3
H2A.Z
H3−5ac
H4−4ac
H4K20 me2
DNA Meth. m5C
0.00
0.25
0.50
0.75
1.00
π̂
TFAP4 − Reverse
H3K4 me1:H2A.Z
H3K4 me1:H3−5ac
H3K4 me1:H4−4ac
H3K4 me1:H4K20 me2
H3K9 me3:H4K20 me3
H2A.Z:H4−4ac
DNA Meth. m5C:H4K20 me2
H3K27 me3:H4K20 me3
DNA Meth. m5C:H4K20 me3
H3K27 me3:DNA Meth. m5C
H2A.Z:H3−5ac
H3K4 me3:H3−5ac
H2A.Z:H3K4 me3
DNA Meth. m5C:H3K9 me3
H3K27 me3:H3K9 me3
H3K9 me2:DNA Meth. m5C
H2A.Z:H4K20 me2
H4−4ac:H3−5ac
H4K20 me2:H3K4 me3
H4K20 me2:H4−4ac
H3K4 me3:H4−4ac
H4K20 me2:H3−5ac
H3K27 me2
H3K4 me1
H3K9 me3
H3K9 me2
H3K27 me3
H3K4 me3
H4K20 me3
H2A.Z
H3−5ac
H4−4ac
H4K20 me2
DNA Meth. m5C
0.00
0.25
0.50
0.75
1.00
π̂
IFI16 − Forward
H3K4 me1:H4−4ac
H3K4 me1:H2A.Z
H3K4 me1:H4K20 me2
H3K27 me3:H3K9 me3
H2A.Z:H3−5ac
DNA Meth. m5C:H3K9 me3
H4−4ac:H3−5ac
H3K27 me3:DNA Meth. m5C
H2A.Z:H4−4ac
H2A.Z:H3K4 me3
DNA Meth. m5C:H4K20 me2
H3K4 me3:H4−4ac
H3K27 me2:DNA Meth. m5C
H4K20 me2:H3K4 me3
H4K20 me2:H4−4ac
H2A.Z:H4K20 me2
H4K20 me2:H3−5ac
H3K9 me2
H4K20 me3
H3K4 me3
H3K9 me3
H3K27 me2
H3K4 me1
H3−5ac
H4−4ac
H3K27 me3
H2A.Z
DNA Meth. m5C
H4K20 me2
0.00
0.25
0.50
0.75
1.00
π̂
IFI16 − Reverse
H3K4 me1:H4−4ac
H3K4 me1:H2A.Z
H3K4 me1:H4K20 me2
H3K27 me3:H3K9 me3
H2A.Z:H3−5ac
DNA Meth. m5C:H3K9 me3
H4−4ac:H3−5ac
H3K27 me3:DNA Meth. m5C
H2A.Z:H4−4ac
H2A.Z:H3K4 me3
DNA Meth. m5C:H4K20 me2
H3K4 me3:H4−4ac
H3K27 me2:DNA Meth. m5C
H4K20 me2:H3K4 me3
H4K20 me2:H4−4ac
H2A.Z:H4K20 me2
H4K20 me2:H3−5ac
H3K9 me2
H4K20 me3
H3K4 me3
H3K9 me3
H3K27 me2
H3K4 me1
H3−5ac
H4−4ac
H3K27 me3
H2A.Z
DNA Meth. m5C
H4K20 me2
0.00
0.25
0.50
0.75
1.00
π̂
RCOR1 (1) − Forward
H2A.Z:H3−5ac
H2A.Z:H4−4ac
H3K9 me3:H4K20 me3
H3K27 me3:H4K20 me3
H3K27 me3:H3K4 me1
H4−4ac:H3−5ac
H3K4 me1:H4−4ac
H3K4 me1:H4K20 me2
H3K4 me1:H2A.Z
H3K27 me3:H3K9 me3
H3K4 me3:H3−5ac
H2A.Z:H3K4 me3
H2A.Z:H4K20 me2
H4K20 me2:H3−5ac
DNA Meth. m5C:H4K20 me3
DNA Meth. m5C:H3K9 me3
H4K20 me2:H4−4ac
H3K4 me1:H3−5ac
H4K20 me2:H3K4 me3
H3K9 me2:DNA Meth. m5C
H3K4 me3:H4−4ac
H3K27 me3:DNA Meth. m5C
H3K27 me2
H4K20 me3
H3K4 me1
H3K9 me2
H3K4 me3
H2A.Z
H3K9 me3
H4−4ac
H3K27 me3
H4K20 me2
DNA Meth. m5C
H3−5ac
0.00
0.25
0.50
0.75
1.00
π̂
RCOR1 (1) − Reverse
H2A.Z:H3−5ac
H2A.Z:H4−4ac
H3K9 me3:H4K20 me3
H3K27 me3:H4K20 me3
H3K27 me3:H3K4 me1
H4−4ac:H3−5ac
H3K4 me1:H4−4ac
H3K4 me1:H4K20 me2
H3K4 me1:H2A.Z
H3K27 me3:H3K9 me3
H3K4 me3:H3−5ac
H2A.Z:H3K4 me3
H2A.Z:H4K20 me2
H4K20 me2:H3−5ac
DNA Meth. m5C:H4K20 me3
DNA Meth. m5C:H3K9 me3
H4K20 me2:H4−4ac
H3K4 me1:H3−5ac
H4K20 me2:H3K4 me3
H3K9 me2:DNA Meth. m5C
H3K4 me3:H4−4ac
H3K27 me3:DNA Meth. m5C
H3K27 me2
H4K20 me3
H3K4 me1
H3K9 me2
H3K4 me3
H2A.Z
H3K9 me3
H4−4ac
H3K27 me3
H4K20 me2
DNA Meth. m5C
H3−5ac
0.00
0.25
0.50
0.75
1.00
π̂

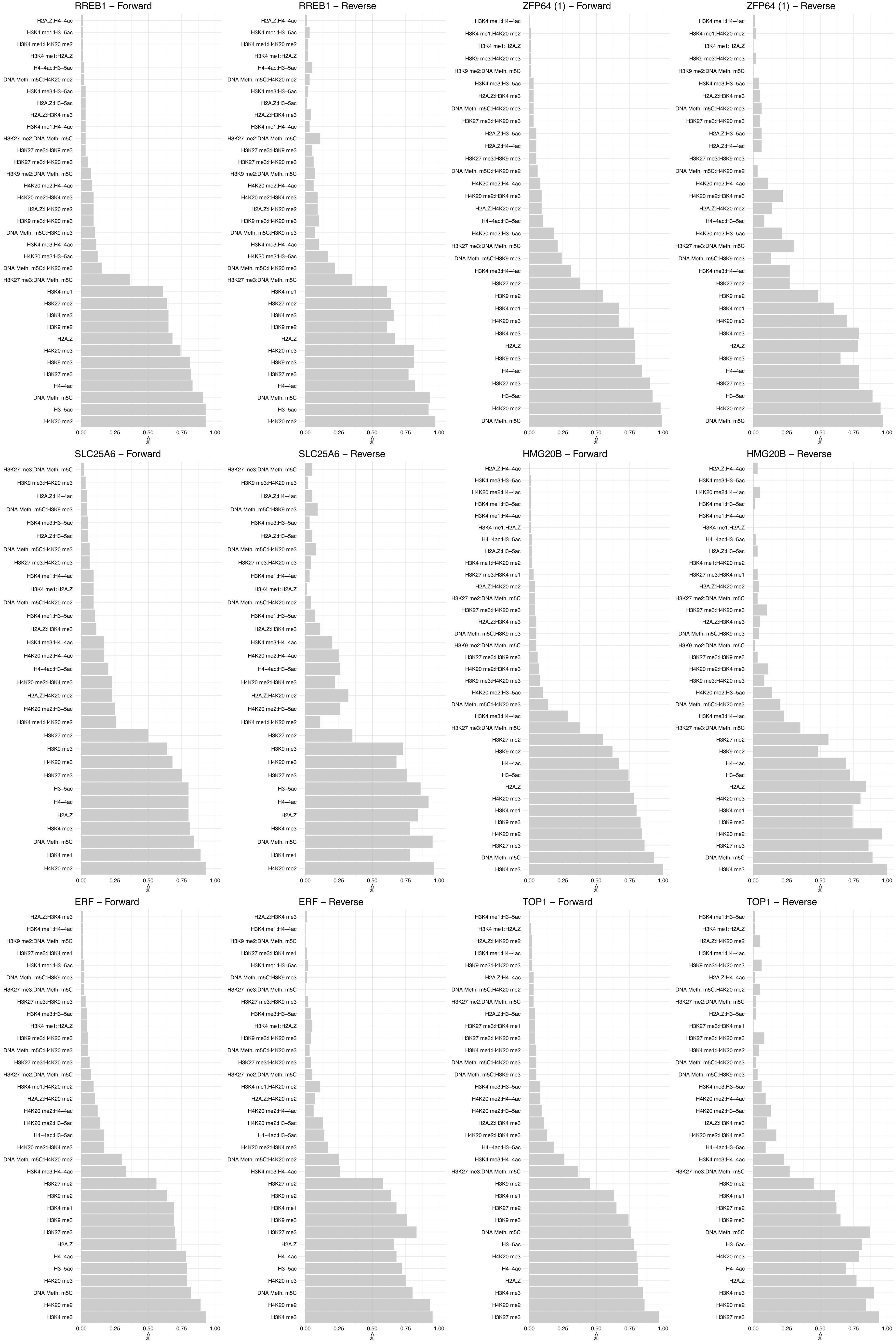
RREB1 – Forward
H2A.Z:H4−4ac
H3K4 me1:H3−5ac
H3K4 me1:H4K20 me2
H3K4 me1:H2A.Z
H4−4ac:H3−5ac
DNA Meth. m5C:H4K20 me2
H3K4 me3:H3−5ac
H2A.Z:H3−5ac
H2A.Z:H3K4 me3
H3K4 me1:H4−4ac
H3K27 me2:DNA Meth. m5C
H3K27 me3:H3K9 me3
H3K27 me3:H4K20 me3
H3K9 me2:DNA Meth. m5C
H4K20 me2:H4−4ac
H4K20 me2:H3K4 me3
H2A.Z:H4K20 me2
H3K9 me3:H4K20 me3
DNA Meth. m5C:H3K9 me3
H3K4 me3:H4−4ac
H4K20 me2:H3−5ac
DNA Meth. m5C:H4K20 me3
H3K27 me3:DNA Meth. m5C
H3K4 me1
H3K27 me2
H3K4 me3
H3K9 me2
H2A.Z
H4K20 me3
H3K9 me3
H3K27 me3
H4−4ac
DNA Meth. m5C
H3−5ac
H4K20 me2
0.00
0.25
0.50
0.75
1.00
π̂
RREB1 – Reverse
H2A.Z:H4−4ac
H3K4 me1:H3−5ac
H3K4 me1:H4K20 me2
H3K4 me1:H2A.Z
H4−4ac:H3−5ac
DNA Meth. m5C:H4K20 me2
H3K4 me3:H3−5ac
H2A.Z:H3−5ac
H2A.Z:H3K4 me3
H3K4 me1:H4−4ac
H3K27 me2:DNA Meth. m5C
H3K27 me3:H3K9 me3
H3K27 me3:H4K20 me3
H3K9 me2:DNA Meth. m5C
H4K20 me2:H4−4ac
H4K20 me2:H3K4 me3
H2A.Z:H4K20 me2
H3K9 me3:H4K20 me3
DNA Meth. m5C:H3K9 me3
H3K4 me3:H4−4ac
H4K20 me2:H3−5ac
DNA Meth. m5C:H4K20 me3
H3K27 me3:DNA Meth. m5C
H3K4 me1
H3K27 me2
H3K4 me3
H3K9 me2
H2A.Z
H4K20 me3
H3K9 me3
H3K27 me3
H4−4ac
DNA Meth. m5C
H3−5ac
H4K20 me2
0.00
0.25
0.50
0.75
1.00
π̂
ZFP64 (1) – Forward
H3K4 me1:H4−4ac
H3K4 me1:H4K20 me2
H3K4 me1:H2A.Z
H3K9 me3:H4K20 me3
H3K9 me2:DNA Meth. m5C
H3K4 me3:H3−5ac
H2A.Z:H3K4 me3
DNA Meth. m5C:H4K20 me3
H3K27 me3:H4K20 me3
H2A.Z:H3−5ac
H2A.Z:H4−4ac
H3K27 me3:H3K9 me3
DNA Meth. m5C:H4K20 me2
H4K20 me2:H4−4ac
H4K20 me2:H3K4 me3
H2A.Z:H4K20 me2
H4−4ac:H3−5ac
H4K20 me2:H3−5ac
H3K27 me3:DNA Meth. m5C
DNA Meth. m5C:H3K9 me3
H3K4 me3:H4−4ac
H3K27 me2
H3K9 me2
H3K4 me1
H4K20 me3
H3K4 me3
H2A.Z
H3K9 me3
H4−4ac
H3K27 me3
H3−5ac
H4K20 me2
DNA Meth. m5C
0.00
0.25
0.50
0.75
1.00
π̂
ZFP64 (1) – Reverse
H3K4 me1:H4−4ac
H3K4 me1:H4K20 me2
H3K4 me1:H2A.Z
H3K9 me3:H4K20 me3
H3K9 me2:DNA Meth. m5C
H3K4 me3:H3−5ac
H2A.Z:H3K4 me3
DNA Meth. m5C:H4K20 me3
H3K27 me3:H4K20 me3
H2A.Z:H3−5ac
H2A.Z:H4−4ac
H3K27 me3:H3K9 me3
DNA Meth. m5C:H4K20 me2
H4K20 me2:H4−4ac
H4K20 me2:H3K4 me3
H2A.Z:H4K20 me2
H4−4ac:H3−5ac
H4K20 me2:H3−5ac
H3K27 me3:DNA Meth. m5C
DNA Meth. m5C:H3K9 me3
H3K4 me3:H4−4ac
H3K27 me2
H3K9 me2
H3K4 me1
H4K20 me3
H3K4 me3
H2A.Z
H3K9 me3
H4−4ac
H3K27 me3
H3−5ac
H4K20 me2
DNA Meth. m5C
0.00
0.25
0.50
0.75
1.00
π̂
SLC25A6 – Forward
H3K27 me3:DNA Meth. m5C
H3K9 me3:H4K20 me3
H2A.Z:H4−4ac
DNA Meth. m5C:H3K9 me3
H3K4 me3:H3−5ac
H2A.Z:H3−5ac
DNA Meth. m5C:H4K20 me3
H3K27 me3:H4K20 me3
H3K4 me1:H4−4ac
H3K4 me1:H2A.Z
DNA Meth. m5C:H4K20 me2
H3K4 me1:H3−5ac
H2A.Z:H3K4 me3
H3K4 me3:H4−4ac
H4K20 me2:H4−4ac
H4−4ac:H3−5ac
H4K20 me2:H3K4 me3
H2A.Z:H4K20 me2
H4K20 me2:H3−5ac
H3K4 me1:H4K20 me2
H3K27 me2
H3K9 me3
H4K20 me3
H3K27 me3
H3−5ac
H4−4ac
H2A.Z
H3K4 me3
DNA Meth. m5C
H3K4 me1
H4K20 me2
0.00
0.25
0.50
0.75
1.00
π̂
SLC25A6 – Reverse
H3K27 me3:DNA Meth. m5C
H3K9 me3:H4K20 me3
H2A.Z:H4−4ac
DNA Meth. m5C:H3K9 me3
H3K4 me3:H3−5ac
H2A.Z:H3−5ac
DNA Meth. m5C:H4K20 me3
H3K27 me3:H4K20 me3
H3K4 me1:H4−4ac
H3K4 me1:H2A.Z
DNA Meth. m5C:H4K20 me2
H3K4 me1:H3−5ac
H2A.Z:H3K4 me3
H3K4 me3:H4−4ac
H4K20 me2:H4−4ac
H4−4ac:H3−5ac
H4K20 me2:H3K4 me3
H2A.Z:H4K20 me2
H4K20 me2:H3−5ac
H3K4 me1:H4K20 me2
H3K27 me2
H3K9 me3
H4K20 me3
H3K27 me3
H3−5ac
H4−4ac
H2A.Z
H3K4 me3
DNA Meth. m5C
H3K4 me1
H4K20 me2
0.00
0.25
0.50
0.75
1.00
π̂
HMG20B – Forward
H2A.Z:H4−4ac
H3K4 me3:H3−5ac
H4K20 me2:H4−4ac
H3K4 me1:H3−5ac
H3K4 me1:H4−4ac
H3K4 me1:H2A.Z
H4−4ac:H3−5ac
H2A.Z:H3−5ac
H3K4 me1:H4K20 me2
H3K27 me3:H3K4 me1
H2A.Z:H4K20 me2
H3K27 me2:DNA Meth. m5C
H3K27 me3:H4K20 me3
H2A.Z:H3K4 me3
DNA Meth. m5C:H3K9 me3
H3K9 me2:DNA Meth. m5C
H3K27 me3:H3K9 me3
H4K20 me2:H3K4 me3
H3K9 me3:H4K20 me3
H4K20 me2:H3−5ac
DNA Meth. m5C:H4K20 me3
H3K4 me3:H4−4ac
H3K27 me3:DNA Meth. m5C
H3K27 me2
H3K9 me2
H4−4ac
H3−5ac
H2A.Z
H4K20 me3
H3K4 me1
H3K9 me3
H4K20 me2
H3K27 me3
DNA Meth. m5C
H3K4 me3
0.00
0.25
0.50
0.75
1.00
π̂
HMG20B – Reverse
H2A.Z:H4−4ac
H3K4 me3:H3−5ac
H4K20 me2:H4−4ac
H3K4 me1:H3−5ac
H3K4 me1:H4−4ac
H3K4 me1:H2A.Z
H4−4ac:H3−5ac
H2A.Z:H3−5ac
H3K4 me1:H4K20 me2
H3K27 me3:H3K4 me1
H2A.Z:H4K20 me2
H3K27 me2:DNA Meth. m5C
H3K27 me3:H4K20 me3
H2A.Z:H3K4 me3
DNA Meth. m5C:H3K9 me3
H3K9 me2:DNA Meth. m5C
H3K27 me3:H3K9 me3
H4K20 me2:H3K4 me3
H3K9 me3:H4K20 me3
H4K20 me2:H3−5ac
DNA Meth. m5C:H4K20 me3
H3K4 me3:H4−4ac
H3K27 me3:DNA Meth. m5C
H3K27 me2
H3K9 me2
H4−4ac
H3−5ac
H2A.Z
H4K20 me3
H3K4 me1
H3K9 me3
H4K20 me2
H3K27 me3
DNA Meth. m5C
H3K4 me3
0.00
0.25
0.50
0.75
1.00
π̂
ERF – Forward
H2A.Z:H3K4 me3
H3K4 me1:H4−4ac
H3K9 me2:DNA Meth. m5C
H3K27 me3:H3K4 me1
H3K4 me1:H3−5ac
DNA Meth. m5C:H3K9 me3
H3K27 me3:DNA Meth. m5C
H3K27 me3:H3K9 me3
H3K4 me3:H3−5ac
H3K4 me1:H2A.Z
H3K9 me3:H4K20 me3
DNA Meth. m5C:H4K20 me3
H3K27 me3:H4K20 me3
H3K27 me2:DNA Meth. m5C
H3K4 me1:H4K20 me2
H2A.Z:H4K20 me2
H4K20 me2:H4−4ac
H4K20 me2:H3−5ac
H4−4ac:H3−5ac
H4K20 me2:H3K4 me3
DNA Meth. m5C:H4K20 me2
H3K4 me3:H4−4ac
H3K27 me2
H3K9 me2
H3K4 me1
H3K9 me3
H3K27 me3
H2A.Z
H4−4ac
H3−5ac
H4K20 me3
DNA Meth. m5C
H4K20 me2
H3K4 me3
0.00
0.25
0.50
0.75
1.00
π̂
ERF – Reverse
H2A.Z:H3K4 me3
H3K4 me1:H4−4ac
H3K9 me2:DNA Meth. m5C
H3K27 me3:H3K4 me1
H3K4 me1:H3−5ac
DNA Meth. m5C:H3K9 me3
H3K27 me3:DNA Meth. m5C
H3K27 me3:H3K9 me3
H3K4 me3:H3−5ac
H3K4 me1:H2A.Z
H3K9 me3:H4K20 me3
DNA Meth. m5C:H4K20 me3
H3K27 me3:H4K20 me3
H3K27 me2:DNA Meth. m5C
H3K4 me1:H4K20 me2
H2A.Z:H4K20 me2
H4K20 me2:H4−4ac
H4K20 me2:H3−5ac
H4−4ac:H3−5ac
H4K20 me2:H3K4 me3
DNA Meth. m5C:H4K20 me2
H3K4 me3:H4−4ac
H3K27 me2
H3K9 me2
H3K4 me1
H3K9 me3
H3K27 me3
H2A.Z
H4−4ac
H3−5ac
H4K20 me3
DNA Meth. m5C
H4K20 me2
H3K4 me3
0.00
0.25
0.50
0.75
1.00
π̂
TOP1 – Forward
H3K4 me1:H3−5ac
H3K4 me1:H2A.Z
H2A.Z:H4K20 me2
H3K4 me1:H4−4ac
H3K9 me3:H4K20 me3
H2A.Z:H4−4ac
DNA Meth. m5C:H4K20 me2
H3K27 me2:DNA Meth. m5C
H2A.Z:H3−5ac
H3K27 me3:H3K4 me1
H3K27 me3:H4K20 me3
H3K4 me1:H4K20 me2
DNA Meth. m5C:H4K20 me3
DNA Meth. m5C:H3K9 me3
H3K4 me3:H3−5ac
H4K20 me2:H4−4ac
H4K20 me2:H3−5ac
H2A.Z:H3K4 me3
H4K20 me2:H3K4 me3
H4−4ac:H3−5ac
H3K4 me3:H4−4ac
H3K27 me3:DNA Meth. m5C
H3K9 me2
H3K4 me1
H3K27 me2
H3K9 me3
DNA Meth. m5C
H3−5ac
H4K20 me3
H4−4ac
H2A.Z
H3K4 me3
H4K20 me2
H3K27 me3
0.00
0.25
0.50
0.75
1.00
π̂
TOP1 – Reverse
H3K4 me1:H3−5ac
H3K4 me1:H2A.Z
H2A.Z:H4K20 me2
H3K4 me1:H4−4ac
H3K9 me3:H4K20 me3
H2A.Z:H4−4ac
DNA Meth. m5C:H4K20 me2
H3K27 me2:DNA Meth. m5C
H2A.Z:H3−5ac
H3K27 me3:H3K4 me1
H3K27 me3:H4K20 me3
H3K4 me1:H4K20 me2
DNA Meth. m5C:H4K20 me3
DNA Meth. m5C:H3K9 me3
H3K4 me3:H3−5ac
H4K20 me2:H4−4ac
H4K20 me2:H3−5ac
H2A.Z:H3K4 me3
H4K20 me2:H3K4 me3
H4−4ac:H3−5ac
H3K4 me3:H4−4ac
H3K27 me3:DNA Meth. m5C
H3K9 me2
H3K4 me1
H3K27 me2
H3K9 me3
DNA Meth. m5C
H3−5ac
H4K20 me3
H4−4ac
H2A.Z
H3K4 me3
H4K20 me2
H3K27 me3
0.00
0.25
0.50
0.75
1.00
π̂

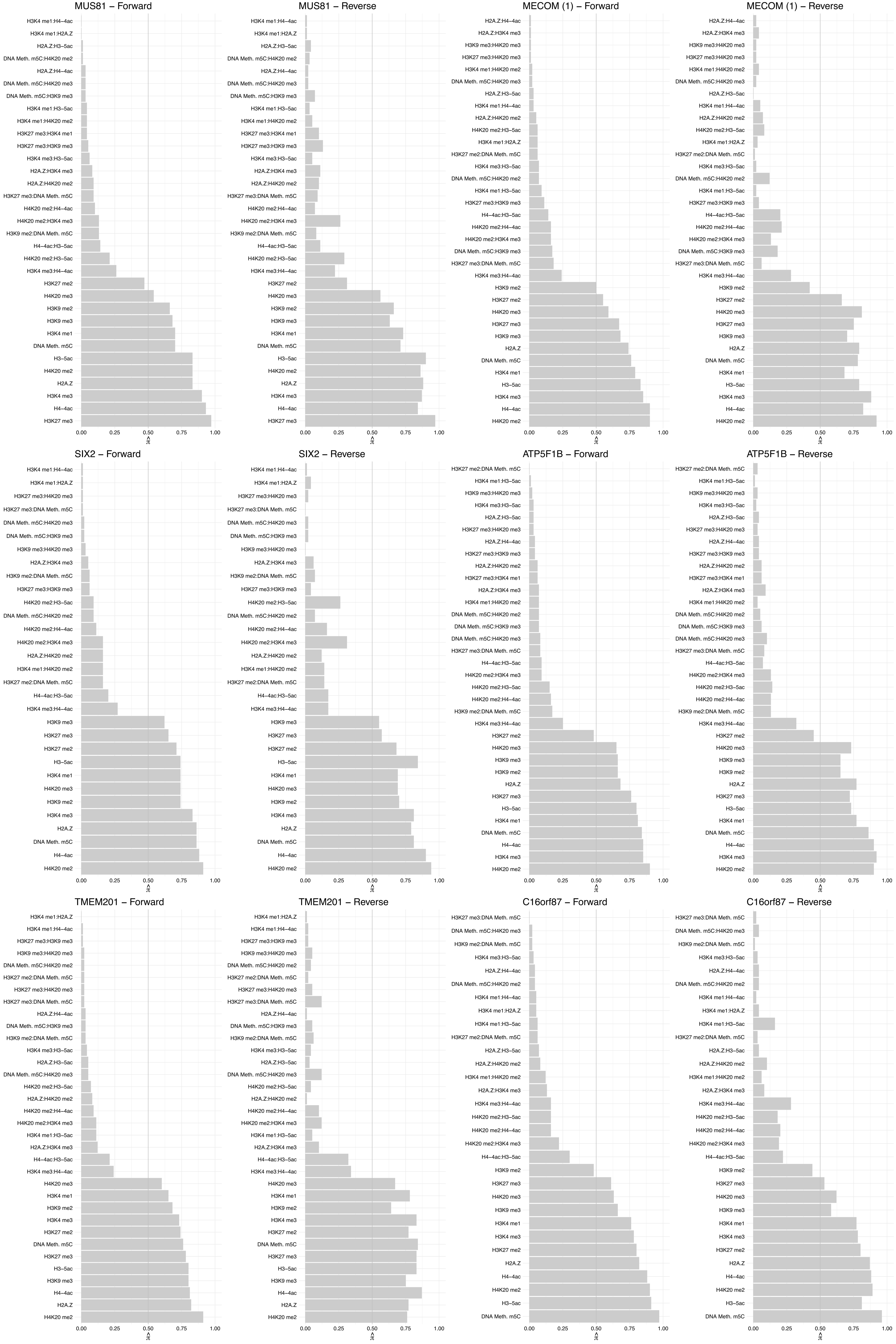
MUS81 – Forward
MUS81 – Reverse
H3K4 me1:H4–4ac
H3K4 me1:H2A.Z
H2A.Z:H3–5ac
DNA Meth. m5C:H4K20 me2
H2A.Z:H4–4ac
DNA Meth. m5C:H4K20 me3
DNA Meth. m5C:H3K9 me3
H3K4 me1:H3–5ac
H3K4 me1:H4K20 me2
H3K27 me3:H3K4 me1
H3K27 me3:H3K9 me3
H3K4 me3:H3–5ac
H2A.Z:H3K4 me3
H2A.Z:H4K20 me2
H3K27 me3:DNA Meth. m5C
H4K20 me2:H4–4ac
H4K20 me2:H3K4 me3
H3K9 me2:DNA Meth. m5C
H4–4ac:H3–5ac
H4K20 me2:H3–5ac
H3K4 me3:H4–4ac
H3K27 me2
H4K20 me3
H3K9 me2
H3K9 me3
H3K4 me1
DNA Meth. m5C
H3–5ac
H4K20 me2
H2A.Z
H3K4 me3
H4–4ac
H3K27 me3
MECOM (1) – Forward
MECOM (1) – Reverse
H2A.Z:H4–4ac
H2A.Z:H3K4 me3
H3K9 me3:H4K20 me3
H3K27 me3:H4K20 me3
H3K4 me1:H4K20 me2
DNA Meth. m5C:H4K20 me3
H2A.Z:H3–5ac
H3K4 me1:H4–4ac
H2A.Z:H4K20 me2
H4K20 me2:H3–5ac
H3K4 me1:H2A.Z
H3K27 me2:DNA Meth. m5C
H3K4 me3:H3–5ac
DNA Meth. m5C:H4K20 me2
H3K4 me1:H3–5ac
H3K27 me3:H3K9 me3
H4–4ac:H3–5ac
H4K20 me2:H4–4ac
H4K20 me2:H3K4 me3
DNA Meth. m5C:H3K9 me3
H3K27 me3:DNA Meth. m5C
H3K4 me3:H4–4ac
H3K9 me2
H3K27 me2
H4K20 me3
H3K27 me3
H3K9 me3
H2A.Z
DNA Meth. m5C
H3K4 me1
H3–5ac
H3K4 me3
H4–4ac
H4K20 me2
SIX2 – Forward
SIX2 – Reverse
H3K4 me1:H4–4ac
H3K4 me1:H2A.Z
H3K27 me3:H4K20 me3
H3K27 me3:DNA Meth. m5C
DNA Meth. m5C:H4K20 me3
DNA Meth. m5C:H3K9 me3
H3K9 me3:H4K20 me3
H2A.Z:H3K4 me3
H3K9 me2:DNA Meth. m5C
H3K27 me3:H3K9 me3
H4K20 me2:H3–5ac
DNA Meth. m5C:H4K20 me2
H4K20 me2:H4–4ac
H4K20 me2:H3K4 me3
H2A.Z:H4K20 me2
H3K4 me1:H4K20 me2
H3K27 me2:DNA Meth. m5C
H4–4ac:H3–5ac
H3K4 me3:H4–4ac
H3K9 me3
H3K27 me3
H3K27 me2
H3–5ac
H3K4 me1
H4K20 me3
H3K9 me2
H3K4 me3
H2A.Z
DNA Meth. m5C
H4–4ac
H4K20 me2
ATP5F1B – Forward
ATP5F1B – Reverse
H3K27 me2:DNA Meth. m5C
H3K4 me1:H3–5ac
H3K9 me3:H4K20 me3
H3K4 me3:H3–5ac
H2A.Z:H3–5ac
H3K27 me3:H4K20 me3
H2A.Z:H4–4ac
H3K27 me3:H3K9 me3
H2A.Z:H4K20 me2
H3K27 me3:H3K4 me1
H2A.Z:H3K4 me3
H3K4 me1:H4K20 me2
DNA Meth. m5C:H4K20 me2
DNA Meth. m5C:H3K9 me3
DNA Meth. m5C:H4K20 me3
H3K27 me3:DNA Meth. m5C
H4–4ac:H3–5ac
H4K20 me2:H3K4 me3
H4K20 me2:H3–5ac
H4K20 me2:H4–4ac
H3K9 me2:DNA Meth. m5C
H3K4 me3:H4–4ac
H3K27 me2
H4K20 me3
H3K9 me3
H3K9 me2
H2A.Z
H3K27 me3
H3–5ac
H3K4 me1
DNA Meth. m5C
H4–4ac
H3K4 me3
H4K20 me2
TMEM201 – Forward
TMEM201 – Reverse
H3K4 me1:H2A.Z
H3K4 me1:H4–4ac
H3K27 me3:H3K9 me3
H3K9 me3:H4K20 me3
DNA Meth. m5C:H4K20 me2
H3K27 me2:DNA Meth. m5C
H3K27 me3:H4K20 me3
H3K27 me3:DNA Meth. m5C
H2A.Z:H4–4ac
DNA Meth. m5C:H3K9 me3
H3K9 me2:DNA Meth. m5C
H3K4 me3:H3–5ac
H2A.Z:H3–5ac
DNA Meth. m5C:H4K20 me3
H4K20 me2:H3–5ac
H2A.Z:H4K20 me2
H4K20 me2:H4–4ac
H4K20 me2:H3K4 me3
H3K4 me1:H3–5ac
H2A.Z:H3K4 me3
H4–4ac:H3–5ac
H3K4 me3:H4–4ac
H4K20 me3
H3K4 me1
H3K9 me2
H3K4 me3
H3K27 me2
DNA Meth. m5C
H3K27 me3
H3–5ac
H3K9 me3
H4–4ac
H2A.Z
H4K20 me2
C16orf87 – Forward
C16orf87 – Reverse
H3K27 me3:DNA Meth. m5C
DNA Meth. m5C:H4K20 me3
H3K9 me2:DNA Meth. m5C
H3K4 me3:H3–5ac
H2A.Z:H4–4ac
DNA Meth. m5C:H4K20 me2
H3K4 me1:H4–4ac
H3K4 me1:H2A.Z
H3K4 me1:H3–5ac
H3K27 me2:DNA Meth. m5C
H2A.Z:H3–5ac
H2A.Z:H4K20 me2
H3K4 me1:H4K20 me2
H2A.Z:H3K4 me3
H3K4 me3:H4–4ac
H4K20 me2:H3–5ac
H4K20 me2:H4–4ac
H4K20 me2:H3K4 me3
H4–4ac:H3–5ac
H3K9 me2
H3K27 me3
H4K20 me3
H3K9 me3
H3K4 me1
H3K4 me3
H3K27 me2
H2A.Z
H4–4ac
H4K20 me2
H3–5ac
DNA Meth. m5C
0.00 0.25 0.50 0.75 1.00
$\hat{\pi}$

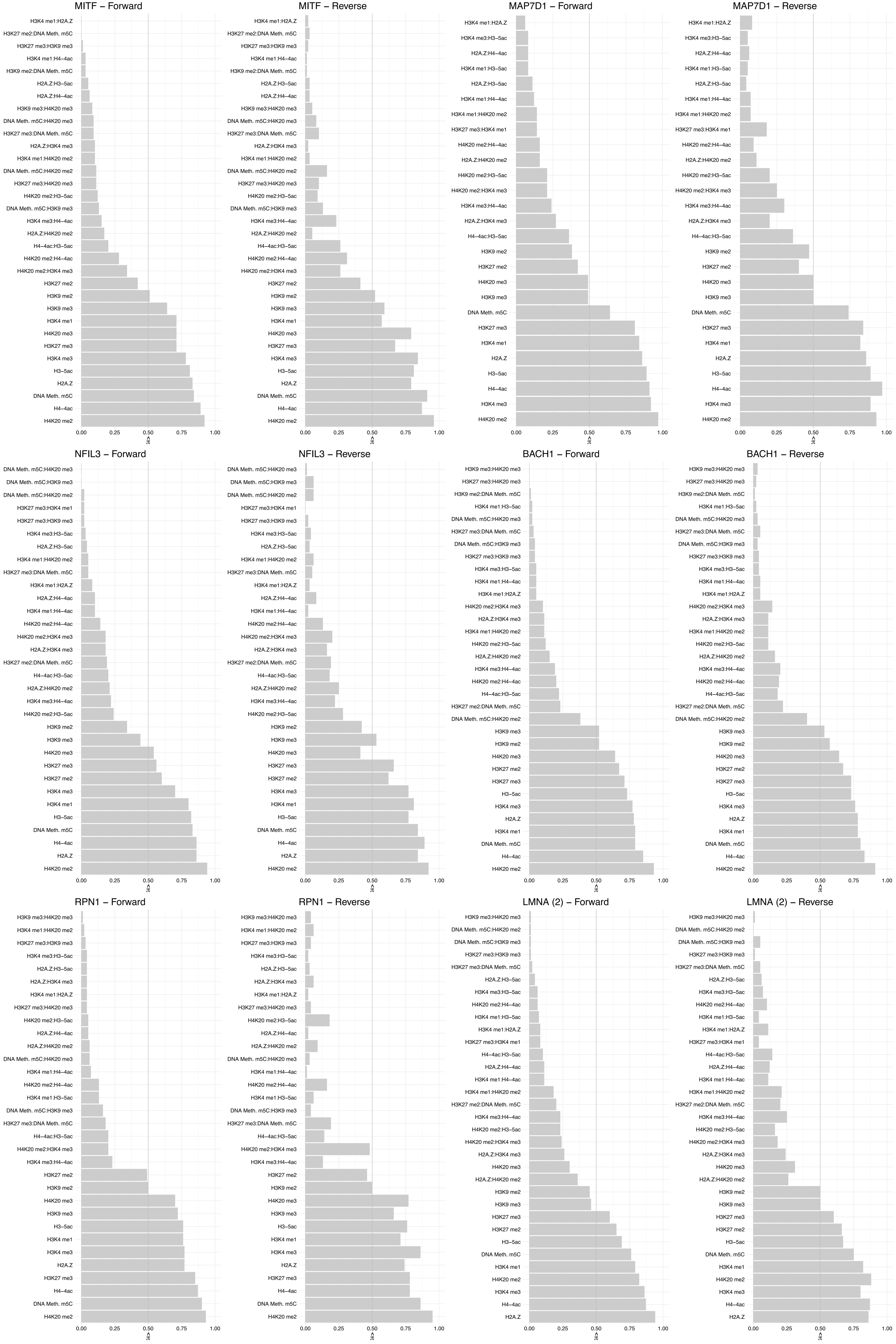

MITF – Forward
H3K4 me1:H2A.Z
H3K27 me2:DNA Meth. m5C
H3K27 me3:H3K9 me3
H3K4 me1:H4–4ac
H3K9 me2:DNA Meth. m5C
H2A.Z:H3–5ac
H2A.Z:H4–4ac
H3K9 me3:H4K20 me3
DNA Meth. m5C:H4K20 me3
H3K27 me3:DNA Meth. m5C
H2A.Z:H3K4 me3
H3K4 me1:H4K20 me2
DNA Meth. m5C:H4K20 me2
H3K27 me3:H4K20 me3
H4K20 me2:H3–5ac
DNA Meth. m5C:H3K9 me3
H3K4 me3:H4–4ac
H2A.Z:H4K20 me2
H4–4ac:H3–5ac
H4K20 me2:H4–4ac
H4K20 me2:H3K4 me3
H3K27 me2
H3K9 me2
H3K9 me3
H3K4 me1
H4K20 me3
H3K27 me3
H3K4 me3
H3–5ac
H2A.Z
DNA Meth. m5C
H4–4ac
H4K20 me2
0.00
0.25
0.50
0.75
1.00
π̂
MITF – Reverse
MAP7D1 – Forward
H3K4 me1:H2A.Z
H3K4 me3:H3–5ac
H2A.Z:H4–4ac
H3K4 me1:H3–5ac
H2A.Z:H3–5ac
H3K4 me1:H4–4ac
H3K4 me1:H4K20 me2
H3K27 me3:H3K4 me1
H4K20 me2:H4–4ac
H2A.Z:H4K20 me2
H4K20 me2:H3–5ac
H4K20 me2:H3K4 me3
H3K4 me3:H4–4ac
H2A.Z:H3K4 me3
H4–4ac:H3–5ac
H3K9 me2
H3K27 me2
H4K20 me3
H3K9 me3
DNA Meth. m5C
H3K27 me3
H3K4 me1
H2A.Z
H3–5ac
H4–4ac
H3K4 me3
H4K20 me2
MAP7D1 – Reverse
NFIL3 – Forward
DNA Meth. m5C:H4K20 me3
DNA Meth. m5C:H3K9 me3
DNA Meth. m5C:H4K20 me2
H3K27 me3:H3K4 me1
H3K27 me3:H3K9 me3
H3K4 me3:H3–5ac
H2A.Z:H3–5ac
H3K4 me1:H4K20 me2
H3K27 me3:DNA Meth. m5C
H3K4 me1:H2A.Z
H2A.Z:H4–4ac
H3K4 me1:H4–4ac
H4K20 me2:H4–4ac
H4K20 me2:H3K4 me3
H2A.Z:H3K4 me3
H3K27 me2:DNA Meth. m5C
H4–4ac:H3–5ac
H2A.Z:H4K20 me2
H3K4 me3:H4–4ac
H4K20 me2:H3–5ac
H3K9 me2
H3K9 me3
H4K20 me3
H3K27 me3
H3K27 me2
H3K4 me3
H3K4 me1
H3–5ac
DNA Meth. m5C
H4–4ac
H2A.Z
H4K20 me2
NFIL3 – Reverse
BACH1 – Forward
H3K9 me3:H4K20 me3
H3K27 me3:H4K20 me3
H3K9 me2:DNA Meth. m5C
H3K4 me1:H3–5ac
DNA Meth. m5C:H4K20 me3
H3K27 me3:DNA Meth. m5C
DNA Meth. m5C:H3K9 me3
H3K27 me3:H3K9 me3
H3K4 me3:H3–5ac
H3K4 me1:H4–4ac
H3K4 me1:H2A.Z
H4K20 me2:H3K4 me3
H2A.Z:H3K4 me3
H3K4 me1:H4K20 me2
H4K20 me2:H3–5ac
H2A.Z:H4K20 me2
H3K4 me3:H4–4ac
H4K20 me2:H4–4ac
H4–4ac:H3–5ac
H3K27 me2:DNA Meth. m5C
DNA Meth. m5C:H4K20 me2
H3K9 me3
H3K9 me2
H4K20 me3
H3K27 me2
H3K27 me3
H3–5ac
H3K4 me3
H2A.Z
H3K4 me1
DNA Meth. m5C
H4–4ac
H4K20 me2
BACH1 – Reverse
RPN1 – Forward
H3K9 me3:H4K20 me3
H3K4 me1:H4K20 me2
H3K27 me3:H3K9 me3
H3K4 me3:H3–5ac
H2A.Z:H3–5ac
H2A.Z:H3K4 me3
H3K4 me1:H2A.Z
H3K27 me3:H4K20 me3
H4K20 me2:H3–5ac
H2A.Z:H4–4ac
H2A.Z:H4K20 me2
DNA Meth. m5C:H4K20 me3
H3K4 me1:H4–4ac
H4K20 me2:H4–4ac
H3K4 me1:H3–5ac
DNA Meth. m5C:H3K9 me3
H3K27 me3:DNA Meth. m5C
H4–4ac:H3–5ac
H4K20 me2:H3K4 me3
H3K4 me3:H4–4ac
H3K27 me2
H3K9 me2
H4K20 me3
H3K9 me3
H3–5ac
H3K4 me1
H3K4 me3
H2A.Z
H3K27 me3
H4–4ac
DNA Meth. m5C
H4K20 me2
RPN1 – Reverse
LMNA (2) – Forward
H3K9 me3:H4K20 me3
DNA Meth. m5C:H4K20 me2
DNA Meth. m5C:H3K9 me3
H3K27 me3:H3K9 me3
H3K27 me3:DNA Meth. m5C
H2A.Z:H3–5ac
H3K4 me3:H3–5ac
H4K20 me2:H4–4ac
H3K4 me1:H3–5ac
H3K4 me1:H2A.Z
H3K27 me3:H3K4 me1
H4–4ac:H3–5ac
H2A.Z:H4–4ac
H3K4 me1:H4–4ac
H3K4 me1:H4K20 me2
H3K27 me2:DNA Meth. m5C
H3K4 me3:H4–4ac
H4K20 me2:H3–5ac
H4K20 me2:H3K4 me3
H2A.Z:H3K4 me3
H4K20 me3
H2A.Z:H4K20 me2
H3K9 me2
H3K9 me3
H3K27 me3
H3K27 me2
H3–5ac
DNA Meth. m5C
H3K4 me1
H4K20 me2
H3K4 me3
H4–4ac
H2A.Z
LMNA (2) – Reverse

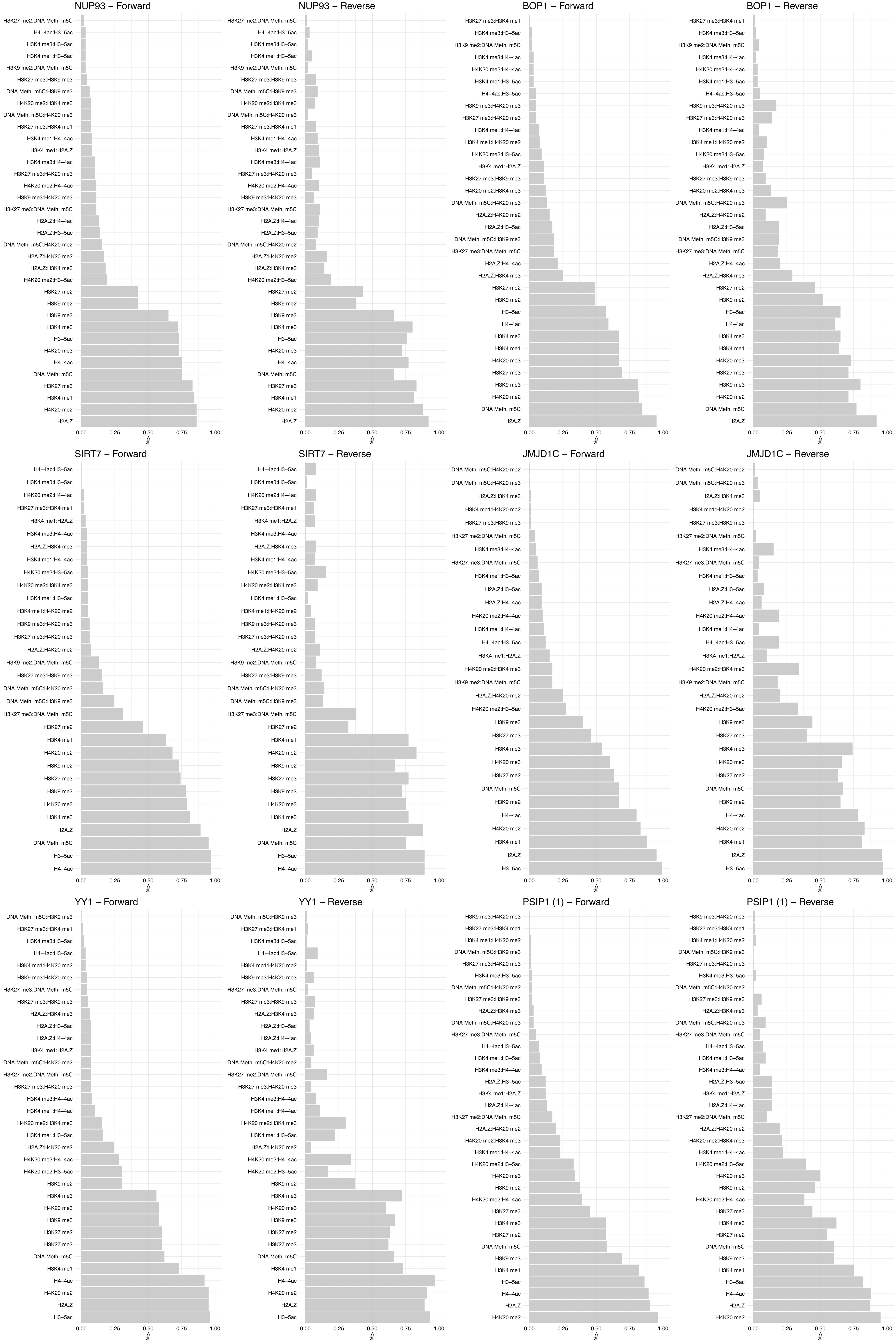
NUP93 – Forward
H3K27 me2:DNA Meth. m5C
H4−4ac:H3−5ac
H3K4 me3:H3−5ac
H3K4 me1:H3−5ac
H3K9 me2:DNA Meth. m5C
H3K27 me3:H3K9 me3
DNA Meth. m5C:H3K9 me3
H4K20 me2:H3K4 me3
DNA Meth. m5C:H4K20 me3
H3K27 me3:H3K4 me1
H3K4 me1:H4−4ac
H3K4 me1:H2A.Z
H3K4 me3:H4−4ac
H3K27 me3:H4K20 me3
H4K20 me2:H4−4ac
H3K9 me3:H4K20 me3
H3K27 me3:DNA Meth. m5C
H2A.Z:H4−4ac
H2A.Z:H3−5ac
DNA Meth. m5C:H4K20 me2
H2A.Z:H4K20 me2
H2A.Z:H3K4 me3
H4K20 me2:H3−5ac
H3K27 me2
H3K9 me2
H3K9 me3
H3K4 me3
H3−5ac
H4K20 me3
H4−4ac
DNA Meth. m5C
H3K27 me3
H3K4 me1
H4K20 me2
H2A.Z
0.00
0.25
0.50
0.75
1.00
π̂
NUP93 – Reverse
H3K27 me2:DNA Meth. m5C
H4−4ac:H3−5ac
H3K4 me3:H3−5ac
H3K4 me1:H3−5ac
H3K9 me2:DNA Meth. m5C
H3K27 me3:H3K9 me3
DNA Meth. m5C:H3K9 me3
H4K20 me2:H3K4 me3
DNA Meth. m5C:H4K20 me3
H3K27 me3:H3K4 me1
H3K4 me1:H4−4ac
H3K4 me1:H2A.Z
H3K4 me3:H4−4ac
H3K27 me3:H4K20 me3
H4K20 me2:H4−4ac
H3K9 me3:H4K20 me3
H3K27 me3:DNA Meth. m5C
H2A.Z:H4−4ac
H2A.Z:H3−5ac
DNA Meth. m5C:H4K20 me2
H2A.Z:H4K20 me2
H2A.Z:H3K4 me3
H4K20 me2:H3−5ac
H3K27 me2
H3K9 me2
H3K9 me3
H3K4 me3
H3−5ac
H4K20 me3
H4−4ac
DNA Meth. m5C
H3K27 me3
H3K4 me1
H4K20 me2
H2A.Z
0.00
0.25
0.50
0.75
1.00
π̂
BOP1 – Forward
H3K27 me3:H3K4 me1
H3K4 me3:H3−5ac
H3K9 me2:DNA Meth. m5C
H3K4 me3:H4−4ac
H4K20 me2:H4−4ac
H3K4 me1:H3−5ac
H4−4ac:H3−5ac
H3K9 me3:H4K20 me3
H3K27 me3:H4K20 me3
H3K4 me1:H4−4ac
H3K4 me1:H4K20 me2
H4K20 me2:H3−5ac
H3K4 me1:H2A.Z
H3K27 me3:H3K9 me3
H4K20 me2:H3K4 me3
DNA Meth. m5C:H4K20 me3
H2A.Z:H4K20 me2
H2A.Z:H3−5ac
DNA Meth. m5C:H3K9 me3
H3K27 me3:DNA Meth. m5C
H2A.Z:H4−4ac
H2A.Z:H3K4 me3
H3K27 me2
H3K9 me2
H3−5ac
H4−4ac
H3K4 me3
H3K4 me1
H4K20 me3
H3K27 me3
H3K9 me3
H4K20 me2
DNA Meth. m5C
H2A.Z
0.00
0.25
0.50
0.75
1.00
π̂
BOP1 – Reverse
H3K27 me3:H3K4 me1
H3K4 me3:H3−5ac
H3K9 me2:DNA Meth. m5C
H3K4 me3:H4−4ac
H4K20 me2:H4−4ac
H3K4 me1:H3−5ac
H4−4ac:H3−5ac
H3K9 me3:H4K20 me3
H3K27 me3:H4K20 me3
H3K4 me1:H4−4ac
H3K4 me1:H4K20 me2
H4K20 me2:H3−5ac
H3K4 me1:H2A.Z
H3K27 me3:H3K9 me3
H4K20 me2:H3K4 me3
DNA Meth. m5C:H4K20 me3
H2A.Z:H4K20 me2
H2A.Z:H3−5ac
DNA Meth. m5C:H3K9 me3
H3K27 me3:DNA Meth. m5C
H2A.Z:H4−4ac
H2A.Z:H3K4 me3
H3K27 me2
H3K9 me2
H3−5ac
H4−4ac
H3K4 me3
H3K4 me1
H4K20 me3
H3K27 me3
H3K9 me3
H4K20 me2
DNA Meth. m5C
H2A.Z
0.00
0.25
0.50
0.75
1.00
π̂
SIRT7 – Forward
H4−4ac:H3−5ac
H3K4 me3:H3−5ac
H4K20 me2:H4−4ac
H3K27 me3:H3K4 me1
H3K4 me1:H2A.Z
H3K4 me3:H4−4ac
H2A.Z:H3K4 me3
H3K4 me1:H4−4ac
H4K20 me2:H3−5ac
H4K20 me2:H3K4 me3
H3K4 me1:H3−5ac
H3K4 me1:H4K20 me2
H3K9 me3:H4K20 me3
H3K27 me3:H4K20 me3
H2A.Z:H4K20 me2
H3K9 me2:DNA Meth. m5C
H3K27 me3:H3K9 me3
DNA Meth. m5C:H4K20 me3
DNA Meth. m5C:H3K9 me3
H3K27 me3:DNA Meth. m5C
H3K27 me2
H3K4 me1
H4K20 me2
H3K9 me2
H3K27 me3
H3K9 me3
H4K20 me3
H3K4 me3
H2A.Z
DNA Meth. m5C
H3−5ac
H4−4ac
0.00
0.25
0.50
0.75
1.00
π̂
SIRT7 – Reverse
H4−4ac:H3−5ac
H3K4 me3:H3−5ac
H4K20 me2:H4−4ac
H3K27 me3:H3K4 me1
H3K4 me1:H2A.Z
H3K4 me3:H4−4ac
H2A.Z:H3K4 me3
H3K4 me1:H4−4ac
H4K20 me2:H3−5ac
H4K20 me2:H3K4 me3
H3K4 me1:H3−5ac
H3K4 me1:H4K20 me2
H3K9 me3:H4K20 me3
H3K27 me3:H4K20 me3
H2A.Z:H4K20 me2
H3K9 me2:DNA Meth. m5C
H3K27 me3:H3K9 me3
DNA Meth. m5C:H4K20 me3
DNA Meth. m5C:H3K9 me3
H3K27 me3:DNA Meth. m5C
H3K27 me2
H3K4 me1
H4K20 me2
H3K9 me2
H3K27 me3
H3K9 me3
H4K20 me3
H3K4 me3
H2A.Z
DNA Meth. m5C
H3−5ac
H4−4ac
0.00
0.25
0.50
0.75
1.00
π̂
JMJD1C – Forward
DNA Meth. m5C:H4K20 me2
DNA Meth. m5C:H4K20 me3
H2A.Z:H3K4 me3
H3K4 me1:H4K20 me2
H3K27 me3:H3K9 me3
H3K27 me2:DNA Meth. m5C
H3K4 me3:H4−4ac
H3K27 me3:DNA Meth. m5C
H3K4 me1:H3−5ac
H2A.Z:H3−5ac
H2A.Z:H4−4ac
H4K20 me2:H4−4ac
H3K4 me1:H4−4ac
H4−4ac:H3−5ac
H3K4 me1:H2A.Z
H4K20 me2:H3K4 me3
H3K9 me2:DNA Meth. m5C
H2A.Z:H4K20 me2
H4K20 me2:H3−5ac
H3K9 me3
H3K27 me3
H3K4 me3
H4K20 me3
H3K27 me2
DNA Meth. m5C
H3K9 me2
H4−4ac
H4K20 me2
H3K4 me1
H2A.Z
H3−5ac
0.00
0.25
0.50
0.75
1.00
π̂
JMJD1C – Reverse
DNA Meth. m5C:H4K20 me2
DNA Meth. m5C:H4K20 me3
H2A.Z:H3K4 me3
H3K4 me1:H4K20 me2
H3K27 me3:H3K9 me3
H3K27 me2:DNA Meth. m5C
H3K4 me3:H4−4ac
H3K27 me3:DNA Meth. m5C
H3K4 me1:H3−5ac
H2A.Z:H3−5ac
H2A.Z:H4−4ac
H4K20 me2:H4−4ac
H3K4 me1:H4−4ac
H4−4ac:H3−5ac
H3K4 me1:H2A.Z
H4K20 me2:H3K4 me3
H3K9 me2:DNA Meth. m5C
H2A.Z:H4K20 me2
H4K20 me2:H3−5ac
H3K9 me3
H3K27 me3
H3K4 me3
H4K20 me3
H3K27 me2
DNA Meth. m5C
H3K9 me2
H4−4ac
H4K20 me2
H3K4 me1
H2A.Z
H3−5ac
0.00
0.25
0.50
0.75
1.00
π̂
YY1 – Forward
DNA Meth. m5C:H3K9 me3
H3K27 me3:H3K4 me1
H3K4 me3:H3−5ac
H4−4ac:H3−5ac
H3K4 me1:H4K20 me2
H3K9 me3:H4K20 me3
H3K27 me3:DNA Meth. m5C
H3K27 me3:H3K9 me3
H2A.Z:H3K4 me3
H2A.Z:H3−5ac
H2A.Z:H4−4ac
H3K4 me1:H2A.Z
DNA Meth. m5C:H4K20 me2
H3K27 me2:DNA Meth. m5C
H3K27 me3:H4K20 me3
H3K4 me3:H4−4ac
H3K4 me1:H4−4ac
H4K20 me2:H3K4 me3
H3K4 me1:H3−5ac
H2A.Z:H4K20 me2
H4K20 me2:H4−4ac
H4K20 me2:H3−5ac
H3K9 me2
H3K4 me3
H4K20 me3
H3K9 me3
H3K27 me2
H3K27 me3
DNA Meth. m5C
H3K4 me1
H4−4ac
H4K20 me2
H2A.Z
H3−5ac
0.00
0.25
0.50
0.75
1.00
π̂
YY1 – Reverse
DNA Meth. m5C:H3K9 me3
H3K27 me3:H3K4 me1
H3K4 me3:H3−5ac
H4−4ac:H3−5ac
H3K4 me1:H4K20 me2
H3K9 me3:H4K20 me3
H3K27 me3:DNA Meth. m5C
H3K27 me3:H3K9 me3
H2A.Z:H3K4 me3
H2A.Z:H3−5ac
H2A.Z:H4−4ac
H3K4 me1:H2A.Z
DNA Meth. m5C:H4K20 me2
H3K27 me2:DNA Meth. m5C
H3K27 me3:H4K20 me3
H3K4 me3:H4−4ac
H3K4 me1:H4−4ac
H4K20 me2:H3K4 me3
H3K4 me1:H3−5ac
H2A.Z:H4K20 me2
H4K20 me2:H4−4ac
H4K20 me2:H3−5ac
H3K9 me2
H3K4 me3
H4K20 me3
H3K9 me3
H3K27 me2
H3K27 me3
DNA Meth. m5C
H3K4 me1
H4−4ac
H4K20 me2
H2A.Z
H3−5ac
0.00
0.25
0.50
0.75
1.00
π̂
PSIP1 (1) – Forward
H3K9 me3:H4K20 me3
H3K27 me3:H3K4 me1
H3K4 me1:H4K20 me2
DNA Meth. m5C:H3K9 me3
H3K27 me3:H4K20 me3
H3K4 me3:H3−5ac
DNA Meth. m5C:H4K20 me2
H3K27 me3:H3K9 me3
H2A.Z:H3K4 me3
DNA Meth. m5C:H4K20 me3
H3K27 me3:DNA Meth. m5C
H4−4ac:H3−5ac
H3K4 me1:H3−5ac
H3K4 me3:H4−4ac
H2A.Z:H3−5ac
H3K4 me1:H2A.Z
H2A.Z:H4−4ac
H3K27 me2:DNA Meth. m5C
H2A.Z:H4K20 me2
H4K20 me2:H3K4 me3
H3K4 me1:H4−4ac
H4K20 me2:H3−5ac
H4K20 me3
H3K9 me2
H4K20 me2:H4−4ac
H3K27 me3
H3K4 me3
H3K27 me2
DNA Meth. m5C
H3K9 me3
H3K4 me1
H3−5ac
H4−4ac
H2A.Z
H4K20 me2
0.00
0.25
0.50
0.75
1.00
π̂
PSIP1 (1) – Reverse
H3K9 me3:H4K20 me3
H3K27 me3:H3K4 me1
H3K4 me1:H4K20 me2
DNA Meth. m5C:H3K9 me3
H3K27 me3:H4K20 me3
H3K4 me3:H3−5ac
DNA Meth. m5C:H4K20 me2
H3K27 me3:H3K9 me3
H2A.Z:H3K4 me3
DNA Meth. m5C:H4K20 me3
H3K27 me3:DNA Meth. m5C
H4−4ac:H3−5ac
H3K4 me1:H3−5ac
H3K4 me3:H4−4ac
H2A.Z:H3−5ac
H3K4 me1:H2A.Z
H2A.Z:H4−4ac
H3K27 me2:DNA Meth. m5C
H2A.Z:H4K20 me2
H4K20 me2:H3K4 me3
H3K4 me1:H4−4ac
H4K20 me2:H3−5ac
H4K20 me3
H3K9 me2
H4K20 me2:H4−4ac
H3K27 me3
H3K4 me3
H3K27 me2
DNA Meth. m5C
H3K9 me3
H3K4 me1
H3−5ac
H4−4ac
H2A.Z
H4K20 me2
0.00
0.25
0.50
0.75
1.00
π̂

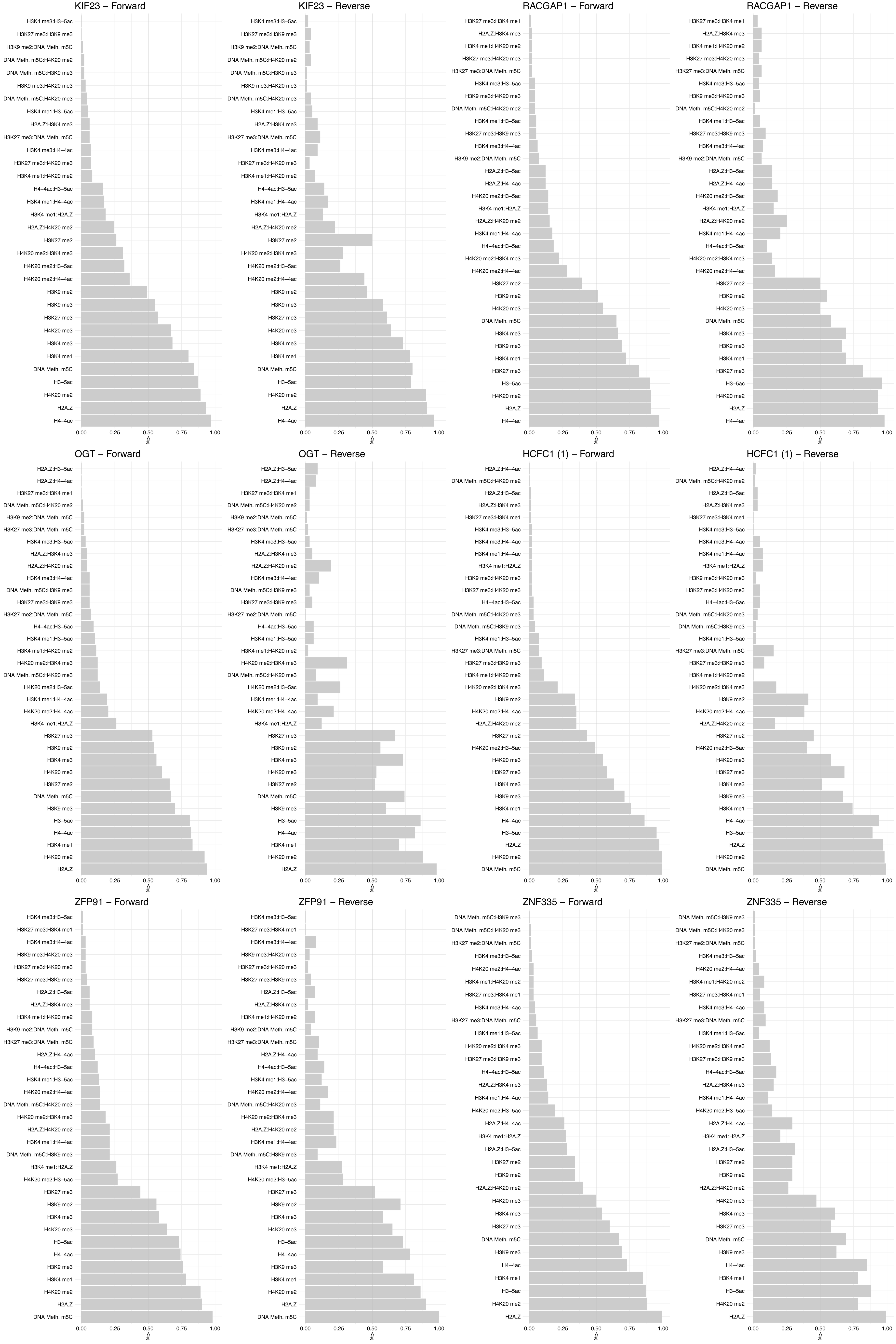
KIF23 – Forward
H3K4 me3:H3–5ac
H3K27 me3:H3K9 me3
H3K9 me2:DNA Meth. m5C
DNA Meth. m5C:H4K20 me2
DNA Meth. m5C:H3K9 me3
H3K9 me3:H4K20 me3
DNA Meth. m5C:H4K20 me3
H3K4 me1:H3–5ac
H2A.Z:H3K4 me3
H3K27 me3:DNA Meth. m5C
H3K4 me3:H4–4ac
H3K27 me3:H4K20 me3
H3K4 me1:H4K20 me2
H4–4ac:H3–5ac
H3K4 me1:H4–4ac
H3K4 me1:H2A.Z
H2A.Z:H4K20 me2
H3K27 me2
H4K20 me2:H3K4 me3
H4K20 me2:H3–5ac
H4K20 me2:H4–4ac
H3K9 me2
H3K9 me3
H3K27 me3
H4K20 me3
H3K4 me3
H3K4 me1
DNA Meth. m5C
H3–5ac
H4K20 me2
H2A.Z
H4–4ac
0.00
0.25
0.50
0.75
1.00
$\hat{\pi}$
KIF23 – Reverse
H3K4 me3:H3–5ac
H3K27 me3:H3K9 me3
H3K9 me2:DNA Meth. m5C
DNA Meth. m5C:H4K20 me2
DNA Meth. m5C:H3K9 me3
H3K9 me3:H4K20 me3
DNA Meth. m5C:H4K20 me3
H3K4 me1:H3–5ac
H2A.Z:H3K4 me3
H3K27 me3:DNA Meth. m5C
H3K4 me3:H4–4ac
H3K27 me3:H4K20 me3
H3K4 me1:H4K20 me2
H4–4ac:H3–5ac
H3K4 me1:H4–4ac
H3K4 me1:H2A.Z
H2A.Z:H4K20 me2
H3K27 me2
H4K20 me2:H3K4 me3
H4K20 me2:H3–5ac
H4K20 me2:H4–4ac
H3K9 me2
H3K9 me3
H3K27 me3
H4K20 me3
H3K4 me3
H3K4 me1
DNA Meth. m5C
H3–5ac
H4K20 me2
H2A.Z
H4–4ac
0.00
0.25
0.50
0.75
1.00
$\hat{\pi}$
RACGAP1 – Forward
H3K27 me3:H3K4 me1
H2A.Z:H3K4 me3
H3K4 me1:H4K20 me2
H3K27 me3:H4K20 me3
H3K27 me3:DNA Meth. m5C
H3K4 me3:H3–5ac
H3K9 me3:H4K20 me3
DNA Meth. m5C:H4K20 me2
H3K4 me1:H3–5ac
H3K27 me3:H3K9 me3
H3K4 me3:H4–4ac
H3K9 me2:DNA Meth. m5C
H2A.Z:H3–5ac
H2A.Z:H4–4ac
H4K20 me2:H3–5ac
H3K4 me1:H2A.Z
H2A.Z:H4K20 me2
H3K4 me1:H4–4ac
H4–4ac:H3–5ac
H4K20 me2:H3K4 me3
H4K20 me2:H4–4ac
H3K27 me2
H3K9 me2
H4K20 me3
DNA Meth. m5C
H3K4 me3
H3K9 me3
H3K4 me1
H3K27 me3
H3–5ac
H4K20 me2
H2A.Z
H4–4ac
0.00
0.25
0.50
0.75
1.00
$\hat{\pi}$
RACGAP1 – Reverse
H3K27 me3:H3K4 me1
H2A.Z:H3K4 me3
H3K4 me1:H4K20 me2
H3K27 me3:H4K20 me3
H3K27 me3:DNA Meth. m5C
H3K4 me3:H3–5ac
H3K9 me3:H4K20 me3
DNA Meth. m5C:H4K20 me2
H3K4 me1:H3–5ac
H3K27 me3:H3K9 me3
H3K4 me3:H4–4ac
H3K9 me2:DNA Meth. m5C
H2A.Z:H3–5ac
H2A.Z:H4–4ac
H4K20 me2:H3–5ac
H3K4 me1:H2A.Z
H2A.Z:H4K20 me2
H3K4 me1:H4–4ac
H4–4ac:H3–5ac
H4K20 me2:H3K4 me3
H4K20 me2:H4–4ac
H3K27 me2
H3K9 me2
H4K20 me3
DNA Meth. m5C
H3K4 me3
H3K9 me3
H3K4 me1
H3K27 me3
H3–5ac
H4K20 me2
H2A.Z
H4–4ac
0.00
0.25
0.50
0.75
1.00
$\hat{\pi}$
OGT – Forward
H2A.Z:H3–5ac
H2A.Z:H4–4ac
H3K27 me3:H3K4 me1
DNA Meth. m5C:H4K20 me2
H3K9 me2:DNA Meth. m5C
H3K27 me3:DNA Meth. m5C
H3K4 me3:H3–5ac
H2A.Z:H3K4 me3
H2A.Z:H4K20 me2
H3K4 me3:H4–4ac
DNA Meth. m5C:H3K9 me3
H3K27 me3:H3K9 me3
H3K27 me2:DNA Meth. m5C
H4–4ac:H3–5ac
H3K4 me1:H3–5ac
H3K4 me1:H4K20 me2
H4K20 me2:H3K4 me3
DNA Meth. m5C:H4K20 me3
H4K20 me2:H3–5ac
H3K4 me1:H4–4ac
H4K20 me2:H4–4ac
H3K4 me1:H2A.Z
H3K27 me3
H3K9 me2
H3K4 me3
H4K20 me3
H3K27 me2
DNA Meth. m5C
H3K9 me3
H3–5ac
H4–4ac
H3K4 me1
H4K20 me2
H2A.Z
0.00
0.25
0.50
0.75
1.00
$\hat{\pi}$
OGT – Reverse
H2A.Z:H3–5ac
H2A.Z:H4–4ac
H3K27 me3:H3K4 me1
DNA Meth. m5C:H4K20 me2
H3K9 me2:DNA Meth. m5C
H3K27 me3:DNA Meth. m5C
H3K4 me3:H3–5ac
H2A.Z:H3K4 me3
H2A.Z:H4K20 me2
H3K4 me3:H4–4ac
DNA Meth. m5C:H3K9 me3
H3K27 me3:H3K9 me3
H3K27 me2:DNA Meth. m5C
H4–4ac:H3–5ac
H3K4 me1:H3–5ac
H3K4 me1:H4K20 me2
H4K20 me2:H3K4 me3
DNA Meth. m5C:H4K20 me3
H4K20 me2:H3–5ac
H3K4 me1:H4–4ac
H4K20 me2:H4–4ac
H3K4 me1:H2A.Z
H3K27 me3
H3K9 me2
H3K4 me3
H4K20 me3
H3K27 me2
DNA Meth. m5C
H3K9 me3
H3–5ac
H4–4ac
H3K4 me1
H4K20 me2
H2A.Z
0.00
0.25
0.50
0.75
1.00
$\hat{\pi}$
HCFC1 (1) – Forward
H2A.Z:H4–4ac
DNA Meth. m5C:H4K20 me2
H2A.Z:H3–5ac
H2A.Z:H3K4 me3
H3K27 me3:H3K4 me1
H3K4 me3:H3–5ac
H3K4 me3:H4–4ac
H3K4 me1:H4–4ac
H3K4 me1:H2A.Z
H3K9 me3:H4K20 me3
H3K27 me3:H4K20 me3
H4–4ac:H3–5ac
DNA Meth. m5C:H4K20 me3
DNA Meth. m5C:H3K9 me3
H3K4 me1:H3–5ac
H3K27 me3:DNA Meth. m5C
H3K27 me3:H3K9 me3
H3K4 me1:H4K20 me2
H4K20 me2:H3K4 me3
H3K9 me2
H4K20 me2:H4–4ac
H2A.Z:H4K20 me2
H3K27 me2
H4K20 me2:H3–5ac
H4K20 me3
H3K27 me3
H3K4 me3
H3K9 me3
H3K4 me1
H4–4ac
H3–5ac
H2A.Z
H4K20 me2
DNA Meth. m5C
0.00
0.25
0.50
0.75
1.00
$\hat{\pi}$
HCFC1 (1) – Reverse
H2A.Z:H4–4ac
DNA Meth. m5C:H4K20 me2
H2A.Z:H3–5ac
H2A.Z:H3K4 me3
H3K27 me3:H3K4 me1
H3K4 me3:H3–5ac
H3K4 me3:H4–4ac
H3K4 me1:H4–4ac
H3K4 me1:H2A.Z
H3K9 me3:H4K20 me3
H3K27 me3:H4K20 me3
H4–4ac:H3–5ac
DNA Meth. m5C:H4K20 me3
DNA Meth. m5C:H3K9 me3
H3K4 me1:H3–5ac
H3K27 me3:DNA Meth. m5C
H3K27 me3:H3K9 me3
H3K4 me1:H4K20 me2
H4K20 me2:H3K4 me3
H3K9 me2
H4K20 me2:H4–4ac
H2A.Z:H4K20 me2
H3K27 me2
H4K20 me2:H3–5ac
H4K20 me3
H3K27 me3
H3K4 me3
H3K9 me3
H3K4 me1
H4–4ac
H3–5ac
H2A.Z
H4K20 me2
DNA Meth. m5C
0.00
0.25
0.50
0.75
1.00
$\hat{\pi}$
ZFP91 – Forward
H3K4 me3:H3–5ac
H3K27 me3:H3K4 me1
H3K4 me3:H4–4ac
H3K9 me3:H4K20 me3
H3K27 me3:H4K20 me3
H3K27 me3:H3K9 me3
H2A.Z:H3–5ac
H2A.Z:H3K4 me3
H3K4 me1:H4K20 me2
H3K9 me2:DNA Meth. m5C
H3K27 me3:DNA Meth. m5C
H2A.Z:H4–4ac
H4–4ac:H3–5ac
H3K4 me1:H3–5ac
H4K20 me2:H4–4ac
DNA Meth. m5C:H4K20 me3
H4K20 me2:H3K4 me3
H2A.Z:H4K20 me2
H3K4 me1:H4–4ac
DNA Meth. m5C:H3K9 me3
H3K4 me1:H2A.Z
H4K20 me2:H3–5ac
H3K27 me3
H3K9 me2
H3K4 me3
H4K20 me3
H3–5ac
H4–4ac
H3K9 me3
H3K4 me1
H4K20 me2
H2A.Z
DNA Meth. m5C
0.00
0.25
0.50
0.75
1.00
$\hat{\pi}$
ZFP91 – Reverse
H3K4 me3:H3–5ac
H3K27 me3:H3K4 me1
H3K4 me3:H4–4ac
H3K9 me3:H4K20 me3
H3K27 me3:H4K20 me3
H3K27 me3:H3K9 me3
H2A.Z:H3–5ac
H2A.Z:H3K4 me3
H3K4 me1:H4K20 me2
H3K9 me2:DNA Meth. m5C
H3K27 me3:DNA Meth. m5C
H2A.Z:H4–4ac
H4–4ac:H3–5ac
H3K4 me1:H3–5ac
H4K20 me2:H4–4ac
DNA Meth. m5C:H4K20 me3
H4K20 me2:H3K4 me3
H2A.Z:H4K20 me2
H3K4 me1:H4–4ac
DNA Meth. m5C:H3K9 me3
H3K4 me1:H2A.Z
H4K20 me2:H3–5ac
H3K27 me3
H3K9 me2
H3K4 me3
H4K20 me3
H3–5ac
H4–4ac
H3K9 me3
H3K4 me1
H4K20 me2
H2A.Z
DNA Meth. m5C
0.00
0.25
0.50
0.75
1.00
$\hat{\pi}$
ZNF335 – Forward
DNA Meth. m5C:H3K9 me3
DNA Meth. m5C:H4K20 me3
H3K27 me2:DNA Meth. m5C
H3K4 me3:H3–5ac
H4K20 me2:H4–4ac
H3K4 me1:H4K20 me2
H3K27 me3:H3K4 me1
H3K4 me3:H4–4ac
H3K27 me3:DNA Meth. m5C
H3K4 me1:H3–5ac
H4K20 me2:H3K4 me3
H3K27 me3:H3K9 me3
H4–4ac:H3–5ac
H2A.Z:H3K4 me3
H3K4 me1:H4–4ac
H4K20 me2:H3–5ac
H2A.Z:H4–4ac
H3K4 me1:H2A.Z
H2A.Z:H3–5ac
H3K27 me2
H3K9 me2
H2A.Z:H4K20 me2
H4K20 me3
H3K4 me3
H3K27 me3
DNA Meth. m5C
H3K9 me3
H4–4ac
H3K4 me1
H3–5ac
H4K20 me2
H2A.Z
0.00
0.25
0.50
0.75
1.00
$\hat{\pi}$
ZNF335 – Reverse
DNA Meth. m5C:H3K9 me3
DNA Meth. m5C:H4K20 me3
H3K27 me2:DNA Meth. m5C
H3K4 me3:H3–5ac
H4K20 me2:H4–4ac
H3K4 me1:H4K20 me2
H3K27 me3:H3K4 me1
H3K4 me3:H4–4ac
H3K27 me3:DNA Meth. m5C
H3K4 me1:H3–5ac
H4K20 me2:H3K4 me3
H3K27 me3:H3K9 me3
H4–4ac:H3–5ac
H2A.Z:H3K4 me3
H3K4 me1:H4–4ac
H4K20 me2:H3–5ac
H2A.Z:H4–4ac
H3K4 me1:H2A.Z
H2A.Z:H3–5ac
H3K27 me2
H3K9 me2
H2A.Z:H4K20 me2
H4K20 me3
H3K4 me3
H3K27 me3
DNA Meth. m5C
H3K9 me3
H4–4ac
H3K4 me1
H3–5ac
H4K20 me2
H2A.Z
0.00
0.25
0.50
0.75
1.00
$\hat{\pi}$

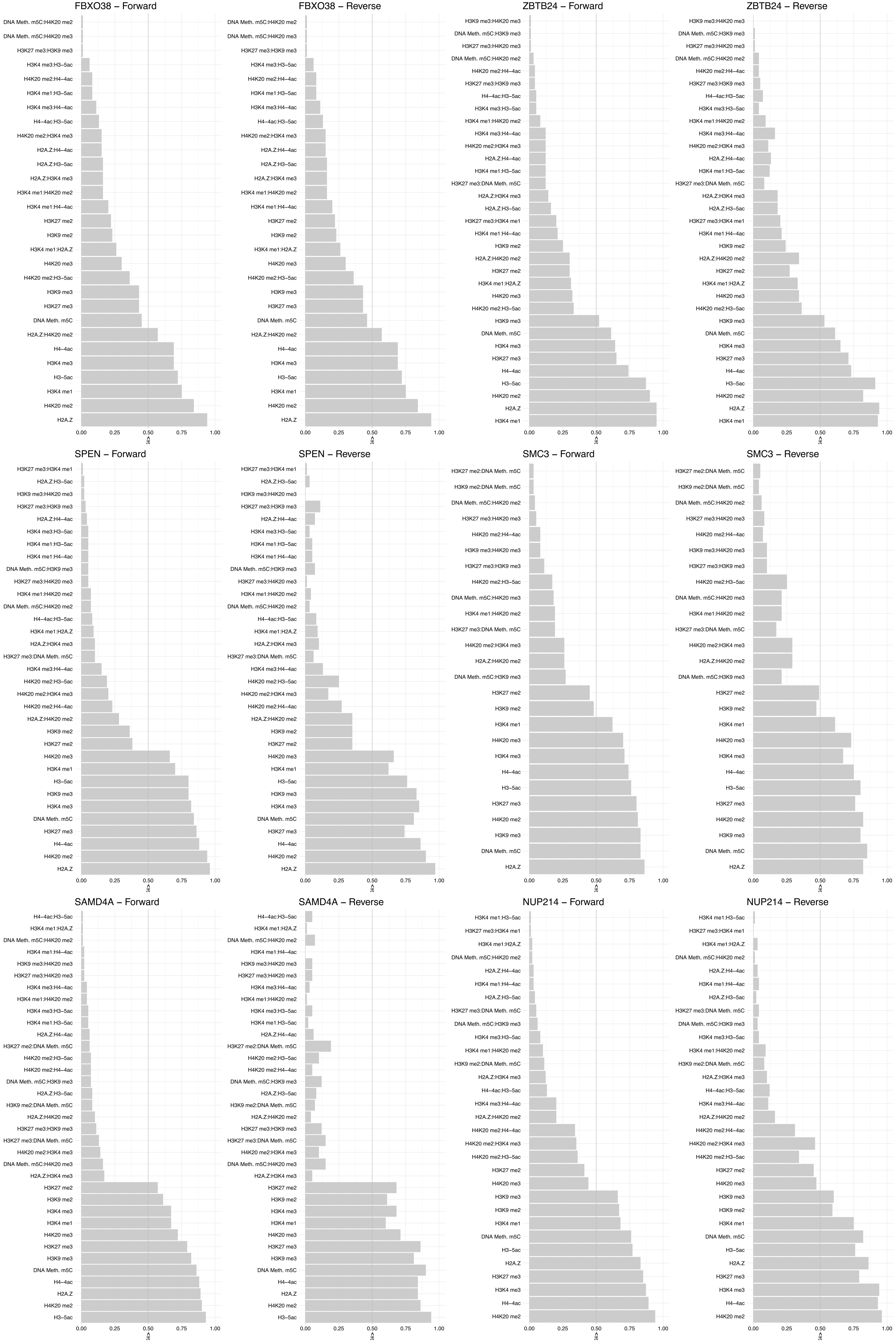
FBXO38 − Forward
DNA Meth. m5C:H4K20 me2
DNA Meth. m5C:H4K20 me3
H3K27 me3:H3K9 me3
H3K4 me3:H3−5ac
H4K20 me2:H4−4ac
H3K4 me1:H3−5ac
H3K4 me3:H4−4ac
H4−4ac:H3−5ac
H4K20 me2:H3K4 me3
H2A.Z:H4−4ac
H2A.Z:H3−5ac
H2A.Z:H3K4 me3
H3K4 me1:H4K20 me2
H3K4 me1:H4−4ac
H3K27 me2
H3K9 me2
H3K4 me1:H2A.Z
H4K20 me3
H4K20 me2:H3−5ac
H3K9 me3
H3K27 me3
DNA Meth. m5C
H2A.Z:H4K20 me2
H4−4ac
H3K4 me3
H3−5ac
H3K4 me1
H4K20 me2
H2A.Z
0.00
0.25
0.50
0.75
1.00
π̂
FBXO38 − Reverse
DNA Meth. m5C:H4K20 me2
DNA Meth. m5C:H4K20 me3
H3K27 me3:H3K9 me3
H3K4 me3:H3−5ac
H4K20 me2:H4−4ac
H3K4 me1:H3−5ac
H3K4 me3:H4−4ac
H4−4ac:H3−5ac
H4K20 me2:H3K4 me3
H2A.Z:H4−4ac
H2A.Z:H3−5ac
H2A.Z:H3K4 me3
H3K4 me1:H4K20 me2
H3K4 me1:H4−4ac
H3K27 me2
H3K9 me2
H3K4 me1:H2A.Z
H4K20 me3
H4K20 me2:H3−5ac
H3K9 me3
H3K27 me3
DNA Meth. m5C
H2A.Z:H4K20 me2
H4−4ac
H3K4 me3
H3−5ac
H3K4 me1
H4K20 me2
H2A.Z
0.00
0.25
0.50
0.75
1.00
π̂
ZBTB24 − Forward
H3K9 me3:H4K20 me3
DNA Meth. m5C:H3K9 me3
H3K27 me3:H4K20 me3
DNA Meth. m5C:H4K20 me2
H4K20 me2:H4−4ac
H3K27 me3:H3K9 me3
H4−4ac:H3−5ac
H3K4 me3:H3−5ac
H3K4 me1:H4K20 me2
H3K4 me3:H4−4ac
H4K20 me2:H3K4 me3
H2A.Z:H4−4ac
H3K4 me1:H3−5ac
H3K27 me3:DNA Meth. m5C
H2A.Z:H3K4 me3
H2A.Z:H3−5ac
H3K27 me3:H3K4 me1
H3K4 me1:H4−4ac
H3K9 me2
H2A.Z:H4K20 me2
H3K27 me2
H3K4 me1:H2A.Z
H4K20 me3
H4K20 me2:H3−5ac
H3K9 me3
DNA Meth. m5C
H3K4 me3
H3K27 me3
H4−4ac
H3−5ac
H4K20 me2
H2A.Z
H3K4 me1
0.00
0.25
0.50
0.75
1.00
π̂
ZBTB24 − Reverse
H3K9 me3:H4K20 me3
DNA Meth. m5C:H3K9 me3
H3K27 me3:H4K20 me3
DNA Meth. m5C:H4K20 me2
H4K20 me2:H4−4ac
H3K27 me3:H3K9 me3
H4−4ac:H3−5ac
H3K4 me3:H3−5ac
H3K4 me1:H4K20 me2
H3K4 me3:H4−4ac
H4K20 me2:H3K4 me3
H2A.Z:H4−4ac
H3K4 me1:H3−5ac
H3K27 me3:DNA Meth. m5C
H2A.Z:H3K4 me3
H2A.Z:H3−5ac
H3K27 me3:H3K4 me1
H3K4 me1:H4−4ac
H3K9 me2
H2A.Z:H4K20 me2
H3K27 me2
H3K4 me1:H2A.Z
H4K20 me3
H4K20 me2:H3−5ac
H3K9 me3
DNA Meth. m5C
H3K4 me3
H3K27 me3
H4−4ac
H3−5ac
H4K20 me2
H2A.Z
H3K4 me1
0.00
0.25
0.50
0.75
1.00
π̂
SPEN − Forward
H3K27 me3:H3K4 me1
H2A.Z:H3−5ac
H3K9 me3:H4K20 me3
H3K27 me3:H3K9 me3
H2A.Z:H4−4ac
H3K4 me3:H3−5ac
H3K4 me1:H3−5ac
H3K4 me1:H4−4ac
DNA Meth. m5C:H3K9 me3
H3K27 me3:H4K20 me3
H3K4 me1:H4K20 me2
DNA Meth. m5C:H4K20 me2
H4−4ac:H3−5ac
H3K4 me1:H2A.Z
H2A.Z:H3K4 me3
H3K27 me3:DNA Meth. m5C
H3K4 me3:H4−4ac
H4K20 me2:H3−5ac
H4K20 me2:H3K4 me3
H4K20 me2:H4−4ac
H2A.Z:H4K20 me2
H3K9 me2
H3K27 me2
H4K20 me3
H3K4 me1
H3−5ac
H3K9 me3
H3K4 me3
DNA Meth. m5C
H3K27 me3
H4−4ac
H4K20 me2
H2A.Z
0.00
0.25
0.50
0.75
1.00
π̂
SPEN − Reverse
H3K27 me3:H3K4 me1
H2A.Z:H3−5ac
H3K9 me3:H4K20 me3
H3K27 me3:H3K9 me3
H2A.Z:H4−4ac
H3K4 me3:H3−5ac
H3K4 me1:H3−5ac
H3K4 me1:H4−4ac
DNA Meth. m5C:H3K9 me3
H3K27 me3:H4K20 me3
H3K4 me1:H4K20 me2
DNA Meth. m5C:H4K20 me2
H4−4ac:H3−5ac
H3K4 me1:H2A.Z
H2A.Z:H3K4 me3
H3K27 me3:DNA Meth. m5C
H3K4 me3:H4−4ac
H4K20 me2:H3−5ac
H4K20 me2:H3K4 me3
H4K20 me2:H4−4ac
H2A.Z:H4K20 me2
H3K9 me2
H3K27 me2
H4K20 me3
H3K4 me1
H3−5ac
H3K9 me3
H3K4 me3
DNA Meth. m5C
H3K27 me3
H4−4ac
H4K20 me2
H2A.Z
0.00
0.25
0.50
0.75
1.00
π̂
SMC3 − Forward
H3K27 me2:DNA Meth. m5C
H3K9 me2:DNA Meth. m5C
DNA Meth. m5C:H4K20 me2
H3K27 me3:H4K20 me3
H4K20 me2:H4−4ac
H3K9 me3:H4K20 me3
H3K27 me3:H3K9 me3
H4K20 me2:H3−5ac
DNA Meth. m5C:H4K20 me3
H3K4 me1:H4K20 me2
H3K27 me3:DNA Meth. m5C
H4K20 me2:H3K4 me3
H2A.Z:H4K20 me2
DNA Meth. m5C:H3K9 me3
H3K27 me2
H3K9 me2
H3K4 me1
H4K20 me3
H3K4 me3
H4−4ac
H3−5ac
H3K27 me3
H4K20 me2
H3K9 me3
DNA Meth. m5C
H2A.Z
0.00
0.25
0.50
0.75
1.00
π̂
SMC3 − Reverse
H3K27 me2:DNA Meth. m5C
H3K9 me2:DNA Meth. m5C
DNA Meth. m5C:H4K20 me2
H3K27 me3:H4K20 me3
H4K20 me2:H4−4ac
H3K9 me3:H4K20 me3
H3K27 me3:H3K9 me3
H4K20 me2:H3−5ac
DNA Meth. m5C:H4K20 me3
H3K4 me1:H4K20 me2
H3K27 me3:DNA Meth. m5C
H4K20 me2:H3K4 me3
H2A.Z:H4K20 me2
DNA Meth. m5C:H3K9 me3
H3K27 me2
H3K9 me2
H3K4 me1
H4K20 me3
H3K4 me3
H4−4ac
H3−5ac
H3K27 me3
H4K20 me2
H3K9 me3
DNA Meth. m5C
H2A.Z
0.00
0.25
0.50
0.75
1.00
π̂
SAMD4A − Forward
H4−4ac:H3−5ac
H3K4 me1:H2A.Z
DNA Meth. m5C:H4K20 me2
H3K4 me1:H4−4ac
H3K9 me3:H4K20 me3
H3K27 me3:H4K20 me3
H3K4 me3:H4−4ac
H3K4 me1:H4K20 me2
H3K4 me3:H3−5ac
H3K4 me1:H3−5ac
H2A.Z:H4−4ac
H3K27 me2:DNA Meth. m5C
H4K20 me2:H3−5ac
H4K20 me2:H4−4ac
DNA Meth. m5C:H3K9 me3
H2A.Z:H3−5ac
H3K9 me2:DNA Meth. m5C
H2A.Z:H4K20 me2
H3K27 me3:H3K9 me3
H3K27 me3:DNA Meth. m5C
H4K20 me2:H3K4 me3
DNA Meth. m5C:H4K20 me3
H2A.Z:H3K4 me3
H3K27 me2
H3K9 me2
H3K4 me3
H3K4 me1
H4K20 me3
H3K27 me3
H3K9 me3
DNA Meth. m5C
H4−4ac
H2A.Z
H4K20 me2
H3−5ac
0.00
0.25
0.50
0.75
1.00
π̂
SAMD4A − Reverse
H4−4ac:H3−5ac
H3K4 me1:H2A.Z
DNA Meth. m5C:H4K20 me2
H3K4 me1:H4−4ac
H3K9 me3:H4K20 me3
H3K27 me3:H4K20 me3
H3K4 me3:H4−4ac
H3K4 me1:H4K20 me2
H3K4 me3:H3−5ac
H3K4 me1:H3−5ac
H2A.Z:H4−4ac
H3K27 me2:DNA Meth. m5C
H4K20 me2:H3−5ac
H4K20 me2:H4−4ac
DNA Meth. m5C:H3K9 me3
H2A.Z:H3−5ac
H3K9 me2:DNA Meth. m5C
H2A.Z:H4K20 me2
H3K27 me3:H3K9 me3
H3K27 me3:DNA Meth. m5C
H4K20 me2:H3K4 me3
DNA Meth. m5C:H4K20 me3
H2A.Z:H3K4 me3
H3K27 me2
H3K9 me2
H3K4 me3
H3K4 me1
H4K20 me3
H3K27 me3
H3K9 me3
DNA Meth. m5C
H4−4ac
H2A.Z
H4K20 me2
H3−5ac
0.00
0.25
0.50
0.75
1.00
π̂
NUP214 − Forward
H3K4 me1:H3−5ac
H3K27 me3:H3K4 me1
H3K4 me1:H2A.Z
DNA Meth. m5C:H4K20 me2
H2A.Z:H4−4ac
H3K4 me1:H4−4ac
H2A.Z:H3−5ac
H3K27 me3:DNA Meth. m5C
DNA Meth. m5C:H3K9 me3
H3K4 me3:H3−5ac
H3K4 me1:H4K20 me2
H3K9 me2:DNA Meth. m5C
H2A.Z:H3K4 me3
H4−4ac:H3−5ac
H3K4 me3:H4−4ac
H2A.Z:H4K20 me2
H4K20 me2:H4−4ac
H4K20 me2:H3K4 me3
H4K20 me2:H3−5ac
H3K27 me2
H4K20 me3
H3K9 me3
H3K9 me2
H3K4 me1
DNA Meth. m5C
H3−5ac
H2A.Z
H3K27 me3
H3K4 me3
H4−4ac
H4K20 me2
0.00
0.25
0.50
0.75
1.00
π̂
NUP214 − Reverse
H3K4 me1:H3−5ac
H3K27 me3:H3K4 me1
H3K4 me1:H2A.Z
DNA Meth. m5C:H4K20 me2
H2A.Z:H4−4ac
H3K4 me1:H4−4ac
H2A.Z:H3−5ac
H3K27 me3:DNA Meth. m5C
DNA Meth. m5C:H3K9 me3
H3K4 me3:H3−5ac
H3K4 me1:H4K20 me2
H3K9 me2:DNA Meth. m5C
H2A.Z:H3K4 me3
H4−4ac:H3−5ac
H3K4 me3:H4−4ac
H2A.Z:H4K20 me2
H4K20 me2:H4−4ac
H4K20 me2:H3K4 me3
H4K20 me2:H3−5ac
H3K27 me2
H4K20 me3
H3K9 me3
H3K9 me2
H3K4 me1
DNA Meth. m5C
H3−5ac
H2A.Z
H3K27 me3
H3K4 me3
H4−4ac
H4K20 me2
0.00
0.25
0.50
0.75
1.00
π̂

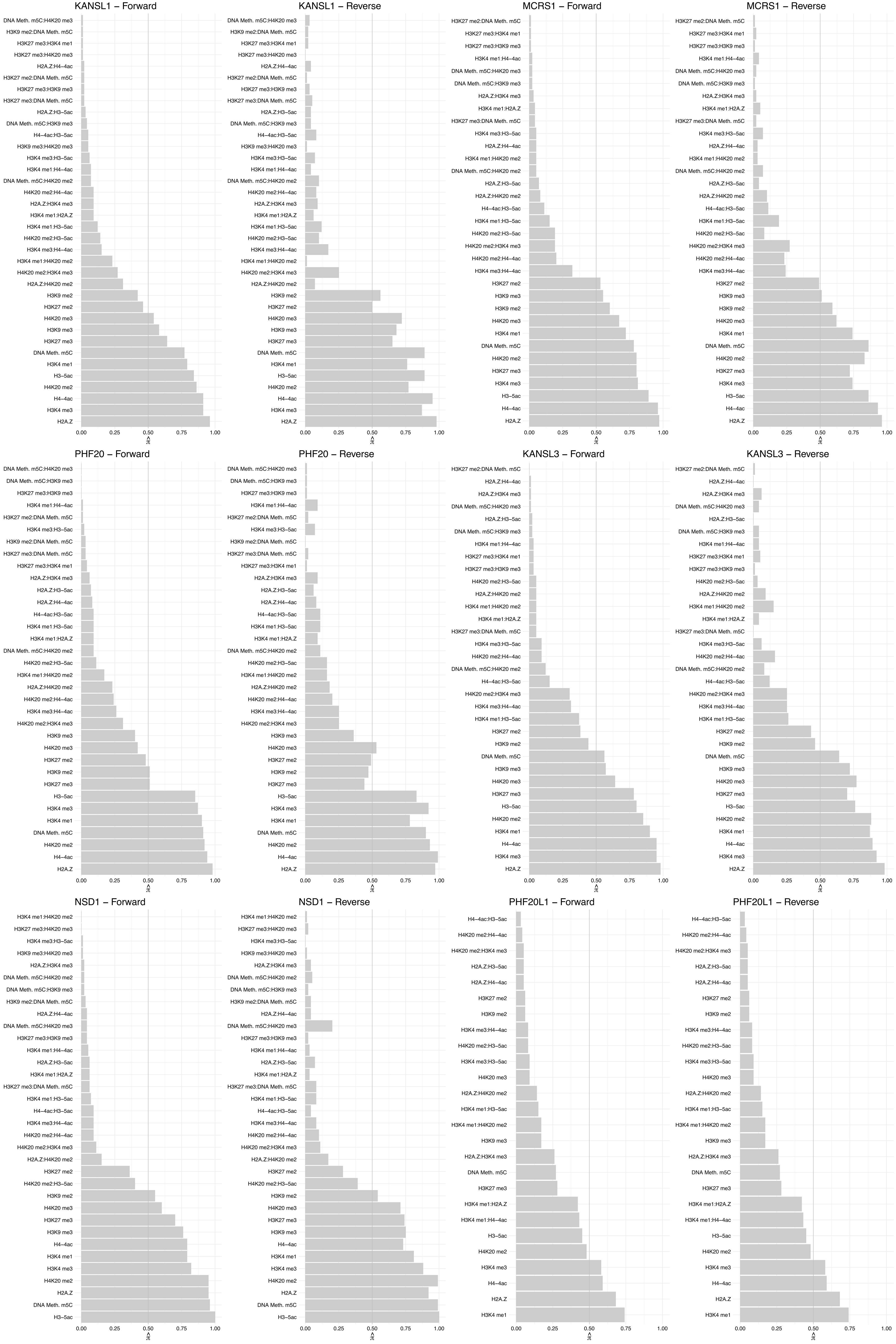
KANSL1 – Forward
DNA Meth. m5C:H4K20 me3
H3K9 me2:DNA Meth. m5C
H3K27 me3:H3K4 me1
H3K27 me3:H4K20 me3
H2A.Z:H4−4ac
H3K27 me2:DNA Meth. m5C
H3K27 me3:H3K9 me3
H3K27 me3:DNA Meth. m5C
H2A.Z:H3−5ac
DNA Meth. m5C:H3K9 me3
H4−4ac:H3−5ac
H3K9 me3:H4K20 me3
H3K4 me3:H3−5ac
H3K4 me1:H4−4ac
DNA Meth. m5C:H4K20 me2
H4K20 me2:H4−4ac
H2A.Z:H3K4 me3
H3K4 me1:H2A.Z
H3K4 me1:H3−5ac
H4K20 me2:H3−5ac
H3K4 me3:H4−4ac
H3K4 me1:H4K20 me2
H4K20 me2:H3K4 me3
H2A.Z:H4K20 me2
H3K9 me2
H3K27 me2
H4K20 me3
H3K9 me3
H3K27 me3
DNA Meth. m5C
H3K4 me1
H3−5ac
H4K20 me2
H4−4ac
H3K4 me3
H2A.Z
0.00 0.25 0.50 0.75 1.00
π̂
KANSL1 – Reverse
DNA Meth. m5C:H4K20 me3
H3K9 me2:DNA Meth. m5C
H3K27 me3:H3K4 me1
H3K27 me3:H4K20 me3
H2A.Z:H4−4ac
H3K27 me2:DNA Meth. m5C
H3K27 me3:H3K9 me3
H3K27 me3:DNA Meth. m5C
H2A.Z:H3−5ac
DNA Meth. m5C:H3K9 me3
H4−4ac:H3−5ac
H3K9 me3:H4K20 me3
H3K4 me3:H3−5ac
H3K4 me1:H4−4ac
DNA Meth. m5C:H4K20 me2
H4K20 me2:H4−4ac
H2A.Z:H3K4 me3
H3K4 me1:H2A.Z
H3K4 me1:H3−5ac
H4K20 me2:H3−5ac
H3K4 me3:H4−4ac
H3K4 me1:H4K20 me2
H4K20 me2:H3K4 me3
H2A.Z:H4K20 me2
H3K9 me2
H3K27 me2
H4K20 me3
H3K9 me3
H3K27 me3
DNA Meth. m5C
H3K4 me1
H3−5ac
H4K20 me2
H4−4ac
H3K4 me3
H2A.Z
0.00 0.25 0.50 0.75 1.00
π̂
MCRS1 – Forward
H3K27 me2:DNA Meth. m5C
H3K27 me3:H3K4 me1
H3K27 me3:H3K9 me3
H3K4 me1:H4−4ac
DNA Meth. m5C:H4K20 me3
DNA Meth. m5C:H3K9 me3
H2A.Z:H3K4 me3
H3K4 me1:H2A.Z
H3K27 me3:DNA Meth. m5C
H3K4 me3:H3−5ac
H2A.Z:H4−4ac
H3K4 me1:H4K20 me2
DNA Meth. m5C:H4K20 me2
H2A.Z:H3−5ac
H2A.Z:H4K20 me2
H4−4ac:H3−5ac
H3K4 me1:H3−5ac
H4K20 me2:H3−5ac
H4K20 me2:H3K4 me3
H4K20 me2:H4−4ac
H3K4 me3:H4−4ac
H3K27 me2
H3K9 me3
H3K9 me2
H4K20 me3
H3K4 me1
DNA Meth. m5C
H4K20 me2
H3K27 me3
H3K4 me3
H3−5ac
H4−4ac
H2A.Z
0.00 0.25 0.50 0.75 1.00
π̂
MCRS1 – Reverse
H3K27 me2:DNA Meth. m5C
H3K27 me3:H3K4 me1
H3K27 me3:H3K9 me3
H3K4 me1:H4−4ac
DNA Meth. m5C:H4K20 me3
DNA Meth. m5C:H3K9 me3
H2A.Z:H3K4 me3
H3K4 me1:H2A.Z
H3K27 me3:DNA Meth. m5C
H3K4 me3:H3−5ac
H2A.Z:H4−4ac
H3K4 me1:H4K20 me2
DNA Meth. m5C:H4K20 me2
H2A.Z:H3−5ac
H2A.Z:H4K20 me2
H4−4ac:H3−5ac
H3K4 me1:H3−5ac
H4K20 me2:H3−5ac
H4K20 me2:H3K4 me3
H4K20 me2:H4−4ac
H3K4 me3:H4−4ac
H3K27 me2
H3K9 me3
H3K9 me2
H4K20 me3
H3K4 me1
DNA Meth. m5C
H4K20 me2
H3K27 me3
H3K4 me3
H3−5ac
H4−4ac
H2A.Z
0.00 0.25 0.50 0.75 1.00
π̂
PHF20 – Forward
DNA Meth. m5C:H4K20 me3
DNA Meth. m5C:H3K9 me3
H3K27 me3:H3K9 me3
H3K4 me1:H4−4ac
H3K27 me2:DNA Meth. m5C
H3K4 me3:H3−5ac
H3K9 me2:DNA Meth. m5C
H3K27 me3:DNA Meth. m5C
H3K27 me3:H3K4 me1
H2A.Z:H3K4 me3
H2A.Z:H3−5ac
H2A.Z:H4−4ac
H4−4ac:H3−5ac
H3K4 me1:H3−5ac
H3K4 me1:H2A.Z
DNA Meth. m5C:H4K20 me2
H4K20 me2:H3−5ac
H3K4 me1:H4K20 me2
H2A.Z:H4K20 me2
H4K20 me2:H4−4ac
H3K4 me3:H4−4ac
H4K20 me2:H3K4 me3
H3K9 me3
H4K20 me3
H3K27 me2
H3K9 me2
H3K27 me3
H3−5ac
H3K4 me3
H3K4 me1
DNA Meth. m5C
H4K20 me2
H4−4ac
H2A.Z
0.00 0.25 0.50 0.75 1.00
π̂
PHF20 – Reverse
DNA Meth. m5C:H4K20 me3
DNA Meth. m5C:H3K9 me3
H3K27 me3:H3K9 me3
H3K4 me1:H4−4ac
H3K27 me2:DNA Meth. m5C
H3K4 me3:H3−5ac
H3K9 me2:DNA Meth. m5C
H3K27 me3:DNA Meth. m5C
H3K27 me3:H3K4 me1
H2A.Z:H3K4 me3
H2A.Z:H3−5ac
H2A.Z:H4−4ac
H4−4ac:H3−5ac
H3K4 me1:H3−5ac
H3K4 me1:H2A.Z
DNA Meth. m5C:H4K20 me2
H4K20 me2:H3−5ac
H3K4 me1:H4K20 me2
H2A.Z:H4K20 me2
H4K20 me2:H4−4ac
H3K4 me3:H4−4ac
H4K20 me2:H3K4 me3
H3K9 me3
H4K20 me3
H3K27 me2
H3K9 me2
H3K27 me3
H3−5ac
H3K4 me3
H3K4 me1
DNA Meth. m5C
H4K20 me2
H4−4ac
H2A.Z
0.00 0.25 0.50 0.75 1.00
π̂
KANSL3 – Forward
H3K27 me2:DNA Meth. m5C
H2A.Z:H4−4ac
H2A.Z:H3K4 me3
DNA Meth. m5C:H4K20 me3
H2A.Z:H3−5ac
DNA Meth. m5C:H3K9 me3
H3K4 me1:H4−4ac
H3K27 me3:H3K4 me1
H3K27 me3:H3K9 me3
H4K20 me2:H3−5ac
H2A.Z:H4K20 me2
H3K4 me1:H4K20 me2
H3K4 me1:H2A.Z
H3K27 me3:DNA Meth. m5C
H3K4 me3:H3−5ac
H4K20 me2:H4−4ac
DNA Meth. m5C:H4K20 me2
H4−4ac:H3−5ac
H4K20 me2:H3K4 me3
H3K4 me3:H4−4ac
H3K4 me1:H3−5ac
H3K27 me2
H3K9 me2
DNA Meth. m5C
H3K9 me3
H4K20 me3
H3K27 me3
H3−5ac
H4K20 me2
H3K4 me1
H4−4ac
H3K4 me3
H2A.Z
0.00 0.25 0.50 0.75 1.00
π̂
KANSL3 – Reverse
H3K27 me2:DNA Meth. m5C
H2A.Z:H4−4ac
H2A.Z:H3K4 me3
DNA Meth. m5C:H4K20 me3
H2A.Z:H3−5ac
DNA Meth. m5C:H3K9 me3
H3K4 me1:H4−4ac
H3K27 me3:H3K4 me1
H3K27 me3:H3K9 me3
H4K20 me2:H3−5ac
H2A.Z:H4K20 me2
H3K4 me1:H4K20 me2
H3K4 me1:H2A.Z
H3K27 me3:DNA Meth. m5C
H3K4 me3:H3−5ac
H4K20 me2:H4−4ac
DNA Meth. m5C:H4K20 me2
H4−4ac:H3−5ac
H4K20 me2:H3K4 me3
H3K4 me3:H4−4ac
H3K4 me1:H3−5ac
H3K27 me2
H3K9 me2
DNA Meth. m5C
H3K9 me3
H4K20 me3
H3K27 me3
H3−5ac
H4K20 me2
H3K4 me1
H4−4ac
H3K4 me3
H2A.Z
0.00 0.25 0.50 0.75 1.00
π̂
NSD1 – Forward
H3K4 me1:H4K20 me2
H3K27 me3:H4K20 me3
H3K4 me3:H3−5ac
H3K9 me3:H4K20 me3
H2A.Z:H3K4 me3
DNA Meth. m5C:H4K20 me2
DNA Meth. m5C:H3K9 me3
H3K9 me2:DNA Meth. m5C
H2A.Z:H4−4ac
DNA Meth. m5C:H4K20 me3
H3K27 me3:H3K9 me3
H3K4 me1:H4−4ac
H2A.Z:H3−5ac
H3K4 me1:H2A.Z
H3K27 me3:DNA Meth. m5C
H3K4 me1:H3−5ac
H4−4ac:H3−5ac
H3K4 me3:H4−4ac
H4K20 me2:H4−4ac
H4K20 me2:H3K4 me3
H2A.Z:H4K20 me2
H3K27 me2
H4K20 me2:H3−5ac
H3K9 me2
H4K20 me3
H3K27 me3
H3K9 me3
H4−4ac
H3K4 me1
H3K4 me3
H4K20 me2
H2A.Z
DNA Meth. m5C
H3−5ac
0.00 0.25 0.50 0.75 1.00
π̂
NSD1 – Reverse
H3K4 me1:H4K20 me2
H3K27 me3:H4K20 me3
H3K4 me3:H3−5ac
H3K9 me3:H4K20 me3
H2A.Z:H3K4 me3
DNA Meth. m5C:H4K20 me2
DNA Meth. m5C:H3K9 me3
H3K9 me2:DNA Meth. m5C
H2A.Z:H4−4ac
DNA Meth. m5C:H4K20 me3
H3K27 me3:H3K9 me3
H3K4 me1:H4−4ac
H2A.Z:H3−5ac
H3K4 me1:H2A.Z
H3K27 me3:DNA Meth. m5C
H3K4 me1:H3−5ac
H4−4ac:H3−5ac
H3K4 me3:H4−4ac
H4K20 me2:H4−4ac
H4K20 me2:H3K4 me3
H2A.Z:H4K20 me2
H3K27 me2
H4K20 me2:H3−5ac
H3K9 me2
H4K20 me3
H3K27 me3
H3K9 me3
H4−4ac
H3K4 me1
H3K4 me3
H4K20 me2
H2A.Z
DNA Meth. m5C
H3−5ac
0.00 0.25 0.50 0.75 1.00
π̂
PHF20L1 – Forward
H4−4ac:H3−5ac
H4K20 me2:H4−4ac
H4K20 me2:H3K4 me3
H2A.Z:H3−5ac
H2A.Z:H4−4ac
H3K27 me2
H3K9 me2
H3K4 me3:H4−4ac
H4K20 me2:H3−5ac
H3K4 me3:H3−5ac
H4K20 me3
H2A.Z:H4K20 me2
H3K4 me1:H3−5ac
H3K4 me1:H4K20 me2
H3K9 me3
H2A.Z:H3K4 me3
DNA Meth. m5C
H3K27 me3
H3K4 me1:H2A.Z
H3K4 me1:H4−4ac
H3−5ac
H4K20 me2
H3K4 me3
H4−4ac
H2A.Z
H3K4 me1
0.00 0.25 0.50 0.75 1.00
π̂
PHF20L1 – Reverse
H4−4ac:H3−5ac
H4K20 me2:H4−4ac
H4K20 me2:H3K4 me3
H2A.Z:H3−5ac
H2A.Z:H4−4ac
H3K27 me2
H3K9 me2
H3K4 me3:H4−4ac
H4K20 me2:H3−5ac
H3K4 me3:H3−5ac
H4K20 me3
H2A.Z:H4K20 me2
H3K4 me1:H3−5ac
H3K4 me1:H4K20 me2
H3K9 me3
H2A.Z:H3K4 me3
DNA Meth. m5C
H3K27 me3
H3K4 me1:H2A.Z
H3K4 me1:H4−4ac
H3−5ac
H4K20 me2
H3K4 me3
H4−4ac
H2A.Z
H3K4 me1
0.00 0.25 0.50 0.75 1.00
π̂

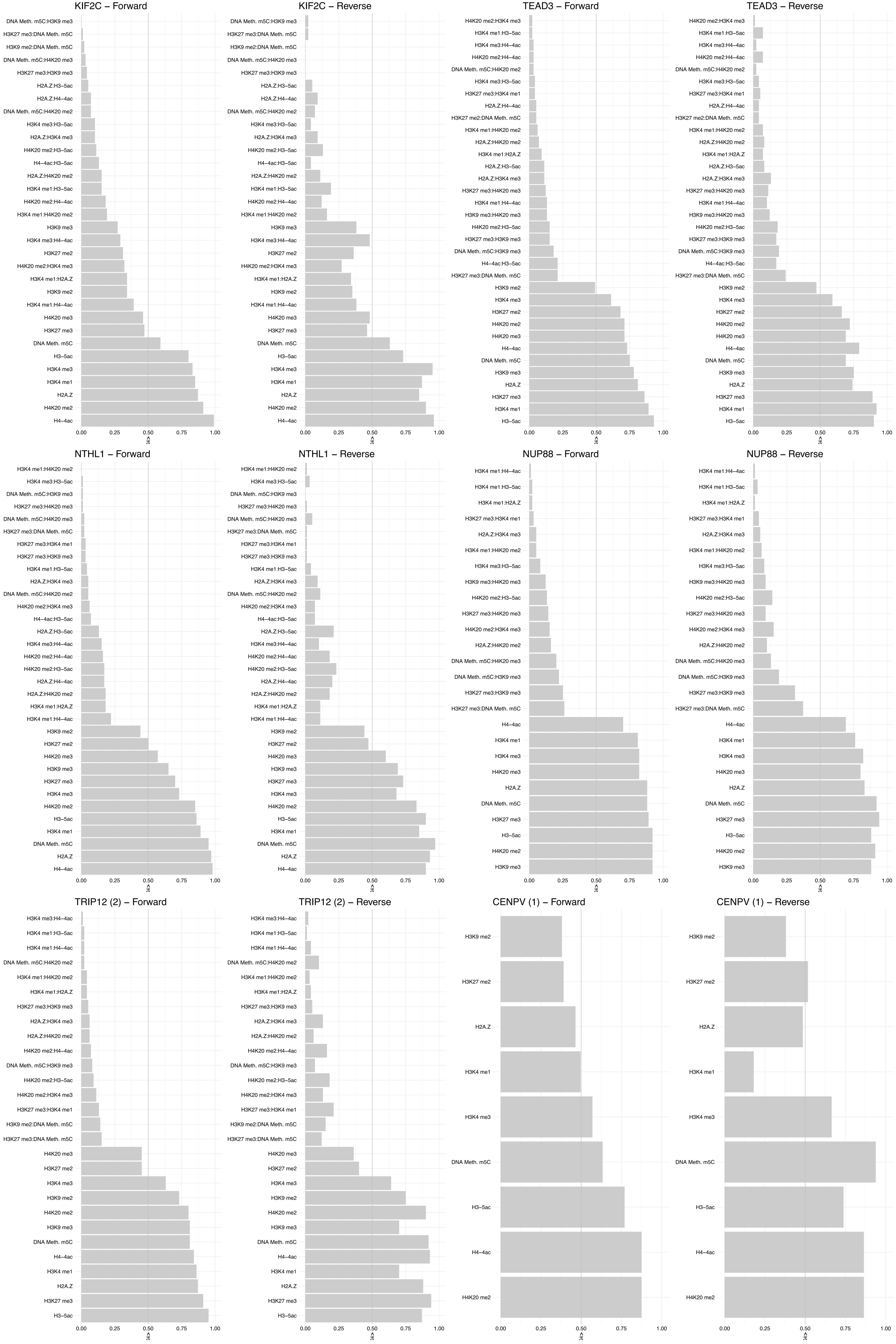
KIF2C – Forward
DNA Meth. m5C:H3K9 me3
H3K27 me3:DNA Meth. m5C
H3K9 me2:DNA Meth. m5C
DNA Meth. m5C:H4K20 me3
H3K27 me3:H3K9 me3
H2A.Z:H3−5ac
H2A.Z:H4−4ac
DNA Meth. m5C:H4K20 me2
H3K4 me3:H3−5ac
H2A.Z:H3K4 me3
H4K20 me2:H3−5ac
H4−4ac:H3−5ac
H2A.Z:H4K20 me2
H3K4 me1:H3−5ac
H4K20 me2:H4−4ac
H3K4 me1:H4K20 me2
H3K9 me3
H3K4 me3:H4−4ac
H3K27 me2
H4K20 me2:H3K4 me3
H3K4 me1:H2A.Z
H3K9 me2
H3K4 me1:H4−4ac
H4K20 me3
H3K27 me3
DNA Meth. m5C
H3−5ac
H3K4 me3
H3K4 me1
H2A.Z
H4K20 me2
H4−4ac
0.00 0.25 0.50 0.75 1.00
$\hat{\pi}$
KIF2C – Reverse
DNA Meth. m5C:H3K9 me3
H3K27 me3:DNA Meth. m5C
H3K9 me2:DNA Meth. m5C
DNA Meth. m5C:H4K20 me3
H3K27 me3:H3K9 me3
H2A.Z:H3−5ac
H2A.Z:H4−4ac
DNA Meth. m5C:H4K20 me2
H3K4 me3:H3−5ac
H2A.Z:H3K4 me3
H4K20 me2:H3−5ac
H4−4ac:H3−5ac
H2A.Z:H4K20 me2
H3K4 me1:H3−5ac
H4K20 me2:H4−4ac
H3K4 me1:H4K20 me2
H3K9 me3
H3K4 me3:H4−4ac
H3K27 me2
H4K20 me2:H3K4 me3
H3K4 me1:H2A.Z
H3K9 me2
H3K4 me1:H4−4ac
H4K20 me3
H3K27 me3
DNA Meth. m5C
H3−5ac
H3K4 me3
H3K4 me1
H2A.Z
H4K20 me2
H4−4ac
0.00 0.25 0.50 0.75 1.00
$\hat{\pi}$
TEAD3 – Forward
H4K20 me2:H3K4 me3
H3K4 me1:H3−5ac
H3K4 me3:H4−4ac
H4K20 me2:H4−4ac
DNA Meth. m5C:H4K20 me2
H3K4 me3:H3−5ac
H3K27 me3:H3K4 me1
H2A.Z:H4−4ac
H3K27 me2:DNA Meth. m5C
H3K4 me1:H4K20 me2
H2A.Z:H4K20 me2
H3K4 me1:H2A.Z
H2A.Z:H3−5ac
H2A.Z:H3K4 me3
H3K27 me3:H4K20 me3
H3K4 me1:H4−4ac
H3K9 me3:H4K20 me3
H4K20 me2:H3−5ac
H3K27 me3:H3K9 me3
DNA Meth. m5C:H3K9 me3
H4−4ac:H3−5ac
H3K27 me3:DNA Meth. m5C
H3K9 me2
H3K4 me3
H3K27 me2
H4K20 me2
H4K20 me3
H4−4ac
DNA Meth. m5C
H3K9 me3
H2A.Z
H3K27 me3
H3K4 me1
H3−5ac
0.00 0.25 0.50 0.75 1.00
$\hat{\pi}$
TEAD3 – Reverse
H4K20 me2:H3K4 me3
H3K4 me1:H3−5ac
H3K4 me3:H4−4ac
H4K20 me2:H4−4ac
DNA Meth. m5C:H4K20 me2
H3K4 me3:H3−5ac
H3K27 me3:H3K4 me1
H2A.Z:H4−4ac
H3K27 me2:DNA Meth. m5C
H3K4 me1:H4K20 me2
H2A.Z:H4K20 me2
H3K4 me1:H2A.Z
H2A.Z:H3−5ac
H2A.Z:H3K4 me3
H3K27 me3:H4K20 me3
H3K4 me1:H4−4ac
H3K9 me3:H4K20 me3
H4K20 me2:H3−5ac
H3K27 me3:H3K9 me3
DNA Meth. m5C:H3K9 me3
H4−4ac:H3−5ac
H3K27 me3:DNA Meth. m5C
H3K9 me2
H3K4 me3
H3K27 me2
H4K20 me2
H4K20 me3
H4−4ac
DNA Meth. m5C
H3K9 me3
H2A.Z
H3K27 me3
H3K4 me1
H3−5ac
0.00 0.25 0.50 0.75 1.00
$\hat{\pi}$
NTHL1 – Forward
H3K4 me1:H4K20 me2
H3K4 me3:H3−5ac
DNA Meth. m5C:H3K9 me3
H3K27 me3:H4K20 me3
DNA Meth. m5C:H4K20 me3
H3K27 me3:DNA Meth. m5C
H3K27 me3:H3K4 me1
H3K27 me3:H3K9 me3
H3K4 me1:H3−5ac
H2A.Z:H3K4 me3
DNA Meth. m5C:H4K20 me2
H4K20 me2:H3K4 me3
H4−4ac:H3−5ac
H2A.Z:H3−5ac
H3K4 me3:H4−4ac
H4K20 me2:H4−4ac
H4K20 me2:H3−5ac
H2A.Z:H4−4ac
H2A.Z:H4K20 me2
H3K4 me1:H2A.Z
H3K4 me1:H4−4ac
H3K9 me2
H3K27 me2
H4K20 me3
H3K9 me3
H3K27 me3
H3K4 me3
H4K20 me2
H3−5ac
H3K4 me1
DNA Meth. m5C
H2A.Z
H4−4ac
0.00 0.25 0.50 0.75 1.00
$\hat{\pi}$
NTHL1 – Reverse
H3K4 me1:H4K20 me2
H3K4 me3:H3−5ac
DNA Meth. m5C:H3K9 me3
H3K27 me3:H4K20 me3
DNA Meth. m5C:H4K20 me3
H3K27 me3:DNA Meth. m5C
H3K27 me3:H3K4 me1
H3K27 me3:H3K9 me3
H3K4 me1:H3−5ac
H2A.Z:H3K4 me3
DNA Meth. m5C:H4K20 me2
H4K20 me2:H3K4 me3
H4−4ac:H3−5ac
H2A.Z:H3−5ac
H3K4 me3:H4−4ac
H4K20 me2:H4−4ac
H4K20 me2:H3−5ac
H2A.Z:H4−4ac
H2A.Z:H4K20 me2
H3K4 me1:H2A.Z
H3K4 me1:H4−4ac
H3K9 me2
H3K27 me2
H4K20 me3
H3K9 me3
H3K27 me3
H3K4 me3
H4K20 me2
H3−5ac
H3K4 me1
DNA Meth. m5C
H2A.Z
H4−4ac
0.00 0.25 0.50 0.75 1.00
$\hat{\pi}$
NUP88 – Forward
H3K4 me1:H4−4ac
H3K4 me1:H3−5ac
H3K4 me1:H2A.Z
H3K27 me3:H3K4 me1
H2A.Z:H3K4 me3
H3K4 me1:H4K20 me2
H3K4 me3:H3−5ac
H3K9 me3:H4K20 me3
H4K20 me2:H3−5ac
H3K27 me3:H4K20 me3
H4K20 me2:H3K4 me3
H2A.Z:H4K20 me2
DNA Meth. m5C:H4K20 me3
DNA Meth. m5C:H3K9 me3
H3K27 me3:H3K9 me3
H3K27 me3:DNA Meth. m5C
H4−4ac
H3K4 me1
H3K4 me3
H4K20 me3
H2A.Z
DNA Meth. m5C
H3K27 me3
H3−5ac
H4K20 me2
H3K9 me3
0.00 0.25 0.50 0.75 1.00
$\hat{\pi}$
NUP88 – Reverse
H3K4 me1:H4−4ac
H3K4 me1:H3−5ac
H3K4 me1:H2A.Z
H3K27 me3:H3K4 me1
H2A.Z:H3K4 me3
H3K4 me1:H4K20 me2
H3K4 me3:H3−5ac
H3K9 me3:H4K20 me3
H4K20 me2:H3−5ac
H3K27 me3:H4K20 me3
H4K20 me2:H3K4 me3
H2A.Z:H4K20 me2
DNA Meth. m5C:H4K20 me3
DNA Meth. m5C:H3K9 me3
H3K27 me3:H3K9 me3
H3K27 me3:DNA Meth. m5C
H4−4ac
H3K4 me1
H3K4 me3
H4K20 me3
H2A.Z
DNA Meth. m5C
H3K27 me3
H3−5ac
H4K20 me2
H3K9 me3
0.00 0.25 0.50 0.75 1.00
$\hat{\pi}$
TRIP12 (2) – Forward
H3K4 me3:H4−4ac
H3K4 me1:H3−5ac
H3K4 me1:H4−4ac
DNA Meth. m5C:H4K20 me2
H3K4 me1:H4K20 me2
H3K4 me1:H2A.Z
H3K27 me3:H3K9 me3
H2A.Z:H3K4 me3
H2A.Z:H4K20 me2
H4K20 me2:H4−4ac
DNA Meth. m5C:H3K9 me3
H4K20 me2:H3−5ac
H4K20 me2:H3K4 me3
H3K27 me3:H3K4 me1
H3K9 me2:DNA Meth. m5C
H3K27 me3:DNA Meth. m5C
H4K20 me3
H3K27 me2
H3K4 me3
H3K9 me2
H4K20 me2
H3K9 me3
DNA Meth. m5C
H4−4ac
H3K4 me1
H2A.Z
H3K27 me3
H3−5ac
0.00 0.25 0.50 0.75 1.00
$\hat{\pi}$
TRIP12 (2) – Reverse
H3K4 me3:H4−4ac
H3K4 me1:H3−5ac
H3K4 me1:H4−4ac
DNA Meth. m5C:H4K20 me2
H3K4 me1:H4K20 me2
H3K4 me1:H2A.Z
H3K27 me3:H3K9 me3
H2A.Z:H3K4 me3
H2A.Z:H4K20 me2
H4K20 me2:H4−4ac
DNA Meth. m5C:H3K9 me3
H4K20 me2:H3−5ac
H4K20 me2:H3K4 me3
H3K27 me3:H3K4 me1
H3K9 me2:DNA Meth. m5C
H3K27 me3:DNA Meth. m5C
H4K20 me3
H3K27 me2
H3K4 me3
H3K9 me2
H4K20 me2
H3K9 me3
DNA Meth. m5C
H4−4ac
H3K4 me1
H2A.Z
H3K27 me3
H3−5ac
0.00 0.25 0.50 0.75 1.00
$\hat{\pi}$
CENPV (1) – Forward
H3K9 me2
H3K27 me2
H2A.Z
H3K4 me1
H3K4 me3
DNA Meth. m5C
H3−5ac
H4−4ac
H4K20 me2
0.00 0.25 0.50 0.75 1.00
$\hat{\pi}$
CENPV (1) – Reverse
H3K9 me2
H3K27 me2
H2A.Z
H3K4 me1
H3K4 me3
DNA Meth. m5C
H3−5ac
H4−4ac
H4K20 me2
0.00 0.25 0.50 0.75 1.00
$\hat{\pi}$

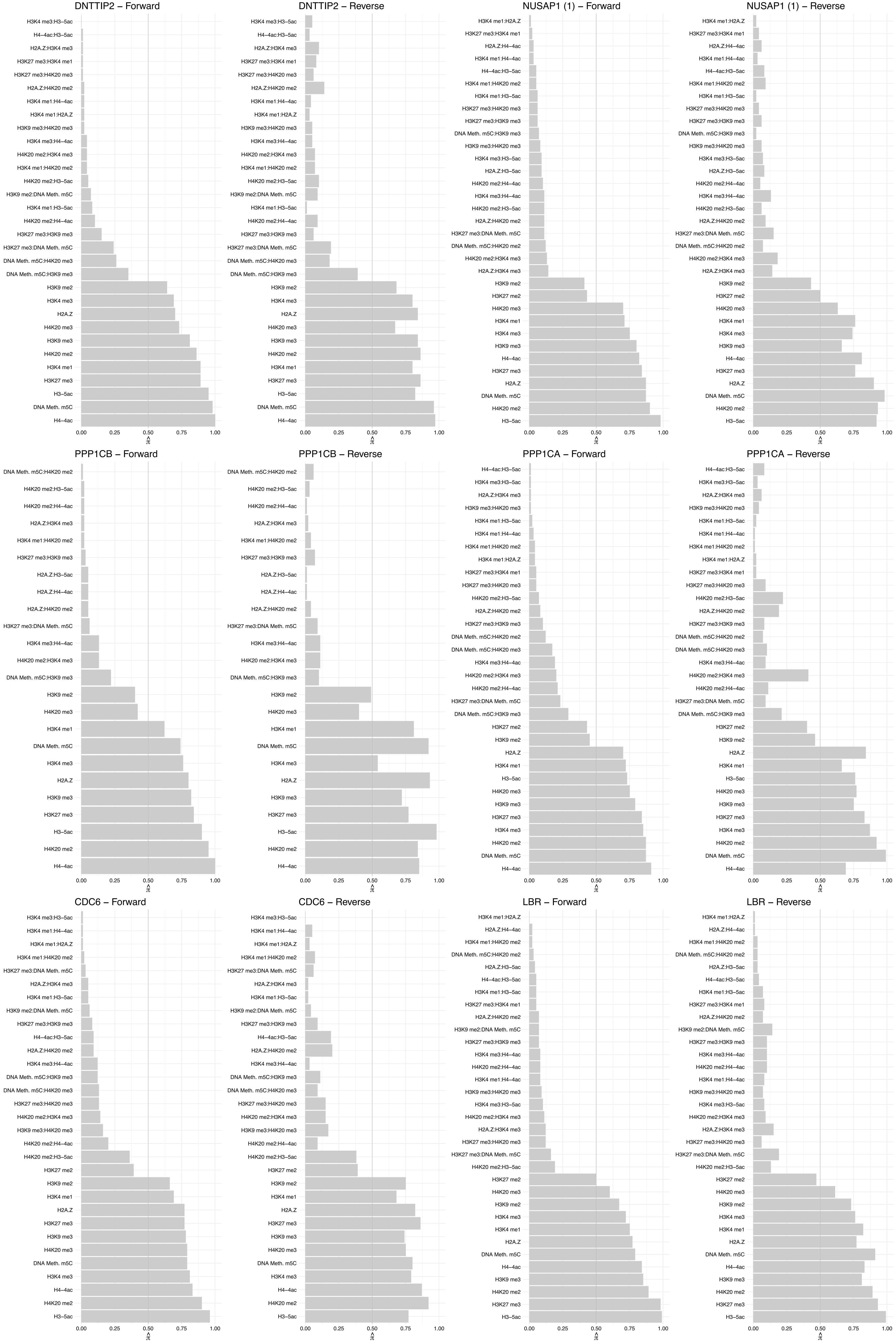
DNTTIP2 – Forward
DNTTIP2 – Reverse
NUSAP1 (1) – Forward
NUSAP1 (1) – Reverse
PPP1CB – Forward
PPP1CB – Reverse
PPP1CA – Forward
PPP1CA – Reverse
CDC6 – Forward
CDC6 – Reverse
LBR – Forward
LBR – Reverse
0.00
0.25
0.50
0.75
1.00
$\hat{\pi}$

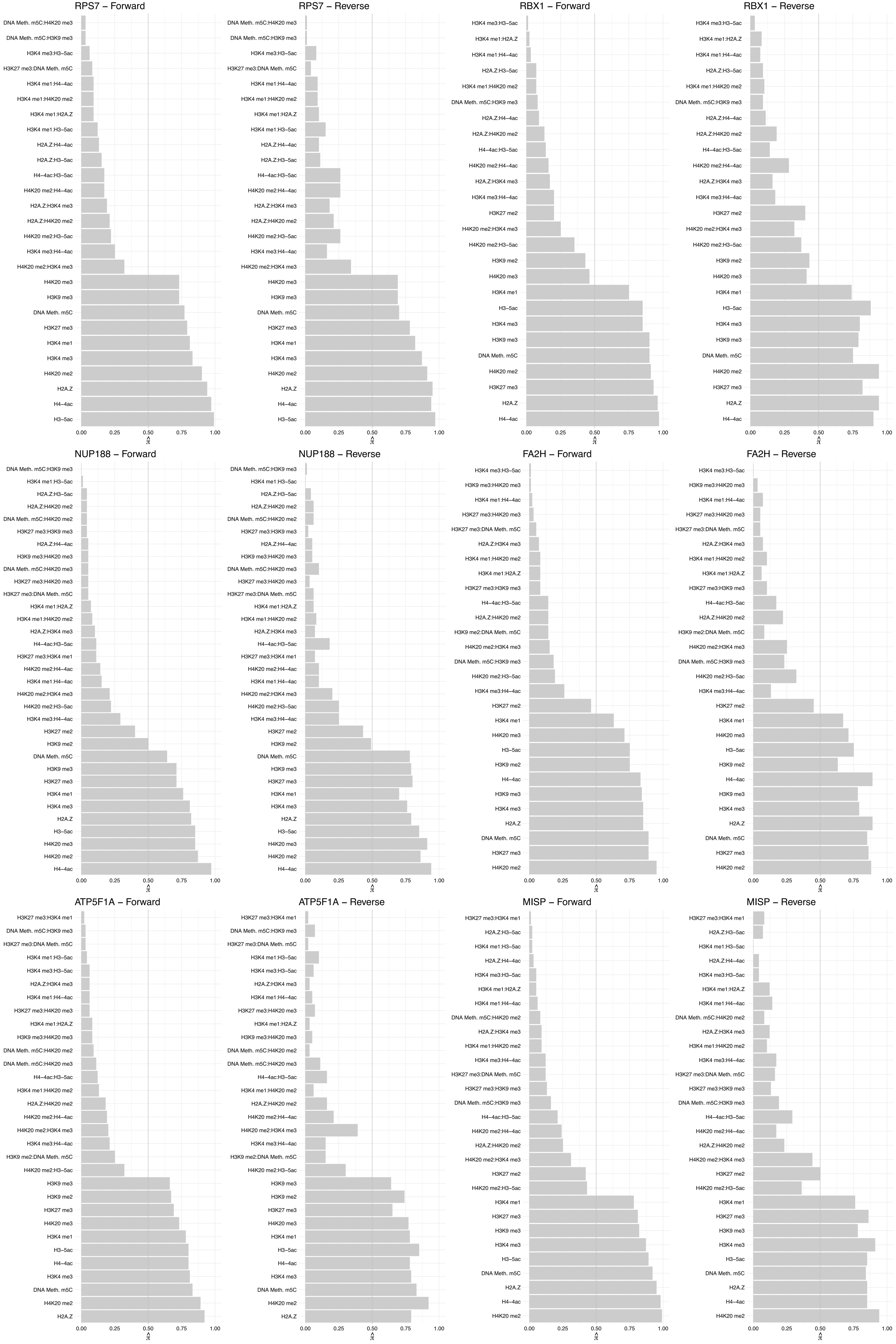
RPS7 – Forward
RPS7 – Reverse
DNA Meth. m5C:H4K20 me3
DNA Meth. m5C:H3K9 me3
H3K4 me3:H3−5ac
H3K27 me3:DNA Meth. m5C
H3K4 me1:H4−4ac
H3K4 me1:H4K20 me2
H3K4 me1:H2A.Z
H3K4 me1:H3−5ac
H2A.Z:H4−4ac
H2A.Z:H3−5ac
H4−4ac:H3−5ac
H4K20 me2:H4−4ac
H2A.Z:H3K4 me3
H2A.Z:H4K20 me2
H4K20 me2:H3−5ac
H3K4 me3:H4−4ac
H4K20 me2:H3K4 me3
H4K20 me3
H3K9 me3
DNA Meth. m5C
H3K27 me3
H3K4 me1
H3K4 me3
H4K20 me2
H2A.Z
H4−4ac
H3−5ac
RBX1 – Forward
RBX1 – Reverse
H3K4 me3:H3−5ac
H3K4 me1:H2A.Z
H3K4 me1:H4−4ac
H2A.Z:H3−5ac
H3K4 me1:H4K20 me2
DNA Meth. m5C:H3K9 me3
H2A.Z:H4−4ac
H2A.Z:H4K20 me2
H4−4ac:H3−5ac
H4K20 me2:H4−4ac
H2A.Z:H3K4 me3
H3K4 me3:H4−4ac
H3K27 me2
H4K20 me2:H3K4 me3
H4K20 me2:H3−5ac
H3K9 me2
H4K20 me3
H3K4 me1
H3−5ac
H3K4 me3
H3K9 me3
DNA Meth. m5C
H4K20 me2
H3K27 me3
H2A.Z
H4−4ac
NUP188 – Forward
NUP188 – Reverse
DNA Meth. m5C:H3K9 me3
H3K4 me1:H3−5ac
H2A.Z:H3−5ac
H2A.Z:H4K20 me2
DNA Meth. m5C:H4K20 me2
H3K27 me3:H3K9 me3
H2A.Z:H4−4ac
H3K9 me3:H4K20 me3
DNA Meth. m5C:H4K20 me3
H3K27 me3:H4K20 me3
H3K27 me3:DNA Meth. m5C
H3K4 me1:H2A.Z
H3K4 me1:H4K20 me2
H2A.Z:H3K4 me3
H4−4ac:H3−5ac
H3K27 me3:H3K4 me1
H4K20 me2:H4−4ac
H3K4 me1:H4−4ac
H4K20 me2:H3K4 me3
H4K20 me2:H3−5ac
H3K4 me3:H4−4ac
H3K27 me2
H3K9 me2
DNA Meth. m5C
H3K9 me3
H3K27 me3
H3K4 me1
H3K4 me3
H2A.Z
H3−5ac
H4K20 me3
H4K20 me2
H4−4ac
FA2H – Forward
FA2H – Reverse
H3K4 me3:H3−5ac
H3K9 me3:H4K20 me3
H3K4 me1:H4−4ac
H3K27 me3:H4K20 me3
H3K27 me3:DNA Meth. m5C
H2A.Z:H3K4 me3
H3K4 me1:H4K20 me2
H3K4 me1:H2A.Z
H3K27 me3:H3K9 me3
H4−4ac:H3−5ac
H2A.Z:H4K20 me2
H3K9 me2:DNA Meth. m5C
H4K20 me2:H3K4 me3
DNA Meth. m5C:H3K9 me3
H4K20 me2:H3−5ac
H3K4 me3:H4−4ac
H3K27 me2
H3K4 me1
H4K20 me3
H3−5ac
H3K9 me2
H4−4ac
H3K9 me3
H3K4 me3
H2A.Z
DNA Meth. m5C
H3K27 me3
H4K20 me2
ATP5F1A – Forward
ATP5F1A – Reverse
H3K27 me3:H3K4 me1
DNA Meth. m5C:H3K9 me3
H3K27 me3:DNA Meth. m5C
H3K4 me1:H3−5ac
H3K4 me3:H3−5ac
H2A.Z:H3K4 me3
H3K4 me1:H4−4ac
H3K27 me3:H4K20 me3
H3K4 me1:H2A.Z
H3K9 me3:H4K20 me3
DNA Meth. m5C:H4K20 me2
DNA Meth. m5C:H4K20 me3
H4−4ac:H3−5ac
H3K4 me1:H4K20 me2
H2A.Z:H4K20 me2
H4K20 me2:H4−4ac
H4K20 me2:H3K4 me3
H3K4 me3:H4−4ac
H3K9 me2:DNA Meth. m5C
H4K20 me2:H3−5ac
H3K9 me3
H3K9 me2
H3K27 me3
H4K20 me3
H3K4 me1
H3−5ac
H4−4ac
H3K4 me3
DNA Meth. m5C
H4K20 me2
H2A.Z
MISP – Forward
MISP – Reverse
H3K27 me3:H3K4 me1
H2A.Z:H3−5ac
H3K4 me1:H3−5ac
H2A.Z:H4−4ac
H3K4 me3:H3−5ac
H3K4 me1:H2A.Z
H3K4 me1:H4−4ac
DNA Meth. m5C:H4K20 me2
H2A.Z:H3K4 me3
H3K4 me1:H4K20 me2
H3K4 me3:H4−4ac
H3K27 me3:DNA Meth. m5C
H3K27 me3:H3K9 me3
DNA Meth. m5C:H3K9 me3
H4−4ac:H3−5ac
H4K20 me2:H4−4ac
H2A.Z:H4K20 me2
H4K20 me2:H3K4 me3
H3K27 me2
H4K20 me2:H3−5ac
H3K4 me1
H3K27 me3
H3K9 me3
H3K4 me3
H3−5ac
DNA Meth. m5C
H2A.Z
H4−4ac
H4K20 me2
0.00
0.25
0.50
0.75
1.00
π̂

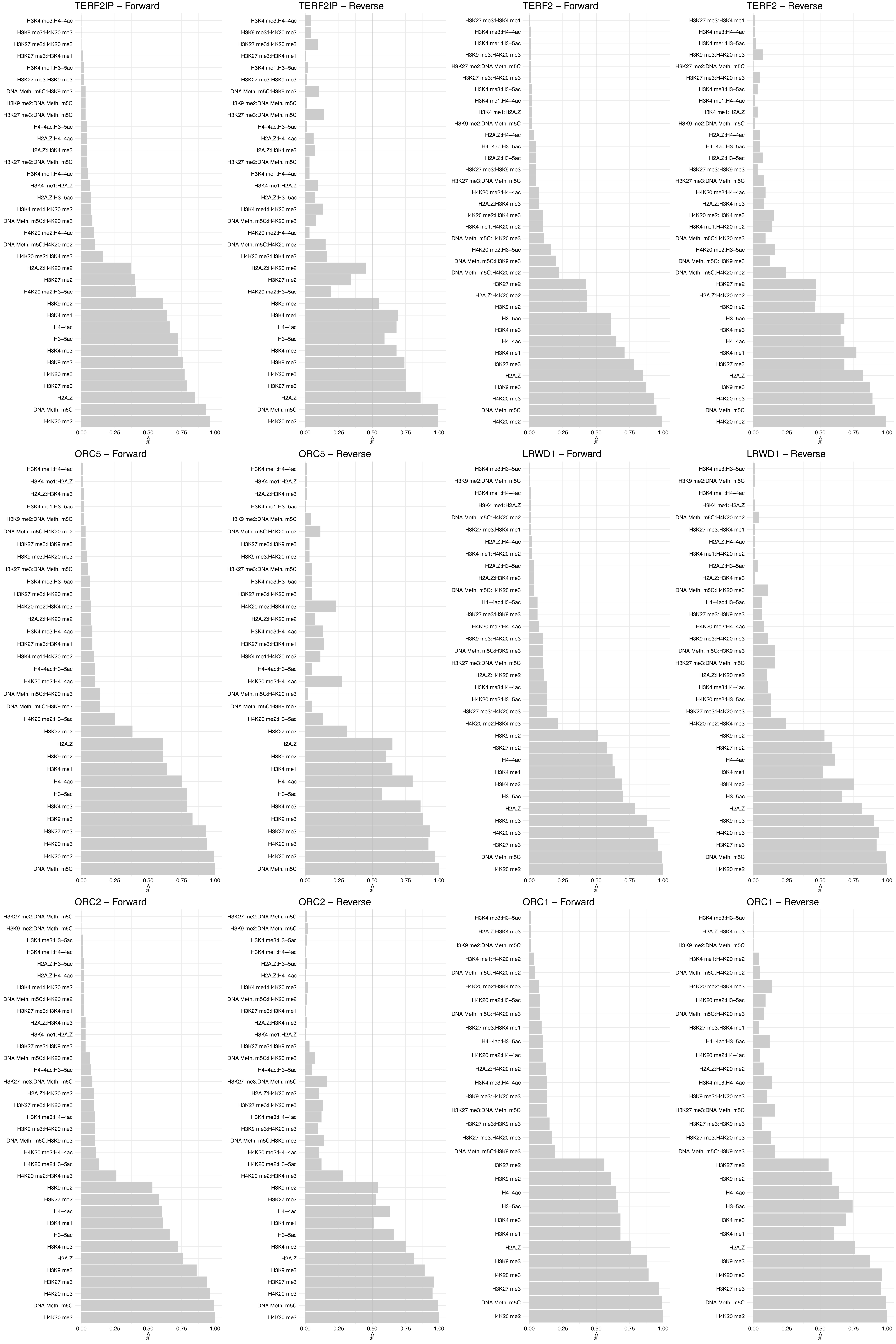
TERF2IP – Forward
TERF2IP – Reverse
TERF2 – Forward
TERF2 – Reverse
ORC5 – Forward
ORC5 – Reverse
LRWD1 – Forward
LRWD1 – Reverse
ORC2 – Forward
ORC2 – Reverse
ORC1 – Forward
ORC1 – Reverse
0.00 0.25 0.50 0.75 1.00
$\hat{\pi}$

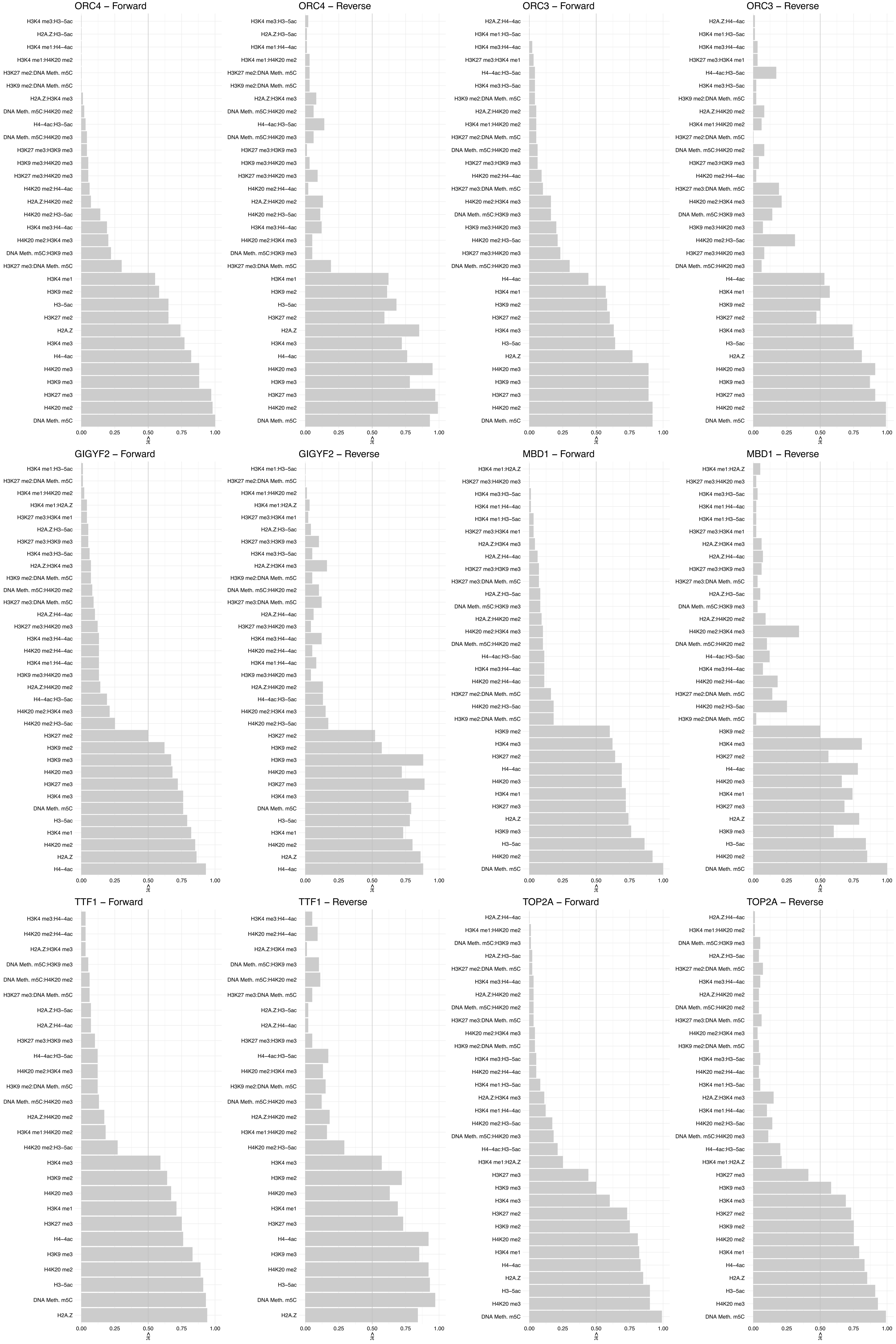
ORC4 – Forward
H3K4 me3:H3–5ac
H2A.Z:H3–5ac
H3K4 me1:H4–4ac
H3K4 me1:H4K20 me2
H3K27 me2:DNA Meth. m5C
H3K9 me2:DNA Meth. m5C
H2A.Z:H3K4 me3
DNA Meth. m5C:H4K20 me2
H4–4ac:H3–5ac
DNA Meth. m5C:H4K20 me3
H3K27 me3:H3K9 me3
H3K9 me3:H4K20 me3
H3K27 me3:H4K20 me3
H4K20 me2:H4–4ac
H2A.Z:H4K20 me2
H4K20 me2:H3–5ac
H3K4 me3:H4–4ac
H4K20 me2:H3K4 me3
DNA Meth. m5C:H3K9 me3
H3K27 me3:DNA Meth. m5C
H3K4 me1
H3K9 me2
H3–5ac
H3K27 me2
H2A.Z
H3K4 me3
H4–4ac
H4K20 me3
H3K9 me3
H3K27 me3
H4K20 me2
DNA Meth. m5C
0.00 0.25 0.50 0.75 1.00
$\hat{\pi}$
ORC4 – Reverse
H3K4 me3:H3–5ac
H2A.Z:H3–5ac
H3K4 me1:H4–4ac
H3K4 me1:H4K20 me2
H3K27 me2:DNA Meth. m5C
H3K9 me2:DNA Meth. m5C
H2A.Z:H3K4 me3
DNA Meth. m5C:H4K20 me2
H4–4ac:H3–5ac
DNA Meth. m5C:H4K20 me3
H3K27 me3:H3K9 me3
H3K9 me3:H4K20 me3
H3K27 me3:H4K20 me3
H4K20 me2:H4–4ac
H2A.Z:H4K20 me2
H4K20 me2:H3–5ac
H3K4 me3:H4–4ac
H4K20 me2:H3K4 me3
DNA Meth. m5C:H3K9 me3
H3K27 me3:DNA Meth. m5C
H3K4 me1
H3K9 me2
H3–5ac
H3K27 me2
H2A.Z
H3K4 me3
H4–4ac
H4K20 me3
H3K9 me3
H3K27 me3
H4K20 me2
DNA Meth. m5C
0.00 0.25 0.50 0.75 1.00
$\hat{\pi}$
ORC3 – Forward
H2A.Z:H4–4ac
H3K4 me1:H3–5ac
H3K4 me3:H4–4ac
H3K27 me3:H3K4 me1
H4–4ac:H3–5ac
H3K4 me3:H3–5ac
H3K9 me2:DNA Meth. m5C
H2A.Z:H4K20 me2
H3K4 me1:H4K20 me2
H3K27 me2:DNA Meth. m5C
DNA Meth. m5C:H4K20 me2
H3K27 me3:H3K9 me3
H4K20 me2:H4–4ac
H3K27 me3:DNA Meth. m5C
H4K20 me2:H3K4 me3
DNA Meth. m5C:H3K9 me3
H3K9 me3:H4K20 me3
H4K20 me2:H3–5ac
H3K27 me3:H4K20 me3
DNA Meth. m5C:H4K20 me3
H4–4ac
H3K4 me1
H3K9 me2
H3K27 me2
H3K4 me3
H3–5ac
H2A.Z
H4K20 me3
H3K9 me3
H3K27 me3
H4K20 me2
DNA Meth. m5C
0.00 0.25 0.50 0.75 1.00
$\hat{\pi}$
ORC3 – Reverse
H2A.Z:H4–4ac
H3K4 me1:H3–5ac
H3K4 me3:H4–4ac
H3K27 me3:H3K4 me1
H4–4ac:H3–5ac
H3K4 me3:H3–5ac
H3K9 me2:DNA Meth. m5C
H2A.Z:H4K20 me2
H3K4 me1:H4K20 me2
H3K27 me2:DNA Meth. m5C
DNA Meth. m5C:H4K20 me2
H3K27 me3:H3K9 me3
H4K20 me2:H4–4ac
H3K27 me3:DNA Meth. m5C
H4K20 me2:H3K4 me3
DNA Meth. m5C:H3K9 me3
H3K9 me3:H4K20 me3
H4K20 me2:H3–5ac
H3K27 me3:H4K20 me3
DNA Meth. m5C:H4K20 me3
H4–4ac
H3K4 me1
H3K9 me2
H3K27 me2
H3K4 me3
H3–5ac
H2A.Z
H4K20 me3
H3K9 me3
H3K27 me3
H4K20 me2
DNA Meth. m5C
0.00 0.25 0.50 0.75 1.00
$\hat{\pi}$
GIGYF2 – Forward
H3K4 me1:H3–5ac
H3K27 me2:DNA Meth. m5C
H3K4 me1:H4K20 me2
H3K4 me1:H2A.Z
H3K27 me3:H3K4 me1
H2A.Z:H3–5ac
H3K27 me3:H3K9 me3
H3K4 me3:H3–5ac
H2A.Z:H3K4 me3
H3K9 me2:DNA Meth. m5C
DNA Meth. m5C:H4K20 me2
H3K27 me3:DNA Meth. m5C
H2A.Z:H4–4ac
H3K27 me3:H4K20 me3
H3K4 me3:H4–4ac
H4K20 me2:H4–4ac
H3K4 me1:H4–4ac
H3K9 me3:H4K20 me3
H2A.Z:H4K20 me2
H4–4ac:H3–5ac
H4K20 me2:H3K4 me3
H4K20 me2:H3–5ac
H3K27 me2
H3K9 me2
H3K9 me3
H4K20 me3
H3K27 me3
H3K4 me3
DNA Meth. m5C
H3–5ac
H3K4 me1
H4K20 me2
H2A.Z
H4–4ac
0.00 0.25 0.50 0.75 1.00
$\hat{\pi}$
GIGYF2 – Reverse
H3K4 me1:H3–5ac
H3K27 me2:DNA Meth. m5C
H3K4 me1:H4K20 me2
H3K4 me1:H2A.Z
H3K27 me3:H3K4 me1
H2A.Z:H3–5ac
H3K27 me3:H3K9 me3
H3K4 me3:H3–5ac
H2A.Z:H3K4 me3
H3K9 me2:DNA Meth. m5C
DNA Meth. m5C:H4K20 me2
H3K27 me3:DNA Meth. m5C
H2A.Z:H4–4ac
H3K27 me3:H4K20 me3
H3K4 me3:H4–4ac
H4K20 me2:H4–4ac
H3K4 me1:H4–4ac
H3K9 me3:H4K20 me3
H2A.Z:H4K20 me2
H4–4ac:H3–5ac
H4K20 me2:H3K4 me3
H4K20 me2:H3–5ac
H3K27 me2
H3K9 me2
H3K9 me3
H4K20 me3
H3K27 me3
H3K4 me3
DNA Meth. m5C
H3–5ac
H3K4 me1
H4K20 me2
H2A.Z
H4–4ac
0.00 0.25 0.50 0.75 1.00
$\hat{\pi}$
MBD1 – Forward
H3K4 me1:H2A.Z
H3K27 me3:H4K20 me3
H3K4 me3:H3–5ac
H3K4 me1:H4–4ac
H3K4 me1:H3–5ac
H3K27 me3:H3K4 me1
H2A.Z:H3K4 me3
H2A.Z:H4–4ac
H3K27 me3:H3K9 me3
H3K27 me3:DNA Meth. m5C
H2A.Z:H3–5ac
DNA Meth. m5C:H3K9 me3
H2A.Z:H4K20 me2
H4K20 me2:H3K4 me3
DNA Meth. m5C:H4K20 me2
H4–4ac:H3–5ac
H3K4 me3:H4–4ac
H4K20 me2:H4–4ac
H3K27 me2:DNA Meth. m5C
H4K20 me2:H3–5ac
H3K9 me2:DNA Meth. m5C
H3K9 me2
H3K4 me3
H3K27 me2
H4–4ac
H4K20 me3
H3K4 me1
H3K27 me3
H2A.Z
H3K9 me3
H3–5ac
H4K20 me2
DNA Meth. m5C
0.00 0.25 0.50 0.75 1.00
$\hat{\pi}$
MBD1 – Reverse
H3K4 me1:H2A.Z
H3K27 me3:H4K20 me3
H3K4 me3:H3–5ac
H3K4 me1:H4–4ac
H3K4 me1:H3–5ac
H3K27 me3:H3K4 me1
H2A.Z:H3K4 me3
H2A.Z:H4–4ac
H3K27 me3:H3K9 me3
H3K27 me3:DNA Meth. m5C
H2A.Z:H3–5ac
DNA Meth. m5C:H3K9 me3
H2A.Z:H4K20 me2
H4K20 me2:H3K4 me3
DNA Meth. m5C:H4K20 me2
H4–4ac:H3–5ac
H3K4 me3:H4–4ac
H4K20 me2:H4–4ac
H3K27 me2:DNA Meth. m5C
H4K20 me2:H3–5ac
H3K9 me2:DNA Meth. m5C
H3K9 me2
H3K4 me3
H3K27 me2
H4–4ac
H4K20 me3
H3K4 me1
H3K27 me3
H2A.Z
H3K9 me3
H3–5ac
H4K20 me2
DNA Meth. m5C
0.00 0.25 0.50 0.75 1.00
$\hat{\pi}$
TTF1 – Forward
H3K4 me3:H4–4ac
H4K20 me2:H4–4ac
H2A.Z:H3K4 me3
DNA Meth. m5C:H3K9 me3
DNA Meth. m5C:H4K20 me2
H3K27 me3:DNA Meth. m5C
H2A.Z:H3–5ac
H2A.Z:H4–4ac
H3K27 me3:H3K9 me3
H4–4ac:H3–5ac
H4K20 me2:H3K4 me3
H3K9 me2:DNA Meth. m5C
DNA Meth. m5C:H4K20 me3
H2A.Z:H4K20 me2
H3K4 me1:H4K20 me2
H4K20 me2:H3–5ac
H3K4 me3
H3K9 me2
H4K20 me3
H3K4 me1
H3K27 me3
H4–4ac
H3K9 me3
H4K20 me2
H3–5ac
DNA Meth. m5C
H2A.Z
0.00 0.25 0.50 0.75 1.00
$\hat{\pi}$
TTF1 – Reverse
H3K4 me3:H4–4ac
H4K20 me2:H4–4ac
H2A.Z:H3K4 me3
DNA Meth. m5C:H3K9 me3
DNA Meth. m5C:H4K20 me2
H3K27 me3:DNA Meth. m5C
H2A.Z:H3–5ac
H2A.Z:H4–4ac
H3K27 me3:H3K9 me3
H4–4ac:H3–5ac
H4K20 me2:H3K4 me3
H3K9 me2:DNA Meth. m5C
DNA Meth. m5C:H4K20 me3
H2A.Z:H4K20 me2
H3K4 me1:H4K20 me2
H4K20 me2:H3–5ac
H3K4 me3
H3K9 me2
H4K20 me3
H3K4 me1
H3K27 me3
H4–4ac
H3K9 me3
H4K20 me2
H3–5ac
DNA Meth. m5C
H2A.Z
0.00 0.25 0.50 0.75 1.00
$\hat{\pi}$
TOP2A – Forward
H2A.Z:H4–4ac
H3K4 me1:H4K20 me2
DNA Meth. m5C:H3K9 me3
H2A.Z:H3–5ac
H3K27 me2:DNA Meth. m5C
H3K4 me3:H4–4ac
H2A.Z:H4K20 me2
DNA Meth. m5C:H4K20 me2
H3K27 me3:DNA Meth. m5C
H4K20 me2:H3K4 me3
H3K9 me2:DNA Meth. m5C
H3K4 me3:H3–5ac
H4K20 me2:H4–4ac
H3K4 me1:H3–5ac
H2A.Z:H3K4 me3
H3K4 me1:H4–4ac
H4K20 me2:H3–5ac
DNA Meth. m5C:H4K20 me3
H4–4ac:H3–5ac
H3K4 me1:H2A.Z
H3K27 me3
H3K9 me3
H3K4 me3
H3K27 me2
H3K9 me2
H4K20 me2
H3K4 me1
H4–4ac
H2A.Z
H3–5ac
H4K20 me3
DNA Meth. m5C
0.00 0.25 0.50 0.75 1.00
$\hat{\pi}$
TOP2A – Reverse
H2A.Z:H4–4ac
H3K4 me1:H4K20 me2
DNA Meth. m5C:H3K9 me3
H2A.Z:H3–5ac
H3K27 me2:DNA Meth. m5C
H3K4 me3:H4–4ac
H2A.Z:H4K20 me2
DNA Meth. m5C:H4K20 me2
H3K27 me3:DNA Meth. m5C
H4K20 me2:H3K4 me3
H3K9 me2:DNA Meth. m5C
H3K4 me3:H3–5ac
H4K20 me2:H4–4ac
H3K4 me1:H3–5ac
H2A.Z:H3K4 me3
H3K4 me1:H4–4ac
H4K20 me2:H3–5ac
DNA Meth. m5C:H4K20 me3
H4–4ac:H3–5ac
H3K4 me1:H2A.Z
H3K27 me3
H3K9 me3
H3K4 me3
H3K27 me2
H3K9 me2
H4K20 me2
H3K4 me1
H4–4ac
H2A.Z
H3–5ac
H4K20 me3
DNA Meth. m5C
0.00 0.25 0.50 0.75 1.00
$\hat{\pi}$

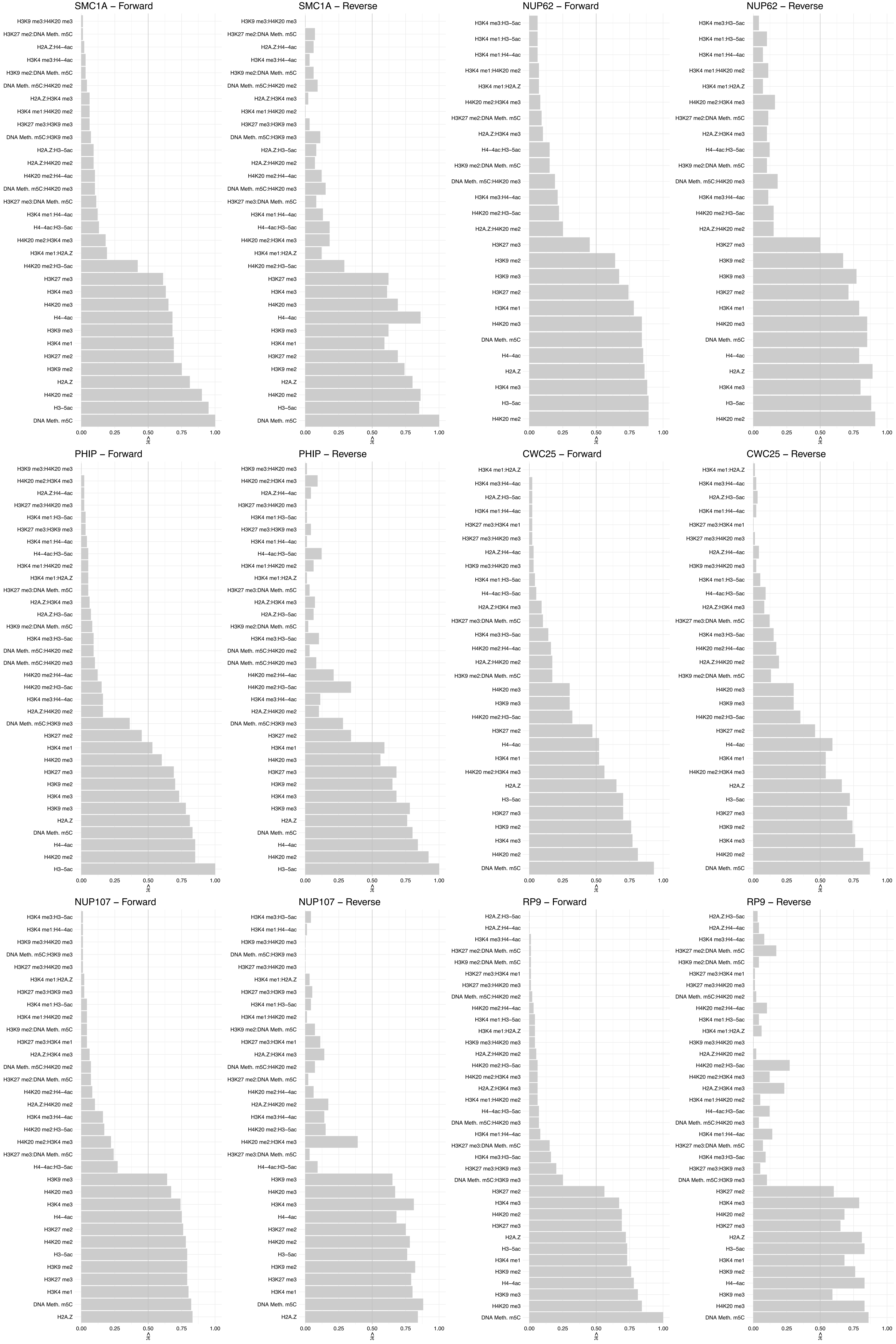
SMC1A – Forward
H3K9 me3:H4K20 me3
H3K27 me2:DNA Meth. m5C
H2A.Z:H4−4ac
H3K4 me3:H4−4ac
H3K9 me2:DNA Meth. m5C
DNA Meth. m5C:H4K20 me2
H2A.Z:H3K4 me3
H3K4 me1:H4K20 me2
H3K27 me3:H3K9 me3
DNA Meth. m5C:H3K9 me3
H2A.Z:H3−5ac
H2A.Z:H4K20 me2
H4K20 me2:H4−4ac
DNA Meth. m5C:H4K20 me3
H3K27 me3:DNA Meth. m5C
H3K4 me1:H4−4ac
H4−4ac:H3−5ac
H4K20 me2:H3K4 me3
H3K4 me1:H2A.Z
H4K20 me2:H3−5ac
H3K27 me3
H3K4 me3
H4K20 me3
H4−4ac
H3K9 me3
H3K4 me1
H3K27 me2
H3K9 me2
H2A.Z
H4K20 me2
H3−5ac
DNA Meth. m5C
0.00 0.25 0.50 0.75 1.00
π̂
SMC1A – Reverse
H3K9 me3:H4K20 me3
H3K27 me2:DNA Meth. m5C
H2A.Z:H4−4ac
H3K4 me3:H4−4ac
H3K9 me2:DNA Meth. m5C
DNA Meth. m5C:H4K20 me2
H2A.Z:H3K4 me3
H3K4 me1:H4K20 me2
H3K27 me3:H3K9 me3
DNA Meth. m5C:H3K9 me3
H2A.Z:H3−5ac
H2A.Z:H4K20 me2
H4K20 me2:H4−4ac
DNA Meth. m5C:H4K20 me3
H3K27 me3:DNA Meth. m5C
H3K4 me1:H4−4ac
H4−4ac:H3−5ac
H4K20 me2:H3K4 me3
H3K4 me1:H2A.Z
H4K20 me2:H3−5ac
H3K27 me3
H3K4 me3
H4K20 me3
H4−4ac
H3K9 me3
H3K4 me1
H3K27 me2
H3K9 me2
H2A.Z
H4K20 me2
H3−5ac
DNA Meth. m5C
0.00 0.25 0.50 0.75 1.00
π̂
NUP62 – Forward
H3K4 me3:H3−5ac
H3K4 me1:H3−5ac
H3K4 me1:H4−4ac
H3K4 me1:H4K20 me2
H3K4 me1:H2A.Z
H4K20 me2:H3K4 me3
H3K27 me2:DNA Meth. m5C
H2A.Z:H3K4 me3
H4−4ac:H3−5ac
H3K9 me2:DNA Meth. m5C
DNA Meth. m5C:H4K20 me3
H3K4 me3:H4−4ac
H4K20 me2:H3−5ac
H2A.Z:H4K20 me2
H3K27 me3
H3K9 me2
H3K9 me3
H3K27 me2
H3K4 me1
H4K20 me3
DNA Meth. m5C
H4−4ac
H2A.Z
H3K4 me3
H3−5ac
H4K20 me2
0.00 0.25 0.50 0.75 1.00
π̂
NUP62 – Reverse
H3K4 me3:H3−5ac
H3K4 me1:H3−5ac
H3K4 me1:H4−4ac
H3K4 me1:H4K20 me2
H3K4 me1:H2A.Z
H4K20 me2:H3K4 me3
H3K27 me2:DNA Meth. m5C
H2A.Z:H3K4 me3
H4−4ac:H3−5ac
H3K9 me2:DNA Meth. m5C
DNA Meth. m5C:H4K20 me3
H3K4 me3:H4−4ac
H4K20 me2:H3−5ac
H2A.Z:H4K20 me2
H3K27 me3
H3K9 me2
H3K9 me3
H3K27 me2
H3K4 me1
H4K20 me3
DNA Meth. m5C
H4−4ac
H2A.Z
H3K4 me3
H3−5ac
H4K20 me2
0.00 0.25 0.50 0.75 1.00
π̂
PHIP – Forward
H3K9 me3:H4K20 me3
H4K20 me2:H3K4 me3
H2A.Z:H4−4ac
H3K27 me3:H4K20 me3
H3K4 me1:H3−5ac
H3K27 me3:H3K9 me3
H3K4 me1:H4−4ac
H4−4ac:H3−5ac
H3K4 me1:H4K20 me2
H3K4 me1:H2A.Z
H3K27 me3:DNA Meth. m5C
H2A.Z:H3K4 me3
H2A.Z:H3−5ac
H3K9 me2:DNA Meth. m5C
H3K4 me3:H3−5ac
DNA Meth. m5C:H4K20 me2
DNA Meth. m5C:H4K20 me3
H4K20 me2:H4−4ac
H4K20 me2:H3−5ac
H3K4 me3:H4−4ac
H2A.Z:H4K20 me2
DNA Meth. m5C:H3K9 me3
H3K27 me2
H3K4 me1
H4K20 me3
H3K27 me3
H3K9 me2
H3K4 me3
H3K9 me3
H2A.Z
DNA Meth. m5C
H4−4ac
H4K20 me2
H3−5ac
0.00 0.25 0.50 0.75 1.00
π̂
PHIP – Reverse
H3K9 me3:H4K20 me3
H4K20 me2:H3K4 me3
H2A.Z:H4−4ac
H3K27 me3:H4K20 me3
H3K4 me1:H3−5ac
H3K27 me3:H3K9 me3
H3K4 me1:H4−4ac
H4−4ac:H3−5ac
H3K4 me1:H4K20 me2
H3K4 me1:H2A.Z
H3K27 me3:DNA Meth. m5C
H2A.Z:H3K4 me3
H2A.Z:H3−5ac
H3K9 me2:DNA Meth. m5C
H3K4 me3:H3−5ac
DNA Meth. m5C:H4K20 me2
DNA Meth. m5C:H4K20 me3
H4K20 me2:H4−4ac
H4K20 me2:H3−5ac
H3K4 me3:H4−4ac
H2A.Z:H4K20 me2
DNA Meth. m5C:H3K9 me3
H3K27 me2
H3K4 me1
H4K20 me3
H3K27 me3
H3K9 me2
H3K4 me3
H3K9 me3
H2A.Z
DNA Meth. m5C
H4−4ac
H4K20 me2
H3−5ac
0.00 0.25 0.50 0.75 1.00
π̂
CWC25 – Forward
H3K4 me1:H2A.Z
H3K4 me3:H4−4ac
H2A.Z:H3−5ac
H3K4 me1:H4−4ac
H3K27 me3:H3K4 me1
H3K27 me3:H4K20 me3
H2A.Z:H4−4ac
H3K9 me3:H4K20 me3
H3K4 me1:H3−5ac
H4−4ac:H3−5ac
H2A.Z:H3K4 me3
H3K27 me3:DNA Meth. m5C
H3K4 me3:H3−5ac
H4K20 me2:H4−4ac
H2A.Z:H4K20 me2
H3K9 me2:DNA Meth. m5C
H4K20 me3
H3K9 me3
H4K20 me2:H3−5ac
H3K27 me2
H4−4ac
H3K4 me1
H4K20 me2:H3K4 me3
H2A.Z
H3−5ac
H3K27 me3
H3K9 me2
H3K4 me3
H4K20 me2
DNA Meth. m5C
0.00 0.25 0.50 0.75 1.00
π̂
CWC25 – Reverse
H3K4 me1:H2A.Z
H3K4 me3:H4−4ac
H2A.Z:H3−5ac
H3K4 me1:H4−4ac
H3K27 me3:H3K4 me1
H3K27 me3:H4K20 me3
H2A.Z:H4−4ac
H3K9 me3:H4K20 me3
H3K4 me1:H3−5ac
H4−4ac:H3−5ac
H2A.Z:H3K4 me3
H3K27 me3:DNA Meth. m5C
H3K4 me3:H3−5ac
H4K20 me2:H4−4ac
H2A.Z:H4K20 me2
H3K9 me2:DNA Meth. m5C
H4K20 me3
H3K9 me3
H4K20 me2:H3−5ac
H3K27 me2
H4−4ac
H3K4 me1
H4K20 me2:H3K4 me3
H2A.Z
H3−5ac
H3K27 me3
H3K9 me2
H3K4 me3
H4K20 me2
DNA Meth. m5C
0.00 0.25 0.50 0.75 1.00
π̂
NUP107 – Forward
H3K4 me3:H3−5ac
H3K4 me1:H4−4ac
H3K9 me3:H4K20 me3
DNA Meth. m5C:H3K9 me3
H3K27 me3:H4K20 me3
H3K4 me1:H2A.Z
H3K27 me3:H3K9 me3
H3K4 me1:H3−5ac
H3K4 me1:H4K20 me2
H3K9 me2:DNA Meth. m5C
H3K27 me3:H3K4 me1
H2A.Z:H3K4 me3
DNA Meth. m5C:H4K20 me2
H3K27 me2:DNA Meth. m5C
H4K20 me2:H4−4ac
H2A.Z:H4K20 me2
H3K4 me3:H4−4ac
H4K20 me2:H3−5ac
H4K20 me2:H3K4 me3
H3K27 me3:DNA Meth. m5C
H4−4ac:H3−5ac
H3K9 me3
H4K20 me3
H3K4 me3
H4−4ac
H3K27 me2
H4K20 me2
H3−5ac
H3K9 me2
H3K27 me3
H3K4 me1
DNA Meth. m5C
H2A.Z
0.00 0.25 0.50 0.75 1.00
π̂
NUP107 – Reverse
H3K4 me3:H3−5ac
H3K4 me1:H4−4ac
H3K9 me3:H4K20 me3
DNA Meth. m5C:H3K9 me3
H3K27 me3:H4K20 me3
H3K4 me1:H2A.Z
H3K27 me3:H3K9 me3
H3K4 me1:H3−5ac
H3K4 me1:H4K20 me2
H3K9 me2:DNA Meth. m5C
H3K27 me3:H3K4 me1
H2A.Z:H3K4 me3
DNA Meth. m5C:H4K20 me2
H3K27 me2:DNA Meth. m5C
H4K20 me2:H4−4ac
H2A.Z:H4K20 me2
H3K4 me3:H4−4ac
H4K20 me2:H3−5ac
H4K20 me2:H3K4 me3
H3K27 me3:DNA Meth. m5C
H4−4ac:H3−5ac
H3K9 me3
H4K20 me3
H3K4 me3
H4−4ac
H3K27 me2
H4K20 me2
H3−5ac
H3K9 me2
H3K27 me3
H3K4 me1
DNA Meth. m5C
H2A.Z
0.00 0.25 0.50 0.75 1.00
π̂
RP9 – Forward
H2A.Z:H3−5ac
H2A.Z:H4−4ac
H3K4 me3:H4−4ac
H3K27 me2:DNA Meth. m5C
H3K9 me2:DNA Meth. m5C
H3K27 me3:H3K4 me1
H3K27 me3:H4K20 me3
DNA Meth. m5C:H4K20 me2
H4K20 me2:H4−4ac
H3K4 me1:H3−5ac
H3K4 me1:H2A.Z
H3K9 me3:H4K20 me3
H2A.Z:H4K20 me2
H4K20 me2:H3−5ac
H4K20 me2:H3K4 me3
H2A.Z:H3K4 me3
H3K4 me1:H4K20 me2
H4−4ac:H3−5ac
DNA Meth. m5C:H4K20 me3
H3K4 me1:H4−4ac
H3K27 me3:DNA Meth. m5C
H3K4 me3:H3−5ac
H3K27 me3:H3K9 me3
DNA Meth. m5C:H3K9 me3
H3K27 me2
H3K4 me3
H4K20 me2
H3K27 me3
H2A.Z
H3−5ac
H3K4 me1
H3K9 me2
H4−4ac
H3K9 me3
H4K20 me3
DNA Meth. m5C
0.00 0.25 0.50 0.75 1.00
π̂
RP9 – Reverse
H2A.Z:H3−5ac
H2A.Z:H4−4ac
H3K4 me3:H4−4ac
H3K27 me2:DNA Meth. m5C
H3K9 me2:DNA Meth. m5C
H3K27 me3:H3K4 me1
H3K27 me3:H4K20 me3
DNA Meth. m5C:H4K20 me2
H4K20 me2:H4−4ac
H3K4 me1:H3−5ac
H3K4 me1:H2A.Z
H3K9 me3:H4K20 me3
H2A.Z:H4K20 me2
H4K20 me2:H3−5ac
H4K20 me2:H3K4 me3
H2A.Z:H3K4 me3
H3K4 me1:H4K20 me2
H4−4ac:H3−5ac
DNA Meth. m5C:H4K20 me3
H3K4 me1:H4−4ac
H3K27 me3:DNA Meth. m5C
H3K4 me3:H3−5ac
H3K27 me3:H3K9 me3
DNA Meth. m5C:H3K9 me3
H3K27 me2
H3K4 me3
H4K20 me2
H3K27 me3
H2A.Z
H3−5ac
H3K4 me1
H3K9 me2
H4−4ac
H3K9 me3
H4K20 me3
DNA Meth. m5C
0.00 0.25 0.50 0.75 1.00
π̂

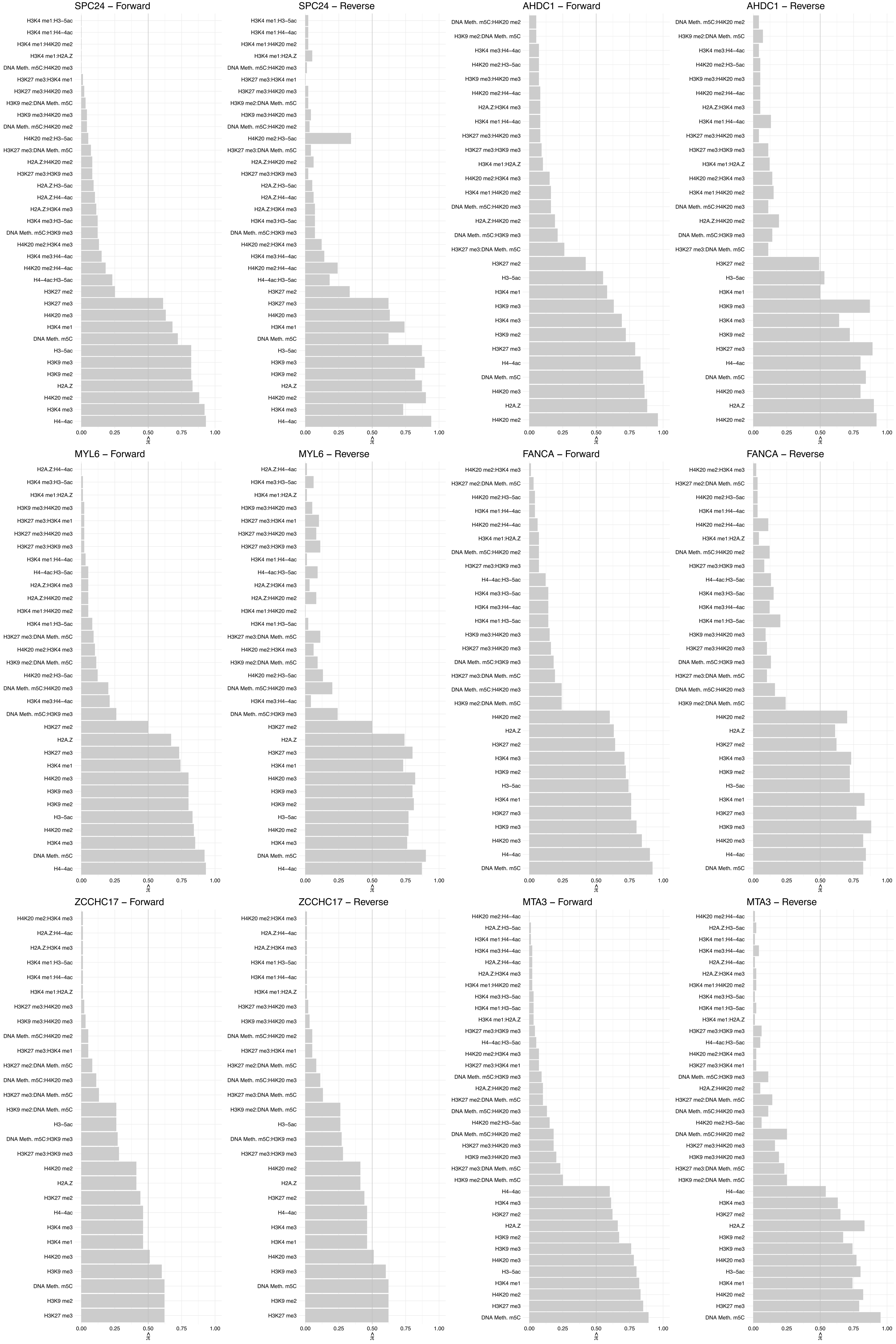
SPC24 – Forward
H3K4 me1:H3−5ac
H3K4 me1:H4−4ac
H3K4 me1:H4K20 me2
H3K4 me1:H2A.Z
DNA Meth. m5C:H4K20 me3
H3K27 me3:H3K4 me1
H3K27 me3:H4K20 me3
H3K9 me2:DNA Meth. m5C
H3K9 me3:H4K20 me3
DNA Meth. m5C:H4K20 me2
H4K20 me2:H3−5ac
H3K27 me3:DNA Meth. m5C
H2A.Z:H4K20 me2
H3K27 me3:H3K9 me3
H2A.Z:H3−5ac
H2A.Z:H4−4ac
H2A.Z:H3K4 me3
H3K4 me3:H3−5ac
DNA Meth. m5C:H3K9 me3
H4K20 me2:H3K4 me3
H3K4 me3:H4−4ac
H4K20 me2:H4−4ac
H4−4ac:H3−5ac
H3K27 me2
H3K27 me3
H4K20 me3
H3K4 me1
DNA Meth. m5C
H3−5ac
H3K9 me3
H3K9 me2
H2A.Z
H4K20 me2
H3K4 me3
H4−4ac
0.00 0.25 0.50 0.75 1.00
$\hat{\pi}$
SPC24 – Reverse
H3K4 me1:H3−5ac
H3K4 me1:H4−4ac
H3K4 me1:H4K20 me2
H3K4 me1:H2A.Z
DNA Meth. m5C:H4K20 me3
H3K27 me3:H3K4 me1
H3K27 me3:H4K20 me3
H3K9 me2:DNA Meth. m5C
H3K9 me3:H4K20 me3
DNA Meth. m5C:H4K20 me2
H4K20 me2:H3−5ac
H3K27 me3:DNA Meth. m5C
H2A.Z:H4K20 me2
H3K27 me3:H3K9 me3
H2A.Z:H3−5ac
H2A.Z:H4−4ac
H2A.Z:H3K4 me3
H3K4 me3:H3−5ac
DNA Meth. m5C:H3K9 me3
H4K20 me2:H3K4 me3
H3K4 me3:H4−4ac
H4K20 me2:H4−4ac
H4−4ac:H3−5ac
H3K27 me2
H3K27 me3
H4K20 me3
H3K4 me1
DNA Meth. m5C
H3−5ac
H3K9 me3
H3K9 me2
H2A.Z
H4K20 me2
H3K4 me3
H4−4ac
0.00 0.25 0.50 0.75 1.00
$\hat{\pi}$
AHDC1 – Forward
DNA Meth. m5C:H4K20 me2
H3K9 me2:DNA Meth. m5C
H3K4 me3:H4−4ac
H4K20 me2:H3−5ac
H3K9 me3:H4K20 me3
H4K20 me2:H4−4ac
H2A.Z:H3K4 me3
H3K4 me1:H4−4ac
H3K27 me3:H4K20 me3
H3K27 me3:H3K9 me3
H3K4 me1:H2A.Z
H4K20 me2:H3K4 me3
H3K4 me1:H4K20 me2
DNA Meth. m5C:H4K20 me3
H2A.Z:H4K20 me2
DNA Meth. m5C:H3K9 me3
H3K27 me3:DNA Meth. m5C
H3K27 me2
H3−5ac
H3K4 me1
H3K9 me3
H3K4 me3
H3K9 me2
H3K27 me3
H4−4ac
DNA Meth. m5C
H4K20 me3
H2A.Z
H4K20 me2
0.00 0.25 0.50 0.75 1.00
$\hat{\pi}$
AHDC1 – Reverse
DNA Meth. m5C:H4K20 me2
H3K9 me2:DNA Meth. m5C
H3K4 me3:H4−4ac
H4K20 me2:H3−5ac
H3K9 me3:H4K20 me3
H4K20 me2:H4−4ac
H2A.Z:H3K4 me3
H3K4 me1:H4−4ac
H3K27 me3:H4K20 me3
H3K27 me3:H3K9 me3
H3K4 me1:H2A.Z
H4K20 me2:H3K4 me3
H3K4 me1:H4K20 me2
DNA Meth. m5C:H4K20 me3
H2A.Z:H4K20 me2
DNA Meth. m5C:H3K9 me3
H3K27 me3:DNA Meth. m5C
H3K27 me2
H3−5ac
H3K4 me1
H3K9 me3
H3K4 me3
H3K9 me2
H3K27 me3
H4−4ac
DNA Meth. m5C
H4K20 me3
H2A.Z
H4K20 me2
0.00 0.25 0.50 0.75 1.00
$\hat{\pi}$
MYL6 – Forward
H2A.Z:H4−4ac
H3K4 me3:H3−5ac
H3K4 me1:H2A.Z
H3K9 me3:H4K20 me3
H3K27 me3:H3K4 me1
H3K27 me3:H4K20 me3
H3K27 me3:H3K9 me3
H3K4 me1:H4−4ac
H4−4ac:H3−5ac
H2A.Z:H3K4 me3
H2A.Z:H4K20 me2
H3K4 me1:H4K20 me2
H3K4 me1:H3−5ac
H3K27 me3:DNA Meth. m5C
H4K20 me2:H3K4 me3
H3K9 me2:DNA Meth. m5C
H4K20 me2:H3−5ac
DNA Meth. m5C:H4K20 me3
H3K4 me3:H4−4ac
DNA Meth. m5C:H3K9 me3
H3K27 me2
H2A.Z
H3K27 me3
H3K4 me1
H4K20 me3
H3K9 me3
H3K9 me2
H3−5ac
H4K20 me2
H3K4 me3
DNA Meth. m5C
H4−4ac
0.00 0.25 0.50 0.75 1.00
$\hat{\pi}$
MYL6 – Reverse
H2A.Z:H4−4ac
H3K4 me3:H3−5ac
H3K4 me1:H2A.Z
H3K9 me3:H4K20 me3
H3K27 me3:H3K4 me1
H3K27 me3:H4K20 me3
H3K27 me3:H3K9 me3
H3K4 me1:H4−4ac
H4−4ac:H3−5ac
H2A.Z:H3K4 me3
H2A.Z:H4K20 me2
H3K4 me1:H4K20 me2
H3K4 me1:H3−5ac
H3K27 me3:DNA Meth. m5C
H4K20 me2:H3K4 me3
H3K9 me2:DNA Meth. m5C
H4K20 me2:H3−5ac
DNA Meth. m5C:H4K20 me3
H3K4 me3:H4−4ac
DNA Meth. m5C:H3K9 me3
H3K27 me2
H2A.Z
H3K27 me3
H3K4 me1
H4K20 me3
H3K9 me3
H3K9 me2
H3−5ac
H4K20 me2
H3K4 me3
DNA Meth. m5C
H4−4ac
0.00 0.25 0.50 0.75 1.00
$\hat{\pi}$
FANCA – Forward
H4K20 me2:H3K4 me3
H3K27 me2:DNA Meth. m5C
H4K20 me2:H3−5ac
H3K4 me1:H4−4ac
H4K20 me2:H4−4ac
H3K4 me1:H2A.Z
DNA Meth. m5C:H4K20 me2
H3K27 me3:H3K9 me3
H4−4ac:H3−5ac
H3K4 me3:H3−5ac
H3K4 me3:H4−4ac
H3K4 me1:H3−5ac
H3K9 me3:H4K20 me3
H3K27 me3:H4K20 me3
DNA Meth. m5C:H3K9 me3
H3K27 me3:DNA Meth. m5C
DNA Meth. m5C:H4K20 me3
H3K9 me2:DNA Meth. m5C
H4K20 me2
H2A.Z
H3K27 me2
H3K4 me3
H3K9 me2
H3−5ac
H3K4 me1
H3K27 me3
H3K9 me3
H4K20 me3
H4−4ac
DNA Meth. m5C
0.00 0.25 0.50 0.75 1.00
$\hat{\pi}$
FANCA – Reverse
H4K20 me2:H3K4 me3
H3K27 me2:DNA Meth. m5C
H4K20 me2:H3−5ac
H3K4 me1:H4−4ac
H4K20 me2:H4−4ac
H3K4 me1:H2A.Z
DNA Meth. m5C:H4K20 me2
H3K27 me3:H3K9 me3
H4−4ac:H3−5ac
H3K4 me3:H3−5ac
H3K4 me3:H4−4ac
H3K4 me1:H3−5ac
H3K9 me3:H4K20 me3
H3K27 me3:H4K20 me3
DNA Meth. m5C:H3K9 me3
H3K27 me3:DNA Meth. m5C
DNA Meth. m5C:H4K20 me3
H3K9 me2:DNA Meth. m5C
H4K20 me2
H2A.Z
H3K27 me2
H3K4 me3
H3K9 me2
H3−5ac
H3K4 me1
H3K27 me3
H3K9 me3
H4K20 me3
H4−4ac
DNA Meth. m5C
0.00 0.25 0.50 0.75 1.00
$\hat{\pi}$
ZCCHC17 – Forward
H4K20 me2:H3K4 me3
H2A.Z:H4−4ac
H2A.Z:H3K4 me3
H3K4 me1:H3−5ac
H3K4 me1:H4−4ac
H3K4 me1:H2A.Z
H3K27 me3:H4K20 me3
H3K9 me3:H4K20 me3
DNA Meth. m5C:H4K20 me2
H3K27 me3:H3K4 me1
H3K27 me2:DNA Meth. m5C
DNA Meth. m5C:H4K20 me3
H3K27 me3:DNA Meth. m5C
H3K9 me2:DNA Meth. m5C
H3−5ac
DNA Meth. m5C:H3K9 me3
H3K27 me3:H3K9 me3
H4K20 me2
H2A.Z
H3K27 me2
H4−4ac
H3K4 me3
H3K4 me1
H4K20 me3
H3K9 me3
DNA Meth. m5C
H3K9 me2
H3K27 me3
0.00 0.25 0.50 0.75 1.00
$\hat{\pi}$
ZCCHC17 – Reverse
H4K20 me2:H3K4 me3
H2A.Z:H4−4ac
H2A.Z:H3K4 me3
H3K4 me1:H3−5ac
H3K4 me1:H4−4ac
H3K4 me1:H2A.Z
H3K27 me3:H4K20 me3
H3K9 me3:H4K20 me3
DNA Meth. m5C:H4K20 me2
H3K27 me3:H3K4 me1
H3K27 me2:DNA Meth. m5C
DNA Meth. m5C:H4K20 me3
H3K27 me3:DNA Meth. m5C
H3K9 me2:DNA Meth. m5C
H3−5ac
DNA Meth. m5C:H3K9 me3
H3K27 me3:H3K9 me3
H4K20 me2
H2A.Z
H3K27 me2
H4−4ac
H3K4 me3
H3K4 me1
H4K20 me3
H3K9 me3
DNA Meth. m5C
H3K9 me2
H3K27 me3
0.00 0.25 0.50 0.75 1.00
$\hat{\pi}$
MTA3 – Forward
H4K20 me2:H4−4ac
H2A.Z:H3−5ac
H3K4 me1:H4−4ac
H3K4 me3:H4−4ac
H2A.Z:H4−4ac
H2A.Z:H3K4 me3
H3K4 me1:H4K20 me2
H3K4 me3:H3−5ac
H3K4 me1:H3−5ac
H3K4 me1:H2A.Z
H3K27 me3:H3K9 me3
H4−4ac:H3−5ac
H4K20 me2:H3K4 me3
H3K27 me3:H3K4 me1
DNA Meth. m5C:H3K9 me3
H2A.Z:H4K20 me2
H3K27 me2:DNA Meth. m5C
DNA Meth. m5C:H4K20 me3
H4K20 me2:H3−5ac
DNA Meth. m5C:H4K20 me2
H3K27 me3:H4K20 me3
H3K9 me3:H4K20 me3
H3K27 me3:DNA Meth. m5C
H3K9 me2:DNA Meth. m5C
H4−4ac
H3K4 me3
H3K27 me2
H2A.Z
H3K9 me2
H3K9 me3
H4K20 me3
H3−5ac
H3K4 me1
H4K20 me2
H3K27 me3
DNA Meth. m5C
0.00 0.25 0.50 0.75 1.00
$\hat{\pi}$
MTA3 – Reverse
H4K20 me2:H4−4ac
H2A.Z:H3−5ac
H3K4 me1:H4−4ac
H3K4 me3:H4−4ac
H2A.Z:H4−4ac
H2A.Z:H3K4 me3
H3K4 me1:H4K20 me2
H3K4 me3:H3−5ac
H3K4 me1:H3−5ac
H3K4 me1:H2A.Z
H3K27 me3:H3K9 me3
H4−4ac:H3−5ac
H4K20 me2:H3K4 me3
H3K27 me3:H3K4 me1
DNA Meth. m5C:H3K9 me3
H2A.Z:H4K20 me2
H3K27 me2:DNA Meth. m5C
DNA Meth. m5C:H4K20 me3
H4K20 me2:H3−5ac
DNA Meth. m5C:H4K20 me2
H3K27 me3:H4K20 me3
H3K9 me3:H4K20 me3
H3K27 me3:DNA Meth. m5C
H3K9 me2:DNA Meth. m5C
H4−4ac
H3K4 me3
H3K27 me2
H2A.Z
H3K9 me2
H3K9 me3
H4K20 me3
H3−5ac
H3K4 me1
H4K20 me2
H3K27 me3
DNA Meth. m5C
0.00 0.25 0.50 0.75 1.00
$\hat{\pi}$

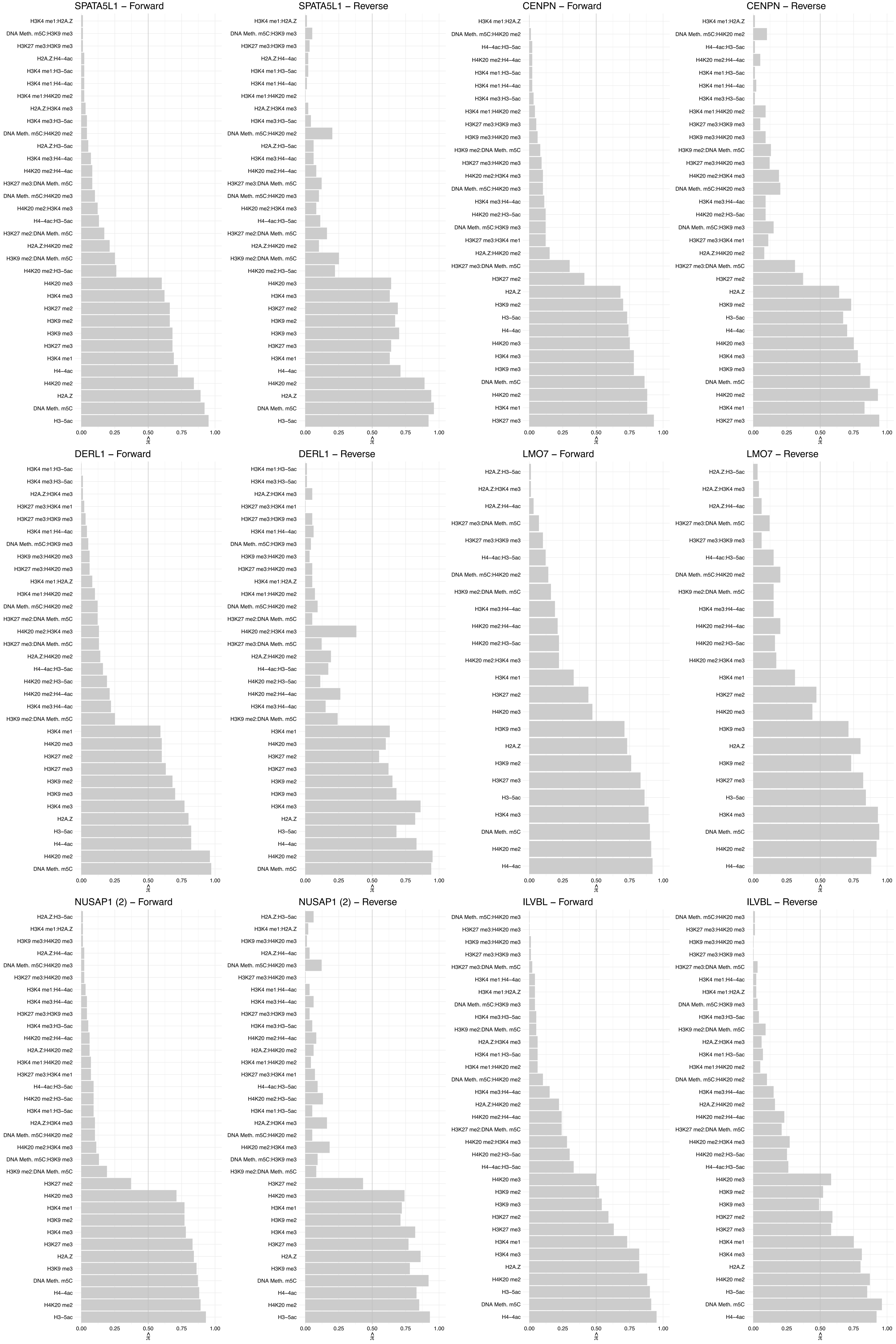
SPATA5L1 − Forward
H3K4 me1:H2A.Z
DNA Meth. m5C:H3K9 me3
H3K27 me3:H3K9 me3
H2A.Z:H4−4ac
H3K4 me1:H3−5ac
H3K4 me1:H4−4ac
H3K4 me1:H4K20 me2
H2A.Z:H3K4 me3
H3K4 me3:H3−5ac
DNA Meth. m5C:H4K20 me2
H2A.Z:H3−5ac
H3K4 me3:H4−4ac
H4K20 me2:H4−4ac
H3K27 me3:DNA Meth. m5C
DNA Meth. m5C:H4K20 me3
H4K20 me2:H3K4 me3
H4−4ac:H3−5ac
H3K27 me2:DNA Meth. m5C
H2A.Z:H4K20 me2
H3K9 me2:DNA Meth. m5C
H4K20 me2:H3−5ac
H4K20 me3
H3K4 me3
H3K27 me2
H3K9 me2
H3K9 me3
H3K27 me3
H3K4 me1
H4−4ac
H4K20 me2
H2A.Z
DNA Meth. m5C
H3−5ac
0.00 0.25 0.50 0.75 1.00
π̂
SPATA5L1 − Reverse
H3K4 me1:H2A.Z
DNA Meth. m5C:H3K9 me3
H3K27 me3:H3K9 me3
H2A.Z:H4−4ac
H3K4 me1:H3−5ac
H3K4 me1:H4−4ac
H3K4 me1:H4K20 me2
H2A.Z:H3K4 me3
H3K4 me3:H3−5ac
DNA Meth. m5C:H4K20 me2
H2A.Z:H3−5ac
H3K4 me3:H4−4ac
H4K20 me2:H4−4ac
H3K27 me3:DNA Meth. m5C
DNA Meth. m5C:H4K20 me3
H4K20 me2:H3K4 me3
H4−4ac:H3−5ac
H3K27 me2:DNA Meth. m5C
H2A.Z:H4K20 me2
H3K9 me2:DNA Meth. m5C
H4K20 me2:H3−5ac
H4K20 me3
H3K4 me3
H3K27 me2
H3K9 me2
H3K9 me3
H3K27 me3
H3K4 me1
H4−4ac
H4K20 me2
H2A.Z
DNA Meth. m5C
H3−5ac
0.00 0.25 0.50 0.75 1.00
π̂
CENPN − Forward
H3K4 me1:H2A.Z
DNA Meth. m5C:H4K20 me2
H4−4ac:H3−5ac
H4K20 me2:H4−4ac
H3K4 me1:H3−5ac
H3K4 me1:H4−4ac
H3K4 me3:H3−5ac
H3K4 me1:H4K20 me2
H3K27 me3:H3K9 me3
H3K9 me3:H4K20 me3
H3K9 me2:DNA Meth. m5C
H3K27 me3:H4K20 me3
H4K20 me2:H3K4 me3
DNA Meth. m5C:H4K20 me3
H3K4 me3:H4−4ac
H4K20 me2:H3−5ac
DNA Meth. m5C:H3K9 me3
H3K27 me3:H3K4 me1
H2A.Z:H4K20 me2
H3K27 me3:DNA Meth. m5C
H3K27 me2
H2A.Z
H3K9 me2
H3−5ac
H4−4ac
H4K20 me3
H3K4 me3
H3K9 me3
DNA Meth. m5C
H4K20 me2
H3K4 me1
H3K27 me3
0.00 0.25 0.50 0.75 1.00
π̂
CENPN − Reverse
H3K4 me1:H2A.Z
DNA Meth. m5C:H4K20 me2
H4−4ac:H3−5ac
H4K20 me2:H4−4ac
H3K4 me1:H3−5ac
H3K4 me1:H4−4ac
H3K4 me3:H3−5ac
H3K4 me1:H4K20 me2
H3K27 me3:H3K9 me3
H3K9 me3:H4K20 me3
H3K9 me2:DNA Meth. m5C
H3K27 me3:H4K20 me3
H4K20 me2:H3K4 me3
DNA Meth. m5C:H4K20 me3
H3K4 me3:H4−4ac
H4K20 me2:H3−5ac
DNA Meth. m5C:H3K9 me3
H3K27 me3:H3K4 me1
H2A.Z:H4K20 me2
H3K27 me3:DNA Meth. m5C
H3K27 me2
H2A.Z
H3K9 me2
H3−5ac
H4−4ac
H4K20 me3
H3K4 me3
H3K9 me3
DNA Meth. m5C
H4K20 me2
H3K4 me1
H3K27 me3
0.00 0.25 0.50 0.75 1.00
π̂
DERL1 − Forward
H3K4 me1:H3−5ac
H3K4 me3:H3−5ac
H2A.Z:H3K4 me3
H3K27 me3:H3K4 me1
H3K27 me3:H3K9 me3
H3K4 me1:H4−4ac
DNA Meth. m5C:H3K9 me3
H3K9 me3:H4K20 me3
H3K27 me3:H4K20 me3
H3K4 me1:H2A.Z
H3K4 me1:H4K20 me2
DNA Meth. m5C:H4K20 me2
H3K27 me2:DNA Meth. m5C
H4K20 me2:H3K4 me3
H3K27 me3:DNA Meth. m5C
H2A.Z:H4K20 me2
H4−4ac:H3−5ac
H4K20 me2:H3−5ac
H4K20 me2:H4−4ac
H3K4 me3:H4−4ac
H3K9 me2:DNA Meth. m5C
H3K4 me1
H4K20 me3
H3K27 me2
H3K27 me3
H3K9 me2
H3K9 me3
H3K4 me3
H2A.Z
H3−5ac
H4−4ac
H4K20 me2
DNA Meth. m5C
0.00 0.25 0.50 0.75 1.00
π̂
DERL1 − Reverse
H3K4 me1:H3−5ac
H3K4 me3:H3−5ac
H2A.Z:H3K4 me3
H3K27 me3:H3K4 me1
H3K27 me3:H3K9 me3
H3K4 me1:H4−4ac
DNA Meth. m5C:H3K9 me3
H3K9 me3:H4K20 me3
H3K27 me3:H4K20 me3
H3K4 me1:H2A.Z
H3K4 me1:H4K20 me2
DNA Meth. m5C:H4K20 me2
H3K27 me2:DNA Meth. m5C
H4K20 me2:H3K4 me3
H3K27 me3:DNA Meth. m5C
H2A.Z:H4K20 me2
H4−4ac:H3−5ac
H4K20 me2:H3−5ac
H4K20 me2:H4−4ac
H3K4 me3:H4−4ac
H3K9 me2:DNA Meth. m5C
H3K4 me1
H4K20 me3
H3K27 me2
H3K27 me3
H3K9 me2
H3K9 me3
H3K4 me3
H2A.Z
H3−5ac
H4−4ac
H4K20 me2
DNA Meth. m5C
0.00 0.25 0.50 0.75 1.00
π̂
LMO7 − Forward
H2A.Z:H3−5ac
H2A.Z:H3K4 me3
H2A.Z:H4−4ac
H3K27 me3:DNA Meth. m5C
H3K27 me3:H3K9 me3
H4−4ac:H3−5ac
DNA Meth. m5C:H4K20 me2
H3K9 me2:DNA Meth. m5C
H3K4 me3:H4−4ac
H4K20 me2:H4−4ac
H4K20 me2:H3−5ac
H4K20 me2:H3K4 me3
H3K4 me1
H3K27 me2
H4K20 me3
H3K9 me3
H2A.Z
H3K9 me2
H3K27 me3
H3−5ac
H3K4 me3
DNA Meth. m5C
H4K20 me2
H4−4ac
0.00 0.25 0.50 0.75 1.00
π̂
LMO7 − Reverse
H2A.Z:H3−5ac
H2A.Z:H3K4 me3
H2A.Z:H4−4ac
H3K27 me3:DNA Meth. m5C
H3K27 me3:H3K9 me3
H4−4ac:H3−5ac
DNA Meth. m5C:H4K20 me2
H3K9 me2:DNA Meth. m5C
H3K4 me3:H4−4ac
H4K20 me2:H4−4ac
H4K20 me2:H3−5ac
H4K20 me2:H3K4 me3
H3K4 me1
H3K27 me2
H4K20 me3
H3K9 me3
H2A.Z
H3K9 me2
H3K27 me3
H3−5ac
H3K4 me3
DNA Meth. m5C
H4K20 me2
H4−4ac
0.00 0.25 0.50 0.75 1.00
π̂
NUSAP1 (2) − Forward
H2A.Z:H3−5ac
H3K4 me1:H2A.Z
H3K9 me3:H4K20 me3
H2A.Z:H4−4ac
DNA Meth. m5C:H4K20 me3
H3K27 me3:H4K20 me3
H3K4 me1:H4−4ac
H3K4 me3:H4−4ac
H3K27 me3:H3K9 me3
H3K4 me3:H3−5ac
H4K20 me2:H4−4ac
H2A.Z:H4K20 me2
H3K4 me1:H4K20 me2
H3K27 me3:H3K4 me1
H4−4ac:H3−5ac
H4K20 me2:H3−5ac
H3K4 me1:H3−5ac
H2A.Z:H3K4 me3
DNA Meth. m5C:H4K20 me2
H4K20 me2:H3K4 me3
DNA Meth. m5C:H3K9 me3
H3K9 me2:DNA Meth. m5C
H3K27 me2
H4K20 me3
H3K4 me1
H3K9 me2
H3K4 me3
H3K27 me3
H2A.Z
H3K9 me3
DNA Meth. m5C
H4−4ac
H4K20 me2
H3−5ac
0.00 0.25 0.50 0.75 1.00
π̂
NUSAP1 (2) − Reverse
H2A.Z:H3−5ac
H3K4 me1:H2A.Z
H3K9 me3:H4K20 me3
H2A.Z:H4−4ac
DNA Meth. m5C:H4K20 me3
H3K27 me3:H4K20 me3
H3K4 me1:H4−4ac
H3K4 me3:H4−4ac
H3K27 me3:H3K9 me3
H3K4 me3:H3−5ac
H4K20 me2:H4−4ac
H2A.Z:H4K20 me2
H3K4 me1:H4K20 me2
H3K27 me3:H3K4 me1
H4−4ac:H3−5ac
H4K20 me2:H3−5ac
H3K4 me1:H3−5ac
H2A.Z:H3K4 me3
DNA Meth. m5C:H4K20 me2
H4K20 me2:H3K4 me3
DNA Meth. m5C:H3K9 me3
H3K9 me2:DNA Meth. m5C
H3K27 me2
H4K20 me3
H3K4 me1
H3K9 me2
H3K4 me3
H3K27 me3
H2A.Z
H3K9 me3
DNA Meth. m5C
H4−4ac
H4K20 me2
H3−5ac
0.00 0.25 0.50 0.75 1.00
π̂
ILVBL − Forward
DNA Meth. m5C:H4K20 me3
H3K27 me3:H4K20 me3
H3K9 me3:H4K20 me3
H3K27 me3:H3K9 me3
H3K27 me3:DNA Meth. m5C
H3K4 me1:H4−4ac
H3K4 me1:H2A.Z
DNA Meth. m5C:H3K9 me3
H3K4 me3:H3−5ac
H3K9 me2:DNA Meth. m5C
H2A.Z:H3K4 me3
H3K4 me1:H3−5ac
H3K4 me1:H4K20 me2
DNA Meth. m5C:H4K20 me2
H3K4 me3:H4−4ac
H2A.Z:H4K20 me2
H4K20 me2:H4−4ac
H3K27 me2:DNA Meth. m5C
H4K20 me2:H3K4 me3
H4K20 me2:H3−5ac
H4−4ac:H3−5ac
H4K20 me3
H3K9 me2
H3K9 me3
H3K27 me2
H3K27 me3
H3K4 me1
H3K4 me3
H2A.Z
H4K20 me2
H3−5ac
DNA Meth. m5C
H4−4ac
0.00 0.25 0.50 0.75 1.00
π̂
ILVBL − Reverse
DNA Meth. m5C:H4K20 me3
H3K27 me3:H4K20 me3
H3K9 me3:H4K20 me3
H3K27 me3:H3K9 me3
H3K27 me3:DNA Meth. m5C
H3K4 me1:H4−4ac
H3K4 me1:H2A.Z
DNA Meth. m5C:H3K9 me3
H3K4 me3:H3−5ac
H3K9 me2:DNA Meth. m5C
H2A.Z:H3K4 me3
H3K4 me1:H3−5ac
H3K4 me1:H4K20 me2
DNA Meth. m5C:H4K20 me2
H3K4 me3:H4−4ac
H2A.Z:H4K20 me2
H4K20 me2:H4−4ac
H3K27 me2:DNA Meth. m5C
H4K20 me2:H3K4 me3
H4K20 me2:H3−5ac
H4−4ac:H3−5ac
H4K20 me3
H3K9 me2
H3K9 me3
H3K27 me2
H3K27 me3
H3K4 me1
H3K4 me3
H2A.Z
H4K20 me2
H3−5ac
DNA Meth. m5C
H4−4ac
0.00 0.25 0.50 0.75 1.00
π̂

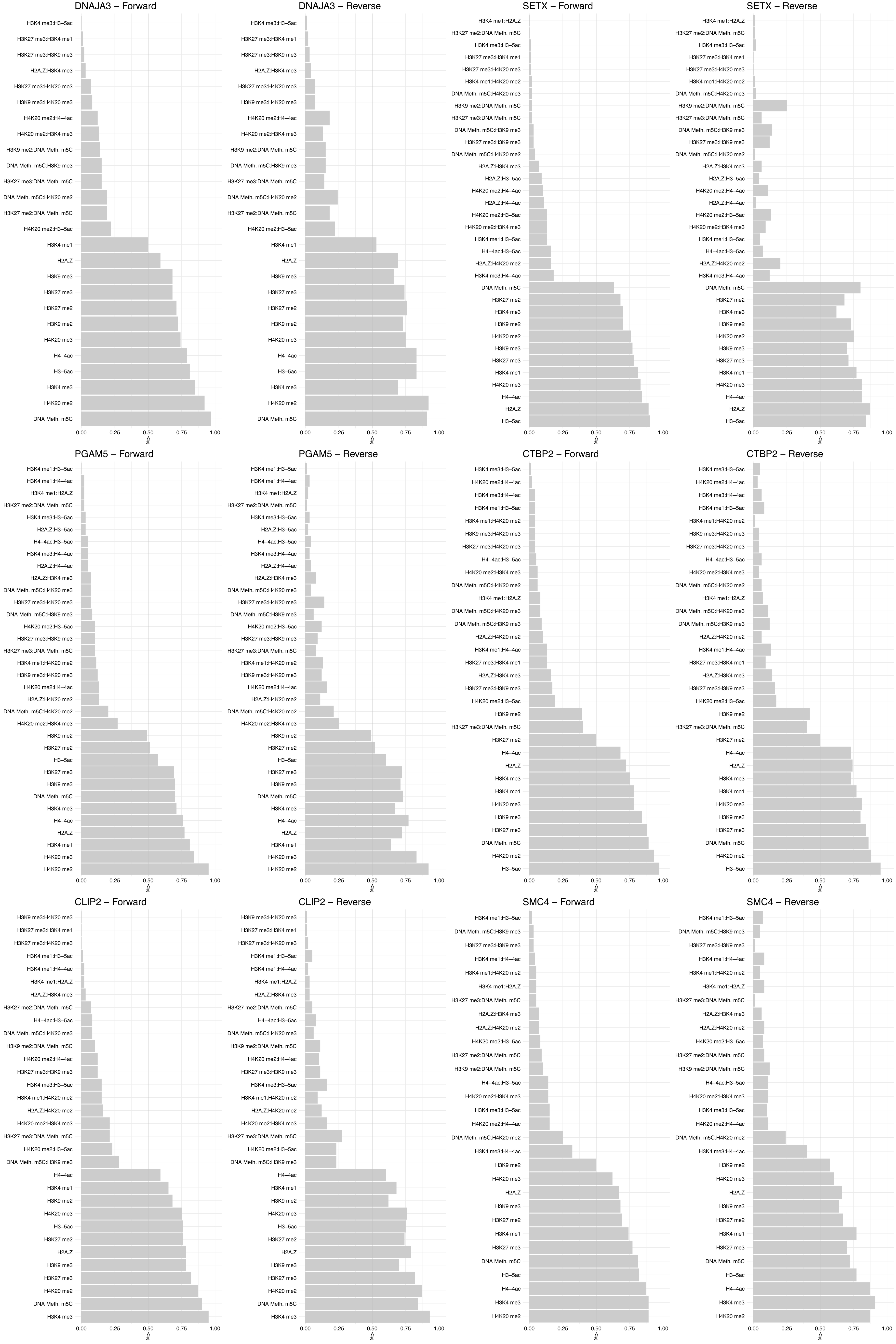

DNAJA3 – Forward
H3K4 me3:H3–5ac
H3K27 me3:H3K4 me1
H3K27 me3:H3K9 me3
H2A.Z:H3K4 me3
H3K27 me3:H4K20 me3
H3K9 me3:H4K20 me3
H4K20 me2:H4–4ac
H4K20 me2:H3K4 me3
H3K9 me2:DNA Meth. m5C
DNA Meth. m5C:H3K9 me3
H3K27 me3:DNA Meth. m5C
DNA Meth. m5C:H4K20 me2
H3K27 me2:DNA Meth. m5C
H4K20 me2:H3–5ac
H3K4 me1
H2A.Z
H3K9 me3
H3K27 me3
H3K27 me2
H3K9 me2
H4K20 me3
H4–4ac
H3–5ac
H3K4 me3
H4K20 me2
DNA Meth. m5C
0.00 0.25 0.50 0.75 1.00
π̂
DNAJA3 – Reverse
H3K4 me3:H3–5ac
H3K27 me3:H3K4 me1
H3K27 me3:H3K9 me3
H2A.Z:H3K4 me3
H3K27 me3:H4K20 me3
H3K9 me3:H4K20 me3
H4K20 me2:H4–4ac
H4K20 me2:H3K4 me3
H3K9 me2:DNA Meth. m5C
DNA Meth. m5C:H3K9 me3
H3K27 me3:DNA Meth. m5C
DNA Meth. m5C:H4K20 me2
H3K27 me2:DNA Meth. m5C
H4K20 me2:H3–5ac
H3K4 me1
H2A.Z
H3K9 me3
H3K27 me3
H3K27 me2
H3K9 me2
H4K20 me3
H4–4ac
H3–5ac
H3K4 me3
H4K20 me2
DNA Meth. m5C
0.00 0.25 0.50 0.75 1.00
π̂
SETX – Forward
H3K4 me1:H2A.Z
H3K27 me2:DNA Meth. m5C
H3K4 me3:H3–5ac
H3K27 me3:H3K4 me1
H3K27 me3:H4K20 me3
H3K4 me1:H4K20 me2
DNA Meth. m5C:H4K20 me3
H3K9 me2:DNA Meth. m5C
H3K27 me3:DNA Meth. m5C
DNA Meth. m5C:H3K9 me3
H3K27 me3:H3K9 me3
DNA Meth. m5C:H4K20 me2
H2A.Z:H3K4 me3
H2A.Z:H3–5ac
H4K20 me2:H4–4ac
H2A.Z:H4–4ac
H4K20 me2:H3–5ac
H4K20 me2:H3K4 me3
H3K4 me1:H3–5ac
H4–4ac:H3–5ac
H2A.Z:H4K20 me2
H3K4 me3:H4–4ac
DNA Meth. m5C
H3K27 me2
H3K4 me3
H3K9 me2
H4K20 me2
H3K9 me3
H3K27 me3
H3K4 me1
H4K20 me3
H4–4ac
H2A.Z
H3–5ac
0.00 0.25 0.50 0.75 1.00
π̂
SETX – Reverse
H3K4 me1:H2A.Z
H3K27 me2:DNA Meth. m5C
H3K4 me3:H3–5ac
H3K27 me3:H3K4 me1
H3K27 me3:H4K20 me3
H3K4 me1:H4K20 me2
DNA Meth. m5C:H4K20 me3
H3K9 me2:DNA Meth. m5C
H3K27 me3:DNA Meth. m5C
DNA Meth. m5C:H3K9 me3
H3K27 me3:H3K9 me3
DNA Meth. m5C:H4K20 me2
H2A.Z:H3K4 me3
H2A.Z:H3–5ac
H4K20 me2:H4–4ac
H2A.Z:H4–4ac
H4K20 me2:H3–5ac
H4K20 me2:H3K4 me3
H3K4 me1:H3–5ac
H4–4ac:H3–5ac
H2A.Z:H4K20 me2
H3K4 me3:H4–4ac
DNA Meth. m5C
H3K27 me2
H3K4 me3
H3K9 me2
H4K20 me2
H3K9 me3
H3K27 me3
H3K4 me1
H4K20 me3
H4–4ac
H2A.Z
H3–5ac
0.00 0.25 0.50 0.75 1.00
π̂
PGAM5 – Forward
H3K4 me1:H3–5ac
H3K4 me1:H4–4ac
H3K4 me1:H2A.Z
H3K27 me2:DNA Meth. m5C
H3K4 me3:H3–5ac
H2A.Z:H3–5ac
H4–4ac:H3–5ac
H3K4 me3:H4–4ac
H2A.Z:H4–4ac
H2A.Z:H3K4 me3
DNA Meth. m5C:H4K20 me3
H3K27 me3:H4K20 me3
DNA Meth. m5C:H3K9 me3
H4K20 me2:H3–5ac
H3K27 me3:H3K9 me3
H3K27 me3:DNA Meth. m5C
H3K4 me1:H4K20 me2
H3K9 me3:H4K20 me3
H4K20 me2:H4–4ac
H2A.Z:H4K20 me2
DNA Meth. m5C:H4K20 me2
H4K20 me2:H3K4 me3
H3K9 me2
H3K27 me2
H3–5ac
H3K27 me3
H3K9 me3
DNA Meth. m5C
H3K4 me3
H4–4ac
H2A.Z
H3K4 me1
H4K20 me3
H4K20 me2
0.00 0.25 0.50 0.75 1.00
π̂
PGAM5 – Reverse
H3K4 me1:H3–5ac
H3K4 me1:H4–4ac
H3K4 me1:H2A.Z
H3K27 me2:DNA Meth. m5C
H3K4 me3:H3–5ac
H2A.Z:H3–5ac
H4–4ac:H3–5ac
H3K4 me3:H4–4ac
H2A.Z:H4–4ac
H2A.Z:H3K4 me3
DNA Meth. m5C:H4K20 me3
H3K27 me3:H4K20 me3
DNA Meth. m5C:H3K9 me3
H4K20 me2:H3–5ac
H3K27 me3:H3K9 me3
H3K27 me3:DNA Meth. m5C
H3K4 me1:H4K20 me2
H3K9 me3:H4K20 me3
H4K20 me2:H4–4ac
H2A.Z:H4K20 me2
DNA Meth. m5C:H4K20 me2
H4K20 me2:H3K4 me3
H3K9 me2
H3K27 me2
H3–5ac
H3K27 me3
H3K9 me3
DNA Meth. m5C
H3K4 me3
H4–4ac
H2A.Z
H3K4 me1
H4K20 me3
H4K20 me2
0.00 0.25 0.50 0.75 1.00
π̂
CTBP2 – Forward
H3K4 me3:H3–5ac
H4K20 me2:H4–4ac
H3K4 me3:H4–4ac
H3K4 me1:H3–5ac
H3K4 me1:H4K20 me2
H3K9 me3:H4K20 me3
H3K27 me3:H4K20 me3
H4–4ac:H3–5ac
H4K20 me2:H3K4 me3
DNA Meth. m5C:H4K20 me2
H3K4 me1:H2A.Z
DNA Meth. m5C:H4K20 me3
DNA Meth. m5C:H3K9 me3
H2A.Z:H4K20 me2
H3K4 me1:H4–4ac
H3K27 me3:H3K4 me1
H2A.Z:H3K4 me3
H3K27 me3:H3K9 me3
H4K20 me2:H3–5ac
H3K9 me2
H3K27 me3:DNA Meth. m5C
H3K27 me2
H4–4ac
H2A.Z
H3K4 me3
H3K4 me1
H4K20 me3
H3K9 me3
H3K27 me3
DNA Meth. m5C
H4K20 me2
H3–5ac
0.00 0.25 0.50 0.75 1.00
π̂
CTBP2 – Reverse
H3K4 me3:H3–5ac
H4K20 me2:H4–4ac
H3K4 me3:H4–4ac
H3K4 me1:H3–5ac
H3K4 me1:H4K20 me2
H3K9 me3:H4K20 me3
H3K27 me3:H4K20 me3
H4–4ac:H3–5ac
H4K20 me2:H3K4 me3
DNA Meth. m5C:H4K20 me2
H3K4 me1:H2A.Z
DNA Meth. m5C:H4K20 me3
DNA Meth. m5C:H3K9 me3
H2A.Z:H4K20 me2
H3K4 me1:H4–4ac
H3K27 me3:H3K4 me1
H2A.Z:H3K4 me3
H3K27 me3:H3K9 me3
H4K20 me2:H3–5ac
H3K9 me2
H3K27 me3:DNA Meth. m5C
H3K27 me2
H4–4ac
H2A.Z
H3K4 me3
H3K4 me1
H4K20 me3
H3K9 me3
H3K27 me3
DNA Meth. m5C
H4K20 me2
H3–5ac
0.00 0.25 0.50 0.75 1.00
π̂
CLIP2 – Forward
H3K9 me3:H4K20 me3
H3K27 me3:H3K4 me1
H3K27 me3:H4K20 me3
H3K4 me1:H3–5ac
H3K4 me1:H4–4ac
H3K4 me1:H2A.Z
H2A.Z:H3K4 me3
H3K27 me2:DNA Meth. m5C
H4–4ac:H3–5ac
DNA Meth. m5C:H4K20 me3
H3K9 me2:DNA Meth. m5C
H4K20 me2:H4–4ac
H3K27 me3:H3K9 me3
H3K4 me3:H3–5ac
H3K4 me1:H4K20 me2
H2A.Z:H4K20 me2
H4K20 me2:H3K4 me3
H3K27 me3:DNA Meth. m5C
H4K20 me2:H3–5ac
DNA Meth. m5C:H3K9 me3
H4–4ac
H3K4 me1
H3K9 me2
H4K20 me3
H3–5ac
H3K27 me2
H2A.Z
H3K9 me3
H3K27 me3
H4K20 me2
DNA Meth. m5C
H3K4 me3
0.00 0.25 0.50 0.75 1.00
π̂
CLIP2 – Reverse
H3K9 me3:H4K20 me3
H3K27 me3:H3K4 me1
H3K27 me3:H4K20 me3
H3K4 me1:H3–5ac
H3K4 me1:H4–4ac
H3K4 me1:H2A.Z
H2A.Z:H3K4 me3
H3K27 me2:DNA Meth. m5C
H4–4ac:H3–5ac
DNA Meth. m5C:H4K20 me3
H3K9 me2:DNA Meth. m5C
H4K20 me2:H4–4ac
H3K27 me3:H3K9 me3
H3K4 me3:H3–5ac
H3K4 me1:H4K20 me2
H2A.Z:H4K20 me2
H4K20 me2:H3K4 me3
H3K27 me3:DNA Meth. m5C
H4K20 me2:H3–5ac
DNA Meth. m5C:H3K9 me3
H4–4ac
H3K4 me1
H3K9 me2
H4K20 me3
H3–5ac
H3K27 me2
H2A.Z
H3K9 me3
H3K27 me3
H4K20 me2
DNA Meth. m5C
H3K4 me3
0.00 0.25 0.50 0.75 1.00
π̂
SMC4 – Forward
H3K4 me1:H3–5ac
DNA Meth. m5C:H3K9 me3
H3K27 me3:H3K9 me3
H3K4 me1:H4–4ac
H3K4 me1:H4K20 me2
H3K4 me1:H2A.Z
H3K27 me3:DNA Meth. m5C
H2A.Z:H3K4 me3
H2A.Z:H4K20 me2
H4K20 me2:H3–5ac
H3K27 me2:DNA Meth. m5C
H3K9 me2:DNA Meth. m5C
H4–4ac:H3–5ac
H4K20 me2:H3K4 me3
H3K4 me3:H3–5ac
H4K20 me2:H4–4ac
DNA Meth. m5C:H4K20 me2
H3K4 me3:H4–4ac
H3K9 me2
H4K20 me3
H2A.Z
H3K9 me3
H3K27 me2
H3K4 me1
H3K27 me3
DNA Meth. m5C
H3–5ac
H4–4ac
H3K4 me3
H4K20 me2
0.00 0.25 0.50 0.75 1.00
π̂
SMC4 – Reverse
H3K4 me1:H3–5ac
DNA Meth. m5C:H3K9 me3
H3K27 me3:H3K9 me3
H3K4 me1:H4–4ac
H3K4 me1:H4K20 me2
H3K4 me1:H2A.Z
H3K27 me3:DNA Meth. m5C
H2A.Z:H3K4 me3
H2A.Z:H4K20 me2
H4K20 me2:H3–5ac
H3K27 me2:DNA Meth. m5C
H3K9 me2:DNA Meth. m5C
H4–4ac:H3–5ac
H4K20 me2:H3K4 me3
H3K4 me3:H3–5ac
H4K20 me2:H4–4ac
DNA Meth. m5C:H4K20 me2
H3K4 me3:H4–4ac
H3K9 me2
H4K20 me3
H2A.Z
H3K9 me3
H3K27 me2
H3K4 me1
H3K27 me3
DNA Meth. m5C
H3–5ac
H4–4ac
H3K4 me3
H4K20 me2
0.00 0.25 0.50 0.75 1.00
π̂

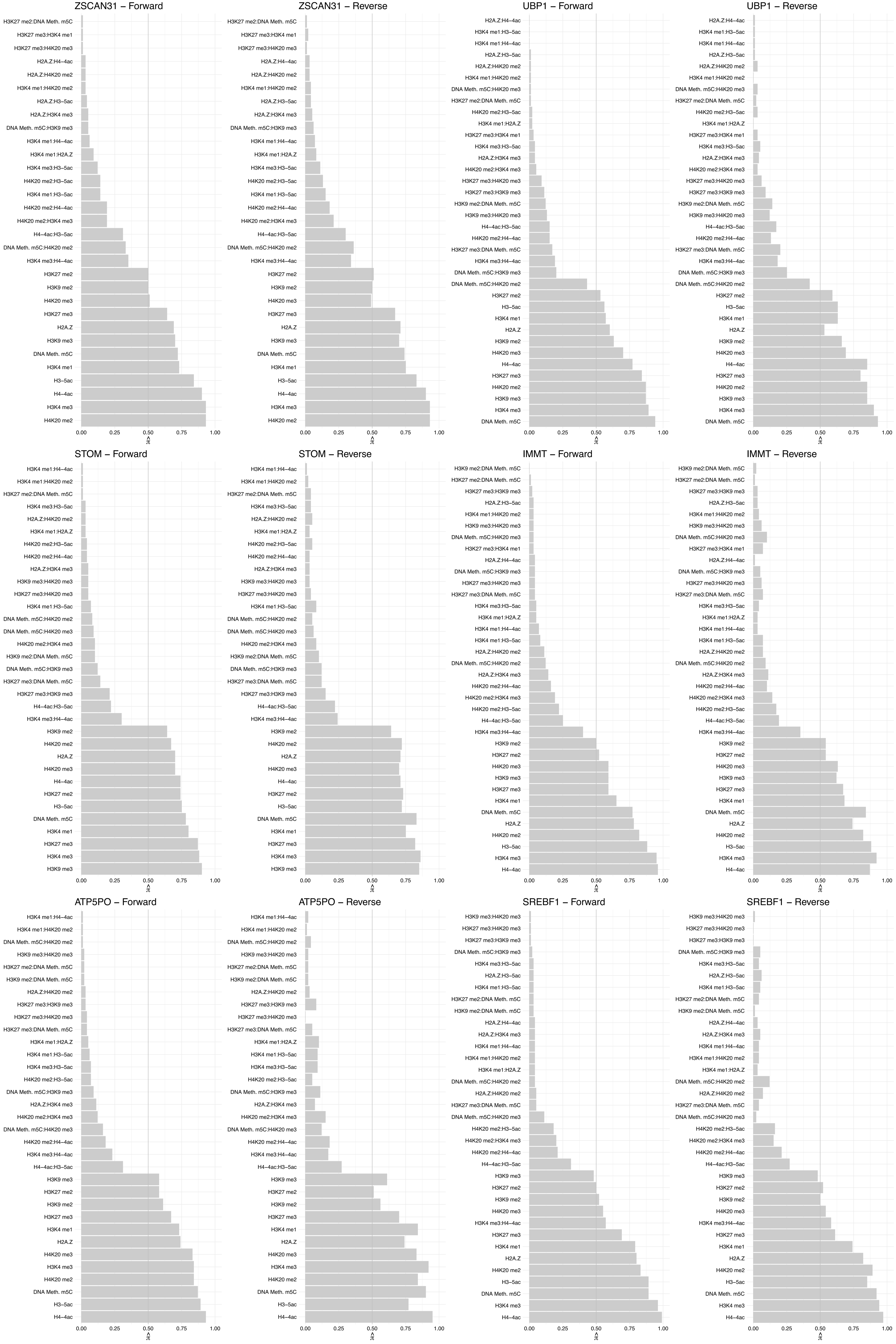
ZSCAN31 − Forward
ZSCAN31 − Reverse
H3K27 me2:DNA Meth. m5C
H3K27 me3:H3K4 me1
H3K27 me3:H4K20 me3
H2A.Z:H4−4ac
H2A.Z:H4K20 me2
H3K4 me1:H4K20 me2
H2A.Z:H3−5ac
H2A.Z:H3K4 me3
DNA Meth. m5C:H3K9 me3
H3K4 me1:H4−4ac
H3K4 me1:H2A.Z
H3K4 me3:H3−5ac
H4K20 me2:H3−5ac
H3K4 me1:H3−5ac
H4K20 me2:H4−4ac
H4K20 me2:H3K4 me3
H4−4ac:H3−5ac
DNA Meth. m5C:H4K20 me2
H3K4 me3:H4−4ac
H3K27 me2
H3K9 me2
H4K20 me3
H3K27 me3
H2A.Z
H3K9 me3
DNA Meth. m5C
H3K4 me1
H3−5ac
H4−4ac
H3K4 me3
H4K20 me2
UBP1 − Forward
UBP1 − Reverse
H2A.Z:H4−4ac
H3K4 me1:H3−5ac
H3K4 me1:H4−4ac
H2A.Z:H3−5ac
H2A.Z:H4K20 me2
H3K4 me1:H4K20 me2
DNA Meth. m5C:H4K20 me3
H3K27 me2:DNA Meth. m5C
H4K20 me2:H3−5ac
H3K4 me1:H2A.Z
H3K27 me3:H3K4 me1
H3K4 me3:H3−5ac
H2A.Z:H3K4 me3
H4K20 me2:H3K4 me3
H3K27 me3:H4K20 me3
H3K27 me3:H3K9 me3
H3K9 me2:DNA Meth. m5C
H3K9 me3:H4K20 me3
H4−4ac:H3−5ac
H4K20 me2:H4−4ac
H3K27 me3:DNA Meth. m5C
H3K4 me3:H4−4ac
DNA Meth. m5C:H3K9 me3
DNA Meth. m5C:H4K20 me2
H3K27 me2
H3−5ac
H3K4 me1
H2A.Z
H3K9 me2
H4K20 me3
H4−4ac
H3K27 me3
H4K20 me2
H3K9 me3
H3K4 me3
DNA Meth. m5C
STOM − Forward
STOM − Reverse
H3K4 me1:H4−4ac
H3K4 me1:H4K20 me2
H3K27 me2:DNA Meth. m5C
H3K4 me3:H3−5ac
H2A.Z:H4K20 me2
H3K4 me1:H2A.Z
H4K20 me2:H3−5ac
H4K20 me2:H4−4ac
H2A.Z:H3K4 me3
H3K9 me3:H4K20 me3
H3K27 me3:H4K20 me3
H3K4 me1:H3−5ac
DNA Meth. m5C:H4K20 me2
DNA Meth. m5C:H4K20 me3
H4K20 me2:H3K4 me3
H3K9 me2:DNA Meth. m5C
DNA Meth. m5C:H3K9 me3
H3K27 me3:DNA Meth. m5C
H3K27 me3:H3K9 me3
H4−4ac:H3−5ac
H3K4 me3:H4−4ac
H3K9 me2
H4K20 me2
H2A.Z
H4K20 me3
H4−4ac
H3K27 me2
H3−5ac
DNA Meth. m5C
H3K4 me1
H3K27 me3
H3K4 me3
H3K9 me3
IMMT − Forward
IMMT − Reverse
H3K9 me2:DNA Meth. m5C
H3K27 me2:DNA Meth. m5C
H3K27 me3:H3K9 me3
H2A.Z:H3−5ac
H3K4 me1:H4K20 me2
H3K9 me3:H4K20 me3
DNA Meth. m5C:H4K20 me3
H3K27 me3:H3K4 me1
H2A.Z:H4−4ac
DNA Meth. m5C:H3K9 me3
H3K27 me3:H4K20 me3
H3K27 me3:DNA Meth. m5C
H3K4 me3:H3−5ac
H3K4 me1:H2A.Z
H3K4 me1:H4−4ac
H3K4 me1:H3−5ac
H2A.Z:H4K20 me2
DNA Meth. m5C:H4K20 me2
H2A.Z:H3K4 me3
H4K20 me2:H4−4ac
H4K20 me2:H3K4 me3
H4K20 me2:H3−5ac
H4−4ac:H3−5ac
H3K4 me3:H4−4ac
H3K9 me2
H3K27 me2
H4K20 me3
H3K9 me3
H3K27 me3
H3K4 me1
DNA Meth. m5C
H2A.Z
H4K20 me2
H3−5ac
H3K4 me3
H4−4ac
ATP5PO − Forward
ATP5PO − Reverse
H3K4 me1:H4−4ac
H3K4 me1:H4K20 me2
DNA Meth. m5C:H4K20 me2
H3K9 me3:H4K20 me3
H3K27 me2:DNA Meth. m5C
H3K9 me2:DNA Meth. m5C
H2A.Z:H4K20 me2
H3K27 me3:H3K9 me3
H3K27 me3:H4K20 me3
H3K27 me3:DNA Meth. m5C
H3K4 me1:H2A.Z
H3K4 me1:H3−5ac
H3K4 me3:H3−5ac
H4K20 me2:H3−5ac
DNA Meth. m5C:H3K9 me3
H2A.Z:H3K4 me3
H4K20 me2:H3K4 me3
DNA Meth. m5C:H4K20 me3
H4K20 me2:H4−4ac
H3K4 me3:H4−4ac
H4−4ac:H3−5ac
H3K9 me3
H3K27 me2
H3K9 me2
H3K27 me3
H3K4 me1
H2A.Z
H4K20 me3
H3K4 me3
H4K20 me2
DNA Meth. m5C
H3−5ac
H4−4ac
SREBF1 − Forward
SREBF1 − Reverse
H3K9 me3:H4K20 me3
H3K27 me3:H4K20 me3
H3K27 me3:H3K9 me3
DNA Meth. m5C:H3K9 me3
H3K4 me3:H3−5ac
H2A.Z:H3−5ac
H3K4 me1:H3−5ac
H3K27 me2:DNA Meth. m5C
H3K9 me2:DNA Meth. m5C
H2A.Z:H4−4ac
H2A.Z:H3K4 me3
H3K4 me1:H4−4ac
H3K4 me1:H4K20 me2
H3K4 me1:H2A.Z
DNA Meth. m5C:H4K20 me2
H2A.Z:H4K20 me2
H3K27 me3:DNA Meth. m5C
DNA Meth. m5C:H4K20 me3
H4K20 me2:H3−5ac
H4K20 me2:H3K4 me3
H4K20 me2:H4−4ac
H4−4ac:H3−5ac
H3K9 me3
H3K27 me2
H3K9 me2
H4K20 me3
H3K4 me3:H4−4ac
H3K27 me3
H3K4 me1
H2A.Z
H4K20 me2
H3−5ac
DNA Meth. m5C
H3K4 me3
H4−4ac
0.00 0.25 0.50 0.75 1.00
π̂

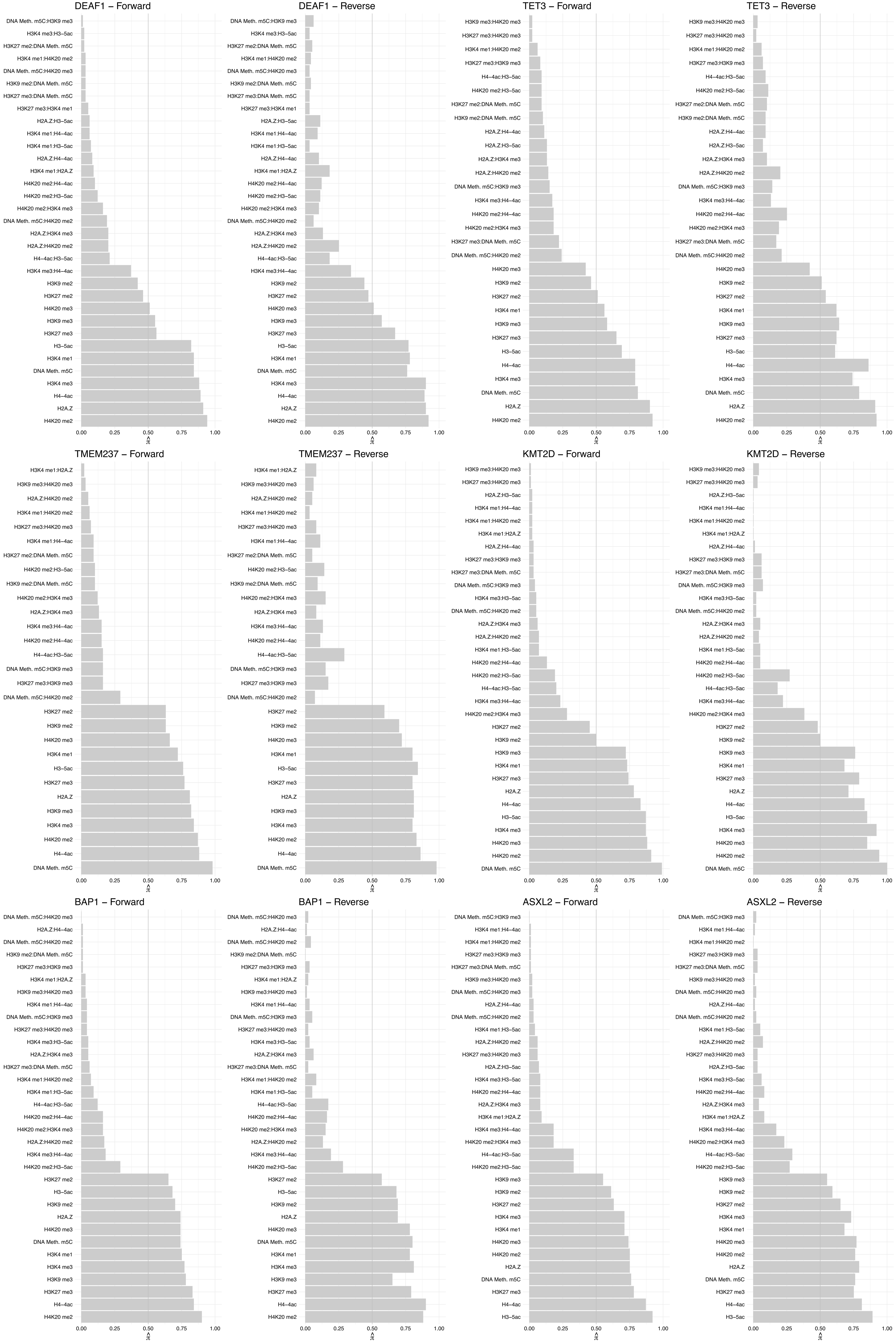
DEAF1 – Forward
DNA Meth. m5C:H3K9 me3
H3K4 me3:H3−5ac
H3K27 me2:DNA Meth. m5C
H3K4 me1:H4K20 me2
DNA Meth. m5C:H4K20 me3
H3K9 me2:DNA Meth. m5C
H3K27 me3:DNA Meth. m5C
H3K27 me3:H3K4 me1
H2A.Z:H3−5ac
H3K4 me1:H4−4ac
H3K4 me1:H3−5ac
H2A.Z:H4−4ac
H3K4 me1:H2A.Z
H4K20 me2:H4−4ac
H4K20 me2:H3−5ac
H4K20 me2:H3K4 me3
DNA Meth. m5C:H4K20 me2
H2A.Z:H3K4 me3
H2A.Z:H4K20 me2
H4−4ac:H3−5ac
H3K4 me3:H4−4ac
H3K9 me2
H3K27 me2
H4K20 me3
H3K9 me3
H3K27 me3
H3−5ac
H3K4 me1
DNA Meth. m5C
H3K4 me3
H4−4ac
H2A.Z
H4K20 me2
0.00
0.25
0.50
0.75
1.00
π̂
DEAF1 – Reverse
DNA Meth. m5C:H3K9 me3
H3K4 me3:H3−5ac
H3K27 me2:DNA Meth. m5C
H3K4 me1:H4K20 me2
DNA Meth. m5C:H4K20 me3
H3K9 me2:DNA Meth. m5C
H3K27 me3:DNA Meth. m5C
H3K27 me3:H3K4 me1
H2A.Z:H3−5ac
H3K4 me1:H4−4ac
H3K4 me1:H3−5ac
H2A.Z:H4−4ac
H3K4 me1:H2A.Z
H4K20 me2:H4−4ac
H4K20 me2:H3−5ac
H4K20 me2:H3K4 me3
DNA Meth. m5C:H4K20 me2
H2A.Z:H3K4 me3
H2A.Z:H4K20 me2
H4−4ac:H3−5ac
H3K4 me3:H4−4ac
H3K9 me2
H3K27 me2
H4K20 me3
H3K9 me3
H3K27 me3
H3−5ac
H3K4 me1
DNA Meth. m5C
H3K4 me3
H4−4ac
H2A.Z
H4K20 me2
0.00
0.25
0.50
0.75
1.00
π̂
TET3 – Forward
H3K9 me3:H4K20 me3
H3K27 me3:H4K20 me3
H3K4 me1:H4K20 me2
H3K27 me3:H3K9 me3
H4−4ac:H3−5ac
H4K20 me2:H3−5ac
H3K27 me2:DNA Meth. m5C
H3K9 me2:DNA Meth. m5C
H2A.Z:H4−4ac
H2A.Z:H3−5ac
H2A.Z:H3K4 me3
H2A.Z:H4K20 me2
DNA Meth. m5C:H3K9 me3
H3K4 me3:H4−4ac
H4K20 me2:H4−4ac
H4K20 me2:H3K4 me3
H3K27 me3:DNA Meth. m5C
DNA Meth. m5C:H4K20 me2
H4K20 me3
H3K9 me2
H3K27 me2
H3K4 me1
H3K9 me3
H3K27 me3
H3−5ac
H4−4ac
H3K4 me3
DNA Meth. m5C
H2A.Z
H4K20 me2
0.00
0.25
0.50
0.75
1.00
π̂
TET3 – Reverse
H3K9 me3:H4K20 me3
H3K27 me3:H4K20 me3
H3K4 me1:H4K20 me2
H3K27 me3:H3K9 me3
H4−4ac:H3−5ac
H4K20 me2:H3−5ac
H3K27 me2:DNA Meth. m5C
H3K9 me2:DNA Meth. m5C
H2A.Z:H4−4ac
H2A.Z:H3−5ac
H2A.Z:H3K4 me3
H2A.Z:H4K20 me2
DNA Meth. m5C:H3K9 me3
H3K4 me3:H4−4ac
H4K20 me2:H4−4ac
H4K20 me2:H3K4 me3
H3K27 me3:DNA Meth. m5C
DNA Meth. m5C:H4K20 me2
H4K20 me3
H3K9 me2
H3K27 me2
H3K4 me1
H3K9 me3
H3K27 me3
H3−5ac
H4−4ac
H3K4 me3
DNA Meth. m5C
H2A.Z
H4K20 me2
0.00
0.25
0.50
0.75
1.00
π̂
TMEM237 – Forward
H3K4 me1:H2A.Z
H3K9 me3:H4K20 me3
H2A.Z:H4K20 me2
H3K4 me1:H4K20 me2
H3K27 me3:H4K20 me3
H3K4 me1:H4−4ac
H3K27 me2:DNA Meth. m5C
H4K20 me2:H3−5ac
H3K9 me2:DNA Meth. m5C
H4K20 me2:H3K4 me3
H2A.Z:H3K4 me3
H3K4 me3:H4−4ac
H4K20 me2:H4−4ac
H4−4ac:H3−5ac
DNA Meth. m5C:H3K9 me3
H3K27 me3:H3K9 me3
DNA Meth. m5C:H4K20 me2
H3K27 me2
H3K9 me2
H4K20 me3
H3K4 me1
H3−5ac
H3K27 me3
H2A.Z
H3K9 me3
H3K4 me3
H4K20 me2
H4−4ac
DNA Meth. m5C
0.00
0.25
0.50
0.75
1.00
π̂
TMEM237 – Reverse
H3K4 me1:H2A.Z
H3K9 me3:H4K20 me3
H2A.Z:H4K20 me2
H3K4 me1:H4K20 me2
H3K27 me3:H4K20 me3
H3K4 me1:H4−4ac
H3K27 me2:DNA Meth. m5C
H4K20 me2:H3−5ac
H3K9 me2:DNA Meth. m5C
H4K20 me2:H3K4 me3
H2A.Z:H3K4 me3
H3K4 me3:H4−4ac
H4K20 me2:H4−4ac
H4−4ac:H3−5ac
DNA Meth. m5C:H3K9 me3
H3K27 me3:H3K9 me3
DNA Meth. m5C:H4K20 me2
H3K27 me2
H3K9 me2
H4K20 me3
H3K4 me1
H3−5ac
H3K27 me3
H2A.Z
H3K9 me3
H3K4 me3
H4K20 me2
H4−4ac
DNA Meth. m5C
0.00
0.25
0.50
0.75
1.00
π̂
KMT2D – Forward
H3K9 me3:H4K20 me3
H3K27 me3:H4K20 me3
H2A.Z:H3−5ac
H3K4 me1:H4−4ac
H3K4 me1:H4K20 me2
H3K4 me1:H2A.Z
H2A.Z:H4−4ac
H3K27 me3:H3K9 me3
H3K27 me3:DNA Meth. m5C
DNA Meth. m5C:H3K9 me3
H3K4 me3:H3−5ac
DNA Meth. m5C:H4K20 me2
H2A.Z:H3K4 me3
H2A.Z:H4K20 me2
H3K4 me1:H3−5ac
H4K20 me2:H4−4ac
H4K20 me2:H3−5ac
H4−4ac:H3−5ac
H3K4 me3:H4−4ac
H4K20 me2:H3K4 me3
H3K27 me2
H3K9 me2
H3K9 me3
H3K4 me1
H3K27 me3
H2A.Z
H4−4ac
H3−5ac
H3K4 me3
H4K20 me3
H4K20 me2
DNA Meth. m5C
0.00
0.25
0.50
0.75
1.00
π̂
KMT2D – Reverse
H3K9 me3:H4K20 me3
H3K27 me3:H4K20 me3
H2A.Z:H3−5ac
H3K4 me1:H4−4ac
H3K4 me1:H4K20 me2
H3K4 me1:H2A.Z
H2A.Z:H4−4ac
H3K27 me3:H3K9 me3
H3K27 me3:DNA Meth. m5C
DNA Meth. m5C:H3K9 me3
H3K4 me3:H3−5ac
DNA Meth. m5C:H4K20 me2
H2A.Z:H3K4 me3
H2A.Z:H4K20 me2
H3K4 me1:H3−5ac
H4K20 me2:H4−4ac
H4K20 me2:H3−5ac
H4−4ac:H3−5ac
H3K4 me3:H4−4ac
H4K20 me2:H3K4 me3
H3K27 me2
H3K9 me2
H3K9 me3
H3K4 me1
H3K27 me3
H2A.Z
H4−4ac
H3−5ac
H3K4 me3
H4K20 me3
H4K20 me2
DNA Meth. m5C
0.00
0.25
0.50
0.75
1.00
π̂
BAP1 – Forward
DNA Meth. m5C:H4K20 me3
H2A.Z:H4−4ac
DNA Meth. m5C:H4K20 me2
H3K9 me2:DNA Meth. m5C
H3K27 me3:H3K9 me3
H3K4 me1:H2A.Z
H3K9 me3:H4K20 me3
H3K4 me1:H4−4ac
DNA Meth. m5C:H3K9 me3
H3K27 me3:H4K20 me3
H3K4 me3:H3−5ac
H2A.Z:H3K4 me3
H3K27 me3:DNA Meth. m5C
H3K4 me1:H4K20 me2
H3K4 me1:H3−5ac
H4−4ac:H3−5ac
H4K20 me2:H4−4ac
H4K20 me2:H3K4 me3
H2A.Z:H4K20 me2
H3K4 me3:H4−4ac
H4K20 me2:H3−5ac
H3K27 me2
H3−5ac
H3K9 me2
H2A.Z
H4K20 me3
DNA Meth. m5C
H3K4 me1
H3K4 me3
H3K9 me3
H3K27 me3
H4−4ac
H4K20 me2
0.00
0.25
0.50
0.75
1.00
π̂
BAP1 – Reverse
DNA Meth. m5C:H4K20 me3
H2A.Z:H4−4ac
DNA Meth. m5C:H4K20 me2
H3K9 me2:DNA Meth. m5C
H3K27 me3:H3K9 me3
H3K4 me1:H2A.Z
H3K9 me3:H4K20 me3
H3K4 me1:H4−4ac
DNA Meth. m5C:H3K9 me3
H3K27 me3:H4K20 me3
H3K4 me3:H3−5ac
H2A.Z:H3K4 me3
H3K27 me3:DNA Meth. m5C
H3K4 me1:H4K20 me2
H3K4 me1:H3−5ac
H4−4ac:H3−5ac
H4K20 me2:H4−4ac
H4K20 me2:H3K4 me3
H2A.Z:H4K20 me2
H3K4 me3:H4−4ac
H4K20 me2:H3−5ac
H3K27 me2
H3−5ac
H3K9 me2
H2A.Z
H4K20 me3
DNA Meth. m5C
H3K4 me1
H3K4 me3
H3K9 me3
H3K27 me3
H4−4ac
H4K20 me2
0.00
0.25
0.50
0.75
1.00
π̂
ASXL2 – Forward
DNA Meth. m5C:H3K9 me3
H3K4 me1:H4−4ac
H3K4 me1:H4K20 me2
H3K27 me3:H3K9 me3
H3K27 me3:DNA Meth. m5C
H3K9 me3:H4K20 me3
DNA Meth. m5C:H4K20 me3
H2A.Z:H4−4ac
DNA Meth. m5C:H4K20 me2
H3K4 me1:H3−5ac
H2A.Z:H4K20 me2
H3K27 me3:H4K20 me3
H2A.Z:H3−5ac
H3K4 me3:H3−5ac
H4K20 me2:H4−4ac
H2A.Z:H3K4 me3
H3K4 me1:H2A.Z
H3K4 me3:H4−4ac
H4K20 me2:H3K4 me3
H4−4ac:H3−5ac
H4K20 me2:H3−5ac
H3K9 me3
H3K9 me2
H3K27 me2
H3K4 me3
H3K4 me1
H4K20 me3
H4K20 me2
H2A.Z
DNA Meth. m5C
H3K27 me3
H4−4ac
H3−5ac
0.00
0.25
0.50
0.75
1.00
π̂
ASXL2 – Reverse
DNA Meth. m5C:H3K9 me3
H3K4 me1:H4−4ac
H3K4 me1:H4K20 me2
H3K27 me3:H3K9 me3
H3K27 me3:DNA Meth. m5C
H3K9 me3:H4K20 me3
DNA Meth. m5C:H4K20 me3
H2A.Z:H4−4ac
DNA Meth. m5C:H4K20 me2
H3K4 me1:H3−5ac
H2A.Z:H4K20 me2
H3K27 me3:H4K20 me3
H2A.Z:H3−5ac
H3K4 me3:H3−5ac
H4K20 me2:H4−4ac
H2A.Z:H3K4 me3
H3K4 me1:H2A.Z
H3K4 me3:H4−4ac
H4K20 me2:H3K4 me3
H4−4ac:H3−5ac
H4K20 me2:H3−5ac
H3K9 me3
H3K9 me2
H3K27 me2
H3K4 me3
H3K4 me1
H4K20 me3
H4K20 me2
H2A.Z
DNA Meth. m5C
H3K27 me3
H4−4ac
H3−5ac
0.00
0.25
0.50
0.75
1.00
π̂

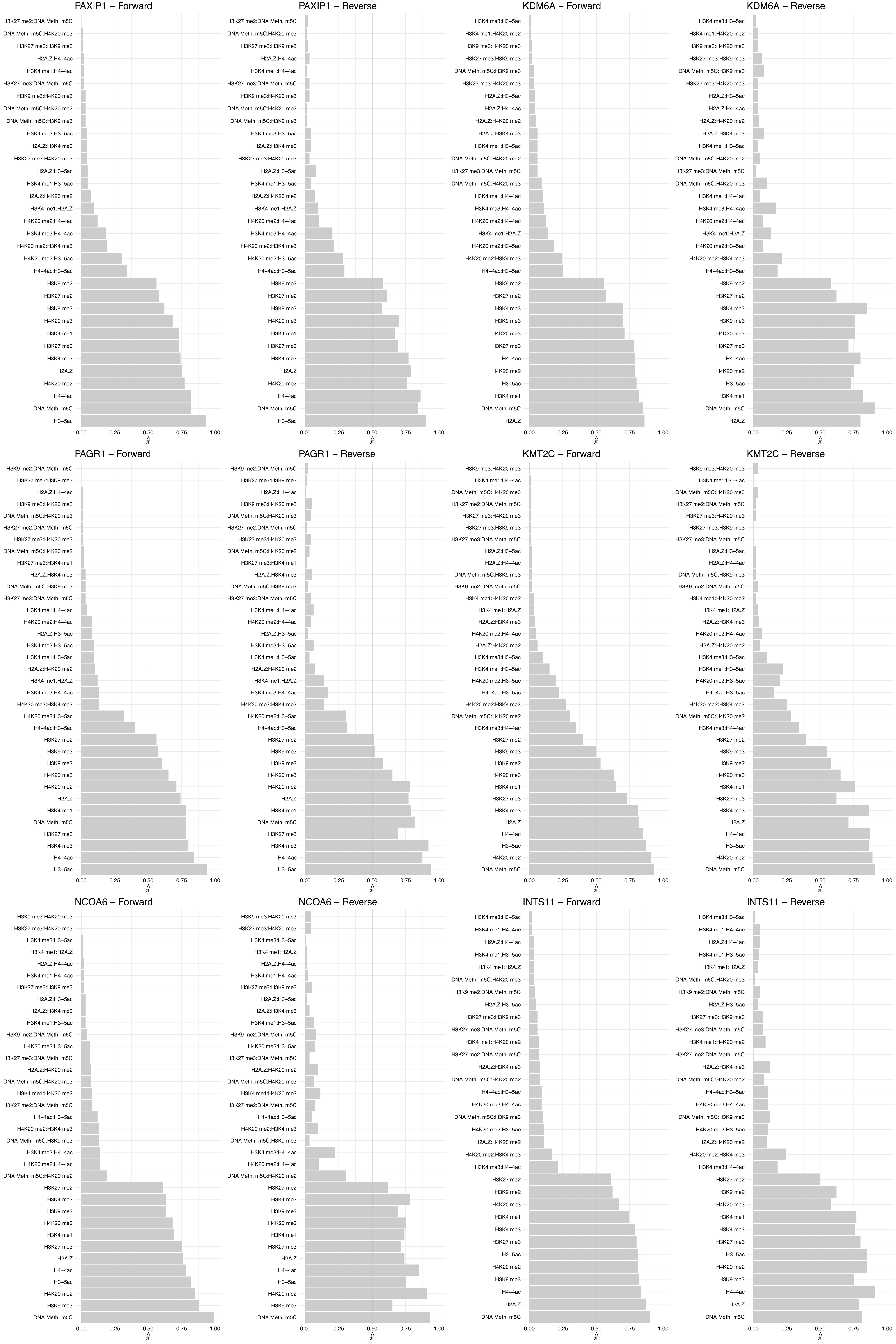

PAXIP1 – Forward
H3K27 me2:DNA Meth. m5C
DNA Meth. m5C:H4K20 me3
H3K27 me3:H3K9 me3
H2A.Z:H4−4ac
H3K4 me1:H4−4ac
H3K27 me3:DNA Meth. m5C
H3K9 me3:H4K20 me3
DNA Meth. m5C:H4K20 me2
DNA Meth. m5C:H3K9 me3
H3K4 me3:H3−5ac
H2A.Z:H3K4 me3
H3K27 me3:H4K20 me3
H2A.Z:H3−5ac
H3K4 me1:H3−5ac
H2A.Z:H4K20 me2
H3K4 me1:H2A.Z
H4K20 me2:H4−4ac
H3K4 me3:H4−4ac
H4K20 me2:H3K4 me3
H4K20 me2:H3−5ac
H4−4ac:H3−5ac
H3K9 me2
H3K27 me2
H3K9 me3
H4K20 me3
H3K4 me1
H3K27 me3
H3K4 me3
H2A.Z
H4K20 me2
H4−4ac
DNA Meth. m5C
H3−5ac
0.00 0.25 0.50 0.75 1.00
π̂
PAXIP1 – Reverse
H3K27 me2:DNA Meth. m5C
DNA Meth. m5C:H4K20 me3
H3K27 me3:H3K9 me3
H2A.Z:H4−4ac
H3K4 me1:H4−4ac
H3K27 me3:DNA Meth. m5C
H3K9 me3:H4K20 me3
DNA Meth. m5C:H4K20 me2
DNA Meth. m5C:H3K9 me3
H3K4 me3:H3−5ac
H2A.Z:H3K4 me3
H3K27 me3:H4K20 me3
H2A.Z:H3−5ac
H3K4 me1:H3−5ac
H2A.Z:H4K20 me2
H3K4 me1:H2A.Z
H4K20 me2:H4−4ac
H3K4 me3:H4−4ac
H4K20 me2:H3K4 me3
H4K20 me2:H3−5ac
H4−4ac:H3−5ac
H3K9 me2
H3K27 me2
H3K9 me3
H4K20 me3
H3K4 me1
H3K27 me3
H3K4 me3
H2A.Z
H4K20 me2
H4−4ac
DNA Meth. m5C
H3−5ac
0.00 0.25 0.50 0.75 1.00
π̂
KDM6A – Forward
H3K4 me3:H3−5ac
H3K4 me1:H4K20 me2
H3K9 me3:H4K20 me3
H3K27 me3:H3K9 me3
DNA Meth. m5C:H3K9 me3
H3K27 me3:H4K20 me3
H2A.Z:H3−5ac
H2A.Z:H4−4ac
H2A.Z:H4K20 me2
H2A.Z:H3K4 me3
H3K4 me1:H3−5ac
DNA Meth. m5C:H4K20 me2
H3K27 me3:DNA Meth. m5C
DNA Meth. m5C:H4K20 me3
H3K4 me1:H4−4ac
H3K4 me3:H4−4ac
H4K20 me2:H4−4ac
H3K4 me1:H2A.Z
H4K20 me2:H3−5ac
H4K20 me2:H3K4 me3
H4−4ac:H3−5ac
H3K9 me2
H3K27 me2
H3K4 me3
H3K9 me3
H4K20 me3
H3K27 me3
H4−4ac
H4K20 me2
H3−5ac
H3K4 me1
DNA Meth. m5C
H2A.Z
0.00 0.25 0.50 0.75 1.00
π̂
KDM6A – Reverse
H3K4 me3:H3−5ac
H3K4 me1:H4K20 me2
H3K9 me3:H4K20 me3
H3K27 me3:H3K9 me3
DNA Meth. m5C:H3K9 me3
H3K27 me3:H4K20 me3
H2A.Z:H3−5ac
H2A.Z:H4−4ac
H2A.Z:H4K20 me2
H2A.Z:H3K4 me3
H3K4 me1:H3−5ac
DNA Meth. m5C:H4K20 me2
H3K27 me3:DNA Meth. m5C
DNA Meth. m5C:H4K20 me3
H3K4 me1:H4−4ac
H3K4 me3:H4−4ac
H4K20 me2:H4−4ac
H3K4 me1:H2A.Z
H4K20 me2:H3−5ac
H4K20 me2:H3K4 me3
H4−4ac:H3−5ac
H3K9 me2
H3K27 me2
H3K4 me3
H3K9 me3
H4K20 me3
H3K27 me3
H4−4ac
H4K20 me2
H3−5ac
H3K4 me1
DNA Meth. m5C
H2A.Z
0.00 0.25 0.50 0.75 1.00
π̂
PAGR1 – Forward
H3K9 me2:DNA Meth. m5C
H3K27 me3:H3K9 me3
H2A.Z:H4−4ac
H3K9 me3:H4K20 me3
DNA Meth. m5C:H4K20 me3
H3K27 me2:DNA Meth. m5C
H3K27 me3:H4K20 me3
DNA Meth. m5C:H4K20 me2
H3K27 me3:H3K4 me1
H2A.Z:H3K4 me3
DNA Meth. m5C:H3K9 me3
H3K27 me3:DNA Meth. m5C
H3K4 me1:H4−4ac
H4K20 me2:H4−4ac
H2A.Z:H3−5ac
H3K4 me3:H3−5ac
H3K4 me1:H3−5ac
H2A.Z:H4K20 me2
H3K4 me1:H2A.Z
H3K4 me3:H4−4ac
H4K20 me2:H3K4 me3
H4K20 me2:H3−5ac
H4−4ac:H3−5ac
H3K27 me2
H3K9 me3
H3K9 me2
H4K20 me3
H4K20 me2
H2A.Z
H3K4 me1
DNA Meth. m5C
H3K27 me3
H3K4 me3
H4−4ac
H3−5ac
0.00 0.25 0.50 0.75 1.00
π̂
PAGR1 – Reverse
H3K9 me2:DNA Meth. m5C
H3K27 me3:H3K9 me3
H2A.Z:H4−4ac
H3K9 me3:H4K20 me3
DNA Meth. m5C:H4K20 me3
H3K27 me2:DNA Meth. m5C
H3K27 me3:H4K20 me3
DNA Meth. m5C:H4K20 me2
H3K27 me3:H3K4 me1
H2A.Z:H3K4 me3
DNA Meth. m5C:H3K9 me3
H3K27 me3:DNA Meth. m5C
H3K4 me1:H4−4ac
H4K20 me2:H4−4ac
H2A.Z:H3−5ac
H3K4 me3:H3−5ac
H3K4 me1:H3−5ac
H2A.Z:H4K20 me2
H3K4 me1:H2A.Z
H3K4 me3:H4−4ac
H4K20 me2:H3K4 me3
H4K20 me2:H3−5ac
H4−4ac:H3−5ac
H3K27 me2
H3K9 me3
H3K9 me2
H4K20 me3
H4K20 me2
H2A.Z
H3K4 me1
DNA Meth. m5C
H3K27 me3
H3K4 me3
H4−4ac
H3−5ac
0.00 0.25 0.50 0.75 1.00
π̂
KMT2C – Forward
H3K9 me3:H4K20 me3
H3K4 me1:H4−4ac
DNA Meth. m5C:H4K20 me3
H3K27 me2:DNA Meth. m5C
H3K27 me3:H4K20 me3
H3K27 me3:H3K9 me3
H3K27 me3:DNA Meth. m5C
H2A.Z:H3−5ac
H2A.Z:H4−4ac
DNA Meth. m5C:H3K9 me3
H3K9 me2:DNA Meth. m5C
H3K4 me1:H4K20 me2
H3K4 me1:H2A.Z
H2A.Z:H3K4 me3
H4K20 me2:H4−4ac
H2A.Z:H4K20 me2
H3K4 me3:H3−5ac
H3K4 me1:H3−5ac
H4K20 me2:H3−5ac
H4−4ac:H3−5ac
H4K20 me2:H3K4 me3
DNA Meth. m5C:H4K20 me2
H3K4 me3:H4−4ac
H3K27 me2
H3K9 me3
H3K9 me2
H4K20 me3
H3K4 me1
H3K27 me3
H3K4 me3
H2A.Z
H4−4ac
H3−5ac
H4K20 me2
DNA Meth. m5C
0.00 0.25 0.50 0.75 1.00
π̂
KMT2C – Reverse
H3K9 me3:H4K20 me3
H3K4 me1:H4−4ac
DNA Meth. m5C:H4K20 me3
H3K27 me2:DNA Meth. m5C
H3K27 me3:H4K20 me3
H3K27 me3:H3K9 me3
H3K27 me3:DNA Meth. m5C
H2A.Z:H3−5ac
H2A.Z:H4−4ac
DNA Meth. m5C:H3K9 me3
H3K9 me2:DNA Meth. m5C
H3K4 me1:H4K20 me2
H3K4 me1:H2A.Z
H2A.Z:H3K4 me3
H4K20 me2:H4−4ac
H2A.Z:H4K20 me2
H3K4 me3:H3−5ac
H3K4 me1:H3−5ac
H4K20 me2:H3−5ac
H4−4ac:H3−5ac
H4K20 me2:H3K4 me3
DNA Meth. m5C:H4K20 me2
H3K4 me3:H4−4ac
H3K27 me2
H3K9 me3
H3K9 me2
H4K20 me3
H3K4 me1
H3K27 me3
H3K4 me3
H2A.Z
H4−4ac
H3−5ac
H4K20 me2
DNA Meth. m5C
0.00 0.25 0.50 0.75 1.00
π̂
NCOA6 – Forward
H3K9 me3:H4K20 me3
H3K27 me3:H4K20 me3
H3K4 me3:H3−5ac
H3K4 me1:H2A.Z
H2A.Z:H4−4ac
H3K4 me1:H4−4ac
H3K27 me3:H3K9 me3
H2A.Z:H3−5ac
H2A.Z:H3K4 me3
H3K4 me1:H3−5ac
H3K9 me2:DNA Meth. m5C
H4K20 me2:H3−5ac
H3K27 me3:DNA Meth. m5C
H2A.Z:H4K20 me2
DNA Meth. m5C:H4K20 me3
H3K4 me1:H4K20 me2
H3K27 me2:DNA Meth. m5C
H4−4ac:H3−5ac
H4K20 me2:H3K4 me3
DNA Meth. m5C:H3K9 me3
H3K4 me3:H4−4ac
H4K20 me2:H4−4ac
DNA Meth. m5C:H4K20 me2
H3K27 me2
H3K4 me3
H3K9 me2
H4K20 me3
H3K4 me1
H3K27 me3
H2A.Z
H4−4ac
H3−5ac
H4K20 me2
H3K9 me3
DNA Meth. m5C
0.00 0.25 0.50 0.75 1.00
π̂
NCOA6 – Reverse
H3K9 me3:H4K20 me3
H3K27 me3:H4K20 me3
H3K4 me3:H3−5ac
H3K4 me1:H2A.Z
H2A.Z:H4−4ac
H3K4 me1:H4−4ac
H3K27 me3:H3K9 me3
H2A.Z:H3−5ac
H2A.Z:H3K4 me3
H3K4 me1:H3−5ac
H3K9 me2:DNA Meth. m5C
H4K20 me2:H3−5ac
H3K27 me3:DNA Meth. m5C
H2A.Z:H4K20 me2
DNA Meth. m5C:H4K20 me3
H3K4 me1:H4K20 me2
H3K27 me2:DNA Meth. m5C
H4−4ac:H3−5ac
H4K20 me2:H3K4 me3
DNA Meth. m5C:H3K9 me3
H3K4 me3:H4−4ac
H4K20 me2:H4−4ac
DNA Meth. m5C:H4K20 me2
H3K27 me2
H3K4 me3
H3K9 me2
H4K20 me3
H3K4 me1
H3K27 me3
H2A.Z
H4−4ac
H3−5ac
H4K20 me2
H3K9 me3
DNA Meth. m5C
0.00 0.25 0.50 0.75 1.00
π̂
INTS11 – Forward
H3K4 me3:H3−5ac
H3K4 me1:H4−4ac
H2A.Z:H4−4ac
H3K4 me1:H3−5ac
H3K4 me1:H2A.Z
DNA Meth. m5C:H4K20 me3
H3K9 me2:DNA Meth. m5C
H2A.Z:H3−5ac
H3K27 me3:H3K9 me3
H3K27 me3:DNA Meth. m5C
H3K4 me1:H4K20 me2
H3K27 me2:DNA Meth. m5C
H2A.Z:H3K4 me3
DNA Meth. m5C:H4K20 me2
H4−4ac:H3−5ac
H4K20 me2:H4−4ac
DNA Meth. m5C:H3K9 me3
H4K20 me2:H3−5ac
H2A.Z:H4K20 me2
H4K20 me2:H3K4 me3
H3K4 me3:H4−4ac
H3K27 me2
H3K9 me2
H4K20 me3
H3K4 me1
H3K4 me3
H3K27 me3
H3−5ac
H4K20 me2
H3K9 me3
H4−4ac
H2A.Z
DNA Meth. m5C
0.00 0.25 0.50 0.75 1.00
π̂
INTS11 – Reverse
H3K4 me3:H3−5ac
H3K4 me1:H4−4ac
H2A.Z:H4−4ac
H3K4 me1:H3−5ac
H3K4 me1:H2A.Z
DNA Meth. m5C:H4K20 me3
H3K9 me2:DNA Meth. m5C
H2A.Z:H3−5ac
H3K27 me3:H3K9 me3
H3K27 me3:DNA Meth. m5C
H3K4 me1:H4K20 me2
H3K27 me2:DNA Meth. m5C
H2A.Z:H3K4 me3
DNA Meth. m5C:H4K20 me2
H4−4ac:H3−5ac
H4K20 me2:H4−4ac
DNA Meth. m5C:H3K9 me3
H4K20 me2:H3−5ac
H2A.Z:H4K20 me2
H4K20 me2:H3K4 me3
H3K4 me3:H4−4ac
H3K27 me2
H3K9 me2
H4K20 me3
H3K4 me1
H3K4 me3
H3K27 me3
H3−5ac
H4K20 me2
H3K9 me3
H4−4ac
H2A.Z
DNA Meth. m5C
0.00 0.25 0.50 0.75 1.00
π̂

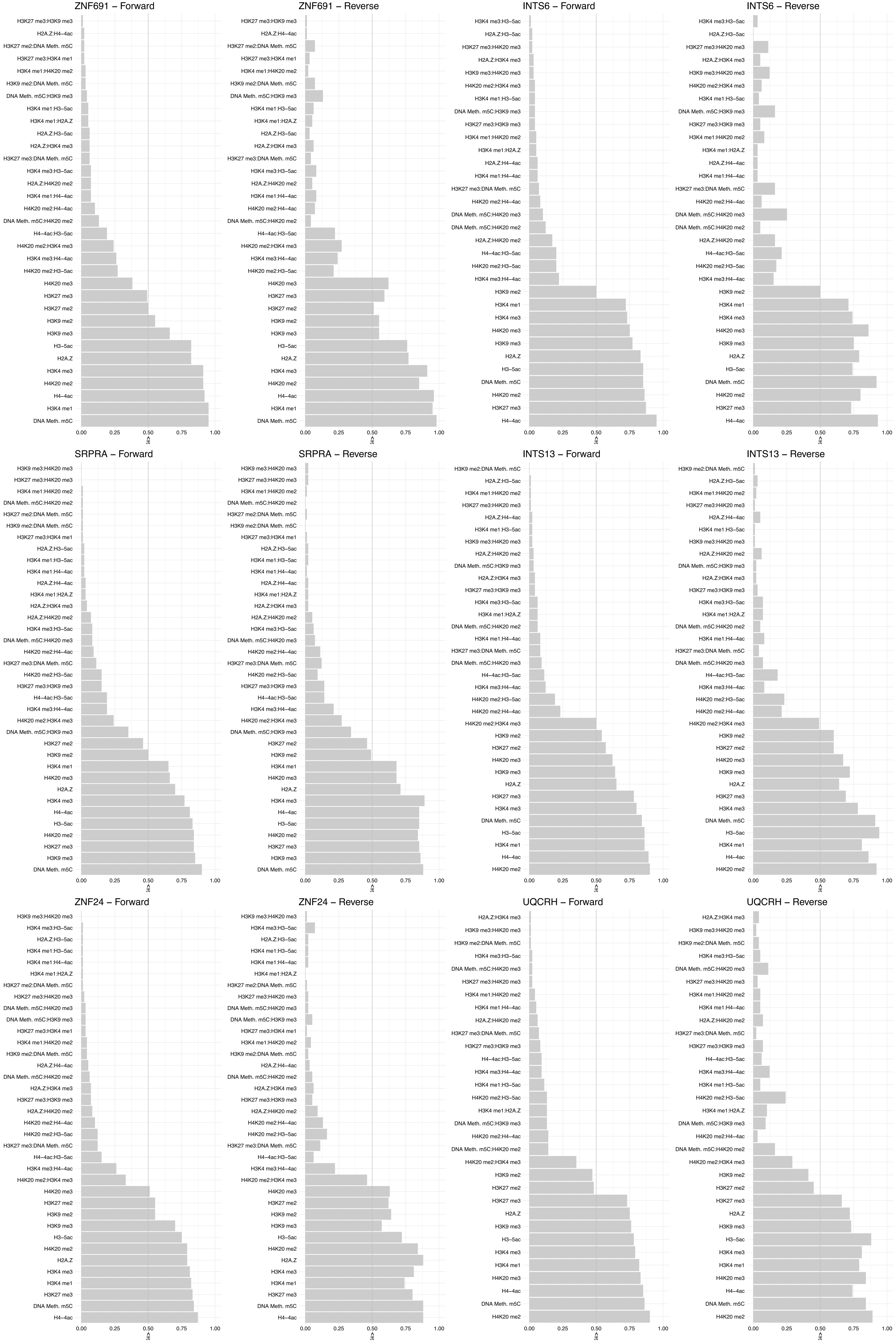
ZNF691 – Forward
H3K27 me3:H3K9 me3
H2A.Z:H4–4ac
H3K27 me2:DNA Meth. m5C
H3K27 me3:H3K4 me1
H3K4 me1:H4K20 me2
H3K9 me2:DNA Meth. m5C
DNA Meth. m5C:H3K9 me3
H3K4 me1:H3–5ac
H3K4 me1:H2A.Z
H2A.Z:H3–5ac
H2A.Z:H3K4 me3
H3K27 me3:DNA Meth. m5C
H3K4 me3:H3–5ac
H2A.Z:H4K20 me2
H3K4 me1:H4–4ac
H4K20 me2:H4–4ac
DNA Meth. m5C:H4K20 me2
H4–4ac:H3–5ac
H4K20 me2:H3K4 me3
H3K4 me3:H4–4ac
H4K20 me2:H3–5ac
H4K20 me3
H3K27 me3
H3K27 me2
H3K9 me2
H3K9 me3
H3–5ac
H2A.Z
H3K4 me3
H4K20 me2
H4–4ac
H3K4 me1
DNA Meth. m5C
0.00
0.25
0.50
0.75
1.00
π̂
ZNF691 – Reverse
H3K27 me3:H3K9 me3
H2A.Z:H4–4ac
H3K27 me2:DNA Meth. m5C
H3K27 me3:H3K4 me1
H3K4 me1:H4K20 me2
H3K9 me2:DNA Meth. m5C
DNA Meth. m5C:H3K9 me3
H3K4 me1:H3–5ac
H3K4 me1:H2A.Z
H2A.Z:H3–5ac
H2A.Z:H3K4 me3
H3K27 me3:DNA Meth. m5C
H3K4 me3:H3–5ac
H2A.Z:H4K20 me2
H3K4 me1:H4–4ac
H4K20 me2:H4–4ac
DNA Meth. m5C:H4K20 me2
H4–4ac:H3–5ac
H4K20 me2:H3K4 me3
H3K4 me3:H4–4ac
H4K20 me2:H3–5ac
H4K20 me3
H3K27 me3
H3K27 me2
H3K9 me2
H3K9 me3
H3–5ac
H2A.Z
H3K4 me3
H4K20 me2
H4–4ac
H3K4 me1
DNA Meth. m5C
0.00
0.25
0.50
0.75
1.00
π̂
INTS6 – Forward
H3K4 me3:H3–5ac
H2A.Z:H3–5ac
H3K27 me3:H4K20 me3
H2A.Z:H3K4 me3
H3K9 me3:H4K20 me3
H4K20 me2:H3K4 me3
H3K4 me1:H3–5ac
DNA Meth. m5C:H3K9 me3
H3K27 me3:H3K9 me3
H3K4 me1:H4K20 me2
H3K4 me1:H2A.Z
H2A.Z:H4–4ac
H3K4 me1:H4–4ac
H3K27 me3:DNA Meth. m5C
H4K20 me2:H4–4ac
DNA Meth. m5C:H4K20 me3
DNA Meth. m5C:H4K20 me2
H2A.Z:H4K20 me2
H4–4ac:H3–5ac
H4K20 me2:H3–5ac
H3K4 me3:H4–4ac
H3K9 me2
H3K4 me1
H3K4 me3
H4K20 me3
H3K9 me3
H2A.Z
H3–5ac
DNA Meth. m5C
H4K20 me2
H3K27 me3
H4–4ac
0.00
0.25
0.50
0.75
1.00
π̂
INTS6 – Reverse
H3K4 me3:H3–5ac
H2A.Z:H3–5ac
H3K27 me3:H4K20 me3
H2A.Z:H3K4 me3
H3K9 me3:H4K20 me3
H4K20 me2:H3K4 me3
H3K4 me1:H3–5ac
DNA Meth. m5C:H3K9 me3
H3K27 me3:H3K9 me3
H3K4 me1:H4K20 me2
H3K4 me1:H2A.Z
H2A.Z:H4–4ac
H3K4 me1:H4–4ac
H3K27 me3:DNA Meth. m5C
H4K20 me2:H4–4ac
DNA Meth. m5C:H4K20 me3
DNA Meth. m5C:H4K20 me2
H2A.Z:H4K20 me2
H4–4ac:H3–5ac
H4K20 me2:H3–5ac
H3K4 me3:H4–4ac
H3K9 me2
H3K4 me1
H3K4 me3
H4K20 me3
H3K9 me3
H2A.Z
H3–5ac
DNA Meth. m5C
H4K20 me2
H3K27 me3
H4–4ac
0.00
0.25
0.50
0.75
1.00
π̂
SRPRA – Forward
H3K9 me3:H4K20 me3
H3K27 me3:H4K20 me3
H3K4 me1:H4K20 me2
DNA Meth. m5C:H4K20 me2
H3K27 me2:DNA Meth. m5C
H3K9 me2:DNA Meth. m5C
H3K27 me3:H3K4 me1
H2A.Z:H3–5ac
H3K4 me1:H3–5ac
H3K4 me1:H4–4ac
H2A.Z:H4–4ac
H3K4 me1:H2A.Z
H2A.Z:H3K4 me3
H2A.Z:H4K20 me2
H3K4 me3:H3–5ac
DNA Meth. m5C:H4K20 me3
H4K20 me2:H4–4ac
H3K27 me3:DNA Meth. m5C
H4K20 me2:H3–5ac
H3K27 me3:H3K9 me3
H4–4ac:H3–5ac
H3K4 me3:H4–4ac
H4K20 me2:H3K4 me3
DNA Meth. m5C:H3K9 me3
H3K27 me2
H3K9 me2
H3K4 me1
H4K20 me3
H2A.Z
H3K4 me3
H4–4ac
H3–5ac
H4K20 me2
H3K27 me3
H3K9 me3
DNA Meth. m5C
0.00
0.25
0.50
0.75
1.00
π̂
SRPRA – Reverse
H3K9 me3:H4K20 me3
H3K27 me3:H4K20 me3
H3K4 me1:H4K20 me2
DNA Meth. m5C:H4K20 me2
H3K27 me2:DNA Meth. m5C
H3K9 me2:DNA Meth. m5C
H3K27 me3:H3K4 me1
H2A.Z:H3–5ac
H3K4 me1:H3–5ac
H3K4 me1:H4–4ac
H2A.Z:H4–4ac
H3K4 me1:H2A.Z
H2A.Z:H3K4 me3
H2A.Z:H4K20 me2
H3K4 me3:H3–5ac
DNA Meth. m5C:H4K20 me3
H4K20 me2:H4–4ac
H3K27 me3:DNA Meth. m5C
H4K20 me2:H3–5ac
H3K27 me3:H3K9 me3
H4–4ac:H3–5ac
H3K4 me3:H4–4ac
H4K20 me2:H3K4 me3
DNA Meth. m5C:H3K9 me3
H3K27 me2
H3K9 me2
H3K4 me1
H4K20 me3
H2A.Z
H3K4 me3
H4–4ac
H3–5ac
H4K20 me2
H3K27 me3
H3K9 me3
DNA Meth. m5C
0.00
0.25
0.50
0.75
1.00
π̂
INTS13 – Forward
H3K9 me2:DNA Meth. m5C
H2A.Z:H3–5ac
H3K4 me1:H4K20 me2
H3K27 me3:H4K20 me3
H2A.Z:H4–4ac
H3K4 me1:H3–5ac
H3K9 me3:H4K20 me3
H2A.Z:H4K20 me2
DNA Meth. m5C:H3K9 me3
H2A.Z:H3K4 me3
H3K27 me3:H3K9 me3
H3K4 me3:H3–5ac
H3K4 me1:H2A.Z
DNA Meth. m5C:H4K20 me2
H3K4 me1:H4–4ac
H3K27 me3:DNA Meth. m5C
DNA Meth. m5C:H4K20 me3
H4–4ac:H3–5ac
H3K4 me3:H4–4ac
H4K20 me2:H3–5ac
H4K20 me2:H4–4ac
H4K20 me2:H3K4 me3
H3K9 me2
H3K27 me2
H4K20 me3
H3K9 me3
H2A.Z
H3K27 me3
H3K4 me3
DNA Meth. m5C
H3–5ac
H3K4 me1
H4–4ac
H4K20 me2
0.00
0.25
0.50
0.75
1.00
π̂
INTS13 – Reverse
H3K9 me2:DNA Meth. m5C
H2A.Z:H3–5ac
H3K4 me1:H4K20 me2
H3K27 me3:H4K20 me3
H2A.Z:H4–4ac
H3K4 me1:H3–5ac
H3K9 me3:H4K20 me3
H2A.Z:H4K20 me2
DNA Meth. m5C:H3K9 me3
H2A.Z:H3K4 me3
H3K27 me3:H3K9 me3
H3K4 me3:H3–5ac
H3K4 me1:H2A.Z
DNA Meth. m5C:H4K20 me2
H3K4 me1:H4–4ac
H3K27 me3:DNA Meth. m5C
DNA Meth. m5C:H4K20 me3
H4–4ac:H3–5ac
H3K4 me3:H4–4ac
H4K20 me2:H3–5ac
H4K20 me2:H4–4ac
H4K20 me2:H3K4 me3
H3K9 me2
H3K27 me2
H4K20 me3
H3K9 me3
H2A.Z
H3K27 me3
H3K4 me3
DNA Meth. m5C
H3–5ac
H3K4 me1
H4–4ac
H4K20 me2
0.00
0.25
0.50
0.75
1.00
π̂
ZNF24 – Forward
H3K9 me3:H4K20 me3
H3K4 me3:H3–5ac
H2A.Z:H3–5ac
H3K4 me1:H3–5ac
H3K4 me1:H4–4ac
H3K4 me1:H2A.Z
H3K27 me2:DNA Meth. m5C
H3K27 me3:H4K20 me3
DNA Meth. m5C:H4K20 me3
DNA Meth. m5C:H3K9 me3
H3K27 me3:H3K4 me1
H3K4 me1:H4K20 me2
H3K9 me2:DNA Meth. m5C
H2A.Z:H4–4ac
DNA Meth. m5C:H4K20 me2
H2A.Z:H3K4 me3
H3K27 me3:H3K9 me3
H2A.Z:H4K20 me2
H4K20 me2:H4–4ac
H4K20 me2:H3–5ac
H3K27 me3:DNA Meth. m5C
H4–4ac:H3–5ac
H3K4 me3:H4–4ac
H4K20 me2:H3K4 me3
H4K20 me3
H3K27 me2
H3K9 me2
H3K9 me3
H3–5ac
H4K20 me2
H2A.Z
H3K4 me3
H3K4 me1
H3K27 me3
DNA Meth. m5C
H4–4ac
0.00
0.25
0.50
0.75
1.00
π̂
ZNF24 – Reverse
H3K9 me3:H4K20 me3
H3K4 me3:H3–5ac
H2A.Z:H3–5ac
H3K4 me1:H3–5ac
H3K4 me1:H4–4ac
H3K4 me1:H2A.Z
H3K27 me2:DNA Meth. m5C
H3K27 me3:H4K20 me3
DNA Meth. m5C:H4K20 me3
DNA Meth. m5C:H3K9 me3
H3K27 me3:H3K4 me1
H3K4 me1:H4K20 me2
H3K9 me2:DNA Meth. m5C
H2A.Z:H4–4ac
DNA Meth. m5C:H4K20 me2
H2A.Z:H3K4 me3
H3K27 me3:H3K9 me3
H2A.Z:H4K20 me2
H4K20 me2:H4–4ac
H4K20 me2:H3–5ac
H3K27 me3:DNA Meth. m5C
H4–4ac:H3–5ac
H3K4 me3:H4–4ac
H4K20 me2:H3K4 me3
H4K20 me3
H3K27 me2
H3K9 me2
H3K9 me3
H3–5ac
H4K20 me2
H2A.Z
H3K4 me3
H3K4 me1
H3K27 me3
DNA Meth. m5C
H4–4ac
0.00
0.25
0.50
0.75
1.00
π̂
UQCRH – Forward
H2A.Z:H3K4 me3
H3K9 me3:H4K20 me3
H3K9 me2:DNA Meth. m5C
H3K4 me3:H3–5ac
DNA Meth. m5C:H4K20 me3
H3K27 me3:H4K20 me3
H3K4 me1:H4K20 me2
H3K4 me1:H4–4ac
H2A.Z:H4K20 me2
H3K27 me3:DNA Meth. m5C
H3K27 me3:H3K9 me3
H4–4ac:H3–5ac
H3K4 me3:H4–4ac
H3K4 me1:H3–5ac
H4K20 me2:H3–5ac
H3K4 me1:H2A.Z
DNA Meth. m5C:H3K9 me3
H4K20 me2:H4–4ac
DNA Meth. m5C:H4K20 me2
H4K20 me2:H3K4 me3
H3K9 me2
H3K27 me2
H3K27 me3
H2A.Z
H3K9 me3
H3–5ac
H3K4 me3
H3K4 me1
H4K20 me3
H4–4ac
DNA Meth. m5C
H4K20 me2
0.00
0.25
0.50
0.75
1.00
π̂
UQCRH – Reverse
H2A.Z:H3K4 me3
H3K9 me3:H4K20 me3
H3K9 me2:DNA Meth. m5C
H3K4 me3:H3–5ac
DNA Meth. m5C:H4K20 me3
H3K27 me3:H4K20 me3
H3K4 me1:H4K20 me2
H3K4 me1:H4–4ac
H2A.Z:H4K20 me2
H3K27 me3:DNA Meth. m5C
H3K27 me3:H3K9 me3
H4–4ac:H3–5ac
H3K4 me3:H4–4ac
H3K4 me1:H3–5ac
H4K20 me2:H3–5ac
H3K4 me1:H2A.Z
DNA Meth. m5C:H3K9 me3
H4K20 me2:H4–4ac
DNA Meth. m5C:H4K20 me2
H4K20 me2:H3K4 me3
H3K9 me2
H3K27 me2
H3K27 me3
H2A.Z
H3K9 me3
H3–5ac
H3K4 me3
H3K4 me1
H4K20 me3
H4–4ac
DNA Meth. m5C
H4K20 me2
0.00
0.25
0.50
0.75
1.00
π̂

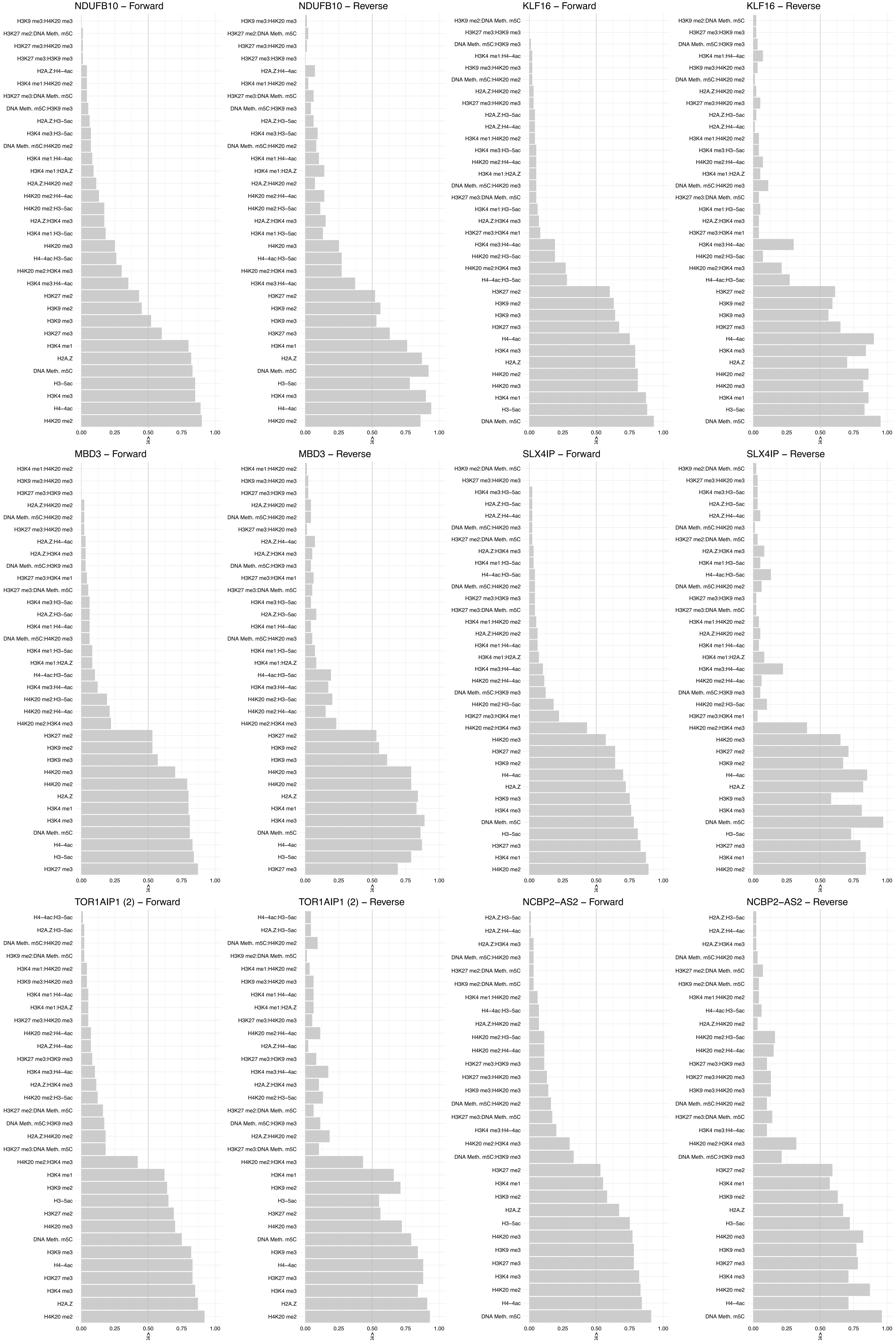
NDUFB10 – Forward
H3K9 me3:H4K20 me3
H3K27 me2:DNA Meth. m5C
H3K27 me3:H4K20 me3
H3K27 me3:H3K9 me3
H2A.Z:H4–4ac
H3K4 me1:H4K20 me2
H3K27 me3:DNA Meth. m5C
DNA Meth. m5C:H3K9 me3
H2A.Z:H3–5ac
H3K4 me3:H3–5ac
DNA Meth. m5C:H4K20 me2
H3K4 me1:H4–4ac
H3K4 me1:H2A.Z
H2A.Z:H4K20 me2
H4K20 me2:H4–4ac
H4K20 me2:H3–5ac
H2A.Z:H3K4 me3
H3K4 me1:H3–5ac
H4K20 me3
H4–4ac:H3–5ac
H4K20 me2:H3K4 me3
H3K4 me3:H4–4ac
H3K27 me2
H3K9 me2
H3K9 me3
H3K27 me3
H3K4 me1
H2A.Z
DNA Meth. m5C
H3–5ac
H3K4 me3
H4–4ac
H4K20 me2
0.00 0.25 0.50 0.75 1.00
$\hat{\pi}$
NDUFB10 – Reverse
H3K9 me3:H4K20 me3
H3K27 me2:DNA Meth. m5C
H3K27 me3:H4K20 me3
H3K27 me3:H3K9 me3
H2A.Z:H4–4ac
H3K4 me1:H4K20 me2
H3K27 me3:DNA Meth. m5C
DNA Meth. m5C:H3K9 me3
H2A.Z:H3–5ac
H3K4 me3:H3–5ac
DNA Meth. m5C:H4K20 me2
H3K4 me1:H4–4ac
H3K4 me1:H2A.Z
H2A.Z:H4K20 me2
H4K20 me2:H4–4ac
H4K20 me2:H3–5ac
H2A.Z:H3K4 me3
H3K4 me1:H3–5ac
H4K20 me3
H4–4ac:H3–5ac
H4K20 me2:H3K4 me3
H3K4 me3:H4–4ac
H3K27 me2
H3K9 me2
H3K9 me3
H3K27 me3
H3K4 me1
H2A.Z
DNA Meth. m5C
H3–5ac
H3K4 me3
H4–4ac
H4K20 me2
0.00 0.25 0.50 0.75 1.00
$\hat{\pi}$
KLF16 – Forward
H3K9 me2:DNA Meth. m5C
H3K27 me3:H3K9 me3
DNA Meth. m5C:H3K9 me3
H3K4 me1:H4–4ac
H3K9 me3:H4K20 me3
DNA Meth. m5C:H4K20 me2
H2A.Z:H4K20 me2
H3K27 me3:H4K20 me3
H2A.Z:H3–5ac
H2A.Z:H4–4ac
H3K4 me1:H4K20 me2
H3K4 me3:H3–5ac
H4K20 me2:H4–4ac
H3K4 me1:H2A.Z
DNA Meth. m5C:H4K20 me3
H3K27 me3:DNA Meth. m5C
H3K4 me1:H3–5ac
H2A.Z:H3K4 me3
H3K27 me3:H3K4 me1
H3K4 me3:H4–4ac
H4K20 me2:H3–5ac
H4K20 me2:H3K4 me3
H4–4ac:H3–5ac
H3K27 me2
H3K9 me2
H3K9 me3
H3K27 me3
H4–4ac
H3K4 me3
H2A.Z
H4K20 me2
H4K20 me3
H3K4 me1
H3–5ac
DNA Meth. m5C
0.00 0.25 0.50 0.75 1.00
$\hat{\pi}$
KLF16 – Reverse
H3K9 me2:DNA Meth. m5C
H3K27 me3:H3K9 me3
DNA Meth. m5C:H3K9 me3
H3K4 me1:H4–4ac
H3K9 me3:H4K20 me3
DNA Meth. m5C:H4K20 me2
H2A.Z:H4K20 me2
H3K27 me3:H4K20 me3
H2A.Z:H3–5ac
H2A.Z:H4–4ac
H3K4 me1:H4K20 me2
H3K4 me3:H3–5ac
H4K20 me2:H4–4ac
H3K4 me1:H2A.Z
DNA Meth. m5C:H4K20 me3
H3K27 me3:DNA Meth. m5C
H3K4 me1:H3–5ac
H2A.Z:H3K4 me3
H3K27 me3:H3K4 me1
H3K4 me3:H4–4ac
H4K20 me2:H3–5ac
H4K20 me2:H3K4 me3
H4–4ac:H3–5ac
H3K27 me2
H3K9 me2
H3K9 me3
H3K27 me3
H4–4ac
H3K4 me3
H2A.Z
H4K20 me2
H4K20 me3
H3K4 me1
H3–5ac
DNA Meth. m5C
0.00 0.25 0.50 0.75 1.00
$\hat{\pi}$
MBD3 – Forward
H3K4 me1:H4K20 me2
H3K9 me3:H4K20 me3
H3K27 me3:H3K9 me3
H2A.Z:H4K20 me2
DNA Meth. m5C:H4K20 me2
H3K27 me3:H4K20 me3
H2A.Z:H4–4ac
H2A.Z:H3K4 me3
DNA Meth. m5C:H3K9 me3
H3K27 me3:H3K4 me1
H3K27 me3:DNA Meth. m5C
H3K4 me3:H3–5ac
H2A.Z:H3–5ac
H3K4 me1:H4–4ac
DNA Meth. m5C:H4K20 me3
H3K4 me1:H3–5ac
H3K4 me1:H2A.Z
H4–4ac:H3–5ac
H3K4 me3:H4–4ac
H4K20 me2:H3–5ac
H4K20 me2:H4–4ac
H4K20 me2:H3K4 me3
H3K27 me2
H3K9 me2
H3K9 me3
H4K20 me3
H4K20 me2
H2A.Z
H3K4 me1
H3K4 me3
DNA Meth. m5C
H4–4ac
H3–5ac
H3K27 me3
0.00 0.25 0.50 0.75 1.00
$\hat{\pi}$
MBD3 – Reverse
H3K4 me1:H4K20 me2
H3K9 me3:H4K20 me3
H3K27 me3:H3K9 me3
H2A.Z:H4K20 me2
DNA Meth. m5C:H4K20 me2
H3K27 me3:H4K20 me3
H2A.Z:H4–4ac
H2A.Z:H3K4 me3
DNA Meth. m5C:H3K9 me3
H3K27 me3:H3K4 me1
H3K27 me3:DNA Meth. m5C
H3K4 me3:H3–5ac
H2A.Z:H3–5ac
H3K4 me1:H4–4ac
DNA Meth. m5C:H4K20 me3
H3K4 me1:H3–5ac
H3K4 me1:H2A.Z
H4–4ac:H3–5ac
H3K4 me3:H4–4ac
H4K20 me2:H3–5ac
H4K20 me2:H4–4ac
H4K20 me2:H3K4 me3
H3K27 me2
H3K9 me2
H3K9 me3
H4K20 me3
H4K20 me2
H2A.Z
H3K4 me1
H3K4 me3
DNA Meth. m5C
H4–4ac
H3–5ac
H3K27 me3
0.00 0.25 0.50 0.75 1.00
$\hat{\pi}$
SLX4IP – Forward
H3K9 me2:DNA Meth. m5C
H3K27 me3:H4K20 me3
H3K4 me3:H3–5ac
H2A.Z:H3–5ac
H2A.Z:H4–4ac
DNA Meth. m5C:H4K20 me3
H3K27 me2:DNA Meth. m5C
H2A.Z:H3K4 me3
H3K4 me1:H3–5ac
H4–4ac:H3–5ac
DNA Meth. m5C:H4K20 me2
H3K27 me3:H3K9 me3
H3K27 me3:DNA Meth. m5C
H3K4 me1:H4K20 me2
H2A.Z:H4K20 me2
H3K4 me1:H4–4ac
H3K4 me1:H2A.Z
H3K4 me3:H4–4ac
H4K20 me2:H4–4ac
DNA Meth. m5C:H3K9 me3
H4K20 me2:H3–5ac
H3K27 me3:H3K4 me1
H4K20 me2:H3K4 me3
H4K20 me3
H3K27 me2
H3K9 me2
H4–4ac
H2A.Z
H3K9 me3
H3K4 me3
DNA Meth. m5C
H3–5ac
H3K27 me3
H3K4 me1
H4K20 me2
0.00 0.25 0.50 0.75 1.00
$\hat{\pi}$
SLX4IP – Reverse
H3K9 me2:DNA Meth. m5C
H3K27 me3:H4K20 me3
H3K4 me3:H3–5ac
H2A.Z:H3–5ac
H2A.Z:H4–4ac
DNA Meth. m5C:H4K20 me3
H3K27 me2:DNA Meth. m5C
H2A.Z:H3K4 me3
H3K4 me1:H3–5ac
H4–4ac:H3–5ac
DNA Meth. m5C:H4K20 me2
H3K27 me3:H3K9 me3
H3K27 me3:DNA Meth. m5C
H3K4 me1:H4K20 me2
H2A.Z:H4K20 me2
H3K4 me1:H4–4ac
H3K4 me1:H2A.Z
H3K4 me3:H4–4ac
H4K20 me2:H4–4ac
DNA Meth. m5C:H3K9 me3
H4K20 me2:H3–5ac
H3K27 me3:H3K4 me1
H4K20 me2:H3K4 me3
H4K20 me3
H3K27 me2
H3K9 me2
H4–4ac
H2A.Z
H3K9 me3
H3K4 me3
DNA Meth. m5C
H3–5ac
H3K27 me3
H3K4 me1
H4K20 me2
0.00 0.25 0.50 0.75 1.00
$\hat{\pi}$
TOR1AIP1 (2) – Forward
H4–4ac:H3–5ac
H2A.Z:H3–5ac
DNA Meth. m5C:H4K20 me2
H3K9 me2:DNA Meth. m5C
H3K4 me1:H4K20 me2
H3K9 me3:H4K20 me3
H3K4 me1:H4–4ac
H3K4 me1:H2A.Z
H3K27 me3:H4K20 me3
H4K20 me2:H4–4ac
H2A.Z:H4–4ac
H3K27 me3:H3K9 me3
H3K4 me3:H4–4ac
H2A.Z:H3K4 me3
H4K20 me2:H3–5ac
H3K27 me2:DNA Meth. m5C
DNA Meth. m5C:H3K9 me3
H2A.Z:H4K20 me2
H3K27 me3:DNA Meth. m5C
H4K20 me2:H3K4 me3
H3K4 me1
H3K9 me2
H3–5ac
H3K27 me2
H4K20 me3
DNA Meth. m5C
H3K9 me3
H4–4ac
H3K27 me3
H3K4 me3
H2A.Z
H4K20 me2
0.00 0.25 0.50 0.75 1.00
$\hat{\pi}$
TOR1AIP1 (2) – Reverse
H4–4ac:H3–5ac
H2A.Z:H3–5ac
DNA Meth. m5C:H4K20 me2
H3K9 me2:DNA Meth. m5C
H3K4 me1:H4K20 me2
H3K9 me3:H4K20 me3
H3K4 me1:H4–4ac
H3K4 me1:H2A.Z
H3K27 me3:H4K20 me3
H4K20 me2:H4–4ac
H2A.Z:H4–4ac
H3K27 me3:H3K9 me3
H3K4 me3:H4–4ac
H2A.Z:H3K4 me3
H4K20 me2:H3–5ac
H3K27 me2:DNA Meth. m5C
DNA Meth. m5C:H3K9 me3
H2A.Z:H4K20 me2
H3K27 me3:DNA Meth. m5C
H4K20 me2:H3K4 me3
H3K4 me1
H3K9 me2
H3–5ac
H3K27 me2
H4K20 me3
DNA Meth. m5C
H3K9 me3
H4–4ac
H3K27 me3
H3K4 me3
H2A.Z
H4K20 me2
0.00 0.25 0.50 0.75 1.00
$\hat{\pi}$
NCBP2–AS2 – Forward
H2A.Z:H3–5ac
H2A.Z:H4–4ac
H2A.Z:H3K4 me3
DNA Meth. m5C:H4K20 me3
H3K27 me2:DNA Meth. m5C
H3K9 me2:DNA Meth. m5C
H3K4 me1:H4K20 me2
H4–4ac:H3–5ac
H2A.Z:H4K20 me2
H4K20 me2:H3–5ac
H4K20 me2:H4–4ac
H3K27 me3:H3K9 me3
H3K27 me3:H4K20 me3
H3K9 me3:H4K20 me3
DNA Meth. m5C:H4K20 me2
H3K27 me3:DNA Meth. m5C
H3K4 me3:H4–4ac
H4K20 me2:H3K4 me3
DNA Meth. m5C:H3K9 me3
H3K27 me2
H3K4 me1
H3K9 me2
H2A.Z
H3–5ac
H4K20 me3
H3K9 me3
H3K27 me3
H3K4 me3
H4K20 me2
H4–4ac
DNA Meth. m5C
0.00 0.25 0.50 0.75 1.00
$\hat{\pi}$
NCBP2–AS2 – Reverse
H2A.Z:H3–5ac
H2A.Z:H4–4ac
H2A.Z:H3K4 me3
DNA Meth. m5C:H4K20 me3
H3K27 me2:DNA Meth. m5C
H3K9 me2:DNA Meth. m5C
H3K4 me1:H4K20 me2
H4–4ac:H3–5ac
H2A.Z:H4K20 me2
H4K20 me2:H3–5ac
H4K20 me2:H4–4ac
H3K27 me3:H3K9 me3
H3K27 me3:H4K20 me3
H3K9 me3:H4K20 me3
DNA Meth. m5C:H4K20 me2
H3K27 me3:DNA Meth. m5C
H3K4 me3:H4–4ac
H4K20 me2:H3K4 me3
DNA Meth. m5C:H3K9 me3
H3K27 me2
H3K4 me1
H3K9 me2
H2A.Z
H3–5ac
H4K20 me3
H3K9 me3
H3K27 me3
H3K4 me3
H4K20 me2
H4–4ac
DNA Meth. m5C
0.00 0.25 0.50 0.75 1.00
$\hat{\pi}$

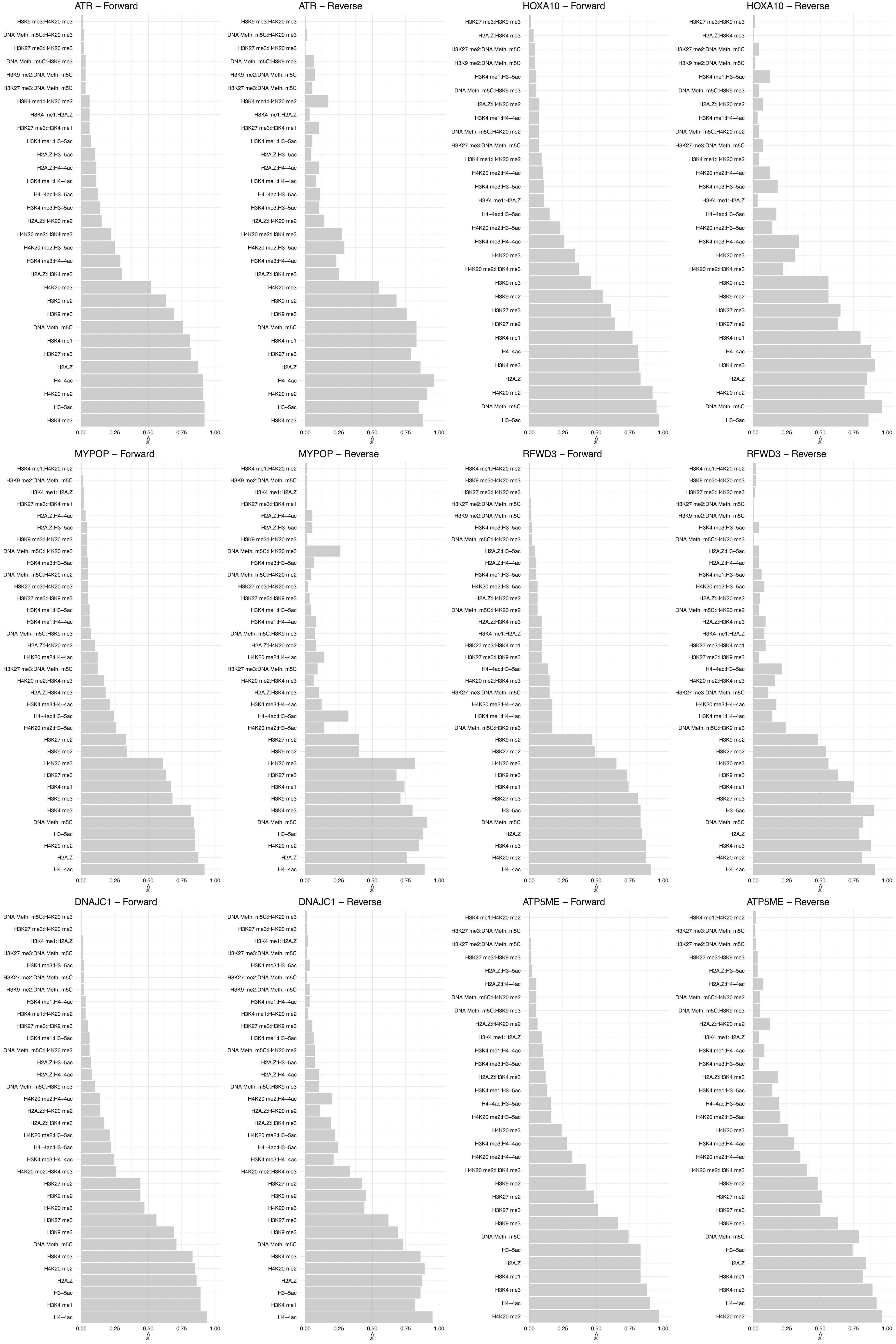
ATR – Forward
ATR – Reverse
HOXA10 – Forward
HOXA10 – Reverse
MYPOP – Forward
MYPOP – Reverse
RFWD3 – Forward
RFWD3 – Reverse
DNAJC1 – Forward
DNAJC1 – Reverse
ATP5ME – Forward
ATP5ME – Reverse
0.00 0.25 0.50 0.75 1.00
$\hat{\pi}$

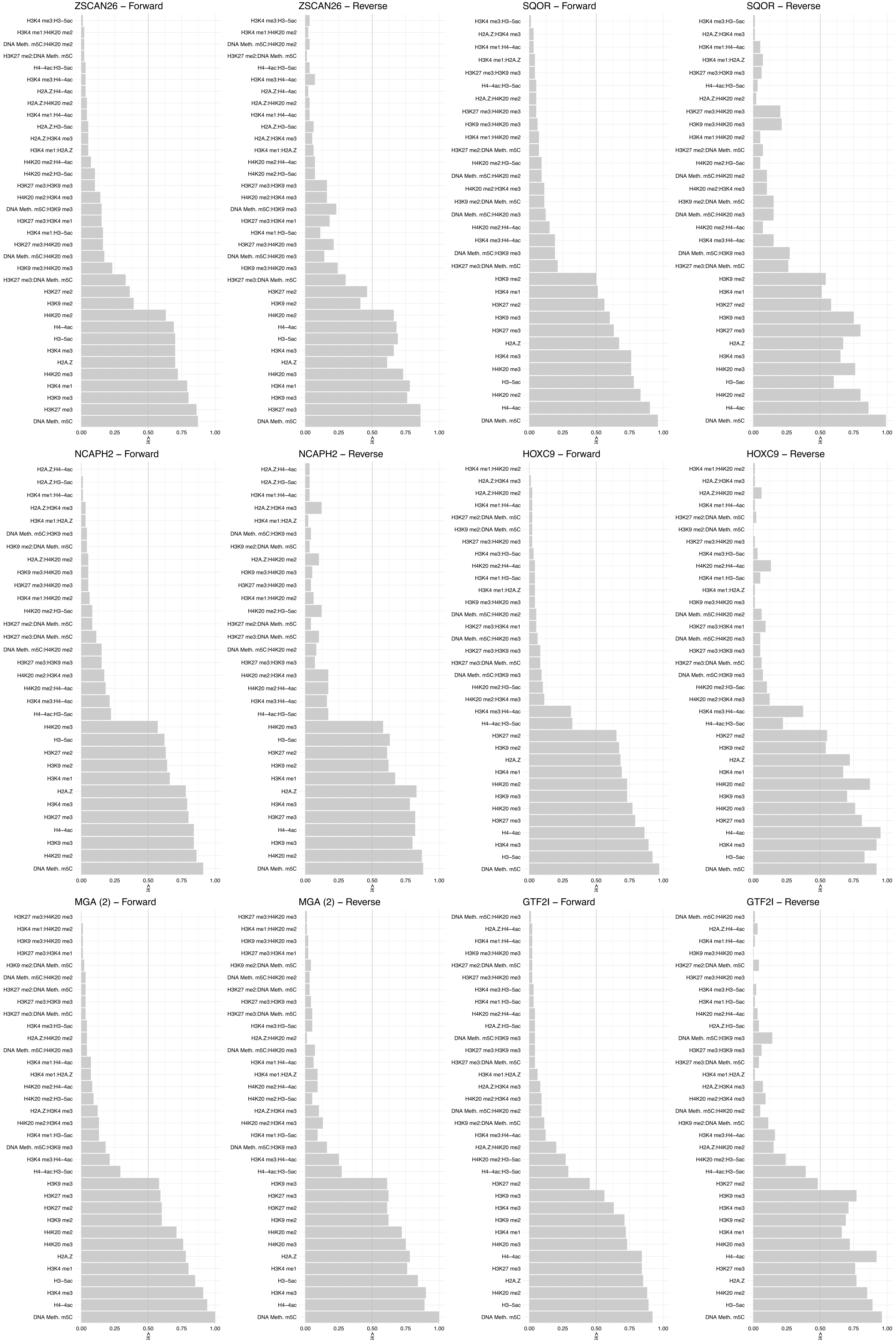
ZSCAN26 – Forward
ZSCAN26 – Reverse
SQOR – Forward
SQOR – Reverse
NCAPH2 – Forward
NCAPH2 – Reverse
HOXC9 – Forward
HOXC9 – Reverse
MGA (2) – Forward
MGA (2) – Reverse
GTF2I – Forward
GTF2I – Reverse
0.00
0.25
0.50
0.75
1.00
$\hat{\pi}$
DNA Meth. m5C
H3K27 me3
H3K9 me3
H3K4 me1
H4K20 me3
H2A.Z
H3K4 me3
H3–5ac
H4–4ac
H4K20 me2
H3K9 me2
H3K27 me2
H3K27 me3:DNA Meth. m5C
H3K9 me3:H4K20 me3
DNA Meth. m5C:H4K20 me3
H3K27 me3:H4K20 me3
H3K4 me1:H3–5ac
H3K27 me3:H3K4 me1
DNA Meth. m5C:H3K9 me3
H4K20 me2:H3K4 me3
H3K27 me3:H3K9 me3
H4K20 me2:H3–5ac
H4K20 me2:H4–4ac
H3K4 me1:H2A.Z
H2A.Z:H3K4 me3
H2A.Z:H3–5ac
H3K4 me1:H4–4ac
H2A.Z:H4K20 me2
H2A.Z:H4–4ac
H3K4 me3:H4–4ac
H4–4ac:H3–5ac
H3K27 me2:DNA Meth. m5C
DNA Meth. m5C:H4K20 me2
H3K4 me1:H4K20 me2
H3K4 me3:H3–5ac
H3K9 me2:DNA Meth. m5C

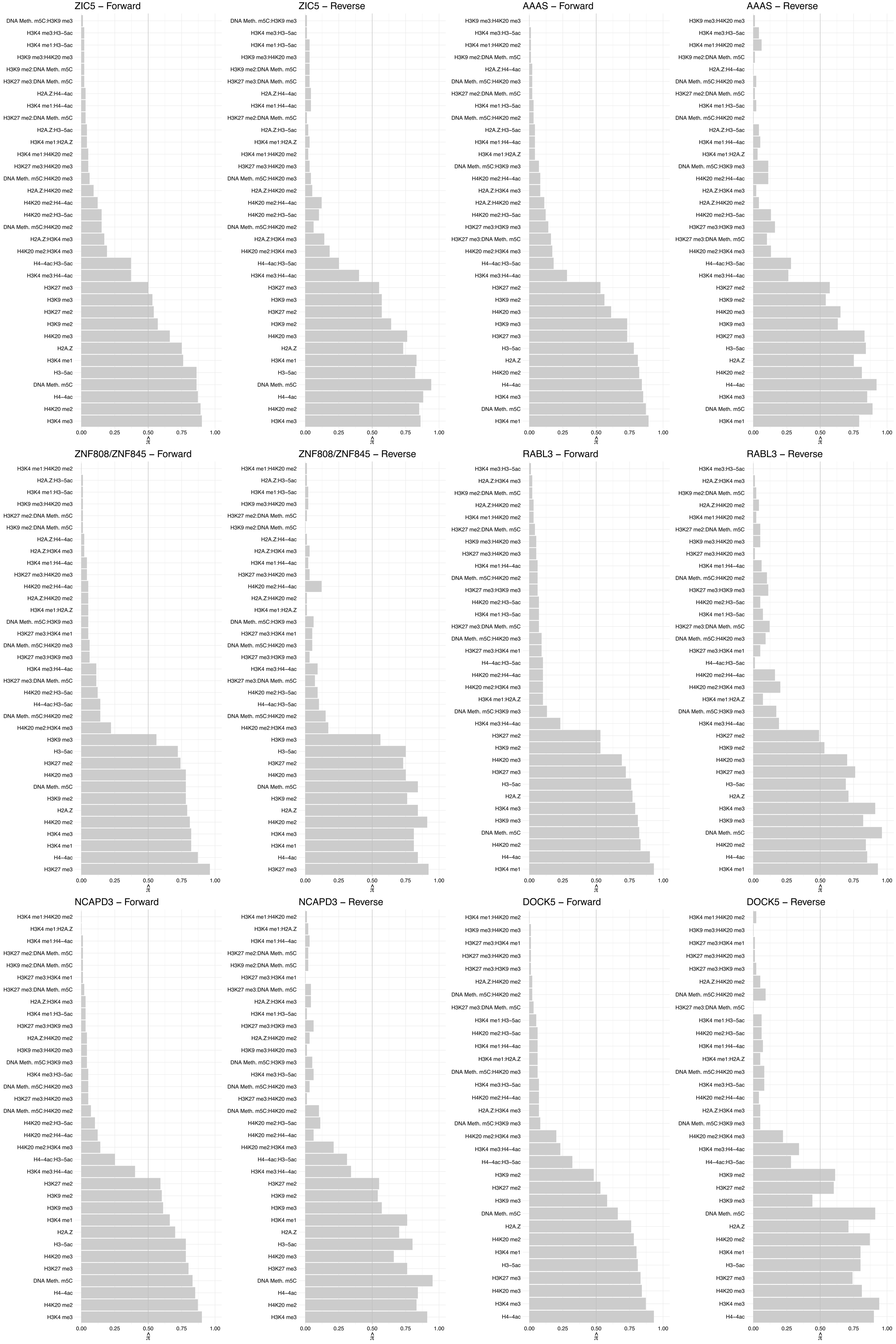
ZIC5 – Forward
DNA Meth. m5C:H3K9 me3
H3K4 me3:H3–5ac
H3K4 me1:H3–5ac
H3K9 me3:H4K20 me3
H3K9 me2:DNA Meth. m5C
H3K27 me3:DNA Meth. m5C
H2A.Z:H4–4ac
H3K4 me1:H4–4ac
H3K27 me2:DNA Meth. m5C
H2A.Z:H3–5ac
H3K4 me1:H2A.Z
H3K4 me1:H4K20 me2
H3K27 me3:H4K20 me3
DNA Meth. m5C:H4K20 me3
H2A.Z:H4K20 me2
H4K20 me2:H4–4ac
H4K20 me2:H3–5ac
DNA Meth. m5C:H4K20 me2
H2A.Z:H3K4 me3
H4K20 me2:H3K4 me3
H4–4ac:H3–5ac
H3K4 me3:H4–4ac
H3K27 me3
H3K9 me3
H3K27 me2
H3K9 me2
H4K20 me3
H2A.Z
H3K4 me1
H3–5ac
DNA Meth. m5C
H4–4ac
H4K20 me2
H3K4 me3
0.00 0.25 0.50 0.75 1.00
$\hat{\pi}$
ZIC5 – Reverse
DNA Meth. m5C:H3K9 me3
H3K4 me3:H3–5ac
H3K4 me1:H3–5ac
H3K9 me3:H4K20 me3
H3K9 me2:DNA Meth. m5C
H3K27 me3:DNA Meth. m5C
H2A.Z:H4–4ac
H3K4 me1:H4–4ac
H3K27 me2:DNA Meth. m5C
H2A.Z:H3–5ac
H3K4 me1:H2A.Z
H3K4 me1:H4K20 me2
H3K27 me3:H4K20 me3
DNA Meth. m5C:H4K20 me3
H2A.Z:H4K20 me2
H4K20 me2:H4–4ac
H4K20 me2:H3–5ac
DNA Meth. m5C:H4K20 me2
H2A.Z:H3K4 me3
H4K20 me2:H3K4 me3
H4–4ac:H3–5ac
H3K4 me3:H4–4ac
H3K27 me3
H3K9 me3
H3K27 me2
H3K9 me2
H4K20 me3
H2A.Z
H3K4 me1
H3–5ac
DNA Meth. m5C
H4–4ac
H4K20 me2
H3K4 me3
0.00 0.25 0.50 0.75 1.00
$\hat{\pi}$
AAAS – Forward
H3K9 me3:H4K20 me3
H3K4 me3:H3–5ac
H3K4 me1:H4K20 me2
H3K9 me2:DNA Meth. m5C
H2A.Z:H4–4ac
DNA Meth. m5C:H4K20 me3
H3K27 me2:DNA Meth. m5C
H3K4 me1:H3–5ac
DNA Meth. m5C:H4K20 me2
H2A.Z:H3–5ac
H3K4 me1:H4–4ac
H3K4 me1:H2A.Z
DNA Meth. m5C:H3K9 me3
H4K20 me2:H4–4ac
H2A.Z:H3K4 me3
H2A.Z:H4K20 me2
H4K20 me2:H3–5ac
H3K27 me3:H3K9 me3
H3K27 me3:DNA Meth. m5C
H4K20 me2:H3K4 me3
H4–4ac:H3–5ac
H3K4 me3:H4–4ac
H3K27 me2
H3K9 me2
H4K20 me3
H3K9 me3
H3K27 me3
H3–5ac
H2A.Z
H4K20 me2
H4–4ac
H3K4 me3
DNA Meth. m5C
H3K4 me1
0.00 0.25 0.50 0.75 1.00
$\hat{\pi}$
AAAS – Reverse
H3K9 me3:H4K20 me3
H3K4 me3:H3–5ac
H3K4 me1:H4K20 me2
H3K9 me2:DNA Meth. m5C
H2A.Z:H4–4ac
DNA Meth. m5C:H4K20 me3
H3K27 me2:DNA Meth. m5C
H3K4 me1:H3–5ac
DNA Meth. m5C:H4K20 me2
H2A.Z:H3–5ac
H3K4 me1:H4–4ac
H3K4 me1:H2A.Z
DNA Meth. m5C:H3K9 me3
H4K20 me2:H4–4ac
H2A.Z:H3K4 me3
H2A.Z:H4K20 me2
H4K20 me2:H3–5ac
H3K27 me3:H3K9 me3
H3K27 me3:DNA Meth. m5C
H4K20 me2:H3K4 me3
H4–4ac:H3–5ac
H3K4 me3:H4–4ac
H3K27 me2
H3K9 me2
H4K20 me3
H3K9 me3
H3K27 me3
H3–5ac
H2A.Z
H4K20 me2
H4–4ac
H3K4 me3
DNA Meth. m5C
H3K4 me1
0.00 0.25 0.50 0.75 1.00
$\hat{\pi}$
ZNF808/ZNF845 – Forward
H3K4 me1:H4K20 me2
H2A.Z:H3–5ac
H3K4 me1:H3–5ac
H3K9 me3:H4K20 me3
H3K27 me2:DNA Meth. m5C
H3K9 me2:DNA Meth. m5C
H2A.Z:H4–4ac
H2A.Z:H3K4 me3
H3K4 me1:H4–4ac
H3K27 me3:H4K20 me3
H4K20 me2:H4–4ac
H2A.Z:H4K20 me2
H3K4 me1:H2A.Z
DNA Meth. m5C:H3K9 me3
H3K27 me3:H3K4 me1
DNA Meth. m5C:H4K20 me3
H3K27 me3:H3K9 me3
H3K4 me3:H4–4ac
H3K27 me3:DNA Meth. m5C
H4K20 me2:H3–5ac
H4–4ac:H3–5ac
DNA Meth. m5C:H4K20 me2
H4K20 me2:H3K4 me3
H3K9 me3
H3–5ac
H3K27 me2
H4K20 me3
DNA Meth. m5C
H3K9 me2
H2A.Z
H4K20 me2
H3K4 me3
H3K4 me1
H4–4ac
H3K27 me3
0.00 0.25 0.50 0.75 1.00
$\hat{\pi}$
ZNF808/ZNF845 – Reverse
H3K4 me1:H4K20 me2
H2A.Z:H3–5ac
H3K4 me1:H3–5ac
H3K9 me3:H4K20 me3
H3K27 me2:DNA Meth. m5C
H3K9 me2:DNA Meth. m5C
H2A.Z:H4–4ac
H2A.Z:H3K4 me3
H3K4 me1:H4–4ac
H3K27 me3:H4K20 me3
H4K20 me2:H4–4ac
H2A.Z:H4K20 me2
H3K4 me1:H2A.Z
DNA Meth. m5C:H3K9 me3
H3K27 me3:H3K4 me1
DNA Meth. m5C:H4K20 me3
H3K27 me3:H3K9 me3
H3K4 me3:H4–4ac
H3K27 me3:DNA Meth. m5C
H4K20 me2:H3–5ac
H4–4ac:H3–5ac
DNA Meth. m5C:H4K20 me2
H4K20 me2:H3K4 me3
H3K9 me3
H3–5ac
H3K27 me2
H4K20 me3
DNA Meth. m5C
H3K9 me2
H2A.Z
H4K20 me2
H3K4 me3
H3K4 me1
H4–4ac
H3K27 me3
0.00 0.25 0.50 0.75 1.00
$\hat{\pi}$
RABL3 – Forward
H3K4 me3:H3–5ac
H2A.Z:H3K4 me3
H3K9 me2:DNA Meth. m5C
H2A.Z:H4K20 me2
H3K4 me1:H4K20 me2
H3K27 me2:DNA Meth. m5C
H3K9 me3:H4K20 me3
H3K27 me3:H4K20 me3
H3K4 me1:H4–4ac
DNA Meth. m5C:H4K20 me2
H3K27 me3:H3K9 me3
H4K20 me2:H3–5ac
H3K4 me1:H3–5ac
H3K27 me3:DNA Meth. m5C
DNA Meth. m5C:H4K20 me3
H3K27 me3:H3K4 me1
H4–4ac:H3–5ac
H4K20 me2:H4–4ac
H4K20 me2:H3K4 me3
H3K4 me1:H2A.Z
DNA Meth. m5C:H3K9 me3
H3K4 me3:H4–4ac
H3K27 me2
H3K9 me2
H4K20 me3
H3K27 me3
H3–5ac
H2A.Z
H3K4 me3
H3K9 me3
DNA Meth. m5C
H4K20 me2
H4–4ac
H3K4 me1
0.00 0.25 0.50 0.75 1.00
$\hat{\pi}$
RABL3 – Reverse
H3K4 me3:H3–5ac
H2A.Z:H3K4 me3
H3K9 me2:DNA Meth. m5C
H2A.Z:H4K20 me2
H3K4 me1:H4K20 me2
H3K27 me2:DNA Meth. m5C
H3K9 me3:H4K20 me3
H3K27 me3:H4K20 me3
H3K4 me1:H4–4ac
DNA Meth. m5C:H4K20 me2
H3K27 me3:H3K9 me3
H4K20 me2:H3–5ac
H3K4 me1:H3–5ac
H3K27 me3:DNA Meth. m5C
DNA Meth. m5C:H4K20 me3
H3K27 me3:H3K4 me1
H4–4ac:H3–5ac
H4K20 me2:H4–4ac
H4K20 me2:H3K4 me3
H3K4 me1:H2A.Z
DNA Meth. m5C:H3K9 me3
H3K4 me3:H4–4ac
H3K27 me2
H3K9 me2
H4K20 me3
H3K27 me3
H3–5ac
H2A.Z
H3K4 me3
H3K9 me3
DNA Meth. m5C
H4K20 me2
H4–4ac
H3K4 me1
0.00 0.25 0.50 0.75 1.00
$\hat{\pi}$
NCAPD3 – Forward
H3K4 me1:H4K20 me2
H3K4 me1:H2A.Z
H3K4 me1:H4–4ac
H3K27 me2:DNA Meth. m5C
H3K9 me2:DNA Meth. m5C
H3K27 me3:H3K4 me1
H3K27 me3:DNA Meth. m5C
H2A.Z:H3K4 me3
H3K4 me1:H3–5ac
H3K27 me3:H3K9 me3
H2A.Z:H4K20 me2
H3K9 me3:H4K20 me3
DNA Meth. m5C:H3K9 me3
H3K4 me3:H3–5ac
DNA Meth. m5C:H4K20 me3
H3K27 me3:H4K20 me3
DNA Meth. m5C:H4K20 me2
H4K20 me2:H3–5ac
H4K20 me2:H4–4ac
H4K20 me2:H3K4 me3
H4–4ac:H3–5ac
H3K4 me3:H4–4ac
H3K27 me2
H3K9 me2
H3K9 me3
H3K4 me1
H2A.Z
H3–5ac
H4K20 me3
H3K27 me3
DNA Meth. m5C
H4–4ac
H4K20 me2
H3K4 me3
0.00 0.25 0.50 0.75 1.00
$\hat{\pi}$
NCAPD3 – Reverse
H3K4 me1:H4K20 me2
H3K4 me1:H2A.Z
H3K4 me1:H4–4ac
H3K27 me2:DNA Meth. m5C
H3K9 me2:DNA Meth. m5C
H3K27 me3:H3K4 me1
H3K27 me3:DNA Meth. m5C
H2A.Z:H3K4 me3
H3K4 me1:H3–5ac
H3K27 me3:H3K9 me3
H2A.Z:H4K20 me2
H3K9 me3:H4K20 me3
DNA Meth. m5C:H3K9 me3
H3K4 me3:H3–5ac
DNA Meth. m5C:H4K20 me3
H3K27 me3:H4K20 me3
DNA Meth. m5C:H4K20 me2
H4K20 me2:H3–5ac
H4K20 me2:H4–4ac
H4K20 me2:H3K4 me3
H4–4ac:H3–5ac
H3K4 me3:H4–4ac
H3K27 me2
H3K9 me2
H3K9 me3
H3K4 me1
H2A.Z
H3–5ac
H4K20 me3
H3K27 me3
DNA Meth. m5C
H4–4ac
H4K20 me2
H3K4 me3
0.00 0.25 0.50 0.75 1.00
$\hat{\pi}$
DOCK5 – Forward
H3K4 me1:H4K20 me2
H3K9 me3:H4K20 me3
H3K27 me3:H3K4 me1
H3K27 me3:H4K20 me3
H3K27 me3:H3K9 me3
H2A.Z:H4K20 me2
DNA Meth. m5C:H4K20 me2
H3K27 me3:DNA Meth. m5C
H3K4 me1:H3–5ac
H4K20 me2:H3–5ac
H3K4 me1:H4–4ac
H3K4 me1:H2A.Z
DNA Meth. m5C:H4K20 me3
H3K4 me3:H3–5ac
H4K20 me2:H4–4ac
H2A.Z:H3K4 me3
DNA Meth. m5C:H3K9 me3
H4K20 me2:H3K4 me3
H3K4 me3:H4–4ac
H4–4ac:H3–5ac
H3K9 me2
H3K27 me2
H3K9 me3
DNA Meth. m5C
H2A.Z
H4K20 me2
H3K4 me1
H3–5ac
H3K27 me3
H4K20 me3
H3K4 me3
H4–4ac
0.00 0.25 0.50 0.75 1.00
$\hat{\pi}$
DOCK5 – Reverse
H3K4 me1:H4K20 me2
H3K9 me3:H4K20 me3
H3K27 me3:H3K4 me1
H3K27 me3:H4K20 me3
H3K27 me3:H3K9 me3
H2A.Z:H4K20 me2
DNA Meth. m5C:H4K20 me2
H3K27 me3:DNA Meth. m5C
H3K4 me1:H3–5ac
H4K20 me2:H3–5ac
H3K4 me1:H4–4ac
H3K4 me1:H2A.Z
DNA Meth. m5C:H4K20 me3
H3K4 me3:H3–5ac
H4K20 me2:H4–4ac
H2A.Z:H3K4 me3
DNA Meth. m5C:H3K9 me3
H4K20 me2:H3K4 me3
H3K4 me3:H4–4ac
H4–4ac:H3–5ac
H3K9 me2
H3K27 me2
H3K9 me3
DNA Meth. m5C
H2A.Z
H4K20 me2
H3K4 me1
H3–5ac
H3K27 me3
H4K20 me3
H3K4 me3
H4–4ac
0.00 0.25 0.50 0.75 1.00
$\hat{\pi}$

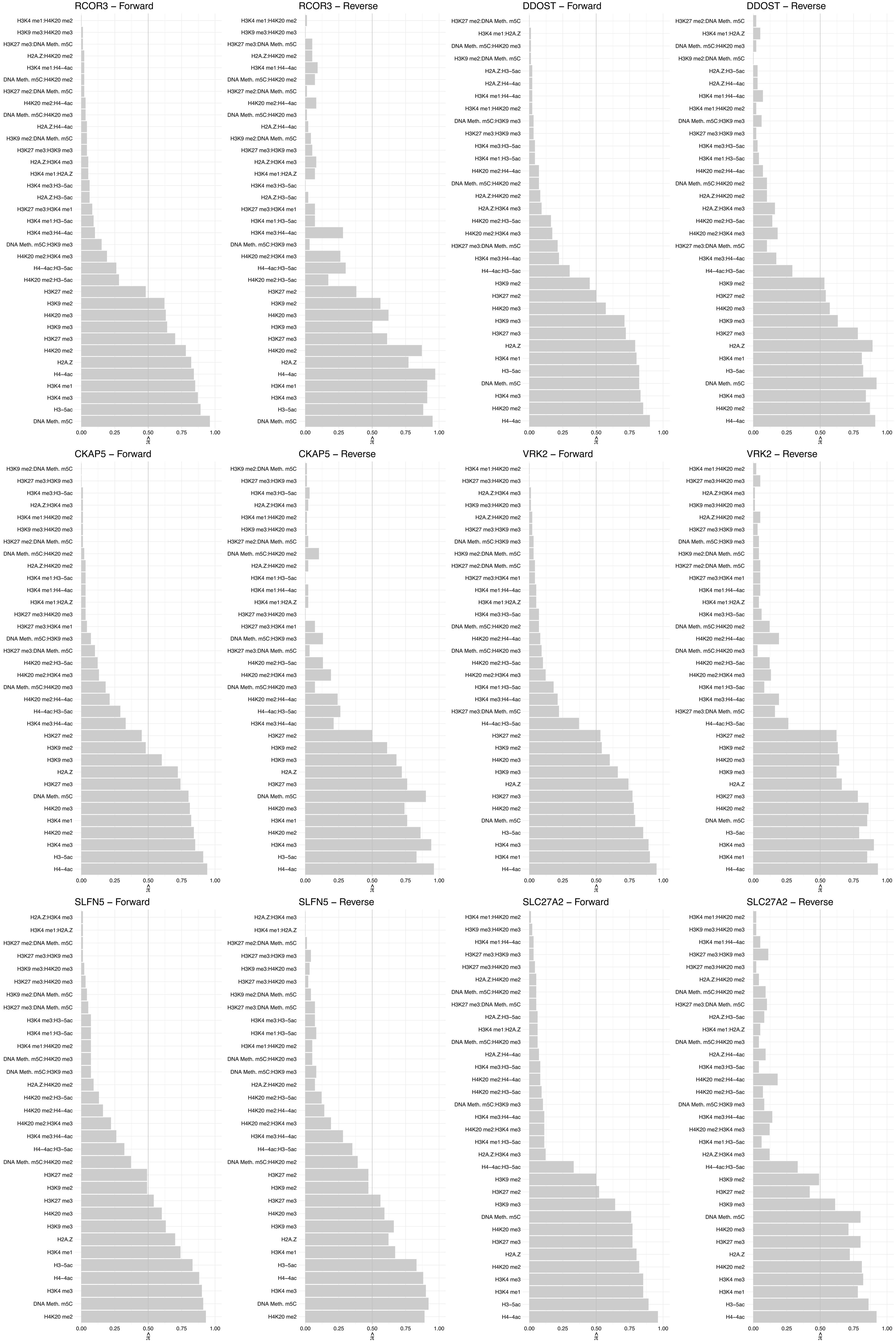
RCOR3 − Forward
RCOR3 − Reverse
H3K4 me1:H4K20 me2
H3K9 me3:H4K20 me3
H3K27 me3:DNA Meth. m5C
H2A.Z:H4K20 me2
H3K4 me1:H4−4ac
DNA Meth. m5C:H4K20 me2
H3K27 me2:DNA Meth. m5C
H4K20 me2:H4−4ac
DNA Meth. m5C:H4K20 me3
H2A.Z:H4−4ac
H3K9 me2:DNA Meth. m5C
H3K27 me3:H3K9 me3
H2A.Z:H3K4 me3
H3K4 me1:H2A.Z
H3K4 me3:H3−5ac
H2A.Z:H3−5ac
H3K27 me3:H3K4 me1
H3K4 me1:H3−5ac
H3K4 me3:H4−4ac
DNA Meth. m5C:H3K9 me3
H4K20 me2:H3K4 me3
H4−4ac:H3−5ac
H4K20 me2:H3−5ac
H3K27 me2
H3K9 me2
H4K20 me3
H3K9 me3
H3K27 me3
H4K20 me2
H2A.Z
H4−4ac
H3K4 me1
H3K4 me3
H3−5ac
DNA Meth. m5C
DDOST − Forward
DDOST − Reverse
H3K27 me2:DNA Meth. m5C
H3K4 me1:H2A.Z
DNA Meth. m5C:H4K20 me3
H3K9 me2:DNA Meth. m5C
H2A.Z:H3−5ac
H2A.Z:H4−4ac
H3K4 me1:H4−4ac
H3K4 me1:H4K20 me2
DNA Meth. m5C:H3K9 me3
H3K27 me3:H3K9 me3
H3K4 me3:H3−5ac
H3K4 me1:H3−5ac
H4K20 me2:H4−4ac
DNA Meth. m5C:H4K20 me2
H2A.Z:H4K20 me2
H2A.Z:H3K4 me3
H4K20 me2:H3−5ac
H4K20 me2:H3K4 me3
H3K27 me3:DNA Meth. m5C
H3K4 me3:H4−4ac
H4−4ac:H3−5ac
H3K9 me2
H3K27 me2
H4K20 me3
H3K9 me3
H3K27 me3
H2A.Z
H3K4 me1
H3−5ac
DNA Meth. m5C
H3K4 me3
H4K20 me2
H4−4ac
CKAP5 − Forward
CKAP5 − Reverse
H3K9 me2:DNA Meth. m5C
H3K27 me3:H3K9 me3
H3K4 me3:H3−5ac
H2A.Z:H3K4 me3
H3K4 me1:H4K20 me2
H3K9 me3:H4K20 me3
H3K27 me2:DNA Meth. m5C
DNA Meth. m5C:H4K20 me2
H2A.Z:H4K20 me2
H3K4 me1:H3−5ac
H3K4 me1:H4−4ac
H3K4 me1:H2A.Z
H3K27 me3:H4K20 me3
H3K27 me3:H3K4 me1
DNA Meth. m5C:H3K9 me3
H3K27 me3:DNA Meth. m5C
H4K20 me2:H3−5ac
H4K20 me2:H3K4 me3
DNA Meth. m5C:H4K20 me3
H4K20 me2:H4−4ac
H4−4ac:H3−5ac
H3K4 me3:H4−4ac
H3K27 me2
H3K9 me2
H3K9 me3
H2A.Z
H3K27 me3
DNA Meth. m5C
H4K20 me3
H3K4 me1
H4K20 me2
H3K4 me3
H3−5ac
H4−4ac
VRK2 − Forward
VRK2 − Reverse
H3K4 me1:H4K20 me2
H3K27 me3:H4K20 me3
H2A.Z:H3K4 me3
H3K9 me3:H4K20 me3
H2A.Z:H4K20 me2
H3K27 me3:H3K9 me3
DNA Meth. m5C:H3K9 me3
H3K9 me2:DNA Meth. m5C
H3K27 me2:DNA Meth. m5C
H3K27 me3:H3K4 me1
H3K4 me1:H4−4ac
H3K4 me1:H2A.Z
H3K4 me3:H3−5ac
DNA Meth. m5C:H4K20 me2
H4K20 me2:H4−4ac
DNA Meth. m5C:H4K20 me3
H4K20 me2:H3−5ac
H4K20 me2:H3K4 me3
H3K4 me1:H3−5ac
H3K4 me3:H4−4ac
H3K27 me3:DNA Meth. m5C
H4−4ac:H3−5ac
H3K27 me2
H3K9 me2
H4K20 me3
H3K9 me3
H2A.Z
H3K27 me3
H4K20 me2
DNA Meth. m5C
H3−5ac
H3K4 me3
H3K4 me1
H4−4ac
SLFN5 − Forward
SLFN5 − Reverse
H2A.Z:H3K4 me3
H3K4 me1:H2A.Z
H3K27 me2:DNA Meth. m5C
H3K27 me3:H3K9 me3
H3K9 me3:H4K20 me3
H3K27 me3:H4K20 me3
H3K9 me2:DNA Meth. m5C
H3K27 me3:DNA Meth. m5C
H3K4 me3:H3−5ac
H3K4 me1:H3−5ac
H3K4 me1:H4K20 me2
DNA Meth. m5C:H4K20 me3
DNA Meth. m5C:H3K9 me3
H2A.Z:H4K20 me2
H4K20 me2:H3−5ac
H4K20 me2:H4−4ac
H4K20 me2:H3K4 me3
H3K4 me3:H4−4ac
H4−4ac:H3−5ac
DNA Meth. m5C:H4K20 me2
H3K27 me2
H3K9 me2
H3K27 me3
H4K20 me3
H3K9 me3
H2A.Z
H3K4 me1
H3−5ac
H4−4ac
H3K4 me3
DNA Meth. m5C
H4K20 me2
SLC27A2 − Forward
SLC27A2 − Reverse
H3K4 me1:H4K20 me2
H3K9 me3:H4K20 me3
H3K4 me1:H4−4ac
H3K27 me3:H3K9 me3
H3K27 me3:H4K20 me3
H2A.Z:H4K20 me2
DNA Meth. m5C:H4K20 me2
H3K27 me3:DNA Meth. m5C
H2A.Z:H3−5ac
H3K4 me1:H2A.Z
DNA Meth. m5C:H4K20 me3
H2A.Z:H4−4ac
H3K4 me3:H3−5ac
H4K20 me2:H4−4ac
H4K20 me2:H3−5ac
DNA Meth. m5C:H3K9 me3
H3K4 me3:H4−4ac
H4K20 me2:H3K4 me3
H3K4 me1:H3−5ac
H2A.Z:H3K4 me3
H4−4ac:H3−5ac
H3K9 me2
H3K27 me2
H3K9 me3
DNA Meth. m5C
H4K20 me3
H3K27 me3
H2A.Z
H4K20 me2
H3K4 me3
H3K4 me1
H3−5ac
H4−4ac
0.00
0.25
0.50
0.75
1.00
π̂

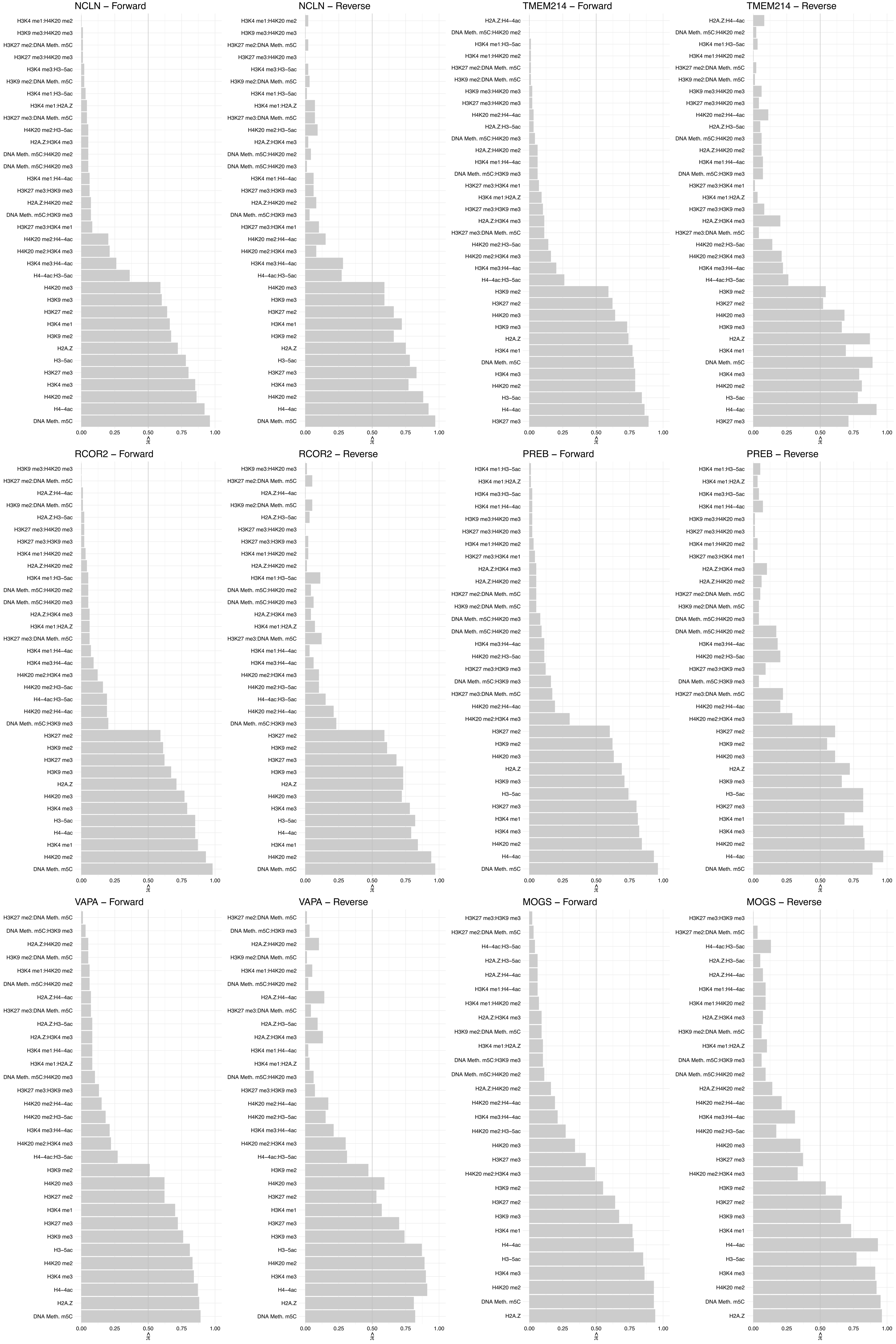
NCLN – Forward
H3K4 me1:H4K20 me2
H3K9 me3:H4K20 me3
H3K27 me2:DNA Meth. m5C
H3K27 me3:H4K20 me3
H3K4 me3:H3–5ac
H3K9 me2:DNA Meth. m5C
H3K4 me1:H3–5ac
H3K4 me1:H2A.Z
H3K27 me3:DNA Meth. m5C
H4K20 me2:H3–5ac
H2A.Z:H3K4 me3
DNA Meth. m5C:H4K20 me2
DNA Meth. m5C:H4K20 me3
H3K4 me1:H4–4ac
H3K27 me3:H3K9 me3
H2A.Z:H4K20 me2
DNA Meth. m5C:H3K9 me3
H3K27 me3:H3K4 me1
H4K20 me2:H4–4ac
H4K20 me2:H3K4 me3
H3K4 me3:H4–4ac
H4–4ac:H3–5ac
H4K20 me3
H3K9 me3
H3K27 me2
H3K4 me1
H3K9 me2
H2A.Z
H3–5ac
H3K27 me3
H3K4 me3
H4K20 me2
H4–4ac
DNA Meth. m5C
0.00
0.25
0.50
0.75
1.00
π̂
NCLN – Reverse
H3K4 me1:H4K20 me2
H3K9 me3:H4K20 me3
H3K27 me2:DNA Meth. m5C
H3K27 me3:H4K20 me3
H3K4 me3:H3–5ac
H3K9 me2:DNA Meth. m5C
H3K4 me1:H3–5ac
H3K4 me1:H2A.Z
H3K27 me3:DNA Meth. m5C
H4K20 me2:H3–5ac
H2A.Z:H3K4 me3
DNA Meth. m5C:H4K20 me2
DNA Meth. m5C:H4K20 me3
H3K4 me1:H4–4ac
H3K27 me3:H3K9 me3
H2A.Z:H4K20 me2
DNA Meth. m5C:H3K9 me3
H3K27 me3:H3K4 me1
H4K20 me2:H4–4ac
H4K20 me2:H3K4 me3
H3K4 me3:H4–4ac
H4–4ac:H3–5ac
H4K20 me3
H3K9 me3
H3K27 me2
H3K4 me1
H3K9 me2
H2A.Z
H3–5ac
H3K27 me3
H3K4 me3
H4K20 me2
H4–4ac
DNA Meth. m5C
0.00
0.25
0.50
0.75
1.00
π̂
TMEM214 – Forward
H2A.Z:H4–4ac
DNA Meth. m5C:H4K20 me2
H3K4 me1:H3–5ac
H3K4 me1:H4K20 me2
H3K27 me2:DNA Meth. m5C
H3K9 me2:DNA Meth. m5C
H3K9 me3:H4K20 me3
H3K27 me3:H4K20 me3
H4K20 me2:H4–4ac
H2A.Z:H3–5ac
DNA Meth. m5C:H4K20 me3
H2A.Z:H4K20 me2
H3K4 me1:H4–4ac
DNA Meth. m5C:H3K9 me3
H3K27 me3:H3K4 me1
H3K4 me1:H2A.Z
H3K27 me3:H3K9 me3
H2A.Z:H3K4 me3
H3K27 me3:DNA Meth. m5C
H4K20 me2:H3–5ac
H4K20 me2:H3K4 me3
H3K4 me3:H4–4ac
H4–4ac:H3–5ac
H3K9 me2
H3K27 me2
H4K20 me3
H3K9 me3
H2A.Z
H3K4 me1
DNA Meth. m5C
H3K4 me3
H4K20 me2
H3–5ac
H4–4ac
H3K27 me3
0.00
0.25
0.50
0.75
1.00
π̂
TMEM214 – Reverse
H2A.Z:H4–4ac
DNA Meth. m5C:H4K20 me2
H3K4 me1:H3–5ac
H3K4 me1:H4K20 me2
H3K27 me2:DNA Meth. m5C
H3K9 me2:DNA Meth. m5C
H3K9 me3:H4K20 me3
H3K27 me3:H4K20 me3
H4K20 me2:H4–4ac
H2A.Z:H3–5ac
DNA Meth. m5C:H4K20 me3
H2A.Z:H4K20 me2
H3K4 me1:H4–4ac
DNA Meth. m5C:H3K9 me3
H3K27 me3:H3K4 me1
H3K4 me1:H2A.Z
H3K27 me3:H3K9 me3
H2A.Z:H3K4 me3
H3K27 me3:DNA Meth. m5C
H4K20 me2:H3–5ac
H4K20 me2:H3K4 me3
H3K4 me3:H4–4ac
H4–4ac:H3–5ac
H3K9 me2
H3K27 me2
H4K20 me3
H3K9 me3
H2A.Z
H3K4 me1
DNA Meth. m5C
H3K4 me3
H4K20 me2
H3–5ac
H4–4ac
H3K27 me3
0.00
0.25
0.50
0.75
1.00
π̂
RCOR2 – Forward
H3K9 me3:H4K20 me3
H3K27 me2:DNA Meth. m5C
H2A.Z:H4–4ac
H3K9 me2:DNA Meth. m5C
H2A.Z:H3–5ac
H3K27 me3:H4K20 me3
H3K27 me3:H3K9 me3
H3K4 me1:H4K20 me2
H2A.Z:H4K20 me2
H3K4 me1:H3–5ac
DNA Meth. m5C:H4K20 me2
DNA Meth. m5C:H4K20 me3
H2A.Z:H3K4 me3
H3K4 me1:H2A.Z
H3K27 me3:DNA Meth. m5C
H3K4 me1:H4–4ac
H3K4 me3:H4–4ac
H4K20 me2:H3K4 me3
H4K20 me2:H3–5ac
H4–4ac:H3–5ac
H4K20 me2:H4–4ac
DNA Meth. m5C:H3K9 me3
H3K27 me2
H3K9 me2
H3K27 me3
H3K9 me3
H2A.Z
H4K20 me3
H3K4 me3
H3–5ac
H4–4ac
H3K4 me1
H4K20 me2
DNA Meth. m5C
0.00
0.25
0.50
0.75
1.00
π̂
RCOR2 – Reverse
H3K9 me3:H4K20 me3
H3K27 me2:DNA Meth. m5C
H2A.Z:H4–4ac
H3K9 me2:DNA Meth. m5C
H2A.Z:H3–5ac
H3K27 me3:H4K20 me3
H3K27 me3:H3K9 me3
H3K4 me1:H4K20 me2
H2A.Z:H4K20 me2
H3K4 me1:H3–5ac
DNA Meth. m5C:H4K20 me2
DNA Meth. m5C:H4K20 me3
H2A.Z:H3K4 me3
H3K4 me1:H2A.Z
H3K27 me3:DNA Meth. m5C
H3K4 me1:H4–4ac
H3K4 me3:H4–4ac
H4K20 me2:H3K4 me3
H4K20 me2:H3–5ac
H4–4ac:H3–5ac
H4K20 me2:H4–4ac
DNA Meth. m5C:H3K9 me3
H3K27 me2
H3K9 me2
H3K27 me3
H3K9 me3
H2A.Z
H4K20 me3
H3K4 me3
H3–5ac
H4–4ac
H3K4 me1
H4K20 me2
DNA Meth. m5C
0.00
0.25
0.50
0.75
1.00
π̂
PREB – Forward
H3K4 me1:H3–5ac
H3K4 me1:H2A.Z
H3K4 me3:H3–5ac
H3K4 me1:H4–4ac
H3K9 me3:H4K20 me3
H3K27 me3:H4K20 me3
H3K4 me1:H4K20 me2
H3K27 me3:H3K4 me1
H2A.Z:H3K4 me3
H2A.Z:H4K20 me2
H3K27 me2:DNA Meth. m5C
H3K9 me2:DNA Meth. m5C
DNA Meth. m5C:H4K20 me3
DNA Meth. m5C:H4K20 me2
H3K4 me3:H4–4ac
H4K20 me2:H3–5ac
H3K27 me3:H3K9 me3
DNA Meth. m5C:H3K9 me3
H3K27 me3:DNA Meth. m5C
H4K20 me2:H4–4ac
H4K20 me2:H3K4 me3
H3K27 me2
H3K9 me2
H4K20 me3
H2A.Z
H3K9 me3
H3–5ac
H3K27 me3
H3K4 me1
H3K4 me3
H4K20 me2
H4–4ac
DNA Meth. m5C
0.00
0.25
0.50
0.75
1.00
π̂
PREB – Reverse
H3K4 me1:H3–5ac
H3K4 me1:H2A.Z
H3K4 me3:H3–5ac
H3K4 me1:H4–4ac
H3K9 me3:H4K20 me3
H3K27 me3:H4K20 me3
H3K4 me1:H4K20 me2
H3K27 me3:H3K4 me1
H2A.Z:H3K4 me3
H2A.Z:H4K20 me2
H3K27 me2:DNA Meth. m5C
H3K9 me2:DNA Meth. m5C
DNA Meth. m5C:H4K20 me3
DNA Meth. m5C:H4K20 me2
H3K4 me3:H4–4ac
H4K20 me2:H3–5ac
H3K27 me3:H3K9 me3
DNA Meth. m5C:H3K9 me3
H3K27 me3:DNA Meth. m5C
H4K20 me2:H4–4ac
H4K20 me2:H3K4 me3
H3K27 me2
H3K9 me2
H4K20 me3
H2A.Z
H3K9 me3
H3–5ac
H3K27 me3
H3K4 me1
H3K4 me3
H4K20 me2
H4–4ac
DNA Meth. m5C
0.00
0.25
0.50
0.75
1.00
π̂
VAPA – Forward
H3K27 me2:DNA Meth. m5C
DNA Meth. m5C:H3K9 me3
H2A.Z:H4K20 me2
H3K9 me2:DNA Meth. m5C
H3K4 me1:H4K20 me2
DNA Meth. m5C:H4K20 me2
H2A.Z:H4–4ac
H3K27 me3:DNA Meth. m5C
H2A.Z:H3–5ac
H2A.Z:H3K4 me3
H3K4 me1:H4–4ac
H3K4 me1:H2A.Z
DNA Meth. m5C:H4K20 me3
H3K27 me3:H3K9 me3
H4K20 me2:H4–4ac
H4K20 me2:H3–5ac
H3K4 me3:H4–4ac
H4K20 me2:H3K4 me3
H4–4ac:H3–5ac
H3K9 me2
H4K20 me3
H3K27 me2
H3K4 me1
H3K27 me3
H3K9 me3
H3–5ac
H4K20 me2
H3K4 me3
H4–4ac
H2A.Z
DNA Meth. m5C
0.00
0.25
0.50
0.75
1.00
π̂
VAPA – Reverse
H3K27 me2:DNA Meth. m5C
DNA Meth. m5C:H3K9 me3
H2A.Z:H4K20 me2
H3K9 me2:DNA Meth. m5C
H3K4 me1:H4K20 me2
DNA Meth. m5C:H4K20 me2
H2A.Z:H4–4ac
H3K27 me3:DNA Meth. m5C
H2A.Z:H3–5ac
H2A.Z:H3K4 me3
H3K4 me1:H4–4ac
H3K4 me1:H2A.Z
DNA Meth. m5C:H4K20 me3
H3K27 me3:H3K9 me3
H4K20 me2:H4–4ac
H4K20 me2:H3–5ac
H3K4 me3:H4–4ac
H4K20 me2:H3K4 me3
H4–4ac:H3–5ac
H3K9 me2
H4K20 me3
H3K27 me2
H3K4 me1
H3K27 me3
H3K9 me3
H3–5ac
H4K20 me2
H3K4 me3
H4–4ac
H2A.Z
DNA Meth. m5C
0.00
0.25
0.50
0.75
1.00
π̂
MOGS – Forward
H3K27 me3:H3K9 me3
H3K27 me2:DNA Meth. m5C
H4–4ac:H3–5ac
H2A.Z:H3–5ac
H2A.Z:H4–4ac
H3K4 me1:H4–4ac
H3K4 me1:H4K20 me2
H2A.Z:H3K4 me3
H3K9 me2:DNA Meth. m5C
H3K4 me1:H2A.Z
DNA Meth. m5C:H3K9 me3
DNA Meth. m5C:H4K20 me2
H2A.Z:H4K20 me2
H4K20 me2:H4–4ac
H3K4 me3:H4–4ac
H4K20 me2:H3–5ac
H4K20 me3
H3K27 me3
H4K20 me2:H3K4 me3
H3K9 me2
H3K27 me2
H3K9 me3
H3K4 me1
H4–4ac
H3–5ac
H3K4 me3
H4K20 me2
DNA Meth. m5C
H2A.Z
0.00
0.25
0.50
0.75
1.00
π̂
MOGS – Reverse
H3K27 me3:H3K9 me3
H3K27 me2:DNA Meth. m5C
H4–4ac:H3–5ac
H2A.Z:H3–5ac
H2A.Z:H4–4ac
H3K4 me1:H4–4ac
H3K4 me1:H4K20 me2
H2A.Z:H3K4 me3
H3K9 me2:DNA Meth. m5C
H3K4 me1:H2A.Z
DNA Meth. m5C:H3K9 me3
DNA Meth. m5C:H4K20 me2
H2A.Z:H4K20 me2
H4K20 me2:H4–4ac
H3K4 me3:H4–4ac
H4K20 me2:H3–5ac
H4K20 me3
H3K27 me3
H4K20 me2:H3K4 me3
H3K9 me2
H3K27 me2
H3K9 me3
H3K4 me1
H4–4ac
H3–5ac
H3K4 me3
H4K20 me2
DNA Meth. m5C
H2A.Z
0.00
0.25
0.50
0.75
1.00
π̂

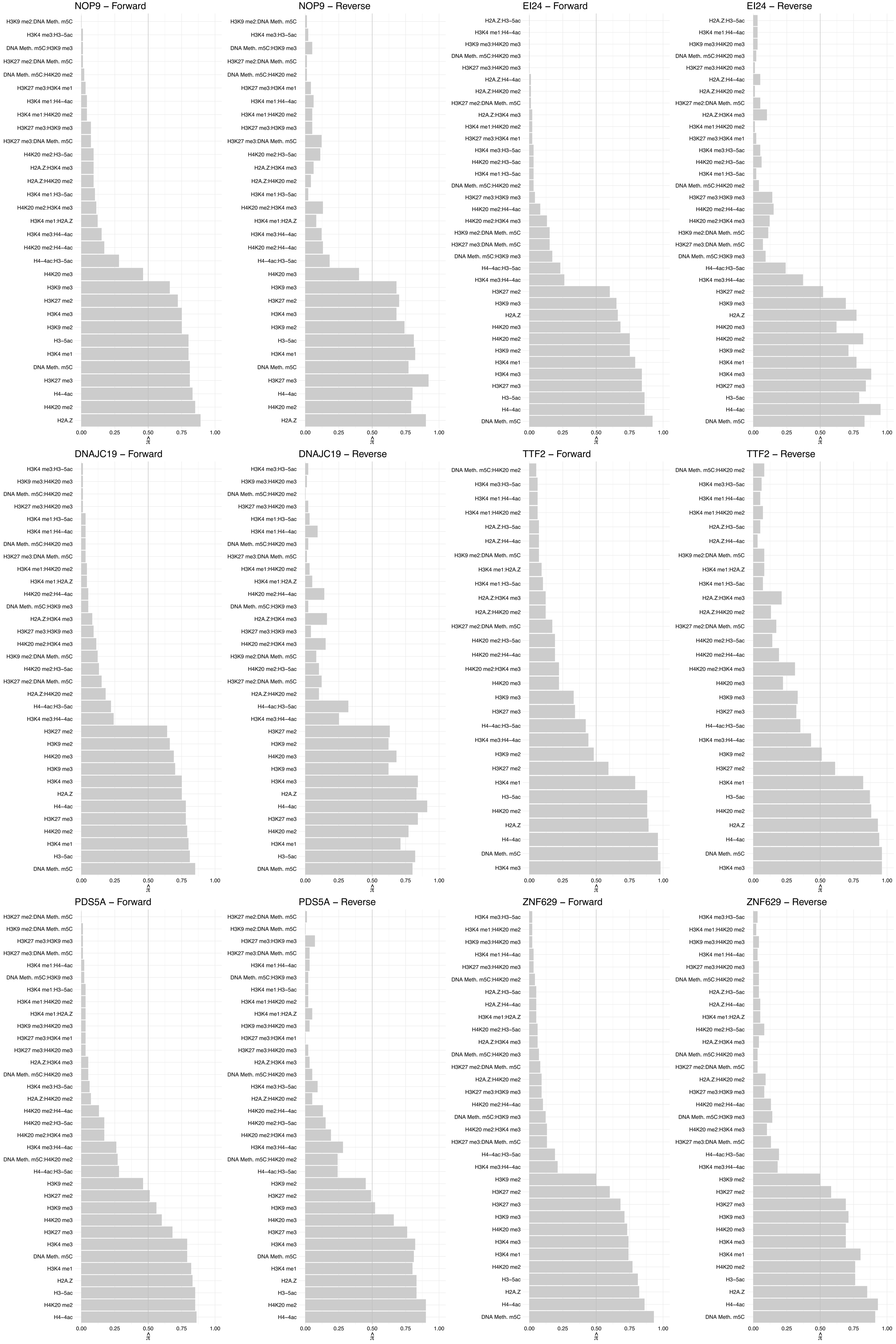
NOP9 – Forward
NOP9 – Reverse
EI24 – Forward
EI24 – Reverse
DNAJC19 – Forward
DNAJC19 – Reverse
TTF2 – Forward
TTF2 – Reverse
PDS5A – Forward
PDS5A – Reverse
ZNF629 – Forward
ZNF629 – Reverse
0.00 0.25 0.50 0.75 1.00
$\hat{\pi}$

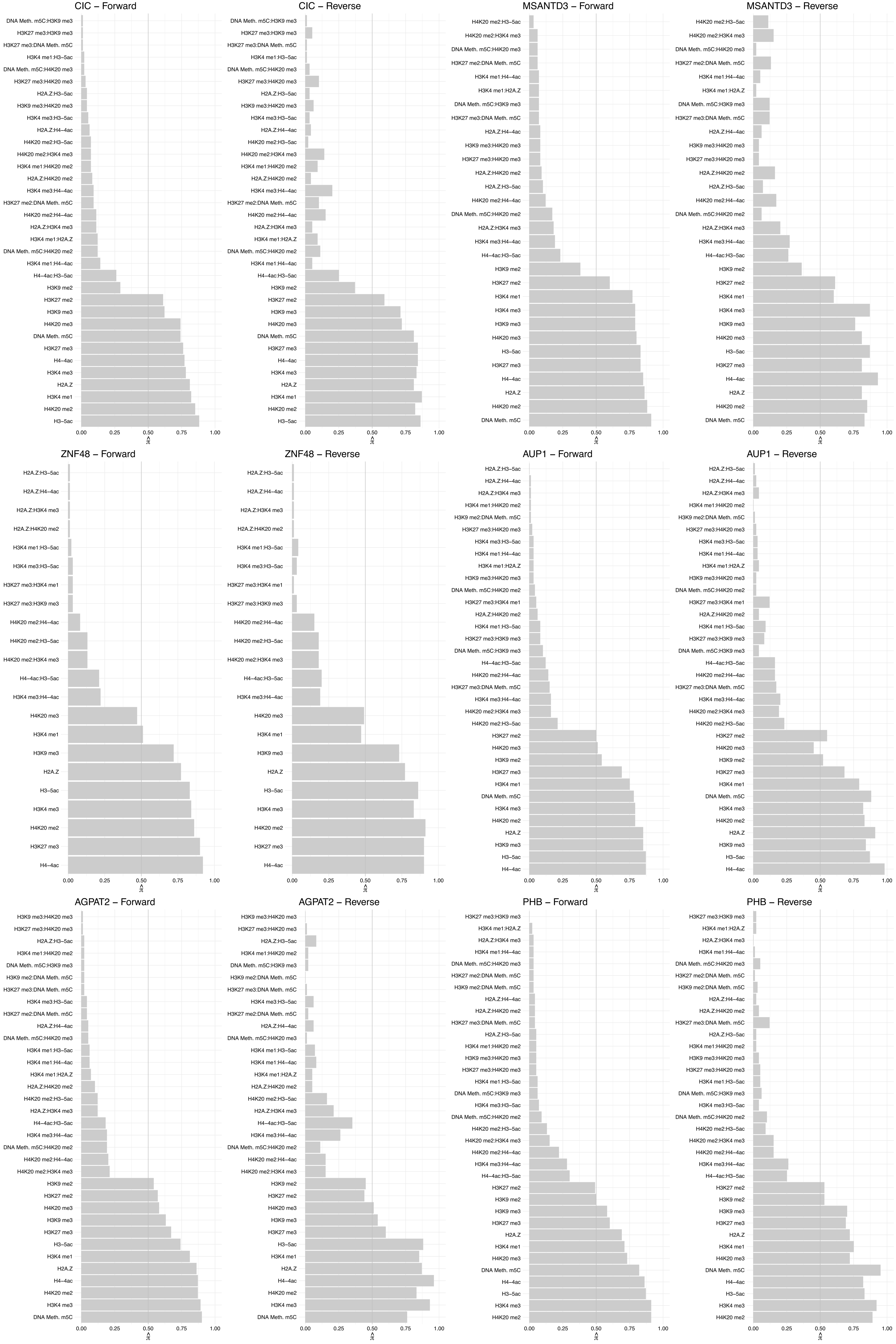
CIC – Forward
DNA Meth. m5C:H3K9 me3
H3K27 me3:H3K9 me3
H3K27 me3:DNA Meth. m5C
H3K4 me1:H3−5ac
DNA Meth. m5C:H4K20 me3
H3K27 me3:H4K20 me3
H2A.Z:H3−5ac
H3K9 me3:H4K20 me3
H3K4 me3:H3−5ac
H2A.Z:H4−4ac
H4K20 me2:H3−5ac
H4K20 me2:H3K4 me3
H3K4 me1:H4K20 me2
H2A.Z:H4K20 me2
H3K4 me3:H4−4ac
H3K27 me2:DNA Meth. m5C
H4K20 me2:H4−4ac
H2A.Z:H3K4 me3
H3K4 me1:H2A.Z
DNA Meth. m5C:H4K20 me2
H3K4 me1:H4−4ac
H4−4ac:H3−5ac
H3K9 me2
H3K27 me2
H3K9 me3
H4K20 me3
DNA Meth. m5C
H3K27 me3
H4−4ac
H3K4 me3
H2A.Z
H3K4 me1
H4K20 me2
H3−5ac
0.00 0.25 0.50 0.75 1.00
π̂
CIC – Reverse
DNA Meth. m5C:H3K9 me3
H3K27 me3:H3K9 me3
H3K27 me3:DNA Meth. m5C
H3K4 me1:H3−5ac
DNA Meth. m5C:H4K20 me3
H3K27 me3:H4K20 me3
H2A.Z:H3−5ac
H3K9 me3:H4K20 me3
H3K4 me3:H3−5ac
H2A.Z:H4−4ac
H4K20 me2:H3−5ac
H4K20 me2:H3K4 me3
H3K4 me1:H4K20 me2
H2A.Z:H4K20 me2
H3K4 me3:H4−4ac
H3K27 me2:DNA Meth. m5C
H4K20 me2:H4−4ac
H2A.Z:H3K4 me3
H3K4 me1:H2A.Z
DNA Meth. m5C:H4K20 me2
H3K4 me1:H4−4ac
H4−4ac:H3−5ac
H3K9 me2
H3K27 me2
H3K9 me3
H4K20 me3
DNA Meth. m5C
H3K27 me3
H4−4ac
H3K4 me3
H2A.Z
H3K4 me1
H4K20 me2
H3−5ac
0.00 0.25 0.50 0.75 1.00
π̂
MSANTD3 – Forward
H4K20 me2:H3−5ac
H4K20 me2:H3K4 me3
DNA Meth. m5C:H4K20 me3
H3K27 me2:DNA Meth. m5C
H3K4 me1:H4−4ac
H3K4 me1:H2A.Z
DNA Meth. m5C:H3K9 me3
H3K27 me3:DNA Meth. m5C
H2A.Z:H4−4ac
H3K9 me3:H4K20 me3
H3K27 me3:H4K20 me3
H2A.Z:H4K20 me2
H2A.Z:H3−5ac
H4K20 me2:H4−4ac
DNA Meth. m5C:H4K20 me2
H2A.Z:H3K4 me3
H3K4 me3:H4−4ac
H4−4ac:H3−5ac
H3K9 me2
H3K27 me2
H3K4 me1
H3K4 me3
H3K9 me3
H4K20 me3
H3−5ac
H3K27 me3
H4−4ac
H2A.Z
H4K20 me2
DNA Meth. m5C
0.00 0.25 0.50 0.75 1.00
π̂
MSANTD3 – Reverse
H4K20 me2:H3−5ac
H4K20 me2:H3K4 me3
DNA Meth. m5C:H4K20 me3
H3K27 me2:DNA Meth. m5C
H3K4 me1:H4−4ac
H3K4 me1:H2A.Z
DNA Meth. m5C:H3K9 me3
H3K27 me3:DNA Meth. m5C
H2A.Z:H4−4ac
H3K9 me3:H4K20 me3
H3K27 me3:H4K20 me3
H2A.Z:H4K20 me2
H2A.Z:H3−5ac
H4K20 me2:H4−4ac
DNA Meth. m5C:H4K20 me2
H2A.Z:H3K4 me3
H3K4 me3:H4−4ac
H4−4ac:H3−5ac
H3K9 me2
H3K27 me2
H3K4 me1
H3K4 me3
H3K9 me3
H4K20 me3
H3−5ac
H3K27 me3
H4−4ac
H2A.Z
H4K20 me2
DNA Meth. m5C
0.00 0.25 0.50 0.75 1.00
π̂
ZNF48 – Forward
H2A.Z:H3−5ac
H2A.Z:H4−4ac
H2A.Z:H3K4 me3
H2A.Z:H4K20 me2
H3K4 me1:H3−5ac
H3K4 me3:H3−5ac
H3K27 me3:H3K4 me1
H3K27 me3:H3K9 me3
H4K20 me2:H4−4ac
H4K20 me2:H3−5ac
H4K20 me2:H3K4 me3
H4−4ac:H3−5ac
H3K4 me3:H4−4ac
H4K20 me3
H3K4 me1
H3K9 me3
H2A.Z
H3−5ac
H3K4 me3
H4K20 me2
H3K27 me3
H4−4ac
0.00 0.25 0.50 0.75 1.00
π̂
ZNF48 – Reverse
H2A.Z:H3−5ac
H2A.Z:H4−4ac
H2A.Z:H3K4 me3
H2A.Z:H4K20 me2
H3K4 me1:H3−5ac
H3K4 me3:H3−5ac
H3K27 me3:H3K4 me1
H3K27 me3:H3K9 me3
H4K20 me2:H4−4ac
H4K20 me2:H3−5ac
H4K20 me2:H3K4 me3
H4−4ac:H3−5ac
H3K4 me3:H4−4ac
H4K20 me3
H3K4 me1
H3K9 me3
H2A.Z
H3−5ac
H3K4 me3
H4K20 me2
H3K27 me3
H4−4ac
0.00 0.25 0.50 0.75 1.00
π̂
AUP1 – Forward
H2A.Z:H3−5ac
H2A.Z:H4−4ac
H2A.Z:H3K4 me3
H3K4 me1:H4K20 me2
H3K9 me2:DNA Meth. m5C
H3K27 me3:H4K20 me3
H3K4 me3:H3−5ac
H3K4 me1:H4−4ac
H3K4 me1:H2A.Z
H3K9 me3:H4K20 me3
DNA Meth. m5C:H4K20 me2
H3K27 me3:H3K4 me1
H2A.Z:H4K20 me2
H3K4 me1:H3−5ac
H3K27 me3:H3K9 me3
DNA Meth. m5C:H3K9 me3
H4−4ac:H3−5ac
H4K20 me2:H4−4ac
H3K27 me3:DNA Meth. m5C
H3K4 me3:H4−4ac
H4K20 me2:H3K4 me3
H4K20 me2:H3−5ac
H3K27 me2
H4K20 me3
H3K9 me2
H3K27 me3
H3K4 me1
DNA Meth. m5C
H3K4 me3
H4K20 me2
H2A.Z
H3K9 me3
H3−5ac
H4−4ac
0.00 0.25 0.50 0.75 1.00
π̂
AUP1 – Reverse
H2A.Z:H3−5ac
H2A.Z:H4−4ac
H2A.Z:H3K4 me3
H3K4 me1:H4K20 me2
H3K9 me2:DNA Meth. m5C
H3K27 me3:H4K20 me3
H3K4 me3:H3−5ac
H3K4 me1:H4−4ac
H3K4 me1:H2A.Z
H3K9 me3:H4K20 me3
DNA Meth. m5C:H4K20 me2
H3K27 me3:H3K4 me1
H2A.Z:H4K20 me2
H3K4 me1:H3−5ac
H3K27 me3:H3K9 me3
DNA Meth. m5C:H3K9 me3
H4−4ac:H3−5ac
H4K20 me2:H4−4ac
H3K27 me3:DNA Meth. m5C
H3K4 me3:H4−4ac
H4K20 me2:H3K4 me3
H4K20 me2:H3−5ac
H3K27 me2
H4K20 me3
H3K9 me2
H3K27 me3
H3K4 me1
DNA Meth. m5C
H3K4 me3
H4K20 me2
H2A.Z
H3K9 me3
H3−5ac
H4−4ac
0.00 0.25 0.50 0.75 1.00
π̂
AGPAT2 – Forward
H3K9 me3:H4K20 me3
H3K27 me3:H4K20 me3
H2A.Z:H3−5ac
H3K4 me1:H4K20 me2
DNA Meth. m5C:H3K9 me3
H3K9 me2:DNA Meth. m5C
H3K27 me3:DNA Meth. m5C
H3K4 me3:H3−5ac
H3K27 me2:DNA Meth. m5C
H2A.Z:H4−4ac
DNA Meth. m5C:H4K20 me3
H3K4 me1:H3−5ac
H3K4 me1:H4−4ac
H3K4 me1:H2A.Z
H2A.Z:H4K20 me2
H4K20 me2:H3−5ac
H2A.Z:H3K4 me3
H4−4ac:H3−5ac
H3K4 me3:H4−4ac
DNA Meth. m5C:H4K20 me2
H4K20 me2:H4−4ac
H4K20 me2:H3K4 me3
H3K9 me2
H3K27 me2
H4K20 me3
H3K9 me3
H3K27 me3
H3−5ac
H3K4 me1
H2A.Z
H4−4ac
H4K20 me2
H3K4 me3
DNA Meth. m5C
0.00 0.25 0.50 0.75 1.00
π̂
AGPAT2 – Reverse
H3K9 me3:H4K20 me3
H3K27 me3:H4K20 me3
H2A.Z:H3−5ac
H3K4 me1:H4K20 me2
DNA Meth. m5C:H3K9 me3
H3K9 me2:DNA Meth. m5C
H3K27 me3:DNA Meth. m5C
H3K4 me3:H3−5ac
H3K27 me2:DNA Meth. m5C
H2A.Z:H4−4ac
DNA Meth. m5C:H4K20 me3
H3K4 me1:H3−5ac
H3K4 me1:H4−4ac
H3K4 me1:H2A.Z
H2A.Z:H4K20 me2
H4K20 me2:H3−5ac
H2A.Z:H3K4 me3
H4−4ac:H3−5ac
H3K4 me3:H4−4ac
DNA Meth. m5C:H4K20 me2
H4K20 me2:H4−4ac
H4K20 me2:H3K4 me3
H3K9 me2
H3K27 me2
H4K20 me3
H3K9 me3
H3K27 me3
H3−5ac
H3K4 me1
H2A.Z
H4−4ac
H4K20 me2
H3K4 me3
DNA Meth. m5C
0.00 0.25 0.50 0.75 1.00
π̂
PHB – Forward
H3K27 me3:H3K9 me3
H3K4 me1:H2A.Z
H2A.Z:H3K4 me3
H3K4 me1:H4−4ac
DNA Meth. m5C:H4K20 me3
H3K27 me2:DNA Meth. m5C
H3K9 me2:DNA Meth. m5C
H2A.Z:H4−4ac
H2A.Z:H4K20 me2
H3K27 me3:DNA Meth. m5C
H2A.Z:H3−5ac
H3K4 me1:H4K20 me2
H3K9 me3:H4K20 me3
H3K27 me3:H4K20 me3
H3K4 me1:H3−5ac
DNA Meth. m5C:H3K9 me3
H3K4 me3:H3−5ac
DNA Meth. m5C:H4K20 me2
H4K20 me2:H3−5ac
H4K20 me2:H3K4 me3
H4K20 me2:H4−4ac
H3K4 me3:H4−4ac
H4−4ac:H3−5ac
H3K27 me2
H3K9 me2
H3K9 me3
H3K27 me3
H2A.Z
H3K4 me1
H4K20 me3
DNA Meth. m5C
H4−4ac
H3−5ac
H3K4 me3
H4K20 me2
0.00 0.25 0.50 0.75 1.00
π̂
PHB – Reverse
H3K27 me3:H3K9 me3
H3K4 me1:H2A.Z
H2A.Z:H3K4 me3
H3K4 me1:H4−4ac
DNA Meth. m5C:H4K20 me3
H3K27 me2:DNA Meth. m5C
H3K9 me2:DNA Meth. m5C
H2A.Z:H4−4ac
H2A.Z:H4K20 me2
H3K27 me3:DNA Meth. m5C
H2A.Z:H3−5ac
H3K4 me1:H4K20 me2
H3K9 me3:H4K20 me3
H3K27 me3:H4K20 me3
H3K4 me1:H3−5ac
DNA Meth. m5C:H3K9 me3
H3K4 me3:H3−5ac
DNA Meth. m5C:H4K20 me2
H4K20 me2:H3−5ac
H4K20 me2:H3K4 me3
H4K20 me2:H4−4ac
H3K4 me3:H4−4ac
H4−4ac:H3−5ac
H3K27 me2
H3K9 me2
H3K9 me3
H3K27 me3
H2A.Z
H3K4 me1
H4K20 me3
DNA Meth. m5C
H4−4ac
H3−5ac
H3K4 me3
H4K20 me2
0.00 0.25 0.50 0.75 1.00
π̂

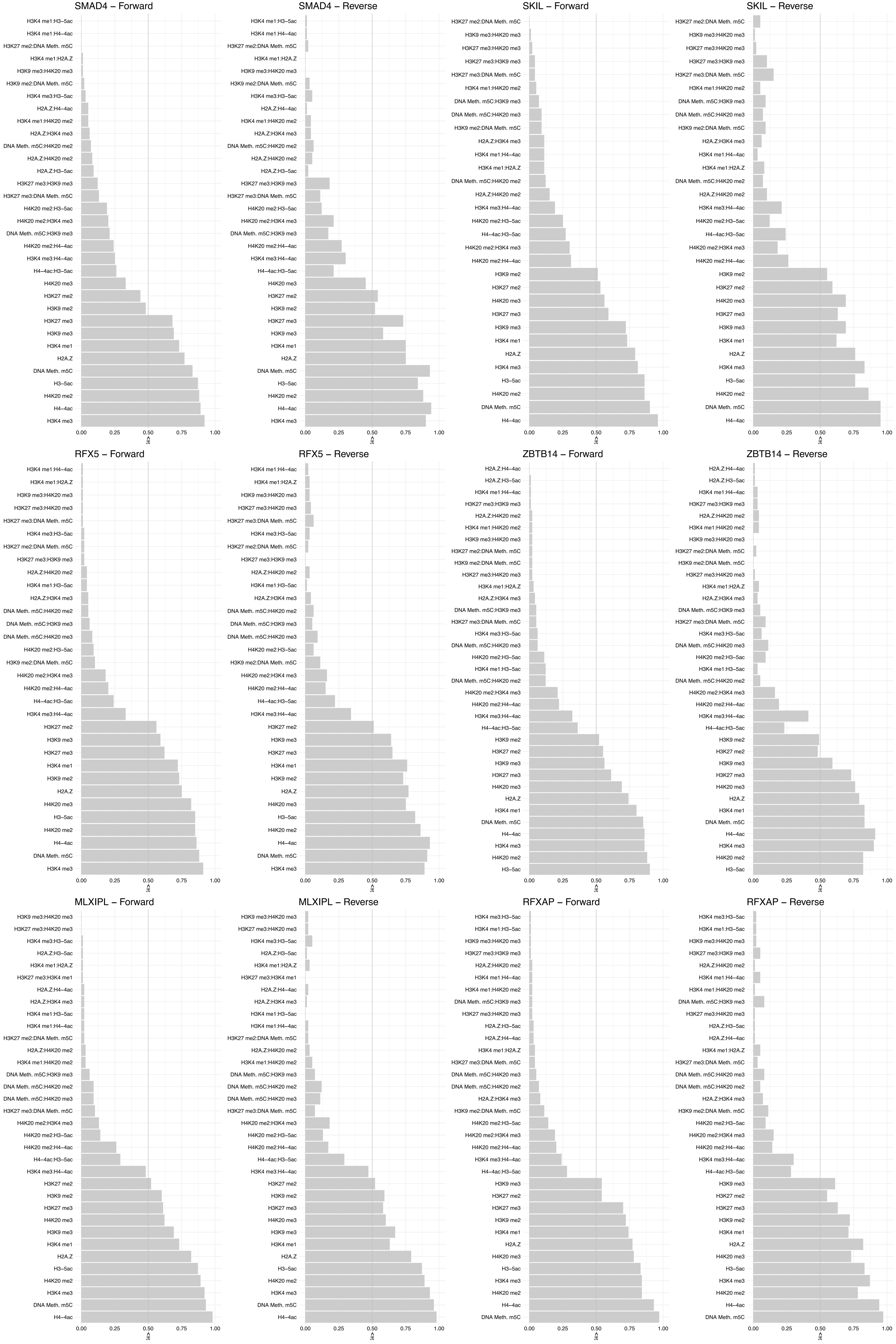
SMAD4 – Forward
H3K4 me1:H3-5ac
H3K4 me1:H4-4ac
H3K27 me2:DNA Meth. m5C
H3K4 me1:H2A.Z
H3K9 me3:H4K20 me3
H3K9 me2:DNA Meth. m5C
H3K4 me3:H3-5ac
H2A.Z:H4-4ac
H3K4 me1:H4K20 me2
H2A.Z:H3K4 me3
DNA Meth. m5C:H4K20 me2
H2A.Z:H4K20 me2
H2A.Z:H3-5ac
H3K27 me3:H3K9 me3
H3K27 me3:DNA Meth. m5C
H4K20 me2:H3-5ac
H4K20 me2:H3K4 me3
DNA Meth. m5C:H3K9 me3
H4K20 me2:H4-4ac
H3K4 me3:H4-4ac
H4-4ac:H3-5ac
H4K20 me3
H3K27 me2
H3K9 me2
H3K27 me3
H3K9 me3
H3K4 me1
H2A.Z
DNA Meth. m5C
H3-5ac
H4K20 me2
H4-4ac
H3K4 me3
0.00 0.25 0.50 0.75 1.00
$\hat{\pi}$
SMAD4 – Reverse
H3K4 me1:H3-5ac
H3K4 me1:H4-4ac
H3K27 me2:DNA Meth. m5C
H3K4 me1:H2A.Z
H3K9 me3:H4K20 me3
H3K9 me2:DNA Meth. m5C
H3K4 me3:H3-5ac
H2A.Z:H4-4ac
H3K4 me1:H4K20 me2
H2A.Z:H3K4 me3
DNA Meth. m5C:H4K20 me2
H2A.Z:H4K20 me2
H2A.Z:H3-5ac
H3K27 me3:H3K9 me3
H3K27 me3:DNA Meth. m5C
H4K20 me2:H3-5ac
H4K20 me2:H3K4 me3
DNA Meth. m5C:H3K9 me3
H4K20 me2:H4-4ac
H3K4 me3:H4-4ac
H4-4ac:H3-5ac
H4K20 me3
H3K27 me2
H3K9 me2
H3K27 me3
H3K9 me3
H3K4 me1
H2A.Z
DNA Meth. m5C
H3-5ac
H4K20 me2
H4-4ac
H3K4 me3
0.00 0.25 0.50 0.75 1.00
$\hat{\pi}$
SKIL – Forward
H3K27 me2:DNA Meth. m5C
H3K9 me3:H4K20 me3
H3K27 me3:H4K20 me3
H3K27 me3:H3K9 me3
H3K27 me3:DNA Meth. m5C
H3K4 me1:H4K20 me2
DNA Meth. m5C:H3K9 me3
DNA Meth. m5C:H4K20 me3
H3K9 me2:DNA Meth. m5C
H2A.Z:H3K4 me3
H3K4 me1:H4-4ac
H3K4 me1:H2A.Z
DNA Meth. m5C:H4K20 me2
H2A.Z:H4K20 me2
H3K4 me3:H4-4ac
H4K20 me2:H3-5ac
H4-4ac:H3-5ac
H4K20 me2:H3K4 me3
H4K20 me2:H4-4ac
H3K9 me2
H3K27 me2
H4K20 me3
H3K27 me3
H3K9 me3
H3K4 me1
H2A.Z
H3K4 me3
H3-5ac
H4K20 me2
DNA Meth. m5C
H4-4ac
0.00 0.25 0.50 0.75 1.00
$\hat{\pi}$
SKIL – Reverse
H3K27 me2:DNA Meth. m5C
H3K9 me3:H4K20 me3
H3K27 me3:H4K20 me3
H3K27 me3:H3K9 me3
H3K27 me3:DNA Meth. m5C
H3K4 me1:H4K20 me2
DNA Meth. m5C:H3K9 me3
DNA Meth. m5C:H4K20 me3
H3K9 me2:DNA Meth. m5C
H2A.Z:H3K4 me3
H3K4 me1:H4-4ac
H3K4 me1:H2A.Z
DNA Meth. m5C:H4K20 me2
H2A.Z:H4K20 me2
H3K4 me3:H4-4ac
H4K20 me2:H3-5ac
H4-4ac:H3-5ac
H4K20 me2:H3K4 me3
H4K20 me2:H4-4ac
H3K9 me2
H3K27 me2
H4K20 me3
H3K27 me3
H3K9 me3
H3K4 me1
H2A.Z
H3K4 me3
H3-5ac
H4K20 me2
DNA Meth. m5C
H4-4ac
0.00 0.25 0.50 0.75 1.00
$\hat{\pi}$
RFX5 – Forward
H3K4 me1:H4-4ac
H3K4 me1:H2A.Z
H3K9 me3:H4K20 me3
H3K27 me3:H4K20 me3
H3K27 me3:DNA Meth. m5C
H3K4 me3:H3-5ac
H3K27 me2:DNA Meth. m5C
H3K27 me3:H3K9 me3
H2A.Z:H4K20 me2
H3K4 me1:H3-5ac
H2A.Z:H3K4 me3
DNA Meth. m5C:H4K20 me2
DNA Meth. m5C:H3K9 me3
DNA Meth. m5C:H4K20 me3
H4K20 me2:H3-5ac
H3K9 me2:DNA Meth. m5C
H4K20 me2:H3K4 me3
H4K20 me2:H4-4ac
H4-4ac:H3-5ac
H3K4 me3:H4-4ac
H3K27 me2
H3K9 me3
H3K27 me3
H3K4 me1
H3K9 me2
H2A.Z
H4K20 me3
H3-5ac
H4K20 me2
H4-4ac
DNA Meth. m5C
H3K4 me3
0.00 0.25 0.50 0.75 1.00
$\hat{\pi}$
RFX5 – Reverse
H3K4 me1:H4-4ac
H3K4 me1:H2A.Z
H3K9 me3:H4K20 me3
H3K27 me3:H4K20 me3
H3K27 me3:DNA Meth. m5C
H3K4 me3:H3-5ac
H3K27 me2:DNA Meth. m5C
H3K27 me3:H3K9 me3
H2A.Z:H4K20 me2
H3K4 me1:H3-5ac
H2A.Z:H3K4 me3
DNA Meth. m5C:H4K20 me2
DNA Meth. m5C:H3K9 me3
DNA Meth. m5C:H4K20 me3
H4K20 me2:H3-5ac
H3K9 me2:DNA Meth. m5C
H4K20 me2:H3K4 me3
H4K20 me2:H4-4ac
H4-4ac:H3-5ac
H3K4 me3:H4-4ac
H3K27 me2
H3K9 me3
H3K27 me3
H3K4 me1
H3K9 me2
H2A.Z
H4K20 me3
H3-5ac
H4K20 me2
H4-4ac
DNA Meth. m5C
H3K4 me3
0.00 0.25 0.50 0.75 1.00
$\hat{\pi}$
ZBTB14 – Forward
H2A.Z:H4-4ac
H2A.Z:H3-5ac
H3K4 me1:H4-4ac
H3K27 me3:H3K9 me3
H2A.Z:H4K20 me2
H3K4 me1:H4K20 me2
H3K9 me3:H4K20 me3
H3K27 me2:DNA Meth. m5C
H3K9 me2:DNA Meth. m5C
H3K27 me3:H4K20 me3
H3K4 me1:H2A.Z
H2A.Z:H3K4 me3
DNA Meth. m5C:H3K9 me3
H3K27 me3:DNA Meth. m5C
H3K4 me3:H3-5ac
DNA Meth. m5C:H4K20 me3
H4K20 me2:H3-5ac
H3K4 me1:H3-5ac
DNA Meth. m5C:H4K20 me2
H4K20 me2:H3K4 me3
H4K20 me2:H4-4ac
H3K4 me3:H4-4ac
H4-4ac:H3-5ac
H3K9 me2
H3K27 me2
H3K9 me3
H3K27 me3
H4K20 me3
H2A.Z
H3K4 me1
DNA Meth. m5C
H4-4ac
H3K4 me3
H4K20 me2
H3-5ac
0.00 0.25 0.50 0.75 1.00
$\hat{\pi}$
ZBTB14 – Reverse
H2A.Z:H4-4ac
H2A.Z:H3-5ac
H3K4 me1:H4-4ac
H3K27 me3:H3K9 me3
H2A.Z:H4K20 me2
H3K4 me1:H4K20 me2
H3K9 me3:H4K20 me3
H3K27 me2:DNA Meth. m5C
H3K9 me2:DNA Meth. m5C
H3K27 me3:H4K20 me3
H3K4 me1:H2A.Z
H2A.Z:H3K4 me3
DNA Meth. m5C:H3K9 me3
H3K27 me3:DNA Meth. m5C
H3K4 me3:H3-5ac
DNA Meth. m5C:H4K20 me3
H4K20 me2:H3-5ac
H3K4 me1:H3-5ac
DNA Meth. m5C:H4K20 me2
H4K20 me2:H3K4 me3
H4K20 me2:H4-4ac
H3K4 me3:H4-4ac
H4-4ac:H3-5ac
H3K9 me2
H3K27 me2
H3K9 me3
H3K27 me3
H4K20 me3
H2A.Z
H3K4 me1
DNA Meth. m5C
H4-4ac
H3K4 me3
H4K20 me2
H3-5ac
0.00 0.25 0.50 0.75 1.00
$\hat{\pi}$
MLXIPL – Forward
H3K9 me3:H4K20 me3
H3K27 me3:H4K20 me3
H3K4 me3:H3-5ac
H2A.Z:H3-5ac
H3K4 me1:H2A.Z
H3K27 me3:H3K4 me1
H2A.Z:H4-4ac
H2A.Z:H3K4 me3
H3K4 me1:H3-5ac
H3K4 me1:H4-4ac
H3K27 me2:DNA Meth. m5C
H2A.Z:H4K20 me2
H3K4 me1:H4K20 me2
DNA Meth. m5C:H3K9 me3
DNA Meth. m5C:H4K20 me2
DNA Meth. m5C:H4K20 me3
H3K27 me3:DNA Meth. m5C
H4K20 me2:H3K4 me3
H4K20 me2:H3-5ac
H4K20 me2:H4-4ac
H4-4ac:H3-5ac
H3K4 me3:H4-4ac
H3K27 me2
H3K9 me2
H3K27 me3
H4K20 me3
H3K9 me3
H3K4 me1
H2A.Z
H3-5ac
H4K20 me2
H3K4 me3
DNA Meth. m5C
H4-4ac
0.00 0.25 0.50 0.75 1.00
$\hat{\pi}$
MLXIPL – Reverse
H3K9 me3:H4K20 me3
H3K27 me3:H4K20 me3
H3K4 me3:H3-5ac
H2A.Z:H3-5ac
H3K4 me1:H2A.Z
H3K27 me3:H3K4 me1
H2A.Z:H4-4ac
H2A.Z:H3K4 me3
H3K4 me1:H3-5ac
H3K4 me1:H4-4ac
H3K27 me2:DNA Meth. m5C
H2A.Z:H4K20 me2
H3K4 me1:H4K20 me2
DNA Meth. m5C:H3K9 me3
DNA Meth. m5C:H4K20 me2
DNA Meth. m5C:H4K20 me3
H3K27 me3:DNA Meth. m5C
H4K20 me2:H3K4 me3
H4K20 me2:H3-5ac
H4K20 me2:H4-4ac
H4-4ac:H3-5ac
H3K4 me3:H4-4ac
H3K27 me2
H3K9 me2
H3K27 me3
H4K20 me3
H3K9 me3
H3K4 me1
H2A.Z
H3-5ac
H4K20 me2
H3K4 me3
DNA Meth. m5C
H4-4ac
0.00 0.25 0.50 0.75 1.00
$\hat{\pi}$
RFXAP – Forward
H3K4 me3:H3-5ac
H3K4 me1:H3-5ac
H3K9 me3:H4K20 me3
H3K27 me3:H3K9 me3
H2A.Z:H4K20 me2
H3K4 me1:H4-4ac
H3K4 me1:H4K20 me2
DNA Meth. m5C:H3K9 me3
H3K27 me3:H4K20 me3
H2A.Z:H3-5ac
H2A.Z:H4-4ac
H3K4 me1:H2A.Z
H3K27 me3:DNA Meth. m5C
DNA Meth. m5C:H4K20 me3
DNA Meth. m5C:H4K20 me2
H2A.Z:H3K4 me3
H3K9 me2:DNA Meth. m5C
H4K20 me2:H3-5ac
H4K20 me2:H3K4 me3
H4K20 me2:H4-4ac
H3K4 me3:H4-4ac
H4-4ac:H3-5ac
H3K9 me3
H3K27 me2
H3K27 me3
H3K9 me2
H3K4 me1
H2A.Z
H4K20 me3
H3-5ac
H3K4 me3
H4K20 me2
H4-4ac
DNA Meth. m5C
0.00 0.25 0.50 0.75 1.00
$\hat{\pi}$
RFXAP – Reverse
H3K4 me3:H3-5ac
H3K4 me1:H3-5ac
H3K9 me3:H4K20 me3
H3K27 me3:H3K9 me3
H2A.Z:H4K20 me2
H3K4 me1:H4-4ac
H3K4 me1:H4K20 me2
DNA Meth. m5C:H3K9 me3
H3K27 me3:H4K20 me3
H2A.Z:H3-5ac
H2A.Z:H4-4ac
H3K4 me1:H2A.Z
H3K27 me3:DNA Meth. m5C
DNA Meth. m5C:H4K20 me3
DNA Meth. m5C:H4K20 me2
H2A.Z:H3K4 me3
H3K9 me2:DNA Meth. m5C
H4K20 me2:H3-5ac
H4K20 me2:H3K4 me3
H4K20 me2:H4-4ac
H3K4 me3:H4-4ac
H4-4ac:H3-5ac
H3K9 me3
H3K27 me2
H3K27 me3
H3K9 me2
H3K4 me1
H2A.Z
H4K20 me3
H3-5ac
H3K4 me3
H4K20 me2
H4-4ac
DNA Meth. m5C
0.00 0.25 0.50 0.75 1.00
$\hat{\pi}$

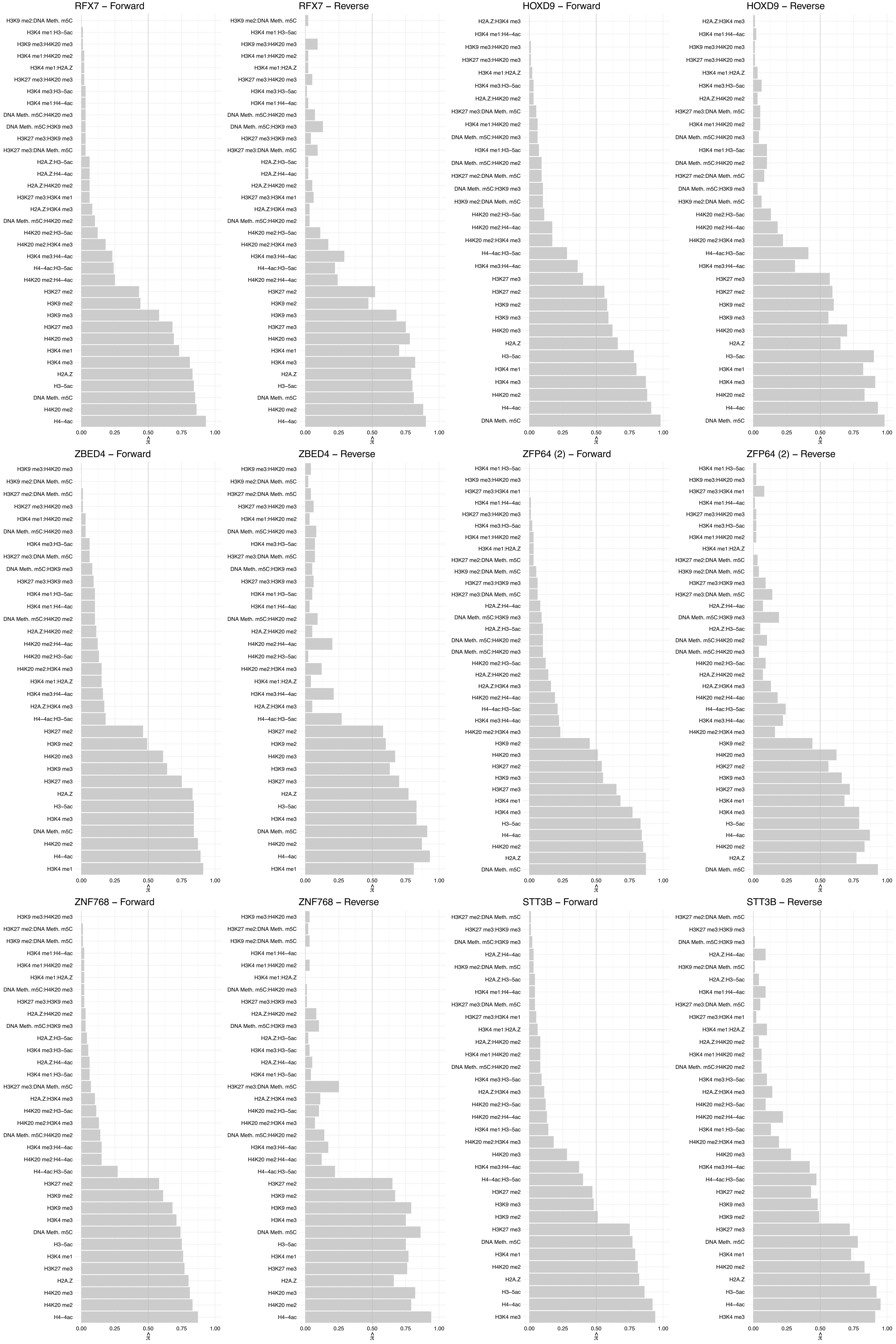
RFX7 – Forward
H3K9 me2:DNA Meth. m5C
H3K4 me1:H3–5ac
H3K9 me3:H4K20 me3
H3K4 me1:H4K20 me2
H3K4 me1:H2A.Z
H3K27 me3:H4K20 me3
H3K4 me3:H3–5ac
H3K4 me1:H4–4ac
DNA Meth. m5C:H4K20 me3
DNA Meth. m5C:H3K9 me3
H3K27 me3:H3K9 me3
H3K27 me3:DNA Meth. m5C
H2A.Z:H3–5ac
H2A.Z:H4–4ac
H2A.Z:H4K20 me2
H3K27 me3:H3K4 me1
H2A.Z:H3K4 me3
DNA Meth. m5C:H4K20 me2
H4K20 me2:H3–5ac
H4K20 me2:H3K4 me3
H3K4 me3:H4–4ac
H4–4ac:H3–5ac
H4K20 me2:H4–4ac
H3K27 me2
H3K9 me2
H3K9 me3
H3K27 me3
H4K20 me3
H3K4 me1
H3K4 me3
H2A.Z
H3–5ac
DNA Meth. m5C
H4K20 me2
H4–4ac
0.00
0.25
0.50
0.75
1.00
π̂
RFX7 – Reverse
H3K9 me2:DNA Meth. m5C
H3K4 me1:H3–5ac
H3K9 me3:H4K20 me3
H3K4 me1:H4K20 me2
H3K4 me1:H2A.Z
H3K27 me3:H4K20 me3
H3K4 me3:H3–5ac
H3K4 me1:H4–4ac
DNA Meth. m5C:H4K20 me3
DNA Meth. m5C:H3K9 me3
H3K27 me3:H3K9 me3
H3K27 me3:DNA Meth. m5C
H2A.Z:H3–5ac
H2A.Z:H4–4ac
H2A.Z:H4K20 me2
H3K27 me3:H3K4 me1
H2A.Z:H3K4 me3
DNA Meth. m5C:H4K20 me2
H4K20 me2:H3–5ac
H4K20 me2:H3K4 me3
H3K4 me3:H4–4ac
H4–4ac:H3–5ac
H4K20 me2:H4–4ac
H3K27 me2
H3K9 me2
H3K9 me3
H3K27 me3
H4K20 me3
H3K4 me1
H3K4 me3
H2A.Z
H3–5ac
DNA Meth. m5C
H4K20 me2
H4–4ac
0.00
0.25
0.50
0.75
1.00
π̂
HOXD9 – Forward
H2A.Z:H3K4 me3
H3K4 me1:H4–4ac
H3K9 me3:H4K20 me3
H3K27 me3:H4K20 me3
H3K4 me1:H2A.Z
H3K4 me3:H3–5ac
H2A.Z:H4K20 me2
H3K27 me3:DNA Meth. m5C
H3K4 me1:H4K20 me2
DNA Meth. m5C:H4K20 me3
H3K4 me1:H3–5ac
DNA Meth. m5C:H4K20 me2
H3K27 me2:DNA Meth. m5C
DNA Meth. m5C:H3K9 me3
H3K9 me2:DNA Meth. m5C
H4K20 me2:H3–5ac
H4K20 me2:H4–4ac
H4K20 me2:H3K4 me3
H4–4ac:H3–5ac
H3K4 me3:H4–4ac
H3K27 me3
H3K27 me2
H3K9 me2
H3K9 me3
H4K20 me3
H2A.Z
H3–5ac
H3K4 me1
H3K4 me3
H4K20 me2
H4–4ac
DNA Meth. m5C
0.00
0.25
0.50
0.75
1.00
π̂
HOXD9 – Reverse
H2A.Z:H3K4 me3
H3K4 me1:H4–4ac
H3K9 me3:H4K20 me3
H3K27 me3:H4K20 me3
H3K4 me1:H2A.Z
H3K4 me3:H3–5ac
H2A.Z:H4K20 me2
H3K27 me3:DNA Meth. m5C
H3K4 me1:H4K20 me2
DNA Meth. m5C:H4K20 me3
H3K4 me1:H3–5ac
DNA Meth. m5C:H4K20 me2
H3K27 me2:DNA Meth. m5C
DNA Meth. m5C:H3K9 me3
H3K9 me2:DNA Meth. m5C
H4K20 me2:H3–5ac
H4K20 me2:H4–4ac
H4K20 me2:H3K4 me3
H4–4ac:H3–5ac
H3K4 me3:H4–4ac
H3K27 me3
H3K27 me2
H3K9 me2
H3K9 me3
H4K20 me3
H2A.Z
H3–5ac
H3K4 me1
H3K4 me3
H4K20 me2
H4–4ac
DNA Meth. m5C
0.00
0.25
0.50
0.75
1.00
π̂
ZBED4 – Forward
H3K9 me3:H4K20 me3
H3K9 me2:DNA Meth. m5C
H3K27 me2:DNA Meth. m5C
H3K27 me3:H4K20 me3
H3K4 me1:H4K20 me2
DNA Meth. m5C:H4K20 me3
H3K4 me3:H3–5ac
H3K27 me3:DNA Meth. m5C
DNA Meth. m5C:H3K9 me3
H3K27 me3:H3K9 me3
H3K4 me1:H3–5ac
H3K4 me1:H4–4ac
DNA Meth. m5C:H4K20 me2
H2A.Z:H4K20 me2
H4K20 me2:H4–4ac
H4K20 me2:H3–5ac
H4K20 me2:H3K4 me3
H3K4 me1:H2A.Z
H3K4 me3:H4–4ac
H2A.Z:H3K4 me3
H4–4ac:H3–5ac
H3K27 me2
H3K9 me2
H4K20 me3
H3K9 me3
H3K27 me3
H2A.Z
H3–5ac
H3K4 me3
DNA Meth. m5C
H4K20 me2
H4–4ac
H3K4 me1
0.00
0.25
0.50
0.75
1.00
π̂
ZBED4 – Reverse
H3K9 me3:H4K20 me3
H3K9 me2:DNA Meth. m5C
H3K27 me2:DNA Meth. m5C
H3K27 me3:H4K20 me3
H3K4 me1:H4K20 me2
DNA Meth. m5C:H4K20 me3
H3K4 me3:H3–5ac
H3K27 me3:DNA Meth. m5C
DNA Meth. m5C:H3K9 me3
H3K27 me3:H3K9 me3
H3K4 me1:H3–5ac
H3K4 me1:H4–4ac
DNA Meth. m5C:H4K20 me2
H2A.Z:H4K20 me2
H4K20 me2:H4–4ac
H4K20 me2:H3–5ac
H4K20 me2:H3K4 me3
H3K4 me1:H2A.Z
H3K4 me3:H4–4ac
H2A.Z:H3K4 me3
H4–4ac:H3–5ac
H3K27 me2
H3K9 me2
H4K20 me3
H3K9 me3
H3K27 me3
H2A.Z
H3–5ac
H3K4 me3
DNA Meth. m5C
H4K20 me2
H4–4ac
H3K4 me1
0.00
0.25
0.50
0.75
1.00
π̂
ZFP64 (2) – Forward
H3K4 me1:H3–5ac
H3K9 me3:H4K20 me3
H3K27 me3:H3K4 me1
H3K4 me1:H4–4ac
H3K27 me3:H4K20 me3
H3K4 me3:H3–5ac
H3K4 me1:H4K20 me2
H3K4 me1:H2A.Z
H3K27 me2:DNA Meth. m5C
H3K9 me2:DNA Meth. m5C
H3K27 me3:H3K9 me3
H3K27 me3:DNA Meth. m5C
H2A.Z:H4–4ac
DNA Meth. m5C:H3K9 me3
H2A.Z:H3–5ac
DNA Meth. m5C:H4K20 me2
DNA Meth. m5C:H4K20 me3
H4K20 me2:H3–5ac
H2A.Z:H4K20 me2
H2A.Z:H3K4 me3
H4K20 me2:H4–4ac
H4–4ac:H3–5ac
H3K4 me3:H4–4ac
H4K20 me2:H3K4 me3
H3K9 me2
H4K20 me3
H3K27 me2
H3K9 me3
H3K27 me3
H3K4 me1
H3K4 me3
H3–5ac
H4–4ac
H4K20 me2
H2A.Z
DNA Meth. m5C
0.00
0.25
0.50
0.75
1.00
π̂
ZFP64 (2) – Reverse
H3K4 me1:H3–5ac
H3K9 me3:H4K20 me3
H3K27 me3:H3K4 me1
H3K4 me1:H4–4ac
H3K27 me3:H4K20 me3
H3K4 me3:H3–5ac
H3K4 me1:H4K20 me2
H3K4 me1:H2A.Z
H3K27 me2:DNA Meth. m5C
H3K9 me2:DNA Meth. m5C
H3K27 me3:H3K9 me3
H3K27 me3:DNA Meth. m5C
H2A.Z:H4–4ac
DNA Meth. m5C:H3K9 me3
H2A.Z:H3–5ac
DNA Meth. m5C:H4K20 me2
DNA Meth. m5C:H4K20 me3
H4K20 me2:H3–5ac
H2A.Z:H4K20 me2
H2A.Z:H3K4 me3
H4K20 me2:H4–4ac
H4–4ac:H3–5ac
H3K4 me3:H4–4ac
H4K20 me2:H3K4 me3
H3K9 me2
H4K20 me3
H3K27 me2
H3K9 me3
H3K27 me3
H3K4 me1
H3K4 me3
H3–5ac
H4–4ac
H4K20 me2
H2A.Z
DNA Meth. m5C
0.00
0.25
0.50
0.75
1.00
π̂
ZNF768 – Forward
H3K9 me3:H4K20 me3
H3K27 me2:DNA Meth. m5C
H3K9 me2:DNA Meth. m5C
H3K4 me1:H4–4ac
H3K4 me1:H4K20 me2
H3K4 me1:H2A.Z
DNA Meth. m5C:H4K20 me3
H3K27 me3:H3K9 me3
H2A.Z:H4K20 me2
DNA Meth. m5C:H3K9 me3
H2A.Z:H3–5ac
H3K4 me3:H3–5ac
H2A.Z:H4–4ac
H3K4 me1:H3–5ac
H3K27 me3:DNA Meth. m5C
H2A.Z:H3K4 me3
H4K20 me2:H3–5ac
H4K20 me2:H3K4 me3
DNA Meth. m5C:H4K20 me2
H3K4 me3:H4–4ac
H4K20 me2:H4–4ac
H4–4ac:H3–5ac
H3K27 me2
H3K9 me2
H3K9 me3
H3K4 me3
DNA Meth. m5C
H3–5ac
H3K4 me1
H3K27 me3
H2A.Z
H4K20 me3
H4K20 me2
H4–4ac
0.00
0.25
0.50
0.75
1.00
π̂
ZNF768 – Reverse
H3K9 me3:H4K20 me3
H3K27 me2:DNA Meth. m5C
H3K9 me2:DNA Meth. m5C
H3K4 me1:H4–4ac
H3K4 me1:H4K20 me2
H3K4 me1:H2A.Z
DNA Meth. m5C:H4K20 me3
H3K27 me3:H3K9 me3
H2A.Z:H4K20 me2
DNA Meth. m5C:H3K9 me3
H2A.Z:H3–5ac
H3K4 me3:H3–5ac
H2A.Z:H4–4ac
H3K4 me1:H3–5ac
H3K27 me3:DNA Meth. m5C
H2A.Z:H3K4 me3
H4K20 me2:H3–5ac
H4K20 me2:H3K4 me3
DNA Meth. m5C:H4K20 me2
H3K4 me3:H4–4ac
H4K20 me2:H4–4ac
H4–4ac:H3–5ac
H3K27 me2
H3K9 me2
H3K9 me3
H3K4 me3
DNA Meth. m5C
H3–5ac
H3K4 me1
H3K27 me3
H2A.Z
H4K20 me3
H4K20 me2
H4–4ac
0.00
0.25
0.50
0.75
1.00
π̂
STT3B – Forward
H3K27 me2:DNA Meth. m5C
H3K27 me3:H3K9 me3
DNA Meth. m5C:H3K9 me3
H2A.Z:H4–4ac
H3K9 me2:DNA Meth. m5C
H2A.Z:H3–5ac
H3K4 me1:H4–4ac
H3K27 me3:DNA Meth. m5C
H3K27 me3:H3K4 me1
H3K4 me1:H2A.Z
H2A.Z:H4K20 me2
H3K4 me1:H4K20 me2
DNA Meth. m5C:H4K20 me2
H3K4 me3:H3–5ac
H2A.Z:H3K4 me3
H4K20 me2:H3–5ac
H4K20 me2:H4–4ac
H3K4 me1:H3–5ac
H4K20 me2:H3K4 me3
H4K20 me3
H3K4 me3:H4–4ac
H4–4ac:H3–5ac
H3K27 me2
H3K9 me3
H3K9 me2
H3K27 me3
DNA Meth. m5C
H3K4 me1
H4K20 me2
H2A.Z
H3–5ac
H4–4ac
H3K4 me3
0.00
0.25
0.50
0.75
1.00
π̂
STT3B – Reverse
H3K27 me2:DNA Meth. m5C
H3K27 me3:H3K9 me3
DNA Meth. m5C:H3K9 me3
H2A.Z:H4–4ac
H3K9 me2:DNA Meth. m5C
H2A.Z:H3–5ac
H3K4 me1:H4–4ac
H3K27 me3:DNA Meth. m5C
H3K27 me3:H3K4 me1
H3K4 me1:H2A.Z
H2A.Z:H4K20 me2
H3K4 me1:H4K20 me2
DNA Meth. m5C:H4K20 me2
H3K4 me3:H3–5ac
H2A.Z:H3K4 me3
H4K20 me2:H3–5ac
H4K20 me2:H4–4ac
H3K4 me1:H3–5ac
H4K20 me2:H3K4 me3
H4K20 me3
H3K4 me3:H4–4ac
H4–4ac:H3–5ac
H3K27 me2
H3K9 me3
H3K9 me2
H3K27 me3
DNA Meth. m5C
H3K4 me1
H4K20 me2
H2A.Z
H3–5ac
H4–4ac
H3K4 me3
0.00
0.25
0.50
0.75
1.00
π̂

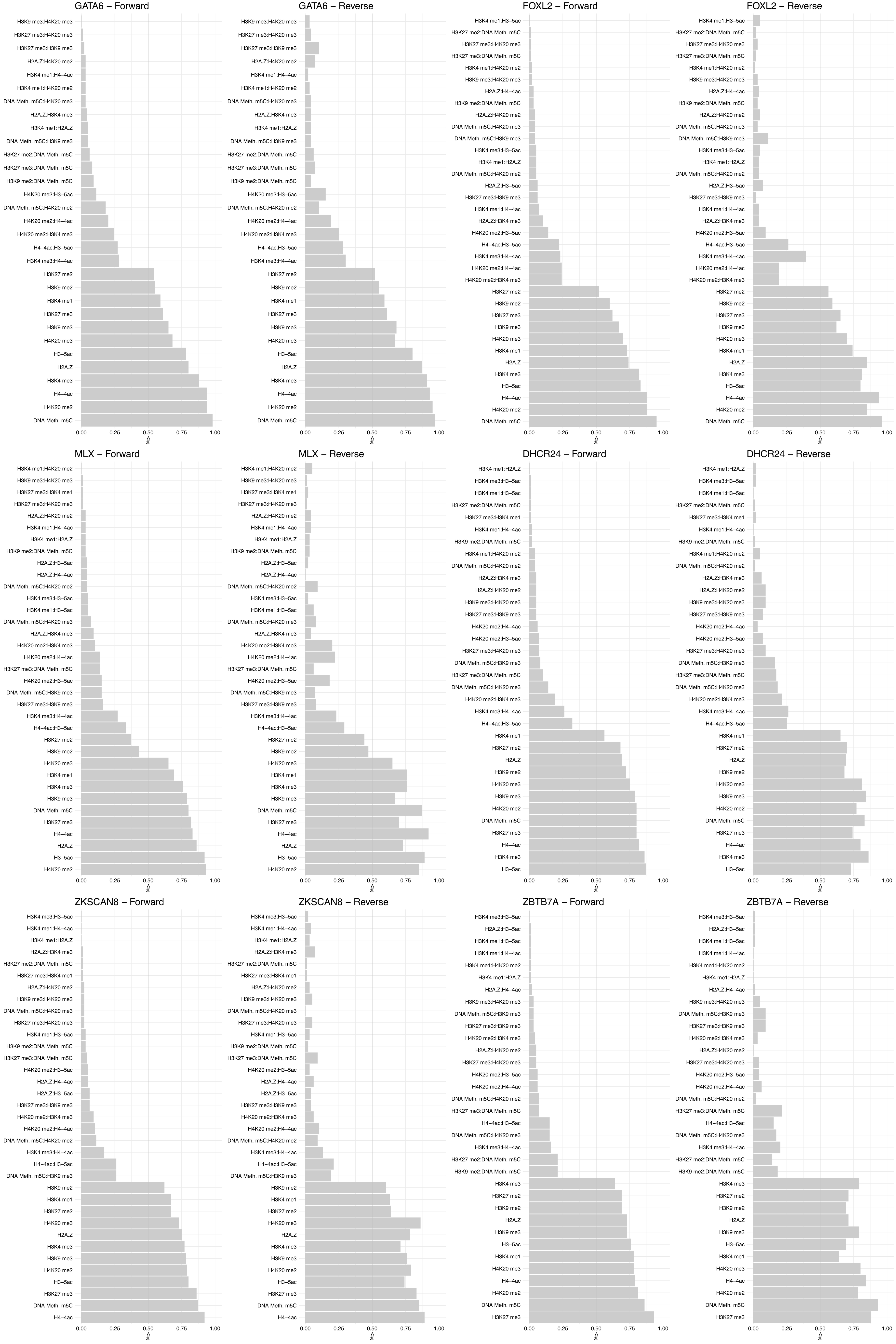
GATA6 – Forward
H3K9 me3:H4K20 me3
H3K27 me3:H4K20 me3
H3K27 me3:H3K9 me3
H2A.Z:H4K20 me2
H3K4 me1:H4−4ac
H3K4 me1:H4K20 me2
DNA Meth. m5C:H4K20 me3
H2A.Z:H3K4 me3
H3K4 me1:H2A.Z
DNA Meth. m5C:H3K9 me3
H3K27 me2:DNA Meth. m5C
H3K27 me3:DNA Meth. m5C
H3K9 me2:DNA Meth. m5C
H4K20 me2:H3−5ac
DNA Meth. m5C:H4K20 me2
H4K20 me2:H4−4ac
H4K20 me2:H3K4 me3
H4−4ac:H3−5ac
H3K4 me3:H4−4ac
H3K27 me2
H3K9 me2
H3K4 me1
H3K27 me3
H3K9 me3
H4K20 me3
H3−5ac
H2A.Z
H3K4 me3
H4−4ac
H4K20 me2
DNA Meth. m5C
0.00
0.25
0.50
0.75
1.00
π̂
GATA6 – Reverse
H3K9 me3:H4K20 me3
H3K27 me3:H4K20 me3
H3K27 me3:H3K9 me3
H2A.Z:H4K20 me2
H3K4 me1:H4−4ac
H3K4 me1:H4K20 me2
DNA Meth. m5C:H4K20 me3
H2A.Z:H3K4 me3
H3K4 me1:H2A.Z
DNA Meth. m5C:H3K9 me3
H3K27 me2:DNA Meth. m5C
H3K27 me3:DNA Meth. m5C
H3K9 me2:DNA Meth. m5C
H4K20 me2:H3−5ac
DNA Meth. m5C:H4K20 me2
H4K20 me2:H4−4ac
H4K20 me2:H3K4 me3
H4−4ac:H3−5ac
H3K4 me3:H4−4ac
H3K27 me2
H3K9 me2
H3K4 me1
H3K27 me3
H3K9 me3
H4K20 me3
H3−5ac
H2A.Z
H3K4 me3
H4−4ac
H4K20 me2
DNA Meth. m5C
0.00
0.25
0.50
0.75
1.00
π̂
FOXL2 – Forward
H3K4 me1:H3−5ac
H3K27 me2:DNA Meth. m5C
H3K27 me3:H4K20 me3
H3K27 me3:DNA Meth. m5C
H3K4 me1:H4K20 me2
H3K9 me3:H4K20 me3
H2A.Z:H4−4ac
H3K9 me2:DNA Meth. m5C
H2A.Z:H4K20 me2
DNA Meth. m5C:H4K20 me3
DNA Meth. m5C:H3K9 me3
H3K4 me3:H3−5ac
H3K4 me1:H2A.Z
DNA Meth. m5C:H4K20 me2
H2A.Z:H3−5ac
H3K27 me3:H3K9 me3
H3K4 me1:H4−4ac
H2A.Z:H3K4 me3
H4K20 me2:H3−5ac
H4−4ac:H3−5ac
H3K4 me3:H4−4ac
H4K20 me2:H4−4ac
H4K20 me2:H3K4 me3
H3K27 me2
H3K9 me2
H3K27 me3
H3K9 me3
H4K20 me3
H3K4 me1
H2A.Z
H3K4 me3
H3−5ac
H4−4ac
H4K20 me2
DNA Meth. m5C
0.00
0.25
0.50
0.75
1.00
π̂
FOXL2 – Reverse
H3K4 me1:H3−5ac
H3K27 me2:DNA Meth. m5C
H3K27 me3:H4K20 me3
H3K27 me3:DNA Meth. m5C
H3K4 me1:H4K20 me2
H3K9 me3:H4K20 me3
H2A.Z:H4−4ac
H3K9 me2:DNA Meth. m5C
H2A.Z:H4K20 me2
DNA Meth. m5C:H4K20 me3
DNA Meth. m5C:H3K9 me3
H3K4 me3:H3−5ac
H3K4 me1:H2A.Z
DNA Meth. m5C:H4K20 me2
H2A.Z:H3−5ac
H3K27 me3:H3K9 me3
H3K4 me1:H4−4ac
H2A.Z:H3K4 me3
H4K20 me2:H3−5ac
H4−4ac:H3−5ac
H3K4 me3:H4−4ac
H4K20 me2:H4−4ac
H4K20 me2:H3K4 me3
H3K27 me2
H3K9 me2
H3K27 me3
H3K9 me3
H4K20 me3
H3K4 me1
H2A.Z
H3K4 me3
H3−5ac
H4−4ac
H4K20 me2
DNA Meth. m5C
0.00
0.25
0.50
0.75
1.00
π̂
MLX – Forward
H3K4 me1:H4K20 me2
H3K9 me3:H4K20 me3
H3K27 me3:H3K4 me1
H3K27 me3:H4K20 me3
H2A.Z:H4K20 me2
H3K4 me1:H4−4ac
H3K4 me1:H2A.Z
H3K9 me2:DNA Meth. m5C
H2A.Z:H3−5ac
H2A.Z:H4−4ac
DNA Meth. m5C:H4K20 me2
H3K4 me3:H3−5ac
H3K4 me1:H3−5ac
DNA Meth. m5C:H4K20 me3
H2A.Z:H3K4 me3
H4K20 me2:H3K4 me3
H4K20 me2:H4−4ac
H3K27 me3:DNA Meth. m5C
H4K20 me2:H3−5ac
DNA Meth. m5C:H3K9 me3
H3K27 me3:H3K9 me3
H3K4 me3:H4−4ac
H4−4ac:H3−5ac
H3K27 me2
H3K9 me2
H4K20 me3
H3K4 me1
H3K4 me3
H3K9 me3
DNA Meth. m5C
H3K27 me3
H4−4ac
H2A.Z
H3−5ac
H4K20 me2
0.00
0.25
0.50
0.75
1.00
π̂
MLX – Reverse
H3K4 me1:H4K20 me2
H3K9 me3:H4K20 me3
H3K27 me3:H3K4 me1
H3K27 me3:H4K20 me3
H2A.Z:H4K20 me2
H3K4 me1:H4−4ac
H3K4 me1:H2A.Z
H3K9 me2:DNA Meth. m5C
H2A.Z:H3−5ac
H2A.Z:H4−4ac
DNA Meth. m5C:H4K20 me2
H3K4 me3:H3−5ac
H3K4 me1:H3−5ac
DNA Meth. m5C:H4K20 me3
H2A.Z:H3K4 me3
H4K20 me2:H3K4 me3
H4K20 me2:H4−4ac
H3K27 me3:DNA Meth. m5C
H4K20 me2:H3−5ac
DNA Meth. m5C:H3K9 me3
H3K27 me3:H3K9 me3
H3K4 me3:H4−4ac
H4−4ac:H3−5ac
H3K27 me2
H3K9 me2
H4K20 me3
H3K4 me1
H3K4 me3
H3K9 me3
DNA Meth. m5C
H3K27 me3
H4−4ac
H2A.Z
H3−5ac
H4K20 me2
0.00
0.25
0.50
0.75
1.00
π̂
DHCR24 – Forward
H3K4 me1:H2A.Z
H3K4 me3:H3−5ac
H3K4 me1:H3−5ac
H3K27 me2:DNA Meth. m5C
H3K27 me3:H3K4 me1
H3K4 me1:H4−4ac
H3K9 me2:DNA Meth. m5C
H3K4 me1:H4K20 me2
DNA Meth. m5C:H4K20 me2
H2A.Z:H3K4 me3
H2A.Z:H4K20 me2
H3K9 me3:H4K20 me3
H3K27 me3:H3K9 me3
H4K20 me2:H4−4ac
H4K20 me2:H3−5ac
H3K27 me3:H4K20 me3
DNA Meth. m5C:H3K9 me3
H3K27 me3:DNA Meth. m5C
DNA Meth. m5C:H4K20 me3
H4K20 me2:H3K4 me3
H3K4 me3:H4−4ac
H4−4ac:H3−5ac
H3K4 me1
H3K27 me2
H2A.Z
H3K9 me2
H4K20 me3
H3K9 me3
H4K20 me2
DNA Meth. m5C
H3K27 me3
H4−4ac
H3K4 me3
H3−5ac
0.00
0.25
0.50
0.75
1.00
π̂
DHCR24 – Reverse
H3K4 me1:H2A.Z
H3K4 me3:H3−5ac
H3K4 me1:H3−5ac
H3K27 me2:DNA Meth. m5C
H3K27 me3:H3K4 me1
H3K4 me1:H4−4ac
H3K9 me2:DNA Meth. m5C
H3K4 me1:H4K20 me2
DNA Meth. m5C:H4K20 me2
H2A.Z:H3K4 me3
H2A.Z:H4K20 me2
H3K9 me3:H4K20 me3
H3K27 me3:H3K9 me3
H4K20 me2:H4−4ac
H4K20 me2:H3−5ac
H3K27 me3:H4K20 me3
DNA Meth. m5C:H3K9 me3
H3K27 me3:DNA Meth. m5C
DNA Meth. m5C:H4K20 me3
H4K20 me2:H3K4 me3
H3K4 me3:H4−4ac
H4−4ac:H3−5ac
H3K4 me1
H3K27 me2
H2A.Z
H3K9 me2
H4K20 me3
H3K9 me3
H4K20 me2
DNA Meth. m5C
H3K27 me3
H4−4ac
H3K4 me3
H3−5ac
0.00
0.25
0.50
0.75
1.00
π̂
ZKSCAN8 – Forward
H3K4 me3:H3−5ac
H3K4 me1:H4−4ac
H3K4 me1:H2A.Z
H2A.Z:H3K4 me3
H3K27 me2:DNA Meth. m5C
H3K27 me3:H3K4 me1
H2A.Z:H4K20 me2
H3K9 me3:H4K20 me3
DNA Meth. m5C:H4K20 me3
H3K27 me3:H4K20 me3
H3K4 me1:H3−5ac
H3K9 me2:DNA Meth. m5C
H3K27 me3:DNA Meth. m5C
H4K20 me2:H3−5ac
H2A.Z:H4−4ac
H2A.Z:H3−5ac
H3K27 me3:H3K9 me3
H4K20 me2:H3K4 me3
H4K20 me2:H4−4ac
DNA Meth. m5C:H4K20 me2
H3K4 me3:H4−4ac
H4−4ac:H3−5ac
DNA Meth. m5C:H3K9 me3
H3K9 me2
H3K4 me1
H3K27 me2
H4K20 me3
H2A.Z
H3K4 me3
H3K9 me3
H4K20 me2
H3−5ac
H3K27 me3
DNA Meth. m5C
H4−4ac
0.00
0.25
0.50
0.75
1.00
π̂
ZKSCAN8 – Reverse
H3K4 me3:H3−5ac
H3K4 me1:H4−4ac
H3K4 me1:H2A.Z
H2A.Z:H3K4 me3
H3K27 me2:DNA Meth. m5C
H3K27 me3:H3K4 me1
H2A.Z:H4K20 me2
H3K9 me3:H4K20 me3
DNA Meth. m5C:H4K20 me3
H3K27 me3:H4K20 me3
H3K4 me1:H3−5ac
H3K9 me2:DNA Meth. m5C
H3K27 me3:DNA Meth. m5C
H4K20 me2:H3−5ac
H2A.Z:H4−4ac
H2A.Z:H3−5ac
H3K27 me3:H3K9 me3
H4K20 me2:H3K4 me3
H4K20 me2:H4−4ac
DNA Meth. m5C:H4K20 me2
H3K4 me3:H4−4ac
H4−4ac:H3−5ac
DNA Meth. m5C:H3K9 me3
H3K9 me2
H3K4 me1
H3K27 me2
H4K20 me3
H2A.Z
H3K4 me3
H3K9 me3
H4K20 me2
H3−5ac
H3K27 me3
DNA Meth. m5C
H4−4ac
0.00
0.25
0.50
0.75
1.00
π̂
ZBTB7A – Forward
H3K4 me3:H3−5ac
H2A.Z:H3−5ac
H3K4 me1:H3−5ac
H3K4 me1:H4−4ac
H3K4 me1:H4K20 me2
H3K4 me1:H2A.Z
H2A.Z:H4−4ac
H3K9 me3:H4K20 me3
DNA Meth. m5C:H3K9 me3
H3K27 me3:H3K9 me3
H4K20 me2:H3K4 me3
H2A.Z:H4K20 me2
H3K27 me3:H4K20 me3
H4K20 me2:H3−5ac
H4K20 me2:H4−4ac
DNA Meth. m5C:H4K20 me2
H3K27 me3:DNA Meth. m5C
H4−4ac:H3−5ac
DNA Meth. m5C:H4K20 me3
H3K4 me3:H4−4ac
H3K27 me2:DNA Meth. m5C
H3K9 me2:DNA Meth. m5C
H3K4 me3
H3K27 me2
H3K9 me2
H2A.Z
H3K9 me3
H3−5ac
H3K4 me1
H4K20 me3
H4−4ac
H4K20 me2
DNA Meth. m5C
H3K27 me3
0.00
0.25
0.50
0.75
1.00
π̂
ZBTB7A – Reverse
H3K4 me3:H3−5ac
H2A.Z:H3−5ac
H3K4 me1:H3−5ac
H3K4 me1:H4−4ac
H3K4 me1:H4K20 me2
H3K4 me1:H2A.Z
H2A.Z:H4−4ac
H3K9 me3:H4K20 me3
DNA Meth. m5C:H3K9 me3
H3K27 me3:H3K9 me3
H4K20 me2:H3K4 me3
H2A.Z:H4K20 me2
H3K27 me3:H4K20 me3
H4K20 me2:H3−5ac
H4K20 me2:H4−4ac
DNA Meth. m5C:H4K20 me2
H3K27 me3:DNA Meth. m5C
H4−4ac:H3−5ac
DNA Meth. m5C:H4K20 me3
H3K4 me3:H4−4ac
H3K27 me2:DNA Meth. m5C
H3K9 me2:DNA Meth. m5C
H3K4 me3
H3K27 me2
H3K9 me2
H2A.Z
H3K9 me3
H3−5ac
H3K4 me1
H4K20 me3
H4−4ac
H4K20 me2
DNA Meth. m5C
H3K27 me3
0.00
0.25
0.50
0.75
1.00
π̂

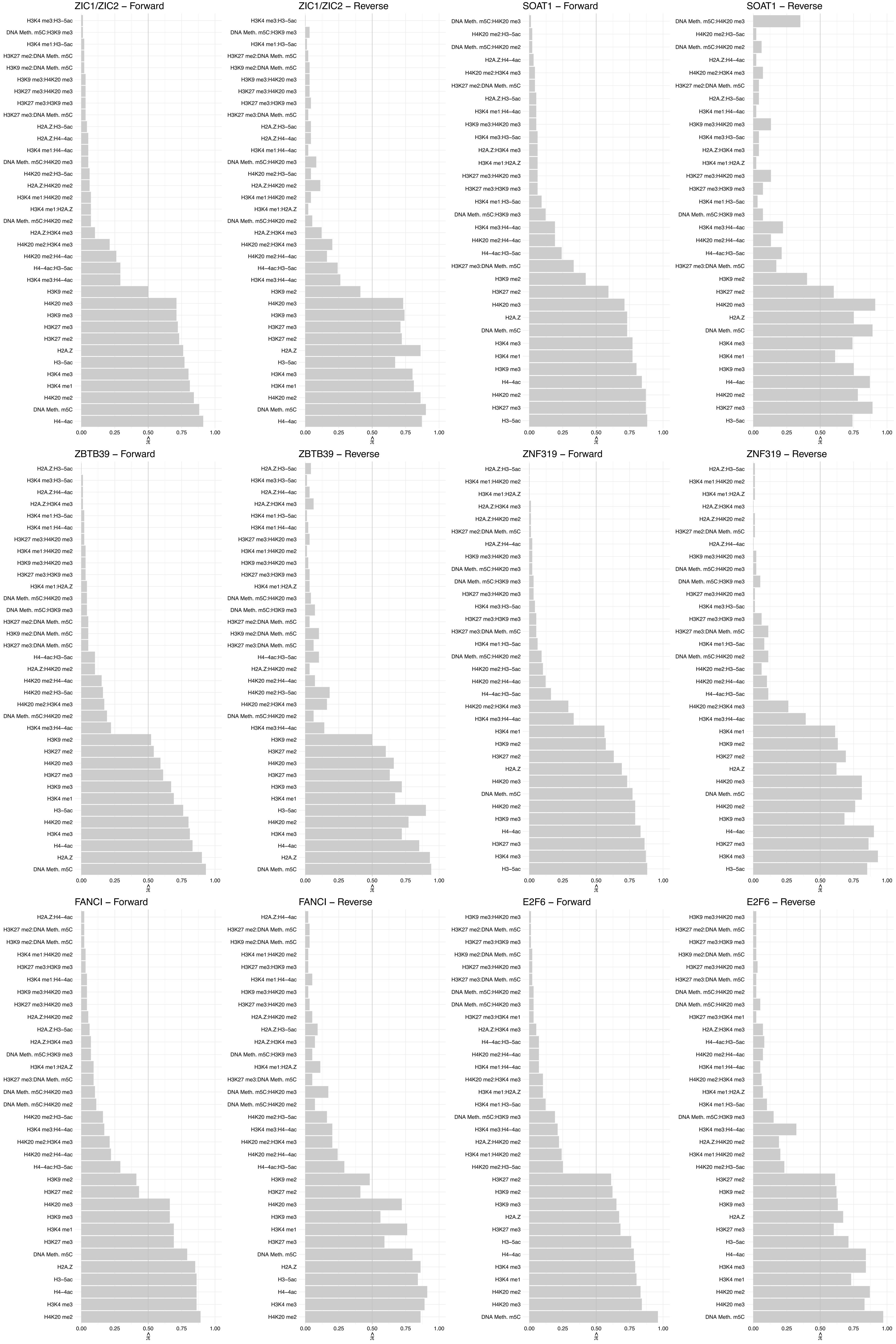
ZIC1/ZIC2 – Forward
ZIC1/ZIC2 – Reverse
SOAT1 – Forward
SOAT1 – Reverse
ZBTB39 – Forward
ZBTB39 – Reverse
ZNF319 – Forward
ZNF319 – Reverse
FANCI – Forward
FANCI – Reverse
E2F6 – Forward
E2F6 – Reverse
0.00
0.25
0.50
0.75
1.00
$\hat{\pi}$

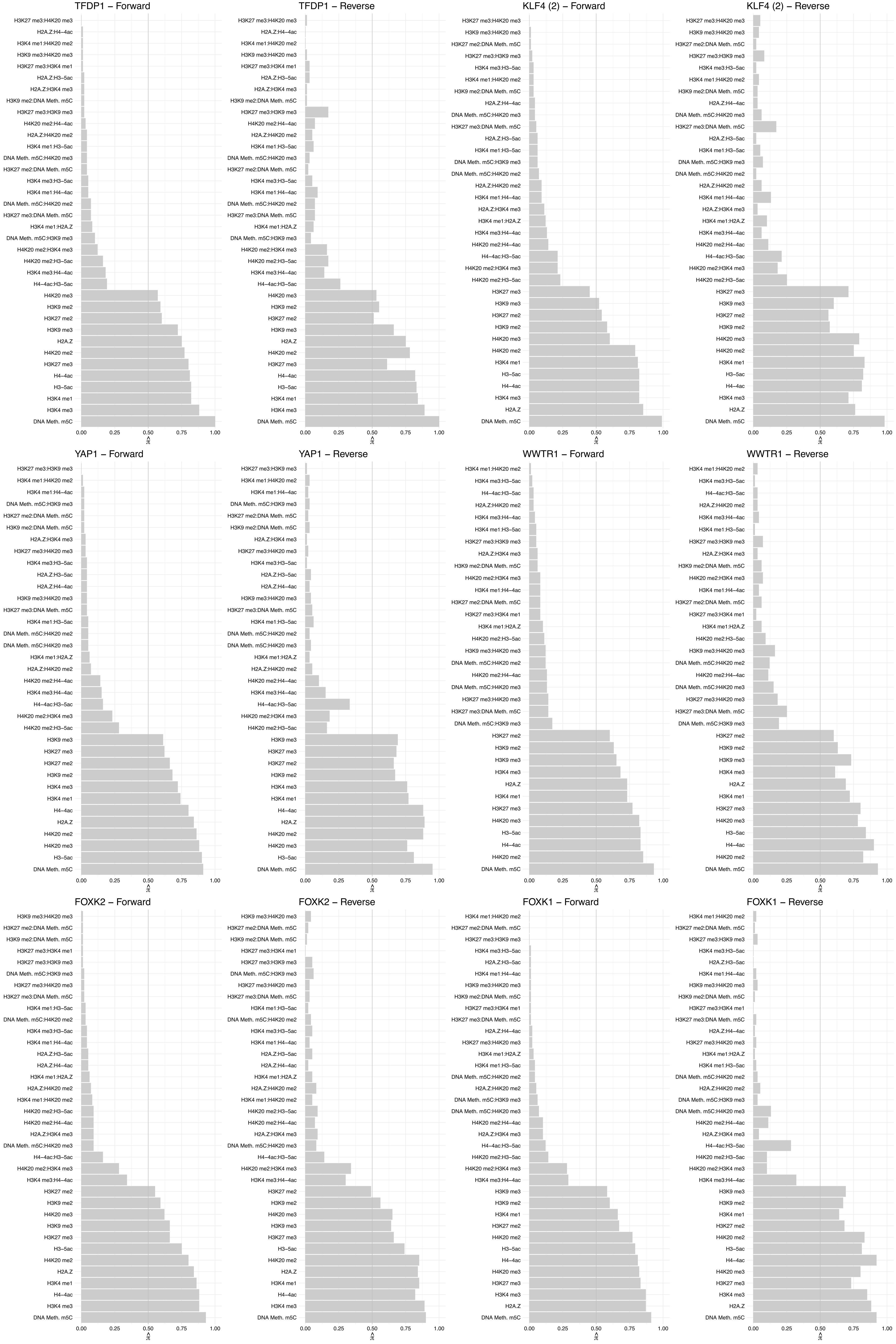
TFDP1 − Forward
H3K27 me3:H4K20 me3
H2A.Z:H4−4ac
H3K4 me1:H4K20 me2
H3K9 me3:H4K20 me3
H3K27 me3:H3K4 me1
H2A.Z:H3−5ac
H2A.Z:H3K4 me3
H3K9 me2:DNA Meth. m5C
H3K27 me3:H3K9 me3
H4K20 me2:H4−4ac
H2A.Z:H4K20 me2
H3K4 me1:H3−5ac
DNA Meth. m5C:H4K20 me3
H3K27 me2:DNA Meth. m5C
H3K4 me3:H3−5ac
H3K4 me1:H4−4ac
DNA Meth. m5C:H4K20 me2
H3K27 me3:DNA Meth. m5C
H3K4 me1:H2A.Z
DNA Meth. m5C:H3K9 me3
H4K20 me2:H3K4 me3
H4K20 me2:H3−5ac
H3K4 me3:H4−4ac
H4−4ac:H3−5ac
H4K20 me3
H3K9 me2
H3K27 me2
H3K9 me3
H2A.Z
H4K20 me2
H3K27 me3
H4−4ac
H3−5ac
H3K4 me1
H3K4 me3
DNA Meth. m5C
0.00 0.25 0.50 0.75 1.00
π̂
TFDP1 − Reverse
H3K27 me3:H4K20 me3
H2A.Z:H4−4ac
H3K4 me1:H4K20 me2
H3K9 me3:H4K20 me3
H3K27 me3:H3K4 me1
H2A.Z:H3−5ac
H2A.Z:H3K4 me3
H3K9 me2:DNA Meth. m5C
H3K27 me3:H3K9 me3
H4K20 me2:H4−4ac
H2A.Z:H4K20 me2
H3K4 me1:H3−5ac
DNA Meth. m5C:H4K20 me3
H3K27 me2:DNA Meth. m5C
H3K4 me3:H3−5ac
H3K4 me1:H4−4ac
DNA Meth. m5C:H4K20 me2
H3K27 me3:DNA Meth. m5C
H3K4 me1:H2A.Z
DNA Meth. m5C:H3K9 me3
H4K20 me2:H3K4 me3
H4K20 me2:H3−5ac
H3K4 me3:H4−4ac
H4−4ac:H3−5ac
H4K20 me3
H3K9 me2
H3K27 me2
H3K9 me3
H2A.Z
H4K20 me2
H3K27 me3
H4−4ac
H3−5ac
H3K4 me1
H3K4 me3
DNA Meth. m5C
0.00 0.25 0.50 0.75 1.00
π̂
KLF4 (2) − Forward
H3K27 me3:H4K20 me3
H3K9 me3:H4K20 me3
H3K27 me2:DNA Meth. m5C
H3K27 me3:H3K9 me3
H3K4 me3:H3−5ac
H3K4 me1:H4K20 me2
H3K9 me2:DNA Meth. m5C
H2A.Z:H4−4ac
DNA Meth. m5C:H4K20 me3
H3K27 me3:DNA Meth. m5C
H2A.Z:H3−5ac
H3K4 me1:H3−5ac
DNA Meth. m5C:H3K9 me3
DNA Meth. m5C:H4K20 me2
H2A.Z:H4K20 me2
H3K4 me1:H4−4ac
H2A.Z:H3K4 me3
H3K4 me1:H2A.Z
H3K4 me3:H4−4ac
H4K20 me2:H4−4ac
H4−4ac:H3−5ac
H4K20 me2:H3K4 me3
H4K20 me2:H3−5ac
H3K27 me3
H3K9 me3
H3K27 me2
H3K9 me2
H4K20 me3
H4K20 me2
H3K4 me1
H3−5ac
H4−4ac
H3K4 me3
H2A.Z
DNA Meth. m5C
0.00 0.25 0.50 0.75 1.00
π̂
KLF4 (2) − Reverse
H3K27 me3:H4K20 me3
H3K9 me3:H4K20 me3
H3K27 me2:DNA Meth. m5C
H3K27 me3:H3K9 me3
H3K4 me3:H3−5ac
H3K4 me1:H4K20 me2
H3K9 me2:DNA Meth. m5C
H2A.Z:H4−4ac
DNA Meth. m5C:H4K20 me3
H3K27 me3:DNA Meth. m5C
H2A.Z:H3−5ac
H3K4 me1:H3−5ac
DNA Meth. m5C:H3K9 me3
DNA Meth. m5C:H4K20 me2
H2A.Z:H4K20 me2
H3K4 me1:H4−4ac
H2A.Z:H3K4 me3
H3K4 me1:H2A.Z
H3K4 me3:H4−4ac
H4K20 me2:H4−4ac
H4−4ac:H3−5ac
H4K20 me2:H3K4 me3
H4K20 me2:H3−5ac
H3K27 me3
H3K9 me3
H3K27 me2
H3K9 me2
H4K20 me3
H4K20 me2
H3K4 me1
H3−5ac
H4−4ac
H3K4 me3
H2A.Z
DNA Meth. m5C
0.00 0.25 0.50 0.75 1.00
π̂
YAP1 − Forward
H3K27 me3:H3K9 me3
H3K4 me1:H4K20 me2
H3K4 me1:H4−4ac
DNA Meth. m5C:H3K9 me3
H3K27 me2:DNA Meth. m5C
H3K9 me2:DNA Meth. m5C
H2A.Z:H3K4 me3
H3K27 me3:H4K20 me3
H3K4 me3:H3−5ac
H2A.Z:H3−5ac
H2A.Z:H4−4ac
H3K9 me3:H4K20 me3
H3K27 me3:DNA Meth. m5C
H3K4 me1:H3−5ac
DNA Meth. m5C:H4K20 me2
DNA Meth. m5C:H4K20 me3
H3K4 me1:H2A.Z
H2A.Z:H4K20 me2
H4K20 me2:H4−4ac
H3K4 me3:H4−4ac
H4−4ac:H3−5ac
H4K20 me2:H3K4 me3
H4K20 me2:H3−5ac
H3K9 me3
H3K27 me3
H3K27 me2
H3K9 me2
H3K4 me3
H3K4 me1
H4−4ac
H2A.Z
H4K20 me2
H4K20 me3
H3−5ac
DNA Meth. m5C
0.00 0.25 0.50 0.75 1.00
π̂
YAP1 − Reverse
H3K27 me3:H3K9 me3
H3K4 me1:H4K20 me2
H3K4 me1:H4−4ac
DNA Meth. m5C:H3K9 me3
H3K27 me2:DNA Meth. m5C
H3K9 me2:DNA Meth. m5C
H2A.Z:H3K4 me3
H3K27 me3:H4K20 me3
H3K4 me3:H3−5ac
H2A.Z:H3−5ac
H2A.Z:H4−4ac
H3K9 me3:H4K20 me3
H3K27 me3:DNA Meth. m5C
H3K4 me1:H3−5ac
DNA Meth. m5C:H4K20 me2
DNA Meth. m5C:H4K20 me3
H3K4 me1:H2A.Z
H2A.Z:H4K20 me2
H4K20 me2:H4−4ac
H3K4 me3:H4−4ac
H4−4ac:H3−5ac
H4K20 me2:H3K4 me3
H4K20 me2:H3−5ac
H3K9 me3
H3K27 me3
H3K27 me2
H3K9 me2
H3K4 me3
H3K4 me1
H4−4ac
H2A.Z
H4K20 me2
H4K20 me3
H3−5ac
DNA Meth. m5C
0.00 0.25 0.50 0.75 1.00
π̂
WWTR1 − Forward
H3K4 me1:H4K20 me2
H3K4 me3:H3−5ac
H4−4ac:H3−5ac
H2A.Z:H4K20 me2
H3K4 me3:H4−4ac
H3K4 me1:H3−5ac
H3K27 me3:H3K9 me3
H2A.Z:H3K4 me3
H3K9 me2:DNA Meth. m5C
H4K20 me2:H3K4 me3
H3K4 me1:H4−4ac
H3K27 me2:DNA Meth. m5C
H3K27 me3:H3K4 me1
H3K4 me1:H2A.Z
H4K20 me2:H3−5ac
H3K9 me3:H4K20 me3
DNA Meth. m5C:H4K20 me2
H4K20 me2:H4−4ac
DNA Meth. m5C:H4K20 me3
H3K27 me3:H4K20 me3
H3K27 me3:DNA Meth. m5C
DNA Meth. m5C:H3K9 me3
H3K27 me2
H3K9 me2
H3K9 me3
H3K4 me3
H2A.Z
H3K4 me1
H3K27 me3
H4K20 me3
H3−5ac
H4−4ac
H4K20 me2
DNA Meth. m5C
0.00 0.25 0.50 0.75 1.00
π̂
WWTR1 − Reverse
H3K4 me1:H4K20 me2
H3K4 me3:H3−5ac
H4−4ac:H3−5ac
H2A.Z:H4K20 me2
H3K4 me3:H4−4ac
H3K4 me1:H3−5ac
H3K27 me3:H3K9 me3
H2A.Z:H3K4 me3
H3K9 me2:DNA Meth. m5C
H4K20 me2:H3K4 me3
H3K4 me1:H4−4ac
H3K27 me2:DNA Meth. m5C
H3K27 me3:H3K4 me1
H3K4 me1:H2A.Z
H4K20 me2:H3−5ac
H3K9 me3:H4K20 me3
DNA Meth. m5C:H4K20 me2
H4K20 me2:H4−4ac
DNA Meth. m5C:H4K20 me3
H3K27 me3:H4K20 me3
H3K27 me3:DNA Meth. m5C
DNA Meth. m5C:H3K9 me3
H3K27 me2
H3K9 me2
H3K9 me3
H3K4 me3
H2A.Z
H3K4 me1
H3K27 me3
H4K20 me3
H3−5ac
H4−4ac
H4K20 me2
DNA Meth. m5C
0.00 0.25 0.50 0.75 1.00
π̂
FOXK2 − Forward
H3K9 me3:H4K20 me3
H3K27 me2:DNA Meth. m5C
H3K9 me2:DNA Meth. m5C
H3K27 me3:H3K4 me1
H3K27 me3:H3K9 me3
DNA Meth. m5C:H3K9 me3
H3K27 me3:H4K20 me3
H3K27 me3:DNA Meth. m5C
H3K4 me1:H3−5ac
DNA Meth. m5C:H4K20 me2
H3K4 me3:H3−5ac
H3K4 me1:H4−4ac
H2A.Z:H3−5ac
H2A.Z:H4−4ac
H3K4 me1:H2A.Z
H2A.Z:H4K20 me2
H3K4 me1:H4K20 me2
H4K20 me2:H3−5ac
H4K20 me2:H4−4ac
H2A.Z:H3K4 me3
DNA Meth. m5C:H4K20 me3
H4−4ac:H3−5ac
H4K20 me2:H3K4 me3
H3K4 me3:H4−4ac
H3K27 me2
H3K9 me2
H4K20 me3
H3K9 me3
H3K27 me3
H3−5ac
H4K20 me2
H2A.Z
H3K4 me1
H4−4ac
H3K4 me3
DNA Meth. m5C
0.00 0.25 0.50 0.75 1.00
π̂
FOXK2 − Reverse
H3K9 me3:H4K20 me3
H3K27 me2:DNA Meth. m5C
H3K9 me2:DNA Meth. m5C
H3K27 me3:H3K4 me1
H3K27 me3:H3K9 me3
DNA Meth. m5C:H3K9 me3
H3K27 me3:H4K20 me3
H3K27 me3:DNA Meth. m5C
H3K4 me1:H3−5ac
DNA Meth. m5C:H4K20 me2
H3K4 me3:H3−5ac
H3K4 me1:H4−4ac
H2A.Z:H3−5ac
H2A.Z:H4−4ac
H3K4 me1:H2A.Z
H2A.Z:H4K20 me2
H3K4 me1:H4K20 me2
H4K20 me2:H3−5ac
H4K20 me2:H4−4ac
H2A.Z:H3K4 me3
DNA Meth. m5C:H4K20 me3
H4−4ac:H3−5ac
H4K20 me2:H3K4 me3
H3K4 me3:H4−4ac
H3K27 me2
H3K9 me2
H4K20 me3
H3K9 me3
H3K27 me3
H3−5ac
H4K20 me2
H2A.Z
H3K4 me1
H4−4ac
H3K4 me3
DNA Meth. m5C
0.00 0.25 0.50 0.75 1.00
π̂
FOXK1 − Forward
H3K4 me1:H4K20 me2
H3K27 me2:DNA Meth. m5C
H3K27 me3:H3K9 me3
H3K4 me3:H3−5ac
H2A.Z:H3−5ac
H3K4 me1:H4−4ac
H3K9 me3:H4K20 me3
H3K9 me2:DNA Meth. m5C
H3K27 me3:H3K4 me1
H3K27 me3:DNA Meth. m5C
H2A.Z:H4−4ac
H3K27 me3:H4K20 me3
H3K4 me1:H2A.Z
H3K4 me1:H3−5ac
DNA Meth. m5C:H4K20 me2
H2A.Z:H4K20 me2
DNA Meth. m5C:H3K9 me3
DNA Meth. m5C:H4K20 me3
H4K20 me2:H4−4ac
H2A.Z:H3K4 me3
H4−4ac:H3−5ac
H4K20 me2:H3−5ac
H4K20 me2:H3K4 me3
H3K4 me3:H4−4ac
H3K9 me3
H3K9 me2
H3K4 me1
H3K27 me2
H4K20 me2
H3−5ac
H4−4ac
H4K20 me3
H3K27 me3
H3K4 me3
H2A.Z
DNA Meth. m5C
0.00 0.25 0.50 0.75 1.00
π̂
FOXK1 − Reverse
H3K4 me1:H4K20 me2
H3K27 me2:DNA Meth. m5C
H3K27 me3:H3K9 me3
H3K4 me3:H3−5ac
H2A.Z:H3−5ac
H3K4 me1:H4−4ac
H3K9 me3:H4K20 me3
H3K9 me2:DNA Meth. m5C
H3K27 me3:H3K4 me1
H3K27 me3:DNA Meth. m5C
H2A.Z:H4−4ac
H3K27 me3:H4K20 me3
H3K4 me1:H2A.Z
H3K4 me1:H3−5ac
DNA Meth. m5C:H4K20 me2
H2A.Z:H4K20 me2
DNA Meth. m5C:H3K9 me3
DNA Meth. m5C:H4K20 me3
H4K20 me2:H4−4ac
H2A.Z:H3K4 me3
H4−4ac:H3−5ac
H4K20 me2:H3−5ac
H4K20 me2:H3K4 me3
H3K4 me3:H4−4ac
H3K9 me3
H3K9 me2
H3K4 me1
H3K27 me2
H4K20 me2
H3−5ac
H4−4ac
H4K20 me3
H3K27 me3
H3K4 me3
H2A.Z
DNA Meth. m5C
0.00 0.25 0.50 0.75 1.00
π̂

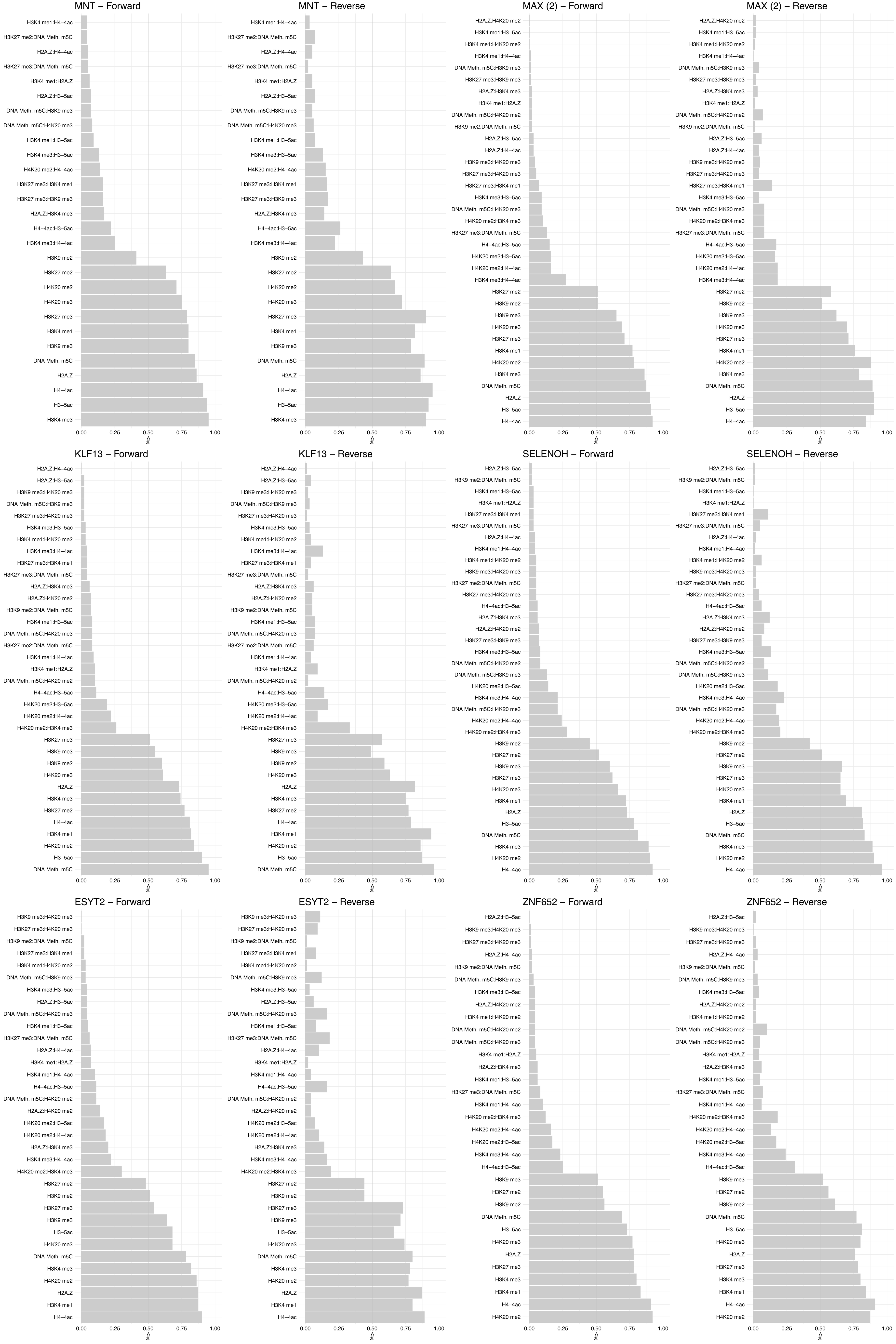
MNT – Forward
H3K4 me1:H4–4ac
H3K27 me2:DNA Meth. m5C
H2A.Z:H4–4ac
H3K27 me3:DNA Meth. m5C
H3K4 me1:H2A.Z
H2A.Z:H3–5ac
DNA Meth. m5C:H3K9 me3
DNA Meth. m5C:H4K20 me3
H3K4 me1:H3–5ac
H3K4 me3:H3–5ac
H4K20 me2:H4–4ac
H3K27 me3:H3K4 me1
H3K27 me3:H3K9 me3
H2A.Z:H3K4 me3
H4–4ac:H3–5ac
H3K4 me3:H4–4ac
H3K9 me2
H3K27 me2
H4K20 me2
H4K20 me3
H3K27 me3
H3K4 me1
H3K9 me3
DNA Meth. m5C
H2A.Z
H4–4ac
H3–5ac
H3K4 me3
0.00
0.25
0.50
0.75
1.00
π̂
MNT – Reverse
H3K4 me1:H4–4ac
H3K27 me2:DNA Meth. m5C
H2A.Z:H4–4ac
H3K27 me3:DNA Meth. m5C
H3K4 me1:H2A.Z
H2A.Z:H3–5ac
DNA Meth. m5C:H3K9 me3
DNA Meth. m5C:H4K20 me3
H3K4 me1:H3–5ac
H3K4 me3:H3–5ac
H4K20 me2:H4–4ac
H3K27 me3:H3K4 me1
H3K27 me3:H3K9 me3
H2A.Z:H3K4 me3
H4–4ac:H3–5ac
H3K4 me3:H4–4ac
H3K9 me2
H3K27 me2
H4K20 me2
H4K20 me3
H3K27 me3
H3K4 me1
H3K9 me3
DNA Meth. m5C
H2A.Z
H4–4ac
H3–5ac
H3K4 me3
0.00
0.25
0.50
0.75
1.00
π̂
MAX (2) – Forward
H2A.Z:H4K20 me2
H3K4 me1:H3–5ac
H3K4 me1:H4K20 me2
H3K4 me1:H4–4ac
DNA Meth. m5C:H3K9 me3
H3K27 me3:H3K9 me3
H2A.Z:H3K4 me3
H3K4 me1:H2A.Z
DNA Meth. m5C:H4K20 me2
H3K9 me2:DNA Meth. m5C
H2A.Z:H3–5ac
H2A.Z:H4–4ac
H3K9 me3:H4K20 me3
H3K27 me3:H4K20 me3
H3K27 me3:H3K4 me1
H3K4 me3:H3–5ac
DNA Meth. m5C:H4K20 me3
H4K20 me2:H3K4 me3
H3K27 me3:DNA Meth. m5C
H4–4ac:H3–5ac
H4K20 me2:H3–5ac
H4K20 me2:H4–4ac
H3K4 me3:H4–4ac
H3K27 me2
H3K9 me2
H3K9 me3
H4K20 me3
H3K27 me3
H3K4 me1
H4K20 me2
H3K4 me3
DNA Meth. m5C
H2A.Z
H3–5ac
H4–4ac
0.00
0.25
0.50
0.75
1.00
π̂
MAX (2) – Reverse
H2A.Z:H4K20 me2
H3K4 me1:H3–5ac
H3K4 me1:H4K20 me2
H3K4 me1:H4–4ac
DNA Meth. m5C:H3K9 me3
H3K27 me3:H3K9 me3
H2A.Z:H3K4 me3
H3K4 me1:H2A.Z
DNA Meth. m5C:H4K20 me2
H3K9 me2:DNA Meth. m5C
H2A.Z:H3–5ac
H2A.Z:H4–4ac
H3K9 me3:H4K20 me3
H3K27 me3:H4K20 me3
H3K27 me3:H3K4 me1
H3K4 me3:H3–5ac
DNA Meth. m5C:H4K20 me3
H4K20 me2:H3K4 me3
H3K27 me3:DNA Meth. m5C
H4–4ac:H3–5ac
H4K20 me2:H3–5ac
H4K20 me2:H4–4ac
H3K4 me3:H4–4ac
H3K27 me2
H3K9 me2
H3K9 me3
H4K20 me3
H3K27 me3
H3K4 me1
H4K20 me2
H3K4 me3
DNA Meth. m5C
H2A.Z
H3–5ac
H4–4ac
0.00
0.25
0.50
0.75
1.00
π̂
KLF13 – Forward
H2A.Z:H4–4ac
H2A.Z:H3–5ac
H3K9 me3:H4K20 me3
DNA Meth. m5C:H3K9 me3
H3K27 me3:H4K20 me3
H3K4 me3:H3–5ac
H3K4 me1:H4K20 me2
H3K4 me3:H4–4ac
H3K27 me3:H3K4 me1
H3K27 me3:DNA Meth. m5C
H2A.Z:H3K4 me3
H2A.Z:H4K20 me2
H3K9 me2:DNA Meth. m5C
H3K4 me1:H3–5ac
DNA Meth. m5C:H4K20 me3
H3K27 me2:DNA Meth. m5C
H3K4 me1:H4–4ac
H3K4 me1:H2A.Z
DNA Meth. m5C:H4K20 me2
H4–4ac:H3–5ac
H4K20 me2:H3–5ac
H4K20 me2:H4–4ac
H4K20 me2:H3K4 me3
H3K27 me3
H3K9 me3
H3K9 me2
H4K20 me3
H2A.Z
H3K4 me3
H3K27 me2
H4–4ac
H3K4 me1
H4K20 me2
H3–5ac
DNA Meth. m5C
0.00
0.25
0.50
0.75
1.00
π̂
KLF13 – Reverse
H2A.Z:H4–4ac
H2A.Z:H3–5ac
H3K9 me3:H4K20 me3
DNA Meth. m5C:H3K9 me3
H3K27 me3:H4K20 me3
H3K4 me3:H3–5ac
H3K4 me1:H4K20 me2
H3K4 me3:H4–4ac
H3K27 me3:H3K4 me1
H3K27 me3:DNA Meth. m5C
H2A.Z:H3K4 me3
H2A.Z:H4K20 me2
H3K9 me2:DNA Meth. m5C
H3K4 me1:H3–5ac
DNA Meth. m5C:H4K20 me3
H3K27 me2:DNA Meth. m5C
H3K4 me1:H4–4ac
H3K4 me1:H2A.Z
DNA Meth. m5C:H4K20 me2
H4–4ac:H3–5ac
H4K20 me2:H3–5ac
H4K20 me2:H4–4ac
H4K20 me2:H3K4 me3
H3K27 me3
H3K9 me3
H3K9 me2
H4K20 me3
H2A.Z
H3K4 me3
H3K27 me2
H4–4ac
H3K4 me1
H4K20 me2
H3–5ac
DNA Meth. m5C
0.00
0.25
0.50
0.75
1.00
π̂
SELENOH – Forward
H2A.Z:H3–5ac
H3K9 me2:DNA Meth. m5C
H3K4 me1:H3–5ac
H3K4 me1:H2A.Z
H3K27 me3:H3K4 me1
H3K27 me3:DNA Meth. m5C
H2A.Z:H4–4ac
H3K4 me1:H4–4ac
H3K4 me1:H4K20 me2
H3K9 me3:H4K20 me3
H3K27 me2:DNA Meth. m5C
H3K27 me3:H4K20 me3
H4–4ac:H3–5ac
H2A.Z:H3K4 me3
H2A.Z:H4K20 me2
H3K27 me3:H3K9 me3
H3K4 me3:H3–5ac
DNA Meth. m5C:H4K20 me2
DNA Meth. m5C:H3K9 me3
H4K20 me2:H3–5ac
H3K4 me3:H4–4ac
DNA Meth. m5C:H4K20 me3
H4K20 me2:H4–4ac
H4K20 me2:H3K4 me3
H3K9 me2
H3K27 me2
H3K9 me3
H3K27 me3
H4K20 me3
H3K4 me1
H2A.Z
H3–5ac
DNA Meth. m5C
H3K4 me3
H4K20 me2
H4–4ac
0.00
0.25
0.50
0.75
1.00
π̂
SELENOH – Reverse
H2A.Z:H3–5ac
H3K9 me2:DNA Meth. m5C
H3K4 me1:H3–5ac
H3K4 me1:H2A.Z
H3K27 me3:H3K4 me1
H3K27 me3:DNA Meth. m5C
H2A.Z:H4–4ac
H3K4 me1:H4–4ac
H3K4 me1:H4K20 me2
H3K9 me3:H4K20 me3
H3K27 me2:DNA Meth. m5C
H3K27 me3:H4K20 me3
H4–4ac:H3–5ac
H2A.Z:H3K4 me3
H2A.Z:H4K20 me2
H3K27 me3:H3K9 me3
H3K4 me3:H3–5ac
DNA Meth. m5C:H4K20 me2
DNA Meth. m5C:H3K9 me3
H4K20 me2:H3–5ac
H3K4 me3:H4–4ac
DNA Meth. m5C:H4K20 me3
H4K20 me2:H4–4ac
H4K20 me2:H3K4 me3
H3K9 me2
H3K27 me2
H3K9 me3
H3K27 me3
H4K20 me3
H3K4 me1
H2A.Z
H3–5ac
DNA Meth. m5C
H3K4 me3
H4K20 me2
H4–4ac
0.00
0.25
0.50
0.75
1.00
π̂
ESYT2 – Forward
H3K9 me3:H4K20 me3
H3K27 me3:H4K20 me3
H3K9 me2:DNA Meth. m5C
H3K27 me3:H3K4 me1
H3K4 me1:H4K20 me2
DNA Meth. m5C:H3K9 me3
H3K4 me3:H3–5ac
H2A.Z:H3–5ac
DNA Meth. m5C:H4K20 me3
H3K4 me1:H3–5ac
H3K27 me3:DNA Meth. m5C
H2A.Z:H4–4ac
H3K4 me1:H2A.Z
H3K4 me1:H4–4ac
H4–4ac:H3–5ac
DNA Meth. m5C:H4K20 me2
H2A.Z:H4K20 me2
H4K20 me2:H3–5ac
H4K20 me2:H4–4ac
H2A.Z:H3K4 me3
H3K4 me3:H4–4ac
H4K20 me2:H3K4 me3
H3K27 me2
H3K9 me2
H3K27 me3
H3K9 me3
H3–5ac
H4K20 me3
DNA Meth. m5C
H3K4 me3
H4K20 me2
H2A.Z
H3K4 me1
H4–4ac
0.00
0.25
0.50
0.75
1.00
π̂
ESYT2 – Reverse
H3K9 me3:H4K20 me3
H3K27 me3:H4K20 me3
H3K9 me2:DNA Meth. m5C
H3K27 me3:H3K4 me1
H3K4 me1:H4K20 me2
DNA Meth. m5C:H3K9 me3
H3K4 me3:H3–5ac
H2A.Z:H3–5ac
DNA Meth. m5C:H4K20 me3
H3K4 me1:H3–5ac
H3K27 me3:DNA Meth. m5C
H2A.Z:H4–4ac
H3K4 me1:H2A.Z
H3K4 me1:H4–4ac
H4–4ac:H3–5ac
DNA Meth. m5C:H4K20 me2
H2A.Z:H4K20 me2
H4K20 me2:H3–5ac
H4K20 me2:H4–4ac
H2A.Z:H3K4 me3
H3K4 me3:H4–4ac
H4K20 me2:H3K4 me3
H3K27 me2
H3K9 me2
H3K27 me3
H3K9 me3
H3–5ac
H4K20 me3
DNA Meth. m5C
H3K4 me3
H4K20 me2
H2A.Z
H3K4 me1
H4–4ac
0.00
0.25
0.50
0.75
1.00
π̂
ZNF652 – Forward
H2A.Z:H3–5ac
H3K9 me3:H4K20 me3
H3K27 me3:H4K20 me3
H2A.Z:H4–4ac
H3K9 me2:DNA Meth. m5C
DNA Meth. m5C:H3K9 me3
H3K4 me3:H3–5ac
H2A.Z:H4K20 me2
H3K4 me1:H4K20 me2
DNA Meth. m5C:H4K20 me2
DNA Meth. m5C:H4K20 me3
H3K4 me1:H2A.Z
H2A.Z:H3K4 me3
H3K4 me1:H3–5ac
H3K27 me3:DNA Meth. m5C
H3K4 me1:H4–4ac
H4K20 me2:H3K4 me3
H4K20 me2:H4–4ac
H4K20 me2:H3–5ac
H3K4 me3:H4–4ac
H4–4ac:H3–5ac
H3K9 me3
H3K27 me2
H3K9 me2
DNA Meth. m5C
H3–5ac
H4K20 me3
H2A.Z
H3K27 me3
H3K4 me3
H3K4 me1
H4–4ac
H4K20 me2
0.00
0.25
0.50
0.75
1.00
π̂
ZNF652 – Reverse
H2A.Z:H3–5ac
H3K9 me3:H4K20 me3
H3K27 me3:H4K20 me3
H2A.Z:H4–4ac
H3K9 me2:DNA Meth. m5C
DNA Meth. m5C:H3K9 me3
H3K4 me3:H3–5ac
H2A.Z:H4K20 me2
H3K4 me1:H4K20 me2
DNA Meth. m5C:H4K20 me2
DNA Meth. m5C:H4K20 me3
H3K4 me1:H2A.Z
H2A.Z:H3K4 me3
H3K4 me1:H3–5ac
H3K27 me3:DNA Meth. m5C
H3K4 me1:H4–4ac
H4K20 me2:H3K4 me3
H4K20 me2:H4–4ac
H4K20 me2:H3–5ac
H3K4 me3:H4–4ac
H4–4ac:H3–5ac
H3K9 me3
H3K27 me2
H3K9 me2
DNA Meth. m5C
H3–5ac
H4K20 me3
H2A.Z
H3K27 me3
H3K4 me3
H3K4 me1
H4–4ac
H4K20 me2
0.00
0.25
0.50
0.75
1.00
π̂

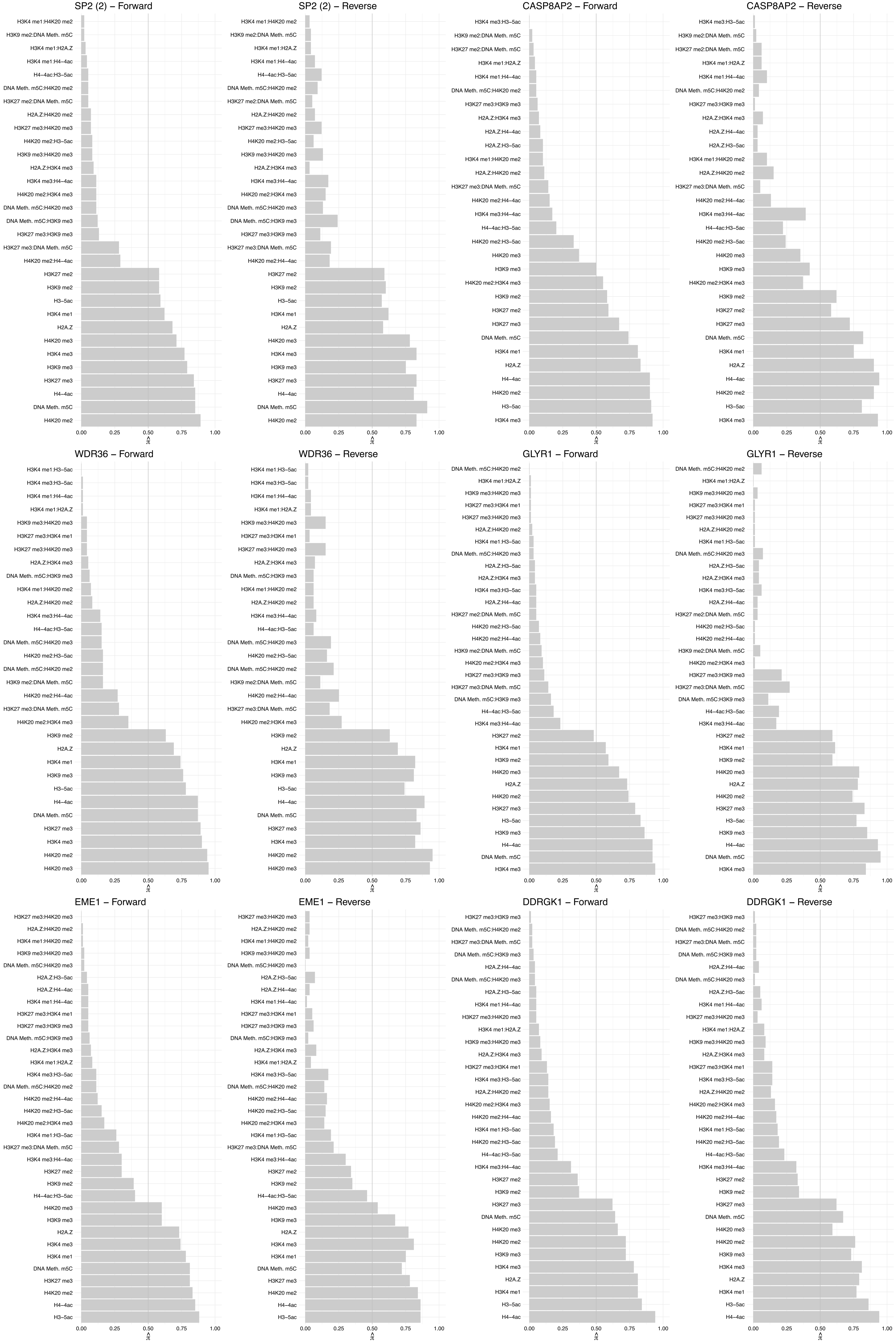
SP2 (2) – Forward
H3K4 me1:H4K20 me2
H3K9 me2:DNA Meth. m5C
H3K4 me1:H2A.Z
H3K4 me1:H4–4ac
H4–4ac:H3–5ac
DNA Meth. m5C:H4K20 me2
H3K27 me2:DNA Meth. m5C
H2A.Z:H4K20 me2
H3K27 me3:H4K20 me3
H4K20 me2:H3–5ac
H3K9 me3:H4K20 me3
H2A.Z:H3K4 me3
H3K4 me3:H4–4ac
H4K20 me2:H3K4 me3
DNA Meth. m5C:H4K20 me3
DNA Meth. m5C:H3K9 me3
H3K27 me3:H3K9 me3
H3K27 me3:DNA Meth. m5C
H4K20 me2:H4–4ac
H3K27 me2
H3K9 me2
H3–5ac
H3K4 me1
H2A.Z
H4K20 me3
H3K4 me3
H3K9 me3
H3K27 me3
H4–4ac
DNA Meth. m5C
H4K20 me2
0.00 0.25 0.50 0.75 1.00
$\hat{\pi}$
SP2 (2) – Reverse
H3K4 me1:H4K20 me2
H3K9 me2:DNA Meth. m5C
H3K4 me1:H2A.Z
H3K4 me1:H4–4ac
H4–4ac:H3–5ac
DNA Meth. m5C:H4K20 me2
H3K27 me2:DNA Meth. m5C
H2A.Z:H4K20 me2
H3K27 me3:H4K20 me3
H4K20 me2:H3–5ac
H3K9 me3:H4K20 me3
H2A.Z:H3K4 me3
H3K4 me3:H4–4ac
H4K20 me2:H3K4 me3
DNA Meth. m5C:H4K20 me3
DNA Meth. m5C:H3K9 me3
H3K27 me3:H3K9 me3
H3K27 me3:DNA Meth. m5C
H4K20 me2:H4–4ac
H3K27 me2
H3K9 me2
H3–5ac
H3K4 me1
H2A.Z
H4K20 me3
H3K4 me3
H3K9 me3
H3K27 me3
H4–4ac
DNA Meth. m5C
H4K20 me2
0.00 0.25 0.50 0.75 1.00
$\hat{\pi}$
CASP8AP2 – Forward
H3K4 me3:H3–5ac
H3K9 me2:DNA Meth. m5C
H3K27 me2:DNA Meth. m5C
H3K4 me1:H2A.Z
H3K4 me1:H4–4ac
DNA Meth. m5C:H4K20 me2
H3K27 me3:H3K9 me3
H2A.Z:H3K4 me3
H2A.Z:H4–4ac
H2A.Z:H3–5ac
H3K4 me1:H4K20 me2
H2A.Z:H4K20 me2
H3K27 me3:DNA Meth. m5C
H4K20 me2:H4–4ac
H3K4 me3:H4–4ac
H4–4ac:H3–5ac
H4K20 me2:H3–5ac
H4K20 me3
H3K9 me3
H4K20 me2:H3K4 me3
H3K9 me2
H3K27 me2
H3K27 me3
DNA Meth. m5C
H3K4 me1
H2A.Z
H4–4ac
H4K20 me2
H3–5ac
H3K4 me3
0.00 0.25 0.50 0.75 1.00
$\hat{\pi}$
CASP8AP2 – Reverse
H3K4 me3:H3–5ac
H3K9 me2:DNA Meth. m5C
H3K27 me2:DNA Meth. m5C
H3K4 me1:H2A.Z
H3K4 me1:H4–4ac
DNA Meth. m5C:H4K20 me2
H3K27 me3:H3K9 me3
H2A.Z:H3K4 me3
H2A.Z:H4–4ac
H2A.Z:H3–5ac
H3K4 me1:H4K20 me2
H2A.Z:H4K20 me2
H3K27 me3:DNA Meth. m5C
H4K20 me2:H4–4ac
H3K4 me3:H4–4ac
H4–4ac:H3–5ac
H4K20 me2:H3–5ac
H4K20 me3
H3K9 me3
H4K20 me2:H3K4 me3
H3K9 me2
H3K27 me2
H3K27 me3
DNA Meth. m5C
H3K4 me1
H2A.Z
H4–4ac
H4K20 me2
H3–5ac
H3K4 me3
0.00 0.25 0.50 0.75 1.00
$\hat{\pi}$
WDR36 – Forward
H3K4 me1:H3–5ac
H3K4 me3:H3–5ac
H3K4 me1:H4–4ac
H3K4 me1:H2A.Z
H3K9 me3:H4K20 me3
H3K27 me3:H3K4 me1
H3K27 me3:H4K20 me3
H2A.Z:H3K4 me3
DNA Meth. m5C:H3K9 me3
H3K4 me1:H4K20 me2
H2A.Z:H4K20 me2
H3K4 me3:H4–4ac
H4–4ac:H3–5ac
DNA Meth. m5C:H4K20 me3
H4K20 me2:H3–5ac
DNA Meth. m5C:H4K20 me2
H3K9 me2:DNA Meth. m5C
H4K20 me2:H4–4ac
H3K27 me3:DNA Meth. m5C
H4K20 me2:H3K4 me3
H3K9 me2
H2A.Z
H3K4 me1
H3K9 me3
H3–5ac
H4–4ac
DNA Meth. m5C
H3K27 me3
H3K4 me3
H4K20 me2
H4K20 me3
0.00 0.25 0.50 0.75 1.00
$\hat{\pi}$
WDR36 – Reverse
H3K4 me1:H3–5ac
H3K4 me3:H3–5ac
H3K4 me1:H4–4ac
H3K4 me1:H2A.Z
H3K9 me3:H4K20 me3
H3K27 me3:H3K4 me1
H3K27 me3:H4K20 me3
H2A.Z:H3K4 me3
DNA Meth. m5C:H3K9 me3
H3K4 me1:H4K20 me2
H2A.Z:H4K20 me2
H3K4 me3:H4–4ac
H4–4ac:H3–5ac
DNA Meth. m5C:H4K20 me3
H4K20 me2:H3–5ac
DNA Meth. m5C:H4K20 me2
H3K9 me2:DNA Meth. m5C
H4K20 me2:H4–4ac
H3K27 me3:DNA Meth. m5C
H4K20 me2:H3K4 me3
H3K9 me2
H2A.Z
H3K4 me1
H3K9 me3
H3–5ac
H4–4ac
DNA Meth. m5C
H3K27 me3
H3K4 me3
H4K20 me2
H4K20 me3
0.00 0.25 0.50 0.75 1.00
$\hat{\pi}$
GLYR1 – Forward
DNA Meth. m5C:H4K20 me2
H3K4 me1:H2A.Z
H3K9 me3:H4K20 me3
H3K27 me3:H3K4 me1
H3K27 me3:H4K20 me3
H2A.Z:H4K20 me2
H3K4 me1:H3–5ac
DNA Meth. m5C:H4K20 me3
H2A.Z:H3–5ac
H2A.Z:H3K4 me3
H3K4 me3:H3–5ac
H2A.Z:H4–4ac
H3K27 me2:DNA Meth. m5C
H4K20 me2:H3–5ac
H4K20 me2:H4–4ac
H3K9 me2:DNA Meth. m5C
H4K20 me2:H3K4 me3
H3K27 me3:H3K9 me3
H3K27 me3:DNA Meth. m5C
DNA Meth. m5C:H3K9 me3
H4–4ac:H3–5ac
H3K4 me3:H4–4ac
H3K27 me2
H3K4 me1
H3K9 me2
H4K20 me3
H2A.Z
H4K20 me2
H3K27 me3
H3–5ac
H3K9 me3
H4–4ac
DNA Meth. m5C
H3K4 me3
0.00 0.25 0.50 0.75 1.00
$\hat{\pi}$
GLYR1 – Reverse
DNA Meth. m5C:H4K20 me2
H3K4 me1:H2A.Z
H3K9 me3:H4K20 me3
H3K27 me3:H3K4 me1
H3K27 me3:H4K20 me3
H2A.Z:H4K20 me2
H3K4 me1:H3–5ac
DNA Meth. m5C:H4K20 me3
H2A.Z:H3–5ac
H2A.Z:H3K4 me3
H3K4 me3:H3–5ac
H2A.Z:H4–4ac
H3K27 me2:DNA Meth. m5C
H4K20 me2:H3–5ac
H4K20 me2:H4–4ac
H3K9 me2:DNA Meth. m5C
H4K20 me2:H3K4 me3
H3K27 me3:H3K9 me3
H3K27 me3:DNA Meth. m5C
DNA Meth. m5C:H3K9 me3
H4–4ac:H3–5ac
H3K4 me3:H4–4ac
H3K27 me2
H3K4 me1
H3K9 me2
H4K20 me3
H2A.Z
H4K20 me2
H3K27 me3
H3–5ac
H3K9 me3
H4–4ac
DNA Meth. m5C
H3K4 me3
0.00 0.25 0.50 0.75 1.00
$\hat{\pi}$
EME1 – Forward
H3K27 me3:H4K20 me3
H2A.Z:H4K20 me2
H3K4 me1:H4K20 me2
H3K9 me3:H4K20 me3
DNA Meth. m5C:H4K20 me3
H2A.Z:H3–5ac
H2A.Z:H4–4ac
H3K4 me1:H4–4ac
H3K27 me3:H3K4 me1
H3K27 me3:H3K9 me3
DNA Meth. m5C:H3K9 me3
H2A.Z:H3K4 me3
H3K4 me1:H2A.Z
H3K4 me3:H3–5ac
DNA Meth. m5C:H4K20 me2
H4K20 me2:H4–4ac
H4K20 me2:H3–5ac
H4K20 me2:H3K4 me3
H3K4 me1:H3–5ac
H3K27 me3:DNA Meth. m5C
H3K4 me3:H4–4ac
H3K27 me2
H3K9 me2
H4–4ac:H3–5ac
H4K20 me3
H3K9 me3
H2A.Z
H3K4 me3
H3K4 me1
DNA Meth. m5C
H3K27 me3
H4K20 me2
H4–4ac
H3–5ac
0.00 0.25 0.50 0.75 1.00
$\hat{\pi}$
EME1 – Reverse
H3K27 me3:H4K20 me3
H2A.Z:H4K20 me2
H3K4 me1:H4K20 me2
H3K9 me3:H4K20 me3
DNA Meth. m5C:H4K20 me3
H2A.Z:H3–5ac
H2A.Z:H4–4ac
H3K4 me1:H4–4ac
H3K27 me3:H3K4 me1
H3K27 me3:H3K9 me3
DNA Meth. m5C:H3K9 me3
H2A.Z:H3K4 me3
H3K4 me1:H2A.Z
H3K4 me3:H3–5ac
DNA Meth. m5C:H4K20 me2
H4K20 me2:H4–4ac
H4K20 me2:H3–5ac
H4K20 me2:H3K4 me3
H3K4 me1:H3–5ac
H3K27 me3:DNA Meth. m5C
H3K4 me3:H4–4ac
H3K27 me2
H3K9 me2
H4–4ac:H3–5ac
H4K20 me3
H3K9 me3
H2A.Z
H3K4 me3
H3K4 me1
DNA Meth. m5C
H3K27 me3
H4K20 me2
H4–4ac
H3–5ac
0.00 0.25 0.50 0.75 1.00
$\hat{\pi}$
DDRGK1 – Forward
H3K27 me3:H3K9 me3
DNA Meth. m5C:H4K20 me2
H3K27 me3:DNA Meth. m5C
DNA Meth. m5C:H3K9 me3
H2A.Z:H4–4ac
DNA Meth. m5C:H4K20 me3
H2A.Z:H3–5ac
H3K4 me1:H4–4ac
H3K27 me3:H4K20 me3
H3K4 me1:H2A.Z
H3K9 me3:H4K20 me3
H2A.Z:H3K4 me3
H3K27 me3:H3K4 me1
H3K4 me3:H3–5ac
H2A.Z:H4K20 me2
H4K20 me2:H3K4 me3
H4K20 me2:H4–4ac
H3K4 me1:H3–5ac
H4K20 me2:H3–5ac
H4–4ac:H3–5ac
H3K4 me3:H4–4ac
H3K27 me2
H3K9 me2
H3K27 me3
DNA Meth. m5C
H4K20 me3
H4K20 me2
H3K9 me3
H3K4 me3
H2A.Z
H3K4 me1
H3–5ac
H4–4ac
0.00 0.25 0.50 0.75 1.00
$\hat{\pi}$
DDRGK1 – Reverse
H3K27 me3:H3K9 me3
DNA Meth. m5C:H4K20 me2
H3K27 me3:DNA Meth. m5C
DNA Meth. m5C:H3K9 me3
H2A.Z:H4–4ac
DNA Meth. m5C:H4K20 me3
H2A.Z:H3–5ac
H3K4 me1:H4–4ac
H3K27 me3:H4K20 me3
H3K4 me1:H2A.Z
H3K9 me3:H4K20 me3
H2A.Z:H3K4 me3
H3K27 me3:H3K4 me1
H3K4 me3:H3–5ac
H2A.Z:H4K20 me2
H4K20 me2:H3K4 me3
H4K20 me2:H4–4ac
H3K4 me1:H3–5ac
H4K20 me2:H3–5ac
H4–4ac:H3–5ac
H3K4 me3:H4–4ac
H3K27 me2
H3K9 me2
H3K27 me3
DNA Meth. m5C
H4K20 me3
H4K20 me2
H3K9 me3
H3K4 me3
H2A.Z
H3K4 me1
H3–5ac
H4–4ac
0.00 0.25 0.50 0.75 1.00
$\hat{\pi}$

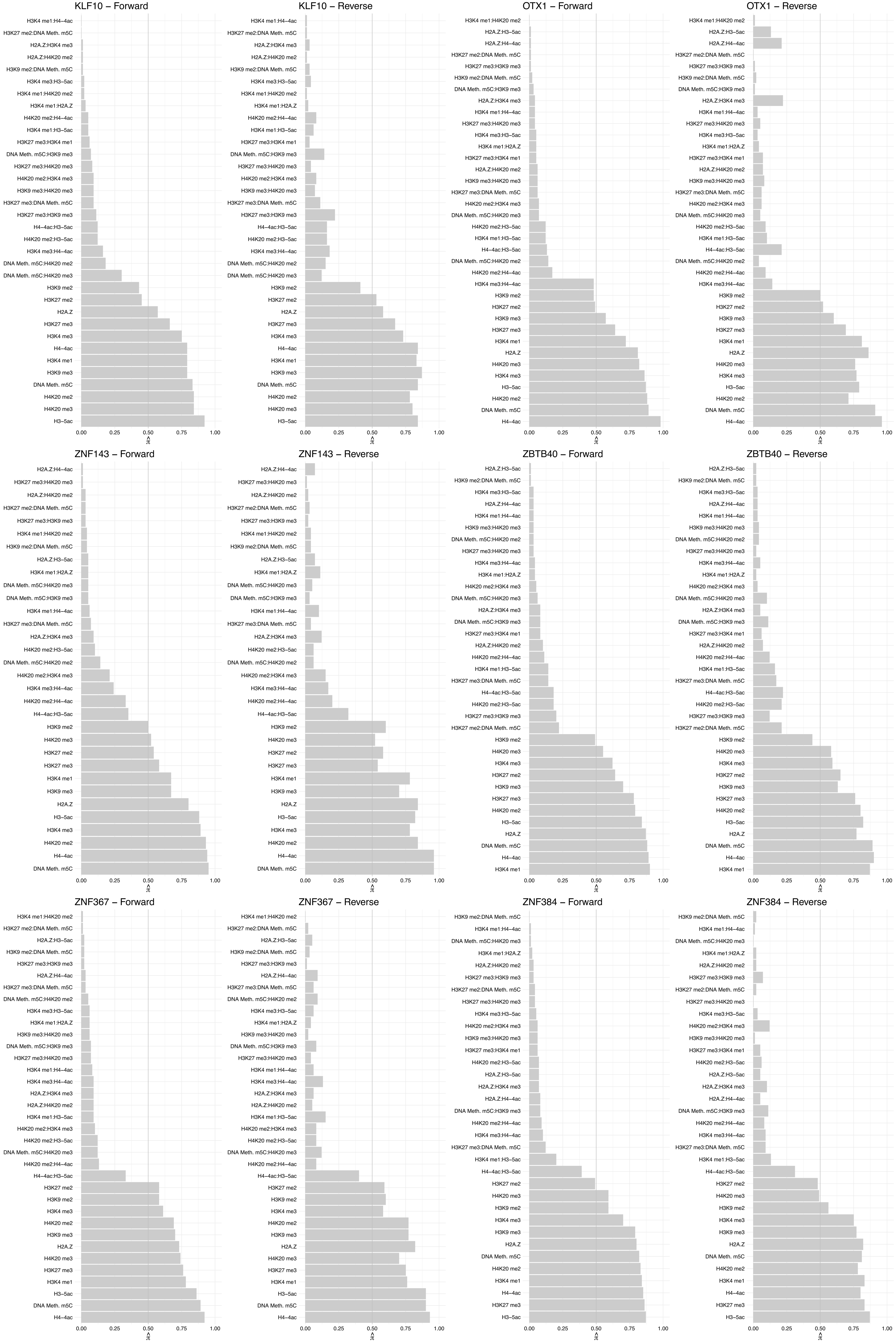

KLF10 − Forward
H3K4 me1:H4−4ac
H3K27 me2:DNA Meth. m5C
H2A.Z:H3K4 me3
H2A.Z:H4K20 me2
H3K9 me2:DNA Meth. m5C
H3K4 me3:H3−5ac
H3K4 me1:H4K20 me2
H3K4 me1:H2A.Z
H4K20 me2:H4−4ac
H3K4 me1:H3−5ac
H3K27 me3:H3K4 me1
DNA Meth. m5C:H3K9 me3
H3K27 me3:H4K20 me3
H4K20 me2:H3K4 me3
H3K9 me3:H4K20 me3
H3K27 me3:DNA Meth. m5C
H3K27 me3:H3K9 me3
H4−4ac:H3−5ac
H4K20 me2:H3−5ac
H3K4 me3:H4−4ac
DNA Meth. m5C:H4K20 me2
DNA Meth. m5C:H4K20 me3
H3K9 me2
H3K27 me2
H2A.Z
H3K27 me3
H3K4 me3
H4−4ac
H3K4 me1
H3K9 me3
DNA Meth. m5C
H4K20 me2
H4K20 me3
H3−5ac
0.00 0.25 0.50 0.75 1.00
π̂
KLF10 − Reverse
H3K4 me1:H4−4ac
H3K27 me2:DNA Meth. m5C
H2A.Z:H3K4 me3
H2A.Z:H4K20 me2
H3K9 me2:DNA Meth. m5C
H3K4 me3:H3−5ac
H3K4 me1:H4K20 me2
H3K4 me1:H2A.Z
H4K20 me2:H4−4ac
H3K4 me1:H3−5ac
H3K27 me3:H3K4 me1
DNA Meth. m5C:H3K9 me3
H3K27 me3:H4K20 me3
H4K20 me2:H3K4 me3
H3K9 me3:H4K20 me3
H3K27 me3:DNA Meth. m5C
H3K27 me3:H3K9 me3
H4−4ac:H3−5ac
H4K20 me2:H3−5ac
H3K4 me3:H4−4ac
DNA Meth. m5C:H4K20 me2
DNA Meth. m5C:H4K20 me3
H3K9 me2
H3K27 me2
H2A.Z
H3K27 me3
H3K4 me3
H4−4ac
H3K4 me1
H3K9 me3
DNA Meth. m5C
H4K20 me2
H4K20 me3
H3−5ac
0.00 0.25 0.50 0.75 1.00
π̂
OTX1 − Forward
H3K4 me1:H4K20 me2
H2A.Z:H3−5ac
H2A.Z:H4−4ac
H3K27 me2:DNA Meth. m5C
H3K27 me3:H3K9 me3
H3K9 me2:DNA Meth. m5C
DNA Meth. m5C:H3K9 me3
H2A.Z:H3K4 me3
H3K4 me1:H4−4ac
H3K27 me3:H4K20 me3
H3K4 me3:H3−5ac
H3K4 me1:H2A.Z
H3K27 me3:H3K4 me1
H2A.Z:H4K20 me2
H3K9 me3:H4K20 me3
H3K27 me3:DNA Meth. m5C
H4K20 me2:H3K4 me3
DNA Meth. m5C:H4K20 me3
H4K20 me2:H3−5ac
H3K4 me1:H3−5ac
H4−4ac:H3−5ac
DNA Meth. m5C:H4K20 me2
H4K20 me2:H4−4ac
H3K4 me3:H4−4ac
H3K9 me2
H3K27 me2
H3K9 me3
H3K27 me3
H3K4 me1
H2A.Z
H4K20 me3
H3K4 me3
H3−5ac
H4K20 me2
DNA Meth. m5C
H4−4ac
0.00 0.25 0.50 0.75 1.00
π̂
OTX1 − Reverse
H3K4 me1:H4K20 me2
H2A.Z:H3−5ac
H2A.Z:H4−4ac
H3K27 me2:DNA Meth. m5C
H3K27 me3:H3K9 me3
H3K9 me2:DNA Meth. m5C
DNA Meth. m5C:H3K9 me3
H2A.Z:H3K4 me3
H3K4 me1:H4−4ac
H3K27 me3:H4K20 me3
H3K4 me3:H3−5ac
H3K4 me1:H2A.Z
H3K27 me3:H3K4 me1
H2A.Z:H4K20 me2
H3K9 me3:H4K20 me3
H3K27 me3:DNA Meth. m5C
H4K20 me2:H3K4 me3
DNA Meth. m5C:H4K20 me3
H4K20 me2:H3−5ac
H3K4 me1:H3−5ac
H4−4ac:H3−5ac
DNA Meth. m5C:H4K20 me2
H4K20 me2:H4−4ac
H3K4 me3:H4−4ac
H3K9 me2
H3K27 me2
H3K9 me3
H3K27 me3
H3K4 me1
H2A.Z
H4K20 me3
H3K4 me3
H3−5ac
H4K20 me2
DNA Meth. m5C
H4−4ac
0.00 0.25 0.50 0.75 1.00
π̂
ZNF143 − Forward
H2A.Z:H4−4ac
H3K27 me3:H4K20 me3
H2A.Z:H4K20 me2
H3K27 me2:DNA Meth. m5C
H3K27 me3:H3K9 me3
H3K4 me1:H4K20 me2
H3K9 me2:DNA Meth. m5C
H2A.Z:H3−5ac
H3K4 me1:H2A.Z
DNA Meth. m5C:H4K20 me3
DNA Meth. m5C:H3K9 me3
H3K4 me1:H4−4ac
H3K27 me3:DNA Meth. m5C
H2A.Z:H3K4 me3
H4K20 me2:H3−5ac
DNA Meth. m5C:H4K20 me2
H4K20 me2:H3K4 me3
H3K4 me3:H4−4ac
H4K20 me2:H4−4ac
H4−4ac:H3−5ac
H3K9 me2
H4K20 me3
H3K27 me2
H3K27 me3
H3K4 me1
H3K9 me3
H2A.Z
H3−5ac
H3K4 me3
H4K20 me2
H4−4ac
DNA Meth. m5C
0.00 0.25 0.50 0.75 1.00
π̂
ZNF143 − Reverse
H2A.Z:H4−4ac
H3K27 me3:H4K20 me3
H2A.Z:H4K20 me2
H3K27 me2:DNA Meth. m5C
H3K27 me3:H3K9 me3
H3K4 me1:H4K20 me2
H3K9 me2:DNA Meth. m5C
H2A.Z:H3−5ac
H3K4 me1:H2A.Z
DNA Meth. m5C:H4K20 me3
DNA Meth. m5C:H3K9 me3
H3K4 me1:H4−4ac
H3K27 me3:DNA Meth. m5C
H2A.Z:H3K4 me3
H4K20 me2:H3−5ac
DNA Meth. m5C:H4K20 me2
H4K20 me2:H3K4 me3
H3K4 me3:H4−4ac
H4K20 me2:H4−4ac
H4−4ac:H3−5ac
H3K9 me2
H4K20 me3
H3K27 me2
H3K27 me3
H3K4 me1
H3K9 me3
H2A.Z
H3−5ac
H3K4 me3
H4K20 me2
H4−4ac
DNA Meth. m5C
0.00 0.25 0.50 0.75 1.00
π̂
ZBTB40 − Forward
H2A.Z:H3−5ac
H3K9 me2:DNA Meth. m5C
H3K4 me3:H3−5ac
H2A.Z:H4−4ac
H3K4 me1:H4−4ac
H3K9 me3:H4K20 me3
DNA Meth. m5C:H4K20 me2
H3K27 me3:H4K20 me3
H3K4 me3:H4−4ac
H3K4 me1:H2A.Z
H4K20 me2:H3K4 me3
DNA Meth. m5C:H4K20 me3
H2A.Z:H3K4 me3
DNA Meth. m5C:H3K9 me3
H3K27 me3:H3K4 me1
H2A.Z:H4K20 me2
H4K20 me2:H4−4ac
H3K4 me1:H3−5ac
H3K27 me3:DNA Meth. m5C
H4−4ac:H3−5ac
H4K20 me2:H3−5ac
H3K27 me3:H3K9 me3
H3K27 me2:DNA Meth. m5C
H3K9 me2
H4K20 me3
H3K4 me3
H3K27 me2
H3K9 me3
H3K27 me3
H4K20 me2
H3−5ac
H2A.Z
DNA Meth. m5C
H4−4ac
H3K4 me1
0.00 0.25 0.50 0.75 1.00
π̂
ZBTB40 − Reverse
H2A.Z:H3−5ac
H3K9 me2:DNA Meth. m5C
H3K4 me3:H3−5ac
H2A.Z:H4−4ac
H3K4 me1:H4−4ac
H3K9 me3:H4K20 me3
DNA Meth. m5C:H4K20 me2
H3K27 me3:H4K20 me3
H3K4 me3:H4−4ac
H3K4 me1:H2A.Z
H4K20 me2:H3K4 me3
DNA Meth. m5C:H4K20 me3
H2A.Z:H3K4 me3
DNA Meth. m5C:H3K9 me3
H3K27 me3:H3K4 me1
H2A.Z:H4K20 me2
H4K20 me2:H4−4ac
H3K4 me1:H3−5ac
H3K27 me3:DNA Meth. m5C
H4−4ac:H3−5ac
H4K20 me2:H3−5ac
H3K27 me3:H3K9 me3
H3K27 me2:DNA Meth. m5C
H3K9 me2
H4K20 me3
H3K4 me3
H3K27 me2
H3K9 me3
H3K27 me3
H4K20 me2
H3−5ac
H2A.Z
DNA Meth. m5C
H4−4ac
H3K4 me1
0.00 0.25 0.50 0.75 1.00
π̂
ZNF367 − Forward
H3K4 me1:H4K20 me2
H3K27 me2:DNA Meth. m5C
H2A.Z:H3−5ac
H3K9 me2:DNA Meth. m5C
H3K27 me3:H3K9 me3
H2A.Z:H4−4ac
H3K27 me3:DNA Meth. m5C
DNA Meth. m5C:H4K20 me2
H3K4 me3:H3−5ac
H3K4 me1:H2A.Z
H3K9 me3:H4K20 me3
DNA Meth. m5C:H3K9 me3
H3K27 me3:H4K20 me3
H3K4 me1:H4−4ac
H3K4 me3:H4−4ac
H2A.Z:H3K4 me3
H2A.Z:H4K20 me2
H3K4 me1:H3−5ac
H4K20 me2:H3K4 me3
H4K20 me2:H3−5ac
DNA Meth. m5C:H4K20 me3
H4K20 me2:H4−4ac
H4−4ac:H3−5ac
H3K27 me2
H3K9 me2
H3K4 me3
H4K20 me2
H3K9 me3
H2A.Z
H4K20 me3
H3K27 me3
H3K4 me1
H3−5ac
DNA Meth. m5C
H4−4ac
0.00 0.25 0.50 0.75 1.00
π̂
ZNF367 − Reverse
H3K4 me1:H4K20 me2
H3K27 me2:DNA Meth. m5C
H2A.Z:H3−5ac
H3K9 me2:DNA Meth. m5C
H3K27 me3:H3K9 me3
H2A.Z:H4−4ac
H3K27 me3:DNA Meth. m5C
DNA Meth. m5C:H4K20 me2
H3K4 me3:H3−5ac
H3K4 me1:H2A.Z
H3K9 me3:H4K20 me3
DNA Meth. m5C:H3K9 me3
H3K27 me3:H4K20 me3
H3K4 me1:H4−4ac
H3K4 me3:H4−4ac
H2A.Z:H3K4 me3
H2A.Z:H4K20 me2
H3K4 me1:H3−5ac
H4K20 me2:H3K4 me3
H4K20 me2:H3−5ac
DNA Meth. m5C:H4K20 me3
H4K20 me2:H4−4ac
H4−4ac:H3−5ac
H3K27 me2
H3K9 me2
H3K4 me3
H4K20 me2
H3K9 me3
H2A.Z
H4K20 me3
H3K27 me3
H3K4 me1
H3−5ac
DNA Meth. m5C
H4−4ac
0.00 0.25 0.50 0.75 1.00
π̂
ZNF384 − Forward
H3K9 me2:DNA Meth. m5C
H3K4 me1:H4−4ac
DNA Meth. m5C:H4K20 me3
H3K4 me1:H2A.Z
H2A.Z:H4K20 me2
H3K27 me3:H3K9 me3
H3K27 me2:DNA Meth. m5C
H3K27 me3:H4K20 me3
H3K4 me3:H3−5ac
H4K20 me2:H3K4 me3
H3K9 me3:H4K20 me3
H3K27 me3:H3K4 me1
H4K20 me2:H3−5ac
H2A.Z:H3−5ac
H2A.Z:H3K4 me3
H2A.Z:H4−4ac
DNA Meth. m5C:H3K9 me3
H4K20 me2:H4−4ac
H3K4 me3:H4−4ac
H3K27 me3:DNA Meth. m5C
H3K4 me1:H3−5ac
H4−4ac:H3−5ac
H3K27 me2
H4K20 me3
H3K9 me2
H3K4 me3
H3K9 me3
H2A.Z
DNA Meth. m5C
H4K20 me2
H3K4 me1
H4−4ac
H3K27 me3
H3−5ac
0.00 0.25 0.50 0.75 1.00
π̂
ZNF384 − Reverse
H3K9 me2:DNA Meth. m5C
H3K4 me1:H4−4ac
DNA Meth. m5C:H4K20 me3
H3K4 me1:H2A.Z
H2A.Z:H4K20 me2
H3K27 me3:H3K9 me3
H3K27 me2:DNA Meth. m5C
H3K27 me3:H4K20 me3
H3K4 me3:H3−5ac
H4K20 me2:H3K4 me3
H3K9 me3:H4K20 me3
H3K27 me3:H3K4 me1
H4K20 me2:H3−5ac
H2A.Z:H3−5ac
H2A.Z:H3K4 me3
H2A.Z:H4−4ac
DNA Meth. m5C:H3K9 me3
H4K20 me2:H4−4ac
H3K4 me3:H4−4ac
H3K27 me3:DNA Meth. m5C
H3K4 me1:H3−5ac
H4−4ac:H3−5ac
H3K27 me2
H4K20 me3
H3K9 me2
H3K4 me3
H3K9 me3
H2A.Z
DNA Meth. m5C
H4K20 me2
H3K4 me1
H4−4ac
H3K27 me3
H3−5ac
0.00 0.25 0.50 0.75 1.00
π̂

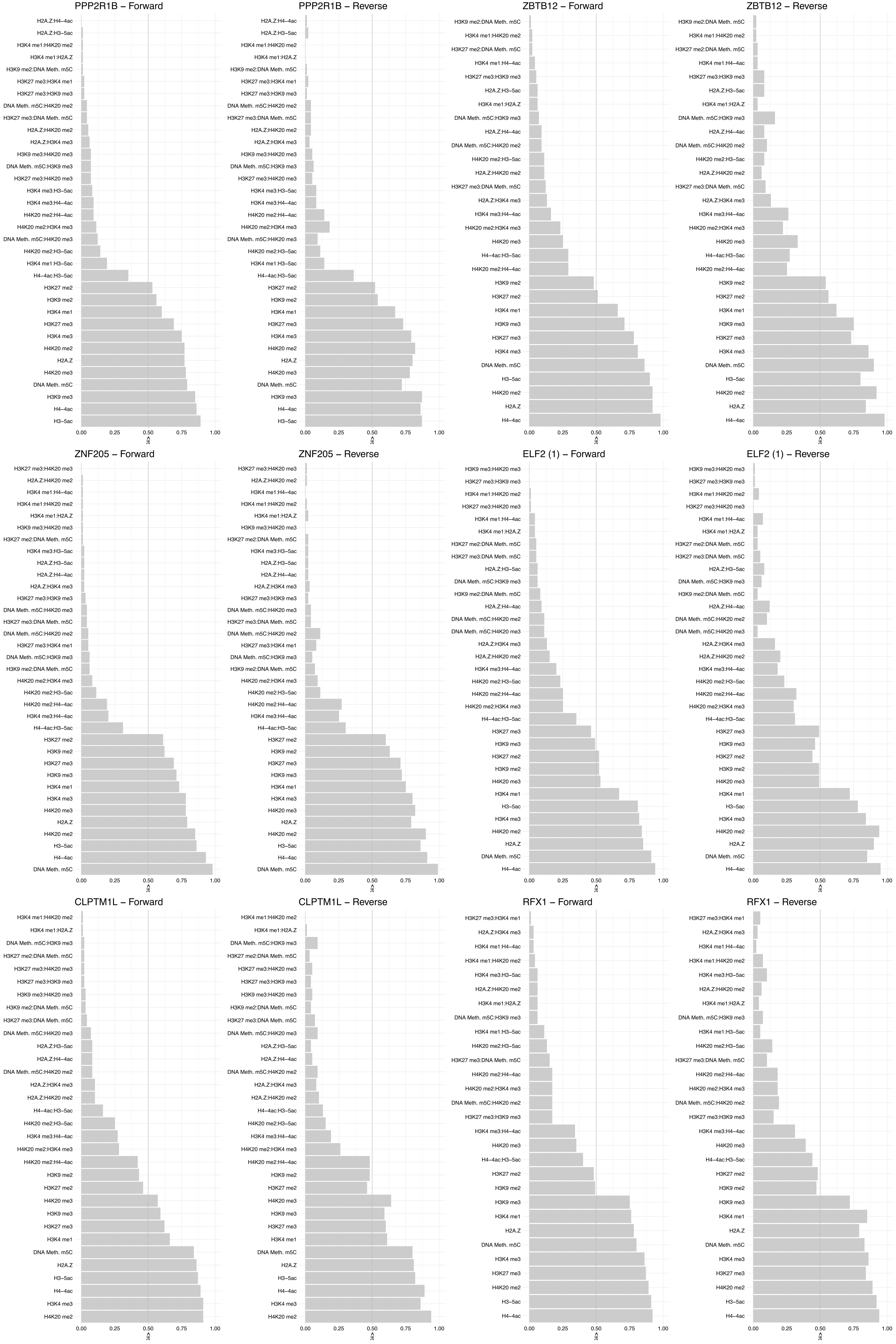
PPP2R1B – Forward
H2A.Z:H4−4ac
H2A.Z:H3−5ac
H3K4 me1:H4K20 me2
H3K4 me1:H2A.Z
H3K9 me2:DNA Meth. m5C
H3K27 me3:H3K4 me1
H3K27 me3:H3K9 me3
DNA Meth. m5C:H4K20 me2
H3K27 me3:DNA Meth. m5C
H2A.Z:H4K20 me2
H2A.Z:H3K4 me3
H3K9 me3:H4K20 me3
DNA Meth. m5C:H3K9 me3
H3K27 me3:H4K20 me3
H3K4 me3:H3−5ac
H3K4 me3:H4−4ac
H4K20 me2:H4−4ac
H4K20 me2:H3K4 me3
DNA Meth. m5C:H4K20 me3
H4K20 me2:H3−5ac
H3K4 me1:H3−5ac
H4−4ac:H3−5ac
H3K27 me2
H3K9 me2
H3K4 me1
H3K27 me3
H3K4 me3
H4K20 me2
H2A.Z
H4K20 me3
DNA Meth. m5C
H3K9 me3
H4−4ac
H3−5ac
0.00 0.25 0.50 0.75 1.00
π̂
PPP2R1B – Reverse
H2A.Z:H4−4ac
H2A.Z:H3−5ac
H3K4 me1:H4K20 me2
H3K4 me1:H2A.Z
H3K9 me2:DNA Meth. m5C
H3K27 me3:H3K4 me1
H3K27 me3:H3K9 me3
DNA Meth. m5C:H4K20 me2
H3K27 me3:DNA Meth. m5C
H2A.Z:H4K20 me2
H2A.Z:H3K4 me3
H3K9 me3:H4K20 me3
DNA Meth. m5C:H3K9 me3
H3K27 me3:H4K20 me3
H3K4 me3:H3−5ac
H3K4 me3:H4−4ac
H4K20 me2:H4−4ac
H4K20 me2:H3K4 me3
DNA Meth. m5C:H4K20 me3
H4K20 me2:H3−5ac
H3K4 me1:H3−5ac
H4−4ac:H3−5ac
H3K27 me2
H3K9 me2
H3K4 me1
H3K27 me3
H3K4 me3
H4K20 me2
H2A.Z
H4K20 me3
DNA Meth. m5C
H3K9 me3
H4−4ac
H3−5ac
0.00 0.25 0.50 0.75 1.00
π̂
ZBTB12 – Forward
H3K9 me2:DNA Meth. m5C
H3K4 me1:H4K20 me2
H3K27 me2:DNA Meth. m5C
H3K4 me1:H4−4ac
H3K27 me3:H3K9 me3
H2A.Z:H3−5ac
H3K4 me1:H2A.Z
DNA Meth. m5C:H3K9 me3
H2A.Z:H4−4ac
DNA Meth. m5C:H4K20 me2
H4K20 me2:H3−5ac
H2A.Z:H4K20 me2
H3K27 me3:DNA Meth. m5C
H2A.Z:H3K4 me3
H3K4 me3:H4−4ac
H4K20 me2:H3K4 me3
H4K20 me3
H4−4ac:H3−5ac
H4K20 me2:H4−4ac
H3K9 me2
H3K27 me2
H3K4 me1
H3K9 me3
H3K27 me3
H3K4 me3
DNA Meth. m5C
H3−5ac
H4K20 me2
H2A.Z
H4−4ac
0.00 0.25 0.50 0.75 1.00
π̂
ZBTB12 – Reverse
H3K9 me2:DNA Meth. m5C
H3K4 me1:H4K20 me2
H3K27 me2:DNA Meth. m5C
H3K4 me1:H4−4ac
H3K27 me3:H3K9 me3
H2A.Z:H3−5ac
H3K4 me1:H2A.Z
DNA Meth. m5C:H3K9 me3
H2A.Z:H4−4ac
DNA Meth. m5C:H4K20 me2
H4K20 me2:H3−5ac
H2A.Z:H4K20 me2
H3K27 me3:DNA Meth. m5C
H2A.Z:H3K4 me3
H3K4 me3:H4−4ac
H4K20 me2:H3K4 me3
H4K20 me3
H4−4ac:H3−5ac
H4K20 me2:H4−4ac
H3K9 me2
H3K27 me2
H3K4 me1
H3K9 me3
H3K27 me3
H3K4 me3
DNA Meth. m5C
H3−5ac
H4K20 me2
H2A.Z
H4−4ac
0.00 0.25 0.50 0.75 1.00
π̂
ZNF205 – Forward
H3K27 me3:H4K20 me3
H2A.Z:H4K20 me2
H3K4 me1:H4−4ac
H3K4 me1:H4K20 me2
H3K4 me1:H2A.Z
H3K9 me3:H4K20 me3
H3K27 me2:DNA Meth. m5C
H3K4 me3:H3−5ac
H2A.Z:H3−5ac
H2A.Z:H4−4ac
H2A.Z:H3K4 me3
H3K27 me3:H3K9 me3
DNA Meth. m5C:H4K20 me3
H3K27 me3:DNA Meth. m5C
DNA Meth. m5C:H4K20 me2
H3K27 me3:H3K4 me1
DNA Meth. m5C:H3K9 me3
H3K9 me2:DNA Meth. m5C
H4K20 me2:H3K4 me3
H4K20 me2:H3−5ac
H4K20 me2:H4−4ac
H3K4 me3:H4−4ac
H4−4ac:H3−5ac
H3K27 me2
H3K9 me2
H3K27 me3
H3K9 me3
H3K4 me1
H3K4 me3
H4K20 me3
H2A.Z
H4K20 me2
H3−5ac
H4−4ac
DNA Meth. m5C
0.00 0.25 0.50 0.75 1.00
π̂
ZNF205 – Reverse
H3K27 me3:H4K20 me3
H2A.Z:H4K20 me2
H3K4 me1:H4−4ac
H3K4 me1:H4K20 me2
H3K4 me1:H2A.Z
H3K9 me3:H4K20 me3
H3K27 me2:DNA Meth. m5C
H3K4 me3:H3−5ac
H2A.Z:H3−5ac
H2A.Z:H4−4ac
H2A.Z:H3K4 me3
H3K27 me3:H3K9 me3
DNA Meth. m5C:H4K20 me3
H3K27 me3:DNA Meth. m5C
DNA Meth. m5C:H4K20 me2
H3K27 me3:H3K4 me1
DNA Meth. m5C:H3K9 me3
H3K9 me2:DNA Meth. m5C
H4K20 me2:H3K4 me3
H4K20 me2:H3−5ac
H4K20 me2:H4−4ac
H3K4 me3:H4−4ac
H4−4ac:H3−5ac
H3K27 me2
H3K9 me2
H3K27 me3
H3K9 me3
H3K4 me1
H3K4 me3
H4K20 me3
H2A.Z
H4K20 me2
H3−5ac
H4−4ac
DNA Meth. m5C
0.00 0.25 0.50 0.75 1.00
π̂
ELF2 (1) – Forward
H3K9 me3:H4K20 me3
H3K27 me3:H3K9 me3
H3K4 me1:H4K20 me2
H3K27 me3:H4K20 me3
H3K4 me1:H4−4ac
H3K4 me1:H2A.Z
H3K27 me2:DNA Meth. m5C
H3K27 me3:DNA Meth. m5C
H2A.Z:H3−5ac
DNA Meth. m5C:H3K9 me3
H3K9 me2:DNA Meth. m5C
H2A.Z:H4−4ac
DNA Meth. m5C:H4K20 me2
DNA Meth. m5C:H4K20 me3
H2A.Z:H3K4 me3
H2A.Z:H4K20 me2
H3K4 me3:H4−4ac
H4K20 me2:H3−5ac
H4K20 me2:H4−4ac
H4K20 me2:H3K4 me3
H4−4ac:H3−5ac
H3K27 me3
H3K9 me3
H3K27 me2
H3K9 me2
H4K20 me3
H3K4 me1
H3−5ac
H3K4 me3
H4K20 me2
H2A.Z
DNA Meth. m5C
H4−4ac
0.00 0.25 0.50 0.75 1.00
π̂
ELF2 (1) – Reverse
H3K9 me3:H4K20 me3
H3K27 me3:H3K9 me3
H3K4 me1:H4K20 me2
H3K27 me3:H4K20 me3
H3K4 me1:H4−4ac
H3K4 me1:H2A.Z
H3K27 me2:DNA Meth. m5C
H3K27 me3:DNA Meth. m5C
H2A.Z:H3−5ac
DNA Meth. m5C:H3K9 me3
H3K9 me2:DNA Meth. m5C
H2A.Z:H4−4ac
DNA Meth. m5C:H4K20 me2
DNA Meth. m5C:H4K20 me3
H2A.Z:H3K4 me3
H2A.Z:H4K20 me2
H3K4 me3:H4−4ac
H4K20 me2:H3−5ac
H4K20 me2:H4−4ac
H4K20 me2:H3K4 me3
H4−4ac:H3−5ac
H3K27 me3
H3K9 me3
H3K27 me2
H3K9 me2
H4K20 me3
H3K4 me1
H3−5ac
H3K4 me3
H4K20 me2
H2A.Z
DNA Meth. m5C
H4−4ac
0.00 0.25 0.50 0.75 1.00
π̂
CLPTM1L – Forward
H3K4 me1:H4K20 me2
H3K4 me1:H2A.Z
DNA Meth. m5C:H3K9 me3
H3K27 me2:DNA Meth. m5C
H3K27 me3:H4K20 me3
H3K27 me3:H3K9 me3
H3K9 me3:H4K20 me3
H3K9 me2:DNA Meth. m5C
H3K27 me3:DNA Meth. m5C
DNA Meth. m5C:H4K20 me3
H2A.Z:H3−5ac
H2A.Z:H4−4ac
DNA Meth. m5C:H4K20 me2
H2A.Z:H3K4 me3
H2A.Z:H4K20 me2
H4−4ac:H3−5ac
H4K20 me2:H3−5ac
H3K4 me3:H4−4ac
H4K20 me2:H3K4 me3
H4K20 me2:H4−4ac
H3K9 me2
H3K27 me2
H4K20 me3
H3K9 me3
H3K27 me3
H3K4 me1
DNA Meth. m5C
H2A.Z
H3−5ac
H4−4ac
H3K4 me3
H4K20 me2
0.00 0.25 0.50 0.75 1.00
π̂
CLPTM1L – Reverse
H3K4 me1:H4K20 me2
H3K4 me1:H2A.Z
DNA Meth. m5C:H3K9 me3
H3K27 me2:DNA Meth. m5C
H3K27 me3:H4K20 me3
H3K27 me3:H3K9 me3
H3K9 me3:H4K20 me3
H3K9 me2:DNA Meth. m5C
H3K27 me3:DNA Meth. m5C
DNA Meth. m5C:H4K20 me3
H2A.Z:H3−5ac
H2A.Z:H4−4ac
DNA Meth. m5C:H4K20 me2
H2A.Z:H3K4 me3
H2A.Z:H4K20 me2
H4−4ac:H3−5ac
H4K20 me2:H3−5ac
H3K4 me3:H4−4ac
H4K20 me2:H3K4 me3
H4K20 me2:H4−4ac
H3K9 me2
H3K27 me2
H4K20 me3
H3K9 me3
H3K27 me3
H3K4 me1
DNA Meth. m5C
H2A.Z
H3−5ac
H4−4ac
H3K4 me3
H4K20 me2
0.00 0.25 0.50 0.75 1.00
π̂
RFX1 – Forward
H3K27 me3:H3K4 me1
H2A.Z:H3K4 me3
H3K4 me1:H4−4ac
H3K4 me1:H4K20 me2
H3K4 me3:H3−5ac
H2A.Z:H4K20 me2
H3K4 me1:H2A.Z
DNA Meth. m5C:H3K9 me3
H3K4 me1:H3−5ac
H4K20 me2:H3−5ac
H3K27 me3:DNA Meth. m5C
H4K20 me2:H4−4ac
H4K20 me2:H3K4 me3
DNA Meth. m5C:H4K20 me2
H3K27 me3:H3K9 me3
H3K4 me3:H4−4ac
H4K20 me3
H4−4ac:H3−5ac
H3K27 me2
H3K9 me2
H3K9 me3
H3K4 me1
H2A.Z
DNA Meth. m5C
H3K4 me3
H3K27 me3
H4K20 me2
H3−5ac
H4−4ac
0.00 0.25 0.50 0.75 1.00
π̂
RFX1 – Reverse
H3K27 me3:H3K4 me1
H2A.Z:H3K4 me3
H3K4 me1:H4−4ac
H3K4 me1:H4K20 me2
H3K4 me3:H3−5ac
H2A.Z:H4K20 me2
H3K4 me1:H2A.Z
DNA Meth. m5C:H3K9 me3
H3K4 me1:H3−5ac
H4K20 me2:H3−5ac
H3K27 me3:DNA Meth. m5C
H4K20 me2:H4−4ac
H4K20 me2:H3K4 me3
DNA Meth. m5C:H4K20 me2
H3K27 me3:H3K9 me3
H3K4 me3:H4−4ac
H4K20 me3
H4−4ac:H3−5ac
H3K27 me2
H3K9 me2
H3K9 me3
H3K4 me1
H2A.Z
DNA Meth. m5C
H3K4 me3
H3K27 me3
H4K20 me2
H3−5ac
H4−4ac
0.00 0.25 0.50 0.75 1.00
π̂

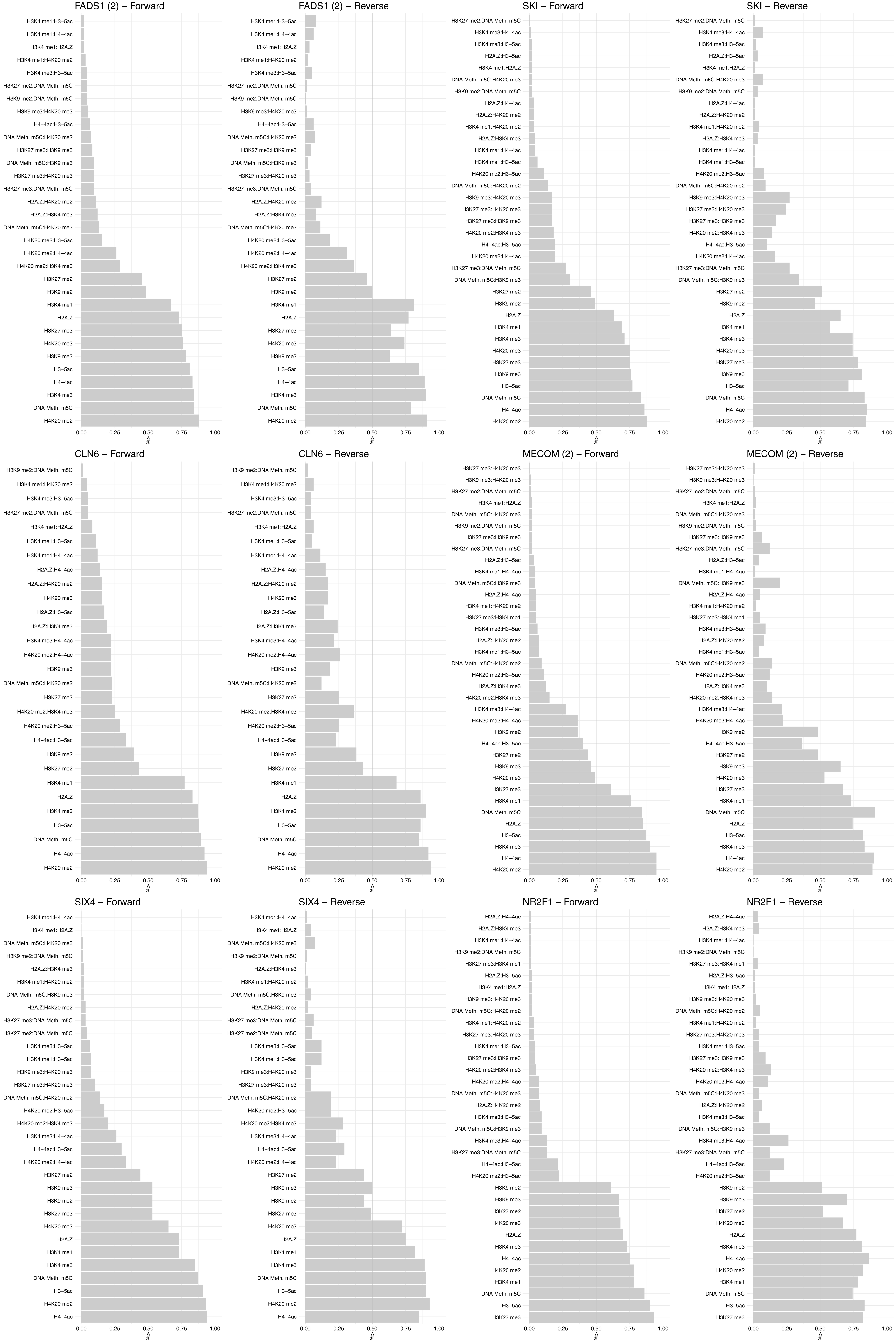
FADS1 (2) – Forward
FADS1 (2) – Reverse
SKI – Forward
SKI – Reverse
CLN6 – Forward
CLN6 – Reverse
MECOM (2) – Forward
MECOM (2) – Reverse
SIX4 – Forward
SIX4 – Reverse
NR2F1 – Forward
NR2F1 – Reverse
0.00
0.25
0.50
0.75
1.00
$\hat{\pi}$

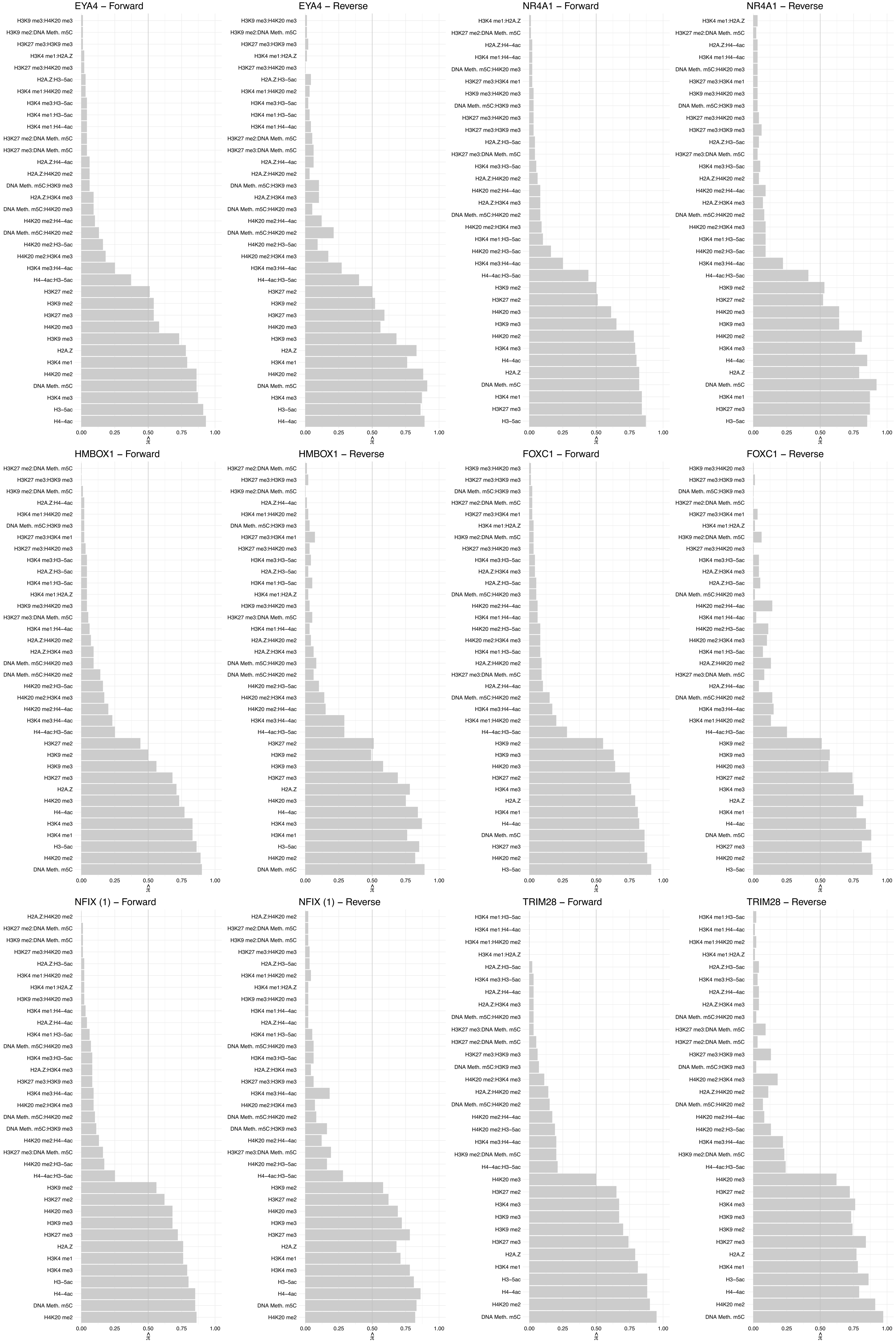
EYA4 – Forward
H3K9 me3:H4K20 me3
H3K9 me2:DNA Meth. m5C
H3K27 me3:H3K9 me3
H3K4 me1:H2A.Z
H3K27 me3:H4K20 me3
H2A.Z:H3−5ac
H3K4 me1:H4K20 me2
H3K4 me3:H3−5ac
H3K4 me1:H3−5ac
H3K4 me1:H4−4ac
H3K27 me2:DNA Meth. m5C
H3K27 me3:DNA Meth. m5C
H2A.Z:H4−4ac
H2A.Z:H4K20 me2
DNA Meth. m5C:H3K9 me3
H2A.Z:H3K4 me3
DNA Meth. m5C:H4K20 me3
H4K20 me2:H4−4ac
DNA Meth. m5C:H4K20 me2
H4K20 me2:H3−5ac
H4K20 me2:H3K4 me3
H3K4 me3:H4−4ac
H4−4ac:H3−5ac
H3K27 me2
H3K9 me2
H3K27 me3
H4K20 me3
H3K9 me3
H2A.Z
H3K4 me1
H4K20 me2
DNA Meth. m5C
H3K4 me3
H3−5ac
H4−4ac
0.00 0.25 0.50 0.75 1.00
$\hat{\pi}$
EYA4 – Reverse
H3K9 me3:H4K20 me3
H3K9 me2:DNA Meth. m5C
H3K27 me3:H3K9 me3
H3K4 me1:H2A.Z
H3K27 me3:H4K20 me3
H2A.Z:H3−5ac
H3K4 me1:H4K20 me2
H3K4 me3:H3−5ac
H3K4 me1:H3−5ac
H3K4 me1:H4−4ac
H3K27 me2:DNA Meth. m5C
H3K27 me3:DNA Meth. m5C
H2A.Z:H4−4ac
H2A.Z:H4K20 me2
DNA Meth. m5C:H3K9 me3
H2A.Z:H3K4 me3
DNA Meth. m5C:H4K20 me3
H4K20 me2:H4−4ac
DNA Meth. m5C:H4K20 me2
H4K20 me2:H3−5ac
H4K20 me2:H3K4 me3
H3K4 me3:H4−4ac
H4−4ac:H3−5ac
H3K27 me2
H3K9 me2
H3K27 me3
H4K20 me3
H3K9 me3
H2A.Z
H3K4 me1
H4K20 me2
DNA Meth. m5C
H3K4 me3
H3−5ac
H4−4ac
0.00 0.25 0.50 0.75 1.00
$\hat{\pi}$
NR4A1 – Forward
H3K4 me1:H2A.Z
H3K27 me2:DNA Meth. m5C
H2A.Z:H4−4ac
H3K4 me1:H4−4ac
DNA Meth. m5C:H4K20 me3
H3K27 me3:H3K4 me1
H3K9 me3:H4K20 me3
DNA Meth. m5C:H3K9 me3
H3K27 me3:H4K20 me3
H3K27 me3:H3K9 me3
H2A.Z:H3−5ac
H3K27 me3:DNA Meth. m5C
H3K4 me3:H3−5ac
H2A.Z:H4K20 me2
H4K20 me2:H4−4ac
H2A.Z:H3K4 me3
DNA Meth. m5C:H4K20 me2
H4K20 me2:H3K4 me3
H3K4 me1:H3−5ac
H4K20 me2:H3−5ac
H3K4 me3:H4−4ac
H4−4ac:H3−5ac
H3K9 me2
H3K27 me2
H4K20 me3
H3K9 me3
H4K20 me2
H3K4 me3
H4−4ac
H2A.Z
DNA Meth. m5C
H3K4 me1
H3K27 me3
H3−5ac
0.00 0.25 0.50 0.75 1.00
$\hat{\pi}$
NR4A1 – Reverse
H3K4 me1:H2A.Z
H3K27 me2:DNA Meth. m5C
H2A.Z:H4−4ac
H3K4 me1:H4−4ac
DNA Meth. m5C:H4K20 me3
H3K27 me3:H3K4 me1
H3K9 me3:H4K20 me3
DNA Meth. m5C:H3K9 me3
H3K27 me3:H4K20 me3
H3K27 me3:H3K9 me3
H2A.Z:H3−5ac
H3K27 me3:DNA Meth. m5C
H3K4 me3:H3−5ac
H2A.Z:H4K20 me2
H4K20 me2:H4−4ac
H2A.Z:H3K4 me3
DNA Meth. m5C:H4K20 me2
H4K20 me2:H3K4 me3
H3K4 me1:H3−5ac
H4K20 me2:H3−5ac
H3K4 me3:H4−4ac
H4−4ac:H3−5ac
H3K9 me2
H3K27 me2
H4K20 me3
H3K9 me3
H4K20 me2
H3K4 me3
H4−4ac
H2A.Z
DNA Meth. m5C
H3K4 me1
H3K27 me3
H3−5ac
0.00 0.25 0.50 0.75 1.00
$\hat{\pi}$
HMBOX1 – Forward
H3K27 me2:DNA Meth. m5C
H3K27 me3:H3K9 me3
H3K9 me2:DNA Meth. m5C
H2A.Z:H4−4ac
H3K4 me1:H4K20 me2
DNA Meth. m5C:H3K9 me3
H3K27 me3:H3K4 me1
H3K27 me3:H4K20 me3
H3K4 me3:H3−5ac
H2A.Z:H3−5ac
H3K4 me1:H3−5ac
H3K4 me1:H2A.Z
H3K9 me3:H4K20 me3
H3K27 me3:DNA Meth. m5C
H3K4 me1:H4−4ac
H2A.Z:H4K20 me2
H2A.Z:H3K4 me3
DNA Meth. m5C:H4K20 me3
DNA Meth. m5C:H4K20 me2
H4K20 me2:H3−5ac
H4K20 me2:H3K4 me3
H4K20 me2:H4−4ac
H3K4 me3:H4−4ac
H4−4ac:H3−5ac
H3K27 me2
H3K9 me2
H3K9 me3
H3K27 me3
H2A.Z
H4K20 me3
H4−4ac
H3K4 me3
H3K4 me1
H3−5ac
H4K20 me2
DNA Meth. m5C
0.00 0.25 0.50 0.75 1.00
$\hat{\pi}$
HMBOX1 – Reverse
H3K27 me2:DNA Meth. m5C
H3K27 me3:H3K9 me3
H3K9 me2:DNA Meth. m5C
H2A.Z:H4−4ac
H3K4 me1:H4K20 me2
DNA Meth. m5C:H3K9 me3
H3K27 me3:H3K4 me1
H3K27 me3:H4K20 me3
H3K4 me3:H3−5ac
H2A.Z:H3−5ac
H3K4 me1:H3−5ac
H3K4 me1:H2A.Z
H3K9 me3:H4K20 me3
H3K27 me3:DNA Meth. m5C
H3K4 me1:H4−4ac
H2A.Z:H4K20 me2
H2A.Z:H3K4 me3
DNA Meth. m5C:H4K20 me3
DNA Meth. m5C:H4K20 me2
H4K20 me2:H3−5ac
H4K20 me2:H3K4 me3
H4K20 me2:H4−4ac
H3K4 me3:H4−4ac
H4−4ac:H3−5ac
H3K27 me2
H3K9 me2
H3K9 me3
H3K27 me3
H2A.Z
H4K20 me3
H4−4ac
H3K4 me3
H3K4 me1
H3−5ac
H4K20 me2
DNA Meth. m5C
0.00 0.25 0.50 0.75 1.00
$\hat{\pi}$
FOXC1 – Forward
H3K9 me3:H4K20 me3
H3K27 me3:H3K9 me3
DNA Meth. m5C:H3K9 me3
H3K27 me2:DNA Meth. m5C
H3K27 me3:H3K4 me1
H3K4 me1:H2A.Z
H3K9 me2:DNA Meth. m5C
H3K27 me3:H4K20 me3
H3K4 me3:H3−5ac
H2A.Z:H3K4 me3
H2A.Z:H3−5ac
DNA Meth. m5C:H4K20 me3
H4K20 me2:H4−4ac
H3K4 me1:H4−4ac
H4K20 me2:H3−5ac
H4K20 me2:H3K4 me3
H3K4 me1:H3−5ac
H2A.Z:H4K20 me2
H3K27 me3:DNA Meth. m5C
H2A.Z:H4−4ac
DNA Meth. m5C:H4K20 me2
H3K4 me3:H4−4ac
H3K4 me1:H4K20 me2
H4−4ac:H3−5ac
H3K9 me2
H3K9 me3
H4K20 me3
H3K27 me2
H3K4 me3
H2A.Z
H3K4 me1
H4−4ac
DNA Meth. m5C
H3K27 me3
H4K20 me2
H3−5ac
0.00 0.25 0.50 0.75 1.00
$\hat{\pi}$
FOXC1 – Reverse
H3K9 me3:H4K20 me3
H3K27 me3:H3K9 me3
DNA Meth. m5C:H3K9 me3
H3K27 me2:DNA Meth. m5C
H3K27 me3:H3K4 me1
H3K4 me1:H2A.Z
H3K9 me2:DNA Meth. m5C
H3K27 me3:H4K20 me3
H3K4 me3:H3−5ac
H2A.Z:H3K4 me3
H2A.Z:H3−5ac
DNA Meth. m5C:H4K20 me3
H4K20 me2:H4−4ac
H3K4 me1:H4−4ac
H4K20 me2:H3−5ac
H4K20 me2:H3K4 me3
H3K4 me1:H3−5ac
H2A.Z:H4K20 me2
H3K27 me3:DNA Meth. m5C
H2A.Z:H4−4ac
DNA Meth. m5C:H4K20 me2
H3K4 me3:H4−4ac
H3K4 me1:H4K20 me2
H4−4ac:H3−5ac
H3K9 me2
H3K9 me3
H4K20 me3
H3K27 me2
H3K4 me3
H2A.Z
H3K4 me1
H4−4ac
DNA Meth. m5C
H3K27 me3
H4K20 me2
H3−5ac
0.00 0.25 0.50 0.75 1.00
$\hat{\pi}$
NFIX (1) – Forward
H2A.Z:H4K20 me2
H3K27 me2:DNA Meth. m5C
H3K9 me2:DNA Meth. m5C
H3K27 me3:H4K20 me3
H2A.Z:H3−5ac
H3K4 me1:H4K20 me2
H3K4 me1:H2A.Z
H3K9 me3:H4K20 me3
H3K4 me1:H4−4ac
H2A.Z:H4−4ac
H3K4 me1:H3−5ac
DNA Meth. m5C:H4K20 me3
H3K4 me3:H3−5ac
H2A.Z:H3K4 me3
H3K27 me3:H3K9 me3
H3K4 me3:H4−4ac
H4K20 me2:H3K4 me3
DNA Meth. m5C:H4K20 me2
DNA Meth. m5C:H3K9 me3
H4K20 me2:H4−4ac
H3K27 me3:DNA Meth. m5C
H4K20 me2:H3−5ac
H4−4ac:H3−5ac
H3K9 me2
H3K27 me2
H4K20 me3
H3K9 me3
H3K27 me3
H2A.Z
H3K4 me1
H3K4 me3
H3−5ac
H4−4ac
DNA Meth. m5C
H4K20 me2
0.00 0.25 0.50 0.75 1.00
$\hat{\pi}$
NFIX (1) – Reverse
H2A.Z:H4K20 me2
H3K27 me2:DNA Meth. m5C
H3K9 me2:DNA Meth. m5C
H3K27 me3:H4K20 me3
H2A.Z:H3−5ac
H3K4 me1:H4K20 me2
H3K4 me1:H2A.Z
H3K9 me3:H4K20 me3
H3K4 me1:H4−4ac
H2A.Z:H4−4ac
H3K4 me1:H3−5ac
DNA Meth. m5C:H4K20 me3
H3K4 me3:H3−5ac
H2A.Z:H3K4 me3
H3K27 me3:H3K9 me3
H3K4 me3:H4−4ac
H4K20 me2:H3K4 me3
DNA Meth. m5C:H4K20 me2
DNA Meth. m5C:H3K9 me3
H4K20 me2:H4−4ac
H3K27 me3:DNA Meth. m5C
H4K20 me2:H3−5ac
H4−4ac:H3−5ac
H3K9 me2
H3K27 me2
H4K20 me3
H3K9 me3
H3K27 me3
H2A.Z
H3K4 me1
H3K4 me3
H3−5ac
H4−4ac
DNA Meth. m5C
H4K20 me2
0.00 0.25 0.50 0.75 1.00
$\hat{\pi}$
TRIM28 – Forward
H3K4 me1:H3−5ac
H3K4 me1:H4−4ac
H3K4 me1:H4K20 me2
H3K4 me1:H2A.Z
H2A.Z:H3−5ac
H3K4 me3:H3−5ac
H2A.Z:H4−4ac
H2A.Z:H3K4 me3
DNA Meth. m5C:H4K20 me3
H3K27 me3:DNA Meth. m5C
H3K27 me2:DNA Meth. m5C
H3K27 me3:H3K9 me3
DNA Meth. m5C:H3K9 me3
H4K20 me2:H3K4 me3
H2A.Z:H4K20 me2
DNA Meth. m5C:H4K20 me2
H4K20 me2:H4−4ac
H4K20 me2:H3−5ac
H3K4 me3:H4−4ac
H3K9 me2:DNA Meth. m5C
H4−4ac:H3−5ac
H4K20 me3
H3K27 me2
H3K4 me3
H3K9 me3
H3K9 me2
H3K27 me3
H2A.Z
H3K4 me1
H3−5ac
H4−4ac
H4K20 me2
DNA Meth. m5C
0.00 0.25 0.50 0.75 1.00
$\hat{\pi}$
TRIM28 – Reverse
H3K4 me1:H3−5ac
H3K4 me1:H4−4ac
H3K4 me1:H4K20 me2
H3K4 me1:H2A.Z
H2A.Z:H3−5ac
H3K4 me3:H3−5ac
H2A.Z:H4−4ac
H2A.Z:H3K4 me3
DNA Meth. m5C:H4K20 me3
H3K27 me3:DNA Meth. m5C
H3K27 me2:DNA Meth. m5C
H3K27 me3:H3K9 me3
DNA Meth. m5C:H3K9 me3
H4K20 me2:H3K4 me3
H2A.Z:H4K20 me2
DNA Meth. m5C:H4K20 me2
H4K20 me2:H4−4ac
H4K20 me2:H3−5ac
H3K4 me3:H4−4ac
H3K9 me2:DNA Meth. m5C
H4−4ac:H3−5ac
H4K20 me3
H3K27 me2
H3K4 me3
H3K9 me3
H3K9 me2
H3K27 me3
H2A.Z
H3K4 me1
H3−5ac
H4−4ac
H4K20 me2
DNA Meth. m5C
0.00 0.25 0.50 0.75 1.00
$\hat{\pi}$

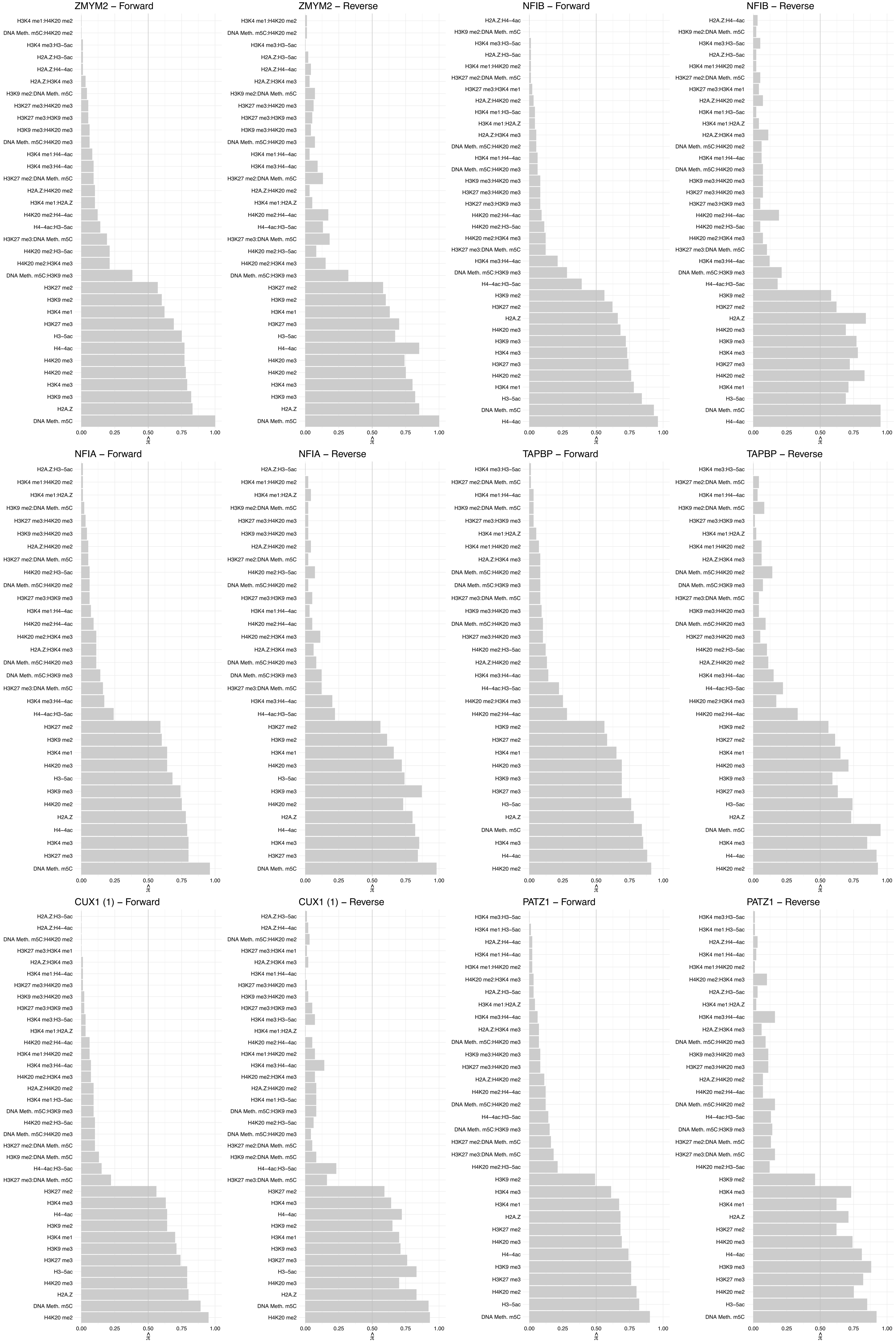
ZMYM2 – Forward
H3K4 me1:H4K20 me2
DNA Meth. m5C:H4K20 me2
H3K4 me3:H3–5ac
H2A.Z:H3–5ac
H2A.Z:H4–4ac
H2A.Z:H3K4 me3
H3K9 me2:DNA Meth. m5C
H3K27 me3:H4K20 me3
H3K27 me3:H3K9 me3
H3K9 me3:H4K20 me3
DNA Meth. m5C:H4K20 me3
H3K4 me1:H4–4ac
H3K4 me3:H4–4ac
H3K27 me2:DNA Meth. m5C
H2A.Z:H4K20 me2
H3K4 me1:H2A.Z
H4K20 me2:H4–4ac
H4–4ac:H3–5ac
H3K27 me3:DNA Meth. m5C
H4K20 me2:H3–5ac
H4K20 me2:H3K4 me3
DNA Meth. m5C:H3K9 me3
H3K27 me2
H3K9 me2
H3K4 me1
H3K27 me3
H3–5ac
H4–4ac
H4K20 me3
H4K20 me2
H3K4 me3
H3K9 me3
H2A.Z
DNA Meth. m5C
0.00 0.25 0.50 0.75 1.00
$\hat{\pi}$
ZMYM2 – Reverse
H3K4 me1:H4K20 me2
DNA Meth. m5C:H4K20 me2
H3K4 me3:H3–5ac
H2A.Z:H3–5ac
H2A.Z:H4–4ac
H2A.Z:H3K4 me3
H3K9 me2:DNA Meth. m5C
H3K27 me3:H4K20 me3
H3K27 me3:H3K9 me3
H3K9 me3:H4K20 me3
DNA Meth. m5C:H4K20 me3
H3K4 me1:H4–4ac
H3K4 me3:H4–4ac
H3K27 me2:DNA Meth. m5C
H2A.Z:H4K20 me2
H3K4 me1:H2A.Z
H4K20 me2:H4–4ac
H4–4ac:H3–5ac
H3K27 me3:DNA Meth. m5C
H4K20 me2:H3–5ac
H4K20 me2:H3K4 me3
DNA Meth. m5C:H3K9 me3
H3K27 me2
H3K9 me2
H3K4 me1
H3K27 me3
H3–5ac
H4–4ac
H4K20 me3
H4K20 me2
H3K4 me3
H3K9 me3
H2A.Z
DNA Meth. m5C
0.00 0.25 0.50 0.75 1.00
$\hat{\pi}$
NFIB – Forward
H2A.Z:H4–4ac
H3K9 me2:DNA Meth. m5C
H3K4 me3:H3–5ac
H2A.Z:H3–5ac
H3K4 me1:H4K20 me2
H3K27 me2:DNA Meth. m5C
H3K27 me3:H3K4 me1
H2A.Z:H4K20 me2
H3K4 me1:H3–5ac
H3K4 me1:H2A.Z
H2A.Z:H3K4 me3
DNA Meth. m5C:H4K20 me2
H3K4 me1:H4–4ac
DNA Meth. m5C:H4K20 me3
H3K9 me3:H4K20 me3
H3K27 me3:H4K20 me3
H3K27 me3:H3K9 me3
H4K20 me2:H4–4ac
H4K20 me2:H3–5ac
H4K20 me2:H3K4 me3
H3K27 me3:DNA Meth. m5C
H3K4 me3:H4–4ac
DNA Meth. m5C:H3K9 me3
H4–4ac:H3–5ac
H3K9 me2
H3K27 me2
H2A.Z
H4K20 me3
H3K9 me3
H3K4 me3
H3K27 me3
H4K20 me2
H3K4 me1
H3–5ac
DNA Meth. m5C
H4–4ac
0.00 0.25 0.50 0.75 1.00
$\hat{\pi}$
NFIB – Reverse
H2A.Z:H4–4ac
H3K9 me2:DNA Meth. m5C
H3K4 me3:H3–5ac
H2A.Z:H3–5ac
H3K4 me1:H4K20 me2
H3K27 me2:DNA Meth. m5C
H3K27 me3:H3K4 me1
H2A.Z:H4K20 me2
H3K4 me1:H3–5ac
H3K4 me1:H2A.Z
H2A.Z:H3K4 me3
DNA Meth. m5C:H4K20 me2
H3K4 me1:H4–4ac
DNA Meth. m5C:H4K20 me3
H3K9 me3:H4K20 me3
H3K27 me3:H4K20 me3
H3K27 me3:H3K9 me3
H4K20 me2:H4–4ac
H4K20 me2:H3–5ac
H4K20 me2:H3K4 me3
H3K27 me3:DNA Meth. m5C
H3K4 me3:H4–4ac
DNA Meth. m5C:H3K9 me3
H4–4ac:H3–5ac
H3K9 me2
H3K27 me2
H2A.Z
H4K20 me3
H3K9 me3
H3K4 me3
H3K27 me3
H4K20 me2
H3K4 me1
H3–5ac
DNA Meth. m5C
H4–4ac
0.00 0.25 0.50 0.75 1.00
$\hat{\pi}$
NFIA – Forward
H2A.Z:H3–5ac
H3K4 me1:H4K20 me2
H3K4 me1:H2A.Z
H3K9 me2:DNA Meth. m5C
H3K27 me3:H4K20 me3
H3K9 me3:H4K20 me3
H2A.Z:H4K20 me2
H3K27 me2:DNA Meth. m5C
H4K20 me2:H3–5ac
DNA Meth. m5C:H4K20 me2
H3K27 me3:H3K9 me3
H3K4 me1:H4–4ac
H4K20 me2:H4–4ac
H4K20 me2:H3K4 me3
H2A.Z:H3K4 me3
DNA Meth. m5C:H4K20 me3
DNA Meth. m5C:H3K9 me3
H3K27 me3:DNA Meth. m5C
H3K4 me3:H4–4ac
H4–4ac:H3–5ac
H3K27 me2
H3K9 me2
H3K4 me1
H4K20 me3
H3–5ac
H3K9 me3
H4K20 me2
H2A.Z
H4–4ac
H3K4 me3
H3K27 me3
DNA Meth. m5C
0.00 0.25 0.50 0.75 1.00
$\hat{\pi}$
NFIA – Reverse
H2A.Z:H3–5ac
H3K4 me1:H4K20 me2
H3K4 me1:H2A.Z
H3K9 me2:DNA Meth. m5C
H3K27 me3:H4K20 me3
H3K9 me3:H4K20 me3
H2A.Z:H4K20 me2
H3K27 me2:DNA Meth. m5C
H4K20 me2:H3–5ac
DNA Meth. m5C:H4K20 me2
H3K27 me3:H3K9 me3
H3K4 me1:H4–4ac
H4K20 me2:H4–4ac
H4K20 me2:H3K4 me3
H2A.Z:H3K4 me3
DNA Meth. m5C:H4K20 me3
DNA Meth. m5C:H3K9 me3
H3K27 me3:DNA Meth. m5C
H3K4 me3:H4–4ac
H4–4ac:H3–5ac
H3K27 me2
H3K9 me2
H3K4 me1
H4K20 me3
H3–5ac
H3K9 me3
H4K20 me2
H2A.Z
H4–4ac
H3K4 me3
H3K27 me3
DNA Meth. m5C
0.00 0.25 0.50 0.75 1.00
$\hat{\pi}$
TAPBP – Forward
H3K4 me3:H3–5ac
H3K27 me2:DNA Meth. m5C
H3K4 me1:H4–4ac
H3K9 me2:DNA Meth. m5C
H3K27 me3:H3K9 me3
H3K4 me1:H2A.Z
H3K4 me1:H4K20 me2
H2A.Z:H3K4 me3
DNA Meth. m5C:H4K20 me2
DNA Meth. m5C:H3K9 me3
H3K27 me3:DNA Meth. m5C
H3K9 me3:H4K20 me3
DNA Meth. m5C:H4K20 me3
H3K27 me3:H4K20 me3
H4K20 me2:H3–5ac
H2A.Z:H4K20 me2
H3K4 me3:H4–4ac
H4–4ac:H3–5ac
H4K20 me2:H3K4 me3
H4K20 me2:H4–4ac
H3K9 me2
H3K27 me2
H3K4 me1
H4K20 me3
H3K9 me3
H3K27 me3
H3–5ac
H2A.Z
DNA Meth. m5C
H3K4 me3
H4–4ac
H4K20 me2
0.00 0.25 0.50 0.75 1.00
$\hat{\pi}$
TAPBP – Reverse
H3K4 me3:H3–5ac
H3K27 me2:DNA Meth. m5C
H3K4 me1:H4–4ac
H3K9 me2:DNA Meth. m5C
H3K27 me3:H3K9 me3
H3K4 me1:H2A.Z
H3K4 me1:H4K20 me2
H2A.Z:H3K4 me3
DNA Meth. m5C:H4K20 me2
DNA Meth. m5C:H3K9 me3
H3K27 me3:DNA Meth. m5C
H3K9 me3:H4K20 me3
DNA Meth. m5C:H4K20 me3
H3K27 me3:H4K20 me3
H4K20 me2:H3–5ac
H2A.Z:H4K20 me2
H3K4 me3:H4–4ac
H4–4ac:H3–5ac
H4K20 me2:H3K4 me3
H4K20 me2:H4–4ac
H3K9 me2
H3K27 me2
H3K4 me1
H4K20 me3
H3K9 me3
H3K27 me3
H3–5ac
H2A.Z
DNA Meth. m5C
H3K4 me3
H4–4ac
H4K20 me2
0.00 0.25 0.50 0.75 1.00
$\hat{\pi}$
CUX1 (1) – Forward
H2A.Z:H3–5ac
H2A.Z:H4–4ac
DNA Meth. m5C:H4K20 me2
H3K27 me3:H3K4 me1
H2A.Z:H3K4 me3
H3K4 me1:H4–4ac
H3K27 me3:H4K20 me3
H3K9 me3:H4K20 me3
H3K27 me3:H3K9 me3
H3K4 me3:H3–5ac
H3K4 me1:H2A.Z
H4K20 me2:H4–4ac
H3K4 me1:H4K20 me2
H3K4 me3:H4–4ac
H4K20 me2:H3K4 me3
H2A.Z:H4K20 me2
H3K4 me1:H3–5ac
DNA Meth. m5C:H3K9 me3
H4K20 me2:H3–5ac
DNA Meth. m5C:H4K20 me3
H3K27 me2:DNA Meth. m5C
H3K9 me2:DNA Meth. m5C
H4–4ac:H3–5ac
H3K27 me3:DNA Meth. m5C
H3K27 me2
H3K4 me3
H4–4ac
H3K9 me2
H3K4 me1
H3K9 me3
H3K27 me3
H3–5ac
H4K20 me3
H2A.Z
DNA Meth. m5C
H4K20 me2
0.00 0.25 0.50 0.75 1.00
$\hat{\pi}$
CUX1 (1) – Reverse
H2A.Z:H3–5ac
H2A.Z:H4–4ac
DNA Meth. m5C:H4K20 me2
H3K27 me3:H3K4 me1
H2A.Z:H3K4 me3
H3K4 me1:H4–4ac
H3K27 me3:H4K20 me3
H3K9 me3:H4K20 me3
H3K27 me3:H3K9 me3
H3K4 me3:H3–5ac
H3K4 me1:H2A.Z
H4K20 me2:H4–4ac
H3K4 me1:H4K20 me2
H3K4 me3:H4–4ac
H4K20 me2:H3K4 me3
H2A.Z:H4K20 me2
H3K4 me1:H3–5ac
DNA Meth. m5C:H3K9 me3
H4K20 me2:H3–5ac
DNA Meth. m5C:H4K20 me3
H3K27 me2:DNA Meth. m5C
H3K9 me2:DNA Meth. m5C
H4–4ac:H3–5ac
H3K27 me3:DNA Meth. m5C
H3K27 me2
H3K4 me3
H4–4ac
H3K9 me2
H3K4 me1
H3K9 me3
H3K27 me3
H3–5ac
H4K20 me3
H2A.Z
DNA Meth. m5C
H4K20 me2
0.00 0.25 0.50 0.75 1.00
$\hat{\pi}$
PATZ1 – Forward
H3K4 me3:H3–5ac
H3K4 me1:H3–5ac
H2A.Z:H4–4ac
H3K4 me1:H4–4ac
H3K4 me1:H4K20 me2
H4K20 me2:H3K4 me3
H2A.Z:H3–5ac
H3K4 me1:H2A.Z
H3K4 me3:H4–4ac
H2A.Z:H3K4 me3
DNA Meth. m5C:H4K20 me3
H3K9 me3:H4K20 me3
H3K27 me3:H4K20 me3
H2A.Z:H4K20 me2
H4K20 me2:H4–4ac
DNA Meth. m5C:H4K20 me2
H4–4ac:H3–5ac
DNA Meth. m5C:H3K9 me3
H3K27 me2:DNA Meth. m5C
H3K27 me3:DNA Meth. m5C
H4K20 me2:H3–5ac
H3K9 me2
H3K4 me3
H3K4 me1
H2A.Z
H3K27 me2
H4K20 me3
H4–4ac
H3K9 me3
H3K27 me3
H4K20 me2
H3–5ac
DNA Meth. m5C
0.00 0.25 0.50 0.75 1.00
$\hat{\pi}$
PATZ1 – Reverse
H3K4 me3:H3–5ac
H3K4 me1:H3–5ac
H2A.Z:H4–4ac
H3K4 me1:H4–4ac
H3K4 me1:H4K20 me2
H4K20 me2:H3K4 me3
H2A.Z:H3–5ac
H3K4 me1:H2A.Z
H3K4 me3:H4–4ac
H2A.Z:H3K4 me3
DNA Meth. m5C:H4K20 me3
H3K9 me3:H4K20 me3
H3K27 me3:H4K20 me3
H2A.Z:H4K20 me2
H4K20 me2:H4–4ac
DNA Meth. m5C:H4K20 me2
H4–4ac:H3–5ac
DNA Meth. m5C:H3K9 me3
H3K27 me2:DNA Meth. m5C
H3K27 me3:DNA Meth. m5C
H4K20 me2:H3–5ac
H3K9 me2
H3K4 me3
H3K4 me1
H2A.Z
H3K27 me2
H4K20 me3
H4–4ac
H3K9 me3
H3K27 me3
H4K20 me2
H3–5ac
DNA Meth. m5C
0.00 0.25 0.50 0.75 1.00
$\hat{\pi}$

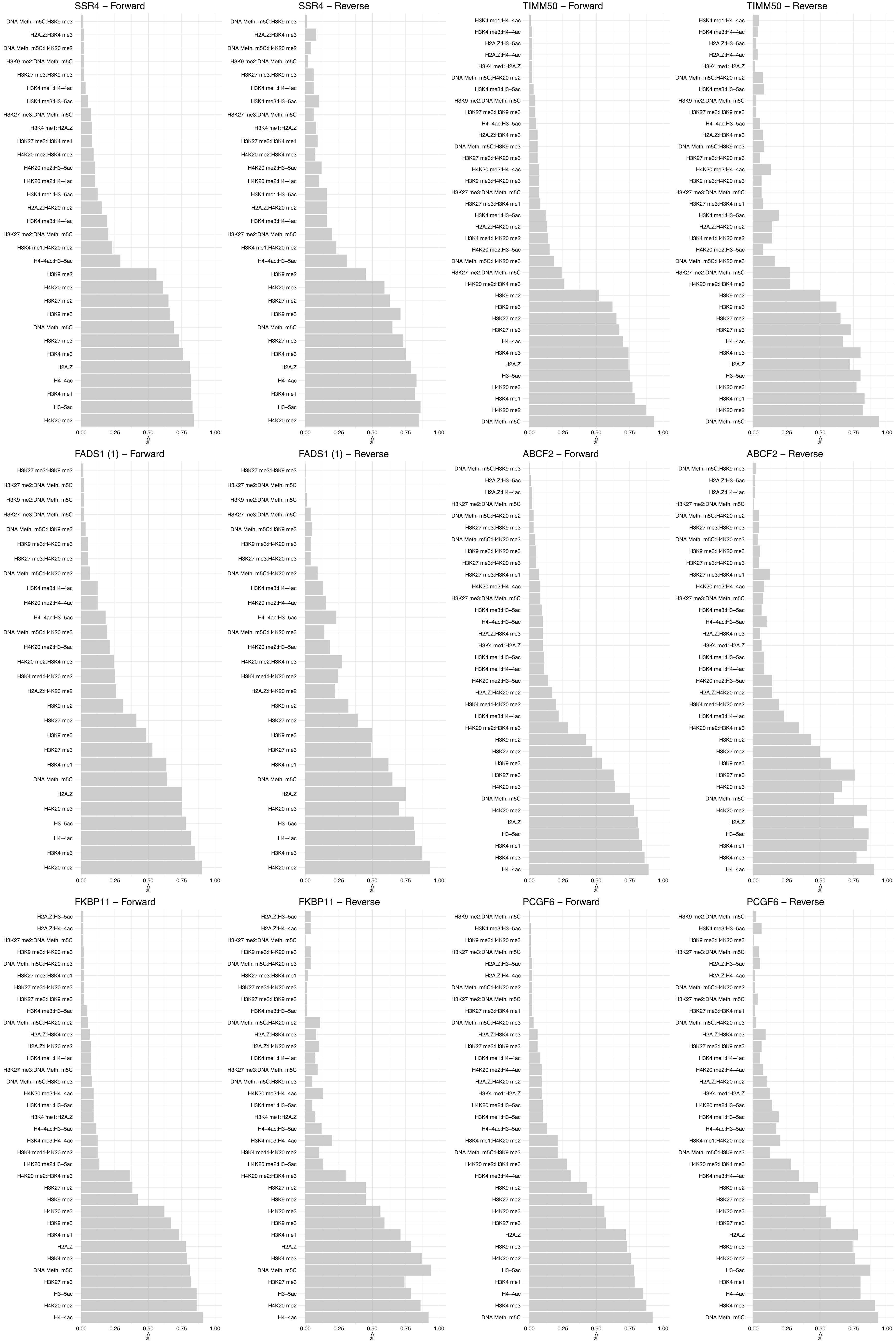
SSR4 − Forward
DNA Meth. m5C:H3K9 me3
H2A.Z:H3K4 me3
DNA Meth. m5C:H4K20 me2
H3K9 me2:DNA Meth. m5C
H3K27 me3:H3K9 me3
H3K4 me1:H4−4ac
H3K4 me3:H3−5ac
H3K27 me3:DNA Meth. m5C
H3K4 me1:H2A.Z
H3K27 me3:H3K4 me1
H4K20 me2:H3K4 me3
H4K20 me2:H3−5ac
H4K20 me2:H4−4ac
H3K4 me1:H3−5ac
H2A.Z:H4K20 me2
H3K4 me3:H4−4ac
H3K27 me2:DNA Meth. m5C
H3K4 me1:H4K20 me2
H4−4ac:H3−5ac
H3K9 me2
H4K20 me3
H3K27 me2
H3K9 me3
DNA Meth. m5C
H3K27 me3
H3K4 me3
H2A.Z
H4−4ac
H3K4 me1
H3−5ac
H4K20 me2
0.00
0.25
0.50
0.75
1.00
$\hat{\pi}$
SSR4 − Reverse
DNA Meth. m5C:H3K9 me3
H2A.Z:H3K4 me3
DNA Meth. m5C:H4K20 me2
H3K9 me2:DNA Meth. m5C
H3K27 me3:H3K9 me3
H3K4 me1:H4−4ac
H3K4 me3:H3−5ac
H3K27 me3:DNA Meth. m5C
H3K4 me1:H2A.Z
H3K27 me3:H3K4 me1
H4K20 me2:H3K4 me3
H4K20 me2:H3−5ac
H4K20 me2:H4−4ac
H3K4 me1:H3−5ac
H2A.Z:H4K20 me2
H3K4 me3:H4−4ac
H3K27 me2:DNA Meth. m5C
H3K4 me1:H4K20 me2
H4−4ac:H3−5ac
H3K9 me2
H4K20 me3
H3K27 me2
H3K9 me3
DNA Meth. m5C
H3K27 me3
H3K4 me3
H2A.Z
H4−4ac
H3K4 me1
H3−5ac
H4K20 me2
0.00
0.25
0.50
0.75
1.00
$\hat{\pi}$
TIMM50 − Forward
H3K4 me1:H4−4ac
H3K4 me3:H4−4ac
H2A.Z:H3−5ac
H2A.Z:H4−4ac
H3K4 me1:H2A.Z
DNA Meth. m5C:H4K20 me2
H3K4 me3:H3−5ac
H3K9 me2:DNA Meth. m5C
H3K27 me3:H3K9 me3
H4−4ac:H3−5ac
H2A.Z:H3K4 me3
DNA Meth. m5C:H3K9 me3
H3K27 me3:H4K20 me3
H4K20 me2:H4−4ac
H3K9 me3:H4K20 me3
H3K27 me3:DNA Meth. m5C
H3K27 me3:H3K4 me1
H3K4 me1:H3−5ac
H2A.Z:H4K20 me2
H3K4 me1:H4K20 me2
H4K20 me2:H3−5ac
DNA Meth. m5C:H4K20 me3
H3K27 me2:DNA Meth. m5C
H4K20 me2:H3K4 me3
H3K9 me2
H3K9 me3
H3K27 me2
H3K27 me3
H4−4ac
H3K4 me3
H2A.Z
H3−5ac
H4K20 me3
H3K4 me1
H4K20 me2
DNA Meth. m5C
0.00
0.25
0.50
0.75
1.00
$\hat{\pi}$
TIMM50 − Reverse
H3K4 me1:H4−4ac
H3K4 me3:H4−4ac
H2A.Z:H3−5ac
H2A.Z:H4−4ac
H3K4 me1:H2A.Z
DNA Meth. m5C:H4K20 me2
H3K4 me3:H3−5ac
H3K9 me2:DNA Meth. m5C
H3K27 me3:H3K9 me3
H4−4ac:H3−5ac
H2A.Z:H3K4 me3
DNA Meth. m5C:H3K9 me3
H3K27 me3:H4K20 me3
H4K20 me2:H4−4ac
H3K9 me3:H4K20 me3
H3K27 me3:DNA Meth. m5C
H3K27 me3:H3K4 me1
H3K4 me1:H3−5ac
H2A.Z:H4K20 me2
H3K4 me1:H4K20 me2
H4K20 me2:H3−5ac
DNA Meth. m5C:H4K20 me3
H3K27 me2:DNA Meth. m5C
H4K20 me2:H3K4 me3
H3K9 me2
H3K9 me3
H3K27 me2
H3K27 me3
H4−4ac
H3K4 me3
H2A.Z
H3−5ac
H4K20 me3
H3K4 me1
H4K20 me2
DNA Meth. m5C
0.00
0.25
0.50
0.75
1.00
$\hat{\pi}$
FADS1 (1) − Forward
H3K27 me3:H3K9 me3
H3K27 me2:DNA Meth. m5C
H3K9 me2:DNA Meth. m5C
H3K27 me3:DNA Meth. m5C
DNA Meth. m5C:H3K9 me3
H3K9 me3:H4K20 me3
H3K27 me3:H4K20 me3
DNA Meth. m5C:H4K20 me2
H3K4 me3:H4−4ac
H4K20 me2:H4−4ac
H4−4ac:H3−5ac
DNA Meth. m5C:H4K20 me3
H4K20 me2:H3−5ac
H4K20 me2:H3K4 me3
H3K4 me1:H4K20 me2
H2A.Z:H4K20 me2
H3K9 me2
H3K27 me2
H3K9 me3
H3K27 me3
H3K4 me1
DNA Meth. m5C
H2A.Z
H4K20 me3
H3−5ac
H4−4ac
H3K4 me3
H4K20 me2
0.00
0.25
0.50
0.75
1.00
$\hat{\pi}$
FADS1 (1) − Reverse
H3K27 me3:H3K9 me3
H3K27 me2:DNA Meth. m5C
H3K9 me2:DNA Meth. m5C
H3K27 me3:DNA Meth. m5C
DNA Meth. m5C:H3K9 me3
H3K9 me3:H4K20 me3
H3K27 me3:H4K20 me3
DNA Meth. m5C:H4K20 me2
H3K4 me3:H4−4ac
H4K20 me2:H4−4ac
H4−4ac:H3−5ac
DNA Meth. m5C:H4K20 me3
H4K20 me2:H3−5ac
H4K20 me2:H3K4 me3
H3K4 me1:H4K20 me2
H2A.Z:H4K20 me2
H3K9 me2
H3K27 me2
H3K9 me3
H3K27 me3
H3K4 me1
DNA Meth. m5C
H2A.Z
H4K20 me3
H3−5ac
H4−4ac
H3K4 me3
H4K20 me2
0.00
0.25
0.50
0.75
1.00
$\hat{\pi}$
ABCF2 − Forward
DNA Meth. m5C:H3K9 me3
H2A.Z:H3−5ac
H2A.Z:H4−4ac
H3K27 me2:DNA Meth. m5C
DNA Meth. m5C:H4K20 me2
H3K27 me3:H3K9 me3
DNA Meth. m5C:H4K20 me3
H3K9 me3:H4K20 me3
H3K27 me3:H4K20 me3
H3K27 me3:H3K4 me1
H4K20 me2:H4−4ac
H3K27 me3:DNA Meth. m5C
H3K4 me3:H3−5ac
H4−4ac:H3−5ac
H2A.Z:H3K4 me3
H3K4 me1:H2A.Z
H3K4 me1:H3−5ac
H3K4 me1:H4−4ac
H4K20 me2:H3−5ac
H2A.Z:H4K20 me2
H3K4 me1:H4K20 me2
H3K4 me3:H4−4ac
H4K20 me2:H3K4 me3
H3K9 me2
H3K27 me2
H3K9 me3
H3K27 me3
H4K20 me3
DNA Meth. m5C
H4K20 me2
H2A.Z
H3−5ac
H3K4 me1
H3K4 me3
H4−4ac
0.00
0.25
0.50
0.75
1.00
$\hat{\pi}$
ABCF2 − Reverse
DNA Meth. m5C:H3K9 me3
H2A.Z:H3−5ac
H2A.Z:H4−4ac
H3K27 me2:DNA Meth. m5C
DNA Meth. m5C:H4K20 me2
H3K27 me3:H3K9 me3
DNA Meth. m5C:H4K20 me3
H3K9 me3:H4K20 me3
H3K27 me3:H4K20 me3
H3K27 me3:H3K4 me1
H4K20 me2:H4−4ac
H3K27 me3:DNA Meth. m5C
H3K4 me3:H3−5ac
H4−4ac:H3−5ac
H2A.Z:H3K4 me3
H3K4 me1:H2A.Z
H3K4 me1:H3−5ac
H3K4 me1:H4−4ac
H4K20 me2:H3−5ac
H2A.Z:H4K20 me2
H3K4 me1:H4K20 me2
H3K4 me3:H4−4ac
H4K20 me2:H3K4 me3
H3K9 me2
H3K27 me2
H3K9 me3
H3K27 me3
H4K20 me3
DNA Meth. m5C
H4K20 me2
H2A.Z
H3−5ac
H3K4 me1
H3K4 me3
H4−4ac
0.00
0.25
0.50
0.75
1.00
$\hat{\pi}$
FKBP11 − Forward
H2A.Z:H3−5ac
H2A.Z:H4−4ac
H3K27 me2:DNA Meth. m5C
H3K9 me3:H4K20 me3
DNA Meth. m5C:H4K20 me3
H3K27 me3:H3K4 me1
H3K27 me3:H4K20 me3
H3K27 me3:H3K9 me3
H3K4 me3:H3−5ac
DNA Meth. m5C:H4K20 me2
H2A.Z:H3K4 me3
H2A.Z:H4K20 me2
H3K4 me1:H4−4ac
H3K27 me3:DNA Meth. m5C
DNA Meth. m5C:H3K9 me3
H4K20 me2:H4−4ac
H3K4 me1:H3−5ac
H3K4 me1:H2A.Z
H4−4ac:H3−5ac
H3K4 me3:H4−4ac
H3K4 me1:H4K20 me2
H4K20 me2:H3−5ac
H4K20 me2:H3K4 me3
H3K27 me2
H3K9 me2
H4K20 me3
H3K9 me3
H3K4 me1
H2A.Z
H3K4 me3
DNA Meth. m5C
H3K27 me3
H3−5ac
H4K20 me2
H4−4ac
0.00
0.25
0.50
0.75
1.00
$\hat{\pi}$
FKBP11 − Reverse
H2A.Z:H3−5ac
H2A.Z:H4−4ac
H3K27 me2:DNA Meth. m5C
H3K9 me3:H4K20 me3
DNA Meth. m5C:H4K20 me3
H3K27 me3:H3K4 me1
H3K27 me3:H4K20 me3
H3K27 me3:H3K9 me3
H3K4 me3:H3−5ac
DNA Meth. m5C:H4K20 me2
H2A.Z:H3K4 me3
H2A.Z:H4K20 me2
H3K4 me1:H4−4ac
H3K27 me3:DNA Meth. m5C
DNA Meth. m5C:H3K9 me3
H4K20 me2:H4−4ac
H3K4 me1:H3−5ac
H3K4 me1:H2A.Z
H4−4ac:H3−5ac
H3K4 me3:H4−4ac
H3K4 me1:H4K20 me2
H4K20 me2:H3−5ac
H4K20 me2:H3K4 me3
H3K27 me2
H3K9 me2
H4K20 me3
H3K9 me3
H3K4 me1
H2A.Z
H3K4 me3
DNA Meth. m5C
H3K27 me3
H3−5ac
H4K20 me2
H4−4ac
0.00
0.25
0.50
0.75
1.00
$\hat{\pi}$
PCGF6 − Forward
H3K9 me2:DNA Meth. m5C
H3K4 me3:H3−5ac
H3K9 me3:H4K20 me3
H3K27 me3:DNA Meth. m5C
H2A.Z:H3−5ac
H2A.Z:H4−4ac
DNA Meth. m5C:H4K20 me2
H3K27 me2:DNA Meth. m5C
H3K27 me3:H3K4 me1
DNA Meth. m5C:H4K20 me3
H2A.Z:H3K4 me3
H3K27 me3:H3K9 me3
H3K4 me1:H4−4ac
H4K20 me2:H4−4ac
H2A.Z:H4K20 me2
H3K4 me1:H2A.Z
H4K20 me2:H3−5ac
H3K4 me1:H3−5ac
H4−4ac:H3−5ac
H3K4 me1:H4K20 me2
DNA Meth. m5C:H3K9 me3
H4K20 me2:H3K4 me3
H3K4 me3:H4−4ac
H3K9 me2
H3K27 me2
H4K20 me3
H3K27 me3
H2A.Z
H3K9 me3
H4K20 me2
H3−5ac
H3K4 me1
H4−4ac
H3K4 me3
DNA Meth. m5C
0.00
0.25
0.50
0.75
1.00
$\hat{\pi}$
PCGF6 − Reverse
H3K9 me2:DNA Meth. m5C
H3K4 me3:H3−5ac
H3K9 me3:H4K20 me3
H3K27 me3:DNA Meth. m5C
H2A.Z:H3−5ac
H2A.Z:H4−4ac
DNA Meth. m5C:H4K20 me2
H3K27 me2:DNA Meth. m5C
H3K27 me3:H3K4 me1
DNA Meth. m5C:H4K20 me3
H2A.Z:H3K4 me3
H3K27 me3:H3K9 me3
H3K4 me1:H4−4ac
H4K20 me2:H4−4ac
H2A.Z:H4K20 me2
H3K4 me1:H2A.Z
H4K20 me2:H3−5ac
H3K4 me1:H3−5ac
H4−4ac:H3−5ac
H3K4 me1:H4K20 me2
DNA Meth. m5C:H3K9 me3
H4K20 me2:H3K4 me3
H3K4 me3:H4−4ac
H3K9 me2
H3K27 me2
H4K20 me3
H3K27 me3
H2A.Z
H3K9 me3
H4K20 me2
H3−5ac
H3K4 me1
H4−4ac
H3K4 me3
DNA Meth. m5C
0.00
0.25
0.50
0.75
1.00
$\hat{\pi}$

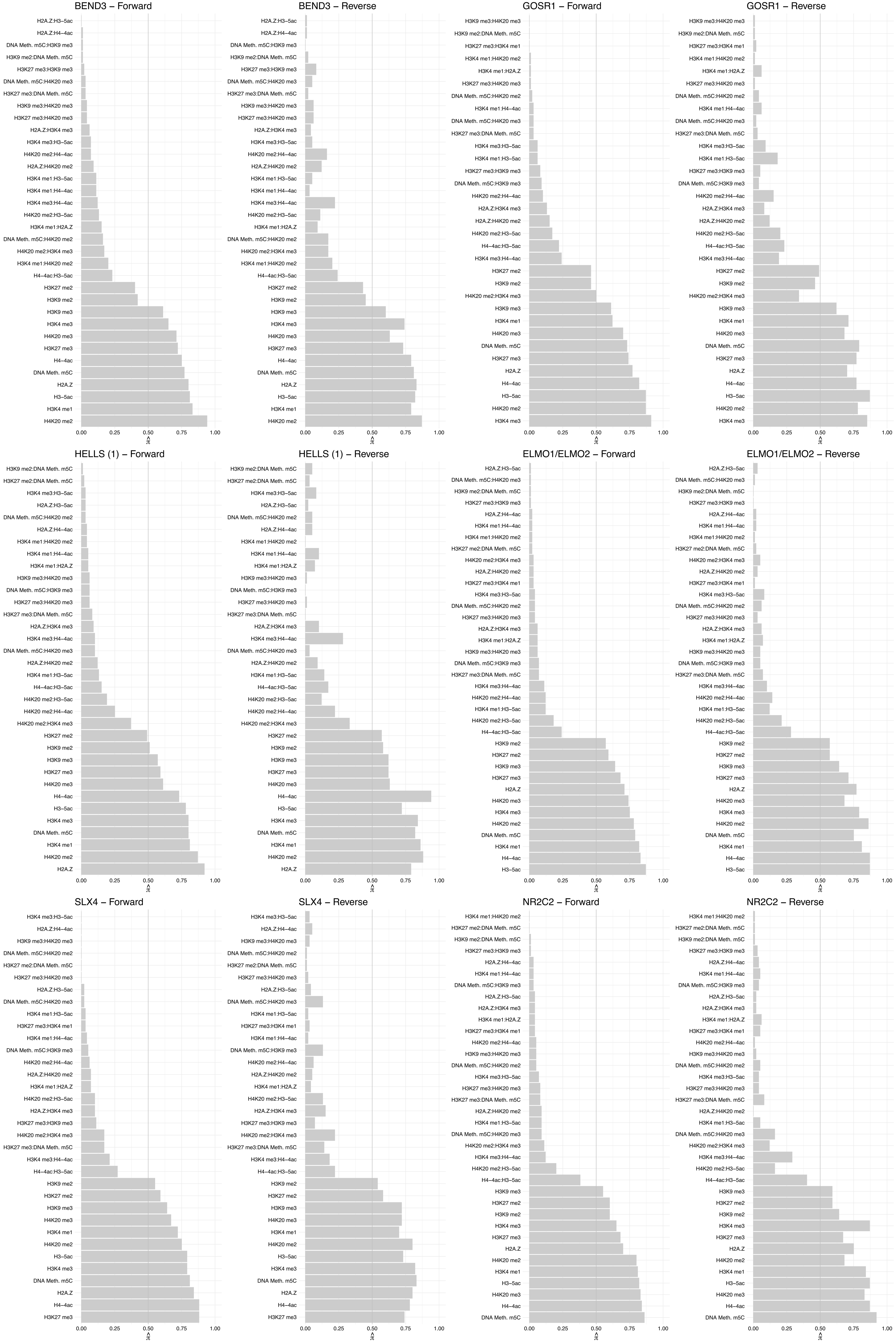

BEND3 – Forward
BEND3 – Reverse
GOSR1 – Forward
GOSR1 – Reverse
HELLS (1) – Forward
HELLS (1) – Reverse
ELMO1/ELMO2 – Forward
ELMO1/ELMO2 – Reverse
SLX4 – Forward
SLX4 – Reverse
NR2C2 – Forward
NR2C2 – Reverse
0.00
0.25
0.50
0.75
1.00
$\hat{\pi}$

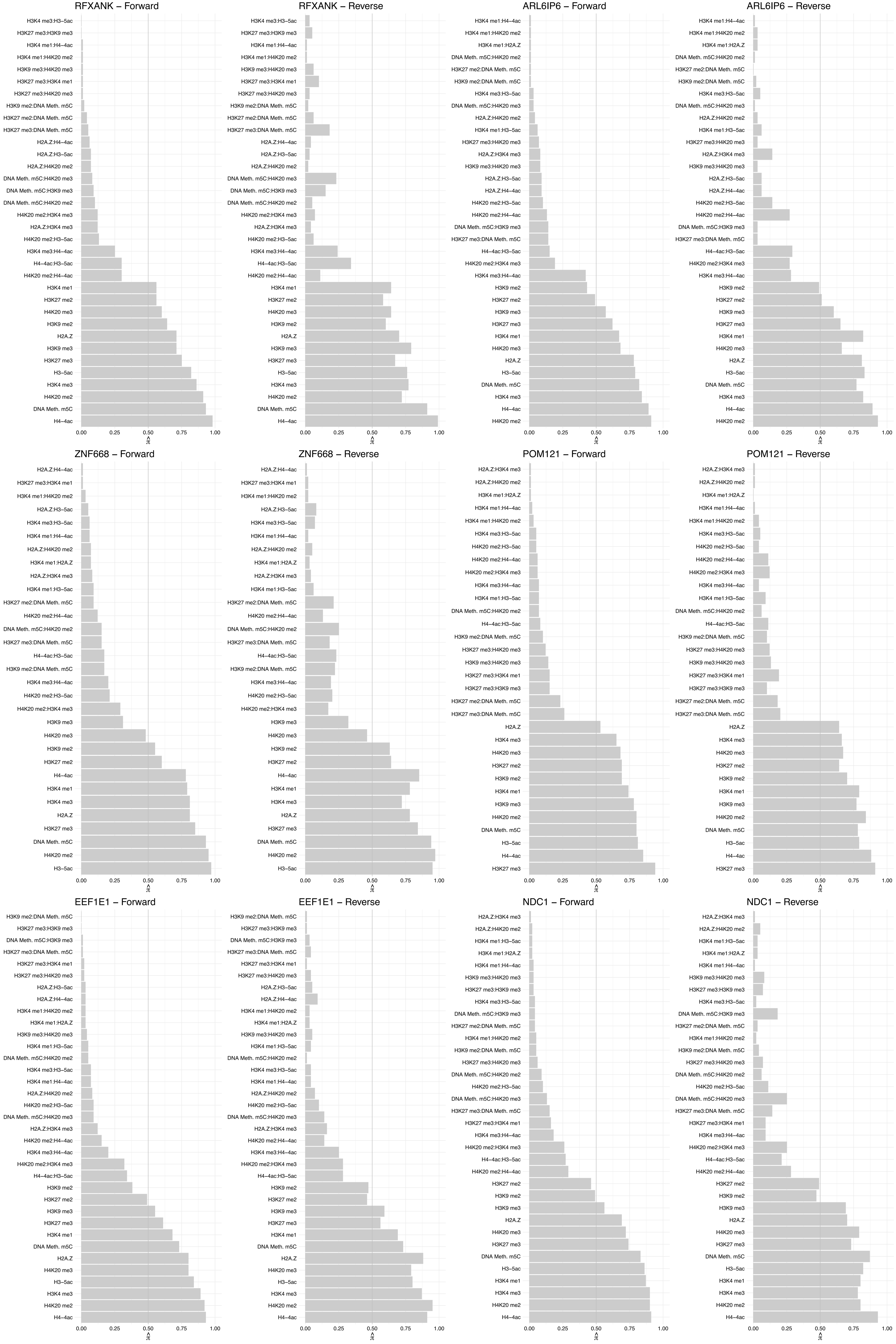
RFXANK – Forward
RFXANK – Reverse
ARL6IP6 – Forward
ARL6IP6 – Reverse
ZNF668 – Forward
ZNF668 – Reverse
POM121 – Forward
POM121 – Reverse
EEF1E1 – Forward
EEF1E1 – Reverse
NDC1 – Forward
NDC1 – Reverse
0.00
0.25
0.50
0.75
1.00
$\hat{\pi}$

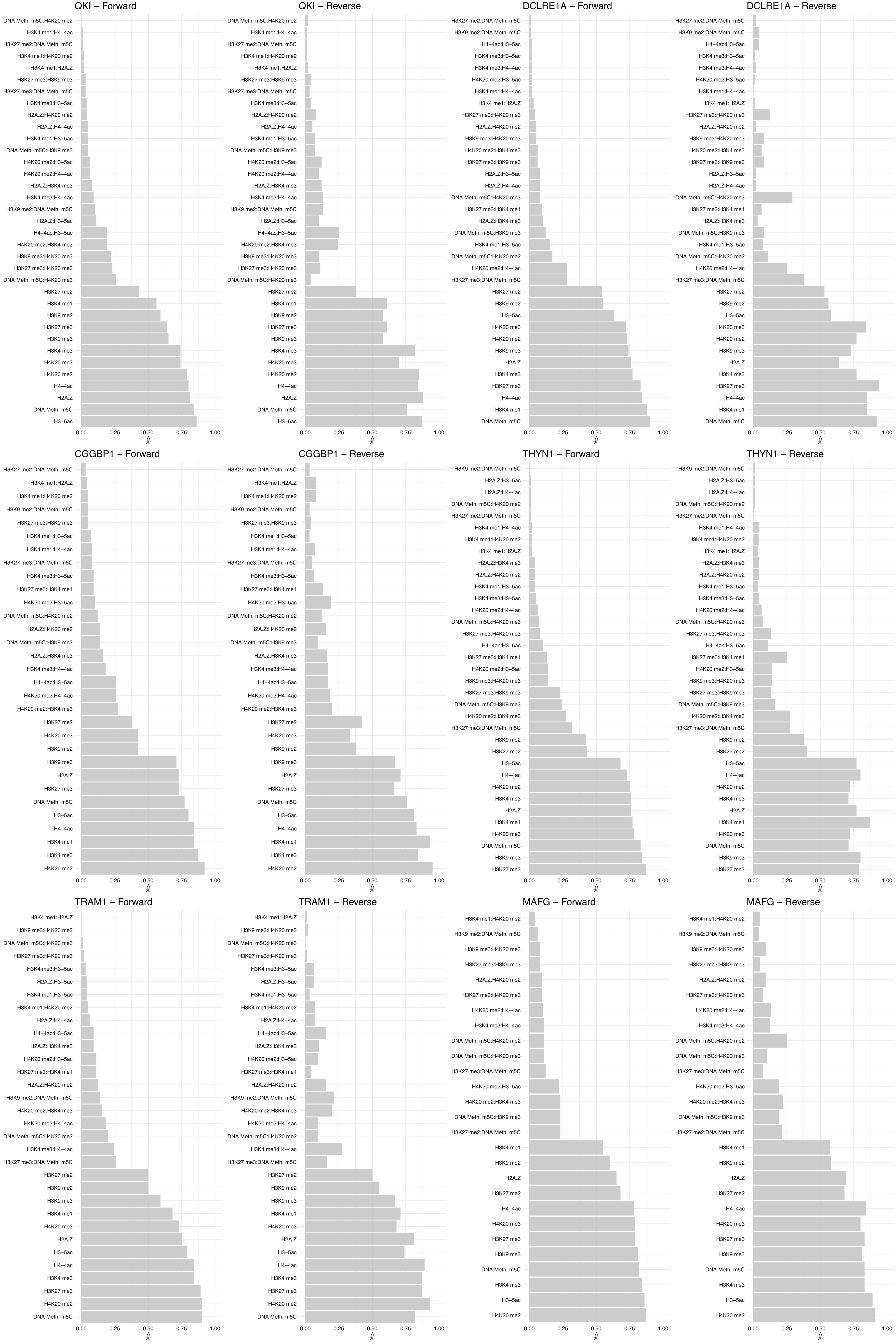
QKI – Forward
DNA Meth. m5C:H4K20 me2
H3K4 me1:H4−4ac
H3K27 me2:DNA Meth. m5C
H3K4 me1:H4K20 me2
H3K4 me1:H2A.Z
H3K27 me3:H3K9 me3
H3K27 me3:DNA Meth. m5C
H3K4 me3:H3−5ac
H2A.Z:H4K20 me2
H2A.Z:H4−4ac
H3K4 me1:H3−5ac
DNA Meth. m5C:H3K9 me3
H4K20 me2:H3−5ac
H4K20 me2:H4−4ac
H2A.Z:H3K4 me3
H3K4 me3:H4−4ac
H3K9 me2:DNA Meth. m5C
H2A.Z:H3−5ac
H4−4ac:H3−5ac
H4K20 me2:H3K4 me3
H3K9 me3:H4K20 me3
H3K27 me3:H4K20 me3
DNA Meth. m5C:H4K20 me3
H3K27 me2
H3K4 me1
H3K9 me2
H3K27 me3
H3K9 me3
H3K4 me3
H4K20 me3
H4K20 me2
H4−4ac
H2A.Z
DNA Meth. m5C
H3−5ac
0.00 0.25 0.50 0.75 1.00
$\hat{\pi}$
QKI – Reverse
DNA Meth. m5C:H4K20 me2
H3K4 me1:H4−4ac
H3K27 me2:DNA Meth. m5C
H3K4 me1:H4K20 me2
H3K4 me1:H2A.Z
H3K27 me3:H3K9 me3
H3K27 me3:DNA Meth. m5C
H3K4 me3:H3−5ac
H2A.Z:H4K20 me2
H2A.Z:H4−4ac
H3K4 me1:H3−5ac
DNA Meth. m5C:H3K9 me3
H4K20 me2:H3−5ac
H4K20 me2:H4−4ac
H2A.Z:H3K4 me3
H3K4 me3:H4−4ac
H3K9 me2:DNA Meth. m5C
H2A.Z:H3−5ac
H4−4ac:H3−5ac
H4K20 me2:H3K4 me3
H3K9 me3:H4K20 me3
H3K27 me3:H4K20 me3
DNA Meth. m5C:H4K20 me3
H3K27 me2
H3K4 me1
H3K9 me2
H3K27 me3
H3K9 me3
H3K4 me3
H4K20 me3
H4K20 me2
H4−4ac
H2A.Z
DNA Meth. m5C
H3−5ac
0.00 0.25 0.50 0.75 1.00
$\hat{\pi}$
DCLRE1A – Forward
H3K27 me2:DNA Meth. m5C
H3K9 me2:DNA Meth. m5C
H4−4ac:H3−5ac
H3K4 me3:H3−5ac
H3K4 me3:H4−4ac
H4K20 me2:H3−5ac
H3K4 me1:H4−4ac
H3K4 me1:H2A.Z
H3K27 me3:H4K20 me3
H2A.Z:H4K20 me2
H3K9 me3:H4K20 me3
H4K20 me2:H3K4 me3
H3K27 me3:H3K9 me3
H2A.Z:H3−5ac
H2A.Z:H4−4ac
DNA Meth. m5C:H4K20 me3
H3K27 me3:H3K4 me1
H2A.Z:H3K4 me3
DNA Meth. m5C:H3K9 me3
H3K4 me1:H3−5ac
DNA Meth. m5C:H4K20 me2
H4K20 me2:H4−4ac
H3K27 me3:DNA Meth. m5C
H3K27 me2
H3K9 me2
H3−5ac
H4K20 me3
H4K20 me2
H3K9 me3
H2A.Z
H3K4 me3
H3K27 me3
H4−4ac
H3K4 me1
DNA Meth. m5C
0.00 0.25 0.50 0.75 1.00
$\hat{\pi}$
DCLRE1A – Reverse
H3K27 me2:DNA Meth. m5C
H3K9 me2:DNA Meth. m5C
H4−4ac:H3−5ac
H3K4 me3:H3−5ac
H3K4 me3:H4−4ac
H4K20 me2:H3−5ac
H3K4 me1:H4−4ac
H3K4 me1:H2A.Z
H3K27 me3:H4K20 me3
H2A.Z:H4K20 me2
H3K9 me3:H4K20 me3
H4K20 me2:H3K4 me3
H3K27 me3:H3K9 me3
H2A.Z:H3−5ac
H2A.Z:H4−4ac
DNA Meth. m5C:H4K20 me3
H3K27 me3:H3K4 me1
H2A.Z:H3K4 me3
DNA Meth. m5C:H3K9 me3
H3K4 me1:H3−5ac
DNA Meth. m5C:H4K20 me2
H4K20 me2:H4−4ac
H3K27 me3:DNA Meth. m5C
H3K27 me2
H3K9 me2
H3−5ac
H4K20 me3
H4K20 me2
H3K9 me3
H2A.Z
H3K4 me3
H3K27 me3
H4−4ac
H3K4 me1
DNA Meth. m5C
0.00 0.25 0.50 0.75 1.00
$\hat{\pi}$
CGGBP1 – Forward
H3K27 me2:DNA Meth. m5C
H3K4 me1:H2A.Z
H3K4 me1:H4K20 me2
H3K9 me2:DNA Meth. m5C
H3K27 me3:H3K9 me3
H3K4 me1:H3−5ac
H3K4 me1:H4−4ac
H3K27 me3:DNA Meth. m5C
H3K4 me3:H3−5ac
H3K27 me3:H3K4 me1
H4K20 me2:H3−5ac
DNA Meth. m5C:H4K20 me2
H2A.Z:H4K20 me2
DNA Meth. m5C:H3K9 me3
H2A.Z:H3K4 me3
H3K4 me3:H4−4ac
H4−4ac:H3−5ac
H4K20 me2:H4−4ac
H4K20 me2:H3K4 me3
H3K27 me2
H4K20 me3
H3K9 me2
H3K9 me3
H2A.Z
H3K27 me3
DNA Meth. m5C
H3−5ac
H4−4ac
H3K4 me1
H3K4 me3
H4K20 me2
0.00 0.25 0.50 0.75 1.00
$\hat{\pi}$
CGGBP1 – Reverse
H3K27 me2:DNA Meth. m5C
H3K4 me1:H2A.Z
H3K4 me1:H4K20 me2
H3K9 me2:DNA Meth. m5C
H3K27 me3:H3K9 me3
H3K4 me1:H3−5ac
H3K4 me1:H4−4ac
H3K27 me3:DNA Meth. m5C
H3K4 me3:H3−5ac
H3K27 me3:H3K4 me1
H4K20 me2:H3−5ac
DNA Meth. m5C:H4K20 me2
H2A.Z:H4K20 me2
DNA Meth. m5C:H3K9 me3
H2A.Z:H3K4 me3
H3K4 me3:H4−4ac
H4−4ac:H3−5ac
H4K20 me2:H4−4ac
H4K20 me2:H3K4 me3
H3K27 me2
H4K20 me3
H3K9 me2
H3K9 me3
H2A.Z
H3K27 me3
DNA Meth. m5C
H3−5ac
H4−4ac
H3K4 me1
H3K4 me3
H4K20 me2
0.00 0.25 0.50 0.75 1.00
$\hat{\pi}$
THYN1 – Forward
H3K9 me2:DNA Meth. m5C
H2A.Z:H3−5ac
H2A.Z:H4−4ac
DNA Meth. m5C:H4K20 me2
H3K27 me2:DNA Meth. m5C
H3K4 me1:H4−4ac
H3K4 me1:H4K20 me2
H3K4 me1:H2A.Z
H2A.Z:H3K4 me3
H2A.Z:H4K20 me2
H3K4 me1:H3−5ac
H3K4 me3:H3−5ac
H4K20 me2:H4−4ac
DNA Meth. m5C:H4K20 me3
H3K27 me3:H4K20 me3
H4−4ac:H3−5ac
H3K27 me3:H3K4 me1
H4K20 me2:H3−5ac
H3K9 me3:H4K20 me3
H3K27 me3:H3K9 me3
DNA Meth. m5C:H3K9 me3
H4K20 me2:H3K4 me3
H3K27 me3:DNA Meth. m5C
H3K9 me2
H3K27 me2
H3−5ac
H4−4ac
H4K20 me2
H3K4 me3
H2A.Z
H3K4 me1
H4K20 me3
DNA Meth. m5C
H3K9 me3
H3K27 me3
0.00 0.25 0.50 0.75 1.00
$\hat{\pi}$
THYN1 – Reverse
H3K9 me2:DNA Meth. m5C
H2A.Z:H3−5ac
H2A.Z:H4−4ac
DNA Meth. m5C:H4K20 me2
H3K27 me2:DNA Meth. m5C
H3K4 me1:H4−4ac
H3K4 me1:H4K20 me2
H3K4 me1:H2A.Z
H2A.Z:H3K4 me3
H2A.Z:H4K20 me2
H3K4 me1:H3−5ac
H3K4 me3:H3−5ac
H4K20 me2:H4−4ac
DNA Meth. m5C:H4K20 me3
H3K27 me3:H4K20 me3
H4−4ac:H3−5ac
H3K27 me3:H3K4 me1
H4K20 me2:H3−5ac
H3K9 me3:H4K20 me3
H3K27 me3:H3K9 me3
DNA Meth. m5C:H3K9 me3
H4K20 me2:H3K4 me3
H3K27 me3:DNA Meth. m5C
H3K9 me2
H3K27 me2
H3−5ac
H4−4ac
H4K20 me2
H3K4 me3
H2A.Z
H3K4 me1
H4K20 me3
DNA Meth. m5C
H3K9 me3
H3K27 me3
0.00 0.25 0.50 0.75 1.00
$\hat{\pi}$
TRAM1 – Forward
H3K4 me1:H2A.Z
H3K9 me3:H4K20 me3
DNA Meth. m5C:H4K20 me3
H3K27 me3:H4K20 me3
H3K4 me3:H3−5ac
H2A.Z:H3−5ac
H3K4 me1:H3−5ac
H3K4 me1:H4K20 me2
H2A.Z:H4−4ac
H4−4ac:H3−5ac
H2A.Z:H3K4 me3
H4K20 me2:H3−5ac
H3K27 me3:H3K4 me1
H2A.Z:H4K20 me2
H3K9 me2:DNA Meth. m5C
H4K20 me2:H3K4 me3
H4K20 me2:H4−4ac
DNA Meth. m5C:H4K20 me2
H3K4 me3:H4−4ac
H3K27 me3:DNA Meth. m5C
H3K27 me2
H3K9 me2
H3K9 me3
H3K4 me1
H4K20 me3
H2A.Z
H3−5ac
H4−4ac
H3K4 me3
H3K27 me3
H4K20 me2
DNA Meth. m5C
0.00 0.25 0.50 0.75 1.00
$\hat{\pi}$
TRAM1 – Reverse
H3K4 me1:H2A.Z
H3K9 me3:H4K20 me3
DNA Meth. m5C:H4K20 me3
H3K27 me3:H4K20 me3
H3K4 me3:H3−5ac
H2A.Z:H3−5ac
H3K4 me1:H3−5ac
H3K4 me1:H4K20 me2
H2A.Z:H4−4ac
H4−4ac:H3−5ac
H2A.Z:H3K4 me3
H4K20 me2:H3−5ac
H3K27 me3:H3K4 me1
H2A.Z:H4K20 me2
H3K9 me2:DNA Meth. m5C
H4K20 me2:H3K4 me3
H4K20 me2:H4−4ac
DNA Meth. m5C:H4K20 me2
H3K4 me3:H4−4ac
H3K27 me3:DNA Meth. m5C
H3K27 me2
H3K9 me2
H3K9 me3
H3K4 me1
H4K20 me3
H2A.Z
H3−5ac
H4−4ac
H3K4 me3
H3K27 me3
H4K20 me2
DNA Meth. m5C
0.00 0.25 0.50 0.75 1.00
$\hat{\pi}$
MAFG – Forward
H3K4 me1:H4K20 me2
H3K9 me2:DNA Meth. m5C
H3K9 me3:H4K20 me3
H3K27 me3:H3K9 me3
H2A.Z:H4K20 me2
H3K27 me3:H4K20 me3
H4K20 me2:H4−4ac
H3K4 me3:H4−4ac
DNA Meth. m5C:H4K20 me2
DNA Meth. m5C:H4K20 me3
H3K27 me3:DNA Meth. m5C
H4K20 me2:H3−5ac
H4K20 me2:H3K4 me3
DNA Meth. m5C:H3K9 me3
H3K27 me2:DNA Meth. m5C
H3K4 me1
H3K9 me2
H2A.Z
H3K27 me2
H4−4ac
H4K20 me3
H3K27 me3
H3K9 me3
DNA Meth. m5C
H3K4 me3
H3−5ac
H4K20 me2
0.00 0.25 0.50 0.75 1.00
$\hat{\pi}$
MAFG – Reverse
H3K4 me1:H4K20 me2
H3K9 me2:DNA Meth. m5C
H3K9 me3:H4K20 me3
H3K27 me3:H3K9 me3
H2A.Z:H4K20 me2
H3K27 me3:H4K20 me3
H4K20 me2:H4−4ac
H3K4 me3:H4−4ac
DNA Meth. m5C:H4K20 me2
DNA Meth. m5C:H4K20 me3
H3K27 me3:DNA Meth. m5C
H4K20 me2:H3−5ac
H4K20 me2:H3K4 me3
DNA Meth. m5C:H3K9 me3
H3K27 me2:DNA Meth. m5C
H3K4 me1
H3K9 me2
H2A.Z
H3K27 me2
H4−4ac
H4K20 me3
H3K27 me3
H3K9 me3
DNA Meth. m5C
H3K4 me3
H3−5ac
H4K20 me2
0.00 0.25 0.50 0.75 1.00
$\hat{\pi}$

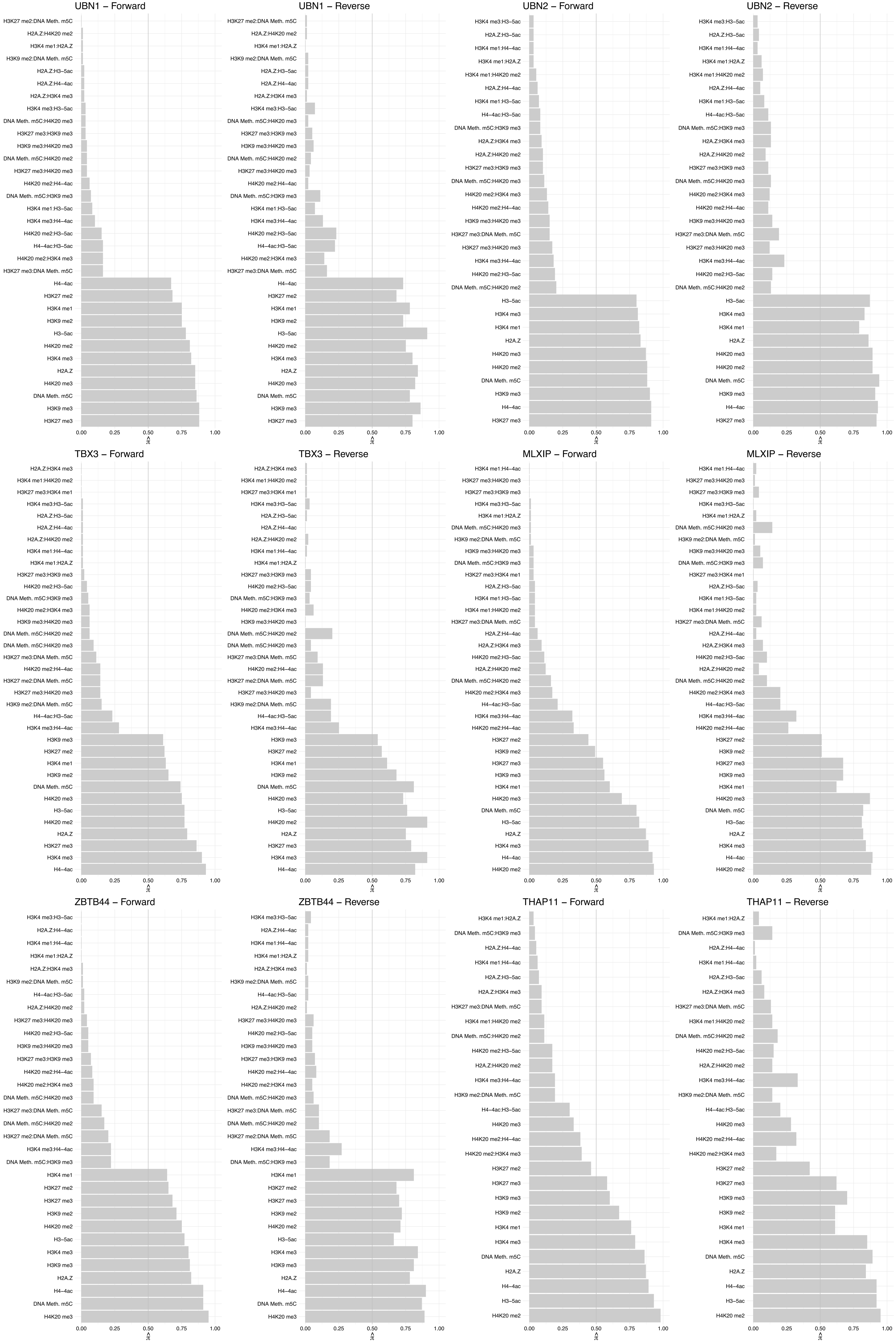
UBN1 – Forward
H3K27 me2:DNA Meth. m5C
H2A.Z:H4K20 me2
H3K4 me1:H2A.Z
H3K9 me2:DNA Meth. m5C
H2A.Z:H3−5ac
H2A.Z:H4−4ac
H2A.Z:H3K4 me3
H3K4 me3:H3−5ac
DNA Meth. m5C:H4K20 me3
H3K27 me3:H3K9 me3
H3K9 me3:H4K20 me3
DNA Meth. m5C:H4K20 me2
H3K27 me3:H4K20 me3
H4K20 me2:H4−4ac
DNA Meth. m5C:H3K9 me3
H3K4 me1:H3−5ac
H3K4 me3:H4−4ac
H4K20 me2:H3−5ac
H4−4ac:H3−5ac
H4K20 me2:H3K4 me3
H3K27 me3:DNA Meth. m5C
H4−4ac
H3K27 me2
H3K4 me1
H3K9 me2
H3−5ac
H4K20 me2
H3K4 me3
H2A.Z
H4K20 me3
DNA Meth. m5C
H3K9 me3
H3K27 me3
0.00 0.25 0.50 0.75 1.00
$\hat{\pi}$
UBN1 – Reverse
H3K27 me2:DNA Meth. m5C
H2A.Z:H4K20 me2
H3K4 me1:H2A.Z
H3K9 me2:DNA Meth. m5C
H2A.Z:H3−5ac
H2A.Z:H4−4ac
H2A.Z:H3K4 me3
H3K4 me3:H3−5ac
DNA Meth. m5C:H4K20 me3
H3K27 me3:H3K9 me3
H3K9 me3:H4K20 me3
DNA Meth. m5C:H4K20 me2
H3K27 me3:H4K20 me3
H4K20 me2:H4−4ac
DNA Meth. m5C:H3K9 me3
H3K4 me1:H3−5ac
H3K4 me3:H4−4ac
H4K20 me2:H3−5ac
H4−4ac:H3−5ac
H4K20 me2:H3K4 me3
H3K27 me3:DNA Meth. m5C
H4−4ac
H3K27 me2
H3K4 me1
H3K9 me2
H3−5ac
H4K20 me2
H3K4 me3
H2A.Z
H4K20 me3
DNA Meth. m5C
H3K9 me3
H3K27 me3
0.00 0.25 0.50 0.75 1.00
$\hat{\pi}$
UBN2 – Forward
H3K4 me3:H3−5ac
H2A.Z:H3−5ac
H3K4 me1:H4−4ac
H3K4 me1:H2A.Z
H3K4 me1:H4K20 me2
H2A.Z:H4−4ac
H3K4 me1:H3−5ac
H4−4ac:H3−5ac
DNA Meth. m5C:H3K9 me3
H2A.Z:H3K4 me3
H2A.Z:H4K20 me2
H3K27 me3:H3K9 me3
DNA Meth. m5C:H4K20 me3
H4K20 me2:H3K4 me3
H4K20 me2:H4−4ac
H3K9 me3:H4K20 me3
H3K27 me3:DNA Meth. m5C
H3K27 me3:H4K20 me3
H3K4 me3:H4−4ac
H4K20 me2:H3−5ac
DNA Meth. m5C:H4K20 me2
H3−5ac
H3K4 me3
H3K4 me1
H2A.Z
H4K20 me3
H4K20 me2
DNA Meth. m5C
H3K9 me3
H4−4ac
H3K27 me3
0.00 0.25 0.50 0.75 1.00
$\hat{\pi}$
UBN2 – Reverse
H3K4 me3:H3−5ac
H2A.Z:H3−5ac
H3K4 me1:H4−4ac
H3K4 me1:H2A.Z
H3K4 me1:H4K20 me2
H2A.Z:H4−4ac
H3K4 me1:H3−5ac
H4−4ac:H3−5ac
DNA Meth. m5C:H3K9 me3
H2A.Z:H3K4 me3
H2A.Z:H4K20 me2
H3K27 me3:H3K9 me3
DNA Meth. m5C:H4K20 me3
H4K20 me2:H3K4 me3
H4K20 me2:H4−4ac
H3K9 me3:H4K20 me3
H3K27 me3:DNA Meth. m5C
H3K27 me3:H4K20 me3
H3K4 me3:H4−4ac
H4K20 me2:H3−5ac
DNA Meth. m5C:H4K20 me2
H3−5ac
H3K4 me3
H3K4 me1
H2A.Z
H4K20 me3
H4K20 me2
DNA Meth. m5C
H3K9 me3
H4−4ac
H3K27 me3
0.00 0.25 0.50 0.75 1.00
$\hat{\pi}$
TBX3 – Forward
H2A.Z:H3K4 me3
H3K4 me1:H4K20 me2
H3K27 me3:H3K4 me1
H3K4 me3:H3−5ac
H2A.Z:H3−5ac
H2A.Z:H4−4ac
H2A.Z:H4K20 me2
H3K4 me1:H4−4ac
H3K4 me1:H2A.Z
H3K27 me3:H3K9 me3
H4K20 me2:H3−5ac
DNA Meth. m5C:H3K9 me3
H4K20 me2:H3K4 me3
H3K9 me3:H4K20 me3
DNA Meth. m5C:H4K20 me2
DNA Meth. m5C:H4K20 me3
H3K27 me3:DNA Meth. m5C
H4K20 me2:H4−4ac
H3K27 me2:DNA Meth. m5C
H3K27 me3:H4K20 me3
H3K9 me2:DNA Meth. m5C
H4−4ac:H3−5ac
H3K4 me3:H4−4ac
H3K9 me3
H3K27 me2
H3K4 me1
H3K9 me2
DNA Meth. m5C
H4K20 me3
H3−5ac
H4K20 me2
H2A.Z
H3K27 me3
H3K4 me3
H4−4ac
0.00 0.25 0.50 0.75 1.00
$\hat{\pi}$
TBX3 – Reverse
H2A.Z:H3K4 me3
H3K4 me1:H4K20 me2
H3K27 me3:H3K4 me1
H3K4 me3:H3−5ac
H2A.Z:H3−5ac
H2A.Z:H4−4ac
H2A.Z:H4K20 me2
H3K4 me1:H4−4ac
H3K4 me1:H2A.Z
H3K27 me3:H3K9 me3
H4K20 me2:H3−5ac
DNA Meth. m5C:H3K9 me3
H4K20 me2:H3K4 me3
H3K9 me3:H4K20 me3
DNA Meth. m5C:H4K20 me2
DNA Meth. m5C:H4K20 me3
H3K27 me3:DNA Meth. m5C
H4K20 me2:H4−4ac
H3K27 me2:DNA Meth. m5C
H3K27 me3:H4K20 me3
H3K9 me2:DNA Meth. m5C
H4−4ac:H3−5ac
H3K4 me3:H4−4ac
H3K9 me3
H3K27 me2
H3K4 me1
H3K9 me2
DNA Meth. m5C
H4K20 me3
H3−5ac
H4K20 me2
H2A.Z
H3K27 me3
H3K4 me3
H4−4ac
0.00 0.25 0.50 0.75 1.00
$\hat{\pi}$
MLXIP – Forward
H3K4 me1:H4−4ac
H3K27 me3:H4K20 me3
H3K27 me3:H3K9 me3
H3K4 me3:H3−5ac
H3K4 me1:H2A.Z
DNA Meth. m5C:H4K20 me3
H3K9 me2:DNA Meth. m5C
H3K9 me3:H4K20 me3
DNA Meth. m5C:H3K9 me3
H3K27 me3:H3K4 me1
H2A.Z:H3−5ac
H3K4 me1:H3−5ac
H3K4 me1:H4K20 me2
H3K27 me3:DNA Meth. m5C
H2A.Z:H4−4ac
H2A.Z:H3K4 me3
H4K20 me2:H3−5ac
H2A.Z:H4K20 me2
DNA Meth. m5C:H4K20 me2
H4K20 me2:H3K4 me3
H4−4ac:H3−5ac
H3K4 me3:H4−4ac
H4K20 me2:H4−4ac
H3K27 me2
H3K9 me2
H3K27 me3
H3K9 me3
H3K4 me1
H4K20 me3
DNA Meth. m5C
H3−5ac
H2A.Z
H3K4 me3
H4−4ac
H4K20 me2
0.00 0.25 0.50 0.75 1.00
$\hat{\pi}$
MLXIP – Reverse
H3K4 me1:H4−4ac
H3K27 me3:H4K20 me3
H3K27 me3:H3K9 me3
H3K4 me3:H3−5ac
H3K4 me1:H2A.Z
DNA Meth. m5C:H4K20 me3
H3K9 me2:DNA Meth. m5C
H3K9 me3:H4K20 me3
DNA Meth. m5C:H3K9 me3
H3K27 me3:H3K4 me1
H2A.Z:H3−5ac
H3K4 me1:H3−5ac
H3K4 me1:H4K20 me2
H3K27 me3:DNA Meth. m5C
H2A.Z:H4−4ac
H2A.Z:H3K4 me3
H4K20 me2:H3−5ac
H2A.Z:H4K20 me2
DNA Meth. m5C:H4K20 me2
H4K20 me2:H3K4 me3
H4−4ac:H3−5ac
H3K4 me3:H4−4ac
H4K20 me2:H4−4ac
H3K27 me2
H3K9 me2
H3K27 me3
H3K9 me3
H3K4 me1
H4K20 me3
DNA Meth. m5C
H3−5ac
H2A.Z
H3K4 me3
H4−4ac
H4K20 me2
0.00 0.25 0.50 0.75 1.00
$\hat{\pi}$
ZBTB44 – Forward
H3K4 me3:H3−5ac
H2A.Z:H4−4ac
H3K4 me1:H4−4ac
H3K4 me1:H2A.Z
H2A.Z:H3K4 me3
H3K9 me2:DNA Meth. m5C
H4−4ac:H3−5ac
H2A.Z:H4K20 me2
H3K27 me3:H4K20 me3
H4K20 me2:H3−5ac
H3K9 me3:H4K20 me3
H3K27 me3:H3K9 me3
H4K20 me2:H4−4ac
H4K20 me2:H3K4 me3
DNA Meth. m5C:H4K20 me3
H3K27 me3:DNA Meth. m5C
DNA Meth. m5C:H4K20 me2
H3K27 me2:DNA Meth. m5C
H3K4 me3:H4−4ac
DNA Meth. m5C:H3K9 me3
H3K4 me1
H3K27 me2
H3K27 me3
H3K9 me2
H4K20 me2
H3−5ac
H3K4 me3
H3K9 me3
H2A.Z
H4−4ac
DNA Meth. m5C
H4K20 me3
0.00 0.25 0.50 0.75 1.00
$\hat{\pi}$
ZBTB44 – Reverse
H3K4 me3:H3−5ac
H2A.Z:H4−4ac
H3K4 me1:H4−4ac
H3K4 me1:H2A.Z
H2A.Z:H3K4 me3
H3K9 me2:DNA Meth. m5C
H4−4ac:H3−5ac
H2A.Z:H4K20 me2
H3K27 me3:H4K20 me3
H4K20 me2:H3−5ac
H3K9 me3:H4K20 me3
H3K27 me3:H3K9 me3
H4K20 me2:H4−4ac
H4K20 me2:H3K4 me3
DNA Meth. m5C:H4K20 me3
H3K27 me3:DNA Meth. m5C
DNA Meth. m5C:H4K20 me2
H3K27 me2:DNA Meth. m5C
H3K4 me3:H4−4ac
DNA Meth. m5C:H3K9 me3
H3K4 me1
H3K27 me2
H3K27 me3
H3K9 me2
H4K20 me2
H3−5ac
H3K4 me3
H3K9 me3
H2A.Z
H4−4ac
DNA Meth. m5C
H4K20 me3
0.00 0.25 0.50 0.75 1.00
$\hat{\pi}$
THAP11 – Forward
H3K4 me1:H2A.Z
DNA Meth. m5C:H3K9 me3
H2A.Z:H4−4ac
H3K4 me1:H4−4ac
H2A.Z:H3−5ac
H2A.Z:H3K4 me3
H3K27 me3:DNA Meth. m5C
H3K4 me1:H4K20 me2
DNA Meth. m5C:H4K20 me2
H4K20 me2:H3−5ac
H2A.Z:H4K20 me2
H3K4 me3:H4−4ac
H3K9 me2:DNA Meth. m5C
H4−4ac:H3−5ac
H4K20 me3
H4K20 me2:H4−4ac
H4K20 me2:H3K4 me3
H3K27 me2
H3K27 me3
H3K9 me3
H3K9 me2
H3K4 me1
H3K4 me3
DNA Meth. m5C
H2A.Z
H4−4ac
H3−5ac
H4K20 me2
0.00 0.25 0.50 0.75 1.00
$\hat{\pi}$
THAP11 – Reverse
H3K4 me1:H2A.Z
DNA Meth. m5C:H3K9 me3
H2A.Z:H4−4ac
H3K4 me1:H4−4ac
H2A.Z:H3−5ac
H2A.Z:H3K4 me3
H3K27 me3:DNA Meth. m5C
H3K4 me1:H4K20 me2
DNA Meth. m5C:H4K20 me2
H4K20 me2:H3−5ac
H2A.Z:H4K20 me2
H3K4 me3:H4−4ac
H3K9 me2:DNA Meth. m5C
H4−4ac:H3−5ac
H4K20 me3
H4K20 me2:H4−4ac
H4K20 me2:H3K4 me3
H3K27 me2
H3K27 me3
H3K9 me3
H3K9 me2
H3K4 me1
H3K4 me3
DNA Meth. m5C
H2A.Z
H4−4ac
H3−5ac
H4K20 me2
0.00 0.25 0.50 0.75 1.00
$\hat{\pi}$

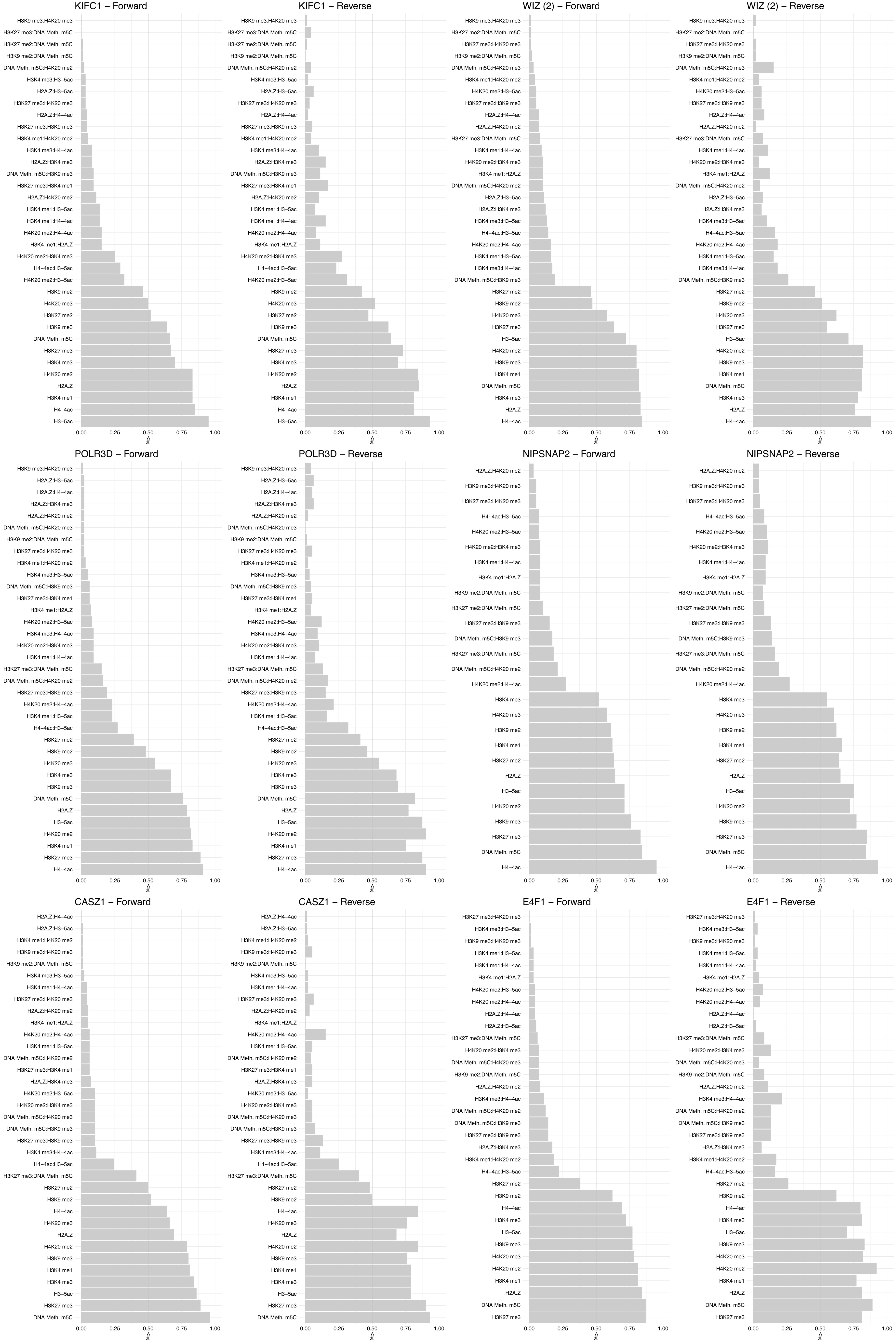

KIFC1 – Forward
KIFC1 – Reverse
WIZ (2) – Forward
WIZ (2) – Reverse
POLR3D – Forward
POLR3D – Reverse
NIPSNAP2 – Forward
NIPSNAP2 – Reverse
CASZ1 – Forward
CASZ1 – Reverse
E4F1 – Forward
E4F1 – Reverse
0.00 0.25 0.50 0.75 1.00
$\hat{\pi}$

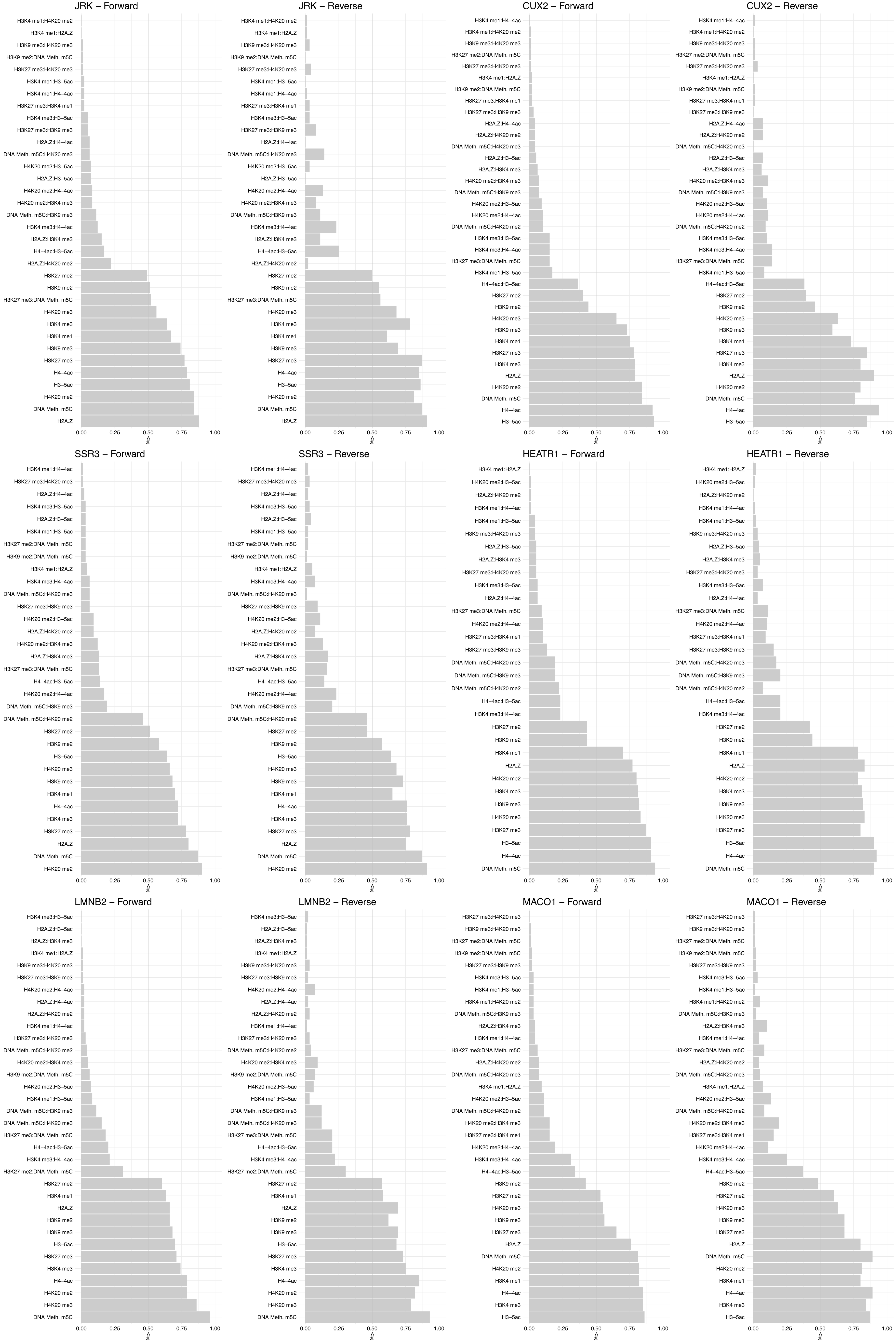
JRK – Forward
H3K4 me1:H4K20 me2
H3K4 me1:H2A.Z
H3K9 me3:H4K20 me3
H3K9 me2:DNA Meth. m5C
H3K27 me3:H4K20 me3
H3K4 me1:H3−5ac
H3K4 me1:H4−4ac
H3K27 me3:H3K4 me1
H3K4 me3:H3−5ac
H3K27 me3:H3K9 me3
H2A.Z:H4−4ac
DNA Meth. m5C:H4K20 me3
H4K20 me2:H3−5ac
H2A.Z:H3−5ac
H4K20 me2:H4−4ac
H4K20 me2:H3K4 me3
DNA Meth. m5C:H3K9 me3
H3K4 me3:H4−4ac
H2A.Z:H3K4 me3
H4−4ac:H3−5ac
H2A.Z:H4K20 me2
H3K27 me2
H3K9 me2
H3K27 me3:DNA Meth. m5C
H4K20 me3
H3K4 me3
H3K4 me1
H3K9 me3
H3K27 me3
H4−4ac
H3−5ac
H4K20 me2
DNA Meth. m5C
H2A.Z
0.00
0.25
0.50
0.75
1.00
$\hat{\pi}$
JRK – Reverse
H3K4 me1:H4K20 me2
H3K4 me1:H2A.Z
H3K9 me3:H4K20 me3
H3K9 me2:DNA Meth. m5C
H3K27 me3:H4K20 me3
H3K4 me1:H3−5ac
H3K4 me1:H4−4ac
H3K27 me3:H3K4 me1
H3K4 me3:H3−5ac
H3K27 me3:H3K9 me3
H2A.Z:H4−4ac
DNA Meth. m5C:H4K20 me3
H4K20 me2:H3−5ac
H2A.Z:H3−5ac
H4K20 me2:H4−4ac
H4K20 me2:H3K4 me3
DNA Meth. m5C:H3K9 me3
H3K4 me3:H4−4ac
H2A.Z:H3K4 me3
H4−4ac:H3−5ac
H2A.Z:H4K20 me2
H3K27 me2
H3K9 me2
H3K27 me3:DNA Meth. m5C
H4K20 me3
H3K4 me3
H3K4 me1
H3K9 me3
H3K27 me3
H4−4ac
H3−5ac
H4K20 me2
DNA Meth. m5C
H2A.Z
0.00
0.25
0.50
0.75
1.00
$\hat{\pi}$
CUX2 – Forward
H3K4 me1:H4−4ac
H3K4 me1:H4K20 me2
H3K9 me3:H4K20 me3
H3K27 me2:DNA Meth. m5C
H3K27 me3:H4K20 me3
H3K4 me1:H2A.Z
H3K9 me2:DNA Meth. m5C
H3K27 me3:H3K4 me1
H3K27 me3:H3K9 me3
H2A.Z:H4−4ac
H2A.Z:H4K20 me2
DNA Meth. m5C:H4K20 me3
H2A.Z:H3−5ac
H2A.Z:H3K4 me3
H4K20 me2:H3K4 me3
DNA Meth. m5C:H3K9 me3
H4K20 me2:H3−5ac
H4K20 me2:H4−4ac
DNA Meth. m5C:H4K20 me2
H3K4 me3:H3−5ac
H3K4 me3:H4−4ac
H3K27 me3:DNA Meth. m5C
H3K4 me1:H3−5ac
H4−4ac:H3−5ac
H3K27 me2
H3K9 me2
H4K20 me3
H3K9 me3
H3K4 me1
H3K27 me3
H3K4 me3
H2A.Z
H4K20 me2
DNA Meth. m5C
H4−4ac
H3−5ac
0.00
0.25
0.50
0.75
1.00
$\hat{\pi}$
CUX2 – Reverse
H3K4 me1:H4−4ac
H3K4 me1:H4K20 me2
H3K9 me3:H4K20 me3
H3K27 me2:DNA Meth. m5C
H3K27 me3:H4K20 me3
H3K4 me1:H2A.Z
H3K9 me2:DNA Meth. m5C
H3K27 me3:H3K4 me1
H3K27 me3:H3K9 me3
H2A.Z:H4−4ac
H2A.Z:H4K20 me2
DNA Meth. m5C:H4K20 me3
H2A.Z:H3−5ac
H2A.Z:H3K4 me3
H4K20 me2:H3K4 me3
DNA Meth. m5C:H3K9 me3
H4K20 me2:H3−5ac
H4K20 me2:H4−4ac
DNA Meth. m5C:H4K20 me2
H3K4 me3:H3−5ac
H3K4 me3:H4−4ac
H3K27 me3:DNA Meth. m5C
H3K4 me1:H3−5ac
H4−4ac:H3−5ac
H3K27 me2
H3K9 me2
H4K20 me3
H3K9 me3
H3K4 me1
H3K27 me3
H3K4 me3
H2A.Z
H4K20 me2
DNA Meth. m5C
H4−4ac
H3−5ac
0.00
0.25
0.50
0.75
1.00
$\hat{\pi}$
SSR3 – Forward
H3K4 me1:H4−4ac
H3K27 me3:H4K20 me3
H2A.Z:H4−4ac
H3K4 me3:H3−5ac
H2A.Z:H3−5ac
H3K4 me1:H3−5ac
H3K27 me2:DNA Meth. m5C
H3K9 me2:DNA Meth. m5C
H3K4 me1:H2A.Z
H3K4 me3:H4−4ac
DNA Meth. m5C:H4K20 me3
H3K27 me3:H3K9 me3
H4K20 me2:H3−5ac
H2A.Z:H4K20 me2
H4K20 me2:H3K4 me3
H2A.Z:H3K4 me3
H3K27 me3:DNA Meth. m5C
H4−4ac:H3−5ac
H4K20 me2:H4−4ac
DNA Meth. m5C:H3K9 me3
DNA Meth. m5C:H4K20 me2
H3K27 me2
H3K9 me2
H3−5ac
H4K20 me3
H3K9 me3
H3K4 me1
H4−4ac
H3K4 me3
H3K27 me3
H2A.Z
DNA Meth. m5C
H4K20 me2
0.00
0.25
0.50
0.75
1.00
$\hat{\pi}$
SSR3 – Reverse
H3K4 me1:H4−4ac
H3K27 me3:H4K20 me3
H2A.Z:H4−4ac
H3K4 me3:H3−5ac
H2A.Z:H3−5ac
H3K4 me1:H3−5ac
H3K27 me2:DNA Meth. m5C
H3K9 me2:DNA Meth. m5C
H3K4 me1:H2A.Z
H3K4 me3:H4−4ac
DNA Meth. m5C:H4K20 me3
H3K27 me3:H3K9 me3
H4K20 me2:H3−5ac
H2A.Z:H4K20 me2
H4K20 me2:H3K4 me3
H2A.Z:H3K4 me3
H3K27 me3:DNA Meth. m5C
H4−4ac:H3−5ac
H4K20 me2:H4−4ac
DNA Meth. m5C:H3K9 me3
DNA Meth. m5C:H4K20 me2
H3K27 me2
H3K9 me2
H3−5ac
H4K20 me3
H3K9 me3
H3K4 me1
H4−4ac
H3K4 me3
H3K27 me3
H2A.Z
DNA Meth. m5C
H4K20 me2
0.00
0.25
0.50
0.75
1.00
$\hat{\pi}$
HEATR1 – Forward
H3K4 me1:H2A.Z
H4K20 me2:H3−5ac
H2A.Z:H4K20 me2
H3K4 me1:H4−4ac
H3K4 me1:H3−5ac
H3K9 me3:H4K20 me3
H2A.Z:H3−5ac
H2A.Z:H3K4 me3
H3K27 me3:H4K20 me3
H3K4 me3:H3−5ac
H2A.Z:H4−4ac
H3K27 me3:DNA Meth. m5C
H4K20 me2:H4−4ac
H3K27 me3:H3K4 me1
H3K27 me3:H3K9 me3
DNA Meth. m5C:H4K20 me3
DNA Meth. m5C:H3K9 me3
DNA Meth. m5C:H4K20 me2
H4−4ac:H3−5ac
H3K4 me3:H4−4ac
H3K27 me2
H3K9 me2
H3K4 me1
H2A.Z
H4K20 me2
H3K4 me3
H3K9 me3
H4K20 me3
H3K27 me3
H3−5ac
H4−4ac
DNA Meth. m5C
0.00
0.25
0.50
0.75
1.00
$\hat{\pi}$
HEATR1 – Reverse
H3K4 me1:H2A.Z
H4K20 me2:H3−5ac
H2A.Z:H4K20 me2
H3K4 me1:H4−4ac
H3K4 me1:H3−5ac
H3K9 me3:H4K20 me3
H2A.Z:H3−5ac
H2A.Z:H3K4 me3
H3K27 me3:H4K20 me3
H3K4 me3:H3−5ac
H2A.Z:H4−4ac
H3K27 me3:DNA Meth. m5C
H4K20 me2:H4−4ac
H3K27 me3:H3K4 me1
H3K27 me3:H3K9 me3
DNA Meth. m5C:H4K20 me3
DNA Meth. m5C:H3K9 me3
DNA Meth. m5C:H4K20 me2
H4−4ac:H3−5ac
H3K4 me3:H4−4ac
H3K27 me2
H3K9 me2
H3K4 me1
H2A.Z
H4K20 me2
H3K4 me3
H3K9 me3
H4K20 me3
H3K27 me3
H3−5ac
H4−4ac
DNA Meth. m5C
0.00
0.25
0.50
0.75
1.00
$\hat{\pi}$
LMNB2 – Forward
H3K4 me3:H3−5ac
H2A.Z:H3−5ac
H2A.Z:H3K4 me3
H3K4 me1:H2A.Z
H3K9 me3:H4K20 me3
H3K27 me3:H3K9 me3
H4K20 me2:H4−4ac
H2A.Z:H4−4ac
H2A.Z:H4K20 me2
H3K4 me1:H4−4ac
H3K27 me3:H4K20 me3
DNA Meth. m5C:H4K20 me2
H4K20 me2:H3K4 me3
H3K9 me2:DNA Meth. m5C
H4K20 me2:H3−5ac
H3K4 me1:H3−5ac
DNA Meth. m5C:H3K9 me3
DNA Meth. m5C:H4K20 me3
H3K27 me3:DNA Meth. m5C
H4−4ac:H3−5ac
H3K4 me3:H4−4ac
H3K27 me2:DNA Meth. m5C
H3K27 me2
H3K4 me1
H2A.Z
H3K9 me2
H3K9 me3
H3−5ac
H3K27 me3
H3K4 me3
H4−4ac
H4K20 me2
H4K20 me3
DNA Meth. m5C
0.00
0.25
0.50
0.75
1.00
$\hat{\pi}$
LMNB2 – Reverse
H3K4 me3:H3−5ac
H2A.Z:H3−5ac
H2A.Z:H3K4 me3
H3K4 me1:H2A.Z
H3K9 me3:H4K20 me3
H3K27 me3:H3K9 me3
H4K20 me2:H4−4ac
H2A.Z:H4−4ac
H2A.Z:H4K20 me2
H3K4 me1:H4−4ac
H3K27 me3:H4K20 me3
DNA Meth. m5C:H4K20 me2
H4K20 me2:H3K4 me3
H3K9 me2:DNA Meth. m5C
H4K20 me2:H3−5ac
H3K4 me1:H3−5ac
DNA Meth. m5C:H3K9 me3
DNA Meth. m5C:H4K20 me3
H3K27 me3:DNA Meth. m5C
H4−4ac:H3−5ac
H3K4 me3:H4−4ac
H3K27 me2:DNA Meth. m5C
H3K27 me2
H3K4 me1
H2A.Z
H3K9 me2
H3K9 me3
H3−5ac
H3K27 me3
H3K4 me3
H4−4ac
H4K20 me2
H4K20 me3
DNA Meth. m5C
0.00
0.25
0.50
0.75
1.00
$\hat{\pi}$
MACO1 – Forward
H3K27 me3:H4K20 me3
H3K9 me3:H4K20 me3
H3K27 me2:DNA Meth. m5C
H3K9 me2:DNA Meth. m5C
H3K27 me3:H3K9 me3
H3K4 me3:H3−5ac
H3K4 me1:H3−5ac
H3K4 me1:H4K20 me2
DNA Meth. m5C:H3K9 me3
H2A.Z:H3K4 me3
H3K4 me1:H4−4ac
H3K27 me3:DNA Meth. m5C
H2A.Z:H4K20 me2
DNA Meth. m5C:H4K20 me3
H3K4 me1:H2A.Z
H4K20 me2:H3−5ac
DNA Meth. m5C:H4K20 me2
H4K20 me2:H3K4 me3
H3K27 me3:H3K4 me1
H4K20 me2:H4−4ac
H3K4 me3:H4−4ac
H4−4ac:H3−5ac
H3K9 me2
H3K27 me2
H4K20 me3
H3K9 me3
H3K27 me3
H2A.Z
DNA Meth. m5C
H4K20 me2
H3K4 me1
H4−4ac
H3K4 me3
H3−5ac
0.00
0.25
0.50
0.75
1.00
$\hat{\pi}$
MACO1 – Reverse
H3K27 me3:H4K20 me3
H3K9 me3:H4K20 me3
H3K27 me2:DNA Meth. m5C
H3K9 me2:DNA Meth. m5C
H3K27 me3:H3K9 me3
H3K4 me3:H3−5ac
H3K4 me1:H3−5ac
H3K4 me1:H4K20 me2
DNA Meth. m5C:H3K9 me3
H2A.Z:H3K4 me3
H3K4 me1:H4−4ac
H3K27 me3:DNA Meth. m5C
H2A.Z:H4K20 me2
DNA Meth. m5C:H4K20 me3
H3K4 me1:H2A.Z
H4K20 me2:H3−5ac
DNA Meth. m5C:H4K20 me2
H4K20 me2:H3K4 me3
H3K27 me3:H3K4 me1
H4K20 me2:H4−4ac
H3K4 me3:H4−4ac
H4−4ac:H3−5ac
H3K9 me2
H3K27 me2
H4K20 me3
H3K9 me3
H3K27 me3
H2A.Z
DNA Meth. m5C
H4K20 me2
H3K4 me1
H4−4ac
H3K4 me3
H3−5ac
0.00
0.25
0.50
0.75
1.00
$\hat{\pi}$

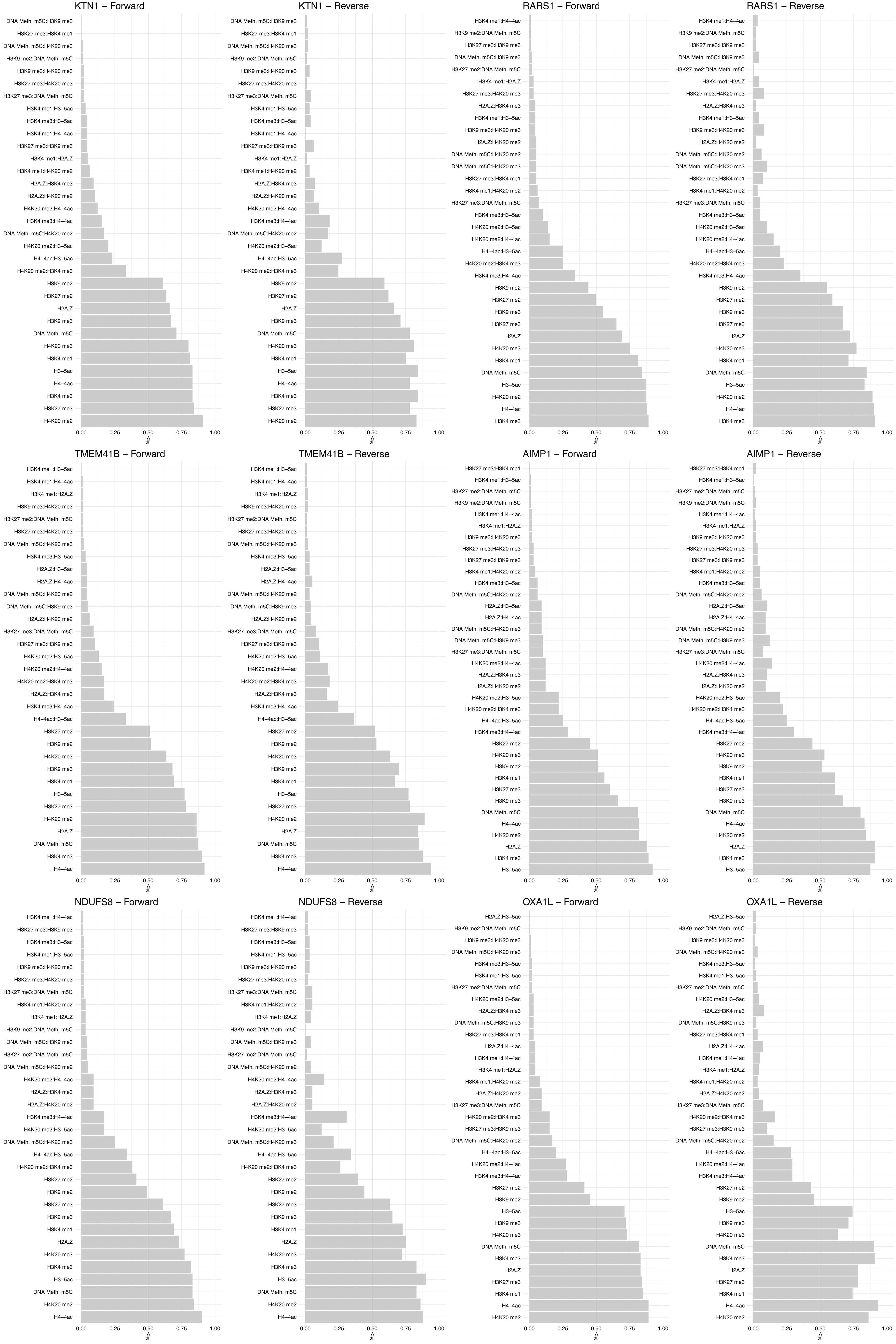
KTN1 – Forward
DNA Meth. m5C:H3K9 me3
H3K27 me3:H3K4 me1
DNA Meth. m5C:H4K20 me3
H3K9 me2:DNA Meth. m5C
H3K9 me3:H4K20 me3
H3K27 me3:H4K20 me3
H3K27 me3:DNA Meth. m5C
H3K4 me1:H3−5ac
H3K4 me3:H3−5ac
H3K4 me1:H4−4ac
H3K27 me3:H3K9 me3
H3K4 me1:H2A.Z
H3K4 me1:H4K20 me2
H2A.Z:H3K4 me3
H2A.Z:H4K20 me2
H4K20 me2:H4−4ac
H3K4 me3:H4−4ac
DNA Meth. m5C:H4K20 me2
H4K20 me2:H3−5ac
H4−4ac:H3−5ac
H4K20 me2:H3K4 me3
H3K9 me2
H3K27 me2
H2A.Z
H3K9 me3
DNA Meth. m5C
H4K20 me3
H3K4 me1
H3−5ac
H4−4ac
H3K4 me3
H3K27 me3
H4K20 me2
0.00
0.25
0.50
0.75
1.00
$\hat{\pi}$
KTN1 – Reverse
DNA Meth. m5C:H3K9 me3
H3K27 me3:H3K4 me1
DNA Meth. m5C:H4K20 me3
H3K9 me2:DNA Meth. m5C
H3K9 me3:H4K20 me3
H3K27 me3:H4K20 me3
H3K27 me3:DNA Meth. m5C
H3K4 me1:H3−5ac
H3K4 me3:H3−5ac
H3K4 me1:H4−4ac
H3K27 me3:H3K9 me3
H3K4 me1:H2A.Z
H3K4 me1:H4K20 me2
H2A.Z:H3K4 me3
H2A.Z:H4K20 me2
H4K20 me2:H4−4ac
H3K4 me3:H4−4ac
DNA Meth. m5C:H4K20 me2
H4K20 me2:H3−5ac
H4−4ac:H3−5ac
H4K20 me2:H3K4 me3
H3K9 me2
H3K27 me2
H2A.Z
H3K9 me3
DNA Meth. m5C
H4K20 me3
H3K4 me1
H3−5ac
H4−4ac
H3K4 me3
H3K27 me3
H4K20 me2
0.00
0.25
0.50
0.75
1.00
$\hat{\pi}$
RARS1 – Forward
H3K4 me1:H4−4ac
H3K9 me2:DNA Meth. m5C
H3K27 me3:H3K9 me3
DNA Meth. m5C:H3K9 me3
H3K27 me2:DNA Meth. m5C
H3K4 me1:H2A.Z
H3K27 me3:H4K20 me3
H2A.Z:H3K4 me3
H3K4 me1:H3−5ac
H3K9 me3:H4K20 me3
H2A.Z:H4K20 me2
DNA Meth. m5C:H4K20 me2
DNA Meth. m5C:H4K20 me3
H3K27 me3:H3K4 me1
H3K4 me1:H4K20 me2
H3K27 me3:DNA Meth. m5C
H3K4 me3:H3−5ac
H4K20 me2:H3−5ac
H4K20 me2:H4−4ac
H4−4ac:H3−5ac
H4K20 me2:H3K4 me3
H3K4 me3:H4−4ac
H3K9 me2
H3K27 me2
H3K9 me3
H3K27 me3
H2A.Z
H4K20 me3
H3K4 me1
DNA Meth. m5C
H3−5ac
H4K20 me2
H4−4ac
H3K4 me3
0.00
0.25
0.50
0.75
1.00
$\hat{\pi}$
RARS1 – Reverse
H3K4 me1:H4−4ac
H3K9 me2:DNA Meth. m5C
H3K27 me3:H3K9 me3
DNA Meth. m5C:H3K9 me3
H3K27 me2:DNA Meth. m5C
H3K4 me1:H2A.Z
H3K27 me3:H4K20 me3
H2A.Z:H3K4 me3
H3K4 me1:H3−5ac
H3K9 me3:H4K20 me3
H2A.Z:H4K20 me2
DNA Meth. m5C:H4K20 me2
DNA Meth. m5C:H4K20 me3
H3K27 me3:H3K4 me1
H3K4 me1:H4K20 me2
H3K27 me3:DNA Meth. m5C
H3K4 me3:H3−5ac
H4K20 me2:H3−5ac
H4K20 me2:H4−4ac
H4−4ac:H3−5ac
H4K20 me2:H3K4 me3
H3K4 me3:H4−4ac
H3K9 me2
H3K27 me2
H3K9 me3
H3K27 me3
H2A.Z
H4K20 me3
H3K4 me1
DNA Meth. m5C
H3−5ac
H4K20 me2
H4−4ac
H3K4 me3
0.00
0.25
0.50
0.75
1.00
$\hat{\pi}$
TMEM41B – Forward
H3K4 me1:H3−5ac
H3K4 me1:H4−4ac
H3K4 me1:H2A.Z
H3K9 me3:H4K20 me3
H3K27 me2:DNA Meth. m5C
H3K27 me3:H4K20 me3
DNA Meth. m5C:H4K20 me3
H3K4 me3:H3−5ac
H2A.Z:H3−5ac
H2A.Z:H4−4ac
DNA Meth. m5C:H4K20 me2
DNA Meth. m5C:H3K9 me3
H2A.Z:H4K20 me2
H3K27 me3:DNA Meth. m5C
H3K27 me3:H3K9 me3
H4K20 me2:H3−5ac
H4K20 me2:H4−4ac
H4K20 me2:H3K4 me3
H2A.Z:H3K4 me3
H3K4 me3:H4−4ac
H4−4ac:H3−5ac
H3K27 me2
H3K9 me2
H4K20 me3
H3K9 me3
H3K4 me1
H3−5ac
H3K27 me3
H4K20 me2
H2A.Z
DNA Meth. m5C
H3K4 me3
H4−4ac
0.00
0.25
0.50
0.75
1.00
$\hat{\pi}$
TMEM41B – Reverse
H3K4 me1:H3−5ac
H3K4 me1:H4−4ac
H3K4 me1:H2A.Z
H3K9 me3:H4K20 me3
H3K27 me2:DNA Meth. m5C
H3K27 me3:H4K20 me3
DNA Meth. m5C:H4K20 me3
H3K4 me3:H3−5ac
H2A.Z:H3−5ac
H2A.Z:H4−4ac
DNA Meth. m5C:H4K20 me2
DNA Meth. m5C:H3K9 me3
H2A.Z:H4K20 me2
H3K27 me3:DNA Meth. m5C
H3K27 me3:H3K9 me3
H4K20 me2:H3−5ac
H4K20 me2:H4−4ac
H4K20 me2:H3K4 me3
H2A.Z:H3K4 me3
H3K4 me3:H4−4ac
H4−4ac:H3−5ac
H3K27 me2
H3K9 me2
H4K20 me3
H3K9 me3
H3K4 me1
H3−5ac
H3K27 me3
H4K20 me2
H2A.Z
DNA Meth. m5C
H3K4 me3
H4−4ac
0.00
0.25
0.50
0.75
1.00
$\hat{\pi}$
AIMP1 – Forward
H3K27 me3:H3K4 me1
H3K4 me1:H3−5ac
H3K27 me2:DNA Meth. m5C
H3K9 me2:DNA Meth. m5C
H3K4 me1:H4−4ac
H3K4 me1:H2A.Z
H3K9 me3:H4K20 me3
H3K27 me3:H4K20 me3
H3K27 me3:H3K9 me3
H3K4 me1:H4K20 me2
H3K4 me3:H3−5ac
DNA Meth. m5C:H4K20 me2
H2A.Z:H3−5ac
H2A.Z:H4−4ac
DNA Meth. m5C:H4K20 me3
DNA Meth. m5C:H3K9 me3
H3K27 me3:DNA Meth. m5C
H4K20 me2:H4−4ac
H2A.Z:H3K4 me3
H2A.Z:H4K20 me2
H4K20 me2:H3−5ac
H4K20 me2:H3K4 me3
H4−4ac:H3−5ac
H3K4 me3:H4−4ac
H3K27 me2
H4K20 me3
H3K9 me2
H3K4 me1
H3K27 me3
H3K9 me3
DNA Meth. m5C
H4−4ac
H4K20 me2
H2A.Z
H3K4 me3
H3−5ac
0.00
0.25
0.50
0.75
1.00
$\hat{\pi}$
AIMP1 – Reverse
H3K27 me3:H3K4 me1
H3K4 me1:H3−5ac
H3K27 me2:DNA Meth. m5C
H3K9 me2:DNA Meth. m5C
H3K4 me1:H4−4ac
H3K4 me1:H2A.Z
H3K9 me3:H4K20 me3
H3K27 me3:H4K20 me3
H3K27 me3:H3K9 me3
H3K4 me1:H4K20 me2
H3K4 me3:H3−5ac
DNA Meth. m5C:H4K20 me2
H2A.Z:H3−5ac
H2A.Z:H4−4ac
DNA Meth. m5C:H4K20 me3
DNA Meth. m5C:H3K9 me3
H3K27 me3:DNA Meth. m5C
H4K20 me2:H4−4ac
H2A.Z:H3K4 me3
H2A.Z:H4K20 me2
H4K20 me2:H3−5ac
H4K20 me2:H3K4 me3
H4−4ac:H3−5ac
H3K4 me3:H4−4ac
H3K27 me2
H4K20 me3
H3K9 me2
H3K4 me1
H3K27 me3
H3K9 me3
DNA Meth. m5C
H4−4ac
H4K20 me2
H2A.Z
H3K4 me3
H3−5ac
0.00
0.25
0.50
0.75
1.00
$\hat{\pi}$
NDUFS8 – Forward
H3K4 me1:H4−4ac
H3K27 me3:H3K9 me3
H3K4 me3:H3−5ac
H3K4 me1:H3−5ac
H3K9 me3:H4K20 me3
H3K27 me3:H4K20 me3
H3K27 me3:DNA Meth. m5C
H3K4 me1:H4K20 me2
H3K4 me1:H2A.Z
H3K9 me2:DNA Meth. m5C
DNA Meth. m5C:H3K9 me3
H3K27 me2:DNA Meth. m5C
DNA Meth. m5C:H4K20 me2
H4K20 me2:H4−4ac
H2A.Z:H3K4 me3
H2A.Z:H4K20 me2
H3K4 me3:H4−4ac
H4K20 me2:H3−5ac
DNA Meth. m5C:H4K20 me3
H4−4ac:H3−5ac
H4K20 me2:H3K4 me3
H3K27 me2
H3K9 me2
H3K27 me3
H3K9 me3
H3K4 me1
H2A.Z
H4K20 me3
H3K4 me3
H3−5ac
DNA Meth. m5C
H4K20 me2
H4−4ac
0.00
0.25
0.50
0.75
1.00
$\hat{\pi}$
NDUFS8 – Reverse
H3K4 me1:H4−4ac
H3K27 me3:H3K9 me3
H3K4 me3:H3−5ac
H3K4 me1:H3−5ac
H3K9 me3:H4K20 me3
H3K27 me3:H4K20 me3
H3K27 me3:DNA Meth. m5C
H3K4 me1:H4K20 me2
H3K4 me1:H2A.Z
H3K9 me2:DNA Meth. m5C
DNA Meth. m5C:H3K9 me3
H3K27 me2:DNA Meth. m5C
DNA Meth. m5C:H4K20 me2
H4K20 me2:H4−4ac
H2A.Z:H3K4 me3
H2A.Z:H4K20 me2
H3K4 me3:H4−4ac
H4K20 me2:H3−5ac
DNA Meth. m5C:H4K20 me3
H4−4ac:H3−5ac
H4K20 me2:H3K4 me3
H3K27 me2
H3K9 me2
H3K27 me3
H3K9 me3
H3K4 me1
H2A.Z
H4K20 me3
H3K4 me3
H3−5ac
DNA Meth. m5C
H4K20 me2
H4−4ac
0.00
0.25
0.50
0.75
1.00
$\hat{\pi}$
OXA1L – Forward
H2A.Z:H3−5ac
H3K9 me2:DNA Meth. m5C
H3K9 me3:H4K20 me3
DNA Meth. m5C:H4K20 me3
H3K4 me3:H3−5ac
H3K4 me1:H3−5ac
H3K27 me2:DNA Meth. m5C
H4K20 me2:H3−5ac
H2A.Z:H3K4 me3
DNA Meth. m5C:H3K9 me3
H3K27 me3:H3K4 me1
H2A.Z:H4−4ac
H3K4 me1:H4−4ac
H3K4 me1:H2A.Z
H3K4 me1:H4K20 me2
H2A.Z:H4K20 me2
H3K27 me3:DNA Meth. m5C
H4K20 me2:H3K4 me3
H3K27 me3:H3K9 me3
DNA Meth. m5C:H4K20 me2
H4−4ac:H3−5ac
H4K20 me2:H4−4ac
H3K4 me3:H4−4ac
H3K27 me2
H3K9 me2
H3−5ac
H3K9 me3
H4K20 me3
DNA Meth. m5C
H3K4 me3
H2A.Z
H3K27 me3
H3K4 me1
H4−4ac
H4K20 me2
0.00
0.25
0.50
0.75
1.00
$\hat{\pi}$
OXA1L – Reverse
H2A.Z:H3−5ac
H3K9 me2:DNA Meth. m5C
H3K9 me3:H4K20 me3
DNA Meth. m5C:H4K20 me3
H3K4 me3:H3−5ac
H3K4 me1:H3−5ac
H3K27 me2:DNA Meth. m5C
H4K20 me2:H3−5ac
H2A.Z:H3K4 me3
DNA Meth. m5C:H3K9 me3
H3K27 me3:H3K4 me1
H2A.Z:H4−4ac
H3K4 me1:H4−4ac
H3K4 me1:H2A.Z
H3K4 me1:H4K20 me2
H2A.Z:H4K20 me2
H3K27 me3:DNA Meth. m5C
H4K20 me2:H3K4 me3
H3K27 me3:H3K9 me3
DNA Meth. m5C:H4K20 me2
H4−4ac:H3−5ac
H4K20 me2:H4−4ac
H3K4 me3:H4−4ac
H3K27 me2
H3K9 me2
H3−5ac
H3K9 me3
H4K20 me3
DNA Meth. m5C
H3K4 me3
H2A.Z
H3K27 me3
H3K4 me1
H4−4ac
H4K20 me2
0.00
0.25
0.50
0.75
1.00
$\hat{\pi}$

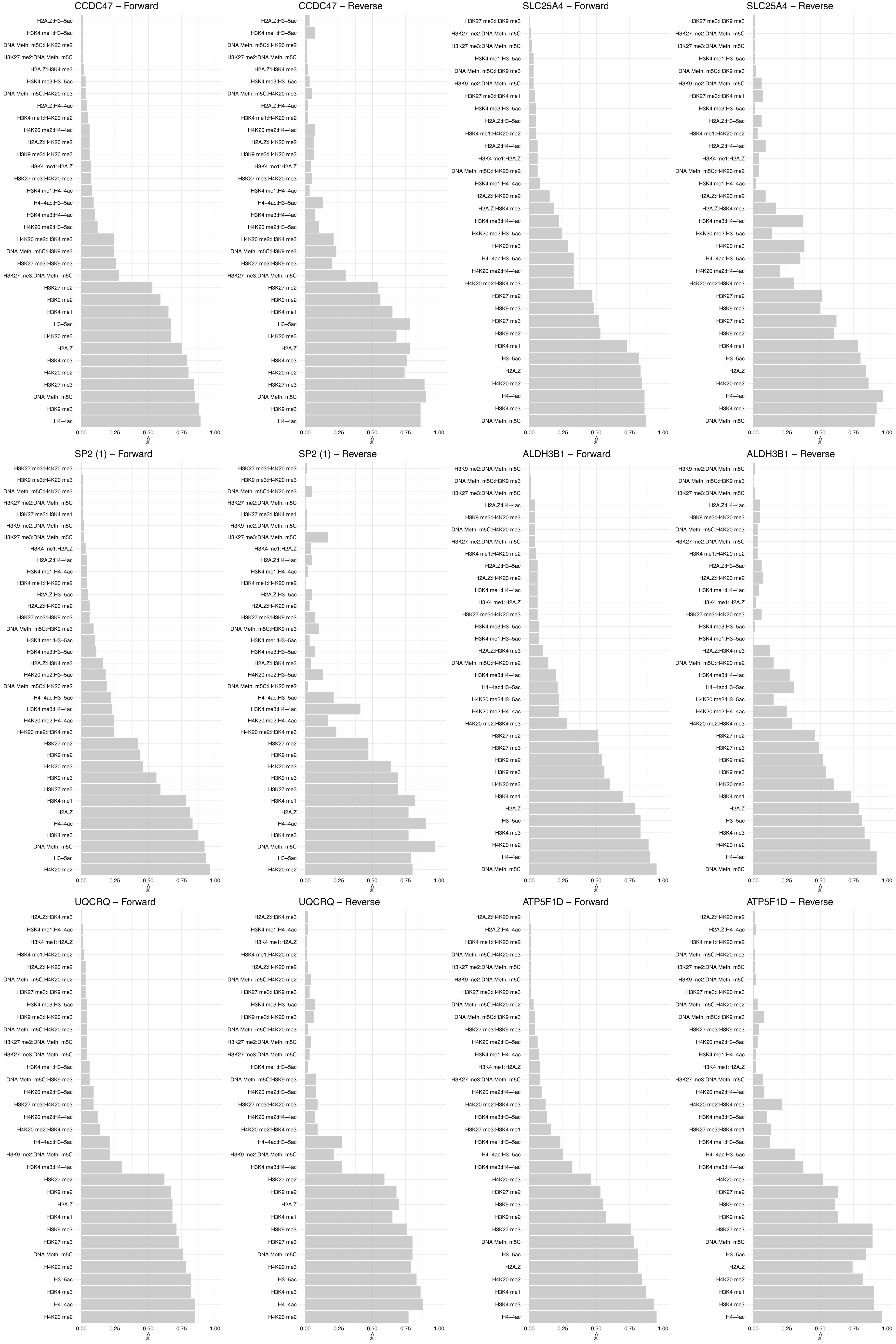

CCDC47 – Forward
H2A.Z:H3–5ac
H3K4 me1:H3–5ac
DNA Meth. m5C:H4K20 me2
H3K27 me2:DNA Meth. m5C
H2A.Z:H3K4 me3
H3K4 me3:H3–5ac
DNA Meth. m5C:H4K20 me3
H2A.Z:H4–4ac
H3K4 me1:H4K20 me2
H4K20 me2:H4–4ac
H2A.Z:H4K20 me2
H3K9 me3:H4K20 me3
H3K4 me1:H2A.Z
H3K27 me3:H4K20 me3
H3K4 me1:H4–4ac
H4–4ac:H3–5ac
H3K4 me3:H4–4ac
H4K20 me2:H3–5ac
H4K20 me2:H3K4 me3
DNA Meth. m5C:H3K9 me3
H3K27 me3:H3K9 me3
H3K27 me3:DNA Meth. m5C
H3K27 me2
H3K9 me2
H3K4 me1
H3–5ac
H4K20 me3
H2A.Z
H3K4 me3
H4K20 me2
H3K27 me3
DNA Meth. m5C
H3K9 me3
H4–4ac
0.00 0.25 0.50 0.75 1.00
π̂
CCDC47 – Reverse
H2A.Z:H3–5ac
H3K4 me1:H3–5ac
DNA Meth. m5C:H4K20 me2
H3K27 me2:DNA Meth. m5C
H2A.Z:H3K4 me3
H3K4 me3:H3–5ac
DNA Meth. m5C:H4K20 me3
H2A.Z:H4–4ac
H3K4 me1:H4K20 me2
H4K20 me2:H4–4ac
H2A.Z:H4K20 me2
H3K9 me3:H4K20 me3
H3K4 me1:H2A.Z
H3K27 me3:H4K20 me3
H3K4 me1:H4–4ac
H4–4ac:H3–5ac
H3K4 me3:H4–4ac
H4K20 me2:H3–5ac
H4K20 me2:H3K4 me3
DNA Meth. m5C:H3K9 me3
H3K27 me3:H3K9 me3
H3K27 me3:DNA Meth. m5C
H3K27 me2
H3K9 me2
H3K4 me1
H3–5ac
H4K20 me3
H2A.Z
H3K4 me3
H4K20 me2
H3K27 me3
DNA Meth. m5C
H3K9 me3
H4–4ac
0.00 0.25 0.50 0.75 1.00
π̂
SLC25A4 – Forward
H3K27 me3:H3K9 me3
H3K27 me2:DNA Meth. m5C
H3K27 me3:DNA Meth. m5C
H3K4 me1:H3–5ac
DNA Meth. m5C:H3K9 me3
H3K9 me2:DNA Meth. m5C
H3K27 me3:H3K4 me1
H3K4 me3:H3–5ac
H2A.Z:H3–5ac
H3K4 me1:H4K20 me2
H2A.Z:H4–4ac
H3K4 me1:H2A.Z
DNA Meth. m5C:H4K20 me2
H3K4 me1:H4–4ac
H2A.Z:H4K20 me2
H2A.Z:H3K4 me3
H3K4 me3:H4–4ac
H4K20 me2:H3–5ac
H4K20 me3
H4–4ac:H3–5ac
H4K20 me2:H4–4ac
H4K20 me2:H3K4 me3
H3K27 me2
H3K9 me3
H3K27 me3
H3K9 me2
H3K4 me1
H3–5ac
H2A.Z
H4K20 me2
H4–4ac
H3K4 me3
DNA Meth. m5C
0.00 0.25 0.50 0.75 1.00
π̂
SLC25A4 – Reverse
H3K27 me3:H3K9 me3
H3K27 me2:DNA Meth. m5C
H3K27 me3:DNA Meth. m5C
H3K4 me1:H3–5ac
DNA Meth. m5C:H3K9 me3
H3K9 me2:DNA Meth. m5C
H3K27 me3:H3K4 me1
H3K4 me3:H3–5ac
H2A.Z:H3–5ac
H3K4 me1:H4K20 me2
H2A.Z:H4–4ac
H3K4 me1:H2A.Z
DNA Meth. m5C:H4K20 me2
H3K4 me1:H4–4ac
H2A.Z:H4K20 me2
H2A.Z:H3K4 me3
H3K4 me3:H4–4ac
H4K20 me2:H3–5ac
H4K20 me3
H4–4ac:H3–5ac
H4K20 me2:H4–4ac
H4K20 me2:H3K4 me3
H3K27 me2
H3K9 me3
H3K27 me3
H3K9 me2
H3K4 me1
H3–5ac
H2A.Z
H4K20 me2
H4–4ac
H3K4 me3
DNA Meth. m5C
0.00 0.25 0.50 0.75 1.00
π̂
SP2 (1) – Forward
H3K27 me3:H4K20 me3
H3K9 me3:H4K20 me3
DNA Meth. m5C:H4K20 me3
H3K27 me2:DNA Meth. m5C
H3K27 me3:H3K4 me1
H3K9 me2:DNA Meth. m5C
H3K27 me3:DNA Meth. m5C
H3K4 me1:H2A.Z
H2A.Z:H4–4ac
H3K4 me1:H4–4ac
H3K4 me1:H4K20 me2
H2A.Z:H3–5ac
H2A.Z:H4K20 me2
H3K27 me3:H3K9 me3
DNA Meth. m5C:H3K9 me3
H3K4 me1:H3–5ac
H3K4 me3:H3–5ac
H2A.Z:H3K4 me3
H4K20 me2:H3–5ac
DNA Meth. m5C:H4K20 me2
H4–4ac:H3–5ac
H3K4 me3:H4–4ac
H4K20 me2:H4–4ac
H4K20 me2:H3K4 me3
H3K27 me2
H3K9 me2
H4K20 me3
H3K9 me3
H3K27 me3
H3K4 me1
H2A.Z
H4–4ac
H3K4 me3
DNA Meth. m5C
H3–5ac
H4K20 me2
0.00 0.25 0.50 0.75 1.00
π̂
SP2 (1) – Reverse
H3K27 me3:H4K20 me3
H3K9 me3:H4K20 me3
DNA Meth. m5C:H4K20 me3
H3K27 me2:DNA Meth. m5C
H3K27 me3:H3K4 me1
H3K9 me2:DNA Meth. m5C
H3K27 me3:DNA Meth. m5C
H3K4 me1:H2A.Z
H2A.Z:H4–4ac
H3K4 me1:H4–4ac
H3K4 me1:H4K20 me2
H2A.Z:H3–5ac
H2A.Z:H4K20 me2
H3K27 me3:H3K9 me3
DNA Meth. m5C:H3K9 me3
H3K4 me1:H3–5ac
H3K4 me3:H3–5ac
H2A.Z:H3K4 me3
H4K20 me2:H3–5ac
DNA Meth. m5C:H4K20 me2
H4–4ac:H3–5ac
H3K4 me3:H4–4ac
H4K20 me2:H4–4ac
H4K20 me2:H3K4 me3
H3K27 me2
H3K9 me2
H4K20 me3
H3K9 me3
H3K27 me3
H3K4 me1
H2A.Z
H4–4ac
H3K4 me3
DNA Meth. m5C
H3–5ac
H4K20 me2
0.00 0.25 0.50 0.75 1.00
π̂
ALDH3B1 – Forward
H3K9 me2:DNA Meth. m5C
DNA Meth. m5C:H3K9 me3
H3K27 me3:DNA Meth. m5C
H2A.Z:H4–4ac
H3K9 me3:H4K20 me3
DNA Meth. m5C:H4K20 me3
H3K27 me2:DNA Meth. m5C
H3K4 me1:H4K20 me2
H2A.Z:H3–5ac
H2A.Z:H4K20 me2
H3K4 me1:H4–4ac
H3K4 me1:H2A.Z
H3K27 me3:H4K20 me3
H3K4 me3:H3–5ac
H3K4 me1:H3–5ac
H2A.Z:H3K4 me3
DNA Meth. m5C:H4K20 me2
H3K4 me3:H4–4ac
H4–4ac:H3–5ac
H4K20 me2:H3–5ac
H4K20 me2:H4–4ac
H4K20 me2:H3K4 me3
H3K27 me2
H3K27 me3
H3K9 me2
H3K9 me3
H4K20 me3
H3K4 me1
H2A.Z
H3–5ac
H3K4 me3
H4K20 me2
H4–4ac
DNA Meth. m5C
0.00 0.25 0.50 0.75 1.00
π̂
ALDH3B1 – Reverse
H3K9 me2:DNA Meth. m5C
DNA Meth. m5C:H3K9 me3
H3K27 me3:DNA Meth. m5C
H2A.Z:H4–4ac
H3K9 me3:H4K20 me3
DNA Meth. m5C:H4K20 me3
H3K27 me2:DNA Meth. m5C
H3K4 me1:H4K20 me2
H2A.Z:H3–5ac
H2A.Z:H4K20 me2
H3K4 me1:H4–4ac
H3K4 me1:H2A.Z
H3K27 me3:H4K20 me3
H3K4 me3:H3–5ac
H3K4 me1:H3–5ac
H2A.Z:H3K4 me3
DNA Meth. m5C:H4K20 me2
H3K4 me3:H4–4ac
H4–4ac:H3–5ac
H4K20 me2:H3–5ac
H4K20 me2:H4–4ac
H4K20 me2:H3K4 me3
H3K27 me2
H3K27 me3
H3K9 me2
H3K9 me3
H4K20 me3
H3K4 me1
H2A.Z
H3–5ac
H3K4 me3
H4K20 me2
H4–4ac
DNA Meth. m5C
0.00 0.25 0.50 0.75 1.00
π̂
UQCRQ – Forward
H2A.Z:H3K4 me3
H3K4 me1:H4–4ac
H3K4 me1:H2A.Z
H3K4 me1:H4K20 me2
H2A.Z:H4K20 me2
DNA Meth. m5C:H4K20 me2
H3K27 me3:H3K9 me3
H3K4 me3:H3–5ac
H3K9 me3:H4K20 me3
DNA Meth. m5C:H4K20 me3
H3K27 me2:DNA Meth. m5C
H3K27 me3:DNA Meth. m5C
H3K4 me1:H3–5ac
DNA Meth. m5C:H3K9 me3
H4K20 me2:H3–5ac
H3K27 me3:H4K20 me3
H4K20 me2:H4–4ac
H4K20 me2:H3K4 me3
H4–4ac:H3–5ac
H3K9 me2:DNA Meth. m5C
H3K4 me3:H4–4ac
H3K27 me2
H3K9 me2
H2A.Z
H3K4 me1
H3K9 me3
H3K27 me3
DNA Meth. m5C
H4K20 me3
H3–5ac
H3K4 me3
H4–4ac
H4K20 me2
0.00 0.25 0.50 0.75 1.00
π̂
UQCRQ – Reverse
H2A.Z:H3K4 me3
H3K4 me1:H4–4ac
H3K4 me1:H2A.Z
H3K4 me1:H4K20 me2
H2A.Z:H4K20 me2
DNA Meth. m5C:H4K20 me2
H3K27 me3:H3K9 me3
H3K4 me3:H3–5ac
H3K9 me3:H4K20 me3
DNA Meth. m5C:H4K20 me3
H3K27 me2:DNA Meth. m5C
H3K27 me3:DNA Meth. m5C
H3K4 me1:H3–5ac
DNA Meth. m5C:H3K9 me3
H4K20 me2:H3–5ac
H3K27 me3:H4K20 me3
H4K20 me2:H4–4ac
H4K20 me2:H3K4 me3
H4–4ac:H3–5ac
H3K9 me2:DNA Meth. m5C
H3K4 me3:H4–4ac
H3K27 me2
H3K9 me2
H2A.Z
H3K4 me1
H3K9 me3
H3K27 me3
DNA Meth. m5C
H4K20 me3
H3–5ac
H3K4 me3
H4–4ac
H4K20 me2
0.00 0.25 0.50 0.75 1.00
π̂
ATP5F1D – Forward
H2A.Z:H4K20 me2
H2A.Z:H4–4ac
H3K4 me1:H4K20 me2
DNA Meth. m5C:H4K20 me3
H3K27 me2:DNA Meth. m5C
H3K9 me2:DNA Meth. m5C
H3K27 me3:H4K20 me3
DNA Meth. m5C:H4K20 me2
DNA Meth. m5C:H3K9 me3
H3K27 me3:H3K9 me3
H4K20 me2:H3–5ac
H3K4 me1:H4–4ac
H3K4 me1:H2A.Z
H3K27 me3:DNA Meth. m5C
H4K20 me2:H4–4ac
H4K20 me2:H3K4 me3
H3K4 me3:H3–5ac
H3K27 me3:H3K4 me1
H3K4 me1:H3–5ac
H4–4ac:H3–5ac
H3K4 me3:H4–4ac
H4K20 me3
H3K27 me2
H3K9 me3
H3K9 me2
H3K27 me3
DNA Meth. m5C
H3–5ac
H2A.Z
H4K20 me2
H3K4 me1
H3K4 me3
H4–4ac
0.00 0.25 0.50 0.75 1.00
π̂
ATP5F1D – Reverse
H2A.Z:H4K20 me2
H2A.Z:H4–4ac
H3K4 me1:H4K20 me2
DNA Meth. m5C:H4K20 me3
H3K27 me2:DNA Meth. m5C
H3K9 me2:DNA Meth. m5C
H3K27 me3:H4K20 me3
DNA Meth. m5C:H4K20 me2
DNA Meth. m5C:H3K9 me3
H3K27 me3:H3K9 me3
H4K20 me2:H3–5ac
H3K4 me1:H4–4ac
H3K4 me1:H2A.Z
H3K27 me3:DNA Meth. m5C
H4K20 me2:H4–4ac
H4K20 me2:H3K4 me3
H3K4 me3:H3–5ac
H3K27 me3:H3K4 me1
H3K4 me1:H3–5ac
H4–4ac:H3–5ac
H3K4 me3:H4–4ac
H4K20 me3
H3K27 me2
H3K9 me3
H3K9 me2
H3K27 me3
DNA Meth. m5C
H3–5ac
H2A.Z
H4K20 me2
H3K4 me1
H3K4 me3
H4–4ac
0.00 0.25 0.50 0.75 1.00
π̂

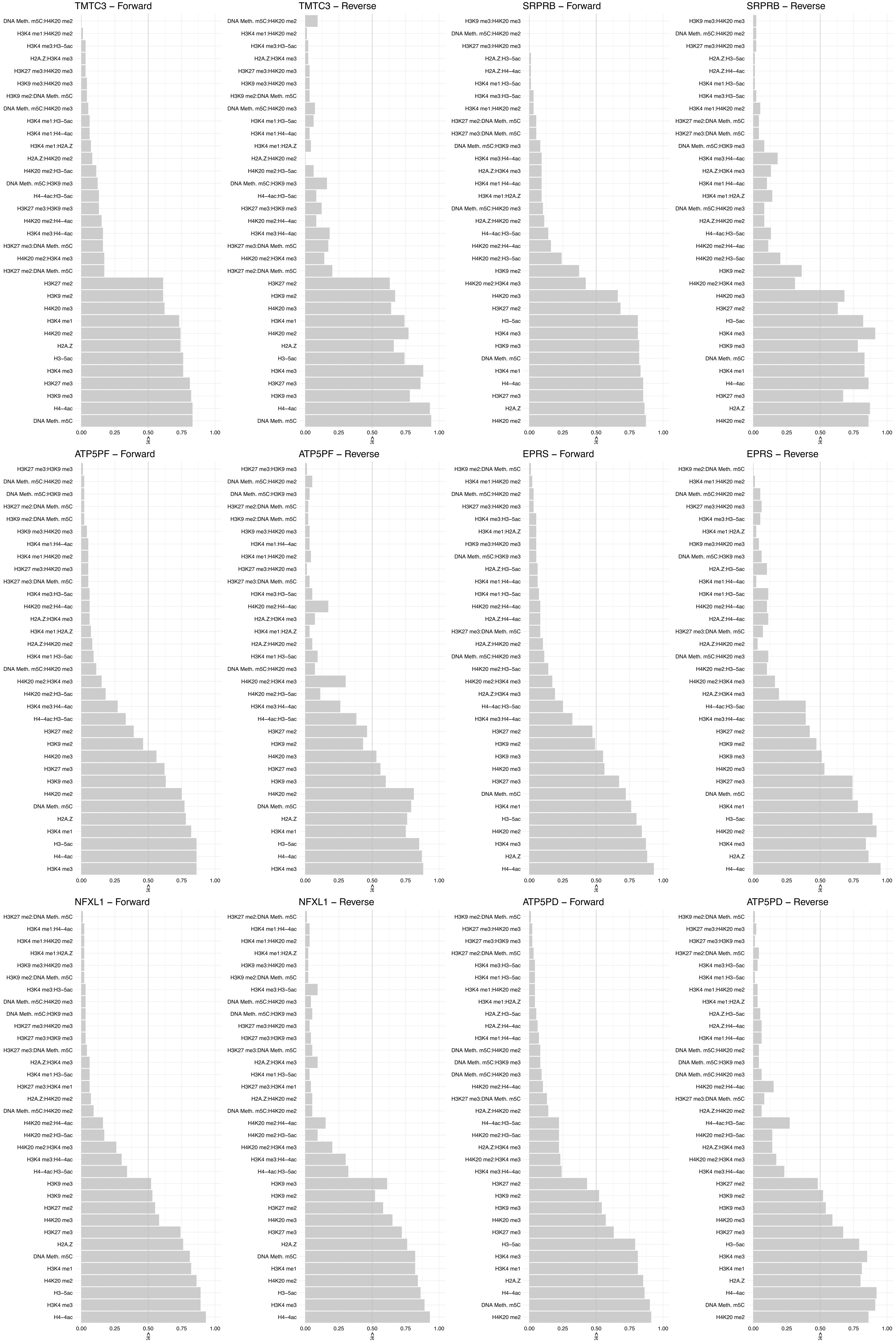
TMTC3 − Forward
DNA Meth. m5C:H4K20 me2
H3K4 me1:H4K20 me2
H3K4 me3:H3−5ac
H2A.Z:H3K4 me3
H3K27 me3:H4K20 me3
H3K9 me3:H4K20 me3
H3K9 me2:DNA Meth. m5C
DNA Meth. m5C:H4K20 me3
H3K4 me1:H3−5ac
H3K4 me1:H4−4ac
H3K4 me1:H2A.Z
H2A.Z:H4K20 me2
H4K20 me2:H3−5ac
DNA Meth. m5C:H3K9 me3
H4−4ac:H3−5ac
H3K27 me3:H3K9 me3
H4K20 me2:H4−4ac
H3K4 me3:H4−4ac
H3K27 me3:DNA Meth. m5C
H4K20 me2:H3K4 me3
H3K27 me2:DNA Meth. m5C
H3K27 me2
H3K9 me2
H4K20 me3
H3K4 me1
H4K20 me2
H2A.Z
H3−5ac
H3K4 me3
H3K27 me3
H3K9 me3
H4−4ac
DNA Meth. m5C
0.00 0.25 0.50 0.75 1.00
π̂
TMTC3 − Reverse
DNA Meth. m5C:H4K20 me2
H3K4 me1:H4K20 me2
H3K4 me3:H3−5ac
H2A.Z:H3K4 me3
H3K27 me3:H4K20 me3
H3K9 me3:H4K20 me3
H3K9 me2:DNA Meth. m5C
DNA Meth. m5C:H4K20 me3
H3K4 me1:H3−5ac
H3K4 me1:H4−4ac
H3K4 me1:H2A.Z
H2A.Z:H4K20 me2
H4K20 me2:H3−5ac
DNA Meth. m5C:H3K9 me3
H4−4ac:H3−5ac
H3K27 me3:H3K9 me3
H4K20 me2:H4−4ac
H3K4 me3:H4−4ac
H3K27 me3:DNA Meth. m5C
H4K20 me2:H3K4 me3
H3K27 me2:DNA Meth. m5C
H3K27 me2
H3K9 me2
H4K20 me3
H3K4 me1
H4K20 me2
H2A.Z
H3−5ac
H3K4 me3
H3K27 me3
H3K9 me3
H4−4ac
DNA Meth. m5C
0.00 0.25 0.50 0.75 1.00
π̂
SRPRB − Forward
H3K9 me3:H4K20 me3
DNA Meth. m5C:H4K20 me2
H3K27 me3:H4K20 me3
H2A.Z:H3−5ac
H2A.Z:H4−4ac
H3K4 me1:H3−5ac
H3K4 me3:H3−5ac
H3K4 me1:H4K20 me2
H3K27 me2:DNA Meth. m5C
H3K27 me3:DNA Meth. m5C
DNA Meth. m5C:H3K9 me3
H3K4 me3:H4−4ac
H2A.Z:H3K4 me3
H3K4 me1:H4−4ac
H3K4 me1:H2A.Z
DNA Meth. m5C:H4K20 me3
H2A.Z:H4K20 me2
H4−4ac:H3−5ac
H4K20 me2:H4−4ac
H4K20 me2:H3−5ac
H3K9 me2
H4K20 me2:H3K4 me3
H4K20 me3
H3K27 me2
H3−5ac
H3K4 me3
H3K9 me3
DNA Meth. m5C
H3K4 me1
H4−4ac
H3K27 me3
H2A.Z
H4K20 me2
0.00 0.25 0.50 0.75 1.00
π̂
SRPRB − Reverse
H3K9 me3:H4K20 me3
DNA Meth. m5C:H4K20 me2
H3K27 me3:H4K20 me3
H2A.Z:H3−5ac
H2A.Z:H4−4ac
H3K4 me1:H3−5ac
H3K4 me3:H3−5ac
H3K4 me1:H4K20 me2
H3K27 me2:DNA Meth. m5C
H3K27 me3:DNA Meth. m5C
DNA Meth. m5C:H3K9 me3
H3K4 me3:H4−4ac
H2A.Z:H3K4 me3
H3K4 me1:H4−4ac
H3K4 me1:H2A.Z
DNA Meth. m5C:H4K20 me3
H2A.Z:H4K20 me2
H4−4ac:H3−5ac
H4K20 me2:H4−4ac
H4K20 me2:H3−5ac
H3K9 me2
H4K20 me2:H3K4 me3
H4K20 me3
H3K27 me2
H3−5ac
H3K4 me3
H3K9 me3
DNA Meth. m5C
H3K4 me1
H4−4ac
H3K27 me3
H2A.Z
H4K20 me2
0.00 0.25 0.50 0.75 1.00
π̂
ATP5PF − Forward
H3K27 me3:H3K9 me3
DNA Meth. m5C:H4K20 me2
DNA Meth. m5C:H3K9 me3
H3K27 me2:DNA Meth. m5C
H3K9 me2:DNA Meth. m5C
H3K9 me3:H4K20 me3
H3K4 me1:H4−4ac
H3K4 me1:H4K20 me2
H3K27 me3:H4K20 me3
H3K27 me3:DNA Meth. m5C
H3K4 me3:H3−5ac
H4K20 me2:H4−4ac
H2A.Z:H3K4 me3
H3K4 me1:H2A.Z
H2A.Z:H4K20 me2
H3K4 me1:H3−5ac
DNA Meth. m5C:H4K20 me3
H4K20 me2:H3K4 me3
H4K20 me2:H3−5ac
H3K4 me3:H4−4ac
H4−4ac:H3−5ac
H3K27 me2
H3K9 me2
H4K20 me3
H3K27 me3
H3K9 me3
H4K20 me2
DNA Meth. m5C
H2A.Z
H3K4 me1
H3−5ac
H4−4ac
H3K4 me3
0.00 0.25 0.50 0.75 1.00
π̂
ATP5PF − Reverse
H3K27 me3:H3K9 me3
DNA Meth. m5C:H4K20 me2
DNA Meth. m5C:H3K9 me3
H3K27 me2:DNA Meth. m5C
H3K9 me2:DNA Meth. m5C
H3K9 me3:H4K20 me3
H3K4 me1:H4−4ac
H3K4 me1:H4K20 me2
H3K27 me3:H4K20 me3
H3K27 me3:DNA Meth. m5C
H3K4 me3:H3−5ac
H4K20 me2:H4−4ac
H2A.Z:H3K4 me3
H3K4 me1:H2A.Z
H2A.Z:H4K20 me2
H3K4 me1:H3−5ac
DNA Meth. m5C:H4K20 me3
H4K20 me2:H3K4 me3
H4K20 me2:H3−5ac
H3K4 me3:H4−4ac
H4−4ac:H3−5ac
H3K27 me2
H3K9 me2
H4K20 me3
H3K27 me3
H3K9 me3
H4K20 me2
DNA Meth. m5C
H2A.Z
H3K4 me1
H3−5ac
H4−4ac
H3K4 me3
0.00 0.25 0.50 0.75 1.00
π̂
EPRS − Forward
H3K9 me2:DNA Meth. m5C
H3K4 me1:H4K20 me2
DNA Meth. m5C:H4K20 me2
H3K27 me3:H4K20 me3
H3K4 me3:H3−5ac
H3K4 me1:H2A.Z
H3K9 me3:H4K20 me3
DNA Meth. m5C:H3K9 me3
H2A.Z:H3−5ac
H3K4 me1:H4−4ac
H3K4 me1:H3−5ac
H4K20 me2:H4−4ac
H2A.Z:H4−4ac
H3K27 me3:DNA Meth. m5C
H2A.Z:H4K20 me2
DNA Meth. m5C:H4K20 me3
H4K20 me2:H3−5ac
H4K20 me2:H3K4 me3
H2A.Z:H3K4 me3
H4−4ac:H3−5ac
H3K4 me3:H4−4ac
H3K27 me2
H3K9 me2
H3K9 me3
H4K20 me3
H3K27 me3
DNA Meth. m5C
H3K4 me1
H3−5ac
H4K20 me2
H3K4 me3
H2A.Z
H4−4ac
0.00 0.25 0.50 0.75 1.00
π̂
EPRS − Reverse
H3K9 me2:DNA Meth. m5C
H3K4 me1:H4K20 me2
DNA Meth. m5C:H4K20 me2
H3K27 me3:H4K20 me3
H3K4 me3:H3−5ac
H3K4 me1:H2A.Z
H3K9 me3:H4K20 me3
DNA Meth. m5C:H3K9 me3
H2A.Z:H3−5ac
H3K4 me1:H4−4ac
H3K4 me1:H3−5ac
H4K20 me2:H4−4ac
H2A.Z:H4−4ac
H3K27 me3:DNA Meth. m5C
H2A.Z:H4K20 me2
DNA Meth. m5C:H4K20 me3
H4K20 me2:H3−5ac
H4K20 me2:H3K4 me3
H2A.Z:H3K4 me3
H4−4ac:H3−5ac
H3K4 me3:H4−4ac
H3K27 me2
H3K9 me2
H3K9 me3
H4K20 me3
H3K27 me3
DNA Meth. m5C
H3K4 me1
H3−5ac
H4K20 me2
H3K4 me3
H2A.Z
H4−4ac
0.00 0.25 0.50 0.75 1.00
π̂
NFXL1 − Forward
H3K27 me2:DNA Meth. m5C
H3K4 me1:H4−4ac
H3K4 me1:H4K20 me2
H3K4 me1:H2A.Z
H3K9 me3:H4K20 me3
H3K9 me2:DNA Meth. m5C
H3K4 me3:H3−5ac
DNA Meth. m5C:H4K20 me3
DNA Meth. m5C:H3K9 me3
H3K27 me3:H4K20 me3
H3K27 me3:H3K9 me3
H3K27 me3:DNA Meth. m5C
H2A.Z:H3K4 me3
H3K4 me1:H3−5ac
H3K27 me3:H3K4 me1
H2A.Z:H4K20 me2
DNA Meth. m5C:H4K20 me2
H4K20 me2:H4−4ac
H4K20 me2:H3−5ac
H4K20 me2:H3K4 me3
H3K4 me3:H4−4ac
H4−4ac:H3−5ac
H3K9 me3
H3K9 me2
H3K27 me2
H4K20 me3
H3K27 me3
H2A.Z
DNA Meth. m5C
H3K4 me1
H4K20 me2
H3−5ac
H3K4 me3
H4−4ac
0.00 0.25 0.50 0.75 1.00
π̂
NFXL1 − Reverse
H3K27 me2:DNA Meth. m5C
H3K4 me1:H4−4ac
H3K4 me1:H4K20 me2
H3K4 me1:H2A.Z
H3K9 me3:H4K20 me3
H3K9 me2:DNA Meth. m5C
H3K4 me3:H3−5ac
DNA Meth. m5C:H4K20 me3
DNA Meth. m5C:H3K9 me3
H3K27 me3:H4K20 me3
H3K27 me3:H3K9 me3
H3K27 me3:DNA Meth. m5C
H2A.Z:H3K4 me3
H3K4 me1:H3−5ac
H3K27 me3:H3K4 me1
H2A.Z:H4K20 me2
DNA Meth. m5C:H4K20 me2
H4K20 me2:H4−4ac
H4K20 me2:H3−5ac
H4K20 me2:H3K4 me3
H3K4 me3:H4−4ac
H4−4ac:H3−5ac
H3K9 me3
H3K9 me2
H3K27 me2
H4K20 me3
H3K27 me3
H2A.Z
DNA Meth. m5C
H3K4 me1
H4K20 me2
H3−5ac
H3K4 me3
H4−4ac
0.00 0.25 0.50 0.75 1.00
π̂
ATP5PD − Forward
H3K9 me2:DNA Meth. m5C
H3K27 me3:H4K20 me3
H3K27 me3:H3K9 me3
H3K27 me2:DNA Meth. m5C
H3K4 me3:H3−5ac
H3K4 me1:H3−5ac
H3K4 me1:H4K20 me2
H3K4 me1:H2A.Z
H2A.Z:H3−5ac
H2A.Z:H4−4ac
H3K4 me1:H4−4ac
DNA Meth. m5C:H4K20 me2
DNA Meth. m5C:H3K9 me3
DNA Meth. m5C:H4K20 me3
H4K20 me2:H4−4ac
H3K27 me3:DNA Meth. m5C
H2A.Z:H4K20 me2
H4−4ac:H3−5ac
H4K20 me2:H3−5ac
H2A.Z:H3K4 me3
H4K20 me2:H3K4 me3
H3K4 me3:H4−4ac
H3K27 me2
H3K9 me2
H3K9 me3
H4K20 me3
H3K27 me3
H3−5ac
H3K4 me3
H3K4 me1
H2A.Z
H4−4ac
DNA Meth. m5C
H4K20 me2
0.00 0.25 0.50 0.75 1.00
π̂
ATP5PD − Reverse
H3K9 me2:DNA Meth. m5C
H3K27 me3:H4K20 me3
H3K27 me3:H3K9 me3
H3K27 me2:DNA Meth. m5C
H3K4 me3:H3−5ac
H3K4 me1:H3−5ac
H3K4 me1:H4K20 me2
H3K4 me1:H2A.Z
H2A.Z:H3−5ac
H2A.Z:H4−4ac
H3K4 me1:H4−4ac
DNA Meth. m5C:H4K20 me2
DNA Meth. m5C:H3K9 me3
DNA Meth. m5C:H4K20 me3
H4K20 me2:H4−4ac
H3K27 me3:DNA Meth. m5C
H2A.Z:H4K20 me2
H4−4ac:H3−5ac
H4K20 me2:H3−5ac
H2A.Z:H3K4 me3
H4K20 me2:H3K4 me3
H3K4 me3:H4−4ac
H3K27 me2
H3K9 me2
H3K9 me3
H4K20 me3
H3K27 me3
H3−5ac
H3K4 me3
H3K4 me1
H2A.Z
H4−4ac
DNA Meth. m5C
H4K20 me2
0.00 0.25 0.50 0.75 1.00
π̂

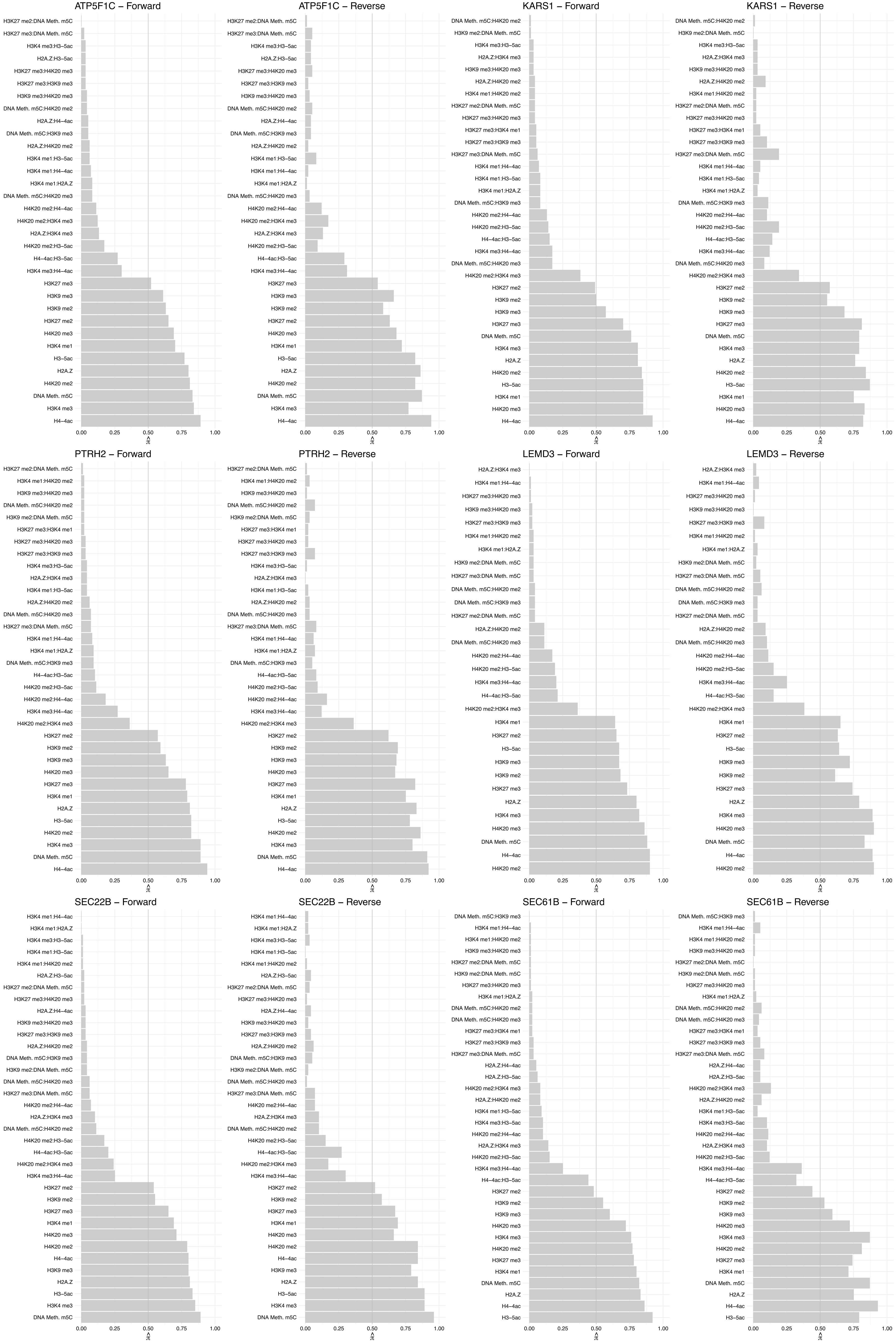
ATP5F1C – Forward
H3K27 me2:DNA Meth. m5C
H3K27 me3:DNA Meth. m5C
H3K4 me3:H3−5ac
H2A.Z:H3−5ac
H3K27 me3:H4K20 me3
H3K27 me3:H3K9 me3
H3K9 me3:H4K20 me3
DNA Meth. m5C:H4K20 me2
H2A.Z:H4−4ac
DNA Meth. m5C:H3K9 me3
H2A.Z:H4K20 me2
H3K4 me1:H3−5ac
H3K4 me1:H4−4ac
H3K4 me1:H2A.Z
DNA Meth. m5C:H4K20 me3
H4K20 me2:H4−4ac
H4K20 me2:H3K4 me3
H2A.Z:H3K4 me3
H4K20 me2:H3−5ac
H4−4ac:H3−5ac
H3K4 me3:H4−4ac
H3K27 me3
H3K9 me3
H3K9 me2
H3K27 me2
H4K20 me3
H3K4 me1
H3−5ac
H2A.Z
H4K20 me2
DNA Meth. m5C
H3K4 me3
H4−4ac
0.00 0.25 0.50 0.75 1.00
π̂
ATP5F1C – Reverse
H3K27 me2:DNA Meth. m5C
H3K27 me3:DNA Meth. m5C
H3K4 me3:H3−5ac
H2A.Z:H3−5ac
H3K27 me3:H4K20 me3
H3K27 me3:H3K9 me3
H3K9 me3:H4K20 me3
DNA Meth. m5C:H4K20 me2
H2A.Z:H4−4ac
DNA Meth. m5C:H3K9 me3
H2A.Z:H4K20 me2
H3K4 me1:H3−5ac
H3K4 me1:H4−4ac
H3K4 me1:H2A.Z
DNA Meth. m5C:H4K20 me3
H4K20 me2:H4−4ac
H4K20 me2:H3K4 me3
H2A.Z:H3K4 me3
H4K20 me2:H3−5ac
H4−4ac:H3−5ac
H3K4 me3:H4−4ac
H3K27 me3
H3K9 me3
H3K9 me2
H3K27 me2
H4K20 me3
H3K4 me1
H3−5ac
H2A.Z
H4K20 me2
DNA Meth. m5C
H3K4 me3
H4−4ac
0.00 0.25 0.50 0.75 1.00
π̂
KARS1 – Forward
DNA Meth. m5C:H4K20 me2
H3K9 me2:DNA Meth. m5C
H3K4 me3:H3−5ac
H2A.Z:H3K4 me3
H3K9 me3:H4K20 me3
H2A.Z:H4K20 me2
H3K4 me1:H4K20 me2
H3K27 me2:DNA Meth. m5C
H3K27 me3:H4K20 me3
H3K27 me3:H3K4 me1
H3K27 me3:H3K9 me3
H3K27 me3:DNA Meth. m5C
H3K4 me1:H4−4ac
H3K4 me1:H3−5ac
H3K4 me1:H2A.Z
DNA Meth. m5C:H3K9 me3
H4K20 me2:H4−4ac
H4K20 me2:H3−5ac
H4−4ac:H3−5ac
H3K4 me3:H4−4ac
DNA Meth. m5C:H4K20 me3
H4K20 me2:H3K4 me3
H3K27 me2
H3K9 me2
H3K9 me3
H3K27 me3
DNA Meth. m5C
H3K4 me3
H2A.Z
H4K20 me2
H3−5ac
H3K4 me1
H4K20 me3
H4−4ac
0.00 0.25 0.50 0.75 1.00
π̂
KARS1 – Reverse
DNA Meth. m5C:H4K20 me2
H3K9 me2:DNA Meth. m5C
H3K4 me3:H3−5ac
H2A.Z:H3K4 me3
H3K9 me3:H4K20 me3
H2A.Z:H4K20 me2
H3K4 me1:H4K20 me2
H3K27 me2:DNA Meth. m5C
H3K27 me3:H4K20 me3
H3K27 me3:H3K4 me1
H3K27 me3:H3K9 me3
H3K27 me3:DNA Meth. m5C
H3K4 me1:H4−4ac
H3K4 me1:H3−5ac
H3K4 me1:H2A.Z
DNA Meth. m5C:H3K9 me3
H4K20 me2:H4−4ac
H4K20 me2:H3−5ac
H4−4ac:H3−5ac
H3K4 me3:H4−4ac
DNA Meth. m5C:H4K20 me3
H4K20 me2:H3K4 me3
H3K27 me2
H3K9 me2
H3K9 me3
H3K27 me3
DNA Meth. m5C
H3K4 me3
H2A.Z
H4K20 me2
H3−5ac
H3K4 me1
H4K20 me3
H4−4ac
0.00 0.25 0.50 0.75 1.00
π̂
PTRH2 – Forward
H3K27 me2:DNA Meth. m5C
H3K4 me1:H4K20 me2
H3K9 me3:H4K20 me3
DNA Meth. m5C:H4K20 me2
H3K9 me2:DNA Meth. m5C
H3K27 me3:H3K4 me1
H3K27 me3:H4K20 me3
H3K27 me3:H3K9 me3
H3K4 me3:H3−5ac
H2A.Z:H3K4 me3
H3K4 me1:H3−5ac
H2A.Z:H4K20 me2
DNA Meth. m5C:H4K20 me3
H3K27 me3:DNA Meth. m5C
H3K4 me1:H4−4ac
H3K4 me1:H2A.Z
DNA Meth. m5C:H3K9 me3
H4−4ac:H3−5ac
H4K20 me2:H3−5ac
H4K20 me2:H4−4ac
H3K4 me3:H4−4ac
H4K20 me2:H3K4 me3
H3K27 me2
H3K9 me2
H3K9 me3
H4K20 me3
H3K27 me3
H3K4 me1
H2A.Z
H3−5ac
H4K20 me2
H3K4 me3
DNA Meth. m5C
H4−4ac
0.00 0.25 0.50 0.75 1.00
π̂
PTRH2 – Reverse
H3K27 me2:DNA Meth. m5C
H3K4 me1:H4K20 me2
H3K9 me3:H4K20 me3
DNA Meth. m5C:H4K20 me2
H3K9 me2:DNA Meth. m5C
H3K27 me3:H3K4 me1
H3K27 me3:H4K20 me3
H3K27 me3:H3K9 me3
H3K4 me3:H3−5ac
H2A.Z:H3K4 me3
H3K4 me1:H3−5ac
H2A.Z:H4K20 me2
DNA Meth. m5C:H4K20 me3
H3K27 me3:DNA Meth. m5C
H3K4 me1:H4−4ac
H3K4 me1:H2A.Z
DNA Meth. m5C:H3K9 me3
H4−4ac:H3−5ac
H4K20 me2:H3−5ac
H4K20 me2:H4−4ac
H3K4 me3:H4−4ac
H4K20 me2:H3K4 me3
H3K27 me2
H3K9 me2
H3K9 me3
H4K20 me3
H3K27 me3
H3K4 me1
H2A.Z
H3−5ac
H4K20 me2
H3K4 me3
DNA Meth. m5C
H4−4ac
0.00 0.25 0.50 0.75 1.00
π̂
LEMD3 – Forward
H2A.Z:H3K4 me3
H3K4 me1:H4−4ac
H3K27 me3:H4K20 me3
H3K9 me3:H4K20 me3
H3K27 me3:H3K9 me3
H3K4 me1:H4K20 me2
H3K4 me1:H2A.Z
H3K9 me2:DNA Meth. m5C
H3K27 me3:DNA Meth. m5C
DNA Meth. m5C:H4K20 me2
DNA Meth. m5C:H3K9 me3
H3K27 me2:DNA Meth. m5C
H2A.Z:H4K20 me2
DNA Meth. m5C:H4K20 me3
H4K20 me2:H4−4ac
H4K20 me2:H3−5ac
H3K4 me3:H4−4ac
H4−4ac:H3−5ac
H4K20 me2:H3K4 me3
H3K4 me1
H3K27 me2
H3−5ac
H3K9 me3
H3K9 me2
H3K27 me3
H2A.Z
H3K4 me3
H4K20 me3
DNA Meth. m5C
H4−4ac
H4K20 me2
0.00 0.25 0.50 0.75 1.00
π̂
LEMD3 – Reverse
H2A.Z:H3K4 me3
H3K4 me1:H4−4ac
H3K27 me3:H4K20 me3
H3K9 me3:H4K20 me3
H3K27 me3:H3K9 me3
H3K4 me1:H4K20 me2
H3K4 me1:H2A.Z
H3K9 me2:DNA Meth. m5C
H3K27 me3:DNA Meth. m5C
DNA Meth. m5C:H4K20 me2
DNA Meth. m5C:H3K9 me3
H3K27 me2:DNA Meth. m5C
H2A.Z:H4K20 me2
DNA Meth. m5C:H4K20 me3
H4K20 me2:H4−4ac
H4K20 me2:H3−5ac
H3K4 me3:H4−4ac
H4−4ac:H3−5ac
H4K20 me2:H3K4 me3
H3K4 me1
H3K27 me2
H3−5ac
H3K9 me3
H3K9 me2
H3K27 me3
H2A.Z
H3K4 me3
H4K20 me3
DNA Meth. m5C
H4−4ac
H4K20 me2
0.00 0.25 0.50 0.75 1.00
π̂
SEC22B – Forward
H3K4 me1:H4−4ac
H3K4 me1:H2A.Z
H3K4 me3:H3−5ac
H3K4 me1:H3−5ac
H3K4 me1:H4K20 me2
H2A.Z:H3−5ac
H3K27 me2:DNA Meth. m5C
H3K27 me3:H4K20 me3
H2A.Z:H4−4ac
H3K9 me3:H4K20 me3
H3K27 me3:H3K9 me3
H2A.Z:H4K20 me2
DNA Meth. m5C:H3K9 me3
H3K9 me2:DNA Meth. m5C
DNA Meth. m5C:H4K20 me3
H3K27 me3:DNA Meth. m5C
H4K20 me2:H4−4ac
H2A.Z:H3K4 me3
DNA Meth. m5C:H4K20 me2
H4K20 me2:H3−5ac
H4−4ac:H3−5ac
H4K20 me2:H3K4 me3
H3K4 me3:H4−4ac
H3K27 me2
H3K9 me2
H3K27 me3
H3K4 me1
H4K20 me3
H4K20 me2
H4−4ac
H3K9 me3
H2A.Z
H3−5ac
H3K4 me3
DNA Meth. m5C
0.00 0.25 0.50 0.75 1.00
π̂
SEC22B – Reverse
H3K4 me1:H4−4ac
H3K4 me1:H2A.Z
H3K4 me3:H3−5ac
H3K4 me1:H3−5ac
H3K4 me1:H4K20 me2
H2A.Z:H3−5ac
H3K27 me2:DNA Meth. m5C
H3K27 me3:H4K20 me3
H2A.Z:H4−4ac
H3K9 me3:H4K20 me3
H3K27 me3:H3K9 me3
H2A.Z:H4K20 me2
DNA Meth. m5C:H3K9 me3
H3K9 me2:DNA Meth. m5C
DNA Meth. m5C:H4K20 me3
H3K27 me3:DNA Meth. m5C
H4K20 me2:H4−4ac
H2A.Z:H3K4 me3
DNA Meth. m5C:H4K20 me2
H4K20 me2:H3−5ac
H4−4ac:H3−5ac
H4K20 me2:H3K4 me3
H3K4 me3:H4−4ac
H3K27 me2
H3K9 me2
H3K27 me3
H3K4 me1
H4K20 me3
H4K20 me2
H4−4ac
H3K9 me3
H2A.Z
H3−5ac
H3K4 me3
DNA Meth. m5C
0.00 0.25 0.50 0.75 1.00
π̂
SEC61B – Forward
DNA Meth. m5C:H3K9 me3
H3K4 me1:H4−4ac
H3K4 me1:H4K20 me2
H3K9 me3:H4K20 me3
H3K27 me2:DNA Meth. m5C
H3K9 me2:DNA Meth. m5C
H3K27 me3:H4K20 me3
H3K4 me1:H2A.Z
DNA Meth. m5C:H4K20 me2
DNA Meth. m5C:H4K20 me3
H3K27 me3:H3K4 me1
H3K27 me3:H3K9 me3
H3K27 me3:DNA Meth. m5C
H2A.Z:H4−4ac
H2A.Z:H3−5ac
H4K20 me2:H3K4 me3
H2A.Z:H4K20 me2
H3K4 me1:H3−5ac
H3K4 me3:H3−5ac
H4K20 me2:H4−4ac
H2A.Z:H3K4 me3
H4K20 me2:H3−5ac
H3K4 me3:H4−4ac
H4−4ac:H3−5ac
H3K27 me2
H3K9 me2
H3K9 me3
H4K20 me3
H3K4 me3
H4K20 me2
H3K27 me3
H3K4 me1
DNA Meth. m5C
H2A.Z
H4−4ac
H3−5ac
0.00 0.25 0.50 0.75 1.00
π̂
SEC61B – Reverse
DNA Meth. m5C:H3K9 me3
H3K4 me1:H4−4ac
H3K4 me1:H4K20 me2
H3K9 me3:H4K20 me3
H3K27 me2:DNA Meth. m5C
H3K9 me2:DNA Meth. m5C
H3K27 me3:H4K20 me3
H3K4 me1:H2A.Z
DNA Meth. m5C:H4K20 me2
DNA Meth. m5C:H4K20 me3
H3K27 me3:H3K4 me1
H3K27 me3:H3K9 me3
H3K27 me3:DNA Meth. m5C
H2A.Z:H4−4ac
H2A.Z:H3−5ac
H4K20 me2:H3K4 me3
H2A.Z:H4K20 me2
H3K4 me1:H3−5ac
H3K4 me3:H3−5ac
H4K20 me2:H4−4ac
H2A.Z:H3K4 me3
H4K20 me2:H3−5ac
H3K4 me3:H4−4ac
H4−4ac:H3−5ac
H3K27 me2
H3K9 me2
H3K9 me3
H4K20 me3
H3K4 me3
H4K20 me2
H3K27 me3
H3K4 me1
DNA Meth. m5C
H2A.Z
H4−4ac
H3−5ac
0.00 0.25 0.50 0.75 1.00
π̂

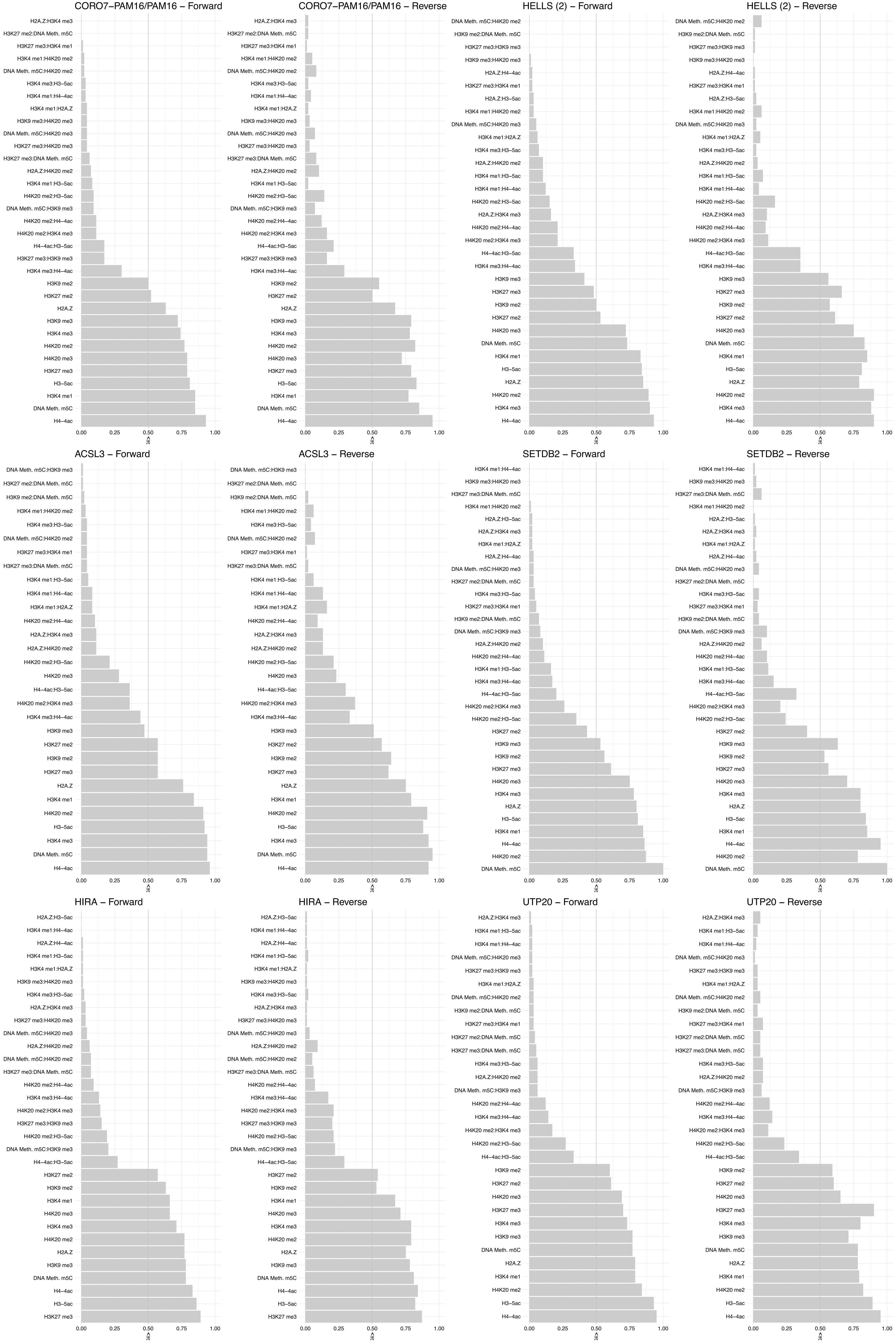
CORO7−PAM16/PAM16 − Forward
H2A.Z:H3K4 me3
H3K27 me2:DNA Meth. m5C
H3K27 me3:H3K4 me1
H3K4 me1:H4K20 me2
DNA Meth. m5C:H4K20 me2
H3K4 me3:H3−5ac
H3K4 me1:H4−4ac
H3K4 me1:H2A.Z
H3K9 me3:H4K20 me3
DNA Meth. m5C:H4K20 me3
H3K27 me3:H4K20 me3
H3K27 me3:DNA Meth. m5C
H2A.Z:H4K20 me2
H3K4 me1:H3−5ac
H4K20 me2:H3−5ac
DNA Meth. m5C:H3K9 me3
H4K20 me2:H4−4ac
H4K20 me2:H3K4 me3
H4−4ac:H3−5ac
H3K27 me3:H3K9 me3
H3K4 me3:H4−4ac
H3K9 me2
H3K27 me2
H2A.Z
H3K9 me3
H3K4 me3
H4K20 me2
H4K20 me3
H3K27 me3
H3−5ac
H3K4 me1
DNA Meth. m5C
H4−4ac
0.00
0.25
0.50
0.75
1.00
π̂
CORO7−PAM16/PAM16 − Reverse
H2A.Z:H3K4 me3
H3K27 me2:DNA Meth. m5C
H3K27 me3:H3K4 me1
H3K4 me1:H4K20 me2
DNA Meth. m5C:H4K20 me2
H3K4 me3:H3−5ac
H3K4 me1:H4−4ac
H3K4 me1:H2A.Z
H3K9 me3:H4K20 me3
DNA Meth. m5C:H4K20 me3
H3K27 me3:H4K20 me3
H3K27 me3:DNA Meth. m5C
H2A.Z:H4K20 me2
H3K4 me1:H3−5ac
H4K20 me2:H3−5ac
DNA Meth. m5C:H3K9 me3
H4K20 me2:H4−4ac
H4K20 me2:H3K4 me3
H4−4ac:H3−5ac
H3K27 me3:H3K9 me3
H3K4 me3:H4−4ac
H3K9 me2
H3K27 me2
H2A.Z
H3K9 me3
H3K4 me3
H4K20 me2
H4K20 me3
H3K27 me3
H3−5ac
H3K4 me1
DNA Meth. m5C
H4−4ac
0.00
0.25
0.50
0.75
1.00
π̂
HELLS (2) − Forward
DNA Meth. m5C:H4K20 me2
H3K9 me2:DNA Meth. m5C
H3K27 me3:H3K9 me3
H3K9 me3:H4K20 me3
H2A.Z:H4−4ac
H3K27 me3:H3K4 me1
H2A.Z:H3−5ac
H3K4 me1:H4K20 me2
DNA Meth. m5C:H4K20 me3
H3K4 me1:H2A.Z
H3K4 me3:H3−5ac
H2A.Z:H4K20 me2
H3K4 me1:H3−5ac
H3K4 me1:H4−4ac
H4K20 me2:H3−5ac
H2A.Z:H3K4 me3
H4K20 me2:H4−4ac
H4K20 me2:H3K4 me3
H4−4ac:H3−5ac
H3K4 me3:H4−4ac
H3K9 me3
H3K27 me3
H3K9 me2
H3K27 me2
H4K20 me3
DNA Meth. m5C
H3K4 me1
H3−5ac
H2A.Z
H4K20 me2
H3K4 me3
H4−4ac
0.00
0.25
0.50
0.75
1.00
π̂
HELLS (2) − Reverse
DNA Meth. m5C:H4K20 me2
H3K9 me2:DNA Meth. m5C
H3K27 me3:H3K9 me3
H3K9 me3:H4K20 me3
H2A.Z:H4−4ac
H3K27 me3:H3K4 me1
H2A.Z:H3−5ac
H3K4 me1:H4K20 me2
DNA Meth. m5C:H4K20 me3
H3K4 me1:H2A.Z
H3K4 me3:H3−5ac
H2A.Z:H4K20 me2
H3K4 me1:H3−5ac
H3K4 me1:H4−4ac
H4K20 me2:H3−5ac
H2A.Z:H3K4 me3
H4K20 me2:H4−4ac
H4K20 me2:H3K4 me3
H4−4ac:H3−5ac
H3K4 me3:H4−4ac
H3K9 me3
H3K27 me3
H3K9 me2
H3K27 me2
H4K20 me3
DNA Meth. m5C
H3K4 me1
H3−5ac
H2A.Z
H4K20 me2
H3K4 me3
H4−4ac
0.00
0.25
0.50
0.75
1.00
π̂
ACSL3 − Forward
DNA Meth. m5C:H3K9 me3
H3K27 me2:DNA Meth. m5C
H3K9 me2:DNA Meth. m5C
H3K4 me1:H4K20 me2
H3K4 me3:H3−5ac
DNA Meth. m5C:H4K20 me2
H3K27 me3:H3K4 me1
H3K27 me3:DNA Meth. m5C
H3K4 me1:H3−5ac
H3K4 me1:H4−4ac
H3K4 me1:H2A.Z
H4K20 me2:H4−4ac
H2A.Z:H3K4 me3
H2A.Z:H4K20 me2
H4K20 me2:H3−5ac
H4K20 me3
H4−4ac:H3−5ac
H4K20 me2:H3K4 me3
H3K4 me3:H4−4ac
H3K9 me3
H3K27 me2
H3K9 me2
H3K27 me3
H2A.Z
H3K4 me1
H4K20 me2
H3−5ac
H3K4 me3
DNA Meth. m5C
H4−4ac
0.00
0.25
0.50
0.75
1.00
π̂
ACSL3 − Reverse
DNA Meth. m5C:H3K9 me3
H3K27 me2:DNA Meth. m5C
H3K9 me2:DNA Meth. m5C
H3K4 me1:H4K20 me2
H3K4 me3:H3−5ac
DNA Meth. m5C:H4K20 me2
H3K27 me3:H3K4 me1
H3K27 me3:DNA Meth. m5C
H3K4 me1:H3−5ac
H3K4 me1:H4−4ac
H3K4 me1:H2A.Z
H4K20 me2:H4−4ac
H2A.Z:H3K4 me3
H2A.Z:H4K20 me2
H4K20 me2:H3−5ac
H4K20 me3
H4−4ac:H3−5ac
H4K20 me2:H3K4 me3
H3K4 me3:H4−4ac
H3K9 me3
H3K27 me2
H3K9 me2
H3K27 me3
H2A.Z
H3K4 me1
H4K20 me2
H3−5ac
H3K4 me3
DNA Meth. m5C
H4−4ac
0.00
0.25
0.50
0.75
1.00
π̂
SETDB2 − Forward
H3K4 me1:H4−4ac
H3K9 me3:H4K20 me3
H3K27 me3:DNA Meth. m5C
H3K4 me1:H4K20 me2
H2A.Z:H3−5ac
H2A.Z:H3K4 me3
H3K4 me1:H2A.Z
H2A.Z:H4−4ac
DNA Meth. m5C:H4K20 me3
H3K27 me2:DNA Meth. m5C
H3K4 me3:H3−5ac
H3K27 me3:H3K4 me1
H3K9 me2:DNA Meth. m5C
DNA Meth. m5C:H3K9 me3
H2A.Z:H4K20 me2
H4K20 me2:H4−4ac
H3K4 me1:H3−5ac
H3K4 me3:H4−4ac
H4−4ac:H3−5ac
H4K20 me2:H3K4 me3
H4K20 me2:H3−5ac
H3K27 me2
H3K9 me3
H3K9 me2
H3K27 me3
H4K20 me3
H3K4 me3
H2A.Z
H3−5ac
H3K4 me1
H4−4ac
H4K20 me2
DNA Meth. m5C
0.00
0.25
0.50
0.75
1.00
π̂
SETDB2 − Reverse
H3K4 me1:H4−4ac
H3K9 me3:H4K20 me3
H3K27 me3:DNA Meth. m5C
H3K4 me1:H4K20 me2
H2A.Z:H3−5ac
H2A.Z:H3K4 me3
H3K4 me1:H2A.Z
H2A.Z:H4−4ac
DNA Meth. m5C:H4K20 me3
H3K27 me2:DNA Meth. m5C
H3K4 me3:H3−5ac
H3K27 me3:H3K4 me1
H3K9 me2:DNA Meth. m5C
DNA Meth. m5C:H3K9 me3
H2A.Z:H4K20 me2
H4K20 me2:H4−4ac
H3K4 me1:H3−5ac
H3K4 me3:H4−4ac
H4−4ac:H3−5ac
H4K20 me2:H3K4 me3
H4K20 me2:H3−5ac
H3K27 me2
H3K9 me3
H3K9 me2
H3K27 me3
H4K20 me3
H3K4 me3
H2A.Z
H3−5ac
H3K4 me1
H4−4ac
H4K20 me2
DNA Meth. m5C
0.00
0.25
0.50
0.75
1.00
π̂
HIRA − Forward
H2A.Z:H3−5ac
H3K4 me1:H4−4ac
H2A.Z:H4−4ac
H3K4 me1:H3−5ac
H3K4 me1:H2A.Z
H3K9 me3:H4K20 me3
H3K4 me3:H3−5ac
H2A.Z:H3K4 me3
H3K27 me3:H4K20 me3
DNA Meth. m5C:H4K20 me3
H2A.Z:H4K20 me2
DNA Meth. m5C:H4K20 me2
H3K27 me3:DNA Meth. m5C
H4K20 me2:H4−4ac
H3K4 me3:H4−4ac
H4K20 me2:H3K4 me3
H3K27 me3:H3K9 me3
H4K20 me2:H3−5ac
DNA Meth. m5C:H3K9 me3
H4−4ac:H3−5ac
H3K27 me2
H3K9 me2
H3K4 me1
H4K20 me3
H3K4 me3
H4K20 me2
H2A.Z
H3K9 me3
DNA Meth. m5C
H4−4ac
H3−5ac
H3K27 me3
0.00
0.25
0.50
0.75
1.00
π̂
HIRA − Reverse
H2A.Z:H3−5ac
H3K4 me1:H4−4ac
H2A.Z:H4−4ac
H3K4 me1:H3−5ac
H3K4 me1:H2A.Z
H3K9 me3:H4K20 me3
H3K4 me3:H3−5ac
H2A.Z:H3K4 me3
H3K27 me3:H4K20 me3
DNA Meth. m5C:H4K20 me3
H2A.Z:H4K20 me2
DNA Meth. m5C:H4K20 me2
H3K27 me3:DNA Meth. m5C
H4K20 me2:H4−4ac
H3K4 me3:H4−4ac
H4K20 me2:H3K4 me3
H3K27 me3:H3K9 me3
H4K20 me2:H3−5ac
DNA Meth. m5C:H3K9 me3
H4−4ac:H3−5ac
H3K27 me2
H3K9 me2
H3K4 me1
H4K20 me3
H3K4 me3
H4K20 me2
H2A.Z
H3K9 me3
DNA Meth. m5C
H4−4ac
H3−5ac
H3K27 me3
0.00
0.25
0.50
0.75
1.00
π̂
UTP20 − Forward
H2A.Z:H3K4 me3
H3K4 me1:H3−5ac
H3K4 me1:H4−4ac
DNA Meth. m5C:H4K20 me3
H3K27 me3:H3K9 me3
H3K4 me1:H2A.Z
DNA Meth. m5C:H4K20 me2
H3K9 me2:DNA Meth. m5C
H3K27 me3:H3K4 me1
H3K27 me2:DNA Meth. m5C
H3K27 me3:DNA Meth. m5C
H3K4 me3:H3−5ac
H2A.Z:H4K20 me2
DNA Meth. m5C:H3K9 me3
H4K20 me2:H4−4ac
H3K4 me3:H4−4ac
H4K20 me2:H3K4 me3
H4K20 me2:H3−5ac
H4−4ac:H3−5ac
H3K9 me2
H3K27 me2
H4K20 me3
H3K27 me3
H3K4 me3
H3K9 me3
DNA Meth. m5C
H2A.Z
H3K4 me1
H4K20 me2
H3−5ac
H4−4ac
0.00
0.25
0.50
0.75
1.00
π̂
UTP20 − Reverse
H2A.Z:H3K4 me3
H3K4 me1:H3−5ac
H3K4 me1:H4−4ac
DNA Meth. m5C:H4K20 me3
H3K27 me3:H3K9 me3
H3K4 me1:H2A.Z
DNA Meth. m5C:H4K20 me2
H3K9 me2:DNA Meth. m5C
H3K27 me3:H3K4 me1
H3K27 me2:DNA Meth. m5C
H3K27 me3:DNA Meth. m5C
H3K4 me3:H3−5ac
H2A.Z:H4K20 me2
DNA Meth. m5C:H3K9 me3
H4K20 me2:H4−4ac
H3K4 me3:H4−4ac
H4K20 me2:H3K4 me3
H4K20 me2:H3−5ac
H4−4ac:H3−5ac
H3K9 me2
H3K27 me2
H4K20 me3
H3K27 me3
H3K4 me3
H3K9 me3
DNA Meth. m5C
H2A.Z
H3K4 me1
H4K20 me2
H3−5ac
H4−4ac
0.00
0.25
0.50
0.75
1.00
π̂

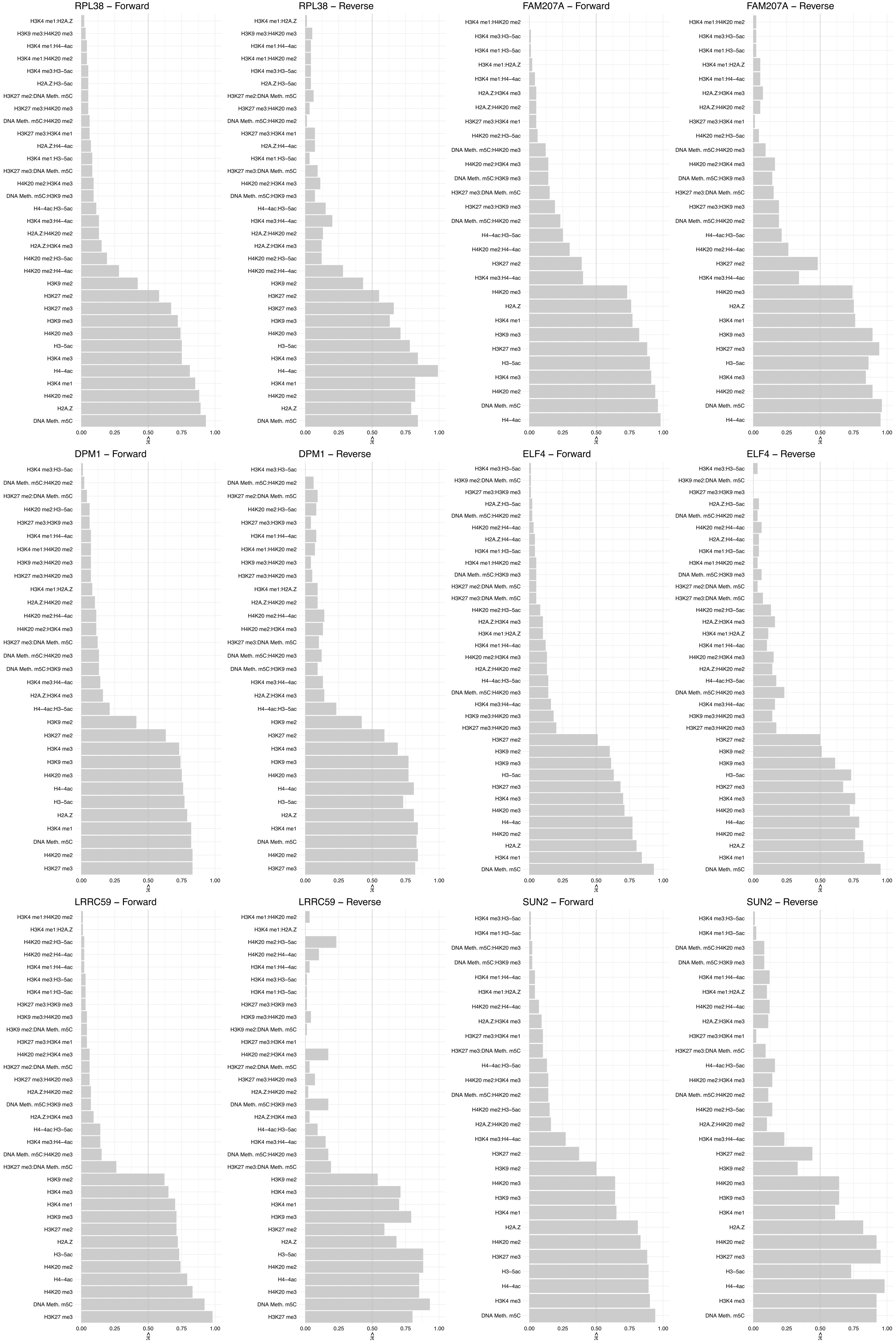

RPL38 – Forward
RPL38 – Reverse
H3K4 me1:H2A.Z
H3K9 me3:H4K20 me3
H3K4 me1:H4−4ac
H3K4 me1:H4K20 me2
H3K4 me3:H3−5ac
H2A.Z:H3−5ac
H3K27 me2:DNA Meth. m5C
H3K27 me3:H4K20 me3
DNA Meth. m5C:H4K20 me2
H3K27 me3:H3K4 me1
H2A.Z:H4−4ac
H3K4 me1:H3−5ac
H3K27 me3:DNA Meth. m5C
H4K20 me2:H3K4 me3
DNA Meth. m5C:H3K9 me3
H4−4ac:H3−5ac
H3K4 me3:H4−4ac
H2A.Z:H4K20 me2
H2A.Z:H3K4 me3
H4K20 me2:H3−5ac
H4K20 me2:H4−4ac
H3K9 me2
H3K27 me2
H3K27 me3
H3K9 me3
H4K20 me3
H3−5ac
H3K4 me3
H4−4ac
H3K4 me1
H4K20 me2
H2A.Z
DNA Meth. m5C
FAM207A – Forward
FAM207A – Reverse
H3K4 me1:H4K20 me2
H3K4 me3:H3−5ac
H3K4 me1:H3−5ac
H3K4 me1:H2A.Z
H3K4 me1:H4−4ac
H2A.Z:H3K4 me3
H2A.Z:H4K20 me2
H3K27 me3:H3K4 me1
H4K20 me2:H3−5ac
DNA Meth. m5C:H4K20 me3
H4K20 me2:H3K4 me3
DNA Meth. m5C:H3K9 me3
H3K27 me3:DNA Meth. m5C
H3K27 me3:H3K9 me3
DNA Meth. m5C:H4K20 me2
H4−4ac:H3−5ac
H4K20 me2:H4−4ac
H3K27 me2
H3K4 me3:H4−4ac
H4K20 me3
H2A.Z
H3K4 me1
H3K9 me3
H3K27 me3
H3−5ac
H3K4 me3
H4K20 me2
DNA Meth. m5C
H4−4ac
DPM1 – Forward
DPM1 – Reverse
H3K4 me3:H3−5ac
DNA Meth. m5C:H4K20 me2
H3K27 me2:DNA Meth. m5C
H4K20 me2:H3−5ac
H3K27 me3:H3K9 me3
H3K4 me1:H4−4ac
H3K4 me1:H4K20 me2
H3K9 me3:H4K20 me3
H3K27 me3:H4K20 me3
H3K4 me1:H2A.Z
H2A.Z:H4K20 me2
H4K20 me2:H4−4ac
H4K20 me2:H3K4 me3
H3K27 me3:DNA Meth. m5C
DNA Meth. m5C:H4K20 me3
DNA Meth. m5C:H3K9 me3
H3K4 me3:H4−4ac
H2A.Z:H3K4 me3
H4−4ac:H3−5ac
H3K9 me2
H3K27 me2
H3K4 me3
H3K9 me3
H4K20 me3
H4−4ac
H3−5ac
H2A.Z
H3K4 me1
DNA Meth. m5C
H4K20 me2
H3K27 me3
ELF4 – Forward
ELF4 – Reverse
H3K4 me3:H3−5ac
H3K9 me2:DNA Meth. m5C
H3K27 me3:H3K9 me3
H2A.Z:H3−5ac
DNA Meth. m5C:H4K20 me2
H4K20 me2:H4−4ac
H2A.Z:H4−4ac
H3K4 me1:H3−5ac
H3K4 me1:H4K20 me2
DNA Meth. m5C:H3K9 me3
H3K27 me2:DNA Meth. m5C
H3K27 me3:DNA Meth. m5C
H4K20 me2:H3−5ac
H2A.Z:H3K4 me3
H3K4 me1:H2A.Z
H3K4 me1:H4−4ac
H4K20 me2:H3K4 me3
H2A.Z:H4K20 me2
H4−4ac:H3−5ac
DNA Meth. m5C:H4K20 me3
H3K4 me3:H4−4ac
H3K9 me3:H4K20 me3
H3K27 me3:H4K20 me3
H3K27 me2
H3K9 me2
H3K9 me3
H3−5ac
H3K27 me3
H3K4 me3
H4K20 me3
H4−4ac
H4K20 me2
H2A.Z
H3K4 me1
DNA Meth. m5C
LRRC59 – Forward
LRRC59 – Reverse
H3K4 me1:H4K20 me2
H3K4 me1:H2A.Z
H4K20 me2:H3−5ac
H4K20 me2:H4−4ac
H3K4 me1:H4−4ac
H3K4 me3:H3−5ac
H3K4 me1:H3−5ac
H3K27 me3:H3K9 me3
H3K9 me3:H4K20 me3
H3K9 me2:DNA Meth. m5C
H3K27 me3:H3K4 me1
H4K20 me2:H3K4 me3
H3K27 me2:DNA Meth. m5C
H3K27 me3:H4K20 me3
H2A.Z:H4K20 me2
DNA Meth. m5C:H3K9 me3
H2A.Z:H3K4 me3
H4−4ac:H3−5ac
H3K4 me3:H4−4ac
DNA Meth. m5C:H4K20 me3
H3K27 me3:DNA Meth. m5C
H3K9 me2
H3K4 me3
H3K4 me1
H3K9 me3
H3K27 me2
H2A.Z
H3−5ac
H4K20 me2
H4−4ac
H4K20 me3
DNA Meth. m5C
H3K27 me3
SUN2 – Forward
SUN2 – Reverse
H3K4 me3:H3−5ac
H3K4 me1:H3−5ac
DNA Meth. m5C:H4K20 me3
DNA Meth. m5C:H3K9 me3
H3K4 me1:H4−4ac
H3K4 me1:H2A.Z
H4K20 me2:H4−4ac
H2A.Z:H3K4 me3
H3K27 me3:H3K4 me1
H3K27 me3:DNA Meth. m5C
H4−4ac:H3−5ac
H4K20 me2:H3K4 me3
DNA Meth. m5C:H4K20 me2
H4K20 me2:H3−5ac
H2A.Z:H4K20 me2
H3K4 me3:H4−4ac
H3K27 me2
H3K9 me2
H4K20 me3
H3K9 me3
H3K4 me1
H2A.Z
H4K20 me2
H3K27 me3
H3−5ac
H4−4ac
H3K4 me3
DNA Meth. m5C
0.00
0.25
0.50
0.75
1.00
π̂

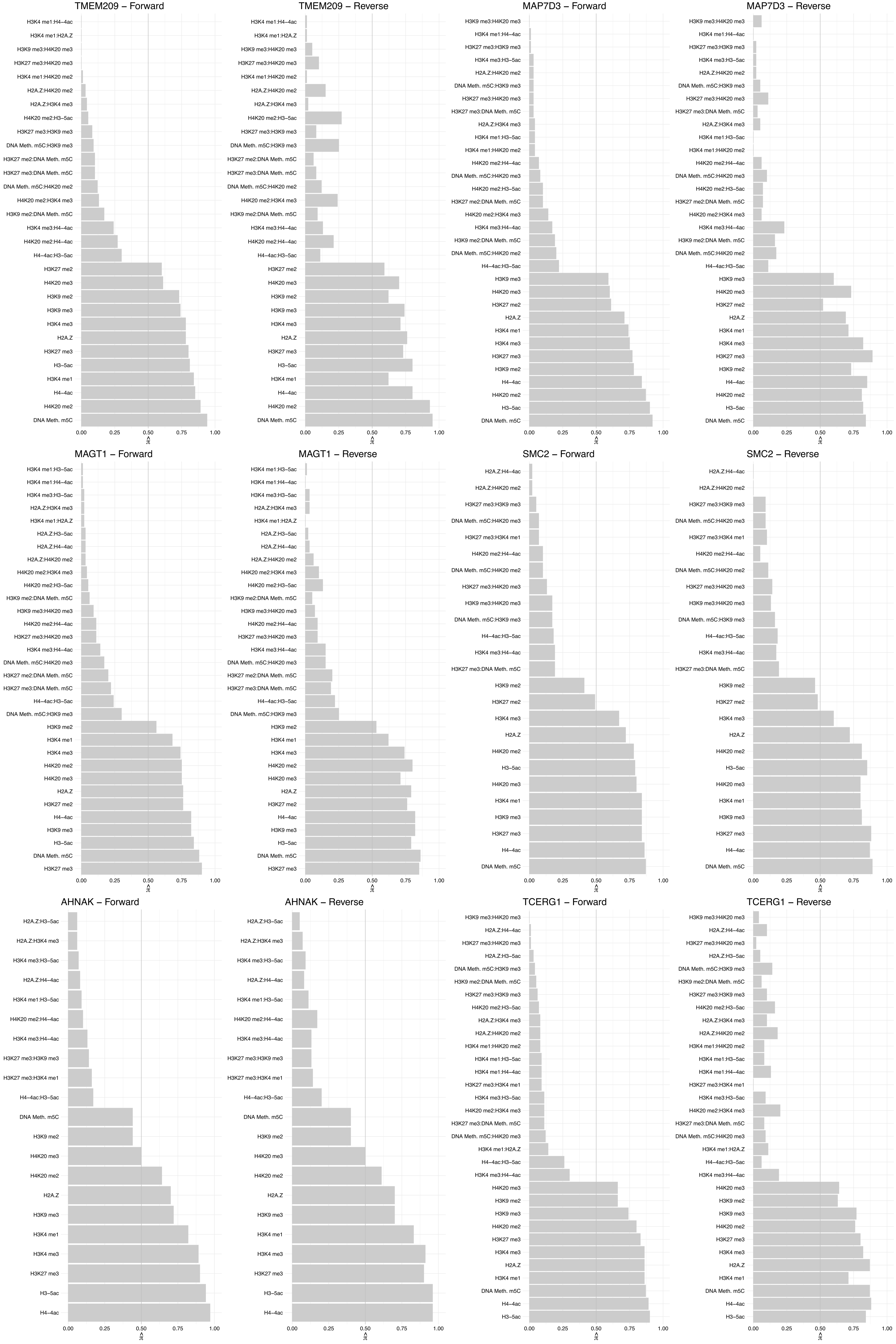
TMEM209 – Forward
H3K4 me1:H4–4ac
H3K4 me1:H2A.Z
H3K9 me3:H4K20 me3
H3K27 me3:H4K20 me3
H3K4 me1:H4K20 me2
H2A.Z:H4K20 me2
H2A.Z:H3K4 me3
H4K20 me2:H3–5ac
H3K27 me3:H3K9 me3
DNA Meth. m5C:H3K9 me3
H3K27 me2:DNA Meth. m5C
H3K27 me3:DNA Meth. m5C
DNA Meth. m5C:H4K20 me2
H4K20 me2:H3K4 me3
H3K9 me2:DNA Meth. m5C
H3K4 me3:H4–4ac
H4K20 me2:H4–4ac
H4–4ac:H3–5ac
H3K27 me2
H4K20 me3
H3K9 me2
H3K9 me3
H3K4 me3
H2A.Z
H3K27 me3
H3–5ac
H3K4 me1
H4–4ac
H4K20 me2
DNA Meth. m5C
0.00 0.25 0.50 0.75 1.00
π̂
TMEM209 – Reverse
MAP7D3 – Forward
H3K9 me3:H4K20 me3
H3K4 me1:H4–4ac
H3K27 me3:H3K9 me3
H3K4 me3:H3–5ac
H2A.Z:H4K20 me2
DNA Meth. m5C:H3K9 me3
H3K27 me3:H4K20 me3
H3K27 me3:DNA Meth. m5C
H2A.Z:H3K4 me3
H3K4 me1:H3–5ac
H3K4 me1:H4K20 me2
H4K20 me2:H4–4ac
DNA Meth. m5C:H4K20 me3
H4K20 me2:H3–5ac
H3K27 me2:DNA Meth. m5C
H4K20 me2:H3K4 me3
H3K4 me3:H4–4ac
H3K9 me2:DNA Meth. m5C
DNA Meth. m5C:H4K20 me2
H4–4ac:H3–5ac
H3K9 me3
H4K20 me3
H3K27 me2
H2A.Z
H3K4 me1
H3K4 me3
H3K27 me3
H3K9 me2
H4–4ac
H4K20 me2
H3–5ac
DNA Meth. m5C
MAP7D3 – Reverse
MAGT1 – Forward
H3K4 me1:H3–5ac
H3K4 me1:H4–4ac
H3K4 me3:H3–5ac
H2A.Z:H3K4 me3
H3K4 me1:H2A.Z
H2A.Z:H3–5ac
H2A.Z:H4–4ac
H2A.Z:H4K20 me2
H4K20 me2:H3K4 me3
H4K20 me2:H3–5ac
H3K9 me2:DNA Meth. m5C
H3K9 me3:H4K20 me3
H4K20 me2:H4–4ac
H3K27 me3:H4K20 me3
H3K4 me3:H4–4ac
DNA Meth. m5C:H4K20 me3
H3K27 me2:DNA Meth. m5C
H3K27 me3:DNA Meth. m5C
H4–4ac:H3–5ac
DNA Meth. m5C:H3K9 me3
H3K9 me2
H3K4 me1
H3K4 me3
H4K20 me2
H4K20 me3
H2A.Z
H3K27 me2
H4–4ac
H3K9 me3
H3–5ac
DNA Meth. m5C
H3K27 me3
MAGT1 – Reverse
SMC2 – Forward
H2A.Z:H4–4ac
H2A.Z:H4K20 me2
H3K27 me3:H3K9 me3
DNA Meth. m5C:H4K20 me3
H3K27 me3:H3K4 me1
H4K20 me2:H4–4ac
DNA Meth. m5C:H4K20 me2
H3K27 me3:H4K20 me3
H3K9 me3:H4K20 me3
DNA Meth. m5C:H3K9 me3
H4–4ac:H3–5ac
H3K4 me3:H4–4ac
H3K27 me3:DNA Meth. m5C
H3K9 me2
H3K27 me2
H3K4 me3
H2A.Z
H4K20 me2
H3–5ac
H4K20 me3
H3K4 me1
H3K9 me3
H3K27 me3
H4–4ac
DNA Meth. m5C
SMC2 – Reverse
AHNAK – Forward
H2A.Z:H3–5ac
H2A.Z:H3K4 me3
H3K4 me3:H3–5ac
H2A.Z:H4–4ac
H3K4 me1:H3–5ac
H4K20 me2:H4–4ac
H3K4 me3:H4–4ac
H3K27 me3:H3K9 me3
H3K27 me3:H3K4 me1
H4–4ac:H3–5ac
DNA Meth. m5C
H3K9 me2
H4K20 me3
H4K20 me2
H2A.Z
H3K9 me3
H3K4 me1
H3K4 me3
H3K27 me3
H3–5ac
H4–4ac
AHNAK – Reverse
TCERG1 – Forward
H3K9 me3:H4K20 me3
H2A.Z:H4–4ac
H3K27 me3:H4K20 me3
H2A.Z:H3–5ac
DNA Meth. m5C:H3K9 me3
H3K9 me2:DNA Meth. m5C
H3K27 me3:H3K9 me3
H4K20 me2:H3–5ac
H2A.Z:H3K4 me3
H2A.Z:H4K20 me2
H3K4 me1:H4K20 me2
H3K4 me1:H3–5ac
H3K4 me1:H4–4ac
H3K27 me3:H3K4 me1
H3K4 me3:H3–5ac
H4K20 me2:H3K4 me3
H3K27 me3:DNA Meth. m5C
DNA Meth. m5C:H4K20 me3
H3K4 me1:H2A.Z
H4–4ac:H3–5ac
H3K4 me3:H4–4ac
H4K20 me3
H3K9 me2
H3K9 me3
H4K20 me2
H3K27 me3
H3K4 me3
H2A.Z
H3K4 me1
DNA Meth. m5C
H4–4ac
H3–5ac
TCERG1 – Reverse

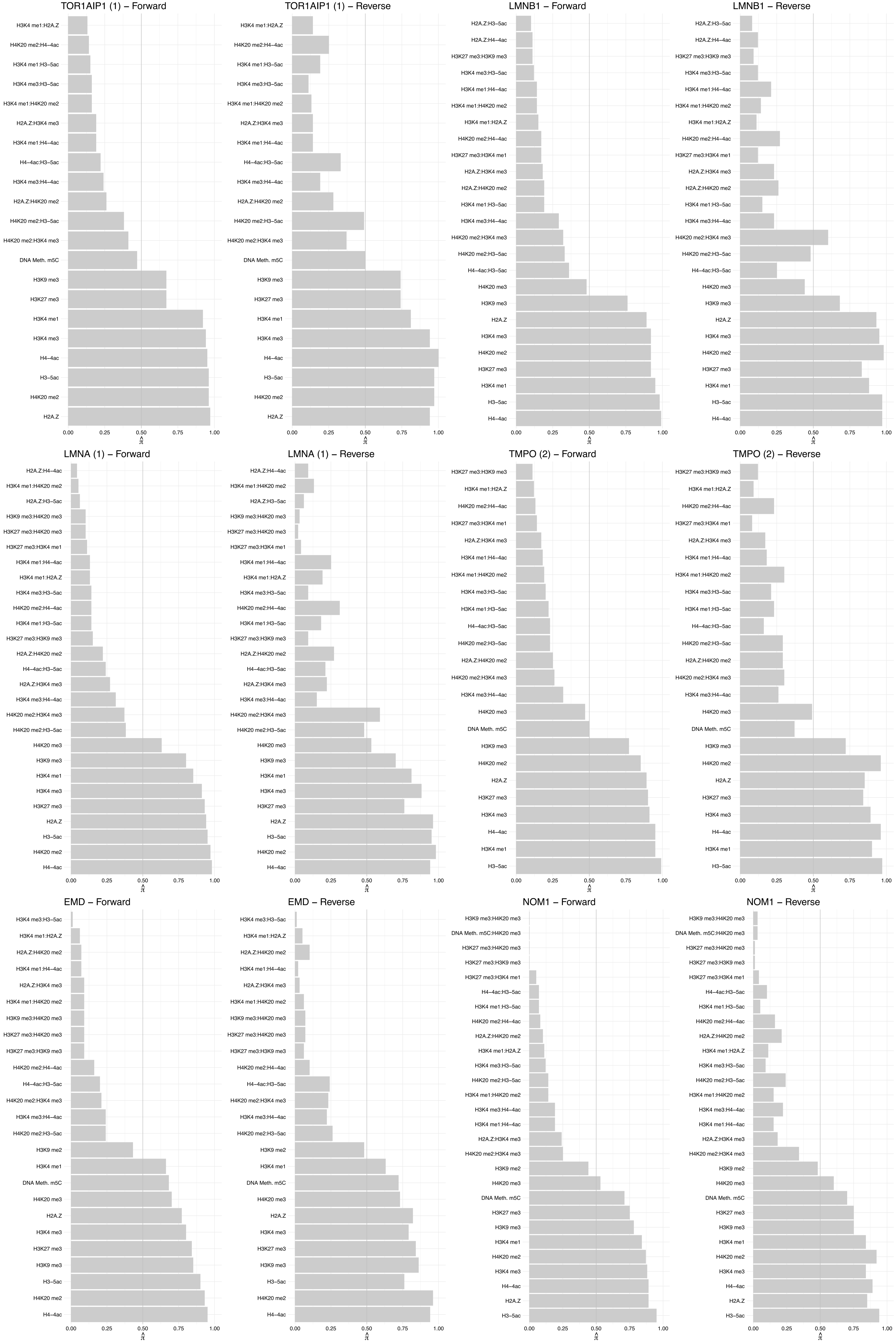

TOR1AIP1 (1) – Forward
H3K4 me1:H2A.Z
H4K20 me2:H4−4ac
H3K4 me1:H3−5ac
H3K4 me3:H3−5ac
H3K4 me1:H4K20 me2
H2A.Z:H3K4 me3
H3K4 me1:H4−4ac
H4−4ac:H3−5ac
H3K4 me3:H4−4ac
H2A.Z:H4K20 me2
H4K20 me2:H3−5ac
H4K20 me2:H3K4 me3
DNA Meth. m5C
H3K9 me3
H3K27 me3
H3K4 me1
H3K4 me3
H4−4ac
H3−5ac
H4K20 me2
H2A.Z
0.00 0.25 0.50 0.75 1.00
π̂
TOR1AIP1 (1) – Reverse
H3K4 me1:H2A.Z
H4K20 me2:H4−4ac
H3K4 me1:H3−5ac
H3K4 me3:H3−5ac
H3K4 me1:H4K20 me2
H2A.Z:H3K4 me3
H3K4 me1:H4−4ac
H4−4ac:H3−5ac
H3K4 me3:H4−4ac
H2A.Z:H4K20 me2
H4K20 me2:H3−5ac
H4K20 me2:H3K4 me3
DNA Meth. m5C
H3K9 me3
H3K27 me3
H3K4 me1
H3K4 me3
H4−4ac
H3−5ac
H4K20 me2
H2A.Z
0.00 0.25 0.50 0.75 1.00
π̂
LMNB1 – Forward
H2A.Z:H3−5ac
H2A.Z:H4−4ac
H3K27 me3:H3K9 me3
H3K4 me3:H3−5ac
H3K4 me1:H4−4ac
H3K4 me1:H4K20 me2
H3K4 me1:H2A.Z
H4K20 me2:H4−4ac
H3K27 me3:H3K4 me1
H2A.Z:H3K4 me3
H2A.Z:H4K20 me2
H3K4 me1:H3−5ac
H3K4 me3:H4−4ac
H4K20 me2:H3K4 me3
H4K20 me2:H3−5ac
H4−4ac:H3−5ac
H4K20 me3
H3K9 me3
H2A.Z
H3K4 me3
H4K20 me2
H3K27 me3
H3K4 me1
H3−5ac
H4−4ac
0.00 0.25 0.50 0.75 1.00
π̂
LMNB1 – Reverse
H2A.Z:H3−5ac
H2A.Z:H4−4ac
H3K27 me3:H3K9 me3
H3K4 me3:H3−5ac
H3K4 me1:H4−4ac
H3K4 me1:H4K20 me2
H3K4 me1:H2A.Z
H4K20 me2:H4−4ac
H3K27 me3:H3K4 me1
H2A.Z:H3K4 me3
H2A.Z:H4K20 me2
H3K4 me1:H3−5ac
H3K4 me3:H4−4ac
H4K20 me2:H3K4 me3
H4K20 me2:H3−5ac
H4−4ac:H3−5ac
H4K20 me3
H3K9 me3
H2A.Z
H3K4 me3
H4K20 me2
H3K27 me3
H3K4 me1
H3−5ac
H4−4ac
0.00 0.25 0.50 0.75 1.00
π̂
LMNA (1) – Forward
H2A.Z:H4−4ac
H3K4 me1:H4K20 me2
H2A.Z:H3−5ac
H3K9 me3:H4K20 me3
H3K27 me3:H4K20 me3
H3K27 me3:H3K4 me1
H3K4 me1:H4−4ac
H3K4 me1:H2A.Z
H3K4 me3:H3−5ac
H4K20 me2:H4−4ac
H3K4 me1:H3−5ac
H3K27 me3:H3K9 me3
H2A.Z:H4K20 me2
H4−4ac:H3−5ac
H2A.Z:H3K4 me3
H3K4 me3:H4−4ac
H4K20 me2:H3K4 me3
H4K20 me2:H3−5ac
H4K20 me3
H3K9 me3
H3K4 me1
H3K4 me3
H3K27 me3
H2A.Z
H3−5ac
H4K20 me2
H4−4ac
0.00 0.25 0.50 0.75 1.00
π̂
LMNA (1) – Reverse
H2A.Z:H4−4ac
H3K4 me1:H4K20 me2
H2A.Z:H3−5ac
H3K9 me3:H4K20 me3
H3K27 me3:H4K20 me3
H3K27 me3:H3K4 me1
H3K4 me1:H4−4ac
H3K4 me1:H2A.Z
H3K4 me3:H3−5ac
H4K20 me2:H4−4ac
H3K4 me1:H3−5ac
H3K27 me3:H3K9 me3
H2A.Z:H4K20 me2
H4−4ac:H3−5ac
H2A.Z:H3K4 me3
H3K4 me3:H4−4ac
H4K20 me2:H3K4 me3
H4K20 me2:H3−5ac
H4K20 me3
H3K9 me3
H3K4 me1
H3K4 me3
H3K27 me3
H2A.Z
H3−5ac
H4K20 me2
H4−4ac
0.00 0.25 0.50 0.75 1.00
π̂
TMPO (2) – Forward
H3K27 me3:H3K9 me3
H3K4 me1:H2A.Z
H4K20 me2:H4−4ac
H3K27 me3:H3K4 me1
H2A.Z:H3K4 me3
H3K4 me1:H4−4ac
H3K4 me1:H4K20 me2
H3K4 me3:H3−5ac
H3K4 me1:H3−5ac
H4−4ac:H3−5ac
H4K20 me2:H3−5ac
H2A.Z:H4K20 me2
H4K20 me2:H3K4 me3
H3K4 me3:H4−4ac
H4K20 me3
DNA Meth. m5C
H3K9 me3
H4K20 me2
H2A.Z
H3K27 me3
H3K4 me3
H4−4ac
H3K4 me1
H3−5ac
0.00 0.25 0.50 0.75 1.00
π̂
TMPO (2) – Reverse
H3K27 me3:H3K9 me3
H3K4 me1:H2A.Z
H4K20 me2:H4−4ac
H3K27 me3:H3K4 me1
H2A.Z:H3K4 me3
H3K4 me1:H4−4ac
H3K4 me1:H4K20 me2
H3K4 me3:H3−5ac
H3K4 me1:H3−5ac
H4−4ac:H3−5ac
H4K20 me2:H3−5ac
H2A.Z:H4K20 me2
H4K20 me2:H3K4 me3
H3K4 me3:H4−4ac
H4K20 me3
DNA Meth. m5C
H3K9 me3
H4K20 me2
H2A.Z
H3K27 me3
H3K4 me3
H4−4ac
H3K4 me1
H3−5ac
0.00 0.25 0.50 0.75 1.00
π̂
EMD – Forward
H3K4 me3:H3−5ac
H3K4 me1:H2A.Z
H2A.Z:H4K20 me2
H3K4 me1:H4−4ac
H2A.Z:H3K4 me3
H3K4 me1:H4K20 me2
H3K9 me3:H4K20 me3
H3K27 me3:H4K20 me3
H3K27 me3:H3K9 me3
H4K20 me2:H4−4ac
H4−4ac:H3−5ac
H4K20 me2:H3K4 me3
H3K4 me3:H4−4ac
H4K20 me2:H3−5ac
H3K9 me2
H3K4 me1
DNA Meth. m5C
H4K20 me3
H2A.Z
H3K4 me3
H3K27 me3
H3K9 me3
H3−5ac
H4K20 me2
H4−4ac
0.00 0.25 0.50 0.75 1.00
π̂
EMD – Reverse
H3K4 me3:H3−5ac
H3K4 me1:H2A.Z
H2A.Z:H4K20 me2
H3K4 me1:H4−4ac
H2A.Z:H3K4 me3
H3K4 me1:H4K20 me2
H3K9 me3:H4K20 me3
H3K27 me3:H4K20 me3
H3K27 me3:H3K9 me3
H4K20 me2:H4−4ac
H4−4ac:H3−5ac
H4K20 me2:H3K4 me3
H3K4 me3:H4−4ac
H4K20 me2:H3−5ac
H3K9 me2
H3K4 me1
DNA Meth. m5C
H4K20 me3
H2A.Z
H3K4 me3
H3K27 me3
H3K9 me3
H3−5ac
H4K20 me2
H4−4ac
0.00 0.25 0.50 0.75 1.00
π̂
NOM1 – Forward
H3K9 me3:H4K20 me3
DNA Meth. m5C:H4K20 me3
H3K27 me3:H4K20 me3
H3K27 me3:H3K9 me3
H3K27 me3:H3K4 me1
H4−4ac:H3−5ac
H3K4 me1:H3−5ac
H4K20 me2:H4−4ac
H2A.Z:H4K20 me2
H3K4 me1:H2A.Z
H3K4 me3:H3−5ac
H4K20 me2:H3−5ac
H3K4 me1:H4K20 me2
H3K4 me3:H4−4ac
H3K4 me1:H4−4ac
H2A.Z:H3K4 me3
H4K20 me2:H3K4 me3
H3K9 me2
H4K20 me3
DNA Meth. m5C
H3K27 me3
H3K9 me3
H3K4 me1
H4K20 me2
H3K4 me3
H4−4ac
H2A.Z
H3−5ac
0.00 0.25 0.50 0.75 1.00
π̂
NOM1 – Reverse
H3K9 me3:H4K20 me3
DNA Meth. m5C:H4K20 me3
H3K27 me3:H4K20 me3
H3K27 me3:H3K9 me3
H3K27 me3:H3K4 me1
H4−4ac:H3−5ac
H3K4 me1:H3−5ac
H4K20 me2:H4−4ac
H2A.Z:H4K20 me2
H3K4 me1:H2A.Z
H3K4 me3:H3−5ac
H4K20 me2:H3−5ac
H3K4 me1:H4K20 me2
H3K4 me3:H4−4ac
H3K4 me1:H4−4ac
H2A.Z:H3K4 me3
H4K20 me2:H3K4 me3
H3K9 me2
H4K20 me3
DNA Meth. m5C
H3K27 me3
H3K9 me3
H3K4 me1
H4K20 me2
H3K4 me3
H4−4ac
H2A.Z
H3−5ac
0.00 0.25 0.50 0.75 1.00
π̂

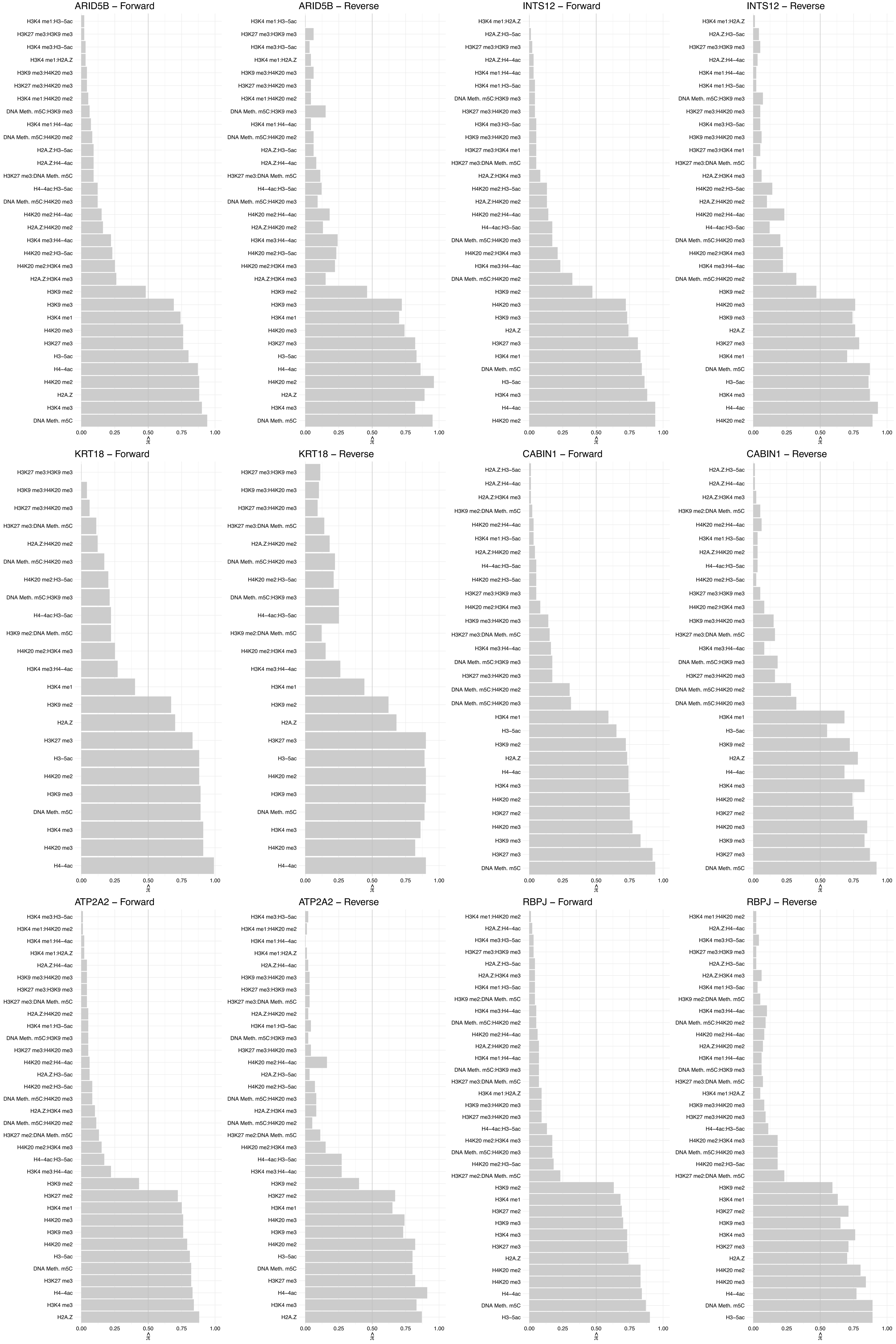

ARID5B – Forward
H3K4 me1:H3–5ac
H3K27 me3:H3K9 me3
H3K4 me3:H3–5ac
H3K4 me1:H2A.Z
H3K9 me3:H4K20 me3
H3K27 me3:H4K20 me3
H3K4 me1:H4K20 me2
DNA Meth. m5C:H3K9 me3
H3K4 me1:H4–4ac
DNA Meth. m5C:H4K20 me2
H2A.Z:H3–5ac
H2A.Z:H4–4ac
H3K27 me3:DNA Meth. m5C
H4–4ac:H3–5ac
DNA Meth. m5C:H4K20 me3
H4K20 me2:H4–4ac
H2A.Z:H4K20 me2
H3K4 me3:H4–4ac
H4K20 me2:H3–5ac
H4K20 me2:H3K4 me3
H2A.Z:H3K4 me3
H3K9 me2
H3K9 me3
H3K4 me1
H4K20 me3
H3K27 me3
H3–5ac
H4–4ac
H4K20 me2
H2A.Z
H3K4 me3
DNA Meth. m5C
0.00 0.25 0.50 0.75 1.00
π̂
ARID5B – Reverse
INTS12 – Forward
H3K4 me1:H2A.Z
H2A.Z:H3–5ac
H3K27 me3:H3K9 me3
H2A.Z:H4–4ac
H3K4 me1:H4–4ac
H3K4 me1:H3–5ac
DNA Meth. m5C:H3K9 me3
H3K27 me3:H4K20 me3
H3K4 me3:H3–5ac
H3K9 me3:H4K20 me3
H3K27 me3:H3K4 me1
H3K27 me3:DNA Meth. m5C
H2A.Z:H3K4 me3
H4K20 me2:H3–5ac
H2A.Z:H4K20 me2
H4K20 me2:H4–4ac
H4–4ac:H3–5ac
DNA Meth. m5C:H4K20 me3
H4K20 me2:H3K4 me3
H3K4 me3:H4–4ac
DNA Meth. m5C:H4K20 me2
H3K9 me2
H4K20 me3
H3K9 me3
H2A.Z
H3K27 me3
H3K4 me1
DNA Meth. m5C
H3–5ac
H3K4 me3
H4–4ac
H4K20 me2
INTS12 – Reverse
KRT18 – Forward
H3K27 me3:H3K9 me3
H3K9 me3:H4K20 me3
H3K27 me3:H4K20 me3
H3K27 me3:DNA Meth. m5C
H2A.Z:H4K20 me2
DNA Meth. m5C:H4K20 me3
H4K20 me2:H3–5ac
DNA Meth. m5C:H3K9 me3
H4–4ac:H3–5ac
H3K9 me2:DNA Meth. m5C
H4K20 me2:H3K4 me3
H3K4 me3:H4–4ac
H3K4 me1
H3K9 me2
H2A.Z
H3K27 me3
H3–5ac
H4K20 me2
H3K9 me3
DNA Meth. m5C
H3K4 me3
H4K20 me3
H4–4ac
KRT18 – Reverse
CABIN1 – Forward
H2A.Z:H3–5ac
H2A.Z:H4–4ac
H2A.Z:H3K4 me3
H3K9 me2:DNA Meth. m5C
H4K20 me2:H4–4ac
H3K4 me1:H3–5ac
H2A.Z:H4K20 me2
H4–4ac:H3–5ac
H4K20 me2:H3–5ac
H3K27 me3:H3K9 me3
H4K20 me2:H3K4 me3
H3K9 me3:H4K20 me3
H3K27 me3:DNA Meth. m5C
H3K4 me3:H4–4ac
DNA Meth. m5C:H3K9 me3
H3K27 me3:H4K20 me3
DNA Meth. m5C:H4K20 me2
DNA Meth. m5C:H4K20 me3
H3K4 me1
H3–5ac
H3K9 me2
H2A.Z
H4–4ac
H3K4 me3
H4K20 me2
H3K27 me2
H4K20 me3
H3K9 me3
H3K27 me3
DNA Meth. m5C
CABIN1 – Reverse
ATP2A2 – Forward
H3K4 me3:H3–5ac
H3K4 me1:H4K20 me2
H3K4 me1:H4–4ac
H3K4 me1:H2A.Z
H2A.Z:H4–4ac
H3K9 me3:H4K20 me3
H3K27 me3:H3K9 me3
H3K27 me3:DNA Meth. m5C
H2A.Z:H4K20 me2
H3K4 me1:H3–5ac
DNA Meth. m5C:H3K9 me3
H3K27 me3:H4K20 me3
H4K20 me2:H4–4ac
H2A.Z:H3–5ac
H4K20 me2:H3–5ac
DNA Meth. m5C:H4K20 me3
H2A.Z:H3K4 me3
DNA Meth. m5C:H4K20 me2
H3K27 me2:DNA Meth. m5C
H4K20 me2:H3K4 me3
H4–4ac:H3–5ac
H3K4 me3:H4–4ac
H3K9 me2
H3K27 me2
H3K4 me1
H4K20 me3
H3K9 me3
H4K20 me2
H3–5ac
DNA Meth. m5C
H3K27 me3
H4–4ac
H3K4 me3
H2A.Z
ATP2A2 – Reverse
RBPJ – Forward
H3K4 me1:H4K20 me2
H2A.Z:H4–4ac
H3K4 me3:H3–5ac
H3K27 me3:H3K9 me3
H2A.Z:H3–5ac
H2A.Z:H3K4 me3
H3K4 me1:H3–5ac
H3K9 me2:DNA Meth. m5C
H3K4 me3:H4–4ac
DNA Meth. m5C:H4K20 me2
H4K20 me2:H4–4ac
H2A.Z:H4K20 me2
H3K4 me1:H4–4ac
DNA Meth. m5C:H3K9 me3
H3K27 me3:DNA Meth. m5C
H3K4 me1:H2A.Z
H3K9 me3:H4K20 me3
H3K27 me3:H4K20 me3
H4–4ac:H3–5ac
H4K20 me2:H3K4 me3
DNA Meth. m5C:H4K20 me3
H4K20 me2:H3–5ac
H3K27 me2:DNA Meth. m5C
H3K9 me2
H3K4 me1
H3K27 me2
H3K9 me3
H3K4 me3
H3K27 me3
H2A.Z
H4K20 me2
H4K20 me3
H4–4ac
DNA Meth. m5C
H3–5ac
RBPJ – Reverse

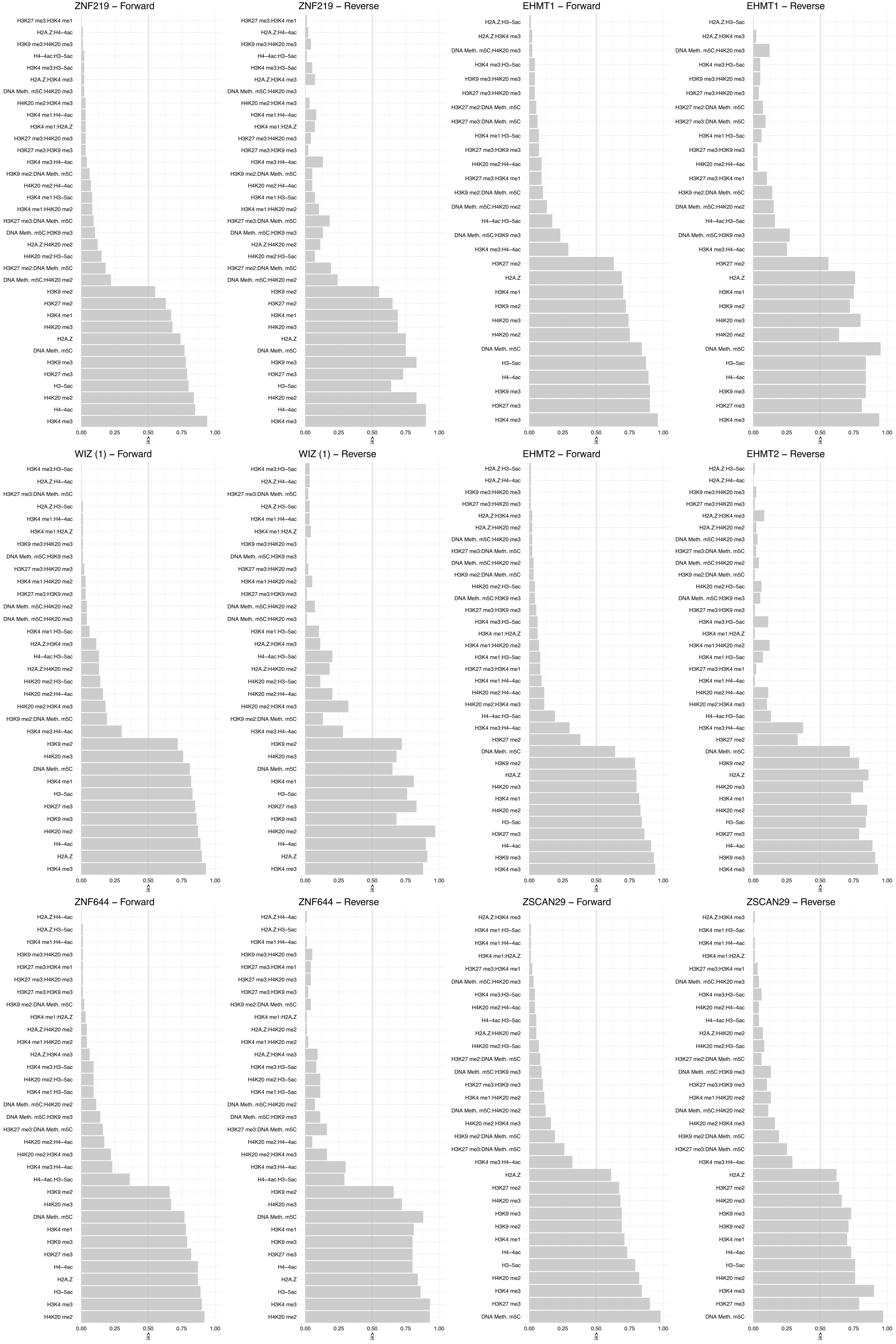
ZNF219 – Forward
H3K27 me3:H3K4 me1
H2A.Z:H4–4ac
H3K9 me3:H4K20 me3
H4–4ac:H3–5ac
H3K4 me3:H3–5ac
H2A.Z:H3K4 me3
DNA Meth. m5C:H4K20 me3
H4K20 me2:H3K4 me3
H3K4 me1:H4–4ac
H3K4 me1:H2A.Z
H3K27 me3:H4K20 me3
H3K27 me3:H3K9 me3
H3K4 me3:H4–4ac
H3K9 me2:DNA Meth. m5C
H4K20 me2:H4–4ac
H3K4 me1:H3–5ac
H3K4 me1:H4K20 me2
H3K27 me3:DNA Meth. m5C
DNA Meth. m5C:H3K9 me3
H2A.Z:H4K20 me2
H4K20 me2:H3–5ac
H3K27 me2:DNA Meth. m5C
DNA Meth. m5C:H4K20 me2
H3K9 me2
H3K27 me2
H3K4 me1
H4K20 me3
H2A.Z
DNA Meth. m5C
H3K9 me3
H3K27 me3
H3–5ac
H4K20 me2
H4–4ac
H3K4 me3
ZNF219 – Reverse
EHMT1 – Forward
H2A.Z:H3–5ac
H2A.Z:H3K4 me3
DNA Meth. m5C:H4K20 me3
H3K4 me3:H3–5ac
H3K9 me3:H4K20 me3
H3K27 me3:H4K20 me3
H3K27 me2:DNA Meth. m5C
H3K27 me3:DNA Meth. m5C
H3K4 me1:H3–5ac
H3K27 me3:H3K9 me3
H4K20 me2:H4–4ac
H3K27 me3:H3K4 me1
H3K9 me2:DNA Meth. m5C
DNA Meth. m5C:H4K20 me2
H4–4ac:H3–5ac
DNA Meth. m5C:H3K9 me3
H3K4 me3:H4–4ac
H3K27 me2
H2A.Z
H3K4 me1
H3K9 me2
H4K20 me3
H4K20 me2
DNA Meth. m5C
H3–5ac
H4–4ac
H3K9 me3
H3K27 me3
H3K4 me3
EHMT1 – Reverse
WIZ (1) – Forward
H3K4 me3:H3–5ac
H2A.Z:H4–4ac
H3K27 me3:DNA Meth. m5C
H2A.Z:H3–5ac
H3K4 me1:H4–4ac
H3K4 me1:H2A.Z
H3K9 me3:H4K20 me3
DNA Meth. m5C:H3K9 me3
H3K27 me3:H4K20 me3
H3K4 me1:H4K20 me2
H3K27 me3:H3K9 me3
DNA Meth. m5C:H4K20 me2
DNA Meth. m5C:H4K20 me3
H3K4 me1:H3–5ac
H2A.Z:H3K4 me3
H4–4ac:H3–5ac
H2A.Z:H4K20 me2
H4K20 me2:H3–5ac
H4K20 me2:H4–4ac
H4K20 me2:H3K4 me3
H3K9 me2:DNA Meth. m5C
H3K4 me3:H4–4ac
H3K9 me2
H4K20 me3
DNA Meth. m5C
H3K4 me1
H3–5ac
H3K27 me3
H3K9 me3
H4K20 me2
H4–4ac
H2A.Z
H3K4 me3
WIZ (1) – Reverse
EHMT2 – Forward
H2A.Z:H3–5ac
H2A.Z:H4–4ac
H3K9 me3:H4K20 me3
H3K27 me3:H4K20 me3
H2A.Z:H3K4 me3
H2A.Z:H4K20 me2
DNA Meth. m5C:H4K20 me3
H3K27 me3:DNA Meth. m5C
DNA Meth. m5C:H4K20 me2
H3K9 me2:DNA Meth. m5C
H4K20 me2:H3–5ac
DNA Meth. m5C:H3K9 me3
H3K27 me3:H3K9 me3
H3K4 me3:H3–5ac
H3K4 me1:H2A.Z
H3K4 me1:H4K20 me2
H3K4 me1:H3–5ac
H3K27 me3:H3K4 me1
H3K4 me1:H4–4ac
H4K20 me2:H4–4ac
H4K20 me2:H3K4 me3
H4–4ac:H3–5ac
H3K4 me3:H4–4ac
H3K27 me2
DNA Meth. m5C
H3K9 me2
H2A.Z
H4K20 me3
H3K4 me1
H4K20 me2
H3–5ac
H3K27 me3
H4–4ac
H3K9 me3
H3K4 me3
EHMT2 – Reverse
ZNF644 – Forward
H2A.Z:H4–4ac
H2A.Z:H3–5ac
H3K4 me1:H4–4ac
H3K9 me3:H4K20 me3
H3K27 me3:H3K4 me1
H3K27 me3:H4K20 me3
H3K27 me3:H3K9 me3
H3K9 me2:DNA Meth. m5C
H3K4 me1:H2A.Z
H2A.Z:H4K20 me2
H3K4 me1:H4K20 me2
H2A.Z:H3K4 me3
H3K4 me3:H3–5ac
H4K20 me2:H3–5ac
H3K4 me1:H3–5ac
DNA Meth. m5C:H4K20 me2
DNA Meth. m5C:H3K9 me3
H3K27 me3:DNA Meth. m5C
H4K20 me2:H4–4ac
H4K20 me2:H3K4 me3
H3K4 me3:H4–4ac
H4–4ac:H3–5ac
H3K9 me2
H4K20 me3
DNA Meth. m5C
H3K4 me1
H3K9 me3
H3K27 me3
H4–4ac
H2A.Z
H3–5ac
H3K4 me3
H4K20 me2
ZNF644 – Reverse
ZSCAN29 – Forward
H2A.Z:H3K4 me3
H3K4 me1:H3–5ac
H3K4 me1:H4–4ac
H3K4 me1:H2A.Z
H3K27 me3:H3K4 me1
DNA Meth. m5C:H4K20 me3
H3K4 me3:H3–5ac
H4K20 me2:H4–4ac
H4–4ac:H3–5ac
H2A.Z:H4K20 me2
H4K20 me2:H3–5ac
H3K27 me2:DNA Meth. m5C
DNA Meth. m5C:H3K9 me3
H3K27 me3:H3K9 me3
H3K4 me1:H4K20 me2
DNA Meth. m5C:H4K20 me2
H4K20 me2:H3K4 me3
H3K9 me2:DNA Meth. m5C
H3K27 me3:DNA Meth. m5C
H3K4 me3:H4–4ac
H2A.Z
H3K27 me2
H4K20 me3
H3K9 me3
H3K9 me2
H3K4 me1
H4–4ac
H3–5ac
H4K20 me2
H3K4 me3
H3K27 me3
DNA Meth. m5C
ZSCAN29 – Reverse
0.00
0.25
0.50
0.75
1.00
π̂

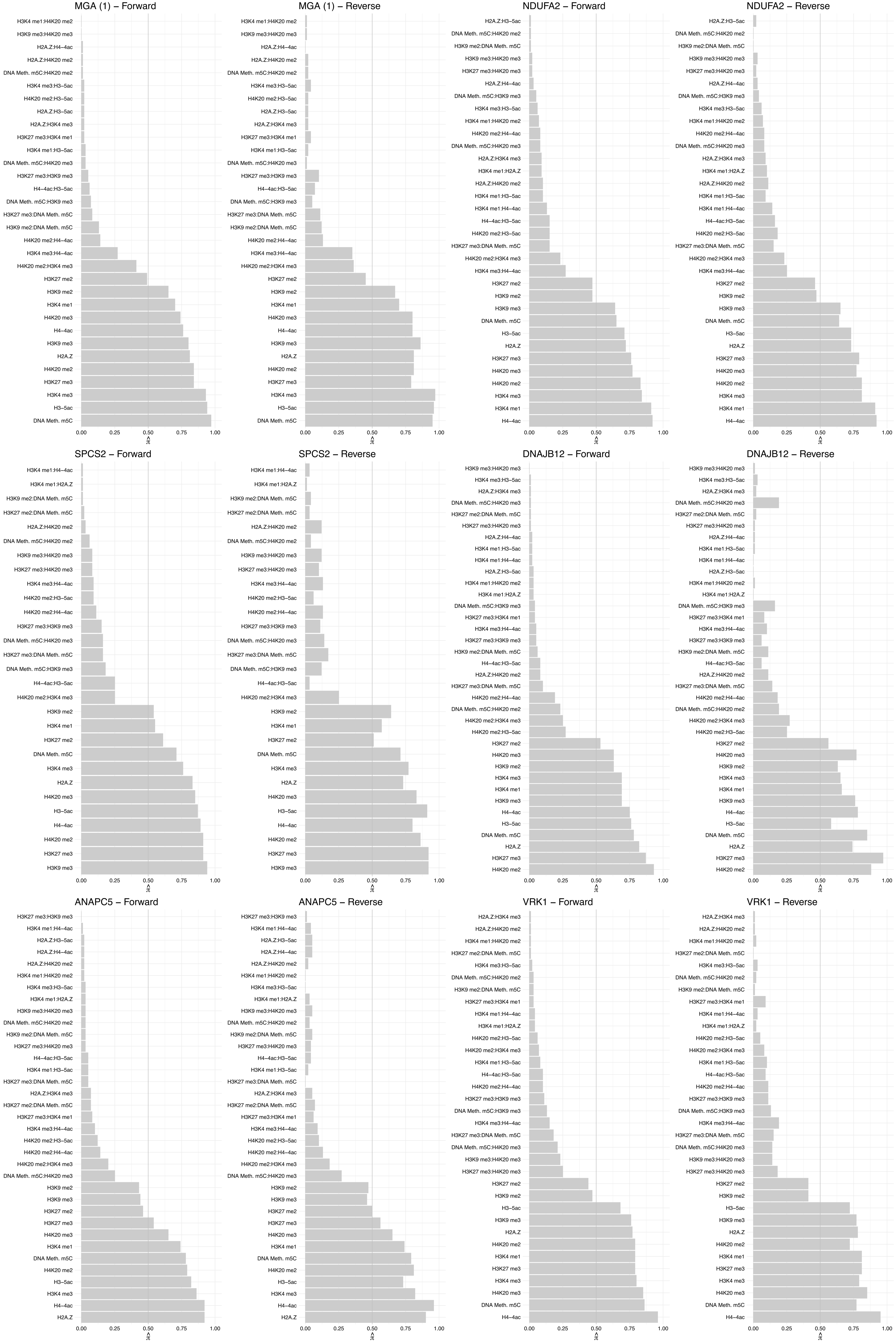
MGA (1) – Forward
H3K4 me1:H4K20 me2
H3K9 me3:H4K20 me3
H2A.Z:H4–4ac
H2A.Z:H4K20 me2
DNA Meth. m5C:H4K20 me2
H3K4 me3:H3–5ac
H4K20 me2:H3–5ac
H2A.Z:H3–5ac
H2A.Z:H3K4 me3
H3K27 me3:H3K4 me1
H3K4 me1:H3–5ac
DNA Meth. m5C:H4K20 me3
H3K27 me3:H3K9 me3
H4–4ac:H3–5ac
DNA Meth. m5C:H3K9 me3
H3K27 me3:DNA Meth. m5C
H3K9 me2:DNA Meth. m5C
H4K20 me2:H4–4ac
H3K4 me3:H4–4ac
H4K20 me2:H3K4 me3
H3K27 me2
H3K9 me2
H3K4 me1
H4K20 me3
H4–4ac
H3K9 me3
H2A.Z
H4K20 me2
H3K27 me3
H3K4 me3
H3–5ac
DNA Meth. m5C
0.00 0.25 0.50 0.75 1.00
$\hat{\pi}$
MGA (1) – Reverse
H3K4 me1:H4K20 me2
H3K9 me3:H4K20 me3
H2A.Z:H4–4ac
H2A.Z:H4K20 me2
DNA Meth. m5C:H4K20 me2
H3K4 me3:H3–5ac
H4K20 me2:H3–5ac
H2A.Z:H3–5ac
H2A.Z:H3K4 me3
H3K27 me3:H3K4 me1
H3K4 me1:H3–5ac
DNA Meth. m5C:H4K20 me3
H3K27 me3:H3K9 me3
H4–4ac:H3–5ac
DNA Meth. m5C:H3K9 me3
H3K27 me3:DNA Meth. m5C
H3K9 me2:DNA Meth. m5C
H4K20 me2:H4–4ac
H3K4 me3:H4–4ac
H4K20 me2:H3K4 me3
H3K27 me2
H3K9 me2
H3K4 me1
H4K20 me3
H4–4ac
H3K9 me3
H2A.Z
H4K20 me2
H3K27 me3
H3K4 me3
H3–5ac
DNA Meth. m5C
0.00 0.25 0.50 0.75 1.00
$\hat{\pi}$
NDUFA2 – Forward
H2A.Z:H3–5ac
DNA Meth. m5C:H4K20 me2
H3K9 me2:DNA Meth. m5C
H3K9 me3:H4K20 me3
H3K27 me3:H4K20 me3
H2A.Z:H4–4ac
DNA Meth. m5C:H3K9 me3
H3K4 me3:H3–5ac
H3K4 me1:H4K20 me2
H4K20 me2:H4–4ac
DNA Meth. m5C:H4K20 me3
H2A.Z:H3K4 me3
H3K4 me1:H2A.Z
H2A.Z:H4K20 me2
H3K4 me1:H3–5ac
H3K4 me1:H4–4ac
H4–4ac:H3–5ac
H4K20 me2:H3–5ac
H3K27 me3:DNA Meth. m5C
H4K20 me2:H3K4 me3
H3K4 me3:H4–4ac
H3K27 me2
H3K9 me2
H3K9 me3
DNA Meth. m5C
H3–5ac
H2A.Z
H3K27 me3
H4K20 me3
H4K20 me2
H3K4 me3
H3K4 me1
H4–4ac
0.00 0.25 0.50 0.75 1.00
$\hat{\pi}$
NDUFA2 – Reverse
H2A.Z:H3–5ac
DNA Meth. m5C:H4K20 me2
H3K9 me2:DNA Meth. m5C
H3K9 me3:H4K20 me3
H3K27 me3:H4K20 me3
H2A.Z:H4–4ac
DNA Meth. m5C:H3K9 me3
H3K4 me3:H3–5ac
H3K4 me1:H4K20 me2
H4K20 me2:H4–4ac
DNA Meth. m5C:H4K20 me3
H2A.Z:H3K4 me3
H3K4 me1:H2A.Z
H2A.Z:H4K20 me2
H3K4 me1:H3–5ac
H3K4 me1:H4–4ac
H4–4ac:H3–5ac
H4K20 me2:H3–5ac
H3K27 me3:DNA Meth. m5C
H4K20 me2:H3K4 me3
H3K4 me3:H4–4ac
H3K27 me2
H3K9 me2
H3K9 me3
DNA Meth. m5C
H3–5ac
H2A.Z
H3K27 me3
H4K20 me3
H4K20 me2
H3K4 me3
H3K4 me1
H4–4ac
0.00 0.25 0.50 0.75 1.00
$\hat{\pi}$
SPCS2 – Forward
H3K4 me1:H4–4ac
H3K4 me1:H2A.Z
H3K9 me2:DNA Meth. m5C
H3K27 me2:DNA Meth. m5C
H2A.Z:H4K20 me2
DNA Meth. m5C:H4K20 me2
H3K9 me3:H4K20 me3
H3K27 me3:H4K20 me3
H3K4 me3:H4–4ac
H4K20 me2:H3–5ac
H4K20 me2:H4–4ac
H3K27 me3:H3K9 me3
DNA Meth. m5C:H4K20 me3
H3K27 me3:DNA Meth. m5C
DNA Meth. m5C:H3K9 me3
H4–4ac:H3–5ac
H4K20 me2:H3K4 me3
H3K9 me2
H3K4 me1
H3K27 me2
DNA Meth. m5C
H3K4 me3
H2A.Z
H4K20 me3
H3–5ac
H4–4ac
H4K20 me2
H3K27 me3
H3K9 me3
0.00 0.25 0.50 0.75 1.00
$\hat{\pi}$
SPCS2 – Reverse
H3K4 me1:H4–4ac
H3K4 me1:H2A.Z
H3K9 me2:DNA Meth. m5C
H3K27 me2:DNA Meth. m5C
H2A.Z:H4K20 me2
DNA Meth. m5C:H4K20 me2
H3K9 me3:H4K20 me3
H3K27 me3:H4K20 me3
H3K4 me3:H4–4ac
H4K20 me2:H3–5ac
H4K20 me2:H4–4ac
H3K27 me3:H3K9 me3
DNA Meth. m5C:H4K20 me3
H3K27 me3:DNA Meth. m5C
DNA Meth. m5C:H3K9 me3
H4–4ac:H3–5ac
H4K20 me2:H3K4 me3
H3K9 me2
H3K4 me1
H3K27 me2
DNA Meth. m5C
H3K4 me3
H2A.Z
H4K20 me3
H3–5ac
H4–4ac
H4K20 me2
H3K27 me3
H3K9 me3
0.00 0.25 0.50 0.75 1.00
$\hat{\pi}$
DNAJB12 – Forward
H3K9 me3:H4K20 me3
H3K4 me3:H3–5ac
H2A.Z:H3K4 me3
DNA Meth. m5C:H4K20 me3
H3K27 me2:DNA Meth. m5C
H3K27 me3:H4K20 me3
H2A.Z:H4–4ac
H3K4 me1:H3–5ac
H3K4 me1:H4–4ac
H2A.Z:H3–5ac
H3K4 me1:H4K20 me2
H3K4 me1:H2A.Z
DNA Meth. m5C:H3K9 me3
H3K27 me3:H3K4 me1
H3K4 me3:H4–4ac
H3K27 me3:H3K9 me3
H3K9 me2:DNA Meth. m5C
H4–4ac:H3–5ac
H2A.Z:H4K20 me2
H3K27 me3:DNA Meth. m5C
H4K20 me2:H4–4ac
DNA Meth. m5C:H4K20 me2
H4K20 me2:H3K4 me3
H4K20 me2:H3–5ac
H3K27 me2
H4K20 me3
H3K9 me2
H3K4 me3
H3K4 me1
H3K9 me3
H4–4ac
H3–5ac
DNA Meth. m5C
H2A.Z
H3K27 me3
H4K20 me2
0.00 0.25 0.50 0.75 1.00
$\hat{\pi}$
DNAJB12 – Reverse
H3K9 me3:H4K20 me3
H3K4 me3:H3–5ac
H2A.Z:H3K4 me3
DNA Meth. m5C:H4K20 me3
H3K27 me2:DNA Meth. m5C
H3K27 me3:H4K20 me3
H2A.Z:H4–4ac
H3K4 me1:H3–5ac
H3K4 me1:H4–4ac
H2A.Z:H3–5ac
H3K4 me1:H4K20 me2
H3K4 me1:H2A.Z
DNA Meth. m5C:H3K9 me3
H3K27 me3:H3K4 me1
H3K4 me3:H4–4ac
H3K27 me3:H3K9 me3
H3K9 me2:DNA Meth. m5C
H4–4ac:H3–5ac
H2A.Z:H4K20 me2
H3K27 me3:DNA Meth. m5C
H4K20 me2:H4–4ac
DNA Meth. m5C:H4K20 me2
H4K20 me2:H3K4 me3
H4K20 me2:H3–5ac
H3K27 me2
H4K20 me3
H3K9 me2
H3K4 me3
H3K4 me1
H3K9 me3
H4–4ac
H3–5ac
DNA Meth. m5C
H2A.Z
H3K27 me3
H4K20 me2
0.00 0.25 0.50 0.75 1.00
$\hat{\pi}$
ANAPC5 – Forward
H3K27 me3:H3K9 me3
H3K4 me1:H4–4ac
H2A.Z:H3–5ac
H2A.Z:H4–4ac
H2A.Z:H4K20 me2
H3K4 me1:H4K20 me2
H3K4 me3:H3–5ac
H3K4 me1:H2A.Z
H3K9 me3:H4K20 me3
DNA Meth. m5C:H4K20 me2
H3K9 me2:DNA Meth. m5C
H3K27 me3:H4K20 me3
H4–4ac:H3–5ac
H3K4 me1:H3–5ac
H3K27 me3:DNA Meth. m5C
H2A.Z:H3K4 me3
H3K27 me2:DNA Meth. m5C
H3K27 me3:H3K4 me1
H3K4 me3:H4–4ac
H4K20 me2:H3–5ac
H4K20 me2:H4–4ac
H4K20 me2:H3K4 me3
DNA Meth. m5C:H4K20 me3
H3K9 me2
H3K9 me3
H3K27 me2
H3K27 me3
H4K20 me3
H3K4 me1
DNA Meth. m5C
H4K20 me2
H3–5ac
H3K4 me3
H4–4ac
H2A.Z
0.00 0.25 0.50 0.75 1.00
$\hat{\pi}$
ANAPC5 – Reverse
H3K27 me3:H3K9 me3
H3K4 me1:H4–4ac
H2A.Z:H3–5ac
H2A.Z:H4–4ac
H2A.Z:H4K20 me2
H3K4 me1:H4K20 me2
H3K4 me3:H3–5ac
H3K4 me1:H2A.Z
H3K9 me3:H4K20 me3
DNA Meth. m5C:H4K20 me2
H3K9 me2:DNA Meth. m5C
H3K27 me3:H4K20 me3
H4–4ac:H3–5ac
H3K4 me1:H3–5ac
H3K27 me3:DNA Meth. m5C
H2A.Z:H3K4 me3
H3K27 me2:DNA Meth. m5C
H3K27 me3:H3K4 me1
H3K4 me3:H4–4ac
H4K20 me2:H3–5ac
H4K20 me2:H4–4ac
H4K20 me2:H3K4 me3
DNA Meth. m5C:H4K20 me3
H3K9 me2
H3K9 me3
H3K27 me2
H3K27 me3
H4K20 me3
H3K4 me1
DNA Meth. m5C
H4K20 me2
H3–5ac
H3K4 me3
H4–4ac
H2A.Z
0.00 0.25 0.50 0.75 1.00
$\hat{\pi}$
VRK1 – Forward
H2A.Z:H3K4 me3
H2A.Z:H4K20 me2
H3K4 me1:H4K20 me2
H3K27 me2:DNA Meth. m5C
H3K4 me3:H3–5ac
DNA Meth. m5C:H4K20 me2
H3K9 me2:DNA Meth. m5C
H3K27 me3:H3K4 me1
H3K4 me1:H4–4ac
H3K4 me1:H2A.Z
H4K20 me2:H3–5ac
H4K20 me2:H3K4 me3
H3K4 me1:H3–5ac
H4–4ac:H3–5ac
H4K20 me2:H4–4ac
H3K27 me3:H3K9 me3
DNA Meth. m5C:H3K9 me3
H3K4 me3:H4–4ac
H3K27 me3:DNA Meth. m5C
DNA Meth. m5C:H4K20 me3
H3K9 me3:H4K20 me3
H3K27 me3:H4K20 me3
H3K27 me2
H3K9 me2
H3–5ac
H3K9 me3
H2A.Z
H4K20 me2
H3K4 me1
H3K27 me3
H3K4 me3
H4K20 me3
DNA Meth. m5C
H4–4ac
0.00 0.25 0.50 0.75 1.00
$\hat{\pi}$
VRK1 – Reverse
H2A.Z:H3K4 me3
H2A.Z:H4K20 me2
H3K4 me1:H4K20 me2
H3K27 me2:DNA Meth. m5C
H3K4 me3:H3–5ac
DNA Meth. m5C:H4K20 me2
H3K9 me2:DNA Meth. m5C
H3K27 me3:H3K4 me1
H3K4 me1:H4–4ac
H3K4 me1:H2A.Z
H4K20 me2:H3–5ac
H4K20 me2:H3K4 me3
H3K4 me1:H3–5ac
H4–4ac:H3–5ac
H4K20 me2:H4–4ac
H3K27 me3:H3K9 me3
DNA Meth. m5C:H3K9 me3
H3K4 me3:H4–4ac
H3K27 me3:DNA Meth. m5C
DNA Meth. m5C:H4K20 me3
H3K9 me3:H4K20 me3
H3K27 me3:H4K20 me3
H3K27 me2
H3K9 me2
H3–5ac
H3K9 me3
H2A.Z
H4K20 me2
H3K4 me1
H3K27 me3
H3K4 me3
H4K20 me3
DNA Meth. m5C
H4–4ac
0.00 0.25 0.50 0.75 1.00
$\hat{\pi}$

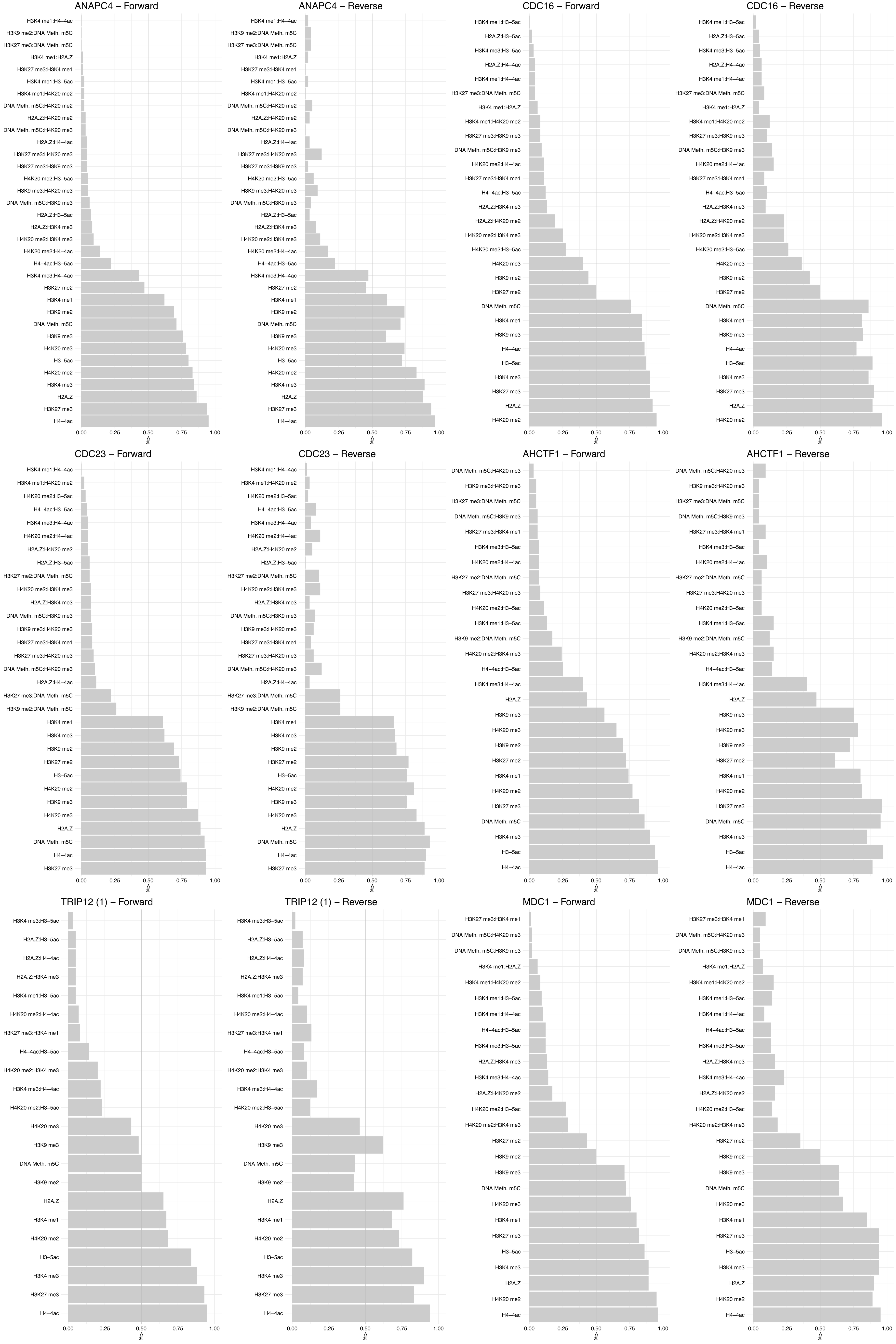
ANAPC4 – Forward
H3K4 me1:H4−4ac
H3K9 me2:DNA Meth. m5C
H3K27 me3:DNA Meth. m5C
H3K4 me1:H2A.Z
H3K27 me3:H3K4 me1
H3K4 me1:H3−5ac
H3K4 me1:H4K20 me2
DNA Meth. m5C:H4K20 me2
H2A.Z:H4K20 me2
DNA Meth. m5C:H4K20 me3
H2A.Z:H4−4ac
H3K27 me3:H4K20 me3
H3K27 me3:H3K9 me3
H4K20 me2:H3−5ac
H3K9 me3:H4K20 me3
DNA Meth. m5C:H3K9 me3
H2A.Z:H3−5ac
H2A.Z:H3K4 me3
H4K20 me2:H3K4 me3
H4K20 me2:H4−4ac
H4−4ac:H3−5ac
H3K4 me3:H4−4ac
H3K27 me2
H3K4 me1
H3K9 me2
DNA Meth. m5C
H3K9 me3
H4K20 me3
H3−5ac
H4K20 me2
H3K4 me3
H2A.Z
H3K27 me3
H4−4ac
0.00 0.25 0.50 0.75 1.00
π̂
ANAPC4 – Reverse
H3K4 me1:H4−4ac
H3K9 me2:DNA Meth. m5C
H3K27 me3:DNA Meth. m5C
H3K4 me1:H2A.Z
H3K27 me3:H3K4 me1
H3K4 me1:H3−5ac
H3K4 me1:H4K20 me2
DNA Meth. m5C:H4K20 me2
H2A.Z:H4K20 me2
DNA Meth. m5C:H4K20 me3
H2A.Z:H4−4ac
H3K27 me3:H4K20 me3
H3K27 me3:H3K9 me3
H4K20 me2:H3−5ac
H3K9 me3:H4K20 me3
DNA Meth. m5C:H3K9 me3
H2A.Z:H3−5ac
H2A.Z:H3K4 me3
H4K20 me2:H3K4 me3
H4K20 me2:H4−4ac
H4−4ac:H3−5ac
H3K4 me3:H4−4ac
H3K27 me2
H3K4 me1
H3K9 me2
DNA Meth. m5C
H3K9 me3
H4K20 me3
H3−5ac
H4K20 me2
H3K4 me3
H2A.Z
H3K27 me3
H4−4ac
0.00 0.25 0.50 0.75 1.00
π̂
CDC16 – Forward
H3K4 me1:H3−5ac
H2A.Z:H3−5ac
H3K4 me3:H3−5ac
H2A.Z:H4−4ac
H3K4 me1:H4−4ac
H3K27 me3:DNA Meth. m5C
H3K4 me1:H2A.Z
H3K4 me1:H4K20 me2
H3K27 me3:H3K9 me3
DNA Meth. m5C:H3K9 me3
H4K20 me2:H4−4ac
H3K27 me3:H3K4 me1
H4−4ac:H3−5ac
H2A.Z:H3K4 me3
H2A.Z:H4K20 me2
H4K20 me2:H3K4 me3
H4K20 me2:H3−5ac
H4K20 me3
H3K9 me2
H3K27 me2
DNA Meth. m5C
H3K4 me1
H3K9 me3
H4−4ac
H3−5ac
H3K4 me3
H3K27 me3
H2A.Z
H4K20 me2
0.00 0.25 0.50 0.75 1.00
π̂
CDC16 – Reverse
H3K4 me1:H3−5ac
H2A.Z:H3−5ac
H3K4 me3:H3−5ac
H2A.Z:H4−4ac
H3K4 me1:H4−4ac
H3K27 me3:DNA Meth. m5C
H3K4 me1:H2A.Z
H3K4 me1:H4K20 me2
H3K27 me3:H3K9 me3
DNA Meth. m5C:H3K9 me3
H4K20 me2:H4−4ac
H3K27 me3:H3K4 me1
H4−4ac:H3−5ac
H2A.Z:H3K4 me3
H2A.Z:H4K20 me2
H4K20 me2:H3K4 me3
H4K20 me2:H3−5ac
H4K20 me3
H3K9 me2
H3K27 me2
DNA Meth. m5C
H3K4 me1
H3K9 me3
H4−4ac
H3−5ac
H3K4 me3
H3K27 me3
H2A.Z
H4K20 me2
0.00 0.25 0.50 0.75 1.00
π̂
CDC23 – Forward
H3K4 me1:H4−4ac
H3K4 me1:H4K20 me2
H4K20 me2:H3−5ac
H4−4ac:H3−5ac
H3K4 me3:H4−4ac
H4K20 me2:H4−4ac
H2A.Z:H4K20 me2
H2A.Z:H3−5ac
H3K27 me2:DNA Meth. m5C
H4K20 me2:H3K4 me3
H2A.Z:H3K4 me3
DNA Meth. m5C:H3K9 me3
H3K9 me3:H4K20 me3
H3K27 me3:H3K4 me1
H3K27 me3:H4K20 me3
DNA Meth. m5C:H4K20 me3
H2A.Z:H4−4ac
H3K27 me3:DNA Meth. m5C
H3K9 me2:DNA Meth. m5C
H3K4 me1
H3K4 me3
H3K9 me2
H3K27 me2
H3−5ac
H4K20 me2
H3K9 me3
H4K20 me3
H2A.Z
DNA Meth. m5C
H4−4ac
H3K27 me3
0.00 0.25 0.50 0.75 1.00
π̂
CDC23 – Reverse
H3K4 me1:H4−4ac
H3K4 me1:H4K20 me2
H4K20 me2:H3−5ac
H4−4ac:H3−5ac
H3K4 me3:H4−4ac
H4K20 me2:H4−4ac
H2A.Z:H4K20 me2
H2A.Z:H3−5ac
H3K27 me2:DNA Meth. m5C
H4K20 me2:H3K4 me3
H2A.Z:H3K4 me3
DNA Meth. m5C:H3K9 me3
H3K9 me3:H4K20 me3
H3K27 me3:H3K4 me1
H3K27 me3:H4K20 me3
DNA Meth. m5C:H4K20 me3
H2A.Z:H4−4ac
H3K27 me3:DNA Meth. m5C
H3K9 me2:DNA Meth. m5C
H3K4 me1
H3K4 me3
H3K9 me2
H3K27 me2
H3−5ac
H4K20 me2
H3K9 me3
H4K20 me3
H2A.Z
DNA Meth. m5C
H4−4ac
H3K27 me3
0.00 0.25 0.50 0.75 1.00
π̂
AHCTF1 – Forward
DNA Meth. m5C:H4K20 me3
H3K9 me3:H4K20 me3
H3K27 me3:DNA Meth. m5C
DNA Meth. m5C:H3K9 me3
H3K27 me3:H3K4 me1
H3K4 me3:H3−5ac
H4K20 me2:H4−4ac
H3K27 me2:DNA Meth. m5C
H3K27 me3:H4K20 me3
H4K20 me2:H3−5ac
H3K4 me1:H3−5ac
H3K9 me2:DNA Meth. m5C
H4K20 me2:H3K4 me3
H4−4ac:H3−5ac
H3K4 me3:H4−4ac
H2A.Z
H3K9 me3
H4K20 me3
H3K9 me2
H3K27 me2
H3K4 me1
H4K20 me2
H3K27 me3
DNA Meth. m5C
H3K4 me3
H3−5ac
H4−4ac
0.00 0.25 0.50 0.75 1.00
π̂
AHCTF1 – Reverse
DNA Meth. m5C:H4K20 me3
H3K9 me3:H4K20 me3
H3K27 me3:DNA Meth. m5C
DNA Meth. m5C:H3K9 me3
H3K27 me3:H3K4 me1
H3K4 me3:H3−5ac
H4K20 me2:H4−4ac
H3K27 me2:DNA Meth. m5C
H3K27 me3:H4K20 me3
H4K20 me2:H3−5ac
H3K4 me1:H3−5ac
H3K9 me2:DNA Meth. m5C
H4K20 me2:H3K4 me3
H4−4ac:H3−5ac
H3K4 me3:H4−4ac
H2A.Z
H3K9 me3
H4K20 me3
H3K9 me2
H3K27 me2
H3K4 me1
H4K20 me2
H3K27 me3
DNA Meth. m5C
H3K4 me3
H3−5ac
H4−4ac
0.00 0.25 0.50 0.75 1.00
π̂
TRIP12 (1) – Forward
H3K4 me3:H3−5ac
H2A.Z:H3−5ac
H2A.Z:H4−4ac
H2A.Z:H3K4 me3
H3K4 me1:H3−5ac
H4K20 me2:H4−4ac
H3K27 me3:H3K4 me1
H4−4ac:H3−5ac
H4K20 me2:H3K4 me3
H3K4 me3:H4−4ac
H4K20 me2:H3−5ac
H4K20 me3
H3K9 me3
DNA Meth. m5C
H3K9 me2
H2A.Z
H3K4 me1
H4K20 me2
H3−5ac
H3K4 me3
H3K27 me3
H4−4ac
0.00 0.25 0.50 0.75 1.00
π̂
TRIP12 (1) – Reverse
H3K4 me3:H3−5ac
H2A.Z:H3−5ac
H2A.Z:H4−4ac
H2A.Z:H3K4 me3
H3K4 me1:H3−5ac
H4K20 me2:H4−4ac
H3K27 me3:H3K4 me1
H4−4ac:H3−5ac
H4K20 me2:H3K4 me3
H3K4 me3:H4−4ac
H4K20 me2:H3−5ac
H4K20 me3
H3K9 me3
DNA Meth. m5C
H3K9 me2
H2A.Z
H3K4 me1
H4K20 me2
H3−5ac
H3K4 me3
H3K27 me3
H4−4ac
0.00 0.25 0.50 0.75 1.00
π̂
MDC1 – Forward
H3K27 me3:H3K4 me1
DNA Meth. m5C:H4K20 me3
DNA Meth. m5C:H3K9 me3
H3K4 me1:H2A.Z
H3K4 me1:H4K20 me2
H3K4 me1:H3−5ac
H3K4 me1:H4−4ac
H4−4ac:H3−5ac
H3K4 me3:H3−5ac
H2A.Z:H3K4 me3
H3K4 me3:H4−4ac
H2A.Z:H4K20 me2
H4K20 me2:H3−5ac
H4K20 me2:H3K4 me3
H3K27 me2
H3K9 me2
H3K9 me3
DNA Meth. m5C
H4K20 me3
H3K4 me1
H3K27 me3
H3−5ac
H3K4 me3
H2A.Z
H4K20 me2
H4−4ac
0.00 0.25 0.50 0.75 1.00
π̂
MDC1 – Reverse
H3K27 me3:H3K4 me1
DNA Meth. m5C:H4K20 me3
DNA Meth. m5C:H3K9 me3
H3K4 me1:H2A.Z
H3K4 me1:H4K20 me2
H3K4 me1:H3−5ac
H3K4 me1:H4−4ac
H4−4ac:H3−5ac
H3K4 me3:H3−5ac
H2A.Z:H3K4 me3
H3K4 me3:H4−4ac
H2A.Z:H4K20 me2
H4K20 me2:H3−5ac
H4K20 me2:H3K4 me3
H3K27 me2
H3K9 me2
H3K9 me3
DNA Meth. m5C
H4K20 me3
H3K4 me1
H3K27 me3
H3−5ac
H3K4 me3
H2A.Z
H4K20 me2
H4−4ac
0.00 0.25 0.50 0.75 1.00
π̂

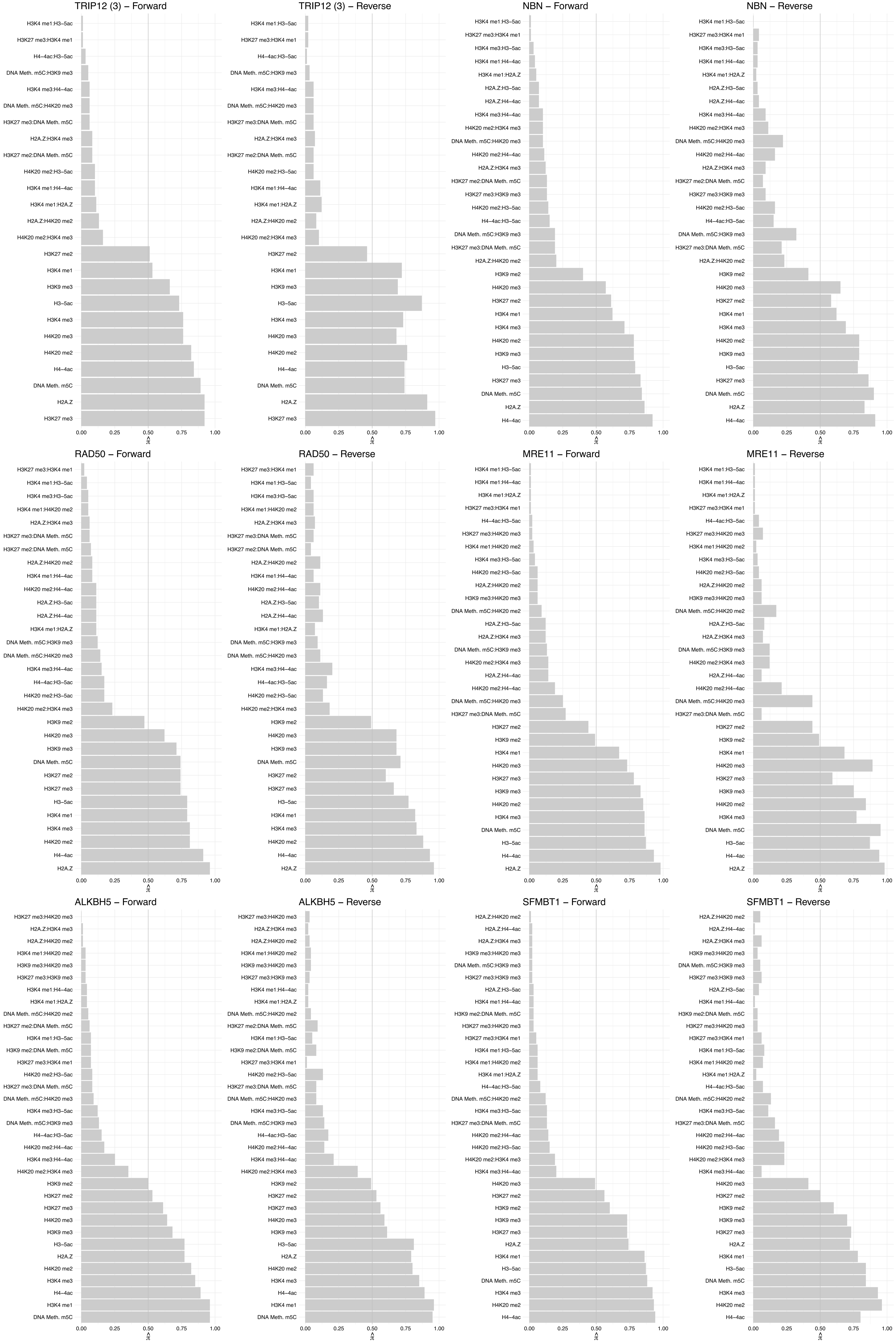

TRIP12 (3) – Forward
TRIP12 (3) – Reverse
NBN – Forward
NBN – Reverse
RAD50 – Forward
RAD50 – Reverse
MRE11 – Forward
MRE11 – Reverse
ALKBH5 – Forward
ALKBH5 – Reverse
SFMBT1 – Forward
SFMBT1 – Reverse
0.00
0.25
0.50
0.75
1.00
$\hat{\pi}$

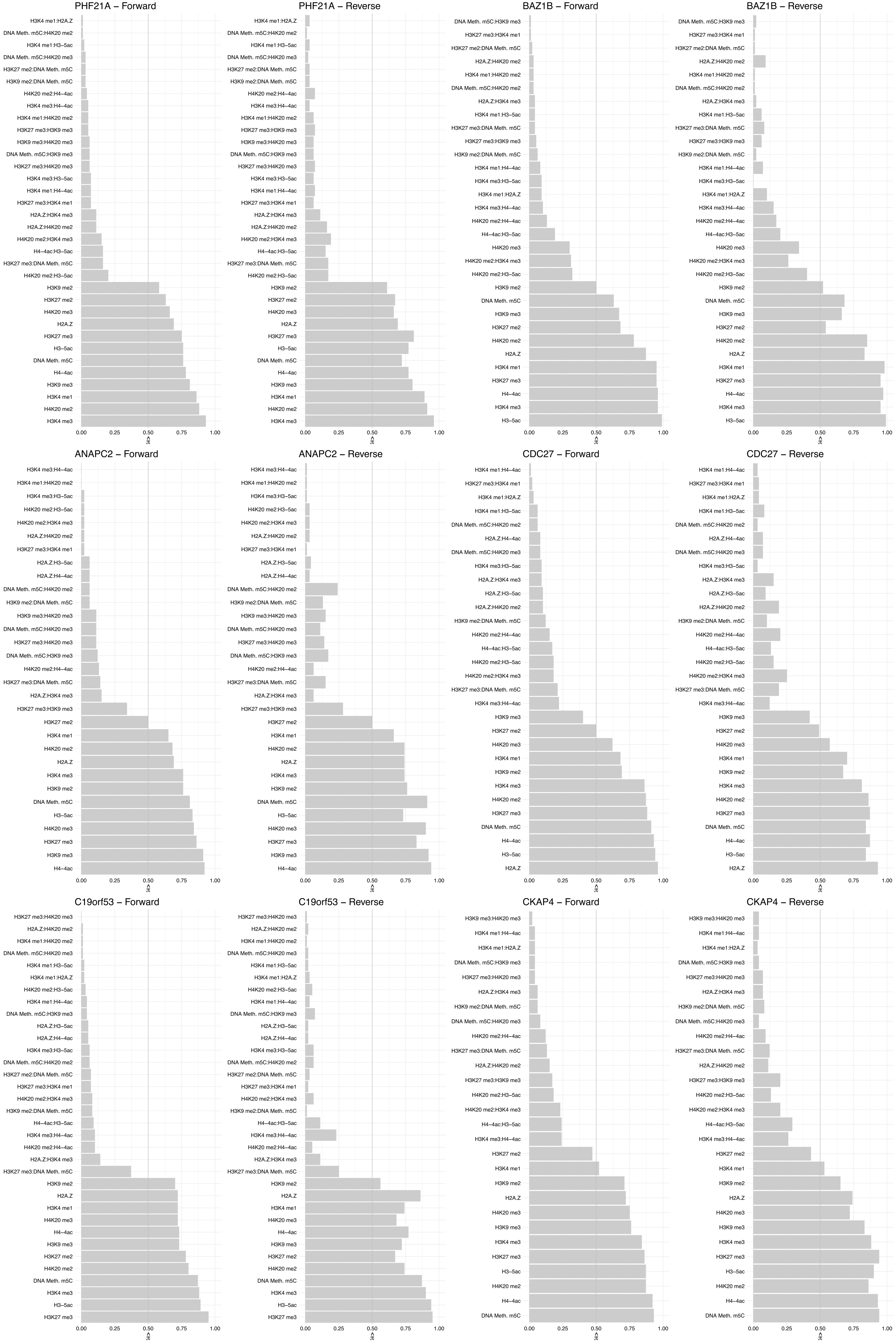
PHF21A – Forward
PHF21A – Reverse
BAZ1B – Forward
BAZ1B – Reverse
ANAPC2 – Forward
ANAPC2 – Reverse
CDC27 – Forward
CDC27 – Reverse
C19orf53 – Forward
C19orf53 – Reverse
CKAP4 – Forward
CKAP4 – Reverse
0.00 0.25 0.50 0.75 1.00
$\hat{\pi}$

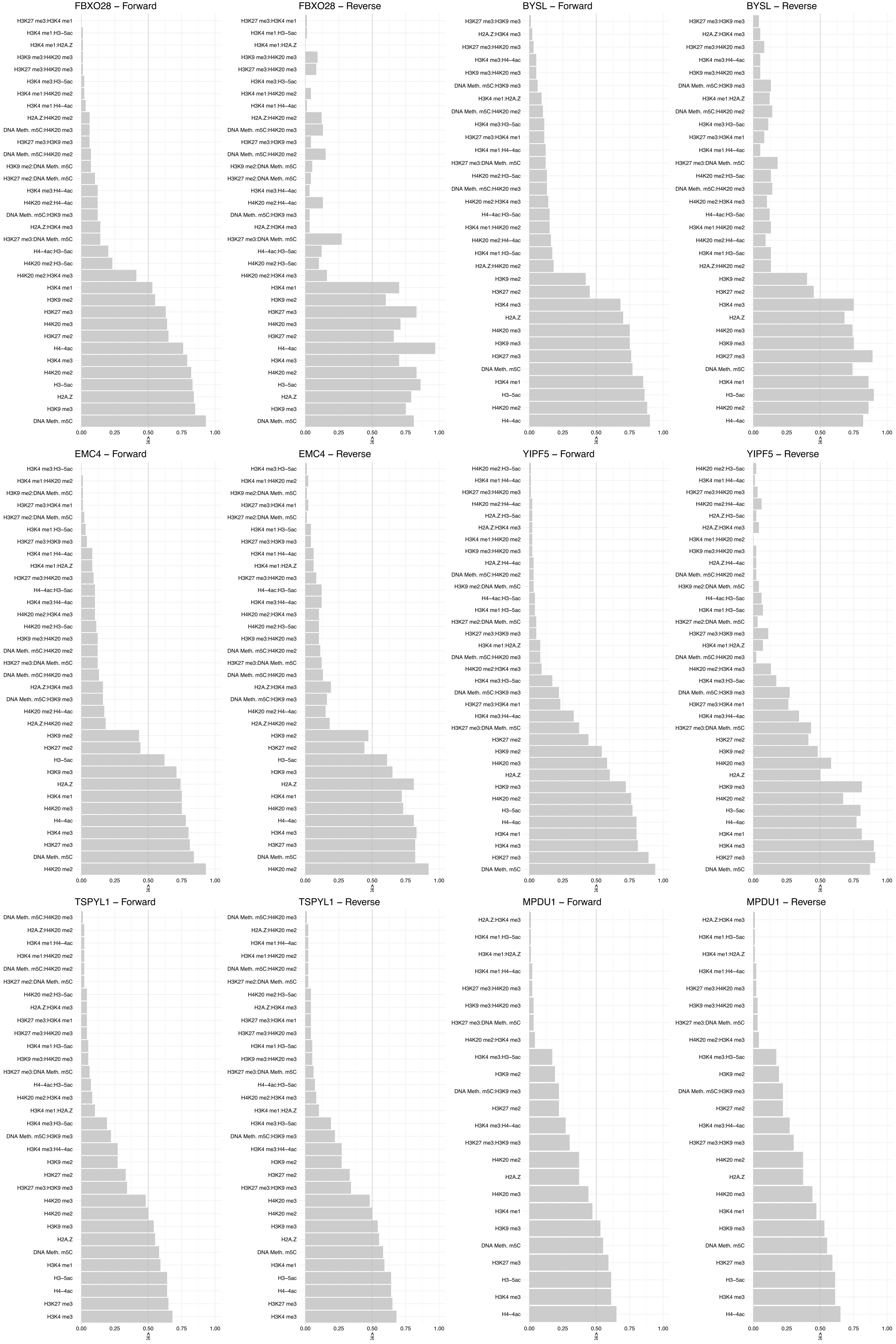
FBXO28 – Forward
H3K27 me3:H3K4 me1
H3K4 me1:H3–5ac
H3K4 me1:H2A.Z
H3K9 me3:H4K20 me3
H3K27 me3:H4K20 me3
H3K4 me3:H3–5ac
H3K4 me1:H4K20 me2
H3K4 me1:H4–4ac
H2A.Z:H4K20 me2
DNA Meth. m5C:H4K20 me3
H3K27 me3:H3K9 me3
DNA Meth. m5C:H4K20 me2
H3K9 me2:DNA Meth. m5C
H3K27 me2:DNA Meth. m5C
H3K4 me3:H4–4ac
H4K20 me2:H4–4ac
DNA Meth. m5C:H3K9 me3
H2A.Z:H3K4 me3
H3K27 me3:DNA Meth. m5C
H4–4ac:H3–5ac
H4K20 me2:H3–5ac
H4K20 me2:H3K4 me3
H3K4 me1
H3K9 me2
H3K27 me3
H4K20 me3
H3K27 me2
H4–4ac
H3K4 me3
H4K20 me2
H3–5ac
H2A.Z
H3K9 me3
DNA Meth. m5C
0.00 0.25 0.50 0.75 1.00
π̂
FBXO28 – Reverse
H3K27 me3:H3K4 me1
H3K4 me1:H3–5ac
H3K4 me1:H2A.Z
H3K9 me3:H4K20 me3
H3K27 me3:H4K20 me3
H3K4 me3:H3–5ac
H3K4 me1:H4K20 me2
H3K4 me1:H4–4ac
H2A.Z:H4K20 me2
DNA Meth. m5C:H4K20 me3
H3K27 me3:H3K9 me3
DNA Meth. m5C:H4K20 me2
H3K9 me2:DNA Meth. m5C
H3K27 me2:DNA Meth. m5C
H3K4 me3:H4–4ac
H4K20 me2:H4–4ac
DNA Meth. m5C:H3K9 me3
H2A.Z:H3K4 me3
H3K27 me3:DNA Meth. m5C
H4–4ac:H3–5ac
H4K20 me2:H3–5ac
H4K20 me2:H3K4 me3
H3K4 me1
H3K9 me2
H3K27 me3
H4K20 me3
H3K27 me2
H4–4ac
H3K4 me3
H4K20 me2
H3–5ac
H2A.Z
H3K9 me3
DNA Meth. m5C
0.00 0.25 0.50 0.75 1.00
π̂
BYSL – Forward
H3K27 me3:H3K9 me3
H2A.Z:H3K4 me3
H3K27 me3:H4K20 me3
H3K4 me3:H4–4ac
H3K9 me3:H4K20 me3
DNA Meth. m5C:H3K9 me3
H3K4 me1:H2A.Z
DNA Meth. m5C:H4K20 me2
H3K4 me3:H3–5ac
H3K27 me3:H3K4 me1
H3K4 me1:H4–4ac
H3K27 me3:DNA Meth. m5C
H4K20 me2:H3–5ac
DNA Meth. m5C:H4K20 me3
H4K20 me2:H3K4 me3
H4–4ac:H3–5ac
H3K4 me1:H4K20 me2
H4K20 me2:H4–4ac
H3K4 me1:H3–5ac
H2A.Z:H4K20 me2
H3K9 me2
H3K27 me2
H3K4 me3
H2A.Z
H4K20 me3
H3K9 me3
H3K27 me3
DNA Meth. m5C
H3K4 me1
H3–5ac
H4K20 me2
H4–4ac
0.00 0.25 0.50 0.75 1.00
π̂
BYSL – Reverse
H3K27 me3:H3K9 me3
H2A.Z:H3K4 me3
H3K27 me3:H4K20 me3
H3K4 me3:H4–4ac
H3K9 me3:H4K20 me3
DNA Meth. m5C:H3K9 me3
H3K4 me1:H2A.Z
DNA Meth. m5C:H4K20 me2
H3K4 me3:H3–5ac
H3K27 me3:H3K4 me1
H3K4 me1:H4–4ac
H3K27 me3:DNA Meth. m5C
H4K20 me2:H3–5ac
DNA Meth. m5C:H4K20 me3
H4K20 me2:H3K4 me3
H4–4ac:H3–5ac
H3K4 me1:H4K20 me2
H4K20 me2:H4–4ac
H3K4 me1:H3–5ac
H2A.Z:H4K20 me2
H3K9 me2
H3K27 me2
H3K4 me3
H2A.Z
H4K20 me3
H3K9 me3
H3K27 me3
DNA Meth. m5C
H3K4 me1
H3–5ac
H4K20 me2
H4–4ac
0.00 0.25 0.50 0.75 1.00
π̂
EMC4 – Forward
H3K4 me3:H3–5ac
H3K4 me1:H4K20 me2
H3K9 me2:DNA Meth. m5C
H3K27 me3:H3K4 me1
H3K27 me2:DNA Meth. m5C
H3K4 me1:H3–5ac
H3K27 me3:H3K9 me3
H3K4 me1:H4–4ac
H3K4 me1:H2A.Z
H3K27 me3:H4K20 me3
H4–4ac:H3–5ac
H3K4 me3:H4–4ac
H4K20 me2:H3K4 me3
H4K20 me2:H3–5ac
H3K9 me3:H4K20 me3
DNA Meth. m5C:H4K20 me2
H3K27 me3:DNA Meth. m5C
DNA Meth. m5C:H4K20 me3
H2A.Z:H3K4 me3
DNA Meth. m5C:H3K9 me3
H4K20 me2:H4–4ac
H2A.Z:H4K20 me2
H3K9 me2
H3K27 me2
H3–5ac
H3K9 me3
H2A.Z
H3K4 me1
H4K20 me3
H4–4ac
H3K4 me3
H3K27 me3
DNA Meth. m5C
H4K20 me2
0.00 0.25 0.50 0.75 1.00
π̂
EMC4 – Reverse
H3K4 me3:H3–5ac
H3K4 me1:H4K20 me2
H3K9 me2:DNA Meth. m5C
H3K27 me3:H3K4 me1
H3K27 me2:DNA Meth. m5C
H3K4 me1:H3–5ac
H3K27 me3:H3K9 me3
H3K4 me1:H4–4ac
H3K4 me1:H2A.Z
H3K27 me3:H4K20 me3
H4–4ac:H3–5ac
H3K4 me3:H4–4ac
H4K20 me2:H3K4 me3
H4K20 me2:H3–5ac
H3K9 me3:H4K20 me3
DNA Meth. m5C:H4K20 me2
H3K27 me3:DNA Meth. m5C
DNA Meth. m5C:H4K20 me3
H2A.Z:H3K4 me3
DNA Meth. m5C:H3K9 me3
H4K20 me2:H4–4ac
H2A.Z:H4K20 me2
H3K9 me2
H3K27 me2
H3–5ac
H3K9 me3
H2A.Z
H3K4 me1
H4K20 me3
H4–4ac
H3K4 me3
H3K27 me3
DNA Meth. m5C
H4K20 me2
0.00 0.25 0.50 0.75 1.00
π̂
YIPF5 – Forward
H4K20 me2:H3–5ac
H3K4 me1:H4–4ac
H3K27 me3:H4K20 me3
H4K20 me2:H4–4ac
H2A.Z:H3–5ac
H2A.Z:H3K4 me3
H3K4 me1:H4K20 me2
H3K9 me3:H4K20 me3
H2A.Z:H4–4ac
DNA Meth. m5C:H4K20 me2
H3K9 me2:DNA Meth. m5C
H4–4ac:H3–5ac
H3K4 me1:H3–5ac
H3K27 me2:DNA Meth. m5C
H3K27 me3:H3K9 me3
H3K4 me1:H2A.Z
DNA Meth. m5C:H4K20 me3
H4K20 me2:H3K4 me3
H3K4 me3:H3–5ac
DNA Meth. m5C:H3K9 me3
H3K27 me3:H3K4 me1
H3K4 me3:H4–4ac
H3K27 me3:DNA Meth. m5C
H3K27 me2
H3K9 me2
H4K20 me3
H2A.Z
H3K9 me3
H4K20 me2
H3–5ac
H4–4ac
H3K4 me1
H3K4 me3
H3K27 me3
DNA Meth. m5C
0.00 0.25 0.50 0.75 1.00
π̂
YIPF5 – Reverse
H4K20 me2:H3–5ac
H3K4 me1:H4–4ac
H3K27 me3:H4K20 me3
H4K20 me2:H4–4ac
H2A.Z:H3–5ac
H2A.Z:H3K4 me3
H3K4 me1:H4K20 me2
H3K9 me3:H4K20 me3
H2A.Z:H4–4ac
DNA Meth. m5C:H4K20 me2
H3K9 me2:DNA Meth. m5C
H4–4ac:H3–5ac
H3K4 me1:H3–5ac
H3K27 me2:DNA Meth. m5C
H3K27 me3:H3K9 me3
H3K4 me1:H2A.Z
DNA Meth. m5C:H4K20 me3
H4K20 me2:H3K4 me3
H3K4 me3:H3–5ac
DNA Meth. m5C:H3K9 me3
H3K27 me3:H3K4 me1
H3K4 me3:H4–4ac
H3K27 me3:DNA Meth. m5C
H3K27 me2
H3K9 me2
H4K20 me3
H2A.Z
H3K9 me3
H4K20 me2
H3–5ac
H4–4ac
H3K4 me1
H3K4 me3
H3K27 me3
DNA Meth. m5C
0.00 0.25 0.50 0.75 1.00
π̂
TSPYL1 – Forward
DNA Meth. m5C:H4K20 me3
H2A.Z:H4K20 me2
H3K4 me1:H4–4ac
H3K4 me1:H4K20 me2
DNA Meth. m5C:H4K20 me2
H3K27 me2:DNA Meth. m5C
H4K20 me2:H3–5ac
H2A.Z:H3K4 me3
H3K27 me3:H3K4 me1
H3K27 me3:H4K20 me3
H3K4 me1:H3–5ac
H3K9 me3:H4K20 me3
H3K27 me3:DNA Meth. m5C
H4–4ac:H3–5ac
H4K20 me2:H3K4 me3
H3K4 me1:H2A.Z
H3K4 me3:H3–5ac
DNA Meth. m5C:H3K9 me3
H3K4 me3:H4–4ac
H3K9 me2
H3K27 me2
H3K27 me3:H3K9 me3
H4K20 me3
H4K20 me2
H3K9 me3
H2A.Z
DNA Meth. m5C
H3K4 me1
H3–5ac
H4–4ac
H3K27 me3
H3K4 me3
0.00 0.25 0.50 0.75 1.00
π̂
TSPYL1 – Reverse
DNA Meth. m5C:H4K20 me3
H2A.Z:H4K20 me2
H3K4 me1:H4–4ac
H3K4 me1:H4K20 me2
DNA Meth. m5C:H4K20 me2
H3K27 me2:DNA Meth. m5C
H4K20 me2:H3–5ac
H2A.Z:H3K4 me3
H3K27 me3:H3K4 me1
H3K27 me3:H4K20 me3
H3K4 me1:H3–5ac
H3K9 me3:H4K20 me3
H3K27 me3:DNA Meth. m5C
H4–4ac:H3–5ac
H4K20 me2:H3K4 me3
H3K4 me1:H2A.Z
H3K4 me3:H3–5ac
DNA Meth. m5C:H3K9 me3
H3K4 me3:H4–4ac
H3K9 me2
H3K27 me2
H3K27 me3:H3K9 me3
H4K20 me3
H4K20 me2
H3K9 me3
H2A.Z
DNA Meth. m5C
H3K4 me1
H3–5ac
H4–4ac
H3K27 me3
H3K4 me3
0.00 0.25 0.50 0.75 1.00
π̂
MPDU1 – Forward
H2A.Z:H3K4 me3
H3K4 me1:H3–5ac
H3K4 me1:H2A.Z
H3K4 me1:H4–4ac
H3K27 me3:H4K20 me3
H3K9 me3:H4K20 me3
H3K27 me3:DNA Meth. m5C
H4K20 me2:H3K4 me3
H3K4 me3:H3–5ac
H3K9 me2
DNA Meth. m5C:H3K9 me3
H3K27 me2
H3K4 me3:H4–4ac
H3K27 me3:H3K9 me3
H4K20 me2
H2A.Z
H4K20 me3
H3K4 me1
H3K9 me3
DNA Meth. m5C
H3K27 me3
H3–5ac
H3K4 me3
H4–4ac
0.00 0.25 0.50 0.75 1.00
π̂
MPDU1 – Reverse
H2A.Z:H3K4 me3
H3K4 me1:H3–5ac
H3K4 me1:H2A.Z
H3K4 me1:H4–4ac
H3K27 me3:H4K20 me3
H3K9 me3:H4K20 me3
H3K27 me3:DNA Meth. m5C
H4K20 me2:H3K4 me3
H3K4 me3:H3–5ac
H3K9 me2
DNA Meth. m5C:H3K9 me3
H3K27 me2
H3K4 me3:H4–4ac
H3K27 me3:H3K9 me3
H4K20 me2
H2A.Z
H4K20 me3
H3K4 me1
H3K9 me3
DNA Meth. m5C
H3K27 me3
H3–5ac
H3K4 me3
H4–4ac
0.00 0.25 0.50 0.75 1.00
π̂

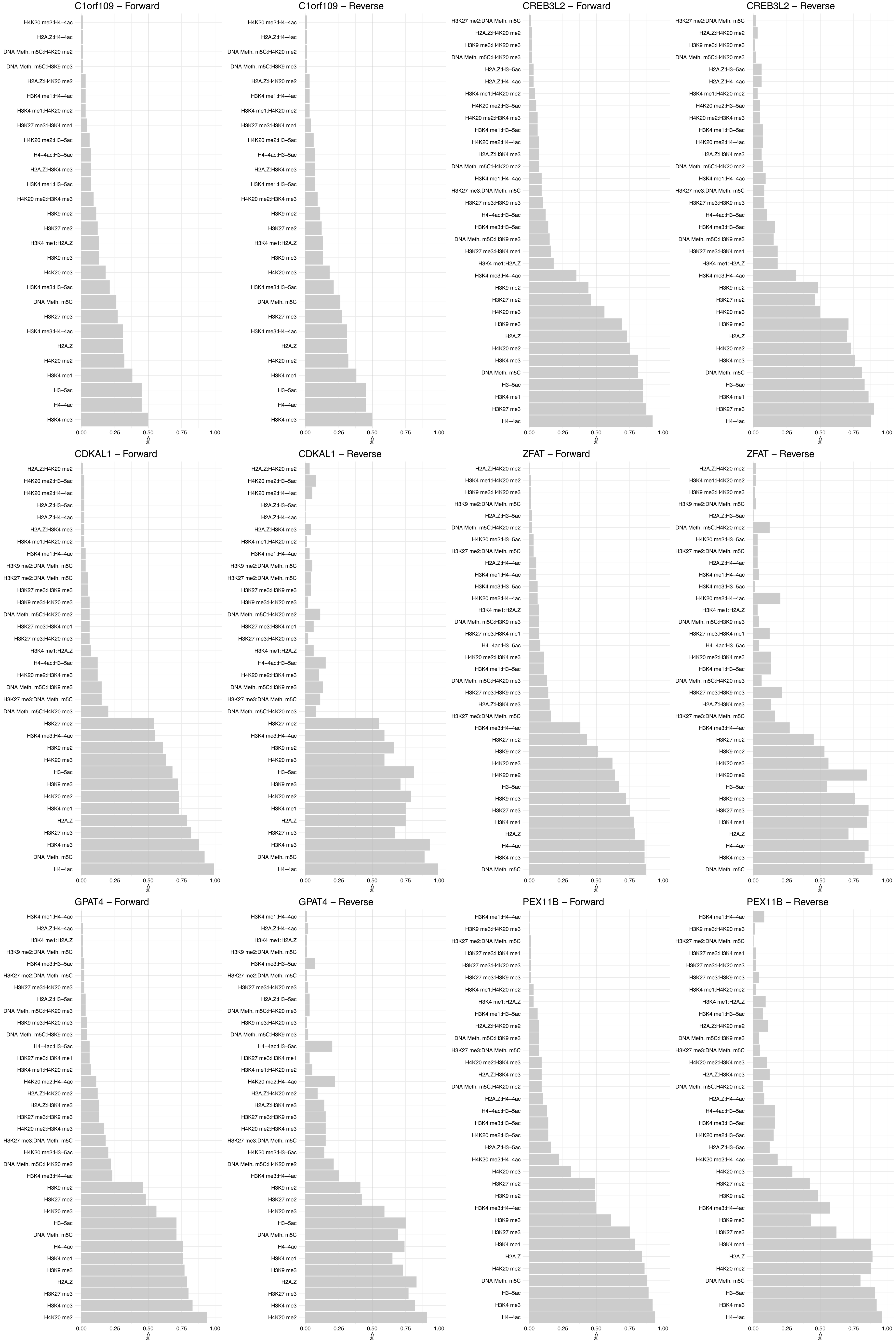
C1orf109 – Forward
H4K20 me2:H4−4ac
H2A.Z:H4−4ac
DNA Meth. m5C:H4K20 me2
DNA Meth. m5C:H3K9 me3
H2A.Z:H4K20 me2
H3K4 me1:H4−4ac
H3K4 me1:H4K20 me2
H3K27 me3:H3K4 me1
H4K20 me2:H3−5ac
H4−4ac:H3−5ac
H2A.Z:H3K4 me3
H3K4 me1:H3−5ac
H4K20 me2:H3K4 me3
H3K9 me2
H3K27 me2
H3K4 me1:H2A.Z
H3K9 me3
H4K20 me3
H3K4 me3:H3−5ac
DNA Meth. m5C
H3K27 me3
H3K4 me3:H4−4ac
H2A.Z
H4K20 me2
H3K4 me1
H3−5ac
H4−4ac
H3K4 me3
0.00
0.25
0.50
0.75
1.00
π̂
C1orf109 – Reverse
H4K20 me2:H4−4ac
H2A.Z:H4−4ac
DNA Meth. m5C:H4K20 me2
DNA Meth. m5C:H3K9 me3
H2A.Z:H4K20 me2
H3K4 me1:H4−4ac
H3K4 me1:H4K20 me2
H3K27 me3:H3K4 me1
H4K20 me2:H3−5ac
H4−4ac:H3−5ac
H2A.Z:H3K4 me3
H3K4 me1:H3−5ac
H4K20 me2:H3K4 me3
H3K9 me2
H3K27 me2
H3K4 me1:H2A.Z
H3K9 me3
H4K20 me3
H3K4 me3:H3−5ac
DNA Meth. m5C
H3K27 me3
H3K4 me3:H4−4ac
H2A.Z
H4K20 me2
H3K4 me1
H3−5ac
H4−4ac
H3K4 me3
0.00
0.25
0.50
0.75
1.00
π̂
CREB3L2 – Forward
H3K27 me2:DNA Meth. m5C
H2A.Z:H4K20 me2
H3K9 me3:H4K20 me3
DNA Meth. m5C:H4K20 me3
H2A.Z:H3−5ac
H2A.Z:H4−4ac
H3K4 me1:H4K20 me2
H4K20 me2:H3−5ac
H4K20 me2:H3K4 me3
H3K4 me1:H3−5ac
H4K20 me2:H4−4ac
H2A.Z:H3K4 me3
DNA Meth. m5C:H4K20 me2
H3K4 me1:H4−4ac
H3K27 me3:DNA Meth. m5C
H3K27 me3:H3K9 me3
H4−4ac:H3−5ac
H3K4 me3:H3−5ac
DNA Meth. m5C:H3K9 me3
H3K27 me3:H3K4 me1
H3K4 me1:H2A.Z
H3K4 me3:H4−4ac
H3K9 me2
H3K27 me2
H4K20 me3
H3K9 me3
H2A.Z
H4K20 me2
H3K4 me3
DNA Meth. m5C
H3−5ac
H3K4 me1
H3K27 me3
H4−4ac
0.00
0.25
0.50
0.75
1.00
π̂
CREB3L2 – Reverse
H3K27 me2:DNA Meth. m5C
H2A.Z:H4K20 me2
H3K9 me3:H4K20 me3
DNA Meth. m5C:H4K20 me3
H2A.Z:H3−5ac
H2A.Z:H4−4ac
H3K4 me1:H4K20 me2
H4K20 me2:H3−5ac
H4K20 me2:H3K4 me3
H3K4 me1:H3−5ac
H4K20 me2:H4−4ac
H2A.Z:H3K4 me3
DNA Meth. m5C:H4K20 me2
H3K4 me1:H4−4ac
H3K27 me3:DNA Meth. m5C
H3K27 me3:H3K9 me3
H4−4ac:H3−5ac
H3K4 me3:H3−5ac
DNA Meth. m5C:H3K9 me3
H3K27 me3:H3K4 me1
H3K4 me1:H2A.Z
H3K4 me3:H4−4ac
H3K9 me2
H3K27 me2
H4K20 me3
H3K9 me3
H2A.Z
H4K20 me2
H3K4 me3
DNA Meth. m5C
H3−5ac
H3K4 me1
H3K27 me3
H4−4ac
0.00
0.25
0.50
0.75
1.00
π̂
CDKAL1 – Forward
H2A.Z:H4K20 me2
H4K20 me2:H3−5ac
H4K20 me2:H4−4ac
H2A.Z:H3−5ac
H2A.Z:H4−4ac
H2A.Z:H3K4 me3
H3K4 me1:H4K20 me2
H3K4 me1:H4−4ac
H3K9 me2:DNA Meth. m5C
H3K27 me2:DNA Meth. m5C
H3K27 me3:H3K9 me3
H3K9 me3:H4K20 me3
DNA Meth. m5C:H4K20 me2
H3K27 me3:H3K4 me1
H3K27 me3:H4K20 me3
H3K4 me1:H2A.Z
H4−4ac:H3−5ac
H4K20 me2:H3K4 me3
DNA Meth. m5C:H3K9 me3
H3K27 me3:DNA Meth. m5C
DNA Meth. m5C:H4K20 me3
H3K27 me2
H3K4 me3:H4−4ac
H3K9 me2
H4K20 me3
H3−5ac
H3K9 me3
H4K20 me2
H3K4 me1
H2A.Z
H3K27 me3
H3K4 me3
DNA Meth. m5C
H4−4ac
0.00
0.25
0.50
0.75
1.00
π̂
CDKAL1 – Reverse
H2A.Z:H4K20 me2
H4K20 me2:H3−5ac
H4K20 me2:H4−4ac
H2A.Z:H3−5ac
H2A.Z:H4−4ac
H2A.Z:H3K4 me3
H3K4 me1:H4K20 me2
H3K4 me1:H4−4ac
H3K9 me2:DNA Meth. m5C
H3K27 me2:DNA Meth. m5C
H3K27 me3:H3K9 me3
H3K9 me3:H4K20 me3
DNA Meth. m5C:H4K20 me2
H3K27 me3:H3K4 me1
H3K27 me3:H4K20 me3
H3K4 me1:H2A.Z
H4−4ac:H3−5ac
H4K20 me2:H3K4 me3
DNA Meth. m5C:H3K9 me3
H3K27 me3:DNA Meth. m5C
DNA Meth. m5C:H4K20 me3
H3K27 me2
H3K4 me3:H4−4ac
H3K9 me2
H4K20 me3
H3−5ac
H3K9 me3
H4K20 me2
H3K4 me1
H2A.Z
H3K27 me3
H3K4 me3
DNA Meth. m5C
H4−4ac
0.00
0.25
0.50
0.75
1.00
π̂
ZFAT – Forward
H2A.Z:H4K20 me2
H3K4 me1:H4K20 me2
H3K9 me3:H4K20 me3
H3K9 me2:DNA Meth. m5C
H2A.Z:H3−5ac
DNA Meth. m5C:H4K20 me2
H4K20 me2:H3−5ac
H3K27 me2:DNA Meth. m5C
H2A.Z:H4−4ac
H3K4 me1:H4−4ac
H3K4 me3:H3−5ac
H4K20 me2:H4−4ac
H3K4 me1:H2A.Z
DNA Meth. m5C:H3K9 me3
H3K27 me3:H3K4 me1
H4−4ac:H3−5ac
H4K20 me2:H3K4 me3
H3K4 me1:H3−5ac
DNA Meth. m5C:H4K20 me3
H3K27 me3:H3K9 me3
H2A.Z:H3K4 me3
H3K27 me3:DNA Meth. m5C
H3K4 me3:H4−4ac
H3K27 me2
H3K9 me2
H4K20 me3
H4K20 me2
H3−5ac
H3K9 me3
H3K27 me3
H3K4 me1
H2A.Z
H4−4ac
H3K4 me3
DNA Meth. m5C
0.00
0.25
0.50
0.75
1.00
π̂
ZFAT – Reverse
H2A.Z:H4K20 me2
H3K4 me1:H4K20 me2
H3K9 me3:H4K20 me3
H3K9 me2:DNA Meth. m5C
H2A.Z:H3−5ac
DNA Meth. m5C:H4K20 me2
H4K20 me2:H3−5ac
H3K27 me2:DNA Meth. m5C
H2A.Z:H4−4ac
H3K4 me1:H4−4ac
H3K4 me3:H3−5ac
H4K20 me2:H4−4ac
H3K4 me1:H2A.Z
DNA Meth. m5C:H3K9 me3
H3K27 me3:H3K4 me1
H4−4ac:H3−5ac
H4K20 me2:H3K4 me3
H3K4 me1:H3−5ac
DNA Meth. m5C:H4K20 me3
H3K27 me3:H3K9 me3
H2A.Z:H3K4 me3
H3K27 me3:DNA Meth. m5C
H3K4 me3:H4−4ac
H3K27 me2
H3K9 me2
H4K20 me3
H4K20 me2
H3−5ac
H3K9 me3
H3K27 me3
H3K4 me1
H2A.Z
H4−4ac
H3K4 me3
DNA Meth. m5C
0.00
0.25
0.50
0.75
1.00
π̂
GPAT4 – Forward
H3K4 me1:H4−4ac
H2A.Z:H4−4ac
H3K4 me1:H2A.Z
H3K9 me2:DNA Meth. m5C
H3K4 me3:H3−5ac
H3K27 me2:DNA Meth. m5C
H3K27 me3:H4K20 me3
H2A.Z:H3−5ac
DNA Meth. m5C:H4K20 me3
H3K9 me3:H4K20 me3
DNA Meth. m5C:H3K9 me3
H4−4ac:H3−5ac
H3K27 me3:H3K4 me1
H3K4 me1:H4K20 me2
H4K20 me2:H4−4ac
H2A.Z:H4K20 me2
H2A.Z:H3K4 me3
H3K27 me3:H3K9 me3
H4K20 me2:H3K4 me3
H3K27 me3:DNA Meth. m5C
H4K20 me2:H3−5ac
DNA Meth. m5C:H4K20 me2
H3K4 me3:H4−4ac
H3K9 me2
H3K27 me2
H4K20 me3
H3−5ac
DNA Meth. m5C
H4−4ac
H3K4 me1
H3K9 me3
H2A.Z
H3K27 me3
H3K4 me3
H4K20 me2
0.00
0.25
0.50
0.75
1.00
π̂
GPAT4 – Reverse
H3K4 me1:H4−4ac
H2A.Z:H4−4ac
H3K4 me1:H2A.Z
H3K9 me2:DNA Meth. m5C
H3K4 me3:H3−5ac
H3K27 me2:DNA Meth. m5C
H3K27 me3:H4K20 me3
H2A.Z:H3−5ac
DNA Meth. m5C:H4K20 me3
H3K9 me3:H4K20 me3
DNA Meth. m5C:H3K9 me3
H4−4ac:H3−5ac
H3K27 me3:H3K4 me1
H3K4 me1:H4K20 me2
H4K20 me2:H4−4ac
H2A.Z:H4K20 me2
H2A.Z:H3K4 me3
H3K27 me3:H3K9 me3
H4K20 me2:H3K4 me3
H3K27 me3:DNA Meth. m5C
H4K20 me2:H3−5ac
DNA Meth. m5C:H4K20 me2
H3K4 me3:H4−4ac
H3K9 me2
H3K27 me2
H4K20 me3
H3−5ac
DNA Meth. m5C
H4−4ac
H3K4 me1
H3K9 me3
H2A.Z
H3K27 me3
H3K4 me3
H4K20 me2
0.00
0.25
0.50
0.75
1.00
π̂
PEX11B – Forward
H3K4 me1:H4−4ac
H3K9 me3:H4K20 me3
H3K27 me2:DNA Meth. m5C
H3K27 me3:H3K4 me1
H3K27 me3:H4K20 me3
H3K27 me3:H3K9 me3
H3K4 me1:H4K20 me2
H3K4 me1:H2A.Z
H3K4 me1:H3−5ac
H2A.Z:H4K20 me2
DNA Meth. m5C:H3K9 me3
H3K27 me3:DNA Meth. m5C
H4K20 me2:H3K4 me3
H2A.Z:H3K4 me3
DNA Meth. m5C:H4K20 me2
H2A.Z:H4−4ac
H4−4ac:H3−5ac
H3K4 me3:H3−5ac
H4K20 me2:H3−5ac
H2A.Z:H3−5ac
H4K20 me2:H4−4ac
H4K20 me3
H3K27 me2
H3K9 me2
H3K4 me3:H4−4ac
H3K9 me3
H3K27 me3
H3K4 me1
H2A.Z
H4K20 me2
DNA Meth. m5C
H3−5ac
H3K4 me3
H4−4ac
0.00
0.25
0.50
0.75
1.00
π̂
PEX11B – Reverse
H3K4 me1:H4−4ac
H3K9 me3:H4K20 me3
H3K27 me2:DNA Meth. m5C
H3K27 me3:H3K4 me1
H3K27 me3:H4K20 me3
H3K27 me3:H3K9 me3
H3K4 me1:H4K20 me2
H3K4 me1:H2A.Z
H3K4 me1:H3−5ac
H2A.Z:H4K20 me2
DNA Meth. m5C:H3K9 me3
H3K27 me3:DNA Meth. m5C
H4K20 me2:H3K4 me3
H2A.Z:H3K4 me3
DNA Meth. m5C:H4K20 me2
H2A.Z:H4−4ac
H4−4ac:H3−5ac
H3K4 me3:H3−5ac
H4K20 me2:H3−5ac
H2A.Z:H3−5ac
H4K20 me2:H4−4ac
H4K20 me3
H3K27 me2
H3K9 me2
H3K4 me3:H4−4ac
H3K9 me3
H3K27 me3
H3K4 me1
H2A.Z
H4K20 me2
DNA Meth. m5C
H3−5ac
H3K4 me3
H4−4ac
0.00
0.25
0.50
0.75
1.00
π̂

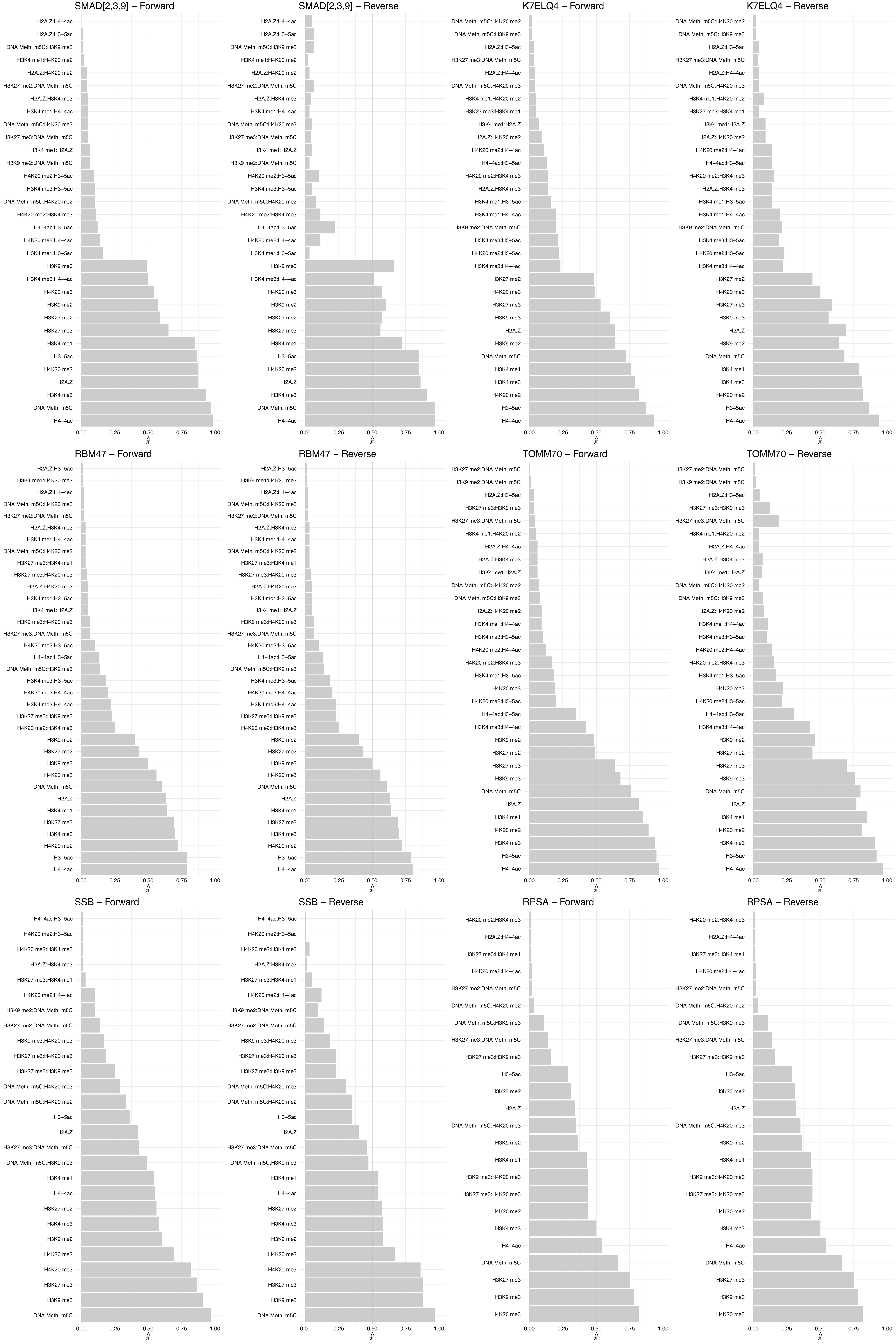
SMAD[2,3,9] – Forward
H2A.Z:H4−4ac
H2A.Z:H3−5ac
DNA Meth. m5C:H3K9 me3
H3K4 me1:H4K20 me2
H2A.Z:H4K20 me2
H3K27 me2:DNA Meth. m5C
H2A.Z:H3K4 me3
H3K4 me1:H4−4ac
DNA Meth. m5C:H4K20 me3
H3K27 me3:DNA Meth. m5C
H3K4 me1:H2A.Z
H3K9 me2:DNA Meth. m5C
H4K20 me2:H3−5ac
H3K4 me3:H3−5ac
DNA Meth. m5C:H4K20 me2
H4K20 me2:H3K4 me3
H4−4ac:H3−5ac
H4K20 me2:H4−4ac
H3K4 me1:H3−5ac
H3K9 me3
H3K4 me3:H4−4ac
H4K20 me3
H3K9 me2
H3K27 me2
H3K27 me3
H3K4 me1
H3−5ac
H4K20 me2
H2A.Z
H3K4 me3
DNA Meth. m5C
H4−4ac
0.00
0.25
0.50
0.75
1.00
$\hat{\pi}$
SMAD[2,3,9] – Reverse
K7ELQ4 – Forward
DNA Meth. m5C:H4K20 me2
DNA Meth. m5C:H3K9 me3
H2A.Z:H3−5ac
H3K27 me3:DNA Meth. m5C
H2A.Z:H4−4ac
DNA Meth. m5C:H4K20 me3
H3K4 me1:H4K20 me2
H3K27 me3:H3K4 me1
H3K4 me1:H2A.Z
H2A.Z:H4K20 me2
H4K20 me2:H4−4ac
H4−4ac:H3−5ac
H4K20 me2:H3K4 me3
H2A.Z:H3K4 me3
H3K4 me1:H3−5ac
H3K4 me1:H4−4ac
H3K9 me2:DNA Meth. m5C
H3K4 me3:H3−5ac
H4K20 me2:H3−5ac
H3K4 me3:H4−4ac
H3K27 me2
H4K20 me3
H3K27 me3
H3K9 me3
H2A.Z
H3K9 me2
DNA Meth. m5C
H3K4 me1
H3K4 me3
H4K20 me2
H3−5ac
H4−4ac
K7ELQ4 – Reverse
RBM47 – Forward
H2A.Z:H3−5ac
H3K4 me1:H4K20 me2
H2A.Z:H4−4ac
DNA Meth. m5C:H4K20 me3
H3K27 me2:DNA Meth. m5C
H2A.Z:H3K4 me3
H3K4 me1:H4−4ac
DNA Meth. m5C:H4K20 me2
H3K27 me3:H3K4 me1
H3K27 me3:H4K20 me3
H2A.Z:H4K20 me2
H3K4 me1:H3−5ac
H3K4 me1:H2A.Z
H3K9 me3:H4K20 me3
H3K27 me3:DNA Meth. m5C
H4K20 me2:H3−5ac
H4−4ac:H3−5ac
DNA Meth. m5C:H3K9 me3
H3K4 me3:H3−5ac
H4K20 me2:H4−4ac
H3K4 me3:H4−4ac
H3K27 me3:H3K9 me3
H4K20 me2:H3K4 me3
H3K9 me2
H3K27 me2
H3K9 me3
H4K20 me3
DNA Meth. m5C
H2A.Z
H3K4 me1
H3K27 me3
H3K4 me3
H4K20 me2
H3−5ac
H4−4ac
RBM47 – Reverse
TOMM70 – Forward
H3K27 me2:DNA Meth. m5C
H3K9 me2:DNA Meth. m5C
H2A.Z:H3−5ac
H3K27 me3:H3K9 me3
H3K27 me3:DNA Meth. m5C
H3K4 me1:H4K20 me2
H2A.Z:H4−4ac
H2A.Z:H3K4 me3
H3K4 me1:H2A.Z
DNA Meth. m5C:H4K20 me2
DNA Meth. m5C:H3K9 me3
H2A.Z:H4K20 me2
H3K4 me1:H4−4ac
H3K4 me3:H3−5ac
H4K20 me2:H4−4ac
H4K20 me2:H3K4 me3
H3K4 me1:H3−5ac
H4K20 me3
H4K20 me2:H3−5ac
H4−4ac:H3−5ac
H3K4 me3:H4−4ac
H3K9 me2
H3K27 me2
H3K27 me3
H3K9 me3
DNA Meth. m5C
H2A.Z
H3K4 me1
H4K20 me2
H3K4 me3
H3−5ac
H4−4ac
TOMM70 – Reverse
SSB – Forward
H4−4ac:H3−5ac
H4K20 me2:H3−5ac
H4K20 me2:H3K4 me3
H2A.Z:H3K4 me3
H3K27 me3:H3K4 me1
H4K20 me2:H4−4ac
H3K9 me2:DNA Meth. m5C
H3K27 me2:DNA Meth. m5C
H3K9 me3:H4K20 me3
H3K27 me3:H4K20 me3
H3K27 me3:H3K9 me3
DNA Meth. m5C:H4K20 me3
DNA Meth. m5C:H4K20 me2
H3−5ac
H2A.Z
H3K27 me3:DNA Meth. m5C
DNA Meth. m5C:H3K9 me3
H3K4 me1
H4−4ac
H3K27 me2
H3K4 me3
H3K9 me2
H4K20 me2
H4K20 me3
H3K27 me3
H3K9 me3
DNA Meth. m5C
SSB – Reverse
RPSA – Forward
H4K20 me2:H3K4 me3
H2A.Z:H4−4ac
H3K27 me3:H3K4 me1
H4K20 me2:H4−4ac
H3K27 me2:DNA Meth. m5C
DNA Meth. m5C:H4K20 me2
DNA Meth. m5C:H3K9 me3
H3K27 me3:DNA Meth. m5C
H3K27 me3:H3K9 me3
H3−5ac
H3K27 me2
H2A.Z
DNA Meth. m5C:H4K20 me3
H3K9 me2
H3K4 me1
H3K9 me3:H4K20 me3
H3K27 me3:H4K20 me3
H4K20 me2
H3K4 me3
H4−4ac
DNA Meth. m5C
H3K27 me3
H3K9 me3
H4K20 me3
RPSA – Reverse

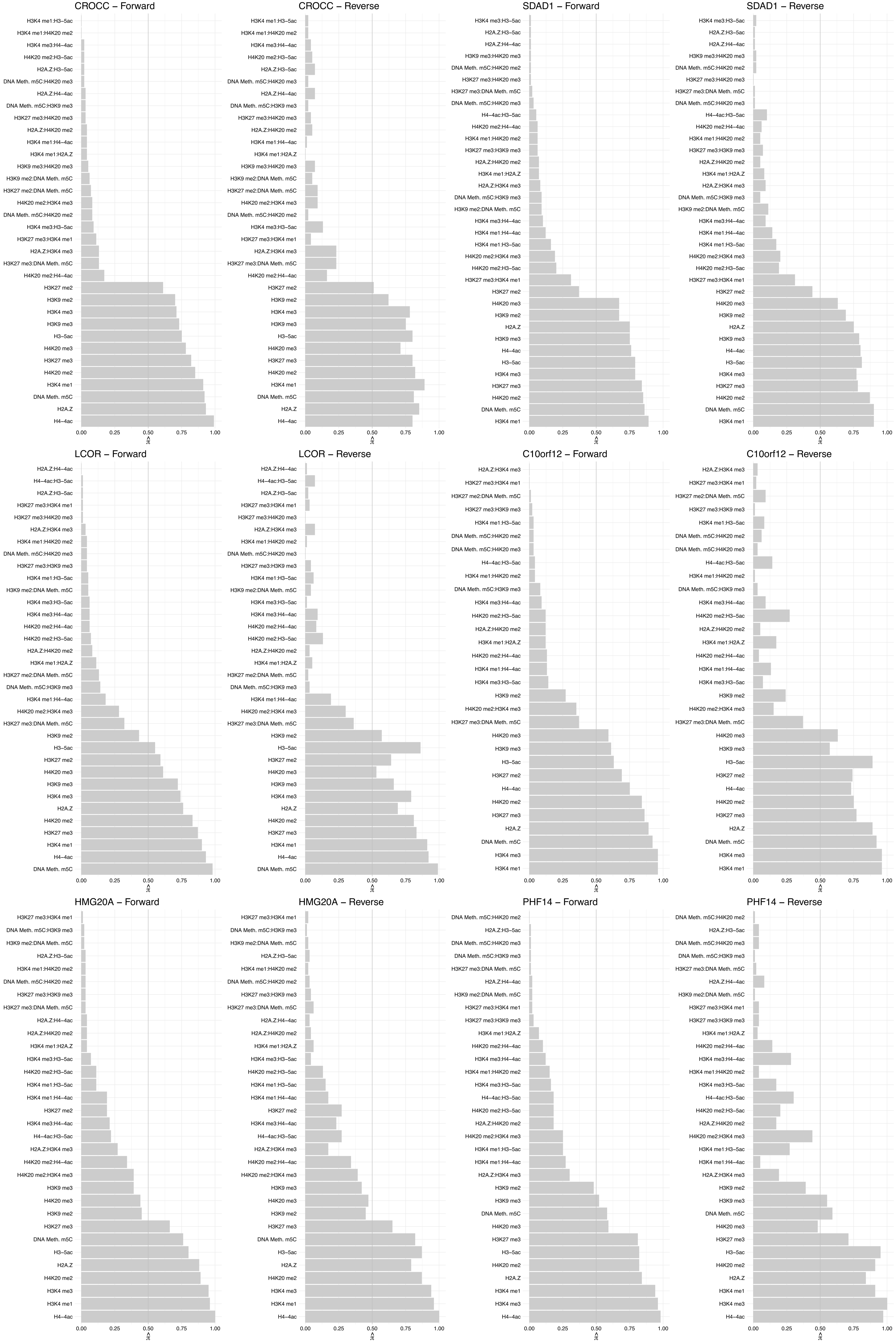
CROCC – Forward
H3K4 me1:H3–5ac
H3K4 me1:H4K20 me2
H3K4 me3:H4–4ac
H4K20 me2:H3–5ac
H2A.Z:H3–5ac
DNA Meth. m5C:H4K20 me3
H2A.Z:H4–4ac
DNA Meth. m5C:H3K9 me3
H3K27 me3:H4K20 me3
H2A.Z:H4K20 me2
H3K4 me1:H4–4ac
H3K4 me1:H2A.Z
H3K9 me3:H4K20 me3
H3K9 me2:DNA Meth. m5C
H3K27 me2:DNA Meth. m5C
H4K20 me2:H3K4 me3
DNA Meth. m5C:H4K20 me2
H3K4 me3:H3–5ac
H3K27 me3:H3K4 me1
H2A.Z:H3K4 me3
H3K27 me3:DNA Meth. m5C
H4K20 me2:H4–4ac
H3K27 me2
H3K9 me2
H3K4 me3
H3K9 me3
H3–5ac
H4K20 me3
H3K27 me3
H4K20 me2
H3K4 me1
DNA Meth. m5C
H2A.Z
H4–4ac
0.00
0.25
0.50
0.75
1.00
$\hat{\pi}$
CROCC – Reverse
H3K4 me1:H3–5ac
H3K4 me1:H4K20 me2
H3K4 me3:H4–4ac
H4K20 me2:H3–5ac
H2A.Z:H3–5ac
DNA Meth. m5C:H4K20 me3
H2A.Z:H4–4ac
DNA Meth. m5C:H3K9 me3
H3K27 me3:H4K20 me3
H2A.Z:H4K20 me2
H3K4 me1:H4–4ac
H3K4 me1:H2A.Z
H3K9 me3:H4K20 me3
H3K9 me2:DNA Meth. m5C
H3K27 me2:DNA Meth. m5C
H4K20 me2:H3K4 me3
DNA Meth. m5C:H4K20 me2
H3K4 me3:H3–5ac
H3K27 me3:H3K4 me1
H2A.Z:H3K4 me3
H3K27 me3:DNA Meth. m5C
H4K20 me2:H4–4ac
H3K27 me2
H3K9 me2
H3K4 me3
H3K9 me3
H3–5ac
H4K20 me3
H3K27 me3
H4K20 me2
H3K4 me1
DNA Meth. m5C
H2A.Z
H4–4ac
0.00
0.25
0.50
0.75
1.00
$\hat{\pi}$
SDAD1 – Forward
H3K4 me3:H3–5ac
H2A.Z:H3–5ac
H2A.Z:H4–4ac
H3K9 me3:H4K20 me3
DNA Meth. m5C:H4K20 me2
H3K27 me3:H4K20 me3
H3K27 me3:DNA Meth. m5C
DNA Meth. m5C:H4K20 me3
H4–4ac:H3–5ac
H4K20 me2:H4–4ac
H3K4 me1:H4K20 me2
H3K27 me3:H3K9 me3
H2A.Z:H4K20 me2
H3K4 me1:H2A.Z
H2A.Z:H3K4 me3
DNA Meth. m5C:H3K9 me3
H3K9 me2:DNA Meth. m5C
H3K4 me3:H4–4ac
H3K4 me1:H4–4ac
H3K4 me1:H3–5ac
H4K20 me2:H3K4 me3
H4K20 me2:H3–5ac
H3K27 me3:H3K4 me1
H3K27 me2
H4K20 me3
H3K9 me2
H2A.Z
H3K9 me3
H4–4ac
H3–5ac
H3K4 me3
H3K27 me3
H4K20 me2
DNA Meth. m5C
H3K4 me1
0.00
0.25
0.50
0.75
1.00
$\hat{\pi}$
SDAD1 – Reverse
H3K4 me3:H3–5ac
H2A.Z:H3–5ac
H2A.Z:H4–4ac
H3K9 me3:H4K20 me3
DNA Meth. m5C:H4K20 me2
H3K27 me3:H4K20 me3
H3K27 me3:DNA Meth. m5C
DNA Meth. m5C:H4K20 me3
H4–4ac:H3–5ac
H4K20 me2:H4–4ac
H3K4 me1:H4K20 me2
H3K27 me3:H3K9 me3
H2A.Z:H4K20 me2
H3K4 me1:H2A.Z
H2A.Z:H3K4 me3
DNA Meth. m5C:H3K9 me3
H3K9 me2:DNA Meth. m5C
H3K4 me3:H4–4ac
H3K4 me1:H4–4ac
H3K4 me1:H3–5ac
H4K20 me2:H3K4 me3
H4K20 me2:H3–5ac
H3K27 me3:H3K4 me1
H3K27 me2
H4K20 me3
H3K9 me2
H2A.Z
H3K9 me3
H4–4ac
H3–5ac
H3K4 me3
H3K27 me3
H4K20 me2
DNA Meth. m5C
H3K4 me1
0.00
0.25
0.50
0.75
1.00
$\hat{\pi}$
LCOR – Forward
H2A.Z:H4–4ac
H4–4ac:H3–5ac
H2A.Z:H3–5ac
H3K27 me3:H3K4 me1
H3K27 me3:H4K20 me3
H2A.Z:H3K4 me3
H3K4 me1:H4K20 me2
DNA Meth. m5C:H4K20 me3
H3K27 me3:H3K9 me3
H3K4 me1:H3–5ac
H3K9 me2:DNA Meth. m5C
H3K4 me3:H3–5ac
H3K4 me3:H4–4ac
H4K20 me2:H4–4ac
H4K20 me2:H3–5ac
H2A.Z:H4K20 me2
H3K4 me1:H2A.Z
H3K27 me2:DNA Meth. m5C
DNA Meth. m5C:H3K9 me3
H3K4 me1:H4–4ac
H4K20 me2:H3K4 me3
H3K27 me3:DNA Meth. m5C
H3K9 me2
H3–5ac
H3K27 me2
H4K20 me3
H3K9 me3
H3K4 me3
H2A.Z
H4K20 me2
H3K27 me3
H3K4 me1
H4–4ac
DNA Meth. m5C
0.00
0.25
0.50
0.75
1.00
$\hat{\pi}$
LCOR – Reverse
H2A.Z:H4–4ac
H4–4ac:H3–5ac
H2A.Z:H3–5ac
H3K27 me3:H3K4 me1
H3K27 me3:H4K20 me3
H2A.Z:H3K4 me3
H3K4 me1:H4K20 me2
DNA Meth. m5C:H4K20 me3
H3K27 me3:H3K9 me3
H3K4 me1:H3–5ac
H3K9 me2:DNA Meth. m5C
H3K4 me3:H3–5ac
H3K4 me3:H4–4ac
H4K20 me2:H4–4ac
H4K20 me2:H3–5ac
H2A.Z:H4K20 me2
H3K4 me1:H2A.Z
H3K27 me2:DNA Meth. m5C
DNA Meth. m5C:H3K9 me3
H3K4 me1:H4–4ac
H4K20 me2:H3K4 me3
H3K27 me3:DNA Meth. m5C
H3K9 me2
H3–5ac
H3K27 me2
H4K20 me3
H3K9 me3
H3K4 me3
H2A.Z
H4K20 me2
H3K27 me3
H3K4 me1
H4–4ac
DNA Meth. m5C
0.00
0.25
0.50
0.75
1.00
$\hat{\pi}$
C10orf12 – Forward
H2A.Z:H3K4 me3
H3K27 me3:H3K4 me1
H3K27 me2:DNA Meth. m5C
H3K27 me3:H3K9 me3
H3K4 me1:H3–5ac
DNA Meth. m5C:H4K20 me2
DNA Meth. m5C:H4K20 me3
H4–4ac:H3–5ac
H3K4 me1:H4K20 me2
DNA Meth. m5C:H3K9 me3
H3K4 me3:H4–4ac
H4K20 me2:H3–5ac
H2A.Z:H4K20 me2
H3K4 me1:H2A.Z
H4K20 me2:H4–4ac
H3K4 me1:H4–4ac
H3K4 me3:H3–5ac
H3K9 me2
H4K20 me2:H3K4 me3
H3K27 me3:DNA Meth. m5C
H4K20 me3
H3K9 me3
H3–5ac
H3K27 me2
H4–4ac
H4K20 me2
H3K27 me3
H2A.Z
DNA Meth. m5C
H3K4 me3
H3K4 me1
0.00
0.25
0.50
0.75
1.00
$\hat{\pi}$
C10orf12 – Reverse
H2A.Z:H3K4 me3
H3K27 me3:H3K4 me1
H3K27 me2:DNA Meth. m5C
H3K27 me3:H3K9 me3
H3K4 me1:H3–5ac
DNA Meth. m5C:H4K20 me2
DNA Meth. m5C:H4K20 me3
H4–4ac:H3–5ac
H3K4 me1:H4K20 me2
DNA Meth. m5C:H3K9 me3
H3K4 me3:H4–4ac
H4K20 me2:H3–5ac
H2A.Z:H4K20 me2
H3K4 me1:H2A.Z
H4K20 me2:H4–4ac
H3K4 me1:H4–4ac
H3K4 me3:H3–5ac
H3K9 me2
H4K20 me2:H3K4 me3
H3K27 me3:DNA Meth. m5C
H4K20 me3
H3K9 me3
H3–5ac
H3K27 me2
H4–4ac
H4K20 me2
H3K27 me3
H2A.Z
DNA Meth. m5C
H3K4 me3
H3K4 me1
0.00
0.25
0.50
0.75
1.00
$\hat{\pi}$
HMG20A – Forward
H3K27 me3:H3K4 me1
DNA Meth. m5C:H3K9 me3
H3K9 me2:DNA Meth. m5C
H2A.Z:H3–5ac
H3K4 me1:H4K20 me2
DNA Meth. m5C:H4K20 me2
H3K27 me3:H3K9 me3
H3K27 me3:DNA Meth. m5C
H2A.Z:H4–4ac
H2A.Z:H4K20 me2
H3K4 me1:H2A.Z
H3K4 me3:H3–5ac
H4K20 me2:H3–5ac
H3K4 me1:H3–5ac
H3K4 me1:H4–4ac
H3K27 me2
H3K4 me3:H4–4ac
H4–4ac:H3–5ac
H2A.Z:H3K4 me3
H4K20 me2:H4–4ac
H4K20 me2:H3K4 me3
H3K9 me3
H4K20 me3
H3K9 me2
H3K27 me3
DNA Meth. m5C
H3–5ac
H2A.Z
H4K20 me2
H3K4 me3
H3K4 me1
H4–4ac
0.00
0.25
0.50
0.75
1.00
$\hat{\pi}$
HMG20A – Reverse
H3K27 me3:H3K4 me1
DNA Meth. m5C:H3K9 me3
H3K9 me2:DNA Meth. m5C
H2A.Z:H3–5ac
H3K4 me1:H4K20 me2
DNA Meth. m5C:H4K20 me2
H3K27 me3:H3K9 me3
H3K27 me3:DNA Meth. m5C
H2A.Z:H4–4ac
H2A.Z:H4K20 me2
H3K4 me1:H2A.Z
H3K4 me3:H3–5ac
H4K20 me2:H3–5ac
H3K4 me1:H3–5ac
H3K4 me1:H4–4ac
H3K27 me2
H3K4 me3:H4–4ac
H4–4ac:H3–5ac
H2A.Z:H3K4 me3
H4K20 me2:H4–4ac
H4K20 me2:H3K4 me3
H3K9 me3
H4K20 me3
H3K9 me2
H3K27 me3
DNA Meth. m5C
H3–5ac
H2A.Z
H4K20 me2
H3K4 me3
H3K4 me1
H4–4ac
0.00
0.25
0.50
0.75
1.00
$\hat{\pi}$
PHF14 – Forward
DNA Meth. m5C:H4K20 me2
H2A.Z:H3–5ac
DNA Meth. m5C:H4K20 me3
DNA Meth. m5C:H3K9 me3
H3K27 me3:DNA Meth. m5C
H2A.Z:H4–4ac
H3K9 me2:DNA Meth. m5C
H3K27 me3:H3K4 me1
H3K27 me3:H3K9 me3
H3K4 me1:H2A.Z
H4K20 me2:H4–4ac
H3K4 me3:H4–4ac
H3K4 me1:H4K20 me2
H3K4 me3:H3–5ac
H4–4ac:H3–5ac
H4K20 me2:H3–5ac
H2A.Z:H4K20 me2
H4K20 me2:H3K4 me3
H3K4 me1:H3–5ac
H3K4 me1:H4–4ac
H2A.Z:H3K4 me3
H3K9 me2
H3K9 me3
DNA Meth. m5C
H4K20 me3
H3K27 me3
H3–5ac
H4K20 me2
H2A.Z
H3K4 me1
H3K4 me3
H4–4ac
0.00
0.25
0.50
0.75
1.00
$\hat{\pi}$
PHF14 – Reverse
DNA Meth. m5C:H4K20 me2
H2A.Z:H3–5ac
DNA Meth. m5C:H4K20 me3
DNA Meth. m5C:H3K9 me3
H3K27 me3:DNA Meth. m5C
H2A.Z:H4–4ac
H3K9 me2:DNA Meth. m5C
H3K27 me3:H3K4 me1
H3K27 me3:H3K9 me3
H3K4 me1:H2A.Z
H4K20 me2:H4–4ac
H3K4 me3:H4–4ac
H3K4 me1:H4K20 me2
H3K4 me3:H3–5ac
H4–4ac:H3–5ac
H4K20 me2:H3–5ac
H2A.Z:H4K20 me2
H4K20 me2:H3K4 me3
H3K4 me1:H3–5ac
H3K4 me1:H4–4ac
H2A.Z:H3K4 me3
H3K9 me2
H3K9 me3
DNA Meth. m5C
H4K20 me3
H3K27 me3
H3–5ac
H4K20 me2
H2A.Z
H3K4 me1
H3K4 me3
H4–4ac
0.00
0.25
0.50
0.75
1.00
$\hat{\pi}$

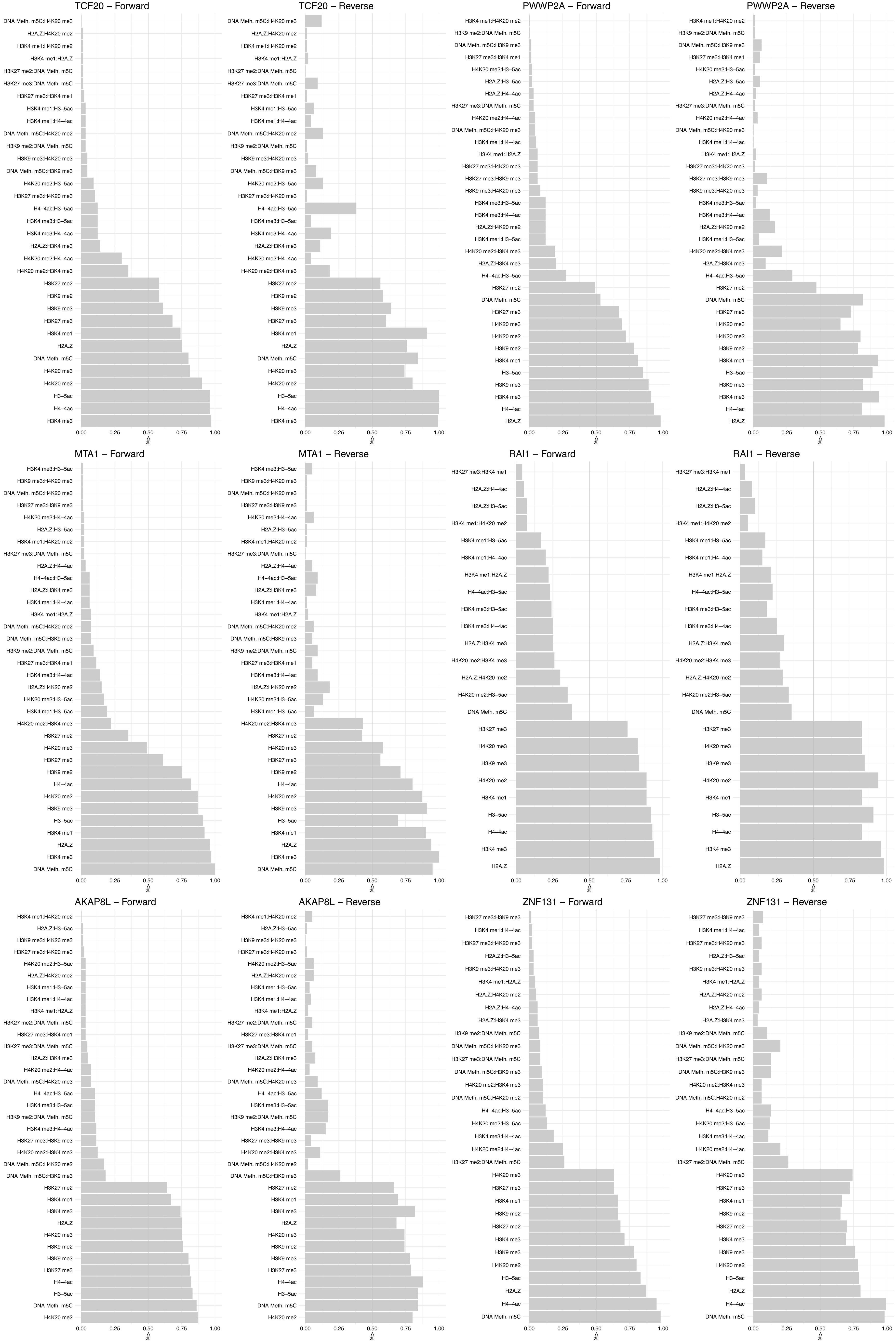
TCF20 – Forward
TCF20 – Reverse
PWWP2A – Forward
PWWP2A – Reverse
MTA1 – Forward
MTA1 – Reverse
RAI1 – Forward
RAI1 – Reverse
AKAP8L – Forward
AKAP8L – Reverse
ZNF131 – Forward
ZNF131 – Reverse
0.00 0.25 0.50 0.75 1.00
$\hat{\pi}$

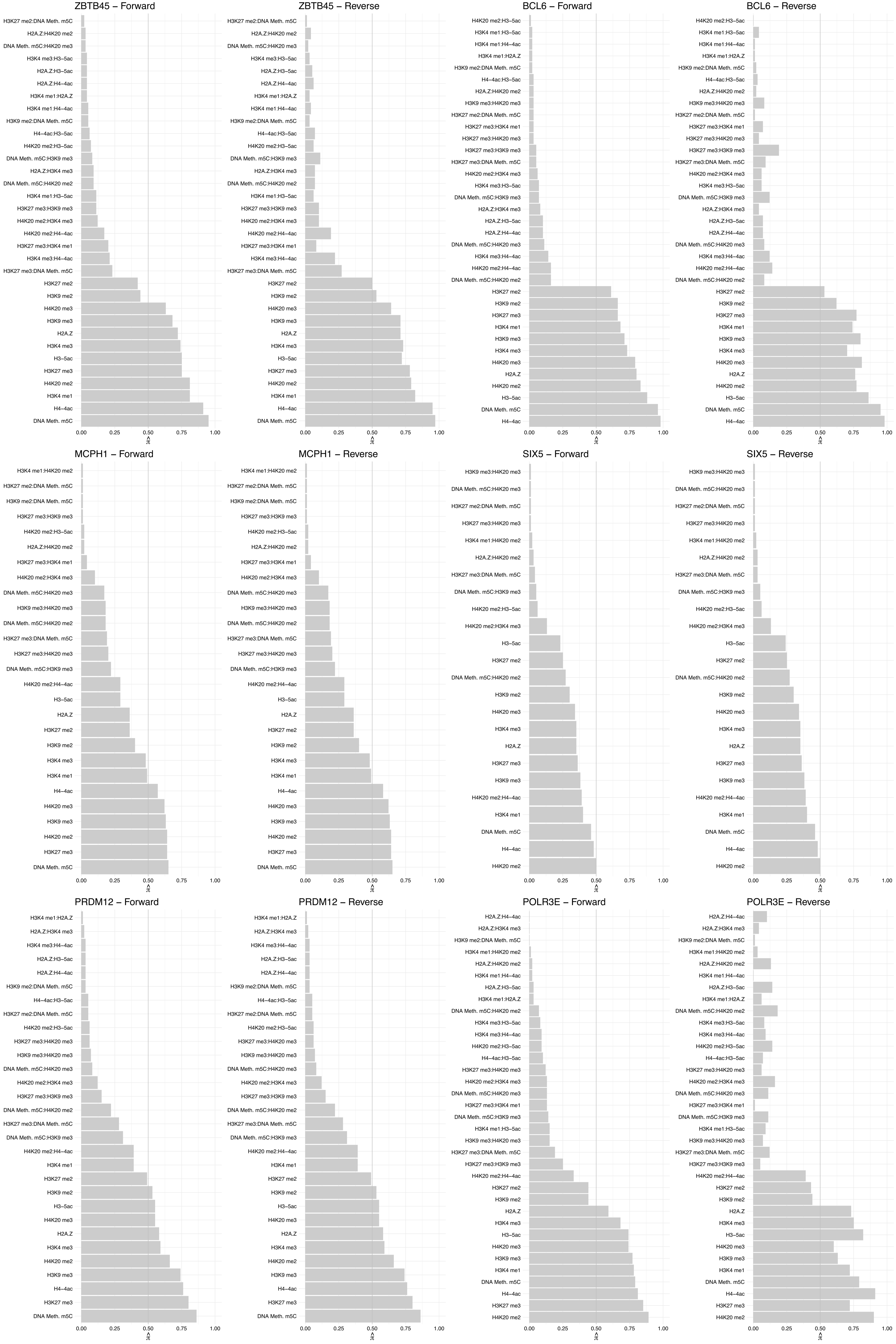
ZBTB45 – Forward
ZBTB45 – Reverse
BCL6 – Forward
BCL6 – Reverse
MCPH1 – Forward
MCPH1 – Reverse
SIX5 – Forward
SIX5 – Reverse
PRDM12 – Forward
PRDM12 – Reverse
POLR3E – Forward
POLR3E – Reverse
0.00
0.25
0.50
0.75
1.00
$\hat{\pi}$

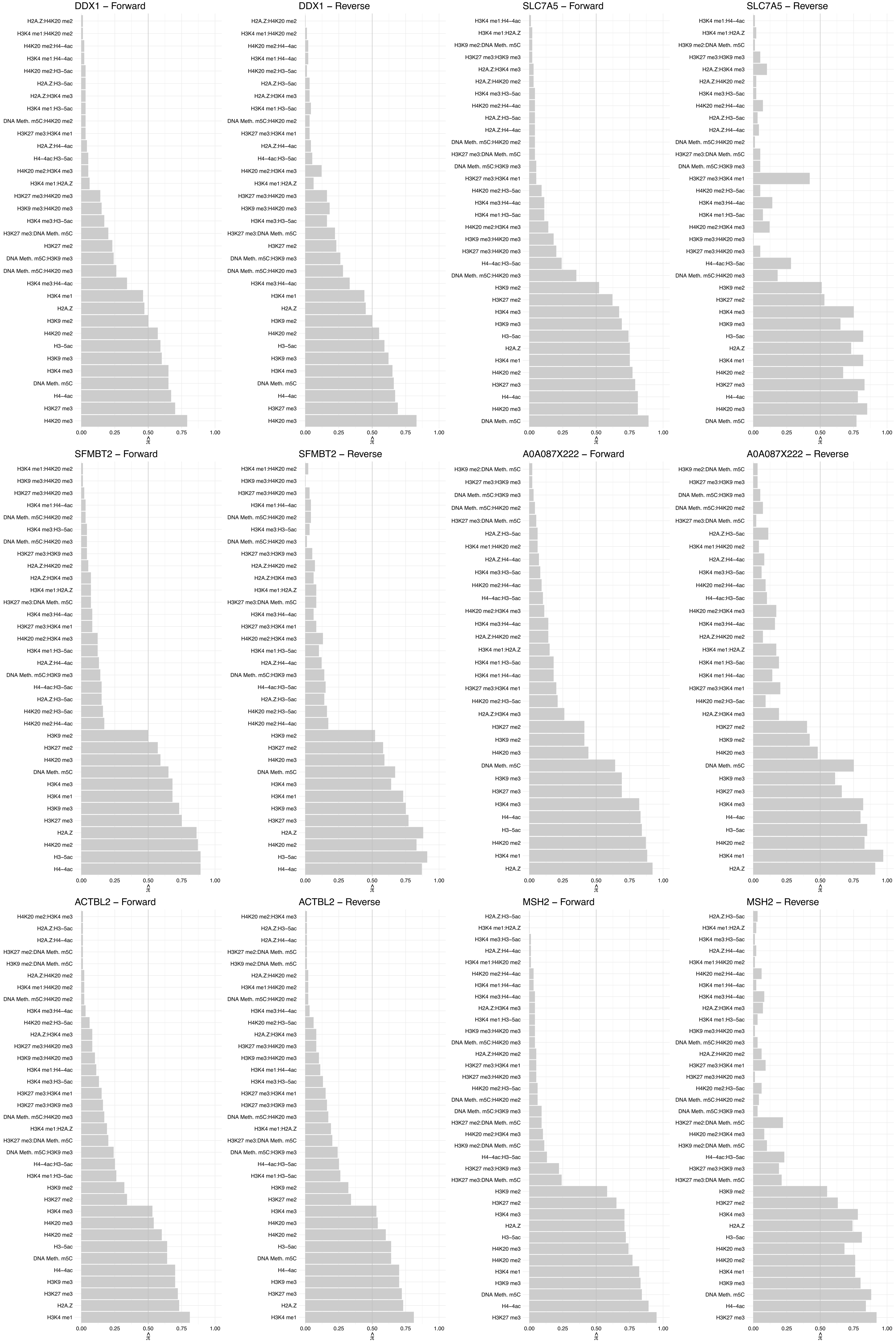

DDX1 – Forward
H2A.Z:H4K20 me2
H3K4 me1:H4K20 me2
H4K20 me2:H4−4ac
H3K4 me1:H4−4ac
H4K20 me2:H3−5ac
H2A.Z:H3−5ac
H2A.Z:H3K4 me3
H3K4 me1:H3−5ac
DNA Meth. m5C:H4K20 me2
H3K27 me3:H3K4 me1
H2A.Z:H4−4ac
H4−4ac:H3−5ac
H4K20 me2:H3K4 me3
H3K4 me1:H2A.Z
H3K27 me3:H4K20 me3
H3K9 me3:H4K20 me3
H3K4 me3:H3−5ac
H3K27 me3:DNA Meth. m5C
H3K27 me2
DNA Meth. m5C:H3K9 me3
DNA Meth. m5C:H4K20 me3
H3K4 me3:H4−4ac
H3K4 me1
H2A.Z
H3K9 me2
H4K20 me2
H3−5ac
H3K9 me3
H3K4 me3
DNA Meth. m5C
H4−4ac
H3K27 me3
H4K20 me3
0.00 0.25 0.50 0.75 1.00
$\hat{\pi}$
DDX1 – Reverse
H2A.Z:H4K20 me2
H3K4 me1:H4K20 me2
H4K20 me2:H4−4ac
H3K4 me1:H4−4ac
H4K20 me2:H3−5ac
H2A.Z:H3−5ac
H2A.Z:H3K4 me3
H3K4 me1:H3−5ac
DNA Meth. m5C:H4K20 me2
H3K27 me3:H3K4 me1
H2A.Z:H4−4ac
H4−4ac:H3−5ac
H4K20 me2:H3K4 me3
H3K4 me1:H2A.Z
H3K27 me3:H4K20 me3
H3K9 me3:H4K20 me3
H3K4 me3:H3−5ac
H3K27 me3:DNA Meth. m5C
H3K27 me2
DNA Meth. m5C:H3K9 me3
DNA Meth. m5C:H4K20 me3
H3K4 me3:H4−4ac
H3K4 me1
H2A.Z
H3K9 me2
H4K20 me2
H3−5ac
H3K9 me3
H3K4 me3
DNA Meth. m5C
H4−4ac
H3K27 me3
H4K20 me3
0.00 0.25 0.50 0.75 1.00
$\hat{\pi}$
SLC7A5 – Forward
H3K4 me1:H4−4ac
H3K4 me1:H2A.Z
H3K9 me2:DNA Meth. m5C
H3K27 me3:H3K9 me3
H2A.Z:H3K4 me3
H2A.Z:H4K20 me2
H3K4 me3:H3−5ac
H4K20 me2:H4−4ac
H2A.Z:H3−5ac
H2A.Z:H4−4ac
DNA Meth. m5C:H4K20 me2
H3K27 me3:DNA Meth. m5C
DNA Meth. m5C:H3K9 me3
H3K27 me3:H3K4 me1
H4K20 me2:H3−5ac
H3K4 me3:H4−4ac
H3K4 me1:H3−5ac
H4K20 me2:H3K4 me3
H3K9 me3:H4K20 me3
H3K27 me3:H4K20 me3
H4−4ac:H3−5ac
DNA Meth. m5C:H4K20 me3
H3K9 me2
H3K27 me2
H3K4 me3
H3K9 me3
H3−5ac
H2A.Z
H3K4 me1
H4K20 me2
H3K27 me3
H4−4ac
H4K20 me3
DNA Meth. m5C
0.00 0.25 0.50 0.75 1.00
$\hat{\pi}$
SLC7A5 – Reverse
H3K4 me1:H4−4ac
H3K4 me1:H2A.Z
H3K9 me2:DNA Meth. m5C
H3K27 me3:H3K9 me3
H2A.Z:H3K4 me3
H2A.Z:H4K20 me2
H3K4 me3:H3−5ac
H4K20 me2:H4−4ac
H2A.Z:H3−5ac
H2A.Z:H4−4ac
DNA Meth. m5C:H4K20 me2
H3K27 me3:DNA Meth. m5C
DNA Meth. m5C:H3K9 me3
H3K27 me3:H3K4 me1
H4K20 me2:H3−5ac
H3K4 me3:H4−4ac
H3K4 me1:H3−5ac
H4K20 me2:H3K4 me3
H3K9 me3:H4K20 me3
H3K27 me3:H4K20 me3
H4−4ac:H3−5ac
DNA Meth. m5C:H4K20 me3
H3K9 me2
H3K27 me2
H3K4 me3
H3K9 me3
H3−5ac
H2A.Z
H3K4 me1
H4K20 me2
H3K27 me3
H4−4ac
H4K20 me3
DNA Meth. m5C
0.00 0.25 0.50 0.75 1.00
$\hat{\pi}$
SFMBT2 – Forward
H3K4 me1:H4K20 me2
H3K9 me3:H4K20 me3
H3K27 me3:H4K20 me3
H3K4 me1:H4−4ac
DNA Meth. m5C:H4K20 me2
H3K4 me3:H3−5ac
DNA Meth. m5C:H4K20 me3
H3K27 me3:H3K9 me3
H2A.Z:H4K20 me2
H2A.Z:H3K4 me3
H3K4 me1:H2A.Z
H3K27 me3:DNA Meth. m5C
H3K4 me3:H4−4ac
H3K27 me3:H3K4 me1
H4K20 me2:H3K4 me3
H3K4 me1:H3−5ac
H2A.Z:H4−4ac
DNA Meth. m5C:H3K9 me3
H4−4ac:H3−5ac
H2A.Z:H3−5ac
H4K20 me2:H3−5ac
H4K20 me2:H4−4ac
H3K9 me2
H3K27 me2
H4K20 me3
DNA Meth. m5C
H3K4 me3
H3K4 me1
H3K9 me3
H3K27 me3
H2A.Z
H4K20 me2
H3−5ac
H4−4ac
0.00 0.25 0.50 0.75 1.00
$\hat{\pi}$
SFMBT2 – Reverse
H3K4 me1:H4K20 me2
H3K9 me3:H4K20 me3
H3K27 me3:H4K20 me3
H3K4 me1:H4−4ac
DNA Meth. m5C:H4K20 me2
H3K4 me3:H3−5ac
DNA Meth. m5C:H4K20 me3
H3K27 me3:H3K9 me3
H2A.Z:H4K20 me2
H2A.Z:H3K4 me3
H3K4 me1:H2A.Z
H3K27 me3:DNA Meth. m5C
H3K4 me3:H4−4ac
H3K27 me3:H3K4 me1
H4K20 me2:H3K4 me3
H3K4 me1:H3−5ac
H2A.Z:H4−4ac
DNA Meth. m5C:H3K9 me3
H4−4ac:H3−5ac
H2A.Z:H3−5ac
H4K20 me2:H3−5ac
H4K20 me2:H4−4ac
H3K9 me2
H3K27 me2
H4K20 me3
DNA Meth. m5C
H3K4 me3
H3K4 me1
H3K9 me3
H3K27 me3
H2A.Z
H4K20 me2
H3−5ac
H4−4ac
0.00 0.25 0.50 0.75 1.00
$\hat{\pi}$
A0A087X222 – Forward
H3K9 me2:DNA Meth. m5C
H3K27 me3:H3K9 me3
DNA Meth. m5C:H3K9 me3
DNA Meth. m5C:H4K20 me2
H3K27 me3:DNA Meth. m5C
H2A.Z:H3−5ac
H3K4 me1:H4K20 me2
H2A.Z:H4−4ac
H3K4 me3:H3−5ac
H4K20 me2:H4−4ac
H4−4ac:H3−5ac
H4K20 me2:H3K4 me3
H3K4 me3:H4−4ac
H2A.Z:H4K20 me2
H3K4 me1:H2A.Z
H3K4 me1:H3−5ac
H3K4 me1:H4−4ac
H3K27 me3:H3K4 me1
H4K20 me2:H3−5ac
H2A.Z:H3K4 me3
H3K27 me2
H3K9 me2
H4K20 me3
DNA Meth. m5C
H3K9 me3
H3K27 me3
H3K4 me3
H4−4ac
H3−5ac
H4K20 me2
H3K4 me1
H2A.Z
0.00 0.25 0.50 0.75 1.00
$\hat{\pi}$
A0A087X222 – Reverse
H3K9 me2:DNA Meth. m5C
H3K27 me3:H3K9 me3
DNA Meth. m5C:H3K9 me3
DNA Meth. m5C:H4K20 me2
H3K27 me3:DNA Meth. m5C
H2A.Z:H3−5ac
H3K4 me1:H4K20 me2
H2A.Z:H4−4ac
H3K4 me3:H3−5ac
H4K20 me2:H4−4ac
H4−4ac:H3−5ac
H4K20 me2:H3K4 me3
H3K4 me3:H4−4ac
H2A.Z:H4K20 me2
H3K4 me1:H2A.Z
H3K4 me1:H3−5ac
H3K4 me1:H4−4ac
H3K27 me3:H3K4 me1
H4K20 me2:H3−5ac
H2A.Z:H3K4 me3
H3K27 me2
H3K9 me2
H4K20 me3
DNA Meth. m5C
H3K9 me3
H3K27 me3
H3K4 me3
H4−4ac
H3−5ac
H4K20 me2
H3K4 me1
H2A.Z
0.00 0.25 0.50 0.75 1.00
$\hat{\pi}$
ACTBL2 – Forward
H4K20 me2:H3K4 me3
H2A.Z:H3−5ac
H2A.Z:H4−4ac
H3K27 me2:DNA Meth. m5C
H3K9 me2:DNA Meth. m5C
H2A.Z:H4K20 me2
H3K4 me1:H4K20 me2
DNA Meth. m5C:H4K20 me2
H3K4 me3:H4−4ac
H4K20 me2:H3−5ac
H2A.Z:H3K4 me3
H3K27 me3:H4K20 me3
H3K9 me3:H4K20 me3
H3K4 me1:H4−4ac
H3K4 me3:H3−5ac
H3K27 me3:H3K4 me1
H3K27 me3:H3K9 me3
DNA Meth. m5C:H4K20 me3
H3K4 me1:H2A.Z
H3K27 me3:DNA Meth. m5C
DNA Meth. m5C:H3K9 me3
H4−4ac:H3−5ac
H3K4 me1:H3−5ac
H3K9 me2
H3K27 me2
H3K4 me3
H4K20 me3
H4K20 me2
H3−5ac
DNA Meth. m5C
H4−4ac
H3K9 me3
H3K27 me3
H2A.Z
H3K4 me1
0.00 0.25 0.50 0.75 1.00
$\hat{\pi}$
ACTBL2 – Reverse
H4K20 me2:H3K4 me3
H2A.Z:H3−5ac
H2A.Z:H4−4ac
H3K27 me2:DNA Meth. m5C
H3K9 me2:DNA Meth. m5C
H2A.Z:H4K20 me2
H3K4 me1:H4K20 me2
DNA Meth. m5C:H4K20 me2
H3K4 me3:H4−4ac
H4K20 me2:H3−5ac
H2A.Z:H3K4 me3
H3K27 me3:H4K20 me3
H3K9 me3:H4K20 me3
H3K4 me1:H4−4ac
H3K4 me3:H3−5ac
H3K27 me3:H3K4 me1
H3K27 me3:H3K9 me3
DNA Meth. m5C:H4K20 me3
H3K4 me1:H2A.Z
H3K27 me3:DNA Meth. m5C
DNA Meth. m5C:H3K9 me3
H4−4ac:H3−5ac
H3K4 me1:H3−5ac
H3K9 me2
H3K27 me2
H3K4 me3
H4K20 me3
H4K20 me2
H3−5ac
DNA Meth. m5C
H4−4ac
H3K9 me3
H3K27 me3
H2A.Z
H3K4 me1
0.00 0.25 0.50 0.75 1.00
$\hat{\pi}$
MSH2 – Forward
H2A.Z:H3−5ac
H3K4 me1:H2A.Z
H3K4 me3:H3−5ac
H2A.Z:H4−4ac
H3K4 me1:H4K20 me2
H4K20 me2:H4−4ac
H3K4 me1:H4−4ac
H3K4 me3:H4−4ac
H2A.Z:H3K4 me3
H3K4 me1:H3−5ac
H3K9 me3:H4K20 me3
DNA Meth. m5C:H4K20 me3
H2A.Z:H4K20 me2
H3K27 me3:H3K4 me1
H3K27 me3:H4K20 me3
H4K20 me2:H3−5ac
DNA Meth. m5C:H4K20 me2
DNA Meth. m5C:H3K9 me3
H3K27 me2:DNA Meth. m5C
H4K20 me2:H3K4 me3
H3K9 me2:DNA Meth. m5C
H4−4ac:H3−5ac
H3K27 me3:H3K9 me3
H3K27 me3:DNA Meth. m5C
H3K9 me2
H3K27 me2
H3K4 me3
H2A.Z
H3−5ac
H4K20 me3
H4K20 me2
H3K4 me1
H3K9 me3
DNA Meth. m5C
H4−4ac
H3K27 me3
0.00 0.25 0.50 0.75 1.00
$\hat{\pi}$
MSH2 – Reverse
H2A.Z:H3−5ac
H3K4 me1:H2A.Z
H3K4 me3:H3−5ac
H2A.Z:H4−4ac
H3K4 me1:H4K20 me2
H4K20 me2:H4−4ac
H3K4 me1:H4−4ac
H3K4 me3:H4−4ac
H2A.Z:H3K4 me3
H3K4 me1:H3−5ac
H3K9 me3:H4K20 me3
DNA Meth. m5C:H4K20 me3
H2A.Z:H4K20 me2
H3K27 me3:H3K4 me1
H3K27 me3:H4K20 me3
H4K20 me2:H3−5ac
DNA Meth. m5C:H4K20 me2
DNA Meth. m5C:H3K9 me3
H3K27 me2:DNA Meth. m5C
H4K20 me2:H3K4 me3
H3K9 me2:DNA Meth. m5C
H4−4ac:H3−5ac
H3K27 me3:H3K9 me3
H3K27 me3:DNA Meth. m5C
H3K9 me2
H3K27 me2
H3K4 me3
H2A.Z
H3−5ac
H4K20 me3
H4K20 me2
H3K4 me1
H3K9 me3
DNA Meth. m5C
H4−4ac
H3K27 me3
0.00 0.25 0.50 0.75 1.00
$\hat{\pi}$

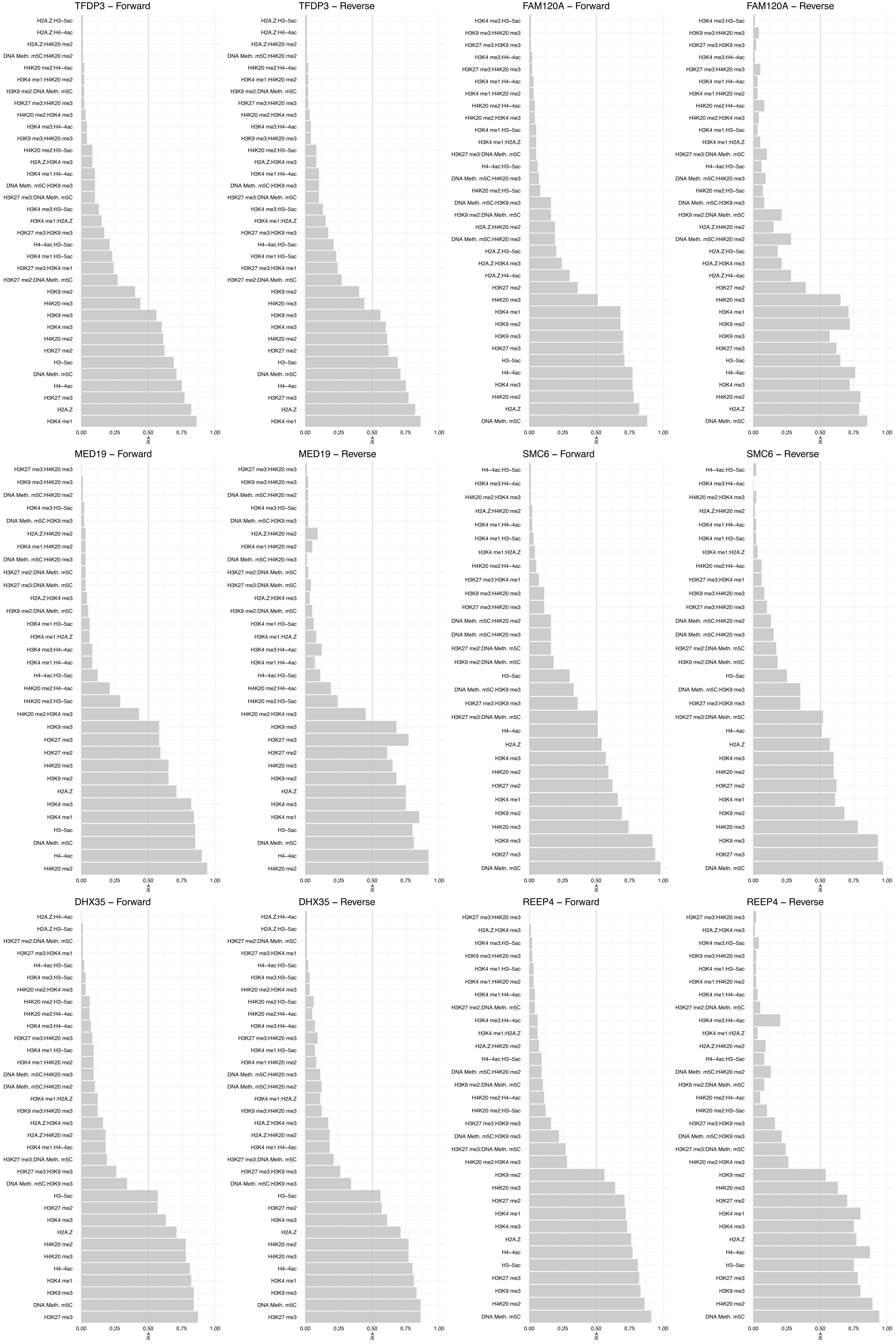
TFDP3 – Forward
H2A.Z:H3–5ac
H2A.Z:H4–4ac
H2A.Z:H4K20 me2
DNA Meth. m5C:H4K20 me2
H4K20 me2:H4–4ac
H3K4 me1:H4K20 me2
H3K9 me2:DNA Meth. m5C
H3K27 me3:H4K20 me3
H4K20 me2:H3K4 me3
H3K4 me3:H4–4ac
H3K9 me3:H4K20 me3
H4K20 me2:H3–5ac
H2A.Z:H3K4 me3
H3K4 me1:H4–4ac
DNA Meth. m5C:H3K9 me3
H3K27 me3:DNA Meth. m5C
H3K4 me3:H3–5ac
H3K4 me1:H2A.Z
H3K27 me3:H3K9 me3
H4–4ac:H3–5ac
H3K4 me1:H3–5ac
H3K27 me3:H3K4 me1
H3K27 me2:DNA Meth. m5C
H3K9 me2
H4K20 me3
H3K9 me3
H3K4 me3
H4K20 me2
H3K27 me2
H3–5ac
DNA Meth. m5C
H4–4ac
H3K27 me3
H2A.Z
H3K4 me1
0.00 0.25 0.50 0.75 1.00
$\hat{\pi}$
TFDP3 – Reverse
H2A.Z:H3–5ac
H2A.Z:H4–4ac
H2A.Z:H4K20 me2
DNA Meth. m5C:H4K20 me2
H4K20 me2:H4–4ac
H3K4 me1:H4K20 me2
H3K9 me2:DNA Meth. m5C
H3K27 me3:H4K20 me3
H4K20 me2:H3K4 me3
H3K4 me3:H4–4ac
H3K9 me3:H4K20 me3
H4K20 me2:H3–5ac
H2A.Z:H3K4 me3
H3K4 me1:H4–4ac
DNA Meth. m5C:H3K9 me3
H3K27 me3:DNA Meth. m5C
H3K4 me3:H3–5ac
H3K4 me1:H2A.Z
H3K27 me3:H3K9 me3
H4–4ac:H3–5ac
H3K4 me1:H3–5ac
H3K27 me3:H3K4 me1
H3K27 me2:DNA Meth. m5C
H3K9 me2
H4K20 me3
H3K9 me3
H3K4 me3
H4K20 me2
H3K27 me2
H3–5ac
DNA Meth. m5C
H4–4ac
H3K27 me3
H2A.Z
H3K4 me1
0.00 0.25 0.50 0.75 1.00
$\hat{\pi}$
FAM120A – Forward
H3K4 me3:H3–5ac
H3K9 me3:H4K20 me3
H3K27 me3:H3K9 me3
H3K4 me3:H4–4ac
H3K27 me3:H4K20 me3
H3K4 me1:H4–4ac
H3K4 me1:H4K20 me2
H4K20 me2:H4–4ac
H4K20 me2:H3K4 me3
H3K4 me1:H3–5ac
H3K4 me1:H2A.Z
H3K27 me3:DNA Meth. m5C
H4–4ac:H3–5ac
DNA Meth. m5C:H4K20 me3
H4K20 me2:H3–5ac
DNA Meth. m5C:H3K9 me3
H3K9 me2:DNA Meth. m5C
H2A.Z:H4K20 me2
DNA Meth. m5C:H4K20 me2
H2A.Z:H3–5ac
H2A.Z:H3K4 me3
H2A.Z:H4–4ac
H3K27 me2
H4K20 me3
H3K4 me1
H3K9 me2
H3K9 me3
H3K27 me3
H3–5ac
H4–4ac
H3K4 me3
H4K20 me2
H2A.Z
DNA Meth. m5C
0.00 0.25 0.50 0.75 1.00
$\hat{\pi}$
FAM120A – Reverse
H3K4 me3:H3–5ac
H3K9 me3:H4K20 me3
H3K27 me3:H3K9 me3
H3K4 me3:H4–4ac
H3K27 me3:H4K20 me3
H3K4 me1:H4–4ac
H3K4 me1:H4K20 me2
H4K20 me2:H4–4ac
H4K20 me2:H3K4 me3
H3K4 me1:H3–5ac
H3K4 me1:H2A.Z
H3K27 me3:DNA Meth. m5C
H4–4ac:H3–5ac
DNA Meth. m5C:H4K20 me3
H4K20 me2:H3–5ac
DNA Meth. m5C:H3K9 me3
H3K9 me2:DNA Meth. m5C
H2A.Z:H4K20 me2
DNA Meth. m5C:H4K20 me2
H2A.Z:H3–5ac
H2A.Z:H3K4 me3
H2A.Z:H4–4ac
H3K27 me2
H4K20 me3
H3K4 me1
H3K9 me2
H3K9 me3
H3K27 me3
H3–5ac
H4–4ac
H3K4 me3
H4K20 me2
H2A.Z
DNA Meth. m5C
0.00 0.25 0.50 0.75 1.00
$\hat{\pi}$
MED19 – Forward
H3K27 me3:H4K20 me3
H3K9 me3:H4K20 me3
DNA Meth. m5C:H4K20 me2
H3K4 me3:H3–5ac
DNA Meth. m5C:H3K9 me3
H2A.Z:H4K20 me2
H3K4 me1:H4K20 me2
DNA Meth. m5C:H4K20 me3
H3K27 me2:DNA Meth. m5C
H3K27 me3:DNA Meth. m5C
H2A.Z:H3K4 me3
H3K9 me2:DNA Meth. m5C
H3K4 me1:H3–5ac
H3K4 me1:H2A.Z
H3K4 me3:H4–4ac
H3K4 me1:H4–4ac
H4–4ac:H3–5ac
H4K20 me2:H4–4ac
H4K20 me2:H3–5ac
H4K20 me2:H3K4 me3
H3K9 me3
H3K27 me3
H3K27 me2
H4K20 me3
H3K9 me2
H2A.Z
H3K4 me3
H3K4 me1
H3–5ac
DNA Meth. m5C
H4–4ac
H4K20 me2
0.00 0.25 0.50 0.75 1.00
$\hat{\pi}$
MED19 – Reverse
H3K27 me3:H4K20 me3
H3K9 me3:H4K20 me3
DNA Meth. m5C:H4K20 me2
H3K4 me3:H3–5ac
DNA Meth. m5C:H3K9 me3
H2A.Z:H4K20 me2
H3K4 me1:H4K20 me2
DNA Meth. m5C:H4K20 me3
H3K27 me2:DNA Meth. m5C
H3K27 me3:DNA Meth. m5C
H2A.Z:H3K4 me3
H3K9 me2:DNA Meth. m5C
H3K4 me1:H3–5ac
H3K4 me1:H2A.Z
H3K4 me3:H4–4ac
H3K4 me1:H4–4ac
H4–4ac:H3–5ac
H4K20 me2:H4–4ac
H4K20 me2:H3–5ac
H4K20 me2:H3K4 me3
H3K9 me3
H3K27 me3
H3K27 me2
H4K20 me3
H3K9 me2
H2A.Z
H3K4 me3
H3K4 me1
H3–5ac
DNA Meth. m5C
H4–4ac
H4K20 me2
0.00 0.25 0.50 0.75 1.00
$\hat{\pi}$
SMC6 – Forward
H4–4ac:H3–5ac
H3K4 me3:H4–4ac
H4K20 me2:H3K4 me3
H2A.Z:H4K20 me2
H3K4 me1:H4–4ac
H3K4 me1:H3–5ac
H3K4 me1:H2A.Z
H4K20 me2:H4–4ac
H3K27 me3:H3K4 me1
H3K9 me3:H4K20 me3
H3K27 me3:H4K20 me3
DNA Meth. m5C:H4K20 me2
DNA Meth. m5C:H4K20 me3
H3K27 me2:DNA Meth. m5C
H3K9 me2:DNA Meth. m5C
H3–5ac
DNA Meth. m5C:H3K9 me3
H3K27 me3:H3K9 me3
H3K27 me3:DNA Meth. m5C
H4–4ac
H2A.Z
H3K4 me3
H4K20 me2
H3K27 me2
H3K4 me1
H3K9 me2
H4K20 me3
H3K9 me3
H3K27 me3
DNA Meth. m5C
0.00 0.25 0.50 0.75 1.00
$\hat{\pi}$
SMC6 – Reverse
H4–4ac:H3–5ac
H3K4 me3:H4–4ac
H4K20 me2:H3K4 me3
H2A.Z:H4K20 me2
H3K4 me1:H4–4ac
H3K4 me1:H3–5ac
H3K4 me1:H2A.Z
H4K20 me2:H4–4ac
H3K27 me3:H3K4 me1
H3K9 me3:H4K20 me3
H3K27 me3:H4K20 me3
DNA Meth. m5C:H4K20 me2
DNA Meth. m5C:H4K20 me3
H3K27 me2:DNA Meth. m5C
H3K9 me2:DNA Meth. m5C
H3–5ac
DNA Meth. m5C:H3K9 me3
H3K27 me3:H3K9 me3
H3K27 me3:DNA Meth. m5C
H4–4ac
H2A.Z
H3K4 me3
H4K20 me2
H3K27 me2
H3K4 me1
H3K9 me2
H4K20 me3
H3K9 me3
H3K27 me3
DNA Meth. m5C
0.00 0.25 0.50 0.75 1.00
$\hat{\pi}$
DHX35 – Forward
H2A.Z:H4–4ac
H2A.Z:H3–5ac
H3K27 me2:DNA Meth. m5C
H3K27 me3:H3K4 me1
H4–4ac:H3–5ac
H3K4 me3:H3–5ac
H4K20 me2:H3K4 me3
H4K20 me2:H3–5ac
H4K20 me2:H4–4ac
H3K4 me3:H4–4ac
H3K27 me3:H4K20 me3
H3K4 me1:H3–5ac
H3K4 me1:H4K20 me2
DNA Meth. m5C:H4K20 me3
DNA Meth. m5C:H4K20 me2
H3K4 me1:H2A.Z
H3K9 me3:H4K20 me3
H2A.Z:H3K4 me3
H2A.Z:H4K20 me2
H3K4 me1:H4–4ac
H3K27 me3:DNA Meth. m5C
H3K27 me3:H3K9 me3
DNA Meth. m5C:H3K9 me3
H3–5ac
H3K27 me2
H3K4 me3
H2A.Z
H4K20 me2
H4K20 me3
H4–4ac
H3K4 me1
H3K9 me3
DNA Meth. m5C
H3K27 me3
0.00 0.25 0.50 0.75 1.00
$\hat{\pi}$
DHX35 – Reverse
H2A.Z:H4–4ac
H2A.Z:H3–5ac
H3K27 me2:DNA Meth. m5C
H3K27 me3:H3K4 me1
H4–4ac:H3–5ac
H3K4 me3:H3–5ac
H4K20 me2:H3K4 me3
H4K20 me2:H3–5ac
H4K20 me2:H4–4ac
H3K4 me3:H4–4ac
H3K27 me3:H4K20 me3
H3K4 me1:H3–5ac
H3K4 me1:H4K20 me2
DNA Meth. m5C:H4K20 me3
DNA Meth. m5C:H4K20 me2
H3K4 me1:H2A.Z
H3K9 me3:H4K20 me3
H2A.Z:H3K4 me3
H2A.Z:H4K20 me2
H3K4 me1:H4–4ac
H3K27 me3:DNA Meth. m5C
H3K27 me3:H3K9 me3
DNA Meth. m5C:H3K9 me3
H3–5ac
H3K27 me2
H3K4 me3
H2A.Z
H4K20 me2
H4K20 me3
H4–4ac
H3K4 me1
H3K9 me3
DNA Meth. m5C
H3K27 me3
0.00 0.25 0.50 0.75 1.00
$\hat{\pi}$
REEP4 – Forward
H3K27 me3:H4K20 me3
H2A.Z:H3K4 me3
H3K4 me3:H3–5ac
H3K9 me3:H4K20 me3
H3K4 me1:H3–5ac
H3K4 me1:H4K20 me2
H3K4 me1:H4–4ac
H3K27 me2:DNA Meth. m5C
H3K4 me3:H4–4ac
H3K4 me1:H2A.Z
H2A.Z:H4K20 me2
H4–4ac:H3–5ac
DNA Meth. m5C:H4K20 me2
H3K9 me2:DNA Meth. m5C
H4K20 me2:H4–4ac
H4K20 me2:H3–5ac
H3K27 me3:H3K9 me3
DNA Meth. m5C:H3K9 me3
H3K27 me3:DNA Meth. m5C
H4K20 me2:H3K4 me3
H3K9 me2
H4K20 me3
H3K27 me2
H3K4 me1
H3K4 me3
H2A.Z
H4–4ac
H3–5ac
H3K27 me3
H3K9 me3
H4K20 me2
DNA Meth. m5C
0.00 0.25 0.50 0.75 1.00
$\hat{\pi}$
REEP4 – Reverse
H3K27 me3:H4K20 me3
H2A.Z:H3K4 me3
H3K4 me3:H3–5ac
H3K9 me3:H4K20 me3
H3K4 me1:H3–5ac
H3K4 me1:H4K20 me2
H3K4 me1:H4–4ac
H3K27 me2:DNA Meth. m5C
H3K4 me3:H4–4ac
H3K4 me1:H2A.Z
H2A.Z:H4K20 me2
H4–4ac:H3–5ac
DNA Meth. m5C:H4K20 me2
H3K9 me2:DNA Meth. m5C
H4K20 me2:H4–4ac
H4K20 me2:H3–5ac
H3K27 me3:H3K9 me3
DNA Meth. m5C:H3K9 me3
H3K27 me3:DNA Meth. m5C
H4K20 me2:H3K4 me3
H3K9 me2
H4K20 me3
H3K27 me2
H3K4 me1
H3K4 me3
H2A.Z
H4–4ac
H3–5ac
H3K27 me3
H3K9 me3
H4K20 me2
DNA Meth. m5C
0.00 0.25 0.50 0.75 1.00
$\hat{\pi}$

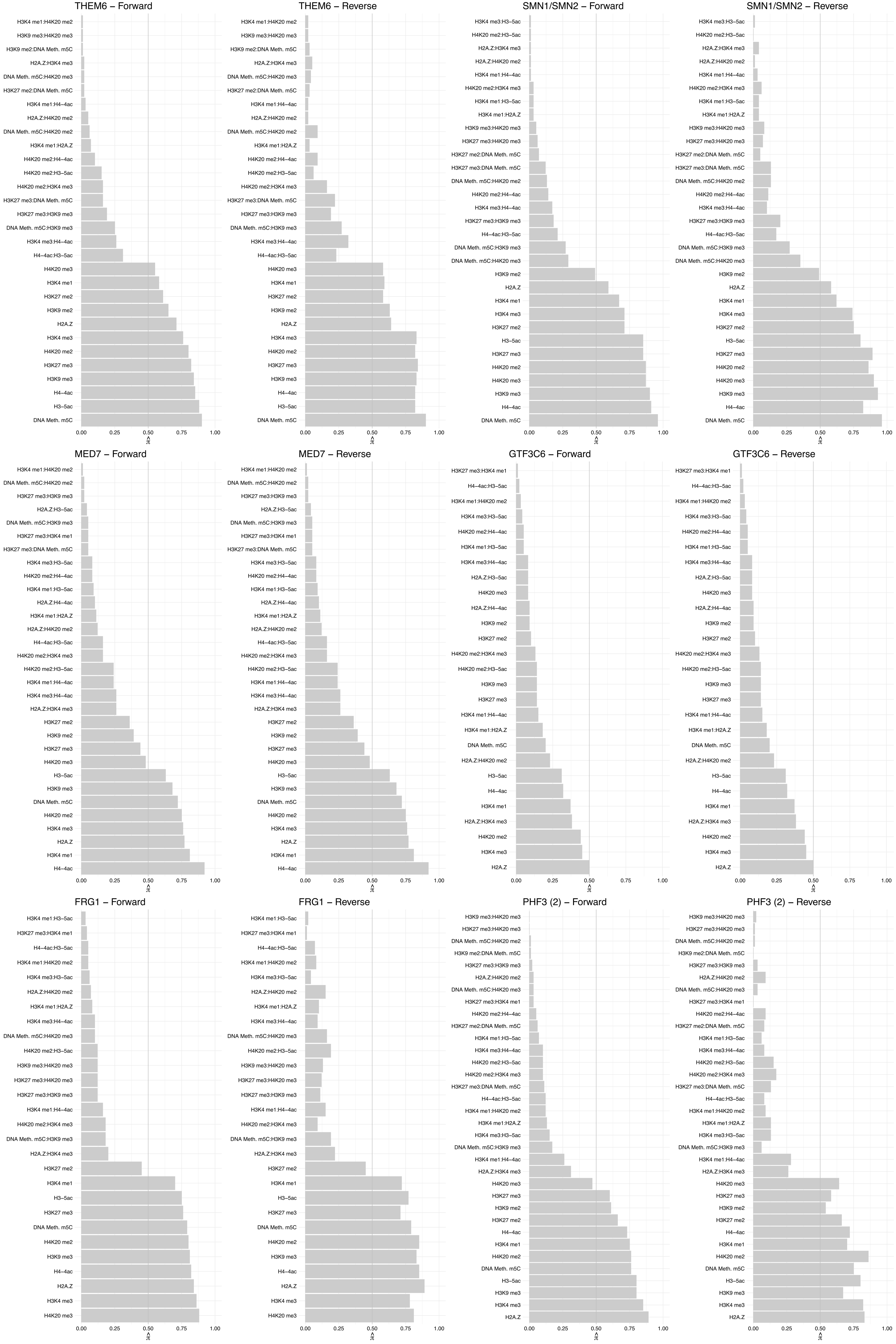
THEM6 – Forward
H3K4 me1:H4K20 me2
H3K9 me3:H4K20 me3
H3K9 me2:DNA Meth. m5C
H2A.Z:H3K4 me3
DNA Meth. m5C:H4K20 me3
H3K27 me2:DNA Meth. m5C
H3K4 me1:H4–4ac
H2A.Z:H4K20 me2
DNA Meth. m5C:H4K20 me2
H3K4 me1:H2A.Z
H4K20 me2:H4–4ac
H4K20 me2:H3–5ac
H4K20 me2:H3K4 me3
H3K27 me3:DNA Meth. m5C
H3K27 me3:H3K9 me3
DNA Meth. m5C:H3K9 me3
H3K4 me3:H4–4ac
H4–4ac:H3–5ac
H4K20 me3
H3K4 me1
H3K27 me2
H3K9 me2
H2A.Z
H3K4 me3
H4K20 me2
H3K27 me3
H3K9 me3
H4–4ac
H3–5ac
DNA Meth. m5C
0.00
0.25
0.50
0.75
1.00
π̂
THEM6 – Reverse
H3K4 me1:H4K20 me2
H3K9 me3:H4K20 me3
H3K9 me2:DNA Meth. m5C
H2A.Z:H3K4 me3
DNA Meth. m5C:H4K20 me3
H3K27 me2:DNA Meth. m5C
H3K4 me1:H4–4ac
H2A.Z:H4K20 me2
DNA Meth. m5C:H4K20 me2
H3K4 me1:H2A.Z
H4K20 me2:H4–4ac
H4K20 me2:H3–5ac
H4K20 me2:H3K4 me3
H3K27 me3:DNA Meth. m5C
H3K27 me3:H3K9 me3
DNA Meth. m5C:H3K9 me3
H3K4 me3:H4–4ac
H4–4ac:H3–5ac
H4K20 me3
H3K4 me1
H3K27 me2
H3K9 me2
H2A.Z
H3K4 me3
H4K20 me2
H3K27 me3
H3K9 me3
H4–4ac
H3–5ac
DNA Meth. m5C
0.00
0.25
0.50
0.75
1.00
π̂
SMN1/SMN2 – Forward
H3K4 me3:H3–5ac
H4K20 me2:H3–5ac
H2A.Z:H3K4 me3
H2A.Z:H4K20 me2
H3K4 me1:H4–4ac
H4K20 me2:H3K4 me3
H3K4 me1:H3–5ac
H3K4 me1:H2A.Z
H3K9 me3:H4K20 me3
H3K27 me3:H4K20 me3
H3K27 me2:DNA Meth. m5C
H3K27 me3:DNA Meth. m5C
DNA Meth. m5C:H4K20 me2
H4K20 me2:H4–4ac
H3K4 me3:H4–4ac
H3K27 me3:H3K9 me3
H4–4ac:H3–5ac
DNA Meth. m5C:H3K9 me3
DNA Meth. m5C:H4K20 me3
H3K9 me2
H2A.Z
H3K4 me1
H3K4 me3
H3K27 me2
H3–5ac
H3K27 me3
H4K20 me2
H4K20 me3
H3K9 me3
H4–4ac
DNA Meth. m5C
0.00
0.25
0.50
0.75
1.00
π̂
SMN1/SMN2 – Reverse
H3K4 me3:H3–5ac
H4K20 me2:H3–5ac
H2A.Z:H3K4 me3
H2A.Z:H4K20 me2
H3K4 me1:H4–4ac
H4K20 me2:H3K4 me3
H3K4 me1:H3–5ac
H3K4 me1:H2A.Z
H3K9 me3:H4K20 me3
H3K27 me3:H4K20 me3
H3K27 me2:DNA Meth. m5C
H3K27 me3:DNA Meth. m5C
DNA Meth. m5C:H4K20 me2
H4K20 me2:H4–4ac
H3K4 me3:H4–4ac
H3K27 me3:H3K9 me3
H4–4ac:H3–5ac
DNA Meth. m5C:H3K9 me3
DNA Meth. m5C:H4K20 me3
H3K9 me2
H2A.Z
H3K4 me1
H3K4 me3
H3K27 me2
H3–5ac
H3K27 me3
H4K20 me2
H4K20 me3
H3K9 me3
H4–4ac
DNA Meth. m5C
0.00
0.25
0.50
0.75
1.00
π̂
MED7 – Forward
H3K4 me1:H4K20 me2
DNA Meth. m5C:H4K20 me2
H3K27 me3:H3K9 me3
H2A.Z:H3–5ac
DNA Meth. m5C:H3K9 me3
H3K27 me3:H3K4 me1
H3K27 me3:DNA Meth. m5C
H3K4 me3:H3–5ac
H4K20 me2:H4–4ac
H3K4 me1:H3–5ac
H2A.Z:H4–4ac
H3K4 me1:H2A.Z
H2A.Z:H4K20 me2
H4–4ac:H3–5ac
H4K20 me2:H3K4 me3
H4K20 me2:H3–5ac
H3K4 me1:H4–4ac
H3K4 me3:H4–4ac
H2A.Z:H3K4 me3
H3K27 me2
H3K9 me2
H3K27 me3
H4K20 me3
H3–5ac
H3K9 me3
DNA Meth. m5C
H4K20 me2
H3K4 me3
H2A.Z
H3K4 me1
H4–4ac
0.00
0.25
0.50
0.75
1.00
π̂
MED7 – Reverse
H3K4 me1:H4K20 me2
DNA Meth. m5C:H4K20 me2
H3K27 me3:H3K9 me3
H2A.Z:H3–5ac
DNA Meth. m5C:H3K9 me3
H3K27 me3:H3K4 me1
H3K27 me3:DNA Meth. m5C
H3K4 me3:H3–5ac
H4K20 me2:H4–4ac
H3K4 me1:H3–5ac
H2A.Z:H4–4ac
H3K4 me1:H2A.Z
H2A.Z:H4K20 me2
H4–4ac:H3–5ac
H4K20 me2:H3K4 me3
H4K20 me2:H3–5ac
H3K4 me1:H4–4ac
H3K4 me3:H4–4ac
H2A.Z:H3K4 me3
H3K27 me2
H3K9 me2
H3K27 me3
H4K20 me3
H3–5ac
H3K9 me3
DNA Meth. m5C
H4K20 me2
H3K4 me3
H2A.Z
H3K4 me1
H4–4ac
0.00
0.25
0.50
0.75
1.00
π̂
GTF3C6 – Forward
H3K27 me3:H3K4 me1
H4–4ac:H3–5ac
H3K4 me1:H4K20 me2
H3K4 me3:H3–5ac
H4K20 me2:H4–4ac
H3K4 me1:H3–5ac
H3K4 me3:H4–4ac
H2A.Z:H3–5ac
H4K20 me3
H2A.Z:H4–4ac
H3K9 me2
H3K27 me2
H4K20 me2:H3K4 me3
H4K20 me2:H3–5ac
H3K9 me3
H3K27 me3
H3K4 me1:H4–4ac
H3K4 me1:H2A.Z
DNA Meth. m5C
H2A.Z:H4K20 me2
H3–5ac
H4–4ac
H3K4 me1
H2A.Z:H3K4 me3
H4K20 me2
H3K4 me3
H2A.Z
0.00
0.25
0.50
0.75
1.00
GTF3C6 – Reverse
H3K27 me3:H3K4 me1
H4–4ac:H3–5ac
H3K4 me1:H4K20 me2
H3K4 me3:H3–5ac
H4K20 me2:H4–4ac
H3K4 me1:H3–5ac
H3K4 me3:H4–4ac
H2A.Z:H3–5ac
H4K20 me3
H2A.Z:H4–4ac
H3K9 me2
H3K27 me2
H4K20 me2:H3K4 me3
H4K20 me2:H3–5ac
H3K9 me3
H3K27 me3
H3K4 me1:H4–4ac
H3K4 me1:H2A.Z
DNA Meth. m5C
H2A.Z:H4K20 me2
H3–5ac
H4–4ac
H3K4 me1
H2A.Z:H3K4 me3
H4K20 me2
H3K4 me3
H2A.Z
0.00
0.25
0.50
0.75
1.00
FRG1 – Forward
H3K4 me1:H3–5ac
H3K27 me3:H3K4 me1
H4–4ac:H3–5ac
H3K4 me1:H4K20 me2
H3K4 me3:H3–5ac
H2A.Z:H4K20 me2
H3K4 me1:H2A.Z
H3K4 me3:H4–4ac
DNA Meth. m5C:H4K20 me2
H4K20 me2:H3–5ac
H3K9 me3:H4K20 me3
H3K27 me3:H4K20 me3
H3K27 me3:H3K9 me3
H3K4 me1:H4–4ac
H4K20 me2:H3K4 me3
DNA Meth. m5C:H3K9 me3
H2A.Z:H3K4 me3
H3K27 me2
H3K4 me1
H3–5ac
H3K27 me3
DNA Meth. m5C
H4K20 me2
H3K9 me3
H4–4ac
H2A.Z
H3K4 me3
H4K20 me3
0.00
0.25
0.50
0.75
1.00
π̂
FRG1 – Reverse
H3K4 me1:H3–5ac
H3K27 me3:H3K4 me1
H4–4ac:H3–5ac
H3K4 me1:H4K20 me2
H3K4 me3:H3–5ac
H2A.Z:H4K20 me2
H3K4 me1:H2A.Z
H3K4 me3:H4–4ac
DNA Meth. m5C:H4K20 me2
H4K20 me2:H3–5ac
H3K9 me3:H4K20 me3
H3K27 me3:H4K20 me3
H3K27 me3:H3K9 me3
H3K4 me1:H4–4ac
H4K20 me2:H3K4 me3
DNA Meth. m5C:H3K9 me3
H2A.Z:H3K4 me3
H3K27 me2
H3K4 me1
H3–5ac
H3K27 me3
DNA Meth. m5C
H4K20 me2
H3K9 me3
H4–4ac
H2A.Z
H3K4 me3
H4K20 me3
0.00
0.25
0.50
0.75
1.00
π̂
PHF3 (2) – Forward
H3K9 me3:H4K20 me3
H3K27 me3:H4K20 me3
DNA Meth. m5C:H4K20 me2
H3K9 me2:DNA Meth. m5C
H3K27 me3:H3K9 me3
H2A.Z:H4K20 me2
DNA Meth. m5C:H4K20 me3
H3K27 me3:H3K4 me1
H4K20 me2:H4–4ac
H3K27 me2:DNA Meth. m5C
H3K4 me1:H3–5ac
H3K4 me3:H4–4ac
H4K20 me2:H3–5ac
H4K20 me2:H3K4 me3
H3K27 me3:DNA Meth. m5C
H4–4ac:H3–5ac
H3K4 me1:H4K20 me2
H3K4 me1:H2A.Z
H3K4 me3:H3–5ac
DNA Meth. m5C:H3K9 me3
H3K4 me1:H4–4ac
H2A.Z:H3K4 me3
H4K20 me3
H3K27 me3
H3K9 me2
H3K27 me2
H4–4ac
H3K4 me1
H4K20 me2
DNA Meth. m5C
H3–5ac
H3K9 me3
H3K4 me3
H2A.Z
0.00
0.25
0.50
0.75
1.00
π̂
PHF3 (2) – Reverse
H3K9 me3:H4K20 me3
H3K27 me3:H4K20 me3
DNA Meth. m5C:H4K20 me2
H3K9 me2:DNA Meth. m5C
H3K27 me3:H3K9 me3
H2A.Z:H4K20 me2
DNA Meth. m5C:H4K20 me3
H3K27 me3:H3K4 me1
H4K20 me2:H4–4ac
H3K27 me2:DNA Meth. m5C
H3K4 me1:H3–5ac
H3K4 me3:H4–4ac
H4K20 me2:H3–5ac
H4K20 me2:H3K4 me3
H3K27 me3:DNA Meth. m5C
H4–4ac:H3–5ac
H3K4 me1:H4K20 me2
H3K4 me1:H2A.Z
H3K4 me3:H3–5ac
DNA Meth. m5C:H3K9 me3
H3K4 me1:H4–4ac
H2A.Z:H3K4 me3
H4K20 me3
H3K27 me3
H3K9 me2
H3K27 me2
H4–4ac
H3K4 me1
H4K20 me2
DNA Meth. m5C
H3–5ac
H3K9 me3
H3K4 me3
H2A.Z
0.00
0.25
0.50
0.75
1.00
π̂

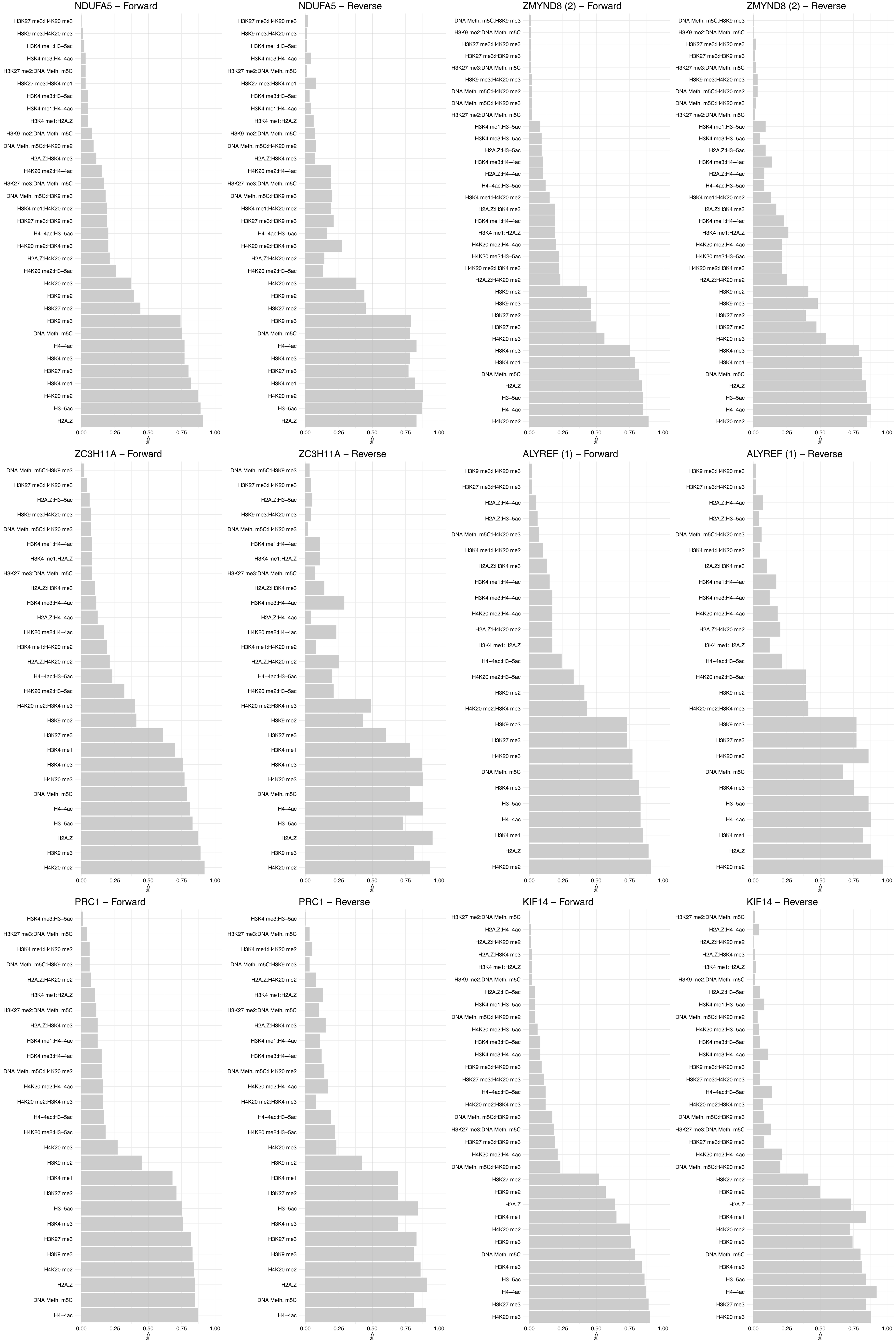
NDUFA5 – Forward
NDUFA5 – Reverse
ZMYND8 (2) – Forward
ZMYND8 (2) – Reverse
ZC3H11A – Forward
ZC3H11A – Reverse
ALYREF (1) – Forward
ALYREF (1) – Reverse
PRC1 – Forward
PRC1 – Reverse
KIF14 – Forward
KIF14 – Reverse
H3K27 me3:H4K20 me3
H3K9 me3:H4K20 me3
H3K4 me1:H3–5ac
H3K4 me3:H4–4ac
H3K27 me2:DNA Meth. m5C
H3K27 me3:H3K4 me1
H3K4 me3:H3–5ac
H3K4 me1:H4–4ac
H3K4 me1:H2A.Z
H3K9 me2:DNA Meth. m5C
DNA Meth. m5C:H4K20 me2
H2A.Z:H3K4 me3
H4K20 me2:H4–4ac
H3K27 me3:DNA Meth. m5C
DNA Meth. m5C:H3K9 me3
H3K4 me1:H4K20 me2
H3K27 me3:H3K9 me3
H4–4ac:H3–5ac
H4K20 me2:H3K4 me3
H2A.Z:H4K20 me2
H4K20 me2:H3–5ac
DNA Meth. m5C:H4K20 me3
H3K9 me2:DNA Meth. m5C
H2A.Z:H3–5ac
H2A.Z:H4–4ac
H4K20 me3
H3K9 me2
H3K27 me2
H3K9 me3
DNA Meth. m5C
H4–4ac
H3K4 me3
H3K27 me3
H3K4 me1
H4K20 me2
H3–5ac
H2A.Z
0.00
0.25
0.50
0.75
1.00
$\hat{\pi}$

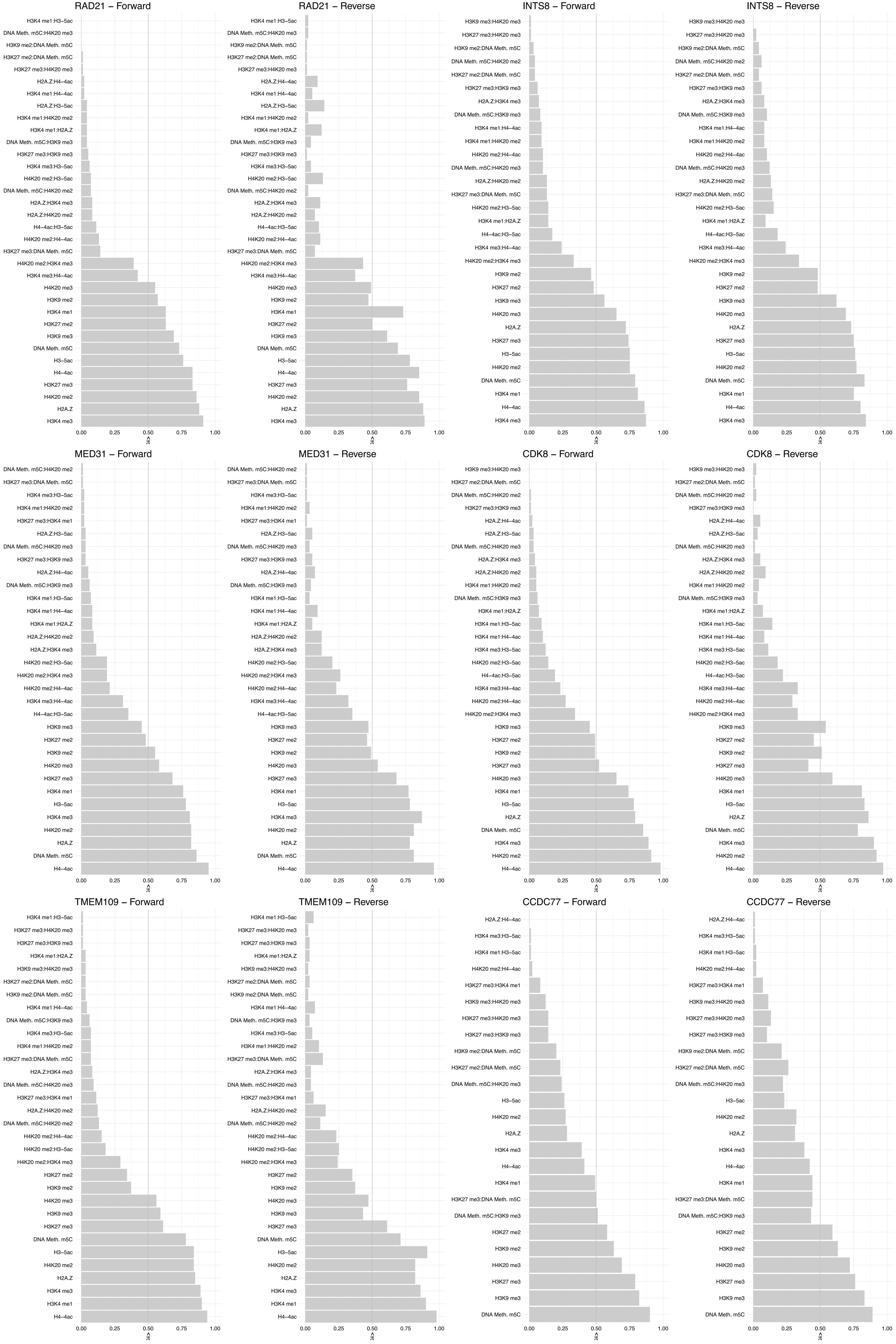
RAD21 – Forward
RAD21 – Reverse
INTS8 – Forward
INTS8 – Reverse
MED31 – Forward
MED31 – Reverse
CDK8 – Forward
CDK8 – Reverse
TMEM109 – Forward
TMEM109 – Reverse
CCDC77 – Forward
CCDC77 – Reverse
0.00
0.25
0.50
0.75
1.00
$\hat{\pi}$

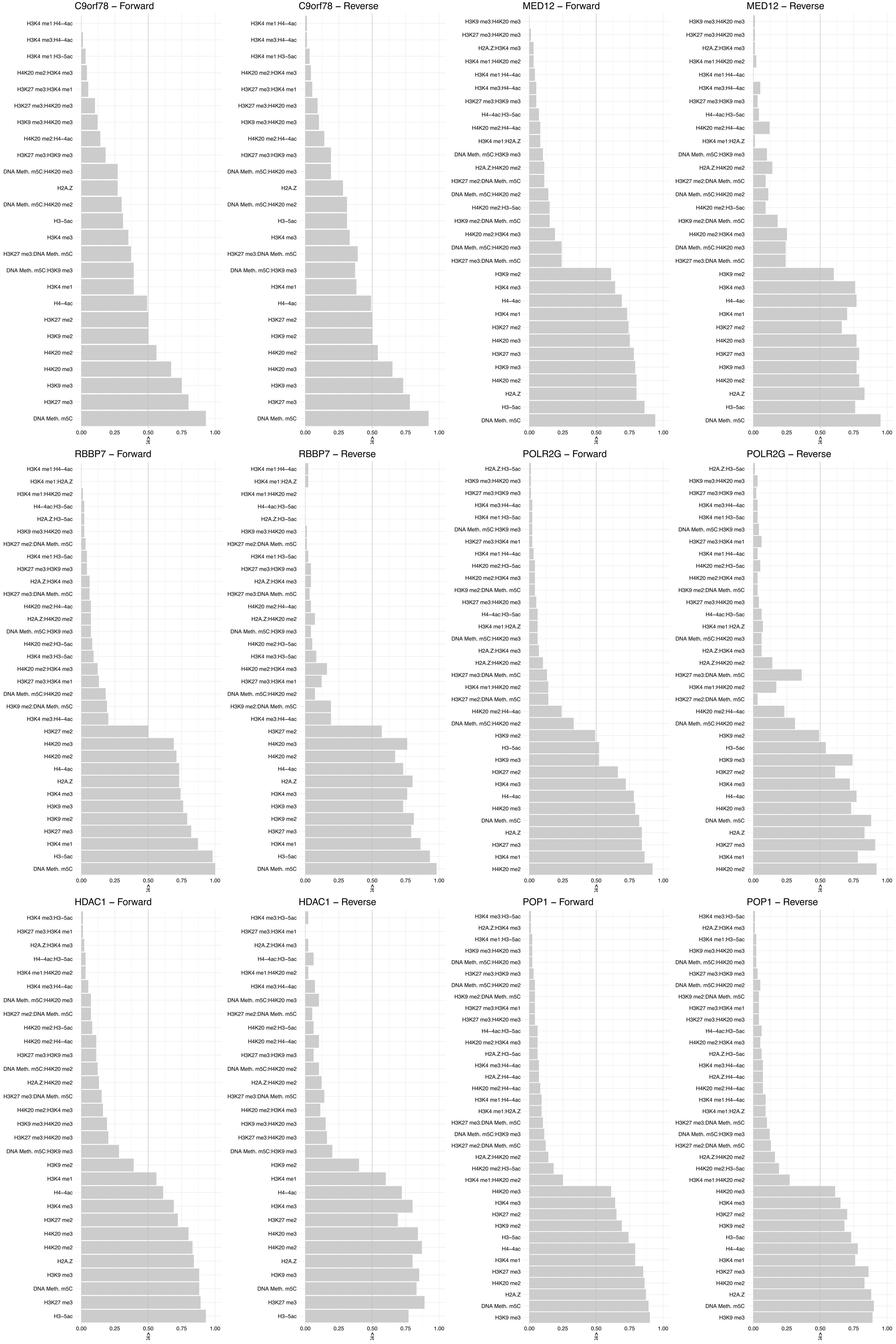
C9orf78 – Forward
H3K4 me1:H4–4ac
H3K4 me3:H4–4ac
H3K4 me1:H3–5ac
H4K20 me2:H3K4 me3
H3K27 me3:H3K4 me1
H3K27 me3:H4K20 me3
H3K9 me3:H4K20 me3
H4K20 me2:H4–4ac
H3K27 me3:H3K9 me3
DNA Meth. m5C:H4K20 me3
H2A.Z
DNA Meth. m5C:H4K20 me2
H3–5ac
H3K4 me3
H3K27 me3:DNA Meth. m5C
DNA Meth. m5C:H3K9 me3
H3K4 me1
H4–4ac
H3K27 me2
H3K9 me2
H4K20 me2
H4K20 me3
H3K9 me3
H3K27 me3
DNA Meth. m5C
0.00 0.25 0.50 0.75 1.00
π̂
C9orf78 – Reverse
H3K4 me1:H4–4ac
H3K4 me3:H4–4ac
H3K4 me1:H3–5ac
H4K20 me2:H3K4 me3
H3K27 me3:H3K4 me1
H3K27 me3:H4K20 me3
H3K9 me3:H4K20 me3
H4K20 me2:H4–4ac
H3K27 me3:H3K9 me3
DNA Meth. m5C:H4K20 me3
H2A.Z
DNA Meth. m5C:H4K20 me2
H3–5ac
H3K4 me3
H3K27 me3:DNA Meth. m5C
DNA Meth. m5C:H3K9 me3
H3K4 me1
H4–4ac
H3K27 me2
H3K9 me2
H4K20 me2
H4K20 me3
H3K9 me3
H3K27 me3
DNA Meth. m5C
0.00 0.25 0.50 0.75 1.00
π̂
MED12 – Forward
H3K9 me3:H4K20 me3
H3K27 me3:H4K20 me3
H2A.Z:H3K4 me3
H3K4 me1:H4K20 me2
H3K4 me1:H4–4ac
H3K4 me3:H4–4ac
H3K27 me3:H3K9 me3
H4–4ac:H3–5ac
H4K20 me2:H4–4ac
H3K4 me1:H2A.Z
DNA Meth. m5C:H3K9 me3
H2A.Z:H4K20 me2
H3K27 me2:DNA Meth. m5C
DNA Meth. m5C:H4K20 me2
H4K20 me2:H3–5ac
H3K9 me2:DNA Meth. m5C
H4K20 me2:H3K4 me3
DNA Meth. m5C:H4K20 me3
H3K27 me3:DNA Meth. m5C
H3K9 me2
H3K4 me3
H4–4ac
H3K4 me1
H3K27 me2
H4K20 me3
H3K27 me3
H3K9 me3
H4K20 me2
H2A.Z
H3–5ac
DNA Meth. m5C
0.00 0.25 0.50 0.75 1.00
π̂
MED12 – Reverse
H3K9 me3:H4K20 me3
H3K27 me3:H4K20 me3
H2A.Z:H3K4 me3
H3K4 me1:H4K20 me2
H3K4 me1:H4–4ac
H3K4 me3:H4–4ac
H3K27 me3:H3K9 me3
H4–4ac:H3–5ac
H4K20 me2:H4–4ac
H3K4 me1:H2A.Z
DNA Meth. m5C:H3K9 me3
H2A.Z:H4K20 me2
H3K27 me2:DNA Meth. m5C
DNA Meth. m5C:H4K20 me2
H4K20 me2:H3–5ac
H3K9 me2:DNA Meth. m5C
H4K20 me2:H3K4 me3
DNA Meth. m5C:H4K20 me3
H3K27 me3:DNA Meth. m5C
H3K9 me2
H3K4 me3
H4–4ac
H3K4 me1
H3K27 me2
H4K20 me3
H3K27 me3
H3K9 me3
H4K20 me2
H2A.Z
H3–5ac
DNA Meth. m5C
0.00 0.25 0.50 0.75 1.00
π̂
RBBP7 – Forward
H3K4 me1:H4–4ac
H3K4 me1:H2A.Z
H3K4 me1:H4K20 me2
H4–4ac:H3–5ac
H2A.Z:H3–5ac
H3K9 me3:H4K20 me3
H3K27 me2:DNA Meth. m5C
H3K4 me1:H3–5ac
H3K27 me3:H3K9 me3
H2A.Z:H3K4 me3
H3K27 me3:DNA Meth. m5C
H4K20 me2:H4–4ac
H2A.Z:H4K20 me2
DNA Meth. m5C:H3K9 me3
H4K20 me2:H3–5ac
H3K4 me3:H3–5ac
H4K20 me2:H3K4 me3
H3K27 me3:H3K4 me1
DNA Meth. m5C:H4K20 me2
H3K9 me2:DNA Meth. m5C
H3K4 me3:H4–4ac
H3K27 me2
H4K20 me3
H4K20 me2
H4–4ac
H2A.Z
H3K4 me3
H3K9 me3
H3K9 me2
H3K27 me3
H3K4 me1
H3–5ac
DNA Meth. m5C
0.00 0.25 0.50 0.75 1.00
π̂
RBBP7 – Reverse
H3K4 me1:H4–4ac
H3K4 me1:H2A.Z
H3K4 me1:H4K20 me2
H4–4ac:H3–5ac
H2A.Z:H3–5ac
H3K9 me3:H4K20 me3
H3K27 me2:DNA Meth. m5C
H3K4 me1:H3–5ac
H3K27 me3:H3K9 me3
H2A.Z:H3K4 me3
H3K27 me3:DNA Meth. m5C
H4K20 me2:H4–4ac
H2A.Z:H4K20 me2
DNA Meth. m5C:H3K9 me3
H4K20 me2:H3–5ac
H3K4 me3:H3–5ac
H4K20 me2:H3K4 me3
H3K27 me3:H3K4 me1
DNA Meth. m5C:H4K20 me2
H3K9 me2:DNA Meth. m5C
H3K4 me3:H4–4ac
H3K27 me2
H4K20 me3
H4K20 me2
H4–4ac
H2A.Z
H3K4 me3
H3K9 me3
H3K9 me2
H3K27 me3
H3K4 me1
H3–5ac
DNA Meth. m5C
0.00 0.25 0.50 0.75 1.00
π̂
POLR2G – Forward
H2A.Z:H3–5ac
H3K9 me3:H4K20 me3
H3K27 me3:H3K9 me3
H3K4 me3:H4–4ac
H3K4 me1:H3–5ac
DNA Meth. m5C:H3K9 me3
H3K27 me3:H3K4 me1
H3K4 me1:H4–4ac
H4K20 me2:H3–5ac
H4K20 me2:H3K4 me3
H3K9 me2:DNA Meth. m5C
H3K27 me3:H4K20 me3
H4–4ac:H3–5ac
H3K4 me1:H2A.Z
DNA Meth. m5C:H4K20 me3
H2A.Z:H3K4 me3
H2A.Z:H4K20 me2
H3K27 me3:DNA Meth. m5C
H3K4 me1:H4K20 me2
H3K27 me2:DNA Meth. m5C
H4K20 me2:H4–4ac
DNA Meth. m5C:H4K20 me2
H3K9 me2
H3–5ac
H3K9 me3
H3K27 me2
H3K4 me3
H4–4ac
H4K20 me3
DNA Meth. m5C
H2A.Z
H3K27 me3
H3K4 me1
H4K20 me2
0.00 0.25 0.50 0.75 1.00
π̂
POLR2G – Reverse
H2A.Z:H3–5ac
H3K9 me3:H4K20 me3
H3K27 me3:H3K9 me3
H3K4 me3:H4–4ac
H3K4 me1:H3–5ac
DNA Meth. m5C:H3K9 me3
H3K27 me3:H3K4 me1
H3K4 me1:H4–4ac
H4K20 me2:H3–5ac
H4K20 me2:H3K4 me3
H3K9 me2:DNA Meth. m5C
H3K27 me3:H4K20 me3
H4–4ac:H3–5ac
H3K4 me1:H2A.Z
DNA Meth. m5C:H4K20 me3
H2A.Z:H3K4 me3
H2A.Z:H4K20 me2
H3K27 me3:DNA Meth. m5C
H3K4 me1:H4K20 me2
H3K27 me2:DNA Meth. m5C
H4K20 me2:H4–4ac
DNA Meth. m5C:H4K20 me2
H3K9 me2
H3–5ac
H3K9 me3
H3K27 me2
H3K4 me3
H4–4ac
H4K20 me3
DNA Meth. m5C
H2A.Z
H3K27 me3
H3K4 me1
H4K20 me2
0.00 0.25 0.50 0.75 1.00
π̂
HDAC1 – Forward
H3K4 me3:H3–5ac
H3K27 me3:H3K4 me1
H2A.Z:H3K4 me3
H4–4ac:H3–5ac
H3K4 me1:H4K20 me2
H3K4 me3:H4–4ac
DNA Meth. m5C:H4K20 me3
H3K27 me2:DNA Meth. m5C
H4K20 me2:H3–5ac
H4K20 me2:H4–4ac
H3K27 me3:H3K9 me3
DNA Meth. m5C:H4K20 me2
H2A.Z:H4K20 me2
H3K27 me3:DNA Meth. m5C
H4K20 me2:H3K4 me3
H3K9 me3:H4K20 me3
H3K27 me3:H4K20 me3
DNA Meth. m5C:H3K9 me3
H3K9 me2
H3K4 me1
H4–4ac
H3K4 me3
H3K27 me2
H4K20 me3
H4K20 me2
H2A.Z
H3K9 me3
DNA Meth. m5C
H3K27 me3
H3–5ac
0.00 0.25 0.50 0.75 1.00
π̂
HDAC1 – Reverse
H3K4 me3:H3–5ac
H3K27 me3:H3K4 me1
H2A.Z:H3K4 me3
H4–4ac:H3–5ac
H3K4 me1:H4K20 me2
H3K4 me3:H4–4ac
DNA Meth. m5C:H4K20 me3
H3K27 me2:DNA Meth. m5C
H4K20 me2:H3–5ac
H4K20 me2:H4–4ac
H3K27 me3:H3K9 me3
DNA Meth. m5C:H4K20 me2
H2A.Z:H4K20 me2
H3K27 me3:DNA Meth. m5C
H4K20 me2:H3K4 me3
H3K9 me3:H4K20 me3
H3K27 me3:H4K20 me3
DNA Meth. m5C:H3K9 me3
H3K9 me2
H3K4 me1
H4–4ac
H3K4 me3
H3K27 me2
H4K20 me3
H4K20 me2
H2A.Z
H3K9 me3
DNA Meth. m5C
H3K27 me3
H3–5ac
0.00 0.25 0.50 0.75 1.00
π̂
POP1 – Forward
H3K4 me3:H3–5ac
H2A.Z:H3K4 me3
H3K4 me1:H3–5ac
H3K9 me3:H4K20 me3
DNA Meth. m5C:H4K20 me3
H3K27 me3:H3K9 me3
DNA Meth. m5C:H4K20 me2
H3K9 me2:DNA Meth. m5C
H3K27 me3:H3K4 me1
H3K27 me3:H4K20 me3
H4–4ac:H3–5ac
H4K20 me2:H3K4 me3
H2A.Z:H3–5ac
H3K4 me3:H4–4ac
H2A.Z:H4–4ac
H4K20 me2:H4–4ac
H3K4 me1:H4–4ac
H3K4 me1:H2A.Z
H3K27 me3:DNA Meth. m5C
DNA Meth. m5C:H3K9 me3
H3K27 me2:DNA Meth. m5C
H2A.Z:H4K20 me2
H4K20 me2:H3–5ac
H3K4 me1:H4K20 me2
H4K20 me3
H3K4 me3
H3K27 me2
H3K9 me2
H3–5ac
H4–4ac
H3K4 me1
H3K27 me3
H4K20 me2
H2A.Z
DNA Meth. m5C
H3K9 me3
0.00 0.25 0.50 0.75 1.00
π̂
POP1 – Reverse
H3K4 me3:H3–5ac
H2A.Z:H3K4 me3
H3K4 me1:H3–5ac
H3K9 me3:H4K20 me3
DNA Meth. m5C:H4K20 me3
H3K27 me3:H3K9 me3
DNA Meth. m5C:H4K20 me2
H3K9 me2:DNA Meth. m5C
H3K27 me3:H3K4 me1
H3K27 me3:H4K20 me3
H4–4ac:H3–5ac
H4K20 me2:H3K4 me3
H2A.Z:H3–5ac
H3K4 me3:H4–4ac
H2A.Z:H4–4ac
H4K20 me2:H4–4ac
H3K4 me1:H4–4ac
H3K4 me1:H2A.Z
H3K27 me3:DNA Meth. m5C
DNA Meth. m5C:H3K9 me3
H3K27 me2:DNA Meth. m5C
H2A.Z:H4K20 me2
H4K20 me2:H3–5ac
H3K4 me1:H4K20 me2
H4K20 me3
H3K4 me3
H3K27 me2
H3K9 me2
H3–5ac
H4–4ac
H3K4 me1
H3K27 me3
H4K20 me2
H2A.Z
DNA Meth. m5C
H3K9 me3
0.00 0.25 0.50 0.75 1.00
π̂

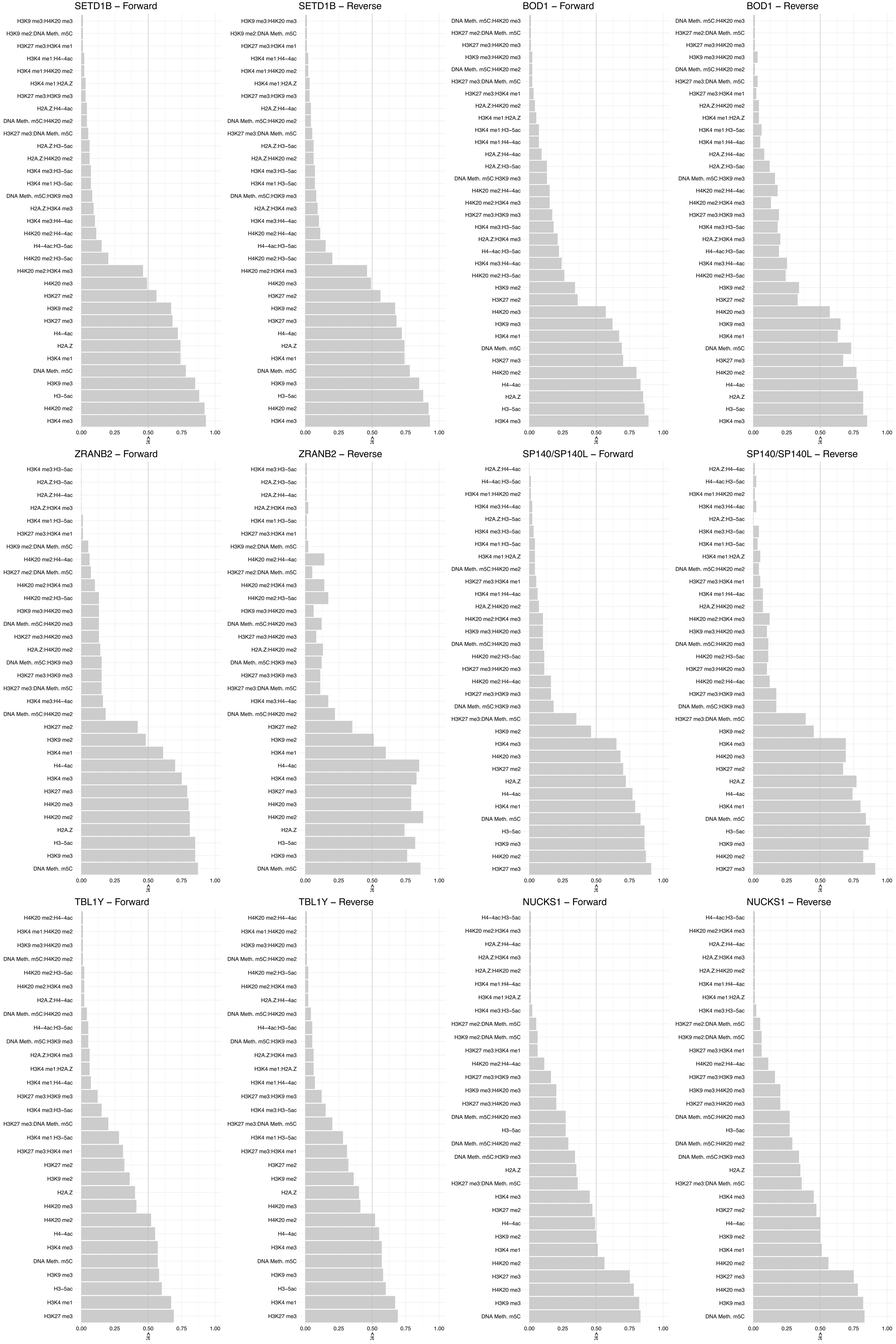
SETD1B – Forward
H3K9 me3:H4K20 me3
H3K9 me2:DNA Meth. m5C
H3K27 me3:H3K4 me1
H3K4 me1:H4–4ac
H3K4 me1:H4K20 me2
H3K4 me1:H2A.Z
H3K27 me3:H3K9 me3
H2A.Z:H4–4ac
DNA Meth. m5C:H4K20 me2
H3K27 me3:DNA Meth. m5C
H2A.Z:H3–5ac
H2A.Z:H4K20 me2
H3K4 me3:H3–5ac
H3K4 me1:H3–5ac
DNA Meth. m5C:H3K9 me3
H2A.Z:H3K4 me3
H3K4 me3:H4–4ac
H4K20 me2:H4–4ac
H4–4ac:H3–5ac
H4K20 me2:H3–5ac
H4K20 me2:H3K4 me3
H4K20 me3
H3K27 me2
H3K9 me2
H3K27 me3
H4–4ac
H2A.Z
H3K4 me1
DNA Meth. m5C
H3K9 me3
H3–5ac
H4K20 me2
H3K4 me3
0.00 0.25 0.50 0.75 1.00
π̂
SETD1B – Reverse
H3K9 me3:H4K20 me3
H3K9 me2:DNA Meth. m5C
H3K27 me3:H3K4 me1
H3K4 me1:H4–4ac
H3K4 me1:H4K20 me2
H3K4 me1:H2A.Z
H3K27 me3:H3K9 me3
H2A.Z:H4–4ac
DNA Meth. m5C:H4K20 me2
H3K27 me3:DNA Meth. m5C
H2A.Z:H3–5ac
H2A.Z:H4K20 me2
H3K4 me3:H3–5ac
H3K4 me1:H3–5ac
DNA Meth. m5C:H3K9 me3
H2A.Z:H3K4 me3
H3K4 me3:H4–4ac
H4K20 me2:H4–4ac
H4–4ac:H3–5ac
H4K20 me2:H3–5ac
H4K20 me2:H3K4 me3
H4K20 me3
H3K27 me2
H3K9 me2
H3K27 me3
H4–4ac
H2A.Z
H3K4 me1
DNA Meth. m5C
H3K9 me3
H3–5ac
H4K20 me2
H3K4 me3
0.00 0.25 0.50 0.75 1.00
π̂
BOD1 – Forward
DNA Meth. m5C:H4K20 me3
H3K27 me2:DNA Meth. m5C
H3K27 me3:H4K20 me3
H3K9 me3:H4K20 me3
DNA Meth. m5C:H4K20 me2
H3K27 me3:DNA Meth. m5C
H3K27 me3:H3K4 me1
H2A.Z:H4K20 me2
H3K4 me1:H2A.Z
H3K4 me1:H3–5ac
H3K4 me1:H4–4ac
H2A.Z:H4–4ac
H2A.Z:H3–5ac
DNA Meth. m5C:H3K9 me3
H4K20 me2:H4–4ac
H4K20 me2:H3K4 me3
H3K27 me3:H3K9 me3
H3K4 me3:H3–5ac
H2A.Z:H3K4 me3
H4–4ac:H3–5ac
H3K4 me3:H4–4ac
H4K20 me2:H3–5ac
H3K9 me2
H3K27 me2
H4K20 me3
H3K9 me3
H3K4 me1
DNA Meth. m5C
H3K27 me3
H4K20 me2
H4–4ac
H2A.Z
H3–5ac
H3K4 me3
0.00 0.25 0.50 0.75 1.00
π̂
BOD1 – Reverse
DNA Meth. m5C:H4K20 me3
H3K27 me2:DNA Meth. m5C
H3K27 me3:H4K20 me3
H3K9 me3:H4K20 me3
DNA Meth. m5C:H4K20 me2
H3K27 me3:DNA Meth. m5C
H3K27 me3:H3K4 me1
H2A.Z:H4K20 me2
H3K4 me1:H2A.Z
H3K4 me1:H3–5ac
H3K4 me1:H4–4ac
H2A.Z:H4–4ac
H2A.Z:H3–5ac
DNA Meth. m5C:H3K9 me3
H4K20 me2:H4–4ac
H4K20 me2:H3K4 me3
H3K27 me3:H3K9 me3
H3K4 me3:H3–5ac
H2A.Z:H3K4 me3
H4–4ac:H3–5ac
H3K4 me3:H4–4ac
H4K20 me2:H3–5ac
H3K9 me2
H3K27 me2
H4K20 me3
H3K9 me3
H3K4 me1
DNA Meth. m5C
H3K27 me3
H4K20 me2
H4–4ac
H2A.Z
H3–5ac
H3K4 me3
0.00 0.25 0.50 0.75 1.00
π̂
ZRANB2 – Forward
H3K4 me3:H3–5ac
H2A.Z:H3–5ac
H2A.Z:H4–4ac
H2A.Z:H3K4 me3
H3K4 me1:H3–5ac
H3K27 me3:H3K4 me1
H3K9 me2:DNA Meth. m5C
H4K20 me2:H4–4ac
H3K27 me2:DNA Meth. m5C
H4K20 me2:H3K4 me3
H4K20 me2:H3–5ac
H3K9 me3:H4K20 me3
DNA Meth. m5C:H4K20 me3
H3K27 me3:H4K20 me3
H2A.Z:H4K20 me2
DNA Meth. m5C:H3K9 me3
H3K27 me3:H3K9 me3
H3K27 me3:DNA Meth. m5C
H3K4 me3:H4–4ac
DNA Meth. m5C:H4K20 me2
H3K27 me2
H3K9 me2
H3K4 me1
H4–4ac
H3K4 me3
H3K27 me3
H4K20 me3
H4K20 me2
H2A.Z
H3–5ac
H3K9 me3
DNA Meth. m5C
0.00 0.25 0.50 0.75 1.00
π̂
ZRANB2 – Reverse
H3K4 me3:H3–5ac
H2A.Z:H3–5ac
H2A.Z:H4–4ac
H2A.Z:H3K4 me3
H3K4 me1:H3–5ac
H3K27 me3:H3K4 me1
H3K9 me2:DNA Meth. m5C
H4K20 me2:H4–4ac
H3K27 me2:DNA Meth. m5C
H4K20 me2:H3K4 me3
H4K20 me2:H3–5ac
H3K9 me3:H4K20 me3
DNA Meth. m5C:H4K20 me3
H3K27 me3:H4K20 me3
H2A.Z:H4K20 me2
DNA Meth. m5C:H3K9 me3
H3K27 me3:H3K9 me3
H3K27 me3:DNA Meth. m5C
H3K4 me3:H4–4ac
DNA Meth. m5C:H4K20 me2
H3K27 me2
H3K9 me2
H3K4 me1
H4–4ac
H3K4 me3
H3K27 me3
H4K20 me3
H4K20 me2
H2A.Z
H3–5ac
H3K9 me3
DNA Meth. m5C
0.00 0.25 0.50 0.75 1.00
π̂
SP140/SP140L – Forward
H2A.Z:H4–4ac
H4–4ac:H3–5ac
H3K4 me1:H4K20 me2
H3K4 me3:H4–4ac
H2A.Z:H3–5ac
H3K4 me3:H3–5ac
H3K4 me1:H3–5ac
H3K4 me1:H2A.Z
DNA Meth. m5C:H4K20 me2
H3K27 me3:H3K4 me1
H3K4 me1:H4–4ac
H2A.Z:H4K20 me2
H4K20 me2:H3K4 me3
H3K9 me3:H4K20 me3
DNA Meth. m5C:H4K20 me3
H4K20 me2:H3–5ac
H3K27 me3:H4K20 me3
H4K20 me2:H4–4ac
H3K27 me3:H3K9 me3
DNA Meth. m5C:H3K9 me3
H3K27 me3:DNA Meth. m5C
H3K9 me2
H3K4 me3
H4K20 me3
H3K27 me2
H2A.Z
H4–4ac
H3K4 me1
DNA Meth. m5C
H3–5ac
H3K9 me3
H4K20 me2
H3K27 me3
0.00 0.25 0.50 0.75 1.00
π̂
SP140/SP140L – Reverse
H2A.Z:H4–4ac
H4–4ac:H3–5ac
H3K4 me1:H4K20 me2
H3K4 me3:H4–4ac
H2A.Z:H3–5ac
H3K4 me3:H3–5ac
H3K4 me1:H3–5ac
H3K4 me1:H2A.Z
DNA Meth. m5C:H4K20 me2
H3K27 me3:H3K4 me1
H3K4 me1:H4–4ac
H2A.Z:H4K20 me2
H4K20 me2:H3K4 me3
H3K9 me3:H4K20 me3
DNA Meth. m5C:H4K20 me3
H4K20 me2:H3–5ac
H3K27 me3:H4K20 me3
H4K20 me2:H4–4ac
H3K27 me3:H3K9 me3
DNA Meth. m5C:H3K9 me3
H3K27 me3:DNA Meth. m5C
H3K9 me2
H3K4 me3
H4K20 me3
H3K27 me2
H2A.Z
H4–4ac
H3K4 me1
DNA Meth. m5C
H3–5ac
H3K9 me3
H4K20 me2
H3K27 me3
0.00 0.25 0.50 0.75 1.00
π̂
TBL1Y – Forward
H4K20 me2:H4–4ac
H3K4 me1:H4K20 me2
H3K9 me3:H4K20 me3
DNA Meth. m5C:H4K20 me2
H4K20 me2:H3–5ac
H4K20 me2:H3K4 me3
H2A.Z:H4–4ac
DNA Meth. m5C:H4K20 me3
H4–4ac:H3–5ac
DNA Meth. m5C:H3K9 me3
H2A.Z:H3K4 me3
H3K4 me1:H2A.Z
H3K4 me1:H4–4ac
H3K27 me3:H3K9 me3
H3K4 me3:H3–5ac
H3K27 me3:DNA Meth. m5C
H3K4 me1:H3–5ac
H3K27 me3:H3K4 me1
H3K27 me2
H3K9 me2
H2A.Z
H4K20 me3
H4K20 me2
H4–4ac
H3K4 me3
DNA Meth. m5C
H3K9 me3
H3–5ac
H3K4 me1
H3K27 me3
0.00 0.25 0.50 0.75 1.00
π̂
TBL1Y – Reverse
H4K20 me2:H4–4ac
H3K4 me1:H4K20 me2
H3K9 me3:H4K20 me3
DNA Meth. m5C:H4K20 me2
H4K20 me2:H3–5ac
H4K20 me2:H3K4 me3
H2A.Z:H4–4ac
DNA Meth. m5C:H4K20 me3
H4–4ac:H3–5ac
DNA Meth. m5C:H3K9 me3
H2A.Z:H3K4 me3
H3K4 me1:H2A.Z
H3K4 me1:H4–4ac
H3K27 me3:H3K9 me3
H3K4 me3:H3–5ac
H3K27 me3:DNA Meth. m5C
H3K4 me1:H3–5ac
H3K27 me3:H3K4 me1
H3K27 me2
H3K9 me2
H2A.Z
H4K20 me3
H4K20 me2
H4–4ac
H3K4 me3
DNA Meth. m5C
H3K9 me3
H3–5ac
H3K4 me1
H3K27 me3
0.00 0.25 0.50 0.75 1.00
π̂
NUCKS1 – Forward
H4–4ac:H3–5ac
H4K20 me2:H3K4 me3
H2A.Z:H4–4ac
H2A.Z:H3K4 me3
H2A.Z:H4K20 me2
H3K4 me1:H4–4ac
H3K4 me1:H2A.Z
H3K4 me3:H3–5ac
H3K27 me2:DNA Meth. m5C
H3K9 me2:DNA Meth. m5C
H3K27 me3:H3K4 me1
H4K20 me2:H4–4ac
H3K27 me3:H3K9 me3
H3K9 me3:H4K20 me3
H3K27 me3:H4K20 me3
DNA Meth. m5C:H4K20 me3
H3–5ac
DNA Meth. m5C:H4K20 me2
DNA Meth. m5C:H3K9 me3
H2A.Z
H3K27 me3:DNA Meth. m5C
H3K4 me3
H3K27 me2
H4–4ac
H3K9 me2
H3K4 me1
H4K20 me2
H3K27 me3
H4K20 me3
H3K9 me3
DNA Meth. m5C
0.00 0.25 0.50 0.75 1.00
π̂
NUCKS1 – Reverse
H4–4ac:H3–5ac
H4K20 me2:H3K4 me3
H2A.Z:H4–4ac
H2A.Z:H3K4 me3
H2A.Z:H4K20 me2
H3K4 me1:H4–4ac
H3K4 me1:H2A.Z
H3K4 me3:H3–5ac
H3K27 me2:DNA Meth. m5C
H3K9 me2:DNA Meth. m5C
H3K27 me3:H3K4 me1
H4K20 me2:H4–4ac
H3K27 me3:H3K9 me3
H3K9 me3:H4K20 me3
H3K27 me3:H4K20 me3
DNA Meth. m5C:H4K20 me3
H3–5ac
DNA Meth. m5C:H4K20 me2
DNA Meth. m5C:H3K9 me3
H2A.Z
H3K27 me3:DNA Meth. m5C
H3K4 me3
H3K27 me2
H4–4ac
H3K9 me2
H3K4 me1
H4K20 me2
H3K27 me3
H4K20 me3
H3K9 me3
DNA Meth. m5C
0.00 0.25 0.50 0.75 1.00
π̂

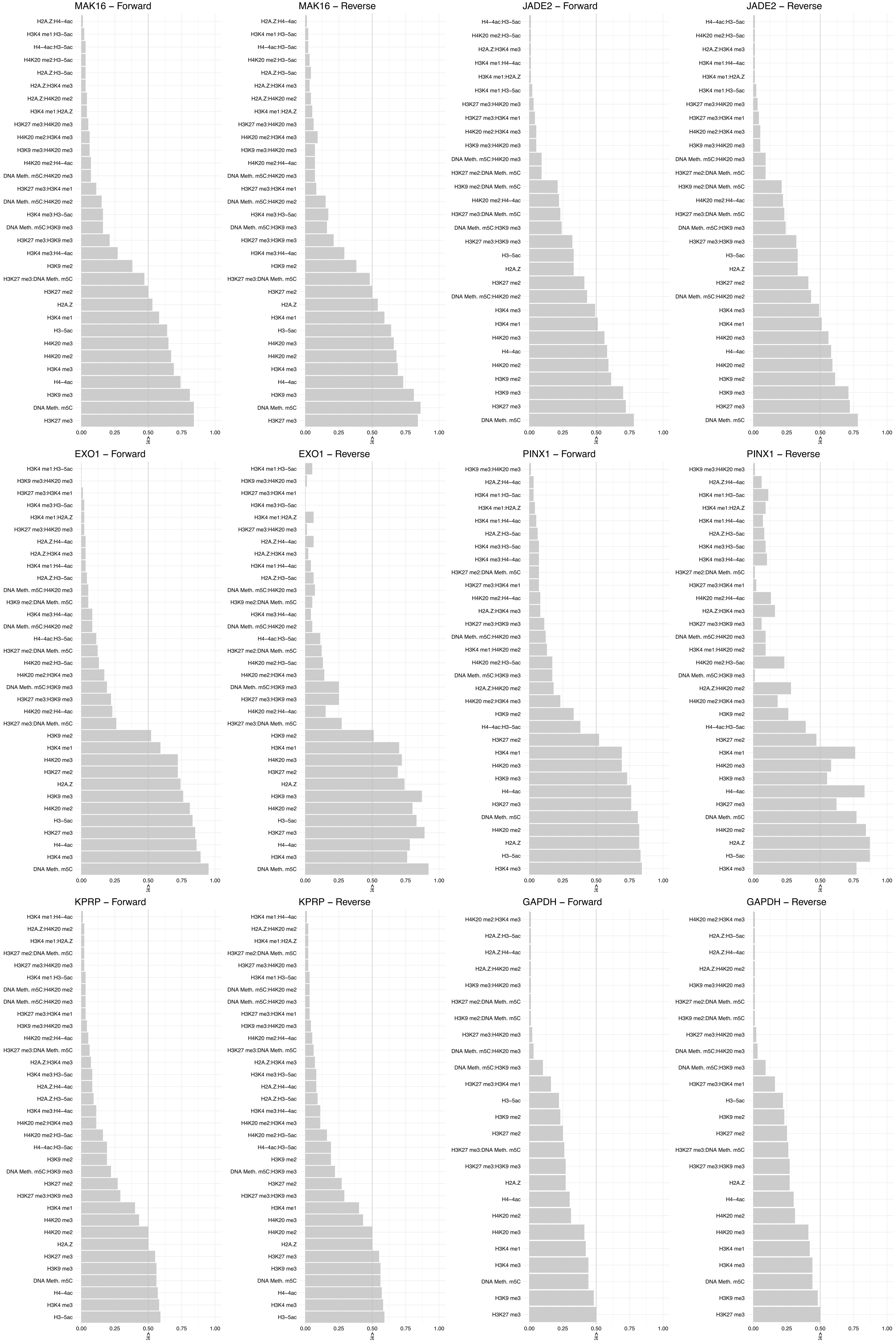
MAK16 – Forward
H2A.Z:H4–4ac
H3K4 me1:H3–5ac
H4–4ac:H3–5ac
H4K20 me2:H3–5ac
H2A.Z:H3–5ac
H2A.Z:H3K4 me3
H2A.Z:H4K20 me2
H3K4 me1:H2A.Z
H3K27 me3:H4K20 me3
H4K20 me2:H3K4 me3
H3K9 me3:H4K20 me3
H4K20 me2:H4–4ac
DNA Meth. m5C:H4K20 me3
H3K27 me3:H3K4 me1
DNA Meth. m5C:H4K20 me2
H3K4 me3:H3–5ac
DNA Meth. m5C:H3K9 me3
H3K27 me3:H3K9 me3
H3K4 me3:H4–4ac
H3K9 me2
H3K27 me3:DNA Meth. m5C
H3K27 me2
H2A.Z
H3K4 me1
H3–5ac
H4K20 me3
H4K20 me2
H3K4 me3
H4–4ac
H3K9 me3
DNA Meth. m5C
H3K27 me3
0.00
0.25
0.50
0.75
1.00
π̂
MAK16 – Reverse
H2A.Z:H4–4ac
H3K4 me1:H3–5ac
H4–4ac:H3–5ac
H4K20 me2:H3–5ac
H2A.Z:H3–5ac
H2A.Z:H3K4 me3
H2A.Z:H4K20 me2
H3K4 me1:H2A.Z
H3K27 me3:H4K20 me3
H4K20 me2:H3K4 me3
H3K9 me3:H4K20 me3
H4K20 me2:H4–4ac
DNA Meth. m5C:H4K20 me3
H3K27 me3:H3K4 me1
DNA Meth. m5C:H4K20 me2
H3K4 me3:H3–5ac
DNA Meth. m5C:H3K9 me3
H3K27 me3:H3K9 me3
H3K4 me3:H4–4ac
H3K9 me2
H3K27 me3:DNA Meth. m5C
H3K27 me2
H2A.Z
H3K4 me1
H3–5ac
H4K20 me3
H4K20 me2
H3K4 me3
H4–4ac
H3K9 me3
DNA Meth. m5C
H3K27 me3
0.00
0.25
0.50
0.75
1.00
π̂
JADE2 – Forward
H4–4ac:H3–5ac
H4K20 me2:H3–5ac
H2A.Z:H3K4 me3
H3K4 me1:H4–4ac
H3K4 me1:H2A.Z
H3K4 me1:H3–5ac
H3K27 me3:H4K20 me3
H3K27 me3:H3K4 me1
H4K20 me2:H3K4 me3
H3K9 me3:H4K20 me3
DNA Meth. m5C:H4K20 me3
H3K27 me2:DNA Meth. m5C
H3K9 me2:DNA Meth. m5C
H4K20 me2:H4–4ac
H3K27 me3:DNA Meth. m5C
DNA Meth. m5C:H3K9 me3
H3K27 me3:H3K9 me3
H3–5ac
H2A.Z
H3K27 me2
DNA Meth. m5C:H4K20 me2
H3K4 me3
H3K4 me1
H4K20 me3
H4–4ac
H4K20 me2
H3K9 me2
H3K9 me3
H3K27 me3
DNA Meth. m5C
0.00
0.25
0.50
0.75
1.00
π̂
JADE2 – Reverse
H4–4ac:H3–5ac
H4K20 me2:H3–5ac
H2A.Z:H3K4 me3
H3K4 me1:H4–4ac
H3K4 me1:H2A.Z
H3K4 me1:H3–5ac
H3K27 me3:H4K20 me3
H3K27 me3:H3K4 me1
H4K20 me2:H3K4 me3
H3K9 me3:H4K20 me3
DNA Meth. m5C:H4K20 me3
H3K27 me2:DNA Meth. m5C
H3K9 me2:DNA Meth. m5C
H4K20 me2:H4–4ac
H3K27 me3:DNA Meth. m5C
DNA Meth. m5C:H3K9 me3
H3K27 me3:H3K9 me3
H3–5ac
H2A.Z
H3K27 me2
DNA Meth. m5C:H4K20 me2
H3K4 me3
H3K4 me1
H4K20 me3
H4–4ac
H4K20 me2
H3K9 me2
H3K9 me3
H3K27 me3
DNA Meth. m5C
0.00
0.25
0.50
0.75
1.00
π̂
EXO1 – Forward
H3K4 me1:H3–5ac
H3K9 me3:H4K20 me3
H3K27 me3:H3K4 me1
H3K4 me3:H3–5ac
H3K4 me1:H2A.Z
H3K27 me3:H4K20 me3
H2A.Z:H4–4ac
H2A.Z:H3K4 me3
H3K4 me1:H4–4ac
H2A.Z:H3–5ac
DNA Meth. m5C:H4K20 me3
H3K9 me2:DNA Meth. m5C
H3K4 me3:H4–4ac
DNA Meth. m5C:H4K20 me2
H4–4ac:H3–5ac
H3K27 me2:DNA Meth. m5C
H4K20 me2:H3–5ac
H4K20 me2:H3K4 me3
DNA Meth. m5C:H3K9 me3
H3K27 me3:H3K9 me3
H4K20 me2:H4–4ac
H3K27 me3:DNA Meth. m5C
H3K9 me2
H3K4 me1
H4K20 me3
H3K27 me2
H2A.Z
H3K9 me3
H4K20 me2
H3–5ac
H3K27 me3
H4–4ac
H3K4 me3
DNA Meth. m5C
0.00
0.25
0.50
0.75
1.00
π̂
EXO1 – Reverse
H3K4 me1:H3–5ac
H3K9 me3:H4K20 me3
H3K27 me3:H3K4 me1
H3K4 me3:H3–5ac
H3K4 me1:H2A.Z
H3K27 me3:H4K20 me3
H2A.Z:H4–4ac
H2A.Z:H3K4 me3
H3K4 me1:H4–4ac
H2A.Z:H3–5ac
DNA Meth. m5C:H4K20 me3
H3K9 me2:DNA Meth. m5C
H3K4 me3:H4–4ac
DNA Meth. m5C:H4K20 me2
H4–4ac:H3–5ac
H3K27 me2:DNA Meth. m5C
H4K20 me2:H3–5ac
H4K20 me2:H3K4 me3
DNA Meth. m5C:H3K9 me3
H3K27 me3:H3K9 me3
H4K20 me2:H4–4ac
H3K27 me3:DNA Meth. m5C
H3K9 me2
H3K4 me1
H4K20 me3
H3K27 me2
H2A.Z
H3K9 me3
H4K20 me2
H3–5ac
H3K27 me3
H4–4ac
H3K4 me3
DNA Meth. m5C
0.00
0.25
0.50
0.75
1.00
π̂
PINX1 – Forward
H3K9 me3:H4K20 me3
H2A.Z:H4–4ac
H3K4 me1:H3–5ac
H3K4 me1:H2A.Z
H3K4 me1:H4–4ac
H2A.Z:H3–5ac
H3K4 me3:H3–5ac
H3K4 me3:H4–4ac
H3K27 me2:DNA Meth. m5C
H3K27 me3:H3K4 me1
H4K20 me2:H4–4ac
H2A.Z:H3K4 me3
H3K27 me3:H3K9 me3
DNA Meth. m5C:H4K20 me3
H3K4 me1:H4K20 me2
H4K20 me2:H3–5ac
DNA Meth. m5C:H3K9 me3
H2A.Z:H4K20 me2
H4K20 me2:H3K4 me3
H3K9 me2
H4–4ac:H3–5ac
H3K27 me2
H3K4 me1
H4K20 me3
H3K9 me3
H4–4ac
H3K27 me3
DNA Meth. m5C
H4K20 me2
H2A.Z
H3–5ac
H3K4 me3
0.00
0.25
0.50
0.75
1.00
π̂
PINX1 – Reverse
H3K9 me3:H4K20 me3
H2A.Z:H4–4ac
H3K4 me1:H3–5ac
H3K4 me1:H2A.Z
H3K4 me1:H4–4ac
H2A.Z:H3–5ac
H3K4 me3:H3–5ac
H3K4 me3:H4–4ac
H3K27 me2:DNA Meth. m5C
H3K27 me3:H3K4 me1
H4K20 me2:H4–4ac
H2A.Z:H3K4 me3
H3K27 me3:H3K9 me3
DNA Meth. m5C:H4K20 me3
H3K4 me1:H4K20 me2
H4K20 me2:H3–5ac
DNA Meth. m5C:H3K9 me3
H2A.Z:H4K20 me2
H4K20 me2:H3K4 me3
H3K9 me2
H4–4ac:H3–5ac
H3K27 me2
H3K4 me1
H4K20 me3
H3K9 me3
H4–4ac
H3K27 me3
DNA Meth. m5C
H4K20 me2
H2A.Z
H3–5ac
H3K4 me3
0.00
0.25
0.50
0.75
1.00
π̂
KPRP – Forward
H3K4 me1:H4–4ac
H2A.Z:H4K20 me2
H3K4 me1:H2A.Z
H3K27 me2:DNA Meth. m5C
H3K27 me3:H4K20 me3
H3K4 me1:H3–5ac
DNA Meth. m5C:H4K20 me2
DNA Meth. m5C:H4K20 me3
H3K27 me3:H3K4 me1
H3K9 me3:H4K20 me3
H4K20 me2:H4–4ac
H3K27 me3:DNA Meth. m5C
H2A.Z:H3K4 me3
H3K4 me3:H3–5ac
H2A.Z:H4–4ac
H2A.Z:H3–5ac
H3K4 me3:H4–4ac
H4K20 me2:H3K4 me3
H4K20 me2:H3–5ac
H4–4ac:H3–5ac
H3K9 me2
DNA Meth. m5C:H3K9 me3
H3K27 me2
H3K27 me3:H3K9 me3
H3K4 me1
H4K20 me3
H4K20 me2
H2A.Z
H3K27 me3
H3K9 me3
DNA Meth. m5C
H4–4ac
H3K4 me3
H3–5ac
0.00
0.25
0.50
0.75
1.00
π̂
KPRP – Reverse
H3K4 me1:H4–4ac
H2A.Z:H4K20 me2
H3K4 me1:H2A.Z
H3K27 me2:DNA Meth. m5C
H3K27 me3:H4K20 me3
H3K4 me1:H3–5ac
DNA Meth. m5C:H4K20 me2
DNA Meth. m5C:H4K20 me3
H3K27 me3:H3K4 me1
H3K9 me3:H4K20 me3
H4K20 me2:H4–4ac
H3K27 me3:DNA Meth. m5C
H2A.Z:H3K4 me3
H3K4 me3:H3–5ac
H2A.Z:H4–4ac
H2A.Z:H3–5ac
H3K4 me3:H4–4ac
H4K20 me2:H3K4 me3
H4K20 me2:H3–5ac
H4–4ac:H3–5ac
H3K9 me2
DNA Meth. m5C:H3K9 me3
H3K27 me2
H3K27 me3:H3K9 me3
H3K4 me1
H4K20 me3
H4K20 me2
H2A.Z
H3K27 me3
H3K9 me3
DNA Meth. m5C
H4–4ac
H3K4 me3
H3–5ac
0.00
0.25
0.50
0.75
1.00
π̂
GAPDH – Forward
H4K20 me2:H3K4 me3
H2A.Z:H3–5ac
H2A.Z:H4–4ac
H2A.Z:H4K20 me2
H3K9 me3:H4K20 me3
H3K27 me2:DNA Meth. m5C
H3K9 me2:DNA Meth. m5C
H3K27 me3:H4K20 me3
DNA Meth. m5C:H4K20 me3
DNA Meth. m5C:H3K9 me3
H3K27 me3:H3K4 me1
H3–5ac
H3K9 me2
H3K27 me2
H3K27 me3:DNA Meth. m5C
H3K27 me3:H3K9 me3
H2A.Z
H4–4ac
H4K20 me2
H4K20 me3
H3K4 me1
H3K4 me3
DNA Meth. m5C
H3K9 me3
H3K27 me3
0.00
0.25
0.50
0.75
1.00
π̂
GAPDH – Reverse
H4K20 me2:H3K4 me3
H2A.Z:H3–5ac
H2A.Z:H4–4ac
H2A.Z:H4K20 me2
H3K9 me3:H4K20 me3
H3K27 me2:DNA Meth. m5C
H3K9 me2:DNA Meth. m5C
H3K27 me3:H4K20 me3
DNA Meth. m5C:H4K20 me3
DNA Meth. m5C:H3K9 me3
H3K27 me3:H3K4 me1
H3–5ac
H3K9 me2
H3K27 me2
H3K27 me3:DNA Meth. m5C
H3K27 me3:H3K9 me3
H2A.Z
H4–4ac
H4K20 me2
H4K20 me3
H3K4 me1
H3K4 me3
DNA Meth. m5C
H3K9 me3
H3K27 me3
0.00
0.25
0.50
0.75
1.00
π̂

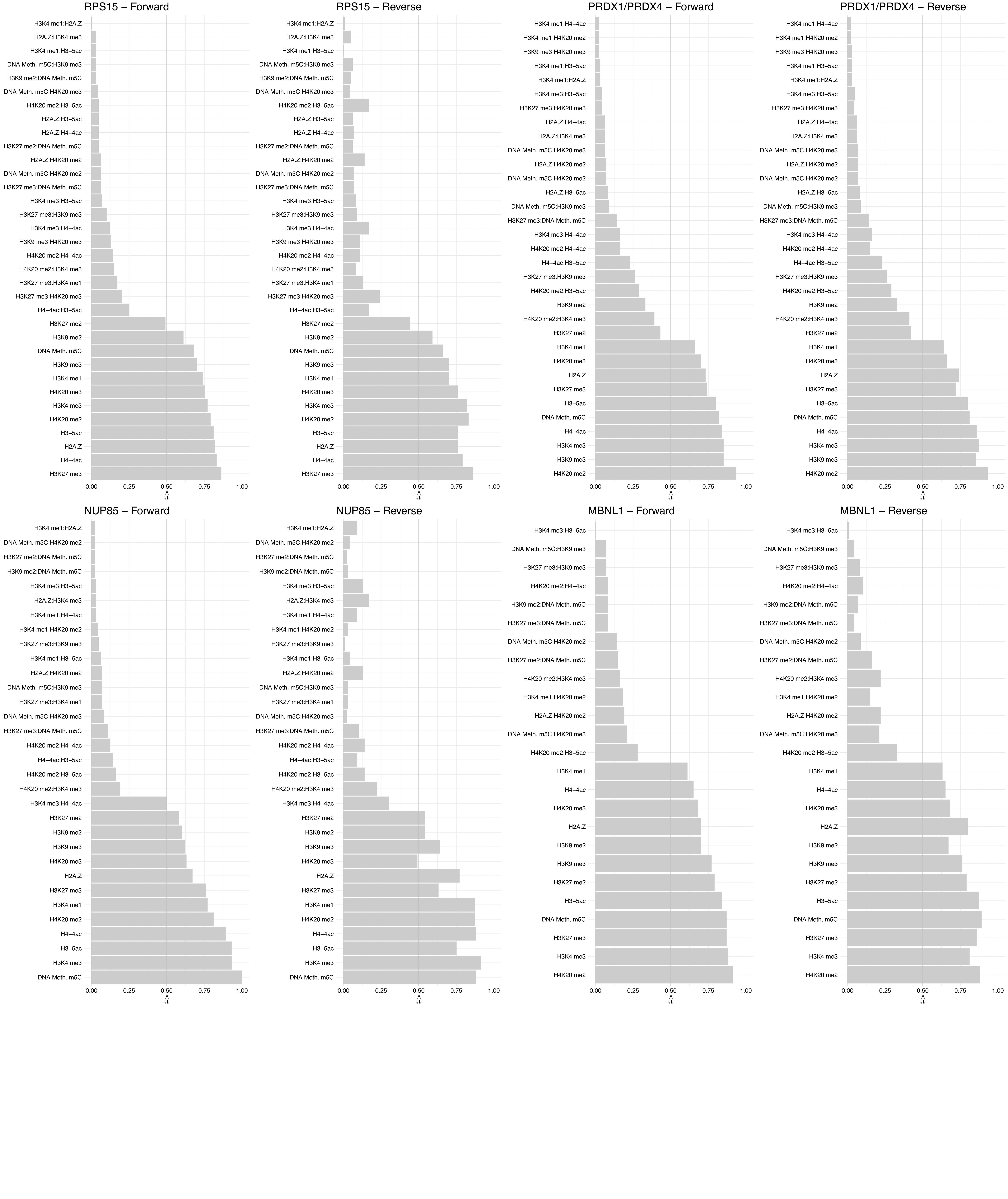
RPS15 – Forward
H3K4 me1:H2A.Z
H2A.Z:H3K4 me3
H3K4 me1:H3−5ac
DNA Meth. m5C:H3K9 me3
H3K9 me2:DNA Meth. m5C
DNA Meth. m5C:H4K20 me3
H4K20 me2:H3−5ac
H2A.Z:H3−5ac
H2A.Z:H4−4ac
H3K27 me2:DNA Meth. m5C
H2A.Z:H4K20 me2
DNA Meth. m5C:H4K20 me2
H3K27 me3:DNA Meth. m5C
H3K4 me3:H3−5ac
H3K27 me3:H3K9 me3
H3K4 me3:H4−4ac
H3K9 me3:H4K20 me3
H4K20 me2:H4−4ac
H4K20 me2:H3K4 me3
H3K27 me3:H3K4 me1
H3K27 me3:H4K20 me3
H4−4ac:H3−5ac
H3K27 me2
H3K9 me2
DNA Meth. m5C
H3K9 me3
H3K4 me1
H4K20 me3
H3K4 me3
H4K20 me2
H3−5ac
H2A.Z
H4−4ac
H3K27 me3
0.00 0.25 0.50 0.75 1.00
$\hat{\pi}$
RPS15 – Reverse
H3K4 me1:H2A.Z
H2A.Z:H3K4 me3
H3K4 me1:H3−5ac
DNA Meth. m5C:H3K9 me3
H3K9 me2:DNA Meth. m5C
DNA Meth. m5C:H4K20 me3
H4K20 me2:H3−5ac
H2A.Z:H3−5ac
H2A.Z:H4−4ac
H3K27 me2:DNA Meth. m5C
H2A.Z:H4K20 me2
DNA Meth. m5C:H4K20 me2
H3K27 me3:DNA Meth. m5C
H3K4 me3:H3−5ac
H3K27 me3:H3K9 me3
H3K4 me3:H4−4ac
H3K9 me3:H4K20 me3
H4K20 me2:H4−4ac
H4K20 me2:H3K4 me3
H3K27 me3:H3K4 me1
H3K27 me3:H4K20 me3
H4−4ac:H3−5ac
H3K27 me2
H3K9 me2
DNA Meth. m5C
H3K9 me3
H3K4 me1
H4K20 me3
H3K4 me3
H4K20 me2
H3−5ac
H2A.Z
H4−4ac
H3K27 me3
0.00 0.25 0.50 0.75 1.00
$\hat{\pi}$
PRDX1/PRDX4 – Forward
H3K4 me1:H4−4ac
H3K4 me1:H4K20 me2
H3K9 me3:H4K20 me3
H3K4 me1:H3−5ac
H3K4 me1:H2A.Z
H3K4 me3:H3−5ac
H3K27 me3:H4K20 me3
H2A.Z:H4−4ac
H2A.Z:H3K4 me3
DNA Meth. m5C:H4K20 me3
H2A.Z:H4K20 me2
DNA Meth. m5C:H4K20 me2
H2A.Z:H3−5ac
DNA Meth. m5C:H3K9 me3
H3K27 me3:DNA Meth. m5C
H3K4 me3:H4−4ac
H4K20 me2:H4−4ac
H4−4ac:H3−5ac
H3K27 me3:H3K9 me3
H4K20 me2:H3−5ac
H3K9 me2
H4K20 me2:H3K4 me3
H3K27 me2
H3K4 me1
H4K20 me3
H2A.Z
H3K27 me3
H3−5ac
DNA Meth. m5C
H4−4ac
H3K4 me3
H3K9 me3
H4K20 me2
0.00 0.25 0.50 0.75 1.00
$\hat{\pi}$
PRDX1/PRDX4 – Reverse
H3K4 me1:H4−4ac
H3K4 me1:H4K20 me2
H3K9 me3:H4K20 me3
H3K4 me1:H3−5ac
H3K4 me1:H2A.Z
H3K4 me3:H3−5ac
H3K27 me3:H4K20 me3
H2A.Z:H4−4ac
H2A.Z:H3K4 me3
DNA Meth. m5C:H4K20 me3
H2A.Z:H4K20 me2
DNA Meth. m5C:H4K20 me2
H2A.Z:H3−5ac
DNA Meth. m5C:H3K9 me3
H3K27 me3:DNA Meth. m5C
H3K4 me3:H4−4ac
H4K20 me2:H4−4ac
H4−4ac:H3−5ac
H3K27 me3:H3K9 me3
H4K20 me2:H3−5ac
H3K9 me2
H4K20 me2:H3K4 me3
H3K27 me2
H3K4 me1
H4K20 me3
H2A.Z
H3K27 me3
H3−5ac
DNA Meth. m5C
H4−4ac
H3K4 me3
H3K9 me3
H4K20 me2
0.00 0.25 0.50 0.75 1.00
$\hat{\pi}$
NUP85 – Forward
H3K4 me1:H2A.Z
DNA Meth. m5C:H4K20 me2
H3K27 me2:DNA Meth. m5C
H3K9 me2:DNA Meth. m5C
H3K4 me3:H3−5ac
H2A.Z:H3K4 me3
H3K4 me1:H4−4ac
H3K4 me1:H4K20 me2
H3K27 me3:H3K9 me3
H3K4 me1:H3−5ac
H2A.Z:H4K20 me2
DNA Meth. m5C:H3K9 me3
H3K27 me3:H3K4 me1
DNA Meth. m5C:H4K20 me3
H3K27 me3:DNA Meth. m5C
H4K20 me2:H4−4ac
H4−4ac:H3−5ac
H4K20 me2:H3−5ac
H4K20 me2:H3K4 me3
H3K4 me3:H4−4ac
H3K27 me2
H3K9 me2
H3K9 me3
H4K20 me3
H2A.Z
H3K27 me3
H3K4 me1
H4K20 me2
H4−4ac
H3−5ac
H3K4 me3
DNA Meth. m5C
0.00 0.25 0.50 0.75 1.00
$\hat{\pi}$
NUP85 – Reverse
H3K4 me1:H2A.Z
DNA Meth. m5C:H4K20 me2
H3K27 me2:DNA Meth. m5C
H3K9 me2:DNA Meth. m5C
H3K4 me3:H3−5ac
H2A.Z:H3K4 me3
H3K4 me1:H4−4ac
H3K4 me1:H4K20 me2
H3K27 me3:H3K9 me3
H3K4 me1:H3−5ac
H2A.Z:H4K20 me2
DNA Meth. m5C:H3K9 me3
H3K27 me3:H3K4 me1
DNA Meth. m5C:H4K20 me3
H3K27 me3:DNA Meth. m5C
H4K20 me2:H4−4ac
H4−4ac:H3−5ac
H4K20 me2:H3−5ac
H4K20 me2:H3K4 me3
H3K4 me3:H4−4ac
H3K27 me2
H3K9 me2
H3K9 me3
H4K20 me3
H2A.Z
H3K27 me3
H3K4 me1
H4K20 me2
H4−4ac
H3−5ac
H3K4 me3
DNA Meth. m5C
0.00 0.25 0.50 0.75 1.00
$\hat{\pi}$
MBNL1 – Forward
H3K4 me3:H3−5ac
DNA Meth. m5C:H3K9 me3
H3K27 me3:H3K9 me3
H4K20 me2:H4−4ac
H3K9 me2:DNA Meth. m5C
H3K27 me3:DNA Meth. m5C
DNA Meth. m5C:H4K20 me2
H3K27 me2:DNA Meth. m5C
H4K20 me2:H3K4 me3
H3K4 me1:H4K20 me2
H2A.Z:H4K20 me2
DNA Meth. m5C:H4K20 me3
H4K20 me2:H3−5ac
H3K4 me1
H4−4ac
H4K20 me3
H2A.Z
H3K9 me2
H3K9 me3
H3K27 me2
H3−5ac
DNA Meth. m5C
H3K27 me3
H3K4 me3
H4K20 me2
0.00 0.25 0.50 0.75 1.00
$\hat{\pi}$
MBNL1 – Reverse
H3K4 me3:H3−5ac
DNA Meth. m5C:H3K9 me3
H3K27 me3:H3K9 me3
H4K20 me2:H4−4ac
H3K9 me2:DNA Meth. m5C
H3K27 me3:DNA Meth. m5C
DNA Meth. m5C:H4K20 me2
H3K27 me2:DNA Meth. m5C
H4K20 me2:H3K4 me3
H3K4 me1:H4K20 me2
H2A.Z:H4K20 me2
DNA Meth. m5C:H4K20 me3
H4K20 me2:H3−5ac
H3K4 me1
H4−4ac
H4K20 me3
H2A.Z
H3K9 me2
H3K9 me3
H3K27 me2
H3−5ac
DNA Meth. m5C
H3K27 me3
H3K4 me3
H4K20 me2
0.00 0.25 0.50 0.75 1.00
$\hat{\pi}$

**Extended Fig. 3c.** Extension of Fig. 3c. Full list of scatter plots for all 55 proteins with robust interaction effects.

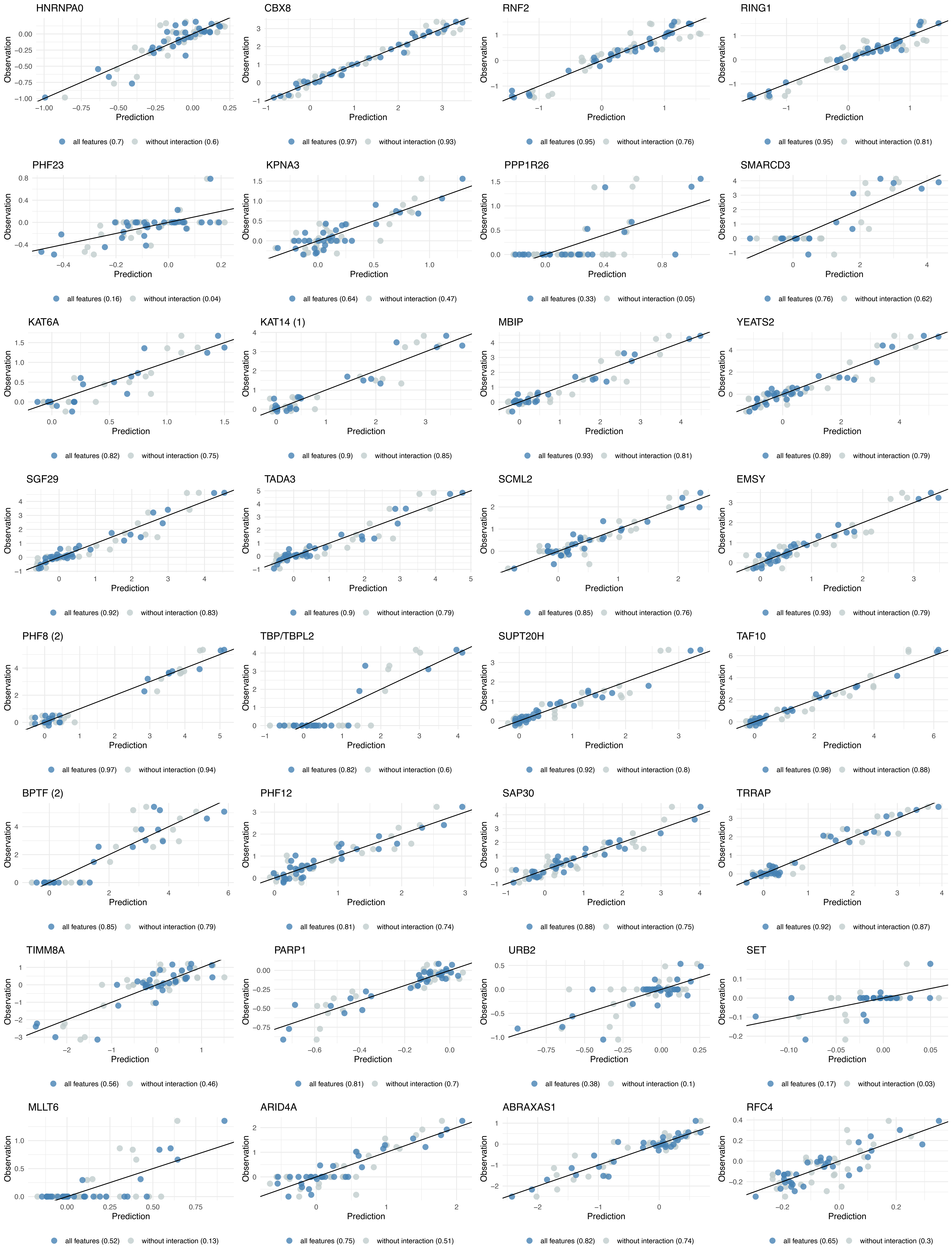
HNRNPA0
all features (0.7) without interaction (0.6)
CBX8
all features (0.97) without interaction (0.93)
RNF2
all features (0.95) without interaction (0.76)
RING1
all features (0.95) without interaction (0.81)
PHF23
all features (0.16) without interaction (0.04)
KPNA3
all features (0.64) without interaction (0.47)
PPP1R26
all features (0.33) without interaction (0.05)
SMARCD3
all features (0.76) without interaction (0.62)
KAT6A
all features (0.82) without interaction (0.75)
KAT14 (1)
all features (0.9) without interaction (0.85)
MBIP
all features (0.93) without interaction (0.81)
YEATS2
all features (0.89) without interaction (0.79)
SGF29
all features (0.92) without interaction (0.83)
TADA3
all features (0.9) without interaction (0.79)
SCML2
all features (0.85) without interaction (0.76)
EMSY
all features (0.93) without interaction (0.79)
PHF8 (2)
all features (0.97) without interaction (0.94)
TBP/TBPL2
all features (0.82) without interaction (0.6)
SUPT20H
all features (0.92) without interaction (0.8)
TAF10
all features (0.98) without interaction (0.88)
BPTF (2)
all features (0.85) without interaction (0.79)
PHF12
all features (0.81) without interaction (0.74)
SAP30
all features (0.88) without interaction (0.75)
TRRAP
all features (0.92) without interaction (0.87)
TIMM8A
all features (0.56) without interaction (0.46)
PARP1
all features (0.81) without interaction (0.7)
URB2
all features (0.38) without interaction (0.1)
SET
all features (0.17) without interaction (0.03)
MLLT6
all features (0.52) without interaction (0.13)
ARID4A
all features (0.75) without interaction (0.51)
ABRAXAS1
all features (0.82) without interaction (0.74)
RFC4
all features (0.65) without interaction (0.3)
Observation
Prediction

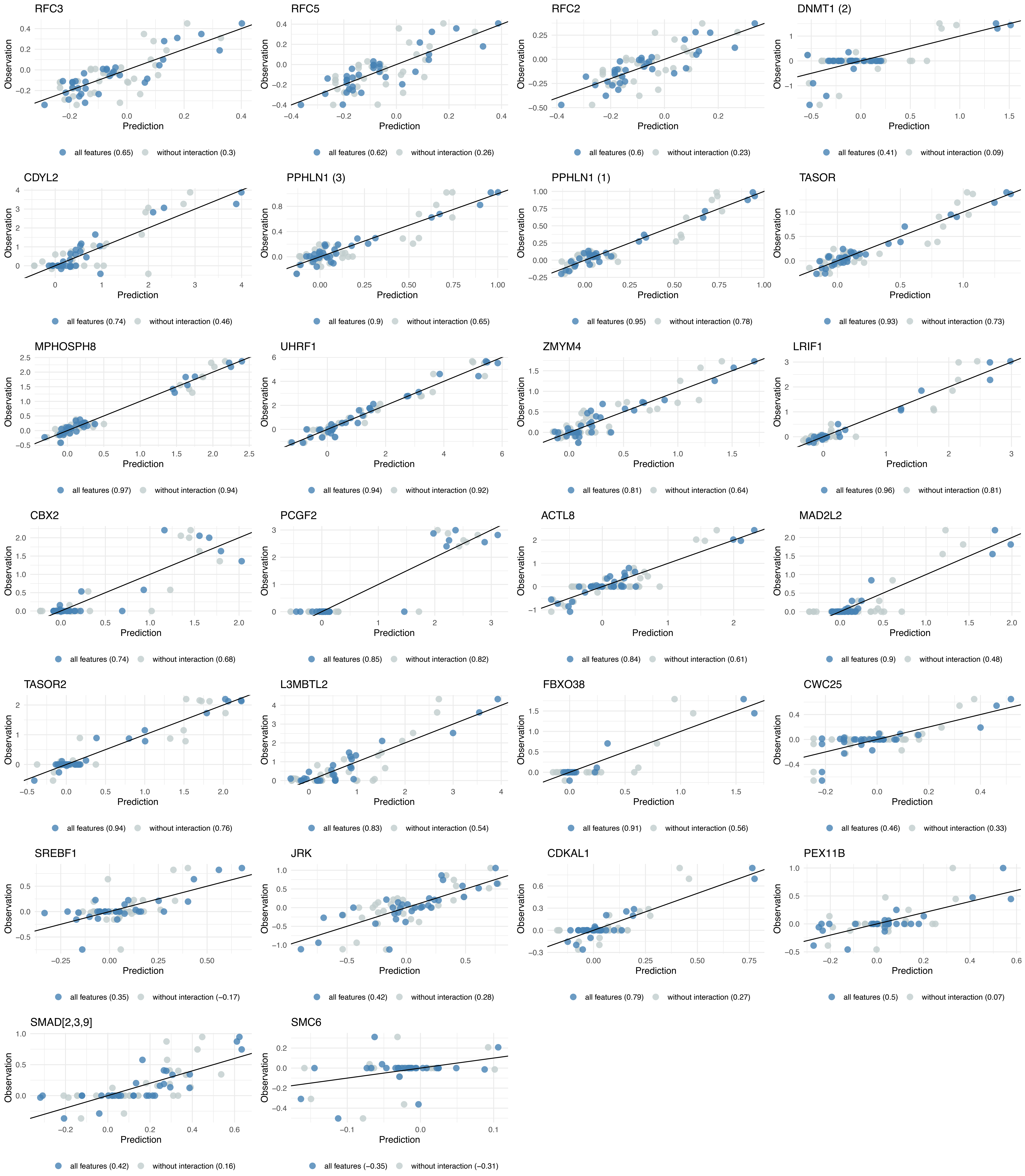
RFC3
all features (0.65) without interaction (0.3)
RFC5
all features (0.62) without interaction (0.26)
RFC2
all features (0.6) without interaction (0.23)
DNMT1 (2)
all features (0.41) without interaction (0.09)
CDYL2
all features (0.74) without interaction (0.46)
PPHLN1 (3)
all features (0.9) without interaction (0.65)
PPHLN1 (1)
all features (0.95) without interaction (0.78)
TASOR
all features (0.93) without interaction (0.73)
MPHOSPH8
all features (0.97) without interaction (0.94)
UHRF1
all features (0.94) without interaction (0.92)
ZMYM4
all features (0.81) without interaction (0.64)
LRIF1
all features (0.96) without interaction (0.81)
CBX2
all features (0.74) without interaction (0.68)
PCGF2
all features (0.85) without interaction (0.82)
ACTL8
all features (0.84) without interaction (0.61)
MAD2L2
all features (0.9) without interaction (0.48)
TASOR2
all features (0.94) without interaction (0.76)
L3MBTL2
all features (0.83) without interaction (0.54)
FBXO38
all features (0.91) without interaction (0.56)
CWC25
all features (0.46) without interaction (0.33)
SREBF1
all features (0.35) without interaction (−0.17)
JRK
all features (0.42) without interaction (0.28)
CDKAL1
all features (0.79) without interaction (0.27)
PEX11B
all features (0.5) without interaction (0.07)
SMAD[2,3,9]
all features (0.42) without interaction (0.16)
SMC6
all features (−0.35) without interaction (−0.31)
Observation
Prediction

**Extended Fig. 5b.** Extension of Fig. 5b. Selection probability heatmaps and model coefficients for all 1915 protein complexes.

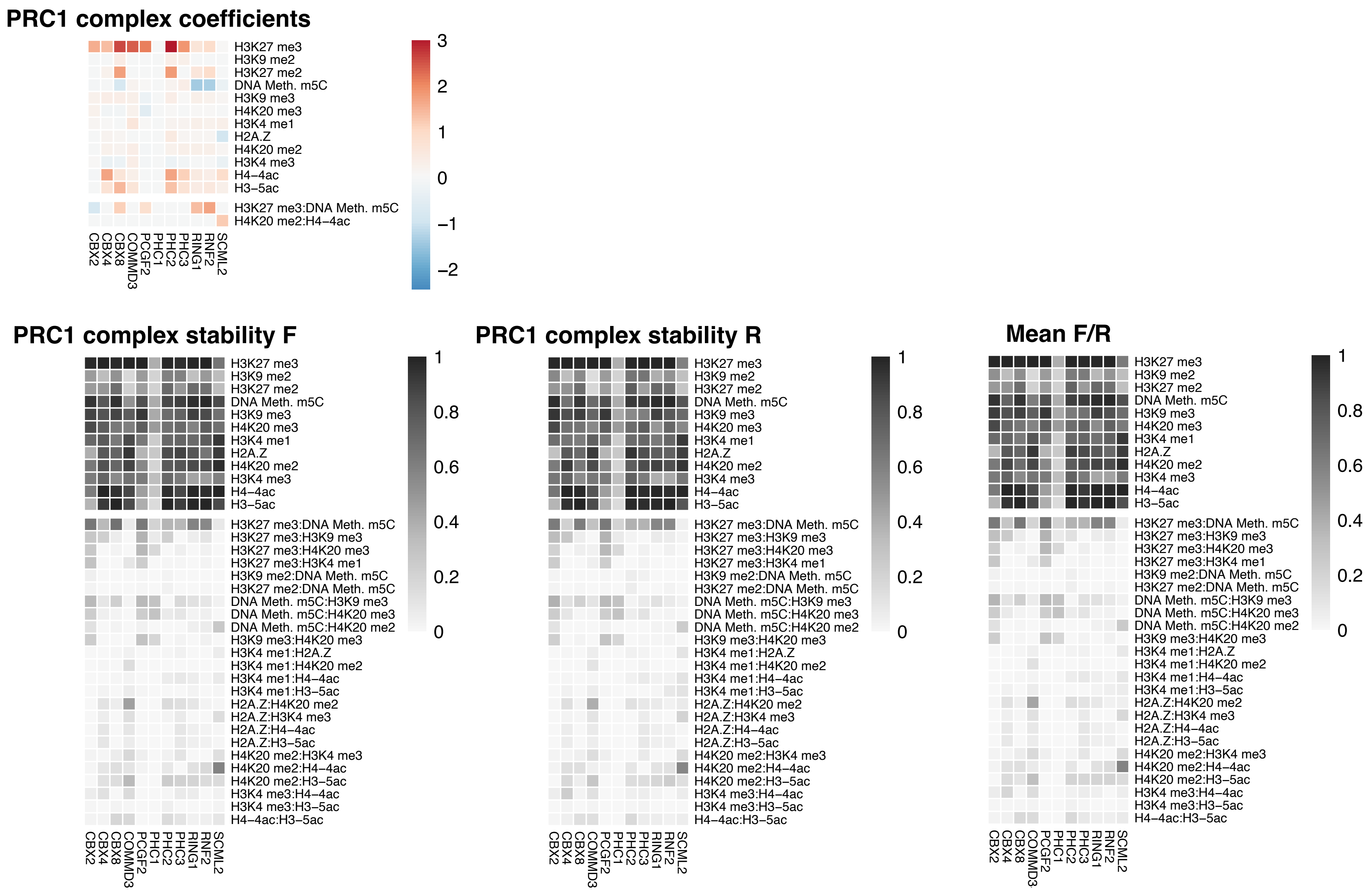
PRC1 complex coefficients
H3K27 me3
H3K9 me2
H3K27 me2
DNA Meth. m5C
H3K9 me3
H4K20 me3
H3K4 me1
H2A.Z
H4K20 me2
H3K4 me3
H4−4ac
H3−5ac
H3K27 me3:DNA Meth. m5C
H4K20 me2:H4−4ac
CBX2 CBX4 CBX8 COMMD3 PCGF2 PHC1 PHC2 PHC3 RING1 RNF2 SCML2
3 2 1 0 −1 −2
PRC1 complex stability F
PRC1 complex stability R
Mean F/R
H3K27 me3:H3K9 me3
H3K27 me3:H4K20 me3
H3K27 me3:H3K4 me1
H3K9 me2:DNA Meth. m5C
H3K27 me2:DNA Meth. m5C
DNA Meth. m5C:H3K9 me3
DNA Meth. m5C:H4K20 me3
DNA Meth. m5C:H4K20 me2
H3K9 me3:H4K20 me3
H3K4 me1:H2A.Z
H3K4 me1:H4K20 me2
H3K4 me1:H4−4ac
H3K4 me1:H3−5ac
H2A.Z:H4K20 me2
H2A.Z:H3K4 me3
H2A.Z:H4−4ac
H2A.Z:H3−5ac
H4K20 me2:H3K4 me3
H4K20 me2:H3−5ac
H3K4 me3:H4−4ac
H3K4 me3:H3−5ac
H4−4ac:H3−5ac
1 0.8 0.6 0.4 0.2 0

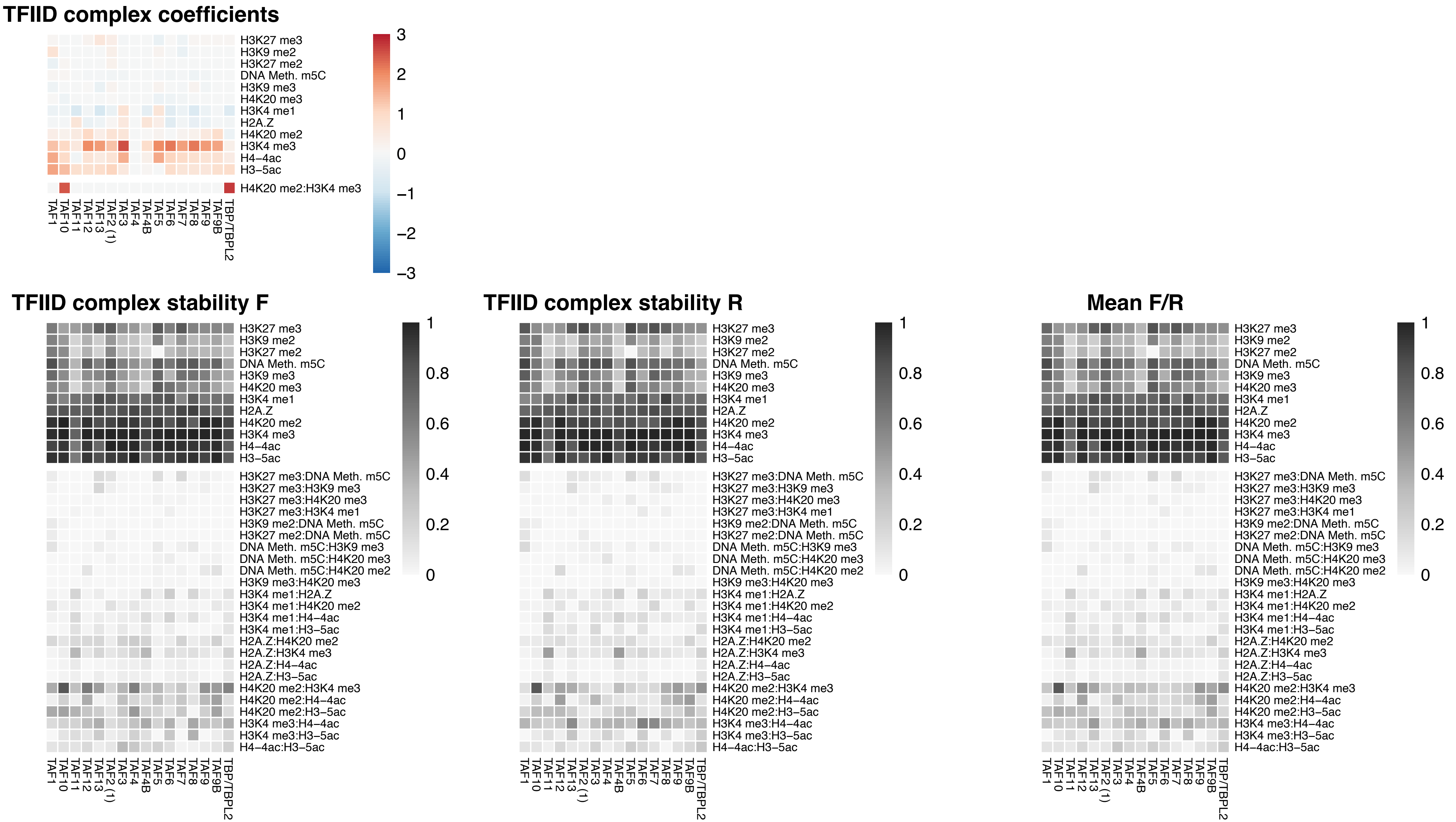
TFIID complex coefficients
H3K27 me3
H3K9 me2
H3K27 me2
DNA Meth. m5C
H3K9 me3
H4K20 me3
H3K4 me1
H2A.Z
H4K20 me2
H3K4 me3
H4−4ac
H3−5ac
H4K20 me2:H3K4 me3
TAF1
TAF10
TAF11
TAF12
TAF13
TAF2 (1)
TAF3
TAF4
TAF4B
TAF5
TAF6
TAF7
TAF8
TAF9
TAF9B
TBP/TBPL2
3
2
1
0
−1
−2
−3
TFIID complex stability F
TFIID complex stability R
Mean F/R
H3K27 me3:DNA Meth. m5C
H3K27 me3:H3K9 me3
H3K27 me3:H4K20 me3
H3K27 me3:H3K4 me1
H3K9 me2:DNA Meth. m5C
H3K27 me2:DNA Meth. m5C
DNA Meth. m5C:H3K9 me3
DNA Meth. m5C:H4K20 me3
DNA Meth. m5C:H4K20 me2
H3K9 me3:H4K20 me3
H3K4 me1:H2A.Z
H3K4 me1:H4K20 me2
H3K4 me1:H4−4ac
H3K4 me1:H3−5ac
H2A.Z:H4K20 me2
H2A.Z:H3K4 me3
H2A.Z:H4−4ac
H2A.Z:H3−5ac
H4K20 me2:H3K4 me3
H4K20 me2:H4−4ac
H4K20 me2:H3−5ac
H3K4 me3:H4−4ac
H3K4 me3:H3−5ac
H4−4ac:H3−5ac
1
0.8
0.6
0.4
0.2
0

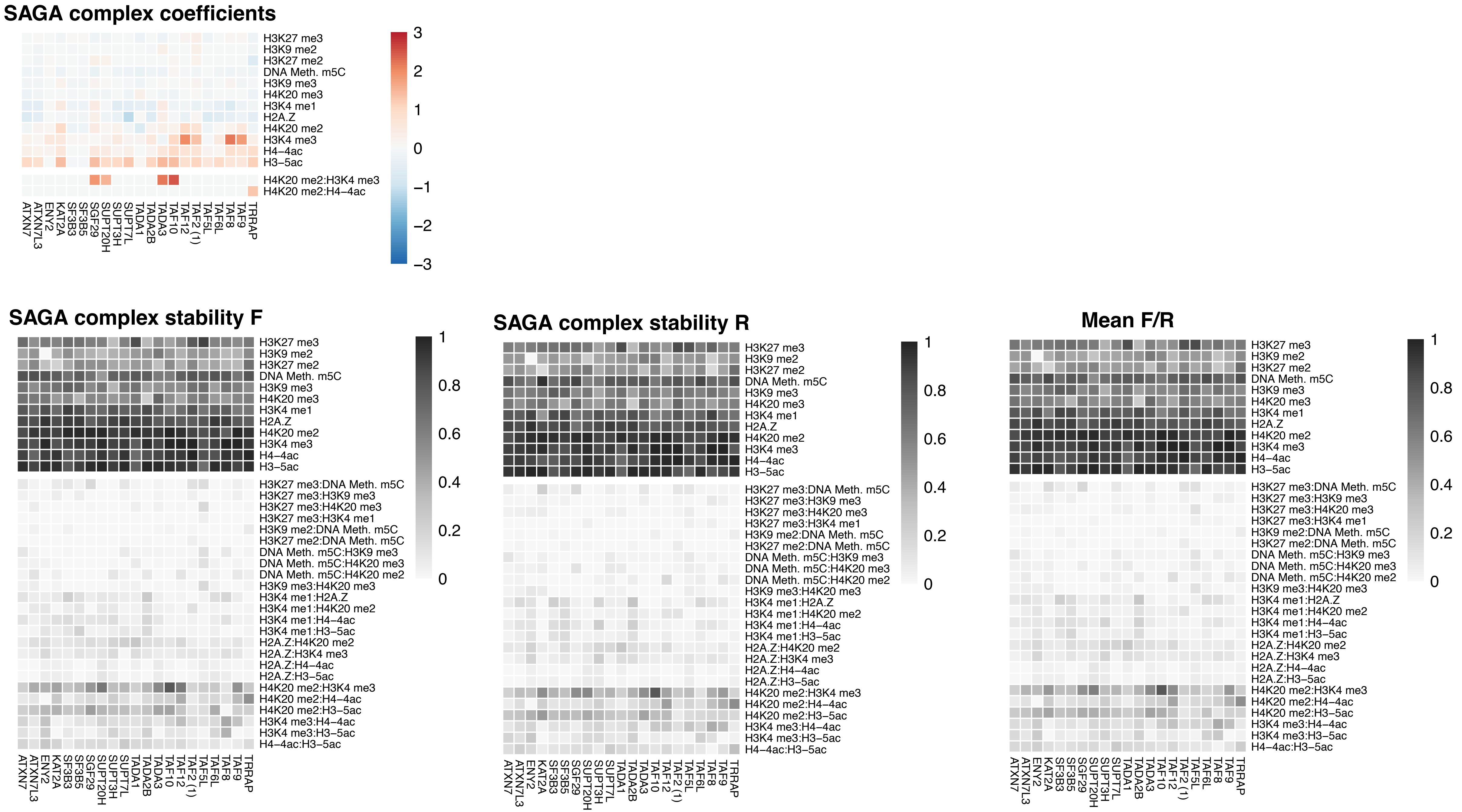

SAGA complex coefficients
H3K27 me3
H3K9 me2
H3K27 me2
DNA Meth. m5C
H3K9 me3
H4K20 me3
H3K4 me1
H2A.Z
H4K20 me2
H3K4 me3
H4−4ac
H3−5ac
H4K20 me2:H3K4 me3
H4K20 me2:H4−4ac
ATXN7
ATXN7L3
ENY2
KAT2A
SF3B3
SF3B5
SGF29
SUPT20H
SUPT3H
SUPT7L
TADA1
TADA2B
TADA3
TAF10
TAF12
TAF2 (1)
TAF5L
TAF6L
TAF8
TAF9
TRRAP
3
2
1
0
−1
−2
−3
SAGA complex stability F
SAGA complex stability R
Mean F/R
H3K27 me3
H3K9 me2
H3K27 me2
DNA Meth. m5C
H3K9 me3
H4K20 me3
H3K4 me1
H2A.Z
H4K20 me2
H3K4 me3
H4−4ac
H3−5ac
H3K27 me3:DNA Meth. m5C
H3K27 me3:H3K9 me3
H3K27 me3:H4K20 me3
H3K27 me3:H3K4 me1
H3K9 me2:DNA Meth. m5C
H3K27 me2:DNA Meth. m5C
DNA Meth. m5C:H3K9 me3
DNA Meth. m5C:H4K20 me3
DNA Meth. m5C:H4K20 me2
H3K9 me3:H4K20 me3
H3K4 me1:H2A.Z
H3K4 me1:H4K20 me2
H3K4 me1:H4−4ac
H3K4 me1:H3−5ac
H2A.Z:H4K20 me2
H2A.Z:H3K4 me3
H2A.Z:H4−4ac
H2A.Z:H3−5ac
H4K20 me2:H3K4 me3
H4K20 me2:H4−4ac
H4K20 me2:H3−5ac
H3K4 me3:H4−4ac
H3K4 me3:H3−5ac
H4−4ac:H3−5ac
1
0.8
0.6
0.4
0.2
0

### HUSH complex coefficients

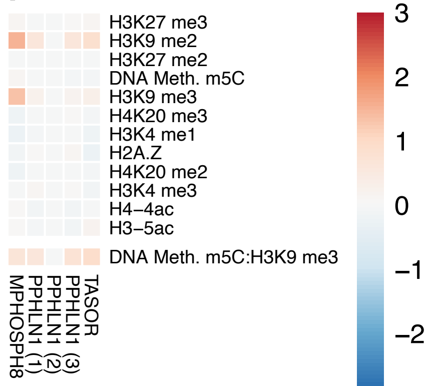


### HUSH complex stability F

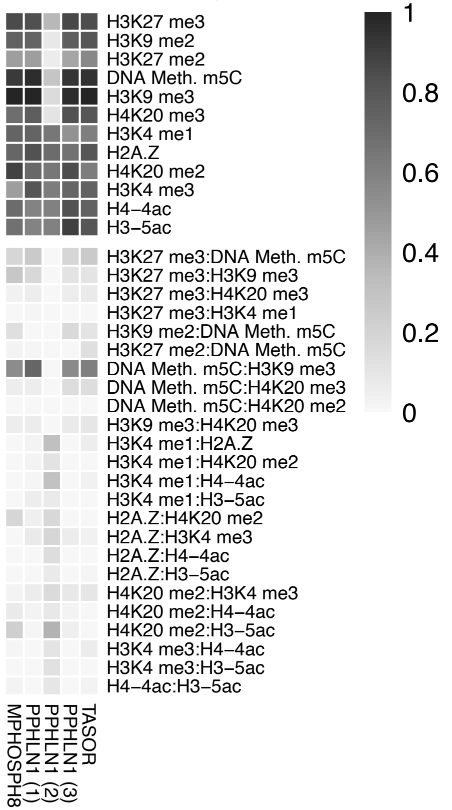


### HUSH complex stability R

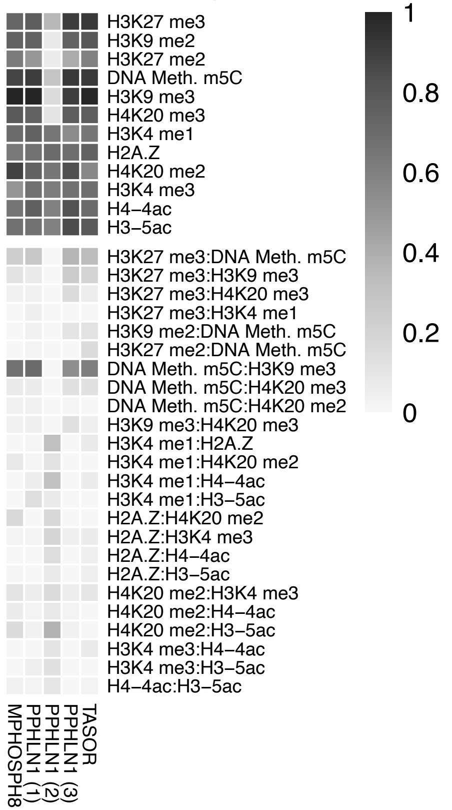


### Mean F/R

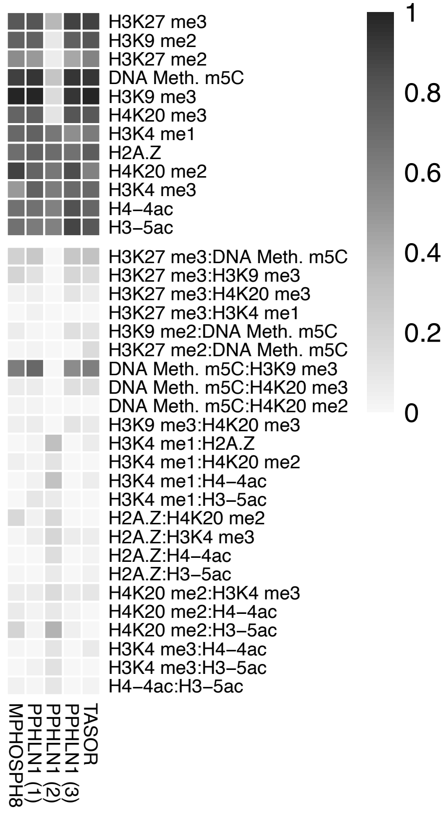

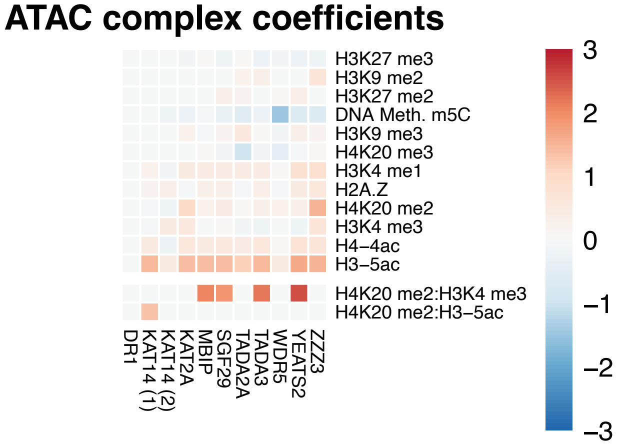
ATAC complex coefficients
H3K27 me3
H3K9 me2
H3K27 me2
DNA Meth. m5C
H3K9 me3
H4K20 me3
H3K4 me1
H2A.Z
H4K20 me2
H3K4 me3
H4−4ac
H3−5ac
H4K20 me2:H3K4 me3
H4K20 me2:H3−5ac
DR1
KAT14 (1)
KAT14 (2)
KAT2A
MBIP
SGF29
TADA2A
TADA3
WDR5
YEATS2
ZZZ3
3
2
1
0
−1
−2
−3

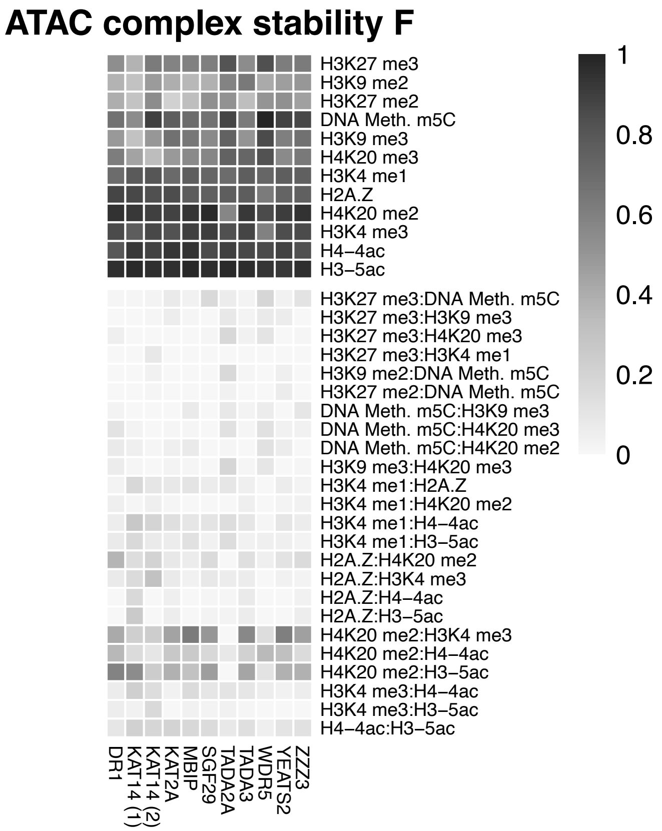
ATAC complex stability F
H3K27 me3
H3K9 me2
H3K27 me2
DNA Meth. m5C
H3K9 me3
H4K20 me3
H3K4 me1
H2A.Z
H4K20 me2
H3K4 me3
H4−4ac
H3−5ac
H3K27 me3:DNA Meth. m5C
H3K27 me3:H3K9 me3
H3K27 me3:H4K20 me3
H3K27 me3:H3K4 me1
H3K9 me2:DNA Meth. m5C
H3K27 me2:DNA Meth. m5C
DNA Meth. m5C:H3K9 me3
DNA Meth. m5C:H4K20 me3
DNA Meth. m5C:H4K20 me2
H3K9 me3:H4K20 me3
H3K4 me1:H2A.Z
H3K4 me1:H4K20 me2
H3K4 me1:H4−4ac
H3K4 me1:H3−5ac
H2A.Z:H4K20 me2
H2A.Z:H3K4 me3
H2A.Z:H4−4ac
H2A.Z:H3−5ac
H4K20 me2:H3K4 me3
H4K20 me2:H4−4ac
H4K20 me2:H3−5ac
H3K4 me3:H4−4ac
H3K4 me3:H3−5ac
H4−4ac:H3−5ac
DR1
KAT14 (1)
KAT14 (2)
KAT2A
MBIP
SGF29
TADA2A
TADA3
WDR5
YEATS2
ZZZ3
1
0.8
0.6
0.4
0.2
0

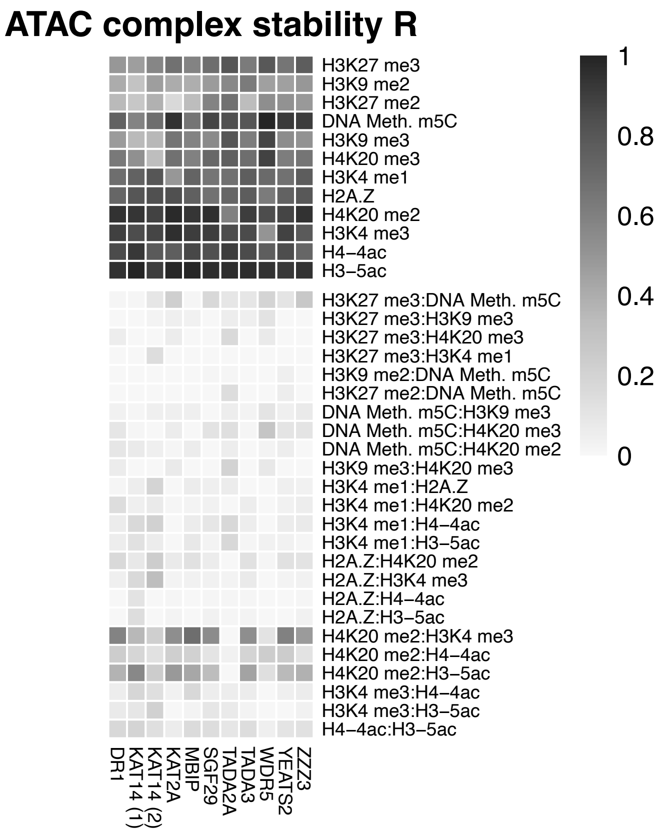
ATAC complex stability R
H3K27 me3
H3K9 me2
H3K27 me2
DNA Meth. m5C
H3K9 me3
H4K20 me3
H3K4 me1
H2A.Z
H4K20 me2
H3K4 me3
H4−4ac
H3−5ac
H3K27 me3:DNA Meth. m5C
H3K27 me3:H3K9 me3
H3K27 me3:H4K20 me3
H3K27 me3:H3K4 me1
H3K9 me2:DNA Meth. m5C
H3K27 me2:DNA Meth. m5C
DNA Meth. m5C:H3K9 me3
DNA Meth. m5C:H4K20 me3
DNA Meth. m5C:H4K20 me2
H3K9 me3:H4K20 me3
H3K4 me1:H2A.Z
H3K4 me1:H4K20 me2
H3K4 me1:H4−4ac
H3K4 me1:H3−5ac
H2A.Z:H4K20 me2
H2A.Z:H3K4 me3
H2A.Z:H4−4ac
H2A.Z:H3−5ac
H4K20 me2:H3K4 me3
H4K20 me2:H4−4ac
H4K20 me2:H3−5ac
H3K4 me3:H4−4ac
H3K4 me3:H3−5ac
H4−4ac:H3−5ac
DR1
KAT14 (1)
KAT14 (2)
KAT2A
MBIP
SGF29
TADA2A
TADA3
WDR5
YEATS2
ZZZ3
1
0.8
0.6
0.4
0.2
0

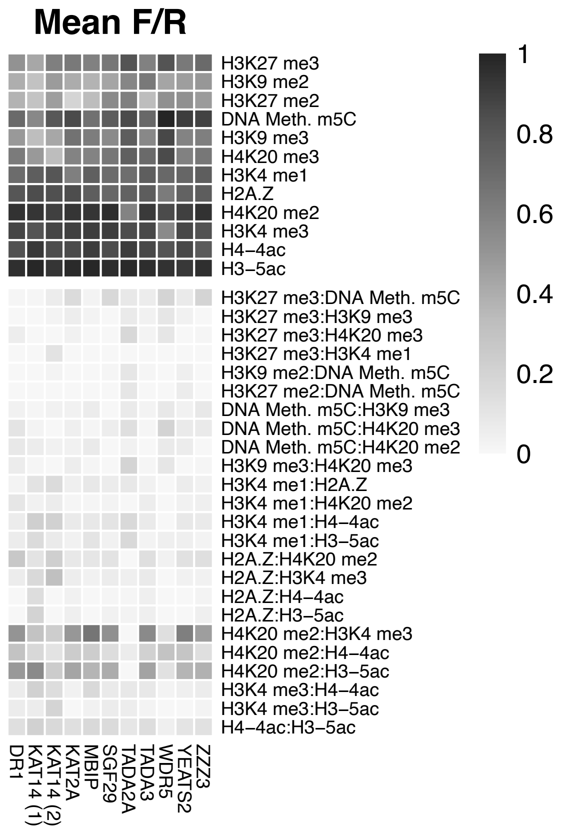
Mean F/R
H3K27 me3
H3K9 me2
H3K27 me2
DNA Meth. m5C
H3K9 me3
H4K20 me3
H3K4 me1
H2A.Z
H4K20 me2
H3K4 me3
H4−4ac
H3−5ac
H3K27 me3:DNA Meth. m5C
H3K27 me3:H3K9 me3
H3K27 me3:H4K20 me3
H3K27 me3:H3K4 me1
H3K9 me2:DNA Meth. m5C
H3K27 me2:DNA Meth. m5C
DNA Meth. m5C:H3K9 me3
DNA Meth. m5C:H4K20 me3
DNA Meth. m5C:H4K20 me2
H3K9 me3:H4K20 me3
H3K4 me1:H2A.Z
H3K4 me1:H4K20 me2
H3K4 me1:H4−4ac
H3K4 me1:H3−5ac
H2A.Z:H4K20 me2
H2A.Z:H3K4 me3
H2A.Z:H4−4ac
H2A.Z:H3−5ac
H4K20 me2:H3K4 me3
H4K20 me2:H4−4ac
H4K20 me2:H3−5ac
H3K4 me3:H4−4ac
H3K4 me3:H3−5ac
H4−4ac:H3−5ac
DR1
KAT14 (1)
KAT14 (2)
KAT2A
MBIP
SGF29
TADA2A
TADA3
WDR5
YEATS2
ZZZ3
1
0.8
0.6
0.4
0.2
0

**Extended Fig. S4.** Extension of Fig. S4. Heatmap of estimated main effects in the linear model for all 1915 proteins.

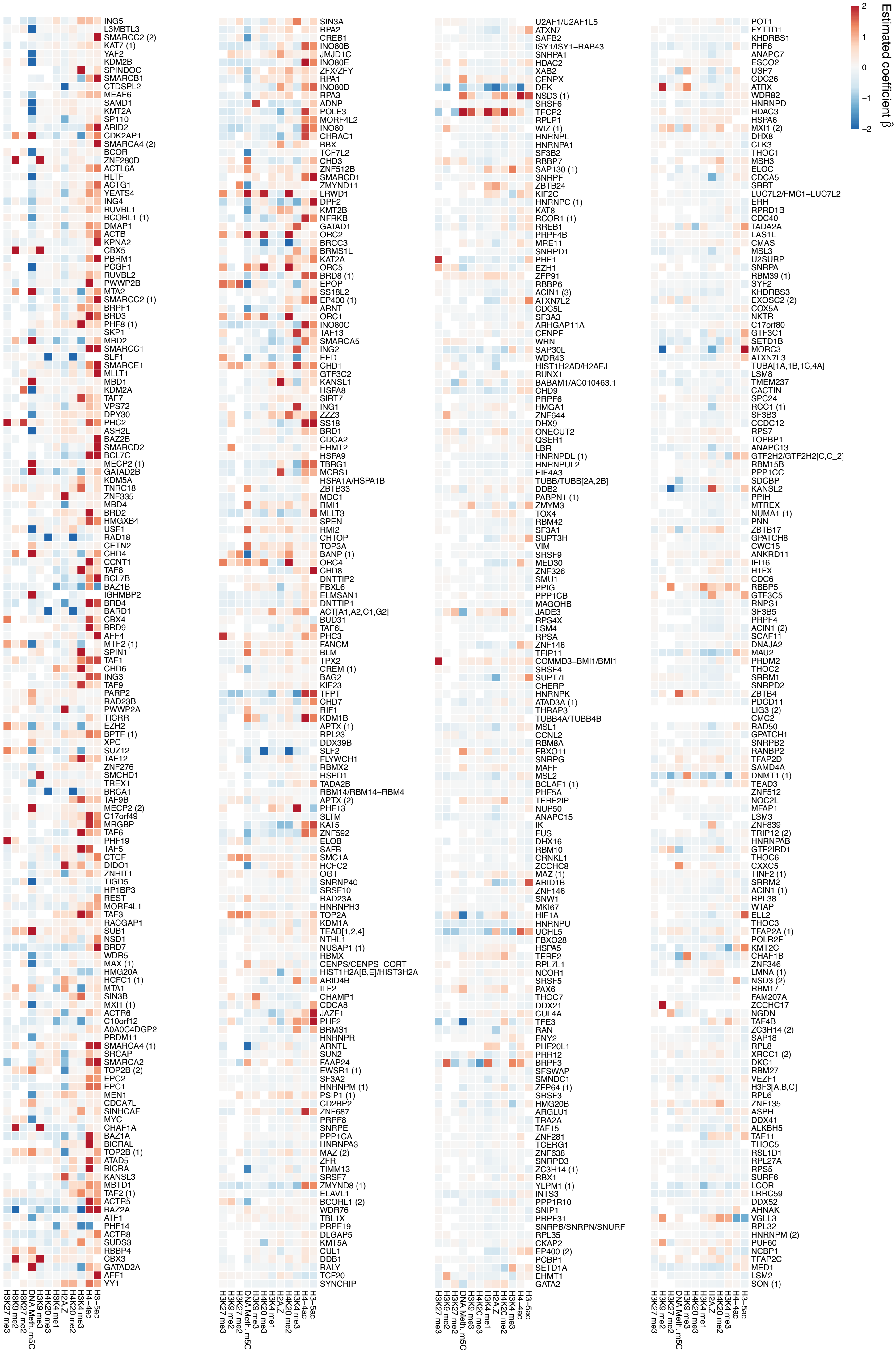

ING5
L3MBTL3
SMARCC2 (2)
KAT7 (1)
YAF2
KDM2B
SPINDOC
SMARCB1
CTDSPL2
MEAF6
SAMD1
KMT2A
SP110
ARID2
CDK2AP1
SMARCA4 (2)
BCOR
ZNF280D
ACTL6A
HLTF
ACTG1
YEATS4
ING4
RUVBL1
BCORL1 (1)
DMAP1
ACTB
KPNA2
CBX5
PBRM1
PCGF1
RUVBL2
PWWP2B
MTA2
SMARCC2 (1)
BRPF1
BRD3
PHF8 (1)
SKP1
MBD2
SMARCC1
SLF1
SMARCE1
MLLT1
MBD1
KDM2A
TAF7
VPS72
DPY30
PHC2
ASH2L
BAZ2B
SMARCD2
BCL7C
MECP2 (1)
GATAD2B
KDM5A
TNRC18
ZNF335
MBD4
BRD2
HMGXB4
USF1
RAD18
CETN2
CHD4
CCNT1
TAF8
BCL7B
BAZ1B
IGHMBP2
BRD4
BARD1
CBX4
BRD9
AFF4
MTF2 (1)
SPIN1
TAF1
CHD6
ING3
TAF9
PARP2
RAD23B
PWWP2A
TICRR
EZH2
BPTF (1)
XPC
SUZ12
TAF12
ZNF276
SMCHD1
TREX1
BRCA1
TAF9B
MECP2 (2)
C17orf49
MRGBP
TAF6
PHF19
TAF5
CTCF
DIDO1
ZNHIT1
TIGD5
HP1BP3
REST
MORF4L1
TAF3
RACGAP1
SUB1
NSD1
BRD7
WDR5
MAX (1)
HMG20A
HCFC1 (1)
MTA1
SIN3B
MXI1 (1)
ACTR6
C10orf12
A0A0C4DGP2
PRDM11
SMARCA4 (1)
SRCAP
SMARCA2
TOP2B (2)
EPC2
EPC1
MEN1
CDCA7L
SINHCAF
MYC
CHAF1A
BAZ1A
BICRAL
TOP2B (1)
ATAD5
BICRA
KANSL3
MBTD1
TAF2 (1)
ACTR5
BAZ2A
ATF1
PHF14
ACTR8
SUDS3
RBBP4
CBX3
GATAD2A
AFF1
YY1
SIN3A
RPA2
CREB1
INO80B
JMJD1C
INO80E
ZFX/ZFY
RPA1
INO80D
RPA3
ADNP
POLE3
MORF4L2
INO80
CHRAC1
BBX
TCF7L2
CHD3
ZNF512B
SMARCD1
ZMYND11
LRWD1
DPF2
KMT2B
NFRKB
GATAD1
ORC2
BRCC3
BRMS1L
KAT2A
ORC5
BRD8 (1)
EPOP
SS18L2
EP400 (1)
ARNT
ORC1
INO80C
TAF13
SMARCA5
ING2
EED
CHD1
GTF3C2
KANSL1
HSPA8
SIRT7
ING1
ZZZ3
SS18
BRD1
CDCA2
EHMT2
HSPA9
TBRG1
MCRS1
HSPA1A/HSPA1B
ZBTB33
MDC1
RMI1
MLLT3
SPEN
RMI2
CHTOP
TOP3A
BANP (1)
ORC4
CHD8
DNTTIP2
FBXL6
ELMSAN1
DNTTIP1
ACT[A1,A2,C1,G2]
BUD31
TAF6L
PHC3
FANCM
BLM
TPX2
CREM (1)
BAG2
KIF23
TFPT
CHD7
RIF1
KDM1B
APTX (1)
RPL23
DDX39B
SLF2
FLYWCH1
RBMX2
HSPD1
TADA2B
RBM14/RBM14−RBM4
APTX (2)
PHF13
SLTM
KAT5
ZNF592
ELOB
SAFB
SMC1A
HCFC2
OGT
SNRNP40
SRSF10
RAD23A
HNRNPH3
TOP2A
KDM1A
TEAD[1,2,4]
NTHL1
NUSAP1 (1)
RBMX
CENPS/CENPS−CORT
HIST1H2A[B,E]/HIST3H2A
ARID4B
ILF2
CHAMP1
CDCA8
JAZF1
PHF2
BRMS1
HNRNPR
ARNTL
SUN2
FAAP24
EWSR1 (1)
SF3A2
HNRNPM (1)
PSIP1 (1)
CD2BP2
ZNF687
PRPF8
SNRPE
PPP1CA
HNRNPA3
MAZ (2)
ZFR
TIMM13
SRSF7
ZMYND8 (1)
ELAVL1
BCORL1 (2)
WDR76
TBL1X
PRPF19
DLGAP5
KMT5A
CUL1
DDB1
RALY
TCF20
SYNCRIP
U2AF1/U2AF1L5
ATXN7
SAFB2
ISY1/ISY1−RAB43
SNRPA1
HDAC2
XAB2
CENPX
DEK
NSD3 (1)
SRSF6
TFCP2
RPLP1
WIZ (1)
HNRNPL
HNRNPA1
SF3B2
RBBP7
SAP130 (1)
SNRPF
ZBTB24
KIF2C
HNRNPC (1)
KAT8
RCOR1 (1)
RREB1
PRPF4B
MRE11
SNRPD1
PHF1
EZH1
ZFP91
RBBP6
ACIN1 (3)
ATXN7L2
CDC5L
SF3A3
ARHGAP11A
CENPF
WRN
SAP30L
WDR43
HIST1H2AD/H2AFJ
RUNX1
BABAM1/AC010463.1
CHD9
PRPF6
HMGA1
ZNF644
DHX9
ONECUT2
QSER1
LBR
HNRNPDL (1)
HNRNPUL2
EIF4A3
TUBB/TUBB[2A,2B]
DDB2
PABPN1 (1)
ZMYM3
TOX4
RBM42
SF3A1
SUPT3H
VIM
SRSF9
MED30
ZNF326
SMU1
PPIG
PPP1CB
MAGOHB
JADE3
RPS4X
LSM4
RPSA
ZNF148
TFIP11
COMMD3−BMI1/BMI1
SRSF4
SUPT7L
CHERP
HNRNPK
ATAD3A (1)
THRAP3
TUBB4A/TUBB4B
MSL1
CCNL2
RBM8A
FBXO11
SNRPG
MAFF
MSL2
BCLAF1 (1)
PHF5A
TERF2IP
NUP50
ANAPC15
IK
FUS
DHX16
RBM10
CRNKL1
ZCCHC8
MAZ (1)
ARID1B
ZNF146
SNW1
MKI67
HIF1A
HNRNPU
UCHL5
FBXO28
HSPA5
TERF2
RPL7L1
NCOR1
SRSF5
PAX6
THOC7
DDX21
CUL4A
TFE3
RAN
ENY2
PHF20L1
PRR12
BRPF3
SFSWAP
SMNDC1
ZFP64 (1)
SRSF3
HMG20B
ARGLU1
TRA2A
TAF15
ZNF281
TCERG1
ZNF638
SNRPD3
ZC3H14 (1)
RBX1
YLPM1 (1)
INTS3
PPP1R10
SNIP1
PRPF31
SNRPB/SNRPN/SNURF
RPL35
CKAP2
EP400 (2)
PCBP1
SETD1A
EHMT1
GATA2
POT1
FYTTD1
KHDRBS1
PHF6
ANAPC7
ESCO2
USP7
CDC26
ATRX
WDR82
HNRNPD
HDAC3
HSPA6
MXI1 (2)
DHX8
CLK3
THOC1
MSH3
ELOC
CDCA5
SRRT
LUC7L2/FMC1−LUC7L2
ERH
RPRD1B
CDC40
TADA2A
LAS1L
CMAS
MSL3
U2SURP
SNRPA
RBM39 (1)
SYF2
KHDRBS3
EXOSC2 (2)
COX5A
NKTR
C17orf80
GTF3C1
SETD1B
MORC3
ATXN7L3
TUBA[1A,1B,1C,4A]
LSM8
TMEM237
CACTIN
SPC24
RCC1 (1)
SF3B3
CCDC12
RPS7
TOPBP1
ANAPC13
GTF2H2/GTF2H2[C,C_2]
RBM15B
PPP1CC
SDCBP
KANSL2
PPIH
MTREX
NUMA1 (1)
PNN
ZBTB17
GPATCH8
CWC15
ANKRD11
IFI16
H1FX
CDC6
RBBP5
GTF3C5
RNPS1
SF3B5
PRPF4
ACIN1 (2)
SCAF11
DNAJA2
MAU2
PRDM2
THOC2
SRRM1
SNRPD2
ZBTB4
PDCD11
LIG3 (2)
CMC2
RAD50
GPATCH1
SNRPB2
RANBP2
TFAP2D
SAMD4A
DNMT1 (1)
TEAD3
ZNF512
NOC2L
MFAP1
LSM3
ZNF839
TRIP12 (2)
HNRNPAB
GTF2IRD1
THOC6
CXXC5
TINF2 (1)
SRRM2
ACIN1 (1)
RPL38
WTAP
ELL2
THOC3
TFAP2A (1)
POLR2F
KMT2C
CHAF1B
ZNF346
LMNA (1)
NSD3 (2)
RBM17
FAM207A
ZCCHC17
NGDN
TAF4B
ZC3H14 (2)
SAP18
RPL8
XRCC1 (2)
DKC1
RBM27
VEZF1
H3F3[A,B,C]
RPL6
ZNF135
ASPH
DDX41
ALKBH5
TAF11
THOC5
RSL1D1
RPL27A
RPS5
SURF6
LCOR
LRRC59
DDX52
AHNAK
VGLL3
RPL32
HNRNPM (2)
PUF60
NCBP1
TFAP2C
MED1
LSM2
SON (1)
H3K27 me3
H3K9 me2
H3K27 me2
DNA Meth. m5C
H3K9 me3
H4K20 me3
H3K4 me1
H2A.Z
H4K20 me2
H3K4 me3
H4−4ac
H3−5ac
Estimated coefficient β
2
1
0
−1
−2

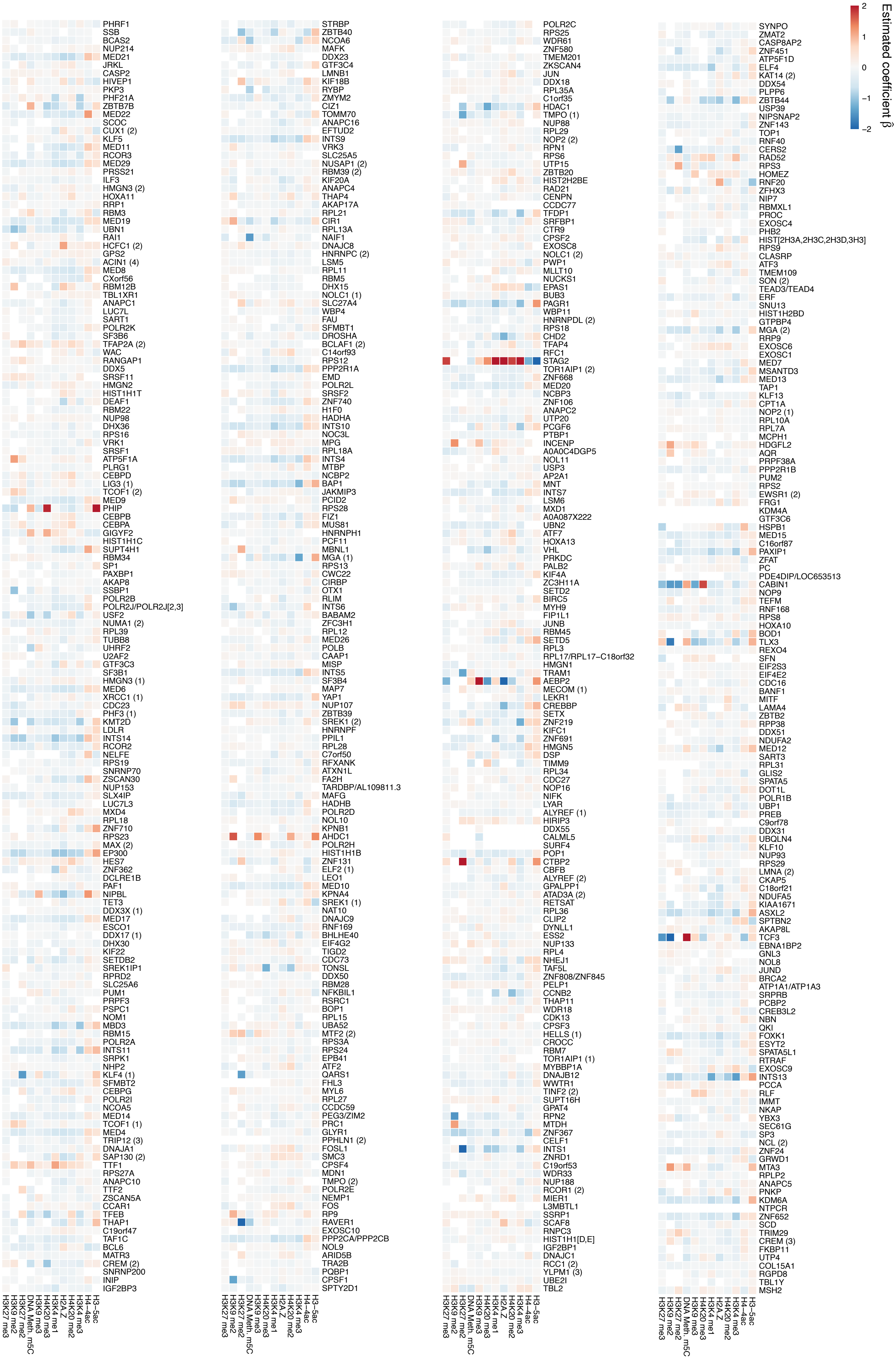
Estimated coefficient β̂
2
1
0
−1
−2
PHRF1
SSB
BCAS2
NUP214
MED21
JRKL
CASP2
HIVEP1
PKP3
PHF21A
ZBTB7B
MED22
SCOC
CUX1 (2)
KLF5
MED11
RCOR3
MED29
PRSS21
ILF3
HMGN3 (2)
HOXA11
RRP1
RBM3
MED19
UBN1
RAI1
HCFC1 (2)
GPS2
ACIN1 (4)
MED8
CXorf56
RBM12B
TBL1XR1
ANAPC1
LUC7L
SART1
POLR2K
SF3B6
TFAP2A (2)
WAC
RANGAP1
DDX5
SRSF11
HMGN2
HIST1H1T
DEAF1
RBM22
NUP98
DHX36
RPS16
VRK1
SRSF1
ATP5F1A
PLRG1
CEBPD
LIG3 (1)
TCOF1 (2)
MED9
PHIP
CEBPB
CEBPA
GIGYF2
HIST1H1C
SUPT4H1
RBM34
SP1
PAXBP1
AKAP8
SSBP1
POLR2B
POLR2J/POLR2J[2,3]
USF2
NUMA1 (2)
RPL39
TUBB8
UHRF2
U2AF2
GTF3C3
SF3B1
HMGN3 (1)
MED6
XRCC1 (1)
CDC23
PHF3 (1)
KMT2D
LDLR
INTS14
RCOR2
NELFE
RPS19
SNRNP70
ZSCAN30
NUP153
SLX4IP
LUC7L3
MXD4
RPL18
ZNF710
RPS23
MAX (2)
EP300
HES7
ZNF362
DCLRE1B
PAF1
NIPBL
TET3
DDX3X (1)
MED17
ESCO1
DDX17 (1)
DHX30
KIF22
SETDB2
SREK1IP1
RPRD2
SLC25A6
PUM1
PRPF3
PSPC1
NOM1
MBD3
RBM15
POLR2A
INTS11
SRPK1
NHP2
KLF4 (1)
SFMBT2
CEBPG
POLR2I
NCOA5
MED14
TCOF1 (1)
MED4
TRIP12 (3)
DNAJA1
SAP130 (2)
TTF1
RPS27A
ANAPC10
TTF2
ZSCAN5A
CCAR1
TFEB
THAP1
C19orf47
TAF1C
BCL6
MATR3
CREM (2)
SNRNP200
INIP
IGF2BP3
H3K27 me3
H3K9 me2
H3K27 me2
DNA Meth. m5C
H3K9 me3
H4K20 me3
H3K4 me1
H2A.Z
H4K20 me2
H3K4 me3
H4–4ac
H3–5ac
STRBP
ZBTB40
NCOA6
MAFK
DDX23
GTF3C4
LMNB1
KIF18B
RYBP
ZMYM2
CIZ1
TOMM70
ANAPC16
EFTUD2
INTS9
VRK3
SLC25A5
NUSAP1 (2)
RBM39 (2)
KIF20A
ANAPC4
THAP4
AKAP17A
RPL21
CIR1
RPL13A
NAIF1
DNAJC8
HNRNPC (2)
LSM5
RPL11
RBM5
DHX15
NOLC1 (1)
SLC27A4
WBP4
FAU
SFMBT1
DROSHA
BCLAF1 (2)
C14orf93
RPS12
PPP2R1A
EMD
POLR2L
SRSF2
ZNF740
H1F0
HADHA
INTS10
NOC3L
MPG
RPL18A
INTS4
MTBP
NCBP2
BAP1
JAKMIP3
PCID2
RPS28
FIZ1
MUS81
HNRNPH1
PCF11
MBNL1
MGA (1)
RPS13
CWC22
CIRBP
OTX1
RLIM
INTS6
BABAM2
ZFC3H1
RPL12
MED26
POLB
CAAP1
MISP
INTS5
SF3B4
MAP7
YAP1
NUP107
ZBTB39
SREK1 (2)
HNRNPF
PPIL1
RPL28
C7orf50
RFXANK
ATXN1L
FA2H
TARDBP/AL109811.3
MAFG
HADHB
POLR2D
NOL10
KPNB1
AHDC1
POLR2H
HIST1H1B
ZNF131
ELF2 (1)
LEO1
MED10
KPNA4
SREK1 (1)
NAT10
DNAJC9
RNF169
BHLHE40
EIF4G2
TIGD2
CDC73
TONSL
DDX50
RBM28
NFKBIL1
RSRC1
BOP1
RPL15
UBA52
MTF2 (2)
RPS3A
RPS24
EPB41
ATF2
QARS1
FHL3
MYL6
RPL27
CCDC59
PEG3/ZIM2
PRC1
GLYR1
PPHLN1 (2)
FOSL1
SMC3
CPSF4
MDN1
TMPO (2)
POLR2E
NEMP1
FOS
RP9
RAVER1
EXOSC10
PPP2CA/PPP2CB
NOL9
ARID5B
TRA2B
PQBP1
CPSF1
SPTY2D1
H3K27 me3
H3K9 me2
H3K27 me2
DNA Meth. m5C
H3K9 me3
H4K20 me3
H3K4 me1
H2A.Z
H4K20 me2
H3K4 me3
H4–4ac
H3–5ac
POLR2C
RPS25
WDR61
ZNF580
TMEM201
ZKSCAN4
JUN
DDX18
RPL35A
C1orf35
HDAC1
TMPO (1)
NUP88
RPL29
NOP2 (2)
RPN1
RPS6
UTP15
ZBTB20
HIST2H2BE
RAD21
CENPN
CCDC77
TFDP1
SRFBP1
CTR9
CPSF2
EXOSC8
NOLC1 (2)
PWP1
MLLT10
NUCKS1
EPAS1
BUB3
PAGR1
WBP11
HNRNPDL (2)
RPS18
CHD2
TFAP4
RFC1
STAG2
TOR1AIP1 (2)
ZNF668
MED20
NCBP3
ZNF106
ANAPC2
UTP20
PCGF6
PTBP1
INCENP
A0A0C4DGP5
NOL11
USP3
AP2A1
MNT
INTS7
LSM6
MXD1
A0A087X222
UBN2
ATF7
HOXA13
VHL
PRKDC
PALB2
KIF4A
ZC3H11A
SETD2
BIRC5
MYH9
FIP1L1
JUNB
RBM45
SETD5
RPL3
RPL17/RPL17–C18orf32
HMGN1
TRAM1
AEBP2
MECOM (1)
LEKR1
CREBBP
SETX
ZNF219
KIFC1
ZNF691
HMGN5
DSP
TIMM9
RPL34
CDC27
NOP16
NIFK
LYAR
ALYREF (1)
HIRIP3
DDX55
CALML5
SURF4
POP1
CTBP2
CBFB
ALYREF (2)
GPALPP1
ATAD3A (2)
RETSAT
RPL36
CLIP2
DYNLL1
ESS2
NUP133
RPL4
NHEJ1
TAF5L
ZNF808/ZNF845
PELP1
CCNB2
THAP11
WDR18
CDK13
CPSF3
HELLS (1)
CROCC
RBM7
TOR1AIP1 (1)
MYBBP1A
DNAJB12
WWTR1
TINF2 (2)
SUPT16H
GPAT4
RPN2
MTDH
ZNF367
CELF1
INTS1
ZNRD1
C19orf53
WDR33
NUP188
RCOR1 (2)
MIER1
L3MBTL1
SSRP1
SCAF8
RNPC3
HIST1H1[D,E]
IGF2BP1
DNAJC1
RCC1 (2)
YLPM1 (3)
UBE2I
TBL2
H3K27 me3
H3K9 me2
H3K27 me2
DNA Meth. m5C
H3K9 me3
H4K20 me3
H3K4 me1
H2A.Z
H4K20 me2
H3K4 me3
H4–4ac
H3–5ac
SYNPO
ZMAT2
CASP8AP2
ZNF451
ATP5F1D
ELF4
KAT14 (2)
DDX54
PLPP6
ZBTB44
USP39
NIPSNAP2
ZNF143
TOP1
RNF40
CERS2
RAD52
RPS3
HOMEZ
RNF20
ZFHX3
NIP7
RBMXL1
PROC
EXOSC4
PHB2
HIST[2H3A,2H3C,2H3D,3H3]
RPS9
CLASRP
ATF3
TMEM109
SON (2)
TEAD3/TEAD4
ERF
SNU13
HIST1H2BD
GTPBP4
MGA (2)
RRP9
EXOSC6
EXOSC1
MED7
MSANTD3
MED13
TAP1
KLF13
CPT1A
NOP2 (1)
RPL10A
RPL7A
MCPH1
HDGFL2
AQR
PRPF38A
PPP2R1B
PUM2
RPS2
EWSR1 (2)
FRG1
KDM4A
GTF3C6
HSPB1
MED15
C16orf87
PAXIP1
ZFAT
PC
PDE4DIP/LOC653513
CABIN1
NOP9
TEFM
RNF168
RPS8
HOXA10
BOD1
TLX3
REXO4
SFN
EIF2S3
EIF4E2
CDC16
BANF1
MITF
LAMA4
ZBTB2
RPP38
DDX51
NDUFA2
MED12
SART3
RPL31
GLIS2
SPATA5
DOT1L
POLR1B
UBP1
PREB
C9orf78
DDX31
UBQLN4
KLF10
NUP93
RPS29
LMNA (2)
CKAP5
C18orf21
NDUFA5
KIAA1671
ASXL2
SPTBN2
AKAP8L
TCF3
EBNA1BP2
GNL3
NOL8
JUND
BRCA2
ATP1A1/ATP1A3
SRPRB
PCBP2
CREB3L2
NBN
QKI
FOXK1
ESYT2
SPATA5L1
RTRAF
EXOSC9
INTS13
PCCA
RLF
IMMT
NKAP
YBX3
SEC61G
SP3
NCL (2)
ZNF24
GRWD1
MTA3
RPLP2
ANAPC5
PNKP
KDM6A
NTPCR
ZNF652
SCD
TRIM29
CREM (3)
FKBP11
UTP4
COL15A1
RGPD8
TBL1Y
MSH2
H3K27 me3
H3K9 me2
H3K27 me2
DNA Meth. m5C
H3K9 me3
H4K20 me3
H3K4 me1
H2A.Z
H4K20 me2
H3K4 me3
H4–4ac
H3–5ac

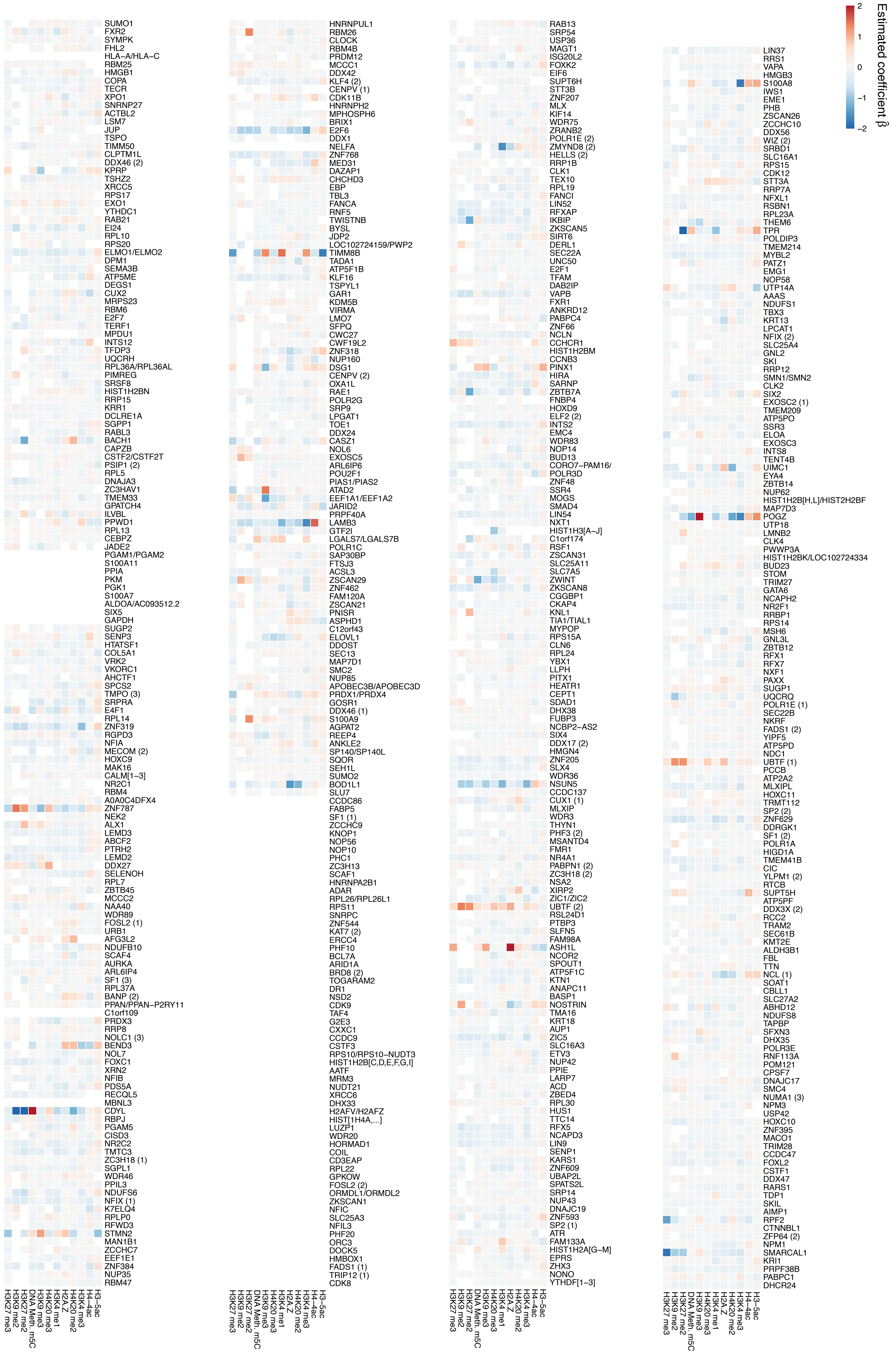
SUMO1
FXR2
SYMPK
FHL2
HLA−A/HLA−C
RBM25
HMGB1
COPA
TECR
XPO1
SNRNP27
ACTBL2
LSM7
JUP
TSPO
TIMM50
CLPTM1L
DDX46 (2)
KPRP
TSHZ2
XRCC5
RPS17
EXO1
YTHDC1
RAB21
EI24
RPL10
RPS20
ELMO1/ELMO2
DPM1
SEMA3B
ATP5ME
DEGS1
CUX2
MRPS23
RBM6
E2F7
TERF1
MPDU1
INTS12
TFDP3
UQCRH
RPL36A/RPL36AL
PIMREG
SRSF8
HIST1H2BN
RRP15
KRR1
DCLRE1A
SGPP1
RABL3
BACH1
CAPZB
CSTF2/CSTF2T
PSIP1 (2)
RPL5
DNAJA3
ZC3HAV1
TMEM33
GPATCH4
ILVBL
PPWD1
RPL13
CEBPZ
JADE2
PGAM1/PGAM2
S100A11
PPIA
PKM
PGK1
S100A7
ALDOA/AC093512.2
SIX5
GAPDH
SUGP2
SENP3
HTATSF1
COL5A1
VRK2
VKORC1
AHCTF1
SPCS2
TMPO (3)
SRPRA
E4F1
RPL14
ZNF319
RGPD3
NFIA
MECOM (2)
HOXC9
MAK16
CALM[1−3]
NR2C1
RBM4
A0A0C4DFX4
ZNF787
NEK2
ALX1
LEMD3
ABCF2
PTRH2
LEMD2
DDX27
SELENOH
RPL7
ZBTB45
MCCC2
NAA40
WDR89
FOSL2 (1)
URB1
AFG3L2
NDUFB10
SCAF4
AURKA
ARL6IP4
SF1 (3)
RPL37A
BANP (2)
PPAN/PPAN−P2RY11
C1orf109
PRDX3
RRP8
NOLC1 (3)
BEND3
NOL7
FOXC1
XRN2
NFIB
PDS5A
RECQL5
MBNL3
CDYL
RBPJ
PGAM5
CISD3
NR2C2
TMTC3
ZC3H18 (1)
SGPL1
WDR46
PPIL3
NDUFS6
NFIX (1)
K7ELQ4
RPLP0
RFWD3
STMN2
MAN1B1
ZCCHC7
EEF1E1
ZNF384
NUP35
RBM47
HNRNPUL1
RBM26
CLOCK
RBM4B
PRDM12
MCCC1
DDX42
KLF4 (2)
CENPV (1)
CDK11B
HNRNPH2
MPHOSPH6
BRIX1
E2F6
DDX1
NELFA
ZNF768
MED31
DAZAP1
CHCHD3
EBP
TBL3
FANCA
RNF5
TWISTNB
BYSL
JDP2
LOC102724159/PWP2
TIMM8B
TADA1
ATP5F1B
KLF16
TSPYL1
GAR1
KDM5B
VIRMA
LMO7
SFPQ
CWC27
CWF19L2
ZNF318
NUP160
DSG1
CENPV (2)
OXA1L
RAE1
POLR2G
SRP9
LPGAT1
TOE1
DDX24
CASZ1
NOL6
EXOSC5
ARL6IP6
POU2F1
PIAS1/PIAS2
ATAD2
EEF1A1/EEF1A2
JARID2
PRPF40A
LAMB3
GTF2I
LGALS7/LGALS7B
POLR1C
SAP30BP
FTSJ3
ACSL3
ZSCAN29
ZNF462
FAM120A
ZSCAN21
PNISR
ASPHD1
C12orf43
ELOVL1
DDOST
SEC13
MAP7D1
SMC2
NUP85
APOBEC3B/APOBEC3D
PRDX1/PRDX4
GOSR1
DDX46 (1)
S100A9
AGPAT2
REEP4
ANKLE2
SP140/SP140L
SQOR
SEH1L
SUMO2
BOD1L1
SLU7
CCDC86
FABP5
SF1 (1)
ZCCHC9
KNOP1
NOP56
NOP10
PHC1
ZC3H13
SCAF1
HNRNPA2B1
ADAR
RPL26/RPL26L1
RPS11
SNRPC
ZNF544
KAT7 (2)
ERCC4
PHF10
BCL7A
ARID1A
BRD8 (2)
TOGARAM2
DR1
NSD2
CDK9
TAF4
G2E3
CXXC1
CCDC9
CSTF3
RPS10/RPS10−NUDT3
HIST1H2B[C,D,E,F,G,I]
AATF
MRM3
NUDT21
XRCC6
DHX33
H2AFV/H2AFZ
HIST[1H4A,...]
LUZP1
WDR20
HORMAD1
COIL
CD3EAP
RPL22
GPKOW
FOSL2 (2)
ORMDL1/ORMDL2
ZKSCAN1
NFIC
SLC25A3
NFIL3
PHF20
ORC3
DOCK5
HMBOX1
FADS1 (1)
TRIP12 (1)
CDK8
RAB13
SRP54
USP36
MAGT1
ISG20L2
FOXK2
EIF6
SUPT6H
STT3B
ZNF207
MLX
KIF14
WDR75
ZRANB2
POLR1E (2)
ZMYND8 (2)
HELLS (2)
RRP1B
CLK1
TEX10
RPL19
FANCI
LIN52
RFXAP
IKBIP
ZKSCAN5
SIRT6
DERL1
SEC22A
UNC50
E2F1
TFAM
DAB2IP
VAPB
FXR1
ANKRD12
PABPC4
ZNF66
NCLN
CCHCR1
HIST1H2BM
CCNB3
PINX1
HIRA
SARNP
ZBTB7A
FNBP4
HOXD9
ELF2 (2)
INTS2
EMC4
WDR83
NOP14
BUD13
CORO7−PAM16/
POLR3D
ZNF48
SSR4
MOGS
SMAD4
LIN54
NXT1
HIST1H3[A−J]
C1orf174
RSF1
ZSCAN31
SLC25A11
SLC7A5
ZWINT
ZKSCAN8
CGGBP1
CKAP4
KNL1
TIA1/TIAL1
MYPOP
RPS15A
CLN6
RPL24
YBX1
LLPH
PITX1
HEATR1
CEPT1
SDAD1
DHX38
FUBP3
NCBP2−AS2
SIX4
DDX17 (2)
HMGN4
ZNF205
SLX4
WDR36
NSUN5
CCDC137
CUX1 (1)
MLXIP
WDR3
THYN1
PHF3 (2)
MSANTD4
FMR1
NR4A1
PABPN1 (2)
ZC3H18 (2)
NSA2
XIRP2
ZIC1/ZIC2
UBTF (2)
RSL24D1
PTBP3
SLFN5
FAM98A
ASH1L
NCOR2
SPOUT1
ATP5F1C
KTN1
ANAPC11
BASP1
NOSTRIN
TMA16
KRT18
AUP1
ZIC5
SLC16A3
ETV3
NUP42
PPIE
LARP7
ACD
ZBED4
RPL30
HUS1
TTC14
RFX5
NCAPD3
LIN9
SENP1
KARS1
ZNF609
UBAP2L
SPATS2L
SRP14
NUP43
DNAJC19
ZNF593
SP2 (1)
ATR
FAM133A
HIST1H2A[G−M]
EPRS
ZHX3
NONO
YTHDF[1−3]
LIN37
RRS1
VAPA
HMGB3
S100A8
IWS1
EME1
PHB
ZSCAN26
ZCCHC10
DDX56
WIZ (2)
SRBD1
SLC16A1
RPS15
CDK12
STT3A
RRP7A
NFXL1
RSBN1
RPL23A
THEM6
TPR
POLDIP3
TMEM214
MYBL2
PATZ1
EMG1
NOP58
UTP14A
AAAS
NDUFS1
TBX3
KRT13
LPCAT1
NFIX (2)
SLC25A4
GNL2
SKI
RRP12
SMN1/SMN2
CLK2
SIX2
EXOSC2 (1)
TMEM209
ATP5PO
SSR3
ELOA
EXOSC3
INTS8
TENT4B
UIMC1
EYA4
ZBTB14
NUP62
HIST1H2B[H,L]/HIST2H2BF
MAP7D3
POGZ
UTP18
LMNB2
CLK4
PWWP3A
HIST1H2BK/LOC102724334
BUD23
STOM
TRIM27
GATA6
NCAPH2
NR2F1
RRBP1
RPS14
MSH6
GNL3L
ZBTB12
RFX1
RFX7
NXF1
PAXX
SUGP1
UQCRQ
POLR1E (1)
SEC22B
NKRF
FADS1 (2)
YIPF5
ATP5PD
NDC1
UBTF (1)
PCCB
ATP2A2
MLXIPL
HOXC11
TRMT112
SP2 (2)
ZNF629
DDRGK1
SF1 (2)
POLR1A
HIGD1A
TMEM41B
CIC
YLPM1 (2)
RTCB
SUPT5H
ATP5PF
DDX3X (2)
RCC2
TRAM2
SEC61B
KMT2E
ALDH3B1
FBL
TTN
NCL (1)
SOAT1
CBLL1
SLC27A2
ABHD12
NDUFS8
TAPBP
SFXN3
DHX35
POLR3E
RNF113A
POM121
CPSF7
DNAJC17
SMC4
NUMA1 (3)
NPM3
USP42
HOXC10
ZNF395
MACO1
TRIM28
CCDC47
FOXL2
CSTF1
DDX47
RARS1
TDP1
SKIL
AIMP1
RPF2
CTNNBL1
ZFP64 (2)
NPM1
SMARCAL1
KRI1
PRPF38B
PABPC1
DHCR24
H3K27 me3
H3K9 me2
H3K27 me2
DNA Meth. m5C
H3K9 me3
H4K20 me3
H3K4 me1
H2A.Z
H4K20 me2
H3K4 me3
H4−4ac
H3−5ac
Estimated coefficient β̂
2
1
0
−1
−2
